## Supplemental Table S2 for "Transcriptomic responses of the human kidney to acute injury at single cell resolution"

**Podocytes**

*This means that positive fold changes imply higher mean value in controls than in AKI and vice versa.

| | baseMean | log2 fold change control/AKI* | lfcSE | stat | pvalue | padj |
|---|---|---|---|---|---|---|
| AL669831.5 | 12.2210027 | -0.403961783 | 0.43214877 | -0.9347748 | 0.34990436 | NA |
| SAMD11 | 48.102533 | -2.068473845 | 0.64228135 | -3.2205105 | 0.00127963 | 0.0545866 |
| NOC2L | 12.6479149 | 0.36560269 | 0.48865539 | 0.74818102 | 0.45435098 | NA |
| HES4 | 23.03128 | 0.570367846 | 0.77026306 | 0.74048448 | 0.45900607 | NA |
| ISG15 | 576.585799 | -1.669597622 | 2.02394467 | -0.8249226 | 0.40941556 | 0.76772515 |
| AGRN | 111.659355 | 0.610607683 | 0.49264955 | 1.23943618 | 0.21518401 | 0.62305077 |
| C1orf159 | 12.7788718 | 0.254827308 | 0.42546806 | 0.59893405 | 0.54921686 | NA |
| SDF4 | 51.4293417 | -0.354406444 | 0.5498265 | -0.6445787 | 0.51920025 | 0.83104823 |
| B3GALT6 | 4.97465606 | -0.21288149 | 0.77972001 | -0.273023 | 0.78483554 | NA |
| UBE2J2 | 22.2247154 | -0.531194327 | 0.46768815 | -1.1357874 | 0.25604553 | NA |
| ACAP3 | 7.91076114 | 0.482434379 | 0.51090963 | 0.94426559 | 0.34503395 | NA |
| INTS11 | 19.0675457 | 0.204104111 | 0.3557585 | 0.57371534 | 0.56616044 | NA |
| DVL1 | 11.8569733 | -0.336006187 | 0.43496176 | -0.7724959 | 0.43982076 | NA |
| MXRA8 | 79.7406274 | -0.142746471 | 0.40220939 | -0.3549059 | 0.72266012 | 0.91509584 |
| AURKAIP1 | 43.2543672 | 0.00657422 | 0.52582306 | 0.01250272 | 0.99002453 | 0.99710744 |
| CCNL2 | 55.3029639 | 0.07928814 | 0.23379733 | 0.33913193 | 0.73451034 | 0.91893089 |
| MRPL20 | 46.4876818 | -0.481406195 | 0.30682234 | -1.5690063 | 0.11664647 | 0.5065504 |
| ANKRD65 | 16.1822483 | -1.745907161 | 0.62292926 | -2.8027374 | 0.00506709 | NA |
| VWA1 | 70.8850489 | -1.786184621 | 0.54370907 | -3.2851845 | 0.00101916 | 0.05167121 |
| ATAD3C | 5.82868676 | -0.966987329 | 0.7494072 | -1.2903363 | 0.19693391 | NA |
| ATAD3B | 9.264103 | -0.180012663 | 0.63690204 | -0.2826379 | 0.77745442 | NA |
| ATAD3A | 5.95827862 | 0.095389785 | 0.64293132 | 0.14836699 | 0.88205315 | NA |
| SSU72 | 39.5444282 | -0.329600673 | 0.38760192 | -0.8503587 | 0.39512568 | 0.7584618 |
| MIB2 | 9.51409055 | 0.255243329 | 0.49889866 | 0.51161357 | 0.60892148 | NA |
| CDK11B | 9.02427123 | 0.168469233 | 0.49028038 | 0.34361814 | 0.73113348 | NA |
| SLC35E2B | 14.235477 | 0.249917694 | 0.51690881 | 0.48348508 | 0.62875135 | NA |
| CDK11A | 4.78124249 | -0.487179429 | 0.57704656 | -0.8442636 | 0.3985221 | NA |
| SLC35E2A | 10.9568293 | 0.008752218 | 0.55178514 | 0.01586164 | 0.98734477 | NA |
| NADK | 11.4037512 | -0.643056116 | 0.49374312 | -1.3024103 | 0.19277617 | NA |
| GNB1 | 155.392165 | -0.220544453 | 0.28481528 | -0.7743421 | 0.43872854 | 0.78335165 |
| PRKCZ | 21.5365712 | 0.080714158 | 0.32054507 | 0.25180284 | 0.80119346 | NA |
| FAAP20 | 19.4240643 | 0.159349407 | 0.42237844 | 0.37726691 | 0.70597526 | NA |
| SKI | 39.465746 | 0.00759542 | 0.31755616 | 0.02391835 | 0.98091774 | 0.99453905 |
| MORN1 | 7.74445844 | -0.150836188 | 0.47788357 | -0.3156338 | 0.75228051 | NA |
| RER1 | 38.6532152 | -0.102657995 | 0.33081456 | -0.3103189 | 0.75631849 | 0.92313949 |
| PEX10 | 10.6140949 | 0.539759357 | 0.5058476 | 1.06703948 | 0.28595401 | NA |
| PANK4 | 3.8199875 | 1.095453489 | 0.7102303 | 1.54239194 | 0.12297838 | NA |
| TNFRSF14 | 24.6732356 | 0.58756527 | 0.48925395 | 1.20094129 | 0.22977398 | NA |
| FAM213B | 5.21454943 | -0.469697732 | 0.77158098 | -0.6087472 | 0.54269204 | NA |
| ARHGEF16 | 20.5426354 | -0.106746555 | 0.44107478 | -0.2420146 | 0.80876882 | NA |
| TPRG1L | 18.2253749 | -0.014170596 | 0.45967685 | -0.0308273 | 0.97540727 | NA |
| WRAP73 | 6.32002732 | -0.46885494 | 0.64136231 | -0.7310298 | 0.46476097 | NA |
| SMIM1 | 7.79233443 | -1.07042863 | 0.68158338 | -1.5705028 | 0.11629817 | NA |
| LRRC47 | 13.8788741 | 0.442983794 | 0.50520062 | 0.87684729 | 0.38056959 | NA |
| CEP104 | 26.122262 | -0.21559529 | 0.34849869 | -0.6186402 | 0.53615343 | NA |
| C1orf174 | 6.00334306 | -0.147728247 | 0.62427005 | -0.2366416 | 0.81293486 | NA |
| NPHP4 | 12.1683926 | -0.532401147 | 0.43531641 | -1.2230211 | 0.22132173 | NA |
| RPL22 | 12.8474251 | -0.72001067 | 0.57554305 | -1.251011 | 0.21093047 | NA |
| RNF207 | 3.99756485 | -0.15363851 | 0.94559973 | -0.1624773 | 0.87092999 | NA |
| ICMT | 24.7736965 | -0.168967867 | 0.46021932 | -0.3671464 | 0.71350982 | NA |
| ACOT7 | 14.1538094 | 0.572242726 | 0.589809 | 0.97021702 | 0.33193833 | NA |
| PLEKHG5 | 5.73621805 | 0.433712978 | 0.58817385 | 0.73738908 | 0.46088577 | NA |
| NOL9 | 15.4076178 | 0.144062037 | 0.49972618 | 0.28828195 | 0.77313092 | NA |
| ZBTB48 | 7.92418947 | 0.063688798 | 0.53678193 | 0.1186493 | 0.90555321 | NA |
| KLHL21 | 12.5252922 | 0.044969946 | 0.53854717 | 0.08350234 | 0.93345212 | NA |
| PHF13 | 9.7206617 | 0.026876522 | 0.51861605 | 0.05182354 | 0.9586693 | NA |
| THAP3 | 7.77959471 | 0.519767146 | 0.60219824 | 0.86311635 | 0.3880735 | NA |
| DNAJC11 | 25.8794764 | 0.232770054 | 0.29577645 | 0.78697966 | 0.43129377 | NA |
| CAMTA1 | 117.945979 | -0.015029361 | 0.24267285 | -0.0619326 | 0.95061651 | 0.98520702 |
| VAMP3 | 28.1995901 | -0.438805651 | 0.39943098 | -1.0985769 | 0.27195265 | NA |
| PER3 | 43.3187237 | -1.559593504 | 0.56859889 | -2.7428712 | 0.00609046 | 0.12229157 |
| PARK7 | 105.601704 | -0.905552033 | 0.42606594 | -2.12538 | 0.03355492 | 0.29047359 |
| ERRFI1 | 63.9620585 | -0.386517005 | 0.47663753 | -0.8109244 | 0.41740908 | 0.77165508 |
| SLC45A1 | 74.3844856 | -0.478645838 | 0.46199256 | -1.0360466 | 0.30018039 | 0.68170865 |
| RERE | 604.968307 | -0.096813972 | 0.25331661 | -0.3821856 | 0.70232367 | 0.90757606 |
| ENO1 | 83.4090663 | -1.223523137 | 0.49158901 | -2.4889148 | 0.01281337 | 0.18904687 |
| GPR157 | 4.99166341 | 0.098305826 | 0.63861014 | 0.15393715 | 0.87765929 | NA |
| H6PD | 26.1647951 | -0.128986401 | 0.35332428 | -0.3650652 | 0.71506273 | NA |
| SPSB1 | 63.2622571 | -1.038941671 | 0.47893363 | -2.1692811 | 0.03006135 | 0.27996878 |
| SLC25A33 | 20.5517187 | 0.640671502 | 0.35019518 | 1.82946977 | 0.06732926 | NA |
| TMEM201 | 4.60937763 | 1.073017904 | 0.61443092 | 1.74636053 | 0.08074832 | NA |
| PIK3CD | 7.59914313 | 0.361176136 | 0.79509831 | 0.45425343 | 0.64964643 | NA |
| CLSTN1 | 63.059107 | -0.060867087 | 0.3769981 | -0.161452 | 0.87173744 | 0.96207879 |
| CTNNBIP1 | 89.0023943 | 0.580676757 | 0.5852918 | 0.99211496 | 0.32114145 | 0.69513974 |
| LZIC | 19.7340143 | 0.045494068 | 0.53094141 | 0.08568566 | 0.9317163 | NA |
| NMNAT1 | 10.7964506 | 0.569346385 | 0.45870056 | 1.24121581 | 0.21452604 | NA |
| UBE4B | 63.6233995 | 0.304713901 | 0.27462093 | 1.10958005 | 0.26718003 | 0.66013596 |
| KIF1B | 139.965804 | 0.164705917 | 0.28439243 | 0.57915014 | 0.56248787 | 0.85386823 |
| PGD | 23.3618586 | -0.340756033 | 0.42983149 | -0.7927666 | 0.42791384 | NA |
| DFFA | 17.5786131 | 0.302952393 | 0.35542921 | 0.85235649 | 0.39401626 | NA |
| PEX14 | 28.8394318 | -0.342502983 | 0.33763872 | -1.0144067 | 0.31038872 | 0.69082013 |
| CASZ1 | 171.502854 | -1.30315762 | 0.46613335 | -2.7956756 | 0.00517914 | 0.11257186 |
| AL139423.1 | 6.89682419 | -1.686678004 | 0.83568228 | -2.0183245 | 0.04355748 | NA |
| SRM | 13.320911 | -0.005948866 | 0.46136801 | -0.012894 | 0.98971238 | NA |
| EXOSC10 | 10.8966615 | -0.077813858 | 0.50837739 | -0.1530632 | 0.87834845 | NA |
| MTOR | 20.2158639 | 0.163789845 | 0.36531403 | 0.44835355 | 0.65389806 | NA |
| UBIAD1 | 4.973932 | -1.035473853 | 0.61613997 | -1.6805822 | 0.0928441 | NA |
| FBXO2 | 36.4529185 | 0.150758149 | 0.7038174 | 0.21420066 | 0.83039058 | 0.95008863 |
| FBXO44 | 7.86079415 | 0.511303461 | 0.50560575 | 1.01126908 | 0.31188766 | NA |
| MAD2L2 | 5.19183553 | -0.369028077 | 0.75630381 | -0.4879363 | 0.62559497 | NA |
| AGTRAP | 24.5349099 | 0.202464169 | 0.58712989 | 0.3448371 | 0.73021684 | NA |
| MTHFR | 12.4434938 | 0.272827445 | 0.40607918 | 0.67185776 | 0.50167425 | NA |
| CLCN6 | 7.30640911 | -0.513217998 | 0.57650825 | -0.890218 | 0.37334886 | NA |
| KIAA2013 | 29.1524218 | 0.662428601 | 0.34384385 | 1.92653907 | 0.0540371 | 0.3615191 |
| PLOD1 | 19.02825 | 0.842478552 | 0.47842796 | 1.76093085 | 0.07825011 | NA |
| MFN2 | 13.1685749 | -0.242809584 | 0.48327581 | -0.5024245 | 0.61536898 | NA |
| MIIP | 12.5764277 | 0.027089196 | 0.65623634 | 0.04127963 | 0.96707297 | NA |
| TNFRSF8 | 8.9067448 | 0.87698115 | 0.69293357 | 1.26560638 | 0.20565404 | NA |
| VPS13D | 91.5978601 | 0.26194595 | 0.22290995 | 1.17512007 | 0.23994668 | 0.64752079 |
| DHRS3 | 139.893866 | 0.747258571 | 0.39287591 | 1.90202184 | 0.0571683 | 0.37377289 |
| PDPN | 27.6830468 | 1.117280916 | 0.59169476 | 1.88827244 | 0.05898939 | NA |
| PRDM2 | 54.6463598 | 0.131499419 | 0.25724949 | 0.51117466 | 0.60922876 | 0.8725395 |
| KAZN | 130.507684 | -0.037606339 | 0.36358303 | -0.1034326 | 0.91761964 | 0.97775374 |
| TMEM51 | 11.141284 | 0.325570541 | 0.62974123 | 0.51699099 | 0.60516245 | NA |
| EFHD2 | 4.42275693 | 0.116326828 | 0.71280268 | 0.16319639 | 0.8703638 | NA |
| CASP9 | 8.30982991 | 0.560798044 | 0.58643928 | 0.9562764 | 0.33893261 | NA |
| DNAJC16 | 30.946139 | 0.303689381 | 0.30554952 | 0.99391216 | 0.32026563 | 0.69496685 |
| DDI2 | 23.2920273 | 0.555667763 | 0.3810682 | 1.45818454 | 0.14478968 | NA |
| PLEKHM2 | 31.9278701 | -0.296300276 | 0.29400564 | -1.0078047 | 0.31354822 | 0.69229808 |
| FBLIM1 | 13.7438073 | -0.799283733 | 0.85646255 | -0.9332384 | 0.3506969 | NA |
| AL450998.2 | 6.26872963 | -1.050067218 | 0.5908196 | -1.777306 | 0.07551791 | NA |

| | | | | | | |
|---|---|---|---|---|---|---|
| SPEN | 63.1413115 | 0.152044719 | 0.26540032 | 0.57288823 | 0.56672038 | 0.85641499 |
| ZBTB17 | 7.38681746 | -0.087991605 | 0.57334678 | -0.1534701 | 0.87802754 | NA |
| EPHA2 | 18.7853984 | 0.477876944 | 0.51240166 | 0.93262176 | 0.35101529 | NA |
| FBXO42 | 36.9729752 | -0.171457513 | 0.31024111 | -0.5526589 | 0.58049699 | 0.86175324 |
| SZRD1 | 28.5561025 | 0.332315386 | 0.40107974 | 0.82855192 | 0.40735801 | 0.76772515 |
| NECAP2 | 20.7084455 | -0.275303462 | 0.40239305 | -0.6841656 | 0.49387063 | NA |
| NBPF1 | 8.49175222 | -1.336510466 | 0.64173661 | -2.0826465 | 0.03728346 | NA |
| CROCC | 15.0714924 | 0.204265049 | 0.48361874 | 0.42236793 | 0.67275648 | NA |
| ATP13A2 | 12.2818467 | -0.667919043 | 0.44870516 | -1.4885477 | 0.13660651 | NA |
| SDHB | 22.3246908 | -0.317548515 | 0.37740485 | -0.8414002 | 0.40012378 | NA |
| RCC2 | 7.64756047 | -0.943239174 | 0.61396709 | -1.5363025 | 0.12446421 | NA |
| ARHGEF10L | 52.0211701 | -0.310682579 | 0.46109971 | -0.6737861 | 0.50044731 | 0.82345406 |
| KLHDC7A | 5.53259563 | -1.470737731 | 1.12263788 | -1.3100731 | 0.1901711 | NA |
| IFFO2 | 9.51712498 | 0.257406613 | 0.5676247 | 0.45348029 | 0.65020293 | NA |
| UBR4 | 54.6860696 | -0.03878833 | 0.2208554 | -0.1756277 | 0.86058643 | 0.95946634 |
| AL035413.1 | 10.2161481 | -1.157276323 | 0.53643287 | -2.1573554 | 0.03097799 | NA |
| EMC1 | 19.4238022 | 0.798931425 | 0.39527194 | 2.02121968 | 0.04325703 | NA |
| MRTO4 | 7.02979189 | -0.531699165 | 0.72035296 | -0.7381092 | 0.46044808 | NA |
| AKR7A2 | 12.208665 | -0.279418225 | 0.50341729 | -0.555043 | 0.57886527 | NA |
| CAPZB | 91.4814484 | -0.051202513 | 0.33114869 | -0.1546209 | 0.87712018 | 0.96359019 |
| NBL1 | 6.87673746 | -0.216319462 | 1.12262322 | -0.1926911 | 0.84720092 | NA |
| TMCO4 | 24.4862261 | 0.030236385 | 0.44296281 | 0.06825942 | 0.94557913 | NA |
| OTUD3 | 7.70891851 | 0.804353063 | 0.4736003 | 1.69837954 | 0.08943615 | NA |
| CAMK2N1 | 12.1397728 | 0.356729699 | 0.57145476 | 0.62424837 | 0.53246449 | NA |
| MUL1 | 7.54281127 | 0.565180193 | 0.52065102 | 1.08552595 | 0.27768878 | NA |
| PINK1 | 20.1388838 | -0.287602045 | 0.45900923 | -0.6265714 | 0.53094023 | NA |
| DDOST | 21.7839367 | -0.468467253 | 0.41113917 | -1.1394372 | 0.25452086 | NA |
| HP1BP3 | 89.2265828 | -0.129779632 | 0.21043921 | -0.6167084 | 0.53742708 | 0.83677705 |
| EIF4G3 | 176.298837 | 0.200732254 | 0.33852886 | 0.59295463 | 0.55321152 | 0.84896932 |
| ECE1 | 8.08574456 | 0.145615709 | 0.72345551 | 0.20127804 | 0.84048117 | NA |
| USP48 | 46.8855083 | 0.438106667 | 0.28978599 | 1.51182832 | 0.13057754 | 0.52613957 |
| HSPG2 | 8.28057351 | 0.482803348 | 0.70657151 | 0.6833043 | 0.49441458 | NA |
| CDC42 | 82.867487 | -0.260315008 | 0.30203592 | -0.8618677 | 0.38876031 | 0.75301425 |
| ZBTB40 | 10.4370573 | 0.282753584 | 0.62511054 | 0.45232573 | 0.65103434 | NA |
| KDM1A | 25.4079449 | -0.126912125 | 0.30453058 | -0.4167467 | 0.67686367 | NA |
| AL031428.1 | 7.34617643 | -0.504845525 | 0.70163388 | -0.7195284 | 0.47181539 | NA |
| LUZP1 | 34.8118878 | 0.145939667 | 0.30054366 | 0.48558558 | 0.62726102 | 0.8806406 |
| LINC01355 | 8.37746869 | 0.346680403 | 0.48270622 | 0.71820166 | 0.47263296 | NA |
| HNRNPR | 64.9481678 | -0.415368705 | 0.22837097 | -1.8188332 | 0.06893688 | 0.40491695 |
| ASAP3 | 7.13099654 | 0.453431844 | 0.75496746 | 0.60059786 | 0.54810786 | NA |
| ID3 | 20.7891561 | -1.02571717 | 1.11898471 | -0.9166499 | 0.35932615 | NA |
| RPL11 | 101.413158 | -0.626215747 | 0.3664727 | -1.7087651 | 0.08749448 | 0.44978149 |
| ELOA | 12.5323722 | 0.045073082 | 0.50875082 | 0.08859559 | 0.92940331 | NA |
| ELOA-AS1 | 9.74930753 | -0.70159169 | 0.47649 | -1.4724164 | 0.14090846 | NA |
| PITHD1 | 20.4624707 | 0.069016361 | 0.39802582 | 0.17339669 | 0.86233962 | NA |
| LYPLA2 | 10.1181569 | 0.782032316 | 0.47363879 | 1.65111544 | 0.09871501 | NA |
| HMGCL | 18.3757661 | -0.523560902 | 0.43903327 | -1.1925313 | 0.233053 | NA |
| FUCA1 | 12.4737631 | -0.225718048 | 0.41545272 | -0.5433062 | 0.58691898 | NA |
| PNRC2 | 45.2312421 | -0.165202694 | 0.44241082 | -0.3734147 | 0.70883984 | 0.91126921 |
| SRSF10 | 66.3776333 | 0.013281183 | 0.25852677 | 0.05137257 | 0.95902865 | 0.98766083 |
| IFNLR1 | 6.4834538 | 0.286084988 | 0.64574013 | 0.44303424 | 0.65774097 | NA |
| STPG1 | 6.60822466 | -0.020256446 | 0.53884299 | -0.0375925 | 0.9700126 | NA |
| NIPAL3 | 28.4705669 | 0.059829395 | 0.32270404 | 0.18540021 | 0.85291515 | NA |
| RCAN3 | 12.0121421 | -0.218777409 | 0.57162647 | -0.3827279 | 0.70192151 | NA |
| SRRM1 | 56.70328 | 0.249749261 | 0.23432149 | 1.06584017 | 0.2864959 | 0.67428533 |
| CLIC4 | 37.0204692 | 0.41047461 | 0.53111914 | 0.77284846 | 0.43961207 | 0.78399202 |
| SYF2 | 17.6931446 | -0.146487852 | 0.36243684 | -0.4041748 | 0.68608415 | NA |
| RSRP1 | 122.654638 | 0.070807389 | 0.31957607 | 0.22156662 | 0.82465127 | 0.94860623 |
| TMEM50A | 59.0617116 | -0.467189015 | 0.45349444 | -1.0301979 | 0.30291712 | 0.68310554 |
| MACO1 | 34.4545101 | 0.148048419 | 0.38342494 | 0.386121 | 0.69940706 | 0.90729979 |
| LDLRAP1 | 7.40867803 | -0.033895746 | 0.54848871 | -0.0617984 | 0.95072335 | NA |
| MAN1C1 | 10.7304857 | 0.975109986 | 0.53203151 | 1.83280497 | 0.06683158 | NA |
| MTFR1L | 21.8660939 | 0.650779233 | 0.34275614 | 1.89866545 | 0.05760848 | NA |
| AL033528.2 | 25.0188007 | 0.767175906 | 0.46336025 | 1.65567914 | 0.09778681 | NA |
| PAQR7 | 49.1961145 | 0.652963388 | 0.43191542 | 1.51178531 | 0.13058849 | 0.52613957 |
| STMN1 | 38.3423636 | -0.812822598 | 0.41345643 | -1.9659208 | 0.04930776 | 0.34686145 |
| PAFAH2 | 9.6786605 | -0.703095828 | 0.45346428 | -1.5504988 | 0.12102184 | NA |
| PDIK1L | 5.0856321 | -1.445129595 | 0.66002945 | -2.1894926 | 0.02856106 | NA |
| ZNF593 | 11.4706942 | -0.956228652 | 0.69405402 | -1.3777438 | 0.16828239 | NA |
| CEP85 | 3.84187604 | -2.296713005 | 0.92082895 | -2.4941798 | 0.01262486 | NA |
| SH3BGRL3 | 41.0964851 | -0.51314444 | 0.35659225 | -1.439023 | 0.15014402 | 0.54834287 |
| DHDDS | 13.8081258 | -0.23536187 | 0.41615949 | -0.5655569 | 0.57169504 | NA |
| HMGN2 | 48.0033246 | -0.547327235 | 0.44125177 | -1.2403967 | 0.2148287 | 0.62305077 |
| ARID1A | 74.231916 | -0.089761364 | 0.25780959 | -0.3481692 | 0.7277131 | 0.91700494 |
| PIGV | 5.2564584 | -0.33690374 | 0.60128618 | -0.5603051 | 0.57527132 | NA |
| ZDHHC18 | 8.03704423 | 0.618679379 | 0.53159039 | 1.16382724 | 0.24449403 | NA |
| GPN2 | 9.4638356 | 0.13209716 | 0.58494067 | 0.22583002 | 0.82133363 | NA |
| NUDC | 27.4711789 | -0.233177635 | 0.35789073 | -0.651533 | 0.51470245 | NA |
| SLC9A1 | 5.38510809 | -0.495103236 | 0.63670254 | -0.7776052 | 0.43680177 | NA |
| WDTC1 | 27.1644267 | 0.227932591 | 0.40869068 | 0.55771419 | 0.57703957 | NA |
| TMEM222 | 16.105494 | 0.103491349 | 0.39013913 | 0.2652678 | 0.79080317 | NA |
| MAP3K6 | 4.0284086 | -0.926827736 | 0.79235884 | -1.1697071 | 0.24211887 | NA |
| WASF2 | 138.379961 | -0.338766453 | 0.24221126 | -1.3986404 | 0.16192085 | 0.56290656 |
| AHDC1 | 44.5368943 | -0.276365642 | 0.29973818 | -0.9220235 | 0.35651632 | 0.72847869 |
| IFI6 | 126.040823 | -2.908995334 | 1.59104423 | -1.828356 | 0.06749613 | 0.40302824 |
| FAM76A | 18.187141 | 0.177051003 | 0.329569 | 0.53721983 | 0.59111577 | NA |
| STX12 | 40.1097829 | -0.045860821 | 0.24529506 | -0.1869619 | 0.85169053 | 0.95532544 |
| PPP1R8 | 14.2724052 | -0.012470243 | 0.39468425 | -0.0315955 | 0.97479464 | NA |
| THEMIS2 | 4.71514989 | -3.333831936 | 1.1428047 | -2.9172368 | 0.00353148 | NA |
| RPA2 | 11.5584277 | -1.098403265 | 0.44784096 | -2.4526637 | 0.01418028 | NA |
| EYA3 | 37.4168585 | 0.138395432 | 0.27515957 | 0.50296428 | 0.61498939 | 0.87553421 |
| DNAJC8 | 23.3431236 | -0.288984285 | 0.34061014 | -0.8484312 | 0.39619785 | NA |
| ATP5IF1 | 89.5001939 | -0.140271323 | 0.43321197 | -0.3237937 | 0.7460942 | 0.92250642 |
| SESN2 | 13.3948309 | -0.09719095 | 0.51818653 | -0.1875598 | 0.85122176 | NA |
| PHACTR4 | 342.631931 | 0.39143684 | 0.27501979 | 1.42330427 | 0.15464797 | 0.55484098 |
| RCC1 | 16.4370484 | -0.868762162 | 0.52846597 | -1.6439321 | 0.10019023 | NA |
| TRNAU1AP | 18.6327006 | -0.140928699 | 0.42018629 | -0.3353958 | 0.73732657 | NA |
| SNHG12 | 4.51011518 | -0.359811289 | 0.6792223 | -0.5297401 | 0.59629214 | NA |
| TAF12 | 9.30337747 | 0.119051953 | 0.51339434 | 0.23189183 | 0.81662203 | NA |
| GMEB1 | 15.7949686 | -0.116151861 | 0.52015225 | -0.2233036 | 0.82329924 | NA |
| YTHDF2 | 33.207824 | 0.048171178 | 0.2866211 | 0.16806571 | 0.86653159 | 0.96089268 |
| EPB41 | 43.7496362 | -0.012834101 | 0.34502555 | -0.0371975 | 0.9703275 | 0.98977467 |
| SRSF4 | 76.7943165 | -0.014739893 | 0.25766537 | -0.0572056 | 0.95438145 | 0.9857324 |
| MECR | 7.7805407 | -0.578713943 | 0.52399218 | -1.1044324 | 0.26940561 | NA |
| SDC3 | 16.2262017 | -0.432458274 | 0.58254862 | -0.7423557 | 0.45787187 | NA |
| PUM1 | 111.402776 | 0.200615105 | 0.23642466 | 0.84853715 | 0.39613889 | 0.75911719 |
| SNRNP40 | 14.4639463 | 0.344563582 | 0.45095945 | 0.76406778 | 0.44482685 | NA |
| ZCCHC17 | 14.1121394 | -0.157104647 | 0.44978995 | -0.3492845 | 0.72687575 | NA |
| SERINC2 | 8.94391186 | -0.867584326 | 0.63238787 | -1.371918 | 0.17008896 | NA |
| TINAGL1 | 6.60025938 | 0.201338973 | 0.85647666 | 0.23507818 | 0.81414805 | NA |
| PEF1 | 16.58359 | -0.338585756 | 0.42657942 | -0.7937227 | 0.4273569 | NA |
| PTP4A2 | 10.3840246 | -0.017761462 | 0.52329364 | -0.0339417 | 0.97292366 | NA |
| KHDRBS1 | 66.8241069 | -0.64610024 | 0.31040326 | -2.0814866 | 0.03738939 | 0.30414888 |

| | | | | | | |
|---|---|---|---|---|---|---|
| TMEM39B | 7.71962577 | 1.014776732 | 0.49352768 | 2.05616984 | 0.03976615 | NA |
| KPNA6 | 72.8651021 | -0.358046479 | 0.32109747 | -1.115071 | 0.26482 | 0.65974932 |
| TXLNA | 26.6034543 | -0.448239972 | 0.34464752 | -1.3005751 | 0.19340395 | NA |
| TMEM234 | 8.00140073 | -0.040921293 | 0.54955479 | -0.0744626 | 0.94064228 | NA |
| EIF3I | 16.0110414 | -0.709638879 | 0.56147529 | -1.2638827 | 0.20627215 | NA |
| HDAC1 | 21.2644297 | -0.151682997 | 0.45067062 | -0.3365717 | 0.73643976 | NA |
| MARCKSL1 | 6.61558513 | 0.181063365 | 0.6823946 | 0.26533528 | 0.79075119 | NA |
| TSSK3 | 5.37495152 | -0.068484521 | 0.77061093 | -0.0888704 | 0.92918489 | NA |
| BSDC1 | 26.7282646 | 0.056845012 | 0.36295134 | 0.15661883 | 0.87554527 | NA |
| ZBTB8A | 8.41840522 | 0.17788458 | 0.51327519 | 0.34656766 | 0.72891615 | NA |
| ZBTB8OS | 29.8020681 | 0.068447345 | 0.28162013 | 0.24304848 | 0.80796784 | 0.94328109 |
| RBBP4 | 38.4263843 | 0.119953628 | 0.25823054 | 0.46452147 | 0.64227418 | 0.88951638 |
| SYNC | 8.22391893 | -0.296323759 | 0.71309089 | -0.4155484 | 0.67774051 | NA |
| KIAA1522 | 11.7233515 | 0.361688664 | 0.43957153 | 0.82282096 | 0.41060982 | NA |
| YARS | 18.2446613 | -0.530426988 | 0.58999488 | -0.8990366 | 0.36863316 | NA |
| S100PBP | 28.6646328 | -0.228269318 | 0.34270981 | -0.6660717 | 0.50536525 | 0.82600448 |
| TMEM54 | 8.27839486 | -0.665403056 | 0.65096052 | -1.0221865 | 0.30669264 | NA |
| RNF19B | 28.6847341 | -0.305909213 | 0.7584659 | -0.4033263 | 0.68670823 | 0.90499319 |
| AL020995.1 | 4.01618727 | -0.672397976 | 0.78587206 | -0.8556074 | 0.39221496 | NA |
| AK2 | 25.9243049 | 0.141454419 | 0.41931898 | 0.33734323 | 0.73585817 | NA |
| ZNF362 | 14.189581 | 0.192048364 | 0.56299463 | 0.34111935 | 0.73301373 | NA |
| AL513327.1 | 5.87852242 | -1.522411642 | 0.93086919 | -1.6354732 | 0.10194988 | NA |
| PHC2 | 33.5205373 | -0.748973273 | 0.51781892 | -1.4464 | 0.14806505 | 0.54691531 |
| CSMD2 | 22.0967081 | -3.019096949 | 0.69372088 | -4.3520342 | 1.35E-05 | NA |
| SMIM12 | 14.3375191 | 0.124564503 | 0.39412154 | 0.31605607 | 0.75195995 | NA |
| ZMYM1 | 16.7552379 | 0.189565712 | 0.3829232 | 0.4950489 | 0.6205656 | NA |
| SFPQ | 91.9793267 | -0.15844616 | 0.24963684 | -0.6347066 | 0.52561976 | 0.83195298 |
| ZMYM4 | 156.132932 | 0.377596831 | 0.17033657 | 2.216769 | 0.02663888 | 0.26417433 |
| KIAA0319L | 97.9773044 | -0.058097864 | 0.26581534 | -0.2185648 | 0.82698911 | 0.949142 |
| NCDN | 5.01188263 | -0.469324451 | 0.70791466 | -0.6629676 | 0.50735133 | NA |
| PSMB2 | 31.2706012 | -0.127967726 | 0.44434679 | -0.2879907 | 0.77335388 | 0.92747586 |
| AGO4 | 33.729653 | -0.208979758 | 0.4024155 | -0.5193134 | 0.60354221 | 0.87047761 |
| AGO1 | 10.3481779 | -0.842541588 | 0.50723112 | -1.6610605 | 0.09670129 | NA |
| AGO3 | 78.1713761 | 0.274420842 | 0.2257241 | 1.21573568 | 0.22408563 | 0.63029911 |
| ADPRHL2 | 19.4782074 | -0.642073179 | 0.49001427 | -1.3103153 | 0.1900892 | NA |
| TRAPPC3 | 14.9038777 | -0.548736504 | 0.48802237 | -1.1244085 | 0.26083977 | NA |
| MAP7D1 | 22.5469038 | 0.377339895 | 0.33705914 | 1.11950648 | 0.26292413 | NA |
| THRAP3 | 75.0348093 | -0.015107075 | 0.22975001 | -0.0657544 | 0.94757336 | 0.98345894 |
| EVA1B | 86.9759889 | 0.478732036 | 0.33531162 | 1.42772279 | 0.15337166 | 0.55384216 |
| STK40 | 31.3949181 | 0.03383531 | 0.43538192 | 0.07771409 | 0.93805549 | 0.97885408 |
| LSM10 | 18.692129 | -0.281627059 | 0.55635305 | -0.5062021 | 0.61271481 | NA |
| OSCP1 | 6.71445582 | 0.224743647 | 0.64908115 | 0.34624892 | 0.72915565 | NA |
| MRPS15 | 11.5558987 | -0.315305651 | 0.46246882 | -0.6817879 | 0.49537307 | NA |
| MEAF6 | 28.3251749 | 0.128202205 | 0.40206259 | 0.31886131 | 0.74983168 | NA |
| SNIP1 | 10.923813 | -0.113999206 | 0.58922288 | -0.1934738 | 0.8465879 | NA |
| DNALI1 | 4.91030896 | -0.863933719 | 0.72882787 | -1.1853742 | 0.23586955 | NA |
| GNL2 | 8.34008605 | -0.040120959 | 0.72451587 | -0.0553762 | 0.95583873 | NA |
| C1orf109 | 5.26250307 | -0.387646945 | 0.70976916 | -0.5461592 | 0.58495652 | NA |
| C1orf122 | 45.8071053 | -0.852336367 | 0.39822307 | -2.1403491 | 0.03232657 | 0.29020358 |
| MTF1 | 11.021096 | 0.04893385 | 0.44373702 | 0.11027669 | 0.91218994 | NA |
| INPP5B | 20.183457 | 0.355669799 | 0.6028426 | 0.58998783 | 0.55519881 | NA |
| SF3A3 | 12.9087344 | 0.096062067 | 0.46299498 | 0.20747971 | 0.83563523 | NA |
| UTP11 | 4.26442033 | -0.468218325 | 0.65782416 | -0.7117682 | 0.47660832 | NA |
| RRAGC | 19.0188211 | -0.304808087 | 0.56200113 | -0.542362 | 0.58756912 | NA |
| AL139260.1 | 17.3962156 | -1.410151306 | 0.70998558 | -1.986169 | 0.04701457 | NA |
| MYCBP | 5.50528546 | 0.698673636 | 0.68566982 | 1.01896513 | 0.30821952 | NA |
| RHBDL2 | 4.18561746 | -2.315852567 | 1.04863904 | -2.2084363 | 0.02721387 | NA |
| AKIRIN1 | 32.8211363 | -0.385294928 | 0.28848949 | -1.3355597 | 0.18169324 | 0.58390314 |
| NDUFS5 | 69.7878377 | -0.897191548 | 0.411545 | -2.180057 | 0.02925324 | 0.27926427 |
| MACF1 | 659.951503 | 0.203393391 | 0.20474287 | 0.99340893 | 0.32051071 | 0.69496685 |
| PABPC4 | 11.6073331 | -0.133447131 | 0.46990027 | -0.2839903 | 0.7764178 | NA |
| PPIE | 16.1871374 | -0.314033536 | 0.35194189 | -0.892288 | 0.37223856 | NA |
| BMP8B | 4.24973186 | -3.598551767 | 1.2168516 | -2.9572643 | 0.00310382 | NA |
| TRIT1 | 16.6701542 | -0.509864481 | 0.36941829 | -1.380182 | 0.16753061 | NA |
| CAP1 | 59.9163941 | -0.327407818 | 0.39592476 | -0.8269445 | 0.4082685 | 0.76772515 |
| PPT1 | 25.698367 | -0.264183396 | 0.39254153 | -0.6730075 | 0.50094252 | NA |
| RLF | 39.836749 | 0.051560465 | 0.24838307 | 0.20758446 | 0.83555344 | 0.95223511 |
| ZMPSTE24 | 20.8167088 | 0.234345686 | 0.36090955 | 0.64931972 | 0.51613174 | NA |
| SMAP2 | 12.9316183 | -0.044022925 | 0.4677717 | -0.094112 | 0.9250202 | NA |
| AL603839.3 | 13.7805477 | -0.218325771 | 0.46128407 | -0.4733 | 0.63599913 | NA |
| ZNF684 | 5.56979583 | 0.125276706 | 0.64729923 | 0.19353755 | 0.846538 | NA |
| NFYC | 25.2960779 | 0.738548332 | 0.32081907 | 2.30207117 | 0.02133116 | NA |
| CTPS1 | 5.61800406 | -1.365734131 | 0.91608039 | -1.4908453 | 0.13600212 | NA |
| SCMH1 | 105.044928 | -0.019100017 | 0.25192439 | -0.0758165 | 0.93956512 | 0.97998296 |
| HIVEP3 | 64.1803364 | 0.441893657 | 0.37824302 | 1.16827974 | 0.24269395 | 0.64918976 |
| FOXJ3 | 68.2760289 | -0.132066535 | 0.26471625 | -0.4988985 | 0.6178509 | 0.87662819 |
| PPCS | 21.4540855 | -0.450994665 | 0.3474395 | -1.2980524 | 0.19426935 | NA |
| CCDC30 | 38.3659936 | 0.25010444 | 0.30592523 | 0.81753454 | 0.41362302 | 0.76946282 |
| PPIH | 7.29667594 | 0.655074796 | 0.52669111 | 1.24375517 | 0.21358968 | NA |
| YBX1 | 47.4667151 | -0.985439402 | 0.40261017 | -2.4476267 | 0.01438006 | 0.19867928 |
| P3H1 | 12.8261345 | 0.251087483 | 0.43695795 | 0.57462619 | 0.56554413 | NA |
| SVBP | 6.95450467 | -0.588212927 | 0.66283478 | -0.8874201 | 0.37485275 | NA |
| ERMAP | 11.1694156 | 0.685610924 | 0.46326364 | 1.47995843 | 0.13888434 | NA |
| SLC2A1 | 6.43208238 | -1.184064762 | 1.02410345 | -1.1561964 | 0.24760081 | NA |
| EBNA1BP2 | 18.5305363 | 0.385231524 | 0.38476575 | 1.00121054 | 0.31672503 | NA |
| ELOVL1 | 16.300821 | -0.240411722 | 0.66456763 | -0.3617566 | 0.71753393 | NA |
| MED8 | 7.98988811 | -0.38052651 | 0.62294694 | -0.610849 | 0.54129955 | NA |
| SZT2 | 15.2802745 | -0.124679293 | 0.43274912 | -0.2881099 | 0.77326264 | NA |
| HYI | 6.41059847 | 0.415906518 | 0.55339361 | 0.75155642 | 0.45231786 | NA |
| PTPRF | 51.6050482 | -0.203539703 | 0.46283465 | -0.4397676 | 0.66010541 | 0.89724783 |
| IPO13 | 6.21566397 | 0.528635676 | 0.67570378 | 0.78234825 | 0.43400994 | NA |
| ATP6V0B | 63.3139883 | -0.221817927 | 0.3908053 | -0.5675919 | 0.57031211 | 0.8573305 |
| B4GALT2 | 16.0790004 | -0.646129148 | 0.58500126 | -1.1044919 | 0.26937982 | NA |
| CCDC24 | 9.28263693 | -0.786606352 | 0.50558295 | -1.5558403 | 0.11974606 | NA |
| DMAP1 | 8.33835496 | -0.184628441 | 0.62174248 | -0.2969532 | 0.76650221 | NA |
| ERI3 | 46.6033715 | -0.152492393 | 0.26346559 | -0.5787943 | 0.56272795 | 0.85388124 |
| RNF220 | 56.2134005 | 0.100200641 | 0.52616686 | 0.1904351 | 0.84896819 | 0.95532127 |
| TMEM53 | 8.85900713 | -0.655705479 | 0.53453579 | -1.2266821 | 0.2199421 | NA |
| RPS8 | 81.2350447 | -1.172239594 | 0.43985479 | -2.6650604 | 0.00769745 | 0.14000382 |
| EIF2B3 | 37.2970042 | 0.45201742 | 0.29661532 | 1.52391801 | 0.12752919 | 0.52301154 |
| HECTD3 | 5.69418522 | -0.006313484 | 0.53226116 | -0.0118616 | 0.99053601 | NA |
| UROD | 8.20865931 | 0.102635058 | 0.62547398 | 0.16409165 | 0.86965899 | NA |
| ZSWIM5 | 94.0712385 | 0.353336417 | 0.40371895 | 0.87520394 | 0.38146295 | 0.74899182 |
| TESK2 | 6.10642201 | -0.192574734 | 0.61994945 | -0.3106297 | 0.75608212 | NA |
| PRDX1 | 148.480159 | -0.647660572 | 0.46165315 | -1.4029159 | 0.16064191 | 0.56283048 |
| AKR1A1 | 11.6324543 | -0.360168617 | 0.55217211 | -0.652276 | 0.51422314 | NA |
| NASP | 32.6282565 | 0.623516522 | 0.32220362 | 1.93516302 | 0.05297029 | 0.35868165 |
| GPBP1L1 | 80.5530052 | 0.306523127 | 0.20465422 | 1.4977611 | 0.13419533 | 0.52741886 |
| TMEM69 | 6.10166035 | -0.529656312 | 0.70426629 | -0.7520682 | 0.45201002 | NA |
| IPP | 24.5339468 | -0.25010135 | 0.37943347 | -0.6591442 | 0.50980321 | NA |
| MAST2 | 24.7761103 | -0.030779685 | 0.33224811 | -0.0926407 | 0.92618904 | NA |
| TSPAN1 | 17.2960885 | -1.21658677 | 0.71340678 | -1.7053199 | 0.08813478 | NA |
| POMGNT1 | 15.0042124 | 0.761439429 | 0.36943147 | 2.06111141 | 0.03929241 | NA |

| | | | | | | |
|---|---|---|---|---|---|---|
| LRRC41 | 25.1422329 | -0.005035004 | 0.36412631 | -0.0138276 | 0.9889675 | NA |
| UQCRH | 29.7015785 | -0.303815079 | 0.4692126 | -0.6474998 | 0.5173085 | 0.83104823 |
| NSUN4 | 13.9858762 | -0.369607752 | 0.41274204 | -0.8954933 | 0.37052342 | NA |
| FAAH | 7.8121169 | 0.561501235 | 0.61326143 | 0.91559848 | 0.35987753 | NA |
| MKNK1 | 16.8200461 | 0.291956137 | 0.44027847 | 0.663117 | 0.50725563 | NA |
| ATPAF1 | 18.6854472 | 0.127926961 | 0.40147103 | 0.31864556 | 0.7499953 | NA |
| EFCAB14-AS | 4.31407159 | -1.570490347 | 1.15539289 | -1.3592695 | 0.1740612 | NA |
| EFCAB14 | 50.8823862 | -0.240954912 | 0.31486221 | -0.765271 | 0.4441102 | 0.78734568 |
| CYP4A11 | 7.72318477 | 1.576772261 | 1.9840791 | 0.7947124 | 0.42678082 | NA |
| CYP4A22-AS: | 6.20460507 | -0.024781678 | 0.70346103 | -0.0352282 | 0.97189776 | NA |
| PDZK1IP1 | 138.51237 | -0.395908692 | 0.50759976 | -0.7799623 | 0.43541305 | 0.78167891 |
| CMPK1 | 35.0365983 | -0.104840399 | 0.34979114 | -0.2997229 | 0.76438856 | 0.92399242 |
| LINC01389 | 6.76538572 | -0.637317777 | 1.00566827 | -0.6337257 | 0.52625988 | NA |
| FOXD2 | 31.5642238 | 0.118774181 | 0.37353198 | 0.31797593 | 0.75050319 | 0.92274989 |
| TRABD2B | 31.201323 | -1.098707762 | 0.71045638 | -1.5464817 | 0.12198828 | 0.51522823 |
| SPATA6 | 76.6483502 | 0.368559299 | 0.33684388 | 1.09415465 | 0.27388716 | 0.66679851 |
| AGBL4 | 47.0902108 | 0.543675552 | 0.49662969 | 1.09473026 | 0.27363483 | 0.6665843 |
| FAF1 | 124.405064 | 0.169690489 | 0.2596917 | 0.65343053 | 0.51347876 | 0.83070544 |
| RNF11 | 69.7228021 | 0.294813636 | 0.32141659 | 0.91723217 | 0.35902099 | 0.72955368 |
| TTC39A | 7.7614123 | -0.576662868 | 0.79733475 | -0.7232381 | 0.46953362 | NA |
| EPS15 | 144.06287 | 0.462064214 | 0.16684959 | 2.76934585 | 0.0056169 | 0.11765106 |
| OSBPL9 | 141.206848 | 0.530410758 | 0.2255957 | 2.35115631 | 0.01871517 | 0.22818497 |
| NRDC | 68.4321465 | 0.032738001 | 0.25813435 | 0.12682543 | 0.89907857 | 0.97097328 |
| TXNDC12 | 20.1076411 | -0.580017502 | 0.33456011 | -1.733672 | 0.08297629 | NA |
| BTF3L4 | 30.8017797 | -0.211218039 | 0.3332812 | -0.6337532 | 0.52624188 | 0.83195298 |
| ZFYVE9 | 127.209638 | 0.525450132 | 0.29027767 | 1.8101638 | 0.07027039 | 0.40663921 |
| CC2D1B | 15.614848 | -0.456645854 | 0.43770518 | -1.0432727 | 0.29682202 | NA |
| PRPF38A | 15.1726366 | 0.445815711 | 0.45061013 | 0.98936016 | 0.32248696 | NA |
| TUT4 | 95.2941005 | 0.496391191 | 0.25160194 | 1.97292274 | 0.04850437 | 0.34339091 |
| GPX7 | 4.97333929 | 0.052730159 | 0.69354487 | 0.07602992 | 0.9393953 | NA |
| COA7 | 4.7717975 | -0.799576929 | 0.75229147 | -1.0628552 | 0.2878476 | NA |
| ZYG11B | 52.8937591 | 0.693277526 | 0.29701586 | 2.33414316 | 0.01958823 | 0.23436534 |
| ECHDC2 | 95.4946037 | 0.333429698 | 0.29413225 | 1.13360468 | 0.25696041 | 0.65588339 |
| SCP2 | 54.3120902 | 0.067981034 | 0.27496445 | 0.24723572 | 0.8047258 | 0.94198206 |
| CPT2 | 5.88710376 | -0.302768956 | 0.58469476 | -0.517824 | 0.6045811 | NA |
| C1orf123 | 10.3515228 | -0.662491719 | 0.51337841 | -1.290455 | 0.19689274 | NA |
| MAGOH | 11.6281211 | -0.516075899 | 0.51950038 | -0.9934081 | 0.32051111 | NA |
| NDC1 | 12.6300039 | 0.156417111 | 0.428543 | 0.36499747 | 0.71511329 | NA |
| YIPF1 | 10.2557011 | -0.225747403 | 0.48185631 | -0.4684953 | 0.63943045 | NA |
| HSPB11 | 26.5984869 | 0.481939564 | 0.39696453 | 1.21406202 | 0.22472404 | NA |
| LRRC42 | 10.3862069 | -0.619321257 | 0.47638197 | -1.3000519 | 0.1935832 | NA |
| TMEM59 | 100.403567 | -0.167341796 | 0.32691377 | -0.5118836 | 0.60873248 | 0.8725395 |
| TCEANC2 | 6.16039867 | -0.250712608 | 0.59412857 | -0.4219838 | 0.67303687 | NA |
| MRPL37 | 9.80523259 | -0.701683935 | 0.54161429 | -1.2955418 | 0.1951334 | NA |
| SSBP3 | 121.117686 | -0.347457048 | 0.31430196 | -1.105488 | 0.26894818 | 0.66147637 |
| ACOT11 | 24.8846061 | 0.596583271 | 0.3839142 | 1.55394949 | 0.12019646 | NA |
| DHCR24 | 6.31386586 | -0.014179535 | 0.54766381 | -0.0258909 | 0.97934432 | NA |
| USP24 | 76.2592174 | -0.08963654 | 0.24531872 | -0.3653881 | 0.71482172 | 0.91376358 |
| PLPP3 | 36.8245411 | 0.130083832 | 0.68752729 | 0.18920534 | 0.84993189 | 0.95532127 |
| PRKAA2 | 47.1315139 | 0.705059685 | 0.32420214 | 2.1747533 | 0.02964861 | 0.27926427 |
| DAB1 | 37.893238 | 0.161844293 | 0.67827373 | 0.23861206 | 0.81140641 | 0.94408044 |
| MYSM1 | 30.449252 | 0.48980092 | 0.35849168 | 1.36628254 | 0.1718503 | 0.57423036 |
| JUN | 23.1031873 | 0.876959064 | 0.55324575 | 1.58511669 | 0.11293982 | NA |
| LINC01135 | 7.7588782 | 0.134621115 | 0.70385333 | 0.19126302 | 0.84831953 | NA |
| FGGY | 51.446051 | -0.060430332 | 0.42671377 | -0.141618 | 0.88738178 | 0.96727237 |
| HOOK1 | 14.5868172 | 0.15107848 | 0.42781853 | 0.35313683 | 0.72398586 | NA |
| NFIA | 37.1323747 | 1.514428146 | 0.53260917 | 2.84341355 | 0.00446331 | 0.10711948 |
| TM2D1 | 57.2606574 | 0.59393726 | 0.2599619 | 2.28470887 | 0.0223299 | 0.24469443 |
| PATJ | 192.421797 | 0.182154613 | 0.2545799 | 0.71551057 | 0.47429361 | 0.80625938 |
| USP1 | 14.980012 | -0.965119963 | 0.41991904 | -2.2983477 | 0.02154201 | NA |
| DOCK7 | 61.699101 | 0.43201122 | 0.28423267 | 1.51992107 | 0.12853081 | 0.52341115 |
| ATG4C | 15.0500457 | 0.132833489 | 0.38168343 | 0.34802005 | 0.72782512 | NA |
| ALG6 | 8.72867747 | 0.74772417 | 0.5301345 | 1.41044238 | 0.1584091 | NA |
| ITGB3BP | 28.3880445 | -0.341868076 | 0.57719346 | -0.5922937 | 0.5536539 | NA |
| EFCAB7 | 12.7685671 | 0.511655303 | 0.51098719 | 1.00130749 | 0.31667817 | NA |
| PGM1 | 14.1200345 | -0.170903427 | 0.40613874 | -0.4208006 | 0.67390069 | NA |
| ROR1 | 18.1307141 | -0.064537297 | 0.74145472 | -0.0870415 | 0.93063856 | NA |
| CACHD1 | 8.29845662 | 0.104377196 | 0.59206572 | 0.17629326 | 0.86006356 | NA |
| RAVER2 | 9.59194105 | 0.215276978 | 0.60206225 | 0.35756598 | 0.72066814 | NA |
| JAK1 | 53.249092 | -0.173788243 | 0.26952007 | -0.6448063 | 0.51905271 | 0.83104823 |
| AK4 | 26.0389851 | 0.081561956 | 0.57878517 | 0.14091922 | 0.88793376 | NA |
| DNAJC6 | 37.0194011 | -0.00184876 | 0.49685694 | -0.0037209 | 0.99703115 | 0.99958475 |
| AL139294.1 | 4.59784863 | -0.378108598 | 0.86823148 | -0.4354928 | 0.66320473 | NA |
| LEPR | 119.984916 | 0.076016668 | 0.472862 | 0.16075867 | 0.87228347 | 0.96207879 |
| PDE4B | 50.1575875 | -0.502587681 | 0.39666838 | -1.2670223 | 0.20514732 | 0.61387077 |
| SGIP1 | 304.686952 | 1.251263229 | 0.21142259 | 5.9183043 | 3.25E-09 | 4.40E-06 |
| WDR78 | 29.0346428 | -0.051495313 | 0.34135002 | -0.1508578 | 0.88008789 | 0.96423929 |
| MIER1 | 61.1041704 | -0.235383128 | 0.23608685 | -0.9970192 | 0.31875518 | 0.69469143 |
| SLC35D1 | 18.469216 | -0.105394224 | 0.37655669 | -0.2798894 | 0.77956236 | NA |
| SERBP1 | 55.2825759 | -0.541241724 | 0.42786798 | -1.2649737 | 0.20588077 | 0.61446093 |
| GADD45A | 119.80671 | -0.945560986 | 0.63755034 | -1.4831158 | 0.13804365 | 0.53418928 |
| GNG12 | 66.9874913 | -0.24499145 | 0.287416 | -0.8523932 | 0.39399588 | 0.75773027 |
| GNG12-AS1 | 16.0367625 | -0.279663565 | 0.65850076 | -0.4246974 | 0.67105727 | NA |
| WLS | 13.7625718 | 0.907140089 | 0.62284444 | 1.45644729 | 0.14526901 | NA |
| LRRC40 | 17.2982427 | 0.107924928 | 0.44498511 | 0.24253604 | 0.80836483 | NA |
| SRSF11 | 108.804636 | 0.130253899 | 0.22606068 | 0.57618997 | 0.56448678 | 0.8549147 |
| ANKRD13C | 47.9548 | 0.480279144 | 0.31781812 | 1.51117611 | 0.13074359 | 0.52613957 |
| HHLA3 | 4.52529117 | -1.351014376 | 0.7742784 | -1.744869 | 0.08100768 | NA |
| PTGER3 | 28.9509106 | 0.013388446 | 0.62103438 | 0.0215583 | 0.9828003 | 0.99552981 |
| ZRANB2-AS1 | 5.20047894 | 0.105372158 | 0.71910262 | 0.14653285 | 0.88350076 | NA |
| ZRANB2 | 103.782052 | -0.042519969 | 0.21767665 | -0.1953355 | 0.84513034 | 0.95509854 |
| NEGR1 | 11.2308061 | 1.173510854 | 0.83995576 | 1.39711031 | 0.1623804 | NA |
| CRYZ | 17.8606217 | 0.11560294 | 0.4467058 | 0.25878988 | 0.79579737 | NA |
| TYW3 | 13.2619833 | 0.100628029 | 0.39664223 | 0.25369974 | 0.79972754 | NA |
| ACADM | 31.103861 | -0.125814496 | 0.32639087 | -0.3854719 | 0.69988785 | 0.90729979 |
| RABGGTB | 11.6464637 | -0.286142835 | 0.55470608 | -0.5158459 | 0.60596207 | NA |
| AC092813.2 | 336.670773 | 0.064552942 | 0.6762934 | 0.09545109 | 0.92395654 | 0.97775374 |
| ST6GALNAC: | 2498.45019 | 0.15824066 | 0.44275958 | 0.35739636 | 0.7207951 | 0.91509584 |
| PIGK | 35.992404 | 0.11105286 | 0.39172445 | 0.28349739 | 0.77679558 | 0.92940498 |
| AC118549.1 | 101.189275 | 0.488695126 | 0.230948 | 2.11603967 | 0.03434143 | 0.29210684 |
| USP33 | 68.2693792 | 0.35220517 | 0.23920855 | 1.47237702 | 0.1409191 | 0.53597868 |
| MIGA1 | 51.4334206 | 0.48069965 | 0.26380228 | 1.82219675 | 0.06842514 | 0.40491695 |
| NEXN | 8.22309274 | 1.123197111 | 0.75474202 | 1.48818681 | 0.13670164 | NA |
| FUBP1 | 49.5871516 | -0.040439577 | 0.28159045 | -0.1436113 | 0.8858074 | 0.96652834 |
| GIPC2 | 12.8195723 | 0.457183477 | 0.43804425 | 1.04369246 | 0.2966277 | NA |
| IFI44L | 51.6199489 | -2.025661561 | 2.37096835 | -0.8543604 | 0.39290531 | 0.75742582 |
| IFI44 | 23.2152533 | 0.775004398 | 1.47199246 | 0.52650025 | 0.59854068 | NA |
| LINC01781 | 9.6889829 | 0.540523428 | 1.10528058 | 0.4890373 | 0.62481529 | NA |
| ADGRL2 | 17.0760782 | 1.116864025 | 0.68106485 | 1.63987912 | 0.1010303 | NA |
| LINC01362 | 6.50425612 | -1.190798004 | 0.8729132 | -1.3641654 | 0.17251551 | NA |
| LINC01725 | 21.1363406 | 0.275434027 | 0.75430767 | 0.36514812 | 0.71500084 | NA |
| TTLL7 | 49.4590234 | 0.007412811 | 0.45683112 | 0.01622659 | 0.98705362 | 0.99585746 |
| PRKACB | 70.9612506 | 0.161550048 | 0.30214715 | 0.53467341 | 0.59287571 | 0.8654248 |

| | | | | | | |
|---|---|---|---|---|---|---|
| RPF1 | 10.7362948 | -0.263944468 | 0.42644511 | -0.6189412 | 0.53595506 | NA |
| GNG5 | 54.5627997 | -0.249009816 | 0.41794315 | -0.5957983 | 0.55130998 | 0.8472507 |
| SPATA1 | 9.64414947 | -0.450485596 | 0.63984873 | -0.7040502 | 0.48140154 | NA |
| CTBS | 20.6605463 | -0.544938305 | 0.46084619 | -1.1824733 | 0.23701797 | NA |
| SSX2IP | 5.26580228 | 0.101330138 | 0.63987194 | 0.15836003 | 0.87417311 | NA |
| WDR63 | 4.52971091 | -0.485336674 | 0.9637792 | -0.5035766 | 0.61455892 | NA |
| MCOLN3 | 7.65573377 | -0.214265875 | 0.91654795 | -0.2337749 | 0.81515976 | NA |
| SYDE2 | 23.5906676 | 0.48132736 | 0.50763133 | 0.94818293 | 0.34303633 | NA |
| C1orf52 | 23.9496989 | 0.388012478 | 0.3156496 | 1.22925065 | 0.21897784 | NA |
| BCL10 | 8.10843665 | -0.222489067 | 0.57879007 | -0.3844037 | 0.70067923 | NA |
| AL078459.1 | 11.725642 | 1.436053424 | 0.72374491 | 1.98419829 | 0.04723374 | NA |
| DDAH1 | 22.753541 | 0.065895671 | 0.54034778 | 0.12195048 | 0.90293824 | NA |
| CYR61 | 34.1735452 | -0.739136413 | 0.89502645 | -0.8258263 | 0.40890261 | 0.76772515 |
| ZNHIT6 | 24.533242 | 0.112469395 | 0.33206245 | 0.33869953 | 0.73483609 | NA |
| AL136382.1 | 6.32707021 | 0.231213823 | 0.61789794 | 0.3741942 | 0.70825984 | NA |
| ODF2L | 72.589042 | 0.376033988 | 0.39637537 | 0.94868151 | 0.34278262 | 0.71725052 |
| SH3GLB1 | 76.2347249 | -0.281001257 | 0.2418764 | -1.1617556 | 0.24533477 | 0.64996527 |
| SELENOF | 55.3571578 | 0.211369003 | 0.2516402 | 0.83996517 | 0.40092791 | 0.76381845 |
| HS2ST1 | 28.763886 | 0.226415649 | 0.42364837 | 0.53444239 | 0.5930355 | 0.8654248 |
| LMO4 | 21.4819248 | -0.270144357 | 0.42389698 | -0.6372878 | 0.52393743 | NA |
| PKN2-AS1 | 9.96812225 | 1.411680695 | 0.72601117 | 1.94443384 | 0.05184316 | NA |
| PKN2 | 96.0010777 | 0.543848491 | 0.17955175 | 3.0289233 | 0.00245427 | 0.07963615 |
| GTF2B | 20.7592 | 0.133373495 | 0.41047028 | 0.3249285 | 0.74523518 | NA |
| KYAT3 | 29.2127421 | 0.224639896 | 0.37461245 | 0.59965946 | 0.54873321 | 0.84550378 |
| GBP3 | 13.3057609 | -2.490978001 | 1.08741292 | -2.2907379 | 0.02197858 | NA |
| GBP1 | 15.2456317 | -1.116703033 | 1.65019308 | -0.6767105 | 0.49858963 | NA |
| GBP2 | 6.48904618 | -0.283197871 | 0.71463862 | -0.3962812 | 0.69189757 | NA |
| GBP4 | 8.19847909 | -0.115563223 | 2.43740364 | -0.0474124 | 0.96218452 | NA |
| LRRC8B | 8.69137781 | -0.075719504 | 0.57291278 | -0.1321659 | 0.89485311 | NA |
| ZNF326 | 23.9537154 | -0.193952997 | 0.32408644 | -0.5984607 | 0.54953258 | NA |
| ZNF644 | 90.8141812 | 0.043442241 | 0.21660811 | 0.20055686 | 0.8410451 | 0.95393706 |
| HFM1 | 71.7352793 | 0.382905297 | 0.76534156 | 0.50030642 | 0.61685933 | 0.87619503 |
| CDC7 | 10.5152089 | -0.876427043 | 0.66021626 | -1.3274848 | 0.18434837 | NA |
| TGFBR3 | 1002.82591 | -0.015965966 | 0.30203167 | -0.0528619 | 0.95784195 | 0.98714493 |
| BTBD8 | 23.640145 | -0.168192313 | 0.45577167 | -0.3690276 | 0.71210717 | NA |
| GLMN | 22.54868 | -0.086412454 | 0.34040842 | -0.2538493 | 0.79961195 | NA |
| RPAP2 | 46.338024 | -0.288202721 | 0.2745511 | -1.0497234 | 0.29384528 | 0.6804998 |
| EVI5 | 110.725965 | 0.517523605 | 0.25647845 | 2.01780543 | 0.04361153 | 0.33029013 |
| RPL5 | 47.8087098 | -1.209108344 | 0.3591195 | -3.3668691 | 0.00076027 | 0.04122345 |
| FAM69A | 5.3864156 | -0.204862235 | 0.72394954 | -0.2829786 | 0.77719323 | NA |
| MTF2 | 40.5662422 | 0.513474204 | 0.30866114 | 1.66355314 | 0.09620176 | 0.47011368 |
| TMED5 | 53.6987738 | 0.026363525 | 0.42751522 | 0.06166687 | 0.95082813 | 0.98520702 |
| CCDC18 | 11.2595629 | -0.690711135 | 0.69868371 | -0.9885891 | 0.3228642 | NA |
| DR1 | 21.5314269 | 0.372063586 | 0.36597508 | 1.01663639 | 0.30932644 | NA |
| FNBP1L | 251.548443 | 0.405376506 | 0.22524099 | 1.79974573 | 0.0719008 | 0.41365905 |
| BCAR3 | 206.687615 | -0.677267479 | 0.34760089 | -1.9484055 | 0.05136646 | 0.35507065 |
| DNTTIP2 | 6.90367559 | -1.579052188 | 0.66463916 | -2.3758037 | 0.01751077 | NA |
| GCLM | 31.7696581 | -0.828494737 | 0.3606585 | -2.2971724 | 0.02160894 | 0.24278631 |
| ARHGAP29 | 412.214199 | -0.308929114 | 0.42199892 | -0.7320614 | 0.4641311 | 0.79869145 |
| ABCD3 | 46.9208013 | 0.20605632 | 0.29728837 | 0.69311935 | 0.48823465 | 0.8169471 |
| F3 | 261.443993 | -0.716118577 | 0.55119049 | -1.2992216 | 0.1938679 | 0.59842328 |
| SLC44A3 | 10.00247 | 0.194643003 | 0.5028707 | 0.38706371 | 0.69870904 | NA |
| CNN3 | 85.320097 | -0.054162549 | 0.37751498 | -0.1434713 | 0.88591801 | 0.96652834 |
| AC105942.1 | 8.1771009 | 0.198940461 | 0.4888574 | 0.40694988 | 0.6840448 | NA |
| ALG14 | 19.8099969 | -0.207432779 | 0.3603764 | -0.5756003 | 0.56488535 | NA |
| PTBP2 | 81.6429558 | 0.307531174 | 0.26857129 | 1.14506348 | 0.25218286 | 0.65211999 |
| DPYD | 129.271407 | 0.319545908 | 0.23894778 | 1.33730436 | 0.18112331 | 0.58390314 |
| DPYD-AS1 | 5.34808674 | -0.675650121 | 0.76731139 | -0.8805423 | 0.37856561 | NA |
| SNX7 | 14.8628823 | 0.012449588 | 0.43500451 | 0.02861945 | 0.9771681 | NA |
| PALMD | 9.79899413 | 0.653941263 | 0.58976161 | 1.10882305 | 0.26750653 | NA |
| AGL | 17.9171567 | 0.03202328 | 0.49357889 | 0.06487976 | 0.94826974 | NA |
| TRMT13 | 17.097212 | 0.268036389 | 0.44477002 | 0.60264042 | 0.54674793 | NA |
| DBT | 21.0792571 | -0.108043557 | 0.35509276 | -0.3042685 | 0.76092331 | NA |
| RTCA | 16.8462928 | -0.085277795 | 0.39952261 | -0.2134492 | 0.83097658 | NA |
| CDC14A | 740.116004 | 0.421600529 | 0.33259897 | 1.26759422 | 0.2049429 | 0.61387077 |
| AL589990.1 | 7.87104852 | 1.014508156 | 0.84757818 | 1.19694936 | 0.23132629 | NA |
| VCAM1 | 18.1254361 | -0.791030281 | 0.70107254 | -1.1283145 | 0.25918714 | NA |
| EXTL2 | 12.9247355 | 0.194322738 | 0.39761265 | 0.48872373 | 0.6250373 | NA |
| SLC30A7 | 34.367935 | -0.30236085 | 0.3004563 | -1.0063389 | 0.31425259 | 0.69309869 |
| DPH5 | 8.98073984 | -0.951750618 | 0.5224068 | -1.8218573 | 0.06847665 | NA |
| AC093157.1 | 8.0896503 | -0.025967339 | 0.55415571 | -0.0468593 | 0.96262538 | NA |
| RNPC3 | 32.6817404 | 0.152161648 | 0.30976967 | 0.49120899 | 0.62327864 | 0.8780438 |
| AMY2B | 7.79299685 | 0.667367349 | 0.63280677 | 1.05461475 | 0.29160156 | NA |
| PRMT6 | 17.295663 | 0.412197636 | 0.4981687 | 0.8274258 | 0.40799576 | NA |
| NTNG1 | 1584.55003 | -0.821484765 | 0.29242971 | -2.80917 | 0.00496694 | 0.11177304 |
| VAV3 | 21.4632364 | 0.828880592 | 0.76168803 | 1.08821533 | 0.27650006 | NA |
| SLC25A24 | 34.802609 | -0.09619537 | 0.31434618 | -0.3060173 | 0.75959147 | 0.92313949 |
| PRPF38B | 30.7124601 | -0.006308887 | 0.37550759 | -0.016801 | 0.98659541 | 0.99585746 |
| STXBP3 | 48.0161434 | 0.503039054 | 0.37985566 | 1.32429 | 0.18540675 | 0.587049 |
| AL449266.1 | 8.70360912 | -0.32796809 | 0.48816661 | -0.6718364 | 0.50168786 | NA |
| GPSM2 | 9.81224327 | -1.875292249 | 0.73902482 | -2.5375227 | 0.01116401 | NA |
| WDR47 | 10.1984925 | 0.040845653 | 0.48642809 | 0.08397059 | 0.93307982 | NA |
| TAF13 | 6.67927044 | 0.109951521 | 0.57502325 | 0.19121231 | 0.84835926 | NA |
| TMEM167B | 17.3276424 | -0.015859507 | 0.4090739 | -0.0387693 | 0.96907433 | NA |
| SARS | 35.1090798 | -1.115839129 | 0.44686565 | -2.4970349 | 0.01252366 | 0.18813325 |
| CELSR2 | 13.153792 | 0.525769051 | 0.59533276 | 0.88315155 | 0.37715439 | NA |
| SORT1 | 197.079229 | 0.14954314 | 0.31869462 | 0.46923647 | 0.63890062 | 0.88715539 |
| PSMA5 | 27.9175304 | -0.602816173 | 0.48459471 | -1.2439595 | 0.21351449 | NA |
| CYB561D1 | 8.38591682 | 0.923461328 | 0.48117013 | 1.91919921 | 0.05495913 | NA |
| AMIGO1 | 4.23040142 | 0.314806531 | 0.69171841 | 0.45510792 | 0.6490316 | NA |
| GNAI3 | 46.6256204 | 0.202312247 | 0.30190585 | 0.67011701 | 0.5027832 | 0.82562294 |
| AMPD2 | 11.4367844 | -0.023749589 | 0.50936398 | -0.046626 | 0.96281133 | NA |
| GSTM4 | 5.70846727 | 0.175626387 | 0.73474959 | 0.2390289 | 0.81108317 | NA |
| GSTM3 | 9.27673317 | -0.540293605 | 0.70285958 | -0.7687078 | 0.44206682 | NA |
| CSF1 | 5.29180149 | -0.104675936 | 0.79082876 | -0.1323623 | 0.89469771 | NA |
| AHCYL1 | 55.9631948 | -0.09095802 | 0.26546132 | -0.3426413 | 0.73186831 | 0.91806182 |
| STRIP1 | 12.3268135 | 0.734932555 | 0.39202074 | 1.87472874 | 0.06083005 | NA |
| KCNC4 | 12.3873407 | -0.549275061 | 0.4285626 | -1.2816682 | 0.19995906 | NA |
| RBM15 | 10.3395006 | 0.14097102 | 0.47748379 | 0.29523729 | 0.76781262 | NA |
| SLC16A4 | 24.796774 | 0.179441666 | 0.38316696 | 0.46831195 | 0.63956153 | NA |
| AL355488.1 | 5.17421206 | 0.424513687 | 0.60945548 | 0.69654585 | 0.48608706 | NA |
| LAMTOR5 | 51.2048797 | -0.435510214 | 0.38097607 | -1.1431432 | 0.25297913 | 0.65272479 |
| LRIF1 | 13.313248 | -0.5002138 | 0.50756637 | -0.9855141 | 0.32437163 | NA |
| DRAM2 | 17.0694197 | -0.229206376 | 0.37437882 | -0.6122312 | 0.54038483 | NA |
| CEPT1 | 19.376248 | 0.30171603 | 0.41078308 | 0.73448993 | 0.4626502 | NA |
| WDR77 | 6.74213161 | -0.57818022 | 0.71486027 | -0.8088017 | 0.4186292 | NA |
| ATP5PB | 28.6090757 | -0.438282633 | 0.42033918 | -1.042688 | 0.2970928 | 0.68103704 |
| RAP1A | 64.7974991 | -0.170365522 | 0.2394832 | -0.7113882 | 0.47684372 | 0.80863008 |
| DDX20 | 5.27152541 | -0.146897998 | 0.64379235 | -0.2281761 | 0.81950938 | NA |
| KCND3 | 56.3266575 | -1.221252553 | 0.8136163 | -1.5010178 | 0.13335096 | 0.52613957 |
| AL445426.1 | 5.45679569 | 1.114890724 | 0.67377246 | 1.65469915 | 0.09798554 | NA |
| CTTNBP2NL | 24.5471629 | 0.561866162 | 0.42430186 | 1.3242133 | 0.18543222 | NA |
| WNT2B | 8.77342807 | -0.210004766 | 0.56382992 | -0.3724612 | 0.7095495 | NA |

| | | | | | | |
|---|---|---|---|---|---|---|
| ST7L | 30.1996529 | -0.063109545 | 0.34949691 | -0.1805725 | 0.85670311 | 0.95618817 |
| CAPZA1 | 51.2260529 | 0.111095355 | 0.38250082 | 0.29044475 | 0.77147601 | 0.92669573 |
| MOV10 | 9.81249855 | 0.566902169 | 0.83566956 | 0.67838078 | 0.49753029 | NA |
| RHOC | 27.0371042 | 0.111403241 | 0.43704175 | 0.25490297 | 0.79879804 | NA |
| SLC16A1-AS1 | 6.89119735 | -0.180826397 | 0.88691317 | -0.2038829 | 0.83844504 | NA |
| LRIG2 | 25.6502357 | 0.17599852 | 0.30485088 | 0.5773266 | 0.56371885 | NA |
| MAGI3 | 109.893429 | -0.40365549 | 0.4996218 | -0.8079221 | 0.41913543 | 0.77304616 |
| PHTF1 | 41.4871203 | 0.314490787 | 0.34739181 | 0.9052913 | 0.36531108 | 0.73398222 |
| RSBN1 | 36.4695539 | 0.052988763 | 0.40752583 | 0.13002553 | 0.89654623 | 0.97097328 |
| HIPK1 | 33.6588193 | 0.078998716 | 0.38280699 | 0.20636696 | 0.83650428 | 0.95252283 |
| TRIM33 | 73.3939897 | 0.23621848 | 0.24986949 | 0.94536743 | 0.34447132 | 0.71842682 |
| BCAS2 | 12.8313895 | -1.015345573 | 0.42504724 | -2.3887829 | 0.01690429 | NA |
| DENND2C | 5.27641645 | 0.119383286 | 0.61462855 | 0.19423648 | 0.84599072 | NA |
| NRAS | 13.5391427 | -0.112579006 | 0.4685867 | -0.2402522 | 0.81013471 | NA |
| CSDE1 | 100.77669 | -0.087030419 | 0.265634 | -0.3276328 | 0.7431893 | 0.92122771 |
| SIKE1 | 17.7366564 | -0.012779727 | 0.43062376 | -0.0296772 | 0.97632446 | NA |
| TSPAN2 | 119.634612 | 1.18493661 | 0.57017503 | 2.07819801 | 0.03769113 | 0.30414888 |
| NGF | 27.8425211 | -1.008752981 | 0.48596627 | -2.0757675 | 0.03791546 | NA |
| VANGL1 | 8.09048333 | -0.13332206 | 0.48798255 | -0.2732107 | 0.78469124 | NA |
| SLC22A15 | 13.1045829 | -0.009684803 | 0.51851648 | -0.0186779 | 0.98509805 | NA |
| ATP1A1 | 167.242481 | -0.081947708 | 0.30118074 | -0.2720881 | 0.78555425 | 0.93615351 |
| LINC01762 | 19.3912131 | 0.535138418 | 0.94209126 | 0.56803246 | 0.57001292 | NA |
| CD58 | 18.2394372 | 0.493604223 | 0.45869182 | 1.07611298 | 0.28187673 | NA |
| IGSF3 | 5.48639799 | -0.728339513 | 0.80381095 | -0.906108 | 0.3648787 | NA |
| TTF2 | 6.58080385 | -0.265345303 | 0.74460772 | -0.3563558 | 0.7215741 | NA |
| MAN1A2 | 101.262104 | 0.099688277 | 0.29458967 | 0.33839705 | 0.735064 | 0.91905659 |
| GDAP2 | 28.8352792 | 0.207648568 | 0.33793881 | 0.61445612 | 0.53891396 | 0.83844842 |
| WDR3 | 7.29414125 | -0.214842628 | 0.53228521 | -0.4036231 | 0.68648986 | NA |
| SPAG17 | 19.2673787 | -0.611660553 | 0.46491122 | -1.3156502 | 0.18829147 | NA |
| WARS2 | 16.3423294 | -0.111914744 | 0.47436903 | -0.2359234 | 0.81349213 | NA |
| AL359915.2 | 45.3647389 | -0.174365515 | 0.35032166 | -0.4977298 | 0.61867453 | 0.87662819 |
| ZNF697 | 6.26329694 | 1.038029476 | 0.92457192 | 1.1227136 | 0.26155915 | NA |
| PHGDH | 4.83928125 | -1.119506027 | 0.8091825 | -1.3835025 | 0.16651083 | NA |
| NOTCH2 | 151.892561 | -0.02755028 | 0.21325774 | -0.1291877 | 0.89720912 | 0.97097328 |
| SEC22B | 43.6613965 | -0.099497375 | 0.28134198 | -0.3536528 | 0.72359911 | 0.91509584 |
| NBPF26 | 14.8703082 | 0.837779258 | 0.54785011 | 1.52921253 | 0.12621177 | NA |
| LINC00623 | 46.5422193 | 0.729868051 | 0.53931084 | 1.35333466 | 0.17594874 | 0.57988078 |
| SRGAP2C | 279.062313 | 0.424695912 | 0.26560164 | 1.59899579 | 0.10982154 | 0.49603135 |
| SRGAP2-AS1 | 16.0660439 | -0.347969039 | 0.48394078 | -0.7190323 | 0.47212103 | NA |
| AC244021.1 | 101.12455 | -0.046846992 | 0.43473431 | -0.1077601 | 0.91418603 | 0.97707273 |
| NBPF15 | 19.9623402 | 0.092224947 | 0.39678217 | 0.23243219 | 0.81620235 | NA |
| FP700111.1 | 24.7077163 | 0.397253484 | 0.4138896 | 0.95980543 | 0.33715315 | NA |
| SRGAP2B | 233.649801 | 0.330106755 | 0.26558708 | 1.24293229 | 0.21389279 | 0.62255988 |
| GPR89A | 8.70474878 | 0.281370028 | 0.4930776 | 0.57064046 | 0.56824339 | NA |
| PDZK1 | 19.4151512 | 1.271377082 | 0.82254376 | 1.54566498 | 0.1221855 | NA |
| RNF115 | 99.2517291 | 0.259947589 | 0.2126206 | 1.22258892 | 0.221485 | 0.62528839 |
| POLR3C | 9.19712858 | -0.104392325 | 0.50976449 | -0.2047854 | 0.83773979 | NA |
| PIAS3 | 9.22639501 | -0.134032245 | 0.48583689 | -0.2758791 | 0.78264092 | NA |
| ITGA10 | 6.98505112 | -2.156833249 | 1.20115779 | -1.7956286 | 0.07255361 | NA |
| PEX11B | 9.38791206 | -0.093498789 | 0.53192377 | -0.1757748 | 0.86047088 | NA |
| RBM8A | 24.5232435 | -0.698782801 | 0.41396331 | -1.6880307 | 0.09140533 | NA |
| LIX1L | 23.8562241 | -0.342938445 | 0.35821556 | -0.9573522 | 0.33838953 | NA |
| POLR3GL | 6.92630603 | -0.71807864 | 0.59792532 | -1.2009504 | 0.22977045 | NA |
| TXNIP | 413.855631 | 0.120601374 | 0.45195034 | 0.26684652 | 0.78958733 | 0.93806861 |
| NBPF12 | 20.037276 | 0.761669605 | 0.41141707 | 1.85133206 | 0.0641218 | NA |
| PRKAB2 | 9.17563045 | 0.184976743 | 0.49118193 | 0.37659517 | 0.70647448 | NA |
| AC242426.2 | 23.6181805 | -0.70899265 | 0.42424762 | -1.6711765 | 0.09468682 | NA |
| FMO5 | 11.2757789 | -0.017977395 | 0.76514672 | -0.0234954 | 0.98125514 | NA |
| CHD1L | 21.1862045 | -0.160812733 | 0.35019119 | -0.4592141 | 0.64608042 | NA |
| BCL9 | 17.232772 | -0.361664754 | 0.37326507 | -0.968922 | 0.33258409 | NA |
| ACP6 | 16.6626204 | -0.018884622 | 0.43113484 | -0.0438021 | 0.96506213 | NA |
| GPR89B | 10.4369087 | 0.864450681 | 0.50136746 | 1.72418584 | 0.0846743 | NA |
| NBPF11 | 14.8829653 | 0.065914234 | 0.51182335 | 0.12878317 | 0.89752922 | NA |
| AC239809.3 | 8.33318568 | 0.940731193 | 0.70410661 | 1.33606356 | 0.1815285 | NA |
| NBPF14 | 23.6186277 | 0.226197989 | 0.30588802 | 0.73947971 | 0.45961575 | NA |
| PDE4DIP | 352.970501 | 0.356780186 | 0.32015139 | 1.11441085 | 0.26510297 | 0.65974932 |
| NBPF9 | 8.42443448 | 0.167518071 | 0.52017802 | 0.32203989 | 0.74742248 | NA |
| AC245297.3 | 229.815261 | -0.041938596 | 0.26572824 | -0.1578251 | 0.87459459 | 0.96229085 |
| NBPF19 | 24.418476 | 0.635588602 | 0.49717042 | 1.27841195 | 0.20110421 | NA |
| HIST2H2AC | 4.51004014 | -1.758810298 | 0.86814938 | -2.0259305 | 0.04277191 | NA |
| BOLA1 | 5.98833222 | -0.902954611 | 0.66258672 | -1.362772 | 0.17295439 | NA |
| SF3B4 | 11.4275652 | 0.246805818 | 0.50103212 | 0.4925948 | 0.62229892 | NA |
| OTUD7B | 45.2126741 | 0.271805152 | 0.35432012 | 0.76711745 | 0.44301169 | 0.78643653 |
| VPS45 | 39.2501007 | -0.10650244 | 0.3445092 | -0.3091425 | 0.75721312 | 0.92313949 |
| PLEKHO1 | 29.5627778 | 0.135464741 | 0.50679195 | 0.26729852 | 0.78923932 | 0.93792988 |
| ANP32E | 15.1393367 | -0.173187868 | 0.47773888 | -0.3625157 | 0.71696666 | NA |
| APH1A | 25.4511937 | 0.266922467 | 0.35559442 | 0.7506374 | 0.45287091 | NA |
| C1orf54 | 5.63174048 | -0.716934989 | 0.71991923 | -0.9958548 | 0.31932072 | NA |
| MRPS21 | 46.3185597 | -0.304879159 | 0.29577862 | -1.0307681 | 0.30264959 | 0.68310554 |
| PRPF3 | 22.0955263 | 0.273054294 | 0.41100217 | 0.66436218 | 0.50645853 | NA |
| RPRD2 | 90.784016 | 0.651359119 | 0.25699899 | 2.53448127 | 0.01126139 | 0.17500466 |
| TARS2 | 7.55999154 | -0.581527978 | 0.57544425 | -1.0105722 | 0.31222121 | NA |
| ECM1 | 42.2162917 | -0.925305107 | 0.9491425 | -0.9748853 | 0.32961714 | 0.70253658 |
| ADAMTSL4 | 8.45550432 | -0.870747172 | 1.150101 | -0.757105 | 0.44898698 | NA |
| MCL1 | 68.8892689 | -0.932156642 | 0.39961005 | -2.3326657 | 0.0196657 | 0.23460022 |
| ENSA | 76.3557839 | -0.420165494 | 0.22558878 | -1.8625283 | 0.06252866 | 0.385435 |
| GOLPH3L | 26.0119701 | 0.339950552 | 0.31958085 | 1.06373882 | 0.28744703 | NA |
| ARNT | 46.9307451 | 0.14466836 | 0.38543247 | 0.37534036 | 0.70740736 | 0.91058211 |
| SETDB1 | 20.9090009 | -0.586991123 | 0.34138531 | -1.7194387 | 0.08553451 | NA |
| CERS2 | 37.6428175 | -0.039832131 | 0.32839736 | -0.1212925 | 0.90345937 | 0.97237083 |
| MINDY1 | 6.54613876 | -0.722464187 | 0.55979159 | -1.2905949 | 0.19684418 | NA |
| PRUNE1 | 9.25613149 | 0.466254388 | 0.48550122 | 0.96035679 | 0.33687568 | NA |
| C1orf56 | 9.01605834 | 0.189515021 | 0.65122697 | 0.29101224 | 0.77104196 | NA |
| CDC42SE1 | 23.8197346 | -0.023018934 | 0.4012581 | -0.0573669 | 0.95425293 | NA |
| GABPB2 | 29.1551497 | -0.080666646 | 0.31883701 | -0.2530028 | 0.80026607 | 0.94085969 |
| SCNM1 | 5.37390043 | -0.265953132 | 0.75830799 | -0.3507191 | 0.72579906 | NA |
| VPS72 | 8.91496684 | -0.177029351 | 0.46139367 | -0.383684 | 0.7012127 | NA |
| PIP5K1A | 94.454942 | 0.458985185 | 0.28726326 | 1.59778588 | 0.11009064 | 0.49667521 |
| PSMD4 | 24.9672184 | -0.252636958 | 0.39480669 | -0.6399004 | 0.52223735 | NA |
| ZNF687 | 3.79280109 | -1.106312634 | 0.78118909 | -1.4161906 | 0.15671971 | NA |
| PI4KB | 24.5640358 | 0.513859566 | 0.3167389 | 1.62234436 | 0.10472963 | NA |
| PSMB4 | 18.8885299 | -0.165945524 | 0.56496133 | -0.293729 | 0.76896499 | NA |
| POGZ | 152.718728 | 0.346365243 | 0.2250274 | 1.53921365 | 0.12375215 | 0.51858498 |
| TUFT1 | 5.05688033 | 0.59762798 | 0.8543798 | 0.69948749 | 0.48424743 | NA |
| SNX27 | 36.0755473 | 0.311798361 | 0.3129392 | 0.99635443 | 0.31907797 | 0.69469143 |
| MRPL9 | 7.3877579 | -0.708873966 | 0.56677858 | -1.250707 | 0.21104138 | NA |
| RORC | 10.4529286 | 0.344963362 | 0.58737929 | 0.58729235 | 0.55700738 | NA |
| THEM4 | 16.3138388 | -0.40629064 | 0.43043878 | -0.9438988 | 0.34522137 | NA |
| S100A10 | 181.798877 | -2.768997122 | 0.42868489 | -6.4592832 | 1.05E-10 | 2.27E-07 |
| S100A11 | 175.541989 | -0.878714887 | 0.41577071 | -2.1134603 | 0.03456139 | 0.29210684 |
| FLG-AS1 | 24.9536563 | 0.965223275 | 0.41982392 | 2.29911456 | 0.02149844 | NA |
| S100A6 | 1132.44508 | -0.878457255 | 0.61326549 | -1.4324257 | 0.15202203 | 0.55086922 |
| S100A4 | 109.838061 | -0.715313239 | 0.96015685 | -0.7449962 | 0.45627399 | 0.79324789 |

| | | | | | | |
|---|---|---|---|---|---|---|
| S100A2 | 7.86529726 | -2.058518103 | 1.03447519 | -1.9899154 | 0.04660026 | NA |
| S100A16 | 72.4638741 | -0.085368569 | 0.4585251 | -0.1861808 | 0.85230298 | 0.95569766 |
| S100A13 | 50.5262761 | -1.056704439 | 0.40796564 | -2.59018 | 0.00959258 | 0.15497902 |
| S100A1 | 5.19285587 | -2.245573891 | 1.00536726 | -2.2335856 | 0.02551035 | NA |
| CHTOP | 24.1659145 | 0.047255661 | 0.32012052 | 0.14761834 | 0.88264398 | NA |
| SNAPIN | 8.38916547 | -0.500747648 | 0.5947907 | -0.8418888 | 0.39985018 | NA |
| ILF2 | 22.6919649 | -0.535742508 | 0.48380715 | -1.1073472 | 0.26814383 | NA |
| NPR1 | 78.7882736 | 0.181738746 | 0.48395567 | 0.37552767 | 0.70726807 | 0.91058211 |
| INTS3 | 41.7984309 | 0.576878898 | 0.26493599 | 2.17742744 | 0.02944869 | 0.27926427 |
| SLC27A3 | 4.96937757 | 0.241376773 | 0.78025503 | 0.30935625 | 0.75705054 | NA |
| GATAD2B | 72.9442497 | 0.071369679 | 0.29263354 | 0.24388755 | 0.80731791 | 0.94328109 |
| DENND4B | 10.3888881 | -0.283991908 | 0.4550117 | -0.624142 | 0.53253434 | NA |
| CRTC2 | 9.22663483 | -0.488535284 | 0.57172682 | -0.8544908 | 0.39283313 | NA |
| SLC39A1 | 31.7259555 | -0.897012111 | 0.35884283 | -2.4997354 | 0.01242861 | 0.18809868 |
| JTB | 28.7772718 | -0.544379703 | 0.48301034 | -1.127056 | 0.2597188 | 0.65595635 |
| RAB13 | 15.934395 | -1.651083551 | 0.55163542 | -2.9930703 | 0.00276186 | NA |
| RPS27 | 23.6138305 | -0.569786072 | 0.50766253 | -1.1223717 | 0.26170442 | NA |
| TPM3 | 60.1684172 | -0.484309379 | 0.32112316 | -1.5081733 | 0.13151017 | 0.52613957 |
| C1orf43 | 40.9304102 | -0.254891767 | 0.28073541 | -0.9079431 | 0.3639083 | 0.7336044 |
| UBAP2L | 29.5331683 | 0.115445737 | 0.31744399 | 0.36367278 | 0.71610237 | 0.91399852 |
| HAX1 | 13.8580493 | -0.851576394 | 0.50293183 | -1.6932243 | 0.09041278 | NA |
| ATP8B2 | 6.18764045 | -0.286462077 | 0.68548206 | -0.4178987 | 0.67602117 | NA |
| IL6R | 35.9384641 | -1.145754465 | 0.53918218 | -2.1249858 | 0.0335878 | 0.29047359 |
| UBE2Q1 | 26.6111113 | -0.021241276 | 0.44820762 | -0.0473916 | 0.96220112 | NA |
| ADAR | 52.7637831 | -0.41466996 | 0.4940275 | -0.8393661 | 0.40126388 | 0.76381845 |
| PMVK | 23.4179286 | -0.606420621 | 0.36687753 | -1.6529239 | 0.09834636 | NA |
| PBXIP1 | 23.5951388 | -0.226993688 | 0.42701099 | -0.5315875 | 0.59501175 | NA |
| PYGO2 | 11.1101658 | 0.279399275 | 0.44477507 | 0.62818107 | 0.52988533 | NA |
| SHC1 | 4.22280341 | -1.533293537 | 0.98506968 | -1.5565331 | 0.11958138 | NA |
| CKS1B | 5.5128193 | -0.968612055 | 0.64899086 | -1.4924895 | 0.13557086 | NA |
| FLAD1 | 5.18713436 | -0.455957151 | 0.78908819 | -0.5778279 | 0.56338034 | NA |
| ZBTB7B | 5.62761983 | -0.354747245 | 0.62307498 | -0.5693492 | 0.56911918 | NA |
| ADAM15 | 12.3389337 | 0.059122502 | 0.56202991 | 0.10519458 | 0.91622141 | NA |
| EFNA1 | 16.589111 | -2.710131181 | 0.7460994 | -3.6323996 | 0.0002808 | NA |
| SLC50A1 | 7.97866112 | -0.989752924 | 0.61612595 | -1.6064133 | 0.10818313 | NA |
| DPM3 | 21.4913576 | -0.211583726 | 0.43507337 | -0.4863173 | 0.62674218 | NA |
| THBS3 | 6.01312349 | 0.310796149 | 0.56779578 | 0.54737312 | 0.58412242 | NA |
| MTX1 | 7.08554307 | -0.567253442 | 0.63449522 | -0.8940232 | 0.37130949 | NA |
| GBA | 14.5163407 | 0.10554211 | 0.53113363 | 0.19871103 | 0.8424888 | NA |
| FAM189B | 6.41935713 | 0.084320719 | 0.57933044 | 0.14554857 | 0.88427777 | NA |
| SCAMP3 | 20.6747637 | 0.272040358 | 0.37710633 | 0.721389 | 0.47067022 | NA |
| CLK2 | 10.374843 | 0.301144841 | 0.43408685 | 0.69374329 | 0.48784321 | NA |
| FDPS | 23.3489291 | -0.278782506 | 0.39436478 | -0.7069153 | 0.4796191 | NA |
| RUSC1 | 8.51011288 | -0.659853367 | 0.58514278 | -1.1276792 | 0.25945541 | NA |
| ASH1L | 169.137242 | 0.100768823 | 0.26264722 | 0.38366605 | 0.70122598 | 0.90757606 |
| MSTO1 | 4.48632793 | 0.117130215 | 0.65592304 | 0.17857311 | 0.85827291 | NA |
| YY1AP1 | 40.8844811 | 0.131726771 | 0.25281096 | 0.52104851 | 0.60233298 | 0.87047761 |
| DAP3 | 33.4325947 | -0.35083217 | 0.33300053 | -1.0535484 | 0.29208973 | 0.67930583 |
| GON4L | 30.3880285 | 0.059421269 | 0.37452041 | 0.15865963 | 0.87393705 | 0.96229085 |
| RIT1 | 8.46551781 | -0.334274396 | 0.57588773 | -0.5804506 | 0.56161077 | NA |
| KHDC4 | 28.0723009 | -0.046981659 | 0.35630673 | -0.1318573 | 0.89509713 | NA |
| ARHGEF2 | 15.0014674 | -0.225669716 | 0.39157813 | -0.5763083 | 0.56440683 | NA |
| SSR2 | 34.8437901 | -0.411928006 | 0.51719515 | -0.7964653 | 0.42576162 | 0.77730788 |
| UBQLN4 | 7.20040921 | 0.020498827 | 0.59272824 | 0.03458385 | 0.97241158 | NA |
| LAMTOR2 | 11.6177217 | -0.30277418 | 0.51439962 | -0.5885972 | 0.55613151 | NA |
| LMNA | 68.5538061 | -0.128047081 | 0.34344891 | -0.3728272 | 0.70927709 | 0.91144677 |
| SLC25A44 | 7.52264655 | 0.091242167 | 0.48329242 | 0.18879288 | 0.85025515 | NA |
| SMG5 | 25.6159276 | 0.494267245 | 0.32847307 | 1.50474208 | 0.1323904 | NA |
| GLMP | 15.2293578 | 0.775440507 | 0.41818338 | 1.85430733 | 0.0636952 | NA |
| CCT3 | 34.5286151 | -0.373475948 | 0.38468711 | -0.9708564 | 0.33161979 | 0.70495276 |
| MEF2D | 27.5721971 | 0.232950312 | 0.52009608 | 0.44789862 | 0.65422637 | NA |
| NAXE | 8.9260302 | -0.410594837 | 0.61139675 | -0.6715686 | 0.5018584 | NA |
| GPATCH4 | 6.31857654 | -1.200423349 | 0.74142063 | -1.6190854 | 0.10542888 | NA |
| NES | 267.046714 | -0.075292138 | 0.59888018 | -0.1257215 | 0.89995235 | 0.97097328 |
| ISG20L2 | 8.80572501 | -0.255769032 | 0.57212761 | -0.4470489 | 0.65483974 | NA |
| RRNAD1 | 7.20787905 | -1.055418009 | 0.67025911 | -1.5746418 | 0.11533915 | NA |
| MRPL24 | 7.32589221 | -0.312751406 | 0.64611638 | -0.4840481 | 0.62835174 | NA |
| HDGF | 33.1583782 | -0.080630679 | 0.38644361 | -0.208648 | 0.83472305 | 0.95179548 |
| PRCC | 22.7142566 | 0.610429899 | 0.33581683 | 1.81774657 | 0.06910288 | NA |
| ARHGEF11 | 37.4781227 | 0.166197971 | 0.28960966 | 0.57386888 | 0.56605653 | 0.85573063 |
| ETV3 | 4.46151413 | -0.051450999 | 0.63758088 | -0.0806972 | 0.93568275 | NA |
| KIRREL1 | 578.826662 | -0.740776711 | 0.32794691 | -2.2588312 | 0.02389389 | 0.25172367 |
| IFI16 | 58.9014235 | -4.160578853 | 1.23600723 | -3.3661444 | 0.00076227 | 0.04122345 |
| DUSP23 | 44.4024775 | -0.377009381 | 0.47540826 | -0.7930224 | 0.4277648 | 0.77844087 |
| SLAMF8 | 21.8155206 | -1.855109218 | 0.72349975 | -2.5640772 | 0.01034505 | NA |
| AL590560.2 | 6.36671658 | 0.106636886 | 0.57727176 | 0.18472562 | 0.85344425 | NA |
| CFAP45 | 9.49803429 | -0.306749278 | 0.76442567 | -0.4012807 | 0.68821346 | NA |
| TAGLN2 | 59.9267045 | -0.431210165 | 0.52010624 | -0.8290809 | 0.40705861 | 0.76772515 |
| IGSF8 | 18.0190587 | 0.169017374 | 0.43555348 | 0.38805194 | 0.6979776 | NA |
| PEA15 | 104.316209 | 0.040914229 | 0.47398959 | 0.08631883 | 0.93121297 | 0.97832857 |
| DCAF8 | 28.030959 | 0.743755627 | 0.29011671 | 2.56364283 | 0.01035801 | NA |
| AL139011.2 | 6.14628026 | 0.630253398 | 0.61753336 | 1.02059814 | 0.30744487 | NA |
| COPA | 56.5235815 | -0.203219907 | 0.2727302 | -0.7451317 | 0.45619213 | 0.79324789 |
| NCSTN | 51.0152527 | 0.542201923 | 0.49144673 | 1.10327711 | 0.26990685 | 0.66267686 |
| F11R | 11.9652626 | 0.030192809 | 0.54369339 | 0.05553279 | 0.95571401 | NA |
| AL591806.3 | 9.53676354 | 0.075415902 | 0.49432903 | 0.15256215 | 0.87874357 | NA |
| USF1 | 5.97201906 | -0.275299892 | 0.76053525 | -0.3619818 | 0.71736566 | NA |
| KLHDC9 | 4.58821104 | -0.353434103 | 0.72054621 | -0.4905086 | 0.62377406 | NA |
| PFDN2 | 26.79264 | -0.906891334 | 0.43659252 | -2.0772031 | 0.03778282 | NA |
| NIT1 | 19.6045047 | -0.873950294 | 0.40533567 | -2.1561149 | 0.03107469 | NA |
| DEDD | 9.06814936 | 0.105883187 | 0.51622595 | 0.20511016 | 0.83748606 | NA |
| UFC1 | 30.5097128 | -0.728970203 | 0.4171908 | -1.7473305 | 0.08058003 | 0.43346499 |
| PPOX | 5.8929274 | 0.900994077 | 0.77985851 | 1.15533018 | 0.24795524 | NA |
| B4GALT3 | 12.4264787 | 0.043613617 | 0.4813508 | 0.09060672 | 0.92780509 | NA |
| ADAMTS4 | 6.10476129 | -3.467906929 | 1.46656816 | -2.3646408 | 0.01804756 | NA |
| NDUFS2 | 21.8902886 | -0.334639591 | 0.51884778 | -0.6449668 | 0.51894871 | NA |
| TOMM40L | 5.58488383 | 0.001746169 | 0.61878394 | 0.00282194 | 0.99774842 | NA |
| SDHC | 38.1578859 | -0.389345134 | 0.42173516 | -0.9231982 | 0.35590393 | 0.72759393 |
| AL592295.4 | 5.27855486 | -1.553343543 | 0.64334411 | -2.4144832 | 0.01575755 | NA |
| DUSP12 | 7.45930933 | -0.658078519 | 0.56358555 | -1.1676639 | 0.24294235 | NA |
| ATF6 | 75.2722595 | -0.396229073 | 0.2769325 | -1.4307785 | 0.1524937 | 0.55175242 |
| NOS1AP | 169.085755 | 0.083532805 | 0.38431247 | 0.21735648 | 0.82793054 | 0.94941654 |
| C1orf226 | 10.341644 | -0.515573053 | 0.46897504 | -1.0993614 | 0.27161046 | NA |
| UHMK1 | 51.0505285 | 0.046020591 | 0.3272173 | 0.14064229 | 0.88815254 | 0.96759245 |
| UAP1 | 10.4366355 | -0.434661739 | 0.57873183 | -0.751059 | 0.45261716 | NA |
| HSD17B7 | 8.30294417 | -0.615482633 | 0.56203549 | -1.0950957 | 0.27347472 | NA |
| PBX1 | 74.022809 | 0.851375258 | 0.3258987 | 2.61239227 | 0.0089911 | 0.14884862 |
| MGST3 | 69.7889092 | -0.34535785 | 0.36540768 | -0.9451302 | 0.34459239 | 0.71842682 |
| ALDH9A1 | 23.6182004 | -0.61568177 | 0.43504529 | -1.4152131 | 0.15700604 | NA |
| TMCO1 | 47.9266687 | -0.492417702 | 0.27706989 | -1.7772328 | 0.07552994 | 0.42138852 |
| UCK2 | 15.9535492 | -0.332520577 | 0.49090418 | -0.6773635 | 0.49817535 | NA |
| POGK | 17.0622745 | -0.444506904 | 0.37829732 | -1.17502 | 0.23998673 | NA |
| TADA1 | 6.28675358 | -1.109374086 | 0.5901234 | -1.8799019 | 0.06012145 | NA |

| | | | | | | |
|---|---|---|---|---|---|---|
| ILDR2 | 57.4352738 | 0.745836698 | 0.47636808 | 1.56567313 | 0.11742516 | 0.50775741 |
| MAEL | 8.71677813 | -0.258666772 | 0.8036371 | -0.3218701 | 0.74755109 | NA |
| POU2F1 | 68.9464951 | -0.061695506 | 0.23974635 | -0.2573366 | 0.79691895 | 0.93934998 |
| CREG1 | 15.9302871 | -0.219454039 | 0.49744749 | -0.4411602 | 0.65909702 | NA |
| MPZL1 | 36.4234293 | 0.292863854 | 0.35835192 | 0.81725208 | 0.41378438 | 0.76946282 |
| MPC2 | 23.1823259 | -0.486288411 | 0.46211587 | -1.0523084 | 0.29265808 | NA |
| DCAF6 | 107.344551 | 0.716031262 | 0.22020413 | 3.2516705 | 0.00114729 | 0.05186671 |
| TIPRL | 17.1094217 | 0.309295923 | 0.37773775 | 0.81881126 | 0.4128941 | NA |
| SFT2D2 | 23.4209205 | -0.280984225 | 0.32083649 | -0.8757864 | 0.38114618 | NA |
| ATP1B1 | 230.00172 | 0.13982351 | 0.54666266 | 0.25577659 | 0.79812335 | 0.94015741 |
| NME7 | 58.6011196 | -0.005349083 | 0.27742525 | -0.0192812 | 0.98461681 | 0.99585746 |
| BLZF1 | 22.8394963 | -0.108271197 | 0.4237927 | -0.2554815 | 0.79835123 | NA |
| SLC19A2 | 7.37282041 | 0.002429588 | 0.5510535 | 0.00440899 | 0.99648215 | NA |
| F5 | 66.8173247 | 0.50468206 | 0.47458135 | 1.06342581 | 0.28758889 | 0.67542591 |
| C1orf112 | 5.2189432 | -0.017061427 | 0.65337632 | -0.0261127 | 0.97916744 | NA |
| SCYL3 | 11.4713554 | -0.075840771 | 0.38979051 | -0.194568 | 0.84573113 | NA |
| KIFAP3 | 44.1518899 | -0.007246513 | 0.25056374 | -0.0289208 | 0.97692773 | 0.99298531 |
| FMO4 | 12.4037638 | 0.675631403 | 0.60082169 | 1.12451233 | 0.26079574 | NA |
| PRRC2C | 122.475978 | -0.155135909 | 0.25330966 | -0.6124358 | 0.54024946 | 0.83956007 |
| VAMP4 | 15.8579865 | 0.046547341 | 0.42830834 | 0.10867718 | 0.91345855 | NA |
| METTL13 | 5.22816228 | -0.036719566 | 0.64666373 | -0.0567831 | 0.95471798 | NA |
| DNM3 | 31.0345472 | 0.938695115 | 0.51741304 | 1.81420848 | 0.06964564 | 0.40663921 |
| PIGC | 26.0049672 | -0.357940403 | 0.3409676 | -1.0497784 | 0.29382003 | NA |
| SUCO | 56.7838897 | -0.129958702 | 0.31259108 | -0.4157467 | 0.67759539 | 0.90499319 |
| AL645568.1 | 15.3363371 | 0.662315861 | 0.55149759 | 1.20094062 | 0.22977423 | NA |
| PRDX6 | 115.425267 | -0.307948652 | 0.35282331 | -0.8728127 | 0.3827652 | 0.74924885 |
| KLHL20 | 23.2063509 | -0.063510091 | 0.32416006 | -0.195922 | 0.84467121 | NA |
| DARS2 | 7.08934149 | -0.240921986 | 0.65374714 | -0.3685247 | 0.71248201 | NA |
| ZBTB37 | 17.7093864 | 0.043418879 | 0.36127079 | 0.12018375 | 0.90433759 | NA |
| RC3H1 | 72.6124611 | 0.165319254 | 0.26502392 | 0.62378995 | 0.53276554 | 0.83658422 |
| RABGAP1L | 680.187164 | 1.209921061 | 0.3721659 | 3.25102616 | 0.00114989 | 0.05186671 |
| CACYBP | 10.38516 | -0.300148158 | 0.64965952 | -0.4620084 | 0.6440753 | NA |
| MRPS14 | 13.2664342 | -0.605767168 | 0.43677364 | -1.3869133 | 0.16546821 | NA |
| KIAA0040 | 85.9532955 | -1.345995165 | 0.58750838 | -2.2910229 | 0.02196209 | 0.24297341 |
| COP1 | 113.311524 | 0.319746796 | 0.20313231 | 1.57408141 | 0.11546863 | 0.50513044 |
| RASAL2 | 119.739164 | 0.026557688 | 0.29257093 | 0.0907735 | 0.92767257 | 0.97775374 |
| RALGPS2 | 31.4581802 | 0.24645701 | 0.5217487 | 0.47236727 | 0.63666466 | 0.88633797 |
| ANGPTL1 | 9.54576408 | -1.380472968 | 0.98245248 | -1.4051295 | 0.15998277 | NA |
| FAM20B | 62.6654127 | 0.208289379 | 0.40713466 | 0.51159825 | 0.60893221 | 0.8725395 |
| TOR3A | 12.6486099 | -0.508083628 | 0.52343203 | -0.9706774 | 0.33170896 | NA |
| ABL2 | 54.4007473 | 0.238045337 | 0.31651942 | 0.75207183 | 0.45200786 | 0.7927748 |
| SOAT1 | 16.4211497 | -0.40594287 | 0.48766897 | -0.8324148 | 0.40517485 | NA |
| AXDND1 | 34.7410553 | -0.475660655 | 0.6478158 | -0.7342529 | 0.4627946 | 0.79793427 |
| AL160286.3 | 11.6891068 | 0.588315006 | 0.92320894 | 0.63725012 | 0.52396194 | NA |
| NPHS2 | 1793.17469 | -0.0913227 | 0.38526281 | -0.23704 | 0.81262575 | 0.94415747 |
| TDRD5 | 29.0759937 | -0.051527853 | 0.53108885 | -0.097023 | 0.9227081 | 0.97775374 |
| TOR1AIP2 | 59.5136265 | -0.092866592 | 0.26794572 | -0.3465873 | 0.72890136 | 0.91700494 |
| TOR1AIP1 | 32.4343367 | 0.082721946 | 0.30823084 | 0.2683766 | 0.78840945 | 0.93749303 |
| CEP350 | 71.3860278 | 0.265468442 | 0.23807368 | 1.11506844 | 0.2648211 | 0.65974932 |
| QSOX1 | 33.0931525 | 0.457194025 | 0.45900411 | 0.99605649 | 0.3192227 | 0.69469143 |
| ACBD6 | 52.8378346 | 0.120694246 | 0.30604275 | 0.39437055 | 0.69330748 | 0.90557575 |
| XPR1 | 51.7660315 | 0.34915442 | 0.30800602 | 1.13359609 | 0.25696402 | 0.65588339 |
| STX6 | 12.2321517 | 0.205016523 | 0.39899399 | 0.51383361 | 0.60736833 | NA |
| MR1 | 14.9232734 | 0.270961309 | 0.44049025 | 0.61513578 | 0.53846506 | NA |
| IER5 | 4.92858781 | -0.566744758 | 1.17166264 | -0.4837098 | 0.6285918 | NA |
| GLUL | 105.4777 | -1.853517666 | 0.48224761 | -3.8434979 | 0.00012129 | 0.01171344 |
| RNASEL | 8.11192851 | -0.283474916 | 0.53109603 | -0.5337545 | 0.59351137 | NA |
| DHX9 | 45.4378873 | -0.049854423 | 0.24210556 | -0.2059202 | 0.83685326 | 0.95264575 |
| LAMC1 | 39.5670976 | 0.62101012 | 0.35683041 | 1.74035086 | 0.08179743 | 0.43779104 |
| SMG7 | 49.8946896 | 0.393442297 | 0.26063522 | 1.50955153 | 0.1311579 | 0.52613957 |
| ARPC5 | 73.781554 | -0.240416213 | 0.38673908 | -0.6216496 | 0.53417227 | 0.83677705 |
| RGL1 | 60.3474846 | -0.610185023 | 0.53231912 | -1.1462767 | 0.25168066 | 0.65183754 |
| COLGALT2 | 28.8202083 | 1.013548987 | 0.45223377 | 2.2412059 | 0.02501274 | 0.25362919 |
| TSEN15 | 52.0739822 | 0.349375776 | 0.355919 | 0.98161598 | 0.32628907 | 0.69764285 |
| C1orf21 | 204.938925 | -1.510912827 | 0.57738274 | -2.6168306 | 0.00887504 | 0.14875902 |
| EDEM3 | 62.6850706 | 0.635501026 | 0.28475035 | 2.23178315 | 0.0256293 | 0.25731925 |
| RNF2 | 16.8249938 | 0.434902199 | 0.47478845 | 0.91599153 | 0.35967134 | NA |
| TRMT1L | 19.7712727 | 0.157866976 | 0.38980255 | 0.40499216 | 0.68548328 | NA |
| SWT1 | 17.7344583 | 0.023384241 | 0.46691964 | 0.05008194 | 0.96005709 | NA |
| IVNS1ABP | 38.4192087 | 0.46006848 | 0.48235011 | 0.95380611 | 0.3401818 | 0.71508205 |
| TPR | 42.3354407 | -0.573528925 | 0.26249055 | -2.1849507 | 0.02889246 | 0.27926427 |
| ODR4 | 25.582664 | -0.25215679 | 0.3363357 | -0.7497176 | 0.45342482 | NA |
| AL596220.1 | 5.76327842 | -0.421198893 | 0.59600388 | -0.7067049 | 0.47974985 | NA |
| ERVMER61-1 | 30.2913104 | -0.084762402 | 0.47660208 | -0.1778473 | 0.85884288 | 0.95804916 |
| AL390957.1 | 7.75398779 | -0.553399192 | 0.73269859 | -0.755289 | 0.45007559 | NA |
| AL136454.1 | 24.5369946 | -0.652462892 | 0.53833658 | -1.2119981 | 0.22551312 | NA |
| UCHL5 | 19.4749739 | 0.132989868 | 0.34518556 | 0.38527066 | 0.7000369 | NA |
| TROVE2 | 76.5815378 | -0.204930121 | 0.35249233 | -0.5813747 | 0.56098795 | 0.85386276 |
| GLRX2 | 10.8211022 | -0.95676704 | 0.53908912 | -1.7747846 | 0.07593347 | NA |
| CDC73 | 70.4549569 | 0.311650241 | 0.25396092 | 1.22715825 | 0.21976312 | 0.62462904 |
| B3GALT2 | 16.3180084 | 2.526715286 | 0.65208167 | 3.87484481 | 0.00010669 | NA |
| LINC01031 | 19.7711403 | 1.222941972 | 0.55611294 | 2.19908924 | 0.02787158 | NA |
| AL357793.1 | 11.0200464 | 1.131835364 | 0.51289076 | 2.20677666 | 0.02732967 | NA |
| AL356272.1 | 6.61413263 | 0.629000412 | 0.57058644 | 1.10237533 | 0.27029853 | NA |
| KCNT2 | 150.738673 | -2.689672722 | 0.62777918 | -4.2844248 | 1.83E-05 | 0.00285812 |
| CFH | 68.3230404 | -1.91102491 | 0.58538476 | -3.2645621 | 0.00109633 | 0.05186671 |
| ZBTB41 | 23.5404092 | -0.183817125 | 0.37187961 | -0.494292 | 0.62109996 | NA |
| CRB1 | 7.58867926 | -0.589066808 | 0.79494346 | -0.7410172 | 0.45868299 | NA |
| DENND1B | 57.3362204 | 0.377392708 | 0.245061 | 1.53999499 | 0.12356157 | 0.51858498 |
| NEK7 | 90.4895756 | 0.38370279 | 0.21232302 | 1.80716531 | 0.07073651 | 0.40760715 |
| MIR181A1HG | 8.29232939 | 1.718570335 | 0.87867427 | 1.95586737 | 0.05048078 | NA |
| LINC00862 | 6.87084582 | 0.739561827 | 0.75678871 | 0.97723687 | 0.3284519 | NA |
| ZNF281 | 9.60664696 | 0.100871367 | 0.45639775 | 0.22101635 | 0.8250797 | NA |
| DDX59 | 10.6523352 | -0.558877322 | 0.44967567 | -1.2428454 | 0.21392482 | NA |
| CAMSAP2 | 135.483303 | 0.517369549 | 0.18456297 | 2.80321421 | 0.00505961 | 0.11214076 |
| TMEM9 | 18.4163195 | -0.161169266 | 0.45641427 | -0.3531206 | 0.72399806 | NA |
| TNNT2 | 133.90212 | 0.819753561 | 0.6524683 | 1.25638833 | 0.20897521 | 0.61736443 |
| TNNI1 | 58.4086274 | 1.423313672 | 0.79333186 | 1.79409619 | 0.07279782 | 0.417635 |
| PHLDA3 | 11.2028237 | -0.595736003 | 0.6056316 | -0.9836607 | 0.32528237 | NA |
| CSRP1 | 108.265857 | 0.15000025 | 0.44016769 | 0.34077979 | 0.73326937 | 0.91884473 |
| AC096677.1 | 7.9561329 | 0.133689699 | 0.52011137 | 0.25704053 | 0.79714749 | NA |
| NAV1 | 15.1112977 | 0.692155087 | 0.84014005 | 0.82385679 | 0.41002093 | NA |
| IPO9-AS1 | 5.0829248 | 0.320013873 | 1.04580901 | 0.30599648 | 0.75960733 | NA |
| IPO9 | 29.285479 | -0.178754165 | 0.27698405 | -0.6453591 | 0.51869452 | 0.83104823 |
| SHISA4 | 7.6398556 | -0.348844234 | 0.62955583 | -0.5541117 | 0.57950242 | NA |
| TIMM17A | 13.792263 | -0.294665588 | 0.50967822 | -0.5781404 | 0.56316931 | NA |
| RNPEP | 15.9832748 | 0.160035074 | 0.36293427 | 0.44094782 | 0.65925077 | NA |
| ELF3 | 12.8497662 | -0.797441511 | 0.66941705 | -1.1912477 | 0.23355636 | NA |
| ARL8A | 25.123684 | 0.008668526 | 0.36486292 | 0.02375831 | 0.98104539 | NA |
| PTPN7 | 33.4410623 | 0.496945166 | 0.3423741 | 1.45146835 | 0.14664949 | 0.54619864 |
| PPP1R12B | 66.9795045 | 0.635566596 | 0.40241355 | 1.57938668 | 0.11424739 | 0.50483797 |
| KDM5B | 49.5822865 | 0.011753959 | 0.3288094 | 0.03574703 | 0.97148407 | 0.99028384 |
| RABIF | 6.3042286 | -0.00537216 | 0.65804963 | -0.0081638 | 0.99348633 | NA |

| | | | | | | |
|---|---|---|---|---|---|---|
| KLHL12 | 14.0563146 | 0.529121064 | 0.43431205 | 1.21829699 | 0.22311113 | NA |
| ADIPOR1 | 15.5281074 | -0.025625195 | 0.38071235 | -0.0673085 | 0.94633607 | NA |
| CYB5R1 | 15.8569553 | -0.140515757 | 0.45029111 | -0.3120554 | 0.75499845 | NA |
| TMEM183A | 7.31313906 | 0.457166702 | 0.5960645 | 0.76697523 | 0.44309624 | NA |
| PPFIA4 | 60.9968032 | 0.77011732 | 0.54559707 | 1.41151294 | 0.15809342 | 0.56051304 |
| ADORA1 | 76.307887 | 0.991865861 | 0.57131089 | 1.73612279 | 0.08254213 | 0.43993543 |
| CHI3L1 | 356.758073 | -1.189486332 | 0.56131985 | -2.1190883 | 0.034083 | 0.29210684 |
| BTG2 | 48.025666 | -1.385628109 | 0.70040254 | -1.9783311 | 0.04789137 | 0.3419103 |
| ATP2B4 | 41.0261954 | -0.69080372 | 0.47240372 | -1.4623164 | 0.14365452 | 0.54150812 |
| SNRPE | 8.73282406 | -0.987711763 | 0.69235849 | -1.4265901 | 0.15369808 | NA |
| SOX13 | 16.1496922 | -0.181809143 | 0.45199323 | -0.4022386 | 0.68750841 | NA |
| PLEKHA6 | 18.6741153 | -0.524906141 | 0.4170579 | -1.258593 | 0.20817739 | NA |
| PPP1R15B | 13.277756 | -0.214751216 | 0.38969946 | -0.5510688 | 0.5815865 | NA |
| MDM4 | 47.4444171 | 0.612744904 | 0.25670111 | 2.38699753 | 0.01698661 | 0.21530523 |
| LRRN2 | 12.6218822 | 0.301022878 | 0.60902 | 0.4942742 | 0.62111254 | NA |
| NFASC | 627.851936 | 0.151315224 | 0.45692129 | 0.33116256 | 0.7405217 | 0.92003444 |
| RBBP5 | 13.7550622 | 0.320232303 | 0.46729818 | 0.68528473 | 0.49316427 | NA |
| DSTYK | 54.5154925 | -0.982598422 | 0.33222888 | -2.9575948 | 0.00310049 | 0.0904424 |
| NUAK2 | 14.0573974 | 0.476458679 | 0.66509153 | 0.71638062 | 0.47375636 | NA |
| CDK18 | 7.48174313 | -1.55199097 | 0.76959682 | -2.0166286 | 0.04373428 | NA |
| ELK4 | 25.3298598 | 0.113225321 | 0.36406799 | 0.31100049 | 0.75580025 | NA |
| NUCKS1 | 139.167057 | -0.174478182 | 0.25993983 | -0.6712253 | 0.50207704 | 0.82522868 |
| RAB29 | 7.64758212 | -0.433663344 | 0.86782026 | -0.4997156 | 0.61727533 | NA |
| SLC41A1 | 9.87758959 | 0.150397904 | 0.50082383 | 0.30030101 | 0.76394756 | NA |
| RHEX | 31.8728281 | 1.241116177 | 0.91396725 | 1.35794382 | 0.17448151 | 0.57784404 |
| SRGAP2 | 372.392626 | 0.234642199 | 0.25607378 | 0.91630699 | 0.3595059 | 0.72980778 |
| RASSF5 | 8.54511806 | -0.065852821 | 0.65577012 | -0.1004206 | 0.92001042 | NA |
| EIF2D | 8.89905044 | 0.191769893 | 0.48548109 | 0.39501002 | 0.6928355 | NA |
| DYRK3 | 8.26724928 | 0.480449965 | 0.56232619 | 0.85439727 | 0.39288491 | NA |
| MAPKAPK2 | 34.9085421 | -0.160572843 | 0.40355805 | -0.3978928 | 0.69070921 | 0.90557575 |
| PIGR | 18.4896086 | 0.03199483 | 1.44832544 | 0.02209091 | 0.98237544 | NA |
| PFKFB2 | 8.74302032 | -1.238872428 | 0.79383212 | -1.5606227 | 0.1186128 | NA |
| CD55 | 46.5053025 | -1.285721138 | 0.4579082 | -2.8078142 | 0.0049879 | 0.11177304 |
| CR1 | 235.815543 | 0.861868669 | 0.48333571 | 1.78316778 | 0.074559 | 0.41891814 |
| CR1L | 5.8940625 | 0.221687337 | 0.58984149 | 0.37584222 | 0.7070342 | NA |
| CD46 | 139.39855 | -0.26084055 | 0.33178167 | -0.7861813 | 0.43176127 | 0.77901411 |
| MIR29B2CH( | 83.3911457 | 0.104014265 | 0.3637566 | 0.28594468 | 0.77492049 | 0.9282568 |
| PLXNA2 | 32.5517492 | -2.051949465 | 0.78012616 | -2.630279 | 0.00853148 | 0.14787902 |
| AL031316.1 | 10.3273697 | -0.509948399 | 0.51674571 | -0.9868459 | 0.32371818 | NA |
| G0S2 | 344.399327 | -1.964634252 | 0.82274503 | -2.3879017 | 0.01694487 | 0.21530523 |
| HSD11B1 | 5.55479525 | -1.24545255 | 0.92791754 | -1.3422017 | 0.17953059 | NA |
| UTP25 | 4.17257153 | -0.88157496 | 0.81079328 | -1.0872993 | 0.27690456 | NA |
| HHAT | 8.70610767 | -0.058048864 | 0.83406308 | -0.0695977 | 0.94451388 | NA |
| RCOR3 | 64.6395788 | 0.344850247 | 0.29537216 | 1.167511 | 0.24300407 | 0.64918976 |
| TRAF5 | 39.3076865 | 0.466313162 | 0.40217795 | 1.15946974 | 0.24626476 | 0.65061974 |
| LINC00467 | 11.0254925 | 0.102944595 | 0.52633607 | 0.19558719 | 0.84493328 | NA |
| SLC30A1 | 20.887884 | 0.129165168 | 0.48004669 | 0.26906793 | 0.78787741 | NA |
| LPGAT1 | 49.7928742 | -0.231337125 | 0.26920553 | -0.8593327 | 0.39015697 | 0.7553588 |
| INTS7 | 13.3894557 | 0.061085688 | 0.41092178 | 0.14865527 | 0.88182566 | NA |
| PPP2R5A | 62.4306585 | 0.068628258 | 0.2997041 | 0.22898672 | 0.81887924 | 0.94694381 |
| AL360091.3 | 9.02604695 | -0.666451972 | 0.95886494 | -0.6950426 | 0.48702861 | NA |
| TMEM206 | 5.75459644 | -0.258472085 | 0.70421262 | -0.367037 | 0.71359142 | NA |
| NENF | 19.9530947 | -0.239125683 | 0.40618159 | -0.5887162 | 0.55605167 | NA |
| ATF3 | 4.78611235 | 0.129765467 | 0.98200393 | 0.13214353 | 0.89487077 | NA |
| NSL1 | 27.3773982 | 0.074078438 | 0.32296101 | 0.2293727 | 0.81857926 | NA |
| TATDN3 | 8.16215227 | -0.350406873 | 0.67069338 | -0.5224546 | 0.60135382 | NA |
| FLVCR1 | 14.5312661 | -0.472509043 | 0.36490116 | -1.294896 | 0.19535612 | NA |
| ANGEL2 | 23.7676849 | 0.126115166 | 0.34743903 | 0.36298503 | 0.71661607 | NA |
| RPS6KC1 | 75.85705 | 0.785718881 | 0.32006536 | 2.45487006 | 0.01409355 | 0.19677241 |
| PROX1 | 4.56899895 | -1.916079123 | 0.93473239 | -2.0498692 | 0.0403772 | NA |
| SMYD2 | 21.3324926 | 0.271079294 | 0.48179642 | 0.56264282 | 0.57367813 | NA |
| PTPN14 | 385.289457 | 0.429403579 | 0.20913192 | 2.0532666 | 0.04004673 | 0.31540889 |
| KCTD3 | 42.115296 | 0.489861078 | 0.29512808 | 1.6598254 | 0.09694958 | 0.47093115 |
| ESRRG | 179.673965 | 1.529090241 | 0.56882823 | 2.68814056 | 0.00718511 | 0.13427683 |
| GPATCH2 | 60.9183069 | 0.304655331 | 0.25055584 | 1.21591791 | 0.2240162 | 0.63029911 |
| SPATA17 | 10.531001 | 0.648023723 | 0.5273877 | 1.22874258 | 0.21916833 | NA |
| RRP15 | 15.278035 | -0.489617217 | 0.51666923 | -0.9476415 | 0.34331198 | NA |
| TGFB2 | 8.55477524 | -1.350176992 | 0.93553475 | -1.4432141 | 0.14896016 | NA |
| LYPLAL1 | 56.6295641 | 0.451629198 | 0.39756359 | 1.13599235 | 0.25595977 | 0.65588339 |
| EPRS | 31.7739253 | -0.351303452 | 0.3020655 | -1.1630042 | 0.24482778 | 0.64996527 |
| BPNT1 | 6.36437816 | 0.145796309 | 0.72759244 | 0.20038184 | 0.84118196 | NA |
| IARS2 | 32.9176508 | 0.255767716 | 0.28784442 | 0.88856235 | 0.37423833 | 0.74189183 |
| RAB3GAP2 | 108.267088 | 0.202420562 | 0.22460369 | 0.90123436 | 0.36746373 | 0.73626927 |
| MARK1 | 45.5351111 | -0.318335089 | 0.41724814 | -0.7629395 | 0.44549948 | 0.78802698 |
| C1orf115 | 26.3413327 | -1.227978924 | 0.54536902 | -2.2516478 | 0.02434454 | NA |
| 2-Mar | 45.3661507 | -0.099660829 | 0.307914 | -0.3236645 | 0.74619205 | 0.92250642 |
| MIA3 | 26.0374899 | -0.448779044 | 0.32068912 | -1.3994209 | 0.16168681 | NA |
| AIDA | 15.6335308 | 0.519818055 | 0.37170316 | 1.3984763 | 0.16197008 | NA |
| BROX | 25.6358368 | 0.431952298 | 0.32724614 | 1.31996149 | 0.18684788 | NA |
| FAM177B | 8.08881513 | -1.035154631 | 0.86711987 | -1.1937849 | 0.23256211 | NA |
| AC093152.1 | 14.9810181 | 0.621221844 | 0.48003568 | 1.29411597 | 0.19562537 | NA |
| DISP1 | 8.28577321 | 0.832428801 | 0.54660085 | 1.52291896 | 0.12777898 | NA |
| SUSD4 | 8.26698636 | -1.292266519 | 0.79221984 | -1.6311969 | 0.10284879 | NA |
| CAPN2 | 314.703878 | -0.306285167 | 0.26453116 | -1.1578415 | 0.24692871 | 0.65061974 |
| TP53BP2 | 24.9288963 | 0.416071297 | 0.31517116 | 1.32014393 | 0.18678697 | NA |
| FBXO28 | 28.5897718 | -0.15651441 | 0.31658607 | -0.4943819 | 0.62103653 | 0.8780438 |
| DEGS1 | 19.1247684 | 0.083213037 | 0.45693151 | 0.18211271 | 0.85549427 | NA |
| NVL | 22.1535173 | 0.412838467 | 0.35125353 | 1.17532902 | 0.23986311 | NA |
| CNIH4 | 25.1145747 | -0.17999346 | 0.33623146 | -0.5353261 | 0.59242441 | NA |
| WDR26 | 107.522186 | 0.342897489 | 0.29288475 | 1.1707591 | 0.24169562 | 0.64918976 |
| CNIH3 | 36.2503687 | 0.309444996 | 0.35690665 | 0.86701942 | 0.38593137 | 0.75125852 |
| DNAH14 | 61.3141807 | -0.156734357 | 0.5120436 | -0.3060957 | 0.75953176 | 0.92313949 |
| LBR | 6.81323564 | -0.413708548 | 0.71011605 | -0.5825929 | 0.56016741 | NA |
| ENAH | 11.8419745 | 0.200771611 | 0.48609126 | 0.41303276 | 0.67958262 | NA |
| SRP9 | 55.742322 | -0.363091855 | 0.32544391 | -1.1156818 | 0.26455835 | 0.65974932 |
| EPHX1 | 35.6939546 | -0.998952197 | 0.52791817 | -1.8922482 | 0.05845792 | 0.37650386 |
| AL591895.1 | 5.85036038 | -0.382741609 | 0.61191628 | -0.6254804 | 0.53165584 | NA |
| TMEM63A | 9.45831068 | -0.435880925 | 0.52400903 | -0.8318195 | 0.40551084 | NA |
| SDE2 | 7.49821373 | -1.165713392 | 0.60334435 | -1.9320864 | 0.05334885 | NA |
| H3F3A | 9.74587383 | -1.068911179 | 0.5795654 | -1.8443323 | 0.06513473 | NA |
| ACBD3 | 56.4977061 | -0.568695695 | 0.25687778 | -2.2138766 | 0.02683727 | 0.26549261 |
| PARP1 | 26.5381026 | -0.28407246 | 0.3428035 | -0.8286743 | 0.40728872 | NA |
| ITPKB | 7.36672378 | -0.579718707 | 0.75917034 | -0.7636214 | 0.4450929 | NA |
| PSEN2 | 10.6044727 | -0.959782114 | 0.54887821 | -1.7486249 | 0.08035588 | NA |
| COQ8A | 11.6855147 | -0.830363365 | 0.67768553 | -1.225293 | 0.22046483 | NA |
| CDC42BPA | 202.832651 | 0.198170996 | 0.24808245 | 0.79881102 | 0.4244 | 0.77714058 |
| ZNF678 | 22.3791879 | 0.610616824 | 0.35300429 | 1.72977169 | 0.08367107 | NA |
| SNAP47 | 13.2791961 | -0.758022975 | 0.51965242 | -1.4587115 | 0.14464452 | NA |
| JMJD4 | 5.22806584 | 0.080889496 | 0.57959764 | 0.13956147 | 0.88900649 | NA |
| ARF1 | 110.745539 | -0.523463467 | 0.3411147 | -1.5345673 | 0.12489015 | 0.52114654 |
| C1orf35 | 7.1050396 | -0.10756875 | 0.49255988 | -0.2183872 | 0.82712747 | NA |
| MRPL55 | 13.9031202 | 0.264435265 | 0.41378426 | 0.63906554 | 0.5227803 | NA |
| GUK1 | 41.4659227 | -0.220065101 | 0.49603541 | -0.443648 | 0.65729712 | 0.89703806 |

| | | | | | | |
|---|---|---|---|---|---|---|
| OBSCN | 21.1866645 | -0.515071881 | 0.48740589 | -1.0567617 | 0.29062035 | NA |
| TRIM11 | 6.80732707 | 0.175918545 | 0.55695084 | 0.3158601 | 0.7521087 | NA |
| RNF187 | 27.3108296 | -0.165144046 | 0.33153952 | -0.4981127 | 0.6184046 | NA |
| RHOU | 7.69956963 | -1.552361197 | 0.85543524 | -1.8147034 | 0.06956951 | NA |
| RAB4A | 22.1284148 | -0.125565791 | 0.39945337 | -0.3143441 | 0.75325974 | NA |
| CCSAP | 4.71934092 | 0.074674681 | 0.73670206 | 0.10136347 | 0.91926193 | NA |
| NUP133 | 19.9349254 | 0.246216223 | 0.33624192 | 0.73225915 | 0.46401041 | NA |
| ABCB10 | 12.541913 | -0.002769167 | 0.46394473 | -0.0059687 | 0.99523766 | NA |
| TAF5L | 7.14713306 | -0.073790093 | 0.51745161 | -0.1426029 | 0.88660382 | NA |
| GALNT2 | 35.7429025 | 0.224803052 | 0.2583465 | 0.870161 | 0.38421242 | 0.75029638 |
| COG2 | 10.7167924 | -0.348858228 | 0.50512616 | -0.6906358 | 0.48979442 | NA |
| C1orf198 | 15.6009875 | 0.292601992 | 0.47095442 | 0.62129577 | 0.53440503 | NA |
| TTC13 | 17.9029493 | 0.076398383 | 0.45001821 | 0.16976731 | 0.86519313 | NA |
| ARV1 | 11.6035627 | -0.048520365 | 0.49902257 | -0.0972308 | 0.92254311 | NA |
| FAM89A | 158.924018 | -0.575019542 | 0.4832347 | -1.1899384 | 0.23407059 | 0.63975088 |
| C1orf131 | 5.4267533 | -1.309545072 | 0.63519797 | -2.061633 | 0.03924269 | NA |
| GNPAT | 22.1196429 | -0.210953796 | 0.34323514 | -0.6146043 | 0.53881607 | NA |
| EXOC8 | 6.35505562 | -0.208950724 | 0.56613143 | -0.3690852 | 0.71206423 | NA |
| SPRTN | 5.94459453 | -0.098133825 | 0.56333201 | -0.1742025 | 0.86170635 | NA |
| EGLN1 | 42.9267718 | -0.141964447 | 0.27356934 | -0.5189341 | 0.60380672 | 0.87047761 |
| DISC1 | 7.48768301 | -0.382687535 | 0.60373928 | -0.6338622 | 0.52617073 | NA |
| SIPA1L2 | 19.6297304 | -0.639272701 | 0.3965933 | -1.61191 | 0.10698153 | NA |
| NTPCR | 10.7487643 | -0.598668214 | 0.46253856 | -1.2943098 | 0.19555842 | NA |
| PCNX2 | 46.0709303 | 0.277964229 | 0.34014397 | 0.81719581 | 0.41381653 | 0.76946282 |
| MAP3K21 | 16.8515464 | 0.197865475 | 0.4802436 | 0.41201064 | 0.68033162 | NA |
| COA6 | 7.88191583 | -1.290286873 | 0.55948472 | -2.3062057 | 0.02109914 | NA |
| TARBP1 | 17.0637084 | -0.195003447 | 0.56794664 | -0.3433482 | 0.73133654 | NA |
| IRF2BP2 | 28.557885 | -0.93014648 | 0.4787682 | -1.9427909 | 0.05204143 | 0.35776279 |
| TOMM20 | 47.1822664 | -0.748173557 | 0.40858297 | -1.8311423 | 0.06707931 | 0.40247583 |
| ARID4B | 122.060297 | 0.36500802 | 0.22349913 | 1.63315184 | 0.10243707 | 0.48317363 |
| B3GALNT2 | 12.4196935 | -0.16518012 | 0.515274 | -0.3205675 | 0.74853814 | NA |
| LYST | 56.9919881 | 0.355459818 | 0.45198088 | 0.78644879 | 0.43160461 | 0.77901411 |
| NID1 | 5.26226662 | -0.597174116 | 0.86004152 | -0.694355 | 0.48745962 | NA |
| GPR137B | 10.4754058 | -0.051617166 | 0.54531283 | -0.0946561 | 0.92458802 | NA |
| ERO1B | 31.1490094 | -0.445059736 | 0.3764318 | -1.1823117 | 0.23708203 | 0.64321384 |
| EDARADD | 9.30325703 | -0.547805129 | 0.62240902 | -0.8801369 | 0.37878517 | NA |
| LGALS8 | 111.027129 | -0.559373921 | 0.42348998 | -1.3208669 | 0.18654573 | 0.58835886 |
| HEATR1 | 17.4368455 | -0.949090892 | 0.36451969 | -2.6036752 | 0.00922301 | NA |
| MTR | 66.6290526 | 0.307602261 | 0.30410152 | 1.01151176 | 0.31177156 | 0.69138624 |
| MT1HL1 | 7.26721492 | 2.290014132 | 2.31052675 | 0.9911221 | 0.32162596 | NA |
| RYR2 | 47.2601797 | -0.909292823 | 0.95777524 | -0.9493802 | 0.34242729 | 0.71725052 |
| FMN2 | 843.162557 | -0.279028134 | 0.21641916 | -1.289295 | 0.19729556 | 0.59987316 |
| RGS7 | 15.7430319 | -0.010759346 | 0.80538912 | -0.0133592 | 0.98934123 | NA |
| FH | 10.3790381 | -0.178669754 | 0.48147461 | -0.3710886 | 0.71057152 | NA |
| OPN3 | 10.1774311 | -0.237396153 | 0.559889 | -0.4240057 | 0.67156163 | NA |
| CEP170 | 38.8971167 | 0.145250255 | 0.34345733 | 0.42290626 | 0.67236365 | 0.90403541 |
| SDCCAG8 | 77.5861423 | 0.257139323 | 0.25817546 | 0.99598668 | 0.31925662 | 0.69469143 |
| AKT3 | 345.714262 | 0.709667301 | 0.26862386 | 2.64186252 | 0.00824515 | 0.144148 |
| ZBTB18 | 9.60335886 | 0.501303305 | 0.60815822 | 0.8242975 | 0.40977054 | NA |
| ADSS | 18.2220476 | -0.421257746 | 0.3939971 | -1.06919 | 0.28498407 | NA |
| CATSPERE | 17.915944 | -0.57807159 | 0.54800827 | -1.0548592 | 0.29148971 | NA |
| DESI2 | 27.1083861 | 0.755989785 | 0.30870711 | 2.44889009 | 0.01432972 | NA |
| COX20 | 31.7644747 | 0.059997415 | 0.3658786 | 0.16398176 | 0.86974551 | 0.96200921 |
| HNRNPU | 143.519946 | -0.076549282 | 0.22949565 | -0.3335544 | 0.73871584 | 0.91993596 |
| EFCAB2 | 55.3182127 | 0.283548118 | 0.45886282 | 0.61793657 | 0.53661715 | 0.83677705 |
| KIF26B | 13.3731484 | 1.385328451 | 0.63531176 | 2.18054906 | 0.02921679 | NA |
| SMYD3 | 139.862297 | 0.4881375 | 0.23434586 | 2.082979 | 0.03725314 | 0.30414888 |
| TFB2M | 5.95236905 | 0.266442394 | 0.65839981 | 0.40468176 | 0.68571145 | NA |
| CNST | 36.4098189 | -0.07082357 | 0.29133737 | -0.2430981 | 0.80792938 | 0.94328109 |
| SCCPDH | 20.4442858 | 0.178888851 | 0.45340107 | 0.39454881 | 0.6931759 | NA |
| AHCTF1 | 49.6459949 | 0.048207289 | 0.29968045 | 0.16086231 | 0.87220185 | 0.96207879 |
| ZNF124 | 57.8580105 | 0.06451486 | 0.31678599 | 0.2036544 | 0.83862358 | 0.95393706 |
| AL390728.6 | 13.4391981 | 0.349876612 | 0.4028475 | 0.86850883 | 0.38511584 | NA |
| ZNF496 | 38.9824289 | -0.515257524 | 0.42993795 | -1.1984463 | 0.23074333 | 0.63682501 |
| SH3BP5L | 6.57883856 | 0.201743967 | 0.53116246 | 0.37981594 | 0.70408205 | NA |
| ZNF672 | 6.56412617 | -0.437172512 | 0.52271007 | -0.8363576 | 0.40295378 | NA |
| ZNF692 | 7.81102502 | 0.320143253 | 0.50017465 | 0.64006293 | 0.52213169 | NA |
| FAM110C | 7.58894502 | -1.153443631 | 0.8030126 | -1.4363954 | 0.15088985 | NA |
| SH3YL1 | 27.7860687 | -0.013269487 | 0.36637863 | -0.036218 | 0.97110856 | NA |
| ACP1 | 18.8758446 | 0.004679326 | 0.38084986 | 0.01228654 | 0.99019701 | NA |
| TMEM18 | 19.6112794 | 1.114640354 | 0.54087154 | 2.0608227 | 0.03931996 | NA |
| EIPR1 | 19.4909314 | 0.26416465 | 0.41601501 | 0.63498826 | 0.52543607 | NA |
| TRAPPC12 | 39.3583306 | -0.145282054 | 0.2905442 | -0.5000343 | 0.61705095 | 0.87619503 |
| ADI1 | 33.6672723 | -1.181760965 | 0.40796216 | -2.8967416 | 0.0037706 | 0.09813187 |
| AC108488.2 | 6.33903239 | -0.506137236 | 0.62208937 | -0.8136086 | 0.41586924 | NA |
| RPS7 | 62.8599933 | -0.682514923 | 0.49807033 | -1.3703184 | 0.17058754 | 0.57276745 |
| NRIR | 3.68258655 | 1.553593145 | 3.69221379 | 0.42077551 | 0.67391902 | NA |
| CMPK2 | 11.5795258 | -2.014311794 | 2.38033827 | -0.8462292 | 0.39742488 | NA |
| RSAD2 | 27.8162473 | -0.943337609 | 1.82527735 | -0.5168188 | 0.60528267 | NA |
| ID2 | 10.4860729 | -1.326628117 | 0.87288206 | -1.5198252 | 0.12855492 | NA |
| KIDINS220 | 78.9812585 | 0.121022056 | 0.27708475 | 0.4367691 | 0.66227881 | 0.89839561 |
| MBOAT2 | 80.7478611 | 0.097297311 | 0.41415479 | 0.23492982 | 0.8142632 | 0.94496468 |
| ASAP2 | 36.9282969 | -0.083856364 | 0.40636139 | -0.2063591 | 0.83651043 | 0.95252283 |
| ITGB1BP1 | 35.2970436 | 0.049298788 | 0.33948465 | 0.14521655 | 0.8845399 | 0.96573186 |
| CPSF3 | 8.56069709 | -0.870859707 | 0.53475991 | -1.628506 | 0.10341764 | NA |
| IAH1 | 28.2967795 | -1.048469079 | 0.39570516 | -2.6496219 | 0.00805819 | NA |
| ADAM17 | 45.0650897 | 0.048268024 | 0.35572658 | 0.13568855 | 0.8920675 | 0.97029385 |
| YWHAQ | 75.4199451 | -0.218857806 | 0.32475093 | -0.6739251 | 0.50035893 | 0.82345406 |
| TAF1B | 27.2852417 | 0.240298552 | 0.41979384 | 0.57242039 | 0.5670372 | NA |
| GRHL1 | 13.5399375 | -0.957239766 | 0.47341692 | -2.0219805 | 0.04317837 | NA |
| AC104794.2 | 5.48369913 | -0.295038472 | 0.66896781 | -0.4410354 | 0.65918738 | NA |
| KLF11 | 11.5362055 | 0.753954879 | 0.55606758 | 1.35586916 | 0.1751408 | NA |
| CYS1 | 26.5799674 | 0.97708727 | 0.42015806 | 2.32552309 | 0.02004401 | NA |
| HPCAL1 | 15.7546984 | 0.139014821 | 0.39435871 | 0.35250856 | 0.7244569 | NA |
| ODC1 | 27.2968531 | 0.282095159 | 0.40597886 | 0.69485184 | 0.48714816 | NA |
| NOL10 | 30.5125419 | 0.116703814 | 0.36198251 | 0.3224018 | 0.74714832 | 0.92250642 |
| PDIA6 | 72.549171 | -0.404608724 | 0.50451302 | -0.8019788 | 0.42256525 | 0.77588259 |
| PQLC3 | 18.1346133 | -0.460385614 | 0.54951987 | -0.8377961 | 0.40214522 | NA |
| ROCK2 | 87.7996343 | 0.246821568 | 0.25884354 | 0.95355505 | 0.34030892 | 0.71508205 |
| E2F6 | 11.3942675 | 0.554330351 | 0.60001577 | 0.92385964 | 0.3555594 | NA |
| LPIN1 | 38.2008252 | -0.179371123 | 0.26365187 | -0.6803332 | 0.4962935 | 0.8234249 |
| MIR3681HG | 20.8200155 | 1.901389561 | 0.56477671 | 3.36662177 | 0.00076095 | NA |
| TRIB2 | 124.920138 | 0.331379707 | 0.42511358 | 0.77950863 | 0.43568016 | 0.78167891 |
| NBAS | 82.4378686 | 0.312882018 | 0.19437864 | 1.60965224 | 0.1074738 | 0.49144726 |
| DDX1 | 31.4327158 | 0.050067002 | 0.29355539 | 0.17055385 | 0.86457459 | 0.96089268 |
| SMC6 | 17.7921343 | -0.804003057 | 0.36838492 | -2.182508 | 0.02907206 | NA |
| GEN1 | 6.68538265 | -0.76855358 | 0.7628353 | -1.0074961 | 0.31369644 | NA |
| KCNS3 | 5.7017511 | -0.625975434 | 0.63716357 | -0.9824407 | 0.32588278 | NA |
| RDH14 | 10.4914034 | -0.735782134 | 0.6005021 | -1.2252782 | 0.22047042 | NA |
| LINC01376 | 43.5540676 | -0.368837191 | 0.34102806 | -1.081545 | 0.27945475 | 0.66958859 |
| TTC32 | 11.1946539 | -0.681064219 | 0.47664279 | -1.4288776 | 0.1530394 | NA |
| WDR35 | 36.1477235 | 0.13017727 | 0.24484748 | 0.53166677 | 0.59495681 | 0.86653654 |
| MATN3 | 5.9482228 | 0.278454885 | 0.76982398 | 0.36171241 | 0.71756696 | NA |

| | | | | | | |
|---|---|---|---|---|---|---|
| LAPTM4A | 103.153495 | -0.171931249 | 0.34299213 | -0.5012688 | 0.61618197 | 0.87600213 |
| SDC1 | 5.27701212 | -0.153684214 | 0.84342448 | -0.1822146 | 0.85541435 | NA |
| PUM2 | 96.9242146 | 0.207698889 | 0.22095202 | 0.94001804 | 0.34720831 | 0.71997796 |
| RHOB | 13.4381568 | -1.082188861 | 0.60682359 | -1.7833665 | 0.07452666 | NA |
| HS1BP3 | 8.86685848 | -0.166934513 | 0.58835612 | -0.2837304 | 0.776617 | NA |
| LDAH | 42.472952 | -0.094277188 | 0.30923899 | -0.3048684 | 0.76046641 | 0.92336765 |
| AC018742.1 | 152.51804 | -0.351481794 | 0.70060939 | -0.5016801 | 0.61589256 | 0.8758977 |
| KLHL29 | 693.284767 | -1.009001269 | 0.43187962 | -2.3363021 | 0.0194755 | 0.23378067 |
| ATAD2B | 67.5476717 | 0.500725497 | 0.23057341 | 2.17165325 | 0.02988183 | 0.27926427 |
| UBXN2A | 28.82251 | -0.250521516 | 0.29639503 | -0.8452285 | 0.3979833 | 0.76034868 |
| FKBP1B | 7.26071812 | 1.247855284 | 0.91096856 | 1.36981158 | 0.17074573 | NA |
| SF3B6 | 27.043742 | -0.567982591 | 0.44247134 | -1.2836596 | 0.19926108 | NA |
| FAM228B | 22.8406876 | -0.896607657 | 0.38928906 | -2.3031926 | 0.02126801 | NA |
| TP53I3 | 8.01476518 | -1.361512143 | 0.75504237 | -1.8032261 | 0.07135271 | NA |
| ITSN2 | 72.0076737 | -0.328813356 | 0.28987198 | -1.1343399 | 0.25665199 | 0.65588339 |
| NCOA1 | 230.505957 | 0.213004493 | 0.31430676 | 0.6776962 | 0.49796434 | 0.82345406 |
| PTRHD1 | 7.19217871 | -0.085518971 | 0.54979721 | -0.1555464 | 0.87639058 | NA |
| DNAJC27 | 9.66051715 | 0.417864282 | 0.4427501 | 0.94379262 | 0.34527563 | NA |
| DNAJC27-AS | 11.2227141 | 1.319900226 | 0.60525343 | 2.18073977 | 0.02920267 | NA |
| DNMT3A | 37.3964433 | 0.468904149 | 0.31047404 | 1.51028457 | 0.13097083 | 0.52613957 |
| DTNB | 37.5373208 | 0.396963406 | 0.41527641 | 0.95590164 | 0.33912193 | 0.71342248 |
| ASXL2 | 46.3125912 | 0.138922668 | 0.27792955 | 0.4998485 | 0.61718176 | 0.87619503 |
| RAB10 | 106.811372 | -0.055247988 | 0.20406242 | -0.2707406 | 0.78659052 | 0.93615351 |
| HADHA | 38.4514185 | -0.075513803 | 0.33724198 | -0.2239158 | 0.82282283 | 0.94740448 |
| HADHB | 38.1465899 | -0.094345577 | 0.41507619 | -0.227297 | 0.8201928 | 0.94728252 |
| SELENOI | 16.4808203 | -0.698286421 | 0.44919758 | -1.5545196 | 0.12006053 | NA |
| MAPRE3 | 12.7262067 | 0.188940908 | 0.46499002 | 0.40633325 | 0.68449776 | NA |
| TMEM214 | 14.2254841 | 0.34995334 | 0.48019965 | 0.72876634 | 0.46614461 | NA |
| AGBL5 | 14.0106643 | -0.131928917 | 0.38571905 | -0.3420337 | 0.73232552 | NA |
| OST4 | 42.8624268 | 0.139970455 | 0.53788372 | 0.26022437 | 0.7946907 | 0.93934998 |
| EMILIN1 | 9.58369997 | -0.110991995 | 0.65617359 | -0.1691504 | 0.86567838 | NA |
| KHK | 4.2234055 | 0.411075766 | 0.83193221 | 0.49412171 | 0.62122022 | NA |
| PREB | 7.37630669 | 0.628180869 | 0.61281673 | 1.02507134 | 0.30532953 | NA |
| ATRAID | 35.0273815 | -0.870876866 | 0.45021423 | -1.934361 | 0.05306875 | 0.35874478 |
| CAD | 6.581632 | -0.641098789 | 0.54260274 | -1.1815252 | 0.23739416 | NA |
| TRIM54 | 12.2728777 | 0.501946528 | 0.77559792 | 0.64717364 | 0.51751957 | NA |
| MPV17 | 8.52148823 | -0.13496989 | 0.53603927 | -0.251791 | 0.80120258 | NA |
| GTF3C2 | 19.0771572 | 0.074168009 | 0.36111203 | 0.20538781 | 0.83726914 | NA |
| EIF2B4 | 6.63249292 | -0.434712285 | 0.66917677 | -0.6496225 | 0.5159361 | NA |
| SNX17 | 20.4479516 | 0.148227829 | 0.46098504 | 0.32154586 | 0.74779677 | NA |
| ZNF513 | 4.0370515 | 0.848435676 | 0.6921432 | 1.22580946 | 0.22027039 | NA |
| PPM1G | 37.7678145 | -0.752852281 | 0.38629489 | -1.9489056 | 0.0513067 | 0.35507065 |
| NRBP1 | 20.3343459 | 0.272432749 | 0.37665394 | 0.72329723 | 0.46949729 | NA |
| IFT172 | 11.456886 | 0.483610849 | 0.44142061 | 1.09557832 | 0.27326335 | NA |
| ZNF512 | 22.854275 | -0.03135037 | 0.36943755 | -0.0848597 | 0.93237291 | NA |
| GPN1 | 6.9823429 | 0.027193269 | 0.52775485 | 0.05152633 | 0.95890612 | NA |
| SUPT7L | 12.3324119 | 0.511363701 | 0.47402971 | 1.07875875 | 0.28069529 | NA |
| SLC4A1AP | 11.3556172 | -0.005343265 | 0.40571531 | -0.01317 | 0.98949218 | NA |
| MRPL33 | 43.7433035 | -0.067962374 | 0.34342389 | -0.1978965 | 0.84312607 | 0.95469552 |
| RBKS | 17.5267072 | -0.289874032 | 0.46206056 | -0.6273507 | 0.53042936 | NA |
| BABAM2 | 117.484481 | -0.110684392 | 0.27855008 | -0.397359 | 0.69110272 | 0.90557575 |
| FOSL2 | 47.4091614 | -1.653283731 | 0.60874385 | -2.7158939 | 0.00660971 | 0.12888932 |
| PPP1CB | 24.8885585 | 0.108445379 | 0.30531994 | 0.35518603 | 0.72245023 | NA |
| TRMT61B | 16.3543553 | 0.307248673 | 0.40598828 | 0.75679197 | 0.4491745 | NA |
| WDR43 | 24.7717233 | -0.599546118 | 0.42700492 | -1.4040731 | 0.16029709 | NA |
| TOGARAM2 | 16.2727135 | 0.298442137 | 0.70435924 | 0.42370728 | 0.67177931 | NA |
| C2orf71 | 7.50967127 | 0.402976807 | 0.840089 | 0.47968347 | 0.63145248 | NA |
| CLIP4 | 200.734805 | 0.367897601 | 0.33556119 | 1.09636517 | 0.272919 | 0.66531666 |
| ALK | 8.73139304 | -0.608347454 | 0.68130695 | -0.8929124 | 0.37190406 | NA |
| YPEL5 | 35.0530984 | -0.495892583 | 0.34272466 | -1.4469124 | 0.14792145 | 0.54691531 |
| LBH | 59.3486416 | 0.713525142 | 0.56558533 | 1.26156939 | 0.20710377 | 0.61629707 |
| LCLAT1 | 47.307656 | 0.61171848 | 0.28570086 | 2.14111529 | 0.03226474 | 0.29020358 |
| GALNT14 | 24.8091689 | 0.33575189 | 0.82161907 | 0.40864666 | 0.68279898 | NA |
| MEMO1 | 7.14976135 | 0.39902529 | 0.55105257 | 0.72411475 | 0.46899529 | NA |
| DPY30 | 20.2743193 | -0.427579871 | 0.4085774 | -1.0465089 | 0.29532615 | NA |
| SPAST | 20.9152424 | 0.124147027 | 0.37872657 | 0.32780121 | 0.74306197 | NA |
| SLC30A6 | 24.1927976 | 0.224729725 | 0.31592443 | 0.71134013 | 0.47687349 | NA |
| YIPF4 | 53.5725176 | 0.206240753 | 0.23837287 | 0.86520228 | 0.38692777 | 0.75125852 |
| BIRC6 | 198.149934 | 0.303563418 | 0.15137671 | 2.00535092 | 0.04492555 | 0.33424083 |
| TTC27 | 24.1000362 | -0.440106393 | 0.3503552 | -1.256172 | 0.20905361 | NA |
| LINC00486 | 38.7613961 | 0.545298483 | 0.35449155 | 1.53825525 | 0.12398622 | 0.51897128 |
| RASGRP3 | 30.1011923 | 1.58495434 | 0.49601736 | 3.19536064 | 0.00139656 | 0.05608372 |
| AC020594.1 | 10.4804508 | 1.594721568 | 0.65393143 | 2.43866786 | 0.01474151 | NA |
| FAM98A | 43.9964414 | 0.864251146 | 0.4869506 | 1.77482304 | 0.07592711 | 0.42186351 |
| LINC01320 | 52.8845758 | 0.039322489 | 0.57992909 | 0.06780569 | 0.94594032 | 0.98249707 |
| AC012593.1 | 23.0579436 | 0.981404913 | 0.89272809 | 1.0993324 | 0.27162311 | NA |
| AC009414.2 | 15.2351794 | -0.734754502 | 0.75611723 | -0.9717468 | 0.33117653 | NA |
| CRIM1 | 1579.77597 | 0.148062708 | 0.32676204 | 0.45312089 | 0.65046169 | 0.89412414 |
| FEZ2 | 247.322963 | -0.029467525 | 0.43638101 | -0.0675271 | 0.94616212 | 0.98249707 |
| STRN | 38.8006258 | 0.404352885 | 0.30228008 | 1.33767624 | 0.181002 | 0.58390314 |
| HEATR5B | 32.1156547 | -0.030928742 | 0.33536035 | -0.0922254 | 0.92651896 | 0.97775374 |
| GPATCH11 | 5.94545987 | -0.645404322 | 0.6024089 | -1.0713725 | 0.28400198 | NA |
| EIF2AK2 | 42.4764592 | 0.033054651 | 0.60488341 | 0.05464632 | 0.95642024 | 0.98641719 |
| CEBPZOS | 32.7146927 | 0.019647267 | 0.34878909 | 0.05632994 | 0.95507897 | 0.9859507 |
| CEBPZ | 21.9029516 | -0.61857572 | 0.44488357 | -1.3904216 | 0.16440089 | NA |
| NDUFAF7 | 10.6822794 | -0.415408265 | 0.49058311 | -0.8467643 | 0.39712651 | NA |
| PRKD3 | 15.2859595 | 0.302169832 | 0.5368635 | 0.56284294 | 0.57354183 | NA |
| AC006369.1 | 52.9008443 | -1.038545591 | 0.40463753 | -2.5666072 | 0.01026989 | 0.1620804 |
| CDC42EP3 | 493.188435 | -0.60791658 | 0.30763673 | -1.9760858 | 0.04814506 | 0.34212397 |
| RMDN2 | 19.8348304 | 1.21787938 | 0.59148409 | 2.05902307 | 0.03949203 | NA |
| CYP1B1 | 62.1750301 | -1.080731465 | 0.87483123 | -1.23536 | 0.21669657 | 0.62433386 |
| CYP1B1-AS1 | 8.65410018 | 0.615524383 | 0.7434723 | 0.82790493 | 0.40772434 | NA |
| ATL2 | 32.5519846 | 0.401385105 | 0.34692559 | 1.15697751 | 0.24728155 | 0.65061974 |
| AC011247.1 | 10.6143488 | -0.782262371 | 0.6560508 | -1.1923808 | 0.23311197 | NA |
| HNRNPLL | 46.1669683 | 0.569539317 | 0.37239809 | 1.52938301 | 0.12616952 | 0.52276701 |
| GALM | 7.28025952 | 0.260628729 | 0.6612795 | 0.39412794 | 0.69348658 | NA |
| SRSF7 | 36.4975555 | -0.307044084 | 0.4113829 | -0.7463706 | 0.4554436 | 0.79324789 |
| GEMIN6 | 5.63362694 | 0.092475322 | 0.69787614 | 0.13250965 | 0.89458119 | NA |
| DHX57 | 18.4729234 | -0.13908163 | 0.37383651 | -0.3720386 | 0.70986407 | NA |
| MORN2 | 7.89946158 | -0.580929301 | 0.54548955 | -1.0649687 | 0.28689009 | NA |
| SOS1 | 117.943467 | 0.270888993 | 0.2211121 | 1.22512061 | 0.22052978 | 0.62462904 |
| MAP4K3 | 171.093224 | 0.481228142 | 0.22485407 | 2.14017977 | 0.03234024 | 0.29020358 |
| AC007388.1 | 124.573713 | 0.296687941 | 0.22176538 | 1.33784609 | 0.18094661 | 0.58390314 |
| TMEM178A | 130.848648 | 0.126217835 | 0.51186762 | 0.24658296 | 0.80523099 | 0.94230147 |
| THUMPD2 | 22.5144779 | -0.235451559 | 0.43573495 | -0.540355 | 0.58895223 | NA |
| SLC8A1-AS1 | 29.3908587 | -0.12516714 | 0.84936072 | -0.1473663 | 0.88284291 | 0.96526488 |
| SLC8A1 | 30.4262713 | 0.790802946 | 1.48842556 | 0.53130164 | 0.59520977 | 0.86653654 |
| PKDCC | 41.3845883 | -2.146073964 | 0.7354006 | -2.918238 | 0.00352016 | 0.09757286 |
| EML4 | 205.759634 | 0.869261981 | 0.31311865 | 2.77614243 | 0.00550081 | 0.11620462 |
| COX7A2L | 42.2512107 | -0.507013842 | 0.36247902 | -1.3987398 | 0.16189101 | 0.56290656 |
| MTA3 | 37.3183485 | -0.027723317 | 0.28256369 | -0.0981135 | 0.92184216 | 0.97775374 |
| ZFP36L2 | 98.3187519 | -0.964924342 | 0.41518339 | -2.3240919 | 0.02012057 | 0.23722175 |
| THADA | 66.9698969 | 0.165524962 | 0.32526883 | 0.50888665 | 0.61083168 | 0.8736013 |

| | | | | | | |
|---|---|---|---|---|---|---|
| PLEKHH2 | 115.700657 | 0.814594064 | 0.32875809 | 2.47779172 | 0.01321983 | 0.19218504 |
| DYNC2LI1 | 31.2589203 | 0.436611741 | 0.31917924 | 1.36792024 | 0.17133704 | 0.57422722 |
| LRPPRC | 45.530626 | 0.278355062 | 0.27803092 | 1.00116583 | 0.31674664 | 0.69424851 |
| PPM1B | 52.2520311 | 0.338601175 | 0.24519712 | 1.38093457 | 0.16729908 | 0.56878882 |
| SLC3A1 | 27.1634497 | 0.958043851 | 0.72787368 | 1.31622269 | 0.18809931 | NA |
| PREPL | 56.1112775 | 0.471241889 | 0.3449261 | 1.36621115 | 0.1718727 | 0.57423036 |
| CAMKMT | 103.512293 | 0.332289638 | 0.24472325 | 1.35781802 | 0.17452144 | 0.57784404 |
| SRBD1 | 34.5012318 | 0.105474939 | 0.27323638 | 0.38602085 | 0.69948123 | 0.90729979 |
| PRKCE | 62.5382057 | 0.615187564 | 0.37457775 | 1.64234944 | 0.10051761 | 0.4819142 |
| EPAS1 | 20.4053913 | -0.352882568 | 0.59490373 | -0.5931759 | 0.55306343 | NA |
| RHOQ | 84.7303581 | -0.800771491 | 0.41080353 | -1.949281 | 0.05126188 | 0.35507065 |
| AC018682.1 | 9.62071328 | -1.530990715 | 0.74527566 | -2.054261 | 0.03995044 | NA |
| PIGF | 18.5473592 | -0.307075317 | 0.44289813 | -0.6933317 | 0.48810141 | NA |
| CRIPT | 11.9000135 | 0.210796259 | 0.43628285 | 0.48316421 | 0.62897915 | NA |
| SOCS5 | 41.2121666 | -0.310521889 | 0.43329162 | -0.716658 | 0.47358517 | 0.8060686 |
| MCFD2 | 34.9378373 | -0.584580733 | 0.43150692 | -1.3547424 | 0.17549964 | 0.57919164 |
| TTC7A | 16.2501536 | -0.962573788 | 0.40844577 | -2.3566746 | 0.0184394 | NA |
| CALM2 | 7.91546698 | -2.02013718 | 0.65200794 | -3.0983322 | 0.00194613 | NA |
| EPCAM | 14.9259318 | -1.359369531 | 0.68391576 | -1.9876272 | 0.04685294 | NA |
| MSH2 | 19.4829968 | 0.217072559 | 0.47044915 | 0.46141556 | 0.64450049 | NA |
| MSH6 | 19.7001205 | 0.05991616 | 0.45442419 | 0.13185073 | 0.89510236 | NA |
| FBXO11 | 104.004819 | 0.018400412 | 0.22825774 | 0.08061243 | 0.93575018 | 0.97832857 |
| FOXN2 | 30.1055709 | 0.356109374 | 0.31448227 | 1.13236709 | 0.25748013 | 0.65588339 |
| PPP1R21 | 56.546179 | 0.416797157 | 0.29266435 | 1.42414734 | 0.15440382 | 0.55470497 |
| STON1-GTF2 | 19.3262289 | -0.231159441 | 0.597906 | -0.386615 | 0.69904123 | NA |
| AC073082.1 | 4.36214159 | -3.830316863 | 1.49071666 | -2.5694466 | 0.01018611 | NA |
| AC009975.1 | 38.8084143 | -0.11798029 | 1.24146601 | -0.095033 | 0.92428858 | 0.97775374 |
| GPR75-ASB3 | 18.6566538 | -0.576397676 | 0.49042803 | -1.1752951 | 0.23987666 | NA |
| ERLEC1 | 31.7049165 | 0.122427944 | 0.31392048 | 0.38999667 | 0.69653901 | 0.90718427 |
| PSME4 | 56.1049064 | -0.082455059 | 0.25134605 | -0.3280539 | 0.74287089 | 0.92122771 |
| ACYP2 | 108.728603 | 0.254896295 | 0.26614256 | 0.95774346 | 0.33819213 | 0.71275062 |
| SPTBN1 | 111.008175 | 0.166609504 | 0.26046257 | 0.63966775 | 0.52238863 | 0.83120735 |
| AC093110.1 | 4.5949472 | -0.621300253 | 0.99383404 | -0.6251549 | 0.53186938 | NA |
| RTN4 | 234.175971 | -0.181079165 | 0.29281238 | -0.6184136 | 0.53630273 | 0.83677705 |
| CLHC1 | 16.508585 | 0.599682818 | 0.57014042 | 1.05181601 | 0.29288397 | NA |
| RPS27A | 115.081956 | -0.565286938 | 0.46711376 | -1.2101697 | 0.22621377 | 0.63327644 |
| MTIF2 | 9.44440672 | 0.242911882 | 0.455017 | 0.53385232 | 0.59344371 | NA |
| AC012358.3 | 15.2387855 | -0.182833697 | 0.48294593 | -0.3785801 | 0.70499973 | NA |
| CCDC88A | 49.5105463 | 0.46009515 | 0.2642775 | 1.74095468 | 0.08169152 | 0.43779104 |
| CFAP36 | 24.3761805 | -0.33774948 | 0.3630509 | -0.9303089 | 0.35221118 | NA |
| PPP4R3B | 58.4666151 | 0.350012323 | 0.25017449 | 1.39907277 | 0.16179116 | 0.56290656 |
| PNPT1 | 33.3375077 | 0.32224609 | 0.75768762 | 0.42530204 | 0.67061651 | 0.90305784 |
| EFEMP1 | 99.4025141 | 0.607642034 | 0.44658625 | 1.36063758 | 0.17362825 | 0.57677001 |
| VRK2 | 49.0471359 | -0.002291709 | 0.3317334 | -0.0069083 | 0.99448803 | 0.99867379 |
| FANCL | 44.9440371 | 0.189528729 | 0.35347131 | 0.53619268 | 0.59182539 | 0.8654248 |
| PAPOLG | 11.9013068 | 0.916972683 | 0.37601329 | 2.43867093 | 0.01474139 | NA |
| REL | 15.0770017 | -0.141712094 | 0.40933943 | -0.346197 | 0.72919465 | NA |
| PUS10 | 10.2151921 | 0.484125642 | 0.63634084 | 0.76079612 | 0.44677885 | NA |
| PEX13 | 18.7092372 | 0.057501863 | 0.34367786 | 0.16731326 | 0.86712357 | NA |
| KIAA1841 | 22.6909347 | 0.610851729 | 0.3983649 | 1.53339747 | 0.12517795 | NA |
| C2orf74 | 12.5800204 | -0.099723822 | 0.54606085 | -0.182624 | 0.85509304 | NA |
| USP34 | 224.770335 | 0.282227386 | 0.20665155 | 1.36571626 | 0.17202804 | 0.57427631 |
| XPO1 | 111.863006 | 0.882002119 | 0.38484653 | 2.29182816 | 0.02191556 | 0.24297341 |
| FAM161A | 8.08391923 | -0.72686488 | 0.52415143 | -1.386746 | 0.16551924 | NA |
| CCT4 | 34.3990981 | 0.391071362 | 0.32319566 | 1.21001426 | 0.22627342 | 0.63327644 |
| COMMD1 | 81.9421945 | 0.182354696 | 0.1904713 | 0.95738673 | 0.33837209 | 0.71275062 |
| AC018462.1 | 4.834145 | 0.012497243 | 0.8184498 | 0.01526941 | 0.98781725 | NA |
| B3GNT2 | 5.04733574 | 0.0494441 | 0.73898879 | 0.06690778 | 0.94665512 | NA |
| EHBP1 | 127.010232 | 0.341385902 | 0.22391886 | 1.52459644 | 0.12735979 | 0.52301154 |
| WDPCP | 91.6641763 | -0.157641023 | 0.26509498 | -0.5946586 | 0.55207167 | 0.84754076 |
| MDH1 | 30.9103125 | -0.487821683 | 0.42833037 | -1.1388912 | 0.25474854 | 0.65520614 |
| UGP2 | 38.2505094 | -0.004528509 | 0.37170063 | -0.0121832 | 0.99027944 | 0.99710744 |
| VPS54 | 69.4222272 | 0.116862008 | 0.21114536 | 0.55346709 | 0.57994361 | 0.86175324 |
| PELI1 | 9.75184076 | 0.122755462 | 0.63093757 | 0.19456039 | 0.84573711 | NA |
| LGALSL | 4.75542139 | -0.594633359 | 0.72151211 | -0.8241488 | 0.409855 | NA |
| AFTPH | 35.5400326 | 0.026555028 | 0.28310246 | 0.09380006 | 0.92526798 | 0.97775374 |
| SERTAD2 | 23.1248775 | 0.740852057 | 0.37355161 | 1.98326558 | 0.04733777 | NA |
| CEP68 | 19.9758748 | -0.510158529 | 0.46976753 | -1.0859808 | 0.27748748 | NA |
| RAB1A | 136.689881 | 0.157952494 | 0.23420727 | 0.67441329 | 0.5000486 | 0.82345406 |
| ACTR2 | 79.909406 | -0.002892635 | 0.25739701 | -0.011238 | 0.99103354 | 0.99717986 |
| SPRED2 | 42.6853386 | 0.044072119 | 0.35000712 | 0.12591778 | 0.89979701 | 0.97097328 |
| MEIS1 | 36.3234057 | -0.286953031 | 0.48086668 | -0.5967414 | 0.55068008 | 0.84700736 |
| ETAA1 | 14.8692926 | -0.078095999 | 0.41573633 | -0.1878498 | 0.85099437 | NA |
| C1D | 23.1227506 | -0.191075639 | 0.35842722 | -0.5330947 | 0.59396805 | NA |
| WDR92 | 7.42400985 | 0.658088815 | 0.52454444 | 1.25459116 | 0.20962721 | NA |
| PNO1 | 8.868555 | -1.377820175 | 0.62524759 | -2.2036393 | 0.02754972 | NA |
| APLF | 22.2096109 | 0.413918392 | 0.45417809 | 0.91135703 | 0.36210729 | NA |
| ARHGAP25 | 8.16507312 | -0.817648484 | 0.7347963 | -1.1127553 | 0.26581353 | NA |
| ANTXR1 | 9.79862163 | 0.608410814 | 0.58487391 | 1.04024269 | 0.29822716 | NA |
| GFPT1 | 49.4586725 | 0.135956623 | 0.26291527 | 0.51711193 | 0.60507803 | 0.87047761 |
| NFU1 | 15.9725284 | 0.043709228 | 0.35472704 | 0.12321933 | 0.90193342 | NA |
| AAK1 | 72.7380361 | 0.088976142 | 0.25621108 | 0.34727672 | 0.72838344 | 0.91700494 |
| ANXA4 | 67.8647807 | 0.244502385 | 0.40865599 | 0.59830857 | 0.54963406 | 0.84615033 |
| GMCL1 | 17.4520476 | -0.232998562 | 0.38682712 | -0.6023325 | 0.54695281 | NA |
| MXD1 | 13.0475335 | -0.241075934 | 0.5155066 | -0.4676486 | 0.64003591 | NA |
| PCBP1 | 65.8434845 | -0.47919405 | 0.38138568 | -1.2564553 | 0.20895095 | 0.61736443 |
| C2orf42 | 9.92618301 | 0.939920099 | 0.51347789 | 1.83049771 | 0.06717555 | NA |
| PCYOX1 | 112.649602 | -0.578637815 | 0.72916156 | -0.793566 | 0.42744814 | 0.77844087 |
| SNRPG | 25.6873478 | -0.284775302 | 0.43102298 | -0.6606963 | 0.50880707 | NA |
| FAM136A | 8.56851267 | 0.453795007 | 0.47345813 | 0.95846914 | 0.33782625 | NA |
| NAGK | 12.8023605 | -0.158238987 | 0.40029803 | -0.3953029 | 0.69261933 | NA |
| MCEE | 15.0065621 | -0.626909326 | 0.44563468 | -1.4067786 | 0.15949305 | NA |
| MPHOSPH10 | 14.5085191 | -0.88931777 | 0.383742 | -2.3174887 | 0.02047712 | NA |
| PAIP2B | 6.10939506 | 0.960059138 | 0.69665556 | 1.37809728 | 0.16817325 | NA |
| ZNF638 | 172.279876 | -0.027161003 | 0.19403726 | -0.1399783 | 0.88867715 | 0.96764417 |
| EXOC6B | 184.969414 | 0.426358019 | 0.2469948 | 1.72618215 | 0.08431465 | 0.44413014 |
| SPR | 4.90848671 | 0.025335811 | 0.66990935 | 0.03781976 | 0.96983139 | NA |
| SFXN5 | 25.6252533 | -0.247087239 | 0.53508776 | -0.4617696 | 0.64424658 | NA |
| SMYD5 | 4.29103744 | -0.344735939 | 0.84569462 | -0.4076364 | 0.68354061 | NA |
| CCT7 | 30.3077649 | -0.365842183 | 0.48024388 | -0.7617842 | 0.44618883 | 0.78855856 |
| ALMS1 | 42.7755104 | 0.498965625 | 0.26486811 | 1.88382671 | 0.05958842 | 0.37905854 |
| NAT8 | 10.10007 | 0.448660886 | 1.82924187 | 0.24527149 | 0.80624622 | NA |
| TPRKB | 6.99595035 | -0.437813723 | 0.59922275 | -0.730636 | 0.4650015 | NA |
| DUSP11 | 8.92179485 | -0.487475279 | 0.47930528 | -1.0170455 | 0.30913178 | NA |
| STAMBP | 19.4633383 | -0.249733359 | 0.3320856 | -0.752015 | 0.45204202 | NA |
| DGUOK | 27.2907758 | 0.038742739 | 0.40199852 | 0.09637533 | 0.92322249 | NA |
| TET3 | 59.9978258 | -0.260788275 | 0.34975907 | -0.7456226 | 0.45589541 | 0.79324789 |
| BOLA3 | 10.0975205 | 0.393045399 | 0.60046691 | 0.6545663 | 0.51274703 | NA |
| MOB1A | 51.3783034 | 0.129578217 | 0.25644627 | 0.50528408 | 0.61335932 | 0.87439243 |
| MTHFD2 | 48.1553513 | -1.510511253 | 0.51146752 | -2.9532887 | 0.00314408 | 0.0904424 |
| AC006030.1 | 11.9904131 | -0.526871322 | 0.55117715 | -0.955902 | 0.33912177 | NA |
| WDR54 | 3.7389453 | -1.226520982 | 0.83795685 | -1.4637042 | 0.14327478 | NA |
| RTKN | 4.37379499 | 0.282659018 | 0.75894517 | 0.37243668 | 0.70956775 | NA |

| | | | | | | |
|---|---|---|---|---|---|---|
| MOGS | 17.4327917 | -0.122637006 | 0.4656964 | -0.2633411 | 0.79228768 | NA |
| CCDC142 | 4.31512765 | -0.453785407 | 0.6504321 | -0.6976676 | 0.48538509 | NA |
| AUP1 | 14.4476401 | 0.473498084 | 0.48851653 | 0.96925704 | 0.33241696 | NA |
| HTRA2 | 7.04300441 | -0.012613238 | 0.60257476 | -0.0209322 | 0.98329971 | NA |
| DOK1 | 6.72418768 | -0.308456688 | 0.55027992 | -0.5605451 | 0.57510772 | NA |
| POLE4 | 14.6581134 | -0.054669258 | 0.5072494 | -0.1077759 | 0.91417346 | NA |
| TACR1 | 10.1092049 | 0.531892299 | 0.60840656 | 0.87423827 | 0.38198851 | NA |
| AC007681.1 | 4.74216236 | -0.764661549 | 0.78428875 | -0.9749745 | 0.3295729 | NA |
| EVA1A | 7.62680058 | 0.529011496 | 0.90739817 | 0.58299819 | 0.55989452 | NA |
| MRPL19 | 35.6585593 | -0.45829565 | 0.35475627 | -1.2918606 | 0.19640543 | 0.59987316 |
| GCFC2 | 31.4169102 | -0.226119567 | 0.31909118 | -0.7086362 | 0.47855025 | 0.81001889 |
| SUCLG1 | 32.7888169 | 0.386247845 | 0.3726726 | 1.03642674 | 0.3000031 | 0.68168772 |
| DNAH6 | 11.1465106 | 0.145939646 | 0.59440796 | 0.24552101 | 0.80605304 | NA |
| TMSB10 | 316.08741 | -0.592006584 | 0.51147862 | -1.1574415 | 0.24709202 | 0.65061974 |
| KCMF1 | 58.7492586 | 0.351746696 | 0.2523793 | 1.39372243 | 0.16340144 | 0.56500958 |
| TCF7L1 | 335.874124 | -0.062349229 | 0.31789828 | -0.1961295 | 0.84450881 | 0.95509854 |
| TGOLN2 | 119.991786 | -0.149437436 | 0.37713762 | -0.3962411 | 0.69192716 | 0.90557575 |
| RETSAT | 5.23891998 | 0.394407671 | 0.54739256 | 0.72052071 | 0.47120446 | NA |
| ELMOD3 | 19.9620449 | 0.305946288 | 0.33352236 | 0.91731867 | 0.35897568 | NA |
| MAT2A | 35.176523 | -0.230185014 | 0.39695985 | -0.5798698 | 0.56200245 | 0.85386823 |
| GGCX | 25.8857176 | 0.254084111 | 0.45559433 | 0.55769945 | 0.57704963 | NA |
| VAMP8 | 52.1987252 | -0.929257776 | 0.43444627 | -2.1389475 | 0.03243992 | 0.29045543 |
| VAMP5 | 207.357802 | -0.708686406 | 0.4256777 | -1.6648427 | 0.09594414 | 0.46942032 |
| RNF181 | 13.8830511 | -0.344340698 | 0.51436589 | -0.669447 | 0.50321038 | NA |
| TMEM150A | 8.79114564 | -0.552200527 | 0.53648109 | -1.029301 | 0.30333825 | NA |
| USP39 | 33.8579472 | -0.022756568 | 0.32019484 | -0.071071 | 0.94334125 | 0.98114803 |
| C2orf68 | 13.4836511 | 0.445554966 | 0.39205366 | 1.13646426 | 0.25576232 | NA |
| ATOH8 | 21.5857465 | -2.034663701 | 0.7406304 | -2.7472052 | 0.00601055 | NA |
| POLR1A | 12.0614665 | -0.349292296 | 0.43668176 | -0.7998784 | 0.42378127 | NA |
| PTCD3 | 29.846325 | 0.215917154 | 0.29068425 | 0.74278931 | 0.45760925 | 0.79434199 |
| IMMT | 38.6970342 | -0.007926288 | 0.3195815 | -0.0248021 | 0.98021283 | 0.99453905 |
| MRPL35 | 10.6311321 | -0.164133669 | 0.49884633 | -0.3290265 | 0.74213565 | NA |
| KDM3A | 19.3837821 | -0.392411248 | 0.45308671 | -0.8660842 | 0.38644398 | NA |
| CHMP3 | 21.0263372 | -0.3152898 | 0.37273905 | -0.8458727 | 0.39762375 | NA |
| RMND5A | 29.3814932 | 0.487217654 | 0.43144143 | 1.12927879 | 0.25878025 | 0.65588339 |
| PLGLB1 | 10.4961815 | 0.909496648 | 0.45601785 | 1.99443212 | 0.04610485 | NA |
| CYTOR | 27.1739804 | -1.007157264 | 0.52993002 | -1.9005477 | 0.05736128 | NA |
| KRCC1 | 44.7998172 | -0.0826816 | 0.26762431 | -0.3089465 | 0.7573622 | 0.92313949 |
| THNSL2 | 13.3071903 | 1.023016855 | 0.67844484 | 1.50788507 | 0.13158394 | NA |
| EIF2AK3 | 38.64134 | 0.472790527 | 0.31632122 | 1.49465322 | 0.13500497 | 0.52779404 |
| RPIA | 15.187676 | 0.098792647 | 0.42366632 | 0.23318504 | 0.81561772 | NA |
| MAL | 13.881838 | 1.466045044 | 0.61708845 | 2.37574538 | 0.01751354 | NA |
| MRPS5 | 27.2614867 | 0.070067776 | 0.31747169 | 0.22070559 | 0.82532168 | NA |
| ZNF514 | 3.14147092 | -1.925637471 | 0.92361725 | -2.0848869 | 0.03707957 | NA |
| AC092835.1 | 7.83335363 | -0.327938701 | 0.48041596 | -0.6826141 | 0.49485074 | NA |
| FAHD2A | 8.18581873 | 0.144793594 | 0.49868814 | 0.29034898 | 0.77154927 | NA |
| ANKRD36C | 33.4140375 | 0.457029774 | 0.38983348 | 1.17237177 | 0.24104783 | 0.64918976 |
| STARD7 | 50.049201 | 0.245869301 | 0.27796985 | 0.88451787 | 0.37641672 | 0.74330391 |
| TMEM127 | 15.4267609 | 0.3168872 | 0.43981035 | 0.72050874 | 0.47121182 | NA |
| CIAO1 | 19.4426138 | -0.091836718 | 0.35861312 | -0.2560886 | 0.79788246 | NA |
| SNRNP200 | 26.1127964 | -0.433774651 | 0.37635748 | -1.1525602 | 0.24909094 | NA |
| KANSL3 | 32.0493112 | 0.089599146 | 0.34525566 | 0.25951536 | 0.79523763 | 0.93934998 |
| LMAN2L | 10.5825162 | 0.111392853 | 0.47289034 | 0.23555747 | 0.81377608 | NA |
| CNNM4 | 6.32074411 | 0.437515378 | 0.61957482 | 0.70615423 | 0.48009223 | NA |
| ANKRD23 | 9.31880458 | -0.534737547 | 0.45609268 | -1.1724318 | 0.24102375 | NA |
| SEMA4C | 4.19086846 | -1.4475127 | 0.84932583 | -1.7043079 | 0.08832359 | NA |
| ANKRD36 | 44.5338816 | 0.538428374 | 0.34310484 | 1.56928236 | 0.11658216 | 0.5065504 |
| AC092683.1 | 34.6134059 | -0.293953589 | 0.42990213 | -0.6837686 | 0.49412128 | 0.82249096 |
| ANKRD36B | 12.8308202 | 0.231399911 | 0.41878787 | 0.55254683 | 0.58057376 | NA |
| COX5B | 56.0842106 | -0.185808994 | 0.55202755 | -0.3365937 | 0.73642321 | 0.91929008 |
| ACTR1B | 19.3622487 | -0.061309687 | 0.57005402 | -0.1075507 | 0.91435214 | NA |
| TMEM131 | 79.3890249 | 0.326858599 | 0.25312454 | 1.2912956 | 0.1966012 | 0.59987316 |
| INPP4A | 27.4062471 | 0.39491708 | 0.30820318 | 1.28135305 | 0.20006969 | NA |
| COA5 | 13.3937414 | -0.131059653 | 0.42373566 | -0.3092958 | 0.75709654 | NA |
| UNC50 | 23.6118411 | -0.133610719 | 0.43502603 | -0.3071327 | 0.75874237 | NA |
| MGAT4A | 26.176736 | 0.775251477 | 0.52600835 | 1.47383872 | 0.14052502 | NA |
| TSGA10 | 19.4168077 | 0.135695655 | 0.54895392 | 0.24718952 | 0.80476156 | NA |
| AC079447.1 | 21.8700453 | 0.319151646 | 0.32107417 | 0.99401222 | 0.32021692 | NA |
| MITD1 | 12.333708 | 0.034236152 | 0.48844381 | 0.0700923 | 0.94412019 | NA |
| TXNDC9 | 14.1766917 | -0.39231613 | 0.50073744 | -0.7834767 | 0.43334721 | NA |
| EIF5B | 47.7350887 | -0.452543156 | 0.3693553 | -1.2252245 | 0.22049066 | 0.62462904 |
| REV1 | 61.6002572 | 0.189720415 | 0.28444422 | 0.66698636 | 0.50478084 | 0.82600448 |
| LONRF2 | 6.62162184 | -0.830118533 | 0.67010217 | -1.238794 | 0.2154218 | NA |
| NPAS2 | 20.3672766 | -0.233807875 | 0.39956934 | -0.5851497 | 0.55844707 | NA |
| RPL31 | 28.6637829 | -0.616088864 | 0.44541242 | -1.3831874 | 0.1666074 | 0.56878882 |
| TBC1D8 | 26.2824594 | 0.073091103 | 0.39622699 | 0.18446775 | 0.85364652 | NA |
| CNOT11 | 14.7469046 | 0.225463983 | 0.44171295 | 0.510431 | 0.60974954 | NA |
| RNF149 | 56.7947757 | 0.45318246 | 0.46632899 | 0.97180846 | 0.33114585 | 0.70431439 |
| MAP4K4 | 39.1186883 | -0.25878093 | 0.41662024 | -0.6211434 | 0.53450525 | 0.83677705 |
| IL1R1 | 11.8323871 | -0.869387157 | 0.57901598 | -1.5014908 | 0.13322867 | NA |
| MFSD9 | 4.33969921 | -0.568829546 | 0.73755375 | -0.7712381 | 0.44056582 | NA |
| TMEM182 | 6.34073796 | 1.131245927 | 0.61350746 | 1.84389922 | 0.06519783 | NA |
| POU3F3 | 8.51747676 | 0.110468215 | 0.77939693 | 0.1417355 | 0.88728893 | NA |
| MRPS9 | 19.9953614 | 0.285721643 | 0.37531758 | 0.76127966 | 0.44649005 | NA |
| TGFBRAP1 | 12.2302366 | -0.090338488 | 0.44202126 | -0.2043759 | 0.83805977 | NA |
| C2orf49 | 10.5494292 | -0.378191587 | 0.47386185 | -0.7981052 | 0.42480947 | NA |
| FHL2 | 6.17855269 | -0.031878488 | 0.88291362 | -0.036106 | 0.97119783 | NA |
| NCK2 | 105.98024 | -0.544534303 | 0.42140887 | -1.2921757 | 0.1962963 | 0.59987316 |
| UXS1 | 18.8982364 | 0.294911028 | 0.35554463 | 0.82946276 | 0.4068426 | NA |
| SULT1C2 | 13.2144943 | -0.484284014 | 1.15639012 | -0.4187895 | 0.67537 | NA |
| GCC2 | 66.4118984 | -0.208257873 | 0.32209891 | -0.646565 | 0.51791353 | 0.83104823 |
| GCC2-AS1 | 6.14295669 | -0.246942598 | 0.72991271 | -0.338318 | 0.73512357 | NA |
| LIMS1 | 84.7702097 | -0.581098278 | 0.29104974 | -1.9965601 | 0.045873 | 0.33590606 |
| RANBP2 | 54.848854 | 0.176010331 | 0.32808211 | 0.53648256 | 0.59162509 | 0.8654248 |
| CCDC138 | 5.35584036 | -0.100105233 | 0.62814433 | -0.1593666 | 0.87338004 | NA |
| 10-Sep | 94.0071353 | -0.650078305 | 0.31309855 | -2.0762738 | 0.03786864 | 0.30414888 |
| SOWAHC | 52.6924885 | -1.22717826 | 0.40229232 | -3.050464 | 0.00228488 | 0.07786717 |
| NPHP1 | 19.6528131 | 0.420957577 | 0.44507742 | 0.94580753 | 0.34424676 | NA |
| SMIM37 | 30.1792974 | 0.470970019 | 0.46877295 | 1.00468686 | 0.31504766 | 0.6938809 |
| MIR4435-2H( | 100.49013 | -0.882788446 | 0.39381342 | -2.2416414 | 0.02498456 | 0.25362919 |
| AC017002.5 | 5.45769659 | -0.96010639 | 1.05417868 | -0.9107625 | 0.36242054 | NA |
| ANAPC1 | 34.2564499 | 0.324571487 | 0.28175059 | 1.15198158 | 0.24932864 | 0.65117642 |
| TMEM87B | 45.8838295 | -0.042483123 | 0.3217401 | -0.1320417 | 0.89495128 | 0.97097328 |
| ZC3H8 | 14.699874 | -0.0795583 | 0.4352473 | -0.1827887 | 0.8549638 | NA |
| ZC3H6 | 33.8541542 | -0.140421094 | 0.37508295 | -0.3743734 | 0.7081265 | 0.9106723 |
| TTL | 13.4563751 | -0.200333085 | 0.57292639 | -0.3496664 | 0.7265891 | NA |
| POLR1B | 7.03634866 | -0.373545832 | 0.72817723 | -0.5129875 | 0.60796005 | NA |
| CHCHD5 | 9.33891905 | -0.260614837 | 0.556662 | -0.4681743 | 0.63965996 | NA |
| SLC20A1 | 20.0865413 | -0.725138308 | 0.58601194 | -1.2374122 | 0.2159341 | NA |
| PSD4 | 6.66443238 | -0.193662713 | 0.66572554 | -0.2909047 | 0.77112418 | NA |
| PAX8 | 107.569831 | 0.122417174 | 0.54607226 | 0.22417761 | 0.8226191 | 0.94740448 |
| CBWD2 | 14.1834609 | 0.443495525 | 0.44517071 | 0.99623699 | 0.31913501 | NA |
| RABL2A | 9.90392224 | -0.450065544 | 0.52814121 | -0.852169 | 0.39412033 | NA |

| | | | | | | |
|---|---|---|---|---|---|---|
| SLC35F5 | 97.85791 | -0.271467742 | 0.26793637 | -1.0131799 | 0.31097424 | 0.69098737 |
| ACTR3 | 94.6666488 | 0.093406764 | 0.23657492 | 0.39482953 | 0.6929687 | 0.90557575 |
| DDX18 | 26.7675092 | -0.889304072 | 0.32805963 | -2.7108001 | 0.00671211 | NA |
| CCDC93 | 31.7600012 | 0.047567349 | 0.37255781 | 0.12767777 | 0.89840399 | 0.97097328 |
| INSIG2 | 27.4604641 | 0.107308526 | 0.40811257 | 0.26293854 | 0.79259795 | NA |
| STEAP3 | 24.0016054 | -3.152325972 | 0.58699199 | -5.3703049 | 7.86E-08 | NA |
| C2orf76 | 10.9619625 | -0.54118758 | 0.5583749 | -0.969219 | 0.33243592 | NA |
| DBI | 46.7099685 | -1.04317278 | 0.37938414 | -2.7496478 | 0.00596593 | 0.12229157 |
| TMEM37 | 8.35027115 | 0.351533175 | 1.019655 | 0.34475698 | 0.73027708 | NA |
| CFAP221 | 15.9263715 | -0.016632713 | 0.65097519 | -0.0255505 | 0.9796159 | NA |
| PTPN4 | 68.2750275 | 0.962428161 | 0.40150031 | 2.39707947 | 0.01652634 | 0.21347398 |
| EPB41L5 | 552.536765 | 0.108503734 | 0.28198918 | 0.38477978 | 0.70040058 | 0.90729979 |
| TMEM185B | 3.33323665 | -0.332226896 | 0.84699902 | -0.39224 | 0.69488089 | NA |
| RALB | 11.1317761 | -0.83975128 | 0.49183247 | -1.7073929 | 0.08774905 | NA |
| GLI2 | 10.0412879 | -2.344784328 | 0.81926091 | -2.8620727 | 0.0042088 | NA |
| TFCP2L1 | 20.6395661 | 0.921271654 | 0.72464231 | 1.27134676 | 0.20360532 | NA |
| CLASP1 | 117.793644 | 0.562000551 | 0.25645729 | 2.19140022 | 0.02842284 | 0.27579678 |
| NIFK-AS1 | 11.5432099 | -0.033537805 | 0.53504062 | -0.0626827 | 0.95001915 | NA |
| NIFK | 8.23380787 | -1.252720734 | 0.65427932 | -1.9146574 | 0.05553621 | NA |
| TSN | 16.253728 | -0.358376344 | 0.42376173 | -0.8457025 | 0.39771874 | NA |
| BIN1 | 5.50009758 | -0.212339533 | 0.86164015 | -0.2464364 | 0.8053444 | NA |
| ERCC3 | 15.4611642 | -0.13647684 | 0.37979151 | -0.3593467 | 0.71933571 | NA |
| MAP3K2 | 122.042228 | -0.210955799 | 0.26886908 | -0.7846042 | 0.43268569 | 0.77964155 |
| IWS1 | 44.4090896 | -0.147308076 | 0.2735174 | -0.5385693 | 0.59018408 | 0.8654248 |
| AC010976.1 | 6.56530645 | -0.717392583 | 0.96205588 | -0.745687 | 0.4558565 | NA |
| LIMS2 | 77.7015595 | 1.091008644 | 0.44382683 | 2.45818542 | 0.01396411 | 0.19677241 |
| WDR33 | 65.405987 | 0.024218243 | 0.27335405 | 0.08859661 | 0.9294025 | 0.97777732 |
| SFT2D3 | 4.94006867 | -0.29536587 | 0.81748831 | -0.361309 | 0.71786848 | NA |
| POLR2D | 7.87053318 | -0.080408698 | 0.47234476 | -0.1702331 | 0.86482684 | NA |
| AMMECR1L | 11.8781581 | -0.217624952 | 0.437072 | -0.4979156 | 0.61854356 | NA |
| SAP130 | 19.806989 | 0.187559236 | 0.41716267 | 0.44960695 | 0.65299387 | NA |
| UGGT1 | 38.0954206 | 0.014989831 | 0.34267472 | 0.04374362 | 0.96510877 | 0.98892249 |
| HS6ST1 | 22.4271228 | -0.571824672 | 0.44617231 | -1.2816229 | 0.19997495 | NA |
| SMPD4 | 16.372058 | 0.03372822 | 0.4193863 | 0.0804228 | 0.935901 | NA |
| MZT2B | 45.7001561 | 0.230487501 | 0.41169122 | 0.55985527 | 0.57557816 | 0.85929076 |
| CCDC115 | 15.1726393 | -0.350295362 | 0.37990966 | -0.9220491 | 0.35650296 | NA |
| IMP4 | 9.90160977 | -0.781337788 | 0.57634048 | -1.3556878 | 0.17519851 | NA |
| PTPN18 | 18.0426448 | -0.409368853 | 0.4560801 | -0.897581 | 0.36940895 | NA |
| FAM168B | 38.2633742 | -0.077120674 | 0.38117379 | -0.2023242 | 0.8396633 | 0.95393706 |
| PLEKHB2 | 37.9842524 | 0.025952613 | 0.29274921 | 0.08865135 | 0.929359 | 0.97777732 |
| MZT2A | 16.144325 | -0.389255029 | 0.40759504 | -0.9550043 | 0.33957551 | NA |
| NCKAP5 | 42.1444236 | 1.408664718 | 0.59362953 | 2.37296941 | 0.01764573 | 0.22021865 |
| MGAT5 | 829.880599 | -0.007786734 | 0.20528625 | -0.0379311 | 0.96974262 | 0.98977467 |
| CCNT2-AS1 | 13.1357778 | 0.138247565 | 0.56558372 | 0.24443342 | 0.80689516 | NA |
| ACMSD | 10.4703751 | 1.207076017 | 1.06452596 | 1.13390942 | 0.25683254 | NA |
| CCNT2 | 43.5656487 | -0.211359776 | 0.26655663 | -0.7929264 | 0.42782073 | 0.77844087 |
| RAB3GAP1 | 102.572759 | 0.178888635 | 0.19881209 | 0.89978752 | 0.36823334 | 0.73626927 |
| ZRANB3 | 28.7389708 | -0.426061498 | 0.32700597 | -1.3029166 | 0.19260324 | 0.59588004 |
| R3HDM1 | 60.3864752 | 0.126943818 | 0.26937277 | 0.47125705 | 0.63745719 | 0.88636488 |
| UBXN4 | 83.0011742 | -0.698040386 | 0.23417396 | -2.9808625 | 0.00287438 | 0.08899601 |
| MCM6 | 46.0760143 | -0.143730059 | 0.35618569 | -0.4035256 | 0.68656157 | 0.90499319 |
| DARS | 48.6909433 | -0.284065521 | 0.30692811 | -0.9255116 | 0.35469986 | 0.72623051 |
| THSD7B | 15.1037138 | -3.654090918 | 1.64534304 | -2.2208687 | 0.02635985 | NA |
| HNMT | 13.3606076 | 0.691626542 | 0.49805215 | 1.38866291 | 0.16493528 | NA |
| AC114763.1 | 16.0964266 | -0.760549887 | 0.54436313 | -1.397137 | 0.16237237 | NA |
| SPOPL | 44.6846299 | 0.344495231 | 0.35917842 | 0.95912006 | 0.33749827 | 0.71222319 |
| LRP1B | 28.1851575 | 1.135497133 | 0.96333084 | 1.1787198 | 0.23850977 | NA |
| GTDC1 | 123.705217 | 0.318905348 | 0.2225075 | 1.43323413 | 0.15179094 | 0.55086922 |
| ZEB2 | 96.8013062 | -0.259006459 | 0.26206139 | -0.9883427 | 0.32298485 | 0.69582324 |
| ACVR2A | 27.7096558 | 0.606479335 | 0.30040524 | 2.01887073 | 0.04350066 | NA |
| ORC4 | 53.138865 | 0.690787766 | 0.33159276 | 2.08324137 | 0.03722923 | 0.30414888 |
| MBD5 | 323.747353 | 0.475312155 | 0.24895624 | 1.9092197 | 0.05623376 | 0.37026642 |
| EPC2 | 57.2658345 | 0.066661428 | 0.28235272 | 0.23609274 | 0.81336071 | 0.94431839 |
| MMADHC | 16.332693 | -0.064048789 | 0.46181445 | -0.1386894 | 0.88969556 | NA |
| AC007364.1 | 16.7182095 | 1.153463158 | 0.56871642 | 2.02818685 | 0.04254118 | NA |
| RND3 | 5.26118264 | -0.58248457 | 1.08611151 | -0.5363027 | 0.59174934 | NA |
| RBM43 | 12.8366081 | 0.336422028 | 0.5849923 | 0.57508796 | 0.56523181 | NA |
| NMI | 10.7314194 | -0.13581079 | 0.99682486 | -0.1362434 | 0.89162888 | NA |
| RIF1 | 59.4645854 | -0.11554593 | 0.24331524 | -0.4748816 | 0.63487136 | 0.88550146 |
| ARL5A | 20.4634004 | 0.048439946 | 0.46365496 | 0.10447412 | 0.9167931 | NA |
| CACNB4 | 7.41483488 | 0.196137395 | 0.57247734 | 0.34261163 | 0.73189065 | NA |
| STAM2 | 39.2987698 | 0.304124762 | 0.286894 | 1.0600597 | 0.28911744 | 0.67724823 |
| FMNL2 | 247.001612 | 0.39153852 | 0.23630212 | 1.65694037 | 0.09753153 | 0.47206192 |
| PRPF40A | 45.6437249 | -0.071334303 | 0.24075989 | -0.2962882 | 0.76701003 | 0.92451492 |
| ARL6IP6 | 27.8359566 | -0.014063362 | 0.38520431 | -0.0365088 | 0.97087663 | NA |
| KCNJ3 | 14.8162189 | 2.41647627 | 0.72979736 | 3.3111606 | 0.0009291 | NA |
| LINC01876 | 22.2103338 | -0.907590015 | 0.87051385 | -1.0425911 | 0.2971377 | NA |
| GPD2 | 26.8874732 | -0.176620544 | 0.36901023 | -0.4786332 | 0.6321996 | NA |
| ACVR1 | 31.1400538 | 0.471782583 | 0.47739791 | 0.98823763 | 0.32303628 | 0.69582324 |
| UPP2 | 13.2365614 | -0.297385904 | 0.6586824 | -0.451486 | 0.65163929 | NA |
| CCDC148 | 120.919091 | 1.132790185 | 0.34405908 | 3.29242931 | 0.00099326 | 0.05099566 |
| PKP4 | 310.356462 | 0.619273021 | 0.35392256 | 1.74974157 | 0.08016292 | 0.43346499 |
| PKP4-AS1 | 8.88909606 | -1.284273863 | 0.81356497 | -1.5785757 | 0.11443342 | NA |
| TANC1 | 397.578 | 0.173748846 | 0.23235345 | 0.74777821 | 0.45459395 | 0.79324789 |
| WDSUB1 | 50.0421164 | 0.362982285 | 0.35940977 | 1.00993995 | 0.31252406 | 0.69213367 |
| BAZ2B | 212.849926 | 0.550833093 | 0.22443972 | 2.45425847 | 0.01411755 | 0.19677241 |
| AC009961.1 | 12.456218 | 0.283878704 | 0.50480961 | 0.56234806 | 0.5738789 | NA |
| 7-Mar | 88.7275309 | 0.374051147 | 0.18788179 | 1.99088558 | 0.04649347 | 0.33771838 |
| CD302 | 5.33113931 | 0.755682849 | 0.69128622 | 1.0931548 | 0.27432584 | NA |
| PLA2R1 | 2663.86674 | 0.466082682 | 0.34586931 | 1.34756877 | 0.17779712 | 0.58063214 |
| ITGB6 | 42.6879592 | -2.346540883 | 0.7484885 | -3.1350393 | 0.00171831 | 0.06518907 |
| LINC02478 | 6.37238102 | 1.141073521 | 0.56110368 | 2.03362331 | 0.04198959 | NA |
| RBMS1 | 543.609738 | 0.570719325 | 0.21817673 | 2.61585795 | 0.00890036 | 0.14875902 |
| TANK | 57.1828682 | -0.015991699 | 0.3520473 | -0.0454249 | 0.96376867 | 0.98892249 |
| PSMD14 | 31.1949888 | 0.026428918 | 0.31882603 | 0.08289448 | 0.93393544 | 0.97832857 |
| DPP4 | 32.0871052 | -0.481285367 | 1.51810905 | -0.3170295 | 0.75122121 | 0.92274989 |
| IFIH1 | 19.1055066 | -0.013449372 | 1.05126862 | -0.0127935 | 0.98979257 | NA |
| GCA | 18.3846716 | -0.073863955 | 0.54132745 | -0.1364497 | 0.8914658 | NA |
| FIGN | 203.170557 | 0.696060672 | 0.31956894 | 2.17812366 | 0.02939683 | 0.27926427 |
| GRB14 | 10.8573803 | 0.0409895 | 0.59800792 | 0.06854341 | 0.94535307 | NA |
| COBLL1 | 102.813405 | 0.790033007 | 0.48068191 | 1.64356718 | 0.10026564 | 0.48127507 |
| AC019197.1 | 90.4472899 | 0.721559714 | 0.92269835 | 0.78201041 | 0.43420846 | 0.78065137 |
| SCN2A | 11.6214156 | 0.20008094 | 1.01114937 | 0.19787476 | 0.84314305 | NA |
| CSRNP3 | 17.0877313 | 1.673975249 | 0.64216184 | 2.60678093 | 0.00913978 | NA |
| TTC21B | 33.230025 | 0.356738687 | 0.29051675 | 1.22794533 | 0.21946749 | 0.62462904 |
| AC010127.1 | 6.80692257 | -1.283782859 | 0.74802224 | -1.7162362 | 0.08611882 | NA |
| SCN9A | 30.9486821 | -0.276084201 | 0.40905888 | -0.6749253 | 0.4997232 | 0.82345406 |
| SCN7A | 33.8138497 | 0.456599231 | 0.72703587 | 0.62802848 | 0.52998528 | 0.834804 |
| STK39 | 33.3786644 | -0.086159141 | 0.37492831 | -0.2298016 | 0.81824591 | 0.94694381 |
| CERS6 | 422.041484 | 0.949410983 | 0.35308054 | 2.68893602 | 0.00716802 | 0.13427683 |
| LRP2 | 65.1423763 | 1.643023746 | 0.9889256 | 1.661423 | 0.09662852 | 0.47093115 |
| FASTKD1 | 23.0781019 | 0.380345932 | 0.3374562 | 1.12709717 | 0.2597014 | NA |
| PPIG | 63.1303441 | 0.16849819 | 0.23346293 | 0.72173423 | 0.47045789 | 0.80448595 |

| | | | | | | |
|---|---|---|---|---|---|---|
| KLHL23 | 23.9718481 | -0.189711954 | 0.44038914 | -0.4307825 | 0.66662649 | NA |
| SSB | 88.0309966 | 0.208633394 | 0.59717874 | 0.34936507 | 0.72681525 | 0.91700494 |
| METTL5 | 16.096895 | -0.628738769 | 0.41703787 | -1.5076299 | 0.13164927 | NA |
| UBR3 | 136.241033 | 0.639269994 | 0.24164407 | 2.64550255 | 0.00815697 | 0.14384634 |
| GORASP2 | 27.0247824 | -0.018325921 | 0.30234512 | -0.0606126 | 0.95166775 | NA |
| TLK1 | 77.2603643 | 0.337632165 | 0.28355267 | 1.19072117 | 0.23376306 | 0.63975088 |
| METTL8 | 16.0861139 | -0.406382512 | 0.42715825 | -0.9513629 | 0.34142019 | NA |
| DCAF17 | 12.7737961 | -0.527487535 | 0.50503415 | -1.0444591 | 0.29627301 | NA |
| DYNC1I2 | 64.7009441 | -0.24221573 | 0.23257538 | -1.0414504 | 0.29766654 | 0.68103704 |
| SLC25A12 | 40.5725878 | 0.459770148 | 0.31975693 | 1.43787393 | 0.15046983 | 0.54834287 |
| HAT1 | 15.2231649 | -0.290916054 | 0.44002339 | -0.6611377 | 0.50852401 | NA |
| METAP1D | 5.07912503 | -0.99213258 | 0.97491025 | -1.0176656 | 0.30883692 | NA |
| ITGA6 | 25.9991271 | 0.744312302 | 0.52221288 | 1.42530439 | 0.15406923 | NA |
| PDK1 | 11.7958609 | -0.498892941 | 0.52553499 | -0.9493049 | 0.34246556 | NA |
| RAPGEF4 | 7.20472698 | -0.411078629 | 0.66664579 | -0.6166373 | 0.53747402 | NA |
| MAP3K20 | 88.3691286 | -0.910621885 | 0.47552939 | -1.9149645 | 0.05549704 | 0.36720388 |
| MAP3K20-AS | 4.54017196 | -2.861020114 | 0.99223531 | -2.8834089 | 0.00393396 | NA |
| SP3 | 47.2741244 | -0.057560763 | 0.28781858 | -0.1999897 | 0.84148861 | 0.95398401 |
| OLA1 | 66.1700929 | 0.457015016 | 0.31634389 | 1.44467789 | 0.14854838 | 0.54724091 |
| CIR1 | 28.7349237 | -0.290586962 | 0.30608182 | -0.9493768 | 0.34242903 | 0.71725052 |
| SCRN3 | 10.5681858 | -0.039237523 | 0.55353733 | -0.0708851 | 0.94348924 | NA |
| GPR155 | 19.7379729 | -0.38631354 | 0.48984389 | -0.7886462 | 0.4303188 | NA |
| AC010894.3 | 87.2314681 | -1.561229357 | 0.48433985 | -3.2234171 | 0.00126671 | 0.0545866 |
| WIPF1 | 26.5768979 | -0.003067683 | 0.38986765 | -0.0078685 | 0.99372189 | NA |
| CHN1 | 42.1560463 | -0.357953567 | 0.41047928 | -0.8720381 | 0.38318759 | 0.74924885 |
| ATF2 | 55.2786423 | 0.125898989 | 0.25511019 | 0.49350827 | 0.6216535 | 0.8780438 |
| ATP5MC3 | 55.1951033 | -0.452274854 | 0.53459704 | -0.8460108 | 0.39754674 | 0.76023081 |
| LNPK | 21.9142581 | 0.277973955 | 0.40040564 | 0.69423088 | 0.48753743 | NA |
| HOXD-AS2 | 4.65740293 | -1.03167454 | 0.8495036 | -1.214444 | 0.22457823 | NA |
| HOXD8 | 7.80812008 | -0.83673048 | 0.61003476 | -1.3716112 | 0.17018452 | NA |
| HOXD3 | 8.52588543 | 0.231251332 | 0.67611299 | 0.3420306 | 0.73232786 | NA |
| HAGLR | 16.3890552 | -0.333972094 | 0.54475885 | -0.6130641 | 0.53983396 | NA |
| HOXD1 | 9.59787792 | -0.016513047 | 0.63147635 | -0.0261499 | 0.97913777 | NA |
| MTX2 | 52.051364 | 0.550663253 | 0.28568624 | 1.92751059 | 0.05391603 | 0.3615191 |
| HNRNPA3 | 99.4499727 | 0.128380414 | 0.22487315 | 0.57090148 | 0.56806643 | 0.85654436 |
| NFE2L2 | 198.240511 | -0.429505818 | 0.21483149 | -1.9992684 | 0.04557932 | 0.33551673 |
| AGPS | 48.5862535 | -0.089609986 | 0.28214558 | -0.3176019 | 0.7507869 | 0.92274989 |
| TTC30A | 4.19825116 | -0.253280025 | 0.64856427 | -0.3905242 | 0.69614898 | NA |
| RBM45 | 12.5667263 | 0.245641315 | 0.42259076 | 0.5812747 | 0.56105533 | NA |
| AC009948.1 | 16.2693828 | -0.006069134 | 0.6366616 | -0.0095327 | 0.99239408 | NA |
| PRKRA | 29.1848081 | -0.495504847 | 0.30044953 | -1.6492116 | 0.0991043 | 0.47796319 |
| PJVK | 9.4049757 | -0.332849282 | 0.54937154 | -0.6058728 | 0.54459921 | NA |
| FKBP7 | 4.49664614 | -1.063326857 | 0.89759981 | -1.1846336 | 0.23616236 | NA |
| PLEKHA3 | 32.1617121 | -0.360493343 | 0.30618335 | -1.1773774 | 0.23904492 | 0.64594683 |
| TTN-AS1 | 46.3343405 | 0.0791323 | 0.34581771 | 0.22882662 | 0.81900367 | 0.94694381 |
| SESTD1 | 37.2522609 | -0.321217582 | 0.36300521 | -0.8848842 | 0.37621909 | 0.74330391 |
| CWC22 | 17.2089164 | -0.067744295 | 0.34792425 | -0.1947099 | 0.84562006 | NA |
| UBE2E3 | 79.2265591 | 0.073722548 | 0.25105119 | 0.29365544 | 0.76902121 | 0.9255638 |
| SSFA2 | 33.2786609 | 0.547249065 | 0.41931752 | 1.30509469 | 0.19186061 | 0.5949439 |
| PDE1A | 70.8795268 | 2.48457482 | 0.83941791 | 2.95987825 | 0.00307761 | 0.0904424 |
| DNAJC10 | 51.850996 | 0.156161418 | 0.2748982 | 0.56806998 | 0.56998745 | 0.8573305 |
| NCKAP1 | 136.902616 | 0.338662377 | 0.22509343 | 1.50454136 | 0.13244203 | 0.52613957 |
| NUP35 | 9.62161953 | -0.099522613 | 0.60478661 | -0.1645582 | 0.86929171 | NA |
| ZNF804A | 367.665875 | 0.332978272 | 0.61023038 | 0.54565994 | 0.58529971 | 0.86235633 |
| FSIP2 | 11.9630366 | 0.391279717 | 0.75803557 | 0.51617593 | 0.60573154 | NA |
| FSIP2-AS1 | 7.53107251 | 0.707603474 | 0.57869133 | 1.22276495 | 0.22141849 | NA |
| LINC01473 | 12.847179 | -0.386285422 | 0.44782865 | -0.8625742 | 0.38837165 | NA |
| ZC3H15 | 34.7366899 | -0.320565432 | 0.29707078 | -1.0790877 | 0.28054862 | 0.66958859 |
| ITGAV | 637.920035 | 0.950472485 | 0.37314179 | 2.54721531 | 0.01085864 | 0.17004884 |
| AC017101.1 | 34.4830407 | 0.263168554 | 0.60630015 | 0.43405656 | 0.66424737 | 0.89968484 |
| FAM171B | 6.97983059 | 0.66168015 | 0.55005312 | 1.20293864 | 0.22900007 | NA |
| AC007319.1 | 106.681482 | -0.125750566 | 0.84631335 | -0.1485863 | 0.88188009 | 0.96464553 |
| CALCRL | 9.74559128 | 0.373596149 | 0.49889057 | 0.7488539 | 0.45394527 | NA |
| TFPI | 116.096145 | -0.917470693 | 0.31037649 | -2.9559929 | 0.00311664 | 0.0904424 |
| LINC01090 | 11.0661106 | -1.319025508 | 0.74944537 | -1.7600022 | 0.07840743 | NA |
| GULP1 | 329.242428 | 0.239515757 | 0.26456378 | 0.90532331 | 0.36529413 | 0.73398222 |
| COL3A1 | 27.2206474 | 1.550766412 | 0.73757254 | 2.10252731 | 0.03550711 | NA |
| WDR75 | 15.6275494 | -0.296656295 | 0.39436373 | -0.7522403 | 0.45190655 | NA |
| SLC40A1 | 15.1041535 | 0.908011077 | 0.59511513 | 1.5257738 | 0.12706621 | NA |
| ASNSD1 | 12.2816795 | -0.084954205 | 0.51139257 | -0.1661233 | 0.86805994 | NA |
| ANKAR | 54.7345311 | 0.658438214 | 0.40633136 | 1.62044646 | 0.10513641 | 0.48679597 |
| OSGEPL1 | 5.07054743 | -0.500158276 | 0.76246965 | -0.6559714 | 0.51184253 | NA |
| ORMDL1 | 35.0736512 | 0.058606808 | 0.3080108 | 0.19027517 | 0.84909351 | 0.95532127 |
| PMS1 | 24.6467515 | -0.36191882 | 0.37348244 | -0.9690384 | 0.33252604 | NA |
| C2orf88 | 14.0954642 | -0.814379021 | 0.51197351 | -1.5906663 | 0.11168469 | NA |
| HIBCH | 67.6542673 | 0.413845809 | 0.30768877 | 1.34501435 | 0.1786206 | 0.58191579 |
| INPP1 | 5.27972461 | -0.176010726 | 0.64102944 | -0.2745751 | 0.78364269 | NA |
| MFSD6 | 125.952637 | 0.069224267 | 0.42376928 | 0.16335367 | 0.87023998 | 0.96207879 |
| NEMP2 | 6.53935328 | -0.091174445 | 0.56078859 | -0.1625826 | 0.87084712 | NA |
| NAB1 | 26.0520639 | -0.423220394 | 0.32425934 | -1.3051911 | 0.1918278 | NA |
| GLS | 383.665036 | 0.464796388 | 0.30397136 | 1.52907955 | 0.12624472 | 0.52276701 |
| STAT1 | 122.970851 | -0.055522109 | 0.5759686 | -0.0963978 | 0.92320463 | 0.97775374 |
| MYO1B | 336.177247 | 0.484030189 | 0.40103189 | 1.20696185 | 0.22744684 | 0.63327644 |
| NABP1 | 9.06942835 | -1.005638556 | 0.98185358 | -1.0242246 | 0.30572922 | NA |
| SLC39A10 | 36.1739002 | 0.429609536 | 0.43036932 | 0.99823457 | 0.31816563 | 0.69469143 |
| DNAH7 | 23.2258733 | 0.765256281 | 0.50962139 | 1.50161727 | 0.13319598 | NA |
| GTF3C3 | 13.507575 | -1.200444006 | 0.44555768 | -2.6942505 | 0.00705471 | NA |
| PGAP1 | 33.1664814 | -1.16711713 | 0.55225327 | -2.113373 | 0.03456886 | 0.29210684 |
| ANKRD44 | 197.100118 | -0.087745672 | 0.29441111 | -0.2980379 | 0.76567423 | 0.9240942 |
| SF3B1 | 62.5449236 | -0.089534154 | 0.23835385 | -0.3756354 | 0.70718794 | 0.91058211 |
| COQ10B | 17.4134243 | -0.722851197 | 0.42449361 | -1.7028553 | 0.08859515 | NA |
| HSPD1 | 36.750388 | -0.764953325 | 0.3885414 | -1.968782 | 0.04897813 | 0.34548746 |
| MOB4 | 15.7681701 | 0.042353313 | 0.413998 | 0.10230318 | 0.91851603 | NA |
| AC011997.1 | 5.78465709 | 0.942808341 | 0.88636371 | 1.06368112 | 0.28747318 | NA |
| PLCL1 | 86.1006576 | 1.947501557 | 0.54733187 | 3.55817313 | 0.00037344 | 0.02657343 |
| FTCDNL1 | 7.44971826 | 0.173146278 | 0.63860926 | 0.27113023 | 0.78629087 | NA |
| C2orf69 | 8.50370028 | -0.395026201 | 0.53320642 | -0.7408504 | 0.45878414 | NA |
| TYW5 | 10.6183194 | -0.006000265 | 0.44191853 | -0.0135778 | 0.98916685 | NA |
| MAIP1 | 7.19879714 | 0.006613433 | 0.53785473 | 0.01229595 | 0.9901895 | NA |
| SPATS2L | 231.522711 | 0.17597898 | 0.37430038 | 0.47015443 | 0.63824469 | 0.886719 |
| KCTD18 | 8.23277861 | -0.499768823 | 0.50320059 | -0.9931801 | 0.32062219 | NA |
| AOX1 | 33.5391043 | -0.061168811 | 0.61651312 | -0.0992174 | 0.92096568 | 0.97775374 |
| BZW1 | 19.5646721 | 0.342293547 | 0.37828347 | 0.90485992 | 0.36553961 | NA |
| CLK1 | 43.144963 | 0.723437986 | 0.38820192 | 1.86356109 | 0.06238337 | 0.38512471 |
| PPIL3 | 7.9828508 | -0.200448939 | 0.51225591 | -0.3913063 | 0.69557088 | NA |
| NIF3L1 | 5.99877666 | -0.423726932 | 0.55728449 | -0.7603422 | 0.44705004 | NA |
| ORC2 | 29.5486018 | -0.200403306 | 0.41668618 | -0.4809454 | 0.6305553 | 0.8822119 |
| AC005037.1 | 10.062001 | -0.922271019 | 0.53818216 | -1.7136782 | 0.08658785 | NA |
| FAM126B | 56.6575345 | 0.655793726 | 0.30188732 | 2.1723129 | 0.02983207 | 0.27926427 |
| NDUFB3 | 31.8203402 | -0.564992768 | 0.48819969 | -1.1572985 | 0.24715043 | 0.65061974 |
| CFLAR | 146.236759 | 0.293133876 | 0.44674206 | 0.65615913 | 0.51172175 | 0.83054958 |
| CFLAR-AS1 | 9.62682601 | -0.113307977 | 0.87425444 | -0.1296053 | 0.89687874 | NA |
| CASP8 | 7.23580157 | 0.34608914 | 0.85608998 | 0.40426725 | 0.68601621 | NA |

| | | | | | | |
|---|---|---|---|---|---|---|
| TRAK2 | 40.5297859 | -0.193036601 | 0.38443996 | -0.5021242 | 0.61558017 | 0.8757605 |
| STRADB | 18.0883673 | 0.018883252 | 0.59127678 | 0.0319364 | 0.97452277 | NA |
| ALS2 | 28.8641398 | 0.282360481 | 0.35096301 | 0.8045306 | 0.4210906 | 0.77473274 |
| SUMO1 | 34.0586494 | -0.168010921 | 0.31172553 | -0.5389707 | 0.58990708 | 0.8654248 |
| NOP58 | 29.4203118 | -0.227596431 | 0.3268101 | -0.696418 | 0.4861671 | 0.81509175 |
| BMPR2 | 214.898827 | -0.810679977 | 0.30173823 | -2.6866996 | 0.00721618 | 0.13427683 |
| FAM117B | 11.0596996 | 0.175649406 | 0.44298775 | 0.39651075 | 0.69172828 | NA |
| ICA1L | 28.533683 | 0.162993395 | 0.28774719 | 0.56644652 | 0.57109028 | 0.8573305 |
| WDR12 | 12.3651063 | -0.654336346 | 0.50575262 | -1.2937874 | 0.19573888 | NA |
| CARF | 25.9393415 | 0.043715197 | 0.35032362 | 0.12478518 | 0.90069362 | NA |
| NBEAL1 | 255.679446 | 0.551898978 | 0.17203205 | 3.20811716 | 0.00133607 | 0.0545866 |
| CYP20A1 | 50.7886426 | -0.12882138 | 0.34403355 | -0.3744442 | 0.70807384 | 0.9106723 |
| ABI2 | 93.3989663 | 0.200241971 | 0.21770525 | 0.91978476 | 0.35768525 | 0.72903084 |
| RAPH1 | 48.3588339 | 0.460061643 | 0.27825687 | 1.6533703 | 0.09825552 | 0.47499929 |
| PARD3B | 1868.53506 | 0.510845899 | 0.26533355 | 1.92529701 | 0.05419221 | 0.3615191 |
| INO80D | 124.917304 | 1.190243887 | 0.69153687 | 1.72115753 | 0.08522224 | 0.44564602 |
| NDUFS1 | 48.1702332 | 0.125413902 | 0.26021829 | 0.48195652 | 0.62983683 | 0.88166074 |
| EEF1B2 | 23.4921814 | -0.79697504 | 0.4287649 | -1.8587693 | 0.06305985 | NA |
| ZDBF2 | 19.2890622 | 0.167588431 | 0.34598615 | 0.48437902 | 0.62811691 | NA |
| FASTKD2 | 14.1785568 | -0.11060524 | 0.52103097 | -0.2122815 | 0.83188742 | NA |
| CPO | 4.50791788 | 0.250140419 | 0.69549828 | 0.35965642 | 0.71910409 | NA |
| KLF7 | 374.10306 | -0.29773822 | 0.27503284 | -1.0825551 | 0.27900593 | 0.66958859 |
| AC007879.1 | 8.0324921 | 0.836329431 | 0.70735197 | 1.18233845 | 0.23707143 | NA |
| AC009226.1 | 5.20280058 | 0.295089998 | 0.87641664 | 0.33670059 | 0.73634262 | NA |
| CREB1 | 51.1708704 | -0.29363996 | 0.27578906 | -1.0647266 | 0.28699966 | 0.67443257 |
| METTL21A | 28.8267675 | -0.530294238 | 0.32617537 | -1.6257949 | 0.1039933 | 0.4862279 |
| CCNYL1 | 20.2817295 | -0.445729776 | 0.41565935 | -1.0723439 | 0.28356559 | NA |
| FZD5 | 5.87740295 | -0.16325922 | 0.70993306 | -0.2299642 | 0.81811955 | NA |
| PLEKHM3 | 28.4773521 | 0.529511447 | 0.30301973 | 1.74744876 | 0.08055953 | NA |
| IDH1 | 8.46936813 | -0.257976734 | 0.51006837 | -0.5057689 | 0.61301887 | NA |
| PIKFYVE | 45.2879479 | 0.167046035 | 0.27250417 | 0.61300359 | 0.53987397 | 0.83929813 |
| PTH2R | 75.7296825 | 1.339822661 | 0.62116735 | 2.15694316 | 0.03101009 | 0.28264481 |
| MAP2 | 24.5855091 | 2.892510022 | 0.59528226 | 4.85905631 | 1.18E-06 | NA |
| RPE | 17.8228591 | -0.390553974 | 0.44331135 | -0.8809925 | 0.37832188 | NA |
| KANSL1L | 124.875869 | 0.27993514 | 0.26666536 | 1.04976191 | 0.29382759 | 0.6804998 |
| AC007038.1 | 19.6171587 | 0.283891775 | 0.51239499 | 0.5540487 | 0.57954552 | NA |
| LANCL1 | 53.9263082 | 0.275688857 | 0.37121742 | 0.74266141 | 0.4576867 | 0.79434199 |
| ERBB4 | 836.620568 | 0.479479729 | 0.2101853 | 2.28122389 | 0.0225352 | 0.24569509 |
| IKZF2 | 41.8526159 | 1.088809882 | 0.47483518 | 2.29302697 | 0.02184645 | 0.24297341 |
| SPAG16 | 148.122045 | 0.667789003 | 0.36061674 | 1.85179701 | 0.06405498 | 0.3906872 |
| AC068051.1 | 7.16805838 | -0.005200332 | 0.82280507 | -0.0063202 | 0.99495721 | NA |
| BARD1 | 9.63875616 | 0.515546376 | 0.56349045 | 0.91491591 | 0.36023578 | NA |
| AC016708.1 | 8.7184807 | -0.20484097 | 0.62704392 | -0.3266772 | 0.74391202 | NA |
| ATIC | 23.9652256 | 0.323701861 | 0.40578801 | 0.79771174 | 0.42503779 | NA |
| PECR | 5.84336542 | -0.385314344 | 0.60494287 | -0.6369434 | 0.52416174 | NA |
| XRCC5 | 81.2042349 | -0.433091221 | 0.23076262 | -1.8767824 | 0.06054792 | 0.38074788 |
| SMARCAL1 | 15.6422993 | -0.142831586 | 0.40587457 | -0.3519107 | 0.72490527 | NA |
| RPL37A | 66.9373594 | -0.15529005 | 0.39029411 | -0.3978796 | 0.69071896 | 0.90557575 |
| IGFBP2 | 1167.78975 | -0.613370574 | 0.47265464 | -1.2977141 | 0.19438561 | 0.59865455 |
| IGFBP5 | 18.7392072 | -0.029130793 | 0.90316117 | -0.0322543 | 0.97426929 | NA |
| TNS1 | 23.6381671 | 1.099638911 | 0.75941896 | 1.44800034 | 0.14761696 | NA |
| ARPC2 | 61.715796 | -0.48487703 | 0.33038296 | -1.4676212 | 0.14220716 | 0.53855483 |
| AAMP | 14.9327103 | 0.290938898 | 0.41378851 | 0.70311015 | 0.48198711 | NA |
| PNKD | 43.4178646 | -0.419704754 | 0.31634715 | -1.3267221 | 0.18460063 | 0.587049 |
| TMBIM1 | 35.1811965 | -0.319855212 | 0.47960596 | -0.6669125 | 0.50482802 | 0.82600448 |
| CTDSP1 | 28.4498787 | -0.088290636 | 0.35912827 | -0.2458471 | 0.80580061 | NA |
| USP37 | 22.7981793 | -0.52899629 | 0.40238167 | -1.314663 | 0.18862319 | NA |
| CNOT9 | 15.0321737 | -0.021994868 | 0.41051767 | -0.0535784 | 0.95727109 | NA |
| ZNF142 | 6.68505588 | -0.223750553 | 0.528383 | -0.4234628 | 0.67195762 | NA |
| RNF25 | 5.73462075 | 0.221836143 | 0.68387279 | 0.32438217 | 0.74564871 | NA |
| STK36 | 7.31967983 | 0.783433714 | 0.60549941 | 1.2938637 | 0.1957125 | NA |
| TTLL4 | 4.230667 | -0.510351903 | 0.7170393 | -0.7117489 | 0.47662031 | NA |
| CYP27A1 | 13.1730378 | 0.855890768 | 0.46939312 | 1.82339861 | 0.06824304 | NA |
| CNPPD1 | 13.8786314 | -0.032439384 | 0.53261608 | -0.0609058 | 0.95143427 | NA |
| RETREG2 | 22.7337811 | -0.062892719 | 0.34034483 | -0.1847912 | 0.85339284 | NA |
| ZFAND2B | 11.9438486 | -0.170969742 | 0.41093386 | -0.4160517 | 0.67737215 | NA |
| ANKZF1 | 9.92670352 | 0.228229736 | 0.51634282 | 0.44201203 | 0.6584805 | NA |
| STK16 | 4.61822305 | -0.809060291 | 0.7175774 | -1.1274885 | 0.25953598 | NA |
| TUBA4A | 5.75649311 | -0.742414208 | 0.74565135 | -0.9956586 | 0.31941603 | NA |
| DNAJB2 | 23.3759363 | 0.697563509 | 0.45545698 | 1.53156837 | 0.12562898 | NA |
| DNPEP | 10.7049407 | -0.442162211 | 0.60013262 | -0.7367742 | 0.4612597 | NA |
| GMPPA | 5.54053156 | 0.840775324 | 0.67991018 | 1.23659765 | 0.21623649 | NA |
| CHPF | 32.3545035 | 0.790906013 | 0.51166217 | 1.54575823 | 0.12216297 | 0.51522823 |
| OBSL1 | 15.3615789 | 0.618591906 | 0.40895811 | 1.51260456 | 0.13038013 | NA |
| EPHA4 | 29.349914 | 1.424002662 | 0.82246632 | 1.73138113 | 0.08338381 | 0.44152054 |
| SGPP2 | 10.4998484 | -1.81373151 | 0.88105519 | -2.0585901 | 0.03953352 | NA |
| FARSB | 17.9887062 | 0.341809924 | 0.36530327 | 0.9356881 | 0.3494338 | NA |
| ACSL3 | 70.6017522 | 0.079419209 | 0.36283305 | 0.21888637 | 0.82673856 | 0.949142 |
| AP1S3 | 53.809792 | -0.747948143 | 0.2737474 | -2.7322566 | 0.00629021 | 0.12568031 |
| WDFY1 | 46.3081704 | 0.053990488 | 0.33968146 | 0.15894447 | 0.87371263 | 0.96229085 |
| MRPL44 | 7.90975681 | -0.548230636 | 0.64462836 | -0.85046 | 0.39506939 | NA |
| SERPINE2 | 9.17622633 | -0.803028713 | 0.77636787 | -1.0343405 | 0.30097702 | NA |
| CUL3 | 90.1461077 | -0.023334565 | 0.19029873 | -0.1226207 | 0.90240746 | 0.97220041 |
| IRS1 | 13.1052569 | 0.159575759 | 0.5547399 | 0.2876587 | 0.773608 | NA |
| RHBDD1 | 58.0915933 | -0.086289208 | 0.34290412 | -0.2516424 | 0.8013175 | 0.94085969 |
| COL4A4 | 600.731885 | -0.15901895 | 0.41039226 | -0.3874804 | 0.69840061 | 0.90729979 |
| COL4A3 | 1303.90159 | 0.406752649 | 0.24067777 | 1.69002996 | 0.09102222 | 0.45906588 |
| AC097662.1 | 149.819499 | -0.334621361 | 0.51534656 | -0.6493133 | 0.51613591 | 0.83104823 |
| MFF | 27.6275975 | 0.293044528 | 0.33866411 | 0.86529549 | 0.38687662 | NA |
| AGFG1 | 77.9189516 | 0.327890869 | 0.23514068 | 1.39444554 | 0.1631831 | 0.56498748 |
| TRIP12 | 123.699673 | -0.031287646 | 0.19308584 | -0.1620401 | 0.87127429 | 0.96207879 |
| FBXO36 | 9.5854807 | 0.24349128 | 0.41543829 | 0.58610697 | 0.55780363 | NA |
| AC009950.1 | 3.98044971 | -0.986274042 | 1.08793659 | -0.9065547 | 0.36464232 | NA |
| SP110 | 21.9546689 | -0.241518662 | 1.04423193 | -0.2312883 | 0.81709081 | NA |
| SP140L | 8.75674871 | -0.839154863 | 0.8207451 | -1.0224305 | 0.30657717 | NA |
| SP100 | 81.8738936 | 0.084632429 | 0.4841454 | 0.17480788 | 0.8612306 | 0.95951831 |
| AC010149.1 | 4.03430244 | -0.258998018 | 0.76623336 | -0.3380145 | 0.73535223 | NA |
| CAB39 | 52.0727765 | 0.320185524 | 0.24928273 | 1.28442722 | 0.19899251 | 0.60367512 |
| ITM2C | 40.6257141 | 0.17908444 | 0.4542627 | 0.394231 | 0.6934105 | 0.90557575 |
| PSMD1 | 79.2570963 | 0.191282107 | 0.18795847 | 1.01768282 | 0.30882871 | 0.6882709 |
| ARMC9 | 55.975054 | -0.425263237 | 0.2381132 | -1.7859709 | 0.07410397 | 0.4186152 |
| B3GNT7 | 10.7275675 | 2.312630188 | 0.60782045 | 3.80479169 | 0.00014192 | NA |
| NCL | 60.6641832 | -1.093364748 | 0.35506241 | -3.0793593 | 0.00207446 | 0.07380722 |
| PTMA | 316.880455 | -1.425426467 | 0.4682463 | -3.0441809 | 0.00233315 | 0.07786717 |
| PDE6D | 13.7320441 | 0.407890918 | 0.40461803 | 1.00808884 | 0.31341181 | NA |
| COPS7B | 11.3907932 | 0.753003718 | 0.45246047 | 1.664242 | 0.09606408 | NA |
| DIS3L2 | 57.9804258 | 0.005861364 | 0.27931005 | 0.02098515 | 0.9832575 | 0.99552981 |
| EIF4E2 | 17.6836856 | 0.022544929 | 0.36270173 | 0.06215832 | 0.95043676 | NA |
| EFHD1 | 23.2766146 | 1.666733464 | 0.4571409 | 3.64599507 | 0.00026636 | NA |
| GIGYF2 | 76.0999685 | 0.16101074 | 0.21041251 | 0.76521469 | 0.44414372 | 0.78734568 |
| ATG16L1 | 8.18242468 | 0.019573823 | 0.46787134 | 0.04183591 | 0.96662951 | NA |
| DGKD | 36.3535591 | -0.706507342 | 0.41201987 | -1.7147409 | 0.08639273 | 0.44638081 |
| USP40 | 51.7037527 | 0.351805985 | 0.26064828 | 1.34973453 | 0.17710115 | 0.58001112 |

| | | | | | | |
|---|---|---|---|---|---|---|
| SH3BP4 | 22.2180048 | 0.020713695 | 0.43600201 | 0.04750825 | 0.96210815 | NA |
| AGAP1 | 117.729876 | 0.136725895 | 0.28771476 | 0.47521334 | 0.63463492 | 0.88547617 |
| IQCA1 | 10.4477375 | 0.420920332 | 0.52714792 | 0.79848618 | 0.42458841 | NA |
| COPS8 | 23.44104 | 0.262862113 | 0.34148071 | 0.76977149 | 0.44143545 | NA |
| RAB17 | 5.98832359 | 0.184057487 | 0.90206429 | 0.20404032 | 0.83832199 | NA |
| LRRFIP1 | 169.35037 | 0.078749291 | 0.26289869 | 0.29954234 | 0.76452627 | 0.92399242 |
| RAMP1 | 103.627863 | -0.543282584 | 0.77884834 | -0.6975461 | 0.48546113 | 0.81450403 |
| ILKAP | 35.8524601 | -1.030703338 | 0.41976286 | -2.455442 | 0.01407115 | 0.19677241 |
| PER2 | 7.35222441 | -1.013946223 | 0.64719785 | -1.5666712 | 0.11719157 | NA |
| TRAF3IP1 | 21.9967632 | 0.062451285 | 0.4023833 | 0.15520347 | 0.87666091 | NA |
| ASB1 | 9.30578255 | 0.012306459 | 0.44158296 | 0.02786896 | 0.97776666 | NA |
| HDAC4 | 50.4192597 | -0.654759329 | 0.33676872 | -1.9442403 | 0.05186648 | 0.35716541 |
| NDUFA10 | 38.0902977 | -0.247325777 | 0.33663176 | -0.7347072 | 0.46251784 | 0.79793427 |
| COPS9 | 25.4463575 | -0.899097443 | 0.59179678 | -1.5192672 | 0.12869525 | NA |
| AC124861.1 | 12.2914266 | 0.718185056 | 0.60128021 | 1.19442656 | 0.23231115 | NA |
| GPC1 | 16.2345747 | -0.288404934 | 0.53847496 | -0.5355958 | 0.59223792 | NA |
| ANKMY1 | 17.4032084 | -0.076564935 | 0.40541847 | -0.1888541 | 0.85020718 | NA |
| RNPEPL1 | 24.8018235 | 0.165455886 | 0.40978388 | 0.40376377 | 0.68638644 | NA |
| CAPN10 | 5.12179068 | 0.163640164 | 0.59761225 | 0.27382331 | 0.78422041 | NA |
| SNED1 | 34.8590004 | -2.168149386 | 0.54782431 | -3.9577458 | 7.57E-05 | 0.00829402 |
| AC005237.1 | 5.00128849 | -1.679065116 | 0.87328742 | -1.9226947 | 0.0545184 | NA |
| MTERF4 | 22.475983 | -0.736165499 | 0.36531193 | -2.0151696 | 0.04388688 | NA |
| PPP1R7 | 21.5240482 | -0.311847479 | 0.31762894 | -0.9817981 | 0.32619934 | NA |
| HDLBP | 84.925135 | -0.392530448 | 0.26060528 | -1.506226 | 0.13200917 | 0.52613957 |
| 2-Sep | 152.732142 | -0.130361844 | 0.28089053 | -0.464102 | 0.6425747 | 0.88951638 |
| FARP2 | 66.4635336 | 0.645333167 | 0.28912842 | 2.23199492 | 0.0256153 | 0.25731925 |
| STK25 | 22.7888448 | 0.315380641 | 0.31334845 | 1.00648541 | 0.31418212 | NA |
| BOK | 19.3549971 | -0.158179738 | 0.6255754 | -0.2528548 | 0.80038042 | NA |
| THAP4 | 13.7905746 | 0.093802326 | 0.40754813 | 0.23016257 | 0.81796544 | NA |
| ATG4B | 32.0722748 | 0.356874315 | 0.40801321 | 0.87466362 | 0.38175696 | 0.74899182 |
| DTYMK | 8.53849958 | -0.238979251 | 0.58803712 | -0.4064016 | 0.68444752 | NA |
| ING5 | 20.6924016 | 0.176568122 | 0.35882189 | 0.49207734 | 0.62266467 | NA |
| D2HGDH | 30.5517391 | 0.716599066 | 0.34893251 | 2.05368958 | 0.04000575 | 0.31540889 |
| LINC01266 | 8.36157035 | 0.027985428 | 0.64730611 | 0.04323368 | 0.96551525 | NA |
| CNTN4 | 166.103712 | 1.200894325 | 1.13728538 | 1.0559305 | 0.29099997 | 0.67792199 |
| TRNT1 | 14.771921 | -0.635163542 | 0.43609582 | -1.456477 | 0.14526079 | NA |
| CRBN | 30.6834806 | -0.295600931 | 0.37339845 | -0.7916501 | 0.42856471 | 0.77844087 |
| AC034195.1 | 39.7439024 | 0.459235847 | 0.44359776 | 1.03525286 | 0.30055084 | 0.68204729 |
| SUMF1 | 130.706004 | 0.060532197 | 0.32719361 | 0.18500421 | 0.85322573 | 0.95569766 |
| ITPR1 | 27.945813 | 1.092585501 | 0.34923683 | 3.12849447 | 0.00175704 | NA |
| AC018816.1 | 6.79896231 | -0.692179919 | 0.67612761 | -1.0237415 | 0.30595737 | NA |
| BHLHE40 | 17.2130101 | -1.791629856 | 0.83450618 | -2.1469342 | 0.03179852 | NA |
| ARL8B | 52.7137506 | 0.257801454 | 0.24959011 | 1.03289932 | 0.30165102 | 0.68219446 |
| AC026202.2 | 6.59884379 | 0.120556209 | 0.66823358 | 0.18041028 | 0.85683048 | NA |
| EDEM1 | 16.6676409 | -0.119095972 | 0.42160975 | -0.2824792 | 0.77757612 | NA |
| LMCD1-AS1 | 41.0749818 | 0.62163747 | 0.48845047 | 1.27267247 | 0.20313429 | 0.61166494 |
| LMCD1 | 10.2964249 | 0.570103349 | 0.63858343 | 0.89276252 | 0.37198436 | NA |
| RAD18 | 11.2267602 | -0.280966848 | 0.52773205 | -0.5324044 | 0.59444596 | NA |
| SRGAP3 | 14.9952516 | -0.400102324 | 0.40018824 | -0.9997853 | 0.31741442 | NA |
| THUMPD3-A | 41.8618228 | 0.042443002 | 0.33349324 | 0.12726795 | 0.89872832 | 0.97097328 |
| THUMPD3 | 9.83455962 | -0.572243012 | 0.44584327 | -1.2835071 | 0.19931447 | NA |
| SETD5 | 126.684034 | -0.134552549 | 0.16141212 | -0.8335963 | 0.40450851 | 0.76739313 |
| MTMR14 | 15.5187608 | -0.168722747 | 0.39699187 | -0.425003 | 0.67083446 | NA |
| OGG1 | 9.51483039 | -0.5966535 | 0.45743067 | -1.3043583 | 0.19211144 | NA |
| CAMK1 | 4.91844251 | -0.217044816 | 0.71407298 | -0.3039533 | 0.76116349 | NA |
| TADA3 | 14.9703677 | -0.164972953 | 0.39818523 | -0.4143121 | 0.67864556 | NA |
| RPUSD3 | 7.20556262 | -0.948811314 | 0.56528231 | -1.6784734 | 0.09325472 | NA |
| JAGN1 | 11.4977909 | -1.250096059 | 0.57234907 | -2.1841497 | 0.02895125 | NA |
| IL17RC | 7.27013497 | -0.095579017 | 0.50817368 | -0.1880834 | 0.8508113 | NA |
| CRELD1 | 29.9646125 | 0.260518472 | 0.49719845 | 0.52397282 | 0.60029745 | 0.86882737 |
| EMC3 | 22.6385602 | -0.189165441 | 0.33894345 | -0.5581033 | 0.57677388 | NA |
| BRK1 | 39.0024497 | -0.498299405 | 0.36764354 | -1.3553874 | 0.17529418 | 0.57898467 |
| VHL | 11.7691067 | -0.365459863 | 0.48331644 | -0.7561503 | 0.44955909 | NA |
| SEC13 | 19.6725499 | -0.52190765 | 0.35734685 | -1.4605072 | 0.14415073 | NA |
| ATG7 | 81.3182152 | 0.245608815 | 0.36774165 | 0.66788415 | 0.50420755 | 0.82600448 |
| VGLL4 | 179.596882 | 0.257548642 | 0.28046296 | 0.91829824 | 0.35846275 | 0.72914991 |
| TAMM41 | 20.6399039 | 0.020306357 | 0.36362122 | 0.05584481 | 0.95546544 | NA |
| PPARG | 12.2112498 | -1.31178224 | 0.73988358 | -1.7729576 | 0.07623573 | NA |
| TSEN2 | 20.5227229 | -0.36784422 | 0.49860499 | -0.7377468 | 0.46066834 | NA |
| MKRN2OS | 4.69464948 | -1.793367625 | 1.07468596 | -1.6687364 | 0.09516962 | NA |
| MKRN2 | 13.3716722 | -0.320086483 | 0.54320635 | -0.5892539 | 0.55569093 | NA |
| RAF1 | 70.2550682 | 0.40342373 | 0.21258128 | 1.89773871 | 0.05773051 | 0.37585064 |
| CAND2 | 21.6047166 | 0.000914565 | 0.42476931 | 0.00215309 | 0.99828209 | NA |
| RPL32 | 94.1279647 | -0.807819327 | 0.38895702 | -2.0768858 | 0.0378121 | 0.30414888 |
| IQSEC1 | 42.4147508 | 0.102617819 | 0.49715093 | 0.2064118 | 0.83646925 | 0.95252283 |
| NUP210 | 9.63685055 | 0.647242316 | 0.42745719 | 1.51416873 | 0.12998305 | NA |
| HDAC11 | 5.73368238 | 0.33213657 | 0.52547206 | 0.63207274 | 0.52733934 | NA |
| XPC | 25.5497431 | -0.479615 | 0.30742908 | -1.5600834 | 0.11874018 | NA |
| LSM3 | 23.8356838 | -0.227698546 | 0.49473609 | -0.4602424 | 0.64534221 | NA |
| SLC6A6 | 138.063407 | -1.831533073 | 0.61953082 | -2.9563228 | 0.00311331 | 0.0904424 |
| CCDC174 | 14.6317804 | -0.092858576 | 0.56453982 | -0.1644854 | 0.86934901 | NA |
| FGD5 | 6.04563543 | 0.191561704 | 0.77138743 | 0.24833397 | 0.80387602 | NA |
| FGD5-AS1 | 49.2864844 | -0.084887282 | 0.2837618 | -0.2991498 | 0.76482576 | 0.92407902 |
| NR2C2 | 47.5819859 | -0.056607248 | 0.31797901 | -0.178022 | 0.85870572 | 0.95804916 |
| MRPS25 | 16.0340482 | 0.026887134 | 0.41890815 | 0.06418384 | 0.94882384 | NA |
| RBSN | 13.0123763 | -0.070168064 | 0.39519888 | -0.1775513 | 0.8590754 | NA |
| CAPN7 | 63.1238263 | 0.017452523 | 0.24529987 | 0.07114771 | 0.9432802 | 0.98114803 |
| SH3BP5-AS1 | 4.87745665 | -0.123203245 | 0.70973779 | -0.1735898 | 0.86218784 | NA |
| SH3BP5 | 73.6476457 | -0.282196766 | 0.4180229 | -0.6750749 | 0.49962819 | 0.82345406 |
| METTL6 | 16.0642792 | -0.259208227 | 0.42854236 | -0.6048602 | 0.54527188 | NA |
| EAF1 | 7.8425962 | 0.132689078 | 0.5584986 | 0.23758176 | 0.8122055 | NA |
| COLQ | 9.89218247 | 0.156085815 | 0.47665149 | 0.32746318 | 0.74331758 | NA |
| HACL1 | 19.2243659 | 0.165547178 | 0.34112789 | 0.48529358 | 0.62746811 | NA |
| BTD | 16.1103662 | -0.19489903 | 0.37476437 | -0.5200575 | 0.60302348 | NA |
| ANKRD28 | 103.184834 | 0.61841158 | 0.21556509 | 2.86879274 | 0.00412042 | 0.10253013 |
| DPH3 | 8.27530463 | 0.019668694 | 0.59629294 | 0.03298495 | 0.97368659 | NA |
| OXNAD1 | 15.1624855 | 0.110181108 | 0.43225832 | 0.25489644 | 0.79880308 | NA |
| RFTN1 | 8.66015457 | 1.314520336 | 0.80654909 | 1.62980822 | 0.10314204 | NA |
| PLCL2 | 44.6125332 | -0.057440459 | 0.3936625 | -0.145913 | 0.88399011 | 0.96552688 |
| TBC1D5 | 268.402748 | 0.276492202 | 0.20376426 | 1.356922 | 0.17480599 | 0.57831412 |
| RAB5A | 49.3403308 | 0.059584041 | 0.2630744 | 0.22649122 | 0.82081939 | 0.94728252 |
| KAT2B | 54.2877477 | 0.56349619 | 0.46399245 | 1.2144512 | 0.22457547 | 0.63080203 |
| ZNF385D | 10.0156974 | 0.564213916 | 0.78483991 | 0.71889045 | 0.47220842 | NA |
| UBE2E2 | 214.786133 | 0.508012632 | 0.30499116 | 1.66566348 | 0.09578046 | 0.46942032 |
| UBE2E1 | 77.7970632 | 0.402116166 | 0.30733891 | 1.30838028 | 0.19074436 | 0.59329687 |
| NKIRAS1 | 9.03758962 | -0.030092002 | 0.56050664 | -0.0536871 | 0.95718442 | NA |
| RPL15 | 152.334105 | -1.363411526 | 0.45131381 | -3.0209834 | 0.00251955 | 0.08046695 |
| NR1D2 | 30.3054682 | -0.167084794 | 0.33727996 | -0.495389 | 0.62032553 | 0.87793351 |
| THRB | 28.120401 | 1.182268986 | 0.6022434 | 1.96310823 | 0.0496336 | NA |
| TOP2B | 55.4684864 | 0.349928438 | 0.25313902 | 1.38235675 | 0.16686219 | 0.56878882 |
| NGLY1 | 28.8812898 | 0.014268332 | 0.35010736 | 0.04075416 | 0.96749188 | 0.98964394 |
| OXSM | 3.84307895 | -0.725928811 | 0.70409638 | -1.0310077 | 0.30253719 | NA |
| NEK10 | 19.6882066 | 1.204698877 | 0.96241327 | 1.251748 | 0.2106617 | NA |

| | | | | | | |
|---|---|---|---|---|---|---|
| SLC4A7 | 34.0936054 | 0.954189577 | 0.81692412 | 1.16802719 | 0.2427958 | 0.64918976 |
| CMC1 | 40.5768372 | 0.372193551 | 0.37811131 | 0.98434916 | 0.32494386 | 0.69623471 |
| AZI2 | 33.8231238 | 0.19730188 | 0.40094797 | 0.49208849 | 0.62265679 | 0.8780438 |
| RBMS3 | 39.4438729 | 0.620113988 | 0.43400643 | 1.4288129 | 0.15305802 | 0.55330064 |
| RBMS3-AS3 | 16.7385931 | 1.270758957 | 0.55122191 | 2.30534915 | 0.02114703 | NA |
| TGFBR2 | 38.0735182 | 1.010008 | 0.4433547 | 2.27810372 | 0.0227204 | 0.2464009 |
| STT3B | 77.136643 | 0.103916118 | 0.30118929 | 0.3450193 | 0.73007986 | 0.91718621 |
| OSBPL10 | 106.049346 | 0.153801302 | 0.28757809 | 0.53481578 | 0.59277725 | 0.8654248 |
| GPD1L | 19.0922931 | 0.930533871 | 0.62845828 | 1.48066132 | 0.13869685 | NA |
| CMTM8 | 5.77650554 | -0.262700262 | 0.69864946 | -0.3760115 | 0.70690831 | NA |
| CMTM7 | 13.7010277 | 0.181145301 | 0.37786127 | 0.47939631 | 0.63165672 | NA |
| CMTM6 | 37.2677378 | 0.188952671 | 0.31246744 | 0.60471156 | 0.54537067 | 0.84428375 |
| DYNC1LI1 | 22.9799532 | -0.386828262 | 0.31723102 | -1.2193898 | 0.22269628 | NA |
| CNOT10 | 25.6317548 | -0.128696545 | 0.31686346 | -0.4061577 | 0.68462671 | NA |
| GLB1 | 28.5768902 | 0.351851527 | 0.28512749 | 1.23401472 | 0.21719742 | 0.62453323 |
| CRTAP | 26.3650132 | -0.28175705 | 0.46503671 | -0.6058813 | 0.54459357 | NA |
| FBXL2 | 63.2925635 | -0.013069621 | 0.28924688 | -0.045185 | 0.96395985 | 0.98892249 |
| UBP1 | 29.5279131 | 0.233619105 | 0.31680993 | 0.73741093 | 0.46087249 | 0.79680257 |
| CLASP2 | 88.3354767 | 0.409368666 | 0.25032689 | 1.63533639 | 0.10197855 | 0.48317363 |
| PDCD6IP | 89.3454058 | 0.063814229 | 0.19287315 | 0.33086114 | 0.74074938 | 0.92003444 |
| LINC01811 | 34.6682238 | -1.591897915 | 1.78665972 | -0.8909911 | 0.37293394 | 0.7407542 |
| STAC | 21.4646641 | 0.499017511 | 0.51453602 | 0.9698398 | 0.33212635 | NA |
| TRANK1 | 17.3621989 | -0.754670963 | 0.83471476 | -0.9041064 | 0.36593898 | NA |
| EPM2AIP1 | 32.8968552 | -0.284310241 | 0.32641646 | -0.8710046 | 0.38375164 | 0.74975754 |
| MLH1 | 10.650378 | -0.213475561 | 0.45468918 | -0.4694978 | 0.63871387 | NA |
| LRRFIP2 | 91.3382009 | 0.468774123 | 0.23770097 | 1.97211704 | 0.04859625 | 0.34339091 |
| GOLGA4 | 233.422779 | 0.037457561 | 0.2561357 | 0.14624107 | 0.88373108 | 0.96552688 |
| ITGA9 | 51.5950212 | 0.021189512 | 0.4017768 | 0.05273951 | 0.95793946 | 0.98714493 |
| CTDSPL | 587.781206 | 0.450491586 | 0.37187014 | 1.21142176 | 0.2257338 | 0.63317863 |
| VILL | 14.5467378 | 0.956171037 | 0.67537337 | 1.41576656 | 0.15684386 | NA |
| PLCD1 | 11.1022304 | 0.741547977 | 0.52689703 | 1.4073869 | 0.1593127 | NA |
| ACAA1 | 19.1733723 | -0.133299362 | 0.43665383 | -0.3052747 | 0.76015695 | NA |
| MYD88 | 5.00480661 | -0.875088385 | 1.21372425 | -0.7209944 | 0.47091296 | NA |
| OXSR1 | 47.4627728 | 0.474091735 | 0.23246899 | 2.03937621 | 0.04141249 | 0.31933285 |
| XYLB | 7.44910141 | -0.352633723 | 0.71929766 | -0.4902473 | 0.62395893 | NA |
| ACVR2B | 14.5930556 | -0.128130066 | 0.52711302 | -0.2430789 | 0.80794426 | NA |
| EXOG | 21.1977786 | 0.474796113 | 0.39019671 | 1.21681218 | 0.22367568 | NA |
| WDR48 | 32.3171572 | -0.220713687 | 0.38656095 | -0.5709674 | 0.56802177 | 0.85654436 |
| GORASP1 | 10.1376291 | -0.1729097 | 0.50036264 | -0.3455688 | 0.72966682 | NA |
| TTC21A | 4.60288742 | -0.821455748 | 0.90303741 | -0.9096586 | 0.36300258 | NA |
| CSRNP1 | 8.89259382 | -1.880640055 | 0.79405359 | -2.3684045 | 0.01786499 | NA |
| SLC25A38 | 9.7580045 | 0.02131768 | 0.46048264 | 0.04629421 | 0.96307575 | NA |
| RPSA | 61.3542276 | -0.551850904 | 0.43480308 | -1.2691973 | 0.20437069 | 0.6137834 |
| MYRIP | 31.7950657 | 0.69988134 | 0.62695184 | 1.11632392 | 0.26428351 | 0.65974932 |
| EIF1B | 27.770972 | -0.439365416 | 0.34437699 | -1.2758269 | 0.20201673 | NA |
| ENTPD3-AS1 | 7.4194874 | -0.125786385 | 0.58405548 | -0.2153672 | 0.82948105 | NA |
| RPL14 | 101.708213 | -1.17509898 | 0.54071355 | -2.1732375 | 0.02976244 | 0.27926427 |
| ZNF621 | 20.3797091 | -0.491607215 | 0.42922387 | -1.1453399 | 0.25206839 | NA |
| CTNNB1 | 85.4996685 | 0.373289465 | 0.23590659 | 1.58236134 | 0.1135671 | 0.50483797 |
| ULK4 | 57.9557292 | 0.348466845 | 0.3726497 | 0.93510566 | 0.34973385 | 0.72238707 |
| TRAK1 | 74.260198 | 0.414541293 | 0.22709802 | 1.82538487 | 0.06794297 | 0.40347978 |
| SEC22C | 29.2082104 | 0.090503629 | 0.35006958 | 0.2585304 | 0.79599759 | 0.93934998 |
| SS18L2 | 8.12999909 | -0.590756692 | 0.56476017 | -1.0460311 | 0.29554668 | NA |
| NKTR | 87.2761344 | -0.118997144 | 0.2062792 | -0.5768742 | 0.56402445 | 0.8549147 |
| AC006059.1 | 14.3907511 | -0.712198368 | 0.49128955 | -1.4496509 | 0.14715588 | NA |
| AC099329.3 | 6.77822333 | -0.342098439 | 0.59092733 | -0.5789179 | 0.56264454 | NA |
| POMGNT2 | 4.72818792 | -0.071459681 | 0.61368276 | -0.116444 | 0.90730066 | NA |
| SNRK | 40.6859792 | -0.180273352 | 0.29618032 | -0.6086608 | 0.5427493 | 0.84207245 |
| ANO10 | 57.1164905 | 0.344537023 | 0.31304709 | 1.10059169 | 0.27107441 | 0.66433703 |
| ABHD5 | 15.2871254 | 0.794809215 | 0.50596871 | 1.57086634 | 0.1162137 | NA |
| TCAIM | 71.7207794 | 0.527893135 | 0.35421829 | 1.49030457 | 0.13614417 | 0.52892793 |
| ZNF445 | 14.0650075 | -0.009696656 | 0.43302518 | -0.0223928 | 0.98213461 | NA |
| AC099669.1 | 20.1418602 | -0.017440621 | 0.41142171 | -0.0423911 | 0.96618692 | NA |
| ZNF197 | 18.1944651 | -0.214048251 | 0.42286391 | -0.5061871 | 0.61272532 | NA |
| KIAA1143 | 15.1621044 | -0.629875029 | 0.4177681 | -1.5077145 | 0.13162761 | NA |
| ZDHHC3 | 22.5771882 | -0.056032709 | 0.3598987 | -0.1556902 | 0.8762772 | NA |
| EXOSC7 | 7.55634814 | 0.450728386 | 0.58592852 | 0.7692549 | 0.441742 | NA |
| LARS2 | 18.7732424 | 0.223940618 | 0.39489967 | 0.56708232 | 0.57065826 | NA |
| LIMD1 | 17.4547538 | 0.576637896 | 0.6200803 | 0.92994068 | 0.3524018 | NA |
| SACM1L | 31.3539516 | 0.387969624 | 0.29071384 | 1.3345413 | 0.18202652 | 0.58390314 |
| LZTFL1 | 8.19027616 | 0.166134702 | 0.51387931 | 0.32329518 | 0.74647171 | NA |
| FYCO1 | 16.1782944 | -0.329476269 | 0.36481026 | -0.9031442 | 0.36644937 | NA |
| LTF | 33.6247452 | -1.674463612 | 1.10204763 | -1.5194113 | 0.128659 | 0.52341115 |
| LRRC2 | 239.674997 | -0.88202555 | 0.44599084 | -1.9776764 | 0.04796523 | 0.3419103 |
| ALS2CL | 549.638089 | -0.563981094 | 0.48986321 | -1.1513032 | 0.24960751 | 0.65148524 |
| TMIE | 8.43223013 | -0.954917999 | 0.618351 | -1.5442976 | 0.12251624 | NA |
| MYL3 | 8.9389579 | 0.614124136 | 0.66559524 | 0.92266906 | 0.35617969 | NA |
| PTH1R | 412.133291 | 0.931183157 | 0.53690621 | 1.73434975 | 0.08285605 | 0.44045101 |
| CCDC12 | 39.4213043 | -0.313049542 | 0.33175351 | -0.9436209 | 0.34536342 | 0.71852078 |
| NBEAL2 | 4.99891797 | -0.499895594 | 0.76619963 | -0.6524352 | 0.51412048 | NA |
| SETD2 | 106.148653 | 0.474003476 | 0.21003116 | 2.25682454 | 0.02401904 | 0.25196701 |
| KIF9-AS1 | 26.4226569 | -0.022027029 | 0.36929761 | -0.0596457 | 0.95243779 | NA |
| KIF9 | 5.43306867 | -0.160937771 | 0.81644688 | -0.1971197 | 0.84373386 | NA |
| KLHL18 | 14.9364997 | 0.301400395 | 0.41479723 | 0.72662105 | 0.46745813 | NA |
| PTPN23 | 15.2848171 | -0.531252868 | 0.4471841 | -1.1879959 | 0.23483503 | NA |
| SCAP | 34.8731611 | 0.621234256 | 0.33388298 | 1.86063472 | 0.06279578 | 0.3863717 |
| ELP6 | 11.4484242 | -0.361209461 | 0.4766328 | -0.7578359 | 0.44854921 | NA |
| SMARCC1 | 65.6131788 | 0.002586602 | 0.26083373 | 0.00991667 | 0.99208777 | 0.99792949 |
| DHX30 | 24.9920159 | 0.488417336 | 0.30771815 | 1.58722306 | 0.11246213 | NA |
| MAP4 | 165.381818 | -0.107843714 | 0.23474347 | -0.4594109 | 0.64593911 | 0.89204257 |
| ZNF589 | 4.31874501 | -0.627046787 | 0.730241 | -0.8586847 | 0.39051449 | NA |
| NME6 | 6.9266103 | -0.663168515 | 0.62762101 | -1.0566385 | 0.29067661 | NA |
| PLXNB1 | 39.6685485 | 0.739321402 | 0.34619911 | 2.13553813 | 0.03271708 | 0.29047359 |
| TMA7 | 63.6530016 | -0.147737434 | 0.40602977 | -0.3638586 | 0.71596359 | 0.91399852 |
| SHISA5 | 32.3052466 | -0.588095878 | 0.47382406 | -1.2411693 | 0.21454321 | 0.62289712 |
| PFKFB4 | 6.73994125 | -1.43863198 | 0.886019 | -1.6237033 | 0.10443914 | NA |
| COL7A1 | 18.3464045 | 1.368022061 | 0.59872432 | 2.28489477 | 0.022319 | NA |
| UQCRC1 | 21.9346605 | -0.100877489 | 0.46011173 | -0.2192456 | 0.82645871 | NA |
| SLC26A6 | 5.9618994 | -0.098716756 | 0.65681542 | -0.150296 | 0.88053106 | NA |
| NCKIPSD | 13.2647845 | 0.139407436 | 0.41370767 | 0.33697088 | 0.73613886 | NA |
| IP6K2 | 63.516508 | 0.339161314 | 0.21955947 | 1.54473552 | 0.12241025 | 0.51557214 |
| PRKAR2A | 63.8398166 | -0.415752291 | 0.33396598 | -1.2448941 | 0.21317067 | 0.62255988 |
| SLC25A20 | 10.1298732 | 0.582243218 | 0.47017359 | 1.23835799 | 0.21558335 | NA |
| ARIH2 | 54.1114097 | 0.282843293 | 0.21558898 | 1.3119562 | 0.18953491 | 0.5922601 |
| P4HTM | 42.5327134 | -0.315494407 | 0.3616957 | -0.8722647 | 0.38306397 | 0.74924885 |
| WDR6 | 29.4864565 | 0.200858777 | 0.36641248 | 0.54817669 | 0.58357059 | 0.86175324 |
| DALRD3 | 7.8962573 | -0.453397219 | 0.52840727 | -0.858045 | 0.39086761 | NA |
| NDUFAF3 | 24.0046032 | -0.518733165 | 0.5162333 | -1.0048425 | 0.31497269 | NA |
| IMPDH2 | 6.31677605 | -0.491647609 | 0.78227554 | -0.6284839 | 0.52968697 | NA |
| QRICH1 | 34.6505547 | 0.304154036 | 0.3252679 | 0.93508776 | 0.34974307 | 0.72238707 |
| QARS | 8.07115481 | 0.817861912 | 0.52247958 | 1.56534714 | 0.11750154 | NA |
| USP19 | 6.19140004 | 0.247298712 | 0.57438934 | 0.43054196 | 0.66680145 | NA |
| LAMB2 | 108.835419 | 0.016317297 | 0.54329217 | 0.03003411 | 0.97603985 | 0.992815 |

| | | | | | | |
|---|---|---|---|---|---|---|
| C3orf62 | 5.14653435 | 0.068941527 | 0.66696857 | 0.10336548 | 0.91767291 | NA |
| USP4 | 25.4333683 | 0.09764636 | 0.31610297 | 0.30890682 | 0.75739241 | NA |
| RHOA | 125.100458 | -0.372978497 | 0.37251442 | -1.0012458 | 0.31670798 | 0.69424851 |
| TCTA | 13.6931123 | -0.846750749 | 0.4009653 | -2.1117806 | 0.03470527 | NA |
| DAG1 | 158.392221 | 0.587970012 | 0.52042168 | 1.12979538 | 0.25856245 | 0.65588339 |
| APEH | 13.438829 | 1.137700635 | 0.45797167 | 2.48421616 | 0.0129837 | NA |
| MST1 | 5.64037638 | 0.701647061 | 0.67185934 | 1.04433624 | 0.29632985 | NA |
| RNF123 | 9.35254691 | 0.615714929 | 0.46847115 | 1.31430702 | 0.1887429 | NA |
| IP6K1 | 47.8926987 | 0.385389179 | 0.26436676 | 1.45778231 | 0.14490056 | 0.54367868 |
| UBA7 | 7.27723386 | 0.710110188 | 0.89508004 | 0.79334826 | 0.42757495 | NA |
| RBM6 | 145.644246 | 0.054564513 | 0.2895568 | 0.18844149 | 0.85053058 | 0.95532127 |
| RBM5 | 45.1202471 | 0.191037783 | 0.2451911 | 0.77913833 | 0.43589823 | 0.78167891 |
| GNAI2 | 54.7708163 | -0.350780878 | 0.42470171 | -0.8259465 | 0.40883446 | 0.76772515 |
| SEMA3B | 180.16163 | 0.551123114 | 0.55581137 | 0.99156502 | 0.32140976 | 0.69513974 |
| IFRD2 | 6.32512375 | -0.128710853 | 0.63647655 | -0.202224 | 0.83974159 | NA |
| HYAL2 | 17.4866925 | 0.159163901 | 0.38564801 | 0.41271807 | 0.67981319 | NA |
| TUSC2 | 11.2931113 | -0.311443126 | 0.56245517 | -0.5537208 | 0.57976994 | NA |
| RASSF1 | 5.08990122 | -0.156804361 | 0.70929101 | -0.221072 | 0.82503639 | NA |
| TMEM115 | 15.2917317 | 0.33038173 | 0.51820741 | 0.63754729 | 0.52376842 | NA |
| HEMK1 | 6.44701106 | 0.765174832 | 0.57612999 | 1.32812881 | 0.18413556 | NA |
| CISH | 3.91686707 | -0.779972828 | 1.07181443 | -0.7277126 | 0.46678956 | NA |
| MAPKAPK3 | 4.57569583 | -1.111203746 | 0.69911897 | -1.5894344 | 0.11196235 | NA |
| DOCK3 | 52.4873297 | 0.654190744 | 0.4195604 | 1.55922899 | 0.11894219 | 0.51050744 |
| MANF | 19.2797646 | -0.691233328 | 0.64964394 | -1.0640187 | 0.28732021 | NA |
| RBM15B | 8.56602982 | -0.236908613 | 0.65925519 | -0.359358 | 0.71932731 | NA |
| DCAF1 | 16.7233205 | 0.00723377 | 0.39376129 | 0.01837095 | 0.98534292 | NA |
| RAD54L2 | 22.3392311 | 0.066372571 | 0.34112613 | 0.194569 | 0.84573037 | NA |
| TEX264 | 24.4514673 | -0.394742591 | 0.42907396 | -0.9199873 | 0.35757939 | NA |
| PCBP4 | 7.60805382 | 0.54436495 | 0.59580773 | 0.91365876 | 0.36089618 | NA |
| RPL29 | 119.914289 | -1.548451034 | 0.41786266 | -3.7056459 | 0.00021085 | 0.01781706 |
| ALAS1 | 5.1619066 | -0.078171881 | 0.64976677 | -0.1203076 | 0.90423948 | NA |
| WDR82 | 30.0085288 | -0.082430048 | 0.30379052 | -0.2713384 | 0.78613074 | 0.93615351 |
| DNAH1 | 3.70556956 | -0.235337611 | 0.64740411 | -0.3635096 | 0.71622424 | NA |
| BAP1 | 12.8910657 | 0.434384987 | 0.45106948 | 0.96301126 | 0.33554187 | NA |
| PHF7 | 5.73299132 | -0.913245398 | 0.5858137 | -1.5589348 | 0.1190118 | NA |
| SEMA3G | 88.7711423 | -0.201860292 | 0.63755824 | -0.3166147 | 0.751536 | 0.92274989 |
| TNNC1 | 24.4750196 | 0.695439242 | 0.81971703 | 0.8483894 | 0.39622114 | NA |
| NISCH | 30.915269 | 0.423054279 | 0.31647131 | 1.33678556 | 0.18129264 | 0.58390314 |
| SMIM4 | 12.9797622 | -0.084759213 | 0.43717206 | -0.1938807 | 0.84626932 | NA |
| PBRM1 | 70.4627522 | -0.061732406 | 0.25839998 | -0.2389025 | 0.81118117 | 0.94408044 |
| GNL3 | 14.9188387 | -0.476766221 | 0.50464095 | -0.9447632 | 0.34477976 | NA |
| GLT8D1 | 18.5810224 | -0.541421425 | 0.46108776 | -1.1742264 | 0.24030435 | NA |
| SPCS1 | 63.4567749 | -0.591466976 | 0.36525991 | -1.6193044 | 0.10538179 | 0.4872104 |
| NEK4 | 17.2300751 | -0.1406579 | 0.4208687 | -0.3342085 | 0.73822223 | NA |
| SFMBT1 | 15.3419876 | 0.557478405 | 0.50238427 | 1.10966533 | 0.26714327 | NA |
| AC096887.1 | 7.24865652 | -0.136369309 | 0.64687174 | -0.2108135 | 0.83303279 | NA |
| PRKCD | 10.6660781 | 0.061116217 | 0.45790061 | 0.13347048 | 0.89382131 | NA |
| TKT | 21.7469154 | -0.392973621 | 0.36294352 | -1.0827404 | 0.27892366 | NA |
| DCP1A | 36.1300151 | 0.325237646 | 0.28084206 | 1.15808026 | 0.24683129 | 0.65061974 |
| CACNA1D | 19.3681711 | 1.001425764 | 0.77366119 | 1.29439835 | 0.19552786 | NA |
| CHDH | 5.4097516 | 0.20023577 | 1.11190484 | 0.18008355 | 0.85708698 | NA |
| IL17RB | 10.1818947 | -1.038057709 | 1.19375155 | -0.869576 | 0.38453215 | NA |
| ACTR8 | 6.67471796 | 0.558626107 | 0.58140751 | 0.96081681 | 0.33664428 | NA |
| SELENOK | 30.930624 | -1.048591549 | 0.54088616 | -1.9386548 | 0.05254339 | 0.35808347 |
| CACNA2D3 | 32.0652758 | -0.878851146 | 0.59006883 | -1.4894045 | 0.1363809 | 0.52934059 |
| ERC2 | 61.7077563 | 0.452429921 | 0.30534532 | 1.48169921 | 0.13842035 | 0.53418928 |
| CCDC66 | 39.8137872 | -0.197026171 | 0.33823804 | -0.5825074 | 0.56022494 | 0.85359593 |
| FAM208A | 67.2047408 | 0.236844888 | 0.24088942 | 0.98320999 | 0.3255041 | 0.69669901 |
| ARHGEF3 | 599.789426 | -0.066856326 | 0.29438277 | -0.2271068 | 0.8203407 | 0.94728252 |
| IL17RD | 56.8399691 | 0.507652512 | 0.55992344 | 0.9066463 | 0.36459387 | 0.73398222 |
| APPL1 | 36.0275355 | -0.094019612 | 0.29526185 | -0.3184279 | 0.75016037 | 0.92274989 |
| PDE12 | 10.4517737 | 0.115740887 | 0.41062049 | 0.28186827 | 0.77804452 | NA |
| ARF4 | 42.2481262 | -0.077374097 | 0.2879238 | -0.2687312 | 0.78813657 | 0.93749303 |
| DENND6A | 54.8569963 | 0.297563361 | 0.2754192 | 1.08040166 | 0.27996336 | 0.66958859 |
| DENND6A-A! | 10.6660413 | -0.348120399 | 0.61561073 | -0.5654879 | 0.57174197 | NA |
| SLMAP | 73.1979596 | 0.427042301 | 0.23377003 | 1.82676244 | 0.0677355 | 0.40302824 |
| FLNB | 15.9670953 | 0.443281105 | 0.56689702 | 0.78194291 | 0.43424813 | NA |
| ABHD6 | 5.86492194 | -0.443822231 | 0.568345 | -0.7809028 | 0.43485964 | NA |
| RPP14 | 5.40209623 | -0.327422343 | 0.55094723 | -0.5942898 | 0.55231827 | NA |
| PXK | 13.1547888 | -0.180839661 | 0.41612561 | -0.4345795 | 0.66386768 | NA |
| PDHB | 16.5386515 | -0.17130602 | 0.50586033 | -0.3386429 | 0.73487874 | NA |
| KCTD6 | 5.07788734 | 0.18687536 | 0.71003323 | 0.26319242 | 0.79240228 | NA |
| ACOX2 | 5.19435106 | -0.149654061 | 0.6520086 | -0.2295277 | 0.81845876 | NA |
| C3orf67 | 6.73706329 | 0.495887009 | 0.68233854 | 0.7267463 | 0.46738139 | NA |
| FHIT | 247.524721 | 0.40371481 | 0.49027329 | 0.82344851 | 0.41025298 | 0.76788553 |
| PTPRG | 428.384323 | 0.963965825 | 0.41903674 | 2.300043274 | 0.02142372 | 0.24262159 |
| THOC7 | 20.5291236 | -0.303156902 | 0.45278718 | -0.6695351 | 0.50315421 | NA |
| ATXN7 | 8.16826177 | -0.324286901 | 0.4983745 | -0.6506892 | 0.51524714 | NA |
| PSMD6-AS2 | 8.98198606 | -0.41317333 | 0.54371937 | -0.7599018 | 0.44731329 | NA |
| PSMD6 | 13.2240198 | -0.616963313 | 0.51272777 | -1.2032961 | 0.22886176 | NA |
| PRICKLE2 | 92.7045362 | 0.89687101 | 0.34199803 | 2.62244495 | 0.00873014 | 0.14875902 |
| ADAMTS9 | 536.04315 | 0.118778403 | 0.30645623 | 0.38758684 | 0.69832181 | 0.90729979 |
| ADAMTS9-A! | 187.44498 | 0.128193866 | 0.3557128 | 0.36038587 | 0.71855859 | 0.91472065 |
| ADAMTS9-A! | 764.269622 | -0.518545046 | 0.56415086 | -0.9191603 | 0.35801176 | 0.72914991 |
| MAGI1 | 88.9503899 | 0.835868686 | 0.50967231 | 1.64001197 | 0.10100268 | 0.48247622 |
| SLC25A26 | 32.9366433 | 0.192107844 | 0.34978022 | 0.54922444 | 0.58285143 | 0.86175324 |
| LRIG1 | 21.3105023 | -1.068026445 | 0.61182542 | -1.7456392 | 0.08087366 | NA |
| SUCLG2 | 89.3857902 | 0.332868208 | 0.22499368 | 1.47945581 | 0.13901853 | 0.53519929 |
| SUCLG2-AS1 | 39.0884487 | 0.996089292 | 0.44366581 | 2.2451342 | 0.02475952 | 0.25362919 |
| FAM19A1 | 10.4074952 | 0.623885919 | 0.78498629 | 0.79477301 | 0.42674555 | NA |
| EOGT | 10.1351742 | 0.366391163 | 0.4892596 | 0.74886862 | 0.4539364 | NA |
| AC109587.1 | 3.6165596 | -1.036054464 | 0.91755876 | -1.1291424 | 0.25883778 | NA |
| TMF1 | 42.6442082 | -0.132166323 | 0.2592534 | -0.5097959 | 0.61019445 | 0.87320729 |
| UBA3 | 26.4123503 | -0.001599608 | 0.32814726 | -0.0048747 | 0.99611059 | NA |
| ARL6IP5 | 45.259024 | -0.424974115 | 0.42305526 | -1.0045357 | 0.31512047 | 0.6938809 |
| FRMD4B | 31.4952417 | 0.805916284 | 0.43092035 | 1.870221 | 0.06145314 | 0.38229128 |
| MITF | 35.2481439 | 1.480750646 | 0.48586567 | 3.0476544 | 0.00230635 | 0.07786717 |
| FOXP1 | 229.550409 | -0.862573429 | 0.32826277 | -2.6276919 | 0.00859663 | 0.14826624 |
| EIF4E3 | 16.7426383 | -0.422356271 | 0.50925495 | -0.8293611 | 0.40690008 | NA |
| RYBP | 50.5297652 | 0.223940723 | 0.31722376 | 0.70593932 | 0.48022588 | 0.81093801 |
| SHQ1 | 19.2871442 | 0.222713015 | 0.43908083 | 0.50722555 | 0.61199657 | NA |
| GXYLT2 | 9.08023426 | -0.119183168 | 0.58789479 | -0.2027287 | 0.83934706 | NA |
| PPP4R2 | 53.4641087 | 0.290172521 | 0.23736753 | 1.22246089 | 0.22153339 | 0.62528839 |
| PDZRN3 | 12.1747632 | 1.848905515 | 0.80847041 | 2.28691798 | 0.02220061 | NA |
| CNTN3 | 40.2709975 | 0.265261629 | 0.65915581 | 0.4024263 | 0.68737032 | 0.90499319 |
| ZNF717 | 24.5077288 | 0.103668216 | 0.35879634 | 0.28893331 | 0.77263241 | NA |
| ROBO2 | 288.393199 | 0.134760873 | 0.37485461 | 0.35950171 | 0.7192198 | 0.91472065 |
| ROBO1 | 155.254925 | 0.214828014 | 0.27826451 | 0.77202807 | 0.4400978 | 0.78399202 |
| GBE1 | 117.768973 | 0.177374089 | 0.36708475 | 0.48319656 | 0.62895618 | 0.88166074 |
| CHMP2B | 18.7471615 | -0.282682675 | 0.38901613 | -0.7266606 | 0.46743392 | NA |
| CGGBP1 | 60.0566033 | -0.000749367 | 0.24486249 | -0.0030604 | 0.99755819 | 0.99961632 |
| ZNF654 | 49.9539389 | -0.194637257 | 0.28567679 | -0.6813198 | 0.49566915 | 0.8234249 |
| C3orf38 | 21.0973172 | -0.131553135 | 0.30996881 | -0.4244077 | 0.67126854 | NA |

| | | | | | | |
|---|---|---|---|---|---|---|
| PROS1 | 24.0561336 | -1.378155378 | 0.54251304 | -2.5403175 | 0.01107519 | NA |
| ARL13B | 64.083959 | 0.629418777 | 0.25933606 | 2.42703915 | 0.01522261 | 0.20310171 |
| NSUN3 | 21.2139191 | 0.306213668 | 0.36305598 | 0.84343375 | 0.3989859 | NA |
| MTRNR2L12 | 47.9241545 | -0.10979764 | 0.47778204 | -0.229807 | 0.81824177 | 0.94694381 |
| ARL6 | 11.5236153 | -0.112647861 | 0.49462927 | -0.227742 | 0.81984682 | NA |
| CRYBG3 | 16.7869535 | -0.661118473 | 0.44667534 | -1.4800872 | 0.13884997 | NA |
| RIOX2 | 15.3217369 | -0.051293382 | 0.46079482 | -0.111315 | 0.91136655 | NA |
| CLDND1 | 25.8925323 | 0.341163511 | 0.32773433 | 1.04097581 | 0.29788678 | NA |
| CPOX | 7.38037508 | 0.976023587 | 0.55986287 | 1.74332615 | 0.08127666 | NA |
| ST3GAL6 | 518.666056 | 0.786790038 | 0.39400542 | 1.99690154 | 0.04583588 | 0.33590606 |
| DCBLD2 | 119.044661 | 0.575860794 | 0.31825042 | 1.8094581 | 0.07037986 | 0.40663921 |
| CMSS1 | 41.0957622 | -0.85817423 | 0.42669755 | -2.0112003 | 0.04430431 | 0.33093605 |
| FILIP1L | 12.2087084 | -0.899175709 | 0.59005786 | -1.5238772 | 0.12753939 | NA |
| TBC1D23 | 26.6769037 | -0.06663628 | 0.31935961 | -0.2086559 | 0.83471684 | NA |
| NIT2 | 11.9217536 | 0.134932573 | 0.43197171 | 0.31236438 | 0.75476362 | NA |
| TOMM70 | 23.116137 | 0.208965723 | 0.35255992 | 0.59270982 | 0.55337537 | NA |
| TMEM45A | 9.9607342 | -0.168398795 | 0.52814041 | -0.3188523 | 0.7498385 | NA |
| TFG | 50.3801921 | -0.182140256 | 0.35140298 | -0.518323 | 0.60423291 | 0.87047761 |
| IMPG2 | 6.1035615 | 1.050769131 | 1.16604918 | 0.9011362 | 0.36751591 | NA |
| SENP7 | 93.6424339 | 0.525990731 | 0.26447739 | 1.9887928 | 0.04672408 | 0.33774256 |
| TRMT10C | 14.0633593 | -0.700900078 | 0.42627208 | -1.6442552 | 0.1001235 | NA |
| PCNP | 56.0285154 | -0.367446974 | 0.26296005 | -1.3973491 | 0.16230863 | 0.56332966 |
| ZBTB11 | 13.9361036 | -0.698164874 | 0.42229134 | -1.6532777 | 0.09827435 | NA |
| RPL24 | 50.1311872 | -0.786805309 | 0.46476464 | -1.6929113 | 0.09047235 | 0.45866472 |
| CEP97 | 10.405204 | 0.188422218 | 0.43168654 | 0.43647925 | 0.66248905 | NA |
| NXPE3 | 15.0832151 | -0.03113861 | 0.44111245 | -0.0705911 | 0.9437232 | NA |
| NFKBIZ | 16.3719841 | -1.668872644 | 0.70734004 | -2.359364 | 0.01830629 | NA |
| ALCAM | 91.9224329 | -0.059489617 | 0.35500163 | -0.1675756 | 0.86691715 | 0.96089268 |
| CBLB | 855.464309 | -0.116720001 | 0.19193767 | -0.6081141 | 0.54311182 | 0.84207245 |
| LINC00882 | 16.1230131 | 0.100398128 | 0.52668035 | 0.19062441 | 0.84881987 | NA |
| DUBR | 23.0191168 | 0.273903663 | 0.49393332 | 0.55453571 | 0.57921227 | NA |
| BBX | 151.219418 | 0.355317611 | 0.22506714 | 1.57871832 | 0.11440068 | 0.50483797 |
| CD47 | 134.737803 | 0.017619581 | 0.31747846 | 0.05549851 | 0.95574132 | 0.98638341 |
| IFT57 | 20.0364968 | -0.533759793 | 0.37974529 | -1.4055732 | 0.15985091 | NA |
| DZIP3 | 14.9779129 | 0.097358927 | 0.51956651 | 0.18738492 | 0.85135885 | NA |
| NECTIN3 | 34.9635356 | -0.295823882 | 0.39567749 | -0.7476389 | 0.454678 | 0.79324789 |
| PHLDB2 | 106.679624 | 0.407807815 | 0.35859623 | 1.13723399 | 0.25544048 | 0.65588339 |
| ABHD10 | 7.46812518 | 0.096550972 | 0.64658152 | 0.14932529 | 0.88129696 | NA |
| CD200 | 62.0788646 | 0.267975997 | 0.47142362 | 0.56843991 | 0.5697363 | 0.8573305 |
| ATG3 | 15.0581973 | 0.0852518 | 0.38745807 | 0.22002845 | 0.825849 | NA |
| SLC35A5 | 13.8509725 | -0.074721436 | 0.3817825 | -0.1957173 | 0.84483145 | NA |
| GTPBP8 | 5.96195983 | -0.897247107 | 0.57778109 | -1.5529188 | 0.12044254 | NA |
| NEPRO | 15.7210656 | -0.093933001 | 0.47477507 | -0.1978474 | 0.84316448 | NA |
| AC078785.1 | 5.18154396 | 0.006373492 | 0.77668398 | 0.00820603 | 0.99345261 | NA |
| CFAP44 | 22.252767 | 0.454361005 | 0.35725794 | 1.271801 | 0.20344383 | NA |
| SPICE1 | 14.8216028 | -0.00305193 | 0.45179688 | -0.0067551 | 0.99461026 | NA |
| USF3 | 25.3047164 | 0.494841221 | 0.30717186 | 1.61095882 | 0.1071887 | NA |
| NAA50 | 28.8755772 | -0.427083876 | 0.29801344 | -1.4331027 | 0.15182848 | 0.55086922 |
| ATP6V1A | 45.041306 | 0.11254947 | 0.27152825 | 0.41450372 | 0.67850524 | 0.90499319 |
| GRAMD1C | 9.53458955 | -0.590577087 | 0.71142046 | -0.8301379 | 0.40646082 | NA |
| ZDHHC23 | 4.03383835 | 0.476621405 | 0.72742031 | 0.65522147 | 0.51232517 | NA |
| CCDC191 | 31.626677 | 0.147951497 | 0.44547377 | 0.33212168 | 0.73979738 | 0.92003444 |
| QTRT2 | 15.107541 | -0.156852592 | 0.43854487 | -0.357666 | 0.72059328 | NA |
| ZBTB20 | 1108.26057 | 0.086378503 | 0.18859498 | 0.45801063 | 0.64694481 | 0.89261695 |
| ZBTB20-AS5 | 20.1159891 | -0.288374995 | 0.590845 | -0.4880722 | 0.62549873 | NA |
| LSAMP | 22.7105253 | 0.164114887 | 0.60261397 | 0.27233834 | 0.78536188 | NA |
| B4GALT4 | 11.8363566 | -0.396330481 | 0.42711758 | -0.9279189 | 0.35344963 | NA |
| ARHGAP31 | 11.3745277 | 0.794412591 | 0.53942302 | 1.47270798 | 0.14082979 | NA |
| TMEM39A | 18.464202 | -0.099850897 | 0.4022818 | -0.2482113 | 0.80397091 | NA |
| POGLUT1 | 27.9219301 | -0.07275818 | 0.48314981 | -0.1505913 | 0.88029809 | NA |
| TIMMDC1 | 28.0417782 | 0.205980335 | 0.38799121 | 0.53088919 | 0.59549557 | NA |
| COX17 | 24.1898973 | -0.060097637 | 0.47027632 | -0.1277922 | 0.89831344 | NA |
| MAATS1 | 13.5822103 | 0.269101677 | 0.47138316 | 0.57087674 | 0.5680832 | NA |
| GSK3B | 109.519361 | 0.259621749 | 0.23080893 | 1.12483408 | 0.26065935 | 0.65595635 |
| LRRC58 | 34.157917 | -0.057944592 | 0.3486902 | -0.1661779 | 0.86801698 | 0.96089268 |
| FSTL1 | 144.898732 | -0.224514761 | 0.54854384 | -0.4092923 | 0.68232518 | 0.90499319 |
| NDUFB4 | 35.2247584 | -0.668123927 | 0.454178 | -1.4710618 | 0.14127439 | 0.53597868 |
| HGD | 8.4442638 | -0.330719503 | 0.80530652 | -0.4106753 | 0.68131064 | NA |
| RABL3 | 14.2586931 | -0.180635693 | 0.39482093 | -0.457513 | 0.64730238 | NA |
| GTF2E1 | 6.11168886 | -0.076381642 | 0.53008511 | -0.1440932 | 0.8854269 | NA |
| GOLGB1 | 123.56588 | -0.144790511 | 0.27091843 | -0.5344432 | 0.59303495 | 0.8654248 |
| IQCB1 | 27.2395603 | -0.511872376 | 0.35356979 | -1.4477266 | 0.14769354 | NA |
| CCDC58 | 4.35943194 | -0.728166527 | 0.86729748 | -0.839581 | 0.40114333 | NA |
| FAM162A | 25.2594478 | -0.513372664 | 0.5172955 | -0.9924166 | 0.32099433 | NA |
| AC083798.2 | 12.3178255 | -0.33669076 | 0.45915999 | -0.7332755 | 0.46339043 | NA |
| KPNA1 | 51.1015534 | 0.108903395 | 0.249073 | 0.43723485 | 0.66194104 | 0.89839561 |
| PARP9 | 43.8609229 | -1.155016131 | 0.73892042 | -1.563113 | 0.11802601 | 0.50926968 |
| DTX3L | 23.9473422 | -1.73017859 | 0.77257644 | -2.2394918 | 0.02512393 | NA |
| PARP14 | 79.6265884 | 0.048172115 | 0.80346038 | 0.05995581 | 0.95219083 | 0.98520702 |
| HSPBAP1 | 12.193086 | 0.419895863 | 0.4631215 | 0.90666459 | 0.36458419 | NA |
| DIRC2 | 21.7317466 | -0.297958359 | 0.3705839 | -0.804024 | 0.42138311 | NA |
| PDIA5 | 10.906781 | -0.202571976 | 0.47026031 | -0.4307656 | 0.6666388 | NA |
| SEC22A | 33.0414181 | 0.176623143 | 0.30163097 | 0.58556038 | 0.55817098 | 0.8520194 |
| ADCY5 | 7.01670188 | 0.912492779 | 0.60437082 | 1.50982269 | 0.13108867 | NA |
| HACD2 | 20.1327695 | 0.385007165 | 0.45966545 | 0.83758125 | 0.40226593 | NA |
| MYLK-AS1 | 9.00776748 | 0.929025411 | 0.58193913 | 1.59643055 | 0.1103927 | NA |
| MYLK | 102.840163 | 0.572840126 | 0.42359886 | 1.35231745 | 0.17627379 | 0.57988078 |
| CCDC14 | 32.0438852 | 0.434745927 | 0.40266671 | 1.07966692 | 0.28029053 | 0.66958859 |
| KALRN | 25.8413177 | 1.956881931 | 0.83840712 | 2.33404736 | 0.01959324 | NA |
| UMPS | 10.1927234 | 0.511683808 | 0.61436634 | 0.83286433 | 0.40492125 | NA |
| ITGB5 | 207.914457 | -0.37863316 | 0.26639278 | -1.4213342 | 0.15521964 | 0.55566712 |
| ITGB5-AS1 | 4.78611432 | -1.830951166 | 0.93455297 | -1.9591732 | 0.0500925 | NA |
| HEG1 | 21.4924281 | 1.126294665 | 0.47241012 | 2.38414592 | 0.01711881 | NA |
| ZNF148 | 151.989748 | 0.45921029 | 0.18049836 | 2.54412446 | 0.0109552 | 0.17090118 |
| SNX4 | 45.0049994 | 0.038640639 | 0.31491859 | 0.1227004 | 0.90234434 | 0.97220041 |
| OSBPL11 | 12.0052334 | -0.251478659 | 0.44472301 | -0.5654726 | 0.57175238 | NA |
| SLC41A3 | 8.30238123 | 0.004207677 | 0.52911654 | 0.00795227 | 0.99365507 | NA |
| ALDH1L1 | 10.3269977 | 0.408577514 | 1.04349457 | 0.39154733 | 0.69539271 | NA |
| AC016924.1 | 5.6310806 | 0.374896363 | 0.85770877 | 0.43709051 | 0.66204571 | NA |
| KLF15 | 9.77523333 | -1.491253038 | 0.6671919 | -2.2351186 | 0.02540956 | NA |
| ZXDC | 39.2490486 | -0.356685018 | 0.29561775 | -1.2065751 | 0.22759583 | 0.63327644 |
| TXNRD3 | 5.30069545 | 0.28161562 | 0.57495079 | 0.48980821 | 0.62426962 | NA |
| CHCHD6 | 19.8580482 | 0.028073161 | 0.43083612 | 0.06515972 | 0.94804683 | NA |
| PLXNA1 | 14.9481401 | -0.171660278 | 0.72772422 | -0.2358864 | 0.81352079 | NA |
| TPRA1 | 11.3950473 | -0.192019192 | 0.571507 | -0.3359875 | 0.73688031 | NA |
| ABTB1 | 8.54572448 | 0.573958036 | 0.63820397 | 0.89933323 | 0.36847519 | NA |
| MGLL | 17.1145747 | -1.20364657 | 0.46715805 | -2.5765297 | 0.00997976 | NA |
| SEC61A1 | 51.9133064 | 0.269482389 | 0.43542474 | 0.61889545 | 0.53598524 | 0.83677705 |
| RUVBL1 | 14.9190571 | -0.319384393 | 0.44171248 | -0.7230595 | 0.46964334 | NA |
| EEFSEC | 29.1660728 | -0.089417796 | 0.33962617 | -0.263283 | 0.79233246 | 0.93928291 |
| RPN1 | 62.8105908 | -0.211180845 | 0.42847271 | -0.4928688 | 0.62210527 | 0.8780438 |
| RAB7A | 88.7392628 | -0.085590847 | 0.20432582 | -0.4188939 | 0.67529364 | 0.90485333 |
| ACAD9 | 14.3355769 | 0.216040723 | 0.41333129 | 0.52268176 | 0.60119574 | NA |

| | | | | | | |
|---|---|---|---|---|---|---|
| CNBP | 85.6793027 | -0.807637627 | 0.36647698 | -2.2037881 | 0.02753925 | 0.26935188 |
| COPG1 | 19.3829701 | -0.135201737 | 0.38378221 | -0.3522877 | 0.72462254 | NA |
| HMCES | 38.2747123 | 1.019942133 | 0.43135744 | 2.36449415 | 0.01805471 | 0.22394469 |
| H1FX | 46.300657 | 0.084322794 | 0.64894368 | 0.12993854 | 0.89661505 | 0.97097328 |
| MBD4 | 15.4747129 | -0.639927315 | 0.38116533 | -1.6788707 | 0.09317724 | NA |
| IFT122 | 16.7288208 | 0.531997982 | 0.39448414 | 1.34859156 | 0.17746819 | NA |
| PLXND1 | 11.7705038 | 0.055184461 | 0.68647471 | 0.08038819 | 0.93592852 | NA |
| TMCC1 | 113.364173 | 0.203484823 | 0.21345379 | 0.95329682 | 0.3404397 | 0.71508205 |
| PIK3R4 | 18.3140743 | 0.389820304 | 0.37291371 | 1.04533649 | 0.29586748 | NA |
| ATP2C1 | 69.0347334 | 0.595840277 | 0.25998811 | 2.29179818 | 0.02191729 | 0.24297341 |
| NEK11 | 48.5128496 | 0.391252075 | 0.33743553 | 1.15948688 | 0.24625778 | 0.65061974 |
| NUDT16 | 8.87769921 | 0.068209361 | 0.55125334 | 0.12373505 | 0.90152506 | NA |
| MRPL3 | 20.9195719 | -0.190175133 | 0.40755456 | -0.466625 | 0.64076822 | NA |
| CPNE4 | 26.4194142 | -0.982193085 | 0.56422856 | -1.7407717 | 0.08172361 | NA |
| DNAJC13 | 44.3130674 | 0.197422215 | 0.29023915 | 0.68020532 | 0.49637446 | 0.8234249 |
| UBA5 | 17.7275283 | 0.092337498 | 0.40513205 | 0.22791951 | 0.81970881 | NA |
| TMEM108 | 5.26214127 | 1.081339652 | 0.81341791 | 1.32937773 | 0.18372338 | NA |
| CDV3 | 64.6217378 | -0.790310863 | 0.35233903 | -2.2430409 | 0.02489417 | 0.25362919 |
| TOPBP1 | 19.1856436 | 0.164035022 | 0.39160977 | 0.41887367 | 0.67530846 | NA |
| SRPRB | 13.0628389 | 0.281591457 | 0.58254999 | 0.48337732 | 0.62882785 | NA |
| RAB6B | 7.13569057 | 0.343632238 | 0.69886928 | 0.49169744 | 0.62293325 | NA |
| RYK | 59.2524042 | 0.204057839 | 0.28199771 | 0.72361524 | 0.46930198 | 0.80448595 |
| AMOTL2 | 79.4912354 | 0.727071253 | 0.45980772 | 1.58125063 | 0.11382074 | 0.50483797 |
| ANAPC13 | 11.8628305 | -0.357269193 | 0.46562088 | -0.7672963 | 0.44290535 | NA |
| CEP63 | 23.0235883 | -0.218297903 | 0.33728145 | -0.6472277 | 0.51748457 | NA |
| PPP2R3A | 86.5356533 | -0.031429655 | 0.31267885 | -0.1005174 | 0.91993359 | 0.97775374 |
| MSL2 | 40.486077 | 0.413604882 | 0.32932752 | 1.25590743 | 0.20914953 | 0.61736443 |
| PCCB | 17.3077096 | -0.227429334 | 0.37577412 | -0.6052288 | 0.54502696 | NA |
| STAG1 | 165.799149 | 0.031563023 | 0.25951491 | 0.12162316 | 0.90319747 | 0.97237083 |
| NCK1 | 19.6800582 | -0.284161047 | 0.44444853 | -0.6393565 | 0.52259106 | NA |
| DBR1 | 4.57472711 | -0.465432097 | 0.67006513 | -0.6946072 | 0.48730147 | NA |
| ARMC8 | 45.1742442 | -0.155307383 | 0.31609227 | -0.4913356 | 0.62318911 | 0.8780438 |
| MRAS | 4.74271894 | -0.272069616 | 0.64238725 | -0.423529 | 0.67190936 | NA |
| CEP70 | 24.4812113 | 0.626960575 | 0.38701496 | 1.61999056 | 0.1052343 | NA |
| FAIM | 7.17549827 | -0.14901608 | 0.62178049 | -0.2396603 | 0.81059364 | NA |
| PIK3CB | 58.4757969 | 0.688108513 | 0.28559387 | 2.40939528 | 0.01597898 | 0.20852711 |
| MRPS22 | 6.99999226 | -0.43515759 | 0.62496989 | -0.6962857 | 0.48624994 | NA |
| COPB2 | 38.8942148 | -0.425244195 | 0.25487399 | -1.6684488 | 0.09522667 | 0.46829657 |
| AC097103.2 | 54.8641219 | -0.06068235 | 0.42474445 | -0.1428679 | 0.88639451 | 0.9665824 |
| NMNAT3 | 12.4973244 | 0.36380477 | 0.50725491 | 0.71720305 | 0.47324882 | NA |
| CLSTN2 | 64.6986882 | 0.765352537 | 0.95071587 | 0.80502763 | 0.42080374 | 0.77473274 |
| SLC25A36 | 67.0231586 | 0.069864287 | 0.26625064 | 0.26240045 | 0.79301273 | 0.93934998 |
| ZBTB38 | 117.900554 | -0.446027944 | 0.26102299 | -1.7087688 | 0.08749378 | 0.44978149 |
| RASA2 | 63.254258 | 0.285717368 | 0.23846717 | 1.19814129 | 0.23086202 | 0.63682501 |
| RNF7 | 32.3414237 | -0.443489302 | 0.44972796 | -0.9861279 | 0.32407034 | 0.69623471 |
| ATP1B3 | 30.3054134 | -0.679005662 | 0.4575185 | -1.4841054 | 0.13778097 | 0.53375321 |
| TFDP2 | 73.8483152 | 0.431966965 | 0.27948201 | 1.54559849 | 0.12220157 | 0.51522823 |
| GK5 | 27.1337876 | 0.013366156 | 0.30573082 | 0.04371871 | 0.96512863 | NA |
| XRN1 | 88.8234889 | 0.467072401 | 0.32903365 | 1.41952775 | 0.15574521 | 0.55613041 |
| ATR | 57.3868666 | -0.090546384 | 0.2863866 | -0.3161684 | 0.75187471 | 0.92274989 |
| PLS1 | 6.03486882 | -0.861406655 | 0.70881533 | -1.2152766 | 0.22426063 | NA |
| TRPC1 | 54.7787233 | -0.20441624 | 0.44172054 | -0.4627728 | 0.64352725 | 0.8902681 |
| PCOLCE2 | 388.062469 | 0.287391587 | 0.5424658 | 0.52978747 | 0.59625929 | 0.86775303 |
| U2SURP | 66.0340326 | 0.072261635 | 0.24899671 | 0.29021121 | 0.77165466 | 0.92669573 |
| CHST2 | 66.8023395 | -4.942796479 | 0.88300795 | -5.5976806 | 2.17E-08 | 1.76E-05 |
| SLC9A9 | 18.6312597 | -0.542286264 | 0.59056439 | -0.9182509 | 0.35848754 | NA |
| C3orf58 | 23.7854297 | -0.444953219 | 0.36868435 | -1.2068677 | 0.22748311 | NA |
| AC107021.1 | 10.7564402 | -0.113412595 | 0.74822025 | -0.1515765 | 0.87952098 | NA |
| LNCSRLR | 5.69789923 | -0.377724206 | 0.75045726 | -0.5033254 | 0.6147355 | NA |
| PLOD2 | 123.134985 | 0.291171897 | 0.34347369 | 0.84772692 | 0.39659007 | 0.75911719 |
| PLSCR4 | 32.9753848 | -0.316774876 | 0.37689294 | -0.8404903 | 0.40063352 | 0.76381845 |
| PLSCR1 | 86.5126067 | -2.40155897 | 0.73746818 | -3.2564916 | 0.00112798 | 0.05186671 |
| GYG1 | 22.4123056 | -0.076976355 | 0.32006303 | -0.2405037 | 0.80993977 | NA |
| HLTF | 26.3759808 | 0.547038373 | 0.33319504 | 1.64179628 | 0.10063223 | NA |
| HPS3 | 28.7213507 | -0.035104117 | 0.28640569 | -0.1225678 | 0.90244935 | 0.97220041 |
| CP | 9.57003718 | -2.951999806 | 1.18001995 | -2.5016525 | 0.01236152 | NA |
| TM4SF1 | 14.2526498 | -0.841603115 | 0.88250798 | -0.9536493 | 0.34026119 | NA |
| WWTR1 | 146.838713 | 0.264895027 | 0.19534395 | 1.35604417 | 0.17508512 | 0.57876547 |
| COMMD2 | 25.4519375 | -0.51846317 | 0.30628577 | -1.6927433 | 0.09050434 | NA |
| ANKUB1 | 23.1403494 | -0.525149868 | 0.57065929 | -0.9202512 | 0.35744153 | NA |
| RNF13 | 74.0259123 | 0.131896115 | 0.28870929 | 0.45684749 | 0.64778067 | 0.89261695 |
| TSC22D2 | 57.8492744 | -0.013320368 | 0.26019191 | -0.0511944 | 0.95917062 | 0.98766083 |
| SERP1 | 49.1969819 | -0.638675954 | 0.32952219 | -1.938188 | 0.05260029 | 0.35808347 |
| EIF2A | 22.1405992 | 0.169716111 | 0.39868144 | 0.42569353 | 0.67033117 | NA |
| SELENOT | 44.2142902 | 0.172758803 | 0.38520814 | 0.4484817 | 0.65380559 | 0.8949739 |
| SIAH2 | 7.74799868 | -1.805318763 | 0.71686986 | -2.5183354 | 0.0117911 | NA |
| MED12L | 8.5631116 | 1.169388936 | 0.62490777 | 1.87129844 | 0.06130373 | NA |
| MBNL1 | 187.6251 | 0.448313949 | 0.23150914 | 1.93648489 | 0.05280834 | 0.35868165 |
| RAP2B | 7.96533323 | -1.184925884 | 0.54107211 | -2.1899593 | 0.02852719 | NA |
| C3orf79 | 9.35343355 | -0.815026571 | 0.91792733 | -0.8878988 | 0.3745952 | NA |
| ARHGEF26 | 241.262989 | -0.056584075 | 0.26738081 | -0.2116235 | 0.83240074 | 0.95050875 |
| DHX36 | 76.5430284 | 0.144568534 | 0.28545747 | 0.50644508 | 0.61254423 | 0.87391153 |
| MME | 1512.20495 | 0.085685061 | 0.38057551 | 0.22514602 | 0.82186568 | 0.94728252 |
| MME-AS1 | 43.3984494 | -0.885835365 | 0.76992659 | -1.1505452 | 0.24991937 | 0.6518054 |
| GMPS | 25.2888789 | -0.022492254 | 0.33872958 | -0.0664018 | 0.94705794 | NA |
| SSR3 | 43.360739 | 0.15866941 | 0.29649801 | 0.53514494 | 0.59254964 | 0.8654248 |
| TIPARP | 33.3803105 | -0.200639448 | 0.44255029 | -0.453371 | 0.65028165 | 0.89412414 |
| LINC00886 | 36.150456 | 0.315010212 | 0.44942463 | 0.70091889 | 0.48335363 | 0.81370025 |
| LEKR1 | 12.933098 | -0.467174697 | 0.5489392 | -0.85105 | 0.39474159 | NA |
| CCNL1 | 37.9073091 | 0.011970858 | 0.36028827 | 0.03322578 | 0.97349454 | 0.99108782 |
| VEPH1 | 5.77821337 | 0.956406978 | 0.81553856 | 1.17273054 | 0.24090388 | NA |
| RSRC1 | 68.5044373 | 0.059498308 | 0.23619939 | 0.25189865 | 0.8011194 | 0.94085969 |
| MLF1 | 7.04968393 | -0.195235986 | 0.78108805 | -0.2499539 | 0.80262302 | NA |
| GFM1 | 18.1506107 | -0.069377985 | 0.37250865 | -0.1862453 | 0.8522524 | NA |
| LXN | 6.67305846 | -0.222603149 | 0.62595726 | -0.3556204 | 0.72212489 | NA |
| RARRES1 | 5.82498225 | -0.327906047 | 0.7110896 | -0.4611318 | 0.64470403 | NA |
| MFSD1 | 14.7720522 | 0.28608846 | 0.43294914 | 0.66078999 | 0.508747 | NA |
| IQCJ-SCHIP1 | 628.491331 | 1.847299809 | 0.80370482 | 2.29848043 | 0.02153446 | 0.24262159 |
| SMC4 | 7.11656326 | -0.179543561 | 0.50407643 | -0.3561832 | 0.72170337 | NA |
| KPNA4 | 59.0264867 | -0.264851034 | 0.33105673 | -0.8000171 | 0.4237009 | 0.77656162 |
| PPM1L | 60.6643913 | -0.316122628 | 0.48446991 | -0.6525124 | 0.51407067 | 0.83070544 |
| B3GALNT1 | 11.7498401 | 0.547180205 | 0.56704664 | 0.96496508 | 0.3345623 | NA |
| NMD3 | 43.5534547 | -0.090254036 | 0.35975788 | -0.2508744 | 0.80191123 | 0.94085969 |
| LINC01322 | 7.46023593 | -0.072166626 | 0.57123533 | -0.1263343 | 0.89946729 | NA |
| SERPINI2 | 39.8229128 | -0.177774787 | 0.59097315 | -0.300817 | 0.76355402 | 0.92336765 |
| WDR49 | 98.1205598 | 0.049113291 | 0.60301781 | 0.08144584 | 0.9350874 | 0.97832857 |
| PDCD10 | 39.1260869 | -0.311778195 | 0.32825967 | -0.9497913 | 0.34221828 | 0.71725052 |
| SERPINI1 | 9.1530639 | -0.317109341 | 0.54827494 | -0.5783765 | 0.56300996 | NA |
| GOLIM4 | 317.272394 | 0.236834548 | 0.39499506 | 0.59958864 | 0.54878042 | 0.84550378 |
| MECOM | 189.268912 | 0.308682176 | 0.52662317 | 0.58615381 | 0.55777216 | 0.85177857 |
| MYNN | 9.86337055 | 0.170645289 | 0.5427961 | 0.31438193 | 0.75323098 | NA |
| SEC62 | 76.1072793 | 0.004955867 | 0.28798949 | 0.0172085 | 0.98627028 | 0.99585746 |
| GPR160 | 6.90922345 | 0.962950187 | 0.76336867 | 1.2614484 | 0.20714734 | NA |

| | | | | | | |
|---|---|---|---|---|---|---|
| PHC3 | 108.057802 | 0.026376605 | 0.21778643 | 0.12111225 | 0.90360213 | 0.97237083 |
| PRKCI | 162.852787 | -0.249382848 | 0.36740011 | -0.6787773 | 0.49727899 | 0.82345406 |
| SKIL | 47.9905946 | 0.290525789 | 0.30388278 | 0.95604557 | 0.33904921 | 0.71342248 |
| AC026316.5 | 89.6523629 | -0.596109376 | 0.53020019 | -1.12431 | 0.26088154 | 0.65595635 |
| SLC7A14 | 10.7797711 | -0.614303231 | 0.91043951 | -0.6747326 | 0.49984566 | NA |
| RPL22L1 | 18.6559911 | -1.363562053 | 0.67300043 | -2.0260939 | 0.04275517 | NA |
| EIF5A2 | 16.2046722 | -0.240819653 | 0.43707517 | -0.5509799 | 0.58164742 | NA |
| TNIK | 74.2879171 | 0.303286429 | 0.52207892 | 0.58092066 | 0.56129393 | 0.85386276 |
| PLD1 | 39.3904158 | 0.363661352 | 0.33833091 | 1.07486884 | 0.28243346 | 0.67148306 |
| FNDC3B | 744.26282 | 0.380472728 | 0.26153792 | 1.45475165 | 0.14573803 | 0.54436111 |
| TNFSF10 | 24.4170354 | -0.651819226 | 0.77410063 | -0.8420342 | 0.3997688 | NA |
| NCEH1 | 15.947072 | 0.398858823 | 0.61353134 | 0.65010342 | 0.51562542 | NA |
| ECT2 | 7.34440833 | -0.259745194 | 0.52149455 | -0.4980784 | 0.61842875 | NA |
| NLGN1 | 55.6585809 | -0.481738814 | 0.59062394 | -0.8156439 | 0.41470383 | 0.77005091 |
| NAALADL2 | 201.136687 | 0.035756753 | 0.21167509 | 0.16892282 | 0.86585735 | 0.96089268 |
| TBL1XR1 | 141.50607 | 0.239806405 | 0.28505741 | 0.84125653 | 0.40020424 | 0.76351288 |
| KCNMB2-AS: | 6.40453817 | -0.52834383 | 0.78012388 | -0.6772563 | 0.49824334 | NA |
| ZMAT3 | 28.3214024 | 0.085634223 | 0.50352161 | 0.1700706 | 0.86495461 | NA |
| PIK3CA | 62.276021 | 0.391187651 | 0.36626601 | 1.06804245 | 0.28550136 | 0.67369331 |
| KCNMB3 | 9.76973976 | -0.403226232 | 0.48743707 | -0.8272375 | 0.40810245 | NA |
| ZNF639 | 9.24005345 | -0.209562271 | 0.46600043 | -0.449704 | 0.65292387 | NA |
| MFN1 | 38.1016327 | 0.42955258 | 0.31061003 | 1.38293209 | 0.16668569 | 0.56878882 |
| GNB4 | 11.56706 | 0.67639412 | 0.74008388 | 0.91394251 | 0.36074705 | NA |
| ACTL6A | 11.4329553 | -0.096849827 | 0.40333626 | -0.2401218 | 0.81023584 | NA |
| MRPL47 | 8.18607013 | -0.857158742 | 0.62397702 | -1.3737024 | 0.16953409 | NA |
| NDUFB5 | 23.0327811 | -0.409897465 | 0.40610579 | -1.0093367 | 0.3128132 | NA |
| USP13 | 16.9442448 | -0.230956471 | 0.45646565 | -0.5059668 | 0.61287995 | NA |
| PEX5L | 20.2239898 | 0.267907072 | 0.88773262 | 0.30178802 | 0.76281366 | NA |
| TTC14 | 72.6314524 | 0.520079099 | 0.26049678 | 1.99648957 | 0.04588066 | 0.33590606 |
| FXR1 | 77.5144273 | 0.697912911 | 0.28049097 | 2.48818316 | 0.01283976 | 0.18904687 |
| DNAJC19 | 19.1950568 | 0.407840151 | 0.38715874 | 1.05341843 | 0.29214926 | NA |
| ATP11B | 65.7361372 | 0.446271927 | 0.23468572 | 1.90157253 | 0.05722706 | 0.37377289 |
| DCUN1D1 | 40.8680388 | 0.398405319 | 0.36777286 | 1.08329179 | 0.27867893 | 0.66958859 |
| MCCC1 | 31.7806408 | 0.28297819 | 0.41813349 | 0.67676519 | 0.49855495 | 0.82345406 |
| LAMP3 | 8.19097929 | 4.765190709 | 2.26507347 | 2.10376872 | 0.03539863 | NA |
| MCF2L2 | 15.8697168 | 0.31721944 | 0.60007421 | 0.52863369 | 0.59705959 | NA |
| B3GNT5 | 8.75123614 | 0.791724808 | 0.50269798 | 1.57495123 | 0.1152677 | NA |
| KLHL24 | 40.2670264 | 0.271874391 | 0.46495612 | 0.58473129 | 0.55872841 | 0.8520194 |
| YEATS2 | 40.4996202 | 0.031301704 | 0.3966321 | 0.07891874 | 0.93709726 | 0.97856407 |
| ABCC5 | 19.8581451 | 0.287908376 | 0.39636798 | 0.72636639 | 0.46761419 | NA |
| EIF2B5 | 12.8171665 | 0.128551595 | 0.45524721 | 0.28237756 | 0.77765402 | NA |
| DVL3 | 19.6097359 | 0.09754959 | 0.34720949 | 0.28095313 | 0.77874635 | NA |
| AP2M1 | 52.57197 | -0.677299905 | 0.41761621 | -1.6218238 | 0.10484108 | 0.4862279 |
| ABCF3 | 6.17790305 | 1.33844533 | 0.54518072 | 2.45504891 | 0.01408654 | NA |
| ALG3 | 5.06740179 | 0.404565803 | 0.77299112 | 0.52337704 | 0.6007119 | NA |
| PSMD2 | 26.7283783 | -0.630041165 | 0.40801144 | -1.5441753 | 0.12254588 | NA |
| EIF4G1 | 36.1099165 | -0.283208098 | 0.34612355 | -0.8182283 | 0.41322682 | 0.76946282 |
| FAM131A | 5.9777893 | -0.609357042 | 0.72945965 | -0.835354 | 0.40351842 | NA |
| POLR2H | 19.5777452 | -0.041866245 | 0.45312715 | -0.092394 | 0.92638498 | NA |
| CHRD | 3.92355122 | 0.16861442 | 0.77275598 | 0.21819879 | 0.82727423 | NA |
| MAGEF1 | 17.3584952 | -0.259237956 | 0.39527405 | -0.6558436 | 0.51192476 | NA |
| VPS8 | 54.7232196 | 0.325949414 | 0.26417472 | 1.23384029 | 0.21726242 | 0.62453323 |
| EHHADH | 16.3370803 | 1.061372678 | 0.61355087 | 1.72988539 | 0.08365075 | NA |
| MAP3K13 | 51.0047609 | 0.662243027 | 0.42107542 | 1.57274205 | 0.11577856 | 0.50513044 |
| TMEM41A | 8.77445651 | -1.371066293 | 0.57151489 | -2.3990036 | 0.01643975 | NA |
| SENP2 | 25.1079998 | -0.153310892 | 0.32559793 | -0.4708595 | 0.63774104 | NA |
| IGF2BP2 | 76.079404 | 0.128016106 | 0.23329697 | 0.54872598 | 0.58319351 | 0.86175324 |
| TRA2B | 46.9562401 | -0.102175271 | 0.28460853 | -0.3590028 | 0.71959297 | 0.91472065 |
| ETV5 | 6.35738323 | -1.673845529 | 0.70860822 | -2.3621593 | 0.01816883 | NA |
| TBCCD1 | 6.28732781 | -0.500206547 | 0.59265389 | -0.8440112 | 0.39866312 | NA |
| AC068631.2 | 6.52884093 | -1.231102281 | 0.62332045 | -1.9750712 | 0.04826007 | NA |
| EIF4A2 | 89.646024 | -0.022124312 | 0.36738764 | -0.0602206 | 0.95197992 | 0.98520702 |
| ST6GAL1 | 200.231654 | -1.745555979 | 0.43337241 | -4.0278429 | 5.63E-05 | 0.00691867 |
| RPL39L | 4.77326288 | -0.442926507 | 0.68089218 | -0.650509 | 0.51536347 | NA |
| RTP4 | 5.92962026 | -0.475980208 | 1.47738255 | -0.322178 | 0.74731782 | NA |
| BCL6 | 31.5699692 | -1.237225332 | 0.72688964 | -1.7020814 | 0.08874011 | 0.45503146 |
| LPP | 255.287689 | 0.296372663 | 0.2698897 | 1.09812513 | 0.27214985 | 0.66496374 |
| TPRG1 | 10.97731 | 0.499398931 | 0.51839519 | 0.96335564 | 0.33536908 | NA |
| P3H2 | 484.348855 | -0.343888748 | 0.31597781 | -1.088332 | 0.27644858 | 0.66941818 |
| P3H2-AS1 | 14.3630589 | -0.36857921 | 0.58488933 | -0.6301691 | 0.52858392 | NA |
| IL1RAP | 99.0023003 | 0.281264389 | 0.38810927 | 0.72470413 | 0.46863356 | 0.80448595 |
| CCDC50 | 65.5764902 | 0.137024679 | 0.28374449 | 0.48291573 | 0.62915558 | 0.88166074 |
| MB21D2 | 34.7140315 | 0.328102842 | 0.40945689 | 0.80131229 | 0.42295088 | 0.77623925 |
| OPA1 | 50.0978266 | 0.223912935 | 0.23279556 | 0.96184365 | 0.33612815 | 0.71116888 |
| HES1 | 19.4056511 | -0.590769351 | 0.50014273 | -1.1812015 | 0.23752268 | NA |
| ATP13A3 | 94.5702956 | 0.57137296 | 0.36301334 | 1.57397237 | 0.11549384 | 0.50513044 |
| LSG1 | 12.7384785 | -0.103578436 | 0.47123659 | -0.2198013 | 0.82602587 | NA |
| FAM43A | 14.8349647 | -0.698307142 | 0.84872045 | -0.8227764 | 0.41063517 | NA |
| XXYLT1 | 8.20069376 | 0.227892185 | 0.50541025 | 0.45090535 | 0.65205777 | NA |
| ACAP2 | 82.4011021 | 0.702569514 | 0.20732821 | 3.38868266 | 0.00070229 | 0.03956247 |
| PPP1R2 | 23.2580622 | 0.093953417 | 0.32334191 | 0.29056987 | 0.77138031 | NA |
| APOD | 226.839796 | -1.934516572 | 1.22968573 | -1.5731796 | 0.11567723 | 0.50513044 |
| MUC20-OT1 | 82.7491501 | 0.24914143 | 0.4126278 | 0.60379215 | 0.54598184 | 0.84440067 |
| TNK2 | 7.90918038 | 0.918473446 | 0.518648 | 1.77089944 | 0.07657743 | NA |
| TFRC | 10.3795241 | 0.842299163 | 0.55043568 | 1.53024086 | 0.12595712 | NA |
| PCYT1A | 16.7153244 | 0.142397443 | 0.38174073 | 0.37302135 | 0.70913255 | NA |
| UBXN7 | 38.9621737 | -0.032109513 | 0.2971064 | -0.1080741 | 0.9139369 | 0.97707273 |
| RNF168 | 19.3016518 | -0.544464494 | 0.34813587 | -1.5639425 | 0.11783107 | NA |
| PIGX | 14.8980609 | -0.160049187 | 0.44115538 | -0.3627955 | 0.71675765 | NA |
| PAK2 | 52.868892 | -0.303820246 | 0.23481858 | -1.2938509 | 0.19571691 | 0.59987316 |
| SENP5 | 52.5573891 | -0.293387923 | 0.28692486 | -1.0225253 | 0.30653235 | 0.68576679 |
| NCBP2 | 17.9774971 | -0.555172917 | 0.41501959 | -1.3377029 | 0.1809933 | NA |
| NCBP2-AS2 | 14.4720734 | -0.812051853 | 0.44637788 | -1.8192027 | 0.06888051 | NA |
| PIGZ | 5.98158032 | -0.148458501 | 0.6208674 | -0.2391147 | 0.81101667 | NA |
| DLG1 | 132.387186 | 0.334288269 | 0.188031 | 1.77783594 | 0.0754308 | 0.42138852 |
| RUBCN | 14.4426174 | 0.273396551 | 0.4401142 | 0.62119457 | 0.53447161 | NA |
| FYTTD1 | 27.7627907 | -0.277095712 | 0.35209042 | -0.7870016 | 0.43128095 | NA |
| LRCH3 | 85.8514991 | 0.006204637 | 0.2474076 | 0.0250786 | 0.97999227 | 0.99453905 |
| IQCG | 13.1103217 | -1.115120035 | 0.47007662 | -2.3722091 | 0.01768208 | NA |
| RPL35A | 67.0965362 | -0.447335571 | 0.40405096 | -1.1071266 | 0.26823919 | 0.66098308 |
| LMLN | 13.1068382 | -0.122238101 | 0.46208597 | -0.2645354 | 0.79136738 | NA |
| ZNF595 | 18.6745574 | 0.361550699 | 0.52139525 | 0.69342921 | 0.48804023 | NA |
| ZNF718 | 50.2237832 | 0.008210783 | 0.25683318 | 0.03196932 | 0.97449651 | 0.9916101 |
| ZNF141 | 24.5092621 | -0.17607398 | 0.33512998 | -0.5253901 | 0.59931202 | NA |
| AC092574.2 | 4.9255863 | 0.090595422 | 1.09431603 | 0.08278726 | 0.9340207 | NA |
| ZNF721 | 90.9990774 | 0.120850176 | 0.27109345 | 0.4457879 | 0.65575047 | 0.89583156 |
| PIGG | 19.7007555 | 0.046307953 | 0.43098777 | 0.10744609 | 0.91443509 | NA |
| ATP5ME | 17.9357806 | 0.149769525 | 0.5569735 | 0.26889884 | 0.78800753 | NA |
| PCGF3 | 24.3718591 | -0.062282278 | 0.31928415 | -0.1950685 | 0.84533933 | NA |
| AC139887.2 | 55.5530822 | 0.027154429 | 0.26607005 | 0.10205744 | 0.91871108 | 0.97775374 |
| GAK | 36.6078307 | -0.20813027 | 0.41342366 | -0.503431 | 0.61466131 | 0.8753744 |
| TMEM175 | 9.38762879 | 1.015825774 | 0.52688232 | 1.92799365 | 0.05385592 | NA |
| IDUA | 6.32036451 | -0.110081759 | 0.6462722 | -0.1703334 | 0.86474792 | NA |

| | | | | | | |
|---|---|---|---|---|---|---|
| FGFRL1 | 8.50824694 | 0.208233232 | 0.50162619 | 0.41511634 | 0.67805674 | NA |
| AC092535.3 | 9.33710372 | 0.58161877 | 0.53944379 | 1.07818235 | 0.28095239 | NA |
| SPON2 | 14.7911415 | 0.836661905 | 0.58688039 | 1.42560889 | 0.15398127 | NA |
| CTBP1-AS | 16.0882165 | 1.055247896 | 0.65723155 | 1.6055953 | 0.10836286 | NA |
| CTBP1 | 33.0587867 | 0.580007383 | 0.39556668 | 1.46626957 | 0.14257486 | 0.53894095 |
| CTBP1-DT | 8.39669528 | -0.429564554 | 0.5659033 | -0.7590777 | 0.4478061 | NA |
| MAEA | 20.9106229 | 0.116233601 | 0.33269559 | 0.34936923 | 0.72681213 | NA |
| UVSSA | 23.2523122 | 0.638332945 | 0.38365329 | 1.66382762 | 0.09614688 | NA |
| SLBP | 27.9786836 | -0.255612354 | 0.32718324 | -0.7812514 | 0.43465467 | NA |
| TMEM129 | 12.8060071 | 0.102512524 | 0.46754441 | 0.2192573 | 0.82644962 | NA |
| LETM1 | 13.1737891 | -0.107322265 | 0.50169753 | -0.2139183 | 0.8306108 | NA |
| NSD2 | 54.2553675 | -0.109363995 | 0.30673694 | -0.35654 | 0.72143619 | 0.91509584 |
| NELFA | 16.8178403 | 0.157014381 | 0.44944256 | 0.34935361 | 0.72682385 | NA |
| C4orf48 | 5.73719697 | 1.581629793 | 0.95211403 | 1.66117687 | 0.09667793 | NA |
| POLN | 6.31791939 | 0.328985921 | 0.61036378 | 0.53899974 | 0.58988703 | NA |
| MXD4 | 60.2169516 | -0.412424932 | 0.38357832 | -1.075204 | 0.28228342 | 0.67148306 |
| ZFYVE28 | 39.4922564 | -0.983696383 | 0.44073313 | -2.2319547 | 0.02561796 | 0.25731925 |
| RNF4 | 21.5742874 | -0.520621315 | 0.32340653 | -1.6098046 | 0.10744052 | NA |
| FAM193A | 48.8718703 | 0.324928131 | 0.29834994 | 1.08908394 | 0.27611687 | 0.66941424 |
| TNIP2 | 12.3107253 | -0.680707525 | 0.49657197 | -1.3708134 | 0.17043312 | NA |
| SH3BP2 | 9.43799042 | 0.004197269 | 0.65267519 | 0.00643087 | 0.99486894 | NA |
| ADD1 | 84.5143779 | 0.341388394 | 0.28891006 | 1.18164245 | 0.2373476 | 0.64350343 |
| MFSD10 | 22.8401063 | 0.203214899 | 0.52922391 | 0.38398662 | 0.70098837 | NA |
| NOP14-AS1 | 5.56218159 | -0.469821817 | 0.62307297 | -0.7540398 | 0.45082532 | NA |
| NOP14 | 7.93162344 | -0.174407556 | 0.73241578 | -0.2381264 | 0.81178304 | NA |
| GRK4 | 7.97793053 | -0.01248828 | 0.59598479 | -0.020954 | 0.98328233 | NA |
| HTT | 49.8975665 | 0.108357969 | 0.31381341 | 0.34529426 | 0.72987316 | 0.91718621 |
| RGS12 | 22.3778656 | 0.29085363 | 0.46874756 | 0.62049097 | 0.53493459 | NA |
| DOK7 | 9.97682321 | -0.068869838 | 0.74600131 | -0.0923187 | 0.92644487 | NA |
| LRPAP1 | 60.7877306 | -0.74412092 | 0.33513699 | -2.2203485 | 0.02639512 | 0.26304323 |
| TMEM128 | 19.3700816 | 0.023134812 | 0.44927841 | 0.05149326 | 0.95893247 | NA |
| STX18 | 39.5503437 | 0.314418842 | 0.3144729 | 0.99982808 | 0.31739371 | 0.69424851 |
| STX18-AS1 | 24.6906166 | 0.391810714 | 0.49098282 | 0.79801309 | 0.42486289 | NA |
| EVC2 | 15.3702105 | -0.174627623 | 0.42385434 | -0.4119991 | 0.68034006 | NA |
| EVC | 26.1039064 | -0.22062614 | 0.40861304 | -0.5399391 | 0.58923906 | NA |
| CRMP1 | 7.12485798 | -0.886959772 | 0.83557748 | -1.0614932 | 0.28846584 | NA |
| WFS1 | 35.7758178 | 0.107878679 | 0.49078157 | 0.21980996 | 0.82601916 | 0.949142 |
| MAN2B2 | 24.9186623 | -0.002269955 | 0.50715974 | -0.0044758 | 0.99642883 | NA |
| MRFAP1 | 62.7651756 | -0.427358656 | 0.30217362 | -1.4142818 | 0.15727917 | 0.55958274 |
| MRFAP1L1 | 10.5752222 | -0.397215004 | 0.49819644 | -0.797306 | 0.42527334 | NA |
| BLOC1S4 | 6.09889461 | -1.243288588 | 0.70258111 | -1.7696015 | 0.07679355 | NA |
| KIAA0232 | 96.3763243 | 0.063767008 | 0.25228533 | 0.25275749 | 0.80045562 | 0.94085969 |
| TBC1D14 | 29.4097525 | -0.138512681 | 0.34963333 | -0.3961655 | 0.6919829 | 0.90557575 |
| TADA2B | 8.51174574 | 0.061000933 | 0.52930621 | 0.11524696 | 0.90824938 | NA |
| GRPEL1 | 11.4178872 | -0.44795787 | 0.48030631 | -0.9326504 | 0.35100051 | NA |
| SORCS2 | 8.98426513 | -0.334944988 | 0.63273259 | -0.5293626 | 0.59655392 | NA |
| AFAP1 | 39.9229675 | 0.731940249 | 0.28244333 | 2.59145879 | 0.009557 | 0.15497902 |
| ABLIM2 | 438.595801 | -1.929441875 | 0.34064094 | -5.6641514 | 1.48E-08 | 1.50E-05 |
| SH3TC1 | 6.16696782 | -0.49578398 | 0.87894668 | -0.564066 | 0.57270918 | NA |
| ACOX3 | 16.1517573 | 0.728060542 | 0.42866075 | 1.69845396 | 0.08942212 | NA |
| TRMT44 | 10.7882801 | 0.662195155 | 0.47756498 | 1.38660744 | 0.1655615 | NA |
| SLC2A9 | 14.7672689 | 0.70785555 | 0.6501339 | 1.08878424 | 0.27624904 | NA |
| WDR1 | 33.8506907 | 0.386419199 | 0.40194326 | 0.96137749 | 0.3363624 | 0.71116888 |
| ZNF518B | 9.50416832 | 0.009321636 | 0.57022479 | 0.0163473 | 0.98695732 | NA |
| AC005699.1 | 8.81454656 | 0.90179973 | 0.73831258 | 1.22143352 | 0.22192192 | NA |
| RAB28 | 52.1432027 | 0.136905593 | 0.23745998 | 0.57654175 | 0.56424906 | 0.8549147 |
| BOD1L1 | 44.1306614 | -0.284770944 | 0.25664357 | -1.109597 | 0.26717271 | 0.66013596 |
| CPEB2-DT | 7.87769175 | 0.055154272 | 0.58451995 | 0.09435824 | 0.92482459 | NA |
| CPEB2 | 60.1816892 | 0.234228977 | 0.37031904 | 0.6325059 | 0.52705636 | 0.83195298 |
| AC098829.1 | 155.290209 | -0.597430755 | 0.51140169 | -1.1682221 | 0.24271719 | 0.64918976 |
| C1QTNF7 | 17.287413 | -0.624028972 | 0.86092694 | -0.7248338 | 0.46855398 | NA |
| CC2D2A | 44.9521285 | -0.518183607 | 0.46571906 | -1.1126528 | 0.26585758 | 0.66013596 |
| FBXL5 | 106.216219 | -0.003493324 | 0.25671192 | -0.013608 | 0.98914276 | 0.99651342 |
| FAM200B | 41.6106317 | -0.072720857 | 0.43343088 | -0.1677796 | 0.86675668 | 0.96089268 |
| BST1 | 33.3657706 | -0.23207598 | 0.57351077 | -0.4046584 | 0.68572859 | 0.90499319 |
| CD38 | 5.94698826 | -1.64187102 | 1.30497477 | -1.258163 | 0.2083328 | NA |
| TAPT1 | 37.0891193 | -0.162308123 | 0.31218335 | -0.5199128 | 0.60312434 | 0.87047761 |
| TAPT1-AS1 | 73.8928991 | 0.557105456 | 0.41965368 | 1.32753621 | 0.18433137 | 0.587049 |
| LDB2 | 18.4329611 | 0.533379331 | 0.64583909 | 0.82587032 | 0.40887766 | NA |
| QDPR | 17.613492 | 0.10885047 | 0.39906852 | 0.27276136 | 0.78503667 | NA |
| LAP3 | 33.4780797 | -1.057275112 | 1.06428767 | -0.993411 | 0.32050969 | 0.69496685 |
| AC006160.1 | 3.71681508 | -0.259564117 | 1.46273593 | -0.1774511 | 0.85915406 | NA |
| MED28 | 19.7674818 | 0.142745775 | 0.35300674 | 0.40437125 | 0.68593974 | NA |
| FAM184B | 25.3273357 | 0.689733207 | 0.56219433 | 1.22685905 | 0.21987557 | NA |
| DCAF16 | 29.7481083 | 0.185627666 | 0.27658874 | 0.6711324 | 0.50213619 | 0.82522868 |
| LCORL | 67.5914647 | -0.066377144 | 0.29481817 | -0.225146 | 0.82186566 | 0.94728252 |
| AC024230.1 | 9.89448845 | 0.538895828 | 0.84918475 | 0.63460375 | 0.52568688 | NA |
| SLIT2 | 13.1800314 | 1.264848072 | 0.68846382 | 1.83720341 | 0.06617987 | NA |
| PACRGL | 26.3031618 | -0.023223043 | 0.28716255 | -0.0808707 | 0.93554476 | NA |
| KCNIP4 | 260.397107 | 0.102355342 | 0.8054667 | 0.12707582 | 0.89888039 | 0.97097328 |
| ADGRA3 | 46.0541977 | -0.095976287 | 0.29438677 | -0.3260211 | 0.74440842 | 0.92221153 |
| PPARGC1A | 28.5630979 | 1.194853257 | 0.65965759 | 1.81132344 | 0.0700908 | 0.40663921 |
| DHX15 | 56.3876477 | 0.164679451 | 0.25377987 | 0.64890666 | 0.51639871 | 0.83104823 |
| CCDC149 | 12.6570187 | 0.479624398 | 0.4207904 | 1.13981783 | 0.2543622 | NA |
| SEPSECS | 9.53840327 | -0.058740352 | 0.51190123 | -0.1147494 | 0.90864377 | NA |
| AC104662.2 | 8.29930995 | 0.161096557 | 0.55099915 | 0.2923717 | 0.77000244 | NA |
| PI4K2B | 4.25666206 | -0.054270678 | 0.86911385 | -0.0624437 | 0.9502095 | NA |
| ZCCHC4 | 9.82925419 | 0.213517685 | 0.51590185 | 0.41387269 | 0.67896735 | NA |
| ANAPC4 | 29.8530566 | 0.550008961 | 0.31089543 | 1.76911238 | 0.07687512 | 0.42346132 |
| SLC34A2 | 5.84947263 | -3.490664298 | 1.11815313 | -3.1218124 | 0.00179741 | NA |
| SEL1L3 | 7.28924338 | -1.961342389 | 0.89719273 | -2.1860881 | 0.02880916 | NA |
| SMIM20 | 10.1697444 | -0.910040232 | 0.54852504 | -1.6590678 | 0.09710213 | NA |
| RBPJ | 58.7297541 | -0.282163707 | 0.24970376 | -1.1299938 | 0.25847883 | 0.65588339 |
| TBC1D19 | 52.1123554 | 0.260214815 | 0.24538618 | 1.0604298 | 0.28894911 | 0.67724823 |
| STIM2 | 44.7092316 | 0.219310267 | 0.32913484 | 0.66632347 | 0.50520437 | 0.82600448 |
| PCDH7 | 24.7734286 | 0.055406352 | 0.85888771 | 0.06450943 | 0.9485646 | NA |
| ARAP2 | 79.288575 | 0.160226994 | 0.33504626 | 0.4782235 | 0.63249114 | 0.88389921 |
| C4orf19 | 44.4278879 | -1.17341006 | 0.49915781 | -2.3507797 | 0.01873412 | 0.22818497 |
| RELL1 | 41.4080574 | -0.319042163 | 0.35076092 | -0.9095716 | 0.36304849 | 0.73259935 |
| PGM2 | 6.60676767 | -0.953221355 | 0.72689998 | -1.3113515 | 0.18973905 | NA |
| TBC1D1 | 216.07506 | -0.459234522 | 0.29575103 | -1.5527741 | 0.12047712 | 0.51168085 |
| KLF3 | 18.3845734 | -1.580044287 | 0.44161548 | -3.5778734 | 0.0003464 | NA |
| FAM114A1 | 50.3869036 | -0.116516564 | 0.43569396 | -0.2674275 | 0.78914 | 0.93792988 |
| KLHL5 | 18.2214589 | 0.352047939 | 0.34038239 | 1.0342719 | 0.30100907 | NA |
| WDR19 | 18.6521026 | -0.655112503 | 0.45144142 | -1.4511573 | 0.14673607 | NA |
| RFC1 | 77.4016351 | -0.17806998 | 0.31796654 | -0.5600274 | 0.57546073 | 0.85929076 |
| RPL9 | 7.67032085 | 0.128224029 | 0.583422 | 0.21977922 | 0.82604311 | NA |
| LIAS | 7.80730851 | -0.171639542 | 0.56430513 | -0.3041609 | 0.76100533 | NA |
| UGDH | 14.513527 | -0.119921136 | 0.49583487 | -0.241857 | 0.80889097 | NA |
| UGDH-AS1 | 20.7083672 | -0.225774103 | 0.36086799 | -0.6256418 | 0.53154991 | NA |
| SMIM14 | 60.3661466 | -0.150013382 | 0.34623566 | -0.4332696 | 0.66481893 | 0.89968484 |
| UBE2K | 126.960683 | 0.338271428 | 0.24491862 | 1.38115847 | 0.16723024 | 0.56878882 |
| PDS5A | 97.2412038 | 0.223793898 | 0.23680313 | 0.94506307 | 0.34462668 | 0.71842682 |

| | | | | | | |
|---|---|---|---|---|---|---|
| N4BP2 | 30.9867811 | 0.655385201 | 0.33999111 | 1.92765395 | 0.05389818 | 0.3615191 |
| RBM47 | 60.6282789 | 0.158193914 | 0.52281341 | 0.30258197 | 0.76220845 | 0.92336765 |
| NSUN7 | 15.4613588 | 0.94600394 | 0.51410256 | 1.84010742 | 0.06575247 | NA |
| APBB2 | 682.37362 | 0.529932759 | 0.31678078 | 1.67286902 | 0.09435308 | 0.46752784 |
| AC131953.1 | 5.38920832 | -0.417826897 | 0.84056977 | -0.4970758 | 0.6191356 | NA |
| UCHL1 | 16.7024119 | 0.19137766 | 0.43782185 | 0.43711308 | 0.66202934 | NA |
| LIMCH1 | 31.6791377 | 1.026748649 | 0.48254756 | 2.12776675 | 0.03335643 | 0.29047359 |
| TMEM33 | 27.5649332 | 0.559314548 | 0.32783699 | 1.70607516 | 0.08799409 | NA |
| SLC30A9 | 61.4316827 | 0.218279073 | 0.26984726 | 0.8088986 | 0.41857347 | 0.77290571 |
| SHISA3 | 21.4346027 | -1.654629967 | 1.09095804 | -1.5166761 | 0.12934848 | NA |
| ATP8A1 | 32.5056856 | 0.16240409 | 0.56693997 | 0.28645729 | 0.7745279 | 0.92806061 |
| KCTD8 | 20.1121928 | 1.8748565 | 0.77735041 | 2.41185503 | 0.01587159 | NA |
| GUF1 | 10.9839679 | 0.381262896 | 0.46494918 | 0.82000982 | 0.41221051 | NA |
| GNPDA2 | 14.6419304 | -0.086386481 | 0.50380724 | -0.1714673 | 0.86385632 | NA |
| GABRA2 | 34.8425907 | 0.497707141 | 0.62928823 | 0.79090489 | 0.4289995 | 0.7784723 |
| COMMD8 | 16.1484474 | -0.321077387 | 0.4152266 | -0.7732582 | 0.43936956 | NA |
| ATP10D | 87.6628303 | -0.240552839 | 0.2674859 | -0.8993103 | 0.36848738 | 0.73626927 |
| NFXL1 | 16.429478 | 0.103074303 | 0.36221314 | 0.28456809 | 0.77597506 | NA |
| TXK | 10.4689898 | -0.229692836 | 0.50553134 | -0.4543592 | 0.64957028 | NA |
| TEC | 13.3231956 | -0.001537025 | 0.46151286 | -0.0033304 | 0.99734273 | NA |
| SLAIN2 | 54.2218718 | -0.065654051 | 0.27600944 | -0.2378689 | 0.81198281 | 0.94415747 |
| FRYL | 94.1550207 | 0.25198306 | 0.21152765 | 1.19125355 | 0.23355406 | 0.63963219 |
| OCIAD1 | 75.8212609 | 0.046249984 | 0.26570816 | 0.17406309 | 0.86181588 | 0.95951831 |
| OCIAD2 | 12.3370762 | -0.066620447 | 0.56306293 | -0.1183179 | 0.90581574 | NA |
| DCUN1D4 | 54.5256335 | 0.122750224 | 0.31169477 | 0.39381547 | 0.69371728 | 0.90557575 |
| LRRC66 | 4.60630223 | -0.13564487 | 0.682666 | -0.1986987 | 0.84249841 | NA |
| SGCB | 78.1236185 | 0.201143258 | 0.42264698 | 0.47591316 | 0.63413625 | 0.88508487 |
| SPATA18 | 38.0452356 | 1.044326894 | 0.55478044 | 1.88241475 | 0.05977972 | 0.37905854 |
| USP46 | 117.309083 | 0.023527724 | 0.44875138 | 0.05242931 | 0.95818662 | 0.98714883 |
| ERVMER34-1 | 14.3102485 | -0.434573117 | 0.60579424 | -0.7173609 | 0.47315142 | NA |
| RASL11B | 157.775027 | -1.389986141 | 0.59484221 | -2.3367308 | 0.01945318 | 0.23378067 |
| SCFD2 | 65.5828819 | 0.097224167 | 0.36190423 | 0.26864612 | 0.78820202 | 0.93749303 |
| AC023154.1 | 110.148596 | -0.429744107 | 0.22694187 | -1.8936308 | 0.05827404 | 0.37650386 |
| AC058822.1 | 33.3830347 | 0.35816864 | 0.56684399 | 0.63186458 | 0.52747537 | 0.83195298 |
| LNX1 | 10.4698336 | -0.977420083 | 0.9251263 | -1.0565261 | 0.29072792 | NA |
| CHIC2 | 49.1117522 | 0.618732232 | 0.34518758 | 1.79245216 | 0.07306057 | 0.41834543 |
| SRD5A3 | 6.92557073 | -0.097144079 | 0.66419875 | -0.1462575 | 0.88371808 | NA |
| TMEM165 | 80.5449822 | -0.060392436 | 0.3087946 | -0.1955748 | 0.844943 | 0.95509854 |
| CLOCK | 57.2427185 | 0.124139224 | 0.31037593 | 0.39996408 | 0.68918297 | 0.90557575 |
| EXOC1 | 45.4711575 | 0.056934755 | 0.33421006 | 0.1703562 | 0.86473002 | 0.96089268 |
| CEP135 | 20.8750555 | 0.140931483 | 0.39052373 | 0.36087815 | 0.71819054 | NA |
| KIAA1211 | 117.074524 | 0.100391126 | 0.41542502 | 0.24165883 | 0.80904453 | 0.94346479 |
| AASDH | 19.8510238 | 0.329815593 | 0.39480398 | 0.83539075 | 0.40349774 | NA |
| PAICS | 12.1845942 | -1.008431209 | 0.46871391 | -2.1514855 | 0.0314379 | NA |
| SRP72 | 29.3807517 | -0.059757225 | 0.31652472 | -0.1887917 | 0.85025611 | 0.95532127 |
| REST | 32.6596276 | -0.244763702 | 0.34552024 | -0.7083918 | 0.47870199 | 0.81001889 |
| NOA1 | 5.41451256 | -0.017209078 | 0.72654712 | -0.0236861 | 0.98110298 | NA |
| POLR2B | 34.293763 | 0.212933092 | 0.31454423 | 0.67695755 | 0.49843289 | 0.82345406 |
| IGFBP7 | 1214.41452 | -0.935332989 | 0.23436919 | -3.9908531 | 6.58E-05 | 0.00764336 |
| AC116049.2 | 15.6941716 | -1.740114993 | 0.99833653 | -1.7430144 | 0.08133109 | NA |
| AC104806.2 | 21.6894214 | 0.277991263 | 0.50352319 | 0.55209228 | 0.58088513 | NA |
| CENPC | 40.9294784 | 0.401205059 | 0.35857518 | 1.11888685 | 0.26318841 | 0.6581333 |
| UBA6 | 50.9036476 | 0.509312139 | 0.38373979 | 1.32723305 | 0.1844316 | 0.587049 |
| UBA6-AS1 | 117.069767 | 0.264463336 | 0.44350954 | 0.59629684 | 0.55097695 | 0.84714272 |
| YTHDC1 | 49.9278003 | -0.032808426 | 0.28566572 | -0.114849 | 0.9085648 | 0.97490445 |
| UGT2A3 | 4.59756613 | 0.927648904 | 1.14596175 | 0.80949377 | 0.41823118 | NA |
| UGT2B7 | 38.0331071 | 0.002717439 | 0.752923 | 0.00360919 | 0.99712029 | 0.99958475 |
| UTP3 | 14.7529845 | -0.788033693 | 0.40670416 | -1.9376091 | 0.05267093 | NA |
| RUFY3 | 113.130409 | 0.049830077 | 0.24615148 | 0.20243663 | 0.8395754 | 0.95393706 |
| GRSF1 | 21.8343565 | -0.48420963 | 0.35898554 | -1.3488277 | 0.17739233 | NA |
| MOB1B | 41.2327469 | 0.816779678 | 0.30770095 | 2.65445939 | 0.00794356 | 0.14193427 |
| DCK | 5.26186335 | 0.659418931 | 0.64185053 | 1.02737148 | 0.30424557 | NA |
| SLC4A4 | 30.6742779 | 1.593389661 | 0.87998178 | 1.81070756 | 0.07018613 | 0.40663921 |
| COX18 | 7.45547593 | -0.008958253 | 0.56085181 | -0.0159726 | 0.98725626 | NA |
| ANKRD17 | 215.195733 | 0.495505444 | 0.24464607 | 2.02539715 | 0.0428266 | 0.32651257 |
| AC053527.1 | 33.071581 | 0.933208534 | 0.46775901 | 1.99506265 | 0.04603605 | 0.33643641 |
| RASSF6 | 6.94838463 | 0.054710911 | 0.62788597 | 0.08713511 | 0.93056412 | NA |
| MTHFD2L | 31.1831391 | 0.077795258 | 0.38641385 | 0.20132627 | 0.84044346 | 0.95393706 |
| BTC | 31.7617923 | -0.169834198 | 0.44429154 | -0.3822585 | 0.70226967 | 0.90757606 |
| RCHY1 | 29.5021281 | 0.036988971 | 0.38342253 | 0.09647052 | 0.92314689 | 0.97775374 |
| THAP6 | 15.1759777 | 0.107142382 | 0.37443556 | 0.28614372 | 0.77476805 | NA |
| CDKL2 | 7.31510412 | 0.381757106 | 0.74321692 | 0.51365502 | 0.6074932 | NA |
| G3BP2 | 47.9132589 | 0.185358015 | 0.25398734 | 0.72979234 | 0.46551713 | 0.80073684 |
| USO1 | 103.836889 | 0.507452786 | 0.22962636 | 2.20990647 | 0.02711165 | 0.26690502 |
| NAAA | 15.8825381 | 0.041879199 | 0.38724553 | 0.10814637 | 0.91387959 | NA |
| SDAD1 | 16.0682762 | 0.18952044 | 0.41637717 | 0.4551653 | 0.64899032 | NA |
| CXCL10 | 13.5009524 | 2.938448058 | 3.7712942 | 0.77916172 | 0.43588446 | NA |
| NUP54 | 23.7620088 | -0.073351173 | 0.34117513 | -0.2149957 | 0.8297707 | NA |
| SCARB2 | 142.353146 | -0.120361927 | 0.46561485 | -0.258501 | 0.79602024 | 0.93934998 |
| SHROOM3 | 135.1734 | 0.786786306 | 0.32583967 | 2.4146425 | 0.01575066 | 0.20674653 |
| 11-Sep | 211.256147 | 0.054141402 | 0.28287525 | 0.19139675 | 0.84821477 | 0.95532127 |
| CCNI | 144.658616 | -0.325606355 | 0.3199559 | -1.0176601 | 0.30883951 | 0.6882709 |
| CCNG2 | 15.8763352 | 0.382233655 | 0.6446203 | 0.59295938 | 0.55320834 | NA |
| CNOT6L | 47.5308057 | 0.370686699 | 0.33357197 | 1.11126452 | 0.26645451 | 0.66013596 |
| MRPL1 | 26.3227565 | 0.48125269 | 0.3464562 | 1.38907223 | 0.16481079 | NA |
| FRAS1 | 70.7733659 | 1.416121018 | 0.68935367 | 2.05427357 | 0.03994922 | 0.31540889 |
| BMP2K | 52.3688921 | 0.341529815 | 0.43407795 | 0.78679374 | 0.43140262 | 0.77901411 |
| PAQR3 | 16.8890943 | 0.395550059 | 0.42119249 | 0.93911945 | 0.34766942 | NA |
| LINC01088 | 12.6871155 | 0.359273866 | 0.71785166 | 0.50048483 | 0.61673373 | NA |
| ANTXR2 | 30.0410501 | 0.339692561 | 0.35685868 | 0.95189658 | 0.34114943 | 0.71620192 |
| CFAP299 | 7.24779561 | 0.98786987 | 0.75093028 | 1.31552808 | 0.18833249 | NA |
| PRKG2 | 21.2947429 | -1.639706767 | 0.92220778 | -1.7780231 | 0.07540005 | NA |
| RASGEF1B | 285.471462 | 1.282894518 | 0.67821545 | 1.8915737 | 0.0585478 | 0.37650386 |
| HNRNPD | 65.1570302 | -0.340244092 | 0.24940729 | -1.3642107 | 0.17250127 | 0.57538252 |
| HNRNPDL | 78.6803591 | 0.099974839 | 0.3036292 | 0.32926622 | 0.74195447 | 0.92085904 |
| ENOPH1 | 22.0807127 | -0.25314395 | 0.39860264 | -0.6350785 | 0.52537724 | NA |
| TMEM150C | 254.074129 | -0.872505108 | 0.415175 | -2.1015358 | 0.03559396 | 0.29549524 |
| SCD5 | 109.195374 | 0.564099034 | 0.41248306 | 1.36756898 | 0.17144703 | 0.57422722 |
| SEC31A | 107.268836 | -0.349383385 | 0.24647809 | -1.4175028 | 0.15633597 | 0.55671527 |
| LIN54 | 17.636963 | -0.301763324 | 0.41536943 | -0.7264938 | 0.46753609 | NA |
| COPS4 | 27.2267813 | -0.094759144 | 0.33039098 | -0.2868091 | 0.77425848 | NA |
| COQ2 | 4.93919568 | -1.210686407 | 0.78893388 | -1.5345854 | 0.1248857 | NA |
| HELQ | 12.3078077 | 0.0728872 | 0.39947895 | 0.18245567 | 0.85522514 | NA |
| MRPS18C | 9.68260111 | -1.11400224 | 0.56910892 | -1.95745 | 0.05029459 | NA |
| ABRAXAS1 | 6.84341774 | -0.411529963 | 0.65948288 | -0.6240192 | 0.53261499 | NA |
| GPAT3 | 25.8950422 | 2.801870421 | 0.88562594 | 3.16371767 | 0.00155768 | NA |
| CDS1 | 43.8322644 | 0.165852798 | 0.50605929 | 0.32773392 | 0.74311285 | 0.92122771 |
| WDFY3 | 163.739178 | 0.699268953 | 0.21676474 | 3.22593493 | 0.00125562 | 0.0545866 |
| ARHGAP24 | 400.932667 | 0.602568443 | 0.33217637 | 1.81400151 | 0.06967749 | 0.40663921 |
| MAPK10 | 308.713535 | 0.468091604 | 0.28961615 | 1.61624825 | 0.10604065 | 0.48930703 |
| PTPN13 | 351.935681 | -0.103767197 | 0.38976516 | -0.26623 | 0.79006204 | 0.93835773 |
| AC093827.5 | 5.39175912 | 0.194565413 | 0.59404342 | 0.32752726 | 0.74326913 | NA |
| AFF1 | 115.155738 | 0.166078339 | 0.36045605 | 0.46074504 | 0.64498153 | 0.89162142 |

| | | | | | | |
|---|---|---|---|---|---|---|
| KLHL8 | 16.5774021 | -0.368148416 | 0.42586457 | -0.864473 | 0.38732812 | NA |
| HSD17B11 | 11.4096093 | -0.197704593 | 0.57748421 | -0.342355 | 0.73208377 | NA |
| NUDT9 | 23.0551954 | -0.486947733 | 0.41366163 | -1.1771644 | 0.23912991 | NA |
| SPP1 | 502.284398 | -2.558490576 | 0.55797716 | -4.5852962 | 4.53E-06 | 0.00096777 |
| PKD2 | 58.95168 | 0.156910055 | 0.34354905 | 0.45673261 | 0.64786325 | 0.89261695 |
| PPM1K | 18.0859525 | 0.702070484 | 0.70028628 | 1.00254782 | 0.31607908 | NA |
| HERC6 | 10.0441701 | -0.322771942 | 1.30772895 | -0.2468187 | 0.80504855 | NA |
| HERC5 | 14.4912751 | -1.075063368 | 1.13902073 | -0.9438488 | 0.34524691 | NA |
| PYURF | 17.3265457 | -1.242473093 | 0.54587875 | -2.2760972 | 0.02284019 | NA |
| HERC3 | 17.7932662 | 0.385214451 | 0.42260621 | 0.91152105 | 0.3620209 | NA |
| NAP1L5 | 4.03339593 | -0.223434492 | 0.73593535 | -0.3036061 | 0.76142797 | NA |
| FAM13A-AS1 | 8.87539759 | 0.268042427 | 0.49654921 | 0.53981039 | 0.5893278 | NA |
| FAM13A | 62.582185 | 0.1924934 | 0.30469092 | 0.63176612 | 0.52753971 | 0.83195298 |
| SNCA | 138.831833 | -0.599048311 | 0.40549667 | -1.4773199 | 0.1395899 | 0.53526806 |
| CCSER1 | 100.867109 | 1.200321561 | 0.67024063 | 1.79088151 | 0.07331231 | 0.41834543 |
| SMARCAD1 | 40.1033519 | 0.435676284 | 0.2605439 | 1.67217996 | 0.09448884 | 0.46752784 |
| PDLIM5 | 553.224582 | 0.06930469 | 0.25596801 | 0.27075528 | 0.78657926 | 0.93615351 |
| BMPR1B | 24.5772088 | 0.049963369 | 0.46680363 | 0.10703295 | 0.91476284 | NA |
| UNC5C | 404.739313 | -0.613963674 | 0.34577152 | -1.775634 | 0.07579326 | 0.42186351 |
| STPG2 | 83.1535284 | 0.911272051 | 0.43445197 | 2.09752083 | 0.0359475 | 0.29695121 |
| RAP1GDS1 | 73.8686424 | 0.44437371 | 0.23099214 | 1.92376121 | 0.05438452 | 0.36220627 |
| TSPAN5 | 76.3927901 | 0.307894086 | 0.43844812 | 0.70223607 | 0.48253195 | 0.81295551 |
| EIF4E | 32.6105421 | 0.396565512 | 0.40613269 | 0.97644322 | 0.32884487 | 0.70199726 |
| METAP1 | 36.2821759 | 0.583160938 | 0.28723141 | 2.03028264 | 0.04232782 | 0.3239276 |
| ADH5 | 43.6439064 | -0.276416548 | 0.41332773 | -0.6687588 | 0.50364936 | 0.82600448 |
| TRMT10A | 6.01005629 | 0.434452579 | 0.64518829 | 0.67337332 | 0.50070982 | NA |
| LAMTOR3 | 16.4921868 | 0.22400754 | 0.44042958 | 0.50861148 | 0.61102458 | NA |
| DNAJB14 | 65.5811063 | 0.239001926 | 0.23480903 | 1.01785662 | 0.3087461 | 0.6882709 |
| H2AFZ | 25.6297784 | -1.014064702 | 0.65290365 | -1.5531613 | 0.1203846 | NA |
| DDIT4L | 17.1750452 | -0.334038445 | 0.73306798 | -0.4556718 | 0.64862597 | NA |
| PPP3CA | 87.0965909 | -0.063586847 | 0.30049862 | -0.2116045 | 0.83241563 | 0.95050875 |
| BANK1 | 19.8265534 | 0.148223555 | 0.68461449 | 0.2165066 | 0.82859288 | NA |
| NFKB1 | 56.3154595 | 0.371695223 | 0.24296249 | 1.5298461 | 0.12605483 | 0.52276701 |
| MANBA | 30.54956 | -0.064125619 | 0.40636907 | -0.1578014 | 0.87461327 | 0.96229085 |
| UBE2D3 | 127.478238 | -0.02579618 | 0.20328545 | -0.1268963 | 0.89902245 | 0.97097328 |
| AC018797.2 | 6.65429808 | 0.316192803 | 0.62920378 | 0.50252845 | 0.61529584 | NA |
| CISD2 | 18.59099 | -0.888028769 | 0.38254454 | -2.3213736 | 0.02026669 | NA |
| SLC9B1 | 6.41985587 | 0.051121483 | 0.67118373 | 0.07616615 | 0.93928691 | NA |
| SLC9B2 | 9.80615598 | -0.569438771 | 0.50737137 | -1.1223313 | 0.26172161 | NA |
| TACR3 | 17.6913554 | 0.623447707 | 0.76519004 | 0.81476192 | 0.4152086 | NA |
| TET2 | 176.768916 | 0.145461741 | 0.40975767 | 0.35499456 | 0.72259367 | 0.91509584 |
| PPA2 | 59.6250433 | 0.290344797 | 0.37760205 | 0.76891742 | 0.44194234 | 0.78620488 |
| ARHGEF38 | 14.4572537 | -0.812354141 | 0.65257242 | -1.244849 | 0.21318725 | NA |
| INTS12 | 241.984336 | -0.734409086 | 0.43383435 | -1.6928329 | 0.09048727 | 0.45866472 |
| GSTCD | 50.6033371 | -0.63175213 | 0.43233286 | -1.4612633 | 0.14394321 | 0.54209252 |
| NPNT | 1143.6162 | -0.000415694 | 0.28508078 | -0.0014582 | 0.99883656 | 0.99961632 |
| AC109361.2 | 14.0268468 | 0.229883842 | 0.76559697 | 0.30026744 | 0.76397317 | NA |
| AC109361.1 | 7.81644584 | -0.02358395 | 0.985463 | -0.0239318 | 0.98090697 | NA |
| TBCK | 101.109479 | 0.111944619 | 0.28164829 | 0.39746244 | 0.69102648 | 0.90557575 |
| AIMP1 | 22.5172528 | 0.062250798 | 0.34341446 | 0.18127017 | 0.85615552 | NA |
| PAPSS1 | 87.3367871 | 0.852820415 | 0.34964363 | 2.43911326 | 0.01472335 | 0.19896332 |
| SGMS2 | 8.75073817 | -0.378623478 | 0.55155553 | -0.6864648 | 0.49242003 | NA |
| CYP2U1 | 8.74734296 | -0.267778656 | 0.58198899 | -0.4601095 | 0.64543763 | NA |
| HADH | 11.1025827 | -0.16720167 | 0.48647679 | -0.3436992 | 0.73107253 | NA |
| RPL34-AS1 | 7.92710018 | -1.34406298 | 0.754976 | -1.7802725 | 0.07503138 | NA |
| RPL34 | 52.971398 | -0.408082876 | 0.47533445 | -0.8585174 | 0.39060685 | 0.75550853 |
| OSTC | 32.1142739 | -0.389031574 | 0.55881644 | -0.6961706 | 0.486322 | 0.81509175 |
| SEC24B-AS1 | 9.34857539 | 0.171627748 | 0.52648186 | 0.32598986 | 0.74443203 | NA |
| SEC24B | 76.2279817 | 0.386377107 | 0.21814893 | 1.7711621 | 0.07653375 | 0.42346132 |
| MCUB | 16.732048 | 0.024836234 | 0.56441144 | 0.04400378 | 0.96490139 | NA |
| CASP6 | 12.8153802 | 0.132630086 | 0.41460469 | 0.31989529 | 0.74904771 | NA |
| PLA2G12A | 10.9924831 | -0.540749759 | 0.44364565 | -1.2188776 | 0.22289064 | NA |
| GAR1 | 10.6255932 | 0.364883043 | 0.42510472 | 0.85833684 | 0.39070649 | NA |
| EGF | 49.5686611 | 4.328939769 | 1.23537714 | 3.50414431 | 0.00045808 | 0.02928507 |
| ELOVL6 | 23.6561021 | 0.41244847 | 0.4056363 | 1.01679378 | 0.30925154 | NA |
| ENPEP | 899.698009 | -1.312032027 | 0.35570677 | -3.6885214 | 0.00022556 | 0.01829751 |
| FAM241A | 6.28416126 | 0.418583219 | 0.6286638 | 0.66583 | 0.50551977 | NA |
| AP1AR | 10.3526016 | -0.260973233 | 0.53337345 | -0.489288 | 0.62463781 | NA |
| ALPK1 | 46.5018636 | -0.873854786 | 0.4098651 | -2.1320547 | 0.03300235 | 0.29047359 |
| LARP7 | 28.3420047 | 0.158860986 | 0.31190169 | 0.50933032 | 0.61052071 | NA |
| ANK2 | 63.5817812 | 0.915602126 | 0.56830133 | 1.61112086 | 0.10715338 | 0.49086361 |
| CAMK2D | 77.110664 | -0.050758599 | 0.23816047 | -0.2131277 | 0.83122734 | 0.95050875 |
| ARSJ | 14.0291663 | 0.929450681 | 0.41571341 | 2.23579672 | 0.02536509 | NA |
| UGT8 | 20.2098765 | 1.965161309 | 0.65490016 | 3.00070365 | 0.00269357 | NA |
| SNHG8 | 10.6660345 | -1.014341241 | 0.75844331 | -1.3373989 | 0.18109247 | NA |
| METTL14 | 17.2094726 | -0.2557588 | 0.3459762 | -0.7392381 | 0.45976241 | NA |
| SEC24D | 31.0134832 | -0.162455955 | 0.30771795 | -0.5279379 | 0.59754247 | 0.86882662 |
| USP53 | 168.854373 | 0.247407396 | 0.31013881 | 0.79773118 | 0.4250265 | 0.77723512 |
| C4orf3 | 50.7585642 | -0.844722219 | 0.44292864 | -1.9071294 | 0.05650384 | 0.37144176 |
| PDE5A | 19.4885334 | -0.665525596 | 0.69478579 | -0.957886 | 0.33812024 | NA |
| AC108866.1 | 19.0595761 | -0.444667608 | 0.51619319 | -0.8614364 | 0.38899773 | NA |
| PRDM5 | 112.791518 | 0.323113255 | 0.34634779 | 0.93291561 | 0.35086354 | 0.72238707 |
| NDNF | 408.369454 | -0.053405177 | 0.57831404 | -0.0923463 | 0.92642288 | 0.97775374 |
| ANXA5 | 84.2137895 | -0.694023524 | 0.44078093 | -1.5745317 | 0.11536459 | 0.50513044 |
| EXOSC9 | 10.8125624 | 0.075290045 | 0.45167207 | 0.16669183 | 0.86761253 | NA |
| BBS7 | 18.5024142 | 0.381924665 | 0.43294059 | 0.88216414 | 0.37768805 | NA |
| KIAA1109 | 115.907443 | 0.522577293 | 0.26363579 | 1.98219403 | 0.04745753 | 0.34008434 |
| SPATA5 | 43.5499007 | 0.087334723 | 0.34750974 | 0.2513159 | 0.80156988 | 0.94085969 |
| SPRY1 | 12.9168856 | -0.338488129 | 0.78799841 | -0.4295543 | 0.66751986 | NA |
| ANKRD50 | 9.00979969 | 0.072654199 | 0.57422044 | 0.12652667 | 0.89931504 | NA |
| FAT4 | 20.4972547 | -0.407551072 | 0.57952654 | -0.7032483 | 0.481901 | NA |
| INTU | 32.45299 | -0.415113574 | 0.31596343 | -1.3138026 | 0.18891264 | 0.59077076 |
| HSPA4L | 13.123739 | -0.345933557 | 0.63847404 | -0.541813 | 0.58794731 | NA |
| MFSD8 | 27.6265508 | 0.150023637 | 0.2948694 | 0.50877994 | 0.61090648 | NA |
| ABHD18 | 57.2022909 | 0.363991393 | 0.31071055 | 1.17148065 | 0.24140563 | 0.64918976 |
| LARP1B | 65.3938457 | 0.120089123 | 0.25551844 | 0.46998222 | 0.63836772 | 0.886719 |
| PGRMC2 | 41.6107212 | -0.431796961 | 0.2779158 | -1.5536971 | 0.12025669 | 0.51128003 |
| AC108062.1 | 17.4099509 | -0.356588765 | 0.42229836 | -0.8444001 | 0.39844588 | NA |
| JADE1 | 71.6661506 | -0.281234203 | 0.28846326 | -0.9749394 | 0.32959031 | 0.70253658 |
| SCLT1 | 47.8758949 | 0.128593266 | 0.44474531 | 0.28913912 | 0.77247492 | 0.92669573 |
| C4orf33 | 7.28678486 | 0.963973999 | 0.69585958 | 1.38529961 | 0.16596087 | NA |
| AC096711.2 | 12.4497357 | 0.052937332 | 0.47773848 | 0.11080818 | 0.91176846 | NA |
| AC109927.1 | 26.6993012 | -0.491156172 | 0.49341084 | -0.9954304 | 0.31952696 | NA |
| NOCT | 5.06114113 | -0.686081442 | 0.72944337 | -0.9405548 | 0.34693307 | NA |
| ELF2 | 166.242 | -0.503859657 | 0.21836404 | -2.3074296 | 0.02103088 | 0.24028525 |
| NDUFC1 | 32.5287724 | -0.489279962 | 0.50150004 | -0.9756329 | 0.32924639 | 0.70248468 |
| NAA15 | 45.069448 | -0.01823994 | 0.30639593 | -0.0595306 | 0.95252948 | 0.98520702 |
| AC097376.2 | 66.434697 | -0.503851643 | 0.33417767 | -1.5077358 | 0.13162216 | 0.52613957 |
| RAB33B | 12.9184342 | 0.264862619 | 0.4763803 | 0.55598986 | 0.57821778 | NA |
| SETD7 | 20.967822 | -0.496331883 | 0.43085216 | -1.1519772 | 0.24933042 | NA |
| MGST2 | 20.4201715 | 0.644890977 | 0.33468064 | 1.92688464 | 0.05399401 | NA |
| MAML3 | 402.04993 | -0.184521107 | 0.26693129 | -0.6912682 | 0.48939702 | 0.81787981 |
| SCOC | 61.0108717 | 0.194212891 | 0.41190918 | 0.47149445 | 0.63728768 | 0.88636488 |

| | | | | | | |
|---|---|---|---|---|---|---|
| SCOC-AS1 | 5.35295308 | -0.277088619 | 0.70225 | -0.3945726 | 0.69315833 | NA |
| ELMOD2 | 19.6714552 | -0.036665584 | 0.38206512 | -0.0959668 | 0.92354691 | NA |
| TBC1D9 | 34.0108703 | 0.502932412 | 0.314102 | 1.60117546 | 0.10933806 | 0.49591853 |
| RNF150 | 127.909894 | 0.621526007 | 0.37982135 | 1.63636405 | 0.10176342 | 0.48317363 |
| ZNF330 | 14.2121979 | -0.300520099 | 0.50647255 | -0.5933591 | 0.55294085 | NA |
| IL15 | 20.1031394 | -0.426351408 | 0.35339315 | -1.2064507 | 0.22764377 | NA |
| INPP4B | 21.3080699 | 1.901274333 | 0.82822915 | 2.29558974 | 0.02169935 | NA |
| AC104596.1 | 12.9254088 | -0.211695135 | 0.815823 | -0.2594866 | 0.79525982 | NA |
| USP38 | 16.5518338 | 0.03463094 | 0.35917682 | 0.09641752 | 0.92318898 | NA |
| GAB1 | 41.5657987 | 0.344627617 | 0.37446275 | 0.9203255 | 0.35740269 | 0.72903084 |
| SMARCA5 | 48.662237 | 0.002924494 | 0.25132354 | 0.01163637 | 0.99071573 | 0.99710744 |
| ANAPC10 | 37.4675586 | 0.546972199 | 0.28399841 | 1.92596921 | 0.05410822 | 0.3615191 |
| ABCE1 | 16.3050145 | 0.378046035 | 0.42230809 | 0.89519012 | 0.37068546 | NA |
| OTUD4 | 41.6962951 | -0.248248321 | 0.2917711 | -0.8508324 | 0.39486243 | 0.75831536 |
| SMAD1 | 45.7195533 | -0.892421948 | 0.30688088 | -2.9080402 | 0.00363702 | 0.09771848 |
| MMAA | 6.78517457 | -0.166945107 | 0.61372323 | -0.2720202 | 0.7856065 | NA |
| ZNF827 | 36.1140191 | 0.656327766 | 0.3958134 | 1.65817468 | 0.09728221 | 0.47141775 |
| LSM6 | 12.191101 | 0.037323589 | 0.50919636 | 0.07329901 | 0.94156818 | NA |
| SLC10A7 | 30.1865026 | 0.006138057 | 0.36870517 | 0.0166476 | 0.98671775 | 0.99585746 |
| TTC29 | 7.50639364 | 2.003712246 | 1.20184584 | 1.66719573 | 0.09547549 | NA |
| EDNRA | 8.17980638 | -0.704835225 | 0.82702005 | -0.8522589 | 0.39407039 | NA |
| TMEM184C | 15.5948714 | -0.247095921 | 0.40979329 | -0.602977 | 0.54652401 | NA |
| PRMT9 | 13.6238626 | 0.187152401 | 0.43795772 | 0.42732983 | 0.66913911 | NA |
| ARHGAP10 | 11.2560278 | 0.556427367 | 0.79383043 | 0.70093983 | 0.48334056 | NA |
| NR3C2 | 187.179273 | 0.99114219 | 0.35349046 | 2.80387252 | 0.00504929 | 0.11214076 |
| DCLK2 | 60.8642308 | -0.012026494 | 0.27375036 | -0.0439323 | 0.96495833 | 0.98892249 |
| LRBA | 208.593941 | 0.49492758 | 0.24296077 | 2.03706791 | 0.04164324 | 0.31989578 |
| RPS3A | 34.4543048 | -0.332350516 | 0.53540401 | -0.6207472 | 0.534766 | 0.83677705 |
| SH3D19 | 91.3399211 | 0.231028255 | 0.27020892 | 0.85499864 | 0.3925519 | 0.75710438 |
| FAM160A1 | 19.7638571 | 1.409162085 | 0.63232103 | 2.22855484 | 0.02584354 | NA |
| GATB | 6.65811706 | 0.124424383 | 0.7197618 | 0.17286883 | 0.86275453 | NA |
| FBXW7 | 109.323407 | 0.746492573 | 0.24103792 | 3.09699227 | 0.00195495 | 0.07143492 |
| ARFIP1 | 46.7948824 | 0.054360894 | 0.2408817 | 0.22567466 | 0.82145447 | 0.94728252 |
| TRIM2 | 33.4131409 | 1.049546848 | 0.48260556 | 2.17475085 | 0.02964879 | 0.27926427 |
| PLRG1 | 8.95459169 | 0.074333859 | 0.47636254 | 0.15604472 | 0.87599777 | NA |
| FGB | 8.09862854 | -3.880273316 | 2.05787857 | -1.8855696 | 0.05935297 | NA |
| MAP9 | 24.2143647 | 0.285200962 | 0.29498107 | 0.96684495 | 0.33362155 | NA |
| GUCY1B1 | 12.7648111 | 0.965814406 | 0.61821015 | 1.56227524 | 0.11822316 | NA |
| CTSO | 16.9967973 | -0.492861265 | 0.65285832 | -0.7549284 | 0.45029196 | NA |
| PDGFC | 71.3435242 | 1.112173184 | 0.38501049 | 2.88868283 | 0.00386859 | 0.09814043 |
| ETFDH | 31.6803397 | 0.161625378 | 0.33333826 | 0.48486897 | 0.6277693 | 0.88104923 |
| PPID | 10.6738111 | -0.835457391 | 0.45688237 | -1.828605 | 0.0674588 | NA |
| FNIP2 | 56.8279005 | 0.155109395 | 0.49418514 | 0.313869 | 0.75362054 | 0.92313949 |
| RAPGEF2 | 144.632666 | 0.773661993 | 0.2357582 | 3.28159098 | 0.00103223 | 0.05168807 |
| NAF1 | 18.5050129 | 0.155561813 | 0.37432831 | 0.41557587 | 0.67772039 | NA |
| TMA16 | 17.5298013 | -0.482187617 | 0.42399426 | -1.1372503 | 0.25543365 | NA |
| TMEM192 | 25.9560119 | -0.20094358 | 0.44174748 | -0.4548834 | 0.64919315 | NA |
| KLHL2 | 29.1195979 | -0.489901401 | 0.3254308 | -1.5053935 | 0.13222294 | 0.52613957 |
| MSMO1 | 23.9773807 | -0.462092115 | 0.56190641 | -0.8223649 | 0.41086924 | NA |
| DDX60 | 43.3059122 | -0.209104798 | 0.67770737 | -0.3085473 | 0.75766589 | 0.92313949 |
| DDX60L | 39.4297543 | 1.114317355 | 1.01179019 | 1.10133244 | 0.270752 | 0.66394807 |
| PALLD | 241.349514 | -0.43728487 | 0.43850066 | -0.9972274 | 0.31865415 | 0.69469143 |
| CBR4 | 111.176847 | 0.155278354 | 0.31628253 | 0.49094824 | 0.62346305 | 0.8780438 |
| SH3RF1 | 326.447446 | 0.197643616 | 0.24639065 | 0.80215549 | 0.42246302 | 0.77588259 |
| AC096741.1 | 6.693841 | -0.6812784 | 0.69354367 | -0.9823151 | 0.32594465 | NA |
| NEK1 | 52.719379 | 0.182264324 | 0.32956516 | 0.55304488 | 0.58023268 | 0.86175324 |
| CLCN3 | 62.9467913 | 0.471828957 | 0.25831713 | 1.82654926 | 0.06776757 | 0.40302824 |
| HPF1 | 14.4096004 | 0.44709955 | 0.41743589 | 1.0710616 | 0.28414173 | NA |
| MFAP3L | 10.2495953 | -0.169348695 | 0.5045763 | -0.3356255 | 0.73715326 | NA |
| GALNT7 | 18.8664743 | 0.484360869 | 0.49205339 | 0.98436649 | 0.32493535 | NA |
| HMGB2 | 9.79204624 | -1.150743302 | 0.71858167 | -1.6014092 | 0.10928631 | NA |
| FBXO8 | 15.2396193 | -0.246963431 | 0.38909958 | -0.6347049 | 0.52562087 | NA |
| CEP44 | 26.0817227 | 0.443207296 | 0.28995988 | 1.52851247 | 0.12638535 | NA |
| HPGD | 180.334912 | 1.34860903 | 0.93591025 | 1.44095979 | 0.14959604 | 0.54834287 |
| WDR17 | 16.3565253 | 1.013671353 | 0.53788901 | 1.88453629 | 0.05949247 | NA |
| SPCS3 | 64.77761 | -0.017161161 | 0.34879987 | -0.0492006 | 0.96075944 | 0.98879479 |
| AGA | 4.17500343 | 0.044772169 | 0.66599839 | 0.06722564 | 0.94640207 | NA |
| AC078881.1 | 25.1910835 | -0.029997638 | 0.4383865 | -0.0684274 | 0.94544543 | NA |
| AC098864.1 | 21.9056728 | -1.945692382 | 0.47388285 | -4.105851 | 4.03E-05 | NA |
| TENM3 | 328.112325 | -1.069043938 | 0.28717937 | -3.7225653 | 0.00019721 | 0.01701871 |
| DCTD | 35.4977232 | 0.059424204 | 0.27231262 | 0.21822053 | 0.82725729 | 0.94918121 |
| WWC2 | 170.909009 | 0.750322281 | 0.23976861 | 3.12935992 | 0.00175188 | 0.06518907 |
| CDKN2AIP | 8.35152592 | -0.021921317 | 0.5401007 | -0.0405875 | 0.96762478 | NA |
| ING2 | 6.26885483 | -1.370506359 | 0.69472051 | -1.9727449 | 0.04852463 | NA |
| RWDD4 | 11.326448 | -0.512663773 | 0.46247996 | -1.1085103 | 0.26764151 | NA |
| TRAPPC11 | 26.9385475 | 0.018438831 | 0.34655653 | 0.05320584 | 0.9575679 | NA |
| STOX2 | 40.4419288 | 0.040555023 | 0.42872494 | 0.0945945 | 0.92463692 | 0.97775374 |
| IRF2 | 27.8634228 | -0.063093562 | 0.40218953 | -0.1568752 | 0.87534321 | NA |
| CASP3 | 11.1567917 | 0.444025817 | 0.48789272 | 0.91008903 | 0.36277556 | NA |
| PRIMPOL | 18.6873039 | 0.291706651 | 0.38081737 | 0.76600142 | 0.44367546 | NA |
| ACSL1 | 36.2467349 | 0.427387401 | 0.37803146 | 1.13056041 | 0.25824016 | 0.65588339 |
| SLC25A4 | 36.1613192 | -0.192959436 | 0.53758816 | -0.3589354 | 0.71964339 | 0.91472065 |
| CFAP97 | 42.5674389 | -0.967005836 | 0.31659109 | -3.0544316 | 0.00225487 | 0.07750653 |
| SNX25 | 291.735552 | 0.838218554 | 0.42956575 | 1.95131609 | 0.05101946 | 0.35507065 |
| LRP2BP | 24.7977734 | 0.626248939 | 0.54913996 | 1.14041771 | 0.25411232 | NA |
| AC112722.1 | 47.3581187 | 0.596784146 | 0.46766608 | 1.27609029 | 0.2019236 | 0.60937659 |
| ANKRD37 | 11.8507086 | 0.59853607 | 0.9726291 | 0.61537956 | 0.53830409 | NA |
| UFSP2 | 19.328326 | -0.150458633 | 0.36540534 | -0.4117582 | 0.68051669 | NA |
| C4orf47 | 8.35525548 | 1.071360669 | 0.78857972 | 1.35859526 | 0.17427488 | NA |
| SORBS2 | 26.0093951 | 1.481883061 | 0.56477828 | 2.62383152 | 0.00869468 | NA |
| TLR3 | 12.5803142 | 0.409711876 | 0.77580816 | 0.52810978 | 0.59742315 | NA |
| CYP4V2 | 10.4565076 | -0.168577193 | 0.4901332 | -0.3439416 | 0.73089021 | NA |
| FAT1 | 312.718706 | 0.281801117 | 0.35323598 | 0.79777015 | 0.42500389 | 0.77723512 |
| LINC01060 | 8.80890755 | 0.10923192 | 1.03442209 | 0.10559705 | 0.91590207 | NA |
| FRG1-DT | 10.876123 | 0.431744579 | 0.75639133 | 0.57079525 | 0.56813844 | NA |
| FRG1 | 15.6454422 | -0.300277493 | 0.42600426 | -0.7048697 | 0.48089133 | NA |
| CCDC127 | 9.6079715 | -0.489760517 | 0.49646497 | -0.9864956 | 0.32388996 | NA |
| SDHA | 30.1320817 | -0.348994083 | 0.29396519 | -1.1871953 | 0.2351506 | 0.64178976 |
| PDCD6 | 15.1743429 | -0.659634209 | 0.44312406 | -1.4885994 | 0.13659289 | NA |
| EXOC3 | 20.199845 | -0.224621118 | 0.365988 | -0.613739 | 0.53938779 | NA |
| SLC9A3 | 8.82855581 | 1.232580973 | 0.5866473 | 2.10105964 | 0.03563573 | NA |
| BRD9 | 12.560167 | -0.909101268 | 0.46059987 | -1.9737332 | 0.04841209 | NA |
| AC116351.1 | 9.3210871 | 0.777249515 | 0.55776847 | 1.39349848 | 0.1634691 | NA |
| SLC12A7 | 23.0577699 | -0.155148353 | 0.41423509 | -0.3745418 | 0.70800127 | NA |
| CLPTM1L | 24.3217395 | 0.027476281 | 0.35936965 | 0.07645688 | 0.93905562 | NA |
| LPCAT1 | 18.4294149 | 0.007223463 | 0.54159022 | 0.01333751 | 0.98935853 | NA |
| MRPL36 | 13.5274873 | -1.624895636 | 0.60483084 | -2.6865291 | 0.00721987 | NA |
| NDUFS6 | 36.3435499 | 0.092505476 | 0.50319876 | 0.18383487 | 0.854143 | 0.95569766 |
| ICE1 | 39.7177887 | -0.266019857 | 0.28587952 | -0.9305313 | 0.35209604 | 0.7234557 |
| MED10 | 19.7452782 | -0.257680044 | 0.36686223 | -0.7023891 | 0.48243651 | NA |
| NSUN2 | 18.0860234 | 0.226795787 | 0.38686886 | 0.58623429 | 0.55771808 | NA |
| SRD5A1 | 5.3085567 | -0.310520652 | 0.63452394 | -0.4893758 | 0.62457568 | NA |
| TENT4A | 27.3352821 | -0.020222362 | 0.35314552 | -0.0572635 | 0.95433527 | NA |

| | | | | | | |
|---|---|---|---|---|---|---|
| MTRR | 13.2202383 | 0.449409334 | 0.55531224 | 0.80929124 | 0.41834764 | NA |
| MIR4458HG | 5.49385944 | -1.020670293 | 0.74940152 | -1.3619806 | 0.17320402 | NA |
| AC021088.1 | 76.5651506 | -0.23816612 | 0.76546989 | -0.3111371 | 0.75569636 | 0.92313949 |
| SEMA5A | 1100.23068 | 0.302984694 | 0.24494887 | 1.23693036 | 0.21611294 | 0.62432626 |
| AC091906.1 | 9.340184 | -0.331968051 | 0.70903979 | -0.4681938 | 0.63964599 | NA |
| SNHG18 | 9.47965764 | -0.246013957 | 0.52893208 | -0.4651145 | 0.6418495 | NA |
| FAM173B | 8.57498569 | -0.123286908 | 0.5295727 | -0.2328045 | 0.81591322 | NA |
| CCT5 | 29.7445213 | -0.448167177 | 0.4280113 | -1.0470919 | 0.29505718 | 0.68103704 |
| CMBL | 8.66616465 | -1.315712263 | 0.93130319 | -1.4127647 | 0.15772492 | NA |
| 6-Mar | 125.255332 | 0.158709285 | 0.23759175 | 0.66799157 | 0.50413897 | 0.82600448 |
| ANKRD33B | 148.140922 | -1.107261186 | 0.48743681 | -2.2715994 | 0.02311072 | 0.24858211 |
| DAP | 26.0592561 | -0.399392354 | 0.36767923 | -1.0862521 | 0.27736745 | NA |
| DNAH5 | 9.03451041 | -1.364594365 | 0.94580723 | -1.4427828 | 0.14908168 | NA |
| TRIO | 129.461008 | -0.357773306 | 0.29583805 | -1.2093553 | 0.22652639 | 0.63327644 |
| OTULIN | 17.7830716 | 0.01664265 | 0.35636099 | 0.04670166 | 0.96275101 | NA |
| ANKH | 126.276945 | -1.613510296 | 0.60177997 | -2.6812296 | 0.00733522 | 0.13572774 |
| LINC02149 | 83.7449693 | 1.310879765 | 0.74504783 | 1.75945719 | 0.07849989 | 0.42784309 |
| FBXL7 | 333.381195 | 0.436457386 | 0.33175112 | 1.31561693 | 0.18830265 | 0.59077076 |
| ZNF622 | 6.41402183 | -0.609539823 | 0.57235473 | -1.0649686 | 0.28689014 | NA |
| RETREG1 | 412.859695 | -0.029268833 | 0.29897575 | -0.097897 | 0.92201407 | 0.97775374 |
| MYO10 | 30.3385586 | -0.825993872 | 0.59476235 | -1.3887797 | 0.16489976 | 0.56632803 |
| LINC02241 | 12.5033451 | 1.038267629 | 0.47365497 | 2.19203363 | 0.02837708 | NA |
| CDH12 | 22.7161386 | -0.774666899 | 0.63964465 | -1.2110895 | 0.22586112 | NA |
| C5orf17 | 17.2090698 | 1.5419813 | 0.93659203 | 1.64637457 | 0.09968666 | NA |
| CDH10 | 20.2176033 | 2.681265146 | 0.95684691 | 2.80218821 | 0.00507573 | NA |
| CDH6 | 29.8654129 | 0.236835829 | 0.79714464 | 0.29710521 | 0.76638618 | 0.92431233 |
| DROSHA | 32.4362473 | 0.496598461 | 0.26544053 | 1.87084643 | 0.06136637 | 0.38229128 |
| C5orf22 | 11.3709198 | -0.039581685 | 0.49457997 | -0.0800309 | 0.93621267 | NA |
| PDZD2 | 56.8715457 | 0.104107368 | 0.52783466 | 0.19723481 | 0.84364379 | 0.95501513 |
| GOLPH3 | 81.6191346 | 0.093252364 | 0.25211189 | 0.36988483 | 0.7114683 | 0.91233495 |
| MTMR12 | 38.0071663 | -0.541362096 | 0.2955065 | -1.8319803 | 0.06695435 | 0.40232126 |
| ZFR | 105.138353 | 0.152547988 | 0.17552213 | 0.86910971 | 0.38478713 | 0.75104967 |
| SUB1 | 86.2805281 | -0.331490624 | 0.40137924 | -0.8258788 | 0.40887282 | 0.76772515 |
| NPR3 | 184.416851 | 0.086191364 | 0.45835218 | 0.18804615 | 0.85084047 | 0.95532127 |
| TARS | 24.9636887 | -0.397884566 | 0.41991343 | -0.9475395 | 0.34336394 | NA |
| C1QTNF3-AN | 32.5476212 | 0.519507671 | 0.34604004 | 1.50129354 | 0.13327966 | 0.52613957 |
| RAI14 | 227.951516 | 0.402947281 | 0.33098254 | 1.21742761 | 0.22344156 | 0.62979776 |
| RAD1 | 10.8353072 | -0.251963041 | 0.54020633 | -0.46642 | 0.64091489 | NA |
| BRIX1 | 9.44173183 | -1.233750017 | 0.58613332 | -2.1048966 | 0.03530031 | NA |
| DNAJC21 | 23.4631364 | -0.468218442 | 0.37011013 | -1.2650787 | 0.20584312 | NA |
| SPEF2 | 24.9106005 | 0.326852243 | 0.49647814 | 0.65834166 | 0.51031861 | NA |
| LMBRD2 | 22.2868814 | 0.532387608 | 0.37497228 | 1.41980525 | 0.15566439 | NA |
| SKP2 | 4.17983132 | -0.291560442 | 0.64254277 | -0.4537604 | 0.65000131 | NA |
| NADK2 | 18.8456966 | 0.343482698 | 0.37837849 | 0.90777544 | 0.36399688 | NA |
| RANBP3L | 12.6589547 | 2.072244943 | 0.97330588 | 2.12907884 | 0.03324774 | NA |
| NIPBL | 182.399036 | 0.251401171 | 0.19000297 | 1.32314336 | 0.18578771 | 0.58779637 |
| CPLANE1 | 86.3852641 | 0.60975884 | 0.28135296 | 2.16723804 | 0.03021671 | 0.27996878 |
| NUP155 | 13.1562472 | 0.029889226 | 0.45498264 | 0.06569311 | 0.94762216 | NA |
| WDR70 | 83.7075395 | 0.21946984 | 0.20114589 | 1.09109781 | 0.27522985 | 0.66766283 |
| EGFLAM | 7.79660161 | -3.273899937 | 0.96399811 | -3.3961684 | 0.00068336 | NA |
| LIFR | 60.1058882 | -0.350480784 | 0.33667299 | -1.0410125 | 0.29786975 | 0.68103704 |
| OSMR-AS1 | 58.6734617 | -2.739289192 | 0.56245662 | -4.8702231 | 1.11E-06 | 0.00028258 |
| OSMR | 82.430421 | -2.002940437 | 0.61355979 | -3.2644584 | 0.00109674 | 0.05186671 |
| RICTOR | 84.6579599 | 0.563562712 | 0.24385289 | 2.31107657 | 0.02082863 | 0.23864663 |
| C9 | 4.18985536 | 0.045628801 | 0.90996162 | 0.05014365 | 0.96000791 | NA |
| DAB2 | 22.0276898 | -1.716786259 | 0.68076671 | -2.5218423 | 0.0116742 | NA |
| TTC33 | 36.3255553 | 0.151649337 | 0.36017935 | 0.4210384 | 0.67372704 | 0.90403541 |
| PTGER4 | 139.08083 | 0.195264966 | 0.39938802 | 0.48891042 | 0.62490512 | 0.87931199 |
| PRKAA1 | 38.3316431 | 0.617993574 | 0.36273274 | 1.703716 | 0.08843416 | 0.45403664 |
| RPL37 | 40.3160364 | -0.530375319 | 0.3324219 | -1.5954885 | 0.11060304 | 0.49679506 |
| CARD6 | 5.14678065 | -0.101878824 | 0.72003831 | -0.1414908 | 0.8874822 | NA |
| OXCT1 | 18.7360617 | -0.303615696 | 0.46554244 | -0.6521762 | 0.51428751 | NA |
| C5orf51 | 8.58011478 | -0.046541345 | 0.45547981 | -0.1021809 | 0.91861307 | NA |
| FBXO4 | 4.96336756 | -1.209660613 | 0.61867538 | -1.9552428 | 0.05055441 | NA |
| GHR | 182.362022 | 0.378808159 | 0.47560738 | 0.79647242 | 0.4257575 | 0.77730788 |
| SELENOP | 88.8246449 | 0.150321945 | 0.40341321 | 0.37262525 | 0.70942738 | 0.91144677 |
| ZNF131 | 48.7498788 | -0.134903094 | 0.28789541 | -0.4685837 | 0.63936724 | 0.8874995 |
| HMGCS1 | 23.6576772 | -0.740255323 | 0.4505541 | -1.6429888 | 0.10038526 | NA |
| CCL28 | 8.88243375 | -1.004869695 | 0.71961324 | -1.3964025 | 0.16259334 | NA |
| TMEM267 | 25.9902478 | 0.004080783 | 0.39165246 | 0.0104194 | 0.99168667 | NA |
| PAIP1 | 17.6484666 | -0.316524175 | 0.47402535 | -0.6677368 | 0.50430161 | NA |
| NNT-AS1 | 19.8108401 | -0.244647909 | 0.38384773 | -0.6373567 | 0.52389255 | NA |
| NNT | 37.9629564 | 0.190093643 | 0.30327521 | 0.62680245 | 0.53078874 | 0.83515914 |
| MRPS30-DT | 13.8650306 | 0.041997298 | 0.57600377 | 0.0729115 | 0.94187654 | NA |
| MRPS30 | 10.4946963 | -0.758885261 | 0.54405235 | -1.3948754 | 0.16305341 | NA |
| PARP8 | 59.1705512 | 0.311264387 | 0.4187143 | 0.74338132 | 0.45725085 | 0.79426529 |
| ITGA1 | 173.680551 | 0.451359847 | 0.3216639 | 1.40320331 | 0.16055622 | 0.56283048 |
| ITGA2 | 24.814167 | -0.247098831 | 0.66457825 | -0.371813 | 0.71003209 | NA |
| MOCS2 | 18.8528543 | -0.391899035 | 0.4607657 | -0.8505386 | 0.39502568 | NA |
| NDUFS4 | 47.6239721 | 0.399912456 | 0.34285567 | 1.16641633 | 0.24344616 | 0.64918976 |
| ARL15 | 157.463783 | 0.572806294 | 0.36907264 | 1.55201505 | 0.12065862 | 0.51191566 |
| SNX18 | 7.58910254 | 0.170721005 | 0.59120185 | 0.2887694 | 0.77275785 | NA |
| GPX8 | 14.1602555 | -0.89233425 | 0.72823598 | -1.2253367 | 0.2204484 | NA |
| DHX29 | 30.2415603 | -0.256395885 | 0.33344522 | -0.7689296 | 0.44193513 | 0.78620488 |
| MTREX | 49.2926826 | 0.285517958 | 0.29506681 | 0.96763834 | 0.33322502 | 0.7076234 |
| PLPP1 | 125.004542 | -0.282671994 | 0.26527027 | -1.0656 | 0.28660451 | 0.67428533 |
| SLC38A9 | 22.18069 | 0.294132617 | 0.3365188 | 0.87404511 | 0.38209369 | NA |
| IL6ST | 92.7888587 | -0.387397121 | 0.28740151 | -1.34793 | 0.17768089 | 0.58063214 |
| AC008914.1 | 3.96148005 | -2.010143962 | 1.2669388 | -1.5866149 | 0.11259989 | NA |
| MAP3K1 | 27.9045525 | 0.969859693 | 0.55194404 | 1.75717033 | 0.07888878 | NA |
| MIER3 | 9.61997072 | -0.160722972 | 0.51923193 | -0.3095398 | 0.75691091 | NA |
| GPBP1 | 88.3431839 | 0.71436518 | 0.31480284 | 2.26924633 | 0.02325335 | 0.2486058 |
| PDE4D | 517.403656 | 0.606027153 | 0.35071835 | 1.72795963 | 0.08399546 | 0.44304225 |
| ELOVL7 | 5.53497647 | 0.042059247 | 0.76055552 | 0.05530069 | 0.95589891 | NA |
| ERCC8 | 18.7284842 | 0.032540939 | 0.35001751 | 0.09296946 | 0.92592782 | NA |
| NDUFAF2 | 32.3820857 | -0.681906057 | 0.40176186 | -1.6972892 | 0.089642 | 0.4567688 |
| SMIM15 | 14.3390206 | -0.491083481 | 0.43337244 | -1.1331673 | 0.257144 | NA |
| ZSWIM6 | 662.589592 | 0.177656582 | 0.32723518 | 0.54290185 | 0.58719739 | 0.86417728 |
| KIF2A | 37.7692147 | 0.262919268 | 0.39974263 | 0.65772136 | 0.51071719 | 0.82958308 |
| DIMT1 | 6.69127139 | -0.44400025 | 0.55154242 | -0.8050156 | 0.42081068 | NA |
| IPO11 | 16.0153725 | 0.055307725 | 0.34470201 | 0.16045083 | 0.87252595 | NA |
| RNF180 | 43.3310492 | 0.376575598 | 0.40437508 | 0.93125322 | 0.35172259 | 0.72305466 |
| SREK1IP1 | 26.0179415 | 0.361607003 | 0.29745227 | 1.21568076 | 0.22410656 | NA |
| CWC27 | 36.3731509 | 0.380995783 | 0.27884573 | 1.36633179 | 0.17183485 | 0.57423036 |
| PPWD1 | 23.602001 | 0.572863935 | 0.39390556 | 1.45431798 | 0.14585817 | NA |
| TRIM23 | 20.0643731 | -0.292990088 | 0.39749147 | -0.7370978 | 0.46106288 | NA |
| TRAPPC13 | 25.7866304 | 0.150144964 | 0.40738039 | 0.36856208 | 0.71245416 | NA |
| SGTB | 7.91114868 | -0.126790391 | 0.58634521 | -0.2162385 | 0.82880187 | NA |
| NLN | 29.5711825 | -0.296421989 | 0.32224948 | -0.9198525 | 0.35764985 | 0.72903084 |
| ERBIN | 240.7125 | 0.626103599 | 0.21676087 | 2.88845312 | 0.00387142 | 0.09814043 |
| SREK1 | 63.7667754 | 0.350287665 | 0.26441319 | 1.32477379 | 0.18524619 | 0.587049 |
| MAST4 | 212.804959 | 0.378798201 | 0.26816952 | 1.41253263 | 0.15779319 | 0.56042835 |
| PIK3R1 | 43.6639355 | -0.224222432 | 0.34531254 | -0.6493319 | 0.5161239 | 0.83104823 |

| | | | | | | |
|---|---|---|---|---|---|---|
| SLC30A5 | 25.4087348 | -0.128620818 | 0.31928403 | -0.4028414 | 0.68706492 | NA |
| MRPS36 | 20.1801702 | -0.742977985 | 0.39846516 | -1.8645996 | 0.06223754 | NA |
| CDK7 | 13.8351144 | -0.39669324 | 0.48215415 | -0.8227519 | 0.4106491 | NA |
| CCDC125 | 15.5597968 | 0.28780575 | 0.50066929 | 0.57484203 | 0.56539814 | NA |
| AK6 | 17.5894523 | -0.544999871 | 0.37647837 | -1.447626 | 0.14772168 | NA |
| RAD17 | 21.9575456 | -0.015182203 | 0.36871417 | -0.0411761 | 0.96715553 | NA |
| MARVELD2 | 6.71245679 | 0.755311129 | 0.65897412 | 1.1461924 | 0.25171555 | NA |
| OCLN | 10.5289308 | 0.405971769 | 0.58321759 | 0.69608972 | 0.48637264 | NA |
| GTF2H2C | 10.8974666 | -0.165716692 | 0.46453051 | -0.3567402 | 0.72128633 | NA |
| NAIP | 38.5170957 | 0.190763144 | 0.3632657 | 0.52513393 | 0.59949009 | 0.86882737 |
| GTF2H2 | 12.492543 | 0.058022872 | 0.48032468 | 0.12079927 | 0.90385003 | NA |
| BDP1 | 65.3206482 | -0.129364998 | 0.24224932 | -0.5340159 | 0.5933305 | 0.8654248 |
| MCCC2 | 20.7619429 | -0.015490624 | 0.3809407 | -0.0406641 | 0.96756365 | NA |
| MAP1B | 169.958689 | -0.644848763 | 0.4312157 | -1.4954204 | 0.13480475 | 0.52779404 |
| MRPS27 | 26.659814 | -0.091533583 | 0.41569823 | -0.2201924 | 0.82572133 | NA |
| PTCD2 | 5.63483904 | -0.63844161 | 0.64008874 | -0.9974267 | 0.31855743 | NA |
| TNPO1 | 124.342284 | 0.420370554 | 0.17223237 | 2.4407174 | 0.01465812 | 0.19896332 |
| FCHO2 | 42.6868243 | 0.217593192 | 0.24853172 | 0.87551479 | 0.38129387 | 0.74899182 |
| FOXD1 | 15.2228363 | -0.312783647 | 0.5969962 | -0.523929 | 0.60032789 | NA |
| BTF3 | 53.0802845 | -0.676318207 | 0.42065935 | -1.6077574 | 0.10788831 | 0.49167977 |
| ANKRA2 | 10.4890127 | 0.307281779 | 0.45554763 | 0.67453271 | 0.49997269 | NA |
| ARHGEF28 | 83.9912541 | 0.357892885 | 0.38591133 | 0.92739667 | 0.35372062 | 0.72532398 |
| ENC1 | 31.8964236 | 1.544611353 | 1.00697869 | 1.5339067 | 0.12505261 | 0.52128816 |
| HEXB | 24.3450179 | -0.090810775 | 0.48537494 | -0.1870941 | 0.85158687 | NA |
| GFM2 | 10.6168493 | -0.210818687 | 0.4620443 | -0.4562738 | 0.64819313 | NA |
| NSA2 | 12.3965379 | -0.836965715 | 0.45946129 | -1.821624 | 0.06851206 | NA |
| FAM169A | 18.2173729 | 0.767040977 | 0.46412638 | 1.65265542 | 0.09840101 | NA |
| HMGCR | 16.0201742 | 0.629622173 | 0.40477198 | 1.55549841 | 0.11982741 | NA |
| COL4A3BP | 51.3866278 | 0.308586185 | 0.25103694 | 1.2292461 | 0.21897955 | 0.62462904 |
| POLK | 75.3946776 | 0.359089249 | 0.26912445 | 1.33428696 | 0.18210983 | 0.58390314 |
| POC5 | 9.26949164 | -0.206245098 | 0.51848226 | -0.3977862 | 0.69078778 | NA |
| IQGAP2 | 803.14777 | 0.925973574 | 0.41086049 | 2.25374208 | 0.0242124 | 0.25196701 |
| AC025188.1 | 7.04632108 | -0.490454015 | 0.66432299 | -0.7382764 | 0.46034648 | NA |
| F2R | 212.849261 | -0.811915839 | 0.49641871 | -1.6355464 | 0.10193455 | 0.48317363 |
| AC008581.2 | 6.83117986 | -1.443599458 | 0.80275802 | -1.7982996 | 0.07212954 | NA |
| ZBED3 | 18.1816367 | 0.178613294 | 0.42278733 | 0.42246605 | 0.67268488 | NA |
| WDR41 | 58.9006402 | 0.070809328 | 0.29406514 | 0.24079471 | 0.80971423 | 0.94346479 |
| TBCA | 93.7783599 | -0.158430118 | 0.38793303 | -0.4083955 | 0.68298331 | 0.90499319 |
| AP3B1 | 97.6724807 | 0.2725687 | 0.23306483 | 1.16949734 | 0.24220331 | 0.64918976 |
| SCAMP1-AS1 | 9.01488246 | -0.490336642 | 0.82717337 | -0.5927858 | 0.5533245 | NA |
| SCAMP1 | 46.2208226 | 0.406446576 | 0.30049607 | 1.35258535 | 0.17618814 | 0.57988078 |
| LHFPL2 | 5.42257773 | -1.031171388 | 0.6453728 | -1.5977918 | 0.11008931 | NA |
| ARSB | 17.7730438 | 0.704180622 | 0.46966485 | 1.49932579 | 0.13378913 | NA |
| DMGDH | 20.1567759 | 0.688846568 | 0.69734729 | 0.98780992 | 0.32324575 | NA |
| BHMT2 | 15.1299442 | 0.943665242 | 0.76940071 | 1.22649386 | 0.22001288 | NA |
| BHMT | 13.4727384 | 3.509710199 | 1.0439446 | 3.36196978 | 0.00077389 | NA |
| JMY | 39.9167366 | 0.190536071 | 0.29750783 | 0.64044052 | 0.52188625 | 0.83108387 |
| HOMER1 | 18.8966241 | 0.408644638 | 0.3751141 | 1.08938758 | 0.275983 | NA |
| TENT2 | 126.127868 | 0.4171207 | 0.20391204 | 2.04559135 | 0.04079659 | 0.31695313 |
| CMYA5 | 133.615562 | 0.462168876 | 0.4128959 | 1.1193351 | 0.26299721 | 0.6581333 |
| MTX3 | 13.4160487 | -0.346923573 | 0.40382474 | -0.8590944 | 0.39028844 | NA |
| THBS4 | 4.88433496 | -0.899525636 | 0.9211315 | -0.9765442 | 0.32879485 | NA |
| AC012636.1 | 8.23782135 | 1.042826918 | 0.76964582 | 1.35494392 | 0.17543543 | NA |
| SERINC5 | 340.863713 | 0.521555631 | 0.34901088 | 1.49438214 | 0.13507577 | 0.52779404 |
| AC010260.1 | 37.3119515 | -0.525078062 | 0.73850565 | -0.7110007 | 0.47708378 | 0.80863008 |
| ZFYVE16 | 50.7793396 | 0.134742739 | 0.33481196 | 0.40244303 | 0.68735801 | 0.90499319 |
| AC008771.1 | 5.25412797 | 0.144612578 | 0.60914705 | 0.23740175 | 0.81234513 | NA |
| FAM151B | 9.73056444 | 0.288795203 | 0.48797111 | 0.59182849 | 0.55396545 | NA |
| DHFR | 85.7485353 | -0.304347361 | 0.33360209 | -0.9123065 | 0.36160739 | 0.73240894 |
| MSH3 | 42.512104 | 0.282189364 | 0.27276489 | 1.03455165 | 0.30087834 | 0.68214788 |
| RASGRF2 | 7.96455265 | -0.61569845 | 0.72319273 | -0.8513615 | 0.39456857 | NA |
| CKMT2-AS1 | 23.4844058 | -0.267572809 | 0.40489424 | -0.6608462 | 0.50871097 | NA |
| ZCCHC9 | 13.4579079 | -0.350318894 | 0.52720401 | -0.6644845 | 0.50638026 | NA |
| SSBP2 | 218.673811 | 0.005022237 | 0.25401024 | 0.01977179 | 0.98422542 | 0.99577746 |
| ATG10 | 57.7731512 | 0.722731357 | 0.31111667 | 2.32302359 | 0.02017789 | 0.23722175 |
| RPS23 | 92.3649733 | -0.977202683 | 0.4811717 | -2.0308815 | 0.04226702 | 0.3239276 |
| ATP6AP1L | 23.5652171 | 0.36803503 | 0.45644125 | 0.80631414 | 0.42006174 | NA |
| TMEM167A | 24.7191196 | -0.530816527 | 0.38521202 | -1.3779854 | 0.16820781 | NA |
| XRCC4 | 20.7193356 | 0.009955596 | 0.57354713 | 0.01735794 | 0.98615106 | NA |
| EDIL3 | 97.7129127 | -1.396435001 | 0.50257878 | -2.7785395 | 0.00546039 | 0.11595461 |
| AC113383.1 | 11.9802523 | -2.004046673 | 0.56721629 | -3.5331261 | 0.00041068 | NA |
| COX7C | 60.1675651 | -0.533355614 | 0.44508639 | -1.1983193 | 0.23079274 | 0.63682501 |
| RASA1 | 81.445815 | 0.18331679 | 0.25405558 | 0.72156174 | 0.47056397 | 0.80448595 |
| CCNH | 97.3406558 | 0.180365576 | 0.37356641 | 0.48282065 | 0.62922309 | 0.88166074 |
| TMEM161B | 21.3187424 | -0.092664451 | 0.41395662 | -0.2238506 | 0.82287353 | NA |
| TMEM161B-. | 57.7385199 | 0.628474394 | 0.41844817 | 1.50191693 | 0.13311856 | 0.52613957 |
| CETN3 | 11.2543055 | 0.00837048 | 0.57295546 | 0.0146093 | 0.98834388 | NA |
| MBLAC2 | 6.19497868 | 0.046063909 | 0.57233415 | 0.08048429 | 0.93585209 | NA |
| LYSMD3 | 16.8989522 | -0.223127822 | 0.45783012 | -0.4873594 | 0.62600365 | NA |
| ADGRV1 | 22.0621499 | 0.513985017 | 0.54947903 | 0.93540424 | 0.34958001 | NA |
| ARRDC3 | 29.6891797 | 0.023006115 | 0.53978386 | 0.04262098 | 0.96600367 | 0.98892249 |
| ARRDC3-AS1 | 10.9099903 | -0.012779496 | 0.48303234 | -0.0264568 | 0.97889298 | NA |
| NR2F1-AS1 | 26.6240506 | 0.454024895 | 0.38530518 | 1.17835141 | 0.23865654 | NA |
| NR2F1 | 21.542368 | -0.453782343 | 0.54462117 | -0.8332073 | 0.40472782 | NA |
| FAM172A | 164.585195 | 0.106704272 | 0.24231848 | 0.44034723 | 0.65968564 | 0.89709157 |
| KIAA0825 | 70.79189 | 0.266656835 | 0.22360852 | 1.19251645 | 0.2330588 | 0.63935677 |
| SLF1 | 11.8223366 | 0.630065724 | 0.4965226 | 1.26895678 | 0.20445648 | NA |
| MCTP1 | 5.56960283 | -1.293322234 | 1.01359783 | -1.2759718 | 0.20196549 | NA |
| TTC37 | 67.4951627 | 0.092219975 | 0.2225625 | 0.41435541 | 0.67861384 | 0.90499319 |
| ARSK | 8.56929534 | 0.036925634 | 0.58920269 | 0.06267051 | 0.95002888 | NA |
| RHOBTB3 | 62.9787269 | -0.708703177 | 0.46575349 | -1.5216272 | 0.12810252 | 0.52341115 |
| GLRX | 18.298139 | -1.250095043 | 0.94308557 | -1.3255372 | 0.18499302 | NA |
| ELL2 | 39.957766 | 0.235031903 | 0.29218852 | 0.80438446 | 0.42117497 | 0.77473274 |
| AC104123.1 | 35.6940283 | -0.269102339 | 0.45541098 | -0.5909 | 0.55458744 | 0.84979473 |
| CAST | 232.980203 | -0.177440168 | 0.20711052 | -0.8567414 | 0.3915878 | 0.75653902 |
| ERAP1 | 46.0071554 | -0.312439479 | 0.28781893 | -1.0855418 | 0.27768176 | 0.66958859 |
| AC009126.1 | 8.28495266 | -0.135576911 | 0.5387338 | -0.2516584 | 0.80130508 | NA |
| ERAP2 | 20.8148663 | 0.003755834 | 0.72340701 | 0.00519187 | 0.99585751 | NA |
| LNPEP | 87.5946371 | 0.107507251 | 0.2607917 | 0.41223417 | 0.68016779 | 0.90499319 |
| LIX1-AS1 | 17.1988673 | -0.251849749 | 0.58423149 | -0.4310787 | 0.66641115 | NA |
| RIOK2 | 6.99705916 | -0.537428445 | 0.55443876 | -0.9693198 | 0.33238567 | NA |
| RGMB | 4.14096014 | -0.821802459 | 0.71367388 | -1.1515098 | 0.24952257 | NA |
| CHD1 | 55.295417 | -0.023492068 | 0.26262317 | -0.0894516 | 0.928723 | 0.97777732 |
| LINC02062 | 19.1041864 | -0.016314125 | 0.3763507 | -0.0433482 | 0.96542397 | NA |
| FAM174A | 20.644419 | -0.349874369 | 0.42061209 | -0.831822 | 0.40550945 | NA |
| SLCO4C1 | 7.26108555 | 2.468772598 | 0.80628318 | 3.06191751 | 0.00219924 | NA |
| PAM | 269.380858 | 0.294557985 | 0.20566947 | 1.4321911 | 0.15208914 | 0.55086922 |
| GIN1 | 6.41720718 | 0.349364033 | 0.53400961 | 0.65422798 | 0.51296494 | NA |
| PPIP5K2 | 44.041249 | 0.69476986 | 0.31599744 | 2.19865658 | 0.02790235 | 0.27202278 |
| C5orf30 | 10.3142997 | 0.372710366 | 0.5903414 | 0.63134715 | 0.52781356 | NA |
| NUDT12 | 9.26246141 | 0.457743999 | 0.43679069 | 1.04797105 | 0.29465194 | NA |
| AC099520.1 | 20.3247921 | -2.901706697 | 1.10700703 | -2.621218 | 0.00876162 | NA |
| EFNA5 | 62.9610409 | 0.033629132 | 0.55096869 | 0.06103638 | 0.95133024 | 0.98520702 |

| | | | | | | |
|---|---|---|---|---|---|---|
| FBXL17 | 253.518329 | 0.260524371 | 0.21125681 | 1.23321172 | 0.21749679 | 0.62454341 |
| FER | 164.888319 | 0.262570802 | 0.17850679 | 1.47092892 | 0.14131034 | 0.53597868 |
| AC008467.1 | 8.17254222 | 0.036671839 | 0.72920564 | 0.05029012 | 0.9598912 | NA |
| PJA2 | 98.3838049 | -0.304181231 | 0.2039303 | -1.4915941 | 0.1358056 | 0.52829049 |
| MAN2A1 | 68.3050744 | -1.276521421 | 0.44585753 | -2.8630702 | 0.00419558 | 0.10376374 |
| TMEM232 | 44.924213 | 0.404755527 | 0.4842473 | 0.83584467 | 0.40324229 | 0.76594067 |
| SLC25A46 | 39.618378 | 0.330683021 | 0.3121088 | 1.05951201 | 0.28936666 | 0.67724823 |
| WDR36 | 15.2167909 | -0.675343775 | 0.5003637 | -1.3497058 | 0.17711038 | NA |
| STARD4 | 14.4902417 | 0.616637119 | 0.57717453 | 1.06837203 | 0.28535273 | NA |
| NREP | 12.2384832 | 0.279944755 | 0.47594121 | 0.58819187 | 0.55640351 | NA |
| EPB41L4A | 113.897483 | -1.103354785 | 0.40133464 | -2.7492139 | 0.00597384 | 0.12229157 |
| EPB41L4A-AS | 4.97384454 | -0.630469461 | 0.71901659 | -0.8768497 | 0.38056829 | NA |
| APC | 43.7331227 | 0.269448268 | 0.34111335 | 0.78990831 | 0.42958132 | 0.77854416 |
| SRP19 | 21.2020666 | -0.055474671 | 0.34869715 | -0.1590913 | 0.87359697 | NA |
| REEP5 | 85.5199447 | -0.517136398 | 0.32403052 | -1.5959497 | 0.11050003 | 0.49679506 |
| DCP2 | 35.8531859 | -0.044028012 | 0.27930518 | -0.1576341 | 0.87474516 | 0.96229085 |
| MCC | 174.275278 | -0.224440445 | 0.31196285 | -0.7194461 | 0.47186612 | 0.80516996 |
| YTHDC2 | 60.7180336 | 0.487044816 | 0.22892975 | 2.12748591 | 0.03337973 | 0.29047359 |
| PGGT1B | 25.0636644 | 0.41111594 | 0.30808516 | 1.33442305 | 0.18206525 | NA |
| CCDC112 | 9.06216184 | -0.094500367 | 0.48400452 | -0.1952469 | 0.84519969 | NA |
| FEM1C | 9.14185428 | -0.191453862 | 0.54091774 | -0.3539427 | 0.72338186 | NA |
| TMED7-TICA | 16.5896992 | 0.224169702 | 0.3942309 | 0.5686254 | 0.56961038 | NA |
| ATG12 | 24.5521394 | 0.221830548 | 0.30196411 | 0.73462554 | 0.46256758 | NA |
| AP3S1 | 52.2089596 | -0.0699474 | 0.24511053 | -0.2853708 | 0.77536004 | 0.9285091 |
| COMMD10 | 79.3234534 | -0.272190392 | 0.27219838 | -0.9999706 | 0.31732472 | 0.69424851 |
| SEMA6A | 9.96551437 | -0.660093666 | 0.72734457 | -0.9075391 | 0.36412176 | NA |
| SEMA6A-AS1 | 8.83045217 | -0.191334645 | 0.48470686 | -0.394743 | 0.69303256 | NA |
| DTWD2 | 23.9387182 | 0.813086865 | 0.36126816 | 2.25064637 | 0.02440794 | NA |
| AC008629.1 | 11.5426357 | 0.3528945 | 0.60303588 | 0.58519652 | 0.55841559 | NA |
| DMXL1 | 95.2821315 | 0.20446373 | 0.24007127 | 0.85167928 | 0.39439212 | 0.75777094 |
| TNFAIP8 | 16.5439862 | -1.190031701 | 0.66903516 | -1.7787282 | 0.07528433 | NA |
| HSD17B4 | 44.8713198 | 0.281299553 | 0.39847591 | 0.70593867 | 0.48022628 | 0.81093801 |
| PRR16 | 10.6938617 | 2.184408721 | 0.72548894 | 3.01094696 | 0.00260434 | NA |
| SRFBP1 | 45.9186349 | 0.612640214 | 0.32678564 | 1.87474643 | 0.06082762 | 0.38132428 |
| LOX | 83.87025 | 0.12988215 | 0.39674276 | 0.32737119 | 0.74338715 | 0.92122771 |
| SNX2 | 26.1154486 | 0.136483058 | 0.40243366 | 0.33914424 | 0.73450107 | NA |
| SNX24 | 57.637923 | 0.37485143 | 0.31319259 | 1.19687198 | 0.23135646 | 0.63691078 |
| PPIC | 12.7256044 | -0.03806244 | 0.560061 | -0.0679612 | 0.94581649 | NA |
| CEP120 | 18.6725126 | 0.557400249 | 0.36715957 | 1.51814168 | 0.12897869 | NA |
| CSNK1G3 | 41.7437888 | 0.13350734 | 0.3026099 | 0.4411863 | 0.65907813 | 0.89709157 |
| ZNF608 | 118.400261 | 0.176917136 | 0.2512717 | 0.70408701 | 0.48137859 | 0.81183849 |
| AC112196.1 | 3.89930628 | -1.339465731 | 0.93458757 | -1.4332159 | 0.15179616 | NA |
| GRAMD2B | 56.7899785 | 0.584724373 | 0.35653873 | 1.6400024 | 0.10100467 | 0.48247622 |
| ALDH7A1 | 55.5638109 | 0.19111651 | 0.38816413 | 0.49236005 | 0.62246483 | 0.8780438 |
| PHAX | 14.8328534 | -0.041695035 | 0.40813323 | -0.1021604 | 0.9186294 | NA |
| C5orf63 | 7.59135238 | -0.896904185 | 0.55008809 | -1.6304737 | 0.10300141 | NA |
| PRRC1 | 23.4161606 | -0.041049444 | 0.49464891 | -0.082987 | 0.93386185 | NA |
| LINC01184 | 35.3046628 | -0.225458848 | 0.44467719 | -0.5070169 | 0.61214297 | 0.87391153 |
| SLC12A2 | 9.56257822 | -0.390104378 | 0.51168098 | -0.7623977 | 0.44582271 | NA |
| SLC27A6 | 9.59750257 | 3.080830011 | 1.36153654 | 2.2627597 | 0.02365051 | NA |
| ISOC1 | 8.26229999 | 0.208126931 | 0.59624347 | 0.34906367 | 0.72704151 | NA |
| ADAMTS19 | 462.107097 | 0.126482686 | 0.48071285 | 0.26311484 | 0.79246206 | 0.93928291 |
| AC008591.1 | 47.4313923 | -0.379007665 | 0.742007 | -0.5107872 | 0.60950008 | 0.87261996 |
| CHSY3 | 13.4792519 | -1.066692024 | 0.7027412 | -1.5179017 | 0.12903919 | NA |
| HINT1 | 141.004287 | -0.701593652 | 0.47647333 | -1.472472 | 0.14089345 | 0.53597868 |
| LYRM7 | 10.0426397 | -0.320394471 | 0.42083282 | -0.7613343 | 0.4464574 | NA |
| CDC42SE2 | 100.793226 | 0.112447053 | 0.20018708 | 0.56170984 | 0.57431373 | 0.85861279 |
| RAPGEF6 | 4.21225876 | 0.331871189 | 0.96579601 | 0.34362452 | 0.73112868 | NA |
| P4HA2 | 18.4203279 | 0.01876911 | 0.54930467 | 0.03416885 | 0.9727425 | NA |
| SLC22A5 | 16.3146235 | 0.430543726 | 0.64439205 | 0.66813941 | 0.50404461 | NA |
| AC116366.3 | 88.4059081 | -0.142081255 | 0.47663757 | -0.2980908 | 0.7656339 | 0.9240942 |
| IRF1 | 23.3096857 | -0.753556121 | 0.62425198 | -1.2071345 | 0.22738034 | NA |
| TH2LCRR | 4.32278797 | -2.691460631 | 1.00695153 | -2.67288 | 0.00752031 | NA |
| KIF3A | 25.3813923 | 0.314041134 | 0.34462046 | 0.91126665 | 0.36215489 | NA |
| 8-Sep | 11.6452069 | -0.008250888 | 0.56064741 | -0.0147167 | 0.98825818 | NA |
| UQCRQ | 31.6609559 | -0.385789395 | 0.4130608 | -0.9339773 | 0.35031562 | 0.72238707 |
| AFF4 | 114.88528 | -0.197120285 | 0.17724274 | -1.1121487 | 0.26607422 | 0.66013596 |
| ZCCHC10 | 12.3345908 | -0.490090102 | 0.63086524 | -0.7768539 | 0.43724499 | NA |
| HSPA4 | 31.0449147 | -0.216876228 | 0.3394002 | -0.6389985 | 0.52282388 | 0.83127153 |
| C5orf15 | 27.5647363 | -0.166350172 | 0.38597848 | -0.430983 | 0.66648072 | NA |
| VDAC1 | 40.2855341 | -0.381832327 | 0.31735401 | -1.2031748 | 0.2289087 | 0.63549192 |
| SKP1 | 134.483616 | -0.467829877 | 0.44140204 | -1.0598725 | 0.28920261 | 0.67724823 |
| CDKL3 | 23.0881215 | 0.305493276 | 0.36892995 | 0.82805226 | 0.40764091 | NA |
| UBE2B | 65.8137835 | -0.171546404 | 0.22814124 | -0.7519307 | 0.45209273 | 0.7927748 |
| CDKN2AIPNL | 4.90616737 | -0.476410468 | 0.63504461 | -0.7502 | 0.45313426 | NA |
| JADE2 | 20.8418577 | -0.298453702 | 0.49191806 | -0.6067143 | 0.54404056 | NA |
| SAR1B | 29.8433394 | -0.426317307 | 0.35975235 | -1.1850299 | 0.23600561 | 0.64267939 |
| SEC24A | 27.8961754 | -0.028305436 | 0.34374383 | -0.0823446 | 0.93437272 | NA |
| CAMLG | 15.946406 | -0.176458258 | 0.43444328 | -0.406171 | 0.68461697 | NA |
| DDX46 | 46.9378287 | -0.165078714 | 0.28451497 | -0.580211 | 0.56177234 | 0.85386823 |
| C5orf24 | 25.7612891 | -0.056784632 | 0.31783147 | -0.1786627 | 0.85820255 | NA |
| TXNDC15 | 36.6774438 | 0.427415245 | 0.3622457 | 1.17990427 | 0.23803829 | 0.64408493 |
| PCBD2 | 19.5407172 | -0.311313373 | 0.40080143 | -0.7767272 | 0.43731973 | NA |
| H2AFY | 30.6750613 | -0.213841565 | 0.28142651 | -0.7598487 | 0.44734503 | 0.78947076 |
| CXCL14 | 90.6430651 | -2.939986858 | 1.36935262 | -2.1469903 | 0.03179405 | 0.28784971 |
| SMAD5 | 61.574808 | -0.160505308 | 0.33286272 | -0.4821967 | 0.62966622 | 0.88166074 |
| SPOCK1 | 878.241187 | -0.486203905 | 0.37206954 | -1.3067555 | 0.19129578 | 0.59410082 |
| KLHL3 | 11.6068996 | 0.537292281 | 0.61945277 | 0.86736601 | 0.3857415 | NA |
| HNRNPA0 | 32.5525726 | -0.595725893 | 0.4298372 | -1.3859338 | 0.16576713 | 0.56743028 |
| AC106791.1 | 9.79036146 | -0.110894144 | 0.51213995 | -0.2165309 | 0.82857391 | NA |
| PKD2L2 | 6.03424943 | -0.303657438 | 0.69250819 | -0.4384893 | 0.66103162 | NA |
| FAM13B | 126.652667 | -0.147905406 | 0.21315829 | -0.6938759 | 0.48776002 | 0.81648973 |
| NME5 | 15.8338443 | 0.004282515 | 0.53712988 | 0.00797296 | 0.99363857 | NA |
| BRD8 | 15.6747193 | -0.14246471 | 0.36867299 | -0.3864257 | 0.69918143 | NA |
| CDC23 | 11.9167998 | 0.024618463 | 0.39500032 | 0.06232517 | 0.95030388 | NA |
| FAM53C | 9.30512128 | 0.063070655 | 0.49276868 | 0.12799242 | 0.89815497 | NA |
| KDM3B | 45.6042467 | 0.191659109 | 0.29053674 | 0.65967253 | 0.509464 | 0.82854289 |
| EGR1 | 17.8308304 | 2.347305284 | 1.38648275 | 1.69299277 | 0.09045684 | NA |
| ETF1 | 27.9134257 | -0.20889037 | 0.3131251 | -0.6671147 | 0.50469885 | NA |
| HSPA9 | 43.3529077 | -0.411276052 | 0.37860987 | -1.0862793 | 0.27735546 | 0.66958859 |
| CTNNA1 | 270.393878 | -0.046694105 | 0.27994807 | -0.1667956 | 0.86753088 | 0.96089268 |
| SIL1 | 117.160359 | 0.227120584 | 0.23983263 | 0.94699617 | 0.34364073 | 0.71842682 |
| PAIP2 | 46.3044218 | 0.035025962 | 0.31205132 | 0.11224424 | 0.91062975 | 0.97557165 |
| DNAJC18 | 8.92725243 | 0.496228895 | 0.57972804 | 0.85596842 | 0.39201525 | NA |
| UBE2D2 | 89.8936437 | 0.094683709 | 0.30687867 | 0.30853793 | 0.75767305 | 0.92313949 |
| CXXC5 | 68.5514752 | -0.427122946 | 0.54531915 | -0.7832532 | 0.43347847 | 0.78003046 |
| PURA | 58.6194856 | -0.242457702 | 0.29750374 | -0.8149736 | 0.41508741 | 0.77017133 |
| IGIP | 9.63262359 | 0.550837667 | 0.60843593 | 0.90533389 | 0.36528853 | NA |
| CYSTM1 | 93.1889286 | -0.244487137 | 0.28217193 | -0.8664474 | 0.38624485 | 0.75125852 |
| PFDN1 | 26.2471211 | 0.112306051 | 0.32037093 | 0.35055007 | 0.72592592 | NA |
| HBEGF | 10.5341415 | 1.224471279 | 0.62038006 | 1.97374377 | 0.04841089 | NA |
| AC008438.1 | 4.38074865 | -0.047273152 | 0.86917882 | -0.0543883 | 0.95662581 | NA |
| SRA1 | 13.8400758 | -0.515757145 | 0.46587821 | -1.1070643 | 0.26826612 | NA |

| | | | | | | |
|---|---|---|---|---|---|---|
| SLC35A4 | 18.3819346 | 0.577559744 | 0.43108155 | 1.33979232 | 0.18031287 | NA |
| NDUFA2 | 26.3762387 | -0.122385479 | 0.41000683 | -0.2984962 | 0.76532448 | NA |
| IK | 10.3978038 | -0.453986625 | 0.55139013 | -0.8233492 | 0.41030944 | NA |
| WDR55 | 5.68645783 | -0.00862236 | 0.66955003 | -0.0128778 | 0.98972525 | NA |
| HARS | 16.5395133 | -0.510908739 | 0.3976231 | -1.2849071 | 0.19882475 | NA |
| ZMAT2 | 13.2402896 | 0.579858651 | 0.41499541 | 1.39726521 | 0.16233383 | NA |
| AC244517.11 | 5.76190757 | -1.504976547 | 1.00081689 | -1.5037481 | 0.13264623 | NA |
| TAF7 | 45.3782962 | -0.513379444 | 0.36590299 | -1.403048 | 0.16060254 | 0.56283048 |
| DIAPH1 | 30.2365884 | -0.435584781 | 0.36791833 | -1.183917 | 0.23644591 | 0.64320899 |
| HDAC3 | 11.9664856 | 0.656899892 | 0.45407967 | 1.4466622 | 0.14799156 | NA |
| DELE1 | 18.1401466 | 0.788392049 | 0.36533974 | 2.1579696 | 0.0309302 | NA |
| RNF14 | 23.3030942 | 0.284531499 | 0.32828967 | 0.86670866 | 0.38610166 | NA |
| GNPDA1 | 13.3478154 | -0.312112341 | 0.57748493 | -0.5404684 | 0.58887407 | NA |
| NDFIP1 | 90.479648 | 0.14429646 | 0.23286337 | 0.61966147 | 0.53548069 | 0.83677705 |
| SPRY4 | 7.19455403 | -1.400176808 | 0.82619586 | -1.6947275 | 0.09012714 | NA |
| SPRY4-AS1 | 67.3978595 | -0.820589542 | 0.53711671 | -1.5277677 | 0.12657023 | 0.52276701 |
| FGF1 | 1026.94526 | -0.494957635 | 0.38620189 | -1.2816033 | 0.19998183 | 0.60620493 |
| ARHGAP26 | 51.4311887 | -1.452072301 | 0.58280233 | -2.4915348 | 0.01271925 | 0.18904687 |
| NR3C1 | 80.2298828 | 0.211365374 | 0.26136645 | 0.80869359 | 0.41869142 | 0.77290571 |
| YIPF5 | 18.4546438 | 0.02328886 | 0.41801808 | 0.05571257 | 0.95557079 | NA |
| KCTD16 | 15.847057 | 1.565766791 | 0.68270141 | 2.29348697 | 0.02181998 | NA |
| PRELID2 | 35.1917255 | 0.407910958 | 0.55234261 | 0.73851076 | 0.46020413 | 0.7966658 |
| SH3RF2 | 5.79569956 | 1.836194558 | 0.81666488 | 2.24840642 | 0.02455029 | NA |
| LARS | 37.4628863 | -0.099763714 | 0.29891189 | -0.3337563 | 0.73856349 | 0.91993596 |
| TCERG1 | 36.8809556 | 0.117962157 | 0.30973956 | 0.38084304 | 0.70331972 | 0.90762481 |
| PPP2R2B | 112.258015 | -0.052916552 | 0.4420299 | -0.1197126 | 0.90471081 | 0.97257012 |
| STK32A | 6.23252016 | 1.371047945 | 1.10121133 | 1.24503618 | 0.21311845 | NA |
| DPYSL3 | 29.0196547 | 0.287667621 | 0.52444371 | 0.54851953 | 0.58333522 | 0.86175324 |
| JAKMIP2 | 43.1920296 | -0.367148571 | 0.59115278 | -0.6210722 | 0.53455211 | 0.83677705 |
| AC091948.1 | 10.3348361 | 0.071060845 | 0.54936794 | 0.12935018 | 0.89708056 | NA |
| FBXO38 | 25.5526716 | 0.524622416 | 0.34087715 | 1.53903662 | 0.12379536 | NA |
| GRPEL2 | 5.12736986 | 0.097468681 | 0.66481315 | 0.14661064 | 0.88343935 | NA |
| CSNK1A1 | 210.29686 | -0.309652612 | 0.19630885 | -1.5773747 | 0.11470934 | 0.50513044 |
| ARHGEF37 | 62.1290605 | -0.495450657 | 0.49131049 | -1.0084268 | 0.31324962 | 0.69213367 |
| PPARGC1B | 13.0248096 | 0.484535768 | 0.80644181 | 0.60083165 | 0.54795212 | NA |
| SLC26A2 | 27.6227052 | -0.329841413 | 0.48410867 | -0.6813376 | 0.49565793 | NA |
| HMGXB3 | 22.2726063 | -0.161315078 | 0.30911042 | -0.5218688 | 0.6017617 | NA |
| PDGFRB | 140.432046 | -0.27199981 | 0.47730788 | -0.5698624 | 0.56877103 | 0.85727808 |
| CAMK2A | 12.1641558 | 1.26916694 | 0.95622685 | 1.32726554 | 0.18442086 | NA |
| TCOF1 | 9.54902772 | -0.491207456 | 0.57828375 | -0.8494229 | 0.39564602 | NA |
| CD74 | 34.1995109 | -1.334245076 | 0.77011269 | -1.7325323 | 0.08317884 | 0.44101093 |
| RPS14 | 109.220928 | -0.207463553 | 0.49871087 | -0.4159997 | 0.67741025 | 0.90499319 |
| NDST1 | 10.4080567 | 0.324474089 | 0.42704236 | 0.7598171 | 0.44736392 | NA |
| SYNPO | 173.332176 | -0.067658449 | 0.60769705 | -0.1113358 | 0.91135005 | 0.97608551 |
| RBM22 | 11.4757664 | -0.437872543 | 0.46530422 | -0.9410457 | 0.34668143 | NA |
| DCTN4 | 36.4361912 | -0.226885294 | 0.30607361 | -0.7412769 | 0.45852557 | 0.79511744 |
| GPX3 | 272.930273 | 0.003368574 | 0.63531535 | 0.00530221 | 0.99576947 | 0.99921845 |
| TNIP1 | 20.349157 | 0.064959747 | 0.47741034 | 0.1360669 | 0.89176839 | NA |
| ANXA6 | 4.98331466 | 1.133929849 | 0.70141843 | 1.61662398 | 0.10595947 | NA |
| GM2A | 4.91162124 | 0.08279724 | 0.61961731 | 0.13362642 | 0.89369799 | NA |
| SLC36A1 | 11.8814441 | 0.471527972 | 0.56969926 | 0.82767876 | 0.40785245 | NA |
| SPARC | 532.069853 | 0.571735679 | 0.60279353 | 0.9484768 | 0.34288678 | 0.71725052 |
| ATOX1 | 57.7815623 | -0.10236254 | 0.37138306 | -0.2756252 | 0.78283593 | 0.93415197 |
| G3BP1 | 58.2797001 | -0.169635475 | 0.26267828 | -0.6457918 | 0.51841422 | 0.83104823 |
| FAM114A2 | 22.0470813 | 0.260585206 | 0.38629246 | 0.6745801 | 0.49994258 | NA |
| MFAP3 | 24.8184063 | -0.113073825 | 0.34791058 | -0.3250083 | 0.74517479 | NA |
| GALNT10 | 34.3559956 | -1.004550452 | 0.55368966 | -1.8142843 | 0.06963397 | 0.40663921 |
| SAP30L-AS1 | 11.675473 | 0.19013565 | 0.42889113 | 0.44331914 | 0.65753491 | NA |
| SAP30L | 10.6256334 | -0.516003131 | 0.44438224 | -1.1611696 | 0.24557295 | NA |
| LARP1 | 86.3769018 | -0.001734829 | 0.31445749 | -0.0055169 | 0.99559818 | 0.99921845 |
| FAXDC2 | 31.323179 | 0.726041508 | 0.58403271 | 1.24315214 | 0.21381178 | 0.62255988 |
| CNOT8 | 18.6046359 | 0.305898109 | 0.39822077 | 0.76816212 | 0.44239087 | NA |
| GEMIN5 | 8.55430135 | 0.157836293 | 0.46795938 | 0.33728631 | 0.73590108 | NA |
| MRPL22 | 13.1702226 | -0.542930951 | 0.44857807 | -1.2103377 | 0.22614934 | NA |
| HAVCR2 | 8.85432912 | -0.93722741 | 0.60092419 | -1.5596433 | 0.11884419 | NA |
| CYFIP2 | 31.7251545 | 0.986593954 | 0.55432613 | 1.77980776 | 0.07510743 | 0.42134955 |
| CLINT1 | 47.4849275 | 0.092223197 | 0.2549076 | 0.36179069 | 0.71750845 | 0.91472065 |
| EBF1 | 9.43243341 | 0.125423488 | 0.88195204 | 0.14221124 | 0.88691316 | NA |
| RNF145 | 116.562887 | 0.132005659 | 0.33958876 | 0.38872211 | 0.69748172 | 0.90729979 |
| UBLCP1 | 20.9665166 | 0.334528984 | 0.34022557 | 0.98325645 | 0.32548124 | NA |
| TTC1 | 32.8127483 | -0.247806292 | 0.34366761 | -0.7210639 | 0.47087023 | 0.80448595 |
| PWWP2A | 36.4735237 | 0.209343191 | 0.34766685 | 0.60213733 | 0.54708273 | 0.84499784 |
| ZBED8 | 6.14678761 | 0.600444551 | 0.63831374 | 0.94067308 | 0.34687242 | NA |
| SLU7 | 9.60852046 | 0.118652213 | 0.51350193 | 0.23106478 | 0.81726447 | NA |
| AC113414.1 | 12.6601534 | -0.651237077 | 0.64842092 | -1.0043431 | 0.31521327 | NA |
| CCNG1 | 19.0256393 | -0.714859026 | 0.409573 | -1.7453764 | 0.08091938 | NA |
| NUDCD2 | 16.3736341 | -1.181158832 | 0.46808243 | -2.5233992 | 0.01162264 | NA |
| MAT2B | 28.6756557 | -0.172084472 | 0.42638209 | -0.4035922 | 0.68651265 | 0.90499319 |
| AC109466.1 | 671.751295 | -0.062107246 | 0.68582947 | -0.0905579 | 0.92784392 | 0.97775374 |
| AC008415.1 | 235.893631 | 0.0019845 | 0.44187511 | 0.00449109 | 0.99641664 | 0.99937337 |
| AC114321.1 | 7.58191514 | 0.588261333 | 0.76895491 | 0.76501408 | 0.44426316 | NA |
| TENM2 | 283.447882 | 0.517925172 | 0.40003833 | 1.29468888 | 0.19542758 | 0.59987316 |
| AC011369.1 | 5.57461716 | 0.43714222 | 0.9202571 | 0.47502184 | 0.63477141 | NA |
| WWC1 | 40.1139661 | 0.013422693 | 0.57213111 | 0.02346087 | 0.98128265 | 0.99453905 |
| RARS | 23.2425643 | -0.104283789 | 0.31635679 | -0.3296398 | 0.74167214 | NA |
| PANK3 | 35.3824643 | 0.128788811 | 0.33015695 | 0.3900836 | 0.69647473 | 0.90718427 |
| SLIT3 | 46.4573325 | 0.06085129 | 0.59346357 | 0.10253585 | 0.91833136 | 0.97775374 |
| KCNIP1 | 15.555647 | 0.274640549 | 1.50701333 | 0.18224162 | 0.85539311 | NA |
| RANBP17 | 38.1701626 | 0.089378299 | 0.53200661 | 0.16800223 | 0.86658152 | 0.96089268 |
| NPM1 | 28.6702781 | -0.688930453 | 0.34612289 | -1.9904215 | 0.04654452 | 0.33771838 |
| FBXW11 | 78.3867157 | 0.118014562 | 0.22312437 | 0.5289183 | 0.59686213 | 0.86831879 |
| STK10 | 5.60426242 | -0.332252498 | 0.60233461 | -0.5516078 | 0.58121706 | NA |
| UBTD2 | 15.6359842 | 0.158261082 | 0.43841613 | 0.36098371 | 0.71811162 | NA |
| DUSP1 | 113.181218 | 0.533543901 | 0.44756446 | 1.19210515 | 0.23322002 | 0.63935677 |
| ERGIC1 | 77.6635826 | -0.25522152 | 0.31371951 | -0.8135341 | 0.41591191 | 0.7708692 |
| RPL26L1 | 6.60907524 | -0.252420249 | 0.64655403 | -0.3904086 | 0.69623443 | NA |
| ATP6V0E1 | 93.3494091 | -0.564509072 | 0.3000561 | -1.8813451 | 0.05992499 | 0.37905854 |
| CREBRF | 66.6784691 | -0.14535069 | 0.34457066 | -0.4218313 | 0.67314816 | 0.90403541 |
| BOD1 | 9.20858956 | -0.079677413 | 0.53932614 | -0.1477351 | 0.88255181 | NA |
| CPEB4 | 527.01446 | 0.531253452 | 0.23427864 | 2.26761366 | 0.02335276 | 0.2486058 |
| SFXN1 | 31.2843754 | 0.348455239 | 0.31759556 | 1.09716659 | 0.27256857 | 0.66531666 |
| THOC3 | 5.83992835 | -0.944251786 | 0.70689458 | -1.3357745 | 0.18162297 | NA |
| AC139491.7 | 29.1990438 | -0.731714467 | 0.74323911 | -0.984494 | 0.32487266 | 0.69623471 |
| SIMC1 | 32.5631119 | -0.078066944 | 0.31067365 | -0.2512828 | 0.80159547 | 0.94085969 |
| KIAA1191 | 26.0135527 | -0.519635508 | 0.41106034 | -1.2641344 | 0.20618179 | NA |
| ARL10 | 6.94028799 | -0.546247686 | 0.60126481 | -0.9084977 | 0.36361534 | NA |
| NOP16 | 3.08800215 | -1.122374527 | 1.00909096 | -1.112263 | 0.26602509 | NA |
| CLTB | 35.6786038 | -0.361658302 | 0.40199768 | -0.8996527 | 0.3683051 | 0.73626927 |
| FAF2 | 31.3901268 | 0.316516889 | 0.31586324 | 1.00206942 | 0.31631007 | 0.69424851 |
| RNF44 | 11.4769974 | 0.504209191 | 0.43089102 | 1.17015478 | 0.24193869 | NA |
| TSPAN17 | 8.06342258 | 0.277375531 | 0.58924618 | 0.47072945 | 0.63783395 | NA |
| UIMC1 | 42.7614226 | -0.142745122 | 0.32424995 | -0.4402318 | 0.65976927 | 0.89709157 |
| ZNF346 | 23.8543791 | 0.250754272 | 0.34216132 | 0.73285394 | 0.46364752 | NA |

| | | | | | | |
|---|---|---|---|---|---|---|
| FGFR4 | 34.7095111 | 1.049290304 | 0.69616573 | 1.50724212 | 0.13174861 | 0.52613957 |
| NSD1 | 119.55647 | 0.083964101 | 0.2782919 | 0.30171234 | 0.76287136 | 0.92336765 |
| PRELID1 | 24.5473065 | -0.198996973 | 0.44660879 | -0.4455733 | 0.65590547 | NA |
| LMAN2 | 43.5123193 | -0.161083898 | 0.33087029 | -0.4868491 | 0.62636528 | 0.88050706 |
| RGS14 | 4.41229044 | -1.459598642 | 0.94642643 | -1.5422209 | 0.12301992 | NA |
| DBN1 | 16.01374 | 0.344485126 | 0.44375054 | 0.77630357 | 0.43756976 | NA |
| PDLIM7 | 30.9573625 | 0.305021601 | 0.54264088 | 0.56210583 | 0.57404392 | 0.85861279 |
| DDX41 | 4.20408888 | 0.037091043 | 0.6702793 | 0.0553367 | 0.95587023 | NA |
| FAM193B | 35.4176951 | 0.303434439 | 0.30264052 | 1.00262332 | 0.31604264 | 0.69424851 |
| TMED9 | 81.1290293 | -0.112347113 | 0.4253288 | -0.2641418 | 0.79167066 | 0.93914731 |
| B4GALT7 | 11.2024408 | 0.841158067 | 0.48968719 | 1.71774571 | 0.08584301 | NA |
| FAM153A | 5.19484753 | -0.643108788 | 0.90091501 | -0.7138396 | 0.47532638 | NA |
| AC106795.3 | 4.15746337 | -0.42346685 | 0.75900361 | -0.5579247 | 0.57689582 | NA |
| RMND5B | 17.2667748 | -0.382677158 | 0.35379481 | -1.0816359 | 0.27941436 | NA |
| NHP2 | 19.1871174 | -0.522543036 | 0.42332231 | -1.2343858 | 0.21705919 | NA |
| HNRNPAB | 23.6843488 | -0.86244876 | 0.5070382 | -1.7009542 | 0.08895159 | NA |
| PHYKPL | 45.1819287 | -0.328731621 | 0.28119406 | -1.1690561 | 0.24238104 | 0.64918976 |
| CLK4 | 26.0336433 | 0.833880015 | 0.4815868 | 1.73152588 | 0.08335801 | NA |
| ZNF354A | 6.50092396 | -0.961444041 | 0.69324124 | -1.3868823 | 0.16547764 | NA |
| ZNF354B | 8.18976832 | -0.380847199 | 0.48610865 | -0.7834611 | 0.43335642 | NA |
| ZFP2 | 6.57243849 | 0.52870174 | 0.55991427 | 0.94425481 | 0.34503945 | NA |
| ZNF354C | 6.00149276 | -0.069621616 | 0.76038443 | -0.0915611 | 0.92704678 | NA |
| RUFY1 | 38.3684392 | 0.177703365 | 0.31127334 | 0.57089169 | 0.56807306 | 0.85654436 |
| HNRNPH1 | 100.399445 | 0.748416833 | 0.2556916 | 2.92702939 | 0.00342217 | 0.09584152 |
| CANX | 191.663234 | -0.156386397 | 0.35510243 | -0.440398 | 0.65964888 | 0.89709157 |
| MAML1 | 14.4214983 | -0.079091979 | 0.4841659 | -0.1633572 | 0.87023721 | NA |
| MGAT4B | 47.1811388 | -0.389501357 | 0.51505689 | -0.7562298 | 0.44951144 | 0.79098412 |
| SQSTM1 | 123.236809 | -0.965746014 | 0.35421505 | -2.7264398 | 0.00640216 | 0.12729006 |
| MRNIP | 15.6304277 | -0.286642638 | 0.42233469 | -0.6787097 | 0.49732183 | NA |
| TBC1D9B | 35.1180811 | -0.497116315 | 0.29965248 | -1.6589762 | 0.0971206 | 0.47119753 |
| RNF130 | 42.9410596 | -0.140013942 | 0.35627666 | -0.3929922 | 0.69432526 | 0.90557575 |
| MAPK9 | 20.273963 | 0.093288496 | 0.39814702 | 0.23430665 | 0.81474692 | NA |
| CNOT6 | 33.4915182 | -0.080287578 | 0.40539173 | -0.1980494 | 0.84300644 | 0.95469552 |
| FLT4 | 27.8403853 | -0.005284023 | 0.55014985 | -0.0096047 | 0.99233668 | NA |
| MGAT1 | 48.6933933 | 0.466491808 | 0.40951467 | 1.13913332 | 0.25464755 | 0.65520614 |
| ZFP62 | 5.46976285 | 0.02470325 | 0.60033251 | 0.04114928 | 0.96717689 | NA |
| TRIM41 | 7.77589654 | 0.090448776 | 0.55125216 | 0.16407877 | 0.86966914 | NA |
| RACK1 | 58.8957939 | -0.549896541 | 0.40670878 | -1.3520646 | 0.17635466 | 0.57988078 |
| TRIM52 | 11.3535552 | -0.385075056 | 0.45964421 | -0.8377677 | 0.4021612 | NA |
| DUSP22 | 15.46952 | -0.808705026 | 0.47817054 | -1.6912481 | 0.09078942 | NA |
| EXOC2 | 58.8441871 | -0.30417248 | 0.22676362 | -1.3413637 | 0.17980241 | 0.58390314 |
| FOXC1 | 305.169043 | -0.69721962 | 0.30140915 | -2.3131999 | 0.02071165 | 0.23840101 |
| GMDS | 442.652513 | 0.152482154 | 0.17552241 | 0.86873327 | 0.38499304 | 0.75104967 |
| GMDS-DT | 155.640984 | -0.335007646 | 0.30895322 | -1.0843313 | 0.27821794 | 0.66958859 |
| WRNIP1 | 19.2960375 | -0.166402362 | 0.38883324 | -0.427953 | 0.66868532 | NA |
| SERPINB1 | 83.189791 | -2.143189991 | 0.66716111 | -3.2124024 | 0.0013163 | 0.0545866 |
| SERPINB9 | 49.157796 | -0.533404489 | 0.37604973 | -1.4184414 | 0.15606195 | 0.55671527 |
| SERPINB6 | 53.4480486 | -0.511299021 | 0.24038541 | -2.1269969 | 0.03342034 | 0.29047359 |
| NQO2 | 16.8529137 | 0.04961642 | 0.38862235 | 0.12767258 | 0.89840809 | NA |
| RIPK1 | 40.8866507 | -0.483906536 | 0.26714932 | -1.811371 | 0.07008344 | 0.40663921 |
| BPHL | 9.33393111 | 0.388308232 | 0.49799966 | 0.77973595 | 0.43554632 | NA |
| TUBB2A | 31.9059681 | -0.303626553 | 0.79342578 | -0.382678 | 0.70195857 | 0.90757606 |
| PSMG4 | 17.6734449 | -0.286233561 | 0.45958537 | -0.6228082 | 0.53341054 | NA |
| SLC22A23 | 21.1361007 | 2.001880294 | 0.6041468 | 3.31356602 | 0.00092114 | NA |
| PXDC1 | 27.4105692 | -0.030400775 | 0.42130714 | -0.0721582 | 0.94247599 | NA |
| FAM50B | 4.99670274 | -1.047207989 | 0.78171895 | -1.3396221 | 0.18036823 | NA |
| PRPF4B | 55.223736 | -0.251597761 | 0.2348088 | -1.0715006 | 0.28394441 | 0.67249058 |
| ECI2 | 15.8960836 | -0.61612045 | 0.40633805 | -1.5162755 | 0.12944969 | NA |
| CDYL | 66.2093307 | -0.078035448 | 0.24665439 | -0.3163757 | 0.75171738 | 0.92274989 |
| LYRM4 | 20.0169972 | -0.361613322 | 0.45857266 | -0.7885628 | 0.4303676 | NA |
| FARS2 | 92.3540872 | 0.064365577 | 0.22054035 | 0.29185397 | 0.77039828 | 0.92621526 |
| RREB1 | 57.1686231 | 0.160770659 | 0.30636842 | 0.5247625 | 0.5997483 | 0.86882737 |
| SSR1 | 57.0654244 | -0.275887687 | 0.30255251 | -0.9118671 | 0.36183867 | 0.73240894 |
| AL139095.4 | 9.13050167 | -0.475711556 | 0.95842638 | -0.4963465 | 0.61964998 | NA |
| RIOK1 | 9.37864327 | -0.853115432 | 0.48144724 | -1.7719812 | 0.07639768 | NA |
| DSP | 9.42914477 | -1.233697388 | 0.71464677 | -1.7263037 | 0.08429279 | NA |
| SNRNP48 | 10.2853931 | 0.235802588 | 0.50328725 | 0.46852486 | 0.6394093 | NA |
| BLOC1S5-TXI | 5.24961957 | -0.427468681 | 0.64831335 | -0.6593551 | 0.5096678 | NA |
| SLC35B3 | 24.4263054 | 0.329939635 | 0.33056779 | 0.99809977 | 0.31823098 | NA |
| GCNT2 | 14.7856778 | 1.515816839 | 0.76979062 | 1.96912876 | 0.04893831 | NA |
| PAK1IP1 | 5.00514756 | -0.898980601 | 0.76283873 | -1.1784674 | 0.23861031 | NA |
| TMEM14C | 33.8014805 | -0.24366774 | 0.48971061 | -0.497575 | 0.61878365 | 0.87662819 |
| AL024498.2 | 8.39428152 | -0.072497053 | 0.48268579 | -0.1501951 | 0.88061067 | NA |
| MAK | 20.440596 | -0.451049003 | 0.61363121 | -0.735049 | 0.46230966 | NA |
| SYCP2L | 6.34243964 | 0.083946251 | 0.6578707 | 0.12760296 | 0.89846319 | NA |
| SMIM13 | 12.1641446 | 0.726806115 | 0.54245084 | 1.33985619 | 0.1802921 | NA |
| AL139807.1 | 38.770153 | -0.571086143 | 0.45901473 | -1.2441565 | 0.21344198 | 0.62255988 |
| NEDD9 | 183.523377 | -0.552501656 | 0.23698434 | -2.3313847 | 0.01973308 | 0.23471374 |
| TMEM170B | 28.2710298 | 0.896671408 | 0.40206873 | 2.23014458 | 0.02573785 | NA |
| HIVEP1 | 54.8041426 | 0.11246034 | 0.3030589 | 0.37108411 | 0.71057489 | 0.91222063 |
| PHACTR1 | 16.5484065 | -2.390382389 | 0.91580731 | -2.6101368 | 0.0090506 | NA |
| TBC1D7 | 6.40003051 | -0.406557914 | 0.62906889 | -0.6462852 | 0.51809467 | NA |
| GFOD1 | 22.3856613 | -0.120475873 | 0.46183698 | -0.2608623 | 0.79419867 | NA |
| SIRT5 | 16.2296226 | 0.454203762 | 0.41236268 | 1.10146669 | 0.27069359 | NA |
| NOL7 | 32.1978195 | -0.532487946 | 0.31776916 | -1.6757068 | 0.09379563 | 0.46635603 |
| RANBP9 | 47.8638166 | 0.08503515 | 0.29335565 | 0.2898705 | 0.77191531 | 0.92669573 |
| MCUR1 | 14.5095686 | 0.21559819 | 0.39492656 | 0.5459197 | 0.58512113 | NA |
| JARID2 | 106.151888 | 0.352539463 | 0.30665175 | 1.14964114 | 0.25029171 | 0.6518054 |
| DTNBP1 | 33.408309 | -0.52594094 | 0.37607286 | -1.3985081 | 0.16196054 | 0.56290656 |
| MYLIP | 75.4976267 | -0.047188449 | 0.47398938 | -0.0995559 | 0.92069689 | 0.97775374 |
| GMPR | 9.11018978 | -0.036418612 | 1.14956375 | -0.0316804 | 0.97472694 | NA |
| ATXN1 | 134.380031 | 0.401801836 | 0.37528261 | 1.07066469 | 0.28432022 | 0.67249058 |
| CAP2 | 37.9443025 | 0.852860624 | 0.43075375 | 1.9799262 | 0.04771182 | 0.34130362 |
| FAM8A1 | 13.108516 | -0.01809674 | 0.52084939 | -0.0347447 | 0.97228334 | NA |
| NUP153 | 40.4931818 | -0.061099611 | 0.27037989 | -0.2259769 | 0.82121939 | 0.94728252 |
| KIF13A | 158.046743 | 0.559708268 | 0.27786146 | 2.01434296 | 0.04397354 | 0.33029013 |
| TPMT | 8.19880808 | 0.04473725 | 0.50822355 | 0.08802672 | 0.92985544 | NA |
| KDM1B | 10.1476046 | -0.328632394 | 0.54071154 | -0.6077777 | 0.54333495 | NA |
| DEK | 42.8879161 | -0.225408222 | 0.25430253 | -0.8863782 | 0.37541376 | 0.74330391 |
| RNF144B | 17.1960207 | -0.222465456 | 0.48537276 | -0.4583394 | 0.64670863 | NA |
| AL022068.1 | 67.1326589 | 0.259578502 | 0.63152752 | 0.41103276 | 0.68104851 | 0.90499319 |
| ID4 | 20.5816974 | -3.374684923 | 0.99558409 | -3.3896533 | 0.00069981 | NA |
| MBOAT1 | 12.7365896 | 0.205237761 | 0.5381012 | 0.38141108 | 0.70289824 | NA |
| E2F3 | 14.9099151 | -0.180924109 | 0.62411592 | -0.2898886 | 0.77190144 | NA |
| CDKAL1 | 138.510461 | 0.359181975 | 0.2488045 | 1.44363133 | 0.14884271 | 0.54750323 |
| SOX4 | 10.4234212 | -0.741738938 | 0.6307446 | -1.1759735 | 0.23960546 | NA |
| CASC15 | 11.8493136 | 0.424782931 | 0.83906036 | 0.50626028 | 0.61267394 | NA |
| DCDC2 | 78.5697799 | 0.049412683 | 0.32260027 | 0.15317 | 0.87826421 | 0.96407027 |
| MRS2 | 26.2530677 | 0.543079928 | 0.31922017 | 1.7012707 | 0.08889217 | NA |
| ALDH5A1 | 5.13887186 | -0.001172474 | 0.63433425 | -0.0018484 | 0.99852523 | NA |
| TDP2 | 16.7202936 | -0.046473066 | 0.43320709 | -0.1072768 | 0.9145694 | NA |
| ACOT13 | 17.4671725 | -0.19222336 | 0.43785988 | -0.4390066 | 0.66065678 | NA |
| C6orf62 | 68.9566737 | 0.066861877 | 0.33258895 | 0.20103457 | 0.84067154 | 0.95393706 |

| | | | | | | |
|---|---|---|---|---|---|---|
| GMNN | 6.31859491 | -1.344219515 | 0.64888458 | -2.0715849 | 0.03830417 | NA |
| RIPOR2 | 47.3779464 | 1.16077157 | 0.39660873 | 2.92674233 | 0.00342533 | 0.09584152 |
| CARMIL1 | 43.8415296 | -0.255354828 | 0.37723327 | -0.6769149 | 0.49845994 | 0.82345406 |
| SLC17A1 | 22.3211088 | 1.667909704 | 1.46619166 | 1.13757959 | 0.25529608 | NA |
| SLC17A3 | 8.0178693 | 1.448791591 | 1.27877893 | 1.13294922 | 0.25723558 | NA |
| TRIM38 | 37.9831622 | -0.403806586 | 0.61954525 | -0.651779 | 0.51454375 | 0.8308079 |
| U91328.2 | 5.38334605 | -1.107248784 | 0.94585942 | -1.1706272 | 0.24174865 | NA |
| HIST1H1C | 6.41527813 | -0.532795575 | 0.62013268 | -0.8591638 | 0.39025013 | NA |
| HIST1H4C | 12.2988227 | -2.695862196 | 0.81590763 | -3.3041267 | 0.00095273 | NA |
| HIST1H2AC | 22.9101893 | 0.361635221 | 0.50441051 | 0.71694624 | 0.47340727 | NA |
| BTN3A2 | 8.95888522 | 0.709674091 | 0.94764182 | 0.74888431 | 0.45392694 | NA |
| BTN2A2 | 4.92843621 | 1.25652433 | 0.59347334 | 2.11723805 | 0.03423965 | NA |
| BTN3A1 | 7.71785932 | 0.524600663 | 1.16989051 | 0.4484186 | 0.65385112 | NA |
| BTN2A1 | 11.5966988 | -0.258266421 | 0.54660978 | -0.4724877 | 0.63657869 | NA |
| HCG11 | 4.86980371 | -0.508019298 | 0.87273617 | -0.5820995 | 0.56049966 | NA |
| HMGN4 | 4.33441743 | -1.293546513 | 1.00635431 | -1.2853788 | 0.19865994 | NA |
| ABT1 | 7.63917699 | -0.329063292 | 0.53230725 | -0.618183 | 0.53645472 | NA |
| ZNF322 | 10.0976791 | 0.109461548 | 0.6305298 | 0.1736025 | 0.86217787 | NA |
| LINC00240 | 11.3901895 | -0.69958391 | 0.63512958 | -1.1014822 | 0.27068686 | NA |
| ZSCAN16-AS | 17.4890089 | -0.47358649 | 0.51899843 | -0.9125008 | 0.36150514 | NA |
| ZKSCAN8 | 14.3098516 | 0.363051474 | 0.40225767 | 0.90253463 | 0.36677294 | NA |
| ZSCAN9 | 8.01433225 | -0.079793127 | 0.62991703 | -0.1266724 | 0.89919966 | NA |
| PGBD1 | 7.04328546 | 0.55626378 | 0.49950615 | 1.11362749 | 0.26543903 | NA |
| ZSCAN31 | 15.2303704 | 0.395794762 | 0.46441774 | 0.85223869 | 0.39408163 | NA |
| ZSCAN23 | 4.89673998 | -0.147949303 | 0.64349369 | -0.2299157 | 0.81815728 | NA |
| TRIM27 | 21.9425488 | -0.43046662 | 0.45907858 | -0.9376752 | 0.34841133 | NA |
| GABBR1 | 21.8654819 | 0.69127998 | 0.44562856 | 1.55124702 | 0.1208425 | NA |
| HLA-F | 20.8203517 | -0.305334661 | 1.06854928 | -0.2857469 | 0.77507197 | NA |
| HLA-A | 481.869791 | -0.312295157 | 0.69594696 | -0.4487341 | 0.65362346 | 0.8949739 |
| ZNRD1 | 7.29497391 | 0.588050854 | 0.59543731 | 0.9875949 | 0.32335109 | NA |
| PPP1R11 | 12.4865102 | -0.641132586 | 0.45519522 | -1.4084783 | 0.15898949 | NA |
| TRIM26 | 14.7461208 | 0.318594689 | 0.55919068 | 0.56974249 | 0.56885237 | NA |
| HCG17 | 33.344494 | -0.255956795 | 0.30997067 | -0.8257452 | 0.40894865 | 0.76772515 |
| HLA-E | 187.882807 | -0.742269285 | 0.68479654 | -1.0839267 | 0.27839728 | 0.66958859 |
| GNL1 | 21.4276825 | -0.174422291 | 0.37957843 | -0.4595158 | 0.6458638 | NA |
| PRR3 | 7.6267751 | -0.151365922 | 0.52819673 | -0.2865711 | 0.77444074 | NA |
| ABCF1 | 19.8177975 | -0.342311427 | 0.40343852 | -0.8484847 | 0.39616806 | NA |
| PPP1R10 | 17.9561701 | 0.377704271 | 0.38018611 | 0.99347205 | 0.32047996 | NA |
| MRPS18B | 19.8209654 | -0.745058737 | 0.37043598 | -2.0113023 | 0.04429354 | NA |
| ATAT1 | 5.00108063 | 0.390634173 | 0.58120308 | 0.67211305 | 0.50151173 | NA |
| C6orf136 | 7.20037249 | 0.252135994 | 0.54640099 | 0.46144864 | 0.64447676 | NA |
| DHX16 | 7.16073266 | -0.44732183 | 0.65704893 | -0.6808044 | 0.49599524 | NA |
| MDC1 | 8.88193844 | -0.58335029 | 0.48071405 | -1.2135079 | 0.2249357 | NA |
| TUBB | 71.9509856 | -0.716870574 | 0.59344701 | -1.2079774 | 0.22705596 | 0.63327644 |
| FLOT1 | 41.0171306 | -1.172135761 | 0.39386419 | -2.9759896 | 0.00292045 | 0.08973738 |
| IER3 | 131.611463 | -1.498940623 | 0.89244217 | -1.6795941 | 0.09303631 | 0.46399004 |
| DDR1 | 30.9264726 | -0.135159654 | 0.33576371 | -0.402544 | 0.68728374 | 0.90499319 |
| POU5F1 | 8.12471145 | -0.31809554 | 1.01113994 | -0.314591 | 0.7530722 | NA |
| HCG27 | 6.29352479 | -1.225219854 | 0.62805403 | -1.9508192 | 0.05107856 | NA |
| HLA-C | 265.830702 | -0.156967055 | 0.80784319 | -0.1943039 | 0.84593796 | 0.95532127 |
| HLA-B | 350.180677 | -0.998315817 | 0.85930041 | -1.1617774 | 0.24532588 | 0.64996527 |
| MICA | 27.9374966 | -0.11839931 | 0.36749826 | -0.3221765 | 0.74731897 | NA |
| NFKBIL1 | 8.76495851 | 0.217384679 | 0.51906015 | 0.4188044 | 0.67535909 | NA |
| AIF1 | 148.510235 | -0.452449383 | 0.65535445 | -0.6903888 | 0.48994971 | 0.81825647 |
| PRRC2A | 43.5494658 | 0.069297567 | 0.35362731 | 0.19596215 | 0.84463979 | 0.95509854 |
| BAG6 | 17.0962879 | -0.284547175 | 0.42439978 | -0.6704697 | 0.50255844 | NA |
| C6orf47 | 8.79296698 | 0.256840065 | 0.46308046 | 0.55463378 | 0.57914518 | NA |
| GPANK1 | 4.98737696 | -0.879866936 | 0.63510185 | -1.385395 | 0.16593172 | NA |
| DDAH2 | 21.2500014 | -0.237881312 | 0.42587233 | -0.5585742 | 0.57645232 | NA |
| CLIC1 | 109.066471 | -0.909496228 | 0.39138329 | -2.3237994 | 0.02013625 | 0.23722175 |
| VARS | 8.3493148 | -0.669602445 | 0.64964735 | -1.0307168 | 0.30267364 | NA |
| LSM2 | 15.4501256 | -0.888998352 | 0.52763142 | -1.6848852 | 0.09201073 | NA |
| HSPA1A | 69.7208083 | 0.511142733 | 0.50329953 | 1.01558356 | 0.30982774 | 0.69009407 |
| HSPA1B | 28.816143 | 0.938748005 | 0.67094016 | 1.39915311 | 0.16176708 | 0.56290656 |
| C6orf48 | 21.7664889 | -0.359513419 | 0.44042996 | -0.8162783 | 0.41434098 | NA |
| NEU1 | 14.3244721 | -0.480682895 | 0.45315752 | -1.0607413 | 0.28880749 | NA |
| EHMT2 | 10.049484 | 0.614931091 | 0.50106604 | 1.2272456 | 0.21973029 | NA |
| NELFE | 9.97793396 | -0.353039745 | 0.5395305 | -0.6543462 | 0.51288878 | NA |
| STK19 | 17.1770669 | -0.368776282 | 0.5041319 | -0.7315075 | 0.4644692 | NA |
| ATF6B | 18.1292467 | 0.059422208 | 0.37429088 | 0.15875943 | 0.87385841 | NA |
| AGPAT1 | 14.5749871 | 0.078246258 | 0.44627212 | 0.17533307 | 0.86081793 | NA |
| RNF5 | 13.9461172 | -0.375866357 | 0.50526278 | -0.7439027 | 0.45693533 | NA |
| PBX2 | 18.6377501 | -0.028496532 | 0.54990028 | -0.0518213 | 0.95867111 | NA |
| HLA-DRA | 13.0217039 | 0.35757139 | 0.66896715 | 0.53451263 | 0.59298692 | NA |
| HLA-DRB1 | 10.9417305 | 1.586904092 | 1.08444594 | 1.46333168 | 0.14337664 | NA |
| PSMB8 | 32.3756236 | -1.592369186 | 0.57971932 | -2.7467934 | 0.0060181 | 0.12229157 |
| PSMB9 | 20.8128689 | -0.489315456 | 0.95907778 | -0.5101937 | 0.60991575 | NA |
| TAP1 | 35.3997552 | -0.857615604 | 1.03170482 | -0.8312606 | 0.40582641 | 0.76772515 |
| BRD2 | 51.9145388 | -0.130919802 | 0.34344529 | -0.3811955 | 0.70305818 | 0.90757606 |
| HLA-DPA1 | 7.22312087 | -0.188546898 | 0.97562674 | -0.1932572 | 0.84675754 | NA |
| HLA-DPB1 | 6.33323482 | -0.532998741 | 0.99266613 | -0.5369366 | 0.59131143 | NA |
| RXRB | 8.44622411 | 0.043517096 | 0.46061758 | 0.09447554 | 0.92473141 | NA |
| SLC39A7 | 24.9912577 | 0.309000184 | 0.41628696 | 0.74227687 | 0.4579196 | NA |
| RING1 | 19.079539 | 0.209220605 | 0.3882851 | 0.53883244 | 0.59000248 | NA |
| VPS52 | 11.171486 | 0.29576533 | 0.39453617 | 0.74965327 | 0.45346356 | NA |
| RPS18 | 50.0862357 | -0.449196732 | 0.48903066 | -0.9185451 | 0.35833354 | 0.72914991 |
| B3GALT4 | 11.1585979 | -0.172299781 | 0.58719391 | -0.2934291 | 0.76919418 | NA |
| WDR46 | 7.79173548 | 0.340807282 | 0.50145254 | 0.67964016 | 0.49673234 | NA |
| PFDN6 | 11.5271116 | 0.353896586 | 0.55270395 | 0.64030045 | 0.52197729 | NA |
| RGL2 | 11.0667365 | -0.059614273 | 0.48133135 | -0.1238529 | 0.90143176 | NA |
| TAPBP | 128.932147 | -0.216681142 | 0.37391361 | -0.5794952 | 0.56225508 | 0.85386823 |
| ZBTB22 | 6.80746335 | 0.382921327 | 0.54668645 | 0.7004405 | 0.48365225 | NA |
| DAXX | 6.405123 | -0.060981034 | 0.61790913 | -0.0986893 | 0.92138495 | NA |
| PHF1 | 16.7744862 | -0.314923225 | 0.42458928 | -0.7417126 | 0.45826147 | NA |
| CUTA | 32.7640077 | -0.469486291 | 0.3754227 | -1.2505538 | 0.21109731 | 0.61954464 |
| SYNGAP1 | 6.77897674 | 0.420262807 | 0.59594966 | 0.7051985 | 0.48068671 | NA |
| ITPR3 | 8.25536212 | 0.183487012 | 0.92304205 | 0.19878511 | 0.84243085 | NA |
| UQCC2 | 15.4752873 | -0.877159101 | 0.42242992 | -2.0764606 | 0.03785137 | NA |
| IP6K3 | 8.98272383 | -2.024620845 | 0.89860913 | -2.2530606 | 0.02425533 | NA |
| LEMD2 | 19.4749123 | -0.378715447 | 0.40317927 | -0.9393227 | 0.34756507 | NA |
| SMIM29 | 11.7950533 | 0.348985926 | 0.6297248 | 0.55418799 | 0.57945019 | NA |
| NUDT3 | 18.138421 | -0.972575367 | 0.6693748 | -1.4529608 | 0.14623463 | NA |
| C6orf106 | 40.0622731 | 0.295913672 | 0.26651793 | 1.11029557 | 0.26687168 | 0.66013596 |
| SNRPC | 20.9535312 | -0.403215582 | 0.43924835 | -0.9179672 | 0.35863603 | NA |
| UHRF1BP1 | 30.1102723 | 0.380112501 | 0.33163187 | 1.14618808 | 0.25171734 | 0.65183754 |
| TAF11 | 15.7359139 | 0.052835406 | 0.36705688 | 0.14394338 | 0.88554518 | NA |
| ANKS1A | 68.0999981 | 0.587568594 | 0.26481884 | 2.21875679 | 0.02650328 | 0.26347374 |
| ZNF76 | 16.0219902 | -0.267320271 | 0.39158764 | -0.6826576 | 0.49482324 | NA |
| PPARD | 12.49691 | -0.380183839 | 0.39875304 | -0.9534318 | 0.34037133 | NA |
| RPL10A | 82.5102461 | -0.21044842 | 0.52028231 | -0.4044889 | 0.68585324 | 0.90499319 |
| TEAD3 | 36.2151987 | 0.350972876 | 0.33846568 | 1.03695263 | 0.29975793 | 0.68168772 |
| FKBP5 | 207.477875 | -0.354589619 | 0.53583006 | -0.6617576 | 0.50812657 | 0.82703105 |
| SRPK1 | 22.9912645 | -0.207953203 | 0.41519683 | -0.5008545 | 0.61647352 | NA |

| | | | | | | |
|---|---|---|---|---|---|---|
| MAPK14 | 35.6131291 | -0.210086089 | 0.30013225 | -0.6999784 | 0.4839408 | 0.81370025 |
| BRPF3 | 10.0003008 | 0.220385127 | 0.63701313 | 0.34596638 | 0.72936798 | NA |
| ETV7 | 4.193243 | -0.327420581 | 1.6728303 | -0.1957285 | 0.84482267 | NA |
| KCTD20 | 50.9967335 | 0.020124614 | 0.2966518 | 0.06783918 | 0.94591366 | 0.98249707 |
| STK38 | 63.9547706 | 0.268850698 | 0.26750965 | 1.00501308 | 0.31489055 | 0.6938809 |
| SRSF3 | 59.6787548 | -0.10194848 | 0.26371091 | -0.3865918 | 0.69905842 | 0.90729979 |
| CDKN1A | 18.4839528 | -1.300477405 | 0.64683417 | -2.0105267 | 0.04437547 | NA |
| PPIL1 | 5.68160945 | 0.117857036 | 0.72638475 | 0.16225153 | 0.87110778 | NA |
| C6orf89 | 35.1770798 | 0.077124279 | 0.25467699 | 0.30283176 | 0.76201808 | 0.92336765 |
| MTCH1 | 32.754109 | -0.341582171 | 0.40893807 | -0.8352907 | 0.40355405 | 0.76594067 |
| TBC1D22B | 15.5412651 | -0.023057427 | 0.38364333 | -0.0601012 | 0.95207503 | NA |
| RNF8 | 14.6431866 | 0.051989443 | 0.37794781 | 0.1375572 | 0.89059038 | NA |
| CMTR1 | 13.0697458 | -0.224548349 | 0.66143781 | -0.3394852 | 0.73424424 | NA |
| CCDC167 | 4.80261324 | -1.412538808 | 0.73402439 | -1.9243759 | 0.05430749 | NA |
| ZFAND3 | 363.796641 | 0.462015856 | 0.17403528 | 2.65472533 | 0.0079373 | 0.14193427 |
| BTBD9 | 180.117282 | -0.162836577 | 0.20873691 | -0.7801044 | 0.43532945 | 0.78167891 |
| GLO1 | 35.6235365 | -0.018909785 | 0.31533643 | -0.059967 | 0.9521819 | 0.98520702 |
| SAYSD1 | 6.5514645 | -0.360128033 | 0.65770294 | -0.5475542 | 0.58399802 | NA |
| DAAM2 | 24.2552077 | -0.59666303 | 0.47551846 | -1.2547631 | 0.20956476 | NA |
| MOCS1 | 8.3379307 | 0.531777683 | 0.61529457 | 0.8642652 | 0.38744222 | NA |
| UNC5CL | 7.14933157 | 0.54857231 | 0.71619545 | 0.76595335 | 0.44370406 | NA |
| OARD1 | 15.4603977 | -0.234382057 | 0.53251712 | -0.4401399 | 0.65983575 | NA |
| NFYA | 6.8299717 | -0.489448357 | 0.61315609 | -0.7982443 | 0.42472873 | NA |
| TFEB | 10.0193492 | -0.922699075 | 0.51093515 | -1.8059025 | 0.07093357 | NA |
| USP49 | 14.4890484 | 0.289376971 | 0.39822035 | 0.72667549 | 0.46742477 | NA |
| MED20 | 3.47029368 | 0.083471356 | 0.69489289 | 0.12012118 | 0.90438715 | NA |
| CCND3 | 32.12759 | -0.198143601 | 0.45194663 | -0.4384226 | 0.66107999 | 0.89797068 |
| TAF8 | 13.4234289 | -0.005789084 | 0.42626514 | -0.0135809 | 0.98916431 | NA |
| MRPS10 | 10.26157 | -0.387994959 | 0.43139934 | -0.899387 | 0.36844656 | NA |
| TRERF1 | 16.397596 | 0.370443353 | 0.68601326 | 0.53999445 | 0.58920086 | NA |
| UBR2 | 101.503772 | 0.148368834 | 0.19945155 | 0.74388408 | 0.45694661 | 0.79407688 |
| TBCC | 4.46719896 | -0.309869764 | 0.6071972 | -0.5103281 | 0.60982165 | NA |
| BICRAL | 57.7268529 | -0.203651912 | 0.27945205 | -0.7287544 | 0.46615191 | 0.80148883 |
| RPL7L1 | 36.6855636 | -0.402619327 | 0.38792416 | -1.0378816 | 0.2993252 | 0.68168772 |
| C6orf226 | 11.475838 | -0.211845965 | 0.43323274 | -0.4889888 | 0.62484961 | NA |
| CNPY3 | 14.505356 | 0.07204409 | 0.49345139 | 0.14600038 | 0.88392109 | NA |
| PEX6 | 10.3868153 | 0.105644659 | 0.43609308 | 0.24225255 | 0.80858448 | NA |
| PPP2R5D | 8.48762337 | 0.464948313 | 0.4866439 | 0.95541793 | 0.33936639 | NA |
| MEA1 | 11.2505307 | -0.718080483 | 0.45133053 | -1.5910301 | 0.11160279 | NA |
| KLHDC3 | 15.4957998 | 0.428845914 | 0.52376231 | 0.81877964 | 0.41291215 | NA |
| RRP36 | 19.8753853 | -0.446418683 | 0.42575402 | -1.0485366 | 0.29439144 | NA |
| CUL7 | 7.4227082 | -0.536889529 | 0.51536609 | -1.0417634 | 0.29752139 | NA |
| KLC4 | 8.99326161 | 0.486623373 | 0.46803453 | 1.03971683 | 0.29847148 | NA |
| MRPL2 | 13.6737038 | -0.284473616 | 0.49879213 | -0.570325 | 0.5684573 | NA |
| PTK7 | 24.2275377 | 0.551548157 | 0.4728791 | 1.16636187 | 0.24346817 | NA |
| SRF | 11.1799599 | 0.198411997 | 0.4506436 | 0.44028584 | 0.6597301 | NA |
| CUL9 | 9.82864752 | -0.278391597 | 0.48147324 | -0.5782078 | 0.56312382 | NA |
| DNPH1 | 15.9303744 | -0.926332695 | 0.82082908 | -1.128533 | 0.2590949 | NA |
| ZNF318 | 17.4075136 | -0.024495899 | 0.42334663 | -0.0578625 | 0.95385814 | NA |
| ABCC10 | 8.7628037 | -0.28344426 | 0.53230917 | -0.5324805 | 0.59439323 | NA |
| TJAP1 | 11.8543731 | 0.441386264 | 0.45105536 | 0.97856339 | 0.32779576 | NA |
| POLR1C | 16.07762 | -1.023347047 | 0.47597641 | -2.1499953 | 0.03155559 | NA |
| YIPF3 | 28.6953961 | -0.142145249 | 0.41316981 | -0.3440359 | 0.73081928 | 0.91722837 |
| XPO5 | 43.8470399 | -0.325348834 | 0.46882671 | -0.6939639 | 0.48770482 | 0.81648973 |
| POLH | 7.63256136 | 0.268366895 | 0.51035287 | 0.52584577 | 0.59899537 | NA |
| GTPBP2 | 11.0390002 | -0.171418896 | 0.73732075 | -0.2324889 | 0.8161583 | NA |
| MAD2L1BP | 4.76727523 | -1.251051229 | 0.72440832 | -1.7269973 | 0.08416814 | NA |
| MRPS18A | 15.8932469 | -0.344290059 | 0.58976302 | -0.583777 | 0.55937038 | NA |
| VEGFA | 1005.34766 | 0.221263535 | 0.44402105 | 0.49831767 | 0.61826016 | 0.87662819 |
| MRPL14 | 16.2273776 | -1.344860995 | 0.55078045 | -2.441737 | 0.01461679 | NA |
| TMEM63B | 12.1413857 | 0.092534975 | 0.5348792 | 0.17300163 | 0.86265014 | NA |
| SLC29A1 | 16.0378829 | 0.36961202 | 0.51090158 | 0.72345053 | 0.46940313 | NA |
| HSP90AB1 | 113.282784 | -0.493474777 | 0.39304321 | -1.255523 | 0.20928898 | 0.61736443 |
| SLC35B2 | 12.9972692 | 0.50724755 | 0.45120222 | 1.12421333 | 0.26092254 | NA |
| CDC5L | 19.4585402 | 0.237768773 | 0.37999771 | 0.62571106 | 0.53150447 | NA |
| SUPT3H | 101.123793 | 0.579597525 | 0.3367496 | 1.72115281 | 0.0852231 | 0.44564602 |
| CLIC5 | 1957.78295 | 0.059866139 | 0.35551796 | 0.16839132 | 0.86627544 | 0.96089268 |
| ENPP4 | 31.9629568 | -0.110498423 | 0.53027609 | -0.208379 | 0.83493302 | 0.95179548 |
| ENPP5 | 10.4132537 | -0.607004601 | 0.61506453 | -0.9868958 | 0.32369372 | NA |
| RCAN2 | 6.48913539 | -0.801516572 | 0.76314018 | -1.0502875 | 0.29358596 | NA |
| CYP39A1 | 10.2876495 | 1.468809571 | 0.66160445 | 2.22007207 | 0.02641388 | NA |
| SLC25A27 | 23.8689803 | 0.964255667 | 0.50811069 | 1.89772757 | 0.05773198 | NA |
| TNFRSF21 | 12.0187274 | 1.59369916 | 1.0948381 | 1.45564824 | 0.14548988 | NA |
| CD2AP | 201.60225 | 0.303473667 | 0.24190547 | 1.25451346 | 0.20965544 | 0.61754717 |
| MUT | 15.7580804 | -0.041121604 | 0.40856997 | -0.1006476 | 0.91983018 | NA |
| PKHD1 | 114.81957 | 1.761775167 | 0.5932293 | 2.96980469 | 0.00297989 | 0.0904424 |
| MCM3 | 8.74866348 | -0.173942145 | 0.52759797 | -0.3296869 | 0.74163654 | NA |
| EFHC1 | 32.3074401 | 0.19876161 | 0.44432071 | 0.44733816 | 0.65463092 | 0.89508505 |
| TRAM2 | 107.022433 | 0.660086418 | 0.38485319 | 1.71516419 | 0.08631513 | 0.44638081 |
| TRAM2-AS1 | 7.14959882 | 0.046313025 | 0.6746435 | 0.06864814 | 0.9452697 | NA |
| TMEM14A | 17.8083971 | 0.194043707 | 0.52892591 | 0.36686368 | 0.7137207 | NA |
| GSTA4 | 25.7626071 | 0.355777218 | 0.46449529 | 0.76594366 | 0.44370983 | NA |
| ICK | 9.86191244 | -0.826520285 | 0.45987712 | -1.7972633 | 0.07229383 | NA |
| FBXO9 | 36.8723091 | 0.136889806 | 0.3559217 | 0.38460651 | 0.70052897 | 0.90729979 |
| ELOVL5 | 52.8281957 | -0.229067534 | 0.30677808 | -0.7466881 | 0.45525187 | 0.79324789 |
| GCLC | 20.717441 | -0.333978133 | 0.49612163 | -0.6731779 | 0.5008341 | NA |
| LRRC1 | 63.9575892 | 1.021094818 | 0.34047881 | 2.99899669 | 0.0027087 | 0.08516665 |
| TINAG | 15.5682773 | 1.08191967 | 1.0882286 | 0.99420257 | 0.32012426 | NA |
| HMGCLL1 | 10.7213953 | -1.742822429 | 0.65689822 | -2.6531088 | 0.00797542 | NA |
| DST | 737.430367 | 0.606438366 | 0.24106008 | 2.5157146 | 0.01187913 | 0.18181798 |
| AL512422.1 | 6.19839618 | 0.121830967 | 0.8043371 | 0.15146755 | 0.87960691 | NA |
| BEND6 | 8.03708124 | 0.290118935 | 0.72138619 | 0.40216868 | 0.68755989 | NA |
| KIAA1586 | 8.67862854 | -0.874212309 | 0.50188838 | -1.7418461 | 0.08153538 | NA |
| ZNF451 | 73.8682511 | 0.359681031 | 0.22317857 | 1.61162886 | 0.10704273 | 0.49086361 |
| ZNF451-AS1 | 10.4332478 | 0.092482043 | 0.7611258 | 0.12150691 | 0.90328954 | NA |
| BAG2 | 6.04687519 | 0.16957143 | 0.6139525 | 0.27619633 | 0.78239727 | NA |
| RAB23 | 5.74698305 | -0.433545511 | 0.7023069 | -0.6173163 | 0.5370261 | NA |
| PRIM2 | 41.0118932 | 0.219744753 | 0.42288296 | 0.51963493 | 0.60331805 | 0.87047761 |
| AL021368.2 | 6.15349435 | 0.592734584 | 0.59114182 | 1.00269439 | 0.31600834 | NA |
| PTP4A1 | 18.5074659 | -0.737456483 | 0.56338594 | -1.3089721 | 0.1905438 | NA |
| PHF3 | 100.088773 | 0.033112451 | 0.26385529 | 0.12549474 | 0.90013189 | 0.97097328 |
| EYS | 16.2364934 | -0.502156528 | 0.71695616 | -0.7004006 | 0.48367716 | NA |
| LMBRD1 | 63.4652797 | 0.284282463 | 0.28956585 | 0.98175411 | 0.326221 | 0.69764285 |
| COL19A1 | 17.9653461 | 0.027643639 | 0.45507055 | 0.06074583 | 0.95156163 | NA |
| FAM135A | 85.9429221 | 0.250531393 | 0.24736568 | 1.01279769 | 0.3111568 | 0.69098737 |
| SDHAF4 | 6.02229845 | 0.269509785 | 0.56785537 | 0.47460991 | 0.63506504 | NA |
| SMAP1 | 62.9504882 | 0.17187798 | 0.22256679 | 0.77225349 | 0.43996431 | 0.78399202 |
| OGFRL1 | 67.9901794 | -0.854315659 | 0.49091684 | -1.7402452 | 0.08181598 | 0.43779104 |
| AL136164.2 | 78.9457087 | 0.764070275 | 0.43935474 | 1.73907369 | 0.0820218 | 0.43831413 |
| LINC00472 | 45.0502794 | -0.08169786 | 0.37890563 | -0.2156153 | 0.82928761 | 0.95008863 |
| AC019205.1 | 16.1166283 | 0.832622224 | 0.46469386 | 1.79176508 | 0.0731706 | NA |
| OOEP | 54.7858951 | -0.558741723 | 0.41993239 | -1.3305516 | 0.18333659 | 0.58584259 |
| MTO1 | 11.1245543 | -0.456829209 | 0.4844852 | -0.9429167 | 0.34572349 | NA |

| | | | | | | |
|---|---|---|---|---|---|---|
| EEF1A1 | 247.095267 | -0.600742183 | 0.3510968 | -1.7110443 | 0.08707294 | 0.448752 |
| SLC17A5 | 62.8952877 | -0.033660195 | 0.46014043 | -0.073152 | 0.94168516 | 0.98114803 |
| COX7A2 | 46.0875155 | -0.230007818 | 0.50994412 | -0.4510451 | 0.65195702 | 0.89449325 |
| TMEM30A | 92.8862835 | -0.213156968 | 0.43164044 | -0.4938299 | 0.6214263 | 0.8780438 |
| SENP6 | 180.291393 | 0.361601972 | 0.20979665 | 1.72358318 | 0.08478312 | 0.44564602 |
| MYO6 | 120.054774 | 0.310274713 | 0.29856024 | 1.03923654 | 0.29869474 | 0.68168772 |
| LINC02540 | 6.38988656 | 0.039337606 | 0.78978311 | 0.04980811 | 0.9602753 | NA |
| MEI4 | 7.57167191 | 0.378885811 | 0.5871734 | 0.64527074 | 0.51875175 | NA |
| IRAK1BP1 | 45.8077554 | 0.099285831 | 0.30995762 | 0.32032067 | 0.74872526 | 0.92274989 |
| PHIP | 194.691547 | -0.119085574 | 0.36881977 | -0.3228828 | 0.74678398 | 0.92250642 |
| HMGN3 | 221.554331 | -0.349345798 | 0.32283207 | -1.0821285 | 0.27919542 | 0.66958859 |
| HMGN3-AS1 | 11.4520941 | -0.511264389 | 0.54965901 | -0.9301483 | 0.35229431 | NA |
| LCA5 | 33.4379848 | -0.515121009 | 0.40665562 | -1.2667254 | 0.20525349 | 0.61387077 |
| SH3BGRL2 | 73.0924327 | 0.612110443 | 0.39805928 | 1.53773691 | 0.12411295 | 0.51897128 |
| ELOVL4 | 16.6605501 | -0.063929127 | 0.64687577 | -0.0988275 | 0.92127522 | NA |
| BCKDHB | 209.697504 | -2.219687871 | 0.50773441 | -4.37175 | 1.23E-05 | 0.00217357 |
| AL359715.1 | 11.3197943 | -2.444805334 | 0.71431031 | -3.4226096 | 0.00062023 | NA |
| TENT5A | 34.4498805 | -2.07941294 | 0.67489202 | -3.0811047 | 0.00206234 | 0.07380722 |
| IBTK | 56.450919 | 0.136024384 | 0.31507648 | 0.43171863 | 0.66594593 | 0.89968484 |
| TPBG | 22.0817711 | -0.149181503 | 0.35322583 | -0.4223403 | 0.67277665 | NA |
| UBE3D | 28.7515589 | 0.330522976 | 0.45644298 | 0.72412764 | 0.46898738 | 0.80448595 |
| DOPEY1 | 48.305195 | 0.138687361 | 0.25723354 | 0.53914961 | 0.58978363 | 0.8654248 |
| PGM3 | 21.8234484 | 0.353722059 | 0.4307415 | 0.82119336 | 0.41153614 | NA |
| ME1 | 35.8866322 | 0.299732115 | 0.29148975 | 1.02827669 | 0.30381969 | 0.68398872 |
| CYB5R4 | 14.8432365 | 0.13295132 | 0.42714408 | 0.31125638 | 0.75560573 | NA |
| CEP162 | 26.3701537 | 0.277744713 | 0.39783059 | 0.69814821 | 0.48508451 | NA |
| NT5E | 45.6737899 | -1.74771882 | 0.50259393 | -3.4773974 | 0.00050631 | 0.03111485 |
| SNX14 | 50.6900807 | 0.386444382 | 0.23574093 | 1.63927572 | 0.10115585 | 0.48247622 |
| SYNCRIP | 21.6174847 | -0.466291247 | 0.36279485 | -1.2852753 | 0.19869611 | NA |
| ZNF292 | 130.794893 | -0.29604154 | 0.24014667 | -1.232753 | 0.21766792 | 0.62454341 |
| SMIM8 | 19.4673835 | 0.388414595 | 0.43555367 | 0.89177207 | 0.37251512 | NA |
| SLC35A1 | 6.68397118 | 0.730893347 | 0.88113193 | 0.82949365 | 0.40682513 | NA |
| RARS2 | 36.837693 | 0.541129219 | 0.25570809 | 2.11619904 | 0.03432788 | 0.29210684 |
| ORC3 | 26.8497691 | 0.099489389 | 0.3659639 | 0.27185575 | 0.78573294 | NA |
| AKIRIN2 | 12.3678721 | -0.163508739 | 0.49518423 | -0.3301978 | 0.74125052 | NA |
| RNGTT | 54.8678816 | 0.367295775 | 0.27614025 | 1.33010591 | 0.18348338 | 0.58584259 |
| PNRC1 | 37.8226833 | 0.103916158 | 0.4474962 | 0.23221685 | 0.81636959 | 0.94632611 |
| PM20D2 | 34.5258931 | 0.140306815 | 0.43084036 | 0.32565848 | 0.74468276 | 0.92226971 |
| UBE2J1 | 23.1288881 | -0.054100194 | 0.33122909 | -0.1633317 | 0.87025731 | NA |
| RRAGD | 11.2317126 | -0.09197203 | 0.620838 | -0.1481418 | 0.8822309 | NA |
| ANKRD6 | 17.4862597 | 0.867266797 | 0.46830227 | 1.85193808 | 0.06403472 | NA |
| LYRM2 | 25.718115 | -0.317007264 | 0.37255788 | -0.850894 | 0.39482824 | NA |
| MDN1 | 24.0897824 | -0.305852745 | 0.39998566 | -0.7646593 | 0.44447446 | NA |
| CASP8AP2 | 21.8127349 | 0.400994682 | 0.34368308 | 1.16675712 | 0.24330847 | NA |
| BACH2 | 36.893576 | 0.188207372 | 0.48288302 | 0.38975769 | 0.69671573 | 0.90718427 |
| MAP3K7 | 36.3865903 | 0.426317948 | 0.30748618 | 1.38646214 | 0.16560583 | 0.56743028 |
| EPHA7 | 49.1353011 | 1.119594282 | 0.4910031 | 2.28021836 | 0.02259474 | 0.24569509 |
| MANEA | 40.3399179 | 0.378595378 | 0.43180486 | 0.87677423 | 0.38060928 | 0.74899182 |
| UFL1 | 39.5122699 | -0.011648492 | 0.29678009 | -0.0392496 | 0.96869141 | 0.98964394 |
| GPR63 | 8.57505946 | 0.105826246 | 0.55735263 | 0.18987305 | 0.84940861 | NA |
| NDUFAF4 | 11.3852415 | -0.505892829 | 0.50505204 | -1.0016648 | 0.31650553 | NA |
| KLHL32 | 7.49133474 | 0.392012265 | 0.79355408 | 0.49399565 | 0.62130925 | NA |
| MMS22L | 15.5796857 | -0.556531443 | 0.39975757 | -1.3921724 | 0.16387021 | NA |
| FBXL4 | 54.524288 | 0.438642572 | 0.25098319 | 1.74769702 | 0.08051651 | 0.43346499 |
| PNISR | 135.997896 | 0.289231475 | 0.2779623 | 1.04054209 | 0.29808812 | 0.68115235 |
| AL513550.1 | 8.93068649 | -0.057358542 | 0.5442779 | -0.1053847 | 0.91607059 | NA |
| USP45 | 33.4195926 | 0.122934539 | 0.33498366 | 0.36698667 | 0.71362896 | 0.91328825 |
| TSTD3 | 8.35673133 | 0.181709759 | 0.49777839 | 0.36504148 | 0.71508044 | NA |
| CCNC | 32.8576338 | 0.296337947 | 0.39458483 | 0.75101201 | 0.45264543 | 0.79291818 |
| SIM1 | 14.0729703 | 1.421309235 | 0.93289705 | 1.52354349 | 0.12762279 | NA |
| ASCC3 | 93.7435872 | -0.023795262 | 0.27705289 | -0.0858871 | 0.93155619 | 0.97832857 |
| GRIK2 | 146.8447 | -0.22612159 | 0.49823131 | -0.4538486 | 0.64993779 | 0.89412414 |
| HACE1 | 32.9856132 | 0.310187598 | 0.36685284 | 0.84553686 | 0.39781116 | 0.76034868 |
| PREP | 18.0940967 | -0.365742448 | 0.376767 | -0.9707391 | 0.33167823 | NA |
| ATG5 | 40.0956757 | 0.028557153 | 0.28590703 | 0.09988266 | 0.92043749 | 0.97775374 |
| CRYBG1 | 12.0843005 | 0.641647557 | 0.67424824 | 0.95164885 | 0.3412751 | NA |
| RTN4IP1 | 5.60485558 | 0.521568155 | 0.62328305 | 0.83680786 | 0.40270058 | NA |
| QRSL1 | 12.9822605 | 0.202659978 | 0.43200173 | 0.46911845 | 0.63898497 | NA |
| LINC02532 | 39.646147 | 0.893503284 | 0.81350373 | 1.0983395 | 0.27205627 | 0.66496374 |
| CD24 | 44.4996645 | -0.933345273 | 0.50189452 | -1.8596443 | 0.06293587 | 0.3863717 |
| C6orf203 | 19.1415398 | 0.156226907 | 0.38625308 | 0.40446773 | 0.6858688 | NA |
| PDSS2 | 73.061529 | 0.161217212 | 0.261176 | 0.61727422 | 0.53705386 | 0.83677705 |
| SOBP | 69.8075423 | -0.023824594 | 0.28435195 | -0.0837856 | 0.93322691 | 0.97832857 |
| SEC63 | 57.0826908 | -0.369286682 | 0.25560495 | -1.4447556 | 0.14852655 | 0.54724091 |
| OSTM1 | 30.4733101 | 0.031883764 | 0.33245209 | 0.09590484 | 0.92359615 | 0.97775374 |
| SNX3 | 47.341811 | -0.153846132 | 0.31133088 | -0.4941563 | 0.62119577 | 0.8780438 |
| AFG1L | 21.8802967 | 0.995010372 | 0.44428239 | 2.23958995 | 0.02511756 | NA |
| FOXO3 | 103.312765 | 0.182724992 | 0.33111311 | 0.55185067 | 0.58105067 | 0.86175324 |
| ARMC2 | 19.970858 | -0.048205903 | 0.33459462 | -0.1440726 | 0.88544317 | NA |
| SESN1 | 22.6111693 | 0.308229279 | 0.46853187 | 0.65786193 | 0.51062685 | NA |
| CEP57L1 | 10.8709598 | -0.088377524 | 0.51158319 | -0.172753 | 0.86284559 | NA |
| CD164 | 107.535006 | -0.112097904 | 0.31398572 | -0.3570159 | 0.72107988 | 0.91509584 |
| PPIL6 | 7.484313 | -0.330202653 | 0.5921096 | -0.5576715 | 0.57706872 | NA |
| MICAL1 | 3.97724366 | 1.232405749 | 0.81755739 | 1.50742414 | 0.13170197 | NA |
| AK9 | 24.3990981 | -0.204306434 | 0.46741118 | -0.4371022 | 0.66203726 | NA |
| FIG4 | 21.9706508 | 0.606689987 | 0.46437928 | 1.3064536 | 0.19139835 | NA |
| WASF1 | 23.0420389 | 1.103220105 | 0.34033733 | 3.2415489 | 0.00118882 | NA |
| CDC40 | 29.3460609 | -0.047119113 | 0.37351623 | -0.1261501 | 0.8996131 | 0.97097328 |
| METTL24 | 25.1365159 | 1.29160999 | 0.48498617 | 2.6631893 | 0.00774039 | NA |
| CDK19 | 36.8594836 | 0.517907322 | 0.35744904 | 1.44889836 | 0.14736597 | 0.54635866 |
| AMD1 | 18.2322187 | -0.996220811 | 0.47269215 | -2.1075468 | 0.03507021 | NA |
| GTF3C6 | 24.1160051 | -0.567727214 | 0.42416186 | -1.3384683 | 0.18074382 | NA |
| RPF2 | 8.67032349 | -1.026315078 | 0.67696668 | -1.5160496 | 0.12950681 | NA |
| SLC16A10 | 121.726798 | 0.349851906 | 0.5102412 | 0.68565985 | 0.49292763 | 0.82174866 |
| MFSD4B | 9.70865141 | 0.486977902 | 0.59198301 | 0.82262142 | 0.41072331 | NA |
| REV3L | 94.3438236 | 0.15824568 | 0.26812753 | 0.59018811 | 0.55506454 | 0.8502046 |
| TRAF3IP2-AS | 44.2284891 | -0.540913997 | 0.35509954 | -1.5232743 | 0.1276901 | 0.52314248 |
| TRAF3IP2 | 12.6955433 | 0.216844031 | 0.50585318 | 0.4286699 | 0.66816347 | NA |
| FYN | 390.612174 | 0.414627151 | 0.27049756 | 1.53283138 | 0.12531741 | 0.52185565 |
| WISP3 | 4.23798247 | 0.280925849 | 1.23277721 | 0.22788047 | 0.81973917 | NA |
| TUBE1 | 17.678942 | 0.157923177 | 0.53144883 | 0.29715594 | 0.76634746 | NA |
| FAM229B | 13.251962 | -0.318488037 | 0.57555218 | -0.5533608 | 0.58001635 | NA |
| HDAC2 | 24.4938891 | -0.705540147 | 0.39734409 | -1.7756402 | 0.07579224 | NA |
| FRK | 10.6193145 | 0.308333886 | 0.67761975 | 0.45502494 | 0.64909129 | NA |
| NT5DC1 | 62.0406132 | 0.221951002 | 0.30922751 | 0.71775955 | 0.47290556 | 0.80580648 |
| TSPYL4 | 11.7502473 | 0.240180394 | 0.52711572 | 0.45565022 | 0.64864152 | NA |
| DSE | 39.3901808 | -0.056531764 | 0.39609968 | -0.1427211 | 0.88651049 | 0.9665824 |
| TSPYL1 | 52.7798883 | 0.012319013 | 0.41459721 | 0.02971321 | 0.97629578 | 0.992815 |
| CALHM5 | 28.3113223 | 1.105306787 | 0.56079604 | 1.9709604 | 0.0487284 | NA |
| RWDD1 | 21.3449059 | -0.496512021 | 0.40491764 | -1.226205 | 0.22012155 | NA |
| ZUP1 | 8.56035433 | 0.470640569 | 0.63573409 | 0.74031042 | 0.45911166 | NA |
| KPNA5 | 43.1291209 | 0.291843975 | 0.27617564 | 1.05673322 | 0.29063336 | 0.67772126 |
| DCBLD1 | 13.5781938 | 0.48850613 | 0.52430633 | 0.93171893 | 0.3514818 | NA |

| | | | | | | |
|---|---|---|---|---|---|---|
| GOPC | 24.5145581 | -0.292747901 | 0.34178157 | -0.8565351 | 0.39170189 | NA |
| NUS1 | 22.9219262 | -0.339601813 | 0.4126443 | -0.8229892 | 0.41051412 | NA |
| SLC35F1 | 5.20689677 | -0.325453902 | 0.85711788 | -0.3797073 | 0.70416271 | NA |
| CEP85L | 116.310633 | 0.700231692 | 0.32942077 | 2.12564526 | 0.03353281 | 0.29047359 |
| MCM9 | 39.9295757 | 0.080056319 | 0.32541625 | 0.24601205 | 0.8056729 | 0.94254667 |
| ASF1A | 8.77767575 | 0.830758306 | 0.51512277 | 1.61273846 | 0.10680134 | NA |
| AL365275.1 | 11.395072 | -0.008561964 | 0.71184679 | -0.0120278 | 0.99040342 | NA |
| FAM184A | 10.6691722 | 0.579964003 | 0.81646966 | 0.71033136 | 0.47749868 | NA |
| MAN1A1 | 75.4346481 | 0.206214309 | 0.3860635 | 0.53414609 | 0.59324046 | 0.8654248 |
| TBC1D32 | 43.4351055 | 0.350227752 | 0.34012776 | 1.0296947 | 0.30315335 | 0.68310554 |
| GJA1 | 146.948436 | -1.60484074 | 0.60442483 | -2.6551535 | 0.00792723 | 0.14193427 |
| HSF2 | 13.9696341 | 0.313521627 | 0.44741115 | 0.70074612 | 0.48346147 | NA |
| SERINC1 | 127.64375 | -0.376029045 | 0.3668839 | -1.0249265 | 0.30539785 | 0.68511459 |
| NKAIN2 | 19.101351 | -0.057117127 | 0.59514173 | -0.0959723 | 0.92354256 | NA |
| RNF217-AS1 | 8.81838815 | 0.037777101 | 0.7717323 | 0.04895104 | 0.96095831 | NA |
| RNF217 | 60.7465034 | -0.115181647 | 0.45904007 | -0.2509185 | 0.80187712 | 0.94085969 |
| HDDC2 | 13.2595611 | -1.326137947 | 0.62469778 | -2.1228472 | 0.03376666 | NA |
| NCOA7 | 39.6533973 | -0.341040081 | 0.33115446 | -1.029852 | 0.30307949 | 0.68310554 |
| HINT3 | 13.7032353 | 0.156139519 | 0.4297076 | 0.36336225 | 0.7163343 | NA |
| TRMT11 | 20.1523983 | -0.19940099 | 0.37188836 | -0.5361851 | 0.59183064 | NA |
| RNF146 | 24.1569979 | 0.388938202 | 0.41801402 | 0.93044296 | 0.35214178 | NA |
| ECHDC1 | 37.1872546 | 0.532321188 | 0.34500154 | 1.54295306 | 0.12284217 | 0.51631899 |
| PTPRK | 83.8575613 | 0.540184173 | 0.41278153 | 1.30864425 | 0.19065489 | 0.59329687 |
| LAMA2 | 7.5341648 | 0.153843433 | 0.78896591 | 0.19499377 | 0.84539783 | NA |
| AL356124.1 | 16.5253311 | -0.590247108 | 0.64782249 | -0.9111248 | 0.36222965 | NA |
| ARHGAP18 | 104.572025 | 0.0188102 | 0.43853412 | 0.04289336 | 0.96578655 | 0.98892249 |
| L3MBTL3 | 36.1421431 | 1.627198685 | 0.4310825 | 3.77468045 | 0.00016021 | 0.01469766 |
| EPB41L2 | 9.11455907 | 0.388452044 | 0.71120463 | 0.54618886 | 0.58493613 | NA |
| AKAP7 | 13.0581112 | 0.559347496 | 0.81923211 | 0.68277048 | 0.49475189 | NA |
| MED23 | 24.002878 | 0.20109782 | 0.35254964 | 0.57040995 | 0.56839968 | NA |
| CTGF | 1932.14715 | -0.49439455 | 0.58977024 | -0.8382833 | 0.40187161 | 0.76453623 |
| STX7 | 87.1321996 | 0.172421327 | 0.25568352 | 0.67435448 | 0.50008598 | 0.82345406 |
| RPS12 | 55.356627 | -0.659242318 | 0.52463378 | -1.2565762 | 0.20890714 | 0.61736443 |
| EYA4 | 25.5509922 | -0.388510919 | 0.62496187 | -0.6216554 | 0.53416849 | NA |
| AL024497.1 | 11.526879 | 1.013944106 | 0.62006735 | 1.63521609 | 0.10200376 | NA |
| TARID | 355.99028 | 0.924424999 | 0.3682959 | 2.5100062 | 0.0120729 | 0.183894 |
| TCF21 | 205.902318 | 0.506355182 | 0.6176737 | 0.81977779 | 0.4123428 | 0.76946282 |
| TBPL1 | 26.802317 | 0.160213319 | 0.45421493 | 0.35272579 | 0.72429402 | NA |
| SLC2A12 | 63.1932686 | -0.227090536 | 0.40893281 | -0.5553248 | 0.57867251 | 0.86175324 |
| SGK1 | 89.6374217 | -0.133157264 | 0.70288167 | -0.1894448 | 0.84974423 | 0.95532127 |
| HBS1L | 28.9235303 | -0.13040518 | 0.31979595 | -0.4077762 | 0.68343798 | 0.90499319 |
| AHI1 | 118.447782 | -0.253626186 | 0.29606265 | -0.8566639 | 0.39163065 | 0.75653902 |
| LINC00271 | 43.4337074 | -0.21629712 | 0.41998154 | -0.5150158 | 0.60654199 | 0.87080602 |
| PDE7B | 322.40724 | -1.098145763 | 0.4156606 | -2.641929 | 0.00824353 | 0.144148 |
| AL138828.1 | 44.739825 | -2.230598181 | 0.49803592 | -4.4787898 | 7.51E-06 | 0.00144987 |
| BCLAF1 | 77.892936 | 0.119300207 | 0.21031105 | 0.567256 | 0.57054026 | 0.8573305 |
| MAP7 | 35.4199925 | -0.152769441 | 0.38629239 | -0.3954762 | 0.69249149 | 0.90557575 |
| MAP3K5 | 77.3282414 | 2.005838436 | 0.61563491 | 3.25816228 | 0.00112136 | 0.05186671 |
| PEX7 | 12.6335189 | 0.314902783 | 0.42698075 | 0.73751048 | 0.46081197 | NA |
| IFNGR1 | 27.6328559 | 0.016617441 | 0.34078978 | 0.04876156 | 0.96110932 | NA |
| TNFAIP3 | 6.21739143 | -0.143338677 | 0.71958214 | -0.1991971 | 0.84210857 | NA |
| PERP | 44.7295887 | -0.472604146 | 0.50168148 | -0.9420403 | 0.34617203 | 0.71908069 |
| ARFGEF3 | 5.48605633 | 0.006529979 | 0.72952816 | 0.00895096 | 0.99285826 | NA |
| HEBP2 | 31.2370961 | -0.892541706 | 0.37893923 | -2.355369 | 0.01850432 | 0.22674781 |
| NHSL1 | 11.3724253 | 0.091612445 | 0.69581118 | 0.1316628 | 0.89525101 | NA |
| CCDC28A | 12.7988425 | -0.097641331 | 0.47339836 | -0.2062562 | 0.83659082 | NA |
| REPS1 | 158.368778 | 0.364865149 | 0.24299086 | 1.50155912 | 0.13321101 | 0.52613957 |
| ABRACL | 39.1290858 | 0.547621392 | 0.36424153 | 1.50345677 | 0.1327213 | 0.52613957 |
| HECA | 14.2731452 | -0.453431084 | 0.4111266 | -1.1028989 | 0.27007107 | NA |
| CITED2 | 96.6765552 | 0.346888462 | 0.50549899 | 0.68622978 | 0.49256822 | 0.82148713 |
| VTA1 | 22.8788472 | -0.404764 | 0.34949712 | -1.1581326 | 0.24680994 | NA |
| HIVEP2 | 192.260954 | 0.230211346 | 0.25247834 | 0.91180631 | 0.36187069 | 0.73240894 |
| AIG1 | 82.5150249 | 0.08650942 | 0.34346731 | 0.2518709 | 0.80114085 | 0.94085969 |
| ADAT2 | 4.70502429 | -0.209825911 | 0.91158432 | -0.2301772 | 0.81795409 | NA |
| PEX3 | 25.0328114 | -0.045182571 | 0.38197414 | -0.118287 | 0.90584027 | NA |
| FUCA2 | 17.6834188 | -0.177419564 | 0.4716909 | -0.3761352 | 0.70681637 | NA |
| PHACTR2 | 41.7544362 | 0.722538906 | 0.38885307 | 1.85812833 | 0.06315079 | 0.3863717 |
| PLAGL1 | 20.7787158 | -0.373272846 | 0.45273551 | -0.8244832 | 0.40966503 | NA |
| SF3B5 | 39.3068573 | -0.620063067 | 0.43031573 | -1.4409491 | 0.14959906 | 0.54834287 |
| STX11 | 18.7894347 | -0.19692314 | 0.51616524 | -0.3815118 | 0.7028235 | NA |
| UTRN | 377.169353 | 0.506594943 | 0.24696746 | 2.05126192 | 0.04024145 | 0.31540889 |
| AL024474.2 | 8.33198033 | -0.146581635 | 0.62906359 | -0.2330156 | 0.81574929 | NA |
| EPM2A | 17.5720404 | -0.033756837 | 0.50024029 | -0.0674812 | 0.94619859 | NA |
| FBXO30 | 8.49054789 | -0.104704313 | 0.56693976 | -0.1846833 | 0.85347743 | NA |
| AL356599.1 | 23.2692164 | -0.274823845 | 0.48478573 | -0.5668976 | 0.57078378 | NA |
| SHPRH | 72.987152 | 0.415452898 | 0.2986363 | 1.39116679 | 0.16417485 | 0.56548642 |
| RAB32 | 5.10184584 | -0.873988402 | 0.75964231 | -1.1505262 | 0.2499272 | NA |
| STXBP5 | 77.9499656 | 0.222173866 | 0.27176099 | 0.81753405 | 0.4136233 | 0.76946282 |
| SAMD5 | 28.2023994 | -0.727111014 | 0.64329264 | -1.1302959 | 0.25835158 | NA |
| AL033504.1 | 18.4084944 | -1.669309536 | 0.77572567 | -2.1519328 | 0.03140265 | NA |
| SASH1 | 217.33603 | 0.452882803 | 0.43396699 | 1.04358813 | 0.29667599 | 0.68103704 |
| UST | 7.52900319 | -0.312074756 | 0.81025438 | -0.3851565 | 0.70012145 | NA |
| TAB2 | 84.448637 | 0.237081633 | 0.278143 | 0.85237317 | 0.39400701 | 0.75773027 |
| PPIL4 | 20.3340884 | -0.257483099 | 0.37052273 | -0.6949185 | 0.48710637 | NA |
| GINM1 | 40.2398632 | -0.427772349 | 0.28006589 | -1.527399 | 0.12666183 | 0.52276701 |
| KATNA1 | 13.6475702 | 0.230720124 | 0.41261117 | 0.55917082 | 0.57604515 | NA |
| LATS1 | 50.9456455 | 0.076779248 | 0.22298517 | 0.34432447 | 0.73060229 | 0.91722837 |
| NUP43 | 15.3138082 | 0.514661545 | 0.44698984 | 1.15139429 | 0.24957006 | NA |
| PCMT1 | 68.5677958 | -0.185150564 | 0.35777877 | -0.5175001 | 0.60480707 | 0.87047761 |
| AL355312.2 | 5.93096255 | 0.643031652 | 0.71928613 | 0.89398589 | 0.37132944 | NA |
| LRP11 | 34.7505241 | -0.76359254 | 0.5207512 | -1.4663289 | 0.1425587 | 0.53894095 |
| AL355312.3 | 5.51918871 | -1.031954398 | 0.75422823 | -1.3682256 | 0.17124146 | NA |
| RAET1E | 21.9558963 | -0.357600272 | 0.72030523 | -0.4964566 | 0.61957231 | NA |
| PLEKHG1 | 148.24973 | 0.222930104 | 0.27994242 | 0.79634271 | 0.42583286 | 0.77730788 |
| MTHFD1L | 36.5086624 | -1.266446333 | 0.73598759 | -1.7207441 | 0.08529726 | 0.44564602 |
| AKAP12 | 19.9039848 | 2.259488704 | 0.95185658 | 2.37377012 | 0.01760751 | NA |
| RMND1 | 20.6919001 | -0.39640034 | 0.37291897 | -1.0629664 | 0.28779718 | NA |
| ARMT1 | 7.97349129 | -0.29515675 | 0.52570774 | -0.5614465 | 0.57449322 | NA |
| SYNE1 | 303.59631 | 0.578058847 | 0.34121942 | 1.69409713 | 0.09024683 | 0.45866472 |
| AL080276.2 | 5.838086 | -0.842326333 | 0.62713322 | -1.3431378 | 0.17922736 | NA |
| MTRF1L | 9.57259638 | -0.876772012 | 0.55100173 | -1.5912328 | 0.1115572 | NA |
| RGS17 | 3.81070239 | -2.442209375 | 1.16713193 | -2.0924878 | 0.0363949 | NA |
| IPCEF1 | 12.3384303 | -1.336580555 | 0.61751991 | -2.1644331 | 0.03043112 | NA |
| CNKSR3 | 184.488074 | -0.773461643 | 0.41986183 | -1.8421814 | 0.06544862 | 0.39680062 |
| SCAF8 | 92.2079603 | 0.516075868 | 0.29072088 | 1.77515931 | 0.07587159 | 0.42186351 |
| TIAM2 | 70.2177596 | 0.222539828 | 0.28151655 | 0.79050355 | 0.42923375 | 0.7784723 |
| TFB1M | 23.0899527 | 0.125074947 | 0.35065865 | 0.35668576 | 0.72132706 | NA |
| ARID1B | 337.553816 | 0.310134127 | 0.19790628 | 1.56707575 | 0.11709699 | 0.50775741 |
| TMEM242 | 11.8817927 | 0.094475389 | 0.50198439 | 0.18820384 | 0.85071686 | NA |
| ZDHHC14 | 25.5133115 | -1.01892678 | 0.38875615 | -2.6209921 | 0.00876743 | NA |
| SNX9 | 60.5527401 | -0.074872608 | 0.26385615 | -0.283763 | 0.77659202 | 0.92940498 |
| SYNJ2 | 101.739148 | 0.242066589 | 0.30695239 | 0.78861281 | 0.43033834 | 0.77892126 |
| SERAC1 | 10.8725631 | -0.939173271 | 0.64835464 | -1.4485487 | 0.14746366 | NA |

| | | | | | | |
|---|---|---|---|---|---|---|
| GTF2H5 | 25.0990358 | -0.778648197 | 0.46624897 | -1.6700266 | 0.09491409 | NA |
| TULP4 | 143.747715 | 0.204581592 | 0.31931561 | 0.64068773 | 0.52172559 | 0.83108387 |
| TMEM181 | 39.6148975 | 0.144543866 | 0.27612483 | 0.5234729 | 0.60064521 | 0.86883629 |
| DYNLT1 | 60.7422643 | -0.689487176 | 0.60951402 | -1.1312081 | 0.25796753 | 0.65588339 |
| SYTL3 | 10.3984129 | -0.365113464 | 0.75277711 | -0.485022 | 0.62766073 | NA |
| EZR | 91.6319351 | 0.254880564 | 0.29784659 | 0.85574445 | 0.39213915 | 0.75666812 |
| RSPH3 | 40.788684 | -0.449603792 | 0.315657 | -1.4243429 | 0.15434725 | 0.55470497 |
| AL035530.2 | 39.4849437 | 0.163616392 | 0.41405507 | 0.39515612 | 0.69272768 | 0.90557575 |
| SOD2.1 | 13.8151105 | -0.146699726 | 0.5493277 | -0.2670532 | 0.78942819 | NA |
| WTAP | 58.311582 | 0.000163405 | 0.30191092 | 0.00054124 | 0.99956816 | 0.99961632 |
| ACAT2 | 5.5521821 | -1.14100539 | 0.65214693 | -1.749614 | 0.08018494 | NA |
| TCP1 | 20.1804545 | -0.332254759 | 0.36782185 | -0.9033035 | 0.36636485 | NA |
| MRPL18 | 15.411542 | -0.433992461 | 0.6345888 | -0.6838956 | 0.49404112 | NA |
| IGF2R | 95.5160284 | -0.199165668 | 0.24517087 | -0.8123545 | 0.41658824 | 0.7711921 |
| SLC22A2 | 7.83644804 | 0.832197779 | 0.9215219 | 0.90306891 | 0.36648932 | NA |
| MAP3K4 | 33.1814266 | 0.114852004 | 0.28326703 | 0.40545489 | 0.68514317 | 0.90499319 |
| AGPAT4 | 21.0749694 | 0.167353814 | 0.41343219 | 0.40479144 | 0.68563082 | NA |
| PRKN | 229.489985 | 0.308847282 | 0.27462344 | 1.12462098 | 0.26074968 | 0.65595635 |
| PACRG | 42.1855436 | 0.974479255 | 0.46489243 | 2.09613922 | 0.03606984 | 0.29723902 |
| QKI | 353.643633 | 0.513166631 | 0.22142973 | 2.31751461 | 0.02047572 | 0.23840101 |
| PDE10A | 327.071045 | -2.432768482 | 0.65271822 | -3.7271343 | 0.00019367 | 0.01701871 |
| LINC00473 | 6.2101527 | -2.849026087 | 1.07318659 | -2.6547351 | 0.00793707 | NA |
| SFT2D1 | 32.7953192 | 0.328679946 | 0.35082239 | 0.93688418 | 0.34881814 | 0.72238707 |
| MPC1 | 23.4480957 | -0.418312898 | 0.39163127 | -1.0681295 | 0.28546212 | NA |
| RPS6KA2 | 62.8573305 | -0.516146271 | 0.52072044 | -0.9912157 | 0.32158027 | 0.69513974 |
| Z94721.1 | 8.33053742 | -0.705715425 | 0.56290502 | -1.2537025 | 0.20995016 | NA |
| FGFR1OP | 30.3598347 | 0.672747946 | 0.37638517 | 1.7873923 | 0.07387411 | 0.41834543 |
| AFDN | 78.5930134 | 0.32943418 | 0.29325377 | 1.12337578 | 0.26127793 | 0.65618779 |
| FRMD1 | 67.7400924 | 1.308232813 | 0.5286218 | 2.47479919 | 0.01333111 | 0.19273104 |
| LINC02519 | 18.3626703 | 0.092430036 | 0.44169796 | 0.20926073 | 0.83424471 | NA |
| WDR27 | 34.6314654 | 0.127080631 | 0.29310226 | 0.43357098 | 0.66460001 | 0.89968484 |
| C6orf120 | 12.6800726 | -0.114480478 | 0.44021944 | -0.2600532 | 0.79482273 | NA |
| PHF10 | 39.5246643 | -0.253932085 | 0.46556176 | -0.5454316 | 0.58545673 | 0.86235633 |
| ERMARD | 19.7876363 | -0.101536532 | 0.39884965 | -0.2545735 | 0.79905257 | NA |
| FAM120B | 55.2511154 | -0.154853908 | 0.27287448 | -0.5674914 | 0.5703804 | 0.8573305 |
| PSMB1 | 62.7825225 | -0.683054901 | 0.35651295 | -1.915933 | 0.05537363 | 0.36698602 |
| TBP | 11.4349313 | -0.414537825 | 0.42006061 | -0.9868524 | 0.323715 | NA |
| PDCD2 | 20.9901311 | -0.179834984 | 0.41554215 | -0.4327719 | 0.66518046 | NA |
| FAM20C | 111.340524 | -0.498556127 | 0.53361256 | -0.9343036 | 0.35014731 | 0.72238707 |
| PDGFA | 27.7354146 | -0.61880564 | 0.46253651 | -1.3378525 | 0.18094451 | NA |
| PRKAR1B | 12.8607413 | -0.390565514 | 0.49543595 | -0.788327 | 0.43050548 | NA |
| DNAAF5 | 15.0926659 | -0.216985918 | 0.36881958 | -0.5883254 | 0.55631392 | NA |
| SUN1 | 39.3540624 | 0.038995003 | 0.32368443 | 0.12047228 | 0.90410903 | 0.97237083 |
| COX19 | 12.9832275 | 0.034076186 | 0.43248884 | 0.0787909 | 0.93719894 | NA |
| C7orf50 | 33.2489341 | -0.07277519 | 0.43224844 | -0.1683643 | 0.86629672 | 0.96089268 |
| ZFAND2A | 8.17469619 | -0.883998899 | 0.53498597 | -1.6523777 | 0.09845757 | NA |
| INTS1 | 22.979834 | 0.190804307 | 0.51365227 | 0.37146591 | 0.71029055 | NA |
| MAFK | 30.8592579 | -0.255965622 | 0.62956885 | -0.4065729 | 0.68432172 | 0.90499319 |
| PSMG3 | 7.84559126 | -0.059552752 | 0.59019949 | -0.1009027 | 0.91962766 | NA |
| MAD1L1 | 32.9751805 | 0.211638925 | 0.42267406 | 0.50071425 | 0.61657224 | 0.87619503 |
| MRM2 | 6.06420483 | -0.715300179 | 0.57183101 | -1.2508944 | 0.21097302 | NA |
| SNX8 | 30.7718683 | -0.26018878 | 0.51332764 | -0.5068669 | 0.61224823 | 0.87391153 |
| EIF3B | 32.4420049 | -0.58807806 | 0.35414946 | -1.6605364 | 0.0968066 | 0.47093115 |
| CHST12 | 21.8711202 | -0.461594631 | 0.3921176 | -1.1771842 | 0.23912201 | NA |
| BRAT1 | 9.48225725 | -0.102238623 | 0.60625012 | -0.168641 | 0.86607903 | NA |
| IQCE | 11.7291302 | 0.353193548 | 0.47348467 | 0.74594505 | 0.45570061 | NA |
| TTYH3 | 8.56197365 | 0.244655927 | 0.61734241 | 0.39630507 | 0.69187999 | NA |
| AMZ1 | 6.65385718 | -0.613691714 | 0.77575591 | -0.7910887 | 0.42889225 | NA |
| GNA12 | 47.7611303 | 0.011941425 | 0.23795075 | 0.05018444 | 0.95997541 | 0.98823865 |
| SDK1 | 24.4652522 | -0.135816618 | 0.50283178 | -0.2701035 | 0.78708064 | NA |
| FOXK1 | 40.9657753 | -0.450880283 | 0.38039921 | -1.1852819 | 0.23590603 | 0.64267939 |
| AP5Z1 | 8.73624782 | -0.233138252 | 0.53709194 | -0.4340751 | 0.66423389 | NA |
| RBAK-RBAKD | 25.9075841 | -0.421140619 | 0.3040884 | -1.3849283 | 0.1660744 | NA |
| WIPI2 | 31.1374322 | -0.175388819 | 0.43282345 | -0.4052202 | 0.68531563 | 0.90499319 |
| TNRC18 | 87.1034381 | 0.50041844 | 0.23506025 | 2.12889439 | 0.033263 | 0.29047359 |
| FBXL18 | 9.89729756 | -0.029811585 | 0.43558295 | -0.0684407 | 0.94543485 | NA |
| ACTB | 833.953826 | 0.135437092 | 0.54412871 | 0.24890635 | 0.80343322 | 0.94126368 |
| RNF216 | 55.2665373 | -0.152416329 | 0.25752908 | -0.5918412 | 0.55395691 | 0.84947041 |
| CCZ1 | 8.40195759 | -0.255505529 | 0.51996076 | -0.4913939 | 0.6231479 | NA |
| PMS2 | 12.0221474 | 0.952451092 | 0.43252897 | 2.20205154 | 0.02766167 | NA |
| AIMP2 | 12.4700804 | -1.005112232 | 0.55730793 | -1.8035132 | 0.07130765 | NA |
| EIF2AK1 | 36.1768651 | -0.086414894 | 0.29564562 | -0.2922921 | 0.77006326 | 0.92621526 |
| USP42 | 21.5973905 | 0.151235055 | 0.34848562 | 0.43397789 | 0.6643045 | NA |
| CYTH3 | 24.5026982 | 0.343561815 | 0.34635869 | 0.99192492 | 0.32123415 | NA |
| FAM220A | 4.93291572 | 0.176562479 | 0.6159367 | 0.28665686 | 0.77437507 | NA |
| RAC1 | 96.9757458 | -0.590663625 | 0.23172044 | -2.5490355 | 0.01080213 | 0.16981952 |
| DAGLB | 5.26310942 | -0.844487174 | 0.63531862 | -1.3292341 | 0.18377075 | NA |
| ZDHHC4 | 11.2621543 | -0.511396839 | 0.51593163 | -0.9912105 | 0.32158281 | NA |
| C7orf26 | 6.54174169 | 0.094280462 | 0.63245543 | 0.14907052 | 0.88149799 | NA |
| ZNF316 | 9.6304604 | 0.912630074 | 0.53686285 | 1.69993149 | 0.08914381 | NA |
| AC073343.2 | 7.10311616 | -0.515821526 | 0.61001249 | -0.8455917 | 0.39778053 | NA |
| ZNF12 | 15.0226097 | -0.370837205 | 0.43355749 | -0.8553357 | 0.39236532 | NA |
| CCZ1B | 12.756549 | -0.664187944 | 0.50279299 | -1.3209968 | 0.18650242 | NA |
| C1GALT1 | 46.63146 | 0.233712154 | 0.38715959 | 0.60365844 | 0.54607075 | 0.84440067 |
| COL28A1 | 11.0384359 | -0.152384635 | 0.57048567 | -0.2671139 | 0.78938149 | NA |
| MIOS | 17.8889864 | 0.248979869 | 0.40706057 | 0.61165313 | 0.54076728 | NA |
| RPA3 | 15.5133444 | -0.467691099 | 0.61063993 | -0.7659032 | 0.44373388 | NA |
| UMAD1 | 112.803449 | 0.120407986 | 0.28872987 | 0.41702642 | 0.67665908 | 0.90499319 |
| GLCCI1 | 29.49266 | -0.224429665 | 0.35261457 | -0.636473 | 0.52446816 | 0.83192915 |
| ICA1 | 17.3292397 | 0.378178943 | 0.6452868 | 0.58606335 | 0.55783295 | NA |
| NDUFA4 | 86.4241406 | -0.702840733 | 0.55311903 | -1.2706862 | 0.20384029 | 0.61333549 |
| PHF14 | 66.8853129 | -0.077776675 | 0.23575495 | -0.3299047 | 0.74147195 | 0.92054185 |
| THSD7A | 926.253334 | 0.306515964 | 0.48051231 | 0.63789409 | 0.52354263 | 0.83178884 |
| TMEM106B | 85.6714864 | 0.228217345 | 0.28135718 | 0.81113033 | 0.41729083 | 0.77165508 |
| SCIN | 14.432736 | 0.316386215 | 0.64593447 | 0.48981163 | 0.6242672 | NA |
| AC011287.1 | 20.2751881 | 1.430549544 | 0.87753311 | 1.63019438 | 0.10306042 | NA |
| ISPD | 21.3594816 | 0.730365786 | 0.4324568 | 1.68887571 | 0.09124325 | NA |
| ANKMY2 | 14.9750821 | 0.959718435 | 0.4618214 | 2.07811598 | 0.03769868 | NA |
| BZW2 | 6.12875062 | -1.513359375 | 0.82159013 | -1.8419883 | 0.06547687 | NA |
| AC073332.1 | 12.6829886 | 0.524664823 | 0.79008287 | 0.66406302 | 0.50664997 | NA |
| SNX13 | 64.8795727 | 0.211258547 | 0.3277068 | 0.6446572 | 0.51914936 | 0.83104823 |
| HDAC9 | 25.9488247 | 2.158360258 | 0.90716077 | 2.37924778 | 0.01734801 | NA |
| TWISTNB | 11.2230803 | -0.408212291 | 0.44583575 | -0.9156114 | 0.35987076 | NA |
| MACC1 | 18.8949653 | 0.672484434 | 0.85913708 | 0.78274405 | 0.43377743 | NA |
| MACC1-AS1 | 7.95300135 | -1.793594417 | 0.77014602 | -2.3289017 | 0.01986427 | NA |
| ITGB8 | 862.471982 | -0.157285214 | 0.47306311 | -0.3324825 | 0.73952494 | 0.92003444 |
| SP4 | 14.5691696 | 0.320516327 | 0.40301134 | 0.79530349 | 0.42643698 | NA |
| RAPGEF5 | 9.495382 | -2.429310399 | 0.78444552 | -3.0968504 | 0.00195589 | NA |
| STEAP1B | 43.5964912 | -0.985632737 | 0.43927213 | -2.2437862 | 0.02484616 | 0.25362919 |
| TOMM7 | 50.6495801 | -0.370852333 | 0.44369325 | -0.8358305 | 0.40325029 | 0.76594067 |
| FAM126A | 73.0162061 | 0.57532269 | 0.40117861 | 1.43408117 | 0.1515491 | 0.55086922 |
| KLHL7 | 22.2180356 | 0.196811433 | 0.35849475 | 0.54899391 | 0.58300963 | NA |
| NUPL2 | 12.9095747 | -0.330467755 | 0.48306565 | -0.6841053 | 0.4939087 | NA |

| | | | | | | |
|---|---|---|---|---|---|---|
| GPNMB | 37.3787889 | -1.217249874 | 0.58716569 | -2.0730943 | 0.0381635 | 0.304707 |
| MALSU1 | 36.7489245 | 0.317959783 | 0.44420421 | 0.71579642 | 0.47411706 | 0.80625938 |
| TRA2A | 105.274546 | 0.383556386 | 0.27821718 | 1.37862222 | 0.16801126 | 0.57025412 |
| CCDC126 | 11.7053305 | 0.09221286 | 0.4824101 | 0.19115035 | 0.8484078 | NA |
| FAM221A | 6.59043289 | -0.075861203 | 0.57683868 | -0.131512 | 0.8953703 | NA |
| MPP6 | 18.4229461 | -0.249337525 | 0.34505804 | -0.7225959 | 0.46992819 | NA |
| OSBPL3 | 23.7487913 | 0.204148848 | 0.60248275 | 0.33884596 | 0.73472577 | NA |
| CYCS | 22.9982511 | -0.743364854 | 0.51007941 | -1.4573512 | 0.14501945 | NA |
| C7orf31 | 5.94211819 | -0.168678676 | 0.56510983 | -0.2984883 | 0.7653305 | NA |
| NFE2L3 | 28.148921 | -0.294380438 | 0.55849383 | -0.527097 | 0.59812621 | NA |
| HNRNPA2B1 | 211.147614 | 0.149467757 | 0.22930177 | 0.65183867 | 0.51450525 | 0.8308079 |
| CBX3 | 41.0616967 | -0.880162667 | 0.30288858 | -2.9058958 | 0.00366203 | 0.09771848 |
| SNX10 | 28.2444033 | -1.400262247 | 0.51668094 | -2.7101101 | 0.00672609 | NA |
| AC004540.1 | 15.7089443 | -1.161084265 | 0.63303725 | -1.8341484 | 0.06663197 | NA |
| SKAP2 | 81.8455259 | 0.046324184 | 0.27095142 | 0.1709686 | 0.86424846 | 0.96089268 |
| HOTAIRM1 | 29.8380994 | 0.085204949 | 0.50915029 | 0.16734735 | 0.86709675 | 0.96089268 |
| HOXA3 | 10.9683568 | -0.055509603 | 0.48673826 | -0.1140441 | 0.90920287 | NA |
| HOXA-AS2 | 9.35357044 | 0.215572966 | 0.57293481 | 0.3762609 | 0.70672295 | NA |
| HOXA7 | 13.0320937 | 0.19518576 | 0.6371139 | 0.30635929 | 0.7593311 | NA |
| HOXA9 | 27.5165296 | -0.050947331 | 0.44136337 | -0.1154317 | 0.90810294 | NA |
| HOXA10-AS | 5.64026687 | -1.232322426 | 0.83498284 | -1.4758656 | 0.13997999 | NA |
| HIBADH | 58.8038157 | 0.2392413 | 0.32102914 | 0.74523236 | 0.45613127 | 0.79324789 |
| TAX1BP1 | 113.845718 | -0.016989976 | 0.24142137 | -0.0703748 | 0.94389537 | 0.98114803 |
| JAZF1 | 137.242121 | 0.558511624 | 0.29529297 | 1.89138136 | 0.05857346 | 0.37650386 |
| CREB5 | 7.35812408 | -0.044251446 | 0.64535548 | -0.0685691 | 0.94533261 | NA |
| CPVL | 8.51945671 | -0.357770873 | 0.68204808 | -0.5245537 | 0.59989345 | NA |
| WIPF3 | 170.698941 | -0.063568499 | 0.30358373 | -0.2093936 | 0.83414096 | 0.95164471 |
| SCRN1 | 26.3693614 | 0.281274689 | 0.46122155 | 0.60984724 | 0.54196301 | NA |
| FKBP14 | 9.60226983 | -1.022245972 | 0.63227354 | -1.616778 | 0.1059262 | NA |
| PLEKHA8 | 14.5159155 | 0.018716671 | 0.39934558 | 0.04686836 | 0.96261815 | NA |
| MTURN | 12.0919602 | -0.357720688 | 0.50743166 | -0.7049633 | 0.48083309 | NA |
| ZNRF2 | 41.3325399 | -0.429427047 | 0.40631816 | -1.0568739 | 0.29056915 | 0.67772126 |
| AC006978.2 | 45.8768616 | -0.567265453 | 0.41212036 | -1.3764558 | 0.16868057 | 0.57156924 |
| NOD1 | 38.2227535 | -0.852425917 | 0.4286231 | -1.988754 | 0.04672836 | 0.33774256 |
| GGCT | 10.4962524 | -1.110561973 | 0.69048227 | -1.6083859 | 0.10775068 | NA |
| AC005154.6 | 24.0793605 | 0.653219236 | 0.37660822 | 1.73447951 | 0.08283304 | NA |
| GARS | 25.0409038 | 0.004609987 | 0.5767613 | 0.00799289 | 0.99362267 | NA |
| AQP1 | 328.842805 | 0.532319149 | 1.27592922 | 0.41720116 | 0.67653128 | 0.90499319 |
| LSM5 | 12.064148 | -0.54567181 | 0.55697858 | -0.9796998 | 0.32723432 | NA |
| AVL9 | 59.8179046 | -0.318340574 | 0.25902466 | -1.2289972 | 0.21907287 | 0.62462904 |
| KBTBD2 | 25.4672674 | 0.087013356 | 0.33682069 | 0.25833733 | 0.79614658 | NA |
| FKBP9 | 47.0143177 | -0.088825299 | 0.41351473 | -0.2148056 | 0.82991885 | 0.95008863 |
| NT5C3A | 32.3365094 | 0.29784176 | 0.65632871 | 0.45379968 | 0.64997301 | 0.89412414 |
| RP9 | 7.10355329 | -0.730253125 | 0.61593954 | -1.1855922 | 0.23578337 | NA |
| BBS9 | 82.2688634 | 0.322937929 | 0.28423698 | 1.13615731 | 0.25589073 | 0.65588339 |
| DPY19L1 | 23.5819315 | 0.080351913 | 0.42753987 | 0.18794016 | 0.85092356 | NA |
| HERPUD2 | 31.5187403 | 0.075310484 | 0.29785792 | 0.25284029 | 0.80039163 | 0.94085969 |
| AC018647.2 | 4.3150129 | -0.334571352 | 0.61052913 | -0.5480029 | 0.58368991 | NA |
| SEPT7-AS1 | 6.68777122 | -0.400675841 | 0.52944077 | -0.7567907 | 0.44917528 | NA |
| 7-Sep | 137.651843 | 0.342589432 | 0.28155193 | 1.21678951 | 0.22368431 | 0.63004415 |
| KIAA0895 | 19.6339292 | 0.187501431 | 0.51264969 | 0.36574962 | 0.71455192 | NA |
| ELMO1 | 21.5884283 | -1.713008653 | 0.67222437 | -2.5482692 | 0.01082589 | NA |
| EPDR1 | 34.9982572 | 0.287969527 | 0.44465746 | 0.64762104 | 0.51723008 | 0.83104823 |
| SFRP4 | 5.36089937 | -0.663509238 | 0.94673666 | -0.7008382 | 0.48340398 | NA |
| STARD3NL | 44.8820204 | 0.661505701 | 0.25213249 | 2.62364324 | 0.00869949 | 0.14875902 |
| VPS41 | 57.7698068 | 0.249297239 | 0.25632802 | 0.97257116 | 0.33076648 | 0.70387662 |
| POU6F2 | 11.3556968 | -0.526309411 | 0.61967511 | -0.8493312 | 0.39569702 | NA |
| YAE1D1 | 9.66995192 | -0.939485381 | 0.49964458 | -1.8803074 | 0.0600662 | NA |
| RALA | 52.0356279 | -0.301940939 | 0.31543192 | -0.9572301 | 0.33845111 | 0.71275062 |
| CDK13 | 119.435427 | -0.039245355 | 0.20725316 | -0.1893595 | 0.84981107 | 0.95532127 |
| MPLKIP | 22.6979575 | -0.20049123 | 0.36169903 | -0.554304 | 0.57937079 | NA |
| SUGCT | 15.5049799 | 1.373323559 | 0.74639327 | 1.83994632 | 0.06577612 | NA |
| GLI3 | 30.0845096 | -0.595293644 | 0.53648065 | -1.1096274 | 0.2671596 | 0.66013596 |
| MRPL32 | 18.7883346 | -0.515243648 | 0.45169841 | -1.1406807 | 0.25400283 | NA |
| STK17A | 8.48502223 | 0.017843317 | 0.51904054 | 0.0343775 | 0.97257612 | NA |
| COA1 | 56.6802175 | -0.126970213 | 0.30941588 | -0.4103545 | 0.68154588 | 0.90499319 |
| BLVRA | 30.6292433 | -0.634074089 | 0.4447601 | -1.4256542 | 0.15396819 | 0.55412156 |
| URGCP | 9.20790915 | -0.491777706 | 0.54636284 | -0.9000936 | 0.36807043 | NA |
| UBE2D4 | 7.28746601 | -0.045875318 | 0.64516436 | -0.0711064 | 0.94331307 | NA |
| DBNL | 18.5518156 | 0.473524444 | 0.4203921 | 1.12638758 | 0.2600015 | NA |
| POLM | 8.78994353 | -0.47859861 | 0.58216671 | -0.8220989 | 0.41102061 | NA |
| POLD2 | 10.6894497 | -0.119171582 | 0.47315193 | -0.2518675 | 0.8011435 | NA |
| YKT6 | 9.35990621 | 0.231621996 | 0.54001723 | 0.42891594 | 0.6679844 | NA |
| NUDCD3 | 48.4865136 | 0.170330113 | 0.28302241 | 0.60182555 | 0.54729027 | 0.84499784 |
| DDX56 | 11.1059583 | 0.443574058 | 0.42475193 | 1.04431323 | 0.29634049 | NA |
| TMED4 | 50.2264901 | -0.34596796 | 0.39476143 | -0.8763976 | 0.38081393 | 0.74899182 |
| OGDH | 46.7167704 | -0.049639975 | 0.26507591 | -0.187267 | 0.85145128 | 0.95532127 |
| ZMIZ2 | 22.0088778 | 0.084805889 | 0.39537991 | 0.21449215 | 0.83016328 | NA |
| PPIA | 142.811642 | -0.530301083 | 0.45587832 | -1.1632514 | 0.24472752 | 0.64996527 |
| H2AFV | 40.6082456 | -0.829667123 | 0.5920836 | -1.4012669 | 0.16113429 | 0.56290656 |
| PURB | 27.3979323 | -0.214247983 | 0.3816185 | -0.5614193 | 0.57451175 | NA |
| SNHG15 | 7.47397689 | -0.163164193 | 0.68794325 | -0.2371768 | 0.81251962 | NA |
| CCM2 | 33.900526 | -0.114842432 | 0.29122974 | -0.3943362 | 0.69333283 | 0.90557575 |
| TBRG4 | 5.98019204 | 0.607137388 | 0.60732795 | 0.99968622 | 0.31746238 | NA |
| RAMP3 | 94.7088204 | -0.786364292 | 0.55874717 | -1.4073705 | 0.15931755 | 0.56239512 |
| IGFBP3 | 7.9209592 | 0.777105229 | 1.23543346 | 0.62901423 | 0.52933974 | NA |
| TNS3 | 315.60481 | 0.808776501 | 0.2788854 | 2.90003168 | 0.00373125 | 0.09813187 |
| HUS1 | 16.8607086 | 0.509885815 | 0.37897163 | 1.34544589 | 0.17848128 | NA |
| UPP1 | 10.7676735 | -0.392751722 | 0.92588334 | -0.4241914 | 0.67142626 | NA |
| GRB10 | 24.8756629 | 0.141751475 | 0.55167307 | 0.25694833 | 0.79721866 | NA |
| COBL | 25.8345212 | -0.276455349 | 0.44643684 | -0.6192485 | 0.53575266 | NA |
| SEC61G | 21.4939669 | -0.18761723 | 0.56466164 | -0.3322649 | 0.73968926 | NA |
| EGFR | 92.153583 | -0.101716467 | 0.25886027 | -0.3929397 | 0.69436405 | 0.90557575 |
| LANCL2 | 20.6686169 | 0.665394597 | 0.37506091 | 1.77409742 | 0.07604704 | NA |
| VOPP1 | 52.8208112 | -1.419418375 | 0.4902403 | -2.8953523 | 0.00378733 | 0.09813187 |
| NIPSNAP2 | 41.9032608 | 0.025675016 | 0.31424017 | 0.08170507 | 0.93488125 | 0.97832857 |
| PSPH | 5.54590584 | -0.231268787 | 0.74422159 | -0.3107526 | 0.75598871 | NA |
| CCT6A | 27.147635 | 0.010124822 | 0.36573633 | 0.02768339 | 0.97791467 | NA |
| SUMF2 | 31.5977174 | -0.320652453 | 0.44565409 | -0.7195097 | 0.47182691 | 0.80516996 |
| PHKG1 | 12.2843288 | 1.433369202 | 0.61192565 | 2.34239111 | 0.01916062 | NA |
| CHCHD2 | 93.6223999 | -0.783723442 | 0.55742014 | -1.4059834 | 0.15972907 | 0.56283048 |
| ZNF727 | 10.1602265 | 0.482945903 | 0.55065292 | 0.8770423 | 0.38046366 | NA |
| ZNF736 | 43.8809564 | -0.245124491 | 0.38191185 | -0.6418353 | 0.52098015 | 0.83108387 |
| ZNF680 | 37.7777404 | 0.070286395 | 0.28230858 | 0.2489701 | 0.80338391 | 0.94126368 |
| ZNF107 | 14.2181817 | 0.34745894 | 0.50328768 | 0.6903784 | 0.48995626 | NA |
| ZNF138 | 14.5034093 | -0.243878728 | 0.52965625 | -0.4604472 | 0.64519528 | NA |
| ZNF117 | 18.7862111 | 0.754275039 | 0.43472804 | 1.73505036 | 0.08273189 | NA |
| ZNF92 | 7.42917347 | -0.322030541 | 0.6382043 | -0.5045885 | 0.6138479 | NA |
| VKORC1L1 | 41.3608161 | 0.060999829 | 0.3172788 | 0.19225939 | 0.84753903 | 0.95532127 |
| GUSB | 23.7162074 | -0.183860665 | 0.44739032 | -0.4109625 | 0.6811 | NA |
| AC068533.4 | 5.96913218 | -0.645822465 | 0.6977896 | -0.9255261 | 0.35469233 | NA |
| CRCP | 7.25429023 | 0.275431269 | 0.4976216 | 0.55349541 | 0.57992422 | NA |
| AC068533.3 | 4.68074485 | -0.476211201 | 0.66848662 | -0.7123721 | 0.4762344 | NA |

| | | | | | | |
|---|---|---|---|---|---|---|
| TPST1 | 42.6107333 | 0.048699267 | 0.49082361 | 0.09921949 | 0.920964 | 0.97775374 |
| AC027644.4 | 20.2326431 | 0.142582266 | 0.31970046 | 0.44598706 | 0.6556066 | NA |
| AC027644.3 | 11.8334617 | 0.091417336 | 0.71541235 | 0.12778272 | 0.89832093 | NA |
| TMEM248 | 56.050897 | 0.465665704 | 0.25706984 | 1.81143654 | 0.0700733 | 0.40663921 |
| SBDS | 73.4365186 | -0.677618942 | 0.33056028 | -2.0499104 | 0.04037318 | 0.31551756 |
| TYW1 | 40.0761313 | 0.124754952 | 0.26101632 | 0.47795843 | 0.63267979 | 0.88389921 |
| AUTS2 | 201.978244 | 0.339801901 | 0.25434875 | 1.33596845 | 0.18155959 | 0.58390314 |
| TYW1B | 20.3916364 | 0.352649656 | 0.31452306 | 1.12122036 | 0.26219408 | NA |
| POM121 | 39.3036379 | 0.489166727 | 0.26241169 | 1.86411944 | 0.06230493 | 0.38512471 |
| NSUN5 | 11.0495752 | 0.578028886 | 0.40711942 | 1.41980181 | 0.15566539 | NA |
| BAZ1B | 47.4399325 | 0.079755312 | 0.2890814 | 0.27589223 | 0.78263084 | 0.93415197 |
| BCL7B | 12.5873261 | 0.369075754 | 0.46064165 | 0.80122098 | 0.42300373 | NA |
| TBL2 | 6.51729243 | -0.378844189 | 0.53748261 | -0.7048492 | 0.48090409 | NA |
| MLXIPL | 6.1240747 | 0.180755317 | 0.87180501 | 0.20733457 | 0.83574858 | NA |
| DNAJC30 | 6.34977811 | -0.871023652 | 0.7359387 | -1.1835546 | 0.23658941 | NA |
| BUD23 | 15.627597 | -0.088968758 | 0.43830831 | -0.2029821 | 0.83914899 | NA |
| CLDN3 | 5.39770497 | -1.242957499 | 0.87183584 | -1.4256784 | 0.1539612 | NA |
| CLDN4 | 8.71883115 | -0.833117241 | 0.59814977 | -1.3928238 | 0.16367308 | NA |
| LIMK1 | 5.38660418 | -0.015560084 | 0.61124915 | -0.0254562 | 0.97969108 | NA |
| EIF4H | 35.1084479 | -0.659265918 | 0.27979819 | -2.3562194 | 0.01846201 | 0.22674781 |
| RFC2 | 4.50802145 | -0.853308634 | 0.66617876 | -1.2809004 | 0.20022866 | NA |
| CLIP2 | 12.9335343 | -0.725794432 | 0.48801343 | -1.4872427 | 0.13695071 | NA |
| GTF2IRD1 | 47.1591328 | -0.751076389 | 0.44639318 | -1.6825445 | 0.09246331 | 0.46324185 |
| GTF2I | 128.688838 | 0.257907302 | 0.2061649 | 1.25097583 | 0.21094329 | 0.61954092 |
| GTF2IRD2 | 12.2975646 | -0.3445811 | 0.47947754 | -0.7186595 | 0.47235073 | NA |
| CASTOR2 | 12.1452862 | 1.012547259 | 0.6164453 | 1.64255816 | 0.10047438 | NA |
| RCC1L | 18.8231511 | -0.242585568 | 0.46623531 | -0.5203072 | 0.60284951 | NA |
| GTF2IRD2B | 15.7763921 | -0.150101297 | 0.44137549 | -0.3400762 | 0.73379915 | NA |
| TRIM73 | 11.7312557 | -0.260833604 | 0.49499163 | -0.5269455 | 0.59823144 | NA |
| POM121C | 29.7393766 | -0.196281497 | 0.28679373 | -0.6843995 | 0.4937229 | 0.82248426 |
| SPDYE5 | 7.70123698 | -0.705817563 | 0.6686306 | -1.0556166 | 0.29114341 | NA |
| HIP1 | 98.9282647 | 1.469735638 | 0.45755613 | 3.21214285 | 0.00131749 | 0.0545866 |
| RHBDD2 | 17.2768551 | -0.020157423 | 0.51313062 | -0.0392832 | 0.96866459 | NA |
| POR | 27.9841846 | -0.074514712 | 0.4046172 | -0.184161 | 0.85388715 | NA |
| TMEM120A | 15.6897305 | 0.405458349 | 0.47078516 | 0.86123859 | 0.38910665 | NA |
| STYXL1 | 9.16847719 | 0.335928062 | 0.60980943 | 0.55087384 | 0.58172016 | NA |
| MDH2 | 21.3190755 | -0.013103828 | 0.48438412 | -0.0270526 | 0.97841782 | NA |
| HSPB1 | 58.6889398 | 0.681471264 | 0.44575061 | 1.52881735 | 0.12630973 | 0.52276701 |
| YWHAG | 52.2851111 | -0.133014446 | 0.32764369 | -0.4059729 | 0.68476254 | 0.90499319 |
| DTX2 | 11.6040026 | 0.713792426 | 0.55486473 | 1.28642602 | 0.19829442 | NA |
| POMZP3 | 9.14589492 | -0.626382297 | 0.54671663 | -1.1457166 | 0.25191245 | NA |
| CCDC146 | 25.8954724 | -0.062154223 | 0.38393142 | -0.1618889 | 0.87139337 | NA |
| GSAP | 32.7349434 | 0.963199417 | 0.44774374 | 2.15122923 | 0.03145811 | 0.28544541 |
| PTPN12 | 65.3173998 | -0.312752564 | 0.25327108 | -1.234853 | 0.21688521 | 0.62433386 |
| APTR | 10.2616265 | 0.021861356 | 0.47958161 | 0.04558422 | 0.96364164 | NA |
| RSBN1L | 43.7439678 | -0.330060439 | 0.26285042 | -1.2556968 | 0.2092259 | 0.61736443 |
| TMEM60 | 6.80210185 | -0.956866174 | 0.79101242 | -1.2096728 | 0.22640449 | NA |
| PHTF2 | 17.4919581 | 0.589846418 | 0.42307526 | 1.39418791 | 0.16326087 | NA |
| MAGI2 | 13369.5996 | 0.345894753 | 0.27562977 | 1.25492521 | 0.20950591 | 0.61754717 |
| AC006355.2 | 67.3195273 | -0.281114558 | 0.62652828 | -0.4486861 | 0.65365808 | 0.8949739 |
| GNAI1 | 25.2039539 | 0.202135992 | 0.55419629 | 0.36473718 | 0.7153076 | NA |
| SEMA3C | 34.0278865 | 1.111389824 | 0.6801786 | 1.63396764 | 0.10226564 | 0.48317363 |
| CACNA2D1 | 78.7007689 | 0.250690461 | 0.19566932 | 1.28119454 | 0.20012534 | 0.60620493 |
| PCLO | 74.709968 | 0.906175362 | 0.76742493 | 1.1808 | 0.23768218 | 0.64355069 |
| AC003984.1 | 11.6113522 | 1.305968554 | 0.63669763 | 2.05115977 | 0.04025139 | NA |
| KIAA1324L | 19.5874381 | 0.21329084 | 0.53018129 | 0.40229794 | 0.68746477 | NA |
| DMTF1 | 67.8406384 | 0.538794066 | 0.40226264 | 1.33940865 | 0.18043767 | 0.58390314 |
| TMEM243 | 14.5155987 | 0.001771524 | 0.4060457 | 0.00436287 | 0.99651895 | NA |
| TP53TG1 | 6.7170326 | 0.499723058 | 0.60635463 | 0.82414322 | 0.40985819 | NA |
| CROT | 15.7957331 | 0.459632763 | 0.45567655 | 1.00868207 | 0.31312713 | NA |
| ABCB1 | 7.60283207 | 0.50324075 | 0.78600979 | 0.64024743 | 0.52201175 | NA |
| SLC25A40 | 15.6280624 | 0.134422837 | 0.55488158 | 0.242255 | 0.80858258 | NA |
| DBF4 | 4.97930779 | -1.073578795 | 0.7380598 | -1.454596 | 0.14578115 | NA |
| SRI | 21.104593 | -0.588217079 | 0.52800893 | -1.1140287 | 0.26526689 | NA |
| CFAP69 | 18.1888288 | 0.559802553 | 0.42405389 | 1.32012125 | 0.18679454 | NA |
| GTPBP10 | 12.9805457 | -0.232251221 | 0.43794109 | -0.5303253 | 0.59588644 | NA |
| CLDN12 | 20.2095414 | 0.730339422 | 0.37551879 | 1.94488117 | 0.05178928 | NA |
| CDK14 | 161.369725 | 0.046216101 | 0.29513914 | 0.15659089 | 0.87556728 | 0.96267305 |
| FZD1 | 9.5890847 | -1.531455118 | 0.83066882 | -1.843641 | 0.06523547 | NA |
| AC079760.1 | 15.9401901 | 0.111432796 | 0.42420656 | 0.26268523 | 0.7927932 | NA |
| MTERF1 | 51.7875234 | 0.130077507 | 0.3085395 | 0.4215911 | 0.6733235 | 0.90403541 |
| AKAP9 | 169.993026 | -0.215684435 | 0.19093398 | -1.1296283 | 0.25863287 | 0.65588339 |
| KRIT1 | 49.445502 | -0.189055726 | 0.24765796 | -0.7633743 | 0.44524019 | 0.78798633 |
| ANKIB1 | 88.1382181 | -0.04445887 | 0.28681796 | -0.1550073 | 0.87681558 | 0.96352317 |
| GATAD1 | 20.4586708 | -0.732260539 | 0.36514435 | -2.0054002 | 0.04492028 | NA |
| AC007566.1 | 15.7268285 | -0.175313381 | 0.47857706 | -0.3663222 | 0.7141247 | NA |
| PEX1 | 16.0377995 | 0.240854539 | 0.46621663 | 0.51661508 | 0.60542489 | NA |
| RBM48 | 6.31218395 | -0.561529885 | 0.56519746 | -0.993511 | 0.320461 | NA |
| FAM133B | 20.273273 | 0.047915773 | 0.30969127 | 0.1547211 | 0.8770412 | NA |
| CDK6 | 157.385195 | 0.517853267 | 0.35499354 | 1.45876813 | 0.14462894 | 0.5436645 |
| SAMD9 | 17.4353199 | -0.063891326 | 0.90803184 | -0.0703624 | 0.94390519 | NA |
| SAMD9L | 28.9419644 | 1.068307683 | 0.8092664 | 1.32009396 | 0.18680365 | 0.58871453 |
| VPS50 | 39.7435809 | 0.443199228 | 0.33285435 | 1.33151102 | 0.18302092 | 0.58584259 |
| GNGT1 | 11.3764886 | 1.137921244 | 0.87943036 | 1.29392991 | 0.19568963 | NA |
| BET1 | 15.8043156 | -0.181660558 | 0.42128418 | -0.4312067 | 0.66631809 | NA |
| AC002074.1 | 6.77642854 | -2.427912485 | 1.02107653 | -2.3777968 | 0.01741642 | NA |
| CASD1 | 32.219728 | 0.194982336 | 0.36942613 | 0.52779791 | 0.59763961 | 0.86882662 |
| SGCE | 40.4802851 | 0.378852472 | 0.37355351 | 1.01418527 | 0.31049435 | 0.69082013 |
| PPP1R9A | 95.4609321 | 0.536983316 | 0.2868883 | 1.87175051 | 0.06124113 | 0.38214464 |
| AC002429.2 | 6.94275643 | -1.130736919 | 0.88047367 | -1.2842371 | 0.19905902 | NA |
| PON2 | 87.1848329 | 0.2911278 | 0.28379045 | 1.02585483 | 0.30496002 | 0.6848669 |
| PDK4 | 102.626026 | 0.927231286 | 0.71899052 | 1.28962936 | 0.19717938 | 0.59987316 |
| DYNC1I1 | 127.248828 | 0.353088279 | 0.61837047 | 0.57099796 | 0.56800103 | 0.85654436 |
| SLC25A13 | 39.0499048 | -0.080285488 | 0.33982876 | -0.2362528 | 0.81323653 | 0.94431839 |
| SEM1 | 66.7793269 | -0.011946144 | 0.3047845 | -0.0391954 | 0.96873462 | 0.98964394 |
| SDHAF3 | 10.6621886 | -0.454171146 | 0.49823191 | -0.9115657 | 0.36199736 | NA |
| ASNS | 14.5171977 | 1.053514695 | 0.7321588 | 1.43891557 | 0.15017444 | NA |
| LMTK2 | 24.4752212 | 0.118508664 | 0.31346237 | 0.37806344 | 0.70538346 | NA |
| TECPR1 | 11.0948479 | -0.012711194 | 0.47417322 | -0.0268071 | 0.97861362 | NA |
| BRI3 | 75.4540115 | -0.089918703 | 0.27126228 | -0.3314825 | 0.74028004 | 0.92003444 |
| BAIAP2L1 | 70.4000967 | -0.52950046 | 0.48492551 | -1.0919212 | 0.27486773 | 0.66766283 |
| TMEM130 | 9.56659448 | 1.241792626 | 1.09941104 | 1.12950714 | 0.25868396 | NA |
| TRRAP | 55.3443968 | 0.251865661 | 0.25696323 | 0.98016225 | 0.32700603 | 0.69880742 |
| SMURF1 | 93.0587276 | 0.052664748 | 0.28979712 | 0.18172971 | 0.85579485 | 0.95615679 |
| ARPC1B | 12.298811 | 0.330375711 | 0.4756568 | 0.69456741 | 0.48732644 | NA |
| PDAP1 | 18.5760145 | -0.291195331 | 0.38409988 | -0.758124 | 0.44837676 | NA |
| BUD31 | 18.482546 | -0.455672731 | 0.42599867 | -1.0696576 | 0.28477344 | NA |
| CPSF4 | 14.1101185 | -0.218356901 | 0.50317288 | -0.43396 | 0.66431749 | NA |
| ZNF789 | 13.758777 | 0.16381082 | 0.42618503 | 0.3843655 | 0.70070757 | NA |
| ZNF394 | 10.5465323 | -0.400777963 | 0.55343169 | -0.7241688 | 0.46896211 | NA |
| ZKSCAN5 | 8.91351041 | 0.403196458 | 0.52274154 | 0.77131131 | 0.44052243 | NA |
| FAM200A | 8.20804835 | -0.49774728 | 0.59546049 | -0.8359031 | 0.40320941 | NA |
| ZSCAN25 | 4.21411838 | -0.778041505 | 0.65473091 | -1.1883378 | 0.23470034 | NA |

| | | | | | | |
|---|---|---|---|---|---|---|
| TRIM4 | 21.9050957 | -0.159292526 | 0.37394138 | -0.4259826 | 0.67012051 | NA |
| ZKSCAN1 | 57.9978905 | -0.149004612 | 0.27188377 | -0.5480453 | 0.58366082 | 0.86175324 |
| ZSCAN21 | 6.12383206 | -0.659334203 | 0.54596564 | -1.2076478 | 0.22718277 | NA |
| ZNF3 | 9.39225582 | 0.082287285 | 0.54214945 | 0.15177971 | 0.87936069 | NA |
| COPS6 | 23.5488555 | -0.110956536 | 0.36008384 | -0.3081408 | 0.75797517 | NA |
| MCM7 | 7.02801745 | 0.447410415 | 0.61521199 | 0.72724593 | 0.46707531 | NA |
| AP4M1 | 11.2152559 | -0.063603259 | 0.44931767 | -0.1415552 | 0.88743135 | NA |
| TAF6 | 11.4247679 | 0.157525492 | 0.5331921 | 0.29543853 | 0.7676589 | NA |
| CNPY4 | 5.40744608 | -0.438687317 | 0.65168756 | -0.6731559 | 0.50084812 | NA |
| LAMTOR4 | 64.687378 | -0.456101826 | 0.39457902 | -1.1559201 | 0.24771383 | 0.65061974 |
| CASTOR3 | 20.0501705 | 0.252900434 | 0.37360712 | 0.67691546 | 0.49845959 | NA |
| SPDYE3 | 13.8244444 | -0.779501738 | 0.49710754 | -1.5680747 | 0.11686371 | NA |
| PILRB | 22.725233 | -0.309891791 | 0.50977528 | -0.6078988 | 0.54325458 | NA |
| MEPCE | 11.086196 | 0.306757433 | 0.5022711 | 0.61074076 | 0.54137121 | NA |
| PPP1R35 | 12.9741236 | -0.483087593 | 0.49706622 | -0.9718777 | 0.33111138 | NA |
| TSC22D4 | 10.9716526 | 0.121348713 | 0.4822848 | 0.25161215 | 0.80134087 | NA |
| LRCH4 | 15.7291677 | 0.031975699 | 0.43607468 | 0.0733262 | 0.94154654 | NA |
| GNB2 | 42.9854881 | -0.804080991 | 0.42547928 | -1.8898241 | 0.05878149 | 0.37677048 |
| GIGYF1 | 15.5497359 | -0.006235964 | 0.41355958 | -0.0150788 | 0.98796935 | NA |
| POP7 | 11.4738976 | -0.550651332 | 0.50937298 | -1.0810376 | 0.2796804 | NA |
| SLC12A9 | 10.63793 | -0.051859641 | 0.46740978 | -0.1109511 | 0.9116551 | NA |
| TRIP6 | 17.1685153 | 0.340460211 | 0.5189417 | 0.6560664 | 0.5117814 | NA |
| SRRT | 12.190782 | -0.113595947 | 0.50384951 | -0.2254561 | 0.82162447 | NA |
| TRIM56 | 37.0574552 | -0.781126385 | 0.51773051 | -1.5087509 | 0.13136245 | 0.52613957 |
| AP1S1 | 11.7784721 | 0.085277922 | 0.50976609 | 0.16728834 | 0.86714318 | NA |
| PLOD3 | 25.3620699 | 0.052779993 | 0.35150512 | 0.15015426 | 0.88064291 | NA |
| ZNHIT1 | 29.0918857 | -0.369042048 | 0.38406259 | -0.9608904 | 0.33660729 | 0.71116888 |
| FIS1 | 60.1560555 | -0.66029413 | 0.38031208 | -1.7361903 | 0.0825302 | 0.43993543 |
| IFT22 | 9.02423126 | -0.741386478 | 0.49849101 | -1.4872615 | 0.13694577 | NA |
| CUX1 | 362.678159 | -0.328953552 | 0.27848077 | -1.1812433 | 0.23750607 | 0.64350343 |
| PRKRIP1 | 49.3896418 | -0.002222951 | 0.27982076 | -0.0079442 | 0.99366152 | 0.99833815 |
| ORAI2 | 12.6611224 | -0.37515236 | 0.64411513 | -0.5824306 | 0.56027667 | NA |
| ALKBH4 | 5.17379186 | -0.27046396 | 0.58452597 | -0.4627065 | 0.64357476 | NA |
| POLR2J | 17.2838032 | -0.515069158 | 0.49937457 | -1.0314285 | 0.30233993 | NA |
| AC105052.3 | 8.45525599 | 0.84188838 | 0.61356522 | 1.37212533 | 0.17002443 | NA |
| FAM185A | 28.4064366 | 0.204102578 | 0.4155651 | 0.49114465 | 0.62332414 | NA |
| FBXL13 | 22.3843272 | -1.17323628 | 0.41577268 | -2.8218215 | 0.00477517 | NA |
| ARMC10 | 29.2985614 | -1.168853468 | 0.416239 | -2.8081306 | 0.004983 | 0.11177304 |
| NAPEPLD | 137.697854 | -0.554779991 | 0.42131271 | -1.3167891 | 0.18790932 | 0.59036421 |
| PMPCB | 23.0088163 | 0.370470445 | 0.33587646 | 1.10299614 | 0.27002884 | NA |
| DNAJC2 | 14.1728666 | -1.13311472 | 0.53007341 | -2.1376562 | 0.03254465 | NA |
| PSMC2 | 22.8839051 | -0.172738581 | 0.38073563 | -0.4536969 | 0.65004698 | NA |
| ORC5 | 18.4174002 | -0.176948474 | 0.36395421 | -0.4861833 | 0.62683718 | NA |
| LHFPL3 | 13.0440066 | -0.687081837 | 0.45178369 | -1.5208204 | 0.12830492 | NA |
| LINC01004 | 68.8360132 | 0.482271633 | 0.25871713 | 1.86408852 | 0.06230927 | 0.38512471 |
| KMT2E-AS1 | 5.85109496 | -0.625927736 | 0.57683618 | -1.0851048 | 0.27787522 | NA |
| KMT2E | 145.846295 | 0.565448275 | 0.25088357 | 2.25382745 | 0.02420702 | 0.25196701 |
| SRPK2 | 110.603604 | 0.179074592 | 0.19950595 | 0.89759024 | 0.36940405 | 0.7377168 |
| PUS7 | 10.6463917 | -0.645296941 | 0.62286426 | -1.0360154 | 0.30019498 | NA |
| RINT1 | 7.32553959 | 0.088642479 | 0.51547246 | 0.17196356 | 0.86346618 | NA |
| EFCAB10 | 4.93893292 | -1.333064142 | 0.67144749 | -1.9853587 | 0.04710457 | NA |
| ATXN7L1 | 57.9210674 | -0.122046582 | 0.38801337 | -0.3145422 | 0.75310926 | 0.92313949 |
| CDHR3 | 15.1655358 | 0.36720353 | 0.66238198 | 0.55436823 | 0.57932686 | NA |
| SYPL1 | 35.2949565 | -0.085571395 | 0.38822444 | -0.2204173 | 0.82554615 | 0.949142 |
| NAMPT | 52.8296991 | -2.637396278 | 0.72088626 | -3.658547 | 0.00025365 | 0.01982419 |
| CCDC71L | 8.49065333 | 0.019140971 | 0.67780649 | 0.02823958 | 0.97747107 | NA |
| PRKAR2B | 92.0682141 | 0.616923243 | 0.34577703 | 1.78416491 | 0.07439687 | 0.41891814 |
| HBP1 | 36.0464018 | 0.576494571 | 0.28756992 | 2.00471094 | 0.04499396 | 0.33424083 |
| COG5 | 96.5318296 | 0.439882823 | 0.28901868 | 1.52198754 | 0.1280122 | 0.52341115 |
| BCAP29 | 37.4763936 | -0.198299738 | 0.32666925 | -0.6070352 | 0.54382754 | 0.84253801 |
| SLC26A4 | 7.26599045 | 0.263000247 | 0.81652472 | 0.3220971 | 0.74737913 | NA |
| CBLL1 | 9.67119673 | -0.021147481 | 0.47108123 | -0.0448914 | 0.9641939 | NA |
| SLC26A3 | 223.230793 | 1.206965571 | 0.61083744 | 1.97591944 | 0.0481639 | 0.34212397 |
| DLD | 23.7053249 | 0.592603743 | 0.344078 | 1.7222948 | 0.08501613 | NA |
| LAMB1 | 25.5079917 | -1.093650676 | 0.75288116 | -1.4526198 | 0.14632935 | NA |
| NRCAM | 9.78478128 | -0.95117075 | 0.70358885 | -1.3518843 | 0.17641232 | NA |
| PNPLA8 | 40.1798972 | -0.588173593 | 0.29407343 | -2.0000909 | 0.04549045 | 0.33547138 |
| DNAJB9 | 48.543369 | -0.418801871 | 0.58478991 | -0.7161578 | 0.4738939 | 0.80625572 |
| IMMP2L | 454.764663 | 0.394107774 | 0.28665112 | 1.37486913 | 0.16917202 | 0.57194586 |
| DOCK4 | 224.076372 | 0.623772803 | 0.21633129 | 2.88341465 | 0.00393389 | 0.09910477 |
| ZNF277 | 77.7809556 | 0.126342426 | 0.271789 | 0.46485482 | 0.64203543 | 0.88951638 |
| AC004112.1 | 4.64653404 | -0.308938028 | 0.79207822 | -0.3900347 | 0.69651086 | NA |
| IFRD1 | 31.8076703 | 0.134037795 | 0.3178558 | 0.42169372 | 0.67324859 | 0.90403541 |
| TMEM168 | 39.9744141 | 0.308613641 | 0.25131671 | 1.22798697 | 0.21945186 | 0.62462904 |
| BMT2 | 32.4785765 | 0.349202494 | 0.26292488 | 1.32814549 | 0.18413005 | 0.587049 |
| SMIM30 | 13.7290215 | -0.464569164 | 0.45772323 | -1.0149565 | 0.31012657 | NA |
| FOXP2 | 39.9758954 | 1.957863338 | 0.79893817 | 2.45058181 | 0.01426255 | 0.19811275 |
| MDFIC | 138.685437 | 0.138720119 | 0.34138945 | 0.40633979 | 0.68449295 | 0.90499319 |
| TES | 10.4894042 | 1.275941679 | 0.46505307 | 2.74364748 | 0.00607607 | NA |
| CAV2 | 5.78111775 | -0.049215672 | 0.66949654 | -0.0735115 | 0.94139912 | NA |
| MET | 13.4572841 | -1.405187417 | 0.80390763 | -1.7479464 | 0.08047331 | NA |
| CAPZA2 | 78.115973 | 0.56758413 | 0.26653049 | 2.12952793 | 0.03321061 | 0.29047359 |
| ST7 | 54.7222154 | 0.481411348 | 0.37089571 | 1.29796958 | 0.19429779 | 0.59865455 |
| CFTR | 8.99404126 | -0.555643426 | 0.49070148 | -1.1323451 | 0.25748937 | NA |
| CTTNBP2 | 182.201707 | 0.046393395 | 0.3044113 | 0.15240366 | 0.87886857 | 0.96423929 |
| LSM8 | 22.0027038 | -0.093017053 | 0.35989575 | -0.2584555 | 0.79605535 | NA |
| TSPAN12 | 12.2023811 | -0.067906471 | 0.47552782 | -0.1428023 | 0.88644631 | NA |
| ING3 | 9.20251401 | -0.131217895 | 0.54376052 | -0.2413156 | 0.80931052 | NA |
| FAM3C | 35.5465332 | -0.339956692 | 0.33587043 | -1.0121662 | 0.3114586 | 0.69107006 |
| AASS | 84.2748344 | 0.492000515 | 0.37107522 | 1.32587812 | 0.18488007 | 0.587049 |
| CADPS2 | 75.7417494 | 0.646362101 | 0.62361657 | 1.03647358 | 0.29998126 | 0.68168772 |
| SLC13A1 | 12.5810099 | 1.318342481 | 1.65298351 | 0.79755332 | 0.42512975 | NA |
| IQUB | 9.13564728 | 0.461622469 | 0.71963874 | 0.64146417 | 0.52122115 | NA |
| NDUFA5 | 33.5516226 | -0.614011333 | 0.37160957 | -1.6523023 | 0.09847295 | 0.47548368 |
| WASL | 230.187336 | -0.261858631 | 0.2306346 | -1.1353831 | 0.25621482 | 0.65588339 |
| AC006333.2 | 5.80532888 | -1.368453172 | 0.76840884 | -1.780892 | 0.07493009 | NA |
| POT1 | 24.2419831 | 0.166060939 | 0.31855216 | 0.52129905 | 0.60215846 | NA |
| POT1-AS1 | 13.8360863 | -0.327344317 | 0.56944519 | -0.5748478 | 0.56539423 | NA |
| ZNF800 | 47.2221641 | 0.087629801 | 0.2910958 | 0.30103424 | 0.76338839 | 0.92336765 |
| GCC1 | 3.47912383 | -0.289952394 | 0.72963008 | -0.3973964 | 0.69107514 | NA |
| ARF5 | 12.9811917 | -0.509675326 | 0.53496566 | -0.9527253 | 0.34072926 | NA |
| SND1 | 90.2335489 | -0.226057188 | 0.31988667 | -0.706679 | 0.47976599 | 0.81093801 |
| LRRC4 | 12.2106421 | -0.100588191 | 0.48130132 | -0.2089921 | 0.83445438 | NA |
| RBM28 | 14.8145606 | -0.51338031 | 0.41032042 | -1.2511693 | 0.21087272 | NA |
| IMPDH1 | 27.3210706 | 0.340444634 | 0.46149062 | 0.73770651 | 0.46069282 | NA |
| HILPDA | 20.3219887 | -1.889930052 | 0.68712262 | -2.750499 | 0.00595046 | NA |
| METTL2B | 17.2490255 | -0.424481615 | 0.50333495 | -0.8433382 | 0.3990393 | NA |
| CALU | 59.9561208 | -0.160275041 | 0.40266478 | -0.3980359 | 0.69060372 | 0.90557575 |
| FLNC | 9.47464473 | 1.329344362 | 0.51968421 | 2.55798491 | 0.01052807 | NA |
| KCP | 7.23986831 | 0.21342376 | 0.63436585 | 0.3364364 | 0.73654181 | NA |
| ATP6V1F | 38.3731165 | -0.564457753 | 0.45432219 | -1.2424173 | 0.21408263 | 0.62255988 |
| TNPO3 | 83.180393 | 0.322980927 | 0.32507832 | 0.99354804 | 0.32044295 | 0.69496685 |
| TSPAN33 | 26.915007 | 1.609433097 | 0.51970901 | 3.09679662 | 0.00195624 | NA |

| | | | | | | |
|---|---|---|---|---|---|---|
| SMO | 21.0137041 | 1.33468599 | 0.64512542 | 2.06887831 | 0.03855751 | NA |
| AHCYL2 | 51.0579391 | 0.748604737 | 0.26599841 | 2.81432034 | 0.00488805 | 0.11177304 |
| NRF1 | 59.2408548 | -0.012015609 | 0.27951653 | -0.0429871 | 0.96571181 | 0.98892249 |
| UBE2H | 106.10731 | 0.536988211 | 0.25198347 | 2.13104537 | 0.0330854 | 0.29047359 |
| ZC3HC1 | 10.2325605 | 0.860811043 | 0.49838697 | 1.72719412 | 0.08413281 | NA |
| KLHDC10 | 58.7286989 | -0.136629145 | 0.24845482 | -0.5499155 | 0.58237736 | 0.86175324 |
| TMEM209 | 10.2555902 | -0.436417439 | 0.55509511 | -0.786203 | 0.43174857 | NA |
| CEP41 | 6.26120513 | -0.057733968 | 0.58708984 | -0.0983392 | 0.92166292 | NA |
| COPG2 | 19.258629 | 0.507208074 | 0.39203289 | 1.2937896 | 0.19573811 | NA |
| AC016831.7 | 199.032634 | -0.394108893 | 0.34141706 | -1.1543327 | 0.24836378 | 0.6507516 |
| LINC00513 | 23.218458 | -1.274873574 | 0.7128053 | -1.7885299 | 0.07369056 | NA |
| AC058791.1 | 4.02082141 | -1.789280375 | 0.98304207 | -1.8201463 | 0.06873673 | NA |
| MKLN1 | 166.758051 | 0.207018759 | 0.29584946 | 0.69974358 | 0.48408745 | 0.81370025 |
| AC008264.2 | 58.8435898 | -0.076415463 | 0.71123797 | -0.1074401 | 0.91443986 | 0.97707273 |
| PODXL | 3571.08563 | -0.160084028 | 0.4142141 | -0.3864765 | 0.69914378 | 0.90729979 |
| AC009518.1 | 7.02137843 | -0.993518717 | 0.83636197 | -1.1879052 | 0.23487077 | NA |
| PLXNA4 | 65.528867 | -2.860756158 | 0.75680116 | -3.7800631 | 0.00015679 | 0.01469766 |
| CHCHD3 | 117.194399 | 0.137993781 | 0.20751911 | 0.66496903 | 0.5060703 | 0.82600448 |
| EXOC4 | 273.019262 | 0.30240052 | 0.24616551 | 1.22844392 | 0.21928037 | 0.62462904 |
| SLC35B4 | 13.7069849 | 0.750754916 | 0.38399559 | 1.95511339 | 0.05056969 | NA |
| AKR1B1 | 15.9812349 | 0.411635036 | 0.41283859 | 0.99708469 | 0.3187234 | NA |
| BPGM | 5.93885555 | -0.172152681 | 0.58520944 | -0.2941728 | 0.76862589 | NA |
| CALD1 | 179.372478 | -0.426786008 | 0.27982748 | -1.5251754 | 0.12721534 | 0.52301154 |
| AGBL3 | 7.61913741 | -0.469120287 | 0.72965441 | -0.6429349 | 0.52026634 | NA |
| CYREN | 11.7802471 | 0.02747016 | 0.42652644 | 0.06440435 | 0.94864826 | NA |
| TMEM140 | 13.4146967 | -0.222541133 | 0.91247757 | -0.2438867 | 0.80731858 | NA |
| WDR91 | 9.12818407 | 0.153897924 | 0.69883316 | 0.22022127 | 0.82569883 | NA |
| CNOT4 | 79.0053874 | -0.007846234 | 0.23171753 | -0.0338612 | 0.97298783 | 0.99106947 |
| NUP205 | 16.8126717 | 0.307713893 | 0.35989845 | 0.855002 | 0.39255004 | NA |
| STMP1 | 44.2637508 | 0.534930882 | 0.34343868 | 1.55757322 | 0.11933445 | 0.51061407 |
| SLC13A4 | 5.79259268 | -0.1242078 | 0.87009209 | -0.1427525 | 0.88648567 | NA |
| FAM180A | 61.5419299 | -0.731443471 | 0.96907333 | -0.7547865 | 0.45037709 | 0.79147725 |
| AC015987.1 | 3.72086887 | -0.866245789 | 1.16871026 | -0.7411981 | 0.45857336 | NA |
| MTPN | 50.6678751 | -0.203848951 | 0.29788634 | -0.6843179 | 0.49377445 | 0.82248426 |
| PTN | 4.52061187 | 1.503175192 | 1.22677843 | 1.22530292 | 0.22046111 | NA |
| CREB3L2 | 92.3300611 | 0.289858912 | 0.36055291 | 0.80392893 | 0.42143802 | 0.77486519 |
| TRIM24 | 26.1878865 | -0.006343312 | 0.40008524 | -0.0158549 | 0.98735015 | NA |
| KIAA1549 | 10.1267233 | 0.851062415 | 0.51182626 | 1.66279551 | 0.09635338 | NA |
| ZC3HAV1 | 34.1748583 | -0.359754557 | 0.47630032 | -0.7553103 | 0.45006279 | 0.79126774 |
| TTC26 | 10.6411432 | -0.108753681 | 0.43800386 | -0.2482939 | 0.80390703 | NA |
| UBN2 | 46.709625 | 0.591622564 | 0.28847669 | 2.05085048 | 0.04028151 | 0.31540889 |
| LUC7L2 | 25.1704518 | 0.498267621 | 0.33907944 | 1.46947165 | 0.1417049 | NA |
| HIPK2 | 351.939342 | -0.935306719 | 0.44330978 | -2.1098265 | 0.0348733 | 0.29312053 |
| TBXAS1 | 67.6975852 | -1.243586029 | 0.46623536 | -2.6672924 | 0.00764651 | 0.13970383 |
| PARP12 | 21.8315551 | -0.274486358 | 0.90405968 | -0.3036153 | 0.76142099 | NA |
| KDM7A | 31.3322695 | 0.175528264 | 0.349363 | 0.50242373 | 0.61536949 | 0.8757605 |
| SLC37A3 | 21.8336167 | 0.82993211 | 0.34886037 | 2.3789808 | 0.01736058 | NA |
| MKRN1 | 23.066804 | -0.000992574 | 0.34319588 | -0.0028922 | 0.9976924 | NA |
| NDUFB2 | 63.1141853 | -0.235209492 | 0.39237143 | -0.5994562 | 0.5488687 | 0.84550378 |
| BRAF | 148.520649 | 0.406426511 | 0.16972249 | 2.39465327 | 0.01663609 | 0.21392089 |
| MRPS33 | 10.4110119 | -0.034014314 | 0.46697574 | -0.0728396 | 0.94193378 | NA |
| TMEM178B | 8.51434963 | 0.368863434 | 0.9508979 | 0.38791066 | 0.69808215 | NA |
| AGK | 19.079063 | 0.250135896 | 0.35144157 | 0.71174248 | 0.47662426 | NA |
| KIAA1147 | 11.7476646 | 0.523848301 | 0.55878971 | 0.93746949 | 0.34851711 | NA |
| SSBP1 | 36.877984 | 0.105081345 | 0.32556303 | 0.32276805 | 0.74687091 | 0.92250642 |
| MGAM | 8.26580967 | 0.044187594 | 0.72602055 | 0.06086273 | 0.95146853 | NA |
| GSTK1 | 26.0425178 | -0.591803475 | 0.48456159 | -1.2213173 | 0.22196589 | NA |
| AC073342.1 | 4.94632565 | -0.175725003 | 0.57262625 | -0.3068756 | 0.75893809 | NA |
| TMEM139 | 8.38333275 | 0.610951635 | 0.58423294 | 1.04573296 | 0.29568434 | NA |
| CASP2 | 10.2774384 | 0.486030116 | 0.46899938 | 1.03631292 | 0.30005618 | NA |
| ZYX | 18.1232479 | 0.006208358 | 0.53422487 | 0.01162125 | 0.9907278 | NA |
| TCAF2 | 19.5814538 | -1.795451702 | 0.68719489 | -2.6127256 | 0.00898234 | NA |
| TCAF1 | 58.3788586 | -0.986288598 | 0.40335384 | -2.4452193 | 0.01447641 | 0.19867928 |
| AC004889.1 | 9.29204966 | 0.819996654 | 0.46684933 | 1.75644817 | 0.07901191 | NA |
| AC074386.1 | 14.4333586 | 0.456427556 | 0.50279755 | 0.90777602 | 0.36399657 | NA |
| TPK1 | 31.5820697 | -0.112602369 | 0.35038653 | -0.3213661 | 0.74793293 | 0.92253893 |
| CNTNAP2 | 21.5433427 | -1.206431568 | 0.59173737 | -2.0387956 | 0.04147043 | NA |
| CUL1 | 45.5410028 | -0.286829241 | 0.26253489 | -1.0925376 | 0.27459687 | 0.66766283 |
| PDIA4 | 26.6203901 | -0.06106113 | 0.41502388 | -0.1471268 | 0.88303195 | NA |
| ZNF786 | 5.56107202 | 0.024430689 | 0.61934067 | 0.03944629 | 0.96853458 | NA |
| ZNF398 | 13.8809496 | -0.677163064 | 0.38879301 | -1.7417059 | 0.08155992 | NA |
| ZNF282 | 10.6329081 | 0.082952297 | 0.4487518 | 0.18485118 | 0.85334577 | NA |
| ZNF212 | 3.49832703 | 0.064816303 | 0.85909595 | 0.07544711 | 0.93985898 | NA |
| ZNF783 | 9.81773608 | 0.805878528 | 0.48520935 | 1.66088828 | 0.09673589 | NA |
| ZNF777 | 5.74542408 | -0.172214303 | 0.64576712 | -0.2666817 | 0.78971421 | NA |
| ZNF746 | 10.1377698 | -0.197757666 | 0.47181811 | -0.4191396 | 0.67511409 | NA |
| ZNF467 | 7.81562685 | -1.711440363 | 0.64693566 | -2.6454568 | 0.00815807 | NA |
| ZNF862 | 20.3172458 | -0.167142328 | 0.3953069 | -0.4228166 | 0.67242906 | NA |
| ATP6V0E2 | 8.19425379 | 0.290341279 | 0.61436323 | 0.47258896 | 0.63650646 | NA |
| AC092666.1 | 6.85584439 | -1.260949855 | 0.72747686 | -1.7333195 | 0.08303889 | NA |
| ACTR3C | 25.0375833 | -0.113406514 | 0.4231118 | -0.2680297 | 0.78867648 | NA |
| LRRC61 | 8.08395764 | -0.342158361 | 0.60454095 | -0.5659805 | 0.57140707 | NA |
| RARRES2 | 118.113867 | 0.190976092 | 0.52275437 | 0.36532663 | 0.7148676 | 0.91376358 |
| AC073111.5 | 4.56325671 | 0.801567488 | 0.87624606 | 0.91477443 | 0.36031006 | NA |
| TMEM176B | 17.3312286 | -1.45041368 | 0.76228313 | -1.9027231 | 0.05707669 | NA |
| TMEM176A | 11.9258669 | -1.831333651 | 0.77456411 | -2.364341 | 0.01806218 | NA |
| KCNH2 | 8.29308679 | -0.414649366 | 0.60823317 | -0.6817276 | 0.49541119 | NA |
| ABCB8 | 11.4902605 | 0.466656069 | 0.47314097 | 0.98629394 | 0.32398889 | NA |
| SLC4A2 | 10.5676682 | 1.031032597 | 0.46245685 | 2.22946767 | 0.0257828 | NA |
| FASTK | 8.70448087 | 0.504590394 | 0.57901535 | 0.8714629 | 0.38350146 | NA |
| TMUB1 | 16.0216266 | 1.139180662 | 0.41077258 | 2.77326365 | 0.00554971 | NA |
| AGAP3 | 23.6510458 | 0.406580639 | 0.37534528 | 1.08321766 | 0.27871182 | NA |
| CHPF2 | 9.66023396 | -0.304956882 | 0.65766013 | -0.4636998 | 0.64286283 | NA |
| SMARCD3 | 11.8489527 | -0.967951106 | 0.53109026 | -1.8225736 | 0.06836799 | NA |
| NUB1 | 32.6792805 | -0.390051034 | 0.61004987 | -0.6393757 | 0.52257858 | 0.83120735 |
| RHEB | 108.150809 | -0.00017731 | 0.30361483 | -0.000584 | 0.99953404 | 0.99961632 |
| PRKAG2 | 27.7189137 | 0.19742973 | 0.62385116 | 0.31646929 | 0.75164633 | NA |
| GALNT11 | 46.9077587 | 0.072474824 | 0.31531589 | 0.22984831 | 0.81820965 | 0.94694381 |
| KMT2C | 187.369228 | 0.44917778 | 0.21272929 | 2.11149944 | 0.03472941 | 0.29285337 |
| ACTR3B | 9.74544597 | 0.475207996 | 0.53057924 | 0.89564001 | 0.37044506 | NA |
| DPP6 | 1174.66166 | 0.12719798 | 0.29928929 | 0.4250001 | 0.6708366 | 0.90305784 |
| AC006019.3 | 6.78666835 | -0.486292385 | 0.59542392 | -0.8167162 | 0.41409061 | NA |
| PAXIP1-AS2 | 4.41460587 | -0.442533431 | 0.72437914 | -0.6109141 | 0.54125645 | NA |
| PAXIP1 | 11.6937491 | 0.092499688 | 0.45151355 | 0.20486581 | 0.83767697 | NA |
| PAXIP1-AS1 | 6.3374154 | -0.493834393 | 0.70232719 | -0.7031401 | 0.48196846 | NA |
| INSIG1 | 14.1040906 | -1.099955842 | 0.48944872 | -2.2473362 | 0.02461855 | NA |
| AC009403.1 | 11.0350817 | -0.685172793 | 0.44829758 | -1.5283883 | 0.12641615 | NA |
| RBM33 | 78.5499148 | 0.388464813 | 0.2133251 | 1.82099909 | 0.068607 | 0.40491695 |
| LINC01006 | 25.534305 | -0.106846329 | 0.38111966 | -0.2803485 | 0.77921013 | NA |
| LMBR1 | 48.9021281 | 0.097992078 | 0.2508967 | 0.39056742 | 0.69611701 | 0.90718427 |
| NOM1 | 12.728447 | -0.537804925 | 0.42620803 | -1.2618367 | 0.20700756 | NA |
| UBE3C | 68.1426972 | 0.002132868 | 0.25902155 | 0.00823433 | 0.99343003 | 0.99833815 |
| DNAJB6 | 54.612002 | 0.028547571 | 0.25429431 | 0.11226193 | 0.91061573 | 0.97557165 |

| | | | | | | |
|---|---|---|---|---|---|---|
| PTPRN2 | 11.2546698 | -0.323329861 | 0.46056074 | -0.7020352 | 0.4826572 | NA |
| ESYT2 | 101.010561 | -0.151589123 | 0.2422436 | -0.6257714 | 0.53146487 | 0.83567299 |
| WDR60 | 64.422196 | -0.399063407 | 0.33036473 | -1.207948 | 0.22706727 | 0.63327644 |
| GTPBP6 | 16.2149442 | -0.370574045 | 0.37357805 | -0.9919588 | 0.32121761 | NA |
| LINC00685 | 8.47309791 | 0.427131447 | 0.53351972 | 0.80059167 | 0.42336807 | NA |
| IL3RA | 6.39002898 | -0.364939021 | 0.62830126 | -0.5808345 | 0.56135203 | NA |
| SLC25A6 | 52.5055711 | -0.654889959 | 0.40127462 | -1.6320244 | 0.10267435 | 0.48317363 |
| LINC00106 | 16.2220342 | 0.826298576 | 0.44565751 | 1.85411117 | 0.06372326 | NA |
| ASMTL-AS1 | 9.65587039 | 0.699818186 | 0.50546864 | 1.38449377 | 0.16620732 | NA |
| ASMTL | 24.0312592 | -0.301648437 | 0.45450692 | -0.6636828 | 0.50689334 | NA |
| AKAP17A | 14.0944952 | -0.251047408 | 0.40368551 | -0.6218886 | 0.53401513 | NA |
| DHRSX | 60.8737168 | -0.357690847 | 0.22917603 | -1.560769 | 0.11857826 | 0.51050744 |
| CD99 | 35.1372255 | -0.371928678 | 0.4719272 | -0.788106 | 0.43063468 | 0.77901411 |
| ARSD | 22.1034568 | -0.109847654 | 0.32688922 | -0.3360394 | 0.73684116 | NA |
| PRKX | 15.6658087 | 0.336346525 | 0.44905081 | 0.74901663 | 0.45384718 | NA |
| PUDP | 20.5275982 | -0.065405569 | 0.49026656 | -0.1334082 | 0.89387058 | NA |
| STS | 10.1609483 | -0.372408327 | 0.49412693 | -0.7536694 | 0.45104779 | NA |
| PNPLA4 | 6.35020283 | -0.145030784 | 0.71048179 | -0.2041302 | 0.83825176 | NA |
| TBL1X | 108.029819 | 0.222068652 | 0.26748034 | 0.83022421 | 0.40641203 | 0.76772515 |
| SHROOM2 | 26.99606 | 0.402692961 | 0.47793234 | 0.84257315 | 0.39946721 | NA |
| WWC3 | 28.3202612 | 0.262299507 | 0.43952214 | 0.59678337 | 0.55065202 | NA |
| MID1 | 13.7568976 | -0.880376274 | 0.67637812 | -1.3016037 | 0.19305189 | NA |
| ARHGAP6 | 62.4124992 | 1.133486088 | 0.5286411 | 2.1441505 | 0.03202083 | 0.28925721 |
| MSL3 | 4.14438242 | 0.729400372 | 0.65025597 | 1.1217127 | 0.26198462 | NA |
| PRPS2 | 6.20508174 | -0.358852236 | 0.62576173 | -0.5734647 | 0.56633012 | NA |
| TMSB4X | 1094.27771 | -0.682604279 | 0.48855928 | -1.397178 | 0.16236005 | 0.56332966 |
| RAB9A | 16.2093141 | -0.658416609 | 0.41762862 | -1.5765601 | 0.11489678 | NA |
| TRAPPC2 | 6.63962212 | 0.322609124 | 0.57455363 | 0.56149523 | 0.57445998 | NA |
| OFD1 | 25.4581566 | 0.198363875 | 0.48308675 | 0.4106175 | 0.68135303 | NA |
| GPM6B | 8.06420615 | -2.313902552 | 0.91589483 | -2.5263846 | 0.01152433 | NA |
| GEMIN8 | 16.4970883 | 0.243204421 | 0.38566095 | 0.63061718 | 0.52829086 | NA |
| MOSPD2 | 24.5105573 | -0.373295772 | 0.37572022 | -0.9935472 | 0.32044336 | NA |
| PIR | 23.3582535 | -0.034340267 | 0.41219416 | -0.0833109 | 0.93360433 | NA |
| BMX | 5.34685546 | -0.698348805 | 0.75338323 | -0.9269503 | 0.35395234 | NA |
| CLTRN | 11.1745266 | 0.4310194 | 1.12924751 | 0.38168727 | 0.70269334 | NA |
| CA5B | 19.9312391 | 0.070887109 | 0.50963184 | 0.13909474 | 0.88937528 | NA |
| ZRSR2 | 9.67695148 | -0.157764866 | 0.56689921 | -0.2782944 | 0.78078639 | NA |
| AP1S2 | 12.990048 | -0.119896875 | 0.50300987 | -0.2383589 | 0.81160275 | NA |
| CTPS2 | 25.2704385 | 0.799534954 | 0.41952658 | 1.90580286 | 0.05667579 | NA |
| SYAP1 | 23.9054407 | -0.316594517 | 0.32409279 | -0.9768638 | 0.32863658 | NA |
| TXLNG | 10.9728302 | -0.937011348 | 0.51150844 | -1.831859 | 0.06697243 | NA |
| RBBP7 | 27.6576471 | -0.197022667 | 0.30511899 | -0.645724 | 0.51845809 | NA |
| REPS2 | 14.985771 | -0.057499579 | 0.53527778 | -0.1074201 | 0.91445573 | NA |
| NHS | 59.2848734 | 0.930268149 | 0.54842428 | 1.69625632 | 0.08983735 | 0.45718982 |
| SCML1 | 15.2275938 | -0.818766186 | 0.42432652 | -1.9295664 | 0.05366059 | NA |
| RAI2 | 5.23197585 | 0.059294952 | 0.61860971 | 0.09585196 | 0.92363815 | NA |
| CDKL5 | 14.1785138 | 0.161670944 | 0.40138762 | 0.4027801 | 0.68711001 | NA |
| PHKA2 | 16.5286331 | 0.195885677 | 0.39545644 | 0.49534072 | 0.62035963 | NA |
| PDHA1 | 16.1165437 | -0.039222787 | 0.41338219 | -0.0948826 | 0.92440806 | NA |
| SH3KBP1 | 18.091164 | 0.082352127 | 0.66128533 | 0.12453343 | 0.90089294 | NA |
| BCLAF3 | 24.3468259 | 0.679309724 | 0.38967275 | 1.7432826 | 0.08128426 | NA |
| EIF1AX | 54.5111513 | -0.165781545 | 0.29202329 | -0.5676997 | 0.57023888 | 0.8573305 |
| RPS6KA3 | 36.2637405 | 0.0682788 | 0.28121545 | 0.2427989 | 0.80816119 | 0.94328109 |
| MBTPS2 | 18.9931856 | -0.37895702 | 0.3560089 | -1.0644594 | 0.28712063 | NA |
| SMS | 25.8760352 | 0.089627674 | 0.3031726 | 0.2956325 | 0.76751075 | NA |
| PRDX4 | 9.67420983 | -0.919040436 | 0.60540455 | -1.51806 | 0.12899927 | NA |
| ACOT9 | 18.9180302 | 0.413730386 | 0.39078529 | 1.05871535 | 0.28972944 | NA |
| SAT1 | 164.425283 | -3.004397267 | 0.56975715 | -5.273119 | 1.34E-07 | 7.77E-05 |
| APOO | 171.230023 | -0.534475128 | 0.31413488 | -1.7014192 | 0.08886429 | 0.4550929 |
| KLHL15 | 10.8602517 | 0.016667153 | 0.58717473 | 0.02838534 | 0.97735482 | NA |
| EIF2S3 | 21.117118 | -0.303497647 | 0.44642566 | -0.6798392 | 0.4966063 | NA |
| ZFX | 45.4147499 | 0.445879752 | 0.3642273 | 1.22417993 | 0.22088436 | 0.62519676 |
| POLA1 | 24.6006939 | -0.20134029 | 0.35906862 | -0.5607293 | 0.5749821 | NA |
| GK | 24.7667982 | 0.758936474 | 0.44816806 | 1.69341938 | 0.09037567 | NA |
| TAB3 | 28.2345814 | 0.206679098 | 0.3656901 | 0.56517553 | 0.57195438 | NA |
| DMD | 149.029331 | 0.869441603 | 0.40276277 | 2.15869405 | 0.03087391 | 0.28264481 |
| AL591501.1 | 14.2276869 | 2.433568139 | 0.61848647 | 3.93471525 | 8.33E-05 | NA |
| TMEM47 | 42.4165363 | 0.67740194 | 0.27699094 | 2.44557433 | 0.01446216 | 0.19867928 |
| AF241726.2 | 9.86464091 | 0.051248606 | 0.5952351 | 0.08609809 | 0.93138844 | NA |
| DYNLT3 | 37.4432003 | -0.262277918 | 0.43033183 | -0.6094783 | 0.54220744 | 0.84163543 |
| RPGR | 11.5451443 | -0.368034097 | 0.52384867 | -0.702558 | 0.48233122 | NA |
| MID1IP1 | 11.6851945 | -0.522383952 | 0.54961966 | -0.9504463 | 0.34188555 | NA |
| BCOR | 19.6414228 | 0.327043745 | 0.43964005 | 0.74388979 | 0.45694315 | NA |
| ATP6AP2 | 61.7312937 | 0.233336226 | 0.36723852 | 0.63538059 | 0.52518023 | 0.83195298 |
| CXorf38 | 23.1599671 | 0.516096782 | 0.39661098 | 1.301267 | 0.19316708 | NA |
| MED14 | 45.58845 | 0.484647946 | 0.36318621 | 1.33443378 | 0.18206173 | 0.58390314 |
| USP9X | 102.237722 | -0.040306943 | 0.27653734 | -0.1457559 | 0.88411411 | 0.96552688 |
| DDX3X | 65.2390618 | -0.235526748 | 0.22496244 | -1.0469603 | 0.29511789 | 0.68103704 |
| CASK | 55.5164549 | 0.405975028 | 0.29799637 | 1.36234889 | 0.1730878 | 0.57564735 |
| MAOA | 37.9970579 | -1.872826563 | 0.47693899 | -3.9267634 | 8.61E-05 | 0.00914516 |
| MAOB | 20.0922767 | 1.240020225 | 0.60845836 | 2.03797056 | 0.04155288 | NA |
| FUNDC1 | 10.789605 | 0.648073647 | 0.60873437 | 1.0646247 | 0.2870458 | NA |
| KDM6A | 93.9166545 | 0.669275718 | 0.48184121 | 1.38899643 | 0.16483384 | 0.56632803 |
| KRBOX4 | 15.4107012 | 0.396546949 | 0.39896433 | 0.99394086 | 0.32025166 | NA |
| ZNF674 | 4.67583681 | 0.685680727 | 0.76031692 | 0.90183542 | 0.3671443 | NA |
| SLC9A7 | 6.92486675 | -0.75369457 | 0.61255889 | -1.2304034 | 0.21854607 | NA |
| RP2 | 11.2409653 | 0.643613814 | 0.51075789 | 1.26011526 | 0.20762779 | NA |
| JADE3 | 10.4309389 | -0.419487833 | 0.54628561 | -0.7678911 | 0.44255191 | NA |
| RGN | 7.88270266 | -0.779857382 | 1.03368251 | -0.7544458 | 0.45058161 | NA |
| NDUFB11 | 45.5812321 | -0.103342418 | 0.43798922 | -0.2359474 | 0.81347349 | 0.94431839 |
| RBM10 | 26.6403016 | 0.526106034 | 0.40536301 | 1.29786393 | 0.1943341 | NA |
| UBA1 | 52.8178119 | 0.380951163 | 0.38670622 | 0.98511774 | 0.32456624 | 0.69623471 |
| CDK16 | 10.9413022 | -0.004267438 | 0.48462772 | -0.0088056 | 0.99297424 | NA |
| USP11 | 22.2185473 | 0.051048281 | 0.35231944 | 0.14489204 | 0.88479611 | NA |
| ZNF41 | 6.18792297 | 0.133539376 | 0.54521106 | 0.24493152 | 0.80650945 | NA |
| ARAF | 14.6519462 | 0.223700588 | 0.43920719 | 0.50932816 | 0.61052222 | NA |
| SYN1 | 9.31501833 | -2.852303427 | 0.78107941 | -3.6517458 | 0.00026046 | NA |
| TIMP1 | 445.439596 | -2.611760907 | 0.52772767 | -4.9490695 | 7.46E-07 | 0.00023577 |
| ELK1 | 13.6378004 | -0.270998432 | 0.54583018 | -0.4964885 | 0.61954977 | NA |
| UXT | 7.1416785 | -1.055814329 | 0.61483011 | -1.7172456 | 0.0859343 | NA |
| ZNF81 | 17.7592965 | 0.027079996 | 0.48530024 | 0.0558005 | 0.95550074 | NA |
| ZNF182 | 7.99620803 | 1.381537075 | 0.50359951 | 2.74332488 | 0.00608205 | NA |
| ZNF630 | 11.6431997 | 0.170781094 | 0.56953475 | 0.29986071 | 0.76428341 | NA |
| EBP | 9.31744279 | -0.44370241 | 0.494613 | -0.8970698 | 0.36968165 | NA |
| TBC1D25 | 5.12516419 | -0.366919573 | 0.58673098 | -0.6253625 | 0.53173315 | NA |
| RBM3 | 37.9072781 | -0.910653646 | 0.59626609 | -1.5272605 | 0.12669625 | 0.52276701 |
| WDR13 | 27.8255647 | 0.044817703 | 0.38381987 | 0.11676754 | 0.90704426 | NA |
| HDAC6 | 12.1289173 | 0.340661529 | 0.41165721 | 0.8275369 | 0.40793281 | NA |
| PCSK1N | 13.5611304 | -0.379981908 | 0.86290242 | -0.4403533 | 0.65968126 | NA |
| TIMM17B | 10.7226423 | -0.540303406 | 0.53676351 | -1.0065949 | 0.31412949 | NA |
| PQBP1 | 6.13903514 | 0.56414023 | 0.56947924 | 0.99062476 | 0.32186884 | NA |
| SLC35A2 | 6.42211503 | 0.031911571 | 0.64626139 | 0.04937874 | 0.96061747 | NA |
| OTUD5 | 28.924868 | 0.500755958 | 0.33396449 | 1.49942876 | 0.13376244 | 0.52673829 |

| | | | | | | |
|---|---|---|---|---|---|---|
| GRIPAP1 | 21.2262783 | -0.09955568 | 0.37323963 | -0.2667339 | 0.78967401 | NA |
| TFE3 | 25.7398291 | 0.310797707 | 0.30256833 | 1.02719839 | 0.30432706 | NA |
| PRAF2 | 5.92988086 | 0.042172122 | 0.76327375 | 0.05525163 | 0.95593799 | NA |
| WDR45 | 14.0769175 | 0.102667078 | 0.43855411 | 0.23410356 | 0.81490458 | NA |
| GPKOW | 3.40022281 | -0.248071003 | 0.75689831 | -0.3277468 | 0.74310311 | NA |
| PLP2 | 23.2253035 | -0.840765658 | 0.55909464 | -1.5037985 | 0.13263327 | NA |
| CCDC22 | 5.35384286 | -1.029610666 | 0.67961689 | -1.5149869 | 0.12977573 | NA |
| PPP1R3F | 7.98781747 | 0.381280383 | 0.53868112 | 0.7078035 | 0.47906729 | NA |
| CLCN5 | 17.8145413 | 0.764013035 | 0.66137951 | 1.15518098 | 0.24801632 | NA |
| SHROOM4 | 10.8927519 | 1.159914969 | 0.74572222 | 1.55542497 | 0.11984489 | NA |
| GSPT2 | 4.38623102 | -0.143511378 | 0.67168415 | -0.213659 | 0.83081297 | NA |
| MAGED1 | 56.0948225 | 0.164391196 | 0.23482733 | 0.70005138 | 0.48389522 | 0.81370025 |
| FAM156A | 6.86769405 | 0.58793177 | 0.51846494 | 1.13398558 | 0.2568006 | NA |
| GPR173 | 32.1644688 | 0.494226076 | 0.43079239 | 1.14724886 | 0.25127878 | 0.65183754 |
| TSPYL2 | 9.13803891 | 0.014805156 | 0.70060416 | 0.02113198 | 0.98314037 | NA |
| KANTR | 25.0679033 | 0.129611165 | 0.41278543 | 0.31399162 | 0.75352741 | NA |
| KDM5C | 16.3540629 | -0.239088789 | 0.45353752 | -0.5271643 | 0.5980795 | NA |
| IQSEC2 | 56.9839342 | -0.683248962 | 0.33001849 | -2.0703354 | 0.03842094 | 0.30615982 |
| SMC1A | 18.5935493 | -1.157186392 | 0.41865438 | -2.7640614 | 0.00570868 | NA |
| HSD17B10 | 16.1104049 | 0.078259718 | 0.42599045 | 0.18371238 | 0.8542391 | NA |
| HUWE1 | 76.304103 | 0.032608412 | 0.27117348 | 0.12024927 | 0.90428569 | 0.97237083 |
| PHF8 | 16.2464496 | 0.157576611 | 0.3963523 | 0.39756704 | 0.69094936 | NA |
| FAM120C | 16.2473075 | 0.173200447 | 0.43982988 | 0.39378964 | 0.69373636 | NA |
| TSR2 | 8.73536083 | -0.905673448 | 0.53796225 | -1.683526 | 0.09227331 | NA |
| FGD1 | 5.93544927 | -0.804359395 | 0.8063715 | -0.9975047 | 0.31851957 | NA |
| MAGED2 | 39.7533314 | -0.653339751 | 0.43165688 | -1.5135627 | 0.13013679 | 0.52613957 |
| APEX2 | 3.56436864 | -0.369256478 | 0.73843977 | -0.5000496 | 0.61704018 | NA |
| FAM104B | 6.44976582 | -0.33902814 | 0.60361749 | -0.5616606 | 0.57434731 | NA |
| MAGEH1 | 8.56215118 | 0.248146352 | 0.82526849 | 0.3006856 | 0.76365425 | NA |
| RRAGB | 22.9698564 | -0.397978115 | 0.35717289 | -1.114245 | 0.26517408 | NA |
| AL445472.1 | 30.1575773 | -0.656952297 | 0.3476526 | -1.8896804 | 0.05880072 | 0.37677048 |
| KLF8 | 10.3750403 | 0.756873873 | 0.41868148 | 1.80775582 | 0.07064451 | NA |
| UBQLN2 | 16.6118652 | -0.505170096 | 0.41623083 | -1.2136777 | 0.2248708 | NA |
| NBDY | 50.9599455 | -0.261788786 | 0.26201349 | -0.9991424 | 0.31772571 | 0.69434024 |
| SPIN3 | 9.71571017 | 0.027514611 | 0.43738389 | 0.06290723 | 0.94984037 | NA |
| FAAH2 | 17.6162467 | -0.440515461 | 0.44325241 | -0.9938253 | 0.32030793 | NA |
| SPIN4 | 5.40381537 | 0.573406393 | 0.68214274 | 0.8405959 | 0.40057435 | NA |
| ARHGEF9 | 17.8355846 | -0.267679482 | 0.35603227 | -0.7518405 | 0.45214699 | NA |
| MTMR8 | 8.51127623 | -0.103536715 | 0.55348805 | -0.1870622 | 0.85161183 | NA |
| ZC4H2 | 27.3019801 | 0.672525259 | 0.42868836 | 1.56879758 | 0.11669511 | NA |
| ZC3H12B | 18.0769599 | -0.20256755 | 0.49552842 | -0.408791 | 0.68269306 | NA |
| LAS1L | 7.93767867 | 0.333219556 | 0.48484118 | 0.68727569 | 0.49190901 | NA |
| MSN | 39.691738 | -0.026596765 | 0.37327824 | -0.0712518 | 0.94319732 | 0.98114803 |
| OPHN1 | 29.8491782 | 0.028656211 | 0.32418692 | 0.0883941 | 0.92956345 | 0.97777732 |
| YIPF6 | 35.3872596 | -0.302012551 | 0.3137095 | -0.9627141 | 0.33569103 | 0.71099887 |
| EFNB1 | 98.4660054 | 0.00109437 | 0.47394689 | 0.00230906 | 0.99815764 | 0.99961632 |
| PJA1 | 9.95771703 | 0.090454511 | 0.54956077 | 0.16459419 | 0.8692634 | NA |
| EDA | 30.381896 | 1.009079989 | 0.3604492 | 2.79950683 | 0.00511807 | 0.11257186 |
| IGBP1 | 15.4175826 | -0.677105456 | 0.38418068 | -1.7624662 | 0.07799056 | NA |
| PDZD11 | 7.6421985 | -0.755889962 | 0.5906764 | -1.2797023 | 0.20064985 | NA |
| SNX12 | 11.0105511 | 0.295272864 | 0.41797184 | 0.70644201 | 0.4799133 | NA |
| MED12 | 7.46336881 | 0.913221852 | 0.51872168 | 1.76052377 | 0.07831904 | NA |
| ZMYM3 | 8.25377889 | 0.620747363 | 0.4690108 | 1.32352466 | 0.18566096 | NA |
| NONO | 28.8397318 | -0.154539936 | 0.31040819 | -0.4978604 | 0.61858247 | 0.87662819 |
| TAF1 | 39.3418206 | -0.014001266 | 0.35916817 | -0.0389825 | 0.96890436 | 0.98964394 |
| OGT | 59.5011632 | 0.743698976 | 0.25557585 | 2.90989538 | 0.0036155 | 0.09771848 |
| NHSL2 | 12.3047094 | 0.784668951 | 0.60524008 | 1.296459 | 0.1948174 | NA |
| PIN4 | 18.8059821 | -0.566213784 | 0.40019172 | -1.4148563 | 0.15711063 | NA |
| RPS4X | 80.3458769 | -0.722560691 | 0.41093534 | -1.7583319 | 0.07869106 | 0.42784309 |
| HDAC8 | 77.9245659 | 0.141772137 | 0.25818081 | 0.54911958 | 0.58292339 | 0.86175324 |
| PHKA1 | 6.41115794 | -0.344655301 | 0.75976671 | -0.4536331 | 0.65009295 | NA |
| NAP1L2 | 6.10249021 | -0.181504097 | 0.67594028 | -0.2685209 | 0.78829838 | NA |
| CHIC1 | 40.8503269 | 0.759890882 | 0.32238733 | 2.35707431 | 0.01841956 | 0.22674781 |
| XIST | 126.618471 | 1.025558693 | 3.7412451 | 0.2741223 | 0.78399064 | 0.93497972 |
| JPX | 126.849851 | 0.033952532 | 0.42530318 | 0.07983136 | 0.93637138 | 0.97834166 |
| FTX | 794.584752 | 0.211903981 | 0.22813353 | 0.92885945 | 0.35296193 | 0.72416887 |
| SLC16A2 | 13.1387689 | -0.093077861 | 0.49320913 | -0.1887189 | 0.85031318 | NA |
| RLIM | 23.8719664 | -0.037620464 | 0.42007305 | -0.089557 | 0.92863929 | NA |
| NEXMIF | 34.9146084 | 0.129713831 | 0.66126409 | 0.1961604 | 0.84448462 | 0.95509854 |
| ABCB7 | 27.576342 | 0.319316392 | 0.28920211 | 1.10412884 | 0.26953725 | NA |
| UPRT | 6.83005756 | -0.113831438 | 0.53097982 | -0.21438 | 0.83025076 | NA |
| ZDHHC15 | 11.0747057 | 0.152043138 | 0.58060809 | 0.26186879 | 0.7934226 | NA |
| PBDC1 | 6.61717445 | -1.007732997 | 0.71929208 | -1.4010067 | 0.16121208 | NA |
| ATRX | 157.810038 | 0.242335685 | 0.23249182 | 1.0423407 | 0.29725375 | 0.68103704 |
| MAGT1 | 59.3341747 | -0.09685015 | 0.44935901 | -0.2155296 | 0.82935447 | 0.95008863 |
| COX7B | 47.0405737 | -0.27036675 | 0.39772981 | -0.6797749 | 0.49664699 | 0.8234249 |
| PGK1 | 71.97979 | -0.290208659 | 0.32527808 | -0.8921864 | 0.37229306 | 0.74056922 |
| TAF9B | 7.77817756 | -0.355640528 | 0.5160824 | -0.6891158 | 0.49075041 | NA |
| BRWD3 | 37.1855173 | 0.328596364 | 0.29963349 | 1.096661 | 0.27278961 | 0.66531666 |
| SH3BGRL | 53.3952309 | 0.132844302 | 0.29639201 | 0.44820473 | 0.65400545 | 0.8949739 |
| RPS6KA6 | 39.575818 | 0.988271703 | 0.39970348 | 2.47251215 | 0.01341672 | 0.19297233 |
| HDX | 13.4922361 | -0.479319533 | 0.45858838 | -1.0452064 | 0.29592756 | NA |
| APOOL | 23.977557 | 0.074145196 | 0.30555533 | 0.24265718 | 0.80827098 | NA |
| CHM | 41.9349474 | 0.151347555 | 0.29885356 | 0.50642714 | 0.61255682 | 0.87391153 |
| DACH2 | 279.679585 | 0.402186424 | 0.7006135 | 0.57404892 | 0.5659347 | 0.85573063 |
| DIAPH2 | 93.8590648 | 0.255934172 | 0.24376176 | 1.0499357 | 0.29374768 | 0.6804998 |
| DIAPH2-AS1 | 7.5521055 | -0.096840255 | 0.82700063 | -0.1170982 | 0.90678227 | NA |
| TSPAN6 | 10.9076083 | -0.676133776 | 0.56944101 | -1.187364 | 0.23508406 | NA |
| SRPX2 | 6.26319471 | -0.792247151 | 0.74657843 | -1.0611707 | 0.28861234 | NA |
| SYTL4 | 34.1991878 | -0.147595602 | 0.37358946 | -0.3950743 | 0.69278804 | 0.90557575 |
| TRMT2B | 15.2060918 | 0.155003508 | 0.44024013 | 0.35208855 | 0.72477186 | NA |
| HNRNPH2 | 23.6664171 | -0.133137086 | 0.32070565 | -0.415138 | 0.67804092 | NA |
| ARMCX4 | 63.7973117 | -0.057474054 | 0.33821739 | -0.1699323 | 0.86506339 | 0.96089268 |
| ARMCX1 | 10.0472193 | -0.738198146 | 0.52791474 | -1.3983283 | 0.16201449 | NA |
| ARMCX6 | 10.0637302 | -0.062744354 | 0.62744132 | -0.1000004 | 0.92034404 | NA |
| ARMCX3 | 44.3566066 | -0.538863165 | 0.40349343 | -1.3354943 | 0.18171463 | 0.58390314 |
| ARMCX2 | 13.5528751 | 0.462725061 | 0.48150817 | 0.96099109 | 0.33655665 | NA |
| ZMAT1 | 50.6677591 | 0.362825725 | 0.36384775 | 0.99719106 | 0.31867178 | 0.69469143 |
| TCEAL2 | 29.8659894 | -1.335575956 | 0.45339114 | -2.9457478 | 0.00322175 | 0.09202406 |
| BEX5 | 5.44006254 | -0.369487236 | 0.66604813 | -0.5547456 | 0.57906871 | NA |
| GPRASP1 | 4.86461704 | -0.423196404 | 0.67530973 | -0.6266701 | 0.53087552 | NA |
| BHLHB9 | 6.8091511 | -0.422657548 | 0.67434889 | -0.6267639 | 0.530814 | NA |
| LINC00630 | 19.2247364 | 0.702495254 | 0.39758 | 1.76692803 | 0.07724028 | NA |
| BEX4 | 66.2103548 | -0.950776927 | 0.47750563 | -1.9911324 | 0.04646634 | 0.33771838 |
| TCEAL8 | 24.0213193 | -0.133058427 | 0.42471769 | -0.3132868 | 0.75406281 | NA |
| BEX2 | 20.9504128 | -1.705020262 | 0.71467896 | -2.3857149 | 0.01704596 | NA |
| TCEAL9 | 57.7001126 | -0.377404511 | 0.36892774 | -1.0229768 | 0.30631883 | 0.68566731 |
| BEX3 | 131.468917 | -0.328835148 | 0.37159614 | -0.8849262 | 0.37619645 | 0.74330391 |
| TCEAL4 | 50.3184741 | -0.327902103 | 0.31357642 | -1.0456848 | 0.29570658 | 0.68103704 |
| TCEAL3 | 30.7941436 | -0.57294154 | 0.3887752 | -1.4737091 | 0.14055994 | 0.53597868 |
| TCEAL1 | 6.98958845 | -0.321649282 | 0.5248982 | -0.6127841 | 0.5400191 | NA |
| MORF4L2 | 47.2865114 | 0.012836976 | 0.33418854 | 0.03841238 | 0.96935889 | 0.98977467 |
| SLC25A53 | 13.6911717 | -0.308680296 | 0.39481233 | -0.7818406 | 0.43430827 | NA |

| | | | | | | |
|---|---|---|---|---|---|---|
| FAM199X | 33.648286 | 0.281770152 | 0.29689736 | 0.94904904 | 0.34259567 | 0.71725052 |
| IL1RAPL2 | 7.72041034 | 2.217416793 | 0.65824437 | 3.36868327 | 0.00075528 | NA |
| RNF128 | 7.34096327 | 1.33477795 | 0.83909822 | 1.59072908 | 0.11167056 | NA |
| TBC1D8B | 13.2350959 | 0.055420058 | 0.53562628 | 0.10346777 | 0.91759173 | NA |
| MORC4 | 11.373668 | -0.560849847 | 0.54600212 | -1.0271935 | 0.30432934 | NA |
| RBM41 | 23.3566322 | 0.270278731 | 0.36631814 | 0.73782513 | 0.46062072 | NA |
| PRPS1 | 8.8899599 | -0.314683163 | 0.59775656 | -0.5264403 | 0.5985823 | NA |
| TSC22D3 | 170.394669 | -0.23185346 | 0.33645822 | -0.6891003 | 0.49076015 | 0.81880839 |
| MID2 | 12.546864 | 0.233729776 | 0.48885553 | 0.47811625 | 0.63256747 | NA |
| PSMD10 | 6.39625472 | 0.144304106 | 0.60534151 | 0.23838462 | 0.8115828 | NA |
| ATG4A | 5.39761117 | 0.068687779 | 0.68327807 | 0.10052683 | 0.91992608 | NA |
| COL4A6 | 12.6515317 | -0.031391899 | 0.68396776 | -0.0458968 | 0.96339254 | NA |
| COL4A5 | 195.267143 | 0.354111063 | 0.24263032 | 1.45946747 | 0.14443649 | 0.54344563 |
| ACSL4 | 100.607704 | -0.278825358 | 0.28869128 | -0.9658254 | 0.33413157 | 0.70906302 |
| TMEM164 | 31.374876 | -0.874682255 | 0.42624195 | -2.0520792 | 0.04016196 | 0.31540889 |
| AMMECR1 | 23.5269939 | -0.251390041 | 0.33460978 | -0.7512932 | 0.45247625 | NA |
| ALG13 | 47.2154358 | 0.452035729 | 0.34212132 | 1.32127321 | 0.18641028 | 0.58835886 |
| AMOT | 19.3626877 | 0.9364018 | 0.68162506 | 1.37377843 | 0.16951048 | NA |
| LRCH2 | 14.5104179 | 0.393955582 | 0.42876565 | 0.91881331 | 0.35819323 | NA |
| PLS3-AS1 | 6.07981968 | -0.509099471 | 0.7766718 | -0.6554886 | 0.51215325 | NA |
| PLS3 | 122.593121 | -0.231829183 | 0.39651337 | -0.5846693 | 0.55877012 | 0.8520194 |
| DANT2 | 99.9466237 | -0.117692241 | 0.37033182 | -0.3178021 | 0.75063503 | 0.92274989 |
| KLHL13 | 25.1572602 | -0.236392804 | 0.6330282 | -0.3734317 | 0.70882716 | NA |
| WDR44 | 30.4422418 | 0.063945564 | 0.31520421 | 0.20287027 | 0.83923643 | 0.95393706 |
| IL13RA1 | 60.0546588 | -1.206280444 | 0.50141296 | -2.4057624 | 0.01613876 | 0.20980383 |
| LONRF3 | 11.2017477 | 0.117303468 | 0.73112442 | 0.16044255 | 0.87253247 | NA |
| PGRMC1 | 54.126031 | -0.591433844 | 0.41114251 | -1.438513 | 0.15028855 | 0.54834287 |
| SLC25A43 | 31.4851148 | -0.153100705 | 0.28335575 | -0.5403127 | 0.58898142 | 0.86533208 |
| SLC25A5 | 28.6211931 | -0.728337404 | 0.44867534 | -1.6233061 | 0.10452399 | 0.4862279 |
| CXorf56 | 6.15192948 | 0.083132314 | 0.53752642 | 0.15465717 | 0.8770916 | NA |
| UBE2A | 18.2345659 | 0.080510578 | 0.42536886 | 0.18927238 | 0.84987934 | NA |
| NKRF | 4.60830307 | -0.140535106 | 0.61240785 | -0.2294796 | 0.81849618 | NA |
| 6-Sep | 8.41433991 | -0.67202854 | 0.59227703 | -1.1346524 | 0.256521 | NA |
| RPL39 | 5.24190269 | 0.494040908 | 0.7629493 | 0.64754094 | 0.51728191 | NA |
| UPF3B | 15.7890136 | -0.481419865 | 0.39693558 | -1.2128413 | 0.2251905 | NA |
| NDUFA1 | 40.9129675 | -0.365104841 | 0.38762582 | -0.9419002 | 0.34624374 | 0.71908069 |
| NKAP | 11.1640351 | -1.147992868 | 0.51343717 | -2.2358975 | 0.02535849 | NA |
| RHOXF1-AS1 | 9.3767426 | 0.401780187 | 0.58349129 | 0.68857958 | 0.49108788 | NA |
| ZBTB33 | 4.63696161 | 0.530188653 | 0.60377997 | 0.87811567 | 0.37988095 | NA |
| LAMP2 | 63.4035984 | -0.083869903 | 0.30253298 | -0.2772257 | 0.78160684 | 0.93415197 |
| CUL4B | 19.9768238 | -0.154484945 | 0.33078212 | -0.4670293 | 0.6404789 | NA |
| MCTS1 | 18.3732624 | 0.039741688 | 0.36459342 | 0.10900275 | 0.91320031 | NA |
| C1GALT1C1 | 12.699059 | -0.286340327 | 0.61450993 | -0.4659653 | 0.64124031 | NA |
| THOC2 | 60.4177809 | 0.255919464 | 0.24518588 | 1.04377732 | 0.29658843 | 0.68103704 |
| XIAP | 68.6248055 | 0.030652206 | 0.24792774 | 0.12363363 | 0.90160537 | 0.97206575 |
| STAG2 | 62.3710165 | 0.029924145 | 0.25958142 | 0.11527845 | 0.90822442 | 0.9747971 |
| TENM1 | 8.75576872 | 0.965684828 | 0.62471363 | 1.54580401 | 0.12215191 | NA |
| SMARCA1 | 32.5726602 | 0.390940175 | 0.33880857 | 1.15386742 | 0.24855453 | 0.65083096 |
| OCRL | 10.8354991 | -0.142067472 | 0.43563907 | -0.3261128 | 0.74433902 | NA |
| APLN | 26.5937421 | -0.771571263 | 0.77381141 | -0.997105 | 0.31871353 | NA |
| ZDHHC9 | 13.3913643 | 0.467758264 | 0.4359782 | 1.07289371 | 0.28331881 | NA |
| UTP14A | 6.1493454 | 0.287125786 | 0.66175547 | 0.43388502 | 0.66437194 | NA |
| BCORL1 | 8.43629769 | -0.266910568 | 0.55436227 | -0.4814732 | 0.63018023 | NA |
| ELF4 | 8.15133475 | 0.365362287 | 0.64391246 | 0.56740987 | 0.57043574 | NA |
| AIFM1 | 13.7118714 | -0.077988964 | 0.38326315 | -0.2034867 | 0.83875462 | NA |
| ZNF280C | 6.61686573 | -0.443925545 | 0.52167119 | -0.8509681 | 0.39478707 | NA |
| SLC25A14 | 7.27100301 | 0.979013416 | 0.65406696 | 1.49680915 | 0.13444293 | NA |
| RBMX2 | 7.45770153 | 0.934798638 | 0.62941008 | 1.48519807 | 0.13749136 | NA |
| ENOX2 | 24.4381064 | -0.065657939 | 0.50984692 | -0.1287797 | 0.89753196 | NA |
| RAP2C | 16.4940189 | 0.827924164 | 0.49077119 | 1.68698609 | 0.09160603 | NA |
| RAP2C-AS1 | 37.3432891 | -0.188287448 | 0.38300252 | -0.4916089 | 0.62299585 | 0.8780438 |
| MBNL3 | 66.3021998 | -0.249564952 | 0.32225011 | -0.7744449 | 0.43866774 | 0.78335165 |
| GPC4 | 18.4020076 | -0.786632117 | 0.38963989 | -2.0188696 | 0.04350078 | NA |
| GPC3 | 11.7055912 | -1.093656143 | 0.71020459 | -1.539917 | 0.12358058 | NA |
| PHF6 | 13.6624004 | 0.042293278 | 0.38980744 | 0.10849787 | 0.91360077 | NA |
| HPRT1 | 25.5135255 | -0.686161127 | 0.34166249 | -2.008301 | 0.04461131 | NA |
| MIR503HG | 39.0707995 | -2.11100052 | 0.60370372 | -3.4967492 | 0.00047096 | 0.02938818 |
| FAM122B | 11.3745601 | 0.725868129 | 0.42425207 | 1.71093597 | 0.08709293 | NA |
| FAM122C | 5.84536813 | -0.569099883 | 0.60241791 | -0.9446928 | 0.34481571 | NA |
| MOSPD1 | 4.94192062 | 0.348424925 | 0.66557133 | 0.52349749 | 0.60062809 | NA |
| RTL8B | 5.80922171 | -1.05909696 | 0.70519639 | -1.5018468 | 0.13313667 | NA |
| RTL8C | 38.156373 | -0.638488888 | 0.46366565 | -1.3770459 | 0.16849805 | 0.57142817 |
| RTL8A | 27.0599898 | -1.014381906 | 0.42274627 | -2.3995053 | 0.01641724 | NA |
| ZNF75D | 17.8780125 | 0.029992483 | 0.43974904 | 0.06820364 | 0.94562353 | NA |
| ZNF449 | 5.36215137 | -1.004021054 | 0.65670035 | -1.5288877 | 0.1262923 | NA |
| INTS6L | 14.1047555 | 0.101611317 | 0.48138053 | 0.21108315 | 0.83282239 | NA |
| MMGT1 | 10.6663551 | -0.901680636 | 0.61566607 | -1.4645612 | 0.14304067 | NA |
| SLC9A6 | 11.4199444 | -0.196229778 | 0.49448828 | -0.396834 | 0.69148985 | NA |
| FHL1 | 45.8703382 | -0.515571384 | 0.48576382 | -1.0613623 | 0.2885253 | 0.67684131 |
| MAP7D3 | 10.1836706 | -0.409858918 | 0.48713081 | -0.8413734 | 0.40013876 | NA |
| HTATSF1 | 12.2283906 | -0.516242662 | 0.48688537 | -1.0602961 | 0.28900991 | NA |
| AL683813.1 | 16.8617337 | -0.120205334 | 0.41876751 | -0.2870455 | 0.77407747 | NA |
| RBMX | 55.8548047 | -0.12467166 | 0.24197925 | -0.5152163 | 0.60640186 | 0.87080602 |
| FGF13 | 75.3884002 | -1.292258828 | 0.82937893 | -1.5581042 | 0.11920854 | 0.51061407 |
| ATP11C | 58.1194405 | -0.13663985 | 0.45753844 | -0.2986412 | 0.76521379 | 0.9240942 |
| LDOC1 | 4.56168965 | -0.361120985 | 0.86177893 | -0.4190413 | 0.67518593 | NA |
| SLITRK2 | 18.9356173 | 0.135275061 | 0.67569221 | 0.20020219 | 0.84132245 | NA |
| FMR1 | 25.4912464 | 0.20172278 | 0.41428175 | 0.48692172 | 0.62631381 | NA |
| IDS | 34.6767545 | -0.168773826 | 0.48627348 | -0.3470759 | 0.72853426 | 0.91700494 |
| TMEM185A | 14.5668611 | -0.014885952 | 0.4185469 | -0.0355658 | 0.97162859 | NA |
| LINC00894 | 98.4509785 | 0.104890213 | 0.32659171 | 0.32116619 | 0.74808446 | 0.92253893 |
| MAMLD1 | 11.1961795 | -1.894194991 | 0.56373273 | -3.360094 | 0.00077916 | NA |
| MTM1 | 19.1067718 | 0.120741807 | 0.36413652 | 0.3315839 | 0.74020348 | NA |
| MTMR1 | 22.5763514 | -0.102350054 | 0.31360065 | -0.3263707 | 0.74414392 | NA |
| CD99L2 | 24.0228614 | -0.0712014 | 0.39829392 | -0.178766 | 0.85812147 | NA |
| VMA21 | 17.1329612 | 0.072655796 | 0.51824013 | 0.14019717 | 0.88850421 | NA |
| CETN2 | 14.801482 | -0.867107808 | 0.56583261 | -1.5324458 | 0.12541246 | NA |
| ZNF185 | 58.6171598 | -0.841445047 | 0.41760601 | -2.0149256 | 0.04391244 | 0.33029013 |
| BGN | 183.501305 | -1.328989743 | 0.51334551 | -2.5888796 | 0.00962887 | 0.15497902 |
| CCNQ | 7.70219857 | -0.44454966 | 0.69880963 | -0.6361527 | 0.52467688 | NA |
| SLC6A8 | 8.59112068 | -0.802427717 | 0.63176479 | -1.2701368 | 0.2040359 | NA |
| BCAP31 | 31.0468434 | -0.075452192 | 0.40241802 | -0.1874971 | 0.85127093 | 0.95532127 |
| IDH3G | 7.97574585 | -0.116408202 | 0.47929899 | -0.2428718 | 0.80810472 | NA |
| SSR4 | 30.7382174 | -0.707783378 | 0.60675167 | -1.1665124 | 0.24340732 | 0.64918976 |
| HCFC1 | 12.8626378 | 0.44109353 | 0.40902045 | 1.07841436 | 0.28084888 | NA |
| TMEM187 | 4.32289698 | -0.01912408 | 0.74180383 | -0.0257805 | 0.97943241 | NA |
| IRAK1 | 9.61095944 | -0.380727611 | 0.62410281 | -0.6100399 | 0.54183538 | NA |
| MECP2 | 56.5417618 | 0.014883104 | 0.27287609 | 0.05454162 | 0.95650365 | 0.98641719 |
| FLNA | 144.621285 | 0.088235834 | 0.43981657 | 0.20061962 | 0.84099602 | 0.95393706 |
| EMD | 17.9720564 | -0.401081296 | 0.36873396 | -1.0877254 | 0.27671634 | NA |
| RPL10 | 152.66321 | -1.22710062 | 0.51400836 | -2.3873165 | 0.01697187 | 0.21530523 |
| DNASE1L1 | 5.66876696 | -0.527449591 | 0.6171919 | -0.8545958 | 0.39277497 | NA |
| TAZ | 13.0440684 | -0.008114165 | 0.45978269 | -0.0176478 | 0.9859198 | NA |

| | | | | | | |
|---|---|---|---|---|---|---|
| ATP6AP1 | 52.3288052 | -0.167070992 | 0.49344616 | -0.33858 | 0.73492617 | 0.91905659 |
| GDI1 | 31.3917929 | -0.039784547 | 0.36681366 | -0.1084598 | 0.91363095 | 0.97707273 |
| FAM50A | 16.561732 | 0.114997608 | 0.55079915 | 0.2087832 | 0.83461749 | NA |
| PLXNA3 | 22.6468852 | -0.097455928 | 0.54701922 | -0.1781581 | 0.85859878 | NA |
| LAGE3 | 3.46932789 | -2.301688671 | 0.89513197 | -2.57134 | 0.01013058 | NA |
| UBL4A | 9.18646364 | -0.392229248 | 0.54737976 | -0.7165578 | 0.47364698 | NA |
| SLC10A3 | 4.61941506 | 0.202605162 | 0.6770422 | 0.29925042 | 0.76474898 | NA |
| FAM3A | 13.4449704 | -0.081353569 | 0.49683644 | -0.1637432 | 0.86993334 | NA |
| G6PD | 5.10847113 | -0.149465662 | 0.61868413 | -0.2415864 | 0.80910067 | NA |
| IKBKG | 6.50277739 | 1.041319798 | 0.52269637 | 1.99220782 | 0.04634826 | NA |
| DKC1 | 8.4969025 | -0.793464458 | 0.49486676 | -1.6033901 | 0.10884856 | NA |
| MPP1 | 4.95444001 | 0.712967224 | 0.61878484 | 1.15220538 | 0.24923669 | NA |
| F8 | 7.50688856 | 0.340207271 | 0.49479079 | 0.68757802 | 0.49171855 | NA |
| F8A1 | 4.67025875 | -0.604280225 | 0.75493158 | -0.8004437 | 0.42345377 | NA |
| FUNDC2 | 21.5717727 | -0.312680477 | 0.40038858 | -0.7809425 | 0.43483629 | NA |
| BRCC3 | 11.6082893 | -0.466912427 | 0.45770187 | -1.0201235 | 0.3076699 | NA |
| VBP1 | 14.54422 | -0.570796822 | 0.49471972 | -1.1537782 | 0.24859112 | NA |
| CLIC2 | 5.43847191 | -0.939228102 | 0.79228268 | -1.1854709 | 0.2358313 | NA |
| TMLHE-AS1 | 9.18117492 | -0.483743865 | 0.71407612 | -0.6774402 | 0.4981267 | NA |
| TMLHE | 23.1410817 | -0.000904244 | 0.35026854 | -0.0025816 | 0.99794021 | NA |
| VAMP7 | 13.6229292 | -0.305831678 | 0.42739251 | -0.7155757 | 0.4742534 | NA |
| ZNF596 | 9.64094305 | -0.424879064 | 0.48950824 | -0.8679712 | 0.38541009 | NA |
| FBXO25 | 27.4190588 | -0.293449218 | 0.4336071 | -0.676763 | 0.49855637 | NA |
| TDRP | 18.0600916 | 0.117282929 | 0.37154917 | 0.31565925 | 0.75226116 | NA |
| ERICH1 | 47.8848209 | -0.237101224 | 0.35756421 | -0.6631011 | 0.5072658 | 0.82657837 |
| AC100810.1 | 7.89146149 | -0.227454831 | 0.70574124 | -0.3222921 | 0.74723141 | NA |
| KBTBD11 | 29.7823483 | -0.797751069 | 0.48875655 | -1.6322054 | 0.10263623 | 0.48317363 |
| AC246817.1 | 15.7084795 | 0.639442837 | 0.52644634 | 1.21464011 | 0.22450338 | NA |
| CSMD1 | 29.5968763 | 2.733126387 | 0.95980292 | 2.84759125 | 0.00440515 | 0.10635282 |
| MCPH1 | 103.247469 | 0.268050781 | 0.23742069 | 1.12901186 | 0.25889283 | 0.65588339 |
| ANGPT2 | 14.4139504 | -0.79500129 | 0.65367476 | -1.2162031 | 0.22390755 | NA |
| MCPH1-AS1 | 11.2445311 | -0.182435883 | 0.55480999 | -0.3288259 | 0.7422873 | NA |
| AGPAT5 | 10.8757191 | -0.393992417 | 0.48784325 | -0.8076209 | 0.41930884 | NA |
| DEFB1 | 38.7193772 | -0.414599989 | 0.84718375 | -0.4893861 | 0.62456834 | 0.87929511 |
| FAM85B | 46.0971767 | 0.514752306 | 0.50155254 | 1.02631782 | 0.3047418 | 0.6848669 |
| MFHAS1 | 42.718005 | -0.27228293 | 0.40221423 | -0.67696 | 0.49843136 | 0.82345406 |
| ERI1 | 14.2720797 | -0.202573712 | 0.4231509 | -0.4787269 | 0.63213293 | NA |
| PPP1R3B | 22.5003778 | -0.42621461 | 0.45289212 | -0.9410952 | 0.34665607 | NA |
| TNKS | 129.191281 | 0.196017875 | 0.23351549 | 0.8394213 | 0.40123294 | 0.76381845 |
| MSRA | 45.8960368 | 0.149369567 | 0.44568909 | 0.33514298 | 0.73751723 | 0.91929008 |
| XKR6 | 131.821225 | 0.392259137 | 0.21568213 | 1.81869091 | 0.0689586 | 0.40491695 |
| MTMR9 | 27.0209859 | 0.136366865 | 0.3039913 | 0.44858805 | 0.65372886 | NA |
| NEIL2 | 4.21036748 | -0.743851223 | 0.73964677 | -1.0056844 | 0.31456741 | NA |
| FDFT1 | 32.6937472 | -0.247694044 | 0.31041784 | -0.7979375 | 0.42490674 | 0.77723512 |
| CTSB | 47.2092315 | -0.814426287 | 0.33404335 | -2.4380856 | 0.01476528 | 0.19896332 |
| FAM66D | 17.3360893 | 0.184914413 | 0.86531448 | 0.2136962 | 0.83078398 | NA |
| AC145124.1 | 6.653755 | 0.467341373 | 0.64962904 | 0.7193973 | 0.47189616 | NA |
| LONRF1 | 37.0176611 | 0.125876447 | 0.32370026 | 0.38886731 | 0.69737431 | 0.90729979 |
| TRMT9B | 11.8222227 | -0.626485261 | 0.83291004 | -0.7521644 | 0.45195221 | NA |
| DLC1 | 124.504947 | -0.277119068 | 0.2621 | -1.0573028 | 0.2903734 | 0.67772126 |
| TUSC3 | 175.986029 | 0.853917635 | 0.32635838 | 2.61650283 | 0.00888356 | 0.14875902 |
| MICU3 | 32.8224484 | 0.400347791 | 0.42383049 | 0.94459413 | 0.34486613 | 0.71842682 |
| ZDHHC2 | 23.6160368 | 0.532697135 | 0.36183713 | 1.47220142 | 0.14096649 | NA |
| CNOT7 | 16.3753261 | 0.406572406 | 0.36425765 | 1.11616711 | 0.26435061 | NA |
| VPS37A | 42.0192612 | -0.68012242 | 0.31932633 | -2.1298664 | 0.03318264 | 0.29047359 |
| MTUS1 | 20.9953956 | 0.757053073 | 0.64569855 | 1.1724559 | 0.24101407 | NA |
| PCM1 | 143.478726 | 0.261217842 | 0.18525551 | 1.4100409 | 0.15852761 | 0.56107153 |
| ASAH1 | 62.5247263 | -0.239405232 | 0.36600063 | -0.6541115 | 0.51303996 | 0.83070544 |
| PSD3 | 135.627801 | 0.283135559 | 0.25181479 | 1.1243802 | 0.26085177 | 0.65595635 |
| SH2D4A | 18.0298867 | -0.433888487 | 0.47549415 | -0.9125002 | 0.36150549 | NA |
| CSGALNACT1 | 28.1629119 | -2.351250351 | 0.91771844 | -2.5620607 | 0.01040531 | NA |
| INTS10 | 37.5551163 | -0.32428095 | 0.43024565 | -0.7537112 | 0.45102268 | 0.79158286 |
| LPL | 41.772861 | -2.964830975 | 0.84665433 | -3.5018199 | 0.00046209 | 0.02928507 |
| ATP6V1B2 | 16.121252 | 0.642341714 | 0.45216904 | 1.42057871 | 0.15543927 | NA |
| XPO7 | 38.3670254 | 0.425591138 | 0.30958132 | 1.37473134 | 0.16921475 | 0.57194586 |
| DMTN | 9.8273275 | 0.918319199 | 0.47770767 | 1.92234553 | 0.05456229 | NA |
| FAM160B2 | 13.6344585 | 0.397706763 | 0.40396341 | 0.98451185 | 0.32486391 | NA |
| BMP1 | 53.1676363 | -1.312398409 | 0.53789567 | -2.4398754 | 0.01469233 | 0.19896332 |
| PHYHIP | 123.2924 | -0.803540101 | 0.34705981 | -2.3152785 | 0.0205977 | 0.23840101 |
| POLR3D | 4.68143952 | -1.14529532 | 0.71415275 | -1.603712 | 0.10877755 | NA |
| SLC39A14 | 56.4591411 | -1.019120748 | 0.49857666 | -2.0440603 | 0.04094759 | 0.31695313 |
| PPP3CC | 61.5152108 | -0.062681685 | 0.24767609 | -0.2530793 | 0.80020696 | 0.94085969 |
| SORBS3 | 23.6653691 | -0.580115925 | 0.31490822 | -1.8421746 | 0.06544962 | NA |
| PDLIM2 | 7.02242106 | -0.040721777 | 0.63594071 | -0.0640339 | 0.94894322 | NA |
| C8orf58 | 17.8442967 | 0.182285291 | 0.52606131 | 0.34650959 | 0.72895978 | NA |
| CCAR2 | 25.5179844 | 1.169848641 | 0.57474782 | 2.03541207 | 0.04180943 | NA |
| BIN3 | 14.3818778 | 0.231584666 | 0.39087746 | 0.59247383 | 0.55353334 | NA |
| PEBP4 | 22.4539308 | -0.673553679 | 0.46730656 | -1.4413529 | 0.149485 | NA |
| RHOBTB2 | 46.6154676 | -1.10260576 | 0.56217403 | -1.9613246 | 0.04984117 | 0.34914643 |
| TNFRSF10B | 13.5954841 | -1.20523365 | 0.53726584 | -2.2432724 | 0.02487925 | NA |
| CHMP7 | 9.29675539 | -0.703710032 | 0.50795705 | -1.3853731 | 0.16593841 | NA |
| R3HCC1 | 6.57193202 | 0.428120772 | 0.6162335 | 0.6947379 | 0.48721957 | NA |
| ENTPD4 | 35.0587985 | 0.168518731 | 0.39875303 | 0.4226143 | 0.67257669 | 0.90403541 |
| SLC25A37 | 111.924678 | -0.320152124 | 0.34736265 | -0.9216654 | 0.35670315 | 0.72849344 |
| ADAM28 | 72.8300667 | -0.054494407 | 0.47156941 | -0.1155597 | 0.90800153 | 0.9747971 |
| AC120193.1 | 13.5758844 | -0.785614871 | 0.66683407 | -1.1781265 | 0.23874619 | NA |
| DOCK5 | 438.753891 | -0.156282838 | 0.28436831 | -0.549579 | 0.5826082 | 0.86175324 |
| KCTD9 | 41.0378584 | 0.640320611 | 0.39125141 | 1.63659629 | 0.10171485 | 0.48317363 |
| PPP2R2A | 79.2064266 | 0.40825584 | 0.29328136 | 1.39202793 | 0.16391394 | 0.56548642 |
| BNIP3L | 162.514298 | -0.102862121 | 0.36416127 | -0.2824631 | 0.77758844 | 0.93007924 |
| PNMA2 | 22.81442 | -0.666515448 | 0.42835546 | -1.5559868 | 0.11971123 | NA |
| DPYSL2 | 93.1714104 | -0.205136645 | 0.31989456 | -0.6412633 | 0.52135162 | 0.83108387 |
| TRIM35 | 12.5358234 | -0.270190682 | 0.54383958 | -0.4968206 | 0.6193156 | NA |
| PTK2B | 8.78365875 | 0.234128918 | 0.85797151 | 0.27288659 | 0.7849404 | NA |
| EPHX2 | 16.4472597 | 0.001521099 | 0.4023251 | 0.00378077 | 0.99698339 | NA |
| CLU | 32.6957282 | -0.348568119 | 0.73786281 | -0.4724023 | 0.63663963 | 0.88633797 |
| CCDC25 | 12.9654139 | -0.563373558 | 0.42468418 | -1.3265706 | 0.18465076 | NA |
| ELP3 | 23.6636546 | 0.148378762 | 0.38300414 | 0.38740772 | 0.69845439 | NA |
| ZNF395 | 10.9183628 | -0.210968396 | 0.44426011 | -0.4748758 | 0.63487547 | NA |
| FBXO16 | 12.6329526 | -0.698732325 | 0.57866393 | -1.2074924 | 0.22724255 | NA |
| FZD3 | 6.79841081 | 0.110781032 | 0.60349259 | 0.18356652 | 0.85435353 | NA |
| EXTL3 | 16.9381446 | 0.347361367 | 0.50650214 | 0.68580435 | 0.49283649 | NA |
| INTS9 | 21.0601193 | 0.175335211 | 0.33842986 | 0.51808434 | 0.60439943 | NA |
| HMBOX1 | 71.7268794 | 0.363867152 | 0.26142475 | 1.3918619 | 0.16396422 | 0.56548642 |
| KIF13B | 38.3401024 | 0.486947921 | 0.44090536 | 1.10442731 | 0.26940782 | 0.66185228 |
| SARAF | 98.1917835 | -0.065169689 | 0.36717888 | -0.1774876 | 0.85912542 | 0.95810083 |
| LEPROTL1 | 25.3075174 | -0.079508635 | 0.36942754 | -0.2152212 | 0.82959487 | NA |
| DCTN6 | 18.1737334 | -0.633855658 | 0.4836557 | -1.3105514 | 0.19000936 | NA |
| RBPMS | 715.09197 | -0.287664767 | 0.2657772 | -1.0823531 | 0.27909568 | 0.66958859 |
| GTF2E2 | 32.9491655 | 0.308070254 | 0.2982376 | 1.03296919 | 0.30161832 | 0.68219446 |
| GSR | 15.510148 | -0.069506608 | 0.4407784 | -0.1576906 | 0.87470061 | NA |
| UBXN8 | 3.35903812 | 0.072434819 | 0.84505138 | 0.08571647 | 0.93169181 | NA |
| PPP2CB | 109.061068 | 0.036657957 | 0.23753393 | 0.15432724 | 0.87735172 | 0.96359019 |

| | | | | | | |
|---|---|---|---|---|---|---|
| PURG | 6.036959 | 0.589800775 | 0.79914491 | 0.73803984 | 0.46049024 | NA |
| WRN | 63.6059265 | 0.199281615 | 0.22155038 | 0.89948667 | 0.36839349 | 0.73626927 |
| NRG1 | 14.126179 | 1.502085829 | 0.92189413 | 1.62934742 | 0.10323949 | NA |
| FUT10 | 21.6593278 | 0.296860607 | 0.44603669 | 0.66555199 | 0.5056975 | NA |
| TTI2 | 7.38372174 | -0.660181714 | 0.48766757 | -1.3537536 | 0.17581501 | NA |
| MAK16 | 4.06923364 | -0.019935347 | 0.74776933 | -0.0266598 | 0.97873111 | NA |
| UNC5D | 141.151901 | 1.172607488 | 1.27405781 | 0.92037228 | 0.35737825 | 0.72903084 |
| ZNF703 | 10.0012391 | -1.306246635 | 0.89514906 | -1.4592504 | 0.1444962 | NA |
| ERLIN2 | 23.3055639 | 0.311310961 | 0.32024819 | 0.97209281 | 0.33100438 | NA |
| PLPBP | 12.7075104 | -0.542833392 | 0.42685388 | -1.2717078 | 0.20347697 | NA |
| RAB11FIP1 | 25.9252444 | -0.344690005 | 0.29491266 | -1.1687867 | 0.24248957 | NA |
| EIF4EBP1 | 9.51965509 | 0.376188311 | 1.00709457 | 0.37353822 | 0.70874791 | NA |
| ASH2L | 17.2285928 | 0.158622113 | 0.50170683 | 0.31616494 | 0.75187731 | NA |
| LSM1 | 18.9778736 | -0.151932693 | 0.4212673 | -0.3606563 | 0.71835642 | NA |
| BAG4 | 15.9770625 | 0.337594491 | 0.41865239 | 0.80638377 | 0.4200216 | NA |
| DDHD2 | 26.9544737 | -0.014118729 | 0.39446037 | -0.0357925 | 0.9714478 | NA |
| PLPP5 | 9.87425764 | -1.120374989 | 0.58358149 | -1.9198261 | 0.05487986 | NA |
| NSD3 | 98.3109764 | -0.347606786 | 0.24255722 | -1.4330919 | 0.15183158 | 0.55086922 |
| FGFR1 | 75.8776997 | -0.196376078 | 0.32204229 | -0.6097835 | 0.54200523 | 0.84163543 |
| TACC1 | 50.1660248 | 0.840727938 | 0.4313951 | 1.94885836 | 0.05131234 | 0.35507065 |
| PLEKHA2 | 72.7538215 | -0.303649751 | 0.34708058 | -0.8748682 | 0.38164564 | 0.74899182 |
| AC108863.1 | 5.57205035 | -1.769981334 | 1.00552075 | -1.7602634 | 0.07836316 | NA |
| TM2D2 | 13.2665895 | -0.438102443 | 0.6170248 | -0.710024 | 0.47768923 | NA |
| ADAM9 | 50.96813 | -0.350441627 | 0.35007929 | -1.001035 | 0.31680988 | 0.69424851 |
| ADAM32 | 10.60601 | -0.056307908 | 0.55204572 | -0.1019986 | 0.91875776 | NA |
| TCIM | 16.0442002 | -1.449960821 | 1.01198879 | -1.4327835 | 0.15191972 | NA |
| SFRP1 | 4.96862757 | -1.191729156 | 0.88922516 | -1.3401883 | 0.18018413 | NA |
| GOLGA7 | 21.7626776 | 0.112419428 | 0.39267195 | 0.2862935 | 0.77465333 | NA |
| GPAT4 | 20.3502549 | 0.574503861 | 0.35045362 | 1.63931498 | 0.10114768 | NA |
| KAT6A | 84.6395208 | -0.311299869 | 0.24979277 | -1.2462325 | 0.21267905 | 0.62255988 |
| AP3M2 | 8.76955146 | 0.211533727 | 0.46071368 | 0.45914358 | 0.64613106 | NA |
| IKBKB | 25.0500089 | 0.352714162 | 0.353022 | 0.99912798 | 0.3177327 | NA |
| POLB | 5.41241529 | -0.128464499 | 0.59853017 | -0.2146333 | 0.83005324 | NA |
| VDAC3 | 23.9005904 | -0.306486145 | 0.33347973 | -0.9190548 | 0.35806691 | NA |
| SLC20A2 | 93.5695786 | 0.255671535 | 0.38360925 | 0.6664895 | 0.50509828 | 0.82600448 |
| SMIM19 | 21.4131402 | -0.527011469 | 0.38337819 | -1.3746517 | 0.16923946 | NA |
| THAP1 | 5.12465516 | -0.379059312 | 0.59826707 | -0.6335955 | 0.52634485 | NA |
| RNF170 | 29.0264099 | -0.231230527 | 0.30336821 | -0.7622108 | 0.44593421 | 0.78845211 |
| HOOK3 | 47.8343586 | 0.113807731 | 0.27674635 | 0.41123481 | 0.68090037 | 0.90499319 |
| FNTA | 23.8367418 | -0.241903001 | 0.35045741 | -0.6902494 | 0.49003739 | NA |
| HGSNAT | 60.8876729 | 0.038090568 | 0.31426919 | 0.12120364 | 0.90352974 | 0.97237083 |
| SPIDR | 199.508834 | -0.432197782 | 0.34061285 | -1.2688828 | 0.20448288 | 0.6137834 |
| CEBPD | 191.68344 | -3.627400987 | 0.56238386 | -6.4500446 | 1.12E-10 | 2.27E-07 |
| PRKDC | 63.8139341 | 0.082414549 | 0.24580316 | 0.33528678 | 0.73740877 | 0.91929008 |
| UBE2V2 | 29.7842581 | -0.169224146 | 0.28610784 | -0.5914698 | 0.55420569 | 0.84953071 |
| PXDNL | 20.3481574 | 1.483445931 | 0.46077659 | 3.21944723 | 0.00128438 | NA |
| PCMTD1 | 180.055469 | 1.042812084 | 0.27096456 | 3.84851832 | 0.00011883 | 0.01171344 |
| AC064807.1 | 12.0104032 | -0.273596686 | 0.47898597 | -0.5711998 | 0.56786422 | NA |
| RB1CC1 | 76.5389521 | 0.108406513 | 0.26480663 | 0.4093799 | 0.68226089 | 0.90499319 |
| ATP6V1H | 47.7412071 | 0.18521051 | 0.361355 | 0.51254447 | 0.60827 | 0.8725395 |
| TCEA1 | 39.3288069 | 0.877337732 | 0.31409456 | 2.79322801 | 0.00521849 | 0.11257186 |
| LYPLA1 | 32.2184127 | -0.185375497 | 0.33827034 | -0.5480099 | 0.5836851 | 0.86175324 |
| MRPL15 | 10.0872834 | -0.348475245 | 0.50085368 | -0.6957626 | 0.48657753 | NA |
| XKR4 | 415.746023 | -3.847746153 | 0.77536318 | -4.9625082 | 6.96E-07 | 0.00023577 |
| AC022679.2 | 5.16483001 | -4.160991572 | 1.33804499 | -3.1097546 | 0.00187243 | NA |
| TMEM68 | 15.8013686 | 0.30783315 | 0.36051451 | 0.85387174 | 0.39317606 | NA |
| TGS1 | 14.5013595 | -0.295834647 | 0.36636039 | -0.8074963 | 0.41938062 | NA |
| LYN | 10.1887379 | -0.071495113 | 0.47079798 | -0.1518594 | 0.87929781 | NA |
| RPS20 | 41.8821055 | -0.451719309 | 0.4474522 | -1.0095365 | 0.31271743 | 0.69213367 |
| PLAG1 | 14.6560419 | -0.11132193 | 0.58771411 | -0.1894151 | 0.84976749 | NA |
| CHCHD7 | 7.42651999 | -0.078680055 | 0.54357558 | -0.1447454 | 0.88491191 | NA |
| IMPAD1 | 64.1496562 | -0.119540913 | 0.33951612 | -0.352092 | 0.72476926 | 0.91549801 |
| FAM110B | 12.447401 | 0.55596467 | 0.67683107 | 0.82142309 | 0.41140531 | NA |
| UBXN2B | 27.325505 | 0.049297686 | 0.32938716 | 0.14966487 | 0.88102903 | NA |
| SDCBP | 28.9869001 | 0.274366639 | 0.35456235 | 0.77381776 | 0.43903857 | 0.78343178 |
| NSMAF | 26.515127 | 0.033357509 | 0.35990824 | 0.09268337 | 0.92615511 | NA |
| TOX | 19.7303667 | 1.350902638 | 0.97962729 | 1.37899654 | 0.16789582 | NA |
| LINC01301 | 10.3740155 | -0.943338739 | 0.92855656 | -1.0159195 | 0.30966771 | NA |
| RAB2A | 140.161583 | -0.059931386 | 0.2504203 | -0.2393232 | 0.81085498 | 0.94408044 |
| CHD7 | 138.81116 | -0.221444195 | 0.29164422 | -0.7592957 | 0.4476757 | 0.78947076 |
| CLVS1 | 5.02416745 | -0.058200757 | 0.705887 | -0.0824505 | 0.93428846 | NA |
| ASPH | 67.8051456 | 0.005429373 | 0.23161423 | 0.02344145 | 0.98129814 | 0.99453905 |
| YTHDF3 | 48.0461211 | -0.066598649 | 0.25870223 | -0.2574336 | 0.79684406 | 0.93934998 |
| CYP7B1 | 36.9262606 | 0.934595174 | 0.75470665 | 1.23835556 | 0.21558425 | 0.62324286 |
| ARMC1 | 19.4668673 | -0.110448865 | 0.32522314 | -0.3396095 | 0.73415063 | NA |
| MTFR1 | 26.2668808 | -0.460725966 | 0.34103053 | -1.3509816 | 0.17670133 | NA |
| PDE7A | 27.6807957 | 0.314442497 | 0.4931386 | 0.63763514 | 0.52371122 | NA |
| ADHFE1 | 15.6011315 | 0.058630203 | 0.532411 | 0.11012207 | 0.91231256 | NA |
| VCPIP1 | 24.5708633 | 0.194827003 | 0.35025272 | 0.55624693 | 0.57804206 | NA |
| SGK3 | 66.2266238 | 0.105714178 | 0.29715959 | 0.35574883 | 0.72202867 | 0.91509584 |
| COPS5 | 5.54977174 | -0.947255835 | 0.71982972 | -1.3159443 | 0.18819273 | NA |
| CSPP1 | 166.401581 | -0.289290177 | 0.27950563 | -1.0350066 | 0.30066582 | 0.68204729 |
| ARFGEF1 | 137.856381 | 0.158367908 | 0.18124981 | 0.87375491 | 0.38225174 | 0.74899182 |
| SULF1 | 348.929933 | -0.312377912 | 0.39213639 | -0.7966053 | 0.42568032 | 0.77730788 |
| NCOA2 | 180.746845 | 0.523769361 | 0.29974005 | 1.74741202 | 0.08056589 | 0.43346499 |
| TRAM1 | 67.7248186 | 0.035873254 | 0.37568108 | 0.09548858 | 0.92392676 | 0.97775374 |
| LACTB2 | 7.59648585 | -1.641086308 | 0.80612882 | -2.0357619 | 0.04177427 | NA |
| TERF1 | 35.2676977 | -0.135874246 | 0.30064592 | -0.4519411 | 0.65131142 | 0.89412414 |
| AC022893.1 | 12.1101408 | -0.921242435 | 0.68842439 | -1.3381897 | 0.1808346 | NA |
| SBSPON | 158.835466 | -0.977003386 | 0.62461064 | -1.5641799 | 0.11777533 | 0.50872923 |
| RPL7 | 72.2586359 | -0.93069631 | 0.48226705 | -1.929836 | 0.05362716 | 0.3615191 |
| RDH10 | 14.2748252 | -1.897207212 | 0.72349289 | -2.6222887 | 0.00873414 | NA |
| RDH10-AS1 | 4.64459498 | -1.494071692 | 0.93024685 | -1.6061024 | 0.10825141 | NA |
| STAU2 | 69.2322568 | 0.487740914 | 0.31377614 | 1.55442319 | 0.12008351 | 0.51128003 |
| AC027018.1 | 6.24136416 | -0.617780145 | 0.79065031 | -0.781357 | 0.43459256 | NA |
| ELOC | 30.4587757 | -0.104998455 | 0.31723212 | -0.330983 | 0.74065729 | 0.92003444 |
| TMEM70 | 9.41278301 | -0.196311236 | 0.5764632 | -0.3405443 | 0.7334467 | NA |
| JPH1 | 34.9104677 | -1.114880009 | 0.51604849 | -2.1604172 | 0.03074039 | 0.28264481 |
| GDAP1 | 20.8635067 | 0.199872878 | 0.46610342 | 0.42881658 | 0.66805671 | NA |
| PEX2 | 21.7291974 | -0.257155054 | 0.34960228 | -0.7355646 | 0.46199573 | NA |
| ZC2HC1A | 46.8286085 | 0.122398463 | 0.30229717 | 0.4048945 | 0.68555506 | 0.90499319 |
| IL7 | 14.4868541 | -0.433240601 | 0.62360356 | -0.6947372 | 0.48722 | NA |
| TPD52 | 57.3002844 | 0.198325215 | 0.27891875 | 0.71105013 | 0.47705317 | 0.80863008 |
| ZBTB10 | 44.8285512 | -0.116499704 | 0.40258425 | -0.2893797 | 0.77229084 | 0.92669573 |
| ZNF704 | 127.705695 | 0.32732884 | 0.30982288 | 1.05650314 | 0.29073841 | 0.67772126 |
| PAG1 | 8.13270996 | 1.209722289 | 0.68125604 | 1.77572339 | 0.07577852 | NA |
| FABP5 | 15.9799592 | -3.198267078 | 1.01205373 | -3.1601752 | 0.00157674 | NA |
| IMPA1 | 13.4425747 | 0.002120623 | 0.41195065 | 0.00514776 | 0.9958927 | NA |
| ZFAND1 | 14.6817433 | -0.010622436 | 0.39534099 | -0.026869 | 0.97856418 | NA |
| CHMP4C | 13.8581972 | -0.174425071 | 0.41185232 | -0.4235136 | 0.67192056 | NA |
| SNX16 | 7.95479168 | 0.242595088 | 0.57343094 | 0.42305894 | 0.67225226 | NA |
| RALYL | 14.4943653 | 1.41222577 | 0.58051793 | 2.43269965 | 0.01498673 | NA |
| LRRCC1 | 8.31595019 | 0.041277036 | 0.51279006 | 0.080495 | 0.93584357 | NA |

| | | | | | | |
|---|---|---|---|---|---|---|
| E2F5 | 10.8058803 | -0.177751452 | 0.46914515 | -0.3788837 | 0.70477422 | NA |
| C8orf59 | 18.1231594 | -1.066241043 | 0.47444466 | -2.2473454 | 0.02461796 | NA |
| CA13 | 9.63704124 | 0.397467254 | 0.47569568 | 0.83554943 | 0.40340843 | NA |
| CA3 | 10.7506973 | 0.573556909 | 0.4061772 | 1.41208543 | 0.15792481 | NA |
| CA2 | 32.759413 | 0.551712225 | 0.45098635 | 1.22334572 | 0.22119915 | 0.62521515 |
| WWP1 | 191.486516 | -0.137130136 | 0.32550119 | -0.4212892 | 0.67354391 | 0.90403541 |
| RMDN1 | 33.9760891 | -0.248961968 | 0.33238533 | -0.7490161 | 0.4538475 | 0.79324789 |
| CPNE3 | 40.8648527 | 0.290276779 | 0.34054095 | 0.85239903 | 0.39399266 | 0.75773027 |
| CNGB3 | 24.8904488 | 2.161091679 | 0.91483628 | 2.36227151 | 0.01816333 | NA |
| MMP16 | 25.2558835 | 1.913573744 | 0.85577243 | 2.23607782 | 0.02534667 | NA |
| AC090568.2 | 9.70812379 | -0.583724704 | 0.94375143 | -0.6185153 | 0.53623571 | NA |
| AF117829.1 | 80.6695522 | 0.035958284 | 0.32982665 | 0.10902177 | 0.91318523 | 0.97707273 |
| RIPK2 | 7.20196778 | -0.489862853 | 0.54281194 | -0.9024541 | 0.3668157 | NA |
| OSGIN2 | 12.4366523 | -0.582443983 | 0.42203493 | -1.3800848 | 0.16756054 | NA |
| NBN | 21.0684988 | -0.917882127 | 0.53343498 | -1.7207011 | 0.08530509 | NA |
| DECR1 | 30.4967757 | -0.192709564 | 0.3802534 | -0.5067925 | 0.61230043 | 0.87391153 |
| TMEM64 | 47.480919 | -0.783282766 | 0.35485705 | -2.2073192 | 0.02729177 | 0.26802763 |
| NECAB1 | 250.945502 | -0.388500319 | 0.38529646 | -1.0083153 | 0.31330311 | 0.69213367 |
| AC103770.1 | 4.42222165 | -1.565503778 | 1.11069417 | -1.4094823 | 0.1586926 | NA |
| PIP4P2 | 17.2175031 | -0.268006917 | 0.423768 | -0.6324378 | 0.52710082 | NA |
| OTUD6B-AS1 | 21.8196499 | -0.502981866 | 0.4177005 | -1.2041687 | 0.22852439 | NA |
| OTUD6B | 11.5621921 | -0.320921323 | 0.44692234 | -0.7180695 | 0.47271441 | NA |
| LRRC69 | 14.7074586 | -0.310829297 | 0.41649752 | -0.7462933 | 0.45549027 | NA |
| RUNX1T1 | 32.6337754 | 0.324502574 | 0.40984296 | 0.79177295 | 0.42849308 | 0.77844087 |
| AC117834.1 | 5.31920674 | -0.973677877 | 0.75251501 | -1.2938983 | 0.19570056 | NA |
| TRIQK | 38.121943 | -0.437382795 | 0.36010482 | -1.2145986 | 0.22451924 | 0.63080203 |
| LINC00535 | 10.1873729 | -2.296308869 | 0.87339022 | -2.62919 | 0.00855885 | NA |
| FAM92A | 11.5749969 | 0.426713933 | 0.46583839 | 0.91601281 | 0.35966017 | NA |
| RBM12B | 18.1350001 | 0.034264105 | 0.45527454 | 0.07526031 | 0.9400076 | NA |
| TMEM67 | 25.2031099 | 0.548178075 | 0.30680007 | 1.78675993 | 0.0739763 | NA |
| PDP1 | 4.62930778 | 0.408522764 | 0.68962127 | 0.59238713 | 0.55359138 | NA |
| VIRMA | 26.8238674 | -0.104478528 | 0.27683884 | -0.3773984 | 0.70587757 | NA |
| ESRP1 | 8.2430641 | 1.278845307 | 0.63034283 | 2.02880916 | 0.04247773 | NA |
| DPY19L4 | 34.1272377 | 0.474542533 | 0.38714019 | 1.22576407 | 0.22028747 | 0.62462904 |
| INTS8 | 25.144642 | 0.643213033 | 0.34177591 | 1.88197299 | 0.05983968 | NA |
| NDUFAF6 | 25.9729765 | 0.240107295 | 0.40353731 | 0.59500643 | 0.55183917 | NA |
| TP53INP1 | 20.6859303 | 0.599422265 | 0.61159764 | 0.9800925 | 0.32704046 | NA |
| PLEKHF2 | 10.6325127 | 0.399933342 | 0.54318164 | 0.73627919 | 0.46156081 | NA |
| C8orf37-AS1 | 14.5880908 | 0.646911375 | 0.74538801 | 0.8678854 | 0.38545707 | NA |
| UQCRB | 36.3474702 | -0.719830012 | 0.40228932 | -1.7893341 | 0.07356102 | 0.41834543 |
| MTERF3 | 7.77944492 | -0.616987624 | 0.59091903 | -1.0441153 | 0.29643203 | NA |
| PTDSS1 | 27.4958403 | -0.034643935 | 0.3878185 | -0.0893303 | 0.92881943 | NA |
| SDC2 | 110.533393 | -0.006013817 | 0.37673774 | -0.0159629 | 0.98726401 | 0.99585746 |
| CPQ | 138.39815 | 0.059357894 | 0.22714957 | 0.26131633 | 0.79384857 | 0.93934998 |
| TSPYL5 | 16.1384419 | -0.686496377 | 0.44070386 | -1.5577272 | 0.11929794 | NA |
| MTDH | 138.782679 | -0.201737969 | 0.20181393 | -0.9996236 | 0.31749269 | 0.69424851 |
| LAPTM4B | 37.1838646 | -0.225562587 | 0.41026805 | -0.5497932 | 0.58246122 | 0.86175324 |
| MATN2 | 8.89645458 | 0.072396204 | 0.94619973 | 0.0765126 | 0.93901129 | NA |
| RPL30 | 78.769612 | -0.955320847 | 0.37966558 | -2.5162166 | 0.01186222 | 0.18181798 |
| RIDA | 5.73879391 | -1.237521322 | 0.68651075 | -1.8026248 | 0.07144716 | NA |
| POP1 | 4.46737677 | 0.683736003 | 0.61184551 | 1.11749779 | 0.26378154 | NA |
| NIPAL2 | 16.2396286 | -0.409889094 | 0.5312615 | -0.7715392 | 0.44038738 | NA |
| STK3 | 122.03985 | -0.322891805 | 0.29104277 | -1.1094308 | 0.26724439 | 0.66013596 |
| AC104986.2 | 19.8791373 | -0.122005604 | 0.3335991 | -0.3657252 | 0.71457012 | NA |
| VPS13B | 178.33244 | 0.462361532 | 0.16205925 | 2.85304009 | 0.00433032 | 0.10580579 |
| COX6C | 65.545992 | -0.570494126 | 0.4116444 | -1.3858906 | 0.1657803 | 0.56743028 |
| POLR2K | 28.6491429 | -0.009441941 | 0.47794588 | -0.0197553 | 0.98423861 | 0.99577746 |
| SPAG1 | 13.0793827 | -0.540586713 | 0.64802847 | -0.8342021 | 0.40416711 | NA |
| RNF19A | 109.993182 | 0.023087605 | 0.28388651 | 0.08132688 | 0.935182 | 0.97832857 |
| ANKRD46 | 16.1533659 | 0.059893959 | 0.35724707 | 0.16765417 | 0.86685535 | NA |
| PABPC1 | 67.2367509 | -0.405948163 | 0.32220245 | -1.2599164 | 0.20769951 | 0.61707831 |
| YWHAZ | 135.146245 | 0.335376738 | 0.23504371 | 1.42686968 | 0.15361746 | 0.55384216 |
| ZNF706 | 36.1593042 | -0.52473352 | 0.31971121 | -1.6412734 | 0.10074067 | 0.4824134 |
| RRM2B | 16.7544448 | -0.279368232 | 0.36933115 | -0.7564167 | 0.44939943 | NA |
| UBR5 | 126.667634 | 0.21079196 | 0.23084886 | 0.91311676 | 0.36118114 | 0.73240575 |
| KLF10 | 36.4332789 | -0.384373973 | 0.59907126 | -0.6416164 | 0.52112225 | 0.83108387 |
| AZIN1 | 23.9135337 | -0.141808807 | 0.3415693 | -0.4151685 | 0.67801857 | NA |
| AZIN1-AS1 | 35.7621208 | -0.021339877 | 0.38740591 | -0.055084 | 0.95607152 | 0.98641719 |
| ATP6V1C1 | 33.3856491 | 0.136918565 | 0.31699488 | 0.43192674 | 0.66579466 | 0.89968484 |
| BAALC-AS1 | 16.0086956 | -0.321537787 | 0.460325 | -0.6985017 | 0.48486351 | NA |
| FZD6 | 28.3021125 | 0.185722692 | 0.2955903 | 0.62831118 | 0.52980011 | NA |
| SLC25A32 | 18.2112379 | 0.14547214 | 0.37542659 | 0.38748492 | 0.69839725 | NA |
| DCAF13 | 8.29853268 | 0.16685862 | 0.52070357 | 0.32044839 | 0.74862845 | NA |
| DPYS | 20.9397019 | 2.378180617 | 1.57184864 | 1.51298322 | 0.13028392 | NA |
| LRP12 | 21.747891 | 0.748987591 | 0.42507022 | 1.76203261 | 0.07806379 | NA |
| ZFPM2 | 159.781915 | -0.123548878 | 0.36760717 | -0.3360894 | 0.73680344 | 0.91929008 |
| OXR1 | 118.233837 | 0.99597664 | 0.37597183 | 2.64907251 | 0.0080713 | 0.1435842 |
| AC090579.1 | 13.6072143 | 0.990282063 | 0.72082319 | 1.37382103 | 0.16949725 | NA |
| EIF3E | 43.3609267 | 0.182456897 | 0.3280478 | 0.55618997 | 0.578081 | 0.86138741 |
| EMC2 | 42.0496261 | 0.358511338 | 0.30669091 | 1.16896628 | 0.24241722 | 0.64918976 |
| NUDCD1 | 46.6166351 | -0.510748358 | 0.34188919 | -1.4939003 | 0.13520168 | 0.52779404 |
| ENY2 | 36.3621657 | -0.562201905 | 0.39606309 | -1.4194756 | 0.15576039 | 0.55613041 |
| EBAG9 | 16.395967 | -0.23084024 | 0.39928402 | -0.5781354 | 0.56317269 | NA |
| SYBU | 14.2076455 | 0.101502823 | 0.41543502 | 0.244329 | 0.80697603 | NA |
| TRPS1 | 120.496364 | 0.927422506 | 0.32309932 | 2.87039447 | 0.0040996 | 0.10253013 |
| EIF3H | 53.3793518 | -0.143701356 | 0.33023946 | -0.4351429 | 0.66345871 | 0.89939456 |
| UTP23 | 11.0269849 | -0.414176945 | 0.56209509 | -0.736845 | 0.46121663 | NA |
| RAD21 | 55.2777834 | -0.081775376 | 0.35987726 | -0.2272313 | 0.82024389 | 0.94728252 |
| MED30 | 10.5933948 | 1.45000366 | 0.5134475 | 2.82405437 | 0.00474203 | NA |
| EXT1 | 106.474646 | 0.667402525 | 0.41692815 | 1.60076149 | 0.10942976 | 0.49591853 |
| SAMD12 | 71.6331466 | 0.402501796 | 0.25496995 | 1.57862443 | 0.11442223 | 0.50483797 |
| AC023590.1 | 6.14961211 | -1.571442595 | 1.08857059 | -1.4435835 | 0.14885616 | NA |
| TNFRSF11B | 13.6527595 | -0.459779152 | 0.71049994 | -0.6471206 | 0.51755389 | NA |
| MAL2 | 164.215464 | 0.480799586 | 0.45972387 | 1.04584428 | 0.29563293 | 0.68103704 |
| MAL2-AS1 | 5.79902909 | 0.035046232 | 0.82248324 | 0.04261027 | 0.96601221 | NA |
| TAF2 | 36.1151875 | 0.168669674 | 0.31574655 | 0.53419324 | 0.59320784 | 0.8654248 |
| DEPTOR | 69.1732524 | 0.34717886 | 0.38115153 | 0.91086834 | 0.36236475 | 0.73259935 |
| AC091563.1 | 7.4173957 | -0.87774154 | 0.65547355 | -1.3390953 | 0.18053965 | NA |
| MRPL13 | 13.9331029 | -0.47436705 | 0.5851713 | -0.8106465 | 0.41756872 | NA |
| MTBP | 5.96237052 | -0.107761886 | 0.67591309 | -0.1594316 | 0.87332887 | NA |
| SNTB1 | 13.0069209 | -0.639558311 | 0.80830522 | -0.7912337 | 0.42880765 | NA |
| ZHX2 | 67.0274428 | -0.184668527 | 0.30116373 | -0.6131831 | 0.53975525 | 0.83929813 |
| DERL1 | 17.7229556 | -0.430748176 | 0.41988506 | -1.0258716 | 0.30495209 | NA |
| C8orf76 | 4.64381449 | 0.644219092 | 0.62771768 | 1.02628796 | 0.30475587 | NA |
| WDYHV1 | 8.70323988 | -0.33449433 | 0.53106317 | -0.6298579 | 0.52878756 | NA |
| FBXO32 | 8.09795858 | -0.681071515 | 0.63772929 | -1.0679634 | 0.28553704 | NA |
| FAM91A1 | 29.3155951 | 0.180395889 | 0.27983279 | 0.64465601 | 0.51915013 | 0.83104823 |
| TMEM65 | 26.278467 | 0.323975071 | 0.42310153 | 0.76571472 | 0.44384607 | NA |
| RNF139 | 11.5393797 | 0.09704417 | 0.46832816 | 0.20721404 | 0.8358427 | NA |
| TATDN1 | 25.3358992 | 0.27398205 | 0.39600835 | 0.69185928 | 0.48902571 | NA |
| NDUFB9 | 47.8458747 | -0.73258874 | 0.33246574 | -2.2035014 | 0.02755943 | 0.26935188 |
| MTSS1 | 776.138615 | -0.468281525 | 0.44887704 | -1.0432289 | 0.29684227 | 0.68103704 |
| SQLE | 22.9420382 | -0.716498953 | 0.49959115 | -1.4341706 | 0.15152358 | NA |

| | | | | | | |
|---|---|---|---|---|---|---|
| WASHC5 | 20.043303 | 0.378335533 | 0.37458059 | 1.0100244 | 0.3124836 | NA |
| NSMCE2 | 102.154003 | 0.21850148 | 0.29749936 | 0.73446034 | 0.46266822 | 0.79793427 |
| FAM84B | 5.88480343 | -0.393597296 | 0.73408384 | -0.5361749 | 0.59183771 | NA |
| PCAT1 | 30.5380253 | 1.225981783 | 0.66077927 | 1.85535751 | 0.06354519 | 0.38816159 |
| MYC | 6.23653876 | -5.061124391 | 1.5253496 | -3.3180094 | 0.00090661 | NA |
| PVT1 | 14.9232911 | -1.38292855 | 0.87157604 | -1.5866987 | 0.1125809 | NA |
| FAM49B | 61.8736282 | 0.511326099 | 0.22389063 | 2.28382089 | 0.02238206 | 0.24469443 |
| ASAP1 | 63.4718902 | -0.354511391 | 0.37292546 | -0.9506227 | 0.34179596 | 0.717188 |
| EFR3A | 63.6417733 | 0.275491729 | 0.21871801 | 1.25957498 | 0.20782272 | 0.61707831 |
| LRRC6 | 19.7959016 | 0.033231335 | 0.44618404 | 0.07447899 | 0.94062926 | NA |
| PHF20L1 | 43.7056059 | 0.326880124 | 0.28619336 | 1.14216528 | 0.25338532 | 0.65294211 |
| NDRG1 | 183.593941 | -0.380650677 | 0.28948899 | -1.3149055 | 0.18854165 | 0.59077076 |
| ST3GAL1 | 186.87957 | -1.258328107 | 0.44769004 | -2.8107127 | 0.00494319 | 0.11177304 |
| ZFAT | 19.2100001 | -0.034546316 | 0.52899868 | -0.0653051 | 0.94793108 | NA |
| AC083843.3 | 6.4049703 | -1.376290096 | 0.75043789 | -1.8339827 | 0.06665657 | NA |
| KHDRBS3 | 14.1892639 | -0.402579292 | 0.50995738 | -0.7894371 | 0.42985656 | NA |
| TRAPPC9 | 127.809167 | -0.00337927 | 0.24557795 | -0.0137605 | 0.98902107 | 0.99651342 |
| CHRAC1 | 16.738475 | 0.055695636 | 0.51958122 | 0.10719332 | 0.91463562 | NA |
| AGO2 | 48.5398193 | -0.444479994 | 0.38504569 | -1.1543565 | 0.24835405 | 0.6507516 |
| PTK2 | 241.941813 | 0.45518495 | 0.18467226 | 2.46482585 | 0.01370799 | 0.19440417 |
| DENND3 | 6.52891255 | -0.239517566 | 0.6987548 | -0.3427777 | 0.7317657 | NA |
| SLC45A4 | 12.2764527 | 0.835838723 | 0.55791458 | 1.4981482 | 0.13409475 | NA |
| PTP4A3 | 4.81182064 | 0.310494412 | 0.8345044 | 0.37207043 | 0.70984041 | NA |
| TSNARE1 | 10.8490942 | 0.350353333 | 0.53625058 | 0.65333884 | 0.51353786 | NA |
| JRK | 11.3412453 | 0.502567124 | 0.50791754 | 0.98946598 | 0.3224352 | NA |
| THEM6 | 6.9721571 | -0.002711544 | 0.70465926 | -0.003848 | 0.99692973 | NA |
| LY6E | 160.332784 | -0.06329651 | 0.89195153 | -0.0709641 | 0.94342635 | 0.98114803 |
| GLI4 | 5.34343687 | 0.177222092 | 0.58566303 | 0.30260078 | 0.76219412 | NA |
| RHPN1 | 14.0720535 | 0.858375165 | 0.41219586 | 2.08244492 | 0.03730185 | NA |
| ZC3H3 | 10.487616 | 0.556410852 | 0.45356463 | 1.22675099 | 0.2199162 | NA |
| GSDMD | 18.9386944 | -0.039679319 | 0.54003971 | -0.0734748 | 0.94142828 | NA |
| EEF1D | 49.7886491 | -0.344405351 | 0.3649824 | -0.9436218 | 0.34536294 | 0.71852078 |
| TIGD5 | 4.34247763 | -0.712464617 | 0.73002391 | -0.975947 | 0.32909075 | NA |
| TSTA3 | 6.86698036 | -0.957971805 | 0.57864887 | -1.6555322 | 0.0978166 | NA |
| ZNF623 | 9.47663987 | 0.12458282 | 0.5102373 | 0.24416643 | 0.80710193 | NA |
| FAM83H | 7.59776949 | -0.128277558 | 0.54596443 | -0.2349559 | 0.81424297 | NA |
| SCRIB | 5.35549919 | 0.440725125 | 0.59724748 | 0.73792714 | 0.46055873 | NA |
| PUF60 | 24.1566977 | -0.417278425 | 0.42648295 | -0.9784176 | 0.32786783 | NA |
| NRBP2 | 17.1088654 | -1.111332783 | 0.45190769 | -2.4592031 | 0.01392458 | NA |
| PLEC | 73.2606914 | -0.359462085 | 0.43412689 | -0.8280116 | 0.40766395 | 0.76772515 |
| PARP10 | 14.5845765 | 0.449391716 | 0.90861924 | 0.4945875 | 0.62089133 | NA |
| GRINA | 24.5407705 | -0.266615018 | 0.58800005 | -0.4534269 | 0.6502414 | NA |
| EXOSC4 | 6.40521368 | -0.076909558 | 0.69351531 | -0.1108981 | 0.91169712 | NA |
| GPAA1 | 25.1174098 | -0.090922871 | 0.61460261 | -0.1479377 | 0.88239197 | NA |
| CYC1 | 18.4805407 | -0.836487953 | 0.68154733 | -1.2273366 | 0.21969612 | NA |
| SHARPIN | 13.4189965 | 0.463269316 | 0.39698032 | 1.16698309 | 0.2432172 | NA |
| MAF1 | 25.2430727 | 0.443272403 | 0.3712832 | 1.19389297 | 0.23251984 | NA |
| HGH1 | 4.75864144 | 0.594258326 | 0.68332206 | 0.86966068 | 0.38448587 | NA |
| MROH1 | 46.9887967 | -0.18247502 | 0.36211045 | -0.5039209 | 0.61431697 | 0.87519131 |
| HSF1 | 21.8470607 | -0.441171021 | 0.36900486 | -1.1955697 | 0.23186452 | NA |
| DGAT1 | 7.57198408 | 0.965619395 | 0.50199064 | 1.92358047 | 0.05440719 | NA |
| SLC52A2 | 8.50787147 | -0.213002359 | 0.59834544 | -0.3559856 | 0.72185135 | NA |
| CPSF1 | 5.46874191 | 0.978888369 | 0.59842156 | 1.63578394 | 0.10188481 | NA |
| VPS28 | 47.8100354 | -0.50076674 | 0.34800164 | -1.4389781 | 0.15015673 | 0.54834287 |
| CYHR1 | 26.2820135 | -0.238059792 | 0.30050458 | -0.7922002 | 0.42824394 | NA |
| KIFC2 | 7.66446772 | 0.974837989 | 0.51792139 | 1.88221225 | 0.0598072 | NA |
| PPP1R16A | 18.5475951 | 0.347895387 | 0.40637624 | 0.85609184 | 0.39194698 | NA |
| LRRC14 | 6.25061813 | -0.542782387 | 0.62596259 | -0.8671163 | 0.38587827 | NA |
| ARHGAP39 | 4.9843388 | 0.519537408 | 0.56434985 | 0.92059458 | 0.35726213 | NA |
| ZNF251 | 23.047593 | 0.475216851 | 0.45684391 | 1.0402171 | 0.29823905 | NA |
| RPL8 | 100.434597 | -0.597169747 | 0.41127862 | -1.4519834 | 0.14650621 | 0.54616653 |
| ZNF7 | 8.36293239 | -0.474315725 | 0.62534695 | -0.7584841 | 0.44816122 | NA |
| COMMD5 | 10.2270335 | 0.621114927 | 0.55408079 | 1.12098259 | 0.26229527 | NA |
| ZNF250 | 16.8361382 | 0.818595066 | 0.60540556 | 1.3521433 | 0.17632948 | NA |
| ZNF16 | 15.642208 | 0.189189104 | 0.4257509 | 0.44436572 | 0.65677819 | NA |
| C8orf33 | 12.3823854 | -0.644358291 | 0.45082686 | -1.4292811 | 0.15292347 | NA |
| WASHC1 | 122.08578 | 0.309030865 | 0.2730147 | 1.13192026 | 0.25766796 | 0.65588339 |
| CBWD1 | 9.44952944 | -0.301589873 | 0.46297962 | -0.6514107 | 0.51478141 | NA |
| DOCK8 | 11.4312123 | 1.236545702 | 0.70305982 | 1.75880581 | 0.0786105 | NA |
| KANK1 | 614.52349 | 0.382219683 | 0.35366291 | 1.08074573 | 0.27981023 | 0.66958859 |
| SMARCA2 | 88.7726429 | 0.696043139 | 0.21025715 | 3.31043746 | 0.0009315 | 0.0490672 |
| VLDLR-AS1 | 11.4314277 | 2.723891965 | 0.67999924 | 4.00572797 | 6.18E-05 | NA |
| VLDLR | 8.94617284 | 2.112376825 | 0.66694206 | 3.1672569 | 0.00153884 | NA |
| PUM3 | 49.9811358 | 0.096555201 | 0.40690414 | 0.23729226 | 0.81243007 | 0.94415747 |
| RFX3 | 95.5141491 | 0.9725943 | 0.43035203 | 2.25999702 | 0.02382144 | 0.25161392 |
| RFX3-AS1 | 62.3882339 | -0.101767248 | 0.4547055 | -0.2238091 | 0.82290582 | 0.94740448 |
| GLIS3 | 80.1347714 | 0.534170775 | 0.56796363 | 0.94050173 | 0.34696026 | 0.71988554 |
| SLC1A1 | 23.3280159 | -1.400274559 | 0.85839318 | -1.6312741 | 0.10283249 | NA |
| SPATA6L | 22.8838725 | 0.312300882 | 0.5219406 | 0.59834564 | 0.54960933 | NA |
| CDC37L1 | 19.5442961 | -0.612758349 | 0.55378144 | -1.1064985 | 0.2685108 | NA |
| AK3 | 64.89062 | 0.040443841 | 0.26762467 | 0.1511215 | 0.87987987 | 0.96423929 |
| RCL1 | 23.9893425 | -0.69156682 | 0.42953964 | -1.6100186 | 0.10739379 | NA |
| JAK2 | 92.1952331 | 0.350179659 | 0.3161506 | 1.10763559 | 0.26801923 | 0.66098308 |
| PLGRKT | 14.1546059 | 0.318387736 | 0.57897111 | 0.54991991 | 0.58237431 | NA |
| CD274 | 6.2552131 | -0.399794179 | 1.27912527 | -0.3125528 | 0.75462045 | NA |
| RIC1 | 74.3015134 | 0.528204254 | 0.27501182 | 1.92066022 | 0.05477456 | 0.36420591 |
| ERMP1 | 38.2979371 | 0.57955884 | 0.41083581 | 1.41068238 | 0.15833829 | 0.56089091 |
| KIAA2026 | 81.7235426 | 0.316694332 | 0.25786402 | 1.22814469 | 0.21939266 | 0.62462904 |
| RANBP6 | 8.29388861 | -0.082865166 | 0.48041722 | -0.1724858 | 0.8630556 | NA |
| UHRF2 | 44.4193211 | 0.128841653 | 0.32466431 | 0.39684575 | 0.69148121 | 0.90557575 |
| GLDC | 8.71583943 | 1.120711452 | 0.72553198 | 1.54467548 | 0.12242478 | NA |
| AL162411.1 | 8.08468361 | 0.6533751 | 0.61572069 | 1.06115502 | 0.28861946 | NA |
| KDM4C | 122.97996 | -0.181437293 | 0.19413702 | -0.9345837 | 0.35000288 | 0.72238707 |
| DMAC1 | 27.1971977 | -0.813812126 | 0.42504701 | -1.9146403 | 0.05553839 | NA |
| PTPRD | 910.861167 | 0.407712621 | 0.35359721 | 1.15304254 | 0.24889292 | 0.65087666 |
| MPDZ | 88.2422259 | -0.12804613 | 0.38096373 | -0.3361111 | 0.73678711 | 0.91929008 |
| NFIB | 97.1371064 | 0.196642658 | 0.52351577 | 0.37561936 | 0.7071999 | 0.91058211 |
| ZDHHC21 | 87.2895934 | 0.717727263 | 0.25374048 | 2.82858797 | 0.00467539 | 0.11154919 |
| FREM1 | 11.3638272 | 1.619873496 | 0.77723394 | 2.08415177 | 0.03714636 | NA |
| TTC39B | 43.364106 | -0.463508095 | 0.5978762 | -0.7752577 | 0.43818743 | 0.78335165 |
| SNAPC3 | 35.4452991 | 0.031528025 | 0.2981131 | 0.10575861 | 0.91577388 | 0.97746812 |
| PSIP1 | 39.4327891 | -0.132913518 | 0.30803451 | -0.431489 | 0.66611281 | 0.89968484 |
| CCDC171 | 30.3383348 | 0.37320942 | 0.50604575 | 0.73750134 | 0.46081753 | 0.79680257 |
| BNC2 | 35.5771569 | 1.589995983 | 0.6927413 | 2.29522331 | 0.02172033 | 0.24297341 |
| CNTLN | 54.174515 | 0.400685032 | 0.3013103 | 1.3298086 | 0.18358134 | 0.58584259 |
| SH3GL2 | 7.47082583 | 2.221481937 | 0.73556029 | 3.02012217 | 0.00252673 | NA |
| RRAGA | 21.3443816 | -0.391592813 | 0.36298463 | -1.0788138 | 0.28067076 | NA |
| HAUS6 | 9.13147802 | 0.157529306 | 0.45407623 | 0.3469226 | 0.72864947 | NA |
| PLIN2 | 21.9338796 | -1.957157587 | 0.92644179 | -2.1125532 | 0.03463903 | NA |
| DENND4C | 81.9064838 | 0.151167719 | 0.2231514 | 0.67742223 | 0.4981381 | 0.82345406 |
| RPS6 | 102.64152 | -0.50904862 | 0.4529007 | -1.123974 | 0.26102405 | 0.65595635 |
| ACER2 | 11.0077779 | -1.486094731 | 0.87960787 | -1.6894969 | 0.09112425 | NA |
| MLLT3 | 87.6663224 | 0.568025945 | 0.43745031 | 1.2984925 | 0.19411815 | 0.59865455 |

| | | | | | | |
|---|---|---|---|---|---|---|
| FOCAD | 58.5485472 | 0.16105936 | 0.28192664 | 0.57128109 | 0.56780911 | 0.85654436 |
| HACD4 | 9.70295665 | -0.170761267 | 0.50945556 | -0.3351838 | 0.73748642 | NA |
| KLHL9 | 30.0145257 | 0.299257965 | 0.34076333 | 0.87819885 | 0.37983582 | 0.7485977 |
| AL391117.1 | 12.350635 | 2.665315699 | 0.97749586 | 2.72667722 | 0.00639756 | NA |
| TUSC1 | 6.7647266 | -0.5588547 | 0.58258609 | -0.9592654 | 0.33742505 | NA |
| CAAP1 | 26.168412 | 0.566761335 | 0.30963812 | 1.83039908 | 0.06719029 | NA |
| PLAA | 21.6422501 | 0.418401838 | 0.30177696 | 1.3864605 | 0.16560633 | NA |
| IFT74 | 30.9761443 | 0.539149779 | 0.31858078 | 1.69234876 | 0.0905795 | 0.45866472 |
| LRRC19 | 3.70249531 | -0.843561257 | 0.9971712 | -0.8459543 | 0.39757824 | NA |
| MOB3B | 16.3164606 | -0.546837242 | 0.49338334 | -1.1083415 | 0.26771435 | NA |
| C9orf72 | 8.1320202 | -1.105424015 | 0.58864999 | -1.877897 | 0.06039526 | NA |
| LINGO2 | 34.2099262 | -0.766980187 | 0.57897702 | -1.3247161 | 0.18526533 | 0.587049 |
| ACO1 | 24.5629327 | 0.375404574 | 0.44469558 | 0.84418328 | 0.398567 | NA |
| DDX58 | 48.3289434 | 0.44570848 | 0.9124945 | 0.48845059 | 0.62523071 | 0.87931199 |
| TOPORS | 16.4011166 | -0.456674938 | 0.38157346 | -1.1968205 | 0.23137653 | NA |
| SMIM27 | 4.5728564 | 0.279825651 | 0.75532784 | 0.37046914 | 0.71103297 | NA |
| NDUFB6 | 21.1480194 | -0.148414285 | 0.38778152 | -0.3827266 | 0.70192252 | NA |
| APTX | 24.6865517 | -0.00183758 | 0.32894892 | -0.0055862 | 0.99554287 | NA |
| DNAJA1 | 58.2008659 | -0.053549423 | 0.42051738 | -0.1273418 | 0.89866991 | 0.97097328 |
| SMU1 | 19.4251017 | 0.195212523 | 0.35179999 | 0.55489633 | 0.57896557 | NA |
| B4GALT1 | 33.3842383 | -0.122168116 | 0.33522948 | -0.3644313 | 0.71553597 | 0.91399852 |
| BAG1 | 24.6241088 | -0.404005104 | 0.34103473 | -1.184645 | 0.23615782 | NA |
| CHMP5 | 35.7491837 | 0.217886252 | 0.41420816 | 0.52603081 | 0.5988668 | 0.86882737 |
| NFX1 | 43.0475558 | -0.278248259 | 0.34123358 | -0.8154188 | 0.41483265 | 0.77005091 |
| AQP3 | 27.2721825 | -0.18222782 | 0.43681381 | -0.417175 | 0.67655038 | NA |
| UBE2R2 | 114.130643 | 0.44529689 | 0.2651882 | 1.67917308 | 0.09311832 | 0.46399004 |
| UBAP2 | 102.188289 | 1.094107197 | 0.31912958 | 3.42841044 | 0.00060713 | 0.0356142 |
| DCAF12 | 12.1007906 | -0.011501003 | 0.40895569 | -0.0281229 | 0.97756416 | NA |
| UBAP1 | 65.809084 | 0.144674151 | 0.24596024 | 0.58820137 | 0.55639714 | 0.85122066 |
| NUDT2 | 8.35352728 | -0.372498483 | 0.55514264 | -0.670996 | 0.50222308 | NA |
| FAM219A | 25.3896716 | 0.587616806 | 0.31802917 | 1.84768209 | 0.06464835 | NA |
| CNTFR | 6.47319706 | -1.879768347 | 0.98851295 | -1.9016123 | 0.05722186 | NA |
| RPP25L | 7.29641972 | -0.110821418 | 0.66136601 | -0.1675644 | 0.86692596 | NA |
| DCTN3 | 17.7474868 | -0.392317163 | 0.5099459 | -0.769331 | 0.44169687 | NA |
| SIGMAR1 | 6.00672526 | 0.600683583 | 0.55369179 | 1.08486995 | 0.27797926 | NA |
| IL11RA | 7.45528036 | -0.276725407 | 0.53285333 | -0.5193275 | 0.60353235 | NA |
| VCP | 65.4271108 | -0.353810426 | 0.33018545 | -1.0715506 | 0.28392192 | 0.67249058 |
| PIGO | 8.34617309 | 0.131330962 | 0.58480368 | 0.22457273 | 0.82231168 | NA |
| STOML2 | 14.5013123 | -0.920981795 | 0.56120928 | -1.6410666 | 0.10078359 | NA |
| FAM214B | 7.48375071 | 0.677562217 | 0.56850175 | 1.19183841 | 0.23332461 | NA |
| UNC13B | 48.9415819 | 0.074318504 | 0.24636754 | 0.30165704 | 0.76291352 | 0.92336765 |
| RUSC2 | 87.1820838 | 0.329958484 | 0.40075368 | 0.82334486 | 0.41031191 | 0.76788553 |
| TESK1 | 9.89028549 | 0.58735715 | 0.43224064 | 1.35886608 | 0.17418903 | NA |
| CCDC107 | 24.9303339 | -0.325027493 | 0.47012658 | -0.6913617 | 0.48933829 | NA |
| TPM2 | 80.5842162 | 1.741856106 | 0.46199673 | 3.77027799 | 0.00016307 | 0.01469766 |
| TLN1 | 77.5740094 | -0.53346847 | 0.33252238 | -1.6043085 | 0.10864608 | 0.49402296 |
| CREB3 | 12.4927719 | -0.249939395 | 0.53317409 | -0.4687763 | 0.63922951 | NA |
| GBA2 | 11.8765448 | -0.239723647 | 0.41482167 | -0.5778957 | 0.56333456 | NA |
| RGP1 | 7.97166561 | -0.422428557 | 0.5213404 | -0.810274 | 0.41778273 | NA |
| HINT2 | 9.86665073 | 0.111259249 | 0.64675358 | 0.17202726 | 0.86341609 | NA |
| TMEM8B | 9.02907576 | 0.120475128 | 0.57335454 | 0.21012327 | 0.83357147 | NA |
| RECK | 29.7374912 | -0.103771895 | 0.34829903 | -0.2979391 | 0.76574965 | 0.9240942 |
| CLTA | 87.3694748 | -0.766589962 | 0.31198856 | -2.4571092 | 0.01400601 | 0.19677241 |
| GNE | 21.4128643 | 0.005887533 | 0.45210238 | 0.01302257 | 0.98960979 | NA |
| RNF38 | 91.7038006 | 0.300701117 | 0.20831897 | 1.44346487 | 0.14888956 | 0.54750323 |
| EBLN3P | 28.344347 | -0.134167466 | 0.34042708 | -0.3941151 | 0.69349603 | NA |
| ZCCHC7 | 136.983334 | 0.123064866 | 0.22275141 | 0.55247626 | 0.58062209 | 0.86175324 |
| GRHPR | 29.0816346 | -0.327197396 | 0.42148804 | -0.7762911 | 0.43757715 | 0.78335165 |
| ZBTB5 | 11.2789508 | 0.047266887 | 0.47017749 | 0.10052988 | 0.91992366 | NA |
| AL138752.2 | 18.8068731 | -0.197031851 | 0.45122855 | -0.4366564 | 0.66236053 | NA |
| TRMT10B | 20.0458399 | 0.03903775 | 0.40057914 | 0.09745328 | 0.92236644 | NA |
| DCAF10 | 70.1485643 | 0.008574988 | 0.23622812 | 0.03629961 | 0.97104346 | 0.99008353 |
| ALDH1B1 | 5.66515563 | -0.401757638 | 0.88632698 | -0.4532838 | 0.65034441 | NA |
| GLIDR | 21.7433704 | 0.691661828 | 0.47834889 | 1.44593589 | 0.14819518 | NA |
| CNTNAP3B | 24.5684898 | 0.74416038 | 0.81737876 | 0.91042295 | 0.3625995 | NA |
| AL512625.3 | 7.70376852 | 0.603847082 | 0.54220641 | 1.11368487 | 0.2654144 | NA |
| CBWD5 | 13.2992827 | -0.227371392 | 0.48551087 | -0.4683137 | 0.63956027 | NA |
| ZNF658 | 6.2705183 | -0.241028618 | 0.69166473 | -0.3484761 | 0.72748267 | NA |
| PGM5 | 11.6729338 | -0.348555401 | 0.62577372 | -0.5569991 | 0.57752804 | NA |
| AL161457.2 | 5.50923614 | 1.265610132 | 0.61760652 | 2.04921758 | 0.04044084 | NA |
| FAM189A2 | 8.9793557 | 0.168460955 | 0.48768318 | 0.34543113 | 0.72977027 | NA |
| APBA1 | 99.6764878 | 0.92997284 | 0.4615466 | 2.01490564 | 0.04391454 | 0.33029013 |
| PTAR1 | 45.485299 | 0.236991181 | 0.29921065 | 0.79205464 | 0.42832882 | 0.77844087 |
| MAMDC2-AS | 20.9722789 | 0.620144357 | 0.39260217 | 1.57957445 | 0.11420435 | NA |
| MAMDC2 | 59.6043229 | -0.339438744 | 0.43197348 | -0.7857861 | 0.43199282 | 0.77905765 |
| SMC5-AS1 | 19.2725477 | 0.576555551 | 0.4510151 | 1.27835089 | 0.20112573 | NA |
| SMC5 | 69.7030796 | 0.148967139 | 0.23224193 | 0.64143086 | 0.52124279 | 0.83108387 |
| KLF9 | 56.8342689 | -0.593925612 | 0.38909845 | -1.5264147 | 0.12690661 | 0.52301154 |
| TRPM3 | 61.4424545 | 2.070384681 | 0.57658278 | 3.59078479 | 0.00032968 | 0.02431268 |
| CEMIP2 | 134.952412 | -1.065343899 | 0.5138436 | -2.0732843 | 0.03814583 | 0.304707 |
| ABHD17B | 14.031442 | -0.200292927 | 0.59047821 | -0.3392046 | 0.7344556 | NA |
| C9orf85 | 22.891618 | -0.092905983 | 0.36106873 | -0.2573083 | 0.79694078 | NA |
| GDA | 21.1064052 | 1.667756175 | 0.94641914 | 1.76217503 | 0.07803973 | NA |
| LINC01504 | 6.57660255 | -0.153043841 | 1.20421222 | -0.1270904 | 0.89886883 | NA |
| ZFAND5 | 107.344768 | -0.41739394 | 0.36353187 | -1.1481633 | 0.25090117 | 0.65183754 |
| TMC1 | 43.5938201 | -0.202034022 | 0.79110843 | -0.255381 | 0.79842888 | 0.94015741 |
| ALDH1A1 | 11.0587646 | -1.15375093 | 0.91745547 | -1.2575552 | 0.20855265 | NA |
| ANXA1 | 80.4381452 | -1.636554807 | 0.7193647 | -2.2750002 | 0.02290592 | 0.24709152 |
| CARNMT1 | 18.0030618 | 0.218808707 | 0.33974361 | 0.64404068 | 0.51954905 | NA |
| NMRK1 | 37.2646752 | -0.649497013 | 0.38607756 | -1.6822967 | 0.09251132 | 0.46324185 |
| OSTF1 | 32.4750595 | -0.689405387 | 0.43090632 | -1.5998962 | 0.10962161 | 0.49598431 |
| PCSK5 | 18.0513284 | 0.686575917 | 0.61871332 | 1.10968342 | 0.26713547 | NA |
| RFK | 14.2446715 | -0.906804651 | 0.45494119 | -1.9932349 | 0.04623573 | NA |
| PRUNE2 | 38.5172137 | -0.654205867 | 0.60928288 | -1.0737309 | 0.2829433 | 0.67190751 |
| PCA3 | 11.8916573 | -1.612454798 | 1.00016375 | -1.6121908 | 0.10692043 | NA |
| VPS13A | 91.3054552 | 0.624606817 | 0.25325035 | 2.46636114 | 0.01364936 | 0.194252 |
| GNAQ | 205.830281 | -0.098966375 | 0.23225329 | -0.426114 | 0.67002479 | 0.90305784 |
| CEP78 | 11.5428842 | 0.558308882 | 0.48073084 | 1.16137522 | 0.24548934 | NA |
| TLE4 | 109.888326 | 0.787927718 | 0.38667388 | 2.03770609 | 0.04157933 | 0.31989578 |
| TLE1 | 21.8846105 | -3.333329103 | 0.80957805 | -4.117366 | 3.83E-05 | NA |
| AL591368.1 | 4.20162757 | 3.360454303 | 1.20511867 | 2.78848414 | 0.00529553 | NA |
| FRMD3 | 553.856403 | 0.045321004 | 0.42719379 | 0.10609003 | 0.91551092 | 0.97744467 |
| UBQLN1 | 52.478604 | -0.097356675 | 0.28928433 | -0.3365432 | 0.73646128 | 0.91929008 |
| GKAP1 | 16.8187738 | 0.137226195 | 0.49805337 | 0.27552508 | 0.78291286 | NA |
| KIF27 | 17.2750331 | 0.171449139 | 0.37173822 | 0.46120934 | 0.64464842 | NA |
| C9orf64 | 5.51598953 | -0.702388503 | 0.65684762 | -1.0693325 | 0.28491987 | NA |
| HNRNPK | 112.812618 | -0.209708768 | 0.25543672 | -0.8209813 | 0.41165693 | 0.7690836 |
| RMI1 | 9.44188891 | 0.346356489 | 0.48560018 | 0.71325445 | 0.47568832 | NA |
| NTRK2 | 26.8823358 | 1.312699895 | 0.49205703 | 2.66777996 | 0.00763542 | NA |
| AGTPBP1 | 40.0285739 | -0.18415779 | 0.27771263 | -0.6631236 | 0.50725142 | 0.82657837 |
| NAA35 | 44.0407616 | 0.026315411 | 0.26825215 | 0.09809954 | 0.92185326 | 0.97775374 |
| GOLM1 | 9.48745586 | -0.662383438 | 0.60713665 | -1.0909956 | 0.2752748 | NA |
| ISCA1 | 14.9110814 | -0.333615397 | 0.45502534 | -0.7331798 | 0.46344876 | NA |

| | | | | | | |
|---|---|---|---|---|---|---|
| TUT7 | 28.4889611 | 0.02553159 | 0.35776276 | 0.07136458 | 0.9431076 | NA |
| AL353613.1 | 5.76682807 | 0.060354892 | 0.69420479 | 0.08694105 | 0.93071837 | NA |
| GAS1 | 101.321877 | -0.346631334 | 0.61423764 | -0.5643277 | 0.57253111 | 0.85816194 |
| AL513318.2 | 18.7463112 | 0.616607852 | 0.54516479 | 1.13104857 | 0.25803465 | NA |
| DAPK1 | 184.532123 | 0.821114614 | 0.25832052 | 3.17866588 | 0.00147955 | 0.05883368 |
| CTSL | 79.6963417 | -0.392167174 | 0.48229196 | -0.8131323 | 0.4161422 | 0.7708692 |
| SPIN1 | 71.5416961 | -0.14593087 | 0.25772719 | -0.5662222 | 0.57124271 | 0.8573305 |
| SECISBP2 | 43.9456552 | -0.275419912 | 0.27612281 | -0.9974544 | 0.31854399 | 0.69469143 |
| SEMA4D | 27.5750562 | 0.078318866 | 0.35087929 | 0.22320744 | 0.82337406 | NA |
| AUH | 61.3202252 | 0.539097366 | 0.2866911 | 1.88041196 | 0.06005195 | 0.37905854 |
| NFIL3 | 8.07961353 | -0.697699675 | 0.94154137 | -0.7410186 | 0.45868216 | NA |
| SPTLC1 | 39.0206877 | -0.528581882 | 0.37064237 | -1.4261237 | 0.15383263 | 0.55412156 |
| IARS | 75.454376 | -0.155195758 | 0.27018961 | -0.5743957 | 0.56570004 | 0.85573063 |
| NOL8 | 17.0687248 | -0.62017217 | 0.42342837 | -1.4646448 | 0.14301785 | NA |
| CENPP | 53.0841297 | 0.560195811 | 0.34288841 | 1.63375545 | 0.10231021 | 0.48317363 |
| OMD | 4.85491194 | 1.130523589 | 0.94831944 | 1.19213373 | 0.23320881 | NA |
| ECM2 | 12.1208191 | 1.117143554 | 0.58433402 | 1.91182357 | 0.05589883 | NA |
| IPPK | 10.7562224 | 0.597414911 | 0.41249258 | 1.44830464 | 0.14753187 | NA |
| BICD2 | 21.9420529 | -0.283796557 | 0.46287316 | -0.6131195 | 0.53979734 | NA |
| ZNF484 | 8.93279181 | -0.557660069 | 0.49846792 | -1.1187482 | 0.26324759 | NA |
| CARD19 | 5.50088252 | -0.356821147 | 0.68056585 | -0.5243007 | 0.60006941 | NA |
| NINJ1 | 23.6382482 | -0.852135549 | 0.42848349 | -1.9887244 | 0.04673164 | NA |
| FAM120AOS | 14.3871823 | -0.563115777 | 0.44852492 | -1.2554838 | 0.20930318 | NA |
| FAM120A | 72.9808892 | -0.048026333 | 0.26043487 | -0.1844082 | 0.85369321 | 0.95569766 |
| PHF2 | 20.7817327 | 0.322390916 | 0.42145228 | 0.76495236 | 0.44429991 | NA |
| PTPDC1 | 22.9132927 | -0.298380615 | 0.38623444 | -0.7725376 | 0.4397961 | NA |
| AL158152.2 | 46.4585545 | -0.032496604 | 0.3348325 | -0.0970533 | 0.92268406 | 0.97775374 |
| ZNF169 | 6.96064076 | 0.080132416 | 0.64503288 | 0.12422997 | 0.90113319 | NA |
| MFSD14B | 32.5228766 | 0.578267391 | 0.2910806 | 1.98662289 | 0.0469642 | 0.33774256 |
| FBP1 | 7.25186375 | 0.699647751 | 0.70913962 | 0.98661495 | 0.32383144 | NA |
| C9orf3 | 334.380961 | 0.338343832 | 0.16692303 | 2.02694513 | 0.04266802 | 0.32591621 |
| FANCC | 61.2181066 | 0.033626532 | 0.25895384 | 0.12985531 | 0.8966809 | 0.97097328 |
| PTCH1 | 9.99099602 | 0.307989371 | 0.47815698 | 0.64411769 | 0.51949912 | NA |
| LINC00476 | 28.0990179 | -0.029701889 | 0.4527276 | -0.0656065 | 0.94769109 | NA |
| ERCC6L2 | 55.6808332 | -0.2701283 | 0.25825532 | -1.0459738 | 0.29557312 | 0.68103704 |
| SLC35D2 | 10.7825446 | 0.290459287 | 0.53561784 | 0.5422883 | 0.58761991 | NA |
| HABP4 | 33.6727658 | 0.687018712 | 0.33054046 | 2.07847083 | 0.03766602 | 0.30414888 |
| CDC14B | 26.0632203 | 0.779200334 | 0.40684037 | 1.91524831 | 0.05546084 | NA |
| AAED1 | 13.312079 | 0.09985035 | 0.4689565 | 0.21292028 | 0.83138914 | NA |
| ZNF510 | 6.83249201 | -0.159127053 | 0.55894652 | -0.284691 | 0.77588087 | NA |
| ZNF782 | 26.938669 | -0.135918794 | 0.3362327 | -0.4042403 | 0.68603605 | NA |
| MFSD14C | 73.8203126 | 0.368222843 | 0.2310806 | 1.59348227 | 0.11105204 | 0.49770949 |
| CTSV | 9.42865796 | 1.754486017 | 0.94643143 | 1.85379095 | 0.06376907 | NA |
| CCDC180 | 8.61052872 | -0.898017295 | 0.64011035 | -1.4029101 | 0.16064364 | NA |
| TDRD7 | 27.5345678 | 0.05723599 | 0.42266888 | 0.13541567 | 0.89228323 | NA |
| TMOD1 | 42.0183948 | 0.498501068 | 0.96749531 | 0.51524908 | 0.60637898 | 0.87080602 |
| TSTD2 | 19.2275082 | -0.487761342 | 0.3989421 | -1.2226369 | 0.22146686 | NA |
| NCBP1 | 19.6823403 | 0.782645679 | 0.44577256 | 1.75570627 | 0.07913858 | NA |
| XPA | 19.3396586 | -0.818085627 | 0.3560198 | -2.2978656 | 0.02156944 | NA |
| TRMO | 6.74879507 | -0.286671076 | 0.68392251 | -0.4191572 | 0.67510122 | NA |
| ANP32B | 26.6071713 | -0.788155735 | 0.36332783 | -2.1692688 | 0.03006228 | NA |
| NANS | 6.43295208 | -0.749265271 | 0.5683835 | -1.318239 | 0.18742367 | NA |
| TRIM14 | 13.2292375 | -0.054908054 | 0.67726351 | -0.0810734 | 0.93538358 | NA |
| CORO2A | 9.78825032 | -0.102827921 | 0.61346278 | -0.1676188 | 0.86688315 | NA |
| ANKS6 | 7.93819129 | 0.481627137 | 0.48335728 | 0.99642056 | 0.31904585 | NA |
| TGFBR1 | 22.1810536 | 0.470511003 | 0.38864662 | 1.21063964 | 0.22603354 | NA |
| ALG2 | 13.6897106 | -0.058917608 | 0.43875795 | -0.1342827 | 0.89317902 | NA |
| SEC61B | 56.9581561 | -0.964475273 | 0.45147571 | -2.1362728 | 0.03265718 | 0.29047359 |
| AL359710.1 | 20.2000463 | 1.041475424 | 0.53641759 | 1.94153855 | 0.05219299 | NA |
| STX17 | 25.3607771 | 0.740990475 | 0.40565621 | 1.82664644 | 0.06775295 | NA |
| ERP44 | 54.4362897 | 0.222153595 | 0.27479912 | 0.80842178 | 0.41884782 | 0.77290571 |
| INVS | 65.7548112 | -0.26895395 | 0.30777904 | -0.873854 | 0.38219777 | 0.74899182 |
| TEX10 | 11.314026 | -0.115888788 | 0.50368143 | -0.2300835 | 0.81802688 | NA |
| MRPL50 | 13.947137 | -0.971003023 | 0.50678466 | -1.9160071 | 0.05536419 | NA |
| ZNF189 | 16.4701765 | -0.546452314 | 0.45674802 | -1.1963978 | 0.23154136 | NA |
| ALDOB | 36.1907977 | 0.704684423 | 1.54216292 | 0.45694551 | 0.64771021 | 0.89261695 |
| TMEM246 | 14.0650651 | 0.415172088 | 0.43504478 | 0.95432035 | 0.33992151 | NA |
| RNF20 | 9.60304422 | -0.232559017 | 0.5587525 | -0.4162111 | 0.6772555 | NA |
| SMC2 | 7.1545125 | 0.547559855 | 0.50991585 | 1.07382396 | 0.28290159 | NA |
| NIPSNAP3A | 19.5058743 | 0.700247665 | 0.37537576 | 1.86545786 | 0.06211725 | NA |
| ABCA1 | 20.3663767 | -1.855553531 | 1.02770776 | -1.8055264 | 0.07099234 | NA |
| SLC44A1 | 66.4872873 | 0.928834195 | 0.2983508 | 3.11322841 | 0.00185053 | 0.06823399 |
| FSD1L | 53.0517871 | 0.6413498 | 0.26953388 | 2.37947751 | 0.0173372 | 0.21770801 |
| FKTN | 20.8962884 | 0.355627059 | 0.37506768 | 0.94816771 | 0.34304409 | NA |
| TMEM38B | 21.3483294 | -0.327811347 | 0.55061409 | -0.5953559 | 0.5516056 | NA |
| LINC01505 | 11.4027043 | -0.410061836 | 0.69653798 | -0.5887143 | 0.55605298 | NA |
| ZNF462 | 109.736497 | 0.280689782 | 0.25376956 | 1.10608136 | 0.2686913 | 0.66139295 |
| AL807761.4 | 5.98347547 | 0.398951912 | 0.82443758 | 0.48390797 | 0.62845119 | NA |
| RAD23B | 121.008685 | 0.265652168 | 0.23031675 | 1.15342098 | 0.24873763 | 0.65087666 |
| ELP1 | 18.8548365 | 0.262625974 | 0.36555333 | 0.71843409 | 0.47248967 | NA |
| FAM206A | 15.6283285 | 0.394121539 | 0.49669599 | 0.79348646 | 0.42749446 | NA |
| CTNNAL1 | 39.7468157 | -0.158136174 | 0.56197141 | -0.2813954 | 0.77840714 | 0.93078401 |
| TMEM245 | 188.015823 | 0.301359064 | 0.29446497 | 1.02341226 | 0.30611296 | 0.68558486 |
| PTPN3 | 40.2959809 | 0.182208505 | 0.33048591 | 0.55133517 | 0.58140393 | 0.86175324 |
| PALM2-AKAF | 7.78277543 | 1.923247274 | 0.69292912 | 2.77553247 | 0.00551114 | NA |
| TXN | 52.949554 | -1.070513213 | 0.47694787 | -2.2445078 | 0.02479975 | 0.25362919 |
| ECPAS | 87.7161549 | 0.123018098 | 0.26174355 | 0.46999476 | 0.63835876 | 0.886719 |
| ZNF483 | 34.0660686 | 0.262274174 | 0.39425873 | 0.66523364 | 0.50590107 | 0.82600448 |
| PTGR1 | 49.4714671 | -0.475819719 | 0.39670434 | -1.1994316 | 0.23036017 | 0.63682501 |
| C9orf84 | 14.4329923 | -0.73094725 | 0.7904961 | -0.924669 | 0.3551381 | NA |
| UGCG | 207.530454 | -0.73909041 | 0.40459035 | -1.8267623 | 0.06773551 | 0.40302824 |
| SUSD1 | 33.997859 | 0.294704672 | 0.31802046 | 0.92668464 | 0.35409029 | 0.72571512 |
| PTBP3 | 301.282896 | 0.099094786 | 0.2969935 | 0.33365978 | 0.7386363 | 0.91993596 |
| HSDL2 | 58.3594786 | -0.156616487 | 0.30190099 | -0.5187677 | 0.60392273 | 0.87047761 |
| C9orf147 | 4.52545377 | -0.874411315 | 0.78834609 | -1.1091719 | 0.26735604 | NA |
| KIAA1958 | 60.4481187 | 0.258617233 | 0.28872756 | 0.89571371 | 0.37040569 | 0.73862609 |
| INIP | 16.6393749 | 0.211169581 | 0.39291223 | 0.53744722 | 0.59095874 | NA |
| SNX30 | 14.4259538 | 0.524431474 | 0.49894723 | 1.05107603 | 0.29322367 | NA |
| SLC31A2 | 12.3610451 | -0.378812103 | 0.64342021 | -0.5887476 | 0.5560306 | NA |
| FKBP15 | 16.3084391 | -0.575874361 | 0.37478243 | -1.5365565 | 0.12440196 | NA |
| SLC31A1 | 7.67979714 | 0.25644087 | 0.59586046 | 0.43037068 | 0.66692602 | NA |
| CDC26 | 10.4465099 | -0.420055661 | 0.60087203 | -0.6990767 | 0.48450407 | NA |
| PRPF4 | 7.09764818 | -0.476632736 | 0.55339882 | -0.8612825 | 0.38908246 | NA |
| HDHD3 | 9.58377541 | -0.86051956 | 0.67073656 | -1.2829472 | 0.19951059 | NA |
| ALAD | 6.3294663 | 0.250645639 | 0.57984541 | 0.43226287 | 0.66555037 | NA |
| POLE3 | 5.29643319 | -0.299033389 | 0.6908139 | -0.4328711 | 0.6651084 | NA |
| RGS3 | 11.4999979 | -0.616787459 | 0.72618373 | -0.8493546 | 0.39568403 | NA |
| ZNF618 | 21.3097794 | 0.687704943 | 0.60960747 | 1.12811108 | 0.25927301 | NA |
| KIF12 | 9.04395857 | 1.107282402 | 0.6988959 | 1.58433095 | 0.11311843 | NA |
| COL27A1 | 12.0847225 | 0.59252959 | 0.49007706 | 1.20905391 | 0.22664214 | NA |
| ATP6V1G1 | 92.1492788 | -1.062430617 | 0.46993672 | -2.2607951 | 0.02377195 | 0.25161392 |
| TMEM268 | 4.57828639 | 1.21059877 | 1.05707063 | 1.14523925 | 0.25211006 | NA |
| PAPPA | 97.4235485 | -2.037228899 | 0.4928705 | -4.1333959 | 3.57E-05 | 0.00483262 |

| | | | | | | |
|---|---|---|---|---|---|---|
| ASTN2 | 44.9069395 | 0.375222429 | 0.49343311 | 0.76043221 | 0.44699628 | 0.78947076 |
| TRIM32 | 4.61182814 | -0.576143926 | 0.68514836 | -0.8409039 | 0.40040177 | NA |
| CDK5RAP2 | 44.8904144 | 0.091283922 | 0.29453841 | 0.30992196 | 0.7566203 | 0.92313949 |
| MEGF9 | 138.24521 | -0.685735437 | 0.33010948 | -2.077297 | 0.03777415 | 0.30414888 |
| FBXW2 | 23.5800182 | -0.196909417 | 0.32800906 | -0.600317 | 0.54829499 | NA |
| PSMD5 | 19.0718846 | -0.33393346 | 0.34087186 | -0.9796451 | 0.32726132 | NA |
| PHF19 | 4.27861558 | -1.140818264 | 0.75816796 | -1.5047039 | 0.13240023 | NA |
| C5 | 10.0879711 | 0.353270579 | 0.62449709 | 0.56568811 | 0.57160582 | NA |
| CNTRL | 30.5117639 | -0.893381901 | 0.33335 | -2.6800117 | 0.00736196 | 0.13572774 |
| RAB14 | 78.6508551 | 0.02319947 | 0.31895376 | 0.07273615 | 0.94201608 | 0.98114803 |
| GSN | 148.130595 | 0.170982669 | 0.35753317 | 0.47822882 | 0.63248735 | 0.88389921 |
| STOM | 13.4649955 | -1.36324754 | 0.5864066 | -2.324748 | 0.02008544 | NA |
| DAB2IP | 58.4621506 | 0.400063237 | 0.34871217 | 1.14725917 | 0.25127452 | 0.65183754 |
| TTLL11 | 12.8533054 | -0.106059424 | 0.51275129 | -0.2068438 | 0.83613185 | NA |
| NDUFA8 | 15.7759874 | -1.065098229 | 0.46445274 | -2.2932327 | 0.02183461 | NA |
| RBM18 | 13.6234878 | 0.395565376 | 0.40869452 | 0.96787542 | 0.33310659 | NA |
| MRRF | 18.8905183 | 0.014739154 | 0.43264741 | 0.03406736 | 0.97282344 | NA |
| PDCL | 11.5349189 | -0.391366939 | 0.54265802 | -0.7212036 | 0.47078423 | NA |
| RC3H2 | 51.2374504 | -0.027359519 | 0.25730968 | -0.1063291 | 0.91532121 | 0.97744467 |
| ZBTB6 | 5.02380954 | -0.152322085 | 0.68946015 | -0.2209295 | 0.82514733 | NA |
| ZBTB26 | 8.53233011 | 0.08327235 | 0.48892404 | 0.17031756 | 0.8647604 | NA |
| RABGAP1 | 73.1746123 | 0.20988413 | 0.22879183 | 0.91735851 | 0.3589548 | 0.72955368 |
| STRBP | 29.9899586 | 0.010697232 | 0.38393499 | 0.02786209 | 0.97777214 | 0.99319905 |
| CRB2 | 180.745447 | 0.408819556 | 0.49667908 | 0.82310605 | 0.41044769 | 0.76788553 |
| DENND1A | 123.579287 | -0.336051578 | 0.29239161 | -1.1493202 | 0.25042397 | 0.6518054 |
| PSMB7 | 30.3013237 | -0.397064574 | 0.34057961 | -1.1658495 | 0.24367529 | 0.64937383 |
| NR6A1 | 28.8837661 | 1.112884918 | 0.70469124 | 1.57925181 | 0.11427831 | 0.50483797 |
| MIR181A2HG | 10.3349824 | 0.461746535 | 0.58317241 | 0.79178391 | 0.42848668 | NA |
| OLFML2A | 18.2597604 | -1.032709508 | 0.61978178 | -1.666247 | 0.09566424 | NA |
| RPL35 | 61.5414926 | -0.313911091 | 0.49714993 | -0.6314214 | 0.52776504 | 0.83195298 |
| ARPC5L | 15.7766384 | -0.640183721 | 0.4669613 | -1.3709567 | 0.17038845 | NA |
| GOLGA1 | 16.4687192 | -0.436851661 | 0.42211054 | -1.0349224 | 0.30070515 | NA |
| SCAI | 16.6193349 | -0.216652651 | 0.41952655 | -0.5164218 | 0.60555985 | NA |
| PPP6C | 33.5809356 | 0.542552358 | 0.3477135 | 1.56034311 | 0.11867882 | 0.51050744 |
| RABEPK | 8.94418258 | 0.440320667 | 0.49917599 | 0.88209504 | 0.37772542 | NA |
| HSPA5 | 145.727941 | -0.811859504 | 0.63538375 | -1.2777467 | 0.20133875 | 0.60910153 |
| GAPVD1 | 50.9557385 | 0.315102358 | 0.21911372 | 1.43807679 | 0.15041227 | 0.54834287 |
| MAPKAP1 | 77.5248121 | 0.084332225 | 0.23925234 | 0.35248235 | 0.72447656 | 0.91541337 |
| PBX3 | 75.273126 | -0.213161236 | 0.38436545 | -0.5545796 | 0.57918225 | 0.86175324 |
| MVB12B | 20.6427671 | 0.025220523 | 0.34949696 | 0.07216235 | 0.9424727 | NA |
| AL161908.1 | 10.7762055 | 0.36820499 | 0.43590679 | 0.84468745 | 0.39828537 | NA |
| LMX1B | 129.315562 | 0.52969092 | 0.50328589 | 1.05246527 | 0.29258614 | 0.67930583 |
| ZBTB43 | 19.5074708 | -0.33067109 | 0.35234324 | -0.9384914 | 0.34799195 | NA |
| ZBTB34 | 5.7637237 | 0.00696457 | 0.58670637 | 0.01187062 | 0.99052884 | NA |
| RALGPS1 | 21.5481607 | 0.523430758 | 0.32722384 | 1.5996107 | 0.10968497 | NA |
| GARNL3 | 11.1318307 | 0.378283818 | 0.50428811 | 0.75013432 | 0.45317381 | NA |
| SLC2A8 | 4.25045589 | 0.392859628 | 0.61352654 | 0.64033029 | 0.5219579 | NA |
| RPL12 | 16.2815146 | -0.552170839 | 0.69755614 | -0.7915791 | 0.42860616 | NA |
| LRSAM1 | 16.0938988 | -0.321151413 | 0.43873476 | -0.7319945 | 0.46417195 | NA |
| FAM129B | 16.7876872 | -1.511749693 | 0.59457693 | -2.5425636 | 0.01100426 | NA |
| STXBP1 | 31.0790556 | -0.023064318 | 0.29323865 | -0.0786537 | 0.93730804 | 0.97856407 |
| PTRH1 | 7.59024535 | -0.528916184 | 0.5480511 | -0.9650855 | 0.33450198 | NA |
| CDK9 | 30.9410728 | 0.927100249 | 0.34378234 | 2.69676522 | 0.00700166 | 0.13332741 |
| FPGS | 19.5044273 | -0.219693737 | 0.39347698 | -0.5583395 | 0.57661259 | NA |
| ENG | 10.3229372 | 0.324335063 | 0.67061115 | 0.48364102 | 0.62864065 | NA |
| AK1 | 4.7451071 | -0.011120816 | 0.71056227 | -0.0156507 | 0.98751304 | NA |
| DPM2 | 10.7257165 | -0.723469399 | 0.63429686 | -1.1405849 | 0.25404272 | NA |
| FAM102A | 10.1475481 | -0.56800319 | 0.56710991 | -1.0015751 | 0.31654883 | NA |
| SLC25A25 | 9.77950834 | -0.07937258 | 0.4754964 | -0.1669257 | 0.86742849 | NA |
| PTGES2 | 10.077051 | 0.008841174 | 0.48085002 | 0.01838655 | 0.98533048 | NA |
| C9orf16 | 20.418946 | 0.05438514 | 0.3966462 | 0.13711247 | 0.8909419 | NA |
| CIZ1 | 12.9467026 | -0.102634084 | 0.44406117 | -0.231126 | 0.81721691 | NA |
| DNM1 | 10.3279782 | -0.528505852 | 0.58844787 | -0.8981354 | 0.36911338 | NA |
| GOLGA2 | 36.9196341 | -0.212742422 | 0.44983709 | -0.4729322 | 0.63626158 | 0.88633797 |
| SWI5 | 16.0424565 | -0.93381384 | 0.51757963 | -1.8041936 | 0.07120097 | NA |
| TRUB2 | 5.69696831 | -0.648383999 | 0.67338039 | -0.9628792 | 0.33560813 | NA |
| COQ4 | 25.3291447 | 0.318773236 | 0.30702593 | 1.0382616 | 0.29914828 | NA |
| SLC27A4 | 20.6148234 | -0.04381327 | 0.42087252 | -0.104101 | 0.91708916 | NA |
| URM1 | 10.4664752 | -0.11933881 | 0.54623523 | -0.2184751 | 0.82705894 | NA |
| ODF2 | 7.57353005 | -0.889484975 | 0.58093112 | -1.5311367 | 0.12573561 | NA |
| GLE1 | 13.626803 | -0.123947477 | 0.41981592 | -0.2952424 | 0.76780868 | NA |
| AL356481.1 | 9.42595653 | 0.575008833 | 0.6981774 | 0.82358557 | 0.41017508 | NA |
| SPTAN1 | 152.910038 | 0.189048701 | 0.29091137 | 0.64984983 | 0.51578923 | 0.83104823 |
| WDR34 | 8.53548504 | -0.15921684 | 0.55358048 | -0.2876128 | 0.77364313 | NA |
| SET | 56.0171833 | -0.981286673 | 0.32248738 | -3.0428685 | 0.00234335 | 0.07786717 |
| ZDHHC12 | 18.6026028 | -0.299277137 | 0.44304999 | -0.6754929 | 0.49936264 | NA |
| ZER1 | 36.6194359 | 0.325356649 | 0.37113874 | 0.87664427 | 0.38067989 | 0.74899182 |
| TBC1D13 | 11.3600499 | 0.614248056 | 0.4943705 | 1.24248524 | 0.21405758 | NA |
| ENDOG | 9.95148956 | -0.499250327 | 0.4871545 | -1.0248296 | 0.30544362 | NA |
| SPOUT1 | 6.56326243 | 0.327296153 | 0.64971648 | 0.50375227 | 0.61443547 | NA |
| KYAT1 | 29.2971755 | 0.164571836 | 0.52412584 | 0.31399298 | 0.75352637 | 0.92313949 |
| LRRC8A | 23.6325453 | -0.511067868 | 0.56455285 | -0.9052613 | 0.36532695 | NA |
| DOLK | 4.06878711 | -0.801913393 | 0.79662378 | -1.00664 | 0.31410779 | NA |
| NUP188 | 7.30923548 | -0.84480123 | 0.57034742 | -1.4812046 | 0.13855206 | NA |
| SH3GLB2 | 21.589482 | 0.198396858 | 0.37195711 | 0.53338638 | 0.59376614 | NA |
| MIGA2 | 6.75299102 | 1.329270131 | 0.58503091 | 2.27213656 | 0.02307826 | NA |
| CRAT | 4.6326204 | -0.316272679 | 0.68741585 | -0.4600893 | 0.64545212 | NA |
| PTPA | 28.7789011 | 0.376684527 | 0.42173275 | 0.89318302 | 0.37175917 | 0.74014106 |
| IER5L | 12.4529634 | -2.162003927 | 0.98034563 | -2.2053487 | 0.02742964 | NA |
| LINC01503 | 14.4157779 | -3.514484625 | 0.9266487 | -3.7926828 | 0.00014903 | NA |
| NTMT1 | 13.1352404 | -0.745316006 | 0.66534548 | -1.120194 | 0.2626311 | NA |
| ASB6 | 5.28033483 | 0.21078969 | 0.62213444 | 0.33881694 | 0.73474764 | NA |
| TOR1B | 10.2560538 | -0.378871457 | 0.4650892 | -0.8146211 | 0.41528925 | NA |
| TOR1A | 7.63733846 | -0.212542175 | 0.5694536 | -0.3732388 | 0.70897073 | NA |
| C9orf78 | 16.1598482 | -0.274137337 | 0.38334345 | -0.715122 | 0.47453366 | NA |
| USP20 | 8.53701128 | 0.13326532 | 0.5369431 | 0.24819263 | 0.80398537 | NA |
| FNBP1 | 201.111763 | -0.279368798 | 0.33988899 | -0.8219413 | 0.41111031 | 0.7685709 |
| GPR107 | 40.7339889 | 0.361979633 | 0.27657666 | 1.30878591 | 0.19060688 | 0.59329687 |
| NCS1 | 14.8256631 | 0.798299066 | 0.47824761 | 1.66921704 | 0.09507437 | NA |
| ASS1 | 25.5082395 | -0.384185618 | 0.50081242 | -0.7671248 | 0.44300733 | NA |
| FUBP3 | 39.9218183 | 0.406106402 | 0.28357307 | 1.43210495 | 0.15211379 | 0.55086922 |
| EXOSC2 | 3.29218785 | -0.308808541 | 0.84295727 | -0.3663395 | 0.71411176 | NA |
| ABL1 | 74.9075882 | -0.065752053 | 0.20622268 | -0.3188401 | 0.74984779 | 0.92274989 |
| AIF1L | 108.556369 | -0.019016188 | 0.45126666 | -0.0421396 | 0.96638743 | 0.98906571 |
| NUP214 | 46.8549524 | 0.40862053 | 0.27936044 | 1.46270008 | 0.14354947 | 0.54150812 |
| AL157938.2 | 8.79817951 | -1.052933087 | 0.72724678 | -1.4478346 | 0.1476633 | NA |
| PRRC2B | 103.327588 | -0.031427955 | 0.24131478 | -0.1302363 | 0.89637944 | 0.97097328 |
| POMT1 | 13.5869794 | 0.765596716 | 0.39143185 | 1.95588766 | 0.05047839 | NA |
| UCK1 | 11.9437385 | 0.243193168 | 0.51094905 | 0.47596364 | 0.63410028 | NA |
| RAPGEF1 | 84.8110028 | 0.071979079 | 0.23452542 | 0.30691376 | 0.75890901 | 0.92313949 |
| MED27 | 102.583733 | -1.210142873 | 0.26828787 | -4.5106134 | 6.46E-06 | 0.00131091 |
| NTNG2 | 6.55108325 | 0.279160294 | 0.58662827 | 0.47587255 | 0.63416518 | NA |
| SETX | 104.260891 | 0.164439359 | 0.25804932 | 0.63724004 | 0.5239685 | 0.83178884 |

| | | | | | | |
|---|---|---|---|---|---|---|
| TTF1 | 12.8973318 | -0.638419821 | 0.40979185 | -1.5579124 | 0.11925401 | NA |
| DDX31 | 11.8497314 | -0.68830029 | 0.54169643 | -1.2706384 | 0.20385731 | NA |
| GTF3C4 | 14.1548809 | 0.079231638 | 0.37382516 | 0.21194838 | 0.8321473 | NA |
| AK8 | 6.7702264 | -0.321979784 | 0.61191881 | -0.5261806 | 0.59876276 | NA |
| SPACA9 | 5.0588554 | -0.06021026 | 0.76049962 | -0.079172 | 0.93689583 | NA |
| TSC1 | 20.5735897 | -0.028940836 | 0.40530705 | -0.0714047 | 0.94307565 | NA |
| GTF3C5 | 10.8208837 | 0.251171171 | 0.42287126 | 0.59396605 | 0.55253482 | NA |
| SURF6 | 6.8372349 | -0.40463519 | 0.57286548 | -0.7063354 | 0.47997956 | NA |
| MED22 | 6.44180865 | 0.035832007 | 0.55606132 | 0.06443895 | 0.94862072 | NA |
| RPL7A | 62.6550103 | -0.591028886 | 0.40529112 | -1.4582823 | 0.14476274 | 0.5436645 |
| SURF1 | 9.74183206 | 0.051151568 | 0.55344636 | 0.09242371 | 0.9263614 | NA |
| SURF2 | 6.1572775 | -0.562062134 | 0.58822817 | -0.9555172 | 0.33931621 | NA |
| SURF4 | 38.2541846 | -0.041162053 | 0.40239107 | -0.1022937 | 0.91852359 | 0.97775374 |
| REXO4 | 7.07892273 | -0.471494999 | 0.53646382 | -0.8788943 | 0.37945858 | NA |
| CACFD1 | 6.77064456 | 0.262188811 | 0.59778097 | 0.43860347 | 0.66094888 | NA |
| VAV2 | 67.1528786 | 0.309303306 | 0.42686232 | 0.72459734 | 0.46869909 | 0.80448595 |
| BRD3OS | 7.51564277 | -0.230085543 | 0.63097762 | -0.3646493 | 0.71537321 | NA |
| BRD3 | 20.157251 | 0.139759894 | 0.36631982 | 0.38152425 | 0.70281428 | NA |
| WDR5 | 9.9977162 | -0.476919099 | 0.58431946 | -0.8161958 | 0.41438815 | NA |
| RXRA | 14.0216335 | -0.117551891 | 0.45244559 | -0.2598144 | 0.79500692 | NA |
| PPP1R26 | 7.52599211 | -0.288591386 | 0.5197017 | -0.555302 | 0.57868812 | NA |
| MRPS2 | 8.43280515 | 0.282681595 | 0.49080392 | 0.57595628 | 0.56464474 | NA |
| CAMSAP1 | 18.0738934 | 0.152233812 | 0.3865075 | 0.39387027 | 0.69367682 | NA |
| UBAC1 | 9.90253985 | -0.134073219 | 0.47011886 | -0.28519 | 0.77549855 | NA |
| NACC2 | 38.5428806 | 0.316801477 | 0.44988699 | 0.70418013 | 0.4813206 | 0.81183849 |
| TMEM250 | 7.12784021 | 0.143408172 | 0.62315192 | 0.23013356 | 0.81798798 | NA |
| QSOX2 | 5.96011709 | 0.049443465 | 0.61275692 | 0.08069018 | 0.93568835 | NA |
| GPSM1 | 47.2326606 | -0.153624576 | 0.4065799 | -0.377846 | 0.70554502 | 0.90991753 |
| SNAPC4 | 4.73992257 | 0.259190144 | 0.61652235 | 0.42040673 | 0.67418835 | NA |
| ENTR1 | 7.83983456 | -0.395917154 | 0.63767177 | -0.6208792 | 0.53467909 | NA |
| PMPCA | 11.3548265 | 0.106204183 | 0.45781432 | 0.23198091 | 0.81655284 | NA |
| INPP5E | 6.41501814 | 0.131510032 | 0.56147941 | 0.23422058 | 0.81481374 | NA |
| SEC16A | 21.4072411 | -0.613446617 | 0.41992129 | -1.460861 | 0.1440536 | NA |
| AGPAT2 | 11.1518632 | -0.278390407 | 0.55815709 | -0.4987671 | 0.61794345 | NA |
| SNHG7 | 17.2644157 | -0.470975302 | 0.6893813 | -0.6831855 | 0.49448964 | NA |
| RABL6 | 34.4993276 | -0.194613376 | 0.37647873 | -0.5169306 | 0.60520461 | 0.87047761 |
| PHPT1 | 53.2486388 | -0.736848484 | 0.58044406 | -1.2694565 | 0.2042783 | 0.6137834 |
| EDF1 | 52.090126 | -0.283798903 | 0.40203337 | -0.7059088 | 0.48024485 | 0.81093801 |
| TRAF2 | 6.54272744 | 0.781247217 | 0.5695481 | 1.37169665 | 0.17015789 | NA |
| FBXW5 | 24.6008171 | -0.069678885 | 0.43845109 | -0.1589205 | 0.87373148 | NA |
| LCNL1 | 11.2669543 | 0.764358716 | 0.61592722 | 1.24098869 | 0.21460993 | NA |
| PAXX | 10.0432929 | 0.008009956 | 0.50263885 | 0.01593581 | 0.9872856 | NA |
| ABCA2 | 9.20151708 | 0.418280753 | 0.72627883 | 0.5759231 | 0.56466716 | NA |
| MAN1B1 | 23.2485589 | 0.223551618 | 0.46610282 | 0.47961868 | 0.63149856 | NA |
| DPP7 | 18.9205376 | 0.260382102 | 0.41437864 | 0.62836757 | 0.52976317 | NA |
| SSNA1 | 22.1052247 | -0.413416571 | 0.37061834 | -1.1154779 | 0.26464568 | NA |
| TPRN | 5.33096981 | 0.249451311 | 0.5991083 | 0.41637098 | 0.67713856 | NA |
| TMEM203 | 13.7531273 | -0.387463911 | 0.47473201 | -0.816174 | 0.41440064 | NA |
| NDOR1 | 6.3229505 | -0.026857699 | 0.58659402 | -0.0457858 | 0.96348095 | NA |
| BX255925.3 | 6.96370757 | 0.802519737 | 0.69361951 | 1.15700282 | 0.24727121 | NA |
| TUBB4B | 53.1917078 | -0.518053792 | 0.53671058 | -0.9652386 | 0.33442529 | 0.70906302 |
| NELFB | 17.5471024 | 0.22877386 | 0.42257907 | 0.54137528 | 0.58824894 | NA |
| EXD3 | 24.5431976 | 0.102818843 | 0.38139552 | 0.26958587 | 0.78747887 | NA |
| NOXA1 | 4.91127808 | -0.488385584 | 0.63991685 | -0.7632016 | 0.44534316 | NA |
| NSMF | 6.15217303 | 0.349809128 | 0.53059049 | 0.6592827 | 0.50971425 | NA |
| PNPLA7 | 12.2968004 | -0.717552317 | 0.68825906 | -1.0425614 | 0.29715148 | NA |
| MRPL41 | 41.694511 | -0.549530801 | 0.32972637 | -1.6666268 | 0.09558863 | 0.46937954 |
| DPH7 | 10.9068995 | -0.031992482 | 0.42187434 | -0.0758341 | 0.93955105 | NA |
| ARRDC1 | 11.0400172 | 0.049524599 | 0.52102536 | 0.09505219 | 0.92427338 | NA |
| ARRDC1-AS1 | 6.5472731 | -0.474513502 | 0.54904626 | -0.8642505 | 0.3874503 | NA |
| EHMT1 | 80.0180813 | -0.073103854 | 0.23923991 | -0.3055671 | 0.75993425 | 0.92313949 |
| AC069287.3 | 6.34914222 | 0.65384823 | 0.88809057 | 0.73624049 | 0.46158436 | NA |
| BET1L | 8.80361654 | -0.371690347 | 0.4793896 | -0.7753409 | 0.43813827 | NA |
| RIC8A | 12.8390321 | -0.793509262 | 0.4926575 | -1.6106712 | 0.1072514 | NA |
| SIRT3 | 7.45456111 | -0.073189011 | 0.5474196 | -0.1336982 | 0.89364124 | NA |
| PSMD13 | 25.4930139 | -0.387980462 | 0.43131655 | -0.899526 | 0.36837254 | NA |
| PGGHG | 10.076328 | -0.860109829 | 0.6131316 | -1.4028144 | 0.1606722 | NA |
| IFITM2 | 304.372133 | -2.837040349 | 0.52958788 | -5.3570719 | 8.46E-08 | 5.72E-05 |
| AC136475.1 | 4.35572969 | -2.243060644 | 0.93543891 | -2.3978697 | 0.01649073 | NA |
| IFITM3 | 693.857483 | -2.490881668 | 0.59674718 | -4.1740988 | 2.99E-05 | 0.00418422 |
| SIGIRR | 16.9579384 | -0.94176765 | 0.50994956 | -1.8467859 | 0.06477818 | NA |
| PTDSS2 | 18.1611556 | -0.914429926 | 0.41950064 | -2.1798058 | 0.02927186 | NA |
| RNH1 | 45.6268754 | -1.037447036 | 0.3717836 | -2.7904594 | 0.00526333 | 0.11257186 |
| HRAS | 17.0776348 | -0.653217378 | 0.44984022 | -1.4521098 | 0.14647109 | NA |
| RASSF7 | 36.808923 | -0.035325014 | 0.45464662 | -0.0776977 | 0.93806849 | 0.97885408 |
| MIR210HG | 5.84604951 | -0.706814432 | 0.63161155 | -1.1190651 | 0.26311238 | NA |
| PHRF1 | 15.6513299 | 0.730441651 | 0.46886402 | 1.55789657 | 0.11925777 | NA |
| IRF7 | 9.99116107 | -1.553046988 | 1.29907471 | -1.1955024 | 0.23189078 | NA |
| DEAF1 | 28.053499 | -0.147947869 | 0.38072026 | -0.3885999 | 0.69757211 | NA |
| EPS8L2 | 26.7610381 | 0.114270792 | 0.39671734 | 0.28804083 | 0.77331548 | NA |
| TALDO1 | 31.3401801 | -0.669478291 | 0.53434257 | -1.2529009 | 0.21024177 | 0.61792799 |
| GATD1 | 13.3764887 | 0.112993027 | 0.56662281 | 0.19941489 | 0.84193821 | NA |
| AP006621.3 | 12.3692387 | -0.713146523 | 0.49771055 | -1.4328539 | 0.15189958 | NA |
| SLC25A22 | 4.34060143 | -1.032750566 | 1.04648714 | -0.9868736 | 0.32370459 | NA |
| RPLP2 | 53.3533526 | -0.044377252 | 0.44256685 | -0.1002724 | 0.92012805 | 0.97775374 |
| PNPLA2 | 16.8212078 | 0.357551363 | 0.4030371 | 0.88714257 | 0.37500215 | NA |
| CD151 | 237.519865 | -0.13897087 | 0.45086266 | -0.3082333 | 0.75790484 | 0.92313949 |
| POLR2L | 99.7348716 | -0.556008233 | 0.41661847 | -1.3345741 | 0.18201577 | 0.58390314 |
| TSPAN4 | 53.8955587 | 0.234295431 | 0.39913359 | 0.58701005 | 0.55719696 | 0.85122066 |
| CHID1 | 11.9528211 | 0.894911233 | 0.56138855 | 1.5941031 | 0.11091294 | NA |
| AP2A2 | 45.6014168 | 0.119062695 | 0.30015369 | 0.39667243 | 0.69160903 | 0.90557575 |
| TOLLIP | 20.5369608 | -0.026940872 | 0.34425312 | -0.0782589 | 0.93762211 | NA |
| MOB2 | 15.5641258 | 0.239162798 | 0.39066692 | 0.61219107 | 0.54041135 | NA |
| DUSP8 | 6.39221684 | 0.168259773 | 0.64105758 | 0.26247217 | 0.79295744 | NA |
| MRPL23 | 7.05566493 | -0.238796241 | 0.69024117 | -0.3459606 | 0.72937234 | NA |
| CD81-AS1 | 5.69493588 | -0.75246682 | 0.78377378 | -0.9600561 | 0.33702697 | NA |
| CD81 | 624.303717 | -0.674548434 | 0.4881673 | -1.3817977 | 0.16703383 | 0.56878882 |
| TSSC4 | 15.6686875 | -0.226742111 | 0.48069872 | -0.4716928 | 0.6371461 | NA |
| KCNQ1 | 80.6362774 | -0.163251251 | 0.37171359 | -0.4391856 | 0.66052706 | 0.89752019 |
| KCNQ1OT1 | 407.061597 | 0.016606367 | 0.36152247 | 0.04593454 | 0.96336243 | 0.98892249 |
| CDKN1C | 1144.20297 | 0.104216052 | 0.48592136 | 0.21447103 | 0.83017976 | 0.95008863 |
| SLC22A18 | 6.41048536 | -0.409703056 | 0.89752004 | -0.4564835 | 0.64804236 | NA |
| NAP1L4 | 47.6621984 | -0.228813744 | 0.35632169 | -0.642155 | 0.52077257 | 0.83108387 |
| CARS | 22.3786409 | -0.478132459 | 0.46660867 | -1.0246969 | 0.30550623 | NA |
| OSBPL5 | 15.5077216 | -0.121504438 | 0.37107392 | -0.32744 | 0.74333513 | NA |
| ZNF195 | 14.9795729 | 0.36146813 | 0.41408551 | 0.87293113 | 0.38270062 | NA |
| AC127526.1 | 10.451039 | 0.844091831 | 0.60216391 | 1.40176423 | 0.16098566 | NA |
| NUP98 | 46.9264745 | 0.025351041 | 0.26960526 | 0.09403022 | 0.92508515 | 0.97775374 |
| PGAP2 | 7.20562432 | -0.109432703 | 0.56278421 | -0.1944488 | 0.8458245 | NA |
| RHOG | 10.7008175 | -0.493397749 | 0.51203345 | -0.9636045 | 0.33524424 | NA |
| STIM1 | 39.8766758 | 0.022108253 | 0.38332754 | 0.05767457 | 0.95400785 | 0.98559751 |
| RRM1 | 8.76545683 | -0.343179736 | 0.44424325 | -0.7725041 | 0.43981591 | NA |
| TRIM21 | 6.88309485 | 0.702801982 | 0.89100153 | 0.78877753 | 0.43024205 | NA |

| | | | | | | |
|---|---|---|---|---|---|---|
| AC104389.4 | 23.6973537 | -0.080125219 | 0.50120102 | -0.1598664 | 0.87298629 | NA |
| TRIM5 | 10.5953957 | -0.149490186 | 0.99469125 | -0.150288 | 0.88053738 | NA |
| TRIM22 | 37.6861249 | -0.893901596 | 0.90805005 | -0.9844189 | 0.32490961 | 0.69623471 |
| FAM160A2 | 10.7547048 | 0.231701081 | 0.45752493 | 0.50642285 | 0.61255984 | NA |
| SMPD1 | 31.478768 | 0.843872439 | 0.47211948 | 1.78741289 | 0.07387078 | 0.41834543 |
| APBB1 | 10.0372016 | 0.238543885 | 0.47576779 | 0.50138721 | 0.61609864 | NA |
| ARFIP2 | 10.2438987 | -0.478424774 | 0.50585561 | -0.9457734 | 0.34426418 | NA |
| AC084337.2 | 8.80968654 | -0.315444899 | 0.50736635 | -0.62173 | 0.53411939 | NA |
| DNHD1 | 12.5821103 | -1.253477225 | 0.47142018 | -2.6589384 | 0.00783873 | NA |
| RRP8 | 4.31501987 | -0.406194573 | 0.6797595 | -0.5975563 | 0.55013603 | NA |
| ILK | 34.7685198 | -0.551906761 | 0.35464489 | -1.5562237 | 0.11965492 | 0.51093023 |
| TAF10 | 11.6226154 | -0.232087824 | 0.60882879 | -0.3812038 | 0.70305206 | NA |
| TPP1 | 30.6698851 | -0.21321674 | 0.37446762 | -0.5693863 | 0.569094 | 0.8573305 |
| MRPL17 | 15.1127558 | -1.369855636 | 0.6950234 | -1.970949 | 0.04872971 | NA |
| AC087280.2 | 8.4004642 | -0.029333558 | 0.80456056 | -0.0364591 | 0.97091629 | NA |
| SYT9 | 35.8521333 | 0.334754885 | 0.30290266 | 1.10515664 | 0.26909172 | 0.66147637 |
| AC107884.1 | 7.47407237 | 0.440404094 | 0.62700129 | 0.70239743 | 0.48243135 | NA |
| PPFIBP2 | 13.6709028 | 1.780323525 | 0.45475125 | 3.91493931 | 9.04E-05 | NA |
| EIF3F | 22.1474072 | -0.159212142 | 0.39604605 | -0.4020041 | 0.687681 | NA |
| TUB | 14.3150289 | -0.007323684 | 0.67166843 | -0.0109037 | 0.99130026 | NA |
| RIC3 | 82.4959506 | -1.21109549 | 0.60949325 | -1.9870532 | 0.04691651 | 0.33774256 |
| STK33 | 106.242132 | 0.449708751 | 0.29959191 | 1.50107106 | 0.13333718 | 0.52613957 |
| TRIM66 | 16.9917424 | 0.411785181 | 0.51996519 | 0.79194759 | 0.42839123 | NA |
| RPL27A | 55.6957001 | -0.207051172 | 0.46422733 | -0.4460125 | 0.65558825 | 0.89583156 |
| ST5 | 41.3248594 | -0.600938157 | 0.42304072 | -1.4205208 | 0.15545611 | 0.5560229 |
| AKIP1 | 13.999755 | -0.111017624 | 0.43105503 | -0.2575486 | 0.7967553 | NA |
| TMEM9B | 39.3081248 | -0.301424831 | 0.44592137 | -0.6759596 | 0.4990663 | 0.82345406 |
| TMEM9B-AS | 9.01708817 | -0.272104835 | 0.56623344 | -0.4805524 | 0.63083466 | NA |
| DENND5A | 37.3800147 | 0.585354237 | 0.4427252 | 1.32216154 | 0.18611436 | 0.58792656 |
| TMEM41B | 35.3749517 | -0.061292992 | 0.32364921 | -0.1893809 | 0.84979426 | 0.95532127 |
| IPO7 | 36.6769066 | -0.445026435 | 0.41258181 | -1.078638 | 0.28074912 | 0.66958859 |
| ZNF143 | 8.41709647 | -0.59601765 | 0.53811348 | -1.1076059 | 0.26803207 | NA |
| WEE1 | 10.5249455 | 0.343366418 | 0.51889133 | 0.66173089 | 0.50814371 | NA |
| SWAP70 | 30.9926734 | -0.087298743 | 0.30445408 | -0.2867386 | 0.77431246 | 0.92806061 |
| SBF2-AS1 | 16.0148921 | -0.32869011 | 0.43365511 | -0.7579528 | 0.44847923 | NA |
| SBF2 | 419.773133 | 0.433049949 | 0.14654505 | 2.95506372 | 0.00312604 | 0.0904424 |
| AC011092.2 | 16.9642505 | -0.207893572 | 0.52933647 | -0.3927437 | 0.69450878 | NA |
| AC100763.1 | 4.09204098 | -0.816650353 | 0.77682945 | -1.0512608 | 0.29313883 | NA |
| ADM | 60.5888775 | 1.343227719 | 0.92656746 | 1.44968152 | 0.14714735 | 0.54635866 |
| AMPD3 | 12.914611 | 0.980061661 | 0.67819988 | 1.44509265 | 0.14843186 | NA |
| MTRNR2L8 | 20.9683874 | -0.493667128 | 0.56868151 | -0.8680907 | 0.38534468 | NA |
| RNF141 | 16.3202985 | 0.279964077 | 0.3583843 | 0.78118399 | 0.43469429 | NA |
| CTR9 | 18.6888436 | -0.291541809 | 0.36196043 | -0.8054522 | 0.4205588 | NA |
| EIF4G2 | 77.0488015 | -0.549149558 | 0.31048366 | -1.7686907 | 0.07694551 | 0.42346132 |
| ZBED5 | 29.7612819 | -0.125298323 | 0.31731723 | -0.3948677 | 0.69294053 | 0.90557575 |
| ZBED5-AS1 | 11.2508526 | -0.040592723 | 0.59152456 | -0.0686239 | 0.94528899 | NA |
| GALNT18 | 23.3778318 | 0.035218862 | 0.66643589 | 0.05284659 | 0.95785414 | NA |
| USP47 | 113.06876 | 0.246224463 | 0.19866995 | 1.23936439 | 0.21521058 | 0.62305077 |
| DKK3 | 6.07305383 | -1.06207971 | 1.07985414 | -0.98354 | 0.32534175 | NA |
| MICAL2 | 38.3854805 | -1.483861053 | 0.5779248 | -2.5675677 | 0.01024148 | 0.1620804 |
| PARVA | 203.76296 | 0.42911664 | 0.23974929 | 1.78985571 | 0.07347711 | 0.41834543 |
| TEAD1 | 347.498595 | 0.413774618 | 0.30752028 | 1.34551977 | 0.17845744 | 0.58191579 |
| ARNTL | 10.9408806 | 1.729160315 | 0.53005566 | 3.26222403 | 0.00110542 | NA |
| BTBD10 | 29.0774935 | 0.090887325 | 0.29380004 | 0.30935096 | 0.75705457 | 0.92313949 |
| FAR1 | 71.0497046 | 0.28065351 | 0.2276937 | 1.23259232 | 0.2177279 | 0.62454341 |
| RRAS2 | 58.0567013 | 0.45690412 | 0.30827509 | 1.48213119 | 0.13830539 | 0.53418928 |
| COPB1 | 41.3122345 | -0.063655466 | 0.29285211 | -0.2173639 | 0.82792479 | 0.94941654 |
| PSMA1 | 11.0018117 | 1.809720628 | 0.62684748 | 2.88701908 | 0.00388911 | NA |
| PDE3B | 10.7613596 | 1.377453745 | 0.56606914 | 2.43336662 | 0.01495915 | NA |
| CYP2R1 | 9.45070457 | 0.171450284 | 0.6028967 | 0.28437755 | 0.77612107 | NA |
| SOX6 | 301.267826 | 0.656511397 | 0.31894539 | 2.0583818 | 0.0395535 | 0.31456664 |
| C11orf58 | 65.9364044 | -0.356598812 | 0.22173173 | -1.6082444 | 0.10778166 | 0.49167977 |
| PLEKHA7 | 99.0216872 | 0.483908975 | 0.46896777 | 1.03185977 | 0.30213782 | 0.68233351 |
| RPS13 | 52.5518802 | -0.918143169 | 0.56383785 | -1.6283816 | 0.103444 | 0.48612129 |
| PIK3C2A | 78.3792199 | -0.087632813 | 0.25173749 | -0.3481119 | 0.72775616 | 0.91700494 |
| NUCB2 | 23.7106281 | -0.769836452 | 0.44833152 | -1.7171143 | 0.0859583 | NA |
| SERGEF | 37.3648868 | -0.133792871 | 0.35740954 | -0.3743405 | 0.708151 | 0.9106723 |
| SAAL1 | 4.52282692 | -0.021518905 | 0.78057068 | -0.0275682 | 0.97800657 | NA |
| SAA1 | 10.3185645 | -0.250668611 | 4.03776998 | -0.062081 | 0.95049836 | NA |
| HPS5 | 56.8817525 | 0.515872208 | 0.49503929 | 1.04208335 | 0.29737304 | 0.68103704 |
| GTF2H1 | 29.0782944 | 0.493262697 | 0.33541211 | 1.47061683 | 0.14139477 | 0.53597868 |
| LDHA | 90.4429268 | -1.816454644 | 0.41400063 | -4.3875649 | 1.15E-05 | 0.0021133 |
| TSG101 | 40.9611355 | -0.117370919 | 0.2794317 | -0.4200344 | 0.67446034 | 0.90403541 |
| UEVLD | 26.1374511 | -0.265754754 | 0.34196767 | -0.7771341 | 0.43707963 | NA |
| SPTY2D1OS | 7.70497833 | -0.393451101 | 0.47195572 | -0.8336611 | 0.40447201 | NA |
| SPTY2D1 | 11.0427268 | -0.146658662 | 0.45139799 | -0.3248988 | 0.74525767 | NA |
| ZDHHC13 | 19.908252 | -0.118722339 | 0.3642821 | -0.3259077 | 0.74449419 | NA |
| NAV2 | 114.933667 | 0.259797854 | 0.83645259 | 0.31059484 | 0.75610865 | 0.92313949 |
| HTATIP2 | 19.78346 | -0.359564231 | 0.39339546 | -0.914002 | 0.3607158 | NA |
| PRMT3 | 20.9938292 | -0.338539564 | 0.38377482 | -0.8821307 | 0.37770611 | NA |
| NELL1 | 106.711905 | 2.097975169 | 1.33057863 | 1.57673896 | 0.1148556 | 0.50513044 |
| ANO5 | 21.6300872 | 0.184669941 | 0.48257014 | 0.38268 | 0.70195704 | NA |
| FANCF | 6.26059689 | 0.227438732 | 0.61841723 | 0.36777554 | 0.71304061 | NA |
| GAS2 | 8.2912436 | 0.239790724 | 0.61589217 | 0.38933881 | 0.69702553 | NA |
| SVIP | 31.0180611 | -0.633070833 | 0.35757968 | -1.7704329 | 0.07665505 | 0.42346132 |
| AC006299.1 | 9.19880049 | -0.36520111 | 0.5122575 | -0.7129249 | 0.47589225 | NA |
| SLC5A12 | 20.4993357 | 2.922209672 | 1.78762916 | 1.6346845 | 0.1021152 | NA |
| BBOX1 | 13.9844246 | 0.745185403 | 1.04017338 | 0.71640499 | 0.47374131 | NA |
| BBOX1-AS1 | 8.51728927 | -0.897025972 | 0.74705224 | -1.200754 | 0.22984664 | NA |
| LGR4 | 210.576488 | 0.240341129 | 0.23254913 | 1.03350689 | 0.30136675 | 0.68219446 |
| LIN7C | 25.7855187 | 0.205768912 | 0.31672735 | 0.64967207 | 0.51590407 | NA |
| BDNF-AS | 60.2362268 | -0.02831286 | 0.30985014 | -0.091376 | 0.92719385 | 0.97775374 |
| METTL15 | 105.800004 | 0.244854027 | 0.24678337 | 0.99218205 | 0.32110873 | 0.69513974 |
| KCNA4 | 6.17339356 | 0.638240687 | 0.87366461 | 0.73053284 | 0.46506455 | NA |
| ARL14EP | 19.2454184 | -0.07412383 | 0.47021323 | -0.1576388 | 0.87474146 | NA |
| MPPED2 | 10.8728787 | 1.233951183 | 0.75114009 | 1.64277103 | 0.10043032 | NA |
| DCDC1 | 16.80383 | 0.44044797 | 0.55027786 | 0.80041013 | 0.42347321 | NA |
| DNAJC24 | 13.9739113 | -0.097656925 | 0.40119016 | -0.243418 | 0.80768157 | NA |
| IMMP1L | 29.3544573 | 0.077988127 | 0.34773564 | 0.22427419 | 0.82254396 | 0.94740448 |
| ELP4 | 54.1973693 | 0.219773441 | 0.28085867 | 0.78250546 | 0.43391758 | 0.78047436 |
| AC131571.1 | 5.96814955 | -1.243499434 | 0.87287911 | -1.4245953 | 0.15427423 | NA |
| RCN1 | 30.1042952 | -0.466299252 | 0.57246794 | -0.8145421 | 0.41533445 | 0.77027733 |
| WT1 | 443.118313 | 0.327687507 | 0.42930154 | 0.76330383 | 0.44528222 | 0.78798633 |
| EIF3M | 110.938547 | 0.137470263 | 0.22799916 | 0.60294196 | 0.5465473 | 0.84449366 |
| CCDC73 | 5.9480653 | 0.003660269 | 0.80984343 | 0.00451972 | 0.99639379 | NA |
| PRRG4 | 4.91382074 | 0.105308747 | 0.8056833 | 0.13070737 | 0.8960068 | NA |
| QSER1 | 32.3762365 | -0.118607821 | 0.32952749 | -0.359933 | 0.71889724 | 0.91472065 |
| TCP11L1 | 15.2334084 | -0.516204559 | 0.43592732 | -1.1841528 | 0.23635258 | NA |
| CSTF3 | 32.0344869 | 0.525337293 | 0.28888068 | 1.81852695 | 0.06898363 | 0.40491695 |
| HIPK3 | 158.884328 | -0.366036364 | 0.34990459 | -1.0461033 | 0.29551332 | 0.68103704 |
| CD59 | 4.79271094 | -0.648006644 | 0.71242097 | -0.9095839 | 0.363042 | NA |
| FBXO3 | 31.4412452 | 0.559562728 | 0.3311019 | 1.69000157 | 0.09102765 | 0.45906588 |
| LMO2 | 25.6047495 | -1.225925186 | 0.77585383 | -1.5800981 | 0.11408439 | NA |

| | | | | | | |
|---|---|---|---|---|---|---|
| CAPRIN1 | 57.0744994 | -0.054553017 | 0.23827169 | -0.228953 | 0.81890545 | 0.94694381 |
| NAT10 | 12.9586649 | -0.053800035 | 0.5009579 | -0.1073943 | 0.91447616 | NA |
| ABTB2 | 21.7605006 | 0.101136708 | 0.83912655 | 0.12052617 | 0.90406635 | NA |
| CAT | 23.2661612 | -0.088817367 | 0.3622387 | -0.2451902 | 0.80630919 | NA |
| APIP | 14.9741792 | 0.600239318 | 0.48768153 | 1.23080181 | 0.218397 | NA |
| PDHX | 27.666731 | 0.210681167 | 0.31053506 | 0.67844566 | 0.49748916 | NA |
| PAMR1 | 31.6845285 | -2.874848815 | 0.83644522 | -3.436984 | 0.00058823 | 0.03508621 |
| FJX1 | 6.29527351 | -2.088181509 | 1.21976126 | -1.7119592 | 0.08690419 | NA |
| TRIM44 | 106.865475 | -0.198348084 | 0.36102037 | -0.5494097 | 0.58272433 | 0.86175324 |
| LDLRAD3 | 6.88716754 | -1.945067782 | 0.83514902 | -2.3290068 | 0.0198587 | NA |
| COMMD9 | 8.82834535 | 0.232361766 | 0.56294897 | 0.41275813 | 0.67978383 | NA |
| TRAF6 | 12.1780104 | 0.052572049 | 0.45263237 | 0.11614735 | 0.90753577 | NA |
| C11orf74 | 8.42539698 | 0.035741145 | 0.62637821 | 0.05706001 | 0.95449739 | NA |
| API5 | 36.4691604 | 0.106408196 | 0.30792413 | 0.34556628 | 0.72966869 | 0.91718621 |
| TTC17 | 136.431629 | 0.047749607 | 0.22830367 | 0.20914954 | 0.83433151 | 0.95164471 |
| HSD17B12 | 69.5336722 | -0.024030721 | 0.2080036 | -0.1155303 | 0.90802479 | 0.9747971 |
| ALKBH3 | 12.617935 | 0.45186533 | 0.46907298 | 0.96331563 | 0.33538915 | NA |
| ACCS | 8.38782542 | 0.815090207 | 0.79709842 | 1.0225716 | 0.30651044 | NA |
| EXT2 | 38.7236514 | 0.102592297 | 0.33985464 | 0.30187111 | 0.76275032 | 0.92336765 |
| TP53I11 | 4.81478691 | 0.189218599 | 0.62393855 | 0.3032648 | 0.76168807 | NA |
| PRDM11 | 10.3450921 | -0.308714571 | 0.49636476 | -0.621951 | 0.53397408 | NA |
| CHST1 | 21.4544723 | -0.07044159 | 0.58201539 | -0.1210305 | 0.90366691 | NA |
| SLC35C1 | 4.22923805 | -0.247485002 | 0.74938638 | -0.3302502 | 0.74121092 | NA |
| CRY2 | 26.0700179 | -0.062592072 | 0.35585385 | -0.1758926 | 0.86037829 | NA |
| MAPK8IP1 | 5.79911289 | 0.950280419 | 0.74607112 | 1.27371291 | 0.20276518 | NA |
| PEX16 | 5.81956956 | -0.016160058 | 0.67273901 | -0.0240213 | 0.98083563 | NA |
| PHF21A | 117.653897 | 0.3166815 | 0.23064348 | 1.37303469 | 0.16974156 | 0.57273205 |
| DGKZ | 7.77314203 | 0.831310351 | 0.54567334 | 1.52345789 | 0.12764419 | NA |
| MDK | 15.9780978 | -0.616002703 | 2.7028639 | -0.2279074 | 0.81971822 | NA |
| AMBRA1 | 154.612028 | -0.095517139 | 0.35227138 | -0.2711465 | 0.78627839 | 0.93615351 |
| ATG13 | 21.1055999 | 0.255869463 | 0.34845977 | 0.73428695 | 0.46277387 | NA |
| ARHGAP1 | 18.1528581 | 0.352496352 | 0.36234243 | 0.9728266 | 0.33063949 | NA |
| CKAP5 | 25.3667067 | 0.277561899 | 0.34819021 | 0.79715595 | 0.42536046 | NA |
| LRP4 | 21.5141606 | -0.60932133 | 0.51221716 | -1.1895762 | 0.23421301 | NA |
| C11orf49 | 79.2201442 | 0.80799308 | 0.22820261 | 3.54068294 | 0.00039909 | 0.02714874 |
| ARFGAP2 | 19.5214655 | 0.492117957 | 0.37290952 | 1.3196712 | 0.18694482 | NA |
| PACSIN3 | 11.7145155 | -0.154140435 | 0.53508842 | -0.2880654 | 0.77329671 | NA |
| DDB2 | 17.642123 | -0.498093089 | 0.43104501 | -1.1555477 | 0.24786619 | NA |
| ACP2 | 7.5843071 | 1.53134758 | 0.60315006 | 2.53891641 | 0.01111964 | NA |
| MADD | 17.7788046 | 0.222864687 | 0.42100532 | 0.52936312 | 0.59655358 | NA |
| SLC39A13 | 18.5593253 | 0.130306348 | 0.51325041 | 0.25388455 | 0.79958475 | NA |
| PSMC3 | 10.5547048 | 0.153723503 | 0.62186378 | 0.24719804 | 0.80475497 | NA |
| CELF1 | 102.155571 | 0.313737107 | 0.26719882 | 1.17417102 | 0.24032653 | 0.64811464 |
| NDUFS3 | 16.7998094 | 0.519110277 | 0.42679391 | 1.21630198 | 0.22386991 | NA |
| KBTBD4 | 5.13066461 | -0.093675915 | 0.66244977 | -0.1414083 | 0.88754737 | NA |
| MTCH2 | 22.823394 | -0.009659901 | 0.42617815 | -0.0226663 | 0.98191642 | NA |
| FNBP4 | 74.5841472 | -0.081021917 | 0.26012705 | -0.3114706 | 0.75544292 | 0.92313949 |
| NUP160 | 31.2927905 | -0.334737117 | 0.29586875 | -1.1313703 | 0.25789926 | 0.65588339 |
| PTPRJ | 50.3987187 | 0.499726612 | 0.49533438 | 1.0088672 | 0.31303833 | 0.69213367 |
| TNKS1BP1 | 24.9961584 | -0.299633589 | 0.42697957 | -0.7017516 | 0.4828341 | NA |
| SSRP1 | 15.5911494 | -0.079406011 | 0.37052286 | -0.214308 | 0.83030688 | NA |
| TIMM10 | 8.5108417 | -0.64545921 | 0.76006853 | -0.8492119 | 0.39576341 | NA |
| UBE2L6 | 35.2325534 | -1.204805156 | 0.89574294 | -1.3450345 | 0.1786141 | 0.58191579 |
| SERPING1 | 73.867081 | -0.642842589 | 0.53918681 | -1.1922446 | 0.23316533 | 0.63935677 |
| CLP1 | 4.58890068 | -0.17801225 | 0.70746225 | -0.2516208 | 0.80133414 | NA |
| ZDHHC5 | 21.7139538 | -0.636056831 | 0.3666942 | -1.7345702 | 0.08281696 | NA |
| MED19 | 7.57935237 | -0.468860261 | 0.61659769 | -0.760399 | 0.44701613 | NA |
| CTNND1 | 108.159263 | 0.153490474 | 0.248818 | 0.61687851 | 0.53731486 | 0.83677705 |
| LPXN | 10.9954778 | 0.38636674 | 0.58218959 | 0.6636442 | 0.50691806 | NA |
| GLYAT | 10.2762169 | 2.89765418 | 1.71395471 | 1.69062471 | 0.0909085 | NA |
| FAM111A | 16.1573145 | -0.298519802 | 0.59014626 | -0.5058404 | 0.61296872 | NA |
| DTX4 | 13.3864973 | 0.688466631 | 0.56958162 | 1.20872339 | 0.22676913 | NA |
| OSBP | 27.9514727 | -0.262759299 | 0.32054349 | -0.8197306 | 0.41236971 | NA |
| PATL1 | 16.2062782 | 0.268358283 | 0.51569513 | 0.52038165 | 0.60279759 | NA |
| STX3 | 17.8504249 | -0.100702724 | 0.52776224 | -0.1908108 | 0.84867384 | NA |
| MRPL16 | 15.1416115 | -0.487514408 | 0.40379408 | -1.2073342 | 0.22730346 | NA |
| PRPF19 | 13.4318473 | -0.061767439 | 0.48365058 | -0.1277109 | 0.89837778 | NA |
| TMEM109 | 30.1397129 | -0.193179499 | 0.52371525 | -0.3688636 | 0.71222939 | 0.91273377 |
| SLC15A3 | 8.72366902 | -1.713829561 | 1.28955035 | -1.3290133 | 0.18384358 | NA |
| VPS37C | 18.7374941 | 0.690600869 | 0.34986804 | 1.97388957 | 0.0483943 | NA |
| DDB1 | 32.0659081 | -0.513712551 | 0.34897462 | -1.4720628 | 0.14100392 | 0.53597868 |
| TKFC | 6.50771841 | -0.104201733 | 0.5792558 | -0.179889 | 0.85723973 | NA |
| CYB561A3 | 7.09943155 | 0.742594463 | 0.5407574 | 1.37324881 | 0.16967501 | NA |
| TMEM138 | 6.43282563 | -0.087018626 | 0.63551961 | -0.1369252 | 0.89108995 | NA |
| TMEM216 | 6.63064214 | -0.403936392 | 0.61351314 | -0.6583989 | 0.51028182 | NA |
| CPSF7 | 34.0413018 | 0.549610688 | 0.4005879 | 1.37201021 | 0.17006026 | 0.57273205 |
| AP003108.2 | 18.1874305 | -0.139287938 | 0.42129343 | -0.3306198 | 0.74093171 | NA |
| TMEM258 | 20.1692317 | -0.467471969 | 0.58873993 | -0.7940212 | 0.42718311 | NA |
| FADS1 | 10.1639794 | -1.440005811 | 0.79196351 | -1.8182729 | 0.06902243 | NA |
| FADS3 | 9.169502 | -0.741983259 | 0.5848619 | -1.2686469 | 0.20456701 | NA |
| RAB3IL1 | 9.17454605 | -1.659550782 | 0.5208265 | -3.1863793 | 0.00144066 | NA |
| FTH1 | 430.253299 | -1.541637119 | 0.51000241 | -3.0228036 | 0.00250445 | 0.08046695 |
| ASRGL1 | 9.37377933 | -0.904266831 | 0.77889475 | -1.1609615 | 0.24565755 | NA |
| AHNAK | 177.507286 | -0.415430145 | 0.42115375 | -0.9864097 | 0.32393211 | 0.69623471 |
| EML3 | 10.3322482 | -0.537415748 | 0.68218188 | -0.7877895 | 0.43081982 | NA |
| B3GAT3 | 23.4643293 | -0.124842267 | 0.40097516 | -0.3113466 | 0.75553712 | NA |
| GANAB | 57.0665752 | 0.123958873 | 0.32238904 | 0.38450089 | 0.70060723 | 0.90729979 |
| UQCC3 | 10.8080285 | -0.777979787 | 0.49663042 | -1.5665166 | 0.11722773 | NA |
| UBXN1 | 17.322695 | -0.803581594 | 0.45621991 | -1.7613909 | 0.07817226 | NA |
| TTC9C | 11.2677866 | -0.391023063 | 0.43807592 | -0.892592 | 0.3720757 | NA |
| POLR2G | 11.180883 | 0.232106145 | 0.56765424 | 0.40888648 | 0.68262297 | NA |
| TAF6L | 4.71462233 | -0.4893593 | 0.67348003 | -0.7266129 | 0.4674631 | NA |
| TMEM223 | 11.2628318 | -0.48389037 | 0.46309991 | -1.0448941 | 0.29607191 | NA |
| TMEM179B | 23.2111209 | -1.066078177 | 0.4846674 | -2.1996078 | 0.02783474 | NA |
| NXF1 | 7.10413739 | 0.426097033 | 0.62796793 | 0.67853311 | 0.49743374 | NA |
| STX5 | 21.3268406 | -0.355725405 | 0.33788138 | -1.0528115 | 0.29242739 | NA |
| WDR74 | 4.01673346 | -1.543923899 | 0.97210764 | -1.5882232 | 0.11223588 | NA |
| SLC3A2 | 67.950794 | -1.246677394 | 0.5418783 | -2.3006594 | 0.02141089 | 0.24262159 |
| SLC22A6 | 8.9226318 | 0.75744452 | 1.70386106 | 0.44454594 | 0.65664792 | NA |
| RARRES3 | 23.1594332 | -0.579816279 | 0.94935122 | -0.61075 | 0.54136509 | NA |
| PLA2G16 | 47.8087218 | -0.261886945 | 0.46263588 | -0.5660757 | 0.5713423 | 0.8573305 |
| ATL3 | 39.4622729 | -0.027591033 | 0.3057438 | -0.0902423 | 0.92809465 | 0.97775374 |
| RTN3 | 59.8335111 | 0.529475382 | 0.28161857 | 1.88011529 | 0.06009237 | 0.37905854 |
| C11orf95 | 7.88224781 | -0.125702426 | 0.61299462 | -0.2050629 | 0.83752302 | NA |
| MARK2 | 29.7143463 | 0.048807834 | 0.3403916 | 0.1433873 | 0.88598431 | 0.96652834 |
| NAA40 | 8.62680279 | 0.012646073 | 0.51114363 | 0.02474074 | 0.98026176 | NA |
| OTUB1 | 16.0082678 | -0.086483186 | 0.39234441 | -0.2204267 | 0.82553885 | NA |
| STIP1 | 17.0374393 | -0.083612682 | 0.44634048 | -0.1873294 | 0.85140239 | NA |
| TRPT1 | 5.44749635 | 0.042029543 | 0.65148186 | 0.06451376 | 0.94856115 | NA |
| NUDT22 | 7.26602781 | 0.395009664 | 0.5568615 | 0.70934992 | 0.47810736 | NA |
| DNAJC4 | 5.11467428 | 0.069369658 | 0.70328815 | 0.09863618 | 0.92142714 | NA |
| VEGFB | 22.2394211 | -1.164810649 | 0.46383208 | -2.5112766 | 0.01202954 | NA |
| FKBP2 | 36.2074854 | -0.188283259 | 0.32569964 | -0.5780886 | 0.56320428 | 0.85428443 |

| | | | | | | |
|---|---|---|---|---|---|---|
| PPP1R14B | 10.4518327 | -0.081472364 | 0.63812348 | -0.1276749 | 0.89840624 | NA |
| BAD | 16.6641278 | -0.295624682 | 0.47557361 | -0.6216171 | 0.53419368 | NA |
| GPR137 | 12.6532968 | -0.111605351 | 0.49771414 | -0.2242358 | 0.82257379 | NA |
| ESRRA | 16.0476245 | 0.516424117 | 0.38096665 | 1.35556252 | 0.1752384 | NA |
| TRMT112 | 43.0219263 | -0.259028477 | 0.35315571 | -0.7334682 | 0.46327293 | 0.79823068 |
| PRDX5 | 29.5978555 | -0.53564338 | 0.46350738 | -1.1556308 | 0.24783222 | 0.65061974 |
| RPS6KA4 | 4.2526954 | -0.355446702 | 0.67366708 | -0.5276296 | 0.59775643 | NA |
| SLC22A12 | 5.3577222 | 1.165504791 | 1.12746839 | 1.03373612 | 0.30125954 | NA |
| SF1 | 62.292163 | -0.23087703 | 0.25505797 | -0.9051943 | 0.36536245 | 0.73398222 |
| MAP4K2 | 5.18319944 | 0.289538282 | 0.59725078 | 0.48478511 | 0.62782879 | NA |
| MEN1 | 4.55622638 | 0.002017458 | 0.64296572 | 0.00313774 | 0.99749645 | NA |
| EHD1 | 38.3290124 | 1.010823835 | 0.40969416 | 2.46726443 | 0.01361498 | 0.194252 |
| ATG2A | 6.99354008 | 0.699558828 | 0.52766872 | 1.32575383 | 0.18492125 | NA |
| PPP2R5B | 6.19052 | -0.365072801 | 0.7347448 | -0.4968702 | 0.61928058 | NA |
| SAC3D1 | 5.95137292 | -0.239023672 | 0.63490599 | -0.376471 | 0.7065668 | NA |
| ZFPL1 | 10.4826765 | 0.088659197 | 0.52006194 | 0.17047815 | 0.86463412 | NA |
| VPS51 | 24.880152 | -0.009294086 | 0.37098855 | -0.0250522 | 0.98001331 | NA |
| TM7SF2 | 8.87322623 | -0.436727521 | 0.63429871 | -0.6885203 | 0.49112521 | NA |
| ZNHIT2 | 4.12363383 | -0.069410593 | 0.7415284 | -0.0936048 | 0.92542312 | NA |
| FAU | 87.6196485 | -0.442191874 | 0.44519185 | -0.9932614 | 0.32058259 | 0.69496685 |
| SYVN1 | 12.0651543 | 0.47315142 | 0.44496737 | 1.06333959 | 0.28762798 | NA |
| MRPL49 | 13.4317219 | -0.393029973 | 0.4915491 | -0.7995742 | 0.42395755 | NA |
| CAPN1 | 21.7046922 | 0.051667938 | 0.38802068 | 0.13315769 | 0.89406867 | NA |
| POLA2 | 44.5117005 | 1.356921023 | 0.64993905 | 2.08776658 | 0.0368189 | 0.30230251 |
| CDC42EP2 | 25.2103455 | 0.143985061 | 0.4872269 | 0.29551953 | 0.76759704 | NA |
| DPF2 | 16.9034775 | -0.131241266 | 0.39143474 | -0.3352826 | 0.7374119 | NA |
| FRMD8 | 12.6059695 | -0.067705257 | 0.44545763 | -0.1519903 | 0.87919456 | NA |
| NEAT1 | 3850.95771 | -0.764788601 | 0.47073001 | -1.6246863 | 0.10422942 | 0.4862279 |
| MALAT1 | 28169.7029 | -0.083719014 | 0.2779205 | -0.3012337 | 0.76323631 | 0.92336765 |
| SCYL1 | 15.5224343 | -0.090108694 | 0.48605344 | -0.1853884 | 0.85292437 | NA |
| LTBP3 | 43.1885239 | -0.120511511 | 0.46020364 | -0.2618656 | 0.79342505 | 0.93934998 |
| SSSCA1 | 11.6770056 | -1.453034485 | 0.63453729 | -2.2899119 | 0.02202642 | NA |
| FAM89B | 11.4939233 | -0.435664783 | 0.71822621 | -0.6065844 | 0.54412678 | NA |
| EHBP1L1 | 6.83244587 | -1.487642997 | 0.95898485 | -1.5512685 | 0.12083735 | NA |
| MAP3K11 | 14.9732866 | -0.200586609 | 0.43469091 | -0.4614465 | 0.64447828 | NA |
| PCNX3 | 9.07190457 | 0.388985151 | 0.48031476 | 0.80985468 | 0.4180237 | NA |
| SIPA1 | 6.83560392 | -0.339464967 | 0.59502476 | -0.5705056 | 0.56833481 | NA |
| RELA | 17.3040856 | -0.117634261 | 0.50353725 | -0.2336158 | 0.81528325 | NA |
| KAT5 | 17.032275 | -0.321196577 | 0.41668237 | -0.7708427 | 0.44080016 | NA |
| RNASEH2C | 16.8268172 | 0.073901938 | 0.59112803 | 0.1250185 | 0.90050891 | NA |
| CFL1 | 110.598009 | -0.312509496 | 0.39859186 | -0.7840338 | 0.43302027 | 0.77989796 |
| MUS81 | 8.2502213 | 0.731247 | 0.53402573 | 1.36931043 | 0.17090226 | NA |
| EFEMP2 | 35.6399697 | -0.149903333 | 0.42312139 | -0.3542797 | 0.72312925 | 0.91509584 |
| FIBP | 5.29394918 | 0.18655969 | 0.76752302 | 0.24306723 | 0.80795332 | NA |
| CCDC85B | 16.9960451 | -0.757023826 | 0.61021993 | -1.2405754 | 0.21476265 | NA |
| C11orf68 | 8.70449772 | 0.097158305 | 0.5337179 | 0.18204056 | 0.8555509 | NA |
| DRAP1 | 16.7143827 | -0.705061613 | 0.48414528 | -1.4563017 | 0.14530922 | NA |
| SART1 | 7.46024293 | -0.863623599 | 0.65713031 | -1.3142349 | 0.18876716 | NA |
| EIF1AD | 5.44134937 | -0.308322488 | 0.57504118 | -0.5361746 | 0.59183787 | NA |
| BANF1 | 25.2586024 | -0.580690758 | 0.51678154 | -1.1236678 | 0.261154 | NA |
| SF3B2 | 32.6475994 | -0.033316271 | 0.37461874 | -0.0889338 | 0.92913452 | 0.97777732 |
| PACS1 | 78.7048856 | -0.359692752 | 0.24089839 | -1.4931306 | 0.13540301 | 0.52807173 |
| KLC2 | 7.93829014 | 0.331195732 | 0.73407757 | 0.45117266 | 0.65186511 | NA |
| RAB1B | 26.9725105 | -0.033427956 | 0.51471808 | -0.0649442 | 0.94821842 | NA |
| YIF1A | 9.97613905 | -0.602226713 | 0.58760537 | -1.0248829 | 0.30541843 | NA |
| BRMS1 | 6.36071497 | -0.316007069 | 0.68269875 | -0.4628792 | 0.64345094 | NA |
| B4GAT1 | 52.476269 | -0.090783545 | 0.60218118 | -0.1507579 | 0.88016673 | 0.96423929 |
| MRPL11 | 8.82380054 | -0.409018523 | 0.5185146 | -0.7888274 | 0.43021289 | NA |
| ZDHHC24 | 12.6979123 | -0.212978115 | 0.43878517 | -0.4853813 | 0.62740589 | NA |
| CTSF | 20.2831217 | -0.191683661 | 0.43926953 | -0.4363691 | 0.66256893 | NA |
| CCS | 15.5044621 | 0.146939048 | 0.36677251 | 0.40062721 | 0.68869461 | NA |
| RBM14-RBM | 14.0721123 | 0.422174593 | 0.3911235 | 1.07938948 | 0.28041414 | NA |
| RBM4 | 11.4245238 | 0.486981164 | 0.55770366 | 0.87318983 | 0.38255962 | NA |
| RBM4B | 10.459693 | -0.184544213 | 0.60162397 | -0.3067434 | 0.75903865 | NA |
| C11orf80 | 27.8299839 | -0.369126353 | 0.53705985 | -0.6873095 | 0.4918877 | NA |
| RCE1 | 7.08604173 | 0.637586207 | 0.54811404 | 1.16323642 | 0.2447336 | NA |
| PC | 29.3950695 | 0.718215471 | 0.43050176 | 1.6683218 | 0.09525186 | 0.46829657 |
| LRFN4 | 23.3467226 | 0.360938108 | 0.45780619 | 0.7884081 | 0.43045804 | NA |
| KDM2A | 114.574552 | -0.023839188 | 0.27552847 | -0.0865217 | 0.93105172 | 0.97832857 |
| GRK2 | 8.68868385 | 0.364041732 | 0.49074973 | 0.7418073 | 0.4582041 | NA |
| ANKRD13D | 15.3250269 | -1.038551493 | 0.42048533 | -2.4698876 | 0.01351555 | NA |
| SSH3 | 11.2158898 | 0.242917105 | 0.56128299 | 0.432789 | 0.66516806 | NA |
| RAD9A | 38.1292982 | 0.164580415 | 0.2895214 | 0.56845682 | 0.56972482 | 0.8573305 |
| POLD4 | 4.59503878 | 0.101479552 | 1.0381986 | 0.0977458 | 0.92213415 | NA |
| PPP1CA | 27.163634 | -0.125848037 | 0.46665764 | -0.2696796 | 0.78740677 | NA |
| RPS6KB2 | 9.4164336 | 0.51731273 | 0.46189961 | 1.11996789 | 0.26272744 | NA |
| CORO1B | 16.6106641 | -0.166313855 | 0.46368468 | -0.3586788 | 0.7198354 | NA |
| TMEM134 | 23.1786809 | 0.369044129 | 0.37673064 | 0.97959681 | 0.32728518 | NA |
| AIP | 11.6741682 | -0.284122314 | 0.44620595 | -0.6367515 | 0.52428672 | NA |
| PITPNM1 | 5.12643786 | 0.299164549 | 0.68064347 | 0.43953195 | 0.66027614 | NA |
| CDK2AP2 | 26.4277112 | -0.679407345 | 0.49328031 | -1.3773251 | 0.16841177 | NA |
| GSTP1 | 154.438342 | -0.445652141 | 0.55288729 | -0.8060452 | 0.4202168 | 0.77437498 |
| NDUFV1 | 19.7885614 | 0.22307236 | 0.38976782 | 0.57232113 | 0.56710443 | NA |
| NUDT8 | 4.58359516 | -0.529402179 | 0.76168484 | -0.695041 | 0.48702962 | NA |
| UNC93B1 | 25.5172028 | -0.111005224 | 0.83416456 | -0.1330735 | 0.89413523 | NA |
| ALDH3B1 | 18.95974 | -0.269194842 | 0.45102087 | -0.5968567 | 0.55060304 | NA |
| NDUFS8 | 26.8655868 | -0.539612269 | 0.51822185 | -1.0412766 | 0.29774721 | NA |
| TCIRG1 | 6.29738502 | 0.965365665 | 0.78693817 | 1.22673635 | 0.2199217 | NA |
| CHKA | 25.4600956 | 0.618122332 | 0.46170545 | 1.33878067 | 0.18064208 | NA |
| KMT5B | 63.5154187 | 0.103999516 | 0.25510473 | 0.40767381 | 0.68351317 | 0.90499319 |
| C11orf24 | 6.16431568 | -0.185542371 | 0.63815703 | -0.2907472 | 0.77124467 | NA |
| LRP5 | 8.4926145 | 1.018954147 | 0.60900405 | 1.67314839 | 0.09429809 | NA |
| PPP6R3 | 140.050498 | 0.602260079 | 0.17020945 | 3.5383469 | 0.00040264 | 0.02714874 |
| CPT1A | 51.9408489 | 0.130471979 | 0.32688888 | 0.39913251 | 0.68979557 | 0.90557575 |
| MRPL21 | 8.01336688 | 0.480772816 | 0.58247514 | 0.82539628 | 0.40914665 | NA |
| IGHMBP2 | 8.26854673 | -0.031770269 | 0.59209224 | -0.0536576 | 0.95720794 | NA |
| MRGPRF | 123.285513 | -0.318068331 | 0.59473571 | -0.5348062 | 0.59278389 | 0.8654248 |
| TPCN2 | 20.5541145 | -0.402268973 | 0.49998338 | -0.8045647 | 0.42107092 | NA |
| CCND1 | 9.36849027 | 0.433903492 | 0.64735155 | 0.67027489 | 0.50268257 | NA |
| ORAOV1 | 11.6630261 | 0.225424693 | 0.41052055 | 0.54911914 | 0.58292369 | NA |
| FADD | 8.31216925 | -0.652544939 | 0.5883417 | -1.1091258 | 0.26737593 | NA |
| PPFIA1 | 53.3240898 | -0.000807975 | 0.32596943 | -0.0024787 | 0.9980223 | 0.99961632 |
| AP002336.2 | 18.1155588 | 0.086924879 | 0.58950758 | 0.14745337 | 0.88277418 | NA |
| AP000487.1 | 4.84965168 | -0.725563187 | 0.65524571 | -1.1073147 | 0.26815791 | NA |
| CTTN | 56.8771591 | 0.193285435 | 0.2576815 | 0.75009434 | 0.45319789 | 0.79324789 |
| SHANK2 | 88.4574251 | 0.180905436 | 0.47160029 | 0.38359908 | 0.70127562 | 0.90757606 |
| DHCR7 | 12.3909501 | -1.363491416 | 0.61477536 | -2.2178693 | 0.02656374 | NA |
| NADSYN1 | 11.470857 | -0.585093767 | 0.46790356 | -1.250458 | 0.21113227 | NA |
| AP002495.2 | 15.4429625 | 0.831202594 | 0.48335021 | 1.71966945 | 0.08549254 | NA |
| RNF121 | 19.4768336 | 0.139373058 | 0.35959582 | 0.38758253 | 0.698325 | NA |
| IL18BP | 10.4619192 | -2.12227745 | 1.30338426 | -1.6282822 | 0.10346505 | NA |
| NUMA1 | 46.7977413 | -0.268086572 | 0.35378337 | -0.7577704 | 0.44858843 | 0.78987065 |
| LAMTOR1 | 30.181185 | -0.694487923 | 0.40036038 | -1.734657 | 0.08280158 | 0.44045101 |

| | | | | | | |
|---|---|---|---|---|---|---|
| ANAPC15 | 4.25829138 | 0.048905081 | 0.714323 | 0.06846354 | 0.94541664 | NA |
| FOLR1 | 6.30363933 | -2.202531267 | 0.8850363 | -2.4886338 | 0.0128235 | NA |
| INPPL1 | 28.3650899 | 0.638243213 | 0.34944307 | 1.82645836 | 0.06778125 | NA |
| CLPB | 8.55973941 | -0.450514692 | 0.59809825 | -0.7532453 | 0.45130252 | NA |
| ARAP1 | 16.2803091 | 0.994295168 | 0.39734641 | 2.50233835 | 0.0123376 | NA |
| ATG16L2 | 6.9341139 | 0.174169927 | 0.62682497 | 0.27786054 | 0.78111941 | NA |
| FCHSD2 | 175.021478 | 1.031616884 | 0.3457132 | 2.98402514 | 0.00284483 | 0.08875882 |
| ARHGEF17 | 51.2075091 | 0.442575996 | 0.33842637 | 1.30774679 | 0.19095921 | 0.59351 |
| FAM168A | 75.7064358 | 0.70326724 | 0.25415614 | 2.76706769 | 0.0056563 | 0.11765106 |
| PLEKHB1 | 5.96212992 | 0.194232458 | 0.70500891 | 0.27550355 | 0.7829294 | NA |
| RAB6A | 75.2345003 | 0.016345932 | 0.19841001 | 0.08238461 | 0.93434087 | 0.97832857 |
| MRPL48 | 17.4675777 | 0.333068558 | 0.45589215 | 0.7305863 | 0.46503188 | NA |
| COA4 | 24.8168521 | -0.424257102 | 0.53834428 | -0.7880777 | 0.43065128 | NA |
| PAAF1 | 17.6690009 | 0.112878427 | 0.43117713 | 0.26179132 | 0.79348233 | NA |
| C2CD3 | 24.7491341 | 0.079454814 | 0.29941769 | 0.26536446 | 0.79072871 | NA |
| PPME1 | 26.4708044 | 0.197151651 | 0.32476442 | 0.60706051 | 0.54381076 | NA |
| P4HA3 | 4.31026262 | -1.871356872 | 0.92215349 | -2.0293334 | 0.04242434 | NA |
| PGM2L1 | 15.680526 | 0.567523604 | 0.64630267 | 0.8781081 | 0.37988506 | NA |
| POLD3 | 14.0913167 | -0.281422375 | 0.40883344 | -0.6883546 | 0.49122951 | NA |
| RNF169 | 37.1994142 | -0.024238715 | 0.30517767 | -0.0794249 | 0.93669464 | 0.97842737 |
| XRRA1 | 37.4525181 | 0.350613206 | 0.51591561 | 0.67959411 | 0.4967615 | 0.8234249 |
| SPCS2 | 48.9062173 | -0.638741605 | 0.4067029 | -1.5705361 | 0.11629044 | 0.50608799 |
| NEU3 | 5.76453602 | 0.752514509 | 0.62609079 | 1.20192555 | 0.22939238 | NA |
| ARRB1 | 62.8375781 | -0.082697693 | 0.3920906 | -0.2109148 | 0.83295379 | 0.95060793 |
| RPS3 | 96.7667331 | -0.38391809 | 0.36945731 | -1.0391406 | 0.29873936 | 0.68168772 |
| SERPINH1 | 19.4312782 | 0.472554668 | 0.44927621 | 1.05181323 | 0.29288525 | NA |
| MAP6 | 101.433396 | 1.159635282 | 0.57557814 | 2.01473129 | 0.04393281 | 0.33029013 |
| UVRAG | 87.009463 | 0.436096885 | 0.34803166 | 1.25303798 | 0.21019188 | 0.61792799 |
| THAP12 | 11.6458309 | -0.105926021 | 0.54744761 | -0.1934907 | 0.84657469 | NA |
| EMSY | 36.9750902 | 0.103130242 | 0.30141032 | 0.34215897 | 0.73223126 | 0.91806182 |
| TSKU | 4.45855016 | -0.804812709 | 0.88387508 | -0.9105503 | 0.36253237 | NA |
| ACER3 | 37.3610328 | 0.236717574 | 0.27458265 | 0.86209952 | 0.38863276 | 0.75301425 |
| PAK1 | 154.728861 | 0.19437562 | 0.34591173 | 0.56192261 | 0.57416875 | 0.85861279 |
| CLNS1A | 41.0174783 | 0.154158706 | 0.31355141 | 0.49165368 | 0.62296418 | 0.8780438 |
| RSF1 | 97.8211117 | 0.142034181 | 0.19949425 | 0.71197129 | 0.47648255 | 0.80863008 |
| AAMDC | 29.1312574 | -0.113606204 | 0.30651845 | -0.3706342 | 0.71091004 | 0.91222063 |
| AP002812.2 | 8.70728701 | 0.39015719 | 0.56964406 | 0.684914 | 0.49339819 | NA |
| INTS4 | 20.7273684 | 0.567994662 | 0.50098196 | 1.13376271 | 0.2568941 | NA |
| ALG8 | 11.5838824 | -0.409827138 | 0.45462116 | -0.9014696 | 0.36733871 | NA |
| KCTD21 | 6.70399652 | 0.152197936 | 0.54566217 | 0.27892338 | 0.78030362 | NA |
| GAB2 | 76.5184074 | 0.174508278 | 0.28096831 | 0.62109594 | 0.5345365 | 0.83677705 |
| AP003086.2 | 20.2070537 | 0.009970199 | 0.6021128 | 0.01655869 | 0.98678868 | NA |
| NARS2 | 21.5599582 | 0.198005342 | 0.42778605 | 0.46286068 | 0.64346423 | NA |
| TENM4 | 11.6267154 | 0.469729163 | 0.77085712 | 0.60935957 | 0.54228613 | NA |
| PRCP | 31.1338841 | 0.474189269 | 0.32144853 | 1.4751639 | 0.14016849 | 0.53597868 |
| RAB30 | 9.61987139 | -0.000255518 | 0.50705531 | -0.0005039 | 0.99959793 | NA |
| RAB30-AS1 | 32.7593904 | 0.37207951 | 0.3680916 | 1.01083401 | 0.31209588 | 0.69172726 |
| PCF11 | 30.9014544 | 0.07598127 | 0.29522013 | 0.25737157 | 0.79689194 | 0.93934998 |
| ANKRD42 | 13.8903944 | 0.597352451 | 0.41138754 | 1.45204313 | 0.14648961 | NA |
| CCDC90B | 20.1963026 | 0.201173771 | 0.41772318 | 0.4815959 | 0.63009304 | NA |
| AP000446.1 | 9.71563727 | 0.64796091 | 0.5248749 | 1.23450543 | 0.21701463 | NA |
| DLG2 | 632.996219 | 1.228144369 | 0.48635644 | 2.525194 | 0.01156344 | 0.17871297 |
| AP001825.1 | 12.4987495 | 1.309408581 | 0.79888279 | 1.63904967 | 0.10120291 | NA |
| TMEM126B | 18.0035664 | -0.585740746 | 0.48715682 | -1.2023659 | 0.2292218 | NA |
| TMEM126A | 7.53411772 | 0.172713603 | 0.5430456 | 0.31804622 | 0.75044987 | NA |
| CREBZF | 45.1165929 | 0.407721459 | 0.29549285 | 1.37980143 | 0.16764779 | 0.56949702 |
| SYTL2 | 11.9712237 | -0.506074824 | 0.61272915 | -0.8259356 | 0.40884062 | NA |
| PICALM | 381.954696 | 0.411563651 | 0.19790346 | 2.07961828 | 0.03756056 | 0.30414888 |
| EED | 13.3355176 | 0.886146426 | 0.40318062 | 2.19788945 | 0.02795698 | NA |
| HIKESHI | 23.237991 | 0.519662277 | 0.35315049 | 1.47150379 | 0.14115492 | NA |
| AP001831.1 | 8.151122 | 0.060263127 | 0.55605704 | 0.1083758 | 0.9136976 | NA |
| ME3 | 20.5783445 | -0.018806586 | 0.50844715 | -0.0369883 | 0.97049435 | NA |
| PRSS23 | 20.1999733 | 0.022691205 | 0.77714386 | 0.0291982 | 0.97670651 | NA |
| TMEM135 | 56.358056 | 0.582584699 | 0.30242902 | 1.92635184 | 0.05406046 | 0.3615191 |
| CTSC | 11.2851776 | -0.639179745 | 0.53552644 | -1.193554 | 0.23265246 | NA |
| NOX4 | 92.6204227 | 1.036917252 | 0.54919875 | 1.88805464 | 0.05901862 | 0.37757019 |
| CHORDC1 | 14.6653423 | 0.128420724 | 0.45159442 | 0.28437181 | 0.77612546 | NA |
| SLC36A4 | 27.1142591 | -0.224489 | 0.41031113 | -0.547119 | 0.584297 | NA |
| SMCO4 | 5.34903743 | -0.796146821 | 0.81776658 | -0.9735624 | 0.33027385 | NA |
| CEP295 | 17.975037 | 0.366251948 | 0.41027258 | 0.89270393 | 0.37201575 | NA |
| TAF1D | 39.7666043 | -0.426025579 | 0.3550288 | -1.1999747 | 0.23014917 | 0.63682501 |
| PANX1 | 8.48889155 | 0.278187443 | 0.49612653 | 0.56071874 | 0.57498929 | NA |
| MRE11 | 17.7780893 | -0.298529991 | 0.3875729 | -0.7702551 | 0.44114861 | NA |
| ANKRD49 | 6.14325177 | -0.227557734 | 0.62529863 | -0.3639185 | 0.71591888 | NA |
| CWC15 | 17.6607877 | -0.233148279 | 0.40085635 | -0.5816255 | 0.56081896 | NA |
| SRSF8 | 15.5069925 | -0.318529503 | 0.40920349 | -0.7784134 | 0.43632531 | NA |
| ENDOD1 | 45.5153117 | -0.778993406 | 0.27250554 | -2.8586333 | 0.0042547 | 0.10458835 |
| AP000787.1 | 5.26240465 | -0.720658725 | 0.71389058 | -1.0094806 | 0.31274418 | NA |
| SESN3 | 22.1401022 | 0.737973117 | 0.49264688 | 1.49797584 | 0.13413953 | NA |
| FAM76B | 9.44398603 | -0.206787838 | 0.43131196 | -0.4794391 | 0.63162625 | NA |
| CEP57 | 22.5531834 | -0.651489797 | 0.35179856 | -1.8518831 | 0.06404262 | NA |
| MTMR2 | 34.9391247 | 0.242185734 | 0.29326859 | 0.82581546 | 0.40890878 | 0.76772515 |
| MAML2 | 458.816644 | 0.68361265 | 0.34395821 | 1.98748754 | 0.0468684 | 0.33774256 |
| CCDC82 | 20.5073314 | -0.019246285 | 0.31792267 | -0.0605376 | 0.95172745 | NA |
| JRKL | 6.26221832 | -0.382415895 | 0.58924314 | -0.6489951 | 0.51634156 | NA |
| ARHGAP42 | 22.0783712 | -0.133159106 | 0.51506041 | -0.258531 | 0.79599709 | NA |
| PGR | 21.4680243 | -0.13742303 | 0.39477094 | -0.3481083 | 0.72775887 | NA |
| TRPC6 | 230.745335 | -0.977786705 | 0.47059973 | -2.077746 | 0.03773276 | 0.30414888 |
| CEP126 | 34.5336098 | -0.379488407 | 0.47297669 | -0.8023406 | 0.42235596 | 0.77588259 |
| CFAP300 | 6.46093263 | -0.662848328 | 0.53679766 | -1.2348197 | 0.21689761 | NA |
| YAP1 | 173.818809 | 0.188021371 | 0.22658392 | 0.82980896 | 0.4066468 | 0.76772515 |
| BIRC3 | 11.4582127 | -0.291401849 | 1.01105987 | -0.2882142 | 0.77318275 | NA |
| BIRC2 | 35.6583788 | 0.516580623 | 0.34110109 | 1.51445023 | 0.12991169 | 0.52613957 |
| TMEM123 | 71.2200568 | 0.103227657 | 0.44111691 | 0.23401428 | 0.81497389 | 0.94551891 |
| MMP7 | 8.95470401 | -1.015719757 | 0.75025383 | -1.3538348 | 0.17578908 | NA |
| DCUN1D5 | 20.0400244 | 0.255369526 | 0.43067247 | 0.5929553 | 0.55321106 | NA |
| DYNC2H1 | 120.325526 | 0.874964298 | 0.32201871 | 2.71712257 | 0.00658522 | 0.12888932 |
| CASP4 | 22.3254127 | -1.488597886 | 0.73442148 | -2.0268986 | 0.04267278 | NA |
| CARD16 | 3.46458777 | -0.687934338 | 1.21514692 | -0.5661326 | 0.57130361 | NA |
| MSANTD4 | 11.1919434 | 0.102735394 | 0.45224357 | 0.22716828 | 0.82029289 | NA |
| KBTBD3 | 7.60769644 | -0.413457719 | 0.66729217 | -0.6196052 | 0.53551772 | NA |
| AASDHPPT | 24.3968154 | 0.252292417 | 0.34806204 | 0.72484899 | 0.46854468 | NA |
| AP000766.1 | 4.56749758 | -0.057428622 | 0.71690505 | -0.0801063 | 0.9361527 | NA |
| CWF19L2 | 35.210811 | -0.194106317 | 0.33427118 | -0.5806852 | 0.56145266 | 0.85386276 |
| ALKBH8 | 8.54654708 | 0.122861588 | 0.47277544 | 0.25987303 | 0.79496171 | NA |
| CUL5 | 80.1674674 | 0.10009108 | 0.22119209 | 0.45250751 | 0.65090342 | 0.89412414 |
| ACAT1 | 20.2677971 | -0.705348008 | 0.40345195 | -1.7482825 | 0.08041511 | NA |
| NPAT | 17.2269246 | 0.134339271 | 0.43668508 | 0.30763421 | 0.75836068 | NA |
| ATM | 64.1330424 | 0.340242227 | 0.32908051 | 1.0339179 | 0.30117455 | 0.68219446 |
| C11orf65 | 12.1029369 | 0.713072978 | 0.85434178 | 0.83464604 | 0.40391703 | NA |
| KDELC2 | 15.8093703 | -0.073740225 | 0.46415081 | -0.1588713 | 0.8737703 | NA |
| EXPH5 | 400.858547 | 0.319177412 | 0.40521114 | 0.78768173 | 0.4308829 | 0.77901411 |
| DDX10 | 41.1762565 | 0.445583815 | 0.35837495 | 1.2433453 | 0.21374062 | 0.62255988 |

| | | | | | | |
|---|---|---|---|---|---|---|
| RDX | 192.751745 | 0.348821473 | 0.29813399 | 1.17001577 | 0.24199462 | 0.64918976 |
| ZC3H12C | 30.9580599 | 0.031748599 | 0.35678207 | 0.08898597 | 0.92909306 | 0.97777732 |
| FDX1 | 24.1835265 | 0.555722407 | 0.37185712 | 1.49445144 | 0.13505767 | NA |
| ARHGAP20 | 15.1177899 | 0.266485997 | 1.04654475 | 0.25463412 | 0.79900571 | NA |
| SIK2 | 64.9894014 | 0.024134432 | 0.29859514 | 0.08082661 | 0.93557985 | 0.97832857 |
| PPP2R1B | 21.5768775 | 0.442097896 | 0.35334748 | 1.25117037 | 0.21087233 | NA |
| C11orf1 | 6.64033721 | -1.389446235 | 0.69409861 | -2.0017995 | 0.0453063 | NA |
| CRYAB | 214.332159 | -0.75042416 | 0.64888593 | -1.1564809 | 0.24748452 | 0.65061974 |
| DIXDC1 | 15.0844602 | -0.057233945 | 0.55904719 | -0.1023777 | 0.91845691 | NA |
| DLAT | 14.5923152 | -0.094153251 | 0.37112743 | -0.2536952 | 0.79973104 | NA |
| NKAPD1 | 8.72792053 | 0.297744595 | 0.54873359 | 0.54260319 | 0.58740304 | NA |
| TIMM8B | 16.9135575 | -0.661052947 | 0.48702865 | -1.3573184 | 0.17468007 | NA |
| IL18 | 6.89669207 | -1.775204404 | 0.82809881 | -2.1437109 | 0.03205606 | NA |
| PTS | 42.778836 | -1.284625144 | 0.31773091 | -4.043123 | 5.27E-05 | 0.00668529 |
| AP002884.1 | 21.336713 | -0.905346492 | 0.51276178 | -1.7656279 | 0.0774583 | NA |
| NCAM1 | 12.9609282 | 1.872458863 | 0.61002509 | 3.06947845 | 0.00214433 | NA |
| TTC12 | 15.8804522 | -0.385659655 | 0.35563623 | -1.0844217 | 0.27817785 | NA |
| ZW10 | 7.4990357 | 0.387744052 | 0.51499859 | 0.75290313 | 0.45150813 | NA |
| USP28 | 16.3716762 | -0.013716489 | 0.38136973 | -0.0359664 | 0.97130917 | NA |
| ZBTB16 | 54.7572777 | -1.099924019 | 0.74570287 | -1.4750165 | 0.14020811 | 0.53597868 |
| AP002518.2 | 6.15200863 | -1.623677825 | 1.38481436 | -1.1724877 | 0.2410013 | NA |
| NNMT | 20.7688057 | -4.480387774 | 1.13621862 | -3.9432445 | 8.04E-05 | NA |
| C11orf71 | 5.91307725 | 0.803173259 | 0.70706413 | 1.13592703 | 0.25598711 | NA |
| REXO2 | 26.7520713 | 0.049582784 | 0.39154656 | 0.12663318 | 0.89923074 | NA |
| CADM1 | 41.4282541 | 1.229329463 | 0.71595159 | 1.71705667 | 0.08596882 | 0.44638081 |
| BUD13 | 8.04541904 | 0.106068677 | 0.52003806 | 0.2039633 | 0.83838218 | NA |
| ZPR1 | 5.97890117 | -0.831155166 | 0.74502366 | -1.1156091 | 0.2645895 | NA |
| SIK3 | 394.000061 | 0.439593154 | 0.28174108 | 1.56027354 | 0.11869525 | 0.51050744 |
| PAFAH1B2 | 49.2959055 | 0.103704774 | 0.29067866 | 0.35676776 | 0.72126567 | 0.91509584 |
| SIDT2 | 11.242352 | -0.239444413 | 0.48752828 | -0.4911395 | 0.62332776 | NA |
| TAGLN | 17.8978792 | 0.871616707 | 0.72014802 | 1.21032994 | 0.22615231 | NA |
| PCSK7 | 24.6312307 | -0.068867602 | 0.28351136 | -0.2429095 | 0.80807551 | NA |
| RNF214 | 16.3788498 | -0.132982984 | 0.43011558 | -0.3091796 | 0.75718487 | NA |
| BACE1 | 52.5859838 | -0.976503299 | 0.30039045 | -3.2507801 | 0.00115089 | 0.05186671 |
| CEP164 | 25.88584 | -0.150471971 | 0.33193117 | -0.4533228 | 0.65031631 | NA |
| DSCAML1 | 17.2441675 | -0.111828727 | 0.85269562 | -0.1311473 | 0.89565879 | NA |
| FXYD2 | 15.3906762 | -0.283944749 | 0.81360627 | -0.3489953 | 0.72709285 | NA |
| UBE4A | 32.9089956 | 0.537216967 | 0.27719537 | 1.93804455 | 0.05261779 | 0.35808347 |
| ATP5MG | 51.6276753 | -0.377433582 | 0.43463833 | -0.8683854 | 0.38518338 | 0.75104967 |
| KMT2A | 89.584226 | -0.247477308 | 0.21889994 | -1.1305499 | 0.25824458 | 0.65588339 |
| TMEM25 | 6.01988113 | -0.553620843 | 0.56364756 | -0.982211 | 0.3259959 | NA |
| IFT46 | 5.36448263 | 0.200977261 | 0.57744452 | 0.34804601 | 0.72780563 | NA |
| ARCN1 | 27.3056621 | -0.370099015 | 0.42015349 | -0.8808662 | 0.37839023 | NA |
| PHLDB1 | 33.634533 | 0.991463454 | 0.41001383 | 2.41812197 | 0.01560085 | 0.20644861 |
| AP002954.1 | 10.5940895 | -0.445363116 | 0.64404293 | -0.6915115 | 0.48924413 | NA |
| DDX6 | 61.0591502 | -0.186762768 | 0.29934553 | -0.6239036 | 0.53269087 | 0.83658422 |
| BCL9L | 22.4487059 | 0.684043733 | 0.49800945 | 1.37355574 | 0.16957965 | NA |
| CCDC84 | 26.1066292 | -0.113622023 | 0.49095462 | -0.2314308 | 0.81698013 | NA |
| RPS25 | 75.1468201 | -0.579723611 | 0.39104331 | -1.4825049 | 0.13820601 | 0.53418928 |
| TRAPPC4 | 28.5963641 | -0.777284114 | 0.45209095 | -1.7193092 | 0.08555809 | 0.44564602 |
| SLC37A4 | 7.99918019 | -0.658072192 | 0.66525905 | -0.9891969 | 0.32256681 | NA |
| HYOU1 | 26.0797356 | 0.050465697 | 0.5797232 | 0.08705137 | 0.93063068 | NA |
| VPS11 | 8.56271984 | -0.024473007 | 0.47535334 | -0.0514838 | 0.95893999 | NA |
| H2AFX | 5.43294293 | -0.045818344 | 0.61298825 | -0.0747459 | 0.9404169 | NA |
| DPAGT1 | 6.23071595 | -0.379259684 | 0.59188119 | -0.64077 | 0.52167215 | NA |
| HINFP | 5.57339781 | -0.163706994 | 0.59356797 | -0.2758016 | 0.78270046 | NA |
| NLRX1 | 6.25063949 | 0.426298073 | 0.65367825 | 0.65215276 | 0.51430263 | NA |
| CBL | 32.3745743 | 0.278547888 | 0.34348166 | 0.81095417 | 0.41739199 | 0.77165508 |
| RNF26 | 6.59431158 | -0.376546141 | 0.53182443 | -0.7080272 | 0.47892839 | NA |
| OAF | 4.88399698 | -0.35954003 | 0.83855413 | -0.4287619 | 0.66809654 | NA |
| TMEM136 | 6.49069155 | 0.047158537 | 0.61669266 | 0.07647008 | 0.93904512 | NA |
| ARHGEF12 | 600.01953 | 0.773077276 | 0.32089034 | 2.40916346 | 0.01598914 | 0.20852711 |
| SC5D | 9.18522006 | -0.126943247 | 0.47331465 | -0.2682005 | 0.78854496 | NA |
| SORL1 | 16.7908451 | 0.930090818 | 0.6569064 | 1.41586505 | 0.15681502 | NA |
| JHY | 30.364722 | 0.656572363 | 0.33711404 | 1.94762687 | 0.05145963 | 0.35507065 |
| HSPA8 | 70.9069654 | -0.356499741 | 0.44141611 | -0.8076274 | 0.41930512 | 0.77304616 |
| VWA5A | 8.43247855 | -1.264949166 | 0.53291661 | -2.3736343 | 0.01761398 | NA |
| TBRG1 | 25.8351857 | 0.295707686 | 0.29493565 | 1.00261764 | 0.31604538 | NA |
| SIAE | 12.4352177 | -0.827578026 | 0.57537196 | -1.4383357 | 0.15033883 | NA |
| MSANTD2 | 22.0633986 | 0.085044142 | 0.33624653 | 0.25292199 | 0.80032849 | NA |
| TMEM218 | 7.04626728 | 0.000898491 | 0.53704332 | 0.00167303 | 0.99866511 | NA |
| PKNOX2 | 16.9222937 | 0.302833499 | 0.55735889 | 0.54333663 | 0.58689805 | NA |
| EI24 | 27.0074416 | 0.024455203 | 0.3298941 | 0.07413046 | 0.94090657 | NA |
| STT3A | 17.2432308 | 0.221454201 | 0.41348784 | 0.53557609 | 0.59225156 | NA |
| PUS3 | 4.50725947 | -1.433400453 | 0.78724395 | -1.8207831 | 0.06863984 | NA |
| CDON | 14.5256454 | -1.083471183 | 0.52478589 | -2.0645966 | 0.03896118 | NA |
| RPUSD4 | 5.49747942 | -0.161643104 | 0.75477682 | -0.2141601 | 0.83042218 | NA |
| FAM118B | 14.281737 | -0.643678209 | 0.44314448 | -1.4525245 | 0.14635582 | NA |
| SRPRA | 27.2415468 | -0.383635288 | 0.4305056 | -0.8911273 | 0.37286089 | NA |
| FOXRED1 | 8.35776922 | -0.12309223 | 0.5570959 | -0.2209534 | 0.82512872 | NA |
| TIRAP | 8.32953348 | -0.130608355 | 0.56257562 | -0.2321614 | 0.81641264 | NA |
| ST3GAL4 | 12.8468643 | 0.774847326 | 0.48247349 | 1.60598942 | 0.10827624 | NA |
| ETS1 | 13.1648269 | -0.639984503 | 0.44588701 | -1.4353065 | 0.15119979 | NA |
| KCNJ1 | 9.09476945 | 1.680510573 | 0.8274108 | 2.03104741 | 0.04225019 | NA |
| ARHGAP32 | 39.7521041 | 0.574981355 | 0.39945894 | 1.43940039 | 0.15003711 | 0.54834287 |
| NFRKB | 13.6856674 | 0.276597726 | 0.3814075 | 0.72520264 | 0.46832772 | NA |
| PRDM10 | 15.4292838 | 0.26012383 | 0.54066673 | 0.48111677 | 0.63043351 | NA |
| APLP2 | 192.861016 | -0.475673155 | 0.32819678 | -1.4493535 | 0.14723888 | 0.54635866 |
| ZBTB44 | 52.3753004 | 0.567123146 | 0.27343609 | 2.07406106 | 0.03807364 | 0.304707 |
| AP003486.1 | 6.21507008 | 1.067347873 | 0.60834999 | 1.7544964 | 0.07934549 | NA |
| SNX19 | 27.3647766 | 0.154928338 | 0.34642907 | 0.44721518 | 0.6547197 | NA |
| NTM | 10.039854 | -0.175260649 | 0.69352555 | -0.2527097 | 0.80049253 | NA |
| OPCML | 9.75431298 | -2.33771181 | 0.83800768 | -2.7896067 | 0.00527721 | NA |
| JAM3 | 23.371099 | 0.266437642 | 0.44095376 | 0.60423034 | 0.54569051 | NA |
| NCAPD3 | 14.0475131 | 0.535460904 | 0.38506813 | 1.39056146 | 0.16435845 | NA |
| VPS26B | 14.6156274 | 0.588400394 | 0.42604793 | 1.3810662 | 0.16725861 | NA |
| THYN1 | 4.97475297 | -0.176097252 | 0.75674213 | -0.2327044 | 0.81599092 | NA |
| ACAD8 | 6.26132845 | 0.614180733 | 0.54241339 | 1.13231116 | 0.25750364 | NA |
| ZMYND11 | 81.4364593 | 0.225341833 | 0.25337819 | 0.88934978 | 0.37381513 | 0.74177796 |
| DIP2C | 80.4187558 | -0.012408243 | 0.26637268 | -0.0465823 | 0.96284617 | 0.98892249 |
| LARP4B | 93.4716026 | 0.460472737 | 0.24587614 | 1.87278332 | 0.06109831 | 0.38184091 |
| GTPBP4 | 13.8004925 | 0.028508474 | 0.46290849 | 0.06158555 | 0.95089288 | NA |
| IDI1 | 15.8838755 | -0.790284029 | 0.45565483 | -1.7343919 | 0.08284858 | NA |
| WDR37 | 29.7158118 | -0.47208792 | 0.3393963 | -1.3909637 | 0.16423644 | 0.56548642 |
| PFKP | 6.74566143 | -1.682691811 | 0.94106409 | -1.7880735 | 0.07376414 | NA |
| PITRM1 | 13.8929155 | -0.344627074 | 0.40487528 | -0.8511932 | 0.39466205 | NA |
| KLF6 | 28.938155 | -0.222006806 | 0.57707529 | -0.3847103 | 0.70045207 | 0.90729979 |
| AKR1C1 | 6.67025074 | -2.062567834 | 0.98974419 | -2.0839403 | 0.0371656 | NA |
| AKR1C3 | 6.46675032 | -0.774200635 | 0.84416759 | -0.9171172 | 0.35908122 | NA |
| NET1 | 4.06258492 | -0.143683255 | 0.77234626 | -0.1860348 | 0.8524175 | NA |
| ASB13 | 5.64094419 | -0.113237264 | 0.5915711 | -0.1914178 | 0.84819824 | NA |
| FAM208B | 65.7739854 | -0.171692131 | 0.33234372 | -0.5166101 | 0.60542834 | 0.87047761 |
| GDI2 | 45.7982777 | -0.59281927 | 0.32153745 | -1.8437021 | 0.06522657 | 0.39604633 |

| | | | | | | |
|---|---|---|---|---|---|---|
| ANKRD16 | 7.849937 | -0.052031601 | 0.52920077 | -0.0983211 | 0.92167733 | NA |
| FBH1 | 32.7617327 | 0.030435196 | 0.27756633 | 0.10965017 | 0.91268682 | 0.97700125 |
| RBM17 | 27.637018 | -0.432079144 | 0.33234243 | -1.3001023 | 0.19356592 | NA |
| PFKFB3 | 33.8666949 | -0.103861073 | 0.47506006 | -0.2186272 | 0.82694043 | 0.949142 |
| LINP1 | 6.12904667 | -3.545433636 | 1.37998512 | -2.5691825 | 0.01019388 | NA |
| LINC00707 | 9.11172782 | -2.5938722 | 1.08950177 | -2.3807875 | 0.01727567 | NA |
| SFMBT2 | 410.825319 | -2.035730074 | 0.47465947 | -4.2888222 | 1.80E-05 | 0.00285812 |
| ITIH5 | 146.238797 | -0.525447221 | 0.56501116 | -0.9299767 | 0.35238314 | 0.723679 |
| KIN | 11.7608264 | -0.312449785 | 0.46200815 | -0.6762863 | 0.49885889 | NA |
| ATP5F1C | 38.4416905 | 0.10772905 | 0.3170134 | 0.33982491 | 0.73398839 | 0.91884473 |
| TAF3 | 86.4926816 | -0.26277641 | 0.2323 | -1.1311942 | 0.25797337 | 0.65588339 |
| USP6NL | 31.6923894 | 0.204311897 | 0.30836625 | 0.66256244 | 0.50761083 | 0.82657837 |
| ECHDC3 | 7.08121698 | -0.855260142 | 0.65156146 | -1.3126316 | 0.18930713 | NA |
| UPF2 | 40.1485682 | -0.169956292 | 0.28919895 | -0.5876795 | 0.55674744 | 0.85122066 |
| DHTKD1 | 21.1871731 | 0.216587304 | 0.45240838 | 0.47874291 | 0.63212154 | NA |
| SEC61A2 | 7.94523304 | 0.704654156 | 0.59877326 | 1.1768297 | 0.23926349 | NA |
| NUDT5 | 21.228129 | -0.393959082 | 0.37678477 | -1.0455812 | 0.29575442 | NA |
| CDC123 | 22.3215197 | -0.165813326 | 0.33384513 | -0.4966774 | 0.61941657 | NA |
| CAMK1D | 9.04325694 | 0.04812908 | 0.57270822 | 0.0840377 | 0.93302646 | NA |
| OPTN | 66.4550011 | -0.079761753 | 0.39371809 | -0.2025859 | 0.83945867 | 0.95393706 |
| PHYH | 9.31814294 | -0.477649216 | 0.54162376 | -0.8818838 | 0.37783965 | NA |
| SEPHS1 | 18.2841587 | 0.158063973 | 0.36246344 | 0.43608253 | 0.66277685 | NA |
| BEND7 | 26.6155309 | -0.128235948 | 0.32735819 | -0.3917298 | 0.6952579 | NA |
| PRPF18 | 4.83852919 | -0.613158486 | 0.62086903 | -0.9875811 | 0.32335787 | NA |
| AL157392.5 | 11.5028416 | -0.021066625 | 0.46704512 | -0.0451062 | 0.96402267 | NA |
| FRMD4A | 35.8761207 | 1.967498711 | 0.55062214 | 3.5732285 | 0.00035261 | 0.02553881 |
| FAM107B | 21.0854177 | 1.04103532 | 0.62641228 | 1.66190121 | 0.09653258 | NA |
| DCLRE1C | 11.886213 | 0.185565155 | 0.55638915 | 0.33351685 | 0.73874417 | NA |
| OLAH | 18.5263398 | 0.453623393 | 0.66718161 | 0.67990992 | 0.4965615 | NA |
| RPP38 | 24.1912731 | -0.105589757 | 0.4197167 | -0.2515739 | 0.80137046 | NA |
| NMT2 | 25.7333722 | -0.409481799 | 0.43908947 | -0.9325703 | 0.35104187 | NA |
| FAM171A1 | 15.3610575 | -0.24223971 | 0.44048469 | -0.549939 | 0.58236121 | NA |
| MINDY3 | 40.0499533 | 0.570169813 | 0.32416238 | 1.75890184 | 0.07859418 | 0.42784309 |
| PTER | 16.0790457 | 0.670059255 | 0.46937557 | 1.42755461 | 0.1534201 | NA |
| RSU1 | 42.0799918 | -0.194231721 | 0.31342231 | -0.6197125 | 0.53544708 | 0.83677705 |
| CUBN | 73.5804559 | 1.73274179 | 0.84268818 | 2.05620754 | 0.03976252 | 0.31540889 |
| TRDMT1 | 12.1901632 | 0.440694469 | 0.49720359 | 0.88634611 | 0.37543105 | NA |
| VIM | 513.986305 | -0.034438299 | 0.46394412 | -0.0742294 | 0.94082784 | 0.98097628 |
| HACD1 | 5.90042064 | -0.226234532 | 0.84022366 | -0.2692551 | 0.78773336 | NA |
| STAM | 19.5157409 | -0.212957727 | 0.37516484 | -0.5676377 | 0.57028097 | NA |
| CACNB2 | 270.032155 | 0.776816387 | 0.45271153 | 1.71591917 | 0.08617684 | 0.44638081 |
| AL450384.2 | 17.8719357 | -0.604192634 | 0.45095694 | -1.3398012 | 0.18031 | NA |
| NSUN6 | 25.2146457 | -0.312126994 | 0.35926062 | -0.8688038 | 0.38495445 | NA |
| ARL5B | 16.4145855 | 0.734129209 | 0.40608664 | 1.80781423 | 0.07063542 | NA |
| MALRD1 | 9.87488751 | 0.844202774 | 0.82794301 | 1.01963875 | 0.30789982 | NA |
| PLXDC2 | 242.393239 | 0.390499851 | 0.28946288 | 1.34904986 | 0.17732095 | 0.58001112 |
| NEBL | 1621.07413 | 0.270576241 | 0.25641755 | 1.05521732 | 0.29132594 | 0.67792199 |
| MLLT10 | 119.674944 | 0.295768673 | 0.18967123 | 1.5593755 | 0.11890753 | 0.51050744 |
| DNAJC1 | 81.095262 | 0.264872191 | 0.34514855 | 0.76741504 | 0.44283479 | 0.78643653 |
| BMI1 | 7.74791606 | 0.595432304 | 0.59057146 | 1.00823075 | 0.3133437 | NA |
| PIP4K2A | 28.9857608 | -0.332171835 | 0.35320481 | -0.940451 | 0.34698625 | 0.71988554 |
| MSRB2 | 23.4407797 | -0.175858964 | 0.36284506 | -0.4846668 | 0.6279127 | NA |
| OTUD1 | 11.8633026 | 0.88669015 | 0.5617391 | 1.57847326 | 0.11445693 | NA |
| KIAA1217 | 78.5893791 | 0.850434681 | 0.4654424 | 1.82715342 | 0.06767671 | 0.40302824 |
| ARHGAP21 | 110.634109 | 0.523766357 | 0.23499092 | 2.22887914 | 0.02582195 | 0.25860202 |
| PDSS1 | 5.70725625 | -0.392049544 | 0.62716361 | -0.6251153 | 0.53189541 | NA |
| ABI1 | 130.348245 | 0.029956979 | 0.32368665 | 0.09254932 | 0.92626161 | 0.97775374 |
| ANKRD26 | 48.6744736 | 0.286388212 | 0.27951984 | 1.02457205 | 0.30556516 | 0.68511459 |
| YME1L1 | 52.1874454 | -0.308064528 | 0.22615731 | -1.3621692 | 0.17314449 | 0.57564735 |
| MASTL | 7.06693007 | 0.295548722 | 0.6066183 | 0.48720706 | 0.6261116 | NA |
| ACBD5 | 18.078396 | -0.271822331 | 0.35466491 | -0.7664201 | 0.44342635 | NA |
| RAB18 | 83.6733728 | -0.584574026 | 0.27753451 | -2.1063111 | 0.03517733 | 0.29357873 |
| MPP7 | 15.9415377 | 1.220199146 | 0.41202695 | 2.9614547 | 0.0030619 | NA |
| WAC-AS1 | 23.1732527 | -0.303486257 | 0.39314459 | -0.7719457 | 0.44014662 | NA |
| WAC | 149.468003 | 0.314516091 | 0.17980444 | 1.74921201 | 0.08025438 | 0.43346499 |
| BAMBI | 70.1005161 | -0.618269431 | 0.85694113 | -0.7214841 | 0.4706117 | 0.80448595 |
| SVIL | 81.4177456 | 0.239986239 | 0.40117976 | 0.59820126 | 0.54970565 | 0.84615033 |
| MTPAP | 9.9612336 | 0.041019316 | 0.48473527 | 0.0846221 | 0.93256183 | NA |
| MAP3K8 | 13.6468191 | -1.457118189 | 0.65720864 | -2.2171318 | 0.02661409 | NA |
| ZNF438 | 35.1135765 | 0.155298771 | 0.32907676 | 0.47192263 | 0.63698201 | 0.88633797 |
| ZEB1 | 49.6843482 | -0.576282484 | 0.37257347 | -1.546762 | 0.12192065 | 0.51522823 |
| ARHGAP12 | 47.4373181 | 0.426386567 | 0.29581581 | 1.44139207 | 0.14947395 | 0.54834287 |
| KIF5B | 80.0862191 | 0.133390305 | 0.27652088 | 0.48238783 | 0.62953047 | 0.88166074 |
| EPC1 | 120.008745 | 0.520072939 | 0.23999451 | 2.16702019 | 0.03023331 | 0.27996878 |
| AL391839.1 | 4.6592533 | 0.018326965 | 0.79472224 | 0.02306084 | 0.98160174 | NA |
| CCDC7 | 57.306251 | 0.145895531 | 0.3787059 | 0.38524757 | 0.700054 | 0.90729979 |
| ITGB1 | 230.797665 | 0.026185999 | 0.36494072 | 0.07175412 | 0.94279759 | 0.98114803 |
| NRP1 | 47.0219275 | -0.485942223 | 0.53379475 | -0.9103541 | 0.36263581 | 0.73259935 |
| PARD3 | 959.484077 | -0.281945811 | 0.38108986 | -0.7398408 | 0.45939663 | 0.79594734 |
| CUL2 | 22.5982068 | -0.216656152 | 0.32411971 | -0.6684449 | 0.50384967 | NA |
| CREM | 42.2029698 | -0.297783882 | 0.34302893 | -0.8681014 | 0.38533885 | 0.75104967 |
| CCNY | 116.370036 | 0.086662889 | 0.19828755 | 0.43705664 | 0.66207027 | 0.89839561 |
| AL117336.3 | 10.5932269 | -0.115568138 | 0.52886178 | -0.2185224 | 0.82702212 | NA |
| FZD8 | 27.8624002 | 1.931878968 | 0.90230122 | 2.1410577 | 0.03226938 | NA |
| ZNF248 | 46.8343666 | 0.12249004 | 0.37936149 | 0.32288475 | 0.74678253 | 0.92250642 |
| ZNF25 | 19.3923776 | -0.637168135 | 0.36988607 | -1.7226065 | 0.08495972 | NA |
| ZNF33A | 46.9348841 | 0.060751591 | 0.34856993 | 0.1742881 | 0.86163905 | 0.95951831 |
| AL117339.5 | 13.2436503 | -0.158053754 | 0.48912781 | -0.3231338 | 0.74659388 | NA |
| LINC00839 | 278.608855 | 0.074490419 | 0.40731479 | 0.18288169 | 0.85489085 | 0.95578763 |
| ZNF33B | 29.0284232 | 0.253951633 | 0.4191875 | 0.60581872 | 0.54463514 | 0.84346702 |
| BMS1 | 18.5441829 | -0.110906435 | 0.34624414 | -0.3203128 | 0.74873121 | NA |
| CSGALNACT2 | 21.8649083 | -0.719738738 | 0.40126671 | -1.7936667 | 0.07286639 | NA |
| HNRNPF | 45.680962 | -1.048458951 | 0.34591753 | -3.0309506 | 0.00243785 | 0.07963615 |
| ZNF487 | 8.78641042 | 0.233749342 | 0.47253851 | 0.49466728 | 0.620835 | NA |
| ZNF32 | 17.2525597 | -0.806458457 | 0.38712504 | -2.0831989 | 0.0372331 | NA |
| CXCL12 | 6.74246043 | -0.984611617 | 1.25602965 | -0.7839079 | 0.43309413 | NA |
| TMEM72-AS | 7.74196469 | 0.936803586 | 0.67884184 | 1.38000272 | 0.16758581 | NA |
| RASSF4 | 21.5809021 | 0.1691461 | 0.35602062 | 0.47510197 | 0.63471429 | NA |
| DEPP1 | 7.64858137 | -2.582521867 | 1.19716909 | -2.1571906 | 0.03099082 | NA |
| ZNF22 | 17.4264411 | -0.973223461 | 0.41079182 | -2.3691403 | 0.01782949 | NA |
| 8-Mar | 27.6027448 | 0.280471578 | 0.35045263 | 0.80031239 | 0.42352983 | NA |
| ZFAND4 | 14.3455831 | 0.20802215 | 0.42373134 | 0.49092935 | 0.62347642 | NA |
| WASHC2C | 14.0060782 | -0.435127569 | 0.45544108 | -0.9553982 | 0.33937639 | NA |
| AGAP4 | 14.8830854 | 0.702820192 | 0.37441356 | 1.8771227 | 0.06050128 | NA |
| TIMM23 | 15.111353 | -0.405850276 | 0.44264329 | -0.9168789 | 0.35920612 | NA |
| NCOA4 | 31.288544 | -0.325009987 | 0.30987239 | -1.0488511 | 0.29424667 | 0.6804998 |
| AL356056.2 | 12.4315475 | 0.365015161 | 0.77181173 | 0.4729329 | 0.63626105 | NA |
| PTPN20 | 7.63180184 | 1.274639185 | 0.61205564 | 2.08255443 | 0.03729186 | NA |
| AGAP9 | 12.0543464 | -0.088265686 | 0.4186819 | -0.210818 | 0.83302928 | NA |
| MAPK8 | 69.5589308 | 0.032122342 | 0.2543453 | 0.12629422 | 0.89949903 | 0.97097328 |
| AL138760.1 | 5.35116141 | -0.772559819 | 1.05894229 | -0.729558 | 0.4656604 | NA |
| ERCC6 | 34.3757298 | -0.564753354 | 0.62799846 | -0.899291 | 0.36849768 | 0.73626927 |
| PARG | 36.7849822 | -0.352144269 | 0.33949985 | -1.0372443 | 0.29962203 | 0.68168772 |

| | | | | | | |
|---|---|---|---|---|---|---|
| TIMM23B | 37.407211 | 0.005716416 | 0.48201925 | 0.01185931 | 0.99053786 | 0.99710744 |
| LINC00843 | 9.61378669 | 0.206632635 | 0.53165317 | 0.38866059 | 0.69752724 | NA |
| AGAP6 | 16.5245598 | 0.256744627 | 0.40764348 | 0.62982641 | 0.52880816 | NA |
| WASHC2A | 17.0284747 | -0.359630438 | 0.4331464 | -0.8302746 | 0.40638357 | NA |
| SGMS1 | 131.239196 | 0.02612504 | 0.27698799 | 0.09431831 | 0.9248563 | 0.97775374 |
| ASAH2B | 6.50692303 | -0.260239753 | 0.59372856 | -0.4383144 | 0.66115842 | NA |
| PRKG1 | 704.232462 | 1.225828883 | 0.4008882 | 3.05778238 | 0.00222981 | 0.07730023 |
| CSTF2T | 5.00666173 | -0.794145242 | 0.6409232 | -1.2390646 | 0.21532158 | NA |
| PCDH15 | 20.2844962 | 2.569866402 | 0.59112626 | 4.34740692 | 1.38E-05 | NA |
| IPMK | 12.6754479 | 0.296598814 | 0.62217258 | 0.4767147 | 0.6335653 | NA |
| CISD1 | 26.2256237 | 0.178473962 | 0.36491243 | 0.4890871 | 0.62478003 | NA |
| UBE2D1 | 14.7933227 | -0.68044391 | 0.44117169 | -1.5423562 | 0.12298705 | NA |
| TFAM | 13.4520667 | -0.178320513 | 0.46417753 | -0.3841645 | 0.70085655 | NA |
| BICC1 | 119.755217 | 0.997916248 | 0.64835055 | 1.53916157 | 0.12376486 | 0.51858498 |
| PHYHIPL | 10.4111741 | 1.989361943 | 0.78983273 | 2.51871298 | 0.01177846 | NA |
| SLC16A9 | 14.3688347 | 3.584957065 | 1.23308054 | 2.90731784 | 0.00364543 | NA |
| CCDC6 | 88.3360104 | -0.345607118 | 0.2687171 | -1.2861374 | 0.19839511 | 0.60276446 |
| ANK3 | 130.432994 | 0.655323003 | 0.43861655 | 1.49406812 | 0.13515781 | 0.52779404 |
| AL592430.1 | 10.1226597 | 0.843838685 | 0.82286619 | 1.02548712 | 0.3051334 | NA |
| RHOBTB1 | 13.2181827 | 1.894324112 | 0.87557219 | 2.16352705 | 0.03050067 | NA |
| CABCOCO1 | 15.5845028 | -0.282891162 | 0.61439525 | -0.4604384 | 0.64520158 | NA |
| ARID5B | 83.4506106 | 0.322912783 | 0.42087319 | 0.76724483 | 0.44293596 | 0.78643653 |
| ADO | 11.8223143 | -0.235443635 | 0.45497926 | -0.5174821 | 0.60481964 | NA |
| NRBF2 | 7.57226087 | 0.089093922 | 0.53289055 | 0.16718991 | 0.86722062 | NA |
| JMJD1C | 260.173867 | 0.267661684 | 0.23995679 | 1.11545786 | 0.26465427 | 0.65974932 |
| REEP3 | 53.9092184 | -0.191224756 | 0.3506549 | -0.545336 | 0.58552242 | 0.86235633 |
| LINC01515 | 37.7975258 | -0.197809464 | 0.48465924 | -0.4081413 | 0.68316992 | 0.90499319 |
| CTNNA3 | 43.740501 | 1.87283994 | 0.51711976 | 3.62167548 | 0.0002927 | 0.02198511 |
| SIRT1 | 23.7687633 | 0.459204739 | 0.31523392 | 1.45671106 | 0.14519616 | NA |
| HERC4 | 194.43301 | 0.072325317 | 0.20462592 | 0.35345139 | 0.72375007 | 0.91509584 |
| PBLD | 15.5329244 | 0.878886618 | 0.40962931 | 2.14556576 | 0.03190764 | NA |
| HNRNPH3 | 53.8146808 | 0.056153991 | 0.25244116 | 0.22244388 | 0.82396836 | 0.94808955 |
| RUFY2 | 43.8943375 | 0.430424503 | 0.30575107 | 1.40776122 | 0.15920179 | 0.56239512 |
| SLC25A16 | 27.8926747 | 0.225190419 | 0.32731655 | 0.68798971 | 0.49145926 | NA |
| TET1 | 11.7178965 | 1.06776152 | 0.48128318 | 2.21857229 | 0.02651584 | NA |
| CCAR1 | 54.7484999 | 0.009185204 | 0.31984354 | 0.0287178 | 0.97708966 | 0.99298531 |
| DDX50 | 28.5895832 | -0.309203043 | 0.30933217 | -0.9995826 | 0.31751257 | 0.69424851 |
| DDX21 | 25.9109567 | -1.355092063 | 0.56833632 | -2.3843137 | 0.01711101 | NA |
| KIF1BP | 21.2146387 | -0.46950677 | 0.3451352 | -1.3603561 | 0.17371727 | NA |
| VPS26A | 36.6878408 | -0.19744565 | 0.29443696 | -0.6705872 | 0.50248356 | 0.82546509 |
| SUPV3L1 | 12.8545954 | 0.300296306 | 0.44468453 | 0.67530189 | 0.49948398 | NA |
| HKDC1 | 8.94136668 | -1.075485227 | 0.89364089 | -1.203487 | 0.22878791 | NA |
| HK1 | 29.4644185 | -0.656211221 | 0.41763726 | -1.5712468 | 0.11612532 | 0.50591226 |
| TSPAN15 | 20.7793877 | -0.179807325 | 0.53864543 | -0.3338139 | 0.73852 | NA |
| H2AFY2 | 10.3841419 | 0.14185566 | 0.47207098 | 0.30049647 | 0.76379849 | NA |
| AIFM2 | 12.0075195 | -0.240573194 | 0.46959895 | -0.512295 | 0.60844456 | NA |
| TYSND1 | 8.84903585 | -0.037062097 | 0.56957872 | -0.0650693 | 0.94811881 | NA |
| SAR1A | 29.9784547 | 0.085553617 | 0.31171833 | 0.27445809 | 0.78373261 | 0.9349469 |
| PPA1 | 78.2397908 | -1.250852336 | 0.46007008 | -2.71883 | 0.00655133 | 0.12888932 |
| LRRC20 | 7.49621537 | -0.281139933 | 0.50942945 | -0.5518722 | 0.58103594 | NA |
| EIF4EBP2 | 25.3149644 | -0.016960834 | 0.32076231 | -0.0528766 | 0.95783019 | NA |
| SGPL1 | 31.3131508 | 0.201156288 | 0.31038155 | 0.64809357 | 0.51692443 | 0.83104823 |
| PCBD1 | 31.2706746 | -0.579854156 | 0.44972285 | -1.2893589 | 0.19727335 | 0.59987316 |
| VSIR | 31.0196989 | 0.161936239 | 0.50114536 | 0.32313227 | 0.74659507 | 0.92250642 |
| PSAP | 423.952518 | -0.543251744 | 0.42062349 | -1.2915393 | 0.19651675 | 0.59987316 |
| CHST3 | 4.90903741 | 0.253642369 | 0.80961431 | 0.3132879 | 0.75406194 | NA |
| SPOCK2 | 640.797703 | 0.59172917 | 0.49042837 | 1.20655574 | 0.22760328 | 0.63327644 |
| ASCC1 | 35.1488801 | 0.367075892 | 0.39169753 | 0.9371412 | 0.34868593 | 0.72238707 |
| ANAPC16 | 115.423617 | -0.226723823 | 0.26715388 | -0.8486638 | 0.39606841 | 0.75911719 |
| DDIT4 | 13.6791997 | -1.745559485 | 0.99083418 | -1.761707 | 0.07811882 | NA |
| DNAJB12 | 20.4055211 | -0.032396897 | 0.42670474 | -0.0759235 | 0.93948 | NA |
| MICU1 | 55.0593281 | 0.428121592 | 0.32566662 | 1.31460079 | 0.1886441 | 0.59077076 |
| MCU | 83.4289436 | 0.658761438 | 0.35115857 | 1.8759657 | 0.06065999 | 0.3808621 |
| P4HA1 | 40.5458809 | 0.401442299 | 0.3944416 | 1.01774837 | 0.30879755 | 0.6882709 |
| ECD | 15.3899634 | -0.246233444 | 0.3747822 | -0.6570041 | 0.51117828 | NA |
| FAM149B1 | 22.1439115 | 0.064456427 | 0.37729401 | 0.17083872 | 0.86435058 | NA |
| DNAJC9 | 19.7830094 | -0.650923596 | 0.3980184 | -1.6354108 | 0.10196295 | NA |
| MRPS16 | 19.3910165 | -0.390465518 | 0.39176311 | -0.9966878 | 0.31891607 | NA |
| CFAP70 | 16.1341961 | 0.454050083 | 0.50043953 | 0.90730259 | 0.36424681 | NA |
| ANXA7 | 46.2012431 | -0.27224798 | 0.28814707 | -0.944823 | 0.34474924 | 0.71842682 |
| PPP3CB | 48.906364 | 0.010416099 | 0.24431482 | 0.04263392 | 0.96599335 | 0.98892249 |
| USP54 | 29.41356 | -0.018354537 | 0.46327783 | -0.0396189 | 0.96839699 | 0.98964394 |
| SEC24C | 23.989127 | 1.074048191 | 0.34415922 | 3.1207887 | 0.00180367 | NA |
| FUT11 | 31.8224467 | -0.14528213 | 0.47227742 | -0.3076203 | 0.75837126 | 0.92313949 |
| CHCHD1 | 15.350442 | -0.960761364 | 0.53766082 | -1.7869284 | 0.07394907 | NA |
| ZSWIM8 | 20.7659861 | -0.226191314 | 0.33652774 | -0.6721327 | 0.5014992 | NA |
| CAMK2G | 32.1969239 | -0.234481164 | 0.33399057 | -0.7020592 | 0.48264223 | 0.81295551 |
| PLAU | 9.19413081 | -2.034871403 | 0.91269194 | -2.2295271 | 0.02577885 | NA |
| VCL | 114.679727 | -0.114991247 | 0.21085754 | -0.5453504 | 0.58551253 | 0.86235633 |
| AP3M1 | 21.073465 | 0.160151862 | 0.40325311 | 0.39714972 | 0.69125706 | NA |
| ADK | 150.810269 | 0.44210965 | 0.2858631 | 1.54657822 | 0.12196499 | 0.51522823 |
| KAT6B | 71.6755872 | 0.533269817 | 0.31087998 | 1.71535593 | 0.08627999 | 0.44638081 |
| SAMD8 | 34.0745098 | -0.017795039 | 0.3028479 | -0.058759 | 0.95314407 | 0.98520702 |
| VDAC2 | 46.5122442 | -0.563388186 | 0.51821894 | -1.0871625 | 0.27696501 | 0.66958859 |
| COMTD1 | 8.74994005 | 0.614524914 | 0.72339974 | 0.84949563 | 0.39560556 | NA |
| LRMDA | 453.25104 | 0.437079351 | 0.31824415 | 1.37340892 | 0.16962526 | 0.57273205 |
| DLG5 | 87.6709237 | -1.76339318 | 0.59450289 | -2.9661642 | 0.00301539 | 0.0904424 |
| POLR3A | 14.4813368 | 0.022588996 | 0.40536746 | 0.05572474 | 0.95556109 | NA |
| RPS24 | 88.7276668 | -0.629372025 | 0.37304959 | -1.6871002 | 0.09158409 | 0.46030369 |
| ZMIZ1-AS1 | 6.15865248 | 0.146278252 | 1.02825146 | 0.14225922 | 0.88687526 | NA |
| ZMIZ1 | 65.5348508 | 0.376798665 | 0.3048766 | 1.2359055 | 0.21649369 | 0.62433386 |
| PPIF | 3.87217054 | -0.454323559 | 1.1417642 | -0.3979136 | 0.69069384 | NA |
| ZCCHC24 | 15.7630679 | -1.232115688 | 0.71627542 | -1.7201703 | 0.08540149 | NA |
| EIF5AL1 | 4.24518242 | -0.461022567 | 0.80029326 | -0.576067 | 0.56456987 | NA |
| NUTM2B-AS | 59.5760294 | -0.103890971 | 0.38367224 | -0.2707805 | 0.78655984 | 0.93615351 |
| TMEM254 | 9.08751506 | -0.535217827 | 0.52465395 | -1.0201349 | 0.30766447 | NA |
| ANXA11 | 51.0320733 | -0.110679998 | 0.37595208 | -0.2943992 | 0.76845287 | 0.92555901 |
| FAM213A | 10.3727805 | -0.608671682 | 0.51230044 | -1.1881147 | 0.23478823 | NA |
| TSPAN14 | 11.9571614 | 0.18121044 | 0.44773823 | 0.40472408 | 0.68568034 | NA |
| NRG3 | 36.7151583 | -0.888532676 | 0.83259461 | -1.0671852 | 0.2858882 | 0.67377254 |
| GHITM | 35.9526141 | 0.051857737 | 0.33098362 | 0.15667766 | 0.8754989 | 0.96267305 |
| CCSER2 | 126.966607 | 0.60293563 | 0.190803 | 3.15999025 | 0.00157774 | 0.06212942 |
| WAPL | 48.4626081 | 0.326002195 | 0.2393162 | 1.36222366 | 0.17312731 | 0.57564735 |
| BMPR1A | 93.9949664 | 0.30382461 | 0.25183262 | 1.20645456 | 0.22764227 | 0.63327644 |
| ADIRF | 21.3850833 | -0.141998833 | 0.79900362 | -0.1777199 | 0.85894297 | NA |
| GLUD1 | 42.1747329 | -0.080767517 | 0.30812058 | -0.2621296 | 0.79322155 | 0.93934998 |
| SHLD2 | 54.064357 | 0.164851974 | 0.31343943 | 0.52594523 | 0.59892626 | 0.86882737 |
| NUTM2A-AS | 68.0811253 | 0.085329461 | 0.24920553 | 0.34240597 | 0.73204539 | 0.91806182 |
| MINPP1 | 11.3218472 | 0.487978761 | 0.44889051 | 1.08707746 | 0.27700258 | NA |
| ATAD1 | 39.4989062 | 0.244165549 | 0.25842759 | 0.94481223 | 0.34475475 | 0.71842682 |
| PTEN | 77.5842686 | 0.294069162 | 0.25736298 | 1.14262417 | 0.25319467 | 0.65286559 |
| RNLS | 34.927059 | 0.473937245 | 0.40992226 | 1.15616371 | 0.2476142 | 0.65061974 |
| LIPA | 22.2639428 | -0.149177524 | 0.56842219 | -0.2624414 | 0.79298115 | NA |

| | | | | | | |
|---|---|---|---|---|---|---|
| IFIT2 | 41.7400546 | 1.356701953 | 1.81793798 | 0.74628616 | 0.45549457 | 0.79324789 |
| IFIT3 | 62.0650885 | -1.080443311 | 1.38811262 | -0.7783542 | 0.43636022 | 0.78209326 |
| IFIT1 | 97.5427548 | -0.622968925 | 0.90117275 | -0.6912869 | 0.48938526 | 0.81787981 |
| IFIT5 | 14.3755494 | -0.174241496 | 0.744286 | -0.2341056 | 0.81490302 | NA |
| SLC16A12 | 16.8436283 | 2.726295988 | 1.00021887 | 2.72569942 | 0.00641654 | NA |
| PANK1 | 9.06746799 | 0.657714024 | 0.51704984 | 1.27205149 | 0.20335482 | NA |
| RPP30 | 11.8119627 | -0.35658907 | 0.46670098 | -0.7640633 | 0.44482954 | NA |
| PCGF5 | 70.1163114 | -0.20556129 | 0.35796933 | -0.5742427 | 0.56580355 | 0.85573063 |
| HECTD2 | 44.6220788 | -0.435589675 | 0.29474398 | -1.4778577 | 0.13944586 | 0.53526806 |
| PPP1R3C | 4.15852797 | -1.118447898 | 0.92362747 | -1.2109297 | 0.22592236 | NA |
| TNKS2 | 68.9808631 | 0.185708868 | 0.23544103 | 0.78877018 | 0.43024634 | 0.77892126 |
| BTAF1 | 54.1817526 | -0.063927327 | 0.27739761 | -0.2304538 | 0.81773918 | 0.94694381 |
| CPEB3 | 56.1845595 | 0.4700681 | 0.33532564 | 1.40182571 | 0.1609673 | 0.56283048 |
| 5-Mar | 21.506862 | -0.062114074 | 0.34595864 | -0.1795419 | 0.85751221 | NA |
| IDE | 25.1129755 | -0.243456528 | 0.32287857 | -0.7540189 | 0.4508379 | NA |
| EXOC6 | 48.9539435 | 0.237562039 | 0.2801335 | 0.84803154 | 0.39642041 | 0.75911719 |
| MYOF | 80.5238637 | -0.4039637 | 0.33681578 | -1.199361 | 0.23038761 | 0.63682501 |
| FRA10AC1 | 22.6572483 | 0.274856079 | 0.42608356 | 0.64507553 | 0.51887824 | NA |
| SLC35G1 | 5.07495787 | 0.568603365 | 0.7614042 | 0.74678254 | 0.45519484 | NA |
| PLCE1 | 846.120946 | 0.544639498 | 0.36518052 | 1.49142538 | 0.13584985 | 0.52829049 |
| PLCE1-AS2 | 6.01234754 | -0.340249793 | 0.87072019 | -0.3907682 | 0.69596855 | NA |
| NOC3L | 10.05047 | -0.421369344 | 0.55023098 | -0.7658045 | 0.44379265 | NA |
| TBC1D12 | 13.4052984 | -0.291797889 | 0.47115198 | -0.6193286 | 0.53569992 | NA |
| PDLIM1 | 9.24099837 | 1.056757217 | 0.6335673 | 1.66794786 | 0.09532608 | NA |
| SORBS1 | 26.4429417 | 0.60406261 | 0.85515161 | 0.70638072 | 0.4799514 | NA |
| ALDH18A1 | 12.0306746 | -0.855229144 | 0.53805858 | -1.5894722 | 0.11195383 | NA |
| TCTN3 | 29.0480619 | 0.197600635 | 0.29436051 | 0.67128786 | 0.50203716 | 0.82522868 |
| ENTPD1-AS1 | 62.0155439 | -0.435878578 | 0.39207263 | -1.1117292 | 0.26625461 | 0.66013596 |
| ZNF518A | 96.3807481 | 0.003861451 | 0.27181376 | 0.01420624 | 0.98866544 | 0.99651342 |
| BLNK | 11.4636562 | -1.017536507 | 0.87024857 | -1.1692481 | 0.24230368 | NA |
| TM9SF3 | 115.02524 | 0.123139824 | 0.25860857 | 0.47616296 | 0.63395829 | 0.88508487 |
| LCOR | 93.2959306 | 0.508040909 | 0.25371598 | 2.0024001 | 0.04524172 | 0.3354231 |
| RRP12 | 12.4152461 | -0.040905937 | 0.46493509 | -0.087982 | 0.92989095 | NA |
| PGAM1 | 67.8117402 | -0.377009407 | 0.41446149 | -0.9096368 | 0.36301411 | 0.73259935 |
| EXOSC1 | 13.3562593 | -0.056728748 | 0.46375819 | -0.122324 | 0.90264244 | NA |
| ZDHHC16 | 8.60197171 | -0.187271093 | 0.49987808 | -0.3746335 | 0.70793303 | NA |
| MMS19 | 21.4023551 | -0.471579226 | 0.32491522 | -1.4513916 | 0.14667084 | NA |
| MORN4 | 8.89933726 | -0.2943397 | 0.47810409 | -0.6156394 | 0.53813257 | NA |
| PI4K2A | 5.87441275 | -0.02921768 | 0.72743658 | -0.0401653 | 0.96796137 | NA |
| AVPI1 | 5.8683208 | -1.3244508 | 0.8001379 | -1.6552782 | 0.09786809 | NA |
| MARVELD1 | 4.39687119 | -0.11052062 | 0.64786142 | -0.170593 | 0.86454381 | NA |
| ZFYVE27 | 14.6605128 | 0.404907863 | 0.41363921 | 0.9788914 | 0.32763364 | NA |
| R3HCC1L | 34.0623766 | -0.157052921 | 0.34109447 | -0.4604382 | 0.64520173 | 0.89162142 |
| PYROXD2 | 5.1002208 | 0.559267215 | 0.55787624 | 1.00249333 | 0.31610538 | NA |
| HPS1 | 14.0081921 | -0.530033603 | 0.44705211 | -1.1856193 | 0.23577268 | NA |
| GOT1 | 14.5585026 | 0.382030888 | 0.52402861 | 0.72902677 | 0.46598529 | NA |
| SLC25A28 | 20.1194998 | -0.263354941 | 0.62064075 | -0.4243275 | 0.67132698 | NA |
| CUTC | 18.388 | -0.271098918 | 0.38656461 | -0.701303 | 0.48311395 | NA |
| COX15 | 11.9003379 | -0.135483459 | 0.49530839 | -0.2735335 | 0.78444311 | NA |
| ABCC2 | 8.35988645 | -1.355950888 | 0.7769607 | -1.7451988 | 0.08095027 | NA |
| DNMBP | 13.7523817 | 0.592026429 | 0.44801725 | 1.32143669 | 0.1863558 | NA |
| ERLIN1 | 13.0699426 | 0.2166562 | 0.43270225 | 0.50070504 | 0.61657872 | NA |
| CHUK | 11.6410978 | 0.39792333 | 0.38041387 | 1.0460274 | 0.29554838 | NA |
| CWF19L1 | 12.2550426 | -0.469559539 | 0.40541365 | -1.1582233 | 0.24677292 | NA |
| BLOC1S2 | 16.8990365 | -0.188475484 | 0.38845744 | -0.4851895 | 0.6275419 | NA |
| SCD | 12.0003389 | -1.491709802 | 0.69687513 | -2.1405697 | 0.03230875 | NA |
| OLMALINC | 14.1050221 | 0.237582631 | 0.40368726 | 0.58853141 | 0.55617565 | NA |
| SEC31B | 8.36093491 | 0.245062424 | 0.55437471 | 0.44205196 | 0.65845161 | NA |
| HIF1AN | 16.0823599 | -0.06900775 | 0.42448068 | -0.1625698 | 0.87085715 | NA |
| PAX2 | 22.1365158 | 1.499755649 | 0.66798393 | 2.24519719 | 0.02475548 | NA |
| SLF2 | 57.6951245 | -0.234730226 | 0.36506514 | -0.6429818 | 0.52023593 | 0.83108387 |
| MRPL43 | 17.2015537 | 0.300043845 | 0.36993993 | 0.81106098 | 0.41733065 | NA |
| SEMA4G | 9.37459942 | 1.98704384 | 0.74583024 | 2.66420391 | 0.00771708 | NA |
| LZTS2 | 46.9079491 | 0.388491194 | 0.28741665 | 1.35166557 | 0.17648232 | 0.57988078 |
| PDZD7 | 8.34110872 | 1.199916935 | 0.58537948 | 2.04981038 | 0.04038294 | NA |
| SFXN3 | 8.20192376 | 0.342106375 | 0.5906181 | 0.57923449 | 0.56243096 | NA |
| BTRC | 59.1493221 | 0.344124037 | 0.21666487 | 1.58827799 | 0.11222349 | 0.50240451 |
| DPCD | 8.14048177 | -0.682702084 | 0.5365325 | -1.2724338 | 0.20321903 | NA |
| POLL | 6.94263256 | -0.200308753 | 0.53918098 | -0.3715056 | 0.71026099 | NA |
| FBXW4 | 56.8237932 | 0.275618312 | 0.22380238 | 1.23152541 | 0.21812642 | 0.62462904 |
| OGA | 94.031372 | 0.430128239 | 0.27459327 | 1.56641945 | 0.11725045 | 0.50775741 |
| ARMH3 | 56.2707138 | 0.218055528 | 0.25249799 | 0.86359313 | 0.38781144 | 0.75225404 |
| HPS6 | 3.40534855 | -0.579283967 | 0.76960226 | -0.7527057 | 0.45162681 | NA |
| LDB1 | 8.36785534 | 0.398561882 | 0.54032839 | 0.73762898 | 0.46073994 | NA |
| PPRC1 | 7.8044019 | -1.4439885 | 0.95492711 | -1.5121453 | 0.13049691 | NA |
| NOLC1 | 10.7650079 | -0.778161116 | 0.53683638 | -1.4495313 | 0.14718927 | NA |
| GBF1 | 73.052138 | 0.182989338 | 0.20410683 | 0.89653708 | 0.36996599 | 0.73847541 |
| NFKB2 | 6.99526776 | -0.311902868 | 0.71882751 | -0.433905 | 0.66435741 | NA |
| FBXL15 | 10.4202441 | -0.48884898 | 0.49432448 | -0.9889233 | 0.32270068 | NA |
| CUEDC2 | 15.7064834 | -0.410879021 | 0.37591793 | -1.0930019 | 0.27439295 | NA |
| ACTR1A | 15.7085427 | -0.043669368 | 0.61088526 | -0.0714854 | 0.94301146 | NA |
| SUFU | 17.7983918 | 0.023822088 | 0.42166795 | 0.0564949 | 0.95494756 | NA |
| TRIM8 | 56.3949467 | 0.221883242 | 0.31705695 | 0.69982141 | 0.48403884 | 0.81370025 |
| ARL3 | 54.0056205 | -0.285968499 | 0.42860891 | -0.6672015 | 0.50464345 | 0.82600448 |
| SFXN2 | 20.9092025 | 0.222180842 | 0.45339127 | 0.49004217 | 0.62410406 | NA |
| WBP1L | 89.0591718 | -0.492156283 | 0.29645387 | -1.6601446 | 0.09688538 | 0.47093115 |
| CNNM2 | 35.6133061 | 0.541421387 | 0.42405443 | 1.27677332 | 0.20168229 | 0.60910153 |
| NT5C2 | 155.010157 | 0.66274845 | 0.25386693 | 2.61061351 | 0.009038 | 0.14901673 |
| PCGF6 | 11.0704496 | -0.390245367 | 0.41086535 | -0.9498133 | 0.34220714 | NA |
| ATP5MD | 17.8343273 | 0.247641337 | 0.4342861 | 0.57022626 | 0.56852425 | NA |
| PDCD11 | 7.7807637 | -0.525894898 | 0.76147474 | -0.6906268 | 0.48980007 | NA |
| SH3PXD2A | 15.3607199 | 0.268355713 | 0.50778965 | 0.52847811 | 0.59716754 | NA |
| STN1 | 46.0055909 | 0.344289009 | 0.53693633 | 0.64121012 | 0.52138618 | 0.83108387 |
| SLK | 204.471566 | 0.056875869 | 0.30901382 | 0.18405607 | 0.85396947 | 0.95569766 |
| CFAP43 | 8.09578958 | 0.5330317 | 0.66014795 | 0.80744279 | 0.41941142 | NA |
| GSTO1 | 23.2131258 | -0.609741358 | 0.51333488 | -1.1878043 | 0.23491054 | NA |
| GSTO2 | 14.4407659 | 0.139888887 | 0.55670257 | 0.25128119 | 0.80159671 | NA |
| XPNPEP1 | 15.2586571 | -0.495565808 | 0.40292306 | -1.2299267 | 0.21872457 | NA |
| ADD3 | 196.628348 | 0.322751675 | 0.32176654 | 1.00306165 | 0.31583111 | 0.69424851 |
| MXI1 | 22.6779702 | -0.72398537 | 0.48242913 | -1.5007082 | 0.13343104 | NA |
| SMNDC1 | 11.6946932 | -0.163119404 | 0.40870956 | -0.3991084 | 0.68981336 | NA |
| DUSP5 | 5.01115741 | -2.072905034 | 1.15692862 | -1.7917311 | 0.07317604 | NA |
| SMC3 | 27.6970924 | -0.27940734 | 0.30018661 | -0.9307788 | 0.35196799 | NA |
| PDCD4 | 26.4301194 | 0.236562063 | 0.35168989 | 0.6726439 | 0.50117386 | NA |
| BBIP1 | 19.9452852 | 0.14292144 | 0.36248411 | 0.39428332 | 0.69337187 | NA |
| SHOC2 | 49.0727394 | 0.215499883 | 0.25991552 | 0.82911509 | 0.40703928 | 0.76772515 |
| GPAM | 8.78034779 | -0.252379278 | 0.59754228 | -0.4223622 | 0.67276066 | NA |
| ACSL5 | 20.4345502 | -0.412710498 | 0.51488799 | -0.8015539 | 0.42281103 | NA |
| AL157786.1 | 23.0722329 | -0.027978639 | 0.65618445 | -0.0426384 | 0.9659898 | NA |
| ZDHHC6 | 412.650936 | -0.019447939 | 0.45055601 | -0.0431643 | 0.96557056 | 0.98892249 |
| VTI1A | 1216.2763 | -0.434337563 | 0.36500225 | -1.1899586 | 0.23406267 | 0.63975088 |
| TCF7L2 | 148.346942 | -0.090570541 | 0.23716189 | -0.3818933 | 0.7025405 | 0.90757606 |
| CASP7 | 12.0232859 | -0.722527753 | 0.61804728 | -1.1690493 | 0.24238376 | NA |

| | | | | | | |
|---|---|---|---|---|---|---|
| NHLRC2 | 54.4077026 | 0.038153482 | 0.2554032 | 0.1493853 | 0.88124961 | 0.9644761 |
| CCDC186 | 42.5875909 | 0.298360192 | 0.25558153 | 1.16737776 | 0.24305785 | 0.64918976 |
| AFAP1L2 | 67.7223061 | 0.160896314 | 0.40945601 | 0.3929514 | 0.69435538 | 0.90557575 |
| ABLIM1 | 43.9755577 | 0.733671987 | 0.45755881 | 1.60344849 | 0.10883567 | 0.49433089 |
| FAM160B1 | 28.0472514 | 0.177853295 | 0.31161838 | 0.5707407 | 0.56817542 | NA |
| TRUB1 | 21.8741441 | 0.30122475 | 0.45355492 | 0.66414173 | 0.5065996 | NA |
| ATRNL1 | 81.1292199 | -0.329083042 | 0.54874936 | -0.5996964 | 0.54870856 | 0.84550378 |
| HSPA12A | 130.875168 | -0.35484031 | 0.34512739 | -1.028143 | 0.30388256 | 0.68398872 |
| ENO4 | 5.44035691 | -1.406579132 | 0.68662442 | -2.0485422 | 0.0405069 | NA |
| SHTN1 | 56.9465209 | 0.243463537 | 0.24563437 | 0.99116234 | 0.32160632 | 0.69513974 |
| PDZD8 | 61.3671501 | -0.076038828 | 0.27545311 | -0.27605 | 0.78250967 | 0.93415197 |
| EMX2OS | 13.457213 | 0.692575196 | 0.74438004 | 0.93040538 | 0.35216123 | NA |
| EMX2 | 6.22725779 | 0.734471771 | 0.65173132 | 1.12695485 | 0.25976157 | NA |
| RAB11FIP2 | 42.3039379 | 0.181006271 | 0.33062947 | 0.54745958 | 0.58406303 | 0.86175324 |
| CASC2 | 21.2855869 | -0.259006183 | 0.43299267 | -0.5981768 | 0.54972195 | NA |
| FAM204A | 33.9376007 | -0.089267508 | 0.27763469 | -0.3215286 | 0.74780981 | 0.92253893 |
| CACUL1 | 39.190095 | 0.201684113 | 0.2965669 | 0.68006279 | 0.4964647 | 0.8234249 |
| EIF3A | 56.1674133 | -0.506221429 | 0.33549374 | -1.5088849 | 0.1313282 | 0.52613957 |
| FAM45A | 14.7366916 | 0.236320338 | 0.58289634 | 0.4054243 | 0.68516565 | NA |
| SFXN4 | 10.0803389 | -0.262879691 | 0.52228329 | -0.5033278 | 0.61473384 | NA |
| PRDX3 | 33.9777403 | 0.123850421 | 0.39480672 | 0.31369887 | 0.75374976 | 0.92313949 |
| GRK5 | 196.155229 | -0.486569997 | 0.40413023 | -1.2039931 | 0.22859225 | 0.63504807 |
| RGS10 | 41.0945823 | -0.10394107 | 0.32788676 | -0.317003 | 0.75124136 | 0.92274989 |
| TIAL1 | 55.1527407 | 0.37360669 | 0.34693527 | 1.07687721 | 0.28153512 | 0.6701329 |
| BAG3 | 15.1153146 | -1.420602778 | 0.64610515 | -2.1987176 | 0.02789801 | NA |
| INPP5F | 20.151442 | 0.304329292 | 0.44355787 | 0.68610956 | 0.49264402 | NA |
| MCMBP | 23.7235905 | -0.557988928 | 0.33614657 | -1.6599572 | 0.09692305 | NA |
| SEC23IP | 24.2651114 | 0.112595569 | 0.34250396 | 0.32874239 | 0.74235041 | NA |
| WDR11 | 42.2920758 | 0.185826521 | 0.24813154 | 0.74890327 | 0.45391551 | 0.79324789 |
| FGFR2 | 22.4778694 | 0.614323978 | 0.50796231 | 1.2093889 | 0.22651347 | NA |
| ATE1 | 60.0207853 | -0.246454046 | 0.25650669 | -0.9608094 | 0.33664799 | 0.71116888 |
| NSMCE4A | 7.31328336 | -0.655134006 | 0.52026288 | -1.2592365 | 0.20794492 | NA |
| TACC2 | 110.109333 | 0.352722195 | 0.32623952 | 1.08117557 | 0.27961902 | 0.66958859 |
| BTBD16 | 6.68719862 | 0.61414556 | 0.83928552 | 0.73174807 | 0.46432235 | NA |
| PLEKHA1 | 79.6572929 | -0.097604102 | 0.24530804 | -0.3978838 | 0.69071582 | 0.90557575 |
| HTRA1 | 1023.00891 | -0.219020807 | 0.4962737 | -0.4413307 | 0.65897362 | 0.89709157 |
| C10orf88 | 8.67195273 | -0.108121095 | 0.50181638 | -0.2154595 | 0.8294091 | NA |
| IKZF5 | 11.8451441 | 0.431572322 | 0.43801281 | 0.98529613 | 0.32447863 | NA |
| ACADSB | 19.9352321 | 0.007219734 | 0.40775485 | 0.01770607 | 0.98587334 | NA |
| BUB3 | 19.2511458 | 0.094810361 | 0.37972506 | 0.2496816 | 0.80283359 | NA |
| OAT | 18.3877003 | 0.267584699 | 0.47304905 | 0.56565952 | 0.57162526 | NA |
| LHPP | 12.3143232 | 0.626024585 | 0.43741902 | 1.43117823 | 0.15237914 | NA |
| ABRAXAS2 | 13.0432733 | -0.404557186 | 0.44941368 | -0.9001889 | 0.36801976 | NA |
| ZRANB1 | 38.218508 | 0.692445055 | 0.42250043 | 1.63892154 | 0.1012296 | 0.48247622 |
| CTBP2 | 113.583448 | -0.144313457 | 0.25799773 | -0.5593594 | 0.57591647 | 0.85929076 |
| EDRF1 | 14.7906374 | 0.327921133 | 0.4419854 | 0.74192752 | 0.45813125 | NA |
| UROS | 12.6293827 | 0.03355463 | 0.50334783 | 0.06666291 | 0.94685006 | NA |
| BCCIP | 16.4691511 | -0.36809914 | 0.39010907 | -0.9435801 | 0.34538429 | NA |
| DHX32 | 26.1861578 | -0.74082602 | 0.35208286 | -2.1041241 | 0.03536762 | NA |
| FANK1 | 13.6879691 | 0.085526612 | 0.47805992 | 0.17890354 | 0.85801344 | NA |
| DOCK1 | 164.816553 | -0.045233373 | 0.23476977 | -0.1926712 | 0.84721648 | 0.95532127 |
| FAM196A | 5.66838843 | -1.506353779 | 0.81558292 | -1.8469658 | 0.06475209 | NA |
| MGMT | 57.4251334 | -0.095353392 | 0.31165722 | -0.305956 | 0.75963816 | 0.92313949 |
| C10orf143 | 6.94865809 | 1.008632933 | 0.55525881 | 1.81650956 | 0.06929224 | NA |
| GLRX3 | 29.7688215 | 0.10862869 | 0.29593454 | 0.36706999 | 0.71356681 | 0.91328825 |
| PPP2R2D | 34.1229003 | 0.10029285 | 0.32773156 | 0.30602133 | 0.7595884 | 0.92313949 |
| BNIP3 | 33.4206128 | -0.559975909 | 0.46373199 | -1.2075421 | 0.22722343 | 0.63327644 |
| STK32C | 7.15151386 | -0.152890978 | 0.54398101 | -0.2810594 | 0.77866484 | NA |
| LRRC27 | 7.33936705 | -0.014221086 | 0.61866174 | -0.0229869 | 0.98166076 | NA |
| INPP5A | 17.8241891 | 0.592170715 | 0.38555837 | 1.53587826 | 0.12456824 | NA |
| TUBGCP2 | 26.6027983 | -0.112107346 | 0.31248862 | -0.3587566 | 0.7197772 | NA |
| FUOM | 13.3434364 | -0.226000309 | 0.72389626 | -0.3121999 | 0.75488864 | NA |
| ECHS1 | 37.8910926 | 0.199753868 | 0.461327 | 0.43299844 | 0.66501591 | 0.89968484 |
| SCART1 | 7.80431195 | 0.449655797 | 0.70205453 | 0.64048557 | 0.52185697 | NA |
| SLC6A12 | 8.45178441 | 1.988915253 | 0.76220762 | 2.60941403 | 0.00906974 | NA |
| KDM5A | 69.5290594 | 0.143545972 | 0.26844095 | 0.53473947 | 0.59283003 | 0.8654248 |
| CCDC77 | 4.5345794 | -0.642382842 | 0.65026141 | -0.987884 | 0.32320946 | NA |
| WNK1 | 120.735315 | 0.066444636 | 0.29321535 | 0.22660695 | 0.82072939 | 0.94728252 |
| RAD52 | 10.9179253 | -0.267280023 | 0.46538459 | -0.5743207 | 0.56575078 | NA |
| ERC1 | 223.4268 | 0.401511038 | 0.18409092 | 2.18104747 | 0.02917991 | 0.27926427 |
| WNT5B | 12.8658662 | -1.826746824 | 0.81672458 | -2.2366742 | 0.02530764 | NA |
| FBXL14 | 6.67785268 | -0.925471291 | 0.7959567 | -1.1627156 | 0.2449449 | NA |
| ADIPOR2 | 37.2400704 | -0.45417424 | 0.35945895 | -1.263494 | 0.20641169 | 0.6151402 |
| DCP1B | 17.1956168 | -0.482607911 | 0.40515579 | -1.1911663 | 0.23358831 | NA |
| CACNA1C | 14.3358376 | 1.025587764 | 0.88709991 | 1.15611303 | 0.24763493 | NA |
| FKBP4 | 13.1082651 | 0.975712391 | 0.5168695 | 1.88773452 | 0.0590616 | NA |
| ITFG2 | 24.5239016 | 0.126436424 | 0.33590927 | 0.37640052 | 0.70661916 | NA |
| NRIP2 | 31.7689974 | 1.135305304 | 0.69873331 | 1.6248049 | 0.10420414 | 0.4862279 |
| TULP3 | 26.4932588 | 0.491280436 | 0.31188067 | 1.57521925 | 0.11520585 | NA |
| TEAD4 | 15.2375769 | -0.57972059 | 0.60274179 | -0.9618059 | 0.33614713 | NA |
| TSPAN9 | 94.5108028 | 0.328995882 | 0.31593956 | 1.04132538 | 0.29772456 | 0.68103704 |
| PARP11 | 17.9439795 | 0.380591678 | 0.47158161 | 0.80705368 | 0.41963556 | NA |
| CCND2 | 8.13172647 | 0.149608316 | 0.7952978 | 0.18811609 | 0.85078564 | NA |
| TIGAR | 9.21761891 | -0.016414869 | 0.45559815 | -0.0360293 | 0.97125902 | NA |
| C12orf4 | 12.4272951 | 0.321512111 | 0.44577885 | 0.72123681 | 0.47076383 | NA |
| AC005833.2 | 5.5420565 | 0.054775112 | 0.60895601 | 0.08994921 | 0.92832757 | NA |
| CD9 | 63.0686782 | -0.47145765 | 0.50598503 | -0.931762 | 0.35145951 | 0.72288022 |
| TNFRSF1A | 81.6669029 | -0.973390219 | 0.33658047 | -2.8919985 | 0.003828 | 0.09814043 |
| SCNN1A | 11.8889153 | -0.349227236 | 0.58695042 | -0.5949859 | 0.55185288 | NA |
| LTBR | 25.6270048 | -0.378918373 | 0.42179399 | -0.8983494 | 0.36899931 | NA |
| TAPBPL | 8.57744936 | -0.737232978 | 0.66374727 | -1.1107134 | 0.26669174 | NA |
| MRPL51 | 25.6783539 | -0.467104842 | 0.49165971 | -0.9500572 | 0.3420832 | NA |
| NCAPD2 | 7.44080633 | -0.106059554 | 0.5536866 | -0.1915516 | 0.84809346 | NA |
| GAPDH | 222.181059 | -1.053852914 | 0.46780931 | -2.2527404 | 0.02427552 | 0.25196701 |
| IFFO1 | 10.1162423 | 0.387159937 | 0.52132388 | 0.74264761 | 0.45769505 | NA |
| CHD4 | 24.5596564 | -0.348373046 | 0.48697712 | -0.7153787 | 0.47437509 | NA |
| AC006064.2 | 5.70520959 | -0.874560849 | 0.69859833 | -1.2518794 | 0.21061382 | NA |
| ING4 | 14.6209872 | -0.079289814 | 0.37438663 | -0.2117859 | 0.83227406 | NA |
| ZNF384 | 11.2219646 | 0.060449828 | 0.44100512 | 0.13707285 | 0.89097321 | NA |
| COPS7A | 19.7642885 | 0.219390693 | 0.33987605 | 0.64550206 | 0.51860188 | NA |
| MLF2 | 31.9518557 | -0.406447512 | 0.36137682 | -1.1247194 | 0.26070797 | 0.65595635 |
| PTMS | 120.990341 | -0.439219382 | 0.50815449 | -0.8643422 | 0.38739992 | 0.75181535 |
| CD4 | 5.1649992 | 0.897160414 | 0.78421955 | 1.14401689 | 0.25261663 | NA |
| USP5 | 12.7638673 | 0.47880537 | 0.42734288 | 1.12042436 | 0.26253297 | NA |
| TPI1 | 43.9347664 | -0.495900687 | 0.43038994 | -1.1522125 | 0.24923374 | 0.65117642 |
| LRRC23 | 16.3843475 | 0.170134854 | 0.40305385 | 0.42211445 | 0.67294148 | NA |
| ATN1 | 39.8631478 | -0.021695746 | 0.36233798 | -0.0598771 | 0.95225352 | 0.98520702 |
| C12orf57 | 24.8029398 | -1.056041639 | 0.72440255 | -1.4578105 | 0.14489279 | NA |
| PTPN6 | 6.10142624 | -0.273697453 | 0.55050411 | -0.497176 | 0.61906492 | NA |
| PHB2 | 19.6898581 | -0.086894384 | 0.47592328 | -0.1825807 | 0.85512706 | NA |
| LPCAT3 | 44.8016436 | 0.05352109 | 0.33567808 | 0.15944172 | 0.87332087 | 0.96229085 |
| C1S | 89.2570249 | -0.920411636 | 0.38783122 | -2.3732273 | 0.01763341 | 0.22021865 |
| C1R | 276.792556 | -2.089187414 | 0.41633067 | -5.0180964 | 5.22E-07 | 0.00023518 |

| | | | | | | |
|---|---|---|---|---|---|---|
| C1RL | 44.6658051 | -2.325694835 | 0.58423309 | -3.9807653 | 6.87E-05 | 0.00773949 |
| RBP5 | 5.65514117 | -0.189513375 | 0.73881935 | -0.2565084 | 0.79755829 | NA |
| CLSTN3 | 6.58210415 | -0.5779073 | 0.59772033 | -0.9668523 | 0.33361786 | NA |
| PEX5 | 8.51514976 | 0.818183045 | 0.50301418 | 1.62656058 | 0.10383046 | NA |
| CD163L1 | 6.98127093 | -0.213713296 | 0.52124295 | -0.4100071 | 0.68180076 | NA |
| SLC2A3 | 16.2701968 | -1.687053255 | 0.50031205 | -3.372002 | 0.00074624 | NA |
| FOXJ2 | 8.21604614 | 0.37187401 | 0.48821246 | 0.76170528 | 0.44623592 | NA |
| NECAP1 | 10.8430162 | 0.533727133 | 0.49161767 | 1.08565491 | 0.2776317 | NA |
| FAM66C | 9.39991332 | 0.495507412 | 0.48744543 | 1.01653926 | 0.30937266 | NA |
| LINC00937 | 7.88017537 | -0.177985407 | 0.50178331 | -0.3547057 | 0.72281007 | NA |
| RIMKLB | 35.4374064 | 0.101759961 | 0.41024784 | 0.24804509 | 0.80409952 | 0.94152069 |
| A2ML1-AS1 | 7.96419172 | 0.590285135 | 0.63483409 | 0.92982583 | 0.35246127 | NA |
| PHC1 | 16.85125 | 0.41393603 | 0.3533226 | 1.17155266 | 0.2413767 | NA |
| M6PR | 29.5095681 | -0.397363801 | 0.30932292 | -1.2846245 | 0.19892354 | 0.60367512 |
| CLEC2D | 13.4651024 | 0.463205287 | 0.60547151 | 0.76503235 | 0.44425228 | NA |
| GABARAPL1 | 35.2848546 | -0.074782654 | 0.32656674 | -0.2289965 | 0.81887161 | 0.94694381 |
| KLRD1 | 17.8390266 | 0.110867001 | 0.56000561 | 0.1979748 | 0.84306478 | NA |
| MAGOHB | 8.68404838 | -0.614175816 | 0.53437634 | -1.149332 | 0.25041911 | NA |
| YBX3 | 152.073791 | -1.335905597 | 0.36640588 | -3.6459721 | 0.00026638 | 0.02038583 |
| PRH1 | 6.83743175 | -0.340715285 | 0.59667835 | -0.57102 | 0.56798607 | NA |
| SMIM10L1 | 9.63453098 | -0.725113486 | 0.55688255 | -1.3020941 | 0.19288422 | NA |
| AC244131.2 | 10.4325543 | -0.232742851 | 0.74311572 | -0.3131987 | 0.75412973 | NA |
| LINC01252 | 24.3629633 | -0.676359119 | 0.55241675 | -1.2243639 | 0.22081499 | NA |
| ETV6 | 103.113651 | -0.600031717 | 0.31130302 | -1.9274844 | 0.05391929 | 0.3615191 |
| BCL2L14 | 20.7112806 | -0.740992698 | 0.40113933 | -1.8472203 | 0.06471523 | NA |
| LRP6 | 50.5503783 | 0.522119448 | 0.33826173 | 1.54353685 | 0.12270057 | 0.51625884 |
| MANSC1 | 22.4151513 | -0.031201069 | 0.47265054 | -0.066013 | 0.94736749 | NA |
| BORCS5 | 26.4624213 | 0.270425632 | 0.33881684 | 0.79814695 | 0.42478522 | NA |
| DUSP16 | 60.5058361 | 0.126427336 | 0.24116618 | 0.52423327 | 0.6001163 | 0.86882737 |
| CREBL2 | 22.6365388 | -0.259055131 | 0.37040438 | -0.6993846 | 0.48431169 | NA |
| CDKN1B | 41.7598067 | -0.30989238 | 0.4832038 | -0.6413285 | 0.52130926 | 0.83108387 |
| APOLD1 | 7.36577731 | 0.209910525 | 0.75101182 | 0.27950362 | 0.77985836 | NA |
| GPRC5A | 46.7581868 | -0.077319964 | 0.44530145 | -0.1736351 | 0.86215224 | 0.95962939 |
| GPRC5D-AS1 | 21.7956371 | 0.913280232 | 0.52041472 | 1.75490854 | 0.07927495 | NA |
| HEBP1 | 46.3828808 | 0.041368086 | 0.41911996 | 0.09870226 | 0.92137468 | 0.97775374 |
| FAM234B | 4.8860752 | 0.378137946 | 0.70235067 | 0.5383891 | 0.59030845 | NA |
| EMP1 | 25.4848417 | 0.424904977 | 0.76640247 | 0.55441494 | 0.57929491 | NA |
| GRIN2B | 14.4714304 | 0.46677996 | 1.05117306 | 0.44405624 | 0.65700192 | NA |
| ATF7IP | 77.5792794 | 0.176206571 | 0.28380754 | 0.62086641 | 0.53468752 | 0.83677705 |
| PLBD1 | 35.1698443 | -0.776045162 | 0.28767776 | -2.6976196 | 0.00698372 | 0.13332741 |
| PLBD1-AS1 | 8.54611735 | -0.26196955 | 0.63024651 | -0.415662 | 0.67765732 | NA |
| AC010168.1 | 10.4696151 | -1.466997493 | 0.65922852 | -2.2253247 | 0.02605945 | NA |
| H2AFJ | 216.214879 | -0.893158374 | 0.37308433 | -2.3939852 | 0.01666642 | 0.21392089 |
| WBP11 | 25.6627484 | -0.358298799 | 0.3660774 | -0.9787515 | 0.32770278 | NA |
| C12orf60 | 7.18023805 | -0.909502857 | 0.71686575 | -1.2687213 | 0.20454048 | NA |
| SMCO3 | 21.6482981 | 0.307739512 | 0.74550776 | 0.41279183 | 0.67975914 | NA |
| MGP | 322.424111 | -0.215784318 | 0.75179562 | -0.2870252 | 0.774093 | 0.92806061 |
| RERG | 647.883893 | -0.261623365 | 0.29568686 | -0.8847988 | 0.37626518 | 0.74330391 |
| PTPRO | 4411.19941 | 0.160085735 | 0.30958151 | 0.51710367 | 0.60508379 | 0.87047761 |
| EPS8 | 36.164971 | -0.830818252 | 0.74207893 | -1.1195821 | 0.26289191 | 0.6581333 |
| STRAP | 34.515296 | -0.381071048 | 0.35789449 | -1.0647581 | 0.28698542 | 0.67443257 |
| DERA | 42.5953452 | 0.318386998 | 0.33746532 | 0.94346584 | 0.34544268 | 0.71852078 |
| MGST1 | 22.694133 | -1.49936824 | 0.86489818 | -1.7335777 | 0.08299303 | NA |
| PIK3C2G | 10.8966854 | -1.133941633 | 1.0327645 | -1.0979673 | 0.27221878 | NA |
| PLEKHA5 | 433.016665 | 0.806661858 | 0.28710813 | 2.80960991 | 0.00496016 | 0.11177304 |
| AEBP2 | 98.371269 | 0.413121734 | 0.32427103 | 1.27400137 | 0.20266293 | 0.610699 |
| AC024901.1 | 36.576821 | -1.293339738 | 0.56140313 | -2.303763 | 0.02123595 | 0.2419467 |
| LINC02398 | 44.8116277 | -1.329079494 | 0.43119973 | -3.0822827 | 0.0020542 | 0.07380722 |
| PDE3A | 155.588982 | -1.109617504 | 0.44630671 | -2.4862218 | 0.01291075 | 0.18904687 |
| PYROXD1 | 13.1884877 | -0.716438339 | 0.46540356 | -1.5393916 | 0.12370872 | NA |
| RECQL | 12.70727 | -1.29057732 | 0.44677368 | -2.8886601 | 0.00386887 | NA |
| GOLT1B | 16.6296381 | 0.144517911 | 0.38208446 | 0.37823551 | 0.70525565 | NA |
| LDHB | 41.0906176 | -0.393205872 | 0.32140873 | -1.2233827 | 0.22118519 | 0.62521515 |
| CMAS | 15.6318364 | -0.231855914 | 0.41680329 | -0.5562718 | 0.57802507 | NA |
| C2CD5 | 48.5810295 | 0.714314347 | 0.2945268 | 2.42529488 | 0.01529596 | 0.20341115 |
| ETNK1 | 63.3653826 | 0.272992072 | 0.33273599 | 0.82044647 | 0.41196163 | 0.76929852 |
| SOX5 | 20.5824853 | 2.000140142 | 0.92147604 | 2.1705829 | 0.02996272 | NA |
| CASC1 | 8.29368412 | -0.097933799 | 0.54181512 | -0.1807513 | 0.85656277 | NA |
| ETFRF1 | 23.5475081 | -0.407356651 | 0.42559313 | -0.9571504 | 0.33849133 | NA |
| KRAS | 26.8502524 | 0.069266838 | 0.31137567 | 0.22245424 | 0.82396029 | NA |
| RASSF8-AS1 | 89.0655042 | -0.277827524 | 0.45729259 | -0.6075487 | 0.54348683 | 0.84233189 |
| RASSF8 | 321.590074 | 0.048196461 | 0.36168105 | 0.1332568 | 0.89399029 | 0.97097328 |
| SSPN | 4.21881465 | -0.154065183 | 0.82681163 | -0.1863365 | 0.85218089 | NA |
| ITPR2 | 48.840171 | 0.782918343 | 0.53791148 | 1.45547804 | 0.14553697 | 0.54436111 |
| INTS13 | 7.53839707 | 0.285181907 | 0.60858845 | 0.46859566 | 0.63935868 | NA |
| FGFR1OP2 | 33.2839801 | 0.921961705 | 0.39719454 | 2.32118422 | 0.0202769 | 0.2375493 |
| TM7SF3 | 34.4739217 | 0.283945551 | 0.33290203 | 0.85294027 | 0.39369243 | 0.75773027 |
| MED21 | 37.828414 | 0.261979829 | 0.44587306 | 0.58756596 | 0.55682367 | 0.85122066 |
| STK38L | 22.6912854 | 0.420300396 | 0.48146027 | 0.87297005 | 0.3826794 | NA |
| PPFIBP1 | 57.1490452 | -0.113808006 | 0.28914876 | -0.3935967 | 0.6938788 | 0.90557575 |
| MRPS35 | 16.296949 | -0.370120151 | 0.45100994 | -0.8206474 | 0.41184712 | NA |
| MANSC4 | 5.40739062 | -0.848654718 | 0.94005543 | -0.9027709 | 0.36664748 | NA |
| KLHL42 | 182.796759 | -0.20920173 | 0.28491215 | -0.7342675 | 0.46278573 | 0.79793427 |
| AC008011.2 | 38.8322475 | 1.077349648 | 0.61340746 | 1.75633608 | 0.07903104 | 0.42911633 |
| CCDC91 | 614.409161 | 1.098608888 | 0.3766472 | 2.91681148 | 0.00353629 | 0.09757286 |
| AC022364.1 | 7.72226207 | -0.169756683 | 0.57958786 | -0.2928921 | 0.76960466 | NA |
| FAR2 | 5.51428303 | -1.661167342 | 0.83483104 | -1.9898246 | 0.04661026 | NA |
| AC009318.1 | 6.80846671 | -0.778772573 | 0.63461031 | -1.2271666 | 0.21975998 | NA |
| ERGIC2 | 29.4398289 | 0.038632951 | 0.28236228 | 0.13682051 | 0.89117267 | 0.96958057 |
| TMTC1 | 36.8426292 | 0.177441859 | 0.88011788 | 0.20161147 | 0.84022048 | 0.95393706 |
| IPO8 | 23.1641688 | 0.38221035 | 0.34503699 | 1.10773733 | 0.26797527 | NA |
| CAPRIN2 | 20.8085039 | -0.050129833 | 0.47736164 | -0.1050144 | 0.9163644 | NA |
| SINHCAF | 39.6482897 | -0.325979959 | 0.41338306 | -0.7885663 | 0.43036552 | 0.77892126 |
| DENND5B | 84.932964 | 0.146062026 | 0.32993457 | 0.44269997 | 0.65798276 | 0.89709157 |
| ETFBKMT | 6.79341165 | 0.715875667 | 0.6380465 | 1.1219804 | 0.26187077 | NA |
| AMN1 | 7.88861012 | -0.417590012 | 0.55490075 | -0.752549 | 0.45172097 | NA |
| KIAA1551 | 19.0547602 | -0.379027763 | 0.36166517 | -1.0480074 | 0.29463521 | NA |
| BICD1 | 166.072433 | -0.534030727 | 0.33501649 | -1.5940431 | 0.11092638 | 0.49769622 |
| FGD4 | 70.6834529 | -0.259527258 | 0.41408956 | -0.6267418 | 0.53082847 | 0.83515914 |
| DNM1L | 49.7948983 | 0.011780035 | 0.25423888 | 0.04633451 | 0.96304363 | 0.98892249 |
| YARS2 | 16.3499385 | -0.126888085 | 0.38428721 | -0.3301908 | 0.74125583 | NA |
| PKP2 | 6.642025 | -0.114153029 | 0.59064646 | -0.1932679 | 0.84674913 | NA |
| ALG10B | 10.3261518 | 0.815633396 | 0.47863026 | 1.70409909 | 0.08836258 | NA |
| CPNE8 | 225.449935 | 0.895335719 | 0.24247636 | 3.69246607 | 0.00022209 | 0.01829751 |
| KIF21A | 60.9782862 | 0.678658877 | 0.41751821 | 1.62545934 | 0.10406472 | 0.4862279 |
| C12orf40 | 6.78941304 | -0.890235535 | 0.94752802 | -0.9395348 | 0.34745624 | NA |
| SLC2A13 | 135.889203 | 1.430227694 | 0.36952308 | 3.87046914 | 0.00010863 | 0.01101469 |
| LRRK2 | 34.1646178 | 0.84266747 | 0.38694306 | 2.17775575 | 0.02942423 | 0.27926427 |
| CNTN1 | 86.8144006 | 0.485475589 | 0.49870489 | 0.97347269 | 0.33031843 | 0.70366153 |
| GXYLT1 | 43.038566 | 0.088098751 | 0.35433854 | 0.24862876 | 0.80364796 | 0.94126368 |
| YAF2 | 56.8950772 | 0.349947117 | 0.30079953 | 1.16338985 | 0.24467137 | 0.64996527 |
| PPHLN1 | 99.0274962 | 0.296505594 | 0.26353937 | 1.12509031 | 0.26055077 | 0.65595635 |
| ZCRB1 | 19.6889461 | -0.092823464 | 0.50680396 | -0.1831546 | 0.85467673 | NA |

| | | | | | | | |
|---|---|---|---|---|---|---|---|
| PRICKLE1 | 56.4116767 | | -0.215107703 | 0.31514757 | -0.6825618 | 0.49488377 | 0.8233177 |
| PUS7L | 13.8805662 | | -0.059431974 | 0.42859851 | -0.1386658 | 0.88971421 | NA |
| IRAK4 | 16.6511123 | | -0.488945817 | 0.37330839 | -1.3097638 | 0.19027575 | NA |
| TWF1 | 10.6363457 | | -0.361673854 | 0.45484028 | -0.7951667 | 0.42651653 | NA |
| TMEM117 | 63.189617 | | 0.377007227 | 0.36759479 | 1.02560548 | 0.30507758 | 0.6848669 |
| ANO6 | 49.6755007 | | 0.322971781 | 0.31541995 | 1.02394215 | 0.3058626 | 0.6854026 |
| AC008124.1 | 10.7386685 | | -0.243229356 | 0.52808209 | -0.46059 | 0.64509275 | NA |
| ARID2 | 115.446031 | | 0.340713485 | 0.22905476 | 1.48747609 | 0.13688911 | 0.5308052 |
| SCAF11 | 120.929707 | | -0.021910611 | 0.22675865 | -0.0966252 | 0.92302401 | 0.97775374 |
| SLC38A1 | 11.7924837 | | -1.095680378 | 0.71919637 | -1.5234787 | 0.12763897 | NA |
| SLC38A2 | 110.362695 | | -0.070215217 | 0.34167461 | -0.2055032 | 0.83717902 | 0.95274919 |
| AC008014.1 | 160.476956 | | -0.31984392 | 0.52593005 | -0.6081492 | 0.54308855 | 0.84207245 |
| SLC38A4 | 5.8492193 | | 0.023910688 | 0.94239099 | 0.02537236 | 0.97975795 | NA |
| AMIGO2 | 71.0492542 | | -0.574977434 | 0.58084117 | -0.9899048 | 0.32222068 | 0.69513974 |
| PCED1B | 216.726221 | | -0.027774062 | 0.34189447 | -0.0812358 | 0.93525445 | 0.97832857 |
| RPAP3 | 13.2820361 | | 0.261388589 | 0.45821076 | 0.57045494 | 0.56836918 | NA |
| RAPGEF3 | 85.612267 | | 0.116148004 | 0.49955648 | 0.23250225 | 0.81614794 | 0.94632611 |
| SLC48A1 | 87.0595011 | | 0.246712196 | 0.35359391 | 0.6977275 | 0.48534763 | 0.81450403 |
| HDAC7 | 28.400159 | | -0.460119276 | 0.49079923 | -0.9374898 | 0.34850666 | NA |
| TMEM106C | 15.4803262 | | -0.218968256 | 0.55404751 | -0.3952157 | 0.69268373 | NA |
| SENP1 | 16.1050083 | | -0.067412748 | 0.36330195 | -0.1855557 | 0.85279319 | NA |
| PFKM | 7.59614243 | | 0.100220145 | 0.52233792 | 0.19186841 | 0.84784528 | NA |
| ASB8 | 12.1431792 | | -0.132152262 | 0.45773321 | -0.2887102 | 0.77280313 | NA |
| ZNF641 | 7.41866873 | | -0.215942383 | 0.6500863 | -0.3321749 | 0.73975716 | NA |
| AC090115.1 | 6.44896543 | | -0.352857813 | 0.57051609 | -0.6184888 | 0.53625317 | NA |
| KANSL2 | 11.8955426 | | 0.338452153 | 0.40493497 | 0.83581854 | 0.403257 | NA |
| CCNT1 | 16.4096157 | | -0.435733795 | 0.36114916 | -1.2065203 | 0.22761693 | NA |
| ADCY6 | 11.3361501 | | -0.081182934 | 0.44695374 | -0.1816361 | 0.85586832 | NA |
| DDX23 | 14.6463974 | | -0.125766384 | 0.4425987 | -0.2841544 | 0.77629203 | NA |
| CCDC65 | 12.863731 | | -0.764845879 | 0.59205642 | -1.2918463 | 0.19641038 | NA |
| DDN | 132.693251 | | 0.607263638 | 1.1720627 | 0.51811532 | 0.60437781 | 0.87047761 |
| DDN-AS1 | 9.797614 | | 0.015296154 | 0.81994897 | 0.01865501 | 0.98511632 | NA |
| PRKAG1 | 24.5456348 | | -0.64206382 | 0.33246457 | -1.9312248 | 0.05345526 | NA |
| KMT2D | 20.8416781 | | 0.041953279 | 0.37029627 | 0.11329652 | 0.90979548 | NA |
| LMBR1L | 16.0070879 | | -0.678342262 | 0.43113677 | -1.5733807 | 0.1156307 | NA |
| TUBA1B | 81.1956413 | | -0.522123525 | 0.64222621 | -0.8129901 | 0.41622376 | 0.7708692 |
| TUBA1A | 30.1966697 | | -0.391156654 | 0.81185562 | -0.4818057 | 0.62994399 | 0.88166074 |
| TUBA1C | 13.7776991 | | -0.530969295 | 0.43825437 | -1.211555 | 0.22568275 | NA |
| SPATS2 | 23.4591663 | | -0.161117565 | 0.44250056 | -0.364107 | 0.71577808 | NA |
| KCNH3 | 9.57694379 | | -0.079527222 | 0.69326897 | -0.1147134 | 0.90867231 | NA |
| MCRS1 | 12.9755663 | | 0.408401561 | 0.38804638 | 1.05245555 | 0.2925906 | NA |
| PRPF40B | 22.5453892 | | 0.243899569 | 0.31349088 | 0.77801169 | 0.43656212 | NA |
| FMNL3 | 9.05501025 | | -0.071667526 | 0.7417418 | -0.0966206 | 0.92302771 | NA |
| TMBIM6 | 165.275694 | | -0.051654028 | 0.31683374 | -0.163032 | 0.87049327 | 0.96207879 |
| NCKAP5L | 11.0949239 | | -0.43131247 | 0.47019665 | -0.9173023 | 0.35898425 | NA |
| AQP2 | 37.7863599 | | 0.487643608 | 0.62949996 | 0.77465232 | 0.43854513 | 0.78335165 |
| SMARCD1 | 18.2507761 | | 0.025701866 | 0.36646657 | 0.07013427 | 0.94408679 | NA |
| GPD1 | 4.92221772 | | 0.704345941 | 0.6676272 | 1.05499886 | 0.29142584 | NA |
| AC074032.1 | 38.9290063 | | -0.527851753 | 0.32762856 | -1.6111286 | 0.10715169 | 0.49086361 |
| CERS5 | 31.6507821 | | 0.399597802 | 0.30616914 | 1.30515374 | 0.1918405 | 0.5949439 |
| LIMA1 | 35.9141993 | | 0.794782475 | 0.37825605 | 2.10117582 | 0.03562554 | 0.29549524 |
| LARP4 | 54.5197141 | | 0.324923944 | 0.27377263 | 1.18683866 | 0.23529127 | 0.64178976 |
| DIP2B | 36.7059971 | | 0.306118464 | 0.44100317 | 0.69414119 | 0.48759367 | 0.81648973 |
| ATF1 | 20.5275058 | | -0.174412638 | 0.36978101 | -0.4716647 | 0.63716616 | NA |
| METTL7A | 8.8996854 | | 1.282199654 | 0.82095294 | 1.56184307 | 0.11832496 | NA |
| SLC11A2 | 30.6663348 | | -0.500760803 | 0.29734781 | -1.6840911 | 0.09216406 | 0.46264534 |
| LETMD1 | 18.3053805 | | -0.406303542 | 0.42105781 | -0.964959 | 0.33456533 | NA |
| CSRNP2 | 15.645057 | | 0.556702177 | 0.37935529 | 1.4674955 | 0.14224131 | NA |
| TFCP2 | 59.2910399 | | 0.552743451 | 0.25928333 | 2.13181251 | 0.03302226 | 0.29047359 |
| POU6F1 | 9.51866222 | | -0.134856613 | 0.51087255 | -0.2639731 | 0.79180064 | NA |
| DAZAP2 | 69.5147414 | | -0.131592997 | 0.41258441 | -0.3189481 | 0.7497659 | 0.92274989 |
| ACVR1B | 25.4435443 | | 0.072393801 | 0.33825236 | 0.21402305 | 0.83052908 | NA |
| ATG101 | 9.38224397 | | -1.076175511 | 0.61098712 | -1.7613718 | 0.07817549 | NA |
| KRT7 | 6.17887053 | | -1.785398505 | 0.88017962 | -2.0284479 | 0.04251456 | NA |
| KRT8 | 14.1610271 | | -1.394777557 | 0.62702326 | -2.2244431 | 0.02611865 | NA |
| KRT18 | 18.2853026 | | -2.298618038 | 0.68138586 | -3.3734454 | 0.00074234 | NA |
| EIF4B | 42.078912 | | -0.269955051 | 0.3092079 | -0.8730535 | 0.3826339 | 0.74924885 |
| AC068888.1 | 24.4416841 | | -0.353897056 | 0.46821184 | -0.7558482 | 0.44974024 | NA |
| TNS2 | 79.8424073 | | -0.315624883 | 0.44213686 | -0.7138624 | 0.47531227 | 0.8073143 |
| SPRYD3 | 8.86117334 | | 0.115952333 | 0.50388374 | 0.23011723 | 0.81800067 | NA |
| CSAD | 21.4922199 | | 0.458973445 | 0.37768614 | 1.2152245 | 0.22428048 | NA |
| ZNF740 | 6.04096781 | | 0.371560764 | 0.55463947 | 0.66991403 | 0.50291259 | NA |
| RARG | 5.80634795 | | 0.39395404 | 0.74693992 | 0.527424 | 0.59789918 | NA |
| MFSD5 | 9.37877307 | | -0.04513493 | 0.49934144 | -0.0903889 | 0.92797817 | NA |
| PFDN5 | 50.5702226 | | -0.212832333 | 0.47210789 | -0.4508129 | 0.65212439 | 0.89449325 |
| C12orf10 | 10.447098 | | -0.042437053 | 0.47387144 | -0.0895539 | 0.92864169 | NA |
| AAAS | 5.07800523 | | -0.335951797 | 0.59895905 | -0.5608928 | 0.57487065 | NA |
| SP1 | 43.0072859 | | -0.119229018 | 0.28499975 | -0.4183478 | 0.67569285 | 0.90499319 |
| PRR13 | 32.4385178 | | -0.53697713 | 0.45402091 | -1.1827145 | 0.2369223 | 0.64321384 |
| PCBP2 | 80.6715886 | | -0.025614056 | 0.22509922 | -0.1137901 | 0.90940421 | 0.97528913 |
| MAP3K12 | 10.930159 | | -0.604791908 | 0.4880195 | -1.2392782 | 0.2152425 | NA |
| ATP5MC2 | 75.3065749 | | 0.040757202 | 0.3153842 | 0.12923032 | 0.8971754 | 0.97097328 |
| CALCOCO1 | 37.8574352 | | 0.319667026 | 0.34416208 | 0.92882698 | 0.35297876 | 0.72416887 |
| HOXC4 | 8.54302121 | | 1.377388059 | 0.56563367 | 2.43512389 | 0.01488669 | NA |
| SMUG1 | 10.359507 | | 0.403386656 | 0.47306119 | 0.85271561 | 0.39381703 | NA |
| LINC02381 | 4.39147542 | | 0.557703404 | 0.71759014 | 0.77718934 | 0.43704707 | NA |
| CBX5 | 41.5422275 | | -0.577006059 | 0.34492222 | -1.6728585 | 0.09435516 | 0.46752784 |
| HNRNPA1 | 31.2420759 | | -0.395713082 | 0.3635554 | -1.0884533 | 0.27639503 | 0.66941818 |
| COPZ1 | 29.2529944 | | -0.17683837 | 0.338729 | -0.5220645 | 0.60162545 | 0.86994397 |
| ZNF385A | 4.76974457 | | -0.399004783 | 0.9843577 | -0.4053453 | 0.6852237 | NA |
| PPP1R1A | 5.34649761 | | -0.29368777 | 0.97787989 | -0.3003311 | 0.76392459 | NA |
| CD63 | 126.129055 | | -0.57044665 | 0.46437276 | -1.228424 | 0.21928784 | 0.62462904 |
| ORMDL2 | 9.81288118 | | -1.503328275 | 0.64060649 | -2.3467266 | 0.01893915 | NA |
| PYM1 | 6.30698422 | | -0.601082416 | 0.56165796 | -1.070193 | 0.28453245 | NA |
| CDK2 | 6.06926928 | | -0.915098841 | 0.80614107 | -1.1351597 | 0.25630842 | NA |
| RAB5B | 32.5270951 | | 0.163918108 | 0.3765844 | 0.43527589 | 0.66336219 | 0.89939456 |
| SUOX | 5.9388183 | | -0.140511674 | 0.65515904 | -0.2144696 | 0.8301809 | NA |
| IKZF4 | 11.0378489 | | -0.344884712 | 0.60108869 | -0.5737668 | 0.56612564 | NA |
| RPS26 | 14.5680772 | | -0.172640374 | 0.44726064 | -0.385995 | 0.69950037 | NA |
| ERBB3 | 8.66552781 | | 0.629879539 | 0.63480849 | 0.99223553 | 0.32108264 | NA |
| PA2G4 | 13.1382913 | | -0.879523467 | 0.45285201 | -1.9421874 | 0.05211442 | NA |
| RPL41 | 177.416993 | | -0.501994647 | 0.4245318 | -1.1824665 | 0.23702064 | 0.64321384 |
| ESYT1 | 34.8917459 | | -0.029723603 | 0.31529032 | -0.0942738 | 0.9248917 | 0.97775374 |
| MYL6 | 200.099441 | | 0.052367755 | 0.52894993 | 0.09900324 | 0.9211357 | 0.97775374 |
| SMARCC2 | 35.6874295 | | -0.087207584 | 0.34124568 | -0.2555566 | 0.79829324 | 0.94015741 |
| RNF41 | 14.1166233 | | -0.260800447 | 0.41547392 | -0.627718 | 0.53018871 | NA |
| NABP2 | 6.58975623 | | -1.286719395 | 0.65928364 | -1.9516932 | 0.05097464 | NA |
| ANKRD52 | 10.3655658 | | 0.27263135 | 0.5024826 | 0.54256874 | 0.58742677 | NA |
| CS | 14.7730118 | | -0.107636267 | 0.39041304 | -0.2756984 | 0.78277969 | NA |
| PAN2 | 13.4280234 | | 0.14120014 | 0.41573727 | 0.33963792 | 0.73412922 | NA |
| STAT2 | 26.4518766 | | 0.434964675 | 0.62416514 | 0.69687435 | 0.48588143 | NA |
| SPRYD4 | 5.4268739 | | -0.240913611 | 0.8685772 | -0.2773658 | 0.78149924 | NA |
| RBMS2 | 62.7889542 | | 0.311587577 | 0.35029537 | 0.88949956 | 0.37373466 | 0.74177796 |

| | | | | | | |
|---|---|---|---|---|---|---|
| BAZ2A | 44.5260002 | 0.038579763 | 0.33384479 | 0.11556198 | 0.9079997 | 0.9747971 |
| ATP5F1B | 58.0732206 | -0.244618194 | 0.40945895 | -0.5974181 | 0.55022827 | 0.84663348 |
| PTGES3 | 78.2447127 | 0.0890289 | 0.24261599 | 0.36695396 | 0.71365336 | 0.91328825 |
| NACA | 47.4245596 | -1.243022772 | 0.44192926 | -2.8127189 | 0.00491246 | 0.11177304 |
| NEMP1 | 19.1157386 | 0.040006929 | 0.44901275 | 0.08909976 | 0.92900263 | NA |
| NAB2 | 12.0499216 | 0.290881209 | 0.60542236 | 0.48045997 | 0.63090036 | NA |
| STAT6 | 31.3709566 | -0.444449003 | 0.41988516 | -1.0585013 | 0.28982694 | 0.67746193 |
| LRP1 | 15.3919123 | -0.838825617 | 0.48282333 | -1.7373345 | 0.08232815 | NA |
| SHMT2 | 5.36385572 | 0.153352339 | 0.74583903 | 0.2056105 | 0.83709517 | NA |
| R3HDM2 | 77.420085 | 0.313623216 | 0.35116903 | 0.89308336 | 0.37181254 | 0.74014106 |
| MARS | 15.1903151 | -0.43750913 | 0.60503512 | -0.7231136 | 0.46961009 | NA |
| DDIT3 | 24.9254614 | -0.981814798 | 1.11587238 | -0.879863 | 0.37893354 | NA |
| MBD6 | 12.2776472 | 0.561561032 | 0.37433648 | 1.50015042 | 0.13357544 | NA |
| DCTN2 | 26.2660772 | -0.444929193 | 0.38163466 | -1.1658511 | 0.24367465 | NA |
| PIP4K2C | 12.7889058 | -0.44416914 | 0.62614995 | -0.7093655 | 0.47809772 | NA |
| DTX3 | 10.8730566 | 0.50354725 | 0.5274734 | 0.95464009 | 0.33975974 | NA |
| OS9 | 58.2469814 | -0.101934945 | 0.32848036 | -0.3103228 | 0.75631549 | 0.92313949 |
| TSPAN31 | 13.3710137 | -0.490180261 | 0.58666516 | -0.8355367 | 0.4034156 | NA |
| CDK4 | 10.7950068 | -0.517958669 | 0.49104078 | -1.054818 | 0.29150855 | NA |
| 9-Mar | 7.42195377 | -0.020000726 | 0.52676598 | -0.0379689 | 0.96971248 | NA |
| TSFM | 7.87138882 | -0.278240457 | 0.56293305 | -0.4942692 | 0.62111609 | NA |
| CTDSP2 | 32.276242 | -0.186814273 | 0.28575818 | -0.6537495 | 0.5132732 | 0.83070544 |
| AC084033.3 | 5.07600636 | -0.755194663 | 0.63576667 | -1.1878488 | 0.23489299 | NA |
| LRIG3 | 8.81734509 | -0.492033524 | 0.50389451 | -0.9764614 | 0.32883588 | NA |
| SLC16A7 | 22.0108941 | -0.390637723 | 0.49068562 | -0.7961059 | 0.42597048 | NA |
| FAM19A2 | 22.0866225 | 1.76433356 | 0.65733071 | 2.68408813 | 0.00727279 | NA |
| USP15 | 74.8796164 | 0.353777635 | 0.31471327 | 1.12412686 | 0.26095921 | 0.65595635 |
| MON2 | 115.064627 | 0.129887135 | 0.21929816 | 0.59228556 | 0.55365938 | 0.84933527 |
| AC048341.1 | 14.8149004 | 0.060270304 | 0.56314424 | 0.10702463 | 0.91476944 | NA |
| PPM1H | 54.186168 | 0.328543269 | 0.3416891 | 0.96152693 | 0.33628729 | 0.71116888 |
| DPY19L2 | 13.4854276 | 1.847223265 | 0.89914508 | 2.05442182 | 0.03993488 | NA |
| RXYLT1 | 13.8239133 | -0.896453463 | 0.47145155 | -1.9014753 | 0.05723979 | NA |
| SRGAP1 | 531.536417 | 0.514651572 | 0.21830997 | 2.35743504 | 0.01840167 | 0.22674781 |
| AC020611.2 | 21.7503917 | -0.38636746 | 0.63701043 | -0.6065324 | 0.54416128 | NA |
| C12orf56 | 16.5852597 | 0.250123956 | 0.43962815 | 0.56894436 | 0.5693939 | NA |
| XPOT | 36.150954 | 0.049518685 | 0.39038162 | 0.12684687 | 0.8990616 | 0.97097328 |
| TBK1 | 39.9385004 | 0.320123948 | 0.26088489 | 1.22706969 | 0.2197964 | 0.62462904 |
| RASSF3 | 6.80535412 | 1.233393447 | 0.7319568 | 1.68506318 | 0.09197638 | NA |
| GNS | 51.3858469 | -0.002516663 | 0.29891506 | -0.0084193 | 0.99328243 | 0.99833815 |
| LEMD3 | 36.3919969 | 0.070546135 | 0.33106003 | 0.21309167 | 0.83125546 | 0.95050875 |
| MSRB3 | 151.942665 | 0.179217139 | 0.34116138 | 0.52531485 | 0.59936433 | 0.86882737 |
| LLPH | 12.5353004 | 0.257608665 | 0.46637354 | 0.55236552 | 0.58069795 | NA |
| IRAK3 | 36.8584619 | -1.058852121 | 0.54304881 | -1.9498286 | 0.05119655 | 0.35507065 |
| HELB | 15.1528531 | -0.841445057 | 0.61231629 | -1.3742 | 0.16937959 | NA |
| GRIP1 | 121.959478 | 1.221779375 | 0.2813226 | 4.34298345 | 1.41E-05 | 0.00237548 |
| CAND1 | 96.0086663 | 0.445589889 | 0.2486683 | 1.79190468 | 0.07314824 | 0.41834543 |
| MDM1 | 18.0199165 | 1.027738855 | 0.36657695 | 2.80361015 | 0.0050534 | NA |
| RAP1B | 97.0161536 | -0.399899247 | 0.21253335 | -1.8815835 | 0.05989259 | 0.37905854 |
| NUP107 | 18.5828008 | -0.588686876 | 0.48194975 | -1.2214694 | 0.22190835 | NA |
| SLC35E3 | 16.2469684 | -0.451410081 | 0.48538622 | -0.9300019 | 0.35237013 | NA |
| MDM2 | 35.687227 | 0.182540333 | 0.50001717 | 0.36506813 | 0.71506055 | 0.91376358 |
| CPM | 16.7322625 | -0.113243964 | 0.77902288 | -0.1453667 | 0.88442138 | NA |
| CPSF6 | 76.89309 | 0.434982326 | 0.25311397 | 1.71852358 | 0.08570116 | 0.44564602 |
| YEATS4 | 6.65291834 | -0.13077444 | 0.60505952 | -0.2161348 | 0.82888265 | NA |
| FRS2 | 78.5895843 | 0.24133832 | 0.21821059 | 1.10598814 | 0.26873165 | 0.66139295 |
| CCT2 | 18.7090451 | -0.458373421 | 0.41707003 | -1.0990323 | 0.271754 | NA |
| RAB3IP | 12.8546106 | 0.6238673 | 0.51611123 | 1.20878459 | 0.22674561 | NA |
| AC025159.1 | 37.8070119 | -0.390815039 | 0.52032523 | -0.7510976 | 0.45259391 | 0.79291818 |
| CNOT2 | 75.609938 | 0.271004255 | 0.20601723 | 1.31544462 | 0.18836052 | 0.59077076 |
| TSPAN8 | 5.95225338 | -0.281580389 | 0.58541698 | -0.4809912 | 0.63052279 | NA |
| ZFC3H1 | 69.3654879 | 0.361405405 | 0.31452428 | 1.14905407 | 0.25053368 | 0.6518054 |
| TMEM19 | 16.957774 | 0.724188463 | 0.48681268 | 1.48761216 | 0.1368532 | NA |
| RAB21 | 64.534825 | -0.05139357 | 0.22116562 | -0.2323759 | 0.81624603 | 0.94632611 |
| TBC1D15 | 46.411227 | 0.019044289 | 0.29481805 | 0.06459675 | 0.94849507 | 0.98391202 |
| TRHDE | 17.5787643 | 0.372347424 | 0.737938 | 0.5045782 | 0.61385513 | NA |
| ATXN7L3B | 35.0067155 | -0.689860223 | 0.31185627 | -2.2121095 | 0.0269591 | 0.26604892 |
| CAPS2 | 42.9925772 | 0.984204079 | 0.4255916 | 2.31255524 | 0.0207471 | 0.23840101 |
| GLIPR1L2 | 8.75005105 | 0.770233559 | 0.89696683 | 0.85870908 | 0.39050104 | NA |
| KRR1 | 18.3022359 | 0.031953463 | 0.4075611 | 0.07840165 | 0.93750856 | NA |
| PHLDA1 | 31.6799897 | -1.871430829 | 1.00693028 | -1.8585506 | 0.06309087 | 0.3863717 |
| NAP1L1 | 75.5614591 | 0.001678153 | 0.30978239 | 0.0054172 | 0.99567772 | 0.99921845 |
| BBS10 | 3.77891713 | -0.422674775 | 0.72751491 | -0.5809843 | 0.56125101 | NA |
| OSBPL8 | 66.1639823 | -0.003861631 | 0.22575593 | -0.0171053 | 0.98635258 | 0.99585746 |
| ZDHHC17 | 52.1635661 | 0.818727905 | 0.26147689 | 3.13116732 | 0.00174113 | 0.06518907 |
| NAV3 | 9.59377122 | 0.482355401 | 0.627516 | 0.76867426 | 0.44208671 | NA |
| SYT1 | 32.8014441 | -0.536937076 | 0.97321766 | -0.5517132 | 0.58114483 | 0.86175324 |
| AC027288.3 | 74.678327 | -1.140597671 | 0.57524458 | -1.9828047 | 0.04738925 | 0.34008434 |
| PAWR | 382.490697 | -0.39582295 | 0.37107624 | -1.066689 | 0.28611229 | 0.67390908 |
| AC073569.2 | 26.9712508 | -0.369006466 | 0.69303036 | -0.5324535 | 0.59441191 | NA |
| PPP1R12A | 107.141704 | 0.064763784 | 0.23483062 | 0.27578935 | 0.78270987 | 0.93415197 |
| PTPRQ | 2479.13882 | 0.883659672 | 0.40661548 | 2.17320715 | 0.02976473 | 0.27926427 |
| AC074031.1 | 20.8715147 | 0.598770112 | 0.52460337 | 1.14137679 | 0.25371316 | NA |
| ACSS3 | 14.8747414 | 1.456516301 | 0.77298891 | 1.88426546 | 0.05952908 | NA |
| CCDC59 | 21.0383638 | 0.406694859 | 0.49303334 | 0.82488307 | 0.40943798 | NA |
| METTL25 | 24.6166735 | 0.435759713 | 0.33461011 | 1.30229094 | 0.19281695 | NA |
| TMTC2 | 26.3826848 | -0.175263405 | 0.60755681 | -0.2884725 | 0.77298511 | NA |
| LRRIQ1 | 13.0998651 | 0.432613627 | 0.49643491 | 0.87144078 | 0.38351353 | NA |
| C12orf29 | 9.87157551 | -0.88816711 | 0.49587671 | -1.7911047 | 0.07327649 | NA |
| CEP290 | 55.4994286 | 0.027699946 | 0.2597546 | 0.1066389 | 0.91507545 | 0.97744467 |
| TMTC3 | 28.9295951 | 0.518342544 | 0.28029822 | 1.84925375 | 0.06442118 | 0.39233079 |
| KITLG | 11.6210952 | -0.483847724 | 0.91266834 | -0.5301463 | 0.59601051 | NA |
| DUSP6 | 31.8772404 | -0.090312867 | 0.88729125 | -0.1017849 | 0.9189274 | 0.97775374 |
| POC1B | 20.3409157 | -0.154114811 | 0.36005965 | -0.4280258 | 0.66863235 | NA |
| POC1B-AS1 | 10.5039204 | 0.099105517 | 0.47362836 | 0.20924743 | 0.83425509 | NA |
| ATP2B1 | 57.4266485 | 0.568135753 | 0.35216281 | 1.61327585 | 0.10668459 | 0.49086361 |
| DCN | 450.288791 | -0.336361529 | 0.52595022 | -0.6395311 | 0.52247748 | 0.83120735 |
| BTG1 | 21.3110149 | -1.115141633 | 0.41565611 | -2.6828467 | 0.00729984 | NA |
| EEA1 | 129.151385 | 0.381695398 | 0.33245658 | 1.14810602 | 0.2509248 | 0.65183754 |
| AC138123.1 | 12.1463401 | 0.753529813 | 0.69818228 | 1.07927376 | 0.28046571 | NA |
| NUDT4 | 16.4200025 | 0.585961808 | 0.43755194 | 1.3391823 | 0.18051133 | NA |
| UBE2N | 36.6696426 | 0.333965471 | 0.33738769 | 0.98985671 | 0.32224416 | 0.69513974 |
| MRPL42 | 28.7593608 | -0.276964355 | 0.30587057 | -0.9054953 | 0.36520307 | 0.73398222 |
| SOCS2 | 13.2061748 | -0.948610475 | 0.68766133 | -1.3794733 | 0.16774886 | NA |
| CRADD | 113.897766 | 0.005475502 | 0.29313658 | 0.01867901 | 0.98509717 | 0.99585746 |
| AC012085.2 | 7.84613638 | -0.960171967 | 0.75925229 | -1.2646284 | 0.20600459 | NA |
| PLXNC1 | 15.4802926 | 0.431606997 | 0.40706458 | 1.0602912 | 0.28901214 | NA |
| CEP83 | 43.9847818 | 0.500198911 | 0.26140217 | 1.91352238 | 0.05568121 | 0.36728373 |
| TMCC3 | 150.975271 | 0.665103271 | 0.36570914 | 1.81866732 | 0.0689622 | 0.40491695 |
| NDUFA12 | 22.6551415 | -0.3487912 | 0.44773934 | -0.779005 | 0.43597675 | NA |
| NR2C1 | 27.8047245 | -0.21665209 | 0.35789982 | -0.6053428 | 0.54495123 | NA |
| FGD6 | 19.9031495 | -1.08906073 | 0.47169113 | -2.3088429 | 0.0209523 | NA |
| VEZT | 110.87252 | 0.069045437 | 0.2140561 | 0.32255766 | 0.74703026 | 0.92250642 |
| METAP2 | 44.3096622 | -0.959226518 | 0.26222453 | -3.6580351 | 0.00025416 | 0.01982419 |

| | | | | | | |
|---|---|---|---|---|---|---|
| NTN4 | 33.8203612 | -0.252506688 | 0.52856847 | -0.477718 | 0.63285091 | 0.88389921 |
| SNRPF | 11.9194524 | -0.611505962 | 0.46100441 | -1.3264645 | 0.1846859 | NA |
| LTA4H | 12.1913552 | -0.139058191 | 0.48271576 | -0.2880747 | 0.77328957 | NA |
| ELK3 | 5.18470608 | -0.923372106 | 1.03137036 | -0.8952866 | 0.37063388 | NA |
| CDK17 | 61.1508156 | 0.748102962 | 0.23308553 | 3.20956421 | 0.00132936 | 0.0545866 |
| CFAP54 | 10.2347701 | 0.700919681 | 0.56281769 | 1.24537607 | 0.21299354 | NA |
| NEDD1 | 18.7754383 | 0.01771029 | 0.3601289 | 0.04917764 | 0.96077773 | NA |
| TMPO | 20.0072524 | -0.372153713 | 0.36069005 | -1.0317826 | 0.30217398 | NA |
| SLC25A3 | 70.9313906 | -0.573781724 | 0.56869541 | -1.0089438 | 0.31300158 | 0.69213367 |
| IKBIP | 12.1916903 | -0.438963176 | 0.54032755 | -0.812402 | 0.416561 | NA |
| APAF1 | 17.5101005 | 0.201393893 | 0.40208314 | 0.50087624 | 0.61645823 | NA |
| ANKS1B | 117.997965 | 0.992299447 | 0.62145343 | 1.59673984 | 0.11032371 | 0.49667521 |
| UHRF1BP1L | 32.115998 | -0.338030914 | 0.36788977 | -0.9188375 | 0.35818058 | 0.72914991 |
| ACTR6 | 10.2731574 | -0.41024955 | 0.50614929 | -0.8105307 | 0.41763522 | NA |
| SCYL2 | 38.0851844 | 0.230696007 | 0.27954112 | 0.82526681 | 0.40922013 | 0.76772515 |
| NR1H4 | 17.2176247 | 1.048598316 | 0.67004295 | 1.5649718 | 0.11758952 | NA |
| UTP20 | 19.8713737 | -0.303524348 | 0.39236972 | -0.7735672 | 0.43918677 | NA |
| ARL1 | 36.0499763 | -0.589204861 | 0.38473602 | -1.5314523 | 0.12565765 | 0.52273582 |
| CHPT1 | 77.5374171 | -0.179585449 | 0.38025429 | -0.4722772 | 0.63672891 | 0.88633797 |
| SYCP3 | 4.16736036 | -1.008672958 | 0.83488013 | -1.208165 | 0.2269838 | NA |
| GNPTAB | 70.9913965 | 0.158581128 | 0.38800359 | 0.40871047 | 0.68275215 | 0.90499319 |
| DRAM1 | 20.833539 | -1.363640499 | 0.45712574 | -2.9830753 | 0.00285368 | NA |
| WASHC3 | 47.6810135 | -0.87668273 | 0.37359468 | -2.3466146 | 0.01894484 | 0.23006068 |
| NUP37 | 9.5587138 | -0.69909784 | 0.49491959 | -1.4125483 | 0.15778857 | NA |
| LINC02456 | 7.86766442 | -0.136585288 | 1.19960336 | -0.1138587 | 0.9093498 | NA |
| IGF1 | 49.7673517 | 0.269394522 | 1.26948451 | 0.2122078 | 0.83194492 | 0.95050875 |
| HSP90B1 | 238.375487 | -0.093695323 | 0.54066485 | -0.1732965 | 0.86241838 | 0.95966226 |
| C12orf73 | 8.54383773 | -0.275695809 | 0.49699855 | -0.5547216 | 0.57908513 | NA |
| TDG | 15.3728135 | -0.203663525 | 0.37433433 | -0.5440685 | 0.58639432 | NA |
| HCFC2 | 19.4926768 | -0.041209434 | 0.38350803 | -0.1074539 | 0.91442889 | NA |
| NFYB | 10.9908604 | 0.415751575 | 0.41125618 | 1.01093088 | 0.31204951 | NA |
| TXNRD1 | 201.380756 | -0.851898117 | 0.31047711 | -2.7438355 | 0.0060726 | 0.12229157 |
| CHST11 | 7.55647107 | -0.430978767 | 0.54399872 | -0.7922422 | 0.42821944 | NA |
| SLC41A2 | 20.8734811 | -0.816302793 | 0.47500134 | -1.7185274 | 0.08570047 | NA |
| C12orf45 | 6.84037287 | -0.199408476 | 0.60853848 | -0.3276842 | 0.74315041 | NA |
| ALDH1L2 | 3.95411351 | -1.489025885 | 1.30081332 | -1.1446884 | 0.25233827 | NA |
| WASHC4 | 62.0926011 | 0.448594323 | 0.33681356 | 1.33187725 | 0.18290053 | 0.58584259 |
| APPL2 | 31.4340693 | 0.002987356 | 0.30837236 | 0.0096875 | 0.99227062 | 0.99792949 |
| C12orf75 | 9.77955327 | -1.051247192 | 0.64856188 | -1.6208896 | 0.10504132 | NA |
| NUAK1 | 7.08997769 | 1.531194474 | 0.60979048 | 2.51101735 | 0.01203838 | NA |
| CKAP4 | 39.5471043 | -0.277157762 | 0.50552177 | -0.5482608 | 0.58351285 | 0.86175324 |
| TCP11L2 | 9.24840466 | 1.269903712 | 0.55881625 | 2.27248887 | 0.023057 | NA |
| POLR3B | 9.92151436 | 0.210689188 | 0.46921853 | 0.44902146 | 0.65341618 | NA |
| RIC8B | 38.464652 | 0.030448647 | 0.37180138 | 0.08189493 | 0.93473027 | 0.97832857 |
| TMEM263 | 24.9103917 | 0.043682604 | 0.36049454 | 0.12117411 | 0.90355313 | NA |
| MTERF2 | 7.76595458 | 0.352425472 | 0.48344604 | 0.72898616 | 0.46601013 | NA |
| CRY1 | 19.7177128 | 0.48989711 | 0.43989499 | 1.11366831 | 0.26542151 | NA |
| PWP1 | 13.8406766 | -0.358479029 | 0.38111472 | -0.9406066 | 0.34690649 | NA |
| PRDM4 | 20.6821656 | -0.598275252 | 0.48862035 | -1.2244174 | 0.22079481 | NA |
| FICD | 9.8386953 | 0.156805976 | 0.66514225 | 0.23574803 | 0.8136282 | NA |
| SART3 | 14.2307045 | 0.0016487 | 0.42001293 | 0.00392536 | 0.99686803 | NA |
| ISCU | 38.9732514 | -0.584995088 | 0.52377225 | -1.1168883 | 0.2640421 | 0.65974932 |
| CORO1C | 18.7665176 | 0.015924674 | 0.40767482 | 0.0390622 | 0.9688408 | NA |
| SSH1 | 23.9303046 | 0.162142057 | 0.32741636 | 0.49521673 | 0.62044714 | NA |
| USP30 | 8.76423079 | 0.013304953 | 0.46110008 | 0.0288548 | 0.97698039 | NA |
| UNG | 20.2115002 | -0.169505189 | 0.48489888 | -0.3495681 | 0.72666284 | NA |
| ACACB | 49.2881382 | 0.170590908 | 0.48652201 | 0.35063349 | 0.72586333 | 0.91659454 |
| KCTD10 | 13.1812122 | -0.68358809 | 0.45822776 | -1.4918086 | 0.13574934 | NA |
| UBE3B | 16.8834645 | -0.238253431 | 0.3802806 | -0.6265201 | 0.53097387 | NA |
| MMAB | 12.8835334 | 0.063839051 | 0.43913785 | 0.1453736 | 0.88441591 | NA |
| MVK | 5.4611499 | -0.283716966 | 0.65296922 | -0.4345028 | 0.66392335 | NA |
| TRPV4 | 9.68727901 | 0.862650953 | 0.62004116 | 1.39128014 | 0.16414049 | NA |
| GLTP | 14.3319934 | 0.036199195 | 0.42164839 | 0.08585161 | 0.93158438 | NA |
| TCHP | 15.627864 | -0.269574043 | 0.39670902 | -0.6795259 | 0.49680473 | NA |
| GIT2 | 32.410232 | 0.249495735 | 0.32635333 | 0.76449574 | 0.44457187 | 0.78741638 |
| ANKRD13A | 20.2184357 | 0.146314548 | 0.40937192 | 0.35741227 | 0.72078319 | NA |
| C12orf76 | 12.1429093 | -0.576275038 | 0.43243795 | -1.332619 | 0.18265686 | NA |
| IFT81 | 17.2309094 | 0.149781907 | 0.36653631 | 0.40864139 | 0.68280285 | NA |
| ATP2A2 | 93.1835487 | 0.006149171 | 0.33103946 | 0.01857534 | 0.98517987 | 0.99585746 |
| ANAPC7 | 20.7509795 | -0.168925516 | 0.38268442 | -0.4414225 | 0.65890715 | NA |
| ARPC3 | 76.3267313 | -0.472084617 | 0.33654279 | -1.4027477 | 0.16069209 | 0.56283048 |
| GPN3 | 8.43017333 | -0.716166894 | 0.62629824 | -1.1434918 | 0.25283446 | NA |
| VPS29 | 39.757057 | -0.538270083 | 0.34640753 | -1.5538637 | 0.12021693 | 0.51128003 |
| PPTC7 | 30.888287 | 0.15027066 | 0.31812939 | 0.47235704 | 0.63667196 | 0.88633797 |
| TCTN1 | 18.2584303 | -0.394183716 | 0.38794403 | -1.016084 | 0.3095894 | NA |
| PPP1CC | 24.8136108 | -0.090589191 | 0.30164528 | -0.300317 | 0.7639354 | NA |
| SH2B3 | 9.66610224 | 0.068135862 | 0.55969988 | 0.12173642 | 0.90310777 | NA |
| ATXN2 | 207.179214 | 0.076631115 | 0.22218919 | 0.34489128 | 0.73017611 | 0.91718621 |
| BRAP | 17.9732363 | -0.171167074 | 0.39093219 | -0.4378434 | 0.66149981 | NA |
| ACAD10 | 13.1990042 | 0.451220937 | 0.48366951 | 0.93291169 | 0.35086557 | NA |
| ALDH2 | 26.8109403 | -0.52670822 | 0.39176381 | -1.3444535 | 0.17880179 | NA |
| MAPKAPK5-A | 5.90816334 | -0.274891278 | 0.57557807 | -0.4775916 | 0.63294088 | NA |
| MAPKAPK5 | 17.1110896 | 0.036613944 | 0.35254279 | 0.10385674 | 0.91728304 | NA |
| TMEM116 | 12.58084 | -0.920984054 | 0.63871797 | -1.441926 | 0.14932325 | NA |
| ERP29 | 42.6069071 | -0.705750989 | 0.37410712 | -1.8864944 | 0.05922836 | 0.37831533 |
| NAA25 | 40.3251652 | -0.052165934 | 0.39093053 | -0.1334404 | 0.89384508 | 0.97097328 |
| TRAFD1 | 24.1329168 | -1.434021372 | 0.58502591 | -2.45121 | 0.01423769 | NA |
| HECTD4 | 116.786322 | -0.222466553 | 0.3959397 | -0.5618698 | 0.57420474 | 0.85861279 |
| RPL6 | 65.3967512 | -0.726988629 | 0.44207288 | -1.6444995 | 0.10007307 | 0.48127507 |
| PTPN11 | 38.3436586 | -0.044967846 | 0.37235406 | -0.1207664 | 0.90387609 | 0.97237083 |
| OAS1 | 49.2362114 | -1.227756369 | 1.55007696 | -0.7920616 | 0.42832478 | 0.77844087 |
| AC004551.1 | 5.08839831 | 0.792527634 | 1.20385424 | 0.65832524 | 0.51032916 | NA |
| OAS3 | 12.3106195 | -0.71358789 | 2.43524198 | -0.2930255 | 0.76950269 | NA |
| OAS2 | 12.5324577 | 2.044567039 | 2.54649766 | 0.80289374 | 0.42203615 | NA |
| DDX54 | 13.0318472 | 0.118588424 | 0.40294115 | 0.29430705 | 0.76852328 | NA |
| TPCN1 | 65.9023033 | -0.119497743 | 0.32626906 | -0.3662552 | 0.71417465 | 0.91349491 |
| SLC8B1 | 10.3189166 | -0.317327343 | 0.50277198 | -0.6311556 | 0.5279388 | NA |
| PLBD2 | 13.8656158 | -0.115486349 | 0.41602887 | -0.2775922 | 0.78132546 | NA |
| SDSL | 9.72316145 | -1.170143241 | 0.77334873 | -1.5130861 | 0.13025778 | NA |
| RBM19 | 12.4704006 | -0.789512177 | 0.57253161 | -1.3789844 | 0.16789956 | NA |
| MED13L | 295.870916 | -0.110289155 | 0.30331328 | -0.3636147 | 0.71614578 | 0.91399852 |
| MAP1LC3B2 | 6.03097835 | -2.31144559 | 0.73151753 | -3.1597952 | 0.0015788 | NA |
| C12orf49 | 11.8963739 | -0.573865223 | 0.41889298 | -1.3699566 | 0.17070044 | NA |
| FBXW8 | 27.3933666 | -0.021385087 | 0.31243754 | -0.068446 | 0.94543064 | NA |
| FBXO21 | 153.139651 | -0.406502193 | 0.38742624 | -1.0492377 | 0.29406875 | 0.6804998 |
| WSB2 | 19.7507388 | -0.00083605 | 0.37567681 | -0.0022255 | 0.99822435 | NA |
| VSIG10 | 8.51011519 | -0.253259531 | 0.5895639 | -0.429571 | 0.66750776 | NA |
| PEBP1 | 187.134968 | -1.006948869 | 0.34789215 | -2.8944283 | 0.0037985 | 0.09813187 |
| TAOK3 | 66.6103352 | 0.479889222 | 0.29642696 | 1.61891219 | 0.10546616 | 0.4872104 |
| SUDS3 | 17.5678819 | 0.178490002 | 0.42097654 | 0.42399038 | 0.67157283 | NA |
| HSPB8 | 58.0436789 | -1.31621875 | 0.47178929 | -2.7898445 | 0.00527334 | 0.11257186 |
| PRKAB1 | 10.8653364 | -0.109753629 | 0.50466343 | -0.2174789 | 0.82783518 | NA |
| CIT | 7.30238677 | -1.104474994 | 0.70369373 | -1.5695393 | 0.11652232 | NA |

| | | | | | | |
|---|---|---|---|---|---|---|
| BICDL1 | 32.2070309 | 1.19793307 | 0.51528857 | 2.324781 | 0.02008367 | 0.23722175 |
| RAB35 | 10.7487094 | 0.016756641 | 0.58261381 | 0.02876115 | 0.97705509 | NA |
| GCN1 | 17.4243337 | 0.100310316 | 0.39701039 | 0.25266421 | 0.80052771 | NA |
| RPLP0 | 92.6378619 | -0.926096699 | 0.56327887 | -1.6441176 | 0.10015192 | 0.48127507 |
| PXN-AS1 | 4.81842777 | -0.636699776 | 0.64988953 | -0.9797046 | 0.32723195 | NA |
| PXN | 13.9842682 | -0.436851188 | 0.51907825 | -0.8415902 | 0.40001735 | NA |
| TRIAP1 | 3.76083376 | -0.835891289 | 0.77985887 | -1.0718494 | 0.28378766 | NA |
| GATC | 9.76832889 | -0.646927163 | 0.55460726 | -1.16646 | 0.24342853 | NA |
| SRSF9 | 42.799664 | -0.158738762 | 0.31038634 | -0.5114232 | 0.60905478 | 0.8725395 |
| DYNLL1 | 200.757614 | -0.637565998 | 0.5568887 | -1.1448715 | 0.25226239 | 0.65211999 |
| COQ5 | 6.24826803 | -0.428358609 | 0.63087879 | -0.6789872 | 0.49714599 | NA |
| RNF10 | 45.1861248 | 0.159883604 | 0.27608072 | 0.57911904 | 0.56250885 | 0.85386823 |
| POP5 | 16.8900193 | -0.611343852 | 0.4292247 | -1.4242979 | 0.15436024 | NA |
| MLEC | 60.5474748 | -0.170956854 | 0.38083259 | -0.4489029 | 0.6535017 | 0.8949739 |
| UNC119B | 10.716828 | 0.05488251 | 0.47103512 | 0.11651469 | 0.90724465 | NA |
| ACADS | 9.1039015 | -0.062472324 | 0.49389818 | -0.1264883 | 0.89934544 | NA |
| SPPL3 | 116.72379 | 0.29921012 | 0.19674685 | 1.52078734 | 0.12831321 | 0.52341115 |
| C12orf43 | 5.44114992 | -0.082679207 | 0.7406073 | -0.111637 | 0.9111112 | NA |
| OASL | 8.06792561 | -14.1645001 | 4.22445231 | -3.3529791 | 0.00079947 | NA |
| P2RX4 | 12.0012503 | 0.329600505 | 0.50733443 | 0.64967107 | 0.51590471 | NA |
| CAMKK2 | 18.4730459 | -0.393886055 | 0.44117953 | -0.8928022 | 0.37196311 | NA |
| ANAPC5 | 42.0926369 | 0.106175086 | 0.2697141 | 0.3936579 | 0.69383363 | 0.90557575 |
| RNF34 | 15.2051555 | -0.827984546 | 0.4368445 | -1.8953759 | 0.05804262 | NA |
| KDM2B | 19.2939814 | 0.116372847 | 0.40996966 | 0.28385722 | 0.7765198 | NA |
| ORAI1 | 3.90071374 | -0.913592847 | 0.7570446 | -1.2067887 | 0.22751354 | NA |
| TMEM120B | 13.6276229 | 0.566241409 | 0.50155902 | 1.12896267 | 0.25891358 | NA |
| SETD1B | 10.6673874 | 0.192047945 | 0.43409153 | 0.44241348 | 0.65819003 | NA |
| WDR66 | 25.3386453 | -0.941635536 | 0.63223108 | -1.4893851 | 0.136386 | NA |
| BCL7A | 6.07206024 | -0.62470493 | 0.86589552 | -0.7214553 | 0.47062942 | NA |
| MLXIP | 72.9736763 | -1.185622602 | 0.37885013 | -3.1295294 | 0.00175087 | 0.06518907 |
| CLIP1 | 90.1523717 | -0.173634897 | 0.30714643 | -0.5653163 | 0.57185861 | 0.85778792 |
| ZCCHC8 | 19.8297142 | 0.355482261 | 0.39937152 | 0.89010419 | 0.37340995 | NA |
| RSRC2 | 44.3005346 | -0.475491537 | 0.265868 | -1.7884497 | 0.07370348 | 0.41834543 |
| DENR | 21.9732283 | -0.210116281 | 0.4116617 | -0.5104101 | 0.6097642 | NA |
| HIP1R | 18.212889 | -0.166236042 | 0.60873052 | -0.2730864 | 0.78478678 | NA |
| VPS37B | 16.4893615 | 0.432950367 | 0.41427263 | 1.04508561 | 0.2959834 | NA |
| ARL6IP4 | 53.2246355 | -0.236443063 | 0.31209034 | -0.757611 | 0.44868392 | 0.78987065 |
| PITPNM2 | 10.5049187 | -0.838153032 | 0.524154 | -1.5990587 | 0.10980755 | NA |
| MPHOSPH9 | 20.1335793 | 0.342707121 | 0.36181732 | 0.94718275 | 0.34354567 | NA |
| C12orf65 | 16.6165554 | -0.634976727 | 0.36921063 | -1.7198224 | 0.08546472 | NA |
| SBNO1 | 29.8107612 | -0.092016132 | 0.29148455 | -0.315681 | 0.75224466 | 0.92289907 |
| KMT5A | 14.8535448 | -0.489837542 | 0.48611448 | -1.0076588 | 0.31361828 | NA |
| RILPL2 | 13.9422911 | -0.286220517 | 0.4911918 | -0.5827062 | 0.56009108 | NA |
| SNRNP35 | 23.9768463 | -0.1930878 | 0.31719303 | -0.6087391 | 0.54269738 | NA |
| RILPL1 | 43.5862868 | 0.171223619 | 0.31525763 | 0.54312283 | 0.58704523 | 0.86417728 |
| TMED2 | 30.4500669 | -0.297557211 | 0.47173011 | -0.6307785 | 0.52818537 | 0.83229209 |
| DDX55 | 11.0259528 | 0.250774032 | 0.60076493 | 0.41742455 | 0.6763679 | NA |
| EIF2B1 | 12.8965503 | -0.195234371 | 0.46742165 | -0.4176836 | 0.67617844 | NA |
| GTF2H3 | 8.70563351 | -0.347362154 | 0.4954397 | -0.7011189 | 0.48322879 | NA |
| TCTN2 | 13.4259682 | 0.452045244 | 0.52073401 | 0.86809242 | 0.38534374 | NA |
| ATP6V0A2 | 11.120605 | -0.345419966 | 0.40858438 | -0.8454067 | 0.39788382 | NA |
| CCDC92 | 34.4493201 | 0.066011443 | 0.27849859 | 0.23702613 | 0.81263652 | 0.94415747 |
| RFLNA | 8.01069654 | 0.211467396 | 0.64736922 | 0.32665655 | 0.74392767 | NA |
| NCOR2 | 82.4187771 | -0.263671408 | 0.35791154 | -0.7366943 | 0.46130825 | 0.79721613 |
| SCARB1 | 4.5468393 | 0.458729562 | 0.80171993 | 0.57218181 | 0.56719881 | NA |
| UBC | 263.202337 | -0.41539536 | 0.32864157 | -1.2639769 | 0.20623831 | 0.61507544 |
| DHX37 | 3.83564102 | -0.153756136 | 0.80276113 | -0.1915341 | 0.84810716 | NA |
| BRI3BP | 10.76409 | -0.382640035 | 0.52430362 | -0.7298062 | 0.46550865 | NA |
| AACS | 11.9325849 | 0.099652282 | 0.4958614 | 0.20096802 | 0.84072358 | NA |
| SLC15A4 | 30.707321 | -0.03862748 | 0.34354367 | -0.1124383 | 0.91047586 | 0.97557165 |
| STX2 | 35.2525872 | 0.327504245 | 0.29506846 | 1.10992632 | 0.26703078 | 0.66013596 |
| RAN | 39.1729672 | -1.086245706 | 0.5126129 | -2.119037 | 0.03408734 | 0.29210684 |
| SFSWAP | 39.4180476 | 0.232795482 | 0.29705814 | 0.78366977 | 0.4332339 | 0.7799364 |
| ULK1 | 27.9451514 | 0.255693044 | 0.40621986 | 0.62944497 | 0.52905779 | NA |
| PUS1 | 4.49976537 | -1.058320215 | 0.73816166 | -1.4337242 | 0.15165099 | NA |
| EP400 | 31.9263891 | -0.180031458 | 0.29122175 | -0.6181937 | 0.53644764 | 0.83677705 |
| DDX51 | 6.5733848 | -0.00378113 | 0.61896497 | -0.0061088 | 0.99512592 | NA |
| FBRSL1 | 18.4305333 | 0.60768944 | 0.38142145 | 1.59322302 | 0.11111017 | NA |
| PGAM5 | 5.00742427 | 0.660600283 | 0.74277086 | 0.88937291 | 0.3738027 | NA |
| ANKLE2 | 30.8051018 | -0.2108401 | 0.2941135 | -0.7168665 | 0.4734565 | 0.8060686 |
| GOLGA3 | 38.4108521 | 0.009621545 | 0.45488914 | 0.02115141 | 0.98312488 | 0.99552981 |
| CHFR | 35.6831537 | 0.099801487 | 0.27210652 | 0.3667736 | 0.7137879 | 0.91328825 |
| ZNF26 | 14.6976896 | 0.01334311 | 0.46022543 | 0.02899255 | 0.97687053 | NA |
| ZNF84 | 29.0858297 | 0.252540428 | 0.30573098 | 0.82602171 | 0.40879178 | 0.76772515 |
| ZNF140 | 11.3125829 | -0.526697566 | 0.50954514 | -1.0336622 | 0.3012941 | NA |
| ZNF891 | 10.760402 | -0.028835017 | 0.46544691 | -0.0619512 | 0.95060166 | NA |
| ZNF268 | 8.42007955 | -0.06210267 | 0.47957673 | -0.1294948 | 0.89696618 | NA |
| MPHOSPH8 | 74.2674897 | -0.148450317 | 0.29252768 | -0.5074744 | 0.61182198 | 0.87391153 |
| PSPC1 | 62.8384381 | 0.215290952 | 0.302579 | 0.71151981 | 0.47676218 | 0.80863008 |
| ZMYM5 | 27.4622353 | 0.096791654 | 0.39512479 | 0.24496477 | 0.80648371 | NA |
| AL355001.2 | 4.36008058 | -0.485980834 | 0.68422667 | -0.7102629 | 0.47754111 | NA |
| ZMYM2 | 173.871598 | 0.367538269 | 0.25356027 | 1.44951045 | 0.14719508 | 0.54635866 |
| GJA3 | 16.8650131 | -0.151426336 | 0.6546187 | -0.2313199 | 0.81706628 | NA |
| CRYL1 | 32.7878335 | 0.331352625 | 0.43604027 | 0.7599129 | 0.44730665 | 0.78947076 |
| IFT88 | 47.1314447 | -0.088971946 | 0.25447739 | -0.3496261 | 0.72661929 | 0.91700494 |
| EEF1AKMT1 | 10.9970311 | -0.318096299 | 0.57862271 | -0.5497473 | 0.58249268 | NA |
| XPO4 | 35.5234004 | 0.393291287 | 0.2809821 | 1.3997023 | 0.16160248 | 0.56290656 |
| LATS2 | 75.039513 | 0.434006108 | 0.24909931 | 1.7423015 | 0.0814557 | 0.43759513 |
| SAP18 | 53.4512255 | -0.403491312 | 0.35547411 | -1.1350793 | 0.25634208 | 0.65588339 |
| MRPL57 | 34.9582647 | -1.266501452 | 0.38761515 | -3.2674199 | 0.00108533 | 0.05186671 |
| ZDHHC20 | 36.4810652 | 0.223100471 | 0.31043842 | 0.71866256 | 0.47234886 | 0.80565474 |
| MICU2 | 72.240708 | 0.118208335 | 0.2035015 | 0.58087205 | 0.56132669 | 0.85386276 |
| FGF9 | 27.0317541 | -0.331713344 | 0.62111287 | -0.5340629 | 0.59329801 | NA |
| SACS | 5.86329095 | -1.113004415 | 0.60417854 | -1.842178 | 0.06544912 | NA |
| TNFRSF19 | 20.5956486 | 1.741428133 | 0.67836474 | 2.56709707 | 0.01025539 | NA |
| MIPEP | 29.4547299 | 0.440034333 | 0.42609691 | 1.03270952 | 0.30173986 | 0.68219446 |
| PARP4 | 39.0352322 | -0.9675958 | 0.42521785 | -2.2755296 | 0.02287418 | 0.24709152 |
| MTMR6 | 40.0314548 | -0.200130497 | 0.385226 | -0.5195145 | 0.60340199 | 0.87047761 |
| NUP58 | 62.0881986 | -0.40300136 | 0.30899172 | -1.3042465 | 0.19214957 | 0.59532067 |
| ATP8A2 | 48.8768553 | -0.169708974 | 0.73430856 | -0.231114 | 0.81722625 | 0.94694381 |
| SHISA2 | 181.537893 | -1.873361654 | 0.69686713 | -2.6882623 | 0.00718249 | 0.13427683 |
| RNF6 | 14.3290475 | -0.064500573 | 0.53597052 | -0.1203435 | 0.90421104 | NA |
| CDK8 | 41.6047674 | 0.145036574 | 0.32878474 | 0.44112927 | 0.65911941 | 0.89709157 |
| WASF3 | 16.8768937 | -0.052613955 | 0.33650371 | -0.1563548 | 0.8757534 | NA |
| USP12 | 41.9800505 | 0.367218965 | 0.33811483 | 1.08607768 | 0.27744463 | 0.66958859 |
| RPL21 | 39.9272112 | -0.256724228 | 0.56822772 | -0.4517981 | 0.65141441 | 0.89412414 |
| GTF3A | 12.4956926 | -0.703030182 | 0.48680886 | -1.4441606 | 0.1486938 | NA |
| MTIF3 | 24.9989528 | -0.283284271 | 0.30609576 | -0.925476 | 0.35471837 | NA |
| LNX2 | 77.2826491 | 0.060212518 | 0.23341006 | 0.25796882 | 0.79643097 | 0.93934998 |
| POLR1D | 25.5247326 | -0.183814683 | 0.41224093 | -0.4458914 | 0.6556757 | NA |
| PAN3-AS1 | 4.84379858 | -3.917187395 | 1.12194372 | -3.4914295 | 0.00048044 | NA |
| PAN3 | 303.563718 | -0.201347331 | 0.25423742 | -0.7919657 | 0.42838065 | 0.77844087 |
| FLT1 | 53.1752746 | -3.180354433 | 0.75479064 | -4.2135584 | 2.51E-05 | 0.0036414 |

| | | | | | | |
|---|---|---|---|---|---|---|
| POMP | 39.7496297 | -0.655722818 | 0.38582276 | -1.6995441 | 0.0892167 | 0.45574679 |
| SLC46A3 | 9.4650308 | -0.288996264 | 0.5623002 | -0.5139537 | 0.60728437 | NA |
| SLC7A1 | 19.8447762 | -0.908515258 | 0.55788383 | -1.6285026 | 0.10341836 | NA |
| UBL3 | 61.6161364 | 0.0084835 | 0.32451796 | 0.02614185 | 0.9791442 | 0.99409483 |
| KATNAL1 | 43.2816061 | 0.356063028 | 0.3262501 | 1.0913806 | 0.27510545 | 0.66766283 |
| HMGB1 | 139.297921 | -0.492267274 | 0.27852336 | -1.7674183 | 0.0771582 | 0.42394701 |
| USPL1 | 17.7028471 | -0.102161358 | 0.43836895 | -0.2330488 | 0.81572351 | NA |
| HSPH1 | 23.0526165 | 0.301436014 | 0.33524524 | 0.89915077 | 0.36857236 | NA |
| B3GLCT | 10.8887644 | -0.604619871 | 0.52940118 | -1.1420826 | 0.2534197 | NA |
| FRY | 747.230957 | 0.885578782 | 0.33302087 | 2.65922911 | 0.00783197 | 0.14181457 |
| FRY-AS1 | 16.9247019 | 1.191988012 | 0.62023276 | 1.92183982 | 0.05462592 | NA |
| N4BP2L1 | 29.2021581 | -0.588596007 | 0.43335178 | -1.3582407 | 0.17438733 | 0.57784404 |
| N4BP2L2 | 280.863089 | 0.025560278 | 0.22041284 | 0.11596547 | 0.90767991 | 0.9747971 |
| PDS5B | 72.8901779 | 0.401964444 | 0.23369412 | 1.72004517 | 0.08542423 | 0.44564602 |
| STARD13 | 82.6780674 | 0.634495562 | 0.36255026 | 1.75008993 | 0.0801028 | 0.43346499 |
| AL139383.1 | 82.9577778 | 0.102649922 | 0.38691218 | 0.26530548 | 0.79077414 | 0.93865377 |
| RFC3 | 11.3197947 | -0.565399057 | 0.64817151 | -0.8722985 | 0.38304554 | NA |
| NBEA | 94.9596666 | 0.506608625 | 0.32013668 | 1.58247602 | 0.11354094 | 0.50483797 |
| SPART | 48.39889 | -0.634325707 | 0.46938296 | -1.3514033 | 0.17656626 | 0.57988078 |
| RFXAP | 9.71049052 | -0.127775581 | 0.49963595 | -0.2557374 | 0.79815364 | NA |
| SMAD9 | 4.97521996 | -0.566559047 | 1.11323009 | -0.5089326 | 0.61079949 | NA |
| ALG5 | 18.2830973 | 0.255672366 | 0.37045731 | 0.69015338 | 0.49009773 | NA |
| EXOSC8 | 7.49271717 | -1.382730678 | 0.62664347 | -2.2065668 | 0.02734434 | NA |
| SUPT20H | 40.4445341 | 0.375251566 | 0.36736803 | 1.02145951 | 0.30703678 | 0.68651664 |
| LINC00571 | 9.22050881 | -0.242434695 | 0.60024407 | -0.4038935 | 0.68629102 | NA |
| UFM1 | 29.8343919 | -0.061897338 | 0.30732749 | -0.2014051 | 0.84038179 | 0.95393706 |
| FREM2 | 10.9478689 | 1.100396756 | 0.67584556 | 1.62817784 | 0.10348718 | NA |
| PROSER1 | 13.2202456 | -0.240970037 | 0.40494316 | -0.5950713 | 0.55179584 | NA |
| NHLRC3 | 31.0298005 | -0.277601304 | 0.41838212 | -0.6635114 | 0.50700308 | 0.82657837 |
| LHFPL6 | 123.226248 | 0.978446876 | 0.37252953 | 2.62649481 | 0.00862693 | 0.14826624 |
| COG6 | 32.7316428 | 0.105259304 | 0.29607648 | 0.3555139 | 0.72220463 | 0.91509584 |
| FOXO1 | 57.1093896 | -0.800243609 | 0.29554272 | -2.7077088 | 0.00677494 | 0.13085322 |
| MRPS31 | 22.9261572 | -0.107024981 | 0.35245275 | -0.3036577 | 0.76138871 | NA |
| ELF1 | 183.890211 | 0.652726101 | 0.22528945 | 2.89727766 | 0.00376416 | 0.09813187 |
| WBP4 | 23.418658 | 0.445922975 | 0.36405401 | 1.22488138 | 0.22061992 | NA |
| KBTBD6 | 7.67823624 | 0.149878764 | 0.55820218 | 0.26850265 | 0.78831243 | NA |
| AL354696.2 | 6.66809659 | -0.928157908 | 0.75151334 | -1.2350518 | 0.21681123 | NA |
| KBTBD7 | 7.27426473 | 0.304245854 | 0.54022918 | 0.56317923 | 0.57331285 | NA |
| MTRF1 | 5.00781182 | -0.030653612 | 0.61400611 | -0.049924 | 0.96018299 | NA |
| NAA16 | 28.6888547 | -0.074649687 | 0.36842416 | -0.2026189 | 0.83943294 | 0.95393706 |
| VWA8 | 87.7175674 | 0.818890939 | 0.29276672 | 2.79707659 | 0.00515673 | 0.11257186 |
| DGKH | 58.7846104 | 0.048920743 | 0.38373475 | 0.12748583 | 0.89855589 | 0.97097328 |
| AKAP11 | 67.4252498 | -0.243182234 | 0.27827929 | -0.8738783 | 0.38218452 | 0.74899182 |
| EPSTI1 | 40.4103284 | -0.072613352 | 1.37057671 | -0.0529801 | 0.95774773 | 0.98714493 |
| DNAJC15 | 24.8701628 | -0.154897788 | 0.41007266 | -0.3777325 | 0.70562929 | NA |
| ENOX1 | 17.8414363 | 0.583289628 | 0.68278413 | 0.85428117 | 0.39294922 | NA |
| CCDC122 | 31.9750704 | -0.307748686 | 0.42705506 | -0.72063 | 0.47113719 | 0.80460313 |
| SMIM2-AS1 | 15.4159587 | 0.640676718 | 0.94825029 | 0.67564094 | 0.49926864 | NA |
| TSC22D1 | 236.852861 | 0.434820182 | 0.30765889 | 1.41331909 | 0.15756192 | 0.56009743 |
| NUFIP1 | 6.46020852 | 0.166181657 | 0.6146554 | 0.27036557 | 0.78687903 | NA |
| AL359706.1 | 4.71376208 | -0.69490973 | 0.84375812 | -0.8235888 | 0.41017325 | NA |
| GPALPP1 | 19.6866673 | 0.142955489 | 0.34709467 | 0.41186311 | 0.68043976 | NA |
| GTF2F2 | 27.5874418 | 0.029480004 | 0.31631319 | 0.09319878 | 0.92574564 | NA |
| TPT1 | 184.053076 | -1.169044572 | 0.41619318 | -2.8088989 | 0.00497113 | 0.11177304 |
| SLC25A30 | 8.50285075 | 0.695782289 | 0.57661705 | 1.20666271 | 0.22756207 | NA |
| COG3 | 27.5767574 | 0.422314584 | 0.3335432 | 1.26614658 | 0.20546061 | NA |
| ZC3H13 | 51.2890725 | -0.030231874 | 0.26028515 | -0.1161491 | 0.90753441 | 0.9747971 |
| AL138686.1 | 25.3236524 | -0.260404645 | 0.64167711 | -0.4058188 | 0.68487573 | NA |
| LRCH1 | 43.9432332 | 0.394945014 | 0.35921212 | 1.09947575 | 0.2715606 | 0.66472529 |
| ESD | 32.9628144 | -0.125260863 | 0.39621846 | -0.3161409 | 0.75189556 | 0.92274989 |
| SUCLA2 | 38.3688625 | -0.106569722 | 0.29894392 | -0.3564873 | 0.72147564 | 0.91509584 |
| NUDT15 | 10.3547564 | -0.180071543 | 0.51686203 | -0.3483938 | 0.72754444 | NA |
| MED4 | 70.9031363 | 0.030209953 | 0.31070556 | 0.09723017 | 0.92254361 | 0.97775374 |
| ITM2B | 1478.05734 | 0.191525646 | 0.43834802 | 0.43692599 | 0.66216502 | 0.89839561 |
| RB1 | 88.7257433 | 0.032077988 | 0.25528064 | 0.12565774 | 0.90000286 | 0.97097328 |
| LPAR6 | 5.19071344 | -0.089238108 | 0.87117689 | -0.102434 | 0.91841222 | NA |
| RCBTB2 | 8.7063706 | 0.230545377 | 0.53028098 | 0.43476079 | 0.66373607 | NA |
| FNDC3A | 112.917288 | 0.215300917 | 0.25538617 | 0.84304061 | 0.39920573 | 0.76232507 |
| CDADC1 | 11.6630567 | 0.194389174 | 0.54278041 | 0.35813594 | 0.72024158 | NA |
| CAB39L | 22.155519 | -0.683135136 | 0.61409419 | -1.1124273 | 0.26595447 | NA |
| SETDB2 | 31.0466981 | 0.113440868 | 0.34087444 | 0.33279371 | 0.73929001 | 0.92003444 |
| PHF11 | 23.7037979 | -0.315272605 | 0.5081524 | -0.6204292 | 0.53497523 | NA |
| RCBTB1 | 9.74851214 | 0.702412908 | 0.54533773 | 1.28803284 | 0.19773453 | NA |
| EBPL | 10.2885367 | -0.998110192 | 0.53234563 | -1.8749289 | 0.06080251 | NA |
| KPNA3 | 48.9408973 | 0.080691002 | 0.27657353 | 0.29175244 | 0.77047591 | 0.92621526 |
| SPRYD7 | 12.7820804 | 0.257234255 | 0.42138551 | 0.61044874 | 0.54156459 | NA |
| DLEU2 | 21.7782745 | 0.331885104 | 0.55169336 | 0.60157531 | 0.54745687 | NA |
| TRIM13 | 14.6752483 | -0.723946675 | 0.39841392 | -1.8170717 | 0.06920613 | NA |
| RNASEH2B | 25.1358865 | -0.469272259 | 0.40906255 | -1.1471895 | 0.25130331 | NA |
| INTS6 | 55.1452346 | 0.501581226 | 0.36617016 | 1.36980366 | 0.1707482 | 0.57283267 |
| INTS6-AS1 | 13.2697119 | 0.774821228 | 0.45095083 | 1.71819447 | 0.08576115 | NA |
| WDFY2 | 25.3168344 | 0.061321483 | 0.43459404 | 0.14110061 | 0.88779047 | NA |
| ALG11 | 5.14880053 | -0.85867948 | 0.67903363 | -1.2645611 | 0.20602872 | NA |
| UTP14C | 10.6213657 | 0.013807935 | 0.6540413 | 0.02111172 | 0.98315654 | NA |
| NEK3 | 12.320664 | -0.332904354 | 0.47450504 | -0.7015823 | 0.48293967 | NA |
| VPS36 | 40.0054824 | -0.000307129 | 0.24907762 | -0.0012331 | 0.99901616 | 0.99961632 |
| CKAP2 | 5.55267838 | 0.421204385 | 0.59096511 | 0.71273985 | 0.47600675 | NA |
| SUGT1 | 32.5077705 | -0.144996604 | 0.38025424 | -0.3813149 | 0.7029696 | 0.90757606 |
| TDRD3 | 70.1133755 | 0.358124445 | 0.29229041 | 1.22523499 | 0.22048669 | 0.62462904 |
| PCDH9 | 67.6198849 | 0.432885893 | 0.64120818 | 0.67510975 | 0.49960604 | 0.82345406 |
| DACH1 | 2206.25696 | 0.413810622 | 0.27402224 | 1.51013516 | 0.13100894 | 0.52613957 |
| MZT1 | 11.4549752 | -0.478771269 | 0.54602782 | -0.8768258 | 0.38058127 | NA |
| DIS3 | 21.0483091 | -0.014750508 | 0.37639747 | -0.0391886 | 0.96873998 | NA |
| PIBF1 | 83.1256012 | -0.013068536 | 0.22137234 | -0.0590342 | 0.95292488 | 0.98520702 |
| KLF12 | 71.3544127 | 0.628449129 | 0.41204163 | 1.52520786 | 0.12720726 | 0.52301154 |
| TBC1D4 | 23.7348982 | 0.700517723 | 0.62415013 | 1.12235452 | 0.26171173 | NA |
| COMMD6 | 26.6727477 | -0.522120885 | 0.51622814 | -1.011415 | 0.31181784 | NA |
| UCHL3 | 12.9959252 | -0.355779914 | 0.48191281 | -0.7382661 | 0.46035273 | NA |
| LMO7 | 56.8487177 | 1.669834948 | 0.33026049 | 5.05611476 | 4.28E-07 | 0.00021694 |
| KCTD12 | 39.0753753 | -1.019746814 | 0.69201313 | -1.4735946 | 0.14059077 | 0.53597868 |
| AC001226.2 | 10.2270173 | -1.667661863 | 0.68033193 | -2.4512474 | 0.01423621 | NA |
| FBXL3 | 50.7376718 | 0.441050796 | 0.28701667 | 1.53667309 | 0.12437338 | 0.51952466 |
| MYCBP2 | 125.908443 | 0.423622896 | 0.20713822 | 2.04512186 | 0.04084285 | 0.31695313 |
| SLAIN1 | 11.0884952 | -0.074150018 | 0.57155628 | -0.1297335 | 0.89677725 | NA |
| RNF219 | 10.1428937 | -0.341501535 | 0.53032024 | -0.6439534 | 0.51960563 | NA |
| RBM26 | 110.671582 | 0.460850596 | 0.22508736 | 2.04742995 | 0.04061589 | 0.31680393 |
| NDFIP2 | 42.5186118 | 0.368893431 | 0.3072051 | 1.20080504 | 0.22982684 | 0.63682501 |
| SPRY2 | 10.2252243 | -0.220149434 | 0.65801826 | -0.3345643 | 0.73795376 | NA |
| GPC5 | 36.152832 | 0.86574635 | 0.97471494 | 0.88820466 | 0.37443067 | 0.74191051 |
| GPC6 | 834.533514 | -0.128433547 | 0.1931415 | -0.6649713 | 0.50606887 | 0.82600448 |
| TGDS | 10.0424426 | 0.482478338 | 0.48380169 | 0.99726469 | 0.31863605 | NA |
| GPR180 | 19.5414084 | -0.071229408 | 0.5255001 | -0.1355459 | 0.89218024 | NA |
| ABCC4 | 46.7394181 | -0.634772497 | 0.54639618 | -1.161744 | 0.24533946 | 0.64996527 |

| | | | | | | |
|---|---|---|---|---|---|---|
| CLDN10 | 30.615728 | 0.711655029 | 0.64715041 | 1.09967485 | 0.27147381 | 0.66472529 |
| CLDN10-AS1 | 20.2782124 | -0.263820373 | 1.05044946 | -0.25115 | 0.80169815 | NA |
| DZIP1 | 7.11082846 | 0.443977649 | 0.70813158 | 0.62697055 | 0.53067854 | NA |
| DNAJC3-DT | 5.3794202 | -0.534531501 | 0.72601928 | -0.7362497 | 0.46157873 | NA |
| DNAJC3 | 83.857796 | -0.224714128 | 0.33653337 | -0.6677321 | 0.50430463 | 0.82600448 |
| UGGT2 | 131.658546 | 0.186611259 | 0.26707727 | 0.69871637 | 0.4847293 | 0.81444161 |
| MBNL2 | 122.60263 | 0.369504609 | 0.24613405 | 1.50123321 | 0.13329525 | 0.52613957 |
| RAP2A | 13.6980618 | -0.43234664 | 0.62100913 | -0.6962001 | 0.4863035 | NA |
| IPO5 | 26.5480868 | -0.250590957 | 0.34506857 | -0.7262063 | 0.46771233 | NA |
| FARP1 | 134.345494 | 0.248923924 | 0.24582636 | 1.01260061 | 0.31125097 | 0.69098737 |
| STK24 | 55.8287056 | 0.174621283 | 0.34393463 | 0.50771649 | 0.61165218 | 0.87391153 |
| DOCK9 | 71.1071229 | 0.077583643 | 0.24808192 | 0.31273397 | 0.75448279 | 0.92313949 |
| UBAC2 | 56.0595492 | 0.260404096 | 0.31507897 | 0.82647246 | 0.40853613 | 0.76772515 |
| TM9SF2 | 77.3988568 | 0.33054154 | 0.32389782 | 1.02051177 | 0.30748581 | 0.68713231 |
| CLYBL | 50.3317132 | 0.270408243 | 0.3564649 | 0.75858308 | 0.448102 | 0.78987065 |
| PCCA | 138.349822 | 0.367159758 | 0.2678582 | 1.37072436 | 0.1704609 | 0.57276745 |
| GGACT | 20.6185449 | -0.780121985 | 0.41075785 | -1.899226 | 0.05753477 | NA |
| TMTC4 | 24.0921534 | 0.435407886 | 0.41235522 | 1.05590486 | 0.29101168 | NA |
| NALCN | 40.3943583 | 0.727219747 | 0.30673053 | 2.37087498 | 0.01774603 | 0.22079113 |
| ITGBL1 | 11.0415473 | 0.359575701 | 0.89940145 | 0.39979444 | 0.68930792 | NA |
| FGF14 | 17.8384589 | 1.958746603 | 0.75511535 | 2.59397004 | 0.00948748 | NA |
| FGF14-AS2 | 4.81707772 | -0.121014934 | 0.6401728 | -0.1890348 | 0.85006555 | NA |
| TPP2 | 26.6151964 | 0.044850285 | 0.31440633 | 0.14265071 | 0.88656605 | NA |
| EFNB2 | 59.6306499 | 0.496797752 | 0.53281653 | 0.93239928 | 0.35113021 | 0.72256933 |
| ARGLU1 | 140.825152 | -0.062900831 | 0.27868607 | -0.225705 | 0.82143089 | 0.94728252 |
| FAM155A | 22.9045511 | -1.406700631 | 0.65438254 | -2.149661 | 0.03158204 | NA |
| LIG4 | 11.5738262 | 0.356016162 | 0.5138327 | 0.69286396 | 0.48839493 | NA |
| ABHD13 | 18.6535413 | 0.691643867 | 0.4575384 | 1.511663 | 0.13061962 | NA |
| TNFSF13B | 23.4664866 | 1.266579045 | 0.56110407 | 2.25729792 | 0.02398947 | NA |
| IRS2 | 21.2529632 | -1.629523168 | 0.65265922 | -2.4967443 | 0.01253393 | NA |
| COL4A1 | 33.2399628 | 0.98429294 | 0.71737724 | 1.37207159 | 0.17004115 | 0.57273205 |
| COL4A2 | 40.8266684 | 0.25488265 | 0.61537534 | 0.41419055 | 0.67873456 | 0.90499319 |
| RAB20 | 9.15964649 | -0.770477651 | 0.60118898 | -1.2815898 | 0.19998659 | NA |
| NAXD | 13.8759588 | -0.061008836 | 0.47485156 | -0.1284798 | 0.89776928 | NA |
| CARS2 | 29.0697382 | -0.138720496 | 0.30897146 | -0.4489751 | 0.65344961 | 0.8949739 |
| ING1 | 11.7169477 | -1.48530892 | 0.72542517 | -2.0475012 | 0.0406089 | NA |
| ANKRD10 | 96.2519008 | 0.042771492 | 0.22567454 | 0.18952733 | 0.84967954 | 0.95532127 |
| ARHGEF7 | 44.3931472 | -0.049055414 | 0.30268233 | -0.162069 | 0.87125154 | 0.96207879 |
| TUBGCP3 | 34.3975582 | -0.127621742 | 0.27735297 | -0.460142 | 0.64541432 | 0.89162142 |
| ATP11A | 67.6240094 | -0.404107701 | 0.30090068 | -1.3429936 | 0.17927404 | 0.58310785 |
| MCF2L | 21.8519668 | 0.695969154 | 0.62531694 | 1.11298625 | 0.26571434 | NA |
| PCID2 | 16.5721555 | -0.284661168 | 0.37240335 | -0.7643894 | 0.44463524 | NA |
| CUL4A | 42.5615207 | -0.241483338 | 0.27903689 | -0.8654172 | 0.38680982 | 0.75125852 |
| LAMP1 | 136.85482 | 0.161187168 | 0.34728372 | 0.46413684 | 0.64254971 | 0.88951638 |
| DCUN1D2 | 38.4646925 | 0.031645389 | 0.31016065 | 0.10202902 | 0.91873364 | 0.97775374 |
| TMCO3 | 91.7424368 | -0.098434716 | 0.40483638 | -0.2431469 | 0.80789159 | 0.94328109 |
| TFDP1 | 18.4626281 | -0.139053755 | 0.61213495 | -0.2271619 | 0.82029783 | NA |
| GAS6 | 47.4646382 | -0.912739674 | 0.52822401 | -1.7279405 | 0.08399889 | 0.44304225 |
| RASA3 | 5.97179548 | -1.795827682 | 0.87664904 | -2.0485138 | 0.04050968 | NA |
| CDC16 | 34.1660378 | -0.4875032 | 0.29156501 | -1.6720223 | 0.09451993 | 0.46752784 |
| UPF3A | 42.9086777 | -0.106358663 | 0.35246552 | -0.3017562 | 0.76283791 | 0.92336765 |
| CHAMP1 | 11.2184597 | -0.418619206 | 0.4640746 | -0.9020515 | 0.36702949 | NA |
| TTC5 | 10.4899022 | -0.005763887 | 0.46579798 | -0.0123742 | 0.99012705 | NA |
| CCNB1IP1 | 12.5365 | -0.312818461 | 0.42950316 | -0.7283263 | 0.46641385 | NA |
| TEP1 | 9.8753449 | -0.112777199 | 0.57925999 | -0.1946918 | 0.84563419 | NA |
| OSGEP | 8.23210032 | 0.335601333 | 0.49880604 | 0.67280928 | 0.50106863 | NA |
| APEX1 | 14.3043137 | -0.6281789 | 0.43352049 | -1.4490178 | 0.14733262 | NA |
| PIP4P1 | 9.20531392 | -0.457904837 | 0.57566159 | -0.795441 | 0.42635701 | NA |
| PNP | 6.40235098 | -0.283629149 | 0.71647819 | -0.3958657 | 0.6922041 | NA |
| METTL17 | 6.13399381 | -0.486327046 | 0.68150558 | -0.7136068 | 0.47547032 | NA |
| NDRG2 | 70.8371941 | -0.86907535 | 0.4939632 | -1.7593929 | 0.0785108 | 0.42784309 |
| ARHGEF40 | 32.0659617 | -0.319952008 | 0.43182351 | -0.7409324 | 0.45873446 | 0.79513972 |
| ZNF219 | 13.1050054 | 0.25547192 | 0.50288686 | 0.50801073 | 0.61144582 | NA |
| HNRNPC | 132.976857 | 0.122875188 | 0.20894083 | 0.58808607 | 0.55647453 | 0.85122066 |
| SUPT16H | 34.3353441 | 0.008608563 | 0.30276796 | 0.02843287 | 0.97731691 | 0.99298531 |
| CHD8 | 41.6442063 | 0.014341973 | 0.38777209 | 0.03698557 | 0.97049651 | 0.98977467 |
| RAB2B | 17.2146964 | 0.350390162 | 0.37884901 | 0.92488076 | 0.35502794 | NA |
| TOX4 | 29.9001986 | -0.029920556 | 0.30163277 | -0.0991953 | 0.9209832 | 0.97775374 |
| METTL3 | 11.1184536 | 0.278964381 | 0.46582098 | 0.59886608 | 0.54926219 | NA |
| DAD1 | 81.1041708 | -0.778114853 | 0.4939844 | -1.575181 | 0.11521467 | 0.50513044 |
| ABHD4 | 7.57483557 | 0.241083368 | 0.56176495 | 0.42915345 | 0.66781156 | NA |
| OXA1L | 8.71751867 | 0.02470553 | 0.57663737 | 0.04284414 | 0.96582578 | NA |
| MRPL52 | 13.7982829 | 0.053678835 | 0.59773803 | 0.08980328 | 0.92844354 | NA |
| MMP14 | 5.90044451 | 0.312728587 | 0.57135613 | 0.54734442 | 0.58414213 | NA |
| LRP10 | 72.6134773 | -0.39534909 | 0.40958867 | -0.9652344 | 0.3344274 | 0.70906302 |
| RBM23 | 28.8370838 | 0.318942391 | 0.30756438 | 1.03699391 | 0.2997387 | 0.68168772 |
| PRMT5 | 9.34617896 | 0.007694292 | 0.54062513 | 0.01423221 | 0.98864472 | NA |
| AL132780.3 | 7.90835642 | -0.46024287 | 0.77985163 | -0.5901672 | 0.55507854 | NA |
| PSMB5 | 39.7297593 | -0.822232679 | 0.40210752 | -2.044808 | 0.04087379 | 0.31695313 |
| ACIN1 | 40.2443927 | 0.19652204 | 0.31529886 | 0.62328814 | 0.53309519 | 0.83677705 |
| C14orf119 | 15.4418371 | -0.889815592 | 0.45461234 | -1.9573063 | 0.05031147 | NA |
| SLC7A8 | 135.813605 | -0.599993009 | 0.49110519 | -1.22172 | 0.22181355 | 0.62564377 |
| HOMEZ | 4.4228851 | -0.932664153 | 0.74970855 | -1.2440356 | 0.21348644 | NA |
| PPP1R3E | 6.37042435 | -0.395556171 | 0.57589249 | -0.6868577 | 0.49217242 | NA |
| BCL2L2-PABI | 15.3958386 | 0.324186736 | 0.36400643 | 0.89060716 | 0.37313996 | NA |
| PABPN1 | 69.3615777 | 1.666410305 | 0.33685128 | 4.94702079 | 7.54E-07 | 0.00023577 |
| SLC22A17 | 4.93233815 | 0.713408556 | 0.63515861 | 1.12319748 | 0.26135363 | NA |
| NGDN | 8.06173157 | -6.95E-05 | 0.48924988 | -0.000142 | 0.99988669 | NA |
| ZFHX2 | 27.8420469 | 0.2257245 | 0.34154025 | 0.6609016 | 0.50867542 | NA |
| THTPA | 5.46821883 | -0.764801036 | 0.79759105 | -0.9588887 | 0.33761482 | NA |
| AP1G2 | 9.02149792 | -0.081354724 | 0.57627689 | -0.141173 | 0.88773329 | NA |
| DHRS4-AS1 | 9.56065078 | 0.446090555 | 0.65320683 | 0.6829239 | 0.49465493 | NA |
| DHRS4 | 4.97530946 | 0.246130096 | 0.60087332 | 0.40962061 | 0.68208428 | NA |
| DHRS4L2 | 14.860957 | 0.017294103 | 0.41326677 | 0.04184731 | 0.96662042 | NA |
| PCK2 | 5.98085178 | -0.120769901 | 0.88892951 | -0.1358599 | 0.89193201 | NA |
| PSME1 | 36.0451448 | -1.055940286 | 0.55061604 | -1.9177434 | 0.05514355 | 0.36605934 |
| EMC9 | 6.47159653 | -0.193052064 | 0.6068254 | -0.3181344 | 0.75038295 | NA |
| PSME2 | 5.41965713 | 0.080454334 | 0.9186658 | 0.08757737 | 0.93021259 | NA |
| AL136295.1 | 9.54165687 | -0.418148933 | 0.44534745 | -0.9389274 | 0.34776801 | NA |
| GMPR2 | 9.00358432 | -0.681106615 | 0.60218333 | -1.1310619 | 0.25802905 | NA |
| TINF2 | 7.84086475 | -0.759378124 | 0.61095935 | -1.2429274 | 0.21389458 | NA |
| DHRS1 | 6.51012311 | -0.018265266 | 0.70881961 | -0.0257686 | 0.97944193 | NA |
| KHNYN | 8.95277989 | -0.102365695 | 0.47474845 | -0.2156209 | 0.82928325 | NA |
| SDR39U1 | 12.0443134 | -0.536464116 | 0.4401411 | -1.2188458 | 0.22290273 | NA |
| NOVA1 | 39.5078894 | 1.014041325 | 0.50302154 | 2.0159004 | 0.04381039 | 0.33029013 |
| PRKD1 | 337.623964 | -0.091518335 | 0.29428882 | -0.3109814 | 0.7558148 | 0.92313949 |
| G2E3 | 20.3087145 | 0.650726987 | 0.33784464 | 1.9261131 | 0.05409025 | NA |
| SCFD1 | 73.8945235 | 0.064346188 | 0.24885896 | 0.25856489 | 0.79597098 | 0.93934998 |
| COCH | 8.32556416 | -1.121620072 | 0.66829184 | -1.6783387 | 0.09328099 | NA |
| STRN3 | 112.670373 | -0.0539009 | 0.27933213 | -0.1929635 | 0.84698757 | 0.95532127 |
| AP4S1 | 16.8965354 | 0.229601064 | 0.35865301 | 0.64017605 | 0.52205815 | NA |
| HECTD1 | 95.7437106 | 0.137459677 | 0.24594059 | 0.55891415 | 0.57622031 | 0.85929076 |
| HEATR5A | 18.5289098 | 0.44504475 | 0.41254575 | 1.07877672 | 0.28068728 | NA |

| | | | | | | |
|---|---|---|---|---|---|---|
| AL139353.1 | 8.0249102 | 0.558188035 | 0.54191903 | 1.0300211 | 0.3030001 | NA |
| NUBPL | 53.5507401 | 0.343910946 | 0.32161799 | 1.06931501 | 0.28492775 | 0.67307335 |
| ARHGAP5 | 155.994514 | 0.139010553 | 0.21538253 | 0.64541238 | 0.51865998 | 0.83104823 |
| AKAP6 | 11.7492335 | 0.663896928 | 0.72194141 | 0.91959945 | 0.35778211 | NA |
| NPAS3 | 1457.9805 | 0.442439257 | 0.23323694 | 1.89695186 | 0.05783429 | 0.3759229 |
| EAPP | 26.2004107 | -0.621498208 | 0.32403668 | -1.9179872 | 0.05511263 | NA |
| SNX6 | 52.4530939 | -0.109137624 | 0.30708668 | -0.3553968 | 0.72229234 | 0.91509584 |
| CFL2 | 15.0604998 | -0.263828894 | 0.51006355 | -0.5172471 | 0.60498368 | NA |
| BAZ1A | 27.2402041 | -0.112620597 | 0.4035188 | -0.2790963 | 0.78017093 | NA |
| SRP54 | 41.6170135 | -0.402953311 | 0.28969134 | -1.3909747 | 0.16423311 | 0.56548642 |
| FAM177A1 | 38.7049374 | -0.169631129 | 0.31388961 | -0.5404165 | 0.58890982 | 0.86533208 |
| PPP2R3C | 19.423271 | -0.016021378 | 0.39105418 | -0.0409697 | 0.96732004 | NA |
| NFKBIA | 57.9034354 | -0.614043917 | 0.35435419 | -1.7328535 | 0.0831217 | 0.44101093 |
| RALGAPA1 | 130.998446 | 0.359875266 | 0.20231781 | 1.77876213 | 0.07527877 | 0.42138852 |
| BRMS1L | 8.70431141 | 0.24072384 | 0.5362282 | 0.44892051 | 0.653489 | NA |
| MBIP | 15.4389698 | 0.174693774 | 0.46912034 | 0.37238585 | 0.70960559 | NA |
| SLC25A21 | 6.10492629 | -0.998990191 | 0.81978603 | -1.2185987 | 0.22299654 | NA |
| MIPOL1 | 51.5718203 | -0.267661075 | 0.30493549 | -0.877763 | 0.38007237 | 0.74870012 |
| SEC23A | 57.2121088 | -0.216809075 | 0.25251071 | -0.8586134 | 0.39055385 | 0.75550853 |
| GEMIN2 | 4.63628728 | 0.182526935 | 0.7191231 | 0.25381876 | 0.79963558 | NA |
| TRAPPC6B | 22.7614001 | 0.107449428 | 0.33379359 | 0.32190381 | 0.74752557 | NA |
| PNN | 47.3833962 | -0.02265882 | 0.27016537 | -0.0838702 | 0.93315964 | 0.97832857 |
| MIA2 | 46.3677628 | -0.039837048 | 0.30255964 | -0.1316668 | 0.89524788 | 0.97097328 |
| FBXO33 | 36.0842465 | 0.152619298 | 0.37163704 | 0.41066762 | 0.68131628 | 0.90499319 |
| AL049875.1 | 9.57973805 | -0.352947005 | 0.53116938 | -0.6644717 | 0.50638848 | NA |
| KLHL28 | 27.5338687 | -0.092621617 | 0.41174622 | -0.2249483 | 0.82201949 | NA |
| TOGARAM1 | 59.131678 | 0.255905519 | 0.25859839 | 0.98958666 | 0.32237619 | 0.69513974 |
| PRPF39 | 16.2514173 | 0.207280956 | 0.37476756 | 0.55309204 | 0.58020038 | NA |
| FKBP3 | 17.3809041 | -0.029117212 | 0.4062497 | -0.0716732 | 0.94286199 | NA |
| FANCM | 6.51635206 | 0.256001848 | 0.6331993 | 0.40429901 | 0.68599286 | NA |
| MIS18BP1 | 15.7677885 | 0.132181673 | 0.38986427 | 0.33904536 | 0.73457556 | NA |
| LINC00871 | 15.4354327 | 2.462282246 | 0.85119822 | 2.89272486 | 0.00381916 | NA |
| MDGA2 | 12.176942 | -1.179390234 | 1.05626784 | -1.1165636 | 0.26418096 | NA |
| RPS29 | 40.3433001 | -0.010709526 | 0.44085329 | -0.0242927 | 0.98061912 | 0.99453905 |
| LRR1 | 5.73333244 | -0.425776228 | 0.63093273 | -0.6748362 | 0.49977986 | NA |
| RPL36AL | 86.3037817 | -1.224587545 | 0.3633763 | -3.370026 | 0.00075161 | 0.04122345 |
| MGAT2 | 11.1417854 | 0.2961622 | 0.53136089 | 0.55736545 | 0.57727777 | NA |
| DNAAF2 | 7.29006895 | -0.694472427 | 0.53462354 | -1.2989934 | 0.19394621 | NA |
| POLE2 | 7.84828 | 0.393909921 | 1.01464243 | 0.38822536 | 0.69784927 | NA |
| KLHDC1 | 27.45682 | 0.130517325 | 0.37860413 | 0.34473296 | 0.73029513 | NA |
| KLHDC2 | 27.8745155 | -0.177428596 | 0.31515076 | -0.5629959 | 0.57343765 | NA |
| NEMF | 75.5532273 | 0.414804733 | 0.23554221 | 1.76106328 | 0.07822769 | 0.4276675 |
| ARF6 | 80.2322054 | 0.415524919 | 0.34983429 | 1.18777641 | 0.23492151 | 0.64164421 |
| LINC01588 | 12.6880986 | 1.036654375 | 0.45159339 | 2.29554815 | 0.02170173 | NA |
| VCPKMT | 10.177316 | 0.558878404 | 0.43031695 | 1.29875991 | 0.19402634 | NA |
| SOS2 | 104.197037 | 0.39169524 | 0.22131342 | 1.76986662 | 0.07674936 | 0.42346132 |
| L2HGDH | 15.8550335 | 0.3044165 | 0.45084137 | 0.67521865 | 0.49953686 | NA |
| ATP5S | 20.1294509 | 0.139978014 | 0.37788674 | 0.3704232 | 0.71106719 | NA |
| CDKL1 | 23.5582975 | 0.878791328 | 0.47845145 | 1.83674087 | 0.06624816 | NA |
| MAP4K5 | 63.6147417 | 0.294816648 | 0.25053976 | 1.17672601 | 0.23930488 | 0.64621878 |
| ATL1 | 20.300507 | -0.156200282 | 0.39751066 | -0.3929461 | 0.69435925 | NA |
| SAV1 | 39.1291009 | 0.321502287 | 0.25961468 | 1.23838255 | 0.21557425 | 0.62324286 |
| NIN | 41.5009429 | -0.532651139 | 0.38562797 | -1.3812565 | 0.16720011 | 0.56878882 |
| PYGL | 5.95779311 | -1.164418308 | 0.74415267 | -1.5647573 | 0.11763983 | NA |
| TMX1 | 19.2437657 | -0.267025102 | 0.42030956 | -0.6353058 | 0.52522899 | NA |
| FRMD6 | 6.17169322 | -1.109213703 | 0.62918438 | -1.762939 | 0.07791077 | NA |
| RTRAF | 26.4788088 | -0.534784736 | 0.38479959 | -1.3897747 | 0.1645973 | NA |
| TXNDC16 | 10.0211464 | -0.161816513 | 0.48716056 | -0.3321626 | 0.73976649 | NA |
| ERO1A | 33.2681129 | -0.255867291 | 0.41364795 | -0.6185629 | 0.53620433 | 0.83677705 |
| AL133453.1 | 8.75907394 | -1.571990908 | 0.79775512 | -1.9705181 | 0.04877902 | NA |
| PSMC6 | 33.4094935 | -0.021485993 | 0.26859854 | -0.079993 | 0.93624285 | 0.97834166 |
| STYX | 11.708272 | 0.099789983 | 0.48709332 | 0.2048683 | 0.83767502 | NA |
| GNPNAT1 | 13.2410546 | 0.294551529 | 0.42027443 | 0.70085522 | 0.48339337 | NA |
| FERMT2 | 263.805302 | 0.289596967 | 0.29372561 | 0.98594389 | 0.32416065 | 0.69623471 |
| DDHD1 | 29.2469682 | 0.150980885 | 0.39570702 | 0.38154715 | 0.70279729 | 0.90757606 |
| CNIH1 | 21.4797325 | 0.491735401 | 0.37549639 | 1.30956093 | 0.19034441 | NA |
| GMFB | 18.9806308 | 0.101401075 | 0.38299363 | 0.26475917 | 0.791195 | NA |
| CGRRF1 | 9.01376335 | -0.153045004 | 0.50109293 | -0.3054224 | 0.76004447 | NA |
| SAMD4A | 75.0420831 | 0.436917768 | 0.3665689 | 1.19191172 | 0.23329586 | 0.63935677 |
| SOCS4 | 16.1850318 | -0.243901456 | 0.41207846 | -0.5918811 | 0.5539302 | NA |
| MAPK1IP1L | 46.9758203 | 0.138903422 | 0.35680225 | 0.38930086 | 0.6970536 | 0.90729979 |
| LGALS3 | 9.37339336 | -0.989939781 | 0.62708356 | -1.5786409 | 0.11441844 | NA |
| FBXO34 | 36.2433519 | -0.070704128 | 0.2917943 | -0.2423081 | 0.80854142 | 0.9434534 |
| ATG14 | 14.5820994 | 0.159201168 | 0.46066839 | 0.34558735 | 0.72965285 | NA |
| KTN1 | 116.27312 | -0.176885325 | 0.27390339 | -0.6457946 | 0.5184124 | 0.83104823 |
| PELI2 | 86.728133 | 0.31423019 | 0.19101521 | 1.64505324 | 0.09995883 | 0.48127507 |
| TMEM260 | 26.1257089 | -0.236156704 | 0.36987095 | -0.6384841 | 0.52315863 | NA |
| EXOC5 | 50.2745862 | 0.088355192 | 0.30077022 | 0.2937631 | 0.76893894 | 0.9255638 |
| AP5M1 | 31.0739705 | 0.268563076 | 0.28762487 | 0.9337269 | 0.35044478 | 0.72238707 |
| NAA30 | 15.96963 | 0.299102096 | 0.38512876 | 0.77662882 | 0.4373778 | NA |
| CCDC198 | 15.5017364 | 0.788600364 | 0.63441245 | 1.24304048 | 0.21385292 | NA |
| ARMH4 | 745.972154 | 0.259086643 | 0.3962534 | 0.65384082 | 0.51321437 | 0.83070544 |
| ACTR10 | 36.9966294 | -0.28167853 | 0.31073551 | -0.9064897 | 0.36467673 | 0.73398222 |
| PSMA3 | 30.9026149 | 0.072212012 | 0.46957407 | 0.15378194 | 0.87778167 | 0.96380142 |
| ARID4A | 35.2502005 | 0.413354469 | 0.34887842 | 1.18480951 | 0.23609277 | 0.64267939 |
| AL139021.1 | 4.83727233 | -0.537556047 | 0.76082373 | -0.7065448 | 0.4798494 | NA |
| TIMM9 | 11.4639133 | -0.450972094 | 0.57188156 | -0.788576 | 0.43035984 | NA |
| KIAA0586 | 26.5677875 | -0.287818552 | 0.41048948 | -0.7011594 | 0.48320355 | NA |
| DAAM1 | 18.2586151 | 0.061617027 | 0.41766354 | 0.14752791 | 0.88271536 | NA |
| GPR135 | 11.8497613 | 0.182405022 | 0.66849469 | 0.27285934 | 0.78496134 | NA |
| L3HYPDH | 12.5087303 | -0.255056742 | 0.57453675 | -0.4439346 | 0.65708987 | NA |
| JKAMP | 24.1786323 | 0.222432905 | 0.38344155 | 0.58009599 | 0.56184989 | NA |
| PCNX4 | 141.072772 | 0.195765322 | 0.31782258 | 0.61595789 | 0.53792231 | 0.83722675 |
| DHRS7 | 25.4963407 | -0.626460045 | 0.36289871 | -1.726267 | 0.0842994 | NA |
| PPM1A | 40.219486 | -0.290459018 | 0.34766381 | -0.8354594 | 0.40345907 | 0.76594067 |
| MNAT1 | 93.4434265 | 0.464480732 | 0.24660159 | 1.88352688 | 0.059629 | 0.37905854 |
| TRMT5 | 5.76236528 | -0.754546645 | 0.67482183 | -1.118142 | 0.26350634 | NA |
| SLC38A6 | 20.0634704 | -0.076393272 | 0.36023257 | -0.2120665 | 0.83205513 | NA |
| HIF1A | 157.939817 | -0.332985275 | 0.35958006 | -0.9260393 | 0.35442554 | 0.72603535 |
| HIF1A-AS2 | 51.3527849 | -0.812887151 | 0.62691373 | -1.2966492 | 0.19475193 | 0.59932765 |
| SNAPC1 | 16.509375 | -0.05206636 | 0.43289678 | -0.1202743 | 0.90426586 | NA |
| PPP2R5E | 90.239061 | 0.311584331 | 0.22655245 | 1.37532978 | 0.16902923 | 0.57194586 |
| WDR89 | 4.93084625 | -0.529937654 | 0.63750755 | -0.8312649 | 0.405824 | NA |
| SGPP1 | 29.6457838 | 0.638666039 | 0.30197645 | 2.11495315 | 0.03443394 | 0.29210684 |
| SYNE2 | 453.11967 | -0.045284109 | 0.24952285 | -0.1814828 | 0.85598862 | 0.95615679 |
| ESR2 | 29.2667458 | -0.181880028 | 0.59526498 | -0.3055446 | 0.75995137 | 0.92313949 |
| MTHFD1 | 9.59869152 | -1.019289195 | 0.50104249 | -2.0343368 | 0.04191765 | NA |
| AL122035.1 | 9.66252966 | -1.516243806 | 0.59117639 | -2.5647909 | 0.0103238 | NA |
| ZBTB25 | 21.433337 | -0.300056311 | 0.44824981 | -0.6693953 | 0.50324335 | NA |
| ZBTB1 | 84.8312834 | -0.281872612 | 0.3113753 | -0.9052504 | 0.36533276 | 0.73398222 |
| AL049869.3 | 9.4783033 | -0.599443087 | 0.58003503 | -1.0334601 | 0.30138861 | NA |
| HSPA2 | 41.1901515 | -1.374485611 | 0.73973227 | -1.8580852 | 0.06315691 | 0.3863717 |
| PLEKHG3 | 55.923763 | 0.178766488 | 0.36812967 | 0.48560739 | 0.62724556 | 0.8806406 |

| | | | | | | |
|---|---|---|---|---|---|---|
| SPTB | 36.7869177 | 0.509364204 | 0.54471167 | 0.93510793 | 0.34973268 | 0.72238707 |
| CHURC1-FNT | 9.8326453 | -0.91522912 | 0.50279031 | -1.8202998 | 0.06871335 | NA |
| FNTB | 4.78396828 | 0.045844613 | 0.8192059 | 0.05596226 | 0.95537187 | NA |
| MAX | 32.3129228 | -0.119650855 | 0.39029324 | -0.3065666 | 0.75917331 | 0.92313949 |
| FUT8 | 42.9135738 | 0.102101677 | 0.28280094 | 0.36103726 | 0.71807159 | 0.91472065 |
| AL391261.2 | 7.77804711 | 0.061421021 | 0.68848921 | 0.08921131 | 0.92891398 | NA |
| AL359232.1 | 74.1638791 | 0.159318572 | 0.56037029 | 0.28430946 | 0.77617324 | 0.92920858 |
| CCDC196 | 6.61403124 | -1.607507617 | 1.01968789 | -1.5764702 | 0.11491747 | NA |
| GPHN | 525.740576 | 0.575249712 | 0.30576942 | 1.88131862 | 0.05992859 | 0.37905854 |
| MPP5 | 282.654123 | 0.77669265 | 0.36392585 | 2.13420577 | 0.03282594 | 0.29047359 |
| ATP6V1D | 43.0733497 | -0.334937621 | 0.32078562 | -1.0441167 | 0.2964314 | 0.68103704 |
| EIF2S1 | 23.9406777 | -0.317777398 | 0.355354 | -0.8942559 | 0.371185 | NA |
| TMEM229B | 14.4455491 | -0.043240855 | 0.47209492 | -0.0915936 | 0.92702097 | NA |
| PLEKHH1 | 6.90909729 | 0.400037882 | 0.64525272 | 0.61997086 | 0.53527697 | NA |
| PIGH | 8.4646471 | -0.200020928 | 0.53773873 | -0.3719667 | 0.70991761 | NA |
| ARG2 | 8.07100275 | -1.021729748 | 0.81077036 | -1.2601962 | 0.20759859 | NA |
| VTI1B | 19.1680096 | 0.280429653 | 0.45792691 | 0.61238955 | 0.54028005 | NA |
| ZFYVE26 | 13.4966428 | 0.346107227 | 0.47438533 | 0.72959091 | 0.46564028 | NA |
| RAD51B | 106.828532 | 0.331289997 | 0.50556274 | 0.65528958 | 0.51228133 | 0.83070544 |
| ZFP36L1 | 157.380002 | -1.00335407 | 0.47335695 | -2.1196564 | 0.03403503 | 0.29210684 |
| ACTN1 | 10.0511249 | 0.519155089 | 0.56617927 | 0.91694471 | 0.35917161 | NA |
| DCAF5 | 56.9652602 | -0.015111688 | 0.33845755 | -0.0446487 | 0.96438733 | 0.98892249 |
| EXD2 | 19.0203766 | 0.552259661 | 0.38365636 | 1.43946438 | 0.15001899 | NA |
| ERH | 38.7484574 | -0.573472197 | 0.30266772 | -1.8947254 | 0.0581288 | 0.37650386 |
| SLC39A9 | 31.461022 | 0.156632034 | 0.26095529 | 0.60022556 | 0.54835592 | 0.84550378 |
| SUSD6 | 107.962696 | -0.526591971 | 0.40235669 | -1.308769 | 0.19061261 | 0.59329687 |
| SRSF5 | 120.250096 | 0.520468102 | 0.27700008 | 1.87894565 | 0.06025192 | 0.37947479 |
| AL357153.2 | 26.8685941 | 0.12523439 | 0.42300312 | 0.2960602 | 0.76718411 | NA |
| MED6 | 13.8146507 | 0.180115516 | 0.41854675 | 0.43033548 | 0.66695162 | NA |
| MAP3K9 | 4.96054 | -0.243498204 | 0.72840954 | -0.3342875 | 0.73816262 | NA |
| PCNX1 | 123.816004 | 0.251702763 | 0.29079978 | 0.86555348 | 0.38673508 | 0.75125852 |
| SIPA1L1 | 115.360037 | 0.866033673 | 0.36364198 | 2.38155585 | 0.01723968 | 0.21715567 |
| DPF3 | 7.66296769 | -0.025190233 | 0.81277062 | -0.030993 | 0.97527509 | NA |
| ZFYVE1 | 15.4530274 | -0.139054622 | 0.40554751 | -0.3428812 | 0.73168783 | NA |
| RBM25 | 66.9748152 | 0.278629454 | 0.26025579 | 1.07059847 | 0.28435001 | 0.67249058 |
| PSEN1 | 37.885621 | 0.096147421 | 0.27954349 | 0.34394441 | 0.73088809 | 0.91722837 |
| PAPLN | 4.1168293 | 0.000892905 | 0.81897529 | 0.00109027 | 0.99913009 | NA |
| NUMB | 95.9713231 | 0.360866519 | 0.18029963 | 2.00148226 | 0.04534044 | 0.3354231 |
| HEATR4 | 5.44800637 | -1.290582957 | 0.87065486 | -1.4823129 | 0.13825705 | NA |
| DNAL1 | 21.7175224 | -0.042581219 | 0.40945345 | -0.1039953 | 0.91717311 | NA |
| PNMA1 | 24.8790416 | -0.463202116 | 0.47159171 | -0.98221 | 0.32599638 | NA |
| ELMSAN1 | 18.5046143 | -0.114696347 | 0.4141723 | -0.2769291 | 0.78183458 | NA |
| AC005480.1 | 10.21345 | -0.347367794 | 0.68417162 | -0.5077203 | 0.61164954 | NA |
| COQ6 | 6.89491571 | 0.566498394 | 0.63163661 | 0.89687391 | 0.3697862 | NA |
| ENTPD5 | 14.0586412 | 0.650067877 | 0.63208153 | 1.02845575 | 0.3037355 | NA |
| BBOF1 | 18.8883271 | 0.42352878 | 0.41996921 | 1.00847579 | 0.31322611 | NA |
| ALDH6A1 | 20.2971778 | 0.232837569 | 0.36634029 | 0.63557728 | 0.52505198 | NA |
| LIN52 | 28.1833832 | 0.482708759 | 0.36201333 | 1.33340052 | 0.1824004 | NA |
| ABCD4 | 26.7253042 | 0.062981764 | 0.31107285 | 0.20246628 | 0.83955221 | NA |
| NPC2 | 76.5387557 | -0.68694933 | 0.4264669 | -1.6107917 | 0.10722514 | 0.49086361 |
| ISCA2 | 13.6892759 | -0.329369868 | 0.5774439 | -0.5703928 | 0.56841128 | NA |
| LTBP2 | 10.891944 | -0.906848724 | 0.77577833 | -1.1689534 | 0.2424224 | NA |
| AREL1 | 25.5503658 | 0.281945623 | 0.29240114 | 0.96424256 | 0.33492433 | NA |
| FCF1 | 11.150063 | -0.496590419 | 0.45302642 | -1.0961622 | 0.27300782 | NA |
| YLPM1 | 56.907615 | -0.114117439 | 0.30791932 | -0.3706082 | 0.71092934 | 0.91222063 |
| DLST | 16.4880777 | -0.36629092 | 0.37747039 | -0.9703832 | 0.33185553 | NA |
| EIF2B2 | 5.56886564 | -0.853648538 | 0.65962619 | -1.2941399 | 0.19561712 | NA |
| MLH3 | 26.3165032 | 0.217503505 | 0.38372909 | 0.56681527 | 0.5708397 | NA |
| ACYP1 | 13.4048642 | -0.028897881 | 0.43566606 | -0.0663303 | 0.94711482 | NA |
| NEK9 | 28.3772543 | 0.144993745 | 0.28543198 | 0.50798002 | 0.61146735 | NA |
| AL049780.2 | 11.5610516 | -1.064237356 | 0.79419723 | -1.3400164 | 0.18024 | NA |
| TMED10 | 85.5993498 | -0.005297842 | 0.32843285 | -0.0161307 | 0.98713015 | 0.99585746 |
| FOS | 14.8558802 | 0.879003323 | 0.80365419 | 1.09375816 | 0.27406106 | NA |
| JDP2 | 14.1468227 | 0.611630146 | 0.46498811 | 1.31536727 | 0.1883865 | NA |
| TTLL5 | 49.3921706 | 0.040487099 | 0.24453874 | 0.16556518 | 0.86849915 | 0.96089268 |
| ERG28 | 12.4222879 | -0.252362083 | 0.45510633 | -0.5545124 | 0.57922825 | NA |
| IFT43 | 36.2313161 | 0.144837206 | 0.26754114 | 0.54136424 | 0.58825655 | 0.86533208 |
| GPATCH2L | 57.535339 | 0.210545027 | 0.23064867 | 0.91283867 | 0.36132739 | 0.73240575 |
| ANGEL1 | 6.41430469 | -0.551748745 | 0.6092123 | -0.9056757 | 0.36510755 | NA |
| IRF2BPL | 46.7127134 | -0.694060829 | 0.46304479 | -1.4989065 | 0.1338979 | 0.52676032 |
| POMT2 | 13.7323683 | 0.61138681 | 0.49879867 | 1.2257186 | 0.22030459 | NA |
| TMED8 | 23.3641984 | 0.128512074 | 0.32504021 | 0.39537285 | 0.69256775 | NA |
| VIPAS39 | 10.5117276 | 0.182022043 | 0.40655192 | 0.44772152 | 0.65435419 | NA |
| AHSA1 | 14.4496708 | 0.296257587 | 0.46241306 | 0.64067738 | 0.52173231 | NA |
| SPTLC2 | 53.1642475 | -0.061477642 | 0.35276131 | -0.1742755 | 0.86164898 | 0.95951831 |
| ALKBH1 | 12.4466283 | 0.560451051 | 0.45467518 | 1.23264053 | 0.21770991 | NA |
| SLIRP | 21.019415 | -1.088294048 | 0.4242385 | -2.5652883 | 0.01030901 | NA |
| SNW1 | 21.8318992 | -0.085730057 | 0.3627038 | -0.2363638 | 0.81315037 | NA |
| ADCK1 | 8.60622243 | 0.616658469 | 0.50473341 | 1.22175085 | 0.22180186 | NA |
| NRXN3 | 48.5678293 | 2.182234581 | 1.03528601 | 2.10785672 | 0.03504338 | 0.29354459 |
| CEP128 | 24.3978128 | 0.038612179 | 0.49356436 | 0.0782313 | 0.93764407 | NA |
| GTF2A1 | 27.1446977 | 0.099024919 | 0.29960304 | 0.33052041 | 0.74100677 | NA |
| STON2 | 60.8186961 | 0.162690666 | 0.38148328 | 0.42646867 | 0.66976637 | 0.90305784 |
| SEL1L | 142.890542 | 0.274207943 | 0.33504092 | 0.8184312 | 0.41311101 | 0.76946282 |
| FLRT2 | 88.3321229 | -1.115378901 | 0.49491016 | -2.2536998 | 0.02421506 | 0.25196701 |
| GALC | 43.7662213 | 0.010810081 | 0.44238528 | 0.0244359 | 0.98050491 | 0.99453905 |
| SPATA7 | 35.4698059 | 0.411219757 | 0.33533783 | 1.22628503 | 0.22009143 | 0.62462904 |
| PTPN21 | 67.7740436 | 0.585236753 | 0.2580087 | 2.26828303 | 0.02331196 | 0.2486058 |
| ZC3H14 | 53.4447987 | 0.154958967 | 0.29021498 | 0.53394544 | 0.59337928 | 0.8654248 |
| EML5 | 8.74437336 | -0.382386576 | 0.50738301 | -0.7536448 | 0.45106252 | NA |
| TTC8 | 20.8579884 | 0.217207833 | 0.39074581 | 0.55588014 | 0.5782928 | NA |
| FOXN3 | 47.6181947 | 0.493223285 | 0.39688409 | 1.24273888 | 0.21396407 | 0.62255988 |
| EFCAB11 | 15.98089 | 0.3277876 | 0.42938067 | 0.76339627 | 0.4452271 | NA |
| TDP1 | 5.26997652 | -0.043127943 | 0.61435238 | -0.0702007 | 0.94403395 | NA |
| PSMC1 | 29.0081894 | -0.057020656 | 0.34418777 | -0.1656673 | 0.86841879 | 0.96089268 |
| NRDE2 | 18.5078803 | 0.139107996 | 0.44897761 | 0.30983281 | 0.7566881 | NA |
| CALM1 | 86.2681104 | -0.018836332 | 0.26729212 | -0.070471 | 0.94381882 | 0.98114803 |
| TTC7B | 23.0725988 | 0.618072716 | 0.42394335 | 1.45791345 | 0.1448644 | NA |
| RPS6KA5 | 15.2453055 | -0.52252834 | 0.47496027 | -1.1001517 | 0.27126604 | NA |
| DGLUCY | 36.4807177 | 0.212803003 | 0.50173734 | 0.42413228 | 0.67146935 | 0.90360971 |
| CCDC88C | 6.07503116 | 0.388433227 | 0.77921368 | 0.49849385 | 0.61813601 | NA |
| PPP4R3A | 50.936865 | 0.29572943 | 0.23810052 | 1.24203603 | 0.21422327 | 0.62255988 |
| TC2N | 7.07739142 | 0.720111823 | 0.85693635 | 0.84033292 | 0.40072175 | NA |
| TRIP11 | 63.2988724 | -0.043718223 | 0.2319885 | -0.18845 | 0.85052394 | 0.95532127 |
| ATXN3 | 48.146496 | 0.356117619 | 0.3327483 | 1.07023122 | 0.28451524 | 0.67249058 |
| NDUFB1 | 22.1342395 | 0.315553518 | 0.42517084 | 0.74218053 | 0.45797796 | NA |
| CPSF2 | 25.3878727 | -0.063077822 | 0.28878747 | -0.218423 | 0.82709955 | NA |
| RIN3 | 18.9429785 | -0.12271514 | 0.40782299 | -0.3009029 | 0.7634885 | NA |
| LGMN | 22.0171194 | 0.358029162 | 0.7252447 | 0.4936667 | 0.62154158 | NA |
| GOLGA5 | 20.2728479 | -0.45665101 | 0.36803454 | -1.240783 | 0.21468591 | NA |
| ITPK1 | 13.2603832 | -0.508596085 | 0.42179961 | -1.2057766 | 0.22790366 | NA |
| MOAP1 | 11.1728642 | -0.897571673 | 0.46700328 | -1.9219815 | 0.05460809 | NA |
| GON7 | 4.95987679 | -0.76302259 | 0.67490474 | -1.1305634 | 0.25823891 | NA |

| | | | | | | |
|---|---|---|---|---|---|---|
| UBR7 | 7.26452756 | 0.109890236 | 0.54781219 | 0.20059838 | 0.84101263 | NA |
| BTBD7 | 78.3192304 | 0.041858667 | 0.21983834 | 0.19040658 | 0.84899054 | 0.95532127 |
| UNC79 | 7.40871759 | 0.037191644 | 0.57039065 | 0.06520381 | 0.94801173 | NA |
| DDX24 | 77.8128222 | -0.89601182 | 0.2671812 | -3.3535737 | 0.00079775 | 0.04257475 |
| IFI27L1 | 6.15691764 | -0.210002329 | 0.79650634 | -0.2636543 | 0.79204631 | NA |
| IFI27 | 55.7616178 | -2.072118654 | 1.8328757 | -1.1305287 | 0.2582535 | 0.65588339 |
| IFI27L2 | 16.2603334 | -0.248095756 | 0.52124785 | -0.475965 | 0.63409928 | NA |
| SERPINA1 | 27.7611827 | -1.536107703 | 0.78798441 | -1.9494138 | 0.05124602 | NA |
| DICER1 | 63.9046486 | -0.236651495 | 0.23028467 | -1.0276476 | 0.30411561 | 0.68413362 |
| DICER1-AS1 | 10.2196213 | 0.06621887 | 0.64112912 | 0.10328477 | 0.91773697 | NA |
| CLMN | 21.8641183 | 0.390201987 | 0.66732299 | 0.58472733 | 0.55873108 | NA |
| SYNE3 | 5.6024087 | -0.276952765 | 0.62843491 | -0.4407024 | 0.65942847 | NA |
| GLRX5 | 22.8102586 | -0.291169198 | 0.40481353 | -0.7192674 | 0.47197615 | NA |
| C14orf132 | 10.8332468 | -0.796092041 | 0.9099466 | -0.8748778 | 0.38164042 | NA |
| ATG2B | 44.8135559 | 0.466463301 | 0.33227757 | 1.40383627 | 0.16036761 | 0.56283048 |
| GSKIP | 6.56663077 | -0.40856824 | 0.56048271 | -0.7289578 | 0.46602748 | NA |
| PAPOLA | 92.9392111 | 0.381210992 | 0.2577503 | 1.47899342 | 0.13914207 | 0.53519929 |
| VRK1 | 10.401638 | 0.569502131 | 0.54625336 | 1.04256042 | 0.29715193 | NA |
| SETD3 | 31.3274456 | -0.020140128 | 0.29276882 | -0.0687919 | 0.94515525 | 0.98220592 |
| CCNK | 19.0074678 | -0.21003299 | 0.3499924 | -0.6001073 | 0.54843473 | NA |
| CCDC85C | 101.756829 | -0.595378966 | 0.52051043 | -1.1438368 | 0.25269134 | 0.65239726 |
| EML1 | 12.584613 | -0.06294484 | 0.54101629 | -0.1163456 | 0.90737868 | NA |
| EVL | 134.837974 | -1.154059121 | 0.50396863 | -2.2899424 | 0.02202466 | 0.24297341 |
| YY1 | 81.8214751 | -0.240195992 | 0.27734758 | -0.8660468 | 0.38646448 | 0.75125852 |
| SLC25A29 | 18.3201188 | -0.454471906 | 0.59251114 | -0.7670268 | 0.4430656 | NA |
| WARS | 35.1880988 | 0.180883898 | 0.82699311 | 0.2187248 | 0.82686443 | 0.949142 |
| WDR25 | 13.9546079 | -0.345821557 | 0.49559696 | -0.6977879 | 0.48530985 | NA |
| PPP2R5C | 55.8684291 | 0.038484065 | 0.23727667 | 0.16219068 | 0.8711557 | 0.96207879 |
| DYNC1H1 | 70.7466094 | -0.103302762 | 0.25775846 | -0.4007735 | 0.68858689 | 0.90557575 |
| HSP90AA1 | 162.340724 | -0.392516452 | 0.30176053 | -1.3007548 | 0.1933424 | 0.59771096 |
| WDR20 | 49.2694657 | -0.013758634 | 0.28822665 | -0.0477355 | 0.96192707 | 0.98892249 |
| MOK | 18.2462064 | 0.263751511 | 0.40051502 | 0.65853088 | 0.51019706 | NA |
| ZNF839 | 7.48703451 | -0.535855211 | 0.5378435 | -0.9963032 | 0.31910284 | NA |
| CINP | 7.48628623 | -0.299466824 | 0.56202355 | -0.5328368 | 0.59414656 | NA |
| TECPR2 | 26.8842725 | 0.117553522 | 0.41615766 | 0.28247353 | 0.77758044 | NA |
| ANKRD9 | 9.7574791 | 0.614293461 | 0.47263329 | 1.29972534 | 0.19369512 | NA |
| RCOR1 | 21.6244716 | 0.079983475 | 0.3657563 | 0.21867969 | 0.82689957 | NA |
| TRAF3 | 44.3704807 | 0.143526992 | 0.31290093 | 0.45869787 | 0.64645114 | 0.89244583 |
| CDC42BPB | 63.4798192 | 0.041311148 | 0.28109653 | 0.14696427 | 0.88316022 | 0.96526488 |
| TNFAIP2 | 23.4427358 | -0.037564865 | 0.58068181 | -0.064691 | 0.94842006 | NA |
| EIF5 | 53.0678212 | -0.737996574 | 0.36541397 | -2.0196179 | 0.04342304 | 0.33029013 |
| MARK3 | 136.786883 | 0.394369085 | 0.2229138 | 1.7691551 | 0.076868 | 0.42346132 |
| CKB | 57.4593055 | -0.047688702 | 0.52604567 | -0.0906551 | 0.92776668 | 0.97775374 |
| BAG5 | 20.3938116 | -0.443208515 | 0.39157778 | -1.1318531 | 0.2576962 | NA |
| KLC1 | 53.5242702 | 0.070947407 | 0.3867156 | 0.18346146 | 0.85443596 | 0.95576179 |
| ZFYVE21 | 26.8111428 | -0.573339801 | 0.42821414 | -1.3389091 | 0.18060027 | NA |
| PPP1R13B | 25.6484079 | -0.368874522 | 0.39413781 | -0.9359024 | 0.34932345 | NA |
| ATP5MPL | 29.7040568 | -0.064365819 | 0.43205809 | -0.1489749 | 0.88157343 | 0.96457022 |
| INF2 | 112.85301 | 0.266949368 | 0.4533082 | 0.58889155 | 0.55593403 | 0.85121496 |
| SIVA1 | 17.3531904 | -0.694069053 | 0.46066589 | -1.5066647 | 0.13189662 | NA |
| AKT1 | 18.6697107 | -1.176289487 | 0.49363409 | -2.3829179 | 0.01717603 | NA |
| CEP170B | 11.1583241 | 0.178057176 | 0.49484399 | 0.35982488 | 0.7189781 | NA |
| NUDT14 | 4.06937886 | -1.245134554 | 0.97344428 | -1.279102 | 0.20086113 | NA |
| BRF1 | 12.948573 | 0.230700883 | 0.36856617 | 0.62594156 | 0.53135327 | NA |
| BTBD6 | 12.1703403 | 0.436675697 | 0.43380804 | 1.00661043 | 0.31412202 | NA |
| PACS2 | 25.4387257 | -0.176259503 | 0.34301428 | -0.5138547 | 0.60735357 | NA |
| MTA1 | 17.0197728 | 0.454631422 | 0.46072708 | 0.98676948 | 0.32375566 | NA |
| CRIP2 | 58.1490432 | 0.449945543 | 0.58284538 | 0.77198097 | 0.4401257 | 0.78399202 |
| NIPA1 | 18.3175011 | 0.601651445 | 0.38075044 | 1.5801727 | 0.11406732 | NA |
| NIPA2 | 20.8370146 | -0.143178248 | 0.41061796 | -0.3486897 | 0.72732228 | NA |
| CYFIP1 | 42.8190748 | -0.16205637 | 0.30868858 | -0.5249834 | 0.59959476 | 0.86882737 |
| TUBGCP5 | 12.3827635 | -0.469268012 | 0.40490685 | -1.158953 | 0.24647534 | NA |
| NDN | 41.3365963 | -0.416872632 | 0.67554713 | -0.6170889 | 0.53717609 | 0.83677705 |
| AC124312.1 | 90.2394424 | 0.377420813 | 0.28946053 | 1.3038766 | 0.19227566 | 0.59532067 |
| PWAR6 | 22.5908457 | 0.851645001 | 0.43130981 | 1.97455512 | 0.04831866 | NA |
| UBE3A | 82.250035 | 0.414386739 | 0.22213906 | 1.86543841 | 0.06211998 | 0.38512471 |
| ATP10A | 592.81694 | 0.179951257 | 0.43231751 | 0.4162479 | 0.67722861 | 0.90499319 |
| HERC2 | 77.1097261 | 0.282801701 | 0.22292952 | 1.26857001 | 0.20459447 | 0.6137834 |
| TJP1 | 619.489146 | 0.324836146 | 0.27217908 | 1.19346479 | 0.23268739 | 0.63935677 |
| FAN1 | 15.3514451 | -0.863543452 | 0.40908398 | -2.1109197 | 0.03477921 | NA |
| MTMR10 | 138.484577 | 0.532672583 | 0.47517275 | 1.12100826 | 0.26228434 | 0.65790062 |
| KLF13 | 34.9055455 | 0.002683532 | 0.38754461 | 0.00692445 | 0.99447513 | 0.99867379 |
| OTUD7A | 21.8831877 | 0.851720438 | 0.47877544 | 1.77895598 | 0.07524698 | NA |
| LINC02256 | 49.4865152 | -0.265831598 | 0.40413821 | -0.657774 | 0.51068338 | 0.82958308 |
| AC123768.3 | 9.31538006 | -0.047809903 | 0.53214173 | -0.0898443 | 0.92841094 | NA |
| GOLGA8N | 5.37136043 | 0.70619396 | 0.54962039 | 1.28487584 | 0.19883567 | NA |
| FMN1 | 10.2587473 | 0.479537553 | 0.61204244 | 0.78350375 | 0.43333135 | NA |
| RYR3 | 393.80975 | -1.313200955 | 0.52338459 | -2.5090554 | 0.01210545 | 0.183894 |
| AVEN | 28.5261963 | 0.460754442 | 0.42891473 | 1.07423319 | 0.28271818 | 0.67176622 |
| EMC7 | 33.250589 | -0.587284553 | 0.47361712 | -1.2399986 | 0.21497592 | 0.62305077 |
| KATNBL1 | 58.9743563 | 0.165185042 | 0.24987074 | 0.66108198 | 0.50855974 | 0.82740405 |
| EMC4 | 14.6115064 | 0.013997591 | 0.41061725 | 0.03408915 | 0.97280606 | NA |
| SLC12A6 | 23.8246964 | 0.761391367 | 0.38197194 | 1.99331755 | 0.04622669 | NA |
| NOP10 | 28.9304883 | -0.607298393 | 0.40391015 | -1.5035482 | 0.13269773 | 0.52613957 |
| LPCAT4 | 6.00385275 | 0.358807513 | 0.67174289 | 0.53414412 | 0.59324183 | NA |
| GOLGA8A | 27.1850693 | 0.384561177 | 0.48901064 | 0.78640656 | 0.43162934 | NA |
| GOLGA8B | 43.7040357 | 0.007160512 | 0.40496585 | 0.01768177 | 0.98589273 | 0.99585746 |
| AQR | 36.1108239 | 0.189632276 | 0.28983671 | 0.65427281 | 0.51293606 | 0.83070544 |
| ZNF770 | 20.7027769 | 0.066118628 | 0.42027109 | 0.15732376 | 0.8749897 | NA |
| AC021231.1 | 33.6404128 | -0.064852978 | 0.42639914 | -0.1520945 | 0.87911238 | 0.96423929 |
| DPH6 | 90.2799733 | -0.028628704 | 0.35427176 | -0.08081 | 0.93559305 | 0.97832857 |
| DPH6-DT | 40.1763004 | 1.465938675 | 0.64559607 | 2.27067473 | 0.02316668 | 0.24858211 |
| C15orf41 | 23.9322013 | 1.087949131 | 0.44961152 | 2.41975366 | 0.01553102 | NA |
| MEIS2 | 32.5971075 | 1.03053292 | 0.39683115 | 2.59690528 | 0.00940679 | 0.15375026 |
| SPRED1 | 54.6344313 | -0.054360866 | 0.35893749 | -0.1514494 | 0.87962123 | 0.96423929 |
| FAM98B | 12.9114356 | -0.307302962 | 0.4266192 | -0.7203214 | 0.4713271 | NA |
| RASGRP1 | 5.95396753 | -0.122399604 | 0.76053745 | -0.1609383 | 0.872142 | NA |
| C15orf53 | 5.36430456 | -0.730071334 | 0.75718163 | -0.9641958 | 0.33494778 | NA |
| AC013652.1 | 54.3731286 | 0.036121835 | 0.58470379 | 0.06177801 | 0.95073962 | 0.98520702 |
| THBS1 | 87.6116443 | 0.134188734 | 0.37217037 | 0.36055728 | 0.71843043 | 0.91472065 |
| FSIP1 | 14.8093132 | -0.418017457 | 0.54007648 | -0.7739968 | 0.4389327 | NA |
| EIF2AK4 | 28.1207142 | -0.00023767 | 0.33012251 | -0.0007199 | 0.99942557 | NA |
| SRP14 | 161.091043 | -0.862593735 | 0.46737083 | -1.8456302 | 0.06494591 | 0.39493348 |
| CCDC9B | 5.08498931 | -0.825941955 | 0.72721514 | -1.1357601 | 0.25605697 | NA |
| IVD | 26.2801805 | -0.230083691 | 0.33789701 | -0.6809285 | 0.49591676 | NA |
| BAHD1 | 8.71439007 | -0.021043932 | 0.45005476 | -0.0467586 | 0.96270562 | NA |
| CHST14 | 5.90376408 | 0.772041535 | 0.66291395 | 1.16461802 | 0.24417365 | NA |
| CCDC32 | 20.4547695 | -0.171578668 | 0.43970218 | -0.3902156 | 0.6963771 | NA |
| RMDN3 | 14.7983819 | -0.094753131 | 0.39489948 | -0.2399424 | 0.81037491 | NA |
| GCHFR | 7.13746036 | -1.206185777 | 0.82585341 | -1.4605325 | 0.14414377 | NA |
| DNAJC17 | 14.8208671 | 0.165688483 | 0.38166567 | 0.43411943 | 0.66420171 | NA |
| ZFYVE19 | 6.4471133 | -0.264284457 | 0.63784133 | -0.414342 | 0.67862365 | NA |
| SPINT1 | 12.8378954 | 0.427824187 | 0.56991204 | 0.75068459 | 0.4528425 | NA |

| | | | | | | |
|---|---|---|---|---|---|---|
| VPS18 | 9.07976024 | -0.238630587 | 0.50887986 | -0.4689331 | 0.63911749 | NA |
| INO80 | 39.5986768 | 0.16581546 | 0.38664942 | 0.42885221 | 0.66803078 | 0.90137487 |
| CHP1 | 26.5916546 | 0.012390989 | 0.32877665 | 0.03768817 | 0.96993631 | NA |
| NDUFAF1 | 6.29533127 | 0.393233905 | 0.64897316 | 0.60593246 | 0.5445596 | NA |
| RTF1 | 39.6546569 | -0.104061982 | 0.30000168 | -0.3468713 | 0.72868799 | 0.91700494 |
| RPAP1 | 6.22619286 | 0.232729855 | 0.5159312 | 0.451087 | 0.65192685 | NA |
| TYRO3 | 326.865688 | -0.555103052 | 0.32408317 | -1.7128414 | 0.08674173 | 0.4476138 |
| MGA | 42.6985976 | -0.015593338 | 0.3299844 | -0.0472548 | 0.96231017 | 0.98892249 |
| MAPKBP1 | 16.2001932 | -0.182988504 | 0.40154419 | -0.455712 | 0.64859709 | NA |
| EHD4 | 60.4463243 | 0.65245389 | 0.33320228 | 1.95813152 | 0.05021458 | 0.35115577 |
| VPS39 | 24.7369025 | 0.50881304 | 0.3114109 | 1.63389607 | 0.10228067 | NA |
| TMEM87A | 45.9800289 | -0.069746678 | 0.28902706 | -0.2413154 | 0.80931068 | 0.94346479 |
| GANC | 16.5454016 | 0.077657144 | 0.53374574 | 0.14549464 | 0.88432035 | NA |
| ZNF106 | 51.8298699 | 0.671013267 | 0.41116825 | 1.63196761 | 0.10268631 | 0.48317363 |
| SNAP23 | 33.0882495 | 0.353370309 | 0.38845596 | 0.90967921 | 0.36299171 | 0.73259935 |
| LRRC57 | 14.8431045 | 0.138842293 | 0.40465295 | 0.3431145 | 0.73151232 | NA |
| HAUS2 | 10.3925948 | 0.196578759 | 0.5230124 | 0.3758587 | 0.70702195 | NA |
| STARD9 | 10.8450577 | 0.04183817 | 0.49797014 | 0.08401743 | 0.93304258 | NA |
| TTBK2 | 49.4190254 | 0.04897959 | 0.2956942 | 0.16564272 | 0.86843813 | 0.96089268 |
| UBR1 | 51.4410506 | -0.020447006 | 0.25186233 | -0.0811833 | 0.93529621 | 0.97832857 |
| AC068724.3 | 12.3950729 | -0.362132297 | 0.54504663 | -0.6644061 | 0.50643044 | NA |
| TMEM62 | 20.6318535 | 0.469345424 | 0.56589971 | 0.82937916 | 0.40688989 | NA |
| CCNDBP1 | 19.4234174 | -0.028396405 | 0.44070962 | -0.0644334 | 0.94862516 | NA |
| ADAL | 9.15586441 | 0.313703584 | 0.52973304 | 0.59219184 | 0.55372214 | NA |
| ZSCAN29 | 4.25519368 | -0.807506852 | 0.69762126 | -1.1575147 | 0.24706215 | NA |
| TUBGCP4 | 10.1791253 | -0.848819776 | 0.47252304 | -1.7963564 | 0.07243786 | NA |
| TP53BP1 | 48.126039 | 0.212450592 | 0.23830772 | 0.89149689 | 0.37266266 | 0.7407542 |
| MAP1A | 8.89413159 | -0.339487132 | 0.71549767 | -0.4744769 | 0.63515985 | NA |
| PPIP5K1 | 12.3580351 | 0.754906818 | 0.47956778 | 1.57413997 | 0.11545509 | NA |
| CATSPER2 | 15.1273703 | 0.443183559 | 0.51948636 | 0.85311875 | 0.39359345 | NA |
| PDIA3 | 128.123653 | -0.055779454 | 0.43425686 | -0.1284481 | 0.8977944 | 0.97097328 |
| SERF2 | 218.409035 | -0.261377771 | 0.40980234 | -0.6378143 | 0.5235946 | 0.83178884 |
| MFAP1 | 13.152698 | -0.424719616 | 0.43654926 | -0.9729019 | 0.33060205 | NA |
| FRMD5 | 6.9791657 | -0.22486877 | 0.51419434 | -0.4373225 | 0.66187746 | NA |
| CASC4 | 108.636578 | 0.202025657 | 0.27915911 | 0.72369357 | 0.46925388 | 0.80448595 |
| AC025043.1 | 4.58760349 | -0.463027119 | 0.66616775 | -0.6950608 | 0.48701717 | NA |
| CTDSPL2 | 45.9333941 | 0.022387 | 0.27479951 | 0.08146666 | 0.93507083 | 0.97832857 |
| EIF3J-DT | 16.3361168 | -0.74255972 | 0.46172785 | -1.6082195 | 0.10778712 | NA |
| EIF3J | 43.9898084 | -0.804451159 | 0.29617382 | -2.7161454 | 0.00660469 | 0.12888932 |
| SPG11 | 57.0025059 | -0.078315333 | 0.25904385 | -0.3023246 | 0.76240461 | 0.92336765 |
| B2M | 930.090397 | -0.362928211 | 0.62462348 | -0.5810352 | 0.56121675 | 0.85386276 |
| TRIM69 | 6.80299668 | -0.379415098 | 0.9290582 | -0.4083868 | 0.68298973 | NA |
| SORD | 7.59299148 | 0.361813959 | 0.84529365 | 0.42803345 | 0.66862677 | NA |
| DUOX1 | 9.08821154 | -0.6089644 | 0.68413372 | -0.8901248 | 0.37339887 | NA |
| GATM | 9.16108085 | 1.508302245 | 0.60567852 | 2.49026866 | 0.01276466 | NA |
| SPATA5L1 | 4.76646556 | -0.889280012 | 0.72712927 | -1.2230013 | 0.22132922 | NA |
| SLC30A4 | 11.980175 | 0.451423833 | 0.40780308 | 1.10696524 | 0.26830896 | NA |
| SQOR | 67.2113865 | -1.374938538 | 0.425141 | -3.2340765 | 0.00122037 | 0.05439352 |
| SEMA6D | 23.7222331 | 0.003518187 | 0.39138453 | 0.00898908 | 0.99282785 | NA |
| MYEF2 | 55.9858325 | 0.233064783 | 0.28796343 | 0.80935548 | 0.4183107 | 0.77290571 |
| SLC12A1 | 67.2814925 | 3.618279167 | 1.28852729 | 2.80807337 | 0.00498389 | 0.11177304 |
| DUT | 28.3335532 | -0.397236239 | 0.41524976 | -0.95662 | 0.33875907 | NA |
| CEP152 | 7.89150605 | 0.67617897 | 0.6588374 | 1.02632147 | 0.30474008 | NA |
| EID1 | 123.757742 | -0.905473187 | 0.32863297 | -2.7552719 | 0.00586434 | 0.1213559 |
| SECISBP2L | 75.5401921 | 0.062759653 | 0.31523378 | 0.19908924 | 0.84219294 | 0.95443827 |
| COPS2 | 23.2106525 | -0.213668592 | 0.39354391 | -0.5429346 | 0.58717486 | NA |
| GALK2 | 44.634349 | 0.221819272 | 0.32641013 | 0.67957227 | 0.49677533 | 0.8234249 |
| FAM227B | 16.1905955 | -0.494458363 | 0.5493251 | -0.9001197 | 0.36805654 | NA |
| DTWD1 | 29.1396581 | 0.286073565 | 0.40256244 | 0.71063154 | 0.47731259 | 0.80867998 |
| SLC27A2 | 9.32091653 | 0.974451126 | 1.08320276 | 0.89960177 | 0.36833221 | NA |
| GABPB1 | 31.1161565 | -0.028906773 | 0.35103907 | -0.0823463 | 0.93437133 | 0.97832857 |
| GABPB1-AS1 | 37.1980299 | -0.264395059 | 0.64675542 | -0.4088022 | 0.6826848 | 0.90499319 |
| USP8 | 43.7229657 | -0.100751667 | 0.26036921 | -0.3869569 | 0.6987881 | 0.90729979 |
| TRPM7 | 79.0797714 | 0.297704699 | 0.29387875 | 1.01301881 | 0.31105118 | 0.69098737 |
| SPPL2A | 76.5790129 | 0.132182765 | 0.24132072 | 0.54774727 | 0.58386545 | 0.86175324 |
| AC021752.1 | 6.41699276 | -0.254779027 | 0.80186263 | -0.317734 | 0.75068671 | NA |
| AP4E1 | 25.0773672 | -0.214078949 | 0.35925262 | -0.5959009 | 0.55124145 | NA |
| DMXL2 | 41.9883592 | 0.77283559 | 0.43342352 | 1.78309565 | 0.07457073 | 0.41891814 |
| LYSMD2 | 8.66412838 | -0.854841753 | 0.76567235 | -1.116459 | 0.26422573 | NA |
| TMOD2 | 31.8899933 | -0.4010028 | 0.50082817 | -0.8006794 | 0.42331727 | 0.77656032 |
| TMOD3 | 148.853171 | -0.153448821 | 0.22230336 | -0.6902677 | 0.49002588 | 0.81825647 |
| AC090971.2 | 6.62744475 | -0.3529674 | 0.61635841 | -0.5726658 | 0.56687098 | NA |
| LEO1 | 6.46548453 | -0.749960019 | 0.62763971 | -1.1948894 | 0.23213026 | NA |
| MAPK6 | 28.1479421 | -0.136503894 | 0.35296778 | -0.3867319 | 0.69895471 | NA |
| GNB5 | 24.4020285 | -0.080527684 | 0.33111303 | -0.243203 | 0.80784814 | NA |
| CERNA1 | 4.29611103 | -0.56356791 | 0.85839895 | -0.6565338 | 0.51148074 | NA |
| MYO5C | 57.2033839 | 0.360620185 | 0.25975627 | 1.38830216 | 0.16504505 | 0.56634749 |
| MYO5A | 40.2355703 | 0.488344965 | 0.24504311 | 1.99289412 | 0.04627304 | 0.33756018 |
| ARPP19 | 64.9615776 | -0.191918482 | 0.2938595 | -0.6530961 | 0.51369435 | 0.83070544 |
| FAM214A | 58.1673952 | 0.386422573 | 0.29041755 | 1.33057584 | 0.18332862 | 0.58584259 |
| WDR72 | 45.3674407 | 1.861239045 | 1.23658143 | 1.5051488 | 0.13228583 | 0.52613957 |
| RSL24D1 | 20.5991876 | -0.7883162 | 0.36882129 | -2.1373934 | 0.03256601 | NA |
| RAB27A | 21.640566 | -0.816309485 | 0.57562736 | -1.4181214 | 0.15615532 | NA |
| PIGB | 17.8019035 | 0.256051474 | 0.4896175 | 0.52296226 | 0.60100052 | NA |
| CCPG1 | 93.114846 | -0.298093614 | 0.19921507 | -1.4963407 | 0.1345649 | 0.52779404 |
| C15orf65 | 4.28622242 | -0.081081389 | 0.84395152 | -0.0960735 | 0.92346219 | NA |
| DNAAF4 | 18.4708493 | 0.269408353 | 0.36034405 | 0.74764203 | 0.45467611 | NA |
| PYGO1 | 16.7263833 | 0.399715979 | 0.51258546 | 0.77980359 | 0.43550649 | NA |
| PRTG | 7.18956489 | -0.780264477 | 0.87906316 | -0.8876091 | 0.37475105 | NA |
| NEDD4 | 24.7689599 | 0.321862913 | 0.34664268 | 0.92851496 | 0.35314051 | NA |
| RFX7 | 57.3264735 | 0.446040158 | 0.24293681 | 1.83603366 | 0.06635268 | 0.39989073 |
| TEX9 | 28.4064272 | 0.356646313 | 0.2969956 | 1.20084712 | 0.22981051 | NA |
| ZNF280D | 38.9678243 | 0.545821855 | 0.2813989 | 1.93967304 | 0.05241944 | 0.35808347 |
| AC090517.4 | 32.8140884 | -0.167113274 | 0.44276774 | -0.3774288 | 0.705855 | 0.91002794 |
| TCF12 | 188.15933 | 0.158560991 | 0.20735523 | 0.76468287 | 0.44446041 | 0.78741638 |
| CGNL1 | 243.589387 | 0.382304902 | 0.25853426 | 1.47873979 | 0.13920987 | 0.53519929 |
| POLR2M | 10.3323235 | 0.223904673 | 0.53979068 | 0.41479907 | 0.678289 | NA |
| ALDH1A2 | 32.379633 | 0.265276702 | 0.65882517 | 0.40265113 | 0.68720489 | 0.90499319 |
| ADAM10 | 67.3619363 | -0.013570224 | 0.31508444 | -0.0430685 | 0.9656469 | 0.98892249 |
| MINDY2 | 62.6733273 | -0.257160996 | 0.29497027 | -0.8718201 | 0.38330655 | 0.74924885 |
| RNF111 | 58.9872813 | -0.019629851 | 0.2704186 | -0.0725906 | 0.9421319 | 0.98114803 |
| SLTM | 76.6962617 | 0.077891965 | 0.26439135 | 0.2946086 | 0.76829289 | 0.92555901 |
| MYO1E | 862.285106 | 0.344774188 | 0.32560332 | 1.058878 | 0.28965534 | 0.67746193 |
| FAM81A | 68.0736669 | 1.046978568 | 0.49444886 | 2.11746582 | 0.03422033 | 0.29210684 |
| GTF2A2 | 79.1971002 | 0.087187586 | 0.41083724 | 0.21221929 | 0.83193596 | 0.95050875 |
| BNIP2 | 46.7598688 | 0.071991126 | 0.27662932 | 0.26024402 | 0.79467555 | 0.93934998 |
| ANXA2 | 421.64187 | 0.174452393 | 0.4287198 | 0.40691471 | 0.68407063 | 0.90499319 |
| ICE2 | 47.2861508 | -0.11874127 | 0.28744297 | -0.4130951 | 0.67953697 | 0.90499319 |
| RORA-AS1 | 16.1812896 | 0.383105226 | 0.64327019 | 0.59555881 | 0.55147 | NA |
| RORA | 236.689875 | 0.848655095 | 0.5108356 | 1.66130765 | 0.09665167 | 0.47093115 |
| AC012404.1 | 5.22880755 | -1.209016536 | 0.89531466 | -1.3503817 | 0.17689358 | NA |
| AC009554.1 | 24.1615682 | -0.150303164 | 0.65653747 | -0.2289331 | 0.81892091 | NA |
| VPS13C | 160.688422 | 0.515530483 | 0.22887141 | 2.25248963 | 0.02429135 | 0.25196701 |

| | | | | | | |
|---|---|---|---|---|---|---|
| TLN2 | 22.1933138 | 1.429021556 | 0.58975023 | 2.42309622 | 0.01538885 | NA |
| TPM1 | 60.9061329 | 1.107056032 | 0.68570106 | 1.61448786 | 0.10642165 | 0.49050705 |
| LACTB | 10.7867956 | -0.116416439 | 0.42656176 | -0.2729181 | 0.78491615 | NA |
| RPS27L | 35.1921767 | -0.666001114 | 0.48893127 | -1.3621569 | 0.17314837 | 0.57564735 |
| RAB8B | 29.6729843 | 0.419037525 | 0.37666878 | 1.11248275 | 0.26593064 | 0.66013596 |
| APH1B | 18.608975 | -0.434197026 | 0.44507523 | -0.9755587 | 0.32928318 | NA |
| CA12 | 32.0702629 | -0.764156045 | 0.55231743 | -1.3835451 | 0.1664978 | 0.56878882 |
| USP3 | 70.2457779 | -0.121362933 | 0.26841497 | -0.4521467 | 0.65116333 | 0.89412414 |
| USP3-AS1 | 10.6261586 | -0.787892987 | 0.69559742 | -1.1326853 | 0.25734642 | NA |
| HERC1 | 67.3318437 | 0.058415401 | 0.24308404 | 0.24030949 | 0.81009034 | 0.94363194 |
| FAM96A | 9.58990017 | 0.102458272 | 0.53221062 | 0.19251452 | 0.8473392 | NA |
| SNX1 | 48.2301158 | 0.248459467 | 0.27217083 | 0.91288059 | 0.36130535 | 0.73240575 |
| PPIB | 89.6979012 | -0.863669046 | 0.54090704 | -1.5967051 | 0.11033145 | 0.49667521 |
| CSNK1G1 | 12.5399897 | -0.124836757 | 0.44087136 | -0.2831591 | 0.77705485 | NA |
| TRIP4 | 24.5269593 | 0.368057501 | 0.34892073 | 1.0548456 | 0.29149595 | NA |
| ZNF609 | 226.265635 | 0.223383377 | 0.22328991 | 1.00041861 | 0.31710797 | 0.69424851 |
| OAZ2 | 33.0366991 | -0.669381976 | 0.45481978 | -1.4717521 | 0.14108783 | 0.53597868 |
| SPG21 | 21.4066934 | -0.086659851 | 0.35664498 | -0.2429863 | 0.808016 | NA |
| MTFMT | 9.43409728 | -0.219460662 | 0.45029042 | -0.4873758 | 0.62599203 | NA |
| UBAP1L | 8.85575002 | 0.913879936 | 0.56603771 | 1.61452129 | 0.1064144 | NA |
| PDCD7 | 15.9449851 | -0.148348556 | 0.44202684 | -0.3356098 | 0.73716511 | NA |
| CLPX | 25.1710457 | 0.180703266 | 0.3265159 | 0.55342869 | 0.5799699 | NA |
| PARP16 | 7.78279665 | -0.194850086 | 0.54646427 | -0.3565651 | 0.72141741 | NA |
| DPP8 | 46.2024829 | 0.348665845 | 0.35411066 | 0.98462398 | 0.3248088 | 0.69623471 |
| HACD3 | 34.075018 | 0.005489682 | 0.38635755 | 0.01420881 | 0.98866339 | 0.99651342 |
| INTS14 | 9.16774213 | -0.120503576 | 0.49296618 | -0.2444459 | 0.80688547 | NA |
| SLC24A1 | 4.78148819 | -0.823952291 | 0.84239723 | -0.9781042 | 0.32802278 | NA |
| DENND4A | 90.690337 | 0.176116595 | 0.25254308 | 0.69737249 | 0.48556971 | 0.81450403 |
| RAB11A | 64.6302762 | -0.128928472 | 0.29833997 | -0.4321529 | 0.66563031 | 0.89968484 |
| MEGF11 | 6.88478965 | -3.564900029 | 1.21651088 | -2.93043 | 0.00338493 | NA |
| DIS3L | 21.2827209 | 0.210980137 | 0.33355158 | 0.63252626 | 0.52704305 | NA |
| TIPIN | 7.70073912 | 0.718718823 | 0.67327382 | 1.06749855 | 0.28574677 | NA |
| MAP2K1 | 20.0850915 | 0.212332872 | 0.37262976 | 0.56982263 | 0.56879801 | NA |
| SNAPC5 | 11.2956331 | -0.938495472 | 0.50249737 | -1.8676625 | 0.06180913 | NA |
| RPL4 | 53.7931453 | -0.621637259 | 0.42852127 | -1.4506567 | 0.14687547 | 0.54635866 |
| SMAD6 | 91.6478452 | 0.155491296 | 0.48000325 | 0.32393801 | 0.74598497 | 0.92250642 |
| AC093334.1 | 7.03262775 | -0.140818994 | 0.72388133 | -0.1945333 | 0.84575836 | NA |
| AC087482.1 | 9.79262762 | 0.124671321 | 0.80489788 | 0.15489086 | 0.87690736 | NA |
| SMAD3 | 54.969002 | -0.126134153 | 0.43299107 | -0.2913089 | 0.77081508 | 0.92634844 |
| AAGAB | 40.3301711 | -0.118303764 | 0.27350033 | -0.4325544 | 0.66533855 | 0.89968484 |
| IQCH | 32.0065142 | 0.251657264 | 0.47345344 | 0.5315354 | 0.59504782 | 0.86653654 |
| IQCH-AS1 | 13.2136771 | 0.358699493 | 0.52403905 | 0.68449 | 0.4936658 | NA |
| C15orf61 | 9.7401047 | -0.200909829 | 0.55323581 | -0.3631541 | 0.71648981 | NA |
| MAP2K5 | 59.9999856 | 0.536474879 | 0.2527761 | 2.12233226 | 0.03380985 | 0.29177178 |
| PIAS1 | 124.090439 | 0.483097269 | 0.23054323 | 2.0954737 | 0.0361289 | 0.29723902 |
| FEM1B | 32.7141066 | 0.129173045 | 0.31223598 | 0.41370326 | 0.67909144 | 0.90499319 |
| CORO2B | 178.180252 | -1.421843054 | 0.43042128 | -3.3033754 | 0.00095528 | 0.04967478 |
| ANP32A | 26.5207617 | -0.68724278 | 0.29695088 | -2.3143315 | 0.02064955 | NA |
| SPESP1 | 3.54398286 | -0.306387705 | 0.70940017 | -0.4318969 | 0.66581638 | NA |
| GLCE | 28.2621205 | -1.074593922 | 0.49102933 | -2.1884516 | 0.02863672 | NA |
| PAQR5 | 15.9937561 | 0.277507918 | 0.55292223 | 0.50189322 | 0.61574264 | NA |
| RPLP1 | 138.779677 | -0.357324339 | 0.47306994 | -0.7553309 | 0.45005047 | 0.79126774 |
| DRAIC | 6.74632922 | 0.543057185 | 0.62623238 | 0.86718158 | 0.38584252 | NA |
| TLE3 | 6.39218323 | -2.364353901 | 0.91441097 | -2.5856579 | 0.00971934 | NA |
| UACA | 329.479932 | 0.756056889 | 0.3833417 | 1.97227928 | 0.04857773 | 0.34339091 |
| LARP6 | 4.65441088 | -0.590065132 | 0.62844296 | -0.9389319 | 0.34776572 | NA |
| LRRC49 | 9.80504047 | 0.257250791 | 0.49236488 | 0.52247998 | 0.60133618 | NA |
| THSD4 | 14.3529657 | 1.114556401 | 0.71158028 | 1.56631154 | 0.1172757 | NA |
| THSD4-AS1 | 49.8256558 | -1.409834623 | 0.35330471 | -3.9904212 | 6.60E-05 | 0.00764336 |
| MYO9A | 159.568699 | 0.276662785 | 0.24720535 | 1.11916182 | 0.26307111 | 0.6581333 |
| SENP8 | 6.2875388 | 0.091975766 | 0.69654593 | 0.13204552 | 0.89494829 | NA |
| PKM | 82.3182831 | -0.773505497 | 0.39717855 | -1.9475007 | 0.05147474 | 0.35507065 |
| PARP6 | 17.2675363 | -0.246966099 | 0.39877888 | -0.6193059 | 0.53571488 | NA |
| AC009690.2 | 5.17506907 | -0.987424057 | 0.84386467 | -1.1701213 | 0.24195214 | NA |
| AC009690.1 | 23.8030589 | -0.514921708 | 0.44368999 | -1.1605439 | 0.24582743 | NA |
| ARIH1 | 147.123791 | 0.323241141 | 0.2487068 | 1.29968761 | 0.19370806 | 0.59838529 |
| BBS4 | 24.8414588 | 0.211553174 | 0.3697293 | 0.57218395 | 0.56719736 | NA |
| ADPGK | 19.2586843 | 0.085193726 | 0.38973653 | 0.21859313 | 0.82696701 | NA |
| NEO1 | 64.8166103 | 0.111832546 | 0.26388383 | 0.42379461 | 0.67171561 | 0.90364129 |
| NPTN | 113.464332 | 0.508225457 | 0.31585924 | 1.60902515 | 0.10761084 | 0.49151979 |
| CD276 | 7.47241462 | -0.136787329 | 0.54540607 | -0.2507991 | 0.80196947 | NA |
| C15orf59 | 15.8487699 | 0.758851981 | 0.84311 | 0.90006284 | 0.36808681 | NA |
| LOXL1-AS1 | 6.74783678 | 0.812820548 | 0.68543524 | 1.18584587 | 0.23568317 | NA |
| LOXL1 | 35.9157472 | -0.785428109 | 0.67151078 | -1.1696433 | 0.24214453 | 0.64918976 |
| PML | 11.6404257 | -0.047775972 | 0.7986539 | -0.0598206 | 0.9522985 | NA |
| UBL7 | 10.2134492 | 0.971220142 | 0.53522211 | 1.81461141 | 0.06958365 | NA |
| ARID3B | 13.2874024 | 0.580857758 | 0.55055567 | 1.05503909 | 0.29140745 | NA |
| CLK3 | 9.7110852 | -0.254448593 | 0.47255107 | -0.5384573 | 0.59026135 | NA |
| EDC3 | 13.9340248 | 0.102491176 | 0.41123435 | 0.24922815 | 0.80318431 | NA |
| CSK | 4.50490437 | 0.695021999 | 0.63072921 | 1.10193406 | 0.27049034 | NA |
| SCAMP2 | 21.4141853 | 0.091999498 | 0.40621057 | 0.22648229 | 0.82082634 | NA |
| MPI | 6.3255146 | -0.297867044 | 0.66115333 | -0.4505264 | 0.65233091 | NA |
| FAM219B | 9.48332462 | -0.646969942 | 0.53341076 | -1.2128926 | 0.2251709 | NA |
| COX5A | 22.9158256 | -0.167570014 | 0.37982058 | -0.441182 | 0.65908123 | NA |
| PPCDC | 27.2352112 | -0.328557682 | 0.32203791 | -1.0202453 | 0.30761212 | NA |
| COMMD4 | 15.6115091 | -0.338144069 | 0.50741269 | -0.6664084 | 0.50515011 | NA |
| MAN2C1 | 20.3744203 | 0.570634022 | 0.39306249 | 1.45176412 | 0.1465672 | NA |
| SIN3A | 31.5168449 | -0.356060529 | 0.28828444 | -1.2351014 | 0.21679275 | 0.62433386 |
| PTPN9 | 48.4619436 | -0.318121812 | 0.24615216 | -1.2923787 | 0.19622602 | 0.59987316 |
| SNUPN | 6.69484252 | 0.308533125 | 0.53419056 | 0.57757128 | 0.5635536 | NA |
| IMP3 | 15.1716278 | -0.449509223 | 0.41368777 | -1.0865906 | 0.2772178 | NA |
| SNX33 | 8.69596052 | -0.228118493 | 0.48158238 | -0.4736853 | 0.63572434 | NA |
| DNM1P35 | 4.98032706 | -0.248053265 | 0.71773773 | -0.3456043 | 0.7296401 | NA |
| UBE2Q2 | 40.1525315 | 0.198182903 | 0.37836351 | 0.52378968 | 0.60042483 | 0.86882737 |
| FBXO22 | 29.9033183 | 0.373979754 | 0.30949561 | 1.20835238 | 0.22691175 | 0.63327644 |
| NRG4 | 13.553091 | -0.838694593 | 0.71722854 | -1.1693547 | 0.24226074 | NA |
| TMEM266 | 116.129964 | -0.607783378 | 0.46102071 | -1.3183429 | 0.18738889 | 0.58964262 |
| ETFA | 39.8566621 | -0.008773978 | 0.26337423 | -0.0333137 | 0.97342441 | 0.99108782 |
| SCAPER | 107.584788 | -0.062558087 | 0.23529147 | -0.2658749 | 0.79033559 | 0.93840783 |
| RCN2 | 75.8341126 | 0.458410006 | 0.37219639 | 1.23163475 | 0.21808555 | 0.62462904 |
| TSPAN3 | 67.1219484 | 0.038760434 | 0.35886558 | 0.10800823 | 0.91398917 | 0.97707273 |
| PEAK1 | 40.7738046 | 0.472429758 | 0.3563599 | 1.32570966 | 0.18493589 | 0.587049 |
| HMG20A | 28.5947104 | 0.262653947 | 0.3430016 | 0.76575138 | 0.44382425 | 0.78734568 |
| LINGO1 | 407.35979 | 1.12193538 | 0.52145042 | 2.15156675 | 0.03143149 | 0.28544541 |
| TBC1D2B | 52.6579874 | -0.751845083 | 0.34676026 | -2.1681985 | 0.03014358 | 0.27996878 |
| IDH3A | 15.1912621 | -0.501749276 | 0.4302702 | -1.166126 | 0.24356349 | NA |
| DNAJA4 | 5.49529851 | 0.505219177 | 0.56990546 | 0.88649646 | 0.37535006 | NA |
| WDR61 | 17.058582 | 0.237811564 | 0.43248798 | 0.54986861 | 0.58240949 | NA |
| IREB2 | 43.9626536 | 0.322537846 | 0.26112406 | 1.23519005 | 0.21675978 | 0.62433386 |
| PSMA4 | 32.7498919 | -0.281718611 | 0.44209294 | -0.6372384 | 0.52396955 | 0.83178884 |
| MORF4L1 | 115.447968 | 0.033631002 | 0.3243655 | 0.10368243 | 0.91742137 | 0.97775374 |
| CTSH | 11.9187659 | -0.604729519 | 0.61570829 | -0.9821689 | 0.32601666 | NA |
| TMED3 | 38.1986679 | 0.584514864 | 0.59037242 | 0.99007821 | 0.3221359 | 0.69513974 |

| | | | | | | |
|---|---|---|---|---|---|---|
| MTHFS | 9.08771142 | 0.293484835 | 0.53887933 | 0.5446207 | 0.58601442 | NA |
| ZFAND6 | 116.614091 | 0.028965539 | 0.22919179 | 0.12638122 | 0.89943016 | 0.97097328 |
| FAH | 9.1348979 | 0.112260648 | 0.44222767 | 0.25385261 | 0.79960943 | NA |
| ARNT2 | 22.2670516 | 0.116034348 | 0.48773718 | 0.23790343 | 0.811956 | NA |
| ABHD17C | 11.6823883 | -0.480784145 | 0.47319639 | -1.0160351 | 0.30961267 | NA |
| CEMIP | 7.02969039 | -1.34715409 | 0.87224284 | -1.5444714 | 0.12247418 | NA |
| AC027808.1 | 9.42519579 | 0.434827705 | 0.44664404 | 0.97354418 | 0.33028292 | NA |
| MESD | 38.0575099 | -0.496003219 | 0.37065286 | -1.338188 | 0.18083516 | 0.58390314 |
| TLNRD1 | 9.07241966 | 0.174142946 | 0.52615574 | 0.33097224 | 0.74066545 | NA |
| AC104041.1 | 17.6381964 | 1.229237014 | 0.85150907 | 1.44359827 | 0.14885201 | NA |
| EFL1 | 19.7469821 | 0.021030837 | 0.38690005 | 0.05435729 | 0.95665051 | NA |
| RPS17 | 6.4333834 | 0.699787991 | 0.81634936 | 0.85721631 | 0.39132535 | NA |
| CPEB1 | 17.0967307 | -1.099006252 | 0.72642898 | -1.5128888 | 0.13030791 | NA |
| AP3B2 | 4.49170098 | -1.114070355 | 0.84430031 | -1.3195191 | 0.18699564 | NA |
| WHAMM | 20.1828384 | -0.160683364 | 0.49063794 | -0.3274989 | 0.74329061 | NA |
| HOMER2 | 6.20511157 | 0.216463238 | 0.60324759 | 0.35882984 | 0.71972239 | NA |
| C15orf40 | 13.3834589 | -0.309115279 | 0.408483 | -0.7567396 | 0.44920586 | NA |
| BTBD1 | 55.7839795 | 0.108674642 | 0.30775691 | 0.35311845 | 0.72399964 | 0.91509584 |
| HDGFL3 | 33.1171916 | -0.403998666 | 0.33554851 | -1.2039948 | 0.22859159 | 0.63504807 |
| WDR73 | 23.3584559 | 0.008249158 | 0.34646247 | 0.02380967 | 0.98100442 | NA |
| SEC11A | 54.6654402 | -0.12460483 | 0.28844209 | -0.4319925 | 0.66574684 | 0.89968484 |
| ZNF592 | 22.6722103 | -0.056419408 | 0.32154953 | -0.175461 | 0.8607174 | NA |
| ALPK3 | 13.8945714 | -0.136556534 | 0.38963347 | -0.3504743 | 0.72598274 | NA |
| SLC28A1 | 10.8024608 | 1.127392097 | 1.01072885 | 1.11542487 | 0.2646684 | NA |
| PDE8A | 107.064582 | 0.372554433 | 0.30869107 | 1.20688441 | 0.22747666 | 0.63327644 |
| AKAP13 | 142.629619 | -0.418767457 | 0.26088181 | -1.6051999 | 0.10844982 | 0.49368402 |
| AC087286.2 | 5.43184719 | -0.587792987 | 0.7740165 | -0.7594063 | 0.44760956 | NA |
| KLHL25 | 4.53406263 | -0.519437052 | 0.72533826 | -0.7161308 | 0.4739106 | NA |
| MRPL46 | 10.1582845 | -0.698719817 | 0.55325738 | -1.26292 | 0.20661793 | NA |
| MRPS11 | 14.5473658 | -0.610172877 | 0.42347261 | -1.4408792 | 0.14961881 | NA |
| DET1 | 11.8055128 | -0.444444251 | 0.48713364 | -0.9123662 | 0.36157599 | NA |
| ISG20 | 16.417041 | -17.57899906 | 2.58424309 | -6.8023783 | 1.03E-11 | NA |
| MFGE8 | 6.80733427 | 0.856218452 | 0.81408454 | 1.05175618 | 0.29291143 | NA |
| ABHD2 | 20.5192384 | 0.254557538 | 0.35962923 | 0.70783328 | 0.4790488 | NA |
| POLG | 10.0806392 | -0.053780924 | 0.44715362 | -0.1202739 | 0.90426615 | NA |
| PEX11A | 4.13775665 | -0.952626862 | 0.77491447 | -1.2293316 | 0.2189475 | NA |
| ANPEP | 16.4346283 | 0.100618595 | 0.56224289 | 0.1789593 | 0.85796966 | NA |
| AP3S2 | 8.79760742 | 0.635791788 | 0.53510415 | 1.18816455 | 0.23476859 | NA |
| ZNF710 | 9.4269413 | -0.90813348 | 0.6154926 | -1.475458 | 0.14008946 | NA |
| IDH2 | 11.3467712 | -0.039114418 | 0.79706536 | -0.049073 | 0.96086109 | NA |
| SEMA4B | 26.8421318 | -1.715552927 | 0.55314089 | -3.1014755 | 0.00192559 | NA |
| CIB1 | 34.6393118 | -1.05473822 | 0.40633468 | -2.5957377 | 0.00943881 | 0.15375026 |
| AC091167.6 | 6.50470313 | 1.022007061 | 0.51413612 | 1.98781417 | 0.04683225 | NA |
| IQGAP1 | 110.542937 | -0.390589356 | 0.25706177 | -1.5194377 | 0.12865235 | 0.52341115 |
| CRTC3 | 47.4270087 | -0.107933231 | 0.30533631 | -0.3534897 | 0.72372137 | 0.91509584 |
| FURIN | 13.9323153 | 0.407747896 | 0.54213044 | 0.75212138 | 0.45197807 | NA |
| MAN2A2 | 13.5376245 | -0.428549925 | 0.59148568 | -0.7245314 | 0.46873958 | NA |
| HDDC3 | 6.13965915 | -0.719753044 | 0.62509658 | -1.1514269 | 0.24955664 | NA |
| UNC45A | 19.3851133 | -0.185083429 | 0.40408723 | -0.4580284 | 0.64693204 | NA |
| SLCO3A1 | 12.7206375 | -1.307883447 | 0.5942029 | -2.2010721 | 0.02773092 | NA |
| FAM174B | 18.0569599 | 0.444658379 | 0.4661838 | 0.95382632 | 0.34017156 | NA |
| AC013394.1 | 69.0923689 | 0.087056327 | 0.2443879 | 0.35622193 | 0.72167437 | 0.91509584 |
| CHD2 | 60.8170078 | 0.079699483 | 0.32814107 | 0.24288177 | 0.80809699 | 0.94328109 |
| RGMA | 8.06004922 | -0.165397796 | 0.79792498 | -0.2072849 | 0.83578737 | NA |
| MCTP2 | 95.8423031 | -0.742798268 | 0.42180283 | -1.7610083 | 0.078237 | 0.4276675 |
| LINC01197 | 17.1772956 | 0.428698835 | 0.50175543 | 0.854398 | 0.3928845 | NA |
| AC012409.2 | 21.9213394 | 0.14452364 | 0.73988852 | 0.19533164 | 0.84513333 | NA |
| NR2F2-AS1 | 172.104491 | 0.640118141 | 0.40017556 | 1.59959329 | 0.10968884 | 0.49598431 |
| NR2F2 | 82.9529235 | -0.017381198 | 0.40004399 | -0.0434482 | 0.96534424 | 0.98892249 |
| AC087477.2 | 14.330243 | -0.434941284 | 0.59053906 | -0.7365157 | 0.46141692 | NA |
| ARRDC4 | 174.942218 | 0.619837496 | 0.57451441 | 1.07888938 | 0.28063704 | 0.66958859 |
| IGF1R | 481.36896 | -0.056967617 | 0.26946584 | -0.2114094 | 0.83256781 | 0.95050875 |
| AC036108.1 | 9.4087403 | -1.819213844 | 0.88654659 | -2.0520228 | 0.04016744 | NA |
| TTC23 | 20.5212894 | 0.089410733 | 0.33642992 | 0.26576332 | 0.79042149 | NA |
| LRRC28 | 37.7004681 | 0.41174672 | 0.27798648 | 1.48117535 | 0.13855985 | 0.53421936 |
| MEF2A | 101.869565 | 0.366465056 | 0.27646536 | 1.32553699 | 0.18499311 | 0.587049 |
| LYSMD4 | 4.75402509 | -0.069739746 | 0.63637183 | -0.1095896 | 0.91273484 | NA |
| AC090825.1 | 34.215159 | 0.603488582 | 0.55220062 | 1.09287921 | 0.27444684 | 0.66766283 |
| LINS1 | 8.03454096 | 0.1586984 | 0.51633575 | 0.30735505 | 0.75857314 | NA |
| ASB7 | 11.4777186 | 0.151857348 | 0.40940357 | 0.37092336 | 0.71069461 | NA |
| AC087762.1 | 10.6320774 | 1.252504016 | 0.89703777 | 1.39626675 | 0.16263418 | NA |
| CHSY1 | 28.9311843 | 0.163425784 | 0.29227394 | 0.55915278 | 0.57605746 | 0.85929076 |
| SELENOS | 61.4020278 | -0.85114332 | 0.38289463 | -2.2229179 | 0.02622134 | 0.26195503 |
| SNRPA1 | 16.1448507 | -0.612055325 | 0.40870468 | -1.4975491 | 0.13425044 | NA |
| PCSK6 | 15.0482464 | -0.095961727 | 1.08983252 | -0.0880518 | 0.9298355 | NA |
| TM2D3 | 31.8735746 | -0.126717444 | 0.30512771 | -0.4152931 | 0.67792733 | 0.90499319 |
| TARSL2 | 50.0965747 | -0.412277245 | 0.36804332 | -1.1201867 | 0.26263419 | 0.6581333 |
| POLR3K | 6.16457172 | -0.770990067 | 0.68009919 | -1.1336436 | 0.25694409 | NA |
| SNRNP25 | 5.12349894 | -1.298309912 | 0.75517946 | -1.7192071 | 0.08557666 | NA |
| RHBDF1 | 4.80337207 | 0.113006359 | 0.71477085 | 0.15810152 | 0.8743768 | NA |
| MPG | 17.7297239 | -0.940900619 | 0.4370254 | -2.1529655 | 0.03132139 | NA |
| NPRL3 | 19.893921 | -0.047123118 | 0.44187575 | -0.1066434 | 0.91507191 | NA |
| LUC7L | 48.42624 | 0.66598242 | 0.34414897 | 1.9351574 | 0.05297098 | 0.35868165 |
| RGS11 | 7.35464661 | 0.674597502 | 0.81505445 | 0.8276717 | 0.40785645 | NA |
| AXIN1 | 12.6494395 | 0.046978011 | 0.41877842 | 0.11217868 | 0.91068173 | NA |
| MRPL28 | 15.0817785 | 0.015126958 | 0.5579968 | 0.0271094 | 0.97837248 | NA |
| TMEM8A | 11.8498235 | 0.38711663 | 0.41955857 | 0.92267603 | 0.35617606 | NA |
| NME4 | 7.80048124 | -0.744274118 | 0.65717114 | -1.1325423 | 0.25740651 | NA |
| RAB11FIP3 | 52.7844119 | 0.800416877 | 0.54992988 | 1.45548897 | 0.14553394 | 0.54436111 |
| CAPN15 | 7.58400149 | 0.298797561 | 0.5219299 | 0.572486 | 0.56699277 | NA |
| PIGQ | 16.5890079 | 0.666164557 | 0.47028121 | 1.41652388 | 0.15662218 | NA |
| RAB40C | 21.7681026 | 0.046208413 | 0.4031724 | 0.11461205 | 0.90875263 | NA |
| METTL26 | 12.1042463 | -0.020838965 | 0.60476851 | -0.0344578 | 0.97251213 | NA |
| MCRIP2 | 10.8329069 | 0.347519898 | 0.46756835 | 0.74324941 | 0.4573307 | NA |
| RHOT2 | 14.1696356 | 0.395040722 | 0.39633247 | 0.99674075 | 0.31889036 | NA |
| STUB1 | 37.6764305 | -0.438363954 | 0.39103225 | -1.121043 | 0.26226957 | 0.65790062 |
| JMJD8 | 26.3324435 | 0.288634379 | 0.51907606 | 0.55605412 | 0.57817386 | NA |
| METRN | 11.6346351 | -0.388113709 | 0.52253922 | -0.7427456 | 0.45763572 | NA |
| FAM173A | 13.1478015 | -0.357812327 | 0.51430081 | -0.6957258 | 0.48660059 | NA |
| NARFL | 5.35154468 | -0.074439938 | 0.74167644 | -0.1003671 | 0.92005286 | NA |
| MSLN | 11.8922984 | -0.481510484 | 0.98752824 | -0.4875916 | 0.62583914 | NA |
| LMF1 | 17.809234 | 0.010466263 | 0.42839289 | 0.02443146 | 0.98050846 | NA |
| UBE2I | 31.2136266 | -0.704459932 | 0.33578643 | -2.0979404 | 0.03591041 | 0.29695121 |
| TSR3 | 16.4660502 | -0.332423505 | 0.3822982 | -0.8695398 | 0.38455194 | NA |
| GNPTG | 12.4531256 | -0.539186057 | 0.40477687 | -1.3320575 | 0.18284131 | NA |
| UNKL | 11.424964 | 0.047488834 | 0.65089305 | 0.0729595 | 0.94183834 | NA |
| C16orf91 | 6.35780573 | -0.888180445 | 0.64225469 | -1.38291 | 0.16669246 | NA |
| CLCN7 | 17.7438015 | 0.301149309 | 0.4574171 | 0.65836915 | 0.51030095 | NA |
| TELO2 | 7.36009621 | 0.549904989 | 0.60573142 | 0.90783634 | 0.3639647 | NA |
| IFT140 | 18.300926 | 0.0172148 | 0.37364078 | 0.04607313 | 0.96325196 | NA |
| JPT2 | 11.8759468 | -0.867154968 | 0.59711519 | -1.4522407 | 0.1464347 | NA |
| MAPK8IP3 | 23.9194532 | 0.351145998 | 0.33723351 | 1.04125475 | 0.29775733 | NA |
| NME3 | 48.6617323 | -0.352332027 | 0.38130573 | -0.9240145 | 0.35547876 | 0.72709121 |

| | | | | | | |
|---|---|---|---|---|---|---|
| MRPS34 | 23.0985985 | -0.380447137 | 0.49358515 | -0.7707832 | 0.44083545 | NA |
| SPSB3 | 20.0367039 | 0.195227644 | 0.33865717 | 0.57647575 | 0.56429366 | NA |
| NUBP2 | 11.1652148 | -0.246033069 | 0.48068658 | -0.5118368 | 0.60876525 | NA |
| HAGH | 30.8206285 | -0.115254875 | 0.29545274 | -0.3900958 | 0.6964657 | 0.90718427 |
| FAHD1 | 7.88797268 | -1.063121097 | 0.68535982 | -1.5511868 | 0.12085693 | NA |
| MSRB1 | 7.34950106 | -0.307968375 | 0.64552186 | -0.4770843 | 0.63330206 | NA |
| NDUFB10 | 30.9099383 | -0.543790532 | 0.34516461 | -1.5754527 | 0.11515198 | 0.50513044 |
| RPS2 | 61.8419026 | -0.46954144 | 0.53254966 | -0.8816857 | 0.37794678 | 0.74559928 |
| SNHG9 | 6.64917447 | -0.359992453 | 0.57744305 | -0.623425 | 0.53300525 | NA |
| GFER | 4.71420777 | 0.305386413 | 0.81150729 | 0.37631999 | 0.70667902 | NA |
| ZNF598 | 6.42565351 | 0.234574143 | 0.66169702 | 0.35450385 | 0.72296132 | NA |
| SLC9A3R2 | 68.9201796 | -0.577572892 | 0.57621241 | -1.0023611 | 0.31616923 | 0.69424851 |
| NTHL1 | 9.92639073 | 0.019551927 | 0.55114673 | 0.03547499 | 0.97170099 | NA |
| TSC2 | 22.901423 | 0.579665891 | 0.33582271 | 1.7261069 | 0.08432818 | NA |
| PKD1 | 33.4362876 | 0.170262372 | 0.4172341 | 0.40807396 | 0.68321937 | 0.90499319 |
| SNHG19 | 6.55487062 | -0.168431176 | 0.7168127 | -0.2349724 | 0.81423018 | NA |
| TRAF7 | 8.21321017 | -0.960552637 | 0.54412891 | -1.7653034 | 0.07751278 | NA |
| MLST8 | 4.6668601 | 0.249584304 | 0.66596134 | 0.37477296 | 0.70782932 | NA |
| PGP | 17.7872172 | -0.40169153 | 0.41430311 | -0.9695595 | 0.33226609 | NA |
| E4F1 | 6.1714115 | -0.353560535 | 0.58866202 | -0.6006172 | 0.54809498 | NA |
| ECI1 | 23.686066 | -0.036396666 | 0.35636755 | -0.1021324 | 0.9186516 | NA |
| RNPS1 | 32.5126565 | -0.129015356 | 0.36534924 | -0.3531289 | 0.72399184 | 0.91509584 |
| ABCA3 | 20.8597073 | -0.239687987 | 0.50384595 | -0.4757168 | 0.63427615 | NA |
| PDPK1 | 50.4826284 | 0.30447424 | 0.23122001 | 1.31681611 | 0.18790027 | 0.59036421 |
| AC093525.4 | 4.91749462 | -0.637240816 | 0.81231665 | -0.7844734 | 0.43276237 | NA |
| KCTD5 | 6.75573773 | -0.815029769 | 0.65820226 | -1.2382664 | 0.21561729 | NA |
| SRRM2 | 146.392668 | -0.111467307 | 0.20176021 | -0.5524742 | 0.58062352 | 0.86175324 |
| ELOB | 88.7404803 | -0.186884697 | 0.39964999 | -0.4676209 | 0.6400557 | 0.88784744 |
| FLYWCH2 | 22.8264083 | -0.161581712 | 0.47960254 | -0.3369075 | 0.7361866 | NA |
| FLYWCH1 | 14.3355193 | -0.141342969 | 0.40277852 | -0.3509198 | 0.72564849 | NA |
| TNFRSF12A | 30.7100536 | -0.824201714 | 0.9904563 | -0.8321434 | 0.40532798 | 0.76772515 |
| HCFC1R1 | 45.32421 | -0.813233981 | 0.41950916 | -1.9385369 | 0.05255775 | 0.35808347 |
| THOC6 | 4.87325276 | -0.290274002 | 0.65343673 | -0.4442266 | 0.65687874 | NA |
| IL32 | 13.8918995 | -0.258243584 | 0.66004502 | -0.3912515 | 0.69561137 | NA |
| ZNF213-AS1 | 12.1632763 | 0.179467974 | 0.43303008 | 0.4144469 | 0.67854685 | NA |
| ZNF205 | 5.99145481 | -0.638858681 | 0.61849661 | -1.0329219 | 0.30164046 | NA |
| ZNF213 | 17.883171 | 0.189269466 | 0.40545277 | 0.46681015 | 0.64063572 | NA |
| ZNF263 | 11.4361628 | -0.306279621 | 0.39807746 | -0.769397 | 0.44165765 | NA |
| TIGD7 | 6.4884761 | -0.591170572 | 0.53509774 | -1.1047899 | 0.26925064 | NA |
| ZNF75A | 19.8873989 | -0.519783715 | 0.40627917 | -1.2793757 | 0.20076477 | NA |
| AC025283.3 | 6.98570474 | -0.543100301 | 0.53364708 | -1.0177144 | 0.30881372 | NA |
| ZSCAN32 | 6.18026361 | -0.242081659 | 0.55562419 | -0.4356932 | 0.66305937 | NA |
| NAA60 | 16.515068 | 0.047461713 | 0.3756271 | 0.12635327 | 0.89945229 | NA |
| CLUAP1 | 24.6249283 | 0.16314484 | 0.29250351 | 0.55775344 | 0.57701276 | NA |
| DNASE1 | 23.7780517 | 0.319102874 | 0.37131386 | 0.85938853 | 0.39012619 | NA |
| TRAP1 | 20.5075515 | -0.310469679 | 0.34350156 | -0.9038378 | 0.36608143 | NA |
| CREBBP | 78.5033121 | 0.245251022 | 0.28444662 | 0.86220404 | 0.38857525 | 0.75301425 |
| ADCY9 | 41.1800518 | -0.308931857 | 0.34188257 | -0.9036198 | 0.36619704 | 0.73456735 |
| GLIS2 | 10.2080852 | 0.115684675 | 0.4909923 | 0.23561403 | 0.81373219 | NA |
| VASN | 145.464219 | -0.081297945 | 0.42504804 | -0.1912677 | 0.84831589 | 0.95532127 |
| DNAJA3 | 14.5011854 | -0.167847309 | 0.406636 | -0.4127704 | 0.67977483 | NA |
| NMRAL1 | 9.70492068 | -0.530994637 | 0.60362319 | -0.879679 | 0.37903324 | NA |
| HMOX2 | 10.3466638 | 0.225234913 | 0.47651462 | 0.47267157 | 0.63644751 | NA |
| CDIP1 | 8.17322512 | 0.609933333 | 0.48647942 | 1.25377005 | 0.2099256 | NA |
| UBALD1 | 6.97672348 | 0.996940779 | 0.59015869 | 1.68927577 | 0.09116659 | NA |
| MGRN1 | 24.0529413 | 0.120318804 | 0.34889099 | 0.34486073 | 0.73019907 | NA |
| NUDT16L1 | 9.80343992 | 0.166676508 | 0.50070751 | 0.33288198 | 0.73922337 | NA |
| ANKS3 | 18.0711513 | -0.104240081 | 0.37828128 | -0.2755624 | 0.78288423 | NA |
| ZNF500 | 8.81241135 | -0.256453501 | 0.64442948 | -0.3979543 | 0.69066385 | NA |
| ROGDI | 5.73592472 | 0.063173439 | 0.61874512 | 0.10209929 | 0.91867786 | NA |
| GLYR1 | 35.238014 | -0.241573958 | 0.3583286 | -0.6741688 | 0.50020401 | 0.82345406 |
| UBN1 | 13.4847237 | -0.520389706 | 0.45151671 | -1.152537 | 0.24910049 | NA |
| ALG1 | 4.96494001 | 0.139377533 | 0.62649856 | 0.22247064 | 0.82394753 | NA |
| RBFOX1 | 562.989914 | -0.172083714 | 0.66040605 | -0.2605726 | 0.79442213 | 0.93934998 |
| METTL22 | 7.03613938 | -0.203529624 | 0.58403523 | -0.3484886 | 0.72747327 | NA |
| ABAT | 12.6840185 | -0.059310477 | 0.49914698 | -0.1188237 | 0.90541505 | NA |
| PMM2 | 20.1686633 | -0.620690802 | 0.33836482 | -1.8343834 | 0.0665971 | NA |
| CARHSP1 | 75.2519598 | -0.997684532 | 0.37898394 | -2.6325246 | 0.00847529 | 0.14753551 |
| USP7 | 34.823268 | 0.401821136 | 0.29551002 | 1.35975468 | 0.17390757 | 0.57722512 |
| C16orf72 | 42.3163963 | 0.222486718 | 0.29816078 | 0.74619713 | 0.45554834 | 0.79324789 |
| AC087190.1 | 5.11276078 | -1.961025324 | 0.81826933 | -2.3965524 | 0.01655013 | NA |
| ATF7IP2 | 30.1570975 | -0.088786009 | 0.32065322 | -0.2768911 | 0.78186376 | 0.93415197 |
| EMP2 | 12.1736429 | -1.00767541 | 0.5851166 | -1.7221788 | 0.08503713 | NA |
| NUBP1 | 6.69172644 | -0.514091626 | 0.57970215 | -0.8868203 | 0.37517566 | NA |
| DEXI | 17.3125786 | -0.467941524 | 0.50931879 | -0.9187596 | 0.35822133 | NA |
| CLEC16A | 181.457293 | -0.460972706 | 0.39931899 | -1.1543972 | 0.24833738 | 0.6507516 |
| AC007014.1 | 8.55123031 | -2.678680717 | 0.92969789 | -2.8812378 | 0.00396117 | NA |
| AC007220.1 | 7.83582292 | -1.17841212 | 0.71180867 | -1.6555181 | 0.09781945 | NA |
| AC099489.1 | 6.06752588 | -0.107620839 | 1.06045547 | -0.1014855 | 0.91916508 | NA |
| LITAF | 91.6081313 | -1.981068624 | 0.62903523 | -3.1493763 | 0.00163619 | 0.06320383 |
| SNN | 3.97050702 | -0.83073671 | 0.88518773 | -0.9384865 | 0.34799446 | NA |
| TXNDC11 | 31.0369206 | -0.221228843 | 0.28574405 | -0.7742203 | 0.43880053 | 0.78335165 |
| ZC3H7A | 44.8433294 | 0.45477854 | 0.25480551 | 1.78480653 | 0.0742927 | 0.41891814 |
| RSL1D1 | 14.9575731 | -0.175079614 | 0.4962093 | -0.3528342 | 0.72421274 | NA |
| AC007216.4 | 15.1564483 | 0.050242972 | 0.48203847 | 0.10423021 | 0.91698666 | NA |
| GSPT1 | 46.6721069 | -0.249059741 | 0.38292119 | -0.6504204 | 0.51542073 | 0.83104823 |
| SNX29 | 95.3849964 | 0.359657201 | 0.453758 | 0.79261897 | 0.42799985 | 0.77844087 |
| CPPED1 | 14.5411661 | -0.341987338 | 0.49958551 | -0.6845421 | 0.49363289 | NA |
| ERCC4 | 9.30083673 | -0.601440878 | 0.45197761 | -1.3306873 | 0.18329192 | NA |
| MKL2 | 35.8835337 | 0.98948991 | 0.42786602 | 2.31261623 | 0.02074375 | 0.23840101 |
| PARN | 39.7581317 | -0.148139709 | 0.32569969 | -0.4548353 | 0.64922775 | 0.89387362 |
| BFAR | 14.0769564 | -0.273500033 | 0.45031044 | -0.6073589 | 0.54361279 | NA |
| NOMO1 | 9.65528664 | 0.413018129 | 0.64399316 | 0.64133931 | 0.52130225 | NA |
| PDXDC1 | 89.1509276 | 0.035860944 | 0.25714167 | 0.13945987 | 0.88908677 | 0.96783036 |
| NTAN1 | 10.5160963 | 0.508302635 | 0.62880294 | 0.80836556 | 0.41888017 | NA |
| RRN3 | 14.3625793 | -0.248899217 | 0.44290363 | -0.5619715 | 0.57413544 | NA |
| C16orf45 | 36.5793426 | -0.129915978 | 0.30879933 | -0.4207133 | 0.67396446 | 0.90403541 |
| MARF1 | 27.5117941 | 0.329032742 | 0.39121297 | 0.84105786 | 0.40031552 | NA |
| NDE1 | 8.90422207 | -0.146088865 | 0.49134485 | -0.2973245 | 0.76621878 | NA |
| FOPNL | 13.2568537 | -0.022756686 | 0.43479825 | -0.0523385 | 0.95825898 | NA |
| ABCC1 | 53.544351 | 0.295497789 | 0.36087217 | 0.81884339 | 0.41287577 | 0.76946282 |
| XYLT1 | 52.0480861 | -0.096702973 | 0.73943981 | -0.1307787 | 0.89595038 | 0.97097328 |
| RPS15A | 18.2854201 | -0.163514792 | 0.62768723 | -0.2605036 | 0.79447533 | NA |
| SMG1 | 89.7016363 | 0.157776368 | 0.26957852 | 0.58527056 | 0.55836581 | 0.8520194 |
| COQ7 | 8.71458121 | -0.512834898 | 0.5381827 | -0.9529011 | 0.34064018 | NA |
| GDE1 | 23.5503776 | 0.074255772 | 0.37322148 | 0.198959 | 0.84229482 | NA |
| CCP110 | 12.6094688 | -0.280530628 | 0.44775529 | -0.6265267 | 0.53096956 | NA |
| VPS35L | 38.304917 | 0.382726655 | 0.30448362 | 1.25696962 | 0.20876464 | 0.61736443 |
| KNOP1 | 7.25651616 | -1.010790215 | 0.60671968 | -1.6659921 | 0.09571498 | NA |
| IQCK | 25.2423479 | -0.207604843 | 0.34676889 | -0.5986836 | 0.54938391 | NA |
| UMOD | 52.7840892 | 1.707382521 | 0.63550388 | 2.68665948 | 0.00721705 | 0.13427683 |
| ACSM2A | 30.5962524 | 2.853556496 | 1.75827327 | 1.62293117 | 0.10460412 | 0.4862279 |
| ACSM2B | 31.7112123 | 2.241505217 | 1.57645503 | 1.42186436 | 0.15506563 | 0.55560618 |

| | | | | | | |
|---|---|---|---|---|---|---|
| ACSM3 | 11.2519729 | -0.810927393 | 0.87239737 | -0.929539 | 0.35260982 | NA |
| THUMPD1 | 22.4126202 | 0.102370063 | 0.3181611 | 0.32175544 | 0.74763798 | NA |
| ERI2 | 10.0212765 | 0.757178173 | 0.54112208 | 1.39927421 | 0.16173077 | NA |
| LYRM1 | 10.7442486 | -0.081196613 | 0.60993599 | -0.1331232 | 0.89409597 | NA |
| TMEM159 | 6.28273449 | 0.36502725 | 0.53298191 | 0.68487737 | 0.49342131 | NA |
| METTL9 | 27.1180745 | -0.18756399 | 0.34477243 | -0.5440226 | 0.58642593 | NA |
| NPIPB4 | 8.56055463 | 1.218598859 | 0.60778967 | 2.00496802 | 0.04496647 | NA |
| UQCRC2 | 28.7623811 | -0.203025401 | 0.32035001 | -0.6337612 | 0.52623669 | 0.83195298 |
| MOSMO | 70.8180544 | -0.257709714 | 0.50065373 | -0.5147464 | 0.60673023 | 0.87080602 |
| EEF2K | 13.2322096 | -0.163960891 | 0.50564523 | -0.3242607 | 0.74574065 | NA |
| POLR3E | 12.8753177 | 0.023383443 | 0.44810507 | 0.05218295 | 0.95838292 | NA |
| CDR2 | 64.8738628 | -0.465565658 | 0.29605638 | -1.5725574 | 0.11582133 | 0.50513044 |
| NPIPB5 | 23.9985224 | 0.175375953 | 0.30454452 | 0.57586312 | 0.56470771 | NA |
| USP31 | 35.8241867 | -0.438865884 | 0.36583706 | -1.1996212 | 0.23028651 | 0.63682501 |
| SCNN1G | 9.31517193 | -0.027411569 | 1.27940163 | -0.0214253 | 0.98290639 | NA |
| SCNN1B | 9.20224446 | -1.282735429 | 0.89573383 | -1.4320498 | 0.15212958 | NA |
| COG7 | 43.0340866 | -0.858163593 | 0.40681835 | -2.1094515 | 0.03490563 | 0.29312053 |
| AC008915.2 | 4.21075254 | -1.292630716 | 0.89548135 | -1.4435038 | 0.14887861 | NA |
| GGA2 | 28.6874693 | -0.276344107 | 0.37029075 | -0.7462895 | 0.45549254 | 0.79324789 |
| EARS2 | 6.12603289 | -0.136246229 | 0.5216613 | -0.2611776 | 0.79395558 | NA |
| UBFD1 | 14.918404 | 0.093536211 | 0.44192106 | 0.21165819 | 0.83237371 | NA |
| NDUFAB1 | 19.9503039 | -0.4105171 | 0.4895037 | -0.8386394 | 0.40167168 | NA |
| PALB2 | 3.99585773 | 0.642096105 | 0.62688549 | 1.02426379 | 0.3057107 | NA |
| DCTN5 | 22.6374085 | -0.345782796 | 0.39774973 | -0.8693476 | 0.38465701 | NA |
| RBBP6 | 49.7045119 | 0.037907303 | 0.30273571 | 0.12521583 | 0.90035269 | 0.97097328 |
| TNRC6A | 153.101569 | 0.590904223 | 0.22596386 | 2.61503864 | 0.00892174 | 0.14875902 |
| AC008731.1 | 4.99735557 | 0.031997774 | 0.79226732 | 0.0403876 | 0.96778412 | NA |
| ARHGAP17 | 31.3439028 | 0.031259911 | 0.30068869 | 0.10396105 | 0.91720026 | 0.97775374 |
| LCMT1 | 22.3622877 | 0.248902176 | 0.30549994 | 0.81473722 | 0.41522274 | NA |
| NSMCE1 | 14.179266 | -0.450346545 | 0.52216781 | -0.8624556 | 0.38843686 | NA |
| IL4R | 33.0088809 | -2.399738618 | 0.760178 | -3.1568115 | 0.00159504 | 0.06220673 |
| GTF3C1 | 47.3875731 | -0.113645669 | 0.31455606 | -0.3612891 | 0.71788336 | 0.91472065 |
| KIAA0556 | 47.5076 | 0.052621102 | 0.26589408 | 0.1979025 | 0.84312135 | 0.95469552 |
| XPO6 | 31.5356812 | 0.053406099 | 0.42548639 | 0.12551776 | 0.90011367 | 0.97097328 |
| NUPR1 | 583.510951 | -1.53687732 | 0.70460877 | -2.1811782 | 0.02917024 | 0.27926427 |
| SGF29 | 10.3623922 | -0.641514341 | 0.51933103 | -1.2352706 | 0.21672982 | NA |
| SULT1A2 | 8.00810219 | -0.550638027 | 0.58084697 | -0.9479916 | 0.34313375 | NA |
| SULT1A1 | 29.1907647 | 0.188379295 | 0.42622382 | 0.44197271 | 0.65850896 | 0.89709157 |
| ATXN2L | 19.7319562 | -0.079746401 | 0.41128056 | -0.1938978 | 0.8462559 | NA |
| TUFM | 31.0517296 | -0.148092364 | 0.43647911 | -0.3392885 | 0.73439237 | 0.91893089 |
| SH2B1 | 20.487604 | 0.367259743 | 0.3436961 | 1.06855954 | 0.28526819 | NA |
| RABEP2 | 9.12810502 | 0.50883687 | 0.50955349 | 0.99859363 | 0.31799159 | NA |
| NFATC2IP | 38.7123675 | -0.122530463 | 0.29682715 | -0.4128007 | 0.67975262 | 0.90499319 |
| SPNS1 | 8.11510001 | 0.175290043 | 0.51218314 | 0.34224095 | 0.73216957 | NA |
| QPRT | 43.0242924 | -1.104639039 | 0.39908728 | -2.7679134 | 0.00564164 | 0.11765106 |
| KIF22 | 7.65847287 | -0.338589958 | 0.57318467 | -0.5907171 | 0.55471002 | NA |
| MAZ | 16.6436627 | -0.071128662 | 0.40578782 | -0.1752854 | 0.86085542 | NA |
| AC009133.1 | 6.79726891 | -0.673036069 | 0.62767062 | -1.0722759 | 0.28359613 | NA |
| MVP | 35.7731564 | -1.026211422 | 0.42456992 | -2.417061 | 0.01564639 | 0.20644861 |
| CDIPT | 23.0127108 | 0.173570999 | 0.47126565 | 0.36830819 | 0.71264344 | NA |
| SEZ6L2 | 18.1072983 | -1.187512337 | 0.55454433 | -2.14142 | 0.03224018 | NA |
| KCTD13 | 12.6515388 | 0.274931066 | 0.406158 | 0.67690668 | 0.49846517 | NA |
| TMEM219 | 45.5915703 | -0.01606006 | 0.49989064 | -0.0321271 | 0.97437065 | 0.9916101 |
| TAOK2 | 8.2668477 | 0.315349035 | 0.57051962 | 0.55274004 | 0.58044143 | NA |
| HIRIP3 | 5.02687385 | -0.578379976 | 0.60603227 | -0.9543716 | 0.33989559 | NA |
| INO80E | 17.0798196 | 1.17119578 | 0.42436008 | 2.75991036 | 0.00578172 | NA |
| PPP4C | 17.3553793 | -0.40004296 | 0.40473356 | -0.9884106 | 0.32295158 | NA |
| YPEL3 | 24.1369749 | 0.383419515 | 0.37786389 | 1.0147027 | 0.31024757 | NA |
| MAPK3 | 14.0298905 | -0.156403659 | 0.4809232 | -0.3252155 | 0.74501801 | NA |
| NPIPB13 | 6.94794304 | -0.259410727 | 0.62183792 | -0.4171677 | 0.67655571 | NA |
| CD2BP2 | 14.4218774 | 0.4888573 | 0.41924279 | 1.16604819 | 0.24359496 | NA |
| TBC1D10B | 8.48214096 | -0.390349922 | 0.58799027 | -0.6638714 | 0.50677263 | NA |
| 1-Sep | 5.7476062 | 0.052482508 | 0.76514756 | 0.06859136 | 0.9453149 | NA |
| DCTPP1 | 10.0170745 | -0.729361627 | 0.60422302 | -1.2071066 | 0.22739108 | NA |
| SEPHS2 | 15.8435534 | -0.914426017 | 0.4916317 | -1.8599818 | 0.0628881 | NA |
| ZNF768 | 7.15936362 | 0.193027943 | 0.54641776 | 0.35326074 | 0.72389297 | NA |
| ZNF688 | 9.2335002 | 0.298834251 | 0.6063419 | 0.49284777 | 0.62212015 | NA |
| ZNF785 | 10.1588574 | 0.11346669 | 0.43338533 | 0.26181479 | 0.79346424 | NA |
| ZNF689 | 3.91840676 | -0.799067038 | 0.67905243 | -1.1767383 | 0.23929996 | NA |
| PRR14 | 11.8995235 | 0.462973377 | 0.42584927 | 1.08717664 | 0.27695875 | NA |
| FBRS | 15.2906997 | -0.100690333 | 0.39658175 | -0.2538955 | 0.79957627 | NA |
| PHKG2 | 7.25624014 | -0.133943265 | 0.58266037 | -0.2298822 | 0.81818329 | NA |
| RNF40 | 13.7721639 | 0.137782198 | 0.45906285 | 0.30013798 | 0.76407191 | NA |
| ZNF629 | 6.67012051 | -0.17679142 | 0.5876739 | -0.3008325 | 0.76354221 | NA |
| BCL7C | 28.9860248 | 0.080068118 | 0.37888355 | 0.21132646 | 0.83263254 | 0.95050875 |
| MIR762HG | 6.24334219 | -0.514445843 | 0.53600432 | -0.9597793 | 0.33716631 | NA |
| FBXL19 | 9.42877985 | 0.670554876 | 0.62192832 | 1.07818675 | 0.28095042 | NA |
| ORAI3 | 8.14719267 | -1.183381413 | 0.66659736 | -1.7752567 | 0.07585552 | NA |
| SETD1A | 7.52069393 | -0.599425474 | 0.51809083 | -1.1569892 | 0.24727679 | NA |
| HSD3B7 | 4.00746187 | -0.813888924 | 0.88418027 | -0.9205011 | 0.35731095 | NA |
| STX4 | 10.2165934 | -0.013465369 | 0.49545842 | -0.0271776 | 0.97831809 | NA |
| BCKDK | 10.6103281 | 0.60053473 | 0.53782776 | 1.11659303 | 0.26416837 | NA |
| KAT8 | 15.7127875 | -0.020590677 | 0.46554959 | -0.0442288 | 0.96472206 | NA |
| FUS | 59.3876112 | -0.200379279 | 0.34793298 | -0.5759134 | 0.56467369 | 0.8549147 |
| AC026471.4 | 9.08822843 | -0.270571694 | 0.65344751 | -0.414068 | 0.67882432 | NA |
| ARMC5 | 4.95083173 | 0.828162835 | 0.70666206 | 1.17193618 | 0.24122268 | NA |
| TGFB1I1 | 21.0428693 | -1.393264587 | 0.5265265 | -2.6461433 | 0.00814153 | NA |
| SLC5A2 | 4.86374473 | 0.821341812 | 0.84577431 | 0.97111227 | 0.33149237 | NA |
| C16orf58 | 12.4013572 | -0.350605325 | 0.45795284 | -0.7655926 | 0.44391872 | NA |
| ZNF720 | 19.0219555 | 0.067819971 | 0.36872684 | 0.18393011 | 0.85406828 | NA |
| ZNF267 | 8.55062077 | 0.514235279 | 0.64484563 | 0.79745485 | 0.42518691 | NA |
| VPS35 | 36.045195 | -0.270074709 | 0.31530895 | -0.8565399 | 0.3916992 | 0.75653902 |
| MYLK3 | 51.9609535 | -0.028900577 | 0.71204931 | -0.0405879 | 0.96762444 | 0.98964394 |
| C16orf87 | 9.09814352 | -0.000157383 | 0.54853274 | -0.0002869 | 0.99977107 | NA |
| GPT2 | 5.05304396 | 0.931564631 | 1.01974514 | 0.91352692 | 0.36096548 | NA |
| DNAJA2 | 18.3062869 | -0.042852147 | 0.37740875 | -0.1135431 | 0.90960002 | NA |
| ITFG1 | 135.800232 | 0.145887466 | 0.19244843 | 0.75806006 | 0.44841503 | 0.78987065 |
| PHKB | 102.932994 | 0.181790775 | 0.24502352 | 0.74193193 | 0.45812858 | 0.79476882 |
| LONP2 | 93.9771091 | 0.278084242 | 0.19789019 | 1.40524521 | 0.15994838 | 0.56283048 |
| SIAH1 | 23.8819105 | -0.150366254 | 0.3734771 | -0.4026117 | 0.68723389 | NA |
| N4BP1 | 59.1503701 | 0.170383505 | 0.36388468 | 0.4682349 | 0.63961662 | 0.88754191 |
| ZNF423 | 20.2549142 | 0.195601484 | 0.64073286 | 0.30527775 | 0.76015462 | NA |
| CNEP1R1 | 9.35014401 | 0.445483493 | 0.60298499 | 0.73879699 | 0.46003028 | NA |
| HEATR3 | 8.91563044 | -0.190609053 | 0.45964606 | -0.4146866 | 0.67837136 | NA |
| TENT4B | 24.9426153 | 0.184395457 | 0.35438494 | 0.52032532 | 0.60283685 | NA |
| ADCY7 | 6.22830172 | 1.597745477 | 0.69260632 | 2.30685951 | 0.02106265 | NA |
| BRD7 | 27.289362 | 0.142116644 | 0.3239021 | 0.43876419 | 0.66083241 | NA |
| NKD1 | 107.349806 | 0.158890536 | 0.42700458 | 0.372105 | 0.70981468 | 0.91165558 |
| CYLD | 27.8149907 | -0.04269605 | 0.35256447 | -0.1211014 | 0.90361072 | NA |
| SALL1 | 17.5325188 | 0.016067116 | 0.51527972 | 0.03118135 | 0.97512491 | NA |
| AC087564.1 | 75.5369968 | 0.328555085 | 0.41057673 | 0.80022822 | 0.42357858 | 0.77656162 |
| HNRNPA1P4 | 5.46075718 | -0.886939105 | 0.70016832 | -1.2667513 | 0.20524424 | NA |
| TOX3 | 10.3594631 | -0.001212552 | 0.65051795 | -0.001864 | 0.99851276 | NA |

| | | | | | | |
|---|---|---|---|---|---|---|
| CHD9 | 229.179895 | 0.606908742 | 0.23217663 | 2.61399579 | 0.00894901 | 0.14875902 |
| RBL2 | 48.0269298 | 0.133916353 | 0.30937364 | 0.43286285 | 0.66511441 | 0.89968484 |
| AKTIP | 12.3803025 | -0.141104014 | 0.44517098 | -0.3169659 | 0.75126949 | NA |
| RPGRIP1L | 10.0995017 | 0.313028533 | 0.53430787 | 0.58585799 | 0.55797095 | NA |
| FTO | 137.144517 | -0.001014582 | 0.20337473 | -0.0049887 | 0.99601958 | 0.99922222 |
| LPCAT2 | 84.5224081 | -0.789679026 | 0.39260179 | -2.0113994 | 0.04428328 | 0.33093605 |
| AC007495.1 | 9.64286072 | -1.135293139 | 0.49200672 | -2.3074748 | 0.02102836 | NA |
| GNAO1 | 18.6297398 | -1.250285808 | 0.82780705 | -1.510359 | 0.13095185 | NA |
| AMFR | 35.2103644 | -0.084892787 | 0.32139462 | -0.2641388 | 0.79167297 | 0.93914731 |
| NUDT21 | 27.2924805 | 0.144358631 | 0.30053144 | 0.48034452 | 0.63098244 | NA |
| OGFOD1 | 12.0060573 | 0.008165003 | 0.44961538 | 0.01815997 | 0.98551123 | NA |
| BBS2 | 24.9918089 | 0.283980411 | 0.34917619 | 0.81328688 | 0.4160536 | NA |
| MT2A | 304.479635 | -1.571804297 | 0.94214914 | -1.6683179 | 0.09525263 | 0.46829657 |
| MT1E | 156.070017 | -0.794983372 | 0.92887098 | -0.8558598 | 0.39207531 | 0.75666812 |
| MT1M | 17.8514696 | -1.143002064 | 1.28562098 | -0.8890661 | 0.37396754 | NA |
| MT1F | 18.562421 | -1.359696971 | 0.94096419 | -1.4450039 | 0.14845678 | NA |
| MT1G | 155.413967 | 0.237623574 | 1.31350013 | 0.18090868 | 0.85643926 | 0.95615679 |
| MT1H | 47.4033832 | -0.076088011 | 1.67297418 | -0.0454807 | 0.96372417 | 0.98892249 |
| MT1X | 69.6828609 | -2.300893737 | 0.94119556 | -2.44465 | 0.01449928 | 0.19867928 |
| NUP93 | 9.71161373 | 0.162144874 | 0.47102133 | 0.34424104 | 0.73066502 | NA |
| SLC12A3 | 5.65615126 | 0.766228485 | 1.02348724 | 0.74864489 | 0.45407127 | NA |
| HERPUD1 | 70.9361847 | -0.350321835 | 0.47798896 | -0.7329078 | 0.46361466 | 0.79842482 |
| NLRC5 | 12.5015609 | 0.058775453 | 0.83466675 | 0.07041787 | 0.94386107 | NA |
| FAM192A | 25.0756612 | -0.13049093 | 0.34417128 | -0.3791453 | 0.70457996 | NA |
| RSPRY1 | 21.8073744 | 0.00169456 | 0.38265198 | 0.00442846 | 0.99646661 | NA |
| ARL2BP | 36.2284778 | -0.110515829 | 0.3436355 | -0.3216077 | 0.7477499 | 0.92253893 |
| CIAPIN1 | 12.2275461 | -0.132970506 | 0.44027805 | -0.3020148 | 0.76264075 | NA |
| COQ9 | 12.1513417 | -0.250598258 | 0.46735625 | -0.5362039 | 0.59181761 | NA |
| POLR2C | 11.0754224 | 0.457653618 | 0.56807463 | 0.80562234 | 0.42046064 | NA |
| DOK4 | 8.04198678 | 1.237540767 | 0.57337427 | 2.15834722 | 0.03090085 | NA |
| ADGRG1 | 11.0797241 | -1.012847394 | 0.52281422 | -1.9372989 | 0.05270882 | NA |
| KATNB1 | 4.78168207 | 1.051261648 | 0.61938085 | 1.69727825 | 0.08964407 | NA |
| KIFC3 | 28.3902007 | -0.205224155 | 0.6372122 | -0.3220656 | 0.74740297 | NA |
| USB1 | 6.59183938 | -0.349301876 | 0.58345736 | -0.5986759 | 0.54938901 | NA |
| MMP15 | 6.40413262 | -0.324001276 | 0.76532797 | -0.4233496 | 0.67204022 | NA |
| CFAP20 | 9.80253019 | -0.070322319 | 0.45327977 | -0.1551411 | 0.87671008 | NA |
| CSNK2A2 | 26.0144234 | 0.1360301 | 0.31220132 | 0.43571276 | 0.66304514 | NA |
| SETD6 | 3.84149604 | -0.972917446 | 0.76197449 | -1.2768373 | 0.2016597 | NA |
| CNOT1 | 53.8231366 | 0.359310665 | 0.25611104 | 1.40294878 | 0.16063211 | 0.56283048 |
| SLC38A7 | 6.2842481 | -0.370416065 | 0.66057871 | -0.5607448 | 0.57497153 | NA |
| GOT2 | 17.1245772 | 0.06899649 | 0.39599973 | 0.17423368 | 0.86168182 | NA |
| AC138305.1 | 15.9577178 | 1.124429448 | 0.7148096 | 1.57304748 | 0.11570782 | NA |
| CDH11 | 17.5327025 | -1.616151371 | 0.76030263 | -2.1256685 | 0.03353087 | NA |
| BEAN1 | 5.82702397 | -1.177089449 | 0.58923803 | -1.9976468 | 0.04575497 | NA |
| AC010542.3 | 12.5386385 | 0.707726412 | 0.56530915 | 1.25192811 | 0.21059606 | NA |
| TK2 | 7.21015844 | -0.866886731 | 0.59992051 | -1.4450027 | 0.14845714 | NA |
| CMTM3 | 18.5042628 | -0.976270187 | 0.54081068 | -1.8051977 | 0.07104375 | NA |
| CMTM4 | 46.4649524 | -1.293650145 | 0.45383614 | -2.8504785 | 0.00436535 | 0.10602312 |
| DYNC1LI2 | 43.6407181 | 0.106249562 | 0.3168435 | 0.33533768 | 0.73737038 | 0.91929008 |
| NAE1 | 21.8002712 | 0.656928314 | 0.32660875 | 2.01136162 | 0.04428727 | NA |
| PDP2 | 8.29896432 | -0.132307616 | 0.64518484 | -0.2050693 | 0.83751797 | NA |
| CDH16 | 19.2412972 | -0.929058869 | 0.48581921 | -1.9123552 | 0.05583066 | NA |
| RRAD | 6.36334213 | -2.326667338 | 1.03996711 | -2.2372509 | 0.02526995 | NA |
| FAM96B | 29.4171498 | -0.4093129 | 0.37922085 | -1.0793523 | 0.2804307 | 0.66958859 |
| CES2 | 11.4903088 | -0.202482382 | 0.46399829 | -0.4363861 | 0.66255666 | NA |
| CBFB | 35.9484725 | 0.000169967 | 0.35346128 | 0.00048087 | 0.99961632 | 0.99961632 |
| C16orf70 | 7.82345473 | -0.791203551 | 0.60165494 | -1.3150454 | 0.18849465 | NA |
| TRADD | 11.6045742 | 0.267437124 | 0.51820027 | 0.51608836 | 0.6057927 | NA |
| NOL3 | 23.2842837 | -0.510051962 | 0.3736199 | -1.3651627 | 0.17220191 | NA |
| KIAA0895L | 4.93960156 | -0.450950481 | 0.67432095 | -0.6687475 | 0.50365653 | NA |
| E2F4 | 9.11905774 | -0.215451527 | 0.51540577 | -0.4180231 | 0.67593021 | NA |
| LRRC29 | 5.74799304 | 0.297515987 | 0.57842429 | 0.51435597 | 0.60700314 | NA |
| TMEM208 | 18.3405204 | -0.598853994 | 0.44359315 | -1.3500073 | 0.17701364 | NA |
| TPPP3 | 133.047938 | 0.083427822 | 0.77503485 | 0.10764396 | 0.91427812 | 0.97707273 |
| ZDHHC1 | 10.6337776 | 0.084796553 | 0.43884028 | 0.19322874 | 0.84677983 | NA |
| HSD11B2 | 11.5837119 | 1.365928833 | 0.9951364 | 1.37260464 | 0.16987529 | NA |
| ATP6V0D1 | 23.4038058 | 0.059218638 | 0.39475108 | 0.15001514 | 0.88075267 | NA |
| AC009061.2 | 8.41184008 | 0.376388817 | 0.60649165 | 0.62060016 | 0.53486273 | NA |
| AC027682.6 | 38.1519344 | 0.082116124 | 0.51134535 | 0.16058838 | 0.8724176 | 0.96207879 |
| RIPOR1 | 399.082513 | -0.073429956 | 0.49099536 | -0.1495533 | 0.88111709 | 0.9644761 |
| AC027682.2 | 8.56018629 | 0.423927641 | 0.6047058 | 0.70104776 | 0.48327321 | NA |
| CTCF | 27.8543566 | -0.163902058 | 0.34767984 | -0.4714166 | 0.63734324 | NA |
| ENKD1 | 8.74781652 | 0.677306348 | 0.54478749 | 1.24324872 | 0.21377619 | NA |
| GFOD2 | 11.8995937 | 0.162531172 | 0.4402964 | 0.36914036 | 0.71202311 | NA |
| RANBP10 | 20.7029977 | 0.485935812 | 0.35507501 | 1.3685441 | 0.17114183 | NA |
| THAP11 | 7.01139103 | 0.00941979 | 0.64172737 | 0.0146788 | 0.98828843 | NA |
| NUTF2 | 8.99698595 | -0.274387996 | 0.56771678 | -0.4833185 | 0.62886964 | NA |
| PSKH1 | 9.89952076 | -0.139666418 | 0.47186284 | -0.2959894 | 0.76723815 | NA |
| SLC12A4 | 9.74941878 | -0.171815574 | 0.66456293 | -0.2585392 | 0.79599079 | NA |
| DUS2 | 5.30724157 | 0.127762157 | 0.74369334 | 0.17179414 | 0.86359937 | NA |
| NFATC3 | 134.442245 | -0.121311723 | 0.23805895 | -0.5095869 | 0.6103409 | 0.87320729 |
| PLA2G15 | 22.8961851 | 0.68893438 | 0.34372713 | 2.00430611 | 0.04503728 | NA |
| SLC7A6 | 12.1191603 | -0.054963221 | 0.4411277 | -0.1245971 | 0.90084254 | NA |
| SLC7A6OS | 13.8432469 | -0.803399142 | 0.45297564 | -1.7736034 | 0.07612877 | NA |
| PRMT7 | 9.79177125 | 0.837087337 | 0.44859888 | 1.86600406 | 0.0620408 | NA |
| ZFP90 | 17.5533251 | -0.545057966 | 0.35355732 | -1.5416396 | 0.1231612 | NA |
| CDH1 | 7.85903603 | 0.564929583 | 0.8287964 | 0.68162649 | 0.49547516 | NA |
| TANGO6 | 35.0937607 | 0.23448799 | 0.35935304 | 0.65252819 | 0.51406049 | 0.83070544 |
| CHTF8 | 20.8835252 | 0.105411442 | 0.38615401 | 0.27297772 | 0.78487034 | NA |
| UTP4 | 6.57393875 | -0.355169317 | 0.76193057 | -0.4661439 | 0.64111247 | NA |
| SNTB2 | 9.71206465 | 0.218524284 | 0.47698608 | 0.45813555 | 0.64685506 | NA |
| VPS4A | 16.9113828 | 0.113314761 | 0.40687198 | 0.27850225 | 0.78062684 | NA |
| COG8 | 10.818386 | 0.232686302 | 0.53679818 | 0.43347074 | 0.66467282 | NA |
| NIP7 | 4.49997335 | -0.797808785 | 0.9783331 | -0.8154777 | 0.41479895 | NA |
| TERF2 | 14.1536839 | -0.185374263 | 0.4116479 | -0.4503224 | 0.65247801 | NA |
| CYB5B | 78.3802579 | -0.121147237 | 0.29625487 | -0.4089291 | 0.68259169 | 0.90499319 |
| NFAT5 | 431.073521 | 0.056870878 | 0.19487188 | 0.29183727 | 0.77041105 | 0.92621526 |
| NQO1 | 113.551878 | -0.693068943 | 0.44497952 | -1.55753 | 0.11934471 | 0.51061407 |
| NOB1 | 11.2831368 | -0.60892039 | 0.46920994 | -1.2977568 | 0.19437092 | NA |
| WWP2 | 50.3281424 | 0.213770811 | 0.38836461 | 0.55043845 | 0.58201868 | 0.86175324 |
| PDPR | 35.4721156 | 0.365470863 | 0.40791152 | 0.89595623 | 0.37027614 | 0.73862609 |
| AC009060.1 | 7.66135357 | 0.223783708 | 0.83049419 | 0.26945849 | 0.78757688 | NA |
| EXOSC6 | 21.4300312 | -0.56871463 | 0.48701363 | -1.1677592 | 0.24290392 | NA |
| AARS | 25.8804753 | -0.776203199 | 0.45410947 | -1.7092865 | 0.08739789 | NA |
| DDX19A | 5.35912978 | 0.246604905 | 0.65575483 | 0.37606266 | 0.70687031 | NA |
| ST3GAL2 | 18.6140149 | 1.024483539 | 0.35507881 | 2.88522861 | 0.0039113 | NA |
| FUK | 5.41321416 | 0.018419813 | 0.56553993 | 0.03257031 | 0.97401724 | NA |
| COG4 | 20.6021988 | 0.542963627 | 0.36933609 | 1.47010715 | 0.14153274 | NA |
| SF3B3 | 28.8547811 | -0.019619329 | 0.3023319 | -0.0648933 | 0.94825892 | 0.98391202 |
| MTSS1L | 38.606302 | 0.441910591 | 0.38203403 | 1.15673097 | 0.24738229 | 0.65061974 |
| VAC14 | 17.740349 | -0.004063176 | 0.38811759 | -0.0104689 | 0.99164715 | NA |
| HYDIN | 20.3679934 | 0.578411101 | 0.49242952 | 1.17460688 | 0.24015203 | NA |
| CMTR2 | 13.1191729 | -0.239243734 | 0.54282327 | -0.4407396 | 0.65940149 | NA |

| | | | | | | |
|---|---|---|---|---|---|---|
| ZNF19 | 7.55367441 | -0.496614426 | 0.59109259 | -0.8401635 | 0.40081672 | NA |
| PHLPP2 | 23.4872645 | -0.333757858 | 0.36195816 | -0.9220896 | 0.35648183 | NA |
| AP1G1 | 62.204028 | 0.417411344 | 0.28348814 | 1.47241199 | 0.14090966 | 0.53597868 |
| IST1 | 47.5150975 | 0.148558164 | 0.23441875 | 0.63372987 | 0.52625713 | 0.83195298 |
| ZNF821 | 6.62481561 | -0.131680914 | 0.63371019 | -0.2077936 | 0.83539014 | NA |
| DHODH | 5.23387616 | -0.13216378 | 0.63201056 | -0.2091164 | 0.83435736 | NA |
| TXNL4B | 10.5493858 | -0.047055588 | 0.63572693 | -0.0740186 | 0.94099562 | NA |
| DHX38 | 10.5563515 | -0.520462727 | 0.48005044 | -1.0841834 | 0.27828348 | NA |
| LINC01572 | 21.3259307 | 0.201869566 | 0.58127025 | 0.34729038 | 0.72837318 | NA |
| AC004943.2 | 13.3701826 | -0.229671257 | 0.4500981 | -0.5102693 | 0.60986278 | NA |
| ZFHX3 | 100.512763 | -0.082929608 | 0.22941189 | -0.3614878 | 0.71773479 | 0.91472065 |
| AC138627.1 | 7.36868544 | 0.912592899 | 0.68250067 | 1.33713113 | 0.18117984 | NA |
| PSMD7 | 26.1662552 | -0.635552629 | 0.31088875 | -2.0443089 | 0.04092304 | NA |
| NPIPB15 | 4.91577861 | -2.445520962 | 1.1336283 | -2.1572512 | 0.0309861 | NA |
| GLG1 | 122.75866 | 0.083224714 | 0.24432865 | 0.3406261 | 0.73338508 | 0.91884473 |
| RFWD3 | 13.5583845 | 0.859275457 | 0.47989386 | 1.79055313 | 0.07336503 | NA |
| WDR59 | 36.8621995 | 0.07327853 | 0.34111413 | 0.21482115 | 0.82990676 | 0.95008863 |
| ZNRF1 | 9.65589463 | -0.139292743 | 0.61747451 | -0.2255846 | 0.82152452 | NA |
| ZFP1 | 4.19236295 | 0.829593732 | 0.80783854 | 1.02693012 | 0.30445337 | NA |
| BCAR1 | 31.8921407 | -0.223456323 | 0.45012852 | -0.4964278 | 0.6195926 | 0.87746773 |
| CFDP1 | 60.2403283 | 0.171129202 | 0.29705364 | 0.57608854 | 0.56455533 | 0.8549147 |
| GABARAPL2 | 74.5206414 | -0.794832406 | 0.54930019 | -1.446991 | 0.14789946 | 0.54691531 |
| ADAT1 | 10.4719828 | 0.011924575 | 0.46028217 | 0.0259071 | 0.97933144 | NA |
| KARS | 19.2795841 | -0.01915182 | 0.46853965 | -0.0408756 | 0.9673951 | NA |
| TERF2IP | 95.7564773 | -0.514428363 | 0.31489074 | -1.6336725 | 0.10232763 | 0.48317363 |
| AC010528.1 | 3.67587954 | -0.018876336 | 1.14447439 | -0.0164935 | 0.98684072 | NA |
| NUDT7 | 7.13010609 | 0.497726171 | 0.60511885 | 0.8225263 | 0.41077742 | NA |
| WWOX | 977.000411 | 1.111089906 | 0.33925575 | 3.27508055 | 0.00105632 | 0.05186671 |
| AC046158.1 | 5.37139846 | 0.01860309 | 0.88632528 | 0.02098901 | 0.98325442 | NA |
| MAF | 27.9427096 | 0.000277904 | 0.48737623 | 0.0005702 | 0.99954504 | NA |
| CDYL2 | 14.3622894 | 0.252205905 | 0.51504308 | 0.48967924 | 0.62436089 | NA |
| CMC2 | 23.2296667 | 0.144351855 | 0.3596039 | 0.40141905 | 0.68811163 | NA |
| ATMIN | 15.0343788 | -0.833194552 | 0.40295827 | -2.0676944 | 0.03866877 | NA |
| GAN | 12.9831125 | -0.311487668 | 0.41762676 | -0.7458518 | 0.45575695 | NA |
| CMIP | 82.0796816 | 1.031873059 | 0.30382505 | 3.39627378 | 0.0006831 | 0.03902329 |
| PLCG2 | 217.288109 | 1.953255867 | 0.45834338 | 4.26155571 | 2.03E-05 | 0.00304964 |
| SDR42E1 | 14.9640602 | 0.644154092 | 0.43446718 | 1.48263004 | 0.13817273 | NA |
| MPHOSPH6 | 15.432266 | -0.640217939 | 0.45968301 | -1.3927379 | 0.16369907 | NA |
| HSBP1 | 50.6163217 | -0.157228071 | 0.38600729 | -0.4073189 | 0.68377377 | 0.90499319 |
| OSGIN1 | 9.27511493 | -1.67532112 | 1.00629976 | -1.6648331 | 0.09594606 | NA |
| MBTPS1 | 46.7611865 | -0.255391905 | 0.39494911 | -0.6466451 | 0.51786167 | 0.83104823 |
| HSDL1 | 10.0777266 | 0.278899636 | 0.53254087 | 0.52371499 | 0.60047679 | NA |
| TAF1C | 7.19183136 | -0.21419421 | 0.5483003 | -0.3906513 | 0.69605502 | NA |
| TLDC1 | 14.7497196 | -0.079674251 | 0.47440253 | -0.1679465 | 0.86662536 | NA |
| COTL1 | 28.8815133 | -2.158223369 | 0.70981026 | -3.0405638 | 0.00236136 | 0.07786717 |
| KLHL36 | 19.8523146 | -0.421149043 | 0.49133504 | -0.8571525 | 0.39136062 | NA |
| USP10 | 44.653784 | -0.045031354 | 0.26918244 | -0.1672893 | 0.86714239 | 0.96089268 |
| ZDHHC7 | 21.0318478 | 0.123061137 | 0.32231356 | 0.38180565 | 0.70260553 | NA |
| KIAA0513 | 5.84898804 | 0.515348246 | 0.68201821 | 0.75562242 | 0.44987561 | NA |
| GSE1 | 30.1502818 | -0.491364277 | 0.4436843 | -1.1074637 | 0.26809348 | 0.66098308 |
| EMC8 | 12.67694 | -0.510521782 | 0.42613689 | -1.198023 | 0.23090808 | NA |
| COX4I1 | 91.8007256 | -0.377942995 | 0.35910587 | -1.0524556 | 0.29259055 | 0.67930583 |
| MTHFSD | 5.76147431 | -0.38018663 | 0.62855754 | -0.6048557 | 0.54527486 | NA |
| FOXC2 | 24.9640907 | -0.727966263 | 0.54958507 | -1.3245743 | 0.18531238 | NA |
| AC009154.1 | 6.28831089 | -0.464010539 | 0.82028747 | -0.5656682 | 0.57161935 | NA |
| FBXO31 | 8.85870237 | -0.501004733 | 0.5240723 | -0.955984 | 0.33908031 | NA |
| MAP1LC3B | 44.2740192 | -0.449582213 | 0.3476343 | -1.293262 | 0.19592047 | 0.59987316 |
| ZCCHC14 | 49.6305483 | 0.21591213 | 0.34133534 | 0.63255134 | 0.52702667 | 0.83195298 |
| KLHDC4 | 19.0159853 | 0.462061237 | 0.37391623 | 1.23573462 | 0.21655722 | NA |
| SLC7A5 | 3.8289226 | -1.382701159 | 1.27069661 | -1.0881442 | 0.27653145 | NA |
| BANP | 20.7773619 | -0.328060392 | 0.41564182 | -0.7892863 | 0.42994469 | NA |
| ZFPM1 | 15.3671556 | -0.647465666 | 0.37340659 | -1.7339428 | 0.08292823 | NA |
| ZC3H18 | 19.3115837 | 0.060431748 | 0.41625313 | 0.14518029 | 0.88456853 | NA |
| CYBA | 48.0986039 | -0.639223698 | 0.43876006 | -1.4568867 | 0.14514766 | 0.54410249 |
| MVD | 10.1332062 | 0.508930685 | 0.59173007 | 0.86007237 | 0.38974915 | NA |
| RNF166 | 8.49549175 | 1.616141871 | 0.59832172 | 2.70112521 | 0.00691053 | NA |
| PIEZO1 | 79.8293497 | 0.357349223 | 0.36175358 | 0.98782499 | 0.32323837 | 0.69588897 |
| APRT | 27.7691019 | -0.978538746 | 0.46455235 | -2.1064122 | 0.03516856 | NA |
| GALNS | 11.8346953 | 0.451459912 | 0.43939035 | 1.02746889 | 0.30419973 | NA |
| TRAPPC2L | 7.35750782 | -0.375373872 | 0.68110516 | -0.5511247 | 0.5815482 | NA |
| ACSF3 | 10.0341204 | -0.131651115 | 0.44401787 | -0.2964996 | 0.76684858 | NA |
| ZNF778 | 5.69149218 | 0.75407802 | 0.63682947 | 1.18411294 | 0.23636836 | NA |
| ANKRD11 | 156.431756 | -0.132375383 | 0.18972617 | -0.6977181 | 0.48535351 | 0.81450403 |
| SPG7 | 41.2785534 | 0.04651199 | 0.36315326 | 0.12807813 | 0.89808714 | 0.97097328 |
| RPL13 | 157.597358 | -0.900862 | 0.40990172 | -2.1977512 | 0.02796684 | 0.27202278 |
| DPEP1 | 6.24938955 | 0.988171412 | 1.00172664 | 0.98646813 | 0.32390345 | NA |
| CHMP1A | 21.2521445 | 0.125255567 | 0.46265644 | 0.27073127 | 0.78659772 | NA |
| SPATA33 | 4.70671817 | -1.434824939 | 0.67422133 | -2.1281215 | 0.03332701 | NA |
| CDK10 | 13.0879897 | 0.141665186 | 0.42445758 | 0.33375582 | 0.73856383 | NA |
| VPS9D1 | 8.25187496 | -0.55350323 | 0.51407646 | -1.0766944 | 0.28161683 | NA |
| ZNF276 | 6.2077268 | -0.277029016 | 0.55600423 | -0.4982498 | 0.61830796 | NA |
| TCF25 | 92.8093783 | -0.566758246 | 0.27610024 | -2.0527264 | 0.04009912 | 0.31540889 |
| DEF8 | 9.96046226 | -0.937153167 | 0.55437657 | -1.6904631 | 0.09093938 | NA |
| GAS8 | 9.9166664 | -0.70667494 | 0.5121896 | -1.3797136 | 0.16767486 | NA |
| FAM157C | 23.5477573 | 1.285165093 | 0.42064311 | 3.05523867 | 0.00224881 | NA |
| RPH3AL | 27.6181265 | 0.174704597 | 0.62090398 | 0.28137136 | 0.77842558 | NA |
| RFLNB | 17.4845461 | 0.615019317 | 1.14559786 | 0.53685446 | 0.59136815 | NA |
| VPS53 | 51.7594938 | -0.229627539 | 0.32001538 | -0.7175516 | 0.47303383 | 0.80580648 |
| GLOD4 | 15.5061008 | 0.08272021 | 0.43636279 | 0.18956752 | 0.84964804 | NA |
| NXN | 89.0570056 | -0.082998452 | 0.37868225 | -0.219177 | 0.82651214 | 0.949142 |
| TIMM22 | 8.42136591 | 0.051884318 | 0.51133449 | 0.10146845 | 0.9191786 | NA |
| ABR | 34.8803385 | 0.601895419 | 0.44797148 | 1.34360211 | 0.17907709 | 0.58293473 |
| YWHAE | 144.178646 | -0.245205346 | 0.2274489 | -1.0780678 | 0.28100349 | 0.66958859 |
| CRK | 41.2140752 | -0.3818799 | 0.32944557 | -1.1591593 | 0.24639125 | 0.65061974 |
| MYO1C | 103.13862 | 0.526118072 | 0.46741905 | 1.12558115 | 0.26034285 | 0.65595635 |
| INPP5K | 7.82535817 | -0.00697549 | 0.64635867 | -0.010792 | 0.99138941 | NA |
| PITPNA | 39.4944436 | -0.038692413 | 0.28951091 | -0.1336475 | 0.89368131 | 0.97097328 |
| SLC43A2 | 14.254269 | 1.876114524 | 0.81589372 | 2.29945944 | 0.02147886 | NA |
| PRPF8 | 46.5978858 | 0.12994745 | 0.29515214 | 0.44027276 | 0.65973957 | 0.89709157 |
| TLCD2 | 6.67981031 | -0.804402925 | 0.76378719 | -1.0531768 | 0.29225998 | NA |
| MIR22HG | 16.1000542 | -0.749449488 | 0.48553199 | -1.5435636 | 0.1226941 | NA |
| WDR81 | 10.1913803 | 0.241904573 | 0.42428813 | 0.57014221 | 0.56858125 | NA |
| SMYD4 | 10.1039241 | -0.410288225 | 0.62659139 | -0.6547939 | 0.51260045 | NA |
| RPA1 | 16.7046828 | -0.159019503 | 0.38220476 | -0.4160584 | 0.67736726 | NA |
| SMG6 | 68.3179977 | 0.051241603 | 0.28284495 | 0.18116499 | 0.85623807 | 0.95615679 |
| SRR | 6.61076748 | -0.161173665 | 0.55505451 | -0.2903745 | 0.77152977 | NA |
| TSR1 | 9.11469733 | -0.797516242 | 0.66531492 | -1.1987049 | 0.23064272 | NA |
| SGSM2 | 9.94920475 | 0.444603325 | 0.51523411 | 0.86291517 | 0.38818411 | NA |
| MNT | 9.31447391 | -1.102792982 | 0.60949931 | -1.8093425 | 0.07039782 | NA |
| METTL16 | 23.8971646 | -0.631595034 | 0.34082041 | -1.8531608 | 0.06385931 | NA |
| PAFAH1B1 | 153.755395 | 0.061839925 | 0.20787745 | 0.29748261 | 0.76609808 | 0.92423969 |
| CLUH | 12.1321025 | 0.796594119 | 0.54724168 | 1.45565322 | 0.14548851 | NA |
| CTNS | 6.45112767 | -0.376091696 | 0.54092939 | -0.6952695 | 0.48688644 | NA |

| | | | | | | |
|---|---|---|---|---|---|---|
| TAX1BP3 | 15.3813352 | 0.126347092 | 0.48793174 | 0.2589442 | 0.7956783 | NA |
| EMC6 | 19.203876 | 0.052010087 | 0.56744073 | 0.0916573 | 0.92697032 | NA |
| ITGAE | 10.4906776 | 0.299831278 | 0.6291127 | 0.47659391 | 0.63365132 | NA |
| NCBP3 | 29.1668875 | 0.529711697 | 0.34061413 | 1.55516656 | 0.11990641 | 0.51128003 |
| CAMKK1 | 4.50700248 | -0.206119041 | 0.9203878 | -0.223948 | 0.82279774 | NA |
| ZZEF1 | 36.8380581 | 0.236805833 | 0.33114866 | 0.71510431 | 0.47454459 | 0.80634808 |
| CYB5D2 | 22.5990445 | -0.249727911 | 0.38652516 | -0.6460845 | 0.51822464 | NA |
| ANKFY1 | 36.1353604 | 0.168662391 | 0.3791123 | 0.44488767 | 0.65640093 | 0.89641824 |
| UBE2G1 | 36.972598 | 0.240181447 | 0.29423236 | 0.81629853 | 0.41432941 | 0.7698214 |
| MYBBP1A | 6.54866305 | -0.386054681 | 0.6804235 | -0.5673741 | 0.57046003 | NA |
| PELP1 | 10.9675385 | -0.168563554 | 0.48804665 | -0.3453841 | 0.72980563 | NA |
| MED11 | 9.75314929 | -0.568642702 | 0.62541468 | -0.9092251 | 0.36323133 | NA |
| CXCL16 | 8.75125841 | 0.739985283 | 0.85173541 | 0.86879713 | 0.3849581 | NA |
| PSMB6 | 28.2971628 | -0.707235324 | 0.65022619 | -1.0876758 | 0.27673824 | NA |
| MINK1 | 17.432106 | 0.173033761 | 0.35316915 | 0.48994585 | 0.62417222 | NA |
| SLC25A11 | 13.8412286 | -0.962489909 | 0.67300608 | -1.4301355 | 0.15267812 | NA |
| RNF167 | 16.3746915 | -0.308329563 | 0.5028668 | -0.6131436 | 0.53978139 | NA |
| PFN1 | 93.0388257 | -0.511440436 | 0.47329779 | -1.0805891 | 0.27987993 | 0.66958859 |
| SPAG7 | 31.4491953 | -0.014699982 | 0.42299273 | -0.0347523 | 0.97227723 | 0.99072304 |
| CAMTA2 | 12.9245865 | 0.109248451 | 0.43699827 | 0.24999744 | 0.80258933 | NA |
| KIF1C | 14.6498633 | -0.297312274 | 0.62163582 | -0.478274 | 0.63245517 | NA |
| ZNF594 | 6.72441542 | -0.456931962 | 0.60548136 | -0.754659 | 0.45045361 | NA |
| AC087500.1 | 14.209397 | -0.041002999 | 0.41282837 | -0.0993221 | 0.9208825 | NA |
| RABEP1 | 65.2339801 | 0.075642764 | 0.23725969 | 0.31881844 | 0.74986419 | 0.92274989 |
| NUP88 | 26.2644853 | 0.386190832 | 0.34133098 | 1.13142625 | 0.25787573 | NA |
| RPAIN | 23.5308117 | -0.331234322 | 0.36254755 | -0.91363 | 0.36091129 | NA |
| C1QBP | 20.5825477 | -1.317246744 | 0.44341409 | -2.9706921 | 0.00297129 | NA |
| DERL2 | 22.7224671 | 0.178706973 | 0.40711984 | 0.43895422 | 0.66069471 | NA |
| MIS12 | 10.0032588 | -0.140095861 | 0.45588783 | -0.3073034 | 0.75861248 | NA |
| NLRP1 | 40.343826 | 0.568090301 | 0.61992371 | 0.91638743 | 0.35946372 | 0.72980778 |
| PITPNM3 | 19.1787329 | -0.219356943 | 0.50513046 | -0.434258 | 0.6641011 | NA |
| KIAA0753 | 17.157255 | -0.139816228 | 0.38285137 | -0.3651971 | 0.71496425 | NA |
| TXNDC17 | 11.04633 | -0.769276267 | 0.63187846 | -1.2174434 | 0.22343555 | NA |
| MED31 | 8.64866405 | -0.349970855 | 0.50713852 | -0.6900893 | 0.49013805 | NA |
| XAF1 | 36.2153798 | -1.668146887 | 1.54779226 | -1.0777589 | 0.28114136 | 0.66958859 |
| ALOX12-AS1 | 7.86133765 | -0.102231884 | 1.04411216 | -0.0979127 | 0.92200158 | NA |
| ACADVL | 50.7986659 | 0.267022093 | 0.28217604 | 0.94629613 | 0.34399757 | 0.71842682 |
| DVL2 | 10.3714521 | 0.008261604 | 0.49726628 | 0.01661404 | 0.98674452 | NA |
| PHF23 | 7.71138615 | -0.492968827 | 0.58177479 | -0.8473534 | 0.39679819 | NA |
| ELP5 | 9.05228804 | -0.352397222 | 0.46153011 | -0.7635411 | 0.44514074 | NA |
| EIF5A | 27.8530261 | -0.227566524 | 0.61590866 | -0.369481 | 0.71176926 | NA |
| GPS2 | 21.576886 | -0.131821077 | 0.53860368 | -0.244746 | 0.80665314 | NA |
| KCTD11 | 5.09308531 | 0.767895413 | 0.62750143 | 1.22373493 | 0.22105224 | NA |
| NLGN2 | 6.21893694 | -0.22680939 | 0.61337462 | -0.369773 | 0.71155162 | NA |
| CHRNB1 | 8.67462102 | -0.394748098 | 0.59351008 | -0.6651077 | 0.50598164 | NA |
| ZBTB4 | 65.1249429 | 0.184543228 | 0.3060411 | 0.60300146 | 0.54650772 | 0.84449366 |
| POLR2A | 55.9588609 | -0.158159354 | 0.38057397 | -0.4155811 | 0.67771655 | 0.90499319 |
| EIF4A1 | 28.913177 | -0.721896342 | 0.73169787 | -0.9866044 | 0.32383661 | 0.69623471 |
| CD68 | 9.44244756 | 1.931234834 | 1.7978808 | 1.0741729 | 0.2827452 | NA |
| MPDU1 | 11.4796194 | 0.326131943 | 0.55332006 | 0.58940922 | 0.55558679 | NA |
| FXR2 | 20.0865936 | 0.04265696 | 0.45451145 | 0.09385233 | 0.92522646 | NA |
| SAT2 | 23.5380321 | -0.637934177 | 0.52584486 | -1.2131604 | 0.22506849 | NA |
| TP53 | 9.56725076 | -0.16102371 | 0.6029988 | -0.2670382 | 0.78943975 | NA |
| KDM6B | 10.5902494 | -0.430915214 | 0.57823453 | -0.7452257 | 0.45613531 | NA |
| NAA38 | 45.920674 | -0.561820225 | 0.36103034 | -1.5561579 | 0.11967054 | 0.51093023 |
| CHD3 | 91.239615 | -0.202559454 | 0.47627325 | -0.4253009 | 0.67061732 | 0.90305784 |
| TRAPPC1 | 15.91345 | -1.118354117 | 0.45146275 | -2.4771792 | 0.01324254 | NA |
| PER1 | 44.0452804 | -1.037540408 | 0.47514949 | -2.1836084 | 0.02899104 | 0.27926427 |
| TMEM107 | 16.5912357 | -1.413925967 | 0.56952632 | -2.482635 | 0.01304147 | NA |
| BORCS6 | 4.4782755 | -0.040303085 | 0.8133204 | -0.0495538 | 0.96047799 | NA |
| CTC1 | 16.3445891 | -0.952358132 | 0.4395473 | -2.1666795 | 0.0302593 | NA |
| SLC25A35 | 3.75940003 | -0.409383362 | 0.88907142 | -0.4604617 | 0.64518484 | NA |
| RANGRF | 7.89081793 | -0.204978672 | 0.60189755 | -0.3405541 | 0.7334393 | NA |
| NDEL1 | 25.4205986 | -0.369209067 | 0.39284481 | -0.9398344 | 0.34730252 | NA |
| MYH10 | 34.2611716 | 0.916110282 | 0.37615496 | 2.4354598 | 0.01487288 | 0.19972267 |
| STX8 | 75.1563447 | 0.039355922 | 0.20295208 | 0.19391732 | 0.84624062 | 0.95532127 |
| USP43 | 5.99293125 | -0.051726125 | 0.69417197 | -0.0745149 | 0.94060072 | NA |
| GAS7 | 30.0232469 | -1.582170575 | 0.78778801 | -2.008371 | 0.04460388 | 0.33256131 |
| SCO1 | 14.9675317 | -0.159768841 | 0.49834491 | -0.3205989 | 0.74851436 | NA |
| TMEM220 | 13.8985836 | -0.447521614 | 0.63768486 | -0.7017912 | 0.4828094 | NA |
| SHISA6 | 73.9307948 | -0.625311946 | 0.73385306 | -0.8520942 | 0.39416181 | 0.75773027 |
| DNAH9 | 29.6810584 | -0.317810175 | 0.64158952 | -0.4953481 | 0.6203544 | 0.87793351 |
| MAP2K4 | 65.1572259 | 0.339669456 | 0.2010139 | 1.68978093 | 0.09106987 | 0.45906588 |
| ARHGAP44 | 20.1773494 | -1.311840851 | 0.45665268 | -2.8727322 | 0.00406939 | NA |
| ELAC2 | 12.5060467 | -0.537173632 | 0.44379053 | -1.2104216 | 0.22611717 | NA |
| HS3ST3A1 | 78.6036352 | -0.936253925 | 0.40973058 | -2.2850477 | 0.02231003 | 0.24469443 |
| COX10 | 18.6421998 | 0.45475182 | 0.44990627 | 1.01077013 | 0.31212646 | NA |
| HS3ST3B1 | 117.903947 | -1.462340744 | 0.4561227 | -3.2060249 | 0.00134582 | 0.0545866 |
| TRIM16 | 5.27058784 | -0.636952301 | 0.61611644 | -1.0338181 | 0.30122123 | NA |
| AC015922.4 | 4.50967966 | 0.336648591 | 0.71796327 | 0.46889389 | 0.63914548 | NA |
| ZSWIM7 | 20.5241791 | -0.078396054 | 0.35412838 | -0.2213775 | 0.82479851 | NA |
| TTC19 | 44.4785163 | -0.407443857 | 0.31436008 | -1.2961056 | 0.19493911 | 0.59944886 |
| NCOR1 | 114.448191 | 0.056942572 | 0.18800714 | 0.30287452 | 0.76198549 | 0.92336765 |
| PIGL | 34.910534 | 0.064435006 | 0.34254922 | 0.18810437 | 0.85079483 | 0.95532127 |
| UBB | 178.627722 | -1.000873707 | 0.3548099 | -2.8208731 | 0.00478931 | 0.11177304 |
| LRRC75A | 5.47192555 | -0.823460269 | 0.82711868 | -0.9955769 | 0.31945575 | NA |
| ZNF624 | 5.89527078 | -0.347187015 | 0.66559349 | -0.5216202 | 0.60193479 | NA |
| AC098850.3 | 19.1577804 | -0.032961743 | 0.69438951 | -0.0474687 | 0.9621397 | NA |
| MPRIP | 75.0652756 | 0.465252611 | 0.25304187 | 1.83863883 | 0.06596833 | 0.3991664 |
| COPS3 | 10.7744519 | -0.331180895 | 0.41797219 | -0.7923515 | 0.42815575 | NA |
| MED9 | 6.66738873 | 0.563017543 | 0.59369988 | 0.94832012 | 0.34296651 | NA |
| RASD1 | 12.782378 | -0.153436722 | 0.78689662 | -0.1949897 | 0.84540103 | NA |
| PEMT | 7.53019226 | -0.08083109 | 0.5787495 | -0.1396651 | 0.88892463 | NA |
| RAI1 | 48.02974 | -0.462456815 | 0.34504196 | -1.3402915 | 0.18015059 | 0.58390314 |
| SMCR5 | 4.3912513 | -0.80815207 | 0.86121828 | -0.9383824 | 0.34804793 | NA |
| SREBF1 | 20.9111632 | -0.129151474 | 0.62401779 | -0.2069676 | 0.83603515 | NA |
| TOM1L2 | 267.079719 | -0.089830704 | 0.36073249 | -0.249023 | 0.80334295 | 0.94126368 |
| ATPAF2 | 5.02691033 | -0.679946738 | 0.65924679 | -1.0313994 | 0.30235356 | NA |
| GID4 | 5.14826175 | -0.662125638 | 0.66877631 | -0.9900555 | 0.32214701 | NA |
| DRG2 | 8.85271272 | 0.146053644 | 0.52483333 | 0.27828576 | 0.780793 | NA |
| ALKBH5 | 35.4287119 | -0.422658154 | 0.31967529 | -1.3221483 | 0.18611876 | 0.58792656 |
| FLII | 20.9353911 | 0.244329872 | 0.38362753 | 0.63689348 | 0.52419423 | NA |
| TOP3A | 8.3617687 | -1.476375629 | 0.58896918 | -2.5067112 | 0.01218602 | NA |
| SMCR8 | 5.59940743 | 0.009508006 | 0.57374099 | 0.01657195 | 0.9867781 | NA |
| SHMT1 | 8.67942425 | 0.9068913 | 0.77061567 | 1.17683993 | 0.2392594 | NA |
| TVP23B | 10.0196093 | -0.046471745 | 0.45759279 | -0.101557 | 0.91910832 | NA |
| PRPSAP2 | 27.6463741 | 0.473711147 | 0.35806352 | 1.32298074 | 0.18584178 | NA |
| SLC5A10 | 5.28219581 | 0.321932103 | 0.85844514 | 0.37501768 | 0.70764732 | NA |
| FAM83G | 8.5701348 | 0.56889145 | 0.66819497 | 0.85138541 | 0.3945553 | NA |
| EPN2 | 97.086787 | 0.112443124 | 0.21325045 | 0.52728201 | 0.59799777 | 0.86882737 |
| B9D1 | 25.0801876 | -0.890726457 | 0.35077972 | -2.5392758 | 0.01110822 | NA |
| MAPK7 | 11.3188576 | -0.115715017 | 0.51563161 | -0.2244141 | 0.82243508 | NA |
| AC004448.2 | 5.64663572 | -1.042767509 | 0.71489491 | -1.4586305 | 0.14466684 | NA |

| | | | | | | |
|---|---|---|---|---|---|---|
| SLC47A1 | 12.7226386 | 1.973562132 | 0.89065074 | 2.21586536 | 0.02670072 | NA |
| ALDH3A2 | 18.6885972 | 0.412642581 | 0.37304642 | 1.10614272 | 0.26866475 | NA |
| SLC47A2 | 9.82922144 | 0.098513255 | 1.03675016 | 0.09502121 | 0.92429798 | NA |
| ULK2 | 23.6602274 | -0.015513177 | 0.35638099 | -0.0435298 | 0.96527925 | NA |
| AKAP10 | 51.3353818 | 0.362883755 | 0.28669549 | 1.26574631 | 0.20560392 | 0.61408653 |
| SPECC1 | 9.39658987 | -0.41586955 | 0.64308747 | -0.6466765 | 0.51784134 | NA |
| CCDC144NL-/ | 18.7771525 | -0.645080108 | 0.7362191 | -0.8762067 | 0.38091768 | NA |
| USP22 | 56.2191788 | -0.153859454 | 0.33319554 | -0.4617692 | 0.64424681 | 0.89091887 |
| DHRS7B | 30.6091033 | -0.629904759 | 0.37490331 | -1.6801793 | 0.09292244 | 0.46399004 |
| TMEM11 | 17.3393867 | -1.027571383 | 0.50957535 | -2.0165249 | 0.04374512 | NA |
| AC087294.1 | 8.52668055 | -1.107289079 | 0.65748602 | -1.6841257 | 0.09215739 | NA |
| NATD1 | 17.1709673 | -1.672449194 | 0.56569572 | -2.9564466 | 0.00311206 | NA |
| MAP2K3 | 27.1875669 | -1.105883889 | 0.44826967 | -2.4670058 | 0.01362481 | NA |
| C17orf51 | 5.16830732 | -0.581518233 | 0.60617617 | -0.9593222 | 0.33739648 | NA |
| WSB1 | 208.327895 | 0.516342357 | 0.31833132 | 1.62202812 | 0.10479733 | 0.4862279 |
| KSR1 | 29.1333765 | -0.788988745 | 0.44384042 | -1.7776406 | 0.0754629 | 0.42138852 |
| LGALS9 | 4.12216035 | -0.332491839 | 1.52378802 | -0.2182008 | 0.82727262 | NA |
| NLK | 443.58649 | 0.964020425 | 0.36125073 | 2.66856325 | 0.00761764 | 0.13970383 |
| IFT20 | 14.2002344 | -0.088586839 | 0.48044679 | -0.1843843 | 0.85371199 | NA |
| TNFAIP1 | 9.89783994 | -0.194039273 | 0.50643436 | -0.3831479 | 0.70161009 | NA |
| POLDIP2 | 13.2512599 | -0.201632218 | 0.49060786 | -0.4109845 | 0.68108391 | NA |
| TMEM199 | 7.71865756 | 0.029721931 | 0.63543688 | 0.04677401 | 0.96269334 | NA |
| SARM1 | 13.230395 | -0.623129216 | 0.40495993 | -1.5387429 | 0.12386708 | NA |
| AC005726.1 | 6.05711005 | -0.477515695 | 0.79121073 | -0.6035253 | 0.54615929 | NA |
| KIAA0100 | 21.3020447 | 0.086549322 | 0.48840081 | 0.17720962 | 0.85934374 | NA |
| SDF2 | 20.2038734 | -0.748439226 | 0.44574137 | -1.6790885 | 0.09313479 | NA |
| SUPT6H | 29.3643157 | 0.358179001 | 0.33223715 | 1.07808233 | 0.28099702 | 0.66958859 |
| RAB34 | 23.7889194 | -0.6448745 | 0.37794758 | -1.7062538 | 0.08796083 | NA |
| RPL23A | 64.606387 | -0.641637593 | 0.42716715 | -1.5020761 | 0.13307744 | 0.52613957 |
| TRAF4 | 14.5350463 | -0.234728716 | 0.49403269 | -0.4751279 | 0.63469581 | NA |
| FAM222B | 72.004984 | 0.288390703 | 0.27605669 | 1.04467927 | 0.29617123 | 0.68103704 |
| ERAL1 | 4.71462512 | -0.033339817 | 0.68098954 | -0.0489579 | 0.96095285 | NA |
| FLOT2 | 23.0671674 | -1.098789006 | 0.46241659 | -2.3761885 | 0.01749252 | NA |
| PHF12 | 23.481429 | -0.213155339 | 0.4191247 | -0.5085726 | 0.61105184 | NA |
| MYO18A | 8.40001379 | -0.068496254 | 0.46786658 | -0.1464013 | 0.88360464 | NA |
| NUFIP2 | 93.1298435 | -0.014591611 | 0.23061608 | -0.0632723 | 0.94954967 | 0.98475414 |
| TAOK1 | 112.058035 | 0.025145369 | 0.27807833 | 0.09042549 | 0.9279491 | 0.97775374 |
| ABHD15-AS1 | 7.21461645 | -0.667363701 | 0.65922157 | -1.0123511 | 0.3113702 | NA |
| GIT1 | 11.3518336 | -0.733627945 | 0.50009209 | -1.4669857 | 0.14237994 | NA |
| SSH2 | 75.2387632 | 0.531036921 | 0.30891811 | 1.71902165 | 0.08561043 | 0.44564602 |
| NSRP1 | 58.358354 | -0.222589809 | 0.24490803 | -0.908871 | 0.36341822 | 0.73298075 |
| BLMH | 17.4472057 | 0.063882289 | 0.38816033 | 0.16457707 | 0.86927688 | NA |
| CPD | 83.5876898 | -0.111678004 | 0.32050994 | -0.3484385 | 0.72751089 | 0.91700494 |
| GOSR1 | 37.3549669 | -0.162724004 | 0.27717105 | -0.5870887 | 0.55714411 | 0.85122066 |
| CRLF3 | 26.7070966 | -0.25715044 | 0.42615164 | -0.6034247 | 0.54622617 | NA |
| ATAD5 | 4.70249567 | -1.48143879 | 0.81573995 | -1.8160675 | 0.06936002 | NA |
| TEFM | 13.544311 | 0.106974524 | 0.39881975 | 0.26822775 | 0.78852402 | NA |
| RNF135 | 12.4096505 | -0.014714236 | 0.42764869 | -0.0344073 | 0.97255236 | NA |
| NF1 | 173.135766 | 0.420341444 | 0.21967702 | 1.91345202 | 0.05569021 | 0.36728373 |
| COPRS | 13.7827429 | -0.571313277 | 0.48102706 | -1.1876947 | 0.23495373 | NA |
| UTP6 | 11.6888565 | 0.36378048 | 0.46414718 | 0.78376105 | 0.43318033 | NA |
| SUZ12 | 57.062352 | 0.336230838 | 0.23719841 | 1.41750882 | 0.15633422 | 0.55671527 |
| LRRC37B | 16.8390543 | -0.008941564 | 0.41772811 | -0.0214052 | 0.98292241 | NA |
| RHOT1 | 33.3409067 | 0.293349453 | 0.26758299 | 1.09629334 | 0.27295043 | 0.66531666 |
| RHBDL3 | 13.279518 | -3.020805522 | 0.8038256 | -3.758036 | 0.00017125 | NA |
| C17orf75 | 8.11443131 | 0.358466297 | 0.48930598 | 0.73260151 | 0.46380151 | NA |
| ZNF207 | 84.3744951 | 0.07497638 | 0.22245233 | 0.33704471 | 0.73608319 | 0.91929008 |
| PSMD11 | 36.1708832 | 0.010312682 | 0.29810579 | 0.03459404 | 0.97240346 | 0.99072304 |
| MYO1D | 431.01897 | 0.138089684 | 0.21552097 | 0.64072506 | 0.52170133 | 0.83108387 |
| AC079336.4 | 8.32394193 | -0.823858959 | 0.67197064 | -1.2260342 | 0.22018582 | NA |
| AC025211.1 | 5.75364718 | 0.181648464 | 0.64019232 | 0.28374046 | 0.77660928 | NA |
| TMEM98 | 60.5628466 | -0.78225405 | 0.46818528 | -1.6708215 | 0.09475693 | 0.46755975 |
| CCL2 | 12.2488209 | -3.333651019 | 1.07772502 | -3.0932297 | 0.00197991 | NA |
| ZNF830 | 6.32543266 | -0.909173715 | 0.56501397 | -1.6091172 | 0.10759071 | NA |
| LIG3 | 8.94332184 | -0.039580723 | 0.49459758 | -0.0800261 | 0.93621648 | NA |
| RFFL | 5.14192702 | -1.168801393 | 0.87679002 | -1.333046 | 0.18251672 | NA |
| SLFN5 | 9.69870328 | 0.36667638 | 1.1628149 | 0.31533512 | 0.75250722 | NA |
| SLFN12 | 4.63834035 | -0.369876396 | 1.0226541 | -0.3616828 | 0.71758908 | NA |
| LINC02001 | 8.19136601 | -0.404075487 | 0.49663617 | -0.8136248 | 0.41585994 | NA |
| AP2B1 | 41.7713791 | -0.04803139 | 0.28682893 | -0.1674566 | 0.86701082 | 0.96089268 |
| TAF15 | 122.194791 | 0.21727004 | 0.26304585 | 0.82597783 | 0.40881667 | 0.76772515 |
| MMP28 | 19.5376737 | 0.647567618 | 0.61407178 | 1.05454711 | 0.2916325 | NA |
| AC015849.3 | 8.53701256 | 0.325885285 | 0.748718 | 0.43525771 | 0.66337537 | NA |
| ZNHIT3 | 9.24063187 | -0.852418884 | 0.63635019 | -1.3395437 | 0.18039374 | NA |
| MYO19 | 17.0237309 | -0.326736958 | 0.4125244 | -0.7920427 | 0.42833575 | NA |
| GGNBP2 | 46.5085104 | -0.230062481 | 0.27183708 | -0.8463249 | 0.39737154 | 0.76023081 |
| AATF | 22.6103414 | -0.518373442 | 0.47680601 | -1.0871789 | 0.27695775 | NA |
| ACACA | 64.2858503 | -0.336982961 | 0.28259011 | -1.1924797 | 0.23307322 | 0.63935677 |
| TADA2A | 16.7380015 | 0.126506913 | 0.37189297 | 0.34017022 | 0.73372834 | NA |
| DUSP14 | 12.3835011 | 0.144030807 | 0.60281095 | 0.23893197 | 0.81115834 | NA |
| SYNRG | 48.1436343 | 0.66278495 | 0.26614307 | 2.49033333 | 0.01276233 | 0.18904687 |
| DDX52 | 19.2720772 | -0.272660422 | 0.36541368 | -0.7461692 | 0.45556523 | NA |
| HNF1B | 12.4250147 | 0.323288398 | 0.51802308 | 0.62408107 | 0.53257435 | NA |
| MRPL45 | 11.3069257 | -0.340032645 | 0.47872252 | -0.7102917 | 0.47752325 | NA |
| SOCS7 | 14.2455002 | -0.229651684 | 0.47744363 | -0.4810027 | 0.63051457 | NA |
| ARHGAP23 | 87.6899486 | 0.254008144 | 0.38320633 | 0.66284956 | 0.50742691 | 0.82657837 |
| MLLT6 | 50.7225127 | -0.477632216 | 0.35654211 | -1.3396236 | 0.18036776 | 0.58390314 |
| CISD3 | 15.4771373 | -0.065172369 | 0.40743068 | -0.1599594 | 0.87291305 | NA |
| PCGF2 | 12.7848325 | 0.457820801 | 0.46521979 | 0.98409571 | 0.32506845 | NA |
| PSMB3 | 24.4235327 | -0.399822212 | 0.48257309 | -0.8285216 | 0.40737519 | NA |
| PIP4K2B | 20.2685783 | 0.123933945 | 0.32525363 | 0.38103785 | 0.70317516 | NA |
| CWC25 | 12.1445574 | -0.596372062 | 0.47195233 | -1.2636278 | 0.20636366 | NA |
| RPL23 | 49.2068338 | -0.338170975 | 0.45241742 | -0.7474756 | 0.45477653 | 0.79324789 |
| LASP1 | 26.6166502 | 0.110256637 | 0.37840359 | 0.29137313 | 0.77076596 | NA |
| RPL19 | 103.595341 | -0.820726338 | 0.36774745 | -2.2317662 | 0.02563042 | 0.25731925 |
| FBXL20 | 68.8623593 | 0.568531407 | 0.30939476 | 1.83755989 | 0.06612728 | 0.3991664 |
| MED1 | 20.0400811 | -0.083818098 | 0.33079055 | -0.2533872 | 0.799969 | NA |
| CDK12 | 39.423547 | -0.260063055 | 0.27855873 | -0.9336023 | 0.3505091 | 0.72238707 |
| STARD3 | 7.06127387 | 0.352379798 | 0.50800483 | 0.69365443 | 0.48789895 | NA |
| PGAP3 | 4.82294523 | 1.148588381 | 0.58173295 | 1.97442552 | 0.04833338 | NA |
| ERBB2 | 14.3046349 | 0.771697819 | 0.41813351 | 1.84557753 | 0.06495357 | NA |
| MIEN1 | 13.0848659 | -0.746087082 | 0.43560226 | -1.7127714 | 0.08675461 | NA |
| GRB7 | 6.3446405 | -1.076348378 | 0.81909825 | -1.314065 | 0.18882432 | NA |
| ORMDL3 | 15.7000843 | -0.310489682 | 0.50969608 | -0.6091663 | 0.54241421 | NA |
| PSMD3 | 27.8997462 | 0.314493001 | 0.37077677 | 0.8482004 | 0.39632638 | NA |
| MED24 | 15.884886 | 0.844446991 | 0.39303165 | 2.14854704 | 0.03167032 | NA |
| THRA | 18.9958781 | -0.83330267 | 0.56607015 | -1.4720838 | 0.14099826 | NA |
| MSL1 | 37.821552 | 0.279167352 | 0.42484144 | 0.65710952 | 0.5111105 | 0.82988959 |
| CASC3 | 30.2549307 | 0.394041238 | 0.40046506 | 0.98395909 | 0.32513563 | 0.69627778 |
| WIPF2 | 40.477389 | 0.258331685 | 0.27420236 | 0.94212059 | 0.34613091 | 0.71908069 |
| RARA | 11.7791384 | -0.18080458 | 0.46440902 | -0.3893218 | 0.69703808 | NA |
| IGFBP4 | 58.5979406 | -2.60972059 | 0.5305891 | -4.9185341 | 8.72E-07 | 0.00023577 |
| KRT10 | 30.4980402 | -0.747583525 | 0.43980159 | -1.69982 | 0.08916479 | 0.45574679 |

| | | | | | | |
|---|---|---|---|---|---|---|
| KRT19 | 17.1567472 | -2.827884054 | 0.7922397 | -3.5694804 | 0.00035769 | NA |
| EIF1 | 177.768525 | -1.013740528 | 0.45024471 | -2.2515323 | 0.02435184 | 0.25196701 |
| JUP | 63.897671 | 0.246447086 | 0.38946875 | 0.63277757 | 0.52687891 | 0.83195298 |
| P3H4 | 7.89764362 | -0.024705345 | 0.6241954 | -0.0395795 | 0.96842837 | NA |
| FKBP10 | 5.56774594 | -0.950845778 | 0.80454431 | -1.1818439 | 0.23726765 | NA |
| NT5C3B | 12.3760833 | -0.24544969 | 0.48981097 | -0.5011111 | 0.61629297 | NA |
| ACLY | 22.8843006 | -0.41863311 | 0.35282507 | -1.1865175 | 0.23541801 | NA |
| CNP | 23.1559202 | -0.36441315 | 0.82864283 | -0.4397711 | 0.66010292 | NA |
| DNAJC7 | 37.0427093 | 0.046978327 | 0.33520364 | 0.14014862 | 0.88854257 | 0.96764417 |
| NKIRAS2 | 13.2795552 | -0.093413764 | 0.52407565 | -0.1782448 | 0.85853072 | NA |
| DHX58 | 7.2441387 | -1.656500503 | 1.16465937 | -1.4223047 | 0.15493781 | NA |
| KAT2A | 5.29618789 | 0.10114302 | 0.6392239 | 0.15822785 | 0.87427726 | NA |
| GHDC | 7.82416481 | -0.391205412 | 0.64881932 | -0.6029497 | 0.54654216 | NA |
| STAT5B | 39.4668475 | -0.164060176 | 0.30386701 | -0.5399078 | 0.58926061 | 0.86533208 |
| STAT5A | 6.51734707 | 0.438159815 | 0.57939197 | 0.75624074 | 0.44950487 | NA |
| STAT3 | 156.561803 | -1.497488995 | 0.31717401 | -4.7213483 | 2.34E-06 | 0.00055898 |
| CAVIN1 | 32.6434686 | 0.205707175 | 0.50413797 | 0.40803746 | 0.68324617 | 0.90499319 |
| ATP6V0A1 | 37.1890983 | -0.141884023 | 0.37637251 | -0.3769776 | 0.70619022 | 0.91017081 |
| NAGLU | 29.4785372 | -0.055323253 | 0.37644024 | -0.1469642 | 0.88316025 | 0.96526488 |
| COASY | 8.13799615 | -0.047464899 | 0.49225487 | -0.0964234 | 0.9231843 | NA |
| MLX | 8.3069792 | -0.396519989 | 0.49591146 | -0.7995782 | 0.42395522 | NA |
| RETREG3 | 30.1630828 | -0.180425673 | 0.33086325 | -0.545318 | 0.58553485 | 0.86235633 |
| TUBG1 | 4.188687 | -0.872859403 | 0.83785052 | -1.0417842 | 0.29751176 | NA |
| TUBG2 | 10.4275978 | 0.1796311 | 0.52658571 | 0.34112414 | 0.73301013 | NA |
| PLEKHH3 | 6.35250001 | 0.277314857 | 0.61250027 | 0.45275875 | 0.65072247 | NA |
| EZH1 | 23.6454077 | -0.023378444 | 0.38502687 | -0.060719 | 0.951583 | NA |
| VPS25 | 10.1897248 | -0.875585455 | 0.54773963 | -1.5985432 | 0.10992214 | NA |
| COA3 | 20.990805 | -0.345494127 | 0.39791514 | -0.8682608 | 0.38525156 | NA |
| BECN1 | 12.6439085 | -0.38052215 | 0.44241966 | -0.8600932 | 0.38973766 | NA |
| PSME3 | 10.8818537 | -0.064392886 | 0.49056753 | -0.131262 | 0.89556803 | NA |
| RUNDC1 | 11.277904 | -0.545600964 | 0.48223035 | -1.1314115 | 0.25788193 | NA |
| RPL27 | 46.9947287 | -0.591440767 | 0.43117006 | -1.3717111 | 0.17015338 | 0.57273205 |
| IFI35 | 24.5775991 | -0.332673859 | 1.41685283 | -0.2347978 | 0.81436571 | NA |
| VAT1 | 9.8592226 | -0.479912342 | 0.64432971 | -0.7448242 | 0.45637802 | NA |
| BRCA1 | 9.47558228 | -0.831446027 | 0.73589245 | -1.1298472 | 0.25854062 | NA |
| NBR1 | 36.3178546 | 0.132216831 | 0.25111366 | 0.52652187 | 0.59852566 | 0.86882737 |
| TMEM106A | 7.19409439 | -0.922098226 | 0.66188245 | -1.393145 | 0.16357595 | NA |
| LINC00854 | 34.5817103 | -0.829118027 | 0.72324523 | -1.1463857 | 0.25163558 | 0.65183754 |
| ARL4D | 6.64808584 | -0.676327256 | 0.87509174 | -0.7728644 | 0.43960263 | NA |
| DHX8 | 13.3889018 | -0.145699045 | 0.40397275 | -0.3606655 | 0.7183495 | NA |
| DUSP3 | 30.1231328 | -0.123861064 | 0.39912028 | -0.3103352 | 0.75630608 | 0.92313949 |
| MPP3 | 12.3158281 | 0.463556907 | 0.62435479 | 0.74245751 | 0.45781019 | NA |
| TMEM101 | 12.1400227 | -0.832654973 | 0.6388116 | -1.3034437 | 0.19242332 | NA |
| LSM12 | 14.2621904 | 0.206994177 | 0.45725031 | 0.45269336 | 0.65076956 | NA |
| G6PC3 | 14.4647146 | -0.35502993 | 0.58278818 | -0.6091921 | 0.54239715 | NA |
| HDAC5 | 15.7501508 | 1.216193109 | 0.40590479 | 2.9962522 | 0.0027332 | NA |
| TMUB2 | 12.9467027 | 0.502147758 | 0.45975644 | 1.09220386 | 0.2747435 | NA |
| ATXN7L3 | 9.41805623 | 0.200675184 | 0.47181497 | 0.42532602 | 0.67059903 | NA |
| UBTF | 22.9181386 | -0.242371848 | 0.47525963 | -0.5099778 | 0.61006704 | NA |
| AC003102.1 | 5.67101716 | -1.470699292 | 0.85821531 | -1.7136717 | 0.08658904 | NA |
| SLC25A39 | 12.4179003 | 0.339872934 | 0.50743811 | 0.66978204 | 0.50299674 | NA |
| GRN | 80.2655993 | -0.224721974 | 0.50140403 | -0.4481854 | 0.65401939 | 0.8949739 |
| GPATCH8 | 138.585511 | 0.295565515 | 0.15522217 | 1.90414495 | 0.05689131 | 0.37278053 |
| FZD2 | 24.2927693 | -0.855304255 | 0.58888145 | -1.4524218 | 0.14638436 | NA |
| CCDC43 | 6.0256336 | -1.780723715 | 0.71414145 | -2.4935168 | 0.01264846 | NA |
| DBF4B | 7.61002962 | 0.207137477 | 0.65826209 | 0.31467326 | 0.75300975 | NA |
| EFTUD2 | 17.91287 | 0.528421103 | 0.38746241 | 1.36379964 | 0.17263063 | NA |
| C1QL1 | 149.535308 | -0.617543051 | 0.53116875 | -1.1626118 | 0.24498705 | 0.64996527 |
| DCAKD | 18.4819704 | 0.088717333 | 0.40120649 | 0.22112637 | 0.82499404 | NA |
| NMT1 | 39.7322194 | -0.203029872 | 0.2765799 | -0.7340731 | 0.46290418 | 0.79793427 |
| PLCD3 | 38.3871255 | -0.391239422 | 0.36156569 | -1.0820701 | 0.27922138 | 0.66958859 |
| ACBD4 | 9.51180861 | 0.278741183 | 0.47136391 | 0.59135029 | 0.55428574 | NA |
| HEXIM1 | 31.457426 | -0.436678185 | 0.48497842 | -0.9004075 | 0.36790345 | 0.73626927 |
| AC138150.1 | 19.9102028 | -0.110988101 | 0.46414925 | -0.2391216 | 0.81101131 | NA |
| MAP3K14 | 13.0452211 | -1.312353028 | 0.52886421 | -2.4814556 | 0.0130847 | NA |
| PLEKHM1 | 15.1139797 | -0.017022964 | 0.41367656 | -0.0411504 | 0.96717598 | NA |
| LINC02210-C | 15.3937664 | 0.223766449 | 0.45031805 | 0.49690758 | 0.61925423 | NA |
| MAPT | 41.7880095 | -1.390925215 | 0.6447023 | -2.157469 | 0.03096914 | 0.28264481 |
| KANSL1 | 197.616816 | 0.697081042 | 0.29248461 | 2.38330842 | 0.01715781 | 0.21679778 |
| ARL17B | 51.7611641 | 1.806936184 | 0.89739245 | 2.01354066 | 0.04405778 | 0.33031121 |
| LRRC37A2 | 10.1443146 | 0.353783183 | 0.58893576 | 0.60071608 | 0.54802911 | NA |
| ARL17A | 11.3410575 | -0.379408206 | 0.46130394 | -0.822469 | 0.41081 | NA |
| NSF | 46.4053706 | 0.804413757 | 0.45652145 | 1.76205031 | 0.0780608 | 0.4276675 |
| WNT3 | 15.0776667 | 1.262480585 | 0.94829151 | 1.3313212 | 0.18308335 | NA |
| AC005670.2 | 7.66679021 | 0.597583267 | 0.56008901 | 1.06694339 | 0.2859974 | NA |
| CDC27 | 69.6365203 | 0.070506964 | 0.27337204 | 0.25791579 | 0.7964719 | 0.93934998 |
| AC068234.1 | 178.724815 | -0.548651992 | 0.41856448 | -1.3107944 | 0.18992723 | 0.59302912 |
| AC068234.2 | 24.0877577 | -1.222564483 | 0.43790674 | -2.7918375 | 0.00524097 | NA |
| EFCAB13 | 86.2712781 | -0.11860303 | 0.40023263 | -0.2963352 | 0.76697408 | 0.92451492 |
| AC040934.1 | 5.6709363 | -0.958875348 | 0.64966499 | -1.4759535 | 0.13995636 | NA |
| NPEPPS | 106.66594 | 0.40611299 | 0.1700207 | 2.38860911 | 0.01691229 | 0.21530523 |
| AC025682.1 | 14.2097228 | 0.47707822 | 0.6115286 | 0.78014049 | 0.43530819 | NA |
| KPNB1 | 76.914669 | -0.128657774 | 0.38899278 | -0.3307459 | 0.74083641 | 0.92003444 |
| MRPL10 | 9.80641162 | 0.25350487 | 0.58744782 | 0.43153598 | 0.6660787 | NA |
| SCRN2 | 10.8781252 | -0.012552058 | 0.44946197 | -0.0279269 | 0.97772049 | NA |
| SP2 | 11.099968 | 0.799506497 | 0.49325826 | 1.62086795 | 0.10504596 | NA |
| PNPO | 6.56928221 | -0.863096006 | 0.6912149 | -1.2486652 | 0.21178754 | NA |
| CDK5RAP3 | 13.767391 | 0.248686598 | 0.46695304 | 0.53257303 | 0.59432918 | NA |
| COPZ2 | 5.53644915 | -0.615003104 | 0.58777583 | -1.0463225 | 0.29541214 | NA |
| NFE2L1 | 105.594068 | 0.012432044 | 0.26239655 | 0.04737884 | 0.96221129 | 0.98892249 |
| CBX1 | 17.3931319 | -0.113414581 | 0.44533462 | -0.2546727 | 0.79897588 | NA |
| SNX11 | 9.49232596 | 0.465393489 | 0.46675276 | 0.99708782 | 0.31872189 | NA |
| SKAP1 | 49.0530799 | 0.678515651 | 0.581414 | 1.16700947 | 0.24320654 | 0.64918976 |
| HOXB2 | 7.0447773 | 0.406143315 | 0.61241401 | 0.66318423 | 0.50721257 | NA |
| HOXB3 | 17.1266679 | -0.224048114 | 0.48292319 | -0.4639415 | 0.64268965 | NA |
| HOXB7 | 10.8554435 | -0.473275063 | 0.4467565 | -1.059358 | 0.28943678 | NA |
| CALCOCO2 | 24.7796603 | -0.334329698 | 0.4366565 | -0.7656584 | 0.44387961 | NA |
| ATP5MC1 | 27.8626189 | -0.025099846 | 0.3563221 | -0.0704415 | 0.9438423 | NA |
| UBE2Z | 40.6160864 | 0.144611151 | 0.34869709 | 0.41471855 | 0.67834795 | 0.90499319 |
| SNF8 | 18.1632469 | 0.241681282 | 0.34474854 | 0.70103643 | 0.48328028 | NA |
| ZNF652 | 37.9860397 | -0.164770792 | 0.36848641 | -0.4471557 | 0.65476266 | 0.89508505 |
| AC091180.5 | 4.80676547 | -0.836751616 | 0.93073032 | -0.8990269 | 0.36863832 | NA |
| PHB | 26.2484237 | -0.247241818 | 0.3779049 | -0.6542435 | 0.51295495 | NA |
| SPOP | 40.6858411 | 0.219602102 | 0.26065792 | 0.84249159 | 0.39951285 | 0.76255252 |
| SLC35B1 | 7.77167031 | 0.313707707 | 0.64810842 | 0.48403584 | 0.62836043 | NA |
| FAM117A | 9.22633008 | 0.318132064 | 0.51074144 | 0.62288281 | 0.53336154 | NA |
| KAT7 | 51.996851 | -0.288672562 | 0.42224008 | -0.6836693 | 0.49418404 | 0.82249096 |
| TAC4 | 5.20149177 | -1.329760335 | 0.70783143 | -1.8786398 | 0.06029369 | NA |
| ITGA3 | 394.312611 | 0.079286269 | 0.42780375 | 0.18533327 | 0.85296764 | 0.95569766 |
| PDK2 | 9.73011286 | 0.103697661 | 0.529587 | 0.19580855 | 0.84476002 | NA |
| SAMD14 | 7.47625561 | 0.727585791 | 0.56971485 | 1.27710519 | 0.20156511 | NA |
| PPP1R9B | 6.89571042 | 0.2615235 | 0.73775115 | 0.35448742 | 0.72297363 | NA |
| XYLT2 | 10.266619 | 0.539216448 | 0.44004343 | 1.22537097 | 0.22043548 | NA |

| | | | | | | |
|---|---|---|---|---|---|---|
| MRPL27 | 16.155681 | -0.964344346 | 0.50198036 | -1.9210798 | 0.05472164 | NA |
| LRRC59 | 11.472976 | 0.008166069 | 0.53214349 | 0.01534561 | 0.98775645 | NA |
| ACSF2 | 13.8441562 | 2.873746423 | 0.70837552 | 4.05681217 | 4.97E-05 | NA |
| RSAD1 | 7.33239939 | -0.076011692 | 0.53914739 | -0.140985 | 0.8878818 | NA |
| SPATA20 | 9.51567043 | -0.595099083 | 0.50996331 | -1.1669449 | 0.24323263 | NA |
| ABCC3 | 11.7380391 | -1.877110074 | 0.72365867 | -2.5939164 | 0.00948896 | NA |
| ANKRD40 | 20.9033347 | -0.309467957 | 0.61270364 | -0.5050859 | 0.61349852 | NA |
| AC005921.2 | 9.32731574 | -0.757820958 | 0.57826101 | -1.3105171 | 0.19002096 | NA |
| LUC7L3 | 129.180563 | 0.509698519 | 0.1824697 | 2.7933323 | 0.00521681 | 0.11257186 |
| TOB1 | 31.3969772 | 0.80450337 | 0.4553158 | 1.76691293 | 0.07724281 | 0.42394701 |
| SPAG9 | 91.9091665 | -0.140395025 | 0.23244959 | -0.6039805 | 0.54585659 | 0.84440067 |
| AC005920.2 | 7.30336248 | -0.029453447 | 0.68299278 | -0.0431241 | 0.96560261 | NA |
| MBTD1 | 35.0548036 | -0.373772466 | 0.29690723 | -1.2588864 | 0.20807137 | 0.61736443 |
| UTP18 | 10.1266323 | -0.757029157 | 0.52658573 | -1.4376181 | 0.15054245 | NA |
| CA10 | 282.176391 | 0.989225295 | 0.36500307 | 2.71018347 | 0.0067246 | 0.13050228 |
| TOM1L1 | 43.8443887 | -0.006785772 | 0.33473973 | -0.0202718 | 0.98382656 | 0.99577746 |
| COX11 | 16.2036612 | -0.206233411 | 0.4403298 | -0.4683612 | 0.63952628 | NA |
| STXBP4 | 38.4813174 | 0.071908765 | 0.37032088 | 0.19417961 | 0.84603525 | 0.95532127 |
| HLF | 6.42459527 | -0.195492598 | 0.91416554 | -0.2138481 | 0.83066549 | NA |
| PCTP | 7.62271567 | -0.038788288 | 0.53585249 | -0.0723861 | 0.94229462 | NA |
| DGKE | 7.36256928 | 0.244787584 | 0.64203485 | 0.38126838 | 0.70300412 | NA |
| TRIM25 | 6.82348696 | -1.40221949 | 0.88957927 | -1.5762727 | 0.11496298 | NA |
| COIL | 7.08992403 | -0.039747301 | 0.50453227 | -0.0787805 | 0.93720722 | NA |
| SCPEP1 | 16.8989126 | -0.196731331 | 0.54751207 | -0.3593187 | 0.71935668 | NA |
| AKAP1 | 13.7656186 | 0.292553943 | 0.41637736 | 0.70261731 | 0.48229427 | NA |
| MSI2 | 444.350888 | -0.069795093 | 0.32180996 | -0.2168829 | 0.82829957 | 0.94957124 |
| MRPS23 | 17.4848472 | -0.041433691 | 0.4021728 | -0.1030246 | 0.91794345 | NA |
| CUEDC1 | 25.2208982 | 0.199194299 | 0.35115739 | 0.56725077 | 0.57054382 | NA |
| VEZF1 | 35.2317724 | -0.009703933 | 0.36244045 | -0.0267739 | 0.97864009 | 0.9938318 |
| AC015813.2 | 12.0960151 | 0.460062315 | 0.4863041 | 0.94603832 | 0.34412904 | NA |
| DYNLL2 | 19.4928335 | 0.400120319 | 0.44511971 | 0.89890497 | 0.36870328 | NA |
| MKS1 | 5.37943862 | 0.548100175 | 0.68035914 | 0.80560419 | 0.42047111 | NA |
| TSPOAP1 | 6.93221834 | -0.128116014 | 0.58751024 | -0.218066 | 0.82737768 | NA |
| SUPT4H1 | 11.9720818 | -1.23226527 | 0.69492552 | -1.7732336 | 0.07619001 | NA |
| MTMR4 | 9.49430435 | 0.662984337 | 0.51249386 | 1.29364348 | 0.1957886 | NA |
| RAD51C | 9.80027611 | -0.014121484 | 0.49219969 | -0.0286906 | 0.97711139 | NA |
| TRIM37 | 20.4341271 | 0.039041045 | 0.348665 | 0.11197294 | 0.91084486 | NA |
| SKA2 | 19.9471384 | 0.152405044 | 0.3758311 | 0.40551473 | 0.6850992 | NA |
| YPEL2 | 21.3599691 | 0.002475167 | 0.50546053 | 0.00489685 | 0.99609289 | NA |
| DHX40 | 18.4697964 | 0.021597014 | 0.36428537 | 0.05928598 | 0.95272433 | NA |
| CLTC | 134.44675 | -0.170358535 | 0.30060396 | -0.5667209 | 0.57090384 | 0.8573305 |
| PTRH2 | 16.6512043 | -1.061995944 | 0.49334952 | -2.1526239 | 0.03134825 | NA |
| VMP1 | 92.8308685 | -0.175411691 | 0.52290139 | -0.3354585 | 0.73727928 | 0.91929008 |
| TUBD1 | 13.4474239 | -0.295045027 | 0.44824147 | -0.6582279 | 0.51039172 | NA |
| RPS6KB1 | 18.0227593 | 0.26996627 | 0.43866822 | 0.61542245 | 0.53827577 | NA |
| HEATR6 | 6.2385946 | 0.009626868 | 0.54044715 | 0.01781278 | 0.98578821 | NA |
| USP32 | 57.2603698 | -0.006362409 | 0.28311866 | -0.0224726 | 0.98207098 | 0.99507367 |
| APPBP2 | 99.2039599 | 0.099170418 | 0.27357163 | 0.36250256 | 0.71697651 | 0.91472065 |
| PPM1D | 23.2803013 | 0.08862433 | 0.37872177 | 0.23400908 | 0.81497793 | NA |
| BCAS3 | 176.675264 | 0.333721178 | 0.21520014 | 1.550748 | 0.12096209 | 0.5126669 |
| NACA2 | 7.84812077 | -0.968407902 | 0.58657196 | -1.6509618 | 0.09874637 | NA |
| INTS2 | 12.814366 | 0.754812497 | 0.41832283 | 1.80437798 | 0.07117207 | NA |
| MED13 | 110.496265 | 0.230488591 | 0.21887555 | 1.05305773 | 0.29231453 | 0.67930583 |
| METTL2A | 8.88581012 | -0.084323078 | 0.45739929 | -0.1843533 | 0.85373629 | NA |
| TLK2 | 57.5181546 | 0.523783269 | 0.27957492 | 1.87349877 | 0.06099954 | 0.38181195 |
| MRC2 | 58.560262 | 0.476041006 | 0.74883685 | 0.63570724 | 0.52496726 | 0.83195298 |
| TANC2 | 58.480972 | 0.244394698 | 0.33122372 | 0.73785386 | 0.46060326 | 0.79680257 |
| CYB561 | 14.6171867 | 0.508378283 | 0.45934979 | 1.10673454 | 0.26840872 | NA |
| DCAF7 | 39.4738028 | -0.169894523 | 0.28550837 | -0.5950597 | 0.55180358 | 0.84744995 |
| TACO1 | 6.940879 | -0.121262355 | 0.63195722 | -0.1918838 | 0.84783322 | NA |
| MAP3K3 | 63.4225966 | 0.178661774 | 0.39904274 | 0.44772591 | 0.65435102 | 0.89508505 |
| CCDC47 | 8.78492023 | -0.51813883 | 0.46369747 | -1.1174071 | 0.26382031 | NA |
| DDX42 | 40.7352034 | -0.212856419 | 0.34427944 | -0.6182664 | 0.53639972 | 0.83677705 |
| FTSJ3 | 8.18603316 | -0.022782464 | 0.47807661 | -0.0476544 | 0.96199166 | NA |
| PSMC5 | 14.9421495 | 0.026131411 | 0.44033554 | 0.05934431 | 0.95267787 | NA |
| SMARCD2 | 9.60024457 | 0.381909348 | 0.48250648 | 0.79151133 | 0.42864566 | NA |
| ERN1 | 11.9213332 | 0.238690001 | 0.44806731 | 0.53271014 | 0.59423424 | NA |
| SNHG25 | 6.25219217 | -0.921492446 | 0.67857392 | -1.3579839 | 0.1744688 | NA |
| TEX2 | 11.2886003 | -0.651751759 | 0.55786671 | -1.168293 | 0.24268861 | NA |
| POLG2 | 9.11472447 | 0.056743116 | 0.54719633 | 0.10369791 | 0.91740909 | NA |
| DDX5 | 220.753364 | 0.103578444 | 0.25638312 | 0.40399868 | 0.68621369 | 0.90499319 |
| CEP95 | 23.9335312 | -0.221208277 | 0.40016306 | -0.5527953 | 0.58040356 | NA |
| SMURF2 | 47.1971462 | 0.453376563 | 0.25942554 | 1.7476173 | 0.08053032 | 0.43346499 |
| LRRC37A3 | 44.8878786 | 0.773179227 | 0.31665036 | 2.44174436 | 0.01461649 | 0.19896332 |
| GNA13 | 29.003652 | -0.500640741 | 0.37435028 | -1.3373591 | 0.18110546 | 0.58390314 |
| CEP112 | 110.762615 | 0.23988957 | 0.2772099 | 0.86537158 | 0.38683488 | 0.75125852 |
| PRKCA | 99.1075875 | -0.600098291 | 0.29991145 | -2.0009182 | 0.0454012 | 0.3354231 |
| HELZ | 79.9239633 | 0.554761096 | 0.25513457 | 2.1743862 | 0.02967614 | 0.27926427 |
| PSMD12 | 22.7247811 | -0.035434332 | 0.43845509 | -0.0808163 | 0.93558802 | NA |
| PITPNC1 | 72.0277495 | -1.092970312 | 0.67759879 | -1.6130051 | 0.1067434 | 0.49086361 |
| NOL11 | 13.3966683 | 0.266898696 | 0.48346329 | 0.55205576 | 0.58091015 | NA |
| BPTF | 138.426044 | 0.043263004 | 0.28734054 | 0.15056352 | 0.88032004 | 0.96423929 |
| KPNA2 | 8.89306641 | -0.511976914 | 0.69403758 | -0.737679 | 0.46070957 | NA |
| AC005332.7 | 8.51956262 | 0.402581842 | 0.52124991 | 0.7723394 | 0.43991344 | NA |
| AMZ2 | 8.11102908 | -0.454877575 | 0.47288674 | -0.9619165 | 0.33609153 | NA |
| ARSG | 9.08086522 | 0.201870263 | 0.68785627 | 0.29347739 | 0.76915728 | NA |
| WIPI1 | 21.2275799 | 0.502071485 | 0.34220411 | 1.46716966 | 0.14232991 | NA |
| PRKAR1A | 94.3688139 | -0.094166116 | 0.29013667 | -0.3245578 | 0.74551578 | 0.92250642 |
| FAM20A | 11.4491707 | -0.754748471 | 0.6691631 | -1.1278991 | 0.25936252 | NA |
| LINC01482 | 9.62384949 | 1.378749266 | 1.00128021 | 1.37698643 | 0.16851644 | NA |
| ABCA10 | 15.7030788 | 0.530866469 | 0.65574263 | 0.80956529 | 0.41819006 | NA |
| ABCA5 | 47.870951 | 0.436139682 | 0.28845819 | 1.51196846 | 0.13054189 | 0.52613957 |
| AC005208.1 | 11.436578 | -0.325278572 | 1.04363421 | -0.3116787 | 0.7552847 | NA |
| KCNJ16 | 34.1392288 | 0.772193027 | 0.64465869 | 1.19783234 | 0.2309823 | 0.63682501 |
| SLC39A11 | 59.5292862 | 0.309307896 | 0.34627006 | 0.89325624 | 0.37171996 | 0.74014106 |
| COG1 | 10.3259974 | 0.21965586 | 0.42581261 | 0.515851 | 0.60595848 | NA |
| FAM104A | 22.790882 | 0.171450893 | 0.3120675 | 0.54940323 | 0.58272876 | NA |
| C17orf80 | 9.34609525 | -0.085237487 | 0.48409826 | -0.1760748 | 0.8602352 | NA |
| CDC42EP4 | 30.9801447 | -1.620450965 | 0.45844622 | -3.5346588 | 0.0004083 | 0.02714874 |
| SDK2 | 7.78683559 | -2.649308818 | 1.01901898 | -2.5998621 | 0.00932612 | NA |
| RPL38 | 32.1320638 | -0.283853805 | 0.50293764 | -0.5643916 | 0.57248762 | 0.85816194 |
| GPRC5C | 13.3410494 | -0.893064582 | 0.61510904 | -1.4518801 | 0.14653494 | NA |
| SLC9A3R1 | 54.1119864 | -0.145215257 | 0.38019676 | -0.3819476 | 0.7025002 | 0.90757606 |
| NAT9 | 17.5945186 | 0.061916557 | 0.40944586 | 0.15122038 | 0.87980188 | NA |
| TMEM104 | 8.40449587 | 0.367813139 | 0.4883714 | 0.75314226 | 0.45136443 | NA |
| MRPL58 | 7.22403215 | 0.067204617 | 0.5788642 | 0.11609738 | 0.90757537 | NA |
| KCTD2 | 24.2265921 | 0.54774028 | 0.47583019 | 1.15112552 | 0.2496806 | NA |
| ATP5PD | 17.1735775 | -0.032627014 | 0.54145699 | -0.0602578 | 0.9519503 | NA |
| SLC16A5 | 11.5235741 | 0.435514905 | 0.52228896 | 0.83385815 | 0.40436092 | NA |
| ARMC7 | 3.9390146 | -1.073616478 | 0.83605071 | -1.2841523 | 0.19908865 | NA |
| NT5C | 29.5028115 | -0.57511224 | 0.38199128 | -1.5055638 | 0.13217919 | 0.52613957 |
| JPT1 | 11.8530077 | -0.404507193 | 0.54899763 | -0.7368105 | 0.46123762 | NA |
| SUMO2 | 15.0864769 | -0.281034144 | 0.42257368 | -0.6650536 | 0.50601621 | NA |

| | | | | | | |
|---|---|---|---|---|---|---|
| NUP85 | 7.47134265 | -0.600236627 | 0.61201776 | -0.9807503 | 0.32671587 | NA |
| GGA3 | 11.8669424 | 0.254019787 | 0.45789049 | 0.55476101 | 0.57905814 | NA |
| MRPS7 | 16.622638 | 0.416757417 | 0.43194881 | 0.96483057 | 0.33462968 | NA |
| MIF4GD | 5.96795492 | -1.043371976 | 0.61399311 | -1.699322 | 0.08925853 | NA |
| GRB2 | 44.2157988 | -0.237745516 | 0.29096573 | -0.8170911 | 0.41387635 | 0.76946282 |
| AC011933.4 | 5.75200536 | 0.011734891 | 0.82097573 | 0.01429383 | 0.98859556 | NA |
| TMEM94 | 12.7660611 | 0.839393301 | 0.44854816 | 1.87135603 | 0.06129575 | NA |
| CASKIN2 | 6.44997787 | 0.451609269 | 0.61099079 | 0.73914252 | 0.45982046 | NA |
| TSEN54 | 3.75668138 | 0.032968218 | 0.77264388 | 0.04266936 | 0.96596511 | NA |
| LLGL2 | 23.2722194 | -0.005097187 | 0.37607493 | -0.0135536 | 0.98918609 | NA |
| MYO15B | 11.4065374 | 0.849193382 | 0.62246075 | 1.36425209 | 0.17248824 | NA |
| RECQL5 | 7.7114343 | 0.672686541 | 0.55669525 | 1.20835689 | 0.22691002 | NA |
| SAP30BP | 20.9111626 | -0.011346396 | 0.35783112 | -0.0317088 | 0.97470428 | NA |
| GALK1 | 6.81450502 | 0.051741967 | 0.642088 | 0.08058392 | 0.93577286 | NA |
| H3F3B | 230.825398 | -1.049032227 | 0.43413952 | -2.4163481 | 0.01567706 | 0.20644861 |
| UNK | 24.5246147 | 0.399798055 | 0.31747975 | 1.2592868 | 0.20792676 | NA |
| WBP2 | 29.9699155 | -0.714440334 | 0.31610968 | -2.2601027 | 0.02381488 | 0.25161392 |
| TRIM47 | 4.13182041 | 0.332431338 | 0.87096565 | 0.38168134 | 0.70269774 | NA |
| TRIM65 | 6.59887571 | -0.50797089 | 0.51319849 | -0.9898137 | 0.3222652 | NA |
| ACOX1 | 19.0905691 | -0.269129172 | 0.37573568 | -0.7162726 | 0.47382304 | NA |
| TEN1 | 6.46807029 | -0.115534868 | 0.57983437 | -0.1992549 | 0.84206332 | NA |
| SRP68 | 18.9359702 | -0.054924312 | 0.39034962 | -0.1407054 | 0.88810265 | NA |
| EXOC7 | 22.6334177 | 0.023257621 | 0.32014303 | 0.07264759 | 0.94208655 | NA |
| UBALD2 | 12.9334135 | -0.344374818 | 0.61768346 | -0.5575264 | 0.57716785 | NA |
| PRPSAP1 | 15.6101639 | 0.824604061 | 0.46627633 | 1.76848794 | 0.07697937 | NA |
| UBE2O | 19.0402072 | 0.26328946 | 0.41887638 | 0.62856124 | 0.52963634 | NA |
| MXRA7 | 46.6887201 | -0.24092063 | 0.45212645 | -0.5328612 | 0.59412968 | 0.86620776 |
| JMJD6 | 6.78252104 | -0.189770744 | 0.53407916 | -0.3553233 | 0.72234744 | NA |
| SRSF2 | 47.0734452 | -0.418641231 | 0.30027239 | -1.3942049 | 0.16325575 | 0.56498748 |
| MFSD11 | 29.8333336 | 0.206641562 | 0.43789609 | 0.47189634 | 0.63700078 | 0.88633797 |
| SEC14L1 | 83.2713271 | 0.039069411 | 0.24452032 | 0.15977981 | 0.87305453 | 0.96229085 |
| 9-Sep | 51.8191087 | 0.473831833 | 0.4459369 | 1.06255353 | 0.28798447 | 0.67596355 |
| TNRC6C | 71.2749312 | -0.295542341 | 0.38119909 | -0.7752966 | 0.43816444 | 0.78335165 |
| SYNGR2 | 31.8305559 | -0.234314276 | 0.47959226 | -0.4885698 | 0.62514633 | 0.87931199 |
| AFMID | 8.9012158 | -0.076711143 | 0.47658272 | -0.1609608 | 0.87212426 | NA |
| SOCS3 | 42.9594446 | -3.659689941 | 0.89689142 | -4.0804158 | 4.50E-05 | 0.00588188 |
| AC061992.1 | 5.83251776 | -2.545294003 | 1.13945079 | -2.2337902 | 0.02549688 | NA |
| PGS1 | 25.1591622 | -0.411523066 | 0.41690218 | -0.9870974 | 0.32359488 | NA |
| CYTH1 | 40.2277775 | 0.074862706 | 0.27600229 | 0.27123944 | 0.78620688 | 0.93615351 |
| USP36 | 20.9347942 | -0.15026906 | 0.39084921 | -0.3844681 | 0.70063152 | NA |
| TIMP2 | 193.580207 | -0.256581239 | 0.43081889 | -0.5955664 | 0.55146495 | 0.8472507 |
| LGALS3BP | 113.254209 | -0.271165447 | 0.67689784 | -0.4006003 | 0.68871446 | 0.90557575 |
| CANT1 | 12.4560935 | 0.288803655 | 0.5945028 | 0.48579023 | 0.6271159 | NA |
| C1QTNF1 | 13.5067796 | -0.034594623 | 0.76018737 | -0.045508 | 0.96370238 | NA |
| CBX4 | 4.1511021 | 0.020114592 | 0.75446103 | 0.02666088 | 0.97873022 | NA |
| TBC1D16 | 19.7436264 | 1.069128044 | 0.62429579 | 1.71253443 | 0.08679822 | NA |
| CCDC40 | 6.08807519 | 0.17167623 | 0.65064805 | 0.26385422 | 0.79189226 | NA |
| GAA | 18.8936283 | 0.308637822 | 0.45815819 | 0.67364903 | 0.50053447 | NA |
| EIF4A3 | 11.7451964 | -0.993089227 | 0.55332661 | -1.7947614 | 0.07269173 | NA |
| SGSH | 13.3314484 | 1.060143374 | 0.47905732 | 2.21297813 | 0.02689915 | NA |
| SLC26A11 | 9.96483554 | -0.204242064 | 0.48701578 | -0.4193746 | 0.67494236 | NA |
| RNF213 | 123.97136 | -0.638590605 | 0.54150539 | -1.1792876 | 0.23828366 | 0.64431903 |
| AC124319.2 | 31.8695213 | 0.340991758 | 0.41738692 | 0.816968 | 0.41394672 | 0.76946282 |
| ENDOV | 13.5357292 | -0.066684383 | 0.37992664 | -0.1755191 | 0.86067176 | NA |
| RPTOR | 48.1247409 | 0.169186237 | 0.27199987 | 0.62200851 | 0.53393627 | 0.83677705 |
| CHMP6 | 6.6739919 | -0.090776732 | 0.59709556 | -0.1520305 | 0.87916289 | NA |
| BAIAP2 | 43.0477416 | 0.077306847 | 0.39471887 | 0.19585293 | 0.84472528 | 0.95509854 |
| NDUFAF8 | 14.4817674 | -0.04892599 | 0.49913975 | -0.0980206 | 0.92191592 | NA |
| SLC38A10 | 52.6635539 | 0.205859861 | 0.28491702 | 0.72252568 | 0.46997134 | 0.80448595 |
| AC110285.2 | 7.00599405 | -0.007245682 | 0.60208992 | -0.0120342 | 0.99039831 | NA |
| BAHCC1 | 4.10765332 | -0.786019773 | 0.83438683 | -0.9420328 | 0.34617584 | NA |
| ACTG1 | 594.228316 | -0.184248225 | 0.48638921 | -0.3788082 | 0.70483029 | 0.90928488 |
| FAAP100 | 5.73963289 | 0.004918321 | 0.6369296 | 0.00772192 | 0.99383886 | NA |
| NPLOC4 | 51.2890599 | 0.166995495 | 0.29880861 | 0.55887109 | 0.5762497 | 0.85929076 |
| OXLD1 | 12.5712362 | -0.435561829 | 0.46351337 | -0.9396964 | 0.34737333 | NA |
| CCDC137 | 9.52797897 | -0.138695101 | 0.47023092 | -0.294951 | 0.76803128 | NA |
| ARL16 | 14.2052415 | 0.283584859 | 0.37531978 | 0.75558196 | 0.44989987 | NA |
| HGS | 17.6283597 | 0.383324183 | 0.39593781 | 0.96814241 | 0.33297325 | NA |
| MCRIP1 | 24.8885744 | -0.008254086 | 0.41856206 | -0.0197201 | 0.98426666 | NA |
| P4HB | 147.605826 | -0.110835827 | 0.45459553 | -0.243812 | 0.80737646 | 0.94328109 |
| ARHGDIA | 68.4173018 | -0.274982939 | 0.4109911 | -0.6690727 | 0.50344908 | 0.82600448 |
| ALYREF | 3.93832764 | -0.239424742 | 0.84811196 | -0.2823032 | 0.77771101 | NA |
| ANAPC11 | 37.1994453 | -0.709459699 | 0.41033512 | -1.7289763 | 0.08381334 | 0.44304225 |
| PCYT2 | 8.50857975 | 0.332034628 | 0.64641604 | 0.51365469 | 0.60749344 | NA |
| SIRT7 | 9.60074531 | 0.227413404 | 0.45798854 | 0.49654825 | 0.61950766 | NA |
| MAFG | 8.13287633 | -0.901231318 | 0.5058024 | -1.7817854 | 0.07478424 | NA |
| ASPSCR1 | 12.6354739 | -0.306691101 | 0.50150608 | -0.6115401 | 0.54084205 | NA |
| CENPX | 10.0015899 | -0.699491633 | 0.59054802 | -1.1844788 | 0.23622357 | NA |
| DCXR | 17.0275688 | -0.612746708 | 0.56607002 | -1.0824574 | 0.27904932 | NA |
| RFNG | 12.1049304 | 0.044765072 | 0.51533922 | 0.08686525 | 0.93077862 | NA |
| GPS1 | 15.2645608 | -0.439261792 | 0.53233424 | -0.8251616 | 0.40927983 | NA |
| DUS1L | 10.2625399 | -0.641962109 | 0.55564723 | -1.1553412 | 0.24795075 | NA |
| FASN | 5.54818242 | -0.816613004 | 0.73749437 | -1.1072803 | 0.26817275 | NA |
| CCDC57 | 32.5948567 | 0.153988624 | 0.28521397 | 0.53990561 | 0.58926213 | 0.86533208 |
| SLC16A3 | 7.85154939 | -0.547233147 | 0.60524199 | -0.904156 | 0.36591271 | NA |
| CSNK1D | 32.4503351 | -0.492408821 | 0.33874967 | -1.4536068 | 0.14605536 | 0.54498669 |
| SECTM1 | 9.13847049 | 0.217851968 | 2.24421183 | 0.09707282 | 0.92266857 | NA |
| OGFOD3 | 17.0633234 | -0.148281457 | 0.44967364 | -0.3297535 | 0.74158623 | NA |
| HEXDC | 12.7306879 | -0.440007929 | 0.45176639 | -0.9739723 | 0.33007031 | NA |
| CYBC1 | 11.8079375 | -0.399809213 | 0.57826143 | -0.6913987 | 0.489315 | NA |
| NARF | 10.8641963 | 0.165546504 | 0.55338225 | 0.29915398 | 0.76482256 | NA |
| FOXK2 | 51.7613939 | -0.171032357 | 0.26379303 | -0.6483581 | 0.51675334 | 0.83104823 |
| WDR45B | 37.504721 | -0.002706007 | 0.29947246 | -0.0090359 | 0.99279048 | 0.99820481 |
| RAB40B | 8.06520729 | 0.265089646 | 0.57975892 | 0.45724117 | 0.64749771 | NA |
| FN3KRP | 8.73391779 | -0.208455093 | 0.51648435 | -0.4036039 | 0.68650402 | NA |
| FN3K | 6.18579284 | 0.524522303 | 0.74469321 | 0.70434683 | 0.48121681 | NA |
| TBCD | 29.9699004 | 0.517431879 | 0.30881008 | 1.67556665 | 0.09382311 | 0.46635603 |
| METRNL | 4.65402609 | -0.885711933 | 0.73890194 | -1.1986867 | 0.23064979 | NA |
| USP14 | 49.6042638 | -0.186229149 | 0.37675906 | -0.4942924 | 0.62109967 | 0.8780438 |
| THOC1 | 27.8033231 | 0.146378214 | 0.32155438 | 0.45522071 | 0.64895046 | NA |
| ENOSF1 | 70.5745258 | -0.121007445 | 0.34884271 | -0.3468825 | 0.72867956 | 0.91700494 |
| YES1 | 43.6237481 | 0.363347292 | 0.24977544 | 1.45469581 | 0.14575349 | 0.54436111 |
| AP005328.1 | 9.36799503 | 0.603463774 | 0.82334741 | 0.73293942 | 0.46359538 | NA |
| METTL4 | 10.4454704 | 0.238362124 | 0.65746111 | 0.36254939 | 0.71694152 | NA |
| SMCHD1 | 79.0697297 | 0.355014569 | 0.3153851 | 1.12565422 | 0.26031191 | 0.65595635 |
| AP001011.1 | 11.070641 | -0.875504903 | 0.74421872 | -1.1764081 | 0.23943185 | NA |
| LPIN2 | 39.3125321 | 0.025306521 | 0.3341658 | 0.07573043 | 0.93963356 | 0.97998296 |
| MYOM1 | 7.52915671 | 1.072003973 | 0.70443455 | 1.52179357 | 0.12806081 | NA |
| MYL12A | 184.761782 | -0.052047048 | 0.5141296 | -0.1012333 | 0.91936524 | 0.97775374 |
| MYL12B | 235.885876 | -0.421453273 | 0.55506122 | -0.7592915 | 0.44767819 | 0.78947076 |
| TGIF1 | 11.1974912 | -0.39361043 | 0.65987091 | -0.5964961 | 0.55084385 | NA |
| DLGAP1 | 19.2948084 | -0.918851488 | 0.53021851 | -1.7329676 | 0.08310143 | NA |
| DLGAP1-AS1 | 11.8558372 | -0.53347733 | 0.42414042 | -1.2577847 | 0.20846963 | NA |

| | | | | | | |
|---|---|---|---|---|---|---|
| LINC00667 | 15.5417612 | -0.441043317 | 0.41942177 | -1.0515508 | 0.29300567 | NA |
| ZBTB14 | 5.64364816 | 0.082457422 | 0.70109587 | 0.11761219 | 0.90637495 | NA |
| EPB41L3 | 25.7714244 | 2.1616977 | 0.77023631 | 2.80653829 | 0.0050077 | NA |
| TMEM200C | 44.3398403 | 0.287853253 | 0.65008015 | 0.44279656 | 0.65791289 | 0.89709157 |
| AP005433.1 | 45.998016 | 0.149263685 | 0.54028016 | 0.2762709 | 0.78234001 | 0.93415197 |
| L3MBTL4 | 46.7472111 | 0.736545561 | 0.59200898 | 1.24414593 | 0.21344586 | 0.62255988 |
| AP001021.2 | 10.0897241 | -0.245641907 | 0.68946455 | -0.3562792 | 0.72163145 | NA |
| ARHGAP28 | 574.190415 | 0.356969584 | 0.34589325 | 1.03202242 | 0.30206162 | 0.68233351 |
| LAMA1 | 11.0274406 | -1.471087352 | 0.90054065 | -1.6335602 | 0.10235123 | NA |
| PTPRM | 116.525026 | 0.613057805 | 0.48352382 | 1.26789576 | 0.20483518 | 0.61387077 |
| RAB12 | 23.0831131 | -0.162109186 | 0.30369346 | -0.5337922 | 0.59348534 | NA |
| NDUFV2 | 33.0596216 | 0.125668335 | 0.34933877 | 0.359732 | 0.71904756 | 0.91472065 |
| NDUFV2-AS1 | 4.69524893 | -0.798035627 | 0.68644719 | -1.1625594 | 0.24500832 | NA |
| ANKRD12 | 217.665839 | -0.029478117 | 0.185625 | -0.1588047 | 0.87382277 | 0.96229085 |
| TWSG1 | 44.6780418 | 0.019282628 | 0.4303136 | 0.04481064 | 0.96425824 | 0.98892249 |
| RALBP1 | 46.6598754 | -0.374515143 | 0.38340696 | -0.9768084 | 0.32866401 | 0.70198063 |
| PPP4R1 | 50.8195558 | 0.116519568 | 0.36438631 | 0.31976933 | 0.7491432 | 0.92274989 |
| RAB31 | 39.8314936 | 0.504049684 | 0.48310194 | 1.04336093 | 0.29678116 | 0.68103704 |
| VAPA | 111.109106 | 0.123280886 | 0.27934295 | 0.4413245 | 0.6589781 | 0.89709157 |
| NAPG | 25.4826139 | -0.042216398 | 0.33933217 | -0.1244102 | 0.90099046 | NA |
| GNAL | 19.2214953 | -0.94441894 | 0.49154999 | -1.921308 | 0.05469289 | NA |
| MPPE1 | 24.7798849 | 0.033943991 | 0.34162041 | 0.09936172 | 0.92085108 | NA |
| IMPA2 | 13.6497151 | -0.731404694 | 0.53215922 | -1.3744095 | 0.16931459 | NA |
| TUBB6 | 29.898099 | -0.375154939 | 0.40593748 | -0.9241693 | 0.3553982 | 0.72709121 |
| AFG3L2 | 30.6801559 | -0.195197215 | 0.34747492 | -0.5617592 | 0.57428013 | 0.85861279 |
| PRELID3A | 5.14277858 | 0.011942188 | 0.56711521 | 0.02105778 | 0.98319956 | NA |
| SPIRE1 | 115.427268 | 0.112321348 | 0.33021348 | 0.34014768 | 0.73374532 | 0.91884473 |
| PSMG2 | 27.9266601 | -0.139400491 | 0.39661069 | -0.3514794 | 0.72522872 | NA |
| CEP76 | 6.18350217 | 0.012762821 | 0.61721934 | 0.02067793 | 0.98350257 | NA |
| PTPN2 | 38.035419 | -0.324508156 | 0.29428446 | -1.1027023 | 0.27015646 | 0.66288844 |
| SEH1L | 19.7955344 | -0.285305268 | 0.47839118 | -0.5963849 | 0.55091814 | NA |
| AP002449.1 | 3.71432087 | -1.301982985 | 1.20586199 | -1.0797114 | 0.2802707 | NA |
| CEP192 | 43.0875579 | 0.313289348 | 0.29249349 | 1.07109851 | 0.28412514 | 0.67249058 |
| LDLRAD4 | 20.3996704 | 1.194608503 | 0.74286812 | 1.60810306 | 0.10781261 | NA |
| FAM210A | 14.8010859 | 0.144991691 | 0.42710389 | 0.3394764 | 0.73425087 | NA |
| RNMT | 25.8040178 | 0.094611065 | 0.30637215 | 0.30881092 | 0.75746536 | NA |
| ZNF519 | 9.25366009 | 0.838466342 | 0.49359176 | 1.6987041 | 0.08937495 | NA |
| ANKRD30B | 7.37290775 | 0.48661245 | 0.79387496 | 0.61295856 | 0.53990374 | NA |
| ROCK1 | 145.940771 | 0.184796336 | 0.20447576 | 0.90375667 | 0.36612444 | 0.73456735 |
| ESCO1 | 29.3189865 | 0.191207037 | 0.37069368 | 0.51580874 | 0.605988 | 0.87080602 |
| SNRPD1 | 18.6491885 | -0.793990171 | 0.48220707 | -1.6465751 | 0.09964541 | NA |
| ABHD3 | 49.8859498 | 0.741486263 | 0.28479799 | 2.60355156 | 0.00922634 | 0.1515062 |
| MIB1 | 298.613101 | 0.722178953 | 0.32098744 | 2.24986673 | 0.02445741 | 0.25241537 |
| RBBP8 | 24.0022839 | 0.323203728 | 0.59289077 | 0.54513199 | 0.58566274 | NA |
| CABLES1 | 42.1361625 | 0.602594215 | 0.4758434 | 1.26637087 | 0.20538033 | 0.61387077 |
| TMEM241 | 60.5388562 | 0.454309753 | 0.38758689 | 1.17214942 | 0.24113707 | 0.64918976 |
| RIOK3 | 61.0551302 | 0.245485332 | 0.28295775 | 0.86756886 | 0.3856304 | 0.75125692 |
| RMC1 | 13.975276 | -0.395697077 | 0.42961635 | -0.9210475 | 0.35702561 | NA |
| NPC1 | 32.6734263 | -0.097658122 | 0.41024402 | -0.2380489 | 0.8118432 | 0.94415747 |
| LAMA3 | 28.349367 | 0.403736612 | 0.96173794 | 0.41979899 | 0.6746323 | NA |
| TTC39C | 6.64386044 | 1.375739909 | 0.59972059 | 2.29396811 | 0.02179233 | NA |
| OSBPL1A | 26.1500428 | 0.529163414 | 0.38033932 | 1.39129294 | 0.16413661 | NA |
| IMPACT | 13.1364475 | 0.250616122 | 0.43712543 | 0.57332771 | 0.56642283 | NA |
| ZNF521 | 69.1964223 | 0.61613638 | 0.34458263 | 1.78806571 | 0.07376541 | 0.41834543 |
| SS18 | 64.3387731 | -0.154555162 | 0.25718973 | -0.6009383 | 0.54788108 | 0.84550378 |
| TAF4B | 10.1706911 | -0.712676986 | 0.47610235 | -1.4968987 | 0.13441962 | NA |
| KCTD1 | 13.7301144 | -0.702296261 | 0.50616764 | -1.3874776 | 0.16529618 | NA |
| CDH2 | 12.5532742 | 0.792139491 | 0.60641918 | 1.30625732 | 0.19146507 | NA |
| DSC2 | 24.2224748 | -0.10692143 | 0.39899023 | -0.2679801 | 0.78871466 | NA |
| DSG2 | 30.5719529 | -0.031723202 | 0.3767991 | -0.0841913 | 0.93290434 | 0.97832857 |
| TRAPPC8 | 65.0731311 | 0.255183638 | 0.23904233 | 1.06752487 | 0.28573489 | 0.67377254 |
| RNF138 | 20.1842681 | 0.395635928 | 0.35835505 | 1.10403335 | 0.26957867 | NA |
| GAREM1 | 148.106151 | 0.306329525 | 0.34584965 | 0.88573033 | 0.37576286 | 0.74330391 |
| DTNA | 49.5906631 | -0.747116892 | 0.44874179 | -1.6649149 | 0.09592974 | 0.46942032 |
| MAPRE2 | 38.9063893 | 0.518876904 | 0.51871662 | 1.000309 | 0.31716099 | 0.69424851 |
| ZNF397 | 16.979088 | 0.113349713 | 0.48964292 | 0.23149464 | 0.81693055 | NA |
| ZSCAN30 | 13.1930608 | -0.475447719 | 0.42251449 | -1.1252815 | 0.26046979 | NA |
| ZNF24 | 29.3377347 | -0.415019565 | 0.33050997 | -1.2556946 | 0.20922673 | 0.61736443 |
| GALNT1 | 31.3262516 | -0.346121681 | 0.33002306 | -1.0487803 | 0.29427925 | 0.6804998 |
| RPRD1A | 62.5897348 | 0.147441164 | 0.22756853 | 0.64789787 | 0.51705101 | 0.83104823 |
| SLC39A6 | 24.0696959 | -0.254077987 | 0.41912381 | -0.6062123 | 0.54437382 | NA |
| ELP2 | 33.4611323 | 0.315844388 | 0.2909686 | 1.08549303 | 0.27770335 | 0.66958859 |
| MOCOS | 4.32890253 | -1.465956334 | 0.89581955 | -1.6364415 | 0.10174721 | NA |
| FHOD3 | 42.2743678 | 2.092537269 | 0.44670942 | 4.68433661 | 2.81E-06 | 0.00063289 |
| TPGS2 | 35.5337083 | -0.036883911 | 0.43863988 | -0.084087 | 0.93298726 | 0.97832857 |
| KIAA1328 | 78.3818734 | 0.495293092 | 0.26008689 | 1.904337 | 0.05686631 | 0.37278053 |
| AC016205.1 | 13.3055133 | -0.935610372 | 0.58933746 | -1.587563 | 0.11238518 | NA |
| PIK3C3 | 63.6425604 | 0.229916527 | 0.24628497 | 0.93353862 | 0.35054194 | 0.72238707 |
| LINC00907 | 5.88148457 | -0.683258209 | 0.60842173 | -1.123001 | 0.26143707 | NA |
| SETBP1 | 62.221254 | -0.114419083 | 0.53356044 | -0.2144445 | 0.83020047 | 0.95008863 |
| EPG5 | 26.5783112 | 0.164796409 | 0.33573939 | 0.49084621 | 0.62353522 | NA |
| ATP5F1A | 41.8535846 | -0.351266298 | 0.47706263 | -0.7363106 | 0.46154167 | 0.79727981 |
| HAUS1 | 7.94300184 | 0.142393137 | 0.49994119 | 0.28481977 | 0.77578222 | NA |
| C18orf25 | 29.4743275 | 0.389604979 | 0.32950928 | 1.18237937 | 0.2370552 | 0.64321384 |
| PIAS2 | 44.8228437 | 0.546988274 | 0.37028875 | 1.47719389 | 0.13962367 | 0.53526806 |
| KATNAL2 | 13.0776664 | 0.387506829 | 0.4597576 | 0.84285029 | 0.39931218 | NA |
| HDHD2 | 7.53412416 | 0.529755396 | 0.54213992 | 0.97715623 | 0.32849181 | NA |
| SMAD2 | 123.811621 | 0.177108261 | 0.23725526 | 0.74648823 | 0.45537254 | 0.79324789 |
| ZBTB7C | 219.53696 | 0.889942046 | 0.41244531 | 2.15772136 | 0.0309495 | 0.28264481 |
| CTIF | 33.0057765 | 0.268237389 | 0.41823381 | 0.64135749 | 0.52129044 | 0.83108387 |
| DYM | 113.018915 | 0.13710784 | 0.20115125 | 0.68161565 | 0.49548201 | 0.8234249 |
| ACAA2 | 14.2653397 | -0.675607311 | 0.45263636 | -1.492605 | 0.1355406 | NA |
| SNHG22 | 8.56045497 | -0.457634359 | 0.62664055 | -0.730298 | 0.46520803 | NA |
| MYO5B | 14.6207684 | 0.932523901 | 0.72353524 | 1.28884379 | 0.1974524 | NA |
| MBD1 | 12.6818868 | -0.253458655 | 0.40907151 | -0.619595 | 0.53552447 | NA |
| CXXC1 | 8.20078583 | 0.364956169 | 0.50424209 | 0.72377173 | 0.46920589 | NA |
| ME2 | 20.4165746 | -0.238761904 | 0.35702499 | -0.668754 | 0.50365239 | NA |
| AC091551.1 | 6.75003047 | -0.810532017 | 0.57341648 | -1.4135136 | 0.15750476 | NA |
| SMAD4 | 44.7678881 | 0.243467351 | 0.2857417 | 0.85205397 | 0.39418414 | 0.75773027 |
| MEX3C | 12.946993 | 0.321141979 | 0.40170008 | 0.7994571 | 0.42402541 | NA |
| MBD2 | 81.3926594 | -0.645989117 | 0.27840365 | -2.3203328 | 0.02032288 | 0.2375493 |
| POLI | 33.1431129 | -0.186553534 | 0.38335148 | -0.4866384 | 0.62651464 | 0.88050706 |
| CCDC68 | 11.6008541 | -0.889967582 | 0.61741339 | -1.4414452 | 0.14945894 | NA |
| TCF4 | 17.7497239 | 0.269217145 | 0.61863187 | 0.4351815 | 0.66343069 | NA |
| AC009271.1 | 25.4444298 | -0.011481798 | 0.50114137 | -0.0229113 | 0.98172103 | NA |
| TXNL1 | 31.446794 | -0.099502387 | 0.27538217 | -0.3613247 | 0.71785671 | 0.91472065 |
| WDR7 | 61.9101814 | 0.025379638 | 0.23038776 | 0.11016053 | 0.91228206 | 0.97682578 |
| FECH | 12.2667486 | 0.530931438 | 0.41759137 | 1.2714138 | 0.20358147 | NA |
| NARS | 28.3786083 | -0.396791636 | 0.42276024 | -0.9385737 | 0.34794967 | NA |
| AC027097.2 | 33.122601 | 0.21001771 | 0.48753428 | 0.43077526 | 0.66663179 | 0.90008606 |
| ATP8B1 | 234.892471 | 0.404557432 | 0.29128807 | 1.38885685 | 0.16487628 | 0.56632803 |
| NEDD4L | 150.346276 | 1.155367885 | 0.33416207 | 3.45750761 | 0.0005452 | 0.03300475 |
| MALT1 | 33.2333394 | 0.005390795 | 0.30480441 | 0.01768608 | 0.98588929 | 0.99585746 |

| | | | | | | |
|---|---|---|---|---|---|---|
| AC104365.1 | 4.29501416 | -1.295613133 | 0.97631108 | -1.3270495 | 0.18449231 | NA |
| ZNF532 | 49.5014852 | 0.503676836 | 0.27644086 | 1.82200575 | 0.06845411 | 0.40491695 |
| SEC11C | 14.5791326 | 0.063925202 | 0.47616048 | 0.13425138 | 0.8932038 | NA |
| LMAN1 | 68.2174518 | 0.114863832 | 0.37113676 | 0.30949193 | 0.75694735 | 0.92313949 |
| CCBE1 | 370.198249 | 0.404842166 | 0.30927469 | 1.30900516 | 0.1905326 | 0.59329687 |
| RNF152 | 19.5720942 | 1.320153736 | 0.86123259 | 1.53286552 | 0.12530899 | NA |
| PIGN | 39.8642709 | 0.350090665 | 0.27741874 | 1.2619575 | 0.20696408 | 0.61629707 |
| KIAA1468 | 50.5645522 | 0.307118962 | 0.29249493 | 1.04999754 | 0.29371924 | 0.6804998 |
| ZCCHC2 | 40.3624201 | 0.180729071 | 0.34979959 | 0.51666461 | 0.6053903 | 0.87047761 |
| PHLPP1 | 45.723125 | 0.241917689 | 0.30395424 | 0.79590168 | 0.42608918 | 0.77742588 |
| BCL2 | 70.3957028 | 0.934610447 | 0.40415036 | 2.31253151 | 0.02074841 | 0.23840101 |
| KDSR | 38.2352523 | -0.694822253 | 0.33001529 | -2.1054244 | 0.03525438 | 0.29361757 |
| VPS4B | 43.6209181 | 0.070134 | 0.27066148 | 0.25912073 | 0.79554209 | 0.93934998 |
| SERPINB8 | 3.98264693 | -1.479332752 | 0.96564025 | -1.5319709 | 0.12552962 | NA |
| DSEL | 3.74631559 | -1.32810844 | 1.00589207 | -1.320329 | 0.18672521 | NA |
| TMX3 | 43.3523591 | 0.897849075 | 0.38305828 | 2.34389682 | 0.01908345 | 0.23105216 |
| CCDC102B | 8.69510545 | -1.185067004 | 0.70436635 | -1.6824583 | 0.09248001 | NA |
| DOK6 | 9.54354802 | 1.272573805 | 0.73814168 | 1.72402377 | 0.08470355 | NA |
| RTTN | 26.320388 | 0.211439915 | 0.42824068 | 0.49374084 | 0.62148922 | NA |
| SOCS6 | 13.1858716 | 0.33274409 | 0.426896 | 0.77945 | 0.43571468 | NA |
| TIMM21 | 5.39770998 | -1.202291344 | 0.6478934 | -1.8556932 | 0.06349731 | NA |
| CYB5A | 117.951105 | 0.135573918 | 0.38895934 | 0.3485555 | 0.72742304 | 0.91700494 |
| CNDP2 | 45.2070145 | -0.438430827 | 0.38803398 | -1.1298774 | 0.25852789 | 0.65588339 |
| LINC00909 | 9.87491493 | -0.773999023 | 0.509989 | -1.5176779 | 0.12909562 | NA |
| ZNF407 | 85.9880643 | 0.050459176 | 0.21141738 | 0.2386709 | 0.81136078 | 0.94408044 |
| ZADH2 | 41.900564 | 0.046573533 | 0.36513282 | 0.1275523 | 0.89850328 | 0.97097328 |
| TSHZ1 | 76.1175273 | 0.542167366 | 0.40138053 | 1.3507565 | 0.17677345 | 0.58001112 |
| ZNF516 | 64.6274759 | 0.523827761 | 0.25953529 | 2.0183296 | 0.04355695 | 0.33029013 |
| ZNF236-DT | 7.37701962 | 0.085462722 | 0.49974392 | 0.17101303 | 0.86421352 | NA |
| ZNF236 | 32.3504892 | 0.306436445 | 0.3431813 | 0.89292874 | 0.37189533 | 0.74014106 |
| MBP | 44.885321 | -0.47441873 | 0.33667283 | -1.4091388 | 0.15879414 | 0.56152488 |
| ATP9B | 74.4042925 | -0.099988435 | 0.22085572 | -0.4527319 | 0.6507418 | 0.89412414 |
| NFATC1 | 11.5337618 | -0.063779768 | 0.56241423 | -0.1134035 | 0.90971063 | NA |
| CTDP1 | 10.6862375 | -0.082196263 | 0.47668307 | -0.1724338 | 0.86309652 | NA |
| PQLC1 | 34.663753 | 0.181862245 | 0.4373712 | 0.41580755 | 0.67755084 | 0.90499319 |
| HSBP1L1 | 34.7696449 | -0.832098836 | 0.38566166 | -2.1575876 | 0.03095991 | 0.28264481 |
| TXNL4A | 32.3986229 | -0.848864766 | 0.40165192 | -2.1134338 | 0.03456365 | 0.29210684 |
| ADNP2 | 16.3725405 | -0.260614574 | 0.35659953 | -0.7308326 | 0.46488138 | NA |
| PARD6G-AS1 | 7.9662146 | -0.818791558 | 0.74843105 | -1.0940107 | 0.2739503 | NA |
| PARD6G | 167.089041 | 0.048134484 | 0.4254905 | 0.11312705 | 0.90992983 | 0.97557165 |
| AC139100.1 | 8.05618363 | 0.167679618 | 0.6888961 | 0.24340335 | 0.80769295 | NA |
| ZCCHC3 | 11.374444 | 0.429336142 | 0.44995882 | 0.95416764 | 0.33999879 | NA |
| NRSN2-AS1 | 14.1508093 | -0.363052014 | 0.39697322 | -0.9145504 | 0.36042771 | NA |
| SOX12 | 7.43751123 | 0.175297726 | 0.49721402 | 0.3525599 | 0.7244184 | NA |
| NRSN2 | 17.5556433 | -0.170593548 | 0.41944697 | -0.4067106 | 0.68422052 | NA |
| RBCK1 | 37.0450347 | -0.360517653 | 0.64643357 | -0.5577026 | 0.57704751 | 0.86016344 |
| TBC1D20 | 14.5790089 | -0.313961086 | 0.40082171 | -0.7832936 | 0.43345471 | NA |
| CSNK2A1 | 63.8255982 | -0.052190585 | 0.20897706 | -0.2497431 | 0.802786 | 0.94126368 |
| PSMF1 | 41.8467476 | -0.043896139 | 0.42825791 | -0.1024993 | 0.91836036 | 0.97775374 |
| SNPH | 5.5833822 | 0.458190866 | 0.70660737 | 0.64843771 | 0.51670188 | NA |
| AL136531.2 | 14.0231882 | -1.018153281 | 0.52233129 | -1.9492481 | 0.0512658 | NA |
| SDCBP2-AS1 | 24.2893088 | 0.0494466 | 0.38650018 | 0.12793422 | 0.89820103 | NA |
| NSFL1C | 23.1142168 | 0.406008476 | 0.35873144 | 1.13178948 | 0.25772295 | NA |
| AL117335.1 | 6.46691183 | -0.641957577 | 0.62776493 | -1.0226082 | 0.30649312 | NA |
| SIRPA | 115.899753 | 0.027888023 | 0.39067979 | 0.07138333 | 0.94309268 | 0.98114803 |
| PDYN-AS1 | 23.8294665 | 0.287767336 | 0.52664321 | 0.54641801 | 0.58477863 | NA |
| STK35 | 13.6503486 | -0.046089939 | 0.46671179 | -0.0987546 | 0.92133311 | NA |
| SNRPB | 34.0175242 | -0.811741398 | 0.38522407 | -2.1071928 | 0.03510087 | 0.29354459 |
| NOP56 | 14.0500362 | -0.97831006 | 0.52277738 | -1.8713703 | 0.06129377 | NA |
| IDH3B | 12.1717857 | 0.363736132 | 0.47160506 | 0.77127275 | 0.44054528 | NA |
| EBF4 | 6.47356243 | -1.094306365 | 0.72505876 | -1.5092658 | 0.13123088 | NA |
| CPXM1 | 11.9717152 | -0.324132209 | 1.24919469 | -0.2594729 | 0.79527036 | NA |
| PCED1A | 12.3577653 | 0.697943476 | 0.47036695 | 1.48382762 | 0.13785466 | NA |
| PTPRA | 94.3609182 | -0.189701532 | 0.22700848 | -0.8356584 | 0.40334713 | 0.76594067 |
| MRPS26 | 21.9333989 | -1.001080732 | 0.49609313 | -2.017929 | 0.04359865 | NA |
| UBOX5-AS1 | 7.0384059 | -0.250299527 | 0.58900765 | -0.4249512 | 0.67087223 | NA |
| UBOX5 | 8.89469842 | 0.833631224 | 0.45475547 | 1.83314173 | 0.0667815 | NA |
| DDRGK1 | 18.2060063 | -0.263846851 | 0.42287628 | -0.6239339 | 0.53267099 | NA |
| ITPA | 19.9341539 | 0.053036201 | 0.3727273 | 0.14229224 | 0.88684918 | NA |
| C20orf194 | 185.127169 | 0.442057937 | 0.2492367 | 1.77364702 | 0.07612156 | 0.42236529 |
| ATRN | 59.7196296 | 0.246012422 | 0.24741602 | 0.99432698 | 0.3200637 | 0.69496685 |
| CENPB | 9.09155474 | -0.991336335 | 0.55383324 | -1.7899546 | 0.07346121 | NA |
| CDC25B | 23.5661994 | -0.729992723 | 0.40739941 | -1.7918355 | 0.07315932 | NA |
| AP5S1 | 4.23753989 | -0.69206333 | 0.68554255 | -1.0095119 | 0.31272922 | NA |
| MAVS | 28.8423122 | -0.237238801 | 0.4154386 | -0.5710562 | 0.56796153 | 0.85654436 |
| PANK2 | 33.6696659 | 0.018191569 | 0.29902582 | 0.06083611 | 0.95148973 | 0.98520702 |
| RNF24 | 43.0509351 | -0.591802703 | 0.54840696 | -1.0791305 | 0.28052953 | 0.66958859 |
| PRNP | 50.3935618 | -0.045853461 | 0.40206915 | -0.1140437 | 0.90920314 | 0.97528913 |
| SLC23A2 | 17.3580801 | -0.285573128 | 0.42863397 | -0.6662401 | 0.50525766 | NA |
| TMEM230 | 35.2623503 | -0.249253714 | 0.31410249 | -0.7935426 | 0.42746175 | 0.77844087 |
| PCNA | 9.75375753 | -0.655610093 | 0.49663965 | -1.3200921 | 0.18680426 | NA |
| CDS2 | 47.9213818 | 0.107387611 | 0.29264743 | 0.36695217 | 0.7136547 | 0.91328825 |
| GPCPD1 | 57.5148139 | 0.277950471 | 0.35332858 | 0.78666286 | 0.43147925 | 0.77901411 |
| SHLD1 | 20.3532708 | 0.181972768 | 0.48436345 | 0.37569467 | 0.7071439 | NA |
| TRMT6 | 6.00436885 | -0.405713445 | 0.56127993 | -0.7228362 | 0.46978054 | NA |
| MCM8 | 7.38033408 | 1.187503408 | 0.69192666 | 1.71622727 | 0.08612045 | NA |
| CRLS1 | 26.5149901 | 0.162554767 | 0.34349656 | 0.4732355 | 0.63604517 | NA |
| BMP2 | 46.0092105 | 0.294813284 | 0.70155345 | 0.42022925 | 0.67431799 | 0.90403541 |
| TMX4 | 41.8708312 | -0.343542675 | 0.44679448 | -0.7689054 | 0.44194949 | 0.78620488 |
| PLCB1 | 88.0039411 | 1.258057788 | 0.89736723 | 1.40194309 | 0.16093224 | 0.56283048 |
| PLCB4 | 57.1799371 | 0.54757084 | 0.47867309 | 1.14393487 | 0.25265065 | 0.65239726 |
| ANKEF1 | 5.24637734 | -0.059966144 | 0.61030169 | -0.0982566 | 0.92172857 | NA |
| MKKS | 9.74508583 | 0.441129586 | 0.50817975 | 0.86805818 | 0.38536249 | NA |
| SLX4IP | 27.0606978 | 0.188996368 | 0.43058291 | 0.43893142 | 0.66071123 | NA |
| BTBD3 | 20.9510207 | 0.580886599 | 0.32697357 | 1.77655523 | 0.07564145 | NA |
| SPTLC3 | 14.4316841 | 0.075180502 | 0.4974045 | 0.1511456 | 0.87986086 | NA |
| TASP1 | 57.7787479 | 0.167894929 | 0.2657386 | 0.63180482 | 0.52751442 | 0.83195298 |
| ESF1 | 18.7170807 | -0.605385909 | 0.40272135 | -1.5032377 | 0.13277776 | NA |
| NDUFAF5 | 12.0153173 | 0.447297737 | 0.43815912 | 1.02085686 | 0.30732226 | NA |
| MACROD2 | 413.470322 | -0.530606223 | 0.4712639 | -1.1259216 | 0.26019869 | 0.65595635 |
| FLRT3 | 256.452741 | -0.834133787 | 0.52703514 | -1.582691 | 0.1134919 | 0.50483797 |
| KIF16B | 48.1163115 | 0.206225319 | 0.33037963 | 0.62420713 | 0.53249157 | 0.83658422 |
| SNRPB2 | 19.9066244 | -0.787004076 | 0.42184202 | -1.865637 | 0.06209217 | NA |
| DSTN | 277.613836 | -0.507469137 | 0.33562483 | -1.5120131 | 0.13053053 | 0.52613957 |
| RRBP1 | 57.5368772 | -0.780036342 | 0.49813295 | -1.56592 | 0.11736735 | 0.50775741 |
| SNX5 | 21.6720799 | 0.177698987 | 0.43681259 | 0.4068083 | 0.68414879 | NA |
| OVOL2 | 7.1609487 | 0.25757834 | 0.74161882 | 0.34731904 | 0.72835165 | NA |
| MGME1 | 13.8472556 | -0.358465277 | 0.4589696 | -0.7810218 | 0.43478965 | NA |
| KAT14 | 8.84461029 | -0.268076962 | 0.52468378 | -0.5109305 | 0.6093997 | NA |
| ZNF133 | 14.3767319 | 0.187206237 | 0.39523928 | 0.47365292 | 0.63574743 | NA |
| DZANK1 | 9.64697911 | 0.323149706 | 0.5338392 | 0.60533154 | 0.54495873 | NA |
| POLR3F | 6.41113134 | 0.754097384 | 0.65248737 | 1.15572717 | 0.24779277 | NA |
| RBBP9 | 5.8799908 | 0.085001592 | 0.60705969 | 0.1400218 | 0.88864277 | NA |

| | | | | | | |
|---|---|---|---|---|---|---|
| SEC23B | 21.2524446 | 0.202688889 | 0.34242229 | 0.59192668 | 0.55389968 | NA |
| DTD1 | 13.6640422 | -1.136711953 | 0.47616553 | -2.3872202 | 0.01697632 | NA |
| SLC24A3 | 14.591447 | -0.336208829 | 0.68927052 | -0.4877749 | 0.62570932 | NA |
| AL121761.1 | 21.2697317 | -1.29466075 | 0.51432786 | -2.5171896 | 0.01182952 | NA |
| RIN2 | 34.1799167 | 0.121185371 | 0.46389472 | 0.26123464 | 0.79391156 | 0.93934998 |
| NAA20 | 25.6419194 | -0.270754036 | 0.41619955 | -0.650539 | 0.51534412 | NA |
| CRNKL1 | 8.25256501 | -0.828173438 | 0.55351304 | -1.4962131 | 0.13459814 | NA |
| RALGAPA2 | 59.9752632 | 0.344378737 | 0.29958328 | 1.14952589 | 0.2503392 | 0.6518054 |
| KIZ | 36.5652123 | 0.488239265 | 0.45487672 | 1.07334415 | 0.28311673 | 0.67192596 |
| KIZ-AS1 | 9.39805989 | -1.769748389 | 0.77380635 | -2.2870688 | 0.0221918 | NA |
| XRN2 | 43.1652316 | -0.214075438 | 0.32182821 | -0.6651854 | 0.5059319 | 0.82600448 |
| AL118508.1 | 22.5091764 | -0.169466784 | 0.97746785 | -0.1733733 | 0.86235805 | NA |
| NXT1 | 7.31123126 | -0.290874957 | 0.71608106 | -0.406204 | 0.68459275 | NA |
| GZF1 | 5.7248854 | 0.146727099 | 0.63216264 | 0.2321034 | 0.8164577 | NA |
| NAPB | 30.9530837 | 0.832054367 | 0.33393603 | 2.49165794 | 0.01271484 | 0.18904687 |
| CST3 | 145.141238 | -0.888876523 | 0.37016737 | -2.4012828 | 0.01633771 | 0.21171163 |
| APMAP | 26.4038875 | -0.286799179 | 0.48394359 | -0.5926294 | 0.55342922 | NA |
| ACSS1 | 17.0872764 | -0.502292469 | 0.48545147 | -1.0346914 | 0.30081305 | NA |
| ENTPD6 | 13.0641497 | 0.67417827 | 0.545544 | 1.23579083 | 0.21653632 | NA |
| PYGB | 19.1198406 | -0.094153762 | 0.41596019 | -0.2263528 | 0.82092702 | NA |
| ABHD12 | 20.4506643 | -0.068771925 | 0.4014118 | -0.1713251 | 0.86396813 | NA |
| NINL | 31.1155327 | 0.647871329 | 0.45045372 | 1.4382639 | 0.15035919 | 0.54834287 |
| ZNF337-AS1 | 9.22867049 | -0.247817912 | 0.55773159 | -0.4443319 | 0.65680267 | NA |
| ZNF337 | 9.49734682 | -0.438124162 | 0.46630553 | -0.9395646 | 0.34744095 | NA |
| FAM182B | 16.7572179 | -0.591124404 | 0.4213973 | -1.4027722 | 0.16068479 | NA |
| HM13 | 47.0642505 | -0.393357884 | 0.39074457 | -1.006688 | 0.31408471 | 0.69309869 |
| ID1 | 31.6865865 | -3.365541888 | 1.46993024 | -2.2895929 | 0.02204492 | 0.24297341 |
| BCL2L1 | 56.3524268 | -0.187087352 | 0.37535235 | -0.4984313 | 0.6181801 | 0.87662819 |
| AL117381.1 | 9.36280694 | -0.43844556 | 0.72398682 | -0.6055988 | 0.54478119 | NA |
| DUSP15 | 6.17274451 | -0.443772379 | 1.30612393 | -0.3397628 | 0.73403513 | NA |
| PDRG1 | 7.16669608 | -0.236203062 | 0.6348642 | -0.3720529 | 0.70985347 | NA |
| TM9SF4 | 35.7513997 | -0.240261289 | 0.26965288 | -0.8910021 | 0.37292803 | 0.7407542 |
| PLAGL2 | 12.1646729 | -0.886730629 | 0.54176068 | -1.6367571 | 0.10168124 | NA |
| POFUT1 | 18.9310037 | -0.600673594 | 0.53975521 | -1.112863 | 0.26576728 | NA |
| KIF3B | 26.1190356 | 0.671885767 | 0.35434694 | 1.89612411 | 0.05794364 | NA |
| ASXL1 | 85.2554988 | -0.140902313 | 0.27900017 | -0.5050259 | 0.61354065 | 0.87439243 |
| NOL4L | 24.6001865 | 0.340085751 | 0.36240451 | 0.93841478 | 0.34803129 | NA |
| FO393400.1 | 7.78782335 | -0.425145628 | 0.54139688 | -0.7852754 | 0.43229214 | NA |
| MAPRE1 | 21.687606 | -0.343112049 | 0.33714899 | -1.0176867 | 0.30882687 | NA |
| AL035071.2 | 8.83300761 | 0.184540538 | 0.71358825 | 0.25860927 | 0.79593673 | NA |
| AL035071.1 | 8.82844284 | -0.102510307 | 0.69077744 | -0.1483985 | 0.88202832 | NA |
| CDK5RAP1 | 18.7694738 | 0.40934234 | 0.37705975 | 1.08561665 | 0.27764863 | NA |
| SNTA1 | 24.187386 | 0.347064287 | 0.50834519 | 0.68273349 | 0.49477526 | NA |
| CBFA2T2 | 66.7261445 | 0.488511519 | 0.27033061 | 1.80708914 | 0.07074838 | 0.40760715 |
| PXMP4 | 7.91290799 | 0.303747788 | 0.62753579 | 0.48403261 | 0.62836273 | NA |
| CHMP4B | 30.7903922 | 0.358533978 | 0.33976783 | 1.05523225 | 0.29131912 | 0.67792199 |
| RALY | 37.1137304 | -0.178222502 | 0.3401873 | -0.5238952 | 0.60035142 | 0.86882737 |
| EIF2S2 | 46.812292 | -0.906311881 | 0.34247834 | -2.6463334 | 0.00813696 | 0.14384634 |
| ASIP | 5.45577511 | 0.784855373 | 1.02931785 | 0.7625005 | 0.44576135 | NA |
| AHCY | 11.3461672 | -0.191008575 | 0.43428213 | -0.439826 | 0.66006313 | NA |
| ITCH | 102.960411 | 0.346585845 | 0.23969367 | 1.44595324 | 0.14819032 | 0.54691531 |
| DYNLRB1 | 104.72848 | -0.62333897 | 0.44153313 | -1.4117604 | 0.15802053 | 0.56051304 |
| MAP1LC3A | 16.2120902 | -0.100690899 | 0.54101113 | -0.1861161 | 0.85235369 | NA |
| PIGU | 15.9784765 | -0.47991537 | 0.46056069 | -1.0420242 | 0.29740047 | NA |
| NCOA6 | 63.1799713 | 0.179393904 | 0.23087439 | 0.77701951 | 0.43714726 | 0.78315781 |
| TP53INP2 | 6.16186823 | 0.569423599 | 0.65378029 | 0.87097089 | 0.38377005 | NA |
| GGT7 | 12.628804 | -0.58077304 | 0.48200864 | -1.2049017 | 0.22824124 | NA |
| ACSS2 | 27.2948505 | 0.043653601 | 0.3310377 | 0.13186897 | 0.89508793 | NA |
| GSS | 15.0882301 | -0.071549126 | 0.39003088 | -0.1834448 | 0.85444905 | NA |
| TRPC4AP | 49.2624227 | 0.01486784 | 0.25757701 | 0.05772192 | 0.95397013 | 0.98559751 |
| EDEM2 | 13.8127986 | 0.363581624 | 0.50018809 | 0.72688981 | 0.46729346 | NA |
| PROCR | 8.35545979 | -0.18869642 | 0.65046059 | -0.2900966 | 0.77174232 | NA |
| MMP24OS | 23.8286885 | 0.346758852 | 0.38768754 | 0.89442866 | 0.37109258 | NA |
| MMP24 | 12.4755015 | -0.424506026 | 0.71130607 | -0.596798 | 0.55064227 | NA |
| UQCC1 | 33.0976717 | 0.430844998 | 0.29426713 | 1.46412888 | 0.14315874 | 0.54064417 |
| CEP250 | 6.59476556 | -0.416403776 | 0.56279958 | -0.7398793 | 0.45937322 | NA |
| ERGIC3 | 20.1364711 | 0.453355467 | 0.4224777 | 1.07308734 | 0.28323193 | NA |
| CPNE1 | 22.8789727 | 0.297765136 | 0.44255057 | 0.67283866 | 0.50104994 | NA |
| ROMO1 | 20.588319 | -0.080769374 | 0.52731023 | -0.1531724 | 0.87826232 | NA |
| RBM39 | 131.308766 | -0.123700951 | 0.21211099 | -0.5831897 | 0.55976558 | 0.85321654 |
| PHF20 | 77.7746908 | -0.164033436 | 0.30454206 | -0.5386233 | 0.59014684 | 0.8654248 |
| SCAND1 | 47.1129108 | -0.816753017 | 0.4834106 | -1.6895637 | 0.09111145 | 0.45906588 |
| NORAD | 104.295634 | -0.492361583 | 0.37328181 | -1.3190077 | 0.18716654 | 0.58940023 |
| EPB41L1 | 20.9457231 | 0.13214366 | 0.51825494 | 0.2549781 | 0.79874002 | NA |
| AAR2 | 9.68862433 | 0.091694579 | 0.46468119 | 0.19732793 | 0.84357092 | NA |
| DLGAP4 | 60.4905423 | 0.305430989 | 0.37023543 | 0.82496425 | 0.40939189 | 0.76772515 |
| DLGAP4-AS1 | 11.8292948 | -0.764403845 | 0.44696934 | -1.710193 | 0.08723019 | NA |
| MYL9 | 450.562133 | -0.115842881 | 0.48041877 | -0.241129 | 0.80945516 | 0.94346479 |
| RAB5IF | 10.370756 | -0.746955932 | 0.62617373 | -1.1928893 | 0.23291274 | NA |
| NDRG3 | 12.7228529 | -0.395962474 | 0.57023155 | -0.6943889 | 0.48743834 | NA |
| SOGA1 | 21.2803575 | 0.155457988 | 0.49125557 | 0.31645033 | 0.75166072 | NA |
| SAMHD1 | 7.76736131 | 0.741302656 | 1.04483341 | 0.70949364 | 0.4780182 | NA |
| RBL1 | 9.50455515 | 0.83807938 | 0.53643496 | 1.56231312 | 0.11821424 | NA |
| MROH8 | 6.83034891 | -0.668995068 | 0.61481649 | -1.0881215 | 0.27654146 | NA |
| RPN2 | 67.6238974 | -0.426667802 | 0.59415602 | -0.7181074 | 0.4726911 | 0.80580648 |
| MANBAL | 15.0834036 | 0.163518094 | 0.42414862 | 0.38552075 | 0.69985164 | NA |
| SRC | 5.70511251 | 0.225052818 | 0.62999341 | 0.35723043 | 0.72091931 | NA |
| BLCAP | 33.5280522 | 0.030631529 | 0.33291349 | 0.09201048 | 0.92668972 | 0.97775374 |
| CTNNBL1 | 23.4201502 | 0.543627297 | 0.43759146 | 1.24231697 | 0.21411964 | NA |
| TTI1 | 8.15367365 | -0.24856444 | 0.5237043 | -0.4746275 | 0.63505253 | NA |
| RPRD1B | 34.5561026 | 0.499086219 | 0.2901061 | 1.72035758 | 0.08536746 | 0.44564602 |
| TGM2 | 9.03983144 | -2.577499338 | 0.91615573 | -2.8133856 | 0.00490228 | NA |
| RALGAPB | 52.4883626 | 0.258835123 | 0.27717005 | 0.93384953 | 0.35038152 | 0.72238707 |
| DHX35 | 7.89656136 | 0.105207616 | 0.56393292 | 0.18656052 | 0.85200523 | NA |
| MAFB | 244.608928 | -0.707460487 | 0.3869668 | -1.8282201 | 0.06751652 | 0.40302824 |
| TOP1 | 62.2717556 | -0.150861503 | 0.3317048 | -0.4548065 | 0.64924844 | 0.89387362 |
| PLCG1 | 6.76480761 | -0.411457811 | 0.56244814 | -0.731548 | 0.4644445 | NA |
| ZHX3 | 27.0009656 | 0.514467671 | 0.43752404 | 1.1758615 | 0.23965022 | NA |
| LPIN3 | 8.19395699 | 0.255883872 | 0.69010031 | 0.37079228 | 0.71079225 | NA |
| CHD6 | 86.1547215 | 0.49879784 | 0.27563903 | 1.80960526 | 0.07035702 | 0.40663921 |
| SRSF6 | 25.8686531 | 0.540727756 | 0.34709284 | 1.55787642 | 0.11926255 | NA |
| L3MBTL1 | 7.86070815 | -0.24701333 | 0.56105936 | -0.4402624 | 0.65974708 | NA |
| SGK2 | 5.51094208 | 0.354220202 | 0.66591234 | 0.53193218 | 0.59477297 | NA |
| IFT52 | 14.257213 | -0.302049377 | 0.38988517 | -0.7747137 | 0.43850887 | NA |
| TOX2 | 7.57116199 | -1.113552708 | 0.8829429 | -1.2611832 | 0.20724287 | NA |
| OSER1 | 17.419108 | -0.146123987 | 0.3876205 | -0.3769769 | 0.70619075 | NA |
| FITM2 | 5.72443501 | -0.218125092 | 0.64377999 | -0.3388193 | 0.73474586 | NA |
| TTPAL | 9.20875092 | -0.606449669 | 0.52512835 | -1.1548599 | 0.2481478 | NA |
| SERINC3 | 68.7440147 | -0.11279468 | 0.33384997 | -0.3378604 | 0.7354684 | 0.91927884 |
| PKIG | 46.0630003 | -0.179437065 | 0.32107101 | -0.5588703 | 0.57625021 | 0.85929076 |
| YWHAB | 91.9248071 | -0.739247159 | 0.4117147 | -1.7955326 | 0.07256889 | 0.41691135 |
| TOMM34 | 7.05221774 | 0.711005156 | 0.62940952 | 1.1296384 | 0.25862862 | NA |
| STK4 | 20.9327724 | -0.252522521 | 0.37653773 | -0.6706433 | 0.50244776 | NA |

| | | | | | | |
|---|---|---|---|---|---|---|
| SLPI | 53.1342668 | -3.546120872 | 1.42592868 | -2.4868852 | 0.0128867 | 0.18904687 |
| SDC4 | 164.019475 | -0.510752373 | 0.48772706 | -1.0472094 | 0.295003 | 0.68103704 |
| PIGT | 34.8982055 | 0.107654704 | 0.55955267 | 0.19239423 | 0.84743341 | 0.95532127 |
| WFDC2 | 95.4349701 | -3.844114995 | 0.77837749 | -4.9386256 | 7.87E-07 | 0.00023577 |
| DNTTIP1 | 8.91552596 | -0.268001773 | 0.49343873 | -0.5431308 | 0.58703975 | NA |
| SNX21 | 15.0108692 | 0.338587329 | 0.37910495 | 0.89312295 | 0.37179133 | NA |
| ACOT8 | 13.2216292 | 0.54849874 | 0.41318127 | 1.32750147 | 0.18434285 | NA |
| CTSA | 41.2335399 | -0.113668215 | 0.43420395 | -0.2617853 | 0.79348696 | 0.93934998 |
| PLTP | 45.8507599 | -0.993642238 | 0.50628334 | -1.9626208 | 0.04969025 | 0.34869143 |
| PCIF1 | 5.14835679 | -1.389669556 | 0.75206702 | -1.8478001 | 0.06463126 | NA |
| ZNF335 | 9.61241177 | 0.403211007 | 0.59641287 | 0.67606021 | 0.49900242 | NA |
| NCOA5 | 24.0284097 | -0.057259531 | 0.43291483 | -0.1322651 | 0.8947746 | NA |
| CD40 | 4.75083224 | 1.092474437 | 0.98485376 | 1.1092758 | 0.26731122 | NA |
| SLC35C2 | 16.3721842 | -0.571296579 | 0.43651502 | -1.3087673 | 0.19061319 | NA |
| ELMO2 | 14.7755955 | 0.092968965 | 0.54267607 | 0.17131576 | 0.86397549 | NA |
| SLC13A3 | 36.9694365 | 1.110437653 | 0.92799137 | 1.19660342 | 0.23146117 | 0.63691078 |
| TP53RK | 5.28643226 | -1.989667995 | 0.85439775 | -2.3287374 | 0.01987298 | NA |
| EYA2 | 40.4751271 | 0.10124594 | 0.62888372 | 0.1609931 | 0.87209884 | 0.96207879 |
| ZMYND8 | 86.913169 | -0.592583532 | 0.3739174 | -1.584798 | 0.11301224 | 0.50426586 |
| NCOA3 | 49.8614001 | 0.184704569 | 0.26689853 | 0.6920404 | 0.48891196 | 0.81774305 |
| SULF2 | 177.838408 | -1.368162713 | 0.42465577 | -3.2218159 | 0.00127381 | 0.0545866 |
| ARFGEF2 | 53.323493 | 0.253318442 | 0.28935199 | 0.87546811 | 0.38131925 | 0.74899182 |
| CSE1L | 21.7661008 | 0.388544557 | 0.36087762 | 1.0766657 | 0.28162964 | NA |
| STAU1 | 48.201717 | -0.228478775 | 0.33138793 | -0.6894601 | 0.49053374 | 0.81876743 |
| DDX27 | 12.1937076 | -0.076528945 | 0.42288756 | -0.1809676 | 0.85639301 | NA |
| ZNFX1 | 29.3982129 | -0.282583113 | 0.67709856 | -0.4173441 | 0.67642671 | 0.90499319 |
| ZFAS1 | 21.0883184 | -1.122915906 | 0.56427737 | -1.990007 | 0.04659017 | NA |
| B4GALT5 | 47.7228794 | -0.10448079 | 0.39596426 | -0.2638642 | 0.79188457 | 0.93914731 |
| SLC9A8 | 17.9072135 | 0.172709134 | 0.38978311 | 0.44309034 | 0.65770039 | NA |
| RNF114 | 36.502194 | 0.088162941 | 0.48431011 | 0.1820382 | 0.85555275 | 0.95615679 |
| CEBPB | 31.0015621 | -3.828544068 | 0.77833087 | -4.9189158 | 8.70E-07 | 0.00023577 |
| PTPN1 | 65.6589021 | -0.791362843 | 0.47073416 | -1.6811247 | 0.09273869 | 0.4638078 |
| PARD6B | 97.7311905 | 0.011352618 | 0.3047832 | 0.03724818 | 0.97028713 | 0.98977467 |
| BCAS4 | 8.91963618 | 0.601906407 | 0.57158806 | 1.05304231 | 0.2923216 | NA |
| ADNP | 89.043994 | 0.019679012 | 0.21698532 | 0.09069283 | 0.92773667 | 0.97775374 |
| DPM1 | 22.4776361 | -0.114774596 | 0.41886376 | -0.2740141 | 0.78407375 | NA |
| ATP9A | 60.4846139 | 0.328583355 | 0.28674406 | 1.14591164 | 0.25183171 | 0.65183754 |
| ZFP64 | 15.4091579 | 0.089789024 | 0.41349055 | 0.21714891 | 0.8280923 | NA |
| TSHZ2 | 33.5886999 | -0.708973794 | 0.85532418 | -0.8288948 | 0.40716392 | 0.76772515 |
| AL109930.1 | 8.95957291 | -0.450001047 | 0.93861562 | -0.4794306 | 0.63163233 | NA |
| ZNF217 | 16.5967192 | -0.79071655 | 0.4471324 | -1.768417 | 0.07699121 | NA |
| CYP24A1 | 22.3998981 | 0.752141699 | 1.0399289 | 0.72326262 | 0.46951855 | NA |
| PFDN4 | 10.802539 | -0.236269351 | 0.47827044 | -0.4940078 | 0.62130064 | NA |
| FAM210B | 16.7165472 | -0.27493566 | 0.39358545 | -0.6985412 | 0.48483879 | NA |
| CSTF1 | 7.09739453 | -0.078663778 | 0.51867219 | -0.1516638 | 0.87945214 | NA |
| RTF2 | 14.1824285 | 0.764942813 | 0.39232794 | 1.94975358 | 0.0512055 | NA |
| BMP7 | 72.8714864 | -0.189181075 | 0.61678976 | -0.3067189 | 0.75905734 | 0.92313949 |
| RAE1 | 11.9913561 | -0.07953176 | 0.40146911 | -0.1981018 | 0.8429654 | NA |
| RBM38 | 10.7734332 | -0.207295458 | 0.55874158 | -0.3710042 | 0.71063442 | NA |
| PCK1 | 50.3917669 | 0.012621699 | 0.87263828 | 0.01446384 | 0.98845993 | 0.99651342 |
| PMEPA1 | 97.8027463 | 0.438749042 | 0.58173181 | 0.75421188 | 0.450722 | 0.79158286 |
| RAB22A | 24.7134257 | -0.07275608 | 0.38347867 | -0.1897265 | 0.84952342 | NA |
| VAPB | 38.9461597 | -0.181125028 | 0.25065013 | -0.7226209 | 0.4699128 | 0.80448595 |
| NPEPL1 | 12.27013 | 0.055043782 | 0.41407687 | 0.13293131 | 0.8942477 | NA |
| GNAS | 199.69877 | -0.501593862 | 0.35144129 | -1.427248 | 0.15350843 | 0.55384216 |
| NELFCD | 17.8304396 | 0.15782689 | 0.35465982 | 0.44500922 | 0.65631309 | NA |
| CTSZ | 37.2782176 | -0.120724112 | 0.32603171 | -0.3702833 | 0.71117138 | 0.91224261 |
| ATP5F1E | 110.497621 | -0.933271904 | 0.49301569 | -1.8929862 | 0.0583597 | 0.37650386 |
| PRELID3B | 9.40904428 | 0.126936081 | 0.53253297 | 0.23836286 | 0.81159967 | NA |
| PHACTR3 | 14.9129823 | -1.549103583 | 0.82949024 | -1.8675368 | 0.06182666 | NA |
| SYCP2 | 5.57833409 | -0.062233161 | 0.60008979 | -0.1037064 | 0.91740234 | NA |
| FAM217B | 14.3773008 | -0.271455234 | 0.45170786 | -0.6009531 | 0.54787123 | NA |
| PPP1R3D | 8.6534944 | -0.443977017 | 0.55975127 | -0.7931684 | 0.42767971 | NA |
| MIR646HG | 16.1595167 | -0.377103059 | 0.99634622 | -0.378486 | 0.70506962 | NA |
| TAF4 | 18.2907793 | 0.038833445 | 0.40583109 | 0.09568869 | 0.92376782 | NA |
| LSM14B | 6.81636544 | -0.188763399 | 0.49588979 | -0.380656 | 0.70345856 | NA |
| PSMA7 | 62.7856945 | -0.459881136 | 0.32316083 | -1.423072 | 0.15471527 | 0.55484098 |
| SS18L1 | 16.9036134 | 0.257758708 | 0.34375133 | 0.74984061 | 0.45335071 | NA |
| MTG2 | 19.1998868 | -0.20315066 | 0.34395859 | -0.5906253 | 0.55477148 | NA |
| OSBPL2 | 14.8572336 | -0.23584009 | 0.4293004 | -0.5493591 | 0.58275902 | NA |
| ADRM1 | 11.8974769 | -0.29281597 | 0.6390017 | -0.4582397 | 0.64678022 | NA |
| AL354836.1 | 11.8708388 | 0.742815772 | 0.66517586 | 1.11672087 | 0.26411369 | NA |
| LAMA5 | 93.1440878 | 0.241764888 | 0.37975957 | 0.6366262 | 0.52436836 | 0.83192915 |
| RPS21 | 6.91999058 | 0.814966639 | 0.76087516 | 1.07109114 | 0.28412845 | NA |
| SLCO4A1 | 6.73711968 | -2.715125565 | 0.99419521 | -2.7309783 | 0.00631466 | NA |
| MRGBP | 5.1544585 | -0.702712673 | 0.64720281 | -1.0857689 | 0.27758126 | NA |
| OGFR | 17.0983148 | -0.560116279 | 0.62465642 | -0.896679 | 0.36989025 | NA |
| TCFL5 | 4.18347396 | -0.210772893 | 0.72379973 | -0.2912033 | 0.77089581 | NA |
| DIDO1 | 45.6429132 | -0.213660075 | 0.27579791 | -0.774698 | 0.43851816 | 0.78335165 |
| GID8 | 18.3703617 | -0.269442143 | 0.37657029 | -0.7155162 | 0.47429013 | NA |
| YTHDF1 | 17.8124561 | 0.22040456 | 0.37620748 | 0.58585905 | 0.55797024 | NA |
| ARFGAP1 | 13.1137614 | 0.781090144 | 0.44753301 | 1.7453241 | 0.08092847 | NA |
| PPDPF | 97.754822 | -0.213845486 | 0.55164605 | -0.3876498 | 0.6982752 | 0.90729979 |
| HELZ2 | 10.4132988 | 0.376594189 | 1.120492 | 0.33609717 | 0.73679759 | NA |
| GMEB2 | 8.09734937 | 0.254200662 | 0.56014642 | 0.4538111 | 0.64996479 | NA |
| MHENCR | 8.60836633 | 0.600416408 | 0.59619349 | 1.00708314 | 0.31389482 | NA |
| ARFRP1 | 13.333845 | 0.266942243 | 0.43504451 | 0.61359754 | 0.53948131 | NA |
| SLC2A4RG | 18.7649182 | -0.476210133 | 0.62380877 | -0.7633912 | 0.44523012 | NA |
| ZBTB46 | 94.6969389 | 0.112689143 | 0.49929305 | 0.2256974 | 0.82143678 | 0.94728252 |
| TPD52L2 | 21.9728213 | -0.248477351 | 0.38862923 | -0.6393687 | 0.52258313 | NA |
| DNAJC5 | 34.3150096 | 0.010048272 | 0.27117867 | 0.03705407 | 0.9704419 | 0.98977467 |
| UCKL1 | 8.9894298 | 0.558617753 | 0.46190666 | 1.20937366 | 0.22651932 | NA |
| ZNF512B | 5.79122706 | -0.45945003 | 0.6004171 | -0.7652181 | 0.44414168 | NA |
| PRPF6 | 32.8343203 | 0.309681932 | 0.28924873 | 1.07064233 | 0.28433028 | 0.67249058 |
| TCEA2 | 14.4547826 | 0.212108946 | 0.4176873 | 0.50781756 | 0.6115813 | NA |
| PCMTD2 | 97.0048958 | 0.770747087 | 0.40604216 | 1.89819475 | 0.05767043 | 0.37585064 |
| MIER2 | 7.12763029 | 0.537897076 | 0.49832434 | 1.07941161 | 0.28040428 | NA |
| TPGS1 | 10.9492981 | -0.529310791 | 0.5241395 | -1.0098662 | 0.31255937 | NA |
| CDC34 | 8.51502019 | 0.157070076 | 0.61095176 | 0.2570908 | 0.79710868 | NA |
| BSG | 74.3648542 | -0.240814447 | 0.36822964 | -0.653979 | 0.51312534 | 0.83070544 |
| RNF126 | 11.0531658 | -0.651576858 | 0.47831067 | -1.3622461 | 0.17312023 | NA |
| PALM | 8.34788195 | -0.33815496 | 0.67433964 | -0.5014609 | 0.61604679 | NA |
| AC006273.1 | 12.0934942 | -0.119908406 | 0.6025112 | -0.1990144 | 0.84225148 | NA |
| LINC01836 | 9.37586859 | -0.474911104 | 0.65064275 | -0.7299107 | 0.46544477 | NA |
| PTBP1 | 56.1775258 | -0.39495082 | 0.32169206 | -1.2277295 | 0.21954854 | 0.62462904 |
| MED16 | 15.701781 | 0.588162482 | 0.3467077 | 1.69642176 | 0.08980604 | NA |
| R3HDM4 | 12.1816485 | 0.090540868 | 0.4949979 | 0.18291162 | 0.85486737 | NA |
| ARID3A | 29.1351793 | 0.078246188 | 0.50251059 | 0.15571052 | 0.8762612 | 0.96317491 |
| WDR18 | 6.35155031 | -0.640666745 | 0.69556649 | -0.9210719 | 0.35701289 | NA |
| TMEM259 | 54.8100014 | 0.117720453 | 0.40661935 | 0.28951021 | 0.77219097 | 0.92669573 |
| CNN2 | 29.2729273 | -0.398563422 | 0.49474334 | -0.8055963 | 0.42047564 | 0.77450009 |
| POLR2E | 15.4640536 | -0.828029915 | 0.42152494 | -1.9643676 | 0.04948748 | NA |
| GPX4 | 45.1634129 | -0.269824587 | 0.32833524 | -0.821796 | 0.41119301 | 0.7685709 |

| | | | | | | |
|---|---|---|---|---|---|---|
| SBNO2 | 28.3698154 | -1.735897421 | 0.62534246 | -2.7759148 | 0.00550466 | NA |
| STK11 | 17.2240192 | 0.214492953 | 0.37810458 | 0.56728472 | 0.57052075 | NA |
| ATP5F1D | 48.3384322 | -0.227159763 | 0.42006015 | -0.5407791 | 0.58865983 | 0.86533208 |
| MIDN | 47.3242899 | -1.195690912 | 0.46217988 | -2.5870683 | 0.00967964 | 0.15518032 |
| CIRBP | 97.2806907 | -0.21151732 | 0.27138299 | -0.7794052 | 0.43574104 | 0.78167891 |
| C19orf24 | 17.0748569 | -0.109478368 | 0.48896758 | -0.223897 | 0.82283746 | NA |
| MUM1 | 49.5252871 | -0.247998961 | 0.32891159 | -0.7539989 | 0.45084991 | 0.79158286 |
| NDUFS7 | 16.8717353 | -0.092968995 | 0.46630658 | -0.1993731 | 0.84197089 | NA |
| GAMT | 4.19264603 | -0.289032016 | 0.85619581 | -0.337577 | 0.73568198 | NA |
| DAZAP1 | 23.2733388 | -0.45572514 | 0.37853964 | -1.2039033 | 0.22862694 | NA |
| RPS15 | 132.134397 | -0.632268529 | 0.37248697 | -1.6974246 | 0.08961642 | 0.4567688 |
| C19orf25 | 14.0903773 | -0.550345199 | 0.45713407 | -1.2039033 | 0.22862696 | NA |
| MBD3 | 17.6889437 | -0.099097572 | 0.5582825 | -0.1775043 | 0.85911225 | NA |
| TCF3 | 9.32443693 | -0.139202442 | 0.48416385 | -0.287511 | 0.77372106 | NA |
| REXO1 | 10.5390716 | -0.005113148 | 0.46366899 | -0.0110276 | 0.99120144 | NA |
| KLF16 | 9.88571803 | -1.294004989 | 0.55223331 | -2.3432215 | 0.01911803 | NA |
| ABHD17A | 22.5989006 | -0.195504765 | 0.50055386 | -0.3905769 | 0.69611002 | NA |
| SCAMP4 | 20.8619223 | 0.164470151 | 0.34651316 | 0.4746433 | 0.63504124 | NA |
| CSNK1G2 | 20.4171513 | -0.407314469 | 0.55672408 | -0.7316272 | 0.46439615 | NA |
| BTBD2 | 16.5689469 | 0.015584958 | 0.43867525 | 0.03552732 | 0.97165926 | NA |
| MKNK2 | 15.3486251 | 0.052355858 | 0.46677881 | 0.11216417 | 0.91069324 | NA |
| MOB3A | 12.8918282 | -0.039957118 | 0.46638555 | -0.085674 | 0.93172557 | NA |
| AP3D1 | 31.7727951 | -0.227642149 | 0.35224241 | -0.6462656 | 0.51810737 | 0.83104823 |
| DOT1L | 14.901923 | -0.538764309 | 0.52280695 | -1.0305225 | 0.30276481 | NA |
| PLEKHJ1 | 19.4678102 | -0.171898852 | 0.39954897 | -0.4302322 | 0.66702671 | NA |
| SF3A2 | 6.8930678 | -0.513387823 | 0.633261 | -0.8107049 | 0.41753514 | NA |
| LSM7 | 23.9553252 | -0.05085094 | 0.39788238 | -0.127804 | 0.89830412 | NA |
| SPPL2B | 12.8992766 | 0.386544641 | 0.41779894 | 0.92519297 | 0.35486554 | NA |
| TIMM13 | 21.6134555 | -0.579527278 | 0.48006709 | -1.2071798 | 0.22736292 | NA |
| GADD45B | 107.757883 | -2.439770012 | 0.90303278 | -2.7017513 | 0.00689753 | 0.13258954 |
| GNG7 | 13.9323938 | 0.444771795 | 0.38198557 | 1.16436807 | 0.24427488 | NA |
| SGTA | 16.8103599 | 0.050037275 | 0.4157301 | 0.12036 | 0.90419798 | NA |
| THOP1 | 7.83324326 | 0.524433146 | 0.62408231 | 0.84032689 | 0.40072513 | NA |
| ZNF555 | 4.26332234 | -0.221028189 | 0.65526525 | -0.337311 | 0.73588246 | NA |
| AES | 68.7439123 | -0.359747438 | 0.32283629 | -1.114334 | 0.26513594 | 0.65974932 |
| GNA11 | 24.4261603 | -0.004717134 | 0.42306314 | -0.01115 | 0.99110381 | NA |
| NCLN | 13.1106092 | 0.39405511 | 0.5578434 | 0.7063902 | 0.47994551 | NA |
| NFIC | 165.135714 | -0.585891085 | 0.36092642 | -1.6232979 | 0.10452574 | 0.4862279 |
| DOHH | 3.45947726 | -0.252008036 | 0.68108552 | -0.3700094 | 0.71137549 | NA |
| FZR1 | 10.0543381 | 0.750154614 | 0.43503579 | 1.72435147 | 0.08464441 | NA |
| MFSD12 | 21.3148838 | -0.134630479 | 0.61851381 | -0.2176677 | 0.82768803 | NA |
| HMG20B | 16.9632589 | -0.220551841 | 0.40474223 | -0.5449193 | 0.58580904 | NA |
| TBXA2R | 13.9815602 | 0.219118246 | 0.81487915 | 0.26889661 | 0.78800924 | NA |
| CACTIN | 5.53829225 | 0.73503697 | 0.64633568 | 1.13723718 | 0.25543915 | NA |
| PIP5K1C | 14.8015818 | 0.285762627 | 0.48923758 | 0.58409787 | 0.55915447 | NA |
| MRPL54 | 21.1809802 | -0.813142085 | 0.43786888 | -1.8570447 | 0.06330479 | NA |
| DAPK3 | 14.5239763 | 0.508741545 | 0.45770329 | 1.11150948 | 0.26634911 | NA |
| EEF2 | 88.1516498 | -0.473799349 | 0.36679662 | -1.2917222 | 0.19645335 | 0.59987316 |
| PIAS4 | 7.85191794 | -0.535487554 | 0.64824948 | -0.8260517 | 0.40877479 | NA |
| ZBTB7A | 43.5253356 | -0.011253193 | 0.38064119 | -0.0295638 | 0.97641495 | 0.992815 |
| MAP2K2 | 31.8051174 | -0.018457633 | 0.27881278 | -0.0662008 | 0.94721795 | 0.98334169 |
| SIRT6 | 5.14930996 | 0.416466238 | 0.80468955 | 0.51754896 | 0.604773 | NA |
| YJU2 | 6.79206404 | 0.860528561 | 0.68625252 | 1.25395322 | 0.20985901 | NA |
| STAP2 | 3.7324849 | -0.476886497 | 0.97899703 | -0.4871174 | 0.62617513 | NA |
| SH3GL1 | 12.142607 | 0.016167652 | 0.51494806 | 0.03139667 | 0.9749532 | NA |
| UBXN6 | 25.9127199 | -0.053998546 | 0.44020804 | -0.122666 | 0.90237161 | NA |
| HDGFL2 | 12.8927897 | 0.582162977 | 0.42459074 | 1.37111556 | 0.17033893 | NA |
| PLIN5 | 7.90460617 | 0.801987803 | 0.62123911 | 1.29094868 | 0.19672148 | NA |
| MYDGF | 43.3560499 | -0.778918535 | 0.58199353 | -1.3383629 | 0.18077817 | 0.58390314 |
| DPP9 | 14.2364361 | -0.723232646 | 0.46186031 | -1.5659121 | 0.1173692 | NA |
| TICAM1 | 6.17337837 | 0.210785776 | 0.62528309 | 0.33710455 | 0.73603809 | NA |
| PLIN3 | 13.2939624 | -0.647471179 | 0.4652389 | -1.3916961 | 0.16401443 | NA |
| KDM4B | 35.6478839 | -0.312046167 | 0.44031322 | -0.7086913 | 0.47851604 | 0.81001889 |
| PTPRS | 18.1733978 | 0.294040677 | 0.41603911 | 0.70676211 | 0.47971433 | NA |
| TINCR | 5.84796743 | -0.921147056 | 0.74606962 | -1.2346664 | 0.2169547 | NA |
| SAFB2 | 23.3108549 | 0.127076037 | 0.34612946 | 0.36713442 | 0.71351875 | NA |
| SAFB | 36.0383396 | -0.222589423 | 0.314843 | -0.7069855 | 0.47957552 | 0.81093801 |
| RPL36 | 71.7169546 | -0.18254996 | 0.45106665 | -0.4047073 | 0.68569269 | 0.90499319 |
| C19orf70 | 29.8344579 | -0.520078784 | 0.44670555 | -1.1642541 | 0.24432104 | 0.64996527 |
| LONP1 | 17.1700071 | -0.417107001 | 0.45340717 | -0.9199391 | 0.35760456 | NA |
| CAPS | 4.8526533 | 0.360390091 | 0.6459145 | 0.55795324 | 0.57687631 | NA |
| RANBP3 | 14.9932423 | 0.025103083 | 0.36217343 | 0.06931233 | 0.94474101 | NA |
| RFX2 | 6.89769662 | 0.637543167 | 0.7719915 | 0.82584222 | 0.4088936 | NA |
| MLLT1 | 17.471753 | -0.33309419 | 0.43723576 | -0.7618183 | 0.44616847 | NA |
| CLPP | 15.6708098 | -1.289296559 | 0.48516998 | -2.657412 | 0.00787431 | NA |
| ALKBH7 | 21.825682 | 0.141817345 | 0.71916046 | 0.19719847 | 0.84367223 | NA |
| GTF2F1 | 14.5188002 | -0.867022993 | 0.41755025 | -2.0764518 | 0.03785218 | NA |
| KHSRP | 17.2908786 | 0.01689912 | 0.50933761 | 0.03317862 | 0.97353215 | NA |
| SLC25A23 | 11.2041261 | -0.485268196 | 0.44659453 | -1.0865968 | 0.27721503 | NA |
| C3 | 8.851895 | -3.181618964 | 1.27372728 | -2.4978808 | 0.01249382 | NA |
| TRIP10 | 26.9782267 | -0.392541305 | 0.38700642 | -1.0143018 | 0.31043877 | NA |
| ZNF557 | 4.93038571 | 0.292050292 | 0.65954229 | 0.44280753 | 0.65790496 | NA |
| INSR | 145.833354 | 0.44156337 | 0.23321928 | 1.89333994 | 0.05831268 | 0.37650386 |
| ZNF358 | 12.7737685 | -0.144228917 | 0.62296129 | -0.2315215 | 0.8169097 | NA |
| PNPLA6 | 18.5198643 | 0.383575798 | 0.51846375 | 0.73983148 | 0.45940226 | NA |
| XAB2 | 5.92574056 | -0.358086516 | 0.61624898 | -0.5810744 | 0.5611903 | NA |
| PET100 | 5.15129388 | -0.475904008 | 0.66825727 | -0.7121569 | 0.47636764 | NA |
| STXBP2 | 8.07788302 | 0.629336589 | 0.59074073 | 1.06533469 | 0.2867245 | NA |
| EVI5L | 6.44600472 | -0.434849057 | 0.57554658 | -0.755541 | 0.44992442 | NA |
| MAP2K7 | 16.8606596 | -0.649081264 | 0.57112872 | -1.1364886 | 0.25575214 | NA |
| SNAPC2 | 3.56699447 | -0.213660486 | 0.8341908 | -0.256129 | 0.7978512 | NA |
| TIMM44 | 6.10702164 | -0.199716705 | 0.63755169 | -0.3132557 | 0.7540864 | NA |
| ELAVL1 | 37.4255578 | 0.384814569 | 0.30187337 | 1.27475493 | 0.202396 | 0.61034808 |
| CERS4 | 22.1607231 | 0.3378449 | 0.44587613 | 0.75771022 | 0.44862448 | NA |
| CD320 | 14.8266346 | 0.056405106 | 0.57828531 | 0.09753854 | 0.92229873 | NA |
| RPS28 | 27.0689765 | -0.192742186 | 0.53129486 | -0.3627782 | 0.71677059 | NA |
| KANK3 | 11.280914 | 0.325136548 | 0.47586577 | 0.68325265 | 0.49444721 | NA |
| ANGPTL4 | 35.988963 | -0.44162585 | 0.67651665 | -0.6527938 | 0.51388924 | 0.83070544 |
| RAB11B-AS1 | 4.49119258 | -0.834319138 | 0.89652321 | -0.9306163 | 0.35205206 | NA |
| RAB11B | 33.5890285 | 0.310595379 | 0.42694142 | 0.72748945 | 0.46692618 | 0.8024799 |
| 2-Mar | 14.4563582 | -0.647482052 | 0.6017159 | -1.0760594 | 0.28190069 | NA |
| HNRNPM | 67.4892566 | 0.252537721 | 0.221607 | 1.13957467 | 0.25446354 | 0.65520614 |
| ZNF414 | 4.39617279 | 0.167338943 | 0.70856945 | 0.23616449 | 0.81330503 | NA |
| ADAMTS10 | 4.96888869 | -1.088575234 | 0.65183036 | -1.6700284 | 0.09491374 | NA |
| ZNF558 | 6.01285994 | 0.253983255 | 0.53647501 | 0.47342979 | 0.63590658 | NA |
| ZNF317 | 7.99944769 | -0.388811431 | 0.55825378 | -0.6964779 | 0.48612958 | NA |
| ZNF699 | 4.00650495 | -0.32230663 | 0.6510887 | -0.4950272 | 0.62058091 | NA |
| ZNF559-ZNF | 7.6611352 | -0.232369734 | 0.49825535 | -0.4663668 | 0.640953 | NA |
| ZNF266 | 12.3563283 | -0.16092476 | 0.40281351 | -0.3995019 | 0.68952343 | NA |
| ZNF426 | 10.9834038 | -0.255607991 | 0.44862767 | -0.5697553 | 0.56884368 | NA |
| ZNF121 | 14.8565922 | 0.195198125 | 0.42235805 | 0.46216267 | 0.64396467 | NA |
| ZNF561 | 8.59955477 | 0.028863095 | 0.62056468 | 0.04651102 | 0.96290295 | NA |
| ZNF562 | 18.2868035 | -0.252174494 | 0.36500719 | -0.6908754 | 0.48964383 | NA |

| | | | | | | |
|---|---|---|---|---|---|---|
| ZNF846 | 40.4883134 | 0.330118656 | 0.3424686 | 0.96393846 | 0.33507678 | 0.71006866 |
| FBXL12 | 8.69944855 | 0.200922422 | 0.47276124 | 0.42499766 | 0.67083838 | NA |
| UBL5 | 31.7955759 | -0.019040704 | 0.48114212 | -0.039574 | 0.96843278 | 0.98964394 |
| PIN1 | 27.8396275 | -0.251627679 | 0.36843646 | -0.6829609 | 0.49463158 | NA |
| C19orf66 | 22.2804696 | -1.026858758 | 0.79735423 | -1.2878326 | 0.19780425 | NA |
| EIF3G | 17.8240066 | -0.277570966 | 0.43354951 | -0.640229 | 0.52202373 | NA |
| DNMT1 | 7.40813912 | -0.404623453 | 0.56935902 | -0.7106649 | 0.47729193 | NA |
| MRPL4 | 13.4033555 | 0.061646701 | 0.49486631 | 0.12457243 | 0.90086205 | NA |
| ICAM1 | 66.7599182 | -1.612537801 | 1.28653298 | -1.2533979 | 0.21006092 | 0.61792799 |
| TYK2 | 11.5678463 | 0.311094843 | 0.43889812 | 0.70880878 | 0.47844315 | NA |
| CDC37 | 31.4660752 | -0.476113379 | 0.31105496 | -1.5306407 | 0.12585822 | 0.52276701 |
| PDE4A | 17.3041979 | 0.328893106 | 0.40968528 | 0.80279454 | 0.4220935 | NA |
| KEAP1 | 14.8162432 | 0.019518206 | 0.43787277 | 0.04457506 | 0.96444602 | NA |
| ATG4D | 6.14358766 | 0.496389028 | 0.51980818 | 0.95494655 | 0.33960473 | NA |
| KRI1 | 6.2144285 | -0.614269999 | 0.52008015 | -1.1811064 | 0.23756046 | NA |
| CDKN2D | 10.3915281 | -1.189094061 | 0.84914216 | -1.4003475 | 0.1614093 | NA |
| AP1M2 | 8.61480941 | -0.724027195 | 0.6195197 | -1.1686912 | 0.24252808 | NA |
| SLC44A2 | 87.6727736 | -0.817674448 | 0.33589544 | -2.4343124 | 0.01492011 | 0.19972267 |
| ILF3-DT | 20.2486717 | -0.409196131 | 0.34837042 | -1.1746007 | 0.24015452 | NA |
| ILF3 | 45.5806985 | -0.065296992 | 0.25581729 | -0.2552486 | 0.79853113 | 0.94015741 |
| QTRT1 | 13.028791 | 0.840392525 | 0.61396094 | 1.36880453 | 0.17106038 | NA |
| DNM2 | 45.8841638 | -0.124291341 | 0.3886873 | -0.3197721 | 0.74914113 | 0.92274989 |
| TMED1 | 14.106342 | -0.502764457 | 0.54617653 | -0.9205164 | 0.35730297 | NA |
| CARM1 | 9.19531367 | -0.387987758 | 0.49741067 | -0.780015 | 0.43538207 | NA |
| YIPF2 | 11.2860189 | -0.580782796 | 0.46637605 | -1.2453101 | 0.21301779 | NA |
| TIMM29 | 8.50492129 | -0.01242292 | 0.60863961 | -0.020411 | 0.98371554 | NA |
| SMARCA4 | 39.0915233 | -0.099820938 | 0.28158298 | -0.3544992 | 0.72296481 | 0.91509584 |
| LDLR | 11.783151 | -1.500132362 | 0.7591854 | -1.9759763 | 0.04815746 | NA |
| KANK2 | 57.62903 | -1.412940455 | 0.4827834 | -2.926655 | 0.00342629 | 0.09584152 |
| DOCK6 | 15.7756797 | 0.270909908 | 0.5007556 | 0.54100226 | 0.58850603 | NA |
| RAB3D | 6.31775946 | 0.235911996 | 0.61979255 | 0.38063058 | 0.70347739 | NA |
| RGL3 | 13.7249461 | 0.779775416 | 0.56703181 | 1.37518814 | 0.16907312 | NA |
| PRKCSH | 27.3770339 | -0.176674574 | 0.5292222 | -0.3338382 | 0.73850167 | NA |
| ECSIT | 8.99430466 | 0.011935897 | 0.51745659 | 0.02306647 | 0.98159725 | NA |
| ELOF1 | 6.10541754 | -0.111521958 | 0.68914589 | -0.1618263 | 0.87144261 | NA |
| ZNF627 | 7.99046409 | -1.016655419 | 0.67232985 | -1.5121379 | 0.13049879 | NA |
| ZNF823 | 5.76560118 | -0.330912279 | 0.61035108 | -0.5421671 | 0.5877034 | NA |
| ZNF441 | 5.71746382 | -0.363232241 | 0.81436972 | -0.4460287 | 0.65557654 | NA |
| ZNF440 | 6.02664385 | 0.075351753 | 0.59737995 | 0.12613706 | 0.89962343 | NA |
| ZNF439 | 20.6624852 | -0.057432635 | 0.50595394 | -0.1135136 | 0.90962341 | NA |
| ZNF69 | 9.60988548 | 0.672784325 | 0.48874079 | 1.37656677 | 0.16864623 | NA |
| ZNF844 | 9.56284536 | -0.386956723 | 0.69654343 | -0.5555385 | 0.57852635 | NA |
| ZNF788P | 6.86901186 | 0.34013274 | 0.58366287 | 0.58275549 | 0.56005791 | NA |
| ZNF136 | 9.30439577 | 0.065912899 | 0.44987456 | 0.14651395 | 0.88351568 | NA |
| ZNF44 | 73.0399994 | -0.000575666 | 0.34787779 | -0.0016548 | 0.99867967 | 0.99961632 |
| ZNF563 | 7.92888884 | -0.660876436 | 0.60587698 | -1.0907766 | 0.2753712 | NA |
| ZNF442 | 7.82993889 | -0.996343183 | 0.73625351 | -1.353261 | 0.17597226 | NA |
| ZNF490 | 9.18242799 | -0.467399974 | 0.50706331 | -0.9217783 | 0.35664421 | NA |
| ZNF791 | 19.139552 | -0.369027559 | 0.39438566 | -0.9357023 | 0.3494265 | NA |
| WDR83 | 4.13087142 | -0.556903946 | 0.70780132 | -0.7868083 | 0.4313941 | NA |
| DHPS | 8.8181006 | 0.210954805 | 0.48378448 | 0.4360512 | 0.66279958 | NA |
| TNPO2 | 24.7201039 | 0.49495767 | 0.31032382 | 1.59497158 | 0.11071859 | NA |
| TRIR | 42.7164472 | -0.308632195 | 0.29603225 | -1.0425628 | 0.29715085 | 0.68103704 |
| ASNA1 | 9.71990445 | -0.180084339 | 0.54045017 | -0.3332117 | 0.73897446 | NA |
| HOOK2 | 13.526773 | 0.166405132 | 0.53842192 | 0.30906084 | 0.75727525 | NA |
| JUNB | 29.7980667 | -2.372216154 | 0.67271313 | -3.5263414 | 0.00042134 | 0.02756401 |
| DNASE2 | 18.207335 | -0.364052846 | 0.53296906 | -0.6830656 | 0.49456537 | NA |
| GCDH | 6.90050841 | -0.26721372 | 0.56744467 | -0.4709071 | 0.63770708 | NA |
| FARSA | 10.5730392 | -0.069179666 | 0.55197339 | -0.1253315 | 0.9002611 | NA |
| CALR | 134.997097 | -0.462860649 | 0.56171695 | -0.8240105 | 0.40993361 | 0.76788553 |
| AC092069.1 | 23.5568221 | 0.002660266 | 0.65092728 | 0.00408689 | 0.99673915 | NA |
| RAD23A | 29.31158 | -0.239380503 | 0.31207761 | -0.7670544 | 0.44304918 | 0.78643653 |
| GADD45GIP1 | 55.6802256 | -0.799175811 | 0.31657497 | -2.5244441 | 0.01158814 | 0.17871297 |
| NFIX | 9.79686325 | -0.802658296 | 0.6475883 | -1.2394577 | 0.21517605 | NA |
| TRMT1 | 3.92863606 | 0.14801339 | 0.68919284 | 0.21476339 | 0.8299518 | NA |
| NACC1 | 8.3970173 | -0.078152822 | 0.5605467 | -0.1394225 | 0.88911629 | NA |
| STX10 | 5.41668397 | 0.459372056 | 0.62480884 | 0.73522016 | 0.46220542 | NA |
| IER2 | 26.5484096 | 0.155954321 | 0.5965495 | 0.26142729 | 0.79376301 | NA |
| CCDC130 | 14.9729217 | 0.410576893 | 0.48903972 | 0.83955735 | 0.40115662 | NA |
| MRI1 | 10.1867537 | 0.330064416 | 0.47008058 | 0.70214434 | 0.48258915 | NA |
| C19orf53 | 34.9861866 | -1.838721535 | 0.51854301 | -3.5459384 | 0.00039122 | 0.02714874 |
| ZSWIM4 | 7.59562897 | 0.037648713 | 0.5181676 | 0.07265741 | 0.94207874 | NA |
| CC2D1A | 7.57098197 | -0.361214452 | 0.58743377 | -0.6149024 | 0.53861916 | NA |
| RFX1 | 13.7980058 | 0.206615808 | 0.47852509 | 0.43177633 | 0.66590399 | NA |
| PRKACA | 11.0428415 | -0.085453619 | 0.42251328 | -0.2022507 | 0.83972072 | NA |
| AC022098.1 | 4.57682387 | -1.41612172 | 0.80826636 | -1.7520483 | 0.0797655 | NA |
| ADGRL1 | 9.52796084 | -0.357101416 | 0.66526259 | -0.5367826 | 0.59141776 | NA |
| DDX39A | 4.68376879 | -0.517104115 | 0.71066031 | -0.7276389 | 0.46683464 | NA |
| PKN1 | 30.929506 | -0.145174897 | 0.42719462 | -0.3398332 | 0.73398218 | 0.91884473 |
| GIPC1 | 34.348674 | 0.200506097 | 0.3967794 | 0.50533394 | 0.61332431 | 0.87439243 |
| DNAJB1 | 11.821591 | 0.730490259 | 0.48522718 | 1.50546032 | 0.13220577 | NA |
| TECR | 13.9157335 | 0.505760943 | 0.45175255 | 1.11955304 | 0.26290428 | NA |
| NDUFB7 | 51.3146233 | 0.044607431 | 0.48711135 | 0.09157543 | 0.92703537 | 0.97775374 |
| ADGRE3 | 11.2185487 | 0.05585191 | 0.66887246 | 0.08350158 | 0.93345272 | NA |
| ZNF333 | 9.37204914 | -0.606313419 | 0.49824304 | -1.2169029 | 0.22364115 | NA |
| SYDE1 | 18.7809376 | -0.148171985 | 0.46943183 | -0.3156411 | 0.75227493 | NA |
| ILVBL | 10.4884048 | -0.345589338 | 0.46097402 | -0.7496937 | 0.45343918 | NA |
| BRD4 | 71.9322318 | -0.264676765 | 0.27612144 | -0.958552 | 0.33778447 | 0.71245649 |
| AKAP8 | 17.9679317 | -0.174659831 | 0.36383186 | -0.4800565 | 0.63118725 | NA |
| AKAP8L | 45.0364624 | -0.129866469 | 0.31709773 | -0.4095471 | 0.68213818 | 0.90499319 |
| WIZ | 10.1769684 | -0.258139118 | 0.48944575 | -0.5274111 | 0.59790814 | NA |
| TPM4 | 79.5768389 | -0.326289805 | 0.28106973 | -1.1608856 | 0.24568842 | 0.65046489 |
| RAB8A | 13.2634977 | -0.517655247 | 0.51327091 | -1.008542 | 0.31319436 | NA |
| FAM32A | 14.080874 | -0.044859999 | 0.48350108 | -0.0927816 | 0.92607708 | NA |
| AP1M1 | 16.6685953 | 0.227740457 | 0.44868734 | 0.50757049 | 0.61175459 | NA |
| EPS15L1 | 22.4705242 | -0.140389164 | 0.32128734 | -0.4369583 | 0.6621416 | NA |
| C19orf44 | 6.17737978 | -0.818425695 | 0.64501614 | -1.2688453 | 0.20449625 | NA |
| SIN3B | 18.4440101 | 0.362602389 | 0.367249 | 0.98734751 | 0.32347231 | NA |
| MYO9B | 26.5731672 | -0.436183873 | 0.32988421 | -1.3222333 | 0.18609049 | NA |
| USE1 | 7.25195729 | -1.276543522 | 0.63035417 | -2.0251211 | 0.04285493 | NA |
| OCEL1 | 10.0363815 | -0.933004836 | 0.54139477 | -1.7233355 | 0.08482787 | NA |
| NR2F6 | 19.8881303 | -0.367378344 | 0.36902859 | -0.9955281 | 0.31947946 | NA |
| ABHD8 | 6.06443109 | -0.612368258 | 0.60180032 | -1.0175605 | 0.30888685 | NA |
| MRPL34 | 17.4866995 | -1.133027511 | 0.48249439 | -2.3482709 | 0.0188608 | NA |
| DDA1 | 11.1115569 | -0.143378546 | 0.48655644 | -0.2946802 | 0.76823819 | NA |
| ANO8 | 5.20589557 | -0.044326742 | 0.63503819 | -0.0698017 | 0.9443515 | NA |
| GTPBP3 | 6.88067957 | 0.52401471 | 0.58729197 | 0.89225588 | 0.3722558 | NA |
| BST2 | 340.028855 | -0.489643952 | 0.63723786 | -0.7683849 | 0.44225854 | 0.78640976 |
| MVB12A | 17.8339309 | -0.509731974 | 0.47500235 | -1.0731146 | 0.28321969 | NA |
| SLC27A1 | 24.1742835 | 0.463880614 | 0.39932528 | 1.16166102 | 0.24537318 | NA |
| AC010618.3 | 13.4455262 | -0.610190742 | 0.42833151 | -1.4245759 | 0.15427984 | NA |
| PGLS | 28.2945206 | -0.430753317 | 0.32324122 | -1.3326064 | 0.18266101 | NA |
| COLGALT1 | 14.3803999 | 0.443599304 | 0.5032546 | 0.88146099 | 0.37806836 | NA |

| | | | | | | |
|---|---|---|---|---|---|---|
| MAP1S | 8.16538525 | -0.131001164 | 0.60018763 | -0.218267 | 0.82722107 | NA |
| RPL18A | 6.46197575 | -1.306178412 | 0.86077747 | -1.5174403 | 0.12915556 | NA |
| CCDC124 | 10.8511341 | -0.628697994 | 0.51222456 | -1.2273874 | 0.219677 | NA |
| ARRDC2 | 9.96012352 | -1.149618365 | 0.75555188 | -1.5215611 | 0.12811908 | NA |
| JUND | 91.6097292 | -0.000648649 | 0.30178303 | -0.0021494 | 0.99828504 | 0.99961632 |
| LSM4 | 23.8598681 | -0.253028979 | 0.41885167 | -0.6041016 | 0.54577607 | NA |
| PGPEP1 | 11.071319 | 0.384655679 | 0.42252861 | 0.910366 | 0.36262952 | NA |
| GDF15 | 21.5866494 | -2.184862259 | 1.03150701 | -2.1181264 | 0.03416436 | NA |
| SSBP4 | 25.6431668 | -0.892692935 | 0.33860838 | -2.6363581 | 0.00838012 | NA |
| ELL | 16.9197862 | -0.034647382 | 0.41771535 | -0.082945 | 0.93389531 | NA |
| FKBP8 | 23.6383441 | -0.223232357 | 0.43710698 | -0.5107042 | 0.60955822 | NA |
| KXD1 | 10.6042942 | 0.116763399 | 0.46324719 | 0.2520542 | 0.80099917 | NA |
| UBA52 | 48.8075575 | -0.428012341 | 0.57005983 | -0.75082 | 0.45276097 | 0.79291818 |
| REX1BD | 19.4044392 | 0.405797207 | 0.37107952 | 1.09355862 | 0.2741486 | NA |
| CRTC1 | 19.9336326 | -0.070109976 | 0.38822112 | -0.1805929 | 0.85668713 | NA |
| UPF1 | 22.0479998 | -0.092175286 | 0.42950463 | -0.2146084 | 0.83007267 | NA |
| COPE | 19.968297 | -0.045866072 | 0.44729319 | -0.1025414 | 0.91832695 | NA |
| DDX49 | 7.35012577 | 0.151621995 | 0.55723584 | 0.27209663 | 0.78554772 | NA |
| SUGP2 | 37.0962628 | 0.288846667 | 0.3279104 | 0.8808707 | 0.37838781 | 0.74610644 |
| ARMC6 | 5.26234695 | 0.348927151 | 0.55910683 | 0.62407958 | 0.53257533 | NA |
| SLC25A42 | 25.9446282 | 0.142981337 | 0.55175412 | 0.25913959 | 0.79552754 | NA |
| TMEM161A | 8.3812872 | -0.0286721 | 0.49434061 | -0.0580007 | 0.95374807 | NA |
| RFXANK | 7.56444186 | -0.535530855 | 0.61624888 | -0.8690172 | 0.38483774 | NA |
| SUGP1 | 10.8132893 | 0.438150744 | 0.47210319 | 0.92808256 | 0.35336474 | NA |
| MAU2 | 25.9833115 | -0.058613262 | 0.31328607 | -0.1870918 | 0.85158864 | NA |
| GATAD2A | 42.9608697 | -0.459997259 | 0.26740122 | -1.7202512 | 0.08538679 | 0.44564602 |
| PBX4 | 6.43998637 | 0.345456347 | 0.61873025 | 0.55833111 | 0.57661831 | NA |
| ATP13A1 | 7.92813146 | 0.098789099 | 0.52886138 | 0.18679583 | 0.85182072 | NA |
| ZNF14 | 7.13375142 | 0.446505308 | 0.51336955 | 0.86975417 | 0.38443476 | NA |
| AC011477.4 | 6.63275127 | 0.288551254 | 0.63629578 | 0.45348604 | 0.65019879 | NA |
| ZNF253 | 19.3351845 | 0.220844139 | 0.41075756 | 0.53765081 | 0.59081814 | NA |
| ZNF682 | 10.2127109 | 1.159810801 | 0.6697409 | 1.73173059 | 0.08332154 | NA |
| ZNF90 | 47.8925719 | -0.258249929 | 0.4591608 | -0.562439 | 0.57381697 | 0.85861279 |
| AC011447.3 | 14.6413671 | 0.891131075 | 0.4832367 | 1.84408816 | 0.06517029 | NA |
| ZNF737 | 16.1940543 | 0.69329556 | 0.50497167 | 1.37293951 | 0.16977115 | NA |
| ZNF430 | 17.3882702 | -0.089954833 | 0.37537528 | -0.2396397 | 0.81060956 | NA |
| ZNF714 | 8.53724082 | -0.129650173 | 0.60205226 | -0.215347 | 0.82949675 | NA |
| ZNF431 | 27.2676072 | 0.490446029 | 0.29712315 | 1.65064899 | 0.09881027 | NA |
| ZNF708 | 30.9926339 | 0.096166423 | 0.31748282 | 0.30290276 | 0.76196396 | 0.92336765 |
| ZNF493 | 9.02814244 | 0.803686286 | 0.48102797 | 1.67076831 | 0.09476745 | NA |
| AC010615.4 | 36.243231 | 0.282176015 | 0.35290254 | 0.79958625 | 0.42395056 | 0.77666823 |
| ZNF429 | 23.9153281 | 0.105191635 | 0.31630033 | 0.33256885 | 0.73945976 | NA |
| AC123912.1 | 5.32743304 | -0.367116519 | 0.71128253 | -0.5161332 | 0.60576139 | NA |
| ZNF100 | 12.3495092 | 0.295742658 | 0.62803637 | 0.47090053 | 0.63771177 | NA |
| ZNF43 | 22.406956 | -0.087801637 | 0.37900798 | -0.2316617 | 0.81680077 | NA |
| ZNF208 | 32.1922336 | 0.320710998 | 0.48408502 | 0.66250965 | 0.50764465 | 0.82657837 |
| ZNF257 | 6.76746936 | 0.451506649 | 0.60340676 | 0.7482625 | 0.45430184 | NA |
| ZNF676 | 3.82240996 | -0.487374215 | 0.7686672 | -0.634051 | 0.52604753 | NA |
| ZNF98 | 15.584419 | 1.209321247 | 0.62603858 | 1.93170401 | 0.05339605 | NA |
| ZNF91 | 94.6825293 | -0.05437876 | 0.29347992 | -0.1852895 | 0.85300194 | 0.95569766 |
| ZNF675 | 9.88092467 | 0.06295557 | 0.4631232 | 0.13593698 | 0.8918711 | NA |
| ZNF254 | 56.5172867 | 0.188692568 | 0.27740302 | 0.68021093 | 0.49637091 | 0.8234249 |
| LINC00662 | 35.3339831 | -0.188206396 | 0.34095865 | -0.5519918 | 0.58095394 | 0.86175324 |
| AC006504.5 | 40.1238093 | -0.421310547 | 0.32666023 | -1.2897516 | 0.19713692 | 0.59987316 |
| AC020905.1 | 6.76328436 | -1.733458435 | 0.81673402 | -2.1224271 | 0.03380189 | NA |
| AC005394.1 | 7.14803986 | 0.54404064 | 1.20500334 | 0.45148476 | 0.65164021 | NA |
| UQCRFS1 | 30.1359199 | -0.70416299 | 0.44616869 | -1.5782439 | 0.1145096 | 0.50483797 |
| POP4 | 5.22658275 | -0.702370611 | 0.92007687 | -0.7633825 | 0.44523529 | NA |
| C19orf12 | 20.7464372 | -0.44704468 | 0.45470772 | -0.9831473 | 0.32553493 | NA |
| URI1 | 86.9538265 | 0.162326738 | 0.21966413 | 0.738977 | 0.45992097 | 0.79651556 |
| TSHZ3 | 54.1247414 | -0.529312454 | 0.33480464 | -1.5809591 | 0.11388739 | 0.50483797 |
| AC025809.1 | 21.3132235 | 0.000952756 | 0.41432416 | 0.00229954 | 0.99816523 | NA |
| ZNF507 | 19.9823358 | -0.150839658 | 0.3530568 | -0.4272391 | 0.6692052 | NA |
| DPY19L3 | 60.3254993 | -0.425256891 | 0.40955744 | -1.0383327 | 0.29911519 | 0.68168772 |
| PDCD5 | 13.1199662 | -0.968083884 | 0.50899763 | -1.9019418 | 0.05717876 | NA |
| ANKRD27 | 20.0193535 | 0.230499418 | 0.35487813 | 0.64951711 | 0.51600419 | NA |
| NUDT19 | 7.93753968 | 0.624676158 | 0.47647795 | 1.31102848 | 0.18984814 | NA |
| CEP89 | 19.7592844 | 0.001528354 | 0.31074437 | 0.00491836 | 0.99607573 | NA |
| RHPN2 | 11.107548 | 0.234676045 | 0.45043778 | 0.52099548 | 0.60236992 | NA |
| GPATCH1 | 6.01737124 | 0.336809565 | 0.63129197 | 0.53352423 | 0.59367074 | NA |
| CEBPG | 7.27908511 | -1.129085375 | 0.65002992 | -1.7369745 | 0.08239168 | NA |
| PEPD | 25.8924655 | -0.103791834 | 0.38241989 | -0.271408 | 0.78607722 | NA |
| LSM14A | 91.2442182 | 0.307875378 | 0.21952714 | 1.40244789 | 0.16078154 | 0.56283048 |
| KIAA0355 | 41.6050112 | 0.51565314 | 0.32659824 | 1.57886076 | 0.114368 | 0.50483797 |
| GPI | 16.0992849 | -1.061974582 | 0.59617405 | -1.7813164 | 0.07486078 | NA |
| UBA2 | 40.8457194 | -0.106494891 | 0.26470036 | -0.4023224 | 0.68744675 | 0.90499319 |
| WTIP | 20.4210781 | 0.067070287 | 0.44750187 | 0.14987711 | 0.88086157 | NA |
| SCGB2B2 | 23.3960078 | 0.001868211 | 0.41668683 | 0.00448349 | 0.99642271 | NA |
| ZNF302 | 29.3331746 | -0.719380034 | 0.37046448 | -1.9418327 | 0.05215736 | 0.35795306 |
| ZNF181 | 7.17271926 | -0.193914847 | 0.4950383 | -0.3917169 | 0.69526743 | NA |
| GRAMD1A | 23.3098496 | -1.029196291 | 0.49105852 | -2.095873 | 0.03609346 | NA |
| HPN | 14.4959309 | 1.132721702 | 0.88335199 | 1.28229937 | 0.19973765 | NA |
| LSR | 19.3303039 | -0.304397475 | 0.50579226 | -0.6018231 | 0.54729188 | NA |
| USF2 | 22.7321373 | -0.449946078 | 0.41282135 | -1.0899293 | 0.2757443 | NA |
| TMEM147 | 17.7570706 | -0.092238035 | 0.4284102 | -0.2153031 | 0.82953103 | NA |
| RBM42 | 12.8463086 | -0.335134005 | 0.68996102 | -0.4857289 | 0.62715939 | NA |
| COX6B1 | 50.631468 | -0.252237831 | 0.39241478 | -0.6427837 | 0.52036445 | 0.83108387 |
| KMT2B | 11.013387 | -0.455912463 | 0.45885096 | -0.993596 | 0.32041961 | NA |
| PROSER3 | 4.48636652 | -0.188028197 | 0.69412151 | -0.2708866 | 0.78647827 | NA |
| PRODH2 | 20.8330446 | 1.425243723 | 0.6619066 | 2.15323992 | 0.03129983 | NA |
| NPHS1 | 1427.54048 | -0.638839987 | 0.60369786 | -1.0582115 | 0.28995905 | 0.67746193 |
| KIRREL2 | 9.33244298 | -0.138827729 | 0.5757374 | -0.2411303 | 0.80945413 | NA |
| APLP1 | 17.6797975 | -0.266149923 | 0.57229554 | -0.4650568 | 0.64189079 | NA |
| SDHAF1 | 7.82696019 | -0.767945917 | 0.55266482 | -1.3895328 | 0.16467078 | NA |
| AC002116.2 | 4.66954103 | -0.611265697 | 0.81252631 | -0.7523026 | 0.45186908 | NA |
| CLIP3 | 34.1581207 | -0.172332733 | 0.35757458 | -0.4819491 | 0.62984214 | 0.88166074 |
| THAP8 | 5.34857929 | 0.149088966 | 0.71946902 | 0.20722083 | 0.8358374 | NA |
| POLR2I | 5.51640642 | 0.002513327 | 0.61455539 | 0.00408967 | 0.99673693 | NA |
| TBCB | 28.8194307 | -0.532161786 | 0.42738086 | -1.2451699 | 0.2130693 | 0.62255988 |
| CAPNS1 | 78.7830493 | -0.027767845 | 0.46901647 | -0.0592044 | 0.95278929 | 0.98520702 |
| COX7A1 | 9.57718459 | 0.666200356 | 0.66958577 | 0.99494401 | 0.3197635 | NA |
| ZNF565 | 24.9786641 | -0.90287187 | 0.35836943 | -2.5193886 | 0.01175588 | NA |
| ZNF146 | 34.4878453 | -0.39422181 | 0.30858768 | -1.2775034 | 0.20142458 | 0.60910153 |
| LINC00665 | 11.3240815 | -0.009327184 | 0.61773455 | -0.015099 | 0.98795319 | NA |
| ZFP14 | 21.5181556 | -0.134507973 | 0.32856102 | -0.4093851 | 0.6822571 | NA |
| ZFP82 | 11.0854837 | 0.12388432 | 0.39890961 | 0.31055737 | 0.75613714 | NA |
| ZNF566 | 14.2634894 | 0.10829508 | 0.37149132 | 0.29151443 | 0.77065791 | NA |
| ZNF260 | 10.7464961 | -0.319693529 | 0.43225981 | -0.7395865 | 0.45955092 | NA |
| ZNF529 | 19.146294 | 0.012820322 | 0.47447479 | 0.02702003 | 0.97844376 | NA |
| ZNF529-AS1 | 16.1052019 | -0.600537132 | 0.44086285 | -1.3621858 | 0.17313924 | NA |
| ZNF382 | 6.91070167 | -0.672876564 | 0.52564302 | -1.2801018 | 0.20050934 | NA |
| ZNF461 | 7.32387559 | 0.211200191 | 0.48894532 | 0.43195053 | 0.66577736 | NA |
| ZNF567 | 8.54621007 | -0.390699183 | 0.50430524 | -0.7747276 | 0.43850064 | NA |

| | | | | | | |
|---|---|---|---|---|---|---|
| AC020928.1 | 21.0021232 | 0.010777488 | 0.36556467 | 0.02948176 | 0.97648037 | NA |
| ZNF790 | 14.3866863 | -0.01928874 | 0.50547586 | -0.0381596 | 0.96956046 | NA |
| ZNF345 | 26.190883 | 0.562324153 | 0.36951876 | 1.52177428 | 0.12806565 | NA |
| ZNF568 | 13.8032322 | -0.63328243 | 0.41725914 | -1.5177197 | 0.12908507 | NA |
| ZNF420 | 27.2707851 | -0.167333289 | 0.28547073 | -0.5861662 | 0.55776384 | NA |
| ZNF383 | 8.99524002 | -0.120178523 | 0.57554529 | -0.2088081 | 0.83459804 | NA |
| HKR1 | 22.0055803 | -0.004292745 | 0.34244349 | -0.0125356 | 0.98999828 | NA |
| ZNF569 | 8.40926924 | -0.338396951 | 0.50938167 | -0.6643289 | 0.50647985 | NA |
| ZNF793 | 11.2872337 | -0.263541899 | 0.45360753 | -0.580991 | 0.56124651 | NA |
| ZNF540 | 7.39779934 | -0.934756335 | 0.68972345 | -1.3552625 | 0.17533394 | NA |
| ZNF571 | 7.16701337 | -1.164078208 | 0.56642402 | -2.0551357 | 0.0398659 | NA |
| ZFP30 | 16.7217173 | -0.727001474 | 0.4628673 | -1.5706477 | 0.1162645 | NA |
| ZNF573 | 12.1118594 | -0.064333812 | 0.46506219 | -0.1383338 | 0.88997663 | NA |
| SIPA1L3 | 43.4522429 | -0.25088904 | 0.34244617 | -0.7326379 | 0.46377931 | 0.79842482 |
| SPINT2 | 7.8171016 | -0.438547271 | 0.53731069 | -0.8161894 | 0.41439184 | NA |
| YIF1B | 6.55623111 | 1.197630055 | 0.63815832 | 1.8766974 | 0.06055958 | NA |
| CATSPERG | 6.16249537 | 0.398729286 | 0.60607286 | 0.65789002 | 0.51060881 | NA |
| PSMD8 | 36.1362566 | -0.614424433 | 0.48110002 | -1.2771241 | 0.20155844 | 0.60910153 |
| EIF3K | 21.5499526 | 0.548564391 | 0.46534891 | 1.17882385 | 0.23846833 | NA |
| ACTN4 | 880.217561 | 0.259416507 | 0.36823431 | 0.70448761 | 0.48112916 | 0.81183849 |
| AC008649.2 | 4.98735851 | 0.391846881 | 0.81702305 | 0.47960321 | 0.63150956 | NA |
| CAPN12 | 10.5549585 | 0.292628048 | 0.55592483 | 0.5263806 | 0.59862379 | NA |
| ECH1 | 32.5271619 | -0.448572463 | 0.49672772 | -0.903055 | 0.36649669 | 0.73480504 |
| HNRNPL | 23.9657245 | 0.210315876 | 0.35361077 | 0.5947666 | 0.55199949 | NA |
| SIRT2 | 13.6866768 | 0.798989223 | 0.48013562 | 1.66409071 | 0.0960943 | NA |
| NFKBIB | 4.45985475 | -1.045702395 | 0.8152875 | -1.282618 | 0.19962595 | NA |
| MRPS12 | 10.5792177 | -1.309297603 | 0.69120851 | -1.8942151 | 0.05819647 | NA |
| FBXO17 | 34.0711037 | 0.274363545 | 0.2940094 | 0.9331795 | 0.3507273 | 0.72238707 |
| FBXO27 | 5.32799354 | 0.683488527 | 0.69897616 | 0.9778424 | 0.32815228 | NA |
| PAK4 | 11.1196468 | 0.452111245 | 0.40940585 | 1.10431066 | 0.2694584 | NA |
| SAMD4B | 39.99198 | 0.151610917 | 0.33126729 | 0.45766945 | 0.64718994 | 0.89261695 |
| PAF1 | 10.0758845 | -0.292417057 | 0.47661103 | -0.613534 | 0.53952333 | NA |
| MED29 | 14.4162448 | -0.729586152 | 0.57215739 | -1.2751494 | 0.20225637 | NA |
| ZFP36 | 17.6363272 | -1.324234746 | 0.8366596 | -1.5827641 | 0.11347525 | NA |
| PLEKHG2 | 8.24797282 | -0.079854782 | 0.55062379 | -0.145026 | 0.88469032 | NA |
| RPS16 | 54.848759 | -0.414390918 | 0.46820575 | -0.8850616 | 0.37612343 | 0.74330391 |
| SUPT5H | 22.9373769 | -0.488433809 | 0.44644447 | -1.0940528 | 0.27393183 | NA |
| TIMM50 | 8.2768954 | -0.245926204 | 0.57665137 | -0.4264729 | 0.66976326 | NA |
| EID2B | 4.91121122 | -0.461573501 | 0.66004025 | -0.6993111 | 0.4843576 | NA |
| EID2 | 7.48384092 | -0.503553965 | 0.54360708 | -0.9263197 | 0.35427984 | NA |
| DYRK1B | 6.00671962 | 0.18559264 | 0.56667008 | 0.32751445 | 0.74327882 | NA |
| FBL | 6.95681484 | -0.452551956 | 0.62108286 | -0.7286499 | 0.46621585 | NA |
| PSMC4 | 13.5634141 | -0.928275856 | 0.56078086 | -1.6553273 | 0.09785813 | NA |
| ZNF546 | 7.69112769 | 0.349207187 | 0.53114977 | 0.65745522 | 0.51088826 | NA |
| ZNF780B | 34.1703304 | 0.748802284 | 0.40750838 | 1.83751383 | 0.06613408 | 0.3991664 |
| ZNF780A | 8.03491725 | 0.248586986 | 0.57108048 | 0.43529239 | 0.66335021 | NA |
| MAP3K10 | 7.47329637 | -0.034623198 | 0.5284113 | -0.0655232 | 0.94775744 | NA |
| AKT2 | 71.9348254 | 0.235389613 | 0.29905186 | 0.7871197 | 0.4312118 | 0.77901411 |
| PLD3 | 128.362248 | -0.500991823 | 0.37133965 | -1.3491471 | 0.17728971 | 0.58001112 |
| PRX | 10.2164273 | -0.18078871 | 0.51428929 | -0.3515312 | 0.7251899 | NA |
| SERTAD1 | 16.7028182 | -0.602297508 | 0.54223955 | -1.1107591 | 0.26667206 | NA |
| SERTAD3 | 9.32920356 | 0.379929795 | 0.55678777 | 0.68236016 | 0.49501124 | NA |
| BLVRB | 40.3395225 | -1.011239335 | 0.45077035 | -2.2433581 | 0.02487373 | 0.25362919 |
| SPTBN4 | 5.50710432 | -0.350339821 | 0.64848897 | -0.5402402 | 0.58903138 | NA |
| SHKBP1 | 9.87980893 | 0.024004779 | 0.48460451 | 0.04953478 | 0.96049312 | NA |
| LTBP4 | 60.6870151 | -0.678266536 | 0.55889223 | -1.2135909 | 0.22490398 | 0.63128756 |
| NUMBL | 8.59768272 | -0.663399944 | 0.58857586 | -1.1271273 | 0.25968865 | NA |
| COQ8B | 9.65771784 | -0.396186066 | 0.55489653 | -0.7139819 | 0.47523838 | NA |
| ITPKC | 7.85879294 | -0.937734179 | 0.71255126 | -1.3160235 | 0.18816617 | NA |
| SNRPA | 11.3168412 | -0.137871374 | 0.45265299 | -0.3045851 | 0.76068214 | NA |
| AC008537.1 | 15.9662511 | 0.449212264 | 0.5492997 | 0.81779084 | 0.41347663 | NA |
| AXL | 51.1411037 | -1.363209158 | 0.55218738 | -2.4687438 | 0.01355883 | 0.194252 |
| HNRNPUL1 | 23.5624986 | -0.474566432 | 0.3265962 | -1.4530678 | 0.14620492 | NA |
| TGFB1 | 20.2017641 | -0.966571181 | 0.51642153 | -1.871671 | 0.06125214 | NA |
| CCDC97 | 6.83456045 | -0.668862449 | 0.58525719 | -1.1428522 | 0.25309998 | NA |
| DMAC2 | 5.86448284 | -1.054134749 | 0.64545904 | -1.6331551 | 0.10243637 | NA |
| RPS19 | 111.924251 | -0.545568137 | 0.40434472 | -1.3492649 | 0.17725189 | 0.58001112 |
| ARHGEF1 | 8.55654614 | -0.41352574 | 0.52575278 | -0.7865403 | 0.43155103 | NA |
| RABAC1 | 79.3846797 | -0.262881118 | 0.4909497 | -0.5354543 | 0.59233576 | 0.8654248 |
| GRIK5 | 16.020224 | 0.033617918 | 0.47537795 | 0.07071829 | 0.94362196 | NA |
| DEDD2 | 6.13497415 | -0.092370071 | 0.54149528 | -0.1705833 | 0.8645514 | NA |
| GSK3A | 13.6098994 | 0.22456009 | 0.44436054 | 0.50535561 | 0.61330909 | NA |
| CIC | 11.4884177 | -0.66319589 | 0.5452676 | -1.216276 | 0.22387981 | NA |
| PAFAH1B3 | 5.44215842 | -0.905262896 | 0.7137617 | -1.2682985 | 0.20469137 | NA |
| MEGF8 | 15.8811126 | 0.009876537 | 0.43007056 | 0.02296492 | 0.98167825 | NA |
| LIPE-AS1 | 19.3732747 | 1.456015703 | 0.37554511 | 3.87707269 | 0.00010572 | NA |
| PHLDB3 | 7.58817775 | -0.51216295 | 0.50746614 | -1.0092554 | 0.31285216 | NA |
| ETHE1 | 25.851453 | -0.033205935 | 0.45408593 | -0.073127 | 0.94170507 | NA |
| XRCC1 | 6.46806943 | -0.084408648 | 0.68061532 | -0.1240181 | 0.90130091 | NA |
| ZNF428 | 30.9171867 | -0.07055549 | 0.42505683 | -0.1659907 | 0.86816425 | 0.96089268 |
| CADM4 | 8.04821414 | 0.142817358 | 0.52904702 | 0.26995211 | 0.7871971 | NA |
| PLAUR | 7.2336195 | -1.209571095 | 0.55934258 | -2.162487 | 0.03058065 | NA |
| SMG9 | 10.2839549 | 0.206134994 | 0.46983811 | 0.43873622 | 0.66085268 | NA |
| ZNF283 | 7.66798389 | 0.213845163 | 0.49844851 | 0.42902157 | 0.66790753 | NA |
| ZNF45 | 4.83364036 | 0.191526585 | 0.59891455 | 0.3197895 | 0.74912791 | NA |
| ZNF155 | 10.272234 | -0.802956323 | 0.47307377 | -1.6973174 | 0.08963667 | NA |
| ZNF224 | 11.9068891 | 0.09419518 | 0.53457655 | 0.17620522 | 0.86013272 | NA |
| ZNF234 | 8.20286987 | -0.088834902 | 0.46663421 | -0.1903737 | 0.84901627 | NA |
| ZNF226 | 13.6103253 | -0.38986187 | 0.4232928 | -0.9210217 | 0.35703908 | NA |
| ZNF227 | 8.57537619 | 0.399853138 | 0.49901018 | 0.80129254 | 0.42296231 | NA |
| ZNF235 | 35.1535421 | 0.111959061 | 0.3694248 | 0.3030632 | 0.7618417 | 0.92336765 |
| ZNF229 | 5.93252652 | 0.048519857 | 0.63700269 | 0.076169 | 0.93928465 | NA |
| ZNF180 | 7.03433532 | 0.230330114 | 0.60891383 | 0.37826389 | 0.70523457 | NA |
| AC243964.2 | 31.6360169 | -0.456382752 | 0.46922153 | -0.9726381 | 0.33073318 | 0.70387662 |
| PVR | 15.2822411 | 0.643306897 | 0.44194078 | 1.4556405 | 0.14549203 | NA |
| CEACAM19 | 51.2212746 | 0.678335696 | 0.36998064 | 1.83343564 | 0.06673781 | 0.40161508 |
| BCL3 | 20.7650406 | -3.209374697 | 0.99124837 | -3.2377099 | 0.00120493 | NA |
| BCAM | 482.880143 | 0.328074237 | 0.50559065 | 0.64889301 | 0.51640753 | 0.83104823 |
| NECTIN2 | 227.484453 | -0.053583388 | 0.38601765 | -0.1388107 | 0.88959972 | 0.96812892 |
| TOMM40 | 9.89217392 | 0.058261442 | 0.44716971 | 0.13028933 | 0.89633752 | NA |
| APOE | 46.2622135 | 0.619216012 | 1.00369133 | 0.61693869 | 0.53727517 | 0.83677705 |
| CLPTM1 | 28.1397676 | -0.166481896 | 0.53966908 | -0.3084889 | 0.75771038 | NA |
| RELB | 17.0065169 | -0.750930768 | 0.56719412 | -1.3239396 | 0.1855231 | NA |
| CLASRP | 20.8777731 | -0.115460972 | 0.36123533 | -0.3196281 | 0.74925025 | NA |
| GEMIN7-AS1 | 5.77470759 | 0.258792025 | 0.61622746 | 0.41996185 | 0.67451332 | NA |
| MARK4 | 28.4995597 | -0.103540233 | 0.29062065 | -0.3562728 | 0.72163627 | NA |
| TRAPPC6A | 14.0776665 | 0.688780002 | 0.50077171 | 1.37543712 | 0.16899597 | NA |
| PPP1R13L | 4.74570395 | 0.00120586 | 0.70866491 | 0.00170159 | 0.99864233 | NA |
| ERCC1 | 11.1315859 | 0.45376096 | 0.43440255 | 1.04456329 | 0.29622485 | NA |
| VASP | 12.7223956 | -0.038737454 | 0.54030546 | -0.0716955 | 0.94284426 | NA |
| OPA3 | 14.4715421 | -1.038966159 | 0.57746673 | -1.7991792 | 0.07199033 | NA |
| EML2 | 14.2207722 | 0.093111648 | 0.37882409 | 0.24579126 | 0.80584383 | NA |
| SNRPD2 | 34.4768296 | -0.805382821 | 0.39446835 | -2.0416919 | 0.04118211 | 0.31816119 |

| | | | | | | |
|---|---|---|---|---|---|---|
| FBXO46 | 8.15356431 | 0.806093789 | 0.48373358 | 1.66640033 | 0.09563371 | NA |
| SIX5 | 7.11963164 | 0.365818928 | 0.53970352 | 0.6778146 | 0.49788925 | NA |
| DM1-AS | 6.71492405 | 0.054254662 | 0.54871969 | 0.098875 | 0.92123752 | NA |
| DMPK | 9.71554558 | 0.17592236 | 0.49088652 | 0.35837684 | 0.72006132 | NA |
| SYMPK | 28.9620007 | -0.01169611 | 0.29252992 | -0.0399826 | 0.96810699 | 0.98964394 |
| MYPOP | 5.44411045 | -0.625831865 | 0.59574877 | -1.0504963 | 0.29349 | NA |
| PPP5C | 20.137698 | -0.225220387 | 0.38659118 | -0.5825803 | 0.5601759 | NA |
| PNMA8A | 4.62111669 | -1.807706802 | 0.96282095 | -1.8775109 | 0.06044811 | NA |
| PPP5D1 | 16.611786 | 1.300787749 | 0.65965241 | 1.97192904 | 0.04861771 | NA |
| CALM3 | 22.402401 | 0.138528564 | 0.42208775 | 0.3281985 | 0.74276159 | NA |
| PRKD2 | 17.3274904 | 0.229250014 | 0.47111736 | 0.48660914 | 0.62653534 | NA |
| STRN4 | 8.93609732 | -0.321937575 | 0.56044923 | -0.5744277 | 0.5656784 | NA |
| FKRP | 11.6836597 | 0.094914921 | 0.46175377 | 0.2055531 | 0.83714001 | NA |
| SLC1A5 | 29.8395446 | -1.038884518 | 0.41988218 | -2.4742286 | 0.01335242 | 0.19273104 |
| AP2S1 | 22.467083 | -0.241137803 | 0.45314879 | -0.5321382 | 0.59463025 | NA |
| ARHGAP35 | 41.6410416 | 0.094417708 | 0.28432744 | 0.33207386 | 0.73983349 | 0.92003444 |
| TMEM160 | 20.8544002 | -0.685738217 | 0.44240287 | -1.5500311 | 0.12113404 | NA |
| ZC3H4 | 17.6540363 | 0.290604725 | 0.49955413 | 0.5817282 | 0.56074977 | NA |
| SAE1 | 26.1162517 | 0.614220971 | 0.39335744 | 1.56148303 | 0.11840983 | NA |
| INAFM1 | 5.97451807 | -0.657905566 | 0.5930594 | -1.1093418 | 0.26728277 | NA |
| DHX34 | 8.80092202 | -0.772956203 | 0.6268534 | -1.2330733 | 0.21754841 | NA |
| NAPA | 38.5568219 | -0.308460794 | 0.39270131 | -0.7854845 | 0.43216956 | 0.77905765 |
| EHD2 | 40.5031949 | -0.438921467 | 0.52296692 | -0.8392911 | 0.401306 | 0.76381845 |
| NOP53 | 6.85764877 | -0.381969814 | 0.54940857 | -0.6952382 | 0.48690605 | NA |
| SELENOW | 59.2830305 | 0.538775691 | 0.41690466 | 1.29232349 | 0.19624514 | 0.59987316 |
| PLA2G4C | 15.6377458 | 0.133498353 | 0.669848 | 0.19929649 | 0.84203083 | NA |
| ZSWIM9 | 6.66022991 | -0.295530422 | 0.60049909 | -0.4921413 | 0.62261944 | NA |
| CARD8 | 11.1464845 | 0.045442692 | 0.44220659 | 0.10276349 | 0.91815068 | NA |
| EMP3 | 37.2127264 | -0.292134918 | 0.60781835 | -0.4806287 | 0.63078044 | 0.88222258 |
| KDELR1 | 31.9628441 | -0.534938488 | 0.33740046 | -1.5854705 | 0.11285947 | 0.50413876 |
| GRWD1 | 6.34451144 | 0.037778465 | 0.81669014 | 0.04625801 | 0.9631046 | NA |
| CYTH2 | 12.4397667 | 0.16239631 | 0.4245364 | 0.38252624 | 0.70207107 | NA |
| RPL18 | 65.8269476 | -0.640309731 | 0.45897074 | -1.3950992 | 0.16298592 | 0.56498748 |
| SPHK2 | 7.62085464 | 0.074663137 | 0.4740954 | 0.15748547 | 0.87486226 | NA |
| DBP | 10.5840753 | -0.818030763 | 0.62109391 | -1.3170806 | 0.18781159 | NA |
| CA11 | 6.80895385 | -0.951865473 | 0.8148842 | -1.1680991 | 0.24276681 | NA |
| BCAT2 | 15.7470075 | -0.479784562 | 0.39647933 | -1.2101124 | 0.22623576 | NA |
| HSD17B14 | 14.0264582 | -1.109057323 | 0.60290298 | -1.8395287 | 0.06583747 | NA |
| PLEKHA4 | 10.7965364 | -1.205240696 | 0.57046784 | -2.1127233 | 0.03462446 | NA |
| PPP1R15A | 15.5154551 | -0.479917021 | 0.50483759 | -0.9506365 | 0.34178895 | NA |
| NUCB1 | 133.855602 | -0.06046613 | 0.39955271 | -0.1513345 | 0.87971182 | 0.96423929 |
| BAX | 16.4377775 | -0.397692352 | 0.49529221 | -0.8029449 | 0.42200658 | NA |
| FTL | 476.231343 | -1.0440883 | 0.62484692 | -1.6709505 | 0.09473145 | 0.46755975 |
| GYS1 | 10.4203853 | 0.291298492 | 0.48905529 | 0.59563509 | 0.55141903 | NA |
| RUVBL2 | 7.72050824 | 0.306010421 | 0.49400447 | 0.61944869 | 0.53562081 | NA |
| SNRNP70 | 76.128102 | 0.434859881 | 0.23325725 | 1.86429312 | 0.06228055 | 0.38512471 |
| TRPM4 | 11.9576597 | -0.320402901 | 0.6168705 | -0.5194006 | 0.60348142 | NA |
| SLC6A16 | 18.3854892 | -0.538709595 | 0.44092729 | -1.2217651 | 0.22179645 | NA |
| TEAD2 | 10.5676057 | -1.021621945 | 0.62024827 | -1.6471178 | 0.09953384 | NA |
| PIH1D1 | 10.4647033 | -0.289742824 | 0.46076512 | -0.6288298 | 0.52946051 | NA |
| RPL13A | 83.8082517 | -0.440090859 | 0.44459511 | -0.9898689 | 0.32223822 | 0.69513974 |
| RPS11 | 71.3426291 | -0.335891842 | 0.42501552 | -0.7903049 | 0.42934974 | 0.7784723 |
| FCGRT | 36.9095188 | -0.3801448 | 0.33549158 | -1.1330979 | 0.25717314 | 0.65588339 |
| NOSIP | 9.41900399 | -0.267970221 | 0.46923433 | -0.5710797 | 0.56794559 | NA |
| PRR12 | 10.739488 | 0.585566879 | 0.48240873 | 1.21383972 | 0.22480894 | NA |
| RRAS | 56.3905873 | -0.040521122 | 0.56185411 | -0.0721204 | 0.94250612 | 0.98114803 |
| SCAF1 | 14.0156976 | -0.136069197 | 0.38442272 | -0.3539572 | 0.72337094 | NA |
| IRF3 | 19.9868252 | -0.026226377 | 0.38423106 | -0.0682568 | 0.94558122 | NA |
| PRMT1 | 24.3952972 | -0.374240042 | 0.41228653 | -0.9077183 | 0.36402706 | NA |
| AP2A1 | 23.0666213 | -0.035418165 | 0.32042796 | -0.1105339 | 0.91198593 | NA |
| MED25 | 21.7735349 | 0.697912665 | 0.4322448 | 1.61462363 | 0.10639223 | NA |
| PTOV1 | 19.3420075 | 0.215498218 | 0.39528841 | 0.54516705 | 0.58563863 | NA |
| PNKP | 9.58408406 | -0.410876767 | 0.64868268 | -0.6334018 | 0.5264713 | NA |
| TBC1D17 | 11.5786815 | 0.102447845 | 0.45175627 | 0.22677681 | 0.8205973 | NA |
| ATF5 | 6.03634092 | -1.329422361 | 0.86980274 | -1.5284182 | 0.12640873 | NA |
| VRK3 | 14.0334293 | 0.102360824 | 0.46809253 | 0.21867647 | 0.82690208 | NA |
| MYH14 | 6.85917893 | 0.002620983 | 0.65522934 | 0.0040001 | 0.99680839 | NA |
| KCNC3 | 7.17038748 | 0.959269263 | 0.59045405 | 1.62462983 | 0.10424146 | NA |
| NR1H2 | 23.3908698 | -0.768517626 | 0.39836521 | -1.9291786 | 0.0537087 | NA |
| NAPSA | 8.76236046 | -1.70245301 | 1.11813161 | -1.5225873 | 0.12786199 | NA |
| EMC10 | 48.5503225 | -0.103645804 | 0.55885467 | -0.1854611 | 0.85286739 | 0.95569766 |
| JOSD2 | 12.8481768 | -0.283249178 | 0.63857583 | -0.4435639 | 0.65735792 | NA |
| CLEC11A | 5.73163201 | -0.360097831 | 0.58910605 | -0.6112615 | 0.54102649 | NA |
| C19orf48 | 6.33921174 | -0.131616747 | 0.84316311 | -0.1560988 | 0.87595515 | NA |
| KLK6 | 64.9318961 | 0.264130525 | 0.54335389 | 0.48611141 | 0.62688818 | 0.8806406 |
| AC011483.1 | 6.59880313 | 1.059250197 | 1.08823265 | 0.97336741 | 0.33037073 | NA |
| KLK7 | 64.2776788 | 0.39900566 | 0.65233224 | 0.61166019 | 0.54076261 | 0.84003567 |
| CTU1 | 7.73910961 | -1.283376937 | 0.65187302 | -1.968753 | 0.04898147 | NA |
| ETFB | 32.7857366 | 0.217869694 | 0.34824853 | 0.62561555 | 0.53156714 | 0.83567299 |
| ZNF175 | 9.27140102 | 0.735192753 | 0.53156183 | 1.3830804 | 0.16664022 | NA |
| SPACA6 | 15.6550028 | -0.694727131 | 0.59085299 | -1.1758037 | 0.23967333 | NA |
| ZNF577 | 13.7322287 | 0.069511426 | 0.4736919 | 0.14674396 | 0.88333412 | NA |
| ZNF350-AS1 | 9.20841679 | 0.349594274 | 0.67466272 | 0.51817636 | 0.60433523 | NA |
| ZNF350 | 5.62367091 | -0.603790973 | 0.65209877 | -0.9259195 | 0.35448781 | NA |
| ZNF615 | 4.84769584 | 0.301152607 | 0.82811483 | 0.36366044 | 0.71611158 | NA |
| ZNF614 | 4.96389528 | -0.645689729 | 0.72636144 | -0.8889372 | 0.37403681 | NA |
| ZNF432 | 17.3029138 | -0.02332244 | 0.38466495 | -0.0606305 | 0.95165346 | NA |
| AC011468.3 | 5.41072921 | -0.345375854 | 0.94666241 | -0.3648353 | 0.71523436 | NA |
| PPP2R1A | 51.5271374 | -0.065651082 | 0.35694548 | -0.1839247 | 0.85407255 | 0.95569766 |
| ZNF766 | 12.7946903 | 0.039208784 | 0.51246525 | 0.07651013 | 0.93901325 | NA |
| ZNF480 | 8.15451364 | -0.220184252 | 0.60532944 | -0.3637428 | 0.71605005 | NA |
| ZNF880 | 7.70429932 | 0.796656926 | 0.6099518 | 1.30609817 | 0.19151918 | NA |
| ZNF528 | 7.67859442 | 0.086587682 | 0.52591325 | 0.16464252 | 0.86922536 | NA |
| ZNF578 | 4.89586117 | 0.625044786 | 0.62052216 | 1.00728842 | 0.31379619 | NA |
| ZNF701 | 8.97297687 | 0.361090925 | 0.53912116 | 0.66977694 | 0.50299999 | NA |
| ZNF83 | 54.3087912 | 0.376704132 | 0.24711172 | 1.52442842 | 0.12740172 | 0.52301154 |
| ZNF611 | 22.0217944 | -0.358473315 | 0.47213915 | -0.7592535 | 0.44770092 | NA |
| ZNF600 | 4.32967138 | -0.015404449 | 0.61385695 | -0.0250945 | 0.97997957 | NA |
| ZNF28 | 4.72466615 | 0.046262904 | 0.62697718 | 0.07378722 | 0.94117969 | NA |
| ZNF160 | 31.1157144 | 0.066476897 | 0.38179074 | 0.17411867 | 0.8617722 | 0.95951831 |
| ZNF415 | 4.94947534 | 0.069096656 | 0.62319239 | 0.11087532 | 0.91171522 | NA |
| ZNF347 | 9.14242951 | -0.423809598 | 0.47421902 | -0.8937001 | 0.37148235 | NA |
| ZNF677 | 10.9165189 | 0.051739042 | 0.44164532 | 0.11715066 | 0.90674066 | NA |
| ZNF525 | 6.06547343 | -0.288856444 | 0.60858302 | -0.4746377 | 0.63504523 | NA |
| ZNF765 | 12.5421019 | -0.564142035 | 0.44166079 | -1.2773197 | 0.20148939 | NA |
| ZNF761 | 9.13486076 | -0.075265909 | 0.52768463 | -0.1426343 | 0.88657903 | NA |
| ZNF331 | 10.7707879 | -0.432457403 | 0.57454951 | -0.7526895 | 0.4516365 | NA |
| MYADM | 31.0071448 | 0.536259896 | 0.49537294 | 1.08253773 | 0.27901366 | 0.66958859 |
| AC008440.2 | 8.33844484 | 0.329897471 | 0.67453406 | 0.48907459 | 0.62478889 | NA |
| NDUFA3 | 12.1733591 | 1.151105827 | 0.56585417 | 2.03428002 | 0.04192337 | NA |
| PRPF31 | 16.2307729 | -0.573631412 | 0.37451713 | -1.531656 | 0.12560734 | NA |
| CNOT3 | 10.5098991 | 0.093586235 | 0.46445568 | 0.20149659 | 0.8403103 | NA |
| LENG1 | 8.85524324 | 0.085401426 | 0.52072844 | 0.16400377 | 0.86972818 | NA |

| | | | | | | |
|---|---|---|---|---|---|---|
| TMC4 | 5.26940985 | -0.993166248 | 0.67134492 | -1.4793681 | 0.13904197 | NA |
| MBOAT7 | 22.1138394 | -0.034022195 | 0.41313063 | -0.0823521 | 0.93436669 | NA |
| TSEN34 | 14.665017 | -0.611915668 | 0.46411512 | -1.3184567 | 0.18735083 | NA |
| RPS9 | 125.608315 | -0.659038618 | 0.40484722 | -1.62787 | 0.10355246 | 0.48612129 |
| LENG8-AS1 | 6.47816893 | -0.167931958 | 0.5791709 | -0.2899523 | 0.7718527 | NA |
| LENG8 | 74.9665441 | 0.749883221 | 0.25804949 | 2.90596669 | 0.0036612 | 0.09771848 |
| AC011476.3 | 4.41414518 | -0.184559253 | 0.91165588 | -0.202444 | 0.83956964 | NA |
| RDH13 | 10.4924201 | 1.355990536 | 0.44801824 | 3.02664136 | 0.00247287 | NA |
| PPP1R12C | 18.8490847 | 0.78514995 | 0.38633667 | 2.03229467 | 0.04212383 | NA |
| PPP6R1 | 12.7434162 | -0.044725046 | 0.4313644 | -0.1036827 | 0.91742112 | NA |
| HSPBP1 | 8.10053301 | 0.201890697 | 0.55223171 | 0.36559056 | 0.71467062 | NA |
| KMT5C | 5.24374906 | 0.591561333 | 0.57684122 | 1.02551848 | 0.30511861 | NA |
| RPL28 | 89.8411367 | -0.499970426 | 0.45799148 | -1.0916588 | 0.2749831 | 0.66766283 |
| UBE2S | 11.4798557 | -0.0791922 | 0.60526455 | -0.130839 | 0.89590268 | NA |
| ISOC2 | 12.5843278 | -0.681766273 | 0.604026 | -1.1287035 | 0.25902292 | NA |
| NAT14 | 9.44305426 | 0.089610468 | 0.60878898 | 0.14719463 | 0.8829784 | NA |
| SSC5D | 9.30184496 | -1.816653331 | 0.62053836 | -2.927544 | 0.00341651 | NA |
| ZNF524 | 6.23339736 | -0.041628479 | 0.60874598 | -0.068384 | 0.94547997 | NA |
| CCDC106 | 9.31893144 | 0.712874045 | 0.52132219 | 1.36743468 | 0.1714891 | NA |
| U2AF2 | 12.6352286 | 0.388554784 | 0.51052755 | 0.76108485 | 0.44660639 | NA |
| EPN1 | 33.0845987 | 0.322431854 | 0.33574165 | 0.96035702 | 0.33687556 | 0.71127916 |
| ZNF787 | 10.0726184 | -0.524218789 | 0.51583768 | -1.0162476 | 0.30951151 | NA |
| ZNF444 | 13.3496584 | 0.268514588 | 0.39633424 | 0.67749531 | 0.49809174 | NA |
| ZSCAN5A | 27.9568644 | 0.162330794 | 0.36823561 | 0.44083405 | 0.65933314 | NA |
| ZNF582 | 5.68645337 | -0.077231395 | 0.5687332 | -0.1357955 | 0.89198297 | NA |
| ZNF583 | 7.10728428 | 0.037569157 | 0.50545562 | 0.07432731 | 0.94074995 | NA |
| ZNF667 | 8.3072958 | -0.293425876 | 0.62555407 | -0.4690656 | 0.63902277 | NA |
| ZNF667-AS1 | 16.2692114 | 0.349319923 | 0.43518731 | 0.80268867 | 0.4221547 | NA |
| ZNF471 | 9.42342716 | -0.562890732 | 0.4685815 | -1.2012654 | 0.22964827 | NA |
| ZFP28 | 5.12632451 | 1.04449799 | 0.57132761 | 1.8281945 | 0.06752037 | NA |
| ZNF470 | 5.26547137 | 0.413239771 | 0.71340849 | 0.57924706 | 0.56242248 | NA |
| ZNF264 | 18.7834956 | -0.215658384 | 0.38395066 | -0.5616825 | 0.57433233 | NA |
| ZNF460-AS1 | 5.48286575 | -0.456370419 | 0.61656679 | -0.74018 | 0.45919077 | NA |
| ZNF460 | 4.03725768 | -0.17060317 | 0.72616131 | -0.2349384 | 0.81425655 | NA |
| AC005261.1 | 23.1257312 | -0.321185152 | 0.35541428 | -0.9036923 | 0.36615857 | NA |
| AC004076.1 | 15.234104 | 0.526472246 | 0.46049072 | 1.14328524 | 0.25292018 | NA |
| ZNF549 | 9.1338541 | -0.314976488 | 0.49123675 | -0.6411908 | 0.52139873 | NA |
| ZNF134 | 8.17219146 | -0.202653257 | 0.55460958 | -0.365398 | 0.7148143 | NA |
| ZNF211 | 4.61795523 | -0.47071808 | 0.74423263 | -0.6324878 | 0.52706816 | NA |
| AC003006.1 | 5.78550657 | -0.721443991 | 0.60491359 | -1.1926397 | 0.2330105 | NA |
| ZNF776 | 5.93080197 | 0.280293358 | 0.61730362 | 0.45406077 | 0.64978509 | NA |
| ZNF586 | 44.1055593 | 0.026410083 | 0.26263618 | 0.10055767 | 0.9199016 | 0.97775374 |
| ZNF552 | 7.65747733 | -0.726697431 | 0.53978281 | -1.3462775 | 0.17821305 | NA |
| ZNF814 | 29.0319385 | 0.075977796 | 0.40491359 | 0.18763953 | 0.85115924 | 0.95532127 |
| ZNF587 | 12.7271394 | 0.234944609 | 0.39831437 | 0.58984718 | 0.55529311 | NA |
| ZNF418 | 6.05746704 | -0.311645002 | 0.60254825 | -0.5172117 | 0.60500839 | NA |
| C19orf18 | 8.3059559 | 0.157390602 | 0.62390372 | 0.25226745 | 0.80083434 | NA |
| ZNF606 | 7.36884577 | 0.031579376 | 0.53755235 | 0.0587466 | 0.95315394 | NA |
| ZNF135 | 11.4044328 | 0.054104113 | 0.44275156 | 0.12219971 | 0.90274086 | NA |
| ZSCAN18 | 20.0486375 | -0.620501703 | 0.47329705 | -1.3110196 | 0.18985114 | NA |
| ZNF329 | 9.8530556 | -0.612982709 | 0.45869345 | -1.3363668 | 0.1814294 | NA |
| ZNF274 | 27.5686041 | -0.284645609 | 0.37891717 | -0.751208 | 0.45252749 | NA |
| AC020915.3 | 5.26896258 | -0.291042466 | 0.58752292 | -0.4953721 | 0.62033748 | NA |
| ERVK3-1 | 4.9527386 | -0.091399 | 0.73005855 | -0.1251941 | 0.90036991 | NA |
| AC010642.2 | 5.01829076 | -0.911529695 | 0.75590825 | -1.2058735 | 0.22786629 | NA |
| A1BG | 11.1022431 | -0.319138332 | 0.57866786 | -0.5515052 | 0.58128741 | NA |
| A1BG-AS1 | 8.08132485 | -0.093028934 | 0.50409078 | -0.184548 | 0.85358359 | NA |
| RPS5 | 42.5242407 | -0.821062843 | 0.54755315 | -1.4995126 | 0.13374071 | 0.52673829 |
| ZNF584 | 5.13183964 | 0.674230173 | 0.89918617 | 0.74982267 | 0.45336152 | NA |
| ZNF324 | 4.97209416 | -1.022981054 | 0.76514276 | -1.3369806 | 0.18122896 | NA |
| ZNF446 | 4.99882076 | -0.794335183 | 0.69699599 | -1.1396553 | 0.25442993 | NA |
| SLC27A5 | 9.5986648 | 0.126645982 | 0.45666602 | 0.27732736 | 0.78152875 | NA |
| ZBTB45 | 3.78896068 | 0.547542639 | 0.69821304 | 0.78420569 | 0.43291943 | NA |
| TRIM28 | 25.7807079 | -0.008669118 | 0.41365904 | -0.0209572 | 0.98327983 | NA |
| CHMP2A | 22.0405065 | -0.404077394 | 0.3905386 | -1.034667 | 0.30082446 | NA |
| UBE2M | 21.1470675 | -0.476491745 | 0.47164363 | -1.0102792 | 0.31236155 | NA |
| MZF1-AS1 | 5.17922139 | 0.025133859 | 0.64548482 | 0.03893795 | 0.96893986 | NA |
| MZF1 | 25.0195963 | 0.059408184 | 0.33698541 | 0.17629304 | 0.86006373 | NA |
| RPS4Y1 | 25.4585301 | -0.406526453 | 1.95734873 | -0.2076924 | 0.83546915 | NA |
| ZFY | 17.5252232 | 0.720399759 | 1.81967838 | 0.395894 | 0.69218322 | NA |
| LINC00278 | 34.2611364 | 0.579756145 | 1.97023131 | 0.29425791 | 0.76856083 | 0.92555901 |
| TTTY15 | 11.7148586 | -0.126536713 | 1.9586624 | -0.0646036 | 0.94848959 | NA |
| USP9Y | 64.6421979 | 0.361759272 | 2.41735896 | 0.14965062 | 0.88104027 | 0.9644761 |
| DDX3Y | 22.5477855 | -0.203069448 | 1.92006608 | -0.1057617 | 0.91577143 | NA |
| UTY | 140.560476 | 0.483988602 | 2.56284255 | 0.18884836 | 0.85021167 | 0.95532127 |
| NLGN4Y | 36.8407533 | -1.822941103 | 2.06661376 | -0.8820909 | 0.37772768 | 0.74552966 |
| TTTY14 | 101.803258 | 0.144254333 | 1.64675733 | 0.08759902 | 0.93019538 | 0.97818835 |
| KDM5D | 14.8091436 | -0.045134545 | 2.05917877 | -0.0219187 | 0.9825128 | NA |
| TTTY10 | 14.148505 | -0.045477325 | 1.97514812 | -0.0230248 | 0.98163052 | NA |
| EIF1AY | 20.3763898 | -0.241375885 | 2.15457857 | -0.1120293 | 0.91080019 | NA |
| AP000547.3 | 5.61963888 | -0.382711524 | 0.58562505 | -0.6535095 | 0.51342788 | NA |
| IL17RA | 11.8497723 | -0.332615453 | 0.46592374 | -0.7138839 | 0.47529898 | NA |
| HDHD5 | 9.78018339 | -0.190152322 | 0.44907168 | -0.4234342 | 0.67197847 | NA |
| ADA2 | 7.51784162 | 0.870476758 | 0.69282398 | 1.25641834 | 0.20896434 | NA |
| CECR2 | 89.842917 | 0.245186563 | 0.38826851 | 0.63148711 | 0.52772207 | 0.83195298 |
| ATP6V1E1 | 54.3261882 | -0.437393585 | 0.29262681 | -1.4947147 | 0.13498892 | 0.52779404 |
| BCL2L13 | 51.5845569 | 0.303237922 | 0.27357477 | 1.10842794 | 0.26767705 | 0.66080225 |
| MICAL3 | 55.603565 | 0.587482016 | 0.27532604 | 2.13376848 | 0.03286174 | 0.29047359 |
| PEX26 | 16.5406047 | 0.13545554 | 0.36332202 | 0.37282502 | 0.70927868 | NA |
| USP18 | 9.24255119 | -1.267767791 | 1.50874482 | -0.8402798 | 0.40075153 | NA |
| DGCR2 | 37.9832343 | -0.128514364 | 0.36988103 | -0.3474478 | 0.7282549 | 0.91700494 |
| SLC25A1 | 30.2642145 | 0.211913046 | 0.430345 | 0.49242596 | 0.62241825 | 0.8780438 |
| CLTCL1 | 7.74303608 | -0.311714644 | 0.48957657 | -0.6367025 | 0.52431862 | NA |
| HIRA | 10.5298297 | -0.205112229 | 0.55735902 | -0.3680074 | 0.71286773 | NA |
| MRPL40 | 23.3726609 | -1.347689448 | 0.44224173 | -3.0474045 | 0.00230827 | NA |
| UFD1 | 11.2505983 | -0.587285934 | 0.62928732 | -0.9332556 | 0.350688 | NA |
| CLDN5 | 103.442978 | 0.177969508 | 0.79948225 | 0.22260595 | 0.8238422 | 0.94808955 |
| GNB1L | 4.90027135 | 0.497888702 | 0.71809405 | 0.69334749 | 0.48809151 | NA |
| TXNRD2 | 52.4125161 | 0.862847645 | 0.70376205 | 1.22605025 | 0.22017976 | 0.62462904 |
| COMT | 108.445311 | -0.582912825 | 0.26830431 | -2.1725809 | 0.02981187 | 0.27926427 |
| TANGO2 | 16.1650679 | -0.385706022 | 0.49496428 | -0.7792603 | 0.43582638 | NA |
| DGCR8 | 17.0264913 | -0.042335696 | 0.42419518 | -0.0998024 | 0.92050121 | NA |
| TRMT2A | 6.48470364 | -0.443761844 | 0.75858118 | -0.5849893 | 0.55855494 | NA |
| RANBP1 | 15.4837724 | 0.09614266 | 0.48181112 | 0.1995443 | 0.841837 | NA |
| ZDHHC8 | 12.9943157 | 0.391023944 | 0.4149828 | 0.94226543 | 0.34605676 | NA |
| DGCR6L | 7.52457198 | 0.148149475 | 0.59151082 | 0.25045945 | 0.80223206 | NA |
| KLHL22 | 22.8957253 | -0.228853663 | 0.44288475 | -0.5167341 | 0.60534177 | NA |
| MED15 | 50.3387929 | 0.105867785 | 0.30676927 | 0.34510557 | 0.730015 | 0.91718621 |
| PI4KA | 59.1918282 | 0.091553299 | 0.30203958 | 0.3031169 | 0.76180078 | 0.92336765 |
| SNAP29 | 16.8569852 | -0.725666931 | 0.44818218 | -1.6191338 | 0.10541847 | NA |
| CRKL | 13.7971653 | -0.063328469 | 0.39534259 | -0.1601863 | 0.87273432 | NA |
| LZTR1 | 8.4222508 | -0.394650149 | 0.54332567 | -0.7263602 | 0.46761798 | NA |
| THAP7 | 10.7749174 | -0.943639396 | 0.4540344 | -2.0783434 | 0.03767774 | NA |

| | | | | | | |
|---|---|---|---|---|---|---|
| GGT2 | 12.5708099 | -0.305469297 | 0.96106874 | -0.3178433 | 0.75060378 | NA |
| HIC2 | 13.4348898 | 0.282952195 | 0.46700367 | 0.60588859 | 0.54458874 | NA |
| UBE2L3 | 52.3075948 | -0.294486813 | 0.27573779 | -1.0679958 | 0.28552239 | 0.67369331 |
| YDJC | 5.68589293 | -0.711038469 | 0.75941396 | -0.9362989 | 0.34911931 | NA |
| SDF2L1 | 15.776307 | -0.858181177 | 0.64739438 | -1.3255926 | 0.18497469 | NA |
| PPIL2 | 15.0046796 | 0.30696722 | 0.4198771 | 0.73108827 | 0.46472524 | NA |
| YPEL1 | 5.79562176 | 0.156959626 | 0.72542962 | 0.21636782 | 0.82870105 | NA |
| MAPK1 | 91.603579 | 0.057829108 | 0.23211682 | 0.24913795 | 0.80325407 | 0.94126368 |
| PPM1F | 9.8743429 | -0.101086846 | 0.49994579 | -0.2021956 | 0.8397638 | NA |
| TOP3B | 5.41251542 | -0.444164848 | 0.56168177 | -0.7907767 | 0.42907432 | NA |
| AC245060.5 | 10.8146312 | 2.037199331 | 0.70533404 | 2.88827591 | 0.0038736 | NA |
| BCR | 21.6800201 | -0.134426085 | 0.44478535 | -0.3022269 | 0.76247912 | NA |
| CHCHD10 | 32.4704728 | -0.240546131 | 0.51746184 | -0.4648577 | 0.64203335 | 0.88951638 |
| SMARCB1 | 15.355804 | -0.196664787 | 0.35468977 | -0.5544699 | 0.57925733 | NA |
| DDT | 40.8468717 | -0.11223638 | 0.61387933 | -0.1828313 | 0.85493036 | 0.95578763 |
| CABIN1 | 41.9256814 | 0.107375947 | 0.29091553 | 0.36909665 | 0.71205568 | 0.91273377 |
| GUCD1 | 12.9204115 | -0.423199993 | 0.47840734 | -0.8846018 | 0.37637143 | NA |
| SNRPD3 | 33.224821 | -0.686203717 | 0.33024191 | -2.077882 | 0.03772023 | 0.30414888 |
| KIAA1671 | 70.2652382 | 0.971568005 | 0.31746121 | 3.06043063 | 0.00221019 | 0.07728042 |
| AL022323.4 | 5.72380729 | 1.106970774 | 0.71094355 | 1.55704454 | 0.11945991 | NA |
| LRP5L | 11.6325704 | 0.631393268 | 0.79844787 | 0.79077582 | 0.42907482 | NA |
| GRK3 | 90.7495899 | -0.250319731 | 0.58539792 | -0.4276061 | 0.66893791 | 0.90229869 |
| HPS4 | 12.4630712 | -0.353363441 | 0.50283584 | -0.7027412 | 0.48221707 | NA |
| SRRD | 6.02700867 | 0.595750715 | 0.61344191 | 0.97116077 | 0.33146823 | NA |
| TFIP11 | 8.60346217 | 0.922537604 | 0.55896569 | 1.6504369 | 0.09885361 | NA |
| TPST2 | 4.9779191 | 0.075347976 | 0.80184826 | 0.09396787 | 0.92513468 | NA |
| MIATNB | 16.5281661 | 0.762202931 | 0.46493348 | 1.6393806 | 0.10113402 | NA |
| PITPNB | 46.4551915 | -0.127962545 | 0.29735079 | -0.430342 | 0.66694685 | 0.90021179 |
| TTC28 | 214.104853 | 0.261770893 | 0.20291687 | 1.29004009 | 0.19703674 | 0.59987316 |
| HSCB | 13.0440995 | -0.772300902 | 0.50311511 | -1.5350382 | 0.12477445 | NA |
| CCDC117 | 10.484378 | -0.334482763 | 0.46892875 | -0.7132912 | 0.47566557 | NA |
| XBP1 | 31.7826282 | -0.97570679 | 0.54603215 | -1.7869036 | 0.07395308 | 0.41834543 |
| Z93930.2 | 37.4710025 | -0.638966082 | 0.37832304 | -1.6889431 | 0.09123034 | 0.4590946 |
| ZNRF3 | 542.654128 | 0.185069153 | 0.29289094 | 0.63187053 | 0.52747148 | 0.83195298 |
| ZNRF3-AS1 | 6.15780148 | -0.238783618 | 0.72432397 | -0.3296641 | 0.74165377 | NA |
| KREMEN1 | 25.8089653 | 0.488274212 | 0.56722438 | 0.86081316 | 0.38934096 | NA |
| RHBDD3 | 9.7405655 | -0.640197839 | 0.53552316 | -1.1954625 | 0.23190639 | NA |
| EWSR1 | 60.1728822 | 0.459424406 | 0.28974621 | 1.58560971 | 0.11282787 | 0.50413876 |
| GAS2L1 | 7.84088779 | -0.092745745 | 0.50151331 | -0.1849318 | 0.85328255 | NA |
| AP1B1 | 15.6503578 | -0.166054982 | 0.39206963 | -0.4235344 | 0.67190539 | NA |
| THOC5 | 5.6960137 | 0.135325755 | 0.59308241 | 0.22817361 | 0.81951128 | NA |
| NIPSNAP1 | 7.32855736 | 0.433983641 | 0.50872546 | 0.85308024 | 0.39361481 | NA |
| NF2 | 16.8631049 | 0.282938272 | 0.41813108 | 0.67667362 | 0.49861306 | NA |
| ZMAT5 | 4.89087585 | -0.224645866 | 0.55351092 | -0.4058562 | 0.68484823 | NA |
| UQCR10 | 82.8191846 | -0.273632966 | 0.48652614 | -0.5624219 | 0.57382859 | 0.85861279 |
| ASCC2 | 23.0966697 | 0.238254252 | 0.3829829 | 0.62210153 | 0.53387511 | NA |
| MTMR3 | 99.8211225 | 0.65553569 | 0.2321618 | 2.82361562 | 0.00474853 | 0.11177304 |
| SF3A1 | 13.4284447 | -0.203800127 | 0.53487542 | -0.3810235 | 0.70318578 | NA |
| RNF215 | 10.3889969 | -0.093343122 | 0.60592691 | -0.1540501 | 0.87757021 | NA |
| PES1 | 7.25978006 | 0.519878755 | 0.54216348 | 0.95889667 | 0.3376108 | NA |
| TCN2 | 8.23203329 | -0.206510474 | 1.21145478 | -0.1704649 | 0.86464456 | NA |
| DUSP18 | 9.28016948 | 0.435556363 | 0.54817984 | 0.79455014 | 0.42687523 | NA |
| OSBP2 | 17.3654894 | 0.34372636 | 0.50408364 | 0.68188359 | 0.49531257 | NA |
| MORC2 | 10.0494133 | -0.696334441 | 0.48793246 | -1.4271123 | 0.15354752 | NA |
| SELENOM | 38.1111761 | -1.46120367 | 0.58503755 | -2.4976237 | 0.01250288 | 0.18813325 |
| RNF185 | 22.4827962 | 0.009697722 | 0.4295305 | 0.02257749 | 0.9819873 | NA |
| LIMK2 | 57.2003708 | -0.549940514 | 0.36192554 | -1.5194852 | 0.1286404 | 0.52341115 |
| PIK3IP1 | 24.7966995 | -1.190912555 | 0.63759527 | -1.867819 | 0.0617873 | NA |
| PIK3IP1-AS1 | 4.91527224 | -0.279065689 | 0.61881482 | -0.450968 | 0.65201261 | NA |
| PATZ1 | 11.5643688 | 0.269481596 | 0.44282454 | 0.60855163 | 0.54282167 | NA |
| DRG1 | 22.8475797 | 0.012956871 | 0.3221487 | 0.04022016 | 0.96791761 | NA |
| EIF4ENIF1 | 17.3773568 | -0.199675385 | 0.36766997 | -0.5430832 | 0.58707252 | NA |
| SFI1 | 23.6793239 | 0.054884439 | 0.46411547 | 0.118256 | 0.90586482 | NA |
| PISD | 34.1457274 | -0.446328685 | 0.40620281 | -1.0987829 | 0.27186279 | 0.66496374 |
| PRR14L | 41.6223896 | 0.120782123 | 0.30139396 | 0.40074501 | 0.68860787 | 0.90557575 |
| YWHAH | 16.2584683 | -0.370217939 | 0.57939825 | -0.6389697 | 0.52284263 | NA |
| RTCB | 25.7462465 | -0.186459864 | 0.31054266 | -0.6004324 | 0.54821812 | NA |
| FBXO7 | 45.7164942 | 0.192841501 | 0.28486174 | 0.67696525 | 0.498428 | 0.82345406 |
| SYN3 | 21.7133589 | 0.283965217 | 0.60253213 | 0.47128643 | 0.6374362 | NA |
| TIMP3 | 268.892417 | 1.127810863 | 0.52195031 | 2.1607629 | 0.03071366 | 0.28264481 |
| LARGE1 | 664.824312 | -0.427574187 | 0.31185086 | -1.3710855 | 0.17034831 | 0.57276745 |
| HMGXB4 | 14.7111263 | 0.109594187 | 0.446024 | 0.24571366 | 0.8059039 | NA |
| TOM1 | 13.0291472 | -0.328386416 | 0.52927382 | -0.6204471 | 0.53496346 | NA |
| HMOX1 | 8.11136458 | -1.036332107 | 0.95680593 | -1.0831163 | 0.2787568 | NA |
| APOL6 | 39.5123495 | -0.680881468 | 0.66742391 | -1.0201634 | 0.30765096 | 0.68713231 |
| RBFOX2 | 124.302449 | 0.449161145 | 0.22851007 | 1.96560766 | 0.04934395 | 0.34686145 |
| APOL2 | 46.8777902 | -0.37513202 | 0.72372825 | -0.5183327 | 0.60422616 | 0.87047761 |
| APOL1 | 103.915922 | 0.027123843 | 0.58978107 | 0.04598968 | 0.96331848 | 0.98892249 |
| MYH9 | 461.955052 | 0.054246968 | 0.38439057 | 0.14112461 | 0.8877715 | 0.96743719 |
| TXN2 | 17.412019 | -0.012622192 | 0.39734758 | -0.0317661 | 0.97465856 | NA |
| EIF3D | 23.1463628 | -0.510306585 | 0.37555057 | -1.3588226 | 0.17420283 | NA |
| IFT27 | 9.92618278 | 0.291540849 | 0.43668015 | 0.66763018 | 0.50436969 | NA |
| TST | 18.9761632 | -0.353294999 | 0.55490465 | -0.6366769 | 0.52433529 | NA |
| MPST | 18.5824919 | -0.430552805 | 0.42825777 | -1.005359 | 0.31472402 | NA |
| C1QTNF6 | 5.21453832 | -2.265673044 | 0.75830967 | -2.9877939 | 0.00280999 | NA |
| CARD10 | 10.4935472 | 0.151560225 | 0.57011798 | 0.2658401 | 0.79036235 | NA |
| CDC42EP1 | 11.8595999 | -0.455611735 | 0.64634497 | -0.7049049 | 0.48086942 | NA |
| GGA1 | 20.1325481 | -0.024804595 | 0.38421091 | -0.0645598 | 0.94852446 | NA |
| LGALS1 | 203.944424 | -1.655485621 | 0.53866403 | -3.0733176 | 0.00211693 | 0.07466323 |
| TRIOBP | 32.8003686 | -0.150991576 | 0.35501932 | -0.4253052 | 0.67061418 | 0.90305784 |
| H1F0 | 29.853755 | -0.578235515 | 0.87754537 | -0.6589238 | 0.50994471 | 0.82899228 |
| ANKRD54 | 4.22309058 | -0.24133718 | 0.71568882 | -0.3372097 | 0.73595886 | NA |
| EIF3L | 22.0829544 | 0.314659123 | 0.3770576 | 0.83451209 | 0.40399248 | NA |
| POLR2F | 12.9893695 | 0.11292515 | 0.45058434 | 0.25061934 | 0.80210843 | NA |
| PICK1 | 8.05395342 | -0.197614922 | 0.53562527 | -0.3689425 | 0.71217059 | NA |
| PLA2G6 | 6.08845139 | 1.086939504 | 0.65220555 | 1.66655972 | 0.09560198 | NA |
| MAFF | 4.68143158 | -1.579149918 | 1.05574815 | -1.4957638 | 0.13471521 | NA |
| TMEM184B | 16.7132048 | 0.036764701 | 0.40508585 | 0.0907578 | 0.92768504 | NA |
| CSNK1E | 12.0915372 | 1.03368186 | 0.52021576 | 1.98702527 | 0.0469196 | NA |
| TPTEP2-CSNI | 41.0295063 | 0.22263105 | 0.29844761 | 0.74596359 | 0.45568941 | 0.79324789 |
| KDELR3 | 16.3810688 | -0.112398888 | 0.71692477 | -0.1567792 | 0.87541887 | NA |
| DDX17 | 375.140183 | 0.524834893 | 0.19124332 | 2.74433058 | 0.00606344 | 0.12229157 |
| FAM227A | 6.19289832 | 0.272887275 | 0.73812863 | 0.36970152 | 0.7116049 | NA |
| CBY1 | 15.2493547 | -0.546280831 | 0.37474254 | -1.4577497 | 0.14490954 | NA |
| TOMM22 | 11.168606 | 0.393899341 | 0.54450756 | 0.72340472 | 0.46943127 | NA |
| JOSD1 | 28.0210208 | 0.073435227 | 0.32178894 | 0.22820929 | 0.81948354 | NA |
| GTPBP1 | 24.8430675 | -0.528758102 | 0.39014867 | -1.3552734 | 0.17533048 | NA |
| SUN2 | 95.5409852 | -0.405805829 | 0.49006392 | -0.8280671 | 0.40763248 | 0.76772515 |
| AL021707.1 | 10.474976 | 0.546868426 | 0.51564328 | 1.06055572 | 0.28889186 | NA |
| CBX6 | 46.4209063 | -0.493523831 | 0.48124162 | -1.0255219 | 0.30511698 | 0.6848669 |
| APOBEC3G | 4.69632547 | -0.308041911 | 1.45318588 | -0.211977 | 0.83212502 | NA |
| CBX7 | 31.2591631 | -0.58265838 | 0.53764582 | -1.0837216 | 0.27848826 | 0.66958859 |
| AL031846.2 | 4.80290582 | -1.01826331 | 0.81080374 | -1.255869 | 0.20916346 | NA |

| | | | | | | |
|---|---|---|---|---|---|---|
| RPL3 | 62.1961521 | -0.369355674 | 0.44430583 | -0.8313095 | 0.40579879 | 0.76772515 |
| TAB1 | 11.889695 | 0.086659891 | 0.41717259 | 0.20773151 | 0.83543861 | NA |
| MIEF1 | 10.1384422 | 0.225953665 | 0.46941632 | 0.48135025 | 0.63026759 | NA |
| ATF4 | 58.7478701 | -0.451846333 | 0.37732814 | -1.1974891 | 0.23111598 | 0.63682501 |
| RPS19BP1 | 32.6531439 | -0.647104159 | 0.35537863 | -1.8208865 | 0.06862411 | 0.40491695 |
| TNRC6B | 204.784699 | 0.17755589 | 0.22441506 | 0.79119417 | 0.42883069 | 0.7784723 |
| MKL1 | 122.484654 | 0.767102012 | 0.29775564 | 2.57628037 | 0.00998696 | 0.15885146 |
| SLC25A17 | 18.6831449 | 0.34314975 | 0.3943043 | 0.87026632 | 0.38415488 | NA |
| ST13 | 60.2518298 | -0.126597657 | 0.27358763 | -0.4627317 | 0.64355672 | 0.8902681 |
| XPNPEP3 | 26.6223955 | 0.406379919 | 0.32584909 | 1.24714149 | 0.21234562 | NA |
| RBX1 | 45.219307 | 0.108748669 | 0.34570829 | 0.31456772 | 0.75308989 | 0.92313949 |
| EP300 | 77.6532747 | 0.14881258 | 0.28314083 | 0.52557795 | 0.59918148 | 0.86882737 |
| L3MBTL2 | 9.01057055 | 0.560608947 | 0.50611792 | 1.10766468 | 0.26800666 | NA |
| RANGAP1 | 44.0947364 | 0.135105434 | 0.67585693 | 0.19990242 | 0.8415569 | 0.95398401 |
| ZC3H7B | 65.0219742 | 0.135783446 | 0.43790421 | 0.31007568 | 0.75650341 | 0.92313949 |
| TEF | 40.6750334 | 0.75071549 | 0.46275742 | 1.6222657 | 0.10474647 | 0.4862279 |
| TOB2 | 20.7119494 | -0.441520839 | 0.40038959 | -1.1027281 | 0.27014527 | NA |
| PHF5A | 7.47063594 | -1.773258712 | 0.59624488 | -2.9740443 | 0.00293903 | NA |
| ACO2 | 26.9690537 | -0.009020152 | 0.30470927 | -0.0296025 | 0.97638408 | NA |
| POLR3H | 6.62129909 | -0.481733805 | 0.58178897 | -0.8280216 | 0.4076583 | NA |
| CSDC2 | 36.3550568 | 0.878613438 | 0.35431889 | 2.47972507 | 0.01314837 | 0.1918338 |
| PMM1 | 11.1255036 | -0.507660597 | 0.51835078 | -0.9793765 | 0.32739396 | NA |
| DESI1 | 14.0097874 | -0.434223782 | 0.48688042 | -0.8918489 | 0.37247391 | NA |
| XRCC6 | 57.5986518 | -0.153959209 | 0.34656855 | -0.4442388 | 0.65686992 | 0.89675678 |
| SNU13 | 49.5683104 | -0.999425616 | 0.34377479 | -2.9072103 | 0.00364668 | 0.09771848 |
| SREBF2 | 74.9983105 | -0.060973956 | 0.27756122 | -0.2196775 | 0.82612232 | 0.949142 |
| WBP2NL | 17.6170214 | -0.324759574 | 0.46270445 | -0.7018726 | 0.48275862 | NA |
| NAGA | 7.54906316 | -0.184547496 | 0.61032648 | -0.302375 | 0.76236618 | NA |
| SMDT1 | 25.3101826 | -0.643649572 | 0.48677703 | -1.3222678 | 0.186079 | NA |
| NDUFA6 | 22.672313 | 0.335606325 | 0.4680425 | 0.71704241 | 0.47334793 | NA |
| TCF20 | 33.640141 | -0.04464984 | 0.28274695 | -0.1579145 | 0.87452418 | 0.96229085 |
| RRP7A | 12.6844063 | -0.172667251 | 0.64078148 | -0.2694635 | 0.78757299 | NA |
| POLDIP3 | 15.3158206 | -0.052651922 | 0.40444761 | -0.1301823 | 0.89642219 | NA |
| CYB5R3 | 85.6392545 | -0.337698523 | 0.37322782 | -0.9048053 | 0.36556852 | 0.73403264 |
| A4GALT | 20.1238163 | -0.069426427 | 0.48592239 | -0.1428755 | 0.88638847 | NA |
| ARFGAP3 | 34.1823203 | -0.853687665 | 0.33121 | -2.5774815 | 0.00995232 | 0.15885146 |
| PACSIN2 | 36.5546876 | -0.132731902 | 0.29064042 | -0.4566877 | 0.64789554 | 0.89261695 |
| TTLL1 | 4.67972434 | -0.105358308 | 0.68524021 | -0.1537538 | 0.87780384 | NA |
| MCAT | 5.11532037 | -0.618301731 | 0.5846122 | -1.0576271 | 0.29022546 | NA |
| TSPO | 48.6828918 | -0.265424707 | 0.51860015 | -0.5118099 | 0.60878404 | 0.8725395 |
| TTLL12 | 6.85301112 | 0.361980947 | 0.5802345 | 0.62385285 | 0.53272423 | NA |
| SCUBE1 | 4.86118496 | -2.172109606 | 0.88671911 | -2.4496028 | 0.01430139 | NA |
| EFCAB6 | 22.197317 | 0.147293722 | 0.43167828 | 0.3412118 | 0.73294414 | NA |
| SAMM50 | 5.35561266 | -0.706167457 | 0.56836045 | -1.2424641 | 0.21406538 | NA |
| RTL6 | 6.44064519 | 0.106183904 | 0.55195249 | 0.1923787 | 0.84744557 | NA |
| PHF21B | 74.733749 | 0.482025814 | 0.42645652 | 1.13030471 | 0.25834785 | 0.65588339 |
| NUP50-DT | 24.8596019 | -0.702400235 | 0.4296563 | -1.6347956 | 0.1020919 | NA |
| NUP50 | 16.4599184 | -0.411360197 | 0.37238217 | -1.1046721 | 0.26930169 | NA |
| KIAA0930 | 13.8775529 | 0.327061868 | 0.66045066 | 0.49521015 | 0.62045179 | NA |
| FAM118A | 15.536712 | 0.751397426 | 0.61606179 | 1.21967867 | 0.22258671 | NA |
| ATXN10 | 132.129149 | 0.296120137 | 0.20447303 | 1.44821119 | 0.147558 | 0.54657099 |
| MIRLET7BHG | 30.6621639 | -0.23252443 | 0.32242045 | -0.7211839 | 0.47079639 | 0.80448595 |
| PPARA | 41.8467183 | 0.226380374 | 0.36705482 | 0.61674814 | 0.53740086 | 0.83677705 |
| TTC38 | 13.1995941 | -0.200794618 | 0.48884154 | -0.410756 | 0.68125142 | NA |
| TRMU | 11.6812885 | -0.32976881 | 0.43398659 | -0.7598595 | 0.4473386 | NA |
| CELSR1 | 9.43971299 | 1.828708058 | 0.81057079 | 2.25607445 | 0.02406597 | NA |
| GRAMD4 | 9.48989511 | -0.250737957 | 0.57034151 | -0.4396278 | 0.66020674 | NA |
| CERK | 35.9116343 | -0.221850916 | 0.47571736 | -0.4663503 | 0.6409648 | 0.88880453 |
| AL118516.1 | 4.08767517 | -0.91426127 | 0.751663 | -1.2163181 | 0.22386379 | NA |
| TBC1D22A | 110.141032 | -0.167502084 | 0.26335909 | -0.6360217 | 0.52476231 | 0.83195298 |
| BRD1 | 21.8884689 | -0.34011588 | 0.36053457 | -0.9433655 | 0.34549398 | NA |
| ZBED4 | 11.0817968 | -0.322686127 | 0.53191353 | -0.6066515 | 0.54408223 | NA |
| ALG12 | 7.30801423 | -0.552517305 | 0.53979839 | -1.0235624 | 0.30604204 | NA |
| CRELD2 | 11.7992063 | -0.346655378 | 0.6598976 | -0.5253169 | 0.59936289 | NA |
| PIM3 | 35.017974 | -0.27243074 | 0.38763607 | -0.7028003 | 0.48218022 | 0.81285244 |
| TRABD | 10.4630064 | 0.23515749 | 0.5212713 | 0.45112303 | 0.65190088 | NA |
| SELENOO | 16.5182235 | -0.66223924 | 0.39675096 | -1.669156 | 0.09508647 | NA |
| TUBGCP6 | 4.47237504 | 0.448264118 | 0.66723477 | 0.67182368 | 0.50169595 | NA |
| PLXNB2 | 134.394687 | -0.102988614 | 0.42769636 | -0.2407984 | 0.80971134 | 0.94346479 |
| DENND6B | 14.5718016 | 0.771542972 | 0.56197091 | 1.37292333 | 0.16977618 | NA |
| PPP6R2 | 59.7835853 | 0.387245656 | 0.25960385 | 1.49167917 | 0.13578327 | 0.52829049 |
| SBF1 | 5.53104172 | -0.037635712 | 0.65258626 | -0.0576716 | 0.95401019 | NA |
| MIOX | 13.2178821 | 2.226713534 | 1.0683194 | 2.08431442 | 0.03713158 | NA |
| LMF2 | 11.4875377 | -0.225369616 | 0.61350039 | -0.3673504 | 0.71335766 | NA |
| NCAPH2 | 6.24684833 | -0.044113207 | 0.69037267 | -0.0638977 | 0.94905171 | NA |
| ARSA | 14.1618211 | 0.484135537 | 0.48680999 | 0.99450617 | 0.3199765 | NA |
| RABL2B | 9.77993668 | -0.519861801 | 0.57120088 | -0.9101208 | 0.36275881 | NA |
| FP671120.1 | 331.338542 | -1.193084978 | 0.51904164 | -2.2986306 | 0.02152593 | 0.24262159 |
| FP236383.1 | 203.727624 | -1.001624156 | 0.55349536 | -1.8096342 | 0.07035253 | 0.40663921 |
| CR381653.1 | 13.6718074 | 0.096899315 | 0.7075791 | 0.13694485 | 0.89107439 | NA |
| AP001347.1 | 12.6990209 | 1.227444063 | 0.4952592 | 2.47838719 | 0.01319778 | NA |
| RBM11 | 6.91261862 | 0.513729547 | 0.58246757 | 0.88198824 | 0.37778317 | NA |
| HSPA13 | 38.0650337 | -0.123747966 | 0.48821992 | -0.2534677 | 0.79990684 | 0.94085969 |
| AF127577.4 | 7.10795278 | 0.589567537 | 0.63814634 | 0.92387514 | 0.35555133 | NA |
| NRIP1 | 77.3845131 | -0.188874655 | 0.26136534 | -0.7226461 | 0.46989732 | 0.80448595 |
| AJ009632.2 | 13.7268562 | 1.08117227 | 0.84025306 | 1.28672221 | 0.19819113 | NA |
| USP25 | 115.138305 | 0.15244907 | 0.28690175 | 0.53136334 | 0.59516702 | 0.86653654 |
| MIR99AHG | 111.595359 | 1.338841378 | 0.34139498 | 3.92167854 | 8.79E-05 | 0.00914516 |
| CXADR | 84.4746334 | -0.149570872 | 0.48866599 | -0.30608 | 0.75954376 | 0.92313949 |
| BTG3 | 52.6025173 | -1.455996141 | 0.49162033 | -2.9616272 | 0.00306018 | 0.0904424 |
| C21orf91 | 7.30249175 | -0.001946233 | 0.64454095 | -0.0030196 | 0.99759074 | NA |
| MRPL39 | 7.7350104 | -0.481161694 | 0.5591695 | -0.8604934 | 0.38951709 | NA |
| ATP5PF | 41.7558131 | -0.294803677 | 0.40606366 | -0.7260036 | 0.46783657 | 0.80370399 |
| GABPA | 17.7787685 | 0.359146785 | 0.5911031 | 0.60758738 | 0.54346117 | NA |
| APP | 193.944556 | 0.16837924 | 0.20342955 | 0.82770294 | 0.40783876 | 0.76772515 |
| ADAMTS1 | 53.2236289 | -2.070559627 | 0.88630039 | -2.3361827 | 0.01948172 | 0.23378067 |
| ADAMTS5 | 29.8121279 | 1.750614798 | 2.0628506 | 0.84863868 | 0.39608238 | 0.75911719 |
| AF165147.1 | 16.9076085 | -1.654294821 | 0.81123008 | -2.0392425 | 0.04142584 | NA |
| N6AMT1 | 6.80691687 | -0.042771147 | 0.5754992 | -0.0743201 | 0.9407557 | NA |
| LTN1 | 28.8367741 | 0.031683524 | 0.37235027 | 0.08509064 | 0.93218933 | 0.97832857 |
| USP16 | 32.1039302 | -0.08377504 | 0.34125284 | -0.2454926 | 0.80607505 | 0.94274522 |
| CCT8 | 20.5094837 | 0.081864992 | 0.32535668 | 0.25161614 | 0.80133778 | NA |
| MAP3K7CL | 9.44713573 | 0.278108782 | 0.61597037 | 0.45149701 | 0.65163138 | NA |
| BACH1 | 321.633547 | -0.048152937 | 0.28816352 | -0.1671028 | 0.86728915 | 0.96089268 |
| GRIK1 | 49.3126004 | -0.620819745 | 0.51829515 | -1.1978112 | 0.23099052 | 0.63682501 |
| GRIK1-AS1 | 18.3735111 | 0.329894988 | 0.5301799 | 0.62223216 | 0.53378922 | NA |
| SOD1 | 78.6985395 | -0.370834059 | 0.29419023 | -1.2605247 | 0.20748014 | 0.61696441 |
| SCAF4 | 29.4641029 | -0.025741149 | 0.31995 | -0.0804537 | 0.93587645 | 0.97832857 |
| URB1 | 10.723348 | -0.419289216 | 0.53095102 | -0.7896947 | 0.42970607 | NA |
| EVA1C | 36.723099 | -1.23200249 | 0.41002287 | -3.0047165 | 0.00265828 | 0.08423439 |
| SYNJ1 | 18.2442192 | 0.885224283 | 0.34789384 | 2.54452419 | 0.01094267 | NA |
| PAXBP1 | 33.2471071 | 0.292709881 | 0.3292733 | 0.88895723 | 0.37402607 | 0.74183361 |
| C21orf62-AS | 8.12639799 | -0.214418198 | 0.57412175 | -0.3734716 | 0.70879744 | NA |

| | | | | | | |
|---|---|---|---|---|---|---|
| IL10RB | 3.73287846 | 0.325655331 | 0.68487188 | 0.47549818 | 0.63443193 | NA |
| IFNAR1 | 54.0509124 | -0.440155744 | 0.35259773 | -1.2483227 | 0.21191291 | 0.62148862 |
| IFNGR2 | 25.7540105 | -1.042493831 | 0.36874935 | -2.8271069 | 0.00469706 | NA |
| TMEM50B | 43.3185744 | 0.1567379 | 0.28451383 | 0.5508973 | 0.58170408 | 0.86175324 |
| GART | 18.2252001 | -0.026599828 | 0.52654282 | -0.0505179 | 0.9597097 | NA |
| SON | 124.415419 | -0.258579128 | 0.24925511 | -1.0374075 | 0.29954597 | 0.68168772 |
| DONSON | 6.38292238 | -0.147327777 | 0.60657851 | -0.2428833 | 0.80809582 | NA |
| AP000311.1 | 15.3532245 | -0.3432225 | 0.59562191 | -0.5762422 | 0.56445146 | NA |
| ITSN1 | 59.2484516 | 0.281093401 | 0.34068533 | 0.8250822 | 0.40932492 | 0.76772515 |
| LINC00649 | 16.582221 | -0.054536857 | 0.39413626 | -0.1383706 | 0.88994756 | NA |
| MRPS6 | 62.7813645 | 0.532528427 | 0.35126623 | 1.51602514 | 0.129513 | 0.52613957 |
| LINC00310 | 7.861009 | 1.227395888 | 0.7053351 | 1.74015995 | 0.08183094 | NA |
| AP000317.1 | 9.11973703 | 0.55432181 | 0.57432358 | 0.96517334 | 0.334458 | NA |
| RCAN1 | 99.2656441 | -0.180628415 | 0.42996899 | -0.4200964 | 0.67441505 | 0.90403541 |
| RUNX1 | 47.8946594 | 0.766419635 | 0.69011683 | 1.11056506 | 0.26675561 | 0.66013596 |
| SETD4 | 11.0622533 | 0.360967389 | 0.43572027 | 0.82843837 | 0.40742228 | NA |
| CBR1 | 19.0142953 | -0.715279995 | 0.5144359 | -1.3904162 | 0.16440254 | NA |
| DOPEY2 | 12.2965516 | 0.454927633 | 0.49687587 | 0.91557602 | 0.35988931 | NA |
| MORC3 | 34.4773049 | 0.142326661 | 0.26494185 | 0.53719961 | 0.59112974 | 0.8654248 |
| HLCS | 57.7363439 | 0.053568949 | 0.32883435 | 0.16290558 | 0.87059278 | 0.96207879 |
| PIGP | 10.1659266 | -0.20182957 | 0.6292655 | -0.3207383 | 0.74840869 | NA |
| TTC3 | 100.329453 | 0.240950546 | 0.29260887 | 0.82345605 | 0.4102487 | 0.76788553 |
| VPS26C | 26.0395532 | -0.160457479 | 0.29675932 | -0.540699 | 0.58871504 | NA |
| DYRK1A | 147.410074 | 0.215961821 | 0.21725308 | 0.99405643 | 0.3201954 | 0.69496685 |
| KCNJ15 | 36.0011126 | 2.182003995 | 0.83378073 | 2.61699978 | 0.00887064 | 0.14875902 |
| ETS2 | 95.0858529 | -1.613182538 | 0.47099272 | -3.4250689 | 0.00061464 | 0.0356142 |
| PSMG1 | 7.84090114 | -0.244118955 | 0.46805454 | -0.5215609 | 0.60197608 | NA |
| BRWD1 | 95.7747796 | 0.013807481 | 0.24269922 | 0.05689133 | 0.95463176 | 0.98573992 |
| HMGN1 | 97.3388654 | 0.180350483 | 0.28757103 | 0.62715109 | 0.5305602 | 0.83515914 |
| WRB | 33.0519402 | 0.30337109 | 0.44953516 | 0.67485508 | 0.49976784 | 0.82345406 |
| LCA5L | 6.0828246 | 0.035425337 | 0.80418186 | 0.0440514 | 0.96486343 | NA |
| SH3BGR | 5.86834692 | 0.519759613 | 1.04766929 | 0.49611038 | 0.61981653 | NA |
| BACE2 | 21.9766368 | -0.452759615 | 0.80643938 | -0.5614304 | 0.57450414 | NA |
| MX1 | 83.7706671 | -2.188259813 | 1.94311073 | -1.1261632 | 0.26009644 | 0.65595635 |
| C2CD2 | 9.46774422 | 0.379898111 | 0.45842546 | 0.82870203 | 0.40727304 | NA |
| ZBTB21 | 13.4225266 | -0.307347744 | 0.4502753 | -0.6825774 | 0.49487392 | NA |
| SLC37A1 | 8.16712435 | -0.566838833 | 0.60681137 | -0.9341269 | 0.35023843 | NA |
| PDE9A | 11.4202007 | 0.551135155 | 0.71129086 | 0.77483796 | 0.43843542 | NA |
| WDR4 | 4.47780585 | -0.56445899 | 0.98140956 | -0.5751513 | 0.56518897 | NA |
| NDUFV3 | 26.9745576 | -0.12619454 | 0.3592394 | -0.3512826 | 0.72537637 | NA |
| PKNOX1 | 31.3028094 | 0.315742637 | 0.27159648 | 1.16254319 | 0.24501489 | 0.64996527 |
| RRP1B | 10.634374 | -0.893536665 | 0.44685684 | -1.9996038 | 0.04554306 | NA |
| PDXK | 46.5241013 | -0.100989407 | 0.30791401 | -0.3279793 | 0.74292735 | 0.92122771 |
| CSTB | 75.7762611 | -0.523893103 | 0.42184816 | -1.2418997 | 0.21427357 | 0.62255988 |
| RRP1 | 4.35888297 | -0.842355394 | 0.73551231 | -1.1452635 | 0.25210003 | NA |
| AGPAT3 | 47.5495847 | 0.382637967 | 0.25350304 | 1.50940188 | 0.13119611 | 0.52613957 |
| TRAPPC10 | 74.0170967 | 0.229246544 | 0.27039411 | 0.84782372 | 0.39653615 | 0.75911719 |
| PFKL | 38.6653692 | -0.033124523 | 0.38909522 | -0.0851322 | 0.93215631 | 0.97832857 |
| C21orf2 | 16.8050717 | 0.44673254 | 0.54677805 | 0.8170272 | 0.41391289 | NA |
| TSPEAR | 16.1711942 | 2.236584813 | 0.81011568 | 2.76082153 | 0.00576562 | NA |
| UBE2G2 | 43.324324 | 0.240187104 | 0.26392682 | 0.9100519 | 0.36279514 | 0.73259935 |
| SUMO3 | 27.1014269 | 0.018305202 | 0.40104357 | 0.04564392 | 0.96359406 | NA |
| PTTG1IP | 81.579178 | -0.426577383 | 0.26306321 | -1.6215775 | 0.10489384 | 0.4862279 |
| FAM207A | 6.79338396 | 0.470169214 | 0.72293604 | 0.65036073 | 0.51545923 | NA |
| ADARB1 | 10.1826029 | -0.211994177 | 0.67672242 | -0.3132661 | 0.75407852 | NA |
| POFUT2 | 17.4732012 | -0.009140281 | 0.45875214 | -0.0199242 | 0.98410382 | NA |
| COL18A1 | 84.2909433 | -0.448376005 | 0.36090858 | -1.2423534 | 0.2141062 | 0.62255988 |
| COL6A1 | 11.3290321 | 0.511378316 | 0.41909254 | 1.22020382 | 0.22238762 | NA |
| FTCD | 6.38623728 | 1.148841906 | 0.84439482 | 1.36055064 | 0.17365574 | NA |
| LSS | 11.3311923 | -0.459267732 | 0.49656748 | -0.9248848 | 0.35502581 | NA |
| MCM3AP | 28.1831875 | 0.123083501 | 0.30524471 | 0.40322894 | 0.68677981 | NA |
| YBEY | 4.85013818 | -0.005973415 | 0.68721836 | -0.0086922 | 0.99306474 | NA |
| PCNT | 16.2699026 | -0.183967931 | 0.38017927 | -0.4838978 | 0.62845837 | NA |
| DIP2A | 31.1277216 | -0.388505548 | 0.29568885 | -1.3138999 | 0.18887989 | 0.59077076 |
| PRMT2 | 89.3690477 | -0.272129405 | 0.28070387 | -0.9694537 | 0.33231887 | 0.70606881 |
| AL592183.1 | 46.6001026 | -0.392842071 | 0.50424252 | -0.7790737 | 0.43593631 | 0.78167891 |

**Proximal tubules**

*This means that positive fold changes imply higher mean value in controls than in AKI and vice versa.

| | baseMean | log2 fold change control/AKI* | lfcSE | stat | pvalue | padj |
|---|---|---|---|---|---|---|
| AL627309.1 | 46.0856473 | 0.328401751 | 0.35916013 | 0.91436027 | 0.36052757 | 0.5586436 |
| AL669831.5 | 349.587562 | -0.039380253 | 0.30188362 | -0.1304485 | 0.89621163 | 0.94618908 |
| NOC2L | 274.350881 | 0.109286339 | 0.32335825 | 0.33797294 | 0.73538358 | 0.85033112 |
| KLHL17 | 55.733763 | 0.01271656 | 0.39229035 | 0.03241619 | 0.97414015 | 0.98702979 |
| PLEKHN1 | 47.4948433 | -4.2041694 | 0.6186955 | -6.7952157 | 1.08E-11 | 2.32E-09 |
| HES4 | 116.388637 | -3.339192016 | 0.90684104 | -3.6822242 | 0.00023121 | 0.00300395 |
| ISG15 | 1169.63657 | -1.43659451 | 1.60885161 | -0.8929316 | 0.37189378 | 0.56932302 |
| AGRN | 499.353221 | -1.299851061 | 0.3727027 | -3.4876352 | 0.00048731 | 0.00526046 |
| C1orf159 | 304.992355 | -0.125810908 | 0.37147089 | -0.3386831 | 0.73484848 | 0.84993403 |
| LINC01342 | 94.1721291 | 0.83000054 | 0.36588475 | 2.26847533 | 0.02330025 | 0.09174958 |
| AL390719.2 | 109.671187 | 0.164844385 | 0.39046863 | 0.42217063 | 0.67290048 | 0.81056744 |
| SDF4 | 473.687245 | -0.385747553 | 0.27579935 | -1.3986529 | 0.1619171 | 0.33813809 |
| B3GALT6 | 61.0958485 | -0.564774304 | 0.46735755 | -1.2084416 | 0.22687745 | 0.41931165 |
| C1QTNF12 | 39.0858734 | 1.636375439 | 0.3641958 | 4.49311996 | 7.02E-06 | 0.0001889 |
| UBE2J2 | 715.129567 | -0.581353441 | 0.33890287 | -1.7153984 | 0.0862722 | 0.22293723 |
| LINC01786 | 36.2560674 | 1.722454827 | 0.49180535 | 3.50231006 | 0.00046124 | 0.0050578 |
| SCNN1D | 51.6699071 | 2.116303532 | 0.42833382 | 4.94078088 | 7.78E-07 | 3.15E-05 |
| ACAP3 | 339.851362 | -0.068597586 | 0.35665184 | -0.1923377 | 0.84747771 | 0.91938085 |
| PUSL1 | 42.683084 | -1.500231457 | 0.51692368 | -2.9022301 | 0.00370516 | 0.02417188 |
| INTS11 | 351.174265 | -0.054298037 | 0.23401573 | -0.2320273 | 0.81651681 | 0.90083699 |
| CPTP | 78.8397927 | -0.898448224 | 0.41301194 | -2.1753565 | 0.02960341 | 0.1080623 |
| DVL1 | 157.554134 | -0.648061612 | 0.29933011 | -2.1650398 | 0.03038464 | 0.11018048 |
| AURKAIP1 | 459.348943 | -0.72928348 | 0.44884904 | -1.6247856 | 0.10420824 | 0.25231575 |
| CCNL2 | 1393.10989 | -0.051534152 | 0.2708949 | -0.1902367 | 0.84912365 | 0.92058356 |
| MRPL20 | 450.607117 | 0.027090282 | 0.29876717 | 0.09067356 | 0.92775198 | 0.96193661 |
| VWA1 | 45.0710487 | -0.059964749 | 0.54542189 | -0.109942 | 0.91245541 | 0.9535692 |
| ATAD3B | 296.151961 | -0.320810353 | 0.41114691 | -0.7802816 | 0.43522517 | 0.62763087 |
| ATAD3A | 172.913781 | -0.36970949 | 0.50217398 | -0.7362179 | 0.46159808 | 0.65000256 |
| SSU72 | 531.394181 | -0.340366886 | 0.38165126 | -0.891827 | 0.37248568 | 0.57001787 |
| MIB2 | 236.600328 | 0.673805156 | 0.30780943 | 2.18903348 | 0.02859441 | 0.10561214 |
| CDK11B | 316.680707 | -0.203888472 | 0.45920434 | -0.4440038 | 0.65703984 | 0.8002528 |
| SLC35E2B | 341.622686 | 0.311030131 | 0.25085195 | 1.23989519 | 0.21501416 | 0.40555926 |
| CDK11A | 189.110388 | -0.609130502 | 0.47316054 | -1.2873654 | 0.19796696 | 0.38408112 |
| SLC35E2A | 270.882248 | 0.668143645 | 0.17105258 | 3.90607175 | 9.38E-05 | 0.00147967 |
| NADK | 495.685555 | -0.135439214 | 0.35163549 | -0.3851693 | 0.70011195 | 0.82770696 |
| GNB1 | 3046.74651 | -0.727555726 | 0.34492501 | -2.1093156 | 0.03491735 | 0.12156255 |
| PRKCZ | 1302.21601 | 0.365016718 | 0.55501048 | 0.65767536 | 0.51074676 | 0.68993848 |
| FAAP20 | 393.375962 | 0.390756847 | 0.25121846 | 1.55544639 | 0.11983979 | 0.27702309 |
| AL590822.1 | 117.713718 | 4.725358359 | 0.84412689 | 5.59792422 | 2.17E-08 | 1.58E-06 |
| SKI | 814.23026 | 0.238846664 | 0.45437498 | 0.52565981 | 0.59912459 | 0.7589133 |
| MORN1 | 383.651251 | 0.807742132 | 0.22714819 | 3.55601402 | 0.00037652 | 0.00438957 |
| RER1 | 542.356114 | -0.362650707 | 0.231771 | -1.5646941 | 0.11765465 | 0.27360212 |
| PEX10 | 137.57292 | -0.565286129 | 0.34807942 | -1.6240148 | 0.10437265 | 0.25237302 |
| PLCH2 | 328.615729 | 3.158308208 | 0.82391914 | 3.83327449 | 0.00012645 | 0.0018711 |
| PANK4 | 64.1561829 | 0.37697215 | 0.21457664 | 1.75681822 | 0.0789488 | 0.20996336 |
| AL139246.5 | 38.6613966 | -1.317988054 | 0.60688261 | -2.1717348 | 0.02987567 | 0.10885324 |
| TNFRSF14-A | 90.456375 | -0.80077351 | 0.39257559 | -2.0397944 | 0.0413708 | 0.13664328 |
| TNFRSF14 | 1183.56461 | 0.434281346 | 0.43732713 | 0.99303546 | 0.32069268 | 0.52113035 |
| FAM213B | 60.594743 | -0.506507905 | 0.45146617 | -1.1219177 | 0.26189742 | 0.45999812 |
| ARHGEF16 | 269.400766 | 0.382341244 | 0.4557997 | 0.8388361 | 0.40156129 | 0.59575718 |
| MEGF6 | 41.4678717 | 0.507953943 | 0.62557469 | 0.81197969 | 0.41680329 | 0.61024596 |
| TPRG1L | 224.454408 | 0.512555763 | 0.27382149 | 1.87186101 | 0.06122584 | 0.17732818 |
| WRAP73 | 121.287539 | 0.129283094 | 0.23847587 | 0.54212234 | 0.58773423 | 0.7503095 |
| SMIM1 | 39.9886605 | -0.085512139 | 0.61743722 | -0.1384953 | 0.889849 | 0.94303079 |
| LRRC47 | 201.055471 | 0.518971975 | 0.29158746 | 1.7798158 | 0.07510611 | 0.20349466 |
| CEP104 | 625.841361 | 0.155260713 | 0.29535197 | 0.52568031 | 0.59911035 | 0.7589133 |
| DFFB | 141.907623 | 1.489039905 | 0.45377505 | 3.28144945 | 0.00103275 | 0.00926827 |
| C1orf174 | 137.493628 | 0.181084261 | 0.36382777 | 0.49771973 | 0.6186816 | 0.77331444 |
| LINC01134 | 83.5001576 | 0.322194272 | 0.47980681 | 0.67150834 | 0.50189675 | 0.68289644 |
| AJAP1 | 264.36015 | 0.507536124 | 0.39208265 | 1.29446207 | 0.19550586 | 0.3808166 |
| NPHP4 | 261.881381 | 0.055247129 | 0.28302781 | 0.19520035 | 0.8452361 | 0.91854292 |
| KCNAB2 | 596.067903 | 2.351223846 | 0.61431072 | 3.8274179 | 0.00012949 | 0.00190245 |
| RPL22 | 333.392476 | -0.218057049 | 0.33919935 | -0.6428581 | 0.52031618 | 0.69607397 |
| RNF207 | 310.621842 | -0.249970754 | 0.38915898 | -0.6423358 | 0.52065516 | 0.69645948 |
| ICMT | 260.501815 | -0.225751935 | 0.23770478 | -0.9497156 | 0.34225679 | 0.54278767 |
| GPR153 | 122.554968 | -0.960272863 | 0.6388947 | -1.5030221 | 0.13283334 | 0.29622814 |
| ACOT7 | 324.982747 | 1.011537874 | 0.49908612 | 2.02678022 | 0.04268489 | 0.13956918 |
| ESPN | 617.738708 | 1.91075601 | 0.33814634 | 5.65067775 | 1.60E-08 | 1.22E-06 |
| TNFRSF25 | 35.3972948 | 0.488223432 | 0.5475762 | 0.8916082 | 0.37260298 | 0.57001787 |
| PLEKHG5 | 62.1323316 | 0.092892533 | 0.36543927 | 0.25419417 | 0.79934555 | 0.89091281 |
| NOL9 | 421.535283 | 0.370484236 | 0.3023249 | 1.22545062 | 0.22040549 | 0.41226774 |
| ZBTB48 | 300.695668 | 0.855918913 | 0.27230143 | 3.14327734 | 0.00167067 | 0.01334602 |
| KLHL21 | 268.513745 | -1.273777571 | 0.54993533 | -2.3162316 | 0.02054563 | 0.08414838 |
| PHF13 | 96.6520075 | -0.469031095 | 0.31400385 | -1.4937113 | 0.1352511 | 0.29973917 |
| THAP3 | 103.520056 | 0.080639771 | 0.23976318 | 0.33633092 | 0.73662133 | 0.85097924 |
| DNAJC11 | 1142.52983 | 0.598165232 | 0.23088982 | 2.59069559 | 0.00957822 | 0.04883308 |
| CAMTA1 | 2616.82794 | 0.173915893 | 0.19802786 | 0.8782395 | 0.37981376 | 0.5763431 |
| VAMP3 | 217.67277 | -0.720360026 | 0.20275076 | -3.5529337 | 0.00038096 | 0.00442527 |
| PER3 | 871.629428 | 0.474663362 | 0.64131939 | 0.74013568 | 0.45921767 | 0.64764858 |
| PARK7 | 605.352895 | -0.515995156 | 0.22750733 | -2.2680375 | 0.02332692 | 0.09180077 |
| ERRFI1 | 3637.64952 | -0.526340902 | 0.88000479 | -0.5981114 | 0.5497656 | 0.71955844 |
| AL034417.2 | 88.3456581 | 0.195458553 | 0.70123309 | 0.2787355 | 0.78044782 | 0.87937511 |
| RERE | 8120.43922 | 0.245885953 | 0.21574502 | 1.13970628 | 0.25440869 | 0.45147332 |
| ENO1 | 3980.72896 | -1.888083682 | 0.38089215 | -4.9570034 | 7.16E-07 | 2.93E-05 |
| SLC2A5 | 239.041087 | 1.332820196 | 0.5921853 | 2.25068099 | 0.02440575 | 0.09469522 |
| GPR157 | 200.588286 | -0.569128121 | 0.33600998 | -1.6937834 | 0.09030645 | 0.22976447 |
| MIR34AHG | 301.231291 | -1.931068549 | 0.64130554 | -3.0111522 | 0.00260258 | 0.01872287 |
| H6PD | 593.437879 | -0.618294412 | 0.29036944 | -2.129337 | 0.03322639 | 0.11768322 |
| SPSB1 | 2310.62964 | -3.832655048 | 0.78934541 | -4.8554853 | 1.20E-06 | 4.49E-05 |
| AL928921.2 | 56.3461281 | -0.876367774 | 0.90139695 | -0.9722329 | 0.33093469 | 0.53067752 |
| SLC25A33 | 460.341772 | -0.382595754 | 0.48780669 | -0.7843184 | 0.43285331 | 0.62552702 |
| TMEM201 | 35.8283037 | -0.502349342 | 0.56736006 | -0.8854154 | 0.37593262 | 0.57260901 |
| PIK3CD | 301.339243 | -0.974462513 | 0.55205669 | -1.7651494 | 0.07753866 | 0.20766363 |
| PIK3CD-AS2 | 45.5075888 | -0.091126329 | 0.32748599 | -0.2782602 | 0.7808126 | 0.87964151 |
| CLSTN1 | 943.278427 | 0.294470156 | 0.22200005 | 1.32644185 | 0.18469339 | 0.36750623 |
| CTNNBIP1 | 276.870366 | 0.342331125 | 0.25493109 | 1.34283786 | 0.17932448 | 0.36062216 |
| LZIC | 254.812571 | 0.404786975 | 0.24164687 | 1.67511783 | 0.09391112 | 0.23588779 |
| NMNAT1 | 513.518751 | 0.522112697 | 0.25093479 | 2.08067079 | 0.03746405 | 0.12780481 |
| UBE4B | 1638.28681 | 0.101511411 | 0.14913804 | 0.68065405 | 0.49609042 | 0.67795727 |
| KIF1B | 3502.52922 | 0.442194705 | 0.1562635 | 2.82980166 | 0.00465769 | 0.02879698 |
| AL139424.2 | 33.0584275 | 0.999562513 | 0.45827977 | 2.18111857 | 0.02917465 | 0.10692593 |
| PGD | 253.506275 | -0.629806378 | 0.47033953 | -1.3390462 | 0.18055562 | 0.36235372 |
| DFFA | 439.742699 | 0.15447767 | 0.28517449 | 0.54169525 | 0.58802846 | 0.75041555 |
| PEX14 | 1386.3095 | 0.239806324 | 0.1973304 | 1.21525283 | 0.22426967 | 0.4165399 |
| CASZ1 | 316.97194 | -1.364803353 | 0.44088394 | -3.0956068 | 0.00196411 | 0.01515025 |
| SRM | 217.695984 | -1.357003895 | 0.63972369 | -2.1212344 | 0.03390208 | 0.11921392 |
| EXOSC10 | 366.709439 | -0.333488866 | 0.30979321 | -1.0764886 | 0.28170879 | 0.4805703 |
| AL109811.1 | 47.9816591 | -0.541946552 | 0.36845585 | -1.4708589 | 0.14132929 | 0.30822602 |
| MTOR | 718.730112 | -0.124818249 | 0.28476577 | -0.438319 | 0.66115505 | 0.80309366 |
| UBIAD1 | 89.7307323 | -0.452150698 | 0.4456231 | -1.0146482 | 0.31027353 | 0.51061584 |
| FBXO2 | 50.3068433 | -0.972415654 | 0.47231123 | -2.0588451 | 0.03950908 | 0.13241191 |

| | | | | | | |
|---|---|---|---|---|---|---|
| FBXO44 | 129.380299 | 0.38494179 | 0.48965061 | 0.78615606 | 0.43177607 | 0.62436486 |
| FBXO6 | 75.9438994 | -0.573348494 | 0.38830018 | -1.47656 | 0.13979363 | 0.30629017 |
| MAD2L2 | 38.6442171 | -0.791661162 | 0.33122123 | -2.3901281 | 0.0168425 | 0.07319599 |
| AGTRAP | 310.777093 | -0.61253805 | 0.36001147 | -1.7014404 | 0.08886032 | 0.22732521 |
| MTHFR | 846.971535 | 0.070606744 | 0.24413537 | 0.28921145 | 0.77241957 | 0.87356065 |
| CLCN6 | 497.942366 | -0.033433602 | 0.33041416 | -0.101187 | 0.91940205 | 0.95690112 |
| NPPA-AS1 | 55.7831464 | -0.658152707 | 0.79564892 | -0.8271898 | 0.40812947 | 0.60198328 |
| KIAA2013 | 278.512354 | 0.356724615 | 0.19697888 | 1.810979 | 0.0701441 | 0.19447112 |
| PLOD1 | 420.995552 | 0.256708311 | 0.37894521 | 0.67742858 | 0.49813407 | 0.68013983 |
| MFN2 | 407.107461 | -0.186761925 | 0.26889926 | -0.6945424 | 0.48734215 | 0.67054796 |
| MIIP | 76.1204639 | 0.360646048 | 0.30756845 | 1.17257164 | 0.24096763 | 0.43618862 |
| VPS13D | 2654.74606 | -0.073968872 | 0.19407302 | -0.3811394 | 0.70309983 | 0.82952224 |
| DHRS3 | 1091.55138 | -0.136832634 | 0.40525188 | -0.3376484 | 0.73562819 | 0.85047703 |
| PRDM2 | 768.799305 | -0.562130823 | 0.26403708 | -2.1289844 | 0.03325555 | 0.11772568 |
| KAZN | 5134.10966 | -2.086189328 | 0.39774595 | -5.2450297 | 1.56E-07 | 8.14E-06 |
| TMEM51-AS | 174.734639 | -1.345160053 | 0.53783447 | -2.501067 | 0.01238198 | 0.05856444 |
| TMEM51 | 1999.34149 | -0.955199111 | 0.3683472 | -2.5932031 | 0.00950866 | 0.04854197 |
| C1orf195 | 73.2613896 | -1.372362882 | 0.41929762 | -3.2730042 | 0.00106411 | 0.00946511 |
| FHAD1 | 90.9601447 | -0.55834917 | 0.60459895 | -0.9235034 | 0.35574494 | 0.55411317 |
| EFHD2 | 101.403181 | -0.578357222 | 0.42697045 | -1.3545603 | 0.17555769 | 0.35574102 |
| CASP9 | 158.210077 | 0.311313376 | 0.22292326 | 1.39650466 | 0.16256258 | 0.33905132 |
| DNAJC16 | 1042.31721 | 0.552284363 | 0.20994952 | 2.63055785 | 0.00852449 | 0.04476825 |
| AGMAT | 578.402442 | 1.912971884 | 0.41631857 | 4.59497131 | 4.33E-06 | 0.00012758 |
| DDI2 | 1302.31918 | 0.467554874 | 0.28696523 | 1.62930844 | 0.10324774 | 0.25095251 |
| PLEKHM2 | 631.313156 | -0.82848527 | 0.42791158 | -1.9361132 | 0.05285383 | 0.1607784 |
| SLC25A34 | 58.7090072 | 2.130861346 | 0.74088016 | 2.87612149 | 0.00402595 | 0.02577471 |
| TMEM82 | 184.48111 | 1.428476482 | 0.51570906 | 2.76992708 | 0.00560688 | 0.0330127 |
| FBLIM1 | 288.236929 | -1.338920361 | 0.59777465 | -2.2398413 | 0.02510123 | 0.09667393 |
| UQCRHL | 69.2741663 | -0.805305522 | 0.39505287 | -2.0384753 | 0.04150242 | 0.13684717 |
| AL450998.2 | 104.074751 | 0.107701398 | 0.36537893 | 0.29476631 | 0.76817241 | 0.87071498 |
| SPEN | 1895.39243 | -0.353910676 | 0.53359031 | -0.6632629 | 0.50716217 | 0.68719571 |
| ZBTB17 | 176.705671 | 0.230157845 | 0.28269258 | 0.81416302 | 0.41555157 | 0.60906387 |
| CLCNKA | 56.9832499 | 2.486862898 | 0.87327364 | 2.84774758 | 0.00440298 | 0.02757044 |
| CLCNKB | 46.3752207 | 1.255974691 | 0.6356702 | 1.97582755 | 0.04817431 | 0.15171981 |
| EPHA2 | 154.669311 | -1.385247759 | 0.49908519 | -2.7755738 | 0.00551044 | 0.03261295 |
| FBXO42 | 1333.44248 | -0.184199308 | 0.3047676 | -0.6043927 | 0.5455826 | 0.71579246 |
| SZRD1 | 443.824992 | -0.105502395 | 0.26250367 | -0.4019083 | 0.68775153 | 0.81938377 |
| SPATA21 | 41.9417663 | -0.888466993 | 0.4163808 | -2.1337848 | 0.0328604 | 0.11668849 |
| NECAP2 | 220.689704 | -0.765148571 | 0.23974476 | -3.1915132 | 0.0014153 | 0.01177152 |
| NBPF1 | 504.833923 | -0.76860115 | 0.28927674 | -2.6569753 | 0.00788452 | 0.04244821 |
| CROCC | 539.634712 | 0.639910412 | 0.36953479 | 1.73166487 | 0.08333325 | 0.21815295 |
| BX284668.5 | 40.8387058 | -0.063434958 | 0.78193349 | -0.0811258 | 0.93534193 | 0.96565906 |
| ATP13A2 | 142.219901 | -0.582428808 | 0.40088775 | -1.4528476 | 0.14626607 | 0.31518141 |
| SDHB | 718.560576 | 0.090514702 | 0.19640422 | 0.46085924 | 0.64489959 | 0.79180937 |
| RCC2 | 334.686431 | -0.469135982 | 0.31886186 | -1.4712828 | 0.14121465 | 0.30819999 |
| ARHGEF10L | 3361.68556 | 0.361205607 | 0.44042536 | 0.82012899 | 0.41214258 | 0.60568653 |
| KLHDC7A | 147.462765 | 0.766888434 | 0.59130704 | 1.29693777 | 0.1946526 | 0.37980117 |
| AL080251.1 | 36.0694289 | -1.426519786 | 1.07784279 | -1.3234952 | 0.18567074 | 0.36878552 |
| IFFO2 | 246.86062 | -0.22585709 | 0.52427286 | -0.4308006 | 0.66661333 | 0.80635371 |
| UBR4 | 2309.15304 | -0.185507547 | 0.1472792 | -1.2595638 | 0.20782677 | 0.39691814 |
| AL035413.1 | 178.15109 | 0.105130147 | 0.32363502 | 0.32484169 | 0.74530089 | 0.85593548 |
| EMC1 | 193.384681 | 0.011818464 | 0.24493174 | 0.04825207 | 0.96151535 | 0.97988733 |
| MRTO4 | 198.078483 | -0.37673694 | 0.45224023 | -0.8330461 | 0.40481876 | 0.5987753 |
| AKR7A3 | 209.630387 | 1.544812652 | 0.31552273 | 4.89604234 | 9.78E-07 | 3.81E-05 |
| AKR7A2 | 318.958717 | 0.226460318 | 0.36111264 | 0.62711822 | 0.53058175 | 0.70458089 |
| PQLC2 | 114.906145 | 0.06268089 | 0.52885283 | 0.11852237 | 0.90565377 | 0.95066597 |
| CAPZB | 1608.7724 | -0.396527344 | 0.23853492 | -1.662345 | 0.09644362 | 0.23978826 |
| NBL1 | 162.154849 | -1.61691241 | 0.46255289 | -3.4956271 | 0.00047295 | 0.00515729 |
| TMCO4 | 593.614145 | -0.041324266 | 0.35954021 | -0.1149364 | 0.90849551 | 0.95197217 |
| RNF186 | 161.281597 | -0.108272826 | 0.89689664 | -0.1207194 | 0.90391329 | 0.94968435 |
| OTUD3 | 373.467373 | -0.135649842 | 0.27551015 | -0.4923588 | 0.62246573 | 0.77596685 |
| CAMK2N1 | 869.899735 | 0.49994516 | 0.35336554 | 1.41481015 | 0.15712417 | 0.33121548 |
| MUL1 | 111.372229 | -0.478479078 | 0.3099087 | -1.5439356 | 0.12260393 | 0.28121354 |
| PINK1 | 330.165637 | -0.120553965 | 0.26420379 | -0.4562916 | 0.64818032 | 0.79355195 |
| PINK1-AS | 36.0856343 | 0.121881913 | 0.28711134 | 0.42451097 | 0.6711932 | 0.80935413 |
| DDOST | 226.376949 | -0.34274596 | 0.2904686 | -1.1799759 | 0.23800979 | 0.43296618 |
| KIF17 | 51.1210025 | -1.39280632 | 0.41824885 | -3.33009 | 0.00086818 | 0.00813963 |
| HP1BP3 | 1726.17342 | -0.053308276 | 0.15386437 | -0.3464628 | 0.72899496 | 0.8460193 |
| EIF4G3 | 4244.25516 | -0.41319135 | 0.30627844 | -1.349071 | 0.17731417 | 0.35789211 |
| ECE1 | 620.350887 | 0.597390155 | 0.34332565 | 1.74001027 | 0.08185721 | 0.21535832 |
| NBPF3 | 163.138918 | -0.5514936 | 0.34712062 | -1.5887664 | 0.11211313 | 0.26513711 |
| ALPL | 426.839268 | 1.318249355 | 0.51126199 | 2.5784224 | 0.00992526 | 0.05016428 |
| RAP1GAP | 448.573902 | 0.015849095 | 0.51647786 | 0.03068688 | 0.97551925 | 0.98773468 |
| USP48 | 1225.38746 | -0.026131227 | 0.16781208 | -0.1557172 | 0.87625595 | 0.93549655 |
| HSPG2 | 192.410573 | -0.682121204 | 0.50198179 | -1.3588565 | 0.17419207 | 0.35398825 |
| CDC42 | 1300.77304 | -0.626406673 | 0.23402614 | -2.6766525 | 0.00743617 | 0.04073845 |
| ZBTB40 | 523.306575 | -0.035480825 | 0.35557358 | -0.0997848 | 0.92051521 | 0.95736677 |
| EPHB2 | 84.6432939 | -2.79293469 | 0.95076166 | -2.9375761 | 0.00330789 | 0.02220434 |
| KDM1A | 638.691869 | 0.301221679 | 0.21604079 | 1.39428149 | 0.16323262 | 0.33982829 |
| AL031428.1 | 154.446868 | -0.448743845 | 0.48311263 | -0.9288597 | 0.3529618 | 0.55184755 |
| LUZP1 | 1239.86512 | 0.110569451 | 0.24861884 | 0.4447348 | 0.65651142 | 0.79982242 |
| LINC01355 | 145.31611 | 0.177278604 | 0.22588253 | 0.78482653 | 0.43255528 | 0.6252278 |
| AL109936.6 | 45.9498545 | 0.630244518 | 0.48500041 | 1.29947214 | 0.19378195 | 0.37885739 |
| HNRNPR | 1000.89849 | -0.28054225 | 0.27934591 | -1.0042826 | 0.3152424 | 0.51532786 |
| ZNF436 | 51.8518305 | 0.172230558 | 0.56197443 | 0.30647401 | 0.75924376 | 0.86508348 |
| ZNF436-AS1 | 37.5212939 | -0.139564391 | 0.56425876 | -0.2473411 | 0.80464424 | 0.89400679 |
| TCEA3 | 655.221299 | 1.068084931 | 0.4965136 | 2.15116956 | 0.03146282 | 0.11296709 |
| ASAP3 | 686.7801 | -0.591846385 | 0.48963597 | -1.2087478 | 0.22675976 | 0.4192899 |
| RPL11 | 1513.20199 | -0.908603888 | 0.4029833 | -2.2546937 | 0.02415256 | 0.0939975 |
| ELOA | 240.925874 | -0.505232732 | 0.31783257 | -1.5896191 | 0.11192069 | 0.26486479 |
| ELOA-AS1 | 282.493417 | 0.957928673 | 0.45304358 | 2.11442942 | 0.03447861 | 0.1204993 |
| PITHD1 | 231.377265 | -0.366513193 | 0.24528377 | -1.4942415 | 0.13511251 | 0.29948045 |
| LYPLA2 | 257.732961 | -0.296843386 | 0.35341848 | -0.8399204 | 0.40095302 | 0.59504797 |
| GALE | 463.588389 | -0.40434437 | 0.30828203 | -1.3116054 | 0.18965331 | 0.37377109 |
| HMGCL | 555.467191 | 0.244854535 | 0.26999829 | 0.9068744 | 0.36447322 | 0.56208758 |
| FUCA1 | 167.520144 | -0.145873466 | 0.21230091 | -0.6871071 | 0.49201524 | 0.67440529 |
| PNRC2 | 277.486022 | -0.698759705 | 0.24384216 | -2.865623 | 0.0041619 | 0.0264107 |
| SRSF10 | 818.991114 | -0.214569312 | 0.19569266 | -1.0964607 | 0.27287722 | 0.47075093 |
| MYOM3 | 1052.78627 | 0.63828196 | 0.37502028 | 1.7019932 | 0.08875664 | 0.22722839 |
| IL22RA1 | 124.329842 | -1.009645815 | 0.81846038 | -1.2335916 | 0.21735514 | 0.40834524 |
| IFNLR1 | 286.49483 | 0.437809112 | 0.40415652 | 1.08326625 | 0.27869026 | 0.47708348 |
| STPG1 | 173.669537 | -0.074385786 | 0.18881859 | -0.3939537 | 0.69361521 | 0.82364929 |
| NIPAL3 | 381.982472 | -0.246531523 | 0.34536232 | -0.7138345 | 0.47532955 | 0.66124102 |
| RCAN3 | 68.5124397 | -1.177792777 | 0.55253328 | -2.1316232 | 0.03303783 | 0.11713646 |
| NCMAP | 132.848202 | -1.926979478 | 0.73405376 | -2.6251204 | 0.00866183 | 0.04535056 |
| SRRM1 | 850.80195 | -0.098337756 | 0.17411346 | -0.5647912 | 0.57221577 | 0.73805981 |
| CLIC4 | 5837.21726 | 1.151039665 | 0.33381842 | 3.44810112 | 0.00056454 | 0.00583134 |
| SYF2 | 259.657139 | -0.603885584 | 0.24300297 | -2.4850955 | 0.01295167 | 0.06048672 |
| RSRP1 | 2485.01335 | 0.11882524 | 0.32282172 | 0.36808316 | 0.71281123 | 0.83565716 |
| RHD | 66.9984504 | 0.095378043 | 1.02548855 | 0.09300742 | 0.92589767 | 0.9608782 |
| TMEM50A | 398.93877 | -0.929490902 | 0.23030725 | -4.0358734 | 5.44E-05 | 0.00097344 |
| MACO1 | 807.712681 | 0.187325322 | 0.23745422 | 0.78889025 | 0.43017616 | 0.62271549 |
| LDLRAP1 | 87.4331411 | -0.070402939 | 0.30119664 | -0.2337441 | 0.81518365 | 0.9000179 |

| | | | | | | |
|---|---|---|---|---|---|---|
| MAN1C1 | 1375.13772 | 2.35410756 | 0.79553146 | 2.95916336 | 0.00308476 | 0.02111703 |
| AL031280.1 | 129.268233 | 1.675160857 | 0.96207745 | 1.74119126 | 0.08165006 | 0.21513103 |
| SELENON | 28.7656928 | 0.114579088 | 0.49844501 | 0.22987308 | 0.8181904 | 0.90181214 |
| AL020996.1 | 33.6388432 | 0.800174048 | 0.56206901 | 1.42362243 | 0.1545558 | 0.3271791 |
| MTFR1L | 278.145694 | 1.083613257 | 0.2751039 | 3.93892358 | 8.18E-05 | 0.00132925 |
| AL033528.2 | 72.8422884 | 1.790496846 | 0.55023305 | 3.25406997 | 0.00113764 | 0.00995129 |
| PAQR7 | 229.256774 | 2.017559684 | 0.53184844 | 3.79348616 | 0.00014855 | 0.00209911 |
| STMN1 | 100.502861 | -0.281650389 | 0.54397394 | -0.5177645 | 0.6046226 | 0.76249535 |
| PAFAH2 | 256.551946 | -0.116074053 | 0.25587933 | -0.4536281 | 0.65009653 | 0.79482956 |
| SLC30A2 | 264.150669 | -0.165594381 | 0.47764462 | -0.3466895 | 0.72882459 | 0.84599052 |
| TRIM63 | 48.0191351 | 1.759667806 | 0.85663188 | 2.05417035 | 0.03995921 | 0.13346025 |
| PDIK1L | 101.595484 | 0.195084343 | 0.31464202 | 0.62001999 | 0.53524462 | 0.70809749 |
| ZNF593 | 156.993159 | -0.941951307 | 0.44753473 | -2.1047558 | 0.03531257 | 0.12247709 |
| CEP85 | 185.160252 | -0.097508425 | 0.31484041 | -0.3097075 | 0.75678343 | 0.86359015 |
| SH3BGRL3 | 367.167034 | -1.907435446 | 0.41385572 | -4.6089383 | 4.05E-06 | 0.0001206 |
| UBXN11 | 201.218861 | 0.834994783 | 0.27468225 | 3.03985708 | 0.0023669 | 0.01749875 |
| CRYBG2 | 47.567375 | -0.943891492 | 0.60092081 | -1.5707419 | 0.11624262 | 0.27158651 |
| DHDDS | 538.158692 | -0.688423421 | 0.45894758 | -1.5000045 | 0.13361324 | 0.29750434 |
| HMGN2 | 366.447533 | 0.455605469 | 0.32575389 | 1.39861866 | 0.16192737 | 0.33813809 |
| RPS6KA1 | 497.089429 | -0.131869976 | 0.24808759 | -0.531546 | 0.59504045 | 0.75552751 |
| ARID1A | 1381.47674 | 0.541686086 | 0.20139137 | 2.68971849 | 0.00715123 | 0.03954092 |
| PIGV | 60.9156049 | 0.517024214 | 0.25485116 | 2.02873011 | 0.04248579 | 0.1390859 |
| ZDHHC18 | 218.301788 | -1.373847068 | 0.44887933 | -3.0612975 | 0.0022038 | 0.01654298 |
| SFN | 98.5810041 | -7.401112064 | 1.61845703 | -4.5729432 | 4.81E-06 | 0.00013907 |
| GPN2 | 226.688273 | -0.157398019 | 0.36877383 | -0.4268145 | 0.66951445 | 0.80805094 |
| GPATCH3 | 103.695509 | 0.086504029 | 0.34480952 | 0.25087483 | 0.80191089 | 0.89234908 |
| NUDC | 565.870192 | -0.264412848 | 0.36386521 | -0.726678 | 0.46742322 | 0.65470775 |
| KDF1 | 66.4227045 | 0.099805788 | 0.2983723 | 0.33450085 | 0.73800165 | 0.85128239 |
| SLC9A1 | 124.508597 | -0.345465644 | 0.39745854 | -0.8691866 | 0.38474507 | 0.58106001 |
| WDTC1 | 518.825515 | 0.504894045 | 0.19219544 | 2.62698247 | 0.00861458 | 0.04515488 |
| TMEM222 | 183.885887 | -0.366827265 | 0.35532892 | -1.0323597 | 0.30190363 | 0.50239081 |
| MAP3K6 | 50.3463611 | -1.17036313 | 0.53983737 | -2.1679921 | 0.03015929 | 0.10959528 |
| WASF2 | 1294.42091 | -0.38638975 | 0.1859001 | -2.0784805 | 0.03766512 | 0.12833497 |
| AHDC1 | 356.090259 | -0.45393746 | 0.43256126 | -1.0494177 | 0.2939859 | 0.49394984 |
| IFI6 | 687.590344 | -1.373756124 | 1.58434025 | -0.867084 | 0.38589597 | 0.58215674 |
| FAM76A | 256.318372 | 0.754542803 | 0.20772652 | 3.63238544 | 0.00028081 | 0.00348153 |
| STX12 | 893.680117 | -0.286891676 | 0.14033919 | -2.0442735 | 0.04092654 | 0.13556792 |
| PPP1R8 | 201.184393 | 0.091812612 | 0.28501224 | 0.32213568 | 0.74734991 | 0.85730398 |
| RPA2 | 189.876139 | 0.035997196 | 0.29239743 | 0.1231105 | 0.90201959 | 0.94861498 |
| SMPDL3B | 92.4768266 | -0.974546248 | 0.28091318 | -3.469208 | 0.000522 | 0.00550807 |
| XKR8 | 40.7080981 | -0.25752896 | 0.39511655 | -0.6517797 | 0.51454328 | 0.69267774 |
| EYA3 | 1771.07906 | 0.205734295 | 0.20732609 | 0.99232226 | 0.32104035 | 0.52141674 |
| DNAJC8 | 431.024046 | -0.178978979 | 0.2849855 | -0.6280284 | 0.52998536 | 0.70421206 |
| ATP5IF1 | 840.422798 | 0.135031974 | 0.36817594 | 0.36675937 | 0.71379852 | 0.83637691 |
| AL353622.1 | 106.818788 | 2.478606335 | 0.53079791 | 4.66958568 | 3.02E-06 | 9.62E-05 |
| SESN2 | 503.232762 | 0.784749734 | 0.33212478 | 2.36281598 | 0.01813667 | 0.07701344 |
| MED18 | 41.3647327 | 0.036374784 | 0.33564872 | 0.10837159 | 0.91370094 | 0.95407167 |
| PHACTR4 | 2747.35211 | 0.411499657 | 0.19622906 | 2.09703728 | 0.03599028 | 0.12406855 |
| RCC1 | 513.770689 | -1.411668184 | 0.42236942 | -3.3422595 | 0.00083099 | 0.00788811 |
| TRNAU1AP | 556.251472 | -0.741561811 | 0.20277068 | -3.6571451 | 0.00025504 | 0.00322792 |
| SNHG12 | 148.178538 | -1.123276412 | 0.4150249 | -2.7065278 | 0.00679909 | 0.03803883 |
| TAF12 | 283.036143 | 0.459949803 | 0.30647088 | 1.50079445 | 0.13340873 | 0.29719381 |
| GMEB1 | 313.096509 | -0.026356963 | 0.4994136 | -0.0527758 | 0.95791053 | 0.9784693 |
| YTHDF2 | 862.383174 | -0.117290512 | 0.24937497 | -0.4703379 | 0.63811359 | 0.78726983 |
| EPB41 | 3554.63023 | -0.632616323 | 0.19903184 | -3.1784679 | 0.00148056 | 0.01222529 |
| TMEM200B | 55.2342624 | -0.322627987 | 0.40829063 | -0.790192 | 0.42941565 | 0.62219636 |
| SRSF4 | 2948.40096 | 0.084343044 | 0.19295322 | 0.43711654 | 0.66202684 | 0.80336802 |
| MECR | 257.224284 | -0.003000352 | 0.22087146 | -0.0135842 | 0.98916175 | 0.99471552 |
| PTPRU | 60.5064737 | -1.943223043 | 0.44275156 | -4.3889694 | 1.14E-05 | 0.00028128 |
| PUM1 | 3640.17902 | 0.014373029 | 0.17518632 | 0.08204424 | 0.93461153 | 0.96509871 |
| SNRNP40 | 270.438114 | 0.063747876 | 0.20763793 | 0.3070146 | 0.75883226 | 0.86482758 |
| ZCCHC17 | 461.701984 | -0.390773611 | 0.20887623 | -1.8708381 | 0.06136752 | 0.17746088 |
| AL451070.1 | 57.2207139 | -0.330150716 | 0.39679182 | -0.8320502 | 0.4053806 | 0.59941229 |
| FABP3 | 123.647917 | -0.512747682 | 0.43693603 | -1.1735074 | 0.24059238 | 0.43581337 |
| SERINC2 | 995.322328 | -1.510038783 | 0.45179079 | -3.3423408 | 0.00083075 | 0.00788811 |
| TINAGL1 | 688.564732 | -0.72508974 | 0.41837724 | -1.7331004 | 0.08307783 | 0.21769219 |
| PEF1 | 198.328778 | -0.254313793 | 0.26843319 | -0.9474007 | 0.34343463 | 0.54383762 |
| PTP4A2 | 349.926813 | -0.999153591 | 0.28377695 | -3.5209118 | 0.00043007 | 0.00482004 |
| KHDRBS1 | 1170.57012 | -0.580958657 | 0.2376135 | -2.4449733 | 0.01448629 | 0.06535996 |
| AL445248.1 | 53.6472295 | -0.672017645 | 0.35070444 | -1.9161937 | 0.05534044 | 0.16589509 |
| TMEM39B | 138.535484 | 0.462520789 | 0.2487697 | 1.85923281 | 0.06299415 | 0.18079162 |
| KPNA6 | 1177.93928 | -0.052796355 | 0.2263056 | -0.2332967 | 0.815531 | 0.90018388 |
| TXLNA | 362.678943 | -0.660341761 | 0.35438815 | -1.8633291 | 0.06241598 | 0.17970023 |
| DCDC2B | 49.6079832 | -0.272959647 | 0.46949085 | -0.581395 | 0.56097425 | 0.72918672 |
| TMEM234 | 146.946688 | 0.11175585 | 0.24651994 | 0.45333391 | 0.65030832 | 0.79482956 |
| EIF3I | 254.366824 | -0.97154724 | 0.39239043 | -2.4759708 | 0.01328745 | 0.06159317 |
| HDAC1 | 365.505325 | 0.062355982 | 0.22802371 | 0.27346272 | 0.78449755 | 0.88176367 |
| MARCKSL1 | 61.995523 | -1.711582414 | 0.76809521 | -2.2283467 | 0.02585741 | 0.0986979 |
| TSSK3 | 61.5652107 | 0.914364831 | 0.39995539 | 2.28616703 | 0.02224449 | 0.08897147 |
| FAM229A | 78.2710494 | 0.201318744 | 0.40392276 | 0.49840901 | 0.61819579 | 0.77299851 |
| BSDC1 | 760.316375 | -0.272442114 | 0.2797706 | -0.9738054 | 0.33015318 | 0.52990746 |
| ZBTB8A | 64.8460079 | -0.913223987 | 0.57017549 | -1.6016542 | 0.10923209 | 0.26052624 |
| ZBTB8OS | 1097.15205 | 0.465337024 | 0.21248932 | 2.18993138 | 0.02852921 | 0.10543271 |
| RBBP4 | 784.010266 | -0.03666558 | 0.21279391 | -0.1723056 | 0.86319729 | 0.92835193 |
| SYNC | 131.264955 | -0.928019516 | 0.37040566 | -2.505414 | 0.01223081 | 0.05800961 |
| KIAA1522 | 781.191298 | -0.376135165 | 0.27205833 | -1.3825534 | 0.16680185 | 0.34469756 |
| YARS | 534.422362 | -0.955773723 | 0.39731736 | -2.4055675 | 0.01614737 | 0.07100802 |
| S100PBP | 917.183343 | 0.061309814 | 0.16340923 | 0.37519187 | 0.70751777 | 0.83240065 |
| TMEM54 | 277.908649 | -1.678891874 | 0.52440412 | -3.2015231 | 0.00136703 | 0.01144612 |
| RNF19B | 1171.02839 | 0.068038075 | 0.45974088 | 0.14799222 | 0.8823489 | 0.93892993 |
| AL020995.1 | 299.860296 | -0.328839217 | 0.66288302 | -0.4960743 | 0.61984201 | 0.77427999 |
| AK2 | 1367.78505 | 0.779544979 | 0.33510547 | 2.32626755 | 0.02000428 | 0.0824493 |
| AZIN2 | 263.024779 | 0.13614127 | 0.45766781 | 0.29746744 | 0.76610967 | 0.86919451 |
| TRIM62 | 89.0110872 | -0.492855153 | 0.37947868 | -1.298769 | 0.19402321 | 0.37905874 |
| ZNF362 | 285.220174 | 1.487045578 | 0.36978368 | 4.02139317 | 5.79E-05 | 0.00101799 |
| AL513327.1 | 358.943949 | -1.868203637 | 0.43539458 | -4.2908289 | 1.78E-05 | 0.00040531 |
| PHC2 | 1843.97616 | -0.87949552 | 0.27944572 | -3.1472857 | 0.00164794 | 0.01323734 |
| AL513327.2 | 54.0141759 | -0.726537247 | 0.50668334 | -1.4339079 | 0.15159855 | 0.3229657 |
| ZSCAN20 | 75.3906492 | -0.143216677 | 0.46147766 | -0.3103437 | 0.75629961 | 0.86345151 |
| SMIM12 | 316.911348 | -0.601164443 | 0.28103436 | -2.1391137 | 0.03242646 | 0.11550663 |
| AC114490.3 | 52.945995 | -0.32726654 | 0.46517699 | -0.7035312 | 0.48172475 | 0.66636403 |
| ZMYM1 | 420.499334 | 0.530832041 | 0.2423332 | 2.19050478 | 0.02848765 | 0.1053075 |
| SFPQ | 1752.1317 | 0.113184083 | 0.22884201 | 0.49459487 | 0.62088612 | 0.77487809 |
| ZMYM4 | 2897.55101 | 0.090588199 | 0.11310117 | 0.80094839 | 0.42316153 | 0.61646031 |
| ZMYM4-AS1 | 57.1202676 | -0.964508324 | 0.70420885 | -1.3696339 | 0.17080121 | 0.34969424 |
| KIAA0319L | 1917.82555 | 0.262145289 | 0.19270639 | 1.36033522 | 0.17372387 | 0.35358992 |
| NCDN | 47.2022463 | 0.535983975 | 0.35000132 | 1.53137702 | 0.12567624 | 0.28601099 |
| PSMB2 | 383.649436 | -0.622086228 | 0.16922053 | -3.6761865 | 0.00023675 | 0.00305814 |
| AGO4 | 640.567217 | -0.819262365 | 0.20267726 | -4.0422017 | 5.30E-05 | 0.00095126 |
| AGO1 | 307.945891 | 0.409637116 | 0.19863839 | 2.06222531 | 0.03918629 | 0.13179715 |
| AGO3 | 2751.00442 | -0.07666479 | 0.18073271 | -0.4241888 | 0.67142813 | 0.80950526 |
| AL138787.2 | 197.335439 | -0.214220678 | 0.7337739 | -0.2919437 | 0.77032966 | 0.8723487 |
| ADPRHL2 | 350.841389 | -1.274769131 | 0.35531526 | -3.5877129 | 0.00033359 | 0.00399466 |

| | | | | | | |
|---|---|---|---|---|---|---|
| TRAPPC3 | 275.585396 | -0.327585683 | 0.28821539 | -1.1366003 | 0.25570543 | 0.45283648 |
| MAP7D1 | 164.201877 | -1.464796574 | 0.51222297 | -2.8596854 | 0.00424061 | 0.02676155 |
| THRAP3 | 1953.20779 | -0.000860854 | 0.30807227 | -0.0027943 | 0.99777045 | 0.99871766 |
| SH3D21 | 71.1340547 | -1.332742277 | 0.50138159 | -2.6581396 | 0.00785733 | 0.04237405 |
| EVA1B | 132.075588 | -1.236390651 | 0.30856343 | -4.0069254 | 6.15E-05 | 0.00106678 |
| STK40 | 901.842831 | -1.385842296 | 0.4845967 | -2.8597848 | 0.00423929 | 0.02676155 |
| LSM10 | 89.7143435 | -0.713739965 | 0.34022572 | -2.0978425 | 0.03591906 | 0.12398454 |
| OSCP1 | 145.704027 | 0.639752922 | 0.28294574 | 2.26104455 | 0.0237565 | 0.09290451 |
| MRPS15 | 211.324248 | -0.750338484 | 0.23544147 | -3.1869427 | 0.00143785 | 0.01191575 |
| LINC01137 | 121.138083 | -0.717415293 | 0.3955991 | -1.8134907 | 0.06975617 | 0.1935522 |
| ZC3H12A | 84.0790897 | -2.475234475 | 0.36278859 | -6.8228014 | 8.93E-12 | 1.94E-09 |
| MEAF6 | 282.996253 | -0.280034623 | 0.24308122 | -1.1520208 | 0.24931253 | 0.44548648 |
| SNIP1 | 163.480897 | -0.898992951 | 0.29180654 | -3.0807842 | 0.00206456 | 0.01576544 |
| DNALI1 | 96.5919636 | 0.190178877 | 0.36789625 | 0.51693616 | 0.60520072 | 0.76287349 |
| GNL2 | 230.520265 | -0.654091366 | 0.47854499 | -1.3668336 | 0.17167747 | 0.35095945 |
| C1orf109 | 113.393968 | -0.207059881 | 0.46466374 | -0.4456123 | 0.65587732 | 0.79912092 |
| YRDC | 55.8047356 | -0.653595569 | 0.43058697 | -1.5179177 | 0.12903516 | 0.29085428 |
| C1orf122 | 329.785595 | -0.821482084 | 0.33452138 | -2.4556938 | 0.01406129 | 0.06403259 |
| MTF1 | 366.462077 | -0.384947332 | 0.24528304 | -1.5694005 | 0.11655465 | 0.27207332 |
| AL929472.2 | 108.6881 | 0.078096391 | 0.51110744 | 0.15279838 | 0.87855727 | 0.93706833 |
| INPP5B | 1060.36055 | 0.58140095 | 0.28431235 | 2.04493736 | 0.04086103 | 0.13544314 |
| SF3A3 | 268.348648 | -0.091681076 | 0.24562471 | -0.3732567 | 0.70895738 | 0.8332772 |
| UTP11 | 82.9612717 | -0.279786238 | 0.30637984 | -0.9132005 | 0.36113708 | 0.55908131 |
| AL354702.2 | 96.6777858 | 2.111201595 | 0.7717308 | 2.73567104 | 0.00622532 | 0.03558403 |
| RRAGC | 430.63201 | -0.96804097 | 0.38159516 | -2.5368271 | 0.01118622 | 0.0545759 |
| AL139260.1 | 1138.55903 | -0.684207363 | 0.53985862 | -1.2673825 | 0.20501856 | 0.39336357 |
| RHBDL2 | 163.558811 | -1.263328201 | 0.44348761 | -2.8486212 | 0.00439091 | 0.02752 |
| AKIRIN1 | 544.076659 | -0.310195967 | 0.22854927 | -1.357239 | 0.1747053 | 0.35466314 |
| NDUFS5 | 766.2847 | -0.945627784 | 0.35619624 | -2.6547944 | 0.00793568 | 0.04260647 |
| MACF1 | 10348.8519 | -0.699255483 | 0.32907983 | -2.124881 | 0.03359654 | 0.11844337 |
| PABPC4 | 306.131541 | -0.761576624 | 0.34644423 | -2.1982661 | 0.02793015 | 0.10400253 |
| PPIE | 196.696517 | 0.201792289 | 0.3112503 | 0.64832801 | 0.51677282 | 0.69403559 |
| TRIT1 | 575.776641 | -0.014195114 | 0.20507784 | -0.0692182 | 0.94481596 | 0.97175601 |
| MFSD2A | 420.076887 | -2.680812438 | 0.7564992 | -3.5437082 | 0.00039454 | 0.00451795 |
| CAP1 | 623.606548 | -0.882520208 | 0.24274752 | -3.6355477 | 0.00027739 | 0.00344402 |
| PPT1 | 124.777995 | -0.418303284 | 0.35969286 | -1.1629457 | 0.24485152 | 0.44021245 |
| RLF | 1293.03874 | -0.635801519 | 0.19397284 | -3.2777864 | 0.00104625 | 0.00936293 |
| AL050341.2 | 31.2656271 | 0.248372416 | 0.57699276 | 0.43046019 | 0.66686092 | 0.80648161 |
| ZMPSTE24 | 432.087279 | -0.105678199 | 0.35082506 | -0.3012276 | 0.76324093 | 0.86790637 |
| SMAP2 | 257.282554 | -0.145167081 | 0.19498141 | -0.7445176 | 0.45656342 | 0.64552736 |
| AL603839.3 | 231.111999 | 0.541148686 | 0.22955164 | 2.35741587 | 0.01840262 | 0.0778293 |
| EXO5 | 69.500029 | 0.013228495 | 0.23444826 | 0.05642394 | 0.95500408 | 0.97731423 |
| ZNF684 | 290.353802 | 1.16744991 | 0.46745042 | 2.49748391 | 0.01250781 | 0.05897647 |
| NFYC | 955.677426 | 1.130349723 | 0.19226919 | 5.87899563 | 4.13E-09 | 3.80E-07 |
| CITED4 | 254.858091 | -2.357118249 | 0.50864814 | -4.634084 | 3.59E-06 | 0.00011046 |
| CTPS1 | 429.231973 | -0.287615204 | 0.40229699 | -0.7149325 | 0.47465074 | 0.66076272 |
| SLFNL1-AS1 | 303.988986 | -0.629598384 | 0.26710201 | -2.3571458 | 0.01841601 | 0.07786191 |
| SLFNL1 | 127.924208 | -0.755426385 | 0.32889039 | -2.2968941 | 0.02162481 | 0.08748858 |
| SCMH1 | 5099.34893 | -0.13708617 | 0.32001897 | -0.4283689 | 0.66838258 | 0.80729015 |
| AC093151.3 | 410.10916 | -1.58591601 | 0.76491889 | -2.0733126 | 0.03814319 | 0.12953758 |
| HIVEP3 | 832.922315 | -0.607199264 | 0.36193001 | -1.6776704 | 0.09341146 | 0.23525917 |
| FOXJ3 | 2338.03373 | -0.063877948 | 0.21477676 | -0.2974156 | 0.76614927 | 0.86919451 |
| RIMKLA | 144.428696 | -2.889587326 | 0.52926809 | -5.459591 | 4.77E-08 | 3.01E-06 |
| ZMYND12 | 87.3992737 | 1.400730736 | 0.58376997 | 2.39945665 | 0.01641942 | 0.07183563 |
| PPCS | 267.847518 | -0.083791427 | 0.26771557 | -0.3129868 | 0.75429073 | 0.86223526 |
| CCDC30 | 2334.27542 | 0.944976004 | 0.37864326 | 2.49568951 | 0.01257126 | 0.05919417 |
| PPIH | 142.055834 | 0.173991095 | 0.41160166 | 0.42271719 | 0.67250162 | 0.81022938 |
| YBX1 | 745.490956 | -1.157517152 | 0.37754746 | -3.0658851 | 0.00217027 | 0.01635898 |
| P3H1 | 174.368104 | -0.50898464 | 0.32686234 | -1.5571835 | 0.11942692 | 0.27649723 |
| SVBP | 77.3892187 | -1.157380033 | 0.397496 | -2.9116772 | 0.00359494 | 0.02370912 |
| ERMAP | 244.030599 | 0.525788983 | 0.30652188 | 1.71533913 | 0.08628307 | 0.22293723 |
| AL512353.1 | 46.1850395 | -1.212679631 | 0.62924928 | -1.9271848 | 0.05395661 | 0.16308249 |
| ZNF691 | 80.2453234 | 0.061998029 | 0.238158 | 0.26032311 | 0.79461455 | 0.88818247 |
| SLC2A1 | 185.625844 | -3.153053302 | 0.52951387 | -5.954619 | 2.61E-09 | 2.63E-07 |
| SLC2A1-AS1 | 55.3860421 | -1.436176801 | 0.64957209 | -2.2109583 | 0.02703873 | 0.10181863 |
| EBNA1BP2 | 1744.11873 | 2.615573977 | 0.50054318 | 5.22547123 | 1.74E-07 | 8.79E-06 |
| CFAP57 | 290.971782 | 1.558164204 | 0.61345731 | 2.53997171 | 0.01108614 | 0.05425126 |
| TMEM125 | 66.0592611 | 0.473300176 | 0.51445144 | 0.92000942 | 0.35756783 | 0.5561453 |
| C1orf210 | 66.6951003 | 0.28731488 | 0.52512822 | 0.54713281 | 0.58428749 | 0.74702254 |
| AL139289.2 | 30.6179952 | 0.009216899 | 0.56947313 | 0.01618496 | 0.98708684 | 0.99360989 |
| ELOVL1 | 152.976063 | -0.444640827 | 0.27828921 | -1.5977653 | 0.11009523 | 0.26171962 |
| MED8 | 142.016401 | -0.243095876 | 0.34781729 | -0.6989183 | 0.48460308 | 0.66827686 |
| SZT2 | 573.959959 | 0.644701992 | 0.2308289 | 2.79298643 | 0.00522239 | 0.03132747 |
| HYI | 134.540691 | 0.17009448 | 0.3018088 | 0.56358357 | 0.57303757 | 0.73835552 |
| PTPRF | 1074.77546 | -1.105703143 | 0.32524275 | -3.3996243 | 0.00067479 | 0.00672675 |
| KDM4A-AS1 | 41.7390119 | -0.779729464 | 0.45069119 | -1.7300748 | 0.08361691 | 0.21835341 |
| AL357079.1 | 48.731595 | -1.965502761 | 0.51404158 | -3.823626 | 0.0001315 | 0.00191757 |
| IPO13 | 177.570386 | 0.041786749 | 0.39782753 | 0.10503735 | 0.91634617 | 0.95516023 |
| DPH2 | 47.1464484 | -0.658064606 | 0.4783534 | -1.3756871 | 0.16891852 | 0.34755609 |
| ATP6V0B | 827.244083 | -0.483830339 | 0.31371908 | -1.5422407 | 0.12301511 | 0.28187364 |
| B4GALT2 | 161.517813 | -0.830127102 | 0.39337652 | -2.1102609 | 0.03483589 | 0.12140745 |
| CCDC24 | 220.381697 | 0.380594186 | 0.47554581 | 0.80033128 | 0.42351889 | 0.61666528 |
| SLC6A9 | 38.1674706 | 0.000194401 | 0.39592117 | 0.00049101 | 0.99960823 | 0.99975658 |
| DMAP1 | 165.145021 | 0.652905917 | 0.30715153 | 2.1256802 | 0.0335299 | 0.11836062 |
| ERI3 | 1695.53504 | 0.923701847 | 0.38899396 | 2.37459175 | 0.01756837 | 0.07530608 |
| RNF220 | 232.476936 | -0.58463759 | 0.34393286 | -1.6998597 | 0.08915733 | 0.22771117 |
| TMEM53 | 196.937998 | 0.770049562 | 0.3650193 | 2.10961327 | 0.03489168 | 0.12150921 |
| ARMH1 | 148.605693 | 0.371061507 | 0.56595995 | 0.65563209 | 0.51206087 | 0.6909646 |
| RPS8 | 1185.06381 | -1.126963538 | 0.42311799 | -2.6634734 | 0.00773385 | 0.04181772 |
| PLK3 | 75.9240628 | -2.640434885 | 0.4503267 | -5.8633763 | 4.54E-09 | 4.14E-07 |
| BTBD19 | 91.7864314 | -0.337399537 | 0.35645693 | -0.9465366 | 0.34387496 | 0.54439007 |
| EIF2B3 | 614.82586 | 0.270182339 | 0.28715324 | 0.94089949 | 0.34675637 | 0.54694932 |
| HECTD3 | 178.9977 | 0.266750204 | 0.27039773 | 0.98651051 | 0.32388266 | 0.52438691 |
| UROD | 139.028246 | 0.256417473 | 0.35606501 | 0.72014229 | 0.47143739 | 0.65777609 |
| ZSWIM5 | 1943.91963 | 0.032127464 | 0.32753337 | 0.09808913 | 0.92186152 | 0.95811091 |
| MUTYH | 135.516618 | 0.295852329 | 0.4205141 | 0.70354913 | 0.4817136 | 0.66636403 |
| TOE1 | 57.6161578 | 0.303640998 | 0.27344417 | 1.11043144 | 0.26681316 | 0.46477327 |
| TESK2 | 375.213983 | -0.010816385 | 0.37495687 | -0.028847 | 0.97698661 | 0.9886052 |
| MMACHC | 52.036798 | 0.262459502 | 0.38009693 | 0.69050677 | 0.48987555 | 0.67270543 |
| PRDX1 | 2167.03618 | -1.681256339 | 0.47091345 | -3.5702024 | 0.00035671 | 0.004198 |
| AKR1A1 | 1390.80443 | 0.678188306 | 0.26711032 | 2.53898209 | 0.01111755 | 0.05434675 |
| NASP | 360.818776 | 0.289682134 | 0.20613079 | 1.40533169 | 0.15992267 | 0.33543382 |
| GPBP1L1 | 2105.34011 | 0.065280915 | 0.12548698 | 0.52022061 | 0.60290982 | 0.76145627 |
| TMEM69 | 140.212227 | -0.221497138 | 0.25127206 | -0.8815032 | 0.3780455 | 0.57462476 |
| IPP | 368.735712 | 0.820304303 | 0.2562193 | 3.20157105 | 0.0013668 | 0.01144612 |
| MAST2 | 2172.63834 | -0.082544285 | 0.30387769 | -0.2716365 | 0.7859015 | 0.88296838 |
| TSPAN1 | 1256.13735 | -0.172113187 | 0.36728567 | -0.4686085 | 0.6393495 | 0.78803844 |
| POMGNT1 | 359.757908 | -0.175421051 | 0.15598132 | -1.1246286 | 0.26074647 | 0.4589178 |
| LURAP1 | 103.366583 | 0.13083613 | 0.54074373 | 0.24195589 | 0.80881434 | 0.8961735 |
| LRRC41 | 730.554073 | -0.767415761 | 0.39667323 | -1.9346296 | 0.05303577 | 0.16111731 |
| UQCRH | 1082.58089 | -0.598117025 | 0.4458281 | -1.3415866 | 0.17973006 | 0.36096117 |
| NSUN4 | 411.804569 | 0.082215902 | 0.19163392 | 0.42902584 | 0.66790442 | 0.80703155 |
| FAAH | 462.574896 | 2.02573137 | 0.43516917 | 4.65504341 | 3.24E-06 | 0.00010137 |

| | | | | | | |
|---|---|---|---|---|---|---|
| MKNK1 | 386.202289 | -0.218383042 | 0.21024179 | -1.0387233 | 0.29893345 | 0.49896259 |
| ATPAF1 | 301.210184 | 0.195313366 | 0.22435499 | 0.87055504 | 0.38399715 | 0.5803781 |
| EFCAB14 | 478.86889 | -0.28565328 | 0.22217196 | -1.2857306 | 0.19853712 | 0.38480603 |
| CYP4A11 | 1795.97109 | 2.102453312 | 0.8911868 | 2.3591612 | 0.0183163 | 0.07760326 |
| CYP4X1 | 75.1851216 | 2.493501764 | 0.89854477 | 2.77504454 | 0.00551941 | 0.03265197 |
| CYP4Z1 | 113.942855 | -0.559983878 | 0.8917092 | -0.6279893 | 0.53001092 | 0.70421206 |
| CYP4A22-AS1 | 1185.76139 | 0.479230821 | 0.69507073 | 0.68947058 | 0.49052718 | 0.67330824 |
| CYP4A22 | 149.635119 | 1.664308945 | 1.02098955 | 1.63009401 | 0.10308163 | 0.25074355 |
| LINC00853 | 108.085522 | -0.698278185 | 0.68361453 | -1.0214502 | 0.3070412 | 0.507123 |
| PDZK1IP1 | 4254.56156 | -0.227523869 | 0.36799221 | -0.6182845 | 0.53638784 | 0.70875054 |
| STIL | 97.3170623 | -0.757940756 | 0.29050082 | -2.609083 | 0.00907852 | 0.04699369 |
| CMPK1 | 977.429976 | -0.226863926 | 0.21128784 | -1.0737197 | 0.28294831 | 0.48202492 |
| LINC01389 | 32.974129 | -1.629925845 | 0.47294254 | -3.4463507 | 0.00056821 | 0.00586483 |
| TRABD2B | 8126.78479 | -0.495231265 | 0.54192483 | -0.9138376 | 0.36080221 | 0.55881535 |
| AL356289.1 | 123.887979 | 0.244167745 | 0.60794235 | 0.40162977 | 0.68795652 | 0.81955676 |
| AL109659.1 | 105.47376 | 0.241280444 | 0.69039959 | 0.34947941 | 0.72672942 | 0.84481763 |
| SLC5A9 | 286.427444 | 1.533411258 | 0.71808241 | 2.13542518 | 0.0327263 | 0.11633281 |
| SPATA6 | 1153.10411 | 0.710241545 | 0.43458421 | 1.6343013 | 0.1021956 | 0.24973128 |
| AGBL4 | 1699.72597 | 0.280664601 | 0.81390727 | 0.34483609 | 0.7302176 | 0.84648502 |
| AC099788.1 | 37.1609144 | -0.083535606 | 0.97032598 | -0.0860902 | 0.93139467 | 0.96354054 |
| FAF1 | 5176.19954 | 0.067186278 | 0.13714032 | 0.48990902 | 0.62419828 | 0.77710134 |
| RNF11 | 923.632826 | 0.272890432 | 0.17801484 | 1.5329645 | 0.1252846 | 0.2853566 |
| TTC39A | 951.605148 | -2.28109404 | 0.43160176 | -5.2851825 | 1.26E-07 | 6.65E-06 |
| TTC39A-AS1 | 52.6960732 | -2.367141694 | 0.33977047 | -6.9668847 | 3.24E-12 | 7.93E-10 |
| EPS15 | 2364.60503 | 0.071696577 | 0.16220923 | 0.44200059 | 0.65848878 | 0.80095001 |
| AC104170.1 | 714.411067 | 1.332772512 | 0.74013639 | 1.80071205 | 0.07174828 | 0.19736176 |
| OSBPL9 | 3812.06768 | -0.50557976 | 0.31932624 | -1.5832703 | 0.11335986 | 0.26693414 |
| NRDC | 1843.8654 | -0.228653463 | 0.12269281 | -1.8636256 | 0.0623743 | 0.17965214 |
| TXNDC12 | 369.280464 | -0.192525829 | 0.16055727 | -1.19911 | 0.23048518 | 0.42369138 |
| BTF3L4 | 509.586242 | -0.067630602 | 0.17064566 | -0.3963218 | 0.69186763 | 0.82235775 |
| ZFYVE9 | 1185.0238 | -0.042544923 | 0.1903357 | -0.2235257 | 0.82312637 | 0.90528711 |
| CC2D1B | 341.290906 | -0.321393162 | 0.31741128 | -1.0125449 | 0.3112776 | 0.51145516 |
| PRPF38A | 214.559281 | 0.042236095 | 0.19860194 | 0.21266709 | 0.83158664 | 0.91086448 |
| TUT4 | 2499.18232 | 0.13455548 | 0.24163427 | 0.55685595 | 0.57762585 | 0.74154889 |
| COA7 | 102.281322 | -0.216386578 | 0.47603418 | -0.454561 | 0.64942509 | 0.79436638 |
| ZYG11B | 1207.36208 | 0.768702955 | 0.19448508 | 3.95250338 | 7.73E-05 | 0.00127412 |
| ZYG11A | 895.927294 | 2.347452015 | 0.46544855 | 5.04341889 | 4.57E-07 | 1.99E-05 |
| ECHDC2 | 2399.52254 | 1.001469977 | 0.33006425 | 3.03416676 | 0.00241201 | 0.01773436 |
| SCP2 | 1408.91928 | 0.481648131 | 0.28864622 | 1.6686452 | 0.09518772 | 0.23793877 |
| SLC1A7 | 22.0615483 | 1.424962052 | 1.56713407 | 0.90927897 | 0.36320289 | 0.56088576 |
| CPT2 | 206.561274 | 0.095190997 | 0.37853788 | 0.2514702 | 0.8014506 | 0.89219475 |
| C1orf123 | 98.7111118 | -0.053498402 | 0.24630351 | -0.2172052 | 0.82804843 | 0.90836814 |
| MAGOH | 210.725937 | -0.12967621 | 0.22904663 | -0.5661564 | 0.57128748 | 0.73727873 |
| LRP8 | 168.784943 | -5.595003721 | 1.01787131 | -5.4967693 | 3.87E-08 | 2.58E-06 |
| GLIS1 | 2461.5801 | 0.992937884 | 0.43870943 | 2.26331559 | 0.02361624 | 0.09269984 |
| NDC1 | 267.549962 | -0.228865108 | 0.36669538 | -0.6241287 | 0.53254307 | 0.70588607 |
| YIPF1 | 307.16964 | -0.252930066 | 0.26530408 | -0.9533591 | 0.34040815 | 0.54098255 |
| DIO1 | 911.637691 | 1.912660532 | 0.78862772 | 2.4253022 | 0.01529565 | 0.06811483 |
| HSPB11 | 362.259224 | 0.322734884 | 0.26707884 | 1.20838807 | 0.22689803 | 0.41931165 |
| LRRC42 | 176.749687 | -0.636882451 | 0.17398785 | -3.6604995 | 0.00025172 | 0.00320667 |
| TMEM59 | 891.093304 | -0.187288684 | 0.26535955 | -0.7057921 | 0.48031742 | 0.66529011 |
| TCEANC2 | 187.79753 | 0.496971291 | 0.3066624 | 1.62058109 | 0.10510751 | 0.25355661 |
| MRPL37 | 168.121 | -0.126816193 | 0.19516964 | -0.6497742 | 0.51583811 | 0.69333133 |
| SSBP3 | 1019.75945 | -0.413969666 | 0.4167997 | -0.9932101 | 0.32060758 | 0.52105682 |
| ACOT11 | 876.387694 | 1.312703245 | 0.57370992 | 2.28809578 | 0.02213194 | 0.08872845 |
| FAM151A | 253.249605 | 3.704785065 | 0.57272721 | 6.468673 | 9.89E-11 | 1.56E-08 |
| MROH7-TTC4 | 44.610276 | 0.117071286 | 0.56531871 | 0.207089 | 0.83594035 | 0.91337062 |
| PARS2 | 37.358971 | -0.242378214 | 0.35225703 | -0.688072 | 0.49140742 | 0.6739091 |
| TTC22 | 261.02799 | -1.336382966 | 0.37384042 | -3.5747418 | 0.00035057 | 0.00413894 |
| DHCR24 | 277.938214 | -0.260219824 | 0.34974885 | -0.7440191 | 0.45686492 | 0.64559252 |
| USP24 | 1877.03681 | -0.259277959 | 0.13697318 | -1.8929104 | 0.05836978 | 0.17150581 |
| AC119674.2 | 126.32144 | -0.37306047 | 0.50390413 | -0.7403402 | 0.45909361 | 0.6475935 |
| PLPP3 | 2402.41493 | -0.391171493 | 0.24881063 | -1.5721655 | 0.11591217 | 0.27122023 |
| PRKAA2 | 2397.23175 | 0.315003652 | 0.17333484 | 1.817313 | 0.0691692 | 0.1926254 |
| FYB2 | 583.417273 | -0.632452684 | 0.73274748 | -0.863125 | 0.38806873 | 0.58387159 |
| DAB1 | 572.441636 | 1.212841488 | 0.62707757 | 1.93411715 | 0.05309873 | 0.16120138 |
| TACSTD2 | 185.114448 | -0.646611402 | 0.55285821 | -1.1695791 | 0.2421704 | 0.43740014 |
| MYSM1 | 1032.58902 | -0.294171298 | 0.19179214 | -1.5338027 | 0.1250782 | 0.28507598 |
| JUN | 1395.99781 | -2.586759481 | 0.40814805 | -6.3377969 | 2.33E-10 | 3.23E-08 |
| LINC01135 | 326.76049 | 0.362022413 | 0.61117122 | 0.59234205 | 0.55362157 | 0.72298883 |
| LINC01358 | 70.375865 | -1.620966671 | 0.57211931 | -2.8332668 | 0.00460749 | 0.0285898 |
| FGGY | 7132.42434 | 1.420939297 | 0.4462508 | 3.18417199 | 0.00145169 | 0.01200862 |
| HOOK1 | 1087.44777 | -0.476171578 | 0.39742117 | -1.1981535 | 0.23085725 | 0.42378291 |
| CYP2J2 | 94.5627704 | -0.772823667 | 1.09468631 | -0.7059773 | 0.48020226 | 0.66519782 |
| LINC01748 | 87.6297963 | -3.247436062 | 1.13742297 | -2.8550822 | 0.00430257 | 0.02710263 |
| AC099792.1 | 122.445537 | -2.993431802 | 1.05876112 | -2.8272967 | 0.00469428 | 0.02895793 |
| NFIA | 7802.80343 | 0.889312707 | 0.40339163 | 2.20458887 | 0.02748296 | 0.10294526 |
| TM2D1 | 975.195387 | 0.068242426 | 0.17636661 | 0.38693507 | 0.69880428 | 0.82658873 |
| AC099791.2 | 39.8192991 | 0.209352889 | 0.54457861 | 0.38443098 | 0.70065904 | 0.82796169 |
| PATJ | 19961.4356 | 0.171965541 | 0.22712395 | 0.75714402 | 0.44896359 | 0.63921457 |
| USP1 | 144.053879 | -0.389807106 | 0.28665773 | -1.3598346 | 0.17388227 | 0.35367328 |
| DOCK7 | 1669.92594 | -0.445352135 | 0.26915391 | -1.6546374 | 0.09799807 | 0.24224699 |
| ANGPTL3 | 108.729228 | 2.639064282 | 0.71333892 | 3.69959385 | 0.00021594 | 0.00284065 |
| AC103923.1 | 93.7689019 | -0.023651943 | 0.70138759 | -0.0337216 | 0.97309912 | 0.98648544 |
| ATG4C | 401.506817 | 0.818300138 | 0.27736473 | 2.95026749 | 0.00317499 | 0.02155502 |
| ALG6 | 189.819982 | 0.523882574 | 0.25904203 | 2.02238443 | 0.04313665 | 0.1405119 |
| ITGB3BP | 595.524387 | -0.536712489 | 0.44702823 | -1.2006233 | 0.22989737 | 0.42303709 |
| EFCAB7 | 317.474511 | 0.316864041 | 0.38138596 | 0.8308225 | 0.40607391 | 0.60009523 |
| PGM1 | 992.156339 | -0.371093567 | 0.26973134 | -1.3757896 | 0.16888679 | 0.34755609 |
| ROR1 | 2800.80739 | 0.533547272 | 0.19228738 | 2.77473888 | 0.0055246 | 0.03266857 |
| ROR1-AS1 | 51.060686 | -1.724847036 | 0.74190698 | -2.3248831 | 0.02007821 | 0.08264625 |
| UBE2U | 55.6752667 | 0.229143871 | 1.17361376 | 0.19524641 | 0.84520005 | 0.91854292 |
| CACHD1 | 1459.76866 | 1.395164791 | 0.41647325 | 3.34995058 | 0.00080826 | 0.00771764 |
| RAVER2 | 419.260516 | -0.499361304 | 0.34644879 | -1.4413712 | 0.14947985 | 0.31978206 |
| JAK1 | 2687.56057 | -0.507566275 | 0.20926831 | -2.425433 | 0.01529014 | 0.06811241 |
| AK4 | 4875.65303 | 0.926660514 | 0.40102494 | 2.31073037 | 0.02084775 | 0.08502236 |
| DNAJC6 | 1750.7989 | -0.073595051 | 0.3793519 | -0.1940021 | 0.84617427 | 0.91896252 |
| AL139294.1 | 164.864986 | -1.170144324 | 0.71578809 | -1.6347636 | 0.10209861 | 0.24959259 |
| LEPR | 923.338322 | -0.239951203 | 0.34626173 | -0.6929764 | 0.48832437 | 0.671495 |
| PDE4B | 99.0275882 | 0.602896083 | 0.62547925 | 0.96389462 | 0.33509876 | 0.53490145 |
| SGIP1 | 140.60563 | 0.228912825 | 0.75117608 | 0.30473924 | 0.76056476 | 0.86630058 |
| WDR78 | 880.844749 | 0.336313474 | 0.37597807 | 0.8945029 | 0.37105288 | 0.56877072 |
| MIER1 | 1204.88914 | -0.437428934 | 0.15906693 | -2.7499678 | 0.00596011 | 0.03447057 |
| SLC35D1 | 694.878746 | 0.900866513 | 0.31716566 | 2.84036584 | 0.00450618 | 0.02805006 |
| IL12RB2 | 135.600976 | 0.264128903 | 0.48970888 | 0.53935902 | 0.58963915 | 0.7515514 |
| SERBP1 | 1042.6223 | -0.31049726 | 0.19253297 | -1.6126966 | 0.10681045 | 0.25646507 |
| LINC01702 | 91.1026936 | 3.565580993 | 0.72953359 | 4.88748025 | 1.02E-06 | 3.93E-05 |
| GADD45A | 1288.66294 | 0.359981542 | 0.35460004 | 1.01517626 | 0.31002182 | 0.51032413 |
| GNG12 | 3364.5216 | -0.190083211 | 0.18492326 | -1.0279032 | 0.30399536 | 0.50421883 |
| GNG12-AS1 | 1170.94321 | -0.011636514 | 0.3548292 | -0.0327947 | 0.97383831 | 0.98684149 |
| WLS | 1208.58879 | 0.857950901 | 0.26340908 | 3.25710446 | 0.00112555 | 0.00988969 |
| LRRC7 | 745.476636 | 0.918088752 | 0.56222624 | 1.6329525 | 0.10247899 | 0.25016553 |
| LRRC40 | 484.620986 | 0.535928363 | 0.21214946 | 2.52618299 | 0.01153094 | 0.05567263 |

| | | | | | | |
|---|---|---|---|---|---|---|
| SRSF11 | 2693.23227 | 0.004370623 | 0.18084143 | 0.02416826 | 0.98071839 | 0.99047407 |
| ANKRD13C | 1191.68163 | 0.185858435 | 0.1201758 | 1.54655456 | 0.1219707 | 0.28025441 |
| HHLA3 | 79.2400854 | 0.336365847 | 0.40315862 | 0.83432632 | 0.40409712 | 0.59815975 |
| CTH | 173.178671 | -0.041648097 | 0.44437183 | -0.0937235 | 0.92532878 | 0.96057873 |
| LINC01788 | 943.075653 | 2.294058166 | 0.70677907 | 3.24579243 | 0.00117124 | 0.01018022 |
| PTGER3 | 963.6654 | 0.429256826 | 0.80655612 | 0.5322095 | 0.59458091 | 0.75522636 |
| AL031429.1 | 103.902371 | 2.316768556 | 0.52491508 | 4.41360638 | 1.02E-05 | 0.00025798 |
| ZRANB2-AS1 | 81.055784 | -0.505350578 | 0.61046666 | -0.8278103 | 0.40777795 | 0.60161981 |
| ZRANB2 | 1616.67585 | -0.057437637 | 0.17739361 | -0.3237864 | 0.74609976 | 0.85651503 |
| NEGR1 | 1235.27281 | 1.580793972 | 0.68767513 | 2.29875113 | 0.02151908 | 0.08724477 |
| LRRIQ3 | 80.3074574 | 0.293343368 | 0.58373536 | 0.50252801 | 0.61529616 | 0.77099044 |
| CRYZ | 1345.9038 | 0.455088961 | 0.30746862 | 1.48011513 | 0.13884252 | 0.30497906 |
| TYW3 | 321.765798 | -0.251391477 | 0.20221034 | -1.2432177 | 0.21378761 | 0.4040798 |
| SLC44A5 | 201.474099 | -1.264383336 | 0.61344167 | -2.0611305 | 0.03929059 | 0.13206379 |
| ACADM | 1402.23263 | 1.454739412 | 0.4319007 | 3.36822654 | 0.00075653 | 0.00736492 |
| RABGGTB | 321.520186 | -0.312568631 | 0.36975583 | -0.8453379 | 0.39792218 | 0.59241006 |
| MSH4 | 41.481184 | 0.445707518 | 0.66699999 | 0.66822718 | 0.50398859 | 0.68461449 |
| ASB17 | 50.6950692 | 1.179260317 | 0.76279483 | 1.54597315 | 0.12211105 | 0.28035345 |
| ST6GALNAC3 | 115.840016 | -0.390959176 | 0.30947622 | -1.2632931 | 0.20648386 | 0.39528971 |
| PIGK | 447.911053 | -0.278162683 | 0.20118573 | -1.3826163 | 0.16678254 | 0.34469756 |
| AK5 | 237.262085 | -3.258762638 | 0.70336681 | -4.6330913 | 3.60E-06 | 0.00011071 |
| AC118549.1 | 3013.56603 | 0.450944176 | 0.17070718 | 2.64162397 | 0.00825096 | 0.04389935 |
| USP33 | 2070.32103 | 0.080910893 | 0.21462453 | 0.3769881 | 0.70618245 | 0.83179881 |
| MIGA1 | 1215.57319 | 0.251359979 | 0.09055795 | 2.77568109 | 0.00550862 | 0.03261295 |
| NEXN | 128.908583 | -0.138178678 | 0.3932926 | -0.3513381 | 0.72533471 | 0.84377177 |
| FUBP1 | 1213.79069 | 0.160183922 | 0.1201482 | 1.33321952 | 0.18245977 | 0.36478648 |
| GIPC2 | 2477.85834 | 1.316751279 | 0.46125271 | 2.85472857 | 0.00430736 | 0.02712036 |
| PTGFR | 302.329535 | 0.869011786 | 0.53179468 | 1.63411148 | 0.10223544 | 0.24973128 |
| IFI44L | 722.513462 | -2.21093813 | 1.73348128 | -1.2754324 | 0.20215625 | 0.38967173 |
| IFI44 | 266.228353 | -1.796684472 | 1.54245858 | -1.1648186 | 0.24409242 | 0.43920163 |
| AL136234.1 | 545.53719 | -0.675875734 | 0.71256096 | -0.9485164 | 0.34286664 | 0.54344046 |
| LINC01781 | 4724.20893 | 0.016973666 | 0.69340896 | 0.02447858 | 0.98047087 | 0.9904425 |
| ADGRL2 | 1709.41547 | -0.27304791 | 0.27467134 | -0.9940896 | 0.32017927 | 0.52060465 |
| LINC01362 | 119.554388 | -1.077534799 | 0.69657925 | -1.5468948 | 0.12188862 | 0.28018235 |
| LINC01725 | 1728.99278 | 0.40318718 | 0.56866902 | 0.70900149 | 0.47832355 | 0.66397743 |
| TTLL7 | 1002.13997 | -1.923239361 | 0.4646678 | -4.1389555 | 3.49E-05 | 0.00068144 |
| AL359504.2 | 41.6333499 | -0.795695601 | 0.66145033 | -1.2029559 | 0.22899337 | 0.42199679 |
| PRKACB | 1175.34483 | -0.150897561 | 0.13200746 | -1.1430988 | 0.25299758 | 0.44966778 |
| RPF1 | 162.725268 | -0.085443679 | 0.30398648 | -0.2810772 | 0.77865116 | 0.87801961 |
| GNG5 | 480.653525 | -0.937783354 | 0.26984622 | -3.4752511 | 0.00051038 | 0.00541883 |
| SPATA1 | 206.352196 | -0.450214297 | 0.58681257 | -0.7672199 | 0.44295077 | 0.63456586 |
| CTBS | 354.216155 | -0.420737999 | 0.34408466 | -1.2227747 | 0.22141482 | 0.4134786 |
| SSX2IP | 707.231447 | 0.555694886 | 0.31412961 | 1.76899875 | 0.07689408 | 0.20663516 |
| MCOLN2 | 185.834977 | 0.356352456 | 0.75324698 | 0.47308846 | 0.63615007 | 0.78575302 |
| WDR63 | 277.295925 | -0.626938655 | 0.49010392 | -1.2791953 | 0.20082828 | 0.38798494 |
| MCOLN3 | 408.900763 | -0.235451856 | 0.29505082 | -0.7980044 | 0.42486793 | 0.61796103 |
| SYDE2 | 395.669997 | 1.431896193 | 0.29011839 | 4.93555822 | 7.99E-07 | 3.18E-05 |
| C1orf52 | 311.450139 | -0.139780186 | 0.18354822 | -0.7615448 | 0.44633176 | 0.63743065 |
| BCL10 | 165.553821 | -0.438738695 | 0.20962414 | -2.0929779 | 0.03635113 | 0.12478461 |
| AL078459.1 | 483.24116 | -0.639633632 | 0.46242493 | -1.3832162 | 0.1665986 | 0.34453334 |
| DDAH1 | 3951.83172 | 0.626596063 | 0.2848129 | 2.20002698 | 0.02780498 | 0.10373513 |
| CYR61 | 210.805818 | -0.551997261 | 0.68046987 | -0.8112001 | 0.41725074 | 0.61047432 |
| ZNHIT6 | 566.26334 | -0.415530206 | 0.27426029 | -1.5150943 | 0.12974852 | 0.2919821 |
| AL136382.1 | 57.5042045 | 0.004485316 | 0.7608777 | 0.00589492 | 0.99529656 | 0.99749068 |
| ODF2L | 900.711621 | -0.184180533 | 0.23993188 | -0.7676368 | 0.44270301 | 0.63434352 |
| AL356270.1 | 40.389895 | 0.092380269 | 0.56524851 | 0.16343302 | 0.87017751 | 0.93290367 |
| SH3GLB1 | 964.464868 | -0.697766446 | 0.14303176 | -4.8784023 | 1.07E-06 | 4.08E-05 |
| SELENOF | 977.121878 | -0.155719178 | 0.14849261 | -1.0486662 | 0.29433179 | 0.49425158 |
| HS2ST1 | 1773.27427 | 0.175261593 | 0.20467882 | 0.85627614 | 0.39184506 | 0.5869326 |
| LMO4 | 250.589451 | 0.447338569 | 0.34350699 | 1.30226919 | 0.19282438 | 0.37749937 |
| PKN2-AS1 | 196.620022 | 0.27440124 | 0.54789117 | 0.50083165 | 0.61648961 | 0.77182636 |
| PKN2 | 2910.11401 | 0.189911762 | 0.12831647 | 1.48002639 | 0.1388662 | 0.30497906 |
| GTF2B | 379.395074 | 0.074917204 | 0.17179834 | 0.43607641 | 0.66278129 | 0.80379726 |
| KYAT3 | 642.199707 | -0.267770993 | 0.33588409 | -0.7972125 | 0.42532763 | 0.61832293 |
| GBP3 | 228.760019 | -1.55439371 | 0.45409532 | -3.423056 | 0.00061921 | 0.00630108 |
| GBP1 | 244.27275 | -1.886561322 | 0.7601972 | -2.4816736 | 0.0130767 | 0.06084261 |
| GBP2 | 675.022963 | -3.893080316 | 0.53131585 | -7.3272429 | 2.35E-13 | 7.49E-11 |
| GBP4 | 45.9770625 | -0.015706692 | 0.77101456 | -0.0203715 | 0.98374705 | 0.99214339 |
| LRRC8B | 432.181723 | -0.467326453 | 0.35134216 | -1.3301178 | 0.18347947 | 0.36586457 |
| LRRC8D | 153.757946 | 1.021662125 | 0.32615407 | 3.13245247 | 0.00173352 | 0.01372699 |
| ZNF326 | 790.538931 | 0.136203807 | 0.16020446 | 0.85018734 | 0.39522094 | 0.59005483 |
| ZNF644 | 2626.67807 | 0.131134985 | 0.15483831 | 0.84691565 | 0.39704214 | 0.59164429 |
| HFM1 | 73.2204755 | 1.175334338 | 0.60726509 | 1.93545515 | 0.05293447 | 0.16094131 |
| CDC7 | 86.6195587 | -2.337331941 | 0.78774501 | -2.9671174 | 0.00300606 | 0.02068478 |
| TGFBR3 | 2442.85205 | 0.784463684 | 0.29366107 | 2.67132337 | 0.00755528 | 0.04112906 |
| BTBD8 | 341.701873 | 0.247628797 | 0.56379085 | 0.43922103 | 0.66050139 | 0.80254344 |
| GLMN | 416.374671 | -0.08747532 | 0.24766203 | -0.3532044 | 0.7239352 | 0.84285907 |
| RPAP2 | 1551.17269 | 0.346358955 | 0.21606577 | 1.60302555 | 0.10892901 | 0.25996684 |
| EVI5 | 2331.44276 | -0.076894848 | 0.22978644 | -0.3346361 | 0.73789957 | 0.85123628 |
| RPL5 | 975.551514 | -0.85273729 | 0.34489114 | -2.4724824 | 0.01341783 | 0.06196774 |
| FAM69A | 902.342407 | 0.410300343 | 0.32252262 | 1.27215988 | 0.20331632 | 0.39134885 |
| MTF2 | 1262.26053 | -0.048322607 | 0.23562779 | -0.2050803 | 0.83750942 | 0.91436098 |
| TMED5 | 985.719819 | -0.6670816 | 0.23633164 | -2.8226504 | 0.00476285 | 0.02924926 |
| CCDC18 | 521.655333 | -0.313199196 | 0.26730924 | -1.1716737 | 0.2413281 | 0.43643228 |
| DR1 | 323.077177 | 0.246831018 | 0.22070097 | 1.1183957 | 0.26339803 | 0.46133361 |
| FNBP1L | 3286.16623 | -0.44404607 | 0.22757008 | -1.9512498 | 0.05102734 | 0.1573841 |
| BCAR3 | 1511.26177 | -2.38096639 | 0.62776869 | -3.7927447 | 0.00014899 | 0.00210105 |
| AL109613.1 | 67.0832788 | 1.108132592 | 0.43093613 | 2.57145438 | 0.01012724 | 0.05095963 |
| DNTTIP2 | 218.540464 | -1.759805981 | 0.65879708 | -2.6712413 | 0.00755713 | 0.04112906 |
| GCLM | 1712.16936 | -2.749054305 | 0.55541676 | -4.9495343 | 7.44E-07 | 3.03E-05 |
| ARHGAP29 | 4503.79132 | 0.066403597 | 0.3604967 | 0.18420029 | 0.85385633 | 0.92301331 |
| AC097059.1 | 199.761129 | 1.43765735 | 0.53442754 | 2.69008846 | 0.00714331 | 0.03954092 |
| ABCD3 | 2978.57461 | 0.718564526 | 0.32408245 | 2.21722749 | 0.02660755 | 0.10077637 |
| SLC44A3 | 1166.96354 | 0.321930995 | 0.31444587 | 1.02380417 | 0.30592778 | 0.50583258 |
| CNN3 | 2217.91844 | -0.421312758 | 0.34519955 | -1.2204904 | 0.22227901 | 0.41454644 |
| AC105942.1 | 428.947737 | 0.319469466 | 0.3256884 | 0.98090527 | 0.32663946 | 0.52677357 |
| ALG14 | 611.092879 | 0.039386828 | 0.23598684 | 0.16690265 | 0.86744665 | 0.93102273 |
| TMEM56-RWDD3 | 49.3374301 | 0.00721157 | 0.39290878 | 0.01835431 | 0.9853562 | 0.99288962 |
| AC092802.1 | 118.497886 | 0.746398068 | 0.45035756 | 1.65734549 | 0.09744965 | 0.24137013 |
| PTBP2 | 2364.79408 | -0.306655945 | 0.17892456 | -1.7138841 | 0.08655002 | 0.22345849 |
| DPYD | 1868.90412 | 0.293684922 | 0.37130369 | 0.79095612 | 0.4289696 | 0.62205719 |
| DPYD-AS1 | 69.4745506 | -0.427446301 | 0.62070967 | -0.6886413 | 0.49104904 | 0.67375209 |
| SNX7 | 714.264119 | 0.088002272 | 0.17886768 | 0.49199649 | 0.62272182 | 0.77610911 |
| LINC01708 | 36.2060125 | 2.724186824 | 0.64525174 | 4.2218977 | 2.42E-05 | 0.00051243 |
| FRRS1 | 190.831351 | -0.368422294 | 0.4882998 | -0.7545002 | 0.45054893 | 0.64075861 |
| AGL | 677.717641 | 0.120528941 | 0.18668561 | 0.64562524 | 0.51852209 | 0.69523449 |
| AC093019.2 | 42.8512851 | -1.040117555 | 0.55112595 | -1.8872593 | 0.05912546 | 0.17307314 |
| SASS6 | 109.855945 | -0.182847622 | 0.27505184 | -0.6647751 | 0.50619434 | 0.68623201 |
| TRMT13 | 443.99103 | 0.245339652 | 0.19915448 | 1.23190627 | 0.2179841 | 0.40911198 |
| LRRC39 | 64.5220094 | 0.008474682 | 0.31795244 | 0.02665393 | 0.97873576 | 0.98916955 |
| DBT | 500.595339 | 0.731706769 | 0.27941546 | 2.61870536 | 0.00882641 | 0.04601226 |
| RTCA | 524.77668 | 0.248840646 | 0.26098621 | 0.9534628 | 0.34035564 | 0.54096182 |
| CDC14A | 677.494108 | 0.489395519 | 0.31889257 | 1.53467206 | 0.1248644 | 0.2847781 |

| | | | | | | |
|---|---|---|---|---|---|---|
| LINC01349 | 254.199806 | 2.582187194 | 0.63256859 | 4.08206675 | 4.46E-05 | 0.00082793 |
| VCAM1 | 2641.79014 | -2.031696927 | 0.85693216 | -2.3708959 | 0.01774503 | 0.07582017 |
| EXTL2 | 327.328774 | 0.799382671 | 0.2771694 | 2.88409422 | 0.00392541 | 0.0252846 |
| SLC30A7 | 873.795934 | -0.639316198 | 0.23254628 | -2.7491999 | 0.00597409 | 0.03450556 |
| DPH5 | 252.886336 | 0.155252265 | 0.24947119 | 0.62232543 | 0.5337279 | 0.70689258 |
| AC093157.1 | 280.945921 | 0.648460337 | 0.29303132 | 2.21293867 | 0.02690187 | 0.10147055 |
| RNPC3 | 962.114711 | 0.289221389 | 0.19100446 | 1.51421275 | 0.12997189 | 0.29229281 |
| AMY2B | 209.531064 | 0.381760798 | 0.41921251 | 0.91066175 | 0.36247362 | 0.56030588 |
| VAV3 | 2961.23021 | 1.55388964 | 0.46823745 | 3.31859323 | 0.00090472 | 0.00839608 |
| SLC25A24 | 578.761588 | -0.772201936 | 0.24044259 | -3.2115855 | 0.00132005 | 0.01114842 |
| FAM102B | 117.447085 | 0.166362862 | 0.33527751 | 0.49619452 | 0.61975718 | 0.77427999 |
| PRPF38B | 794.933367 | 0.09239556 | 0.37395944 | 0.24707374 | 0.80485115 | 0.89408337 |
| STXBP3 | 694.24714 | -0.037268883 | 0.18280959 | -0.2038672 | 0.83845726 | 0.91454366 |
| AL449266.1 | 172.467499 | -0.15784603 | 0.24854073 | -0.6350912 | 0.52536893 | 0.70084961 |
| GPSM2 | 155.315495 | -1.504410696 | 0.46710433 | -3.2207167 | 0.0012787 | 0.01087249 |
| WDR47 | 302.599032 | 0.406459299 | 0.2405021 | 1.69004471 | 0.0910194 | 0.23112318 |
| TAF13 | 242.418573 | -0.813364069 | 0.46615032 | -1.7448536 | 0.08101036 | 0.21382803 |
| AL356488.3 | 33.7633969 | 0.154102497 | 0.5347297 | 0.28818765 | 0.7732031 | 0.87420952 |
| TMEM167B | 165.050576 | -0.441402743 | 0.20328979 | -2.1712982 | 0.02990864 | 0.10891544 |
| KIAA1324 | 43.1885848 | 1.667905628 | 0.40562461 | 4.11194388 | 3.92E-05 | 0.00074382 |
| SARS | 419.852218 | -0.69540106 | 0.23161615 | -3.0023859 | 0.00267872 | 0.01904422 |
| CELSR2 | 60.4937478 | 0.30074097 | 0.53661178 | 0.56044422 | 0.57517648 | 0.73976511 |
| SORT1 | 730.781208 | -0.057780517 | 0.34892281 | -0.1655968 | 0.86847423 | 0.93161498 |
| PSMA5 | 949.01968 | -0.931820236 | 0.46238009 | -2.0152689 | 0.04387648 | 0.14207769 |
| SYPL2 | 486.189949 | -0.023922145 | 0.46267496 | -0.051704 | 0.95876455 | 0.97890396 |
| ATXN7L2 | 73.6542635 | -0.431134147 | 0.45384621 | -0.9499565 | 0.34213437 | 0.54270009 |
| CYB561D1 | 104.109539 | -0.44723 | 0.37298033 | -1.1990713 | 0.23050023 | 0.42369138 |
| GNAI3 | 906.139675 | -0.585616072 | 0.2917841 | -2.0070185 | 0.0447477 | 0.14414963 |
| AMPD2 | 159.450547 | -0.352394276 | 0.42082403 | -0.8373911 | 0.40237279 | 0.5965737 |
| GSTM4 | 124.917348 | 0.751710749 | 0.48625826 | 1.54590844 | 0.12212668 | 0.28035345 |
| GSTM3 | 92.0635042 | -0.363181873 | 0.48142463 | -0.7543899 | 0.45061514 | 0.64075861 |
| CSF1 | 136.049431 | -3.377538386 | 0.53884873 | -6.2680641 | 3.66E-10 | 4.86E-08 |
| AHCYL1 | 2884.39594 | -0.972708929 | 0.38520861 | -2.5251484 | 0.01156494 | 0.05579749 |
| STRIP1 | 293.485339 | -0.130477432 | 0.26561185 | -0.4912335 | 0.62326132 | 0.77635796 |
| KCNC4 | 44.5943306 | -0.450289982 | 0.46457665 | -0.969248 | 0.33242147 | 0.53186659 |
| RBM15-AS1 | 54.8738691 | -1.352424842 | 0.55821049 | -2.4227865 | 0.01540198 | 0.0684771 |
| RBM15 | 337.544251 | -0.083440567 | 0.26035155 | -0.3204919 | 0.74859546 | 0.85823001 |
| SLC16A4 | 2064.43858 | 0.181648051 | 0.33642664 | 0.53993361 | 0.58924282 | 0.75126513 |
| AL355488.1 | 78.7915547 | -0.003140359 | 0.26162522 | -0.0120033 | 0.990423 | 0.99518064 |
| LAMTOR5 | 932.771473 | -0.265936362 | 0.25541621 | -1.0411883 | 0.29778818 | 0.49814285 |
| LRIF1 | 205.144773 | -0.309937299 | 0.23339205 | -1.3279686 | 0.18418849 | 0.36695809 |
| DRAM2 | 527.3859 | -0.593689356 | 0.26306973 | -2.2567756 | 0.0240221 | 0.09363008 |
| CEPT1 | 1157.43388 | -0.045539679 | 0.25805013 | -0.1764761 | 0.85991993 | 0.92628466 |
| DENND2D | 468.891559 | -1.056011749 | 0.51171273 | -2.0636808 | 0.03904799 | 0.13150633 |
| CHI3L2 | 100.22535 | -3.74669434 | 0.79429265 | -4.71702 | 2.39E-06 | 7.98E-05 |
| OVGP1 | 101.45782 | -1.582202334 | 0.53416918 | -2.9619873 | 0.0030566 | 0.02099041 |
| WDR77 | 218.162366 | -0.535599029 | 0.27779643 | -1.928027 | 0.05385176 | 0.16298214 |
| ATP5PB | 598.037895 | -0.269817349 | 0.26176705 | -1.0307537 | 0.30265636 | 0.50284408 |
| RAP1A | 1430.73862 | -0.13493751 | 0.16834765 | -0.8015408 | 0.42281863 | 0.61622276 |
| INKA2 | 81.4325611 | -0.918504841 | 0.47187089 | -1.9465173 | 0.05159263 | 0.15834756 |
| DDX20 | 117.215377 | 0.221686442 | 0.33279734 | 0.66613045 | 0.50532772 | 0.68537856 |
| KCND3 | 143.070086 | -1.792148464 | 0.61616868 | -2.9085355 | 0.00363126 | 0.02382656 |
| AL445426.1 | 37.8423947 | 0.503402544 | 0.60813588 | 0.82777972 | 0.40779526 | 0.60161981 |
| CTTNBP2NL | 1351.07195 | -0.320680901 | 0.29157602 | -1.0998192 | 0.27141091 | 0.46966663 |
| WNT2B | 591.769593 | -0.248523174 | 0.48517912 | -0.5122297 | 0.60849023 | 0.76540108 |
| ST7L | 722.666041 | 0.348664653 | 0.17096965 | 2.03933652 | 0.04141645 | 0.13668488 |
| CAPZA1 | 1188.35584 | -0.511008401 | 0.16078497 | -3.17821 | 0.00148187 | 0.01222551 |
| MOV10 | 501.896749 | 0.331590337 | 0.35035822 | 0.94643231 | 0.34392813 | 0.54439007 |
| AL603832.1 | 95.1174404 | -0.425859591 | 0.532312 | -0.8000188 | 0.42369992 | 0.61678577 |
| RHOC | 748.07572 | -1.092122823 | 0.43210488 | -2.5274485 | 0.01148947 | 0.05558791 |
| SLC16A1 | 298.506254 | 0.13786514 | 0.56401611 | 0.24443475 | 0.80689413 | 0.89531977 |
| SLC16A1-AS1 | 499.947993 | -0.213534761 | 0.34948947 | -0.6109905 | 0.54120584 | 0.71231293 |
| AL357055.3 | 92.1075152 | -0.502797817 | 0.54792894 | -0.9176333 | 0.3588109 | 0.5573055 |
| LRIG2 | 734.238755 | 0.705910744 | 0.32449035 | 2.17544449 | 0.02959682 | 0.1080623 |
| MAGI3 | 3240.82421 | 0.358472725 | 0.28437847 | 1.26054806 | 0.20747172 | 0.39646102 |
| PHTF1 | 405.042207 | -0.741782022 | 0.3015727 | -2.4597121 | 0.01390485 | 0.06355699 |
| RSBN1 | 444.405478 | 0.137335833 | 0.12569501 | 1.09261169 | 0.27456433 | 0.47243952 |
| AP4B1 | 68.1652171 | 0.003594439 | 0.37515785 | 0.00958114 | 0.99235547 | 0.99626489 |
| HIPK1 | 555.166653 | -0.467951998 | 0.28043067 | -1.6686905 | 0.09517873 | 0.23793877 |
| SYT6 | 185.49192 | 1.679217126 | 0.5802901 | 2.89375454 | 0.00380666 | 0.0246238 |
| AL121999.1 | 622.382957 | 2.719847691 | 0.67967411 | 4.00169384 | 6.29E-05 | 0.00108297 |
| TRIM33 | 2202.90892 | -0.058461003 | 0.27132978 | -0.2154611 | 0.82940786 | 0.90944441 |
| BCAS2 | 208.99617 | -0.309753059 | 0.23610807 | -1.3119122 | 0.18954976 | 0.37367446 |
| DENND2C | 116.441083 | -0.553580687 | 0.55482262 | -0.9977616 | 0.31839499 | 0.51856466 |
| NRAS | 228.75009 | -0.782881719 | 0.39403368 | -1.9868396 | 0.04694017 | 0.14921358 |
| CSDE1 | 1271.97525 | -0.179842186 | 0.14265775 | -1.2606548 | 0.20743323 | 0.39644274 |
| SIKE1 | 309.811587 | -0.241904814 | 0.27692627 | -0.8735351 | 0.38237148 | 0.57906281 |
| VANGL1 | 227.027048 | 0.459743859 | 0.2576329 | 1.78449204 | 0.07434375 | 0.20222857 |
| SLC22A15 | 121.943929 | -2.046050733 | 0.6803404 | -3.0073927 | 0.00263499 | 0.01884831 |
| ATP1A1 | 5195.73519 | -0.934444734 | 0.2318639 | -4.0301432 | 5.57E-05 | 0.00098844 |
| LINC01762 | 791.859252 | 0.163300655 | 0.65631898 | 0.24881294 | 0.80350548 | 0.89315921 |
| AL390066.1 | 103.283869 | 0.091808095 | 0.3178185 | 0.28886958 | 0.77268119 | 0.87376019 |
| CD58 | 1148.18871 | -0.704054306 | 0.2106247 | -3.3426959 | 0.00082969 | 0.00788811 |
| IGSF3 | 835.928619 | -0.75767999 | 0.50688751 | -1.4947695 | 0.13497461 | 0.29936987 |
| PTGFRN | 122.111994 | -1.628173084 | 0.54584441 | -2.982852 | 0.00285576 | 0.0199612 |
| TTF2 | 259.373537 | -0.758088131 | 0.19847227 | -3.8196174 | 0.00013366 | 0.00193459 |
| MAN1A2 | 1990.80207 | -0.25138259 | 0.24075354 | -1.0441491 | 0.29641642 | 0.49657539 |
| TENT5C | 121.092029 | 1.043293011 | 0.49390699 | 2.11232689 | 0.03465842 | 0.12088116 |
| GDAP2 | 768.776011 | -0.003491952 | 0.15939308 | -0.0219078 | 0.9825215 | 0.99141801 |
| WDR3 | 163.740946 | -0.294385768 | 0.33609605 | -0.8758977 | 0.38108563 | 0.57750588 |
| SPAG17 | 408.004982 | -0.864476398 | 0.60685858 | -1.4245105 | 0.15429876 | 0.32695817 |
| WARS2 | 438.502332 | 0.31602066 | 0.2434757 | 1.29795566 | 0.19430257 | 0.37933419 |
| AL359915.2 | 1070.55326 | -0.153081396 | 0.35802552 | -0.4275712 | 0.66896335 | 0.80745691 |
| AL359915.1 | 345.969257 | 2.220729775 | 0.85288545 | 2.60378432 | 0.00922008 | 0.04753847 |
| LINC01780 | 80.5779907 | 2.598797176 | 0.87314712 | 2.97635657 | 0.00291695 | 0.02026493 |
| HAO2 | 960.243653 | 3.49293146 | 0.901176 | 3.87597037 | 0.0001062 | 0.00163338 |
| ZNF697 | 729.067575 | 0.456160789 | 0.48458535 | 0.94134252 | 0.34652936 | 0.54670612 |
| PHGDH | 110.715221 | 1.010448317 | 0.47534379 | 2.12572109 | 0.03352649 | 0.11836062 |
| HMGCS2 | 66.6389811 | 2.342331193 | 1.12700595 | 2.07836631 | 0.03767563 | 0.12833497 |
| NOTCH2 | 688.290654 | -1.103042913 | 0.23545529 | -4.6847234 | 2.80E-06 | 9.06E-05 |
| SEC22B | 749.720667 | -0.58271177 | 0.25388282 | -2.2951997 | 0.02172168 | 0.08763551 |
| NBPF26 | 119.181524 | -0.786782091 | 0.59012523 | -1.333246 | 0.18245109 | 0.36478648 |
| LINC00623 | 623.090176 | -0.728662561 | 0.45215621 | -1.6115284 | 0.10706461 | 0.25683139 |
| SRGAP2C | 64.4388036 | -1.244551587 | 0.44556515 | -2.7931978 | 0.00521898 | 0.03132072 |
| AC239800.3 | 53.9808278 | -1.196494967 | 0.61517118 | -1.9449789 | 0.05177752 | 0.1587111 |
| AC245595.1 | 43.8202746 | -0.131044326 | 0.46778318 | -0.280139 | 0.77937084 | 0.87852271 |
| NBPF15 | 306.913239 | -1.589287585 | 0.46523953 | -3.4160631 | 0.00063534 | 0.00643171 |
| FP700111.1 | 1506.25798 | -0.317552225 | 0.23860911 | -1.3308471 | 0.18323934 | 0.36558246 |
| SRGAP2B | 119.914048 | -1.212034367 | 0.48133367 | -2.5180752 | 0.01179981 | 0.05659203 |
| NBPF20 | 43.2916966 | -0.394905541 | 0.41090826 | -0.9610553 | 0.33652438 | 0.53661467 |
| GPR89A | 416.040493 | 0.818840116 | 0.23911937 | 3.42439897 | 0.00061616 | 0.00627934 |
| PDZK1 | 5995.49376 | 1.198019328 | 0.53773299 | 2.22790744 | 0.02588669 | 0.09872811 |
| RNF115 | 1709.02961 | -0.218527612 | 0.31238607 | -0.6995434 | 0.4842125 | 0.66811967 |

| | | | | | | |
|---|---|---|---|---|---|---|
| POLR3C | 277.990393 | -0.490481673 | 0.2845073 | -1.7239687 | 0.08471349 | 0.22016835 |
| PIAS3 | 64.2642165 | -0.516448722 | 0.28710165 | -1.7988358 | 0.07204466 | 0.19797838 |
| PEX11B | 101.339472 | 0.448480529 | 0.43270112 | 1.03646722 | 0.29998423 | 0.50029003 |
| RBM8A | 235.273096 | -0.52812114 | 0.24213544 | -2.1810981 | 0.02917616 | 0.10692593 |
| LIX1L | 105.307901 | -0.370543622 | 0.31377722 | -1.1809131 | 0.23763726 | 0.43246069 |
| TXNIP | 2748.02646 | -0.258042335 | 0.74950835 | -0.3442821 | 0.73063412 | 0.84678203 |
| NBPF12 | 597.531059 | 0.087210573 | 0.24889889 | 0.35038554 | 0.72604937 | 0.84424489 |
| PRKAB2 | 275.180153 | 0.168487048 | 0.23409616 | 0.71973436 | 0.47168857 | 0.65792563 |
| AC242426.2 | 494.519757 | -0.048871518 | 0.32378074 | -0.1509402 | 0.88002292 | 0.9374737 |
| FMO5 | 1203.28857 | 1.191772408 | 0.63258545 | 1.88397064 | 0.05956895 | 0.17383682 |
| CHD1L | 651.100285 | -0.142155493 | 0.26301136 | -0.5404918 | 0.58885789 | 0.75097461 |
| LINC00624 | 99.3902225 | -0.861113896 | 0.46771749 | -1.8410983 | 0.06560715 | 0.18599321 |
| BCL9 | 615.874431 | 0.398538318 | 0.22484124 | 1.77253212 | 0.07630627 | 0.20548723 |
| ACP6 | 439.433686 | 0.034967303 | 0.37765172 | 0.0925914 | 0.92622817 | 0.96090956 |
| GPR89B | 928.081787 | 1.57080819 | 0.32225015 | 4.87449946 | 1.09E-06 | 4.14E-05 |
| NBPF11 | 476.770658 | -0.078292777 | 0.25028038 | -0.3128203 | 0.75441721 | 0.86224896 |
| AC239809.3 | 164.098346 | -0.847021662 | 0.4503134 | -1.8809604 | 0.05997731 | 0.1745825 |
| NBPF14 | 214.871435 | -0.489755683 | 0.25971854 | -1.8857171 | 0.05933309 | 0.17344394 |
| PDE4DIP | 1016.5124 | -0.031720181 | 0.38160753 | -0.0831225 | 0.93375412 | 0.96473023 |
| NBPF9 | 311.533944 | -0.633548406 | 0.26028488 | -2.4340576 | 0.01493062 | 0.06688038 |
| AC245297.3 | 1379.73988 | -0.884117436 | 0.33143624 | -2.667534 | 0.00764102 | 0.04147937 |
| NBPF19 | 286.408268 | -0.861369031 | 0.64183536 | -1.3420405 | 0.17958287 | 0.36086531 |
| HIST2H2BE | 43.8846224 | -0.970054447 | 0.65030582 | -1.4916896 | 0.13578053 | 0.30037826 |
| BOLA1 | 56.0409246 | -0.276465868 | 0.39850583 | -0.6937561 | 0.48783515 | 0.67102422 |
| SF3B4 | 187.349801 | -0.193408286 | 0.34875942 | -0.5545607 | 0.57919515 | 0.74258983 |
| MTMR11 | 214.95422 | 0.606032277 | 0.44697218 | 1.35586129 | 0.17514331 | 0.3551841 |
| OTUD7B | 1619.86152 | -0.765288462 | 0.3886032 | -1.9693313 | 0.04891506 | 0.15328731 |
| VPS45 | 715.678598 | 0.180209345 | 0.11861452 | 1.51928576 | 0.12869058 | 0.29046822 |
| ANP32E | 225.601349 | 0.118513418 | 0.32054183 | 0.3697284 | 0.71158487 | 0.83507652 |
| APH1A | 268.233626 | -0.071534537 | 0.29027786 | -0.2464347 | 0.80534575 | 0.89441535 |
| C1orf54 | 128.941881 | -0.842585542 | 0.44411374 | -1.8972292 | 0.0577977 | 0.17059281 |
| CIART | 46.9645589 | -1.767451192 | 0.71746831 | -2.4634554 | 0.0137605 | 0.06310311 |
| MRPS21 | 646.494704 | -0.276210882 | 0.23642732 | -1.1682697 | 0.24269798 | 0.43783354 |
| PRPF3 | 526.281931 | 0.150426246 | 0.25967217 | 0.5792929 | 0.56239155 | 0.73014439 |
| RPRD2 | 3003.85019 | 0.710021093 | 0.20494216 | 3.46449502 | 0.00053123 | 0.00556693 |
| TARS2 | 151.289156 | 0.455621852 | 0.3952415 | 1.15276825 | 0.24900552 | 0.44519533 |
| ADAMTSL4 | 50.6253622 | -1.378458401 | 0.46783062 | -2.9464903 | 0.00321402 | 0.02173648 |
| MCL1 | 686.936169 | -0.839633256 | 0.2863145 | -2.9325558 | 0.00336184 | 0.02248941 |
| ENSA | 1319.00386 | -0.113000764 | 0.25066804 | -0.4507984 | 0.65213482 | 0.79625975 |
| GOLPH3L | 562.753375 | 0.100652685 | 0.21594163 | 0.46611061 | 0.64113633 | 0.78933685 |
| CTSS | 48.7421593 | -1.549647938 | 0.55430285 | -2.7956702 | 0.00517922 | 0.03113667 |
| CTSK | 39.583229 | -0.022582226 | 0.38035982 | -0.0593707 | 0.95265686 | 0.97608518 |
| ARNT | 786.514977 | 0.100759535 | 0.1617296 | 0.62301235 | 0.53327641 | 0.7065165 |
| SETDB1 | 686.814838 | 0.113379189 | 0.14522592 | 0.78070903 | 0.43497365 | 0.62740017 |
| CERS2 | 721.545356 | -0.195930558 | 0.22048284 | -0.8886431 | 0.37419491 | 0.57131832 |
| ANXA9 | 72.2969556 | 1.542716943 | 0.42222633 | 3.65376776 | 0.00025842 | 0.00326166 |
| MINDY1 | 198.411152 | -0.224473095 | 0.42234039 | -0.5314981 | 0.59507369 | 0.75552751 |
| PRUNE1 | 289.845158 | 0.38876079 | 0.33649412 | 1.15532714 | 0.24795649 | 0.44427968 |
| BNIPL | 40.0978333 | 0.856416917 | 0.43512854 | 1.96819295 | 0.04904584 | 0.15352166 |
| C1orf56 | 87.1330891 | -0.66395399 | 0.4472642 | -1.4844783 | 0.13768209 | 0.30335382 |
| CDC42SE1 | 245.624475 | -0.211134668 | 0.19338942 | -1.0917591 | 0.27493898 | 0.47266884 |
| GABPB2 | 415.535516 | 1.233045752 | 0.32416687 | 3.80373767 | 0.00014253 | 0.0020266 |
| SEMA6C | 170.644003 | 2.162108651 | 0.49679351 | 4.35212743 | 1.35E-05 | 0.00032252 |
| SCNM1 | 139.792694 | 0.131246751 | 0.23986501 | 0.54716922 | 0.58426248 | 0.74702254 |
| LYSMD1 | 55.3838912 | 0.642464571 | 0.28472168 | 2.25646523 | 0.02404151 | 0.09367168 |
| VPS72 | 132.823935 | 0.069801734 | 0.21497253 | 0.32470072 | 0.74540759 | 0.85593548 |
| PIP5K1A | 1949.32706 | 0.13989777 | 0.22461906 | 0.62282234 | 0.53340127 | 0.70661363 |
| PSMD4 | 496.86117 | -0.117059586 | 0.15931903 | -0.7347496 | 0.46249203 | 0.65092701 |
| ZNF687 | 63.093206 | 0.226306876 | 0.32504597 | 0.69623036 | 0.48628458 | 0.66963057 |
| PI4KB | 560.97418 | 0.368374696 | 0.20649037 | 1.78397999 | 0.07442692 | 0.20232093 |
| RFX5 | 51.6026537 | -0.213598298 | 0.52795275 | -0.4045784 | 0.68578741 | 0.81810131 |
| SELENBP1 | 194.168681 | 1.849950575 | 0.62091667 | 2.97938625 | 0.00288826 | 0.02011659 |
| PSMB4 | 253.183039 | -0.683493565 | 0.43224269 | -1.5812727 | 0.1138157 | 0.26754791 |
| POGZ | 2108.83886 | 0.579956208 | 0.26014096 | 2.22939209 | 0.02578783 | 0.09854301 |
| CGN | 550.852361 | -0.002502608 | 0.52701506 | -0.0047486 | 0.99621114 | 0.99803144 |
| TUFT1 | 583.671164 | -0.615728781 | 0.35975614 | -1.7115171 | 0.0869857 | 0.22424543 |
| SNX27 | 1153.76653 | 0.463879295 | 0.32488201 | 1.42783927 | 0.15333813 | 0.32545768 |
| MRPL9 | 128.831969 | -0.19923514 | 0.29066256 | -0.6854517 | 0.49305894 | 0.67529565 |
| TDRKH | 55.3094908 | -0.291808554 | 0.41354325 | -0.7056301 | 0.48041822 | 0.66534319 |
| RORC | 405.008342 | 1.368700941 | 0.61677294 | 2.21913262 | 0.0264777 | 0.10036701 |
| AL450992.2 | 43.991019 | -0.455463485 | 0.25187128 | -1.8083185 | 0.07055695 | 0.19526026 |
| THEM4 | 489.88728 | 0.503753748 | 0.22423092 | 2.24658466 | 0.02466658 | 0.09542897 |
| S100A10 | 1391.94551 | -2.558309411 | 0.53670933 | -4.7666573 | 1.87E-06 | 6.42E-05 |
| S100A11 | 2343.0759 | -2.303997112 | 0.47472391 | -4.8533412 | 1.21E-06 | 4.51E-05 |
| FLG-AS1 | 601.86541 | 0.958149255 | 0.46328873 | 2.06814713 | 0.03862619 | 0.1307282 |
| S100A6 | 1772.16522 | -1.636949586 | 0.72031028 | -2.2725617 | 0.0230526 | 0.0911662 |
| S100A4 | 50.4720323 | 1.255630612 | 0.75266677 | 1.66824239 | 0.09526762 | 0.23797755 |
| S100A2 | 152.454395 | -0.478886662 | 0.92839823 | -0.5158203 | 0.60597992 | 0.7635748 |
| S100A16 | 448.204917 | -2.163895154 | 0.54007911 | -4.0066263 | 6.16E-05 | 0.00106678 |
| S100A14 | 76.2942827 | -1.827803561 | 0.66118265 | -2.7644458 | 0.00570196 | 0.0333718 |
| S100A13 | 451.981492 | -0.851898555 | 0.27929753 | -3.0501471 | 0.00228729 | 0.01702059 |
| S100A1 | 503.896455 | -2.021528286 | 0.57369907 | -3.5236737 | 0.00042561 | 0.00478181 |
| AL162258.2 | 40.5904501 | 0.208079038 | 0.39410554 | 0.52797795 | 0.59751464 | 0.75757406 |
| CHTOP | 449.332052 | 0.479857473 | 0.2445207 | 1.96244108 | 0.04971115 | 0.15482636 |
| SNAPIN | 61.293505 | -0.755327593 | 0.39327534 | -1.9206075 | 0.0547812 | 0.16467727 |
| ILF2 | 413.854297 | -0.54148158 | 0.29582963 | -1.8303832 | 0.06719266 | 0.18880889 |
| NPR1 | 274.663503 | -2.37654207 | 0.5130789 | -4.6319232 | 3.62E-06 | 0.00011109 |
| INTS3 | 834.880989 | -0.327020613 | 0.20606474 | -1.58698 | 0.11251718 | 0.26577166 |
| SLC27A3 | 98.5622501 | -0.764024663 | 0.3752575 | -2.036001 | 0.04175025 | 0.13739993 |
| GATAD2B | 2338.63479 | 0.416550404 | 0.26260834 | 1.58620403 | 0.11269303 | 0.26591198 |
| DENND4B | 282.310738 | 0.032150099 | 0.21136513 | 0.15210692 | 0.8791026 | 0.93706833 |
| CRTC2 | 166.537569 | 0.068742185 | 0.33143141 | 0.20740999 | 0.83568968 | 0.91316952 |
| SLC39A1 | 341.353767 | -1.653354788 | 0.53484019 | -3.0913062 | 0.00199278 | 0.0153326 |
| CREB3L4 | 38.3119058 | 0.582848226 | 0.37351694 | 1.56043316 | 0.11865755 | 0.27517534 |
| JTB | 322.555366 | -0.099489744 | 0.32144754 | -0.3095054 | 0.75693711 | 0.86367577 |
| RAB13 | 405.460785 | -1.486660682 | 0.30635679 | -4.85271 | 1.22E-06 | 4.51E-05 |
| RPS27 | 385.258839 | -0.680424731 | 0.39554232 | -1.7202324 | 0.0853902 | 0.22117023 |
| NUP210L | 61.4518182 | -0.865323306 | 0.46150871 | -1.874988 | 0.06079437 | 0.17639891 |
| TPM3 | 711.519953 | -1.190209246 | 0.34470021 | -3.4528823 | 0.00055463 | 0.0057637 |
| C1orf43 | 626.265702 | -0.105776286 | 0.2569469 | -0.4116659 | 0.68058429 | 0.81442647 |
| UBAP2L | 802.798748 | 0.058967626 | 0.23062009 | 0.25569162 | 0.79818897 | 0.89012761 |
| HAX1 | 130.008442 | -0.129457088 | 0.27697396 | -0.467398 | 0.64021512 | 0.78873168 |
| IL6R | 1965.51257 | -0.956032531 | 0.49058647 | -1.9487544 | 0.05132476 | 0.15790414 |
| UBE2Q1 | 191.46344 | -0.333225805 | 0.33906696 | -0.9827729 | 0.32571925 | 0.52586969 |
| ADAR | 1050.77668 | -0.206590884 | 0.32798495 | -0.6298792 | 0.52877364 | 0.70313352 |
| PMVK | 247.885023 | 0.128756567 | 0.29880695 | 0.43090219 | 0.66653949 | 0.80635371 |
| PBXIP1 | 114.550998 | -0.159028715 | 0.39793074 | -0.3996392 | 0.6894223 | 0.82043055 |
| PYGO2 | 87.2108963 | 0.354602642 | 0.24388779 | 1.45395818 | 0.1459579 | 0.31471527 |
| SHC1 | 52.2514846 | -0.718999859 | 0.30393028 | -2.3656737 | 0.0179973 | 0.07663529 |
| CKS1B | 50.4785552 | -0.383763975 | 0.4524763 | -0.8481416 | 0.39635911 | 0.59104595 |
| FLAD1 | 125.650885 | -0.237203508 | 0.3581493 | -0.6623034 | 0.50777678 | 0.68768861 |
| ZBTB7B | 189.419589 | 0.259620356 | 0.26202891 | 0.99080808 | 0.3217793 | 0.5221533 |
| ADAM15 | 84.4568997 | -1.178719707 | 0.42759751 | -2.7566103 | 0.00584039 | 0.03399331 |

| | | | | | | |
|---|---|---|---|---|---|---|
| EFNA1 | 1188.62686 | -1.730492492 | 0.50082912 | -3.4552554 | 0.00054977 | 0.00572903 |
| SLC50A1 | 140.52751 | -1.229907761 | 0.27824224 | -4.4202769 | 9.86E-06 | 0.00025255 |
| DPM3 | 211.667458 | -0.603205758 | 0.3448207 | -1.7493316 | 0.08023371 | 0.21239155 |
| THBS3 | 206.882359 | 1.035451295 | 0.4060989 | 2.54975151 | 0.01077997 | 0.05320889 |
| MTX1 | 120.542661 | -0.757923413 | 0.44576454 | -1.7002775 | 0.08907874 | 0.22768161 |
| GBA | 224.963813 | 0.303396075 | 0.24510288 | 1.23783153 | 0.21577853 | 0.40633004 |
| FAM189B | 74.0287283 | 0.246806479 | 0.33128195 | 0.7450043 | 0.45626912 | 0.64534938 |
| SCAMP3 | 298.822643 | -0.126790699 | 0.35146451 | -0.3607497 | 0.7182866 | 0.83920512 |
| CLK2 | 331.952819 | 0.388035166 | 0.19970753 | 1.94301716 | 0.05201409 | 0.15901947 |
| HCN3 | 118.069119 | 0.238964987 | 0.33730219 | 0.70845963 | 0.47865988 | 0.66413513 |
| PKLR | 196.527223 | 1.826977927 | 0.78152076 | 2.33772156 | 0.0194017 | 0.0806854 |
| FDPS | 915.954016 | 0.053906702 | 0.42584716 | 0.12658697 | 0.89926731 | 0.94743919 |
| RUSC1-AS1 | 43.6532536 | 0.475882406 | 0.41055443 | 1.15912134 | 0.24640673 | 0.44202291 |
| RUSC1 | 142.79788 | -0.440287032 | 0.30493812 | -1.443857 | 0.14877921 | 0.31889237 |
| ASH1L | 5566.71047 | 0.34025796 | 0.14022879 | 2.42644862 | 0.01524741 | 0.06796626 |
| MSTO1 | 119.905576 | -0.424161013 | 0.47957214 | -0.8844572 | 0.37644948 | 0.57294604 |
| YY1AP1 | 811.394966 | 0.209198823 | 0.17302978 | 1.20903365 | 0.22664992 | 0.41919989 |
| DAP3 | 919.391008 | -0.346026873 | 0.27634869 | -1.2521387 | 0.21051934 | 0.39989367 |
| GON4L | 1863.82906 | 0.609018834 | 0.1982324 | 3.07224674 | 0.00212454 | 0.01610679 |
| RIT1 | 584.859779 | -0.425868851 | 0.27886393 | -1.5271565 | 0.12672211 | 0.28781772 |
| KHDC4 | 995.724033 | -0.210286169 | 0.22389032 | -0.9392374 | 0.34760885 | 0.54753787 |
| ARHGEF2 | 168.322503 | 0.143810113 | 0.42347111 | 0.3395984 | 0.73415898 | 0.84943603 |
| SSR2 | 423.863603 | -0.833779364 | 0.28791435 | -2.8959285 | 0.00378039 | 0.02452331 |
| UBQLN4 | 252.322803 | 0.320792848 | 0.20141888 | 1.59266522 | 0.11123531 | 0.26374372 |
| LAMTOR2 | 161.248564 | -0.494208963 | 0.33935976 | -1.4562981 | 0.14531024 | 0.31371357 |
| RAB25 | 61.3915291 | -0.674165033 | 0.6347721 | -1.0620584 | 0.28820918 | 0.48801522 |
| MEX3A | 60.5664447 | -0.491564916 | 0.45966183 | -1.0694056 | 0.28488696 | 0.4843043 |
| LMNA | 815.543738 | -1.110483436 | 0.31687411 | -3.504494 | 0.00045748 | 0.0050326 |
| SLC25A44 | 433.815819 | 1.526332204 | 0.29216597 | 5.22419574 | 1.75E-07 | 8.79E-06 |
| PAQR6 | 59.3688213 | 0.372027153 | 0.64442845 | 0.57729784 | 0.56373827 | 0.73104641 |
| SMG5 | 856.437205 | -0.606850678 | 0.14268372 | -4.2531179 | 2.11E-05 | 0.00046048 |
| TMEM79 | 58.7168895 | 0.639178025 | 0.43086791 | 1.48346632 | 0.13795056 | 0.30371689 |
| GLMP | 224.351725 | 0.022961025 | 0.26961824 | 0.08516125 | 0.9321332 | 0.96391408 |
| CCT3 | 703.3649 | -0.56024891 | 0.27666052 | -2.0250411 | 0.04286315 | 0.13988694 |
| RHBG | 33.526269 | 0.673029119 | 0.81400707 | 0.82680992 | 0.40834481 | 0.6021713 |
| MEF2D | 261.045869 | -0.196348049 | 0.27546968 | -0.7127755 | 0.47598466 | 0.66169592 |
| NAXE | 158.485441 | -0.634384928 | 0.37328106 | -1.6994833 | 0.08922816 | 0.22776623 |
| GPATCH4 | 129.175567 | -1.169309381 | 0.52805988 | -2.21435 | 0.02680471 | 0.10124337 |
| ISG20L2 | 147.601819 | -0.617999844 | 0.49466152 | -1.2493388 | 0.21154117 | 0.40103556 |
| RRNAD1 | 200.079193 | 0.499770139 | 0.29602576 | 1.68826572 | 0.09136023 | 0.23173107 |
| MRPL24 | 128.941295 | 0.219953262 | 0.23633322 | 0.93069125 | 0.3520133 | 0.55108727 |
| HDGF | 737.203458 | -0.381250052 | 0.31326285 | -1.2170292 | 0.22359309 | 0.41588945 |
| PRCC | 398.228393 | 0.399668592 | 0.28836568 | 1.38597836 | 0.16575351 | 0.34319991 |
| ARHGEF11 | 939.211336 | 0.186642685 | 0.21233646 | 0.87899497 | 0.379404 | 0.57597637 |
| ETV3 | 112.87517 | -0.264539516 | 0.27121732 | -0.9753784 | 0.32937259 | 0.5289719 |
| KIRREL1 | 211.860597 | -1.298201049 | 0.52894239 | -2.4543335 | 0.0141146 | 0.06416877 |
| IFI16 | 201.719627 | -3.19183274 | 0.68801706 | -4.6391767 | 3.50E-06 | 0.00010897 |
| DUSP23 | 1398.98015 | -1.559484275 | 0.61661345 | -2.5291117 | 0.01143516 | 0.05544457 |
| TAGLN2 | 446.403978 | -1.643539853 | 0.54605021 | -3.0098695 | 0.0026136 | 0.01874653 |
| IGSF8 | 122.273493 | 0.05894788 | 0.33160969 | 0.17776284 | 0.85890923 | 0.92584688 |
| PEA15 | 157.048185 | -2.316301223 | 0.76812053 | -3.015544 | 0.00256519 | 0.01855205 |
| DCAF8 | 292.395268 | 0.655620458 | 0.26550063 | 2.46937439 | 0.01353495 | 0.06233992 |
| AL139011.2 | 99.9780342 | -0.255058624 | 0.42365743 | -0.6020398 | 0.54714766 | 0.71747374 |
| COPA | 1407.88292 | -0.720635873 | 0.24164951 | -2.9821533 | 0.00286229 | 0.01999661 |
| NCSTN | 229.98766 | 0.156842439 | 0.21210652 | 0.73945129 | 0.45963301 | 0.64816765 |
| F11R | 378.89746 | -0.64055185 | 0.313826 | -2.0411051 | 0.04124038 | 0.13640973 |
| AL591806.3 | 163.052141 | 0.354620574 | 0.3630217 | 0.97685778 | 0.32863957 | 0.52846816 |
| USF1 | 126.984504 | 0.15932743 | 0.38626415 | 0.41248309 | 0.67998537 | 0.81435096 |
| KLHDC9 | 36.632511 | -0.001268081 | 0.58650555 | -0.0021621 | 0.9982749 | 0.99914962 |
| PFDN2 | 450.26642 | -0.667896508 | 0.35771001 | -1.8671451 | 0.06188132 | 0.1784948 |
| NIT1 | 434.64129 | 0.404552264 | 0.27429046 | 1.47490462 | 0.14023819 | 0.30697512 |
| DEDD | 188.118704 | 0.123434274 | 0.18276859 | 0.67535823 | 0.4994482 | 0.68105217 |
| UFC1 | 194.719062 | -0.795788843 | 0.26527138 | -2.9999047 | 0.00270064 | 0.01918015 |
| USP21 | 54.3006501 | 0.513042996 | 0.32336458 | 1.5865776 | 0.11260834 | 0.26584957 |
| PPOX | 90.9121221 | 0.376669946 | 0.30150955 | 1.2492803 | 0.21156257 | 0.40103556 |
| B4GALT3 | 115.126391 | 0.230584178 | 0.28338862 | 0.81366773 | 0.41583533 | 0.60934946 |
| NDUFS2 | 504.866348 | 0.105639182 | 0.26570726 | 0.39757733 | 0.69094178 | 0.82139973 |
| TOMM40L | 59.6259414 | 0.225611354 | 0.33872392 | 0.66606265 | 0.50537106 | 0.68537856 |
| NR1I3 | 163.492312 | 2.660286418 | 0.86934779 | 3.06009453 | 0.00221267 | 0.01660049 |
| SDHC | 1104.37419 | 0.646135874 | 0.26109484 | 2.47471715 | 0.01333417 | 0.061768 |
| AL592295.4 | 141.953867 | 0.122140329 | 0.43620707 | 0.28000538 | 0.77947337 | 0.87856603 |
| AL592295.3 | 40.5227463 | -0.752533951 | 0.68847398 | -1.0930463 | 0.27437346 | 0.47233234 |
| DUSP12 | 153.675603 | -0.051177256 | 0.25379105 | -0.2016511 | 0.84018946 | 0.91531369 |
| ATF6 | 1437.13338 | -0.115504461 | 0.13384122 | -0.8629962 | 0.38813958 | 0.58387159 |
| NOS1AP | 629.099 | -0.902680075 | 0.4708188 | -1.9172558 | 0.05520544 | 0.16565258 |
| UHMK1 | 581.00285 | -0.157414257 | 0.15529715 | -1.0136326 | 0.31075808 | 0.51104512 |
| UAP1 | 213.255338 | -0.006096604 | 0.28335776 | -0.0215156 | 0.98283438 | 0.99158773 |
| DDR2 | 227.114087 | 1.464738995 | 0.67452149 | 2.17152311 | 0.02989165 | 0.10888251 |
| HSD17B7 | 250.574579 | 0.251824111 | 0.41300631 | 0.6097343 | 0.54203783 | 0.71302423 |
| RGS5 | 98.1928975 | 0.628561489 | 0.44785086 | 1.40350627 | 0.16046592 | 0.3363674 |
| NUF2 | 56.8657807 | 0.147141098 | 0.60568 | 0.24293537 | 0.80805546 | 0.89590037 |
| PBX1 | 5440.92206 | 1.626575194 | 0.24543459 | 6.62732652 | 3.42E-11 | 6.42E-09 |
| MGST3 | 2357.54235 | -0.489679156 | 0.23914439 | -2.0476297 | 0.0405963 | 0.13473448 |
| ALDH9A1 | 417.807707 | -0.113706343 | 0.26952137 | -0.4218825 | 0.6731108 | 0.81072605 |
| TMCO1 | 587.135396 | -0.072959212 | 0.15831831 | -0.4608387 | 0.6449143 | 0.79180937 |
| AL451074.2 | 49.4120077 | 1.13506843 | 0.48952649 | 2.31870687 | 0.02041094 | 0.08368911 |
| UCK2 | 364.376775 | -0.609963533 | 0.45067524 | -1.3534436 | 0.17591395 | 0.35616729 |
| FAM78B | 197.59303 | 1.303847091 | 0.36702796 | 3.55244622 | 0.00038167 | 0.00442897 |
| POGK | 216.580987 | -0.591311868 | 0.34463457 | -1.7157648 | 0.08620509 | 0.22281976 |
| TADA1 | 219.008456 | 0.288457119 | 0.24179379 | 1.1929881 | 0.23287403 | 0.42678341 |
| ILDR2 | 321.149742 | 0.805436587 | 0.2797186 | 2.87945314 | 0.00398365 | 0.0255783 |
| MAEL | 76.8991702 | 0.622255411 | 0.78490654 | 0.79277643 | 0.42790809 | 0.62093332 |
| POU2F1 | 2335.83814 | -0.052714597 | 0.28853774 | -0.1826957 | 0.85503681 | 0.92334276 |
| CREG1 | 289.854536 | -0.621477022 | 0.37400757 | -1.6616696 | 0.09657903 | 0.23999435 |
| MPZL1 | 908.429204 | -1.337493047 | 0.26807607 | -4.9892296 | 6.06E-07 | 2.50E-05 |
| ADCY10 | 132.129861 | 1.407579457 | 0.53728472 | 2.61980178 | 0.00879809 | 0.04594111 |
| MPC2 | 543.785329 | 0.060069449 | 0.32698964 | 0.18370444 | 0.85424533 | 0.92322796 |
| DCAF6 | 2773.23775 | -0.074154605 | 0.23918086 | -0.3100357 | 0.75653381 | 0.86351 |
| TIPRL | 346.294664 | -0.098130022 | 0.17989891 | -0.5454731 | 0.58542815 | 0.74799251 |
| SFT2D2 | 1256.79019 | -1.102470925 | 0.2003325 | -5.5032057 | 3.73E-08 | 2.51E-06 |
| TBX19 | 262.908021 | -1.091865705 | 0.33904933 | -3.2203741 | 0.00128023 | 0.01087249 |
| AL023755.1 | 103.087748 | -0.576865255 | 0.68939369 | -0.8367719 | 0.40272081 | 0.59683145 |
| LINC00970 | 384.774825 | 0.987238541 | 0.65394552 | 1.50966481 | 0.13112897 | 0.29412287 |
| ATP1B1 | 4335.04536 | 1.077368646 | 0.30334866 | 3.55158527 | 0.00038292 | 0.00442897 |
| NME7 | 1670.52835 | 0.033678611 | 0.31329352 | 0.10749859 | 0.91439345 | 0.9543495 |
| BLZF1 | 494.503306 | -0.627012875 | 0.30470174 | -2.0577922 | 0.03961008 | 0.13258635 |
| CCDC181 | 79.9407568 | 0.29929428 | 0.37901921 | 0.78965465 | 0.42972949 | 0.6224561 |
| SLC19A2 | 570.028884 | -0.304890798 | 0.39927016 | -0.7636203 | 0.44509355 | 0.63643828 |
| C1orf112 | 100.983915 | -0.203351654 | 0.27799148 | -0.7315032 | 0.46447185 | 0.65217329 |
| SCYL3 | 338.424521 | 0.296106659 | 0.25148334 | 1.17744045 | 0.23901975 | 0.43403362 |
| KIFAP3 | 901.388971 | 0.236812755 | 0.23171165 | 1.02201491 | 0.30677385 | 0.50680356 |
| GORAB | 130.743819 | -0.037750662 | 0.29901653 | -0.1262494 | 0.8995345 | 0.94743919 |
| MROH9 | 68.6228414 | -0.93537942 | 0.63965525 | -1.4623181 | 0.14365407 | 0.31126563 |

| | | | | | | |
|---|---|---|---|---|---|---|
| FMO2 | 109.79018 | 2.767270841 | 1.02161314 | 2.70872674 | 0.00675419 | 0.03787496 |
| FMO1 | 1218.26929 | 2.499551135 | 0.64615687 | 3.86833486 | 0.00010958 | 0.00166892 |
| FMO4 | 1260.35729 | 2.073406678 | 0.6888055 | 3.01014827 | 0.0026112 | 0.01873913 |
| PRRC2C | 2856.65936 | -0.368479731 | 0.23209732 | -1.5876087 | 0.11237485 | 0.26548122 |
| MYOCOS | 687.541224 | -2.794123515 | 0.83310752 | -3.353857 | 0.00079694 | 0.00764957 |
| VAMP4 | 215.083097 | -0.129357465 | 0.23873576 | -0.5418437 | 0.58792619 | 0.75041555 |
| METTL13 | 141.45454 | -0.046431032 | 0.27417792 | -0.1693464 | 0.86552422 | 0.92955312 |
| DNM3 | 448.233332 | -0.531537735 | 0.27219178 | -1.952806 | 0.05084259 | 0.15713627 |
| PIGC | 222.907507 | -0.240997895 | 0.35003931 | -0.6884881 | 0.49114546 | 0.67375209 |
| SUCO | 1384.42593 | -1.167240558 | 0.22051654 | -5.293211 | 1.20E-07 | 6.44E-06 |
| AL031599.1 | 191.054736 | -2.457225989 | 0.81154053 | -3.0278537 | 0.00246297 | 0.01799537 |
| AL645568.1 | 272.36452 | 0.283821781 | 0.25662331 | 1.10598598 | 0.26873259 | 0.46679984 |
| PRDX6 | 842.272322 | -0.739698891 | 0.21885459 | -3.3798646 | 0.00072522 | 0.00714118 |
| KLHL20 | 1039.08808 | 0.581892175 | 0.21710551 | 2.68022762 | 0.00735721 | 0.0404512 |
| CENPL | 49.2371814 | -0.718870358 | 0.40062749 | -1.794361 | 0.07275556 | 0.19907103 |
| DARS2 | 135.32936 | -0.031129345 | 0.22589289 | -0.1378058 | 0.89039392 | 0.94326785 |
| ZBTB37 | 301.67256 | 0.611054551 | 0.30331665 | 2.01457637 | 0.04394906 | 0.1422455 |
| RC3H1 | 2058.84163 | -0.096866181 | 0.27225863 | -0.3557874 | 0.72199981 | 0.84225118 |
| AL121983.2 | 61.8937479 | 1.00578487 | 0.49320877 | 2.03926801 | 0.04142329 | 0.13668488 |
| RABGAP1L | 10702.2175 | 0.37748203 | 0.25369557 | 1.48793309 | 0.13676854 | 0.30178467 |
| Z99127.1 | 34.699176 | -0.930807552 | 0.47151858 | -1.9740634 | 0.04837454 | 0.15215288 |
| CACYBP | 235.603114 | -0.579873046 | 0.46605462 | -1.2442169 | 0.21341973 | 0.40349576 |
| MRPS14 | 159.457793 | -0.399781083 | 0.29148691 | -1.3715233 | 0.17021189 | 0.34924222 |
| KIAA0040 | 114.264838 | -0.470239035 | 0.39949854 | -1.1770732 | 0.23916629 | 0.43415782 |
| AL513329.1 | 43.5780979 | -0.211226025 | 0.73328075 | -0.2880561 | 0.77330376 | 0.87420952 |
| COP1 | 3127.98395 | -0.129145605 | 0.18549089 | -0.6962369 | 0.48628048 | 0.66963057 |
| AL359265.3 | 59.9739484 | 0.304028002 | 0.61935485 | 0.49087853 | 0.62351236 | 0.77652952 |
| PAPPA2 | 124.744655 | -0.135229906 | 0.47932559 | -0.2821254 | 0.77784739 | 0.87758633 |
| AL122019.1 | 52.9297298 | 0.948056832 | 0.76614026 | 1.23744552 | 0.21592173 | 0.40653696 |
| SEC16B | 772.996215 | 1.074259164 | 0.67101936 | 1.60093616 | 0.10939106 | 0.26077541 |
| RASAL2 | 4594.60371 | -0.70594426 | 0.40582049 | -1.7395481 | 0.0819384 | 0.21543891 |
| RALGPS2 | 1797.59609 | -0.476616965 | 0.28755082 | -1.6575051 | 0.09741739 | 0.24133385 |
| ANGPTL1 | 97.3746932 | -1.854974085 | 0.63510827 | -2.9207211 | 0.00349222 | 0.02319182 |
| FAM20B | 337.156195 | 0.050406161 | 0.28620839 | 0.17611699 | 0.86020203 | 0.92629345 |
| TOR3A | 114.431879 | -0.076804889 | 0.34215397 | -0.2244746 | 0.822388 | 0.90483804 |
| ABL2 | 1951.55655 | -0.230223898 | 0.29968265 | -0.7682257 | 0.44235313 | 0.63404423 |
| SOAT1 | 770.645397 | -1.39128435 | 0.24953575 | -5.575491 | 2.47E-08 | 1.76E-06 |
| AXDND1 | 80.5712967 | 0.790127697 | 0.43306831 | 1.82448747 | 0.06807841 | 0.19052689 |
| NPHS2 | 37.0079044 | -0.339329774 | 0.6937126 | -0.4891504 | 0.62473525 | 0.77762859 |
| TOR1AIP2 | 985.396659 | -0.344758184 | 0.29883998 | -1.1536548 | 0.24864172 | 0.44486777 |
| TOR1AIP1 | 348.358271 | 0.112824569 | 0.17524739 | 0.64380171 | 0.51970402 | 0.69597507 |
| CEP350 | 1967.31417 | 0.259776743 | 0.09634062 | 2.69644037 | 0.00700849 | 0.03898759 |
| QSOX1 | 151.556643 | -0.751524485 | 0.41847176 | -1.7958786 | 0.07251382 | 0.19878939 |
| LHX4 | 50.1668779 | 0.832433393 | 0.56880325 | 1.46348213 | 0.1433355 | 0.3110032 |
| ACBD6 | 2262.67364 | -0.085609527 | 0.24131011 | -0.3547698 | 0.72276209 | 0.84228022 |
| XPR1 | 4137.18341 | 0.138481233 | 0.21660978 | 0.63931201 | 0.52261998 | 0.69847426 |
| KIAA1614 | 132.554039 | -0.27171796 | 0.47019769 | -0.5778803 | 0.56334497 | 0.73081294 |
| AL162431.2 | 36.0914709 | -0.040343016 | 0.31169112 | -0.1294327 | 0.89701529 | 0.9465267 |
| STX6 | 375.332345 | -0.321789017 | 0.19321314 | -1.6654614 | 0.09582074 | 0.23882914 |
| MR1 | 106.029111 | -0.642847776 | 0.41754447 | -1.5395912 | 0.12366004 | 0.28302023 |
| CACNA1E | 2536.22493 | 0.873294662 | 0.7880009 | 1.1082407 | 0.26775788 | 0.465756 |
| GLUL | 222.943555 | -0.784332865 | 0.23007005 | -3.4091046 | 0.00065176 | 0.0065641 |
| RNASEL | 101.422751 | -0.044803373 | 0.52569219 | -0.0852274 | 0.93208062 | 0.96391408 |
| NPL | 2031.18879 | 1.20880804 | 0.45672021 | 2.64671457 | 0.00812779 | 0.04343378 |
| DHX9 | 1366.71548 | -0.301079436 | 0.18522286 | -1.6254983 | 0.10405643 | 0.25221955 |
| LAMC1 | 2298.14788 | -1.855760571 | 0.3642817 | -5.0943009 | 3.50E-07 | 1.57E-05 |
| LAMC2 | 100.375988 | -2.856943413 | 0.83470263 | -3.4227081 | 0.00062001 | 0.00630447 |
| SMG7 | 1560.81692 | 0.081631063 | 0.19860816 | 0.41101565 | 0.68106106 | 0.81452787 |
| ARPC5 | 468.809993 | -1.313121623 | 0.36616656 | -3.586132 | 0.00033562 | 0.00400726 |
| RGL1 | 2905.79457 | 1.561830155 | 0.39614877 | 3.94253439 | 8.06E-05 | 0.00131406 |
| TSEN15 | 438.745151 | 0.475997393 | 0.27519827 | 1.72965256 | 0.08369237 | 0.21850514 |
| C1orf21 | 105.062347 | -0.646492437 | 0.34378116 | -1.8805348 | 0.06003523 | 0.17467691 |
| EDEM3 | 903.982097 | -0.047184912 | 0.26311931 | -0.179329 | 0.85767941 | 0.92517602 |
| FAM129A | 67.8672094 | -2.493439617 | 0.54842804 | -4.5465211 | 5.45E-06 | 0.00015414 |
| RNF2 | 310.345176 | -0.271141886 | 0.26952052 | -1.0060157 | 0.314408 | 0.51489517 |
| TRMT1L | 389.999121 | 0.550573612 | 0.33251684 | 1.65577662 | 0.09776706 | 0.24193774 |
| SWT1 | 624.025605 | -0.018431562 | 0.20995519 | -0.0877881 | 0.93004512 | 0.96282888 |
| IVNS1ABP | 1060.41574 | -0.072276289 | 0.3383616 | -0.2136067 | 0.83085381 | 0.9106114 |
| HMCN1 | 101.398684 | 0.835326508 | 0.49093599 | 1.70149782 | 0.08884955 | 0.22732521 |
| TPR | 1153.71686 | -0.318089867 | 0.21704562 | -1.4655438 | 0.14277261 | 0.31024713 |
| ODR4 | 726.393837 | -0.076178863 | 0.14356918 | -0.5306074 | 0.59569089 | 0.75582107 |
| AL596220.1 | 235.48205 | 0.564197163 | 0.37008583 | 1.52450356 | 0.12738297 | 0.28860138 |
| ERVMER61-1 | 96.571292 | 0.47886342 | 0.37438165 | 1.27907822 | 0.20086951 | 0.38800992 |
| BRINP3 | 128.444378 | 1.691721178 | 0.63887848 | 2.64795454 | 0.00809804 | 0.04334239 |
| AL390957.1 | 136.679617 | -0.901418075 | 0.47716305 | -1.8891196 | 0.0588758 | 0.17258568 |
| AL136454.1 | 364.439638 | -0.921778869 | 0.40262389 | -2.2894292 | 0.02205443 | 0.08849533 |
| UCHL5 | 465.219963 | -0.234197845 | 0.15628963 | -1.4984861 | 0.134007 | 0.29804218 |
| TROVE2 | 637.946853 | -0.284194112 | 0.16032274 | -1.7726376 | 0.07628878 | 0.2054805 |
| GLRX2 | 144.95907 | -0.703462167 | 0.39217106 | -1.7937636 | 0.07285091 | 0.19915585 |
| CDC73 | 1473.4658 | -0.336037807 | 0.23632992 | -1.4219013 | 0.15505492 | 0.32773135 |
| LINC01031 | 33.7152003 | -0.495506491 | 0.66577496 | -0.7442552 | 0.45672208 | 0.64552736 |
| KCNT2 | 1213.61521 | -2.601319858 | 0.88192545 | -2.9495916 | 0.00318194 | 0.02159153 |
| ZBTB41 | 306.108459 | -0.087637671 | 0.13548607 | -0.646839 | 0.51773615 | 0.69445243 |
| CRB1 | 81.9279552 | -0.661326541 | 0.65542047 | -1.0090111 | 0.3129693 | 0.51332576 |
| DENND1B | 1737.2304 | 0.572896107 | 0.24751448 | 2.31459634 | 0.02063503 | 0.08440597 |
| NEK7 | 1840.20583 | 0.035999639 | 0.17868548 | 0.2014693 | 0.84033163 | 0.91539582 |
| MIR181A1HG | 510.034584 | 1.338701392 | 0.41214743 | 3.24811293 | 0.00116173 | 0.01011681 |
| LINC00862 | 745.072642 | 0.403984774 | 0.43737093 | 0.92366627 | 0.3556601 | 0.55404398 |
| AC104461.1 | 214.442312 | -0.039875561 | 0.63383603 | -0.0629115 | 0.949837 | 0.97435384 |
| ZNF281 | 289.582692 | -0.186836248 | 0.22356643 | -0.835708 | 0.4033192 | 0.59752446 |
| DDX59 | 157.167748 | 0.181681532 | 0.32111032 | 0.56579164 | 0.57153543 | 0.73745987 |
| CAMSAP2 | 1176.41888 | -0.168868471 | 0.22788924 | -0.7410112 | 0.45868667 | 0.64743262 |
| INAVA | 586.723919 | -0.017025026 | 0.34283978 | -0.0496588 | 0.96039426 | 0.97961929 |
| TMEM9 | 213.660831 | -0.070358375 | 0.37740103 | -0.1864287 | 0.8521086 | 0.92213818 |
| LAD1 | 263.227693 | 0.798060528 | 0.47323956 | 1.68637745 | 0.09172312 | 0.23235055 |
| CSRP1 | 128.095302 | -0.774851721 | 0.51425136 | -1.5067568 | 0.13187301 | 0.29526028 |
| AC096677.1 | 122.254747 | -0.895439167 | 0.55420568 | -1.6157163 | 0.10615566 | 0.25550055 |
| NAV1 | 217.587978 | -1.854876921 | 0.54948216 | -3.3756818 | 0.00073633 | 0.00721435 |
| IPO9-AS1 | 70.492363 | -1.371683241 | 0.61913175 | -2.2154949 | 0.02672611 | 0.10108577 |
| IPO9 | 711.710087 | -0.323542912 | 0.23794559 | -1.3597349 | 0.17391384 | 0.35368502 |
| TIMM17A | 302.326373 | -0.275043249 | 0.20934838 | -1.3138064 | 0.18891135 | 0.37289861 |
| RNPEP | 412.061182 | -0.144712476 | 0.23631815 | -0.6123629 | 0.54029766 | 0.71182814 |
| ELF3-AS1 | 34.306494 | -0.757030671 | 0.56572463 | -1.3381611 | 0.18084394 | 0.362773 |
| ELF3 | 1644.10815 | -1.072882777 | 0.49238365 | -2.1789569 | 0.02933487 | 0.10732805 |
| ARL8A | 335.974362 | -0.183615563 | 0.28410387 | -0.6462973 | 0.51808683 | 0.69478683 |
| PPP1R12B | 2259.1508 | 0.189643291 | 0.20292811 | 0.93453435 | 0.35002832 | 0.5493439 |
| KDM5B | 1273.3609 | -0.75197288 | 0.22664405 | -3.3178585 | 0.0009071 | 0.0084125 |
| RABIF | 126.007046 | -0.811391365 | 0.41269451 | -1.9660823 | 0.04928911 | 0.15396641 |
| KLHL12 | 298.189645 | 0.190488622 | 0.24669658 | 0.77215752 | 0.44002114 | 0.63195408 |
| ADIPOR1 | 177.692528 | -0.391931074 | 0.22156985 | -1.7688827 | 0.07691345 | 0.20663516 |
| CYB5R1 | 155.625627 | -0.001523537 | 0.25161774 | -0.006055 | 0.99516887 | 0.99749068 |
| TMEM183A | 122.77472 | -0.05659165 | 0.2054564 | -0.2754436 | 0.78297545 | 0.8809896 |
| PPFIA4 | 95.6206128 | -3.9901129 | 0.90739261 | -4.397339 | 1.10E-05 | 0.00027411 |

| | | | | | | |
|---|---|---|---|---|---|---|
| BTG2 | 134.338863 | -0.804134334 | 0.57553486 | -1.3971948 | 0.16235499 | 0.33882458 |
| ATP2B4 | 549.361241 | -1.117201122 | 0.50850805 | -2.1970176 | 0.02801919 | 0.10415702 |
| SNRPE | 135.068962 | -0.466862824 | 0.24055018 | -1.9408126 | 0.05228101 | 0.15953225 |
| SOX13 | 405.587082 | 0.654966734 | 0.41550691 | 1.57630767 | 0.11495492 | 0.26944033 |
| ETNK2 | 398.186604 | 0.935489065 | 0.64633248 | 1.44738055 | 0.14779037 | 0.31746789 |
| REN | 56.6626062 | -0.176540792 | 0.76086303 | -0.232027 | 0.81651702 | 0.90083699 |
| GOLT1A | 100.511383 | -0.948836231 | 0.53307984 | -1.7799139 | 0.07509005 | 0.20349136 |
| PLEKHA6 | 3150.89834 | -0.143932877 | 0.3221257 | -0.4468221 | 0.65500353 | 0.79869525 |
| AL606489.1 | 326.205766 | -0.275927937 | 0.41383219 | -0.6667629 | 0.50492362 | 0.6850618 |
| PPP1R15B | 436.998586 | -0.49516933 | 0.35322473 | -1.4018535 | 0.160959 | 0.33698869 |
| PIK3C2B | 327.064917 | 0.193327344 | 0.38721682 | 0.49927414 | 0.61758627 | 0.77258875 |
| MDM4 | 1657.41039 | 0.351573804 | 0.21659145 | 1.62321181 | 0.10454413 | 0.25263333 |
| AL512306.3 | 61.0041681 | 0.910560853 | 0.50529696 | 1.80203115 | 0.0715405 | 0.19702747 |
| LRRN2 | 410.965905 | 1.348030675 | 0.84854536 | 1.58863714 | 0.11214234 | 0.26516043 |
| NFASC | 156.229282 | 3.089780271 | 0.41523834 | 7.44098019 | 9.99E-14 | 3.60E-11 |
| RBBP5 | 438.165047 | 0.107516144 | 0.21678714 | 0.49595259 | 0.61992786 | 0.77431667 |
| DSTYK | 618.399588 | 0.09195943 | 0.19770575 | 0.46513281 | 0.64183635 | 0.78982207 |
| NUAK2 | 312.619961 | -0.078832336 | 0.55769039 | -0.141355 | 0.88758949 | 0.94209263 |
| LEMD1 | 145.762004 | -2.701075504 | 0.58968114 | -4.5805696 | 4.64E-06 | 0.00013495 |
| CDK18 | 805.380196 | -0.627771337 | 0.35972254 | -1.7451543 | 0.08095801 | 0.21373102 |
| MFSD4A | 106.895908 | 0.015424821 | 0.50112709 | 0.03078026 | 0.97544478 | 0.98773468 |
| ELK4 | 928.461729 | -0.936161505 | 0.2897 | -3.2314861 | 0.00123148 | 0.01057879 |
| SLC45A3 | 75.9992819 | 0.340418135 | 0.52347013 | 0.65031053 | 0.51549166 | 0.69315508 |
| NUCKS1 | 1364.74334 | 0.649022841 | 0.34154501 | 1.9002557 | 0.05739957 | 0.16993001 |
| RAB29 | 787.800975 | 0.844682588 | 0.29402595 | 2.87281647 | 0.0040683 | 0.0259731 |
| SLC41A1 | 393.746606 | 1.331277848 | 0.2746447 | 4.84727295 | 1.25E-06 | 4.59E-05 |
| PM20D1 | 86.6191097 | 2.306491496 | 0.6277311 | 3.67433047 | 0.00023847 | 0.00306926 |
| SLC26A9 | 100.978984 | 3.369148364 | 0.53422359 | 6.30662601 | 2.85E-10 | 3.91E-08 |
| RHEX | 9090.91604 | -0.349191893 | 0.72044278 | -0.4846907 | 0.62789579 | 0.78007501 |
| AC244035.1 | 63.279735 | 0.398265402 | 0.66723083 | 0.59689298 | 0.55057883 | 0.720309 |
| SRGAP2 | 230.973984 | -1.655369767 | 0.37849382 | -4.3735715 | 1.22E-05 | 0.00029865 |
| IKBKE | 67.1404867 | -0.761197712 | 0.56702475 | -1.3424418 | 0.1794528 | 0.36082727 |
| EIF2D | 187.547512 | 0.242127631 | 0.24279672 | 0.99724426 | 0.31864596 | 0.51878355 |
| DYRK3 | 140.93217 | -0.641801692 | 0.43819979 | -1.4646326 | 0.14302119 | 0.31058166 |
| MAPKAPK2 | 1233.31567 | -0.327289641 | 0.33293198 | -0.9830526 | 0.32558156 | 0.52577125 |
| PIGR | 2391.42353 | -2.093872967 | 0.66524892 | -3.147503 | 0.00164671 | 0.01323734 |
| FCAMR | 219.142892 | 1.796491329 | 0.83713274 | 2.14600534 | 0.03187255 | 0.11403735 |
| PFKFB2 | 1411.76052 | -0.468775989 | 0.51208753 | -0.9154216 | 0.35997034 | 0.55846014 |
| YOD1 | 366.392856 | -1.425805934 | 0.46774481 | -3.0482561 | 0.00230174 | 0.01710022 |
| CD55 | 92.7705793 | -0.667277506 | 0.3430984 | -1.9448575 | 0.05179213 | 0.1587111 |
| CR1L | 245.236458 | -2.355448829 | 0.74176293 | -3.1754739 | 0.00149592 | 0.01232246 |
| CD46 | 1344.07314 | -0.324365638 | 0.2132677 | -1.5209318 | 0.12827695 | 0.28995582 |
| MIR29B2CHG | 1793.75202 | 0.644666514 | 0.37803392 | 1.70531394 | 0.08813589 | 0.22623195 |
| PLXNA2 | 480.34528 | -0.286188774 | 0.19367608 | -1.4776671 | 0.1394969 | 0.30573777 |
| LAMB3 | 110.409746 | -1.459834657 | 0.78055243 | -1.8702583 | 0.06144796 | 0.17761855 |
| AL031316.1 | 159.218094 | 1.130228978 | 0.6871743 | 1.64474861 | 0.10002166 | 0.2460337 |
| G0S2 | 1076.44596 | 0.408262661 | 0.8097687 | 0.50417194 | 0.61414055 | 0.76975835 |
| HSD11B1 | 57.7202443 | -0.688763438 | 1.01942103 | -0.6756418 | 0.49926811 | 0.68099623 |
| IRF6 | 129.610152 | -0.187882839 | 0.45075534 | -0.4168178 | 0.6768117 | 0.81285008 |
| UTP25 | 201.677427 | -0.939486664 | 0.42103509 | -2.2313738 | 0.02565638 | 0.09820497 |
| SYT14 | 177.647455 | -1.477599633 | 0.68600577 | -2.1539172 | 0.03124667 | 0.11235521 |
| HHAT | 219.590451 | -0.587861628 | 0.55909968 | -1.0514433 | 0.29305502 | 0.49326436 |
| RCOR3 | 1877.81223 | 0.565142738 | 0.28347029 | 1.99365777 | 0.04618947 | 0.1475458 |
| TRAF5 | 136.284676 | -2.112686152 | 0.28988453 | -7.2880265 | 3.15E-13 | 9.58E-11 |
| LINC00467 | 338.586222 | 0.435704154 | 0.46128263 | 0.94454924 | 0.34488905 | 0.54539422 |
| SLC30A1 | 369.267625 | -0.593020842 | 0.40964964 | -1.4476293 | 0.14772074 | 0.3174178 |
| LPGAT1 | 4247.55879 | -1.540224186 | 0.35158083 | -4.3808537 | 1.18E-05 | 0.00029056 |
| INTS7 | 346.271067 | -0.663759457 | 0.25744519 | -2.5782554 | 0.00993006 | 0.05017002 |
| PPP2R5A | 1080.15574 | -0.431046756 | 0.2126435 | -2.0270864 | 0.04265358 | 0.13953522 |
| AL360091.3 | 172.615564 | -0.072981464 | 0.47082748 | -0.1550068 | 0.87681595 | 0.93580287 |
| TMEM206 | 518.462699 | 0.532376073 | 0.34284621 | 1.55281304 | 0.12046781 | 0.27789502 |
| NENF | 286.262117 | -0.192220951 | 0.2455448 | -0.7828345 | 0.43372429 | 0.62599614 |
| ATF3 | 338.879041 | -2.249107819 | 0.63172232 | -3.5602792 | 0.00037046 | 0.00432899 |
| NSL1 | 481.282099 | 0.126359736 | 0.22149487 | 0.57048607 | 0.56834807 | 0.73472607 |
| TATDN3 | 206.6395 | 0.478956671 | 0.36828686 | 1.30049896 | 0.19343001 | 0.37838522 |
| FLVCR1-DT | 67.165561 | 0.250927836 | 0.4279232 | 0.58638521 | 0.55761668 | 0.7262747 |
| FLVCR1 | 412.565636 | -0.685627591 | 0.33722519 | -2.0331446 | 0.04203791 | 0.13799537 |
| ANGEL2 | 397.677608 | 0.31445628 | 0.19453752 | 1.61642996 | 0.10600139 | 0.25531227 |
| RPS6KC1 | 1803.43175 | -0.466761484 | 0.29056007 | -1.6064199 | 0.10818168 | 0.25869614 |
| AL592402.1 | 45.2564375 | -0.168659302 | 1.13841156 | -0.1481532 | 0.88222187 | 0.93886764 |
| AC096639.1 | 53.4935163 | 0.809732506 | 1.33109464 | 0.60832076 | 0.54297475 | 0.71356452 |
| SMYD2 | 493.974466 | -1.190860271 | 0.21756931 | -5.4734755 | 4.41E-08 | 2.81E-06 |
| PTPN14 | 3632.43134 | -0.509209106 | 0.3637014 | -1.4000746 | 0.16149097 | 0.3376383 |
| KCTD3 | 1034.71496 | -0.224006374 | 0.36367093 | -0.615959 | 0.5379216 | 0.70986341 |
| USH2A | 107.270029 | 2.265121882 | 0.45862578 | 4.93893275 | 7.86E-07 | 3.16E-05 |
| ESRRG | 11121.6418 | 1.300131867 | 0.42413171 | 3.06539653 | 0.00217382 | 0.01635898 |
| GPATCH2 | 2102.32566 | -0.084673739 | 0.21240302 | -0.3986466 | 0.69015363 | 0.82084207 |
| SPATA17 | 581.989039 | 0.833340964 | 0.39531018 | 2.10806855 | 0.03502506 | 0.12175716 |
| RRP15 | 255.216928 | -0.163036059 | 0.29560411 | -0.5515352 | 0.58126687 | 0.7443409 |
| TGFB2 | 71.7901765 | -0.090877595 | 0.49560823 | -0.1833658 | 0.85451102 | 0.92322796 |
| LYPLAL1 | 2332.34739 | 1.081956324 | 0.36443802 | 2.96883497 | 0.00298931 | 0.02061091 |
| AC096642.1 | 82.7509529 | 1.085816966 | 0.52601835 | 2.06421878 | 0.03899698 | 0.13143142 |
| AL513283.1 | 662.484853 | 3.237319999 | 0.9298294 | 3.48162794 | 0.00049838 | 0.00534107 |
| ZC3H11B | 103.713305 | 0.540593651 | 0.47015096 | 1.14982994 | 0.25021392 | 0.44645694 |
| SLC30A10 | 46.4497281 | 1.088252616 | 0.92786223 | 1.17286013 | 0.2408519 | 0.43616818 |
| EPRS | 475.512452 | -0.419341681 | 0.28984358 | -1.4467862 | 0.14795682 | 0.31767609 |
| BPNT1 | 311.445789 | -0.223356963 | 0.29386669 | -0.7600622 | 0.44721741 | 0.63794453 |
| IARS2 | 568.191906 | -0.40230498 | 0.1644914 | -2.4457509 | 0.01445508 | 0.06528364 |
| RAB3GAP2 | 1963.56718 | -0.352217609 | 0.18077235 | -1.9484042 | 0.05136662 | 0.15790414 |
| MARK1 | 101.012229 | -1.690878103 | 0.7763532 | -2.1779753 | 0.02940787 | 0.10754582 |
| C1orf115 | 240.584331 | 0.327709926 | 0.27872757 | 1.1757356 | 0.23970055 | 0.43488755 |
| 2-Mar | 2100.67857 | 2.001098663 | 0.52545342 | 3.80832742 | 0.00013991 | 0.00200494 |
| 1-Mar | 303.600179 | 2.079361032 | 0.43630844 | 4.7658052 | 1.88E-06 | 6.43E-05 |
| DUSP10 | 53.2875211 | -0.711037901 | 0.59740688 | -1.1902071 | 0.23396501 | 0.42816534 |
| AL513314.2 | 178.897658 | 0.732180572 | 0.254301 | 2.87918876 | 0.003987 | 0.0255783 |
| TAF1A | 79.3147752 | -0.384162116 | 0.29639289 | -1.2961246 | 0.19493256 | 0.38013098 |
| MIA3 | 625.983373 | 0.048648142 | 0.27883325 | 0.17447038 | 0.86149581 | 0.92717675 |
| AIDA | 269.125471 | -0.846340325 | 0.17427467 | -4.8563589 | 1.20E-06 | 4.49E-05 |
| BROX | 386.175129 | -0.235746496 | 0.21587831 | -1.0920342 | 0.27481807 | 0.47259765 |
| FAM177B | 562.708535 | -2.465677323 | 0.58592192 | -4.2082012 | 2.57E-05 | 0.00053138 |
| AC093152.1 | 1085.0348 | -0.225397305 | 0.15695148 | -1.4360955 | 0.15097518 | 0.32213835 |
| DISP1 | 141.272429 | -0.22886776 | 0.38360096 | -0.5966298 | 0.55075458 | 0.72034285 |
| CAPN2 | 721.0313 | -2.267027475 | 0.46044258 | -4.9235834 | 8.50E-07 | 3.37E-05 |
| TP53BP2 | 748.443339 | -1.333916649 | 0.30281886 | -4.4049985 | 1.06E-05 | 0.00026654 |
| FBXO28 | 1546.76126 | 0.354136281 | 0.19933746 | 1.77656663 | 0.07563957 | 0.20439939 |
| DEGS1 | 452.170494 | -1.157962886 | 0.3457945 | -3.3487025 | 0.00081191 | 0.00773911 |
| NVL | 787.121942 | 0.229982677 | 0.24977901 | 0.92074462 | 0.35718378 | 0.55591212 |
| CNIH4 | 523.938249 | -0.733419688 | 0.24587089 | -2.9829464 | 0.00285488 | 0.0199612 |
| WDR26 | 1563.88043 | 0.038071965 | 0.18138646 | 0.20989419 | 0.83375026 | 0.91221382 |
| CNIH3 | 497.454815 | 0.259153898 | 0.36578995 | 0.70847737 | 0.47864886 | 0.66413513 |
| DNAH14 | 2326.41985 | -0.15825382 | 0.30471599 | -0.5193486 | 0.60351768 | 0.76169806 |
| LBR | 762.273881 | -0.048576487 | 0.31706906 | -0.1532048 | 0.87823681 | 0.93702747 |
| ENAH | 994.886056 | -1.405196164 | 0.41560232 | -3.3811076 | 0.00072194 | 0.00711918 |

| | | | | | | |
|---|---|---|---|---|---|---|
| SRP9 | 460.043575 | -0.351563129 | 0.27837466 | -1.2629135 | 0.20662025 | 0.39538515 |
| EPHX1 | 1081.55951 | -0.592794789 | 0.48109734 | -1.2321722 | 0.21788475 | 0.4090601 |
| AL591895.1 | 385.934399 | 0.217725149 | 0.47360424 | 0.4597196 | 0.6457175 | 0.79225766 |
| TMEM63A | 1440.95175 | 0.063665911 | 0.48099756 | 0.13236223 | 0.89469779 | 0.94546469 |
| AL117348.2 | 36.0128857 | 0.264734465 | 0.66121715 | 0.40037447 | 0.68888072 | 0.82001633 |
| SDE2 | 118.398148 | -0.19572607 | 0.23444097 | -0.8348629 | 0.4037949 | 0.59790609 |
| H3F3A | 150.773092 | -0.30519128 | 0.30436347 | -1.0027198 | 0.31599608 | 0.515945 |
| ACBD3 | 1054.75625 | -0.569955561 | 0.24336387 | -2.3419892 | 0.01918127 | 0.07996279 |
| LIN9 | 115.629195 | 0.49969769 | 0.20470122 | 2.44110756 | 0.01464229 | 0.06591196 |
| PARP1 | 234.270592 | -0.311088273 | 0.27319257 | -1.1387142 | 0.25482236 | 0.45191488 |
| STUM | 229.406585 | -2.178556866 | 0.72466656 | -3.0062887 | 0.00264458 | 0.01887063 |
| ITPKB | 106.000192 | -1.024508813 | 0.49896639 | -2.0532622 | 0.04004716 | 0.1336563 |
| PSEN2 | 69.0849272 | 0.638759461 | 0.38288525 | 1.6682791 | 0.09526033 | 0.23797755 |
| COQ8A | 516.386223 | 0.410180271 | 0.32874924 | 1.24769954 | 0.21214111 | 0.40179884 |
| CDC42BPA | 5067.6545 | 0.304491462 | 0.23233585 | 1.31056598 | 0.19000444 | 0.37414033 |
| ZNF678 | 267.408368 | -0.167644425 | 0.28591132 | -0.5863511 | 0.55763957 | 0.7262747 |
| SNAP47 | 190.153188 | 0.301495838 | 0.19974417 | 1.50940996 | 0.13119405 | 0.29422069 |
| JMJD4 | 76.7112612 | -0.501943248 | 0.46723622 | -1.0742815 | 0.28269652 | 0.48175669 |
| ARF1 | 886.298841 | -0.827081144 | 0.34378959 | -2.4057772 | 0.01613811 | 0.07099005 |
| C1orf35 | 100.373998 | -0.819091783 | 0.23963258 | -3.4181153 | 0.00063056 | 0.00638813 |
| MRPL55 | 157.705977 | 0.367787324 | 0.34998211 | 1.05087463 | 0.29331617 | 0.49326436 |
| GUK1 | 677.736614 | -0.838219711 | 0.32921685 | -2.5461021 | 0.01089333 | 0.05367179 |
| IBA57 | 60.3837331 | -0.244733255 | 0.49800947 | -0.4914229 | 0.62312738 | 0.77626165 |
| OBSCN | 433.919316 | 0.579956367 | 0.39909088 | 1.45319374 | 0.14616997 | 0.31507341 |
| TRIM11 | 108.719592 | -0.534017451 | 0.41786614 | -1.277963 | 0.20126248 | 0.3883312 |
| RNF187 | 233.198494 | -0.728705117 | 0.43090691 | -1.6910964 | 0.09081839 | 0.23086939 |
| RHOU | 89.1242293 | -1.505999284 | 0.61422418 | -2.4518723 | 0.01421151 | 0.06448101 |
| RAB4A | 299.769234 | -0.059748933 | 0.1685854 | -0.3544135 | 0.72302906 | 0.84233055 |
| NUP133 | 489.048258 | 0.043349385 | 0.1469086 | 0.29507726 | 0.76793486 | 0.87064377 |
| ABCB10 | 309.370066 | 0.286237466 | 0.21152929 | 1.35318124 | 0.17599774 | 0.35623169 |
| TAF5L | 167.039653 | -0.308085235 | 0.33951346 | -0.9074316 | 0.36417862 | 0.56199352 |
| URB2 | 94.864164 | -0.287035082 | 0.54964961 | -0.5222147 | 0.60152088 | 0.76075353 |
| GALNT2 | 3329.88693 | -0.407316228 | 0.30381874 | -1.3406554 | 0.18003237 | 0.36146239 |
| PGBD5 | 164.217971 | -0.804366695 | 0.32304312 | -2.489967 | 0.0127755 | 0.05992941 |
| COG2 | 364.82481 | 0.602387189 | 0.20037072 | 3.0063634 | 0.00264393 | 0.01887063 |
| AGT | 602.929024 | -1.297435417 | 0.61886017 | -2.096492 | 0.03603857 | 0.12408456 |
| C1orf198 | 100.834385 | -1.5226899 | 0.43371117 | -3.5108385 | 0.0004467 | 0.00494659 |
| TTC13 | 217.690911 | 0.148146638 | 0.20765074 | 0.71344141 | 0.47557266 | 0.66132438 |
| ARV1 | 143.278041 | 0.323456058 | 0.23399344 | 1.38232961 | 0.16687052 | 0.34469756 |
| FAM89A | 87.2001458 | -2.03952855 | 0.4201142 | -4.8547004 | 1.21E-06 | 4.49E-05 |
| C1orf131 | 147.180644 | -0.013477212 | 0.29233448 | -0.046102 | 0.96322893 | 0.98083195 |
| GNPAT | 527.105803 | -0.164963128 | 0.15954482 | -1.033961 | 0.30115438 | 0.50169217 |
| EXOC8 | 109.435429 | -0.406935667 | 0.26767649 | -1.5202518 | 0.1284477 | 0.2901504 |
| SPRTN | 164.109125 | 0.17025852 | 0.28810999 | 0.59094972 | 0.55455411 | 0.72406869 |
| EGLN1 | 1659.76543 | -1.109686694 | 0.28383449 | -3.909626 | 9.24E-05 | 0.00146557 |
| TSNAX-DISC1 | 78.2260334 | -1.022624847 | 0.49223072 | -2.0775315 | 0.03775253 | 0.12846919 |
| DISC1 | 229.973558 | -1.549605388 | 0.3713037 | -4.1734175 | 3.00E-05 | 0.00060396 |
| SIPA1L2 | 151.80491 | -1.693794402 | 0.66594582 | -2.5434418 | 0.01097664 | 0.05392715 |
| NTPCR | 244.895185 | 0.045744276 | 0.23493277 | 0.1947122 | 0.84561826 | 0.91865716 |
| PCNX2 | 1173.07447 | -0.924438898 | 0.35072431 | -2.6357993 | 0.00839394 | 0.04441884 |
| MAP3K21 | 1581.2042 | -0.068213649 | 0.32339704 | -0.2109285 | 0.83294307 | 0.91175754 |
| KCNK1 | 571.38169 | -0.856758109 | 0.27675408 | -3.095738 | 0.00196324 | 0.01515025 |
| SLC35F3 | 349.182891 | -1.481746301 | 0.85854144 | -1.7258879 | 0.08436758 | 0.21956074 |
| AL355472.1 | 76.3447158 | 0.789959881 | 0.62740895 | 1.25908291 | 0.20800039 | 0.39708375 |
| COA6 | 132.348583 | -0.882289646 | 0.33944962 | -2.599177 | 0.00934476 | 0.04797325 |
| TARBP1 | 497.443829 | -0.107822602 | 0.26506825 | -0.406773 | 0.68417474 | 0.81689199 |
| IRF2BP2 | 676.052778 | -0.605539391 | 0.37617961 | -1.6097082 | 0.10746158 | 0.25751328 |
| TOMM20 | 484.633852 | -0.753399171 | 0.29647193 | -2.5412159 | 0.01104677 | 0.0540972 |
| ARID4B | 2969.59164 | 0.078715634 | 0.1696302 | 0.46404256 | 0.64261725 | 0.7904123 |
| TBCE | 42.7308475 | 0.033029409 | 0.50586882 | 0.06529244 | 0.94794116 | 0.97344198 |
| B3GALNT2 | 611.051576 | -0.252328931 | 0.21924388 | -1.1509053 | 0.24977122 | 0.44590902 |
| GNG4 | 136.364975 | -1.986475155 | 0.68897159 | -2.8832468 | 0.00393599 | 0.02532892 |
| LYST | 916.452352 | -0.181390327 | 0.20558767 | -0.8823016 | 0.37761375 | 0.57414883 |
| NID1 | 932.175656 | -0.091712517 | 0.40699855 | -0.2253387 | 0.82171581 | 0.90424362 |
| GPR137B | 1824.42309 | 0.057821827 | 0.43971586 | 0.13149816 | 0.89538124 | 0.94589524 |
| ERO1B | 479.408336 | -0.667611922 | 0.2800624 | -2.383797 | 0.01713505 | 0.07399815 |
| EDARADD | 71.8766589 | -1.146944146 | 0.37414139 | -3.0655367 | 0.0021728 | 0.01635898 |
| LGALS8 | 455.962752 | -1.003055575 | 0.23510531 | -4.2664097 | 1.99E-05 | 0.00044201 |
| AL359921.1 | 74.4681684 | -0.202555491 | 0.52252703 | -0.387646 | 0.69827805 | 0.82618017 |
| HEATR1 | 513.139441 | -0.276436953 | 0.35572726 | -0.7771037 | 0.43709761 | 0.62895831 |
| MTR | 2838.62144 | 0.489658336 | 0.33035411 | 1.48222264 | 0.13828106 | 0.30419616 |
| MT1HL1 | 564.572199 | -0.425754114 | 0.94360384 | -0.4512001 | 0.65184536 | 0.79603534 |
| RYR2 | 107.223147 | 2.153124364 | 0.45752548 | 4.70602068 | 2.53E-06 | 8.30E-05 |
| LINC01139 | 245.879254 | -0.390054371 | 0.78000633 | -0.5000656 | 0.61702885 | 0.77210284 |
| CHRM3 | 649.730079 | -0.295236295 | 0.31508518 | -0.9370047 | 0.34875613 | 0.54865148 |
| CHRM3-AS1 | 122.259647 | 2.350120348 | 0.94442749 | 2.4884074 | 0.01283166 | 0.06013328 |
| AL356361.2 | 97.9174348 | 2.73884307 | 0.88641552 | 3.08979595 | 0.00200294 | 0.01538188 |
| FMN2 | 40.4837887 | -0.228261541 | 0.48653345 | -0.469159 | 0.63895599 | 0.78803333 |
| RGS7 | 48.3072591 | 0.151629628 | 0.40595729 | 0.37351128 | 0.70876795 | 0.83319745 |
| FH | 166.644627 | 0.090479528 | 0.2226619 | 0.40635389 | 0.6844826 | 0.81708281 |
| KMO | 1024.25548 | 1.220021939 | 0.71090764 | 1.7161469 | 0.08613516 | 0.222723 |
| OPN3 | 201.383309 | -0.90781598 | 0.2651752 | -3.4234573 | 0.0006183 | 0.00629646 |
| PLD5 | 40.3595193 | 0.160041944 | 0.41448263 | 0.38612461 | 0.69940438 | 0.82708445 |
| CEP170 | 948.61047 | -0.346663687 | 0.23733215 | -1.4606689 | 0.14410634 | 0.31185122 |
| SDCCAG8 | 2386.63355 | 0.27253318 | 0.2638997 | 1.032715 | 0.30173729 | 0.50234486 |
| AKT3 | 2182.21679 | -0.688066177 | 0.41012197 | -1.6777111 | 0.09340351 | 0.23525917 |
| ZBTB18 | 344.326841 | 0.454704119 | 0.29173995 | 1.55859393 | 0.11909252 | 0.27592988 |
| AL645465.1 | 141.388951 | -0.490018158 | 0.47701786 | -1.0272533 | 0.30430122 | 0.50454298 |
| ADSS | 542.840847 | -0.356062166 | 0.24997138 | -1.4244117 | 0.15432733 | 0.32695817 |
| CATSPERE | 633.818418 | 0.585310663 | 0.38621489 | 1.51550517 | 0.12964452 | 0.2918961 |
| DESI2 | 546.855744 | -0.294056467 | 0.18126069 | -1.6222848 | 0.10474238 | 0.25294288 |
| COX20 | 459.766575 | -0.052201516 | 0.21583132 | -0.2418626 | 0.80888666 | 0.8961735 |
| HNRNPU | 2545.27847 | -0.463942019 | 0.13046896 | -3.555957 | 0.00037661 | 0.00438957 |
| EFCAB2 | 1441.63741 | 0.0366612 | 0.29186064 | 0.125612 | 0.90003906 | 0.94743919 |
| KIF26B | 880.083438 | -1.619590859 | 0.6076529 | -2.6653224 | 0.00769146 | 0.04169133 |
| SMYD3 | 5813.31607 | 0.053850479 | 0.19937714 | 0.27009355 | 0.78708829 | 0.88351603 |
| TFB2M | 180.085818 | 0.12652063 | 0.32762277 | 0.38617776 | 0.69936502 | 0.82708445 |
| CNST | 542.386109 | 0.157942657 | 0.27398378 | 0.57646718 | 0.56429945 | 0.73149731 |
| SCCPDH | 301.015904 | -0.139158105 | 0.21130857 | -0.658554 | 0.5101822 | 0.68954593 |
| AHCTF1 | 1263.24536 | 0.184959792 | 0.2616793 | 0.70681859 | 0.47967922 | 0.66481737 |
| ZNF669 | 40.1253836 | -0.956005838 | 0.37563515 | -2.5450383 | 0.01092658 | 0.05375829 |
| ZNF124 | 575.977504 | 0.018314207 | 0.21256739 | 0.08615718 | 0.93134147 | 0.96354054 |
| AL390728.6 | 233.703152 | -0.27565661 | 0.36581277 | -0.7535456 | 0.4511221 | 0.64095339 |
| ZNF496 | 263.616097 | -0.348080937 | 0.30966656 | -1.1240508 | 0.26099149 | 0.45911323 |
| OR2T2 | 72.9789911 | 4.169830273 | 0.72298321 | 5.76753406 | 8.04E-09 | 6.85E-07 |
| OR2G6 | 210.020064 | 3.402237392 | 0.9083671 | 3.74544321 | 0.00018008 | 0.00246008 |
| AC098483.1 | 45.3515049 | 3.645934942 | 0.78934213 | 4.61895396 | 3.86E-06 | 0.00011619 |
| OR2T10 | 179.39909 | 3.347611365 | 0.70223656 | 4.76707075 | 1.87E-06 | 6.42E-05 |
| OR2T11 | 365.17913 | 3.853736031 | 0.67082318 | 5.74478659 | 9.20E-09 | 7.55E-07 |
| LYPD8 | 75.99108 | -0.214050462 | 0.44776221 | -0.4780449 | 0.63261821 | 0.78342389 |
| SH3BP5L | 200.239732 | 0.3915155 | 0.34134427 | 1.14698132 | 0.25138934 | 0.44773827 |
| ZNF672 | 112.132548 | 0.238285325 | 0.26264695 | 0.90724575 | 0.36427686 | 0.56199352 |
| ZNF692 | 224.69694 | 0.545206147 | 0.22678258 | 2.40409178 | 0.0162127 | 0.07115834 |

| | | | | | | |
|---|---|---|---|---|---|---|
| PGBD2 | 74.2329414 | 0.521541123 | 0.34636611 | 1.50575104 | 0.1321311 | 0.2954593 |
| FAM110C | 540.853312 | -1.381670822 | 0.48693341 | -2.8374944 | 0.00454691 | 0.02825229 |
| SH3YL1 | 1688.13033 | 0.948579584 | 0.41364502 | 2.29322135 | 0.02183526 | 0.08797648 |
| ACP1 | 319.995608 | -0.434764416 | 0.2583013 | -1.6831678 | 0.09234262 | 0.23361762 |
| ALKAL2 | 221.118262 | 0.368749254 | 0.75973203 | 0.48536753 | 0.62741566 | 0.77969053 |
| TMEM18 | 90.224853 | -0.141009836 | 0.30261629 | -0.4659691 | 0.64123763 | 0.78935287 |
| LINC01115 | 78.5497134 | 1.350751348 | 1.02116777 | 1.32275164 | 0.18591798 | 0.36911614 |
| SNTG2 | 336.191243 | 0.064780344 | 0.77080836 | 0.08404209 | 0.93302297 | 0.96462105 |
| PXDN | 265.178886 | -2.148701618 | 0.67730439 | -3.1724312 | 0.00151168 | 0.01242245 |
| MYT1L | 44.1864045 | 0.164000551 | 0.85918616 | 0.19087895 | 0.84862043 | 0.92025635 |
| LINC01250 | 49.800428 | 0.918736502 | 0.36235489 | 2.53546045 | 0.01122996 | 0.05474007 |
| EIPR1 | 884.948346 | 0.301164026 | 0.19211992 | 1.56758352 | 0.11697836 | 0.27255182 |
| TRAPPC12 | 1060.14603 | 0.325115638 | 0.19164179 | 1.69647566 | 0.08979584 | 0.22873658 |
| ADI1 | 650.444844 | 0.00926486 | 0.3451727 | 0.02684123 | 0.97858637 | 0.98916955 |
| AC108488.2 | 118.895011 | -0.645504282 | 0.4302581 | -1.5002722 | 0.1335439 | 0.29739825 |
| RNASEH1 | 51.5374478 | -0.674120013 | 0.43299602 | -1.5568734 | 0.11950053 | 0.27650132 |
| RNASEH1-AS | 37.0663151 | -0.30742148 | 0.30593214 | -1.0048682 | 0.31496032 | 0.51523844 |
| RPS7 | 1238.25132 | -0.69903442 | 0.41376079 | -1.6894651 | 0.09113034 | 0.23131917 |
| COLEC11 | 190.107519 | 1.867871141 | 0.48976731 | 3.81379298 | 0.00013685 | 0.0019682 |
| ALLC | 220.033231 | 2.378196704 | 0.61735255 | 3.85225055 | 0.00011704 | 0.00176289 |
| CMPK2 | 42.4547763 | -0.632751463 | 1.50974185 | -0.4191124 | 0.67513402 | 0.81161744 |
| RSAD2 | 68.9912924 | -2.785616458 | 1.38260667 | -2.014757 | 0.04393012 | 0.14221779 |
| RNF144A | 59.3926982 | -0.989268419 | 0.42363627 | -2.3351835 | 0.01953384 | 0.08103823 |
| LINC00299 | 54.4684492 | 0.847899182 | 0.75477607 | 1.12337847 | 0.26127679 | 0.45940659 |
| ID2 | 444.944092 | -0.434447668 | 0.40729328 | -1.0666704 | 0.28612071 | 0.48573899 |
| KIDINS220 | 1467.59125 | 0.021374992 | 0.18721488 | 0.11417357 | 0.9091002 | 0.9523146 |
| MBOAT2 | 684.060819 | -0.878783282 | 0.54752009 | -1.6050247 | 0.10848836 | 0.25926944 |
| ASAP2 | 2225.0095 | -0.422319992 | 0.30661859 | -1.3773464 | 0.16840519 | 0.34690862 |
| ITGB1BP1 | 492.290995 | -0.671223237 | 0.30601928 | -2.1934018 | 0.02827844 | 0.10478849 |
| CPSF3 | 262.431662 | 0.164568869 | 0.23366246 | 0.70430171 | 0.4812449 | 0.66596909 |
| IAH1 | 308.181454 | -0.143966474 | 0.21191334 | -0.6793648 | 0.49690672 | 0.67880211 |
| ADAM17 | 1003.16784 | -0.746235246 | 0.30786947 | -2.423869 | 0.01535614 | 0.06833983 |
| AC080162.1 | 40.0951476 | -0.774189093 | 0.40790039 | -1.8979857 | 0.05769797 | 0.17045895 |
| YWHAQ | 959.82674 | -0.74434807 | 0.30494242 | -2.4409463 | 0.01464883 | 0.06591975 |
| TAF1B | 848.454837 | 0.678787258 | 0.24678979 | 2.75046738 | 0.00595103 | 0.03444712 |
| AC010969.2 | 42.3178782 | 0.51288458 | 0.49670312 | 1.03257774 | 0.30180155 | 0.50234486 |
| GRHL1 | 314.001089 | 0.745866693 | 0.27946483 | 2.66891076 | 0.00760977 | 0.04135887 |
| KLF11 | 196.067716 | 0.31893041 | 0.30669063 | 1.03990922 | 0.29838208 | 0.49840643 |
| CYS1 | 774.774431 | 1.116567863 | 0.49827524 | 2.24086562 | 0.02503478 | 0.09647223 |
| C2orf48 | 42.016326 | 0.920260529 | 0.51588472 | 1.78384916 | 0.07444818 | 0.20232093 |
| HPCAL1 | 2182.43184 | 0.992540355 | 0.3743666 | 2.6512524 | 0.00801939 | 0.04298857 |
| ODC1 | 428.595053 | -0.669953343 | 0.31362517 | -2.1361594 | 0.03266642 | 0.11623393 |
| NOL10 | 962.155409 | 0.003263439 | 0.26976515 | 0.01209733 | 0.99034796 | 0.99518064 |
| ATP6V1C2 | 36.2810835 | -0.040016145 | 0.40242965 | -0.0994364 | 0.92079181 | 0.9574582 |
| PDIA6 | 771.262651 | -0.744258008 | 0.3918474 | -1.8993568 | 0.05751758 | 0.1701324 |
| PQLC3 | 94.4962421 | -0.152108148 | 0.35244612 | -0.4315784 | 0.66604783 | 0.80617161 |
| ROCK2 | 2495.02451 | 0.132167273 | 0.18254332 | 0.72403237 | 0.46904586 | 0.65617047 |
| E2F6 | 267.892048 | -0.368179896 | 0.31409373 | -1.1721975 | 0.24111776 | 0.43618862 |
| GREB1 | 302.527001 | 1.980175736 | 0.39215284 | 5.04949986 | 4.43E-07 | 1.94E-05 |
| LPIN1 | 645.671889 | -1.651131284 | 0.65894904 | -2.5057041 | 0.01222079 | 0.05798211 |
| MIR3681HG | 63.5155984 | 0.592828365 | 0.38523517 | 1.53887394 | 0.12383507 | 0.28332621 |
| TRIB2 | 45.1207268 | 0.348518476 | 0.35170026 | 0.99095315 | 0.32170846 | 0.52210014 |
| LINC00276 | 49.8062312 | 0.465153126 | 0.34845056 | 1.33491857 | 0.181903 | 0.36399188 |
| FAM84A | 50.2078631 | -1.512594531 | 0.59255801 | -2.5526522 | 0.01069062 | 0.05292029 |
| NBAS | 2409.30176 | -0.083989436 | 0.15716689 | -0.5343965 | 0.59306723 | 0.75436977 |
| DDX1 | 646.959883 | -0.025707662 | 0.16453165 | -0.1562475 | 0.87583792 | 0.93528875 |
| MYCN | 102.405751 | 4.579954097 | 0.59213225 | 7.73468111 | 1.04E-14 | 4.74E-12 |
| VSNL1 | 104.533773 | 1.621586198 | 0.58306738 | 2.78113004 | 0.005417 | 0.03215715 |
| SMC6 | 738.485355 | 0.057944329 | 0.19063824 | 0.30394914 | 0.76116664 | 0.86655408 |
| GEN1 | 337.253914 | 0.374835402 | 0.41260027 | 0.90847104 | 0.36362941 | 0.56136995 |
| KCNS3 | 179.479463 | 0.646426508 | 0.61126251 | 1.05752684 | 0.29027121 | 0.49029544 |
| RDH14 | 138.654353 | -0.23319406 | 0.32743636 | -0.7121813 | 0.4763525 | 0.6620617 |
| LINC01376 | 1112.00827 | 0.312776152 | 0.49037658 | 0.63782849 | 0.52358533 | 0.69915092 |
| AC010096.1 | 31.820599 | 1.87419608 | 0.57983548 | 3.23228941 | 0.00122803 | 0.01055995 |
| LINC00954 | 59.4316434 | -1.060116488 | 0.57524772 | -1.8428869 | 0.06534552 | 0.18555855 |
| TTC32 | 145.259248 | 0.028528133 | 0.22816581 | 0.12503246 | 0.90049785 | 0.94768577 |
| WDR35 | 696.633495 | 0.215391911 | 0.26711986 | 0.80634931 | 0.42004147 | 0.61361061 |
| LAPTM4A | 2069.25579 | -0.798354917 | 0.35729509 | -2.2344413 | 0.02545405 | 0.097676 |
| AC098828.3 | 110.770067 | -0.354953111 | 1.0044838 | -0.3533687 | 0.72381207 | 0.84282267 |
| SDC1 | 669.153478 | 0.543066319 | 0.40792822 | 1.3312791 | 0.18309719 | 0.3655277 |
| PUM2 | 3527.24479 | -0.166908174 | 0.2199218 | -0.7589433 | 0.44788647 | 0.63841127 |
| RHOB | 934.979229 | -0.303528439 | 0.40749468 | -0.7448648 | 0.45635346 | 0.64540207 |
| HS1BP3 | 367.951001 | 0.276325792 | 0.36742702 | 0.75205627 | 0.45201722 | 0.64143458 |
| GDF7 | 164.521338 | 0.238276873 | 0.497397 | 0.47904767 | 0.63190471 | 0.78299746 |
| LDAH | 1152.39332 | 0.645906071 | 0.39296014 | 1.6436936 | 0.10023951 | 0.2462335 |
| AC018742.1 | 91.5813583 | 2.125614976 | 0.68975991 | 3.08167369 | 0.0020584 | 0.01573361 |
| AC018467.1 | 1482.91744 | 0.704266218 | 0.51555015 | 1.36604795 | 0.17192391 | 0.3513466 |
| KLHL29 | 5457.37321 | -3.70571123 | 0.89559481 | -4.1377096 | 3.51E-05 | 0.00068203 |
| AC009242.1 | 42.5934294 | -0.6980142 | 0.57489142 | -1.214167 | 0.22468395 | 0.41685746 |
| ATAD2B | 1587.76391 | 0.40164322 | 0.14542341 | 2.76188827 | 0.00574681 | 0.03354837 |
| UBXN2A | 731.186933 | -0.075732644 | 0.19981452 | -0.3790147 | 0.70467694 | 0.83052509 |
| WDCP | 37.7182106 | -0.224480594 | 0.33435222 | -0.6713896 | 0.50197239 | 0.68293156 |
| SF3B6 | 248.028581 | -0.742437033 | 0.26470903 | -2.8047287 | 0.00503589 | 0.03043519 |
| FAM228B | 335.072959 | -0.216509886 | 0.30596854 | -0.7076214 | 0.47918039 | 0.6645225 |
| TP53I3 | 230.921993 | 0.405410938 | 0.47592506 | 0.85183776 | 0.39430415 | 0.58900686 |
| ITSN2 | 2387.57745 | -0.183473177 | 0.17347444 | -1.0576381 | 0.29022046 | 0.49027014 |
| NCOA1 | 3662.45466 | 0.006811683 | 0.11905448 | 0.05721484 | 0.95437406 | 0.97689532 |
| PTRHD1 | 61.8720702 | -0.819749399 | 0.46647608 | -1.7573235 | 0.07886268 | 0.20989723 |
| CENPO | 57.9619364 | -0.104998044 | 0.39089921 | -0.2686064 | 0.78823256 | 0.88421655 |
| ADCY3 | 51.7091135 | -1.020296818 | 0.45950292 | -2.220436 | 0.02638919 | 0.10017076 |
| DNAJC27 | 224.856206 | 0.293030917 | 0.34489676 | 0.84961923 | 0.39553682 | 0.59027092 |
| DNAJC27-AS | 177.741901 | 1.26995991 | 0.59218495 | 2.14453258 | 0.03199024 | 0.11430923 |
| DNMT3A | 784.5073 | 0.60802734 | 0.24853672 | 2.44642864 | 0.01442794 | 0.0651825 |
| DTNB | 1196.16442 | -0.195953313 | 0.21757095 | -0.900641 | 0.36777923 | 0.56521471 |
| ASXL2 | 1681.73013 | 0.056747428 | 0.16561262 | 0.34265158 | 0.73186059 | 0.84761805 |
| KIF3C | 50.0212314 | -1.044125364 | 0.37616687 | -2.7756973 | 0.00550835 | 0.03261295 |
| RAB10 | 4166.21434 | -0.725369066 | 0.44491115 | -1.6303684 | 0.10302367 | 0.25064703 |
| HADHA | 1369.63058 | 0.092404866 | 0.18455157 | 0.50069944 | 0.61658267 | 0.77182636 |
| HADHB | 933.943766 | 0.010371349 | 0.23312562 | 0.04448824 | 0.96451522 | 0.98155603 |
| ADGRF3 | 49.2149641 | 0.046664952 | 0.52950302 | 0.08812972 | 0.92977358 | 0.96271389 |
| SELENOI | 582.258149 | -0.615493418 | 0.29693537 | -2.0728194 | 0.03818909 | 0.12960084 |
| KCNK3 | 80.6018307 | -4.338712927 | 0.55669106 | -7.7937535 | 6.50E-15 | 3.18E-12 |
| CENPA | 151.493388 | -0.890889749 | 0.71820535 | -1.2404388 | 0.21481315 | 0.40529166 |
| AC013472.2 | 39.6579224 | -0.883158921 | 0.66220194 | -1.3336701 | 0.182312 | 0.36465061 |
| MAPRE3 | 220.00195 | -0.485857173 | 0.21804504 | -2.2282422 | 0.02586437 | 0.0986979 |
| TMEM214 | 206.6134 | -0.709354587 | 0.38407706 | -1.8469069 | 0.06476063 | 0.18443256 |
| AGBL5 | 211.866079 | -0.393827829 | 0.26790579 | -1.4700236 | 0.14155536 | 0.30857178 |
| OST4 | 341.208755 | -0.803934099 | 0.47067969 | -1.708028 | 0.08763114 | 0.22540065 |
| KHK | 922.773511 | 2.163455077 | 0.49385765 | 4.38072607 | 1.18E-05 | 0.00029056 |
| CGREF1 | 76.6554471 | 0.090721421 | 0.4126481 | 0.21985178 | 0.82598659 | 0.9066142 |
| PREB | 118.709504 | -0.066740134 | 0.31224723 | -0.2137413 | 0.83074879 | 0.91060165 |
| SLC5A6 | 73.5380355 | 0.092518413 | 0.35923086 | 0.25754584 | 0.79675744 | 0.88911002 |
| ATRAID | 199.321741 | -0.015931403 | 0.31215762 | -0.0510364 | 0.95929651 | 0.97915536 |

| | | | | | | |
|---|---|---|---|---|---|---|
| CAD | 133.855842 | -0.319959381 | 0.38821308 | -0.824185 | 0.40983448 | 0.60365368 |
| MPV17 | 302.396809 | -0.345066111 | 0.3655045 | -0.9440817 | 0.3451279 | 0.5454262 |
| GTF3C2 | 646.308258 | 0.408495182 | 0.22810292 | 1.79083717 | 0.07331943 | 0.20019301 |
| AC074117.1 | 105.916768 | 0.56252635 | 0.2896885 | 1.94183181 | 0.05215746 | 0.15936745 |
| EIF2B4 | 180.370253 | 0.144552601 | 0.26137081 | 0.55305563 | 0.58022532 | 0.74366782 |
| SNX17 | 155.171027 | -0.427497225 | 0.34513603 | -1.238634 | 0.21548107 | 0.40616048 |
| ZNF513 | 117.217677 | 0.219229481 | 0.30382506 | 0.72156485 | 0.47056206 | 0.65738999 |
| PPM1G | 889.571633 | -1.349979034 | 0.44168341 | -3.0564404 | 0.00223982 | 0.01674003 |
| NRBP1 | 559.585165 | -0.968793042 | 0.47134369 | -2.0553856 | 0.03984177 | 0.13313291 |
| KRTCAP3 | 74.7331043 | 0.116316992 | 0.35079134 | 0.33158456 | 0.74020298 | 0.85230954 |
| IFT172 | 463.017736 | 0.865645428 | 0.30337837 | 2.85335254 | 0.00432606 | 0.02721308 |
| ZNF512 | 420.485613 | -0.102928334 | 0.21783677 | -0.4725021 | 0.63656843 | 0.78600518 |
| GPN1 | 143.426053 | -0.85156911 | 0.50084744 | -1.7002565 | 0.08908269 | 0.22768161 |
| SUPT7L | 212.239889 | 0.033319739 | 0.20815698 | 0.16007024 | 0.87282574 | 0.9342357 |
| SLC4A1AP | 266.36717 | 0.04738417 | 0.1887599 | 0.25102879 | 0.80179185 | 0.89234824 |
| MRPL33 | 657.745471 | -0.433201296 | 0.24961903 | -1.7354498 | 0.08266116 | 0.21689062 |
| RBKS | 1134.44462 | 0.456385068 | 0.55556346 | 0.82148144 | 0.41137209 | 0.60487849 |
| BABAM2 | 4535.21319 | -0.188015777 | 0.34799891 | -0.5402769 | 0.58900607 | 0.75109371 |
| FOSL2 | 1007.52666 | -1.67485057 | 0.40642959 | -4.1208874 | 3.77E-05 | 0.0007195 |
| PPP1CB | 761.015531 | -1.08228645 | 0.27237684 | -3.9734892 | 7.08E-05 | 0.00118683 |
| SPDYA | 86.8915893 | -1.346687827 | 0.42202606 | -3.1910063 | 0.00141778 | 0.01178504 |
| TRMT61B | 337.357517 | 0.598971689 | 0.19886685 | 3.0119233 | 0.00259598 | 0.01870827 |
| WDR43 | 926.148368 | -1.059709319 | 0.42614042 | -2.4867609 | 0.0128912 | 0.06031539 |
| CLIP4 | 1160.82901 | -0.75628821 | 0.17601949 | -4.2966164 | 1.73E-05 | 0.00039751 |
| ALK | 120.2557 | 0.450156985 | 0.41532168 | 1.08387547 | 0.27842002 | 0.47668128 |
| AC016907.2 | 49.2942262 | 1.822069285 | 0.68984768 | 2.64126318 | 0.00825975 | 0.04389935 |
| YPEL5 | 475.159439 | -0.185609039 | 0.27000585 | -0.687426 | 0.49181433 | 0.67419732 |
| LBH | 137.429014 | 0.604533472 | 0.49944554 | 1.2104092 | 0.22612191 | 0.41873184 |
| LCLAT1 | 1872.2951 | 1.000817104 | 0.2369195 | 4.22429189 | 2.40E-05 | 0.00050914 |
| CAPN13 | 285.071162 | 1.834758793 | 0.53982452 | 3.39880594 | 0.00067681 | 0.00673582 |
| GALNT14 | 5171.12102 | 0.738484617 | 0.26542375 | 2.78228542 | 0.00539775 | 0.03208494 |
| MEMO1 | 109.62442 | -0.353267384 | 0.31021357 | -1.1387877 | 0.25479173 | 0.45191488 |
| DPY30 | 338.293645 | 0.112359556 | 0.20951249 | 0.53629049 | 0.5917578 | 0.75320124 |
| SPAST | 612.492062 | 0.488091962 | 0.16044105 | 3.04218881 | 0.00234865 | 0.01739216 |
| SLC30A6 | 638.21832 | 0.261874796 | 0.1839567 | 1.42356761 | 0.15457168 | 0.3271791 |
| YIPF4 | 953.325426 | 0.525863664 | 0.20484146 | 2.56717398 | 0.01025312 | 0.05144197 |
| AL133245.1 | 61.0290134 | 1.19630823 | 0.62316333 | 1.91973462 | 0.05489143 | 0.16489082 |
| BIRC6 | 5960.49356 | -0.010247957 | 0.15399552 | -0.0665471 | 0.94694225 | 0.97277505 |
| TTC27 | 692.949985 | 0.056957855 | 0.2514845 | 0.22648654 | 0.82082302 | 0.90384168 |
| LINC00486 | 1083.97624 | 0.639555091 | 0.18651962 | 3.42888908 | 0.00060606 | 0.00618557 |
| LTBP1 | 110.865735 | 1.43147132 | 0.55310723 | 2.5880539 | 0.00965199 | 0.04910906 |
| RASGRP3 | 471.97576 | 0.607735357 | 0.40706501 | 1.49296878 | 0.13544536 | 0.29987878 |
| AC020594.1 | 61.3187637 | -0.076992634 | 0.29714609 | -0.259107 | 0.79555268 | 0.88843413 |
| FAM98A | 293.455207 | 0.25858019 | 0.23669911 | 1.09244262 | 0.2746386 | 0.47250801 |
| LINC01317 | 111.031696 | 0.65090132 | 0.54837093 | 1.1869727 | 0.23523839 | 0.42963526 |
| AC068599.1 | 445.524677 | 1.867120438 | 0.58831735 | 3.17366206 | 0.00150529 | 0.01238475 |
| LINC01320 | 12172.2991 | -0.139014489 | 0.52700173 | -0.2637837 | 0.79194656 | 0.88678699 |
| AC073218.2 | 100.317178 | -0.414390413 | 0.6729343 | -0.6157962 | 0.53802901 | 0.70986341 |
| AC073218.1 | 63.3448147 | -1.606315499 | 0.98081719 | -1.6377318 | 0.10147766 | 0.24829601 |
| AC012593.1 | 5898.81359 | -1.047028311 | 1.25090109 | -0.8370193 | 0.40258174 | 0.59681894 |
| CRIM1 | 2579.6943 | -0.238247573 | 0.2603956 | -0.9149447 | 0.36022067 | 0.55851021 |
| FEZ2 | 731.638192 | -0.279650243 | 0.23457433 | -1.1921605 | 0.23319834 | 0.42710552 |
| STRN | 1176.14002 | 0.136663041 | 0.18146566 | 0.75310692 | 0.45138567 | 0.64108831 |
| HEATR5B | 886.243031 | -0.204673707 | 0.19556031 | -1.0466015 | 0.29528342 | 0.49516146 |
| GPATCH11 | 90.1175265 | -0.540249223 | 0.35361316 | -1.5277973 | 0.12656288 | 0.28755136 |
| EIF2AK2 | 813.603837 | -0.238637914 | 0.51946298 | -0.4593935 | 0.64595162 | 0.79225766 |
| CEBPZOS | 593.822675 | -0.031593306 | 0.1586185 | -0.1991779 | 0.84212355 | 0.91676495 |
| CEBPZ | 576.07915 | -0.455807227 | 0.22056402 | -2.066553 | 0.0387763 | 0.13093592 |
| NDUFAF7 | 388.347486 | -0.144746193 | 0.12238037 | -1.1827566 | 0.23690561 | 0.43136764 |
| PRKD3 | 781.254461 | -0.289466756 | 0.20721149 | -1.3969629 | 0.16242473 | 0.33890627 |
| QPCT | 61.5204117 | -1.916721506 | 0.50364234 | -3.8057196 | 0.00014139 | 0.00202093 |
| CDC42EP3 | 119.809576 | -0.286545968 | 0.33910894 | -0.8449968 | 0.39811263 | 0.59255618 |
| RMDN2 | 451.675249 | 1.450780193 | 0.31101274 | 4.66469698 | 3.09E-06 | 9.83E-05 |
| RMDN2-AS1 | 50.1234804 | 0.259424353 | 0.59136852 | 0.43868475 | 0.66088998 | 0.80294442 |
| CYP1B1 | 70.5183605 | -0.305690518 | 0.52755231 | -0.5794506 | 0.56228515 | 0.73014439 |
| CYP1B1-AS1 | 105.727962 | 0.746914303 | 0.53052215 | 1.40788524 | 0.15916505 | 0.33410037 |
| ATL2 | 1289.49207 | 1.061090088 | 0.15672521 | 6.77038529 | 1.28E-11 | 2.59E-09 |
| AC011247.1 | 94.9889006 | -0.101357869 | 0.74732542 | -0.1356275 | 0.89211577 | 0.94368197 |
| HNRNPLL | 621.336964 | -0.033612999 | 0.18336184 | -0.1833151 | 0.85455077 | 0.92322796 |
| GALM | 1885.04255 | 2.010287482 | 0.42088231 | 4.77636491 | 1.78E-06 | 6.19E-05 |
| SRSF7 | 913.144009 | 0.026898223 | 0.2444506 | 0.11003541 | 0.91238129 | 0.9535692 |
| GEMIN6 | 74.2831944 | 0.439714635 | 0.26266306 | 1.67406345 | 0.09411813 | 0.23621021 |
| DHX57 | 748.341995 | 0.320504601 | 0.21793006 | 1.47067643 | 0.14137864 | 0.30828461 |
| MORN2 | 141.759404 | -0.064844161 | 0.32368274 | -0.2003325 | 0.84122057 | 0.91607296 |
| ARHGEF33 | 62.59857 | 0.862549563 | 0.53552252 | 1.61066907 | 0.10725187 | 0.25714844 |
| SOS1 | 3013.98115 | 0.303806081 | 0.16304555 | 1.86332028 | 0.06241722 | 0.17970023 |
| CDKL4 | 34.5275756 | -0.759764802 | 0.42959641 | -1.7685548 | 0.0769682 | 0.20666074 |
| MAP4K3 | 2711.86459 | 0.169853984 | 0.24675727 | 0.68834441 | 0.49123592 | 0.67380875 |
| AC007388.1 | 2171.0566 | 0.785240884 | 0.35650675 | 2.20259753 | 0.02762313 | 0.10333679 |
| TMEM178A | 96.7039891 | 0.866666597 | 0.48879157 | 1.77308007 | 0.07621544 | 0.2054356 |
| THUMPD2 | 523.399007 | 0.024917508 | 0.18759595 | 0.13282541 | 0.89433146 | 0.94518003 |
| SLC8A1 | 187.324948 | 1.197495198 | 0.60787572 | 1.96996713 | 0.04884214 | 0.15318465 |
| PKDCC | 216.783222 | -0.690072161 | 0.67286035 | -1.0255801 | 0.30508957 | 0.50529195 |
| EML4 | 3006.86371 | 0.450286291 | 0.21283285 | 2.11568044 | 0.034372 | 0.12031025 |
| COX7A2L | 717.062874 | -0.273562174 | 0.30350012 | -0.9013577 | 0.36739817 | 0.56469238 |
| MTA3 | 1349.91314 | 0.622369658 | 0.26004737 | 2.39329341 | 0.01669788 | 0.07266389 |
| OXER1 | 76.1858302 | 2.737361575 | 0.52197617 | 5.24422705 | 1.57E-07 | 8.15E-06 |
| HAAO | 513.905959 | 2.488820237 | 0.44993314 | 5.53153352 | 3.17E-08 | 2.20E-06 |
| ZFP36L2 | 1012.50355 | -0.822732435 | 0.34939546 | -2.3547313 | 0.0185361 | 0.07815267 |
| THADA | 1844.61886 | 0.320025219 | 0.20828916 | 1.53644682 | 0.12442883 | 0.28426229 |
| PLEKHH2 | 475.911505 | -0.579158275 | 0.49555474 | -1.168707 | 0.24252172 | 0.43768864 |
| DYNC2LI1 | 743.973535 | 0.512468617 | 0.43615495 | 1.17496917 | 0.24000705 | 0.43509809 |
| LRPPRC | 1826.49566 | -0.003807395 | 0.19978589 | -0.0190574 | 0.98479533 | 0.99254336 |
| PPM1B | 1609.87095 | 0.029826998 | 0.18808831 | 0.15857976 | 0.87399998 | 0.93468316 |
| SLC3A1 | 5402.6267 | 0.924572658 | 0.46474216 | 1.98943144 | 0.0466536 | 0.14854377 |
| PREPL | 1250.71224 | 0.285329188 | 0.31404123 | 0.90857237 | 0.3635759 | 0.56136995 |
| CAMKMT | 5225.66081 | 0.149028318 | 0.11544136 | 1.29094389 | 0.19672314 | 0.38248 |
| LINC01833 | 51.63066 | -0.744175093 | 0.69798389 | -1.066178 | 0.28634317 | 0.48599627 |
| SRBD1 | 1079.98362 | 0.05347622 | 0.17862899 | 0.29937033 | 0.7646575 | 0.86865244 |
| PRKCE | 3188.28711 | 0.474004689 | 0.32455567 | 1.46047266 | 0.14416022 | 0.31188802 |
| EPAS1 | 1133.34113 | -0.988472601 | 0.35437697 | -2.7893251 | 0.0052818 | 0.03152282 |
| ATP6V1E2 | 89.724896 | -1.135257656 | 0.39881803 | -2.8465555 | 0.0044195 | 0.02764861 |
| RHOQ | 1198.00013 | -1.838945906 | 0.43871816 | -4.1916339 | 2.77E-05 | 0.00056575 |
| AC018682.1 | 229.151392 | -2.69948657 | 0.44517497 | -6.0638777 | 1.33E-09 | 1.48E-07 |
| PIGF | 364.718772 | -0.045461393 | 0.2492272 | -0.1824094 | 0.85526142 | 0.92351255 |
| CRIPT | 136.826558 | 0.012571735 | 0.28153201 | 0.04465473 | 0.96438252 | 0.98149623 |
| SOCS5 | 608.67223 | 0.333007978 | 0.21227719 | 1.56874123 | 0.11670825 | 0.27215379 |
| MCFD2 | 483.131673 | -0.613711608 | 0.32200307 | -1.9059185 | 0.05666078 | 0.16850711 |
| TTC7A | 860.553726 | -0.058716151 | 0.37683409 | -0.1558143 | 0.87617938 | 0.93548767 |
| AC106869.1 | 460.598096 | 0.383308912 | 0.57814498 | 0.66299791 | 0.5073319 | 0.68735774 |
| EPCAM | 1002.78525 | -0.625585752 | 0.3273642 | -1.9109779 | 0.05600742 | 0.16710188 |
| MSH2 | 284.126832 | 0.039166452 | 0.22991425 | 0.17035243 | 0.86473298 | 0.92922135 |
| MSH6 | 215.848422 | -0.187307146 | 0.17661486 | -1.0605401 | 0.28889893 | 0.48876052 |

| | | | | | | |
|---|---|---|---|---|---|---|
| FBXO11 | 2374.71339 | 0.042588766 | 0.14793926 | 0.28788008 | 0.77343854 | 0.87422228 |
| FOXN2 | 396.353604 | -0.123733823 | 0.32333156 | -0.382684 | 0.70195406 | 0.82874111 |
| PPP1R21 | 3534.97643 | 1.235986901 | 0.3397366 | 3.63807401 | 0.00027468 | 0.00342905 |
| NRXN1 | 67.9078289 | 0.078963718 | 0.42445073 | 0.18603742 | 0.85241542 | 0.92213818 |
| AC007402.1 | 101.984745 | 0.754843122 | 0.37977156 | 1.98762415 | 0.04685327 | 0.14907563 |
| ASB3 | 64.3513578 | 0.471597243 | 0.31305051 | 1.50645738 | 0.1319498 | 0.29538395 |
| GPR75-ASB3 | 351.449309 | -1.224441996 | 0.35130332 | -3.4854267 | 0.00049135 | 0.00529479 |
| ERLEC1 | 268.955005 | -0.349457108 | 0.23625092 | -1.4791778 | 0.13909281 | 0.30514868 |
| PSME4 | 1711.50206 | -0.559343346 | 0.33256676 | -1.6818979 | 0.09258863 | 0.23406721 |
| ACYP2 | 2413.9961 | 0.465341163 | 0.21011785 | 2.21466743 | 0.0267829 | 0.10118887 |
| C2orf73 | 46.4341221 | 1.236006367 | 0.739239 | 1.67199833 | 0.09452465 | 0.23686461 |
| SPTBN1 | 5278.30272 | -0.63710348 | 0.27050599 | -2.3552287 | 0.01851131 | 0.07812023 |
| AC092839.1 | 121.148893 | -0.740574748 | 0.52139787 | -1.420364 | 0.15550175 | 0.32857445 |
| AC093110.1 | 127.028884 | -1.528986253 | 0.54102892 | -2.8260712 | 0.00471228 | 0.02902976 |
| EML6 | 310.3743 | 0.709066104 | 0.45264368 | 1.5664995 | 0.11723173 | 0.2729102 |
| RTN4 | 3161.12072 | -1.033361506 | 0.40224291 | -2.5689987 | 0.01019928 | 0.05119062 |
| CLHC1 | 295.444493 | 0.25973458 | 0.30722879 | 0.84541093 | 0.39788144 | 0.59241006 |
| RPS27A | 1810.17792 | -0.528484873 | 0.47384037 | -1.1153226 | 0.2647122 | 0.4624535 |
| MTIF2 | 318.466172 | 0.513028798 | 0.27213888 | 1.88517271 | 0.05940652 | 0.17358457 |
| AC012358.3 | 273.004837 | 0.460887816 | 0.47468211 | 0.97093993 | 0.33157819 | 0.53138575 |
| CCDC88A | 518.902938 | -0.420138588 | 0.26486098 | -1.5862608 | 0.11268016 | 0.26591198 |
| CFAP36 | 598.79062 | -0.01012498 | 0.1801355 | -0.0562076 | 0.95517644 | 0.97735171 |
| PPP4R3B | 1652.62561 | 0.295207338 | 0.11487633 | 2.56978388 | 0.0101762 | 0.05114967 |
| PNPT1 | 549.302373 | -0.148137484 | 0.45812901 | -0.3233532 | 0.74642776 | 0.85674806 |
| EFEMP1 | 84.6929649 | 0.860490117 | 0.48436523 | 1.77653154 | 0.07564535 | 0.20439939 |
| AC011306.1 | 35.9825756 | 2.321513097 | 0.82500299 | 2.81394506 | 0.00489376 | 0.02975987 |
| CCDC85A | 70.9877584 | 1.203454773 | 0.61447615 | 1.95850526 | 0.05017075 | 0.15567187 |
| VRK2 | 812.706949 | -0.560104501 | 0.19180183 | -2.920225 | 0.00349779 | 0.02321413 |
| FANCL | 369.884493 | 0.115422482 | 0.29410621 | 0.3924517 | 0.69472449 | 0.82425245 |
| BCL11A | 63.2462878 | -1.317212661 | 0.73559991 | -1.7906645 | 0.07334714 | 0.20019301 |
| PAPOLG | 307.302464 | -0.061314253 | 0.21504739 | -0.2851197 | 0.77555241 | 0.87601952 |
| REL | 440.20918 | -0.378696677 | 0.25470543 | -1.4868025 | 0.13706697 | 0.30224855 |
| PUS10 | 645.393222 | 0.400001707 | 0.45266196 | 0.88366539 | 0.37687687 | 0.57334642 |
| PEX13 | 687.862325 | 0.170453826 | 0.16886323 | 1.00941946 | 0.31277351 | 0.51306684 |
| KIAA1841 | 172.552904 | -1.270079387 | 0.28022467 | -4.5323611 | 5.83E-06 | 0.00016316 |
| C2orf74 | 213.85587 | 1.361919254 | 0.50062405 | 2.72044313 | 0.00651945 | 0.0368252 |
| USP34 | 5032.14203 | -0.048643333 | 0.16219663 | -0.2999035 | 0.76425078 | 0.86833493 |
| AC016727.1 | 162.804002 | 0.493292858 | 0.31293977 | 1.57631885 | 0.11495234 | 0.26944033 |
| XPO1 | 2356.08852 | 0.639819322 | 0.31235038 | 2.04840258 | 0.04052057 | 0.13458091 |
| FAM161A | 131.100471 | 0.123858779 | 0.26427713 | 0.46867005 | 0.63930549 | 0.78803844 |
| CCT4 | 510.042312 | -0.294316231 | 0.21930553 | -1.3420375 | 0.17958385 | 0.36086531 |
| COMMD1 | 2458.77787 | 0.722451147 | 0.17458082 | 4.13820456 | 3.50E-05 | 0.00068152 |
| AC018462.1 | 177.325386 | 0.31818392 | 0.54127169 | 0.58784512 | 0.55663626 | 0.72562693 |
| B3GNT2 | 396.825788 | 0.422830685 | 0.27870955 | 1.51710151 | 0.12924106 | 0.29117475 |
| EHBP1 | 4777.43276 | 0.242289751 | 0.22117246 | 1.09547885 | 0.2733069 | 0.47113178 |
| AC092567.1 | 69.1202258 | -0.253286426 | 0.6094063 | -0.4156282 | 0.67768211 | 0.81339787 |
| WDPCP | 3518.16714 | 0.100560298 | 0.28301969 | 0.35531203 | 0.72235585 | 0.84228022 |
| MDH1 | 911.704885 | -0.093074796 | 0.45559145 | -0.2042944 | 0.83812342 | 0.91451662 |
| UGP2 | 1558.1501 | -0.493678018 | 0.34206674 | -1.4432213 | 0.14895814 | 0.31912616 |
| VPS54 | 1639.53155 | 0.062370276 | 0.17063408 | 0.36552063 | 0.71472281 | 0.8368246 |
| PELI1 | 852.341333 | 0.329043844 | 0.2450037 | 1.34301583 | 0.17926685 | 0.36056127 |
| AC012368.1 | 73.4566646 | 0.929958472 | 0.62802685 | 1.48076229 | 0.13866993 | 0.30470483 |
| AC008074.1 | 302.140959 | -0.325637755 | 0.34270015 | -0.9502119 | 0.34200461 | 0.54263886 |
| LGALSL | 69.2211484 | -0.593520711 | 0.36432724 | -1.6290868 | 0.10329464 | 0.25095251 |
| AFTPH | 2266.00215 | 0.543654309 | 0.14952432 | 3.63589224 | 0.00027702 | 0.00344295 |
| SERTAD2 | 838.961026 | 0.314219232 | 0.32807454 | 0.95776781 | 0.33817986 | 0.53836156 |
| SLC1A4 | 69.3653463 | -2.207975947 | 0.75984288 | -2.9058323 | 0.00366278 | 0.02398743 |
| CEP68 | 241.29251 | 0.120651715 | 0.27578351 | 0.43748705 | 0.66175816 | 0.8032633 |
| RAB1A | 2979.46585 | -0.031099528 | 0.19736743 | -0.1575717 | 0.87479428 | 0.93490877 |
| ACTR2 | 1313.73396 | -0.601402038 | 0.2379183 | -2.5277671 | 0.01147905 | 0.05558791 |
| SPRED2 | 1668.55661 | -1.084484076 | 0.51033441 | -2.125046 | 0.03358278 | 0.11842764 |
| MEIS1 | 479.573617 | 1.038049319 | 0.44766037 | 2.31883231 | 0.02040413 | 0.08368911 |
| LINC01798 | 61.3386057 | 1.7407267 | 0.62806546 | 2.77156892 | 0.00557869 | 0.03286766 |
| LINC01829 | 69.3765364 | -0.398158012 | 0.60231169 | -0.6610498 | 0.50858039 | 0.68836885 |
| ETAA1 | 206.40848 | 0.272188533 | 0.20184464 | 1.34850514 | 0.17749597 | 0.35810086 |
| C1D | 279.673891 | 0.138233551 | 0.16810202 | 0.82231938 | 0.41089515 | 0.6045054 |
| WDR92 | 77.0441851 | 0.43935471 | 0.26908454 | 1.63277573 | 0.10251617 | 0.25021175 |
| AC017083.3 | 38.9047602 | -0.136823005 | 0.397814 | -0.3439371 | 0.73089357 | 0.84689653 |
| PNO1 | 144.514235 | -0.738270537 | 0.52469177 | -1.4070557 | 0.15941088 | 0.33446272 |
| APLF | 475.349305 | -0.10181119 | 0.31987978 | -0.3182795 | 0.7502729 | 0.85921887 |
| ARHGAP25 | 41.9458708 | -0.711321527 | 0.51985909 | -1.3682968 | 0.1712192 | 0.35023154 |
| ANTXR1 | 95.6202755 | -1.059212701 | 0.65197279 | -1.6246272 | 0.10424203 | 0.25231575 |
| GFPT1 | 1190.90941 | -1.101766606 | 0.34599813 | -3.1843137 | 0.00145098 | 0.01200862 |
| NFU1 | 351.282029 | 0.456857317 | 0.18667094 | 2.44739388 | 0.01438935 | 0.06507252 |
| AAK1 | 4556.36925 | -0.338401838 | 0.23025388 | -1.4696901 | 0.14164571 | 0.30862148 |
| ANXA4 | 4009.64451 | -0.773498316 | 0.28685363 | -2.6964913 | 0.00700742 | 0.03898759 |
| GMCL1 | 678.218999 | -0.368899936 | 0.25531077 | -1.4449055 | 0.14848443 | 0.31840991 |
| MXD1 | 441.441185 | -1.097645089 | 0.36178348 | -3.0339834 | 0.00241348 | 0.01773436 |
| PCBP1 | 624.576842 | -1.120343123 | 0.38009086 | -2.9475666 | 0.00320286 | 0.02171195 |
| C2orf42 | 288.04754 | 0.716261555 | 0.24114157 | 2.97029483 | 0.00297514 | 0.02055456 |
| PCYOX1 | 405.660247 | -0.19408188 | 0.46516166 | -0.4172353 | 0.67650628 | 0.81279385 |
| SNRPG | 270.369279 | -0.216556008 | 0.28778945 | -0.7524807 | 0.45176203 | 0.64135703 |
| FAM136A | 264.473911 | -0.083380813 | 0.2929906 | -0.2845853 | 0.77596189 | 0.87633761 |
| TGFA | 587.870628 | -1.038014989 | 0.24659367 | -4.2094146 | 2.56E-05 | 0.00052933 |
| NAGK | 239.004559 | -0.244106812 | 0.30864613 | -0.7908954 | 0.42900502 | 0.62205719 |
| MCEE | 324.623793 | 0.106506184 | 0.2487501 | 0.42816539 | 0.66853071 | 0.80729015 |
| MPHOSPH10 | 264.198478 | -0.461985948 | 0.35234494 | -1.3111752 | 0.18979858 | 0.37389611 |
| PAIP2B | 566.138781 | -0.830240193 | 0.50568573 | -1.6418106 | 0.10062926 | 0.24683702 |
| ZNF638 | 4405.63614 | 0.045532119 | 0.15711836 | 0.28979502 | 0.77197306 | 0.87329455 |
| DYSF | 392.976347 | -1.041297496 | 0.56768328 | -1.8342931 | 0.0666105 | 0.1877107 |
| EXOC6B | 4271.53182 | -0.321415282 | 0.27419021 | -1.1722347 | 0.24110284 | 0.43618862 |
| SPR | 117.504533 | -0.676146732 | 0.36543174 | -1.8502682 | 0.0642749 | 0.18343727 |
| EMX1 | 311.395661 | 0.821526011 | 0.27121631 | 3.02904352 | 0.00245329 | 0.01795371 |
| SFXN5 | 2069.9028 | -0.007016372 | 0.41956829 | -0.0167228 | 0.98665773 | 0.99360989 |
| RAB11FIP5 | 380.390415 | 0.289145645 | 0.33653488 | 0.85918477 | 0.39023858 | 0.58561422 |
| SMYD5 | 98.8139585 | 0.001888958 | 0.25274231 | 0.00747385 | 0.99403679 | 0.99701904 |
| PRADC1 | 61.1739277 | -0.035564676 | 0.39171982 | -0.0907911 | 0.92765858 | 0.96193661 |
| CCT7 | 364.921908 | -0.45065296 | 0.26868837 | -1.6772328 | 0.09349696 | 0.23540831 |
| ALMS1 | 988.71152 | 0.104385818 | 0.26760747 | 0.39007064 | 0.69648431 | 0.82526889 |
| NAT8 | 1512.20796 | 1.010543153 | 0.86451215 | 1.16891723 | 0.24243698 | 0.43759332 |
| TPRKB | 162.299657 | 0.505711972 | 0.1701588 | 2.97200003 | 0.00295867 | 0.02049239 |
| DUSP11 | 216.961804 | -0.026454807 | 0.20964153 | -0.1261907 | 0.89958099 | 0.94743919 |
| STAMBP | 571.767255 | -0.054713656 | 0.19928534 | -0.2745493 | 0.7836625 | 0.88133085 |
| DGUOK | 445.335251 | -0.302475889 | 0.22764733 | -1.3287039 | 0.18394569 | 0.36652763 |
| DGUOK-AS1 | 52.9114808 | -0.235100393 | 0.34200102 | -0.687426 | 0.49181431 | 0.67419732 |
| TET3 | 717.574303 | 0.004905612 | 0.31363807 | 0.015641 | 0.9875208 | 0.99360989 |
| BOLA3 | 95.3247256 | -0.306728822 | 0.38800458 | -0.7905289 | 0.42921897 | 0.62214435 |
| MOB1A | 672.286459 | -0.014923386 | 0.11707614 | -0.1274674 | 0.89857051 | 0.94743919 |
| MTHFD2 | 62.0970416 | -3.864984622 | 0.85131606 | -4.5400114 | 5.63E-06 | 0.00015832 |
| AC006030.1 | 119.340089 | -0.471133061 | 0.46533384 | -1.0124625 | 0.31131697 | 0.51145516 |
| DCTN1 | 130.771555 | 0.125887032 | 0.32007365 | 0.39330646 | 0.69409315 | 0.82385996 |
| WDR54 | 56.0018452 | -1.200086876 | 0.53061203 | -2.2617031 | 0.02371575 | 0.09290451 |
| RTKN | 692.592307 | 0.812236594 | 0.24370533 | 3.33286347 | 0.00085957 | 0.00808658 |

| | | | | | | |
|---|---|---|---|---|---|---|
| MOGS | 311.125561 | -0.011800117 | 0.27535291 | -0.0428545 | 0.9658175 | 0.98244605 |
| CCDC142 | 168.020451 | 0.490944108 | 0.29264255 | 1.67762381 | 0.09342056 | 0.23525917 |
| TTC31 | 161.399448 | 0.282065828 | 0.30697616 | 0.91885254 | 0.35817271 | 0.55662963 |
| PCGF1 | 44.415134 | -0.010166095 | 0.43567676 | -0.023334 | 0.98138383 | 0.99060551 |
| AUP1 | 267.595791 | -0.13591804 | 0.37802408 | -0.3595486 | 0.71918471 | 0.83996803 |
| HTRA2 | 84.8379542 | -0.695840102 | 0.34102458 | -2.0404397 | 0.04130655 | 0.13656268 |
| HK2 | 59.6558956 | -4.648233427 | 0.60995962 | -7.6205592 | 2.53E-14 | 1.05E-11 |
| POLE4 | 144.972747 | -0.286435962 | 0.35973523 | -0.7962411 | 0.42589193 | 0.61876852 |
| TACR1 | 72.1881316 | -0.870038794 | 0.55757586 | -1.5603954 | 0.11866648 | 0.27517534 |
| AC007681.1 | 114.240639 | -0.137141655 | 0.37880567 | -0.362037 | 0.71732441 | 0.8386157 |
| EVA1A | 1962.41683 | 0.313948353 | 0.47923775 | 0.65509938 | 0.51240377 | 0.6911936 |
| AC007099.1 | 319.313515 | -0.443143355 | 0.67548508 | -0.6560372 | 0.51180017 | 0.69074882 |
| MRPL19 | 479.54077 | 0.019533069 | 0.24406449 | 0.08003241 | 0.93621148 | 0.96587767 |
| GCFC2 | 789.839745 | 0.869580184 | 0.26711975 | 3.25539458 | 0.00113235 | 0.00992377 |
| LRRTM4 | 144.885363 | -0.552102688 | 0.78022518 | -0.7076197 | 0.47918147 | 0.6645225 |
| REG1A | 268.01289 | -0.873794099 | 1.40084893 | -0.6237604 | 0.53278494 | 0.70613839 |
| CTNNA2 | 59.7486835 | 0.349310465 | 0.32696068 | 1.0683562 | 0.28535987 | 0.48492781 |
| SUCLG1 | 1517.1642 | 1.519364513 | 0.43043413 | 3.52984208 | 0.00041581 | 0.00470642 |
| DNAH6 | 486.344998 | 1.635255375 | 0.67227542 | 2.43241882 | 0.01499835 | 0.06705232 |
| TRABD2A | 36.2368181 | 0.254745196 | 0.70828753 | 0.35966353 | 0.71909877 | 0.83993922 |
| TMSB10 | 2919.45128 | -1.755225328 | 0.4445247 | -3.948544 | 7.86E-05 | 0.00129072 |
| KCMF1 | 1591.87373 | -0.560039145 | 0.37395138 | -1.4976256 | 0.13423056 | 0.29834576 |
| TCF7L1 | 87.1767317 | 0.107292691 | 0.50407621 | 0.21285014 | 0.83144385 | 0.91085365 |
| TGOLN2 | 1204.50911 | 0.050331914 | 0.26607509 | 0.18916432 | 0.84996403 | 0.92105755 |
| RETSAT | 425.324341 | -0.013661668 | 0.26902017 | -0.0507831 | 0.95949839 | 0.97928849 |
| ELMOD3 | 512.663474 | 0.409006226 | 0.23769708 | 1.72070359 | 0.08530463 | 0.22099234 |
| CAPG | 127.524974 | -0.571893375 | 0.43566264 | -1.3126978 | 0.18928481 | 0.37331323 |
| SH2D6 | 30.8401744 | 1.704904983 | 0.50465594 | 3.37835115 | 0.00072922 | 0.00716003 |
| MAT2A | 1490.78279 | 0.540324929 | 0.26128451 | 2.0679562 | 0.03864414 | 0.13073699 |
| GGCX | 263.396744 | -0.104651239 | 0.23565385 | -0.4440888 | 0.65697837 | 0.80024905 |
| VAMP8 | 712.407872 | -0.506177911 | 0.25593512 | -1.9777587 | 0.04795594 | 0.15122358 |
| VAMP5 | 154.257434 | -1.023731832 | 0.48829542 | -2.0965419 | 0.03603414 | 0.12408456 |
| RNF181 | 121.701524 | -0.562303624 | 0.41046187 | -1.369929 | 0.17070907 | 0.34969424 |
| TMEM150A | 312.264689 | 1.084154566 | 0.27195368 | 3.9865412 | 6.70E-05 | 0.00114157 |
| USP39 | 563.492428 | 0.037419685 | 0.21634757 | 0.17296097 | 0.86268211 | 0.9280163 |
| C2orf68 | 230.253923 | 0.065832588 | 0.19978718 | 0.32951357 | 0.74176754 | 0.85342854 |
| ST3GAL5 | 124.219238 | -0.170830908 | 0.56590017 | -0.3018746 | 0.76274763 | 0.86770535 |
| POLR1A | 460.263772 | -0.147539931 | 0.36911317 | -0.3997146 | 0.68936673 | 0.82043055 |
| PTCD3 | 902.477188 | 0.024323423 | 0.15516768 | 0.15675573 | 0.87543736 | 0.93520536 |
| IMMT | 909.162822 | 0.055923182 | 0.18253817 | 0.30636432 | 0.75932727 | 0.86510672 |
| MRPL35 | 159.489779 | -0.022487922 | 0.22795442 | -0.098651 | 0.92141541 | 0.95782201 |
| REEP1 | 263.167577 | 0.023799784 | 0.48586291 | 0.04898457 | 0.96093159 | 0.97978927 |
| KDM3A | 1218.29048 | -0.9871595 | 0.32404113 | -3.0464018 | 0.00231598 | 0.01718741 |
| CHMP3 | 187.022382 | -0.791065418 | 0.30504857 | -2.5932442 | 0.00950752 | 0.04854197 |
| RNF103-CHM | 324.852413 | -3.707918403 | 0.83568039 | -4.4370054 | 9.12E-06 | 0.00023591 |
| AC015971.1 | 50.8943891 | -0.922100934 | 0.43870941 | -2.101849 | 0.03556651 | 0.12301542 |
| RMND5A | 746.193914 | 0.124424231 | 0.26094748 | 0.47681714 | 0.63349234 | 0.78395238 |
| PLGLB1 | 185.804253 | 1.108501297 | 0.37116308 | 2.98656134 | 0.00282134 | 0.0198014 |
| CYTOR | 192.360365 | -3.173125578 | 0.67389642 | -4.7086251 | 2.49E-06 | 8.22E-05 |
| KRCC1 | 798.54192 | 0.155586846 | 0.25679026 | 0.60589076 | 0.5445873 | 0.71493708 |
| FABP1 | 409.813577 | 0.667174413 | 0.87134284 | 0.76568531 | 0.44386357 | 0.63517875 |
| THNSL2 | 415.580607 | 1.499621557 | 0.79148537 | 1.89469271 | 0.05813312 | 0.17121573 |
| EIF2AK3 | 1194.61403 | -1.779316981 | 0.38602497 | -4.6093313 | 4.04E-06 | 0.0001206 |
| RPIA | 211.001472 | -0.487166599 | 0.38591078 | -1.2623814 | 0.20681158 | 0.39558558 |
| AL845331.2 | 287.573761 | 0.582207982 | 0.81442003 | 0.71487434 | 0.47468669 | 0.66076272 |
| ANKRD20A8I | 72.1550427 | 0.956864987 | 0.71961944 | 1.32968196 | 0.18362308 | 0.36602053 |
| MAL | 218.184344 | 0.14811376 | 0.59267252 | 0.24990826 | 0.80265829 | 0.89272007 |
| MRPS5 | 719.027364 | -0.33000673 | 0.21959114 | -1.5028235 | 0.13288456 | 0.29626686 |
| ZNF514 | 75.4664904 | 0.119531931 | 0.30184763 | 0.39600089 | 0.69210437 | 0.82256172 |
| ZNF2 | 88.4783391 | 0.08361692 | 0.34634528 | 0.24142647 | 0.8092246 | 0.89639903 |
| AC092835.1 | 108.435183 | 0.31390226 | 0.2974392 | 1.05534933 | 0.29126559 | 0.49143001 |
| KCNIP3 | 146.218124 | -1.649473596 | 0.49651466 | -3.3221045 | 0.00089341 | 0.00831929 |
| FAHD2A | 154.707279 | -0.106903976 | 0.38972922 | -0.2743032 | 0.78385162 | 0.88133085 |
| LINC00342 | 51.5622565 | -0.321126205 | 0.48533911 | -0.6616533 | 0.50819347 | 0.68818493 |
| ANKRD36C | 553.647111 | -0.730375296 | 0.26065654 | -2.8020602 | 0.00507774 | 0.03065035 |
| STARD7 | 567.531962 | 0.39916843 | 0.19595814 | 2.03700863 | 0.04164918 | 0.13719906 |
| TMEM127 | 253.086232 | -0.318044658 | 0.29146984 | -1.0911752 | 0.2751958 | 0.47286769 |
| CIAO1 | 392.103126 | -0.458183484 | 0.27762394 | -1.6503746 | 0.09886635 | 0.24379742 |
| SNRNP200 | 515.729151 | -0.371248667 | 0.28124454 | -1.3200209 | 0.18682805 | 0.37015656 |
| NEURL3 | 91.1921714 | -2.06711857 | 0.60151055 | -3.4365458 | 0.00058918 | 0.00604033 |
| ARID5A | 92.967575 | -3.459682828 | 0.61724158 | -5.605071 | 2.08E-08 | 1.55E-06 |
| KANSL3 | 724.514139 | 0.248487416 | 0.20639975 | 1.20391336 | 0.22862307 | 0.42159779 |
| LMAN2L | 559.545087 | 0.727618934 | 0.17491758 | 4.15978166 | 3.19E-05 | 0.00063281 |
| CNNM4 | 204.858611 | -0.381291857 | 0.41141159 | -0.9267893 | 0.35403595 | 0.55245569 |
| CNNM3 | 101.063871 | 0.329419537 | 0.25774584 | 1.27807894 | 0.20122159 | 0.38830696 |
| ANKRD23 | 229.744169 | -0.109505121 | 0.32246869 | -0.3395837 | 0.73417003 | 0.84943603 |
| SEMA4C | 41.308073 | -0.69033714 | 0.52840615 | -1.3064517 | 0.19139898 | 0.37575277 |
| ANKRD36 | 711.981142 | -0.2808654 | 0.27895342 | -1.0068541 | 0.31400489 | 0.51459409 |
| AC092683.1 | 701.01058 | -0.473476124 | 0.32822747 | -1.4425244 | 0.1491545 | 0.31943447 |
| ANKRD36B | 253.841821 | -0.93743051 | 0.48029881 | -1.9517652 | 0.05096608 | 0.15730514 |
| COX5B | 701.038136 | -0.552841638 | 0.42335614 | -1.3058548 | 0.19160195 | 0.37593437 |
| ACTR1B | 210.170432 | -0.596225792 | 0.32880213 | -1.813327 | 0.0697814 | 0.193583 |
| C2orf92 | 107.396341 | 0.77017406 | 0.30612892 | 2.51584873 | 0.01187461 | 0.05683147 |
| TMEM131 | 1999.3868 | 0.23823712 | 0.12321213 | 1.93355249 | 0.05316817 | 0.16130504 |
| INPP4A | 522.177973 | -0.214429693 | 0.2734716 | -0.7841022 | 0.43298013 | 0.62552702 |
| COA5 | 186.798424 | 0.173770148 | 0.25509246 | 0.68120456 | 0.49574206 | 0.6776839 |
| UNC50 | 241.06705 | -0.185151026 | 0.20356588 | -0.9095386 | 0.36306588 | 0.56080054 |
| MGAT4A | 1412.71124 | 1.058268175 | 0.39844963 | 2.65596477 | 0.00790819 | 0.04251897 |
| KIAA1211L | 119.266233 | 0.989241935 | 0.76350779 | 1.29565402 | 0.19509471 | 0.38023079 |
| TSGA10 | 1328.85972 | 0.229701007 | 0.31616118 | 0.72653134 | 0.4675131 | 0.65476674 |
| AC079447.1 | 636.046909 | 0.509101824 | 0.26449237 | 1.92482609 | 0.05425112 | 0.16379297 |
| MITD1 | 420.268747 | -0.116601367 | 0.18637253 | -0.625636 | 0.53155371 | 0.70522693 |
| LYG1 | 338.723116 | 0.199447103 | 0.45647209 | 0.43693165 | 0.66216092 | 0.80342035 |
| TXNDC9 | 350.422177 | -0.509819555 | 0.23685145 | -2.1524865 | 0.03135906 | 0.11267067 |
| EIF5B | 1203.40273 | -0.854887558 | 0.35852867 | -2.3844329 | 0.01710547 | 0.07395389 |
| REV1 | 1274.76664 | 0.386075685 | 0.22173869 | 1.74112906 | 0.08166096 | 0.21513103 |
| AFF3 | 643.195416 | -0.535598078 | 0.56384552 | -0.9499022 | 0.34216197 | 0.54270009 |
| PDCL3 | 49.6958286 | -0.704416836 | 0.41328101 | -1.70445 | 0.08829704 | 0.22639124 |
| NPAS2 | 3983.87 | -1.050417127 | 0.40123599 | -2.6179534 | 0.00884589 | 0.04605492 |
| RPL31 | 637.632971 | -0.241749708 | 0.29559251 | -0.8178479 | 0.41344406 | 0.60717813 |
| TBC1D8 | 1457.13437 | -1.00289787 | 0.23593694 | -4.2507031 | 2.13E-05 | 0.00046365 |
| CNOT11 | 458.791596 | -0.334586189 | 0.18270511 | -1.8312908 | 0.06705715 | 0.18854236 |
| RNF149 | 2372.77318 | -0.352853097 | 0.19436645 | -1.8154012 | 0.06946227 | 0.19305585 |
| MAP4K4 | 2279.03323 | -1.087639869 | 0.46237273 | -2.3523011 | 0.01865767 | 0.07852031 |
| IL1R1 | 1049.70552 | -1.205285924 | 0.36736714 | -3.2808757 | 0.00103485 | 0.00927713 |
| IL1RL2 | 118.909902 | -2.279688145 | 0.52111002 | -4.3746772 | 1.22E-05 | 0.00029767 |
| IL18R1 | 137.808233 | -0.219137843 | 0.5034185 | -0.4352995 | 0.66334501 | 0.80414523 |
| SLC9A2 | 125.259972 | 0.193596966 | 0.59949 | 0.32293611 | 0.74674363 | 0.85681039 |
| MFSD9 | 164.620499 | -0.486638966 | 0.24217614 | -2.0094422 | 0.04449026 | 0.14362411 |
| TMEM182 | 226.453368 | -0.984607928 | 0.25682252 | -3.8338068 | 0.00012618 | 0.0018711 |
| POU3F3 | 654.514452 | -0.392836944 | 0.40702133 | -0.9651508 | 0.3344693 | 0.53408905 |
| AC010884.1 | 63.2719708 | -0.009695772 | 0.66482267 | -0.014584 | 0.98836407 | 0.99423941 |
| MRPS9 | 484.584958 | 0.256054002 | 0.18154033 | 1.41045247 | 0.15840612 | 0.33317059 |

| | | | | | | |
|---|---|---|---|---|---|---|
| TGFBRAP1 | 361.394288 | -0.103718498 | 0.23996947 | -0.4322154 | 0.66558487 | 0.80579154 |
| AC012360.3 | 63.7814507 | -0.334879859 | 0.41946807 | -0.7983441 | 0.42467084 | 0.61773991 |
| C2orf49 | 190.516231 | -0.275619927 | 0.18049279 | -1.5270412 | 0.12675077 | 0.28783511 |
| FHL2 | 307.116464 | -1.981207094 | 0.4654781 | -4.2562842 | 2.08E-05 | 0.00045585 |
| NCK2 | 812.063765 | -0.523837961 | 0.23046286 | -2.2729821 | 0.02302726 | 0.0911662 |
| UXS1 | 552.970765 | -1.768853279 | 0.32708665 | -5.4079042 | 6.38E-08 | 3.83E-06 |
| RGPD3 | 45.9449722 | 1.138483107 | 0.65367557 | 1.74166385 | 0.08156728 | 0.21500822 |
| ST6GAL2 | 57.9363685 | 0.806303385 | 0.56668135 | 1.42285146 | 0.15477922 | 0.32735053 |
| LINC01789 | 284.893126 | 3.403766562 | 0.82499889 | 4.12578319 | 3.69E-05 | 0.00070732 |
| LINC01885 | 89.1647772 | 5.130872813 | 0.53571502 | 9.57761605 | 9.93E-22 | 2.72E-18 |
| LINC01886 | 108.070872 | 4.557088742 | 0.71635864 | 6.36146264 | 2.00E-10 | 2.80E-08 |
| LINC01594 | 203.38469 | -0.963122382 | 0.56729857 | -1.6977346 | 0.08955787 | 0.22830012 |
| SULT1C2 | 1964.94113 | 0.711943308 | 0.39403125 | 1.80681941 | 0.07079044 | 0.19563413 |
| SULT1C4 | 430.374982 | -1.283287648 | 0.57626091 | -2.2269212 | 0.02595254 | 0.09892421 |
| GCC2 | 2405.29494 | -0.687971959 | 0.17896662 | -3.8441357 | 0.00012098 | 0.00180637 |
| GCC2-AS1 | 343.341603 | -0.075976529 | 0.34949473 | -0.2173896 | 0.82790471 | 0.90828317 |
| LIMS1 | 2476.47104 | -0.659625666 | 0.29082444 | -2.2681233 | 0.02332169 | 0.09180077 |
| RANBP2 | 1223.96039 | 0.012058322 | 0.20703893 | 0.05824181 | 0.95355602 | 0.97656847 |
| CCDC138 | 457.416293 | 0.447771108 | 0.29385174 | 1.52379939 | 0.12755883 | 0.28889633 |
| EDAR | 162.775237 | 0.024436954 | 0.56148327 | 0.04352214 | 0.96528531 | 0.98197757 |
| SH3RF3 | 852.178952 | -0.690051398 | 0.65135273 | -1.0594128 | 0.28941184 | 0.48926592 |
| 10-Sep | 1612.19244 | -0.71609588 | 0.3243767 | -2.2076058 | 0.02727176 | 0.10238676 |
| SOWAHC | 162.680505 | -0.480460227 | 0.46488511 | -1.0335032 | 0.30136849 | 0.50186587 |
| MALL | 30.8570872 | 0.355544189 | 0.47005661 | 0.7563859 | 0.44941786 | 0.63975186 |
| NPHP1 | 353.116078 | 0.519936091 | 0.51948289 | 1.00087241 | 0.3168885 | 0.51693918 |
| SMIM37 | 294.081053 | 0.130062686 | 0.3096144 | 0.42007957 | 0.67442733 | 0.81105241 |
| LINC01106 | 44.8154295 | 1.485620552 | 0.35370139 | 4.2002112 | 2.67E-05 | 0.000548 |
| AC112229.3 | 38.1724242 | 1.888356132 | 0.50404094 | 3.74643405 | 0.00017937 | 0.00245507 |
| MIR4435-2H | 1496.74911 | -3.70573991 | 0.57170432 | -6.481917 | 9.06E-11 | 1.47E-08 |
| BCL2L11 | 385.482673 | -0.951406467 | 0.37420495 | -2.5424743 | 0.01100707 | 0.05399925 |
| AC017002.5 | 101.864592 | -3.327144065 | 0.5847754 | -5.6896102 | 1.27E-08 | 1.01E-06 |
| ANAPC1 | 1056.42954 | 0.062615893 | 0.1802025 | 0.34747515 | 0.72823438 | 0.84570945 |
| MERTK | 414.562622 | -0.118271606 | 0.35448817 | -0.3336405 | 0.73865086 | 0.8516729 |
| TMEM87B | 688.014765 | -1.00654374 | 0.14863356 | -6.7719815 | 1.27E-11 | 2.59E-09 |
| FBLN7 | 58.501123 | 0.071974432 | 0.64703939 | 0.11123655 | 0.91142876 | 0.95319043 |
| ZC3H8 | 556.14464 | 0.541563847 | 0.26007411 | 2.08234434 | 0.03731103 | 0.12750493 |
| ZC3H6 | 599.972016 | 0.714694138 | 0.55827411 | 1.28018499 | 0.20048008 | 0.3875854 |
| RGPD8 | 71.024619 | 0.787899748 | 0.38451896 | 2.04905303 | 0.04045693 | 0.1344883 |
| TTL | 327.552966 | -1.112882342 | 0.5417951 | -2.054065 | 0.0399694 | 0.13346178 |
| POLR1B | 210.058385 | 0.392368714 | 0.43263271 | 0.90693261 | 0.36444243 | 0.56208758 |
| CHCHD5 | 109.535033 | -0.600301067 | 0.33615582 | -1.7857822 | 0.07413454 | 0.2017796 |
| SLC20A1 | 832.146649 | -1.791297631 | 0.55024932 | -3.2554291 | 0.00113221 | 0.00992377 |
| PSD4 | 135.770561 | -1.058489353 | 0.35645382 | -2.9694992 | 0.00298286 | 0.02058711 |
| PAX8 | 24108.095 | -0.768043506 | 0.19466509 | -3.9454609 | 7.96E-05 | 0.00130309 |
| CBWD2 | 351.433453 | -0.122303755 | 0.21399253 | -0.5715328 | 0.56763852 | 0.73443659 |
| RABL2A | 261.390755 | 0.535726496 | 0.21562926 | 2.48447954 | 0.0129741 | 0.06054853 |
| AC017074.1 | 175.790873 | 0.066119918 | 0.47963165 | 0.13785562 | 0.89035453 | 0.94326785 |
| SLC35F5 | 2785.97505 | -0.775959961 | 0.26010292 | -2.9832805 | 0.00285177 | 0.01995362 |
| ACTR3 | 1696.14575 | -0.866836877 | 0.24441656 | -3.5465555 | 0.0003903 | 0.00447688 |
| LINC01191 | 116.248142 | -3.826940205 | 0.59852907 | -6.3939086 | 1.62E-10 | 2.36E-08 |
| DPP10 | 222.785836 | -1.825561761 | 0.7674429 | -2.3787591 | 0.01737102 | 0.07475813 |
| DDX18 | 460.44092 | -1.02279117 | 0.36556963 | -2.7978012 | 0.00514518 | 0.03098144 |
| AC009404.1 | 88.2854224 | -0.664215835 | 0.44853711 | -1.4808492 | 0.13864676 | 0.30470276 |
| CCDC93 | 1338.32113 | -0.333125416 | 0.27515405 | -1.2106869 | 0.22601541 | 0.41859117 |
| AC009303.2 | 57.5527526 | -0.536714565 | 0.46863064 | -1.1452827 | 0.25209207 | 0.44852362 |
| INSIG2 | 765.749623 | -1.940789597 | 0.45998004 | -4.2192909 | 2.45E-05 | 0.00051521 |
| C2orf76 | 192.344413 | -0.367118554 | 0.29218234 | -1.2564707 | 0.20894536 | 0.39816684 |
| DBI | 1504.16157 | -1.100488174 | 0.50218019 | -2.1914209 | 0.02842135 | 0.10514747 |
| TMEM37 | 959.396773 | -0.05919222 | 0.53336468 | -0.1109789 | 0.91163309 | 0.95319043 |
| SCTR | 312.903304 | -1.790237619 | 0.93092848 | -1.9230668 | 0.05447167 | 0.16420566 |
| AC013275.1 | 82.7746207 | -1.254220752 | 0.90582986 | -1.3846096 | 0.16617186 | 0.34380645 |
| CFAP221 | 837.800917 | -1.690630426 | 0.52783378 | -3.2029599 | 0.00136023 | 0.01141044 |
| TMEM177 | 87.7516066 | 0.0934628 | 0.40551436 | 0.23047963 | 0.81771909 | 0.90160299 |
| PTPN4 | 2101.86245 | 0.649686972 | 0.2067037 | 3.14308337 | 0.00167178 | 0.01334602 |
| EPB41L5 | 3579.62292 | 0.580925201 | 0.28115704 | 2.06619478 | 0.03881009 | 0.13096256 |
| AC012363.1 | 34.0298321 | -1.027426466 | 0.44091084 | -2.3302363 | 0.01979366 | 0.08184366 |
| RALB | 692.715955 | -0.555019087 | 0.23310663 | -2.3809665 | 0.01726728 | 0.07440511 |
| TFCP2L1 | 1582.54524 | 0.360359678 | 0.55265803 | 0.65204821 | 0.51437007 | 0.69258061 |
| CLASP1 | 3755.24604 | 0.012852141 | 0.22932497 | 0.05604336 | 0.95530727 | 0.97737295 |
| NIFK-AS1 | 229.359849 | -0.472958357 | 0.41713125 | -1.1338358 | 0.25686342 | 0.45406588 |
| NIFK | 130.468077 | -0.681430364 | 0.41949482 | -1.6244071 | 0.10428896 | 0.25233693 |
| TSN | 247.901008 | -0.452668108 | 0.24008392 | -1.8854578 | 0.05936805 | 0.17350914 |
| CNTNAP5 | 81.6720055 | 0.967793132 | 0.68180485 | 1.41945769 | 0.15576562 | 0.32903058 |
| GYPC | 66.0307772 | -2.021253579 | 0.49802361 | -4.0585497 | 4.94E-05 | 0.00089621 |
| BIN1 | 943.688456 | 0.401499127 | 0.36050333 | 1.11371821 | 0.2654001 | 0.46324196 |
| ERCC3 | 393.184117 | -0.078005719 | 0.3218353 | -0.2423778 | 0.80848747 | 0.89601696 |
| MAP3K2 | 2243.09533 | 0.33288121 | 0.08544136 | 3.8960196 | 9.78E-05 | 0.00152835 |
| PROC | 278.89292 | 0.980825012 | 0.38927002 | 2.5196521 | 0.01174709 | 0.05639836 |
| IWS1 | 1390.89207 | 0.066804063 | 0.1706462 | 0.391477 | 0.69544469 | 0.82453597 |
| AC010976.1 | 160.051418 | 0.277389104 | 0.57400275 | 0.48325397 | 0.62891542 | 0.78084635 |
| MYO7B | 1459.52521 | 0.061504314 | 0.58996344 | 0.10425106 | 0.91697011 | 0.95564119 |
| WDR33 | 2102.27387 | 0.353528682 | 0.26153693 | 1.35173522 | 0.17646003 | 0.3567986 |
| SFT2D3 | 47.2626844 | 0.008684097 | 0.31947988 | 0.02718198 | 0.97831459 | 0.98906609 |
| POLR2D | 173.855158 | -0.152017903 | 0.37647613 | -0.4037916 | 0.68636597 | 0.8183732 |
| AMMECR1L | 247.949589 | -0.030995407 | 0.25687187 | -0.1206649 | 0.9039565 | 0.94968435 |
| SAP130 | 545.989188 | 0.228494019 | 0.23028894 | 0.99220578 | 0.32109716 | 0.52141674 |
| UGGT1 | 662.687382 | -0.117130739 | 0.26271823 | -0.4458417 | 0.65571161 | 0.79908554 |
| HS6ST1 | 266.368723 | -0.219512018 | 0.36345581 | -0.6039579 | 0.54587162 | 0.71607448 |
| SMPD4 | 401.376502 | -0.187044679 | 0.3180097 | -0.5881729 | 0.55641626 | 0.72549978 |
| MZT2B | 521.713181 | -0.580004946 | 0.32935815 | -1.7610159 | 0.07823571 | 0.20883678 |
| CCDC115 | 88.1622448 | -0.041423083 | 0.34637719 | -0.1195895 | 0.90480832 | 0.95028792 |
| IMP4 | 168.966126 | -0.544777733 | 0.42113874 | -1.2935826 | 0.19580964 | 0.38113643 |
| PTPN18 | 412.70712 | -0.024528505 | 0.2615772 | -0.0937716 | 0.92529061 | 0.96057873 |
| FAM168B | 944.115259 | 0.076730508 | 0.27980289 | 0.27423057 | 0.78390743 | 0.88133085 |
| PLEKHB2 | 1092.60059 | -0.692368789 | 0.28030556 | -2.4700501 | 0.01350941 | 0.06226895 |
| MZT2A | 280.243837 | -0.256304946 | 0.19610337 | -1.306989 | 0.19121645 | 0.3755558 |
| ANKRD30BL | 111.146266 | -0.708866231 | 0.41981371 | -1.6885257 | 0.09131036 | 0.23164744 |
| GPR39 | 706.618162 | -0.009499632 | 0.20599197 | -0.0461165 | 0.96321738 | 0.98083195 |
| LYPD1 | 88.8548904 | -3.385127425 | 0.86821152 | -3.8989663 | 9.66E-05 | 0.00151505 |
| NCKAP5 | 11487.7455 | -0.028454953 | 0.61149135 | -0.0465337 | 0.96288488 | 0.98077306 |
| AC016909.2 | 57.2690633 | -1.499135085 | 0.9651984 | -1.5531885 | 0.1203781 | 0.27778159 |
| AC010890.1 | 83.3202424 | -1.787122924 | 1.01022266 | -1.7690387 | 0.07688742 | 0.20663516 |
| MGAT5 | 1419.7427 | -0.572155625 | 0.35693408 | -1.6029728 | 0.10894065 | 0.25996684 |
| TMEM163 | 467.420647 | -0.16524765 | 0.54722001 | -0.3019766 | 0.76266988 | 0.86770535 |
| CCNT2-AS1 | 828.002546 | -0.698093198 | 0.44379755 | -1.5729992 | 0.115719 | 0.27086072 |
| ACMSD | 3562.54782 | 0.754188312 | 0.59592439 | 1.26557719 | 0.2056645 | 0.3943821 |
| CCNT2 | 1207.98011 | -0.249515532 | 0.10944003 | -2.2799294 | 0.02261188 | 0.09006311 |
| MAP3K19 | 48.6856907 | -0.874584107 | 0.48206133 | -1.814259 | 0.06963786 | 0.19331655 |
| RAB3GAP1 | 2145.39508 | -0.22363819 | 0.14961124 | -1.4947954 | 0.13496784 | 0.29936987 |
| ZRANB3 | 826.025297 | -0.012606743 | 0.23258352 | -0.0542031 | 0.95677337 | 0.97808644 |
| R3HDM1 | 1713.61483 | 0.113383867 | 0.18502478 | 0.61280369 | 0.54000615 | 0.71158088 |
| UBXN4 | 1589.91549 | -0.616678765 | 0.26063272 | -2.3660835 | 0.01797739 | 0.0765743 |
| MCM6 | 40.1509879 | -0.596877478 | 0.43799316 | -1.3627553 | 0.17295966 | 0.35258151 |

| | | | | | | |
|---|---|---|---|---|---|---|
| DARS | 1636.34661 | -0.982429808 | 0.2200351 | -4.4648777 | 8.01E-06 | 0.00021159 |
| DARS-AS1 | 75.3475654 | -1.270042983 | 0.56356425 | -2.2535904 | 0.02422195 | 0.09419154 |
| THSD7B | 54.6191229 | 0.267493933 | 0.49464526 | 0.54077933 | 0.58865969 | 0.75086157 |
| HNMT | 1018.73453 | 0.635993524 | 0.55914009 | 1.13744934 | 0.25535049 | 0.45232479 |
| AC114763.1 | 370.263024 | 0.435668014 | 0.34336013 | 1.26883693 | 0.20449923 | 0.39280703 |
| SPOPL | 808.018653 | 0.009302519 | 0.17973445 | 0.05175702 | 0.9587223 | 0.97890396 |
| LRP1B | 1061.77258 | 0.364730371 | 0.79140024 | 0.46086715 | 0.64489392 | 0.79180937 |
| KYNU | 1278.31016 | 0.799271736 | 0.3708586 | 2.15519265 | 0.03114676 | 0.11211362 |
| ARHGAP15 | 68.2200971 | 0.749944643 | 0.40396349 | 1.85646638 | 0.06338711 | 0.18169116 |
| GTDC1 | 1558.29032 | -0.275445283 | 0.18042271 | -1.5266664 | 0.12684399 | 0.28790372 |
| ZEB2 | 1913.59314 | 0.177724653 | 0.27394247 | 0.64876635 | 0.51648941 | 0.69386001 |
| TEX41 | 113.883683 | 2.132551986 | 0.64398883 | 3.31147358 | 0.00092806 | 0.00856051 |
| ACVR2A | 790.152837 | 0.702986569 | 0.13714196 | 5.12597715 | 2.96E-07 | 1.36E-05 |
| ORC4 | 1663.85535 | 0.487394169 | 0.25898451 | 1.88194332 | 0.05984371 | 0.17441585 |
| MBD5 | 7014.11487 | 0.587779399 | 0.36142157 | 1.62629863 | 0.10388614 | 0.25194259 |
| EPC2 | 1296.59699 | 0.409123498 | 0.18928603 | 2.16140356 | 0.03066418 | 0.1108991 |
| AC105402.3 | 76.6534889 | 0.072111471 | 0.98687952 | 0.07307019 | 0.94175027 | 0.96947585 |
| MMADHC | 283.480796 | -0.061867046 | 0.24928774 | -0.2481752 | 0.80399882 | 0.89349861 |
| AC007364.1 | 3464.86552 | 0.580457441 | 0.33730332 | 1.72087674 | 0.08527319 | 0.22099234 |
| LINC01931 | 138.880078 | -0.433089697 | 0.69228804 | -0.6255918 | 0.53158274 | 0.70522693 |
| RND3 | 366.001166 | -3.722036461 | 0.99402796 | -3.7443981 | 0.00018083 | 0.00246008 |
| RBM43 | 66.1881635 | -0.000746908 | 0.54118743 | -0.0013801 | 0.99889882 | 0.99926333 |
| NMI | 206.562515 | -0.784199028 | 0.50598172 | -1.5498564 | 0.12117598 | 0.2791528 |
| RIF1 | 1129.93879 | -0.498869169 | 0.2421281 | -2.0603523 | 0.03936488 | 0.13218382 |
| NEB | 143.288764 | 0.650059776 | 0.44011532 | 1.47702148 | 0.13966988 | 0.30606795 |
| ARL5A | 408.223027 | 0.593946489 | 0.24126304 | 2.46182134 | 0.01382335 | 0.06332183 |
| CACNB4 | 108.353248 | 0.433148043 | 0.3114586 | 1.39070826 | 0.16431391 | 0.34125012 |
| STAM2 | 744.757114 | 0.248560164 | 0.2120389 | 1.17223853 | 0.2411013 | 0.43618862 |
| FMNL2 | 10205.8796 | -1.741719643 | 0.51187912 | -3.4025995 | 0.00066748 | 0.00668278 |
| PRPF40A | 1115.10316 | -0.030407208 | 0.14513052 | -0.2095163 | 0.83404522 | 0.91224528 |
| ARL6IP6 | 357.932473 | -0.278238663 | 0.23720632 | -1.1729817 | 0.24080316 | 0.43613755 |
| GALNT13 | 176.121077 | -0.978378845 | 0.60204467 | -1.6250934 | 0.10414265 | 0.25231575 |
| KCNJ3 | 1916.11924 | 1.428986719 | 0.59620135 | 2.396819 | 0.01653809 | 0.0721246 |
| LINC01876 | 363.663497 | -1.054355715 | 0.9239139 | -1.141184 | 0.25379338 | 0.45069836 |
| GPD2 | 572.994493 | -0.578329627 | 0.20475014 | -2.8245628 | 0.00473452 | 0.02911441 |
| ACVR1C | 88.6794044 | 0.121275901 | 0.43051925 | 0.28169681 | 0.778176 | 0.87771268 |
| AC019186.1 | 120.487357 | 1.312567729 | 0.50969839 | 2.57518517 | 0.01001864 | 0.05052448 |
| ACVR1 | 2530.94788 | 0.626107083 | 0.30628995 | 2.04416463 | 0.04093729 | 0.13557076 |
| UPP2 | 1313.74315 | 1.274077154 | 0.75355705 | 1.69075076 | 0.09088442 | 0.2308828 |
| CCDC148 | 1354.20394 | 0.627015325 | 0.55859034 | 1.12249582 | 0.26165168 | 0.45993814 |
| PKP4 | 7267.5131 | 0.147888889 | 0.24088008 | 0.61395234 | 0.53924682 | 0.71085365 |
| PKP4-AS1 | 171.802574 | -0.412060066 | 0.63518749 | -0.6487219 | 0.51651812 | 0.69386001 |
| TANC1 | 1830.55001 | -0.680610171 | 0.17944285 | -3.7929076 | 0.00014889 | 0.00210105 |
| WDSUB1 | 879.913736 | 0.571870759 | 0.30888292 | 1.85141596 | 0.06410974 | 0.18311151 |
| BAZ2B | 4694.38398 | 0.141901133 | 0.28411681 | 0.49944646 | 0.6174649 | 0.77250742 |
| AC009961.1 | 152.143027 | 0.517105711 | 0.27832035 | 1.85795149 | 0.0631759 | 0.18116153 |
| 7-Mar | 1958.28628 | -0.098327844 | 0.19643529 | -0.500561 | 0.61668012 | 0.77187785 |
| PLA2R1 | 766.036213 | -0.479320771 | 0.33355508 | -1.4370064 | 0.15071619 | 0.3217861 |
| ITGB6 | 1353.26023 | -2.939609877 | 0.66407164 | -4.4266457 | 9.57E-06 | 0.00024613 |
| LINC02478 | 47.9668805 | -1.047095817 | 0.51761413 | -2.0229274 | 0.04308063 | 0.14039615 |
| RBMS1 | 3958.98079 | -0.448235664 | 0.24060806 | -1.8629288 | 0.06247229 | 0.17972542 |
| TANK | 1740.27907 | -0.864700528 | 0.18540685 | -4.6638003 | 3.10E-06 | 9.85E-05 |
| AC009299.3 | 39.4491764 | -0.503106004 | 0.54731971 | -0.9192178 | 0.35798168 | 0.55650968 |
| PSMD14 | 883.176179 | -0.88322794 | 0.3423873 | -2.5796165 | 0.00989101 | 0.05004653 |
| DPP4 | 2268.10711 | -0.888258892 | 0.62303654 | -1.4256931 | 0.15395695 | 0.32636683 |
| IFIH1 | 269.448192 | -0.530801758 | 0.58942493 | -0.9005418 | 0.36783202 | 0.56523245 |
| GCA | 293.465089 | -0.694587731 | 0.38554207 | -1.8015874 | 0.07161035 | 0.19714061 |
| KCNH7 | 59.5780943 | 0.13325429 | 0.54977051 | 0.24238166 | 0.80848444 | 0.89601696 |
| FIGN | 2032.42352 | 0.529799277 | 0.29143866 | 1.81787577 | 0.06908312 | 0.19255756 |
| AC016766.1 | 49.8927568 | 1.214718564 | 0.49031321 | 2.47743391 | 0.01323309 | 0.06138274 |
| GRB14 | 636.914936 | 0.597311071 | 0.40862794 | 1.461748 | 0.14381028 | 0.31145639 |
| COBLL1 | 3830.00011 | 0.185049621 | 0.32955812 | 0.5615083 | 0.57445108 | 0.73927339 |
| AC019197.1 | 1165.91562 | -0.117672986 | 0.8541417 | -0.1377675 | 0.89042415 | 0.94326785 |
| SLC38A11 | 137.685135 | 1.348687511 | 0.59222157 | 2.27733602 | 0.02276617 | 0.09045097 |
| SCN2A | 153.352766 | -0.088891213 | 0.92537732 | -0.0960594 | 0.92347338 | 0.95941131 |
| CSRNP3 | 763.654199 | 0.990296445 | 0.50361388 | 1.96638037 | 0.04925469 | 0.15396641 |
| GALNT3 | 239.742159 | 0.168936973 | 0.38139608 | 0.44294365 | 0.65780649 | 0.80061744 |
| AC009495.1 | 176.48803 | 1.821023902 | 0.36336867 | 5.01150504 | 5.40E-07 | 2.28E-05 |
| TTC21B | 1407.32421 | 1.149930614 | 0.28669796 | 4.01094806 | 6.05E-05 | 0.00105329 |
| AC010127.1 | 266.040719 | -0.236186858 | 0.76182485 | -0.3100278 | 0.75653984 | 0.86351 |
| SCN9A | 909.133677 | 0.720089462 | 0.67380241 | 1.06869528 | 0.285207 | 0.48472812 |
| AC073050.1 | 151.679185 | 0.451287481 | 0.74621066 | 0.60477223 | 0.54533035 | 0.71570692 |
| STK39 | 1193.33591 | -3.419905814 | 0.5599122 | -6.1079323 | 1.01E-09 | 1.17E-07 |
| CERS6 | 1434.25322 | 0.493786012 | 0.22867486 | 2.15933668 | 0.03082406 | 0.11127346 |
| NOSTRIN | 1956.92291 | -0.940428249 | 0.69152478 | -1.3599343 | 0.17385073 | 0.3536616 |
| SPC25 | 296.990968 | -1.318654544 | 0.74144377 | -1.7784957 | 0.07532248 | 0.20395995 |
| LRP2 | 28560.0978 | 1.1527556 | 0.77910701 | 1.47958571 | 0.13898384 | 0.30510114 |
| FASTKD1 | 528.352367 | 0.367036893 | 0.24614226 | 1.49115757 | 0.13592013 | 0.30059869 |
| PPIG | 1274.06825 | 0.305496688 | 0.14204665 | 2.15067852 | 0.03150158 | 0.11304387 |
| CCDC173 | 65.1015603 | 0.223410241 | 0.5428282 | 0.41156712 | 0.68065674 | 0.81442647 |
| KLHL23 | 507.67902 | 0.096589665 | 0.24632644 | 0.39212057 | 0.69496913 | 0.82432865 |
| SSB | 327.563425 | -0.457560091 | 0.23017259 | -1.9879 | 0.04682275 | 0.14901311 |
| METTL5 | 202.972881 | -0.336379379 | 0.2012007 | -1.6718599 | 0.09455196 | 0.23688973 |
| UBR3 | 2485.86988 | 0.34182495 | 0.20140184 | 1.69722852 | 0.08965347 | 0.22841637 |
| MYO3B | 113.483427 | 1.243117115 | 0.51941127 | 2.39331949 | 0.0166967 | 0.07266389 |
| AC007405.1 | 48.7932414 | -0.609065505 | 0.64837286 | -0.9393754 | 0.34753805 | 0.54753787 |
| GORASP2 | 556.714004 | -0.306583986 | 0.25069622 | -1.2229302 | 0.22135605 | 0.4134786 |
| TLK1 | 1927.3013 | 0.093267408 | 0.15629274 | 0.59674817 | 0.55067553 | 0.720309 |
| METTL8 | 644.450059 | -0.19738771 | 0.34859774 | -0.5662335 | 0.57123507 | 0.73727873 |
| DCAF17 | 281.068062 | 0.410560426 | 0.25222717 | 1.62774066 | 0.10357989 | 0.25155378 |
| DYNC1I2 | 1155.90779 | -0.360800909 | 0.13538531 | -2.6649931 | 0.00769899 | 0.04169133 |
| SLC25A12 | 1072.188 | 0.28285384 | 0.30450076 | 0.92891014 | 0.35293566 | 0.55184755 |
| HAT1 | 328.192805 | -0.062387064 | 0.18740186 | -0.3329053 | 0.7392058 | 0.85193361 |
| METAP1D | 195.189284 | 0.775164284 | 0.35280226 | 2.19716361 | 0.02800876 | 0.10415702 |
| ITGA6 | 3531.23003 | -0.483765969 | 0.36742763 | -1.3166293 | 0.18796293 | 0.37161516 |
| AC078883.1 | 179.074214 | -2.424789385 | 0.37897254 | -6.3983248 | 1.57E-10 | 2.32E-08 |
| PDK1 | 698.928137 | -3.7329842 | 0.54618339 | -6.8346718 | 8.22E-12 | 1.82E-09 |
| RAPGEF4 | 98.8083293 | -0.887133936 | 0.56469678 | -1.5709917 | 0.11618459 | 0.27158651 |
| MAP3K20 | 410.454082 | -2.688993198 | 0.72553453 | -3.7062236 | 0.00021037 | 0.00278214 |
| SP3 | 1099.04279 | 0.097547561 | 0.15780864 | 0.61813829 | 0.53648419 | 0.70877964 |
| OLA1 | 2114.14478 | -0.474431442 | 0.20643478 | -2.2982147 | 0.02154957 | 0.08731304 |
| CIR1 | 690.00002 | 0.045270255 | 0.26695829 | 0.16957801 | 0.86534202 | 0.92952143 |
| SCRN3 | 232.031387 | 0.618512459 | 0.36146208 | 1.71114065 | 0.08705515 | 0.22434009 |
| GPR155 | 2319.62689 | 1.483116333 | 0.44998282 | 3.29593994 | 0.00098093 | 0.00891028 |
| AC010894.3 | 596.454503 | -0.896714653 | 0.31191301 | -2.874887 | 0.00404172 | 0.02582745 |
| WIPF1 | 1801.148 | 0.582669463 | 0.25302464 | 2.30281712 | 0.02128913 | 0.08651354 |
| CHN1 | 161.872699 | -0.500479947 | 0.40643519 | -1.2313893 | 0.21817729 | 0.4093288 |
| ATF2 | 1088.3254 | -0.207744656 | 0.16578828 | -1.253072 | 0.2101795 | 0.39979605 |
| ATP5MC3 | 866.241948 | -0.857808861 | 0.51323141 | -1.6713881 | 0.09464505 | 0.23698747 |
| LNPK | 633.245889 | -0.755231244 | 0.37898162 | -1.9927912 | 0.04628431 | 0.14777989 |
| HOXD-AS2 | 62.9565938 | 0.225271216 | 0.43721572 | 0.51524043 | 0.60638502 | 0.76359388 |
| HOXD8 | 49.2657279 | 0.541579765 | 0.41718906 | 1.29816386 | 0.19423103 | 0.37930258 |
| HOXD3 | 137.073534 | 0.958944521 | 0.46137047 | 2.07846966 | 0.03766612 | 0.12833497 |

| | | | | | | |
|---|---|---|---|---|---|---|
| HAGLR | 74.6815064 | 0.000193683 | 0.39696572 | 0.00048791 | 0.99961071 | 0.99975658 |
| MTX2 | 1554.73465 | 0.148379913 | 0.26150469 | 0.56740824 | 0.57043686 | 0.73646497 |
| AC079305.3 | 101.848232 | 0.328174463 | 0.4595159 | 0.71417434 | 0.47511938 | 0.66116359 |
| HNRNPA3 | 1389.24353 | -0.232698867 | 0.23283657 | -0.9994086 | 0.31759679 | 0.51761699 |
| NFE2L2 | 3303.20939 | -0.572844132 | 0.22869602 | -2.5048278 | 0.0122511 | 0.05806858 |
| AGPS | 1619.30718 | -0.147107689 | 0.36032525 | -0.4082636 | 0.68308015 | 0.81587484 |
| TTC30A | 38.426706 | 0.177858071 | 0.43189901 | 0.41180477 | 0.68048253 | 0.81442647 |
| PDE11A | 29.1521355 | -0.779220569 | 0.85242451 | -0.9141227 | 0.36065239 | 0.55871211 |
| RBM45 | 272.644018 | 0.475358529 | 0.29820217 | 1.5940814 | 0.1109178 | 0.26312744 |
| OSBPL6 | 1045.81485 | 1.159517295 | 0.31286795 | 3.70609169 | 0.00021048 | 0.00278214 |
| AC009948.1 | 149.870584 | -0.607784457 | 0.59636051 | -1.0191561 | 0.30812886 | 0.50824576 |
| PRKRA | 581.313063 | -0.189506013 | 0.18571944 | -1.0203887 | 0.30754416 | 0.5077145 |
| PJVK | 147.693823 | 0.527225177 | 0.32803191 | 1.60723747 | 0.10800228 | 0.25838981 |
| PLEKHA3 | 351.933244 | -0.234802788 | 0.21941656 | -1.0701234 | 0.28456378 | 0.48393495 |
| TTN-AS1 | 1655.0378 | 0.231694167 | 0.26817296 | 0.86397288 | 0.38760279 | 0.5837081 |
| TTN | 125.520772 | -0.711376307 | 0.6795766 | -1.0467934 | 0.29519487 | 0.49508665 |
| SESTD1 | 3067.31489 | -0.970155825 | 0.4380442 | -2.2147442 | 0.02677763 | 0.10118887 |
| ZNF385B | 1665.66781 | 2.866302497 | 0.72112181 | 3.97478268 | 7.04E-05 | 0.00118185 |
| CWC22 | 538.867236 | 0.483754893 | 0.16782645 | 2.88247113 | 0.00394569 | 0.02536755 |
| UBE2E3 | 1422.26235 | 0.416673592 | 0.18983857 | 2.19488372 | 0.02817194 | 0.10450684 |
| LINC01934 | 411.394896 | 3.231515434 | 0.72038721 | 4.48580344 | 7.26E-06 | 0.00019333 |
| CERKL | 231.895597 | 1.080310221 | 0.61793094 | 1.74827015 | 0.08041726 | 0.21275418 |
| SSFA2 | 1327.98231 | 0.654313195 | 0.26603281 | 2.45952067 | 0.01391227 | 0.06355699 |
| PDE1A | 622.790192 | -0.229719837 | 0.69235417 | -0.3317953 | 0.74004386 | 0.85230954 |
| DNAJC10 | 586.215373 | -0.691449695 | 0.27313853 | -2.5314982 | 0.01135764 | 0.05518581 |
| NCKAP1 | 2592.5588 | -0.161791389 | 0.18366841 | -0.8808885 | 0.37837816 | 0.57505593 |
| NUP35 | 270.286381 | -0.07932118 | 0.29585388 | -0.2681093 | 0.78861518 | 0.88450603 |
| FSIP2 | 196.818114 | 0.201597713 | 0.448914 | 0.44907869 | 0.65337489 | 0.79727674 |
| FSIP2-AS1 | 107.737598 | 0.569464878 | 0.61484872 | 0.92618697 | 0.35434882 | 0.55281804 |
| LINC01473 | 283.170384 | 0.144895927 | 0.60242322 | 0.24052182 | 0.80992575 | 0.89677969 |
| AC017071.1 | 46.252871 | 0.472544879 | 0.47998177 | 0.98450589 | 0.32486684 | 0.52511199 |
| ZC3H15 | 617.153388 | -0.2114716 | 0.21773598 | -0.9712295 | 0.33143403 | 0.53127895 |
| ITGAV | 3772.07272 | -0.351139506 | 0.58101062 | -0.6043599 | 0.54560441 | 0.71579246 |
| AC017101.1 | 288.377084 | -1.543548426 | 0.5374784 | -2.8718334 | 0.00408098 | 0.0260419 |
| AC007319.1 | 1617.2341 | -1.226709958 | 0.47289804 | -2.5940263 | 0.00948592 | 0.04849815 |
| CALCRL | 498.366714 | 0.495028917 | 0.46884681 | 1.05584362 | 0.29103966 | 0.49116975 |
| TFPI | 5187.01178 | -1.431992126 | 0.36628987 | -3.9094505 | 9.25E-05 | 0.00146557 |
| LINC01090 | 175.933916 | -2.434189829 | 0.54232938 | -4.4883975 | 7.18E-06 | 0.00019174 |
| GULP1 | 4346.16845 | -2.086300463 | 0.57381194 | -3.635861 | 0.00027705 | 0.00344295 |
| COL5A2 | 351.294997 | -0.327383572 | 0.57665836 | -0.5677254 | 0.57022147 | 0.7363548 |
| WDR75 | 600.617958 | -0.287763428 | 0.22205102 | -1.2959338 | 0.19499829 | 0.38020506 |
| SLC40A1 | 280.493709 | 0.400889705 | 0.40363185 | 0.99320633 | 0.32060941 | 0.52105682 |
| ASNSD1 | 159.271183 | -0.281606011 | 0.23957236 | -1.1754528 | 0.2398136 | 0.43490488 |
| ANKAR | 846.259554 | 0.263202047 | 0.29429491 | 0.89434794 | 0.37113576 | 0.56877072 |
| OSGEPL1 | 93.6513586 | 0.890970449 | 0.39797979 | 2.23873288 | 0.0251733 | 0.09678834 |
| ORMDL1 | 558.941957 | -0.122376602 | 0.26807108 | -0.456508 | 0.64802469 | 0.79355195 |
| PMS1 | 545.37641 | 0.029915696 | 0.19066088 | 0.15690526 | 0.87531951 | 0.93515234 |
| C2orf88 | 326.51368 | 0.268407132 | 0.33038441 | 0.8124086 | 0.41655721 | 0.60995083 |
| HIBCH | 3882.62367 | 1.285672906 | 0.32904407 | 3.90729703 | 9.33E-05 | 0.00147559 |
| INPP1 | 277.087494 | -1.051926849 | 0.18966653 | -5.5461913 | 2.92E-08 | 2.05E-06 |
| MFSD6 | 276.851971 | -1.141202054 | 0.31318913 | -3.6438112 | 0.00026863 | 0.00336882 |
| NEMP2 | 84.2016427 | 0.916816052 | 0.29776926 | 3.07894793 | 0.00207733 | 0.01582766 |
| AC108047.1 | 68.169644 | 0.730448919 | 0.44213253 | 1.65210401 | 0.09851335 | 0.24316991 |
| NAB1 | 335.851379 | 0.612738044 | 0.34792423 | 1.76112495 | 0.07821726 | 0.20883678 |
| GLS | 9199.61983 | -0.128940713 | 0.3718544 | -0.3467505 | 0.72877874 | 0.84599052 |
| STAT1 | 2187.6981 | -2.312409732 | 0.87846952 | -2.6323164 | 0.00848048 | 0.04465693 |
| STAT4 | 39.0481799 | -2.153180824 | 0.60675927 | -3.5486575 | 0.0003872 | 0.00445248 |
| MYO1B | 582.833499 | -2.545855262 | 0.57702967 | -4.4120006 | 1.02E-05 | 0.00025902 |
| NABP1 | 1664.28652 | -0.625606415 | 0.49453695 | -1.2650347 | 0.20585888 | 0.39458924 |
| AC098617.1 | 119.378889 | -1.122919481 | 0.73414123 | -1.5295687 | 0.12612351 | 0.28669568 |
| TMEFF2 | 36.9993506 | -0.594170352 | 0.48638945 | -1.2215938 | 0.22186126 | 0.41408665 |
| SLC39A10 | 642.10863 | 0.471148598 | 0.29718647 | 1.58536354 | 0.11288376 | 0.26617886 |
| DNAH7 | 1552.73341 | 0.709924383 | 0.42624956 | 1.66551348 | 0.09581036 | 0.23882914 |
| STK17B | 76.6036612 | -0.832420012 | 0.31234502 | -2.6650657 | 0.00769733 | 0.04169133 |
| CCDC150 | 168.068771 | 1.657446792 | 0.47209623 | 3.51082402 | 0.00044672 | 0.00494659 |
| GTF3C3 | 413.072133 | 0.230618388 | 0.16359493 | 1.40969156 | 0.15863078 | 0.33343844 |
| PGAP1 | 225.380338 | -0.066864931 | 0.27488245 | -0.2432492 | 0.80781237 | 0.89588422 |
| ANKRD44 | 654.500371 | -0.933557197 | 0.40913823 | -2.2817647 | 0.02250323 | 0.08977061 |
| SF3B1 | 1348.31616 | 0.10736174 | 0.12994274 | 0.82622348 | 0.40867733 | 0.60246722 |
| COQ10B | 326.30195 | -0.529142464 | 0.34210225 | -1.5467378 | 0.12192649 | 0.28022241 |
| HSPD1 | 1898.46251 | -0.706494325 | 0.4715617 | -1.4982012 | 0.13408097 | 0.29810025 |
| MOB4 | 100.131808 | -0.323191882 | 0.28028888 | -1.1530671 | 0.24888285 | 0.44512491 |
| AC011997.1 | 170.037903 | -0.443020735 | 0.38554888 | -1.149065 | 0.25052917 | 0.44684494 |
| PLCL1 | 5579.79301 | 1.976113117 | 0.59118316 | 3.34264111 | 0.00082985 | 0.00788811 |
| SATB2 | 1519.62954 | 0.969470046 | 0.36500322 | 2.65605887 | 0.00790598 | 0.04251897 |
| SATB2-AS1 | 45.919325 | 0.191164058 | 0.41567578 | 0.45988741 | 0.64559704 | 0.79225766 |
| FTCDNL1 | 559.695126 | 1.386322655 | 0.5212075 | 2.65982866 | 0.00781804 | 0.04220634 |
| C2orf69 | 137.406006 | -0.642446088 | 0.30400091 | -2.1133031 | 0.03457483 | 0.12069351 |
| TYW5 | 408.862401 | 0.082834237 | 0.16825347 | 0.49231814 | 0.62249446 | 0.77596685 |
| MAIP1 | 188.379247 | -0.335971387 | 0.23799327 | -1.4116844 | 0.1580429 | 0.33266186 |
| SPATS2L | 6242.10385 | -0.743922239 | 0.45048032 | -1.6513979 | 0.09865736 | 0.24335009 |
| KCTD18 | 135.585943 | -0.009870706 | 0.26417311 | -0.0373645 | 0.97019435 | 0.9852064 |
| SGO2 | 71.6231384 | 0.123595641 | 0.34090341 | 0.36255325 | 0.71693864 | 0.83851445 |
| AOX1 | 3002.00098 | 0.003112151 | 0.48787606 | 0.00637898 | 0.99491035 | 0.99749068 |
| BZW1 | 248.380624 | -1.026067629 | 0.43866627 | -2.3390621 | 0.01933222 | 0.08044528 |
| CLK1 | 1083.88758 | 0.287420871 | 0.27071244 | 1.06172021 | 0.28836272 | 0.4881546 |
| PPIL3 | 194.878424 | 0.211818559 | 0.27634288 | 0.7665063 | 0.4433751 | 0.63504102 |
| NIF3L1 | 113.87941 | -0.14066619 | 0.24995187 | -0.5627731 | 0.5735894 | 0.73871933 |
| ORC2 | 1282.96186 | 0.4069504 | 0.19155408 | 2.12446736 | 0.03363108 | 0.11853465 |
| AC005037.1 | 378.201021 | 0.70034528 | 0.25229184 | 2.77593318 | 0.00550435 | 0.03261295 |
| FAM126B | 1284.88454 | 0.463015511 | 0.26198747 | 1.76731931 | 0.07717477 | 0.20709409 |
| NDUFB3 | 330.040813 | -0.153429787 | 0.40279532 | -0.3809125 | 0.70326815 | 0.82957801 |
| CFLAR | 3459.57526 | 0.168202203 | 0.26406343 | 0.63697651 | 0.52414015 | 0.69968728 |
| CFLAR-AS1 | 294.883534 | -0.424922167 | 0.54769199 | -0.7758415 | 0.43784258 | 0.62974903 |
| CASP10 | 288.414109 | 0.020077112 | 0.41047794 | 0.04891155 | 0.96098978 | 0.97978927 |
| CASP8 | 620.236681 | -0.191748143 | 0.24071468 | -0.7965785 | 0.42569586 | 0.61870566 |
| ALS2CR12 | 84.3215962 | 0.358566789 | 0.67170331 | 0.53381722 | 0.593468 | 0.75444301 |
| TRAK2 | 943.638099 | -0.511526397 | 0.24583451 | -2.0807754 | 0.03745447 | 0.12780481 |
| STRADB | 680.448949 | 0.092213812 | 0.43208332 | 0.21341674 | 0.83100193 | 0.91066065 |
| C2CD6 | 53.2832245 | -0.12327908 | 0.50716122 | -0.2430767 | 0.80794597 | 0.89590037 |
| TMEM237 | 39.5732906 | 0.031560827 | 0.40755213 | 0.07743998 | 0.93827354 | 0.96676805 |
| ALS2 | 1040.86602 | 0.431802798 | 0.22498188 | 1.9192781 | 0.05494915 | 0.16502804 |
| SUMO1 | 493.64422 | -0.015106888 | 0.15157232 | -0.0996679 | 0.92060802 | 0.95736677 |
| NOP58 | 784.976811 | -0.510520892 | 0.34956621 | -1.4604412 | 0.14416886 | 0.31188802 |
| BMPR2 | 2594.60095 | 0.136605296 | 0.16881384 | 0.80920672 | 0.41839625 | 0.6115331 |
| FAM117B | 670.910305 | 0.918419857 | 0.2221273 | 4.13465556 | 3.55E-05 | 0.0006892 |
| ICA1L | 325.615908 | -0.62678356 | 0.36803172 | -1.7030694 | 0.08855508 | 0.2269255 |
| WDR12 | 653.546783 | 0.627425187 | 0.33260199 | 1.88641442 | 0.05923913 | 0.17320622 |
| CARF | 487.753626 | 0.94950052 | 0.41930261 | 2.26447557 | 0.02354488 | 0.09249919 |
| NBEAL1 | 4442.4369 | 0.285321924 | 0.21346649 | 1.33661224 | 0.18134924 | 0.36362698 |
| CYP20A1 | 1004.28155 | 0.471055406 | 0.30524427 | 1.543208 | 0.12278032 | 0.2814768 |
| ABI2 | 1432.21659 | 0.588557828 | 0.19199502 | 3.06548489 | 0.00217317 | 0.01635898 |
| RAPH1 | 717.718786 | -0.529608592 | 0.22292285 | -2.3757483 | 0.0175134 | 0.07518204 |

| | | | | | | |
|---|---|---|---|---|---|---|
| PARD3B | 5738.12534 | 0.416790852 | 0.24014347 | 1.73559102 | 0.08263618 | 0.21686656 |
| NRP2 | 56.6483636 | -1.375257593 | 0.71026941 | -1.9362478 | 0.05283736 | 0.16076396 |
| AC007679.1 | 63.6743324 | -1.185218294 | 0.65005087 | -1.8232701 | 0.0682625 | 0.19083704 |
| INO80D | 2733.78059 | 0.190101079 | 0.56954576 | 0.33377665 | 0.7385481 | 0.85162605 |
| NDUFS1 | 1531.40034 | 0.057253075 | 0.16987012 | 0.33704029 | 0.73608653 | 0.8507199 |
| EEF1B2 | 443.037937 | -0.500552453 | 0.40496966 | -1.2360246 | 0.21644943 | 0.4072577 |
| ZDBF2 | 370.542974 | -0.153683638 | 0.26620858 | -0.5773054 | 0.56373319 | 0.73104641 |
| MDH1B | 67.4451766 | 1.623647225 | 0.39915609 | 4.06769996 | 4.75E-05 | 0.00087102 |
| FASTKD2 | 472.634495 | -0.27692127 | 0.23351191 | -1.1858979 | 0.23566264 | 0.42995285 |
| CPO | 89.5052736 | 0.447030776 | 0.34719056 | 1.28756603 | 0.19789707 | 0.38405425 |
| KLF7 | 847.415197 | -0.514167531 | 0.7100874 | -0.7240905 | 0.46901018 | 0.65617047 |
| AC009226.1 | 81.7765897 | -0.731291974 | 0.50903935 | -1.4366119 | 0.15082831 | 0.32197534 |
| CREB1 | 1013.61653 | -0.033696766 | 0.17263623 | -0.1951894 | 0.84524466 | 0.91854292 |
| METTL21A | 246.821309 | 0.045699511 | 0.26624485 | 0.17164468 | 0.86371688 | 0.92861929 |
| CCNYL1 | 733.580121 | -0.165205991 | 0.23381062 | -0.7065804 | 0.4798273 | 0.6649472 |
| FZD5 | 257.420537 | -0.374313517 | 0.26696236 | -1.4021209 | 0.16087913 | 0.33687295 |
| PLEKHM3 | 985.401506 | 0.264778797 | 0.28270451 | 0.93659204 | 0.34896845 | 0.54884167 |
| IDH1 | 878.378023 | 1.347387153 | 0.50537633 | 2.66610658 | 0.00767354 | 0.04162294 |
| PIKFYVE | 1440.2018 | 0.180605258 | 0.13397879 | 1.34801381 | 0.17765394 | 0.35833014 |
| PTH2R | 26606.0205 | 0.676656001 | 0.42935949 | 1.5759661 | 0.11503362 | 0.26957869 |
| MAP2 | 106.897525 | -1.123495933 | 0.42899436 | -2.6189061 | 0.00882122 | 0.04601226 |
| RPE | 463.170867 | -0.175649113 | 0.27219197 | -0.6453133 | 0.51872414 | 0.69543739 |
| KANSL1L | 6648.08244 | 0.767306318 | 0.40541886 | 1.89262609 | 0.05840761 | 0.17158018 |
| AC007038.2 | 45.4622316 | 0.122139449 | 0.41956699 | 0.29110833 | 0.77096847 | 0.87268354 |
| AC007038.1 | 501.541322 | -0.016299888 | 0.28822324 | -0.056553 | 0.95490128 | 0.97731423 |
| ACADL | 1023.17891 | 1.457548661 | 0.4874818 | 2.98995506 | 0.00279018 | 0.01963299 |
| LANCL1-AS1 | 101.070216 | 0.076996483 | 0.60799391 | 0.12664022 | 0.89922516 | 0.94743919 |
| LANCL1 | 474.359414 | 0.250769025 | 0.32976566 | 0.76044615 | 0.44698795 | 0.63774996 |
| ERBB4 | 777.222653 | 1.305498612 | 0.46247544 | 2.82284963 | 0.00475989 | 0.0292442 |
| IKZF2 | 280.284842 | 0.878680643 | 0.44398589 | 1.97907334 | 0.04780775 | 0.15092143 |
| SPAG16 | 2987.12365 | 0.247919371 | 0.39087583 | 0.63426631 | 0.52590704 | 0.70084961 |
| AC068051.1 | 140.827787 | -0.543083359 | 0.72268656 | -0.7514784 | 0.45236478 | 0.64164525 |
| BARD1 | 80.7880941 | -1.02943339 | 0.46546448 | -2.2116261 | 0.02699251 | 0.10167254 |
| AC016708.1 | 115.378186 | -0.065154304 | 0.46128925 | -0.1412439 | 0.88767725 | 0.94209263 |
| ATIC | 374.856708 | -0.084119986 | 0.28131638 | -0.2990227 | 0.76492272 | 0.86874558 |
| FN1 | 69.9016414 | 0.922915326 | 0.43747605 | 2.10963622 | 0.0348897 | 0.12150921 |
| MREG | 517.736125 | 1.048391568 | 0.34194297 | 3.06598371 | 0.00216955 | 0.01635898 |
| PECR | 950.780885 | 1.536244092 | 0.41691956 | 3.68474939 | 0.00022893 | 0.00297997 |
| TMEM169 | 90.8208189 | 0.126734608 | 0.43645605 | 0.29037198 | 0.77153168 | 0.87306073 |
| XRCC5 | 1922.43652 | -0.264564248 | 0.22199785 | -1.1917424 | 0.23336228 | 0.4272905 |
| SMARCAL1 | 282.447133 | 0.157042431 | 0.3336839 | 0.47063232 | 0.63790332 | 0.7871047 |
| RPL37A | 1095.52163 | -0.95505809 | 0.33914111 | -2.8161083 | 0.00486093 | 0.02961278 |
| IGFBP2 | 65.2053536 | 0.377252393 | 0.51717132 | 0.72945343 | 0.46572435 | 0.65353026 |
| IGFBP5 | 230.520783 | 0.290279669 | 0.62132469 | 0.46719481 | 0.64036051 | 0.78883989 |
| DIRC3 | 1126.12668 | 0.619034425 | 0.67209846 | 0.92104723 | 0.35702577 | 0.55584812 |
| TNS1 | 4332.74175 | 0.412472137 | 0.41803241 | 0.98669895 | 0.32379025 | 0.52429922 |
| ARPC2 | 1028.25992 | -1.077956224 | 0.34448489 | -3.1291829 | 0.00175293 | 0.01385665 |
| AAMP | 241.863761 | -0.314148007 | 0.29907833 | -1.050387 | 0.2935402 | 0.49350614 |
| PNKD | 725.058665 | -0.138244085 | 0.18139497 | -0.7621164 | 0.44599053 | 0.6372179 |
| TMBIM1 | 680.438922 | -1.144453505 | 0.28367753 | -4.0343467 | 5.48E-05 | 0.00097724 |
| CATIP-AS2 | 101.971192 | -0.124535513 | 0.45225991 | -0.2753627 | 0.78303759 | 0.8809896 |
| CATIP | 85.8031008 | 1.57844192 | 0.65755449 | 2.40047318 | 0.01637389 | 0.07168218 |
| CTDSP1 | 349.356446 | 0.310990421 | 0.25955915 | 1.19814854 | 0.2308592 | 0.42378291 |
| VIL1 | 277.1 | 1.31712099 | 0.61432214 | 2.1440233 | 0.03203102 | 0.11442512 |
| USP37 | 804.802205 | -0.637003895 | 0.19471055 | -3.2715428 | 0.00106962 | 0.00949213 |
| CNOT9 | 419.87617 | -0.386269294 | 0.16792949 | -2.3001873 | 0.02143761 | 0.08698795 |
| PLCD4 | 273.533974 | 0.005635481 | 0.78305165 | 0.00719682 | 0.99425782 | 0.99709481 |
| AC012510.1 | 54.1017779 | -1.977128366 | 0.98952186 | -1.9980644 | 0.04570968 | 0.1465254 |
| ZNF142 | 266.870247 | -0.811904915 | 0.40555132 | -2.0019782 | 0.04528707 | 0.14557924 |
| BCS1L | 100.210354 | -0.175848031 | 0.28430475 | -0.6185195 | 0.53623295 | 0.70869254 |
| RNF25 | 105.30067 | -0.43386319 | 0.36585253 | -1.1858964 | 0.23566322 | 0.42995285 |
| STK36 | 294.869177 | 0.989174877 | 0.24319332 | 4.06744271 | 4.75E-05 | 0.00087102 |
| TTLL4 | 237.444187 | -0.429287438 | 0.25128957 | -1.7083377 | 0.08757371 | 0.22533749 |
| CYP27A1 | 687.836756 | 0.727735721 | 0.40806386 | 1.78338684 | 0.07452335 | 0.20235571 |
| IHH | 56.7051815 | 1.149659653 | 0.63411905 | 1.81300286 | 0.06983138 | 0.19368246 |
| CNPPD1 | 361.44639 | 0.11069557 | 0.21929292 | 0.50478405 | 0.61371052 | 0.76957315 |
| RETREG2 | 340.75159 | -0.543280122 | 0.29512175 | -1.8408678 | 0.06564093 | 0.18605051 |
| ZFAND2B | 290.082477 | 0.568217778 | 0.22737979 | 2.49898101 | 0.0124551 | 0.0588291 |
| ATG9A | 40.1904095 | 0.317777826 | 0.44380465 | 0.71603086 | 0.47397229 | 0.66023762 |
| ANKZF1 | 483.380365 | -0.02259347 | 0.34620182 | -0.065261 | 0.94796621 | 0.97344198 |
| GLB1L | 210.482321 | -0.313759765 | 0.32712105 | -0.9591549 | 0.3374807 | 0.53770173 |
| STK16 | 104.133913 | -0.146706838 | 0.29029719 | -0.5053677 | 0.61330057 | 0.76933839 |
| TUBA4A | 513.895496 | -1.379568851 | 0.46268581 | -2.9816537 | 0.00286696 | 0.02001389 |
| DNAJB2 | 337.40664 | -0.289909564 | 0.52518374 | -0.5520155 | 0.58093773 | 0.74419752 |
| DNPEP | 260.93848 | -0.397642135 | 0.34219874 | -1.162021 | 0.24522692 | 0.44071395 |
| GMPPA | 120.371449 | 0.097744174 | 0.42235852 | 0.23142465 | 0.81698492 | 0.90111634 |
| CHPF | 118.34471 | -1.506703607 | 0.58641605 | -2.5693424 | 0.01018917 | 0.05117135 |
| OBSL1 | 210.8021 | 0.594082169 | 0.37364647 | 1.58995794 | 0.11184429 | 0.26477541 |
| STK11IP | 103.955632 | 0.44949159 | 0.27154782 | 1.6552944 | 0.09786479 | 0.24200482 |
| EPHA4 | 275.777787 | 1.744351444 | 0.47357942 | 3.68333452 | 0.0002302 | 0.00299373 |
| SGPP2 | 836.58782 | -3.923133966 | 0.65308368 | -6.0070923 | 1.89E-09 | 2.01E-07 |
| FARSB | 488.751882 | -0.404746649 | 0.31530585 | -1.2836636 | 0.19925968 | 0.38577668 |
| MOGAT1 | 1325.71415 | -0.2553998 | 0.54223072 | -0.4710168 | 0.63762871 | 0.7869599 |
| ACSL3 | 894.771095 | -0.862222714 | 0.34384567 | -2.5075863 | 0.01215589 | 0.05778572 |
| AP1S3 | 413.104123 | -0.524578893 | 0.35661428 | -1.470998 | 0.14129166 | 0.30819999 |
| WDFY1 | 970.769611 | -0.369434369 | 0.25223551 | -1.4646406 | 0.143019 | 0.31058166 |
| MRPL44 | 164.581323 | -0.288660991 | 0.22227272 | -1.2986794 | 0.19405399 | 0.37906484 |
| SERPINE2 | 2925.31163 | -2.654523618 | 0.62330152 | -4.2588114 | 2.06E-05 | 0.0004532 |
| CUL3 | 2087.38012 | 0.02767596 | 0.16524717 | 0.1674822 | 0.86699065 | 0.93082485 |
| DOCK10 | 56.6798344 | 0.079363679 | 0.38826159 | 0.20440775 | 0.83803487 | 0.91449279 |
| IRS1 | 284.385883 | -0.207244116 | 0.30861981 | -0.6715192 | 0.50188986 | 0.68289644 |
| RHBDD1 | 872.311502 | -0.106699033 | 0.25472266 | -0.4188832 | 0.67530152 | 0.8117476 |
| COL4A4 | 1224.17998 | 0.120763386 | 0.29407497 | 0.4106551 | 0.68132546 | 0.81457974 |
| COL4A3 | 1778.57323 | 0.193985337 | 0.32706426 | 0.59311078 | 0.55310701 | 0.72252338 |
| AC097662.1 | 265.933196 | -0.056174568 | 0.42108876 | -0.1334032 | 0.89387455 | 0.94488613 |
| MFF | 807.091761 | 0.06279272 | 0.18122357 | 0.34649312 | 0.72897215 | 0.8460193 |
| AGFG1 | 2302.41428 | -0.471891477 | 0.18830931 | -2.5059381 | 0.0122127 | 0.05798211 |
| SLC19A3 | 332.581738 | -0.450858089 | 0.50984295 | -0.8843078 | 0.37653009 | 0.57294604 |
| DNER | 143.252624 | -1.648481274 | 0.49174424 | -3.3523143 | 0.00080139 | 0.00767084 |
| TRIP12 | 3419.80334 | 0.053046534 | 0.12749544 | 0.41606612 | 0.67736162 | 0.81315429 |
| FBXO36 | 477.457503 | 0.987069963 | 0.2780087 | 3.55050021 | 0.0003845 | 0.00443631 |
| AC009950.1 | 32.091932 | -1.31722185 | 0.80724682 | -1.6317461 | 0.10273298 | 0.25037114 |
| SP110 | 256.577515 | -0.698765376 | 0.59365325 | -1.1770598 | 0.23917164 | 0.43415782 |
| SP140L | 214.040984 | -1.119666708 | 0.40701435 | -2.7509268 | 0.00594269 | 0.03442792 |
| SP100 | 1405.7413 | -0.374698732 | 0.3969138 | -0.9440305 | 0.34515407 | 0.5454262 |
| AC010149.1 | 59.8710491 | -0.08564936 | 0.40463048 | -0.211673 | 0.83236213 | 0.91150088 |
| CAB39 | 1566.30113 | 0.004806675 | 0.23899852 | 0.02011174 | 0.98395424 | 0.99214482 |
| ITM2C | 217.636416 | -1.413090654 | 0.63293838 | -2.232588 | 0.02557612 | 0.09797986 |
| C2orf72 | 50.0307977 | 0.828073842 | 0.75920527 | 1.0907114 | 0.27539989 | 0.47286813 |
| PSMD1 | 1424.24863 | -0.41796081 | 0.29648317 | -1.4097286 | 0.15861984 | 0.33343844 |
| ARMC9 | 1150.33143 | -0.278374422 | 0.29449958 | -0.9452456 | 0.34453352 | 0.5451369 |
| NCL | 737.677169 | -1.148135467 | 0.4177416 | -2.7484346 | 0.00598806 | 0.03455929 |
| PTMA | 1717.30546 | -1.051773802 | 0.40532287 | -2.5949036 | 0.00946174 | 0.04844682 |

| | | | | | | |
|---|---|---|---|---|---|---|
| PDE6D | 351.254776 | -0.417239175 | 0.27591347 | -1.5122102 | 0.13048039 | 0.29300371 |
| COPS7B | 237.891284 | 0.421125153 | 0.18425046 | 2.28561252 | 0.02227694 | 0.08904928 |
| DIS3L2 | 2021.73326 | 0.331484056 | 0.20910722 | 1.58523487 | 0.11291298 | 0.26620197 |
| AC019130.1 | 113.349723 | -0.055264757 | 0.51629924 | -0.1070402 | 0.91475711 | 0.95437477 |
| TIGD1 | 74.0263178 | 0.128968941 | 0.28369051 | 0.4546114 | 0.64938883 | 0.79436638 |
| EIF4E2 | 724.12027 | 0.143397646 | 0.22595332 | 0.63463393 | 0.52566719 | 0.70084961 |
| EFHD1 | 762.736503 | 1.800385325 | 0.30191355 | 5.96324793 | 2.47E-09 | 2.53E-07 |
| GIGYF2 | 3158.06396 | 0.161409369 | 0.13812265 | 1.16859451 | 0.24256704 | 0.4377128 |
| SNORC | 93.3500574 | -0.391914604 | 0.32698372 | -1.1985753 | 0.23069313 | 0.42378291 |
| NGEF | 729.566642 | 1.235504946 | 0.4433627 | 2.78666865 | 0.00532529 | 0.03175022 |
| INPP5D | 47.4887881 | -0.807148601 | 0.57985267 | -1.3919891 | 0.1639257 | 0.34080823 |
| ATG16L1 | 368.371783 | -0.415251833 | 0.4205871 | -0.9873147 | 0.32348838 | 0.52401995 |
| DGKD | 621.842695 | -1.022930689 | 0.43355085 | -2.3594249 | 0.01830328 | 0.07757672 |
| USP40 | 1806.18559 | 0.218306641 | 0.35890897 | 0.60825073 | 0.54302119 | 0.71356452 |
| AC114812.2 | 1826.7982 | -1.522853734 | 0.847242 | -1.7974247 | 0.07226822 | 0.19835412 |
| MROH2A | 84.6989784 | 1.805423525 | 0.73857729 | 2.44446119 | 0.01450687 | 0.06543128 |
| ARL4C | 70.3448409 | -2.334163242 | 0.97771032 | -2.3873771 | 0.01696907 | 0.07355949 |
| SH3BP4 | 274.789786 | -2.104483174 | 0.6504772 | -3.2352912 | 0.00121519 | 0.0104793 |
| AGAP1 | 12316.7966 | -0.479441988 | 0.2937364 | -1.6322185 | 0.10263346 | 0.25027051 |
| IQCA1 | 630.282023 | -0.25725671 | 0.35118886 | -0.7325309 | 0.46384456 | 0.65164168 |
| ACKR3 | 76.8233358 | 1.228168616 | 0.55891321 | 2.19742276 | 0.02799027 | 0.10415702 |
| AC105760.2 | 114.582585 | 0.63686203 | 0.42238079 | 1.5077912 | 0.13160798 | 0.29485952 |
| COPS8 | 174.459337 | -0.446207065 | 0.24167842 | -1.8462843 | 0.06485094 | 0.18457471 |
| RAB17 | 1104.44426 | 0.258214588 | 0.36754123 | 0.70254592 | 0.48233878 | 0.66670759 |
| LRRFIP1 | 4287.11084 | -1.886219971 | 0.38904594 | -4.8483219 | 1.25E-06 | 4.58E-05 |
| RBM44 | 41.3444801 | -0.72752177 | 0.624877 | -1.164264 | 0.24431706 | 0.43943757 |
| UBE2F-SCLY | 56.2034636 | -0.610166221 | 0.41474501 | -1.471184 | 0.14124137 | 0.30819999 |
| ILKAP | 672.247718 | -0.104950846 | 0.23686564 | -0.4430818 | 0.65770659 | 0.80056693 |
| PER2 | 262.111792 | 0.656342024 | 0.47261496 | 1.38874577 | 0.16491007 | 0.34222897 |
| TRAF3IP1 | 739.244621 | 0.285450204 | 0.20668575 | 1.38108317 | 0.16725339 | 0.34521039 |
| ASB1 | 147.0338 | 0.058441404 | 0.29165911 | 0.20037572 | 0.84118675 | 0.91607296 |
| HDAC4 | 1378.35299 | -0.624420047 | 0.35595923 | -1.7541898 | 0.07939799 | 0.21087159 |
| AC062017.1 | 38.2815628 | 0.468849521 | 0.70593648 | 0.66415256 | 0.50659267 | 0.68655979 |
| NDUFA10 | 1072.21866 | 0.067624148 | 0.21491179 | 0.31466001 | 0.75301981 | 0.8613901 |
| COPS9 | 263.878109 | -0.537215671 | 0.41721961 | -1.2876089 | 0.19788215 | 0.38405425 |
| GPC1 | 62.7225884 | -0.895453785 | 0.41159162 | -2.175588 | 0.02958608 | 0.1080623 |
| ANKMY1 | 369.522573 | 0.179036564 | 0.28500097 | 0.62819634 | 0.52987533 | 0.70418859 |
| RNPEPL1 | 280.576324 | -0.821341999 | 0.38190481 | -2.1506458 | 0.03150416 | 0.11304387 |
| CAPN10-DT | 45.7777781 | -0.499690917 | 0.40439401 | -1.2356536 | 0.21658735 | 0.40740535 |
| CAPN10 | 137.046484 | -0.321579438 | 0.30880756 | -1.0413587 | 0.2977091 | 0.49812308 |
| GPR35 | 82.2678791 | 1.44086287 | 0.52449503 | 2.74714305 | 0.00601169 | 0.03466649 |
| AGXT | 97.0974199 | 2.643908514 | 0.74401614 | 3.5535634 | 0.00038005 | 0.00441844 |
| C2orf54 | 95.2513219 | 3.061071318 | 1.01479965 | 3.01642922 | 0.00255771 | 0.01853967 |
| SNED1 | 171.337825 | -0.165500905 | 0.4161922 | -0.397655 | 0.69088452 | 0.82139973 |
| AC005237.1 | 89.6525942 | -0.871526523 | 0.69487549 | -1.2542197 | 0.20976216 | 0.39933472 |
| MTERF4 | 456.152168 | 0.19940813 | 0.24428939 | 0.81627832 | 0.41434097 | 0.60793873 |
| PASK | 82.239881 | 1.152207827 | 0.3430436 | 3.3587795 | 0.00078288 | 0.00754632 |
| PPP1R7 | 302.606951 | -0.150597954 | 0.20247284 | -0.7437934 | 0.45700149 | 0.64571894 |
| ANO7 | 58.6335687 | -1.899802952 | 0.37065669 | -5.1255056 | 2.97E-07 | 1.36E-05 |
| HDLBP | 1703.21243 | -0.445304001 | 0.31406584 | -1.4178683 | 0.15622921 | 0.32985733 |
| 2-Sep | 2047.92483 | -0.055331462 | 0.15997831 | -0.3458685 | 0.72944153 | 0.84632259 |
| FARP2 | 1219.16528 | 0.261451158 | 0.1867317 | 1.40014343 | 0.16147037 | 0.3376383 |
| STK25 | 416.640186 | 0.17234413 | 0.22387828 | 0.76981176 | 0.44141156 | 0.63333363 |
| BOK | 219.971421 | 0.486882508 | 0.41008625 | 1.18726854 | 0.23512171 | 0.42961829 |
| THAP4 | 422.914249 | 0.025976857 | 0.25752866 | 0.10086977 | 0.91965384 | 0.95690112 |
| ATG4B | 1190.14895 | 0.076239495 | 0.24710589 | 0.30852965 | 0.75767934 | 0.86418282 |
| DTYMK | 100.228803 | -0.425043414 | 0.27047817 | -1.5714518 | 0.11607773 | 0.27150936 |
| ING5 | 548.545488 | 0.7490537 | 0.2938157 | 2.54939986 | 0.01079085 | 0.0532434 |
| D2HGDH | 1365.4283 | 2.325671955 | 0.59750166 | 3.89232716 | 9.93E-05 | 0.00154476 |
| CHL1 | 945.679092 | 1.03343057 | 0.76021697 | 1.35938899 | 0.17402336 | 0.3537824 |
| LINC01266 | 522.528736 | 0.183127163 | 0.57635827 | 0.31773147 | 0.75068864 | 0.85940776 |
| CNTN6 | 78.8563161 | -1.296304261 | 0.69911868 | -1.8541977 | 0.06371088 | 0.18227615 |
| CNTN4 | 7063.50703 | 0.873737555 | 0.43184672 | 2.02325854 | 0.0430465 | 0.14036697 |
| CNTN4-AS2 | 51.1501539 | -0.236374146 | 0.57220832 | -0.4130911 | 0.6795399 | 0.81423861 |
| TRNT1 | 458.762687 | -0.191612507 | 0.27535092 | -0.6958848 | 0.486501 | 0.66974606 |
| CRBN | 748.48576 | 0.207647515 | 0.33542997 | 0.61904879 | 0.53588422 | 0.70860168 |
| AC024060.1 | 46.5195786 | 0.110102336 | 0.33082793 | 0.33280847 | 0.73927886 | 0.85193361 |
| AC034195.1 | 2299.8238 | 2.076160845 | 0.67480791 | 3.0766694 | 0.00209327 | 0.01590493 |
| SUMF1 | 4252.47688 | 0.291181043 | 0.37291316 | 0.78082801 | 0.43490366 | 0.62736523 |
| LRRN1 | 141.655669 | -0.992523364 | 0.40971445 | -2.4224758 | 0.01541515 | 0.06849477 |
| SETMAR | 77.1514541 | 0.562861448 | 0.32056681 | 1.75583198 | 0.0791171 | 0.21028856 |
| ITPR1 | 897.891293 | 0.581025349 | 0.38883467 | 1.49427351 | 0.13510415 | 0.29948045 |
| EGOT | 475.196035 | 0.368824695 | 0.53260484 | 0.69249219 | 0.48862829 | 0.67177813 |
| AC018816.1 | 130.022857 | 0.039374625 | 0.30709367 | 0.12821699 | 0.89797726 | 0.94717771 |
| BHLHE40-AS | 89.9174326 | -0.287904046 | 0.5044294 | -0.5707519 | 0.56816782 | 0.73463601 |
| BHLHE40 | 1074.47463 | -1.316920637 | 0.45947946 | -2.8661142 | 0.00415544 | 0.02638196 |
| ARL8B | 1397.5288 | -0.454642604 | 0.38366963 | -1.1849846 | 0.23602353 | 0.43043834 |
| AC026202.2 | 209.805023 | -1.180269273 | 0.41374804 | -2.8526281 | 0.00433593 | 0.02723966 |
| EDEM1 | 494.03986 | -0.60815547 | 0.3757969 | -1.6183089 | 0.10559604 | 0.25457008 |
| AC069277.1 | 68.9640156 | 0.365320278 | 0.90384898 | 0.40418287 | 0.68607825 | 0.81810131 |
| GRM7-AS3 | 73.4839319 | 1.596444787 | 0.51246856 | 3.11520533 | 0.00183817 | 0.01434023 |
| LMCD1-AS1 | 1220.59016 | -0.079322909 | 0.36006831 | -0.2202996 | 0.82563782 | 0.90652176 |
| LMCD1 | 938.982591 | -2.983680164 | 0.88121921 | -3.3858547 | 0.00070957 | 0.0070132 |
| RAD18 | 224.842899 | -0.045925185 | 0.27396955 | -0.1676288 | 0.86687532 | 0.93077393 |
| SRGAP3 | 860.753894 | 0.072005058 | 0.51604543 | 0.1395324 | 0.88902945 | 0.94267382 |
| THUMPD3-A | 1008.99406 | -0.14886944 | 0.24966132 | -0.5962855 | 0.55098448 | 0.72050604 |
| THUMPD3 | 213.366718 | -0.479560195 | 0.21990554 | -2.1807555 | 0.02920151 | 0.10696555 |
| SETD5 | 3483.14372 | -0.158557219 | 0.1380322 | -1.1486973 | 0.25068081 | 0.44702114 |
| MTMR14 | 418.665299 | 0.1636092 | 0.26128555 | 0.6261701 | 0.53120338 | 0.70499658 |
| BRPF1 | 82.0211943 | -0.243310898 | 0.43862528 | -0.5547124 | 0.57909137 | 0.74258983 |
| OGG1 | 682.150795 | 0.319333433 | 0.35399715 | 0.90207911 | 0.36701485 | 0.56442063 |
| CAMK1 | 273.518353 | 0.119779572 | 0.39868075 | 0.30043982 | 0.7638417 | 0.86815739 |
| TADA3 | 256.523841 | -0.106082166 | 0.20023604 | -0.5297856 | 0.59626062 | 0.75640391 |
| TTLL3 | 53.8961076 | -0.383300262 | 0.39503432 | -0.9702961 | 0.33189892 | 0.53146477 |
| RPUSD3 | 132.071977 | -0.037094751 | 0.23744619 | -0.1562238 | 0.8758566 | 0.93528875 |
| JAGN1 | 137.985389 | -0.926520515 | 0.32515019 | -2.8495155 | 0.00437859 | 0.02746787 |
| IL17RE | 70.5861419 | 1.848640694 | 0.58337792 | 3.16885615 | 0.0015304 | 0.01255369 |
| IL17RC | 218.202317 | 0.610022669 | 0.37903193 | 1.60942292 | 0.1075239 | 0.25757254 |
| CRELD1 | 215.603156 | 0.167240408 | 0.4620637 | 0.36194232 | 0.71739513 | 0.8386157 |
| PRRT3-AS1 | 53.5517088 | -0.473283189 | 0.32848189 | -1.4408197 | 0.14963562 | 0.31999867 |
| EMC3 | 465.398995 | -0.280974095 | 0.17800278 | -1.5784815 | 0.11445503 | 0.26859016 |
| FANCD2 | 56.8563201 | -0.006516843 | 0.30698311 | -0.0212287 | 0.98306324 | 0.99169805 |
| BRK1 | 405.421457 | -0.563338196 | 0.20221498 | -2.7858381 | 0.00533895 | 0.03180401 |
| VHL | 278.364565 | -0.183708424 | 0.20940498 | -0.8772878 | 0.38033036 | 0.57674392 |
| IRAK2 | 519.113818 | -1.933310392 | 0.39318831 | -4.9170089 | 8.79E-07 | 3.46E-05 |
| GHRL | 44.1451237 | 0.152389115 | 0.45588449 | 0.33427133 | 0.73817482 | 0.85141049 |
| SEC13 | 550.855881 | -0.207654438 | 0.2956744 | -0.7023078 | 0.48248723 | 0.66670759 |
| HRH1 | 598.964793 | -3.678886836 | 0.46463168 | -7.9178562 | 2.42E-15 | 1.38E-12 |
| ATG7 | 1907.99977 | -0.32687793 | 0.21692716 | -1.5068557 | 0.13184764 | 0.29525169 |
| VGLL4 | 1997.24826 | -0.476671105 | 0.17577838 | -2.7117732 | 0.00669244 | 0.03761101 |
| TAMM41 | 636.282317 | 0.256684031 | 0.28784704 | 0.8917376 | 0.3725336 | 0.57001787 |
| SYN2 | 826.322302 | 0.558880662 | 0.35973157 | 1.5536047 | 0.12027874 | 0.27766015 |
| PPARG | 3720.93858 | -0.109076084 | 0.3465279 | -0.3147685 | 0.7529374 | 0.8613901 |

| | | | | | | |
|---|---|---|---|---|---|---|
| TSEN2 | 710.71586 | 0.593854086 | 0.36703996 | 1.61795485 | 0.10567233 | 0.25465025 |
| MKRN2OS | 108.338416 | 0.191986639 | 0.42977273 | 0.44671667 | 0.65507965 | 0.79871702 |
| MKRN2 | 187.441871 | 0.086837983 | 0.20393007 | 0.42582237 | 0.67023729 | 0.80863854 |
| RAF1 | 1311.76105 | 0.222499074 | 0.13034468 | 1.70700536 | 0.08782106 | 0.22582532 |
| RPL32 | 1292.43407 | -0.879824907 | 0.4501433 | -1.954544 | 0.05063691 | 0.15671262 |
| IQSEC1 | 223.480658 | 0.004691528 | 0.29159373 | 0.01608926 | 0.98716318 | 0.99360989 |
| NUP210 | 551.958693 | 1.348229963 | 0.32657064 | 4.12844817 | 3.65E-05 | 0.0007031 |
| HDAC11 | 180.988712 | -0.012788555 | 0.36983197 | -0.0345794 | 0.97241516 | 0.98615674 |
| CHCHD4 | 55.2618548 | -0.22718846 | 0.40726072 | -0.5578453 | 0.57695005 | 0.74130618 |
| TMEM43 | 184.001841 | -0.564382909 | 0.24616413 | -2.2927098 | 0.02186472 | 0.08804338 |
| AC093495.1 | 45.3888765 | 0.748229771 | 0.66991437 | 1.11690359 | 0.26403555 | 0.46207385 |
| XPC | 1144.68364 | 1.394981082 | 0.30189431 | 4.62075978 | 3.82E-06 | 0.00011569 |
| LSM3 | 261.283006 | -0.263754193 | 0.26450207 | -0.9971725 | 0.31868078 | 0.51878355 |
| AC093496.1 | 55.2907784 | 2.089688652 | 0.79031858 | 2.64410922 | 0.00819062 | 0.04369947 |
| SLC6A6 | 2797.41307 | -0.916800683 | 0.29659391 | -3.0910974 | 0.00199418 | 0.0153326 |
| CCDC174 | 367.427933 | 0.058815589 | 0.18698 | 0.31455551 | 0.75309916 | 0.8613901 |
| C3orf20 | 67.2972749 | -0.529719372 | 0.78244016 | -0.6770094 | 0.49839997 | 0.68036733 |
| FGD5 | 979.87623 | -0.056296938 | 0.32411553 | -0.173694 | 0.86210592 | 0.92768769 |
| FGD5-AS1 | 388.737452 | 0.06593767 | 0.25239944 | 0.26124333 | 0.79390487 | 0.88760636 |
| NR2C2 | 1101.77569 | -0.157401336 | 0.37347468 | -0.4214512 | 0.67342567 | 0.81072605 |
| MRPS25 | 690.556855 | -0.210928452 | 0.14576648 | -1.4470299 | 0.14788856 | 0.31757928 |
| RBSN | 341.358245 | -0.072512313 | 0.16442589 | -0.441003 | 0.65921082 | 0.80168598 |
| CAPN7 | 1455.11383 | -0.124392988 | 0.13789605 | -0.9020779 | 0.36701547 | 0.56442063 |
| SH3BP5-AS1 | 176.16196 | 0.59661908 | 0.26515857 | 2.25004636 | 0.024446 | 0.09481646 |
| SH3BP5 | 838.223236 | 0.306881412 | 0.52055118 | 0.58953168 | 0.55550466 | 0.72476748 |
| METTL6 | 554.073931 | 0.835553545 | 0.25823099 | 3.23568269 | 0.00121352 | 0.0104793 |
| EAF1 | 220.633405 | 0.381165606 | 0.22050368 | 1.72861338 | 0.08387831 | 0.21878593 |
| COLQ | 252.673884 | 0.772709716 | 0.46001904 | 1.67973421 | 0.09300904 | 0.23469615 |
| HACL1 | 714.606236 | 0.482589756 | 0.22819429 | 2.11481959 | 0.03444533 | 0.12041369 |
| BTD | 361.879433 | 1.54800658 | 0.28422037 | 5.44650123 | 5.14E-08 | 3.21E-06 |
| ANKRD28 | 2039.44722 | -0.293140748 | 0.14351316 | -2.0426053 | 0.04109153 | 0.135983 |
| DPH3 | 96.6874651 | -0.855525546 | 0.38945389 | -2.1967313 | 0.02803964 | 0.10415702 |
| OXNAD1 | 1026.66691 | 0.375695896 | 0.23573144 | 1.59374538 | 0.11099308 | 0.26321826 |
| RFTN1 | 218.936041 | -1.345076022 | 0.39153816 | -3.4353638 | 0.00059176 | 0.00605768 |
| PLCL2 | 2239.83865 | 1.514261742 | 0.24788029 | 6.10884281 | 1.00E-09 | 1.17E-07 |
| TBC1D5 | 5659.0214 | 0.51145353 | 0.26193856 | 1.95257063 | 0.05087049 | 0.15715165 |
| KCNH8 | 88.0402131 | -2.148965804 | 0.70917836 | -3.0302191 | 0.00244376 | 0.01791266 |
| EFHB | 121.175259 | -0.206285568 | 0.49632418 | -0.4156267 | 0.67768321 | 0.81339787 |
| RAB5A | 1106.29181 | -0.423291678 | 0.17524814 | -2.4153848 | 0.0157186 | 0.06963632 |
| KAT2B | 825.993325 | -0.272524121 | 0.32819232 | -0.8303793 | 0.40632435 | 0.60022783 |
| ZNF385D | 603.173393 | -0.356068124 | 0.66792643 | -0.5330948 | 0.59396794 | 0.75475281 |
| UBE2E2 | 10106.5281 | 0.237558515 | 0.29080824 | 0.8168906 | 0.41399095 | 0.60768622 |
| UBE2E1 | 1805.82255 | -0.209738785 | 0.22598003 | -0.9281297 | 0.35334028 | 0.55189396 |
| NKIRAS1 | 316.503603 | -0.037535964 | 0.26127586 | -0.1436641 | 0.88576571 | 0.94094134 |
| RPL15 | 1630.30154 | -1.462182272 | 0.39857146 | -3.6685574 | 0.00024392 | 0.00313057 |
| NR1D2 | 546.340971 | -0.914505113 | 0.54435471 | -1.6799802 | 0.09296117 | 0.23469615 |
| THRB | 3858.98972 | 0.254401307 | 0.29171221 | 0.87209689 | 0.38315552 | 0.57980931 |
| RARB | 123.65639 | 0.224629252 | 0.6219353 | 0.36117785 | 0.7179665 | 0.83896074 |
| TOP2B | 1013.4815 | 0.169232485 | 0.15233968 | 1.11088904 | 0.26661611 | 0.4645363 |
| NGLY1 | 1202.07275 | -0.629758507 | 0.25719764 | -2.4485392 | 0.01434368 | 0.06492333 |
| OXSM | 165.421722 | -0.525117211 | 0.32648528 | -1.6083948 | 0.10774875 | 0.25793326 |
| NEK10 | 387.773079 | -0.470649664 | 0.51315011 | -0.9171774 | 0.35904971 | 0.55755006 |
| SLC4A7 | 1206.7573 | -0.911224188 | 0.3645971 | -2.4992634 | 0.01244518 | 0.0588025 |
| LINC01967 | 63.9269808 | -0.620742329 | 0.64836211 | -0.9574007 | 0.33836505 | 0.53854735 |
| CMC1 | 1192.7834 | 0.116096834 | 0.22856007 | 0.5079489 | 0.61148918 | 0.76769445 |
| AZI2 | 508.651488 | -0.294475095 | 0.29034293 | -1.014232 | 0.31047205 | 0.51075857 |
| RBMS3 | 417.023597 | 0.013362702 | 0.57106013 | 0.02339982 | 0.98133135 | 0.99060551 |
| RBMS3-AS3 | 167.167612 | 0.447375935 | 0.61149463 | 0.73161057 | 0.46440629 | 0.65214805 |
| TGFBR2 | 810.93191 | 0.055512071 | 0.26763482 | 0.20741722 | 0.83568404 | 0.91316952 |
| STT3B | 1339.79378 | -0.080718338 | 0.19959206 | -0.4044166 | 0.68590642 | 0.81810131 |
| OSBPL10 | 6061.49349 | 0.172317027 | 0.18730279 | 0.9199918 | 0.35757705 | 0.5561453 |
| OSBPL10-AS | 110.961541 | -0.947856139 | 0.40553808 | -2.3372802 | 0.01942462 | 0.08075621 |
| ZNF860 | 113.951807 | -1.222623381 | 0.39250938 | -3.1148896 | 0.00184014 | 0.01434742 |
| GPD1L | 341.031168 | 0.714250244 | 0.21825816 | 3.27250195 | 0.001066 | 0.00946966 |
| CMTM8 | 1061.60512 | -0.214810553 | 0.33864591 | -0.634322 | 0.5258707 | 0.70084961 |
| CMTM7 | 722.699104 | -0.88329886 | 0.39530704 | -2.2344628 | 0.02545264 | 0.097676 |
| CMTM6 | 544.165312 | -0.565829053 | 0.38363609 | -1.4749109 | 0.1402365 | 0.30697512 |
| DYNC1LI1 | 624.645607 | 0.079116702 | 0.15898051 | 0.49765031 | 0.61873054 | 0.77331444 |
| CNOT10 | 789.141512 | 0.161002843 | 0.16947803 | 0.94999243 | 0.3421161 | 0.54270009 |
| TRIM71 | 574.615613 | 0.920904905 | 0.3255904 | 2.8284154 | 0.00467791 | 0.02889593 |
| GLB1 | 828.838603 | -0.256193563 | 0.18575601 | -1.3791939 | 0.16783498 | 0.34604603 |
| CRTAP | 452.803368 | -0.291156216 | 0.27129962 | -1.0731906 | 0.28318558 | 0.48236918 |
| FBXL2 | 1336.43392 | -0.751738797 | 0.22648729 | -3.3191213 | 0.00090301 | 0.0083859 |
| UBP1 | 574.570305 | 0.201900041 | 0.19695981 | 1.02508242 | 0.3053243 | 0.5053224 |
| CLASP2 | 2403.67699 | 0.288103601 | 0.17313377 | 1.66405204 | 0.09610203 | 0.23941693 |
| PDCD6IP | 1690.39737 | -0.071415112 | 0.1719887 | -0.4152314 | 0.67797249 | 0.81346017 |
| ARPP21 | 266.418743 | 1.10022617 | 0.50491798 | 2.1790196 | 0.02933021 | 0.10732805 |
| STAC | 133.61047 | -0.535991893 | 0.55811547 | -0.9603602 | 0.33687395 | 0.53698468 |
| TRANK1 | 159.542517 | -1.820486138 | 0.75582824 | -2.4085977 | 0.01601394 | 0.07057977 |
| EPM2AIP1 | 404.808022 | -0.115854889 | 0.14861524 | -0.7795626 | 0.43564837 | 0.62792337 |
| MLH1 | 242.18711 | 0.162918294 | 0.20531703 | 0.79349625 | 0.42748875 | 0.6204562 |
| LRRFIP2 | 3894.91588 | -0.858651809 | 0.23846824 | -3.6006967 | 0.00031737 | 0.00383609 |
| GOLGA4 | 8425.45236 | 0.051978695 | 0.33470693 | 0.15529614 | 0.87658786 | 0.93570514 |
| ITGA9 | 179.007516 | -0.966413223 | 0.44003572 | -2.1962154 | 0.02807653 | 0.10426578 |
| CTDSPL | 3262.53084 | -1.069558559 | 0.30401479 | -3.5181136 | 0.00043463 | 0.00485528 |
| PLCD1 | 100.647186 | -0.427409469 | 0.29816326 | -1.4334746 | 0.15172224 | 0.32307857 |
| DLEC1 | 40.0150924 | 0.511898181 | 0.44346496 | 1.15431484 | 0.24837112 | 0.44467383 |
| ACAA1 | 1061.26796 | 0.898768433 | 0.45145342 | 1.99083316 | 0.04649923 | 0.14825889 |
| MYD88 | 82.6605236 | -0.691097535 | 0.36339498 | -1.9017806 | 0.05719984 | 0.16955843 |
| OXSR1 | 1760.85425 | -0.426721017 | 0.20344788 | -2.0974463 | 0.03595408 | 0.12403257 |
| SLC22A13 | 66.1360814 | 4.090036762 | 0.4421445 | 9.25045267 | 2.24E-20 | 5.11E-17 |
| SLC22A14 | 213.57319 | 3.294660643 | 0.53208203 | 6.1920164 | 5.94E-10 | 7.47E-08 |
| XYLB | 2300.12343 | 1.447707636 | 0.30254614 | 4.78508058 | 1.71E-06 | 6.02E-05 |
| ACVR2B | 518.364203 | 0.799934466 | 0.24963395 | 3.20442983 | 0.0013533 | 0.01136625 |
| EXOG | 286.314103 | -0.240128581 | 0.23268452 | -1.0319921 | 0.30207583 | 0.50243337 |
| SCN11A | 66.2011917 | 0.615536219 | 0.71635524 | 0.85926114 | 0.39019646 | 0.58561422 |
| WDR48 | 661.808986 | -0.506817468 | 0.39258679 | -1.2909692 | 0.19671434 | 0.38248 |
| GORASP1 | 149.457965 | -0.026612656 | 0.25163076 | -0.1057607 | 0.91577218 | 0.95496466 |
| TTC21A | 196.008812 | 0.350811415 | 0.36660296 | 0.95692467 | 0.33860528 | 0.53886712 |
| CSRNP1 | 152.153558 | -1.791434517 | 0.46555974 | -3.8479154 | 0.00011913 | 0.00178953 |
| SLC25A38 | 193.915369 | 0.284615349 | 0.27715644 | 1.02691227 | 0.30446177 | 0.50474813 |
| RPSA | 936.581701 | -0.780059101 | 0.35552435 | -2.1941088 | 0.02822759 | 0.10465867 |
| MYRIP | 1151.54884 | 0.508118327 | 0.60187512 | 0.84422551 | 0.3985434 | 0.59275547 |
| EIF1B | 263.461788 | -0.773883564 | 0.38460314 | -2.0121613 | 0.04420293 | 0.1429322 |
| ENTPD3-AS1 | 452.194712 | 0.498420027 | 0.47111673 | 1.05795443 | 0.29007622 | 0.49020771 |
| RPL14 | 1714.62118 | -1.309546459 | 0.4902156 | -2.6713684 | 0.00755427 | 0.04112906 |
| ZNF620 | 53.287897 | -0.039957054 | 0.29514954 | -0.135379 | 0.89231221 | 0.9437441 |
| ZNF621 | 466.455539 | -0.146320398 | 0.26295636 | -0.5564437 | 0.5779076 | 0.74184112 |
| CTNNB1 | 1901.79326 | -0.048475194 | 0.20146489 | -0.2406136 | 0.8098546 | 0.89677969 |
| ULK4 | 3079.49469 | 0.940975101 | 0.33413098 | 2.8161863 | 0.00485975 | 0.02961278 |
| TRAK1 | 2003.86541 | 0.044203648 | 0.3077235 | 0.14364729 | 0.88577899 | 0.94094134 |
| SEC22C | 261.118648 | -0.132713516 | 0.29076869 | -0.456423 | 0.64808583 | 0.79355195 |
| SS18L2 | 162.096542 | -0.122564352 | 0.24932599 | -0.4915827 | 0.62301435 | 0.77619139 |

| | | | | | | |
|---|---|---|---|---|---|---|
| NKTR | 2518.17132 | -0.124957971 | 0.16567078 | -0.7542547 | 0.45069628 | 0.64075861 |
| AC006059.1 | 267.223961 | -1.229067802 | 0.46238676 | -2.6580947 | 0.00785838 | 0.04237405 |
| CCDC13 | 76.8365893 | 0.119253256 | 0.49424459 | 0.24128389 | 0.8093351 | 0.89644899 |
| AC099329.3 | 111.662702 | -0.644227442 | 0.39486569 | -1.6315103 | 0.10278269 | 0.25038257 |
| AC092042.3 | 31.4158872 | 0.213132282 | 0.71237257 | 0.29918654 | 0.76479772 | 0.86873973 |
| CYP8B1 | 161.296152 | 0.272216439 | 0.83729492 | 0.32511417 | 0.74509467 | 0.85586295 |
| POMGNT2 | 59.8655442 | -0.14591126 | 0.35415206 | -0.4120017 | 0.68033816 | 0.81442647 |
| SNRK | 1490.6806 | -0.460308451 | 0.27078743 | -1.6998885 | 0.08915189 | 0.22771117 |
| ANO10 | 3240.14664 | -0.738433312 | 0.38371085 | -1.9244525 | 0.05429789 | 0.16386199 |
| ABHD5 | 845.765379 | -1.907397561 | 0.6923566 | -2.7549352 | 0.00587038 | 0.0341099 |
| AC006058.1 | 97.1332199 | -0.377869386 | 0.56103202 | -0.6735255 | 0.50061302 | 0.68175883 |
| TCAIM | 1384.25488 | 0.691723718 | 0.37040154 | 1.8674969 | 0.06183223 | 0.17846586 |
| ZNF445 | 291.813422 | 0.299052291 | 0.16297953 | 1.83490711 | 0.06651945 | 0.18764811 |
| ZNF852 | 92.6915444 | 0.443974939 | 0.20986967 | 2.11547931 | 0.03438912 | 0.12031025 |
| ZKSCAN7 | 53.2390997 | 0.682157526 | 0.37824475 | 1.80348179 | 0.07131258 | 0.19663681 |
| AC099669.1 | 243.747142 | -0.162247831 | 0.30998329 | -0.5234083 | 0.60069014 | 0.76020679 |
| ZNF197 | 235.882289 | 0.034390343 | 0.33266323 | 0.10337885 | 0.9176623 | 0.95587788 |
| ZNF35 | 71.8968105 | -0.656146172 | 0.44605339 | -1.4710037 | 0.14129012 | 0.30819999 |
| ZNF502 | 58.7159256 | 0.452636691 | 0.31637534 | 1.43069524 | 0.15251758 | 0.32411759 |
| KIAA1143 | 98.8427014 | -0.495117229 | 0.25081223 | -1.9740554 | 0.04837545 | 0.15215288 |
| ZDHHC3 | 365.363296 | -0.335740069 | 0.18727427 | -1.792772 | 0.07300938 | 0.19946973 |
| EXOSC7 | 191.831027 | -0.540489416 | 0.31276632 | -1.7280934 | 0.08397147 | 0.2189456 |
| CDCP1 | 74.5158699 | -3.224280929 | 0.89424732 | -3.6055807 | 0.00031146 | 0.00377798 |
| LARS2 | 620.136015 | 0.00757609 | 0.13172225 | 0.05751565 | 0.95413444 | 0.97682198 |
| LIMD1 | 442.406897 | -0.64441435 | 0.33102796 | -1.9467067 | 0.05156991 | 0.15831326 |
| SACM1L | 750.371071 | 0.118169721 | 0.1278176 | 0.92451841 | 0.35521647 | 0.55366752 |
| SLC6A20 | 687.59852 | -0.870841115 | 0.61233308 | -1.422169 | 0.1549772 | 0.32761759 |
| LZTFL1 | 289.101578 | -0.015205447 | 0.19627411 | -0.0774705 | 0.93824928 | 0.96676805 |
| FYCO1 | 357.104451 | 0.074269705 | 0.19092948 | 0.38899024 | 0.69728336 | 0.82550208 |
| LINC02009 | 45.5198346 | -2.832477693 | 0.6254466 | -4.5287283 | 5.93E-06 | 0.00016532 |
| LTF | 3062.65712 | -3.466718061 | 1.31626115 | -2.6337616 | 0.00844448 | 0.04456863 |
| LRRC2 | 68.7867773 | -0.743414896 | 0.5007192 | -1.4846942 | 0.13762485 | 0.30328357 |
| ALS2CL | 201.222131 | -2.568590625 | 0.30196775 | -8.5061753 | 1.80E-17 | 1.45E-14 |
| MYL3 | 80.475868 | 1.156472058 | 0.45624245 | 2.53477521 | 0.01125195 | 0.05482776 |
| PTH1R | 3724.88245 | 2.674034994 | 0.65221438 | 4.0999326 | 4.13E-05 | 0.0007768 |
| CCDC12 | 677.869658 | 0.098164123 | 0.32602628 | 0.30109267 | 0.76334383 | 0.86795138 |
| NBEAL2 | 91.488126 | -0.10277768 | 0.53325135 | -0.1927378 | 0.84716434 | 0.91938085 |
| SETD2 | 3831.05771 | 0.420347689 | 0.19875705 | 2.11488193 | 0.03444001 | 0.12041369 |
| KIF9-AS1 | 1305.80586 | 0.760593385 | 0.31188559 | 2.43869359 | 0.01474046 | 0.06626681 |
| KIF9 | 196.97186 | 0.317638207 | 0.2288003 | 1.38827704 | 0.1650527 | 0.34236945 |
| KLHL18 | 395.2063 | 0.083290419 | 0.3597051 | 0.23155195 | 0.81688603 | 0.90111634 |
| AC099778.1 | 46.4971587 | -0.190449395 | 0.43703517 | -0.4357759 | 0.66299933 | 0.80379726 |
| PTPN23 | 305.098067 | -0.250642692 | 0.33369163 | -0.7511207 | 0.45258001 | 0.64185352 |
| SCAP | 732.931918 | 0.63393541 | 0.19685151 | 3.22037367 | 0.00128024 | 0.01087249 |
| ELP6 | 181.910099 | 0.317169445 | 0.32335961 | 0.9808567 | 0.32666341 | 0.52677357 |
| SMARCC1 | 3524.72928 | 0.486910346 | 0.20099854 | 2.42245712 | 0.01541595 | 0.06849477 |
| DHX30 | 457.672287 | 0.509921804 | 0.22545106 | 2.26178489 | 0.0237107 | 0.09290451 |
| MAP4 | 4329.67832 | -0.623748093 | 0.31068852 | -2.0076316 | 0.04468245 | 0.14397329 |
| ZNF589 | 144.741079 | 0.653555983 | 0.26055117 | 2.50835947 | 0.01212932 | 0.05774804 |
| NME6 | 123.333518 | -0.41142428 | 0.41761232 | -0.9851823 | 0.32453452 | 0.52488433 |
| PLXNB1 | 803.362527 | 0.299608361 | 0.23727597 | 1.26269996 | 0.20669703 | 0.39542166 |
| CCDC51 | 55.405037 | -0.591270138 | 0.31225223 | -1.8935658 | 0.05828267 | 0.17132329 |
| TMA7 | 421.76211 | -0.370206917 | 0.34955662 | -1.0590757 | 0.28956531 | 0.48946501 |
| ATRIP | 49.4554297 | 0.074257874 | 0.30486631 | 0.24357521 | 0.80755983 | 0.8957856 |
| SHISA5 | 589.264541 | -1.050695432 | 0.38427731 | -2.7342115 | 0.00625299 | 0.03569494 |
| PFKFB4 | 343.721902 | -4.233030078 | 0.77144673 | -5.4871321 | 4.09E-08 | 2.67E-06 |
| UQCRC1 | 430.859446 | 0.28460682 | 0.28971115 | 0.98238132 | 0.32591203 | 0.52605701 |
| SLC26A6 | 285.883429 | 2.049569613 | 0.58939802 | 3.4773948 | 0.00050631 | 0.00538821 |
| NCKIPSD | 165.865021 | -0.520034818 | 0.31261899 | -1.663478 | 0.09621679 | 0.23950313 |
| IP6K2 | 1126.7147 | 0.428264811 | 0.27262312 | 1.57090424 | 0.11620489 | 0.27158651 |
| PRKAR2A | 1309.6042 | 0.078108467 | 0.29646141 | 0.26346926 | 0.79218891 | 0.88678699 |
| SLC25A20 | 387.681763 | 1.836642537 | 0.52459336 | 3.50107848 | 0.00046338 | 0.00507312 |
| ARIH2 | 1140.94957 | 0.033181047 | 0.24312564 | 0.13647696 | 0.89144424 | 0.94341489 |
| P4HTM | 243.404014 | -0.143577799 | 0.27991845 | -0.5129272 | 0.60800221 | 0.76485741 |
| WDR6 | 840.561395 | 0.552663331 | 0.36325553 | 1.52141754 | 0.12815509 | 0.28987159 |
| DALRD3 | 165.591133 | 0.308671525 | 0.35909307 | 0.85958641 | 0.39001707 | 0.5854741 |
| NDUFAF3 | 152.639414 | -0.541806893 | 0.43446517 | -1.2470663 | 0.21237317 | 0.4021828 |
| IMPDH2 | 117.113396 | -0.641090796 | 0.42200088 | -1.5191693 | 0.12871988 | 0.29046822 |
| QRICH1 | 778.819692 | 0.322666253 | 0.25668708 | 1.25704126 | 0.2087387 | 0.39797925 |
| QARS | 175.051867 | 0.234862291 | 0.29519589 | 0.79561504 | 0.42625582 | 0.61912563 |
| USP19 | 126.374835 | 0.534166951 | 0.27478395 | 1.94395255 | 0.05190117 | 0.15893865 |
| LAMB2 | 407.090426 | 0.892494303 | 0.34432973 | 2.59197574 | 0.00954265 | 0.04869736 |
| CCDC71 | 36.9871273 | -0.096601545 | 0.36815844 | -0.2623912 | 0.79301984 | 0.8871958 |
| KLHDC8B | 61.528979 | 1.266779066 | 0.46486806 | 2.72502924 | 0.00642958 | 0.03651123 |
| C3orf62 | 133.820785 | -0.180776945 | 0.3864793 | -0.4677532 | 0.63996106 | 0.78856044 |
| USP4 | 693.561532 | -0.093863221 | 0.27267111 | -0.344236 | 0.73066879 | 0.84678203 |
| RHOA | 1390.08408 | -0.878003244 | 0.39808586 | -2.2055625 | 0.02741465 | 0.1028387 |
| TCTA | 152.412926 | -0.432221815 | 0.26628465 | -1.6231571 | 0.10455584 | 0.25263333 |
| DAG1 | 552.434131 | -0.159016135 | 0.36151682 | -0.4398582 | 0.66003982 | 0.8020537 |
| BSN | 55.2034659 | 0.423775821 | 0.35216189 | 1.20335515 | 0.22883892 | 0.42176886 |
| APEH | 242.439301 | 0.297351549 | 0.361836 | 0.82178543 | 0.41119903 | 0.60468888 |
| MST1 | 1002.72867 | 1.416448101 | 0.47532261 | 2.97997206 | 0.00288275 | 0.02010881 |
| RNF123 | 257.804841 | 0.137527359 | 0.26734874 | 0.51441184 | 0.60696408 | 0.763902 |
| GMPPB | 70.8485772 | -0.78004446 | 0.45386158 | -1.7186836 | 0.085672 | 0.22181547 |
| IP6K1 | 1063.13619 | 0.019038481 | 0.3818324 | 0.04986083 | 0.96023329 | 0.9796009 |
| UBA7 | 45.4225422 | -0.062697211 | 0.96201291 | -0.0651729 | 0.9480363 | 0.97344198 |
| MON1A | 89.7476356 | 0.651034948 | 0.40396236 | 1.61162279 | 0.10704405 | 0.25682702 |
| RBM6 | 5059.70299 | 0.486152584 | 0.20363063 | 2.38742367 | 0.01696693 | 0.07355949 |
| RBM5 | 1293.41975 | 0.290013179 | 0.11021914 | 2.63124142 | 0.00850736 | 0.044727 |
| SLC38A3 | 72.5307557 | -2.498491701 | 1.10627832 | -2.2584658 | 0.02391664 | 0.09342415 |
| GNAI2 | 419.611518 | -0.625599162 | 0.26081972 | -2.3985884 | 0.0164584 | 0.07193729 |
| SEMA3B | 757.334586 | 0.135705073 | 0.41990683 | 0.32317901 | 0.74655968 | 0.85675598 |
| IFRD2 | 244.93505 | -0.740671992 | 0.37607104 | -1.9695002 | 0.04889568 | 0.15326161 |
| HYAL1 | 222.029318 | 0.840835958 | 0.43459668 | 1.93475006 | 0.05302097 | 0.16110807 |
| HYAL2 | 97.8707309 | -0.610504165 | 0.36713108 | -1.6629051 | 0.09633143 | 0.23963972 |
| TUSC2 | 134.96183 | -0.512609398 | 0.37591292 | -1.3636387 | 0.17268131 | 0.35223399 |
| RASSF1 | 182.809782 | -1.534187227 | 0.46527877 | -3.2973506 | 0.00097602 | 0.0088833 |
| NPRL2 | 59.8128758 | 0.419948451 | 0.34797894 | 1.20682147 | 0.22750091 | 0.42009362 |
| TMEM115 | 110.184553 | -0.07215712 | 0.40007866 | -0.1803573 | 0.85687205 | 0.92453042 |
| HEMK1 | 203.525732 | 1.033590541 | 0.23197412 | 4.45562873 | 8.36E-06 | 0.00021923 |
| CISH | 283.649189 | 1.256738656 | 0.51676268 | 2.43194547 | 0.01501797 | 0.06711813 |
| MAPKAPK3 | 270.702399 | 0.897776932 | 0.39382101 | 2.27965728 | 0.02262802 | 0.0900849 |
| DOCK3 | 208.776905 | -1.07202387 | 0.40025579 | -2.678347 | 0.00739865 | 0.04061407 |
| MANF | 259.204188 | -1.556608254 | 0.61578215 | -2.5278554 | 0.01147616 | 0.05558791 |
| RBM15B | 93.676652 | -0.921704386 | 0.4608919 | -1.9998277 | 0.04551887 | 0.14611878 |
| DCAF1 | 643.452098 | 0.3694635 | 0.27229232 | 1.35686346 | 0.1748246 | 0.35485277 |
| RAD54L2 | 661.072681 | 0.093381319 | 0.22641564 | 0.41243316 | 0.68002196 | 0.81435096 |
| TEX264 | 334.679741 | 0.158514484 | 0.30482501 | 0.52001798 | 0.60305104 | 0.7614943 |
| RRP9 | 55.1241532 | -0.201542917 | 0.48224562 | -0.4179259 | 0.67600132 | 0.81237508 |
| PARP3 | 101.154813 | 0.519467424 | 0.50077442 | 1.0373282 | 0.29958293 | 0.49986405 |
| PCBP4 | 68.1642553 | 0.433837436 | 0.41472484 | 1.04608501 | 0.29552179 | 0.49544966 |
| ABHD14B | 65.094088 | 0.322812807 | 0.30872117 | 1.04564518 | 0.29572488 | 0.4955986 |
| ABHD14A | 51.1448819 | 0.491996379 | 0.52619881 | 0.93500094 | 0.34978781 | 0.54932878 |

| | | | | | | |
|---|---|---|---|---|---|---|
| ACY1 | 67.9693294 | 3.080230958 | 0.65025363 | 4.73696847 | 2.17E-06 | 7.34E-05 |
| RPL29 | 1849.69239 | -1.395581958 | 0.42422166 | -3.289747 | 0.00100277 | 0.00906067 |
| POC1A | 59.0202787 | 0.154330805 | 0.42463304 | 0.36344512 | 0.7162724 | 0.83792317 |
| ALAS1 | 134.861656 | -0.257765755 | 0.26153522 | -0.9855872 | 0.32433573 | 0.52483341 |
| WDR82 | 596.107978 | 0.073502455 | 0.19946407 | 0.36849973 | 0.71250065 | 0.83543596 |
| GLYCTK | 210.463758 | 1.614771411 | 0.41473048 | 3.89354412 | 9.88E-05 | 0.00153877 |
| DNAH1 | 292.353543 | 0.792856151 | 0.34274073 | 2.31328255 | 0.02070711 | 0.08462503 |
| BAP1 | 161.492385 | 0.025632538 | 0.34589567 | 0.07410482 | 0.94092698 | 0.96884661 |
| PHF7 | 154.049732 | 0.947661959 | 0.28622015 | 3.31095472 | 0.00092978 | 0.00856537 |
| NISCH | 436.287551 | 0.43731891 | 0.22675231 | 1.92861945 | 0.05377812 | 0.16279521 |
| NT5DC2 | 57.0027584 | -0.514002863 | 0.59955708 | -0.8573043 | 0.39127673 | 0.58646585 |
| SMIM4 | 519.448953 | -0.202237047 | 0.30735166 | -0.6579989 | 0.51053885 | 0.6897256 |
| PBRM1 | 1924.21637 | 0.027783282 | 0.21181427 | 0.13116814 | 0.8956423 | 0.94595231 |
| GNL3 | 340.650463 | -1.323048342 | 0.39692069 | -3.3332814 | 0.00085828 | 0.00808514 |
| GLT8D1 | 169.666756 | 0.45209028 | 0.24507889 | 1.84467245 | 0.0650852 | 0.18504933 |
| SPCS1 | 417.474787 | -0.773465353 | 0.3258422 | -2.3737421 | 0.01760884 | 0.07540368 |
| NEK4 | 449.993388 | 0.257775844 | 0.19461173 | 1.32456475 | 0.18531555 | 0.36840033 |
| SFMBT1 | 593.344565 | 0.769498615 | 0.27054395 | 2.84426468 | 0.00445141 | 0.0277848 |
| AC096887.1 | 76.7928959 | 0.363604945 | 0.43350526 | 0.83875556 | 0.40160649 | 0.59575976 |
| RFT1 | 80.9341145 | 0.688423724 | 0.36788903 | 1.87128092 | 0.06130615 | 0.17738359 |
| PRKCD | 193.059858 | 0.372146558 | 0.3040765 | 1.22385832 | 0.22100568 | 0.41322123 |
| TKT | 1512.13218 | -1.518886344 | 0.60438206 | -2.5131228 | 0.01196677 | 0.05719263 |
| DCP1A | 1527.98685 | -0.334893999 | 0.14862508 | -2.2532805 | 0.02424147 | 0.09423647 |
| CACNA1D | 3954.09971 | -0.233116647 | 0.55286696 | -0.4216505 | 0.67328012 | 0.81072605 |
| CHDH | 2083.19206 | 0.336854836 | 0.37827683 | 0.89049821 | 0.37319843 | 0.57072753 |
| IL17RB | 3330.53206 | -1.164034577 | 0.6205649 | -1.8757661 | 0.06068741 | 0.17616313 |
| AC012467.2 | 47.7634969 | -0.480322511 | 0.47422099 | -1.0128664 | 0.31112398 | 0.51127879 |
| ACTR8 | 187.951049 | -0.005103237 | 0.21313933 | -0.0239432 | 0.98089792 | 0.99047407 |
| SELENOK | 352.838014 | -1.31530262 | 0.35983112 | -3.6553331 | 0.00025685 | 0.00324481 |
| CACNA2D3 | 1401.27656 | -2.169608617 | 0.35426154 | -6.1243132 | 9.11E-10 | 1.09E-07 |
| WNT5A | 42.9055637 | 0.281693994 | 0.79649203 | 0.35366832 | 0.72358747 | 0.84266894 |
| ERC2 | 747.332276 | -0.763605439 | 0.41467051 | -1.8414751 | 0.06555196 | 0.18587521 |
| CCDC66 | 1063.11847 | 0.537646696 | 0.16169265 | 3.3251152 | 0.00088382 | 0.0082524 |
| FAM208A | 1058.29304 | 0.178116502 | 0.16428106 | 1.08421814 | 0.27826809 | 0.47653932 |
| ARHGEF3 | 4952.47195 | -0.993255416 | 0.57692229 | -1.7216451 | 0.08513383 | 0.22086814 |
| SPATA12 | 103.614888 | -1.569149335 | 0.46466745 | -3.3769298 | 0.000733 | 0.00718684 |
| IL17RD | 429.692432 | 0.766331973 | 0.33168644 | 2.31041089 | 0.02086542 | 0.08506909 |
| HESX1 | 42.2329296 | 0.829145719 | 0.34628023 | 2.3944356 | 0.01664597 | 0.07254889 |
| APPL1 | 670.872083 | -0.125309082 | 0.14591055 | -0.8588076 | 0.39044669 | 0.58579825 |
| ASB14 | 126.078071 | -0.302815601 | 0.45285018 | -0.6686883 | 0.50369436 | 0.68439121 |
| DNAH12 | 146.079551 | -0.226022558 | 0.59416651 | -0.3804027 | 0.70364649 | 0.82988147 |
| PDE12 | 184.974179 | -0.344247368 | 0.27736411 | -1.2411388 | 0.21455446 | 0.40485931 |
| ARF4 | 870.582855 | -1.455966939 | 0.5722596 | -2.544242 | 0.01095152 | 0.05384903 |
| ARF4-AS1 | 36.6221053 | -0.26841725 | 0.40589781 | -0.6612927 | 0.50842463 | 0.68823852 |
| DENND6A | 767.706324 | -0.01554278 | 0.18920879 | -0.0821462 | 0.93453047 | 0.96509871 |
| DENND6A-A! | 139.450367 | -0.797283397 | 0.48097079 | -1.6576545 | 0.09738722 | 0.24130272 |
| SLMAP | 3200.40457 | -0.466399271 | 0.27048044 | -1.7243364 | 0.08464712 | 0.22010729 |
| FLNB | 2126.4039 | -0.56390295 | 0.27504874 | -2.0501928 | 0.04034562 | 0.13434242 |
| ABHD6 | 1312.03066 | 1.919998527 | 0.35762264 | 5.36878344 | 7.93E-08 | 4.58E-06 |
| RPP14 | 125.038086 | 0.368326864 | 0.31690059 | 1.16227888 | 0.2451222 | 0.44058353 |
| PXK | 435.389248 | 0.122383118 | 0.38442026 | 0.31835762 | 0.75021368 | 0.85921887 |
| PDHB | 190.994817 | -0.320886001 | 0.23585569 | -1.3605184 | 0.17366594 | 0.35354806 |
| KCTD6 | 326.399333 | 0.559382127 | 0.39940656 | 1.40053315 | 0.16135372 | 0.33760884 |
| ACOX2 | 390.424533 | 2.273391588 | 0.54863305 | 4.14373796 | 3.42E-05 | 0.00067004 |
| FAM107A | 77.7118154 | 0.513631785 | 0.77998009 | 0.6585191 | 0.51020463 | 0.68954593 |
| FAM3D-AS1 | 83.2038549 | 0.499336919 | 0.9207001 | 0.5423448 | 0.587581 | 0.75018375 |
| C3orf67 | 151.789729 | -1.679551963 | 0.52451242 | -3.2021205 | 0.0013642 | 0.01143675 |
| C3orf67-AS1 | 42.8486715 | -0.906647482 | 0.5242036 | -1.7295713 | 0.0837069 | 0.21850514 |
| FHIT | 25978.1286 | 0.670599741 | 0.39459879 | 1.69944702 | 0.08923499 | 0.22776623 |
| PTPRG | 15441.0433 | 0.601269853 | 0.21892434 | 2.74647325 | 0.00602398 | 0.034679 |
| C3orf14 | 39.9173877 | -0.534026556 | 0.38474887 | -1.3879873 | 0.16514091 | 0.34242025 |
| THOC7 | 344.006096 | -0.216850312 | 0.13441867 | -1.6132455 | 0.10669118 | 0.25642924 |
| ATXN7 | 322.111928 | -0.14934002 | 0.1827522 | -0.8171722 | 0.41383003 | 0.60754543 |
| PSMD6-AS2 | 147.35834 | 0.024853895 | 0.2845644 | 0.08734014 | 0.93040115 | 0.96299996 |
| PSMD6 | 195.004522 | -0.095686501 | 0.25521776 | -0.374921 | 0.70771921 | 0.8325358 |
| PSMD6-AS1 | 49.3559232 | 0.069926637 | 0.28615014 | 0.24437044 | 0.80694393 | 0.89531977 |
| PRICKLE2 | 2985.29559 | 0.043332033 | 0.2959033 | 0.14643984 | 0.88357418 | 0.93957729 |
| ADAMTS9 | 4568.58979 | -0.491344292 | 0.28036247 | -1.7525323 | 0.07968232 | 0.21135017 |
| ADAMTS9-A! | 10338.4623 | 0.980088911 | 0.71576553 | 1.36928766 | 0.17090937 | 0.3498066 |
| ADAMTS9-A! | 5194.17393 | -0.24601699 | 0.51208565 | -0.4804216 | 0.63092766 | 0.78242337 |
| MAGI1 | 11315.2007 | 0.443210969 | 0.2398852 | 1.84759612 | 0.06466079 | 0.18422479 |
| MAGI1-AS1 | 140.931894 | -0.324327086 | 0.63945066 | -0.5071964 | 0.612017 | 0.76821585 |
| SLC25A26 | 1214.78975 | -0.247338855 | 0.32304693 | -0.7656437 | 0.44388832 | 0.63517875 |
| LRIG1 | 549.235095 | -0.30641075 | 0.31821896 | -0.9628928 | 0.33560131 | 0.53551655 |
| SUCLG2 | 3487.47785 | 0.111630767 | 0.2359912 | 0.47302935 | 0.63619224 | 0.78575302 |
| SUCLG2-AS1 | 1961.42348 | 0.528883339 | 0.36819144 | 1.43643571 | 0.1508784 | 0.32203211 |
| FAM19A1 | 603.523886 | 0.42276355 | 0.53086847 | 0.79636214 | 0.42582158 | 0.61876852 |
| EOGT | 170.195378 | -0.552338052 | 0.41816116 | -1.3208736 | 0.1865435 | 0.36982235 |
| AC109587.1 | 85.0500474 | -1.222535115 | 0.55949099 | -2.1850846 | 0.02888265 | 0.10630893 |
| TMF1 | 835.666011 | -0.420245954 | 0.12507495 | -3.359953 | 0.00077956 | 0.00753023 |
| UBA3 | 706.105457 | -0.251522226 | 0.14010674 | -1.7952186 | 0.07261888 | 0.19887875 |
| ARL6IP5 | 720.118238 | -0.893302009 | 0.23535839 | -3.7954968 | 0.00014735 | 0.00208431 |
| FRMD4B | 6125.94378 | 1.885186988 | 0.52267231 | 3.60682393 | 0.00030997 | 0.00377224 |
| MITF | 3186.7022 | -0.250405992 | 0.4985265 | -0.5022922 | 0.61546197 | 0.77111743 |
| FOXP1 | 3926.83379 | 0.252492383 | 0.34324443 | 0.73560519 | 0.461971 | 0.65039406 |
| EIF4E3 | 329.2626 | -0.664315094 | 0.33643674 | -1.9745617 | 0.04831791 | 0.15207661 |
| LINC00870 | 51.3347161 | 2.794413625 | 0.71979676 | 3.88222589 | 0.0001035 | 0.00159588 |
| RYBP | 1067.61371 | -0.365688421 | 0.27674935 | -1.3213705 | 0.18637786 | 0.36965438 |
| SHQ1 | 451.38981 | -0.33974112 | 0.28592571 | -1.1882146 | 0.23474886 | 0.42919869 |
| GXYLT2 | 461.355972 | -1.768080451 | 0.44461215 | -3.9766804 | 6.99E-05 | 0.00117534 |
| PPP4R2 | 1148.86267 | -0.008416331 | 0.24038525 | -0.0350118 | 0.9720703 | 0.98609884 |
| PDZRN3 | 3724.60821 | 1.594711611 | 0.48469135 | 3.29015903 | 0.00100131 | 0.00905338 |
| CNTN3 | 1478.95679 | 0.998499102 | 0.21149761 | 4.72108918 | 2.35E-06 | 7.84E-05 |
| LINC02018 | 99.7767616 | 1.454935202 | 0.55561042 | 2.61862476 | 0.0088285 | 0.04601226 |
| ZNF717 | 494.343503 | 0.47594747 | 0.32008947 | 1.48692012 | 0.13703591 | 0.30222867 |
| ROBO2 | 518.409634 | 0.438389877 | 0.60557463 | 0.72392378 | 0.46911253 | 0.65617047 |
| ROBO1 | 120.043173 | -0.317266846 | 0.45480183 | -0.6975936 | 0.48543139 | 0.66906064 |
| LINC02027 | 4256.85471 | 0.367763029 | 0.63124173 | 0.58260253 | 0.5601609 | 0.72861316 |
| GBE1 | 6323.18678 | -2.007356206 | 0.49415404 | -4.0622074 | 4.86E-05 | 0.00088605 |
| CADM2 | 69.4631503 | -0.181728497 | 0.38054237 | -0.4775513 | 0.63296961 | 0.7836086 |
| CHMP2B | 338.173968 | -0.856022476 | 0.29358249 | -2.9157818 | 0.00354799 | 0.02344852 |
| CGGBP1 | 732.592584 | -0.516518956 | 0.2817405 | -1.8333145 | 0.06675581 | 0.18804396 |
| ZNF654 | 1116.95149 | -0.575945966 | 0.24652995 | -2.336211 | 0.01948025 | 0.08091336 |
| C3orf38 | 192.429806 | -0.209362612 | 0.19062441 | -1.0982991 | 0.27207391 | 0.47033143 |
| PROS1 | 1494.76901 | -0.175095807 | 0.26555169 | -0.6593662 | 0.50966064 | 0.68915039 |
| ARL13B | 568.696642 | 0.052928855 | 0.26740076 | 0.19793831 | 0.84309333 | 0.9175292 |
| DHFR2 | 61.6350811 | 0.37856527 | 0.35682362 | 1.06093108 | 0.28872123 | 0.48858047 |
| NSUN3 | 441.174161 | 0.256366408 | 0.20378626 | 1.25801614 | 0.20838592 | 0.39759824 |
| MTRNR2L12 | 1219.43139 | -0.567668866 | 0.57988031 | -0.9789414 | 0.32760891 | 0.52749153 |
| EPHA6 | 389.571594 | 1.249001631 | 0.78957374 | 1.58186825 | 0.11367965 | 0.26731977 |
| ARL6 | 291.588471 | 0.125137089 | 0.28558699 | 0.43817503 | 0.66125941 | 0.80310852 |
| CRYBG3 | 604.44451 | -1.046573419 | 0.28772192 | -3.6374477 | 0.00027535 | 0.00343115 |
| RIOX2 | 233.844595 | -1.355145099 | 0.24654869 | -5.4964603 | 3.87E-08 | 2.58E-06 |

| | | | | | | |
|---|---|---|---|---|---|---|
| CLDND1 | 215.543066 | -0.33901302 | 0.19284196 | -1.7579837 | 0.07875028 | 0.20970788 |
| CPOX | 126.207016 | 0.361551415 | 0.27447911 | 1.31722743 | 0.1877624 | 0.37136212 |
| ST3GAL6-AS | 122.527983 | 0.058417929 | 0.51584666 | 0.11324669 | 0.90983498 | 0.95264747 |
| ST3GAL6 | 1357.67916 | 0.125970364 | 0.36324922 | 0.34678771 | 0.72875081 | 0.84599052 |
| DCBLD2 | 814.45699 | -1.105080108 | 0.64676418 | -1.7086291 | 0.08751967 | 0.22528303 |
| COL8A1 | 108.865298 | 0.245930659 | 0.48042822 | 0.51189886 | 0.60872179 | 0.76555185 |
| CMSS1 | 1174.2788 | -0.505716487 | 0.41257058 | -1.2257696 | 0.22028538 | 0.41209932 |
| FILIP1L | 262.577827 | -0.991555051 | 0.3342684 | -2.966344 | 0.00301363 | 0.02072647 |
| TBC1D23 | 904.054535 | -0.248233226 | 0.2207906 | -1.1242925 | 0.26088895 | 0.45905068 |
| NIT2 | 756.691029 | 1.081018659 | 0.3690883 | 2.92888901 | 0.00340176 | 0.02267894 |
| TOMM70 | 800.803938 | 0.202215029 | 0.20727382 | 0.97559367 | 0.32926586 | 0.52891681 |
| LNP1 | 120.038859 | 1.743275451 | 0.5577949 | 3.12529832 | 0.00177625 | 0.01399022 |
| TFG | 1384.0571 | -0.494951788 | 0.18068744 | -2.7392706 | 0.00615757 | 0.03528502 |
| ABI3BP | 767.98569 | 0.770872699 | 0.71914886 | 1.07192369 | 0.28375431 | 0.48299079 |
| SENP7 | 1020.99425 | 0.068752707 | 0.27175818 | 0.25299222 | 0.80027422 | 0.89131054 |
| TRMT10C | 203.666823 | -0.867564675 | 0.29418506 | -2.949044 | 0.00318759 | 0.02161912 |
| PCNP | 837.008216 | -0.361172594 | 0.146586 | -2.4638956 | 0.01374361 | 0.06305902 |
| ZBTB11 | 534.802201 | -0.25320316 | 0.35465443 | -0.7139433 | 0.47526223 | 0.66124102 |
| ZBTB11-AS1 | 35.0926165 | 0.184578664 | 0.46379602 | 0.3979738 | 0.6906495 | 0.8212201 |
| RPL24 | 705.874854 | -0.5966227 | 0.36393543 | -1.6393641 | 0.10113745 | 0.24777319 |
| CEP97 | 177.602342 | 0.268377099 | 0.22855485 | 1.17423496 | 0.24030092 | 0.43557323 |
| NFKBIZ | 1492.52806 | -1.591779918 | 0.57951378 | -2.7467508 | 0.00601889 | 0.03467496 |
| ALCAM | 2592.91093 | 0.449382001 | 0.34251406 | 1.31201037 | 0.18951664 | 0.37366291 |
| CBLB | 1894.61294 | -0.251637351 | 0.29523821 | -0.8523197 | 0.39403666 | 0.58879979 |
| LINC00882 | 183.682694 | 0.452058353 | 0.53394656 | 0.84663594 | 0.39719807 | 0.59178195 |
| DUBR | 79.8032053 | -1.478086736 | 0.31520085 | -4.6893489 | 2.74E-06 | 8.92E-05 |
| AC063944.1 | 109.887832 | 1.347287958 | 0.50325503 | 2.67714751 | 0.00742519 | 0.04069457 |
| BBX | 4205.00792 | -0.377921185 | 0.35558704 | -1.0628092 | 0.28786848 | 0.48756075 |
| CD47 | 318.170146 | -1.341850228 | 0.35257347 | -3.805874 | 0.0001413 | 0.00202093 |
| IFT57 | 541.849862 | 0.466277918 | 0.3905893 | 1.19378056 | 0.23256382 | 0.42628407 |
| HHLA2 | 704.862001 | -0.64983111 | 0.54484856 | -1.1926821 | 0.23299391 | 0.42690116 |
| CIP2A | 86.3230475 | 0.544767992 | 0.53776729 | 1.01301809 | 0.31105152 | 0.51127879 |
| DZIP3 | 787.301517 | 0.546716715 | 0.55370524 | 0.98737861 | 0.32345707 | 0.52401995 |
| C3orf85 | 242.507071 | 2.480235126 | 0.7523885 | 3.29648198 | 0.00097904 | 0.008899 |
| NECTIN3-AS | 72.0359152 | 0.539777792 | 0.51565717 | 1.04677647 | 0.29520269 | 0.49508665 |
| NECTIN3 | 1289.59537 | 0.503760232 | 0.15089984 | 3.33837493 | 0.0008427 | 0.00797547 |
| PLCXD2 | 112.364947 | -0.933145391 | 0.41941434 | -2.2248772 | 0.02608949 | 0.09936332 |
| PHLDB2 | 4911.75405 | 0.350257222 | 0.33226609 | 1.05414676 | 0.29181573 | 0.49181338 |
| ABHD10 | 263.768374 | 1.359601786 | 0.38096448 | 3.5688413 | 0.00035856 | 0.0042088 |
| C3orf52 | 356.103511 | -0.307069437 | 0.5750773 | -0.533962 | 0.59336781 | 0.75440985 |
| AC128688.2 | 58.1006294 | -0.145338102 | 0.57699026 | -0.25189 | 0.80112605 | 0.89204182 |
| SLC9C1 | 502.774179 | -0.579257567 | 0.54476321 | -1.0633199 | 0.2876369 | 0.48746773 |
| AC112487.1 | 66.7213287 | -0.419408695 | 0.71520652 | -0.5864162 | 0.55759586 | 0.7262747 |
| CD200 | 166.115176 | 0.127428783 | 0.53805831 | 0.23683081 | 0.81278805 | 0.89861746 |
| AC092894.1 | 443.876643 | 0.858465808 | 0.4050875 | 2.1192108 | 0.03407266 | 0.11956833 |
| ATG3 | 302.591643 | 0.022782596 | 0.17589218 | 0.12952592 | 0.89694152 | 0.94652224 |
| SLC35A5 | 189.847825 | -0.288994836 | 0.29802566 | -0.9696978 | 0.33219713 | 0.53166443 |
| GTPBP8 | 149.265065 | -0.162365944 | 0.30915369 | -0.5251949 | 0.5994477 | 0.75918226 |
| NEPRO | 345.201409 | -0.338463086 | 0.14294234 | -2.3678295 | 0.01789278 | 0.0762613 |
| AC078785.1 | 87.4438608 | -0.057495441 | 0.46980232 | -0.1223822 | 0.90259634 | 0.94883718 |
| CFAP44 | 639.770851 | 0.631381482 | 0.21877912 | 2.88593118 | 0.00390257 | 0.02517299 |
| SPICE1 | 573.357492 | 0.259071682 | 0.18563213 | 1.39561872 | 0.16282934 | 0.33934952 |
| USF3 | 619.59658 | 0.677276233 | 0.17759738 | 3.81354861 | 0.00013699 | 0.0019682 |
| NAA50 | 1115.77926 | -0.187968345 | 0.28176796 | -0.6671033 | 0.50470614 | 0.6850618 |
| ATP6V1A | 1643.81248 | 0.019357926 | 0.23091792 | 0.08383033 | 0.93319133 | 0.96462605 |
| GRAMD1C | 1142.13624 | 1.010260743 | 0.49979425 | 2.02135326 | 0.04324321 | 0.14072523 |
| ZDHHC23 | 178.759353 | -0.331871018 | 0.44451309 | -0.7465945 | 0.45530838 | 0.64465572 |
| CCDC191 | 564.373575 | 0.346924324 | 0.49140474 | 0.70598489 | 0.48019753 | 0.66519782 |
| QTRT2 | 275.369208 | -0.62011974 | 0.21297622 | -2.9116854 | 0.00359484 | 0.02370912 |
| ZBTB20 | 40075.2076 | 0.710982568 | 0.3096308 | 2.29622688 | 0.02166291 | 0.08753273 |
| ZBTB20-AS1 | 163.446153 | -0.470358572 | 0.70401577 | -0.668108 | 0.50406465 | 0.68462289 |
| ZBTB20-AS5 | 691.022884 | -0.411434757 | 0.63928343 | -0.6435874 | 0.51984303 | 0.69597507 |
| ZBTB20-AS2 | 199.974058 | -0.067008401 | 0.71305031 | -0.0939743 | 0.92512958 | 0.96051743 |
| LSAMP | 737.500445 | -1.927230618 | 0.60055135 | -3.2091021 | 0.0013315 | 0.01123132 |
| IGSF11 | 1148.16777 | 2.059908663 | 0.47734132 | 4.31537891 | 1.59E-05 | 0.00037021 |
| UPK1B | 42.8427545 | -0.107794291 | 0.58382313 | -0.1846352 | 0.85351519 | 0.92286287 |
| B4GALT4 | 643.275353 | 1.048602232 | 0.32851292 | 3.19196648 | 0.00141308 | 0.011767 |
| B4GALT4-AS | 98.6673133 | 0.196173391 | 0.61466412 | 0.31915543 | 0.74960865 | 0.85874516 |
| ARHGAP31 | 836.980967 | -0.003341631 | 0.39441437 | -0.0084724 | 0.99324009 | 0.99654041 |
| TMEM39A | 363.635562 | -0.48138236 | 0.23135916 | -2.0806713 | 0.03746401 | 0.12780481 |
| POGLUT1 | 198.339962 | -0.011726483 | 0.29962289 | -0.0391375 | 0.96878078 | 0.98432195 |
| TIMMDC1 | 519.881699 | 0.051635733 | 0.14188952 | 0.36391506 | 0.71592144 | 0.83776419 |
| PLA1A | 202.990557 | -1.248451499 | 0.64526027 | -1.934803 | 0.05301448 | 0.16110807 |
| COX17 | 248.286113 | -0.79610287 | 0.40745451 | -1.9538448 | 0.05071957 | 0.15693301 |
| MAATS1 | 400.391972 | -0.24293768 | 0.54445055 | -0.4462071 | 0.65544769 | 0.79904496 |
| GSK3B | 3599.54437 | -0.156166332 | 0.26169595 | -0.5967472 | 0.55067616 | 0.720309 |
| AC092910.3 | 140.921485 | 0.495393968 | 0.29848664 | 1.65968555 | 0.09697773 | 0.24076684 |
| GPR156 | 64.839182 | -1.332623919 | 0.90663838 | -1.4698516 | 0.14160194 | 0.30858563 |
| LRRC58 | 582.678711 | -0.715770206 | 0.3403846 | -2.1028278 | 0.03548082 | 0.12281204 |
| FSTL1 | 128.892287 | -0.281223671 | 0.39633827 | -0.7095547 | 0.47798034 | 0.66359531 |
| NDUFB4 | 458.921051 | -1.159319052 | 0.41960799 | -2.7628622 | 0.00572969 | 0.03350551 |
| HGD | 2085.67872 | 0.881009787 | 0.42579475 | 2.06909498 | 0.03853717 | 0.13052361 |
| RABL3 | 335.862036 | -0.640969407 | 0.22270291 | -2.8781367 | 0.00400032 | 0.02563457 |
| GTF2E1 | 152.638532 | -0.367983882 | 0.23165531 | -1.5884975 | 0.11217389 | 0.26518928 |
| STXBP5L | 37.0966587 | -0.008478623 | 0.31002327 | -0.0273483 | 0.9781819 | 0.98906609 |
| GOLGB1 | 3545.04499 | -0.236330922 | 0.39410808 | -0.5996602 | 0.54873273 | 0.71869915 |
| IQCB1 | 1164.2201 | -0.15815946 | 0.16189269 | -0.9769401 | 0.32859881 | 0.52846816 |
| EAF2 | 870.972654 | 2.784501482 | 0.51612258 | 5.39503909 | 6.85E-08 | 4.07E-06 |
| SLC15A2 | 697.664111 | 3.096421399 | 0.59973098 | 5.16301724 | 2.43E-07 | 1.16E-05 |
| ILDR1 | 110.285709 | -0.737183523 | 0.52574882 | -1.4021591 | 0.16086773 | 0.33687295 |
| CASR | 195.122331 | 1.991457453 | 0.5496671 | 3.62302468 | 0.00029118 | 0.00358274 |
| CCDC58 | 388.26603 | 0.092161462 | 0.38463267 | 0.23960903 | 0.81063336 | 0.89723445 |
| FAM162A | 929.990988 | -1.097798585 | 0.51178546 | -2.1450367 | 0.03194991 | 0.1142247 |
| WDR5B | 40.4259677 | 0.04309579 | 0.32056893 | 0.13443533 | 0.89305835 | 0.94424181 |
| AC083798.2 | 260.354454 | 0.459023389 | 0.25974201 | 1.76722815 | 0.07719003 | 0.20709409 |
| KPNA1 | 1398.81799 | -0.219414366 | 0.27214132 | -0.8062516 | 0.42009782 | 0.61362753 |
| PARP9 | 705.074993 | -0.821452394 | 0.58574332 | -1.4024102 | 0.16079279 | 0.33687295 |
| DTX3L | 285.581061 | -1.198189241 | 0.48905093 | -2.4500296 | 0.01428445 | 0.06472626 |
| PARP14 | 1892.83564 | -1.789120636 | 0.72378159 | -2.4719068 | 0.01343945 | 0.06204601 |
| HSPBAP1 | 527.10744 | -0.470324923 | 0.23115717 | -2.0346543 | 0.04188567 | 0.13771018 |
| DIRC2 | 641.92865 | -0.831047122 | 0.26327987 | -3.1565159 | 0.00159666 | 0.01294996 |
| LINC02035 | 52.9169483 | -0.63679985 | 0.36289886 | -1.7547585 | 0.07930063 | 0.2107355 |
| PDIA5 | 788.865703 | -0.80102401 | 0.31233798 | -2.5646065 | 0.01032929 | 0.05174837 |
| SEC22A | 859.145127 | -0.311161535 | 0.19428794 | -1.6015484 | 0.10925551 | 0.26053676 |
| ADCY5 | 261.01624 | -0.757938727 | 0.67377522 | -1.1249133 | 0.26062576 | 0.4589178 |
| HACD2 | 542.409653 | -0.180427111 | 0.18607212 | -0.9696622 | 0.33221487 | 0.53166443 |
| MYLK-AS1 | 152.607775 | -0.033952013 | 0.24851002 | -0.1366223 | 0.89132934 | 0.94341489 |
| MYLK | 954.221282 | 1.268945933 | 0.38043014 | 3.33555572 | 0.00085129 | 0.00802638 |
| CCDC14 | 1103.99626 | 0.381569681 | 0.25124777 | 1.51869877 | 0.12883834 | 0.29060179 |
| KALRN | 784.014221 | -1.50572275 | 0.52297932 | -2.8791248 | 0.0039878 | 0.0255783 |
| UMPS | 251.261582 | -0.275672221 | 0.22278786 | -1.2373754 | 0.21594774 | 0.40653696 |
| ITGB5 | 1068.62578 | -0.254221766 | 0.30596987 | -0.8308719 | 0.406046 | 0.60009523 |
| ITGB5-AS1 | 49.3245525 | -1.27192274 | 0.70016353 | -1.8166081 | 0.06927714 | 0.19280848 |

| | | | | | | |
|---|---|---|---|---|---|---|
| MUC13 | 70.4311024 | 1.419018387 | 0.82967106 | 1.71033853 | 0.08720329 | 0.22463738 |
| HEG1 | 176.877282 | 0.098907807 | 0.62702726 | 0.15774084 | 0.87466102 | 0.93490877 |
| SLC12A8 | 411.03809 | -1.866323492 | 0.45928523 | -4.0635391 | 4.83E-05 | 0.00088218 |
| ZNF148 | 3493.90973 | 0.197831539 | 0.13952686 | 1.41787423 | 0.15622749 | 0.32985733 |
| SNX4 | 1582.2001 | -0.056051053 | 0.17951186 | -0.3122415 | 0.75485699 | 0.86259482 |
| OSBPL11 | 490.694987 | 0.312975917 | 0.21238668 | 1.47361371 | 0.14058563 | 0.30748479 |
| AC092902.3 | 65.7683168 | 0.09240737 | 0.51754136 | 0.1785507 | 0.85829051 | 0.92547105 |
| AC092903.2 | 53.9398298 | 1.394056587 | 0.43244049 | 3.22369578 | 0.00126548 | 0.01080742 |
| SLC41A3 | 161.402671 | -0.253466707 | 0.24856239 | -1.0197307 | 0.30785618 | 0.50791823 |
| ALDH1L1 | 2370.5691 | 0.570772012 | 0.50715479 | 1.12543945 | 0.26040286 | 0.45872536 |
| ALDH1L1-AS2 | 77.4907328 | 0.205136712 | 0.52428732 | 0.39126774 | 0.69559934 | 0.824648 |
| AC016924.1 | 67.6337056 | 0.475795185 | 0.87364239 | 0.54461092 | 0.58602114 | 0.74861061 |
| AC063919.1 | 70.4327484 | 1.449485583 | 0.9683505 | 1.49686047 | 0.13442957 | 0.29873964 |
| KLF15 | 306.996006 | 1.094828682 | 0.53920433 | 2.03045234 | 0.04231058 | 0.13871111 |
| ZXDC | 1565.02028 | 0.21999561 | 0.2255512 | 0.97536883 | 0.32937734 | 0.5289719 |
| CHST13 | 55.8357082 | 0.439135267 | 0.38024562 | 1.15487266 | 0.24814258 | 0.44443882 |
| TXNRD3NB | 148.30025 | 0.085116767 | 0.44439396 | 0.19153448 | 0.84810687 | 0.91986577 |
| TXNRD3 | 204.203629 | 0.369507877 | 0.28289726 | 1.30615572 | 0.19149961 | 0.37589647 |
| CHCHD6 | 334.548871 | -0.175399289 | 0.32919725 | -0.5328091 | 0.59416573 | 0.75493416 |
| PLXNA1 | 131.084622 | -1.103921694 | 0.44899447 | -2.4586532 | 0.01394593 | 0.06365528 |
| TPRA1 | 160.668179 | -0.060943912 | 0.3746568 | -0.162666 | 0.87078144 | 0.93335949 |
| ABTB1 | 109.42903 | 0.201813342 | 0.38987273 | 0.51763903 | 0.60471015 | 0.7625356 |
| MGLL | 1960.97662 | -1.650699292 | 0.59972519 | -2.7524261 | 0.00591555 | 0.03434326 |
| SEC61A1 | 813.538222 | -0.813926302 | 0.37654157 | -2.1615842 | 0.03065024 | 0.11087961 |
| RUVBL1 | 594.518195 | 0.234365236 | 0.25502896 | 0.918975 | 0.35810865 | 0.55659318 |
| EEFSEC | 1294.92874 | 0.724696822 | 0.28393878 | 2.55229959 | 0.01070145 | 0.05295477 |
| RPN1 | 512.372765 | -0.325701342 | 0.32239141 | -1.0102668 | 0.31236747 | 0.51264618 |
| RAB7A | 3089.92216 | -0.585846347 | 0.40318935 | -1.4530303 | 0.14621534 | 0.31512164 |
| ACAD9 | 385.311614 | 0.079926099 | 0.24889397 | 0.32112509 | 0.7481156 | 0.85789513 |
| KIAA1257 | 102.69677 | 1.217707937 | 0.44859583 | 2.71448789 | 0.00663783 | 0.03736541 |
| CNBP | 866.568221 | -0.996473248 | 0.44797946 | -2.2243726 | 0.02612338 | 0.09938197 |
| COPG1 | 376.613633 | -0.674171914 | 0.31598073 | -2.1335855 | 0.03287672 | 0.1167162 |
| HMCES | 163.0282 | 0.196244914 | 0.22508474 | 0.87187126 | 0.38327861 | 0.57986755 |
| H1FX | 168.869783 | -1.493035694 | 0.49673115 | -3.0057219 | 0.00264951 | 0.01888552 |
| EFCAB12 | 60.0942113 | 0.194926519 | 0.54850692 | 0.35537659 | 0.72230749 | 0.84228022 |
| MBD4 | 722.552929 | -0.28256774 | 0.29274878 | -0.9652226 | 0.33443333 | 0.53408905 |
| IFT122 | 605.754012 | -0.241097659 | 0.30691995 | -0.7855392 | 0.43213748 | 0.62475566 |
| TMCC1 | 3236.94723 | -0.007332905 | 0.19651146 | -0.0373154 | 0.97023352 | 0.9852064 |
| TMCC1-AS1 | 58.7994658 | -0.089428675 | 0.35812216 | -0.2497156 | 0.80280732 | 0.89272007 |
| ALG1L2 | 44.8234945 | 0.040403796 | 0.54077856 | 0.07471412 | 0.94044217 | 0.96849292 |
| LINC02021 | 64.6909068 | 1.502717095 | 0.58850549 | 2.55344616 | 0.01066628 | 0.05283354 |
| PIK3R4 | 406.626166 | 0.208575384 | 0.1544479 | 1.35045785 | 0.17686917 | 0.35735135 |
| ATP2C1 | 834.274255 | -0.239293114 | 0.21643832 | -1.1055949 | 0.26890188 | 0.46697556 |
| AC055733.2 | 37.3991792 | -1.008362383 | 0.70092861 | -1.4386093 | 0.15026126 | 0.3209648 |
| ASTE1 | 64.1175408 | 0.376834553 | 0.36675922 | 1.02747124 | 0.30419862 | 0.50443388 |
| NEK11 | 1190.01576 | -0.612239725 | 0.29671938 | -2.0633628 | 0.03907818 | 0.13157567 |
| NUDT16 | 213.405388 | 0.363161557 | 0.30592564 | 1.18709095 | 0.23519175 | 0.42961829 |
| MRPL3 | 498.769929 | -0.088715539 | 0.26721942 | -0.3319951 | 0.73989296 | 0.85230954 |
| CPNE4 | 500.929668 | 1.676602833 | 0.64168478 | 2.61281378 | 0.00898002 | 0.04660907 |
| DNAJC13 | 1497.66691 | -0.14634369 | 0.2162335 | -0.6767855 | 0.49854208 | 0.68045095 |
| NPHP3-ACAD | 83.4506954 | -0.540011766 | 0.49192201 | -1.0977589 | 0.27230979 | 0.47033143 |
| ACKR4 | 148.900244 | -1.041688584 | 1.01269856 | -1.0286265 | 0.30365522 | 0.50383756 |
| UBA5 | 754.957454 | -0.703019723 | 0.53478345 | -1.3145877 | 0.18864852 | 0.37263859 |
| CDV3 | 618.043317 | -1.458684948 | 0.48805095 | -2.9887964 | 0.00280079 | 0.0196946 |
| TOPBP1 | 348.234702 | -0.032431034 | 0.28881127 | -0.1122914 | 0.91059233 | 0.95300367 |
| TF | 61.2530108 | -0.054779067 | 0.44259585 | -0.1237677 | 0.90149921 | 0.94833069 |
| SRPRB | 250.898045 | -0.452923701 | 0.4128672 | -1.0970203 | 0.27263252 | 0.47062144 |
| RAB6B | 78.024151 | -2.826538508 | 0.65589732 | -4.3094223 | 1.64E-05 | 0.00037806 |
| SLCO2A1 | 297.576546 | -1.270345424 | 0.73917423 | -1.7186008 | 0.08568709 | 0.22181547 |
| RYK | 1667.27542 | -0.450372681 | 0.20758884 | -2.1695419 | 0.03004157 | 0.10932258 |
| AMOTL2 | 324.970489 | 0.982367686 | 0.60131326 | 1.63370367 | 0.10232109 | 0.24986908 |
| ANAPC13 | 126.900926 | 0.029678045 | 0.28229576 | 0.10513103 | 0.91627183 | 0.95515537 |
| CEP63 | 561.170713 | 0.325298751 | 0.16062065 | 2.02526106 | 0.04284057 | 0.13987985 |
| EPHB1 | 168.910916 | -0.609864589 | 0.41923884 | -1.4546949 | 0.14575375 | 0.31442346 |
| PPP2R3A | 11531.2546 | -1.157458566 | 0.20716548 | -5.5871208 | 2.31E-08 | 1.67E-06 |
| MSL2 | 751.935055 | 0.44932283 | 0.38812325 | 1.15768078 | 0.24699433 | 0.4429031 |
| PCCB | 359.612774 | 0.485202068 | 0.19608818 | 2.47440758 | 0.01334573 | 0.0617798 |
| STAG1 | 4778.29124 | -0.204633533 | 0.44438761 | -0.4604843 | 0.64516861 | 0.79191602 |
| SLC35G2 | 136.148738 | -1.848294685 | 0.34562557 | -5.3476792 | 8.91E-08 | 4.94E-06 |
| NCK1-DT | 89.7057329 | -0.358710066 | 0.27562973 | -1.3014201 | 0.1931147 | 0.37787626 |
| NCK1 | 838.469512 | -0.112775195 | 0.18553359 | -0.6078425 | 0.54329197 | 0.71379134 |
| IL20RB | 82.7089307 | -0.642065141 | 0.53492258 | -1.2002955 | 0.23002461 | 0.42313439 |
| DZIP1L | 184.945407 | 1.148357415 | 0.22682991 | 5.06263675 | 4.13E-07 | 1.82E-05 |
| DBR1 | 89.6741945 | 0.150606529 | 0.21838574 | 0.68963538 | 0.49042351 | 0.67330079 |
| ARMC8 | 1450.85419 | -0.091878714 | 0.13614409 | -0.6748638 | 0.49976231 | 0.68139272 |
| NME9 | 133.516299 | -0.869198732 | 0.45466052 | -1.9117533 | 0.05590785 | 0.16695618 |
| MRAS | 633.323443 | -2.279774367 | 0.5347463 | -4.2632822 | 2.01E-05 | 0.00044752 |
| ESYT3 | 65.2972535 | -1.118611889 | 0.28679582 | -3.9003773 | 9.60E-05 | 0.00150797 |
| CEP70 | 792.309035 | 0.385249606 | 0.42520879 | 0.90602455 | 0.36492286 | 0.5624016 |
| FAIM | 143.269647 | -0.522219901 | 0.41695988 | -1.2524464 | 0.21040725 | 0.39989367 |
| PIK3CB | 2643.29257 | 0.999849408 | 0.20438077 | 4.89209142 | 9.98E-07 | 3.87E-05 |
| MRPS22 | 141.658677 | -0.05711891 | 0.24279237 | -0.2352583 | 0.81400829 | 0.8993248 |
| COPB2 | 761.336224 | -0.576082959 | 0.22340197 | -2.5786835 | 0.00991776 | 0.05014488 |
| AC097103.2 | 863.55437 | 0.410009339 | 0.39327129 | 1.04256109 | 0.29715162 | 0.49762219 |
| NMNAT3 | 515.220956 | 1.266251959 | 0.37215825 | 3.40245572 | 0.00066783 | 0.00668278 |
| CLSTN2 | 1026.64254 | 0.713693601 | 0.4599743 | 1.5515945 | 0.12075928 | 0.27837129 |
| SLC25A36 | 1782.4663 | 0.028265051 | 0.41473723 | 0.06815171 | 0.94566487 | 0.97196767 |
| PXYLP1 | 326.454739 | 0.622890768 | 0.25072402 | 2.48436814 | 0.01297816 | 0.06054853 |
| AC117383.1 | 373.274337 | -1.991592001 | 0.41579696 | -4.7898186 | 1.67E-06 | 5.93E-05 |
| ZBTB38 | 3649.34611 | -1.316404691 | 0.42222581 | -3.1177741 | 0.00182222 | 0.01422394 |
| RASA2 | 1545.52008 | -0.996825411 | 0.23823722 | -4.1841717 | 2.86E-05 | 0.00058119 |
| RNF7 | 349.052689 | -1.214482067 | 0.30829859 | -3.9393046 | 8.17E-05 | 0.00132872 |
| ATP1B3 | 799.535823 | -1.432293566 | 0.62823744 | -2.2798602 | 0.02261598 | 0.09006311 |
| TFDP2 | 3089.91708 | 0.882497131 | 0.29203452 | 3.02189318 | 0.00251199 | 0.01826157 |
| GK5 | 473.490841 | -0.172194913 | 0.303899 | -0.5666189 | 0.57097314 | 0.73701185 |
| XRN1 | 1952.39142 | 0.025688389 | 0.28510618 | 0.09010113 | 0.92820685 | 0.96208463 |
| ATR | 1987.29668 | -0.325184772 | 0.23247838 | -1.3987743 | 0.16188069 | 0.33813809 |
| PLS1 | 1202.52481 | -0.524215826 | 0.35437831 | -1.4792548 | 0.13907222 | 0.30514868 |
| TRPC1 | 556.405975 | -1.178762999 | 0.35145582 | -3.3539436 | 0.00079669 | 0.00764957 |
| PCOLCE2 | 55.3291263 | -1.169820947 | 0.42370436 | -2.7609367 | 0.00576358 | 0.0336318 |
| U2SURP | 1231.31863 | 0.040399724 | 0.19819738 | 0.20383581 | 0.83848181 | 0.91454366 |
| SLC9A9 | 269.264161 | -0.463537287 | 0.4715132 | -0.9830844 | 0.32556589 | 0.52577125 |
| C3orf58 | 155.303001 | -0.880020398 | 0.37266483 | -2.3614259 | 0.01820481 | 0.07727882 |
| AC107021.1 | 265.678053 | -3.654549985 | 0.67200554 | -5.4382736 | 5.38E-08 | 3.31E-06 |
| LNCSRLR | 101.411255 | -2.821889666 | 0.58113219 | -4.8558481 | 1.20E-06 | 4.49E-05 |
| PLOD2 | 2804.68775 | -3.07972294 | 0.58805162 | -5.2371643 | 1.63E-07 | 8.40E-06 |
| PLSCR4 | 1058.76678 | -0.088533733 | 0.44995441 | -0.1967616 | 0.84401413 | 0.91794824 |
| PLSCR2 | 188.078968 | -2.060155825 | 0.59458005 | -3.4648923 | 0.00053044 | 0.00556296 |
| PLSCR1 | 1548.03626 | -2.033409957 | 0.54372569 | -3.7397717 | 0.00018419 | 0.00249472 |
| GYG1 | 148.778583 | -0.895327677 | 0.35900825 | -2.4938917 | 0.01263511 | 0.05945398 |
| HLTF | 284.818781 | 0.87727086 | 0.31140243 | 2.81716123 | 0.00484502 | 0.02956843 |
| HPS3 | 670.125315 | -0.016955879 | 0.2814124 | -0.0602528 | 0.95195431 | 0.97565526 |
| CP | 123.467684 | -2.477179571 | 0.71539634 | -3.4626674 | 0.00053485 | 0.00560059 |

| | | | | | | |
|---|---|---|---|---|---|---|
| TM4SF18 | 156.277092 | -0.072237116 | 0.69489371 | -0.1039542 | 0.9172057 | 0.95576543 |
| TM4SF1 | 108.187753 | -1.225709897 | 0.65073114 | -1.8835888 | 0.05962062 | 0.17395057 |
| WWTR1 | 4138.79346 | 0.254688375 | 0.24708015 | 1.03079252 | 0.30263813 | 0.50284408 |
| COMMD2 | 249.557271 | -0.655034569 | 0.20828289 | -3.1449274 | 0.00166128 | 0.01329308 |
| ANKUB1 | 690.308311 | -1.018920564 | 0.49304845 | -2.0665729 | 0.03877442 | 0.13093592 |
| RNF13 | 2101.95005 | 0.02065248 | 0.15642305 | 0.13202964 | 0.89496084 | 0.94552399 |
| PFN2 | 38.9131872 | -0.912625234 | 0.3227156 | -2.8279552 | 0.00468464 | 0.02892447 |
| AC117386.2 | 53.1001187 | -0.409484875 | 0.71865316 | -0.5697949 | 0.56881685 | 0.73498987 |
| TSC22D2 | 1598.42591 | -0.602377882 | 0.26307246 | -2.2897793 | 0.02203411 | 0.08847748 |
| SERP1 | 848.054441 | -0.43701904 | 0.21539619 | -2.028908 | 0.04246766 | 0.13905979 |
| EIF2A | 485.339404 | -0.538348753 | 0.24095344 | -2.2342439 | 0.02546703 | 0.09769845 |
| SELENOT | 453.281012 | -0.129345276 | 0.20640249 | -0.6266653 | 0.53087866 | 0.70477034 |
| ERICH6 | 64.9135769 | 1.303162207 | 0.37944179 | 3.43441927 | 0.00059382 | 0.0060743 |
| ERICH6-AS1 | 129.878717 | 0.809266262 | 0.51849851 | 1.56078802 | 0.11857378 | 0.27505343 |
| SIAH2 | 284.361986 | -0.433797146 | 0.31306438 | -1.3856484 | 0.1658543 | 0.34332794 |
| MED12L | 66.8722899 | -0.127604349 | 0.33570132 | -0.3801127 | 0.70386173 | 0.82999249 |
| SUCNR1 | 283.314973 | 1.170337599 | 0.51815091 | 2.25868099 | 0.02390324 | 0.09339842 |
| MBNL1 | 4963.29055 | 0.284448005 | 0.27392509 | 1.03841529 | 0.29907675 | 0.499141 |
| RAP2B | 230.991304 | -2.845950497 | 0.35377587 | -8.0445014 | 8.66E-16 | 5.50E-13 |
| ARHGEF26 | 48.0951163 | 0.327390895 | 0.38476242 | 0.85089104 | 0.39482988 | 0.58959943 |
| DHX36 | 1180.14218 | 0.002935184 | 0.18567134 | 0.01580849 | 0.98738717 | 0.99360989 |
| MME | 4119.00671 | 2.219786147 | 0.33941036 | 6.54012482 | 6.15E-11 | 1.05E-08 |
| MME-AS1 | 96.5770793 | 0.533271408 | 0.85538804 | 0.62342631 | 0.53300442 | 0.70622441 |
| DWORF | 98.1209545 | 4.589534998 | 0.78350331 | 5.85770978 | 4.69E-09 | 4.26E-07 |
| PLCH1 | 964.962073 | 0.06610916 | 0.58475989 | 0.11305351 | 0.90998813 | 0.95270509 |
| GMPS | 529.76795 | -0.001645525 | 0.24650978 | -0.0066753 | 0.99467393 | 0.99736617 |
| KCNAB1 | 91.3738118 | -0.928520925 | 0.50504833 | -1.8384794 | 0.0659918 | 0.18656221 |
| AC084036.1 | 35.4624085 | -0.993788587 | 0.84530617 | -1.1756552 | 0.23973269 | 0.4348883 |
| SSR3 | 581.637749 | -0.917570682 | 0.35807624 | -2.5625009 | 0.01039213 | 0.05191143 |
| TIPARP | 432.078843 | -0.534594481 | 0.41550532 | -1.2866128 | 0.19822927 | 0.38448119 |
| LINC00886 | 823.835544 | 1.708794439 | 0.48353715 | 3.53394656 | 0.0004094 | 0.00465701 |
| LEKR1 | 623.26714 | 1.497115844 | 0.51344369 | 2.91583257 | 0.00354741 | 0.02344852 |
| CCNL1 | 2119.62415 | -0.494975337 | 0.29068981 | -1.7027613 | 0.08861276 | 0.22698842 |
| VEPH1 | 332.958338 | 1.791598992 | 0.57413449 | 3.12052144 | 0.00180531 | 0.01416451 |
| PQLC2L | 51.1854174 | -3.531374096 | 0.79846062 | -4.4227279 | 9.75E-06 | 0.00025017 |
| RSRC1 | 1931.84002 | -0.153613345 | 0.21087557 | -0.7284549 | 0.46633516 | 0.65425344 |
| MLF1 | 128.884132 | -0.054088753 | 0.28799556 | -0.1878111 | 0.85102476 | 0.92167424 |
| GFM1 | 451.973902 | 0.142827229 | 0.30371766 | 0.47026318 | 0.638167 | 0.78726983 |
| LXN | 84.7767084 | -0.470845801 | 0.48225139 | -0.9763493 | 0.3288914 | 0.52869297 |
| RARRES1 | 476.176368 | -1.548356322 | 0.6414744 | -2.4137461 | 0.01578946 | 0.06977854 |
| MFSD1 | 321.501142 | -0.374027987 | 0.26579402 | -1.40721 | 0.15936514 | 0.33441793 |
| IQCJ-SCHIP1 | 2810.08453 | 0.289811184 | 0.5647842 | 0.51313614 | 0.6078561 | 0.76481398 |
| AC079594.2 | 42.9322005 | 0.004829771 | 0.53379925 | 0.00904792 | 0.99278091 | 0.99629729 |
| SMC4 | 253.155635 | -0.031027092 | 0.28328472 | -0.1095262 | 0.91278516 | 0.95362395 |
| KPNA4 | 1313.31142 | -0.458493722 | 0.31676313 | -1.447434 | 0.14777541 | 0.31746789 |
| PPM1L | 1304.23566 | 0.309633754 | 0.23922882 | 1.29429952 | 0.19556198 | 0.3808166 |
| B3GALNT1 | 244.02512 | 0.608141491 | 0.37386536 | 1.62663236 | 0.1038152 | 0.2519021 |
| NMD3 | 945.057306 | 0.113414709 | 0.20953075 | 0.54127954 | 0.58831492 | 0.75056149 |
| LINC01322 | 816.915967 | -1.263232659 | 0.4552921 | -2.7745543 | 0.00552774 | 0.03267302 |
| PDCD10 | 796.432117 | -0.479888486 | 0.22603379 | -2.1230829 | 0.0337469 | 0.11879014 |
| SERPINI1 | 278.500101 | 0.437287792 | 0.53572973 | 0.81624702 | 0.41435887 | 0.60793873 |
| GOLIM4 | 1016.96883 | 0.138151941 | 0.43103584 | 0.32051149 | 0.74858062 | 0.85823001 |
| LINC02082 | 40.25392 | 0.909840504 | 0.69033405 | 1.31797135 | 0.18751324 | 0.37110077 |
| MECOM | 1447.91401 | 0.544867385 | 0.44911986 | 1.21318923 | 0.22505748 | 0.41726807 |
| MYNN | 213.999063 | 0.027984423 | 0.17017078 | 0.16444905 | 0.86937765 | 0.93243814 |
| LRRC34 | 56.0144482 | -0.107764236 | 0.50095763 | -0.2151165 | 0.82967651 | 0.90964453 |
| SEC62 | 1092.19925 | 0.06505606 | 0.1864888 | 0.34884701 | 0.72720417 | 0.84508584 |
| GPR160 | 1300.30354 | 0.280939686 | 0.40062034 | 0.70126166 | 0.48313974 | 0.66724397 |
| PHC3 | 2299.58639 | 0.058252324 | 0.14795191 | 0.39372472 | 0.69378429 | 0.82370734 |
| AC008040.5 | 53.8216538 | -0.20271898 | 0.58318583 | -0.3476061 | 0.72813599 | 0.84566684 |
| PRKCI | 1086.56098 | -0.518478378 | 0.22533988 | -2.3008727 | 0.02139883 | 0.08688202 |
| SKIL | 970.566279 | 0.760889345 | 0.31196114 | 2.43905165 | 0.01472586 | 0.0662229 |
| AC026316.5 | 153.505762 | 3.337672573 | 0.61070523 | 5.46527592 | 4.62E-08 | 2.93E-06 |
| RPL22L1 | 179.301001 | -1.188428728 | 0.47885766 | -2.4817996 | 0.01307208 | 0.06084261 |
| EIF5A2 | 92.3297054 | -0.423978866 | 0.28931726 | -1.4654461 | 0.14279924 | 0.31024713 |
| SLC2A2 | 1073.41613 | 0.475524905 | 0.77949808 | 0.61003986 | 0.5418354 | 0.71282636 |
| TNIK | 5360.82807 | -0.59139864 | 0.38056014 | -1.5540215 | 0.12017929 | 0.27760322 |
| PLD1 | 1375.50301 | 1.077156655 | 0.52543034 | 2.05004657 | 0.04035989 | 0.13434242 |
| FNDC3B | 9269.30461 | -1.331666948 | 0.25595411 | -5.2027566 | 1.96E-07 | 9.54E-06 |
| TNFSF10 | 4238.50631 | -0.612021735 | 0.60724203 | -1.0078712 | 0.31351632 | 0.51385486 |
| LINC02068 | 386.2547 | 1.621751649 | 0.54818037 | 2.95842707 | 0.00309213 | 0.02111703 |
| NCEH1 | 2058.72251 | -0.660347031 | 0.37397888 | -1.7657335 | 0.07744057 | 0.20760375 |
| ECT2 | 219.220277 | -1.338498605 | 0.34817303 | -3.8443489 | 0.00012087 | 0.00180637 |
| SPATA16 | 51.6449297 | 2.8700698 | 0.56636037 | 5.06756821 | 4.03E-07 | 1.79E-05 |
| NLGN1 | 19470.8264 | 0.77454584 | 0.51982447 | 1.49001419 | 0.1362205 | 0.30106005 |
| NAALADL2 | 4428.11365 | 0.902904524 | 0.52167545 | 1.73077825 | 0.08349132 | 0.21823333 |
| TBL1XR1 | 5836.89595 | -0.419317476 | 0.39261689 | -1.0680067 | 0.28551748 | 0.4851355 |
| LINC00501 | 157.269167 | 1.033707889 | 0.64860548 | 1.59373905 | 0.11099449 | 0.26321826 |
| LINC00578 | 383.836256 | -0.12881068 | 0.81183452 | -0.1586662 | 0.87393189 | 0.93468316 |
| LINC02015 | 305.736803 | 0.475122531 | 0.76048257 | 0.62476453 | 0.53212562 | 0.70560925 |
| KCNMB2-AS1 | 317.723489 | -1.249049876 | 0.71812966 | -1.7393097 | 0.0819803 | 0.21547536 |
| ZMAT3 | 307.871754 | -1.161939605 | 0.33127328 | -3.5074957 | 0.00045235 | 0.00498818 |
| PIK3CA | 1079.15252 | -0.26683517 | 0.19431035 | -1.3732422 | 0.16967707 | 0.34859523 |
| KCNMB3 | 115.283536 | -0.23813293 | 0.4803898 | -0.4957077 | 0.62010064 | 0.77432081 |
| ZNF639 | 201.299318 | -0.328991928 | 0.23702716 | -1.3879925 | 0.16513932 | 0.34242025 |
| MFN1 | 899.924918 | -0.55851406 | 0.24387271 | -2.2901868 | 0.02201049 | 0.08843004 |
| GNB4 | 39.1292401 | -1.339494313 | 0.44818861 | -2.9886844 | 0.00280181 | 0.0196946 |
| ACTL6A | 159.446187 | -0.3681213 | 0.30454016 | -1.2087775 | 0.22674833 | 0.4192899 |
| MRPL47 | 159.917053 | -0.4633067 | 0.25631201 | -1.8075887 | 0.07067054 | 0.19541679 |
| NDUFB5 | 359.569516 | 0.284441023 | 0.36269233 | 0.78424879 | 0.43289414 | 0.62552702 |
| USP13 | 235.307411 | -0.115381317 | 0.4362206 | -0.2645022 | 0.79139295 | 0.88653344 |
| TTC14 | 1070.81556 | 0.517782223 | 0.1937013 | 2.67309624 | 0.00751547 | 0.04102099 |
| FXR1 | 1245.02997 | -0.005123146 | 0.2111233 | -0.0242661 | 0.98064033 | 0.99047407 |
| DNAJC19 | 188.604724 | 0.320014313 | 0.34604982 | 0.92476369 | 0.35508884 | 0.55359449 |
| ATP11B | 1496.83329 | -0.328208503 | 0.14504837 | -2.2627521 | 0.02365097 | 0.09278303 |
| DCUN1D1 | 773.343625 | 0.388182225 | 0.15358861 | 2.52741548 | 0.01149055 | 0.05558791 |
| MCCC1 | 1067.4268 | 0.256455727 | 0.24865288 | 1.03138049 | 0.30236243 | 0.50269269 |
| MCF2L2 | 182.593688 | -0.715332753 | 0.39419444 | -1.8146698 | 0.06957467 | 0.19324417 |
| B3GNT5 | 276.712759 | -0.64849853 | 0.29653192 | -2.1869434 | 0.02874665 | 0.10602395 |
| KLHL6 | 37.6113482 | 1.856303285 | 0.58783897 | 3.15784321 | 0.00158941 | 0.01291408 |
| KLHL24 | 1747.94703 | -0.219919758 | 0.31770493 | -0.6922139 | 0.48880303 | 0.67188359 |
| YEATS2 | 874.421091 | -1.460619381 | 0.38626933 | -3.7813496 | 0.00015598 | 0.00217503 |
| ABCC5 | 514.746136 | -0.398092574 | 0.43045779 | -0.9248121 | 0.35506366 | 0.55359449 |
| EIF2B5 | 187.398392 | -0.022293626 | 0.2570106 | -0.086742 | 0.93087655 | 0.96334654 |
| DVL3 | 359.213934 | -0.175206976 | 0.26508991 | -0.6609342 | 0.50865454 | 0.68840124 |
| AP2M1 | 605.280876 | -0.800501715 | 0.31680028 | -2.526834 | 0.01150959 | 0.05564794 |
| ABCF3 | 128.797011 | -0.068422189 | 0.3469062 | -0.1972354 | 0.84364332 | 0.91781904 |
| ALG3 | 134.095906 | -0.439698292 | 0.46919195 | -0.9371395 | 0.34868682 | 0.54865148 |
| PSMD2 | 364.483751 | -0.901645419 | 0.4392853 | -2.0525281 | 0.04011837 | 0.13382878 |
| EIF4G1 | 916.921493 | -0.465118333 | 0.39908578 | -1.1654596 | 0.24383302 | 0.43900266 |
| FAM131A | 101.793445 | -0.198143629 | 0.33534215 | -0.59087 | 0.5546075 | 0.72406944 |
| CLCN2 | 66.0377383 | 0.683343457 | 0.36816933 | 1.85605753 | 0.06344536 | 0.1817441 |
| POLR2H | 551.449567 | 0.262913423 | 0.23986734 | 1.09607844 | 0.27304445 | 0.47082907 |

| | | | | | | |
|---|---|---|---|---|---|---|
| THPO | 45.9245799 | 0.668634362 | 0.52260458 | 1.27942691 | 0.20074676 | 0.38798494 |
| CHRD | 158.159993 | 1.195894363 | 0.49686262 | 2.4068914 | 0.01608895 | 0.07082824 |
| MAGEF1 | 134.188415 | -0.668618569 | 0.30393121 | -2.199901 | 0.02781392 | 0.10374025 |
| VPS8 | 1924.8817 | -0.116980568 | 0.17846047 | -0.6554985 | 0.51214686 | 0.69101261 |
| EHHADH | 2780.09181 | 2.028230007 | 0.57242491 | 3.54322455 | 0.00039527 | 0.00452246 |
| MAP3K13 | 7193.7714 | 1.010077593 | 0.45226141 | 2.23339328 | 0.02552302 | 0.09785848 |
| TMEM41A | 432.1447 | -0.580895088 | 0.30159019 | -1.9261073 | 0.05409097 | 0.16341744 |
| LIPH | 56.3005232 | -0.519491549 | 0.64789297 | -0.8018169 | 0.42265886 | 0.61619214 |
| SENP2 | 826.350402 | 0.463229818 | 0.16405798 | 2.82357388 | 0.00474915 | 0.02919129 |
| IGF2BP2 | 3416.15559 | -0.116456971 | 0.27005566 | -0.4312332 | 0.6662988 | 0.80629979 |
| IGF2BP2-AS1 | 74.672417 | -0.653923914 | 0.49951955 | -1.3091057 | 0.19049854 | 0.37457516 |
| TRA2B | 851.619929 | -0.025155731 | 0.13038632 | -0.1929323 | 0.847012 | 0.91938085 |
| ETV5 | 524.766264 | -1.077491707 | 0.39227413 | -2.7467825 | 0.0060183 | 0.03467496 |
| DGKG | 502.805227 | -0.306277146 | 0.50785076 | -0.6030849 | 0.54645218 | 0.71676747 |
| CRYGS | 82.3378027 | 0.784739116 | 0.37463764 | 2.09466172 | 0.03620107 | 0.12445651 |
| TBCCD1 | 142.573955 | 0.261694423 | 0.31128224 | 0.84069822 | 0.40051702 | 0.59474837 |
| DNAJB11 | 148.319728 | -0.213129377 | 0.48097794 | -0.4431167 | 0.6576813 | 0.80056693 |
| AC068631.1 | 45.9260202 | 0.707231086 | 0.42500552 | 1.66405152 | 0.09610213 | 0.23941693 |
| AC068631.2 | 63.9606209 | -0.775352557 | 0.39441684 | -1.9658201 | 0.04931939 | 0.15398584 |
| KNG1 | 63.4956694 | 2.151466392 | 0.58315093 | 3.68938171 | 0.0002248 | 0.00293181 |
| EIF4A2 | 931.816998 | -0.103958363 | 0.23743046 | -0.4378476 | 0.66149674 | 0.80318508 |
| RFC4 | 40.9602639 | -0.333427832 | 0.37766487 | -0.882867 | 0.37730816 | 0.57374783 |
| ST6GAL1 | 531.879024 | -2.517353133 | 0.7992972 | -3.1494582 | 0.00163574 | 0.01318884 |
| MASP1 | 240.264838 | -0.033556387 | 0.5216361 | -0.0643291 | 0.94870817 | 0.97378635 |
| AC072022.1 | 127.516136 | -2.997711224 | 0.82981828 | -3.6124912 | 0.00030327 | 0.00370491 |
| BCL6 | 1965.63258 | -2.021503607 | 0.56839943 | -3.5564842 | 0.00037585 | 0.00438824 |
| LPP | 19857.6756 | -0.263640193 | 0.29807757 | -0.8844684 | 0.37644341 | 0.57294604 |
| TPRG1 | 444.130846 | -0.854649332 | 0.38082074 | -2.24423 | 0.02481761 | 0.09576997 |
| P3H2 | 4330.08482 | -0.322006089 | 0.50840099 | -0.6333703 | 0.52649185 | 0.7009757 |
| P3H2-AS1 | 61.3596825 | -0.328289698 | 0.63711056 | -0.515279 | 0.60635807 | 0.76359388 |
| CLDN1 | 569.469014 | -3.818281574 | 0.43960333 | -8.6857431 | 3.76E-18 | 3.97E-15 |
| CLDN16 | 46.0261815 | -0.450934291 | 0.62016279 | -0.7271225 | 0.46715094 | 0.65464135 |
| IL1RAP | 204.329053 | -0.626024072 | 0.37302174 | -1.6782509 | 0.09329813 | 0.23516686 |
| GMNC | 64.9133262 | 0.856769922 | 0.7829835 | 1.09423752 | 0.27385082 | 0.47170685 |
| CCDC50 | 1710.17455 | -0.698877512 | 0.37518879 | -1.8627356 | 0.06249948 | 0.17972542 |
| FGF12 | 80.7340238 | -0.319785606 | 0.3288324 | -0.9724881 | 0.33080777 | 0.53061146 |
| MB21D2 | 932.345784 | 0.596206228 | 0.37589004 | 1.58611873 | 0.11271238 | 0.26591198 |
| ATP13A4 | 75.9042682 | 1.316194269 | 0.63042566 | 2.08778664 | 0.03681709 | 0.12613363 |
| OPA1 | 1335.69662 | 0.043510145 | 0.10611127 | 0.41004265 | 0.68177466 | 0.81468059 |
| OPA1-AS1 | 46.5175168 | -1.329356581 | 0.69405233 | -1.915355 | 0.05544725 | 0.16608729 |
| HES1 | 225.112653 | -0.452979818 | 0.60324105 | -0.7509101 | 0.45270674 | 0.64196682 |
| LINC00887 | 84.885371 | -2.866416197 | 0.84820958 | -3.3793726 | 0.00072651 | 0.00714884 |
| CPN2 | 128.897902 | -0.184530606 | 0.51902711 | -0.3555317 | 0.72219128 | 0.84228022 |
| ATP13A3 | 3280.5479 | -0.773378344 | 0.36284291 | -2.1314413 | 0.03305281 | 0.11715925 |
| LINC00884 | 343.2482 | 0.409543732 | 0.37748334 | 1.08493193 | 0.27795181 | 0.47611665 |
| TMEM44-AS | 141.682658 | 0.768841798 | 0.50546371 | 1.52106231 | 0.1282447 | 0.28992962 |
| TMEM44 | 73.4519557 | -2.071549223 | 0.41409413 | -5.0026047 | 5.66E-07 | 2.37E-05 |
| LSG1 | 388.448302 | -0.115049861 | 0.37372145 | -0.3078492 | 0.75819706 | 0.86448741 |
| XXYLT1 | 305.648254 | -0.186998656 | 0.29329344 | -0.6375821 | 0.52374574 | 0.69929698 |
| ACAP2 | 1761.1823 | -0.204698781 | 0.16855971 | -1.2143993 | 0.2245953 | 0.4167494 |
| PPP1R2 | 424.756272 | -0.302546502 | 0.17727079 | -1.7066912 | 0.08787945 | 0.22586981 |
| MUC20-OT1 | 5103.92728 | 0.039712562 | 0.42470193 | 0.0935069 | 0.92550087 | 0.96065575 |
| MUC20 | 59.0411979 | 0.678043806 | 0.64493507 | 1.05133654 | 0.29310405 | 0.49326436 |
| MUC4 | 147.763281 | -0.331336909 | 0.99169516 | -0.3341117 | 0.73829531 | 0.85147781 |
| TNK2 | 125.113421 | 0.399920321 | 0.39337064 | 1.01665016 | 0.30931988 | 0.50959707 |
| TFRC | 456.935276 | -0.612794899 | 0.27724526 | -2.2102989 | 0.02708442 | 0.1019347 |
| SLC51A | 182.919057 | -0.052271502 | 0.6673054 | -0.0783322 | 0.9375638 | 0.96641041 |
| PCYT1A | 437.176809 | -0.244117272 | 0.20570702 | -1.1867231 | 0.23533686 | 0.42967809 |
| UBXN7 | 693.407935 | -0.048315122 | 0.17825189 | -0.2710497 | 0.7863528 | 0.88319714 |
| RNF168 | 410.84872 | -0.447528461 | 0.32117432 | -1.393413 | 0.16349494 | 0.3402651 |
| WDR53 | 109.962703 | -0.009880289 | 0.38035539 | -0.0259765 | 0.97927611 | 0.98952729 |
| FBXO45 | 80.615839 | -0.354223857 | 0.49295627 | -0.7185706 | 0.47240556 | 0.65845669 |
| PIGX | 173.85845 | -0.436149001 | 0.1929544 | -2.2603735 | 0.02379808 | 0.09304058 |
| PAK2 | 1945.66098 | -1.196717787 | 0.34406073 | -3.4782167 | 0.00050476 | 0.00538667 |
| SENP5 | 1510.74053 | -0.253207915 | 0.36413353 | -0.695371 | 0.4868228 | 0.67003516 |
| NCBP2 | 312.164362 | -0.390853476 | 0.30316713 | -1.2892343 | 0.19731663 | 0.38330769 |
| NCBP2-AS2 | 129.74506 | -0.799538771 | 0.40470732 | -1.9755975 | 0.04820038 | 0.15174154 |
| PIGZ | 71.7251858 | -0.337070071 | 0.43376278 | -0.7770839 | 0.43710928 | 0.62895831 |
| DLG1 | 4807.264 | 0.134596383 | 0.15578869 | 0.86396761 | 0.38760568 | 0.5837081 |
| BDH1 | 475.438703 | 1.427179456 | 0.37179444 | 3.8386251 | 0.00012373 | 0.00184337 |
| AC024560.1 | 75.5725725 | 3.605519958 | 2.46023048 | 1.46552121 | 0.14277877 | 0.31024713 |
| RUBCN | 381.659936 | -0.381021115 | 0.24361842 | -1.5640078 | 0.11781573 | 0.2737583 |
| FYTTD1 | 423.970127 | -0.413475065 | 0.26043846 | -1.5876114 | 0.11237424 | 0.26548122 |
| LRCH3 | 1699.90817 | -0.294263352 | 0.24977024 | -1.1781362 | 0.23874232 | 0.43383194 |
| IQCG | 295.587536 | -0.226340146 | 0.20427791 | -1.1080011 | 0.26786135 | 0.46575866 |
| RPL35A | 821.949836 | -0.893603216 | 0.41646827 | -2.1456694 | 0.03189937 | 0.1141035 |
| LMLN | 486.772843 | 0.406830689 | 0.26755547 | 1.52054708 | 0.12837354 | 0.29007848 |
| ZNF595 | 323.280907 | 0.099010505 | 0.16253065 | 0.60918051 | 0.54240479 | 0.71323317 |
| ZNF718 | 699.370612 | 0.176487784 | 0.18183448 | 0.97059579 | 0.3317496 | 0.53141196 |
| ZNF141 | 560.932928 | 0.087811134 | 0.15517787 | 0.56587407 | 0.57147939 | 0.73745697 |
| AC092574.2 | 88.9210979 | -0.743321398 | 0.60832487 | -1.2219152 | 0.2217397 | 0.41391613 |
| ZNF721 | 2268.204 | 0.033417695 | 0.17126797 | 0.19511935 | 0.84529951 | 0.91854292 |
| PIGG | 392.114178 | -0.182894934 | 0.25297442 | -0.722978 | 0.46969341 | 0.65667969 |
| PDE6B | 36.3726736 | -0.203840928 | 0.5774572 | -0.3529975 | 0.72409034 | 0.84296809 |
| ATP5ME | 327.412474 | -0.206885561 | 0.40599015 | -0.5095827 | 0.61034384 | 0.76674759 |
| MYL5 | 50.0510436 | 1.170394469 | 0.39228178 | 2.9835555 | 0.0028492 | 0.01994588 |
| PCGF3 | 605.83182 | -0.125838583 | 0.21651327 | -0.581205 | 0.56110233 | 0.72927619 |
| AC139887.2 | 1270.08218 | -0.123305111 | 0.37165224 | -0.3317755 | 0.74005878 | 0.85230954 |
| GAK | 1492.49666 | -0.407356191 | 0.31898941 | -1.277021 | 0.20159482 | 0.38891769 |
| TMEM175 | 126.816809 | 0.554015599 | 0.20762268 | 2.66837714 | 0.00762186 | 0.04140821 |
| DGKQ | 78.966077 | -0.58704749 | 0.39687594 | -1.4791713 | 0.13909455 | 0.30514868 |
| SLC26A1 | 54.2752312 | 0.961437867 | 0.32503272 | 2.95797254 | 0.0030967 | 0.02111763 |
| IDUA | 96.9238394 | 0.052409886 | 0.37468718 | 0.13987638 | 0.88875766 | 0.94260301 |
| FGFRL1 | 138.637862 | -0.358787446 | 0.3192972 | -1.1236786 | 0.26114938 | 0.45929649 |
| RNF212 | 89.1988757 | 0.280837792 | 0.48182369 | 0.58286423 | 0.5599847 | 0.72854361 |
| AC092535.3 | 235.352202 | 0.287433232 | 0.48078401 | 0.59784275 | 0.54994486 | 0.71955844 |
| SPON2 | 612.09032 | -1.708693082 | 0.46737076 | -3.6559692 | 0.00025621 | 0.00323976 |
| CTBP1-AS | 163.263319 | 0.152570495 | 0.43934433 | 0.34726861 | 0.72838953 | 0.84581797 |
| CTBP1 | 539.829812 | 0.050776496 | 0.29148252 | 0.17420083 | 0.86170764 | 0.92733191 |
| CTBP1-DT | 119.732533 | 0.361220792 | 0.41412187 | 0.87225722 | 0.38306807 | 0.57974098 |
| MAEA | 613.530738 | -0.016731005 | 0.25725115 | -0.0650376 | 0.94814404 | 0.97344198 |
| UVSSA | 765.267735 | 0.94653492 | 0.25423916 | 3.72300996 | 0.00019686 | 0.00263772 |
| FAM53A | 69.1935981 | 0.558805879 | 0.50903585 | 1.09777314 | 0.27230356 | 0.47033143 |
| SLBP | 446.006467 | -0.493279265 | 0.35687479 | -1.3822194 | 0.16690433 | 0.34469756 |
| TMEM129 | 133.871915 | 0.191402075 | 0.3423861 | 0.55902409 | 0.57614528 | 0.74061928 |
| TACC3 | 36.6351891 | 0.167001781 | 0.3531819 | 0.4728492 | 0.63632077 | 0.78577016 |
| FGFR3 | 560.281653 | 2.16048442 | 0.68421757 | 3.15759857 | 0.00159074 | 0.01291726 |
| LETM1 | 430.041472 | 0.054698735 | 0.36478245 | 0.14994892 | 0.88080491 | 0.93801515 |
| NSD2 | 1110.03702 | -0.376816262 | 0.2542251 | -1.482215 | 0.13828309 | 0.30419616 |
| NELFA | 370.965279 | 0.006578906 | 0.24018843 | 0.02739061 | 0.97814819 | 0.98906609 |
| POLN | 236.47619 | 0.535730558 | 0.35000881 | 1.53062021 | 0.12586328 | 0.28619908 |
| MXD4 | 293.518115 | -0.139282673 | 0.32215543 | -0.4323462 | 0.66548982 | 0.80579154 |
| ZFYVE28 | 699.046942 | -0.099956799 | 0.30130503 | -0.3317462 | 0.74008091 | 0.85230954 |

| | | | | | | |
|---|---|---|---|---|---|---|
| RNF4 | 794.858693 | 0.295437749 | 0.27977697 | 1.05597593 | 0.29097921 | 0.49112819 |
| FAM193A | 1469.47425 | 0.156872855 | 0.22260948 | 0.7046998 | 0.48099708 | 0.66589506 |
| TNIP2 | 352.964476 | -0.839384703 | 0.34987159 | -2.3991222 | 0.01643443 | 0.07187834 |
| SH3BP2 | 1791.45234 | 0.075945039 | 0.25522202 | 0.2975646 | 0.7660355 | 0.86919451 |
| ADD1 | 2465.77806 | 0.487162628 | 0.1844809 | 2.64072122 | 0.00827298 | 0.04393556 |
| MFSD10 | 311.794672 | -0.914527021 | 0.34004117 | -2.6894597 | 0.00715678 | 0.03955563 |
| NOP14-AS1 | 82.3435288 | -0.395086952 | 0.36035492 | -1.0963828 | 0.27291127 | 0.47075093 |
| NOP14 | 256.706066 | -0.225966827 | 0.39997494 | -0.5649525 | 0.5721061 | 0.73798779 |
| GRK4 | 253.221229 | 0.109133251 | 0.42884137 | 0.25448396 | 0.79912169 | 0.89075067 |
| HTT | 1204.83953 | -0.0060531 | 0.12763828 | -0.0474239 | 0.96217541 | 0.98034181 |
| RGS12 | 1268.37252 | -0.272722547 | 0.25161825 | -1.0838743 | 0.27842054 | 0.47668128 |
| LRPAP1 | 579.911543 | 0.2498651 | 0.27788376 | 0.89917129 | 0.36856143 | 0.56603603 |
| ADRA2C | 62.5359779 | 0.316120404 | 0.76104574 | 0.41537635 | 0.67786641 | 0.81346017 |
| TMEM128 | 135.891897 | 0.145041844 | 0.22897559 | 0.63343802 | 0.52644764 | 0.7009757 |
| LYAR | 53.728218 | -0.800847306 | 0.54670203 | -1.4648698 | 0.14295643 | 0.31053944 |
| ZBTB49 | 113.796658 | 0.035743756 | 0.22909138 | 0.15602401 | 0.8760141 | 0.93538406 |
| STX18 | 1547.11053 | -0.687996257 | 0.19920214 | -3.4537594 | 0.00055283 | 0.00574935 |
| STX18-AS1 | 488.121821 | 0.15269737 | 0.43070775 | 0.35452664 | 0.72294424 | 0.84233055 |
| STK32B | 1597.60838 | 0.721195551 | 0.36635643 | 1.96856254 | 0.04900335 | 0.15345874 |
| EVC2 | 843.002294 | -0.604291031 | 0.28037729 | -2.1552781 | 0.03114008 | 0.11211362 |
| EVC | 2389.66225 | -0.619499893 | 0.50482865 | -1.2271488 | 0.21976666 | 0.41140966 |
| CRMP1 | 517.139839 | -1.059645898 | 0.55057361 | -1.9246217 | 0.05427671 | 0.16383413 |
| WFS1 | 56.5003954 | -0.080601063 | 0.36635561 | -0.2200077 | 0.82586514 | 0.9066142 |
| MAN2B2 | 150.350899 | -0.111632385 | 0.34025593 | -0.3280836 | 0.74284847 | 0.8540591 |
| MRFAP1 | 371.985192 | -1.093637508 | 0.35620244 | -3.0702696 | 0.00213866 | 0.01618695 |
| LINC02482 | 64.6039078 | 0.519109348 | 0.38962473 | 1.33233162 | 0.18275124 | 0.36499654 |
| AC093323.1 | 47.2738993 | 0.185764643 | 0.38969699 | 0.47668997 | 0.63358291 | 0.78395238 |
| MRFAP1L1 | 138.466178 | -0.093114921 | 0.26937706 | -0.3456676 | 0.72959253 | 0.84639784 |
| BLOC1S4 | 113.994982 | -1.153349073 | 0.35893131 | -3.2132863 | 0.00131225 | 0.01109628 |
| KIAA0232 | 2067.94949 | 0.072954477 | 0.25719944 | 0.28364944 | 0.77667904 | 0.87664193 |
| TBC1D14 | 2255.61679 | 0.12394481 | 0.33255282 | 0.37270714 | 0.70936642 | 0.83354355 |
| TADA2B | 152.272492 | -0.245737727 | 0.44073176 | -0.5575676 | 0.57713972 | 0.74138147 |
| GRPEL1 | 307.192584 | -1.14909558 | 0.37811204 | -3.0390346 | 0.00237338 | 0.01751851 |
| LINC02447 | 42.995618 | 1.489728112 | 0.70940664 | 2.09996357 | 0.03573205 | 0.12342541 |
| SORCS2 | 609.758873 | 0.074584726 | 0.82265111 | 0.09066386 | 0.92775969 | 0.96193661 |
| AFAP1 | 419.084895 | -0.808090845 | 0.31527386 | -2.5631394 | 0.01037304 | 0.0518386 |
| ABLIM2 | 553.952309 | -2.670761128 | 0.35482829 | -7.5269115 | 5.20E-14 | 2.09E-11 |
| SH3TC1 | 821.411894 | 0.301077258 | 0.34875802 | 0.86328412 | 0.38798127 | 0.58387159 |
| AC104825.1 | 41.4688355 | 0.683727032 | 0.36976908 | 1.84906493 | 0.06444844 | 0.18381081 |
| ACOX3 | 318.075127 | -0.056044495 | 0.20846073 | -0.2688492 | 0.78804575 | 0.88421655 |
| TRMT44 | 252.295614 | -0.007324494 | 0.25840578 | -0.0283449 | 0.97738704 | 0.98871175 |
| SLC2A9 | 6259.52001 | 0.770807586 | 0.46578114 | 1.65487076 | 0.09795072 | 0.24217361 |
| WDR1 | 569.759496 | -0.53487953 | 0.29446522 | -1.8164438 | 0.06930232 | 0.19283941 |
| ZNF518B | 58.2993923 | -0.135269722 | 0.35969221 | -0.3760708 | 0.70686429 | 0.83210663 |
| CLNK | 39.3952987 | 0.364102651 | 0.39585607 | 0.91978545 | 0.35768489 | 0.55624991 |
| HS3ST1 | 795.49822 | -2.451299553 | 0.77948743 | -3.1447583 | 0.00166224 | 0.01329308 |
| AC005699.1 | 3240.51175 | 0.054866536 | 0.7892054 | 0.06952124 | 0.94457473 | 0.9715808 |
| LINC02360 | 30.7802109 | -1.277662023 | 0.73424167 | -1.740111 | 0.08183952 | 0.21535832 |
| RAB28 | 1350.73029 | 0.284815497 | 0.17021654 | 1.67325397 | 0.09427731 | 0.23650422 |
| BOD1L1 | 1485.01877 | -0.897748699 | 0.5288265 | -1.6976243 | 0.0895787 | 0.22831075 |
| LINC01182 | 325.23723 | -0.811265806 | 0.45939392 | -1.7659481 | 0.07740456 | 0.20754779 |
| LINC00504 | 243.145526 | 0.812553688 | 0.4836795 | 1.67994238 | 0.09296853 | 0.23469615 |
| CPEB2-DT | 372.82197 | 1.186189802 | 0.45353075 | 2.61545615 | 0.00891084 | 0.04635326 |
| CPEB2 | 2770.85158 | -0.046752357 | 0.29667806 | -0.1575862 | 0.87478291 | 0.93490877 |
| AC098829.1 | 2783.03107 | 2.741356268 | 0.5415137 | 5.06239505 | 4.14E-07 | 1.82E-05 |
| C1QTNF7 | 240.532249 | 2.782370067 | 0.77612862 | 3.58493423 | 0.00033716 | 0.00402007 |
| CC2D2A | 799.371917 | 0.803822389 | 0.4167477 | 1.92879863 | 0.05375587 | 0.16276378 |
| FBXL5 | 1949.99556 | 0.510224292 | 0.1888921 | 2.70114152 | 0.00691019 | 0.03855027 |
| FAM200B | 316.016418 | 0.324111079 | 0.3732006 | 0.86846345 | 0.38514067 | 0.58152841 |
| BST1 | 54.6643831 | 1.399056947 | 0.56610269 | 2.47138368 | 0.01345913 | 0.06211594 |
| PROM1 | 704.917036 | -0.478203599 | 0.502457 | -0.9517304 | 0.34123373 | 0.54195003 |
| TAPT1 | 875.778842 | 0.555618571 | 0.19498689 | 2.84951756 | 0.00437856 | 0.02746787 |
| TAPT1-AS1 | 1951.3141 | 1.599197182 | 0.59473377 | 2.68892952 | 0.00716816 | 0.03960254 |
| AC097515.1 | 371.405269 | 1.466402083 | 0.87791982 | 1.67031436 | 0.09485718 | 0.23735075 |
| LDB2 | 129.602536 | 1.11001985 | 0.41913306 | 2.648371 | 0.00808807 | 0.04330593 |
| QDPR | 546.132548 | 0.659067284 | 0.30138341 | 2.18680676 | 0.02875663 | 0.10602395 |
| LAP3 | 659.763612 | -0.622099842 | 0.7148677 | -0.8702307 | 0.38417433 | 0.58045387 |
| AC006160.1 | 115.686856 | -0.944098677 | 0.61063269 | -1.5460991 | 0.12208063 | 0.28034162 |
| MED28 | 391.688834 | -0.169663516 | 0.28869417 | -0.5876929 | 0.55673844 | 0.72562693 |
| FAM184B | 84.5379353 | -1.411356999 | 0.49658849 | -2.8421058 | 0.00448166 | 0.02791011 |
| DCAF16 | 278.320739 | 0.351661829 | 0.19084594 | 1.84264769 | 0.06538047 | 0.18561931 |
| LCORL | 1763.5313 | -0.094473429 | 0.18147633 | -0.5205827 | 0.60265753 | 0.76134809 |
| AC024230.1 | 203.702515 | -0.282002863 | 0.59159865 | -0.4766794 | 0.63359047 | 0.78395238 |
| SLIT2 | 5295.40395 | 1.550557278 | 0.50342174 | 3.08003641 | 0.00206975 | 0.01578748 |
| PACRGL | 628.295712 | 0.237061118 | 0.25102372 | 0.94437737 | 0.34497684 | 0.5454262 |
| KCNIP4 | 11765.2597 | -1.024633642 | 0.59850142 | -1.7119987 | 0.08689691 | 0.22405869 |
| AC096719.1 | 46.904406 | 0.406027996 | 0.56692787 | 0.71618987 | 0.47387412 | 0.66016796 |
| ADGRA3 | 1601.93997 | 0.071580067 | 0.18267545 | 0.39184285 | 0.69517434 | 0.82442931 |
| AC097512.1 | 106.698985 | 1.098303806 | 0.77590161 | 1.41551944 | 0.15691625 | 0.33084927 |
| AC092440.1 | 44.6248822 | -0.59917638 | 0.81568319 | -0.73457 | 0.46260143 | 0.65094732 |
| AC093607.1 | 421.026726 | 0.104754958 | 0.64538139 | 0.16231481 | 0.87105795 | 0.93338059 |
| PPARGC1A | 2992.55629 | 1.132376154 | 0.42124404 | 2.68817135 | 0.00718445 | 0.03967658 |
| DHX15 | 1173.4304 | -0.342984856 | 0.24428059 | -1.404061 | 0.16030068 | 0.33607241 |
| SOD3 | 209.243479 | -0.825691502 | 0.6391749 | -1.2918084 | 0.19642351 | 0.38211425 |
| CCDC149 | 293.775052 | -1.149072119 | 0.43029032 | -2.6704578 | 0.00757479 | 0.04118511 |
| SEPSECS | 700.173237 | 0.646648423 | 0.49819473 | 1.29798326 | 0.19429309 | 0.37933419 |
| AC104662.2 | 175.62672 | 0.245073733 | 0.51092956 | 0.47966246 | 0.63146742 | 0.78273865 |
| PI4K2B | 83.3046206 | -0.413531526 | 0.33607109 | -1.2304883 | 0.21851432 | 0.40973677 |
| ZCCHC4 | 266.61575 | -0.024114599 | 0.18366392 | -0.1312974 | 0.89554003 | 0.94595231 |
| ANAPC4 | 544.257767 | -0.05792894 | 0.22656597 | -0.2556824 | 0.79819606 | 0.89012761 |
| SLC34A2 | 1185.42705 | -3.740448125 | 0.87583562 | -4.2707194 | 1.95E-05 | 0.00043426 |
| SEL1L3 | 2347.32757 | -1.250728775 | 0.29186808 | -4.2852537 | 1.83E-05 | 0.00041355 |
| AC092436.4 | 35.0876815 | -2.257419726 | 0.53493992 | -4.21995 | 2.44E-05 | 0.00051521 |
| SMIM20 | 268.556926 | -1.028988615 | 0.41209818 | -2.4969502 | 0.01252666 | 0.059045 |
| RBPJ | 1801.78128 | -0.181586629 | 0.23858832 | -0.7610877 | 0.44660471 | 0.63757779 |
| TBC1D19 | 2629.63536 | 1.228161328 | 0.34158061 | 3.59552416 | 0.00032374 | 0.00390281 |
| STIM2 | 541.169151 | 0.231924611 | 0.22616003 | 1.02548894 | 0.30513254 | 0.50529195 |
| LINC02506 | 82.7947805 | -1.190539754 | 1.21073203 | -0.9833223 | 0.32544886 | 0.52575488 |
| LINC02484 | 73.5336419 | 3.813294607 | 0.70965728 | 5.37343128 | 7.73E-08 | 4.51E-06 |
| ARAP2 | 1161.61486 | -0.057172211 | 0.22074123 | -0.2590011 | 0.79563443 | 0.88843413 |
| LINC02505 | 94.53735 | -0.704776538 | 1.24732735 | -0.5650293 | 0.57205381 | 0.73798779 |
| C4orf19 | 3441.49791 | -0.086534852 | 0.32559361 | -0.2657756 | 0.790412 | 0.8860758 |
| RELL1 | 272.622364 | -1.808833147 | 0.34630049 | -5.2233052 | 1.76E-07 | 8.79E-06 |
| PGM2 | 136.829752 | -0.59980023 | 0.30928842 | -1.9392909 | 0.05246592 | 0.15984672 |
| TBC1D1 | 2319.34538 | -0.171943534 | 0.2215367 | -0.7761402 | 0.43766622 | 0.62956143 |
| KLF3 | 630.98763 | -0.264294035 | 0.26571556 | -0.9946502 | 0.31990643 | 0.520351 |
| TLR1 | 32.0848293 | -0.056324664 | 0.44348393 | -0.127005 | 0.89893646 | 0.94743919 |
| FAM114A1 | 332.971638 | -1.388570694 | 0.36332034 | -3.8218909 | 0.00013243 | 0.00191957 |
| KLHL5 | 670.495497 | 0.154198932 | 0.23414243 | 0.65856894 | 0.51017261 | 0.68954593 |
| WDR19 | 1034.53475 | -0.131741971 | 0.14030226 | -0.9389868 | 0.34773753 | 0.54767761 |
| RFC1 | 869.239845 | 0.020501176 | 0.25341848 | 0.0808985 | 0.93552267 | 0.96567582 |
| RPL9 | 151.845317 | -0.531349033 | 0.36840195 | -1.4423079 | 0.14921554 | 0.31943447 |
| LIAS | 293.209145 | 0.999986752 | 0.37897139 | 2.63868667 | 0.00832279 | 0.04416885 |

| | | | | | | |
|---|---|---|---|---|---|---|
| UGDH | 1301.8115 | -1.459410073 | 0.57108773 | -2.5554919 | 0.01060378 | 0.05265984 |
| UGDH-AS1 | 521.981037 | 0.840928137 | 0.37924596 | 2.21736876 | 0.0265979 | 0.10076768 |
| SMIM14 | 1167.32275 | 0.08352872 | 0.25906872 | 0.32241916 | 0.74713517 | 0.85712938 |
| AC108471.2 | 41.8942781 | 0.899853481 | 0.60862209 | 1.4785094 | 0.13927148 | 0.30543908 |
| UBE2K | 3131.90132 | -0.083923965 | 0.28473358 | -0.2947456 | 0.76818824 | 0.87071498 |
| PDS5A | 3218.14831 | 0.166542975 | 0.19819921 | 0.84028072 | 0.40075101 | 0.5949339 |
| N4BP2 | 494.36348 | -0.218731781 | 0.24536885 | -0.8914407 | 0.3726928 | 0.57001787 |
| RBM47 | 12402.0157 | 0.222343468 | 0.19818002 | 1.12192677 | 0.26189358 | 0.45999812 |
| NSUN7 | 723.598343 | 0.449109629 | 0.42069906 | 1.06753181 | 0.28573176 | 0.48536166 |
| APBB2 | 2136.9911 | 0.637970677 | 0.23044278 | 2.7684559 | 0.00563226 | 0.03312299 |
| UCHL1 | 142.908161 | -2.795662639 | 0.52737441 | -5.3010965 | 1.15E-07 | 6.21E-06 |
| LIMCH1 | 2837.18871 | -0.140268148 | 0.36614142 | -0.3830983 | 0.7016469 | 0.82861543 |
| TMEM33 | 516.301985 | -0.286534011 | 0.37315039 | -0.7678781 | 0.44255961 | 0.63420435 |
| SLC30A9 | 1971.46987 | 0.342218005 | 0.16954051 | 2.01850286 | 0.04353892 | 0.14131848 |
| ATP8A1 | 346.989391 | 0.079970929 | 0.39027713 | 0.20490806 | 0.83764396 | 0.91436098 |
| KCTD8 | 1233.42158 | 0.455572279 | 0.33216564 | 1.37152138 | 0.17021248 | 0.34924222 |
| GUF1 | 285.441732 | 0.442565813 | 0.21660533 | 2.04318983 | 0.04103365 | 0.13582426 |
| GNPDA2 | 236.08104 | 0.081622112 | 0.20156276 | 0.4049464 | 0.68551691 | 0.81807246 |
| COMMD8 | 123.164632 | -0.231051245 | 0.35725519 | -0.6467401 | 0.51780019 | 0.69447038 |
| ATP10D | 1006.73935 | 0.552568764 | 0.53895762 | 1.02525456 | 0.30524309 | 0.50531003 |
| CORIN | 42.4008055 | -0.623659953 | 0.59915831 | -1.0408934 | 0.29792501 | 0.49825013 |
| AC107068.2 | 102.197452 | -1.165762786 | 0.54750654 | -2.1292217 | 0.03323593 | 0.1176866 |
| NFXL1 | 418.909048 | -0.326187441 | 0.210499 | -1.5495914 | 0.12123963 | 0.27925249 |
| NIPAL1 | 154.011401 | -0.56434596 | 0.31749899 | -1.7774733 | 0.07549041 | 0.20417265 |
| TXK | 611.627479 | 0.011043209 | 0.36041506 | 0.03064025 | 0.97555644 | 0.98773468 |
| TEC | 908.230776 | -0.397082095 | 0.20303889 | -1.9556948 | 0.05050112 | 0.15643364 |
| SLAIN2 | 896.949156 | -0.012542825 | 0.20182329 | -0.0621476 | 0.95044532 | 0.97465176 |
| FRYL | 3419.06573 | -0.150417517 | 0.18105856 | -0.8307672 | 0.40610515 | 0.60009523 |
| OCIAD1 | 1381.26308 | 0.195582593 | 0.12590335 | 1.55343438 | 0.1203194 | 0.27769287 |
| OCIAD2 | 1125.83086 | -0.084924088 | 0.43649555 | -0.1945589 | 0.84573829 | 0.91865716 |
| CWH43 | 51.1999199 | 1.259627489 | 1.07996959 | 1.16635459 | 0.24347111 | 0.43865122 |
| DCUN1D4 | 2245.31548 | -0.117706605 | 0.23766422 | -0.4952643 | 0.62041356 | 0.77455476 |
| LRRC66 | 173.041278 | 0.717484525 | 0.36922053 | 1.94324112 | 0.05198703 | 0.15901947 |
| SGCB | 406.947325 | 0.069686533 | 0.26984918 | 0.25824253 | 0.79621974 | 0.88867422 |
| SPATA18 | 333.908287 | 0.209633949 | 0.49713405 | 0.42168495 | 0.67325499 | 0.81072605 |
| USP46 | 325.037199 | -0.194561568 | 0.29352431 | -0.6628465 | 0.50742885 | 0.68742116 |
| USP46-AS1 | 34.2383204 | -0.125295675 | 0.41422167 | -0.3024846 | 0.76228268 | 0.86753642 |
| SCFD2 | 1900.98593 | 0.183748678 | 0.49685619 | 0.36982266 | 0.71151463 | 0.83506559 |
| AC058822.1 | 575.652969 | -0.762241866 | 0.46835749 | -1.6274788 | 0.10363546 | 0.25157313 |
| LNX1 | 273.569178 | -0.395852867 | 0.51677415 | -0.7660075 | 0.44367185 | 0.63517875 |
| CHIC2 | 1575.82581 | -0.224932917 | 0.21453916 | -1.0484469 | 0.29443276 | 0.49433977 |
| SRD5A3 | 57.910255 | -0.05800044 | 0.4434673 | -0.1307885 | 0.89594259 | 0.9460654 |
| TMEM165 | 1758.89745 | -0.693254103 | 0.17413324 | -3.9811705 | 6.86E-05 | 0.00115761 |
| CLOCK | 1346.42109 | 0.095084171 | 0.13928121 | 0.68267768 | 0.49481054 | 0.6769747 |
| EXOC1 | 1443.58111 | 0.108242143 | 0.1955716 | 0.55346556 | 0.57994465 | 0.74348124 |
| CEP135 | 933.848071 | 1.341468321 | 0.27779461 | 4.82899334 | 1.37E-06 | 4.99E-05 |
| KIAA1211 | 159.51755 | -0.676166059 | 0.6569693 | -1.0292202 | 0.30337622 | 0.50367949 |
| AASDH | 530.893636 | 0.419368783 | 0.35001332 | 1.19815091 | 0.23085827 | 0.42378291 |
| PPAT | 220.291261 | -0.250986658 | 0.45018261 | -0.5575219 | 0.57717091 | 0.74138147 |
| PAICS | 258.766874 | -0.319379049 | 0.24792945 | -1.2881852 | 0.1976815 | 0.38390766 |
| SRP72 | 544.496663 | -0.336387984 | 0.2374105 | -1.4169044 | 0.15651088 | 0.33019688 |
| THEGL | 36.8261954 | 1.799929898 | 0.42358575 | 4.24926924 | 2.14E-05 | 0.00046588 |
| REST | 665.83206 | -0.054531285 | 0.26105814 | -0.2088856 | 0.83453755 | 0.91256531 |
| NOA1 | 119.639733 | -0.21991702 | 0.31733691 | -0.693008 | 0.48830453 | 0.671495 |
| POLR2B | 833.940299 | -0.24909188 | 0.30014374 | -0.8299086 | 0.40659044 | 0.60035928 |
| IGFBP7 | 3061.5653 | -0.808670684 | 0.3431007 | -2.3569485 | 0.0184258 | 0.07786855 |
| AC093725.2 | 131.641391 | 2.541098124 | 0.84393838 | 3.01099959 | 0.00260389 | 0.01872287 |
| ADGRL3 | 77.8469524 | 1.261217311 | 0.49902055 | 2.52738551 | 0.01149153 | 0.05558791 |
| AC116049.2 | 420.829237 | 0.717165501 | 0.6081561 | 1.17924576 | 0.23830033 | 0.43332218 |
| AC104806.2 | 217.282778 | 0.850761188 | 0.47688744 | 1.7839874 | 0.07442571 | 0.20232093 |
| CENPC | 630.634926 | 0.660022649 | 0.29860765 | 2.210334 | 0.02708199 | 0.1019347 |
| UBA6 | 1419.17843 | -0.876016073 | 0.21761831 | -4.0254705 | 5.69E-05 | 0.00100568 |
| UBA6-AS1 | 2649.63999 | 0.605301739 | 0.54268038 | 1.1153927 | 0.26468218 | 0.4624535 |
| TMPRSS11D | 38.7569161 | -0.983864278 | 0.87407811 | -1.1256022 | 0.26033392 | 0.45866286 |
| YTHDC1 | 1551.42254 | -0.025032594 | 0.20366391 | -0.1229113 | 0.90217734 | 0.94861498 |
| UGT2A3 | 1314.29841 | 0.505722414 | 0.5556637 | 0.91012318 | 0.36275755 | 0.56045575 |
| UGT2B7 | 9519.49082 | -0.369622321 | 0.49074191 | -0.7531909 | 0.45133523 | 0.64108831 |
| AC111000.4 | 1094.71389 | 0.78450419 | 0.63969167 | 1.22637863 | 0.22005623 | 0.41183925 |
| ENAM | 54.3619782 | -0.191125175 | 0.48305139 | -0.3956622 | 0.69235424 | 0.82270698 |
| UTP3 | 223.618386 | -0.755344539 | 0.29392495 | -2.5698551 | 0.0101741 | 0.05114967 |
| RUFY3 | 8565.02741 | -0.191486339 | 0.25935659 | -0.738313 | 0.46032428 | 0.64907266 |
| GRSF1 | 583.243243 | -0.277588049 | 0.17819133 | -1.5578089 | 0.11927855 | 0.27622083 |
| MOB1B | 1472.80915 | 0.206346355 | 0.23500035 | 0.87806828 | 0.37990666 | 0.57641971 |
| SLC4A4 | 11456.5227 | 1.426087245 | 0.52014327 | 2.74172008 | 0.00611184 | 0.0350523 |
| GC | 253.44758 | 2.609502204 | 1.09371187 | 2.38591377 | 0.01703675 | 0.07375953 |
| ADAMTS3 | 206.098305 | -0.947659313 | 0.52373008 | -1.8094422 | 0.07038233 | 0.19497385 |
| COX18 | 116.132127 | -0.021247889 | 0.27388788 | -0.0775788 | 0.93816312 | 0.96676805 |
| ANKRD17 | 6456.05823 | 0.24638349 | 0.24377236 | 1.01071134 | 0.31215461 | 0.51235819 |
| AC053527.1 | 953.345462 | 0.764309984 | 0.60012237 | 1.27359023 | 0.20280868 | 0.39065466 |
| ALB | 135.243771 | 1.558015165 | 1.4771918 | 1.0547142 | 0.29155605 | 0.4916955 |
| AFM | 1164.15911 | 6.630795968 | 0.74553103 | 8.89405765 | 5.89E-19 | 8.08E-16 |
| LINC02499 | 995.674344 | 6.528660937 | 0.99311305 | 6.57393533 | 4.90E-11 | 8.72E-09 |
| RASSF6 | 1852.95854 | -1.716057058 | 0.32393623 | -5.297515 | 1.17E-07 | 6.31E-06 |
| CXCL1 | 93.2469523 | -2.891191403 | 0.9965168 | -2.9012972 | 0.00371621 | 0.02419863 |
| CXCL2 | 194.534403 | -3.723283534 | 0.64168619 | -5.8023433 | 6.54E-09 | 5.71E-07 |
| MTHFD2L | 652.24343 | -0.57018558 | 0.24012239 | -2.3745623 | 0.01756977 | 0.07530608 |
| BTC | 91.2417506 | -0.243938459 | 0.64941928 | -0.3756255 | 0.70719531 | 0.83227665 |
| PARM1 | 1258.65148 | 1.373063422 | 0.40846255 | 3.36154058 | 0.00077509 | 0.00750824 |
| AC110760.1 | 48.2624824 | 1.703298684 | 0.81704368 | 2.08470945 | 0.03709568 | 0.12683226 |
| RCHY1 | 465.099074 | 0.20806589 | 0.17918238 | 1.16119615 | 0.24556214 | 0.44114289 |
| THAP6 | 314.46466 | 0.172844595 | 0.19327903 | 0.89427493 | 0.37117481 | 0.56877072 |
| CDKL2 | 991.664825 | 0.745516254 | 0.49821525 | 1.49637383 | 0.13455627 | 0.29897272 |
| G3BP2 | 837.469225 | -0.149977381 | 0.17514258 | -0.8563159 | 0.39182305 | 0.5869326 |
| USO1 | 1499.98906 | -0.257899722 | 0.21242975 | -1.2140471 | 0.22472975 | 0.416886 |
| NAAA | 143.874257 | 0.288024248 | 0.25382121 | 1.13475246 | 0.25647905 | 0.45375428 |
| SDAD1 | 567.103231 | 0.075812632 | 0.24363517 | 0.31117277 | 0.75566928 | 0.8629475 |
| NUP54 | 602.085393 | -0.085189479 | 0.15858102 | -0.5371984 | 0.59113055 | 0.75268244 |
| SCARB2 | 1089.40453 | -0.786195281 | 0.21879778 | -3.5932507 | 0.00032658 | 0.0039199 |
| FAM47E-STB | 41.8563274 | 0.78079767 | 0.50099619 | 1.55849024 | 0.11911708 | 0.2759371 |
| CCDC158 | 232.76111 | 0.676981901 | 0.66520679 | 1.01770142 | 0.30881987 | 0.50906843 |
| SHROOM3 | 2162.17475 | 1.25734614 | 0.51089139 | 2.46108304 | 0.01385183 | 0.06339467 |
| 11-Sep | 2219.83032 | -0.53396179 | 0.35051832 | -1.5233492 | 0.12767138 | 0.28906368 |
| CCNI | 1418.43146 | -0.316289786 | 0.23050487 | -1.372161 | 0.17001333 | 0.34901494 |
| CCNG2 | 649.151776 | 0.062792199 | 0.44946152 | 0.13970539 | 0.88889276 | 0.94267336 |
| CXCL13 | 46.8262897 | 0.941560134 | 0.58829032 | 1.60050252 | 0.10948715 | 0.26077541 |
| CNOT6L | 965.004862 | -0.362305208 | 0.23093203 | -1.5688825 | 0.11667533 | 0.27215379 |
| MRPL1 | 593.221703 | 0.37657685 | 0.15263917 | 2.46710502 | 0.01362104 | 0.06265221 |
| FRAS1 | 3560.89671 | 1.137299042 | 0.41561568 | 2.73641997 | 0.00621117 | 0.03551794 |
| ANXA3 | 110.527486 | -1.228562712 | 0.37053799 | -3.3156187 | 0.0009144 | 0.00845732 |
| LINC01094 | 55.480208 | -3.611379641 | 0.83163442 | -4.3425086 | 1.41E-05 | 0.00033405 |
| BMP2K | 2948.10431 | -0.574224056 | 0.28201669 | -2.036135 | 0.0417368 | 0.13739993 |
| PAQR3 | 183.846864 | -1.502958586 | 0.38586424 | -3.8950451 | 9.82E-05 | 0.00153276 |
| LINC01088 | 108.831537 | 0.584011589 | 0.56721723 | 1.02960834 | 0.3031939 | 0.50355977 |

| | | | | | | |
|---|---|---|---|---|---|---|
| ANTXR2 | 302.186618 | 0.749045217 | 0.3685139 | 2.03261047 | 0.04209189 | 0.13812103 |
| FGF5 | 116.17945 | -0.028790316 | 0.71463177 | -0.0402869 | 0.96786438 | 0.98372512 |
| CFAP299 | 539.960972 | 0.155886596 | 0.6799065 | 0.22927652 | 0.818654 | 0.90192453 |
| PRKG2 | 178.68665 | -3.612794934 | 0.67504017 | -5.3519703 | 8.70E-08 | 4.87E-06 |
| RASGEF1B | 5671.61988 | 0.46017418 | 0.5428302 | 0.84773135 | 0.3965876 | 0.59132234 |
| HNRNPD | 1500.28582 | -0.286800815 | 0.20877034 | -1.3737622 | 0.16951551 | 0.34855751 |
| AC124016.1 | 39.342769 | -0.260906234 | 0.46278763 | -0.563771 | 0.57291001 | 0.73835552 |
| HNRNPDL | 1406.44702 | -0.299324967 | 0.19959112 | -1.4996908 | 0.13369451 | 0.29763693 |
| ENOPH1 | 275.029267 | -0.588606296 | 0.28922286 | -2.0351306 | 0.04183774 | 0.13765274 |
| TMEM150C | 474.537504 | 0.242350181 | 0.54178204 | 0.44732044 | 0.65464371 | 0.79832752 |
| SCD5 | 575.610474 | 1.475569783 | 0.39426361 | 3.74259692 | 0.00018213 | 0.00247416 |
| SEC31A | 3466.10847 | -0.351062255 | 0.25257351 | -1.3899409 | 0.16454682 | 0.34163031 |
| THAP9 | 138.829544 | -0.184806526 | 0.62554218 | -0.2954342 | 0.76766225 | 0.87040668 |
| LIN54 | 608.072574 | -0.723253717 | 0.35964128 | -2.011042 | 0.04432102 | 0.14321269 |
| COPS4 | 299.135466 | 0.354111395 | 0.16794515 | 2.10849439 | 0.03498825 | 0.12170384 |
| COQ2 | 39.9293923 | 0.224887945 | 0.35901325 | 0.6264057 | 0.53104887 | 0.70485977 |
| HELQ | 436.946579 | 0.46193946 | 0.24319002 | 1.8995001 | 0.05749875 | 0.17011341 |
| MRPS18C | 181.905096 | -0.339167247 | 0.2975415 | -1.139899 | 0.2543284 | 0.45144773 |
| ABRAXAS1 | 145.757278 | 0.135725178 | 0.31696891 | 0.42819713 | 0.6685076 | 0.80729015 |
| GPAT3 | 4296.28069 | 1.874485013 | 0.53763897 | 3.48651257 | 0.00048936 | 0.00527749 |
| CDS1 | 526.927483 | -0.494982677 | 0.29922715 | -1.6542038 | 0.09808612 | 0.24237722 |
| WDFY3 | 9226.06813 | 1.161284637 | 0.29454262 | 3.942671 | 8.06E-05 | 0.00131406 |
| WDFY3-AS1 | 52.4251733 | -0.526183247 | 0.70457904 | -0.7468051 | 0.45518119 | 0.64460881 |
| ARHGAP24 | 4161.25654 | 1.653880545 | 0.43223382 | 3.82635616 | 0.00013005 | 0.00190658 |
| MAPK10 | 5729.03379 | 1.007580606 | 0.55513993 | 1.81500294 | 0.06952346 | 0.19318022 |
| AC104827.1 | 93.534376 | 0.49071259 | 0.631197 | 0.77743175 | 0.43690409 | 0.62879508 |
| PTPN13 | 2478.83265 | 0.429184424 | 0.336228 | 1.27646843 | 0.20178998 | 0.38918465 |
| AC093827.5 | 231.705987 | -0.118530968 | 0.578707 | -0.2048203 | 0.83771249 | 0.91436098 |
| AFF1 | 8850.28798 | 0.204640761 | 0.39761156 | 0.51467507 | 0.6067801 | 0.7638107 |
| KLHL8 | 653.232386 | 0.536223653 | 0.11509613 | 4.65891976 | 3.18E-06 | 9.99E-05 |
| HSD17B11 | 403.375223 | 0.266568604 | 0.34946156 | 0.76279808 | 0.44558383 | 0.63688501 |
| NUDT9 | 631.86611 | 0.173019835 | 0.21138223 | 0.81851647 | 0.41306234 | 0.60684304 |
| SPARCL1 | 114.638545 | 1.508332451 | 0.52662058 | 2.864173 | 0.004181 | 0.02648356 |
| SPP1 | 39409.4048 | -1.983030663 | 0.65766355 | -3.0152662 | 0.00256754 | 0.01855205 |
| PKD2 | 2924.60436 | -0.060275453 | 0.24554799 | -0.2454732 | 0.80609005 | 0.89466205 |
| ABCG2 | 84.7608591 | 1.047921727 | 0.79546224 | 1.31737457 | 0.1877131 | 0.37136212 |
| PPM1K | 558.748865 | 0.619528654 | 0.41416776 | 1.49583988 | 0.13469539 | 0.29911927 |
| HERC6 | 228.557337 | -1.034574226 | 0.88765794 | -1.16551 | 0.24381261 | 0.43900266 |
| HERC5 | 280.860956 | -1.00355915 | 0.67485797 | -1.4870672 | 0.13699707 | 0.30219163 |
| PYURF | 135.34643 | -0.612405282 | 0.29919112 | -2.0468698 | 0.04067087 | 0.13494931 |
| AC098582.1 | 47.0440055 | -0.894692095 | 0.78653226 | -1.1375148 | 0.25532314 | 0.45232479 |
| HERC3 | 989.530917 | -0.239110071 | 0.36754828 | -0.6505542 | 0.51533432 | 0.69315508 |
| FAM13A-AS1 | 403.027714 | 0.065643117 | 0.25051097 | 0.26203689 | 0.793293 | 0.88726528 |
| FAM13A | 3347.54177 | -0.013828977 | 0.31498075 | -0.0439042 | 0.96498077 | 0.98181349 |
| TIGD2 | 72.9569896 | 0.275721342 | 0.26927551 | 1.02393767 | 0.30586472 | 0.50583258 |
| GPRIN3 | 279.203272 | -0.991911854 | 0.65990813 | -1.503106 | 0.13281172 | 0.29622814 |
| CCSER1 | 6726.41308 | 0.841904126 | 0.32708614 | 2.57395229 | 0.01005441 | 0.05066759 |
| GRID2 | 55.7428825 | 0.760571487 | 0.34754944 | 2.18838356 | 0.02864167 | 0.10570582 |
| SMARCAD1 | 910.446061 | 0.335267105 | 0.1723008 | 1.94582439 | 0.05167584 | 0.15855449 |
| PDLIM5 | 4042.90279 | -0.394663276 | 0.1941185 | -2.0331049 | 0.04204193 | 0.13799537 |
| BMPR1B | 117.420377 | 0.081259918 | 0.54647941 | 0.14869713 | 0.88179263 | 0.93867887 |
| STPG2 | 3280.97694 | 2.071636227 | 0.5293564 | 3.91349994 | 9.10E-05 | 0.00145666 |
| STPG2-AS1 | 58.2804961 | 0.861957699 | 0.83629777 | 1.03068277 | 0.30268961 | 0.50284408 |
| RAP1GDS1 | 1545.8566 | -0.416451638 | 0.26574741 | -1.5670957 | 0.11709231 | 0.27273936 |
| TSPAN5 | 73.414871 | -0.644294186 | 0.47623396 | -1.3528942 | 0.17608942 | 0.35636464 |
| EIF4E | 928.056239 | -0.745396826 | 0.49833554 | -1.495773 | 0.13471283 | 0.29911927 |
| AC019131.1 | 50.4982658 | -0.569957627 | 0.77852007 | -0.732104 | 0.46410511 | 0.65185867 |
| METAP1 | 527.126318 | 0.026442283 | 0.17921351 | 0.14754626 | 0.88270087 | 0.93903973 |
| ADH5 | 635.305351 | 0.251279569 | 0.22692937 | 1.10730297 | 0.26816296 | 0.46607875 |
| AP002026.1 | 133.560012 | 1.152950276 | 0.56283048 | 2.04848586 | 0.04051242 | 0.13458091 |
| ADH6 | 371.876398 | 2.298448207 | 0.75809112 | 3.0318891 | 0.00243028 | 0.01782481 |
| TRMT10A | 135.530473 | 0.378843462 | 0.32985455 | 1.1485167 | 0.25075532 | 0.44706631 |
| MTTP | 240.232308 | 2.180355967 | 0.73004994 | 2.98658467 | 0.00282113 | 0.0198014 |
| LAMTOR3 | 252.261788 | -0.148026129 | 0.28633474 | -0.5169688 | 0.60517793 | 0.76287349 |
| DNAJB14 | 1198.61093 | -0.133220686 | 0.13266282 | -1.0042051 | 0.31527974 | 0.51532786 |
| H2AFZ | 365.420887 | -1.262500438 | 0.55007541 | -2.2951407 | 0.02172506 | 0.08763551 |
| DDIT4L | 163.647417 | -2.019169755 | 1.08771021 | -1.856349 | 0.06340383 | 0.18169208 |
| EMCN | 97.8868851 | 1.475949017 | 0.49231914 | 2.99795172 | 0.00271801 | 0.01926356 |
| PPP3CA | 4044.15392 | -0.007449401 | 0.22601518 | -0.0329597 | 0.97370669 | 0.98680944 |
| BANK1 | 622.005683 | 0.59641391 | 0.70961062 | 0.84048053 | 0.40063902 | 0.59483958 |
| AP002075.1 | 144.091213 | 0.310209559 | 0.61464582 | 0.50469644 | 0.61377206 | 0.76957315 |
| SLC39A8 | 684.521112 | 0.609256578 | 0.44853376 | 1.35832938 | 0.17435919 | 0.35411786 |
| NFKB1 | 1518.6112 | -0.572898106 | 0.2668066 | -2.1472411 | 0.03177409 | 0.11380389 |
| MANBA | 1072.51583 | -0.469021404 | 0.2476003 | -1.8942683 | 0.0581894 | 0.1712176 |
| UBE2D3 | 3126.61103 | -0.512439842 | 0.21246746 | -2.411851 | 0.01587177 | 0.0700658 |
| AC018797.2 | 92.0627741 | 0.071512025 | 0.32754358 | 0.21832827 | 0.82717335 | 0.90755343 |
| CISD2 | 446.503297 | -0.756444933 | 0.29293908 | -2.5822603 | 0.00981555 | 0.04975658 |
| SLC9B1 | 104.074225 | -0.475197137 | 0.4840472 | -0.9817165 | 0.32623952 | 0.52644376 |
| SLC9B2 | 192.594144 | -0.459322573 | 0.31578011 | -1.4545646 | 0.14578984 | 0.31445182 |
| BDH2 | 312.252825 | 1.148286193 | 0.57309418 | 2.00366055 | 0.04510644 | 0.1451688 |
| CXXC4 | 35.7084358 | 1.114417242 | 0.84549718 | 1.31806145 | 0.18748308 | 0.37109467 |
| CXXC4-AS1 | 119.013613 | -0.672474886 | 0.95238644 | -0.7060946 | 0.48012933 | 0.66519782 |
| AC004053.1 | 2800.43719 | 1.064027203 | 0.75144152 | 1.41598139 | 0.15678095 | 0.33061485 |
| AC096577.1 | 6024.61401 | 1.383727393 | 0.93951488 | 1.47281051 | 0.14080214 | 0.30766378 |
| TET2 | 2017.41333 | 0.609211388 | 0.40533565 | 1.50298002 | 0.1328442 | 0.29622814 |
| PPA2 | 1946.16935 | 0.165760814 | 0.2573597 | 0.64408223 | 0.51952211 | 0.69597507 |
| ARHGEF38 | 647.416733 | -2.501974009 | 0.66954126 | -3.7368481 | 0.00018634 | 0.00251643 |
| INTS12 | 375.523389 | -0.449824343 | 0.17708806 | -2.5401167 | 0.01108155 | 0.05424814 |
| GSTCD | 591.678746 | -0.221844348 | 0.21848732 | -1.0153649 | 0.30993194 | 0.51023746 |
| NPNT | 427.985326 | -1.313054255 | 0.72897804 | -1.8012261 | 0.07166726 | 0.19725764 |
| AC109361.1 | 40.1774076 | 0.092216792 | 0.55926282 | 0.1648899 | 0.86903064 | 0.9321389 |
| TBCK | 3072.46916 | 0.561091429 | 0.42538024 | 1.31903499 | 0.18715741 | 0.37066416 |
| AIMP1 | 470.735888 | -0.157938389 | 0.1814753 | -0.8703024 | 0.38413516 | 0.58045387 |
| PAPSS1 | 464.439752 | -0.544799827 | 0.28270571 | -1.9270917 | 0.05396821 | 0.16308249 |
| SGMS2 | 1088.88152 | -0.644559406 | 0.45991043 | -1.401489 | 0.16106789 | 0.3371137 |
| AC096564.1 | 156.650781 | -1.063174393 | 0.54527165 | -1.9498068 | 0.05119915 | 0.15766945 |
| CYP2U1 | 39.8688468 | 0.107986716 | 0.41970518 | 0.25729183 | 0.7969535 | 0.88912922 |
| HADH | 417.852438 | 1.462939369 | 0.4090934 | 3.57605221 | 0.00034882 | 0.00412181 |
| RPL34-AS1 | 318.332864 | 1.014054944 | 0.40439256 | 2.50760039 | 0.0121554 | 0.05778572 |
| RPL34 | 893.500173 | -0.387996455 | 0.40949742 | -0.9474943 | 0.34338698 | 0.54383762 |
| OSTC | 461.791906 | -0.826107232 | 0.32230597 | -2.5631149 | 0.01037377 | 0.0518386 |
| COL25A1 | 59.4136845 | -0.781401296 | 0.34023223 | -2.2966704 | 0.02163758 | 0.08748858 |
| SEC24B-AS1 | 81.0027316 | 0.123445882 | 0.5794524 | 0.21303886 | 0.83129665 | 0.91080925 |
| SEC24B | 1706.9293 | -0.084237857 | 0.15040368 | -0.5600784 | 0.57542594 | 0.73990275 |
| MCUB | 138.041983 | -3.665310215 | 0.66458574 | -5.5151803 | 3.48E-08 | 2.38E-06 |
| CASP6 | 377.809469 | -0.083157254 | 0.39439402 | -0.2108482 | 0.83300576 | 0.91175754 |
| PLA2G12A | 403.656558 | -0.136413949 | 0.37988424 | -0.3590935 | 0.71952517 | 0.84029407 |
| AC126283.2 | 251.032467 | 0.781929168 | 0.66352877 | 1.17844048 | 0.23862105 | 0.4338025 |
| CFI | 35.2176447 | -0.119698982 | 0.65974639 | -0.1814318 | 0.85602865 | 0.92383161 |
| GAR1 | 171.286935 | -0.743526214 | 0.53098717 | -1.4002715 | 0.16143202 | 0.3376268 |
| RRH | 34.9301358 | 0.16139451 | 0.69162139 | 0.23335674 | 0.81548441 | 0.90018388 |
| EGF | 939.01532 | 0.106264127 | 0.59776128 | 0.17777018 | 0.85890347 | 0.92584688 |
| ELOVL6 | 778.801224 | -0.999355052 | 0.4139211 | -2.4143612 | 0.01576283 | 0.06974211 |

| | | | | | | |
|---|---|---|---|---|---|---|
| ENPEP | 5436.94734 | 1.744659343 | 0.50391868 | 3.46218428 | 0.00053581 | 0.00560455 |
| PANCR | 86.1170591 | 0.625706073 | 0.95565313 | 0.65474182 | 0.51263399 | 0.6911936 |
| PITX2 | 27.961719 | -0.247766333 | 0.84343909 | -0.2937572 | 0.76894342 | 0.87121073 |
| FAM241A | 175.533851 | -1.172199049 | 0.51268776 | -2.28638 | 0.02223204 | 0.08897147 |
| AP1AR | 922.910444 | -0.57210052 | 0.25472525 | -2.2459513 | 0.02470713 | 0.095502 |
| TIFA | 66.9766492 | -2.460641756 | 0.60414967 | -4.0729009 | 4.64E-05 | 0.00085427 |
| ALPK1 | 1470.09585 | -0.858018238 | 0.44387298 | -1.9330265 | 0.05323293 | 0.16146577 |
| ZGRF1 | 74.2542533 | 0.378190098 | 0.42706463 | 0.88555705 | 0.37585626 | 0.57255633 |
| LARP7 | 509.393792 | 0.290699413 | 0.1614489 | 1.8005661 | 0.0717713 | 0.19738547 |
| ANK2 | 3462.1345 | 1.47436284 | 0.64199511 | 2.29653282 | 0.02164544 | 0.08749454 |
| CAMK2D | 3057.39529 | -0.170516827 | 0.31508696 | -0.5411739 | 0.58838775 | 0.75058454 |
| ARSJ | 855.41168 | 0.337000344 | 0.33221942 | 1.01439086 | 0.31039629 | 0.51075653 |
| UGT8 | 277.98967 | -0.231828479 | 0.38772313 | -0.5979227 | 0.54989148 | 0.71955844 |
| SNHG8 | 268.138212 | -0.047800478 | 0.43599183 | -0.1096362 | 0.91269792 | 0.95360549 |
| PRSS12 | 75.8509212 | 1.375264232 | 0.72585234 | 1.89468871 | 0.05813365 | 0.17121573 |
| METTL14 | 377.003499 | 0.312814086 | 0.14715087 | 2.12580517 | 0.03351949 | 0.11836062 |
| SEC24D | 835.082337 | -2.432470209 | 0.34033231 | -7.1473385 | 8.85E-13 | 2.43E-10 |
| USP53 | 1910.52985 | -0.59267254 | 0.20146175 | -2.9418614 | 0.00326246 | 0.02200715 |
| C4orf3 | 633.011489 | -1.101978948 | 0.4314819 | -2.5539401 | 0.01065116 | 0.05278214 |
| PDE5A | 462.941452 | -0.757425477 | 0.45778525 | -1.6545432 | 0.09801719 | 0.24225057 |
| AC108866.1 | 318.831347 | 0.008312778 | 0.46574067 | 0.01784851 | 0.9857597 | 0.99299282 |
| PRDM5 | 94.6527862 | -0.89129978 | 0.75315311 | -1.1834244 | 0.23664098 | 0.43127748 |
| QRFPR | 127.58335 | -2.498289371 | 0.76327033 | -3.2731383 | 0.0010636 | 0.00946511 |
| ANXA5 | 1005.24684 | -2.091656781 | 0.48954166 | -4.2726839 | 1.93E-05 | 0.00043186 |
| EXOSC9 | 93.4491621 | 0.095740584 | 0.26955882 | 0.35517511 | 0.72245841 | 0.84228022 |
| BBS7 | 231.896495 | -0.482653944 | 0.25720993 | -1.8764981 | 0.06058692 | 0.17598324 |
| KIAA1109 | 2945.20335 | 0.329769767 | 0.27093852 | 1.21713872 | 0.22355144 | 0.41588214 |
| BBS12 | 117.233344 | -0.078948693 | 0.350924 | -0.2249738 | 0.82199968 | 0.9044834 |
| FGF2 | 110.16324 | -2.053531529 | 0.80184312 | -2.5610141 | 0.01043671 | 0.05209327 |
| NUDT6 | 104.768271 | 0.876229748 | 0.3581133 | 2.44679477 | 0.01441329 | 0.0651378 |
| SPATA5 | 1509.66249 | -0.046570809 | 0.37750436 | -0.123365 | 0.90181811 | 0.9485283 |
| SPRY1 | 125.340293 | -1.37115555 | 0.40835315 | -3.357769 | 0.00078574 | 0.00756332 |
| LINC01091 | 225.006667 | 1.012111208 | 0.66503914 | 1.52188217 | 0.12803861 | 0.28975156 |
| ANKRD50 | 267.183575 | -0.711559113 | 0.38052352 | -1.8699478 | 0.06149107 | 0.17770569 |
| FAT4 | 375.139639 | 0.691644611 | 0.33707138 | 2.05192328 | 0.04017712 | 0.13395957 |
| AC093772.1 | 72.9295402 | 0.76065803 | 1.01535936 | 0.74915154 | 0.45376587 | 0.64306956 |
| INTU | 641.64948 | 0.394405497 | 0.32422562 | 1.21645384 | 0.22381209 | 0.41608467 |
| HSPA4L | 1213.50785 | -0.246115606 | 0.49230208 | -0.499928 | 0.61712576 | 0.77215361 |
| MFSD8 | 569.988588 | 0.022860618 | 0.1500592 | 0.152344 | 0.87891562 | 0.93706833 |
| ABHD18 | 1507.26799 | 0.30007783 | 0.1272081 | 2.35895224 | 0.01832661 | 0.07760361 |
| LARP1B | 3106.58161 | 0.092626862 | 0.23636744 | 0.39187658 | 0.69514941 | 0.82442931 |
| PGRMC2 | 903.12024 | -0.14977213 | 0.16115857 | -0.9293463 | 0.35270963 | 0.55164205 |
| AC108062.1 | 1049.45611 | 0.96371675 | 0.30229708 | 3.18797903 | 0.00143271 | 0.0118875 |
| AC078850.2 | 279.923835 | 3.755862973 | 0.75458331 | 4.9774 | 6.44E-07 | 2.64E-05 |
| JADE1 | 912.114095 | 1.058904365 | 0.25062702 | 4.22502084 | 2.39E-05 | 0.00050858 |
| SCLT1 | 796.740237 | -0.183278867 | 0.27570462 | -0.6647653 | 0.50620062 | 0.68623201 |
| C4orf33 | 94.9402994 | 0.321289301 | 0.58173901 | 0.55229114 | 0.5807489 | 0.74402516 |
| SNHG27 | 65.3981764 | -0.169218601 | 0.85871857 | -0.1970594 | 0.84378102 | 0.91781904 |
| AC096711.2 | 665.969718 | -0.352925361 | 0.35070779 | -1.0063231 | 0.31426015 | 0.51482776 |
| SLC7A11 | 188.426371 | -6.359402582 | 1.06047533 | -5.9967473 | 2.01E-09 | 2.11E-07 |
| AC109927.1 | 101.226419 | -0.626578652 | 0.48750895 | -1.2852659 | 0.19869937 | 0.38501162 |
| NOCT | 536.314108 | -0.4001124 | 0.37942957 | -1.0545103 | 0.29164934 | 0.49171433 |
| ELF2 | 2751.77996 | -0.324311508 | 0.14588312 | -2.2230913 | 0.02620964 | 0.09965481 |
| NDUFC1 | 420.797727 | -0.104153398 | 0.32094106 | -0.324525 | 0.74554059 | 0.8560165 |
| NAA15 | 1181.3968 | -0.220487894 | 0.26519495 | -0.8314181 | 0.40573746 | 0.59987524 |
| AC097376.2 | 1692.79454 | -0.74082877 | 0.26760182 | -2.7683996 | 0.00563323 | 0.03312299 |
| RAB33B | 120.036232 | -0.25927932 | 0.24591712 | -1.0543362 | 0.29172903 | 0.49178819 |
| SETD7 | 260.377334 | -2.608210539 | 0.46580877 | -5.5993161 | 2.15E-08 | 1.58E-06 |
| MGST2 | 647.818489 | 0.792924008 | 0.39457377 | 2.00957101 | 0.04447661 | 0.14361388 |
| MAML3 | 1628.75552 | -0.43559337 | 0.48969459 | -0.8895205 | 0.37372342 | 0.571065 |
| SCOC | 587.71849 | -0.356900333 | 0.31096786 | -1.1477081 | 0.25108909 | 0.44731974 |
| SCOC-AS1 | 117.261285 | -0.287669668 | 0.52463612 | -0.5483223 | 0.58347064 | 0.74632624 |
| ELMOD2 | 370.740686 | 0.264071638 | 0.23586776 | 1.11957498 | 0.26289492 | 0.46086465 |
| TBC1D9 | 257.571517 | -1.013427339 | 0.3403765 | -2.9773717 | 0.00290731 | 0.02021748 |
| RNF150 | 478.087173 | 1.661560105 | 0.802729 | 2.06988921 | 0.03846272 | 0.13033585 |
| ZNF330 | 330.77925 | -0.101179117 | 0.27916027 | -0.362441 | 0.71702254 | 0.83851445 |
| AC107220.1 | 154.309589 | 4.437959059 | 0.86400365 | 5.13650501 | 2.80E-07 | 1.30E-05 |
| LINC02432 | 90.3756193 | 1.08686358 | 0.63592214 | 1.70911424 | 0.08742979 | 0.22509394 |
| IL15 | 782.020747 | -0.381827932 | 0.2619293 | -1.4577519 | 0.14490895 | 0.31304443 |
| INPP4B | 602.079774 | -1.227230219 | 0.5054458 | -2.4280154 | 0.0151817 | 0.06778356 |
| AC104596.1 | 137.021531 | 0.521573268 | 0.69252271 | 0.75314969 | 0.45135996 | 0.64108831 |
| USP38 | 396.323872 | -0.184672436 | 0.26665325 | -0.6925565 | 0.48858793 | 0.67177813 |
| GAB1 | 2533.39678 | -0.200728523 | 0.28796841 | -0.6970505 | 0.4857712 | 0.66931729 |
| SMARCA5 | 587.389987 | -0.36768727 | 0.17021409 | -2.1601459 | 0.03076138 | 0.111135 |
| GYPA | 63.5850798 | 2.116777668 | 0.72433263 | 2.92238341 | 0.00347364 | 0.02311317 |
| ANAPC10 | 1006.05025 | 0.334970474 | 0.12863923 | 2.60395279 | 0.00921554 | 0.04753847 |
| ABCE1 | 367.32604 | -0.454283937 | 0.27464212 | -1.6540942 | 0.09810837 | 0.24238851 |
| OTUD4 | 866.75638 | -0.1083879 | 0.12004419 | -0.9029 | 0.36657897 | 0.56400246 |
| SMAD1 | 728.503716 | -0.030970496 | 0.27964106 | -0.1107509 | 0.91181389 | 0.95319043 |
| MMAA | 308.585948 | 0.09739806 | 0.33042495 | 0.29476606 | 0.7681726 | 0.87071498 |
| C4orf51 | 59.8874105 | -0.271034897 | 0.70907737 | -0.382236 | 0.70228632 | 0.82899058 |
| ZNF827 | 2245.43009 | 0.430867437 | 0.38076592 | 1.13158088 | 0.25781068 | 0.45521203 |
| LSM6 | 187.054122 | 0.513607753 | 0.21277942 | 2.41380371 | 0.01578696 | 0.06977854 |
| SLC10A7 | 773.210434 | -0.332873134 | 0.21208473 | -1.569529 | 0.11652474 | 0.27204983 |
| TMEM184C | 364.247443 | -0.110695518 | 0.23052686 | -0.4801849 | 0.63109592 | 0.78256123 |
| PRMT9 | 476.878572 | 0.735797183 | 0.20888069 | 3.52257153 | 0.00042738 | 0.00479781 |
| ARHGAP10 | 3430.21794 | 0.507005112 | 0.20086345 | 2.52412831 | 0.01159856 | 0.055881 |
| NR3C2 | 2020.53878 | 0.831563639 | 0.39657085 | 2.09688544 | 0.03600372 | 0.12406855 |
| AC069272.1 | 46.3066823 | -0.403037901 | 0.44644602 | -0.9027696 | 0.36664819 | 0.56404564 |
| DCLK2 | 257.418404 | -1.237741599 | 0.5331145 | -2.3217181 | 0.02024812 | 0.08322068 |
| LRBA | 9843.65995 | 0.072883723 | 0.17377544 | 0.41941325 | 0.67491414 | 0.81142427 |
| RPS3A | 506.177448 | -0.578441474 | 0.41702299 | -1.3870733 | 0.1654194 | 0.342767 |
| SH3D19 | 4382.41339 | -0.099882774 | 0.43547908 | -0.229363 | 0.81858682 | 0.90192453 |
| PRSS48 | 114.480359 | -0.951751231 | 0.52749433 | -1.8042871 | 0.07118631 | 0.19638678 |
| FAM160A1 | 3392.0811 | -0.384837961 | 0.24876315 | -1.5470055 | 0.12186193 | 0.28016797 |
| AC092611.2 | 49.5143928 | -0.321938107 | 0.40601672 | -0.7929183 | 0.4278254 | 0.62087906 |
| GATB | 272.00636 | 0.229098586 | 0.2425511 | 0.94453741 | 0.34489509 | 0.54539422 |
| AC079340.1 | 51.3730877 | 1.800796595 | 0.90154145 | 1.99746401 | 0.0457748 | 0.14663127 |
| FBXW7 | 2721.40231 | -0.298730688 | 0.20187775 | -1.4797604 | 0.13893721 | 0.30504762 |
| TIGD4 | 59.2206833 | 0.526960237 | 0.48020491 | 1.09736538 | 0.2724817 | 0.47050768 |
| ARFIP1 | 1291.4898 | -0.027981276 | 0.1412706 | -0.1980686 | 0.84299135 | 0.91749107 |
| FHDC1 | 68.7410482 | -2.759766068 | 0.60856154 | -4.5349006 | 5.76E-06 | 0.00016187 |
| TRIM2 | 926.423775 | -0.60204253 | 0.35773452 | -1.682931 | 0.09238844 | 0.23365755 |
| TMEM131L | 906.601357 | 0.431992724 | 0.39209306 | 1.10176069 | 0.27056573 | 0.46873665 |
| TLR2 | 162.003238 | -1.11515342 | 0.50623803 | -2.2028243 | 0.02760714 | 0.10330631 |
| PLRG1 | 242.987324 | 0.057193062 | 0.23840164 | 0.23990213 | 0.81040613 | 0.89720029 |
| FGB | 1704.49405 | -5.170973134 | 1.39445401 | -3.7082421 | 0.0002087 | 0.00276663 |
| FGA | 695.921094 | -5.857787763 | 1.24349413 | -4.7107482 | 2.47E-06 | 8.17E-05 |
| LRAT | 441.111268 | -1.672575878 | 0.47616165 | -3.512622 | 0.00044371 | 0.00492861 |
| MAP9 | 257.536424 | -0.193898557 | 0.31466902 | -0.6161984 | 0.53776356 | 0.70986341 |
| GUCY1A1 | 660.368904 | 0.239297014 | 0.33311122 | 0.71836971 | 0.47252936 | 0.65856227 |
| GUCY1B1 | 333.571572 | 0.477433586 | 0.43720965 | 1.09200148 | 0.27483245 | 0.47259765 |
| CTSO | 181.120491 | -0.552508224 | 0.42496341 | -1.3001313 | 0.19355598 | 0.3785776 |

| | | | | | | |
|---|---|---|---|---|---|---|
| PDGFC | 4059.38908 | 0.317514373 | 0.42717345 | 0.74329145 | 0.45730525 | 0.64588182 |
| GLRB | 82.3188641 | -1.011415744 | 0.44163618 | -2.290156 | 0.02201228 | 0.08843004 |
| TMEM144 | 186.027258 | -1.76086277 | 0.51680236 | -3.4072267 | 0.00065627 | 0.0065949 |
| ETFDH | 887.41252 | 0.248013088 | 0.34749147 | 0.71372426 | 0.4753977 | 0.66124102 |
| PPID | 471.122438 | -0.520000672 | 0.42415808 | -1.2259596 | 0.22021387 | 0.41207091 |
| FNIP2 | 12343.7348 | -0.508859654 | 0.47849623 | -1.0634559 | 0.28757524 | 0.48746773 |
| C4orf45 | 287.238151 | -1.170565407 | 0.50614238 | -2.3127196 | 0.02073806 | 0.08469587 |
| RAPGEF2 | 4994.5022 | 0.10559017 | 0.25564204 | 0.41303915 | 0.67957793 | 0.81423861 |
| FSTL5 | 58.2825082 | 0.65029341 | 0.547769 | 1.18716725 | 0.23516166 | 0.42961829 |
| NAF1 | 299.803426 | -0.071220831 | 0.13919152 | -0.5116751 | 0.60887845 | 0.76560837 |
| TMA16 | 222.542552 | -0.157767316 | 0.35971749 | -0.4385867 | 0.66096102 | 0.80295955 |
| 1-Mar | 149.184671 | -0.281170328 | 0.50478054 | -0.557015 | 0.57751718 | 0.74154889 |
| TMEM192 | 1057.65084 | 0.91032785 | 0.35976046 | 2.53037214 | 0.01139416 | 0.05526531 |
| KLHL2 | 953.595589 | -0.695287096 | 0.39116009 | -1.7775001 | 0.075486 | 0.20417265 |
| GK3P | 91.1643928 | 1.280020684 | 0.48583433 | 2.63468551 | 0.00842153 | 0.04450615 |
| MSMO1 | 615.35334 | -0.627287315 | 0.51563406 | -1.2165358 | 0.22378087 | 0.41608307 |
| CPE | 372.708405 | 0.060955561 | 0.70751371 | 0.0861546 | 0.93134352 | 0.96354054 |
| TLL1 | 720.976282 | 1.717009501 | 0.49521106 | 3.46722768 | 0.00052586 | 0.00552339 |
| DDX60 | 331.162589 | -1.224698847 | 1.05543903 | -1.1603691 | 0.24589856 | 0.44147485 |
| DDX60L | 485.49979 | -1.96903185 | 0.82418637 | -2.3890614 | 0.01689148 | 0.07336072 |
| PALLD | 1030.09399 | -0.924984411 | 0.3645006 | -2.537676 | 0.01115913 | 0.05449168 |
| CBR4 | 1912.00915 | 0.908605019 | 0.34917786 | 2.60212665 | 0.00926476 | 0.04768761 |
| SH3RF1 | 4044.23446 | 0.315275678 | 0.41924918 | 0.75200071 | 0.45205063 | 0.64143458 |
| AC096741.1 | 73.8101608 | -0.597579107 | 0.53633966 | -1.1141803 | 0.26520183 | 0.4630717 |
| NEK1 | 1496.10619 | -0.151253819 | 0.2319464 | -0.6521068 | 0.51433229 | 0.69258061 |
| CLCN3 | 1039.39366 | -0.236282631 | 0.19352903 | -1.2209157 | 0.22211795 | 0.41444934 |
| HPF1 | 135.309322 | 0.17470489 | 0.33481536 | 0.52179472 | 0.60181326 | 0.76091269 |
| LINC02275 | 948.758611 | 1.761752436 | 0.66810356 | 2.63694516 | 0.00836564 | 0.04432461 |
| AC084866.2 | 54.0554021 | 0.715949123 | 0.97545495 | 0.73396432 | 0.46297049 | 0.65120457 |
| MFAP3L | 823.981353 | 0.084660523 | 0.23437332 | 0.36122082 | 0.71793438 | 0.83896074 |
| AADAT | 48.9474394 | 0.065318559 | 0.5543781 | 0.11782312 | 0.90620781 | 0.95081066 |
| GALNTL6 | 46.3857959 | -0.094347187 | 0.48048407 | -0.1963586 | 0.84432948 | 0.91821836 |
| AC105285.1 | 142.745022 | -0.784491424 | 0.36025336 | -2.1776103 | 0.02943507 | 0.10759105 |
| GALNT7 | 1237.31269 | -1.051997674 | 0.26861132 | -3.9164309 | 8.99E-05 | 0.00144413 |
| HMGB2 | 122.173879 | -0.352477845 | 0.4971795 | -0.7089549 | 0.47835246 | 0.66397743 |
| SAP30 | 83.9991619 | -1.274005463 | 0.43995665 | -2.8957523 | 0.00378251 | 0.02452547 |
| FBXO8 | 213.577237 | 0.143077407 | 0.26581093 | 0.53826759 | 0.59039232 | 0.75230153 |
| CEP44 | 381.834534 | -0.094560285 | 0.31483985 | -0.3003441 | 0.76391471 | 0.86816839 |
| HPGD | 157.940014 | 2.584584904 | 0.55835112 | 4.62895979 | 3.68E-06 | 0.00011244 |
| GPM6A | 1478.25268 | 1.305160337 | 0.53859293 | 2.42327789 | 0.01538116 | 0.06842892 |
| WDR17 | 192.121512 | 0.263193137 | 0.3555275 | 0.74028912 | 0.45912458 | 0.6475935 |
| SPATA4 | 37.883637 | 0.273856995 | 0.63356432 | 0.43224813 | 0.66556108 | 0.80579154 |
| AC019163.1 | 44.1201128 | 0.687457421 | 0.43440828 | 1.58251456 | 0.11353215 | 0.26719688 |
| SPCS3 | 672.546592 | 0.004685437 | 0.34614766 | 0.01353595 | 0.98920021 | 0.99471552 |
| VEGFC | 56.7112258 | 1.05169074 | 0.69315951 | 1.51724203 | 0.12920559 | 0.29114269 |
| AGA | 97.8430616 | 0.282943305 | 0.29359351 | 0.96372465 | 0.33518399 | 0.53497519 |
| AC078881.1 | 485.426502 | 0.40138435 | 0.39931024 | 1.00519423 | 0.31480333 | 0.51522499 |
| AC098864.1 | 118.453574 | -1.253595667 | 0.6825077 | -1.8367495 | 0.06624689 | 0.18714416 |
| TENM3 | 1204.59199 | -1.925898394 | 0.6319988 | -3.0473134 | 0.00230897 | 0.01714465 |
| DCTD | 275.396247 | -0.527288038 | 0.31900348 | -1.6529225 | 0.09834663 | 0.24284584 |
| WWC2 | 3056.06276 | 0.56238205 | 0.16163648 | 3.47930149 | 0.00050272 | 0.00537086 |
| CDKN2AIP | 121.16156 | 0.074444076 | 0.2755577 | 0.27015785 | 0.78703882 | 0.88351603 |
| AC107214.1 | 92.3933068 | 0.150198764 | 0.28736078 | 0.52268358 | 0.60119446 | 0.76048104 |
| ING2 | 97.2604837 | -0.842148893 | 0.27256271 | -3.0897436 | 0.00200329 | 0.01538188 |
| RWDD4 | 328.32006 | 0.400365149 | 0.20121836 | 1.98970483 | 0.04662346 | 0.14848229 |
| TRAPPC11 | 614.698017 | 0.158270407 | 0.23113182 | 0.68476253 | 0.49349378 | 0.67582369 |
| STOX2 | 7428.82466 | -0.789759741 | 0.47463256 | -1.6639392 | 0.09612457 | 0.23942931 |
| ENPP6 | 926.733165 | 2.74874617 | 0.65275344 | 4.21100222 | 2.54E-05 | 0.00052679 |
| IRF2 | 725.616333 | 0.154419219 | 0.25039946 | 0.61669151 | 0.53743822 | 0.70967393 |
| LINC02427 | 63.1148131 | 0.48945503 | 0.74016501 | 0.66127826 | 0.50843388 | 0.68823852 |
| CASP3 | 239.214105 | -0.376756938 | 0.23912536 | -1.5755624 | 0.11512668 | 0.26975067 |
| PRIMPOL | 192.510022 | 0.337388646 | 0.44412508 | 0.75967032 | 0.44745167 | 0.63814588 |
| ACSL1 | 2805.53983 | 0.921194347 | 0.37973211 | 2.42590583 | 0.01527023 | 0.06804586 |
| SLC25A4 | 204.255903 | -0.540058186 | 0.42480485 | -1.2713089 | 0.20361878 | 0.39166469 |
| CFAP97 | 814.410532 | -0.469712379 | 0.24342534 | -1.9295953 | 0.05365701 | 0.16250035 |
| SNX25 | 1145.90373 | -0.818995478 | 0.29854024 | -2.7433337 | 0.00608189 | 0.03495363 |
| LRP2BP | 133.85291 | -1.060001247 | 0.32009594 | -3.3115111 | 0.00092794 | 0.00856051 |
| AC112722.1 | 182.596099 | -0.515835986 | 0.3555468 | -1.4508245 | 0.14682875 | 0.3161454 |
| ANKRD37 | 294.319284 | -2.121679712 | 0.84152133 | -2.5212429 | 0.01169411 | 0.05624251 |
| UFSP2 | 446.287678 | -0.163008695 | 0.33560143 | -0.4857211 | 0.62716492 | 0.77952027 |
| C4orf47 | 415.240446 | -1.472732956 | 0.58637958 | -2.5115693 | 0.01201957 | 0.05736301 |
| CCDC110 | 43.6854526 | -0.483287192 | 0.51572779 | -0.9370974 | 0.34870844 | 0.54865148 |
| SORBS2 | 5000.55755 | 1.397238606 | 0.46214594 | 3.02337092 | 0.00249976 | 0.01818692 |
| TLR3 | 63.3458988 | -0.330780736 | 0.66938875 | -0.4941534 | 0.62119783 | 0.77507036 |
| FAM149A | 922.698192 | -0.521453621 | 0.29973157 | -1.7397354 | 0.08190549 | 0.21540264 |
| CYP4V2 | 563.804086 | 1.623700782 | 0.36077598 | 4.50057889 | 6.78E-06 | 0.00018467 |
| KLKB1 | 765.968361 | 2.544447004 | 0.3989968 | 6.37711127 | 1.80E-10 | 2.58E-08 |
| F11 | 238.056144 | 3.220503949 | 0.70465031 | 4.57035767 | 4.87E-06 | 0.00014036 |
| F11-AS1 | 313.179133 | 0.77226692 | 0.71407044 | 1.08149963 | 0.27947493 | 0.47799091 |
| AC018709.1 | 284.118789 | 0.947736505 | 0.67043885 | 1.41360618 | 0.15747757 | 0.33177901 |
| FAT1 | 3408.68727 | 0.156653645 | 0.26192234 | 0.59809196 | 0.54977857 | 0.71955844 |
| LINC01060 | 3987.43247 | 0.54781803 | 0.65377407 | 0.83793172 | 0.40206905 | 0.59625235 |
| FRG1-DT | 166.722889 | 0.933518702 | 0.4770455 | 1.9568756 | 0.0503621 | 0.15610883 |
| FRG1 | 137.801381 | 0.140392659 | 0.19392812 | 0.72394173 | 0.46910151 | 0.65617047 |
| CCDC127 | 87.5832061 | -0.518209011 | 0.22974208 | -2.2556121 | 0.02409493 | 0.09382649 |
| SDHA | 945.262257 | -0.343795192 | 0.2067955 | -1.6624888 | 0.09641481 | 0.23976012 |
| PDCD6 | 552.27946 | -0.606949002 | 0.27238367 | -2.228287 | 0.02586138 | 0.0986979 |
| AHRR | 30.3315438 | -0.925950367 | 0.84420999 | -1.0968247 | 0.27271803 | 0.47062144 |
| EXOC3-AS1 | 46.5303457 | 0.303980134 | 0.36523439 | 0.83228783 | 0.4052465 | 0.59927865 |
| EXOC3 | 504.551571 | -0.049158967 | 0.20609204 | -0.2385292 | 0.81147068 | 0.89779874 |
| SLC9A3 | 602.062874 | 0.818752811 | 0.61286141 | 1.33595101 | 0.18156529 | 0.36379409 |
| CEP72 | 67.143408 | 0.197021426 | 0.34010781 | 0.57929109 | 0.56239277 | 0.73014439 |
| ZDHHC11B | 110.940428 | 1.637107553 | 0.70322858 | 2.32798781 | 0.01991275 | 0.08221207 |
| ZDHHC11 | 220.747303 | 1.139665818 | 0.49381597 | 2.30787556 | 0.02100606 | 0.08551531 |
| BRD9 | 364.894677 | -0.014084669 | 0.37929902 | -0.0371334 | 0.97037863 | 0.98525777 |
| TRIP13 | 42.9073963 | 0.731125608 | 0.66028608 | 1.10728612 | 0.26817024 | 0.46607875 |
| SLC12A7 | 1355.25998 | 0.16029455 | 0.3358319 | 0.47730591 | 0.6331443 | 0.78375408 |
| AC114291.1 | 237.118788 | 7.395032981 | 0.9472587 | 7.80677229 | 5.87E-15 | 2.98E-12 |
| SLC6A19 | 587.331679 | 2.974832877 | 0.68272229 | 4.35731034 | 1.32E-05 | 0.00031775 |
| SLC6A18 | 305.037899 | 3.427111239 | 0.90954493 | 3.76794057 | 0.0001646 | 0.00227437 |
| CLPTM1L | 918.24449 | 0.170709691 | 0.39486383 | 0.43232546 | 0.66550488 | 0.80579154 |
| LPCAT1 | 435.92261 | -2.309589966 | 0.62853599 | -3.6745548 | 0.00023826 | 0.00306926 |
| MRPL36 | 129.654329 | -0.976031121 | 0.36996473 | -2.6381735 | 0.00833539 | 0.04421565 |
| NDUFS6 | 521.690272 | -0.42578901 | 0.30058895 | -1.4165158 | 0.15662454 | 0.33033582 |
| AC010451.1 | 67.5463287 | 2.07922923 | 1.246232 | 1.66841265 | 0.09523384 | 0.23797755 |
| LINC01020 | 597.262894 | 3.535986478 | 1.0585187 | 3.34050449 | 0.00083626 | 0.00792935 |
| ADAMTS16 | 224.164832 | 0.784673369 | 0.88385471 | 0.88778547 | 0.37465616 | 0.5715597 |
| ICE1 | 930.018578 | -0.236210301 | 0.30516435 | -0.7740429 | 0.43890542 | 0.63061599 |
| LINC02145 | 43.4694492 | 1.024482601 | 0.58244688 | 1.75892881 | 0.0785896 | 0.20949091 |
| MED10 | 264.124072 | -0.593594511 | 0.26896982 | -2.2069186 | 0.02731975 | 0.10253881 |
| NSUN2 | 482.357362 | -0.103894156 | 0.27959382 | -0.3715896 | 0.71019843 | 0.83402073 |
| SRD5A1 | 236.740878 | -0.572414108 | 0.35675202 | -1.6045154 | 0.1086005 | 0.25942611 |
| TENT4A | 771.038679 | -0.632243513 | 0.31660065 | -1.9969748 | 0.04582792 | 0.14673284 |

| | | | | | | |
|---|---|---|---|---|---|---|
| LINC02196 | 41.6044224 | -0.974090161 | 0.84978383 | -1.1462799 | 0.25167934 | 0.4480801 |
| ADCY2 | 230.891952 | -0.2474516 | 0.70168101 | -0.3526554 | 0.7243468 | 0.84312343 |
| C5orf49 | 44.4670618 | -0.243436583 | 0.43610316 | -0.5582087 | 0.57670188 | 0.74112625 |
| MTRR | 784.61938 | -0.457232463 | 0.25574729 | -1.7878292 | 0.07380357 | 0.20107842 |
| FASTKD3 | 111.488292 | 0.676409952 | 0.27202307 | 2.48659038 | 0.01289738 | 0.06031539 |
| MIR4458HG | 112.791336 | -0.328649469 | 0.41281624 | -0.7961156 | 0.42596482 | 0.61876852 |
| SEMA5A | 3653.6777 | 0.025742539 | 0.52292533 | 0.04922794 | 0.96073764 | 0.97967794 |
| SNHG18 | 189.674623 | 0.036414625 | 0.34324445 | 0.10608948 | 0.91551136 | 0.95487096 |
| LINC02112 | 100.073385 | 0.138129575 | 0.65482582 | 0.21094094 | 0.83293336 | 0.91175754 |
| FAM173B | 234.377001 | 0.367852371 | 0.30342534 | 1.21233238 | 0.22538517 | 0.41770613 |
| CCT5 | 498.197971 | -0.877411756 | 0.28051289 | -3.1278839 | 0.0017607 | 0.013902 |
| CMBL | 1890.93239 | 0.109233682 | 0.6201352 | 0.17614495 | 0.86018007 | 0.92629345 |
| 6-Mar | 2777.13741 | -0.316265701 | 0.25955662 | -1.2184844 | 0.22303994 | 0.41521234 |
| ANKRD33B | 1829.21823 | 0.254266284 | 0.62899298 | 0.40424344 | 0.68603372 | 0.81810131 |
| DAP | 804.998302 | -0.219087696 | 0.28171372 | -0.7776962 | 0.43674812 | 0.62868147 |
| CTNND2 | 80.4508761 | 1.925072039 | 0.39839047 | 4.83212372 | 1.35E-06 | 4.92E-05 |
| LINC02220 | 484.052546 | 2.659524736 | 0.77548392 | 3.4295034 | 0.00060469 | 0.00617619 |
| DNAH5 | 644.759778 | 0.527842177 | 1.02483532 | 0.51505073 | 0.60651757 | 0.7636205 |
| TRIO | 8390.33215 | -1.295635921 | 0.3471538 | -3.7321669 | 0.00018984 | 0.00255863 |
| OTULINL | 67.686221 | 0.427679136 | 0.5074478 | 0.84280419 | 0.39933796 | 0.5934221 |
| OTULIN | 807.673482 | -0.16923392 | 0.26767893 | -0.6322273 | 0.52723838 | 0.70176703 |
| ANKH | 965.502813 | 0.419842022 | 0.3643716 | 1.15223586 | 0.24922417 | 0.44538665 |
| FBXL7 | 2202.59832 | 1.718012916 | 0.5494717 | 3.12666316 | 0.00176802 | 0.01394075 |
| ZNF622 | 143.670983 | -0.920106143 | 0.32270275 | -2.8512497 | 0.00435478 | 0.02734352 |
| RETREG1 | 5024.21037 | -1.351498957 | 0.588373 | -2.2970105 | 0.02161817 | 0.08748753 |
| MYO10 | 1403.60054 | -1.66680498 | 0.42781385 | -3.8960987 | 9.78E-05 | 0.00152835 |
| LINC02241 | 300.510186 | 0.464320517 | 0.37593011 | 1.23512458 | 0.21678414 | 0.40766364 |
| AC093274.1 | 34.873755 | -0.17162984 | 0.81221369 | -0.2113112 | 0.83264445 | 0.91165888 |
| CDH12 | 102.546975 | 0.028437548 | 0.45786192 | 0.06210944 | 0.95047568 | 0.97465176 |
| C5orf17 | 351.851642 | -0.224653119 | 0.57728822 | -0.3891524 | 0.69716338 | 0.82550208 |
| CDH9 | 5803.99795 | 1.873189104 | 0.64728418 | 2.89392075 | 0.00380464 | 0.0246238 |
| AC113386.1 | 868.261776 | -1.834954792 | 0.67757965 | -2.708102 | 0.00676692 | 0.03790527 |
| CDH6 | 12711.0776 | -0.684453795 | 0.72851271 | -0.9395221 | 0.34746275 | 0.54753787 |
| DROSHA | 569.651847 | 0.045647642 | 0.19721873 | 0.23145693 | 0.81695984 | 0.90111634 |
| C5orf22 | 242.553659 | -0.453148022 | 0.22935424 | -1.975756 | 0.04818241 | 0.15171981 |
| PDZD2 | 6969.48147 | 2.688142528 | 0.70131474 | 3.83300445 | 0.00012659 | 0.0018711 |
| AC025178.1 | 112.435683 | -0.509697492 | 0.49739026 | -1.0247436 | 0.30548418 | 0.50546495 |
| GOLPH3 | 1691.86518 | -0.257416431 | 0.25243099 | -1.0197497 | 0.30784718 | 0.50791823 |
| MTMR12 | 955.755578 | -0.014586601 | 0.25018993 | -0.0583021 | 0.95350799 | 0.97656847 |
| ZFR | 2494.20158 | 0.142088866 | 0.13857212 | 1.02537842 | 0.30518467 | 0.50529195 |
| SUB1 | 726.700145 | -0.486704866 | 0.19497215 | -2.4962789 | 0.01255039 | 0.05913653 |
| LINC02061 | 411.389506 | 8.281171833 | 0.72374246 | 11.4421529 | 2.57E-30 | 3.53E-26 |
| NPR3 | 2175.47995 | 3.92639373 | 0.38102231 | 10.3048919 | 6.70E-25 | 3.06E-21 |
| AC010343.3 | 51.1977832 | -0.129467633 | 0.42576325 | -0.3040836 | 0.76106418 | 0.8665094 |
| TARS | 391.643407 | -0.713790576 | 0.23846368 | -2.9932884 | 0.00275989 | 0.01948035 |
| ADAMTS12 | 84.5876492 | -0.844018291 | 0.50041673 | -1.6866309 | 0.09167435 | 0.23229747 |
| SLC45A2 | 182.177539 | 0.891777786 | 0.56996555 | 1.56461699 | 0.11767274 | 0.27360212 |
| AMACR | 142.703883 | 0.055734769 | 0.64343792 | 0.08662027 | 0.93097335 | 0.96337398 |
| C1QTNF3-AM | 743.223066 | 0.184498731 | 0.31134999 | 0.59257664 | 0.55346452 | 0.72285261 |
| RAI14 | 2077.28383 | 0.460994595 | 0.30478925 | 1.5125028 | 0.130406 | 0.29297692 |
| TTC23L | 54.0097281 | -0.364050095 | 0.5088566 | -0.7154277 | 0.47434481 | 0.66055515 |
| RAD1 | 168.192598 | 0.164109401 | 0.20134029 | 0.81508476 | 0.41502379 | 0.60853959 |
| BRIX1 | 163.670673 | -0.42948201 | 0.36755292 | -1.1684903 | 0.24260906 | 0.437731 |
| DNAJC21 | 435.385018 | -0.304928108 | 0.16006275 | -1.9050535 | 0.05677312 | 0.16873139 |
| AGXT2 | 3551.75536 | 1.749229676 | 0.72847154 | 2.40123269 | 0.01633994 | 0.07155642 |
| PRLR | 2753.13893 | 1.696111139 | 0.49219328 | 3.4460266 | 0.00056889 | 0.00586745 |
| SPEF2 | 1580.68987 | 1.396136508 | 0.5238335 | 2.66522953 | 0.00769358 | 0.04169133 |
| AC137810.1 | 64.5052193 | 0.564857907 | 0.80638274 | 0.70048363 | 0.48362533 | 0.66764552 |
| UGT3A1 | 819.080281 | 0.727946445 | 0.45034408 | 1.61642283 | 0.10600293 | 0.25531227 |
| LMBRD2 | 382.083422 | 0.44639217 | 0.2531127 | 1.76361032 | 0.07779761 | 0.20811316 |
| SKP2 | 94.9604458 | 0.939769714 | 0.26058222 | 3.60642295 | 0.00031045 | 0.00377243 |
| NADK2 | 386.194679 | -0.065023276 | 0.32132957 | -0.202357 | 0.83963767 | 0.91500346 |
| NIPBL | 4193.759 | -0.231916418 | 0.18957205 | -1.2233682 | 0.22119065 | 0.41345428 |
| CPLANE1 | 1591.08598 | 0.410630169 | 0.12750508 | 3.22050044 | 0.00127967 | 0.01087249 |
| NUP155 | 257.604312 | 0.195315135 | 0.25901464 | 0.75406987 | 0.45080727 | 0.64075861 |
| WDR70 | 2557.02733 | 0.330412462 | 0.16772979 | 1.96990927 | 0.04884877 | 0.15318465 |
| GDNF-AS1 | 211.287357 | 3.389625366 | 0.63126573 | 5.36956972 | 7.89E-08 | 4.58E-06 |
| LINC02107 | 86.9163502 | 3.987738009 | 0.69167586 | 5.76532771 | 8.15E-09 | 6.90E-07 |
| EGFLAM | 39.1264815 | -0.508950071 | 0.72334771 | -0.7036036 | 0.48167965 | 0.66636403 |
| LIFR | 1841.57574 | 0.041274997 | 0.527008 | 0.07831949 | 0.93757392 | 0.96641041 |
| LIFR-AS1 | 59.0485525 | 0.217306022 | 0.64530924 | 0.33674711 | 0.73630754 | 0.85083186 |
| OSMR-AS1 | 796.752069 | -1.643659911 | 0.40472672 | -4.0611599 | 4.88E-05 | 0.00088885 |
| OSMR | 1668.35639 | -2.448940598 | 0.52433521 | -4.6705629 | 3.00E-06 | 9.60E-05 |
| RICTOR | 2905.32461 | -0.202664693 | 0.19216971 | -1.0546131 | 0.2916023 | 0.4916955 |
| FYB1 | 142.716537 | -0.636352546 | 0.74382877 | -0.8555095 | 0.39226917 | 0.58726848 |
| C9 | 393.264612 | -0.163716705 | 0.41742559 | -0.3922057 | 0.69490622 | 0.82432536 |
| DAB2 | 1168.75778 | -0.178166263 | 0.3072317 | -0.5799085 | 0.56197635 | 0.73014439 |
| TTC33 | 387.635954 | 0.064241225 | 0.2167933 | 0.29632477 | 0.76698207 | 0.86985134 |
| PRKAA1 | 793.405485 | 0.021160938 | 0.19464675 | 0.10871457 | 0.91342889 | 0.95399223 |
| RPL37 | 730.064196 | -0.557002852 | 0.33871788 | -1.6444448 | 0.10008436 | 0.2460337 |
| C6 | 123.212209 | -0.171387977 | 0.84682349 | -0.2023893 | 0.83961243 | 0.91500346 |
| PLCXD3 | 678.3573 | 1.76560917 | 0.75875411 | 2.32698465 | 0.01996608 | 0.08238263 |
| OXCT1 | 291.154137 | 0.557981919 | 0.50677178 | 1.10105168 | 0.27087417 | 0.46903388 |
| C5orf51 | 189.387832 | -0.114540907 | 0.20421248 | -0.5608908 | 0.57487197 | 0.7396699 |
| FBXO4 | 118.576109 | -0.019078116 | 0.39325742 | -0.048513 | 0.96130737 | 0.97988733 |
| GHR | 2961.64046 | 1.162705886 | 0.40736685 | 2.85419861 | 0.00431455 | 0.02715317 |
| SELENOP | 773.618812 | 1.083852158 | 0.5077909 | 2.13444581 | 0.03280631 | 0.11654204 |
| ZNF131 | 1028.61975 | -0.11968843 | 0.23899839 | -0.5007918 | 0.61651767 | 0.77182636 |
| NIM1K | 57.863161 | -1.547662666 | 0.62595598 | -2.4724785 | 0.01341798 | 0.06196774 |
| HMGCS1 | 1019.85057 | -0.287281721 | 0.4237528 | -0.6779465 | 0.49780563 | 0.67975909 |
| CCL28 | 96.6417246 | -3.996398706 | 0.72803852 | -5.4892682 | 4.04E-08 | 2.65E-06 |
| TMEM267 | 376.682732 | -0.565418728 | 0.25909612 | -2.182274 | 0.02908931 | 0.10681148 |
| C5orf34 | 40.4968554 | -0.950845728 | 0.54686356 | -1.7387257 | 0.08208302 | 0.21570398 |
| PAIP1 | 430.660779 | -0.059859094 | 0.17832506 | -0.335674 | 0.73711668 | 0.85110831 |
| NNT-AS1 | 460.975862 | -0.437742137 | 0.32234428 | -1.3579957 | 0.17446505 | 0.35428037 |
| NNT | 1263.24219 | -0.134150034 | 0.25969711 | -0.5165634 | 0.60546094 | 0.76301233 |
| MRPS30-DT | 91.8877268 | -0.166659944 | 0.39665847 | -0.4201598 | 0.67436872 | 0.81105241 |
| MRPS30 | 213.341467 | -0.340408741 | 0.18878223 | -1.8031821 | 0.07135962 | 0.19668739 |
| AC093297.2 | 66.1015724 | 0.102484645 | 0.29422615 | 0.3483193 | 0.7276004 | 0.84525966 |
| PARP8 | 1099.16314 | -1.369748267 | 0.3405908 | -4.0216831 | 5.78E-05 | 0.00101799 |
| AC022126.1 | 89.8226931 | -0.487160408 | 0.62716654 | -0.776764 | 0.437298 | 0.62909779 |
| ITGA1 | 757.378166 | 0.917847125 | 0.22179359 | 4.13829414 | 3.50E-05 | 0.00068152 |
| ITGA2 | 569.622922 | -1.15243487 | 0.35231408 | -3.2710441 | 0.00107151 | 0.00949213 |
| MOCS2 | 181.013624 | 0.268631856 | 0.38649137 | 0.69505266 | 0.4870223 | 0.67024244 |
| NDUFS4 | 1104.39681 | 0.327998033 | 0.19130811 | 1.71450144 | 0.08643667 | 0.22324995 |
| ARL15 | 4604.99673 | 0.600832841 | 0.32836065 | 1.8297955 | 0.06728052 | 0.18892497 |
| SNX18 | 109.16692 | 0.433350777 | 0.32440757 | 1.33582202 | 0.18160745 | 0.36381342 |
| AC026704.1 | 96.3785619 | -1.08966862 | 0.57289889 | -1.9020261 | 0.05716775 | 0.16949995 |
| DHX29 | 738.04336 | 0.27511087 | 0.19032757 | 1.44545987 | 0.14832876 | 0.31826835 |
| MTREX | 1178.38098 | 0.131344411 | 0.15460339 | 0.8495571 | 0.39557137 | 0.59027092 |
| PLPP1 | 2388.0744 | 0.628962303 | 0.36373954 | 1.72915572 | 0.08378123 | 0.21857428 |
| SLC38A9 | 1101.8729 | -0.001872828 | 0.19884948 | -0.0094183 | 0.99248538 | 0.99626489 |
| IL6ST | 1858.63722 | -0.25153105 | 0.25702776 | -0.9786143 | 0.32777058 | 0.52756592 |

| | | | | | | |
|---|---|---|---|---|---|---|
| AC008892.1 | 174.846657 | 0.172945727 | 0.41831701 | 0.41343221 | 0.67928998 | 0.81417094 |
| ANKRD55 | 79.4715964 | 0.135061455 | 0.61229575 | 0.22058205 | 0.82541788 | 0.90642548 |
| LINC01948 | 294.680253 | 0.008522063 | 0.73600269 | 0.01157885 | 0.99076162 | 0.99526343 |
| C5orf67 | 1219.51229 | 1.149045034 | 0.93055417 | 1.2347965 | 0.21690625 | 0.40780368 |
| MAP3K1 | 2212.33813 | -0.236000871 | 0.43400821 | -0.5437705 | 0.5865994 | 0.74927946 |
| AC008937.2 | 73.1730621 | -1.476778851 | 0.64243111 | -2.298735 | 0.02151999 | 0.08724477 |
| SETD9 | 70.7144183 | -0.39315373 | 0.29959258 | -1.3122946 | 0.18942074 | 0.37352757 |
| MIER3 | 263.261527 | 0.167361232 | 0.21864448 | 0.76544917 | 0.44400412 | 0.63521183 |
| GPBP1 | 2446.68368 | 0.03561571 | 0.28191174 | 0.12633638 | 0.89946566 | 0.94743919 |
| PLK2 | 62.5907572 | -1.072997912 | 0.61633657 | -1.7409285 | 0.08169611 | 0.21518228 |
| PDE4D | 32494.8809 | 1.048190554 | 0.5250462 | 1.99637775 | 0.04589282 | 0.14683774 |
| AC092343.1 | 55.1769965 | 0.36537908 | 0.79747029 | 0.45817266 | 0.6468284 | 0.79280318 |
| DEPDC1B | 69.707541 | -0.004744402 | 0.67633396 | -0.0070149 | 0.99440298 | 0.99716743 |
| ELOVL7 | 1354.48072 | -0.201754374 | 0.36744599 | -0.5490722 | 0.58295591 | 0.74580704 |
| ERCC8 | 512.867268 | 0.311258833 | 0.20647397 | 1.50749669 | 0.13168339 | 0.29498027 |
| NDUFAF2 | 1324.94132 | -0.67412786 | 0.3482999 | -1.9354811 | 0.05293129 | 0.16094131 |
| SMIM15 | 154.647914 | -0.758605119 | 0.33519517 | -2.2631744 | 0.02362494 | 0.09270744 |
| SMIM15-AS1 | 77.0394506 | 0.814713889 | 0.34476711 | 2.36308474 | 0.01812352 | 0.07698657 |
| ZSWIM6 | 4551.22929 | -0.564064143 | 0.31482048 | -1.7917009 | 0.07318089 | 0.19989846 |
| KIF2A | 789.894043 | -1.026688334 | 0.29567636 | -3.4723382 | 0.00051595 | 0.00545684 |
| DIMT1 | 156.723846 | -0.24134115 | 0.23667112 | -1.0197322 | 0.30785551 | 0.50791823 |
| IPO11 | 435.389315 | -0.009386831 | 0.2254513 | -0.0416357 | 0.96678908 | 0.9830686 |
| RNF180 | 1847.35045 | 0.026923051 | 0.34821863 | 0.07731651 | 0.93837175 | 0.96679657 |
| SREK1IP1 | 472.188511 | -0.318064777 | 0.20278386 | -1.5684916 | 0.11676646 | 0.27223752 |
| CWC27 | 965.573509 | 0.277801319 | 0.16333929 | 1.70076242 | 0.08898761 | 0.22760835 |
| ADAMTS6 | 64.6437737 | 1.539263369 | 0.59459111 | 2.58877629 | 0.00963176 | 0.04906079 |
| PPWD1 | 425.284279 | 0.192383963 | 0.31909689 | 0.6029014 | 0.54657429 | 0.71685903 |
| TRIM23 | 409.473627 | 0.270449904 | 0.25065839 | 1.07895813 | 0.28060639 | 0.47928621 |
| TRAPPC13 | 463.330107 | 0.293223937 | 0.22261617 | 1.31717266 | 0.18778076 | 0.37136212 |
| SGTB | 118.498192 | -1.42341505 | 0.37027341 | -3.844227 | 0.00012093 | 0.00180637 |
| NLN | 551.625987 | -1.860801492 | 0.363222 | -5.1230419 | 3.01E-07 | 1.37E-05 |
| ERBIN | 4604.88631 | 0.3831297 | 0.16328136 | 2.34643868 | 0.01895378 | 0.07937082 |
| SREK1 | 1561.58375 | 0.241330658 | 0.2295732 | 1.05121441 | 0.29316013 | 0.49326436 |
| MAST4 | 23826.4955 | 0.213953503 | 0.29255344 | 0.73133135 | 0.46457678 | 0.65225381 |
| AC112206.2 | 67.7920275 | 0.45479362 | 0.45083476 | 1.00878118 | 0.31307959 | 0.51338461 |
| PIK3R1 | 1557.2406 | -0.760009732 | 0.55483601 | -1.3697917 | 0.17075195 | 0.34969424 |
| SLC30A5 | 580.41505 | -0.29403103 | 0.2914274 | -1.0089341 | 0.31300625 | 0.51332576 |
| MRPS36 | 476.809371 | 0.014259753 | 0.30874808 | 0.04618572 | 0.96316222 | 0.98083195 |
| CDK7 | 501.153296 | -0.064311616 | 0.2336603 | -0.2752355 | 0.7831353 | 0.88102721 |
| CCDC125 | 611.769513 | 0.651843883 | 0.38894355 | 1.67593444 | 0.09375104 | 0.23571272 |
| AK6 | 246.112359 | -0.123122998 | 0.22049709 | -0.5583883 | 0.57657925 | 0.74103815 |
| RAD17 | 460.062802 | 0.305506459 | 0.12262331 | 2.49142247 | 0.01272327 | 0.05974577 |
| MARVELD2 | 232.465336 | -0.155902174 | 0.23112853 | -0.6745259 | 0.49997704 | 0.68139272 |
| OCLN | 692.704875 | -0.539626276 | 0.30556095 | -1.7660184 | 0.07739276 | 0.20754779 |
| GTF2H2C | 490.973641 | 0.038711809 | 0.30239791 | 0.12801613 | 0.89813621 | 0.94727247 |
| SMN2 | 35.790054 | 0.718303259 | 0.89448199 | 0.80303826 | 0.42195262 | 0.61561518 |
| NAIP | 853.709259 | 0.870868211 | 0.29436047 | 2.95850942 | 0.00309131 | 0.02111703 |
| GTF2H2 | 485.39459 | 0.400184877 | 0.46197462 | 0.86624863 | 0.38635383 | 0.58239877 |
| LINC02197 | 232.682604 | 1.546218301 | 0.85905783 | 1.79990014 | 0.07187641 | 0.19759525 |
| BDP1 | 1733.47632 | -0.109643439 | 0.14692535 | -0.7462527 | 0.45551476 | 0.64475342 |
| MCCC2 | 710.466683 | 0.260988325 | 0.17460528 | 1.49473326 | 0.13498407 | 0.29936987 |
| MAP1B | 711.593788 | -5.891622724 | 1.29945292 | -4.5339255 | 5.79E-06 | 0.00016229 |
| MRPS27 | 641.408961 | -0.055383783 | 0.23644148 | -0.2342389 | 0.81479955 | 0.8996663 |
| PTCD2 | 151.841292 | 0.811937437 | 0.30223894 | 2.68640913 | 0.00722246 | 0.03982885 |
| TNPO1 | 1743.49942 | -0.55393308 | 0.25397538 | -2.1810503 | 0.0291797 | 0.10692593 |
| FCHO2 | 2051.59055 | -0.430091281 | 0.23343225 | -1.8424672 | 0.06540683 | 0.18562958 |
| TMEM171 | 257.605136 | -1.426233826 | 0.42434213 | -3.3610469 | 0.00077648 | 0.0075115 |
| AC116345.3 | 313.024696 | 1.02369524 | 0.98397325 | 1.04036897 | 0.29816851 | 0.49833608 |
| TMEM174 | 135.528009 | 1.340979767 | 0.70800542 | 1.89402471 | 0.05822173 | 0.1712176 |
| BTF3 | 441.524111 | -0.930991787 | 0.33591611 | -2.7715009 | 0.00557985 | 0.03286766 |
| ANKRA2 | 173.371812 | 0.114986302 | 0.42932659 | 0.26782944 | 0.78883061 | 0.88460289 |
| UTP15 | 64.4488064 | -0.061504641 | 0.35439735 | -0.1735471 | 0.86222139 | 0.92773912 |
| ARHGEF28 | 6169.63034 | 0.365642237 | 0.27909547 | 1.31009733 | 0.19016291 | 0.3741919 |
| ENC1 | 40.2407839 | 0.417425976 | 0.57855221 | 0.72150096 | 0.47060135 | 0.65738999 |
| HEXB | 779.144603 | -0.096264323 | 0.40863507 | -0.2355753 | 0.81376225 | 0.89924534 |
| GFM2 | 350.193076 | 0.147833839 | 0.22735868 | 0.650223 | 0.51554819 | 0.69315508 |
| NSA2 | 258.029638 | -0.521676182 | 0.25247883 | -2.0662175 | 0.03880795 | 0.13096256 |
| FAM169A | 161.667857 | -0.376825911 | 0.75151473 | -0.5014219 | 0.61607426 | 0.7715916 |
| ANKRD31 | 58.9315544 | 0.805988195 | 0.55053426 | 1.46401098 | 0.14319095 | 0.31083232 |
| HMGCR | 373.988299 | -0.266094613 | 0.40483826 | -0.6572862 | 0.51099689 | 0.69007235 |
| COL4A3BP | 2470.43289 | 0.020736205 | 0.1894901 | 0.1094316 | 0.91286017 | 0.95362963 |
| POLK | 1270.51985 | 0.079181202 | 0.18370624 | 0.43102074 | 0.66645329 | 0.80635371 |
| ANKDD1B | 380.725602 | 0.020730972 | 0.52411461 | 0.03955427 | 0.96844848 | 0.98409989 |
| POC5 | 290.740371 | -0.338388436 | 0.15124886 | -2.2372958 | 0.02526701 | 0.09712141 |
| IQGAP2 | 2432.46414 | 1.343800818 | 0.44936567 | 2.99043946 | 0.00278576 | 0.01961195 |
| F2R | 34.8656457 | -0.277036143 | 0.42280537 | -0.6552333 | 0.51231758 | 0.6911749 |
| F2RL1 | 229.123784 | 0.080165996 | 0.24182152 | 0.33150894 | 0.74026009 | 0.85230954 |
| AC008581.2 | 95.454631 | -0.651394645 | 0.54332264 | -1.1989094 | 0.23056316 | 0.42375024 |
| ZBED3 | 269.180692 | -0.568981241 | 0.30660639 | -1.8557383 | 0.06349086 | 0.18179846 |
| AC022414.1 | 93.2361393 | -0.965827523 | 0.89946443 | -1.0737807 | 0.28292099 | 0.48202492 |
| WDR41 | 593.039976 | -0.081193739 | 0.23237997 | -0.3494008 | 0.72678846 | 0.84481763 |
| TBCA | 2217.92869 | -0.335612201 | 0.33360945 | -1.0060033 | 0.31441399 | 0.51489517 |
| AP3B1 | 2693.4681 | -0.144116341 | 0.20618457 | -0.6989676 | 0.48457225 | 0.66827686 |
| SCAMP1-AS1 | 58.7666404 | -0.389319354 | 0.49124403 | -0.7925172 | 0.42805915 | 0.62108678 |
| SCAMP1 | 968.021471 | 0.051324485 | 0.17487672 | 0.29348953 | 0.769148 | 0.87130169 |
| LHFPL2 | 1067.40245 | -0.750787317 | 0.59158214 | -1.2691176 | 0.20439912 | 0.39266976 |
| ARSB | 2730.43365 | 1.123991433 | 0.46533034 | 2.41546992 | 0.01571492 | 0.06963632 |
| DMGDH | 2795.03921 | 2.055971439 | 0.53193894 | 3.86505157 | 0.00011107 | 0.00168778 |
| BHMT2 | 4748.02455 | 1.268850702 | 0.43050083 | 2.94738267 | 0.00320476 | 0.02171413 |
| BHMT | 5143.08933 | 3.694591178 | 0.4603775 | 8.02513409 | 1.01E-15 | 6.04E-13 |
| JMY | 886.046233 | 0.219699351 | 0.21144019 | 1.03906144 | 0.29877616 | 0.49882154 |
| HOMER1 | 833.183665 | -1.355285852 | 0.5931637 | -2.2848429 | 0.02232204 | 0.08915159 |
| TENT2 | 3220.50671 | 0.277925069 | 0.17279704 | 1.60839018 | 0.10774975 | 0.25793326 |
| CMYA5 | 951.830446 | 1.030295497 | 0.24117296 | 4.27201919 | 1.94E-05 | 0.00043244 |
| MTX3 | 185.377571 | 0.389335389 | 0.17912064 | 2.17359312 | 0.02973571 | 0.10846027 |
| THBS4 | 178.840006 | 0.871842021 | 0.44365061 | 1.96515458 | 0.04939635 | 0.15398584 |
| SERINC5 | 523.31779 | -0.682848247 | 0.31960066 | -2.1365671 | 0.03263322 | 0.11618274 |
| AC010260.1 | 38.6312796 | -1.046068848 | 0.47117456 | -2.2201301 | 0.02640994 | 0.10018511 |
| ZFYVE16 | 964.341107 | -0.099378118 | 0.12899152 | -0.7704237 | 0.44104862 | 0.63296549 |
| AC008771.1 | 40.0567898 | 0.962191614 | 0.42398771 | 2.26938564 | 0.02324488 | 0.09163579 |
| FAM151B | 355.37143 | 0.372177086 | 0.27689206 | 1.34412335 | 0.1789085 | 0.36010262 |
| DHFR | 2313.86062 | -0.3592739 | 0.47439711 | -0.7573273 | 0.44885377 | 0.639148 |
| MSH3 | 2216.89296 | 1.387484531 | 0.2879845 | 4.81791395 | 1.45E-06 | 5.19E-05 |
| RASGRF2 | 1194.99172 | 1.003299567 | 0.49807723 | 2.0143454 | 0.04397328 | 0.14229024 |
| CKMT2-AS1 | 760.982895 | 0.737347529 | 0.2989687 | 2.46630346 | 0.01365156 | 0.06275049 |
| CKMT2 | 41.817502 | -0.030542497 | 0.54568546 | -0.0559709 | 0.955365 | 0.97737295 |
| ZCCHC9 | 290.67651 | 0.176208755 | 0.27392779 | 0.64326718 | 0.52005075 | 0.69605855 |
| ACOT12 | 96.1825126 | 2.672833162 | 0.60164277 | 4.44255841 | 8.89E-06 | 0.00023121 |
| SSBP2 | 1296.10688 | -0.947450505 | 0.35201653 | -2.6914944 | 0.00711327 | 0.03945835 |
| ATG10 | 1044.06867 | 0.169451954 | 0.25204605 | 0.67230552 | 0.50138922 | 0.68247686 |
| RPS23 | 1539.02861 | -0.965350941 | 0.52464409 | -1.8400111 | 0.06576661 | 0.18625267 |
| ATP6AP1L | 433.549879 | 0.685947407 | 0.4933638 | 1.39034806 | 0.16442321 | 0.34142538 |
| TMEM167A | 372.212993 | -0.424211022 | 0.2195615 | -1.9320829 | 0.05334927 | 0.16174707 |

| | | | | | | |
|---|---|---|---|---|---|---|
| XRCC4 | 671.08015 | -0.149110868 | 0.25622963 | -0.5819423 | 0.56060553 | 0.72893225 |
| VCAN | 224.757582 | -2.373217014 | 0.87831162 | -2.7020217 | 0.00689193 | 0.03846891 |
| EDIL3 | 397.430999 | -1.441427642 | 0.51842024 | -2.7804232 | 0.00542881 | 0.03219935 |
| AC113383.1 | 107.115483 | -1.304963784 | 0.53581192 | -2.4354885 | 0.0148717 | 0.06670365 |
| COX7C | 882.629765 | -1.010264643 | 0.36393583 | -2.7759417 | 0.00550421 | 0.03261295 |
| RASA1 | 2278.03426 | -0.117682226 | 0.18547562 | -0.6344889 | 0.52576179 | 0.70084961 |
| CCNH | 2067.66611 | 0.073072512 | 0.32382099 | 0.22565712 | 0.82146811 | 0.90404363 |
| TMEM161B | 673.114412 | 0.038194426 | 0.18112262 | 0.21087607 | 0.83298398 | 0.91175754 |
| TMEM161B-/ | 1753.54454 | -0.226038625 | 0.37904988 | -0.5963295 | 0.55095513 | 0.72050604 |
| MEF2C | 64.173642 | 0.16074038 | 0.46463573 | 0.34594925 | 0.72938086 | 0.84632259 |
| MEF2C-AS1 | 63.5052081 | -0.704414639 | 0.62058769 | -1.1350767 | 0.25634318 | 0.45367281 |
| CETN3 | 88.1919286 | 0.408237602 | 0.39762481 | 1.02669047 | 0.30456624 | 0.50478491 |
| MBLAC2 | 78.1630024 | 0.398114994 | 0.28082888 | 1.41764264 | 0.15629513 | 0.32994568 |
| POLR3G | 43.1954978 | -0.136388225 | 0.53396672 | -0.2554246 | 0.79839519 | 0.89012761 |
| LYSMD3 | 280.389451 | -0.645465095 | 0.27714499 | -2.3289798 | 0.01986014 | 0.082069 |
| ADGRV1 | 742.479611 | 3.641830856 | 0.42297407 | 8.61005701 | 7.30E-18 | 6.67E-15 |
| LUCAT1 | 290.165013 | -8.299953759 | 1.08737652 | -7.6330081 | 2.29E-14 | 9.82E-12 |
| ARRDC3 | 1046.14159 | 0.343584568 | 0.35766966 | 0.96061982 | 0.33674336 | 0.53683894 |
| ARRDC3-AS1 | 151.703156 | -0.503425547 | 0.26764087 | -1.8809741 | 0.05997544 | 0.1745825 |
| AC114316.2 | 40.5070308 | 0.643059379 | 0.79644036 | 0.80741687 | 0.41942635 | 0.61277734 |
| AC124854.1 | 526.934173 | 1.992235212 | 0.90613189 | 2.19861506 | 0.02790531 | 0.10396793 |
| NR2F1-AS1 | 247.603187 | 0.232742941 | 0.37136699 | 0.62671952 | 0.53084311 | 0.70477034 |
| NR2F1 | 199.602664 | 1.590429736 | 0.37046845 | 4.29302341 | 1.76E-05 | 0.00040199 |
| FAM172A | 3087.30091 | 0.347175609 | 0.22956447 | 1.51232295 | 0.13045173 | 0.29298735 |
| KIAA0825 | 2158.91171 | 1.284005686 | 0.46775661 | 2.74502949 | 0.00605055 | 0.0348027 |
| SLF1 | 408.72359 | 0.798753209 | 0.29491381 | 2.70842928 | 0.00676025 | 0.03788339 |
| MCTP1 | 126.55212 | -0.469024523 | 0.45616879 | -1.028182 | 0.30386424 | 0.50406234 |
| TTC37 | 1411.52141 | -0.266550648 | 0.22836895 | -1.167193 | 0.24313243 | 0.4382342 |
| ARSK | 173.939601 | 0.220423292 | 0.22198966 | 0.99294395 | 0.32073727 | 0.52114103 |
| RHOBTB3 | 436.492371 | -0.819369736 | 0.49973815 | -1.6395981 | 0.10108875 | 0.24769817 |
| GLRX | 1275.49849 | -1.280289237 | 0.53959133 | -2.3727016 | 0.01765852 | 0.07554583 |
| AC008592.4 | 44.0258071 | 0.101573204 | 0.80263163 | 0.12655021 | 0.8992964 | 0.94743919 |
| LINC01554 | 57.2671179 | -0.341114414 | 1.07442411 | -0.3174858 | 0.750875 | 0.85954933 |
| ELL2 | 2417.18136 | -0.611585661 | 0.32483279 | -1.8827707 | 0.05973145 | 0.17419978 |
| AC104123.1 | 1016.30836 | 0.702713775 | 0.47711891 | 1.47282732 | 0.1407976 | 0.30766378 |
| CAST | 2822.51853 | -1.16577388 | 0.24150147 | -4.8271916 | 1.38E-06 | 5.02E-05 |
| ERAP1 | 470.722787 | -1.059679147 | 0.32244396 | -3.2863979 | 0.00101478 | 0.00914922 |
| AC009126.1 | 72.7600131 | -0.053911261 | 0.53419655 | -0.1009203 | 0.91961375 | 0.95690112 |
| ERAP2 | 158.695212 | -0.861093363 | 0.49038591 | -1.7559504 | 0.07909687 | 0.21028635 |
| LNPEP | 1178.48651 | -0.425377918 | 0.1636217 | -2.5997647 | 0.00932877 | 0.04792709 |
| LIX1-AS1 | 610.278515 | 0.144290763 | 0.38068794 | 0.37902635 | 0.70466829 | 0.83052509 |
| RIOK2 | 149.907594 | 0.105336203 | 0.28316629 | 0.37199414 | 0.70989721 | 0.83388131 |
| LINC01340 | 57.3676224 | -0.100694365 | 0.73387009 | -0.1372101 | 0.89086475 | 0.94331061 |
| RGMB | 71.0335327 | -0.608555719 | 0.25542925 | -2.3824825 | 0.01719635 | 0.07419957 |
| CHD1 | 2105.61222 | 0.025131783 | 0.18329237 | 0.13711309 | 0.89094141 | 0.94331329 |
| LINC02062 | 337.42887 | 0.153643591 | 0.44481549 | 0.34540971 | 0.72978638 | 0.84643611 |
| AC021086.1 | 58.4770537 | 0.544966115 | 0.58687865 | 0.92858399 | 0.35310472 | 0.55189396 |
| FAM174A | 198.737492 | -0.188142253 | 0.23088058 | -0.8148899 | 0.41513534 | 0.60858397 |
| SLCO4C1 | 1658.34077 | 1.451060267 | 0.41850788 | 3.46722326 | 0.00052586 | 0.00552339 |
| PAM | 6839.63356 | -0.842141158 | 0.31495544 | -2.6738422 | 0.00749877 | 0.04096679 |
| GIN1 | 182.597358 | 0.547395585 | 0.14818757 | 3.69393737 | 0.00022081 | 0.00288799 |
| PPIP5K2 | 1403.43792 | 0.4827203 | 0.19939166 | 2.42096539 | 0.01547935 | 0.06868743 |
| C5orf30 | 68.9302952 | -0.631280365 | 0.31382462 | -2.0115706 | 0.04426522 | 0.14309987 |
| NUDT12 | 230.595177 | 0.056962943 | 0.41823148 | 0.13619956 | 0.89166353 | 0.94341489 |
| AC099520.1 | 86.6484965 | 1.991140215 | 0.48562458 | 4.10016356 | 4.13E-05 | 0.0007768 |
| EFNA5 | 456.462642 | -0.499560277 | 0.63401122 | -0.787936 | 0.43073413 | 0.6233843 |
| FBXL17 | 4967.97571 | 0.062587143 | 0.14729914 | 0.42489821 | 0.67091088 | 0.80916633 |
| FER | 2442.47664 | -0.191563958 | 0.26560227 | -0.7212437 | 0.4707596 | 0.65738999 |
| AC008467.1 | 130.157478 | -0.743111537 | 0.65503489 | -1.134461 | 0.25660122 | 0.45384415 |
| PJA2 | 1298.3936 | -0.626133163 | 0.1371784 | -4.5643713 | 5.01E-06 | 0.00014277 |
| MAN2A1 | 2318.32861 | -0.202205302 | 0.27940346 | -0.7237036 | 0.4692477 | 0.65619364 |
| TMEM232 | 1446.94143 | 1.147338047 | 0.55029027 | 2.08496882 | 0.03707213 | 0.12678336 |
| SLC25A46 | 701.484349 | 0.024258526 | 0.17219628 | 0.14087717 | 0.88796698 | 0.94209263 |
| WDR36 | 397.204162 | -0.308002787 | 0.2164237 | -1.4231472 | 0.15469348 | 0.32727019 |
| STARD4 | 170.676562 | -1.217508783 | 0.48979992 | -2.4857268 | 0.01292872 | 0.06043786 |
| STARD4-AS1 | 136.30134 | -0.140534472 | 0.43088196 | -0.3261554 | 0.74430679 | 0.85510127 |
| NREP | 845.397286 | 1.108622875 | 0.2391798 | 4.63510236 | 3.57E-06 | 0.00011039 |
| EPB41L4A | 4146.77076 | 0.269781189 | 0.34197927 | 0.78888171 | 0.43018115 | 0.62271549 |
| EPB41L4A-A! | 81.3087643 | -0.950172795 | 0.45521467 | -2.0873071 | 0.03686039 | 0.12621666 |
| AC010261.2 | 67.5381633 | -0.168607547 | 0.68530818 | -0.2460317 | 0.80565768 | 0.89453456 |
| AC104126.1 | 120.289923 | 0.637741634 | 0.71011683 | 0.89807988 | 0.36914296 | 0.56667517 |
| APC | 941.762666 | 0.056412251 | 0.1760175 | 0.32049228 | 0.74859518 | 0.85823001 |
| SRP19 | 317.150149 | 0.044142792 | 0.21584803 | 0.20450867 | 0.83795602 | 0.91447955 |
| REEP5 | 983.063558 | -0.590993893 | 0.27250565 | -2.16874 | 0.03010243 | 0.10948005 |
| DCP2 | 606.059759 | -0.130098025 | 0.13118016 | -0.9917508 | 0.32131912 | 0.52171538 |
| MCC | 500.225931 | -0.008957981 | 0.51850015 | -0.0172767 | 0.98621586 | 0.99331772 |
| YTHDC2 | 1082.08625 | 0.391962363 | 0.21530138 | 1.82052881 | 0.06867851 | 0.19172635 |
| KCNN2 | 58.2759166 | 4.336642426 | 0.79008711 | 5.48881554 | 4.05E-08 | 2.65E-06 |
| PGGT1B | 371.075441 | -0.227408336 | 0.15438664 | -1.4729794 | 0.14075659 | 0.30766378 |
| CCDC112 | 112.667072 | -0.510382572 | 0.42412146 | -1.2033877 | 0.22882631 | 0.42176886 |
| FEM1C | 430.837935 | -1.844026047 | 0.40127177 | -4.5954542 | 4.32E-06 | 0.00012758 |
| TMED7-TICAI | 109.158746 | -0.930763236 | 0.33649167 | -2.7660811 | 0.00567344 | 0.03324749 |
| ATG12 | 453.571411 | -0.210546375 | 0.18030416 | -1.1677289 | 0.24291615 | 0.43805442 |
| AP3S1 | 782.231778 | -0.405107229 | 0.25468001 | -1.5906518 | 0.11168795 | 0.26454238 |
| COMMD10 | 3371.6389 | 0.328057201 | 0.15901988 | 2.06299488 | 0.03911311 | 0.13162864 |
| SEMA6A | 2735.6112 | -1.532651448 | 0.58232622 | -2.6319465 | 0.00848972 | 0.04467482 |
| SEMA6A-AS1 | 436.468528 | -0.391236906 | 0.3376356 | -1.1587549 | 0.2465561 | 0.44223299 |
| LINC00992 | 71.8825571 | -0.064011838 | 0.75238038 | -0.0850791 | 0.93219851 | 0.96391408 |
| LINC02147 | 1969.71062 | 1.964555184 | 0.76025607 | 2.58407038 | 0.00976419 | 0.04960614 |
| AC093206.1 | 101.105281 | 1.476621218 | 0.35331023 | 4.17938987 | 2.92E-05 | 0.00059005 |
| DTWD2 | 932.409848 | 1.308291525 | 0.1756968 | 7.44630255 | 9.60E-14 | 3.56E-11 |
| AC008629.1 | 309.65876 | 1.138318513 | 0.50344741 | 2.26104751 | 0.02375631 | 0.09290451 |
| DMXL1 | 4722.88224 | 0.712634027 | 0.28874957 | 2.4680003 | 0.01358702 | 0.06255871 |
| TNFAIP8 | 1110.79186 | -2.175518696 | 0.29288975 | -7.4277733 | 1.10E-13 | 3.88E-11 |
| HSD17B4 | 1019.88981 | -0.060011018 | 0.29105542 | -0.2061842 | 0.83664706 | 0.91377858 |
| SRFBP1 | 544.074321 | -0.0098388 | 0.29781638 | -0.0330365 | 0.97364551 | 0.98680944 |
| AC119150.1 | 67.1693049 | -0.360230658 | 0.46690871 | -0.7715227 | 0.44039719 | 0.63242789 |
| SNX2 | 908.682691 | 0.233438587 | 0.24053195 | 0.97050969 | 0.3317925 | 0.53141196 |
| SNX24 | 1542.99929 | -0.169357297 | 0.18507497 | -0.915074 | 0.36015278 | 0.55850369 |
| PPIC | 203.585073 | -0.284731938 | 0.33735984 | -0.8440007 | 0.39866902 | 0.59287797 |
| PRDM6 | 77.3096404 | 0.085473926 | 0.42267651 | 0.20222067 | 0.83974422 | 0.91504682 |
| CEP120 | 344.522272 | 0.059583177 | 0.23855176 | 0.24977044 | 0.80276488 | 0.89272007 |
| CSNK1G3 | 1072.68845 | -0.301947438 | 0.21994987 | -1.3728012 | 0.16981416 | 0.34878317 |
| ZNF608 | 1898.72711 | 0.299412375 | 0.49997537 | 0.59885425 | 0.54927008 | 0.71915608 |
| AC112196.1 | 62.3235863 | -0.8377354 | 0.80924866 | -1.0352015 | 0.30057483 | 0.50097024 |
| LINC02240 | 77.8540418 | 0.443885257 | 0.51631486 | 0.85971815 | 0.38994443 | 0.58542916 |
| AC010235.1 | 206.452002 | 0.435322657 | 0.69957577 | 0.62226663 | 0.53376656 | 0.70689258 |
| GRAMD2B | 4017.54333 | -0.278385696 | 0.31645954 | -0.8796881 | 0.37902829 | 0.57578863 |
| AC093535.1 | 131.64729 | -1.537763783 | 0.64436037 | -2.3864965 | 0.01700977 | 0.073666 |
| ALDH7A1 | 969.52808 | 1.863338995 | 0.45655283 | 4.08132174 | 4.48E-05 | 0.00082835 |
| PHAX | 250.624539 | 0.263989719 | 0.20099626 | 1.31340615 | 0.18904613 | 0.37309941 |
| LMNB1 | 57.8986348 | 0.267585856 | 0.48556499 | 0.55108144 | 0.58157785 | 0.74455393 |
| 3-Mar | 141.641 | -1.109106138 | 0.42475268 | -2.6111811 | 0.00902301 | 0.04677701 |

| | | | | | | |
|---|---|---|---|---|---|---|
| C5orf63 | 106.345565 | -0.217956347 | 0.31923793 | -0.6827395 | 0.49477146 | 0.6769747 |
| PRRC1 | 436.08106 | -0.735790568 | 0.31016162 | -2.3722811 | 0.01767863 | 0.07560719 |
| CTXN3 | 42.4281391 | 3.791440314 | 0.63333158 | 5.9865013 | 2.14E-09 | 2.23E-07 |
| CCDC192 | 57.4677893 | 0.420855369 | 0.78858284 | 0.53368568 | 0.59355902 | 0.75444301 |
| LINC01184 | 1043.11341 | 0.402987177 | 0.26228655 | 1.53643856 | 0.12443086 | 0.28426229 |
| SLC12A2 | 387.081099 | -0.078983138 | 0.23792423 | -0.3319676 | 0.73991372 | 0.85230954 |
| FBN2 | 51.6765736 | 0.475203882 | 0.62447981 | 0.76095956 | 0.44668122 | 0.63757779 |
| SLC27A6 | 176.946403 | 1.558690893 | 1.38218245 | 1.12770271 | 0.25944549 | 0.45761798 |
| ISOC1 | 188.782285 | 0.211618438 | 0.21137441 | 1.00115449 | 0.31675212 | 0.51681007 |
| CHSY3 | 391.673588 | -0.344714322 | 0.53567005 | -0.6435199 | 0.51988682 | 0.69597507 |
| HINT1 | 1988.33436 | -1.353679995 | 0.35396557 | -3.8243267 | 0.00013113 | 0.00191473 |
| LYRM7 | 225.363128 | -0.117473072 | 0.26483666 | -0.443568 | 0.65735493 | 0.80028102 |
| CDC42SE2 | 3612.83397 | -0.972873122 | 0.33839198 | -2.8749887 | 0.00404042 | 0.02582745 |
| RAPGEF6 | 36.6916438 | -0.723230432 | 0.35747178 | -2.0231819 | 0.0430544 | 0.14036697 |
| P4HA2-AS1 | 41.018065 | 0.752096713 | 0.6764328 | 1.11185725 | 0.26619953 | 0.46404642 |
| P4HA2 | 1456.5108 | 2.243764942 | 0.40838006 | 5.49430589 | 3.92E-08 | 2.60E-06 |
| SLC22A4 | 502.095141 | -0.315643405 | 0.24615136 | -1.2823143 | 0.19973242 | 0.38650408 |
| SLC22A5 | 2466.74445 | 0.204135433 | 0.47555652 | 0.42925588 | 0.66773702 | 0.80690041 |
| AC116366.3 | 630.123451 | -0.31950004 | 0.48365343 | -0.6605971 | 0.50887074 | 0.68848991 |
| IRF1 | 249.818177 | -2.065206292 | 0.63674006 | -3.2434056 | 0.0011811 | 0.0102529 |
| TH2LCRR | 89.1973561 | -1.091555078 | 0.60788857 | -1.79565 | 0.07255021 | 0.19879007 |
| AC004039.1 | 42.4343813 | -0.467152719 | 0.65927023 | -0.7085906 | 0.47857854 | 0.66413513 |
| KIF3A | 386.233723 | -0.19828831 | 0.25436323 | -0.7795479 | 0.43565705 | 0.62792337 |
| 8-Sep | 238.81941 | 0.281607869 | 0.4142931 | 0.67973102 | 0.49667479 | 0.67856373 |
| SHROOM1 | 78.290114 | -0.047038927 | 0.42699636 | -0.1101624 | 0.91228061 | 0.9535692 |
| UQCRQ | 436.8733 | -0.510365713 | 0.37373942 | -1.3655656 | 0.17207534 | 0.35140594 |
| LEAP2 | 116.287095 | 0.117115089 | 0.86010925 | 0.13616304 | 0.89169239 | 0.94341489 |
| AFF4 | 3429.01346 | -0.246555328 | 0.25092243 | -0.9825958 | 0.3258064 | 0.52594845 |
| ZCCHC10 | 530.232209 | 0.215533217 | 0.21488226 | 1.00302939 | 0.31584668 | 0.51590706 |
| HSPA4 | 932.259084 | -0.09159062 | 0.3836999 | -0.2387038 | 0.81133528 | 0.8977214 |
| FSTL4 | 141.551716 | 1.123869301 | 0.58734015 | 1.91348966 | 0.0556854 | 0.16647322 |
| C5orf15 | 856.624406 | -0.253359506 | 0.30006501 | -0.8443487 | 0.39847458 | 0.59275547 |
| VDAC1 | 900.700866 | -0.604931484 | 0.18097939 | -3.3425436 | 0.00083014 | 0.00788811 |
| AC008608.2 | 34.7663062 | 0.006646507 | 0.47318763 | 0.01404624 | 0.98879309 | 0.99458453 |
| TCF7 | 95.7070091 | -1.483555545 | 0.35361642 | -4.1953808 | 2.72E-05 | 0.00055731 |
| SKP1 | 746.493272 | -0.571853265 | 0.22749972 | -2.5136438 | 0.0119491 | 0.05712814 |
| CDKL3 | 416.692731 | 0.448337725 | 0.26819643 | 1.67167671 | 0.09458809 | 0.23693694 |
| UBE2B | 1266.1839 | -0.592169189 | 0.23901931 | -2.4774952 | 0.01323082 | 0.06138274 |
| CDKN2AIPNL | 75.9370128 | 0.553965601 | 0.31315333 | 1.76899157 | 0.07689528 | 0.20663516 |
| LINC01843 | 59.4765065 | 1.887920395 | 0.54022007 | 3.49472465 | 0.00047455 | 0.00516654 |
| JADE2 | 454.948129 | 0.162303666 | 0.43938816 | 0.36938562 | 0.71184032 | 0.83509031 |
| SAR1B | 729.400401 | 0.322073454 | 0.19306953 | 1.66817344 | 0.0952813 | 0.23797755 |
| SEC24A | 729.452095 | -0.471285876 | 0.23342069 | -2.0190407 | 0.04348298 | 0.14123609 |
| CAMLG | 240.255744 | -0.32928197 | 0.26376581 | -1.2483876 | 0.21188914 | 0.4015283 |
| DDX46 | 1186.86383 | 0.077189496 | 0.30579863 | 0.25241937 | 0.80071693 | 0.89172494 |
| C5orf24 | 395.8419 | 0.433175338 | 0.22214558 | 1.94996156 | 0.0511807 | 0.15766945 |
| TXNDC15 | 221.386188 | 0.03250547 | 0.38472851 | 0.08448937 | 0.93266735 | 0.96432612 |
| PCBD2 | 1114.70402 | 0.378585379 | 0.44616731 | 0.84852783 | 0.39614408 | 0.59081295 |
| C5orf66 | 93.5475039 | -1.532890241 | 0.55020867 | -2.7860162 | 0.00533602 | 0.03180037 |
| H2AFY | 697.271319 | -0.736657981 | 0.33624762 | -2.1908199 | 0.02846483 | 0.10525152 |
| AC034206.1 | 51.6064929 | -0.914640917 | 0.71868005 | -1.2726677 | 0.20313597 | 0.39106527 |
| CXCL14 | 2158.56323 | -0.683384274 | 0.43228846 | -1.5808525 | 0.11391178 | 0.26768193 |
| SLC25A48 | 2228.16504 | 0.903360462 | 0.55375243 | 1.63134355 | 0.10281785 | 0.25038257 |
| LECT2 | 62.9645902 | 0.355292545 | 0.93006468 | 0.38200842 | 0.70245512 | 0.82904704 |
| TGFBI | 55.5992961 | -1.655318046 | 0.66378725 | -2.4937479 | 0.01264023 | 0.05945766 |
| SMAD5 | 1027.37257 | 0.087852437 | 0.19315957 | 0.45481794 | 0.64924022 | 0.79435292 |
| SMIM32 | 239.260345 | 0.615001217 | 0.45632616 | 1.3477229 | 0.17774752 | 0.3584501 |
| TRPC7 | 168.919155 | 0.407715049 | 0.82710749 | 0.49294082 | 0.6220544 | 0.77562992 |
| TRPC7-AS1 | 589.713399 | 2.166253988 | 0.36658163 | 5.90933591 | 3.43E-09 | 3.25E-07 |
| SPOCK1 | 672.75305 | -0.709583574 | 0.81713055 | -0.8683846 | 0.38518383 | 0.58152841 |
| KLHL3 | 37.9765211 | 0.119483369 | 0.64199854 | 0.18611159 | 0.85235725 | 0.92213818 |
| HNRNPA0 | 287.438316 | -1.022568239 | 0.33422045 | -3.0595621 | 0.00221661 | 0.01660517 |
| AC106791.1 | 76.9170933 | 0.089969903 | 0.71282333 | 0.12621627 | 0.89956073 | 0.94743919 |
| PKD2L2 | 52.0951095 | -1.104703922 | 0.43247609 | -2.5543699 | 0.01063802 | 0.05275517 |
| FAM13B | 1490.68744 | -0.254728705 | 0.17672118 | -1.4414158 | 0.14946725 | 0.31978206 |
| NME5 | 457.659686 | 0.850486005 | 0.52710777 | 1.61349549 | 0.1066369 | 0.25638871 |
| BRD8 | 415.184935 | 0.542707111 | 0.2743045 | 1.97848418 | 0.04787411 | 0.15106134 |
| CDC23 | 160.667566 | 0.109014883 | 0.18910267 | 0.57648517 | 0.56428729 | 0.73149731 |
| FAM53C | 214.884505 | -0.998974912 | 0.31741865 | -3.1471841 | 0.00164851 | 0.01323734 |
| KDM3B | 1216.90463 | 0.316080513 | 0.15430569 | 2.04840483 | 0.04052035 | 0.13458091 |
| EGR1 | 120.24955 | 0.544560794 | 0.7221926 | 0.75403818 | 0.4508263 | 0.64075861 |
| ETF1 | 645.726846 | -0.559930382 | 0.4358205 | -1.2847729 | 0.19887165 | 0.38523653 |
| HSPA9 | 1562.22632 | -0.981430895 | 0.39500251 | -2.4846194 | 0.012969 | 0.06054704 |
| CTNNA1 | 4249.51965 | -0.18935893 | 0.17563808 | -1.07812 | 0.2809802 | 0.47950643 |
| SIL1 | 1732.31219 | 0.15619255 | 0.21884974 | 0.71369768 | 0.47541414 | 0.66124102 |
| AC011405.1 | 53.3319416 | -0.64013682 | 0.58049535 | -1.1027424 | 0.27013906 | 0.46841191 |
| MATR3 | 50.4405332 | -2.355722172 | 0.68990731 | -3.4145488 | 0.00063888 | 0.00645803 |
| PAIP2 | 919.982543 | -0.084219077 | 0.2255846 | -0.373337 | 0.70889764 | 0.8332772 |
| AC135457.1 | 50.4758213 | 2.17952995 | 0.60459355 | 3.60495071 | 0.00031221 | 0.00378381 |
| SLC23A1 | 224.4675 | 2.862773042 | 0.53404024 | 5.36059421 | 8.29E-08 | 4.70E-06 |
| PROB1 | 40.6903127 | 1.198633289 | 0.48367669 | 2.47817049 | 0.0132058 | 0.06129764 |
| SPATA24 | 66.8297311 | 0.858158748 | 0.37127394 | 2.31138965 | 0.02081134 | 0.08489913 |
| DNAJC18 | 89.0070627 | -0.206655026 | 0.4149395 | -0.4980365 | 0.6184583 | 0.77318569 |
| UBE2D2 | 5802.29542 | -0.999098696 | 0.40505179 | -2.4665949 | 0.01364046 | 0.06272048 |
| CXXC5 | 551.919858 | 0.22528655 | 0.32795757 | 0.68693811 | 0.49212173 | 0.67441637 |
| AC008667.1 | 35.1647291 | 0.207083262 | 0.51643703 | 0.40098453 | 0.68843151 | 0.8199051 |
| NRG2 | 316.891582 | -0.023962637 | 0.47699286 | -0.0502369 | 0.95993362 | 0.97951389 |
| PURA | 916.820463 | -0.037458605 | 0.2309614 | -0.1621856 | 0.87115973 | 0.93338059 |
| IGIP | 129.137877 | 0.025926928 | 0.38784195 | 0.06684921 | 0.94670175 | 0.97260087 |
| CYSTM1 | 2498.39207 | -0.444137964 | 0.22290815 | -1.9924707 | 0.04631944 | 0.14782317 |
| PFDN1 | 630.44304 | -0.038856259 | 0.20881223 | -0.1860823 | 0.85238022 | 0.92213818 |
| AC008438.1 | 102.522648 | 0.786113112 | 0.36721793 | 2.14072641 | 0.03229611 | 0.11512333 |
| ANKHD1-EIF4 | 167.685391 | 0.276164113 | 0.37106312 | 0.74425103 | 0.45672462 | 0.64552736 |
| SRA1 | 237.348694 | -0.707233353 | 0.30904102 | -2.2884773 | 0.02210974 | 0.08869131 |
| APBB3 | 110.841379 | 0.489249198 | 0.3010727 | 1.62502014 | 0.10415827 | 0.25231575 |
| SLC35A4 | 171.061645 | -0.649156992 | 0.40954778 | -1.585058 | 0.11295316 | 0.26625089 |
| NDUFA2 | 290.132733 | -0.546898037 | 0.30268712 | -1.8068097 | 0.07079195 | 0.19563413 |
| TMCO6 | 33.9308017 | 0.075381887 | 0.33389185 | 0.22576738 | 0.82138235 | 0.90404363 |
| IK | 186.721911 | -0.394193134 | 0.33067689 | -1.1920795 | 0.23323009 | 0.42710552 |
| WDR55 | 168.95333 | -0.758071474 | 0.33999175 | -2.2296761 | 0.02576896 | 0.09849835 |
| HARS | 357.906919 | -0.34953108 | 0.25387349 | -1.3767923 | 0.16857646 | 0.34711458 |
| HARS2 | 94.3282716 | -0.182447767 | 0.24457312 | -0.7459845 | 0.45567676 | 0.64490647 |
| ZMAT2 | 148.378268 | -0.275731715 | 0.27354623 | -1.0079894 | 0.31345953 | 0.51382322 |
| TAF7 | 378.335655 | -0.701334879 | 0.22952262 | -3.0556243 | 0.00224593 | 0.01675825 |
| DIAPH1 | 2497.74581 | -0.017706908 | 0.16118376 | -0.1098554 | 0.91252405 | 0.9535692 |
| AC008781.2 | 62.9084365 | -0.896316206 | 0.64709292 | -1.385143 | 0.16600875 | 0.34352083 |
| HDAC3 | 389.337557 | -0.365035775 | 0.25866009 | -1.4112567 | 0.15816894 | 0.33282494 |
| RELL2 | 75.5160753 | -0.236138951 | 0.54629836 | -0.4322527 | 0.66555775 | 0.80579154 |
| FCHSD1 | 49.0285937 | -0.399532096 | 0.39429707 | -1.0132769 | 0.31092794 | 0.51126311 |
| PCDH1 | 141.460991 | -0.936571653 | 0.44419883 | -2.1084514 | 0.03499196 | 0.12170384 |
| DELE1 | 359.521178 | 0.036144046 | 0.19290931 | 0.18736289 | 0.85137612 | 0.92193178 |
| RNF14 | 578.89764 | -0.307304292 | 0.26285971 | -1.169081 | 0.24237099 | 0.43756794 |
| AC005740.3 | 39.7486025 | -0.285242907 | 0.76347529 | -0.3736112 | 0.70869361 | 0.83318152 |

| | | | | | | |
|---|---|---|---|---|---|---|
| GNPDA1 | 580.149024 | -0.147755954 | 0.43860748 | -0.3368751 | 0.73621103 | 0.85079206 |
| NDFIP1 | 1174.98116 | -0.513731221 | 0.16778255 | -3.061887 | 0.00219946 | 0.01652854 |
| SPRY4 | 185.819646 | -0.92017038 | 0.48202304 | -1.9089759 | 0.0562652 | 0.16762161 |
| SPRY4-AS1 | 287.950183 | -1.730400769 | 0.51329661 | -3.3711517 | 0.00074855 | 0.00731262 |
| FGF1 | 77.1916829 | -0.023669484 | 0.53574085 | -0.0441808 | 0.96476025 | 0.981662 |
| ARHGAP26 | 6765.74469 | -1.237869265 | 0.42662112 | -2.9015659 | 0.00371303 | 0.02418938 |
| ARHGAP26-A | 60.7653593 | -1.674160748 | 0.58893398 | -2.8426968 | 0.00447336 | 0.02789643 |
| NR3C1 | 2266.51397 | 0.052126842 | 0.23782826 | 0.2191785 | 0.826511 | 0.90704454 |
| YIPF5 | 222.5753 | -0.714761554 | 0.32372014 | -2.2079614 | 0.02724697 | 0.10236573 |
| KCTD16 | 964.158432 | 0.641060003 | 0.54482412 | 1.1766366 | 0.23934058 | 0.43434945 |
| PRELID2 | 674.603631 | -2.003572506 | 0.51846647 | -3.8644206 | 0.00011135 | 0.00169028 |
| SH3RF2 | 335.545842 | -2.374538685 | 0.7337931 | -3.2359785 | 0.00121227 | 0.010477 |
| PLAC8L1 | 37.525497 | -0.018195091 | 0.48797393 | -0.037287 | 0.97025616 | 0.9852064 |
| LARS | 854.55078 | 0.229745439 | 0.21018655 | 1.09305492 | 0.27436969 | 0.47233234 |
| AC091959.3 | 37.2638605 | -0.071668745 | 0.32345142 | -0.221575 | 0.82464475 | 0.90637524 |
| TCERG1 | 1106.71506 | -0.418186083 | 0.23861799 | -1.7525338 | 0.07968207 | 0.21135017 |
| PPP2R2B | 1717.99296 | 0.970915881 | 1.1963585 | 0.81155931 | 0.41704455 | 0.61036016 |
| STK32A | 294.551024 | -0.06731849 | 0.6373324 | -0.1056254 | 0.91587957 | 0.95496466 |
| AC091948.1 | 297.23549 | 1.267786037 | 0.61761233 | 2.05272139 | 0.04009961 | 0.13379876 |
| FBXO38 | 706.789139 | -0.095705101 | 0.14863827 | -0.6438793 | 0.51965372 | 0.69597507 |
| ABLIM3 | 3092.45053 | -1.284501717 | 0.49582525 | -2.5906339 | 0.00957993 | 0.04883308 |
| AC012613.1 | 79.9515244 | -1.610360512 | 0.70232825 | -2.2928887 | 0.02185441 | 0.08802775 |
| AFAP1L1 | 147.677652 | -2.874411984 | 0.37091087 | -7.7496029 | 9.22E-15 | 4.36E-12 |
| GRPEL2 | 63.4449739 | -0.242202458 | 0.38214101 | -0.6338039 | 0.52620882 | 0.7009757 |
| CSNK1A1 | 3558.00555 | -0.424219685 | 0.16605714 | -2.5546609 | 0.01062913 | 0.05274777 |
| ARHGEF37 | 230.33244 | 0.162663316 | 0.38606001 | 0.42134205 | 0.67350533 | 0.81072605 |
| PPARGC1B | 757.086561 | 1.157099211 | 0.4441579 | 2.60515281 | 0.00918333 | 0.04744662 |
| SLC26A2 | 102.38723 | -0.816577222 | 0.48581548 | -1.6808382 | 0.09279434 | 0.23445752 |
| TIGD6 | 41.8060466 | 0.044078997 | 0.31372028 | 0.14050413 | 0.88826169 | 0.94229571 |
| HMGXB3 | 484.27879 | -0.404138409 | 0.27885562 | -1.4492748 | 0.14726086 | 0.31672755 |
| CAMK2A | 142.522827 | 2.591743918 | 1.0022001 | 2.58605434 | 0.00970816 | 0.04937654 |
| ARSI | 150.109899 | 2.823488081 | 0.72100668 | 3.91603595 | 9.00E-05 | 0.0014448 |
| TCOF1 | 180.029969 | -0.519308331 | 0.37960903 | -1.3680084 | 0.17130946 | 0.35036388 |
| CD74 | 1610.83763 | -1.022487244 | 0.52670301 | -1.9412975 | 0.0522222 | 0.15948419 |
| RPS14 | 1220.36668 | -0.782855553 | 0.40124483 | -1.951067 | 0.05104908 | 0.1573841 |
| NDST1 | 659.393021 | 0.783036882 | 0.26981026 | 2.9021761 | 0.0037058 | 0.02417188 |
| SYNPO | 116.846132 | 1.553784874 | 0.39308936 | 3.95275229 | 7.73E-05 | 0.00127412 |
| RBM22 | 175.082214 | -0.166887201 | 0.28883961 | -0.577785 | 0.56340927 | 0.73082718 |
| DCTN4 | 703.513199 | -0.04884814 | 0.16839911 | -0.2900736 | 0.77175992 | 0.87317484 |
| SMIM3 | 54.8396014 | -0.011801209 | 0.40730717 | -0.0289737 | 0.97688554 | 0.9886052 |
| ZNF300 | 101.582983 | -0.065356827 | 0.36681796 | -0.1781724 | 0.85858761 | 0.92564577 |
| GPX3 | 23397.173 | 0.0281863 | 0.52661441 | 0.0535236 | 0.95731473 | 0.97834441 |
| TNIP1 | 1206.7936 | -0.805458529 | 0.24356351 | -3.3069754 | 0.00094309 | 0.00865838 |
| ANXA6 | 363.763153 | 0.246070055 | 0.49806068 | 0.49405638 | 0.62126636 | 0.77507036 |
| GM2A | 54.8091362 | -0.163020003 | 0.3925817 | -0.4152512 | 0.67795806 | 0.81346017 |
| SLC36A2 | 3283.23669 | 2.90340986 | 0.88764401 | 3.27091697 | 0.00107199 | 0.00949213 |
| SLC36A1 | 658.903686 | 0.023556003 | 0.47169074 | 0.04993951 | 0.96017059 | 0.9796009 |
| SPARC | 205.88475 | -0.112329368 | 0.58194225 | -0.1930249 | 0.84693944 | 0.91937902 |
| ATOX1 | 525.132951 | 0.190313527 | 0.24711517 | 0.77014101 | 0.44121625 | 0.63313978 |
| G3BP1 | 1249.21772 | -0.209573178 | 0.22800627 | -0.9191553 | 0.35801433 | 0.55650968 |
| LINC01933 | 70.8646117 | -0.63841288 | 0.30209806 | -2.1132637 | 0.0345782 | 0.12069351 |
| LINC01470 | 56.1799899 | 0.193536893 | 0.32570779 | 0.59420407 | 0.55237563 | 0.72184315 |
| FAM114A2 | 189.612991 | 0.314067583 | 0.19532446 | 1.60792756 | 0.10785105 | 0.25813066 |
| MFAP3 | 293.216627 | -0.154736008 | 0.20999778 | -0.7368459 | 0.46121604 | 0.64973158 |
| GALNT10 | 407.839521 | -0.62320451 | 0.34437737 | -1.8096558 | 0.07034918 | 0.1949214 |
| SAP30L-AS1 | 236.853902 | 0.213060368 | 0.3292134 | 0.64718012 | 0.51751538 | 0.69429219 |
| SAP30L | 162.705698 | -0.106585443 | 0.20370932 | -0.5232232 | 0.60081893 | 0.76021647 |
| LARP1 | 2789.09525 | 0.749399319 | 0.25755022 | 2.90972109 | 0.00361751 | 0.0237819 |
| FAXDC2 | 210.788452 | 0.990359106 | 0.49889179 | 1.98511808 | 0.04713134 | 0.14971709 |
| CNOT8 | 290.34271 | -0.159991367 | 0.16447352 | -0.9727485 | 0.33067832 | 0.53050185 |
| GEMIN5 | 203.276069 | 0.016569366 | 0.36021951 | 0.04599797 | 0.96331187 | 0.98084354 |
| MRPL22 | 236.253578 | -0.392830079 | 0.16773048 | -2.3420315 | 0.01917909 | 0.07996279 |
| SGCD | 895.285892 | 0.183586031 | 0.762468 | 0.24077867 | 0.80972666 | 0.89673777 |
| TIMD4 | 613.869686 | -1.072395721 | 0.40196449 | -2.6678867 | 0.007633 | 0.04145226 |
| HAVCR1 | 1650.90001 | -2.084040199 | 0.4114242 | -5.0654293 | 4.07E-07 | 1.80E-05 |
| HAVCR2 | 1067.86905 | -0.265035783 | 0.38011568 | -0.6972503 | 0.48564615 | 0.66928934 |
| AC011377.1 | 42.6662485 | 2.106652923 | 0.52898513 | 3.9824426 | 6.82E-05 | 0.00115285 |
| CYFIP2 | 1538.46632 | 0.534544001 | 0.49073031 | 1.08928263 | 0.27602927 | 0.47377075 |
| THG1L | 52.6119274 | -0.503277514 | 0.48801811 | -1.0312681 | 0.30241511 | 0.50269269 |
| LSM11 | 53.0288344 | -0.408999582 | 0.35530177 | -1.151133 | 0.24967753 | 0.44586762 |
| CLINT1 | 2016.1904 | 0.063403498 | 0.18789569 | 0.33743988 | 0.73578533 | 0.85058695 |
| EBF1 | 53.015983 | 0.255342108 | 0.53073753 | 0.48110807 | 0.63043969 | 0.78203742 |
| RNF145 | 583.800307 | -1.313055871 | 0.1471077 | -8.9258131 | 4.42E-19 | 6.74E-16 |
| UBLCP1 | 248.535434 | 0.309501526 | 0.2771288 | 1.11681472 | 0.26407355 | 0.46207385 |
| AC008691.1 | 38.8802262 | -0.423396483 | 0.46398443 | -0.912523 | 0.36149345 | 0.55944346 |
| ADRA1B | 443.746732 | 1.643791096 | 0.55183939 | 2.97874914 | 0.00289428 | 0.02014822 |
| TTC1 | 796.564172 | -0.306928822 | 0.2147318 | -1.429359 | 0.15290108 | 0.32481583 |
| PWWP2A | 555.480477 | 0.254058523 | 0.26408654 | 0.96202754 | 0.33603577 | 0.53602261 |
| ZBED8 | 38.1768035 | 0.344458886 | 0.44344625 | 0.77677709 | 0.43729029 | 0.62909779 |
| SLU7 | 182.359386 | -0.461848941 | 0.2504572 | -1.8440234 | 0.06517973 | 0.18520981 |
| MIR3142HG | 153.892217 | -2.919744561 | 0.7709225 | -3.7873386 | 0.00015227 | 0.00213671 |
| GABRB2 | 217.692089 | 0.657237556 | 0.94764472 | 0.69354848 | 0.48796541 | 0.67113605 |
| AC113414.1 | 2915.11328 | 0.668946549 | 0.71584802 | 0.93448124 | 0.35005571 | 0.5493439 |
| CCNG1 | 465.368417 | -0.387223956 | 0.33801913 | -1.1455682 | 0.25197387 | 0.44840158 |
| NUDCD2 | 121.866922 | -0.63544726 | 0.33581677 | -1.892244 | 0.05845848 | 0.17167378 |
| MAT2B | 263.137179 | 0.032732726 | 0.26304726 | 0.12443668 | 0.90096954 | 0.94785397 |
| AC109466.1 | 144.173265 | 2.291002646 | 0.82127955 | 2.78955276 | 0.00527809 | 0.03152282 |
| AC008415.1 | 201.740735 | 1.937060513 | 0.97739719 | 1.98185603 | 0.04749536 | 0.15042597 |
| TENM2 | 795.374251 | 1.107489305 | 0.80806793 | 1.37053986 | 0.17051844 | 0.34958066 |
| AC008637.1 | 54.9227007 | 1.262013123 | 0.61388692 | 2.05577457 | 0.03980425 | 0.13307998 |
| WWC1 | 8445.37487 | -0.208970655 | 0.42121533 | -0.4961136 | 0.61981426 | 0.77427999 |
| RARS | 700.988437 | -0.744291882 | 0.2622085 | -2.8385498 | 0.0045319 | 0.02818458 |
| PANK3 | 573.107253 | -0.494096462 | 0.24571928 | -2.0108168 | 0.04434481 | 0.14325579 |
| SLIT3 | 994.448792 | 1.373123369 | 0.87411878 | 1.57086589 | 0.1162138 | 0.27158651 |
| SPDL1 | 56.7013511 | -0.02004759 | 0.35561831 | -0.0563739 | 0.95504395 | 0.97731423 |
| FAM196B | 72.6961378 | 2.207138921 | 0.84729305 | 2.60492979 | 0.00918931 | 0.04745963 |
| KCNIP1 | 85.0011568 | -0.899608068 | 0.47157941 | -1.9076492 | 0.05643658 | 0.16787677 |
| AC008514.1 | 68.9381086 | -1.243812736 | 0.72418129 | -1.7175433 | 0.08587995 | 0.22227274 |
| RANBP17 | 1351.30732 | 0.273976579 | 0.31277345 | 0.87595855 | 0.38105257 | 0.57750588 |
| NPM1 | 632.02161 | -0.934169657 | 0.3510048 | -2.6614156 | 0.00778128 | 0.04204102 |
| FBXW11 | 2233.09362 | 0.058450884 | 0.22337232 | 0.2616747 | 0.79357225 | 0.88739587 |
| STK10 | 363.747357 | -1.555619525 | 0.68797949 | -2.2611423 | 0.02375045 | 0.09290451 |
| UBTD2 | 1028.14437 | -0.028282357 | 0.30976012 | -0.0913041 | 0.92725099 | 0.96170016 |
| SH3PXD2B | 265.843321 | -1.593904357 | 0.50082715 | -3.1825438 | 0.00145987 | 0.01206178 |
| DUSP1 | 648.014886 | -0.906876752 | 0.71667519 | -1.2653944 | 0.20572999 | 0.39445251 |
| ERGIC1 | 1508.393 | -0.47145817 | 0.26435968 | -1.7833966 | 0.07452176 | 0.20235571 |
| RPL26L1 | 80.6797901 | -0.264286603 | 0.26877322 | -0.983307 | 0.32545635 | 0.52575488 |
| ATP6V0E1 | 1462.3435 | -0.551631881 | 0.23256734 | -2.3719232 | 0.01769577 | 0.07565686 |
| CREBRF | 1335.51933 | -0.029709643 | 0.29067535 | -0.102209 | 0.91859077 | 0.95655425 |
| BNIP1 | 97.7461793 | 0.48453394 | 0.22948884 | 2.1113617 | 0.03474123 | 0.12110837 |
| STC2 | 97.2499455 | -0.595861014 | 0.60957432 | -0.9775035 | 0.32831996 | 0.5282489 |
| BOD1 | 185.099066 | -0.154909082 | 0.17792767 | -0.8706295 | 0.38395647 | 0.5803781 |
| CPEB4 | 4589.16731 | 1.267771493 | 0.32461883 | 3.90541572 | 9.41E-05 | 0.00148199 |

| | | | | | | |
|---|---|---|---|---|---|---|
| SFXN1 | 1405.89144 | 0.506884599 | 0.30433929 | 1.66552467 | 0.09580813 | 0.23882914 |
| THOC3 | 260.955825 | -0.669935741 | 0.53289421 | -1.2571646 | 0.20869404 | 0.39796455 |
| SIMC1 | 356.141732 | -0.351659046 | 0.17284246 | -2.034564 | 0.04189476 | 0.13771018 |
| KIAA1191 | 792.505465 | 0.064664479 | 0.41566869 | 0.15556736 | 0.87637406 | 0.93554978 |
| AC138956.1 | 66.3990155 | -0.539959019 | 0.7510522 | -0.7189367 | 0.4721799 | 0.65827602 |
| AC138956.2 | 65.4108793 | -0.843320811 | 0.95485666 | -0.883191 | 0.37713308 | 0.57360814 |
| ARL10 | 206.638752 | -0.296941058 | 0.22802592 | -1.302225 | 0.19283947 | 0.37749937 |
| NOP16 | 84.9302724 | -0.885337717 | 0.58685511 | -1.5086138 | 0.1313975 | 0.29451893 |
| CLTB | 323.425441 | -0.053988837 | 0.41553597 | -0.1299258 | 0.89662514 | 0.94627667 |
| FAF2 | 960.371675 | -0.086718896 | 0.30790621 | -0.2816406 | 0.77821909 | 0.87771268 |
| RNF44 | 201.32679 | -0.330714726 | 0.23114059 | -1.4307947 | 0.15248907 | 0.32411759 |
| CDHR2 | 857.724238 | -0.731618099 | 0.69022227 | -1.0599746 | 0.28915614 | 0.48901458 |
| TSPAN17 | 128.501705 | -0.241768984 | 0.35114772 | -0.6885108 | 0.4911312 | 0.67375209 |
| UIMC1 | 1277.4749 | -0.143922621 | 0.21447796 | -0.6710369 | 0.50219703 | 0.68316938 |
| ZNF346 | 510.703381 | 0.469706311 | 0.28034115 | 1.67548117 | 0.09383986 | 0.23579523 |
| FGFR4 | 443.042402 | 0.599343888 | 0.4316146 | 1.38860892 | 0.1649517 | 0.34226355 |
| NSD1 | 2158.89527 | 0.693058059 | 0.12084715 | 5.73499724 | 9.75E-09 | 7.96E-07 |
| RAB24 | 46.3795938 | 0.253189807 | 0.6275779 | 0.40343965 | 0.68662483 | 0.81853944 |
| MXD3 | 50.3419898 | -0.407583825 | 0.39204805 | -1.0396272 | 0.29851312 | 0.49856456 |
| PRELID1 | 554.20951 | -1.062940242 | 0.57594192 | -1.8455685 | 0.06495489 | 0.18475548 |
| LMAN2 | 465.537443 | -0.436031428 | 0.34937312 | -1.2480394 | 0.21201662 | 0.40167407 |
| RGS14 | 406.367267 | -0.862757571 | 0.44402064 | -1.9430574 | 0.05200923 | 0.15901947 |
| SLC34A1 | 1914.91928 | 4.866941727 | 0.75144082 | 6.47681305 | 9.37E-11 | 1.49E-08 |
| F12 | 118.954939 | -1.841817764 | 0.40944787 | -4.4982961 | 6.85E-06 | 0.0001863 |
| GRK6 | 292.711621 | 1.404261219 | 0.37532328 | 3.74147115 | 0.00018295 | 0.00248036 |
| PRR7-AS1 | 42.6053467 | -1.978004857 | 0.50832138 | -3.8912486 | 9.97E-05 | 0.00154813 |
| PRR7 | 105.670328 | -2.527964836 | 0.42603441 | -5.9337105 | 2.96E-09 | 2.92E-07 |
| DBN1 | 110.958794 | -0.821008284 | 0.30955165 | -2.6522497 | 0.00799574 | 0.04289533 |
| PDLIM7 | 121.00994 | -1.510892601 | 0.51976944 | -2.9068515 | 0.00365086 | 0.02392084 |
| DDX41 | 117.215174 | -0.849879069 | 0.39287758 | -2.163216 | 0.03052457 | 0.1105125 |
| FAM193B | 837.621788 | -0.189975425 | 0.22816523 | -0.8326222 | 0.40505785 | 0.59906431 |
| TMED9 | 365.28736 | -1.422811811 | 0.3976157 | -3.5783592 | 0.00034576 | 0.0040962 |
| B4GALT7 | 137.351208 | -0.174448463 | 0.40297536 | -0.4329011 | 0.66508664 | 0.80579154 |
| AC106795.3 | 54.9923837 | -1.11321093 | 0.5141108 | -2.1653133 | 0.0303637 | 0.11016286 |
| RMND5B | 370.668729 | 0.058164875 | 0.28037298 | 0.20745535 | 0.83565426 | 0.91316952 |
| NHP2 | 155.273886 | -1.422112088 | 0.48886132 | -2.9090297 | 0.00362552 | 0.02380032 |
| HNRNPAB | 429.156883 | -1.244527493 | 0.39706047 | -3.1343525 | 0.00172234 | 0.01367 |
| PHYKPL | 2038.70742 | -0.291992262 | 0.25539341 | -1.1433038 | 0.25291247 | 0.4495748 |
| COL23A1 | 680.336603 | -1.19251338 | 0.60126414 | -1.9833436 | 0.04732906 | 0.15010169 |
| CLK4 | 641.312536 | 0.924927939 | 0.32720019 | 2.8267952 | 0.00470164 | 0.02899027 |
| ZNF354A | 330.787875 | 0.150943876 | 0.2819437 | 0.53536886 | 0.59239482 | 0.75373209 |
| ZNF354B | 280.133754 | 0.02350338 | 0.29853541 | 0.07872895 | 0.93724822 | 0.96636538 |
| ZFP2 | 109.559972 | 0.320949211 | 0.33760796 | 0.95065653 | 0.34177876 | 0.54246891 |
| ZNF354C | 89.5549727 | 0.566089234 | 0.36678312 | 1.54338956 | 0.12273629 | 0.28142293 |
| ADAMTS2 | 43.0325295 | -0.304458533 | 0.35550509 | -0.8564112 | 0.39177039 | 0.5869326 |
| RUFY1 | 894.518053 | 0.275361076 | 0.16610616 | 1.65774152 | 0.09736965 | 0.24130272 |
| AC136604.2 | 86.7538769 | 0.211969683 | 0.27035055 | 0.78405495 | 0.43300787 | 0.62552702 |
| HNRNPH1 | 2973.12477 | 0.697684594 | 0.22200976 | 3.14258525 | 0.00167463 | 0.01336097 |
| CANX | 1867.0221 | -0.511695062 | 0.29119454 | -1.7572275 | 0.07887904 | 0.20990001 |
| MAML1 | 672.995794 | 0.111023213 | 0.29314341 | 0.37873344 | 0.70488582 | 0.83069984 |
| LTC4S | 201.961849 | -0.292157706 | 0.61033963 | -0.4786805 | 0.63216591 | 0.78317951 |
| MGAT4B | 744.666354 | -1.643528946 | 0.42025743 | -3.9107672 | 9.20E-05 | 0.00146557 |
| SQSTM1 | 6947.98662 | -2.817962381 | 0.62194431 | -4.5308918 | 5.87E-06 | 0.00016397 |
| MRNIP | 435.200061 | 0.359352567 | 0.21565778 | 1.66630929 | 0.09565183 | 0.23872476 |
| TBC1D9B | 562.221052 | -0.843113935 | 0.25805965 | -3.2671281 | 0.00108645 | 0.00960149 |
| RNF130 | 1612.72864 | -0.547555298 | 0.36740105 | -1.4903477 | 0.13613284 | 0.30096337 |
| MAPK9 | 442.66376 | -0.063567864 | 0.15341295 | -0.4143579 | 0.67861203 | 0.81376982 |
| CNOT6 | 628.298637 | -0.241815611 | 0.25844608 | -0.9356521 | 0.34945236 | 0.54909516 |
| MGAT1 | 396.772337 | -0.523638201 | 0.42280602 | -1.2384833 | 0.21553691 | 0.40620987 |
| LINC00847 | 41.4590742 | 0.344439174 | 0.41231865 | 0.83537133 | 0.40350867 | 0.59761138 |
| ZFP62 | 119.029881 | -0.032377694 | 0.25472543 | -0.1271082 | 0.89885475 | 0.94743919 |
| TRIM41 | 213.869178 | 0.829550421 | 0.25133189 | 3.3006175 | 0.00096472 | 0.00881564 |
| RACK1 | 1122.10965 | -1.123619053 | 0.39833969 | -2.820756 | 0.00479106 | 0.02935677 |
| TRIM52 | 233.901436 | 0.387289491 | 0.23963787 | 1.61614478 | 0.10606302 | 0.25536725 |
| AC138035.1 | 88.8406736 | 0.661957908 | 0.33711779 | 1.96358046 | 0.04957877 | 0.15448423 |
| DUSP22 | 330.760189 | -0.806369181 | 0.19149836 | -4.2108413 | 2.54E-05 | 0.00052679 |
| EXOC2 | 1219.76014 | -0.462299633 | 0.24070796 | -1.920583 | 0.0547843 | 0.16467727 |
| AL356130.2 | 59.2070095 | 0.953397234 | 1.05104971 | 0.90709053 | 0.36435893 | 0.56203779 |
| AL033381.1 | 99.8838334 | -0.660575716 | 0.83613763 | -0.7900323 | 0.42950892 | 0.62220237 |
| FOXQ1 | 63.8213261 | -0.683466663 | 0.55542197 | -1.2305359 | 0.2184965 | 0.40973677 |
| FOXC1 | 168.565389 | -0.076840777 | 0.5161966 | -0.1488595 | 0.88166449 | 0.93863884 |
| GMDS | 3497.77564 | -1.266281635 | 0.33238631 | -3.8096684 | 0.00013915 | 0.00199725 |
| GMDS-DT | 5648.10637 | -0.228553308 | 0.38157026 | -0.5989809 | 0.5491856 | 0.71911416 |
| MYLK4 | 416.157055 | 0.437650557 | 0.84305617 | 0.51912384 | 0.60367438 | 0.76169806 |
| WRNIP1 | 416.412379 | -0.18196429 | 0.20832424 | -0.8734667 | 0.38240874 | 0.57906281 |
| SERPINB1 | 292.875245 | -1.106420115 | 0.3263133 | -3.3906682 | 0.00069722 | 0.00691524 |
| SERPINB9 | 83.1631166 | -0.87555806 | 0.38991568 | -2.2455061 | 0.02473566 | 0.09553442 |
| SERPINB6 | 800.290051 | -0.477781097 | 0.22819363 | -2.093753 | 0.03628199 | 0.12461396 |
| NQO2 | 2733.8401 | 0.450161377 | 0.42702581 | 1.05417837 | 0.29180126 | 0.49181338 |
| RIPK1 | 835.218371 | -0.33870621 | 0.2959777 | -1.144364 | 0.25247272 | 0.44896778 |
| BPHL | 562.655934 | 1.154014818 | 0.40222228 | 2.86909724 | 0.00411645 | 0.02620725 |
| TUBB2A | 121.753546 | -0.607396901 | 0.32043308 | -1.8955499 | 0.05801959 | 0.17099 |
| PSMG4 | 282.733014 | -0.135462935 | 0.26815364 | -0.5051691 | 0.61344006 | 0.76943841 |
| SLC22A23 | 1500.57976 | 0.544507303 | 0.40916082 | 1.33079044 | 0.18325797 | 0.36558246 |
| PXDC1 | 295.864369 | -0.727756929 | 0.31777632 | -2.2901547 | 0.02201235 | 0.08843004 |
| PRPF4B | 1704.44201 | -0.132757137 | 0.22780722 | -0.5827609 | 0.56005428 | 0.72854361 |
| FAM217A | 51.0048441 | -0.488929267 | 0.62703517 | -0.7797478 | 0.43553936 | 0.62792337 |
| ECI2 | 684.964079 | 0.0972778 | 0.33056272 | 0.29427941 | 0.76854441 | 0.87097463 |
| CDYL | 1567.34877 | -0.704157348 | 0.42583861 | -1.653578 | 0.09821329 | 0.24256029 |
| LYRM4 | 990.104008 | -0.272339282 | 0.34772993 | -0.783192 | 0.4335144 | 0.62582221 |
| FARS2 | 2420.78221 | 0.654714081 | 0.23614262 | 2.77253667 | 0.00556213 | 0.03281965 |
| F13A1 | 165.276595 | 2.577859159 | 0.65900813 | 3.9117259 | 9.16E-05 | 0.00146557 |
| RREB1 | 2680.39036 | 0.185123947 | 0.16752247 | 1.10506935 | 0.26912954 | 0.46717453 |
| SSR1 | 600.167868 | -0.364235989 | 0.33459855 | -1.0885761 | 0.27634085 | 0.47404858 |
| AL139095.4 | 103.790377 | -0.52308537 | 0.47649989 | -1.097766 | 0.27230668 | 0.47033143 |
| RIOK1 | 171.506967 | -0.206072372 | 0.37984181 | -0.5425216 | 0.58745926 | 0.75016807 |
| DSP | 1254.52525 | -0.99755014 | 0.3945129 | -2.5285616 | 0.0114531 | 0.0555119 |
| SNRNP48 | 248.852175 | -0.024835004 | 0.15600621 | -0.1591924 | 0.87351729 | 0.9345535 |
| BLOC1S5-TXI | 227.944424 | 0.577119066 | 0.495702 | 1.16424599 | 0.24432434 | 0.43943757 |
| SLC35B3 | 335.26658 | 0.48213032 | 0.24214521 | 1.99107928 | 0.04647217 | 0.14820709 |
| GCNT2 | 2627.01115 | 0.398637395 | 0.41606232 | 0.95811943 | 0.33800254 | 0.53822035 |
| C6orf52 | 35.077885 | 0.577787224 | 0.33990649 | 1.69984169 | 0.08916071 | 0.22771117 |
| PAK1IP1 | 110.773254 | -0.481830652 | 0.46288867 | -1.0409212 | 0.2979121 | 0.49825013 |
| TMEM14C | 252.211372 | 0.03423113 | 0.24876782 | 0.13760272 | 0.8905544 | 0.94326785 |
| AL024498.2 | 42.590427 | -0.665536315 | 0.52540663 | -1.2667071 | 0.20526004 | 0.39375871 |
| MAK | 276.138776 | -0.927883694 | 0.43360202 | -2.1399432 | 0.03235936 | 0.11529756 |
| SMIM13 | 168.482958 | 0.241453183 | 0.20161585 | 1.19759026 | 0.23107657 | 0.42401159 |
| AL139807.1 | 378.925439 | -0.860938728 | 0.6520039 | -1.3204503 | 0.18668473 | 0.36999531 |
| NEDD9 | 1339.12343 | -0.6642381 | 0.54136012 | -1.2269801 | 0.21983006 | 0.41147217 |
| TMEM170B | 243.770116 | 0.250839259 | 0.29606059 | 0.8472565 | 0.39685217 | 0.59158811 |
| AL157373.2 | 42.1871532 | -1.209948711 | 0.55288594 | -2.1884237 | 0.02863875 | 0.10570582 |
| HIVEP1 | 1296.46675 | -0.608868003 | 0.38651832 | -1.575263 | 0.11519575 | 0.26983097 |
| EDN1 | 55.01333 | 0.109679819 | 0.53774387 | 0.20396294 | 0.83838246 | 0.91454366 |

| | | | | | | |
|---|---|---|---|---|---|---|
| PHACTR1 | 236.04646 | -1.107826586 | 0.4790281 | -2.3126547 | 0.02074163 | 0.08469587 |
| TBC1D7 | 429.84421 | -0.201847915 | 0.61262431 | -0.3294807 | 0.74179234 | 0.85342854 |
| GFOD1 | 118.336247 | -1.661761342 | 0.55056674 | -3.0182741 | 0.00254219 | 0.01843692 |
| SIRT5 | 363.120368 | 0.644131797 | 0.4932737 | 1.30583042 | 0.19161023 | 0.37593437 |
| NOL7 | 298.719231 | -0.925202539 | 0.30017637 | -3.0821964 | 0.00205479 | 0.01571709 |
| RANBP9 | 1178.84935 | 0.196273586 | 0.14456088 | 1.35772271 | 0.17455169 | 0.35440379 |
| MCUR1 | 503.47591 | 0.379505462 | 0.24669841 | 1.53833766 | 0.12396608 | 0.28346879 |
| CD83 | 143.337044 | 1.815051691 | 0.49761709 | 3.64748666 | 0.00026482 | 0.00332405 |
| JARID2 | 1500.59175 | 0.617278975 | 0.30279586 | 2.03859779 | 0.04149019 | 0.13683975 |
| DTNBP1 | 1970.2853 | 0.114451038 | 0.2117189 | 0.54058016 | 0.588797 | 0.75096682 |
| MYLIP | 282.663974 | 0.321378999 | 0.33744947 | 0.95237665 | 0.340906 | 0.54164814 |
| GMPR | 172.105359 | 0.394300414 | 0.44458189 | 0.88690165 | 0.37513185 | 0.57196133 |
| ATXN1 | 5176.14817 | -0.224349093 | 0.2232831 | -1.0047742 | 0.3150056 | 0.51523844 |
| CAP2 | 729.785333 | -0.131850615 | 0.286205 | -0.4606859 | 0.64502396 | 0.79180937 |
| FAM8A1 | 192.782067 | -0.099540526 | 0.25277497 | -0.3937911 | 0.69373529 | 0.82370734 |
| NUP153 | 1217.02966 | -0.204216948 | 0.21719233 | -0.9402586 | 0.34708494 | 0.5471809 |
| KIF13A | 1656.23698 | -0.102569467 | 0.29257239 | -0.3505781 | 0.72590491 | 0.84414851 |
| TPMT | 698.735937 | 0.415200581 | 0.26550775 | 1.56379837 | 0.11786493 | 0.27381461 |
| KDM1B | 488.644337 | -0.458928248 | 0.32064336 | -1.4312732 | 0.15235193 | 0.32404718 |
| DEK | 575.658683 | -0.647962039 | 0.28859514 | -2.2452285 | 0.02475347 | 0.09556779 |
| RNF144B | 732.938808 | -0.817956647 | 0.60567377 | -1.3504905 | 0.17685871 | 0.35735135 |
| MIR548A1HG | 147.381631 | -0.15475852 | 0.77814093 | -0.1988824 | 0.84235475 | 0.91687823 |
| AL022068.1 | 191.495134 | 0.36434093 | 0.55530913 | 0.65610469 | 0.51175676 | 0.69074882 |
| ID4 | 196.729704 | -0.177848795 | 0.37102717 | -0.4793417 | 0.63169556 | 0.78287984 |
| MBOAT1 | 101.161217 | -1.19593088 | 0.74820394 | -1.5984023 | 0.10995348 | 0.2615641 |
| E2F3 | 1731.97213 | -0.074631585 | 0.13971591 | -0.5341667 | 0.59322622 | 0.75436977 |
| CDKAL1 | 5263.6133 | -0.180674492 | 0.2465634 | -0.7327709 | 0.46369815 | 0.65162094 |
| AL513188.1 | 89.1266452 | -1.089391105 | 0.53139999 | -2.0500397 | 0.04036055 | 0.13434242 |
| AL512380.2 | 41.581552 | -0.579867947 | 0.36449219 | -1.5908926 | 0.11163374 | 0.26445968 |
| SOX4 | 725.790961 | -1.32348722 | 0.5936834 | -2.2292812 | 0.0257952 | 0.09854371 |
| CASC15 | 941.791151 | -0.681291073 | 0.60704129 | -1.1223142 | 0.26172886 | 0.45993814 |
| NBAT1 | 38.2815713 | -0.603769228 | 0.60203785 | -1.0028759 | 0.31592076 | 0.51590706 |
| DCDC2 | 3335.83619 | -1.085310933 | 0.37857801 | -2.8668092 | 0.00414633 | 0.02636073 |
| MRS2 | 465.942226 | 0.118500651 | 0.18095592 | 0.65485923 | 0.51255839 | 0.6911936 |
| GPLD1 | 54.4810898 | 0.664296823 | 0.37773338 | 1.75863944 | 0.07863877 | 0.20950469 |
| ALDH5A1 | 187.981733 | 1.068136011 | 0.41777568 | 2.55672137 | 0.01056638 | 0.05255203 |
| KIAA0319 | 158.667402 | -2.956830429 | 0.6744744 | -4.3839031 | 1.17E-05 | 0.00028738 |
| TDP2 | 969.02598 | -0.480197277 | 0.40916134 | -1.1736135 | 0.24054986 | 0.43579395 |
| ACOT13 | 578.45825 | 1.236187325 | 0.47391548 | 2.60845525 | 0.00909519 | 0.04704444 |
| C6orf62 | 1057.89166 | -0.008640349 | 0.34863243 | -0.0247836 | 0.98022761 | 0.99026974 |
| GMNN | 101.056861 | -0.402991039 | 0.29273726 | -1.3766305 | 0.16862652 | 0.34715586 |
| RIPOR2 | 878.233231 | -0.738784166 | 0.57015408 | -1.2957623 | 0.19505739 | 0.38021212 |
| CARMIL1 | 2735.0425 | -0.477972365 | 0.30202044 | -1.5825828 | 0.11351658 | 0.26719688 |
| SCGN | 52.6664303 | 0.29441635 | 0.89802416 | 0.32784903 | 0.74302581 | 0.8540591 |
| SLC17A4 | 149.546277 | -0.137566168 | 0.90247452 | -0.1524322 | 0.87884607 | 0.93706833 |
| SLC17A1 | 9114.39695 | 1.283222044 | 0.70500138 | 1.82016954 | 0.06873319 | 0.19183992 |
| SLC17A3 | 3636.94126 | -0.07489239 | 0.66653767 | -0.1123603 | 0.91053771 | 0.95300367 |
| TRIM38 | 2785.24234 | -0.971218996 | 0.3683829 | -2.6364389 | 0.00837813 | 0.04437366 |
| U91328.2 | 298.395535 | -1.238570339 | 0.63811972 | -1.9409686 | 0.05226208 | 0.15953225 |
| U91328.1 | 43.1424166 | -0.422169631 | 0.4103677 | -1.0287594 | 0.30359275 | 0.50379489 |
| HIST1H1C | 213.262288 | -1.495713108 | 0.47403146 | -3.1553035 | 0.00160331 | 0.01299622 |
| HFE | 90.0852222 | -0.248289914 | 0.59986137 | -0.4139122 | 0.67893844 | 0.8139067 |
| HIST1H4C | 215.723248 | -2.360669661 | 0.62520512 | -3.7758323 | 0.00015947 | 0.00222146 |
| HIST1H2BC | 97.6844802 | -2.566527666 | 0.85980866 | -2.9849986 | 0.0028358 | 0.01988248 |
| HIST1H2AC | 812.377896 | -1.043245019 | 0.66040462 | -1.5797058 | 0.11417426 | 0.26811487 |
| HIST1H1E | 59.9639872 | -2.500184897 | 0.67970454 | -3.6783408 | 0.00023476 | 0.00303853 |
| HIST1H2BD | 172.854486 | -0.822086598 | 0.54579684 | -1.5062136 | 0.13201236 | 0.29538611 |
| HIST1H2BE | 55.0507494 | -0.345530853 | 0.4208389 | -0.8210526 | 0.41161633 | 0.6051727 |
| HIST1H4H | 50.0982303 | -1.131727576 | 0.75299179 | -1.5029747 | 0.13284558 | 0.29622814 |
| BTN3A2 | 85.1445983 | -1.226462156 | 0.66428413 | -1.8462915 | 0.06484989 | 0.18457471 |
| BTN2A2 | 76.979068 | -0.649850248 | 0.51304811 | -1.2666458 | 0.20528196 | 0.39375871 |
| BTN3A1 | 76.7045643 | -1.040544853 | 0.81030927 | -1.284133 | 0.19909543 | 0.38556105 |
| BTN3A3 | 43.7166865 | -0.287607492 | 0.66140369 | -0.4348441 | 0.66367559 | 0.80442661 |
| BTN2A1 | 155.620394 | -0.283521616 | 0.22780095 | -1.2446025 | 0.21327792 | 0.40328327 |
| HCG11 | 62.4172229 | -0.431918115 | 0.45639126 | -0.9463768 | 0.34395641 | 0.54439007 |
| HMGN4 | 169.940829 | -0.269418489 | 0.37186773 | -0.7245009 | 0.4687583 | 0.65617047 |
| ABT1 | 55.4475975 | -0.563890666 | 0.40367284 | -1.3969002 | 0.16244359 | 0.33890627 |
| ZNF322 | 169.373125 | 0.105807263 | 0.19127025 | 0.55318202 | 0.58013878 | 0.74366055 |
| AL513548.1 | 26.7447624 | 0.9606196 | 0.7538183 | 1.2743384 | 0.20254351 | 0.39019873 |
| LINC00240 | 167.651991 | -0.028688341 | 0.49656552 | -0.0577735 | 0.95392903 | 0.97680451 |
| ZNF391 | 66.3760875 | 0.613435623 | 0.38254121 | 1.60358051 | 0.10880655 | 0.25978251 |
| ZNF184 | 109.007871 | 0.532141315 | 0.3905432 | 1.36256711 | 0.17301898 | 0.35259425 |
| HIST1H2BN | 120.961524 | -1.417603656 | 0.30170669 | -4.6986152 | 2.62E-06 | 8.57E-05 |
| AL121944.1 | 135.162619 | -1.378696359 | 0.63662996 | -2.1656165 | 0.0303405 | 0.11013699 |
| ZNF165 | 125.409052 | -2.928038548 | 0.98046526 | -2.9863766 | 0.00282305 | 0.01980323 |
| ZSCAN16-AS | 182.615972 | 0.343236632 | 0.22804012 | 1.505159 | 0.1322832 | 0.29558723 |
| ZSCAN16 | 42.4754257 | 0.173641156 | 0.33239367 | 0.52239609 | 0.60139458 | 0.76066397 |
| ZKSCAN8 | 266.227401 | 0.162977058 | 0.16847125 | 0.96738795 | 0.33335013 | 0.53275938 |
| ZSCAN9 | 118.256673 | -0.323884946 | 0.28576276 | -1.133405 | 0.25704421 | 0.4542683 |
| ZKSCAN4 | 34.4900694 | 0.095941678 | 0.45782529 | 0.20955959 | 0.83401142 | 0.91224528 |
| PGBD1 | 124.717527 | 0.402333452 | 0.28218921 | 1.42575773 | 0.15393829 | 0.32636683 |
| ZSCAN31 | 190.292067 | -1.549098972 | 0.85208037 | -1.8180198 | 0.06906111 | 0.19255756 |
| ZKSCAN3 | 78.1372799 | 0.226127115 | 0.40054448 | 0.56454933 | 0.57238034 | 0.73806372 |
| ZSCAN12 | 99.9579943 | -0.266720901 | 0.25919955 | -1.0290176 | 0.30347139 | 0.50371548 |
| TRIM27 | 535.365228 | -0.471029675 | 0.31857029 | -1.4785738 | 0.13925426 | 0.30543908 |
| GABBR1 | 194.63791 | -0.748586204 | 0.52912516 | -1.4147621 | 0.15713827 | 0.33121548 |
| HLA-F | 152.062732 | -2.070984523 | 0.8086955 | -2.5608953 | 0.01044028 | 0.05209327 |
| HLA-A | 2524.81373 | -1.202969692 | 0.4446812 | -2.7052407 | 0.00682549 | 0.03817095 |
| ZNRD1 | 65.9778292 | -0.157693835 | 0.24230242 | -0.6508141 | 0.51516648 | 0.69315508 |
| PPP1R11 | 142.434328 | -0.431220888 | 0.28738082 | -1.5005208 | 0.13347956 | 0.29730327 |
| TRIM10 | 50.6116244 | 1.699212862 | 0.47267324 | 3.59489964 | 0.00032452 | 0.00390874 |
| TRIM15 | 159.62392 | 0.608577862 | 0.43859629 | 1.38755815 | 0.16527163 | 0.34261003 |
| TRIM26 | 641.513886 | 0.35144102 | 0.23334149 | 1.50612318 | 0.13203555 | 0.29538611 |
| HCG17 | 1010.84592 | 0.204989277 | 0.32615735 | 0.6284981 | 0.52967769 | 0.70399419 |
| HLA-E | 1129.69435 | -1.617550712 | 0.4347581 | -3.7205764 | 0.00019877 | 0.00265548 |
| GNL1 | 344.867871 | -0.264932855 | 0.33723942 | -0.7855928 | 0.43210609 | 0.62475566 |
| PRR3 | 147.172264 | -0.099391057 | 0.26846959 | -0.3702135 | 0.71122345 | 0.83493832 |
| ABCF1 | 496.65018 | -0.34079767 | 0.38225497 | -0.8915454 | 0.37263664 | 0.57001787 |
| PPP1R10 | 722.938407 | 0.836660003 | 0.30461116 | 2.74664923 | 0.00602075 | 0.03467496 |
| MRPS18B | 299.333793 | -0.324039916 | 0.26679885 | -1.2145477 | 0.22453867 | 0.41674812 |
| ATAT1 | 187.508133 | -0.025594453 | 0.31178567 | -0.0820899 | 0.93457523 | 0.96509871 |
| C6orf136 | 173.999917 | 0.39203667 | 0.22782723 | 1.72076303 | 0.08529383 | 0.22099234 |
| DHX16 | 242.472331 | 0.115954593 | 0.27621812 | 0.41979357 | 0.67463626 | 0.81123249 |
| PPP1R18 | 85.1779819 | -1.846753005 | 0.50462328 | -3.6596667 | 0.00025254 | 0.00321114 |
| MDC1 | 202.344703 | -0.018926354 | 0.34562643 | -0.0547596 | 0.95633001 | 0.97772921 |
| TUBB | 623.618083 | -1.322322443 | 0.62378817 | -2.1198261 | 0.03402071 | 0.11944721 |
| FLOT1 | 981.449513 | -0.655783361 | 0.34329268 | -1.9102748 | 0.05609784 | 0.16726845 |
| IER3 | 551.258172 | -2.924683677 | 0.63079035 | -4.6365384 | 3.54E-06 | 0.00011012 |
| DDR1 | 266.099061 | -1.067529917 | 0.4233799 | -2.5214469 | 0.01168733 | 0.05622965 |
| VARS2 | 153.250036 | 0.640883924 | 0.39512374 | 1.62198283 | 0.10480702 | 0.25296529 |
| PSORS1C1 | 56.1594481 | -2.936400386 | 0.88558327 | -3.3157812 | 0.00091387 | 0.00845732 |
| CCHCR1 | 159.987941 | -0.080371362 | 0.4277662 | -0.1878862 | 0.85096587 | 0.92167424 |
| POU5F1 | 1303.22896 | -1.067627046 | 0.46945409 | -2.2741884 | 0.02295466 | 0.09098888 |

| | | | | | | |
|---|---|---|---|---|---|---|
| HCG27 | 87.580403 | -0.276653815 | 0.44504813 | -0.6216267 | 0.53418734 | 0.70733681 |
| HLA-C | 1039.41639 | -1.485020163 | 0.67593469 | -2.1969876 | 0.02802133 | 0.10415702 |
| HLA-B | 1551.10761 | -2.095823059 | 0.58486772 | -3.5834138 | 0.00033913 | 0.00403165 |
| MICA | 337.376234 | 0.308398014 | 0.37718188 | 0.81763742 | 0.41356426 | 0.60725498 |
| MCCD1 | 37.6520225 | -0.428175183 | 1.00445173 | -0.4262775 | 0.66990564 | 0.80830956 |
| NFKBIL1 | 109.174712 | 0.288369849 | 0.25653437 | 1.12409828 | 0.26097134 | 0.45911323 |
| PRRC2A | 415.556705 | 0.198201076 | 0.21734265 | 0.91192907 | 0.36180606 | 0.55961134 |
| BAG6 | 348.798499 | -0.07426477 | 0.2473399 | -0.3002539 | 0.7639835 | 0.86817458 |
| APOM | 196.410927 | 0.495705745 | 0.66955966 | 0.7403459 | 0.45909014 | 0.6475935 |
| C6orf47 | 88.4247632 | -0.730940773 | 0.35833792 | -2.0398086 | 0.04136939 | 0.13664328 |
| GPANK1 | 107.779141 | 0.103406617 | 0.27016694 | 0.38275082 | 0.70190453 | 0.82874111 |
| DDAH2 | 178.171181 | -0.317611444 | 0.42490139 | -0.7474945 | 0.45476513 | 0.64414855 |
| CLIC1 | 1233.35449 | -2.305795163 | 0.40768695 | -5.6557983 | 1.55E-08 | 1.19E-06 |
| VARS | 169.130509 | -0.25898477 | 0.44710597 | -0.579247 | 0.56242255 | 0.73014439 |
| LSM2 | 136.093334 | -0.3675304 | 0.21008104 | -1.7494696 | 0.08020988 | 0.21236949 |
| HSPA1A | 1223.14096 | -1.112591654 | 0.659133 | -1.6879623 | 0.09141846 | 0.23179483 |
| HSPA1B | 521.591473 | -1.287872804 | 0.73180107 | -1.7598673 | 0.0784303 | 0.20920143 |
| C6orf48 | 196.648097 | -0.671266651 | 0.36686487 | -1.8297382 | 0.0672891 | 0.18892497 |
| NEU1 | 384.648544 | -1.420532304 | 0.42203027 | -3.3659489 | 0.00076281 | 0.00741022 |
| SLC44A4 | 991.460705 | -0.101194301 | 0.38889188 | -0.2602119 | 0.79470031 | 0.8882059 |
| EHMT2-AS1 | 80.5361746 | 1.334786739 | 0.50692297 | 2.6331155 | 0.00846056 | 0.04460341 |
| EHMT2 | 182.506755 | 0.589672979 | 0.2379171 | 2.47848086 | 0.01319432 | 0.06128585 |
| C2 | 129.288875 | 0.426904282 | 0.41582039 | 1.02665549 | 0.30458271 | 0.50478491 |
| CFB | 59.7536168 | -0.20138734 | 0.78859203 | -0.2553758 | 0.79843284 | 0.89012761 |
| NELFE | 181.242434 | -0.396020452 | 0.3456555 | -1.1457085 | 0.25191577 | 0.44838456 |
| SKIV2L | 145.157072 | 0.088453332 | 0.24816882 | 0.35642403 | 0.72152303 | 0.84205332 |
| DXO | 54.3070808 | 0.29148723 | 0.40082467 | 0.72721878 | 0.46709194 | 0.65464135 |
| STK19 | 394.753963 | 0.652282327 | 0.3823839 | 1.70583106 | 0.08803954 | 0.22615404 |
| CYP21A2 | 106.844092 | -0.044930065 | 0.71435259 | -0.0628962 | 0.94984916 | 0.97435384 |
| TNXB | 60.7355082 | -0.493417277 | 0.42762226 | -1.1538625 | 0.24855656 | 0.44483154 |
| ATF6B | 356.834034 | 0.274002743 | 0.24820459 | 1.10393907 | 0.26961957 | 0.46774781 |
| AGPAT1 | 146.893952 | 0.255869797 | 0.28805769 | 0.88825886 | 0.37440152 | 0.57141985 |
| RNF5 | 193.459563 | -0.386015155 | 0.41595299 | -0.9280259 | 0.35339413 | 0.55189396 |
| PBX2 | 192.754794 | 0.320873644 | 0.28167964 | 1.13914391 | 0.25464313 | 0.45183086 |
| HLA-DRA | 462.023092 | -0.613268033 | 0.57629254 | -1.064161 | 0.28725578 | 0.48700247 |
| HLA-DRB5 | 125.695562 | -1.093519616 | 0.89071962 | -1.2276811 | 0.21956671 | 0.41120384 |
| HLA-DRB1 | 443.398801 | 0.394006614 | 0.79332543 | 0.49665194 | 0.61943452 | 0.77398259 |
| HLA-DQB1 | 108.775315 | -0.482315642 | 0.90681373 | -0.5318795 | 0.59480945 | 0.75540194 |
| PSMB8 | 217.895483 | -1.962762886 | 0.63231056 | -3.1041121 | 0.00190851 | 0.01478798 |
| PSMB9 | 136.556215 | -2.325429522 | 0.82810877 | -2.8081209 | 0.00498315 | 0.03021436 |
| TAP1 | 224.222992 | -2.028971057 | 0.66691126 | -3.0423404 | 0.00234746 | 0.01739216 |
| HLA-DMA | 82.2401146 | 1.05626126 | 0.53966008 | 1.95727144 | 0.05031556 | 0.15603517 |
| BRD2 | 835.746331 | -0.296776281 | 0.41156388 | -0.7210941 | 0.47085163 | 0.65738999 |
| HLA-DPA1 | 107.800798 | -0.806586617 | 0.65382386 | -1.2336451 | 0.21733517 | 0.40834524 |
| HLA-DPB1 | 97.2368952 | -0.501339505 | 0.53536822 | -0.9364387 | 0.34904737 | 0.54885767 |
| RXRB | 137.429205 | 0.285372939 | 0.33084541 | 0.86255673 | 0.38838123 | 0.58397778 |
| SLC39A7 | 234.435663 | -0.757053731 | 0.5010256 | -1.5110081 | 0.13078639 | 0.29359468 |
| HSD17B8 | 39.3465979 | 1.381728838 | 0.44998438 | 3.07061513 | 0.00213618 | 0.01617716 |
| RING1 | 237.169069 | 0.383474801 | 0.27132507 | 1.41334082 | 0.15755554 | 0.33183985 |
| HCG25 | 85.2696181 | 0.870645732 | 0.50493 | 1.72428995 | 0.08465551 | 0.22010729 |
| VPS52 | 261.057165 | 1.181831628 | 0.21240567 | 5.56403058 | 2.64E-08 | 1.86E-06 |
| RPS18 | 784.707384 | -0.986904499 | 0.43643993 | -2.2612608 | 0.02374311 | 0.09290451 |
| WDR46 | 170.845827 | 0.278957856 | 0.45009468 | 0.61977594 | 0.53540531 | 0.70816292 |
| PFDN6 | 135.78922 | -0.651672108 | 0.36689237 | -1.7761942 | 0.07570091 | 0.20449988 |
| RGL2 | 178.848023 | 0.060209327 | 0.35694147 | 0.16868123 | 0.86604738 | 0.93003067 |
| TAPBP | 946.250225 | -0.770895542 | 0.3817774 | -2.0192278 | 0.04346355 | 0.14123609 |
| ZBTB22 | 43.6124437 | 0.546890015 | 0.33516888 | 1.63168496 | 0.10274587 | 0.25037114 |
| DAXX | 96.8646015 | -0.622099385 | 0.41274399 | -1.5072282 | 0.13175217 | 0.29508612 |
| PHF1 | 273.477511 | -0.256826508 | 0.24233564 | -1.0597967 | 0.2892371 | 0.48903082 |
| CUTA | 270.759706 | -0.338507897 | 0.34910069 | -0.9696569 | 0.33221754 | 0.53166443 |
| SYNGAP1 | 175.336749 | 0.191466522 | 0.22143863 | 0.86464824 | 0.38723188 | 0.58340157 |
| SYNGAP1-AS | 62.7782541 | -0.089674793 | 0.43931923 | -0.2041222 | 0.83825803 | 0.9145179 |
| BAK1 | 40.5014072 | -0.57800029 | 0.51533803 | -1.1215945 | 0.2620349 | 0.46002317 |
| ITPR3 | 227.224101 | -2.950108515 | 0.58698768 | -5.025844 | 5.01E-07 | 2.15E-05 |
| UQCC2 | 203.898398 | 0.079712319 | 0.17582841 | 0.45335289 | 0.65029466 | 0.79482956 |
| IP6K3 | 100.334775 | 2.13843883 | 0.71136177 | 3.00611997 | 0.00264605 | 0.01887063 |
| LEMD2 | 312.387202 | -0.245300811 | 0.28296281 | -0.8669012 | 0.38599613 | 0.58224375 |
| HMGA1 | 141.992887 | -3.628478207 | 0.93178849 | -3.8941007 | 9.86E-05 | 0.00153699 |
| SMIM29 | 95.9190553 | -1.116039077 | 0.4730029 | -2.3594762 | 0.01830076 | 0.07757672 |
| NUDT3 | 222.162949 | -0.649435964 | 0.48995553 | -1.3254998 | 0.18500543 | 0.36794391 |
| C6orf106 | 2331.00542 | -0.12195828 | 0.21330673 | -0.5717507 | 0.56749086 | 0.73431485 |
| SNRPC | 201.007683 | -0.032258234 | 0.28388816 | -0.1136301 | 0.90953104 | 0.95254752 |
| UHRF1BP1 | 510.209691 | -0.169781307 | 0.14520492 | -1.1692532 | 0.24230165 | 0.43756794 |
| TAF11 | 300.899202 | 0.000326914 | 0.16582243 | 0.00197147 | 0.99842699 | 0.99920455 |
| ANKS1A | 1951.85951 | -0.053463108 | 0.1491372 | -0.3584827 | 0.71998211 | 0.84054124 |
| SCUBE3 | 39.1342441 | -0.532302117 | 0.56527748 | -0.9416652 | 0.34636409 | 0.54664509 |
| ZNF76 | 390.768437 | 0.523117513 | 0.23254276 | 2.24955404 | 0.02447727 | 0.09490085 |
| PPARD | 679.767798 | 0.079826118 | 0.43517616 | 0.18343403 | 0.85445747 | 0.92322796 |
| RPL10A | 828.945309 | -0.882551909 | 0.44906905 | -1.9652922 | 0.04938042 | 0.15398584 |
| TEAD3 | 278.77501 | 0.057522383 | 0.24685627 | 0.23301974 | 0.81574608 | 0.90027629 |
| FKBP5 | 4293.02525 | -1.4113574 | 0.92364979 | -1.5280222 | 0.12650702 | 0.28751977 |
| SRPK1 | 1826.92848 | -0.075595741 | 0.28087531 | -0.2691434 | 0.78781932 | 0.88414669 |
| SLC26A8 | 62.317966 | -0.564005787 | 0.64784009 | -0.8705941 | 0.3839758 | 0.5803781 |
| MAPK14 | 1982.88855 | -0.470650804 | 0.18418053 | -2.5553776 | 0.01060727 | 0.05265984 |
| MAPK13 | 165.198839 | -1.240764718 | 0.35676619 | -3.4778091 | 0.00050553 | 0.00538667 |
| Z84485.1 | 68.4608844 | 1.167770396 | 0.53439809 | 2.1852069 | 0.02887368 | 0.10630447 |
| BRPF3 | 364.154816 | 0.375310079 | 0.21937713 | 1.71079859 | 0.0871183 | 0.22446063 |
| PNPLA1 | 390.983384 | 3.428669397 | 0.63450431 | 5.40369756 | 6.53E-08 | 3.91E-06 |
| ETV7 | 33.8016337 | -3.142516156 | 1.05901698 | -2.9673898 | 0.0030034 | 0.02067684 |
| KCTD20 | 935.562295 | 0.132517961 | 0.12628662 | 1.04934288 | 0.29402034 | 0.49394984 |
| STK38 | 1725.59577 | 0.242000086 | 0.24891576 | 0.97221682 | 0.33094269 | 0.53067752 |
| SRSF3 | 801.724186 | -0.174540626 | 0.23733943 | -0.7354051 | 0.46209281 | 0.65049874 |
| CDKN1A | 339.47933 | -3.955505166 | 0.44899144 | -8.8097563 | 1.25E-18 | 1.56E-15 |
| PPIL1 | 73.6260458 | -0.38811782 | 0.38493434 | -1.0082702 | 0.31332477 | 0.51371402 |
| C6orf89 | 973.408561 | -0.226811845 | 0.22258632 | -1.0189838 | 0.30821068 | 0.50831955 |
| MTCH1 | 706.182692 | -0.867415438 | 0.25885028 | -3.3510315 | 0.00080511 | 0.00769572 |
| PIM1 | 219.824087 | -0.991777651 | 0.55076087 | -1.800741 | 0.07174372 | 0.19736176 |
| TMEM217 | 233.996769 | -1.707582758 | 0.66744397 | -2.5583912 | 0.01051577 | 0.05235732 |
| TBC1D22B | 833.09497 | -0.161081702 | 0.19896933 | -0.8095805 | 0.41818129 | 0.6112841 |
| RNF8 | 271.400761 | -0.375500482 | 0.22546452 | -1.6654526 | 0.09582249 | 0.23882914 |
| CMTR1 | 228.372818 | 0.232214673 | 0.30201681 | 0.76887997 | 0.44196457 | 0.63361661 |
| CCDC167 | 59.9510918 | -0.385118294 | 0.40316668 | -0.9552334 | 0.33945966 | 0.53985075 |
| ZFAND3 | 10805.3529 | 0.136651892 | 0.22091457 | 0.61857347 | 0.53619738 | 0.70869254 |
| BTBD9 | 4727.35465 | 0.038066541 | 0.26197903 | 0.14530377 | 0.88447104 | 0.94002757 |
| GLO1 | 319.64136 | -0.659958364 | 0.18249253 | -3.6163583 | 0.00029878 | 0.00366965 |
| SAYSD1 | 113.941313 | -0.511858459 | 0.2229209 | -2.2961439 | 0.02166766 | 0.08753273 |
| KCNK5 | 2187.92368 | 0.23513618 | 0.54663968 | 0.4301484 | 0.6670877 | 0.80661354 |
| KIF6 | 160.047805 | 0.650280931 | 0.66294582 | 0.98089604 | 0.32664401 | 0.52677357 |
| AL590999.1 | 98.6663282 | 3.152528834 | 0.74731658 | 4.21846498 | 2.46E-05 | 0.00051631 |
| MOCS1 | 327.006181 | 1.118123733 | 0.3737229 | 2.99185236 | 0.0027729 | 0.01955154 |
| UNC5CL | 1426.75758 | 0.250821746 | 0.65311996 | 0.38403626 | 0.70095157 | 0.82798546 |
| OARD1 | 406.564401 | -0.036806496 | 0.3258076 | -0.11297 | 0.91005431 | 0.95270509 |
| NFYA | 248.199408 | -0.305372918 | 0.24578623 | -1.242433 | 0.21407686 | 0.40445105 |

| | | | | | | |
|---|---|---|---|---|---|---|
| FOXP4 | 190.806791 | 0.702816222 | 0.36212755 | 1.94079741 | 0.05228285 | 0.15953225 |
| TFEB | 222.080321 | 0.341114359 | 0.40032673 | 0.85208989 | 0.39416421 | 0.58886199 |
| FRS3 | 41.3265687 | 0.754919045 | 0.42458697 | 1.77800803 | 0.07540254 | 0.20409608 |
| USP49 | 386.886697 | -0.235905392 | 0.23446172 | -1.0061574 | 0.31433988 | 0.51489517 |
| MED20 | 87.3965901 | -0.031100325 | 0.26792159 | -0.11608 | 0.90758918 | 0.95167724 |
| CCND3 | 568.90344 | -0.995643301 | 0.53203851 | -1.8713745 | 0.06129319 | 0.17738359 |
| TAF8 | 340.430019 | 0.58166951 | 0.22692987 | 2.56321263 | 0.01037085 | 0.0518386 |
| C6orf132 | 92.0991055 | -1.194121549 | 0.33626257 | -3.5511581 | 0.00038354 | 0.00442897 |
| MRPS10 | 138.798428 | -0.171395901 | 0.2139872 | -0.8009633 | 0.42315287 | 0.61646031 |
| TRERF1 | 204.904326 | 0.198374872 | 0.61439679 | 0.32287746 | 0.74678805 | 0.85681039 |
| UBR2 | 2253.26252 | 0.12701381 | 0.21393981 | 0.59368945 | 0.55271984 | 0.72222411 |
| TBCC | 58.4577252 | -0.240326093 | 0.49284903 | -0.4876262 | 0.62581464 | 0.77848801 |
| BICRAL | 1112.0258 | 0.799557446 | 0.26416882 | 3.0266912 | 0.00247246 | 0.01803622 |
| RPL7L1 | 498.443184 | -0.462272843 | 0.29387602 | -1.5730199 | 0.1157142 | 0.27086072 |
| C6orf226 | 59.8462437 | -0.31145121 | 0.42968008 | -0.7248444 | 0.46854748 | 0.65594733 |
| CNPY3 | 127.431516 | -0.545830856 | 0.38989486 | -1.3999437 | 0.16153018 | 0.33766878 |
| AL035587.1 | 48.8949776 | -0.561192405 | 0.67061864 | -0.8368279 | 0.4026893 | 0.59683145 |
| PEX6 | 308.158363 | -0.393553336 | 0.25539661 | -1.5409497 | 0.12332902 | 0.28240409 |
| PPP2R5D | 127.758393 | 0.700618771 | 0.19232327 | 3.6429225 | 0.00026956 | 0.00337738 |
| MEA1 | 69.9328841 | -1.108151331 | 0.36671773 | -3.0218101 | 0.00251268 | 0.01826157 |
| KLHDC3 | 224.428486 | 0.044118825 | 0.25196099 | 0.17510181 | 0.86099964 | 0.92686107 |
| RRP36 | 290.796882 | -0.22251671 | 0.27792422 | -0.8006381 | 0.42334121 | 0.61654251 |
| CUL7 | 191.660277 | 0.219436605 | 0.3043407 | 0.72102287 | 0.47089544 | 0.65738999 |
| KLC4 | 416.875832 | 0.858173657 | 0.2833307 | 3.02887634 | 0.00245465 | 0.01795406 |
| MRPL2 | 155.402227 | 0.40322983 | 0.2461004 | 1.63847693 | 0.10132224 | 0.24804858 |
| PTK7 | 86.6608301 | -0.490395447 | 0.55171225 | -0.8888609 | 0.37407784 | 0.57127053 |
| SRF | 74.5028729 | 0.136825715 | 0.37619566 | 0.36370892 | 0.71607538 | 0.83776419 |
| CUL9 | 282.367195 | 0.659074552 | 0.1668791 | 3.94941339 | 7.83E-05 | 0.00128759 |
| DNPH1 | 418.792747 | 0.097551574 | 0.50343552 | 0.19377173 | 0.84635462 | 0.91896252 |
| SLC22A7 | 2742.33725 | 1.776432221 | 0.91846076 | 1.93414057 | 0.05309585 | 0.16120138 |
| CRIP3 | 92.6730367 | 0.632361099 | 0.77914303 | 0.81161106 | 0.41701485 | 0.61036016 |
| ZNF318 | 217.562469 | 0.822413773 | 0.23409189 | 3.51320919 | 0.00044273 | 0.00492172 |
| ABCC10 | 273.409828 | 0.06563319 | 0.15681101 | 0.41854963 | 0.67554531 | 0.81196944 |
| TJAP1 | 304.83536 | 0.04128284 | 0.23490853 | 0.17574007 | 0.86049816 | 0.92646675 |
| POLR1C | 367.34359 | -0.579930809 | 0.25999026 | -2.2305867 | 0.02570852 | 0.09832854 |
| YIPF3 | 259.670405 | -0.78648319 | 0.33666503 | -2.3361 | 0.01948603 | 0.08091336 |
| XPO5 | 999.415882 | -0.845432764 | 0.27373493 | -3.0885089 | 0.00201164 | 0.01542142 |
| POLH | 297.177343 | -0.574439901 | 0.26810068 | -2.1426275 | 0.03214302 | 0.11473551 |
| GTPBP2 | 348.332558 | -0.989822725 | 0.3442603 | -2.875216 | 0.00403751 | 0.02582462 |
| MAD2L1BP | 71.7961573 | -0.577755134 | 0.30080451 | -1.9206997 | 0.05476958 | 0.16467727 |
| MRPS18A | 129.488131 | -0.657020453 | 0.29381797 | -2.236148 | 0.02534208 | 0.09732808 |
| VEGFA | 1409.38033 | -0.084995225 | 0.43520267 | -0.1953003 | 0.84515784 | 0.91854292 |
| AL157371.2 | 82.0917685 | -5.988822539 | 1.34321909 | -4.4585597 | 8.25E-06 | 0.00021667 |
| MRPL14 | 333.803719 | -0.980565206 | 0.24528894 | -3.9975925 | 6.40E-05 | 0.00109692 |
| TMEM63B | 441.650687 | 0.082133512 | 0.31200224 | 0.26324655 | 0.79236055 | 0.88689255 |
| SLC29A1 | 44.0518008 | -1.487685237 | 0.35772529 | -4.1587365 | 3.20E-05 | 0.00063479 |
| HSP90AB1 | 1670.27642 | -1.314011366 | 0.40116937 | -3.2754529 | 0.00105493 | 0.00941089 |
| SLC35B2 | 111.188013 | -1.155473033 | 0.41787277 | -2.7651312 | 0.00568999 | 0.03333022 |
| NFKBIE | 67.6998657 | -1.561451998 | 0.444554 | -3.5124012 | 0.00044408 | 0.00492872 |
| AARS2 | 60.6393498 | 0.20095995 | 0.46854972 | 0.42889696 | 0.66799821 | 0.80707373 |
| CDC5L | 300.292812 | -0.159984696 | 0.22614223 | -0.7074517 | 0.47928584 | 0.66453277 |
| SUPT3H | 2270.60988 | -0.03809519 | 0.33095384 | -0.1151073 | 0.90836011 | 0.95197217 |
| RUNX2 | 193.121022 | -0.590545383 | 0.492981 | -1.197907 | 0.23095323 | 0.42389875 |
| CLIC5 | 168.799106 | 0.628104021 | 0.41363296 | 1.51850574 | 0.12888695 | 0.29066362 |
| AL035701.1 | 48.2800549 | 0.608094452 | 0.422188 | 1.44034043 | 0.14977111 | 0.32014622 |
| ENPP4 | 243.957979 | -0.232477684 | 0.37859306 | -0.6140569 | 0.53917772 | 0.71083091 |
| ENPP5 | 156.160326 | 0.002598318 | 0.44180074 | 0.0058812 | 0.99530751 | 0.99749068 |
| RCAN2 | 184.484894 | -0.193020334 | 0.45129546 | -0.4277028 | 0.66886751 | 0.80741232 |
| CYP39A1 | 112.140877 | -0.341271761 | 0.70177611 | -0.4862972 | 0.62675646 | 0.77912423 |
| SLC25A27 | 688.551583 | 0.576377588 | 0.58621572 | 0.98321756 | 0.32550037 | 0.52576404 |
| TDRD6 | 81.4527938 | 1.387306267 | 0.4579122 | 3.02963381 | 0.0024485 | 0.01793781 |
| ADGRF5 | 61.4456845 | 0.792045915 | 0.50675266 | 1.56298323 | 0.11805654 | 0.27411802 |
| ADGRF1 | 84.0927129 | -0.826072887 | 0.53338937 | -1.5487239 | 0.12144809 | 0.27946713 |
| TNFRSF21 | 1275.54313 | 1.097919594 | 0.44790948 | 2.45120868 | 0.01423774 | 0.06457865 |
| AL355353.1 | 52.3843379 | 1.282073075 | 0.63225184 | 2.02778859 | 0.04258183 | 0.13933376 |
| CD2AP | 3407.14282 | -0.189348106 | 0.14316796 | -1.3225592 | 0.18598201 | 0.36918977 |
| PTCHD4 | 233.7046 | -2.973230626 | 1.06736716 | -2.7855744 | 0.0053433 | 0.03181606 |
| MUT | 299.841194 | 0.150872368 | 0.4008479 | 0.37638308 | 0.70663212 | 0.83210663 |
| CENPQ | 36.2704655 | -0.231387143 | 0.39920847 | -0.5796148 | 0.5621744 | 0.73014439 |
| AL138826.1 | 87.7295551 | 3.020043182 | 0.66695773 | 4.52808782 | 5.95E-06 | 0.00016542 |
| AL355997.1 | 77.6466858 | 1.125978698 | 0.8746339 | 1.28737143 | 0.19796486 | 0.38408112 |
| PKHD1 | 22581.5413 | 1.420246691 | 0.52992403 | 2.68009491 | 0.00736013 | 0.0404512 |
| MCM3 | 103.440681 | -0.597899351 | 0.35070226 | -1.7048631 | 0.08821996 | 0.22636297 |
| EFHC1 | 439.47466 | 0.097310166 | 0.3148016 | 0.30911585 | 0.7572334 | 0.86387001 |
| TRAM2 | 185.628035 | 0.561512346 | 0.49163908 | 1.14212309 | 0.25340286 | 0.4503297 |
| TRAM2-AS1 | 42.0579371 | 0.665478482 | 0.51758321 | 1.28574201 | 0.19853311 | 0.38480603 |
| TMEM14A | 118.616979 | 0.486908443 | 0.33759233 | 1.44229713 | 0.14921857 | 0.31943447 |
| GSTA2 | 194.758708 | 3.171470092 | 1.09565422 | 2.89459031 | 0.00379654 | 0.02459316 |
| GSTA1 | 320.814215 | 1.90308813 | 0.77778243 | 2.44681297 | 0.01441256 | 0.0651378 |
| GSTA4 | 174.278577 | 0.580457721 | 0.52021944 | 1.11579397 | 0.26451033 | 0.46242769 |
| ICK | 186.739566 | 0.073070959 | 0.33091428 | 0.22081537 | 0.8252362 | 0.90642548 |
| FBXO9 | 876.573609 | 0.348221951 | 0.13413834 | 2.59599128 | 0.00943185 | 0.04832984 |
| GCM1 | 73.7776752 | 1.794172051 | 0.6995147 | 2.56488111 | 0.01032112 | 0.05174526 |
| ELOVL5 | 1398.41483 | 0.02614692 | 0.43458231 | 0.06016563 | 0.95202372 | 0.97565526 |
| GCLC | 2706.65015 | -1.713603757 | 0.41523877 | -4.1267913 | 3.68E-05 | 0.00070521 |
| LINC01564 | 79.424012 | -2.311648321 | 0.73333424 | -3.1522438 | 0.00162021 | 0.01308211 |
| LRRC1 | 839.150832 | 0.095007812 | 0.40922027 | 0.2321679 | 0.81640761 | 0.90083699 |
| MLIP | 127.47927 | 1.587707745 | 0.52361632 | 3.03219683 | 0.00242781 | 0.0178243 |
| TINAG | 8766.74434 | 1.060526174 | 0.50754934 | 2.08950359 | 0.03666242 | 0.12582058 |
| DST | 2722.20741 | -0.760979557 | 0.43106491 | -1.7653479 | 0.07750531 | 0.20765144 |
| KIAA1586 | 135.876202 | -0.162253609 | 0.19113444 | -0.8488978 | 0.39593815 | 0.59067525 |
| ZNF451 | 1382.9969 | 0.055371867 | 0.13334719 | 0.41524583 | 0.67796195 | 0.81346017 |
| ZNF451-AS1 | 136.909529 | -0.775367262 | 0.56052112 | -1.3832971 | 0.16657377 | 0.34453334 |
| BAG2 | 141.756801 | -0.219822407 | 0.29292383 | -0.7504422 | 0.45298841 | 0.64216694 |
| RAB23 | 144.375207 | 0.684033879 | 0.29721121 | 2.30150766 | 0.02136295 | 0.08678779 |
| PRIM2 | 744.090471 | -0.144217529 | 0.3459877 | -0.4168285 | 0.67680387 | 0.81285008 |
| AL021368.2 | 137.754094 | 0.259877845 | 0.35027341 | 0.74192855 | 0.45813063 | 0.64698089 |
| AL445250.1 | 84.0340075 | -0.086062005 | 0.53554164 | -0.1607009 | 0.87232901 | 0.93392281 |
| KHDRBS2 | 2851.61049 | 0.56693237 | 0.56046708 | 1.01153555 | 0.31176018 | 0.51189467 |
| LGSN | 265.661932 | 4.485816174 | 0.81297784 | 5.51775944 | 3.43E-08 | 2.36E-06 |
| PTP4A1 | 836.528888 | -0.608740732 | 0.29083079 | -2.0931096 | 0.03633937 | 0.12477549 |
| PHF3 | 2747.62565 | 0.143155787 | 0.13128667 | 1.09040613 | 0.27553428 | 0.47298039 |
| EYS | 288.551353 | -0.319933926 | 0.51610409 | -0.6199019 | 0.53532235 | 0.70813196 |
| ADGRB3 | 107.075307 | 0.594650248 | 0.5278782 | 1.12649139 | 0.25995758 | 0.45823541 |
| LMBRD1 | 1666.87163 | 0.015449438 | 0.29450901 | 0.05245829 | 0.95816353 | 0.97865481 |
| COL19A1 | 1680.66617 | 1.341301999 | 0.44844724 | 2.99099177 | 0.00278073 | 0.01959664 |
| FAM135A | 4147.69625 | -1.281852746 | 0.35582811 | -3.6024494 | 0.00031523 | 0.00381705 |
| SDHAF4 | 129.149964 | 0.927728083 | 0.36726939 | 2.52601528 | 0.01153645 | 0.0556796 |
| SMAP1 | 2085.47546 | -0.251283398 | 0.20962588 | -1.1987232 | 0.2306356 | 0.42378291 |
| OGFRL1 | 494.543265 | -2.896311308 | 0.55437508 | -5.2244616 | 1.75E-07 | 8.79E-06 |
| AL136164.2 | 888.947744 | 0.763963395 | 0.420781 | 1.81558435 | 0.06943416 | 0.19304949 |
| LINC00472 | 355.144301 | 0.599576396 | 0.37225426 | 1.61066363 | 0.10725306 | 0.25714844 |
| KCNQ5 | 123.62221 | -1.083927072 | 0.71790293 | -1.5098519 | 0.13108123 | 0.2940639 |

| | | | | | | |
|---|---|---|---|---|---|---|
| AC019205.1 | 60.1081766 | -0.525012464 | 0.45380941 | -1.1569008 | 0.2473129 | 0.44330037 |
| OOEP | 740.725854 | -1.0048465 | 0.3894935 | -2.57988 | 0.00988346 | 0.05003642 |
| DDX43 | 43.9428921 | -0.078358534 | 0.7933675 | -0.098767 | 0.92132326 | 0.95779886 |
| MTO1 | 407.517345 | 0.849525077 | 0.18332281 | 4.6340391 | 3.59E-06 | 0.00011046 |
| EEF1A1 | 4224.10332 | -0.81212674 | 0.43649516 | -1.860563 | 0.06280592 | 0.18032691 |
| AL121972.1 | 113.594256 | 0.212431416 | 0.52446781 | 0.40504185 | 0.68544675 | 0.81805995 |
| SLC17A5 | 1331.74237 | -0.181919538 | 0.34751985 | -0.5234796 | 0.60064057 | 0.76020679 |
| CD109 | 346.770029 | 1.191535627 | 0.25077746 | 4.7513666 | 2.02E-06 | 6.86E-05 |
| AL357507.1 | 67.0424209 | -0.18051028 | 0.36322819 | -0.4969611 | 0.61921648 | 0.77378068 |
| COL12A1 | 80.6976594 | 0.067507528 | 0.52297861 | 0.12908277 | 0.89729216 | 0.94667361 |
| COX7A2 | 475.368824 | -0.594205252 | 0.33872683 | -1.7542314 | 0.07939086 | 0.21087159 |
| TMEM30A | 597.378268 | -0.653079687 | 0.24592627 | -2.6555914 | 0.00791695 | 0.04252688 |
| AL080250.1 | 68.2717089 | 0.253629324 | 0.39977809 | 0.63442528 | 0.52580332 | 0.70084961 |
| FILIP1 | 40.1418142 | 0.240254857 | 0.50186068 | 0.47872819 | 0.63213201 | 0.78317951 |
| SENP6 | 2839.86093 | 0.019425488 | 0.1605927 | 0.12096121 | 0.90372176 | 0.9495833 |
| MYO6 | 7840.35059 | -0.091509813 | 0.19972065 | -0.458189 | 0.64681664 | 0.79280318 |
| MEI4 | 478.00601 | 0.703945227 | 0.46594177 | 1.51080087 | 0.13083919 | 0.29366511 |
| IRAK1BP1 | 712.946932 | 0.640499498 | 0.50650583 | 1.26454515 | 0.20603442 | 0.39476011 |
| PHIP | 2581.78103 | 0.243289171 | 0.19944528 | 1.21982915 | 0.22252965 | 0.41472082 |
| HMGN3 | 2804.9954 | 0.150775014 | 0.30225257 | 0.49883782 | 0.61789364 | 0.7728322 |
| HMGN3-AS1 | 66.8393876 | -0.415717651 | 0.27606524 | -1.5058674 | 0.13210121 | 0.2954593 |
| LCA5 | 170.281883 | -0.359042942 | 0.36171711 | -0.992607 | 0.32090151 | 0.52128431 |
| SH3BGRL2 | 353.617408 | 0.571060684 | 0.27352539 | 2.08777947 | 0.03681773 | 0.12613363 |
| BCKDHB | 2283.5848 | 0.356670239 | 0.49475441 | 0.72090361 | 0.47096882 | 0.65738999 |
| AL359715.1 | 109.121139 | 0.541357435 | 0.78719826 | 0.68770151 | 0.49164077 | 0.67409423 |
| TENT5A | 230.346515 | -0.42013137 | 0.47377533 | -0.8867734 | 0.3752009 | 0.57200298 |
| IBTK | 1720.14833 | -0.496818329 | 0.19084486 | -2.6032575 | 0.00923426 | 0.04756631 |
| TPBG | 50.8109287 | -2.294996197 | 0.6386976 | -3.5932438 | 0.00032659 | 0.0039199 |
| UBE3D | 1051.58033 | 0.155562448 | 0.23189965 | 0.67081796 | 0.5023365 | 0.6832913 |
| DOPEY1 | 1638.28821 | 0.226922694 | 0.14709981 | 1.54264434 | 0.12291709 | 0.28169614 |
| PGM3 | 680.15571 | -1.063056257 | 0.31350122 | -3.3909159 | 0.00069659 | 0.00691399 |
| RWDD2A | 97.8185153 | -0.16086381 | 0.34315373 | -0.4687806 | 0.63922647 | 0.78803844 |
| ME1 | 605.391499 | -2.077349554 | 0.48755855 | -4.2607181 | 2.04E-05 | 0.00045123 |
| CYB5R4 | 397.722986 | -0.266203158 | 0.28437656 | -0.9360939 | 0.34922486 | 0.54887281 |
| CEP162 | 757.143265 | 0.538873001 | 0.21490313 | 2.50751583 | 0.01215831 | 0.05778572 |
| NT5E | 383.097954 | 0.364764154 | 0.60342835 | 0.60448627 | 0.5455204 | 0.71579246 |
| SNX14 | 1524.18526 | 0.354200076 | 0.16202117 | 2.18613448 | 0.02880577 | 0.10612074 |
| AL589666.1 | 89.1581783 | -0.223362811 | 0.34371467 | -0.6498495 | 0.51578944 | 0.69333133 |
| SYNCRIP | 635.295602 | -0.127298054 | 0.34026515 | -0.3741143 | 0.70831929 | 0.83295577 |
| ZNF292 | 5339.19599 | -1.142314795 | 0.34342284 | -3.3262633 | 0.00088019 | 0.00822969 |
| SMIM8 | 427.28418 | 0.585714921 | 0.33337525 | 1.75692384 | 0.07893079 | 0.20995621 |
| SLC35A1 | 66.6350843 | -0.315293297 | 0.38764615 | -0.8133534 | 0.41601549 | 0.60944659 |
| RARS2 | 1004.49446 | -0.007745339 | 0.21263846 | -0.0364249 | 0.97094354 | 0.98568139 |
| ORC3 | 560.461824 | 0.197534046 | 0.31148769 | 0.63416324 | 0.52597429 | 0.70084961 |
| AKIRIN2 | 429.847372 | -0.492954484 | 0.27250589 | -1.8089682 | 0.07045595 | 0.19509893 |
| AL139042.1 | 90.7288083 | 1.244458627 | 0.85097549 | 1.46239069 | 0.14363418 | 0.31126563 |
| RNGTT | 1813.03397 | -0.231757071 | 0.20648517 | -1.1223909 | 0.26169628 | 0.45993814 |
| PNRC1 | 954.618828 | -1.126764786 | 0.38428791 | -2.9320849 | 0.00336695 | 0.02249309 |
| SRSF12 | 170.069812 | -1.728038341 | 0.33786722 | -5.1145486 | 3.14E-07 | 1.43E-05 |
| PM20D2 | 444.368229 | -0.839698813 | 0.16718873 | -5.0224608 | 5.10E-07 | 2.18E-05 |
| GABRR2 | 44.7261594 | 0.514359134 | 0.51714887 | 0.99460554 | 0.31992815 | 0.520351 |
| UBE2J1 | 452.695348 | -1.154969858 | 0.18148647 | -6.3639448 | 1.97E-10 | 2.78E-08 |
| RRAGD | 383.414381 | -1.477971825 | 0.31714332 | -4.6602647 | 3.16E-06 | 9.95E-05 |
| ANKRD6 | 1540.64329 | -0.990857705 | 0.3025294 | -3.2752443 | 0.00105571 | 0.00941089 |
| LYRM2 | 332.246023 | -0.56136902 | 0.23914098 | -2.3474397 | 0.01890293 | 0.07923624 |
| MDN1 | 1265.94263 | 0.007284047 | 0.20698809 | 0.03519066 | 0.97192771 | 0.98607792 |
| CASP8AP2 | 425.838839 | 0.34688295 | 0.17999591 | 1.9271713 | 0.05395829 | 0.16308249 |
| BACH2 | 656.539003 | -2.459694244 | 0.51423723 | -4.7831898 | 1.73E-06 | 6.06E-05 |
| MAP3K7 | 1128.13645 | -0.315483781 | 0.16635896 | -1.8964039 | 0.05790665 | 0.17081613 |
| EPHA7 | 2378.40233 | 1.136130973 | 0.56571923 | 2.00829478 | 0.04461198 | 0.1438815 |
| MANEA | 178.02256 | 0.480869182 | 0.2732585 | 1.75975929 | 0.07844863 | 0.20920143 |
| UFL1-AS1 | 344.193203 | -0.411267152 | 0.47131988 | -0.8725861 | 0.38288875 | 0.57966159 |
| UFL1 | 609.768283 | -0.185067404 | 0.33538096 | -0.5518125 | 0.58107682 | 0.74422219 |
| NDUFAF4 | 172.098419 | -0.861400648 | 0.29269386 | -2.943009 | 0.00325039 | 0.02193653 |
| KLHL32 | 477.820037 | 1.278316939 | 0.59337528 | 2.15431444 | 0.03121552 | 0.11231721 |
| MMS22L | 280.668115 | 0.012929425 | 0.23141239 | 0.05587179 | 0.95544394 | 0.97737295 |
| AL589740.1 | 72.1423928 | 0.468514509 | 0.41934365 | 1.11725671 | 0.26388458 | 0.46194967 |
| FBXL4 | 1167.4085 | 0.260010195 | 0.24999909 | 1.04004458 | 0.29831919 | 0.49840643 |
| COQ3 | 67.5352112 | 1.215238902 | 0.24990752 | 4.86275436 | 1.16E-06 | 4.37E-05 |
| PNISR | 3277.07657 | 0.642803278 | 0.25755135 | 2.49582566 | 0.01256643 | 0.05919179 |
| AL513550.1 | 132.694703 | 7.03E-05 | 0.44875315 | 0.00015676 | 0.99987493 | 0.99987493 |
| USP45 | 880.351599 | -0.055548077 | 0.20106456 | -0.2762699 | 0.7823408 | 0.88078402 |
| TSTD3 | 145.247909 | -0.534407254 | 0.2723849 | -1.9619562 | 0.04976758 | 0.15496689 |
| CCNC | 983.580297 | -0.612958705 | 0.21724734 | -2.8214786 | 0.00478028 | 0.02930382 |
| SIM1 | 346.339575 | 0.382409473 | 0.45155143 | 0.8468791 | 0.39706251 | 0.59164429 |
| ASCC3 | 1785.54906 | -0.978567872 | 0.21109336 | -4.6357113 | 3.56E-06 | 0.00011031 |
| HACE1 | 811.122063 | -0.05744348 | 0.51400289 | -0.1117571 | 0.91101599 | 0.95310114 |
| PREP | 490.173435 | -0.236926356 | 0.241722 | -0.9801605 | 0.3270069 | 0.52691716 |
| ATG5 | 1106.66402 | -0.21670726 | 0.18086527 | -1.1981696 | 0.230851 | 0.42378291 |
| CRYBG1 | 701.607665 | 0.605235209 | 0.54961621 | 1.10119607 | 0.27081133 | 0.46898433 |
| RTN4IP1 | 216.835347 | 0.234878521 | 0.35693284 | 0.65804682 | 0.51050805 | 0.6897256 |
| QRSL1 | 459.88964 | 0.048473825 | 0.18357252 | 0.26405818 | 0.79173509 | 0.88677176 |
| LINC02532 | 7470.54892 | 0.94611209 | 0.66929081 | 1.41360389 | 0.15747824 | 0.33177901 |
| CD24 | 2212.26994 | -0.552309954 | 0.42577415 | -1.29719 | 0.19456584 | 0.37973999 |
| C6orf203 | 306.190547 | 0.686132016 | 0.24385108 | 2.8137338 | 0.00489698 | 0.02976623 |
| BEND3 | 54.0439452 | -0.258658016 | 0.40056772 | -0.6457286 | 0.51845517 | 0.69521277 |
| PDSS2 | 2756.74974 | 0.37067335 | 0.16416304 | 2.25795867 | 0.02394824 | 0.09347027 |
| SOBP | 1504.85028 | 0.516622907 | 0.53579013 | 0.96422624 | 0.33493251 | 0.53471101 |
| SEC63 | 1358.44064 | -0.048577714 | 0.20651243 | -0.235229 | 0.81403099 | 0.8993248 |
| OSTM1 | 611.366297 | -0.054368996 | 0.28528881 | -0.1905753 | 0.84885835 | 0.92036873 |
| SNX3 | 1003.87914 | -0.346084973 | 0.19883712 | -1.7405451 | 0.08176336 | 0.21527667 |
| AFG1L | 908.836547 | 0.600186715 | 0.29561105 | 2.03032569 | 0.04232344 | 0.1387201 |
| FOXO3 | 3686.67575 | 0.120966293 | 0.21437975 | 0.56426176 | 0.572576 | 0.73810771 |
| ARMC2 | 453.835923 | 0.585425485 | 0.39266038 | 1.49092068 | 0.13598232 | 0.30067909 |
| SESN1 | 1017.40001 | 0.293763489 | 0.23920005 | 1.22810798 | 0.21940644 | 0.41095983 |
| CEP57L1 | 473.446604 | 0.638884115 | 0.2202776 | 2.90035892 | 0.00372736 | 0.02424816 |
| CD164 | 1346.79916 | 0.362342927 | 0.23672487 | 1.53065 | 0.12585592 | 0.28619908 |
| PPIL6 | 224.371637 | 0.137014197 | 0.40397547 | 0.33916464 | 0.7344857 | 0.84972954 |
| SMPD2 | 60.3179534 | -0.44722232 | 0.31968543 | -1.3989449 | 0.16182949 | 0.33808823 |
| MICAL1 | 285.639036 | 0.195170749 | 0.21843488 | 0.89349627 | 0.37159147 | 0.56909545 |
| ZBTB24 | 123.165357 | -0.341402273 | 0.25828373 | -1.321811 | 0.1862311 | 0.36941673 |
| AK9 | 688.409342 | 0.671264113 | 0.24285836 | 2.76401484 | 0.00570949 | 0.03340164 |
| FIG4 | 516.023309 | -0.068506346 | 0.19614342 | -0.3492666 | 0.72688917 | 0.84486304 |
| WASF1 | 864.390852 | -0.514295991 | 0.26170657 | -1.9651627 | 0.04939541 | 0.15398584 |
| CDC40 | 420.604404 | 0.072229147 | 0.17689962 | 0.40830585 | 0.68304915 | 0.81587484 |
| METTL24 | 546.096827 | 0.247704547 | 0.46499358 | 0.5327053 | 0.59423759 | 0.75495548 |
| DDO | 102.251089 | 0.984578159 | 0.74182686 | 1.32723445 | 0.18443114 | 0.36733474 |
| CDK19 | 1596.4868 | 0.445244134 | 0.28987715 | 1.53597525 | 0.12454445 | 0.28443348 |
| AMD1 | 679.713068 | -0.983258999 | 0.24262837 | -4.052531 | 5.07E-05 | 0.00091621 |
| GTF3C6 | 361.761188 | -0.857902049 | 0.32466606 | -2.6424137 | 0.00823174 | 0.04383548 |
| RPF2 | 359.58498 | -0.420991562 | 0.29229585 | -1.4402927 | 0.14978462 | 0.32014622 |
| SLC16A10 | 2178.2781 | 0.725282388 | 0.45196606 | 1.60472755 | 0.10855378 | 0.2593597 |
| MFSD4B | 863.715776 | 1.186934475 | 0.29285478 | 4.05297975 | 5.06E-05 | 0.00091566 |

| | | | | | | |
|---|---|---|---|---|---|---|
| AL080317.1 | 93.9658819 | 1.259522652 | 0.39876415 | 3.15856543 | 0.00158548 | 0.01289008 |
| AL080317.2 | 172.729887 | 1.13253583 | 0.31193998 | 3.630621 | 0.00028274 | 0.00349776 |
| REV3L | 3027.25763 | 0.02993484 | 0.20139384 | 0.14863831 | 0.88183904 | 0.93867887 |
| AL080317.3 | 326.932338 | 1.394363604 | 0.32629537 | 4.2733171 | 1.93E-05 | 0.00043134 |
| TRAF3IP2-AS | 873.108318 | -0.534865742 | 0.22532637 | -2.3737379 | 0.01760904 | 0.07540368 |
| TRAF3IP2 | 296.042814 | -1.173758552 | 0.20952518 | -5.6019928 | 2.12E-08 | 1.56E-06 |
| FYN | 168.5767 | 0.578381963 | 0.5521146 | 1.04757592 | 0.29483404 | 0.49487664 |
| TUBE1 | 310.568671 | 0.434426479 | 0.3408081 | 1.2746953 | 0.20241711 | 0.39001003 |
| FAM229B | 83.9290342 | 0.409851814 | 0.4756944 | 0.86158638 | 0.38891517 | 0.58446006 |
| LAMA4 | 38.4200739 | -1.174183687 | 0.68807004 | -1.7064886 | 0.08791714 | 0.22592431 |
| MARCKS | 193.373361 | -1.575800501 | 0.57468211 | -2.7420386 | 0.00610592 | 0.0350523 |
| LINC01268 | 124.061201 | -0.693545616 | 0.59165293 | -1.172217 | 0.24110995 | 0.43618862 |
| HDAC2 | 428.384907 | -0.386395346 | 0.21460782 | -1.8004719 | 0.07178616 | 0.19738675 |
| HDAC2-AS2 | 64.1732247 | 0.4365871 | 0.48340653 | 0.90314689 | 0.36644794 | 0.56392747 |
| FRK | 1786.24063 | 0.246335655 | 0.30648909 | 0.80373385 | 0.4215507 | 0.61529075 |
| NT5DC1 | 3173.49048 | 0.369488627 | 0.31518207 | 1.17230219 | 0.24107575 | 0.43618862 |
| COL10A1 | 122.67719 | -1.053780343 | 0.84578825 | -1.2459151 | 0.21279557 | 0.40264894 |
| TSPYL4 | 88.4164847 | -0.128331312 | 0.30429087 | -0.4217389 | 0.67321557 | 0.81072605 |
| DSE | 889.46262 | 1.222341334 | 0.3242424 | 3.76983806 | 0.00016335 | 0.00226463 |
| TSPYL1 | 279.874764 | -0.136676825 | 0.29390698 | -0.4650343 | 0.6419069 | 0.78982207 |
| RWDD1 | 206.94771 | -0.186106863 | 0.27130211 | -0.6859765 | 0.49272794 | 0.67504467 |
| ZUP1 | 351.843753 | -0.391633944 | 0.19787879 | -1.9791608 | 0.04779791 | 0.15092143 |
| KPNA5 | 640.085072 | 0.013717696 | 0.21253495 | 0.06454325 | 0.94853767 | 0.97368425 |
| DCBLD1 | 311.488563 | -0.27400281 | 0.29522425 | -0.9281176 | 0.35334659 | 0.55189396 |
| GOPC | 419.353343 | 0.024354772 | 0.20430339 | 0.11920885 | 0.9051099 | 0.950459 |
| NUS1 | 531.102806 | -0.565864847 | 0.27498697 | -2.0577879 | 0.0396105 | 0.13258635 |
| SLC35F1 | 100.873481 | -0.380623387 | 0.31512707 | -1.207841 | 0.22710844 | 0.41948194 |
| CEP85L | 591.516114 | -0.70267698 | 0.21489578 | -3.26985 | 0.00107605 | 0.00952185 |
| MCM9 | 1007.7768 | 0.220365917 | 0.28689401 | 0.76810915 | 0.44242234 | 0.63407392 |
| ASF1A | 55.907176 | 0.143014355 | 0.35382912 | 0.40419046 | 0.68607267 | 0.81810131 |
| AL365275.1 | 417.779589 | 1.032781295 | 0.54324503 | 1.90113346 | 0.05728453 | 0.1696994 |
| FAM184A | 483.694554 | 0.555445622 | 0.37155938 | 1.49490405 | 0.13493949 | 0.29936987 |
| MAN1A1 | 3052.64868 | -0.188324181 | 0.24770833 | -0.7602658 | 0.4470957 | 0.6378373 |
| TBC1D32 | 619.943065 | 0.214725984 | 0.29658099 | 0.72400453 | 0.46906295 | 0.65617047 |
| HSF2 | 196.32524 | 0.036592114 | 0.21577372 | 0.16958559 | 0.86533605 | 0.92952143 |
| SERINC1 | 888.545552 | -0.469463761 | 0.20759642 | -2.2614252 | 0.02373294 | 0.09290451 |
| PKIB | 274.743246 | -0.231689303 | 0.59149106 | -0.3917038 | 0.69527708 | 0.82442931 |
| SMPDL3A | 246.852329 | -0.391554804 | 0.48118874 | -0.813724 | 0.41580311 | 0.60934946 |
| NKAIN2 | 136.021195 | -0.544398614 | 0.46454889 | -1.1718866 | 0.24124259 | 0.43635685 |
| RNF217-AS1 | 109.269677 | -1.542892459 | 0.57190151 | -2.697829 | 0.00697933 | 0.03887268 |
| RNF217 | 1233.7844 | -0.873714197 | 0.21587752 | -4.0472681 | 5.18E-05 | 0.00093581 |
| TPD52L1 | 917.29409 | -1.34679838 | 0.36731771 | -3.6665762 | 0.00024582 | 0.00315196 |
| HDDC2 | 231.089132 | -0.517286487 | 0.28796241 | -1.7963681 | 0.07243599 | 0.19873501 |
| AL365259.1 | 270.512482 | 2.264820344 | 0.67026457 | 3.37899457 | 0.00072751 | 0.00715354 |
| HEY2 | 52.6405599 | 0.311683565 | 0.46683606 | 0.667651 | 0.50435639 | 0.68481556 |
| NCOA7 | 2973.55668 | 1.05989212 | 0.31106007 | 3.4073551 | 0.00065596 | 0.0065949 |
| NCOA7-AS1 | 48.7249637 | -0.351552318 | 0.62343741 | -0.5638935 | 0.5728266 | 0.73835552 |
| HINT3 | 200.69045 | -0.420864229 | 0.32049217 | -1.313181 | 0.18912196 | 0.37309941 |
| TRMT11 | 1163.40456 | -0.09878107 | 0.21060708 | -0.4690302 | 0.63904808 | 0.78803844 |
| AC020559.1 | 312.607338 | -0.049539875 | 0.34879451 | -0.1420317 | 0.88705497 | 0.94159858 |
| RNF146 | 578.726538 | 0.896641047 | 0.28855721 | 3.10732503 | 0.00188789 | 0.01465304 |
| ECHDC1 | 1373.47328 | 0.370151744 | 0.2523312 | 1.46692816 | 0.1423956 | 0.30981214 |
| PTPRK | 9428.54767 | -0.105492685 | 0.24155374 | -0.4367255 | 0.66231042 | 0.80353061 |
| AL590006.1 | 55.3213979 | -0.886728908 | 0.71127063 | -1.2466829 | 0.2125138 | 0.40233794 |
| LAMA2 | 82.6733089 | 0.128207175 | 0.52207378 | 0.2455729 | 0.80601287 | 0.89466205 |
| AL356124.1 | 73.0823236 | 0.410081478 | 0.38126007 | 1.07559513 | 0.28210837 | 0.48105285 |
| ARHGAP18 | 768.400509 | 0.210984759 | 0.24983879 | 0.84448361 | 0.39839922 | 0.59273397 |
| L3MBTL3 | 354.707538 | -0.138610292 | 0.30284807 | -0.4576892 | 0.64717574 | 0.79303039 |
| SAMD3 | 240.619468 | -0.108688046 | 0.55826214 | -0.19469 | 0.84563566 | 0.91865716 |
| TMEM200A | 641.856261 | 0.504307227 | 0.33878217 | 1.48858844 | 0.13659578 | 0.30154909 |
| SMLR1 | 38.8558743 | 1.032383208 | 0.73969535 | 1.39568703 | 0.16280876 | 0.33934952 |
| EPB41L2 | 1484.70869 | -1.1336452 | 0.35136253 | -3.226426 | 0.00125347 | 0.01072489 |
| AKAP7 | 409.450232 | 0.309991198 | 0.41117592 | 0.7539138 | 0.45090099 | 0.64079832 |
| MED23 | 730.482381 | -0.044603237 | 0.20613238 | -0.2163815 | 0.82869037 | 0.90892685 |
| ENPP3 | 1289.60249 | -0.851949829 | 0.56940945 | -1.496199 | 0.13460181 | 0.29902546 |
| ENPP1 | 1321.60336 | 0.12221475 | 0.60285417 | 0.20272689 | 0.83934851 | 0.91490443 |
| CTGF | 125.351976 | -0.284831403 | 0.59272343 | -0.4805469 | 0.63083857 | 0.78238366 |
| STX7 | 2126.92058 | 0.261382845 | 0.36745252 | 0.71133775 | 0.47687496 | 0.66253042 |
| VNN1 | 426.972797 | -1.436037901 | 0.73037406 | -1.9661677 | 0.04927924 | 0.15396641 |
| SLC18B1 | 242.203701 | -0.160741398 | 0.37774354 | -0.4255305 | 0.67044999 | 0.80882398 |
| RPS12 | 737.628433 | -0.597792034 | 0.44136098 | -1.3544288 | 0.1755996 | 0.35574102 |
| TBPL1 | 252.787915 | 0.187631788 | 0.22634445 | 0.8289657 | 0.40712382 | 0.60095264 |
| SGK1 | 662.139713 | 0.992593449 | 0.74250247 | 1.33682174 | 0.18128083 | 0.36354299 |
| AL078590.2 | 371.296635 | 2.958390931 | 0.83882439 | 3.52682989 | 0.00042057 | 0.00475244 |
| AL078590.3 | 1295.33061 | 2.716213818 | 0.84772156 | 3.20413441 | 0.00135469 | 0.01137095 |
| ALDH8A1 | 1271.42436 | 1.137078359 | 0.45818304 | 2.481712 | 0.01307529 | 0.06084261 |
| HBS1L | 608.70207 | -0.041677362 | 0.14740717 | -0.2827363 | 0.77737897 | 0.87735969 |
| AHI1 | 2260.60275 | -0.43297592 | 0.13847889 | -3.1266564 | 0.00176806 | 0.01394075 |
| LINC00271 | 988.006198 | 0.15207415 | 0.63623813 | 0.23902081 | 0.81108945 | 0.89759431 |
| PDE7B | 2053.31445 | 0.622632972 | 0.37432377 | 1.6633541 | 0.09624157 | 0.23950313 |
| AL138828.1 | 4054.989 | -3.897589967 | 0.89933864 | -4.3338402 | 1.47E-05 | 0.00034451 |
| BCLAF1 | 1259.42792 | -0.137761779 | 0.1359024 | -1.0136817 | 0.31073466 | 0.51104512 |
| AL023284.4 | 67.8620887 | -0.024522017 | 0.3763176 | -0.0651631 | 0.94804415 | 0.97344198 |
| MAP7 | 5129.17552 | 0.140761722 | 0.21860524 | 0.64390828 | 0.51963491 | 0.69597507 |
| MAP3K5 | 2247.86527 | 0.347842386 | 0.42678349 | 0.81503242 | 0.41505375 | 0.60853959 |
| PEX7 | 298.426645 | 0.522892733 | 0.24701503 | 2.11684583 | 0.03427293 | 0.12014811 |
| IFNGR1 | 362.125171 | -0.360277384 | 0.23031122 | -1.5643067 | 0.11774556 | 0.27368804 |
| TNFAIP3 | 177.344465 | -2.242291684 | 0.50469245 | -4.4428873 | 8.88E-06 | 0.00023121 |
| PERP | 245.939364 | -2.20917215 | 0.50175718 | -4.402871 | 1.07E-05 | 0.00026824 |
| ARFGEF3 | 247.0556 | -3.577697795 | 0.44345126 | -8.0678489 | 7.15E-16 | 5.16E-13 |
| HEBP2 | 376.639295 | -0.568137264 | 0.23208383 | -2.447983 | 0.01436584 | 0.06498766 |
| NHSL1 | 1290.08018 | 0.072905711 | 0.29013432 | 0.25128262 | 0.80159561 | 0.89227474 |
| CCDC28A | 194.84548 | 0.243284056 | 0.36304563 | 0.67011977 | 0.50278144 | 0.68363952 |
| REPS1 | 1587.5206 | 0.55169974 | 0.18544735 | 2.97496696 | 0.0029302 | 0.02033634 |
| ABRACL | 318.066173 | -0.700308451 | 0.45834336 | -1.5279123 | 0.12653431 | 0.28753412 |
| HECA | 164.339529 | -0.8741664 | 0.27020502 | -3.2351967 | 0.00121559 | 0.0104793 |
| AL031772.1 | 68.9443498 | -0.050740306 | 0.35623384 | -0.1424354 | 0.88673611 | 0.94155177 |
| CITED2 | 205.461134 | 0.096033139 | 0.46829433 | 0.20507004 | 0.8375174 | 0.91436098 |
| FILNC1 | 68.6723578 | 0.069882781 | 1.01641859 | 0.06875394 | 0.94518548 | 0.97196767 |
| VTA1 | 647.1746 | 0.130373568 | 0.19406645 | 0.67179858 | 0.50171193 | 0.68278052 |
| ADGRG6 | 745.617304 | 0.074115211 | 0.39764485 | 0.18638544 | 0.85214251 | 0.92213818 |
| HIVEP2 | 2616.07984 | -1.104465852 | 0.3159824 | -3.4953398 | 0.00047346 | 0.00515874 |
| LINC01277 | 524.395214 | 1.38241737 | 0.38583929 | 3.58288392 | 0.00033982 | 0.00403634 |
| AL023581.2 | 57.6606394 | 1.401949566 | 0.4337226 | 3.23236454 | 0.0012277 | 0.01055995 |
| AIG1 | 3868.64132 | 0.994348558 | 0.39645126 | 2.50812307 | 0.01213744 | 0.05775145 |
| ADAT2 | 149.947693 | -0.248354296 | 0.32900404 | -0.754867 | 0.45032879 | 0.64071587 |
| PEX3 | 640.969262 | -0.300159719 | 0.31921635 | -0.9403018 | 0.34706277 | 0.5471809 |
| FUCA2 | 314.236924 | -0.492649771 | 0.25815826 | -1.9083247 | 0.05634927 | 0.16770632 |
| PHACTR2 | 1254.29247 | -0.602066421 | 0.35366511 | -1.7023631 | 0.08868733 | 0.22709456 |
| LTV1 | 39.5977137 | 0.304156754 | 0.44209609 | 0.68798789 | 0.49146041 | 0.67391434 |
| PLAGL1 | 1376.61616 | -1.028999222 | 0.28575642 | -3.6009662 | 0.00031704 | 0.0038355 |
| SF3B5 | 328.278226 | -0.930760798 | 0.35923821 | -2.5909293 | 0.00957172 | 0.04882751 |
| UTRN | 1120.59387 | -0.425867053 | 0.45927035 | -0.9272688 | 0.353787 | 0.55219294 |

| | | | | | | |
|---|---|---|---|---|---|---|
| EPM2A | 603.947454 | 0.407372513 | 0.27948813 | 1.45756644 | 0.14496009 | 0.31310556 |
| FBXO30 | 239.540168 | -0.73214755 | 0.36137571 | -2.026001 | 0.04276468 | 0.13970212 |
| AL356599.1 | 530.439537 | 1.067616022 | 0.50902413 | 2.09737803 | 0.03596012 | 0.12403257 |
| SHPRH | 1748.5787 | 0.443962107 | 0.16871229 | 2.63147467 | 0.00850152 | 0.04471617 |
| RAB32 | 211.269003 | -0.739371325 | 0.28623929 | -2.5830532 | 0.00979302 | 0.04967911 |
| STXBP5-AS1 | 91.1132808 | -0.966828879 | 0.22241654 | -4.3469288 | 1.38E-05 | 0.00032796 |
| STXBP5 | 1214.94837 | -0.82420406 | 0.24929916 | -3.3060844 | 0.0009461 | 0.00868014 |
| SAMD5 | 10117.8356 | -0.374105502 | 0.32510707 | -1.1507147 | 0.24984961 | 0.44598106 |
| AL033504.1 | 5656.39176 | -0.464816335 | 0.36652385 | -1.2681749 | 0.20473551 | 0.39304057 |
| AL513164.1 | 246.940417 | -1.601641628 | 0.57414439 | -2.7896147 | 0.00527708 | 0.03152282 |
| SASH1 | 6918.8703 | -0.262726649 | 0.53761651 | -0.4886878 | 0.62506271 | 0.77775369 |
| UST | 205.303561 | -1.035075518 | 0.58628895 | -1.7654699 | 0.07748482 | 0.20765144 |
| TAB2 | 1418.67905 | -0.017151329 | 0.21677268 | -0.0791213 | 0.93693617 | 0.96611631 |
| ZC3H12D | 79.357498 | 0.961200213 | 0.52348237 | 1.83616539 | 0.0663332 | 0.1872383 |
| PPIL4 | 561.523748 | 0.21880296 | 0.14447005 | 1.51452128 | 0.12989368 | 0.29221282 |
| GINM1 | 843.740034 | 0.270906971 | 0.1937829 | 1.39799217 | 0.16211542 | 0.33842766 |
| AL078581.3 | 56.5513819 | -0.438367733 | 0.71268121 | -0.6150965 | 0.53849098 | 0.71019877 |
| KATNA1 | 362.894656 | 0.48661367 | 0.2199607 | 2.21227551 | 0.02694763 | 0.101591 |
| LATS1 | 909.320539 | 0.199517383 | 0.21684845 | 0.9200775 | 0.35753226 | 0.5561453 |
| NUP43 | 278.827535 | 0.17303187 | 0.21134818 | 0.81870529 | 0.41295458 | 0.60674975 |
| PCMT1 | 700.314376 | -0.263125577 | 0.31197475 | -0.8434195 | 0.39899389 | 0.59316804 |
| LRP11 | 152.365824 | 0.050087169 | 0.41024718 | 0.12209022 | 0.90282757 | 0.94900747 |
| PPP1R14C | 739.142048 | 0.23820574 | 0.45492624 | 0.52361398 | 0.60054706 | 0.76020679 |
| IYD | 495.075132 | 2.213325985 | 0.70180317 | 3.1537703 | 0.00161176 | 0.01304155 |
| PLEKHG1 | 867.233218 | -0.489564438 | 0.51983434 | -0.9417701 | 0.34631036 | 0.54664509 |
| MTHFD1L | 1295.76345 | -2.230271235 | 0.33852443 | -6.5882136 | 4.45E-11 | 8.03E-09 |
| AL138733.1 | 120.800728 | 1.132860525 | 0.59151247 | 1.91519299 | 0.0554679 | 0.16611284 |
| AKAP12 | 3270.96053 | -1.858251248 | 0.53752534 | -3.4570486 | 0.00054613 | 0.0057056 |
| ZBTB2 | 162.363104 | -0.970742623 | 0.38453308 | -2.524471 | 0.01158726 | 0.05584618 |
| RMND1 | 793.341611 | 0.505331541 | 0.346616 | 1.45790022 | 0.14486805 | 0.31300541 |
| ARMT1 | 259.34226 | 0.537472852 | 0.21333827 | 2.51934569 | 0.01175732 | 0.05641571 |
| CCDC170 | 91.8988638 | -0.946912368 | 0.69333431 | -1.3657371 | 0.1720215 | 0.35134835 |
| ESR1 | 657.368525 | 2.37362088 | 0.54457221 | 4.35868898 | 1.31E-05 | 0.00031687 |
| SYNE1 | 2930.85122 | 1.017747389 | 0.33536856 | 3.03471315 | 0.00240765 | 0.01771422 |
| FBXO5 | 39.8119276 | 0.119710648 | 0.52571921 | 0.22770834 | 0.81987299 | 0.90294063 |
| AL080276.2 | 79.1756664 | 0.145180755 | 0.36260268 | 0.40038523 | 0.6888728 | 0.82001633 |
| MTRF1L | 245.421106 | -0.354043522 | 0.2400465 | -1.4748956 | 0.14024062 | 0.30697512 |
| RGS17 | 91.6340457 | -3.831531887 | 0.87219267 | -4.3929879 | 1.12E-05 | 0.00027762 |
| OPRM1 | 68.3865272 | -0.93238484 | 0.74795737 | -1.2465748 | 0.21255343 | 0.4023574 |
| IPCEF1 | 596.349157 | -0.28396649 | 0.52000688 | -0.5460822 | 0.58500945 | 0.74766662 |
| CNKSR3 | 6002.13599 | 0.316221968 | 0.23981268 | 1.3186207 | 0.18729595 | 0.37080981 |
| SCAF8 | 2067.83288 | 0.370418616 | 0.20110209 | 1.84194313 | 0.06548347 | 0.18571941 |
| TIAM2 | 1018.19409 | 0.117150743 | 0.23394146 | 0.50076948 | 0.61653336 | 0.77182636 |
| TFB1M | 610.721304 | 0.033835801 | 0.25978934 | 0.13024323 | 0.896374 | 0.94620317 |
| CLDN20 | 40.4249095 | -0.3289666 | 0.51929151 | -0.6334912 | 0.52641293 | 0.7009757 |
| ARID1B | 6215.04062 | 0.174382758 | 0.15665476 | 1.11316604 | 0.26563712 | 0.46352448 |
| TMEM242 | 349.054356 | 0.162758805 | 0.18191582 | 0.89469296 | 0.37095124 | 0.5686868 |
| ZDHHC14 | 2771.92912 | -1.795052689 | 0.45037784 | -3.9856594 | 6.73E-05 | 0.00114298 |
| SNX9 | 2296.90618 | -0.888556102 | 0.36602903 | -2.4275564 | 0.01520092 | 0.06782522 |
| SYNJ2 | 309.723004 | -1.659012911 | 0.53061956 | -3.1265581 | 0.00176866 | 0.01394075 |
| SERAC1 | 365.545752 | -0.219587498 | 0.27116223 | -0.8098012 | 0.41805444 | 0.61116384 |
| GTF2H5 | 230.594235 | 0.196891075 | 0.28022443 | 0.70261923 | 0.48229308 | 0.66670759 |
| TULP4 | 3001.13881 | -0.102226558 | 0.28098509 | -0.3638149 | 0.71599625 | 0.83776419 |
| TMEM181 | 523.660104 | -0.850048219 | 0.40364046 | -2.105954 | 0.03520835 | 0.12220838 |
| DYNLT1 | 832.499559 | -1.518350499 | 0.24193915 | -6.2757536 | 3.48E-10 | 4.68E-08 |
| SYTL3 | 649.157873 | -2.981642497 | 0.63674585 | -4.6826257 | 2.83E-06 | 9.11E-05 |
| EZR | 3234.1595 | 0.064939807 | 0.18835221 | 0.34477857 | 0.73026084 | 0.84648502 |
| C6orf99 | 53.2235687 | 0.362606817 | 0.4628659 | 0.78339496 | 0.43339521 | 0.62578197 |
| RSPH3 | 772.38802 | -0.251703997 | 0.17241245 | -1.4598946 | 0.14431905 | 0.31211441 |
| AL035530.2 | 345.785712 | 0.21903454 | 0.4369151 | 0.5013206 | 0.61614551 | 0.7715916 |
| SOD2 | 219.017536 | -1.851832598 | 0.51769191 | -3.5770939 | 0.00034744 | 0.00410897 |
| SOD2.1 | 259.978272 | -1.428653579 | 0.32963201 | -4.3340863 | 1.46E-05 | 0.00034451 |
| WTAP | 1952.78105 | -1.125565729 | 0.23344772 | -4.8214896 | 1.42E-06 | 5.13E-05 |
| ACAT2 | 222.195118 | -0.938883238 | 0.4361643 | -2.1525908 | 0.03135085 | 0.11267067 |
| TCP1 | 464.035777 | -0.79713055 | 0.29203138 | -2.7296058 | 0.00634101 | 0.03609478 |
| MRPL18 | 233.853604 | -0.612519062 | 0.29495413 | -2.0766587 | 0.03783307 | 0.12871131 |
| IGF2R | 3550.41858 | -0.307830775 | 0.30636166 | -1.0047954 | 0.31499539 | 0.51523844 |
| SLC22A1 | 156.521061 | 0.288324038 | 0.54517189 | 0.52886813 | 0.59689693 | 0.75700095 |
| SLC22A2 | 1943.99109 | 0.219238373 | 0.35652302 | 0.61493468 | 0.53859786 | 0.71027139 |
| SLC22A3 | 773.748278 | -0.69539375 | 0.55463348 | -1.2537897 | 0.20991845 | 0.39949496 |
| LPA | 55.9055522 | 2.017796307 | 0.85787047 | 2.35209904 | 0.0186678 | 0.07853885 |
| PLG | 2192.50727 | 2.315506983 | 0.70751679 | 3.27272372 | 0.00106517 | 0.00946837 |
| MAP3K4 | 866.526816 | 0.280045208 | 0.12334161 | 2.27048452 | 0.0231782 | 0.09147815 |
| AGPAT4 | 221.490587 | -1.844122418 | 0.58585882 | -3.147725 | 0.00164546 | 0.01323734 |
| PRKN | 8046.35494 | 1.496311099 | 0.49056463 | 3.05018138 | 0.00228703 | 0.01702059 |
| PACRG | 628.338809 | 0.617190506 | 0.56039421 | 1.1013506 | 0.2707441 | 0.46898433 |
| QKI | 3127.89285 | 0.116490736 | 0.16615978 | 0.70107661 | 0.48325521 | 0.6673362 |
| PDE10A | 2792.94768 | 0.913551746 | 0.37165634 | 2.45805506 | 0.01396918 | 0.06372047 |
| SFT2D1 | 437.751921 | -0.182409812 | 0.1374683 | -1.3269227 | 0.18453425 | 0.36743332 |
| MPC1 | 1286.21981 | -0.056007936 | 0.29005114 | -0.1930968 | 0.84688319 | 0.91937902 |
| AL022069.1 | 31.8332974 | 0.251513672 | 0.60716013 | 0.41424603 | 0.67869394 | 0.81376982 |
| RPS6KA2 | 682.549049 | -0.71114384 | 0.25418778 | -2.7977106 | 0.00514662 | 0.03098144 |
| AL159163.1 | 40.4810893 | -0.589665189 | 0.45557556 | -1.2943302 | 0.1955514 | 0.3808166 |
| Z94721.1 | 600.862003 | 1.164224523 | 0.36066232 | 3.2280182 | 0.00124651 | 0.0106787 |
| FGFR1OP | 1462.00001 | 0.685487605 | 0.24776327 | 2.76670395 | 0.00566262 | 0.03324749 |
| CCR6 | 38.1541797 | 0.674032468 | 0.55145046 | 1.22229014 | 0.22159793 | 0.41376417 |
| LINC01558 | 81.559118 | 1.079234461 | 0.68309717 | 1.57991354 | 0.11412667 | 0.26804905 |
| AFDN | 4289.66591 | -0.182546338 | 0.30798742 | -0.5927071 | 0.55337717 | 0.7228074 |
| FRMD1 | 155.045182 | 2.47631547 | 0.70276053 | 3.52369743 | 0.00042557 | 0.00478181 |
| AL606970.3 | 34.4471572 | 1.711928458 | 0.86628564 | 1.97617087 | 0.04813543 | 0.15165818 |
| DACT2 | 107.621318 | 1.274549923 | 1.09324611 | 1.16583989 | 0.24367919 | 0.43890532 |
| BX322234.1 | 67.1500179 | -0.823800095 | 0.65469441 | -1.2582971 | 0.20828432 | 0.39745973 |
| THBS2 | 165.647383 | -3.509820731 | 0.96133225 | -3.6509966 | 0.00026122 | 0.0032861 |
| WDR27 | 1104.71604 | 0.436182886 | 0.24723398 | 1.76425133 | 0.07768967 | 0.20798679 |
| C6orf120 | 80.6372505 | -0.932638957 | 0.26913199 | -3.4653589 | 0.00052952 | 0.00555757 |
| PHF10 | 385.149921 | -0.122723342 | 0.18956663 | -0.6473889 | 0.51738025 | 0.69425481 |
| TCTE3 | 59.3305233 | -0.363546932 | 0.37087155 | -0.9802503 | 0.3269626 | 0.52691716 |
| ERMARD | 251.037875 | 0.560487148 | 0.34106238 | 1.6433567 | 0.10030915 | 0.2463163 |
| DLL1 | 45.2308015 | 0.311539707 | 0.45375109 | 0.68658723 | 0.49234288 | 0.67462692 |
| FAM120B | 1363.09916 | -0.094133212 | 0.24949 | -0.3773025 | 0.70594878 | 0.83159504 |
| PSMB1 | 577.158936 | -0.762163556 | 0.26553582 | -2.8702853 | 0.00410102 | 0.02613325 |
| TBP | 204.82165 | -0.010621036 | 0.19904537 | -0.0533599 | 0.95744517 | 0.97835851 |
| PDCD2 | 227.088584 | -0.65521842 | 0.26528112 | -2.4699022 | 0.013515 | 0.06226895 |
| AC093627.5 | 51.0831596 | 0.650548196 | 0.35180347 | 1.84918075 | 0.06443172 | 0.18380136 |
| AC093627.3 | 95.5934845 | 2.460239818 | 0.52456772 | 4.69003281 | 2.73E-06 | 8.91E-05 |
| FAM20C | 2196.1461 | 1.725366844 | 0.39214967 | 4.39976615 | 1.08E-05 | 0.00027155 |
| PDGFA | 168.029586 | -0.495751569 | 0.29229036 | -1.6960928 | 0.08986831 | 0.22886597 |
| PRKAR1B | 808.636302 | -0.716943642 | 0.48858295 | -1.4673939 | 0.14226894 | 0.30963485 |
| DNAAF5 | 559.665381 | -0.078930147 | 0.40469606 | -0.1950356 | 0.84536506 | 0.91854292 |
| SUN1 | 880.553318 | -0.394061534 | 0.19387767 | -2.0325267 | 0.04210036 | 0.13812103 |
| AC073957.3 | 45.7858658 | -0.662656665 | 0.79520663 | -0.8333138 | 0.40466777 | 0.59871533 |
| COX19 | 334.324019 | -0.474983364 | 0.23033238 | -2.062165 | 0.03919203 | 0.13179715 |
| C7orf50 | 773.910442 | 0.159430856 | 0.27378552 | 0.58232026 | 0.56035098 | 0.72874874 |

| | | | | | | |
|---|---|---|---|---|---|---|
| GPR146 | 60.239053 | -1.231743756 | 0.58139121 | -2.1186144 | 0.03412307 | 0.11965335 |
| GPER1 | 93.5023491 | 0.657593866 | 0.52948614 | 1.24194727 | 0.21425602 | 0.40451891 |
| ZFAND2A | 103.983136 | -1.273106339 | 0.75350886 | -1.6895705 | 0.09111015 | 0.23131078 |
| MICALL2 | 142.882183 | -2.014588575 | 0.44251621 | -4.5525758 | 5.30E-06 | 0.00015018 |
| INTS1 | 376.771736 | 0.594800366 | 0.35560274 | 1.67265404 | 0.09439542 | 0.23672884 |
| MAFK | 133.523077 | -0.397621406 | 0.37335503 | -1.0649954 | 0.28687799 | 0.48654252 |
| TMEM184A | 201.925086 | 0.776280962 | 0.34666838 | 2.23926092 | 0.02513894 | 0.09676482 |
| PSMG3 | 119.206598 | -0.545498929 | 0.26783483 | -2.036699 | 0.04168022 | 0.13726832 |
| PSMG3-AS1 | 96.2795107 | 0.726156123 | 0.3512825 | 2.06715711 | 0.03871935 | 0.13084964 |
| MAD1L1 | 593.702424 | -0.064581106 | 0.3435248 | -0.1879955 | 0.8508802 | 0.92167424 |
| MRM2 | 93.0091271 | -0.336006909 | 0.39664015 | -0.8471329 | 0.39692107 | 0.5916211 |
| SNX8 | 489.387569 | 0.334145565 | 0.45851358 | 0.72875827 | 0.46614954 | 0.65405996 |
| EIF3B | 858.148234 | -0.673454685 | 0.36991977 | -1.8205426 | 0.06867642 | 0.19172635 |
| CHST12 | 303.106317 | 0.355344113 | 0.35581654 | 0.99867226 | 0.31795348 | 0.51803024 |
| BRAT1 | 182.17885 | 0.610809191 | 0.27245339 | 2.24188512 | 0.0249688 | 0.0962721 |
| IQCE | 467.497777 | 0.025897923 | 0.21054705 | 0.12300302 | 0.90210471 | 0.94861498 |
| TTYH3 | 1470.83498 | 0.328183548 | 0.3405759 | 0.96361353 | 0.33523972 | 0.53500185 |
| AMZ1 | 236.023389 | -2.304275628 | 0.78402755 | -2.9390238 | 0.00329248 | 0.02215604 |
| GNA12 | 1174.02867 | -0.495323954 | 0.26280555 | -1.8847545 | 0.05946299 | 0.17371252 |
| SDK1 | 6392.38234 | -1.055162265 | 0.47789315 | -2.207946 | 0.02724804 | 0.10236573 |
| AC017000.1 | 42.9983127 | 1.428239874 | 0.87508675 | 1.63211232 | 0.10265583 | 0.25027051 |
| FOXK1 | 1486.24626 | -0.132371401 | 0.24096532 | -0.549338 | 0.58277353 | 0.74571292 |
| AP5Z1 | 427.499205 | -0.59634777 | 0.49858398 | -1.1960829 | 0.23166421 | 0.42480553 |
| RBAK-RBAKD | 671.196332 | -0.371741783 | 0.33374921 | -1.1138357 | 0.26534968 | 0.46321295 |
| WIPI2 | 879.10402 | -0.148171384 | 0.28181966 | -0.5257667 | 0.59905033 | 0.7589133 |
| TNRC18 | 1669.6591 | 1.174785781 | 0.41928354 | 2.80188865 | 0.00508044 | 0.03065035 |
| FBXL18 | 275.042503 | 0.000647764 | 0.41922862 | 0.00154513 | 0.99876716 | 0.99920455 |
| ACTB | 3009.79746 | -0.085005722 | 0.49970896 | -0.1701105 | 0.86492327 | 0.92933317 |
| RNF216 | 1523.19563 | -0.343533488 | 0.31276568 | -1.0983733 | 0.2720415 | 0.47033143 |
| CCZ1 | 223.136392 | -0.092988283 | 0.27271397 | -0.3409737 | 0.73312341 | 0.84871134 |
| PMS2 | 461.445263 | 0.73272984 | 0.31377685 | 2.33519407 | 0.01953328 | 0.08103823 |
| AIMP2 | 252.11327 | -0.260234649 | 0.29305722 | -0.8879995 | 0.37454104 | 0.5715597 |
| EIF2AK1 | 530.66224 | 0.1011998 | 0.15611026 | 0.64825849 | 0.51681778 | 0.69403559 |
| USP42 | 577.22446 | 0.342798704 | 0.20800297 | 1.64804715 | 0.099343 | 0.24460112 |
| CYTH3 | 371.364885 | -0.105138741 | 0.31057612 | -0.3385281 | 0.73496526 | 0.84999737 |
| FAM220A | 43.5031863 | 0.277806425 | 0.38930367 | 0.71359826 | 0.47547563 | 0.66125654 |
| RAC1 | 1260.13791 | -0.688918233 | 0.25037568 | -2.7515382 | 0.00593161 | 0.0343928 |
| DAGLB | 269.745494 | -0.717908422 | 0.52012594 | -1.3802588 | 0.16750696 | 0.34557766 |
| ZDHHC4 | 125.591427 | 0.024946413 | 0.20462229 | 0.12191445 | 0.90296677 | 0.94908102 |
| C7orf26 | 123.186271 | 0.79235885 | 0.31918775 | 2.4824225 | 0.01304925 | 0.06079742 |
| ZNF316 | 316.86685 | 0.273992121 | 0.42823753 | 0.63981342 | 0.52229391 | 0.69817462 |
| AC073343.2 | 165.773038 | -0.633543527 | 0.50621417 | -1.2515326 | 0.21074023 | 0.40007037 |
| ZNF12 | 317.264681 | -0.20052655 | 0.16477541 | -1.2169689 | 0.22361605 | 0.41588945 |
| CCZ1B | 318.125077 | -0.163912884 | 0.29043594 | -0.5643685 | 0.5725034 | 0.73810771 |
| C1GALT1 | 2062.46875 | 0.677053443 | 0.28304577 | 2.39202812 | 0.01675556 | 0.07284124 |
| COL28A1 | 405.340543 | -0.332416463 | 0.44237108 | -0.7514426 | 0.45238635 | 0.64164525 |
| AC004982.1 | 37.6381091 | 0.101061585 | 0.37214502 | 0.27156506 | 0.78595647 | 0.88296838 |
| AC004982.2 | 109.014732 | -0.226601904 | 0.38203989 | -0.5931368 | 0.55308962 | 0.72252338 |
| MIOS | 456.524021 | -0.073829665 | 0.18464678 | -0.3998427 | 0.68927239 | 0.82041131 |
| RPA3 | 226.922652 | -0.601423771 | 0.49895183 | -1.2053744 | 0.2280588 | 0.42089699 |
| UMAD1 | 2168.01489 | 0.187626301 | 0.18962065 | 0.98948244 | 0.32242716 | 0.52286645 |
| GLCCI1 | 530.167237 | 0.489927037 | 0.23479553 | 2.0866114 | 0.03692328 | 0.12633733 |
| ICA1 | 470.913279 | -1.433171155 | 0.57622401 | -2.4871771 | 0.01287613 | 0.0602777 |
| NDUFA4 | 762.898956 | -0.667131719 | 0.39802344 | -1.6761116 | 0.09371633 | 0.23571272 |
| PHF14 | 1566.46341 | 0.08032171 | 0.31740848 | 0.25305471 | 0.80022594 | 0.89131054 |
| THSD7A | 4343.92825 | 0.45018656 | 0.52171133 | 0.86290354 | 0.3881905 | 0.58387159 |
| TMEM106B | 711.270733 | 0.166446587 | 0.2522855 | 0.65975486 | 0.50941116 | 0.6889789 |
| SCIN | 340.869536 | -0.535130533 | 0.54448038 | -0.9828279 | 0.32569213 | 0.52586969 |
| ARL4A | 49.3797358 | -1.497466933 | 0.57643293 | -2.5978164 | 0.00938186 | 0.04810969 |
| AC011287.1 | 352.205475 | 0.775740041 | 0.47897053 | 1.61959868 | 0.10531851 | 0.25397727 |
| ETV1 | 144.80894 | -1.628860045 | 0.62082637 | -2.6236966 | 0.00869812 | 0.04550579 |
| DGKB | 1371.20676 | 3.475331762 | 0.57580362 | 6.03561989 | 1.58E-09 | 1.74E-07 |
| AGMO | 1864.23358 | 1.451096641 | 0.69846602 | 2.07754794 | 0.03775102 | 0.12846919 |
| ISPD | 1291.10518 | 0.812359012 | 0.31951459 | 2.5424786 | 0.01100693 | 0.05399925 |
| SOSTDC1 | 113.722154 | 1.179675748 | 0.75354064 | 1.56551046 | 0.11746327 | 0.27335636 |
| ANKMY2 | 403.527348 | 0.37413407 | 0.2161621 | 1.73080325 | 0.08348686 | 0.21823333 |
| BZW2 | 277.334863 | -0.454165126 | 0.31323343 | -1.4499255 | 0.14707931 | 0.31648604 |
| AHR | 49.5491255 | -1.091017487 | 0.5001986 | -2.1811686 | 0.02917095 | 0.10692593 |
| AC073332.1 | 221.360286 | -0.860422505 | 0.3996864 | -2.152744 | 0.0313388 | 0.11265695 |
| AC019117.1 | 179.014605 | -0.957514017 | 0.68511381 | -1.3975985 | 0.16223368 | 0.33862297 |
| SNX13 | 1677.94393 | -0.09491404 | 0.14801279 | -0.6412557 | 0.52135658 | 0.69719362 |
| HDAC9 | 300.602421 | -2.286965005 | 0.70195735 | -3.2579828 | 0.00112207 | 0.00987178 |
| TWISTNB | 101.973075 | -0.292944013 | 0.2914723 | -1.0050492 | 0.31487314 | 0.51522797 |
| MACC1 | 3159.23651 | -0.824191189 | 0.45345806 | -1.8175687 | 0.06913007 | 0.19255756 |
| MACC1-AS1 | 61.043518 | -1.677997927 | 0.79777787 | -2.1033398 | 0.03543608 | 0.12278117 |
| ITGB8 | 6966.14898 | -0.3685451 | 0.52490633 | -0.7021159 | 0.48260687 | 0.66677677 |
| SP4 | 157.056795 | 0.381680141 | 0.34123177 | 1.11853637 | 0.26333798 | 0.46128738 |
| DNAH11 | 269.864615 | 2.568846099 | 1.38837677 | 1.85025142 | 0.06427732 | 0.18343727 |
| RAPGEF5 | 1933.81464 | -4.967069908 | 0.8618919 | -5.7629848 | 8.26E-09 | 6.95E-07 |
| STEAP1B | 800.099793 | -0.933021562 | 0.41528122 | -2.2467222 | 0.02465778 | 0.09542186 |
| TOMM7 | 354.952334 | -0.050542526 | 0.34026048 | -0.1485407 | 0.88191608 | 0.93868797 |
| FAM126A | 476.526213 | 0.912063309 | 0.2524199 | 3.61327818 | 0.00030235 | 0.00369698 |
| KLHL7 | 555.227854 | 0.119558516 | 0.19672159 | 0.60775493 | 0.54335003 | 0.71379134 |
| NUPL2 | 559.773218 | -0.08890559 | 0.20907526 | -0.4252325 | 0.67066721 | 0.80901483 |
| MALSU1 | 249.759399 | 0.1903207 | 0.2081466 | 0.9143589 | 0.36052829 | 0.5586436 |
| TRA2A | 1636.08216 | 0.080328468 | 0.25802591 | 0.31131938 | 0.75555783 | 0.86293197 |
| CCDC126 | 242.442388 | -0.097145311 | 0.2431319 | -0.3995581 | 0.68948205 | 0.82043055 |
| FAM221A | 157.770107 | 0.17487157 | 0.24434922 | 0.71566248 | 0.47419978 | 0.66042028 |
| AC004485.1 | 2400.09979 | 1.630393288 | 0.85316049 | 1.9110042 | 0.05600404 | 0.16710188 |
| NPY | 117.76815 | 0.410940492 | 1.09871396 | 0.37401954 | 0.70838978 | 0.8329672 |
| MPP6 | 2517.06921 | -0.272219778 | 0.23076341 | -1.1796488 | 0.23813994 | 0.43308798 |
| GSDME | 124.524798 | -1.051744665 | 0.45201343 | -2.3267996 | 0.01997593 | 0.08239847 |
| OSBPL3 | 298.710889 | -1.417812641 | 0.6051065 | -2.3430795 | 0.01912531 | 0.07989959 |
| CYCS | 329.741124 | -0.815546211 | 0.36256297 | -2.2493919 | 0.02448757 | 0.09490085 |
| NFE2L3 | 201.491485 | -1.955035754 | 0.6129456 | -3.1895746 | 0.00142482 | 0.01182923 |
| HNRNPA2B1 | 4189.23778 | -0.432050436 | 0.15922345 | -2.7134849 | 0.00665796 | 0.03744796 |
| CBX3 | 897.061121 | -1.18808921 | 0.26524386 | -4.4792336 | 7.49E-06 | 0.00019861 |
| SNX10 | 3398.73025 | -1.5429462 | 0.38511589 | -4.0064465 | 6.16E-05 | 0.00106678 |
| AC004540.1 | 370.155337 | -0.472764345 | 0.61937116 | -0.7632973 | 0.4452861 | 0.63664719 |
| SKAP2 | 2464.38998 | -0.273427986 | 0.35473666 | -0.7707914 | 0.44083056 | 0.63271879 |
| HOTAIRM1 | 193.523292 | -1.126945529 | 0.36281499 | -3.1061162 | 0.00189562 | 0.01469643 |
| HOXA3 | 314.919867 | 1.360933252 | 0.39130152 | 3.47796564 | 0.00050523 | 0.00538667 |
| HOXA-AS2 | 172.556343 | 0.480563388 | 0.47836409 | 1.00459753 | 0.31509069 | 0.51523844 |
| HOXA7 | 45.7366127 | -0.128546288 | 0.54281427 | -0.2368145 | 0.8128007 | 0.89861746 |
| HOXA9 | 227.302774 | 0.10900302 | 0.35351446 | 0.30834105 | 0.75782283 | 0.86418282 |
| HOXA10-AS | 59.7044915 | -0.250048461 | 0.36192236 | -0.6908898 | 0.4896348 | 0.67262219 |
| HIBADH | 2727.99384 | 0.311014114 | 0.27164834 | 1.14491448 | 0.25224458 | 0.44873673 |
| TAX1BP1 | 2890.1717 | -0.268854932 | 0.20134918 | -1.3352671 | 0.18178896 | 0.36397622 |
| JAZF1 | 817.541131 | -1.619064458 | 0.41016737 | -3.9473264 | 7.90E-05 | 0.00129575 |
| CREB5 | 3580.16822 | -1.247982541 | 0.54445197 | -2.2921812 | 0.02189519 | 0.08811432 |
| AC005162.3 | 396.292289 | 0.78849293 | 0.45011918 | 1.75174257 | 0.07981808 | 0.21149728 |
| CPVL | 663.331393 | -0.335143004 | 0.3748714 | -0.8940213 | 0.37131051 | 0.56879226 |
| AC007255.1 | 68.4114987 | -0.242373206 | 0.44324819 | -0.5468115 | 0.58450824 | 0.74723507 |

| | | | | | | |
|---|---|---|---|---|---|---|
| WIPF3 | 207.277009 | -1.574695688 | 0.39767218 | -3.9597834 | 7.50E-05 | 0.00124639 |
| SCRN1 | 182.997667 | -1.516687864 | 0.47644532 | -3.1833408 | 0.00145586 | 0.01203588 |
| FKBP14 | 142.027812 | -0.377584196 | 0.28049123 | -1.3461533 | 0.17825309 | 0.3592582 |
| PLEKHA8 | 865.40907 | 0.168020739 | 0.24287709 | 0.69179327 | 0.48906717 | 0.67217925 |
| MTURN | 240.183093 | -0.198626113 | 0.31158623 | -0.6374676 | 0.52382034 | 0.69932847 |
| ZNRF2 | 1409.42777 | 0.828094842 | 0.28924163 | 2.86298632 | 0.00419669 | 0.0265577 |
| AC006978.2 | 1349.07091 | 0.727617104 | 0.29294796 | 2.48377595 | 0.01299976 | 0.06060805 |
| NOD1 | 194.391814 | -1.282624437 | 0.45998659 | -2.7883953 | 0.00529699 | 0.03159521 |
| GGCT | 96.7505481 | -0.391711356 | 0.42984831 | -0.9112781 | 0.36214886 | 0.5599915 |
| AC005154.6 | 384.865648 | 0.897722656 | 0.39779531 | 2.25674517 | 0.024024 | 0.09363008 |
| GARS | 636.612963 | -1.185734481 | 0.39462891 | -3.0046822 | 0.00265858 | 0.01894034 |
| AQP1 | 2348.00421 | -0.178522206 | 0.78862907 | -0.2263703 | 0.82091342 | 0.9038686 |
| PDE1C | 58.553244 | -0.001995633 | 0.54283146 | -0.0036763 | 0.99706671 | 0.99859663 |
| LSM5 | 177.547441 | -0.252433965 | 0.27868914 | -0.9057905 | 0.36504676 | 0.56248732 |
| AVL9 | 2633.55172 | -0.311545501 | 0.14688935 | -2.1209537 | 0.0339257 | 0.11926606 |
| KBTBD2 | 448.087977 | -0.470128834 | 0.16897241 | -2.7822817 | 0.00539782 | 0.03208494 |
| FKBP9 | 263.89254 | 0.100068678 | 0.182192 | 0.54924848 | 0.58283494 | 0.74572188 |
| NT5C3A | 977.033646 | -0.033982175 | 0.1625148 | -0.209102 | 0.83436858 | 0.91245334 |
| RP9 | 136.450034 | -0.167992394 | 0.20054717 | -0.8376702 | 0.40221594 | 0.59640566 |
| BBS9 | 1799.50468 | 0.545109765 | 0.27620804 | 1.97354777 | 0.04843319 | 0.15222969 |
| DPY19L1 | 363.535343 | -0.836693601 | 0.35582011 | -2.3514512 | 0.01870034 | 0.07860338 |
| HERPUD2 | 603.068634 | -0.083391138 | 0.16579732 | -0.5029703 | 0.61498512 | 0.77067115 |
| AC018647.2 | 53.4753271 | -0.11011776 | 0.40679357 | -0.2706969 | 0.78662416 | 0.88335715 |
| SEPT7-AS1 | 119.50875 | 0.332826749 | 0.45545842 | 0.73075112 | 0.46493119 | 0.65256722 |
| 7-Sep | 1377.46901 | -0.391327225 | 0.17193944 | -2.2759596 | 0.02284843 | 0.09064642 |
| EEPD1 | 285.167624 | -0.038387602 | 0.48243998 | -0.0795697 | 0.9365795 | 0.96596653 |
| KIAA0895 | 539.347483 | 0.698975969 | 0.23092024 | 3.0269151 | 0.00247063 | 0.01803247 |
| AC006960.2 | 86.2520486 | 2.031537449 | 0.89018668 | 2.28214766 | 0.02248062 | 0.08970652 |
| AOAH | 596.855643 | 0.930603616 | 0.92690115 | 1.00399446 | 0.31538128 | 0.51543237 |
| AC007349.1 | 45.9073869 | 1.984198778 | 0.68151243 | 2.9114638 | 0.0035974 | 0.02370912 |
| ELMO1 | 133.994402 | 1.319690758 | 0.45359368 | 2.90941166 | 0.0036211 | 0.02379404 |
| EPDR1 | 382.782841 | -0.670196525 | 0.275254 | -2.4348294 | 0.01489881 | 0.06678156 |
| SFRP4 | 105.708681 | -1.300869744 | 0.72506587 | -1.7941401 | 0.07279081 | 0.19911068 |
| STARD3NL | 373.8148 | -0.248502031 | 0.19074289 | -1.3028115 | 0.19263912 | 0.37726881 |
| VPS41 | 1457.47237 | 0.072406265 | 0.19985132 | 0.36230066 | 0.71712737 | 0.8385655 |
| YAE1D1 | 176.163222 | 0.478343026 | 0.24619983 | 1.94290558 | 0.05202757 | 0.15901947 |
| RALA | 890.128381 | -0.646135938 | 0.35158005 | -1.837806 | 0.06609099 | 0.1867469 |
| LINC00265 | 91.9356038 | 0.383087575 | 0.26265704 | 1.45850869 | 0.14470038 | 0.31269244 |
| CDK13 | 2979.14243 | 0.12381807 | 0.1646304 | 0.75209724 | 0.45199258 | 0.64143458 |
| MPLKIP | 310.581487 | 0.65330498 | 0.2717722 | 2.40386975 | 0.01622255 | 0.07116846 |
| SUGCT | 6832.90618 | 1.917139391 | 0.46763042 | 4.09968926 | 4.14E-05 | 0.0007768 |
| AC004988.1 | 59.9332321 | -0.071956562 | 0.64900247 | -0.1108726 | 0.91171741 | 0.95319043 |
| LINC01449 | 39.6477645 | 1.25948422 | 0.84432449 | 1.49170637 | 0.13577614 | 0.30037826 |
| GLI3 | 327.380453 | 0.711560961 | 0.54712717 | 1.30054035 | 0.19341583 | 0.37838522 |
| MRPL32 | 237.742896 | -0.722650633 | 0.25604106 | -2.8224014 | 0.00476655 | 0.02925888 |
| AC005537.1 | 65.4034577 | -0.48891994 | 0.66296206 | -0.737478 | 0.46083172 | 0.64952394 |
| HECW1 | 1734.24975 | 2.76011051 | 0.81482438 | 3.3873686 | 0.00070567 | 0.00699286 |
| STK17A | 333.946654 | -0.893634899 | 0.38530739 | -2.3192779 | 0.02037997 | 0.08363722 |
| COA1 | 1680.56026 | 0.272318149 | 0.14654118 | 1.85830458 | 0.06312577 | 0.18105566 |
| BLVRA | 363.301673 | -0.693364209 | 0.22746199 | -3.048264 | 0.00230168 | 0.01710022 |
| URGCP | 590.098196 | 1.420944158 | 0.31487032 | 4.51279164 | 6.40E-06 | 0.00017504 |
| UBE2D4 | 203.947359 | 0.386273251 | 0.42140145 | 0.9166396 | 0.35933153 | 0.55773494 |
| SPDYE1 | 98.9060398 | 0.022092626 | 0.27528329 | 0.08025415 | 0.93603512 | 0.96587767 |
| AC004951.1 | 117.706694 | 0.65742613 | 0.33451705 | 1.96529931 | 0.0493796 | 0.15398584 |
| DBNL | 336.404916 | 0.158734954 | 0.19448004 | 0.81620179 | 0.41438474 | 0.60793873 |
| POLM | 254.797707 | 0.290745833 | 0.35581082 | 0.81713601 | 0.41385071 | 0.60754543 |
| AEBP1 | 96.4984729 | -1.511259098 | 0.762629 | -1.9816439 | 0.04751911 | 0.15042597 |
| POLD2 | 178.33652 | -0.355493328 | 0.30467079 | -1.1668113 | 0.24328658 | 0.43837638 |
| YKT6 | 214.31469 | -0.479903694 | 0.50426466 | -0.9516901 | 0.34125416 | 0.54195003 |
| CAMK2B | 42.4166651 | 2.255657297 | 0.38670461 | 5.83302411 | 5.44E-09 | 4.84E-07 |
| NUDCD3 | 1567.5185 | -0.08116775 | 0.24225993 | -0.3350441 | 0.73759185 | 0.85118107 |
| DDX56 | 395.830958 | 0.110499035 | 0.18233076 | 0.60603617 | 0.54449074 | 0.71487878 |
| TMED4 | 564.378514 | 0.006693617 | 0.2885205 | 0.0231998 | 0.9814909 | 0.99060551 |
| OGDH | 2321.34662 | 0.669051713 | 0.29263956 | 2.28626541 | 0.02223874 | 0.08897147 |
| ZMIZ2 | 768.877312 | 0.153058918 | 0.40492365 | 0.37799451 | 0.70543467 | 0.83113229 |
| PPIA | 1494.08596 | -1.116105519 | 0.3233417 | -3.4517834 | 0.00055689 | 0.0057803 |
| H2AFV | 520.744732 | 0.312610492 | 0.21687486 | 1.44143256 | 0.14946252 | 0.31978206 |
| PURB | 354.217259 | -0.271173047 | 0.14371394 | -1.8868945 | 0.05917451 | 0.17312808 |
| SNHG15 | 98.2152866 | -0.364075586 | 0.40269009 | -0.9041086 | 0.3659378 | 0.56343439 |
| CCM2 | 684.772178 | -0.073576267 | 0.24790449 | -0.2967928 | 0.76662471 | 0.8695271 |
| TBRG4 | 248.951293 | -0.144568821 | 0.24673445 | -0.5859288 | 0.55792336 | 0.72653007 |
| IGFBP3 | 94.1439648 | -0.978735688 | 0.63482542 | -1.5417399 | 0.12313681 | 0.28205819 |
| AC004870.4 | 184.563166 | 0.83239085 | 1.0879255 | 0.76511751 | 0.44420157 | 0.63542799 |
| AC004870.3 | 118.264342 | 0.291143917 | 0.92471591 | 0.31484688 | 0.75287792 | 0.8613901 |
| TNS3 | 6493.4902 | 0.176408578 | 0.38851705 | 0.4540562 | 0.64978838 | 0.79473983 |
| HUS1 | 283.306457 | 0.018820444 | 0.24402813 | 0.07712408 | 0.93852484 | 0.96688162 |
| UPP1 | 327.091559 | -2.782059149 | 0.66684005 | -4.1720037 | 3.02E-05 | 0.0006065 |
| ABCA13 | 214.088602 | -1.535672778 | 0.77273844 | -1.9873125 | 0.04688777 | 0.14915078 |
| ZPBP | 55.9714485 | 1.141143397 | 0.49536893 | 2.30362327 | 0.0212438 | 0.08638053 |
| FIGNL1 | 45.6978769 | -0.159843787 | 0.50155958 | -0.3186935 | 0.74995893 | 0.85900284 |
| DDC | 195.887495 | 2.68424995 | 0.5295464 | 5.0689608 | 4.00E-07 | 1.78E-05 |
| GRB10 | 2933.1657 | 1.005466866 | 0.30368033 | 3.31093842 | 0.00092984 | 0.00856537 |
| COBL | 3664.95625 | 0.471324884 | 0.31292753 | 1.50617905 | 0.13202122 | 0.29538611 |
| SEC61G | 245.270861 | -1.219620212 | 0.4423798 | -2.7569528 | 0.00583428 | 0.03397215 |
| EGFR | 3145.70396 | -0.590730437 | 0.28608455 | -2.0648806 | 0.0389343 | 0.13131704 |
| LANCL2 | 566.918749 | 0.230828534 | 0.33402291 | 0.691056 | 0.48953035 | 0.67261352 |
| VOPP1 | 850.349426 | -1.793664116 | 0.44367161 | -4.0427742 | 5.28E-05 | 0.00095018 |
| NIPSNAP2 | 807.315237 | 0.460776498 | 0.20643729 | 2.23204101 | 0.02561225 | 0.09806345 |
| PSPH | 308.435915 | 0.874791099 | 0.36653483 | 2.38665204 | 0.01700258 | 0.07365812 |
| CCT6A | 701.441197 | -0.799657548 | 0.43402151 | -1.8424376 | 0.06541117 | 0.18562958 |
| SUMF2 | 549.507811 | -0.012761785 | 0.19711861 | -0.0647417 | 0.94837969 | 0.973595 |
| PHKG1 | 127.422214 | -0.266029865 | 0.28587349 | -0.930586 | 0.35206776 | 0.55108727 |
| CHCHD2 | 720.933697 | -1.021823226 | 0.3945581 | -2.5897915 | 0.00960341 | 0.04893453 |
| ZNF727 | 106.372838 | -0.131210271 | 0.51157025 | -0.2564853 | 0.7975761 | 0.889679 |
| ZNF736 | 618.106844 | 0.044312567 | 0.2599976 | 0.17043452 | 0.86466842 | 0.92922135 |
| ZNF680 | 450.76234 | 0.656400423 | 0.3414263 | 1.9225245 | 0.05453979 | 0.16433874 |
| ZNF107 | 146.981446 | 0.164598883 | 0.5410922 | 0.30419748 | 0.76097745 | 0.8665094 |
| ZNF138 | 331.838975 | 0.334490837 | 0.20763379 | 1.61096536 | 0.10718727 | 0.25708066 |
| ZNF273 | 43.817072 | -1.057972615 | 0.45594 | -2.3204207 | 0.02031813 | 0.08343338 |
| ZNF117 | 174.474805 | -0.185104068 | 0.40283757 | -0.4595005 | 0.64587479 | 0.79225766 |
| ZNF92 | 74.6486455 | -0.107844264 | 0.24990519 | -0.4315407 | 0.66607525 | 0.80617161 |
| VKORC1L1 | 904.654813 | -0.421653621 | 0.42733117 | -0.9867139 | 0.3237829 | 0.52429922 |
| GUSB | 233.509394 | 0.398135747 | 0.20954996 | 1.89995624 | 0.05743886 | 0.17000961 |
| ASL | 343.645315 | 0.45877156 | 0.34347468 | 1.33567795 | 0.18165456 | 0.36381342 |
| AC068533.4 | 301.06544 | 0.149807798 | 0.47834664 | 0.31317832 | 0.75414519 | 0.86214079 |
| CRCP | 136.940631 | -0.286735714 | 0.24260682 | -1.1818947 | 0.23724749 | 0.43180871 |
| AC068533.3 | 142.495126 | -0.275497006 | 0.38687074 | -0.7121164 | 0.47639268 | 0.6620617 |
| TPST1 | 1303.19276 | -0.756978963 | 0.3206216 | -2.3609731 | 0.01822705 | 0.07732535 |
| AC008267.5 | 35.7320784 | 0.245836619 | 0.51755048 | 0.47500027 | 0.63478678 | 0.78468826 |
| AC027644.4 | 901.86036 | -0.062434147 | 0.45043292 | -0.1386092 | 0.88975897 | 0.94300829 |
| AC027644.3 | 120.030835 | -0.051612748 | 0.3498831 | -0.1475143 | 0.88272612 | 0.93903973 |
| AC073335.2 | 48.2003172 | 1.322661177 | 0.4471461 | 2.95800677 | 0.00309635 | 0.02111763 |
| TMEM248 | 597.434186 | -0.252173064 | 0.24029066 | -1.0494501 | 0.293971 | 0.49394984 |

| | | | | | | |
|---|---|---|---|---|---|---|
| SBDS | 318.034546 | -0.787189266 | 0.26805982 | -2.9366179 | 0.00331813 | 0.02225125 |
| TYW1 | 1734.65076 | 0.455157539 | 0.18511879 | 2.4587322 | 0.01394286 | 0.06365528 |
| AC006480.2 | 184.43194 | 0.112253888 | 0.34915185 | 0.32150449 | 0.74782811 | 0.85769676 |
| LINC01372 | 121.547574 | 0.940597553 | 0.39608743 | 2.37472207 | 0.01756217 | 0.07530608 |
| AUTS2 | 2824.84468 | 0.780703059 | 0.26370106 | 2.96056098 | 0.00307079 | 0.02105621 |
| GALNT17 | 55.3507759 | 0.594470348 | 0.47893644 | 1.24123016 | 0.21452074 | 0.4048514 |
| CALN1 | 113.858787 | 0.231661175 | 0.4113867 | 0.56312267 | 0.57335136 | 0.73865654 |
| TYW1B | 1028.87491 | 0.867005371 | 0.17777603 | 4.87695313 | 1.08E-06 | 4.10E-05 |
| POM121 | 1373.27661 | 0.315409871 | 0.20765194 | 1.51893533 | 0.12877877 | 0.29051525 |
| NSUN5 | 214.976059 | 0.316634009 | 0.28128422 | 1.12567283 | 0.26030403 | 0.45866286 |
| BAZ1B | 931.572787 | 0.256052714 | 0.28533718 | 0.89736891 | 0.3695221 | 0.56695645 |
| BCL7B | 219.274287 | -0.313378619 | 0.33479512 | -0.936031 | 0.34925722 | 0.54887281 |
| TBL2 | 112.39694 | -0.117807112 | 0.38406577 | -0.3067368 | 0.7590437 | 0.86492743 |
| MLXIPL | 2781.17768 | 1.517276765 | 0.58292554 | 2.60286547 | 0.00924482 | 0.04760285 |
| VPS37D | 70.5123591 | 0.583712355 | 0.32891066 | 1.77468361 | 0.07595014 | 0.20503394 |
| DNAJC30 | 75.787939 | -0.197827175 | 0.30751338 | -0.6433124 | 0.5200214 | 0.69605855 |
| BUD23 | 332.845449 | -0.107789922 | 0.26197228 | -0.4114555 | 0.6807386 | 0.81442647 |
| STX1A | 48.6871025 | -0.382049886 | 0.42825032 | -0.8921182 | 0.3723296 | 0.56984389 |
| ABHD11 | 307.778878 | -0.716617835 | 0.45122834 | -1.588149 | 0.11225265 | 0.26532973 |
| CLDN3 | 593.552215 | -2.036622173 | 0.49732192 | -4.0951788 | 4.22E-05 | 0.00078884 |
| CLDN4 | 905.837712 | -2.252946689 | 0.6074537 | -3.7088369 | 0.00020821 | 0.00276281 |
| METTL27 | 82.9617887 | -0.87343485 | 0.59680833 | -1.4635098 | 0.14332793 | 0.3110032 |
| LIMK1 | 101.160302 | -1.102812411 | 0.47386984 | -2.3272475 | 0.0199521 | 0.0823497 |
| EIF4H | 479.21217 | -0.603309544 | 0.18650275 | -3.2348561 | 0.00121704 | 0.01048521 |
| LAT2 | 33.6221214 | -0.027753956 | 0.36257569 | -0.0765467 | 0.9389842 | 0.96728216 |
| RFC2 | 94.341876 | -0.507652289 | 0.36148674 | -1.4043455 | 0.16021599 | 0.33594623 |
| CLIP2 | 217.897517 | -0.819380292 | 0.55759672 | -1.4694855 | 0.14170115 | 0.30869321 |
| GTF2IRD1 | 1777.18485 | -0.667809823 | 0.36166894 | -1.8464672 | 0.0648244 | 0.18457471 |
| GTF2I | 4399.56202 | 0.259123536 | 0.16122915 | 1.6071755 | 0.10801587 | 0.25838981 |
| GTF2IRD2 | 258.054678 | 0.071758578 | 0.43736008 | 0.16407208 | 0.8696744 | 0.93268344 |
| CASTOR2 | 510.29023 | -0.928549622 | 0.39070457 | -2.3766029 | 0.01747289 | 0.07506735 |
| RCC1L | 389.777654 | -0.098900547 | 0.29616325 | -0.3339393 | 0.73842536 | 0.85155616 |
| GTF2IRD2B | 469.859541 | 0.369574701 | 0.38858916 | 0.95106796 | 0.34156988 | 0.54220015 |
| AC211486.5 | 108.253414 | -1.875153353 | 0.53021341 | -3.5366012 | 0.00040531 | 0.00462967 |
| TRIM73 | 369.041804 | 0.278652815 | 0.41588423 | 0.67002496 | 0.50284188 | 0.68363952 |
| POM121C | 1094.99161 | 0.157593629 | 0.1857321 | 0.84849967 | 0.39615976 | 0.59081295 |
| SPDYE5 | 270.298688 | -0.356529203 | 0.39794675 | -0.8959219 | 0.37029448 | 0.56780697 |
| HIP1 | 802.47712 | 0.195160806 | 0.31941071 | 0.6110027 | 0.54119779 | 0.71231293 |
| RHBDD2 | 228.706613 | 0.220589114 | 0.26066732 | 0.84624766 | 0.3974146 | 0.59186337 |
| POR | 1914.66049 | -0.489164584 | 0.48045054 | -1.0181372 | 0.30861274 | 0.5088602 |
| TMEM120A | 240.271279 | 0.640693795 | 0.22628219 | 2.83139293 | 0.00463457 | 0.02869292 |
| STYXL1 | 551.447719 | 0.825507566 | 0.23707872 | 3.48199775 | 0.00049769 | 0.00533788 |
| MDH2 | 347.317538 | -0.497759822 | 0.24683398 | -2.0165774 | 0.04373963 | 0.14170153 |
| HSPB1 | 1389.31608 | -0.713478403 | 0.34269561 | -2.0819596 | 0.03734616 | 0.12759268 |
| YWHAG | 786.427953 | -0.958384071 | 0.32810791 | -2.9209417 | 0.00348975 | 0.02318798 |
| ZP3 | 69.0045774 | 0.313692601 | 0.44794721 | 0.70028922 | 0.4837467 | 0.66774583 |
| DTX2 | 379.404227 | 0.116233488 | 0.28859233 | 0.40276014 | 0.68712469 | 0.81885047 |
| SPDYE16 | 81.7873209 | 0.955780501 | 0.43479973 | 2.19820857 | 0.02793425 | 0.10400253 |
| POMZP3 | 216.946188 | -0.131365475 | 0.31843112 | -0.4125397 | 0.67994389 | 0.81435096 |
| CCDC146 | 910.197888 | -0.156427014 | 0.37585768 | -0.4161868 | 0.6772733 | 0.81315429 |
| FGL2 | 89.0120628 | 2.239624411 | 0.66782199 | 3.35362482 | 0.0007976 | 0.00765064 |
| GSAP | 1144.51681 | 0.560807409 | 0.48503412 | 1.1562226 | 0.24759012 | 0.44373924 |
| PTPN12 | 1971.96673 | -0.731460693 | 0.23117466 | -3.1641041 | 0.00155561 | 0.01271483 |
| APTR | 433.497983 | -0.064609073 | 0.36629481 | -0.1763854 | 0.85999114 | 0.92628466 |
| RSBN1L | 893.367043 | 0.280986516 | 0.14141241 | 1.98700042 | 0.04692235 | 0.14921358 |
| TMEM60 | 102.8382 | 0.101109309 | 0.27496019 | 0.36772345 | 0.71307945 | 0.83590011 |
| PHTF2 | 538.948862 | 0.417220057 | 0.14170298 | 2.94432806 | 0.00323657 | 0.021854 |
| MAGI2 | 8652.56736 | -0.129418862 | 0.26475947 | -0.4888167 | 0.62497144 | 0.77771949 |
| GNAI1 | 1048.40545 | -1.289925235 | 0.44319732 | -2.9104987 | 0.00360852 | 0.02374558 |
| SEMA3C | 305.609328 | -3.337244227 | 0.85758586 | -3.8914404 | 9.97E-05 | 0.00154813 |
| CACNA2D1 | 3936.78431 | 0.75106627 | 0.2302536 | 3.26190897 | 0.00110665 | 0.00977247 |
| PCLO | 4551.8173 | 1.268674354 | 0.53126537 | 2.38802382 | 0.01693924 | 0.07349991 |
| SEMA3E | 82.1260536 | -1.639904502 | 0.89916951 | -1.8237991 | 0.06818246 | 0.19072999 |
| SEMA3A | 44.719959 | -0.275500956 | 0.41113977 | -0.6700907 | 0.50279994 | 0.68363952 |
| AC003984.1 | 655.657675 | 3.229475654 | 0.70548796 | 4.57764815 | 4.70E-06 | 0.00013627 |
| SEMA3D | 291.389609 | 1.319080601 | 0.59363062 | 2.2220562 | 0.02627951 | 0.09983736 |
| KIAA1324L | 150.944279 | 0.39790035 | 0.56192951 | 0.70809656 | 0.4788853 | 0.66431341 |
| DMTF1 | 1430.56262 | 0.049834236 | 0.2724638 | 0.18290223 | 0.85487473 | 0.92324046 |
| TMEM243 | 394.612085 | -1.524624388 | 0.30880206 | -4.9372222 | 7.92E-07 | 3.18E-05 |
| TP53TG1 | 51.8418413 | 0.158124428 | 0.46157664 | 0.34257459 | 0.73191852 | 0.84761805 |
| CROT | 229.761578 | 0.668553854 | 0.31306676 | 2.13549934 | 0.03272024 | 0.11633281 |
| ABCB4 | 52.0631391 | 0.985995317 | 0.51837254 | 1.90209789 | 0.05715836 | 0.16949995 |
| ABCB1 | 1626.86938 | -0.279355039 | 0.46200469 | -0.6046584 | 0.54540596 | 0.71573763 |
| RUNDC3B | 793.355153 | 0.849181958 | 0.43312146 | 1.96060931 | 0.04992462 | 0.15526052 |
| SLC25A40 | 286.423563 | 0.106714524 | 0.24859032 | 0.42927868 | 0.66772043 | 0.80690041 |
| DBF4 | 156.398678 | -0.532743242 | 0.30596664 | -1.7411808 | 0.08165189 | 0.21513103 |
| ADAM22 | 510.341428 | 0.571408026 | 0.38723373 | 1.47561534 | 0.14004718 | 0.30674764 |
| SRI | 234.10625 | -0.798661022 | 0.29786163 | -2.6813155 | 0.00733333 | 0.04035247 |
| AC003991.1 | 52.4921855 | 0.133937575 | 0.37055216 | 0.36145404 | 0.71776005 | 0.83887594 |
| ZNF804B | 5293.73859 | 3.2183434 | 0.70055035 | 4.59402157 | 4.35E-06 | 0.00012789 |
| STEAP1 | 98.4174148 | 0.908431599 | 0.42726367 | 2.12616158 | 0.03348981 | 0.11834101 |
| STEAP2 | 380.667775 | 0.660941723 | 0.40242062 | 1.64241515 | 0.100504 | 0.24661803 |
| CFAP69 | 1333.60947 | 1.001482551 | 0.35901011 | 2.78956645 | 0.00527787 | 0.03152282 |
| AC002064.1 | 85.1063307 | 1.415312681 | 0.46769001 | 3.02617685 | 0.00247667 | 0.01804772 |
| GTPBP10 | 547.590617 | 0.105604749 | 0.22315855 | 0.47322744 | 0.63605092 | 0.78572008 |
| CLDN12 | 843.893863 | -0.49273482 | 0.21180395 | -2.3263722 | 0.0199987 | 0.0824493 |
| CDK14 | 5562.65457 | 1.166454765 | 0.31540534 | 3.69827208 | 0.00021707 | 0.00285274 |
| FZD1 | 179.493097 | -0.06268512 | 0.44987182 | -0.13934 | 0.88918152 | 0.94267382 |
| AC079760.2 | 207.167759 | 1.713228828 | 0.49035687 | 3.49384076 | 0.00047612 | 0.00517956 |
| AC079760.1 | 126.382826 | 0.286348726 | 0.53061107 | 0.53965841 | 0.58943263 | 0.75135804 |
| MTERF1 | 745.178871 | -0.009213624 | 0.43521297 | -0.0211704 | 0.98310974 | 0.99169805 |
| AKAP9 | 3155.59121 | -0.003226266 | 0.19775239 | -0.0163147 | 0.98698335 | 0.99360989 |
| KRIT1 | 1615.69551 | 0.520371347 | 0.19320047 | 2.69342697 | 0.00707216 | 0.03927802 |
| ANKIB1 | 5158.00926 | 0.789812507 | 0.18735489 | 4.21559586 | 2.49E-05 | 0.00052085 |
| GATAD1 | 404.913799 | -0.089013822 | 0.38473744 | -0.2313625 | 0.81703318 | 0.90111634 |
| AC007566.1 | 364.194149 | 0.397877372 | 0.49765814 | 0.79949938 | 0.42400091 | 0.61709284 |
| PEX1 | 454.471466 | 0.372857177 | 0.23330365 | 1.59816264 | 0.11000679 | 0.2616455 |
| RBM48 | 156.428647 | -0.245326382 | 0.30588362 | -0.8020252 | 0.42253836 | 0.61619214 |
| FAM133B | 713.552951 | 0.052642199 | 0.1270844 | 0.41423021 | 0.67870552 | 0.81376982 |
| CDK6 | 1243.38239 | -0.935339929 | 0.46960144 | -1.991774 | 0.04639586 | 0.14799815 |
| SAMD9 | 67.0945228 | -1.055823452 | 0.8255608 | -1.2789167 | 0.20092641 | 0.38801046 |
| SAMD9L | 202.516088 | -0.809501862 | 0.99902898 | -0.8102887 | 0.41777428 | 0.61081943 |
| VPS50 | 1454.18793 | 0.256235047 | 0.15095892 | 1.69738262 | 0.08962435 | 0.22838463 |
| GNGT1 | 1066.2612 | 0.253327572 | 0.41485031 | 0.61064814 | 0.54143254 | 0.71250152 |
| GNG11 | 67.1420017 | 0.305159796 | 0.44189096 | 0.69057715 | 0.48983131 | 0.67270543 |
| BET1 | 315.249092 | -1.158688998 | 0.30294466 | -3.8247547 | 0.0001309 | 0.00191473 |
| CASD1 | 556.410239 | 0.250179596 | 0.28180914 | 0.88776254 | 0.3746685 | 0.5715597 |
| SGCE | 551.333407 | 0.104498752 | 0.35505084 | 0.29432053 | 0.76851298 | 0.87097463 |
| PEG10 | 39.8505926 | -0.221353332 | 0.39827856 | -0.5557752 | 0.57836456 | 0.74211119 |
| PPP1R9A | 3570.89226 | -0.150860804 | 0.21416721 | -0.7044066 | 0.48117957 | 0.66594592 |
| AC002429.2 | 270.578046 | -0.687172045 | 0.47024421 | -1.4613089 | 0.14393069 | 0.3116187 |
| PON2 | 781.978651 | -0.837435416 | 0.27843598 | -3.0076408 | 0.00263284 | 0.01884831 |
| PDK4 | 2626.9637 | -0.02495506 | 1.32494205 | -0.0188348 | 0.98497287 | 0.99257632 |

| | | | | | | |
|---|---|---|---|---|---|---|
| AC002451.1 | 46.2887132 | 1.045357327 | 0.94846021 | 1.10216255 | 0.27039101 | 0.46867091 |
| DYNC1I1 | 415.572275 | -2.365220273 | 0.82551825 | -2.8651339 | 0.00416833 | 0.02643929 |
| SLC25A13 | 2724.94701 | 0.121489195 | 0.23089409 | 0.52616849 | 0.59877115 | 0.75878837 |
| SEM1 | 704.891533 | -0.106866876 | 0.19728642 | -0.5416839 | 0.58803629 | 0.75041555 |
| SDHAF3 | 318.207674 | 0.407343035 | 0.21814884 | 1.86727117 | 0.06186373 | 0.17848161 |
| ASNS | 110.574528 | -1.064013741 | 0.45427857 | -2.3422054 | 0.01917016 | 0.07996279 |
| LMTK2 | 2018.42922 | -0.498568644 | 0.19985974 | -2.4945926 | 0.01261018 | 0.05935705 |
| TECPR1 | 232.806675 | 0.419584176 | 0.26073187 | 1.60925542 | 0.1075605 | 0.25757529 |
| BRI3 | 1507.05083 | -0.434531965 | 0.31144251 | -1.3952237 | 0.16294839 | 0.33942014 |
| BAIAP2L1 | 1148.7863 | -0.890190333 | 0.46516625 | -1.9137036 | 0.05565804 | 0.16645501 |
| TRRAP | 1458.31354 | 0.188181185 | 0.14457722 | 1.30159639 | 0.19305439 | 0.37781219 |
| SMURF1 | 2181.80986 | -0.636336451 | 0.2771683 | -2.2958486 | 0.02168454 | 0.08757513 |
| AC004834.1 | 63.7311296 | 0.870690431 | 0.46515635 | 1.87182318 | 0.06123107 | 0.17732818 |
| ARPC1B | 182.311691 | -1.201873859 | 0.36095508 | -3.3297048 | 0.00086938 | 0.00814532 |
| PDAP1 | 242.849615 | -0.752153983 | 0.31601262 | -2.3801391 | 0.01730611 | 0.07452554 |
| BUD31 | 158.908791 | -0.756523222 | 0.34897273 | -2.1678577 | 0.03016951 | 0.10959616 |
| PTCD1 | 40.4418984 | -0.122182009 | 0.39283924 | -0.3110229 | 0.75578319 | 0.86300568 |
| CPSF4 | 196.587394 | 0.28969154 | 0.22304153 | 1.2988233 | 0.19400458 | 0.37905874 |
| ZNF789 | 432.534202 | 0.699957027 | 0.23716604 | 2.95133754 | 0.00316401 | 0.02149609 |
| ZNF394 | 224.426615 | -0.111931978 | 0.26235561 | -0.4266422 | 0.66963994 | 0.80813125 |
| ZKSCAN5 | 201.899091 | 0.485044873 | 0.28878465 | 1.67960754 | 0.0930337 | 0.23469615 |
| FAM200A | 146.295161 | 0.205581603 | 0.2670732 | 0.7697575 | 0.44144375 | 0.63333363 |
| ZSCAN25 | 161.684068 | 0.40452967 | 0.2660062 | 1.5207528 | 0.12832188 | 0.29000957 |
| CYP3A5 | 1055.44088 | 0.363753779 | 0.61804262 | 0.58855776 | 0.55615797 | 0.72533372 |
| TRIM4 | 523.73803 | 0.02590015 | 0.2866987 | 0.09033926 | 0.92801762 | 0.96208348 |
| AZGP1 | 725.934277 | 0.808273027 | 0.51425865 | 1.57172472 | 0.11601441 | 0.27141312 |
| ZKSCAN1 | 2562.20043 | 0.295341685 | 0.31356161 | 0.94189365 | 0.34624709 | 0.54664509 |
| ZSCAN21 | 254.057469 | 1.028686897 | 0.21990304 | 4.67791115 | 2.90E-06 | 9.28E-05 |
| ZNF3 | 304.93961 | 0.260970025 | 0.30718987 | 0.8495398 | 0.39558099 | 0.59027092 |
| COPS6 | 273.354249 | -0.37802808 | 0.19095295 | -1.9796923 | 0.04773812 | 0.15084057 |
| MCM7 | 137.011861 | 0.071074689 | 0.41973372 | 0.1693328 | 0.86553487 | 0.92955312 |
| AP4M1 | 151.886567 | -0.120626231 | 0.22024218 | -0.5476981 | 0.58389919 | 0.74666538 |
| TAF6 | 141.499356 | -0.545960776 | 0.29746441 | -1.8353818 | 0.06644913 | 0.18748832 |
| AC073842.2 | 47.8561499 | 1.084503156 | 0.4203804 | 2.57981378 | 0.00988536 | 0.05003642 |
| CNPY4 | 37.1425663 | -0.074474826 | 0.36043214 | -0.2066265 | 0.83630158 | 0.91369247 |
| LAMTOR4 | 458.27774 | -0.065411153 | 0.26187363 | -0.2497814 | 0.80275643 | 0.89272007 |
| C7orf43 | 77.7779054 | 0.336302959 | 0.39347628 | 0.85469692 | 0.39271896 | 0.58759947 |
| STAG3 | 189.00933 | -0.645917205 | 0.41773936 | -1.5462206 | 0.1220513 | 0.28032125 |
| CASTOR3 | 1069.46664 | -0.830083014 | 0.26621995 | -3.1180346 | 0.00182061 | 0.01421946 |
| SPDYE3 | 399.407634 | -0.433883128 | 0.48826889 | -0.8886151 | 0.37420996 | 0.57131832 |
| PILRB | 1278.04377 | -0.058673926 | 0.3856204 | -0.1521546 | 0.87906498 | 0.93706833 |
| PILRA | 242.4405 | 0.230913608 | 0.33465419 | 0.69000663 | 0.49019002 | 0.67304764 |
| ZCWPW1 | 85.4998837 | 0.619294756 | 0.52384201 | 1.18221668 | 0.23711974 | 0.43169083 |
| MEPCE | 109.184042 | -0.547394891 | 0.50352938 | -1.0871161 | 0.27698551 | 0.47487684 |
| PPP1R35 | 230.781464 | -0.796946499 | 0.27053497 | -2.945817 | 0.00322103 | 0.02177054 |
| TSC22D4 | 181.363558 | -0.426809204 | 0.24662793 | -1.7305793 | 0.08352682 | 0.21827952 |
| AGFG2 | 85.6139831 | 0.603817931 | 0.40690861 | 1.48391534 | 0.13783138 | 0.3035435 |
| LRCH4 | 108.931313 | 0.175564939 | 0.2415053 | 0.72696101 | 0.46724984 | 0.65466556 |
| MOSPD3 | 52.5194856 | -0.189063366 | 0.3552763 | -0.5321587 | 0.5946161 | 0.75522636 |
| GNB2 | 421.808337 | -1.073623254 | 0.36530087 | -2.939011 | 0.00329261 | 0.02215604 |
| GIGYF1 | 562.533613 | 0.336820687 | 0.40557157 | 0.83048396 | 0.40626521 | 0.60022783 |
| POP7 | 70.2966361 | -1.494273421 | 0.41151138 | -3.6311837 | 0.00028212 | 0.00349329 |
| EPHB4 | 160.17148 | 0.737607757 | 0.31642815 | 2.33104342 | 0.01975107 | 0.08176621 |
| SLC12A9 | 363.283594 | 0.868095109 | 0.30344979 | 2.86075371 | 0.00422635 | 0.02670844 |
| TRIP6 | 326.606838 | 0.300396013 | 0.38006956 | 0.79037114 | 0.42931105 | 0.62217874 |
| SRRT | 294.187725 | 0.008833854 | 0.41853914 | 0.0211064 | 0.98316078 | 0.99169805 |
| TRIM56 | 861.488403 | -1.417696154 | 0.40133062 | -3.5324894 | 0.00041167 | 0.00467887 |
| AP1S1 | 264.708958 | -1.406991942 | 0.42746327 | -3.291492 | 0.00099657 | 0.00901653 |
| MOGAT3 | 99.3705725 | 0.075762977 | 0.94049843 | 0.0805562 | 0.9357949 | 0.96587767 |
| PLOD3 | 408.148577 | -1.0965664 | 0.28207102 | -3.8875542 | 0.00010126 | 0.00156832 |
| ZNHIT1 | 369.479796 | -0.204526477 | 0.26840578 | -0.7620047 | 0.44605718 | 0.6372179 |
| CLDN15 | 92.5625567 | -0.097420365 | 0.39019614 | -0.2496702 | 0.80284237 | 0.89272007 |
| FIS1 | 517.402877 | -0.059789056 | 0.28948515 | -0.2065358 | 0.83637238 | 0.91369699 |
| IFT22 | 132.667775 | -0.561669901 | 0.31296929 | -1.7946486 | 0.0727097 | 0.19904772 |
| COL26A1 | 73.5508648 | 0.513264676 | 0.33805018 | 1.51830912 | 0.12893649 | 0.29068151 |
| CUX1 | 11541.5162 | 0.782892735 | 0.20428153 | 3.83242062 | 0.00012689 | 0.00187219 |
| SH2B2 | 223.146393 | -0.14730386 | 0.37821564 | -0.3894706 | 0.69692808 | 0.82544394 |
| PRKRIP1 | 2504.98148 | 0.073518155 | 0.20946527 | 0.35098017 | 0.72560322 | 0.84394089 |
| ORAI2 | 344.296317 | -1.642258012 | 0.70760969 | -2.3208529 | 0.02029478 | 0.08337291 |
| ALKBH4 | 108.230409 | 0.121865568 | 0.28797683 | 0.42317838 | 0.67216512 | 0.80998957 |
| LRWD1 | 48.9396034 | -0.470929685 | 0.52135281 | -0.9032841 | 0.36637515 | 0.56391756 |
| POLR2J | 137.478961 | -0.342787316 | 0.2833937 | -1.2095799 | 0.22644014 | 0.4189815 |
| AC093668.2 | 116.847292 | 0.920572115 | 0.48706817 | 1.89002725 | 0.05875432 | 0.17230325 |
| SPDYE2 | 127.126978 | -0.039661309 | 0.58317558 | -0.0680092 | 0.94577831 | 0.97196767 |
| AC105052.1 | 120.920078 | 0.599466296 | 0.46368677 | 1.29282596 | 0.19607126 | 0.38159148 |
| AC105052.3 | 367.461086 | 0.858200687 | 0.41579859 | 2.06398173 | 0.03901945 | 0.13144251 |
| FAM185A | 780.433236 | 0.64025775 | 0.23869969 | 2.68227302 | 0.00731238 | 0.04026948 |
| FBXL13 | 635.96666 | -0.201266866 | 0.27940689 | -0.7203361 | 0.47131809 | 0.65767659 |
| LRRC17 | 81.2796492 | 0.755519933 | 0.4719731 | 1.60076907 | 0.10942808 | 0.26077541 |
| ARMC10 | 461.485968 | -0.390955834 | 0.12334891 | -3.1695119 | 0.00152695 | 0.0125329 |
| NAPEPLD | 1492.60047 | 0.85218306 | 0.40641087 | 2.09685104 | 0.03600676 | 0.12406855 |
| PMPCB | 518.239818 | 0.421358534 | 0.24048631 | 1.75211026 | 0.07975485 | 0.21145278 |
| DNAJC2 | 482.786548 | -0.388147957 | 0.39984244 | -0.9707523 | 0.33167166 | 0.53141196 |
| PSMC2 | 344.08299 | -0.169729273 | 0.15759426 | -1.0770016 | 0.28147954 | 0.48029877 |
| SLC26A5 | 69.4285735 | -0.63187414 | 0.46615483 | -1.3555027 | 0.17525745 | 0.35536299 |
| RELN | 95.5504652 | -0.404971572 | 0.52058216 | -0.7779206 | 0.43661584 | 0.62857823 |
| ORC5 | 541.959571 | 0.143739153 | 0.18985652 | 0.7570936 | 0.44899379 | 0.63921457 |
| LHFPL3 | 872.840308 | 0.027717533 | 0.50267966 | 0.05513956 | 0.95602728 | 0.97756553 |
| LINC01004 | 1290.22713 | 0.628443055 | 0.2087459 | 3.01056473 | 0.00260762 | 0.01873307 |
| KMT2E-AS1 | 156.147085 | -0.001534447 | 0.43801677 | -0.0035032 | 0.99720488 | 0.99866204 |
| KMT2E | 2565.14907 | 0.283994022 | 0.21304813 | 1.33300408 | 0.18253046 | 0.36485995 |
| SRPK2 | 5705.19266 | 0.041396071 | 0.26461146 | 0.15644096 | 0.87568545 | 0.93525172 |
| PUS7 | 508.329619 | -0.855161984 | 0.44955004 | -1.902262 | 0.05713691 | 0.16949995 |
| RINT1 | 246.285909 | -0.047852865 | 0.28711656 | -0.166667 | 0.86763205 | 0.93103202 |
| EFCAB10 | 86.4319213 | -0.862140441 | 0.41102788 | -2.097523 | 0.0359473 | 0.12403257 |
| ATXN7L1 | 2975.71966 | 1.132493313 | 0.33119674 | 3.41939753 | 0.0006276 | 0.00636751 |
| CDHR3 | 3075.6182 | 1.365181782 | 0.46658464 | 2.92590385 | 0.00343457 | 0.02286433 |
| SYPL1 | 696.370337 | -0.140635068 | 0.25244355 | -0.5570951 | 0.57746244 | 0.74154889 |
| NAMPT | 4444.01825 | -3.87734151 | 0.53265904 | -7.2792184 | 3.36E-13 | 1.00E-10 |
| CCDC71L | 85.1162672 | -0.67455442 | 0.36691597 | -1.8384439 | 0.06599703 | 0.18656221 |
| HBP1 | 861.647395 | -0.717022766 | 0.33113492 | -2.1653493 | 0.03036094 | 0.11016286 |
| COG5 | 2892.9834 | -0.600860438 | 0.31730977 | -1.8936084 | 0.05827701 | 0.17132329 |
| DUS4L | 104.405131 | 0.075899163 | 0.23453784 | 0.32361159 | 0.74623211 | 0.85659522 |
| BCAP29 | 427.593321 | -0.480481909 | 0.20238896 | -2.3740519 | 0.01759408 | 0.07538668 |
| CBLL1 | 208.179111 | -0.076355232 | 0.26508081 | -0.2880451 | 0.77331221 | 0.87420952 |
| SLC26A3 | 3679.53769 | 0.738021207 | 0.68618525 | 1.07554222 | 0.28213204 | 0.48105285 |
| DLD | 536.675753 | 0.033989586 | 0.27714975 | 0.12263979 | 0.90239235 | 0.9486955 |
| LAMB1 | 3638.93801 | -0.767925653 | 0.53874127 | -1.4254071 | 0.15403955 | 0.32649143 |
| NRCAM | 2680.40787 | -3.145337792 | 0.53207896 | -5.9114117 | 3.39E-09 | 3.25E-07 |
| PNPLA8 | 1003.02233 | -0.708825554 | 0.24293832 | -2.9177181 | 0.00352603 | 0.02335973 |
| DNAJB9 | 292.317594 | -0.750792416 | 0.40581131 | -1.8501023 | 0.06429881 | 0.18346041 |
| IMMP2L | 11940.4183 | 0.438860162 | 0.32387262 | 1.35503938 | 0.17540501 | 0.35550443 |
| LRRN3 | 54.0480564 | -0.813513803 | 0.72709833 | -1.1188498 | 0.26320423 | 0.46128738 |

| | | | | | | |
|---|---|---|---|---|---|---|
| DOCK4 | 2884.41745 | -0.456204582 | 0.38073956 | -1.1982064 | 0.23083668 | 0.42378291 |
| ZNF277 | 1521.9484 | -0.292346466 | 0.20383751 | -1.4342133 | 0.15151141 | 0.32288044 |
| AC004112.1 | 66.6775081 | -0.086531935 | 0.35436091 | -0.2441915 | 0.80708248 | 0.89532855 |
| IFRD1 | 1533.17918 | -0.291701765 | 0.49849626 | -0.5851634 | 0.55843786 | 0.72696559 |
| LSMEM1 | 282.931222 | 0.251417862 | 0.51962065 | 0.48384886 | 0.62849313 | 0.78055893 |
| TMEM168 | 418.01898 | 0.143315705 | 0.240915 | 0.59488079 | 0.55192316 | 0.72159584 |
| BMT2 | 975.189799 | 0.049385271 | 0.31550599 | 0.15652721 | 0.87561748 | 0.93525172 |
| SMIM30 | 120.570167 | -0.463629986 | 0.3645335 | -1.2718447 | 0.20342832 | 0.39145524 |
| FOXP2 | 4464.00798 | -0.470635634 | 0.3820534 | -1.2318583 | 0.21800204 | 0.40911198 |
| MDFIC | 329.290796 | -0.668803385 | 0.50545954 | -1.3231591 | 0.18578248 | 0.36895399 |
| TFEC | 1055.47267 | 1.42129968 | 0.72713317 | 1.95466215 | 0.05062296 | 0.15670481 |
| TES | 513.501957 | -2.046944694 | 0.37387577 | -5.4749327 | 4.38E-08 | 2.80E-06 |
| AC002066.1 | 74.2605016 | -0.020737478 | 0.53036989 | -0.0391 | 0.96881064 | 0.98432195 |
| CAV2 | 105.567457 | -1.266637145 | 0.35657938 | -3.5521884 | 0.00038204 | 0.00442897 |
| CAV1 | 48.0319972 | -0.251701117 | 0.54203459 | -0.4643636 | 0.64238728 | 0.79027127 |
| AC006159.2 | 360.863614 | 2.234127716 | 0.91149187 | 2.45106708 | 0.01424334 | 0.06458269 |
| LINC01510 | 603.952263 | 2.175691565 | 0.865056 | 2.51508753 | 0.01190028 | 0.05691458 |
| MET | 3692.52168 | -1.573212709 | 0.35454336 | -4.4372929 | 9.11E-06 | 0.00023591 |
| CAPZA2 | 1458.19998 | -0.577480415 | 0.19423058 | -2.9731693 | 0.00294742 | 0.02043513 |
| ST7 | 4111.99111 | 0.937834219 | 0.24328907 | 3.85481437 | 0.00011582 | 0.00174836 |
| ST7-AS2 | 193.08772 | -0.126145807 | 0.61589078 | -0.2048185 | 0.83771396 | 0.91436098 |
| CFTR | 52.7107398 | -0.401116573 | 0.61977129 | -0.647201 | 0.51750189 | 0.69429219 |
| CTTNBP2 | 773.790397 | 0.499806749 | 0.39745399 | 1.25752102 | 0.20856503 | 0.39782924 |
| LSM8 | 382.866258 | -0.248199765 | 0.18120128 | -1.3697462 | 0.17076614 | 0.34969424 |
| ANKRD7 | 112.696076 | -0.418617601 | 0.59166107 | -0.7075294 | 0.47923755 | 0.66453277 |
| KCND2 | 118.16257 | 0.525551739 | 0.97753865 | 0.53762758 | 0.59083419 | 0.75259958 |
| TSPAN12 | 1042.56115 | -0.167349601 | 0.60969197 | -0.2744822 | 0.78371407 | 0.88133085 |
| ING3 | 234.99424 | 0.307047604 | 0.17815222 | 1.7235126 | 0.08479587 | 0.22025715 |
| FAM3C | 812.111461 | -1.362455503 | 0.37321193 | -3.6506216 | 0.00026161 | 0.00328675 |
| AASS | 1047.54857 | 1.448643172 | 0.59925101 | 2.417423 | 0.01563084 | 0.06931476 |
| CADPS2 | 2217.45356 | -0.111618229 | 0.4053018 | -0.2753953 | 0.78301252 | 0.8809896 |
| AC004594.1 | 100.951214 | -0.89167104 | 0.65286982 | -1.3657716 | 0.17201067 | 0.35134835 |
| AC073311.1 | 51.1603469 | 1.6310525 | 0.82960559 | 1.96605776 | 0.04929194 | 0.15396641 |
| SLC13A1 | 5815.62769 | 1.615782815 | 0.84105448 | 1.92113929 | 0.05471415 | 0.16461081 |
| IQUB | 373.12703 | 0.858600432 | 0.47975383 | 1.78966874 | 0.07350718 | 0.2005117 |
| NDUFA5 | 571.712069 | -0.026456001 | 0.30803195 | -0.0858872 | 0.93155609 | 0.96354054 |
| ASB15 | 74.0847587 | 3.393956483 | 0.84144099 | 4.03350504 | 5.50E-05 | 0.0009782 |
| WASL | 1501.64197 | 0.259203907 | 0.18397889 | 1.40887853 | 0.15887109 | 0.33373885 |
| AC006333.2 | 37.6355162 | -0.483558797 | 0.42456623 | -1.1389478 | 0.25472491 | 0.45191488 |
| POT1 | 476.571725 | 0.091356985 | 0.27208011 | 0.33577238 | 0.73704252 | 0.85110831 |
| POT1-AS1 | 133.860666 | 0.259316973 | 0.341828 | 0.75861829 | 0.44808093 | 0.63857822 |
| AC073486.1 | 430.906438 | 0.626481025 | 0.73495003 | 0.85241309 | 0.39398486 | 0.58879979 |
| GRM8 | 188.737874 | -1.819955412 | 0.81944997 | -2.2209476 | 0.02635451 | 0.10006685 |
| ZNF800 | 796.246862 | -0.502549342 | 0.23504538 | -2.1380949 | 0.03250904 | 0.11577069 |
| GCC1 | 50.9148789 | -0.41253458 | 0.33816467 | -1.2199222 | 0.22249438 | 0.41472082 |
| ARF5 | 132.227412 | -0.796541286 | 0.47708871 | -1.6695874 | 0.09500103 | 0.2376673 |
| SND1 | 3562.43786 | -0.789057847 | 0.34220824 | -2.3057827 | 0.02112278 | 0.08593942 |
| LRRC4 | 46.0303825 | -0.608436697 | 0.41759036 | -1.457018 | 0.1451114 | 0.31338301 |
| RBM28 | 420.707695 | -0.127046978 | 0.27737832 | -0.4580278 | 0.64693248 | 0.79280318 |
| IMPDH1 | 45.9959833 | -1.424401176 | 0.44284604 | -3.2164704 | 0.00129778 | 0.01098744 |
| HILPDA | 844.249631 | -5.233304408 | 0.81661791 | -6.4085105 | 1.47E-10 | 2.21E-08 |
| METTL2B | 379.6742 | -0.266571347 | 0.28522044 | -0.9346152 | 0.34998666 | 0.5493439 |
| AC090114.2 | 57.3439425 | -0.309784879 | 0.49679572 | -0.6235659 | 0.5329127 | 0.70617115 |
| FAM71F2 | 69.674642 | 0.330236994 | 0.50258647 | 0.65707497 | 0.51113271 | 0.69018777 |
| CALU | 686.851206 | -0.89356985 | 0.37402459 | -2.3890671 | 0.01689122 | 0.07336072 |
| KCP | 585.905955 | 0.007467487 | 0.41221775 | 0.01811539 | 0.9855468 | 0.99298265 |
| ATP6V1F | 552.356732 | -1.627497689 | 0.38914655 | -4.1822231 | 2.89E-05 | 0.00058513 |
| IRF5 | 146.166998 | 0.381821709 | 0.33894145 | 1.1265123 | 0.25994874 | 0.45823541 |
| TNPO3 | 1676.39366 | -0.338568597 | 0.22340966 | -1.5154609 | 0.12965573 | 0.2918961 |
| TSPAN33 | 488.29114 | 0.805412295 | 0.43110652 | 1.86824427 | 0.06172803 | 0.17827773 |
| SMO | 261.553779 | 1.507392654 | 0.36505453 | 4.12922601 | 3.64E-05 | 0.00070171 |
| AHCYL2 | 3466.4135 | 0.416892204 | 0.25071658 | 1.66280267 | 0.09635194 | 0.23964727 |
| STRIP2 | 424.310617 | 1.835509017 | 0.66449152 | 2.76227607 | 0.00573999 | 0.03352283 |
| AC078846.1 | 130.103419 | 0.123149681 | 0.26798716 | 0.45953576 | 0.64584948 | 0.79225766 |
| NRF1 | 1681.40464 | 0.358875212 | 0.17606434 | 2.03831859 | 0.04151808 | 0.13686588 |
| UBE2H | 5118.15321 | -0.423871266 | 0.41106694 | -1.031149 | 0.30247095 | 0.50272455 |
| AC073320.1 | 59.9810906 | -0.383239868 | 0.40115986 | -0.9553296 | 0.33941107 | 0.5398361 |
| ZC3HC1 | 680.08998 | 0.270255535 | 0.24867429 | 1.08678517 | 0.27713176 | 0.47500877 |
| KLHDC10 | 2120.20355 | 0.228953951 | 0.16773404 | 1.36498207 | 0.17225869 | 0.35167607 |
| TMEM209 | 198.349966 | 0.284018542 | 0.25531186 | 1.11243773 | 0.26594999 | 0.46378836 |
| AC087071.2 | 41.5121099 | -0.676532234 | 0.49569183 | -1.3648242 | 0.1723083 | 0.35167607 |
| CEP41 | 55.409526 | 0.019761711 | 0.43077043 | 0.04587527 | 0.96340966 | 0.98087026 |
| MEST | 92.8852807 | 0.037335225 | 0.44685742 | 0.08355064 | 0.93341371 | 0.96466122 |
| COPG2 | 244.358506 | 0.007873119 | 0.27300515 | 0.02883872 | 0.97699322 | 0.9886052 |
| AC016831.7 | 11482.6546 | -1.069801759 | 0.34155104 | -3.1321871 | 0.00173509 | 0.01373148 |
| LINC00513 | 2101.55629 | -1.687222238 | 0.54263331 | -3.109323 | 0.00187517 | 0.01458268 |
| AC058791.1 | 205.005773 | -1.831769592 | 0.56546042 | -3.2394302 | 0.00119769 | 0.01038375 |
| MKLN1 | 7210.6206 | -0.865438824 | 0.29794727 | -2.9046711 | 0.00367639 | 0.02403067 |
| PODXL | 115.702366 | -0.224097538 | 0.41701934 | -0.5373792 | 0.59100568 | 0.75259958 |
| PLXNA4 | 77.6855821 | 0.487093797 | 0.43925536 | 1.10890803 | 0.26746986 | 0.4655504 |
| CHCHD3 | 3787.64982 | -0.1635934 | 0.20118885 | -0.8131335 | 0.4161415 | 0.60947233 |
| EXOC4 | 8126.36774 | 0.279196786 | 0.25813186 | 1.08160529 | 0.27942795 | 0.47799091 |
| LRGUK | 436.210545 | 0.780452359 | 0.35151267 | 2.2202681 | 0.02640057 | 0.10018511 |
| SLC35B4 | 645.279796 | 0.865394324 | 0.34121111 | 2.53624307 | 0.01120489 | 0.05463731 |
| AKR1B1 | 709.82031 | -2.921676693 | 0.72333463 | -4.0391771 | 5.36E-05 | 0.00096109 |
| AKR1B10 | 669.531608 | -6.317294232 | 1.57040901 | -4.0227063 | 5.75E-05 | 0.00101494 |
| BPGM | 194.511254 | 0.065074191 | 0.36765804 | 0.17699651 | 0.85951113 | 0.92627715 |
| CALD1 | 2569.07895 | -0.20510433 | 0.23758938 | -0.8632723 | 0.38798777 | 0.58387159 |
| AGBL3 | 918.571848 | 0.673477738 | 0.46283799 | 1.45510472 | 0.14564027 | 0.31432707 |
| CYREN | 428.283131 | -0.219959621 | 0.26181465 | -0.8401349 | 0.40083275 | 0.5949339 |
| TMEM140 | 772.14507 | -1.549656451 | 0.44761026 | -3.4620665 | 0.00053604 | 0.00560455 |
| AC083862.2 | 73.1102495 | -1.58488266 | 0.67549307 | -2.3462604 | 0.01896285 | 0.07937082 |
| WDR91 | 766.242603 | -0.128420638 | 0.51319914 | -0.2502355 | 0.80240524 | 0.89272007 |
| AC009542.1 | 102.187234 | -1.166646092 | 0.68132669 | -1.7123153 | 0.08683859 | 0.22395042 |
| STRA8 | 97.032126 | -1.535694036 | 0.7562821 | -2.0305836 | 0.04229725 | 0.13870059 |
| CNOT4 | 2076.46275 | 0.14505754 | 0.15003225 | 0.96684242 | 0.33362281 | 0.53297994 |
| NUP205 | 400.53695 | 0.047142434 | 0.1818749 | 0.25920252 | 0.79547899 | 0.88843413 |
| STMP1 | 462.83879 | -0.488750467 | 0.22913922 | -2.1329848 | 0.03292598 | 0.11683053 |
| SLC13A4 | 71.8605986 | -1.471424478 | 0.58236663 | -2.5266291 | 0.0115163 | 0.05566079 |
| MTPN | 549.043425 | -0.547792228 | 0.25072426 | -2.1848393 | 0.02890063 | 0.10631802 |
| AC009264.1 | 39.4169815 | 0.795871008 | 0.49426027 | 1.61022654 | 0.10734841 | 0.25733205 |
| PTN | 75.2682888 | -0.065494765 | 0.78183601 | -0.0837705 | 0.93323893 | 0.96462605 |
| DGKI | 201.092884 | 0.90260592 | 0.44589341 | 2.02426386 | 0.04294301 | 0.14011422 |
| CREB3L2 | 939.352728 | -0.154642707 | 0.35321505 | -0.4378146 | 0.66152068 | 0.80318508 |
| TRIM24 | 1039.07511 | -0.592238065 | 0.33404126 | -1.7729488 | 0.07623719 | 0.2054356 |
| ATP6V0A4 | 68.5595635 | 0.835797951 | 0.43809808 | 1.90778731 | 0.05641871 | 0.16786006 |
| TMEM213 | 81.9098794 | 0.662795074 | 0.45951732 | 1.44237234 | 0.14919736 | 0.31943447 |
| KIAA1549 | 2257.89747 | 1.600457958 | 0.32422006 | 4.93633237 | 7.96E-07 | 3.18E-05 |
| ZC3HAV1L | 58.5868142 | -1.619604167 | 0.40027019 | -4.0462772 | 5.20E-05 | 0.00093854 |
| ZC3HAV1 | 1237.98201 | -1.200600866 | 0.43035205 | -2.789811 | 0.00527388 | 0.03152282 |
| TTC26 | 343.637346 | -0.154101788 | 0.32328183 | -0.4766794 | 0.63359043 | 0.78395238 |
| UBN2 | 1393.24408 | 0.392612258 | 0.24358063 | 1.61183697 | 0.10699742 | 0.25680505 |
| LUC7L2 | 259.855349 | 0.382197216 | 0.26918776 | 1.4198165 | 0.15566111 | 0.32886049 |

| | | | | | | |
|---|---|---|---|---|---|---|
| HIPK2 | 5610.26912 | -0.154908956 | 0.2239767 | -0.6916298 | 0.48916986 | 0.67225298 |
| TBXAS1 | 212.767548 | -0.765476409 | 0.32106416 | -2.3841852 | 0.01711699 | 0.07395389 |
| PARP12 | 326.02547 | -1.283402116 | 0.6311721 | -2.0333632 | 0.04201584 | 0.13799537 |
| KDM7A | 744.578404 | -0.743130383 | 0.19919829 | -3.7306062 | 0.00019102 | 0.00257201 |
| SLC37A3 | 520.068292 | -0.468714533 | 0.27880635 | -1.6811473 | 0.09273431 | 0.23434905 |
| MKRN1 | 457.843539 | -0.199162407 | 0.18821338 | -1.0581735 | 0.28997636 | 0.49009938 |
| DENND2A | 103.982298 | -0.246520987 | 0.54045541 | -0.4561357 | 0.64829243 | 0.79361832 |
| NDUFB2 | 674.722744 | -0.426507967 | 0.30781631 | -1.3855925 | 0.16587137 | 0.34332794 |
| BRAF | 3469.18403 | 0.143972424 | 0.17175242 | 0.83825557 | 0.40188719 | 0.59604714 |
| MRPS33 | 175.729288 | -0.019542194 | 0.2067482 | -0.0945217 | 0.92469473 | 0.96021142 |
| TMEM178B | 2769.63839 | -1.168500061 | 0.42602416 | -2.7428023 | 0.00609173 | 0.03499556 |
| AGK | 427.964041 | 0.124244728 | 0.19393712 | 0.64064438 | 0.52175376 | 0.69758864 |
| AC004918.1 | 249.981127 | 0.96935513 | 0.42338767 | 2.2895214 | 0.02204908 | 0.08849533 |
| KIAA1147 | 642.801135 | 0.721569258 | 0.42500025 | 1.69780901 | 0.08954381 | 0.22830012 |
| WEE2-AS1 | 68.5429388 | 0.695067978 | 0.37796077 | 1.83899504 | 0.06591591 | 0.18652136 |
| SSBP1 | 691.175285 | -0.477147369 | 0.22149977 | -2.1541664 | 0.03122713 | 0.11231721 |
| MGAM | 799.094055 | 1.415643096 | 0.65129352 | 2.17358694 | 0.02973617 | 0.10846027 |
| MGAM2 | 73.2019611 | 1.177418407 | 0.80105541 | 1.46983392 | 0.14160674 | 0.30858563 |
| GSTK1 | 686.877541 | 0.196211514 | 0.42685277 | 0.45967024 | 0.64575294 | 0.79225766 |
| AC073342.1 | 112.444194 | 0.337575214 | 0.58586027 | 0.57620431 | 0.56447709 | 0.73162067 |
| TMEM139 | 249.18546 | 1.405414415 | 0.60442568 | 2.32520631 | 0.02006093 | 0.08262475 |
| CASP2 | 131.061915 | -0.08609809 | 0.23643345 | -0.3641536 | 0.71574332 | 0.83766168 |
| ZYX | 172.60435 | -1.52837417 | 0.44021342 | -3.4718937 | 0.0005168 | 0.00546167 |
| EPHA1 | 52.952537 | -0.155239515 | 0.47391553 | -0.3275679 | 0.74323839 | 0.85423182 |
| EPHA1-AS1 | 42.3787894 | -2.976840745 | 0.48819892 | -6.097598 | 1.08E-09 | 1.23E-07 |
| TCAF2 | 362.045351 | -3.28672579 | 0.63687561 | -5.1607029 | 2.46E-07 | 1.16E-05 |
| TCAF1 | 540.068222 | -0.145183498 | 0.35586089 | -0.4079782 | 0.68328965 | 0.81604817 |
| ARHGEF35 | 46.8466974 | 0.7548574 | 0.46507326 | 1.62309354 | 0.10456941 | 0.25263333 |
| AC004889.1 | 356.347111 | 0.286586043 | 0.34854428 | 0.82223713 | 0.41094195 | 0.6045054 |
| AC074386.1 | 740.731709 | -0.086052869 | 0.30915733 | -0.2783465 | 0.78074636 | 0.87963919 |
| ARHGEF5 | 171.900294 | -0.516769567 | 0.33464335 | -1.5442397 | 0.12253026 | 0.2810916 |
| TPK1 | 1124.42962 | -1.452136932 | 0.42095391 | -3.449634 | 0.00056135 | 0.00581586 |
| CNTNAP2 | 163.177658 | 0.504359703 | 0.27749212 | 1.81756405 | 0.06913079 | 0.19255756 |
| CUL1 | 1320.67294 | -0.28906743 | 0.20855395 | -1.3860559 | 0.16572984 | 0.34319991 |
| EZH2 | 192.572959 | -0.960975181 | 0.36474613 | -2.6346412 | 0.00842263 | 0.04450615 |
| PDIA4 | 390.509488 | -1.571298028 | 0.4980746 | -3.1547443 | 0.00160639 | 0.01300577 |
| ZNF786 | 195.879599 | 0.285332991 | 0.4505075 | 0.63335902 | 0.52649922 | 0.7009757 |
| ZNF425 | 117.584747 | 0.344453078 | 0.33641119 | 1.02390494 | 0.30588018 | 0.50583258 |
| ZNF398 | 488.732392 | 0.065677602 | 0.31842302 | 0.20625896 | 0.83658863 | 0.91377858 |
| ZNF282 | 247.515793 | 0.300990944 | 0.30443388 | 0.98869069 | 0.3228145 | 0.52321371 |
| ZNF212 | 201.629677 | 0.520165941 | 0.37631616 | 1.38225779 | 0.16689256 | 0.34469756 |
| ZNF783 | 129.461128 | 0.230238272 | 0.25901782 | 0.8888897 | 0.37406236 | 0.57127053 |
| ZNF777 | 151.267511 | -0.202652581 | 0.35181275 | -0.576024 | 0.56459898 | 0.731678 |
| ZNF746 | 386.790912 | -1.137825345 | 0.38289461 | -2.9716411 | 0.00296213 | 0.020506 |
| KRBA1 | 60.362926 | -0.377865378 | 0.36097958 | -1.0467777 | 0.29520212 | 0.49508665 |
| ZNF467 | 59.967202 | 0.588462914 | 0.63695904 | 0.92386304 | 0.35555763 | 0.55394731 |
| ZNF862 | 377.429933 | 0.429709813 | 0.32885858 | 1.30667054 | 0.19132463 | 0.37566061 |
| ATP6V0E2 | 93.4671305 | 0.310910472 | 0.52293755 | 0.59454608 | 0.55214693 | 0.72175072 |
| AC092666.1 | 114.895414 | -1.882209335 | 0.56630132 | -3.3236888 | 0.00088835 | 0.00828344 |
| ACTR3C | 3107.63207 | -0.614672973 | 0.3242631 | -1.8955995 | 0.05801302 | 0.17099 |
| LRRC61 | 237.68908 | -1.848258945 | 0.53169678 | -3.4761522 | 0.00050866 | 0.00540904 |
| RARRES2 | 412.5641 | -1.471241563 | 0.68122209 | -2.1597091 | 0.03079519 | 0.11120031 |
| AC073111.5 | 40.7907 | -0.203132784 | 0.61958906 | -0.3278508 | 0.74302446 | 0.8540591 |
| TMEM176B | 1775.27924 | -0.088554981 | 0.29466646 | -0.3005262 | 0.76377584 | 0.86815453 |
| TMEM176A | 876.27799 | -0.51868255 | 0.39653401 | -1.3080405 | 0.19085957 | 0.37512362 |
| AOC1 | 306.743057 | 2.070375348 | 0.65393486 | 3.16602691 | 0.00154536 | 0.01264616 |
| ABCB8 | 235.678689 | -0.171464396 | 0.29642206 | -0.5784468 | 0.5629625 | 0.73052419 |
| SLC4A2 | 378.045301 | -0.214050047 | 0.44267528 | -0.4835374 | 0.62871422 | 0.78066726 |
| FASTK | 237.8074 | 0.883034136 | 0.37925419 | 2.32834381 | 0.01989385 | 0.0821588 |
| TMUB1 | 251.657298 | 0.002239339 | 0.37034972 | 0.00604655 | 0.99517558 | 0.99749068 |
| AGAP3 | 450.747671 | -0.138774134 | 0.31373353 | -0.4423312 | 0.65824955 | 0.80094333 |
| CHPF2 | 117.665787 | -1.01407695 | 0.45015285 | -2.2527392 | 0.0242756 | 0.09431566 |
| SMARCD3 | 72.7480754 | -1.845315142 | 0.48667742 | -3.7916597 | 0.00014964 | 0.00210809 |
| NUB1 | 438.136162 | -0.543711751 | 0.38477275 | -1.4130724 | 0.15763444 | 0.33190403 |
| RHEB | 1137.66957 | -1.064533118 | 0.30265511 | -3.5173143 | 0.00043594 | 0.00486596 |
| PRKAG2 | 2983.58495 | 0.516330285 | 0.49581103 | 1.04138523 | 0.29769679 | 0.49812308 |
| GALNT11 | 2399.87956 | 0.635694873 | 0.25914323 | 2.4530638 | 0.01416452 | 0.0643104 |
| KMT2C | 5430.98428 | -0.225929222 | 0.23189608 | -0.9742692 | 0.32992287 | 0.52966184 |
| ACTR3B | 689.596031 | 0.122803718 | 0.26398417 | 0.46519349 | 0.6417929 | 0.78982207 |
| DPP6 | 3317.8435 | -1.135758221 | 0.48188952 | -2.3568851 | 0.01842895 | 0.07786855 |
| PAXIP1-AS2 | 163.398695 | 0.084207051 | 0.3386512 | 0.24865422 | 0.80362826 | 0.89315921 |
| PAXIP1 | 338.516884 | 0.085995535 | 0.22348631 | 0.38479107 | 0.70039222 | 0.82796169 |
| PAXIP1-AS1 | 122.639779 | -0.52491002 | 0.49858246 | -1.0528048 | 0.29243045 | 0.49248608 |
| INSIG1 | 263.982712 | -1.333287053 | 0.59369456 | -2.2457458 | 0.0247203 | 0.095502 |
| AC009403.1 | 517.405239 | -0.074288267 | 0.39567226 | -0.187752 | 0.85107105 | 0.92167424 |
| RBM33 | 3556.12461 | 0.070734269 | 0.16728181 | 0.42284496 | 0.67240838 | 0.81018826 |
| SHH | 98.2637853 | 0.700104417 | 0.53443646 | 1.30998626 | 0.19020048 | 0.37420382 |
| AC021218.1 | 102.697213 | -0.151290596 | 0.74881167 | -0.2020409 | 0.83988474 | 0.91512719 |
| LINC01006 | 673.676821 | 0.21799202 | 0.28532312 | 0.76401807 | 0.44485647 | 0.63616564 |
| RNF32 | 60.3000545 | -0.215657085 | 0.41311644 | -0.5220249 | 0.60165297 | 0.76078523 |
| LMBR1 | 2263.6565 | -0.181006872 | 0.15364757 | -1.1780653 | 0.23877056 | 0.43383194 |
| NOM1 | 302.193365 | -0.567299186 | 0.4530071 | -1.2522965 | 0.21046186 | 0.39989367 |
| UBE3C | 2483.51712 | -0.242258793 | 0.27769883 | -0.8723796 | 0.38300133 | 0.57970398 |
| DNAJB6 | 1092.41521 | -0.621452526 | 0.37354822 | -1.6636474 | 0.09618292 | 0.23950313 |
| PTPRN2 | 51.6639391 | 0.223870651 | 0.50608104 | 0.44236127 | 0.6582278 | 0.80094333 |
| NCAPG2 | 74.1796148 | 0.17490189 | 0.35325236 | 0.49511882 | 0.62051625 | 0.77455476 |
| ESYT2 | 1957.99137 | -0.450006428 | 0.23974872 | -1.876992 | 0.06051919 | 0.17586104 |
| WDR60 | 2051.7658 | 0.186598784 | 0.20194423 | 0.92401145 | 0.35548036 | 0.55390875 |
| PLCXD1 | 37.9240356 | -0.192136865 | 0.40195848 | -0.4780018 | 0.63264894 | 0.78342389 |
| GTPBP6 | 401.672518 | 0.258038858 | 0.32629039 | 0.79082581 | 0.42904565 | 0.62205719 |
| LINC00685 | 212.547079 | 0.857324716 | 0.46156571 | 1.85742721 | 0.0632504 | 0.18133721 |
| PPP2R3B | 124.911573 | 0.365547067 | 0.35780631 | 1.02163393 | 0.30695419 | 0.50704039 |
| IL3RA | 184.063705 | 0.328379368 | 0.52054102 | 0.63084244 | 0.52814355 | 0.70263648 |
| SLC25A6 | 955.609976 | -1.021781937 | 0.40065501 | -2.5502787 | 0.01076368 | 0.05314762 |
| LINC00106 | 290.531182 | 0.847987534 | 0.3166406 | 2.67807586 | 0.00740464 | 0.04061645 |
| ASMTL-AS1 | 191.765069 | 1.34884223 | 0.27896685 | 4.83513453 | 1.33E-06 | 4.86E-05 |
| ASMTL | 241.62204 | 0.096858457 | 0.34140017 | 0.28370945 | 0.77663305 | 0.87664193 |
| AKAP17A | 504.648676 | -0.367527665 | 0.21215378 | -1.7323645 | 0.08320869 | 0.21795175 |
| DHRSX | 3130.68762 | 0.337174562 | 0.25532136 | 1.32058894 | 0.18663846 | 0.36995711 |
| CD99 | 520.360671 | -0.535512471 | 0.32996055 | -1.6229591 | 0.10459815 | 0.2526391 |
| GYG2 | 72.415746 | 1.046321852 | 0.66975618 | 1.56224292 | 0.11823077 | 0.27444356 |
| ARSD | 384.235161 | 0.267849322 | 0.45999313 | 0.58228984 | 0.56037147 | 0.72874874 |
| ARSE | 346.201379 | 0.18825839 | 0.26530806 | 0.70958412 | 0.47796207 | 0.66359531 |
| ARSF | 534.965689 | 3.292358471 | 0.53460397 | 6.15849984 | 7.34E-10 | 8.91E-08 |
| PRKX | 676.533919 | -0.75853051 | 0.38144453 | -1.9885736 | 0.04674829 | 0.14881069 |
| BX890604.1 | 172.984345 | -0.382492386 | 0.39362852 | -0.971709 | 0.33119532 | 0.5310205 |
| AC097347.1 | 41.9339669 | 0.207669885 | 0.56322222 | 0.36871749 | 0.71233831 | 0.83538853 |
| PUDP | 355.611355 | -0.086165699 | 0.34511773 | -0.2496704 | 0.80284221 | 0.89272007 |
| STS | 109.11428 | -0.944637162 | 0.48899122 | -1.931808 | 0.05338321 | 0.16177838 |
| PNPLA4 | 102.199643 | 0.439634255 | 0.3978534 | 1.10501569 | 0.26915279 | 0.46717453 |
| AC078960.1 | 37.8676199 | 0.018610244 | 0.54618899 | 0.0340729 | 0.97281902 | 0.98642035 |
| ANOS1 | 37.3374419 | 0.013365606 | 0.3679256 | 0.03632693 | 0.97102168 | 0.98568139 |
| TBL1X | 2456.49684 | 0.422913584 | 0.24201949 | 1.74743609 | 0.08056172 | 0.21301303 |

| | | | | | | |
|---|---|---|---|---|---|---|
| GPR143 | 159.865377 | 0.180470809 | 0.40003811 | 0.45113404 | 0.65189295 | 0.79603534 |
| SHROOM2 | 574.37073 | 0.649167891 | 0.38407481 | 1.69021212 | 0.09098738 | 0.23108468 |
| WWC3 | 559.750899 | -0.567645805 | 0.19117589 | -2.9692333 | 0.00298544 | 0.02059457 |
| CLCN4 | 182.699921 | 0.2298474 | 0.34184978 | 0.67236375 | 0.50135216 | 0.68247686 |
| MID1 | 550.292155 | -1.467798737 | 0.54926677 | -2.6722875 | 0.00753361 | 0.04107354 |
| AC073529.1 | 167.417948 | 0.463218843 | 0.50684525 | 0.91392558 | 0.36075595 | 0.55880685 |
| HCCS | 56.437189 | -0.23753277 | 0.30495128 | -0.7789204 | 0.43602661 | 0.62830455 |
| ARHGAP6 | 7785.66486 | 0.333172094 | 0.45439563 | 0.73322029 | 0.46342408 | 0.65150297 |
| MSL3 | 92.775576 | -0.305546385 | 0.31216533 | -0.9787967 | 0.32768045 | 0.52749441 |
| PRPS2 | 57.5006885 | -0.433223283 | 0.42655148 | -1.0156413 | 0.30980025 | 0.51008192 |
| TMSB4X | 970.757877 | -0.587792993 | 0.4807013 | -1.2227822 | 0.22141197 | 0.4134786 |
| TCEANC | 44.2372036 | 1.01572376 | 0.45071703 | 2.25357308 | 0.02422304 | 0.09419154 |
| RAB9A | 583.47573 | -1.440443724 | 0.30720955 | -4.6887986 | 2.75E-06 | 8.93E-05 |
| TRAPPC2 | 230.913703 | -0.248184884 | 0.30234067 | -0.8208783 | 0.41171561 | 0.60522897 |
| OFD1 | 961.675548 | -0.490303581 | 0.26484291 | -1.8512996 | 0.06412646 | 0.18312113 |
| GPM6B | 78.5701286 | -1.230455944 | 0.39121439 | -3.1452216 | 0.00165961 | 0.01329148 |
| GEMIN8 | 306.558712 | 0.092526516 | 0.24275806 | 0.38114703 | 0.70309415 | 0.82952224 |
| MOSPD2 | 217.952006 | -0.233362151 | 0.16813435 | -1.3879505 | 0.16515213 | 0.34242025 |
| ASB9 | 115.52637 | 3.260415605 | 0.48985116 | 6.65593118 | 2.82E-11 | 5.43E-09 |
| PIGA | 113.25882 | -1.108244722 | 0.34792695 | -3.1852799 | 0.00144614 | 0.01197719 |
| VEGFD | 110.799048 | -0.614938874 | 0.63434066 | -0.9694142 | 0.33233855 | 0.53179599 |
| PIR | 920.545934 | -0.636828662 | 0.50471985 | -1.2617468 | 0.2070399 | 0.39591183 |
| BMX | 280.954808 | -1.490010966 | 0.66746482 | -2.2323438 | 0.02559225 | 0.09801424 |
| ACE2 | 769.007071 | 0.894832018 | 0.51610764 | 1.73380889 | 0.082952 | 0.21748719 |
| CLTRN | 2181.73894 | -0.58284843 | 0.59055162 | -0.9869559 | 0.32366424 | 0.52421904 |
| CA5B | 774.799683 | -0.568529279 | 0.31654164 | -1.7960647 | 0.07248422 | 0.19878777 |
| ZRSR2 | 351.974027 | 0.097358578 | 0.27816225 | 0.35000644 | 0.72633386 | 0.84450405 |
| AP1S2 | 149.786233 | -1.394216593 | 0.31553012 | -4.4186482 | 9.93E-06 | 0.00025352 |
| CTPS2 | 372.719321 | 0.060362921 | 0.24405686 | 0.24733139 | 0.80465177 | 0.89400679 |
| SYAP1 | 491.591876 | -0.23698962 | 0.19040516 | -1.2446597 | 0.21325688 | 0.40328327 |
| TXLNG | 415.002966 | -0.007270275 | 0.21487509 | -0.0338349 | 0.97300881 | 0.98646685 |
| RBBP7 | 316.349429 | -0.125685916 | 0.18581925 | -0.676388 | 0.4987943 | 0.6806345 |
| REPS2 | 2044.43907 | -0.263319244 | 0.66990378 | -0.3930702 | 0.6942676 | 0.82399567 |
| NHS | 15128.0254 | 0.229384007 | 0.3888352 | 0.58992604 | 0.55524024 | 0.72468843 |
| NHS-AS1 | 78.6859977 | -0.25636774 | 0.72842228 | -0.3519493 | 0.72487626 | 0.84347804 |
| SCML1 | 421.525253 | -0.27319045 | 0.26260411 | -1.0403129 | 0.29819454 | 0.49833608 |
| RAI2 | 88.5874367 | -1.407270896 | 0.53102975 | -2.6500792 | 0.00804729 | 0.04312127 |
| SCML2 | 109.654273 | 0.747939995 | 0.28475981 | 2.62656448 | 0.00862516 | 0.04519309 |
| CDKL5 | 1026.26444 | 0.825363671 | 0.28630789 | 2.8827835 | 0.00394178 | 0.0253543 |
| PHKA2 | 769.665397 | -0.551757957 | 0.38337025 | -1.43923 | 0.15008537 | 0.32063906 |
| ADGRG2 | 76.6227529 | -0.71696059 | 0.59668367 | -1.2015757 | 0.22952796 | 0.42266884 |
| PDHA1 | 553.763266 | -0.125767119 | 0.32971356 | -0.3814436 | 0.70287413 | 0.82939874 |
| MAP3K15 | 140.582151 | -0.113252112 | 0.49383106 | -0.2293337 | 0.81860955 | 0.90192453 |
| SH3KBP1 | 1161.42647 | 0.049591269 | 0.39779787 | 0.12466449 | 0.90078917 | 0.94780971 |
| BCLAF3 | 628.956787 | 0.36565185 | 0.25107404 | 1.45635068 | 0.1452957 | 0.31371357 |
| MAP7D2 | 336.788984 | 1.029306281 | 0.47022952 | 2.18894441 | 0.02860088 | 0.10561214 |
| EIF1AX | 763.764941 | 0.479766667 | 0.30288616 | 1.58398347 | 0.11319748 | 0.26668922 |
| RPS6KA3 | 1804.32464 | -0.519343776 | 0.31445728 | -1.6515559 | 0.09862511 | 0.24335009 |
| MBTPS2 | 370.557593 | 0.318172657 | 0.25426356 | 1.25134981 | 0.21080688 | 0.40007037 |
| SMS | 1073.34907 | -0.94993406 | 0.45370223 | -2.093739 | 0.03628324 | 0.12461396 |
| PHEX | 89.7513925 | -2.167869313 | 1.04780273 | -2.068967 | 0.03854918 | 0.13053202 |
| PTCHD1-AS | 137.751948 | -0.636236042 | 0.55869772 | -1.138784 | 0.25479325 | 0.45191488 |
| PRDX4 | 140.853646 | -1.820253389 | 0.35270027 | -5.1609072 | 2.46E-07 | 1.16E-05 |
| ACOT9 | 521.039085 | 0.692959169 | 0.47780217 | 1.45030562 | 0.14697331 | 0.31635729 |
| SAT1 | 3363.53962 | -0.403597968 | 0.46825503 | -0.8619191 | 0.38873202 | 0.58431296 |
| APOO | 2080.79485 | -0.402638615 | 0.36976919 | -1.0888917 | 0.27620163 | 0.47394788 |
| KLHL15 | 104.315955 | 0.296208326 | 0.26248969 | 1.12845701 | 0.25912696 | 0.45712397 |
| EIF2S3 | 604.818085 | -0.540196543 | 0.32772282 | -1.6483336 | 0.09928423 | 0.24450253 |
| ZFX | 887.41171 | 0.022407554 | 0.32962107 | 0.06797974 | 0.94580176 | 0.97196767 |
| PDK3 | 635.609895 | -0.56581807 | 0.48576113 | -1.1648072 | 0.24409703 | 0.43920163 |
| POLA1 | 554.282486 | -0.001731631 | 0.28534808 | -0.0060685 | 0.99515808 | 0.99749068 |
| IL1RAPL1 | 106.408848 | -0.30279716 | 0.53719051 | -0.5636681 | 0.57298003 | 0.73835552 |
| GK | 2571.02636 | 1.530687593 | 0.40077174 | 3.8193501 | 0.0001338 | 0.00193465 |
| GK-AS1 | 146.433668 | 1.090454002 | 0.55716666 | 1.95714151 | 0.05033083 | 0.15604721 |
| TAB3 | 742.738684 | -0.076780612 | 0.21948037 | -0.3498291 | 0.72646698 | 0.84458719 |
| DMD | 257.89896 | 0.198374696 | 0.22385612 | 0.88617052 | 0.37552564 | 0.57224764 |
| TMEM47 | 178.293335 | -0.496329836 | 0.45308643 | -1.0954418 | 0.27332311 | 0.47113178 |
| AF241726.2 | 448.559665 | 0.496881012 | 0.32707061 | 1.51918575 | 0.12871575 | 0.29046822 |
| DYNLT3 | 168.890126 | -0.106994138 | 0.24878967 | -0.4300586 | 0.66715301 | 0.80662135 |
| RPGR | 293.016993 | -0.091160189 | 0.31885974 | -0.2858943 | 0.77495906 | 0.87561595 |
| MID1IP1 | 141.373065 | 0.173806306 | 0.47703151 | 0.36434974 | 0.71559686 | 0.83756179 |
| BCOR | 790.166143 | 0.644951269 | 0.20019775 | 3.221571 | 0.0012749 | 0.01087249 |
| ATP6AP2 | 422.631598 | -0.313623929 | 0.29181859 | -1.0747223 | 0.2824991 | 0.4816188 |
| CXorf38 | 369.552794 | -0.715492115 | 0.17995646 | -3.975918 | 7.01E-05 | 0.00117766 |
| MED14 | 788.480075 | -0.065129358 | 0.18526451 | -0.3515479 | 0.72517731 | 0.84366028 |
| USP9X | 2134.61094 | -0.14641146 | 0.230906 | -0.6340739 | 0.52603262 | 0.70084961 |
| DDX3X | 1196.85001 | -0.623663959 | 0.28917024 | -2.1567363 | 0.03102621 | 0.11185595 |
| CASK | 1875.5267 | -0.859695395 | 0.23255138 | -3.6967976 | 0.00021834 | 0.00286386 |
| Z93403.1 | 1546.23844 | 1.197786508 | 0.62518877 | 1.91587977 | 0.0553804 | 0.16595959 |
| MAOA | 1666.25696 | -0.327656245 | 0.5929386 | -0.5525973 | 0.58053922 | 0.74389558 |
| MAOB | 1142.17605 | 1.239151165 | 0.45814413 | 2.7047191 | 0.00683622 | 0.03821535 |
| EFHC2 | 61.7632206 | -0.751890079 | 0.81995824 | -0.9169858 | 0.35915007 | 0.55757957 |
| FUNDC1 | 61.0526292 | -0.05367192 | 0.3543371 | -0.1514714 | 0.87960391 | 0.93717301 |
| KDM6A | 2683.41945 | 0.152185396 | 0.37168623 | 0.40944588 | 0.68221247 | 0.81512259 |
| MIR222HG | 176.987196 | -1.575750385 | 0.60663744 | -2.5975159 | 0.00939008 | 0.0481338 |
| KRBOX4 | 367.731725 | 0.164171889 | 0.23358621 | 0.70283211 | 0.48216039 | 0.66670759 |
| ZNF674 | 120.931548 | 0.087545078 | 0.30728101 | 0.28490234 | 0.77571896 | 0.87613545 |
| SLC9A7 | 340.070584 | -0.496678907 | 0.29994384 | -1.6559063 | 0.09774079 | 0.2419164 |
| RP2 | 168.498141 | -1.019918461 | 0.36797178 | -2.7717301 | 0.00557592 | 0.03286766 |
| JADE3 | 509.459493 | -0.680180768 | 0.28687839 | -2.3709724 | 0.01774135 | 0.07582017 |
| RGN | 1256.22768 | -0.003308894 | 0.77591695 | -0.0042645 | 0.99659744 | 0.99819957 |
| NDUFB11 | 307.157166 | -0.352895805 | 0.42616698 | -0.8280693 | 0.40763124 | 0.60157207 |
| RBM10 | 412.534475 | 0.463773919 | 0.22085695 | 2.09988372 | 0.03573907 | 0.12342541 |
| UBA1 | 625.724701 | -0.159288273 | 0.22138994 | -0.7194919 | 0.47183787 | 0.65801724 |
| CDK16 | 240.061109 | -0.01320971 | 0.3105522 | -0.0425362 | 0.96607126 | 0.98263125 |
| USP11 | 136.51244 | 0.049813731 | 0.36378996 | 0.13692992 | 0.89108619 | 0.94333086 |
| ZNF41 | 215.98715 | 0.053431649 | 0.20621762 | 0.25910321 | 0.79555561 | 0.88843413 |
| ARAF | 188.72808 | 0.464573922 | 0.30200864 | 1.53828023 | 0.12398011 | 0.28346879 |
| TIMP1 | 291.455508 | -2.485030124 | 0.54420206 | -4.5663739 | 4.96E-06 | 0.00014171 |
| ELK1 | 116.331792 | -0.469139555 | 0.241359 | -1.9437417 | 0.0519266 | 0.15895347 |
| UXT | 139.707301 | -0.244253144 | 0.20192718 | -1.20961 | 0.22642857 | 0.4189815 |
| ZNF81 | 657.310432 | 0.799922032 | 0.2602904 | 3.07319067 | 0.00211783 | 0.01606481 |
| ZNF182 | 98.7258675 | 0.305251871 | 0.23227985 | 1.31415561 | 0.18879384 | 0.37272032 |
| ZNF630 | 121.192919 | 0.389499917 | 0.62713484 | 0.62107843 | 0.53454802 | 0.70751736 |
| FTSJ1 | 51.784061 | -0.71132812 | 0.40295789 | -1.7652667 | 0.07751896 | 0.20765144 |
| EBP | 232.132702 | 0.58395541 | 0.29180441 | 2.00118775 | 0.04537216 | 0.14575022 |
| TBC1D25 | 162.085929 | -0.080177099 | 0.25635362 | -0.3127598 | 0.75446319 | 0.86224896 |
| RBM3 | 498.471653 | -0.34486173 | 0.3265628 | -1.0560349 | 0.29095225 | 0.49112819 |
| WDR13 | 976.737928 | -1.040071142 | 0.26020117 | -3.997181 | 6.41E-05 | 0.00109692 |
| WAS | 39.4611475 | 1.636368123 | 0.65897891 | 2.4831874 | 0.01302126 | 0.06068764 |
| GLOD5 | 46.8539201 | 1.468547243 | 0.5510735 | 2.66488451 | 0.00770148 | 0.04169133 |
| HDAC6 | 566.743277 | 1.214393567 | 0.38865973 | 3.12456749 | 0.00178067 | 0.01400321 |
| PCSK1N | 1058.55394 | -0.301011595 | 0.65056688 | -0.4626912 | 0.6435857 | 0.79117952 |

| | | | | | | |
|---|---|---|---|---|---|---|
| TIMM17B | 107.334456 | 0.001552512 | 0.35641525 | 0.00435591 | 0.9965245 | 0.99819945 |
| PQBP1 | 130.397865 | 0.722508963 | 0.38878843 | 1.85836024 | 0.06311787 | 0.18105566 |
| SLC35A2 | 81.5082839 | -0.536363834 | 0.42428884 | -1.2641479 | 0.20617696 | 0.3949228 |
| OTUD5 | 508.714533 | 0.057451611 | 0.22147325 | 0.25940655 | 0.79532158 | 0.88840738 |
| GRIPAP1 | 437.288764 | 0.077696575 | 0.19970515 | 0.38905645 | 0.69723439 | 0.82550208 |
| TFE3 | 345.726212 | -1.12136196 | 0.31915963 | -3.5134831 | 0.00044227 | 0.00492064 |
| CCDC120 | 243.169408 | -0.760289788 | 0.3807948 | -1.9965866 | 0.04587011 | 0.14679934 |
| PRAF2 | 37.3521858 | 0.023048266 | 0.55320662 | 0.04166303 | 0.96676732 | 0.9830686 |
| WDR45 | 336.660811 | 0.100393886 | 0.21501802 | 0.46690917 | 0.64056486 | 0.78888372 |
| GPKOW | 61.0801564 | -0.11980568 | 0.25146609 | -0.4764288 | 0.63376895 | 0.78396092 |
| MAGIX | 75.6957896 | 0.244719796 | 0.37264159 | 0.65671628 | 0.51136337 | 0.69043121 |
| PLP2 | 90.4474604 | -2.527669859 | 0.41901697 | -6.0323807 | 1.62E-09 | 1.74E-07 |
| PRICKLE3 | 37.4133527 | 0.007528398 | 0.33803164 | 0.02227128 | 0.98223156 | 0.99119841 |
| CCDC22 | 57.7280601 | 0.179770472 | 0.35044058 | 0.51298418 | 0.60796239 | 0.76485741 |
| PPP1R3F | 60.6843934 | 0.023206182 | 0.36564043 | 0.06346722 | 0.94939446 | 0.97405313 |
| CLCN5 | 1798.76693 | 0.938670099 | 0.23896215 | 3.92811209 | 8.56E-05 | 0.00138225 |
| SHROOM4 | 2250.24606 | 0.800552035 | 0.31630327 | 2.53096351 | 0.01137497 | 0.05523061 |
| GSPT2 | 44.1386183 | -0.41139245 | 0.48614834 | -0.8462282 | 0.39742544 | 0.59186337 |
| MAGED1 | 1387.45531 | 0.319406838 | 0.16251585 | 1.9653889 | 0.04936924 | 0.15398584 |
| FAM156A | 189.485888 | -0.001715108 | 0.23426688 | -0.0073212 | 0.9941586 | 0.99706827 |
| GPR173 | 45.8506908 | -0.622520336 | 0.68100538 | -0.9141196 | 0.36065403 | 0.55871211 |
| TSPYL2 | 95.4234321 | -1.291993972 | 0.50599965 | -2.5533495 | 0.01066924 | 0.05283354 |
| KANTR | 953.019414 | 0.321561108 | 0.23615111 | 1.3616752 | 0.17330041 | 0.35296118 |
| KDM5C | 474.910158 | -0.359652487 | 0.37841311 | -0.9504229 | 0.34189741 | 0.5425642 |
| IQSEC2 | 444.77769 | 0.774650429 | 0.28447655 | 2.72307303 | 0.00646778 | 0.0366338 |
| SMC1A | 393.404421 | -0.17094492 | 0.29393124 | -0.5815813 | 0.56084873 | 0.72909271 |
| HSD17B10 | 183.990856 | -0.40843929 | 0.29827846 | -1.3693221 | 0.17089861 | 0.3498066 |
| HUWE1 | 2007.28954 | 0.062495261 | 0.16635561 | 0.37567269 | 0.70716024 | 0.83227665 |
| PHF8 | 728.43985 | 0.169282225 | 0.26635674 | 0.635547 | 0.52507172 | 0.70065792 |
| FAM120C | 348.936993 | 0.536482033 | 0.25042499 | 2.14228632 | 0.03217045 | 0.11477365 |
| WNK3 | 116.841135 | 0.207652318 | 0.32093114 | 0.64703075 | 0.51761204 | 0.69435391 |
| TSR2 | 88.212752 | -0.614437958 | 0.29863771 | -2.0574694 | 0.0396411 | 0.13265637 |
| FGD1 | 100.20102 | 0.043045851 | 0.34422836 | 0.12505027 | 0.90048375 | 0.94768577 |
| GNL3L | 140.081529 | -0.152478183 | 0.46439147 | -0.3283398 | 0.74265478 | 0.8540591 |
| MAGED2 | 235.260737 | -0.547674621 | 0.35747838 | -1.5320496 | 0.12551019 | 0.28572798 |
| PFKFB1 | 121.520458 | -0.841446317 | 0.90243758 | -0.932415 | 0.35112211 | 0.55041523 |
| APEX2 | 78.4591706 | -0.873888949 | 0.31946635 | -2.7354648 | 0.00622923 | 0.03559149 |
| FAM104B | 150.172801 | 0.594395095 | 0.32754384 | 1.81470393 | 0.06956942 | 0.19324417 |
| USP51 | 61.080741 | 0.59575055 | 0.44700343 | 1.33276507 | 0.18260891 | 0.36487178 |
| RRAGB | 850.174851 | -0.022774153 | 0.20528217 | -0.1109407 | 0.91166335 | 0.95319043 |
| AL445472.1 | 920.271365 | 0.04964328 | 0.39541949 | 0.12554586 | 0.90009142 | 0.94743919 |
| KLF8 | 412.805717 | 0.665694069 | 0.54812438 | 1.21449455 | 0.22455893 | 0.41674812 |
| UBQLN2 | 211.263099 | -0.34490708 | 0.25433502 | -1.3561132 | 0.17506315 | 0.35517926 |
| NBDY | 695.628783 | 0.157266916 | 0.2076292 | 0.75744121 | 0.44878557 | 0.639148 |
| SPIN3 | 261.679282 | 0.175425278 | 0.20210028 | 0.86801106 | 0.38538828 | 0.5816469 |
| SPIN2B | 35.3392442 | 0.060192914 | 0.37402967 | 0.16093085 | 0.87214786 | 0.93380181 |
| FAAH2 | 1003.67114 | -0.661246073 | 0.32350344 | -2.0440156 | 0.040952 | 0.13558675 |
| ZXDB | 34.4424077 | -0.555164736 | 0.39760053 | -1.3962877 | 0.16262788 | 0.33908432 |
| SPIN4 | 39.0212146 | 0.249668088 | 0.35878947 | 0.69586236 | 0.48651503 | 0.66974606 |
| ARHGEF9 | 597.303712 | 0.85332113 | 0.24568273 | 3.47326454 | 0.00051417 | 0.00544645 |
| AMER1 | 50.5469812 | 0.47479189 | 0.40555201 | 1.17072997 | 0.24170734 | 0.43681951 |
| MTMR8 | 300.198721 | 0.43572368 | 0.20554281 | 2.11986822 | 0.03401716 | 0.11944721 |
| ZC4H2 | 536.203723 | 0.288116286 | 0.19734219 | 1.45998324 | 0.14429468 | 0.31211096 |
| ZC3H12B | 257.310749 | 0.177131623 | 0.40462744 | 0.43776473 | 0.66155684 | 0.80318508 |
| LAS1L | 146.710962 | -0.052083157 | 0.3308431 | -0.1574255 | 0.87490948 | 0.93490877 |
| MSN | 1989.14284 | -0.357369739 | 0.311519 | -1.1471844 | 0.25130541 | 0.44764695 |
| AR | 863.365883 | 0.384677203 | 0.34113604 | 1.12763578 | 0.25947377 | 0.45761798 |
| OPHN1 | 1148.98168 | 0.966918739 | 0.22502866 | 4.29686921 | 1.73E-05 | 0.00039751 |
| YIPF6 | 468.566823 | -0.160897989 | 0.23079479 | -0.6971474 | 0.48571055 | 0.6693108 |
| STARD8 | 97.1481158 | 1.049573648 | 0.29564899 | 3.55006678 | 0.00038513 | 0.00443989 |
| EFNB1 | 92.0947165 | -0.452891896 | 0.32799662 | -1.3807822 | 0.16734593 | 0.34532068 |
| EDA | 1983.52573 | 0.996562972 | 0.25036741 | 3.98040207 | 6.88E-05 | 0.00115993 |
| IGBP1 | 429.461697 | 0.003923342 | 0.24457662 | 0.01604136 | 0.98720139 | 0.99360989 |
| PDZD11 | 43.5889115 | -0.620123185 | 0.47837622 | -1.2963086 | 0.1948692 | 0.38011558 |
| DLG3 | 304.300429 | -0.37182776 | 0.3897504 | -0.9540151 | 0.34007601 | 0.54064276 |
| TEX11 | 46.1741274 | -0.484614295 | 0.36744101 | -1.3188901 | 0.18720587 | 0.37070657 |
| SNX12 | 174.882116 | -0.246053689 | 0.31260767 | -0.7871006 | 0.43122296 | 0.62376247 |
| MED12 | 95.4396835 | 0.324760905 | 0.25122891 | 1.29268922 | 0.19611857 | 0.38162936 |
| GJB1 | 182.972153 | 1.476534149 | 0.69300635 | 2.1306214 | 0.03312035 | 0.1173683 |
| ZMYM3 | 235.596071 | 0.561326426 | 0.39432377 | 1.4235166 | 0.15458645 | 0.3271791 |
| NONO | 608.841581 | -0.291071003 | 0.2347741 | -1.2397918 | 0.21505242 | 0.40557561 |
| TAF1 | 1172.62688 | 0.497398089 | 0.17346901 | 2.86735999 | 0.00413912 | 0.0263271 |
| OGT | 1560.93252 | -0.311957542 | 0.35744236 | -0.8727492 | 0.38279978 | 0.57959092 |
| GCNA | 154.861989 | 0.163342535 | 0.36270432 | 0.45034626 | 0.65246079 | 0.7965868 |
| NHSL2 | 149.868538 | -0.9688757 | 0.39355427 | -2.4618605 | 0.01382184 | 0.06332183 |
| PIN4 | 432.923165 | -0.029100802 | 0.24319791 | -0.1196589 | 0.90475333 | 0.95028792 |
| RPS4X | 1060.31262 | -1.026460744 | 0.46345652 | -2.214794 | 0.0267742 | 0.10118887 |
| HDAC8 | 2499.8784 | 0.245736877 | 0.1847609 | 1.33002638 | 0.18350958 | 0.36587139 |
| PHKA1 | 242.712959 | -1.463210775 | 0.56204135 | -2.6033863 | 0.00923079 | 0.04756631 |
| CHIC1 | 722.82629 | 0.433328758 | 0.26367446 | 1.6434233 | 0.10029538 | 0.2463163 |
| TSIX | 76.7787359 | -4.420515496 | 2.56058174 | -1.7263716 | 0.08428059 | 0.219376 |
| XIST | 2979.19543 | -2.751017815 | 3.68647313 | -0.7462465 | 0.4555185 | 0.64475342 |
| JPX | 2227.692 | -0.032548548 | 0.32530992 | -0.100054 | 0.92030148 | 0.95724483 |
| FTX | 17051.7946 | 0.454247681 | 0.20949179 | 2.16833162 | 0.03013346 | 0.10953047 |
| SLC16A2 | 282.830985 | -0.02634337 | 0.48836283 | -0.0539422 | 0.95698121 | 0.97810315 |
| RLIM | 308.009097 | -0.422168011 | 0.18008668 | -2.3442489 | 0.01906544 | 0.07969808 |
| NEXMIF | 62.1593684 | -0.123248671 | 0.34267079 | -0.3596708 | 0.71909334 | 0.83993922 |
| ABCB7 | 676.724113 | 0.210206091 | 0.20275911 | 1.03672824 | 0.29986253 | 0.50014793 |
| UPRT | 162.510508 | -0.041476772 | 0.19204245 | -0.2159771 | 0.8290056 | 0.90919985 |
| ZDHHC15 | 106.949613 | 0.154402137 | 0.30801215 | 0.50128586 | 0.61616996 | 0.7715916 |
| PBDC1 | 119.713025 | -1.350965618 | 0.52856743 | -2.5559002 | 0.01059135 | 0.052638 |
| AC233296.1 | 55.4832454 | -0.272492149 | 0.61643114 | -0.442048 | 0.65845448 | 0.80095001 |
| ATRX | 3361.1121 | 0.316645593 | 0.20441696 | 1.54901823 | 0.12137734 | 0.27946713 |
| MAGT1 | 737.664825 | -0.071939027 | 0.29045957 | -0.2476731 | 0.80438734 | 0.8938579 |
| COX7B | 682.663021 | -0.494147496 | 0.44776817 | -1.1035789 | 0.26977586 | 0.46795971 |
| PGK1 | 2032.16226 | -1.693749287 | 0.43701132 | -3.8757561 | 0.00010629 | 0.00163338 |
| TAF9B | 48.2474143 | -0.028115846 | 0.34707704 | -0.0810075 | 0.93543598 | 0.96565906 |
| BRWD3 | 1035.98598 | -0.206374776 | 0.3151809 | -0.654782 | 0.51260813 | 0.6911936 |
| HMGN5 | 121.993292 | 0.260350115 | 0.49797098 | 0.52282186 | 0.60109822 | 0.76042947 |
| SH3BGRL | 488.400684 | 0.19530374 | 0.28351879 | 0.68885642 | 0.49091363 | 0.67363631 |
| RPS6KA6 | 409.877545 | -0.267888754 | 0.3043442 | -0.8802164 | 0.37874209 | 0.57554521 |
| HDX | 442.703369 | -1.074573902 | 0.30112245 | -3.5685613 | 0.00035895 | 0.0042088 |
| APOOL | 706.43619 | -0.032939456 | 0.22697825 | -0.1451216 | 0.88461483 | 0.94010044 |
| ZNF711 | 96.8901534 | 1.296643144 | 0.40438422 | 3.20646323 | 0.00134377 | 0.01130701 |
| POF1B | 54.7018536 | -0.823207294 | 0.70647097 | -1.1652387 | 0.2439224 | 0.43900266 |
| CHM | 1029.36053 | -0.228433394 | 0.2109853 | -1.0826981 | 0.27894244 | 0.47745554 |
| DACH2 | 51.7103472 | -0.418928612 | 0.59590275 | -0.7030151 | 0.48204636 | 0.66670759 |
| KLHL4 | 200.60887 | 0.780866722 | 0.42873687 | 1.82131928 | 0.06855834 | 0.19150787 |
| PCDH11X | 100.291267 | -0.312462534 | 0.44839249 | -0.6968505 | 0.48589635 | 0.66931729 |
| DIAPH2 | 2906.02584 | -0.34621559 | 0.21710178 | -1.5947156 | 0.11077586 | 0.26297275 |
| DIAPH2-AS1 | 152.251531 | -0.890472172 | 0.53739218 | -1.6570248 | 0.09751446 | 0.24144341 |
| PCDH19 | 35.24869 | 0.471069858 | 0.45522369 | 1.03480964 | 0.30075782 | 0.50121428 |
| TSPAN6 | 82.7420692 | -0.471125683 | 0.41900408 | -1.124394 | 0.26084591 | 0.45903388 |

| | | | | | | |
|---|---|---|---|---|---|---|
| SYTL4 | 183.620886 | 0.044132978 | 0.43451415 | 0.10156856 | 0.91909913 | 0.95679351 |
| CSTF2 | 43.9537852 | -0.804251795 | 0.3753833 | -2.1424816 | 0.03215475 | 0.11474751 |
| TRMT2B | 322.852893 | 0.594167719 | 0.22082376 | 2.6906875 | 0.0071305 | 0.03953225 |
| TIMM8A | 56.8581548 | -0.587605729 | 0.31670609 | -1.8553661 | 0.06354397 | 0.18191253 |
| GLA | 133.857584 | -2.518693702 | 0.71768743 | -3.5094578 | 0.00044902 | 0.0049635 |
| HNRNPH2 | 153.692056 | -0.727737139 | 0.28616285 | -2.5430874 | 0.01098777 | 0.0539432 |
| ARMCX4 | 838.950139 | 0.178664842 | 0.43971652 | 0.40631824 | 0.68450879 | 0.81708281 |
| ARMCX1 | 67.6448316 | -1.251994236 | 0.57245743 | -2.1870521 | 0.02873872 | 0.10602395 |
| ARMCX6 | 109.383175 | -0.686385731 | 0.37373395 | -1.8365624 | 0.06627452 | 0.18714974 |
| ARMCX3 | 342.370525 | -0.911691656 | 0.23543308 | -3.8724025 | 0.00010777 | 0.00164679 |
| ARMCX2 | 63.7450801 | -0.925245258 | 0.36855598 | -2.5104606 | 0.01205738 | 0.05746539 |
| NXF5 | 42.4942196 | 1.020554537 | 0.75378289 | 1.35391046 | 0.17576495 | 0.35597422 |
| ZMAT1 | 964.490528 | 1.407200397 | 0.59760908 | 2.35471724 | 0.01853681 | 0.07815267 |
| ARMCX5 | 66.3139439 | -0.290932682 | 0.3800401 | -0.7655315 | 0.44395509 | 0.63520798 |
| AL035427.1 | 49.6542309 | -0.972605898 | 0.76265369 | -1.2752917 | 0.20220603 | 0.38971288 |
| GPRASP1 | 74.8315173 | 0.658505134 | 0.28902071 | 2.27840122 | 0.02270268 | 0.09027725 |
| GPRASP2 | 43.404387 | 0.642151373 | 0.50155877 | 1.28031133 | 0.20043566 | 0.38755419 |
| BHLHB9 | 206.599374 | -0.303293379 | 0.48735291 | -0.622328 | 0.53372619 | 0.70689258 |
| LINC00630 | 548.025358 | 0.166486522 | 0.17162417 | 0.97006452 | 0.33201433 | 0.53158748 |
| Z68871.1 | 80.0777745 | 0.3341588 | 0.24375243 | 1.37089425 | 0.17040792 | 0.34945862 |
| BEX4 | 210.814659 | -0.425515013 | 0.29422226 | -1.4462366 | 0.14811084 | 0.31795376 |
| TCEAL8 | 104.682605 | -1.089778691 | 0.25468973 | -4.2788481 | 1.88E-05 | 0.00042353 |
| BEX2 | 207.889391 | -1.404221837 | 0.33190203 | -4.2308323 | 2.33E-05 | 0.0004971 |
| TCEAL9 | 286.367418 | -1.347954664 | 0.39807219 | -3.3862066 | 0.00070866 | 0.0070132 |
| BEX3 | 332.739866 | 0.042334003 | 0.39399874 | 0.10744705 | 0.91443433 | 0.9543495 |
| TCEAL4 | 355.820637 | -0.4100051 | 0.23247128 | -1.7636807 | 0.07778576 | 0.20811316 |
| TCEAL3 | 105.285187 | -1.025229192 | 0.23642566 | -4.3363703 | 1.45E-05 | 0.00034174 |
| TCEAL1 | 55.602251 | -0.426728294 | 0.34575264 | -1.2342011 | 0.21712798 | 0.40803033 |
| MORF4L2 | 1309.18122 | -1.005612704 | 0.27380889 | -3.672681 | 0.00024002 | 0.00308625 |
| SLC25A53 | 447.131781 | 0.340799969 | 0.19154336 | 1.77923145 | 0.07520182 | 0.20371372 |
| FAM199X | 882.564826 | -0.404111795 | 0.27072813 | -1.4926849 | 0.13551969 | 0.29999488 |
| IL1RAPL2 | 671.759116 | 0.488660011 | 0.798432 | 0.61202458 | 0.5405215 | 0.71198617 |
| NRK | 59.7095574 | 0.295809463 | 0.69378312 | 0.42637166 | 0.66983705 | 0.80829795 |
| MUM1L1 | 125.183887 | 0.81694885 | 0.53215707 | 1.53516489 | 0.12474333 | 0.28469145 |
| CXorf57 | 55.275571 | 0.01015304 | 0.37713888 | 0.02692122 | 0.97852257 | 0.98916955 |
| RNF128 | 541.413281 | 0.067220068 | 0.36040347 | 0.18651338 | 0.85204219 | 0.92213818 |
| TBC1D8B | 589.695198 | 0.25314594 | 0.57549915 | 0.43987196 | 0.66002985 | 0.8020537 |
| MORC4 | 438.077898 | -0.681929378 | 0.37234948 | -1.8314229 | 0.06703744 | 0.18852734 |
| CLDN2 | 949.84913 | -0.502404392 | 0.64186254 | -0.7827289 | 0.4337863 | 0.62601693 |
| RBM41 | 646.050649 | 0.193772941 | 0.16016837 | 1.20980775 | 0.22635267 | 0.41893277 |
| NUP62CL | 75.0482203 | -0.289910616 | 0.43141934 | -0.6719926 | 0.50158839 | 0.68268018 |
| PRPS1 | 70.4094595 | -0.087919692 | 0.38878374 | -0.2261404 | 0.82109226 | 0.9039929 |
| TSC22D3 | 328.760041 | 0.370697057 | 0.68667497 | 0.53984355 | 0.58930493 | 0.75126513 |
| MID2 | 261.968157 | -1.172162372 | 0.38425994 | -3.0504412 | 0.00228505 | 0.01702059 |
| PSMD10 | 50.1254225 | -0.354368647 | 0.415153 | -0.8535857 | 0.3933346 | 0.58819959 |
| ATG4A | 132.057214 | -0.170437796 | 0.24419145 | -0.6979679 | 0.48519726 | 0.66893973 |
| COL4A5 | 131.979402 | -0.076625854 | 0.82601871 | -0.0927653 | 0.92609004 | 0.96090956 |
| ACSL4 | 1690.5163 | -2.231689264 | 0.58487951 | -3.8156393 | 0.00013583 | 0.00195982 |
| TMEM164 | 667.654671 | -0.491087442 | 0.30719174 | -1.5986349 | 0.10990174 | 0.2615318 |
| AMMECR1 | 555.339846 | -0.246256196 | 0.19009076 | -1.2954664 | 0.19515937 | 0.38030275 |
| CHRDL1 | 47.771285 | 1.024271256 | 0.57419822 | 1.78382869 | 0.07445151 | 0.20232093 |
| PAK3 | 60.6084751 | -0.938181345 | 0.55858034 | -1.6795818 | 0.09303872 | 0.23469615 |
| ALG13 | 919.653389 | 0.043056428 | 0.32100792 | 0.13412887 | 0.89330067 | 0.94442517 |
| ALG13-AS1 | 65.3326599 | -1.338203771 | 0.65258557 | -2.0506181 | 0.04030415 | 0.13431778 |
| TRPC5 | 123.178879 | 1.688775326 | 0.56141241 | 3.00808334 | 0.00262901 | 0.0188472 |
| AMOT | 358.526754 | 0.06036426 | 0.43947287 | 0.13735606 | 0.89074936 | 0.94331061 |
| PLS3 | 501.712745 | -0.051644725 | 0.34388266 | -0.1501812 | 0.88062162 | 0.93789282 |
| DANT2 | 3962.58052 | 0.169083107 | 0.23255498 | 0.72706723 | 0.46718477 | 0.65464135 |
| KLHL13 | 2395.81182 | -0.468915722 | 0.54968167 | -0.8530678 | 0.39362171 | 0.58850053 |
| WDR44 | 876.060984 | -0.268020607 | 0.22107807 | -1.2123346 | 0.22538431 | 0.41770613 |
| DOCK11 | 705.152514 | 0.200616353 | 0.34692693 | 0.57826688 | 0.56308395 | 0.73061262 |
| IL13RA1 | 549.737751 | 0.138067713 | 0.38234114 | 0.36111132 | 0.71801623 | 0.83896074 |
| LINC01285 | 119.595786 | 1.3729247 | 0.48074117 | 2.85585008 | 0.00429218 | 0.0270496 |
| LONRF3 | 359.68578 | 0.696315908 | 0.79215085 | 0.87901933 | 0.3793908 | 0.57597637 |
| PGRMC1 | 562.236664 | -0.496670253 | 0.31547476 | -1.5743581 | 0.11540468 | 0.27026344 |
| SLC25A43 | 657.296076 | -0.271568175 | 0.22533922 | -1.2051527 | 0.22814435 | 0.4209982 |
| SLC25A5 | 551.605569 | -0.798622173 | 0.49309942 | -1.6195967 | 0.10531895 | 0.25397727 |
| CXorf56 | 91.1455561 | 0.473720315 | 0.27874398 | 1.6994818 | 0.08922844 | 0.22776623 |
| UBE2A | 245.15281 | -0.410034225 | 0.25209232 | -1.6265241 | 0.10383822 | 0.25191335 |
| NKRF | 102.824119 | 0.141723526 | 0.2652507 | 0.5343003 | 0.59313379 | 0.75436977 |
| 6-Sep | 118.475334 | -1.329198804 | 0.56450742 | -2.3546171 | 0.0185418 | 0.07815267 |
| RPL39 | 70.288484 | -0.394006671 | 0.43331573 | -0.9092831 | 0.36320071 | 0.56088576 |
| UPF3B | 565.964386 | -0.009926138 | 0.3082853 | -0.0321979 | 0.97431423 | 0.98703942 |
| NDUFA1 | 441.974275 | -0.847007087 | 0.37797689 | -2.2408965 | 0.02503278 | 0.09647223 |
| NKAP | 191.688107 | -0.023825039 | 0.2302988 | -0.1034527 | 0.91760367 | 0.95587788 |
| RHOXF1-AS1 | 224.472875 | 0.156307751 | 0.50472833 | 0.3096869 | 0.75679907 | 0.86359015 |
| ZBTB33 | 64.9832245 | -0.785166006 | 0.38682702 | -2.02976 | 0.04238094 | 0.138859 |
| LAMP2 | 1311.55463 | -1.046567209 | 0.27384142 | -3.8218003 | 0.00013248 | 0.00191957 |
| CUL4B | 288.848437 | 0.350934979 | 0.16677679 | 2.10421951 | 0.0353593 | 0.12254614 |
| MCTS1 | 277.178169 | 0.014482562 | 0.17462219 | 0.08293655 | 0.93390199 | 0.96479108 |
| C1GALT1C1 | 88.126458 | -0.644092105 | 0.32330781 | -1.9921947 | 0.0463497 | 0.14788532 |
| GLUD2 | 40.7902476 | 0.089854988 | 0.37365375 | 0.24047661 | 0.80996079 | 0.89677969 |
| GRIA3 | 549.5404 | 1.396350823 | 0.43005263 | 3.24693012 | 0.00116657 | 0.01015249 |
| THOC2 | 2360.68313 | 0.182826284 | 0.16949952 | 1.07862417 | 0.2807553 | 0.47942107 |
| XIAP | 1712.61066 | -0.139287015 | 0.18890651 | -0.7373331 | 0.46091983 | 0.64958134 |
| STAG2 | 1861.74637 | 0.211440809 | 0.1975294 | 1.07042704 | 0.28442714 | 0.48376259 |
| TENM1 | 1892.10258 | 1.329206535 | 0.56563842 | 2.34992264 | 0.01877731 | 0.07885436 |
| AL442647.1 | 54.9767202 | -0.68000205 | 0.66740196 | -1.0188793 | 0.30826027 | 0.50834017 |
| SMARCA1 | 790.093066 | -0.557140093 | 0.40311478 | -1.382088 | 0.16694469 | 0.34472895 |
| OCRL | 281.227258 | 0.186808715 | 0.18666721 | 1.00075805 | 0.3169438 | 0.51693918 |
| XPNPEP2 | 724.97939 | 2.063367796 | 0.80277251 | 2.57030202 | 0.01016099 | 0.05111071 |
| ZDHHC9 | 575.612016 | 0.165090827 | 0.17632637 | 0.93627987 | 0.34912911 | 0.54887281 |
| AL034405.1 | 48.7131984 | -0.625235601 | 0.3650643 | -1.7126725 | 0.0867728 | 0.22382288 |
| UTP14A | 146.322151 | -0.308899101 | 0.44736795 | -0.6904811 | 0.48989171 | 0.67270543 |
| BCORL1 | 277.948191 | 0.509940251 | 0.3352193 | 1.5212139 | 0.12820617 | 0.28992962 |
| ELF4 | 62.0762573 | -2.462896831 | 0.47137577 | -5.2249118 | 1.74E-07 | 8.79E-06 |
| AIFM1 | 1037.54302 | 0.120186221 | 0.41201939 | 0.2917004 | 0.7705157 | 0.87241523 |
| ZNF280C | 272.652492 | -0.230373765 | 0.26960858 | -0.8544749 | 0.3928419 | 0.58771927 |
| SLC25A14 | 115.465766 | 0.736154364 | 0.2294825 | 3.20788892 | 0.00133713 | 0.01126494 |
| RBMX2 | 107.18585 | 0.230452777 | 0.28843009 | 0.79899007 | 0.42429617 | 0.61739147 |
| ENOX2 | 1083.75694 | -0.316984775 | 0.26072851 | -1.2157657 | 0.2240742 | 0.41634609 |
| STK26 | 56.3545533 | -1.197675617 | 0.51406268 | -2.3298241 | 0.01981545 | 0.08190904 |
| RAP2C | 298.765036 | -0.417704241 | 0.23367057 | -1.7875774 | 0.07384421 | 0.20114916 |
| RAP2C-AS1 | 1100.44156 | 0.674550957 | 0.30928417 | 2.18100709 | 0.02918289 | 0.10692593 |
| MBNL3 | 242.072807 | 0.4425018 | 0.37904768 | 1.16740407 | 0.24304723 | 0.4381755 |
| HS6ST2 | 57.3281683 | 1.05916889 | 0.50134164 | 2.1126689 | 0.03462912 | 0.1208097 |
| GPC4 | 1337.30381 | -0.483760237 | 0.43568776 | -1.110337 | 0.26685384 | 0.46477327 |
| GPC3 | 455.515605 | 2.403876876 | 0.85088155 | 2.82516041 | 0.0047257 | 0.02909332 |
| PHF6 | 263.71883 | 0.053040294 | 0.21086874 | 0.25153227 | 0.80140262 | 0.89219475 |
| HPRT1 | 209.823638 | -1.525355253 | 0.29417474 | -5.1852013 | 2.16E-07 | 1.04E-05 |
| MIR503HG | 72.5967338 | 0.656606273 | 0.59467133 | 1.10414988 | 0.26952813 | 0.46764836 |
| FAM122B | 152.326769 | 0.370598405 | 0.21082707 | 1.75783119 | 0.07877624 | 0.20970788 |
| FAM122C | 115.547159 | 0.137692852 | 0.52520592 | 0.26216927 | 0.79319095 | 0.88726528 |

| | | | | | | |
|---|---|---|---|---|---|---|
| MOSPD1 | 116.296667 | -0.347620018 | 0.21784112 | -1.5957502 | 0.11054457 | 0.26251463 |
| RTL8C | 294.571634 | -0.513334843 | 0.47705066 | -1.0760594 | 0.2819007 | 0.48077801 |
| RTL8A | 160.508206 | -1.076436396 | 0.38561967 | -2.7914458 | 0.00524731 | 0.03140827 |
| ZNF75D | 1196.76406 | -0.326561804 | 0.22675057 | -1.4401808 | 0.14981627 | 0.32016395 |
| ZNF449 | 124.308638 | -0.143921311 | 0.29147575 | -0.4937677 | 0.62147024 | 0.7751836 |
| SMIM10L2A | 111.280798 | -1.641901411 | 0.45773781 | -3.586991 | 0.00033452 | 0.00399757 |
| INTS6L | 317.074123 | 0.035028614 | 0.34476768 | 0.10160063 | 0.91907368 | 0.95679351 |
| MMGT1 | 98.3513132 | -0.414749913 | 0.2741278 | -1.5129801 | 0.13028471 | 0.29285373 |
| SLC9A6 | 176.828389 | -0.130557608 | 0.17199797 | -0.7590648 | 0.44781378 | 0.63841127 |
| FHL1 | 150.098447 | -1.588285492 | 0.56075453 | -2.8324078 | 0.00461989 | 0.02863562 |
| MAP7D3 | 246.490238 | -0.61436032 | 0.19420058 | -3.1635349 | 0.00155866 | 0.01272692 |
| HTATSF1 | 161.846283 | -0.34172148 | 0.33134871 | -1.0313047 | 0.30239796 | 0.50269269 |
| AL683813.1 | 194.463131 | 0.320663527 | 0.43944378 | 0.72970319 | 0.46557163 | 0.65338286 |
| RBMX | 985.732634 | 0.331689378 | 0.16626379 | 1.99495862 | 0.04604739 | 0.1471736 |
| FGF13 | 2548.99216 | -1.180993248 | 0.6127077 | -1.9274986 | 0.05391752 | 0.16308249 |
| ATP11C | 1145.78598 | 0.838209322 | 0.16435424 | 5.10001652 | 3.40E-07 | 1.53E-05 |
| LDOC1 | 56.8274147 | -0.707354328 | 0.56419198 | -1.2537476 | 0.20993377 | 0.39949496 |
| SLITRK4 | 77.8259073 | -0.147592536 | 0.68153159 | -0.2165601 | 0.8285512 | 0.90884693 |
| AL589669.1 | 4118.48874 | 1.278997974 | 0.52330442 | 2.44408023 | 0.0145222 | 0.06547887 |
| FMR1 | 1072.05908 | -0.093290643 | 0.3974266 | -0.2347368 | 0.81441303 | 0.89947803 |
| AFF2 | 449.231412 | -0.5201688 | 0.73722739 | -0.7055744 | 0.48045282 | 0.66534319 |
| IDS | 360.888099 | 0.195186569 | 0.3155838 | 0.61849362 | 0.53625 | 0.70869254 |
| LINC00893 | 92.9268531 | 0.535113919 | 0.35945126 | 1.48869675 | 0.13656724 | 0.30154199 |
| TMEM185A | 198.006969 | -0.214923878 | 0.17304704 | -1.2419968 | 0.21423773 | 0.40451891 |
| CXorf40B | 37.7473224 | -0.530555193 | 0.33533808 | -1.5821501 | 0.1136153 | 0.26730603 |
| LINC00894 | 1735.77632 | 0.580946563 | 0.47867867 | 1.21364622 | 0.22488285 | 0.41705712 |
| MAMLD1 | 205.081502 | -2.57212823 | 0.71750065 | -3.5848445 | 0.00033728 | 0.00402007 |
| MTM1 | 930.312247 | 0.290753104 | 0.17751173 | 1.63793745 | 0.10143474 | 0.24823532 |
| MTMR1 | 535.59515 | -0.129276541 | 0.17785759 | -0.7268542 | 0.46731525 | 0.65466556 |
| CD99L2 | 403.555566 | 0.216509088 | 0.17126175 | 1.2641999 | 0.20615829 | 0.3949228 |
| HMGB3 | 70.3588643 | -1.002143377 | 0.50991708 | -1.9653066 | 0.04937877 | 0.15398584 |
| VMA21 | 170.92047 | -0.46672241 | 0.29566974 | -1.5785261 | 0.11444479 | 0.26859016 |
| GABRE | 47.3764001 | -0.213761496 | 0.37296604 | -0.5731393 | 0.56655038 | 0.73330557 |
| CETN2 | 89.6595586 | -0.495552202 | 0.43433803 | -1.1409367 | 0.25389626 | 0.45078869 |
| NSDHL | 87.389574 | 0.044721311 | 0.3029378 | 0.14762539 | 0.88263842 | 0.93903973 |
| ZNF185 | 54.036895 | -0.101568535 | 0.66788344 | -0.1520752 | 0.87912759 | 0.93706833 |
| PNMA3 | 59.3771066 | 0.290930162 | 1.02295986 | 0.28440037 | 0.77610358 | 0.8764136 |
| PNMA6A | 56.4873849 | 0.10819842 | 0.40837051 | 0.2649516 | 0.79104675 | 0.88629048 |
| ZNF275 | 60.4017978 | 0.732080505 | 0.47184563 | 1.55152545 | 0.12077581 | 0.27837129 |
| BGN | 50.3731454 | 0.578623711 | 0.51815659 | 1.11669661 | 0.26412407 | 0.46207385 |
| CCNQ | 71.593691 | -0.681790193 | 0.48717711 | -1.3994709 | 0.16167181 | 0.33791332 |
| SLC6A8 | 882.62806 | -0.755755977 | 0.50266635 | -1.5034943 | 0.13271163 | 0.29620664 |
| BCAP31 | 499.465555 | -0.673393893 | 0.36432207 | -1.8483478 | 0.06455205 | 0.18402972 |
| ABCD1 | 67.0563134 | 0.112097008 | 0.35235826 | 0.31813362 | 0.75038358 | 0.85927383 |
| IDH3G | 189.296464 | -0.007622365 | 0.28034005 | -0.0271897 | 0.97830842 | 0.98906609 |
| SSR4 | 231.79031 | -0.670494443 | 0.49064685 | -1.366552 | 0.17176576 | 0.35108758 |
| RENBP | 391.784042 | -0.142690869 | 0.48619339 | -0.2934858 | 0.76915083 | 0.87130169 |
| HCFC1 | 160.65226 | -0.445119539 | 0.3591503 | -1.2393684 | 0.2152091 | 0.40575944 |
| TMEM187 | 50.4116139 | 0.257913592 | 0.34507116 | 0.74742145 | 0.4548092 | 0.64414855 |
| IRAK1 | 227.215219 | -1.515235865 | 0.5231349 | -2.8964534 | 0.00377407 | 0.02450551 |
| MECP2 | 992.168897 | 0.411845055 | 0.26592971 | 1.54869892 | 0.12145411 | 0.27946713 |
| FLNA | 226.645891 | -0.994926074 | 0.66722391 | -1.4911427 | 0.13592403 | 0.30059869 |
| EMD | 192.233223 | -0.693455905 | 0.2863694 | -2.4215433 | 0.01545476 | 0.06862958 |
| RPL10 | 1827.4246 | -0.903507778 | 0.49138869 | -1.8386825 | 0.0659619 | 0.18656221 |
| AC245140.2 | 38.9078265 | -0.225131385 | 0.4415841 | -0.5098267 | 0.61017285 | 0.76667332 |
| DNASE1L1 | 154.28049 | -0.194783529 | 0.38133062 | -0.5107996 | 0.60949138 | 0.76602773 |
| TAZ | 281.81284 | 0.295877432 | 0.22684963 | 1.3042888 | 0.19213513 | 0.37658332 |
| AC244090.1 | 48.282648 | 0.474417496 | 0.38887884 | 1.21996223 | 0.22247919 | 0.41472082 |
| ATP6AP1 | 292.459113 | 0.182636651 | 0.34096461 | 0.53564694 | 0.59220258 | 0.75355744 |
| GDI1 | 272.265628 | -0.253629109 | 0.22764763 | -1.1141303 | 0.26522331 | 0.4630717 |
| FAM50A | 349.24435 | -0.483733178 | 0.47216292 | -1.0245048 | 0.3055969 | 0.50559043 |
| PLXNA3 | 170.160722 | -1.371265639 | 0.32014222 | -4.2833014 | 1.84E-05 | 0.0004165 |
| UBL4A | 86.928556 | -0.966722767 | 0.41119594 | -2.3510027 | 0.0187229 | 0.07867405 |
| SLC10A3 | 54.0817536 | -0.783498473 | 0.44503469 | -1.7605335 | 0.0783174 | 0.20901412 |
| FAM3A | 189.739993 | 0.234388644 | 0.18681548 | 1.25465324 | 0.20960467 | 0.39920122 |
| G6PD | 81.794744 | -1.35907732 | 0.59355396 | -2.2897283 | 0.02203707 | 0.08847748 |
| IKBKG | 123.802413 | 0.014089538 | 0.23901361 | 0.05894869 | 0.95299298 | 0.97628361 |
| GAB3 | 45.7342135 | -2.137617121 | 0.5321105 | -4.0172428 | 5.89E-05 | 0.00103211 |
| DKC1 | 185.858518 | -1.054654006 | 0.4190468 | -2.5167929 | 0.01184284 | 0.05671903 |
| MPP1 | 281.201401 | 0.302611347 | 0.24014753 | 1.26010601 | 0.20763112 | 0.39665502 |
| F8 | 164.556097 | 0.654239874 | 0.34477415 | 1.8975897 | 0.05775015 | 0.17049836 |
| F8A1 | 31.1366937 | -0.546922952 | 0.53752652 | -1.0174809 | 0.30892473 | 0.50906843 |
| FUNDC2 | 259.72454 | -0.516635506 | 0.30414062 | -1.6986731 | 0.08938079 | 0.22801357 |
| MTCP1 | 83.8060186 | -1.287009679 | 0.64870278 | -1.9839744 | 0.0472587 | 0.14994791 |
| BRCC3 | 380.951902 | -0.342061576 | 0.14058824 | -2.4330739 | 0.01497125 | 0.06695297 |
| VBP1 | 139.915476 | -0.228030447 | 0.20366105 | -1.1196566 | 0.26286011 | 0.46086257 |
| TMLHE-AS1 | 250.314367 | -0.315967358 | 0.57209372 | -0.5523 | 0.58074283 | 0.74402516 |
| TMLHE | 657.907807 | 0.050412277 | 0.26052365 | 0.19350365 | 0.84656455 | 0.91911764 |
| VAMP7 | 302.629335 | -0.313302202 | 0.22344592 | -1.4021388 | 0.1608738 | 0.33687295 |
| ZNF596 | 170.057092 | 0.233920356 | 0.2613955 | 0.89489052 | 0.37084561 | 0.56858846 |
| FBXO25 | 503.082835 | -0.275834779 | 0.20795637 | -1.326407 | 0.18470493 | 0.36750623 |
| TDRP | 702.039351 | -0.142896959 | 0.33935261 | -0.421087 | 0.67369158 | 0.81073146 |
| ERICH1 | 1613.47765 | -1.121159403 | 0.45776863 | -2.4491836 | 0.01431805 | 0.06483563 |
| DLGAP2 | 443.709984 | 1.513955636 | 0.70699836 | 2.14138493 | 0.03224301 | 0.11497267 |
| AC100810.1 | 93.4527392 | 0.061248206 | 0.38396306 | 0.15951588 | 0.87326245 | 0.93448422 |
| AC019257.8 | 36.4324886 | 0.510263025 | 0.50701373 | 1.00640868 | 0.31421902 | 0.51482191 |
| KBTBD11 | 397.905574 | -0.453261002 | 0.32562948 | -1.3919532 | 0.16393656 | 0.34080823 |
| MYOM2 | 191.361397 | 2.054707423 | 0.30904754 | 6.64851564 | 2.96E-11 | 5.64E-09 |
| AC246817.2 | 203.082388 | 0.797628233 | 0.62759096 | 1.27093645 | 0.20375126 | 0.39180955 |
| AC246817.1 | 221.008026 | 0.308239598 | 0.37391993 | 0.82434653 | 0.40974269 | 0.60360018 |
| CSMD1 | 1451.5634 | 2.197401315 | 0.79694317 | 2.75728736 | 0.00582831 | 0.03395183 |
| MCPH1 | 1438.40861 | -0.194570852 | 0.16007174 | -1.2155228 | 0.22416675 | 0.41640515 |
| ANGPT2 | 192.998728 | -1.335235464 | 0.54817053 | -2.4358031 | 0.01485877 | 0.06669933 |
| MCPH1-AS1 | 437.769588 | 0.05835135 | 0.51004645 | 0.11440399 | 0.90891755 | 0.95219603 |
| AGPAT5 | 1073.6451 | -0.672518822 | 0.42038122 | -1.5997832 | 0.10964668 | 0.26106081 |
| AF233439.1 | 127.30488 | -1.119350125 | 0.63798597 | -1.7545059 | 0.07934387 | 0.21080953 |
| DEFB1 | 919.297699 | -1.745862986 | 0.434999 | -4.0134873 | 5.98E-05 | 0.00104467 |
| FAM85B | 2874.71787 | 1.10336066 | 0.42546412 | 2.59331071 | 0.00950568 | 0.04854197 |
| PRAG1 | 839.803687 | -0.302128627 | 0.45485864 | -0.6642253 | 0.50654611 | 0.68655979 |
| MFHAS1 | 2284.69422 | -0.370155703 | 0.32110682 | -1.1527494 | 0.24901324 | 0.44519533 |
| ERI1 | 620.603967 | -0.910048839 | 0.30094252 | -3.0239956 | 0.0024946 | 0.0181687 |
| PPP1R3B | 295.674969 | -0.950595941 | 0.33564995 | -2.8321051 | 0.00462426 | 0.02864202 |
| AC022784.3 | 86.9878925 | 0.636374477 | 0.69852676 | 0.91102376 | 0.36228286 | 0.560096 |
| TNKS | 2613.71558 | -0.147033912 | 0.16251449 | -0.9047434 | 0.36560135 | 0.56313043 |
| AC034111.1 | 62.1487844 | 1.606465616 | 0.50629303 | 3.1729957 | 0.00150875 | 0.01240576 |
| MSRA | 8932.64267 | 1.264378121 | 0.22950041 | 5.509263 | 3.60E-08 | 2.43E-06 |
| AC104964.1 | 232.465975 | 0.762834911 | 0.38236589 | 1.99503912 | 0.04603861 | 0.1471736 |
| PRSS51 | 142.783718 | -0.074822272 | 0.83642533 | -0.0894548 | 0.92872046 | 0.96224497 |
| AC104964.4 | 91.210918 | 1.045584297 | 0.41785303 | 2.50227765 | 0.01233971 | 0.05840484 |
| PRSS55 | 134.044608 | 0.562180165 | 0.74151805 | 0.75814764 | 0.4483626 | 0.63884679 |
| PINX1 | 40.1519397 | 0.040573809 | 0.41475675 | 0.09782556 | 0.92207081 | 0.95821263 |
| XKR6 | 1056.79049 | 0.022890659 | 0.46388728 | 0.0493453 | 0.96064412 | 0.97967485 |
| MTMR9 | 421.44971 | -0.184892558 | 0.2106856 | -0.8775757 | 0.38017403 | 0.57657954 |

| | | | | | | |
|---|---|---|---|---|---|---|
| NEIL2 | 85.7085459 | 0.395106357 | 0.37831406 | 1.0443872 | 0.29630628 | 0.49645156 |
| FDFT1 | 884.809589 | -0.303353074 | 0.34266288 | -0.8852814 | 0.37600488 | 0.5726357 |
| CTSB | 3907.7653 | -1.142250345 | 0.50175476 | -2.2765112 | 0.02281543 | 0.09054172 |
| FAM66D | 135.1254 | 0.157485565 | 0.81733432 | 0.19268194 | 0.84720807 | 0.91938085 |
| FAM86B1 | 39.4356014 | -0.408233147 | 0.5242365 | -0.7787194 | 0.43614501 | 0.62836238 |
| AC145124.1 | 310.948325 | 0.705198609 | 0.62139076 | 1.13487141 | 0.2564292 | 0.45375428 |
| LONRF1 | 575.117662 | -0.704610646 | 0.30131612 | -2.3384433 | 0.01936427 | 0.08055417 |
| TRMT9B | 1146.43789 | 1.048274357 | 0.55261148 | 1.8969464 | 0.05783501 | 0.1706662 |
| DLC1 | 956.197666 | -1.379906464 | 0.62038935 | -2.2242588 | 0.02613104 | 0.0993835 |
| SGCZ | 42.9068719 | 0.090087998 | 0.39295418 | 0.22925827 | 0.81866818 | 0.90192453 |
| TUSC3 | 1042.72455 | -2.413003468 | 0.40065556 | -6.0226382 | 1.72E-09 | 1.84E-07 |
| MSR1 | 190.92106 | -0.999018052 | 0.66102621 | -1.5113138 | 0.13070851 | 0.29346791 |
| AC068992.1 | 70.9787779 | -2.567392802 | 0.75849053 | -3.3848713 | 0.00071212 | 0.00702734 |
| MICU3 | 337.29903 | -0.099191541 | 0.43271329 | -0.2292316 | 0.81868895 | 0.90192453 |
| ZDHHC2 | 734.167541 | -0.394453995 | 0.21612483 | -1.8251212 | 0.06798275 | 0.1903267 |
| CNOT7 | 356.279781 | -0.203981501 | 0.17772799 | -1.1477173 | 0.25108527 | 0.44731974 |
| VPS37A | 726.551604 | -0.667210125 | 0.22292525 | -2.9929769 | 0.00276271 | 0.01948967 |
| SLC7A2 | 251.320089 | -0.9207605 | 0.56009337 | -1.6439411 | 0.10018837 | 0.24615892 |
| MTUS1 | 2096.13885 | 0.090298125 | 0.38268665 | 0.23595839 | 0.81346496 | 0.89903065 |
| AC027117.2 | 470.354601 | 0.260440151 | 0.41477172 | 0.62791203 | 0.53006157 | 0.70421206 |
| PCM1 | 2354.97601 | 0.112231324 | 0.1941442 | 0.57808229 | 0.56320856 | 0.73070513 |
| ASAH1 | 1494.83222 | -0.131313334 | 0.37325528 | -0.3518057 | 0.72498398 | 0.84350696 |
| NAT1 | 337.212316 | 2.677870177 | 0.63045554 | 4.24751627 | 2.16E-05 | 0.0004688 |
| PSD3 | 334.147619 | -2.433215848 | 0.67479915 | -3.6058372 | 0.00031115 | 0.0037776 |
| SH2D4A | 190.758392 | -0.722723304 | 0.29006257 | -2.4916118 | 0.01271649 | 0.05974577 |
| CSGALNACT1 | 80.7544535 | -0.646538052 | 0.57104231 | -1.1322069 | 0.25754746 | 0.45486445 |
| INTS10 | 1137.3823 | 0.003406304 | 0.27557941 | 0.01236052 | 0.99013799 | 0.99507452 |
| ATP6V1B2 | 1382.90276 | 0.552582303 | 0.30989575 | 1.78312323 | 0.07456625 | 0.20243207 |
| AC018541.1 | 83.5793408 | 4.090376717 | 0.56350161 | 7.25885541 | 3.90E-13 | 1.11E-10 |
| XPO7 | 1163.15236 | 0.616784338 | 0.17241134 | 3.57740003 | 0.00034703 | 0.00410771 |
| NPM2 | 91.7495523 | -0.82116371 | 0.33159996 | -2.4763685 | 0.01327265 | 0.0615454 |
| DMTN | 488.473126 | 0.608916106 | 0.38404762 | 1.58552241 | 0.11284769 | 0.26617886 |
| FAM160B2 | 208.450786 | -0.43143793 | 0.26054009 | -1.6559368 | 0.09773461 | 0.2419164 |
| NUDT18 | 33.4054822 | -0.626410585 | 0.45494168 | -1.376903 | 0.16854223 | 0.34711458 |
| BMP1 | 316.086051 | -1.461404672 | 0.40005009 | -3.6530542 | 0.00025914 | 0.00326473 |
| PHYHIP | 59.2656636 | 0.186620462 | 0.52670041 | 0.35431995 | 0.72309912 | 0.84233055 |
| POLR3D | 97.7528277 | -0.918674186 | 0.44928721 | -2.044737 | 0.04088079 | 0.13544913 |
| PIWIL2 | 66.6479861 | -0.141060433 | 0.4920451 | -0.2866819 | 0.77435588 | 0.87502853 |
| SLC39A14 | 3956.46155 | -1.004077779 | 0.39766889 | -2.5249091 | 0.01157282 | 0.05580386 |
| PPP3CC | 1262.04378 | 0.011896401 | 0.19935723 | 0.05967379 | 0.95241545 | 0.97591078 |
| SORBS3 | 151.075278 | 0.017211902 | 0.25302355 | 0.0680249 | 0.94576581 | 0.97196767 |
| CCAR2 | 190.427958 | 0.028737409 | 0.22618582 | 0.12705221 | 0.89889908 | 0.94743919 |
| AC037459.3 | 36.2904663 | -0.277609618 | 0.52618362 | -0.5275908 | 0.5977834 | 0.75777463 |
| BIN3 | 626.079699 | -0.586501952 | 0.32155454 | -1.8239579 | 0.06815844 | 0.19070171 |
| PEBP4 | 81.8266588 | 0.004769724 | 0.60383594 | 0.00789904 | 0.99369754 | 0.99682467 |
| RHOBTB2 | 178.839852 | 0.049060367 | 0.50042777 | 0.09803686 | 0.92190303 | 0.95811091 |
| TNFRSF10B | 1272.72464 | -1.243888794 | 0.21135634 | -5.8852683 | 3.97E-09 | 3.68E-07 |
| TNFRSF10D | 573.948533 | -0.79020955 | 0.17574662 | -4.4963 | 6.91E-06 | 0.00018731 |
| TNFRSF10A | 450.573538 | -1.161309866 | 0.31412438 | -3.6969747 | 0.00021818 | 0.00286386 |
| CHMP7 | 175.855264 | 0.096107396 | 0.259241 | 0.37072606 | 0.71084158 | 0.83470449 |
| R3HCC1 | 79.025108 | 0.263476379 | 0.28172065 | 0.93523986 | 0.3496647 | 0.54925232 |
| LOXL2 | 55.6611553 | -0.26227397 | 0.28947787 | -0.9060242 | 0.36492302 | 0.5624016 |
| ENTPD4 | 394.452961 | -0.465808429 | 0.18069198 | -2.5779142 | 0.00993987 | 0.05018261 |
| SLC25A37 | 1828.48757 | -1.208418487 | 0.3921461 | -3.0815517 | 0.00205925 | 0.01573361 |
| ADAM28 | 293.695928 | 0.742172558 | 0.6923854 | 1.07190671 | 0.28376194 | 0.48299079 |
| AC120193.1 | 101.574124 | 0.208329083 | 0.58670482 | 0.3550833 | 0.72252719 | 0.84228022 |
| DOCK5 | 1868.7757 | -1.265769817 | 0.21859249 | -5.7905456 | 7.02E-09 | 6.01E-07 |
| KCTD9 | 346.319927 | -1.004707653 | 0.29806398 | -3.3707785 | 0.00074956 | 0.00731262 |
| PPP2R2A | 2236.33408 | 0.03256029 | 0.23352898 | 0.13942719 | 0.88911258 | 0.94267382 |
| BNIP3L | 4064.39332 | -1.383612151 | 0.54247801 | -2.5505405 | 0.0107556 | 0.05312686 |
| PNMA2 | 173.28363 | -3.640685109 | 0.56118272 | -6.4875218 | 8.73E-11 | 1.44E-08 |
| DPYSL2 | 1138.06828 | -1.570622508 | 0.3254804 | -4.8255518 | 1.40E-06 | 5.05E-05 |
| ADRA1A | 87.0609826 | 1.329771051 | 0.78718455 | 1.68927484 | 0.09116677 | 0.23136881 |
| AC090150.1 | 251.856291 | 2.623570421 | 0.63341896 | 4.14191964 | 3.44E-05 | 0.00067441 |
| TRIM35 | 273.34435 | -0.724821182 | 0.41358833 | -1.7525185 | 0.0796847 | 0.21135017 |
| PTK2B | 652.178467 | 0.986812458 | 0.36115463 | 2.73238213 | 0.00628782 | 0.03582174 |
| EPHX2 | 1558.336 | 1.35904618 | 0.61537721 | 2.20847661 | 0.02721107 | 0.10229898 |
| CLU | 3427.02292 | -2.824654994 | 0.61078758 | -4.6246111 | 3.75E-06 | 0.00011432 |
| CCDC25 | 213.609322 | 0.052968385 | 0.20218885 | 0.2619748 | 0.79334087 | 0.88726528 |
| NUGGC | 637.314034 | 3.070108499 | 0.66666044 | 4.60520577 | 4.12E-06 | 0.00012225 |
| ELP3 | 854.911235 | 0.387895099 | 0.16276261 | 2.38319542 | 0.01716308 | 0.07409586 |
| ZNF395 | 359.847449 | -0.67527661 | 0.33810299 | -1.9972512 | 0.0457979 | 0.14667098 |
| FBXO16 | 483.598503 | -0.102509094 | 0.39620293 | -0.2587288 | 0.79584452 | 0.88843413 |
| FZD3 | 350.865083 | 0.068445853 | 0.20559609 | 0.33291419 | 0.73919906 | 0.85193361 |
| EXTL3 | 361.50796 | -0.222345546 | 0.41895233 | -0.530718 | 0.5956142 | 0.75582107 |
| INTS9 | 668.682559 | -0.048327517 | 0.18336992 | -0.263552 | 0.79212512 | 0.88678699 |
| HMBOX1 | 1792.51013 | -0.343448573 | 0.14946271 | -2.297888 | 0.02156816 | 0.08736253 |
| KIF13B | 2921.93252 | 0.600123585 | 0.27827081 | 2.15661711 | 0.03103551 | 0.11186004 |
| AC108449.1 | 119.772402 | -0.407746009 | 0.62427488 | -0.6531514 | 0.51365868 | 0.69196393 |
| SARAF | 1053.67885 | -0.989168276 | 0.25300712 | -3.909646 | 9.24E-05 | 0.00146557 |
| LEPROTL1 | 619.101179 | -0.498754293 | 0.27469894 | -1.8156397 | 0.06942566 | 0.19304949 |
| AC026979.2 | 42.5172392 | 0.021260226 | 0.44962266 | 0.0472846 | 0.9622864 | 0.98038202 |
| DCTN6 | 165.686488 | -0.632421746 | 0.25894224 | -2.4423275 | 0.0145929 | 0.06577603 |
| RBPMS-AS1 | 167.16097 | -1.3417968 | 0.41269055 | -3.2513388 | 0.00114863 | 0.01002718 |
| RBPMS | 19595.144 | -0.027668414 | 0.28815098 | -0.0960205 | 0.92350425 | 0.95941131 |
| GTF2E2 | 1132.60457 | 0.090126723 | 0.27442522 | 0.32841997 | 0.74259414 | 0.8540591 |
| SMIM18 | 41.1788982 | -0.736246664 | 0.3576211 | -2.0587338 | 0.03951974 | 0.13241191 |
| GSR | 1123.90827 | -0.462558802 | 0.35791404 | -1.292374 | 0.19622765 | 0.38178742 |
| UBXN8 | 97.1800606 | -0.210376299 | 0.2986282 | -0.7044757 | 0.48113661 | 0.66594592 |
| PPP2CB | 1654.99055 | -0.593635135 | 0.21160157 | -2.8054383 | 0.00502482 | 0.03038165 |
| WRN | 781.746119 | -0.46450167 | 0.28059135 | -1.6554383 | 0.09783562 | 0.24200482 |
| NRG1 | 5056.61883 | 0.133175855 | 0.45669198 | 0.29160979 | 0.77058499 | 0.87242161 |
| AC090204.1 | 75.6410271 | 0.754612017 | 0.49407298 | 1.52732904 | 0.12667921 | 0.28776797 |
| FUT10 | 721.916067 | 0.58189889 | 0.24256576 | 2.39893252 | 0.01644294 | 0.07189265 |
| TTI2 | 191.840014 | 0.040616569 | 0.27208026 | 0.14928157 | 0.88133146 | 0.93835717 |
| MAK16 | 83.6839674 | -0.371054484 | 0.4484272 | -0.8274576 | 0.40797776 | 0.60182427 |
| RNF122 | 143.706186 | -0.529241051 | 0.35163274 | -1.5050961 | 0.13229938 | 0.29558723 |
| AF279873.3 | 62.9788336 | 2.889405356 | 0.52743507 | 5.47821999 | 4.30E-08 | 2.78E-06 |
| UNC5D | 2862.89273 | -0.325111838 | 0.55648151 | -0.5842276 | 0.55906722 | 0.72767395 |
| LINC01605 | 140.972973 | 0.090574678 | 0.72594454 | 0.12476804 | 0.90070719 | 0.94779621 |
| ERLIN2 | 274.576889 | 0.233483359 | 0.21503414 | 1.08579669 | 0.27756895 | 0.47552026 |
| PLPBP | 216.951506 | -0.039338058 | 0.16514628 | -0.2382013 | 0.81172497 | 0.89793513 |
| RAB11FIP1 | 352.637573 | -0.306383075 | 0.23095603 | -1.3265862 | 0.18464561 | 0.3674949 |
| EIF4EBP1 | 187.4773 | -1.129350146 | 0.31818419 | -3.5493597 | 0.00038617 | 0.00444435 |
| ASH2L | 473.783196 | 0.657710932 | 0.30645366 | 2.14620025 | 0.03185701 | 0.11401148 |
| LSM1 | 188.106758 | -0.442847008 | 0.24064191 | -1.8402738 | 0.06572805 | 0.1862204 |
| BAG4 | 545.842365 | 0.651415628 | 0.14508608 | 4.48985629 | 7.13E-06 | 0.0001908 |
| DDHD2 | 501.808409 | -0.092673583 | 0.2298601 | -0.4031739 | 0.68682033 | 0.81863011 |
| PLPP5 | 164.037126 | -1.071767619 | 0.47461177 | -2.2581986 | 0.02393328 | 0.09346253 |
| NSD3 | 1550.86606 | 0.019563912 | 0.15816941 | 0.1236896 | 0.90156105 | 0.94833069 |
| FGFR1 | 1429.41258 | -1.332667392 | 0.81119704 | -1.6428405 | 0.10041593 | 0.24644604 |
| TACC1 | 1802.31599 | 0.469491199 | 0.32487922 | 1.44512536 | 0.14842268 | 0.3183724 |
| PLEKHA2 | 1232.31586 | 0.175758514 | 0.41082988 | 0.42781337 | 0.668787 | 0.80738625 |

| | | | | | | |
|---|---|---|---|---|---|---|
| TM2D2 | 141.676276 | -0.353252472 | 0.34925524 | -1.011445 | 0.31180349 | 0.51190447 |
| ADAM9 | 1878.0024 | -0.91015222 | 0.33576808 | -2.7106574 | 0.006715 | 0.03772232 |
| ADAM32 | 158.487022 | -1.24969153 | 0.48803609 | -2.5606539 | 0.01044754 | 0.05209327 |
| IDO2 | 225.073026 | 5.40173179 | 0.92243265 | 5.85596334 | 4.74E-09 | 4.28E-07 |
| TCIM | 479.505168 | -2.81659668 | 0.91055248 | -3.0932832 | 0.00197955 | 0.01525223 |
| ZMAT4 | 41.6912347 | 0.280139932 | 0.43220703 | 0.64816144 | 0.51688054 | 0.6940373 |
| SFRP1 | 65.1318685 | 0.079881109 | 0.44699865 | 0.17870548 | 0.85816897 | 0.92545981 |
| GOLGA7 | 310.84301 | -0.340432692 | 0.21980869 | -1.5487681 | 0.12143748 | 0.27946713 |
| AC009630.1 | 69.633646 | 0.24992561 | 0.40530757 | 0.61663198 | 0.5374775 | 0.70967393 |
| GPAT4 | 1072.39784 | -0.059204283 | 0.28500354 | -0.2077317 | 0.83543843 | 0.91316952 |
| KAT6A | 2077.44639 | 0.089590781 | 0.17342256 | 0.51660395 | 0.60543266 | 0.76301233 |
| AP3M2 | 276.791753 | 0.208821191 | 0.18139372 | 1.15120409 | 0.24964828 | 0.44586762 |
| PLAT | 49.8795171 | -0.113921376 | 0.54916453 | -0.2074449 | 0.83566243 | 0.91316952 |
| IKBKB | 791.493094 | -0.173613642 | 0.22631596 | -0.7671295 | 0.44300453 | 0.63457657 |
| POLB | 173.996711 | 0.226648795 | 0.1931124 | 1.17366254 | 0.24053022 | 0.43579395 |
| VDAC3 | 564.526422 | -0.759966525 | 0.3482327 | -2.1823525 | 0.02908352 | 0.10681148 |
| SLC20A2 | 1686.68865 | -0.238791103 | 0.40899388 | -0.5838501 | 0.55932119 | 0.72786628 |
| SMIM19 | 277.308565 | -0.002804578 | 0.22144906 | -0.0126647 | 0.98989533 | 0.99504952 |
| THAP1 | 53.8279207 | -0.390840757 | 0.35828112 | -1.0908773 | 0.27532686 | 0.47286769 |
| RNF170 | 594.400461 | 0.497211738 | 0.32394941 | 1.53484377 | 0.12482221 | 0.2847781 |
| HOOK3 | 1186.09759 | -0.024064323 | 0.14345487 | -0.1677484 | 0.86678122 | 0.93074581 |
| FNTA | 293.721439 | -0.803937533 | 0.23075618 | -3.4839263 | 0.00049412 | 0.0053162 |
| POMK | 97.6544345 | 0.117218137 | 0.22481756 | 0.52139227 | 0.60209353 | 0.7609161 |
| HGSNAT | 936.262405 | -0.33359236 | 0.33334009 | -1.0007568 | 0.3169444 | 0.51693918 |
| SPIDR | 3771.06646 | -0.554416797 | 0.29815971 | -1.8594625 | 0.06296161 | 0.18073609 |
| CEBPD | 1871.51568 | -1.617456546 | 0.44462798 | -3.637775 | 0.000275 | 0.00342991 |
| PRKDC | 1297.72888 | -0.265981849 | 0.33508131 | -0.793783 | 0.42732177 | 0.62027951 |
| MCM4 | 55.8661904 | -0.562578354 | 0.50291275 | -1.1186401 | 0.26329372 | 0.46128738 |
| UBE2V2 | 578.151447 | 0.201326562 | 0.24496368 | 0.8218629 | 0.41115493 | 0.60468888 |
| SNTG1 | 121.693399 | 1.812537069 | 0.47248174 | 3.83620551 | 0.00012495 | 0.00185758 |
| PXDNL | 218.622639 | 0.834359207 | 0.24054727 | 3.46858729 | 0.0005232 | 0.0055147 |
| PCMTD1 | 2339.84848 | 0.170629682 | 0.21932936 | 0.77796099 | 0.43659201 | 0.62857823 |
| AC064807.1 | 237.227135 | 0.557478142 | 0.36070422 | 1.5455271 | 0.12221882 | 0.28051797 |
| AC064807.2 | 96.5530858 | 1.260495982 | 0.6734748 | 1.87163051 | 0.06125774 | 0.17733048 |
| ST18 | 707.881735 | 0.919865271 | 0.84051681 | 1.09440437 | 0.27377767 | 0.47167804 |
| RB1CC1 | 2336.44427 | -0.474105802 | 0.26604823 | -1.7820295 | 0.07474442 | 0.20287559 |
| ATP6V1H | 1654.55447 | -0.101537759 | 0.29167222 | -0.3481228 | 0.72774794 | 0.84535941 |
| TCEA1 | 608.27576 | 0.179592811 | 0.22838664 | 0.78635427 | 0.43165997 | 0.62432871 |
| LYPLA1 | 964.951338 | -0.027038385 | 0.1996035 | -0.1354605 | 0.89224781 | 0.9437441 |
| MRPL15 | 211.881035 | -0.217248227 | 0.25123783 | -0.8647114 | 0.38719719 | 0.58340157 |
| RP1 | 69.8857143 | -0.326383912 | 0.79464581 | -0.4107288 | 0.68127141 | 0.81457974 |
| XKR4 | 1043.57503 | -4.119237243 | 0.78755126 | -5.230437 | 1.69E-07 | 8.68E-06 |
| TMEM68 | 370.852292 | 0.614881864 | 0.23702779 | 2.59413409 | 0.00948295 | 0.04849815 |
| TGS1 | 403.172445 | -0.029326534 | 0.2267293 | -0.129346 | 0.89708386 | 0.9465267 |
| LYN | 368.056662 | -1.24692112 | 0.37378875 | -3.3358979 | 0.00085024 | 0.00802638 |
| RPS20 | 711.91207 | -0.726521698 | 0.42871731 | -1.6946404 | 0.09014365 | 0.2294701 |
| PLAG1 | 138.294791 | -0.021273633 | 0.44459526 | -0.0478494 | 0.96183624 | 0.98006908 |
| CHCHD7 | 103.765646 | -0.047726261 | 0.30000001 | -0.1590875 | 0.87359991 | 0.9345535 |
| IMPAD1 | 494.474841 | -0.453839953 | 0.23722385 | -1.9131296 | 0.05573147 | 0.16655041 |
| FAM110B | 735.071524 | -1.062461497 | 0.41652375 | -2.5507825 | 0.01074814 | 0.05312686 |
| AC027698.1 | 45.5302305 | -1.63783323 | 0.53351429 | -3.0698957 | 0.00214134 | 0.01619828 |
| UBXN2B | 593.446209 | 0.223911578 | 0.22933727 | 0.97634185 | 0.32889509 | 0.52869297 |
| SDCBP | 750.106013 | -1.436069957 | 0.41644948 | -3.4483653 | 0.00056399 | 0.00583134 |
| NSMAF | 503.843886 | -0.906500545 | 0.42590292 | -2.1284206 | 0.03330223 | 0.11783006 |
| TOX | 1209.02782 | -2.774270949 | 0.84487373 | -3.2836516 | 0.00102472 | 0.00922244 |
| LINC01301 | 198.520491 | 0.177452801 | 0.61644725 | 0.28786372 | 0.77345106 | 0.87422228 |
| RAB2A | 2114.19103 | -0.315333129 | 0.22160865 | -1.4229279 | 0.15475705 | 0.32735053 |
| CHD7 | 1834.73183 | -0.286905299 | 0.16671975 | -1.7208836 | 0.08527194 | 0.22099234 |
| CLVS1 | 74.2762112 | -0.733495219 | 0.31925719 | -2.2975057 | 0.02158994 | 0.08739908 |
| ASPH | 2214.11556 | -0.95853146 | 0.32184315 | -2.9782565 | 0.00289893 | 0.02017039 |
| NKAIN3 | 346.597885 | 0.504173527 | 0.79438999 | 0.63466752 | 0.52564528 | 0.70084961 |
| GGH | 209.529712 | 0.802216368 | 0.39858921 | 2.01263944 | 0.04415257 | 0.14280305 |
| TTPA | 71.1598532 | 2.318858917 | 0.74815803 | 3.09942397 | 0.00193897 | 0.01497324 |
| YTHDF3 | 1024.15182 | -0.058429455 | 0.20684538 | -0.2824789 | 0.77757632 | 0.87743794 |
| AC022639.1 | 40.9278496 | 0.17871327 | 0.39598156 | 0.45131716 | 0.65176098 | 0.79601879 |
| CYP7B1 | 766.119522 | 0.285892045 | 0.3790489 | 0.75423526 | 0.45070797 | 0.64075861 |
| ARMC1 | 404.917007 | 0.556205725 | 0.21687024 | 2.56469367 | 0.01032669 | 0.05174837 |
| MTFR1 | 1659.42535 | 0.219042194 | 0.21361061 | 1.02542748 | 0.30516153 | 0.50529195 |
| PDE7A | 3247.77209 | 0.511774762 | 0.3980242 | 1.28578806 | 0.19851704 | 0.38480603 |
| TRIM55 | 274.064432 | -0.429386012 | 0.48454384 | -0.8861654 | 0.37552838 | 0.57224764 |
| RRS1-AS1 | 57.132738 | 0.485394735 | 0.63667671 | 0.76238809 | 0.44582841 | 0.63715672 |
| ADHFE1 | 638.40067 | 1.342039243 | 0.62113273 | 2.16063198 | 0.03072378 | 0.11102843 |
| VXN | 234.319092 | 2.510907621 | 0.54946952 | 4.56969411 | 4.88E-06 | 0.00014036 |
| VCPIP1 | 396.014605 | -0.35059998 | 0.20086128 | -1.7454831 | 0.08090081 | 0.21362738 |
| SGK3 | 2548.3093 | 1.152988302 | 0.31968259 | 3.60666589 | 0.00031016 | 0.00377224 |
| MCMDC2 | 40.8716912 | 0.406709585 | 0.64350068 | 0.63202666 | 0.52736945 | 0.70181098 |
| PPP1R42 | 95.4701078 | 0.255581602 | 0.49581603 | 0.51547668 | 0.60621996 | 0.76359388 |
| COPS5 | 90.8670755 | -0.317897269 | 0.38541838 | -0.8248109 | 0.40947898 | 0.60345429 |
| CSPP1 | 2417.49731 | 0.682681703 | 0.181929 | 3.75246226 | 0.00017511 | 0.00240499 |
| ARFGEF1 | 3194.08892 | -0.000442062 | 0.15437331 | -0.0028636 | 0.99771519 | 0.99871766 |
| C8orf34 | 70.4215982 | 0.304223482 | 0.44583943 | 0.6823611 | 0.49501065 | 0.67697759 |
| SULF1 | 44.6205274 | -0.894473948 | 0.5715968 | -1.5648687 | 0.1176137 | 0.2735671 |
| SLCO5A1 | 105.141156 | -2.563043775 | 0.74254582 | -3.4516978 | 0.00055707 | 0.0057803 |
| NCOA2 | 6073.30755 | 0.210240386 | 0.24769213 | 0.8487972 | 0.39599415 | 0.59069451 |
| TRAM1 | 1203.43708 | -1.177154472 | 0.31711612 | -3.7120613 | 0.00020558 | 0.00273845 |
| LACTB2-AS1 | 166.368895 | -1.337437204 | 0.71043703 | -1.8825556 | 0.05976062 | 0.17421072 |
| LACTB2 | 1217.33737 | -0.328700744 | 0.363515 | -0.9042288 | 0.36587408 | 0.56342388 |
| XKR9 | 452.457916 | 0.303242556 | 0.55650136 | 0.54490892 | 0.58581616 | 0.7484185 |
| MSC-AS1 | 618.810222 | -1.150629517 | 0.71253277 | -1.6148443 | 0.10634441 | 0.25582008 |
| TERF1 | 611.362184 | 0.227727898 | 0.13500935 | 1.68675651 | 0.09165018 | 0.23229457 |
| AC022893.1 | 149.580669 | 0.101089545 | 0.33590912 | 0.30094314 | 0.76345786 | 0.86800903 |
| SBSPON | 104.755469 | -0.946732375 | 0.68469849 | -1.3826997 | 0.16675697 | 0.34469756 |
| RPL7 | 1057.18253 | -0.608943334 | 0.50625843 | -1.202831 | 0.22904173 | 0.42202916 |
| RDH10 | 663.768722 | -0.33964441 | 0.4858762 | -0.6990349 | 0.48453024 | 0.66827686 |
| RDH10-AS1 | 177.256903 | -0.704323468 | 0.51366805 | -1.3711646 | 0.17032364 | 0.34933803 |
| STAU2 | 2552.02629 | 0.221976577 | 0.22709887 | 0.97744466 | 0.32834906 | 0.5282489 |
| AC027018.1 | 291.198227 | -0.389284751 | 0.52039639 | -0.7480543 | 0.45442741 | 0.64387394 |
| AC022826.2 | 36.266817 | 0.931423852 | 0.47001807 | 1.9816767 | 0.04751544 | 0.15042597 |
| ELOC | 444.690205 | -0.617027291 | 0.31604876 | -1.9523168 | 0.05090061 | 0.15720924 |
| TMEM70 | 101.177513 | -0.724470769 | 0.3485604 | -2.0784655 | 0.0376665 | 0.12833497 |
| JPH1 | 147.128434 | -1.16685013 | 0.50277799 | -2.3208059 | 0.02029732 | 0.08337291 |
| GDAP1 | 743.714542 | 0.538510544 | 0.39571941 | 1.36083935 | 0.17356446 | 0.35339395 |
| MIR2052HG | 172.780328 | 0.165499915 | 0.38175317 | 0.43352598 | 0.6646327 | 0.80542131 |
| CASC9 | 300.726265 | 1.658014555 | 0.71906428 | 2.30579462 | 0.02112211 | 0.08593942 |
| HNF4G | 2477.09919 | 0.681347134 | 0.18975785 | 3.59061373 | 0.0003299 | 0.00395621 |
| PEX2 | 297.43842 | 0.117551765 | 0.17957005 | 0.65462901 | 0.51270664 | 0.69122356 |
| PKIA-AS1 | 49.4925594 | -0.686794155 | 0.99198516 | -0.6923432 | 0.48872184 | 0.67183936 |
| ZC2HC1A | 330.98787 | -0.672521717 | 0.38238098 | -1.758774 | 0.0786159 | 0.20949091 |
| IL7 | 742.732253 | -0.709988758 | 0.59555548 | -1.1921455 | 0.23320422 | 0.42710552 |
| AC083837.1 | 81.0791661 | -1.294279787 | 0.73883288 | -1.7517896 | 0.07980999 | 0.21149728 |
| TPD52 | 2472.33103 | 1.631137147 | 0.4425192 | 3.68602567 | 0.00022778 | 0.0029679 |
| AC034114.2 | 345.020676 | 4.362897783 | 0.71645231 | 6.08958578 | 1.13E-09 | 1.28E-07 |
| ZBTB10 | 556.176948 | 0.143351693 | 0.36660426 | 0.39102571 | 0.69577823 | 0.82478874 |

| | | | | | | |
|---|---|---|---|---|---|---|
| ZNF704 | 1814.60611 | 1.012382768 | 0.29189888 | 3.46826534 | 0.00052383 | 0.0055147 |
| PAG1 | 79.5238606 | -1.943977308 | 0.4605809 | -4.2207076 | 2.44E-05 | 0.00051435 |
| IMPA1 | 412.17854 | 0.990231497 | 0.28393022 | 3.48758752 | 0.0004874 | 0.00526046 |
| ZFAND1 | 429.918869 | -0.039653865 | 0.21315072 | -0.1860367 | 0.85241595 | 0.92213818 |
| CHMP4C | 185.580497 | -0.791880098 | 0.22650376 | -3.4961013 | 0.00047211 | 0.00515223 |
| SNX16 | 283.014324 | -0.56903552 | 0.33911534 | -1.6779999 | 0.09334711 | 0.23524708 |
| RALYL | 4557.65745 | 2.544911888 | 0.64586198 | 3.94033395 | 8.14E-05 | 0.0013246 |
| LRRCC1 | 43.9394372 | 0.480090353 | 0.50611883 | 0.9485724 | 0.34283813 | 0.54344046 |
| E2F5 | 292.437301 | -0.151692693 | 0.25805629 | -0.5878279 | 0.55664779 | 0.72562693 |
| C8orf59 | 262.472797 | -0.277969849 | 0.32430611 | -0.8571218 | 0.39137755 | 0.58655282 |
| CA13 | 649.645079 | 0.796663259 | 0.26521356 | 3.00385569 | 0.00266582 | 0.018982 |
| CA1 | 204.019271 | -0.366430464 | 0.51834911 | -0.7069183 | 0.47961726 | 0.66481737 |
| CA3 | 930.081806 | 0.510022129 | 0.3278723 | 1.55555116 | 0.11981486 | 0.27702309 |
| CA2 | 486.12606 | 0.18470452 | 0.45622306 | 0.40485573 | 0.68558356 | 0.81808078 |
| AC084128.1 | 141.829805 | 6.572824192 | 1.056005 | 6.22423586 | 4.84E-10 | 6.20E-08 |
| SLC7A13 | 982.474556 | 6.267329919 | 1.01271599 | 6.18863531 | 6.07E-10 | 7.56E-08 |
| WWP1 | 3092.84256 | 0.401429569 | 0.21172777 | 1.89597028 | 0.05796397 | 0.17093636 |
| RMDN1 | 783.30861 | 0.509896098 | 0.2945239 | 1.73125544 | 0.08340622 | 0.21821893 |
| CPNE3 | 613.23601 | 0.02992861 | 0.16246264 | 0.18421841 | 0.85384212 | 0.92301331 |
| CNGB3 | 77.641552 | -0.904213385 | 0.40433927 | -2.236274 | 0.02533383 | 0.09732366 |
| CNBD1 | 45.1459322 | 0.372533914 | 0.35122446 | 1.0606719 | 0.28883903 | 0.4887195 |
| AF117829.1 | 1479.26411 | -0.025420163 | 0.40515334 | -0.0627421 | 0.94997189 | 0.97435384 |
| RIPK2 | 287.611193 | -0.926299622 | 0.30281275 | -3.0589849 | 0.00222088 | 0.0166257 |
| OSGIN2 | 413.100355 | -0.439924627 | 0.33544995 | -1.3114464 | 0.18970699 | 0.3738167 |
| NBN | 424.696735 | -0.126665713 | 0.26583801 | -0.4764771 | 0.63373455 | 0.78396092 |
| DECR1 | 1049.99419 | 0.145345851 | 0.32371662 | 0.44899101 | 0.65343814 | 0.79728295 |
| CALB1 | 44.1589696 | 2.860890728 | 0.74390833 | 3.84575707 | 0.00012018 | 0.00180035 |
| LINC00534 | 111.100083 | 0.532128179 | 0.61360587 | 0.86721494 | 0.38582425 | 0.5821126 |
| AB015752.1 | 57.1318015 | -0.036124591 | 0.51316427 | -0.0703958 | 0.94387866 | 0.97115634 |
| TMEM64 | 482.780566 | -0.941556376 | 0.32589822 | -2.8891117 | 0.00386332 | 0.02496676 |
| NECAB1 | 203.931147 | -1.45627398 | 0.67282719 | -2.1644101 | 0.03043289 | 0.11032626 |
| PIP4P2 | 153.503503 | -0.966241797 | 0.29101823 | -3.3202106 | 0.0008995 | 0.00836458 |
| OTUD6B-AS1 | 207.771376 | 0.224457814 | 0.38515627 | 0.58277076 | 0.56004763 | 0.72854361 |
| OTUD6B | 292.488372 | -0.258790582 | 0.22566198 | -1.1468063 | 0.25146169 | 0.44780893 |
| LRRC69 | 610.182163 | -0.434711297 | 0.19200465 | -2.2640665 | 0.02357002 | 0.09257144 |
| SLC26A7 | 150.073594 | 0.160036126 | 0.3876756 | 0.41280938 | 0.67974628 | 0.81423861 |
| AC117834.1 | 85.8950126 | -1.380818548 | 0.69759902 | -1.9793872 | 0.04777243 | 0.1509142 |
| TRIQK | 510.400311 | -0.423793221 | 0.26775845 | -1.5827445 | 0.11347971 | 0.26717044 |
| LINC00535 | 315.316725 | 0.543738216 | 0.32787624 | 1.65836419 | 0.09724398 | 0.24120942 |
| FAM92A | 627.101436 | 0.260744419 | 0.28381792 | 0.91870317 | 0.35825085 | 0.55668795 |
| AC120053.1 | 32.809899 | 0.26267463 | 0.43665215 | 0.60156495 | 0.54746377 | 0.71775092 |
| RBM12B | 419.799401 | -0.124936272 | 0.19727793 | -0.6333008 | 0.52653723 | 0.7009757 |
| TMEM67 | 719.646438 | 0.39698962 | 0.24729847 | 1.60530558 | 0.10842657 | 0.25923653 |
| PDP1 | 233.806335 | -0.819050778 | 0.35429885 | -2.3117512 | 0.0207914 | 0.08486759 |
| GEM | 359.199341 | -3.805364131 | 0.71477989 | -5.3238265 | 1.02E-07 | 5.57E-06 |
| RAD54B | 48.134076 | -0.255824082 | 0.54417898 | -0.4701102 | 0.6382763 | 0.7873338 |
| VIRMA | 540.875447 | 0.06092044 | 0.15854554 | 0.38424569 | 0.70079636 | 0.82796169 |
| AC023632.2 | 48.2745078 | 0.680971697 | 0.3791734 | 1.79593743 | 0.07250447 | 0.19878939 |
| ESRP1 | 59.9595897 | -1.150454766 | 0.76492745 | -1.5040051 | 0.13258006 | 0.29597311 |
| DPY19L4 | 419.364674 | 0.392284392 | 0.25413365 | 1.54361451 | 0.12268175 | 0.28134494 |
| INTS8 | 568.774087 | 0.385670804 | 0.18779515 | 2.05367823 | 0.04000684 | 0.13355427 |
| NDUFAF6 | 681.9126 | 0.408034471 | 0.25125399 | 1.62399198 | 0.10437752 | 0.25237302 |
| TP53INP1 | 559.383829 | 0.348398602 | 0.39356936 | 0.88522796 | 0.3760337 | 0.5726357 |
| PLEKHF2 | 571.236968 | 0.912186826 | 0.47667822 | 1.91363226 | 0.05566716 | 0.16645501 |
| C8orf37-AS1 | 774.206654 | 1.927371982 | 0.69985257 | 2.75396858 | 0.00588774 | 0.03419631 |
| C8orf37 | 58.4875666 | 0.787319807 | 0.48329052 | 1.62908185 | 0.10329569 | 0.25095251 |
| UQCRB | 760.112438 | -0.58574466 | 0.46344808 | -1.2638841 | 0.20627163 | 0.39493857 |
| MTERF3 | 164.593006 | 0.062352655 | 0.22434868 | 0.27792744 | 0.78106806 | 0.87978469 |
| PTDSS1 | 277.860054 | 0.083269656 | 0.3347104 | 0.24878121 | 0.80353003 | 0.89315921 |
| SDC2 | 848.488272 | -0.1746107 | 0.41864359 | -0.4170868 | 0.67661495 | 0.81279385 |
| CPQ | 2310.16834 | 0.570989334 | 0.46161802 | 1.23693033 | 0.21611295 | 0.40679212 |
| MTDH | 2334.73066 | -0.424327242 | 0.36242366 | -1.1708045 | 0.24167738 | 0.43681951 |
| LAPTM4B | 897.716677 | -1.311052316 | 0.34790682 | -3.7684007 | 0.0001643 | 0.00227247 |
| MATN2 | 113.706089 | 0.529203488 | 0.51308834 | 1.03140814 | 0.30234946 | 0.50269269 |
| RPL30 | 858.565288 | -0.589583326 | 0.39331449 | -1.4990125 | 0.13387039 | 0.29778667 |
| ERICH5 | 133.905157 | -0.558193307 | 0.26812249 | -2.0818593 | 0.03735532 | 0.12759268 |
| RIDA | 816.338699 | 1.540685874 | 0.45819414 | 3.36251764 | 0.00077235 | 0.00749761 |
| POP1 | 415.16807 | 0.808781126 | 0.30604672 | 2.64267206 | 0.00822547 | 0.04381907 |
| NIPAL2 | 180.162047 | -0.470268674 | 0.5079409 | -0.9258335 | 0.35453254 | 0.55297878 |
| AP003438.1 | 35.679791 | 0.701069433 | 0.58381277 | 1.20084636 | 0.22981081 | 0.42293458 |
| STK3 | 5068.6543 | -0.04420626 | 0.24550834 | -0.1800601 | 0.85710537 | 0.92470235 |
| AC104986.2 | 289.178671 | 0.044719464 | 0.25227466 | 0.17726499 | 0.85930025 | 0.9261227 |
| VPS13B | 3968.7802 | 0.072524092 | 0.1270561 | 0.5708037 | 0.56813272 | 0.73463601 |
| COX6C | 667.937906 | -0.203755026 | 0.29939743 | -0.6805503 | 0.49615605 | 0.67797937 |
| POLR2K | 157.069364 | -0.597078266 | 0.3674548 | -1.6249026 | 0.10418331 | 0.25231575 |
| SPAG1 | 689.397644 | -0.647564522 | 0.25401981 | -2.5492678 | 0.01079494 | 0.05324441 |
| RNF19A | 4478.21654 | -0.302697636 | 0.33625831 | -0.9001938 | 0.36801714 | 0.56539016 |
| AP000424.1 | 189.245031 | -2.312600955 | 0.79173622 | -2.9209235 | 0.00348995 | 0.02318798 |
| ANKRD46 | 501.601037 | 0.35062627 | 0.33692052 | 1.04067949 | 0.29802433 | 0.49833608 |
| PABPC1 | 1551.88672 | -0.76276983 | 0.43019579 | -1.773076 | 0.07621611 | 0.2054356 |
| YWHAZ | 1850.93007 | -0.779019007 | 0.23657443 | -3.2929129 | 0.00099155 | 0.00897702 |
| AP003469.4 | 49.6280418 | -0.768340195 | 0.39404002 | -1.949904 | 0.05118757 | 0.15766945 |
| ZNF706 | 606.669489 | -0.491564056 | 0.20902339 | -2.351718 | 0.01868694 | 0.07858789 |
| GRHL2 | 93.9949685 | 0.12376279 | 0.78522848 | 0.15761373 | 0.87476118 | 0.93490877 |
| NCALD | 333.269547 | 1.666345713 | 0.43314511 | 3.84708425 | 0.00011953 | 0.00179259 |
| RRM2B | 299.694632 | -0.266987585 | 0.25979984 | -1.0276665 | 0.30410675 | 0.50434255 |
| UBR5-AS1 | 58.3835686 | -0.059018134 | 0.50052878 | -0.1179116 | 0.90613773 | 0.95080991 |
| UBR5 | 2840.04183 | 0.286573776 | 0.18164394 | 1.57766771 | 0.11464197 | 0.26884473 |
| KLF10 | 364.528707 | -0.130175852 | 0.3662162 | -0.3554618 | 0.72224369 | 0.84228022 |
| AZIN1 | 575.643353 | -0.230885524 | 0.22200676 | -1.0399932 | 0.29834306 | 0.49840643 |
| AZIN1-AS1 | 642.081866 | 0.591015446 | 0.51627078 | 1.14477803 | 0.25230112 | 0.44873698 |
| AP003354.2 | 96.6625946 | 0.084063881 | 0.54938506 | 0.1530145 | 0.87838684 | 0.93705514 |
| ATP6V1C1 | 1915.69365 | -0.240334849 | 0.35747418 | -0.6723139 | 0.5013839 | 0.68247686 |
| BAALC-AS1 | 167.724807 | 0.182246631 | 0.52994751 | 0.34389562 | 0.73092478 | 0.84689653 |
| FZD6 | 538.162146 | 0.01260852 | 0.20806894 | 0.0605978 | 0.95167952 | 0.97544838 |
| SLC25A32 | 373.282488 | 0.023985066 | 0.24140608 | 0.09935568 | 0.92085587 | 0.9574582 |
| DCAF13 | 284.095 | -0.150452826 | 0.36588476 | -0.4112028 | 0.68092385 | 0.81446268 |
| RIMS2 | 1376.92047 | -4.395997614 | 0.84484568 | -5.2033143 | 1.96E-07 | 9.54E-06 |
| DPYS | 6905.69449 | 1.232628399 | 0.63629273 | 1.93720334 | 0.05272049 | 0.16051116 |
| LRP12 | 379.63418 | 0.39479885 | 0.25859333 | 1.52671705 | 0.12683139 | 0.28790372 |
| ZFPM2 | 110.39233 | -0.738667269 | 0.49103422 | -1.5043091 | 0.13250179 | 0.29584656 |
| AC027031.2 | 41.1781871 | -1.68456847 | 0.51530171 | -3.2690915 | 0.00107893 | 0.00954126 |
| OXR1 | 12335.573 | 0.634227861 | 0.34599413 | 1.83305958 | 0.06679371 | 0.18811206 |
| AC090579.1 | 1032.84964 | -0.376226157 | 0.62779103 | -0.5992856 | 0.54898242 | 0.71891678 |
| ANGPT1 | 120.573322 | 2.996563411 | 0.41894761 | 7.15259701 | 8.52E-13 | 2.38E-10 |
| EIF3E | 1512.85871 | -0.040636407 | 0.1838983 | -0.2209722 | 0.8251141 | 0.90642548 |
| EMC2 | 757.64781 | 0.173626503 | 0.39768182 | 0.43659653 | 0.66240398 | 0.803573 |
| NUDCD1 | 415.915073 | -0.343211003 | 0.29786378 | -1.1522415 | 0.24922185 | 0.44538665 |
| ENY2 | 497.704584 | -0.496365208 | 0.3053874 | -1.6253624 | 0.10408535 | 0.25224504 |
| EBAG9 | 179.595948 | 0.362460595 | 0.28570253 | 1.26866427 | 0.20456083 | 0.39281526 |
| LINC01609 | 23.6686049 | 1.633269316 | 0.88418821 | 1.84719644 | 0.06471867 | 0.18435139 |
| TRPS1 | 8619.06085 | 0.44397566 | 0.27804714 | 1.596764 | 0.11031832 | 0.26206816 |
| EIF3H | 1061.4195 | -0.29281956 | 0.24912454 | -1.1753943 | 0.23983702 | 0.43490488 |

| | | | | | | |
|---|---|---|---|---|---|---|
| UTP23 | 187.349311 | -0.136899148 | 0.30188149 | -0.4534864 | 0.65019854 | 0.79482956 |
| RAD21 | 637.038503 | -0.224200699 | 0.18435972 | -1.2161046 | 0.22394509 | 0.41621903 |
| SLC30A8 | 1350.98031 | 4.607531731 | 0.85901497 | 5.36373858 | 8.15E-08 | 4.66E-06 |
| MED30 | 85.0243183 | -0.031758917 | 0.21831998 | -0.1454696 | 0.88434013 | 0.94002757 |
| EXT1 | 6197.52766 | -0.861408784 | 0.30283556 | -2.844477 | 0.00444844 | 0.02777893 |
| SAMD12 | 4402.85341 | -0.212617041 | 0.34711943 | -0.6125184 | 0.54019482 | 0.71176107 |
| AC023590.1 | 300.905875 | -0.955824113 | 0.50716319 | -1.884648 | 0.05947737 | 0.17371753 |
| SAMD12-AS1 | 284.499907 | -1.033598868 | 0.4429038 | -2.3336871 | 0.01961211 | 0.08126458 |
| TNFRSF11B | 943.665342 | -1.275599902 | 0.64729085 | -1.970675 | 0.04876106 | 0.15301462 |
| COLEC10 | 162.164002 | -1.049027038 | 0.62487355 | -1.6787829 | 0.09319436 | 0.23499174 |
| MAL2 | 279.57998 | -1.152072264 | 0.38248604 | -3.0120635 | 0.00259478 | 0.01870827 |
| ENPP2 | 1093.6931 | 1.097628238 | 0.40275248 | 2.72531714 | 0.00642398 | 0.03651123 |
| TAF2 | 1219.86325 | 0.189890303 | 0.2406103 | 0.78920271 | 0.42999354 | 0.62265243 |
| DEPTOR | 2810.8666 | 0.217132475 | 0.54351639 | 0.39949573 | 0.68952797 | 0.82043055 |
| AC091563.1 | 149.653451 | 0.042376009 | 0.43147505 | 0.09821196 | 0.92176399 | 0.95811091 |
| COL14A1 | 154.384208 | 0.883965263 | 0.55445839 | 1.59428603 | 0.11087198 | 0.26306427 |
| MRPL13 | 244.930957 | -0.165774785 | 0.27130919 | -0.611018 | 0.54118769 | 0.71231293 |
| MTBP | 90.2134552 | 0.004898858 | 0.39548462 | 0.01238698 | 0.99011688 | 0.99507452 |
| SNTB1 | 1701.27827 | -1.214689694 | 0.55192434 | -2.2008265 | 0.02774831 | 0.10358009 |
| ZHX2 | 3542.15629 | -0.415226491 | 0.28590478 | -1.4523244 | 0.14641143 | 0.31539548 |
| AC104316.1 | 127.513017 | -1.787929339 | 0.4838445 | -3.6952561 | 0.00021967 | 0.00287579 |
| DERL1 | 379.28463 | -0.957057839 | 0.23725256 | -4.03392 | 5.49E-05 | 0.00097774 |
| TBC1D31 | 126.69692 | 0.245844355 | 0.23507527 | 1.04581121 | 0.29564821 | 0.49559129 |
| C8orf76 | 86.4987244 | 0.204403731 | 0.27419791 | 0.74546056 | 0.45599334 | 0.64515904 |
| ZHX1-C8orf7( | 42.6677379 | 0.764130667 | 0.45853237 | 1.66647049 | 0.09561974 | 0.23869237 |
| ATAD2 | 88.0906772 | -0.134570153 | 0.27266666 | -0.4935336 | 0.62163561 | 0.77531932 |
| WDYHV1 | 139.99058 | -0.041233977 | 0.39574879 | -0.1041923 | 0.91701674 | 0.95564119 |
| FBXO32 | 375.063927 | -1.926754104 | 0.53757257 | -3.5841749 | 0.00033815 | 0.0040234 |
| FAM91A1 | 980.837734 | 0.155972541 | 0.18245961 | 0.85483324 | 0.39264348 | 0.58757307 |
| FER1L6 | 330.569552 | 1.980331537 | 0.95519472 | 2.07322288 | 0.03815154 | 0.12953758 |
| TMEM65 | 1227.17538 | -0.51298299 | 0.25546501 | -2.0080362 | 0.04463944 | 0.1438823 |
| TRMT12 | 99.5773997 | 0.296307222 | 0.29164042 | 1.01600191 | 0.30962848 | 0.50986035 |
| RNF139-AS1 | 47.5350184 | -0.044979328 | 0.35817067 | -0.1255807 | 0.90006383 | 0.94743919 |
| RNF139 | 149.25482 | 0.090869639 | 0.19297491 | 0.47088835 | 0.63772047 | 0.78700229 |
| TATDN1 | 721.669729 | -0.150318655 | 0.15132363 | -0.9933588 | 0.32053515 | 0.52105682 |
| NDUFB9 | 385.167561 | -0.372005252 | 0.27498945 | -1.3527983 | 0.17612008 | 0.35637407 |
| MTSS1 | 1776.21713 | 1.613160701 | 0.38577325 | 4.1816292 | 2.89E-05 | 0.00058513 |
| SQLE | 184.746074 | -2.217336554 | 0.59910498 | -3.7010818 | 0.00021468 | 0.00283219 |
| WASHC5 | 745.694603 | 0.13142344 | 0.27091436 | 0.48511064 | 0.62759786 | 0.77977555 |
| NSMCE2 | 4383.32404 | -0.002377782 | 0.27205369 | -0.0087401 | 0.99302648 | 0.99644319 |
| TRIB1 | 242.343446 | -0.830248125 | 0.38861044 | -2.1364535 | 0.03264247 | 0.11618548 |
| AC091114.1 | 43.372765 | 1.620857243 | 0.58022414 | 2.79350191 | 0.00521407 | 0.03130499 |
| FAM84B | 329.777823 | 0.326699249 | 0.44521433 | 0.73380219 | 0.46306931 | 0.65120457 |
| PCAT1 | 4734.64844 | 0.187238752 | 0.26426417 | 0.70852872 | 0.47861698 | 0.66413513 |
| MYC | 154.629714 | -3.688615803 | 0.5763993 | -6.3994106 | 1.56E-10 | 2.32E-08 |
| PVT1 | 801.237391 | -2.786482522 | 0.46809169 | -5.9528562 | 2.64E-09 | 2.64E-07 |
| CCDC26 | 261.143969 | -0.276995841 | 0.75540854 | -0.3666835 | 0.71385512 | 0.83637691 |
| AC103718.1 | 44.5797015 | -0.228506311 | 0.83395191 | -0.2740042 | 0.78408141 | 0.88144049 |
| FAM49B | 1810.33898 | -0.874461989 | 0.26314371 | -3.3231347 | 0.00089012 | 0.00829427 |
| ASAP1 | 2275.31567 | -1.070541724 | 0.31045477 | -3.4483018 | 0.00056412 | 0.00583134 |
| ADCY8 | 179.412715 | -0.250510997 | 0.85794111 | -0.2919909 | 0.77029358 | 0.8723487 |
| EFR3A | 1880.73741 | 0.151234826 | 0.23250473 | 0.65045913 | 0.5153957 | 0.69315508 |
| KCNQ3 | 1115.1577 | -1.83060885 | 0.41445484 | -4.4169078 | 1.00E-05 | 0.00025478 |
| LRRC6 | 1175.19962 | 0.116183785 | 0.40912672 | 0.28397995 | 0.77642575 | 0.87658111 |
| PHF20L1 | 1078.9634 | 0.295879668 | 0.12396886 | 2.38672567 | 0.01699917 | 0.07365812 |
| TG | 258.019478 | 0.477178916 | 0.48894171 | 0.97594234 | 0.32909304 | 0.52891681 |
| SLA | 61.3481978 | -0.383960842 | 0.77067576 | -0.4982132 | 0.61833377 | 0.77310052 |
| NDRG1 | 6655.33802 | -1.948391193 | 0.62772182 | -3.1039086 | 0.00190982 | 0.0147898 |
| ST3GAL1 | 1576.10521 | -0.213208641 | 0.36983627 | -0.5764947 | 0.56428087 | 0.73149731 |
| AC105180.1 | 71.2126509 | 0.088737144 | 0.61832995 | 0.14351099 | 0.88588663 | 0.94094134 |
| ZFAT | 1537.74844 | 1.236845543 | 0.46417257 | 2.66462437 | 0.00770744 | 0.04169133 |
| AC083843.3 | 249.120531 | -0.397069794 | 0.43479362 | -0.9132374 | 0.36111769 | 0.55908131 |
| KHDRBS3 | 2309.59342 | -1.421644449 | 0.5110661 | -2.7817233 | 0.00540711 | 0.03212627 |
| LINC02055 | 106.114394 | -0.144290338 | 1.03612775 | -0.1392592 | 0.88924532 | 0.94267382 |
| AC046195.1 | 483.443161 | 0.817131537 | 0.95661385 | 0.85419162 | 0.39299883 | 0.58788987 |
| FAM135B | 555.767628 | -1.345758348 | 0.79046514 | -1.7024892 | 0.08866371 | 0.2270765 |
| TRAPPC9 | 3583.41582 | -0.066907441 | 0.14678287 | -0.4558259 | 0.64851515 | 0.79376872 |
| AC107375.1 | 68.9768138 | 0.748234783 | 0.33001941 | 2.26724476 | 0.02337528 | 0.09196468 |
| CHRAC1 | 108.069513 | -0.697626407 | 0.38066126 | -1.8326698 | 0.06685168 | 0.18817227 |
| AGO2 | 980.010734 | -0.7955079 | 0.39006855 | -2.0394054 | 0.04140958 | 0.13668488 |
| PTK2 | 6494.74976 | -0.128800226 | 0.20737878 | -0.6210868 | 0.5345425 | 0.70751736 |
| DENND3 | 283.834085 | -1.125031073 | 0.40257678 | -2.7945751 | 0.0051968 | 0.03122862 |
| SLC45A4 | 738.968212 | -0.069627298 | 0.43133658 | -0.1614222 | 0.87176089 | 0.93353404 |
| PTP4A3 | 70.0920085 | -1.853550381 | 0.67096894 | -2.762498 | 0.00573609 | 0.03352283 |
| TSNARE1 | 437.372674 | 0.147785544 | 0.28202474 | 0.52401624 | 0.60026725 | 0.76000953 |
| JRK | 156.525837 | 0.450362339 | 0.36567616 | 1.2315879 | 0.21810306 | 0.40924554 |
| THEM6 | 92.1498667 | -0.16459827 | 0.3876125 | -0.4246464 | 0.67109443 | 0.8093165 |
| LY6E-DT | 252.164804 | -0.556193054 | 0.75743479 | -0.7343115 | 0.46275893 | 0.6511021 |
| LY6E | 812.932338 | -0.705514896 | 0.65939161 | -1.0699482 | 0.2846426 | 0.48398841 |
| AC138696.1 | 60.6914751 | 0.406783505 | 0.45557135 | 0.89290845 | 0.3719062 | 0.56932302 |
| GLI4 | 141.193957 | 0.257415944 | 0.30472664 | 0.84474383 | 0.39825388 | 0.59258207 |
| ZNF696 | 48.7582261 | 0.259393159 | 0.38762002 | 0.66919443 | 0.50337146 | 0.68404435 |
| TOP1MT | 231.97118 | 0.602384357 | 0.22792578 | 2.64289697 | 0.00822 | 0.043807 |
| RHPN1 | 313.091057 | -0.037566466 | 0.4689725 | -0.0801038 | 0.93615472 | 0.96587767 |
| ZC3H3 | 650.020284 | 0.447855024 | 0.39564605 | 1.13195879 | 0.25765176 | 0.45499004 |
| AC067930.4 | 59.6868089 | 1.163035171 | 0.70040185 | 1.66052555 | 0.09680877 | 0.24043447 |
| GSDMD | 381.877369 | -0.267712265 | 0.53535061 | -0.500069 | 0.61702646 | 0.77210284 |
| MROH6 | 57.509643 | 0.036612942 | 0.62569411 | 0.05851572 | 0.95333784 | 0.97649094 |
| NAPRT | 249.045267 | 1.146248292 | 0.30613321 | 3.74427949 | 0.00018091 | 0.00246008 |
| EEF1D | 1133.12635 | -0.647661623 | 0.42634496 | -1.5191023 | 0.12873674 | 0.29046822 |
| TIGD5 | 44.3407404 | -1.087754009 | 0.31109318 | -3.4965536 | 0.00047131 | 0.0051476 |
| PYCR3 | 41.1475771 | -0.128201356 | 0.60974521 | -0.210254 | 0.83346945 | 0.91197939 |
| TSTA3 | 178.262924 | -0.973282558 | 0.51684432 | -1.8831252 | 0.0596834 | 0.1740967 |
| ZNF623 | 250.310317 | -0.090908136 | 0.35874326 | -0.2534072 | 0.79995354 | 0.89126832 |
| ZNF707 | 85.2942149 | 0.650953942 | 0.33462335 | 1.94533331 | 0.05173488 | 0.15864204 |
| FAM83H | 145.464302 | -0.487856916 | 0.39746506 | -1.2274209 | 0.21966444 | 0.41133067 |
| IQANK1 | 117.00664 | 0.37619292 | 0.55773297 | 0.67450365 | 0.49999117 | 0.68139272 |
| SCRIB | 152.157188 | -0.051276724 | 0.19306739 | -0.2655898 | 0.79055516 | 0.88610185 |
| PUF60 | 392.658162 | -0.546995981 | 0.39886061 | -1.3713963 | 0.17025144 | 0.34924222 |
| NRBP2 | 826.060198 | -1.339308472 | 0.58361169 | -2.2948623 | 0.02174102 | 0.08764991 |
| EPPK1 | 149.547099 | 0.031597699 | 0.65554807 | 0.04820043 | 0.96155651 | 0.97988733 |
| PLEC | 561.345786 | -0.225972304 | 0.27885209 | -0.8103662 | 0.41772975 | 0.61081943 |
| PARP10 | 512.965975 | 1.137831468 | 0.46340575 | 2.45536759 | 0.01407406 | 0.06404818 |
| GRINA | 504.037369 | 0.50562051 | 0.33152781 | 1.52512245 | 0.12722856 | 0.28832864 |
| SPATC1 | 113.743325 | 0.015254488 | 0.84906827 | 0.01796615 | 0.98566586 | 0.99298265 |
| OPLAH | 117.461087 | 1.270480336 | 0.43726825 | 2.90549411 | 0.00366674 | 0.02399044 |
| EXOSC4 | 66.317074 | -1.083296243 | 0.47343822 | -2.288147 | 0.02212896 | 0.08872845 |
| GPAA1 | 300.003347 | 0.472910755 | 0.39077562 | 1.21018489 | 0.22620795 | 0.41880492 |
| CYC1 | 347.581551 | -0.533479485 | 0.41662499 | -1.2804788 | 0.20037678 | 0.387495 |
| SHARPIN | 316.635385 | 0.132240318 | 0.1661004 | 0.79614689 | 0.42594667 | 0.61876852 |
| MAF1 | 249.0201 | -0.871935883 | 0.41931771 | -2.0794158 | 0.03757915 | 0.12813367 |
| HGH1 | 39.42282 | 0.228404352 | 0.49976763 | 0.4570211 | 0.64765588 | 0.79339712 |
| MROH1 | 1188.51041 | -1.052855842 | 0.52618811 | -2.0009115 | 0.04540193 | 0.14577752 |

| | | | | | | |
|---|---|---|---|---|---|---|
| BOP1 | 54.9833818 | -0.056898353 | 0.56404006 | -0.1008764 | 0.91964855 | 0.95690112 |
| SCX | 85.9243052 | -0.514699997 | 0.64107903 | -0.8028651 | 0.42205269 | 0.61569564 |
| HSF1 | 456.768472 | -0.51524766 | 0.28440594 | -1.8116628 | 0.07003832 | 0.19421712 |
| DGAT1 | 110.688518 | 0.528389372 | 0.32654024 | 1.61814474 | 0.10563141 | 0.2545964 |
| SLC52A2 | 129.185885 | -0.966048684 | 0.48110072 | -2.0079968 | 0.04464364 | 0.1438823 |
| FBXL6 | 55.1619011 | -0.565102335 | 0.40614409 | -1.3913839 | 0.16410905 | 0.34097966 |
| ADCK5 | 76.8876255 | 0.581439717 | 0.27652709 | 2.10265012 | 0.03549637 | 0.12283481 |
| CPSF1 | 161.484403 | 0.536698567 | 0.34210745 | 1.56880115 | 0.11669428 | 0.27215379 |
| SLC39A4 | 133.800117 | 0.097304476 | 0.40594764 | 0.23969711 | 0.81056507 | 0.89723132 |
| VPS28 | 504.505498 | -0.581593747 | 0.35623804 | -1.6325987 | 0.10255343 | 0.25025567 |
| TONSL | 34.9996981 | 0.299763626 | 0.42024161 | 0.71331257 | 0.47565236 | 0.66135476 |
| CYHR1 | 276.560801 | 0.748836292 | 0.17377542 | 4.3092188 | 1.64E-05 | 0.00037806 |
| KIFC2 | 274.845033 | 0.676970243 | 0.31928119 | 2.12029479 | 0.03398119 | 0.11939568 |
| PPP1R16A | 776.858476 | 0.279010497 | 0.25010485 | 1.11557412 | 0.26460448 | 0.46244212 |
| GPT | 356.53452 | 2.064267525 | 0.59946649 | 3.4435078 | 0.00057422 | 0.00590903 |
| MFSD3 | 105.32283 | 0.745713281 | 0.32848844 | 2.27013555 | 0.02319936 | 0.0915353 |
| LRRC14 | 71.4598788 | 0.436609395 | 0.25823484 | 1.69074549 | 0.09088542 | 0.2308828 |
| ARHGAP39 | 197.524458 | 0.033458294 | 0.38348211 | 0.08724864 | 0.93047388 | 0.96300252 |
| ZNF251 | 780.192564 | 0.605295135 | 0.27144324 | 2.22991421 | 0.02575314 | 0.09846535 |
| ZNF34 | 53.5719117 | 0.605859873 | 0.31958102 | 1.8957943 | 0.05798725 | 0.17096822 |
| RPL8 | 1528.61949 | -1.016460579 | 0.46636715 | -2.1795287 | 0.02929242 | 0.1072636 |
| ZNF517 | 40.7104769 | 0.542775136 | 0.40623011 | 1.33612729 | 0.18150767 | 0.36373182 |
| ZNF7 | 263.786743 | -0.226215898 | 0.24678456 | -0.9166534 | 0.35932431 | 0.55773494 |
| COMMD5 | 102.317446 | -0.895504242 | 0.30919417 | -2.896252 | 0.00377649 | 0.02450963 |
| ZNF250 | 316.156609 | -0.768178504 | 0.50064831 | -1.5343675 | 0.12493926 | 0.28485404 |
| ZNF16 | 184.515639 | 0.507837328 | 0.36899299 | 1.37627906 | 0.16873525 | 0.34732754 |
| C8orf33 | 127.919731 | -0.675134609 | 0.43699932 | -1.5449328 | 0.12236251 | 0.28080075 |
| WASHC1 | 1218.62503 | 0.639029225 | 0.14221555 | 4.49338493 | 7.01E-06 | 0.0001889 |
| CBWD1 | 427.84323 | 0.410995309 | 0.30450235 | 1.34972785 | 0.1771033 | 0.35762447 |
| C9orf66 | 125.333893 | 1.977213234 | 0.72991585 | 2.70882353 | 0.00675222 | 0.03787496 |
| DOCK8 | 3073.93559 | 0.347772233 | 0.40431862 | 0.86014401 | 0.38970967 | 0.58526903 |
| AL161725.1 | 55.4646595 | 0.271268969 | 0.65970512 | 0.4111973 | 0.68092786 | 0.81446268 |
| KANK1 | 6864.97893 | 0.350865821 | 0.1581169 | 2.21902795 | 0.02648482 | 0.10036701 |
| SMARCA2 | 1207.89593 | 0.316145146 | 0.21288315 | 1.48506423 | 0.13752681 | 0.30315293 |
| VLDLR-AS1 | 416.644835 | -0.467304007 | 0.51773604 | -0.9025912 | 0.36674288 | 0.56412801 |
| VLDLR | 361.570424 | -1.095791524 | 0.33688673 | -3.2527002 | 0.00114314 | 0.00998663 |
| KCNV2 | 65.0834766 | -0.48515727 | 0.55984812 | -0.8665873 | 0.38616817 | 0.58237187 |
| PUM3 | 3727.95689 | 0.567376572 | 0.36999571 | 1.53346797 | 0.12516059 | 0.28521633 |
| RFX3 | 1729.08438 | 0.296700783 | 0.28916657 | 1.02605492 | 0.3048657 | 0.5051123 |
| RFX3-AS1 | 1793.71957 | 0.488416081 | 0.39567877 | 1.23437526 | 0.2170631 | 0.40796434 |
| GLIS3 | 11029.8095 | -0.516688425 | 0.24730182 | -2.089303 | 0.03668046 | 0.12582058 |
| SLC1A1 | 4135.80457 | 0.64081604 | 0.31133429 | 2.05828933 | 0.03956237 | 0.13252233 |
| SPATA6L | 568.399911 | -0.716805496 | 0.37996073 | -1.8865252 | 0.05922422 | 0.17320622 |
| PLPP6 | 39.1961302 | -0.162522993 | 0.3043294 | -0.5340365 | 0.59331631 | 0.75440985 |
| CDC37L1 | 543.924032 | 0.285792294 | 0.17520995 | 1.63114196 | 0.10286037 | 0.25038257 |
| AK3 | 1540.61908 | 0.447498635 | 0.36435065 | 1.22820868 | 0.21936864 | 0.41094519 |
| RCL1 | 677.608529 | 0.335969295 | 0.26278316 | 1.27850391 | 0.20107181 | 0.38812445 |
| JAK2 | 1375.57499 | -0.015727851 | 0.17427588 | -0.0902469 | 0.92809105 | 0.96208348 |
| PLGRKT | 348.811793 | -0.128093414 | 0.25345682 | -0.5053856 | 0.61328806 | 0.76933839 |
| RIC1 | 1445.9105 | -0.013661067 | 0.17947474 | -0.0761169 | 0.93932606 | 0.96748891 |
| ERMP1 | 384.077203 | 0.229906218 | 0.31938517 | 0.71983998 | 0.47162352 | 0.65790186 |
| KIAA2026 | 1969.72703 | 0.181613723 | 0.15914977 | 1.1411498 | 0.25380759 | 0.45069836 |
| RANBP6 | 159.855357 | -0.292609362 | 0.25732667 | -1.1371125 | 0.25549127 | 0.45251569 |
| UHRF2 | 880.17458 | -0.161691353 | 0.2550045 | -0.6340726 | 0.52603347 | 0.70084961 |
| AL133480.1 | 79.2098039 | -0.694256427 | 0.45357696 | -1.5306254 | 0.125862 | 0.28619908 |
| GLDC | 2406.96976 | 1.631073662 | 0.37046074 | 4.40282459 | 1.07E-05 | 0.00026824 |
| AL162411.1 | 511.810263 | 0.873083183 | 0.241458 | 3.61588008 | 0.00029933 | 0.00367264 |
| AL354707.1 | 40.2944122 | -1.087772341 | 0.78191186 | -1.39117 | 0.16417387 | 0.34106264 |
| KDM4C | 4958.76641 | -0.330006796 | 0.14969332 | -2.2045526 | 0.0274855 | 0.10294526 |
| DMAC1 | 151.183704 | 0.089259658 | 0.28946901 | 0.30835652 | 0.75781106 | 0.86418282 |
| PTPRD | 8095.60519 | 1.201390094 | 0.39284797 | 3.05815525 | 0.00222704 | 0.01665361 |
| LURAP1L-AS | 58.6436791 | -2.689063322 | 0.72812173 | -3.6931508 | 0.00022149 | 0.00289143 |
| LURAP1L | 329.704857 | -2.593217054 | 0.72286646 | -3.5874082 | 0.00033398 | 0.00399466 |
| MPDZ | 1311.29474 | 0.18561265 | 0.30875586 | 0.60116316 | 0.54773132 | 0.71803301 |
| NFIB | 9869.48894 | 0.95496793 | 0.35281857 | 2.7066827 | 0.00679592 | 0.0380366 |
| AL136366.1 | 43.0748469 | -0.13948475 | 0.46675633 | -0.2988385 | 0.76506329 | 0.8688254 |
| ZDHHC21 | 1527.3096 | 0.400757962 | 0.2103392 | 1.90529376 | 0.0567419 | 0.16867516 |
| FREM1 | 637.188704 | 3.124232059 | 0.75387437 | 4.14423435 | 3.41E-05 | 0.00066954 |
| TTC39B | 2249.862 | 0.587096505 | 0.5141045 | 1.14197892 | 0.25346278 | 0.45036201 |
| SNAPC3 | 847.364448 | 0.036580231 | 0.23062298 | 0.15861486 | 0.87397232 | 0.93468316 |
| PSIP1 | 475.1466 | 0.620990851 | 0.26306526 | 2.36059617 | 0.01824559 | 0.07738003 |
| CCDC171 | 815.261446 | 1.084254718 | 0.48742563 | 2.22445159 | 0.02611808 | 0.09938197 |
| BNC2 | 11483.2511 | 0.903601577 | 0.43644935 | 2.07034692 | 0.03841987 | 0.13022283 |
| AL162725.2 | 561.438807 | 2.214618625 | 1.0238646 | 2.1629995 | 0.03054122 | 0.11054356 |
| CNTLN | 980.585464 | 0.520364194 | 0.30459577 | 1.7083763 | 0.08756654 | 0.22533749 |
| SH3GL2 | 1720.73296 | 0.741494247 | 0.45168597 | 1.64161452 | 0.10066992 | 0.24689257 |
| ADAMTSL1 | 126.783115 | 0.48213691 | 0.41151157 | 1.17162419 | 0.24134797 | 0.43643228 |
| SAXO1 | 90.9155684 | 0.574500132 | 0.42907649 | 1.33892243 | 0.18059593 | 0.36235848 |
| RRAGA | 182.730032 | -1.558219035 | 0.31778136 | -4.9034312 | 9.42E-07 | 3.68E-05 |
| HAUS6 | 322.477931 | -0.619518952 | 0.23533386 | -2.6325109 | 0.00847563 | 0.04464852 |
| PLIN2 | 3559.12082 | -1.740869101 | 0.55338721 | -3.1458427 | 0.00165609 | 0.01328264 |
| DENND4C | 2861.43455 | -0.244076143 | 0.18836448 | -1.2957652 | 0.19505637 | 0.38021212 |
| RPS6 | 1710.19055 | -0.908475015 | 0.48217799 | -1.8841072 | 0.05955048 | 0.17383682 |
| ACER2 | 186.653901 | -0.336695102 | 0.43963534 | -0.7658509 | 0.44376505 | 0.63517875 |
| MLLT3 | 1729.58156 | -0.715008019 | 0.37816997 | -1.8907054 | 0.05866367 | 0.17214793 |
| FOCAD | 1532.85137 | -0.496851297 | 0.25508945 | -1.9477532 | 0.0514445 | 0.15796366 |
| KLHL9 | 210.857112 | -0.244459719 | 0.26534939 | -0.9212748 | 0.35690696 | 0.55573369 |
| DMRTA1 | 133.742343 | 0.480253234 | 0.3754242 | 1.27922822 | 0.2008167 | 0.38798494 |
| LINC01239 | 990.554327 | 0.870218277 | 0.71317242 | 1.22020743 | 0.22238625 | 0.41467125 |
| AL391117.1 | 4151.85723 | 2.582080266 | 0.67103608 | 3.84790082 | 0.00011913 | 0.00178953 |
| TUSC1 | 55.5460305 | -0.78862354 | 0.40036584 | -1.9697573 | 0.04886619 | 0.15320423 |
| CAAP1 | 1175.69741 | 0.25759699 | 0.25712011 | 1.00185469 | 0.31641378 | 0.51644245 |
| PLAA | 402.446922 | -0.353803559 | 0.25764386 | -1.3732272 | 0.16968172 | 0.34859523 |
| IFT74 | 908.471231 | 0.402265607 | 0.16542378 | 2.43172783 | 0.01502699 | 0.06713658 |
| LRRC19 | 690.409294 | 0.628689581 | 0.52202213 | 1.20433512 | 0.22846008 | 0.42141062 |
| MOB3B | 1513.008 | 0.221848735 | 0.40447384 | 0.54848723 | 0.58335739 | 0.74625103 |
| C9orf72 | 335.954291 | -1.113642286 | 0.25931658 | -4.294528 | 1.75E-05 | 0.00039994 |
| LINGO2 | 955.001384 | 1.007578058 | 0.77206322 | 1.30504606 | 0.19187717 | 0.37620659 |
| ACO1 | 2281.64867 | 0.31421402 | 0.31050584 | 1.01194237 | 0.31156561 | 0.5116978 |
| DDX58 | 1394.29408 | -0.833992666 | 0.4694555 | -1.7765106 | 0.0756488 | 0.20439939 |
| TOPORS | 330.897845 | -0.086935525 | 0.24467925 | -0.355304 | 0.72236183 | 0.84228022 |
| SMIM27 | 54.8454967 | -0.106745029 | 0.40263172 | -0.2651183 | 0.79091835 | 0.88629048 |
| NDUFB6 | 325.730635 | 0.297714311 | 0.13839931 | 2.15112563 | 0.03146628 | 0.11296709 |
| APTX | 991.531704 | 0.288584185 | 0.29171706 | 0.98926056 | 0.32253567 | 0.52288545 |
| DNAJA1 | 583.656227 | -0.963262852 | 0.35377367 | -2.7228223 | 0.00647269 | 0.03664647 |
| SMU1 | 347.297112 | 0.16485001 | 0.14081382 | 1.17069485 | 0.24172145 | 0.43681951 |
| B4GALT1 | 1497.83458 | 0.33530644 | 0.37743703 | 0.88837716 | 0.37433791 | 0.57138638 |
| BAG1 | 293.749706 | -0.356117712 | 0.31214588 | -1.1408695 | 0.25392423 | 0.45078869 |
| CHMP5 | 300.033242 | -0.360856681 | 0.31099058 | -1.160346 | 0.24590796 | 0.44147485 |
| NFX1 | 1762.80065 | -0.300671511 | 0.27683349 | -1.0861096 | 0.27743051 | 0.47540192 |
| AQP7 | 710.850235 | 1.582607859 | 0.5618697 | 2.81668128 | 0.00485227 | 0.0295863 |
| AQP3 | 484.458713 | -0.452807599 | 0.60776591 | -0.7450362 | 0.45624984 | 0.64534938 |
| NOL6 | 56.4323295 | -0.866307458 | 0.48661123 | -1.7802866 | 0.07502906 | 0.2033663 |

| | | | | | | |
|---|---|---|---|---|---|---|
| PRSS3 | 53.831721 | -0.285760179 | 0.59817412 | -0.4777207 | 0.63284898 | 0.78360082 |
| UBE2R2-AS1 | 53.9035795 | -1.01939082 | 0.57959211 | -1.7588073 | 0.07861025 | 0.20949091 |
| UBE2R2 | 2995.49191 | 0.224717712 | 0.30987971 | 0.72517724 | 0.4683433 | 0.65572846 |
| UBAP2 | 1651.13917 | -0.252656147 | 0.32335574 | -0.7813566 | 0.43459278 | 0.62698276 |
| DCAF12 | 275.980247 | 0.142559223 | 0.25674368 | 0.55525894 | 0.57871756 | 0.74239416 |
| UBAP1 | 2317.21896 | -0.681342643 | 0.29978125 | -2.2727994 | 0.02303827 | 0.0911662 |
| KIF24 | 98.609248 | 0.177217701 | 0.42561053 | 0.41638467 | 0.67712854 | 0.81308812 |
| NUDT2 | 108.529917 | -0.02226915 | 0.47426176 | -0.0469554 | 0.96254878 | 0.98057644 |
| MYORG | 74.1761612 | 0.967483288 | 0.44537186 | 2.17230449 | 0.0298327 | 0.10875449 |
| C9orf24 | 51.5152258 | -2.320660037 | 0.75176131 | -3.0869639 | 0.00202212 | 0.01548448 |
| FAM219A | 983.152262 | -1.923894526 | 0.42165138 | -4.5627612 | 5.05E-06 | 0.00014357 |
| DNAI1 | 72.6487444 | -1.180977263 | 0.52748862 | -2.2388677 | 0.02516452 | 0.09678637 |
| RPP25L | 63.7205691 | -0.226264944 | 0.43392778 | -0.5214346 | 0.60206408 | 0.7609161 |
| DCTN3 | 103.799369 | -0.502563603 | 0.43785006 | -1.1477984 | 0.25105179 | 0.44731974 |
| SIGMAR1 | 78.0267708 | 0.009051291 | 0.36489755 | 0.02480502 | 0.98021049 | 0.99026974 |
| IL11RA | 78.0543311 | 0.944633214 | 0.57872562 | 1.63226437 | 0.10262381 | 0.25027051 |
| AL162231.2 | 57.1044234 | 0.356739144 | 0.46386253 | 0.76906221 | 0.44185638 | 0.63352776 |
| DNAJB5 | 53.7116247 | -0.308556641 | 0.68368505 | -0.451314 | 0.65176325 | 0.79601879 |
| VCP | 1010.16157 | -1.187403515 | 0.36538175 | -3.2497614 | 0.00115502 | 0.01007115 |
| PIGO | 72.5071548 | 0.436805325 | 0.35191213 | 1.24123408 | 0.21451929 | 0.4048514 |
| STOML2 | 244.180738 | -0.494378148 | 0.32777475 | -1.5082862 | 0.13148128 | 0.29462383 |
| FAM214B | 150.913565 | -0.659971358 | 0.34157856 | -1.9321217 | 0.05334448 | 0.16174707 |
| UNC13B | 1715.95148 | 0.169299393 | 0.17929855 | 0.94423181 | 0.3450512 | 0.5454262 |
| RUSC2 | 1670.92495 | -0.275394562 | 0.43648454 | -0.6309377 | 0.52808124 | 0.70262178 |
| TESK1 | 153.731301 | -0.491684376 | 0.26829309 | -1.832639 | 0.06685627 | 0.18817227 |
| CD72 | 128.306441 | 0.169436951 | 0.43598397 | 0.38863115 | 0.69754902 | 0.82567395 |
| CCDC107 | 215.703379 | -0.477843562 | 0.34808732 | -1.3727692 | 0.1698241 | 0.34878317 |
| TLN1 | 543.804549 | -0.587337412 | 0.23818922 | -2.4658438 | 0.01366909 | 0.06278895 |
| CREB3 | 147.437885 | -1.250085963 | 0.3549185 | -3.5221775 | 0.00042802 | 0.00480101 |
| GBA2 | 452.519635 | 0.311751812 | 0.27502584 | 1.13353645 | 0.25698904 | 0.45422937 |
| RGP1 | 200.792135 | -0.606003959 | 0.19343212 | -3.1329025 | 0.00173087 | 0.01372295 |
| AL133410.1 | 122.46767 | -0.49060346 | 0.35994153 | -1.3630088 | 0.17287975 | 0.35252347 |
| NPR2 | 48.4763173 | -0.172284281 | 0.41006759 | -0.4201363 | 0.67438588 | 0.81105241 |
| HINT2 | 116.832125 | 0.785653719 | 0.34364594 | 2.28623016 | 0.0222408 | 0.08897147 |
| TMEM8B | 107.046848 | 0.668999144 | 0.49848348 | 1.34206883 | 0.17957368 | 0.36086531 |
| HRCT1 | 57.4129935 | -2.14304801 | 0.54219467 | -3.9525435 | 7.73E-05 | 0.00127412 |
| SPAAR | 60.0700801 | -0.403015112 | 0.70565319 | -0.5711235 | 0.56791594 | 0.73451775 |
| RECK | 188.535743 | -1.299910859 | 0.35818733 | -3.6291369 | 0.00028437 | 0.00350843 |
| CLTA | 1139.54039 | -0.501880338 | 0.3370335 | -1.4891111 | 0.13645811 | 0.30143937 |
| GNE | 303.843263 | -0.7268562 | 0.26672895 | -2.7250743 | 0.0064287 | 0.03651123 |
| RNF38 | 2486.17713 | 0.305183859 | 0.19659286 | 1.5523649 | 0.12057494 | 0.27804856 |
| MELK | 39.0819021 | -0.835147324 | 0.39178904 | -2.131625 | 0.03303768 | 0.11713646 |
| EBLN3P | 598.576504 | -0.105628679 | 0.24235349 | -0.4358455 | 0.66294883 | 0.80379726 |
| ZCCHC7 | 4567.14287 | -0.128546267 | 0.26735473 | -0.4808079 | 0.63065304 | 0.78222434 |
| GRHPR | 617.639811 | 0.81374467 | 0.37938542 | 2.14490235 | 0.03196066 | 0.1142333 |
| ZBTB5 | 438.879871 | -0.275334216 | 0.25123639 | -1.0959169 | 0.27311512 | 0.47089169 |
| POLR1E | 48.2255378 | -1.020079743 | 0.52229333 | -1.9530783 | 0.05081032 | 0.15707196 |
| AL138752.2 | 547.275108 | -0.193961804 | 0.21419505 | -0.9055382 | 0.36518033 | 0.56260838 |
| TRMT10B | 413.262697 | 0.15200119 | 0.32869846 | 0.46243353 | 0.64377046 | 0.79133367 |
| DCAF10 | 1343.10193 | -0.032212897 | 0.16820477 | -0.19151 | 0.84812604 | 0.91986577 |
| AL135785.1 | 135.124861 | 3.602866552 | 0.98028424 | 3.67532844 | 0.00023754 | 0.00306304 |
| ALDH1B1 | 246.72733 | 0.07054315 | 0.49518359 | 0.14245858 | 0.8867178 | 0.94155177 |
| FAM95C | 219.575212 | -0.625664756 | 0.50650492 | -1.235259 | 0.21673413 | 0.40762551 |
| ANKRD18A | 212.530772 | -1.127803634 | 0.39220763 | -2.875527 | 0.00403354 | 0.02581124 |
| CNTNAP3 | 648.157417 | -0.46433479 | 0.59059829 | -0.7862109 | 0.43174397 | 0.62436486 |
| GLIDR | 661.269814 | 0.14599303 | 0.57551462 | 0.25367388 | 0.79974751 | 0.89115837 |
| FAM95B1 | 57.8428376 | -0.556207267 | 1.01553166 | -0.5477006 | 0.58389752 | 0.74666538 |
| BMS1P14 | 453.390174 | -0.473115983 | 0.5588985 | -0.846515 | 0.39726549 | 0.59181807 |
| FP325318.1 | 54.2657075 | -1.227232438 | 0.66755507 | -1.8383988 | 0.06600366 | 0.18656221 |
| CBWD6 | 225.413537 | -0.19602102 | 0.32987685 | -0.5942248 | 0.55236174 | 0.72184315 |
| CNTNAP3B | 8081.29492 | -0.205840731 | 0.57492254 | -0.3580321 | 0.72031928 | 0.84086326 |
| BX088651.4 | 62.1471244 | -0.943798718 | 0.77860001 | -1.212174 | 0.22544577 | 0.41776195 |
| BX088645.1 | 39.7884771 | -0.650011854 | 0.62138319 | -1.0460725 | 0.29552757 | 0.49544966 |
| FAM27C | 43.2313973 | -0.322962089 | 0.56258795 | -0.5740651 | 0.56592377 | 0.7327021 |
| AL391987.2 | 52.0117276 | -0.861810522 | 0.36687511 | -2.3490569 | 0.01882103 | 0.07898954 |
| AL590399.3 | 37.9673452 | -0.292520438 | 0.7134103 | -0.4100311 | 0.6817831 | 0.81468059 |
| AL590399.1 | 37.9924033 | -0.787533397 | 0.84433983 | -0.9327209 | 0.3509641 | 0.55041523 |
| AL512625.3 | 160.507572 | 0.696455451 | 0.39471597 | 1.76444712 | 0.07765673 | 0.20793921 |
| AL512625.1 | 58.604305 | -0.55492794 | 0.52594396 | -1.0551085 | 0.29137571 | 0.4915553 |
| CBWD5 | 776.13082 | -0.127137099 | 0.26426324 | -0.4811002 | 0.63044528 | 0.78203742 |
| ZNF658 | 168.236889 | 0.38230037 | 0.38975848 | 0.98086479 | 0.32665942 | 0.52677357 |
| CBWD3 | 153.651171 | -0.048334242 | 0.25514494 | -0.1894384 | 0.84974925 | 0.92090248 |
| PGM5 | 234.254676 | -0.822107986 | 0.62496824 | -1.3154396 | 0.18836219 | 0.37213746 |
| AL161457.2 | 107.772508 | 0.300273977 | 0.44414789 | 0.67606756 | 0.49899775 | 0.68080506 |
| TMEM252 | 320.451092 | -1.676370163 | 0.6128627 | -2.7353111 | 0.00623213 | 0.03559328 |
| AL353616.2 | 2663.89834 | -1.656197337 | 0.60589089 | -2.7334911 | 0.00626668 | 0.03572476 |
| PIP5K1B | 265.372465 | -0.225738479 | 0.46657461 | -0.4838208 | 0.62851309 | 0.78055893 |
| APBA1 | 254.14782 | -0.011635203 | 0.34636442 | -0.0335924 | 0.9732022 | 0.98651698 |
| PTAR1 | 1732.25758 | 0.381439188 | 0.25543796 | 1.49327524 | 0.13536515 | 0.2998465 |
| MAMDC2-AS | 379.518364 | -0.188418707 | 0.51343472 | -0.3669769 | 0.71363621 | 0.8362667 |
| MAMDC2 | 773.203153 | -0.616296787 | 0.45526512 | -1.3537097 | 0.17582902 | 0.35604793 |
| SMC5-AS1 | 574.512844 | 0.071240207 | 0.369931 | 0.19257701 | 0.84729025 | 0.91938085 |
| SMC5 | 1342.05369 | -0.519795461 | 0.18072554 | -2.8761595 | 0.00402546 | 0.02577471 |
| KLF9 | 677.321967 | 1.277531625 | 0.45344556 | 2.81738698 | 0.00484162 | 0.02956843 |
| TRPM3 | 14627.1632 | 1.021280846 | 0.28493007 | 3.5843211 | 0.00033796 | 0.0040234 |
| CEMIP2 | 2281.96181 | -1.233228544 | 0.31539928 | -3.9100551 | 9.23E-05 | 0.00146557 |
| ABHD17B | 701.858028 | 0.411975887 | 0.2243061 | 1.83666821 | 0.06625889 | 0.18714416 |
| C9orf85 | 963.893461 | 0.089824966 | 0.19349475 | 0.4642243 | 0.64248705 | 0.79032308 |
| GDA | 9363.20413 | -0.611342182 | 0.47122375 | -1.2973501 | 0.19451075 | 0.37968654 |
| AL135924.2 | 79.3502583 | -0.306086846 | 0.64447055 | -0.4749431 | 0.63482752 | 0.78468826 |
| LINC01504 | 168.96671 | 0.573247683 | 1.12647028 | 0.50888842 | 0.61083044 | 0.76707767 |
| ZFAND5 | 1794.36692 | -0.589626485 | 0.31479867 | -1.8730272 | 0.06106463 | 0.17703317 |
| TMC1 | 293.837462 | 2.723503945 | 0.72259238 | 3.76907371 | 0.00016385 | 0.00226864 |
| ALDH1A1 | 2560.75532 | -0.052527709 | 0.56207206 | -0.0934537 | 0.92554314 | 0.96065575 |
| ANXA1 | 171.361066 | -3.108870341 | 0.95317922 | -3.2615801 | 0.00110793 | 0.00977247 |
| RORB | 147.10068 | 2.40979161 | 0.52934182 | 4.55243006 | 5.30E-06 | 0.00015018 |
| C9orf41-AS1 | 72.4996752 | -0.328640497 | 0.35076858 | -0.9369154 | 0.34880205 | 0.54866074 |
| CARNMT1 | 619.003287 | 0.063189036 | 0.21191524 | 0.2981807 | 0.76556525 | 0.86903543 |
| NMRK1 | 575.479874 | 0.639942544 | 0.40411902 | 1.58354966 | 0.11329624 | 0.26683016 |
| OSTF1 | 323.302808 | -0.289379509 | 0.18030043 | -1.6049851 | 0.10849708 | 0.25926944 |
| PCSK5 | 4967.71538 | -0.372043506 | 0.48886147 | -0.7610408 | 0.44663272 | 0.63757779 |
| RFK | 227.596171 | -0.752526327 | 0.25904501 | -2.9050022 | 0.0036725 | 0.02401671 |
| GCNT1 | 219.553952 | -0.234855917 | 0.43762784 | -0.5366567 | 0.59150477 | 0.75294909 |
| PRUNE2 | 10183.1905 | -0.344479709 | 0.31505631 | -1.0933909 | 0.27422219 | 0.47226581 |
| PCA3 | 979.265284 | -1.174496056 | 0.65088649 | -1.804456 | 0.07115984 | 0.19638678 |
| VPS13A | 5069.42945 | 0.065558219 | 0.2253761 | 0.29088363 | 0.77114032 | 0.87272401 |
| GNA14 | 224.992341 | 1.81719968 | 0.36330202 | 5.00189804 | 5.68E-07 | 2.37E-05 |
| GNAQ | 5408.60988 | -0.447046172 | 0.17452475 | -2.5615058 | 0.01042195 | 0.05204141 |
| CEP78 | 270.912946 | 0.20495592 | 0.27266345 | 0.75168095 | 0.45224295 | 0.64164104 |
| PSAT1 | 924.64581 | 0.975012103 | 0.54447155 | 1.7907494 | 0.07333352 | 0.20019301 |
| TLE4 | 853.705509 | 0.43311984 | 0.29406329 | 1.47287964 | 0.14078349 | 0.30766378 |
| AL161782.1 | 32.5147311 | 1.753390813 | 0.38367886 | 4.56994378 | 4.88E-06 | 0.00014036 |
| LINC01507 | 46.2374288 | 1.510159094 | 0.4764285 | 3.16974969 | 0.0015257 | 0.01253015 |

| | | | | | | |
|---|---|---|---|---|---|---|
| TLE1 | 572.631202 | -2.875951041 | 0.35730348 | -8.049043 | 8.34E-16 | 5.50E-13 |
| AL591368.1 | 91.8034953 | 1.995556958 | 0.89253058 | 2.23584155 | 0.02536215 | 0.09737788 |
| RASEF | 847.368247 | -1.278444463 | 0.43385974 | -2.9466769 | 0.00321209 | 0.02173648 |
| AL137847.1 | 57.5226449 | 0.41380163 | 0.4182178 | 0.98944051 | 0.32244766 | 0.52286645 |
| FRMD3 | 2630.98286 | 0.915231402 | 0.4166181 | 2.19681143 | 0.02803392 | 0.10415702 |
| IDNK | 340.909585 | 1.668890647 | 0.47798233 | 3.49153208 | 0.00048026 | 0.00520692 |
| UBQLN1 | 1051.40976 | -0.284953227 | 0.15572204 | -1.8298837 | 0.06726733 | 0.18892497 |
| GKAP1 | 555.576094 | 0.602594062 | 0.27316896 | 2.20593902 | 0.02738827 | 0.10276786 |
| KIF27 | 695.975792 | 0.394720092 | 0.20750766 | 1.90219526 | 0.05714563 | 0.16949995 |
| C9orf64 | 112.314062 | 1.079905078 | 0.20070922 | 5.38044569 | 7.43E-08 | 4.37E-06 |
| HNRNPK | 1316.81221 | -0.076015266 | 0.17391935 | -0.4370719 | 0.66205918 | 0.80336802 |
| AL354733.3 | 91.1750105 | -0.704153204 | 0.67001939 | -1.0509445 | 0.29328407 | 0.49326436 |
| RMI1 | 156.515275 | 0.153654943 | 0.30443369 | 0.50472385 | 0.61375281 | 0.76957315 |
| NTRK2 | 84.2315193 | 0.508597834 | 0.48274431 | 1.05355532 | 0.29208656 | 0.49214879 |
| AGTPBP1 | 1150.22366 | -0.454475356 | 0.22018576 | -2.0640543 | 0.03901257 | 0.13144251 |
| NAA35 | 973.414039 | -0.181058672 | 0.16199598 | -1.1176739 | 0.2637063 | 0.46169654 |
| GOLM1 | 422.780098 | -1.446516292 | 0.48835419 | -2.9620229 | 0.00305625 | 0.02099041 |
| C9orf153 | 95.8174612 | -0.865697733 | 0.706574 | -1.2252046 | 0.22049813 | 0.41238476 |
| ISCA1 | 320.966111 | -0.302913389 | 0.30302278 | -0.999639 | 0.31748524 | 0.51755172 |
| TUT7 | 1091.65113 | -0.718039444 | 0.2941045 | -2.4414432 | 0.01462869 | 0.06588377 |
| AL513318.2 | 87.151293 | 0.002437518 | 0.45699186 | 0.00533383 | 0.99574424 | 0.9977776 |
| DAPK1 | 2891.79545 | 0.040597977 | 0.37374051 | 0.10862611 | 0.91349905 | 0.95399223 |
| CTSL | 1597.32539 | -1.859681263 | 0.68199313 | -2.7268328 | 0.00639454 | 0.03636929 |
| AL772337.2 | 37.9465276 | 2.793558219 | 0.73421484 | 3.80482396 | 0.00014191 | 0.00202193 |
| CDK20 | 51.4103736 | 0.259268036 | 0.39381955 | 0.65834222 | 0.51031825 | 0.6896315 |
| SPIN1 | 1859.17965 | 0.748987693 | 0.20241476 | 3.70026225 | 0.00021538 | 0.00283862 |
| NXNL2 | 476.862058 | 2.276374985 | 0.60795448 | 3.74431811 | 0.00018088 | 0.00246008 |
| SHC3 | 234.745745 | -1.14920086 | 0.46697498 | -2.4609474 | 0.01385707 | 0.06339748 |
| CKS2 | 139.176948 | -1.18510926 | 0.5576885 | -2.125038 | 0.03358344 | 0.11842764 |
| SECISBP2 | 1132.26519 | 0.236890691 | 0.25892218 | 0.91491078 | 0.36023847 | 0.55851021 |
| SEMA4D | 2594.14943 | -0.472279931 | 0.33008943 | -1.4307636 | 0.15249799 | 0.32411759 |
| GADD45G | 135.03212 | -0.314913796 | 0.54123455 | -0.5818435 | 0.56067211 | 0.72893225 |
| LINC01508 | 302.637061 | 0.425570341 | 0.68939836 | 0.61730686 | 0.53703234 | 0.70939003 |
| SYK | 61.058452 | -1.056338836 | 0.89890351 | -1.1751415 | 0.2399381 | 0.43503063 |
| AL158071.1 | 73.7343811 | -0.988720985 | 0.64783152 | -1.5262008 | 0.12695986 | 0.28802364 |
| AL158071.4 | 280.975003 | -0.168593515 | 0.50155185 | -0.3361437 | 0.73676247 | 0.85107056 |
| AUH | 3040.14549 | -0.211319701 | 0.32753137 | -0.6451892 | 0.51880458 | 0.6954772 |
| NFIL3 | 838.552579 | -1.818047391 | 0.43768431 | -4.1537869 | 3.27E-05 | 0.00064588 |
| SPTLC1 | 857.132631 | -0.318035259 | 0.23008326 | -1.3822616 | 0.16689138 | 0.34469756 |
| IARS | 1023.61092 | -1.466612382 | 0.5332885 | -2.7501294 | 0.00595717 | 0.03446812 |
| NOL8 | 405.369013 | -0.216921285 | 0.28636414 | -0.7575016 | 0.44874942 | 0.639148 |
| CENPP | 672.291464 | 0.339477284 | 0.3340288 | 1.01631143 | 0.30948111 | 0.50977712 |
| ECM2 | 126.644516 | 1.765270616 | 0.64383636 | 2.74180013 | 0.00611035 | 0.0350523 |
| IPPK | 281.196873 | 0.142367847 | 0.25648626 | 0.55507006 | 0.57884674 | 0.74245057 |
| BICD2 | 255.33142 | -0.080574154 | 0.26348453 | -0.3058022 | 0.75975524 | 0.86552236 |
| ZNF484 | 159.20643 | 0.092414997 | 0.24299325 | 0.3803192 | 0.70370848 | 0.82988318 |
| SUSD3 | 191.918418 | 1.26920644 | 0.5864989 | 2.1640389 | 0.03046136 | 0.11040029 |
| CARD19 | 249.87853 | -1.027204271 | 0.29138868 | -3.5252031 | 0.00042316 | 0.00476622 |
| NINJ1 | 282.888494 | -0.493120917 | 0.27736632 | -1.7778688 | 0.07542541 | 0.20411769 |
| FAM120AOS | 148.125083 | -0.460243056 | 0.27095518 | -1.6985948 | 0.08939556 | 0.22801357 |
| FAM120A | 1517.43398 | -0.211583648 | 0.13537682 | -1.5629237 | 0.11807053 | 0.27411802 |
| PHF2 | 302.22456 | 0.19666931 | 0.29500206 | 0.66667097 | 0.50498232 | 0.6850618 |
| PTPDC1 | 670.209189 | -0.350023793 | 0.24444778 | -1.431896 | 0.15217359 | 0.32378818 |
| AL158152.2 | 660.91662 | -0.070034719 | 0.19103566 | -0.3666055 | 0.71391332 | 0.83637691 |
| ZNF169 | 214.849765 | 0.520051433 | 0.22824886 | 2.27844044 | 0.02270035 | 0.09027725 |
| MFSD14B | 637.25874 | 0.014376385 | 0.25990761 | 0.05531345 | 0.95588875 | 0.97756553 |
| PCAT7 | 100.664646 | 0.098610946 | 0.36678927 | 0.26884905 | 0.78804585 | 0.88421655 |
| FBP1 | 1424.40738 | 2.326324181 | 0.50422251 | 4.61368565 | 3.96E-06 | 0.00011891 |
| C9orf3 | 7292.8079 | 0.927319487 | 0.21689835 | 4.27536439 | 1.91E-05 | 0.00042879 |
| AL353768.1 | 68.9826019 | -0.74508433 | 0.59759375 | -1.2468074 | 0.21246812 | 0.40230701 |
| FANCC | 3097.7116 | 0.650984742 | 0.29958186 | 2.17297783 | 0.02978199 | 0.10859848 |
| PTCH1 | 189.539807 | -0.720058445 | 0.5081245 | -1.4170906 | 0.15645644 | 0.33013369 |
| LINC00476 | 626.198258 | 0.577334349 | 0.41419603 | 1.39386741 | 0.16335765 | 0.34003695 |
| ERCC6L2 | 968.596755 | -0.050326478 | 0.19826244 | -0.2538377 | 0.79962096 | 0.8910898 |
| AL160269.1 | 149.004058 | 0.38370875 | 0.34260764 | 1.11996552 | 0.26272845 | 0.46074961 |
| SLC35D2 | 786.40334 | 0.06363223 | 0.23078792 | 0.27571733 | 0.78276518 | 0.88097236 |
| ZNF367 | 84.9020535 | 0.183720754 | 0.42660931 | 0.43065341 | 0.6667204 | 0.80638282 |
| HABP4 | 882.720988 | 0.511449085 | 0.38788093 | 1.31857239 | 0.18731211 | 0.37080981 |
| CDC14B | 2363.18463 | -0.021405258 | 0.29628536 | -0.0722454 | 0.9424066 | 0.96993297 |
| AAED1 | 125.187114 | -0.776242785 | 0.38124542 | -2.0360711 | 0.04174321 | 0.13739993 |
| ZNF510 | 127.177164 | -0.119738436 | 0.21912525 | -0.5464383 | 0.58476467 | 0.74749318 |
| ZNF782 | 759.191192 | -0.018418474 | 0.2054988 | -0.0896281 | 0.92858273 | 0.9622077 |
| MFSD14C | 1408.12227 | -0.133292571 | 0.17708256 | -0.7527143 | 0.45162164 | 0.64124856 |
| AL590705.1 | 245.414908 | -0.377739376 | 0.57295049 | -0.659288 | 0.50971085 | 0.68915039 |
| CCDC180 | 197.873021 | 0.507239001 | 0.5627305 | 0.90138885 | 0.36738161 | 0.56469238 |
| TDRD7 | 416.295602 | -0.55783897 | 0.299478 | -1.8627044 | 0.06250387 | 0.17972542 |
| TSTD2 | 353.469361 | -0.19583303 | 0.1459699 | -1.3415988 | 0.17972613 | 0.36096117 |
| NCBP1 | 318.47941 | -0.088129373 | 0.21315895 | -0.4134444 | 0.67928105 | 0.81417094 |
| XPA | 342.619917 | 0.199712342 | 0.18784489 | 1.06317686 | 0.28770175 | 0.48751736 |
| TRMO | 112.542443 | 0.088788845 | 0.39570772 | 0.22437987 | 0.82246174 | 0.90484654 |
| ANP32B | 436.828488 | -0.415402774 | 0.35512065 | -1.1697511 | 0.24210115 | 0.43733268 |
| NANS | 219.334025 | -0.955444377 | 0.30637464 | -3.1185492 | 0.00181744 | 0.01420275 |
| TRIM14 | 883.937193 | -0.257797757 | 0.45302223 | -0.569062 | 0.56931403 | 0.73547716 |
| CORO2A | 151.490693 | -1.013419615 | 0.49447732 | -2.0494764 | 0.04041555 | 0.13439494 |
| TBC1D2 | 128.33554 | -0.612704881 | 0.37506581 | -1.633593 | 0.10234434 | 0.24988135 |
| ANKS6 | 390.565223 | 0.063567744 | 0.22543882 | 0.28197337 | 0.77796393 | 0.87758633 |
| GALNT12 | 71.6527156 | -1.119160238 | 0.58863274 | -1.9012878 | 0.05726433 | 0.16967621 |
| TGFBR1 | 615.151283 | -0.434362231 | 0.22944009 | -1.8931401 | 0.05833924 | 0.17145282 |
| ALG2 | 237.059851 | -0.522860239 | 0.3457098 | -1.5124253 | 0.13042571 | 0.29297692 |
| SEC61B | 616.740545 | -1.064475041 | 0.31776724 | -3.3498577 | 0.00080853 | 0.00771764 |
| AL359710.1 | 344.18154 | 0.477805685 | 0.52851051 | 0.9040609 | 0.36596311 | 0.56343439 |
| STX17-AS1 | 32.861897 | 0.316551123 | 0.56591623 | 0.55936039 | 0.57591579 | 0.74039371 |
| STX17 | 516.421552 | 0.110299224 | 0.3144847 | 0.35073002 | 0.7257909 | 0.84408756 |
| ERP44 | 1326.11882 | -0.153341666 | 0.24433905 | -0.6275774 | 0.53028082 | 0.70431775 |
| INVS | 3458.40222 | 0.556932403 | 0.23165066 | 2.40419083 | 0.01620831 | 0.07115834 |
| TEX10 | 356.133602 | -0.485972263 | 0.17666845 | -2.7507586 | 0.00594574 | 0.03443105 |
| MSANTD3 | 83.1669289 | -2.566957954 | 0.68399407 | -3.752895 | 0.0001748 | 0.00240325 |
| PLPPR1 | 1650.89083 | 4.07382363 | 0.76812893 | 5.30356748 | 1.14E-07 | 6.20E-06 |
| MRPL50 | 145.61362 | -0.420160824 | 0.34945657 | -1.2023263 | 0.22923714 | 0.42233245 |
| ZNF189 | 656.579189 | 0.536337018 | 0.38018049 | 1.41074314 | 0.15832036 | 0.33304132 |
| ALDOB | 6114.26512 | 1.579006491 | 0.54409278 | 2.90209053 | 0.00370681 | 0.02417188 |
| TMEM246-A! | 1473.96328 | 3.548326322 | 0.743107 | 4.77498707 | 1.80E-06 | 6.22E-05 |
| TMEM246 | 217.582888 | 0.420653226 | 0.54228204 | 0.77570931 | 0.43792064 | 0.62979522 |
| RNF20 | 300.226651 | 1.152957234 | 0.31095092 | 3.70784315 | 0.00020903 | 0.00276831 |
| LINC01492 | 36.6442489 | -0.494596016 | 0.44522331 | -1.1108942 | 0.26661387 | 0.4645363 |
| SMC2 | 94.3842154 | 0.025269913 | 0.27256031 | 0.09271311 | 0.92613148 | 0.96090956 |
| NIPSNAP3A | 125.336292 | -1.028346633 | 0.40804346 | -2.520189 | 0.01172919 | 0.05633214 |
| ABCA1 | 1095.66449 | -0.613679109 | 0.64077764 | -0.9577099 | 0.33820905 | 0.53836156 |
| SLC44A1 | 1301.81773 | 0.495309533 | 0.16092965 | 3.07780164 | 0.00208534 | 0.01585342 |
| FSD1L | 983.163117 | -1.046817064 | 0.20637206 | -5.0724746 | 3.93E-07 | 1.75E-05 |
| FKTN | 435.919263 | 0.045760986 | 0.24471653 | 0.1869959 | 0.85166385 | 0.92197716 |
| TMEM38B | 762.650273 | -0.455150037 | 0.43214753 | -1.0532284 | 0.29223634 | 0.49231264 |
| LINC01505 | 726.376567 | 0.15557577 | 0.36950461 | 0.42103878 | 0.67372676 | 0.81073146 |

| | | | | | | |
|---|---|---|---|---|---|---|
| ZNF462 | 2507.94681 | 0.10820539 | 0.2794484 | 0.38721062 | 0.69860029 | 0.82649009 |
| AL807761.4 | 65.1151987 | -0.624969165 | 0.46330868 | -1.3489261 | 0.17736071 | 0.35793334 |
| RAD23B | 1646.59732 | -0.364221207 | 0.17898401 | -2.0349371 | 0.0418572 | 0.13765274 |
| ELP1 | 369.456994 | 0.242754375 | 0.16190017 | 1.49940774 | 0.13376789 | 0.29775194 |
| FAM206A | 140.208003 | -0.494889583 | 0.18185694 | -2.7213126 | 0.00650232 | 0.03678388 |
| CTNNAL1 | 489.670781 | -1.785308236 | 0.42036741 | -4.2470187 | 2.17E-05 | 0.0004691 |
| TMEM245 | 2367.12639 | -0.091232348 | 0.3247999 | -0.2808879 | 0.77879641 | 0.87801961 |
| EPB41L4B | 61.0500979 | -0.885094705 | 0.38843876 | -2.2785952 | 0.02269114 | 0.09027725 |
| PTPN3 | 3049.54471 | -0.762642704 | 0.22695624 | -3.3603073 | 0.00077856 | 0.00752588 |
| PALM2 | 721.186178 | -1.724829524 | 0.48834309 | -3.5320036 | 0.00041242 | 0.00468359 |
| PALM2-AKAF | 218.164578 | -0.71303541 | 0.42461892 | -1.6792361 | 0.09310604 | 0.23481223 |
| TXN | 1368.54493 | -0.783248179 | 0.43791678 | -1.7885777 | 0.07368286 | 0.20082937 |
| LPAR1 | 197.552313 | -3.612236374 | 0.71899929 | -5.0239777 | 5.06E-07 | 2.17E-05 |
| ECPAS | 2268.38768 | -0.465478633 | 0.25041068 | -1.858861 | 0.06304685 | 0.18090499 |
| ZNF483 | 362.069208 | -0.98069571 | 0.39382287 | -2.4901949 | 0.01276731 | 0.05992941 |
| PTGR1 | 6060.75995 | -0.922073694 | 0.60057891 | -1.5353081 | 0.12470816 | 0.28465857 |
| C9orf84 | 218.034511 | -0.996809572 | 0.65489091 | -1.5221002 | 0.12798398 | 0.28967577 |
| UGCG | 5572.54848 | -0.847382982 | 0.41403542 | -2.0466437 | 0.04069308 | 0.13499034 |
| SUSD1 | 572.814068 | -1.078824805 | 0.27888144 | -3.8683995 | 0.00010955 | 0.00166892 |
| PTBP3 | 2290.35144 | -0.897396316 | 0.29938783 | -2.9974375 | 0.0027226 | 0.01928612 |
| HSDL2 | 2608.13237 | -0.001524389 | 0.29429607 | -0.0051798 | 0.99586715 | 0.9977776 |
| C9orf147 | 288.2993 | 0.587216746 | 0.50580033 | 1.16096552 | 0.24565592 | 0.44119572 |
| KIAA1958 | 3563.07465 | 0.659837082 | 0.25419343 | 2.59580699 | 0.00943691 | 0.04833769 |
| INIP | 353.003251 | 0.175118487 | 0.26992229 | 0.64877371 | 0.51648465 | 0.69386001 |
| SNX30 | 1643.22668 | 1.149407199 | 0.26400262 | 4.3537719 | 1.34E-05 | 0.00032153 |
| SLC31A2 | 106.672363 | 0.0242312 | 0.391978 | 0.06181776 | 0.95070796 | 0.97481703 |
| FKBP15 | 400.178016 | 0.09915808 | 0.16337438 | 0.60693776 | 0.54389222 | 0.71416138 |
| SLC31A1 | 563.231652 | 0.352711394 | 0.19350877 | 1.82271533 | 0.06834652 | 0.19099403 |
| CDC26 | 188.278751 | -0.112915171 | 0.25654453 | -0.4401387 | 0.65983666 | 0.8020537 |
| PRPF4 | 340.583343 | -0.094133791 | 0.44472673 | -0.2116666 | 0.83236715 | 0.91150088 |
| RNF183 | 341.626351 | 0.562613005 | 0.46530404 | 1.20912986 | 0.22661296 | 0.4191881 |
| WDR31 | 223.502079 | 0.760749384 | 0.40846521 | 1.8624582 | 0.06253853 | 0.17978516 |
| BSPRY | 227.603343 | -1.061424363 | 0.3258341 | -3.2575607 | 0.00112374 | 0.00988014 |
| HDHD3 | 1337.66745 | 0.807536545 | 0.27977832 | 2.88634422 | 0.00389746 | 0.02515181 |
| ALAD | 349.089993 | 2.080462052 | 0.44407488 | 4.68493523 | 2.80E-06 | 9.06E-05 |
| POLE3 | 75.1756511 | -0.991738608 | 0.65919126 | -1.5044778 | 0.13245839 | 0.29579784 |
| RGS3 | 1032.05286 | -0.100744238 | 0.54982898 | -0.1832283 | 0.85461888 | 0.92322796 |
| ZNF618 | 1073.81875 | 1.124016752 | 0.43545102 | 2.58127022 | 0.00984375 | 0.04987876 |
| AMBP | 42.3177493 | -1.276873571 | 0.58843938 | -2.1699322 | 0.03001198 | 0.10926274 |
| KIF12 | 1563.95002 | 1.49280054 | 0.40057659 | 3.72662948 | 0.00019406 | 0.00260779 |
| COL27A1 | 2992.14505 | -0.496518381 | 0.38680384 | -1.2836439 | 0.1992666 | 0.38577668 |
| AKNA | 43.4260787 | -1.934260124 | 0.50493951 | -3.8306769 | 0.00012779 | 0.00188348 |
| WHRN | 229.834471 | -0.337592254 | 0.35581558 | -0.9487844 | 0.3427303 | 0.54335001 |
| ATP6V1G1 | 566.375149 | -0.907561689 | 0.23213333 | -3.9096569 | 9.24E-05 | 0.00146557 |
| TMEM268 | 136.159738 | -0.354086317 | 0.52491325 | -0.6745616 | 0.49995435 | 0.68139272 |
| TNFSF15 | 558.492263 | -2.858359661 | 0.82245113 | -3.4754158 | 0.00051006 | 0.00541883 |
| TNC | 346.184214 | -3.16665481 | 0.83672836 | -3.7845673 | 0.00015398 | 0.00215803 |
| PAPPA | 3002.48646 | -0.27275702 | 0.58182085 | -0.468799 | 0.63921333 | 0.78803844 |
| AL137024.1 | 138.338358 | -0.567806246 | 0.54417874 | -1.0434186 | 0.29675444 | 0.49702018 |
| ASTN2 | 5979.49587 | 0.106086455 | 0.37206369 | 0.28512983 | 0.77554468 | 0.87601952 |
| ASTN2-AS1 | 121.141261 | -1.426611621 | 0.73219773 | -1.9483967 | 0.05136752 | 0.15790414 |
| AL133284.1 | 43.8216948 | -0.657739497 | 0.69420625 | -0.9474699 | 0.34339941 | 0.54383762 |
| TRIM32 | 77.9558134 | -0.484624943 | 0.26812246 | -1.8074761 | 0.07068807 | 0.19542586 |
| BRINP1 | 245.505101 | 2.512300162 | 0.60239835 | 4.17049645 | 3.04E-05 | 0.00060818 |
| CDK5RAP2 | 630.994831 | 0.109448402 | 0.27643617 | 0.39592649 | 0.69215926 | 0.82256172 |
| MEGF9 | 800.516014 | -0.364029555 | 0.2782749 | -1.3081652 | 0.19081727 | 0.37509426 |
| FBXW2 | 524.549375 | 0.307979898 | 0.25176926 | 1.2232625 | 0.22123057 | 0.4134582 |
| PSMD5 | 375.433476 | -0.360101929 | 0.21508939 | -1.6741966 | 0.09409197 | 0.23621021 |
| TRAF1 | 77.6329834 | -0.831250669 | 0.41098895 | -2.0225621 | 0.04311832 | 0.14048556 |
| C5 | 111.097307 | 1.408815145 | 0.43923474 | 3.20743106 | 0.00133926 | 0.01127596 |
| CNTRL | 155.983221 | -0.498771545 | 0.49016758 | -1.0175531 | 0.30889038 | 0.50906843 |
| RAB14 | 631.799675 | -0.335291631 | 0.15791018 | -2.123306 | 0.03372822 | 0.11878539 |
| GSN | 404.245867 | 0.251321665 | 0.3404342 | 0.7382386 | 0.46036947 | 0.64907266 |
| STOM | 1237.85764 | -0.785656939 | 0.29401481 | -2.6721679 | 0.00753629 | 0.04107354 |
| AL359644.1 | 272.612736 | 0.282459965 | 0.50721151 | 0.55688793 | 0.577604 | 0.74154889 |
| DAB2IP | 1256.41124 | -1.097359753 | 0.37782069 | -2.9044459 | 0.00367904 | 0.0240365 |
| TTLL11 | 321.523383 | 0.052859471 | 0.24851421 | 0.21270201 | 0.8315594 | 0.91086448 |
| NDUFA8 | 204.870147 | 0.134286295 | 0.20564729 | 0.65299326 | 0.51376063 | 0.69196393 |
| RBM18 | 122.220794 | 0.027142771 | 0.21517569 | 0.12614237 | 0.89961923 | 0.94743919 |
| MRRF | 870.24926 | 0.004213831 | 0.17634156 | 0.02389585 | 0.98093569 | 0.99047407 |
| PDCL | 127.210496 | -0.056225653 | 0.19589604 | -0.2870178 | 0.77409868 | 0.87481001 |
| RC3H2 | 869.085064 | -0.164373323 | 0.16971384 | -0.9685323 | 0.33277862 | 0.53225164 |
| ZBTB6 | 68.6014278 | -0.290838593 | 0.2895242 | -1.0045398 | 0.31511848 | 0.51523844 |
| ZBTB26 | 143.883162 | 0.153866527 | 0.18529743 | 0.83037595 | 0.40632626 | 0.60022783 |
| RABGAP1 | 2262.95279 | 0.099984427 | 0.15811196 | 0.63236474 | 0.52714857 | 0.70172142 |
| STRBP | 1690.83799 | -0.052788343 | 0.20184942 | -0.2615234 | 0.79368893 | 0.88743732 |
| DENND1A | 4421.68923 | 0.921780169 | 0.28482485 | 3.23630525 | 0.00121088 | 0.01047161 |
| AL390774.2 | 53.3502068 | -0.300703727 | 0.84388733 | -0.3563316 | 0.72159224 | 0.8420624 |
| NEK6 | 1332.76371 | -1.30565079 | 0.32909233 | -3.96743 | 7.27E-05 | 0.00121444 |
| PSMB7 | 1030.3634 | -0.634643899 | 0.32396026 | -1.9590177 | 0.05011071 | 0.15561112 |
| NR6A1 | 4889.17955 | 0.256491019 | 0.28102659 | 0.91269306 | 0.36140399 | 0.55943134 |
| MIR181A2HG | 321.859854 | -0.226911764 | 0.44421829 | -0.5108114 | 0.60948313 | 0.76602773 |
| RPL35 | 822.107387 | -0.751967385 | 0.44847436 | -1.6767233 | 0.09359661 | 0.23561593 |
| ARPC5L | 261.890196 | -0.641701107 | 0.43996522 | -1.4585269 | 0.14469537 | 0.31269244 |
| GOLGA1 | 382.469152 | 0.435220422 | 0.39267007 | 1.1083616 | 0.26770568 | 0.4657243 |
| SCAI | 406.935326 | 0.386497506 | 0.16080095 | 2.40357728 | 0.01623554 | 0.07119017 |
| PPP6C | 576.043908 | 0.412302692 | 0.20315756 | 2.02947252 | 0.04241018 | 0.13890476 |
| RABEPK | 246.145542 | 0.153193827 | 0.38110922 | 0.40196831 | 0.68770735 | 0.81938377 |
| HSPA5 | 998.299777 | -1.891612808 | 0.5434323 | -3.4808619 | 0.0004998 | 0.00534801 |
| GAPVD1 | 2002.85319 | 0.291845029 | 0.18419703 | 1.58441768 | 0.1130987 | 0.26653225 |
| MAPKAP1 | 2361.80634 | 0.105371309 | 0.29672378 | 0.35511583 | 0.72250282 | 0.84228022 |
| PBX3 | 411.069606 | -1.138490867 | 0.70351837 | -1.6182816 | 0.10560192 | 0.25457008 |
| MVB12B | 117.761948 | -0.461874006 | 0.4330882 | -1.0664664 | 0.28621286 | 0.48583526 |
| ZBTB43 | 792.06997 | -1.13163906 | 0.46087015 | -2.4554401 | 0.01407122 | 0.06404818 |
| ZBTB34 | 205.267267 | -0.227245848 | 0.38837766 | -0.5851157 | 0.55846995 | 0.72696559 |
| RALGPS1 | 876.49964 | -0.043850202 | 0.24168979 | -0.1814318 | 0.85602869 | 0.92383161 |
| GARNL3 | 627.714788 | 0.383688391 | 0.26877488 | 1.42754558 | 0.1534227 | 0.32558676 |
| SLC2A8 | 108.616473 | 0.091722717 | 0.28045956 | 0.32704436 | 0.74363433 | 0.85454358 |
| ZNF79 | 85.7962132 | 0.151616892 | 0.35983697 | 0.42134885 | 0.67350036 | 0.81072605 |
| RPL12 | 375.344095 | -0.712925681 | 0.46415531 | -1.5359637 | 0.12454729 | 0.28443348 |
| LRSAM1 | 646.682826 | -0.630815856 | 0.24399503 | -2.5853635 | 0.00972764 | 0.0494389 |
| FAM129B | 415.306657 | -2.69448459 | 0.51733055 | -5.208439 | 1.90E-07 | 9.32E-06 |
| STXBP1 | 339.89063 | -0.862942817 | 0.36483375 | -2.3653042 | 0.01801527 | 0.07668797 |
| PTRH1 | 138.392791 | -0.4857733 | 0.23428572 | -2.0734225 | 0.03813297 | 0.12953758 |
| TOR2A | 32.8877757 | -0.41758341 | 0.40092594 | -1.0415475 | 0.29762152 | 0.49810722 |
| CDK9 | 249.481898 | -0.077178231 | 0.22939839 | -0.3364375 | 0.73654094 | 0.85095808 |
| FPGS | 304.108479 | -0.347680264 | 0.42284658 | -0.8222374 | 0.41094181 | 0.6045054 |
| ENG | 65.0073926 | 0.339483349 | 0.44312612 | 0.76610999 | 0.44361086 | 0.63517875 |
| ST6GALNAC4 | 113.07688 | -1.834365011 | 0.65968629 | -2.7806626 | 0.00542481 | 0.03218954 |
| DPM2 | 201.260165 | -1.262452232 | 0.63621201 | -1.9843263 | 0.04721948 | 0.14989269 |
| FAM102A | 912.396003 | -0.364801352 | 0.77064557 | -0.4733711 | 0.63594843 | 0.78566428 |
| NAIF1 | 33.0564484 | -0.262323611 | 0.42618697 | -0.615513 | 0.53821601 | 0.70990443 |
| SLC25A25 | 253.861213 | -0.168348468 | 0.53238649 | -0.3162148 | 0.7518395 | 0.8605113 |

| | | | | | | |
|---|---|---|---|---|---|---|
| SLC25A25-AS | 146.585613 | 0.801473136 | 0.39099763 | 2.04981583 | 0.04038241 | 0.13437462 |
| PTGES2 | 281.744948 | -0.496792315 | 0.26210086 | -1.8954242 | 0.05803622 | 0.17100226 |
| LCN2 | 318.599256 | -9.365280284 | 1.06604862 | -8.7850405 | 1.56E-18 | 1.79E-15 |
| C9orf16 | 204.58211 | -1.203743944 | 0.34471086 | -3.4920395 | 0.00047935 | 0.00520223 |
| CIZ1 | 295.283216 | -0.023855836 | 0.26743785 | -0.0892014 | 0.92892184 | 0.96234084 |
| DNM1 | 1065.5596 | 0.481700165 | 0.28615189 | 1.68337232 | 0.09230304 | 0.23357387 |
| GOLGA2 | 649.917673 | -0.419678602 | 0.34540403 | -1.2150368 | 0.22435206 | 0.41663645 |
| SWI5 | 113.217261 | 0.179192485 | 0.20600242 | 0.86985621 | 0.38437899 | 0.58069909 |
| TRUB2 | 179.898638 | -0.050154796 | 0.30031197 | -0.167009 | 0.86736298 | 0.93100583 |
| COQ4 | 626.065274 | 0.862544681 | 0.20809044 | 4.14504707 | 3.40E-05 | 0.00066813 |
| SLC27A4 | 374.149352 | -0.386378358 | 0.32362879 | -1.1938937 | 0.23251957 | 0.42625996 |
| URM1 | 239.810153 | -0.867675536 | 0.25994612 | -3.3379053 | 0.00084413 | 0.0079796 |
| ODF2 | 223.940045 | 0.029963269 | 0.41256291 | 0.07262715 | 0.94210282 | 0.96969312 |
| GLE1 | 357.024905 | 0.349243039 | 0.27065936 | 1.29034162 | 0.19693207 | 0.38266911 |
| SPTAN1 | 1060.114 | -0.255954473 | 0.18876883 | -1.3559149 | 0.17512625 | 0.3551841 |
| WDR34 | 221.29865 | 0.945088694 | 0.40836297 | 2.31433495 | 0.02064936 | 0.08443936 |
| SET | 707.821414 | -1.204460321 | 0.35204131 | -3.4213608 | 0.00062309 | 0.00633109 |
| PKN3 | 53.808514 | -1.593953732 | 0.43540847 | -3.6608239 | 0.00025141 | 0.00320667 |
| ZDHHC12 | 101.388109 | -1.123600848 | 0.55718203 | -2.0165777 | 0.0437396 | 0.14170153 |
| ZER1 | 1100.42735 | 0.758563033 | 0.32482712 | 2.33528234 | 0.01952868 | 0.08103823 |
| TBC1D13 | 450.503288 | 1.582238971 | 0.22812208 | 6.93593097 | 4.04E-12 | 9.54E-10 |
| ENDOG | 155.44274 | -0.020062627 | 0.34955814 | -0.0573942 | 0.95423114 | 0.97682198 |
| SPOUT1 | 117.310602 | 0.378632531 | 0.3050041 | 1.24140145 | 0.21445748 | 0.40484351 |
| KYAT1 | 447.855139 | 0.482770906 | 0.58561719 | 0.82437968 | 0.40972386 | 0.60360018 |
| LRRC8A | 274.063395 | -0.530970867 | 0.46507691 | -1.141684 | 0.2535854 | 0.4504205 |
| PHYHD1 | 233.882007 | 1.379236021 | 0.58411459 | 2.36124221 | 0.01821383 | 0.07729318 |
| DOLK | 34.4537229 | -0.762676059 | 0.56412488 | -1.3519632 | 0.1763871 | 0.35675637 |
| NUP188 | 268.72839 | -0.142539072 | 0.29984422 | -0.4753771 | 0.63451822 | 0.78453036 |
| SH3GLB2 | 203.305703 | 0.033448514 | 0.37881924 | 0.08829677 | 0.92964081 | 0.96271389 |
| MIGA2 | 235.766679 | 0.69788183 | 0.18776639 | 3.71675582 | 0.0002018 | 0.0026907 |
| DOLPP1 | 35.06599 | 0.627174897 | 0.40107509 | 1.56373434 | 0.11787997 | 0.27381461 |
| CRAT | 199.482487 | 0.586888315 | 0.29496139 | 1.98971233 | 0.04662263 | 0.14848229 |
| PTPA | 379.92238 | -0.027274689 | 0.30618575 | -0.0890789 | 0.92901921 | 0.96236897 |
| IER5L | 434.052395 | -2.029788103 | 0.43045574 | -4.7154397 | 2.41E-06 | 8.02E-05 |
| LINC01503 | 74.8215729 | -1.650722346 | 0.4120718 | -4.0059095 | 6.18E-05 | 0.00106785 |
| NTMT1 | 202.520696 | -0.554143471 | 0.20709267 | -2.6758237 | 0.00745458 | 0.04080671 |
| ASB6 | 86.4254791 | -0.011544274 | 0.34090184 | -0.0338639 | 0.97298566 | 0.98646685 |
| TOR1B | 121.700628 | -0.148516303 | 0.27983165 | -0.5307345 | 0.5956028 | 0.75582107 |
| TOR1A | 97.8772405 | -0.341806627 | 0.30719206 | -1.1126805 | 0.26584566 | 0.46372441 |
| C9orf78 | 321.889721 | 0.070040974 | 0.31852537 | 0.21989135 | 0.82595577 | 0.9066142 |
| USP20 | 300.661143 | 0.749814192 | 0.24935381 | 3.00702925 | 0.00263814 | 0.01885351 |
| FNBP1 | 2031.53147 | 0.167472895 | 0.23287867 | 0.71914226 | 0.47205327 | 0.65816643 |
| GPR107 | 754.643761 | -0.095018672 | 0.22386744 | -0.4244417 | 0.67124373 | 0.80935413 |
| NCS1 | 607.774851 | -0.828072724 | 0.4642636 | -1.7836262 | 0.07448442 | 0.20233016 |
| HMCN2 | 51.3367104 | 2.946532746 | 0.55583398 | 5.30110218 | 1.15E-07 | 6.21E-06 |
| ASS1 | 1965.72479 | 1.947990043 | 0.37647229 | 5.17432514 | 2.29E-07 | 1.09E-05 |
| FUBP3 | 873.277365 | -0.272155742 | 0.27658097 | -0.9840003 | 0.32511538 | 0.52545178 |
| EXOSC2 | 72.9627868 | -0.182395192 | 0.39818525 | -0.4580662 | 0.6469049 | 0.79280318 |
| ABL1 | 3286.11753 | 0.774034696 | 0.23023866 | 3.36187982 | 0.00077414 | 0.00750432 |
| AIF1L | 718.140957 | 0.165892546 | 0.32767055 | 0.50627847 | 0.61266118 | 0.76888361 |
| NUP214 | 1291.51337 | 0.251302128 | 0.2045348 | 1.22865218 | 0.21920224 | 0.41074574 |
| AL157938.2 | 285.835846 | -0.497175039 | 0.66734159 | -0.7450083 | 0.45626669 | 0.64534938 |
| AL157938.3 | 74.435211 | 0.007157384 | 0.456577 | 0.01567618 | 0.98749273 | 0.99360989 |
| PLPP7 | 52.3764447 | 1.069184876 | 0.34054383 | 3.1396395 | 0.00169156 | 0.01346469 |
| PRRC2B | 2646.90604 | -0.144678006 | 0.27388478 | -0.5282441 | 0.59732996 | 0.75748003 |
| POMT1 | 243.433026 | 0.30186496 | 0.18931155 | 1.59454065 | 0.11081499 | 0.26301207 |
| UCK1 | 134.051277 | 0.116030634 | 0.2373264 | 0.4889074 | 0.62490726 | 0.77771949 |
| RAPGEF1 | 2518.14643 | -0.827893589 | 0.37465579 | -2.2097445 | 0.0271229 | 0.10202347 |
| MED27 | 1008.48255 | -0.217923421 | 0.2016514 | -1.0806938 | 0.27983333 | 0.47832342 |
| SETX | 2427.19771 | -0.449666048 | 0.21329568 | -2.1081817 | 0.03501527 | 0.12175402 |
| TTF1 | 305.585634 | -0.186838703 | 0.24774566 | -0.7541553 | 0.45075598 | 0.64075861 |
| DDX31 | 602.286427 | -0.718028821 | 0.31539636 | -2.276592 | 0.0228106 | 0.09054172 |
| GTF3C4 | 196.55389 | -0.33539466 | 0.30394532 | -1.1034704 | 0.26982295 | 0.46798218 |
| AK8 | 194.769988 | 0.614206879 | 0.53158235 | 1.15543129 | 0.24791386 | 0.44426137 |
| SPACA9 | 37.8337737 | 0.54893738 | 0.38276901 | 1.43412181 | 0.15153751 | 0.32288584 |
| TSC1 | 751.897972 | 0.07416924 | 0.18796964 | 0.39458096 | 0.69315217 | 0.82317075 |
| GTF3C5 | 223.788938 | -0.368639652 | 0.42620788 | -0.8649292 | 0.38707764 | 0.5832974 |
| RALGDS | 118.9911 | -0.801869533 | 0.43047236 | -1.8627666 | 0.06249511 | 0.17972542 |
| ABO | 854.833952 | -0.252400504 | 0.46876242 | -0.5384401 | 0.59027322 | 0.75221969 |
| SURF6 | 208.157566 | -0.124105242 | 0.42778371 | -0.2901121 | 0.77173046 | 0.87317484 |
| MED22 | 96.8679203 | -0.065036921 | 0.35538751 | -0.1830028 | 0.8547958 | 0.92322796 |
| RPL7A | 877.688548 | -0.900130701 | 0.44802509 | -2.0091078 | 0.0445257 | 0.14370469 |
| SURF1 | 141.914992 | -0.103775013 | 0.22964635 | -0.4518905 | 0.65134784 | 0.79573901 |
| SURF2 | 70.5026497 | -0.668941728 | 0.37440886 | -1.786661 | 0.07399229 | 0.20147245 |
| SURF4 | 675.599759 | -0.672121208 | 0.37442124 | -1.7950937 | 0.07263877 | 0.19889326 |
| REXO4 | 178.629605 | -0.10868098 | 0.35703371 | -0.3043998 | 0.76082334 | 0.8665094 |
| ADAMTS13 | 69.9052991 | 1.185178723 | 0.5116755 | 2.31627022 | 0.02054352 | 0.08414838 |
| CACFD1 | 145.143575 | 0.569602215 | 0.34066161 | 1.67204696 | 0.09451506 | 0.23686461 |
| ADAMTSL2 | 53.0202591 | 3.38293015 | 0.6857376 | 4.93327208 | 8.09E-07 | 3.21E-05 |
| DBH-AS1 | 84.5854709 | 3.158537789 | 0.84893497 | 3.72058863 | 0.00019876 | 0.00265548 |
| SARDH | 245.547205 | 3.100668472 | 0.6759573 | 4.58707746 | 4.49E-06 | 0.00013109 |
| VAV2 | 1372.03088 | -0.427204027 | 0.23320838 | -1.8318554 | 0.06697297 | 0.18842333 |
| BRD3OS | 80.5555597 | 0.227430584 | 0.31836389 | 0.71437304 | 0.47499654 | 0.6611115 |
| BRD3 | 222.844874 | 0.154708665 | 0.37712144 | 0.41023566 | 0.68163308 | 0.81468059 |
| WDR5 | 290.771158 | 0.213818565 | 0.28482186 | 0.75070982 | 0.45282731 | 0.64207138 |
| RXRA | 476.294929 | 0.611124037 | 0.32011194 | 1.90909477 | 0.05624987 | 0.16761238 |
| PPP1R26-AS1 | 396.742705 | 0.735851649 | 0.44750068 | 1.64435872 | 0.10010213 | 0.2460337 |
| PPP1R26 | 192.811409 | 0.319605302 | 0.2440553 | 1.30956099 | 0.19034439 | 0.37443321 |
| C9orf116 | 53.9633374 | 0.035155593 | 0.55244742 | 0.06363609 | 0.94926 | 0.97398808 |
| MRPS2 | 153.276702 | -0.173198254 | 0.39364401 | -0.439987 | 0.65994651 | 0.8020537 |
| CAMSAP1 | 594.508221 | -0.772000952 | 0.32412922 | -2.3817691 | 0.0172297 | 0.07428987 |
| AL355574.1 | 53.6272095 | -0.162983921 | 0.33765683 | -0.482691 | 0.62931514 | 0.7812011 |
| UBAC1 | 277.918338 | 0.038231462 | 0.36869415 | 0.10369424 | 0.91741199 | 0.95587788 |
| NACC2 | 603.060557 | -0.250260956 | 0.3499925 | -0.7150466 | 0.47458023 | 0.66074873 |
| TMEM250 | 113.443822 | -0.19088845 | 0.4918216 | -0.3881254 | 0.69792325 | 0.82590296 |
| QSOX2 | 149.46271 | 0.077496508 | 0.24217655 | 0.32000005 | 0.74896829 | 0.85857831 |
| SNAPC4 | 289.971784 | -0.664811316 | 0.42099645 | -1.5791376 | 0.1143045 | 0.2683288 |
| ENTR1 | 271.322169 | -0.297958257 | 0.31825384 | -0.9362283 | 0.34915565 | 0.54887281 |
| PMPCA | 268.852743 | -0.20658407 | 0.3018567 | -0.6843779 | 0.49373654 | 0.67602105 |
| INPP5E | 111.643879 | -0.2395892 | 0.27230023 | -0.8798715 | 0.37892894 | 0.57574455 |
| SEC16A | 556.755427 | -0.262967328 | 0.35882078 | -0.7328654 | 0.46364054 | 0.65160677 |
| NOTCH1 | 343.828278 | 1.411423418 | 0.32522812 | 4.33979518 | 1.43E-05 | 0.00033762 |
| EGFL7 | 95.4269555 | -0.368450896 | 0.49548592 | -0.7436153 | 0.45710927 | 0.64580465 |
| AGPAT2 | 216.240498 | -0.3425979 | 0.3969282 | -0.8631231 | 0.38806979 | 0.58387159 |
| SNHG7 | 256.789616 | -0.138189461 | 0.28032856 | -0.4929553 | 0.62204415 | 0.77562992 |
| RABL6 | 696.013545 | 0.572027506 | 0.32185626 | 1.77727629 | 0.07552279 | 0.20417967 |
| AJM1 | 84.8238129 | -0.442835754 | 0.66417069 | -0.6667499 | 0.50493189 | 0.6850618 |
| PHPT1 | 532.453514 | -0.31832848 | 0.37769539 | -0.842818 | 0.39933023 | 0.5934221 |
| MAMDC4 | 33.3479925 | 1.585691895 | 0.58208411 | 2.72416281 | 0.00644647 | 0.03657359 |
| EDF1 | 794.165944 | -0.31817813 | 0.37201139 | -0.8552914 | 0.39238986 | 0.58736138 |
| TRAF2 | 161.23424 | -0.021796098 | 0.31822952 | -0.0684918 | 0.94539418 | 0.97196767 |
| FBXW5 | 541.474984 | -0.442738241 | 0.39485888 | -1.1212569 | 0.26217853 | 0.46007129 |
| LCN12 | 61.0319491 | 1.189168137 | 0.53038035 | 2.24210445 | 0.02495462 | 0.09624452 |

| | | | | | | |
|---|---|---|---|---|---|---|
| PAXX | 133.241615 | 0.111698028 | 0.3604634 | 0.30987342 | 0.75665722 | 0.86357206 |
| ABCA2 | 50.3908223 | 0.412711139 | 0.61945333 | 0.66625058 | 0.50525095 | 0.68535128 |
| ENTPD2 | 94.3757322 | 0.926580911 | 0.68159139 | 1.35943753 | 0.17400798 | 0.3537824 |
| UAP1L1 | 156.110969 | 0.524811459 | 0.42211359 | 1.24329439 | 0.21375937 | 0.4040798 |
| MAN1B1 | 283.379606 | 0.17027996 | 0.29734109 | 0.5726755 | 0.56686443 | 0.73357352 |
| DPP7 | 278.335216 | -0.045666641 | 0.33548566 | -0.136121 | 0.89172563 | 0.94341489 |
| ANAPC2 | 52.0340624 | 0.392581589 | 0.31340058 | 1.25265113 | 0.2103327 | 0.39989367 |
| SSNA1 | 323.843061 | -0.725063155 | 0.33206778 | -2.1834794 | 0.02900052 | 0.10659967 |
| TPRN | 296.684444 | 0.723092184 | 0.26749381 | 2.70321095 | 0.00686731 | 0.03835942 |
| TMEM203 | 90.2575751 | -0.296665388 | 0.32507604 | -0.9126031 | 0.36145134 | 0.55944145 |
| NDOR1 | 98.9803478 | 0.144438853 | 0.28790202 | 0.50169448 | 0.61588245 | 0.7715135 |
| BX255925.3 | 85.2179289 | 0.224355867 | 0.45628337 | 0.49170292 | 0.62292937 | 0.77619139 |
| RNF208 | 57.4729212 | 0.767556725 | 0.52466054 | 1.46295874 | 0.14347867 | 0.31108227 |
| SLC34A3 | 109.13737 | 3.265527396 | 0.53238779 | 6.13373839 | 8.58E-10 | 1.03E-07 |
| TUBB4B | 862.993708 | -0.724139385 | 0.30086661 | -2.4068453 | 0.01609099 | 0.07082824 |
| STPG3-AS1 | 92.5324375 | 3.098782636 | 0.94614168 | 3.27517824 | 0.00105595 | 0.00941089 |
| NELFB | 379.960979 | -0.000382107 | 0.21032305 | -0.0018168 | 0.99855043 | 0.99920455 |
| EXD3 | 753.449139 | -0.141420058 | 0.18843687 | -0.7504904 | 0.45295941 | 0.64216694 |
| NOXA1 | 214.935153 | -0.202136682 | 0.30631474 | -0.6598986 | 0.50931888 | 0.68896021 |
| ENTPD8 | 37.8549162 | 2.191729793 | 0.52615623 | 4.16554941 | 3.11E-05 | 0.00061972 |
| NSMF | 408.302539 | 0.977129332 | 0.36825934 | 2.65337282 | 0.00796918 | 0.04276959 |
| PNPLA7 | 579.016955 | 0.446927667 | 0.56279529 | 0.79412119 | 0.42712488 | 0.62012506 |
| MRPL41 | 614.701517 | -0.447558135 | 0.38002868 | -1.1776957 | 0.23891795 | 0.43402823 |
| DPH7 | 217.644614 | -0.171812822 | 0.25884698 | -0.6637621 | 0.50684256 | 0.68683055 |
| ZMYND19 | 77.1616822 | -0.272141844 | 0.51999867 | -0.523351 | 0.60072999 | 0.76020679 |
| ARRDC1 | 290.156275 | 0.010649564 | 0.26601142 | 0.04003423 | 0.96806583 | 0.98378398 |
| ARRDC1-AS1 | 73.195377 | 0.212532946 | 0.24191904 | 0.87852923 | 0.37965658 | 0.57616838 |
| EHMT1 | 2426.6436 | -0.139267433 | 0.24487964 | -0.5687179 | 0.56954761 | 0.73565672 |
| LINC01001 | 68.113813 | 0.375199299 | 0.47550974 | 0.7890465 | 0.43008483 | 0.62270759 |
| AC069287.3 | 60.873162 | -0.218465324 | 0.59814391 | -0.3652387 | 0.71493321 | 0.83692796 |
| BET1L | 166.848717 | -0.245490145 | 0.30920596 | -0.7939373 | 0.42723194 | 0.6202148 |
| RIC8A | 154.228621 | -0.747886055 | 0.41129192 | -1.8183825 | 0.06900568 | 0.19248289 |
| SIRT3 | 217.703441 | 0.202104802 | 0.35575117 | 0.56810721 | 0.56996218 | 0.73612282 |
| PSMD13 | 374.085498 | -0.142226951 | 0.26572261 | -0.535246 | 0.5924798 | 0.75377024 |
| PGGHG | 322.558106 | -0.575180106 | 0.59302897 | -0.9699022 | 0.33209524 | 0.53165493 |
| IFITM2 | 489.615024 | -2.147180633 | 0.46223896 | -4.6451746 | 3.40E-06 | 0.00010609 |
| IFITM3 | 1734.79042 | -2.504042759 | 0.64124344 | -3.9049799 | 9.42E-05 | 0.00148296 |
| SIGIRR | 664.161007 | 1.081015204 | 0.39765147 | 2.71849921 | 0.00655788 | 0.03699131 |
| ANO9 | 329.469034 | 0.536719967 | 0.51584369 | 1.04047015 | 0.29812152 | 0.49833608 |
| PTDSS2 | 379.336846 | 0.166999739 | 0.21662628 | 0.77091171 | 0.44075926 | 0.63268271 |
| RNH1 | 446.526366 | -0.630684269 | 0.27387357 | -2.30283 | 0.02128841 | 0.08651354 |
| HRAS | 192.665938 | -0.376605136 | 0.41904479 | -0.8987229 | 0.36880029 | 0.56627598 |
| LMNTD2 | 67.1580016 | 1.78192536 | 0.4202121 | 4.24053798 | 2.23E-05 | 0.00048133 |
| AP006284.1 | 64.0650848 | -0.886095408 | 0.69303081 | -1.2785801 | 0.20104496 | 0.38812445 |
| RASSF7 | 427.699685 | -0.712063495 | 0.37735039 | -1.8870087 | 0.05915915 | 0.17312008 |
| MIR210HG | 200.662461 | -3.673599592 | 0.67467156 | -5.4450192 | 5.18E-08 | 3.21E-06 |
| PHRF1 | 445.563481 | 0.006965474 | 0.26603979 | 0.02618208 | 0.97911211 | 0.98943451 |
| IRF7 | 142.248192 | -0.602427018 | 0.7517177 | -0.8014006 | 0.42289977 | 0.61623726 |
| CDHR5 | 833.012885 | 1.079674422 | 0.45459341 | 2.37503316 | 0.01754737 | 0.07528071 |
| DEAF1 | 971.376484 | 0.131458009 | 0.28808201 | 0.45632148 | 0.64815882 | 0.79355195 |
| AC131934.1 | 134.387562 | -0.748554225 | 0.53916393 | -1.3883611 | 0.16502712 | 0.3423682 |
| EPS8L2 | 1671.61648 | 0.613489969 | 0.32142194 | 1.90867486 | 0.05630405 | 0.16770091 |
| AP006621.4 | 247.017322 | 0.22526833 | 0.60252134 | 0.3738761 | 0.7084965 | 0.83302123 |
| TALDO1 | 869.185418 | -1.61548697 | 0.39061649 | -4.1357367 | 3.54E-05 | 0.00068694 |
| GATD1 | 282.986363 | -0.079007943 | 0.38988543 | -0.202644 | 0.8394133 | 0.91490443 |
| AP006621.3 | 569.537002 | 1.170539705 | 0.3293083 | 3.55454053 | 0.00037864 | 0.00440953 |
| SLC25A22 | 85.2560503 | -1.304319388 | 0.40753754 | -3.2004889 | 0.00137195 | 0.01147362 |
| PIDD1 | 134.108908 | 0.807224355 | 0.23403863 | 3.44910732 | 0.00056244 | 0.00582282 |
| RPLP2 | 976.433692 | -0.999039217 | 0.45908394 | -2.176158 | 0.02954345 | 0.10795841 |
| PNPLA2 | 398.803033 | -0.411127461 | 0.18912778 | -2.1738079 | 0.02971956 | 0.10845741 |
| CD151 | 795.95258 | -1.063891533 | 0.3628165 | -2.932313 | 0.00336448 | 0.02249309 |
| POLR2L | 577.753112 | -0.989419883 | 0.40002381 | -2.4734025 | 0.01338333 | 0.06189114 |
| TSPAN4 | 265.876786 | -0.956071765 | 0.26600332 | -3.59421 | 0.00032538 | 0.00391357 |
| AP006623.1 | 35.1885661 | -1.255930257 | 0.3599069 | -3.4895976 | 0.00048375 | 0.00523373 |
| CHID1 | 173.690265 | 0.252331774 | 0.30628202 | 0.82385436 | 0.41002232 | 0.60386547 |
| AP2A2 | 1703.48734 | 0.279251059 | 0.25246936 | 1.10607902 | 0.26869232 | 0.46678905 |
| TOLLIP | 507.373729 | -0.185152264 | 0.29079926 | -0.6367013 | 0.52431944 | 0.69979031 |
| MOB2 | 249.326619 | -0.22862425 | 0.28114215 | -0.8131981 | 0.41610451 | 0.60947233 |
| DUSP8 | 92.6214755 | -1.478553082 | 0.64649163 | -2.2870413 | 0.02219341 | 0.08889687 |
| KRTAP5-AS1 | 94.200644 | -1.288188361 | 0.8806297 | -1.4628037 | 0.14352111 | 0.31111893 |
| MRPL23 | 140.445043 | 0.345063379 | 0.45793865 | 0.75351442 | 0.45114085 | 0.64095339 |
| CD81-AS1 | 81.1517891 | -0.513517806 | 0.42889501 | -1.1973042 | 0.231188 | 0.42415927 |
| CD81 | 705.622721 | -1.191733624 | 0.35604968 | -3.3470993 | 0.00081662 | 0.0077786 |
| TSSC4 | 129.829872 | -0.309507116 | 0.49798327 | -0.6215211 | 0.5342568 | 0.70733681 |
| KCNQ1 | 753.526609 | -0.148198751 | 0.40935793 | -0.3620273 | 0.71733162 | 0.8386157 |
| KCNQ1OT1 | 4552.53356 | 0.197145777 | 0.41427732 | 0.47587876 | 0.63416075 | 0.78430402 |
| CDKN1C | 416.76303 | -1.686609828 | 0.58902192 | -2.8634076 | 0.00419111 | 0.02653467 |
| SLC22A18AS | 94.1964345 | 0.279008643 | 0.5490431 | 0.50817257 | 0.61133233 | 0.76763761 |
| SLC22A18 | 742.324124 | 0.504363859 | 0.6220843 | 0.81076449 | 0.41750093 | 0.61055001 |
| NAP1L4 | 2006.04817 | -0.238969058 | 0.15510737 | -1.5406686 | 0.12339746 | 0.2825136 |
| CARS | 637.771879 | -0.545523035 | 0.31135284 | -1.7521055 | 0.07975566 | 0.21145278 |
| OSBPL5 | 133.355226 | 0.582213416 | 0.42956364 | 1.35536009 | 0.17530286 | 0.35540249 |
| ZNF195 | 536.688847 | 0.486958839 | 0.20813382 | 2.33964296 | 0.01930218 | 0.08034468 |
| AC127526.1 | 548.15805 | 0.902193636 | 0.35392709 | 2.54909463 | 0.0108003 | 0.05325169 |
| NUP98 | 1179.65408 | -0.11276801 | 0.28128184 | -0.4009075 | 0.6884882 | 0.8199051 |
| PGAP2 | 174.527877 | 0.636212245 | 0.32771572 | 1.94135406 | 0.05221535 | 0.15948419 |
| RHOG | 101.105287 | -0.974833858 | 0.3759092 | -2.5932695 | 0.00950682 | 0.04854197 |
| STIM1 | 1482.21815 | -0.802973988 | 0.30400784 | -2.6412937 | 0.00825901 | 0.04389935 |
| RRM1 | 99.991728 | 0.424366326 | 0.35972323 | 1.17970232 | 0.23811863 | 0.43308798 |
| TRIM21 | 51.0661938 | -1.310068252 | 0.58041841 | -2.2571101 | 0.0240012 | 0.09359682 |
| AC104389.4 | 496.135716 | 0.068353605 | 0.61035712 | 0.11198953 | 0.91083171 | 0.95310114 |
| TRIM5 | 110.787842 | -1.132874263 | 0.63969478 | -1.7709606 | 0.07656725 | 0.20594727 |
| TRIM22 | 264.219808 | -2.345671346 | 1.08890483 | -2.1541564 | 0.03122791 | 0.11231721 |
| FAM160A2 | 437.015251 | 0.822497739 | 0.21244109 | 3.87165085 | 0.0001081 | 0.00165004 |
| CAVIN3 | 63.1079376 | -2.253551222 | 0.55704338 | -4.0455578 | 5.22E-05 | 0.00094019 |
| SMPD1 | 355.719147 | 0.259486503 | 0.4155995 | 0.62436673 | 0.53238677 | 0.7058154 |
| APBB1 | 222.896721 | -0.275098495 | 0.30931961 | -0.8893665 | 0.37380616 | 0.57108348 |
| TRIM3 | 77.4502978 | -0.023565297 | 0.36867282 | -0.0639193 | 0.94903451 | 0.97382962 |
| ARFIP2 | 176.482236 | -0.578734683 | 0.29506867 | -1.9613559 | 0.04983752 | 0.1551439 |
| AC084337.2 | 314.810186 | -0.136663425 | 0.49298384 | -0.2772168 | 0.7816136 | 0.8801822 |
| DNHD1 | 482.904277 | -0.17171355 | 0.39957341 | -0.4297422 | 0.6673832 | 0.80675734 |
| RRP8 | 107.504143 | 0.131280077 | 0.25760702 | 0.50961374 | 0.6103221 | 0.76674759 |
| ILK | 376.004627 | -0.731167712 | 0.25567808 | -2.85972 | 0.00424015 | 0.02676155 |
| TAF10 | 162.523437 | -0.344374786 | 0.33544177 | -1.0266306 | 0.30459444 | 0.50478491 |
| TPP1 | 486.137403 | 0.095207798 | 0.33484895 | 0.28433059 | 0.77615705 | 0.8764136 |
| MRPL17 | 163.583347 | -1.340717317 | 0.3958175 | -3.3872108 | 0.00070607 | 0.00699286 |
| AC087280.2 | 325.439699 | -0.137715629 | 0.46174395 | -0.2982511 | 0.76551154 | 0.86903543 |
| ZNF215 | 168.960842 | -0.215226536 | 0.42053943 | -0.5117868 | 0.60880021 | 0.76558023 |
| ZNF214 | 119.029312 | 0.588384386 | 0.48237018 | 1.21977769 | 0.22254916 | 0.41472082 |
| NLRP14 | 42.0271318 | 0.45435012 | 0.41099163 | 1.10549725 | 0.26894419 | 0.46698986 |
| SYT9 | 532.999418 | -0.626150131 | 0.41586576 | -1.5056544 | 0.1321559 | 0.2954593 |
| AC107884.1 | 266.734271 | 0.364111874 | 0.34927273 | 1.04248583 | 0.29718649 | 0.49762219 |
| PPFIBP2 | 1043.36807 | 0.494127214 | 0.36146431 | 1.36701522 | 0.17162054 | 0.35089539 |

| | | | | | | |
|---|---|---|---|---|---|---|
| EIF3F | 295.471601 | -0.276872138 | 0.24799153 | -1.116458 | 0.26422612 | 0.46213441 |
| TUB | 685.030698 | 1.627573301 | 0.72168997 | 2.25522505 | 0.02411921 | 0.09389434 |
| RIC3 | 639.450082 | 0.092461403 | 0.3638388 | 0.25412739 | 0.79939715 | 0.89091281 |
| STK33 | 2731.86517 | 0.460774171 | 0.28275707 | 1.62957612 | 0.10319111 | 0.25087631 |
| TRIM66 | 936.10225 | 0.740687766 | 0.51942179 | 1.42598515 | 0.15387263 | 0.32633949 |
| RPL27A | 915.591799 | -0.994599397 | 0.40739219 | -2.4413806 | 0.01463123 | 0.06588377 |
| ST5 | 268.972535 | -0.979498603 | 0.53602826 | -1.8273264 | 0.0676507 | 0.18961187 |
| AKIP1 | 71.8652989 | -0.403392006 | 0.29855506 | -1.3511478 | 0.17664809 | 0.3570735 |
| TMEM9B | 180.728357 | -0.021050163 | 0.18235824 | -0.115433 | 0.9081019 | 0.95196591 |
| TMEM9B-AS | 160.190394 | 0.651440216 | 0.58556809 | 1.11249268 | 0.26592637 | 0.46378836 |
| DENND5A | 1928.92905 | -1.289121251 | 0.29352055 | -4.3919285 | 1.12E-05 | 0.00027834 |
| TMEM41B | 1082.50286 | -0.354710229 | 0.33881307 | -1.0469201 | 0.29513641 | 0.49508665 |
| IPO7 | 1031.73324 | -0.454390207 | 0.33721782 | -1.347468 | 0.17782957 | 0.35856279 |
| AC132192.2 | 37.1900977 | 0.40727192 | 0.57239243 | 0.71152569 | 0.47675853 | 0.66243581 |
| ZNF143 | 406.140996 | -0.091882174 | 0.44064352 | -0.2085182 | 0.83482441 | 0.91279955 |
| WEE1 | 129.743034 | -0.364968738 | 0.28210353 | -1.2937404 | 0.19575511 | 0.3810844 |
| SWAP70 | 654.349462 | -0.592341167 | 0.24918664 | -2.3770984 | 0.01744943 | 0.07500136 |
| SBF2-AS1 | 272.680466 | -1.359127407 | 0.27186723 | -4.9992322 | 5.76E-07 | 2.39E-05 |
| SBF2 | 5483.62978 | -0.107213268 | 0.25898652 | -0.4139724 | 0.67889433 | 0.8139067 |
| AC011092.2 | 172.786124 | -0.818665714 | 0.51789416 | -1.5807587 | 0.11393322 | 0.26768643 |
| AC100763.1 | 51.296928 | -0.67782065 | 0.65830229 | -1.0296495 | 0.30317455 | 0.50355977 |
| ADM | 572.32879 | -2.087296997 | 0.90379534 | -2.3094797 | 0.02091698 | 0.08522859 |
| AMPD3 | 151.1313 | -2.597438038 | 0.52129669 | -4.9826483 | 6.27E-07 | 2.58E-05 |
| MTRNR2L8 | 644.370252 | -0.299890513 | 0.71377795 | -0.4201454 | 0.67437925 | 0.81105241 |
| RNF141 | 132.0346 | -0.166027904 | 0.17311259 | -0.9590747 | 0.33752111 | 0.53770361 |
| CTR9 | 374.80347 | -0.295984235 | 0.23901441 | -1.2383531 | 0.21558516 | 0.40624496 |
| EIF4G2 | 1103.1344 | -0.32158352 | 0.25253078 | -1.2734429 | 0.20286093 | 0.39070041 |
| ZBED5 | 452.093679 | -0.154068813 | 0.19762039 | -0.77962 | 0.43561457 | 0.62792337 |
| ZBED5-AS1 | 177.226645 | 1.276032526 | 0.56300614 | 2.26646288 | 0.02342306 | 0.09212621 |
| GALNT18 | 2922.28258 | 0.291197971 | 0.46959606 | 0.6201031 | 0.53518991 | 0.70809345 |
| USP47 | 2640.37226 | -0.29838301 | 0.16992908 | -1.7559267 | 0.07910093 | 0.21028635 |
| MICAL2 | 177.394658 | -1.785417288 | 0.56767588 | -3.145135 | 0.0016601 | 0.01329148 |
| PARVA | 883.501516 | -0.610022568 | 0.30437603 | -2.0041741 | 0.04505141 | 0.14502576 |
| TEAD1 | 5628.78968 | -0.346272072 | 0.29805218 | -1.1617834 | 0.24532346 | 0.4408227 |
| LINC00958 | 622.320019 | -2.005599988 | 0.51337398 | -3.9067036 | 9.36E-05 | 0.00147751 |
| ARNTL | 485.855332 | 0.518506374 | 0.42552038 | 1.21852301 | 0.22302529 | 0.41521234 |
| BTBD10 | 603.273366 | -0.302459089 | 0.30694436 | -0.9853874 | 0.32443384 | 0.52483341 |
| FAR1 | 960.913081 | -0.271415919 | 0.21777089 | -1.246337 | 0.2126407 | 0.40246701 |
| SPON1 | 1043.40986 | -1.308359239 | 0.50284381 | -2.6019198 | 0.00927035 | 0.04769848 |
| SPON1-AS1 | 364.752407 | -1.640363942 | 0.88107759 | -1.8617701 | 0.0626355 | 0.17998842 |
| RRAS2 | 4706.78437 | -1.64888474 | 0.6864393 | -2.4020838 | 0.01630197 | 0.07143578 |
| COPB1 | 881.782474 | -0.44853923 | 0.21625367 | -2.0741346 | 0.03806681 | 0.12937807 |
| AC018523.2 | 40.7372568 | -1.115780803 | 0.6536988 | -1.7068729 | 0.08784567 | 0.22582532 |
| PSMA1 | 278.561494 | -0.315357915 | 0.36892329 | -0.8548062 | 0.39265845 | 0.58757307 |
| PDE3B | 833.375621 | -0.266969217 | 0.28470032 | -0.9377201 | 0.34838827 | 0.54845045 |
| CYP2R1 | 155.532038 | -0.162525764 | 0.25781924 | -0.6303865 | 0.52844176 | 0.70282856 |
| CALCB | 300.11736 | 0.593650733 | 0.53937971 | 1.10061747 | 0.27106318 | 0.46924261 |
| INSC | 75.6103776 | 0.040209726 | 0.80836012 | 0.04974234 | 0.96032771 | 0.97961929 |
| SOX6 | 6953.51457 | 0.44394225 | 0.44268241 | 1.00284593 | 0.3159352 | 0.51590706 |
| C11orf58 | 927.6463 | -0.303392922 | 0.15974706 | -1.8992082 | 0.05753711 | 0.17015344 |
| PLEKHA7 | 9116.40571 | 1.012566897 | 0.20118879 | 5.03291909 | 4.83E-07 | 2.09E-05 |
| RPS13 | 737.307011 | -0.930338149 | 0.46331674 | -2.007996 | 0.04464372 | 0.1438823 |
| PIK3C2A | 2204.93729 | -0.17485549 | 0.12130268 | -1.4414809 | 0.14944887 | 0.31978206 |
| NUCB2 | 325.64351 | -0.628619964 | 0.35542091 | -1.7686634 | 0.07695006 | 0.20666074 |
| NCR3LG1 | 194.338897 | -0.173379627 | 0.48111384 | -0.3603713 | 0.71856947 | 0.83946406 |
| AC124798.1 | 88.3220098 | 0.222445583 | 0.44350087 | 0.50156741 | 0.61597185 | 0.77155497 |
| USH1C | 1467.71416 | -0.024468779 | 0.35482625 | -0.0689599 | 0.94502155 | 0.97189453 |
| KCNC1 | 76.3582729 | -1.849158983 | 0.61400952 | -3.0116129 | 0.00259864 | 0.01871756 |
| SERGEF | 1031.46285 | -0.333545177 | 0.20050837 | -1.6634975 | 0.09621289 | 0.23950313 |
| SAAL1 | 91.2154687 | 0.520101409 | 0.24762441 | 2.10036406 | 0.03569683 | 0.12334168 |
| SAA1 | 2344.04605 | -6.915818945 | 1.66566858 | -4.1519778 | 3.30E-05 | 0.00065008 |
| HPS5 | 260.285311 | -0.189958733 | 0.17923562 | -1.0598269 | 0.28922336 | 0.48903082 |
| GTF2H1 | 693.563744 | 0.07200008 | 0.27705551 | 0.25987601 | 0.79495942 | 0.88829145 |
| LDHA | 1844.11059 | -2.644734761 | 0.46788344 | -5.6525505 | 1.58E-08 | 1.21E-06 |
| LDHC | 45.0331712 | 0.855499194 | 0.84450534 | 1.01301809 | 0.31105152 | 0.51127879 |
| TSG101 | 907.123546 | -0.45069554 | 0.28559957 | -1.578068 | 0.11454999 | 0.26867501 |
| UEVLD | 675.885441 | -0.095443827 | 0.15201555 | -0.6278557 | 0.5300985 | 0.70421206 |
| SPTY2D1OS | 159.52607 | 0.116394121 | 0.2697796 | 0.43144153 | 0.66614736 | 0.8061877 |
| SPTY2D1 | 224.768042 | -0.937042849 | 0.44631591 | -2.0995058 | 0.03577234 | 0.12350917 |
| ZDHHC13 | 372.529973 | -0.891800265 | 0.23436246 | -3.8052181 | 0.00014168 | 0.00202181 |
| NAV2 | 1059.81906 | -4.038999815 | 1.23829582 | -3.2617406 | 0.0011073 | 0.00977247 |
| HTATIP2 | 273.923979 | -2.015726729 | 0.3948072 | -5.1055977 | 3.30E-07 | 1.49E-05 |
| PRMT3 | 757.276872 | -0.184174468 | 0.25726476 | -0.7158947 | 0.4740564 | 0.66028768 |
| NELL1 | 233.099172 | 2.999480102 | 0.48361698 | 6.20218108 | 5.57E-10 | 7.07E-08 |
| ANO5 | 463.590819 | 0.604192938 | 0.36042061 | 1.67635515 | 0.09366865 | 0.23571272 |
| FANCF | 50.3177534 | -0.406411476 | 0.44995021 | -0.9032366 | 0.36640036 | 0.56391756 |
| GAS2 | 406.480114 | 0.832353259 | 0.41935632 | 1.98483537 | 0.04716279 | 0.14978229 |
| SVIP | 253.541949 | -0.169509484 | 0.27087017 | -0.6257961 | 0.53144871 | 0.70522693 |
| AC006299.1 | 144.513109 | 0.374785065 | 0.55622642 | 0.67379946 | 0.50043882 | 0.68158931 |
| AC090092.1 | 31.0066912 | 0.183910418 | 0.58242475 | 0.31576683 | 0.7521795 | 0.86075508 |
| LUZP2 | 191.363913 | 0.418106975 | 0.71610489 | 0.58386276 | 0.55931265 | 0.72786628 |
| ANO3 | 32.0692392 | 1.363796148 | 0.43131917 | 3.16191872 | 0.00156733 | 0.01278015 |
| SLC5A12 | 7706.45061 | 2.614609524 | 0.93354496 | 2.80073229 | 0.00509868 | 0.03073333 |
| BBOX1 | 3351.26472 | 0.838684606 | 0.71043178 | 1.180528 | 0.23779028 | 0.43268172 |
| BBOX1-AS1 | 784.774643 | 0.561525383 | 0.46230434 | 1.21462279 | 0.22450999 | 0.41674812 |
| CCDC34 | 53.7897761 | 0.132765973 | 0.3808246 | 0.34862762 | 0.72736889 | 0.84519832 |
| LGR4 | 4604.60104 | -0.021139008 | 0.26508959 | -0.0797429 | 0.93644175 | 0.96589714 |
| LIN7C | 292.666742 | -0.231783073 | 0.26500138 | -0.8746486 | 0.38176516 | 0.57844716 |
| BDNF-AS | 914.68378 | 0.464377506 | 0.25887979 | 1.79379587 | 0.07284576 | 0.19915585 |
| KIF18A | 36.7613607 | -0.647267393 | 0.52074381 | -1.242967 | 0.21387998 | 0.40419867 |
| METTL15 | 2380.68796 | 0.132894874 | 0.28979539 | 0.45858174 | 0.64653455 | 0.79274077 |
| AC090833.1 | 61.0521265 | 0.852675193 | 0.65409405 | 1.30359723 | 0.19237094 | 0.3768513 |
| ARL14EP | 249.040456 | 0.126601532 | 0.18536637 | 0.68298005 | 0.49461945 | 0.67689185 |
| MPPED2 | 370.871954 | -0.244162261 | 0.7324145 | -0.3333662 | 0.73885785 | 0.8518399 |
| DCDC1 | 1246.79967 | 0.305232409 | 0.38542439 | 0.79193849 | 0.42839654 | 0.62144475 |
| DNAJC24 | 513.259742 | -0.156712232 | 0.37179407 | -0.4215028 | 0.67338799 | 0.81072605 |
| IMMP1L | 1220.74383 | 0.226456384 | 0.28762006 | 0.7873456 | 0.43107959 | 0.62368671 |
| ELP4 | 2177.72399 | 0.535862439 | 0.2440234 | 2.19594695 | 0.02809574 | 0.10430886 |
| AC131571.1 | 199.437594 | -0.571399933 | 0.69418724 | -0.8231208 | 0.41043932 | 0.60421993 |
| RCN1 | 204.885815 | -1.630561409 | 0.47184245 | -3.4557328 | 0.0005488 | 0.00572903 |
| AL078612.1 | 41.2349013 | -0.827663714 | 0.44280059 | -1.8691568 | 0.06160101 | 0.17794837 |
| WT1 | 46.4926616 | 0.317919364 | 0.47134378 | 0.67449573 | 0.4999962 | 0.68139272 |
| EIF3M | 467.807329 | -0.166687891 | 0.24908586 | -0.6691985 | 0.50336885 | 0.68404435 |
| CCDC73 | 471.299512 | -0.027584919 | 0.43118247 | -0.063975 | 0.94899009 | 0.97382962 |
| PRRG4 | 554.821264 | -0.529485556 | 0.45877609 | -1.1541263 | 0.24844839 | 0.44469601 |
| QSER1 | 891.395333 | -0.297896117 | 0.23790304 | -1.2521745 | 0.21050628 | 0.39989367 |
| DEPDC7 | 377.908293 | 1.160746488 | 0.57411337 | 2.0218071 | 0.04319628 | 0.1406059 |
| TCP11L1 | 994.606396 | 0.885594836 | 0.30539623 | 2.89982241 | 0.00373374 | 0.02427818 |
| CSTF3 | 908.939604 | 0.098284908 | 0.172329 | 0.57033296 | 0.56845189 | 0.73472607 |
| HIPK3 | 2446.02387 | 0.38822152 | 0.23849834 | 1.6277745 | 0.10357271 | 0.25155378 |
| KIAA1549L | 158.846387 | -0.44405837 | 0.72107463 | -0.6158286 | 0.53800766 | 0.70986341 |
| AL049629.1 | 453.238614 | -0.084587458 | 0.55776689 | -0.1516538 | 0.87946001 | 0.93716208 |
| CD59 | 167.663631 | -0.785094331 | 0.27361898 | -2.8692978 | 0.00411384 | 0.02620281 |

| | | | | | | |
|---|---|---|---|---|---|---|
| FBXO3 | 632.501766 | 0.061707004 | 0.23746191 | 0.25986064 | 0.79497127 | 0.88829145 |
| LMO2 | 71.245885 | 1.263406718 | 0.34349936 | 3.67804675 | 0.00023503 | 0.00303916 |
| CAPRIN1 | 1051.44648 | -0.015385451 | 0.21929508 | -0.0701587 | 0.94406737 | 0.9712047 |
| NAT10 | 361.045648 | -0.265578292 | 0.26876576 | -0.9881403 | 0.32308392 | 0.52358848 |
| ABTB2 | 3080.32094 | -2.822566148 | 0.48367719 | -5.8356403 | 5.36E-09 | 4.80E-07 |
| CAT | 1411.31003 | 0.531288394 | 0.46252591 | 1.14866732 | 0.25069319 | 0.44702114 |
| APIP | 251.632757 | -0.175766154 | 0.34763066 | -0.5056118 | 0.61312921 | 0.76933839 |
| PDHX | 726.204589 | -0.036878993 | 0.19733999 | -0.1868805 | 0.85175434 | 0.92197716 |
| CD44 | 171.540738 | -3.642664611 | 0.90906984 | -4.0070239 | 6.15E-05 | 0.00106678 |
| FJX1 | 55.5648003 | -2.716856324 | 0.75146833 | -3.615397 | 0.00029989 | 0.00367342 |
| TRIM44 | 3133.33101 | 0.033217141 | 0.20389446 | 0.16291341 | 0.87058661 | 0.93322364 |
| LDLRAD3 | 619.505072 | -1.465640033 | 0.35411483 | -4.1388836 | 3.49E-05 | 0.00068144 |
| COMMD9 | 177.78023 | -0.056515501 | 0.21739759 | -0.2599638 | 0.79489171 | 0.88829145 |
| PRR5L | 64.9582041 | -1.274659528 | 0.42813596 | -2.9772307 | 0.00290865 | 0.02021748 |
| TRAF6 | 385.478725 | -0.486860333 | 0.2290967 | -2.1251303 | 0.03357574 | 0.11842764 |
| RAG1 | 86.0403491 | 0.23350043 | 0.500104 | 0.46690375 | 0.64056875 | 0.78888372 |
| C11orf74 | 144.71231 | 0.829620134 | 0.26805178 | 3.0949995 | 0.00196813 | 0.01517277 |
| LRRC4C | 308.152108 | -0.587868512 | 0.6832975 | -0.8603405 | 0.38960137 | 0.58523463 |
| API5 | 735.327663 | -0.044869868 | 0.24354018 | -0.1842401 | 0.8538251 | 0.92301331 |
| TTC17 | 3795.67336 | 0.171934325 | 0.13545544 | 1.26930544 | 0.20433215 | 0.39259612 |
| AC068205.1 | 94.7649511 | 0.559800404 | 0.27098525 | 2.0657966 | 0.03884769 | 0.13105717 |
| HSD17B12 | 2125.35988 | -0.496342002 | 0.18122884 | -2.7387584 | 0.00616717 | 0.03532527 |
| AC087521.2 | 43.9652132 | -1.216574956 | 0.578884 | -2.1015868 | 0.03558949 | 0.12306386 |
| ALKBH3 | 221.275249 | 0.211290676 | 0.17860743 | 1.18298932 | 0.23681338 | 0.43136226 |
| C11orf96 | 122.424404 | -3.168047243 | 0.98389795 | -3.2198942 | 0.00128238 | 0.01088395 |
| ACCS | 213.028644 | 0.582849941 | 0.62234637 | 0.93653625 | 0.34899716 | 0.54884167 |
| EXT2 | 691.098409 | -0.36001139 | 0.18728491 | -1.9222659 | 0.05457231 | 0.16434269 |
| CD82 | 53.6673483 | -0.414411746 | 0.50008336 | -0.8286853 | 0.40728249 | 0.60112212 |
| TSPAN18 | 970.576345 | 1.039501657 | 0.43766506 | 2.37510769 | 0.01754383 | 0.07528071 |
| TP53I11 | 67.0888288 | 0.147227283 | 0.44401926 | 0.3315786 | 0.74020748 | 0.85230954 |
| PRDM11 | 146.194862 | -0.044029286 | 0.33665318 | -0.1307853 | 0.89594515 | 0.9460654 |
| SLC35C1 | 78.2569735 | -1.702528479 | 0.44134158 | -3.8576208 | 0.0001145 | 0.00173223 |
| CRY2 | 532.230679 | 0.52578484 | 0.36018938 | 1.45974552 | 0.14436003 | 0.31215379 |
| MAPK8IP1 | 65.4201086 | 0.410568248 | 0.44061 | 0.93181782 | 0.35143068 | 0.55058411 |
| PEX16 | 113.436882 | -0.217330993 | 0.2732849 | -0.7952543 | 0.4264656 | 0.61929907 |
| LARGE2 | 149.612591 | 0.347563336 | 0.5399238 | 0.64372665 | 0.51975271 | 0.69597507 |
| PHF21A | 2173.03425 | -0.113939303 | 0.35173676 | -0.3239335 | 0.74598842 | 0.85645894 |
| DGKZ | 530.21695 | 0.174089321 | 0.17840901 | 0.97578773 | 0.32916966 | 0.52891681 |
| MDK | 98.1823859 | -2.61661681 | 0.82630157 | -3.1666608 | 0.001542 | 0.01262617 |
| AMBRA1 | 2964.27607 | -0.11347619 | 0.24560708 | -0.4620233 | 0.64406463 | 0.79148232 |
| HARBI1 | 55.4745417 | 0.384027261 | 0.4645578 | 0.82665119 | 0.40843479 | 0.60217443 |
| ATG13 | 568.45308 | 0.137654056 | 0.2334246 | 0.5897153 | 0.55538154 | 0.72476748 |
| ARHGAP1 | 336.386447 | -0.094131795 | 0.29935569 | -0.314448 | 0.75318081 | 0.8613901 |
| ZNF408 | 38.1355022 | 0.210144885 | 0.35005337 | 0.60032243 | 0.54829138 | 0.71849234 |
| CKAP5 | 636.714433 | -0.192774835 | 0.18914629 | -1.0191838 | 0.30811572 | 0.50824576 |
| LRP4 | 473.72074 | -0.77880507 | 0.45769235 | -1.7015908 | 0.08883211 | 0.22732521 |
| C11orf49 | 2557.57796 | 0.111854866 | 0.19018686 | 0.58813141 | 0.55644409 | 0.72549978 |
| ARFGAP2 | 297.419392 | 0.121496353 | 0.22600585 | 0.53758057 | 0.59086665 | 0.75259958 |
| PACSIN3 | 85.0241715 | -0.210410631 | 0.28919671 | -0.7275692 | 0.46687732 | 0.65461205 |
| DDB2 | 236.866625 | -0.542065231 | 0.52573867 | -1.0310545 | 0.30251525 | 0.50273722 |
| ACP2 | 145.883369 | 0.637529275 | 0.26812794 | 2.37770552 | 0.01742073 | 0.07491801 |
| NR1H3 | 234.473278 | 0.868145808 | 0.28614987 | 3.03388498 | 0.00241426 | 0.01773436 |
| MADD | 366.445601 | 0.683178296 | 0.1482084 | 4.60957881 | 4.03E-06 | 0.0001206 |
| SLC39A13 | 207.359797 | -0.897891012 | 0.37660521 | -2.3841705 | 0.01711767 | 0.07395389 |
| PSMC3 | 146.015427 | -0.339356314 | 0.38231086 | -0.887645 | 0.37473175 | 0.5715597 |
| CELF1 | 3394.59489 | 0.097604004 | 0.32709641 | 0.29839522 | 0.76540154 | 0.86903543 |
| NDUFS3 | 215.973305 | 0.033433959 | 0.25178067 | 0.13279001 | 0.89435945 | 0.94518003 |
| KBTBD4 | 43.4008723 | -0.319172307 | 0.33153201 | -0.9627194 | 0.33568834 | 0.53553074 |
| MTCH2 | 1243.30986 | 0.429961851 | 0.16389056 | 2.62346937 | 0.00870393 | 0.04551878 |
| AGBL2 | 49.6411955 | 0.87815115 | 0.50720213 | 1.73136328 | 0.08338699 | 0.21821721 |
| FNBP4 | 2088.48482 | -0.70132028 | 0.2597943 | -2.6995214 | 0.00694393 | 0.03870697 |
| NUP160 | 959.89341 | 0.159486811 | 0.18587182 | 0.85804729 | 0.39086635 | 0.58629953 |
| PTPRJ | 6656.38778 | 0.241583097 | 0.30279364 | 0.79784732 | 0.4249591 | 0.61802805 |
| FOLH1 | 288.546767 | 3.384404033 | 0.60475898 | 5.59628574 | 2.19E-08 | 1.59E-06 |
| TNKS1BP1 | 349.925117 | -1.826713027 | 0.33884803 | -5.3909507 | 7.01E-08 | 4.14E-06 |
| SSRP1 | 174.305686 | -0.624294936 | 0.32450227 | -1.9238538 | 0.05437291 | 0.16401617 |
| TIMM10 | 104.066228 | -1.155059625 | 0.48032361 | -2.4047529 | 0.0161834 | 0.07109804 |
| UBE2L6 | 85.7045469 | -2.397190631 | 0.89389408 | -2.681739 | 0.00732406 | 0.04031761 |
| SERPING1 | 467.022232 | -1.792481804 | 0.50924915 | -3.5198523 | 0.00043179 | 0.00483143 |
| CLP1 | 36.0730843 | -0.843625942 | 0.38589612 | -2.1861478 | 0.0288048 | 0.10612074 |
| ZDHHC5 | 671.031559 | -0.617331998 | 0.23574649 | -2.6186265 | 0.00882846 | 0.04601226 |
| MED19 | 119.404853 | -0.599969576 | 0.26366628 | -2.2754884 | 0.02287665 | 0.09073211 |
| CTNND1 | 1724.37053 | 0.052601692 | 0.28139539 | 0.1869316 | 0.85171426 | 0.92197716 |
| OR9Q1 | 418.287191 | 0.986658218 | 0.89540744 | 1.10190978 | 0.2705009 | 0.46869989 |
| LPXN | 403.470237 | 0.864109067 | 0.33826343 | 2.55454475 | 0.01063268 | 0.05274777 |
| GLYAT | 3866.30172 | 2.219906498 | 0.70239913 | 3.16046305 | 0.00157519 | 0.01282892 |
| GLYATL2 | 89.7497538 | -2.03859684 | 0.51074091 | -3.99145 | 6.57E-05 | 0.00112098 |
| GLYATL1 | 2231.83138 | 1.029535153 | 0.71683262 | 1.4362281 | 0.15093745 | 0.32210798 |
| AP001636.3 | 169.466653 | -0.7112055 | 0.76688158 | -0.9273994 | 0.35371921 | 0.55215001 |
| FAM111A-DT | 53.3736272 | 0.291087164 | 0.40375079 | 0.72095751 | 0.47093566 | 0.65738999 |
| FAM111A | 275.735097 | 0.726046813 | 0.40110634 | 1.81011056 | 0.07027864 | 0.19476534 |
| DTX4 | 296.527857 | 0.267106129 | 0.5519745 | 0.48391027 | 0.62844955 | 0.78055893 |
| OSBP | 898.414382 | -0.279454479 | 0.2204768 | -1.2675006 | 0.20497634 | 0.39334944 |
| AP000442.2 | 47.9520771 | -0.709404804 | 0.44149416 | -1.6068272 | 0.10809229 | 0.25852748 |
| PATL1 | 616.704826 | -0.417901173 | 0.27394923 | -1.5254694 | 0.12714206 | 0.28824615 |
| STX3 | 2570.90303 | 0.366419251 | 0.38441015 | 0.95319868 | 0.34048941 | 0.54104897 |
| MRPL16 | 164.160858 | -0.204662766 | 0.41486968 | -0.4933182 | 0.62178777 | 0.77543854 |
| GIF | 36.2889918 | 0.363792883 | 0.72157547 | 0.5041647 | 0.61414564 | 0.76975835 |
| AP002347.1 | 83.2960226 | 1.39800952 | 0.68362097 | 2.04500679 | 0.04085419 | 0.13544314 |
| AP000790.3 | 60.8806435 | 1.671141742 | 0.8735265 | 1.91309793 | 0.05573552 | 0.16655041 |
| PRPF19 | 252.142559 | -1.012202187 | 0.49784216 | -2.0331789 | 0.04203445 | 0.13799537 |
| TMEM109 | 89.2633983 | -0.027237616 | 0.33595516 | -0.0810752 | 0.93538219 | 0.96565906 |
| TMEM132A | 46.6933929 | -3.000646525 | 0.68908877 | -4.3545137 | 1.33E-05 | 0.00032127 |
| VPS37C | 285.049107 | -0.234241249 | 0.28623889 | -0.8183418 | 0.41316204 | 0.60685942 |
| VWCE | 40.5722667 | -0.383277005 | 0.66188203 | -0.5790715 | 0.56254095 | 0.73018454 |
| DDB1 | 596.123731 | -0.144241144 | 0.27577626 | -0.5230368 | 0.60094861 | 0.76031038 |
| TKFC | 258.015563 | 2.318898541 | 0.47533333 | 4.87846818 | 1.07E-06 | 4.08E-05 |
| CYB561A3 | 93.7856822 | 0.372505535 | 0.22844676 | 1.63060108 | 0.10297452 | 0.25057194 |
| TMEM138 | 160.582539 | 0.299559283 | 0.23919012 | 1.25238989 | 0.21042783 | 0.39989367 |
| TMEM216 | 48.1310628 | 0.002047755 | 0.39704863 | 0.00515744 | 0.99588498 | 0.9977776 |
| CPSF7 | 709.197627 | 0.417835827 | 0.2354491 | 1.77463339 | 0.07595844 | 0.20503394 |
| AP003108.2 | 448.347571 | 0.374683481 | 0.34946831 | 1.07215295 | 0.28365134 | 0.48292248 |
| DAGLA | 61.6146559 | -0.800143922 | 0.49168841 | -1.6273394 | 0.10366503 | 0.25158226 |
| MYRF | 176.452704 | -0.741327485 | 0.56895614 | -1.3029607 | 0.19258819 | 0.37722296 |
| TMEM258 | 193.462794 | -1.037479695 | 0.2679155 | -3.8724138 | 0.00010776 | 0.00164679 |
| FADS2 | 139.735045 | -0.741517442 | 0.7635805 | -0.9711058 | 0.3314956 | 0.5313155 |
| FADS1 | 178.184143 | -3.257272241 | 0.68950306 | -4.7240867 | 2.31E-06 | 7.75E-05 |
| FADS3 | 463.483424 | -1.683288699 | 0.4277596 | -3.9351278 | 8.32E-05 | 0.00134884 |
| RAB3IL1 | 105.068989 | 0.384999701 | 0.54883096 | 0.70149049 | 0.48299697 | 0.66711401 |
| FTH1 | 15342.8017 | -2.128983197 | 0.60334499 | -3.5286333 | 0.00041771 | 0.00472407 |
| AP003733.3 | 32.8081771 | -3.267477424 | 0.93635227 | -3.4895814 | 0.00048378 | 0.00523373 |
| INCENP | 69.3802384 | 0.087793499 | 0.35694394 | 0.24595879 | 0.80571413 | 0.89453456 |
| ASRGL1 | 1140.91285 | 0.769957506 | 0.40959576 | 1.87979851 | 0.06013554 | 0.17489453 |

| | | | | | | |
|---|---|---|---|---|---|---|
| AHNAK | 1378.90304 | -1.313784411 | 0.44338308 | -2.963091 | 0.00304566 | 0.02093627 |
| MTA2 | 66.4566751 | -0.254308612 | 0.44263922 | -0.574528 | 0.56561058 | 0.73247225 |
| EML3 | 127.225628 | 0.567462061 | 0.31814235 | 1.78367342 | 0.07447675 | 0.20233016 |
| B3GAT3 | 206.113577 | -0.192831196 | 0.29483676 | -0.654027 | 0.51309443 | 0.69167833 |
| GANAB | 697.341463 | 0.180622614 | 0.19166346 | 0.94239462 | 0.34599064 | 0.5463646 |
| INTS5 | 52.2594372 | -0.977521865 | 0.35761966 | -2.733412 | 0.00626819 | 0.03572476 |
| CSKMT | 66.9075398 | -0.347695819 | 0.35422383 | -0.981571 | 0.32631126 | 0.52645345 |
| UQCC3 | 181.006836 | -0.504760139 | 0.35182848 | -1.4346768 | 0.15137923 | 0.32276202 |
| UBXN1 | 223.805852 | -0.629598757 | 0.32321002 | -1.9479556 | 0.05142028 | 0.15796366 |
| TTC9C | 268.186155 | 0.155982589 | 0.23388199 | 0.66692861 | 0.50481773 | 0.6850618 |
| POLR2G | 79.203531 | 0.120625029 | 0.33888408 | 0.35594776 | 0.72187969 | 0.84218273 |
| TAF6L | 170.433231 | 0.629607616 | 0.34600331 | 1.81965778 | 0.06881114 | 0.19197929 |
| TMEM223 | 165.512155 | -0.769012407 | 0.23709499 | -3.2434781 | 0.0011808 | 0.0102529 |
| TMEM179B | 239.525132 | -0.748213423 | 0.30537168 | -2.450173 | 0.01427876 | 0.06472188 |
| NXF1 | 311.046073 | 0.608269949 | 0.38275652 | 1.58918245 | 0.11201921 | 0.2649607 |
| STX5 | 526.220319 | -0.098097554 | 0.25716071 | -0.381464 | 0.70285897 | 0.82939874 |
| AP001160.1 | 42.8246393 | -1.043817983 | 0.36351077 | -2.8714912 | 0.0040854 | 0.02605798 |
| WDR74 | 108.922235 | -0.098077873 | 0.42589487 | -0.2302866 | 0.81786909 | 0.90160299 |
| SLC3A2 | 1276.16021 | -0.602642737 | 0.51694906 | -1.1657681 | 0.24370821 | 0.43890532 |
| SLC22A6 | 4506.83278 | 1.352406372 | 0.88598383 | 1.52644589 | 0.12689886 | 0.28796083 |
| SLC22A8 | 1422.81261 | 2.090729172 | 0.88823884 | 2.35379167 | 0.01858302 | 0.07827827 |
| SLC22A24 | 789.514516 | 1.400709746 | 0.65953518 | 2.12378322 | 0.03368827 | 0.1187057 |
| SLC22A10 | 134.471515 | 1.404860709 | 0.66260553 | 2.12020672 | 0.03398862 | 0.11939568 |
| HRASLS5 | 21.8123328 | 2.881724027 | 1.21256983 | 2.37654273 | 0.01747574 | 0.07506735 |
| RARRES3 | 830.49524 | -1.48014871 | 0.74279621 | -1.9926713 | 0.04629745 | 0.14778741 |
| HRASLS2 | 35.0841468 | 1.296148656 | 0.74223361 | 1.74628127 | 0.08076209 | 0.21341931 |
| PLA2G16 | 1079.66941 | -0.334470448 | 0.22412129 | -1.4923636 | 0.13560386 | 0.30013275 |
| ATL3 | 475.06032 | -0.57645376 | 0.22408595 | -2.5724672 | 0.01009765 | 0.05082942 |
| RTN3 | 1077.99863 | -0.569008654 | 0.3412798 | -1.667279 | 0.09545894 | 0.23833436 |
| C11orf95 | 88.3345298 | 0.800123452 | 0.27932555 | 2.86448354 | 0.0041769 | 0.02648138 |
| SPINDOC | 82.0252383 | -0.477996905 | 0.34618754 | -1.3807455 | 0.16735721 | 0.34532068 |
| MARK2 | 842.255005 | -0.122143163 | 0.27735363 | -0.4403878 | 0.65965623 | 0.8020214 |
| NAA40 | 204.998015 | -0.1637912 | 0.27052746 | -0.6054513 | 0.54487917 | 0.71518326 |
| OTUB1 | 261.414162 | 0.001834899 | 0.19960026 | 0.00919287 | 0.99266525 | 0.99629729 |
| MACROD1 | 880.880747 | 0.810862723 | 0.34561089 | 2.34617241 | 0.01896733 | 0.07937082 |
| AP000721.2 | 137.022909 | 0.223644339 | 0.60452622 | 0.36994978 | 0.71141991 | 0.83502592 |
| FLRT1 | 43.9393594 | -0.149336215 | 0.73526295 | -0.2031059 | 0.83905229 | 0.91480192 |
| STIP1 | 377.056297 | -0.844479294 | 0.44892766 | -1.8811033 | 0.05995787 | 0.1745825 |
| TRPT1 | 107.934522 | 0.050835802 | 0.29066514 | 0.17489473 | 0.86116235 | 0.92696343 |
| NUDT22 | 209.334141 | 0.377578182 | 0.20270489 | 1.86269891 | 0.06250464 | 0.17972542 |
| AP001453.1 | 78.7566357 | -0.179296699 | 0.43526322 | -0.4119271 | 0.68039289 | 0.81442647 |
| DNAJC4 | 109.374295 | 0.754900519 | 0.35092738 | 2.15115882 | 0.03146367 | 0.11296709 |
| VEGFB | 424.732892 | -0.59577704 | 0.31756588 | -1.8760738 | 0.06064514 | 0.17607773 |
| FKBP2 | 448.597859 | -0.098017482 | 0.34301479 | -0.2857529 | 0.77506736 | 0.87561595 |
| PPP1R14B | 244.068023 | -1.52754229 | 0.38662313 | -3.9509852 | 7.78E-05 | 0.00128069 |
| PLCB3 | 79.148777 | -0.033567743 | 0.31601587 | -0.1062217 | 0.91540645 | 0.9548342 |
| BAD | 217.996965 | 0.011572238 | 0.28886198 | 0.04006148 | 0.96804411 | 0.98378398 |
| GPR137 | 203.04154 | 0.61317615 | 0.24786476 | 2.47383354 | 0.0133672 | 0.06183738 |
| ESRRA | 773.659343 | 0.482306284 | 0.1764222 | 2.73381849 | 0.00626045 | 0.03571038 |
| TRMT112 | 276.466265 | -0.981170669 | 0.47454792 | -2.0675903 | 0.03867856 | 0.13077629 |
| PRDX5 | 406.399114 | -0.591094803 | 0.37379751 | -1.5813235 | 0.11380408 | 0.26754791 |
| CCDC88B | 39.7289538 | -1.673801984 | 0.5088517 | -3.2893709 | 0.00100412 | 0.00906379 |
| RPS6KA4 | 107.059608 | -0.894763686 | 0.55479058 | -1.6127954 | 0.10678897 | 0.25646507 |
| SLC22A11 | 1191.50263 | 0.107684914 | 0.29383008 | 0.36648704 | 0.71400168 | 0.83638578 |
| SLC22A12 | 1310.83071 | 0.865054869 | 0.35541366 | 2.43393819 | 0.01493555 | 0.0668806 |
| NRXN2 | 110.326715 | 0.820482503 | 0.49972023 | 1.64188371 | 0.10061411 | 0.24683702 |
| SF1 | 1463.51524 | 0.058041757 | 0.22282133 | 0.26048564 | 0.79448919 | 0.88811477 |
| MAP4K2 | 57.6380239 | 0.899400675 | 0.47274837 | 1.90249343 | 0.05710668 | 0.16947024 |
| MEN1 | 132.617335 | 0.519950584 | 0.24028564 | 2.16388537 | 0.03047315 | 0.11041381 |
| CDC42BPG | 33.2959389 | 0.312699439 | 0.40211163 | 0.77764337 | 0.43677929 | 0.62868147 |
| EHD1 | 359.240084 | 0.206003697 | 0.38028306 | 0.54171147 | 0.58801729 | 0.75041555 |
| MIR194-2HG | 142.738751 | 1.594034213 | 0.64238653 | 2.48142535 | 0.01308581 | 0.06086434 |
| ATG2A | 296.24587 | -0.780615372 | 0.36544325 | -2.1360782 | 0.03267304 | 0.11623393 |
| PPP2R5B | 176.764398 | -0.046421275 | 0.32677771 | -0.1420577 | 0.88703447 | 0.94159858 |
| MAJIN | 155.434491 | 2.172973315 | 0.63146785 | 3.44114639 | 0.00057926 | 0.00595638 |
| BATF2 | 60.1527129 | -0.672029756 | 0.59685272 | -1.1259558 | 0.26018425 | 0.45851703 |
| SAC3D1 | 41.6624745 | -0.172711249 | 0.46010594 | -0.3753728 | 0.70738324 | 0.83235488 |
| ZFPL1 | 103.32521 | -0.082698269 | 0.29977752 | -0.2758655 | 0.78265139 | 0.88095163 |
| VPS51 | 501.845766 | 0.118522736 | 0.28776459 | 0.41187394 | 0.68043182 | 0.81442647 |
| TM7SF2 | 116.871407 | 0.689020695 | 0.3599913 | 1.91399265 | 0.0556211 | 0.16642618 |
| ZNHIT2 | 49.5223137 | -0.413009849 | 0.45218223 | -0.9133704 | 0.36104778 | 0.55906936 |
| FAU | 821.847892 | -0.773696195 | 0.40667744 | -1.9024812 | 0.05710827 | 0.16947024 |
| SYVN1 | 174.883224 | -0.886475842 | 0.35600356 | -2.4900758 | 0.01277159 | 0.05992941 |
| MRPL49 | 92.9740576 | -0.755038942 | 0.36690662 | -2.0578504 | 0.03960449 | 0.13258635 |
| AP003068.2 | 33.6194177 | -0.866314194 | 0.44724007 | -1.9370228 | 0.05274256 | 0.16051116 |
| CAPN1 | 463.535461 | -0.973561904 | 0.24824425 | -3.9217904 | 8.79E-05 | 0.00141569 |
| POLA2 | 98.040375 | 0.04342847 | 0.28384901 | 0.15299849 | 0.87839946 | 0.93705514 |
| DPF2 | 271.072442 | -0.003401289 | 0.144453 | -0.023546 | 0.98121475 | 0.99060551 |
| SLC25A45 | 222.720926 | -0.177629207 | 0.35223723 | -0.5042886 | 0.61405862 | 0.76975835 |
| FRMD8 | 230.858078 | -0.413851302 | 0.29658538 | -1.3953867 | 0.16289926 | 0.33939202 |
| NEAT1 | 226385.829 | -1.368636408 | 0.3512399 | -3.8965859 | 9.76E-05 | 0.00152826 |
| MALAT1 | 756554.886 | -0.889718032 | 0.37857613 | -2.3501694 | 0.01876487 | 0.07882625 |
| SCYL1 | 292.917751 | -0.709291524 | 0.32067478 | -2.2118719 | 0.02697552 | 0.10163645 |
| LTBP3 | 357.152883 | 0.162540519 | 0.35270279 | 0.46084274 | 0.64491143 | 0.79180937 |
| SSSCA1 | 106.744814 | -1.501072933 | 0.55203298 | -2.7191726 | 0.00654455 | 0.03694649 |
| FAM89B | 83.6286612 | -1.242581267 | 0.40360322 | -3.0787199 | 0.00207892 | 0.01583098 |
| EHBP1L1 | 70.0865995 | -1.59323711 | 0.48652725 | -3.274713 | 0.00105769 | 0.00942028 |
| MAP3K11 | 269.636968 | 0.005630725 | 0.29170619 | 0.01930273 | 0.98459961 | 0.99241906 |
| PCNX3 | 181.530035 | -0.056453273 | 0.34765345 | -0.1623838 | 0.87100366 | 0.93338059 |
| SIPA1 | 60.8630061 | -1.358670597 | 0.43558204 | -3.1192071 | 0.00181338 | 0.0141918 |
| RELA | 197.458911 | -1.14926006 | 0.25610228 | -4.4875043 | 7.21E-06 | 0.00019217 |
| KAT5 | 204.446713 | -0.215632196 | 0.2840924 | -0.7590214 | 0.44783978 | 0.63841127 |
| RNASEH2C | 180.865099 | 0.20179533 | 0.39649396 | 0.50894931 | 0.61078775 | 0.76707767 |
| AP001266.2 | 58.3632364 | -1.393655058 | 0.79128043 | -1.7612657 | 0.07819345 | 0.20883678 |
| AP5B1 | 103.957413 | 0.246727072 | 0.24113997 | 1.02316953 | 0.3062277 | 0.50620635 |
| OVOL1 | 40.3401826 | 0.761963545 | 0.5584562 | 1.36441058 | 0.17243838 | 0.35183282 |
| CFL1 | 807.49068 | -1.16100346 | 0.39257144 | -2.9574323 | 0.00310213 | 0.02114414 |
| SNX32 | 37.6454327 | -0.753079853 | 0.43585205 | -1.7278337 | 0.08401805 | 0.21898162 |
| MUS81 | 145.492451 | -0.457707689 | 0.23591598 | -1.9401301 | 0.05236388 | 0.15964229 |
| EFEMP2 | 72.9605886 | -0.311961457 | 0.54287415 | -0.5746478 | 0.56552949 | 0.73247225 |
| FIBP | 71.4912453 | -0.217803289 | 0.38386154 | -0.5674007 | 0.57044201 | 0.73646497 |
| CCDC85B | 66.941997 | -0.604767249 | 0.44668363 | -1.3539051 | 0.17576665 | 0.35597422 |
| FOSL1 | 153.59293 | -9.887407284 | 1.62463923 | -6.0859095 | 1.16E-09 | 1.30E-07 |
| C11orf68 | 66.5531512 | -0.308822079 | 0.49064471 | -0.629421 | 0.52907348 | 0.70339575 |
| DRAP1 | 167.623653 | -0.442794253 | 0.36271834 | -1.2207661 | 0.2221746 | 0.41444934 |
| SART1 | 161.369445 | -0.178887137 | 0.45609322 | -0.3922162 | 0.6948985 | 0.82432536 |
| EIF1AD | 75.3996286 | -0.315324001 | 0.30044809 | -1.0495124 | 0.29394235 | 0.49394984 |
| BANF1 | 177.205605 | -0.850609024 | 0.31623649 | -2.6897878 | 0.00714975 | 0.03954092 |
| SF3B2 | 528.752129 | -0.301392995 | 0.19410971 | -1.5526941 | 0.12049623 | 0.27791383 |
| PACS1 | 3359.83771 | -0.89252322 | 0.24592966 | -3.6291808 | 0.00028432 | 0.00350843 |
| KLC2 | 143.437994 | -0.203674441 | 0.34524036 | -0.5899497 | 0.55522439 | 0.72468843 |
| RAB1B | 249.342353 | -0.154245281 | 0.33421637 | -0.4615132 | 0.64443042 | 0.79162826 |
| YIF1A | 174.800729 | -0.68062477 | 0.52772315 | -1.2897383 | 0.19714153 | 0.38302181 |

| | | | | | | |
|---|---|---|---|---|---|---|
| BRMS1 | 132.882328 | -0.099819422 | 0.41141076 | -0.2426272 | 0.80829424 | 0.89601696 |
| B4GAT1 | 70.9908958 | 0.357073551 | 0.52171554 | 0.68442192 | 0.49370878 | 0.67602105 |
| AP001107.9 | 51.1130175 | 2.465446774 | 0.52902295 | 4.6603777 | 3.16E-06 | 9.95E-05 |
| SLC29A2 | 51.1470086 | 0.079015131 | 0.45834122 | 0.17239368 | 0.86312804 | 0.92835028 |
| MRPL11 | 88.6662507 | -0.152228367 | 0.27624911 | -0.5510547 | 0.58159618 | 0.74455393 |
| AP002748.3 | 43.0253172 | 0.182681097 | 0.43849614 | 0.41660822 | 0.67696499 | 0.81296296 |
| DPP3 | 92.2502674 | -0.397927764 | 0.45239226 | -0.8796078 | 0.37907181 | 0.57579093 |
| ZDHHC24 | 158.334592 | -0.487773256 | 0.26508586 | -1.8400576 | 0.06575978 | 0.18625267 |
| CTSF | 117.815165 | 0.59577645 | 0.39303951 | 1.51581824 | 0.12956532 | 0.29180937 |
| CCS | 449.239595 | 0.578883651 | 0.31908034 | 1.81422536 | 0.06964304 | 0.19331655 |
| RBM14-RBM | 390.338951 | 0.470169456 | 0.30929539 | 1.52013081 | 0.1284781 | 0.29017126 |
| RBM4 | 222.936664 | 0.974699431 | 0.37937029 | 2.5692561 | 0.01019171 | 0.05117135 |
| RBM4B | 185.367604 | -0.11988506 | 0.36912879 | -0.3247784 | 0.74534879 | 0.85593548 |
| AP001157.1 | 46.8358757 | 0.047538795 | 0.4807458 | 0.09888551 | 0.92122917 | 0.95777368 |
| SPTBN2 | 69.0804161 | 0.207459665 | 0.42727664 | 0.48553945 | 0.62729373 | 0.77960968 |
| C11orf80 | 755.125567 | -1.193634037 | 0.2223974 | -5.3671222 | 8.00E-08 | 4.61E-06 |
| RCE1 | 53.2760292 | -0.464456607 | 0.32609977 | -1.4242776 | 0.15436612 | 0.32695817 |
| PC | 1311.606 | 0.189419082 | 0.4692846 | 0.4036337 | 0.6864821 | 0.81844047 |
| RHOD | 107.433983 | -1.136545346 | 0.38599862 | -2.9444285 | 0.00323552 | 0.021854 |
| KDM2A | 3559.59798 | -0.03457302 | 0.2189683 | -0.1578905 | 0.87454306 | 0.93490877 |
| GRK2 | 190.265775 | -0.333266485 | 0.28528427 | -1.1681909 | 0.24272978 | 0.43783354 |
| ANKRD13D | 282.94549 | -0.916184184 | 0.3429884 | -2.6711813 | 0.00755848 | 0.04112906 |
| SSH3 | 165.07122 | -0.704375908 | 0.25604173 | -2.75102 | 0.005941 | 0.03442792 |
| RAD9A | 1417.28198 | 0.460405989 | 0.20267245 | 2.27167528 | 0.02310613 | 0.09131812 |
| POLD4 | 62.1713141 | 0.167700421 | 0.43098406 | 0.3891105 | 0.69719441 | 0.82550208 |
| CLCF1 | 124.523116 | -1.047098853 | 0.33129944 | -3.1605814 | 0.00157455 | 0.01282892 |
| PPP1CA | 376.623979 | -0.455114903 | 0.31993794 | -1.42251 | 0.15487826 | 0.32750946 |
| RPS6KB2 | 184.398023 | 0.399825502 | 0.30579419 | 1.30749868 | 0.1910434 | 0.37532348 |
| CORO1B | 187.023772 | -0.836531192 | 0.35506849 | -2.3559714 | 0.01847434 | 0.07801226 |
| TMEM134 | 220.55617 | 0.130057667 | 0.3459475 | 0.37594626 | 0.70695685 | 0.83213891 |
| AIP | 107.949705 | -0.167364798 | 0.35968609 | -0.465308 | 0.64171094 | 0.78982207 |
| PITPNM1 | 180.011074 | -0.757288893 | 0.49254859 | -1.5374907 | 0.12417318 | 0.28386068 |
| CDK2AP2 | 351.608854 | -0.833032695 | 0.49515263 | -1.6823756 | 0.09249604 | 0.23387625 |
| GSTP1 | 1547.46912 | -1.638793983 | 0.60142252 | -2.724863 | 0.00643282 | 0.03651123 |
| NDUFV1 | 539.989518 | 0.349402235 | 0.37357757 | 0.93528698 | 0.34964042 | 0.54925232 |
| NUDT8 | 162.957238 | 0.071737765 | 0.38840891 | 0.1846965 | 0.85346709 | 0.92286287 |
| ACY3 | 257.948993 | 1.736669321 | 0.64473361 | 2.69362305 | 0.007068 | 0.03927083 |
| AP003716.1 | 534.020739 | 1.301161234 | 0.51612217 | 2.52103339 | 0.01170107 | 0.05625627 |
| UNC93B1 | 169.332106 | -0.503111757 | 0.43570889 | -1.154697 | 0.24821452 | 0.4445096 |
| ALDH3B1 | 140.127762 | 0.959256688 | 0.52499037 | 1.82718913 | 0.06767134 | 0.18961187 |
| NDUFS8 | 514.045407 | -0.164686613 | 0.26922711 | -0.6117014 | 0.54073531 | 0.71213095 |
| TCIRG1 | 39.2250951 | 0.79504171 | 0.62189029 | 1.2784276 | 0.2010987 | 0.38812445 |
| CHKA | 1751.67571 | -0.49534994 | 0.49515824 | -1.0003872 | 0.31712319 | 0.51716924 |
| KMT5B | 1257.7086 | 0.510164417 | 0.19831247 | 2.57252822 | 0.01009587 | 0.05082942 |
| C11orf24 | 116.150024 | -0.397361486 | 0.39047971 | -1.0176239 | 0.30885672 | 0.50906843 |
| LRP5 | 999.390773 | 0.661675869 | 0.31594875 | 2.09425066 | 0.03623766 | 0.12455104 |
| PPP6R3 | 3976.76055 | 0.032876348 | 0.18372337 | 0.17894483 | 0.85798102 | 0.92542853 |
| TESMIN | 99.1621902 | 1.216560981 | 0.40960086 | 2.97011331 | 0.0029769 | 0.02055635 |
| CPT1A | 1403.77867 | 0.034755072 | 0.4212355 | 0.08250746 | 0.93424318 | 0.96493643 |
| MRPL21 | 128.410091 | -0.402185904 | 0.24475217 | -1.6432373 | 0.10033384 | 0.24633279 |
| IGHMBP2 | 302.106281 | 0.142642744 | 0.30669435 | 0.4650974 | 0.64186171 | 0.78982207 |
| TPCN2 | 770.226752 | 0.016188986 | 0.43227412 | 0.03745074 | 0.97012561 | 0.9852064 |
| AP003071.2 | 47.5936677 | -3.424239476 | 1.09139328 | -3.1374936 | 0.00170399 | 0.0135479 |
| CCND1 | 289.553952 | -0.760992464 | 0.61260105 | -1.2422317 | 0.21415107 | 0.40445105 |
| ORAOV1 | 339.151602 | -0.081804136 | 0.30869875 | -0.2649967 | 0.79101204 | 0.88629048 |
| ANO1 | 85.1356531 | -0.653798218 | 0.84406023 | -0.7745872 | 0.43858365 | 0.63046712 |
| FADD | 86.6106494 | -0.711659972 | 0.25723503 | -2.7665749 | 0.00566486 | 0.03324749 |
| PPFIA1 | 2139.92749 | -0.425718487 | 0.35877306 | -1.1865955 | 0.23538723 | 0.42967809 |
| AP002336.2 | 718.415504 | -0.802091715 | 0.42940796 | -1.8679014 | 0.06177582 | 0.17834059 |
| AP000487.1 | 208.209048 | 0.805146721 | 0.51978628 | 1.54899572 | 0.12138275 | 0.27946713 |
| CTTN | 2818.56939 | -1.244446108 | 0.30493766 | -4.0809853 | 4.48E-05 | 0.00082843 |
| SHANK2 | 5720.10986 | 0.118864217 | 0.34757661 | 0.34197991 | 0.73236601 | 0.84799299 |
| SHANK2-AS1 | 41.6732525 | 0.031081402 | 0.69345257 | 0.04482124 | 0.9642498 | 0.98143402 |
| DHCR7 | 61.7309105 | -0.875396471 | 0.47169364 | -1.8558581 | 0.06347379 | 0.18178754 |
| NADSYN1 | 588.740484 | -0.279947728 | 0.25068032 | -1.1167519 | 0.26410041 | 0.46207385 |
| AP003498.1 | 83.7902762 | 1.486458991 | 0.76005188 | 1.95573358 | 0.05049655 | 0.15643364 |
| FAM86C1 | 52.8641534 | 0.305508342 | 0.46957214 | 0.65061002 | 0.51529826 | 0.69315508 |
| AP002495.2 | 411.990886 | 0.689701617 | 0.28612689 | 2.41047463 | 0.01593178 | 0.07030808 |
| RNF121 | 933.885194 | 0.40876334 | 0.17675258 | 2.31263017 | 0.02074298 | 0.08469587 |
| IL18BP | 88.4689386 | 0.210732299 | 0.31889343 | 0.66082359 | 0.50872545 | 0.68842924 |
| NUMA1 | 807.473208 | 0.586689119 | 0.29544005 | 1.98581444 | 0.04705393 | 0.14954052 |
| AP002490.1 | 37.3273115 | -0.007301712 | 0.46543606 | -0.0156879 | 0.98748338 | 0.99360989 |
| LAMTOR1 | 454.409841 | -0.744080094 | 0.29097748 | -2.5571741 | 0.01055264 | 0.05250272 |
| ANAPC15 | 69.5744675 | -0.047590086 | 0.35643156 | -0.1335182 | 0.8937836 | 0.94486286 |
| FOLR1 | 170.588787 | 0.257039175 | 0.5353948 | 0.48009277 | 0.63116143 | 0.78257166 |
| INPPL1 | 478.370288 | 0.731436344 | 0.22325539 | 3.27623155 | 0.00105202 | 0.00940644 |
| CLPB | 246.971682 | 0.090465466 | 0.40334149 | 0.22429001 | 0.82253165 | 0.90485083 |
| ARAP1 | 428.832187 | -0.164117853 | 0.27895151 | -0.5883383 | 0.55630525 | 0.72545677 |
| ATG16L2 | 152.409741 | 0.136538724 | 0.45574889 | 0.29959201 | 0.76448838 | 0.8685323 |
| FCHSD2 | 1503.20632 | -0.716670372 | 0.34042567 | -2.1052183 | 0.03527231 | 0.12236839 |
| P2RY6 | 209.07746 | 0.030584476 | 1.02655475 | 0.02979332 | 0.97623188 | 0.98827256 |
| ARHGEF17 | 401.723369 | 0.8157156 | 0.24269808 | 3.3610303 | 0.00077652 | 0.0075115 |
| FAM168A | 1285.33989 | 0.48006158 | 0.24410049 | 1.9666555 | 0.04922294 | 0.15396641 |
| RAB6A | 2053.45735 | -0.512073003 | 0.2345807 | -2.1829289 | 0.02904104 | 0.10672 |
| MRPL48 | 315.132615 | 0.582870355 | 0.18427667 | 3.16301759 | 0.00156143 | 0.01273959 |
| COA4 | 324.315488 | -0.090115268 | 0.21339319 | -0.4222968 | 0.67280838 | 0.81052772 |
| PAAF1 | 650.781866 | 0.635729269 | 0.40475871 | 1.57063763 | 0.11626684 | 0.27158651 |
| C2CD3 | 491.054604 | -0.094815213 | 0.1660381 | -0.5710449 | 0.56796919 | 0.73451775 |
| PPME1 | 866.597452 | -0.919667957 | 0.26228258 | -3.5064012 | 0.00045421 | 0.00500471 |
| P4HA3 | 123.406747 | -1.867752544 | 0.69761072 | -2.6773564 | 0.00742056 | 0.04068547 |
| PGM2L1 | 223.50364 | -1.811199379 | 0.41566267 | -4.357378 | 1.32E-05 | 0.00031775 |
| KCNE3 | 134.095227 | 2.048273363 | 0.628203 | 3.26052784 | 0.00111205 | 0.00979619 |
| POLD3 | 570.514328 | 1.128697907 | 0.39087184 | 2.88764193 | 0.00388141 | 0.02507189 |
| RNF169 | 1079.66314 | -0.415902918 | 0.44833631 | -0.9276583 | 0.35358483 | 0.5520031 |
| XRRA1 | 947.060849 | 0.189563636 | 0.46605148 | 0.40674399 | 0.68419603 | 0.81689199 |
| SPCS2 | 694.591222 | -0.634688934 | 0.26749691 | -2.3726963 | 0.01765878 | 0.07554583 |
| NEU3 | 320.587461 | 0.484687467 | 0.2025554 | 2.39286377 | 0.01671745 | 0.07269862 |
| SLCO2B1 | 301.511487 | 1.637290775 | 0.73316268 | 2.23318892 | 0.02553649 | 0.09788273 |
| ARRB1 | 80.0994734 | -1.754294869 | 0.47913469 | -3.6613814 | 0.00025086 | 0.00320459 |
| RPS3 | 1304.36916 | -0.867127929 | 0.3727611 | -2.3262297 | 0.0200063 | 0.0824493 |
| SERPINH1 | 259.624296 | -0.47979594 | 0.47662455 | -1.0066538 | 0.31410115 | 0.51469031 |
| UVRAG | 1930.4968 | -0.037084801 | 0.20661016 | -0.1794917 | 0.85755166 | 0.92511102 |
| THAP12 | 255.115948 | 0.183706809 | 0.20542701 | 0.89426803 | 0.3711785 | 0.56877072 |
| GVQW3 | 142.792371 | 1.27209231 | 0.53923498 | 2.35906859 | 0.01832087 | 0.07760326 |
| AP002360.1 | 53.7974363 | 0.598536307 | 0.52424099 | 1.14171978 | 0.25357051 | 0.4504205 |
| EMSY | 1059.29629 | 0.112532535 | 0.14963498 | 0.75204697 | 0.45202281 | 0.64143458 |
| TSKU | 418.269099 | -0.364594601 | 0.65289367 | -0.5584288 | 0.57655164 | 0.74103815 |
| AP003119.2 | 61.9769746 | -1.300596462 | 0.68298058 | -1.9042949 | 0.05687178 | 0.16891474 |
| ACER3 | 1532.29738 | 0.113113662 | 0.23001979 | 0.49175621 | 0.6228917 | 0.77619139 |
| CAPN5 | 198.743903 | 0.13451149 | 0.58408304 | 0.23029515 | 0.81786243 | 0.90160299 |
| MYO7A | 287.301264 | 2.34978515 | 0.6418894 | 3.66073211 | 0.0002515 | 0.00320667 |
| PAK1 | 1079.30684 | 0.186469799 | 0.20964229 | 0.88946654 | 0.3737524 | 0.571065 |
| CLNS1A | 766.562078 | 0.186205114 | 0.23786112 | 0.78283123 | 0.43372622 | 0.62599614 |

| | | | | | | |
|---|---|---|---|---|---|---|
| AQP11 | 153.184074 | 0.567902329 | 0.5203973 | 1.09128608 | 0.27514702 | 0.47286769 |
| RSF1 | 1982.31791 | -0.086869098 | 0.19491976 | -0.4456659 | 0.65583857 | 0.79912092 |
| AAMDC | 495.802999 | 0.108690409 | 0.34860583 | 0.31178597 | 0.75520319 | 0.86270273 |
| AP002812.2 | 212.177962 | -0.260827113 | 0.28735873 | -0.9076707 | 0.36405221 | 0.56189625 |
| INTS4 | 528.379753 | 0.034537139 | 0.2155545 | 0.16022463 | 0.87270413 | 0.93417849 |
| ALG8 | 318.854035 | -0.045453208 | 0.33020363 | -0.1376521 | 0.89051541 | 0.94326785 |
| KCTD21-AS1 | 68.9190834 | 0.606629329 | 0.70551506 | 0.85983895 | 0.38987782 | 0.58542916 |
| KCTD21 | 126.118156 | -0.065526123 | 0.32091785 | -0.2041835 | 0.83821012 | 0.9145179 |
| USP35 | 111.070105 | -1.010289641 | 0.50848285 | -1.9868706 | 0.04693673 | 0.14921358 |
| GAB2 | 1956.71644 | -1.550047255 | 0.41893689 | -3.6999541 | 0.00021564 | 0.00283934 |
| AP003086.2 | 484.040876 | -1.943689224 | 0.52025861 | -3.7360059 | 0.00018697 | 0.00252239 |
| NARS2 | 796.980275 | 0.149929566 | 0.20165194 | 0.74350669 | 0.45717497 | 0.64583091 |
| AP003110.1 | 46.3356346 | 0.064250679 | 0.33129537 | 0.19393775 | 0.84622463 | 0.91896252 |
| TENM4 | 1512.77708 | 1.054317939 | 0.43426206 | 2.42783803 | 0.01518912 | 0.06779464 |
| AP001978.1 | 130.150257 | 2.826685776 | 0.51924096 | 5.44388055 | 5.21E-08 | 3.22E-06 |
| MIR4300HG | 112.524716 | 1.560023976 | 0.58621368 | 2.66118657 | 0.00778658 | 0.04205305 |
| PRCP | 517.072089 | 0.306513173 | 0.39099581 | 0.78392956 | 0.43308144 | 0.62552702 |
| RAB30 | 385.601689 | 0.266921599 | 0.31156338 | 0.85671686 | 0.39160138 | 0.58682412 |
| RAB30-AS1 | 1272.08486 | 0.35415504 | 0.21192466 | 1.67113656 | 0.09469471 | 0.23698747 |
| AP001767.3 | 98.9859352 | -0.360685908 | 0.63153855 | -0.5711225 | 0.56791657 | 0.73451775 |
| PCF11 | 761.61436 | 0.41331838 | 0.23523038 | 1.7570791 | 0.07890433 | 0.20992656 |
| PCF11-AS1 | 65.239637 | 0.146290405 | 0.31752627 | 0.46071906 | 0.64500018 | 0.79180937 |
| ANKRD42 | 550.513109 | 0.032082791 | 0.28712767 | 0.11173702 | 0.91103192 | 0.95310114 |
| CCDC90B | 280.533555 | -0.449927291 | 0.26708443 | -1.6845882 | 0.09206804 | 0.23313812 |
| AP000446.1 | 289.527793 | 0.147884367 | 0.22976846 | 0.64362345 | 0.51981964 | 0.69597507 |
| DLG2 | 9777.77217 | 1.369260028 | 0.54775867 | 2.49975056 | 0.01242808 | 0.05874195 |
| AP001825.1 | 75.1681594 | 0.163067708 | 0.69122798 | 0.23591017 | 0.81350238 | 0.89903065 |
| TMEM126B | 231.060594 | -0.231920387 | 0.19818109 | -1.1702448 | 0.24190248 | 0.43703142 |
| TMEM126A | 87.8514386 | -0.162732413 | 0.25971899 | -0.6265711 | 0.53094042 | 0.70478407 |
| CREBZF | 906.42982 | 0.198139612 | 0.2131016 | 0.92978943 | 0.35248012 | 0.55147186 |
| SYTL2 | 897.254522 | -0.421809788 | 0.38764774 | -1.0881265 | 0.27653926 | 0.4742304 |
| PICALM | 7195.32535 | 0.255859 | 0.20447224 | 1.25131412 | 0.2108199 | 0.40007037 |
| EED | 324.994621 | 0.368793691 | 0.16400559 | 2.24866535 | 0.0245338 | 0.09502253 |
| HIKESHI | 389.951816 | 0.633500232 | 0.21589047 | 2.93435941 | 0.00334237 | 0.02238542 |
| AP001831.1 | 101.701509 | -0.862728177 | 0.3808803 | -2.26509 | 0.02350716 | 0.09240395 |
| ME3 | 311.211074 | -0.66428786 | 0.3110515 | -2.1356202 | 0.03271038 | 0.11633281 |
| PRSS23 | 373.459088 | -0.508322332 | 0.7038087 | -0.722245 | 0.47014384 | 0.65697438 |
| AP001528.2 | 523.429148 | 0.547848398 | 0.82820147 | 0.6614917 | 0.50829703 | 0.6881892 |
| FZD4 | 444.783333 | 0.478487915 | 0.44245248 | 1.08144477 | 0.27949932 | 0.47799091 |
| AP001528.1 | 231.400012 | 0.961668788 | 0.53359898 | 1.8022313 | 0.07150902 | 0.19698034 |
| TMEM135 | 2299.34766 | 0.066408454 | 0.33636129 | 0.19743191 | 0.84348956 | 0.91781467 |
| RAB38 | 178.442259 | -0.474230534 | 0.31994261 | -1.4822362 | 0.13827744 | 0.30419616 |
| CTSC | 823.11204 | 0.362974815 | 0.36002038 | 1.0082063 | 0.31335543 | 0.51371402 |
| GRM5-AS1 | 49.5426551 | -2.223415933 | 0.68483715 | -3.2466345 | 0.00116778 | 0.01015659 |
| GRM5 | 342.592239 | 0.639919231 | 0.56623936 | 1.13012142 | 0.25842507 | 0.45600314 |
| NOX4 | 8286.65092 | 2.126634483 | 0.43713387 | 4.86495016 | 1.14E-06 | 4.33E-05 |
| AP003400.1 | 1551.23636 | 0.831348455 | 0.75027129 | 1.1080638 | 0.26783426 | 0.46575866 |
| CHORDC1 | 509.675855 | -0.446199384 | 0.53464464 | -0.8345719 | 0.4039588 | 0.59801958 |
| SLC36A4 | 669.48356 | -1.290232086 | 0.28971958 | -4.4533824 | 8.45E-06 | 0.00022111 |
| SMCO4 | 161.860443 | -0.879204168 | 0.37965817 | -2.3157783 | 0.02057038 | 0.08417692 |
| CEP295 | 591.578127 | -0.043358207 | 0.16181563 | -0.2679482 | 0.78873918 | 0.88457274 |
| TAF1D | 2277.11566 | -0.166468828 | 0.29036166 | -0.5733154 | 0.56643113 | 0.73322046 |
| AP001273.2 | 190.19133 | 1.299598452 | 0.34763853 | 3.73836142 | 0.00018522 | 0.00250381 |
| VSTM5 | 98.4890874 | 1.591867692 | 0.48542695 | 3.27931459 | 0.0010406 | 0.00932251 |
| PANX1 | 155.100813 | -1.139374525 | 0.42389292 | -2.6878829 | 0.00719066 | 0.03969488 |
| MRE11 | 417.516896 | -0.044728989 | 0.18493909 | -0.2418579 | 0.80889024 | 0.8961735 |
| ANKRD49 | 125.722162 | 0.138759746 | 0.28095651 | 0.49388335 | 0.62138856 | 0.77515226 |
| PIWIL4 | 342.165829 | -2.306443551 | 0.7085371 | -3.2552192 | 0.00113305 | 0.00992377 |
| AP000943.3 | 85.0067923 | -2.183108235 | 0.78921529 | -2.7661758 | 0.00567179 | 0.03324749 |
| AMOTL1 | 351.517799 | -0.771252667 | 0.51806557 | -1.4887163 | 0.13656208 | 0.30154199 |
| CWC15 | 274.986986 | -0.203835444 | 0.19120215 | -1.066073 | 0.28639066 | 0.48601669 |
| SRSF8 | 268.594402 | -0.341840239 | 0.36725294 | -0.9308033 | 0.35195534 | 0.55108727 |
| ENDOD1 | 766.144151 | -1.484100386 | 0.48365104 | -3.0685355 | 0.00215111 | 0.01625426 |
| AP000787.1 | 281.790253 | -1.232699635 | 0.46261634 | -2.6646263 | 0.00770739 | 0.04169133 |
| SESN3 | 559.776369 | 0.376448504 | 0.42810406 | 0.87933878 | 0.37921761 | 0.57594857 |
| FAM76B | 245.190265 | 0.034160974 | 0.16457434 | 0.20757169 | 0.83556341 | 0.91316952 |
| CEP57 | 626.399455 | -0.124358986 | 0.32854778 | -0.3785111 | 0.70505094 | 0.83082301 |
| MTMR2 | 1200.17575 | -0.079116438 | 0.20920963 | -0.3781682 | 0.70530562 | 0.83105168 |
| MAML2 | 17426.5191 | 0.263706412 | 0.43440506 | 0.60705189 | 0.54381648 | 0.71416138 |
| CCDC82 | 619.785101 | -0.816121286 | 0.26842286 | -3.0404314 | 0.0023624 | 0.0174751 |
| JRKL | 164.444952 | -0.732414308 | 0.24414262 | -2.9999445 | 0.00270029 | 0.01918015 |
| CNTN5 | 289.103382 | 0.495628593 | 0.37275472 | 1.32963734 | 0.18363779 | 0.36602053 |
| ARHGAP42 | 3399.94643 | 0.232689696 | 0.29449954 | 0.79011904 | 0.42945825 | 0.62219636 |
| TRPC6 | 156.313474 | 0.679793878 | 0.61991714 | 1.0965883 | 0.2728214 | 0.47074056 |
| CEP126 | 510.925945 | 0.328750474 | 0.43975677 | 0.74757342 | 0.4547175 | 0.64414855 |
| YAP1 | 3197.20352 | -0.447160771 | 0.22804321 | -1.9608599 | 0.04989537 | 0.15522376 |
| BIRC3 | 1035.12984 | -3.149326388 | 0.58006022 | -5.4293094 | 5.66E-08 | 3.46E-06 |
| BIRC2 | 831.276639 | 0.183441277 | 0.18085658 | 1.0142914 | 0.31044373 | 0.51075857 |
| TMEM123 | 783.55562 | -0.932995796 | 0.2358734 | -3.9554938 | 7.64E-05 | 0.00126716 |
| MMP7 | 607.410288 | -2.576580591 | 0.75821764 | -3.3982071 | 0.00067829 | 0.00674208 |
| DCUN1D5 | 335.701802 | -0.410168402 | 0.19406649 | -2.1135458 | 0.03455408 | 0.12067077 |
| DYNC2H1 | 1390.0051 | 0.838710394 | 0.42660129 | 1.96602871 | 0.04929529 | 0.15396641 |
| PDGFD | 91.388428 | -0.045309577 | 0.52709856 | -0.0859604 | 0.93149794 | 0.96354054 |
| CASP4 | 582.310632 | -1.385660581 | 0.45996939 | -3.0125061 | 0.002591 | 0.01869204 |
| GRIA4 | 565.009121 | 0.952564512 | 0.97885124 | 0.97314533 | 0.33048108 | 0.53036984 |
| MSANTD4 | 223.584262 | 0.453865066 | 0.23678002 | 1.91682164 | 0.05526059 | 0.16578178 |
| KBTBD3 | 175.803326 | 0.59974192 | 0.35090632 | 1.70912259 | 0.08742824 | 0.22509394 |
| AASDHPPT | 452.520325 | 0.138129765 | 0.46955571 | 0.29417119 | 0.76862709 | 0.87099633 |
| GUCY1A2 | 106.856904 | 0.83419923 | 0.38818961 | 2.14894786 | 0.03163853 | 0.11340726 |
| AP000766.1 | 117.650105 | 1.078682545 | 0.23607656 | 4.56920649 | 4.90E-06 | 0.00014039 |
| CWF19L2 | 1211.45256 | 0.692662388 | 0.17377 | 3.98608718 | 6.72E-05 | 0.00114234 |
| ALKBH8 | 180.375663 | 0.342513333 | 0.24877906 | 1.37677718 | 0.16858115 | 0.34711458 |
| ELMOD1 | 288.003764 | 1.254927884 | 0.52338597 | 2.39771022 | 0.01649791 | 0.07206002 |
| SLC35F2 | 464.403295 | 0.10627221 | 0.33240869 | 0.31970347 | 0.74919313 | 0.85862795 |
| RAB39A | 53.9681902 | -0.484691973 | 0.45632855 | -1.0621557 | 0.28816501 | 0.48800071 |
| CUL5 | 2277.01431 | 0.439735181 | 0.15980137 | 2.75176109 | 0.00592757 | 0.03438395 |
| AP002433.1 | 51.2688414 | -0.256447487 | 0.5801187 | -0.4420604 | 0.65844552 | 0.80095001 |
| ACAT1 | 1814.68721 | 0.127846424 | 0.29226489 | 0.43743339 | 0.66179707 | 0.8032633 |
| NPAT | 488.912309 | -0.086421483 | 0.19041643 | -0.4538552 | 0.64993306 | 0.79477446 |
| ATM | 1607.09665 | -0.129452978 | 0.17468808 | -0.7410522 | 0.45866182 | 0.64743262 |
| C11orf65 | 348.30893 | 0.877945458 | 0.53822212 | 1.63119543 | 0.10284909 | 0.25038257 |
| KDELC2 | 173.123434 | 0.342352168 | 0.30753932 | 1.11319803 | 0.26562339 | 0.46352448 |
| EXPH5 | 434.032938 | 0.454505116 | 0.33888102 | 1.34119379 | 0.17985754 | 0.36116427 |
| DDX10 | 1603.05831 | 0.054050931 | 0.29680125 | 0.18211153 | 0.8554952 | 0.92354672 |
| AP003049.2 | 395.020867 | 4.933871197 | 0.91088284 | 5.41658159 | 6.07E-08 | 3.67E-06 |
| AP002444.1 | 142.301069 | 6.209077463 | 1.23039057 | 5.04642801 | 4.50E-07 | 1.97E-05 |
| RDX | 3917.22058 | 2.071012722 | 0.61513137 | 3.3667812 | 0.00076051 | 0.00739838 |
| ZC3H12C | 2249.67372 | 0.355103094 | 0.3121244 | 1.13769731 | 0.2552469 | 0.45232479 |
| AP000924.1 | 120.811311 | 3.63394112 | 0.62753827 | 5.79078804 | 7.01E-09 | 6.01E-07 |
| FDX1 | 516.12995 | 0.337623198 | 0.22790545 | 1.48141784 | 0.13849526 | 0.30456515 |
| ARHGAP20 | 43.7119292 | 0.877964717 | 0.52786901 | 1.66322457 | 0.09626749 | 0.23952414 |
| COLCA1 | 42.1301405 | 0.62929605 | 0.73347321 | 0.85796732 | 0.39091051 | 0.5863016 |
| COLCA2 | 70.9482368 | -0.065631607 | 0.51072388 | -0.128507 | 0.89774774 | 0.94708137 |

| | | | | | | |
|---|---|---|---|---|---|---|
| SIK2 | 3997.15952 | 0.112071309 | 0.32233477 | 0.34768606 | 0.72807596 | 0.84566684 |
| PPP2R1B | 927.440645 | 0.209061655 | 0.44034444 | 0.47476846 | 0.63495201 | 0.78468826 |
| C11orf1 | 83.1327147 | 0.005960282 | 0.47475802 | 0.01255436 | 0.98998333 | 0.99506501 |
| CRYAB | 10227.3117 | -2.294144384 | 0.60291379 | -3.8050952 | 0.00014175 | 0.00202181 |
| DIXDC1 | 961.976479 | -0.188378828 | 0.56285952 | -0.3346818 | 0.73786514 | 0.85123628 |
| DLAT | 276.004444 | 0.253490153 | 0.20342409 | 1.24611668 | 0.21272156 | 0.40256447 |
| PIH1D2 | 62.5536854 | 1.001094145 | 0.41931223 | 2.38746709 | 0.01696492 | 0.07355949 |
| NKAPD1 | 182.555534 | -0.047769071 | 0.2069035 | -0.2308761 | 0.81741107 | 0.90138805 |
| TIMM8B | 206.444212 | -0.791525227 | 0.26246033 | -3.01579 | 0.00256311 | 0.01855205 |
| AP002884.3 | 176.737439 | 0.836970333 | 0.35307579 | 2.37051181 | 0.01776348 | 0.07587534 |
| IL18 | 294.408778 | -1.415888529 | 0.34820364 | -4.0662658 | 4.78E-05 | 0.00087426 |
| BCO2 | 93.7647563 | -0.143473056 | 0.41587925 | -0.3449873 | 0.73010393 | 0.84648502 |
| PTS | 463.96721 | -0.67250683 | 0.25427896 | -2.6447601 | 0.00817489 | 0.04364805 |
| AP002884.1 | 293.535827 | -0.379238203 | 0.54902677 | -0.6907463 | 0.489725 | 0.67267868 |
| AP003063.1 | 72.1216356 | -0.385666665 | 0.86106421 | -0.4478954 | 0.65422872 | 0.79789243 |
| TTC12 | 466.198167 | 0.560311406 | 0.24386432 | 2.29763584 | 0.02158252 | 0.08739488 |
| ZW10 | 210.987983 | 0.147853316 | 0.22828167 | 0.64767933 | 0.51719238 | 0.69425481 |
| USP28 | 299.643676 | -0.145957967 | 0.30275344 | -0.4821018 | 0.62973366 | 0.78157908 |
| ZBTB16 | 3224.86093 | -0.882166822 | 0.67670468 | -1.3036216 | 0.19236264 | 0.3768513 |
| AP002518.2 | 302.015636 | -2.716414202 | 0.78619887 | -3.4551235 | 0.00055004 | 0.00572903 |
| NNMT | 2023.53946 | -4.584067913 | 0.81512955 | -5.6237293 | 1.87E-08 | 1.40E-06 |
| C11orf71 | 92.0603899 | 0.035919889 | 0.48349693 | 0.07429186 | 0.94077815 | 0.96876614 |
| AP002373.1 | 63.4158494 | -0.777968769 | 0.63856721 | -1.2183037 | 0.2231086 | 0.41522737 |
| REXO2 | 354.492256 | -0.316819304 | 0.33411744 | -0.9482274 | 0.34301372 | 0.54361071 |
| CADM1 | 1487.98821 | -1.042855391 | 0.40359064 | -2.5839435 | 0.00976778 | 0.04960614 |
| AP000462.1 | 50.5270586 | -1.7088489 | 0.54788993 | -3.1189639 | 0.00181488 | 0.0141918 |
| BUD13 | 117.965299 | 0.258000905 | 0.21767666 | 1.18524834 | 0.23591927 | 0.43030545 |
| ZPR1 | 100.64847 | -0.825706983 | 0.42959985 | -1.9220374 | 0.05460105 | 0.16434269 |
| SIK3 | 6860.79689 | 0.119088805 | 0.32235677 | 0.36943168 | 0.71180599 | 0.83509031 |
| PAFAH1B2 | 708.623621 | -0.445155046 | 0.30600811 | -1.4547165 | 0.14574777 | 0.31442346 |
| SIDT2 | 154.828107 | -0.058762067 | 0.30958037 | -0.189812 | 0.84945646 | 0.92079871 |
| TAGLN | 89.6369479 | 0.6972265 | 0.52293377 | 1.33329791 | 0.18243406 | 0.36478648 |
| PCSK7 | 522.953461 | 0.152758065 | 0.23413732 | 0.65242937 | 0.51412422 | 0.69238561 |
| RNF214 | 473.620669 | 0.564995516 | 0.15536776 | 3.63650429 | 0.00027636 | 0.00344061 |
| BACE1 | 223.555478 | 0.503785231 | 0.25715733 | 1.9590545 | 0.05010641 | 0.15561112 |
| CEP164 | 577.692077 | 1.230345817 | 0.30802384 | 3.99432005 | 6.49E-05 | 0.00110887 |
| AP000892.1 | 92.6758229 | 1.346215839 | 0.73229888 | 1.83834206 | 0.06601202 | 0.18656221 |
| DSCAML1 | 4320.77526 | -1.285011333 | 0.69331609 | -1.8534278 | 0.06382106 | 0.18248315 |
| FXYD2 | 840.930332 | -0.079384749 | 0.59598576 | -0.1331991 | 0.89403595 | 0.94498386 |
| AP000757.2 | 61.0197046 | -1.586089721 | 0.65145248 | -2.4346975 | 0.01490424 | 0.06678407 |
| AP000757.1 | 36.1679612 | -0.969933964 | 0.76480725 | -1.2682071 | 0.20472401 | 0.39304057 |
| SMIM35 | 397.5428 | 4.70283599 | 0.67049093 | 7.01401878 | 2.32E-12 | 5.77E-10 |
| TMPRSS4 | 80.8186312 | 3.517295346 | 0.56175934 | 6.26121383 | 3.82E-10 | 4.99E-08 |
| MPZL3 | 241.631659 | -0.340037402 | 0.34252518 | -0.992737 | 0.32083816 | 0.52124318 |
| MPZL2 | 127.823473 | -1.041982883 | 0.43767266 | -2.3807356 | 0.01727811 | 0.07442836 |
| UBE4A | 748.481706 | -0.00891626 | 0.16676263 | -0.0534668 | 0.95736 | 0.97834441 |
| ATP5MG | 830.063414 | -0.271281262 | 0.32112761 | -0.8447771 | 0.39823529 | 0.59258207 |
| KMT2A | 1934.71385 | -0.322532873 | 0.17613172 | -1.8312027 | 0.06707029 | 0.18854236 |
| AP001267.3 | 96.8071038 | 0.788243695 | 0.34031804 | 2.31619722 | 0.0205475 | 0.08414838 |
| TMEM25 | 151.081348 | 0.675376217 | 0.36256974 | 1.86274846 | 0.06249766 | 0.17972542 |
| IFT46 | 162.742896 | 1.248800956 | 0.37898566 | 3.29511405 | 0.00098382 | 0.00891877 |
| ARCN1 | 412.675831 | -0.51580261 | 0.2393089 | -2.1553842 | 0.03113178 | 0.11211362 |
| PHLDB1 | 552.045764 | 0.552085547 | 0.43303807 | 1.27491228 | 0.20234029 | 0.38991683 |
| TREH | 156.387842 | 2.426221792 | 0.5957671 | 4.07243332 | 4.65E-05 | 0.00085484 |
| AP002954.1 | 104.017551 | -0.370900727 | 0.34637583 | -1.0708043 | 0.28425744 | 0.48359398 |
| DDX6 | 1041.17448 | 0.065321362 | 0.14579951 | 0.44802181 | 0.65413745 | 0.79785212 |
| BCL9L | 338.415964 | -0.337269753 | 0.51049301 | -0.6606746 | 0.50882103 | 0.68848991 |
| UPK2 | 37.6064777 | 1.288716447 | 0.47506332 | 2.71272566 | 0.00667323 | 0.03751846 |
| CCDC84 | 715.301259 | 0.455820706 | 0.27078267 | 1.68334519 | 0.09230829 | 0.23357387 |
| RPS25 | 977.670335 | -0.767664594 | 0.40226649 | -1.9083483 | 0.05634621 | 0.16770632 |
| TRAPPC4 | 256.199293 | -0.233447627 | 0.323107 | -0.7225087 | 0.46998176 | 0.65686103 |
| SLC37A4 | 288.494132 | 0.981217528 | 0.550954 | 1.78094274 | 0.0749218 | 0.20315592 |
| HYOU1 | 193.120082 | -1.222016988 | 0.53855498 | -2.2690663 | 0.02326429 | 0.09168593 |
| VPS11 | 175.004575 | 0.131520495 | 0.17048944 | 0.77142899 | 0.4404527 | 0.63244135 |
| HMBS | 65.6242798 | -0.182641537 | 0.30786071 | -0.5932603 | 0.55300696 | 0.72252338 |
| H2AFX | 63.4863326 | -1.33389855 | 0.39093719 | -3.4120533 | 0.00064475 | 0.00649828 |
| DPAGT1 | 88.1864055 | -0.205589892 | 0.32392755 | -0.6346786 | 0.52563804 | 0.70084961 |
| C2CD2L | 124.709387 | 0.675569436 | 0.35328779 | 1.91223544 | 0.055846 | 0.16684419 |
| HINFP | 223.204071 | -0.208847165 | 0.25734881 | -0.8115334 | 0.41705941 | 0.61036016 |
| NLRX1 | 138.861156 | -0.085159244 | 0.35994103 | -0.2365922 | 0.81297317 | 0.8986644 |
| PDZD3 | 552.304947 | 1.631753426 | 0.84084174 | 1.94061897 | 0.05230451 | 0.15953225 |
| CBL | 813.799347 | -0.461116163 | 0.32669926 | -1.4114392 | 0.15811514 | 0.33276281 |
| MCAM | 85.7923617 | -1.506038453 | 0.74368468 | -2.0251035 | 0.04285674 | 0.13988694 |
| RNF26 | 52.6422728 | -0.704053097 | 0.42194197 | -1.6686017 | 0.09519634 | 0.23793877 |
| USP2 | 344.860118 | 2.482715723 | 0.54865076 | 4.52512947 | 6.04E-06 | 0.00016714 |
| USP2-AS1 | 270.267405 | 0.148452557 | 0.79335421 | 0.18712015 | 0.85156643 | 0.92197716 |
| THY1 | 209.408101 | 3.268150206 | 0.59975593 | 5.44913367 | 5.06E-08 | 3.18E-06 |
| NECTIN1 | 76.1783123 | 0.639016918 | 0.36255384 | 1.76254349 | 0.07797751 | 0.20849257 |
| TRIM29 | 80.2288485 | 0.498625497 | 0.60600205 | 0.82281157 | 0.41061516 | 0.60441387 |
| OAF | 1329.23977 | 0.027920778 | 0.25573936 | 0.1091767 | 0.91306234 | 0.95369546 |
| TMEM136 | 39.944672 | -1.284620103 | 0.53413714 | -2.405038 | 0.01617079 | 0.0710654 |
| ARHGEF12 | 6459.53032 | -0.250628314 | 0.2281545 | -1.0985026 | 0.27198508 | 0.47033143 |
| TECTA | 41.2060756 | 0.429058578 | 0.38768555 | 1.106718 | 0.26841587 | 0.46636789 |
| SC5D | 191.9122 | -0.475525201 | 0.28368638 | -1.6762356 | 0.09369206 | 0.23571272 |
| SORL1 | 150.010084 | -0.333700971 | 0.5335145 | -0.6254769 | 0.53165814 | 0.70525869 |
| AP000755.1 | 43.1650239 | 0.676819684 | 0.73908042 | 0.91575918 | 0.35979322 | 0.5583251 |
| UBASH3B | 611.697915 | -4.587292832 | 0.70782362 | -6.4808417 | 9.12E-11 | 1.47E-08 |
| JHY | 744.98955 | 0.879032185 | 0.35762854 | 2.45794755 | 0.01397336 | 0.06372047 |
| HSPA8 | 879.261903 | -0.639496768 | 0.42833353 | -1.4929879 | 0.13544036 | 0.29987878 |
| GRAMD1B | 3144.07685 | 2.719053698 | 0.78695909 | 3.45513982 | 0.00055001 | 0.00572903 |
| ZNF202 | 120.598081 | -0.157257559 | 0.30508522 | -0.5154545 | 0.60623543 | 0.76359388 |
| VWA5A | 130.167289 | -0.958342052 | 0.44580813 | -2.1496738 | 0.03158103 | 0.11326038 |
| TBRG1 | 680.50025 | 0.017629844 | 0.17486115 | 0.10082196 | 0.91969179 | 0.95690112 |
| SIAE | 308.738815 | 0.453733033 | 0.34344681 | 1.32111588 | 0.18646273 | 0.3697157 |
| SPA17 | 77.2062059 | -0.220388021 | 0.25608106 | -0.8606182 | 0.38944836 | 0.5850689 |
| MSANTD2 | 555.023579 | 0.428968955 | 0.20243794 | 2.11901464 | 0.03408923 | 0.11959587 |
| AP000866.1 | 55.0801733 | 1.269030145 | 0.44864833 | 2.8285632 | 0.00467575 | 0.02889561 |
| TMEM218 | 64.1702394 | 0.514322603 | 0.40852692 | 1.25896871 | 0.20804163 | 0.39710718 |
| PKNOX2 | 109.195088 | -1.387889038 | 0.90282366 | -1.5372759 | 0.12422576 | 0.28388839 |
| EI24 | 269.586059 | -0.484551595 | 0.34972294 | -1.3855299 | 0.16589049 | 0.34332794 |
| STT3A-AS1 | 69.3462323 | -0.065633473 | 0.44794336 | -0.1465218 | 0.88350947 | 0.93957729 |
| STT3A | 329.181968 | -0.713510879 | 0.29467219 | -2.4213716 | 0.01546206 | 0.06863293 |
| CHEK1 | 90.8274495 | -0.201551946 | 0.28699892 | -0.7022742 | 0.48250816 | 0.66670759 |
| HYLS1 | 91.5233449 | 0.020254134 | 0.2695926 | 0.07512867 | 0.94011234 | 0.968226 |
| PUS3 | 79.8239845 | -0.096206019 | 0.34337168 | -0.2801804 | 0.77933909 | 0.87852271 |
| CDON | 277.102957 | -1.388545713 | 0.6173024 | -2.2493768 | 0.02448853 | 0.09490085 |
| RPUSD4 | 208.385784 | -0.225434577 | 0.30225844 | -0.7458339 | 0.45576778 | 0.64490647 |
| FAM118B | 415.545318 | -0.476751473 | 0.21182853 | -2.2506481 | 0.02440784 | 0.09469522 |
| SRPRA | 255.932341 | -0.852098563 | 0.44302498 | -1.9233646 | 0.05443428 | 0.16412904 |
| FOXRED1 | 250.616876 | -0.117876169 | 0.39556491 | -0.2979945 | 0.76570736 | 0.86912477 |
| TIRAP | 118.820577 | 0.490057383 | 0.30968487 | 1.58243889 | 0.11354941 | 0.26719688 |
| DCPS | 108.87046 | 0.20457326 | 0.46387673 | 0.44100781 | 0.65920734 | 0.80168598 |
| ST3GAL4 | 85.4036532 | 0.897080878 | 0.54795395 | 1.63714648 | 0.10159987 | 0.24855066 |

| | | | | | | | |
|---|---|---|---|---|---|---|---|
| ETS1 | 1072.87922 | | -3.304185934 | 0.56067239 | -5.893256 | 3.79E-09 | 3.53E-07 |
| KCNJ1 | 53.2559838 | | 1.080793307 | 0.48757644 | 2.21666434 | 0.02664604 | 0.1008664 |
| ARHGAP32 | 3515.32206 | | 0.565183583 | 0.24914183 | 2.26852142 | 0.02329745 | 0.09174958 |
| BARX2 | 181.486965 | | -0.595583053 | 0.55346897 | -1.0760911 | 0.2818865 | 0.48077801 |
| TMEM45B | 105.594113 | | -1.412422356 | 0.53480872 | -2.640986 | 0.00826651 | 0.04391825 |
| NFRKB | 406.016889 | | 0.172600274 | 0.15735191 | 1.0969061 | 0.27268244 | 0.47062144 |
| PRDM10 | 514.700663 | | -0.098368464 | 0.29759583 | -0.3305438 | 0.74098908 | 0.85293394 |
| APLP2 | 1678.91669 | | -0.762502349 | 0.19725511 | -3.8655645 | 0.00011083 | 0.00168611 |
| ST14 | 672.608691 | | -0.741331514 | 0.42079307 | -1.7617484 | 0.07811182 | 0.2087093 |
| ZBTB44 | 1658.9041 | | 0.794065092 | 0.3214939 | 2.46992276 | 0.01351422 | 0.06226895 |
| AP002986.1 | 297.730475 | | 1.593769616 | 0.64741719 | 2.4617351 | 0.01382667 | 0.06332183 |
| ADAMTS15 | 142.379024 | | 1.921318173 | 0.71075754 | 2.70319773 | 0.00686759 | 0.03835942 |
| AP003486.1 | 40.4444916 | | 0.27279935 | 0.37760049 | 0.72245496 | 0.47001481 | 0.65686103 |
| SNX19 | 674.590358 | | 0.278471689 | 0.4910634 | 0.56707889 | 0.57066058 | 0.73667777 |
| NTM | 75.1206538 | | -0.712361472 | 0.4226246 | -1.6855656 | 0.09187951 | 0.23270369 |
| OPCML | 248.925576 | | 3.309895902 | 0.60778623 | 5.44582245 | 5.16E-08 | 3.21E-06 |
| NCAPD3 | 160.311364 | | 0.105044017 | 0.27795996 | 0.37791061 | 0.705497 | 0.83113428 |
| VPS26B | 173.081565 | | 0.28232748 | 0.24540999 | 1.1504319 | 0.24996603 | 0.44613077 |
| THYN1 | 55.9532095 | | 0.144359445 | 0.34827349 | 0.41450024 | 0.67850779 | 0.81376982 |
| ACAD8 | 192.691728 | | 0.493966246 | 0.2563256 | 1.92710458 | 0.0539666 | 0.16308249 |
| GLB1L3 | 37.0852148 | | -0.09698681 | 0.82606324 | -0.1174085 | 0.90653638 | 0.9510826 |
| GLB1L2 | 136.561018 | | 0.855046305 | 0.30925704 | 2.76484022 | 0.00569507 | 0.03334571 |
| B3GAT1 | 99.7170133 | | -2.777141438 | 0.79144627 | -3.508945 | 0.00044989 | 0.00496731 |
| AP004550.1 | 67.3735689 | | -1.312735037 | 0.81380905 | -1.613075 | 0.10672821 | 0.25646507 |
| ZMYND11 | 1476.07007 | | 0.060341233 | 0.18254644 | 0.33055278 | 0.74098232 | 0.85293394 |
| DIP2C | 7505.82089 | | 0.26510212 | 0.17377912 | 1.52551196 | 0.12713145 | 0.28824615 |
| PRR26 | 135.959165 | | -0.397069812 | 0.48221794 | -0.823424 | 0.41026693 | 0.60416081 |
| LARP4B | 2845.11362 | | -0.029304734 | 0.20137198 | -0.1455254 | 0.88429608 | 0.94002757 |
| GTPBP4 | 395.447342 | | -0.566096355 | 0.47036516 | -1.2035253 | 0.22877312 | 0.42176102 |
| IDI1 | 246.139601 | | -0.837810898 | 0.40822442 | -2.0523292 | 0.04013768 | 0.13386063 |
| WDR37 | 1004.91156 | | 0.077190084 | 0.19819583 | 0.38946371 | 0.69693315 | 0.82544394 |
| ADARB2 | 47.7394343 | | -0.158708252 | 0.5866334 | -0.2705408 | 0.78674426 | 0.88341964 |
| PFKP | 931.212185 | | -3.726220698 | 0.57840015 | -6.4422886 | 1.18E-10 | 1.79E-08 |
| PITRM1 | 508.366551 | | -1.157915711 | 0.39130022 | -2.9591492 | 0.0030849 | 0.02111703 |
| PITRM1-AS1 | 54.4470854 | | -1.744492222 | 0.52902492 | -3.2975615 | 0.00097528 | 0.0088833 |
| KLF6 | 4050.81408 | | -2.525935044 | 0.40974975 | -6.1645799 | 7.07E-10 | 8.73E-08 |
| AKR1E2 | 55.7836412 | | -1.073618401 | 0.46785924 | -2.2947466 | 0.02174765 | 0.08764991 |
| AKR1C1 | 2299.19227 | | -5.473024871 | 0.78722836 | -6.9522709 | 3.59E-12 | 8.64E-10 |
| AKR1C2 | 443.447589 | | -3.92093598 | 0.97833623 | -4.0077592 | 6.13E-05 | 0.00106625 |
| AKR1C3 | 1038.21619 | | -1.270659391 | 0.38122406 | -3.3331039 | 0.00085883 | 0.00808514 |
| AL355303.1 | 57.550117 | | -2.846224872 | 0.8279333 | -3.4377466 | 0.00058658 | 0.00602262 |
| TUBAL3 | 86.7666229 | | -0.02690699 | 0.53800198 | -0.0500128 | 0.96011218 | 0.9796009 |
| NET1 | 745.825146 | | 0.100612903 | 0.39086329 | 0.257412 | 0.79686074 | 0.88911002 |
| ASB13 | 671.830989 | | 1.123738473 | 0.39882513 | 2.81762203 | 0.00483807 | 0.02956843 |
| FAM208B | 2360.2601 | | -0.339524831 | 0.23624692 | -1.4371609 | 0.15067229 | 0.3217425 |
| AL365356.1 | 119.884739 | | -0.628810874 | 0.54769986 | -1.1480939 | 0.2509298 | 0.44721035 |
| GDI2 | 1457.04506 | | -0.211229526 | 0.26448752 | -0.798637 | 0.42450091 | 0.61755827 |
| ANKRD16 | 194.599332 | | 0.254293555 | 0.25394103 | 1.00138823 | 0.31663915 | 0.51668724 |
| FBH1 | 861.33551 | | 0.147048107 | 0.23849618 | 0.61656377 | 0.5375225 | 0.70967393 |
| IL15RA | 64.8626814 | | -3.068416326 | 0.59699256 | -5.1397899 | 2.75E-07 | 1.28E-05 |
| RBM17 | 608.564855 | | -0.169097361 | 0.18844113 | -0.8973485 | 0.369533 | 0.56695645 |
| PFKFB3 | 635.263989 | | -1.113334718 | 0.59820448 | -1.8611274 | 0.06272619 | 0.1801735 |
| AL137145.2 | 71.2012334 | | -1.970898815 | 0.62805899 | -3.1380792 | 0.00170059 | 0.01352872 |
| PRKCQ | 153.454352 | | -0.533940849 | 0.61911539 | -0.8624254 | 0.38845346 | 0.58402233 |
| SFMBT2 | 248.6789 | | 0.016028152 | 0.64155252 | 0.02498338 | 0.98006822 | 0.99025469 |
| ITIH5 | 252.336442 | | -0.512255081 | 0.74991749 | -0.6830819 | 0.49455508 | 0.67687134 |
| KIN | 272.534956 | | 0.115003012 | 0.18929845 | 0.60752221 | 0.54350441 | 0.71392572 |
| ATP5F1C | 1042.25949 | | -0.073902326 | 0.27193762 | -0.271762 | 0.78580499 | 0.8829436 |
| TAF3 | 1162.0481 | | 0.225498464 | 0.2052947 | 1.09841349 | 0.27202398 | 0.47033143 |
| CELF2 | 197.704329 | | -2.230113895 | 0.71425904 | -3.1222761 | 0.00179459 | 0.0140965 |
| USP6NL | 1081.30639 | | -0.141144269 | 0.20146069 | -0.7006045 | 0.48354985 | 0.66763205 |
| ECHDC3 | 293.986579 | | 0.282778835 | 0.36085335 | 0.78363921 | 0.43325184 | 0.62564086 |
| PROSER2 | 369.82713 | | -1.931388231 | 0.41017179 | -4.7087301 | 2.49E-06 | 8.22E-05 |
| UPF2 | 1238.52677 | | 0.058673235 | 0.23896478 | 0.24553089 | 0.80604539 | 0.89466205 |
| DHTKD1 | 1510.64944 | | 1.320735838 | 0.29277154 | 4.51114829 | 6.45E-06 | 0.00017605 |
| SEC61A2 | 225.24127 | | 0.108130185 | 0.20498637 | 0.5274994 | 0.59784683 | 0.75778496 |
| NUDT5 | 628.211439 | | -0.063244064 | 0.24490398 | -0.2582402 | 0.7962215 | 0.88867422 |
| CDC123 | 514.73745 | | -0.258748834 | 0.28765489 | -0.8995113 | 0.36838036 | 0.56582134 |
| CAMK1D | 179.023339 | | -0.559648547 | 0.47822927 | -1.1702515 | 0.24189976 | 0.43703142 |
| CCDC3 | 73.5992395 | | 0.530829485 | 0.46096639 | 1.15155789 | 0.24950279 | 0.44575166 |
| OPTN | 1311.87806 | | -0.058953216 | 0.28624056 | -0.2059569 | 0.83682458 | 0.91389966 |
| PHYH | 356.918532 | | 1.075359303 | 0.39699988 | 2.70871444 | 0.00675444 | 0.03787496 |
| SEPHS1 | 369.843007 | | 0.512107456 | 0.26348071 | 1.94362406 | 0.0519408 | 0.15895347 |
| BEND7 | 2355.82619 | | -0.202959238 | 0.18606445 | -1.0908008 | 0.27536056 | 0.47286769 |
| PRPF18 | 207.810322 | | 0.045873785 | 0.23834354 | 0.19246917 | 0.84737471 | 0.91938085 |
| AL157392.5 | 430.439229 | | 0.330296239 | 0.18988239 | 1.739478 | 0.08195072 | 0.21543891 |
| FRMD4A | 1904.64843 | | 1.900561379 | 0.52025722 | 3.65311869 | 0.00025907 | 0.00326473 |
| AL157392.2 | 61.5205518 | | 1.144436544 | 0.9640805 | 1.18707572 | 0.23519776 | 0.42961829 |
| FAM107B | 2485.24277 | | 0.138281051 | 0.36859499 | 0.37515716 | 0.70754359 | 0.83240065 |
| CDNF | 91.6993162 | | 1.051875435 | 0.57653964 | 1.82446334 | 0.06808205 | 0.19052689 |
| HSPA14.1 | 136.796765 | | 0.000402093 | 0.24994879 | 0.0016087 | 0.99871644 | 0.99920455 |
| SUV39H2 | 77.5088925 | | -0.284698724 | 0.39394403 | -0.7226883 | 0.46987143 | 0.65683764 |
| DCLRE1C | 259.059421 | | -0.346706431 | 0.19312775 | -1.7952181 | 0.07261897 | 0.19887875 |
| RPP38 | 455.692779 | | 0.223057006 | 0.16076344 | 1.38748587 | 0.16529366 | 0.34261003 |
| NMT2 | 477.856541 | | 0.099969341 | 0.21818028 | 0.45819605 | 0.6468116 | 0.79280318 |
| FAM171A1 | 979.669369 | | -0.557124083 | 0.56585221 | -0.9845752 | 0.32483275 | 0.52511199 |
| MINDY3 | 710.730078 | | 0.16101375 | 0.18600519 | 0.86564118 | 0.38668697 | 0.58283684 |
| PTER | 2642.87787 | | 1.60547953 | 0.27614663 | 5.81386614 | 6.10E-09 | 5.36E-07 |
| C1QL3 | 164.504682 | | 2.641406988 | 0.69813822 | 3.78350146 | 0.00015464 | 0.00216508 |
| RSU1 | 1807.32281 | | 0.522425953 | 0.31525681 | 1.65714406 | 0.09749035 | 0.24142733 |
| CUBN | 26731.6181 | | 2.632077711 | 0.76726846 | 3.4304521 | 0.00060258 | 0.00615922 |
| TRDMT1 | 137.887535 | | 0.500314082 | 0.27525948 | 1.81760888 | 0.06912393 | 0.19255756 |
| VIM-AS1 | 46.7155367 | | -1.49929268 | 0.50005645 | -2.9982469 | 0.00271538 | 0.01925487 |
| VIM | 1726.14357 | | -2.521412463 | 0.60193268 | -4.1888612 | 2.80E-05 | 0.00057185 |
| HACD1 | 82.3734991 | | -1.854337549 | 0.42998797 | -4.3125336 | 1.61E-05 | 0.00037409 |
| STAM | 658.341439 | | -0.813223552 | 0.41496365 | -1.9597465 | 0.05002543 | 0.15545195 |
| CACNB2 | 920.45281 | | 2.771884288 | 0.65606818 | 4.22499422 | 2.39E-05 | 0.00050858 |
| AL450384.2 | 271.386496 | | 0.656463784 | 0.26977396 | 2.43338458 | 0.0149584 | 0.06691738 |
| NSUN6 | 801.580495 | | 0.622183398 | 0.19985602 | 3.11315808 | 0.00185097 | 0.01442366 |
| ARL5B | 357.919785 | | -1.362600689 | 0.35088931 | -3.8832779 | 0.00010306 | 0.00159078 |
| MALRD1 | 61.6194593 | | 0.58331702 | 0.44429672 | 1.31289968 | 0.18921675 | 0.3732327 |
| PLXDC2 | 6766.20629 | | 1.160121834 | 0.35572423 | 3.26129551 | 0.00110904 | 0.00977599 |
| NEBL | 2009.13016 | | 1.062526023 | 0.39403165 | 2.69654991 | 0.00700619 | 0.03898759 |
| CASC10 | 43.611589 | | 1.623230298 | 0.5575835 | 2.91118781 | 0.00360058 | 0.02370912 |
| MLLT10 | 4126.159 | | 0.371843792 | 0.18966159 | 1.96056453 | 0.04992984 | 0.15526052 |
| DNAJC1 | 3336.00644 | | -0.185020374 | 0.25673932 | -0.7206546 | 0.47112205 | 0.6575369 |
| BMI1 | 88.6135568 | | -0.334221054 | 0.29324986 | -1.1397143 | 0.25440536 | 0.45147332 |
| PIP4K2A | 1991.56579 | | 0.137887405 | 0.34418147 | 0.40062414 | 0.68869687 | 0.82001112 |
| ARMC3 | 108.48932 | | -0.744640686 | 0.80223709 | -0.9282053 | 0.35330111 | 0.55189396 |
| MSRB2 | 350.554805 | | 0.07873675 | 0.39020542 | 0.20178282 | 0.84008651 | 0.91527429 |
| C10orf67 | 425.464449 | | 1.342090086 | 0.71113453 | 1.88725202 | 0.05912644 | 0.17307314 |
| OTUD1 | 117.273035 | | 0.480321773 | 0.29563974 | 1.62468607 | 0.10422947 | 0.25231575 |
| KIAA1217 | 4487.88353 | | 0.357875509 | 0.28566958 | 1.25276031 | 0.21029295 | 0.39989367 |

| | | | | | | |
|---|---|---|---|---|---|---|
| ARHGAP21 | 1574.87367 | -0.522755816 | 0.20764637 | -2.5175293 | 0.01181811 | 0.05665575 |
| THNSL1 | 50.63888 | 0.93100163 | 0.57663153 | 1.61455207 | 0.10640773 | 0.25592749 |
| MYO3A | 232.822145 | -1.454205763 | 0.76619793 | -1.8979505 | 0.0577026 | 0.17045895 |
| APBB1IP | 738.301586 | -0.479092596 | 0.42359923 | -1.1310044 | 0.25805324 | 0.45540566 |
| PDSS1 | 572.14369 | 0.934700729 | 0.2656652 | 3.51834091 | 0.00043425 | 0.00485507 |
| ABI1 | 1881.78045 | -0.060990424 | 0.20550562 | -0.2967823 | 0.76663275 | 0.8695271 |
| ANKRD26 | 1482.63396 | 0.732117863 | 0.27708016 | 2.64226011 | 0.00823548 | 0.04383833 |
| YME1L1 | 937.050312 | -0.401823238 | 0.15672416 | -2.5638882 | 0.01035069 | 0.05180591 |
| MASTL | 203.728832 | 0.280909746 | 0.24999815 | 1.12364731 | 0.26116268 | 0.45929649 |
| ACBD5 | 550.79748 | -0.168740879 | 0.18086943 | -0.9329431 | 0.35084936 | 0.55033368 |
| RAB18 | 1202.43908 | -1.141210715 | 0.31811355 | -3.587432 | 0.00033395 | 0.00399466 |
| MPP7 | 856.423235 | 0.301441711 | 0.33744834 | 0.89329736 | 0.37169795 | 0.56913135 |
| AL358234.1 | 55.7170268 | -1.125071451 | 0.83366347 | -1.3495511 | 0.17716002 | 0.35765809 |
| WAC-AS1 | 339.294271 | 0.091344894 | 0.38553374 | 0.236931 | 0.81271032 | 0.89861746 |
| WAC | 3551.90003 | -0.184160842 | 0.20465846 | -0.8998448 | 0.36820287 | 0.56561209 |
| BAMBI | 104.081913 | 0.586514181 | 0.4950125 | 1.18484722 | 0.23607786 | 0.43048014 |
| SVIL | 4400.04594 | -0.31983666 | 0.46290506 | -0.6909336 | 0.48960727 | 0.67262219 |
| MTPAP | 268.430815 | 0.150451949 | 0.18774384 | 0.80136822 | 0.4229185 | 0.61623726 |
| MAP3K8 | 289.597266 | -1.457222229 | 0.36266554 | -4.0180885 | 5.87E-05 | 0.00102973 |
| ZNF438 | 985.388772 | 0.140638889 | 0.20818801 | 0.67553789 | 0.49933409 | 0.68099623 |
| ZEB1-AS1 | 52.6201164 | -1.086206603 | 0.39511552 | -2.7490861 | 0.00597617 | 0.03450556 |
| ZEB1 | 1854.04769 | -1.908471861 | 0.57648004 | -3.3105602 | 0.00093109 | 0.0085712 |
| ARHGAP12 | 1769.81763 | -0.254355391 | 0.20547776 | -1.2378731 | 0.21576312 | 0.40633004 |
| KIF5B | 1145.19494 | -0.13665179 | 0.18541878 | -0.73699 | 0.46112841 | 0.64969737 |
| EPC1 | 1716.40635 | 0.527109351 | 0.17248097 | 3.05604356 | 0.00224279 | 0.01675223 |
| AL391839.1 | 50.4397818 | 0.449461192 | 0.56952258 | 0.78918942 | 0.43000131 | 0.62265243 |
| CCDC7 | 1121.14694 | 0.356546909 | 0.51167149 | 0.69682778 | 0.48591058 | 0.66931729 |
| ITGB1 | 1495.75692 | -0.774486234 | 0.3498424 | -2.2138146 | 0.02684153 | 0.10132045 |
| ITGB1-DT | 63.040933 | -2.149601766 | 0.7936096 | -2.7086388 | 0.00675598 | 0.03787496 |
| NRP1 | 9428.85451 | 0.424330613 | 0.46963817 | 0.90352667 | 0.36624644 | 0.56380727 |
| PARD3 | 20127.0892 | -0.237202633 | 0.26648924 | -0.8901021 | 0.37341107 | 0.57086578 |
| CUL2 | 633.727916 | 0.115410391 | 0.16750817 | 0.68898365 | 0.49083356 | 0.67359387 |
| CREM | 981.189372 | -0.790568996 | 0.35484716 | -2.2279141 | 0.02588624 | 0.09872811 |
| CCNY | 3884.38921 | -0.4462408 | 0.25246967 | -1.7675026 | 0.07714409 | 0.20709246 |
| AL117336.3 | 388.2425 | 0.126020979 | 0.2849634 | 0.44223567 | 0.65831868 | 0.80095001 |
| FZD8 | 58.41376 | -0.13991367 | 0.5699713 | -0.2454749 | 0.80608871 | 0.89466205 |
| ZNF248 | 956.746614 | 0.064285035 | 0.36572714 | 0.17577322 | 0.86047211 | 0.92646675 |
| ZNF25 | 95.5766726 | 0.58858091 | 0.40144239 | 1.46616532 | 0.14260325 | 0.31006707 |
| ZNF33A | 1935.07993 | 0.191753294 | 0.19460528 | 0.98534476 | 0.32445475 | 0.52483341 |
| AL117339.5 | 95.6847358 | -0.518505654 | 0.5861924 | -0.8845315 | 0.37640936 | 0.57294604 |
| ZNF33B | 1242.25257 | 0.538124615 | 0.29800815 | 1.80573794 | 0.07095928 | 0.19589907 |
| BMS1 | 617.210273 | -0.018745829 | 0.20967035 | -0.0894062 | 0.9287591 | 0.96224497 |
| CSGALNACT2 | 417.847889 | -0.693312028 | 0.37464482 | -1.8505849 | 0.0642293 | 0.18337659 |
| RASGEF1A | 42.3623777 | -0.96084405 | 0.58385859 | -1.6456794 | 0.09982979 | 0.2456673 |
| FXYD4 | 64.1962239 | 1.432625456 | 0.71088426 | 2.01527245 | 0.04387611 | 0.14207769 |
| HNRNPF | 953.333231 | -0.57677025 | 0.35707388 | -1.6152687 | 0.10625253 | 0.25564392 |
| ZNF487 | 276.083247 | 0.486374017 | 0.19889955 | 2.44532483 | 0.01447217 | 0.06531777 |
| ZNF239 | 43.5506223 | 0.210641011 | 0.40490017 | 0.52022949 | 0.60290363 | 0.76145627 |
| ZNF32 | 168.729518 | 0.222727647 | 0.20139848 | 1.10590532 | 0.2687675 | 0.46680133 |
| CXCL12 | 95.4034369 | 1.588926945 | 0.86787951 | 1.83081515 | 0.06712814 | 0.18866627 |
| TMEM72-AS | 746.12226 | 0.101626557 | 0.54759089 | 0.18558847 | 0.85276749 | 0.92227269 |
| RASSF4 | 3296.46383 | 0.60376691 | 0.55144637 | 1.09487874 | 0.27356977 | 0.47143837 |
| DEPP1 | 224.022211 | -1.970666987 | 0.41140674 | -4.7900697 | 1.67E-06 | 5.93E-05 |
| ZNF22 | 184.022503 | -1.204734384 | 0.2734697 | -4.405367 | 1.06E-05 | 0.00026654 |
| ALOX5 | 111.893103 | -3.345300809 | 0.77520997 | -4.315348 | 1.59E-05 | 0.00037021 |
| 8-Mar | 908.203261 | -0.047459736 | 0.22029652 | -0.2154357 | 0.82942764 | 0.90944441 |
| ZFAND4 | 356.782423 | 0.425061016 | 0.21653085 | 1.96305058 | 0.0496403 | 0.15464081 |
| WASHC2C | 568.08278 | 0.032119833 | 0.29721492 | 0.10806938 | 0.91394066 | 0.9541766 |
| AGAP4 | 625.75773 | 0.924528615 | 0.26891538 | 3.43799089 | 0.00058605 | 0.0060217 |
| TIMM23 | 421.100843 | 0.214350887 | 0.26807753 | 0.79958542 | 0.42395104 | 0.61708579 |
| NCOA4 | 503.495383 | 0.464023791 | 0.15930617 | 2.91277969 | 0.00358227 | 0.02366371 |
| AL356056.2 | 61.4957509 | -0.114991625 | 0.4226169 | -0.2720942 | 0.78554956 | 0.88272895 |
| PTPN20 | 887.919417 | -0.34492138 | 0.57677829 | -0.5980138 | 0.54983072 | 0.71955844 |
| AGAP9 | 513.454925 | 0.423692741 | 0.4342683 | 0.97564742 | 0.32923922 | 0.52891681 |
| BX547991.1 | 155.377641 | 1.883358458 | 0.70006214 | 2.69027325 | 0.00713935 | 0.03954092 |
| MAPK8 | 3383.54934 | 0.110583122 | 0.16513924 | 0.66963563 | 0.50309009 | 0.68384133 |
| ARHGAP22 | 334.899951 | -4.022742516 | 0.47018982 | -8.5555713 | 1.17E-17 | 1.00E-14 |
| WDFY4 | 213.437229 | 2.808064482 | 1.01653077 | 2.76239988 | 0.00573782 | 0.03352283 |
| VSTM4 | 352.232874 | 1.300128269 | 0.54900306 | 2.36816217 | 0.0178767 | 0.07621645 |
| AL138760.1 | 47.9490812 | -0.962788557 | 0.63213681 | -1.5230699 | 0.12774122 | 0.28917405 |
| ERCC6 | 535.341865 | 0.18329686 | 0.16287867 | 1.12535831 | 0.26043723 | 0.45872695 |
| OGDHL | 559.766315 | 1.625418201 | 0.52114367 | 3.11894455 | 0.001815 | 0.0141918 |
| PARG | 1721.51029 | 0.594632532 | 0.19747248 | 3.01121715 | 0.00260203 | 0.01872287 |
| TIMM23B | 1573.40839 | 0.269196124 | 0.5201739 | 0.51751179 | 0.60479894 | 0.76257742 |
| LINC00843 | 359.112009 | 0.441000901 | 0.21331651 | 2.06735471 | 0.03870074 | 0.13081901 |
| AGAP6 | 522.993442 | 0.078129363 | 0.22012533 | 0.35493127 | 0.72264108 | 0.84228022 |
| WASHC2A | 538.292201 | -0.022965071 | 0.3833103 | -0.0599125 | 0.95222534 | 0.97578893 |
| SGMS1 | 2567.51818 | -0.154556714 | 0.15674189 | -0.9860588 | 0.32410427 | 0.52462178 |
| SGMS1-AS1 | 67.9325654 | 0.923469114 | 0.40553652 | 2.27715402 | 0.02277703 | 0.09046791 |
| ASAH2B | 140.98003 | -0.272469475 | 0.39224447 | -0.694642 | 0.4872797 | 0.67052935 |
| A1CF | 1435.92261 | 1.535766082 | 0.62797171 | 2.44559755 | 0.01446123 | 0.06528989 |
| PRKG1 | 684.732596 | -1.002579564 | 0.55831485 | -1.7957243 | 0.07253837 | 0.19879007 |
| CSTF2T | 45.0824175 | -0.262762217 | 0.35667594 | -0.7366973 | 0.46130643 | 0.6497711 |
| PCDH15 | 2408.0398 | 3.62456419 | 0.88881892 | 4.07795569 | 4.54E-05 | 0.00083704 |
| IPMK | 639.566419 | -1.067305348 | 0.25317852 | -4.2156237 | 2.49E-05 | 0.00052085 |
| CISD1 | 844.066221 | 0.028014873 | 0.24170229 | 0.11590653 | 0.90772662 | 0.95171808 |
| UBE2D1 | 289.881874 | -0.747703849 | 0.33303008 | -2.2451541 | 0.02475824 | 0.09556779 |
| TFAM | 207.263846 | -0.274867811 | 0.22471154 | -1.2232029 | 0.22125307 | 0.4134582 |
| BICC1 | 16000.8726 | 0.676669783 | 0.35405824 | 1.91118214 | 0.05598118 | 0.16710188 |
| PHYHIPL | 2013.91517 | 0.985429421 | 0.42286864 | 2.33034407 | 0.01978797 | 0.08184366 |
| FAM13C | 42.0283611 | 0.082406953 | 0.70315546 | 0.11719592 | 0.9067048 | 0.95109493 |
| SLC16A9 | 5878.08188 | 2.706930532 | 0.81317671 | 3.3288343 | 0.0008721 | 0.00816524 |
| CCDC6 | 1324.24352 | -0.147030074 | 0.21631024 | -0.6797185 | 0.49668271 | 0.67856373 |
| ANK3 | 14429.0274 | 0.759953534 | 0.51662862 | 1.47098613 | 0.14129487 | 0.30819999 |
| AL592430.1 | 66.5418822 | -0.194407535 | 0.87849039 | -0.2212973 | 0.82486097 | 0.90642548 |
| RHOBTB1 | 5413.04013 | 1.619690797 | 0.59150892 | 2.73823565 | 0.00617698 | 0.03535193 |
| TMEM26 | 148.302034 | 2.359684861 | 0.56111416 | 4.20535609 | 2.61E-05 | 0.00053649 |
| AC022398.1 | 43.3710859 | 1.735488408 | 0.5842578 | 2.97041546 | 0.00297397 | 0.02055456 |
| ARID5B | 3401.98538 | 0.154948889 | 0.40502624 | 0.38256506 | 0.70204228 | 0.82877388 |
| RTKN2 | 69.9086858 | -1.163260069 | 0.53232003 | -2.1852645 | 0.02886946 | 0.10630447 |
| AC024597.1 | 130.519978 | -5.893953492 | 1.66290569 | -3.5443703 | 0.00039355 | 0.00451038 |
| ZNF365 | 333.362726 | -0.600163424 | 1.30576954 | -0.4596243 | 0.64578591 | 0.79225766 |
| ADO | 57.3439239 | -0.844017346 | 0.3738165 | -2.2578387 | 0.02395572 | 0.09347027 |
| NRBF2 | 183.530332 | -0.403997558 | 0.29024389 | -1.3919244 | 0.16394529 | 0.34080823 |
| JMJD1C | 7635.53377 | -0.076152611 | 0.23308228 | -0.3267199 | 0.74387977 | 0.85468232 |
| REEP3 | 892.863523 | -0.878970216 | 0.3210496 | -2.737802 | 0.00618513 | 0.0353838 |
| LINC01515 | 1817.53663 | 1.081504294 | 0.62141306 | 1.74039518 | 0.08178965 | 0.21530454 |
| CTNNA3 | 199.267311 | 1.755483346 | 0.43008346 | 4.08172718 | 4.47E-05 | 0.00082802 |
| DNAJC12 | 200.459196 | 1.02948399 | 0.38945359 | 2.64340611 | 0.00820765 | 0.04375818 |
| SIRT1 | 531.436341 | 0.286971936 | 0.23925159 | 1.19945675 | 0.23035039 | 0.42352956 |
| HERC4 | 4870.22281 | -0.477007212 | 0.27478695 | -1.7359166 | 0.08257859 | 0.21675694 |
| AC016395.1 | 253.827818 | 2.637800755 | 0.65870092 | 4.00454998 | 6.21E-05 | 0.00107266 |
| PBLD | 1794.24304 | 2.041837688 | 0.42326274 | 4.82404312 | 1.41E-06 | 5.07E-05 |

| | | | | | | |
|---|---|---|---|---|---|---|
| HNRNPH3 | 1222.60256 | 0.126093807 | 0.23293174 | 0.54133372 | 0.58827758 | 0.75056149 |
| RUFY2 | 870.256382 | 0.15271152 | 0.19327707 | 0.7901171 | 0.42945939 | 0.62219636 |
| DNA2 | 363.221151 | -0.548017553 | 0.40805094 | -1.3430126 | 0.1792679 | 0.36056127 |
| SLC25A16 | 1988.21532 | 0.247738575 | 0.28692591 | 0.8634235 | 0.38790466 | 0.58387159 |
| TET1 | 161.13643 | 0.627691548 | 0.45819443 | 1.369924 | 0.17071063 | 0.34969424 |
| CCAR1 | 1638.68205 | 0.232927373 | 0.25470977 | 0.9144815 | 0.3604639 | 0.5586436 |
| STOX1 | 85.5159164 | 0.982899492 | 0.55361932 | 1.77540678 | 0.07583075 | 0.20481025 |
| DDX50 | 587.789725 | -0.099616293 | 0.25567094 | -0.389627 | 0.69681241 | 0.82544394 |
| DDX21 | 846.503937 | -1.613472153 | 0.50426782 | -3.1996334 | 0.00137603 | 0.01150072 |
| KIF1BP | 248.427405 | -0.42772913 | 0.24845011 | -1.7215896 | 0.08514389 | 0.22086814 |
| VPS26A | 657.111187 | -0.284537968 | 0.20567377 | -1.3834432 | 0.16652902 | 0.3444934 |
| SUPV3L1 | 376.215107 | 0.04719976 | 0.27446871 | 0.17196773 | 0.8634629 | 0.92856473 |
| AL596223.1 | 94.8192311 | -3.161552405 | 0.78657499 | -4.0193909 | 5.83E-05 | 0.00102537 |
| HKDC1 | 3185.55428 | -3.187480715 | 0.7216763 | -4.4167734 | 1.00E-05 | 0.00025478 |
| HK1 | 421.323911 | -2.696050217 | 0.40704656 | -6.6234442 | 3.51E-11 | 6.50E-09 |
| TSPAN15 | 132.288862 | -2.024336053 | 0.67314703 | -3.0072717 | 0.00263604 | 0.01884831 |
| FAM241B | 57.043768 | -0.877583032 | 0.44536619 | -1.9704752 | 0.04878393 | 0.15305136 |
| H2AFY2 | 225.812074 | -0.568054926 | 0.55035387 | -1.032163 | 0.30199574 | 0.50241408 |
| AIFM2 | 283.97517 | -0.70116499 | 0.25682668 | -2.7301096 | 0.00633133 | 0.03605464 |
| TYSND1 | 113.541358 | 0.558195683 | 0.28460872 | 1.96127401 | 0.04984707 | 0.1551439 |
| SAR1A | 453.772711 | -0.222704945 | 0.20421254 | -1.0905547 | 0.27546887 | 0.47292733 |
| PPA1 | 1027.07085 | -1.235381008 | 0.30029872 | -4.1138404 | 3.89E-05 | 0.00073876 |
| LRRC20 | 53.4011224 | 0.213265675 | 0.29576873 | 0.72105552 | 0.47087535 | 0.65738999 |
| EIF4EBP2 | 861.939303 | -0.020150441 | 0.21884749 | -0.0920753 | 0.92663824 | 0.96120551 |
| SGPL1 | 580.194268 | -0.034946581 | 0.18675887 | -0.1871214 | 0.85156545 | 0.92197716 |
| PCBD1 | 473.419693 | -1.054077499 | 0.48770425 | -2.1613047 | 0.03067181 | 0.1108991 |
| SLC29A3 | 61.8764404 | 1.331232788 | 0.52761482 | 2.52311483 | 0.01163204 | 0.05598328 |
| CDH23 | 38.1920675 | -0.303692927 | 0.62291658 | -0.4875339 | 0.62588005 | 0.77848801 |
| PSAP | 1534.03572 | -0.743626672 | 0.42225485 | -1.761085 | 0.07822402 | 0.20883678 |
| CHST3 | 163.295242 | 0.047129073 | 0.23192273 | 0.20321023 | 0.83897071 | 0.91478574 |
| ASCC1 | 564.21686 | -0.138684916 | 0.18868752 | -0.7349978 | 0.46234081 | 0.65078102 |
| ANAPC16 | 1186.80408 | 0.225186611 | 0.24105149 | 0.93418467 | 0.35020864 | 0.54948602 |
| DDIT4 | 456.295453 | -2.864679436 | 0.66947856 | -4.2789711 | 1.88E-05 | 0.00042353 |
| DNAJB12 | 290.194304 | 0.417537172 | 0.35103146 | 1.18945799 | 0.23425949 | 0.42856658 |
| MICU1 | 3007.92158 | 0.769556243 | 0.23337444 | 3.2975173 | 0.00097544 | 0.0088833 |
| MCU | 2395.66614 | -0.967760186 | 0.47198935 | -2.0503856 | 0.04032681 | 0.13432798 |
| PLA2G12B | 173.581829 | 1.368124591 | 0.68645698 | 1.99302307 | 0.04625892 | 0.14773323 |
| P4HA1 | 3017.72125 | -0.497316981 | 0.57523288 | -0.8645489 | 0.3872864 | 0.58341958 |
| NUDT13 | 147.625298 | 1.140505426 | 0.48141595 | 2.36906446 | 0.01783315 | 0.07610178 |
| ECD | 382.727527 | 0.035235748 | 0.14260406 | 0.24708796 | 0.80484015 | 0.89408337 |
| FAM149B1 | 618.441025 | 0.579817257 | 0.32550211 | 1.78130105 | 0.07486328 | 0.20307758 |
| DNAJC9 | 458.08701 | -0.26114725 | 0.23726622 | -1.1006508 | 0.27104869 | 0.46924261 |
| DNAJC9-AS1 | 34.0742663 | -0.323195125 | 0.49150802 | -0.6575582 | 0.51082207 | 0.6899686 |
| MRPS16 | 296.876451 | -0.71077273 | 0.23719061 | -2.9966309 | 0.00272981 | 0.01930728 |
| CFAP70 | 616.724061 | 1.303793573 | 0.49055975 | 2.65776711 | 0.00786602 | 0.04238775 |
| ANXA7 | 1016.74823 | -0.252493864 | 0.14657666 | -1.7226062 | 0.08495977 | 0.22059928 |
| MSS51 | 49.716906 | 0.041082884 | 0.31624771 | 0.12990729 | 0.89663977 | 0.94627667 |
| PPP3CB | 1034.25425 | 0.526041855 | 0.18975921 | 2.7721545 | 0.00556866 | 0.03284407 |
| USP54 | 885.374514 | 0.551109044 | 0.2581928 | 2.13448648 | 0.03280298 | 0.11654204 |
| AGAP5 | 198.225807 | 0.371326136 | 0.20944338 | 1.77291899 | 0.07624213 | 0.2054356 |
| AC022400.6 | 90.1059484 | 0.757363479 | 0.26970953 | 2.80807092 | 0.00498392 | 0.03021436 |
| SEC24C | 403.94134 | -0.100739491 | 0.22297964 | -0.4517879 | 0.65142182 | 0.7957436 |
| FUT11 | 291.844575 | -2.385062691 | 0.38062015 | -6.2662544 | 3.70E-10 | 4.87E-08 |
| CHCHD1 | 197.990611 | -0.740861838 | 0.37994828 | -1.9499018 | 0.05118782 | 0.15766945 |
| ZSWIM8 | 937.486525 | 0.234768506 | 0.19376254 | 1.21162998 | 0.22565405 | 0.41803488 |
| CAMK2G | 1064.7538 | 1.472101727 | 0.34598892 | 4.2547655 | 2.09E-05 | 0.00045822 |
| VCL | 1065.28031 | -0.638381715 | 0.29569944 | -2.1588871 | 0.03085893 | 0.1113407 |
| AP3M1 | 298.432099 | -0.287234616 | 0.2873523 | -0.9995904 | 0.31750875 | 0.51755172 |
| ADK | 4146.04288 | -0.412824286 | 0.34880314 | -1.1835452 | 0.23659314 | 0.43124763 |
| KAT6B | 1940.22971 | 0.280429898 | 0.28456774 | 0.98545919 | 0.32439857 | 0.52483341 |
| SAMD8 | 636.990433 | 0.146998869 | 0.24966444 | 0.58878576 | 0.556005 | 0.72527222 |
| VDAC2 | 409.628375 | -0.516039369 | 0.23131478 | -2.2308967 | 0.02568797 | 0.09829845 |
| COMTD1 | 230.878589 | -0.304591887 | 0.37034657 | -0.8224509 | 0.41082029 | 0.6045054 |
| AC010997.3 | 161.027709 | 1.008838933 | 0.51325261 | 1.96557975 | 0.04934718 | 0.15398584 |
| ZNF503-AS2 | 33.6829649 | -0.182849418 | 0.34437336 | -0.5309627 | 0.59544461 | 0.75582107 |
| AC010997.5 | 61.2938661 | 0.549522223 | 0.41592214 | 1.32121416 | 0.18642996 | 0.36970421 |
| LRMDA | 20315.2866 | 0.334119302 | 0.27880406 | 1.19840185 | 0.23076061 | 0.42378291 |
| KCNMA1 | 882.072567 | -3.501268773 | 0.91323586 | -3.8339151 | 0.00012612 | 0.0018711 |
| DLG5 | 615.204602 | -0.595938444 | 0.30199849 | -1.973316 | 0.04845957 | 0.15224617 |
| POLR3A | 510.683618 | 0.08841172 | 0.27154936 | 0.3255825 | 0.74474025 | 0.85552754 |
| RPS24 | 1717.31698 | -0.488951185 | 0.41283737 | -1.1843676 | 0.2362676 | 0.43076882 |
| ZMIZ1-AS1 | 82.6613536 | -0.775424129 | 0.4092958 | -1.8945323 | 0.05815438 | 0.1712176 |
| ZMIZ1 | 1163.74034 | -0.638164056 | 0.36840743 | -1.7322236 | 0.08323376 | 0.21797575 |
| PPIF | 174.0791 | -1.530099965 | 0.47726586 | -3.2059699 | 0.00134608 | 0.01131541 |
| ZCCHC24 | 98.1593075 | -0.5813435 | 0.51564319 | -1.1274143 | 0.25956736 | 0.45772416 |
| EIF5AL1 | 56.6035352 | -1.52031107 | 0.53628666 | -2.8348851 | 0.00458422 | 0.02845827 |
| NUTM2B-AS | 2238.99674 | -0.653228561 | 0.28833433 | -2.2655247 | 0.0234805 | 0.09232564 |
| AL135925.1 | 42.2350032 | -0.352786538 | 0.49039412 | -0.7193939 | 0.47189826 | 0.65801724 |
| TMEM254-A | 112.427057 | 1.896424936 | 0.41410627 | 4.57956106 | 4.66E-06 | 0.00013531 |
| TMEM254 | 214.535956 | 1.006413726 | 0.29481586 | 3.41370277 | 0.00064086 | 0.00647335 |
| ANXA11 | 1084.72493 | -1.248026321 | 0.23931256 | -5.2150472 | 1.84E-07 | 9.09E-06 |
| FAM213A | 540.833609 | -0.111667104 | 0.29147336 | -0.3831126 | 0.70163631 | 0.82861543 |
| TSPAN14 | 598.439581 | -0.562347036 | 0.30738363 | -1.8294632 | 0.06733025 | 0.18900178 |
| NRG3 | 5693.89764 | -1.926566033 | 0.62969579 | -3.0595187 | 0.00221693 | 0.01660517 |
| GHITM | 606.156683 | -0.516339016 | 0.28982271 | -1.7815685 | 0.07481962 | 0.20299931 |
| CCSER2 | 2122.48923 | -0.321342573 | 0.18777012 | -1.7113616 | 0.08701439 | 0.22427722 |
| GRID1 | 144.190491 | -2.102381686 | 0.65578587 | -3.2058966 | 0.00134642 | 0.01131541 |
| WAPL | 1970.73428 | 0.052236658 | 0.20138263 | 0.25939008 | 0.79533428 | 0.88840738 |
| BMPR1A | 2328.77947 | -0.198618315 | 0.17484408 | -1.1359739 | 0.25596748 | 0.45318346 |
| MMRN2 | 66.1801421 | 1.178696709 | 0.51421047 | 2.29224564 | 0.02189147 | 0.08811432 |
| ADIRF | 643.204507 | 0.389457683 | 0.73398125 | 0.53060985 | 0.59568917 | 0.75582107 |
| GLUD1 | 1726.99113 | 0.586357263 | 0.32097049 | 1.82682607 | 0.06772592 | 0.18972394 |
| SHLD2 | 2754.67434 | 0.830893174 | 0.22824384 | 3.64037496 | 0.00027224 | 0.00340165 |
| NUTM2A-AS | 2335.58074 | -0.151260969 | 0.22646014 | -0.6679364 | 0.50417419 | 0.68463598 |
| LINC00864 | 178.24176 | 1.981261433 | 0.77954625 | 2.54155726 | 0.01103599 | 0.05408304 |
| MINPP1 | 400.432593 | 0.602852341 | 0.38492812 | 1.56614263 | 0.11731523 | 0.27305822 |
| PAPSS2 | 594.147371 | -0.51722789 | 0.3558028 | -1.4536926 | 0.14603156 | 0.31482457 |
| ATAD1 | 942.722509 | -0.156329563 | 0.18959437 | -0.8245475 | 0.40962854 | 0.60360018 |
| PTEN | 2093.99254 | 0.190372439 | 0.1595845 | 1.1929256 | 0.23289851 | 0.42678341 |
| RNLS | 938.867064 | 0.376977672 | 0.34392425 | 1.09610669 | 0.27303209 | 0.47082907 |
| STAMBPL1 | 55.6081507 | -0.869618858 | 0.50354945 | -1.7269781 | 0.0841716 | 0.21918352 |
| ACTA2 | 85.3843037 | -0.66359157 | 0.47516139 | -1.3965604 | 0.16254582 | 0.33905132 |
| FAS | 231.760319 | -2.098570311 | 0.53392266 | -3.9304762 | 8.48E-05 | 0.00137034 |
| LIPA | 1088.07015 | 1.355027744 | 0.46114712 | 2.93838494 | 0.00329927 | 0.02218092 |
| IFIT2 | 297.018533 | -2.187741959 | 0.87077209 | -2.5124162 | 0.01199076 | 0.05728731 |
| IFIT3 | 485.130392 | -2.45498799 | 1.02102831 | -2.404427 | 0.01619784 | 0.07113868 |
| IFIT1 | 466.807986 | 0.563726503 | 1.30835877 | 0.43086538 | 0.66656625 | 0.80635371 |
| IFIT5 | 115.19387 | -0.72473666 | 0.43716544 | -1.6578087 | 0.09735609 | 0.24130272 |
| SLC16A12 | 3582.5593 | 2.31553799 | 0.54988601 | 4.21094183 | 2.54E-05 | 0.00052679 |
| SLC16A12-AS | 129.442211 | 1.913750648 | 0.86902538 | 2.20218038 | 0.02765257 | 0.10339164 |
| PANK1 | 1183.24663 | 2.212404867 | 0.69935782 | 3.16348054 | 0.00155895 | 0.01272692 |
| AL157400.3 | 49.7493206 | 1.59592825 | 0.87874207 | 1.81615096 | 0.06934722 | 0.19292517 |
| KIF20B | 148.776885 | -0.160328634 | 0.42707654 | -0.3754096 | 0.70735586 | 0.83235488 |

| | | | | | | |
|---|---|---|---|---|---|---|
| RPP30 | 186.822272 | 0.228239825 | 0.23996588 | 0.95113448 | 0.34153612 | 0.54220015 |
| PCGF5 | 2239.96079 | -0.652560851 | 0.25857659 | -2.5236656 | 0.01161383 | 0.05593493 |
| HECTD2 | 361.068854 | -0.234121997 | 0.2887028 | -0.8109447 | 0.41739744 | 0.61047432 |
| PPP1R3C | 130.171306 | -3.557943884 | 0.86681566 | -4.1046142 | 4.05E-05 | 0.00076463 |
| TNKS2 | 1579.50695 | -0.132823589 | 0.1941103 | -0.6842686 | 0.49380555 | 0.67604801 |
| BTAF1 | 1283.68796 | -0.338458011 | 0.20820472 | -1.625602 | 0.10403435 | 0.25221062 |
| CPEB3 | 1251.9135 | 0.637297476 | 0.28948271 | 2.20150448 | 0.02770033 | 0.10345734 |
| 5-Mar | 636.868811 | -0.241525745 | 0.30236278 | -0.7987946 | 0.42440955 | 0.61749089 |
| IDE | 778.695826 | -0.253880879 | 0.22316743 | -1.1376251 | 0.25527705 | 0.45232479 |
| EXOC6 | 554.933025 | -0.137982559 | 0.26763247 | -0.5155673 | 0.60615663 | 0.76359388 |
| MYOF | 423.027001 | -2.353505437 | 0.54382701 | -4.327673 | 1.51E-05 | 0.00035309 |
| RBP4 | 705.526669 | -0.146741949 | 0.81078156 | -0.1809883 | 0.85637679 | 0.92413451 |
| FRA10AC1 | 532.874077 | 0.38433849 | 0.16703165 | 2.30099199 | 0.02139208 | 0.08688038 |
| PLCE1 | 159.599218 | -1.067400226 | 0.54665679 | -1.9525967 | 0.05086741 | 0.15715165 |
| NOC3L | 261.749884 | 0.083165584 | 0.35398622 | 0.23494018 | 0.81425517 | 0.89947803 |
| TBC1D12 | 787.678664 | 0.132018293 | 0.16662694 | 0.79229862 | 0.42818658 | 0.62120591 |
| HELLS | 103.137702 | -0.527813551 | 0.32208814 | -1.638724 | 0.10127076 | 0.24796683 |
| CYP2C9 | 89.7447396 | 1.725226327 | 0.84620715 | 2.03877541 | 0.04147245 | 0.13681416 |
| AL157834.1 | 48.256949 | 0.367614825 | 0.48648188 | 0.75565985 | 0.44985316 | 0.64023853 |
| ACSM6 | 41.7008595 | -1.019881219 | 0.71709401 | -1.422242 | 0.15495601 | 0.32761759 |
| PDLIM1 | 227.708927 | -0.4520613 | 0.41423667 | -1.0913116 | 0.27513577 | 0.47286769 |
| SORBS1 | 1877.47851 | -0.31853603 | 0.4815175 | -0.6615253 | 0.50827547 | 0.6881892 |
| ALDH18A1 | 506.544117 | -1.087834052 | 0.25983292 | -4.1866676 | 2.83E-05 | 0.00057569 |
| TCTN3 | 419.440934 | 0.129967558 | 0.17550548 | 0.74053277 | 0.45897678 | 0.6475935 |
| ENTPD1 | 55.0540709 | -1.115821307 | 0.6513487 | -1.7130936 | 0.08669532 | 0.22374933 |
| ENTPD1-AS1 | 2213.4146 | 0.100346775 | 0.39766116 | 0.25234241 | 0.80077641 | 0.89172494 |
| AL365273.2 | 260.837882 | -1.184343497 | 0.58703827 | -2.0174894 | 0.04364446 | 0.1415605 |
| CC2D2B | 134.241425 | 0.100355889 | 0.63806705 | 0.1572811 | 0.87502331 | 0.93490877 |
| CCNJ | 67.0102415 | -0.430970152 | 0.26583624 | -1.6211866 | 0.10497762 | 0.25333244 |
| ZNF518A | 1651.21112 | 0.354699166 | 0.36637133 | 0.96814117 | 0.33297387 | 0.53250179 |
| BLNK | 449.107813 | -1.992784722 | 0.68454018 | -2.9111289 | 0.00360125 | 0.02370912 |
| TLL2 | 49.1324531 | -3.11323493 | 0.45958496 | -6.7740139 | 1.25E-11 | 2.59E-09 |
| TM9SF3 | 1732.01565 | -0.299255617 | 0.16934483 | -1.7671376 | 0.07720519 | 0.20709422 |
| PIK3AP1 | 863.59313 | -0.722138174 | 0.33734486 | -2.1406526 | 0.03230206 | 0.11512333 |
| LCOR | 3181.49037 | 0.00904876 | 0.19952952 | 0.04535048 | 0.96382795 | 0.98122324 |
| RRP12 | 382.149339 | -0.942831534 | 0.44329729 | -2.1268606 | 0.03343167 | 0.11819651 |
| PGAM1 | 646.236954 | -1.614968536 | 0.38336867 | -4.212573 | 2.52E-05 | 0.00052594 |
| EXOSC1 | 256.808776 | -0.277454574 | 0.25429384 | -1.0910786 | 0.27523829 | 0.47286769 |
| ZDHHC16 | 234.919029 | -0.022159587 | 0.21485346 | -0.1031381 | 0.91785333 | 0.95600423 |
| MMS19 | 606.532467 | -0.076595535 | 0.17815035 | -0.4299488 | 0.66723287 | 0.80664676 |
| UBTD1 | 371.991852 | -0.037546477 | 0.35127936 | -0.106885 | 0.91488025 | 0.9544306 |
| AL355315.1 | 198.237205 | 1.238574943 | 0.50468922 | 2.45413394 | 0.01412244 | 0.0641831 |
| MORN4 | 185.544877 | 0.075551113 | 0.35656009 | 0.21188887 | 0.83219374 | 0.91145661 |
| PI4K2A | 70.1487133 | -1.022711249 | 0.48426295 | -2.1118924 | 0.03469568 | 0.12098032 |
| ZFYVE27 | 477.564788 | -0.513735809 | 0.28098119 | -1.8283637 | 0.06749499 | 0.18930915 |
| R3HCC1L | 1330.52959 | -0.180377805 | 0.17444097 | -1.0340335 | 0.30112051 | 0.50169217 |
| LOXL4 | 39.1751974 | -0.95684174 | 0.48924564 | -1.9557491 | 0.05049471 | 0.15643364 |
| PYROXD2 | 125.972191 | 0.51508421 | 0.29494353 | 1.74638248 | 0.08074451 | 0.21341401 |
| HPS1 | 394.690258 | -0.184355755 | 0.23489264 | -0.7848511 | 0.43254086 | 0.6252278 |
| HPSE2 | 47.1954354 | -0.075608664 | 0.33852221 | -0.2233492 | 0.82326374 | 0.90529291 |
| CNNM1 | 166.058301 | -2.235372512 | 0.72925067 | -3.0653006 | 0.00217451 | 0.01635898 |
| GOT1 | 1177.80788 | 0.15921155 | 0.19895492 | 0.80023931 | 0.42357216 | 0.61666528 |
| AL391684.1 | 58.5619341 | 0.503641288 | 0.51681348 | 0.97451268 | 0.32980204 | 0.52952988 |
| SLC25A28 | 281.198103 | -1.012128235 | 0.35947526 | -2.8155713 | 0.00486906 | 0.02963596 |
| ENTPD7 | 219.707652 | -0.607434274 | 0.44234069 | -1.3732272 | 0.16968172 | 0.34859523 |
| CUTC | 512.719859 | -0.930712876 | 0.20800604 | -4.4744512 | 7.66E-06 | 0.00020271 |
| COX15 | 324.975057 | -0.319359893 | 0.1775978 | -1.7982198 | 0.07214218 | 0.1981272 |
| ABCC2 | 5082.84063 | -1.591891656 | 0.65739707 | -2.4215071 | 0.0154563 | 0.06862958 |
| DNMBP | 887.843697 | 0.54528319 | 0.30743799 | 1.77363633 | 0.07612332 | 0.2054356 |
| DNMBP-AS1 | 59.9607224 | -0.526757739 | 0.55564747 | -0.9480071 | 0.34312584 | 0.54372554 |
| ERLIN1 | 259.900979 | 0.214487495 | 0.26292364 | 0.81577866 | 0.41462674 | 0.60822868 |
| CHUK | 305.866202 | 0.217732315 | 0.27643869 | 0.78763328 | 0.43091124 | 0.62350896 |
| CWF19L1 | 321.373899 | 0.113154259 | 0.19880851 | 0.56916205 | 0.56924617 | 0.73547528 |
| BLOC1S2 | 168.658716 | -1.41877524 | 0.3401146 | -4.1714624 | 3.03E-05 | 0.0006065 |
| SCD | 188.400655 | -2.573915526 | 0.60766604 | -4.2357403 | 2.28E-05 | 0.00048865 |
| OLMALINC | 104.25867 | -0.303622435 | 0.43627559 | -0.6959418 | 0.48646526 | 0.66974606 |
| SEC31B | 158.709246 | 0.645833946 | 0.4728179 | 1.36592534 | 0.1719624 | 0.3513466 |
| HIF1AN | 316.318543 | -0.114226174 | 0.19883216 | -0.5744854 | 0.56563937 | 0.73247225 |
| PAX2 | 2919.99677 | 1.109617171 | 0.538794 | 2.05944605 | 0.03945153 | 0.13234222 |
| SLF2 | 810.973541 | 0.548215858 | 0.20381894 | 2.68971984 | 0.0071512 | 0.03954092 |
| MRPL43 | 187.166812 | -0.534391675 | 0.30980049 | -1.7249543 | 0.08453571 | 0.21995653 |
| SEMA4G | 221.203055 | 1.378245541 | 0.5232157 | 2.63418232 | 0.00843402 | 0.04453201 |
| TWNK | 34.0183497 | -0.77594872 | 0.62974061 | -1.232172 | 0.21788486 | 0.4090601 |
| LZTS2 | 405.396249 | -0.842883457 | 0.25938527 | -3.2495424 | 0.00115591 | 0.01007249 |
| PDZD7 | 50.4001689 | 0.967301988 | 0.69632617 | 1.38915071 | 0.16478693 | 0.342077 |
| SFXN3 | 196.821619 | 0.091216728 | 0.36527713 | 0.24971924 | 0.80280448 | 0.89272007 |
| BTRC | 1948.99953 | 0.399725147 | 0.19236597 | 2.07794109 | 0.03771479 | 0.12840451 |
| DPCD | 195.243567 | -0.283567807 | 0.32260418 | -0.8789961 | 0.3794034 | 0.57597637 |
| POLL | 249.213165 | 0.389105727 | 0.24174068 | 1.60959973 | 0.10748527 | 0.25752501 |
| FBXW4 | 1033.50213 | 0.296821124 | 0.27210061 | 1.09085065 | 0.27533861 | 0.47286769 |
| AC010789.1 | 47.6506735 | 0.313373662 | 0.63591916 | 0.49278852 | 0.62216202 | 0.77569355 |
| NPM3 | 44.3459471 | -0.957066543 | 0.49407058 | -1.9371049 | 0.05273252 | 0.16051116 |
| OGA | 3510.89944 | 0.247730498 | 0.262379 | 0.94417046 | 0.34508255 | 0.5454262 |
| KCNIP2 | 34.3955008 | -0.042240078 | 0.30354033 | -0.139158 | 0.88932526 | 0.94267382 |
| ARMH3 | 1646.63362 | 0.24023371 | 0.14607916 | 1.64454469 | 0.10006374 | 0.2460337 |
| HPS6 | 35.4582653 | -0.674124389 | 0.50797678 | -1.3270772 | 0.18448315 | 0.36738494 |
| LDB1 | 116.951558 | -0.15912935 | 0.33002965 | -0.4821668 | 0.62968744 | 0.78157908 |
| PPRC1 | 224.419532 | -1.525976611 | 0.69644174 | -2.1911045 | 0.02844423 | 0.10520375 |
| NOLC1 | 166.857749 | -1.301918821 | 0.57515105 | -2.263612 | 0.02359799 | 0.09265472 |
| GBF1 | 2456.3777 | 0.368032725 | 0.23920405 | 1.53857233 | 0.12390874 | 0.28341943 |
| NFKB2 | 221.063953 | -1.248244403 | 0.51223249 | -2.4368708 | 0.01481497 | 0.06655811 |
| FBXL15 | 221.748344 | -0.533432457 | 0.35148388 | -1.5176584 | 0.12910054 | 0.29095382 |
| CUEDC2 | 185.731453 | -0.028783499 | 0.22764542 | -0.1264401 | 0.8993836 | 0.94743919 |
| MFSD13A | 35.4015718 | 0.067832482 | 0.44618037 | 0.15202928 | 0.87916384 | 0.93706833 |
| ACTR1A | 288.495622 | -0.206551135 | 0.31388999 | -0.6580367 | 0.51051455 | 0.6897256 |
| SUFU | 1050.53329 | 0.956066837 | 0.31501035 | 3.03503312 | 0.00240509 | 0.01771422 |
| TRIM8 | 467.866323 | 0.086188954 | 0.28543411 | 0.30195744 | 0.76268451 | 0.86770535 |
| ARL3 | 431.091761 | 0.367239752 | 0.34324788 | 1.06989665 | 0.28466583 | 0.48398841 |
| SFXN2 | 962.666715 | 2.50023769 | 0.43864731 | 5.69988148 | 1.20E-08 | 9.55E-07 |
| WBP1L | 1282.54772 | 0.919572294 | 0.18615514 | 4.93981694 | 7.82E-07 | 3.15E-05 |
| CYP17A1 | 118.211218 | 1.937278808 | 0.6983993 | 2.77388422 | 0.00553914 | 0.03271218 |
| BORCS7-ASN | 104.914477 | -0.398418304 | 0.45831063 | -0.8693194 | 0.38467247 | 0.58101439 |
| CNNM2 | 1749.6912 | 1.215954552 | 0.27966299 | 4.34792803 | 1.37E-05 | 0.00032704 |
| NT5C2 | 3996.71176 | 0.012252035 | 0.18482733 | 0.06628908 | 0.94714767 | 0.97291316 |
| PCGF6 | 307.993413 | 0.616898758 | 0.26303859 | 2.34527857 | 0.01901287 | 0.07950256 |
| TAF5 | 53.2626372 | -0.024813757 | 0.44450386 | -0.0558235 | 0.95548242 | 0.97737295 |
| ATP5MD | 422.560021 | -0.473559101 | 0.31051623 | -1.5250704 | 0.12724155 | 0.28832864 |
| PDCD11 | 365.477042 | -0.492095738 | 0.41831379 | -1.1763794 | 0.23944329 | 0.43447832 |
| SH3PXD2A | 234.627681 | 0.628840081 | 0.41894441 | 1.50101077 | 0.13335277 | 0.29711744 |
| STN1 | 131.217451 | -0.018639404 | 0.39896156 | -0.0467198 | 0.96273655 | 0.98069485 |
| SLK | 862.15732 | -0.113128651 | 0.14078461 | -0.8035584 | 0.42165206 | 0.61537317 |
| CFAP43 | 599.82446 | 0.250558914 | 0.51989866 | 0.48193798 | 0.62985 | 0.78158193 |
| GSTO1 | 576.175314 | -1.352294482 | 0.38539814 | -3.5088246 | 0.00045009 | 0.00496731 |

| | | | | | | |
|---|---|---|---|---|---|---|
| GSTO2 | 296.594121 | 0.332512175 | 0.44070707 | 0.75449703 | 0.45055083 | 0.64075861 |
| ITPRIP | 39.3909375 | -1.341714899 | 0.55086612 | -2.4356461 | 0.01486522 | 0.06669933 |
| SORCS3 | 96.8073819 | 1.811034987 | 0.61616705 | 2.93919481 | 0.00329066 | 0.02215604 |
| SORCS1 | 2607.66016 | 2.103887868 | 0.59531059 | 3.53410122 | 0.00040916 | 0.00465701 |
| LINC01435 | 160.181692 | 0.278116872 | 0.87481905 | 0.3179136 | 0.75055048 | 0.85939315 |
| XPNPEP1 | 650.460335 | -0.970816017 | 0.23494715 | -4.1320613 | 3.60E-05 | 0.00069408 |
| ADD3 | 2987.39952 | 1.126650411 | 0.33445474 | 3.36861846 | 0.00075546 | 0.00735969 |
| MXI1 | 1124.72012 | -1.259861479 | 0.39254962 | -3.2094324 | 0.00132997 | 0.01122533 |
| SMNDC1 | 311.505429 | -0.289309646 | 0.22506166 | -1.2854684 | 0.19862866 | 0.38492903 |
| DUSP5 | 117.82813 | -3.483751081 | 0.82814201 | -4.2067074 | 2.59E-05 | 0.0005341 |
| SMC3 | 427.712538 | -0.037131869 | 0.19004793 | -0.1953816 | 0.84509422 | 0.91854292 |
| RBM20 | 706.475062 | 0.080315948 | 0.351943 | 0.22820726 | 0.81948512 | 0.90265851 |
| PDCD4 | 482.428848 | -0.533780885 | 0.38172036 | -1.3983558 | 0.16200625 | 0.33825128 |
| BBIP1 | 268.51342 | -0.120567625 | 0.20971086 | -0.5749231 | 0.56534328 | 0.73243481 |
| SHOC2 | 1423.68328 | -0.241238532 | 0.17537059 | -1.3755929 | 0.16894771 | 0.34755609 |
| GPAM | 82.6800416 | 0.325062359 | 0.25187369 | 1.29057687 | 0.19685044 | 0.38261897 |
| ZDHHC6 | 220.791317 | -0.260183008 | 0.17152078 | -1.5169183 | 0.12928731 | 0.29123109 |
| VTI1A | 3762.09216 | -0.112122405 | 0.17619224 | -0.636364 | 0.52453918 | 0.70001544 |
| TCF7L2 | 2949.72115 | 0.448033422 | 0.28049871 | 1.59727444 | 0.11020455 | 0.26188864 |
| HABP2 | 138.614284 | -1.267680206 | 0.76979357 | -1.6467794 | 0.09960339 | 0.24519823 |
| CASP7 | 282.973018 | -0.8506094 | 0.34740606 | -2.4484588 | 0.01434689 | 0.06492333 |
| DCLRE1A | 48.1580852 | 0.420746615 | 0.33622553 | 1.25138211 | 0.2107951 | 0.40007037 |
| NHLRC2 | 697.587423 | 0.249046982 | 0.1850592 | 1.34576928 | 0.17837693 | 0.3594319 |
| CCDC186 | 1020.10736 | 0.016968642 | 0.20478325 | 0.08286147 | 0.93396169 | 0.96479108 |
| ABLIM1 | 6473.53285 | -0.676385885 | 0.22464655 | -3.0108893 | 0.00260484 | 0.01872287 |
| FAM160B1 | 574.877657 | -0.350162896 | 0.19272625 | -1.8168926 | 0.06923356 | 0.19272631 |
| TRUB1 | 293.844247 | 0.073468887 | 0.21881841 | 0.33575276 | 0.73705731 | 0.85110831 |
| ATRNL1 | 767.054254 | -3.32869925 | 0.66501316 | -5.0054638 | 5.57E-07 | 2.34E-05 |
| GFRA1 | 102.425644 | 0.049601499 | 0.63147675 | 0.07854842 | 0.93739182 | 0.96641041 |
| HSPA12A | 1371.9963 | 0.499163661 | 0.35197514 | 1.41817875 | 0.15613859 | 0.32976758 |
| ENO4 | 175.518694 | -0.426857231 | 0.36081141 | -1.1830481 | 0.2367901 | 0.43136226 |
| SHTN1 | 2431.88507 | -0.345037188 | 0.26793316 | -1.2877734 | 0.19782486 | 0.38402285 |
| PDZD8 | 2429.44456 | 0.195876058 | 0.24330235 | 0.8050726 | 0.42077779 | 0.61442433 |
| AC005871.2 | 105.860187 | -1.418455135 | 0.7479832 | -1.896373 | 0.05791074 | 0.17081613 |
| EMX2OS | 2048.02756 | -0.350228045 | 0.35897829 | -0.9756246 | 0.32925053 | 0.52891681 |
| EMX2 | 718.253721 | 0.610330728 | 0.2849924 | 2.14156846 | 0.03222822 | 0.11494985 |
| RAB11FIP2 | 824.053683 | -0.155594855 | 0.22414376 | -0.6941744 | 0.48757285 | 0.67073074 |
| AC022395.1 | 58.909423 | -1.188189017 | 0.67415029 | -1.7624987 | 0.07798507 | 0.20849257 |
| CASC2 | 737.670961 | 0.78020545 | 0.57316888 | 1.36121391 | 0.1734461 | 0.35320542 |
| FAM204A | 654.105516 | 0.183424359 | 0.18289977 | 1.00286817 | 0.31592447 | 0.51590706 |
| CACUL1 | 1162.07397 | -0.341366396 | 0.15791942 | -2.1616492 | 0.03064522 | 0.11087961 |
| EIF3A | 1735.29531 | -0.589186047 | 0.41101653 | -1.4334851 | 0.15171927 | 0.32307857 |
| FAM45A | 301.609216 | -0.248840518 | 0.17734801 | -1.4031199 | 0.16058109 | 0.33650588 |
| SFXN4 | 471.710965 | 1.016699329 | 0.29830154 | 3.40829395 | 0.0006537 | 0.0065788 |
| PRDX3 | 388.808992 | -0.180501194 | 0.36833741 | -0.4900431 | 0.62410343 | 0.77705383 |
| GRK5 | 182.745154 | -0.642024193 | 0.38203539 | -1.6805359 | 0.09285309 | 0.23456272 |
| TIAL1 | 895.205558 | -0.115024538 | 0.21926882 | -0.5245823 | 0.59987361 | 0.75965147 |
| BAG3 | 638.39538 | 0.13533093 | 0.4399669 | 0.30759343 | 0.75839172 | 0.86454385 |
| INPP5F | 1122.4165 | 0.373195519 | 0.33665271 | 1.10854749 | 0.26762544 | 0.46564381 |
| MCMBP | 403.024766 | -0.170802453 | 0.17953093 | -0.9513818 | 0.34141061 | 0.54213568 |
| SEC23IP | 751.616048 | -0.321295798 | 0.33429677 | -0.9611095 | 0.33649712 | 0.53661467 |
| PLPP4 | 367.488959 | 1.634510491 | 0.44410368 | 3.68047051 | 0.0002328 | 0.00302182 |
| WDR11-AS1 | 162.27554 | 0.190773351 | 0.25886393 | 0.73696381 | 0.46114436 | 0.64969737 |
| WDR11 | 1057.55122 | 0.186423188 | 0.24512521 | 0.76052229 | 0.44694245 | 0.63774996 |
| FGFR2 | 1376.9248 | 1.000227203 | 0.65260462 | 1.53266951 | 0.1253573 | 0.28542734 |
| ATE1 | 2624.83931 | -0.152418882 | 0.15911115 | -0.9579397 | 0.33809318 | 0.53830215 |
| NSMCE4A | 179.865246 | 0.331751436 | 0.27251719 | 1.21735967 | 0.2234674 | 0.41578222 |
| TACC2 | 1267.78517 | -0.154168327 | 0.33560681 | -0.4593719 | 0.64596714 | 0.79225766 |
| PLEKHA1 | 9048.84854 | -1.9460785 | 0.34217548 | -5.6873698 | 1.29E-08 | 1.01E-06 |
| HTRA1 | 157.448501 | -0.34493463 | 0.350263 | -0.9847875 | 0.32472845 | 0.52501213 |
| C10orf88 | 222.154352 | -0.147255972 | 0.20632919 | -0.7136943 | 0.47541623 | 0.66124102 |
| PSTK | 114.122809 | -0.702035304 | 0.34990458 | -2.0063621 | 0.04481763 | 0.14430707 |
| IKZF5 | 307.284647 | -0.169418732 | 0.32374207 | -0.5233139 | 0.60075581 | 0.76020679 |
| ACADSB | 1354.62475 | -0.87803065 | 0.73380969 | -1.1965373 | 0.23148696 | 0.4245373 |
| BUB3 | 364.296509 | -0.415548943 | 0.25179661 | -1.6503357 | 0.09887429 | 0.24379742 |
| CHST15 | 167.490914 | -1.416541322 | 0.45564204 | -3.1088907 | 0.00187791 | 0.01458387 |
| OAT | 314.395866 | -0.007238593 | 0.31215581 | -0.023189 | 0.98149948 | 0.99060551 |
| LHPP | 545.0663 | 0.626665009 | 0.32704485 | 1.91614396 | 0.05534677 | 0.16589509 |
| ABRAXAS2 | 312.492809 | -0.24600904 | 0.17733482 | -1.3872574 | 0.16536329 | 0.34270254 |
| ZRANB1 | 745.234476 | -0.18176025 | 0.31419975 | -0.5784863 | 0.56293585 | 0.73052419 |
| CTBP2 | 4166.77319 | -1.488612605 | 0.2783246 | -5.3484765 | 8.87E-08 | 4.94E-06 |
| EDRF1 | 421.835066 | -0.010337732 | 0.20431569 | -0.0505969 | 0.95964677 | 0.97933252 |
| UROS | 283.838947 | 0.472324375 | 0.2849006 | 1.65785671 | 0.09734639 | 0.24130272 |
| BCCIP | 428.827644 | -0.197837808 | 0.27497972 | -0.7194633 | 0.47185552 | 0.65801724 |
| DHX32 | 1281.51456 | -0.786913146 | 0.26258463 | -2.9967982 | 0.00272831 | 0.01930665 |
| FANK1 | 751.145664 | -0.212792167 | 0.23423592 | -0.9084523 | 0.3636393 | 0.56136995 |
| ADAM12 | 72.5632285 | -1.571646087 | 0.2462473 | -6.382389 | 1.74E-10 | 2.52E-08 |
| DOCK1 | 3821.49734 | -0.262787127 | 0.20826911 | -1.2617672 | 0.20703258 | 0.39591183 |
| AL359094.1 | 48.7003324 | -1.030717533 | 0.55241317 | -1.8658453 | 0.06206301 | 0.1789812 |
| FAM196A | 139.690709 | -0.753337323 | 0.49386734 | -1.525384 | 0.12716335 | 0.28824675 |
| CLRN3 | 350.784755 | 1.108250113 | 0.61302683 | 1.80783298 | 0.0706325 | 0.19541679 |
| PTPRE | 326.762651 | -1.584663277 | 0.37733423 | -4.1996276 | 2.67E-05 | 0.0005486 |
| MGMT | 1748.83434 | 0.537570874 | 0.33512984 | 1.60406748 | 0.10869918 | 0.25957136 |
| AL157832.3 | 187.825834 | 0.692640107 | 0.41194161 | 1.68140361 | 0.09268455 | 0.23426647 |
| C10orf143 | 145.024947 | 1.194460391 | 0.35434963 | 3.37085268 | 0.00074936 | 0.00731262 |
| GLRX3 | 820.018175 | -0.946262091 | 0.37194764 | -2.5440734 | 0.01095681 | 0.05384903 |
| PPP2R2D | 1872.47685 | -0.198592028 | 0.33086671 | -0.6002176 | 0.54836122 | 0.71851517 |
| BNIP3 | 1385.91487 | -1.911626109 | 0.54014822 | -3.539077 | 0.00040153 | 0.00459029 |
| AL162274.2 | 34.8581091 | -0.39081719 | 0.52001273 | -0.7515531 | 0.45231985 | 0.64164525 |
| STK32C | 160.121569 | -0.538833913 | 0.30286222 | -1.7791388 | 0.07521701 | 0.2037146 |
| LRRC27 | 219.110452 | 0.279779258 | 0.45215333 | 0.61877075 | 0.53606739 | 0.70869254 |
| PWWP2B | 48.0977224 | -1.229563041 | 0.51018543 | -2.4100317 | 0.01595114 | 0.07037085 |
| INPP5A | 636.705614 | -0.211816826 | 0.21506725 | -0.9848865 | 0.32467984 | 0.52501213 |
| CFAP46 | 253.947774 | 1.200415137 | 0.4920651 | 2.43954535 | 0.01470576 | 0.06615419 |
| ADAM8 | 48.9622562 | 1.859515321 | 0.32058773 | 5.80033216 | 6.62E-09 | 5.74E-07 |
| TUBGCP2 | 1018.46707 | 0.638388674 | 0.1911167 | 3.34030816 | 0.00083685 | 0.00792935 |
| AL360181.1 | 58.7850469 | 1.136643272 | 0.66590246 | 1.70692157 | 0.08783663 | 0.22582532 |
| PRAP1 | 41.0233969 | 1.031106909 | 0.47310498 | 2.17944632 | 0.02929853 | 0.1072636 |
| FUOM | 148.924106 | 0.583218077 | 0.44667226 | 1.30569575 | 0.19165604 | 0.37593437 |
| ECHS1 | 1065.73071 | 0.083697029 | 0.37801853 | 0.22140986 | 0.82477332 | 0.90642548 |
| PAOX | 110.099784 | -0.220037876 | 0.32985983 | -0.6670648 | 0.50473076 | 0.6850618 |
| SCART1 | 97.3878269 | -0.07249773 | 0.47437929 | -0.1528265 | 0.87853509 | 0.93706833 |
| IQSEC3 | 370.071878 | 2.074829119 | 0.87264803 | 2.37762425 | 0.01742457 | 0.07491801 |
| AC026369.1 | 76.0857582 | 2.499235961 | 1.09849615 | 2.27514312 | 0.02289735 | 0.09078794 |
| SLC6A12 | 1441.67524 | 1.462071885 | 0.55497543 | 2.6344804 | 0.00842662 | 0.04451007 |
| SLC6A13 | 3642.26087 | 2.172787886 | 0.63920097 | 3.39922496 | 0.00067577 | 0.00673168 |
| KDM5A | 1429.25259 | 0.033344331 | 0.11927192 | 0.27956565 | 0.77981076 | 0.87887404 |
| CCDC77 | 97.9690508 | 0.054067927 | 0.3411205 | 0.15850096 | 0.87406206 | 0.93468316 |
| B4GALNT3 | 69.1871558 | -0.542920013 | 0.40365489 | -1.3450104 | 0.17862188 | 0.35963134 |
| WNK1 | 2131.97635 | -0.190647969 | 0.13983525 | -1.3633756 | 0.17276416 | 0.35234018 |
| RAD52 | 374.599134 | 0.653911183 | 0.25684239 | 2.54596284 | 0.01089768 | 0.05367391 |
| ERC1 | 5172.51626 | -0.190171355 | 0.21468604 | -0.8858115 | 0.37571914 | 0.57247468 |
| WNT5B | 105.534464 | -1.77852046 | 0.52698133 | -3.3749212 | 0.00073837 | 0.00722399 |

| | | | | | | |
|---|---|---|---|---|---|---|
| FBXL14 | 112.365517 | -0.15537609 | 0.30433096 | -0.5105497 | 0.60966638 | 0.76617741 |
| ADIPOR2 | 1023.42203 | -0.223558271 | 0.30479786 | -0.733464 | 0.46327545 | 0.65136082 |
| DCP1B | 456.813666 | 0.230853542 | 0.35901917 | 0.64301174 | 0.52021648 | 0.69607397 |
| CACNA1C | 221.656505 | 1.393556104 | 0.63513759 | 2.19410113 | 0.02822814 | 0.10465867 |
| FKBP4 | 366.626071 | -0.835906071 | 0.51803044 | -1.6136234 | 0.10660913 | 0.2563669 |
| ITFG2-AS1 | 65.4482417 | 0.762540717 | 0.35106887 | 2.17205447 | 0.02985155 | 0.10879427 |
| ITFG2 | 852.805403 | 0.492391229 | 0.36417775 | 1.35206291 | 0.17635519 | 0.35674448 |
| TULP3 | 1887.44115 | 0.525951076 | 0.24882818 | 2.11371184 | 0.03453989 | 0.12065194 |
| TEAD4 | 593.105195 | -0.714756043 | 0.49958614 | -1.4306963 | 0.15251727 | 0.32411759 |
| TSPAN9 | 3798.83614 | 0.239113513 | 0.29734993 | 0.80414855 | 0.42131119 | 0.61500665 |
| CRACR2A | 41.8058594 | -0.602124104 | 0.5395839 | -1.1159045 | 0.26446302 | 0.46242769 |
| PARP11 | 291.805923 | 0.287548451 | 0.21114486 | 1.36185388 | 0.173244 | 0.35289873 |
| CCND2 | 70.7434565 | -2.430061472 | 0.84911962 | -2.86186 | 0.00421163 | 0.02662768 |
| TIGAR | 188.418636 | -0.402136474 | 0.48053009 | -0.8368601 | 0.40267119 | 0.59683145 |
| C12orf4 | 276.400705 | 0.387657283 | 0.13534741 | 2.86416469 | 0.00418111 | 0.02648356 |
| AKAP3 | 36.8931918 | 0.787968081 | 0.48031089 | 1.64053761 | 0.10089344 | 0.247308 |
| AC005833.2 | 46.2553362 | -0.971761789 | 0.3723801 | -2.6095965 | 0.00906491 | 0.04695378 |
| CD9 | 344.775203 | -0.743819405 | 0.50713678 | -1.4667037 | 0.14245667 | 0.30981487 |
| PLEKHG6 | 52.8247093 | -0.747974157 | 0.45059106 | -1.6599845 | 0.09691758 | 0.2406611 |
| TNFRSF1A | 1190.68667 | -0.483326242 | 0.38130268 | -1.2675658 | 0.20495304 | 0.39334944 |
| SCNN1A | 807.388314 | -0.578305913 | 0.43402122 | -1.332437 | 0.18271665 | 0.36498063 |
| LTBR | 927.293301 | -0.809981497 | 0.36059025 | -2.2462657 | 0.02468699 | 0.09545406 |
| AC005840.2 | 51.6780929 | 0.384059807 | 0.63958471 | 0.60048309 | 0.54818433 | 0.71842074 |
| TAPBPL | 122.984678 | 0.15929052 | 0.47868843 | 0.33276451 | 0.73931205 | 0.85193361 |
| MRPL51 | 259.749104 | -0.679043561 | 0.33368702 | -2.0349714 | 0.04185375 | 0.13765274 |
| NCAPD2 | 207.910329 | 0.536055172 | 0.26492615 | 2.02341362 | 0.04303052 | 0.14036634 |
| GAPDH | 3368.42093 | -1.775969056 | 0.4396444 | -4.039558 | 5.36E-05 | 0.00096078 |
| IFFO1 | 73.1987519 | 0.437725708 | 0.26226401 | 1.669027 | 0.09511203 | 0.23785816 |
| CHD4 | 323.00774 | 0.090028026 | 0.35531132 | 0.25337787 | 0.79997623 | 0.89126832 |
| AC006064.2 | 93.2440131 | -0.338508787 | 0.42228194 | -0.801618 | 0.42277399 | 0.61622276 |
| ING4 | 256.299258 | -0.083073924 | 0.29461453 | -0.281975 | 0.7779627 | 0.87758633 |
| ZNF384 | 287.035206 | 0.457958437 | 0.16073935 | 2.84907492 | 0.00438466 | 0.02749335 |
| COPS7A | 201.858192 | 0.282088771 | 0.23186806 | 1.21659173 | 0.22375959 | 0.41608307 |
| MLF2 | 428.669167 | -0.350622444 | 0.26559771 | -1.320126 | 0.18679295 | 0.37015627 |
| PTMS | 619.180195 | 0.407228938 | 0.30032911 | 1.35594228 | 0.17511753 | 0.3551841 |
| USP5 | 187.053156 | -0.064436682 | 0.36518464 | -0.1764496 | 0.85994074 | 0.92628466 |
| TPI1 | 994.437151 | -1.062682008 | 0.41262597 | -2.5754123 | 0.01001207 | 0.05050989 |
| SPSB2 | 117.577413 | -0.034577327 | 0.33413459 | -0.1034832 | 0.91757946 | 0.95587788 |
| LRRC23 | 485.233823 | -0.766398331 | 0.43825621 | -1.7487449 | 0.08033512 | 0.21261661 |
| ENO2 | 81.9527808 | -4.682107513 | 0.64039455 | -7.3112857 | 2.65E-13 | 8.24E-11 |
| ATN1 | 387.147028 | -0.603985129 | 0.27094979 | -2.2291404 | 0.02580457 | 0.09855201 |
| C12orf57 | 143.992137 | -0.901266431 | 0.52273047 | -1.7241513 | 0.08468053 | 0.22012441 |
| PTPN6 | 153.727556 | -0.086272777 | 0.34635304 | -0.2490891 | 0.80329184 | 0.89307496 |
| PHB2 | 374.082293 | -0.494720518 | 0.30821734 | -1.6051028 | 0.10847118 | 0.25926944 |
| EMG1.1 | 121.318742 | 0.312187988 | 0.40117517 | 0.77818372 | 0.43646071 | 0.62857823 |
| LPCAT3 | 1843.66104 | 0.31278949 | 0.24837472 | 1.25934509 | 0.20790571 | 0.3970136 |
| C1S | 581.383155 | -0.957223562 | 0.38476826 | -2.4877924 | 0.01285387 | 0.06019407 |
| C1R | 503.624502 | -3.492575703 | 0.57249404 | -6.1006324 | 1.06E-09 | 1.22E-07 |
| C1RL | 849.207085 | -1.513647923 | 0.43181488 | -3.5053167 | 0.00045606 | 0.0050211 |
| C1RL-AS1 | 170.135443 | -1.233513055 | 0.37399754 | -3.298185 | 0.00097312 | 0.00887462 |
| AC233309.1 | 50.1441686 | 1.808642079 | 0.69247837 | 2.61183911 | 0.00900566 | 0.04670474 |
| RBP5 | 929.284099 | 0.337358516 | 0.49530969 | 0.68110623 | 0.49580428 | 0.67770136 |
| AC018653.3 | 45.5156532 | 0.460434332 | 0.72107639 | 0.63853752 | 0.52312384 | 0.69880698 |
| CLSTN3 | 768.008686 | -0.082725857 | 0.43379141 | -0.1907042 | 0.84875732 | 0.92033199 |
| PEX5 | 322.046635 | 0.382956857 | 0.27029259 | 1.41682339 | 0.15653457 | 0.33019688 |
| CD163L1 | 78.2380641 | -0.065668364 | 0.28008436 | -0.2344592 | 0.81462848 | 0.89954987 |
| SLC2A14 | 57.5880618 | 0.904992311 | 0.72319065 | 1.25138829 | 0.21079285 | 0.40007037 |
| FOXJ2 | 97.3222862 | 0.372797111 | 0.38160221 | 0.97692597 | 0.3286058 | 0.52846816 |
| NECAP1 | 152.827972 | 0.092827044 | 0.22309528 | 0.416087 | 0.67734634 | 0.81315429 |
| LINC00937 | 678.4464 | 0.814524295 | 0.32157725 | 2.532904 | 0.01131219 | 0.05507176 |
| RIMKLB | 755.947366 | -0.742059587 | 0.32934842 | -2.2531141 | 0.02425195 | 0.0942505 |
| A2ML1-AS1 | 95.5676735 | -0.832932479 | 0.59815195 | -1.3925099 | 0.16376806 | 0.34068429 |
| AC006581.2 | 61.0815664 | -0.07722817 | 0.40322405 | -0.1915267 | 0.84811297 | 0.91986577 |
| PHC1 | 183.709087 | 0.281589422 | 0.37404254 | 0.75282726 | 0.45155372 | 0.64124856 |
| M6PR | 459.488046 | -0.340838006 | 0.30133584 | -1.1310902 | 0.25801715 | 0.4554006 |
| KLRG1 | 150.810869 | 0.718093969 | 0.30588938 | 2.34756097 | 0.01889678 | 0.07923468 |
| A2M | 74.4381194 | 0.444522364 | 0.39830189 | 1.11604382 | 0.26440338 | 0.46238544 |
| CLEC2D | 158.012069 | -0.923276032 | 0.47260691 | -1.9535813 | 0.05075075 | 0.15695984 |
| CLEC7A | 90.6712727 | -1.425638373 | 1.20255721 | -1.1855057 | 0.23581758 | 0.43017721 |
| GABARAPL1 | 1302.68198 | -0.189394972 | 0.5280048 | -0.3586993 | 0.71982003 | 0.84047498 |
| KLRD1 | 704.112933 | -0.620787133 | 0.69160708 | -0.8976009 | 0.36939837 | 0.56695645 |
| AC022075.1 | 66.3167776 | -1.408383745 | 0.50370806 | -2.7960318 | 0.00517343 | 0.0311155 |
| EIF2S3B | 78.055464 | -0.660590111 | 0.34053022 | -1.9398869 | 0.05239344 | 0.15967814 |
| MAGOHB | 95.5735019 | -0.139249922 | 0.36038736 | -0.3863896 | 0.69920816 | 0.82699511 |
| YBX3 | 1990.75702 | -0.632284457 | 0.33606476 | -1.8814363 | 0.05991259 | 0.17457947 |
| PRH1 | 122.225166 | 0.430614321 | 0.41389547 | 1.04039389 | 0.29815694 | 0.49833608 |
| SMIM10L1 | 152.351518 | -0.844583214 | 0.27469498 | -3.074622 | 0.0021077 | 0.01599678 |
| AC244131.2 | 362.481226 | -0.471726631 | 0.60314884 | -0.7821065 | 0.43415198 | 0.62641276 |
| PRB2 | 53.9589146 | -1.691144144 | 0.94945818 | -1.7811676 | 0.07488508 | 0.2030965 |
| LINC01252 | 158.845772 | -0.01756261 | 0.78492352 | -0.0223749 | 0.98214888 | 0.99118794 |
| ETV6 | 4139.98575 | -1.068075525 | 0.40862448 | -2.6138314 | 0.00895332 | 0.04650366 |
| BCL2L14 | 295.006475 | -0.305468374 | 0.40419303 | -0.7557487 | 0.44979985 | 0.64022914 |
| LRP6 | 1371.38888 | 0.814856048 | 0.27521278 | 2.96082198 | 0.00306819 | 0.02105621 |
| MANSC1 | 71.5048423 | -0.081516888 | 0.29858919 | -0.2730068 | 0.78484796 | 0.88201288 |
| BORCS5 | 454.933934 | 0.391215071 | 0.18610816 | 2.10208446 | 0.03554588 | 0.1229751 |
| DUSP16 | 1161.27986 | -0.425331638 | 0.30375243 | -1.4002576 | 0.1614362 | 0.3376268 |
| CREBL2 | 314.158534 | 0.127124502 | 0.20363362 | 0.62428052 | 0.53244337 | 0.70582218 |
| CDKN1B | 141.609108 | 0.201899976 | 0.29082351 | 0.6942354 | 0.4875346 | 0.67073074 |
| APOLD1 | 68.4920544 | 0.494151171 | 0.51015229 | 0.96863463 | 0.33272752 | 0.53223201 |
| GPRC5D-AS1 | 234.570911 | -0.018922005 | 0.35067076 | -0.0539595 | 0.95696746 | 0.97810315 |
| HEBP1 | 187.528843 | -0.48130144 | 0.18668745 | -2.5781136 | 0.00993413 | 0.05017214 |
| AC023790.2 | 34.967279 | 0.513838217 | 0.48467176 | 1.06017774 | 0.28906374 | 0.48896403 |
| FAM234B | 41.4739093 | -0.65258585 | 0.3307863 | -1.9728321 | 0.04851469 | 0.15235992 |
| EMP1 | 110.428743 | -1.718927184 | 0.69079435 | -2.4883342 | 0.01283431 | 0.06013328 |
| ATF7IP | 2111.87401 | 0.213304774 | 0.2876618 | 0.74151235 | 0.45838285 | 0.64719928 |
| PLBD1 | 123.417169 | 0.17755971 | 0.31945437 | 0.55582182 | 0.57833267 | 0.74211119 |
| PLBD1-AS1 | 140.028626 | 0.88079782 | 0.41429496 | 2.12601623 | 0.03350191 | 0.11835326 |
| AC010168.1 | 320.345953 | -0.046511806 | 0.51756893 | -0.0898659 | 0.92839377 | 0.96208463 |
| HIST4H4 | 34.3125455 | -0.781889446 | 0.43574345 | -1.7943803 | 0.07275249 | 0.19907103 |
| H2AFJ | 455.438564 | -0.959340371 | 0.39466957 | -2.4307432 | 0.01506789 | 0.06729735 |
| WBP11 | 270.711651 | -0.076399992 | 0.1994402 | -0.3830722 | 0.70166625 | 0.82861543 |
| C12orf60 | 129.298085 | 0.083940495 | 0.4694268 | 0.17881488 | 0.85808306 | 0.92545981 |
| MGP | 144.196706 | -0.11175632 | 0.43728219 | -0.2555703 | 0.79828269 | 0.89012761 |
| RERG | 1628.80644 | 0.687806898 | 0.47039816 | 1.46218025 | 0.14369182 | 0.31129821 |
| RERG-AS1 | 49.7441161 | 2.646417679 | 0.84023156 | 3.149629 | 0.00163478 | 0.01318884 |
| PTPRO | 156.842781 | -1.319626575 | 0.64951353 | -2.0317153 | 0.04218248 | 0.13835734 |
| EPS8 | 6430.75123 | -0.640143924 | 0.24447581 | -2.6184347 | 0.00883342 | 0.04602041 |
| STRAP | 543.86833 | -0.588166988 | 0.204855 | -2.8711381 | 0.00408997 | 0.02607497 |
| DERA | 2278.12059 | -0.223723824 | 0.34157935 | -0.6549688 | 0.51248783 | 0.6911936 |
| MGST1 | 5284.81452 | -1.091372312 | 0.36729364 | -2.971389 | 0.00296456 | 0.02051248 |
| AC007529.2 | 392.361283 | -0.442011968 | 0.53395419 | -0.8278088 | 0.4077788 | 0.60161981 |
| RERGL | 216.705229 | 1.924629217 | 0.60768335 | 3.16715803 | 0.00153937 | 0.01261213 |
| PIK3C2G | 111.32543 | 1.706377009 | 0.51116053 | 3.33824092 | 0.00084311 | 0.00797547 |

| | | | | | | |
|---|---|---|---|---|---|---|
| PLCZ1 | 145.280318 | 2.840177948 | 0.71315241 | 3.98256797 | 6.82E-05 | 0.00115285 |
| PLEKHA5 | 17480.2 | 1.319845229 | 0.31309643 | 4.21545923 | 2.49E-05 | 0.00052085 |
| AEBP2 | 4350.6856 | 0.371392943 | 0.21777671 | 1.7053841 | 0.08812281 | 0.22623195 |
| AC024901.1 | 377.945317 | 0.310943331 | 0.46758635 | 0.6649966 | 0.50605267 | 0.68616717 |
| LINC02398 | 106.795003 | 0.065323024 | 0.55042233 | 0.11867801 | 0.90553046 | 0.95066597 |
| PDE3A | 1183.55412 | 2.455740343 | 0.64177515 | 3.82648091 | 0.00012999 | 0.00190658 |
| PYROXD1 | 329.732174 | -0.13261612 | 0.30653851 | -0.4326247 | 0.66528746 | 0.80579154 |
| RECQL | 327.271352 | -0.773484519 | 0.25046571 | -3.0881853 | 0.00201383 | 0.0154296 |
| GOLT1B | 302.016864 | -0.820722429 | 0.2003958 | -4.0955072 | 4.21E-05 | 0.0007888 |
| SPX | 227.911676 | -0.01186993 | 0.5905151 | -0.020101 | 0.98396282 | 0.99214482 |
| LDHB | 1907.31666 | -0.399830857 | 0.37715667 | -1.0601188 | 0.28909058 | 0.48896403 |
| CMAS | 327.459126 | -0.22068143 | 0.24510125 | -0.9003685 | 0.3679242 | 0.56531073 |
| C2CD5 | 1177.80542 | 0.724706649 | 0.21405421 | 3.38562198 | 0.00071017 | 0.0070132 |
| ETNK1 | 2171.62185 | -0.366962626 | 0.31022998 | -1.1828729 | 0.23685954 | 0.43136764 |
| AC084816.1 | 203.667869 | -1.38407206 | 0.67127921 | -2.0618426 | 0.03922272 | 0.13186801 |
| SOX5 | 1991.62143 | 1.40112782 | 0.70790974 | 1.97924641 | 0.04778827 | 0.15092143 |
| BCAT1 | 104.884182 | -3.27673568 | 1.25410944 | -2.6127988 | 0.00898041 | 0.04660907 |
| LRMP | 447.530802 | 1.234215785 | 0.73646394 | 1.67586723 | 0.0937642 | 0.23571272 |
| CASC1 | 356.276509 | 1.282475046 | 0.66082929 | 1.9407055 | 0.05229401 | 0.15953225 |
| ETFRF1 | 316.50195 | 0.107896947 | 0.38411464 | 0.28089778 | 0.77878881 | 0.87801961 |
| KRAS | 640.692603 | -0.866802275 | 0.26600953 | -3.2585384 | 0.00111988 | 0.0098588 |
| RASSF8-AS1 | 1193.41525 | 0.930416843 | 0.35724271 | 2.60443897 | 0.00920248 | 0.04750976 |
| RASSF8 | 2516.25921 | 0.096997939 | 0.39686437 | 0.2444108 | 0.80691268 | 0.89531977 |
| BHLHE41 | 150.320913 | -3.404948508 | 0.61010599 | -5.5809131 | 2.39E-08 | 1.72E-06 |
| SSPN | 63.4201692 | -1.47673724 | 0.52670388 | -2.8037334 | 0.00505147 | 0.03051584 |
| AC022509.2 | 42.4001945 | 0.345171006 | 0.49148198 | 0.70230654 | 0.48248802 | 0.66670759 |
| AC024145.1 | 123.629244 | -1.081633062 | 0.63861028 | -1.6937295 | 0.0903167 | 0.22976447 |
| ITPR2 | 4762.59677 | 0.58514077 | 0.21194616 | 2.76079916 | 0.00576601 | 0.0336318 |
| AC023051.1 | 93.2081429 | -0.875685609 | 0.65654423 | -1.3337801 | 0.18227594 | 0.36463169 |
| INTS13 | 247.940882 | 0.05920596 | 0.28118472 | 0.21055895 | 0.83323144 | 0.91179175 |
| FGFR1OP2 | 798.62459 | 0.378820114 | 0.30261966 | 1.25180273 | 0.21064176 | 0.40006465 |
| TM7SF3 | 2030.19 | 1.433304457 | 0.44503357 | 3.22066593 | 0.00127893 | 0.01087249 |
| MED21 | 255.073465 | 0.636522482 | 0.17938722 | 3.54831575 | 0.0003877 | 0.00445452 |
| STK38L | 726.267534 | 0.91824153 | 0.19396133 | 4.73414732 | 2.20E-06 | 7.43E-05 |
| ARNTL2 | 174.725942 | -3.439666409 | 0.56225034 | -6.1176778 | 9.49E-10 | 1.12E-07 |
| PPFIBP1 | 4515.53879 | -0.191156646 | 0.32800025 | -0.5827942 | 0.56003185 | 0.72854361 |
| REP15 | 39.7877537 | 0.558247704 | 0.51775628 | 1.07820558 | 0.28094202 | 0.47950097 |
| MRPS35 | 443.631333 | -0.331551651 | 0.18217784 | -1.819934 | 0.06876906 | 0.19190095 |
| KLHL42 | 642.787289 | -0.904377406 | 0.25689408 | -3.5204292 | 0.00043085 | 0.00482488 |
| CCDC91 | 3470.55253 | -0.260963305 | 0.20851663 | -1.2515227 | 0.21074383 | 0.40007037 |
| AC022364.1 | 59.3408895 | 0.452901466 | 0.34871103 | 1.29878733 | 0.19401692 | 0.37905874 |
| FAR2 | 149.461783 | -1.245138633 | 0.46400757 | -2.6834447 | 0.0072868 | 0.04014478 |
| AC009318.1 | 117.737078 | 0.271981846 | 0.50068171 | 0.54322305 | 0.58697624 | 0.74969096 |
| ERGIC2 | 500.836722 | -0.187199747 | 0.15491321 | -1.208417 | 0.22688691 | 0.41931165 |
| TMTC1 | 256.252082 | -1.001460309 | 0.52045162 | -1.924214 | 0.05432776 | 0.16391605 |
| IPO8 | 656.256346 | 0.600512383 | 0.19314113 | 3.10918958 | 0.00187601 | 0.01458268 |
| CAPRIN2 | 378.82749 | 0.346505707 | 0.21855938 | 1.58540761 | 0.11287375 | 0.26617886 |
| AC010198.1 | 61.9828068 | 0.250993998 | 0.46698289 | 0.53748007 | 0.59093605 | 0.75259958 |
| DDX11-AS1 | 38.1426633 | -0.12957666 | 0.38493928 | -0.3366158 | 0.73640651 | 0.85087449 |
| DDX11 | 157.530948 | -0.259617573 | 0.49268166 | -0.5269479 | 0.59822976 | 0.75820022 |
| SINHCAF | 777.802126 | -0.610531598 | 0.24309842 | -2.5114585 | 0.01202334 | 0.05736301 |
| DENND5B | 4426.48122 | 1.018858996 | 0.2084528 | 4.88772044 | 1.02E-06 | 3.93E-05 |
| DENND5B-A! | 110.842567 | -0.588915393 | 0.69581666 | -0.8463658 | 0.39734873 | 0.59186337 |
| ETFBKMT | 115.756752 | 1.467290062 | 0.44705496 | 3.28212459 | 0.00103028 | 0.00926644 |
| AMN1 | 309.593117 | -0.571542764 | 0.33550682 | -1.7035206 | 0.08847069 | 0.22675164 |
| KIAA1551 | 548.284793 | -0.267625198 | 0.24061816 | -1.1122402 | 0.26603487 | 0.46387736 |
| BICD1 | 1217.94686 | -1.702780919 | 0.41615901 | -4.0916593 | 4.28E-05 | 0.00079982 |
| FGD4 | 7429.49725 | 0.413361435 | 0.27591747 | 1.49813432 | 0.13409836 | 0.29810025 |
| DNM1L | 924.063469 | -0.398160454 | 0.13889764 | -2.8665747 | 0.0041494 | 0.02636578 |
| YARS2 | 384.072965 | -0.210429952 | 0.16823675 | -1.2507966 | 0.2110087 | 0.40034114 |
| PKP2 | 379.004877 | 0.420119183 | 0.31455639 | 1.33559257 | 0.18168248 | 0.36381618 |
| ALG10B | 87.66147 | 0.544745399 | 0.3433024 | 1.58678006 | 0.11256247 | 0.26583283 |
| CPNE8 | 2539.24077 | -0.334324249 | 0.19359108 | -1.7269611 | 0.08417466 | 0.21918352 |
| KIF21A | 3516.54525 | 0.258764472 | 0.23828222 | 1.08595793 | 0.2774976 | 0.47545746 |
| C12orf40 | 148.825461 | -0.962464984 | 0.51590447 | -1.8655876 | 0.06209909 | 0.17900992 |
| SLC2A13 | 2391.10411 | -0.429483457 | 0.31352962 | -1.3698338 | 0.1707388 | 0.34969424 |
| LINC02471 | 1784.18744 | -3.128351092 | 1.21118302 | -2.5828888 | 0.00979769 | 0.04968439 |
| AC079630.1 | 344.8417 | -1.440732742 | 0.77734033 | -1.8534131 | 0.06382317 | 0.18248315 |
| LRRK2 | 4365.74714 | -0.03281245 | 0.57103453 | -0.0574614 | 0.95417764 | 0.97682198 |
| AC090630.1 | 67.4808164 | 0.596463577 | 0.79244713 | 0.75268564 | 0.45163884 | 0.64124856 |
| GXYLT1 | 635.283413 | -0.658348499 | 0.22959654 | -2.8674147 | 0.0041384 | 0.0263271 |
| YAF2 | 1559.73061 | -0.101161558 | 0.24957988 | -0.4053274 | 0.68523689 | 0.8178807 |
| PPHLN1 | 1811.67835 | -0.100675872 | 0.1327336 | -0.7584807 | 0.44816326 | 0.63862916 |
| ZCRB1 | 191.91396 | -0.060914218 | 0.22635242 | -0.2691123 | 0.78784327 | 0.88414669 |
| PRICKLE1 | 675.11175 | -0.672523958 | 0.53635126 | -1.2538872 | 0.20988303 | 0.39949496 |
| PUS7L | 309.064963 | -0.103526876 | 0.27940327 | -0.3705285 | 0.71098874 | 0.83480577 |
| IRAK4 | 293.395186 | -0.077934161 | 0.34835571 | -0.2237201 | 0.82297513 | 0.9051934 |
| TWF1 | 202.919906 | -0.51795793 | 0.26875308 | -1.9272632 | 0.05394684 | 0.16308249 |
| TMEM117 | 1621.03981 | 0.25361724 | 0.25875514 | 0.98014379 | 0.32701515 | 0.52691716 |
| ANO6 | 2493.23917 | -1.172277168 | 0.45449141 | -2.5793165 | 0.0098996 | 0.05007154 |
| AC008124.1 | 122.007816 | 0.285604581 | 0.24834032 | 1.15005322 | 0.25012195 | 0.44635094 |
| ARID2 | 2261.33253 | 0.266176366 | 0.15409963 | 1.72730043 | 0.08411372 | 0.21910809 |
| SCAF11 | 2833.33202 | -0.130189923 | 0.17097775 | -0.7614437 | 0.44639212 | 0.63743065 |
| SLC38A1 | 609.786643 | -3.151921912 | 0.52901163 | -5.9581335 | 2.55E-09 | 2.59E-07 |
| SLC38A2 | 1958.09088 | -1.282231175 | 0.42211436 | -3.0376393 | 0.00238439 | 0.01759034 |
| AC008014.1 | 5458.9177 | 0.329476641 | 0.51246402 | 0.64292638 | 0.52027187 | 0.69607397 |
| SLC38A4 | 364.714721 | -0.626670234 | 0.86152987 | -0.7273923 | 0.46698564 | 0.65463001 |
| PCED1B | 109.82826 | -1.815080801 | 0.75439773 | -2.4059998 | 0.01612827 | 0.07096958 |
| RPAP3 | 236.598361 | 0.259476847 | 0.27203298 | 0.95384333 | 0.34016295 | 0.54071826 |
| AC004241.1 | 41.0619646 | 0.123122564 | 0.47564159 | 0.25885576 | 0.79574654 | 0.88843413 |
| RAPGEF3 | 830.649028 | -0.507634307 | 0.33993758 | -1.4933162 | 0.13535443 | 0.2998465 |
| SLC48A1 | 317.782196 | -1.029840936 | 0.37876816 | -2.7189216 | 0.00654951 | 0.03695931 |
| HDAC7 | 332.282569 | -1.709257675 | 0.52012128 | -3.2862675 | 0.00101525 | 0.00914922 |
| VDR | 798.050877 | 0.244126624 | 0.50717787 | 0.48134321 | 0.63027259 | 0.78196473 |
| TMEM106C | 195.829879 | -0.757669863 | 0.349362 | -2.1687243 | 0.03010362 | 0.10948005 |
| SENP1 | 285.715079 | 0.063701846 | 0.23974751 | 0.26570389 | 0.79046727 | 0.8860758 |
| PFKM | 140.916455 | 0.414781597 | 0.36985759 | 1.12146298 | 0.26209084 | 0.46004343 |
| ASB8 | 210.640339 | 0.146801603 | 0.2411683 | 0.60871019 | 0.54271655 | 0.71350621 |
| ZNF641 | 134.601571 | 0.547501403 | 0.26119174 | 2.09616658 | 0.03606742 | 0.12412153 |
| AC090115.1 | 68.4517581 | 0.738531372 | 0.51122857 | 1.4446207 | 0.14856446 | 0.31853168 |
| KANSL2 | 378.325492 | 0.640696555 | 0.28292789 | 2.26452242 | 0.023542 | 0.09249919 |
| CCNT1 | 484.346574 | 0.117333648 | 0.20012579 | 0.58629949 | 0.55767427 | 0.7262747 |
| ADCY6 | 336.864419 | 0.573227153 | 0.21427802 | 2.67515612 | 0.00746945 | 0.04085831 |
| DDX23 | 224.494515 | -0.291785676 | 0.24767284 | -1.1781093 | 0.23875304 | 0.43383194 |
| AC073610.3 | 112.022465 | -0.703506414 | 0.42242281 | -1.6654082 | 0.09583135 | 0.23882914 |
| CCDC65 | 237.245569 | -0.74589666 | 0.3480895 | -2.1428301 | 0.03212675 | 0.11470729 |
| PRKAG1 | 324.779821 | -0.348760512 | 0.38340914 | -0.9096302 | 0.36301759 | 0.56078915 |
| KMT2D | 409.061959 | 0.072304016 | 0.3562154 | 0.20297835 | 0.83915195 | 0.91483781 |
| LMBR1L | 352.614049 | -0.049626561 | 0.31541197 | -0.1573389 | 0.87497779 | 0.93490877 |
| AC011603.2 | 57.7109298 | 0.359227586 | 0.30728652 | 1.16903137 | 0.24239099 | 0.43756794 |
| TUBA1B | 688.983854 | -0.973359001 | 0.66933657 | -1.4542146 | 0.14588682 | 0.3146115 |
| TUBA1A | 262.40946 | -0.967125149 | 0.70648661 | -1.3689221 | 0.17102362 | 0.34998817 |
| TUBA1C | 751.692944 | -1.409793647 | 0.51887854 | -2.7170013 | 0.00658764 | 0.03712859 |

| | | | | | | |
|---|---|---|---|---|---|---|
| AC010173.1 | 59.4532781 | -0.10555454 | 0.52680318 | -0.2003681 | 0.84119272 | 0.91607296 |
| DNAJC22 | 402.146401 | 1.47471262 | 0.42226513 | 3.49238549 | 0.00047873 | 0.00519961 |
| SPATS2 | 1230.7759 | -0.328469981 | 0.27080477 | -1.2129402 | 0.22515269 | 0.41738815 |
| MCRS1 | 158.967704 | 0.169643109 | 0.34831596 | 0.487038 | 0.6262314 | 0.77885435 |
| PRPF40B | 649.518932 | 0.179023329 | 0.25730958 | 0.69575074 | 0.48658494 | 0.66977503 |
| FMNL3 | 53.5645486 | -1.798712019 | 0.79539633 | -2.2614035 | 0.02373428 | 0.09290451 |
| TMBIM6 | 2903.49682 | 0.010576197 | 0.33466065 | 0.03160275 | 0.97478885 | 0.98732216 |
| NCKAP5L | 224.562254 | 0.164204717 | 0.24082104 | 0.68185369 | 0.49533147 | 0.67732527 |
| BCDIN3D | 119.893768 | 0.045606362 | 0.31388727 | 0.14529535 | 0.88447768 | 0.94002757 |
| AQP2 | 277.821911 | 1.02962468 | 0.73527242 | 1.40033089 | 0.16141425 | 0.3376268 |
| RACGAP1 | 91.3346142 | -0.148132034 | 0.26283502 | -0.5635932 | 0.57303101 | 0.73835552 |
| SMARCD1 | 266.311634 | -0.12843602 | 0.27048984 | -0.4748275 | 0.63490992 | 0.78468826 |
| GPD1 | 497.449079 | 1.417912472 | 0.58727647 | 2.41438665 | 0.01576173 | 0.06974211 |
| AC074032.1 | 829.886197 | 0.51636479 | 0.19214478 | 2.68737354 | 0.00720164 | 0.03973947 |
| CERS5 | 462.741618 | -0.407360463 | 0.18776655 | -2.1695049 | 0.03004437 | 0.10932258 |
| LIMA1 | 1372.02901 | -0.448314432 | 0.25499418 | -1.7581359 | 0.07872438 | 0.20969202 |
| FAM186A | 35.8915797 | -0.447546472 | 0.71643888 | -0.624682 | 0.53217979 | 0.70560925 |
| LARP4 | 3024.01407 | 0.657243611 | 0.18262896 | 3.59879188 | 0.0003197 | 0.00386089 |
| DIP2B | 5117.9522 | -0.411447073 | 0.36726028 | -1.1203147 | 0.2625797 | 0.46060659 |
| ATF1 | 651.416652 | 0.367116541 | 0.22885314 | 1.60415777 | 0.10867928 | 0.25956907 |
| METTL7A | 712.538419 | 1.157989756 | 0.47045401 | 2.46143029 | 0.01383843 | 0.06335449 |
| HIGD1C | 34.7541172 | -0.002039325 | 0.66820714 | -0.0030519 | 0.99756491 | 0.99870545 |
| SLC11A2 | 1320.69954 | -0.482024973 | 0.24219071 | -1.9902703 | 0.04656117 | 0.14842184 |
| LETMD1 | 637.038243 | 0.227695259 | 0.20609107 | 1.10482834 | 0.26923398 | 0.46725628 |
| CSRNP2 | 234.24471 | -0.128682287 | 0.2482543 | -0.5183487 | 0.60421502 | 0.76216808 |
| TFCP2 | 1323.56074 | 0.46087968 | 0.18593267 | 2.47874504 | 0.01318455 | 0.06126124 |
| POU6F1 | 157.591335 | 0.276450456 | 0.26041491 | 1.06157691 | 0.2884278 | 0.48820448 |
| DAZAP2 | 762.852949 | -0.530135159 | 0.28345055 | -1.8702915 | 0.06144335 | 0.17761855 |
| SMAGP | 122.105299 | -1.104604027 | 0.37394094 | -2.9539532 | 0.00313732 | 0.02133095 |
| GALNT6 | 46.1607296 | -0.222677664 | 0.66894893 | -0.3328769 | 0.73922719 | 0.85193361 |
| SLC4A8 | 48.4559435 | 1.165667777 | 0.44398206 | 2.62548394 | 0.00865259 | 0.04531946 |
| SCN8A | 478.668492 | 0.608443628 | 0.36843731 | 1.65141697 | 0.09865346 | 0.24335009 |
| AC068987.2 | 59.8517098 | 1.487446126 | 0.65093526 | 2.28509071 | 0.02230751 | 0.08911951 |
| AC068987.3 | 78.6220313 | 0.208449055 | 0.4510325 | 0.46215973 | 0.64396678 | 0.79148232 |
| FIGNL2 | 613.10501 | 0.759681814 | 0.42354222 | 1.79363892 | 0.07287082 | 0.19917056 |
| AC068987.1 | 30.2674146 | 0.599124748 | 0.45904238 | 1.30516218 | 0.19183763 | 0.37618289 |
| ACVR1B | 1487.12342 | 0.423233609 | 0.23686453 | 1.78681713 | 0.07396705 | 0.20144374 |
| NR4A1 | 80.2688947 | 1.051412811 | 0.71147858 | 1.47778561 | 0.13946517 | 0.30571711 |
| ATG101 | 143.634723 | -0.82356118 | 0.42552753 | -1.9353887 | 0.05294261 | 0.16094131 |
| AC078864.2 | 190.985247 | -1.927588232 | 0.78442787 | -2.4573174 | 0.0139979 | 0.0637651 |
| AC078864.1 | 497.895365 | -0.69907819 | 0.48144386 | -1.4520451 | 0.14648907 | 0.31551314 |
| KRT7 | 196.889243 | -1.094218717 | 0.60681106 | -1.803228 | 0.07135241 | 0.19668739 |
| KRT8 | 1239.42274 | -1.672650612 | 0.43695536 | -3.8279668 | 0.00012921 | 0.00190025 |
| KRT18 | 1203.79182 | -2.170406913 | 0.58519147 | -3.7088834 | 0.00020818 | 0.00276281 |
| EIF4B | 1137.37053 | -0.523899136 | 0.33759966 | -1.5518355 | 0.12070159 | 0.27829381 |
| AC068888.1 | 370.329545 | -0.612213948 | 0.39079719 | -1.5665771 | 0.11721356 | 0.2729102 |
| TNS2 | 190.303694 | -0.437476838 | 0.52362623 | -0.8354754 | 0.40345009 | 0.59761138 |
| SPRYD3 | 82.5278084 | -0.740660551 | 0.42790077 | -1.7309166 | 0.08346664 | 0.21823333 |
| IGFBP6 | 48.1440295 | -1.134532496 | 0.67884377 | -1.6712719 | 0.09466799 | 0.23698747 |
| CSAD | 730.183053 | 0.705814599 | 0.26825766 | 2.63110693 | 0.00851073 | 0.044727 |
| ZNF740 | 142.149393 | 0.37744214 | 0.21978828 | 1.71729873 | 0.0859246 | 0.22234634 |
| MFSD5 | 85.8672199 | -0.722244786 | 0.41065573 | -1.7587598 | 0.07861831 | 0.20949091 |
| ESPL1 | 76.0185205 | 2.494196013 | 0.82707108 | 3.01569729 | 0.00256389 | 0.01855205 |
| PFDN5 | 653.672054 | 0.012661124 | 0.35122218 | 0.03604876 | 0.97124348 | 0.98569779 |
| C12orf10 | 88.8412999 | -0.292907822 | 0.3878916 | -0.755128 | 0.45017216 | 0.64055951 |
| AAAS | 119.167986 | 0.956555044 | 0.24791435 | 3.85840938 | 0.00011413 | 0.00172856 |
| SP1 | 982.30113 | 0.279890616 | 0.30048185 | 0.93147262 | 0.35160914 | 0.55067488 |
| PRR13 | 740.247508 | -0.050143781 | 0.30081671 | -0.1666921 | 0.86761229 | 0.93103202 |
| PCBP2 | 2080.64419 | 0.117268217 | 0.20232789 | 0.57959493 | 0.56218781 | 0.73014439 |
| MAP3K12 | 163.401789 | -0.024700519 | 0.34916909 | -0.0707409 | 0.94360401 | 0.97094663 |
| TARBP2 | 78.4756223 | 0.761337369 | 0.34724465 | 2.19251001 | 0.0283427 | 0.10494149 |
| ATP5MC2 | 771.343663 | -0.001340007 | 0.22022488 | -0.0060847 | 0.99514512 | 0.99749068 |
| CALCOCO1 | 406.641111 | 0.291348954 | 0.31686216 | 0.91948169 | 0.35784368 | 0.55643373 |
| HOXC4 | 151.045437 | 0.920712314 | 0.46493106 | 1.98032006 | 0.04766758 | 0.15068715 |
| AC023794.1 | 89.7769466 | 0.001342957 | 0.44881212 | 0.00299225 | 0.99761253 | 0.99870545 |
| SMUG1 | 209.268084 | 0.439792527 | 0.29697345 | 1.48091529 | 0.13862915 | 0.30470276 |
| LINC02381 | 70.1095863 | -0.215052461 | 0.43995391 | -0.4888068 | 0.62497849 | 0.77771949 |
| CBX5 | 432.921968 | -0.327951868 | 0.31148489 | -1.052866 | 0.29240239 | 0.49248608 |
| AC078778.1 | 36.5546143 | -0.491926548 | 0.60647758 | -0.8111207 | 0.41729634 | 0.61047432 |
| HNRNPA1 | 475.196765 | -0.29788842 | 0.30388082 | -0.9802804 | 0.32694772 | 0.52691716 |
| COPZ1 | 424.52494 | -0.256583155 | 0.2441657 | -1.0508567 | 0.29332442 | 0.49326436 |
| PPP1R1A | 332.851509 | -1.341439634 | 0.52860325 | -2.537706 | 0.01115817 | 0.05449168 |
| METTL7B | 115.17685 | -0.659620695 | 0.68952784 | -0.9566266 | 0.33875574 | 0.53901002 |
| ITGA7 | 166.873419 | -0.502591563 | 0.61969029 | -0.8110367 | 0.4173446 | 0.61047432 |
| CD63 | 1072.65783 | -1.579844787 | 0.45229366 | -3.4929625 | 0.00047769 | 0.0051925 |
| AC009779.2 | 30.2441295 | -0.917080848 | 0.4915665 | -1.8656292 | 0.06209325 | 0.17900992 |
| ORMDL2 | 131.388866 | -1.594412831 | 0.33441877 | -4.7677133 | 1.86E-06 | 6.42E-05 |
| PYM1 | 161.947228 | 0.384372462 | 0.3012911 | 1.27575113 | 0.20204351 | 0.38956398 |
| CDK2 | 39.3734871 | -0.799847402 | 0.44455358 | -1.7992148 | 0.0719847 | 0.19785328 |
| RAB5B | 432.982612 | 0.061511634 | 0.183601 | 0.33502886 | 0.73760332 | 0.85118107 |
| SUOX | 227.616491 | 0.536183458 | 0.62596335 | 0.85657324 | 0.39168078 | 0.58687893 |
| IKZF4 | 171.758384 | 0.257754975 | 0.36725218 | 0.70184736 | 0.48277436 | 0.66687778 |
| RPS26 | 207.635004 | -0.704081568 | 0.33792441 | -2.0835475 | 0.03720135 | 0.1271618 |
| ERBB3 | 819.760806 | 0.210669574 | 0.45201495 | 0.46606772 | 0.64116703 | 0.78933685 |
| PA2G4 | 227.545562 | -0.600808563 | 0.44494052 | -1.3503121 | 0.1769159 | 0.35735135 |
| RPL41 | 2298.80001 | -0.8699336 | 0.37031705 | -2.3491589 | 0.01881587 | 0.07898954 |
| ESYT1 | 275.864522 | -0.936820768 | 0.25659944 | -3.6509073 | 0.00026132 | 0.0032861 |
| AC034102.5 | 46.1663937 | 0.231809416 | 0.526482 | 0.44029885 | 0.65972068 | 0.8020214 |
| MYL6B | 105.748893 | -0.009406571 | 0.32901192 | -0.0285904 | 0.9771913 | 0.98865966 |
| MYL6 | 1208.48617 | -0.437207406 | 0.40081456 | -1.0907972 | 0.27536212 | 0.47286769 |
| SMARCC2 | 656.657976 | -0.139555774 | 0.25143936 | -0.5550276 | 0.57887581 | 0.74245057 |
| RNF41 | 224.204001 | 0.101607627 | 0.23479849 | 0.43274396 | 0.66520079 | 0.80579154 |
| NABP2 | 77.7063333 | 0.13278089 | 0.29929848 | 0.44364037 | 0.65730261 | 0.80028102 |
| SLC39A5 | 678.283301 | 2.294973611 | 0.45792777 | 5.01164981 | 5.40E-07 | 2.28E-05 |
| ANKRD52 | 257.929981 | -0.376564001 | 0.28178543 | -1.3363501 | 0.18143486 | 0.36369225 |
| COQ10A | 76.4455431 | 0.409560769 | 0.28242496 | 1.45015782 | 0.14701451 | 0.31639629 |
| CS | 406.42627 | -0.005193343 | 0.26770827 | -0.0193993 | 0.9845226 | 0.99241906 |
| AC073896.1 | 108.248221 | 0.001495968 | 0.46944457 | 0.00318668 | 0.9974574 | 0.99870545 |
| AC073896.2 | 106.730564 | 0.702291981 | 0.26998445 | 2.60123123 | 0.00928898 | 0.04774056 |
| PAN2 | 311.453133 | 0.748367895 | 0.36251276 | 2.0643905 | 0.03898071 | 0.13143142 |
| STAT2 | 644.911832 | -0.316620242 | 0.48899622 | -0.6474902 | 0.51731476 | 0.69425481 |
| TIMELESS | 66.9955594 | -0.367291658 | 0.39298074 | -0.9346302 | 0.34997892 | 0.5493439 |
| SPRYD4 | 92.6342985 | -0.012264518 | 0.34425621 | -0.0356261 | 0.97158047 | 0.9858938 |
| GLS2 | 50.5768607 | -0.215774301 | 0.37226041 | -0.5796327 | 0.56216235 | 0.73014439 |
| RBMS2 | 525.094451 | 0.446087968 | 0.29950632 | 1.48941088 | 0.1363792 | 0.3013622 |
| BAZ2A | 830.208875 | -0.045423227 | 0.17209568 | -0.2639417 | 0.79182484 | 0.88678699 |
| ATP5F1B | 1058.32436 | -0.20828648 | 0.36236777 | -0.5747931 | 0.56543126 | 0.73247225 |
| PTGES3 | 1625.40488 | -0.387703575 | 0.24942165 | -1.5544102 | 0.12008659 | 0.27748262 |
| NACA | 720.918928 | -0.816313351 | 0.35482682 | -2.3005965 | 0.02141445 | 0.0869197 |
| NEMP1 | 148.860579 | -0.191635661 | 0.25844467 | -0.7414959 | 0.45839282 | 0.64719928 |
| NAB2 | 116.319779 | -0.878245569 | 0.28991335 | -3.0293382 | 0.0024509 | 0.01794578 |
| STAT6 | 958.015858 | -0.222772298 | 0.18980385 | -1.1736975 | 0.24051623 | 0.43579395 |
| LRP1 | 85.799188 | 0.471060673 | 0.46836169 | 1.00576261 | 0.31452978 | 0.51502325 |

| | | | | | | |
|---|---|---|---|---|---|---|
| SHMT2 | 144.865647 | -0.237398311 | 0.25459551 | -0.9324528 | 0.35110255 | 0.55041523 |
| R3HDM2 | 2953.60811 | 0.75167454 | 0.15730929 | 4.77832258 | 1.77E-06 | 6.18E-05 |
| MARS | 289.244283 | -0.329516504 | 0.37026563 | -0.8899462 | 0.37349476 | 0.57092591 |
| DDIT3 | 424.540287 | -1.51048504 | 0.48866303 | -3.0910565 | 0.00199446 | 0.0153326 |
| MBD6 | 209.825315 | 0.571277053 | 0.39046931 | 1.46305237 | 0.14345305 | 0.3110808 |
| DCTN2 | 631.189554 | -0.527665491 | 0.25423072 | -2.0755379 | 0.03793671 | 0.12896788 |
| PIP4K2C | 173.979052 | -0.632978961 | 0.27670559 | -2.2875539 | 0.02216351 | 0.08880306 |
| DTX3 | 58.1957488 | -0.647895445 | 0.64812705 | -0.9996427 | 0.31748347 | 0.51755172 |
| OS9 | 529.313266 | 0.132955218 | 0.21249637 | 0.62568229 | 0.53152335 | 0.70522693 |
| AC025165.2 | 47.6791186 | -0.479457712 | 0.49677348 | -0.9651435 | 0.33447292 | 0.53408905 |
| TSPAN31 | 130.293569 | -0.308338184 | 0.27566058 | -1.1185429 | 0.26333519 | 0.46128738 |
| CDK4 | 225.322415 | -0.174625372 | 0.44743092 | -0.3902845 | 0.69632615 | 0.82516803 |
| 9-Mar | 268.242968 | -0.540866497 | 0.27685944 | -1.953578 | 0.05075115 | 0.15695984 |
| CYP27B1 | 123.159647 | 2.440933242 | 0.63684903 | 3.83282873 | 0.00012668 | 0.0018711 |
| TSFM | 250.316518 | 0.061830254 | 0.30217398 | 0.20461806 | 0.83787054 | 0.91445907 |
| AVIL | 227.563325 | 1.664609214 | 0.42110233 | 3.95298034 | 7.72E-05 | 0.00127412 |
| CTDSP2 | 403.12391 | 0.072439933 | 0.18321315 | 0.3953861 | 0.69255797 | 0.82275022 |
| AC084033.3 | 43.2077898 | 0.173839584 | 0.27877739 | 0.62357848 | 0.53290445 | 0.70617115 |
| ATP23 | 52.476887 | 1.15030283 | 0.40823932 | 2.81771692 | 0.00483664 | 0.02956843 |
| LRIG3 | 543.410772 | 1.229432096 | 0.42277933 | 2.90797586 | 0.00363776 | 0.02384641 |
| AC068305.2 | 105.467712 | 1.339778003 | 0.69966844 | 1.91487558 | 0.05550837 | 0.16619244 |
| SLC16A7 | 430.098348 | -1.305649361 | 0.5032068 | -2.5946577 | 0.00946852 | 0.0484634 |
| FAM19A2 | 485.889549 | 0.387717369 | 0.50082259 | 0.7741611 | 0.43883553 | 0.63061599 |
| USP15 | 2160.25055 | 0.248339308 | 0.20307207 | 1.22291219 | 0.22136286 | 0.4134786 |
| MON2 | 2570.90296 | 0.169505161 | 0.14121509 | 1.20033321 | 0.23000996 | 0.42313439 |
| AC048341.1 | 192.901192 | 0.003677172 | 0.32624422 | 0.01127122 | 0.99100706 | 0.99543703 |
| PPM1H | 1322.45854 | 0.79520668 | 0.35830668 | 2.2193465 | 0.02646316 | 0.1003404 |
| DPY19L2 | 908.249718 | 0.100464107 | 0.36621556 | 0.27433052 | 0.78383063 | 0.88133085 |
| RXYLT1 | 205.308149 | 0.05865136 | 0.40753745 | 0.14391649 | 0.88556641 | 0.94094134 |
| SRGAP1 | 3069.4751 | -0.124901419 | 0.39050826 | -0.3198432 | 0.74908718 | 0.85857831 |
| AC020611.2 | 98.332744 | -0.971788535 | 0.4927248 | -1.9722745 | 0.04857828 | 0.15251088 |
| C12orf66 | 120.578946 | 0.560728068 | 0.31592473 | 1.7748787 | 0.07591792 | 0.2050053 |
| C12orf56 | 50.9643902 | 0.037606673 | 0.37438846 | 0.10044827 | 0.91998845 | 0.95701657 |
| XPOT | 357.860345 | -0.747855714 | 0.42986607 | -1.7397412 | 0.08190447 | 0.21540264 |
| TBK1 | 902.888732 | -0.6678068 | 0.20750183 | -3.2183176 | 0.00128945 | 0.01093042 |
| RASSF3 | 225.739022 | -1.448007771 | 0.45918349 | -3.1534404 | 0.00161358 | 0.01304859 |
| GNS | 1128.37559 | -0.076129612 | 0.20337284 | -0.3743352 | 0.70815495 | 0.83287619 |
| TBC1D30 | 266.801034 | -0.495089912 | 0.3985484 | -1.2422329 | 0.21415066 | 0.40445105 |
| LINC02389 | 41.6267889 | -0.607592791 | 0.48890112 | -1.2427723 | 0.21395173 | 0.40427853 |
| LEMD3 | 835.457966 | -0.013259142 | 0.18406079 | -0.0720368 | 0.94257266 | 0.97003104 |
| MSRB3 | 57.6930441 | -1.313801159 | 0.59836575 | -2.195649 | 0.02811708 | 0.10435981 |
| HMGA2 | 217.24325 | -4.620966419 | 1.05907164 | -4.3632236 | 1.28E-05 | 0.00031092 |
| LLPH | 100.159308 | -0.735113675 | 0.33508906 | -2.193786 | 0.0282508 | 0.10471437 |
| HELB | 412.780064 | 0.485604955 | 0.44025389 | 1.10301116 | 0.27002232 | 0.46826871 |
| GRIP1 | 6605.47529 | -2.088104023 | 0.3081161 | -6.7770039 | 1.23E-11 | 2.59E-09 |
| CAND1 | 857.417465 | 0.235351934 | 0.13240098 | 1.77756937 | 0.07547461 | 0.20417265 |
| DYRK2 | 205.513676 | -0.275203917 | 0.25618765 | -1.0742279 | 0.28272056 | 0.48175669 |
| MDM1 | 176.13802 | 1.242224219 | 0.22449233 | 5.53348184 | 3.14E-08 | 2.18E-06 |
| LINC02384 | 196.163089 | 0.846959206 | 0.85158664 | 0.9945661 | 0.31994734 | 0.520351 |
| RAP1B | 2759.86176 | -0.727578556 | 0.20839307 | -3.4913759 | 0.00048054 | 0.00520692 |
| NUP107 | 610.808546 | -0.340211042 | 0.27057341 | -1.2573706 | 0.20861948 | 0.39787772 |
| SLC35E3 | 613.217698 | -0.233805972 | 0.38935795 | -0.6004911 | 0.54817901 | 0.71842074 |
| MDM2 | 1149.40765 | -0.706423802 | 0.30396198 | -2.3240532 | 0.02012264 | 0.08277943 |
| CPM | 2990.45755 | -0.089920844 | 0.49052523 | -0.1833154 | 0.85455053 | 0.92322796 |
| CPSF6 | 1199.05989 | 0.0237235 | 0.17937448 | 0.13225683 | 0.89478115 | 0.9454799 |
| YEATS4 | 92.9477885 | -0.12301082 | 0.2267807 | -0.5424219 | 0.58752789 | 0.75018375 |
| FRS2 | 1418.85277 | 0.135494634 | 0.18155935 | 0.74628288 | 0.45549655 | 0.64475342 |
| CCT2 | 495.910376 | -0.616862249 | 0.27924939 | -2.2090013 | 0.02717455 | 0.10218974 |
| BEST3 | 43.0229789 | -1.665222279 | 0.55453018 | -3.0029426 | 0.00267383 | 0.01901929 |
| RAB3IP | 1701.37259 | 0.122235139 | 0.232333 | 0.52612043 | 0.59880453 | 0.75878837 |
| AC025263.2 | 40.3728931 | 0.621073823 | 0.68609005 | 0.9052366 | 0.36534006 | 0.56279121 |
| MYRFL | 382.393386 | 0.247304418 | 0.68229318 | 0.36246063 | 0.71700784 | 0.83851445 |
| AC025159.1 | 543.240224 | -1.141345101 | 0.68940485 | -1.6555513 | 0.09781272 | 0.24200482 |
| CNOT2 | 2083.92975 | -0.238775419 | 0.21032587 | -1.1352641 | 0.25626467 | 0.45359244 |
| KCNMB4 | 318.781638 | 0.794438244 | 0.52228585 | 1.52107939 | 0.12823991 | 0.28992962 |
| TSPAN8 | 54.8992965 | -0.513390889 | 0.57694522 | -0.8898434 | 0.37354998 | 0.57094666 |
| ZFC3H1 | 2120.03745 | -0.30645339 | 0.23637358 | -1.296479 | 0.19481049 | 0.38005515 |
| THAP2 | 87.9342543 | 0.217675165 | 0.41759945 | 0.52125348 | 0.6021902 | 0.76094319 |
| TMEM19 | 170.693196 | 1.364236483 | 0.36610721 | 3.72633052 | 0.00019429 | 0.00260833 |
| RAB21 | 1436.36243 | 0.313019893 | 0.19083412 | 1.64027217 | 0.10094859 | 0.24739895 |
| TBC1D15 | 1404.08737 | -0.31391537 | 0.23711958 | -1.3238695 | 0.1855464 | 0.36868553 |
| TRHDE | 2719.95966 | 1.0378693 | 0.4566153 | 2.27296217 | 0.02302846 | 0.0911662 |
| AC090502.1 | 56.1677339 | 0.506409455 | 0.94292821 | 0.53706045 | 0.59122586 | 0.75273387 |
| ATXN7L3B | 372.926413 | -0.129733872 | 0.26680451 | -0.4862507 | 0.62678945 | 0.77912423 |
| CAPS2 | 949.743202 | 0.421860295 | 0.45242492 | 0.93244265 | 0.35110781 | 0.55041523 |
| GLIPR1L1 | 69.7246559 | -0.737593573 | 0.7915353 | -0.9318518 | 0.35141313 | 0.55058411 |
| KRR1 | 344.037042 | 0.111501308 | 0.22680523 | 0.49161701 | 0.62299012 | 0.77619139 |
| AC078923.1 | 436.164697 | 1.747602694 | 0.48971696 | 3.56859743 | 0.0003589 | 0.0042088 |
| PHLDA1 | 118.706242 | -0.723481308 | 0.5720857 | -1.264638 | 0.20600113 | 0.3947515 |
| NAP1L1 | 873.330847 | -0.517984788 | 0.27297324 | -1.8975662 | 0.05775326 | 0.17049836 |
| BBS10 | 56.8577019 | 0.122274903 | 0.30697683 | 0.39831965 | 0.69039458 | 0.82103405 |
| OSBPL8 | 3814.29571 | 0.479710245 | 0.25406942 | 1.888107 | 0.05901159 | 0.1728728 |
| AC107032.2 | 89.0687009 | 0.737698454 | 0.40105505 | 1.8393945 | 0.06585718 | 0.18643214 |
| ZDHHC17 | 993.101378 | 0.283624467 | 0.16632796 | 1.70521216 | 0.08815486 | 0.22623829 |
| CSRP2 | 232.998303 | -0.942411917 | 0.56585747 | -1.6654581 | 0.09582139 | 0.23882914 |
| NAV3 | 545.525932 | 0.109287385 | 0.85565111 | 0.12772424 | 0.89836721 | 0.94743919 |
| AC128707.1 | 1017.11755 | 3.416131288 | 0.92492005 | 3.69343415 | 0.00022125 | 0.00289096 |
| AC079362.1 | 58.7596323 | 1.126901371 | 1.04247298 | 1.08098856 | 0.2797022 | 0.4782088 |
| SYT1 | 116.939904 | 1.715624906 | 0.75069031 | 2.28539636 | 0.0222896 | 0.08907391 |
| AC027288.3 | 2966.45303 | 3.335919988 | 0.51622291 | 6.46216953 | 1.03E-10 | 1.61E-08 |
| PAWR | 2797.61163 | 0.132370412 | 0.30560946 | 0.43313584 | 0.66491608 | 0.80569349 |
| AC073569.2 | 288.431854 | 1.435944723 | 0.39430754 | 3.64168717 | 0.00027086 | 0.00338744 |
| PPP1R12A | 2266.66069 | 0.372244208 | 0.17643309 | 2.10983214 | 0.03487282 | 0.12150526 |
| OTOGL | 646.602331 | 3.952093236 | 0.72857881 | 5.4243867 | 5.82E-08 | 3.54E-06 |
| PTPRQ | 51.2250562 | 0.690342333 | 0.37856495 | 1.823577 | 0.06821605 | 0.19078501 |
| LIN7A | 883.866463 | 0.985149626 | 0.44747004 | 2.20159908 | 0.02769364 | 0.10345734 |
| ACSS3 | 1836.10726 | 0.610795314 | 0.4905925 | 1.24501559 | 0.21312602 | 0.40310727 |
| PPFIA2 | 57.2861665 | 0.243266724 | 0.56303762 | 0.43206122 | 0.66569692 | 0.80585601 |
| CCDC59 | 190.220203 | -0.239268487 | 0.2049997 | -1.167165 | 0.24314373 | 0.4382342 |
| METTL25 | 764.580343 | 0.525684706 | 0.25433885 | 2.06686751 | 0.03874664 | 0.13090959 |
| TMTC2 | 3637.06556 | -0.235931206 | 0.51111424 | -0.4616017 | 0.64436697 | 0.79162826 |
| LRRIQ1 | 371.48722 | 0.208190769 | 0.5653737 | 0.36823568 | 0.71269751 | 0.83559531 |
| MGAT4C | 40.1770723 | 0.565549446 | 0.39514302 | 1.43125251 | 0.15235786 | 0.32404718 |
| C12orf29 | 153.669326 | -0.945777801 | 0.28003463 | -3.3773601 | 0.00073185 | 0.00718074 |
| CEP290 | 1047.18769 | 0.110093932 | 0.17606148 | 0.62531525 | 0.53176417 | 0.70533109 |
| TMTC3 | 599.501658 | 0.41718079 | 0.21219929 | 1.96598578 | 0.04930025 | 0.15396641 |
| KITLG | 208.372478 | -0.110902644 | 0.51639637 | -0.2147626 | 0.82995239 | 0.90987422 |
| DUSP6 | 331.623674 | -1.441506194 | 0.39413432 | -3.6573983 | 0.00025479 | 0.00322771 |
| POC1B | 777.325046 | -0.515530072 | 0.25646457 | -2.0101415 | 0.04441622 | 0.14345266 |
| POC1B-AS1 | 298.097044 | 0.457973534 | 0.25221662 | 1.81579446 | 0.06940191 | 0.19303814 |
| AC025034.1 | 141.482397 | 0.639772444 | 0.34324632 | 1.86388724 | 0.06233754 | 0.179584 |
| ATP2B1 | 1190.95032 | -0.090680409 | 0.13977682 | -0.6487514 | 0.51649906 | 0.69386001 |
| DCN | 41.8644208 | 0.066199908 | 0.4531178 | 0.14609867 | 0.8838435 | 0.93979077 |

| | | | | | | |
|---|---|---|---|---|---|---|
| BTG1 | 816.812368 | -2.040888777 | 0.35499899 | -5.7489988 | 8.98E-09 | 7.41E-07 |
| AC025164.1 | 48.4566849 | -1.051212313 | 0.56777485 | -1.8514598 | 0.06410343 | 0.18311151 |
| EEA1 | 1692.36076 | -0.282740316 | 0.31900068 | -0.8863314 | 0.37543897 | 0.57223863 |
| AC138123.1 | 2495.26586 | 1.72729404 | 0.55556394 | 3.10908235 | 0.00187669 | 0.01458268 |
| AC138123.2 | 96.6749927 | 1.386186701 | 0.73906763 | 1.87558844 | 0.06071182 | 0.17619667 |
| AC124947.1 | 208.185891 | 0.768884439 | 0.50968332 | 1.50855326 | 0.13141299 | 0.29451893 |
| NUDT4 | 826.831634 | -0.254101112 | 0.35625807 | -0.7132501 | 0.475691 | 0.66135476 |
| UBE2N | 512.405437 | -0.239666543 | 0.22790141 | -1.0516238 | 0.29297219 | 0.49326436 |
| MRPL42 | 424.413972 | 0.29921956 | 0.19691389 | 1.51954518 | 0.12862532 | 0.2904559 |
| SOCS2 | 64.1150814 | 1.525521795 | 0.49876605 | 3.05859191 | 0.0022238 | 0.01663843 |
| CRADD | 3693.48942 | 0.53781027 | 0.33107004 | 1.62446071 | 0.10427752 | 0.25233693 |
| AC012085.2 | 224.044342 | -0.164748453 | 0.6717106 | -0.245267 | 0.80624969 | 0.89476678 |
| PLXNC1 | 154.368229 | 0.997203816 | 0.4094233 | 2.43563032 | 0.01486587 | 0.06669933 |
| CEP83 | 662.544291 | 0.048655181 | 0.23923357 | 0.20337941 | 0.8388385 | 0.91471434 |
| TMCC3 | 716.286376 | -0.181579397 | 0.46615626 | -0.3895247 | 0.69688801 | 0.82544394 |
| NDUFA12 | 280.848738 | -0.057937287 | 0.24039396 | -0.2410097 | 0.80954756 | 0.89661187 |
| NR2C1 | 793.518073 | 0.122104151 | 0.16371156 | 0.74584929 | 0.45575846 | 0.64490647 |
| FGD6 | 1218.2341 | -0.063854178 | 0.19050227 | -0.3351885 | 0.73748287 | 0.85118107 |
| VEZT | 1740.26347 | -0.282754104 | 0.14547135 | -1.9437098 | 0.05193045 | 0.15895347 |
| METAP2 | 557.740002 | -0.002241306 | 0.22433224 | -0.009991 | 0.99202846 | 0.99609195 |
| NTN4 | 2478.40974 | 2.247433573 | 0.36064665 | 6.23167746 | 4.61E-10 | 5.97E-08 |
| SNRPF | 149.921373 | -0.409063357 | 0.19706058 | -2.0758254 | 0.0379101 | 0.12894139 |
| CCDC38 | 45.4264117 | 0.535331827 | 0.42622879 | 1.25597294 | 0.20912578 | 0.39839987 |
| AMDHD1 | 185.439761 | 0.698256183 | 0.39551398 | 1.76543996 | 0.07748985 | 0.20765144 |
| LTA4H | 305.502347 | 0.055339993 | 0.22657947 | 0.24424099 | 0.80704418 | 0.89532855 |
| ELK3 | 131.40186 | -2.058365518 | 0.60310647 | -3.4129389 | 0.00064266 | 0.00648674 |
| CDK17 | 1120.93955 | 0.062897667 | 0.16251949 | 0.38701615 | 0.69874425 | 0.82658873 |
| CFAP54 | 136.322767 | -2.116976553 | 0.75573246 | -2.8012249 | 0.0050909 | 0.03069996 |
| NEDD1 | 355.129646 | 0.061947953 | 0.35067247 | 0.17665474 | 0.8597796 | 0.92628466 |
| TMPO | 523.570684 | 0.781749186 | 0.38310876 | 2.04054115 | 0.04129646 | 0.13656225 |
| SLC25A3 | 1111.53806 | -0.714916349 | 0.35696723 | -2.0027507 | 0.04520406 | 0.1454147 |
| IKBIP | 255.28213 | -1.740981251 | 0.34702565 | -5.0168661 | 5.25E-07 | 2.24E-05 |
| APAF1 | 231.686196 | -0.752942507 | 0.22792412 | -3.3034788 | 0.00095493 | 0.00874365 |
| ANKS1B | 6012.56994 | 2.023384035 | 0.6521933 | 3.10242995 | 0.00191939 | 0.01485549 |
| AC069437.1 | 91.3353371 | 0.712539884 | 0.75969843 | 0.93792465 | 0.34828313 | 0.5484109 |
| UHRF1BP1L | 1146.83208 | 0.336004539 | 0.22768546 | 1.47574002 | 0.1400137 | 0.30672331 |
| ACTR6 | 366.035373 | 0.311032732 | 0.27138835 | 1.14607988 | 0.2517621 | 0.44816924 |
| DEPDC4 | 227.471717 | 1.490722194 | 0.528673 | 2.81974341 | 0.00480621 | 0.02942326 |
| SCYL2 | 1274.52246 | 0.250714498 | 0.20023043 | 1.25212983 | 0.21052256 | 0.39989367 |
| NR1H4 | 4936.31053 | -0.947925891 | 0.32508259 | -2.915954 | 0.00354603 | 0.02344852 |
| GAS2L3 | 981.976321 | -1.514322286 | 0.4180474 | -3.6223698 | 0.00029192 | 0.00358861 |
| ANO4 | 77.8166655 | -0.622373981 | 0.49103127 | -1.2674834 | 0.20498249 | 0.39334944 |
| SLC5A8 | 2279.37358 | 0.48466249 | 0.92183311 | 0.52575947 | 0.59905534 | 0.7589133 |
| UTP20 | 415.325833 | 0.008885352 | 0.30556712 | 0.02907823 | 0.9768022 | 0.9886052 |
| ARL1 | 336.6473 | -0.71336065 | 0.27588568 | -2.585711 | 0.00971784 | 0.04940741 |
| CHPT1 | 2337.59045 | 0.181346681 | 0.40449148 | 0.44833251 | 0.65391324 | 0.79768212 |
| SYCP3 | 86.0117386 | -0.517494384 | 0.69067898 | -0.7492546 | 0.45370378 | 0.64304805 |
| GNPTAB | 550.054765 | -0.155218796 | 0.25097074 | -0.6184737 | 0.53626315 | 0.70869254 |
| DRAM1 | 1318.27101 | -0.502247552 | 0.35848883 | -1.4010131 | 0.16121015 | 0.33735993 |
| WASHC3 | 553.075458 | 0.217098299 | 0.33306513 | 0.65181937 | 0.51451771 | 0.69267774 |
| NUP37 | 153.681027 | 0.380488125 | 0.31194244 | 1.21973825 | 0.22256412 | 0.41472082 |
| PARPBP | 40.4153578 | 0.028921123 | 0.54580684 | 0.05298784 | 0.9577416 | 0.97844258 |
| PAH | 2578.46979 | 2.691201341 | 0.50259757 | 5.35458488 | 8.58E-08 | 4.82E-06 |
| NT5DC3 | 268.154365 | -1.51361376 | 0.2763844 | -5.4764805 | 4.34E-08 | 2.79E-06 |
| HSP90B1 | 2005.52606 | -1.213100866 | 0.49162474 | -2.4675342 | 0.01360472 | 0.06259816 |
| C12orf73 | 166.424414 | 0.159242751 | 0.29402117 | 0.54160301 | 0.58809202 | 0.7504168 |
| TDG | 547.627166 | -0.479151216 | 0.28831405 | -1.6619072 | 0.09653137 | 0.23992921 |
| HCFC2 | 340.336258 | 0.210846213 | 0.22041416 | 0.95659104 | 0.33877371 | 0.53901002 |
| NFYB | 205.853244 | 0.809781577 | 0.20308483 | 3.9874055 | 6.68E-05 | 0.00113884 |
| TXNRD1 | 6274.4615 | -3.165653023 | 0.53358343 | -5.9328173 | 2.98E-09 | 2.92E-07 |
| CHST11 | 1959.11936 | -1.824155225 | 0.61134566 | -2.9838361 | 0.00284659 | 0.01994588 |
| SLC41A2 | 1690.61964 | -1.404401626 | 0.3551866 | -3.9539825 | 7.69E-05 | 0.00127238 |
| C12orf45 | 191.352337 | -0.425942292 | 0.35879415 | -1.1871495 | 0.23516866 | 0.42961829 |
| WASHC4 | 2286.16589 | -0.113511321 | 0.19680611 | -0.5767672 | 0.5640967 | 0.731442 |
| APPL2 | 984.712138 | -0.18166072 | 0.1962847 | -0.9254961 | 0.35470792 | 0.55318938 |
| C12orf75 | 1497.28305 | -0.340329822 | 0.37535068 | -0.9066983 | 0.36456636 | 0.56216797 |
| AC011313.1 | 205.154394 | -1.136486671 | 0.48903948 | -2.323916 | 0.02013 | 0.08278484 |
| NUAK1 | 138.733318 | -0.454059572 | 0.52117885 | -0.8712164 | 0.38363599 | 0.58015209 |
| CKAP4 | 130.348997 | -0.612688378 | 0.42110542 | -1.4549525 | 0.14568242 | 0.31436853 |
| TCP11L2 | 289.867472 | -0.540829796 | 0.36009413 | -1.5019123 | 0.13311976 | 0.29669473 |
| POLR3B | 313.790998 | 0.949709025 | 0.23401602 | 4.0583077 | 4.94E-05 | 0.00089621 |
| RIC8B | 946.843673 | -0.152918386 | 0.17238727 | -0.8870631 | 0.37504492 | 0.5718924 |
| TMEM263 | 560.83721 | -0.508280347 | 0.29363614 | -1.730987 | 0.08345408 | 0.21823333 |
| MTERF2 | 206.808296 | 0.664679379 | 0.27728919 | 2.39706196 | 0.01652713 | 0.07210819 |
| CRY1 | 865.153966 | -0.382706338 | 0.61901536 | -0.6182501 | 0.53641046 | 0.70875054 |
| BTBD11 | 1293.09112 | -3.603412449 | 0.77981259 | -4.6208698 | 3.82E-06 | 0.00011569 |
| PWP1 | 356.530104 | -0.551336038 | 0.27278851 | -2.0211117 | 0.04326821 | 0.14077316 |
| PRDM4 | 586.377888 | -0.537855224 | 0.32836232 | -1.6379931 | 0.10142313 | 0.24823532 |
| AC007622.2 | 47.1601164 | -1.258549418 | 0.61186297 | -2.0569139 | 0.03969451 | 0.13280271 |
| FICD | 59.526588 | -1.670815686 | 0.49636436 | -3.3661073 | 0.00076237 | 0.00741022 |
| SART3 | 391.781921 | -0.480834734 | 0.15949178 | -3.0147933 | 0.00257154 | 0.01857121 |
| ISCU | 661.814874 | -0.609837254 | 0.35108654 | -1.7369998 | 0.08238721 | 0.21646185 |
| AC007569.1 | 31.6714438 | 0.094499423 | 0.59510676 | 0.15879407 | 0.87383112 | 0.93468316 |
| CORO1C | 1293.06085 | -1.101148042 | 0.36459939 | -3.0201588 | 0.00252642 | 0.01834227 |
| SSH1 | 944.527398 | -0.127683278 | 0.26611818 | -0.4797992 | 0.63137021 | 0.78268893 |
| DAO | 266.731937 | 4.081979185 | 0.54760598 | 7.45422688 | 9.04E-14 | 3.44E-11 |
| USP30 | 318.847686 | 0.960709278 | 0.37120916 | 2.58805384 | 0.00965199 | 0.04910906 |
| ALKBH2 | 54.3047881 | 0.254061756 | 0.40391175 | 0.62900314 | 0.529347 | 0.7036229 |
| UNG | 122.003433 | 0.092647426 | 0.34217214 | 0.27076262 | 0.78657361 | 0.88335715 |
| ACACB | 1672.97488 | 0.970508497 | 0.51712357 | 1.87674387 | 0.06055321 | 0.1759226 |
| KCTD10 | 203.68687 | -0.337708941 | 0.27001611 | -1.2506992 | 0.21104422 | 0.40034114 |
| UBE3B | 393.046115 | 0.215288868 | 0.27983218 | 0.76934993 | 0.4416856 | 0.63341542 |
| MMAB | 340.39645 | 0.295107987 | 0.28589889 | 1.03221102 | 0.30197327 | 0.50241408 |
| MVK | 440.632619 | -0.547450589 | 0.30923245 | -1.770353 | 0.07666836 | 0.20617877 |
| FAM222A | 52.5504444 | 0.805123778 | 0.64814101 | 1.24220465 | 0.21416106 | 0.40445105 |
| TRPV4 | 1305.86276 | 2.45892611 | 0.61413959 | 4.00385539 | 6.23E-05 | 0.00107446 |
| GLTP | 199.60502 | -0.268017594 | 0.33476343 | -0.8006179 | 0.42335289 | 0.61654251 |
| TCHP | 417.511616 | -0.573510467 | 0.27098696 | -2.1163766 | 0.03431279 | 0.12022632 |
| GIT2 | 1049.21302 | -0.589013143 | 0.20073394 | -2.9342977 | 0.00334303 | 0.02238542 |
| ANKRD13A | 656.872787 | -1.015574962 | 0.34638807 | -2.9318994 | 0.00336896 | 0.02249309 |
| C12orf76 | 196.255888 | 0.124195551 | 0.29218265 | 0.42506136 | 0.67079195 | 0.80909409 |
| IFT81 | 599.963881 | 0.502543505 | 0.26245298 | 1.91479443 | 0.05551873 | 0.16619244 |
| ATP2A2 | 1798.08908 | -0.1270161 | 0.30891306 | -0.411171 | 0.68094713 | 0.81446268 |
| ANAPC7 | 520.920714 | 0.045850192 | 0.40313396 | 0.11373438 | 0.90944836 | 0.95253371 |
| ARPC3 | 999.585527 | -0.853913501 | 0.21362223 | -3.9973064 | 6.41E-05 | 0.00109692 |
| GPN3 | 99.8167349 | -0.186457616 | 0.21836573 | -0.8538776 | 0.39317279 | 0.58802176 |
| FAM216A | 53.1524951 | -0.225559534 | 0.32905966 | -0.685467 | 0.49304929 | 0.67529565 |
| VPS29 | 550.60217 | -0.480446187 | 0.16240682 | -2.9582882 | 0.00309353 | 0.02111703 |
| PPTC7 | 676.985907 | -0.264726767 | 0.23680138 | -1.1179274 | 0.26359798 | 0.46156588 |
| TCTN1 | 416.560641 | 0.876259914 | 0.39692201 | 2.20763751 | 0.02726955 | 0.10238676 |
| HVCN1 | 84.7946719 | 0.171199537 | 0.62068176 | 0.27582498 | 0.78268249 | 0.88095163 |
| PPP1CC | 295.124382 | 0.145439994 | 0.20428651 | 0.71194126 | 0.47650115 | 0.6621453 |
| PHETA1 | 87.7297446 | 0.288054387 | 0.48432403 | 0.59475552 | 0.55200691 | 0.7216365 |

| | | | | | | |
|---|---|---|---|---|---|---|
| SH2B3 | 136.408811 | -0.643753606 | 0.39343539 | -1.6362372 | 0.10178996 | 0.24897127 |
| ATXN2 | 4502.42405 | 0.45081096 | 0.21160362 | 2.13045011 | 0.03313447 | 0.117388 |
| BRAP | 420.10534 | -0.06520449 | 0.19768166 | -0.3298459 | 0.74151639 | 0.85325429 |
| ACAD10 | 547.434517 | 0.709269305 | 0.16419585 | 4.3196544 | 1.56E-05 | 0.00036429 |
| ALDH2 | 1352.03006 | 0.197944077 | 0.4097354 | 0.48310221 | 0.62902316 | 0.78090938 |
| MAPKAPK5-A | 107.767181 | 0.175048954 | 0.30083646 | 0.58187413 | 0.56065147 | 0.72893225 |
| MAPKAPK5 | 442.982252 | 0.174422283 | 0.23018292 | 0.75775512 | 0.44859759 | 0.63911519 |
| TMEM116 | 399.582346 | 0.243534243 | 0.28180566 | 0.86419218 | 0.38748233 | 0.58365058 |
| ERP29 | 551.540718 | -0.521558674 | 0.30237794 | -1.7248569 | 0.08455327 | 0.21996046 |
| NAA25 | 1032.56125 | -0.281211128 | 0.34672479 | -0.81105 | 0.41733698 | 0.61047432 |
| TRAFD1 | 251.982901 | -0.298411329 | 0.32559152 | -0.9165206 | 0.35939391 | 0.55776861 |
| HECTD4 | 2357.54081 | 0.131344535 | 0.14947035 | 0.87873303 | 0.37954604 | 0.5760644 |
| RPL6 | 1222.83798 | -0.93498683 | 0.40071002 | -2.3333253 | 0.01963108 | 0.08131859 |
| PTPN11 | 1016.15253 | -0.472528015 | 0.35354935 | -1.3365263 | 0.18137731 | 0.36363006 |
| OAS1 | 239.191691 | -0.393242787 | 1.02717575 | -0.3828389 | 0.70183925 | 0.82874111 |
| AC004551.1 | 39.9019536 | -0.970271192 | 1.02674083 | -0.9450011 | 0.34465832 | 0.54521392 |
| OAS3 | 67.6494297 | -2.368231716 | 1.12705042 | -2.1012651 | 0.0356177 | 0.12313035 |
| RASAL1 | 361.314895 | -2.120697699 | 0.35144824 | -6.0341679 | 1.60E-09 | 1.74E-07 |
| DDX54 | 252.664617 | -0.316706377 | 0.37216122 | -0.8509924 | 0.39477356 | 0.58957955 |
| RITA1 | 84.4659537 | -0.034314932 | 0.25984588 | -0.1320588 | 0.89493779 | 0.94552399 |
| IQCD | 95.5043286 | 0.342295707 | 0.47095939 | 0.72680514 | 0.46734534 | 0.65466556 |
| TPCN1 | 3367.17947 | 0.909175596 | 0.60418576 | 1.5047948 | 0.13237684 | 0.2956689 |
| SLC8B1 | 219.630259 | -0.221041475 | 0.24896091 | -0.8878561 | 0.37461814 | 0.5715597 |
| PLBD2 | 228.872224 | -0.534272909 | 0.38299509 | -1.3949863 | 0.16301997 | 0.33943715 |
| SDSL | 173.801347 | 0.331476157 | 0.39261264 | 0.84428295 | 0.39851131 | 0.59275547 |
| RBM19 | 459.934931 | -0.435375967 | 0.4088574 | -1.0648602 | 0.28693921 | 0.48658614 |
| MED13L | 6728.81563 | -0.238035925 | 0.18244923 | -1.3046694 | 0.19200546 | 0.37640429 |
| MAP1LC3B2 | 55.9978245 | -2.404508184 | 0.33768548 | -7.1205555 | 1.07E-12 | 2.89E-10 |
| C12orf49 | 441.479169 | -1.520194194 | 0.52761328 | -2.881266 | 0.00396081 | 0.02545282 |
| RNFT2 | 68.5029697 | -0.393111677 | 0.62039036 | -0.6336521 | 0.52630787 | 0.7009757 |
| FBXW8 | 698.330805 | -0.101586614 | 0.32553639 | -0.3120592 | 0.75499556 | 0.86260933 |
| TESC | 267.211498 | -3.666694678 | 0.84524928 | -4.3380039 | 1.44E-05 | 0.0003398 |
| FBXO21 | 1197.23945 | 0.86045306 | 0.32744583 | 2.62777224 | 0.0085946 | 0.04506742 |
| KSR2 | 73.2770398 | 0.705681418 | 0.45188796 | 1.56162918 | 0.11837537 | 0.27473268 |
| RFC5 | 72.0518066 | -0.098755907 | 0.45184274 | -0.2185626 | 0.82699082 | 0.9074258 |
| WSB2 | 364.308377 | -0.145275697 | 0.27429433 | -0.5296343 | 0.59636548 | 0.75646693 |
| VSIG10 | 471.316284 | 0.779651131 | 0.33694389 | 2.31389011 | 0.02067375 | 0.0845139 |
| PEBP1 | 2717.36478 | -0.165576135 | 0.38646151 | -0.4284415 | 0.66832974 | 0.80729015 |
| TAOK3 | 2328.07099 | 0.171970285 | 0.18530238 | 0.92805224 | 0.35338047 | 0.55189396 |
| SUDS3 | 477.357207 | 0.0696245 | 0.14648518 | 0.47530063 | 0.63457271 | 0.78453036 |
| HSPB8 | 537.88419 | -2.413868802 | 0.32798399 | -7.3597153 | 1.84E-13 | 6.03E-11 |
| CCDC60 | 56.9469285 | -1.663850996 | 0.57496037 | -2.8938534 | 0.00380546 | 0.0246238 |
| AC002070.1 | 411.974944 | -0.640738242 | 0.47479207 | -1.3495134 | 0.17717213 | 0.35765809 |
| PRKAB1 | 725.019919 | -0.08636882 | 0.40548237 | -0.2130027 | 0.83132489 | 0.91080925 |
| CIT | 755.557272 | 0.115420148 | 0.68077306 | 0.16954277 | 0.86536974 | 0.92952143 |
| BICDL1 | 6504.15053 | 1.094931511 | 0.37750809 | 2.90041864 | 0.00372665 | 0.02424816 |
| RAB35 | 258.212499 | -0.477384686 | 0.45176785 | -1.0567035 | 0.29064691 | 0.4907486 |
| GCN1 | 512.742919 | -0.283000746 | 0.3866913 | -0.7318519 | 0.46425899 | 0.65200799 |
| RPLP0 | 1810.59811 | -1.262484371 | 0.49596671 | -2.5455023 | 0.01091207 | 0.05370617 |
| PXN-AS1 | 80.0401621 | 0.51259065 | 0.22453006 | 2.28294885 | 0.02243338 | 0.08954408 |
| PXN | 536.496848 | 0.209638038 | 0.32827132 | 0.63861211 | 0.5230753 | 0.69880698 |
| SIRT4 | 85.285249 | 0.899363659 | 0.35679 | 2.52070868 | 0.01171188 | 0.05627745 |
| TRIAP1 | 41.6615779 | -0.680260345 | 0.41581746 | -1.6359591 | 0.10184815 | 0.24906915 |
| GATC | 148.540897 | -0.148348462 | 0.26459166 | -0.5606695 | 0.5750229 | 0.73973147 |
| SRSF9 | 393.384445 | -0.342392192 | 0.27729349 | -1.2347646 | 0.21691812 | 0.40780368 |
| DYNLL1 | 622.804024 | -0.734570701 | 0.39221425 | -1.8728812 | 0.06108479 | 0.17704536 |
| NRAV | 44.1256867 | -0.189639584 | 0.29897914 | -0.6342904 | 0.52589135 | 0.70084961 |
| COQ5 | 215.588097 | 1.343909426 | 0.22786954 | 5.89771423 | 3.69E-09 | 3.46E-07 |
| RNF10 | 1060.8794 | -0.096128577 | 0.22430158 | -0.4285684 | 0.66823733 | 0.80729015 |
| POP5 | 167.463404 | 0.126473185 | 0.28496914 | 0.44381362 | 0.65717734 | 0.80027901 |
| CABP1 | 287.638514 | -1.479437543 | 0.80953013 | -1.8275262 | 0.06762069 | 0.18958412 |
| MLEC | 1147.54199 | -0.34267743 | 0.39096011 | -0.8765023 | 0.38075703 | 0.57723861 |
| UNC119B | 202.247799 | 0.160457161 | 0.22047825 | 0.72776865 | 0.46675522 | 0.65457476 |
| ACADS | 281.539938 | 0.57706341 | 0.27392681 | 2.10663359 | 0.03514935 | 0.12206541 |
| SPPL3 | 2897.93625 | -0.045531177 | 0.17194964 | -0.2647937 | 0.79116841 | 0.88635434 |
| HNF1A | 431.524156 | 1.60564789 | 0.5018854 | 3.19923209 | 0.00137794 | 0.01150972 |
| C12orf43 | 89.5784865 | -0.217383017 | 0.36932568 | -0.5885944 | 0.55613341 | 0.72533372 |
| OASL | 43.7835562 | -1.869992693 | 1.42399311 | -1.3132035 | 0.18911439 | 0.37309941 |
| P2RX4 | 632.149953 | -0.634461919 | 0.35295501 | -1.7975717 | 0.07224491 | 0.19832987 |
| CAMKK2 | 528.872469 | 0.498856722 | 0.25610643 | 1.94784927 | 0.051433 | 0.15796366 |
| ANAPC5 | 934.393502 | 0.273372534 | 0.22115063 | 1.23613724 | 0.21640755 | 0.40723481 |
| RNF34 | 422.792203 | -0.364034361 | 0.42454123 | -0.857477 | 0.3911813 | 0.58646585 |
| KDM2B | 311.902506 | 0.195844365 | 0.28860987 | 0.67857819 | 0.49740517 | 0.67927993 |
| ORAI1 | 126.086108 | -0.871285167 | 0.38392015 | -2.2694437 | 0.02324136 | 0.09163579 |
| TMEM120B | 846.854864 | 0.297226353 | 0.43971229 | 0.67595644 | 0.4990683 | 0.68080506 |
| RHOF | 215.108819 | -2.331493535 | 0.70747952 | -3.2954926 | 0.00098249 | 0.00891266 |
| SETD1B | 235.311801 | 0.082932535 | 0.43354588 | 0.19128894 | 0.84829922 | 0.91998081 |
| HPD | 675.862575 | 1.322095766 | 0.76803029 | 1.72141096 | 0.08517628 | 0.22091036 |
| WDR66 | 335.197075 | -0.376806772 | 0.43367782 | -0.8688634 | 0.38492187 | 0.58126298 |
| BCL7A | 676.13996 | 0.15435868 | 0.33321808 | 0.46323621 | 0.64319507 | 0.79092642 |
| MLXIP | 1177.89442 | -0.891760735 | 0.37225318 | -2.3955758 | 0.01659428 | 0.07234663 |
| LRRC43 | 77.1928619 | 1.003972057 | 0.34503616 | 2.90975894 | 0.00361708 | 0.0237819 |
| B3GNT4 | 38.0710333 | 0.862973662 | 0.69701648 | 1.2380965 | 0.21568028 | 0.40631248 |
| CLIP1 | 2550.72778 | -0.551075045 | 0.40903102 | -1.3472696 | 0.17789342 | 0.3586388 |
| ZCCHC8 | 478.802321 | 0.044083811 | 0.2338647 | 0.18850135 | 0.85048366 | 0.92140211 |
| RSRC2 | 793.755456 | -0.209044801 | 0.17361032 | -1.2041036 | 0.22854956 | 0.42151894 |
| KNTC1 | 126.59704 | 0.043115073 | 0.30753224 | 0.14019692 | 0.88850441 | 0.94248026 |
| DENR | 801.592464 | -0.213269358 | 0.24766746 | -0.8611117 | 0.3891765 | 0.58478867 |
| CCDC62 | 47.0561893 | -0.259188956 | 0.33782873 | -0.76722 | 0.44295073 | 0.63456586 |
| HIP1R | 977.246652 | 0.275684033 | 0.35319416 | 0.78054529 | 0.43506998 | 0.62747309 |
| VPS37B | 686.160467 | -0.532988685 | 0.29169949 | -1.8271842 | 0.06767208 | 0.18961187 |
| ABCB9 | 100.454438 | 0.839445214 | 0.39804189 | 2.10893687 | 0.03495003 | 0.12161972 |
| OGFOD2 | 76.6085758 | 0.372370435 | 0.26424234 | 1.40920048 | 0.1587759 | 0.33362875 |
| ARL6IP4 | 686.575537 | 0.017247742 | 0.22196558 | 0.07770458 | 0.93806305 | 0.96676805 |
| PITPNM2 | 1066.92782 | -1.243218301 | 0.37045584 | -3.355915 | 0.00079103 | 0.00760887 |
| MPHOSPH9 | 744.466554 | 0.592392292 | 0.15059557 | 3.93366338 | 8.37E-05 | 0.00135388 |
| C12orf65 | 355.042899 | -0.28687427 | 0.27786671 | -1.0324168 | 0.30187689 | 0.50239081 |
| AC068768.1 | 43.8765831 | -0.507073864 | 0.2714425 | -1.868071 | 0.06175218 | 0.1783099 |
| SBNO1 | 1129.05262 | -0.112006947 | 0.22828302 | -0.4906495 | 0.62367438 | 0.77659018 |
| AC137767.1 | 79.3915626 | 0.493155264 | 0.52112591 | 0.9463265 | 0.34398208 | 0.54439007 |
| KMT5A | 192.292302 | -0.582367795 | 0.3209963 | -1.8142508 | 0.06963913 | 0.19331655 |
| RILPL2 | 294.000049 | 0.146323428 | 0.32947988 | 0.44410429 | 0.65696719 | 0.80024905 |
| SNRNP35 | 279.946325 | 0.676534648 | 0.16371239 | 4.13245856 | 3.59E-05 | 0.00069386 |
| RILPL1 | 805.759025 | -0.5703624 | 0.18399294 | -3.0999145 | 0.00193577 | 0.01495689 |
| AC055713.1 | 64.5259572 | 0.801097303 | 0.31655814 | 2.53064823 | 0.0113852 | 0.05524138 |
| TMED2 | 386.523861 | -0.782552813 | 0.33262311 | -2.3526712 | 0.01863911 | 0.07846629 |
| DDX55 | 239.442957 | 0.219785696 | 0.31196569 | 0.70451881 | 0.48110974 | 0.66594592 |
| EIF2B1 | 241.017377 | 0.235972177 | 0.20631663 | 1.14373804 | 0.25273229 | 0.44931278 |
| GTF2H3 | 256.594479 | 0.091383303 | 0.19039722 | 0.47996134 | 0.63125489 | 0.78261674 |
| TCTN2 | 162.030758 | 0.518399433 | 0.33790507 | 1.534157 | 0.12499103 | 0.28492468 |
| ATP6V0A2 | 263.548476 | -0.054662569 | 0.35986944 | -0.1518956 | 0.87926931 | 0.93710788 |
| DNAH10 | 103.838649 | 0.950730121 | 0.70830004 | 1.34227032 | 0.17950836 | 0.36086531 |
| CCDC92 | 981.645019 | -0.103372493 | 0.19252112 | -0.5369411 | 0.59130833 | 0.75276895 |

| | | | | | | |
|---|---|---|---|---|---|---|
| RFLNA | 79.9062137 | -0.145661659 | 0.62346367 | -0.233633 | 0.81526994 | 0.90004068 |
| NCOR2 | 1026.67123 | 0.083534155 | 0.32703103 | 0.25543189 | 0.79838954 | 0.89012761 |
| SCARB1 | 674.166304 | -1.284628745 | 0.61946915 | -2.0737574 | 0.03810185 | 0.12946505 |
| UBC | 3030.32256 | -1.929906716 | 0.33106849 | -5.8293277 | 5.57E-09 | 4.92E-07 |
| DHX37 | 149.98151 | -0.547344439 | 0.46941218 | -1.166021 | 0.24360597 | 0.43883651 |
| BRI3BP | 176.44957 | -0.058233893 | 0.40668814 | -0.1431905 | 0.8861397 | 0.94113722 |
| AACS | 288.396594 | -0.910776575 | 0.22677776 | -4.0161635 | 5.92E-05 | 0.0010342 |
| TMEM132B | 49.2427822 | 0.077002337 | 0.43423886 | 0.17732715 | 0.85925143 | 0.9261227 |
| LINC00939 | 175.860534 | 1.092603927 | 0.65777025 | 1.66107227 | 0.09669893 | 0.24024873 |
| LINC02405 | 37.3938267 | -0.379347305 | 0.67223683 | -0.564306 | 0.57254587 | 0.73810771 |
| SLC15A4 | 668.893947 | -1.670024113 | 0.3256473 | -5.1283217 | 2.92E-07 | 1.35E-05 |
| GLT1D1 | 95.6678832 | -0.304635464 | 0.73438058 | -0.4148196 | 0.67827397 | 0.81367944 |
| TMEM132D | 411.686644 | -3.2154127 | 0.77794466 | -4.1332152 | 3.58E-05 | 0.00069256 |
| STX2 | 451.537396 | -0.406643165 | 0.28888653 | -1.4076224 | 0.1592429 | 0.33421259 |
| RAN | 416.20518 | -1.573011535 | 0.41370254 | -3.8022767 | 0.00014337 | 0.00203648 |
| SFSWAP | 979.734171 | 0.18348487 | 0.26539419 | 0.69136731 | 0.48933474 | 0.67241216 |
| ULK1 | 353.966798 | 0.17682419 | 0.27495453 | 0.64310338 | 0.52015702 | 0.69607397 |
| PUS1 | 244.356584 | -0.416899649 | 0.46825145 | -0.8903328 | 0.37328719 | 0.57079959 |
| EP400 | 904.707824 | -0.008844924 | 0.27525654 | -0.0321334 | 0.97436568 | 0.98703942 |
| DDX51 | 112.496199 | -0.343888747 | 0.4280741 | -0.8033393 | 0.42177865 | 0.61549238 |
| NOC4L | 43.4644136 | 0.146534481 | 0.49005951 | 0.29901365 | 0.76492963 | 0.86874558 |
| GALNT9 | 76.2200355 | 0.881554595 | 0.72316258 | 1.21902685 | 0.222834 | 0.41499805 |
| FBRSL1 | 527.712655 | 0.226243259 | 0.28467533 | 0.79474135 | 0.42676397 | 0.61966672 |
| POLE | 160.964982 | -0.158579344 | 0.31128801 | -0.5094297 | 0.6104511 | 0.76681206 |
| AC135586.2 | 52.8367942 | 1.327565502 | 0.46621332 | 2.84754951 | 0.00440572 | 0.027575 |
| PGAM5 | 88.5402967 | -0.509224543 | 0.49773721 | -1.0230791 | 0.30627045 | 0.50621597 |
| ANKLE2 | 956.793546 | -1.329135929 | 0.48004712 | -2.7687614 | 0.00562698 | 0.03311681 |
| GOLGA3 | 805.777672 | -1.155517819 | 0.51397625 | -2.248193 | 0.02456389 | 0.0951122 |
| CHFR | 878.498049 | -0.451372424 | 0.3012382 | -1.4983904 | 0.13403186 | 0.2980491 |
| ZNF26 | 545.723437 | 0.102821747 | 0.2305721 | 0.44594184 | 0.65563926 | 0.79908554 |
| ZNF84 | 582.789969 | -0.032556421 | 0.2592822 | -0.1255637 | 0.90007733 | 0.94743919 |
| ZNF140 | 311.386561 | 0.192946751 | 0.22833395 | 0.84501999 | 0.39809968 | 0.59255618 |
| ZNF891 | 224.578616 | 0.601972403 | 0.17850782 | 3.37224662 | 0.00074558 | 0.00728931 |
| ZNF268 | 95.9186592 | 0.030734032 | 0.26636747 | 0.11538208 | 0.90814228 | 0.95196591 |
| MPHOSPH8 | 1349.54378 | 0.260401909 | 0.25304583 | 1.02907016 | 0.3034467 | 0.50371548 |
| PSPC1 | 2006.51128 | 0.15573584 | 0.2076483 | 0.74999813 | 0.45325583 | 0.64247959 |
| ZMYM5 | 763.396733 | 0.13075095 | 0.23314295 | 0.5608188 | 0.57492107 | 0.7396699 |
| AL355001.1 | 48.0548159 | 0.309083876 | 0.35427578 | 0.87243862 | 0.38296914 | 0.57970398 |
| ZMYM2 | 3679.81397 | 0.309292963 | 0.21671083 | 1.42721507 | 0.15351791 | 0.32573839 |
| GJB2 | 67.1011074 | -0.162809713 | 0.49740376 | -0.327319 | 0.74342661 | 0.8543765 |
| CRYL1 | 4113.6519 | 1.294469956 | 0.44974872 | 2.87820711 | 0.00399942 | 0.02563457 |
| IFT88 | 3105.47865 | 0.730410225 | 0.26872003 | 2.71810856 | 0.00656563 | 0.03701978 |
| EEF1AKMT1 | 199.695776 | 0.177695015 | 0.3731447 | 0.47620941 | 0.6339252 | 0.78408344 |
| XPO4 | 1080.1728 | 0.222905931 | 0.19906026 | 1.11979121 | 0.26280275 | 0.46082094 |
| LATS2 | 910.961367 | 0.541811863 | 0.43105027 | 1.25695747 | 0.20876904 | 0.39797925 |
| SAP18 | 457.602542 | -0.525233935 | 0.28817836 | -1.8226002 | 0.06836397 | 0.19100386 |
| MRPL57 | 331.03475 | -0.599251213 | 0.2909015 | -2.0599798 | 0.03940047 | 0.13226584 |
| ZDHHC20 | 954.582591 | 0.117009724 | 0.19939414 | 0.58682628 | 0.55732039 | 0.7261364 |
| MICU2 | 1814.22081 | 0.380816469 | 0.1625575 | 2.34265707 | 0.01914697 | 0.07991704 |
| FGF9 | 59.7750294 | 1.363580831 | 0.67281297 | 2.02668631 | 0.0426945 | 0.13956918 |
| AL136962.1 | 954.741018 | -0.543264 | 0.66795905 | -0.8133193 | 0.416035 | 0.60944659 |
| LINC00621 | 2060.13853 | -2.567650155 | 0.74324581 | -3.4546446 | 0.00055102 | 0.00573486 |
| AL157931.1 | 840.405101 | -1.804636377 | 0.71064271 | -2.5394426 | 0.01110293 | 0.05431399 |
| SACS | 204.766738 | -2.335695765 | 0.61075861 | -3.8242535 | 0.00013117 | 0.00191473 |
| TNFRSF19 | 682.62117 | 1.08139392 | 0.62286544 | 1.73615976 | 0.08253559 | 0.21668845 |
| MIPEP | 1120.86726 | -0.087673426 | 0.22162245 | -0.3955981 | 0.69240152 | 0.82270698 |
| PARP4 | 2448.42612 | -0.87123437 | 0.2179141 | -3.9980634 | 6.39E-05 | 0.00109692 |
| CENPJ | 109.459288 | -0.012512192 | 0.36490423 | -0.034289 | 0.97264672 | 0.9863186 |
| PABPC3 | 47.7856562 | -0.990481848 | 0.53491794 | -1.851652 | 0.06407582 | 0.18309093 |
| MTMR6 | 480.757034 | -0.175573202 | 0.22178536 | -0.7916357 | 0.42857315 | 0.62163515 |
| NUP58 | 1385.20395 | -0.253026218 | 0.26002887 | -0.9730697 | 0.33051865 | 0.53036984 |
| ATP8A2 | 36.9668263 | 0.193297287 | 0.4469211 | 0.43250876 | 0.66537169 | 0.80579154 |
| RNF6 | 201.033077 | -0.900321634 | 0.37156674 | -2.4230415 | 0.01539117 | 0.06845125 |
| CDK8 | 1917.66729 | -0.111148542 | 0.1576706 | -0.7049415 | 0.48084667 | 0.66575407 |
| WASF3 | 250.125966 | -0.647268985 | 0.40121326 | -1.6132792 | 0.10668387 | 0.25642924 |
| USP12 | 1234.42373 | -0.707604126 | 0.28770861 | -2.4594472 | 0.01391512 | 0.06355699 |
| LINC02340 | 393.212865 | 0.433231727 | 0.43793299 | 0.98926487 | 0.32253357 | 0.52288545 |
| RPL21 | 557.386136 | -0.3088854 | 0.39952569 | -0.7731303 | 0.43944529 | 0.63125934 |
| GTF3A | 270.366845 | -0.51009337 | 0.23173176 | -2.2012234 | 0.02772022 | 0.1035034 |
| MTIF3 | 499.425335 | -0.046706761 | 0.17475573 | -0.2672688 | 0.78926218 | 0.88501446 |
| LNX2 | 964.337602 | -0.298427626 | 0.24699372 | -1.2082397 | 0.22695507 | 0.41931165 |
| POLR1D | 585.751145 | -0.024582162 | 0.18884686 | -0.1301698 | 0.89643207 | 0.94620317 |
| FLT3 | 102.268401 | 2.057971684 | 0.676416 | 3.04246451 | 0.00234649 | 0.01739216 |
| PAN3 | 4542.40439 | 0.25041642 | 0.16739495 | 1.4959616 | 0.13466366 | 0.29911438 |
| FLT1 | 79.759731 | 0.376672092 | 0.4151927 | 0.90722235 | 0.36428923 | 0.56199352 |
| POMP | 375.450062 | -0.777537698 | 0.25676163 | -3.0282473 | 0.00245977 | 0.01798188 |
| SLC46A3 | 93.8143852 | 0.6936552 | 0.48946733 | 1.41716343 | 0.15643515 | 0.33013369 |
| MTUS2 | 39.3471829 | -1.173815001 | 0.38992569 | -3.0103557 | 0.00260942 | 0.01873615 |
| SLC7A1 | 94.8725478 | -2.419148053 | 0.63818124 | -3.7906913 | 0.00015023 | 0.00211198 |
| UBL3 | 751.006576 | -0.061493034 | 0.15250162 | -0.4032287 | 0.68677996 | 0.81863011 |
| KATNAL1 | 724.995202 | -0.174614142 | 0.24049968 | -0.7260473 | 0.46780978 | 0.65504839 |
| HMGB1 | 1888.37567 | 0.508547546 | 0.27078635 | 1.87803984 | 0.06037572 | 0.17548132 |
| USPL1 | 592.726875 | -0.273470548 | 0.31014275 | -0.881757 | 0.37790822 | 0.57453282 |
| HSPH1 | 1238.83278 | -1.752984192 | 0.84630215 | -2.0713455 | 0.03832652 | 0.12998945 |
| B3GLCT | 318.78934 | -0.17796246 | 0.20541979 | -0.8663355 | 0.3863062 | 0.58239101 |
| FRY | 446.582587 | 0.081094784 | 0.58301477 | 0.13909559 | 0.88937461 | 0.94267382 |
| BRCA2 | 55.3573018 | -1.288987245 | 0.68238755 | -1.8889372 | 0.05890024 | 0.1726204 |
| N4BP2L1 | 718.540426 | 0.198372506 | 0.44467003 | 0.44611171 | 0.65551655 | 0.79904496 |
| N4BP2L2 | 8281.9215 | 0.375170364 | 0.19185293 | 1.95551021 | 0.05052288 | 0.15646568 |
| PDS5B | 1508.42119 | 0.155394 | 0.18422089 | 0.84351997 | 0.3989377 | 0.59314883 |
| KL | 1350.79293 | 1.975223236 | 0.36838244 | 5.36188223 | 8.24E-08 | 4.68E-06 |
| STARD13 | 2528.86761 | -0.945116495 | 0.27808507 | -3.3986597 | 0.00067717 | 0.00673582 |
| AL139383.1 | 398.246531 | -0.000499711 | 0.71576828 | -0.0006981 | 0.99944296 | 0.9997347 |
| RFC3 | 182.10248 | 0.107091724 | 0.45219202 | 0.23682798 | 0.81279024 | 0.89861746 |
| LINC02343 | 68.740201 | 0.241417176 | 0.65766339 | 0.36708319 | 0.71355696 | 0.8362667 |
| NBEA | 3564.18584 | 0.014627258 | 0.22433336 | 0.06520322 | 0.94801219 | 0.97344198 |
| DCLK1 | 206.559073 | -1.798445706 | 0.87487917 | -2.0556504 | 0.03981623 | 0.13307998 |
| SPART | 1680.64722 | -0.116137835 | 0.14568452 | -0.7971872 | 0.4253423 | 0.61832293 |
| SPART-AS1 | 155.371739 | -0.145369553 | 0.41995027 | -0.346159 | 0.72922325 | 0.84621258 |
| RFXAP | 134.972097 | 0.399297511 | 0.17085028 | 2.33711941 | 0.01943298 | 0.08076647 |
| SMAD9 | 674.086434 | -1.21519138 | 0.65506605 | -1.8550669 | 0.06358667 | 0.18199675 |
| ALG5 | 346.387722 | -0.132118515 | 0.15981882 | -0.8266768 | 0.40842027 | 0.60217443 |
| EXOSC8 | 147.346843 | -0.242435178 | 0.27941421 | -0.8676551 | 0.38558315 | 0.58181288 |
| SUPT20H | 1197.19613 | 0.418763309 | 0.2393088 | 1.7498868 | 0.08013785 | 0.21221976 |
| LINC02334 | 381.665294 | 2.390134627 | 0.47435879 | 5.0386642 | 4.69E-07 | 2.03E-05 |
| LINC00571 | 494.752968 | -0.479716219 | 0.22675589 | -2.1155623 | 0.03438205 | 0.12031025 |
| UFM1 | 661.069226 | -0.941149211 | 0.23736967 | -3.9649094 | 7.34E-05 | 0.00122436 |
| FREM2 | 2824.7952 | 1.776207322 | 0.5642916 | 3.14767633 | 0.00164574 | 0.01323734 |
| FREM2-AS1 | 50.7512988 | 0.100213006 | 0.85860389 | 0.11671623 | 0.90708493 | 0.95122652 |
| PROSER1 | 328.235022 | 0.220632936 | 0.14074359 | 1.56762329 | 0.11696907 | 0.27255182 |
| NHLRC3 | 628.477237 | 0.260730965 | 0.46183166 | 0.56455845 | 0.57237414 | 0.73806372 |
| AL354809.1 | 98.9486012 | 0.739608963 | 0.31682653 | 2.33442879 | 0.01957328 | 0.08115274 |
| LHFPL6 | 458.067885 | -0.52216521 | 0.67490802 | -0.7736835 | 0.43911797 | 0.63085526 |

| | | | | | | |
|---|---|---|---|---|---|---|
| COG6 | 955.226929 | 0.086331514 | 0.20110767 | 0.42928008 | 0.66771941 | 0.80690041 |
| LINC00598 | 312.810698 | 0.192208725 | 0.57239945 | 0.33579474 | 0.73702566 | 0.85110831 |
| FOXO1 | 3415.3339 | 0.422828664 | 0.26005803 | 1.62590121 | 0.10397067 | 0.25210084 |
| MRPS31 | 744.015208 | 0.47843391 | 0.30242637 | 1.58198477 | 0.11365305 | 0.26731977 |
| SLC25A15 | 82.584135 | 0.159333546 | 0.4170768 | 0.38202448 | 0.70244321 | 0.82904704 |
| ELF1 | 2473.69742 | -0.359816025 | 0.25073621 | -1.4350382 | 0.15127623 | 0.32263003 |
| WBP4 | 488.300266 | -0.202253191 | 0.17323805 | -1.1674871 | 0.2430137 | 0.4381727 |
| KBTBD6 | 64.1961932 | 0.439807728 | 0.33537899 | 1.31137531 | 0.189731 | 0.3738167 |
| AL354696.2 | 232.871417 | 0.261904326 | 0.39048243 | 0.67071988 | 0.502399 | 0.6833085 |
| KBTBD7 | 74.3867473 | 0.792988636 | 0.42763393 | 1.85436322 | 0.06368721 | 0.18224648 |
| MTRF1 | 268.376829 | -0.20902363 | 0.34203467 | -0.6111182 | 0.5411213 | 0.71231293 |
| NAA16 | 740.820233 | 0.086552722 | 0.21034475 | 0.41148031 | 0.68072038 | 0.81442647 |
| VWA8 | 5250.3988 | 1.059236423 | 0.27448765 | 3.85895842 | 0.00011387 | 0.00172659 |
| DGKH | 4382.04871 | -0.007527443 | 0.34554894 | -0.021784 | 0.98262025 | 0.99144466 |
| AKAP11 | 853.471416 | 0.439430468 | 0.16026639 | 2.74187541 | 0.00610895 | 0.0350523 |
| LINC02341 | 68.8966434 | 1.345747877 | 0.26068082 | 5.16243534 | 2.44E-07 | 1.16E-05 |
| EPSTI1 | 1749.62796 | -0.725070956 | 0.83282703 | -0.8706141 | 0.38396489 | 0.5803781 |
| DNAJC15 | 1059.5324 | -0.00267383 | 0.29634184 | -0.0090228 | 0.99280095 | 0.99629729 |
| ENOX1 | 466.657923 | 0.027046859 | 0.43070142 | 0.06279724 | 0.94992797 | 0.97435384 |
| CCDC122 | 673.643966 | -0.516693016 | 0.49041737 | -1.0535781 | 0.29207611 | 0.49214879 |
| LINC00390 | 338.755161 | -1.916683811 | 0.73296345 | -2.6149787 | 0.0089233 | 0.04636532 |
| SMIM2-AS1 | 8782.19225 | -0.511429138 | 0.29342831 | -1.7429441 | 0.08134338 | 0.21450053 |
| SMIM2 | 81.3316526 | -1.77682643 | 0.69564818 | -2.5542027 | 0.01064313 | 0.05276143 |
| SERP2 | 85.615118 | -0.718500921 | 0.37376363 | -1.9223404 | 0.05456294 | 0.16434269 |
| TSC22D1 | 3757.53919 | 0.128058398 | 0.13065195 | 0.98014913 | 0.32701251 | 0.52691716 |
| TSC22D1-AS | 33.1158803 | -0.368715151 | 0.47935655 | -0.7691877 | 0.44178191 | 0.63348725 |
| NUFIP1 | 198.097739 | 0.469329249 | 0.26489706 | 1.771742 | 0.07643739 | 0.20563835 |
| AL359706.1 | 113.155824 | -0.612800655 | 0.57142559 | -1.0724068 | 0.28353738 | 0.48284841 |
| GPALPP1 | 468.235081 | 0.190135019 | 0.13709516 | 1.38688349 | 0.16547729 | 0.34277681 |
| GTF2F2 | 1040.99605 | -0.60984806 | 0.39868459 | -1.5296504 | 0.12610328 | 0.28669568 |
| TPT1 | 3986.19992 | -1.015648529 | 0.45600795 | -2.2272606 | 0.02592986 | 0.09886527 |
| AL138963.1 | 89.567377 | 0.092787938 | 0.3521994 | 0.26345286 | 0.79220155 | 0.88678699 |
| SLC25A30 | 649.454086 | 0.595124183 | 0.45178268 | 1.31727977 | 0.18774487 | 0.37136212 |
| SLC25A30-AS | 40.0147491 | 0.593794041 | 0.6604397 | 0.89908896 | 0.36860528 | 0.56603995 |
| COG3 | 768.979026 | -0.012813143 | 0.15387169 | -0.0832716 | 0.93363557 | 0.96473023 |
| ZC3H13 | 914.758103 | 0.305658846 | 0.19148707 | 1.59623751 | 0.11043578 | 0.26230171 |
| AL138686.1 | 206.988561 | -1.409619446 | 0.69789909 | -2.0198041 | 0.04340371 | 0.14111354 |
| LRCH1 | 3919.68243 | -0.308592146 | 0.31015252 | -0.994969 | 0.31975133 | 0.52029864 |
| ESD | 435.626001 | 0.04964229 | 0.23665386 | 0.20976751 | 0.83384914 | 0.91224528 |
| SUCLA2 | 1168.27247 | 0.392119391 | 0.34204774 | 1.14638792 | 0.25163468 | 0.44805879 |
| NUDT15 | 54.5311047 | -0.544367598 | 0.37184574 | -1.4639608 | 0.14320465 | 0.31083232 |
| MED4 | 616.484561 | -0.141743667 | 0.22169532 | -0.6393625 | 0.52258716 | 0.69847426 |
| ITM2B | 4987.08996 | -0.234512825 | 0.33513128 | -0.6997641 | 0.48407466 | 0.66799672 |
| RB1 | 2189.61817 | -0.347908591 | 0.17798347 | -1.9547242 | 0.05061563 | 0.15670481 |
| LPAR6 | 339.460564 | 0.167866545 | 0.625135 | 0.26852847 | 0.78829256 | 0.88421655 |
| RCBTB2 | 398.749753 | 0.229621248 | 0.28634914 | 0.80189257 | 0.4226151 | 0.61619214 |
| FNDC3A | 3941.24123 | 0.355247937 | 0.19393775 | 1.83176271 | 0.06698678 | 0.18842352 |
| CDADC1 | 406.243944 | 1.388412221 | 0.29207941 | 4.75354359 | 2.00E-06 | 6.80E-05 |
| CAB39L | 289.818907 | -0.196043338 | 0.39389442 | -0.4977053 | 0.61869177 | 0.77331444 |
| SETDB2 | 614.966146 | 0.217779104 | 0.25216047 | 0.86365284 | 0.38777863 | 0.58384025 |
| PHF11 | 357.585271 | -0.644004348 | 0.36736508 | -1.7530364 | 0.07959576 | 0.21127403 |
| RCBTB1 | 341.333991 | 0.301098188 | 0.29127877 | 1.0337114 | 0.3012711 | 0.50176464 |
| EBPL | 188.928314 | 0.008271256 | 0.34438127 | 0.02401773 | 0.98083847 | 0.99047407 |
| KPNA3 | 1048.84084 | -0.048631723 | 0.26761745 | -0.181721 | 0.85580165 | 0.92378521 |
| SPRYD7 | 256.844403 | 0.105760889 | 0.22597244 | 0.46802561 | 0.63976627 | 0.78846218 |
| DLEU2 | 3191.18444 | -0.087539875 | 0.32584666 | -0.2686536 | 0.78819627 | 0.88421655 |
| TRIM13 | 806.209138 | 0.440144621 | 0.39346699 | 1.11863164 | 0.26329732 | 0.46128738 |
| RNASEH2B | 247.768875 | -0.075580137 | 0.31076428 | -0.2432073 | 0.80784482 | 0.89588422 |
| FAM124A | 469.83333 | 0.424688131 | 0.3581096 | 1.18591664 | 0.23565522 | 0.42995285 |
| INTS6 | 1496.56913 | 0.298972011 | 0.23732263 | 1.25977033 | 0.20775223 | 0.39683107 |
| INTS6-AS1 | 176.879751 | 0.048114577 | 0.41855309 | 0.11495454 | 0.90848116 | 0.95197217 |
| WDFY2 | 1050.94593 | 0.093242589 | 0.15196264 | 0.61358889 | 0.53948702 | 0.71103352 |
| ATP7B | 358.081313 | 0.382157899 | 0.22551471 | 1.69460298 | 0.09015076 | 0.2294701 |
| ALG11 | 134.162073 | 0.302670734 | 0.26215611 | 1.1545439 | 0.24827725 | 0.44456385 |
| UTP14C | 159.491243 | -0.100384614 | 0.32237434 | -0.3113915 | 0.75550305 | 0.86293197 |
| NEK5 | 135.006171 | 0.368235835 | 0.58117672 | 0.6336039 | 0.52633935 | 0.7009757 |
| NEK3 | 291.606729 | -0.492774818 | 0.4556449 | -1.0814887 | 0.27947978 | 0.47799091 |
| VPS36 | 765.32849 | -0.190925209 | 0.12735217 | -1.4991909 | 0.13382411 | 0.29777495 |
| CKAP2 | 104.190553 | 0.629454729 | 0.29191118 | 2.15632279 | 0.03105847 | 0.11191336 |
| SUGT1 | 560.196761 | 0.015073498 | 0.13745469 | 0.10966158 | 0.91267777 | 0.95360549 |
| DIAPH3 | 297.026774 | 0.08379853 | 0.42537556 | 0.19699893 | 0.84382838 | 0.91781904 |
| TDRD3 | 1503.3097 | 0.394194145 | 0.21331106 | 1.84797797 | 0.06460553 | 0.18410561 |
| PCDH9 | 200.230754 | 1.399192547 | 0.47317085 | 2.95705564 | 0.00310592 | 0.02115947 |
| KLHL1 | 55.9092068 | 2.147673755 | 0.48663992 | 4.41327077 | 1.02E-05 | 0.00025798 |
| DACH1 | 1659.55399 | 3.073272211 | 0.53702664 | 5.72275558 | 1.05E-08 | 8.40E-07 |
| MZT1 | 92.4399324 | -0.322297778 | 0.31834787 | -1.0124075 | 0.31134325 | 0.51145516 |
| BORA | 69.5413704 | -0.448832099 | 0.27302427 | -1.6439275 | 0.10019119 | 0.24615892 |
| DIS3 | 415.489116 | -0.289448045 | 0.27128898 | -1.0669363 | 0.28600062 | 0.48564716 |
| PIBF1 | 2369.57916 | 0.045898481 | 0.18424246 | 0.24912 | 0.80326796 | 0.89307496 |
| KLF5 | 228.968202 | -2.148024121 | 0.45437087 | -4.7274688 | 2.27E-06 | 7.64E-05 |
| KLF12 | 2137.02961 | 1.138995549 | 0.25209666 | 4.5180906 | 6.24E-06 | 0.00017141 |
| TBC1D4 | 2324.54956 | 0.917224819 | 0.41644807 | 2.20249505 | 0.02763036 | 0.10333679 |
| COMMD6 | 328.4879 | -0.532343958 | 0.37803745 | -1.4081778 | 0.15907842 | 0.33408049 |
| UCHL3 | 503.641154 | -0.987743816 | 0.38754607 | -2.5487133 | 0.01081212 | 0.05329078 |
| LMO7-AS1 | 63.3118801 | -0.809848498 | 0.48826196 | -1.6586352 | 0.09718931 | 0.24111745 |
| LMO7 | 1546.59699 | 0.918756915 | 0.38879964 | 2.36306012 | 0.01812473 | 0.07698657 |
| KCTD12 | 79.6657499 | -0.239911621 | 0.63035009 | -0.3806006 | 0.70349964 | 0.82977967 |
| AC001226.2 | 177.6733 | -0.643104263 | 0.54020845 | -1.1904743 | 0.23386004 | 0.42803038 |
| FBXL3 | 504.406605 | 0.068352615 | 0.19898977 | 0.34349814 | 0.73122374 | 0.84717131 |
| MYCBP2 | 2573.22122 | 0.152766165 | 0.14572081 | 1.04834829 | 0.29447817 | 0.49435545 |
| SLAIN1 | 271.022462 | 0.073204383 | 0.33966011 | 0.21552246 | 0.82936 | 0.90944441 |
| RNF219 | 277.686507 | -0.472344477 | 0.32331979 | -1.4609204 | 0.14403729 | 0.31175101 |
| RBM26 | 2364.37057 | 0.33021781 | 0.1610435 | 2.05048826 | 0.04031681 | 0.1343273 |
| RBM26-AS1 | 69.1580614 | 0.705580493 | 0.45483256 | 1.55129725 | 0.12083047 | 0.27840363 |
| NDFIP2 | 1673.85357 | -0.107758686 | 0.24723562 | -0.4358542 | 0.66294251 | 0.80379726 |
| SPRY2 | 102.435824 | -0.367329056 | 0.36646847 | -1.0023483 | 0.31617539 | 0.51617629 |
| MIR4500HG | 103.548971 | 1.70769947 | 0.89715145 | 1.90346844 | 0.05697945 | 0.16916121 |
| LINC01040 | 90.0927886 | 0.331657513 | 0.89645401 | 0.36996601 | 0.71140782 | 0.83502592 |
| LINC00410 | 28.3215836 | 2.395723272 | 0.95646819 | 2.50476 | 0.01225345 | 0.05806858 |
| GPC5 | 242.2481 | 1.53873437 | 0.66032988 | 2.330251 | 0.01979289 | 0.08184366 |
| GPC6 | 6341.02923 | 0.643969604 | 0.43906788 | 1.46667436 | 0.14246466 | 0.30981487 |
| TGDS | 256.842877 | -0.191039803 | 0.22042533 | -0.8666871 | 0.38611345 | 0.58235663 |
| GPR180 | 306.115857 | -0.836934671 | 0.33383901 | -2.5070008 | 0.01217604 | 0.05782987 |
| ABCC4 | 5392.9986 | 0.034196412 | 0.2858234 | 0.11964175 | 0.90476695 | 0.95028792 |
| CLDN10 | 11288.8556 | 0.816460954 | 0.72055294 | 1.13310336 | 0.25717085 | 0.45431639 |
| CLDN10-AS1 | 1632.29892 | -0.14226223 | 0.64467173 | -0.2206739 | 0.82534635 | 0.90642548 |
| DZIP1 | 1159.56658 | 0.799921558 | 0.46225085 | 1.73049235 | 0.08354235 | 0.21827952 |
| DNAJC3-DT | 188.877654 | -0.713919063 | 0.4657063 | -1.5329813 | 0.12528046 | 0.2853566 |
| DNAJC3 | 1595.22686 | -0.580002608 | 0.39926067 | -1.4526916 | 0.14630941 | 0.31522526 |
| UGGT2 | 2304.26964 | -0.353177149 | 0.26924326 | -1.31174 | 0.18960789 | 0.3737353 |
| MBNL2 | 3395.24846 | -0.070182839 | 0.48473575 | -0.1447858 | 0.88488002 | 0.94030936 |
| RAP2A | 276.741055 | 0.114015945 | 0.19000136 | 0.60007963 | 0.54845317 | 0.71856697 |
| IPO5 | 514.085484 | -0.310798992 | 0.27037444 | -1.1495132 | 0.25034442 | 0.44663166 |

| | | | | | | |
|---|---|---|---|---|---|---|
| FARP1 | 12883.1384 | 0.85727322 | 0.32388379 | 2.64685438 | 0.00812443 | 0.04343276 |
| STK24 | 1925.94805 | -0.293561939 | 0.22064803 | -1.3304535 | 0.1833689 | 0.3657505 |
| SLC15A1 | 1428.16852 | -1.87976942 | 0.66138534 | -2.8421698 | 0.00448076 | 0.02791011 |
| DOCK9 | 1080.10237 | -0.364686131 | 0.29716289 | -1.2272264 | 0.21973753 | 0.41140966 |
| UBAC2 | 2543.5212 | 0.091557431 | 0.18494091 | 0.49506316 | 0.62055554 | 0.77455476 |
| TM9SF2 | 1205.48428 | -0.447907805 | 0.16863677 | -2.6560506 | 0.00790617 | 0.04251897 |
| CLYBL | 2490.95906 | 0.3893964 | 0.39137504 | 0.99494439 | 0.31976331 | 0.52029864 |
| AL355338.1 | 137.340094 | -0.301242474 | 0.4172857 | -0.7219094 | 0.47035018 | 0.65719571 |
| PCCA | 9011.80891 | -0.024270473 | 0.21204816 | -0.1144574 | 0.90887525 | 0.95219603 |
| PCCA-AS1 | 70.1688338 | -0.581197684 | 0.62597956 | -0.9284611 | 0.35316842 | 0.55189396 |
| GGACT | 407.688848 | 0.690697624 | 0.3835178 | 1.80095326 | 0.07171025 | 0.19733635 |
| TMTC4 | 451.327764 | 0.584681304 | 0.32206996 | 1.815386 | 0.06946461 | 0.19305585 |
| FGF14 | 616.664624 | 0.855260693 | 0.65079833 | 1.31417161 | 0.18878846 | 0.37272032 |
| TPP2 | 909.304686 | -0.014885211 | 0.13850065 | -0.1074739 | 0.91441299 | 0.9543495 |
| TEX30 | 180.662249 | 0.003754007 | 0.29093997 | 0.01290303 | 0.98970516 | 0.99493132 |
| KDELC1 | 55.139045 | 0.346470614 | 0.35820793 | 0.96723323 | 0.33342745 | 0.53275938 |
| SLC10A2 | 75.2829662 | 1.418788157 | 0.68894811 | 2.05935415 | 0.03946032 | 0.13234222 |
| ARGLU1 | 1322.71018 | -0.339489092 | 0.19473045 | -1.7433796 | 0.08126733 | 0.21441175 |
| FAM155A | 97.6784045 | -1.094741221 | 0.62117049 | -1.7623845 | 0.07800436 | 0.20850347 |
| LIG4 | 188.080242 | 0.10896126 | 0.22459694 | 0.48514134 | 0.62757609 | 0.77977555 |
| ABHD13 | 254.466256 | 0.007499885 | 0.22907745 | 0.03273952 | 0.97388231 | 0.98684149 |
| TNFSF13B | 300.640868 | 0.242798106 | 0.38239097 | 0.63494727 | 0.5254628 | 0.70084961 |
| MYO16 | 58.5289713 | -1.262655958 | 0.67482961 | -1.8710737 | 0.06133486 | 0.17740387 |
| IRS2 | 368.960032 | -0.4948894 | 0.52345181 | -0.9454345 | 0.3444371 | 0.54504726 |
| COL4A1 | 3123.45513 | -0.549095828 | 0.33973752 | -1.6162354 | 0.10604342 | 0.25536492 |
| COL4A2 | 4051.24932 | -0.117551281 | 0.26367792 | -0.4458139 | 0.65573168 | 0.79908554 |
| RAB20 | 938.035455 | 0.263387582 | 0.38588333 | 0.68255755 | 0.49488647 | 0.67697759 |
| NAXD | 325.81016 | -0.024902421 | 0.1409335 | -0.1766963 | 0.85974699 | 0.92628466 |
| CARS2 | 720.438474 | -0.159894991 | 0.30879509 | -0.5178029 | 0.6045958 | 0.76249535 |
| ING1 | 79.3995103 | -0.491876864 | 0.21240416 | -2.3157591 | 0.02057143 | 0.08417692 |
| ANKRD10 | 2551.5775 | -0.281416453 | 0.21078526 | -1.335086 | 0.1818482 | 0.3639885 |
| ARHGEF7 | 2433.00383 | -0.043378965 | 0.24052785 | -0.180349 | 0.85687856 | 0.92453042 |
| TUBGCP3 | 1114.18752 | 0.431845444 | 0.38909492 | 1.10987171 | 0.26705431 | 0.46506333 |
| ATP11AUN | 64.0039897 | 3.769687073 | 0.48141069 | 7.83050136 | 4.86E-15 | 2.56E-12 |
| ATP11A | 3647.7655 | 0.038871403 | 0.22634168 | 0.17173772 | 0.86364373 | 0.92861348 |
| MCF2L | 1945.21726 | 1.261864563 | 0.43826049 | 2.87925697 | 0.00398613 | 0.0255783 |
| F10 | 34.3590305 | -0.080080069 | 0.52768717 | -0.1517567 | 0.87937883 | 0.93715173 |
| PROZ | 161.487937 | 3.331623241 | 0.60784048 | 5.48108156 | 4.23E-08 | 2.75E-06 |
| PCID2 | 484.748029 | -0.190121955 | 0.18821504 | -1.0101316 | 0.31243225 | 0.51269111 |
| CUL4A | 1483.20443 | -0.480202654 | 0.31824789 | -1.508895 | 0.13132562 | 0.29441941 |
| LAMP1 | 1088.72926 | -0.104414724 | 0.24638855 | -0.4237807 | 0.67172572 | 0.80972162 |
| GRTP1 | 1084.93558 | 0.575631682 | 0.37504381 | 1.53483852 | 0.1248235 | 0.2847781 |
| GRTP1-AS1 | 73.3305293 | -0.06884798 | 0.5942569 | -0.1158556 | 0.907767 | 0.95171808 |
| ADPRHL1 | 46.4843981 | -0.252179399 | 0.69735844 | -0.3616209 | 0.71763533 | 0.8388017 |
| DCUN1D2 | 406.724011 | -0.055364231 | 0.26914892 | -0.2057011 | 0.83702439 | 0.91404504 |
| TMCO3 | 662.881647 | -1.513175858 | 0.31400294 | -4.8189863 | 1.44E-06 | 5.18E-05 |
| TFDP1 | 334.872054 | -0.314911225 | 0.42749448 | -0.7366439 | 0.4613389 | 0.6497711 |
| GAS6 | 67.8948824 | -0.831320859 | 0.62797661 | -1.3238086 | 0.18556661 | 0.36868553 |
| RASA3 | 149.941221 | -3.605784443 | 0.68215431 | -5.285878 | 1.25E-07 | 6.65E-06 |
| CDC16 | 690.379508 | -0.182434218 | 0.1885509 | -0.9675595 | 0.33326441 | 0.53275938 |
| UPF3A | 764.177274 | -0.075930453 | 0.30780393 | -0.2466845 | 0.80515242 | 0.89434556 |
| CHAMP1 | 170.036302 | 0.114745212 | 0.28102407 | 0.40831098 | 0.68304538 | 0.81587484 |
| TTC5 | 207.278474 | 0.476774884 | 0.26425206 | 1.80424281 | 0.07119325 | 0.19638678 |
| CCNB1IP1 | 333.776216 | 0.231313128 | 0.24540945 | 0.94256 | 0.34590601 | 0.54629954 |
| PARP2 | 73.9775757 | 0.350863386 | 0.30725708 | 1.14192125 | 0.25348675 | 0.45036201 |
| TEP1 | 377.336284 | -0.094561831 | 0.31965553 | -0.2958242 | 0.76736436 | 0.87015054 |
| OSGEP | 210.009451 | 0.218812578 | 0.41747733 | 0.52413044 | 0.60018782 | 0.75997916 |
| APEX1 | 237.344068 | -0.336010821 | 0.36487851 | -0.9208841 | 0.35711094 | 0.55586188 |
| PIP4P1 | 128.066036 | -0.045215066 | 0.32423834 | -0.1394501 | 0.88909449 | 0.94267382 |
| PNP | 329.021051 | 0.061146603 | 0.37810219 | 0.16171978 | 0.87152653 | 0.93353404 |
| METTL17 | 310.733048 | 0.373900688 | 0.23487031 | 1.59194532 | 0.11139699 | 0.26403572 |
| NDRG2 | 121.616686 | 0.064607626 | 0.4441448 | 0.14546523 | 0.88434357 | 0.94002757 |
| ARHGEF40 | 208.00028 | -0.663566138 | 0.35794346 | -1.8538295 | 0.06376356 | 0.18238879 |
| ZNF219 | 99.8353616 | -0.389718916 | 0.41384141 | -0.9417108 | 0.34634073 | 0.54664509 |
| HNRNPC | 5430.35001 | -0.075705879 | 0.23944437 | -0.3161731 | 0.75187109 | 0.8605113 |
| SUPT16H | 548.368894 | -0.210412206 | 0.253235 | -0.830897 | 0.40603181 | 0.60009523 |
| CHD8 | 992.599314 | 0.05066652 | 0.24178347 | 0.20955328 | 0.83401635 | 0.91224528 |
| RAB2B | 332.61511 | 0.071440375 | 0.27106139 | 0.26355792 | 0.79212059 | 0.88678699 |
| TOX4 | 463.549305 | -0.412609674 | 0.19683996 | -2.0961682 | 0.03606727 | 0.12412153 |
| METTL3 | 210.634484 | -0.324508508 | 0.2588517 | -1.2536464 | 0.20997055 | 0.39950948 |
| DAD1 | 687.245464 | -0.960690473 | 0.36408218 | -2.6386638 | 0.00832335 | 0.04416885 |
| ABHD4 | 156.83824 | -1.113124684 | 0.33840991 | -3.2892793 | 0.00100444 | 0.00906379 |
| OXA1L | 352.898687 | 0.199195953 | 0.21763324 | 0.91528276 | 0.36004321 | 0.55846014 |
| SLC7A7 | 1355.93171 | 2.156992733 | 0.4067549 | 5.30292991 | 1.14E-07 | 6.20E-06 |
| MRPL52 | 244.681154 | -0.330106264 | 0.27759422 | -1.1891684 | 0.23437342 | 0.42868381 |
| MMP14 | 233.07767 | -1.227133919 | 0.46318991 | -2.6493106 | 0.00806562 | 0.04320259 |
| LRP10 | 583.938383 | -0.980242837 | 0.3561614 | -2.7522433 | 0.00591885 | 0.03434789 |
| RBM23 | 569.664691 | 0.242758841 | 0.21924281 | 1.1072602 | 0.26818145 | 0.46607875 |
| PRMT5 | 118.219893 | -0.315158968 | 0.31219048 | -1.0095086 | 0.31273079 | 0.51305816 |
| AL132780.1 | 96.1052386 | 0.144211317 | 0.61427857 | 0.23476534 | 0.81439087 | 0.89947803 |
| AL132780.3 | 129.772408 | 1.051785499 | 0.52055953 | 2.02049032 | 0.04333255 | 0.14091561 |
| AL132780.2 | 99.9583118 | 1.445662303 | 0.33934322 | 4.26017734 | 2.04E-05 | 0.00045159 |
| C14orf93 | 89.1810981 | 0.401323896 | 0.27914048 | 1.437713 | 0.1505155 | 0.32145778 |
| PSMB5 | 417.072361 | -1.009761157 | 0.2855576 | -3.5361032 | 0.00040608 | 0.00463454 |
| ACIN1 | 1351.08674 | -0.138741945 | 0.18582787 | -0.7466154 | 0.45529576 | 0.64465572 |
| C14orf119 | 200.247095 | -0.868532355 | 0.27724279 | -3.13275 | 0.00173177 | 0.01372295 |
| SLC7A8 | 2595.46987 | 2.962454423 | 0.48977396 | 6.04861567 | 1.46E-09 | 1.61E-07 |
| RNF212B | 2093.50705 | 5.518206441 | 0.7498258 | 7.35931792 | 1.85E-13 | 6.03E-11 |
| HOMEZ | 165.67121 | 0.559146126 | 0.24461412 | 2.2858293 | 0.02226425 | 0.08902451 |
| PPP1R3E | 185.518086 | 0.120367031 | 0.43982526 | 0.27367012 | 0.78433814 | 0.88165679 |
| BCL2L2 | 50.0676902 | 0.129324477 | 0.58701497 | 0.22030865 | 0.82563078 | 0.90652176 |
| BCL2L2-PABI | 360.787456 | 0.002871345 | 0.24201851 | 0.01186416 | 0.99053399 | 0.99518064 |
| PABPN1 | 831.579347 | 1.130347055 | 0.31652699 | 3.57109212 | 0.0003555 | 0.00418984 |
| SLC22A17 | 137.120599 | -0.453237414 | 0.52311991 | -0.8664121 | 0.38626422 | 0.58239101 |
| NGDN | 232.294179 | -0.196529808 | 0.28161331 | -0.6978712 | 0.48525778 | 0.66895588 |
| ZFHX2 | 181.390348 | 0.146996144 | 0.49692245 | 0.29581305 | 0.76737286 | 0.87015054 |
| THTPA | 42.7871746 | 0.410263892 | 0.40193193 | 1.02072979 | 0.30738248 | 0.50760144 |
| AP1G2 | 265.849926 | -0.48016234 | 0.34074896 | -1.4091381 | 0.15879433 | 0.33362875 |
| AL135999.1 | 102.113093 | 0.982906291 | 0.34921903 | 2.81458401 | 0.00488404 | 0.02971396 |
| LINC00596 | 81.5831007 | 2.294122518 | 0.78376146 | 2.92706726 | 0.00342175 | 0.02280112 |
| DHRS4-AS1 | 377.962957 | 1.200571101 | 0.44924552 | 2.67241643 | 0.00753071 | 0.04107354 |
| DHRS4 | 198.697137 | 1.942899139 | 0.48815698 | 3.98007038 | 6.89E-05 | 0.00116013 |
| DHRS4L2 | 294.400022 | 0.788437429 | 0.51245892 | 1.53853782 | 0.12391717 | 0.28341943 |
| NRL | 144.156346 | 1.075218202 | 0.48021633 | 2.23902884 | 0.02515404 | 0.09678637 |
| PCK2 | 358.174736 | 1.865182482 | 0.33144157 | 5.62748498 | 1.83E-08 | 1.38E-06 |
| DCAF11 | 65.4323766 | 0.07891084 | 0.42863506 | 0.18409796 | 0.85393661 | 0.92302729 |
| PSME1 | 316.382608 | -1.121158592 | 0.44542397 | -2.5170594 | 0.01183389 | 0.05669594 |
| EMC9 | 76.5836556 | -0.33143539 | 0.45180144 | -0.7335864 | 0.46320085 | 0.65132274 |
| PSME2 | 87.9528354 | -0.172211705 | 0.46996938 | -0.3664318 | 0.71404293 | 0.83638578 |
| REC8 | 127.874352 | 0.893906951 | 0.64079149 | 1.3950044 | 0.16301451 | 0.33943715 |
| IPO4 | 106.494386 | -1.07725108 | 0.50780253 | -2.1213976 | 0.03388836 | 0.11919623 |
| AL136295.1 | 240.41058 | 0.03526536 | 0.19787643 | 0.17821911 | 0.85855091 | 0.92564577 |
| GMPR2 | 119.262835 | -0.236082357 | 0.32008102 | -0.7375706 | 0.46077541 | 0.64951136 |

| | | | | | | |
|---|---|---|---|---|---|---|
| TINF2 | 81.4910318 | -0.643832009 | 0.3357842 | -1.9173982 | 0.05518737 | 0.16563461 |
| RABGGTA | 51.1447354 | 0.112650425 | 0.44864329 | 0.2510913 | 0.80174353 | 0.89234824 |
| DHRS1 | 238.128261 | -0.163965075 | 0.2118212 | -0.774073 | 0.43888762 | 0.63061599 |
| CIDEB | 124.709464 | -0.483689655 | 0.50009458 | -0.9671964 | 0.33344588 | 0.53275938 |
| LTB4R | 176.314995 | -0.457252116 | 0.5142784 | -0.889114 | 0.37394183 | 0.57122708 |
| KHNYN | 189.33842 | -0.588226704 | 0.20318635 | -2.8950109 | 0.00379145 | 0.02457185 |
| SDR39U1 | 201.30915 | 0.164563719 | 0.30162184 | 0.54559616 | 0.58534357 | 0.74795416 |
| STXBP6 | 209.585735 | 0.679858797 | 0.41678356 | 1.63120349 | 0.10284739 | 0.25038257 |
| NOVA1 | 1252.92669 | -0.544882032 | 0.53219339 | -1.0238422 | 0.30590983 | 0.50583258 |
| LINC02588 | 61.4472629 | 0.596397635 | 0.55705866 | 1.07061909 | 0.28434074 | 0.48367566 |
| LINC02294 | 471.842844 | 1.946045928 | 0.78860364 | 2.46771106 | 0.013598 | 0.06258826 |
| AL110292.1 | 3251.57548 | 1.456982509 | 0.80748399 | 1.80434848 | 0.07117669 | 0.19638678 |
| PRKD1 | 9092.21027 | -1.400468961 | 0.46709182 | -2.9982734 | 0.00271514 | 0.01925487 |
| G2E3-AS1 | 43.8712023 | -0.348415024 | 0.44144811 | -0.7892548 | 0.42996312 | 0.62265243 |
| G2E3 | 344.332287 | 0.023052086 | 0.16801401 | 0.13720336 | 0.89087006 | 0.94331061 |
| SCFD1 | 1557.11344 | -0.402974926 | 0.16417853 | -2.4544923 | 0.01410837 | 0.06416172 |
| COCH | 181.370823 | -0.259393998 | 0.2974753 | -0.871985 | 0.38321656 | 0.57983766 |
| AL049830.3 | 41.1087088 | 0.032900904 | 0.54240527 | 0.06065742 | 0.95163204 | 0.97544838 |
| STRN3 | 5256.86035 | 0.118420888 | 0.20421799 | 0.57987489 | 0.56199899 | 0.73014439 |
| AP4S1 | 192.407526 | 0.713005113 | 0.40883385 | 1.74399728 | 0.08115956 | 0.21418062 |
| HECTD1 | 2515.23968 | -0.145223142 | 0.29853304 | -0.4864558 | 0.62664401 | 0.77912423 |
| HEATR5A | 348.439144 | -0.040876608 | 0.23200357 | -0.1761896 | 0.86014502 | 0.92629345 |
| AL139353.1 | 112.703193 | 0.74003487 | 0.33422751 | 2.21416502 | 0.02681743 | 0.1012635 |
| NUBPL | 1420.06411 | 0.755622926 | 0.28651006 | 2.6373347 | 0.00835603 | 0.04429086 |
| ARHGAP5 | 3191.76356 | -0.55103742 | 0.29973694 | -1.8384035 | 0.06600298 | 0.18656221 |
| AKAP6 | 1018.89582 | 0.611933378 | 0.3996504 | 1.53117167 | 0.12572697 | 0.28603309 |
| NPAS3 | 907.835409 | -0.49563845 | 0.5445877 | -0.9101169 | 0.36276088 | 0.56045575 |
| EGLN3 | 405.434803 | -1.79941711 | 0.43767482 | -4.1113106 | 3.93E-05 | 0.00074483 |
| EAPP | 430.776569 | 0.02055768 | 0.27839128 | 0.07384456 | 0.94113407 | 0.96898706 |
| SNX6 | 1162.14785 | -0.208644353 | 0.26485077 | -0.7877808 | 0.43082493 | 0.62344989 |
| CFL2 | 147.526505 | -0.433212418 | 0.31402791 | -1.3795348 | 0.16772993 | 0.34593352 |
| BAZ1A | 1187.10306 | -1.022778137 | 0.29380784 | -3.4811124 | 0.00049934 | 0.00534718 |
| SRP54 | 1298.84919 | -0.418885268 | 0.25043604 | -1.6726237 | 0.09440139 | 0.23672884 |
| FAM177A1 | 1336.65595 | 0.751762968 | 0.2503414 | 3.00295106 | 0.00267375 | 0.01901929 |
| PPP2R3C | 489.876772 | -0.134182912 | 0.21478818 | -0.6247221 | 0.53215349 | 0.70560925 |
| NFKBIA | 697.123642 | -1.399491709 | 0.37207124 | -3.7613541 | 0.000169 | 0.00232807 |
| RALGAPA1 | 2857.33217 | -0.096290051 | 0.17185 | -0.5603145 | 0.57526492 | 0.73976511 |
| BRMS1L | 208.736276 | 0.43437384 | 0.19769624 | 2.19717801 | 0.02800774 | 0.10415702 |
| PTCSC3 | 57.9881131 | 3.504767114 | 0.58616721 | 5.97912516 | 2.24E-09 | 2.31E-07 |
| MBIP | 365.726636 | -0.08433347 | 0.22604248 | -0.3730868 | 0.70908383 | 0.83335437 |
| SLC25A21 | 737.073284 | 1.171819263 | 0.45277106 | 2.58810551 | 0.00965054 | 0.04910906 |
| MIPOL1 | 1058.92817 | 0.533666716 | 0.30169564 | 1.76889102 | 0.07691206 | 0.20663516 |
| SEC23A | 1607.47663 | -0.113888733 | 0.20641377 | -0.5517497 | 0.58111987 | 0.74422219 |
| GEMIN2 | 111.032089 | 0.293904274 | 0.33850892 | 0.86823199 | 0.38526735 | 0.58152841 |
| TRAPPC6B | 524.447026 | 0.112843456 | 0.1656271 | 0.68131033 | 0.49567515 | 0.67766001 |
| PNN | 1028.41671 | -0.35606566 | 0.30710515 | -1.1594259 | 0.24628264 | 0.44197383 |
| MIA2 | 5706.79762 | 0.711148231 | 0.23242836 | 3.05964486 | 0.002216 | 0.01660517 |
| FBXO33 | 534.721739 | -0.015456783 | 0.15385724 | -0.1004619 | 0.91997767 | 0.95701657 |
| AL049875.1 | 212.321472 | 0.629850064 | 0.26146209 | 2.40895369 | 0.01599833 | 0.07053364 |
| LINC02307 | 134.211855 | 2.932878629 | 0.6678324 | 4.39163875 | 1.12E-05 | 0.00027834 |
| C14orf28 | 78.6317875 | -0.342966944 | 0.31234091 | -1.0980532 | 0.27218124 | 0.47033143 |
| KLHL28 | 636.08581 | -0.2474214 | 0.20837617 | -1.1873786 | 0.23507833 | 0.42961829 |
| TOGARAM1 | 1362.53499 | 0.408633363 | 0.18385443 | 2.22259186 | 0.02624333 | 0.09975523 |
| PRPF39 | 672.307812 | -0.115776127 | 0.24388305 | -0.4747199 | 0.63498666 | 0.78468826 |
| FKBP3 | 309.553176 | -0.106752391 | 0.27353669 | -0.3902672 | 0.69633899 | 0.82516803 |
| FANCM | 289.594772 | 0.590424979 | 0.35809443 | 1.64879688 | 0.09918925 | 0.24439818 |
| MIS18BP1 | 263.331711 | -0.705871839 | 0.2377969 | -2.9683811 | 0.00299373 | 0.02062593 |
| LINC02303 | 168.821145 | 1.610942006 | 0.78175527 | 2.06067304 | 0.03933424 | 0.1321563 |
| LINC00871 | 5447.27901 | 1.966385119 | 0.72210594 | 2.72312551 | 0.00646675 | 0.0366338 |
| MDGA2 | 47.3599462 | 0.37286505 | 0.39083106 | 0.95403126 | 0.34006782 | 0.54064276 |
| RPS29 | 691.094886 | -0.376901654 | 0.24889788 | -1.5142823 | 0.12995425 | 0.29229281 |
| LRR1 | 86.6829218 | -0.615069578 | 0.44824437 | -1.3721747 | 0.17000906 | 0.34901494 |
| RPL36AL | 777.510484 | -1.39954317 | 0.41659286 | -3.3594987 | 0.00078084 | 0.00753731 |
| MGAT2 | 68.0875399 | -0.997144544 | 0.47148164 | -2.114917 | 0.03443702 | 0.12041369 |
| DNAAF2 | 199.13324 | 0.027266172 | 0.17050018 | 0.15991873 | 0.8729451 | 0.9342905 |
| POLE2 | 140.421332 | -0.98321132 | 0.41897919 | -2.346683 | 0.01894136 | 0.0793488 |
| KLHDC1 | 395.849715 | 1.11581576 | 0.47026658 | 2.37273028 | 0.01765715 | 0.07554583 |
| KLHDC2 | 873.211632 | 0.578228112 | 0.36382011 | 1.58932424 | 0.11198721 | 0.26494075 |
| NEMF | 1792.31973 | 0.154143796 | 0.12380365 | 1.24506662 | 0.21310726 | 0.40310727 |
| AL627171.2 | 51.0429817 | -0.831893465 | 0.67513097 | -1.2321957 | 0.217876 | 0.4090601 |
| ARF6 | 254.25841 | -0.642360909 | 0.32624451 | -1.9689555 | 0.0489582 | 0.1533524 |
| LINC01588 | 174.135565 | -0.114483205 | 0.39240386 | -0.2917484 | 0.77047899 | 0.87241523 |
| VCPKMT | 244.947376 | 0.511952796 | 0.27127323 | 1.88722194 | 0.05913048 | 0.17307314 |
| SOS2 | 1714.69744 | 0.347702723 | 0.16814377 | 2.06788946 | 0.03865042 | 0.13073699 |
| L2HGDH | 1371.03635 | 1.315218336 | 0.32176646 | 4.08749352 | 4.36E-05 | 0.000811 |
| ATP5S | 316.451502 | 0.429716058 | 0.2003077 | 2.14527975 | 0.03193049 | 0.11418502 |
| AL359397.2 | 53.998829 | 1.631849299 | 0.34309241 | 4.75629671 | 1.97E-06 | 6.72E-05 |
| CDKL1 | 1199.48676 | 1.579614133 | 0.29255565 | 5.39936291 | 6.69E-08 | 3.99E-06 |
| MAP4K5 | 1356.7127 | -0.305836412 | 0.21858043 | -1.3991939 | 0.16175484 | 0.33798377 |
| ATL1 | 207.159376 | 1.510978699 | 0.42976952 | 3.51578839 | 0.00043845 | 0.00488605 |
| SAV1 | 439.185512 | -0.903854556 | 0.20656164 | -4.3757135 | 1.21E-05 | 0.00029679 |
| NIN | 506.129993 | 0.037799958 | 0.26783997 | 0.14112889 | 0.88776812 | 0.94209263 |
| PYGL | 194.461986 | -1.026576782 | 0.36778363 | -2.7912519 | 0.00525046 | 0.03141337 |
| TRIM9 | 148.617864 | -3.122753812 | 0.68063672 | -4.5879891 | 4.48E-06 | 0.00013108 |
| TMX1 | 261.051887 | -0.157601244 | 0.22540516 | -0.6991909 | 0.48443271 | 0.66827686 |
| FRMD6 | 78.0065243 | -0.702929018 | 0.51869703 | -1.3551823 | 0.17535949 | 0.35546474 |
| RTRAF | 284.832162 | -0.607420873 | 0.26751198 | -2.2706305 | 0.02316935 | 0.09147815 |
| NID2 | 173.7876 | -1.763348816 | 0.75270449 | -2.3426841 | 0.01914559 | 0.07991704 |
| TXNDC16 | 184.873279 | 0.714592385 | 0.29054496 | 2.45948985 | 0.01391346 | 0.06355699 |
| GPR137C | 53.9082437 | -0.058986291 | 0.54910935 | -0.1074218 | 0.91445439 | 0.9543495 |
| ERO1A | 1243.77386 | -2.680597704 | 0.45251808 | -5.9237362 | 3.15E-09 | 3.04E-07 |
| AL133453.1 | 103.248443 | -3.382165149 | 0.64758524 | -5.2227335 | 1.76E-07 | 8.79E-06 |
| PSMC6 | 741.936001 | -0.51259957 | 0.1903478 | -2.6929629 | 0.00708201 | 0.0393168 |
| STYX | 205.390457 | 0.26480227 | 0.20610452 | 1.28479602 | 0.19886357 | 0.38523653 |
| GNPNAT1 | 142.936804 | -0.66317953 | 0.36987174 | -1.7929986 | 0.07297314 | 0.19941046 |
| FERMT2 | 1834.69219 | 0.278525989 | 0.3037238 | 0.91703708 | 0.35912321 | 0.55757957 |
| DDHD1 | 517.745438 | -0.337747273 | 0.23727196 | -1.4234606 | 0.15460268 | 0.3271791 |
| CNIH1 | 244.908678 | -0.418597099 | 0.25954012 | -1.6128416 | 0.10677893 | 0.25646507 |
| GMFB | 365.407623 | -0.447992527 | 0.27985631 | -1.6007948 | 0.10942237 | 0.26077541 |
| CGRRF1 | 242.763736 | 0.303977656 | 0.27952712 | 1.08747106 | 0.27682868 | 0.47466734 |
| SAMD4A | 6067.86374 | -1.328082034 | 0.44742147 | -2.968302 | 0.0029945 | 0.02062593 |
| GCH1 | 710.230193 | 1.095977614 | 0.2423261 | 4.52273856 | 6.10E-06 | 0.00016802 |
| WDHD1 | 120.343568 | -0.556179695 | 0.49921747 | -1.114103 | 0.26523499 | 0.4630717 |
| SOCS4 | 285.959626 | -0.367231984 | 0.20723892 | -1.7720223 | 0.07639086 | 0.20555349 |
| MAPK1IP1L | 1199.19279 | -0.408653303 | 0.25532929 | -1.6004952 | 0.10948876 | 0.26077541 |
| LGALS3 | 210.435932 | -1.50099867 | 0.3907333 | -3.8414915 | 0.00012229 | 0.00182396 |
| FBXO34 | 2926.34247 | 0.686253399 | 0.25923102 | 2.64726571 | 0.00811456 | 0.04340196 |
| AL158801.3 | 156.258995 | 0.163235213 | 0.5204192 | 0.31366101 | 0.75377852 | 0.86193728 |
| ATG14 | 532.692257 | 0.262904647 | 0.26836394 | 0.97965713 | 0.3272554 | 0.52707579 |
| KTN1-AS1 | 123.23685 | 0.020206395 | 0.28252831 | 0.0715199 | 0.94298399 | 0.9703815 |
| KTN1 | 2754.06686 | -0.014034336 | 0.16885932 | -0.0831126 | 0.93376201 | 0.96473023 |
| AL355773.1 | 44.0193767 | -0.63521272 | 0.6057176 | -1.0486945 | 0.29431874 | 0.49425158 |
| PELI2 | 2468.00292 | 0.693914221 | 0.23795079 | 2.91620892 | 0.00354313 | 0.02344852 |

| | | | | | | |
|---|---|---|---|---|---|---|
| LINC02284 | 45.3038622 | 1.180534392 | 0.46051073 | 2.56353287 | 0.01036129 | 0.05183292 |
| TMEM260 | 768.73864 | 0.296058328 | 0.22362172 | 1.32392473 | 0.18552805 | 0.36868553 |
| EXOC5 | 1119.23124 | -0.055046387 | 0.14910238 | -0.3691852 | 0.71198972 | 0.83516404 |
| AP5M1 | 728.200452 | 0.287275861 | 0.22426709 | 1.28095418 | 0.20020976 | 0.38728128 |
| NAA30 | 318.306392 | -0.216488508 | 0.14249894 | -1.5192289 | 0.12870489 | 0.29046822 |
| CCDC198 | 3324.70208 | 0.904685662 | 0.44979733 | 2.01131844 | 0.04429183 | 0.14315212 |
| SLC35F4 | 68.5777868 | -1.135372848 | 0.57721409 | -1.9669874 | 0.04918466 | 0.15388591 |
| ARMH4 | 846.631333 | -0.031180738 | 0.24604735 | -0.1267266 | 0.89915681 | 0.94743919 |
| ACTR10 | 834.733768 | -0.307675592 | 0.17247134 | -1.7839231 | 0.07443617 | 0.20232093 |
| PSMA3 | 724.38466 | -0.416066646 | 0.28148729 | -1.478101 | 0.13938074 | 0.30562979 |
| ARID4A | 1239.27479 | 0.399136358 | 0.27120355 | 1.47172249 | 0.14109583 | 0.30805998 |
| AL139021.2 | 114.800934 | 1.860176977 | 0.54297676 | 3.42588694 | 0.0006128 | 0.00624969 |
| AL139021.1 | 184.387044 | 0.532524311 | 0.40784562 | 1.30570071 | 0.19165436 | 0.37593437 |
| TIMM9 | 301.68597 | -0.119011937 | 0.16343479 | -0.7281922 | 0.46649596 | 0.65434508 |
| KIAA0586 | 684.08039 | 0.258894602 | 0.26531838 | 0.97578841 | 0.32916933 | 0.52891681 |
| DAAM1 | 907.144722 | -0.558440268 | 0.32198827 | -1.7343497 | 0.08285605 | 0.21731876 |
| GPR135 | 873.367736 | 0.249544636 | 0.37431913 | 0.66666279 | 0.50498755 | 0.6850618 |
| L3HYPDH | 700.835878 | -0.083228278 | 0.32893804 | -0.2530211 | 0.80025188 | 0.89131054 |
| JKAMP | 473.64966 | 0.246324006 | 0.26283634 | 0.93717636 | 0.34866785 | 0.54865148 |
| RTN1 | 113.775595 | 1.260079531 | 0.65618779 | 1.92030324 | 0.05481961 | 0.16471117 |
| LRRC9 | 48.8740599 | -0.038351644 | 0.58986133 | -0.0650181 | 0.94815961 | 0.97344198 |
| AL157911.1 | 299.930943 | 0.811600217 | 0.36137789 | 2.24584913 | 0.02471367 | 0.095502 |
| PCNX4 | 2118.05137 | -0.491474738 | 0.33334904 | -1.4743547 | 0.14038611 | 0.30714643 |
| DHRS7 | 206.012855 | -0.867874725 | 0.24683505 | -3.516011 | 0.00043808 | 0.00488593 |
| AL157756.1 | 45.4740165 | 1.931051135 | 0.45053245 | 4.28615327 | 1.82E-05 | 0.00041309 |
| PPM1A | 1242.814 | 0.283769864 | 0.28742455 | 0.9872847 | 0.32350309 | 0.52401995 |
| MNAT1 | 2444.19632 | 0.360654318 | 0.22091922 | 1.6325167 | 0.10257068 | 0.25025567 |
| TRMT5 | 88.796547 | 0.363647213 | 0.23918395 | 1.52036629 | 0.12841894 | 0.29013325 |
| SLC38A6 | 758.131492 | 0.340132125 | 0.25725387 | 1.32216523 | 0.18611314 | 0.36923618 |
| PRKCH | 153.367006 | -0.116790572 | 0.44496752 | -0.2624699 | 0.7929592 | 0.8871958 |
| LINC01303 | 53.2036591 | -0.066721498 | 0.71995585 | -0.0926744 | 0.92616221 | 0.96090956 |
| HIF1A | 10042.6488 | -1.58888361 | 0.52498529 | -3.0265298 | 0.00247378 | 0.01803626 |
| HIF1A-AS2 | 1280.62121 | -2.713940657 | 0.47910484 | -5.6646071 | 1.47E-08 | 1.15E-06 |
| SNAPC1 | 555.118516 | -0.749550333 | 0.36407932 | -2.0587556 | 0.03951766 | 0.13241191 |
| PPP2R5E | 2826.39044 | -0.28985377 | 0.30859675 | -0.9392638 | 0.3475953 | 0.54753787 |
| WDR89 | 340.047003 | 0.824748555 | 0.19243452 | 4.28586599 | 1.82E-05 | 0.00041309 |
| SGPP1 | 1357.52949 | 0.972593327 | 0.27520972 | 3.53400788 | 0.00040931 | 0.00465701 |
| SYNE2 | 29226.1449 | 0.547194686 | 0.35218668 | 1.55370635 | 0.12025448 | 0.27766015 |
| ESR2 | 1316.73007 | -0.21883643 | 0.56902798 | -0.3845794 | 0.70054907 | 0.82796169 |
| MTHFD1 | 572.96885 | 0.691352275 | 0.2460593 | 2.80969775 | 0.0049588 | 0.03010201 |
| AL122035.1 | 141.502424 | -0.830990389 | 0.33271789 | -2.4975825 | 0.01250434 | 0.05897647 |
| ZBTB25 | 383.395526 | -0.49304483 | 0.26909437 | -1.8322376 | 0.06691603 | 0.18830179 |
| ZBTB1 | 776.03098 | -0.503034268 | 0.22286749 | -2.2571003 | 0.02400181 | 0.09359682 |
| AL049869.3 | 72.0055336 | -1.974048817 | 0.43640874 | -4.5233943 | 6.09E-06 | 0.00016784 |
| PLEKHG3 | 652.186392 | -0.410089243 | 0.36165199 | -1.1339333 | 0.25682252 | 0.45405215 |
| CHURC1-FNT | 230.23488 | -0.326842787 | 0.37268268 | -0.8770002 | 0.38048653 | 0.57691691 |
| GPX2 | 186.495828 | -2.20725982 | 1.54361717 | -1.429927 | 0.15273798 | 0.32453566 |
| RAB15 | 188.667959 | -0.689727228 | 0.54056054 | -1.2759482 | 0.20197384 | 0.38948445 |
| FNTB | 38.9044827 | -0.391680464 | 0.44689188 | -0.8764546 | 0.38078292 | 0.57723861 |
| MAX | 986.414366 | -0.92335314 | 0.24194043 | -3.8164483 | 0.00013539 | 0.00195547 |
| AL139022.1 | 56.2321659 | -1.750201325 | 0.68595113 | -2.5514956 | 0.01072617 | 0.05303881 |
| FUT8 | 2343.45 | -1.236155401 | 0.28138798 | -4.3930641 | 1.12E-05 | 0.00027762 |
| AL391261.2 | 45.9157524 | 1.419421353 | 0.73218891 | 1.9385999 | 0.05255008 | 0.16006753 |
| AL359232.1 | 1226.71971 | 1.324461875 | 0.50868141 | 2.60371592 | 0.00922192 | 0.04753847 |
| CCDC196 | 54.1377637 | -0.122150437 | 0.75370611 | -0.1620664 | 0.87125356 | 0.93338059 |
| GPHN | 11741.4727 | 1.281587294 | 0.27359924 | 4.68417713 | 2.81E-06 | 9.07E-05 |
| FAM71D | 35.0617373 | 1.350141451 | 0.33519873 | 4.02788358 | 5.63E-05 | 0.0009967 |
| MPP5 | 1644.48889 | 0.803315136 | 0.21014884 | 3.82260086 | 0.00013205 | 0.00191957 |
| ATP6V1D | 624.665944 | -0.556700613 | 0.36581859 | -1.5217942 | 0.12806065 | 0.2897536 |
| EIF2S1 | 400.157701 | -0.608119041 | 0.28915386 | -2.1030985 | 0.03545716 | 0.12281204 |
| PLEKHH1 | 498.609017 | -0.692345039 | 0.23021115 | -3.0074349 | 0.00263463 | 0.01884831 |
| PIGH | 191.984035 | -0.121238946 | 0.28640907 | -0.4233069 | 0.67207134 | 0.80998957 |
| ARG2 | 655.798049 | -2.078432126 | 0.71273248 | -2.9161462 | 0.00354384 | 0.02344852 |
| VTI1B | 238.859508 | 0.030525729 | 0.22223987 | 0.13735487 | 0.8907503 | 0.94331061 |
| RDH11 | 125.821189 | -0.221592099 | 0.30118857 | -0.7357255 | 0.46189779 | 0.65035778 |
| RDH12 | 73.0442937 | 2.954978076 | 0.52187408 | 5.66224337 | 1.49E-08 | 1.16E-06 |
| ZFYVE26 | 473.52775 | -0.164473397 | 0.25736311 | -0.6390714 | 0.5227765 | 0.69854722 |
| RAD51B | 2609.23428 | -0.145226951 | 0.49932893 | -0.2908443 | 0.77117044 | 0.87272401 |
| ZFP36L1 | 2148.17295 | -0.771294224 | 0.44515881 | -1.7326271 | 0.08316196 | 0.21787099 |
| ACTN1 | 525.63292 | -1.234658288 | 0.35936394 | -3.4356766 | 0.00059108 | 0.00605522 |
| DCAF5 | 1972.33075 | 0.191196945 | 0.20117895 | 0.95038244 | 0.34191796 | 0.5425642 |
| AL359317.2 | 120.517711 | -0.203368879 | 0.3932889 | -0.5170979 | 0.60508779 | 0.76287144 |
| EXD2 | 371.844786 | 0.2233729 | 0.16321919 | 1.36854555 | 0.17114138 | 0.3501246 |
| ERH | 290.521875 | -0.641109523 | 0.23227783 | -2.7600978 | 0.00577841 | 0.03368976 |
| SLC39A9 | 602.20981 | -0.070158206 | 0.19707855 | -0.3559911 | 0.72184724 | 0.84218273 |
| SUSD6 | 1269.48808 | -0.36865777 | 0.19196274 | -1.9204653 | 0.05479916 | 0.16468582 |
| SRSF5 | 1652.24204 | 0.491513104 | 0.2230691 | 2.20341186 | 0.02756573 | 0.10317955 |
| SYNJ2BP-CO | 31.2404581 | -0.293191767 | 0.40502658 | -0.7238828 | 0.46913771 | 0.65617047 |
| AL357153.2 | 409.928119 | -0.629185133 | 0.40995926 | -1.5347504 | 0.12484515 | 0.2847781 |
| MED6 | 319.532058 | -0.146290663 | 0.23308256 | -0.6276345 | 0.53024338 | 0.70431775 |
| TTC9 | 60.5532418 | -3.1907677 | 0.65259685 | -4.8893397 | 1.01E-06 | 3.92E-05 |
| MAP3K9 | 513.342952 | -0.641355744 | 0.33894737 | -1.8921986 | 0.05846452 | 0.17167378 |
| PCNX1 | 3131.11011 | -0.117444585 | 0.24628575 | -0.4768631 | 0.63345961 | 0.78395238 |
| AC005476.2 | 97.2388025 | 0.826486311 | 0.48872439 | 1.6911092 | 0.09081595 | 0.23086939 |
| SIPA1L1 | 7608.08673 | 0.471443673 | 0.27657397 | 1.70458442 | 0.08827196 | 0.22639124 |
| RGS6 | 153.140353 | 0.642667258 | 0.5595964 | 1.1484478 | 0.25078375 | 0.44706631 |
| DPF3 | 1563.16378 | 1.620347686 | 0.66047072 | 2.45332251 | 0.01415434 | 0.06428545 |
| DCAF4 | 201.739594 | -0.801663302 | 0.30320183 | -2.6439923 | 0.00819345 | 0.04369947 |
| ZFYVE1 | 796.716635 | 0.069758123 | 0.26774667 | 0.26053778 | 0.79444898 | 0.88811477 |
| RBM25 | 1393.54539 | 0.104597832 | 0.24718739 | 0.42315197 | 0.67218438 | 0.80998957 |
| PSEN1 | 1528.20456 | -0.276962284 | 0.15659763 | -1.7686238 | 0.07695668 | 0.20666074 |
| PAPLN | 1236.06082 | 0.246345105 | 0.42890783 | 0.57435441 | 0.565728 | 0.73251782 |
| AC004846.2 | 41.2512103 | 0.66538408 | 0.39419457 | 1.68795851 | 0.09141919 | 0.23179483 |
| NUMB | 6051.6674 | 0.488483562 | 0.20411555 | 2.39317177 | 0.01670342 | 0.07266389 |
| HEATR4 | 218.101562 | -1.190412014 | 0.53776625 | -2.2136235 | 0.02685469 | 0.10132045 |
| AC005225.3 | 54.6114977 | -0.021297856 | 0.40001944 | -0.0532421 | 0.95753905 | 0.9783815 |
| ACOT2 | 73.4521155 | 0.066445343 | 0.54109005 | 0.12279905 | 0.90226623 | 0.94863567 |
| ACOT4 | 63.0432451 | -0.262206855 | 0.56642429 | -0.462916 | 0.64342459 | 0.79112136 |
| ACOT6 | 94.5742963 | 1.229572861 | 0.68643378 | 1.79124761 | 0.07325357 | 0.20005712 |
| AC005225.2 | 131.067068 | 0.694701605 | 0.45842379 | 1.51541352 | 0.12966772 | 0.2918961 |
| DNAL1 | 445.468957 | 0.273801211 | 0.26140052 | 1.04743945 | 0.29489695 | 0.49487664 |
| PNMA1 | 104.445256 | -1.224975978 | 0.31581309 | -3.8788005 | 0.00010497 | 0.0016167 |
| ELMSAN1 | 636.90841 | -0.787891056 | 0.39418739 | -1.9987728 | 0.04563294 | 0.14631361 |
| AC005520.2 | 48.0215558 | -0.411230694 | 0.74040468 | -0.5554134 | 0.57861192 | 0.74232811 |
| LINC02274 | 357.865495 | -0.344068097 | 0.56240635 | -0.6117785 | 0.54068434 | 0.71213095 |
| AC005480.1 | 254.008457 | 0.095258067 | 0.3926827 | 0.24258279 | 0.80832861 | 0.89601696 |
| FAM161B | 48.6670087 | -0.109440014 | 0.33172557 | -0.3299113 | 0.74146698 | 0.85325429 |
| COQ6 | 161.626406 | 0.455480075 | 0.27119006 | 1.67956035 | 0.09304289 | 0.23469615 |
| ENTPD5 | 1709.58905 | 1.217927109 | 0.43208838 | 2.81869908 | 0.00482187 | 0.02950597 |
| BBOF1 | 659.742644 | 0.667922656 | 0.36184674 | 1.84587169 | 0.06491084 | 0.1846685 |
| ALDH6A1 | 1571.13859 | 1.630599132 | 0.41226149 | 3.95525451 | 7.65E-05 | 0.00126716 |
| LIN52 | 2570.91294 | 1.514651086 | 0.2941143 | 5.14987222 | 2.61E-07 | 1.22E-05 |
| ABCD4 | 998.862784 | 0.02578076 | 0.18912616 | 0.13631514 | 0.89157216 | 0.94341489 |

| | | | | | | |
|---|---|---|---|---|---|---|
| NPC2 | 545.776669 | -0.777238894 | 0.28959904 | -2.6838449 | 0.00727809 | 0.04011288 |
| ISCA2 | 90.7743966 | -0.838193875 | 0.33074273 | -2.5342776 | 0.01126794 | 0.05488618 |
| AREL1 | 490.681631 | 0.170527841 | 0.22108442 | 0.77132455 | 0.44051459 | 0.63246396 |
| FCF1 | 227.726628 | 0.076343176 | 0.22646237 | 0.33711197 | 0.7360325 | 0.8507199 |
| YLPM1 | 1031.71594 | 0.274791655 | 0.14067295 | 1.95340786 | 0.05077129 | 0.15698669 |
| DLST | 671.020328 | 0.016515815 | 0.26863757 | 0.06147992 | 0.95097701 | 0.97501996 |
| RPS6KL1 | 41.8961641 | -0.085088495 | 0.83856062 | -0.1014697 | 0.91917761 | 0.95680191 |
| EIF2B2 | 162.685684 | -0.31904112 | 0.24170445 | -1.3199638 | 0.1868471 | 0.37015656 |
| AL049780.1 | 45.9408692 | -0.032248383 | 0.35055417 | -0.0919926 | 0.92670393 | 0.96120551 |
| MLH3 | 1021.38473 | 0.38335408 | 0.24585506 | 1.55926863 | 0.11893281 | 0.27565304 |
| ACYP1 | 299.560271 | 0.237007717 | 0.3299711 | 0.7182681 | 0.47259199 | 0.6585826 |
| ZC2HC1C | 125.001353 | -0.102777031 | 0.55613801 | -0.1848049 | 0.85338206 | 0.92286287 |
| NEK9 | 688.187619 | 0.17921854 | 0.17294438 | 1.03627848 | 0.30007224 | 0.50031507 |
| AL049780.2 | 91.1903065 | 0.404453431 | 0.4157226 | 0.97289257 | 0.3306067 | 0.53044903 |
| TMED10 | 751.165061 | -0.408001815 | 0.25491359 | -1.6005495 | 0.10947674 | 0.26077541 |
| FOS | 100.334284 | 0.007200507 | 0.76325508 | 0.00943395 | 0.99247291 | 0.99626489 |
| JDP2 | 233.377594 | 2.796931365 | 0.50723405 | 5.51408437 | 3.51E-08 | 2.38E-06 |
| FLVCR2 | 648.084344 | 1.021472316 | 0.32358759 | 3.15671038 | 0.0015956 | 0.012949 |
| TTLL5 | 1341.62018 | -0.283627697 | 0.14577678 | -1.9456301 | 0.05169919 | 0.15856808 |
| ERG28 | 108.999483 | -0.297099079 | 0.34466533 | -0.8619929 | 0.38869141 | 0.58431296 |
| AF107885.2 | 60.8896849 | -0.838474463 | 0.55435953 | -1.5125102 | 0.13040411 | 0.29297692 |
| IFT43 | 1605.23844 | -0.365836495 | 0.27626855 | -1.3242061 | 0.18543461 | 0.36858356 |
| GPATCH2L | 2429.43883 | -0.884722415 | 0.29090579 | -3.0412678 | 0.00235584 | 0.01743603 |
| ESRRB | 192.412666 | 0.413727347 | 0.67657794 | 0.61149991 | 0.54086868 | 0.71216974 |
| VASH1-AS1 | 43.8082463 | 0.570765017 | 0.37232553 | 1.53297309 | 0.12528249 | 0.2853566 |
| ANGEL1 | 190.824078 | 0.803847475 | 0.27420332 | 2.9315746 | 0.00337248 | 0.02249471 |
| IRF2BPL | 166.42571 | -0.638694235 | 0.43124262 | -1.4810554 | 0.1385918 | 0.30467968 |
| POMT2 | 334.857313 | -0.526027404 | 0.36676537 | -1.4342341 | 0.15150547 | 0.32288044 |
| GSTZ1 | 99.0953161 | -0.44208005 | 0.25081566 | -1.7625696 | 0.07797312 | 0.20849257 |
| TMED8 | 621.388375 | -0.184198564 | 0.17080224 | -1.0784318 | 0.2808411 | 0.47944813 |
| SAMD15 | 52.8022632 | 0.150642273 | 0.39206687 | 0.38422597 | 0.70081097 | 0.82796169 |
| NOXRED1 | 146.730004 | 0.018804312 | 0.36746272 | 0.05117339 | 0.95918736 | 0.97911685 |
| VIPAS39 | 249.887032 | 0.046116352 | 0.28972408 | 0.15917335 | 0.8735323 | 0.9345535 |
| AHSA1 | 241.15774 | -0.559493011 | 0.33311252 | -1.6795917 | 0.09303679 | 0.23469615 |
| SPTLC2 | 1267.58198 | -0.307964959 | 0.23244072 | -1.3249183 | 0.18519826 | 0.36822056 |
| ALKBH1 | 372.168373 | 0.251834491 | 0.22128046 | 1.13807831 | 0.25508778 | 0.45226856 |
| SLIRP | 380.405137 | -0.391020667 | 0.28472502 | -1.3733274 | 0.16965059 | 0.34859523 |
| SNW1 | 441.279256 | -0.318222069 | 0.22221901 | -1.43202 | 0.15213809 | 0.32376289 |
| ADCK1 | 294.32934 | 0.521445635 | 0.35500674 | 1.46883305 | 0.14187808 | 0.30898043 |
| NRXN3 | 198.672606 | 1.010502826 | 0.33690501 | 2.99937012 | 0.00270538 | 0.01920389 |
| CEP128 | 393.367664 | 0.053734479 | 0.3591773 | 0.14960433 | 0.88107679 | 0.93815891 |
| TSHR | 68.1548141 | 0.306698439 | 0.59016539 | 0.51968219 | 0.6032851 | 0.76169806 |
| AC007262.2 | 339.485757 | 0.980604398 | 0.39262424 | 2.49756456 | 0.01250497 | 0.05897647 |
| GTF2A1 | 373.153183 | 0.328615181 | 0.29643036 | 1.10857466 | 0.26761372 | 0.46564381 |
| STON2 | 3030.62556 | 0.3058409 | 0.59358253 | 0.5152458 | 0.60638127 | 0.76359388 |
| SEL1L | 755.914576 | -0.470715227 | 0.27196324 | -1.7308046 | 0.08348662 | 0.21823333 |
| GALC | 174.765502 | 0.73679116 | 0.35171358 | 2.09486126 | 0.03618333 | 0.12442671 |
| KCNK10 | 361.614326 | 3.227944803 | 0.43071002 | 7.4944735 | 6.66E-14 | 2.61E-11 |
| SPATA7 | 826.735608 | 0.991685736 | 0.25714829 | 3.85647409 | 0.00011503 | 0.00173845 |
| PTPN21 | 543.228546 | 0.176590224 | 0.27143158 | 0.65058836 | 0.51531225 | 0.69315508 |
| ZC3H14 | 1279.83759 | 0.278408546 | 0.13583583 | 2.04959573 | 0.0404039 | 0.13438879 |
| EML5 | 526.244337 | -3.135521751 | 0.99075758 | -3.1647719 | 0.00155205 | 0.01269325 |
| TTC8 | 528.811151 | 0.159787366 | 0.28692936 | 0.55688747 | 0.57760432 | 0.74154889 |
| FOXN3 | 1601.44661 | 0.570008888 | 0.20178127 | 2.82488509 | 0.00472976 | 0.0290982 |
| EFCAB11 | 601.605352 | 0.786701501 | 0.45166333 | 1.7417874 | 0.08154565 | 0.21499255 |
| TDP1 | 315.947836 | -0.386627562 | 0.21166931 | -1.8265641 | 0.06776534 | 0.18979557 |
| KCNK13 | 73.7388753 | 1.354240902 | 0.50597779 | 2.6764829 | 0.00743994 | 0.04074279 |
| PSMC1 | 370.610555 | -0.429363543 | 0.31256092 | -1.3736956 | 0.16953622 | 0.34855751 |
| NRDE2 | 469.145312 | 0.335447306 | 0.20526561 | 1.63421096 | 0.10221456 | 0.24973128 |
| CALM1 | 1419.9246 | -0.021197076 | 0.25469881 | -0.0832241 | 0.93367336 | 0.96473023 |
| TTC7B | 2136.67736 | -0.130151761 | 0.32002057 | -0.4066981 | 0.68422973 | 0.81689199 |
| RPS6KA5 | 778.619632 | -0.969272952 | 0.21890421 | -4.4278406 | 9.52E-06 | 0.00024523 |
| DGLUCY | 1811.32719 | 0.53319917 | 0.35826273 | 1.48829091 | 0.13667419 | 0.30162504 |
| GPR68 | 31.6114291 | 0.687148791 | 0.68725678 | 0.99984288 | 0.31738655 | 0.51753717 |
| CCDC88C | 688.95688 | 0.138928333 | 0.22802621 | 0.60926476 | 0.54234896 | 0.71322817 |
| PPP4R3A | 1520.47188 | 0.394847346 | 0.1584795 | 2.4914727 | 0.01272147 | 0.05974577 |
| CATSPERB | 488.832303 | -0.10020644 | 0.39574198 | -0.2532115 | 0.80010474 | 0.89131054 |
| TC2N | 444.092142 | -0.634353908 | 0.34173432 | -1.856278 | 0.06341394 | 0.18169208 |
| FBLN5 | 314.083398 | 0.913757888 | 0.23706856 | 3.85440356 | 0.00011601 | 0.00174937 |
| TRIP11 | 1289.77301 | 0.397272818 | 0.12263508 | 3.23947118 | 0.00119752 | 0.01038375 |
| ATXN3 | 701.142179 | 0.072859817 | 0.23957487 | 0.30412128 | 0.7610355 | 0.8665094 |
| NDUFB1 | 394.393558 | 0.565348206 | 0.22562154 | 2.50573689 | 0.01221965 | 0.05798211 |
| CPSF2 | 460.257435 | 0.432481537 | 0.1951963 | 2.21562363 | 0.02671729 | 0.10108029 |
| RIN3 | 454.048846 | 0.655944637 | 0.41076482 | 1.59688613 | 0.11029109 | 0.26204888 |
| LGMN | 1462.15578 | 0.250643276 | 0.54502027 | 0.45987881 | 0.64560321 | 0.79225766 |
| GOLGA5 | 450.372568 | -0.223372935 | 0.2588667 | -0.8628879 | 0.38819911 | 0.58387159 |
| ITPK1 | 729.315547 | 0.294472716 | 0.31705847 | 0.92876469 | 0.35301105 | 0.55186159 |
| MOAP1 | 155.37637 | -1.238940616 | 0.3852789 | -3.215698 | 0.00130128 | 0.01101025 |
| AL110118.2 | 70.1538273 | -0.041339138 | 0.40701026 | -0.1015678 | 0.91909974 | 0.95679351 |
| GON7 | 78.9924916 | -0.375893713 | 0.24140073 | -1.5571358 | 0.11943825 | 0.27649723 |
| UBR7 | 85.9155215 | 0.017056358 | 0.32115083 | 0.05311011 | 0.95764417 | 0.97841597 |
| BTBD7 | 3609.81349 | 0.096878215 | 0.25717201 | 0.37670591 | 0.70639217 | 0.83197435 |
| UNC79 | 1127.53282 | 1.330999111 | 0.76159898 | 1.74763773 | 0.08052678 | 0.21296171 |
| DDX24 | 885.381101 | -0.410378547 | 0.20571166 | -1.9949212 | 0.04605148 | 0.1471736 |
| IFI27L1 | 108.786193 | -0.6325402 | 0.31902945 | -1.9827016 | 0.04740077 | 0.15022483 |
| IFI27 | 631.52743 | -1.394823733 | 1.34730279 | -1.0352712 | 0.30054229 | 0.50097024 |
| IFI27L2 | 60.1510806 | -1.292993788 | 0.65366542 | -1.9780667 | 0.04792118 | 0.15117507 |
| PPP4R4 | 76.9131948 | -1.875953707 | 0.69093414 | -2.7150977 | 0.00662562 | 0.03732733 |
| SERPINA6 | 51.1464132 | 0.75659091 | 0.71963049 | 1.05136028 | 0.29309315 | 0.49326436 |
| SERPINA1 | 3810.63588 | -2.538968977 | 0.81256976 | -3.1246166 | 0.00178037 | 0.01400321 |
| DICER1 | 1629.23956 | -0.124032285 | 0.15450942 | -0.802749 | 0.42211983 | 0.61572805 |
| DICER1-AS1 | 191.548718 | 0.668912069 | 0.44292436 | 1.51021738 | 0.13098797 | 0.29390277 |
| CLMN | 2994.16927 | 0.038533349 | 0.4032081 | 0.0955669 | 0.92386455 | 0.95964015 |
| SYNE3 | 50.0728879 | 1.614626235 | 0.86202961 | 1.87305194 | 0.06106121 | 0.17703317 |
| SNHG10 | 48.6657044 | 0.233466749 | 0.32064695 | 0.72811155 | 0.46654531 | 0.65434733 |
| GLRX5 | 245.172336 | -0.350652464 | 0.31438869 | -1.1153469 | 0.26470178 | 0.4624535 |
| ATG2B | 1066.65349 | 0.504286625 | 0.17616043 | 2.8626555 | 0.00420107 | 0.02657318 |
| GSKIP | 198.254012 | -0.636732409 | 0.24064759 | -2.6459122 | 0.00814709 | 0.04351997 |
| AK7 | 135.460372 | -0.512100987 | 0.5102515 | -1.0036246 | 0.31555957 | 0.51566226 |
| PAPOLA | 1580.41634 | -0.096919128 | 0.19802902 | -0.4894188 | 0.62454522 | 0.77746266 |
| VRK1 | 181.810955 | 0.241048177 | 0.30809392 | 0.78238537 | 0.43398813 | 0.62624226 |
| LINC01550 | 311.502059 | 1.182730303 | 0.60089771 | 1.96827229 | 0.04903672 | 0.15352166 |
| SETD3 | 3115.6951 | 0.54953836 | 0.28258092 | 1.94471145 | 0.05180972 | 0.1587295 |
| CCNK | 451.505785 | -0.511255153 | 0.19846194 | -2.5760866 | 0.00999256 | 0.05043006 |
| CCDC85C | 417.091827 | 0.121172217 | 0.30256576 | 0.40048225 | 0.68880135 | 0.82001633 |
| AL160313.1 | 47.2067684 | 0.398092877 | 0.56242137 | 0.70781962 | 0.47905728 | 0.66448474 |
| CYP46A1 | 111.024878 | 1.081515948 | 0.60484241 | 1.78809543 | 0.07376061 | 0.20100134 |
| EML1 | 207.295553 | -0.489577651 | 0.37814261 | -1.2946905 | 0.19542701 | 0.38077015 |
| EVL | 1490.97705 | -1.018369888 | 0.34421471 | -2.958531 | 0.00309109 | 0.02111703 |
| AL133523.1 | 78.0989979 | 0.764685486 | 0.23343491 | 3.27579745 | 0.00105364 | 0.00941089 |
| YY1 | 1225.30406 | -0.239215896 | 0.22449901 | -1.0655543 | 0.28662515 | 0.48629422 |
| SLC25A29 | 1062.51233 | -0.552972631 | 0.38152203 | -1.4493858 | 0.14722986 | 0.31671057 |
| WARS | 662.192578 | -1.112572042 | 0.53998398 | -2.0603797 | 0.03936226 | 0.13218382 |

| | | | | | | |
|---|---|---|---|---|---|---|
| AL157871.4 | 129.028215 | -1.181247212 | 0.6567655 | -1.7985829 | 0.07208468 | 0.19802023 |
| WDR25 | 520.015738 | 0.334174726 | 0.37057704 | 0.90176856 | 0.36717983 | 0.56454671 |
| MEG3 | 162.926399 | -0.876361179 | 0.96882248 | -0.9045632 | 0.36569683 | 0.5632142 |
| MEG8 | 117.522372 | -0.034861987 | 0.99242923 | -0.0351279 | 0.97197773 | 0.98607792 |
| PPP2R5C | 1444.9259 | -0.007141576 | 0.2138002 | -0.033403 | 0.97335319 | 0.98659707 |
| AL118558.3 | 33.3537952 | -0.023312673 | 0.56595275 | -0.0411919 | 0.96714291 | 0.98328281 |
| DYNC1H1 | 1517.48975 | -0.568311603 | 0.26249003 | -2.1650788 | 0.03038165 | 0.11018048 |
| HSP90AA1 | 3222.33418 | -0.825987276 | 0.38895269 | -2.1236189 | 0.03370202 | 0.11872363 |
| WDR20 | 1536.21266 | 0.092246324 | 0.22496382 | 0.41004959 | 0.68176957 | 0.81468059 |
| MOK | 1051.75746 | 0.999315883 | 0.29757978 | 3.35814449 | 0.00078468 | 0.00755836 |
| ZNF839 | 315.208437 | 0.907958578 | 0.25481659 | 3.56318469 | 0.00036638 | 0.00428499 |
| CINP | 129.410262 | -0.141204987 | 0.25586646 | -0.5518699 | 0.58103752 | 0.74422219 |
| TECPR2 | 633.569586 | 0.352403114 | 0.27562429 | 1.27856336 | 0.20105086 | 0.38812445 |
| ANKRD9 | 131.949911 | -0.509781443 | 0.34005458 | -1.4991165 | 0.1338434 | 0.29777495 |
| RCOR1 | 1128.80239 | 0.194552895 | 0.2215886 | 0.87799145 | 0.37994836 | 0.57641971 |
| TRAF3 | 1244.90304 | -0.366107957 | 0.31094877 | -1.17739 | 0.23903989 | 0.43403362 |
| AMN | 1254.78135 | 1.499161684 | 0.42790098 | 3.5035248 | 0.00045914 | 0.00504653 |
| CDC42BPB | 1575.95514 | -0.479293753 | 0.31906267 | -1.5021931 | 0.13304724 | 0.29658132 |
| AL117209.1 | 50.7284778 | -0.894207184 | 0.42870092 | -2.0858532 | 0.03699192 | 0.12654062 |
| EXOC3L4 | 348.646426 | 0.725281307 | 0.61321934 | 1.18274368 | 0.23691075 | 0.43136764 |
| TNFAIP2 | 649.691943 | -1.503000383 | 0.57021759 | -2.6358366 | 0.00839301 | 0.04441884 |
| AL161669.3 | 108.232603 | -0.026983305 | 0.5191027 | -0.0519807 | 0.9585441 | 0.9788247 |
| EIF5 | 1029.84488 | -0.680441349 | 0.37329341 | -1.8228057 | 0.06833282 | 0.19099403 |
| MARK3 | 3223.09474 | -0.152000298 | 0.171069 | -0.8885321 | 0.37425458 | 0.57132281 |
| CKB | 562.834787 | -1.377862135 | 0.50649319 | -2.7203962 | 0.00652037 | 0.0368252 |
| TRMT61A | 62.5676423 | -0.598157279 | 0.59637816 | -1.0029832 | 0.31586896 | 0.51590706 |
| BAG5 | 320.017664 | -1.39032765 | 0.24163272 | -5.7538881 | 8.72E-09 | 7.25E-07 |
| KLC1 | 723.938025 | -0.46553499 | 0.26661914 | -1.7460674 | 0.08079924 | 0.21347632 |
| AL049840.1 | 81.1619088 | -2.533146037 | 0.65372595 | -3.8749357 | 0.00010665 | 0.00163706 |
| ZFYVE21 | 238.88069 | -0.247815673 | 0.2266926 | -1.0931793 | 0.27431506 | 0.47233234 |
| PPP1R13B | 2468.46581 | 0.115187498 | 0.23268232 | 0.4950419 | 0.62057054 | 0.77455476 |
| AL132712.1 | 92.597408 | 1.564253611 | 0.54919321 | 2.84827558 | 0.00439568 | 0.02753731 |
| ATP5MPL | 473.851834 | -0.219195335 | 0.27388881 | -0.8003077 | 0.42353251 | 0.61666528 |
| TDRD9 | 262.949531 | -0.971100614 | 0.52710752 | -1.8423198 | 0.06542839 | 0.18564002 |
| ASPG | 211.897432 | 2.479427306 | 0.50408796 | 4.91864022 | 8.71E-07 | 3.44E-05 |
| INF2 | 325.837435 | -1.035239787 | 0.26443777 | -3.9148711 | 9.05E-05 | 0.0014501 |
| ADSSL1 | 60.9173009 | -0.192690157 | 0.6486714 | -0.2970536 | 0.76642561 | 0.86943605 |
| SIVA1 | 221.300672 | 0.166200996 | 0.27242116 | 0.61008842 | 0.54180324 | 0.71282636 |
| AKT1 | 269.391649 | -0.506552214 | 0.24698118 | -2.050975 | 0.04026939 | 0.13423455 |
| AL583810.1 | 60.075253 | 0.325034886 | 0.30070126 | 1.08092294 | 0.27973139 | 0.4782088 |
| CEP170B | 243.162514 | -0.314615026 | 0.33461718 | -0.9402238 | 0.34710279 | 0.5471809 |
| AHNAK2 | 80.0945731 | -3.576717556 | 0.84524041 | -4.2315979 | 2.32E-05 | 0.00049618 |
| CLBA1 | 133.469766 | 0.33107285 | 0.23163242 | 1.42930275 | 0.15291724 | 0.32481583 |
| NUDT14 | 116.236141 | 0.221102226 | 0.46108633 | 0.47952458 | 0.63156549 | 0.78278942 |
| BRF1 | 407.65407 | 0.086017416 | 0.24942795 | 0.34485878 | 0.73020054 | 0.84648502 |
| BTBD6 | 71.0629083 | -0.267236113 | 0.41088952 | -0.6503843 | 0.51544399 | 0.69315508 |
| PACS2 | 728.440781 | -0.125467258 | 0.26452042 | -0.4743197 | 0.63527191 | 0.78496999 |
| TEX22 | 95.8267085 | 0.245805896 | 0.3860254 | 0.63676094 | 0.52428057 | 0.69979031 |
| MTA1 | 282.868248 | 0.358806006 | 0.32389268 | 1.10779288 | 0.26795128 | 0.46585593 |
| CRIP2 | 108.300308 | -0.203189802 | 0.65867185 | -0.3084841 | 0.75771399 | 0.86418282 |
| AC087386.1 | 152.480964 | 0.240326687 | 0.40318642 | 0.59606841 | 0.55112953 | 0.72062696 |
| AC026495.1 | 89.3075214 | 5.111153872 | 0.76783913 | 6.65654259 | 2.80E-11 | 5.43E-09 |
| NIPA1 | 466.592591 | -0.383557735 | 0.20800237 | -1.8440066 | 0.06518218 | 0.18520981 |
| NIPA2 | 442.852919 | -0.685271716 | 0.35772389 | -1.9156443 | 0.05541039 | 0.16601317 |
| CYFIP1 | 1067.37423 | -0.316107589 | 0.28502685 | -1.1090449 | 0.2674108 | 0.46550671 |
| TUBGCP5 | 451.492329 | 0.059957808 | 0.30660699 | 0.19555265 | 0.84496033 | 0.91854292 |
| PWRN1 | 46.813334 | 1.188850488 | 0.65937776 | 1.80298845 | 0.07139003 | 0.19673163 |
| AC124312.1 | 2058.61192 | 0.507566662 | 0.24878638 | 2.04017064 | 0.04133333 | 0.13661828 |
| PWAR6 | 429.578663 | 0.387365191 | 0.37514476 | 1.03257523 | 0.30180273 | 0.50234486 |
| UBE3A | 2266.11686 | 0.035692647 | 0.13896751 | 0.25684167 | 0.797301 | 0.88944452 |
| ATP10A | 121.830238 | -1.15139086 | 0.78157936 | -1.4731592 | 0.1407081 | 0.30765448 |
| GABRB3 | 873.276014 | 0.659655716 | 0.47270568 | 1.39548931 | 0.16286834 | 0.33937919 |
| GABRG3 | 75.3058465 | -0.027648521 | 0.53809432 | -0.0513823 | 0.95902089 | 0.97901984 |
| HERC2 | 2330.28149 | 0.042622346 | 0.15984341 | 0.26665064 | 0.78973816 | 0.88547575 |
| GOLGA8M | 219.655755 | -0.839531738 | 1.00768432 | -0.8331297 | 0.40477158 | 0.59877014 |
| APBA2 | 73.7602826 | -2.110798602 | 0.75134757 | -2.8093504 | 0.00496416 | 0.03012117 |
| FAM189A1 | 1356.45728 | -0.566729411 | 0.42042582 | -1.3479891 | 0.1776619 | 0.35833014 |
| AC022613.1 | 43.956903 | 1.487444805 | 0.43711145 | 3.40289598 | 0.00066676 | 0.00668278 |
| TJP1 | 2264.05989 | -0.805285903 | 0.27660461 | -2.9113249 | 0.003599 | 0.02370912 |
| AC091057.2 | 43.7943325 | 1.044438664 | 0.35901404 | 2.90918614 | 0.00362371 | 0.02379981 |
| AC091057.6 | 65.933933 | -0.761525376 | 0.28892714 | -2.6357004 | 0.00839638 | 0.04441884 |
| FAN1 | 437.035413 | 0.052478958 | 0.28215904 | 0.1859907 | 0.85245205 | 0.92213818 |
| MTMR10 | 1836.67248 | -0.188422539 | 0.22388825 | -0.8415919 | 0.40001642 | 0.59423156 |
| LINC02352 | 38.0599499 | 1.02497727 | 0.49740725 | 2.06064 | 0.0393374 | 0.1321563 |
| KLF13 | 815.506514 | 0.202906937 | 0.33938438 | 0.59786764 | 0.54992826 | 0.71955844 |
| OTUD7A | 1536.67821 | -0.374329043 | 0.36466765 | -1.0264937 | 0.30465893 | 0.50483075 |
| CHRNA7 | 82.1994472 | 0.57147007 | 0.34196449 | 1.67113862 | 0.0946943 | 0.23698747 |
| LINC02256 | 1369.08323 | -0.925562438 | 0.58196382 | -1.5904123 | 0.11174189 | 0.2645787 |
| AC123768.3 | 207.070494 | -0.166682931 | 0.55840744 | -0.298497 | 0.76532389 | 0.86903543 |
| GOLGA8N | 147.347542 | -0.342846993 | 0.61779013 | -0.5549571 | 0.57892403 | 0.74245057 |
| FMN1 | 484.255051 | -0.889051428 | 0.48459936 | -1.8346112 | 0.06656332 | 0.1877107 |
| RYR3 | 75.0409893 | 0.024252391 | 0.36182491 | 0.06702797 | 0.94655943 | 0.97252756 |
| AC010809.2 | 58.2341333 | 1.626623392 | 0.45257893 | 3.59412089 | 0.00032549 | 0.00391357 |
| AC010809.1 | 107.770887 | 0.616276224 | 0.41421318 | 1.48782378 | 0.13679737 | 0.30179971 |
| AVEN | 1272.40349 | 0.723544928 | 0.3337725 | 2.16777874 | 0.03017553 | 0.10959616 |
| CHRM5 | 142.762211 | 0.322561718 | 0.46069493 | 0.70016337 | 0.48382529 | 0.66778705 |
| EMC7 | 236.430892 | -0.512492683 | 0.27005796 | -1.8977137 | 0.05773381 | 0.17049836 |
| PGBD4 | 48.2655318 | 0.636231526 | 0.32792377 | 1.9401812 | 0.05235767 | 0.15964229 |
| KATNBL1 | 1198.49257 | -0.510819008 | 0.21250428 | -2.4038058 | 0.01622539 | 0.07116846 |
| EMC4 | 111.513078 | -0.84939885 | 0.34557528 | -2.4579271 | 0.01397415 | 0.06372047 |
| SLC12A6 | 2415.43344 | 1.800775119 | 0.30307957 | 5.94159184 | 2.82E-09 | 2.80E-07 |
| NOP10 | 383.459122 | -1.599895868 | 0.42965619 | -3.7236654 | 0.00019635 | 0.00263345 |
| LPCAT4 | 110.160245 | -0.93191921 | 0.35389543 | -2.633318 | 0.00845551 | 0.04459397 |
| GOLGA8A | 464.598913 | -0.30756669 | 0.65248025 | -0.4713808 | 0.6373688 | 0.78678081 |
| GOLGA8B | 896.011056 | 0.10290942 | 0.32219404 | 0.319402 | 0.74942169 | 0.85872756 |
| AQR | 786.177521 | 0.163861243 | 0.17571449 | 0.93254256 | 0.3510562 | 0.55041523 |
| ZNF770 | 466.73721 | 0.030271161 | 0.25482851 | 0.11879032 | 0.90544147 | 0.95066597 |
| AC021231.1 | 633.407329 | 0.400383636 | 0.6131329 | 0.65301281 | 0.51374802 | 0.69196393 |
| DPH6 | 2465.19745 | 0.52265703 | 0.48105753 | 1.08647511 | 0.27726885 | 0.47518431 |
| DPH6-DT | 1699.3874 | 1.712571386 | 0.60035347 | 2.85260514 | 0.00433625 | 0.02723966 |
| C15orf41 | 408.422395 | 0.234663307 | 0.24757716 | 0.94783909 | 0.34321138 | 0.54379822 |
| MEIS2 | 143.233682 | -0.784718757 | 0.52130599 | -1.505294 | 0.13224852 | 0.29556994 |
| SPRED1 | 1304.98641 | -0.421007068 | 0.33328829 | -1.2631919 | 0.20652022 | 0.3953041 |
| FAM98B | 266.363145 | 0.20311986 | 0.18556248 | 1.09461708 | 0.27368443 | 0.47157668 |
| RASGRP1 | 471.908559 | 0.257864484 | 0.2309638 | 1.11647141 | 0.2642204 | 0.46213441 |
| C15orf53 | 232.0553 | 1.137233936 | 0.69716638 | 1.63122316 | 0.10284324 | 0.25038257 |
| AC013652.1 | 731.194052 | 0.994675724 | 0.59050617 | 1.68444596 | 0.09209551 | 0.23316461 |
| THBS1 | 484.958462 | 2.15038094 | 0.50315722 | 4.27377542 | 1.92E-05 | 0.00043115 |
| FSIP1 | 205.27267 | 0.758900532 | 0.37990461 | 1.99760813 | 0.04575916 | 0.14661543 |
| GPR176 | 226.110332 | -1.265495007 | 0.46211383 | -2.738492 | 0.00617217 | 0.03533914 |
| EIF2AK4 | 707.299822 | -0.483466164 | 0.25575316 | -1.8903624 | 0.05870951 | 0.1722087 |
| SRP14 | 981.745714 | -1.344548907 | 0.3439429 | -3.9092213 | 9.26E-05 | 0.00146557 |
| SRP14-AS1 | 168.297708 | 0.181972782 | 0.37081229 | 0.49074097 | 0.62360966 | 0.77658015 |

| | | | | | | |
|---|---|---|---|---|---|---|
| BMF | 59.9378141 | 1.332101473 | 0.90504954 | 1.47185476 | 0.1410601 | 0.30805998 |
| CCDC9B | 58.9436498 | -0.863623278 | 0.49073961 | -1.7598402 | 0.07843491 | 0.20920143 |
| KNSTRN | 69.9983654 | 0.396549951 | 0.3771986 | 1.05130283 | 0.29311953 | 0.49326436 |
| IVD | 474.57345 | 0.866465364 | 0.39172124 | 2.21194379 | 0.02697055 | 0.10163645 |
| BAHD1 | 211.218688 | 0.24991271 | 0.25699145 | 0.97245534 | 0.33082407 | 0.53061146 |
| AC091045.1 | 63.7070784 | -1.226232608 | 0.54585353 | -2.2464499 | 0.02467521 | 0.0954354 |
| CCDC32 | 413.871515 | -0.3203475 | 0.34914518 | -0.9175195 | 0.3588705 | 0.55733491 |
| RPUSD2 | 42.9910622 | -0.367285529 | 0.32377881 | -1.1343717 | 0.25663865 | 0.45384415 |
| KNL1 | 164.531749 | 1.773802704 | 0.54974859 | 3.22657072 | 0.00125283 | 0.01072489 |
| RMDN3 | 319.45111 | -0.164262959 | 0.26047398 | -0.630631 | 0.52828185 | 0.70268407 |
| GCHFR | 289.152563 | -0.960101616 | 0.54808493 | -1.7517388 | 0.07981874 | 0.21149728 |
| DNAJC17 | 498.188196 | 0.20328831 | 0.1883722 | 1.07918426 | 0.2805056 | 0.47921202 |
| ZFYVE19 | 263.975243 | 0.379576911 | 0.28989331 | 1.30936762 | 0.19040985 | 0.3744545 |
| SPINT1-AS1 | 112.23622 | -0.007452305 | 0.3944209 | -0.0188943 | 0.98492543 | 0.99257632 |
| SPINT1 | 510.763813 | -1.236619109 | 0.29202371 | -4.2346531 | 2.29E-05 | 0.00049025 |
| VPS18 | 155.261635 | -0.252104715 | 0.3967297 | -0.6354571 | 0.52513032 | 0.70066789 |
| INO80 | 1196.36707 | -0.132401136 | 0.31237632 | -0.4238514 | 0.6716742 | 0.80972162 |
| AC020661.3 | 38.8015373 | -0.67834542 | 0.49597744 | -1.3676941 | 0.17140785 | 0.35051281 |
| INO80-AS1 | 38.8990621 | -1.407959985 | 0.48598793 | -2.8971089 | 0.00376619 | 0.02446595 |
| CHP1 | 763.055623 | 0.107273207 | 0.34106354 | 0.31452558 | 0.75312189 | 0.8613901 |
| OIP5 | 44.0402506 | -0.57779839 | 0.40508673 | -1.4263573 | 0.15376525 | 0.32616221 |
| NUSAP1 | 131.225655 | -0.219616737 | 0.54712197 | -0.4014036 | 0.68812299 | 0.81968383 |
| NDUFAF1 | 148.576076 | 0.533841767 | 0.21467909 | 2.48669662 | 0.01289353 | 0.06031539 |
| RTF1 | 692.05586 | -0.262528237 | 0.29704068 | -0.8838124 | 0.37679749 | 0.57328928 |
| RPAP1 | 144.021283 | 0.19509986 | 0.24161358 | 0.80748713 | 0.41938589 | 0.61277734 |
| MGA | 1500.49729 | 0.139758995 | 0.24619133 | 0.56768448 | 0.57024923 | 0.7363548 |
| MAPKBP1 | 474.046807 | -0.115019323 | 0.46661273 | -0.2464985 | 0.80529639 | 0.89441535 |
| AC020659.1 | 70.8355079 | 0.271525031 | 0.79952677 | 0.33960718 | 0.73415237 | 0.84943603 |
| EHD4 | 202.745127 | -1.83078978 | 0.25204365 | -7.2637808 | 3.76E-13 | 1.10E-10 |
| VPS39 | 686.120829 | 0.283792865 | 0.10510102 | 2.70019131 | 0.00692996 | 0.03864482 |
| TMEM87A | 1199.01451 | -0.395413265 | 0.23221629 | -1.7027801 | 0.08860923 | 0.22698842 |
| GANC | 929.6239 | 0.737376687 | 0.41065671 | 1.79560364 | 0.07255758 | 0.19879007 |
| ZNF106 | 798.937189 | 0.0698262 | 0.18387495 | 0.37974829 | 0.70413227 | 0.83024009 |
| SNAP23 | 843.033459 | -0.484789336 | 0.37311006 | -1.2993199 | 0.19383416 | 0.37890542 |
| LRRC57 | 99.0846381 | -0.377369073 | 0.2630532 | -1.4345732 | 0.15140875 | 0.32276202 |
| HAUS2 | 168.449388 | -0.083467009 | 0.24507198 | -0.3405816 | 0.73341857 | 0.84892488 |
| STARD9 | 167.55029 | -0.752986001 | 0.26731922 | -2.8168046 | 0.0048504 | 0.0295863 |
| CDAN1 | 95.4160469 | 0.270111012 | 0.27731622 | 0.97401807 | 0.33004757 | 0.52979998 |
| TTBK2 | 1276.7141 | 0.031062246 | 0.21620539 | 0.14367008 | 0.885761 | 0.94094134 |
| UBR1 | 1252.84708 | -0.009411712 | 0.13515709 | -0.0696354 | 0.9444839 | 0.97156028 |
| AC068724.3 | 202.539658 | -0.664947117 | 0.31837011 | -2.0885978 | 0.03674394 | 0.1260068 |
| TMEM62 | 330.122659 | -0.407320319 | 0.23622676 | -1.7242768 | 0.08465788 | 0.22010729 |
| CCNDBP1 | 143.296982 | -0.631920823 | 0.24009885 | -2.6319194 | 0.0084904 | 0.04467482 |
| ADAL | 265.045628 | 0.795844754 | 0.34675135 | 2.29514537 | 0.02172479 | 0.08763551 |
| ZSCAN29 | 115.862965 | -0.367654917 | 0.29250864 | -1.2569027 | 0.20878885 | 0.39797925 |
| TUBGCP4 | 271.110495 | -0.188785054 | 0.23122483 | -0.8164567 | 0.41423899 | 0.60792011 |
| TP53BP1 | 953.998385 | 0.118484416 | 0.23231866 | 0.51000817 | 0.61004574 | 0.76658388 |
| PPIP5K1 | 240.58254 | -0.484326516 | 0.24891339 | -1.9457632 | 0.05168319 | 0.15855449 |
| AC011330.3 | 43.4649305 | -0.189812087 | 0.55678245 | -0.3409089 | 0.73317217 | 0.84871134 |
| CATSPER2 | 297.524691 | -0.01308007 | 0.49170818 | -0.0266013 | 0.97877775 | 0.98916955 |
| PDIA3 | 899.074482 | -0.883673436 | 0.32255528 | -2.7396031 | 0.00615134 | 0.03526409 |
| AC018512.1 | 48.0709718 | 0.390783525 | 0.45294444 | 0.8627626 | 0.38826801 | 0.58387159 |
| SERF2 | 2639.52162 | -0.793277632 | 0.35983969 | -2.2045307 | 0.02748704 | 0.10294526 |
| MFAP1 | 184.577796 | -0.322459525 | 0.20494294 | -1.5734112 | 0.11562363 | 0.27072994 |
| FRMD5 | 1223.89978 | -0.445653813 | 0.5413152 | -0.8232797 | 0.41034896 | 0.60419441 |
| CASC4 | 2429.14538 | 0.144577731 | 0.2045499 | 0.70680911 | 0.47968511 | 0.66481737 |
| AC025043.1 | 83.4352055 | -0.526817418 | 0.39148986 | -1.3456732 | 0.17840793 | 0.3594319 |
| CTDSPL2 | 1416.47519 | 0.187641506 | 0.19515997 | 0.96147537 | 0.3363132 | 0.53640273 |
| EIF3J-DT | 300.301526 | -0.29113746 | 0.28803135 | -1.0107839 | 0.31211986 | 0.51235819 |
| EIF3J | 672.520903 | -0.495970006 | 0.37396004 | -1.3262647 | 0.18475203 | 0.3675466 |
| SPG11 | 2096.16799 | -0.087116575 | 0.14824047 | -0.5876707 | 0.55675337 | 0.72562693 |
| PATL2 | 72.6886919 | -1.144200252 | 0.5970727 | -1.91635 | 0.05532057 | 0.16588908 |
| B2M | 4508.29682 | -1.66874509 | 0.4615296 | -3.6156838 | 0.00029956 | 0.00367264 |
| TRIM69 | 125.595411 | -0.688303114 | 0.40478688 | -1.7004086 | 0.08905409 | 0.22768161 |
| SORD | 1217.28468 | 0.952411697 | 0.61532402 | 1.54782142 | 0.1216653 | 0.27976283 |
| DUOX1 | 138.718288 | -0.910001472 | 0.44565885 | -2.0419239 | 0.04115908 | 0.13617366 |
| SHF | 33.8251199 | 0.152127059 | 0.43838951 | 0.34701346 | 0.72858121 | 0.84596888 |
| AC051619.5 | 62.6413644 | 0.832345181 | 0.82437575 | 1.00966723 | 0.31265475 | 0.51299481 |
| SLC28A2 | 44.6669606 | 0.650115486 | 0.89371422 | 0.72743106 | 0.46696193 | 0.65463001 |
| GATM | 2023.82093 | 2.092074663 | 0.48084462 | 4.35083302 | 1.36E-05 | 0.00032386 |
| AC025580.3 | 34.7404994 | 0.783313362 | 0.76414536 | 1.02508423 | 0.30532345 | 0.5053224 |
| SPATA5L1 | 206.964216 | -0.161663438 | 0.28309551 | -0.5710562 | 0.56796157 | 0.73451775 |
| AC025580.2 | 98.2420421 | -1.628241868 | 0.44505273 | -3.658537 | 0.00025366 | 0.00321936 |
| SLC30A4 | 418.537842 | 0.974853536 | 0.30268169 | 3.22072185 | 0.00127868 | 0.01087249 |
| SQOR | 356.584496 | -1.814701342 | 0.4034888 | -4.4975259 | 6.87E-06 | 0.0001866 |
| AC068722.1 | 49.8349928 | -1.581586156 | 0.54732706 | -2.8896546 | 0.00385665 | 0.02493545 |
| SLC24A5 | 47.3229876 | -1.610935539 | 0.68759375 | -2.3428595 | 0.01913659 | 0.07991704 |
| MYEF2 | 1283.24713 | -0.716245291 | 0.27534053 | -2.6013072 | 0.00928693 | 0.04774056 |
| SLC12A1 | 429.875388 | 3.202786581 | 0.69232117 | 4.62615722 | 3.73E-06 | 0.00011372 |
| DUT | 285.498565 | 0.458051189 | 0.28924014 | 1.58363632 | 0.1132765 | 0.26682953 |
| FBN1 | 151.167249 | 0.361589887 | 0.25898556 | 1.39617782 | 0.16266096 | 0.33910171 |
| CEP152 | 533.308957 | 0.674640461 | 0.23858197 | 2.82770938 | 0.00468823 | 0.02893365 |
| EID1 | 368.235057 | -0.78932636 | 0.3346939 | -2.358353 | 0.01835623 | 0.07770031 |
| SECISBP2L | 1748.62143 | 0.003647585 | 0.23123027 | 0.01577469 | 0.98741414 | 0.99360989 |
| COPS2 | 489.344714 | -0.068406412 | 0.17411447 | -0.3928818 | 0.69440676 | 0.82408948 |
| GALK2 | 2117.95956 | 0.588468758 | 0.24855933 | 2.36751827 | 0.01790784 | 0.07630174 |
| FAM227B | 679.171536 | 0.621425048 | 0.60015819 | 1.03543542 | 0.30046561 | 0.50091001 |
| FGF7 | 47.4084701 | 0.10646306 | 0.49851698 | 0.21355955 | 0.83089055 | 0.9106114 |
| DTWD1 | 433.614059 | 0.360019042 | 0.26375544 | 1.36497295 | 0.17226155 | 0.35167607 |
| ATP8B4 | 1548.48789 | 0.72794757 | 0.56906309 | 1.27920363 | 0.20082536 | 0.38798494 |
| SLC27A2 | 2491.30948 | 0.387202996 | 0.49393473 | 0.7839153 | 0.43308981 | 0.62552702 |
| GABPB1 | 635.528709 | -0.207515399 | 0.26965423 | -0.7695611 | 0.44156029 | 0.63336825 |
| GABPB1-AS1 | 900.439037 | -0.446537245 | 0.34583838 | -1.2911732 | 0.19664363 | 0.38248 |
| USP8 | 1596.96471 | 0.249905769 | 0.11273557 | 2.21674284 | 0.02664067 | 0.1008664 |
| USP50 | 188.93192 | 1.084500982 | 0.485662 | 2.23303652 | 0.02554654 | 0.09789387 |
| TRPM7 | 3245.59846 | -0.047303852 | 0.29939221 | -0.1579996 | 0.87445711 | 0.93490877 |
| SPPL2A | 1288.4891 | -0.395988347 | 0.11696125 | -3.3856371 | 0.00071013 | 0.0070132 |
| AC021752.1 | 83.538112 | -0.525115989 | 0.42015518 | -1.2498144 | 0.21136735 | 0.40077636 |
| AP4E1 | 645.878169 | -0.132109619 | 0.14207936 | -0.9298298 | 0.35245923 | 0.55147186 |
| AC066613.2 | 48.8355417 | 0.517697764 | 0.67869451 | 0.76278466 | 0.44559183 | 0.63688501 |
| DMXL2 | 1753.55207 | 1.196716782 | 0.23798237 | 5.02859435 | 4.94E-07 | 2.13E-05 |
| LYSMD2 | 42.6470652 | -0.560075422 | 0.43401782 | -1.2904434 | 0.19689676 | 0.38265475 |
| TMOD2 | 53.6845798 | -0.556401992 | 0.53660051 | -1.0369017 | 0.29978167 | 0.50007391 |
| TMOD3 | 1644.0365 | -0.502510037 | 0.26074605 | -1.927201 | 0.05395459 | 0.16308249 |
| AC090971.2 | 74.3082787 | -0.783093024 | 0.49948936 | -1.5677872 | 0.1169308 | 0.27253367 |
| LEO1 | 210.652769 | 0.240525935 | 0.22443713 | 1.07168515 | 0.28386147 | 0.48310023 |
| MAPK6 | 1437.62832 | -0.067238974 | 0.36642425 | -0.1835003 | 0.85440545 | 0.92322796 |
| AC023906.5 | 40.0612263 | -1.383645899 | 0.52826665 | -2.6192187 | 0.00881314 | 0.04600218 |
| GNB5 | 679.011936 | 0.911836999 | 0.27622854 | 3.3010239 | 0.00096333 | 0.00880875 |
| MYO5C | 256.465105 | -1.387568457 | 0.50557481 | -2.7445364 | 0.00605964 | 0.03484041 |
| MYO5A | 751.650478 | -0.114028674 | 0.2034261 | -0.560541 | 0.57511047 | 0.73976511 |
| ARPP19 | 779.832859 | -0.699011818 | 0.2694535 | -2.5941835 | 0.00948159 | 0.04849815 |
| FAM214A | 1412.28524 | 0.415916793 | 0.23857797 | 1.74331601 | 0.08127843 | 0.21441175 |

| | | | | | | |
|---|---|---|---|---|---|---|
| WDR72 | 13887.7322 | 1.231071294 | 0.65448788 | 1.88096883 | 0.05997616 | 0.1745825 |
| UNC13C | 49.0455802 | 1.136396403 | 0.67378967 | 1.68657439 | 0.09168522 | 0.23229747 |
| RSL24D1 | 205.026096 | -0.587427236 | 0.24945575 | -2.3548354 | 0.01853091 | 0.07815267 |
| RAB27A | 429.682424 | -0.663400377 | 0.30336902 | -2.1867769 | 0.02875881 | 0.10602395 |
| PIGB | 403.762703 | 0.171887499 | 0.20707029 | 0.83009252 | 0.40648648 | 0.60029924 |
| CCPG1 | 1004.22987 | 0.155548359 | 0.14706596 | 1.0576775 | 0.29020249 | 0.49027014 |
| C15orf65 | 48.2891457 | 0.097347733 | 0.43132672 | 0.22569372 | 0.82143964 | 0.90404363 |
| DNAAF4 | 501.165165 | 0.69614412 | 0.26704495 | 2.60684249 | 0.00913814 | 0.04723094 |
| PRTG | 186.544191 | -0.041050111 | 0.48164033 | -0.0852298 | 0.9320787 | 0.96391408 |
| NEDD4 | 2323.75974 | -0.432988168 | 0.2540779 | -1.7041551 | 0.08835211 | 0.22649006 |
| RFX7 | 2013.04152 | 0.348823404 | 0.23795121 | 1.46594505 | 0.14266326 | 0.31014834 |
| TEX9 | 794.805702 | 0.326678997 | 0.3530415 | 0.92532746 | 0.3547956 | 0.55326317 |
| MNS1 | 76.0030944 | 0.144829628 | 0.53587099 | 0.27026958 | 0.78695287 | 0.88351603 |
| AC090518.1 | 109.455961 | 0.945946531 | 0.52836683 | 1.79032155 | 0.07340223 | 0.20030349 |
| ZNF280D | 602.10318 | 0.789936692 | 0.39069602 | 2.02187035 | 0.04318975 | 0.1406059 |
| AC090517.4 | 450.647448 | 0.163107239 | 0.40987399 | 0.39794484 | 0.69067085 | 0.8212201 |
| TCF12 | 6778.63492 | 0.2600213 | 0.1703113 | 1.52674132 | 0.12682535 | 0.28790372 |
| CGNL1 | 10185.7216 | 0.91781563 | 0.50813323 | 1.80624995 | 0.07087931 | 0.19578819 |
| AC025271.4 | 32.3558874 | 0.733640622 | 0.63618965 | 1.15317912 | 0.24883687 | 0.44510075 |
| POLR2M | 105.889501 | -0.300989418 | 0.31262331 | -0.9627862 | 0.33565482 | 0.53553074 |
| ALDH1A2 | 615.233575 | -2.381096516 | 0.50504242 | -4.7146465 | 2.42E-06 | 8.04E-05 |
| LIPC | 413.791314 | -0.955298361 | 0.27946991 | -3.4182513 | 0.00063025 | 0.00638813 |
| ADAM10 | 2290.26013 | -0.676464334 | 0.18065229 | -3.7445655 | 0.00018071 | 0.00246008 |
| MINDY2 | 1035.25042 | -0.937928079 | 0.22053605 | -4.2529467 | 2.11E-05 | 0.00046048 |
| AC090515.4 | 48.497314 | -1.070143558 | 0.32279618 | -3.31523 | 0.00091568 | 0.00846026 |
| RNF111 | 1872.94745 | 0.06525486 | 0.1613487 | 0.40443376 | 0.68589378 | 0.81810131 |
| SLTM | 2146.09083 | -0.072876972 | 0.18383663 | -0.3964225 | 0.69179338 | 0.82234081 |
| MYO1E | 2201.54965 | -1.849861728 | 0.32529791 | -5.6866696 | 1.30E-08 | 1.01E-06 |
| FAM81A | 94.5752218 | -0.849332819 | 0.44101464 | -1.9258608 | 0.05412175 | 0.16345401 |
| GCNT3 | 282.8605 | -3.09326748 | 0.43734854 | -7.0727742 | 1.52E-12 | 3.93E-10 |
| GTF2A2 | 334.730227 | -0.326435728 | 0.18162579 | -1.7972983 | 0.07228828 | 0.19836946 |
| BNIP2 | 629.621772 | -0.187262291 | 0.15497777 | -1.2083171 | 0.22692532 | 0.41931165 |
| ANXA2 | 1361.03122 | -3.21756703 | 0.52234837 | -6.1598106 | 7.28E-10 | 8.91E-08 |
| ICE2 | 677.37293 | 0.380567245 | 0.30059693 | 1.26603835 | 0.20549935 | 0.39412056 |
| RORA-AS1 | 537.391926 | -0.443112212 | 0.71355125 | -0.6209956 | 0.53460249 | 0.70752113 |
| RORA | 12493.627 | 1.407873933 | 0.44626349 | 3.15480417 | 0.00160606 | 0.01300577 |
| AC012404.1 | 173.740435 | 1.227666534 | 0.90819308 | 1.35176821 | 0.17644948 | 0.3567986 |
| AC009554.1 | 266.607806 | -1.233399226 | 0.48248726 | -2.5563353 | 0.01057811 | 0.05259129 |
| VPS13C | 2698.89666 | -0.637786305 | 0.16165313 | -3.9454003 | 7.97E-05 | 0.00130309 |
| TLN2 | 3015.23894 | 1.066361994 | 0.37851327 | 2.8172381 | 0.00484386 | 0.02956843 |
| AC103740.1 | 57.4192613 | -0.331932839 | 0.64510305 | -0.5145423 | 0.60687286 | 0.76385733 |
| TPM1 | 2398.30365 | -1.03062808 | 0.39650987 | -2.5992495 | 0.00934278 | 0.04797325 |
| LACTB | 438.940432 | -0.508864933 | 0.29473275 | -1.72653 | 0.08425211 | 0.21934354 |
| RPS27L | 800.136667 | -0.117100774 | 0.37548933 | -0.3118618 | 0.75514556 | 0.86270273 |
| RAB8B | 869.267828 | -0.52242156 | 0.20080755 | -2.6016032 | 0.00927892 | 0.04772461 |
| APH1B | 188.524623 | -0.909734034 | 0.44382904 | -2.0497398 | 0.04038983 | 0.13437462 |
| CA12 | 3669.42564 | -1.294243846 | 0.36961618 | -3.5015887 | 0.00046249 | 0.00506746 |
| LINC02568 | 44.252757 | 0.661177446 | 0.6284508 | 1.05207511 | 0.29276509 | 0.49298908 |
| USP3 | 1492.27682 | -0.369287129 | 0.12697479 | -2.9083499 | 0.00363342 | 0.02382929 |
| USP3-AS1 | 318.410331 | -0.008943249 | 0.43353133 | -0.0206288 | 0.98354173 | 0.99200931 |
| HERC1 | 1878.75584 | 0.253188493 | 0.12125881 | 2.08800089 | 0.03679775 | 0.12612824 |
| AC073167.1 | 60.3627699 | -0.871058628 | 0.64700885 | -1.3462855 | 0.17821047 | 0.35922513 |
| DAPK2 | 60.1021984 | -1.644126522 | 0.60728408 | -2.7073434 | 0.00678241 | 0.03797649 |
| FAM96A | 423.040036 | 0.082772804 | 0.27444113 | 0.30160496 | 0.76295323 | 0.8677952 |
| SNX1 | 1749.84533 | -0.277601683 | 0.17738195 | -1.564994 | 0.11758432 | 0.2735671 |
| PPIB | 838.526446 | -1.234239659 | 0.40639322 | -3.0370577 | 0.002389 | 0.01761484 |
| CSNK1G1 | 299.039175 | -0.511559064 | 0.22079078 | -2.3169403 | 0.02050698 | 0.08405778 |
| TRIP4 | 1415.52554 | 0.858198854 | 0.17980831 | 4.77285433 | 1.82E-06 | 6.27E-05 |
| ZNF609 | 3492.78454 | 0.112865198 | 0.25300306 | 0.4461021 | 0.65552349 | 0.79904496 |
| OAZ2 | 255.515129 | -0.085933502 | 0.23547566 | -0.3649358 | 0.71515931 | 0.83712116 |
| AC100830.1 | 74.8994807 | -0.04740128 | 0.47571406 | -0.0996424 | 0.92062825 | 0.95736677 |
| RBPMS2 | 227.48378 | 1.177947984 | 0.36875072 | 3.19442901 | 0.00140108 | 0.01168161 |
| AC069368.1 | 51.5290421 | -2.089199731 | 0.61838583 | -3.3784728 | 0.0007289 | 0.00716003 |
| ANKDD1A | 56.5548603 | -1.395946859 | 0.49277823 | -2.8328095 | 0.00461409 | 0.02861778 |
| SPG21 | 594.44776 | 0.302546608 | 0.29239778 | 1.03470898 | 0.30080484 | 0.5012317 |
| MTFMT | 477.34833 | 0.379150562 | 0.21852112 | 1.73507511 | 0.0827275 | 0.21702313 |
| SLC51B | 104.285758 | 1.111491202 | 0.65823606 | 1.68859056 | 0.09129792 | 0.23164744 |
| UBAP1L | 228.182782 | 0.105730416 | 0.4214728 | 0.25085941 | 0.80192281 | 0.89234908 |
| PDCD7 | 331.883986 | -0.440979376 | 0.25143121 | -1.7538769 | 0.07945161 | 0.21097311 |
| CLPX | 703.690447 | -0.329334079 | 0.28610221 | -1.1511064 | 0.24968847 | 0.44586762 |
| PARP16 | 199.057003 | 0.452310167 | 0.30910796 | 1.46327571 | 0.14339195 | 0.31104186 |
| DPP8 | 970.741883 | 0.219171877 | 0.16689666 | 1.31321904 | 0.18910915 | 0.37309941 |
| HACD3 | 538.358448 | -0.115175639 | 0.19665432 | -0.5856756 | 0.55809354 | 0.72661361 |
| INTS14 | 230.394808 | -0.266146725 | 0.22410772 | -1.1875839 | 0.23499738 | 0.42959577 |
| SLC24A1 | 158.664376 | 0.280617969 | 0.32384713 | 0.86651368 | 0.38620853 | 0.58237187 |
| DENND4A | 3788.50774 | -0.139406697 | 0.30007951 | -0.4645659 | 0.64224238 | 0.79016392 |
| RAB11A | 1517.98387 | -0.796490608 | 0.3941897 | -2.0205769 | 0.04332358 | 0.14091561 |
| MEGF11 | 1749.02386 | -4.204510992 | 0.84838588 | -4.9558946 | 7.20E-07 | 2.94E-05 |
| DIS3L | 287.699208 | 0.327191223 | 0.3049259 | 1.0730188 | 0.28326268 | 0.48244055 |
| TIPIN | 180.03008 | 1.395419631 | 0.54061755 | 2.58115859 | 0.00984693 | 0.04987876 |
| MAP2K1 | 1100.19398 | -1.210413019 | 0.37594125 | -3.2196866 | 0.00128331 | 0.01088509 |
| SNAPC5 | 120.117447 | -0.470431566 | 0.27767568 | -1.6941763 | 0.09023179 | 0.2296337 |
| RPL4 | 977.127899 | -0.949137085 | 0.44599872 | -2.1281162 | 0.03332745 | 0.11788888 |
| ZWILCH | 60.8730861 | -0.321448075 | 0.41502375 | -0.7745293 | 0.43861782 | 0.63046712 |
| SMAD6 | 121.961828 | 0.631010793 | 0.54035459 | 1.16777169 | 0.24289887 | 0.43805442 |
| AC093334.1 | 37.7477643 | -1.215902723 | 0.61479562 | -1.9777348 | 0.04795863 | 0.15122358 |
| AC087482.1 | 3312.63144 | -0.358761764 | 0.62885125 | -0.5705034 | 0.56833633 | 0.73472607 |
| SMAD3 | 4877.93916 | -1.411616406 | 0.40157942 | -3.5151612 | 0.00043949 | 0.00489362 |
| AAGAB | 809.378593 | -0.191993019 | 0.20361403 | -0.9429263 | 0.34571862 | 0.54612944 |
| IQCH | 721.530377 | 0.426317406 | 0.40362835 | 1.05621274 | 0.29087103 | 0.49106654 |
| IQCH-AS1 | 502.407748 | 0.750662197 | 0.3241576 | 2.31573219 | 0.0205729 | 0.08417692 |
| C15orf61 | 82.3559656 | -0.076996154 | 0.37960241 | -0.2028337 | 0.83926503 | 0.91488833 |
| MAP2K5 | 1643.26217 | 0.359979165 | 0.23388424 | 1.53913389 | 0.12377161 | 0.2832283 |
| PIAS1 | 2559.18937 | 0.210432395 | 0.16684804 | 1.26122182 | 0.20722894 | 0.39612324 |
| FEM1B | 347.614879 | -0.211952446 | 0.17798464 | -1.1908469 | 0.23371371 | 0.42781969 |
| CORO2B | 128.923287 | -2.860172903 | 0.67503771 | -4.2370565 | 2.26E-05 | 0.00048732 |
| ANP32A | 757.047334 | -0.032458345 | 0.27811133 | -0.1167099 | 0.90708994 | 0.95122652 |
| SPESP1 | 185.344798 | 1.067213553 | 0.43427396 | 2.45746616 | 0.0139921 | 0.0637651 |
| NOX5 | 45.565607 | 0.38680962 | 0.80825089 | 0.47857618 | 0.63224017 | 0.78320072 |
| GLCE | 1228.53238 | 0.946081897 | 0.3279989 | 2.88440567 | 0.00392153 | 0.02527148 |
| AC026992.2 | 280.786052 | 1.26631668 | 0.53429755 | 2.37005892 | 0.01778525 | 0.07594468 |
| AC026992.1 | 181.676585 | 0.363954416 | 0.70654791 | 0.5151164 | 0.60647168 | 0.7636205 |
| PAQR5 | 3756.18689 | -0.809447999 | 0.48346016 | -1.6742807 | 0.09407545 | 0.23621021 |
| AC027237.3 | 80.6855331 | -1.238264939 | 0.62451279 | -1.9827696 | 0.04739318 | 0.15022483 |
| KIF23 | 371.110235 | -0.671500082 | 0.60512985 | -1.1096793 | 0.26713724 | 0.46514864 |
| RPLP1 | 1998.7212 | -0.61410334 | 0.43611503 | -1.4081224 | 0.15909483 | 0.33408049 |
| DRAIC | 1541.84973 | 1.642993352 | 0.65428604 | 2.51112395 | 0.01203474 | 0.05739743 |
| TLE3 | 403.73753 | -0.322523944 | 0.31038826 | -1.0390984 | 0.29875898 | 0.49882154 |
| UACA | 1837.69547 | -0.35691389 | 0.2850065 | -1.2523009 | 0.21046025 | 0.39989367 |
| AC009269.2 | 42.2372021 | -1.247232936 | 0.49866409 | -2.5011485 | 0.01237913 | 0.05856444 |
| LARP6 | 119.062143 | -0.037355682 | 0.25282822 | -0.1477512 | 0.88253909 | 0.93903973 |
| LRRC49 | 158.289186 | -0.405071092 | 0.52242638 | -0.7753649 | 0.43812405 | 0.63002165 |
| THSD4 | 309.777775 | -0.274035688 | 0.47155256 | -0.581135 | 0.56114948 | 0.72927619 |

| | | | | | | | |
|---|---|---|---|---|---|---|---|
| THSD4-AS1 | 349.474203 | | -0.295176532 | 0.26546469 | -1.1119239 | 0.26617088 | 0.46404642 |
| MYO9A | 6505.51831 | | 0.380510325 | 0.26586371 | 1.43122325 | 0.15236624 | 0.32404718 |
| AC022872.1 | 142.593861 | | -0.3828816 | 0.61735298 | -0.6201988 | 0.53512689 | 0.70807841 |
| SENP8 | 171.81723 | | 0.604108975 | 0.36770152 | 1.64293302 | 0.10039679 | 0.2464432 |
| GRAMD2A | 146.062068 | | 1.073485238 | 0.58188632 | 1.8448367 | 0.0650613 | 0.18501975 |
| PKM | 2594.28565 | | -2.304417986 | 0.29161475 | -7.9022683 | 2.74E-15 | 1.50E-12 |
| PARP6 | 529.92872 | | 0.121552587 | 0.31552571 | 0.3852383 | 0.70006087 | 0.82770696 |
| AC009690.3 | 38.7011845 | | -0.455135627 | 0.69020216 | -0.6594237 | 0.50962376 | 0.68915039 |
| AC009690.2 | 294.298071 | | -0.969138277 | 0.52831907 | -1.8343806 | 0.06659751 | 0.1877107 |
| AC009690.1 | 1392.74083 | | -0.221337419 | 0.52260967 | -0.4235234 | 0.67191344 | 0.80987667 |
| TMEM202-A! | 167.788767 | | 0.565270696 | 0.50698095 | 1.11497423 | 0.26486147 | 0.4626553 |
| ARIH1 | 3451.80089 | | -0.238795658 | 0.22771955 | -1.0486393 | 0.29434418 | 0.49425158 |
| BBS4 | 503.333109 | | 0.259187177 | 0.27727076 | 0.93478006 | 0.34990166 | 0.5493439 |
| ADPGK | 535.202247 | | -0.426866849 | 0.27488373 | -1.5528996 | 0.12044711 | 0.27789405 |
| NEO1 | 2538.99678 | | 0.707081208 | 0.34395364 | 2.05574567 | 0.03980704 | 0.13307998 |
| NPTN | 912.953867 | | 0.08038691 | 0.17572667 | 0.45745424 | 0.64734458 | 0.79309548 |
| CD276 | 171.188979 | | -1.208987455 | 0.47883327 | -2.524861 | 0.01157441 | 0.05580386 |
| LOXL1-AS1 | 44.4870786 | | -0.276258032 | 0.56772857 | -0.4866023 | 0.62654019 | 0.77912423 |
| LOXL1 | 68.7340846 | | 0.074076249 | 0.47038763 | 0.15747916 | 0.87486723 | 0.93490877 |
| STOML1 | 50.1925085 | | -0.557430598 | 0.37294226 | -1.4946834 | 0.13499709 | 0.29936987 |
| PML | 355.026663 | | -1.293231546 | 0.27065725 | -4.7781154 | 1.77E-06 | 6.18E-05 |
| STRA6 | 97.2275096 | | 6.825359533 | 0.8487069 | 8.04206907 | 8.83E-16 | 5.50E-13 |
| UBL7 | 50.5113851 | | 0.234259845 | 0.2860971 | 0.81881238 | 0.41289346 | 0.60672499 |
| UBL7-AS1 | 42.0739266 | | -0.19461713 | 0.42008516 | -0.4632802 | 0.64316354 | 0.79092642 |
| AC012435.2 | 65.90043 | | 0.283302615 | 0.3251793 | 0.8712197 | 0.38363421 | 0.58015209 |
| ARID3B | 295.877193 | | -0.157270715 | 0.3855584 | -0.4079037 | 0.68334435 | 0.81604817 |
| CLK3 | 245.308213 | | 0.154970358 | 0.18434202 | 0.84066757 | 0.40053419 | 0.59474837 |
| EDC3 | 350.613666 | | 0.222566981 | 0.23284192 | 0.95587162 | 0.3391371 | 0.53952556 |
| CSK | 109.185157 | | -0.122462309 | 0.44035196 | -0.278101 | 0.78093485 | 0.87970694 |
| ULK3 | 288.487153 | | 1.239343978 | 0.29804767 | 4.1582073 | 3.21E-05 | 0.00063534 |
| SCAMP2 | 595.179244 | | 0.545742482 | 0.23044598 | 2.36820133 | 0.01787481 | 0.07621645 |
| MPI | 115.970469 | | 0.036513574 | 0.30307806 | 0.12047581 | 0.90410624 | 0.94976888 |
| FAM219B | 199.967655 | | -0.005792173 | 0.21730191 | -0.026655 | 0.97873494 | 0.98916955 |
| COX5A | 454.1419 | | -0.118879962 | 0.24943486 | -0.4765972 | 0.63364896 | 0.78395399 |
| SCAMP5 | 344.719867 | | 1.200715059 | 0.34419699 | 3.48845311 | 0.00048582 | 0.00525173 |
| PPCDC | 778.449906 | | 0.187208433 | 0.40542257 | 0.46176125 | 0.64425254 | 0.79162826 |
| C15orf39 | 39.5070272 | | -0.610917433 | 0.44292625 | -1.3792757 | 0.16780975 | 0.34604603 |
| COMMD4 | 213.849866 | | 0.390093284 | 0.28989377 | 1.34564219 | 0.17841794 | 0.3594319 |
| NEIL1 | 310.83396 | | 0.514546406 | 0.35847167 | 1.4353893 | 0.15117619 | 0.32246687 |
| MAN2C1 | 520.547061 | | 0.705083256 | 0.21769211 | 3.23890125 | 0.00119991 | 0.01038988 |
| SIN3A | 655.621097 | | 0.002263393 | 0.21378654 | 0.01058716 | 0.99155282 | 0.99576632 |
| PTPN9 | 1118.89013 | | -0.585406834 | 0.23414049 | -2.5002375 | 0.01241101 | 0.05868149 |
| SNUPN | 188.830579 | | 0.278940144 | 0.32591349 | 0.85587175 | 0.39206872 | 0.5871393 |
| IMP3 | 154.343168 | | -0.644304031 | 0.35622554 | -1.8086969 | 0.0704981 | 0.19517622 |
| SNX33 | 105.660998 | | 0.163527166 | 0.26542185 | 0.61610288 | 0.53782662 | 0.70986341 |
| UBE2Q2 | 493.149424 | | -1.320819408 | 0.21005633 | -6.2879297 | 3.22E-10 | 4.37E-08 |
| FBXO22 | 636.40129 | | 0.285542502 | 0.22576973 | 1.26475104 | 0.20596059 | 0.39472899 |
| NRG4 | 45.6241412 | | 0.041853787 | 0.47322499 | 0.08844374 | 0.929524 | 0.96267363 |
| TMEM266 | 90.2404502 | | -0.067125892 | 0.53091851 | -0.1264335 | 0.89938877 | 0.94743919 |
| ETFA | 1681.41722 | | 0.510230354 | 0.22096216 | 2.30912997 | 0.02093637 | 0.08528227 |
| SCAPER | 2911.62174 | | -0.021430145 | 0.29217622 | -0.0733466 | 0.94153028 | 0.96932218 |
| RCN2 | 370.092602 | | 0.189982187 | 0.22659627 | 0.8384171 | 0.40179649 | 0.59597711 |
| TSPAN3 | 1015.88084 | | -0.591122689 | 0.23088425 | -2.5602556 | 0.01045952 | 0.05213406 |
| PEAK1 | 1612.32006 | | -0.566165281 | 0.2688677 | -2.1057393 | 0.035227 | 0.12224216 |
| HMG20A | 764.71727 | | 0.174938682 | 0.1632942 | 1.07130984 | 0.28403014 | 0.48332726 |
| LINGO1 | 6132.91213 | | 0.922666441 | 0.62612261 | 1.47361942 | 0.14058409 | 0.30748479 |
| TBC1D2B | 531.237077 | | -0.370860502 | 0.28322294 | -1.3094296 | 0.19038887 | 0.3744545 |
| CIB2 | 60.259546 | | -1.447093371 | 0.54946478 | -2.6336417 | 0.00844746 | 0.04456863 |
| IDH3A | 608.007147 | | -0.279872587 | 0.24047015 | -1.1638558 | 0.24448244 | 0.43966424 |
| DNAJA4 | 298.445295 | | -1.807825587 | 0.8197279 | -2.2053972 | 0.02742624 | 0.10285402 |
| WDR61 | 343.145017 | | -0.317322496 | 0.16871161 | -1.8808575 | 0.05999131 | 0.17458616 |
| IREB2 | 1004.53783 | | -0.03375175 | 0.12993837 | -0.259752 | 0.79505508 | 0.8882949 |
| HYKK | 288.471516 | | 1.592345402 | 0.49852963 | 3.1940838 | 0.00140275 | 0.01168848 |
| PSMA4 | 405.927937 | | -0.540547079 | 0.27983762 | -1.9316455 | 0.05340327 | 0.16180342 |
| MORF4L1 | 1100.08931 | | -0.372004699 | 0.12173346 | -3.0558952 | 0.0022439 | 0.01675223 |
| CTSH | 1456.12903 | | 1.225491223 | 0.34379115 | 3.56463867 | 0.00036436 | 0.00426495 |
| RASGRF1 | 142.468315 | | -0.048231706 | 0.6021025 | -0.0801055 | 0.93615337 | 0.96587767 |
| TMED3 | 269.789428 | | -2.138619272 | 0.57111324 | -3.7446501 | 0.00018065 | 0.00246008 |
| MTHFS | 143.060841 | | 0.23416125 | 0.41789124 | 0.56034017 | 0.57524743 | 0.73976511 |
| ZFAND6 | 2820.61675 | | -0.099947881 | 0.17923009 | -0.5576512 | 0.57708257 | 0.74138147 |
| FAH | 451.145904 | | 0.726607217 | 0.30674568 | 2.36876102 | 0.01784778 | 0.07614054 |
| CTXND1 | 223.787483 | | 0.020737644 | 0.72974243 | 0.02841776 | 0.97732896 | 0.98871175 |
| AC016705.2 | 139.783571 | | -0.367565541 | 0.65656087 | -0.5598347 | 0.57559221 | 0.74004713 |
| ARNT2 | 409.918537 | | -0.651581076 | 0.70073813 | -0.9298496 | 0.35244896 | 0.55147186 |
| ABHD17C | 159.258878 | | 0.023890492 | 0.43021403 | 0.05553164 | 0.95571492 | 0.97753783 |
| CEMIP | 147.904094 | | -1.565908536 | 0.79638671 | -1.9662665 | 0.04926783 | 0.15396641 |
| AC027808.1 | 265.070838 | | -0.312979264 | 0.3832203 | -0.8167085 | 0.41409505 | 0.60777394 |
| MESD | 589.736857 | | -0.238421323 | 0.22063924 | -1.0805935 | 0.27987798 | 0.47834009 |
| TLNRD1 | 140.876346 | | -0.223836125 | 0.54223292 | -0.4128044 | 0.67974994 | 0.81423861 |
| CFAP161 | 119.148113 | | -0.203442978 | 0.97602371 | -0.2084406 | 0.83488495 | 0.91279955 |
| AC104041.1 | 89.386059 | | -0.413504572 | 0.62678753 | -0.6597205 | 0.50943322 | 0.6889789 |
| EFL1 | 1008.40806 | | -1.243637585 | 0.39716856 | -3.1312589 | 0.00174059 | 0.013767 |
| SAXO2 | 77.3644717 | | 1.144211092 | 0.66487242 | 1.72094836 | 0.0852602 | 0.22099234 |
| GOLGA6L10 | 45.2639688 | | 0.467956961 | 0.73353841 | 0.63794473 | 0.52350966 | 0.6991222 |
| GOLGA6L9 | 55.4733619 | | -0.073170411 | 0.30703238 | -0.238315 | 0.81163681 | 0.89791007 |
| RPS17 | 99.0336222 | | -0.196219031 | 0.57349363 | -0.3421468 | 0.73224039 | 0.84791916 |
| CPEB1 | 106.896302 | | 0.57839672 | 0.55617708 | 1.03995065 | 0.29836283 | 0.49840643 |
| AP3B2 | 63.5023768 | | -1.702094218 | 0.55755431 | -3.0527864 | 0.00226727 | 0.01689913 |
| AC105339.3 | 42.7979802 | | -0.29960603 | 0.62999707 | -0.4755673 | 0.63438265 | 0.78445331 |
| SNHG21 | 67.4688629 | | -0.114155516 | 0.31104199 | -0.36701 | 0.71361157 | 0.8362667 |
| WHAMM | 885.615431 | | -0.691897919 | 0.24321582 | -2.84479 | 0.00444407 | 0.0277643 |
| HOMER2 | 510.97858 | | 0.056068275 | 0.63978243 | 0.08763647 | 0.93016561 | 0.96282888 |
| AC022558.2 | 125.481424 | | -0.181974738 | 0.52652 | -0.3456179 | 0.72962989 | 0.84639784 |
| C15orf40 | 347.757791 | | 0.279289169 | 0.1925648 | 1.45036457 | 0.14695688 | 0.31635729 |
| BTBD1 | 658.37475 | | -0.037522597 | 0.22095306 | -0.1698216 | 0.86515046 | 0.92950442 |
| HDGFL3 | 46.5605333 | | -0.709464588 | 0.68919343 | -1.0294129 | 0.3032857 | 0.50363546 |
| ADAMTSL3 | 384.316682 | | 1.044174126 | 0.71450949 | 1.46138595 | 0.14390955 | 0.3116187 |
| GOLGA6L4 | 64.2060317 | | -0.482568913 | 0.39821295 | -1.2118363 | 0.22557505 | 0.41794501 |
| AC048382.5 | 197.050609 | | 0.500350483 | 0.30710557 | 1.62924586 | 0.10326098 | 0.25095251 |
| ZSCAN2 | 187.976822 | | 0.751077578 | 0.26202315 | 2.86645505 | 0.00415097 | 0.02636578 |
| WDR73 | 701.281179 | | -0.002169911 | 0.13181592 | -0.0164617 | 0.98686607 | 0.99360989 |
| NMB | 68.6611267 | | -1.523428884 | 0.50894969 | -2.9932799 | 0.00275996 | 0.01948035 |
| SEC11A | 1137.04867 | | -0.236919611 | 0.21591347 | -1.0972896 | 0.27251481 | 0.47050768 |
| ZNF592 | 936.870428 | | -0.077749317 | 0.18457396 | -0.4212367 | 0.67358228 | 0.81073146 |
| ALPK3 | 605.278203 | | -1.526636693 | 0.70064787 | -2.1788929 | 0.02933962 | 0.10732805 |
| SLC28A1 | 5646.85107 | | 1.119345663 | 0.58945743 | 1.89894234 | 0.05757206 | 0.17022006 |
| PDE8A | 4644.04416 | | 0.02318862 | 0.27639594 | 0.08389638 | 0.93313882 | 0.96462605 |
| AKAP13 | 5753.3566 | | -0.33263427 | 0.23290532 | -1.4281952 | 0.15323568 | 0.32534099 |
| AC087286.2 | 180.202557 | | -0.680949634 | 0.48874345 | -1.393266 | 0.16353937 | 0.3402651 |
| AC021739.2 | 39.1279572 | | 0.013777528 | 0.49020762 | 0.02810549 | 0.97757801 | 0.98883195 |
| KLHL25 | 170.182862 | | -0.152306762 | 0.45175339 | -0.3371458 | 0.73600699 | 0.8507199 |
| AGBL1 | 58.3879388 | | 1.245766702 | 0.48684118 | 2.55887699 | 0.01050109 | 0.05230321 |
| MRPL46 | 144.855909 | | -0.227427816 | 0.32462916 | -0.7005773 | 0.48356686 | 0.66763205 |

| | | | | | | |
|---|---|---|---|---|---|---|
| MRPS11 | 249.572408 | -0.546985176 | 0.30037698 | -1.8209957 | 0.06860751 | 0.1916062 |
| DET1 | 186.446052 | -0.039068842 | 0.37738325 | -0.1035256 | 0.9175458 | 0.95587788 |
| AEN | 103.843599 | -2.413074397 | 0.69927251 | -3.4508355 | 0.00055885 | 0.00579441 |
| ISG20 | 204.05735 | -4.159391125 | 0.87072955 | -4.7769036 | 1.78E-06 | 6.19E-05 |
| ABHD2 | 584.2243 | -0.46701508 | 0.43774378 | -1.0668686 | 0.28603119 | 0.48564716 |
| FANCI | 72.1151986 | -0.766059453 | 0.32719403 | -2.3413002 | 0.01921671 | 0.08008618 |
| POLG | 314.445399 | -0.453807252 | 0.2328269 | -1.9491187 | 0.05128126 | 0.15784789 |
| PEX11A | 165.345149 | 0.055843396 | 0.39612255 | 0.14097505 | 0.88788965 | 0.94209263 |
| WDR93 | 100.944776 | 0.797671206 | 0.50152113 | 1.59050369 | 0.11172132 | 0.26457569 |
| MESP1 | 41.6817923 | -1.415275225 | 0.39029552 | -3.6261631 | 0.00028766 | 0.00354268 |
| ANPEP | 6432.01137 | 0.218164627 | 0.42092684 | 0.51829583 | 0.60425188 | 0.76216808 |
| AP3S2 | 73.9280379 | 0.11588258 | 0.32928588 | 0.35192089 | 0.72489759 | 0.84347804 |
| ZNF710 | 198.95113 | -1.045937163 | 0.23664791 | -4.4198031 | 9.88E-06 | 0.00025264 |
| ZNF710-AS1 | 69.655697 | 0.643044611 | 0.52939877 | 1.21466965 | 0.22449211 | 0.41674812 |
| IDH2 | 558.738898 | -0.072963104 | 0.37648546 | -0.1938006 | 0.84633203 | 0.91896252 |
| SEMA4B | 1317.48402 | -1.467549084 | 0.38464479 | -3.8153359 | 0.000136 | 0.00196017 |
| CIB1 | 777.809627 | -0.949153624 | 0.35489896 | -2.6744334 | 0.00748556 | 0.04091094 |
| AC091167.6 | 72.749895 | 0.265876186 | 0.31576119 | 0.84201666 | 0.39977863 | 0.59394815 |
| ZNF774 | 36.3532025 | 0.190920077 | 0.29220516 | 0.6533768 | 0.51351339 | 0.69196393 |
| IQGAP1 | 1599.18959 | -1.259933216 | 0.26371214 | -4.7776837 | 1.77E-06 | 6.18E-05 |
| CRTC3 | 1463.25149 | -0.522770773 | 0.16619617 | -3.1455044 | 0.00165801 | 0.01329024 |
| CRTC3-AS1 | 177.191737 | 0.44976532 | 0.35229225 | 1.2766824 | 0.2017144 | 0.38909361 |
| FURIN | 151.274426 | -0.644962828 | 0.3513597 | -1.8356198 | 0.06641391 | 0.18742751 |
| MAN2A2 | 212.65093 | 0.185849362 | 0.25238304 | 0.73637817 | 0.46150059 | 0.64993204 |
| HDDC3 | 84.94468 | 0.271396335 | 0.31066643 | 0.87359403 | 0.38233938 | 0.57906281 |
| UNC45A | 313.785439 | -0.32053031 | 0.24447783 | -1.3110813 | 0.1898303 | 0.37390486 |
| RCCD1 | 102.573511 | 0.158013415 | 0.40702605 | 0.3882145 | 0.6978573 | 0.82589622 |
| AC104035.1 | 301.389551 | -2.259267002 | 0.8216212 | -2.7497672 | 0.00596376 | 0.03447714 |
| SLCO3A1 | 1271.04124 | -2.723049482 | 0.46082109 | -5.9091251 | 3.44E-09 | 3.25E-07 |
| FAM174B | 303.599476 | 0.654426151 | 0.34225658 | 1.91209222 | 0.05586437 | 0.16686269 |
| AC013394.1 | 1238.7126 | -0.150856771 | 0.22332574 | -0.675501 | 0.49935749 | 0.68099623 |
| CHD2 | 2445.56792 | -0.007883273 | 0.26643721 | -0.0295877 | 0.97639585 | 0.98836555 |
| MCTP2 | 1711.31473 | -0.613281558 | 0.21887736 | -2.8019415 | 0.00507961 | 0.03065035 |
| AC012409.2 | 270.526039 | 0.118491855 | 0.69248868 | 0.17111017 | 0.86413714 | 0.92885255 |
| NR2F2-AS1 | 2158.95166 | 0.574804654 | 0.61665298 | 0.93213635 | 0.35126607 | 0.55051498 |
| NR2F2 | 640.862494 | 0.435069121 | 0.23324609 | 1.86527939 | 0.06214225 | 0.17909669 |
| AC087477.2 | 155.969654 | 0.062047939 | 0.54676194 | 0.11348255 | 0.909648 | 0.95259723 |
| LINC00923 | 155.003451 | -1.141657782 | 0.89198527 | -1.2799065 | 0.20057801 | 0.38772003 |
| ARRDC4 | 107.275344 | -0.65934669 | 0.47686692 | -1.3826639 | 0.16676794 | 0.34469756 |
| FAM169B | 182.789231 | -1.251301352 | 0.63062358 | -1.9842286 | 0.04723036 | 0.14989269 |
| IGF1R | 11548.4031 | 0.945754781 | 0.3059266 | 3.09144343 | 0.00199186 | 0.0153326 |
| AC036108.1 | 90.3334844 | -0.193208425 | 0.49471876 | -0.3905419 | 0.69613585 | 0.82508568 |
| SYNM | 43.9640904 | -1.459599784 | 0.57481319 | -2.5392594 | 0.01110874 | 0.05432306 |
| TTC23 | 724.097548 | 0.432337824 | 0.18702629 | 2.311642 | 0.02079742 | 0.08486759 |
| LRRC28 | 1702.36179 | 1.257364234 | 0.36093765 | 3.48360512 | 0.00049471 | 0.00531674 |
| MEF2A | 3590.56718 | -0.150962769 | 0.24872292 | -0.6069516 | 0.54388305 | 0.71416138 |
| LYSMD4 | 139.609612 | 0.364091919 | 0.35683648 | 1.02033267 | 0.30757071 | 0.5077145 |
| AC090825.1 | 186.298401 | -0.280698876 | 0.55684659 | -0.5040865 | 0.61420056 | 0.76975835 |
| ADAMTS17 | 212.662509 | -0.577731454 | 0.71450293 | -0.8085781 | 0.41875784 | 0.61199634 |
| AC027020.2 | 44.6556192 | 0.984490023 | 0.44007993 | 2.23707094 | 0.0252817 | 0.09715064 |
| LINS1 | 261.162641 | 0.490804642 | 0.17807972 | 2.75609513 | 0.0058496 | 0.03403245 |
| ASB7 | 481.626869 | 0.36045136 | 0.15440633 | 2.3344338 | 0.01957302 | 0.08115274 |
| AC087762.1 | 6166.09991 | 1.099206508 | 0.63312747 | 1.73615355 | 0.08253669 | 0.21668845 |
| LRRK1 | 85.0855802 | -1.85124169 | 0.62956105 | -2.9405277 | 0.00327654 | 0.02208038 |
| CHSY1 | 557.573936 | -0.769420086 | 0.25412705 | -3.0276985 | 0.00246424 | 0.01799537 |
| SELENOS | 526.327075 | -1.454651232 | 0.35821691 | -4.0608112 | 4.89E-05 | 0.000889 |
| SNRPA1 | 319.794306 | -0.531440836 | 0.17811732 | -2.9836561 | 0.00284827 | 0.01994588 |
| PCSK6 | 44.0522895 | -0.973090595 | 0.42837335 | -2.2715946 | 0.023111 | 0.09131812 |
| TM2D3 | 357.175039 | -0.60141659 | 0.25243633 | -2.3824486 | 0.01719793 | 0.07419957 |
| TARSL2 | 732.287058 | -0.356453954 | 0.28953933 | -1.2311072 | 0.21828277 | 0.40947064 |
| OR4F15 | 27.992529 | -1.379232022 | 0.84405271 | -1.6340591 | 0.10224644 | 0.24973128 |
| POLR3K | 38.3364201 | -0.242845467 | 0.33523661 | -0.7244002 | 0.46882008 | 0.65617047 |
| SNRNP25 | 65.7208523 | -0.653411549 | 0.49038475 | -1.3324467 | 0.18271344 | 0.36498063 |
| RHBDF1 | 63.1494442 | -1.687648361 | 0.35179658 | -4.7972278 | 1.61E-06 | 5.74E-05 |
| MPG | 231.05559 | -1.147570834 | 0.20046637 | -5.7245056 | 1.04E-08 | 8.40E-07 |
| NPRL3 | 619.888068 | 0.245244956 | 0.20086147 | 1.22096568 | 0.22209901 | 0.41444934 |
| LUC7L | 1234.29109 | 0.246839182 | 0.21815965 | 1.13146124 | 0.25786101 | 0.45523177 |
| FAM234A | 198.43821 | 0.946114598 | 0.37163355 | 2.54582667 | 0.01090193 | 0.05367556 |
| AXIN1 | 401.091122 | -0.882705288 | 0.46981725 | -1.8788269 | 0.06026813 | 0.17520579 |
| MRPL28 | 151.486121 | -0.211222754 | 0.40732583 | -0.5185597 | 0.60406783 | 0.76207618 |
| TMEM8A | 284.1446 | -0.797267595 | 0.33260796 | -2.397019 | 0.01652907 | 0.07210819 |
| NME4 | 99.8248027 | -1.026705421 | 0.50107918 | -2.0489884 | 0.04046325 | 0.1344883 |
| DECR2 | 278.194102 | 0.708870609 | 0.34946208 | 2.02846216 | 0.0425131 | 0.13914209 |
| RAB11FIP3 | 9281.46405 | 1.775840568 | 0.46685301 | 3.80385373 | 0.00014246 | 0.0020266 |
| CAPN15 | 505.161212 | 0.346764164 | 0.3584527 | 0.9673917 | 0.33334826 | 0.53275938 |
| PIGQ | 240.808586 | 0.617369739 | 0.30846633 | 2.00141697 | 0.04534747 | 0.14573923 |
| RAB40C | 583.231598 | -0.22132911 | 0.2974434 | -0.744105 | 0.45681298 | 0.64558567 |
| METTL26 | 154.006893 | -0.20811813 | 0.38844099 | -0.535778 | 0.59211201 | 0.75351214 |
| MCRIP2 | 571.135892 | 0.997443804 | 0.3603116 | 2.76828116 | 0.00563528 | 0.03312299 |
| WDR90 | 168.586769 | 0.414252686 | 0.30121571 | 1.37526921 | 0.169048 | 0.34771022 |
| RHOT2 | 316.482298 | 0.204997286 | 0.22815364 | 0.89850545 | 0.36891614 | 0.5663904 |
| RHBDL1 | 36.0577917 | 1.318839674 | 0.48940659 | 2.69477303 | 0.00704366 | 0.03916732 |
| STUB1 | 403.729067 | -0.286882029 | 0.2875754 | -0.9975889 | 0.31847874 | 0.51857782 |
| JMJD8 | 174.998119 | -0.162100963 | 0.38544758 | -0.4205526 | 0.67408185 | 0.8109927 |
| WDR24 | 44.3883982 | 0.403992367 | 0.35017637 | 1.15368255 | 0.24863034 | 0.44486777 |
| METRN | 207.253407 | -0.453006719 | 0.39932284 | -1.1344373 | 0.25661117 | 0.45384415 |
| FAM173A | 250.36366 | -0.56230593 | 0.30932147 | -1.8178691 | 0.06908415 | 0.19255756 |
| HAGHL | 67.1191758 | -1.956805889 | 0.62875241 | -3.1122042 | 0.00185696 | 0.01445392 |
| NARFL | 178.041335 | -0.242732777 | 0.21624907 | -1.1224686 | 0.26166327 | 0.45993814 |
| RPUSD1 | 44.8438218 | -0.593803507 | 0.52707009 | -1.126612 | 0.25990655 | 0.45823541 |
| LMF1 | 522.031504 | 0.3093118 | 0.33821871 | 0.9145319 | 0.36043743 | 0.5586436 |
| CACNA1H | 96.9651743 | 1.645475669 | 0.92793449 | 1.77326707 | 0.07618446 | 0.2054356 |
| AC120498.4 | 88.36923 | -0.983858259 | 0.94273207 | -1.0436245 | 0.29665917 | 0.49692132 |
| UBE2I | 797.498483 | -0.836924848 | 0.31493782 | -2.6574289 | 0.00787392 | 0.04240779 |
| TSR3 | 181.575764 | -0.538375811 | 0.33602031 | -1.6022121 | 0.10910871 | 0.26027726 |
| GNPTG | 411.883149 | 0.318241294 | 0.35892106 | 0.88666097 | 0.37526146 | 0.57203167 |
| UNKL | 264.213354 | -0.318939031 | 0.27435719 | -1.1624956 | 0.24503421 | 0.44048313 |
| C16orf91 | 47.0437143 | -1.030269792 | 0.46622931 | -2.209792 | 0.0271196 | 0.10202347 |
| CCDC154 | 26.5404311 | 0.907868263 | 0.61858784 | 1.46764646 | 0.14220028 | 0.30958374 |
| CLCN7 | 689.022698 | -0.108572509 | 0.38267779 | -0.2837178 | 0.77662663 | 0.87664193 |
| TELO2 | 113.575035 | 0.40699788 | 0.48688458 | 0.83592272 | 0.40319838 | 0.59741003 |
| IFT140 | 435.108409 | 0.589739849 | 0.3586466 | 1.64434808 | 0.10010432 | 0.2460337 |
| AL133297.2 | 35.0286639 | -0.33910284 | 0.55677014 | -0.6090536 | 0.54248893 | 0.71327538 |
| CRAMP1 | 74.3540012 | 0.14706472 | 0.32531805 | 0.45206443 | 0.65122257 | 0.79571294 |
| JPT2 | 379.268678 | -1.176240423 | 0.28751776 | -4.0910183 | 4.29E-05 | 0.00080094 |
| MAPK8IP3 | 824.146906 | -0.39345188 | 0.2345289 | -1.6776264 | 0.09342006 | 0.23525917 |
| NME3 | 488.464965 | 0.28314108 | 0.43314511 | 0.65368643 | 0.51331384 | 0.69190607 |
| MRPS34 | 280.961536 | -0.743955664 | 0.31001329 | -2.3997541 | 0.01640609 | 0.07180021 |
| EME2 | 143.077852 | 0.709836242 | 0.28047448 | 2.53084079 | 0.01137895 | 0.05523061 |
| SPSB3 | 355.251939 | 0.312193541 | 0.21885821 | 1.42646487 | 0.1537342 | 0.32614683 |
| NUBP2 | 115.973359 | 0.025330834 | 0.28724166 | 0.08818649 | 0.92972845 | 0.96271389 |
| HAGH | 1150.68682 | 0.801968848 | 0.36123795 | 2.22005703 | 0.0264149 | 0.10018511 |
| FAHD1 | 441.769746 | 0.044207145 | 0.40179879 | 0.11002309 | 0.91239106 | 0.9535692 |

| | | | | | | |
|---|---|---|---|---|---|---|
| MEIOB | 50.5951512 | 0.838674256 | 0.56869804 | 1.47472684 | 0.140286 | 0.30702542 |
| MSRB1 | 341.410516 | -0.565374012 | 0.31677762 | -1.7847663 | 0.07429924 | 0.20214761 |
| NDUFB10 | 358.43501 | -0.51244553 | 0.38623317 | -1.3267776 | 0.18458226 | 0.36747552 |
| RPS2 | 909.993092 | -0.975307688 | 0.42762234 | -2.2807688 | 0.02256213 | 0.08995321 |
| SNHG9 | 120.770084 | 0.003374363 | 0.40378543 | 0.00835682 | 0.9933323 | 0.99654041 |
| TBL3 | 55.139896 | -0.200971698 | 0.49652499 | -0.4047565 | 0.68565653 | 0.81809663 |
| GFER | 72.0868253 | -0.023374685 | 0.25999747 | -0.0899035 | 0.92836389 | 0.96208463 |
| ZNF598 | 131.534851 | -0.819547976 | 0.47259675 | -1.734138 | 0.0828936 | 0.21737566 |
| SLC9A3R2 | 58.7439037 | 0.514626226 | 0.2975636 | 1.7294663 | 0.08372567 | 0.21851253 |
| TSC2 | 427.937255 | 0.232796661 | 0.21210964 | 1.09752984 | 0.27240984 | 0.47044497 |
| PKD1 | 338.78679 | 0.08089156 | 0.33437656 | 0.24191755 | 0.80884405 | 0.8961735 |
| SNHG19 | 155.74785 | -0.383623078 | 0.38954834 | -0.9847894 | 0.32472753 | 0.52501213 |
| TRAF7 | 182.664714 | -0.556684781 | 0.30115487 | -1.8485 | 0.06453005 | 0.18400527 |
| MLST8 | 81.1580834 | 0.20405383 | 0.45517698 | 0.44829559 | 0.65393988 | 0.79768212 |
| BRICD5 | 36.2864212 | -0.124792559 | 0.56272822 | -0.2217635 | 0.82449802 | 0.90635932 |
| PGP | 299.013826 | -0.877067978 | 0.37196459 | -2.3579341 | 0.01837696 | 0.07774473 |
| E4F1 | 169.318787 | -0.011404488 | 0.29813255 | -0.0382531 | 0.9694859 | 0.98478903 |
| ECI1 | 738.489369 | 0.076955361 | 0.14005367 | 0.54947051 | 0.58268259 | 0.74567914 |
| RNPS1 | 473.295851 | -0.457359627 | 0.27448764 | -1.6662303 | 0.09566755 | 0.23872476 |
| ABCA3 | 255.572198 | -1.494552414 | 0.40439293 | -3.6957926 | 0.0002192 | 0.00287247 |
| TBC1D24 | 41.9332126 | 0.552434965 | 0.43514746 | 1.26953509 | 0.20425028 | 0.39249385 |
| AMDHD2 | 370.048971 | 0.664166411 | 0.46268166 | 1.43547167 | 0.15115273 | 0.32246687 |
| PDPK1 | 1895.60559 | -0.184564959 | 0.19829953 | -0.9307382 | 0.35198898 | 0.55108727 |
| AC093525.6 | 200.702606 | 0.447707981 | 0.38181242 | 1.17258622 | 0.24096178 | 0.43618862 |
| AC093525.7 | 157.05409 | 0.347407402 | 0.36206963 | 0.9595044 | 0.3373047 | 0.53748379 |
| AC093525.4 | 556.182563 | 0.041830489 | 0.37437062 | 0.1117355 | 0.91103313 | 0.95310114 |
| KCTD5 | 144.232319 | -0.163270777 | 0.37041652 | -0.4407762 | 0.65937505 | 0.80181456 |
| SRRM2-AS1 | 45.4000426 | 0.606221622 | 0.377077 | 1.60768653 | 0.10790385 | 0.25821195 |
| SRRM2 | 4093.09226 | 0.528683682 | 0.20876857 | 2.53239121 | 0.01132875 | 0.0550845 |
| ELOB | 446.714361 | -0.683540048 | 0.37850026 | -1.805917 | 0.0709313 | 0.19586128 |
| FLYWCH2 | 98.3269008 | 0.372162343 | 0.30252784 | 1.23017554 | 0.21863138 | 0.40984049 |
| FLYWCH1 | 318.854934 | 0.436334724 | 0.32319601 | 1.35006224 | 0.17699602 | 0.3574605 |
| TNFRSF12A | 557.422622 | -2.99469648 | 0.60606961 | -4.9411758 | 7.77E-07 | 3.15E-05 |
| HCFC1R1 | 236.717957 | -0.518615777 | 0.35794333 | -1.4488768 | 0.147372 | 0.31681745 |
| THOC6 | 66.9426693 | -0.692890332 | 0.54423283 | -1.2731506 | 0.20296462 | 0.39079028 |
| MMP25-AS1 | 130.057394 | -0.261117135 | 0.49271466 | -0.5299561 | 0.59614238 | 0.75632392 |
| IL32 | 1760.60923 | -1.15020001 | 0.45794015 | -2.5116819 | 0.01201573 | 0.05736301 |
| ZNF213-AS1 | 195.729856 | 0.469193109 | 0.15882744 | 2.95410611 | 0.00313576 | 0.02133095 |
| ZNF205 | 127.392775 | 0.885207433 | 0.35280576 | 2.5090504 | 0.01210562 | 0.05765523 |
| ZNF213 | 268.88516 | -0.07793365 | 0.35444049 | -0.219878 | 0.8259662 | 0.9066142 |
| ZNF200 | 58.6102941 | -0.063118691 | 0.2594019 | -0.2433239 | 0.80775446 | 0.89588422 |
| ZNF263 | 136.626004 | -0.07225243 | 0.27266973 | -0.2649815 | 0.79102372 | 0.88629048 |
| TIGD7 | 111.96233 | 0.143748004 | 0.27687131 | 0.51918707 | 0.6036303 | 0.76169806 |
| ZNF75A | 444.374607 | -0.1301684 | 0.501248 | -0.2596886 | 0.79510397 | 0.8882949 |
| AC025283.3 | 121.911521 | -0.179492886 | 0.39427346 | -0.4552497 | 0.64892959 | 0.79404373 |
| ZSCAN32 | 125.638869 | -0.023666922 | 0.26042196 | -0.0908791 | 0.92758863 | 0.96193661 |
| NAA60 | 204.003426 | 0.376925599 | 0.27399895 | 1.37564616 | 0.16893121 | 0.34755609 |
| CLUAP1 | 485.410034 | 0.289464029 | 0.30545865 | 0.94763737 | 0.34331409 | 0.54383762 |
| SLX4 | 60.7809813 | 0.011498743 | 0.37864326 | 0.03036827 | 0.97577335 | 0.98788132 |
| DNASE1 | 580.781143 | 0.176025249 | 0.26088426 | 0.67472545 | 0.49985021 | 0.68139272 |
| TRAP1 | 870.610522 | 0.364619065 | 0.18606643 | 1.95961773 | 0.05004049 | 0.1554635 |
| CREBBP | 1836.2939 | 0.464471274 | 0.16527086 | 2.81036401 | 0.00494855 | 0.03005307 |
| ADCY9 | 3132.56358 | 0.371863531 | 0.24734323 | 1.50343118 | 0.13272789 | 0.29620664 |
| LINC01569 | 70.9054426 | 0.914150729 | 0.61260493 | 1.49223535 | 0.13563746 | 0.30015865 |
| TFAP4 | 53.3359876 | 0.292534238 | 0.47390809 | 0.61728053 | 0.53704971 | 0.70939003 |
| GLIS2 | 377.627014 | 1.017089449 | 0.4052279 | 2.5099196 | 0.01207587 | 0.0575335 |
| GLIS2-AS1 | 40.9743237 | 0.69448801 | 0.53002246 | 1.3102992 | 0.19009464 | 0.3741919 |
| PAM16 | 53.882526 | -0.105075657 | 0.43894659 | -0.2393814 | 0.81080984 | 0.89735732 |
| VASN | 176.105157 | -0.631690525 | 0.58365429 | -1.0823025 | 0.27911812 | 0.47769659 |
| DNAJA3 | 412.70461 | -0.03387886 | 0.1651141 | -0.2051845 | 0.83742796 | 0.91436098 |
| NMRAL1 | 369.003966 | 0.562099832 | 0.20656689 | 2.7211516 | 0.00650549 | 0.03678662 |
| HMOX2 | 742.124151 | 0.417785479 | 0.30654816 | 1.36287059 | 0.1729233 | 0.35255982 |
| CDIP1 | 396.28936 | 0.246083279 | 0.17515973 | 1.40490782 | 0.16004869 | 0.33564679 |
| UBALD1 | 188.020339 | 0.187003602 | 0.27736041 | 0.674226 | 0.50016764 | 0.68142311 |
| MGRN1 | 813.224283 | 0.07366118 | 0.21462238 | 0.34321294 | 0.73143827 | 0.84727663 |
| NUDT16L1 | 145.623312 | 0.197898989 | 0.33564726 | 0.58960407 | 0.55545612 | 0.72476748 |
| ANKS3 | 571.646425 | 0.233710084 | 0.21753431 | 1.07435966 | 0.28266152 | 0.48175669 |
| ZNF500 | 180.497523 | 0.747487847 | 0.31076171 | 2.40534091 | 0.01615739 | 0.07102927 |
| ROGDI | 136.885494 | 1.63079284 | 0.44897133 | 3.63228726 | 0.00028092 | 0.00348153 |
| GLYR1 | 1085.77142 | -0.016975307 | 0.27238892 | -0.0623201 | 0.95030791 | 0.97462556 |
| UBN1 | 652.349425 | 0.637762034 | 0.35970725 | 1.77300302 | 0.07622821 | 0.2054356 |
| PPL | 280.221646 | -0.348245008 | 0.27130897 | -1.2835735 | 0.19929121 | 0.38577668 |
| NAGPA | 136.00168 | 0.156154634 | 0.28766232 | 0.54284007 | 0.58723992 | 0.74995785 |
| ALG1 | 179.637724 | 0.996906017 | 0.34227392 | 2.91259707 | 0.00358437 | 0.02366615 |
| EEF2KMT | 47.0351242 | -0.630554616 | 0.49656955 | -1.2698213 | 0.20414828 | 0.39237089 |
| AC026458.2 | 80.8532995 | 1.467243602 | 0.37398994 | 3.92321676 | 8.74E-05 | 0.00140899 |
| RBFOX1 | 430.059598 | 0.36735718 | 0.48064826 | 0.76429524 | 0.44469132 | 0.63606218 |
| METTL22 | 208.987082 | -0.431238184 | 0.30278625 | -1.424233 | 0.15437902 | 0.32695817 |
| ABAT | 1284.56387 | 1.646731283 | 0.44381882 | 3.71036832 | 0.00020696 | 0.00275351 |
| TMEM186 | 71.9173161 | -0.067133419 | 0.26194426 | -0.256289 | 0.79772773 | 0.88977573 |
| PMM2 | 1209.73718 | -0.546949222 | 0.34855611 | -1.5691856 | 0.1166047 | 0.2720878 |
| AC022167.2 | 76.0236817 | 0.057364771 | 0.40205668 | 0.14267832 | 0.88654424 | 0.94149391 |
| CARHSP1 | 1515.33702 | -1.138666387 | 0.38993827 | -2.9201196 | 0.00349897 | 0.02321413 |
| USP7 | 576.017471 | -0.117666593 | 0.18297385 | -0.6430787 | 0.52017302 | 0.69607397 |
| C16orf72 | 642.573392 | -0.148372716 | 0.26033109 | -0.5699385 | 0.5687194 | 0.73493324 |
| AC087190.1 | 69.3713337 | -0.862751827 | 0.57043343 | -1.5124496 | 0.13041952 | 0.29297692 |
| ATF7IP2 | 432.091478 | 0.648407744 | 0.46907696 | 1.38230567 | 0.16687786 | 0.34469756 |
| NUBP1 | 160.226139 | -0.20490231 | 0.29731354 | -0.6891792 | 0.49071049 | 0.67349241 |
| AC133065.1 | 93.0232129 | 2.770448925 | 0.73491233 | 3.76976793 | 0.0001634 | 0.00226463 |
| CIITA | 266.322237 | 0.325622836 | 0.70976823 | 0.45877348 | 0.64639684 | 0.79264283 |
| DEXI | 206.291289 | -0.080351108 | 0.25744046 | -0.3121153 | 0.75495289 | 0.86260933 |
| CLEC16A | 1375.08288 | -0.055117101 | 0.20096377 | -0.2742639 | 0.78388185 | 0.88133085 |
| AC007014.1 | 58.1946974 | -1.273111574 | 0.48092286 | -2.6472262 | 0.00811551 | 0.04340196 |
| AC007220.1 | 86.8319622 | 0.884281603 | 0.35483476 | 2.49209408 | 0.01269924 | 0.05971474 |
| SOCS1 | 74.9113704 | -2.095341799 | 0.58239277 | -3.5978156 | 0.0003209 | 0.003872 |
| AC099489.1 | 59.6239287 | -1.197664842 | 0.44842036 | -2.6708529 | 0.00756588 | 0.04115297 |
| LITAF | 814.132879 | -1.410704056 | 0.32745721 | -4.3080562 | 1.65E-05 | 0.00037909 |
| TXNDC11 | 1275.9373 | 0.576362003 | 0.15160201 | 3.80180973 | 0.00014364 | 0.00203814 |
| ZC3H7A | 1169.53526 | 0.306085058 | 0.16726268 | 1.82996622 | 0.06725499 | 0.18892497 |
| RSL1D1 | 250.533328 | -0.591448036 | 0.40196226 | -1.4714019 | 0.14118246 | 0.30819999 |
| AC007216.4 | 387.644291 | 0.282785404 | 0.25896269 | 1.09199285 | 0.27483624 | 0.47259765 |
| GSPT1 | 1077.64886 | -0.347753671 | 0.25850112 | -1.3452695 | 0.17853822 | 0.35951571 |
| NPIPB2 | 233.572485 | 0.646659848 | 0.70271626 | 0.92022895 | 0.35745313 | 0.5561453 |
| SNX29 | 17676.6552 | 0.582264961 | 0.28855594 | 2.01785815 | 0.04360604 | 0.14150283 |
| AC010333.2 | 130.153576 | -0.652017365 | 0.67384238 | -0.9676111 | 0.33323863 | 0.53275938 |
| CPPED1 | 1443.51487 | 0.091407434 | 0.41315762 | 0.22124107 | 0.82490473 | 0.90642548 |
| AC109597.1 | 77.3291659 | -0.430537458 | 0.63250493 | -0.6806863 | 0.49606999 | 0.67795727 |
| SHISA9 | 934.432354 | -2.314878485 | 0.64865465 | -3.568738 | 0.0003587 | 0.0042088 |
| U91319.1 | 427.827615 | -1.856767327 | 0.72148667 | -2.5735296 | 0.0100667 | 0.05071088 |
| ERCC4 | 225.454511 | 0.268226825 | 0.22347608 | 1.20024847 | 0.23004286 | 0.42313439 |
| MKL2 | 2510.25732 | -0.138439939 | 0.18242379 | -0.7588919 | 0.44791721 | 0.63841127 |
| AC130650.1 | 48.3642537 | -0.934054507 | 0.64623071 | -1.4453886 | 0.14834876 | 0.31826835 |
| MIR193BHG | 359.169983 | -0.37506025 | 0.56099473 | -0.6685629 | 0.50377436 | 0.68443207 |

| | | | | | | |
|---|---|---|---|---|---|---|
| PARN | 1035.86892 | 0.424805665 | 0.22114331 | 1.92095191 | 0.05473777 | 0.16464574 |
| BFAR | 495.728009 | -0.369630279 | 0.2691592 | -1.3732775 | 0.16966609 | 0.34859523 |
| NOMO1 | 200.273771 | 0.435128132 | 0.32646097 | 1.33286419 | 0.18257638 | 0.36485995 |
| NPIPA1 | 99.7876864 | 0.57394065 | 0.36940936 | 1.5536711 | 0.12026289 | 0.27766015 |
| PDXDC1 | 2709.04468 | 0.579381294 | 0.18575547 | 3.1190537 | 0.00181433 | 0.0141918 |
| NTAN1 | 154.842331 | -0.392460857 | 0.32450433 | -1.2094164 | 0.22650291 | 0.41904108 |
| RRN3 | 484.589525 | -0.046736168 | 0.27948685 | -0.1672214 | 0.86719589 | 0.9309351 |
| NPIPA5 | 32.7511581 | 0.779481252 | 0.58953817 | 1.32218963 | 0.18610501 | 0.36923618 |
| C16orf45 | 434.728704 | -0.269884823 | 0.48035581 | -0.5618436 | 0.5742226 | 0.73911815 |
| MARF1 | 669.405741 | 0.183442739 | 0.18903046 | 0.97044011 | 0.33182716 | 0.53141196 |
| NDE1 | 170.170857 | -0.221121502 | 0.3978867 | -0.5557399 | 0.5783887 | 0.74211119 |
| MYH11 | 39.5205356 | -0.038693787 | 0.36062056 | -0.1072978 | 0.91455274 | 0.95437477 |
| FOPNL | 292.979782 | 0.081821627 | 0.21743872 | 0.37629741 | 0.70669581 | 0.83210663 |
| ABCC1 | 545.727068 | -1.055979424 | 0.38719064 | -2.7272855 | 0.00638578 | 0.03633451 |
| ABCC6 | 1830.93793 | 1.550646394 | 0.35617873 | 4.35356262 | 1.34E-05 | 0.00032153 |
| NOMO3 | 72.0583577 | 0.267744375 | 0.23085316 | 1.15980381 | 0.24612869 | 0.4417554 |
| XYLT1 | 94.6168603 | -0.325119265 | 0.78755713 | -0.4128199 | 0.67973856 | 0.81423861 |
| NOMO2 | 118.831259 | -0.01718083 | 0.40784043 | -0.0421264 | 0.96639797 | 0.98289063 |
| RPS15A | 235.324775 | -0.575351249 | 0.50058598 | -1.1493555 | 0.25040942 | 0.44668947 |
| SMG1 | 3422.625 | 0.07085409 | 0.20332808 | 0.34847173 | 0.72748594 | 0.84519832 |
| AC092287.1 | 132.853159 | 0.605879353 | 0.51490042 | 1.17669229 | 0.23931835 | 0.43434945 |
| TMC7 | 63.0146854 | -0.248112867 | 0.33945589 | -0.7309134 | 0.46483204 | 0.65254534 |
| COQ7 | 131.517486 | 0.655415907 | 0.26054744 | 2.51553386 | 0.01188523 | 0.05686241 |
| SYT17 | 52.5157775 | -1.06147271 | 0.9008982 | -1.1782382 | 0.23870164 | 0.43383194 |
| GDE1 | 178.14038 | -0.635264491 | 0.3068262 | -2.0704376 | 0.03841139 | 0.13022283 |
| CCP110 | 227.873432 | 0.194227055 | 0.21508908 | 0.90300752 | 0.3665219 | 0.56397797 |
| VPS35L | 1264.85931 | 0.375772492 | 0.22750274 | 1.65172736 | 0.09859014 | 0.24331564 |
| KNOP1 | 191.258875 | -1.021153202 | 0.54525197 | -1.8728097 | 0.06109466 | 0.17704536 |
| IQCK | 773.421272 | 0.289163538 | 0.35871145 | 0.80611738 | 0.42017517 | 0.61367513 |
| UMOD | 733.450335 | 1.691179571 | 0.55780173 | 3.03186506 | 0.00243048 | 0.01782481 |
| ACSM5 | 461.794675 | 0.249468193 | 0.74933158 | 0.33292097 | 0.73919394 | 0.85193361 |
| AC137056.1 | 64.443814 | 2.906677275 | 0.85430083 | 3.40240484 | 0.00066796 | 0.00668278 |
| ACSM2A | 10082.0474 | 2.066272231 | 0.74004102 | 2.79210501 | 0.00523664 | 0.03137175 |
| ACSM2B | 10403.044 | 1.927580014 | 0.75014538 | 2.56960858 | 0.01018135 | 0.0511568 |
| ACSM3 | 1142.26674 | -1.513803326 | 0.57824669 | -2.6179196 | 0.00884677 | 0.04605492 |
| ACSM1 | 84.8119458 | -1.317259068 | 0.5975567 | -2.2044085 | 0.02749563 | 0.10294526 |
| THUMPD1 | 511.44976 | -0.021322517 | 0.17126067 | -0.1245033 | 0.9009168 | 0.94785397 |
| ERI2 | 213.09975 | 0.031566072 | 0.3669338 | 0.08602661 | 0.93144526 | 0.96354054 |
| REXO5 | 37.1370034 | -0.413256032 | 0.55996796 | -0.7379994 | 0.4605148 | 0.64921077 |
| LYRM1 | 336.558723 | 0.134577085 | 0.26307306 | 0.51155782 | 0.60896051 | 0.76564132 |
| DNAH3 | 53.0676061 | -1.275176116 | 0.47824831 | -2.6663474 | 0.00766804 | 0.0416096 |
| TMEM159 | 266.352599 | -0.602836929 | 0.34807661 | -1.7319088 | 0.08328981 | 0.21808088 |
| ANKS4B | 294.760999 | 1.155807822 | 0.46167013 | 2.50353606 | 0.01229591 | 0.05823777 |
| CRYM | 704.938055 | -0.258433501 | 0.64145028 | -0.4028894 | 0.68702961 | 0.81880835 |
| NPIPB3 | 118.858191 | 1.098112029 | 0.56260789 | 1.95182478 | 0.05095901 | 0.15730514 |
| METTL9 | 1503.12135 | 0.565852724 | 0.2856178 | 1.98115356 | 0.04757406 | 0.15049689 |
| IGSF6 | 66.0790891 | 0.160320882 | 0.58000878 | 0.27641113 | 0.78223231 | 0.88073421 |
| NPIPB4 | 421.821356 | 1.246877612 | 0.43913516 | 2.83939368 | 0.00451994 | 0.0281229 |
| UQCRC2 | 997.490888 | 0.049516578 | 0.15736594 | 0.3146588 | 0.75302073 | 0.8613901 |
| MOSMO | 969.404217 | 0.070621988 | 0.25320649 | 0.27891065 | 0.78031339 | 0.87935343 |
| EEF2K | 537.378642 | -0.480969909 | 0.54886035 | -0.8763065 | 0.38086342 | 0.57729679 |
| POLR3E | 329.50378 | 0.075354479 | 0.2333904 | 0.3228688 | 0.74679461 | 0.85681039 |
| CDR2 | 2882.12719 | 0.106823285 | 0.26008751 | 0.41072055 | 0.68127746 | 0.81457974 |
| AC092338.2 | 80.5253148 | -0.570011585 | 0.49246818 | -1.1574587 | 0.247085 | 0.44300772 |
| NPIPB5 | 1100.89857 | 0.242732581 | 0.30310872 | 0.80081029 | 0.42324148 | 0.61651127 |
| USP31 | 1263.96331 | -0.422334714 | 0.28752083 | -1.4688839 | 0.14186429 | 0.30898043 |
| COG7 | 672.688343 | 0.234663561 | 0.17263063 | 1.35933911 | 0.17403916 | 0.3537824 |
| AC008915.2 | 46.7516877 | 0.455639991 | 0.317987 | 1.43288876 | 0.15188963 | 0.32333455 |
| GGA2 | 1186.56931 | -0.198723059 | 0.39568616 | -0.5022239 | 0.61551001 | 0.77111743 |
| EARS2 | 199.276975 | -0.273224315 | 0.37390493 | -0.730732 | 0.46494284 | 0.65256722 |
| UBFD1 | 222.352688 | -0.153757982 | 0.32087808 | -0.4791788 | 0.63181142 | 0.78295264 |
| NDUFAB1 | 290.488348 | 0.061994259 | 0.19779095 | 0.31343324 | 0.75395153 | 0.8620632 |
| PALB2 | 239.451218 | 0.379073129 | 0.34055474 | 1.11310484 | 0.2656634 | 0.46352448 |
| DCTN5 | 412.039961 | -0.424721359 | 0.23614562 | -1.798557 | 0.07208878 | 0.19802023 |
| PRKCB | 58.5445895 | -1.63965347 | 0.7750689 | -2.1154938 | 0.03438788 | 0.12031025 |
| RBBP6 | 835.674807 | -0.014337287 | 0.23410582 | -0.0612428 | 0.95116587 | 0.97514066 |
| TNRC6A | 6488.77254 | 0.994348071 | 0.25571473 | 3.88850533 | 0.00010086 | 0.00156395 |
| AC008731.1 | 150.618948 | -0.577867398 | 0.57036916 | -1.0131463 | 0.31099029 | 0.51127879 |
| SLC5A11 | 1499.93759 | 1.292091204 | 0.78316111 | 1.64984086 | 0.09897549 | 0.24400306 |
| ARHGAP17 | 932.213894 | -0.073634054 | 0.16594271 | -0.4437318 | 0.65723651 | 0.80027901 |
| LCMT1 | 559.022329 | 0.172487003 | 0.13575513 | 1.27057448 | 0.20388007 | 0.39194728 |
| ZKSCAN2 | 47.0984465 | 0.330214879 | 0.33237479 | 0.99350159 | 0.32046558 | 0.52100838 |
| AC008741.2 | 75.5288884 | -0.234664721 | 0.37755415 | -0.6215392 | 0.53424488 | 0.70733681 |
| HS3ST4 | 39.855669 | -0.304860613 | 0.54971877 | -0.5545756 | 0.57918499 | 0.74258983 |
| KDM8 | 51.2675958 | 0.767439249 | 0.33414523 | 2.29672362 | 0.02163454 | 0.08748858 |
| NSMCE1 | 263.722409 | 0.294875971 | 0.27890099 | 1.05727831 | 0.29038459 | 0.49042651 |
| AC106739.1 | 135.211098 | 0.709905708 | 0.71781314 | 0.988984 | 0.32267097 | 0.52304293 |
| IL4R | 787.656176 | -1.887131055 | 0.42092386 | -4.4833074 | 7.35E-06 | 0.00019523 |
| GTF3C1 | 746.285267 | 0.201602917 | 0.23460643 | 0.85932392 | 0.39016183 | 0.58561422 |
| KIAA0556 | 1010.44706 | -0.221033518 | 0.2339058 | -0.9449681 | 0.34467516 | 0.54521392 |
| XPO6 | 925.467045 | -0.606539846 | 0.42458122 | -1.4285602 | 0.15313067 | 0.3252188 |
| NPIPB6 | 25.7786369 | 1.132122164 | 0.99910056 | 1.13314136 | 0.2571549 | 0.45431639 |
| NUPR1 | 594.979643 | -1.397183408 | 0.69881746 | -1.9993539 | 0.04557008 | 0.14614624 |
| SGF29 | 201.50777 | 0.092615946 | 0.35396151 | 0.26165542 | 0.79358712 | 0.88739587 |
| SULT1A2 | 49.3758978 | 0.888685227 | 0.79976662 | 1.11118069 | 0.26649058 | 0.46443565 |
| SULT1A1 | 195.101185 | 1.30896313 | 0.58123065 | 2.25205456 | 0.02431882 | 0.09443006 |
| ATXN2L | 549.676011 | -0.25266163 | 0.20719686 | -1.2194279 | 0.22268183 | 0.41477101 |
| TUFM | 580.522005 | -0.452657875 | 0.39760506 | -1.1384611 | 0.25492801 | 0.45204375 |
| SH2B1 | 463.261123 | 0.994814839 | 0.26028454 | 3.82202809 | 0.00013236 | 0.00191957 |
| AC133550.2 | 84.4169845 | -0.003151045 | 0.26892613 | -0.0117171 | 0.99065129 | 0.99522554 |
| ATP2A1 | 38.837713 | 0.14261367 | 0.29602985 | 0.48175435 | 0.62998046 | 0.78167305 |
| RABEP2 | 251.742575 | 0.032301199 | 0.34228513 | 0.09436927 | 0.92481582 | 0.96026441 |
| NFATC2IP | 887.024997 | -0.050377861 | 0.30397444 | -0.1657306 | 0.86836898 | 0.93161498 |
| AC109460.2 | 70.5482352 | -0.225495889 | 0.44703432 | -0.5044263 | 0.61396181 | 0.76974065 |
| SPNS1 | 305.375802 | -0.249368473 | 0.45680155 | -0.5459011 | 0.58513391 | 0.74775597 |
| LAT | 87.4366932 | 0.926531515 | 0.45721273 | 2.02647794 | 0.04271583 | 0.1396056 |
| AC009093.2 | 101.987982 | 0.439051749 | 0.92851029 | 0.47285609 | 0.63631585 | 0.78577016 |
| NPIPB11 | 91.7916112 | 1.36508634 | 0.32104113 | 4.25206058 | 2.12E-05 | 0.00046158 |
| NPIPB12 | 29.9437403 | 0.883508223 | 0.59154685 | 1.4935558 | 0.13529176 | 0.29978082 |
| QPRT | 698.524524 | 0.619660388 | 0.31640136 | 1.95846308 | 0.0501757 | 0.15567187 |
| KIF22 | 161.265777 | 0.04837094 | 0.24795164 | 0.19508215 | 0.84532863 | 0.91854292 |
| MAZ | 206.476603 | -0.420667793 | 0.38176997 | -1.1018881 | 0.27051031 | 0.46869989 |
| AC009133.1 | 846.755505 | -0.627673161 | 0.45098054 | -1.3917965 | 0.16398402 | 0.34082319 |
| MVP | 1657.26701 | -0.306734887 | 0.2849064 | -1.0766163 | 0.2816517 | 0.48053272 |
| CDIPT | 232.266814 | -0.08169396 | 0.24121049 | -0.3386833 | 0.73484833 | 0.84993403 |
| SEZ6L2 | 86.8830396 | -1.411258833 | 0.66299119 | -2.1286238 | 0.0332854 | 0.11780091 |
| ASPHD1 | 173.131179 | -1.532191289 | 0.48704282 | -3.1459068 | 0.00165573 | 0.01328264 |
| KCTD13 | 424.672713 | -0.197270021 | 0.30470231 | -0.6474188 | 0.5173609 | 0.69425481 |
| TMEM219 | 347.266933 | -0.053928825 | 0.32973379 | -0.1635526 | 0.87008335 | 0.93290304 |
| TAOK2 | 171.51953 | 0.327064578 | 0.33751889 | 0.969026 | 0.33253221 | 0.53198168 |
| HIRIP3 | 39.3174444 | 0.204235594 | 0.33568264 | 0.6084187 | 0.54290981 | 0.71356452 |
| INO80E | 271.71542 | 0.961154538 | 0.27669114 | 3.4737453 | 0.00051325 | 0.0054409 |
| PPP4C | 370.326311 | -0.69558302 | 0.35081449 | -1.9827659 | 0.04739358 | 0.15022483 |

| | | | | | | |
|---|---|---|---|---|---|---|
| YPEL3 | 326.061955 | -0.530721875 | 0.34559645 | -1.5356693 | 0.1246195 | 0.28455098 |
| MAPK3 | 103.94163 | -0.032006683 | 0.33688678 | -0.0950072 | 0.92430908 | 0.95992984 |
| NPIPB13 | 282.250511 | 0.685084874 | 0.34325613 | 1.99584162 | 0.04595117 | 0.14699011 |
| CD2BP2 | 245.950803 | 0.257655 | 0.36866162 | 0.69889292 | 0.48461895 | 0.66827686 |
| TBC1D10B | 176.885244 | 0.040748925 | 0.35448872 | 0.11495126 | 0.90848376 | 0.95197217 |
| ZNF48 | 63.9088051 | 0.408155407 | 0.30668763 | 1.33085057 | 0.18323818 | 0.36558246 |
| 1-Sep | 58.7569937 | 0.040449042 | 0.3153638 | 0.12826152 | 0.89794201 | 0.94717771 |
| ZNF771 | 166.385295 | -0.708267051 | 0.45032617 | -1.5727868 | 0.11576819 | 0.27092959 |
| DCTPP1 | 80.151496 | -0.979130129 | 0.35811816 | -2.734098 | 0.00625514 | 0.03569494 |
| SEPHS2 | 851.8433 | -1.059009856 | 0.37739282 | -2.8061208 | 0.00501419 | 0.03037717 |
| ITGAL | 116.225426 | -0.289861454 | 0.70401956 | -0.4117236 | 0.68054204 | 0.81442647 |
| ZNF768 | 153.101136 | 0.873074807 | 0.21355881 | 4.08821723 | 4.35E-05 | 0.00080957 |
| AC002310.4 | 83.2792021 | 1.030706121 | 0.31408836 | 3.28158015 | 0.00103227 | 0.00926827 |
| AC002310.1 | 38.169492 | 0.165213977 | 0.37696613 | 0.43827274 | 0.66118858 | 0.80309366 |
| ZNF688 | 84.3736365 | 0.182195136 | 0.29862229 | 0.61011901 | 0.54178297 | 0.71282636 |
| ZNF785 | 209.617089 | 0.334909691 | 0.321861 | 1.04054138 | 0.29808845 | 0.49833608 |
| AC093249.2 | 98.6755367 | 0.054576159 | 0.60633954 | 0.09000924 | 0.92827987 | 0.96208463 |
| ZNF689 | 61.6305445 | -0.181073312 | 0.60377334 | -0.2999028 | 0.7642513 | 0.86833493 |
| PRR14 | 159.85929 | 0.135834938 | 0.35329324 | 0.38448213 | 0.70062114 | 0.82796169 |
| FBRS | 238.425974 | 0.155118171 | 0.19087816 | 0.81265542 | 0.41641565 | 0.60980868 |
| SRCAP | 59.7372801 | 0.128533756 | 0.2983832 | 0.43076741 | 0.6666375 | 0.80635371 |
| PHKG2 | 213.041829 | -0.202744448 | 0.30148776 | -0.6724799 | 0.50127825 | 0.68247686 |
| CCDC189 | 55.7863735 | 0.318691418 | 0.34478322 | 0.92432405 | 0.35531762 | 0.55376221 |
| RNF40 | 283.861843 | 0.096153369 | 0.22113473 | 0.43481805 | 0.66369451 | 0.80442661 |
| ZNF629 | 72.9857082 | -0.435829733 | 0.42173573 | -1.0334191 | 0.30140783 | 0.50187041 |
| BCL7C | 361.155785 | 0.321297082 | 0.32150549 | 0.99935177 | 0.31762432 | 0.51761699 |
| MIR762HG | 111.879837 | 0.0375267 | 0.30808184 | 0.12180757 | 0.90305142 | 0.94909721 |
| FBXL19-AS1 | 76.0219262 | -1.585507305 | 0.70530711 | -2.2479673 | 0.02457828 | 0.09514104 |
| FBXL19 | 170.519151 | 0.258972856 | 0.39213276 | 0.66042138 | 0.50898345 | 0.68857444 |
| ORAI3 | 130.758391 | -0.447518671 | 0.31718183 | -1.4109215 | 0.15826776 | 0.33298176 |
| SETD1A | 231.938844 | -0.208228244 | 0.44119367 | -0.4719656 | 0.63695133 | 0.78640713 |
| HSD3B7 | 237.143251 | -1.090742937 | 0.58887321 | -1.8522543 | 0.06398931 | 0.18288188 |
| STX1B | 39.354035 | 0.363281466 | 0.46868162 | 0.77511353 | 0.43827258 | 0.63010302 |
| STX4 | 290.274288 | -0.113241199 | 0.30439911 | -0.3720155 | 0.70988128 | 0.83388131 |
| AC135050.3 | 96.1772196 | -0.844615884 | 0.53905608 | -1.5668423 | 0.11715155 | 0.27281623 |
| ZNF668 | 77.4328767 | -0.680959734 | 0.54470039 | -1.2501547 | 0.21124305 | 0.40059609 |
| ZNF646 | 56.3955608 | -0.012213268 | 0.38542754 | -0.0316876 | 0.9747212 | 0.98732216 |
| AC135050.2 | 33.3955655 | -1.377542324 | 0.57009386 | -2.416343 | 0.01567728 | 0.06949824 |
| BCKDK | 203.131739 | -0.384213335 | 0.33333916 | -1.1526198 | 0.24906645 | 0.4452209 |
| KAT8 | 328.237087 | 0.225004522 | 0.34685688 | 0.64869558 | 0.51653516 | 0.69386001 |
| PRSS8 | 390.412158 | -0.80088485 | 0.42788953 | -1.8717094 | 0.06124682 | 0.17733048 |
| FUS | 1433.86313 | 0.263873325 | 0.21105035 | 1.2502861 | 0.21119505 | 0.40056048 |
| AC026471.4 | 147.174186 | 0.293516691 | 0.35209256 | 0.83363503 | 0.40448668 | 0.59854247 |
| AC026471.1 | 47.3684935 | -0.449290173 | 0.39549271 | -1.1360264 | 0.2559455 | 0.45318346 |
| ARMC5 | 79.0742914 | 0.673222239 | 0.45010453 | 1.49570198 | 0.13473133 | 0.29911927 |
| SLC5A2 | 1105.74342 | 3.211526347 | 0.62671472 | 5.12438313 | 2.99E-07 | 1.37E-05 |
| AC026471.3 | 30.3930699 | -0.724728603 | 1.06164039 | -0.6826498 | 0.49482816 | 0.6769747 |
| C16orf58 | 295.054824 | 0.568347782 | 0.2538571 | 2.23884922 | 0.02516573 | 0.09678637 |
| ZNF720 | 546.198928 | 0.104071809 | 0.31731463 | 0.32797671 | 0.74292927 | 0.8540591 |
| ZNF267 | 399.570643 | -0.958099222 | 0.24448361 | -3.9188689 | 8.90E-05 | 0.00143128 |
| AC136944.1 | 34.8499526 | 0.723114108 | 0.35899978 | 2.01424668 | 0.04398364 | 0.14229024 |
| VPS35 | 621.482546 | -0.628764681 | 0.19031667 | -3.3037815 | 0.0009539 | 0.00874006 |
| MYLK3 | 36.5204607 | 0.143921044 | 0.4069257 | 0.35367892 | 0.72357952 | 0.84266894 |
| C16orf87 | 645.263142 | 0.298124753 | 0.32459257 | 0.91845834 | 0.35837896 | 0.5568239 |
| GPT2 | 1047.04882 | 0.621399554 | 0.51582077 | 1.20468114 | 0.22832642 | 0.42122075 |
| DNAJA2 | 263.828141 | -0.141932797 | 0.21279178 | -0.6670032 | 0.50477009 | 0.6850618 |
| ITFG1 | 3174.62958 | -0.014638505 | 0.21829266 | -0.0670591 | 0.94653467 | 0.97252756 |
| PHKB | 2264.28474 | -0.042163531 | 0.18400035 | -0.2291492 | 0.81875296 | 0.90192453 |
| LINC02133 | 48.1994826 | -0.284286394 | 0.53137795 | -0.5349985 | 0.59265091 | 0.75391796 |
| LONP2 | 1682.45567 | 0.229451959 | 0.12550692 | 1.8282016 | 0.0675193 | 0.1893386 |
| SIAH1 | 467.194981 | 0.09847386 | 0.18896493 | 0.5211224 | 0.60228151 | 0.76094319 |
| N4BP1 | 1124.45211 | 0.042085872 | 0.21879716 | 0.19235109 | 0.8474672 | 0.91938085 |
| AC026470.2 | 43.2192247 | -1.030469411 | 0.70363544 | -1.4644933 | 0.1430592 | 0.310615 |
| AC007611.1 | 75.6210256 | 0.066262581 | 0.49091305 | 0.13497824 | 0.89262908 | 0.94393356 |
| CNEP1R1 | 114.245658 | -0.075858692 | 0.19279552 | -0.3934671 | 0.69397453 | 0.82385996 |
| HEATR3 | 225.177625 | 0.383567091 | 0.30739194 | 1.24781114 | 0.21210023 | 0.40177692 |
| TENT4B | 1295.45634 | 0.952256919 | 0.28004673 | 3.4003501 | 0.000673 | 0.00672359 |
| BRD7 | 356.245676 | -0.186636978 | 0.1870728 | -0.9976703 | 0.31843925 | 0.51857513 |
| CYLD | 498.700214 | -0.796270183 | 0.42349207 | -1.8802481 | 0.06007427 | 0.17475339 |
| AC025810.1 | 499.927208 | 2.342366669 | 0.7553988 | 3.10083451 | 0.00192976 | 0.01492733 |
| SALL1 | 776.564543 | 0.441747269 | 0.49275757 | 0.89647993 | 0.3699965 | 0.56741351 |
| AC087564.1 | 1741.5529 | 0.146948327 | 0.51181943 | 0.28710971 | 0.77402832 | 0.87480262 |
| AC007344.1 | 235.265994 | 0.255234203 | 0.58489777 | 0.43637404 | 0.66256537 | 0.80369766 |
| HNRNPA1P4 | 61.187868 | -0.927772067 | 0.3958743 | -2.3436027 | 0.0190985 | 0.07981195 |
| TOX3 | 1477.59608 | -0.384528315 | 0.50325892 | -0.7640765 | 0.44482165 | 0.63616564 |
| AC007906.2 | 87.4263636 | 0.233237321 | 0.63808459 | 0.36552727 | 0.71471785 | 0.8368246 |
| CHD9 | 6370.43603 | 0.619592086 | 0.19616295 | 3.1585581 | 0.00158552 | 0.01289008 |
| AC007342.4 | 45.955225 | 1.182112132 | 0.6842526 | 1.72759611 | 0.08406066 | 0.2190115 |
| RBL2 | 1132.66811 | 0.254633274 | 0.32350549 | 0.78710649 | 0.43121953 | 0.62376247 |
| AKTIP | 165.006729 | 0.418309272 | 0.23714731 | 1.76392163 | 0.07774517 | 0.20807328 |
| RPGRIP1L | 288.010223 | 0.280805586 | 0.28657142 | 0.97987994 | 0.32714539 | 0.52705245 |
| FTO | 4762.58537 | 0.595237813 | 0.21857276 | 2.72329365 | 0.00646346 | 0.0366338 |
| IRX3 | 131.851079 | -0.405483381 | 0.5600691 | -0.7239881 | 0.46907302 | 0.65617047 |
| CRNDE | 265.744413 | 0.386520578 | 0.41583186 | 0.92951169 | 0.35262397 | 0.55163396 |
| IRX5 | 68.5523579 | 0.71161278 | 0.53447592 | 1.3314216 | 0.18305033 | 0.36548738 |
| LPCAT2 | 115.084431 | 0.683579798 | 0.34901205 | 1.95861374 | 0.05015804 | 0.15567187 |
| AMFR | 791.64393 | 0.403575674 | 0.23677564 | 1.70446454 | 0.08829434 | 0.22639124 |
| NUDT21 | 311.414489 | 0.004931812 | 0.15824395 | 0.03116588 | 0.97513725 | 0.98760206 |
| OGFOD1 | 270.992707 | -0.319588138 | 0.25939401 | -1.2320567 | 0.21792789 | 0.40908486 |
| BBS2 | 567.684838 | 0.36601135 | 0.25890284 | 1.41370158 | 0.15744954 | 0.33177901 |
| MT2A | 7396.0749 | -1.620373998 | 0.7687435 | -2.1078214 | 0.03504644 | 0.12176973 |
| MT1E | 2891.13849 | -1.351570025 | 0.85638681 | -1.5782238 | 0.11451421 | 0.26863705 |
| MT1M | 449.639 | -1.776893859 | 1.38449868 | -1.2834204 | 0.19934483 | 0.38582597 |
| MT1F | 782.116318 | 0.046789821 | 0.66581245 | 0.07027478 | 0.94397496 | 0.97118252 |
| MT1G | 12204.5379 | 0.078506376 | 0.84138833 | 0.09330576 | 0.92566066 | 0.96070498 |
| MT1H | 4341.43822 | 0.464822296 | 0.94435747 | 0.49221011 | 0.62257082 | 0.77599147 |
| MT1X | 2884.98461 | -0.509836665 | 0.75284108 | -0.6772169 | 0.49826838 | 0.68025544 |
| NUP93 | 376.541378 | -0.638868145 | 0.39930894 | -1.5999345 | 0.10961312 | 0.26102624 |
| SLC12A3 | 87.2118433 | 2.042630641 | 0.46784419 | 4.36604894 | 1.27E-05 | 0.00030856 |
| HERPUD1 | 1031.91132 | -0.220644123 | 0.46203122 | -0.4775524 | 0.63296881 | 0.7836086 |
| NLRC5 | 243.312826 | -2.811510019 | 0.66343936 | -4.2377799 | 2.26E-05 | 0.00048651 |
| CPNE2 | 50.1796028 | -2.260349902 | 0.34423575 | -6.5662847 | 5.16E-11 | 9.07E-09 |
| FAM192A | 578.50957 | -0.071514052 | 0.27056574 | -0.264313 | 0.79153871 | 0.88662427 |
| RSPRY1 | 1012.93903 | 0.164428614 | 0.17053188 | 0.9642104 | 0.33494045 | 0.53471101 |
| ARL2BP | 372.170795 | -0.54184839 | 0.21901878 | -2.4739815 | 0.01336166 | 0.06183265 |
| PLLP | 334.979927 | 0.678675205 | 0.39454468 | 1.7201479 | 0.08540556 | 0.22117023 |
| CX3CL1 | 258.739156 | 0.307460579 | 0.53607378 | 0.57354153 | 0.56627809 | 0.73309159 |
| CIAPIN1 | 377.937335 | 0.176433448 | 0.14657562 | 1.20370259 | 0.22870455 | 0.42169132 |
| COQ9 | 593.907778 | 0.495249775 | 0.24802388 | 1.99678259 | 0.04584881 | 0.14676543 |
| POLR2C | 194.415592 | -0.140218695 | 0.26939513 | -0.5204945 | 0.60271893 | 0.76135548 |
| DOK4 | 396.178115 | -0.248781853 | 0.22253847 | -1.1179274 | 0.26359799 | 0.46156588 |
| CCDC102A | 36.274841 | -0.219796002 | 0.41009956 | -0.5359577 | 0.59198781 | 0.75342404 |
| ADGRG5 | 301.586087 | 2.62705088 | 0.6433648 | 4.08329904 | 4.44E-05 | 0.00082466 |

| | | | | | | | |
|---|---|---|---|---|---|---|---|
| ADGRG1 | 2009.33578 | | -0.287884965 | 0.59518893 | -0.4836867 | 0.62860824 | 0.78060637 |
| KATNB1 | 44.6144747 | | -0.638565065 | 0.36841304 | -1.7332857 | 0.0830449 | 0.21766795 |
| KIFC3 | 3316.69993 | | 1.61926029 | 0.30794515 | 5.2582751 | 1.45E-07 | 7.64E-06 |
| AC092118.1 | 60.2694614 | | 0.662775662 | 0.33969313 | 1.95110118 | 0.05104501 | 0.1573841 |
| USB1 | 145.553438 | | -0.917916517 | 0.28605862 | -3.2088406 | 0.00133271 | 0.01123463 |
| MMP15 | 89.0083901 | | -0.465647997 | 0.40459751 | -1.1508919 | 0.24977671 | 0.44590902 |
| CFAP20 | 102.471965 | | 0.280072629 | 0.17207763 | 1.6275946 | 0.10361088 | 0.25157313 |
| CSNK2A2 | 520.555367 | | -0.244889704 | 0.22912148 | -1.0688204 | 0.28515062 | 0.48469241 |
| CCDC113 | 43.4666789 | | -1.120428448 | 0.47594932 | -2.3540919 | 0.01856802 | 0.07823911 |
| SETD6 | 136.061 | | -0.110886241 | 0.32600497 | -0.3401367 | 0.73375362 | 0.84924097 |
| CNOT1 | 1601.85166 | | 0.118695848 | 0.18856953 | 0.62945402 | 0.52905186 | 0.70339575 |
| AC009118.3 | 37.4920418 | | -0.008801044 | 0.48953168 | -0.0179785 | 0.98565601 | 0.99298265 |
| SLC38A7 | 320.046299 | | -0.08240556 | 0.46664323 | -0.1765922 | 0.85982871 | 0.92628466 |
| GOT2 | 430.147948 | | 0.223480192 | 0.17231215 | 1.29694967 | 0.19464851 | 0.37980117 |
| AC092378.1 | 102.211192 | | 0.494167412 | 0.34171023 | 1.44615926 | 0.14813254 | 0.31795376 |
| AC138305.1 | 307.65244 | | 0.608034659 | 0.50331902 | 1.20805022 | 0.22702795 | 0.41938976 |
| CDH11 | 85.5915398 | | 2.08944384 | 0.59185165 | 3.5303506 | 0.00041501 | 0.00470127 |
| BEAN1 | 103.744505 | | -1.854438749 | 0.67253573 | -2.7573832 | 0.0058266 | 0.03395183 |
| AC010542.3 | 233.231101 | | -0.981031203 | 0.56101873 | -1.7486603 | 0.08034975 | 0.21261661 |
| TK2 | 207.711772 | | -0.163069417 | 0.39093993 | -0.4171214 | 0.6765896 | 0.81279385 |
| CMTM3 | 59.3381514 | | -2.069164632 | 0.4439606 | -4.6606942 | 3.15E-06 | 9.95E-05 |
| CMTM4 | 1377.0835 | | -0.104137874 | 0.243389 | -0.427866 | 0.66874869 | 0.80738625 |
| DYNC1LI2 | 1163.40671 | | -0.221791574 | 0.2210684 | -1.0032713 | 0.31573 | 0.51587926 |
| AC018557.1 | 51.0809818 | | -0.668885376 | 0.51027388 | -1.310836 | 0.18991318 | 0.37401435 |
| NAE1 | 285.475713 | | -0.201680813 | 0.16794083 | -1.200904 | 0.22978844 | 0.42293458 |
| PDP2 | 222.420271 | | 0.103728587 | 0.33794988 | 0.30693482 | 0.75889299 | 0.86482758 |
| CDH16 | 932.508471 | | 0.52719575 | 0.40296326 | 1.30829731 | 0.19077249 | 0.37506002 |
| RRAD | 383.438895 | | -1.988888465 | 0.62312448 | -3.1917996 | 0.00141389 | 0.011767 |
| FAM96B | 487.403835 | | -1.121335218 | 0.3397779 | -3.3002006 | 0.00096616 | 0.00881723 |
| CES2 | 731.259165 | | 0.049552891 | 0.34851284 | 0.14218383 | 0.8869348 | 0.94159858 |
| CES3 | 215.284936 | | 1.104478593 | 0.78157468 | 1.41314531 | 0.157613 | 0.33190403 |
| CES4A | 135.928402 | | -0.475482365 | 0.75337894 | -0.6311331 | 0.52795352 | 0.70252003 |
| CBFB | 1086.26554 | | -0.506887333 | 0.16318828 | -3.1061503 | 0.0018954 | 0.01469643 |
| C16orf70 | 719.825952 | | 0.252285853 | 0.29580017 | 0.85289286 | 0.39371872 | 0.58851718 |
| TRADD | 190.986772 | | -0.214685986 | 0.36327279 | -0.5909773 | 0.55453561 | 0.72406869 |
| FBXL8 | 33.8396923 | | -1.186565547 | 0.54127886 | -2.192152 | 0.02836853 | 0.10500877 |
| HSF4 | 132.322004 | | -0.094501082 | 0.64169464 | -0.147268 | 0.8829205 | 0.93917362 |
| NOL3 | 291.622407 | | -0.833774351 | 0.4625946 | -1.8023867 | 0.07148459 | 0.19695261 |
| KIAA0895L | 144.141417 | | 0.580922257 | 0.4415155 | 1.31574602 | 0.1882593 | 0.37204012 |
| E2F4 | 216.916548 | | -0.001735371 | 0.32830834 | -0.0052858 | 0.99578256 | 0.9977776 |
| ELMO3 | 146.803095 | | 0.687585707 | 0.30052174 | 2.28797329 | 0.02213908 | 0.08873109 |
| LRRC29 | 135.526646 | | 0.897724281 | 0.41771028 | 2.14915537 | 0.03162209 | 0.11337796 |
| TMEM208 | 228.150166 | | -0.874185256 | 0.37227544 | -2.3482217 | 0.01886329 | 0.07914268 |
| FHOD1 | 84.0229745 | | 0.24400457 | 0.28962509 | 0.84248422 | 0.39951697 | 0.59362375 |
| LRRC36 | 163.790507 | | 0.405147592 | 0.64239986 | 0.63067821 | 0.52825095 | 0.70268407 |
| ZDHHC1 | 204.996986 | | 0.337171387 | 0.30669002 | 1.0993882 | 0.27159878 | 0.46987308 |
| HSD11B2 | 198.945831 | | 3.085024795 | 0.68135964 | 4.52774805 | 5.96E-06 | 0.00016542 |
| ATP6V0D1 | 600.168152 | | 0.017387921 | 0.38456205 | 0.04521486 | 0.96393605 | 0.98126041 |
| AC009061.2 | 133.870426 | | 1.266620439 | 0.45259518 | 2.79857255 | 0.0051329 | 0.03092603 |
| AC027682.6 | 70.970767 | | -0.593861438 | 0.59492335 | -0.998215 | 0.3181751 | 0.51832969 |
| RIPOR1 | 738.958131 | | -0.519951185 | 0.39495052 | -1.316497 | 0.18800728 | 0.37164924 |
| AC027682.4 | 75.7828226 | | 0.099515811 | 0.32986342 | 0.30168792 | 0.76288998 | 0.8677952 |
| CTCF | 713.102848 | | 0.166402047 | 0.3801152 | 0.43776741 | 0.66155489 | 0.80318508 |
| ENKD1 | 146.902932 | | 0.74017496 | 0.37408805 | 1.9786116 | 0.04785975 | 0.1510508 |
| C16orf86 | 35.8692429 | | 0.694156343 | 0.53469958 | 1.29821748 | 0.19421261 | 0.37930258 |
| GFOD2 | 417.080133 | | 0.021717763 | 0.31807603 | 0.06827853 | 0.94556392 | 0.97196767 |
| RANBP10 | 775.501658 | | 0.398635798 | 0.23337253 | 1.70815222 | 0.0876081 | 0.22538368 |
| TSNAXIP1 | 58.4262854 | | 1.492391812 | 0.42913439 | 3.4776794 | 0.00050577 | 0.00538667 |
| CENPT | 150.036118 | | -0.577953683 | 0.47834296 | -1.2082412 | 0.22695449 | 0.41931165 |
| THAP11 | 37.3041232 | | -0.38617813 | 0.35335255 | -1.0928975 | 0.2744388 | 0.47234212 |
| NUTF2 | 169.200616 | | -0.847125695 | 0.40281908 | -2.102993 | 0.03546639 | 0.12281204 |
| PSKH1 | 238.183688 | | 0.697284509 | 0.24178505 | 2.88390251 | 0.0039278 | 0.02528812 |
| LCAT | 71.8535142 | | -0.034463717 | 0.50848566 | -0.0677772 | 0.94596302 | 0.97206051 |
| SLC12A4 | 96.0329895 | | -0.789489971 | 0.31605944 | -2.4979161 | 0.01249257 | 0.05897554 |
| DUS2 | 191.347879 | | 0.341733877 | 0.21679137 | 1.57632601 | 0.11495069 | 0.26944033 |
| DDX28 | 34.1595623 | | -0.858202852 | 0.45310228 | -1.8940599 | 0.05821706 | 0.1712176 |
| NFATC3 | 2909.72291 | | 1.106309185 | 0.28824037 | 3.83814797 | 0.00012397 | 0.00184495 |
| AC020978.5 | 57.7357561 | | 0.046635063 | 0.413013 | 0.11291427 | 0.91009852 | 0.95270509 |
| ESRP2 | 117.189354 | | -0.39694614 | 0.3590922 | -1.1054157 | 0.26897952 | 0.46699205 |
| PLA2G15 | 410.305694 | | 0.505594006 | 0.40088361 | 1.261199 | 0.20723716 | 0.39612324 |
| SLC7A6 | 200.338629 | | -0.968804372 | 0.3843464 | -2.5206542 | 0.01171369 | 0.05627745 |
| SLC7A6OS | 245.882736 | | -0.906168626 | 0.35400216 | -2.5597828 | 0.01047376 | 0.05218605 |
| PRMT7 | 330.748804 | | 1.022033804 | 0.33151228 | 3.08294403 | 0.00204964 | 0.01568642 |
| ZFP90 | 294.567848 | | 0.377647101 | 0.38903849 | 0.97071912 | 0.33168817 | 0.53141196 |
| CDH1 | 390.322478 | | -0.939533052 | 0.54379022 | -1.7277491 | 0.08403322 | 0.21898162 |
| TANGO6 | 1328.00103 | | 1.126564345 | 0.26940063 | 4.18174354 | 2.89E-05 | 0.00058513 |
| AC009137.1 | 38.0760497 | | 0.840997046 | 0.55110201 | 1.5260279 | 0.12700292 | 0.28807363 |
| CHTF8 | 478.396065 | | 0.395233229 | 0.33220448 | 1.18972878 | 0.23415301 | 0.42845219 |
| UTP4 | 894.27061 | | 0.639302927 | 0.19783953 | 3.23142164 | 0.00123176 | 0.01057879 |
| SNTB2 | 1371.96653 | | 0.176779733 | 0.39360476 | 0.44913006 | 0.65333783 | 0.79727674 |
| VPS4A | 351.411425 | | 0.176045123 | 0.26579933 | 0.66232342 | 0.50776397 | 0.68768861 |
| COG8 | 228.403491 | | 0.755107427 | 0.33443187 | 2.25788121 | 0.02395307 | 0.09347027 |
| AC026464.3 | 37.2808148 | | 0.290176489 | 0.35635914 | 0.81428104 | 0.41548397 | 0.60902991 |
| NIP7 | 92.9599854 | | -0.841787471 | 0.35431377 | -2.3758249 | 0.01750977 | 0.07518204 |
| TERF2 | 310.367721 | | 0.202788624 | 0.18570867 | 1.09197179 | 0.2748455 | 0.47259765 |
| CYB5B | 711.848109 | | -0.011018533 | 0.20040143 | -0.0549823 | 0.95615256 | 0.9776207 |
| NFAT5 | 5166.67675 | | -0.142337453 | 0.13332598 | -1.0675898 | 0.28570561 | 0.48536166 |
| NQO1 | 539.690905 | | -3.262262561 | 0.88643623 | -3.6801999 | 0.00023305 | 0.00302217 |
| AC092115.3 | 74.7283936 | | -0.438050579 | 0.5758732 | -0.7606719 | 0.44685304 | 0.63774996 |
| NOB1 | 123.356126 | | -1.044382887 | 0.32654531 | -3.1982787 | 0.00138251 | 0.01153632 |
| WWP2 | 1467.59861 | | 0.014645055 | 0.30197027 | 0.04849833 | 0.96131909 | 0.97988733 |
| CLEC18A | 87.7911623 | | -0.109711269 | 0.86493239 | -0.1268438 | 0.89906407 | 0.94743919 |
| PDPR | 970.729933 | | 0.5087596 | 0.36077607 | 1.4101811 | 0.15848621 | 0.33328458 |
| AC009060.1 | 839.411974 | | 1.566804292 | 0.50215089 | 3.12018622 | 0.00180737 | 0.01417253 |
| CLEC18C | 87.7321142 | | 0.54321366 | 0.88420866 | 0.61435008 | 0.53898402 | 0.71071226 |
| EXOSC6 | 238.053059 | | -0.462867228 | 0.37957145 | -1.2194469 | 0.22267462 | 0.41477101 |
| AARS | 475.144358 | | -0.268925068 | 0.34555516 | -0.7782406 | 0.43642716 | 0.62857823 |
| AC012184.2 | 62.9382561 | | 0.887530989 | 0.35709709 | 2.48540526 | 0.0129404 | 0.0604547 |
| DDX19A | 85.9665841 | | -0.360902514 | 0.30647162 | -1.177605 | 0.23895412 | 0.43402823 |
| ST3GAL2 | 413.876609 | | 0.050195363 | 0.34974691 | 0.1435191 | 0.88588023 | 0.94094134 |
| FUK | 209.271992 | | 0.338887946 | 0.20044049 | 1.69071605 | 0.09089105 | 0.2308828 |
| COG4 | 420.099941 | | 0.52342727 | 0.18998235 | 2.75513635 | 0.00586677 | 0.03410339 |
| SF3B3 | 589.818806 | | -0.196114119 | 0.19507685 | -1.0053172 | 0.31474411 | 0.51518958 |
| IL34 | 52.3076088 | | 0.503383409 | 0.57279033 | 0.87882665 | 0.37949527 | 0.57605113 |
| MTSS1L | 315.738838 | | -1.471637157 | 0.45485644 | -3.2353882 | 0.00121477 | 0.0104793 |
| VAC14 | 672.488996 | | -0.6729343 | 0.44548557 | -1.5105636 | 0.13089967 | 0.29375274 |
| HYDIN | 1309.59804 | | -0.480500806 | 0.47833706 | -1.0045235 | 0.31512637 | 0.51523844 |
| CMTR2 | 369.076832 | | 0.030827573 | 0.32466735 | 0.09495126 | 0.92435354 | 0.95992984 |
| ZNF19 | 337.607248 | | -0.094895133 | 0.45327292 | -0.2093554 | 0.83417081 | 0.91230984 |
| MARVELD3 | 552.166495 | | 0.371259792 | 0.3669093 | 1.01185713 | 0.31160637 | 0.51170343 |
| PHLPP2 | 525.411811 | | -0.805999387 | 0.1908121 | -4.2240477 | 2.40E-05 | 0.00050914 |
| AP1G1 | 1881.34454 | | 0.204989632 | 0.18823516 | 1.08900822 | 0.27615026 | 0.47391907 |
| IST1 | 1119.29626 | | 0.172922789 | 0.14033763 | 1.23219114 | 0.21787769 | 0.4090601 |

| | | | | | | |
|---|---|---|---|---|---|---|
| ZNF821 | 318.096906 | 0.459541231 | 0.36093642 | 1.27319164 | 0.20295005 | 0.39079028 |
| PKD1L3 | 56.8203612 | -1.031592309 | 0.68085331 | -1.5151462 | 0.12973538 | 0.2919821 |
| DHODH | 123.651455 | 0.560475736 | 0.30692212 | 1.82611712 | 0.06783262 | 0.1899452 |
| TXNL4B | 311.677998 | -0.148383933 | 0.30511039 | -0.4863287 | 0.62673415 | 0.77912423 |
| HP | 467.364406 | -7.990698194 | 1.79446466 | -4.4529705 | 8.47E-06 | 0.00022111 |
| DHX38 | 241.909062 | -0.235714971 | 0.29814787 | -0.7905975 | 0.42917889 | 0.62214435 |
| PMFBP1 | 52.1715499 | 0.983955821 | 0.64000872 | 1.53741003 | 0.12419293 | 0.28386068 |
| LINC01572 | 757.356245 | 0.841061494 | 0.51463307 | 1.63429351 | 0.10219724 | 0.24973128 |
| AC004943.2 | 304.348742 | 0.36996201 | 0.33015875 | 1.12055793 | 0.26247608 | 0.46048377 |
| ZFHX3 | 1329.46149 | -0.712341654 | 0.24297935 | -2.9316963 | 0.00337116 | 0.02249471 |
| AC138627.1 | 405.678729 | 1.083881937 | 0.61163563 | 1.77210399 | 0.0763773 | 0.20555349 |
| PSMD7 | 463.524434 | -0.89201223 | 0.3246455 | -2.7476501 | 0.0060024 | 0.0346275 |
| NPIPB15 | 231.682418 | -1.029430551 | 0.85896161 | -1.1984593 | 0.23073825 | 0.42378291 |
| CLEC18B | 105.631307 | 1.170430325 | 0.96949973 | 1.20725183 | 0.22733517 | 0.41984414 |
| AC009053.3 | 639.08693 | 2.060883526 | 0.56211321 | 3.66631401 | 0.00024607 | 0.00315225 |
| GLG1 | 3922.60348 | 0.55348368 | 0.17415478 | 3.17811364 | 0.00148237 | 0.01222551 |
| RFWD3 | 344.948852 | -0.192967282 | 0.28515545 | -0.6767091 | 0.49859056 | 0.68045095 |
| MLKL | 65.9087022 | -2.642141473 | 0.47901206 | -5.5158141 | 3.47E-08 | 2.38E-06 |
| FA2H | 63.617937 | -1.293202831 | 0.48261335 | -2.6795836 | 0.00737138 | 0.04048057 |
| WDR59 | 1809.11435 | 0.386139571 | 0.20946306 | 1.84347333 | 0.06525993 | 0.18536341 |
| ZNRF1 | 762.81559 | 0.049615394 | 0.29869375 | 0.16610791 | 0.86807203 | 0.93140222 |
| LDHD | 115.52811 | 0.836535629 | 0.56357474 | 1.48433841 | 0.13771916 | 0.30335382 |
| ZFP1 | 102.274924 | 0.143401866 | 0.23086374 | 0.6211537 | 0.5344985 | 0.70751736 |
| BCAR1 | 1173.36621 | -1.115490559 | 0.42421606 | -2.629534 | 0.0085502 | 0.04486889 |
| CFDP1 | 2906.98239 | 0.499604847 | 0.22691164 | 2.20175944 | 0.02768231 | 0.10344639 |
| AC009163.4 | 42.5332214 | -0.371289622 | 0.39440154 | -0.9414 | 0.3464999 | 0.54670612 |
| TMEM231 | 37.7950929 | 0.62180537 | 0.45599523 | 1.36362254 | 0.17268639 | 0.35223399 |
| GABARAPL2 | 282.119335 | -0.437570819 | 0.3276017 | -1.3356793 | 0.18165411 | 0.36381342 |
| ADAT1 | 148.643309 | -0.174077632 | 0.28671423 | -0.6071468 | 0.54375348 | 0.71416138 |
| KARS | 279.824225 | -0.580908361 | 0.39218326 | -1.4812166 | 0.13854886 | 0.30463413 |
| TERF2IP | 760.558972 | -0.3564482 | 0.40362629 | -0.8831144 | 0.37717445 | 0.57360814 |
| AC010528.1 | 44.1561718 | -0.756901057 | 0.63247512 | -1.1967286 | 0.23141236 | 0.42445728 |
| MON1B | 94.657578 | -0.14704204 | 0.42416783 | -0.3466601 | 0.72884673 | 0.84599052 |
| SYCE1L | 51.7760142 | -0.515641822 | 0.5252748 | -0.9816611 | 0.32626685 | 0.52644376 |
| NUDT7 | 102.016083 | 0.922993753 | 0.49325834 | 1.87121773 | 0.06131491 | 0.17738359 |
| VAT1L | 49.1073938 | 0.603129083 | 0.4160261 | 1.44973856 | 0.14713144 | 0.31654852 |
| WWOX | 14136.0206 | 0.420357976 | 0.26297926 | 1.59844532 | 0.1099439 | 0.2615641 |
| AC046158.1 | 120.866108 | 0.61066222 | 0.66354868 | 0.92029754 | 0.3574173 | 0.5561453 |
| MAF | 1670.7581 | 1.486645982 | 0.3889236 | 3.82246279 | 0.00013213 | 0.00191957 |
| AC009159.3 | 27.7693967 | 1.980711537 | 0.63401501 | 3.12407672 | 0.00178364 | 0.01401855 |
| LINC01229 | 59.7168462 | 1.155721808 | 0.8698094 | 1.32870697 | 0.18394466 | 0.36652763 |
| MAFTRR | 136.286709 | 1.34282202 | 0.68793886 | 1.95194964 | 0.05094418 | 0.15730514 |
| AC105411.1 | 94.3954321 | -0.021485878 | 0.60860764 | -0.0353033 | 0.97183787 | 0.98607792 |
| DYNLRB2 | 67.9039978 | 0.756545294 | 0.49650904 | 1.52372915 | 0.12757639 | 0.28889633 |
| CDYL2 | 1139.00694 | -0.098510305 | 0.30847155 | -0.3193497 | 0.74946133 | 0.85872756 |
| CMC2 | 734.477846 | 0.659331411 | 0.19795685 | 3.33068245 | 0.00086633 | 0.00813345 |
| CENPN | 58.8051883 | 0.673181248 | 0.40070278 | 1.68000143 | 0.09295704 | 0.23469615 |
| ATMIN | 189.272133 | -0.506990058 | 0.3406419 | -1.4883373 | 0.13666195 | 0.30162504 |
| BCO1 | 181.552438 | 0.294663432 | 0.33052743 | 0.89149463 | 0.37266387 | 0.57001787 |
| GAN | 525.823645 | -0.501432953 | 0.25198309 | -1.9899468 | 0.04659679 | 0.14846635 |
| CMIP | 5426.20022 | 0.131174587 | 0.35591098 | 0.3685601 | 0.71245564 | 0.83543596 |
| PLCG2 | 1411.13828 | -0.354737813 | 0.31301762 | -1.1332839 | 0.25709507 | 0.45429962 |
| SDR42E1 | 38.9819074 | -0.659887047 | 0.3439874 | -1.9183465 | 0.05506709 | 0.1653098 |
| AC092142.1 | 50.2048757 | 0.124083546 | 0.37770705 | 0.32851795 | 0.74252007 | 0.85405074 |
| MPHOSPH6 | 210.667015 | -0.081542289 | 0.2477965 | -0.3290696 | 0.7421031 | 0.85370947 |
| CDH13 | 58.010117 | 0.428444942 | 0.39702664 | 1.07913399 | 0.280528 | 0.47921202 |
| HSBP1 | 385.651874 | -0.852972912 | 0.29058403 | -2.9353743 | 0.00333146 | 0.02232971 |
| MLYCD | 122.250028 | 0.062456822 | 0.41511596 | 0.15045633 | 0.8804046 | 0.93773455 |
| AC009119.2 | 165.390799 | 0.636225292 | 0.50489817 | 1.26010615 | 0.20763107 | 0.39665502 |
| OSGIN1 | 381.056983 | -2.189418706 | 0.71359201 | -3.0681659 | 0.00215377 | 0.01626541 |
| MBTPS1 | 680.579017 | 0.024217185 | 0.18534862 | 0.13065748 | 0.89604627 | 0.94608737 |
| HSDL1 | 177.646066 | -0.079378796 | 0.4336477 | -0.183049 | 0.85475954 | 0.92322796 |
| DNAAF1 | 135.460618 | 0.775709595 | 0.68187969 | 1.13760478 | 0.25528555 | 0.45232479 |
| TAF1C | 220.39908 | 0.364410444 | 0.2209208 | 1.6495072 | 0.09904377 | 0.24412748 |
| ATP2C2 | 248.035105 | 1.418494385 | 0.4633368 | 3.06147574 | 0.00220249 | 0.0165422 |
| ATP2C2-AS1 | 38.7206601 | -0.313858852 | 0.43133791 | -0.7276403 | 0.4668338 | 0.65461205 |
| TLDC1 | 365.224204 | -0.673501422 | 0.28752844 | -2.3423819 | 0.0191611 | 0.07995164 |
| COTL1 | 792.545586 | 1.041455732 | 0.51841957 | 2.00890514 | 0.04454719 | 0.14374021 |
| KLHL36 | 813.364598 | -0.101780108 | 0.18522261 | -0.5495015 | 0.58266131 | 0.74567914 |
| USP10 | 1219.55546 | -0.392589802 | 0.26669041 | -1.4720807 | 0.14099909 | 0.30799594 |
| ZDHHC7 | 584.169739 | -0.293956853 | 0.22436791 | -1.3101555 | 0.19014324 | 0.3741919 |
| KIAA0513 | 149.819391 | -0.619625794 | 0.47887184 | -1.2939282 | 0.19569021 | 0.38101218 |
| GSE1 | 1906.21595 | 0.734792517 | 0.2656273 | 2.76625372 | 0.00567044 | 0.03324749 |
| EMC8 | 197.145293 | -0.306498517 | 0.32625793 | -0.9394362 | 0.34750682 | 0.54753787 |
| COX4I1 | 1344.77056 | -0.563879778 | 0.39068544 | -1.4433089 | 0.14893347 | 0.31912319 |
| IRF8 | 108.452649 | -2.158179172 | 0.79973617 | -2.6986139 | 0.00696289 | 0.03879688 |
| MTHFSD | 132.506536 | 0.195864139 | 0.31806689 | 0.61579543 | 0.53802955 | 0.70986341 |
| C16orf95 | 46.4535466 | 0.254806755 | 0.36520452 | 0.69770975 | 0.48535873 | 0.66902776 |
| AC010531.1 | 46.1928761 | -0.531647517 | 0.36240592 | -1.4669946 | 0.14237754 | 0.30981214 |
| FBXO31 | 461.927355 | -0.070825552 | 0.42365089 | -0.167179 | 0.86722917 | 0.9309351 |
| MAP1LC3B | 604.993381 | -1.649654546 | 0.43420923 | -3.7992157 | 0.00014515 | 0.00205541 |
| ZCCHC14 | 1877.57243 | -0.077456761 | 0.34777043 | -0.2227238 | 0.82375045 | 0.90561016 |
| KLHDC4 | 624.666359 | 0.161810992 | 0.46841328 | 0.34544493 | 0.72975991 | 0.84643611 |
| SLC7A5 | 108.919021 | -2.179937206 | 0.5800887 | -3.7579377 | 0.00017132 | 0.00235771 |
| BANP | 893.222299 | -0.246920263 | 0.46304316 | -0.5332554 | 0.5938568 | 0.75469239 |
| ZFPM1 | 74.9534006 | -1.02370925 | 0.36783711 | -2.7830505 | 0.00538504 | 0.0320368 |
| ZC3H18 | 424.538658 | -0.196020374 | 0.3759535 | -0.5213953 | 0.60209145 | 0.7609161 |
| CYBA | 1165.97861 | -0.71716308 | 0.46829078 | -1.5314482 | 0.12565866 | 0.28601099 |
| MVD | 241.360381 | 0.487851707 | 0.37977525 | 1.28458004 | 0.19893908 | 0.3853127 |
| SNAI3-AS1 | 83.9403999 | 0.261910481 | 0.3073732 | 0.85209276 | 0.39416261 | 0.58886199 |
| SNAI3 | 35.5512131 | -0.705534517 | 0.74744082 | -0.9439336 | 0.34520357 | 0.54544154 |
| RNF166 | 73.6917267 | 0.907808961 | 0.31775958 | 2.85690511 | 0.00427794 | 0.02697226 |
| PIEZO1 | 433.846736 | 0.425732153 | 0.53387561 | 0.79743698 | 0.42519729 | 0.61830885 |
| APRT | 370.416583 | -1.348269537 | 0.38712773 | -3.4827511 | 0.00049629 | 0.00532705 |
| GALNS | 350.436303 | 0.463440161 | 0.34274065 | 1.35215988 | 0.17632418 | 0.35673439 |
| TRAPPC2L | 123.947315 | 0.119360066 | 0.3638588 | 0.32803952 | 0.74288178 | 0.8540591 |
| ACSF3 | 318.611464 | -0.098201354 | 0.31967215 | -0.307194 | 0.75869575 | 0.86481851 |
| AC009113.1 | 35.4103522 | 0.617217185 | 0.476873 | 1.29430096 | 0.19556148 | 0.3808166 |
| ZNF778 | 143.177889 | -0.410517127 | 0.34594152 | -1.1866662 | 0.2353593 | 0.42967809 |
| ANKRD11 | 3535.33183 | -0.268671783 | 0.43354474 | -0.6197095 | 0.53544908 | 0.70816292 |
| AC137932.3 | 36.9609665 | 0.360342587 | 0.50018291 | 0.72042164 | 0.47126543 | 0.65767005 |
| SPG7 | 1838.94939 | -0.170374232 | 0.43411994 | -0.3924589 | 0.69471918 | 0.82425245 |
| RPL13 | 2942.10796 | -0.797427463 | 0.40231003 | -1.9821217 | 0.04746562 | 0.15039556 |
| CPNE7 | 129.726875 | -1.269839092 | 0.55684283 | -2.2804264 | 0.02258241 | 0.09000788 |
| DPEP1 | 1122.38846 | 1.605000557 | 0.42784876 | 3.75132688 | 0.0001759 | 0.00241349 |
| CHMP1A | 357.777697 | 0.189925233 | 0.32779564 | 0.57940134 | 0.5623184 | 0.73014439 |
| SPATA33 | 175.09842 | 0.359925321 | 0.25524734 | 1.41010409 | 0.15850895 | 0.33328458 |
| CDK10 | 851.401238 | 0.509117823 | 0.18759752 | 2.71388358 | 0.00664995 | 0.03741827 |
| LINC02166 | 96.7370275 | 1.094423686 | 0.43593838 | 2.5105009 | 0.012056 | 0.05746539 |
| SPATA2L | 130.904737 | -0.228908854 | 0.25116352 | -0.9113937 | 0.36208796 | 0.55998416 |
| VPS9D1 | 281.089249 | 0.026262272 | 0.27534425 | 0.09537977 | 0.92401318 | 0.95972181 |
| ZNF276 | 235.225164 | 0.24810418 | 0.21809161 | 1.13761453 | 0.25528148 | 0.45232479 |
| FANCA | 55.0170461 | -0.218555032 | 0.45346486 | -0.4819669 | 0.62982949 | 0.78158193 |

| | | | | | | |
|---|---|---|---|---|---|---|
| SPIRE2 | 61.4232178 | 0.194263581 | 0.42807203 | 0.45381049 | 0.64996523 | 0.79477446 |
| TCF25 | 1856.1636 | -0.386027072 | 0.31266827 | -1.2346218 | 0.21697128 | 0.40784768 |
| MC1R | 53.416976 | -0.208691115 | 0.65350567 | -0.3193409 | 0.74946799 | 0.85872756 |
| DEF8 | 220.77869 | -0.187179242 | 0.28749743 | -0.651064 | 0.51500515 | 0.69315508 |
| DBNDD1 | 94.5226909 | -0.980925752 | 0.29446356 | -3.3312297 | 0.00086463 | 0.00812304 |
| GAS8 | 668.970271 | 0.377395792 | 0.23205871 | 1.62629446 | 0.10388703 | 0.25194259 |
| FAM157C | 357.472215 | 0.73767127 | 0.3369179 | 2.18946892 | 0.02856278 | 0.10552829 |
| AC133919.2 | 41.9301775 | 0.639077415 | 0.7967949 | 0.80206013 | 0.42251818 | 0.61619214 |
| LINC02193 | 135.756843 | -0.441847056 | 0.35644083 | -1.2396084 | 0.21512026 | 0.40564774 |
| AC240565.1 | 31.9920801 | 0.302584108 | 0.44340614 | 0.68240848 | 0.4949807 | 0.67697759 |
| RPH3AL | 605.098156 | 0.109611212 | 0.31414548 | 0.34891864 | 0.72715039 | 0.84508584 |
| AC129507.2 | 84.2079489 | 0.62364093 | 0.42357594 | 1.47232379 | 0.14093346 | 0.30790165 |
| C17orf97 | 69.505705 | 1.130383689 | 0.47317116 | 2.38895304 | 0.01689646 | 0.07336072 |
| VPS53 | 1962.35743 | -0.485938088 | 0.28799798 | -1.6872968 | 0.09154629 | 0.23207416 |
| AC015853.3 | 82.7829325 | -1.24598616 | 0.46113828 | -2.7019795 | 0.0068928 | 0.03846891 |
| FAM57A | 80.5508147 | -3.126445306 | 0.44013776 | -7.1033335 | 1.22E-12 | 3.21E-10 |
| GEMIN4 | 35.9118815 | -0.812243402 | 0.50832756 | -1.597874 | 0.11007102 | 0.26171962 |
| GLOD4 | 276.005018 | -0.291581648 | 0.14964138 | -1.9485362 | 0.05135084 | 0.15790414 |
| MRM3 | 67.073451 | -0.168759157 | 0.25993256 | -0.6492421 | 0.51618192 | 0.69372542 |
| NXN | 183.360582 | -0.578932118 | 0.37317855 | -1.5513542 | 0.12081682 | 0.27840363 |
| TIMM22 | 105.296866 | -0.470009983 | 0.29093376 | -1.6155223 | 0.10619763 | 0.25555669 |
| ABR | 354.122786 | -0.407928849 | 0.56653009 | -0.720048 | 0.47149545 | 0.65779014 |
| YWHAE | 1955.9532 | -0.061203753 | 0.2141824 | -0.2857553 | 0.77506556 | 0.87561595 |
| CRK | 1059.30029 | -0.515345431 | 0.46762621 | -1.1020457 | 0.27044182 | 0.46869972 |
| MYO1C | 473.781727 | -1.414535542 | 0.30619249 | -4.619759 | 3.84E-06 | 0.00011599 |
| INPP5K | 135.905841 | -0.138323519 | 0.38963416 | -0.3550087 | 0.72258307 | 0.84228022 |
| PITPNA | 1012.02 | -0.557339343 | 0.29948002 | -1.8610234 | 0.06274087 | 0.18017789 |
| SLC43A2 | 994.641325 | 2.002790993 | 0.5448323 | 3.67597698 | 0.00023694 | 0.00305814 |
| SCARF1 | 62.394289 | 1.36656221 | 0.33290598 | 4.10494942 | 4.04E-05 | 0.00076457 |
| RILP | 65.8659526 | 0.462439428 | 0.41118697 | 1.12464515 | 0.26073943 | 0.4589178 |
| PRPF8 | 724.898068 | -0.018393914 | 0.2200536 | -0.0835883 | 0.93338374 | 0.96466122 |
| TLCD2 | 51.3554156 | -0.831689972 | 0.51569044 | -1.6127698 | 0.10679453 | 0.25646507 |
| MIR22HG | 444.069659 | -0.838526986 | 0.74549636 | -1.1247902 | 0.26067795 | 0.4589178 |
| WDR81 | 393.153679 | 0.099363743 | 0.54698162 | 0.18165828 | 0.85585091 | 0.92378521 |
| SERPINF2 | 1557.41118 | 1.822724941 | 0.49660505 | 3.67037131 | 0.0002422 | 0.00311135 |
| SMYD4 | 230.250252 | -0.150078921 | 0.29634528 | -0.5064326 | 0.61255298 | 0.76881821 |
| RPA1 | 371.631725 | -0.202574368 | 0.19920538 | -1.0169121 | 0.30919524 | 0.50945296 |
| RTN4RL1 | 76.5821061 | 0.73992518 | 0.49684317 | 1.489253 | 0.13642076 | 0.30140544 |
| DPH1 | 103.808 | 0.893088201 | 0.35262018 | 2.53272003 | 0.01131813 | 0.05507193 |
| SMG6 | 1994.03857 | -0.165762918 | 0.35332179 | -0.4691557 | 0.63895837 | 0.78803333 |
| AC090617.4 | 43.9354273 | -0.628787568 | 0.28642613 | -2.195287 | 0.02814302 | 0.10442781 |
| SRR | 90.0838374 | 0.099125596 | 0.38421558 | 0.25799473 | 0.79641097 | 0.88881332 |
| TSR1 | 242.198254 | -0.937169283 | 0.44280979 | -2.116415 | 0.03430953 | 0.12022632 |
| SGSM2 | 290.722336 | 0.218792002 | 0.24470475 | 0.89410605 | 0.37126515 | 0.56878634 |
| MNT | 179.442644 | -0.861662287 | 0.47115197 | -1.8288415 | 0.06742335 | 0.18918565 |
| METTL16 | 753.596311 | 0.120427458 | 0.23445928 | 0.51363912 | 0.60750432 | 0.76448285 |
| PAFAH1B1 | 3168.66362 | -0.021296843 | 0.18528509 | -0.1149409 | 0.90849193 | 0.95197217 |
| CLUH | 1069.90109 | 0.724338709 | 0.3095747 | 2.33978654 | 0.01929476 | 0.08033819 |
| RAP1GAP2 | 308.613957 | -1.293397457 | 0.54044529 | -2.393207 | 0.01670181 | 0.07266389 |
| SPATA22 | 411.399499 | 0.886061475 | 0.86563607 | 1.02359584 | 0.30602621 | 0.5059343 |
| ASPA | 1465.54452 | 1.974561339 | 0.88037408 | 2.24286627 | 0.02490544 | 0.09608186 |
| TRPV1 | 313.786754 | 1.41049172 | 0.37294176 | 3.78206967 | 0.00015553 | 0.00217314 |
| CTNS | 319.683675 | 0.188257768 | 0.23884355 | 0.78820538 | 0.43057659 | 0.6232221 |
| TAX1BP3 | 160.9543 | -1.214302981 | 0.59540708 | -2.03945 | 0.04140513 | 0.13668488 |
| EMC6 | 174.921709 | -1.078795394 | 0.42402498 | -2.5441789 | 0.0109535 | 0.05384903 |
| ITGAE | 165.309229 | -0.053949859 | 0.25137694 | -0.2146174 | 0.83006565 | 0.90992561 |
| NCBP3 | 529.900308 | 0.120795369 | 0.16836878 | 0.71744517 | 0.47309946 | 0.65922275 |
| CAMKK1 | 52.0406686 | -1.602907176 | 0.5648065 | -2.8379758 | 0.00454006 | 0.02822251 |
| ZZEF1 | 1187.13058 | 0.178054037 | 0.22361608 | 0.79624879 | 0.42588744 | 0.61876852 |
| CYB5D2 | 300.527689 | 0.255277456 | 0.30262788 | 0.84353583 | 0.39892884 | 0.59314883 |
| ANKFY1 | 852.346338 | -0.209817104 | 0.2047122 | -1.024937 | 0.30539294 | 0.50537498 |
| UBE2G1 | 1336.18965 | 0.153791811 | 0.28789507 | 0.53419397 | 0.59320734 | 0.75436977 |
| SPNS3 | 136.92427 | 1.493609511 | 0.4390111 | 3.40221352 | 0.00066842 | 0.00668278 |
| AC127521.1 | 156.92058 | 2.580793143 | 0.39066931 | 6.60608102 | 3.95E-11 | 7.21E-09 |
| SPNS2 | 967.704801 | 0.062132299 | 0.39406191 | 0.15767142 | 0.87471573 | 0.93490877 |
| MYBBP1A | 132.929825 | -0.877707112 | 0.50695001 | -1.7313484 | 0.08338964 | 0.21821721 |
| GGT6 | 111.191335 | 2.004601355 | 0.42358739 | 4.73243872 | 2.22E-06 | 7.47E-05 |
| SMTNL2 | 719.336903 | 1.180605322 | 0.34518509 | 3.42020952 | 0.00062573 | 0.00635324 |
| PELP1 | 231.713469 | 0.042085985 | 0.35691021 | 0.11791757 | 0.90613297 | 0.95080991 |
| AC091153.2 | 36.9794287 | -0.772821768 | 0.44729261 | -1.7277767 | 0.08402826 | 0.21898162 |
| ARRB2 | 82.6492219 | 0.449260558 | 0.39895877 | 1.12608268 | 0.26013052 | 0.4584813 |
| MED11 | 120.163041 | -0.420292364 | 0.32949814 | -1.2755531 | 0.20211355 | 0.38964423 |
| CXCL16 | 123.590638 | -1.671107496 | 0.43691278 | -3.824808 | 0.00013087 | 0.00191473 |
| TM4SF5 | 68.1376804 | 0.912750348 | 0.62436781 | 1.46187926 | 0.1437743 | 0.31142767 |
| GLTPD2 | 75.8621615 | 0.624696279 | 0.72860981 | 0.85738109 | 0.3912343 | 0.58646585 |
| PSMB6 | 248.1047 | -0.757954424 | 0.40121762 | -1.8891354 | 0.05887368 | 0.17258568 |
| PLD2 | 90.574717 | -0.750201132 | 0.41723603 | -1.7980258 | 0.07217293 | 0.19817194 |
| MINK1 | 761.889196 | -0.592505236 | 0.25058868 | -2.3644533 | 0.0180567 | 0.0768252 |
| SLC25A11 | 200.844929 | 0.381707153 | 0.29193141 | 1.30752341 | 0.19103501 | 0.37532348 |
| RNF167 | 184.797577 | -0.098410544 | 0.23910069 | -0.4115862 | 0.68064275 | 0.81442647 |
| PFN1 | 998.702688 | -1.418567101 | 0.43813739 | -3.2377221 | 0.00120488 | 0.01042633 |
| SPAG7 | 337.547362 | -0.445203985 | 0.48819367 | -0.9119413 | 0.3617996 | 0.55961134 |
| CAMTA2 | 240.135134 | 0.198952465 | 0.18220229 | 1.09193176 | 0.2748631 | 0.47259765 |
| KIF1C | 518.003412 | 0.077311064 | 0.33478016 | 0.23093084 | 0.81736853 | 0.90138805 |
| ZFP3 | 81.2623118 | 0.177246794 | 0.30699249 | 0.57736524 | 0.56369275 | 0.73104641 |
| ZNF232 | 85.6374153 | -0.524629397 | 0.69099428 | -0.7592384 | 0.44770995 | 0.6383814 |
| USP6 | 37.6393599 | -0.666037729 | 0.52497566 | -1.2687021 | 0.20454733 | 0.39281526 |
| ZNF594 | 234.600604 | 0.48153729 | 0.19959975 | 2.41251445 | 0.01584291 | 0.06996094 |
| AC087500.1 | 358.218178 | 1.067508853 | 0.3226316 | 3.30875477 | 0.00093712 | 0.00860931 |
| SCIMP | 62.7010303 | 0.65431714 | 0.50120414 | 1.3054903 | 0.19172595 | 0.37601768 |
| RABEP1 | 1582.26792 | -0.046746902 | 0.26166758 | -0.17865 | 0.85821257 | 0.92545981 |
| NUP88 | 726.419732 | 0.008823701 | 0.17890197 | 0.04932143 | 0.96066314 | 0.97967485 |
| RPAIN | 388.400343 | -0.065261504 | 0.23099966 | -0.2825177 | 0.77754654 | 0.87743794 |
| C1QBP | 245.070551 | -1.173841165 | 0.31115213 | -3.7725635 | 0.00016158 | 0.00224546 |
| DHX33 | 94.4813358 | -0.184637115 | 0.38610903 | -0.4781994 | 0.63250828 | 0.78339125 |
| DERL2 | 244.398287 | -0.524970259 | 0.27546545 | -1.9057572 | 0.05668172 | 0.16853283 |
| MIS12 | 149.52225 | 0.026486162 | 0.28624968 | 0.09252818 | 0.9262784 | 0.96090956 |
| NLRP1 | 42.7006553 | 0.102758266 | 0.46004204 | 0.22336712 | 0.82324979 | 0.90529291 |
| WSCD1 | 87.7493372 | -1.43696021 | 0.65979718 | -2.1778817 | 0.02941484 | 0.10754582 |
| PITPNM3 | 60.3766305 | 0.236607052 | 0.61579594 | 0.38422964 | 0.70080825 | 0.82796169 |
| KIAA0753 | 603.74059 | 0.018354265 | 0.22223378 | 0.0825899 | 0.93417764 | 0.96493643 |
| TXNDC17 | 165.227451 | -0.590703683 | 0.42750021 | -1.3817623 | 0.1670447 | 0.34488351 |
| MED31 | 129.545978 | -0.461596841 | 0.25798928 | -1.7892094 | 0.0735811 | 0.20063182 |
| XAF1 | 246.955528 | -1.913398094 | 1.65046243 | -1.1593103 | 0.24632972 | 0.44200046 |
| ALOX12-AS1 | 459.494849 | 0.194936631 | 0.62007915 | 0.31437379 | 0.75323716 | 0.8613901 |
| SLC16A13 | 272.614924 | 0.845285541 | 0.50495847 | 1.6739704 | 0.09413642 | 0.23621021 |
| ACADVL | 2185.32489 | -0.12165914 | 0.43113829 | -0.2821812 | 0.77780454 | 0.87758633 |
| DVL2 | 201.722434 | -0.004363531 | 0.21711831 | -0.0200975 | 0.98396561 | 0.99214482 |
| PHF23 | 132.892469 | -0.562921239 | 0.3210283 | -1.7534942 | 0.07951722 | 0.21110644 |
| ELP5 | 159.403567 | -0.441779423 | 0.20208565 | -2.1861 | 0.02880829 | 0.10612074 |
| EIF5A | 336.2276 | -1.275859352 | 0.56168586 | -2.2714821 | 0.02311781 | 0.09131869 |
| GPS2 | 264.028743 | -1.136949615 | 0.40877662 | -2.781347 | 0.00541338 | 0.03214959 |
| NEURL4 | 38.0787395 | 0.759548436 | 0.55340179 | 1.3725081 | 0.16990532 | 0.34889771 |

| | | | | | | |
|---|---|---|---|---|---|---|
| KCTD11 | 38.5714271 | -0.50783185 | 0.37277187 | -1.3623127 | 0.17309922 | 0.35265622 |
| TNK1 | 42.3619368 | -0.230767828 | 0.29436752 | -0.7839446 | 0.43307262 | 0.62552702 |
| NLGN2 | 85.8859944 | -0.502634437 | 0.59729276 | -0.8415211 | 0.40005609 | 0.59423156 |
| TMEM102 | 33.5958287 | -0.93921693 | 0.58780251 | -1.5978444 | 0.11007762 | 0.26171962 |
| CHRNB1 | 45.1420096 | -0.572364389 | 0.47064732 | -1.2161216 | 0.22393859 | 0.41621903 |
| ZBTB4 | 433.942949 | -0.070978418 | 0.29725065 | -0.2387831 | 0.81127381 | 0.8977214 |
| POLR2A | 759.861613 | -0.003222014 | 0.32238102 | -0.0099944 | 0.99202573 | 0.99609195 |
| EIF4A1 | 382.037947 | -1.557050379 | 0.45712688 | -3.4061668 | 0.00065882 | 0.00661122 |
| CD68 | 116.805311 | -1.894457249 | 0.6718129 | -2.8199179 | 0.00480359 | 0.0294204 |
| MPDU1 | 246.688595 | -0.285024429 | 0.30234477 | -0.9427133 | 0.34582758 | 0.5462386 |
| FXR2 | 260.971573 | 0.269239335 | 0.26803994 | 1.00447471 | 0.31514986 | 0.51523844 |
| SHBG | 69.8165196 | 0.784834457 | 0.49990654 | 1.56996236 | 0.11642387 | 0.27190696 |
| SAT2 | 1469.77389 | 0.963924639 | 0.51037398 | 1.88866336 | 0.05893695 | 0.17269106 |
| TP53 | 249.97279 | -0.428999896 | 0.36342592 | -1.1804329 | 0.2378281 | 0.43269309 |
| WRAP53 | 59.6873924 | 0.267167093 | 0.57742267 | 0.46268896 | 0.64358734 | 0.79117952 |
| DNAH2 | 37.4615833 | 0.087305304 | 0.4695888 | 0.18591863 | 0.85250858 | 0.92213818 |
| KDM6B | 540.65256 | -0.792038404 | 0.64459016 | -1.2287473 | 0.21916658 | 0.41073506 |
| TMEM88 | 43.8088685 | 0.133496504 | 0.77956203 | 0.17124552 | 0.86403072 | 0.92885255 |
| NAA38 | 391.070131 | -0.697640007 | 0.2844356 | -2.4527169 | 0.01417819 | 0.06435113 |
| CYB5D1 | 124.209985 | 0.064077682 | 0.29041279 | 0.22064346 | 0.82537006 | 0.90642548 |
| CHD3 | 147.788889 | -0.028857982 | 0.49184577 | -0.0586728 | 0.9532127 | 0.97643573 |
| TRAPPC1 | 99.3803651 | -1.386443877 | 0.47458322 | -2.9213925 | 0.0034847 | 0.02317557 |
| CNTROB | 169.146205 | 0.120198984 | 0.30451248 | 0.39472596 | 0.69304514 | 0.82311496 |
| PER1 | 499.200244 | -0.497134086 | 0.54326957 | -0.9150781 | 0.36015063 | 0.55850369 |
| TMEM107 | 79.7598712 | -0.669955041 | 0.31406147 | -2.1331971 | 0.03290856 | 0.11679896 |
| BORCS6 | 37.4969205 | -0.330184833 | 0.45464064 | -0.7262545 | 0.46768274 | 0.6549374 |
| CTC1 | 222.876714 | -0.169930338 | 0.17801652 | -0.9545762 | 0.33979203 | 0.54031664 |
| PFAS | 69.5561962 | 0.096270982 | 0.35653104 | 0.27002132 | 0.78714385 | 0.88351603 |
| SLC25A35 | 27.4755031 | 1.399141157 | 0.49435709 | 2.8302237 | 0.00465155 | 0.028772 |
| RANGRF | 70.0602031 | -0.061819051 | 0.30343115 | -0.2037334 | 0.83856186 | 0.9145582 |
| AC135178.3 | 36.6936129 | -0.005553622 | 0.40141618 | -0.0138351 | 0.98896156 | 0.99462148 |
| NDEL1 | 725.633914 | -0.343834497 | 0.36687172 | -0.9372063 | 0.34865244 | 0.54865148 |
| MYH10 | 1080.35311 | 0.116601733 | 0.25804057 | 0.45187364 | 0.65136001 | 0.79573901 |
| STX8 | 1709.084 | 0.246149022 | 0.24981521 | 0.98532441 | 0.32446475 | 0.52483341 |
| AC005695.2 | 69.5509031 | -0.598304592 | 0.52108785 | -1.1481837 | 0.25089272 | 0.44720241 |
| AC118755.2 | 45.6017984 | 0.313189134 | 0.3406504 | 0.91938578 | 0.35789383 | 0.55644858 |
| USP43 | 316.857399 | -0.881174454 | 0.29035717 | -3.0347949 | 0.00240699 | 0.01771422 |
| GAS7 | 42.177951 | 0.292055164 | 0.33555562 | 0.8703629 | 0.38410211 | 0.58045387 |
| MYHAS | 182.262464 | 0.761481148 | 0.82681733 | 0.9209787 | 0.35706155 | 0.55584812 |
| MYH3 | 70.9675537 | 0.06825993 | 0.4226909 | 0.161489 | 0.87170828 | 0.93353404 |
| SCO1 | 271.217508 | -0.25896011 | 0.27607162 | -0.9380179 | 0.34823523 | 0.54839847 |
| ADPRM | 65.0520122 | -0.244924228 | 0.27290228 | -0.8974796 | 0.36946305 | 0.56695645 |
| TMEM220 | 347.130746 | 0.343782513 | 0.32996059 | 1.04188962 | 0.29746286 | 0.49796328 |
| TMEM220-A: | 390.569263 | 0.263522421 | 0.39764274 | 0.66271152 | 0.50751533 | 0.68747037 |
| AC015908.3 | 39.90272 | -0.667560426 | 0.68479976 | -0.9748257 | 0.32964671 | 0.52934249 |
| AC015908.2 | 66.7613348 | -0.025481727 | 0.73493154 | -0.0346722 | 0.97234109 | 0.98615459 |
| SHISA6 | 123.701132 | 0.201242044 | 0.58630425 | 0.34323825 | 0.73141923 | 0.84727663 |
| DNAH9 | 90.0047916 | -1.38829266 | 0.58585285 | -2.3696952 | 0.01780276 | 0.07599576 |
| AC005410.2 | 33.6888166 | 0.72276585 | 0.5151059 | 1.40314031 | 0.160575 | 0.33650588 |
| ZNF18 | 87.5069547 | 0.744668761 | 0.35834384 | 2.07808443 | 0.03770158 | 0.12839145 |
| MAP2K4 | 2245.18482 | -0.210608952 | 0.15926838 | -1.3223525 | 0.18605079 | 0.36923618 |
| ARHGAP44 | 142.768493 | -0.282760996 | 0.36586252 | -0.7728613 | 0.43960445 | 0.6314218 |
| ELAC2 | 253.422879 | -0.170345802 | 0.25335164 | -0.672369 | 0.50134878 | 0.68247686 |
| COX10 | 525.229612 | 0.030596753 | 0.17663602 | 0.17321922 | 0.86247912 | 0.92794358 |
| PMP22 | 139.047898 | -3.179682952 | 1.18955682 | -2.672998 | 0.00751767 | 0.04102099 |
| TRIM16 | 239.191479 | 0.292814647 | 0.31247663 | 0.93707695 | 0.34871898 | 0.54865148 |
| ZSWIM7 | 548.293989 | -0.160138271 | 0.18094018 | -0.8850343 | 0.37613812 | 0.57273108 |
| TTC19 | 1220.06028 | -0.393260813 | 0.18985361 | -2.0713897 | 0.0383224 | 0.12998945 |
| NCOR1 | 3808.12477 | 0.345617218 | 0.11928711 | 2.89735585 | 0.00376323 | 0.02445829 |
| PIGL | 1242.77782 | 0.625148017 | 0.36466399 | 1.71431246 | 0.08647136 | 0.22329745 |
| CENPV | 156.542543 | 0.778606277 | 0.3759047 | 2.07128634 | 0.03833205 | 0.12998945 |
| UBB | 1107.16256 | -1.211974098 | 0.40316543 | -3.0061458 | 0.00264582 | 0.01887063 |
| LRRC75A | 98.9446096 | -0.893680062 | 0.33653203 | -2.6555572 | 0.00791775 | 0.04252688 |
| ZNF287 | 30.2566753 | 0.369617094 | 0.37329076 | 0.99015871 | 0.32209655 | 0.52244289 |
| ZNF624 | 74.0368049 | 0.462255899 | 0.31437301 | 1.47040582 | 0.14145188 | 0.30839524 |
| AC098850.3 | 485.093754 | 0.562239632 | 0.65445008 | 0.85910239 | 0.39028403 | 0.5856183 |
| MPRIP | 1114.12621 | 0.043313468 | 0.27673419 | 0.1565165 | 0.87562591 | 0.93525172 |
| COPS3 | 179.430841 | -0.119737642 | 0.27697149 | -0.4323104 | 0.66551586 | 0.80579154 |
| NT5M | 41.0172838 | -0.426115777 | 0.48340385 | -0.8814902 | 0.37805253 | 0.57462476 |
| MED9 | 211.535506 | -0.452250843 | 0.28094683 | -1.6097382 | 0.10745502 | 0.25751328 |
| RASD1 | 425.678684 | -1.844752478 | 0.68671229 | -2.6863543 | 0.00722365 | 0.03982885 |
| PEMT | 147.338414 | -0.092137731 | 0.44677479 | -0.2062286 | 0.83661236 | 0.91377858 |
| RAI1 | 563.346999 | -0.803053886 | 0.32571808 | -2.4654876 | 0.01368269 | 0.06283036 |
| SMCR5 | 51.1760443 | -1.059885255 | 0.45183432 | -2.3457387 | 0.01898941 | 0.07942871 |
| SREBF1 | 513.703231 | -0.423573827 | 0.41068576 | -1.0313818 | 0.3023618 | 0.50269269 |
| TOM1L2 | 1335.88712 | 0.400317746 | 0.25188219 | 1.58930548 | 0.11199144 | 0.26494075 |
| DRC3 | 197.863679 | 1.060660539 | 0.40255317 | 2.63483343 | 0.00841786 | 0.04450615 |
| ATPAF2 | 204.913022 | 0.276158341 | 0.1772051 | 1.55841079 | 0.1191359 | 0.2759371 |
| GID4 | 156.429797 | 0.044852328 | 0.23303153 | 0.19247322 | 0.84737154 | 0.91938085 |
| DRG2 | 177.918697 | 0.075048725 | 0.29412861 | 0.25515615 | 0.7986025 | 0.89024434 |
| MYO15A | 38.569526 | -1.562006954 | 0.65036043 | -2.4017558 | 0.0163166 | 0.07147701 |
| ALKBH5 | 727.451457 | -0.51104286 | 0.32822744 | -1.5569779 | 0.11947574 | 0.27650132 |
| FLII | 457.981829 | -1.27763546 | 0.3433879 | -3.720677 | 0.00019869 | 0.00265548 |
| MIEF2 | 99.5522208 | -0.848071519 | 0.35437376 | -2.3931555 | 0.01670416 | 0.07266389 |
| TOP3A | 291.254543 | -0.278371933 | 0.31966275 | -0.8708301 | 0.38384695 | 0.5803781 |
| SMCR8 | 161.874954 | -0.209545154 | 0.40353751 | -0.5192706 | 0.60357207 | 0.76169806 |
| SHMT1 | 2441.45866 | 1.050964281 | 0.31793719 | 3.30557205 | 0.00094783 | 0.00869021 |
| TRIM16L | 226.250551 | -0.990730179 | 0.50219657 | -1.9727936 | 0.04851908 | 0.15235992 |
| TVP23B | 305.34552 | -0.422379772 | 0.21319929 | -1.98115 | 0.04757445 | 0.15049689 |
| PRPSAP2 | 1222.38988 | 0.902945976 | 0.2278539 | 3.96282865 | 7.41E-05 | 0.00123208 |
| SLC5A10 | 2005.74551 | 1.342795664 | 0.28053859 | 4.78649178 | 1.70E-06 | 6.01E-05 |
| FAM83G | 195.131891 | -0.277523555 | 0.44246928 | -0.6272154 | 0.53051804 | 0.70456455 |
| AC007952.6 | 48.7525403 | 2.65180607 | 0.57740767 | 4.59260624 | 4.38E-06 | 0.00012848 |
| EPN2 | 1074.42654 | -0.39668599 | 0.19648131 | -2.0189502 | 0.04349239 | 0.14123609 |
| EPN2-AS1 | 37.3821698 | -2.059875343 | 0.58001002 | -3.5514479 | 0.00038312 | 0.00442897 |
| B9D1 | 247.987519 | 0.168757628 | 0.27865182 | 0.60562185 | 0.54476589 | 0.71510305 |
| MAPK7 | 63.295559 | -0.860863864 | 0.40508927 | -2.1251214 | 0.03357649 | 0.11842764 |
| AC004448.2 | 94.3148579 | -1.168008345 | 0.30554872 | -3.8226583 | 0.00013202 | 0.00191957 |
| SLC47A1 | 3042.66162 | 1.845521037 | 0.47193385 | 3.91055029 | 9.21E-05 | 0.00146557 |
| AC025627.1 | 314.118745 | 0.687354551 | 0.72942507 | 0.94232373 | 0.34602692 | 0.5463646 |
| ALDH3A2 | 881.980409 | 1.344299824 | 0.36037232 | 3.7303082 | 0.00019125 | 0.00257253 |
| SLC47A2 | 4577.18637 | -0.366258613 | 0.57412717 | -0.6379399 | 0.52351282 | 0.6991222 |
| AC005722.2 | 44.0567854 | 0.238383891 | 0.48083751 | 0.49576808 | 0.62005805 | 0.77432081 |
| ULK2 | 409.114818 | 0.055164128 | 0.33304522 | 0.16563555 | 0.86844377 | 0.93161498 |
| AKAP10 | 1322.29373 | -0.065166906 | 0.19400303 | -0.3359066 | 0.73694126 | 0.85110831 |
| SPECC1 | 100.52196 | -2.072290945 | 0.42728218 | -4.8499353 | 1.24E-06 | 4.55E-05 |
| CCDC144NL-/ | 205.489435 | -1.046531748 | 0.39929638 | -2.6209397 | 0.00876878 | 0.04584043 |
| USP22 | 1197.82926 | -0.048568225 | 0.21909633 | -0.2216752 | 0.82456673 | 0.90636216 |
| DHRS7B | 919.82682 | -0.124416408 | 0.30298735 | -0.4106324 | 0.68134214 | 0.81457974 |
| TMEM11 | 125.251814 | -0.669995333 | 0.31764925 | -2.10923 | 0.03492473 | 0.12156255 |
| AC087294.1 | 130.005811 | -0.364201892 | 0.32472535 | -1.121569 | 0.26204573 | 0.46002317 |
| NATD1 | 61.4338038 | -0.881062323 | 0.43639737 | -2.0189451 | 0.04349293 | 0.14123609 |
| MAP2K3 | 708.44348 | -0.973311715 | 0.59390336 | -1.6388385 | 0.10124689 | 0.24795268 |

| | | | | | | |
|---|---|---|---|---|---|---|
| C17orf51 | 48.1936817 | -0.480241203 | 0.51862762 | -0.9259846 | 0.35445397 | 0.55291915 |
| WSB1 | 3599.39403 | -1.234428803 | 0.44953765 | -2.7459965 | 0.00603274 | 0.03471485 |
| KSR1 | 896.325369 | -0.09139965 | 0.29641024 | -0.3083552 | 0.75781204 | 0.86418282 |
| LGALS9 | 36.4532077 | -0.289218603 | 0.78349767 | -0.3691378 | 0.71202502 | 0.83516404 |
| AC005697.1 | 63.2093425 | 0.769884707 | 0.71174218 | 1.08169043 | 0.27939011 | 0.47799091 |
| NLK | 2164.74802 | 0.157158952 | 0.17588882 | 0.89351305 | 0.37158249 | 0.56909545 |
| IFT20 | 143.069876 | -0.142091538 | 0.31313201 | -0.4537752 | 0.64999064 | 0.79477446 |
| TNFAIP1 | 374.272213 | -1.003892061 | 0.26219523 | -3.8287961 | 0.00012877 | 0.00189589 |
| POLDIP2 | 306.055429 | -0.25549119 | 0.18029275 | -1.4170908 | 0.15645639 | 0.33013369 |
| TMEM199 | 101.314543 | -0.15172996 | 0.22513551 | -0.6739495 | 0.50034343 | 0.68152712 |
| SARM1 | 202.876553 | -0.213028071 | 0.28763663 | -0.7406152 | 0.45892677 | 0.6475935 |
| SLC46A1 | 46.8208141 | 0.835905029 | 0.34724456 | 2.40725159 | 0.01607309 | 0.07079495 |
| AC005726.1 | 1779.56735 | 1.70173902 | 0.55268487 | 3.07904037 | 0.00207669 | 0.01582766 |
| SLC13A2 | 258.039873 | 3.235319076 | 0.85559994 | 3.78134561 | 0.00015598 | 0.00217503 |
| AC005726.5 | 123.364914 | 1.704064905 | 0.67045511 | 2.54165399 | 0.01103293 | 0.05408304 |
| KIAA0100 | 1227.90486 | 0.086593556 | 0.1980901 | 0.43714227 | 0.66200817 | 0.80336802 |
| SDF2 | 292.296534 | -0.203193373 | 0.24368433 | -0.8338385 | 0.404372 | 0.59843738 |
| SUPT6H | 991.17857 | -0.130748059 | 0.31292298 | -0.4178282 | 0.6760727 | 0.81238963 |
| PROCA1 | 34.2550549 | 0.596284568 | 0.70018086 | 0.85161507 | 0.39442778 | 0.58912733 |
| RAB34 | 125.022494 | -1.219165268 | 0.4936005 | -2.4699433 | 0.01351345 | 0.06226895 |
| RPL23A | 983.632006 | -1.019486838 | 0.41074112 | -2.4820667 | 0.01306228 | 0.06083748 |
| TLCD1 | 49.3804728 | -0.71336737 | 0.50978445 | -1.399351 | 0.16170776 | 0.33793692 |
| NEK8 | 107.769832 | -0.638498925 | 0.38778533 | -1.6465268 | 0.09965535 | 0.24528208 |
| TRAF4 | 564.235753 | -0.068602159 | 0.27584913 | -0.2486945 | 0.80359711 | 0.89315921 |
| FAM222B | 2199.66418 | 0.181904627 | 0.25884739 | 0.70274855 | 0.48221247 | 0.66670759 |
| ERAL1 | 117.712296 | 0.150476241 | 0.29775897 | 0.50536259 | 0.61330419 | 0.76933839 |
| FLOT2 | 919.411799 | -0.502326013 | 0.38998796 | -1.2880552 | 0.19772675 | 0.38391783 |
| DHRS13 | 88.5961178 | -1.800647488 | 0.52262488 | -3.4453918 | 0.00057023 | 0.00587682 |
| PHF12 | 486.376239 | 0.269586945 | 0.2191476 | 1.23016154 | 0.21863662 | 0.40984049 |
| PIPOX | 389.656961 | 1.690954151 | 0.5489356 | 3.08042354 | 0.00206706 | 0.01577575 |
| MYO18A | 560.284142 | -0.613397724 | 0.46018808 | -1.3329283 | 0.18255532 | 0.36485995 |
| AC024619.3 | 59.4643843 | -0.965672062 | 0.67360671 | -1.4335844 | 0.1516909 | 0.32307857 |
| NUFIP2 | 1722.94858 | -0.49828272 | 0.31517765 | -1.5809583 | 0.11388758 | 0.26767096 |
| TAOK1 | 2544.40286 | -0.00249522 | 0.19251169 | -0.0129614 | 0.98965859 | 0.99493132 |
| ABHD15-AS1 | 344.12343 | 0.460110425 | 0.49868618 | 0.92264522 | 0.35619212 | 0.55474667 |
| ABHD15 | 79.6249711 | -0.139324421 | 0.33977693 | -0.4100467 | 0.68177166 | 0.81468059 |
| GIT1 | 204.453408 | -0.517588426 | 0.37923621 | -1.3648181 | 0.17231024 | 0.35167607 |
| SSH2 | 5660.60514 | 0.161540438 | 0.1182664 | 1.36590309 | 0.17196938 | 0.3513466 |
| AC023389.1 | 158.049329 | -0.75615994 | 0.30050465 | -2.5163003 | 0.01185941 | 0.05677852 |
| EFCAB5 | 177.465772 | -1.971842636 | 0.37786839 | -5.2183318 | 1.81E-07 | 8.97E-06 |
| NSRP1 | 1677.5216 | 0.083699981 | 0.15863207 | 0.52763594 | 0.59775205 | 0.75777463 |
| AC104984.3 | 64.6733736 | -0.632822136 | 0.61885015 | -1.0225773 | 0.30650773 | 0.50646126 |
| BLMH | 602.082126 | 0.32168719 | 0.13255291 | 2.42685878 | 0.01523018 | 0.06792146 |
| CPD | 2246.83329 | -1.905572459 | 0.35572806 | -5.3568236 | 8.47E-08 | 4.78E-06 |
| GOSR1 | 954.149971 | 0.046590579 | 0.24594801 | 0.18943264 | 0.84975375 | 0.92090248 |
| CRLF3 | 612.315036 | 0.008891045 | 0.23357145 | 0.03806563 | 0.96963535 | 0.98486786 |
| ATAD5 | 165.899608 | 0.363225288 | 0.38734098 | 0.93774042 | 0.34837783 | 0.54845045 |
| AC130324.2 | 97.1243906 | -0.745238142 | 0.53554172 | -1.3915594 | 0.16405585 | 0.34092078 |
| TEFM | 320.872322 | -0.022299376 | 0.18671635 | -0.1194292 | 0.90493536 | 0.95034853 |
| ADAP2 | 331.415911 | 0.426997738 | 0.2602342 | 1.64082101 | 0.10083458 | 0.24725215 |
| RNF135 | 341.071634 | 0.584378821 | 0.24504742 | 2.38475809 | 0.01709036 | 0.07395389 |
| NF1 | 5096.45813 | 0.067457678 | 0.14990425 | 0.45000511 | 0.65270676 | 0.79674517 |
| OMG | 91.6002333 | -0.4778339 | 0.5083746 | -0.9399248 | 0.34725613 | 0.54729675 |
| RAB11FIP4 | 208.719854 | 0.672078306 | 0.32945465 | 2.03997212 | 0.0413531 | 0.13664328 |
| COPRS | 151.331486 | -0.127323538 | 0.35705943 | -0.3565892 | 0.72139936 | 0.84198067 |
| UTP6 | 392.227821 | 0.048505848 | 0.28222444 | 0.17186977 | 0.86353992 | 0.9285747 |
| AC004253.1 | 94.7596212 | -0.442616779 | 0.45567689 | -0.9713391 | 0.33137944 | 0.53125357 |
| SUZ12 | 1072.26809 | 0.13256364 | 0.15600294 | 0.84975093 | 0.39546358 | 0.59027092 |
| LRRC37B | 424.127199 | -0.1308115 | 0.17656172 | -0.7408825 | 0.45876466 | 0.64747603 |
| RHOT1 | 917.749767 | 0.526733918 | 0.1359894 | 3.8733453 | 0.00010735 | 0.00164594 |
| C17orf75 | 153.075591 | 0.939662649 | 0.39105777 | 2.40287423 | 0.01626678 | 0.07130437 |
| ZNF207 | 2606.11137 | 0.18190154 | 0.15034054 | 1.20993009 | 0.22630572 | 0.41890243 |
| PSMD11 | 937.469281 | 0.112635431 | 0.28504687 | 0.39514706 | 0.69273436 | 0.82281714 |
| MYO1D | 1275.61259 | -1.463029851 | 0.3351371 | -4.3654667 | 1.27E-05 | 0.00030884 |
| TMEM98 | 134.619041 | -0.121204587 | 0.45726787 | -0.2650626 | 0.79096128 | 0.88629048 |
| ASIC2 | 79.4232455 | -0.560796311 | 0.47260096 | -1.186617 | 0.23537874 | 0.42967809 |
| CCL2 | 197.922087 | -2.38098256 | 0.72560035 | -3.2813966 | 0.00103294 | 0.00926827 |
| TMEM132E | 86.2756693 | 3.53796786 | 0.354177 | 9.98926475 | 1.70E-23 | 5.82E-20 |
| CCT6B | 116.523054 | 0.908365654 | 0.45869245 | 1.98033703 | 0.04766567 | 0.15068715 |
| ZNF830 | 124.757143 | -0.605077198 | 0.37801 | -1.600691 | 0.10944538 | 0.26077541 |
| LIG3 | 128.532473 | -0.365144493 | 0.18400134 | -1.9844664 | 0.04720387 | 0.14987802 |
| RFFL | 407.670696 | -1.313736994 | 0.43680004 | -3.0076393 | 0.00263285 | 0.01884831 |
| SLFN5 | 184.275912 | -2.527134971 | 0.54996337 | -4.5950969 | 4.33E-06 | 0.00012758 |
| SLFN12 | 43.7443062 | 0.871779738 | 0.40444713 | 2.15548506 | 0.0311239 | 0.11211362 |
| SLFN13 | 74.1719567 | -0.770099119 | 0.69990001 | -1.1002988 | 0.27120197 | 0.46942359 |
| LINC02001 | 172.278893 | 0.322640276 | 0.19569277 | 1.64870821 | 0.09920743 | 0.24439903 |
| AP2B1 | 1192.42579 | -0.320694395 | 0.1502509 | -2.1343926 | 0.03281066 | 0.11654204 |
| TAF15 | 2139.74296 | -0.12462377 | 0.14784698 | -0.842924 | 0.39927093 | 0.5934221 |
| AC015849.3 | 118.844041 | -0.718324287 | 0.47060139 | -1.5263964 | 0.12691117 | 0.28796083 |
| CCL15-CCL14 | 49.6843934 | 1.695688255 | 0.71904186 | 2.35826084 | 0.01836079 | 0.07770031 |
| ZNHIT3 | 122.923428 | -0.122542178 | 0.26265462 | -0.4665525 | 0.64082006 | 0.78905143 |
| MYO19 | 554.649487 | -0.332864571 | 0.37449812 | -0.8888284 | 0.3740953 | 0.57127053 |
| GGNBP2 | 1279.29032 | 0.045489132 | 0.20618531 | 0.22062256 | 0.82538633 | 0.90642548 |
| DHRS11 | 58.5480998 | 1.635020504 | 0.36141722 | 4.52391422 | 6.07E-06 | 0.00016776 |
| MRM1 | 71.4008072 | 0.950120148 | 0.2865481 | 3.31574396 | 0.000914 | 0.00845732 |
| LHX1-DT | 212.789974 | 1.510580606 | 0.56837448 | 2.65772068 | 0.00786711 | 0.04238775 |
| LHX1 | 141.856464 | 1.955382716 | 0.56197466 | 3.47948554 | 0.00050238 | 0.00537086 |
| AATF | 1169.23903 | 0.018225614 | 0.26734365 | 0.06817298 | 0.94564794 | 0.97196767 |
| ACACA | 1648 | -0.321697314 | 0.24925413 | -1.2906398 | 0.19682859 | 0.38261897 |
| TADA2A | 389.61154 | 0.197205165 | 0.23036112 | 0.85606964 | 0.39195926 | 0.58703951 |
| DUSP14 | 91.8808954 | -1.187384383 | 0.45015777 | -2.6377072 | 0.00834686 | 0.04425936 |
| SYNRG | 1511.79847 | 0.123364434 | 0.21911862 | 0.56300297 | 0.57343286 | 0.73865654 |
| DDX52 | 839.907648 | -0.581846833 | 0.28723985 | -2.025648 | 0.04280087 | 0.13978353 |
| AC243585.2 | 40.8153612 | -1.66834146 | 0.64204354 | -2.5984865 | 0.00936357 | 0.04803387 |
| HNF1B | 2180.0556 | 1.358243824 | 0.40508734 | 3.3529654 | 0.00079951 | 0.00766353 |
| MRPL45 | 335.689885 | 0.351071101 | 0.16988838 | 2.06648094 | 0.03878309 | 0.13093592 |
| SOCS7 | 426.755143 | 0.963720498 | 0.24433226 | 3.94430313 | 8.00E-05 | 0.00130751 |
| MLLT6 | 816.023301 | -0.162253655 | 0.2642261 | -0.6140713 | 0.53916824 | 0.71083091 |
| CISD3 | 217.929964 | 0.222309454 | 0.24651848 | 0.9017963 | 0.36716508 | 0.56454671 |
| PCGF2 | 247.636062 | 0.622297597 | 0.31535779 | 1.97330658 | 0.04846064 | 0.15224617 |
| PSMB3 | 351.240724 | -0.58537224 | 0.30454124 | -1.9221444 | 0.05458759 | 0.16434269 |
| PIP4K2B | 357.168788 | 0.532885329 | 0.27886633 | 1.91089876 | 0.05601759 | 0.16710188 |
| CWC25 | 472.757411 | -0.655403399 | 0.38550137 | -1.7001325 | 0.08910601 | 0.22769874 |
| RPL23 | 776.234169 | -0.680045151 | 0.44789732 | -1.5183059 | 0.1289373 | 0.29068151 |
| LASP1 | 835.813423 | -0.222315201 | 0.34029285 | -0.6533055 | 0.51355932 | 0.69196393 |
| RPL19 | 1335.69659 | -1.06380531 | 0.47629155 | -2.2335171 | 0.02551487 | 0.09785458 |
| FBXL20 | 2347.70973 | -0.152812058 | 0.25430529 | -0.6009 | 0.54790657 | 0.71819407 |
| MED1 | 387.88355 | 0.138802688 | 0.24711944 | 0.5616826 | 0.57433229 | 0.73918993 |
| CDK12 | 1217.17447 | 0.071029824 | 0.180884 | 0.39268163 | 0.69455464 | 0.82419362 |
| STARD3 | 278.629722 | -0.280022481 | 0.29223541 | -0.9582086 | 0.33795758 | 0.53821129 |
| PGAP3 | 90.7340036 | 0.857875977 | 0.35975994 | 2.38457892 | 0.01709868 | 0.07395389 |
| ERBB2 | 671.636682 | 0.972091207 | 0.41248915 | 2.35664674 | 0.01844078 | 0.07789454 |
| MIEN1 | 165.338242 | -1.018915588 | 0.34104415 | -2.9876354 | 0.00281145 | 0.01975218 |

| | | | | | | |
|---|---|---|---|---|---|---|
| GRB7 | 585.010206 | 0.060503083 | 0.37982478 | 0.15929209 | 0.87343875 | 0.9345535 |
| GSDMB | 112.741677 | 0.167121144 | 0.45130476 | 0.37030663 | 0.71115404 | 0.83492834 |
| ORMDL3 | 99.6458026 | -0.015485957 | 0.29291348 | -0.0528687 | 0.95783652 | 0.97846662 |
| AC090844.2 | 80.071232 | 0.583960543 | 0.58667675 | 0.99537018 | 0.31955625 | 0.5200852 |
| PSMD3 | 285.683489 | -0.651911846 | 0.33242653 | -1.9610704 | 0.04987081 | 0.15518255 |
| MED24 | 343.762204 | 0.296805323 | 0.13584329 | 2.18490974 | 0.02889547 | 0.10631802 |
| THRA | 218.187931 | 0.398315493 | 0.32304433 | 1.2330057 | 0.21757364 | 0.40869972 |
| NR1D1 | 129.52235 | -2.068420498 | 0.69054038 | -2.995365 | 0.00274117 | 0.0193776 |
| MSL1 | 459.240078 | -0.256615432 | 0.22113747 | -1.1604339 | 0.24587217 | 0.44147485 |
| CASC3 | 513.612928 | 0.10275375 | 0.20659041 | 0.49737908 | 0.61892175 | 0.7734829 |
| RAPGEFL1 | 54.7406602 | -0.561536142 | 0.42862577 | -1.3100849 | 0.19016713 | 0.3741919 |
| WIPF2 | 1385.92837 | -0.060728636 | 0.2923191 | -0.2077477 | 0.83542593 | 0.91316952 |
| RARA | 214.479573 | 0.134098992 | 0.45637481 | 0.29383522 | 0.76888382 | 0.87121073 |
| IGFBP4 | 1553.46925 | 0.608932898 | 0.19175915 | 3.1755089 | 0.00149574 | 0.01232246 |
| KRT10 | 309.391261 | -1.184921513 | 0.43102433 | -2.7490827 | 0.00597623 | 0.03450556 |
| TMEM99 | 98.524381 | 0.050216625 | 0.3632264 | 0.13825158 | 0.89004159 | 0.94316197 |
| KRT19 | 317.138667 | -0.789164621 | 0.58193127 | -1.3561131 | 0.17506319 | 0.35517926 |
| EIF1 | 1443.74457 | -1.342561116 | 0.3519302 | -3.8148506 | 0.00013627 | 0.00196196 |
| JUP | 828.619347 | 0.448899308 | 0.26082153 | 1.72109757 | 0.08523312 | 0.22099234 |
| P3H4 | 105.871255 | 0.220734278 | 0.30500805 | 0.72369985 | 0.46925002 | 0.65619364 |
| NT5C3B | 204.120586 | 0.027235372 | 0.25786755 | 0.10561768 | 0.9158857 | 0.95496466 |
| KLHL11 | 43.4691393 | -0.250109145 | 0.45226446 | -0.5530153 | 0.58025296 | 0.74366782 |
| ACLY | 1539.83557 | -0.11859274 | 0.32459706 | -0.3653537 | 0.71484739 | 0.83689897 |
| TTC25 | 178.422005 | 1.287041854 | 0.41263327 | 3.11909372 | 0.00181408 | 0.0141918 |
| CNP | 269.182118 | -0.693429029 | 0.47511952 | -1.4594834 | 0.14443212 | 0.31226042 |
| DNAJC7 | 971.904882 | 0.006391389 | 0.20668374 | 0.03092352 | 0.97533053 | 0.98772483 |
| NKIRAS2 | 123.383149 | -0.300142205 | 0.2363706 | -1.269795 | 0.20415767 | 0.39237089 |
| DHX58 | 72.0395767 | -0.123517144 | 0.89590732 | -0.1378682 | 0.89034457 | 0.94326785 |
| KAT2A | 172.597713 | 0.690768822 | 0.37687922 | 1.83286525 | 0.06682261 | 0.1881548 |
| GHDC | 134.113418 | 0.848207744 | 0.36480912 | 2.32507278 | 0.02006807 | 0.08262932 |
| STAT5B | 1557.0962 | 0.474807003 | 0.20389451 | 2.32868953 | 0.01987552 | 0.08210782 |
| STAT5A | 163.198856 | 0.215653764 | 0.3350651 | 0.6436175 | 0.5198235 | 0.69597507 |
| STAT3 | 3180.76963 | -0.759220584 | 0.29385047 | -2.583697 | 0.00977476 | 0.04962321 |
| CAVIN1 | 81.9989455 | -0.694100088 | 0.51025882 | -1.3602902 | 0.1737381 | 0.35358992 |
| ATP6V0A1 | 2618.41244 | 0.066834329 | 0.19041952 | 0.35098465 | 0.72559986 | 0.84394089 |
| NAGLU | 490.86124 | 0.162716627 | 0.41813818 | 0.38914559 | 0.69716845 | 0.82550208 |
| AC067852.2 | 34.3895387 | -0.122010201 | 0.40971335 | -0.2977941 | 0.76586035 | 0.86915448 |
| COASY | 276.980953 | -0.239489406 | 0.27815881 | -0.8609808 | 0.3892486 | 0.58483289 |
| MLX | 260.889958 | -0.673124126 | 0.35753982 | -1.882655 | 0.05974714 | 0.17420848 |
| RETREG3 | 552.232598 | -0.042380895 | 0.26131772 | -0.1621815 | 0.87116294 | 0.93338059 |
| TUBG1 | 105.076916 | -0.48546606 | 0.27579222 | -1.7602602 | 0.0783637 | 0.20909699 |
| TUBG2 | 352.373669 | -0.004391104 | 0.22722021 | -0.0193253 | 0.98458158 | 0.99241906 |
| PLEKHH3 | 180.905171 | 0.540365926 | 0.47203585 | 1.14475611 | 0.2523102 | 0.44873698 |
| EZH1 | 771.667368 | 0.008155764 | 0.22042583 | 0.03700004 | 0.97048497 | 0.98529276 |
| VPS25 | 173.581674 | -1.034305976 | 0.30281835 | -3.4155987 | 0.00063642 | 0.00643794 |
| WNK4 | 54.1793573 | -0.950990558 | 0.60685954 | -1.5670687 | 0.11709865 | 0.27273936 |
| COA3 | 342.789303 | -0.44365792 | 0.34157763 | -1.2988495 | 0.19399559 | 0.37905874 |
| BECN1 | 278.483471 | -0.2600693 | 0.20456288 | -1.2713416 | 0.20360714 | 0.39166469 |
| PSME3 | 216.173749 | -0.941262508 | 0.31893515 | -2.9512661 | 0.00316474 | 0.02149609 |
| LINC00671 | 3173.97867 | 2.711898187 | 0.88104146 | 3.07805965 | 0.00208353 | 0.01585342 |
| G6PC | 991.956576 | 3.02057703 | 1.38331464 | 2.18357917 | 0.02899319 | 0.10659967 |
| RUNDC1 | 282.119372 | -0.73303572 | 0.33246703 | -2.2048374 | 0.02746551 | 0.10294526 |
| RPL27 | 560.81908 | -1.097327301 | 0.39114058 | -2.805455 | 0.00502456 | 0.03038165 |
| IFI35 | 217.598673 | -0.730833839 | 0.78646326 | -0.9292663 | 0.35275107 | 0.55164392 |
| VAT1 | 312.450271 | -1.255802126 | 0.42118721 | -2.9815771 | 0.00286768 | 0.02001389 |
| BRCA1 | 244.594576 | -0.060431922 | 0.40774658 | -0.1482095 | 0.88217742 | 0.93886764 |
| NBR1 | 1109.70281 | 0.611752591 | 0.39721869 | 1.54009013 | 0.12353838 | 0.28278902 |
| TMEM106A | 919.302544 | 1.844297369 | 0.50136646 | 3.67854161 | 0.00023457 | 0.00303853 |
| LINC00854 | 1059.72943 | 0.869341637 | 0.52517905 | 1.65532429 | 0.09785873 | 0.24200482 |
| LINC00910 | 37.185328 | 0.246795186 | 0.54009162 | 0.4569506 | 0.64770656 | 0.79339712 |
| ARL4D | 74.3214885 | -0.866357398 | 0.54191107 | -1.5987077 | 0.10988557 | 0.2615318 |
| DHX8 | 339.498283 | -0.031787658 | 0.23548746 | -0.1349866 | 0.89262245 | 0.94393356 |
| DUSP3 | 502.571308 | -0.762305149 | 0.32571442 | -2.3404096 | 0.0192626 | 0.08025303 |
| MPP3 | 109.655506 | -1.625445064 | 0.41832228 | -3.8856287 | 0.00010207 | 0.00157902 |
| PYY | 72.61378 | 0.959125481 | 0.65168232 | 1.47176846 | 0.14108341 | 0.30805998 |
| NAGS | 93.9956079 | 0.426679803 | 0.64895841 | 0.65748405 | 0.51086972 | 0.6899686 |
| TMEM101 | 139.30211 | -0.171198784 | 0.22603182 | -0.7574101 | 0.44880421 | 0.639148 |
| LSM12 | 377.736492 | -0.03700509 | 0.36415213 | -0.1016199 | 0.91905841 | 0.95679351 |
| G6PC3 | 117.002422 | -0.550169319 | 0.40560444 | -1.3564184 | 0.17496609 | 0.35508738 |
| HDAC5 | 227.550555 | 0.105334955 | 0.31907951 | 0.33012134 | 0.74130828 | 0.85315807 |
| ASB16-AS1 | 85.0577132 | 0.07244661 | 0.25512057 | 0.28397009 | 0.7764333 | 0.87658111 |
| TMUB2 | 256.443956 | 0.189635882 | 0.24902576 | 0.76151111 | 0.44635184 | 0.63743065 |
| ATXN7L3 | 124.921234 | -0.297401961 | 0.35875436 | -0.8289849 | 0.40711293 | 0.60095264 |
| AC004596.1 | 35.8003358 | -0.52197799 | 0.39314556 | -1.3276965 | 0.18427838 | 0.36708383 |
| UBTF | 358.244475 | 0.301578166 | 0.29801066 | 1.01197105 | 0.31155189 | 0.5116978 |
| AC003102.1 | 48.0335691 | 0.923708327 | 0.5338966 | 1.73012588 | 0.08360779 | 0.21835341 |
| SLC25A39 | 411.195023 | -0.362372248 | 0.39980329 | -0.9063764 | 0.36473668 | 0.56229577 |
| GRN | 485.653376 | -0.856292182 | 0.44960669 | -1.9045361 | 0.0568404 | 0.1688581 |
| GPATCH8 | 2806.10891 | 0.25101007 | 0.18203779 | 1.37888988 | 0.16792871 | 0.34613516 |
| CCDC43 | 107.586065 | 0.047089929 | 0.3441469 | 0.1368309 | 0.89116446 | 0.94333086 |
| DBF4B | 102.671908 | 0.257170823 | 0.46929308 | 0.5479962 | 0.58369452 | 0.74654294 |
| EFTUD2 | 449.459014 | -0.115191176 | 0.32296241 | -0.3566705 | 0.72133846 | 0.84198067 |
| DCAKD | 467.573307 | 0.30722658 | 0.30111293 | 1.02030351 | 0.30758454 | 0.5077145 |
| NMT1 | 880.750296 | -0.078822621 | 0.20784326 | -0.3792407 | 0.70450914 | 0.83047014 |
| PLCD3 | 59.9673717 | -0.806839616 | 0.47304633 | -1.7056249 | 0.08807794 | 0.22621028 |
| ACBD4 | 360.591724 | 2.079422056 | 0.49239334 | 4.22309139 | 2.41E-05 | 0.00051051 |
| HEXIM1 | 270.441174 | 0.188117891 | 0.30791394 | 0.61094309 | 0.54123726 | 0.71231293 |
| AC138150.1 | 399.684345 | 0.798863591 | 0.40475289 | 1.97370696 | 0.04841507 | 0.15222969 |
| HEXIM2 | 78.4709662 | 1.631996819 | 0.3503287 | 4.65847305 | 3.19E-06 | 9.99E-05 |
| MAP3K14 | 1468.64204 | 0.175762303 | 0.29814977 | 0.58951011 | 0.55551913 | 0.72476748 |
| ARHGAP27 | 453.584658 | 0.028186161 | 0.22397451 | 0.1258454 | 0.8998543 | 0.94743919 |
| PLEKHM1 | 700.51749 | -1.657318079 | 0.64380081 | -2.5742715 | 0.01004514 | 0.05063948 |
| AC091132.2 | 41.1174889 | -1.918499564 | 0.62537829 | -3.0677425 | 0.00215682 | 0.01627951 |
| LINC02210-C | 181.940124 | 0.317092954 | 0.41412283 | 0.76569784 | 0.44385611 | 0.63517875 |
| MAPT-AS1 | 87.7115908 | 3.82147471 | 0.8756499 | 4.36415821 | 1.28E-05 | 0.00031014 |
| MAPT | 1567.26226 | 1.058062867 | 0.42521972 | 2.48827328 | 0.01283651 | 0.06013328 |
| KANSL1 | 6397.28642 | 0.278283428 | 0.29579614 | 0.94079467 | 0.34681009 | 0.54697111 |
| KANSL1-AS1 | 45.5420892 | 0.171946476 | 0.52264501 | 0.32899286 | 0.74216109 | 0.85370947 |
| ARL17B | 1210.32615 | 1.441418463 | 0.94485729 | 1.52554092 | 0.12712424 | 0.28824615 |
| LRRC37A | 37.2518276 | 0.434610426 | 0.46839498 | 0.92787165 | 0.35347415 | 0.55195605 |
| LRRC37A2 | 237.746049 | 0.649915972 | 0.25754811 | 2.52347405 | 0.01162016 | 0.05594576 |
| ARL17A | 318.82773 | -0.217747035 | 0.54615172 | -0.3986933 | 0.6901192 | 0.82084207 |
| NSF | 1207.39875 | 0.120675385 | 0.35209519 | 0.34273511 | 0.73179775 | 0.84761805 |
| WNT3 | 42.0012397 | -0.154672662 | 0.42641077 | -0.3627316 | 0.71680539 | 0.83847512 |
| AC005670.2 | 143.22945 | -0.071409601 | 0.48610966 | -0.1469002 | 0.8832108 | 0.93940952 |
| CDC27 | 1277.74646 | -0.624636256 | 0.17148211 | -3.6425738 | 0.00026993 | 0.00337888 |
| AC068234.1 | 1586.88973 | -2.970228292 | 0.64746333 | -4.587485 | 4.49E-06 | 0.00013109 |
| AC068234.2 | 244.905486 | -3.065516193 | 0.47030579 | -6.5181341 | 7.12E-11 | 1.19E-08 |
| EFCAB13 | 933.920796 | -1.975539975 | 0.49016201 | -4.0303817 | 5.57E-05 | 0.00098844 |
| AC040934.1 | 122.298265 | -0.268541518 | 0.3909185 | -0.6869501 | 0.49211415 | 0.67441637 |
| NPEPPS | 3114.86947 | -0.355357132 | 0.18494936 | -1.9213753 | 0.05468441 | 0.16455746 |
| AC025682.1 | 228.337485 | -0.224432358 | 0.35022684 | -0.64082 | 0.52163964 | 0.69750411 |
| KPNB1 | 1340.23378 | -0.588236037 | 0.32565013 | -1.8063436 | 0.07086468 | 0.19578819 |

| | | | | | | |
|---|---|---|---|---|---|---|
| TBKBP1 | 105.562709 | 0.77022607 | 0.56223741 | 1.3699303 | 0.17070866 | 0.34969424 |
| OSBPL7 | 44.3709585 | -0.231662987 | 0.40627429 | -0.5702133 | 0.56853306 | 0.73476171 |
| MRPL10 | 97.807902 | -0.549043168 | 0.29288403 | -1.8746094 | 0.06084647 | 0.17647537 |
| SCRN2 | 584.017347 | 1.260483973 | 0.54818392 | 2.29938152 | 0.02148328 | 0.08714749 |
| SP2 | 151.686925 | 0.151908946 | 0.23605646 | 0.64352801 | 0.51988155 | 0.69597507 |
| SP2-AS1 | 103.47997 | -0.261313119 | 0.31500949 | -0.8295405 | 0.40679865 | 0.60060202 |
| PNPO | 264.748264 | -0.043319262 | 0.38941426 | -0.1112421 | 0.91142436 | 0.95319043 |
| AC018521.2 | 68.8477499 | -2.101549604 | 0.74242158 | -2.8306688 | 0.00464508 | 0.02874497 |
| PRR15L | 82.7023326 | -0.53209929 | 0.95725742 | -0.5558581 | 0.57830786 | 0.74211119 |
| CDK5RAP3 | 460.341265 | 0.007694363 | 0.31840298 | 0.02416549 | 0.98072061 | 0.99047407 |
| COPZ2 | 51.617347 | 0.299971541 | 0.49855123 | 0.60168649 | 0.54738285 | 0.71771348 |
| NFE2L1 | 938.459176 | 0.508551098 | 0.19996098 | 2.54325166 | 0.01098261 | 0.05393717 |
| CBX1 | 287.643499 | 0.09589054 | 0.29669243 | 0.32319847 | 0.74654494 | 0.85675598 |
| SNX11 | 104.643308 | 0.068742669 | 0.30450758 | 0.22575028 | 0.82139565 | 0.90404363 |
| SKAP1 | 247.893027 | -0.609649651 | 0.71084385 | -0.8576421 | 0.39109009 | 0.58644261 |
| HOXB2 | 101.538076 | -0.383876261 | 0.47648832 | -0.8056362 | 0.42045262 | 0.61401492 |
| HOXB-AS1 | 83.7493785 | 0.00719672 | 0.47681164 | 0.01509342 | 0.98795765 | 0.99390352 |
| HOXB3 | 209.096283 | -0.619799028 | 0.56955751 | -1.0882115 | 0.27650175 | 0.47422542 |
| HOXB-AS3 | 46.7507971 | -0.221658875 | 0.53480099 | -0.4144698 | 0.67853006 | 0.81376982 |
| HOXB7 | 92.1553394 | -0.439416918 | 0.42688575 | -1.0293549 | 0.30331296 | 0.50363546 |
| TTLL6 | 113.728093 | 2.102993345 | 0.47867617 | 4.39335288 | 1.12E-05 | 0.00027762 |
| CALCOCO2 | 705.673858 | -0.249201777 | 0.20668738 | -1.2056942 | 0.22793542 | 0.42078261 |
| ATP5MC1 | 390.205316 | -0.465371327 | 0.35753811 | -1.3015992 | 0.19305342 | 0.37781219 |
| UBE2Z | 804.452183 | -0.844712444 | 0.2967752 | -2.846304 | 0.00442299 | 0.02765784 |
| SNF8 | 323.439427 | 0.170086738 | 0.24087398 | 0.70612333 | 0.48011145 | 0.66519782 |
| AC091133.1 | 55.1122591 | -0.822175454 | 0.33222151 | -2.4747809 | 0.01333179 | 0.061768 |
| IGF2BP1 | 118.447918 | 1.339334102 | 0.631241 | 2.12174765 | 0.03385893 | 0.11912331 |
| ZNF652 | 1015.64585 | 0.019421393 | 0.19874641 | 0.09771947 | 0.92215506 | 0.95822753 |
| AC091180.5 | 64.3274086 | -1.109455216 | 0.47265008 | -2.3473078 | 0.01890963 | 0.07924007 |
| PHB | 494.807117 | -0.364071508 | 0.36476083 | -0.9981102 | 0.31822592 | 0.51835088 |
| SPOP | 923.811296 | 0.324638492 | 0.14417224 | 2.25174065 | 0.02433867 | 0.09448034 |
| SLC35B1 | 146.770044 | -0.153327368 | 0.30759783 | -0.498467 | 0.61815492 | 0.77299851 |
| FAM117A | 225.109953 | 0.656760701 | 0.33501769 | 1.9603762 | 0.04995184 | 0.15529367 |
| KAT7 | 319.684809 | 0.297188652 | 0.14696737 | 2.02214034 | 0.04316186 | 0.1405606 |
| TAC4 | 55.948862 | -0.293885305 | 0.37492482 | -0.7838513 | 0.43312738 | 0.62552702 |
| ITGA3 | 2130.94635 | -3.905120229 | 0.59567745 | -6.555763 | 5.54E-11 | 9.60E-09 |
| PDK2 | 248.28961 | 1.219567502 | 0.43149242 | 2.82639382 | 0.00470753 | 0.02901357 |
| PPP1R9B | 130.174363 | -0.528670061 | 0.44059207 | -1.1999082 | 0.23017498 | 0.4233206 |
| AC002401.1 | 122.798684 | 4.319427434 | 0.66057269 | 6.53891317 | 6.20E-11 | 1.05E-08 |
| XYLT2 | 133.200002 | -0.362761084 | 0.25665532 | -1.4134173 | 0.15753305 | 0.33183985 |
| MRPL27 | 285.190211 | -0.557704149 | 0.28959317 | -1.9258194 | 0.05412692 | 0.16345401 |
| LRRC59 | 233.191241 | -1.53041685 | 0.48624377 | -3.1474272 | 0.00164714 | 0.01323734 |
| ACSF2 | 3895.23294 | 3.135143626 | 0.52889191 | 5.92775876 | 3.07E-09 | 2.99E-07 |
| CHAD | 286.56268 | 1.224143426 | 0.84841113 | 1.44286582 | 0.14905827 | 0.31929079 |
| AC021491.4 | 90.5416383 | 4.69526519 | 0.63494384 | 7.39477244 | 1.42E-13 | 4.85E-11 |
| RSAD1 | 521.692179 | 2.658232771 | 0.29198554 | 9.10398765 | 8.71E-20 | 1.49E-16 |
| SPATA20 | 234.668936 | 0.100824154 | 0.42956516 | 0.23471213 | 0.81443217 | 0.89947803 |
| ABCC3 | 3965.2186 | -1.989425898 | 0.37840291 | -5.2574276 | 1.46E-07 | 7.64E-06 |
| ANKRD40 | 223.357757 | -0.351013918 | 0.27795845 | -1.2628288 | 0.20665073 | 0.39538827 |
| AC005921.2 | 49.1678699 | -1.200560191 | 0.35518622 | -3.3800866 | 0.00072463 | 0.00714055 |
| LUC7L3 | 2322.90379 | 0.261040585 | 0.21732134 | 1.20117324 | 0.22968401 | 0.42281476 |
| TOB1 | 540.719099 | 0.134663786 | 0.46245664 | 0.29119224 | 0.7709043 | 0.87268354 |
| SPAG9 | 6858.42601 | -0.267046053 | 0.40730336 | -0.6556441 | 0.51205313 | 0.6909646 |
| AC005920.2 | 450.405158 | -0.879784489 | 0.59952051 | -1.4674802 | 0.14224547 | 0.30963294 |
| MBTD1 | 1266.61172 | 0.000336298 | 0.20740287 | 0.00162147 | 0.99870625 | 0.99920455 |
| UTP18 | 316.883949 | -0.254069272 | 0.27747275 | -0.9156549 | 0.35984794 | 0.55834681 |
| TOM1L1 | 2012.8205 | 0.183344072 | 0.26806591 | 0.68395148 | 0.49400581 | 0.6761871 |
| COX11 | 339.900584 | 0.393712164 | 0.19869243 | 1.98151568 | 0.04753347 | 0.15043669 |
| STXBP4 | 1104.49622 | -0.159321625 | 0.26979424 | -0.5905301 | 0.5548353 | 0.72429786 |
| HLF | 586.844604 | -0.124611323 | 0.349884 | -0.3561504 | 0.72172794 | 0.84214905 |
| MMD | 133.588887 | -1.593663582 | 0.4894244 | -3.2561997 | 0.00112914 | 0.0099149 |
| PCTP | 472.537959 | 0.653364582 | 0.46940402 | 1.39190239 | 0.16395196 | 0.34080823 |
| ANKFN1 | 129.225156 | -0.071487866 | 0.87620918 | -0.0815877 | 0.93497461 | 0.9654009 |
| C17orf67 | 44.7138222 | -2.287691847 | 0.65873858 | -3.4728372 | 0.00051499 | 0.00545092 |
| DGKE | 72.1316176 | -2.149100575 | 0.5098804 | -4.2149111 | 2.50E-05 | 0.00052132 |
| TRIM25 | 341.786858 | -1.265842944 | 0.49531547 | -2.5556297 | 0.01059958 | 0.05265984 |
| COIL | 184.132154 | 0.085074876 | 0.18031238 | 0.47181939 | 0.63705571 | 0.78646515 |
| SCPEP1 | 199.449309 | 0.604222773 | 0.22948994 | 2.63289436 | 0.00846607 | 0.0446153 |
| AC007114.1 | 48.857182 | 1.262835584 | 0.49491383 | 2.55162719 | 0.01072212 | 0.05303792 |
| AKAP1 | 522.735604 | 0.6580871 | 0.24815273 | 2.65194387 | 0.00800298 | 0.04291741 |
| AC007114.2 | 70.7318635 | 0.66793346 | 0.55580907 | 1.20173185 | 0.22946744 | 0.42264313 |
| MSI2 | 4894.64816 | -0.056290368 | 0.37201905 | -0.1513104 | 0.87973083 | 0.93723539 |
| AC015845.2 | 535.60524 | 0.046919338 | 0.94819994 | 0.04948254 | 0.96053475 | 0.97967485 |
| MRPS23 | 298.863677 | -0.23572723 | 0.22197995 | -1.0619303 | 0.28826734 | 0.48805341 |
| CUEDC1 | 998.619773 | -0.915658883 | 0.44462344 | -2.0594031 | 0.03945564 | 0.13234222 |
| VEZF1 | 435.241098 | -0.5185256 | 0.22930084 | -2.2613332 | 0.02373863 | 0.09290451 |
| AC015813.2 | 324.883012 | -0.090307909 | 0.23501692 | -0.3842613 | 0.70078478 | 0.82796169 |
| SRSF1 | 145.90362 | -0.693472292 | 0.28136746 | -2.4646499 | 0.01371472 | 0.06295634 |
| DYNLL2 | 365.799459 | 0.46523264 | 0.2709422 | 1.71709184 | 0.08596239 | 0.22236088 |
| MKS1 | 149.440318 | 0.604054863 | 0.34714266 | 1.74007673 | 0.08184555 | 0.21535832 |
| TSPOAP1 | 187.667586 | 1.225491313 | 0.70206307 | 1.7455573 | 0.08088791 | 0.21362738 |
| TSPOAP1-AS | 80.9973108 | 0.100153132 | 0.49229488 | 0.20344135 | 0.83879009 | 0.91471434 |
| SUPT4H1 | 122.709179 | -0.65276679 | 0.30449687 | -2.1437553 | 0.03205249 | 0.114472 |
| RNF43 | 274.440949 | 0.978687618 | 0.45341078 | 2.15850101 | 0.0308889 | 0.11141951 |
| MTMR4 | 446.193767 | 1.113336015 | 0.40876062 | 2.72368709 | 0.00645576 | 0.03661115 |
| SEPT4-AS1 | 56.9530506 | 0.656206195 | 0.5217352 | 1.257738 | 0.20848652 | 0.39773483 |
| TEX14 | 42.9631465 | -0.375470599 | 0.38286808 | -0.9806788 | 0.32675117 | 0.52685311 |
| RAD51C | 88.5813811 | 0.332682266 | 0.19930546 | 1.66920797 | 0.09507617 | 0.23781188 |
| PPM1E | 187.942279 | -0.414613215 | 0.60456279 | -0.6858067 | 0.49283501 | 0.67512387 |
| TRIM37 | 573.643328 | -0.106431282 | 0.28633154 | -0.3717065 | 0.71011142 | 0.83398999 |
| SKA2 | 139.397532 | 0.96984895 | 0.29212542 | 3.31997453 | 0.00090026 | 0.00836598 |
| GDPD1 | 84.5775001 | 0.273506941 | 0.44962034 | 0.60830642 | 0.54298426 | 0.71356452 |
| YPEL2 | 2050.46983 | -0.123330094 | 0.46978271 | -0.2625258 | 0.79291608 | 0.8871958 |
| LINC01476 | 37.925048 | -0.522890958 | 0.59786244 | -0.8746008 | 0.38179116 | 0.57844716 |
| DHX40 | 433.061954 | -0.12920992 | 0.32672563 | -0.3954692 | 0.69249666 | 0.8227487 |
| CLTC | 2676.70866 | -0.064672118 | 0.18273989 | -0.3539026 | 0.7234119 | 0.84260765 |
| PTRH2 | 389.897034 | -1.040180094 | 0.33419861 | -3.1124609 | 0.00185535 | 0.01444956 |
| VMP1 | 7690.24768 | -0.917936175 | 0.32716013 | -2.8057703 | 0.00501965 | 0.03037717 |
| TUBD1 | 306.127736 | 0.237795245 | 0.25910925 | 0.91774124 | 0.35875435 | 0.55728081 |
| RPS6KB1 | 561.827965 | 0.269296639 | 0.20185128 | 1.33413389 | 0.18215998 | 0.36445291 |
| AC005702.1 | 42.4115984 | -0.523826547 | 0.57436853 | -0.9120043 | 0.36176648 | 0.55961134 |
| RNFT1 | 186.848013 | 0.207384753 | 0.2217608 | 0.93517318 | 0.34969906 | 0.54925232 |
| RNFT1-DT | 40.8279811 | -0.441474877 | 0.54440421 | -0.8109321 | 0.41740464 | 0.61047432 |
| HEATR6 | 132.192036 | -0.231546312 | 0.2264334 | -1.0225802 | 0.30650638 | 0.50646126 |
| AC025048.4 | 38.0972876 | -0.130629376 | 0.47585885 | -0.2745129 | 0.78369052 | 0.88133085 |
| CA4 | 77.4248484 | 0.876944308 | 0.66173719 | 1.32521539 | 0.18509972 | 0.36807804 |
| USP32 | 2714.20228 | -0.381436208 | 0.22146479 | -1.7223334 | 0.08500913 | 0.22066909 |
| APPBP2 | 2131.17902 | 0.181521839 | 0.116413 | 1.55929185 | 0.11892732 | 0.27565304 |
| AC011921.1 | 252.792706 | 0.732187159 | 0.41292763 | 1.77316099 | 0.07620203 | 0.2054356 |
| PPM1D | 514.016398 | 0.198221452 | 0.21300966 | 0.93057495 | 0.35207348 | 0.55108727 |
| BCAS3 | 5789.56387 | 0.637309065 | 0.30162978 | 2.11288511 | 0.0346106 | 0.12077585 |
| AC005746.2 | 75.0453985 | 0.468456104 | 0.52388135 | 0.89420268 | 0.37121346 | 0.56877072 |
| TBX2 | 365.049269 | 1.24454607 | 0.42037338 | 2.960573 | 0.00307067 | 0.02105621 |

| | | | | | | |
|---|---|---|---|---|---|---|
| NACA2 | 85.903464 | -0.935570546 | 0.4266442 | -2.1928589 | 0.02831754 | 0.10490502 |
| BRIP1 | 78.2091194 | 0.920095647 | 0.63917604 | 1.43950272 | 0.15000814 | 0.32052402 |
| INTS2 | 298.751666 | 0.178427546 | 0.24065013 | 0.74143963 | 0.45842692 | 0.64719928 |
| MED13 | 3962.01797 | -0.318794148 | 0.28599108 | -1.1146996 | 0.26497916 | 0.4628019 |
| EFCAB3 | 85.3916023 | -1.610886204 | 0.35677611 | -4.515118 | 6.33E-06 | 0.00017348 |
| METTL2A | 156.709115 | -0.368249242 | 0.26851478 | -1.37143 | 0.17024094 | 0.34924222 |
| TLK2 | 1601.11606 | 0.084100314 | 0.20580494 | 0.40864089 | 0.68280322 | 0.81575732 |
| TANC2 | 1710.37038 | -1.121723109 | 0.27191927 | -4.1252064 | 3.70E-05 | 0.0007081 |
| AC015923.1 | 94.0198176 | -2.025510436 | 0.64331764 | -3.1485386 | 0.00164089 | 0.01322263 |
| CYB561 | 202.711828 | -0.117768704 | 0.33789335 | -0.3485381 | 0.72743611 | 0.84519832 |
| ACE | 64.1160079 | 0.826505249 | 0.92120553 | 0.89719962 | 0.36961241 | 0.56701481 |
| KCNH6 | 296.23321 | 1.699302066 | 0.84244909 | 2.01709764 | 0.04368533 | 0.14164143 |
| DCAF7 | 1178.77438 | 0.018502165 | 0.19342395 | 0.09565602 | 0.92379377 | 0.95963935 |
| TACO1 | 122.135701 | 0.354826713 | 0.31917758 | 1.11169058 | 0.26627121 | 0.46411235 |
| MAP3K3 | 466.889649 | -0.08092367 | 0.26591888 | -0.3043171 | 0.7608863 | 0.8665094 |
| CCDC47 | 295.410818 | -0.693794446 | 0.25936289 | -2.6749951 | 0.00747303 | 0.04085875 |
| DDX42 | 825.747769 | 0.189028769 | 0.25953016 | 0.72834991 | 0.46639942 | 0.65427662 |
| FTSJ3 | 237.906758 | 0.35327866 | 0.32140523 | 1.09916897 | 0.27169437 | 0.46991984 |
| PSMC5 | 207.880165 | -0.088699175 | 0.34459911 | -0.2573982 | 0.79687143 | 0.88911002 |
| SMARCD2 | 259.663847 | 0.441975058 | 0.22491997 | 1.96503257 | 0.04941047 | 0.15399483 |
| ERN1 | 617.878729 | -0.60111494 | 0.33919502 | -1.7721809 | 0.07636453 | 0.20555349 |
| SNHG25 | 441.421678 | 0.114344644 | 0.43387615 | 0.26354213 | 0.79213276 | 0.88678699 |
| TEX2 | 1016.78585 | 0.011538822 | 0.22169545 | 0.05204808 | 0.95849039 | 0.9788247 |
| PECAM1 | 69.0272302 | -0.291255951 | 0.42747804 | -0.6813355 | 0.49565925 | 0.67766001 |
| POLG2 | 316.282535 | 0.316599874 | 0.21204462 | 1.49308141 | 0.13541588 | 0.29987878 |
| DDX5 | 3231.58966 | -0.003287808 | 0.31030759 | -0.0105953 | 0.99154632 | 0.99576632 |
| CEP95 | 1107.63126 | 0.14877537 | 0.18669617 | 0.79688498 | 0.42551784 | 0.61851251 |
| SMURF2 | 1326.47464 | 0.257705955 | 0.16783737 | 1.53545037 | 0.12467324 | 0.28462626 |
| LRRC37A3 | 1501.54651 | 0.761205461 | 0.25834631 | 2.94645382 | 0.0032144 | 0.02173648 |
| AC103810.2 | 46.4887954 | -0.251906097 | 0.6638815 | -0.3794444 | 0.70435791 | 0.8304347 |
| GNA13 | 1353.56934 | -1.516704761 | 0.43293218 | -3.5033311 | 0.00045948 | 0.00504653 |
| RGS9 | 213.271428 | -1.779519971 | 0.35653107 | -4.9912059 | 6.00E-07 | 2.48E-05 |
| AC004805.1 | 52.380106 | -0.189211758 | 0.44971016 | -0.4207416 | 0.6739438 | 0.8108978 |
| CEP112 | 1998.6103 | -0.687456097 | 0.42642637 | -1.6121332 | 0.10693296 | 0.2566953 |
| PRKCA | 5732.19558 | 0.006514882 | 0.27678474 | 0.02353772 | 0.98122135 | 0.99060551 |
| PRKCA-AS1 | 200.217428 | -0.758843089 | 0.39861971 | -1.9036768 | 0.05695229 | 0.16911722 |
| HELZ | 3400.16321 | 0.056828191 | 0.22599317 | 0.25145977 | 0.80145866 | 0.89219475 |
| AC007448.3 | 76.0169523 | 2.597899504 | 0.48415712 | 5.36581911 | 8.06E-08 | 4.62E-06 |
| PSMD12 | 615.965534 | -0.762447626 | 0.33546349 | -2.2728185 | 0.02303712 | 0.0911662 |
| PITPNC1 | 5310.04325 | -0.165294845 | 0.56422607 | -0.2929585 | 0.76955384 | 0.87161415 |
| NOL11 | 371.545693 | 0.229042062 | 0.21319389 | 1.0743369 | 0.28267171 | 0.48175669 |
| BPTF | 2694.06895 | 0.27382938 | 0.2112628 | 1.2961552 | 0.19492202 | 0.38013098 |
| KPNA2 | 111.702418 | -0.742082469 | 0.47444973 | -1.5640908 | 0.11779624 | 0.2737583 |
| AC005332.7 | 114.145526 | 0.111595292 | 0.19859497 | 0.56192405 | 0.57416776 | 0.73911704 |
| AC005332.6 | 41.8657177 | 0.094142702 | 0.32345053 | 0.2910575 | 0.77100734 | 0.87268354 |
| AMZ2 | 92.1685882 | 0.582935493 | 0.30510959 | 1.91057743 | 0.05605891 | 0.16718874 |
| ARSG | 272.797853 | -0.181647462 | 0.40121215 | -0.4527467 | 0.65073117 | 0.79525429 |
| WIPI1 | 791.482982 | -0.041334178 | 0.38292611 | -0.107943 | 0.91404094 | 0.95420862 |
| PRKAR1A | 1199.56579 | -0.309084912 | 0.27703918 | -1.1156722 | 0.26456248 | 0.46242769 |
| FAM20A | 1917.99379 | 0.150944224 | 0.76412696 | 0.19753815 | 0.84340643 | 0.91779707 |
| AC079210.1 | 93.9799914 | -0.700354978 | 0.65895319 | -1.0628296 | 0.28785922 | 0.48756075 |
| LINC01482 | 281.881191 | 0.443750278 | 0.68209233 | 0.65057216 | 0.51532271 | 0.69315508 |
| ABCA9 | 43.0819671 | -0.241665932 | 0.48485261 | -0.4984317 | 0.61817977 | 0.77299851 |
| ABCA6 | 63.7633807 | -0.442099762 | 0.55137708 | -0.8018102 | 0.42266276 | 0.61619214 |
| ABCA10 | 248.303654 | 0.427361036 | 0.37113372 | 1.15150151 | 0.24952597 | 0.44575166 |
| ABCA5 | 959.104818 | 0.274729158 | 0.30641337 | 0.89659652 | 0.36993426 | 0.56738154 |
| MAP2K6 | 109.12878 | 1.77408315 | 0.65456106 | 2.7103402 | 0.00672142 | 0.03774295 |
| AC005208.1 | 1714.27071 | 0.758027868 | 0.71011213 | 1.0674763 | 0.28575681 | 0.48536166 |
| KCNJ16 | 3604.70965 | 1.034095479 | 0.39459473 | 2.62065203 | 0.00877618 | 0.04586164 |
| SOX9 | 685.364073 | -1.16580556 | 0.8055796 | -1.4471637 | 0.14785108 | 0.31754853 |
| LINC00511 | 515.590489 | -1.176816987 | 0.83672792 | -1.4064512 | 0.15959018 | 0.33478767 |
| SLC39A11 | 2915.34344 | 0.191850901 | 0.19869701 | 0.96554498 | 0.33427192 | 0.53395469 |
| SSTR2 | 327.373667 | 0.405701787 | 0.7776317 | 0.52171457 | 0.60186908 | 0.76091306 |
| COG1 | 217.976789 | -0.259388989 | 0.16753598 | -1.5482584 | 0.12156009 | 0.27961472 |
| FAM104A | 264.165119 | -0.247410239 | 0.18927596 | -1.3071403 | 0.19116506 | 0.37550867 |
| C17orf80 | 148.001903 | 0.107234422 | 0.26138448 | 0.41025551 | 0.68161852 | 0.81468059 |
| CDC42EP4 | 222.479658 | -0.525452202 | 0.30509109 | -1.7222797 | 0.08501886 | 0.22066909 |
| SDK2 | 64.0041298 | -0.164790717 | 0.69980351 | -0.2354814 | 0.8138351 | 0.89925335 |
| RPL38 | 458.929932 | -0.926339425 | 0.40367882 | -2.2947437 | 0.02174782 | 0.08764991 |
| GPRC5C | 860.950006 | -0.56337766 | 0.2141305 | -2.6310014 | 0.00851337 | 0.044727 |
| SLC9A3R1 | 1026.9807 | 0.078197508 | 0.18923014 | 0.41324025 | 0.6794306 | 0.81421186 |
| NAT9 | 223.025014 | -1.084850065 | 0.39864285 | -2.7213584 | 0.00650142 | 0.03678388 |
| TMEM104 | 145.34013 | -0.663443412 | 0.53046494 | -1.2506829 | 0.2110502 | 0.40034114 |
| FDXR | 53.3899821 | -0.424980872 | 0.44975497 | -0.9449165 | 0.34470153 | 0.54521392 |
| CDR2L | 70.7926832 | -1.14712952 | 0.50070101 | -2.2910469 | 0.0219607 | 0.08832609 |
| MRPL58 | 95.1098999 | -0.058953359 | 0.30416775 | -0.1938186 | 0.84631794 | 0.91896252 |
| KCTD2 | 196.016044 | -0.386550293 | 0.25516431 | -1.5149074 | 0.12979587 | 0.2920407 |
| ATP5PD | 290.683921 | -0.354747047 | 0.38076603 | -0.9316667 | 0.35150878 | 0.55063839 |
| SLC16A5 | 107.99816 | 0.055755781 | 0.52608175 | 0.10598311 | 0.91559575 | 0.95488632 |
| ARMC7 | 136.515707 | -0.136701271 | 0.47189547 | -0.2896855 | 0.77205686 | 0.87329455 |
| NT5C | 306.425493 | -0.382744741 | 0.27410456 | -1.3963457 | 0.1626104 | 0.33908432 |
| JPT1 | 279.319088 | -1.118290263 | 0.21000124 | -5.3251603 | 1.01E-07 | 5.55E-06 |
| SUMO2 | 225.465337 | 0.009325442 | 0.33431739 | 0.02789398 | 0.97774671 | 0.98892962 |
| NUP85 | 263.747282 | 0.142199776 | 0.21714533 | 0.65485994 | 0.51255794 | 0.6911936 |
| GGA3 | 371.437115 | -0.117455904 | 0.27447418 | -0.4279306 | 0.66870164 | 0.80738625 |
| MRPS7 | 197.84874 | -0.519832959 | 0.25554821 | -2.0341874 | 0.0419327 | 0.13780185 |
| MIF4GD | 40.2059575 | -0.807632209 | 0.3525943 | -2.2905424 | 0.02198989 | 0.08841755 |
| SLC25A19 | 50.8512385 | 0.272940665 | 0.28746971 | 0.94945887 | 0.34238728 | 0.5429318 |
| GRB2 | 1256.89343 | 0.260902192 | 0.23447037 | 1.1127299 | 0.26582445 | 0.46372441 |
| AC011933.4 | 150.092786 | -0.703603969 | 0.44961253 | -1.5649118 | 0.11760359 | 0.2735671 |
| TMEM94 | 288.175948 | 0.439188145 | 0.17130089 | 2.56384046 | 0.01035211 | 0.05180591 |
| CASKIN2 | 70.0285071 | 0.091775259 | 0.26595325 | 0.34508041 | 0.73003392 | 0.84648502 |
| TSEN54 | 177.184217 | -0.112120623 | 0.23671424 | -0.4736539 | 0.63574673 | 0.78548589 |
| LLGL2 | 1853.38113 | 0.66706882 | 0.34187563 | 1.95120318 | 0.05103288 | 0.1573841 |
| MYO15B | 548.437545 | 0.483122807 | 0.2466703 | 1.95857714 | 0.05016233 | 0.15567187 |
| RECQL5 | 232.039696 | 0.919778299 | 0.21966836 | 4.1871224 | 2.83E-05 | 0.0005754 |
| SMIM5 | 74.4564775 | 1.893994657 | 0.6722343 | 2.8174621 | 0.00484048 | 0.02956843 |
| SAP30BP | 470.271528 | -0.241630048 | 0.38540725 | -0.6269473 | 0.53069376 | 0.70466138 |
| ITGB4 | 134.995906 | -1.21287935 | 0.45612354 | -2.6591027 | 0.00783491 | 0.04228073 |
| GALK1 | 104.616391 | 0.264226855 | 0.41049907 | 0.64367224 | 0.519788 | 0.69597507 |
| H3F3B | 1270.93773 | -1.145238717 | 0.32223648 | -3.5540319 | 0.00037937 | 0.00441432 |
| UNK | 611.47868 | 0.548357349 | 0.24128696 | 2.27263568 | 0.02304814 | 0.0911662 |
| UNC13D | 35.3626163 | 1.432124805 | 0.48747607 | 2.93783611 | 0.00330512 | 0.02219659 |
| WBP2 | 525.379995 | -0.801358978 | 0.23270544 | -3.4436624 | 0.00057389 | 0.00590903 |
| TRIM47 | 288.656625 | -0.766840518 | 0.32876005 | -2.3325234 | 0.01967317 | 0.08146833 |
| TRIM65 | 157.028428 | 0.31703689 | 0.30164052 | 1.0510421 | 0.29323925 | 0.49326436 |
| FBF1 | 48.5657611 | -0.368711578 | 0.39457548 | -0.9344513 | 0.35007113 | 0.5493439 |
| ACOX1 | 1466.42198 | 0.830012425 | 0.34449122 | 2.40938628 | 0.01597938 | 0.07047275 |
| TEN1 | 166.829014 | 0.227954364 | 0.32076167 | 0.71066584 | 0.47729133 | 0.66284015 |
| CDK3 | 109.29965 | 1.286721443 | 0.38699647 | 3.32489195 | 0.00088453 | 0.00825339 |
| EVPL | 296.521245 | 0.102065952 | 0.3895294 | 0.26202374 | 0.79330314 | 0.88726528 |
| SRP68 | 294.085516 | 0.057021205 | 0.17015256 | 0.33511811 | 0.73753599 | 0.85118107 |
| EXOC7 | 566.450839 | 0.083221854 | 0.27618664 | 0.30132469 | 0.76316691 | 0.86789421 |

| | | | | | | |
|---|---|---|---|---|---|---|
| RNF157-AS1 | 46.5118646 | 0.21053878 | 0.57349596 | 0.36711467 | 0.71353348 | 0.8362667 |
| RNF157 | 795.411189 | -0.000642382 | 0.352014 | -0.0018249 | 0.99854396 | 0.99920455 |
| UBALD2 | 280.32663 | -0.404989111 | 0.32064473 | -1.2630462 | 0.20657257 | 0.39534909 |
| QRICH2 | 46.5891955 | 0.439399444 | 0.54139923 | 0.81159968 | 0.41702137 | 0.61036016 |
| PRPSAP1 | 335.123076 | -0.091807364 | 0.2075076 | -0.4424289 | 0.65817885 | 0.80094333 |
| SPHK1 | 84.7283367 | -5.353170653 | 1.06784423 | -5.0130633 | 5.36E-07 | 2.27E-05 |
| UBE2O | 817.897013 | -1.063119435 | 0.3206857 | -3.3151445 | 0.00091596 | 0.00846026 |
| RHBDF2 | 172.720796 | -1.398265639 | 0.37122473 | -3.7666285 | 0.00016547 | 0.00228404 |
| AC015802.4 | 54.6357089 | 0.827162116 | 0.40383067 | 2.04828951 | 0.04053164 | 0.13458508 |
| ST6GALNAC2 | 208.878268 | 0.847387216 | 0.60127223 | 1.40932372 | 0.15873947 | 0.33361574 |
| MXRA7 | 183.283602 | -0.722984923 | 0.45256278 | -1.5975351 | 0.11014648 | 0.26179605 |
| JMJD6 | 241.433523 | -1.025947836 | 0.36096694 | -2.842221 | 0.00448004 | 0.02791011 |
| METTL23 | 122.871321 | -0.278436356 | 0.31954635 | -0.8713489 | 0.38356369 | 0.58015209 |
| AC005837.2 | 81.2711817 | -0.308863199 | 0.74482126 | -0.414681 | 0.67837547 | 0.81372998 |
| SRSF2 | 869.935478 | -0.102343775 | 0.26020966 | -0.3933127 | 0.6940885 | 0.82385996 |
| MFSD11 | 507.303166 | -0.21181154 | 0.36978912 | -0.5727901 | 0.56678684 | 0.73354237 |
| SEC14L1 | 684.421787 | -0.102868366 | 0.26654926 | -0.3859263 | 0.69955127 | 0.82718678 |
| 9-Sep | 1605.1949 | -0.625329716 | 0.42156066 | -1.4833683 | 0.13797658 | 0.30371689 |
| TNRC6C | 824.891309 | -0.015928485 | 0.38260288 | -0.0416319 | 0.96679214 | 0.9830686 |
| TMC6 | 80.2708213 | -1.411808844 | 0.46746253 | -3.020154 | 0.00252646 | 0.01834227 |
| SYNGR2 | 269.155488 | -0.616763519 | 0.38656254 | -1.5955077 | 0.11059874 | 0.26259778 |
| TK1 | 45.2712253 | 0.015428057 | 0.37535345 | 0.04110275 | 0.96721399 | 0.98328281 |
| AFMID | 1595.91408 | 2.349102511 | 0.34869937 | 6.73675574 | 1.62E-11 | 3.22E-09 |
| SOCS3 | 243.924121 | -3.581858113 | 0.6659501 | -5.3785683 | 7.51E-08 | 4.40E-06 |
| PGS1 | 516.19049 | -1.105636606 | 0.32977662 | -3.3526834 | 0.00080032 | 0.00766598 |
| DNAH17 | 45.5573944 | -1.736994421 | 0.53102787 | -3.2710042 | 0.00107166 | 0.00949213 |
| CYTH1 | 1052.98572 | 0.318867929 | 0.2722838 | 1.17108668 | 0.24156394 | 0.43676519 |
| USP36 | 752.527074 | -0.832626492 | 0.53790046 | -1.5479193 | 0.12164173 | 0.27975558 |
| TIMP2 | 81.1933288 | -0.593349401 | 0.56645122 | -1.0474854 | 0.29487575 | 0.49487664 |
| LGALS3BP | 268.459852 | -0.992092719 | 0.68832788 | -1.4413084 | 0.14949759 | 0.31978206 |
| CANT1 | 108.956225 | -0.515625164 | 0.37422966 | -1.3778308 | 0.16825552 | 0.34675664 |
| ENGASE | 151.134397 | 0.628159486 | 0.49981401 | 1.25678648 | 0.20883096 | 0.39800417 |
| CBX4 | 105.423875 | -1.071964042 | 0.29527806 | -3.6303546 | 0.00028303 | 0.00349822 |
| TBC1D16 | 1048.73056 | -0.388396347 | 0.19407114 | -2.0013091 | 0.04535909 | 0.14574238 |
| CCDC40 | 406.160261 | 0.103906049 | 0.39492713 | 0.26310182 | 0.7924721 | 0.88694497 |
| GAA | 245.592093 | 0.408430008 | 0.28732952 | 1.42146903 | 0.15518045 | 0.32794611 |
| EIF4A3 | 199.221005 | -1.217596455 | 0.29259456 | -4.1613776 | 3.16E-05 | 0.00063023 |
| CARD14 | 95.6318462 | -1.251723134 | 0.44827885 | -2.7922869 | 0.00523369 | 0.03137037 |
| AC087741.1 | 271.549514 | 0.000103105 | 0.27907718 | 0.00036945 | 0.99970522 | 0.99977816 |
| SGSH | 162.022345 | -0.212935602 | 0.21732448 | -0.9798049 | 0.32718241 | 0.52705245 |
| SLC26A11 | 561.136177 | -0.296117374 | 0.2879169 | -1.0284821 | 0.30372311 | 0.50388921 |
| RNF213 | 9020.87145 | -1.346179354 | 0.75227167 | -1.7894856 | 0.07353664 | 0.20055048 |
| AC124319.2 | 1764.03623 | -0.396213401 | 0.39034819 | -1.0150256 | 0.31009363 | 0.51038106 |
| ENDOV | 416.298288 | -0.028332574 | 0.16134382 | -0.1756037 | 0.86060528 | 0.92650931 |
| RPTOR | 1463.23897 | 0.003301146 | 0.21636778 | 0.0152571 | 0.98782707 | 0.9938451 |
| CHMP6 | 103.240731 | -0.101156692 | 0.36727745 | -0.2754231 | 0.7829912 | 0.8809896 |
| BAIAP2-DT | 58.3444658 | -0.066661735 | 0.54212549 | -0.1229637 | 0.90213587 | 0.94861498 |
| BAIAP2 | 566.060986 | 1.328148782 | 0.57521077 | 2.30897759 | 0.02094482 | 0.08529136 |
| CEP131 | 32.8200076 | 0.345299015 | 0.40484259 | 0.85292167 | 0.39370274 | 0.58851718 |
| TEPSIN | 130.938776 | 0.302087447 | 0.27986561 | 1.07940181 | 0.28040864 | 0.47912756 |
| NDUFAF8 | 223.962196 | -0.416366785 | 0.26713313 | -1.558649 | 0.11907948 | 0.27592988 |
| SLC38A10 | 598.083661 | 0.796865385 | 0.29351446 | 2.71491012 | 0.00662938 | 0.03733314 |
| LINC00482 | 64.4492809 | 0.634154944 | 0.89178979 | 0.71110362 | 0.47702003 | 0.66266479 |
| AC110285.6 | 68.5813398 | 0.762392494 | 0.62467436 | 1.22046388 | 0.22228907 | 0.41454644 |
| AC110285.2 | 33.2796141 | 1.149694468 | 0.68000303 | 1.69071962 | 0.09089036 | 0.2308828 |
| BAHCC1 | 79.7153417 | 1.366328034 | 0.44224698 | 3.08951354 | 0.00200485 | 0.01538188 |
| ACTG1 | 1638.41629 | -0.253458399 | 0.43791676 | -0.5787821 | 0.56273618 | 0.73036879 |
| FAAP100 | 121.129008 | -0.105999489 | 0.43060621 | -0.2461634 | 0.80555575 | 0.89453456 |
| NPLOC4 | 2564.33663 | 0.470159427 | 0.22102699 | 2.12715841 | 0.03340692 | 0.1181395 |
| TSPAN10 | 38.9207957 | 1.06532868 | 0.43771919 | 2.43381765 | 0.01494052 | 0.06688103 |
| OXLD1 | 134.739903 | 0.030545344 | 0.22412056 | 0.13628979 | 0.89159219 | 0.94341489 |
| CCDC137 | 239.384297 | -0.176069608 | 0.35273376 | -0.4991572 | 0.61766862 | 0.77262127 |
| ARL16 | 368.260026 | 0.950864828 | 0.25965835 | 3.66198442 | 0.00025027 | 0.00320004 |
| HGS | 561.475739 | -1.034149107 | 0.45269748 | -2.2844154 | 0.02234713 | 0.08922578 |
| AC139530.1 | 42.7904923 | -0.285423408 | 0.40824754 | -0.699143 | 0.48446267 | 0.66827686 |
| SLC25A10 | 198.837392 | 1.497608432 | 0.32979349 | 4.54104916 | 5.60E-06 | 0.00015787 |
| MCRIP1 | 274.496544 | -0.146943143 | 0.46238029 | -0.3177972 | 0.75063881 | 0.85940776 |
| P4HB | 1341.47088 | -1.271675963 | 0.48732498 | -2.609503 | 0.00906739 | 0.04695378 |
| ARHGDIA | 652.234078 | -1.334305319 | 0.59963538 | -2.2251945 | 0.02606819 | 0.09932591 |
| ALYREF | 38.0520034 | -0.295121922 | 0.43764637 | -0.6743388 | 0.50009595 | 0.68141606 |
| ANAPC11 | 304.272331 | -0.460098097 | 0.27477746 | -1.674439 | 0.09404436 | 0.23617919 |
| PCYT2 | 179.586402 | 0.845852895 | 0.37084024 | 2.28090912 | 0.02255383 | 0.08994626 |
| SIRT7 | 324.828022 | -0.630953794 | 0.42208367 | -1.4948548 | 0.13495234 | 0.29936987 |
| MAFG | 382.066197 | -1.91216387 | 0.36612143 | -5.2227587 | 1.76E-07 | 8.79E-06 |
| ASPSCR1 | 440.594152 | 1.321948758 | 0.43139068 | 3.06438875 | 0.00218115 | 0.01639992 |
| CENPX | 126.979318 | -0.767209231 | 0.36860077 | -2.0814097 | 0.03739642 | 0.12770123 |
| LRRC45 | 57.6744694 | 1.218728091 | 0.37258541 | 3.27100327 | 0.00107167 | 0.00949213 |
| RAC3 | 63.917514 | 0.00327969 | 0.36386579 | 0.00901346 | 0.9928084 | 0.99629729 |
| DCXR | 457.055268 | 1.393119811 | 0.43701828 | 3.18778385 | 0.00143368 | 0.01188833 |
| RFNG | 319.357227 | 0.644630919 | 0.18430368 | 3.49765616 | 0.00046937 | 0.00513046 |
| GPS1 | 288.490208 | 0.682475575 | 0.31127337 | 2.19252798 | 0.0283414 | 0.10494149 |
| DUS1L | 293.036684 | -0.140496033 | 0.30486669 | -0.4608442 | 0.64491042 | 0.79180937 |
| FASN | 122.672342 | -0.475277575 | 0.41144668 | -1.1551377 | 0.24803404 | 0.44436055 |
| CCDC57 | 1177.10338 | 0.238697975 | 0.2622217 | 0.9102907 | 0.36266922 | 0.56044048 |
| SLC16A3 | 329.743793 | -1.385459619 | 0.45947727 | -3.0152952 | 0.00256729 | 0.01855205 |
| CSNK1D | 1182.65667 | -0.467097166 | 0.32315928 | -1.4454085 | 0.14834318 | 0.31826835 |
| AC132872.1 | 58.5296445 | 0.166632379 | 0.41719456 | 0.39941168 | 0.68958989 | 0.820433 |
| SECTM1 | 116.463089 | -1.736401858 | 0.59095021 | -2.9383217 | 0.00329994 | 0.02218092 |
| OGFOD3 | 192.370126 | 0.194337532 | 0.29889505 | 0.65018651 | 0.51557175 | 0.69315508 |
| HEXDC | 501.097549 | 0.511379615 | 0.24407395 | 2.09518309 | 0.03615472 | 0.12435953 |
| CYBC1 | 244.278421 | -0.673214598 | 0.40172149 | -1.6758242 | 0.09377263 | 0.23571272 |
| NARF | 406.930847 | -0.413690972 | 0.33887959 | -1.220761 | 0.22217654 | 0.41444934 |
| FOXK2 | 1258.84846 | -0.126813967 | 0.2565037 | -0.4943943 | 0.62102773 | 0.77498426 |
| WDR45B | 805.223551 | -0.789310548 | 0.33815495 | -2.3341683 | 0.01958691 | 0.0811847 |
| RAB40B | 718.367635 | 0.120864956 | 0.37860157 | 0.3192405 | 0.74954414 | 0.85874303 |
| FN3KRP | 191.867318 | 0.272193027 | 0.23183887 | 1.1740612 | 0.24037051 | 0.43564176 |
| FN3K | 968.102125 | 1.781092167 | 0.55436992 | 3.21282254 | 0.00131437 | 0.01110736 |
| TBCD | 900.987665 | 0.25550001 | 0.22028858 | 1.15984231 | 0.24611301 | 0.4417554 |
| B3GNTL1 | 60.8716401 | 0.162590406 | 0.34773573 | 0.46756888 | 0.64009292 | 0.78865202 |
| METRNL | 98.6448979 | -0.072346079 | 0.33967322 | -0.2129873 | 0.83133687 | 0.91080925 |
| USP14 | 975.380692 | -0.534501356 | 0.25612033 | -2.086915 | 0.03689582 | 0.12630644 |
| THOC1 | 638.757199 | 0.014924006 | 0.23337281 | 0.06394921 | 0.94901067 | 0.97382962 |
| TYMS | 154.453104 | -0.014534049 | 0.45104758 | -0.0322229 | 0.97429432 | 0.98703942 |
| ENOSF1 | 5237.46818 | 0.209272952 | 0.26443111 | 0.79140822 | 0.42870581 | 0.62176178 |
| YES1 | 1624.21453 | -0.391065824 | 0.19363974 | -2.0195536 | 0.04342971 | 0.14116459 |
| LINC00470 | 108.772706 | 1.089877422 | 1.09932997 | 0.99140154 | 0.32148955 | 0.52188291 |
| AP005230.1 | 91.5109251 | 1.716553467 | 0.98838732 | 1.73672146 | 0.08243635 | 0.21654945 |
| METTL4 | 210.729732 | 0.181362155 | 0.19408687 | 0.93443803 | 0.35007799 | 0.5493439 |
| SMCHD1 | 1782.59172 | -0.173313577 | 0.16334529 | -1.0610259 | 0.28867814 | 0.48856788 |
| AP001011.1 | 237.173732 | -1.010725159 | 0.51572961 | -1.9597967 | 0.05001956 | 0.15545195 |
| LPIN2 | 2355.17714 | -0.536351667 | 0.23543952 | -2.2780868 | 0.0227214 | 0.09032549 |
| AP000919.2 | 161.729233 | -1.088219865 | 0.50676856 | -2.1473705 | 0.0317638 | 0.11379675 |
| AP000919.3 | 55.054458 | -2.059991158 | 0.64021261 | -3.2176673 | 0.00129238 | 0.01094846 |

| | | | | | | |
|---|---|---|---|---|---|---|
| MYOM1 | 108.699894 | 0.301471038 | 0.66029193 | 0.45657235 | 0.64797846 | 0.79355195 |
| MYL12A | 675.085121 | -1.502384914 | 0.3511824 | -4.2780758 | 1.89E-05 | 0.0004243 |
| MYL12B | 1405.99908 | -1.275380441 | 0.28731275 | -4.4389971 | 9.04E-06 | 0.00023463 |
| TGIF1 | 475.038595 | -0.76981796 | 0.39005862 | -1.9735956 | 0.04842775 | 0.15222969 |
| DLGAP1 | 4245.74962 | -2.211521663 | 0.68402436 | -3.2331037 | 0.00122453 | 0.01054312 |
| DLGAP1-AS1 | 91.3280992 | -1.435470489 | 0.4349656 | -3.3001931 | 0.00096618 | 0.00881723 |
| DLGAP1-AS2 | 43.2849116 | -2.681874909 | 0.61982307 | -4.3268395 | 1.51E-05 | 0.00035382 |
| DLGAP1-AS3 | 59.2652172 | -1.737945585 | 0.91252447 | -1.9045468 | 0.056839 | 0.1688581 |
| LINC00667 | 216.984693 | 0.201341359 | 0.24496284 | 0.82192614 | 0.41111894 | 0.60468888 |
| ZBTB14 | 36.4617365 | 0.267882098 | 0.29397871 | 0.91122959 | 0.36217441 | 0.5599915 |
| EPB41L3 | 7223.81996 | 1.090966759 | 0.29094324 | 3.74975804 | 0.00017701 | 0.00242621 |
| AP005059.1 | 40.9723501 | -0.373855221 | 0.88939174 | -0.4203493 | 0.67423028 | 0.81105241 |
| L3MBTL4 | 9525.91991 | 0.320378142 | 0.27492942 | 1.16531051 | 0.24389333 | 0.43900266 |
| AP001021.2 | 419.824934 | -0.432319281 | 0.62823342 | -0.6881507 | 0.49135787 | 0.67390858 |
| ARHGAP28 | 518.376266 | 1.959192014 | 0.3703418 | 5.29022649 | 1.22E-07 | 6.52E-06 |
| LAMA1 | 1302.68669 | -1.873775358 | 0.37458372 | -5.0022872 | 5.67E-07 | 2.37E-05 |
| PTPRM | 13720.0895 | 0.295417877 | 0.3003403 | 0.98361053 | 0.32530705 | 0.52569956 |
| RAB12 | 620.5318 | -0.279518844 | 0.1672533 | -1.6712306 | 0.09467613 | 0.23698747 |
| MTCL1 | 549.587611 | -1.084044289 | 0.50829987 | -2.1326866 | 0.03295045 | 0.11688711 |
| NDUFV2 | 878.650602 | -0.382111352 | 0.24327288 | -1.5707109 | 0.11624982 | 0.27158651 |
| AP005899.1 | 57.9392318 | -1.280016167 | 0.53688954 | -2.3841332 | 0.0171194 | 0.07395389 |
| NDUFV2-AS1 | 91.7777637 | -0.52684782 | 0.38466526 | -1.3696267 | 0.17080345 | 0.34969424 |
| ANKRD12 | 4614.26242 | -0.514249495 | 0.31823597 | -1.6159377 | 0.10610779 | 0.25543018 |
| TWSG1 | 406.030051 | -0.041308971 | 0.26891098 | -0.1536158 | 0.87791269 | 0.93675457 |
| RALBP1 | 549.476917 | -0.308994833 | 0.16826311 | -1.836379 | 0.06630163 | 0.18718773 |
| PPP4R1 | 884.54323 | -0.663503598 | 0.2115525 | -3.1363543 | 0.00171062 | 0.01359277 |
| RAB31 | 246.521818 | -2.385806187 | 0.75945476 | -3.1414724 | 0.00168101 | 0.01340405 |
| VAPA | 1530.45205 | -0.12828028 | 0.22781833 | -0.5630815 | 0.57337941 | 0.73865654 |
| NAPG | 656.042356 | -0.390895352 | 0.27458358 | -1.4235933 | 0.15456424 | 0.3271791 |
| PIEZO2 | 35.8788056 | 0.697275056 | 0.42032081 | 1.65891156 | 0.09713361 | 0.24102289 |
| LINC01928 | 76.0935205 | 2.236746724 | 1.8643267 | 1.19976114 | 0.23023212 | 0.42336889 |
| LINC01255 | 58.9608923 | 2.681448509 | 2.41498217 | 1.11033884 | 0.26685304 | 0.46477327 |
| GNAL | 233.319848 | -0.835665642 | 0.26510761 | -3.1521752 | 0.00162059 | 0.01308211 |
| MPPE1 | 479.162678 | 0.24969705 | 0.23908228 | 1.04439799 | 0.29630129 | 0.49645156 |
| IMPA2 | 1777.89783 | 0.677126271 | 0.42677231 | 1.58662187 | 0.11259831 | 0.26584957 |
| TUBB6 | 158.946389 | -2.544830512 | 0.5525441 | -4.6056605 | 4.11E-06 | 0.00012225 |
| AFG3L2 | 1200.55504 | -0.223949497 | 0.2562905 | -0.8738112 | 0.3822211 | 0.57897056 |
| SPIRE1 | 882.249782 | -2.049176332 | 0.45610946 | -4.4927293 | 7.03E-06 | 0.0001889 |
| PSMG2 | 315.967594 | -0.340859903 | 0.1755898 | -1.9412284 | 0.05223058 | 0.15948419 |
| CEP76 | 108.807058 | 0.114396143 | 0.31258657 | 0.36596627 | 0.71439025 | 0.83664962 |
| PTPN2 | 1216.60982 | -0.464158208 | 0.21105482 | -2.1992306 | 0.02786153 | 0.1038613 |
| SEH1L | 384.0194 | -0.293761907 | 0.30028717 | -0.9782699 | 0.32794084 | 0.52777798 |
| CEP192 | 1053.96696 | -0.000984451 | 0.22346157 | -0.0044055 | 0.99648496 | 0.99819945 |
| LDLRAD4 | 805.321433 | 1.392247869 | 0.58907793 | 2.36343581 | 0.01810636 | 0.07695625 |
| FAM210A | 658.734216 | -1.237845996 | 0.38803626 | -3.1900267 | 0.0014226 | 0.0118179 |
| RNMT | 1029.78348 | -0.935422324 | 0.31605497 | -2.9596824 | 0.00307956 | 0.02110579 |
| ZNF519 | 69.9148315 | 0.843000876 | 0.43593809 | 1.93376285 | 0.05314229 | 0.16129792 |
| ANKRD30B | 61.6228078 | -0.200073812 | 0.58128662 | -0.3441913 | 0.73070241 | 0.84678203 |
| ROCK1 | 2022.59855 | -0.146684118 | 0.1076588 | -1.3624907 | 0.17304308 | 0.35259425 |
| GREB1L | 286.215179 | -0.346874635 | 0.49370089 | -0.7026008 | 0.48230457 | 0.66670759 |
| ESCO1 | 937.660094 | 0.165334997 | 0.21672125 | 0.76289243 | 0.44552755 | 0.63688501 |
| SNRPD1 | 206.693516 | -0.755223176 | 0.42600937 | -1.7727854 | 0.07626428 | 0.20545488 |
| ABHD3 | 654.974628 | -0.862801166 | 0.41725255 | -2.0678152 | 0.0386574 | 0.13073699 |
| MIB1 | 1620.53102 | -0.008928431 | 0.24878649 | -0.0358879 | 0.97137173 | 0.98575496 |
| CTAGE1 | 81.9061769 | 1.260512093 | 0.29926201 | 4.21206856 | 2.53E-05 | 0.00052632 |
| AC090912.1 | 123.694784 | -0.678117163 | 0.41611587 | -1.6296354 | 0.10317857 | 0.25087631 |
| RBBP8 | 1249.00235 | -1.212007658 | 0.29536978 | -4.1033571 | 4.07E-05 | 0.00076774 |
| CABLES1 | 1437.43438 | -0.49710109 | 0.33740211 | -1.4733194 | 0.14066493 | 0.30760915 |
| TMEM241 | 911.646209 | -0.358343581 | 0.24303985 | -1.4744231 | 0.1403677 | 0.30714643 |
| RIOK3 | 2084.79345 | -0.226424446 | 0.26462736 | -0.855635 | 0.39219968 | 0.58726848 |
| RMC1 | 687.887056 | -0.704625162 | 0.21293709 | -3.3090766 | 0.00093604 | 0.00860518 |
| NPC1 | 3231.08884 | -2.196695213 | 0.70053392 | -3.1357442 | 0.00171419 | 0.01361318 |
| ANKRD29 | 94.6748567 | -0.889978428 | 0.58489918 | -1.5215929 | 0.12811112 | 0.28981996 |
| LAMA3 | 126.930105 | -2.205993678 | 0.39832967 | -5.5381103 | 3.06E-08 | 2.14E-06 |
| TTC39C | 232.543491 | -2.250865456 | 0.47037392 | -4.7852684 | 1.71E-06 | 6.02E-05 |
| AC090772.3 | 34.0109733 | 1.594099118 | 0.38663225 | 4.12303713 | 3.74E-05 | 0.00071381 |
| OSBPL1A | 1802.73292 | 0.43523826 | 0.24856154 | 1.75102821 | 0.07994105 | 0.21173952 |
| IMPACT | 162.420538 | 0.62981408 | 0.37063459 | 1.69928576 | 0.08926535 | 0.22776623 |
| ZNF521 | 886.403657 | 1.243658544 | 0.2767831 | 4.49326044 | 7.01E-06 | 0.0001889 |
| SS18 | 1331.21046 | -0.279468772 | 0.14832939 | -1.8841092 | 0.05955021 | 0.17383682 |
| TAF4B | 480.322769 | -0.536496266 | 0.32083612 | -1.6721817 | 0.09448851 | 0.23686064 |
| KCTD1 | 692.622305 | -0.791594428 | 0.43157396 | -1.8342034 | 0.06662381 | 0.1877107 |
| AQP4-AS1 | 570.815423 | 3.086546405 | 0.93984128 | 3.28411452 | 0.00102303 | 0.00921335 |
| CHST9 | 172.550206 | -0.017742061 | 0.61904454 | -0.0286604 | 0.97713544 | 0.98865966 |
| CDH2 | 5097.60835 | 0.115968241 | 0.22830718 | 0.5079483 | 0.6114896 | 0.76769445 |
| DSC2 | 311.967248 | -1.154226632 | 0.16768111 | -6.8834623 | 5.84E-12 | 1.33E-09 |
| DSG2 | 448.298612 | -1.368644688 | 0.39289587 | -3.4834794 | 0.00049494 | 0.00531674 |
| B4GALT6 | 242.074318 | -0.780478018 | 0.33266696 | -2.3461242 | 0.01896978 | 0.07937082 |
| TRAPPC8 | 1545.00994 | 0.177270706 | 0.15348131 | 1.15499862 | 0.24809099 | 0.4444045 |
| RNF125 | 51.3118855 | 0.00497481 | 0.65783744 | 0.00756237 | 0.99396616 | 0.99701904 |
| RNF138 | 509.478512 | 0.153599358 | 0.17502906 | 0.87756487 | 0.3801799 | 0.57657954 |
| GAREM1 | 4579.99477 | 0.142420733 | 0.43579848 | 0.32680411 | 0.74381605 | 0.85468075 |
| AC025887.2 | 79.1296741 | 1.215551247 | 0.38699067 | 3.14103504 | 0.00168352 | 0.01341628 |
| KLHL14 | 105.935999 | -1.164205125 | 0.55324269 | -2.1043299 | 0.03534968 | 0.12254377 |
| CCDC178 | 350.193217 | 1.473401422 | 0.66237286 | 2.22442902 | 0.02611959 | 0.09938197 |
| ASXL3 | 407.139968 | 0.936643816 | 0.48209312 | 1.94286909 | 0.05203198 | 0.15901947 |
| NOL4 | 510.597162 | 2.830294105 | 0.65073221 | 4.34939909 | 1.37E-05 | 0.00032542 |
| DTNA | 2252.14023 | -2.429096025 | 0.6781077 | -3.5821685 | 0.00034075 | 0.00404391 |
| MAPRE2 | 561.051285 | -1.310098706 | 0.31386694 | -4.1740576 | 2.99E-05 | 0.00060315 |
| ZNF397 | 390.414708 | 0.190910638 | 0.29229293 | 0.65314833 | 0.51366066 | 0.69196393 |
| ZSCAN30 | 429.200523 | 0.323388555 | 0.30576081 | 1.05765206 | 0.2902141 | 0.49027014 |
| ZNF24 | 555.913699 | -0.119272098 | 0.17932927 | -0.6651011 | 0.50598583 | 0.68614442 |
| ZNF396 | 59.6531042 | 0.196156013 | 0.32038728 | 0.61224656 | 0.54037464 | 0.71186114 |
| GALNT1 | 697.610586 | -0.12191969 | 0.23355403 | -0.5220192 | 0.60165696 | 0.76078523 |
| C18orf21 | 61.993047 | 0.094824788 | 0.31549135 | 0.30056224 | 0.76374833 | 0.86815453 |
| RPRD1A | 1235.91035 | -0.010484529 | 0.21768865 | -0.048163 | 0.96158637 | 0.97988733 |
| SLC39A6 | 194.010478 | -0.963127764 | 0.3081833 | -3.1251783 | 0.00177697 | 0.01399022 |
| ELP2 | 750.280304 | -0.164594702 | 0.13977652 | -1.1775562 | 0.23897359 | 0.43402823 |
| MOCOS | 82.2038769 | -2.306415314 | 0.44529782 | -5.1794893 | 2.22E-07 | 1.07E-05 |
| FHOD3 | 4200.41623 | 0.410804669 | 0.2483771 | 1.65395547 | 0.09813656 | 0.24241446 |
| TPGS2 | 401.028609 | -1.140348632 | 0.2195956 | -5.1929484 | 2.07E-07 | 1.00E-05 |
| KIAA1328 | 1386.67343 | 0.106172294 | 0.1971363 | 0.53857304 | 0.59018149 | 0.75217273 |
| AC016493.1 | 35.7491453 | -1.376437985 | 0.69641862 | -1.976452 | 0.0481036 | 0.15161097 |
| CELF4 | 130.859119 | -1.451019902 | 0.72637566 | -1.9976164 | 0.04575827 | 0.14661543 |
| AC016205.1 | 281.79787 | 0.757936166 | 0.3651664 | 2.07559122 | 0.03793177 | 0.12896788 |
| PIK3C3 | 1701.20436 | -0.13742316 | 0.23851215 | -0.5761684 | 0.56450137 | 0.73162067 |
| LINC00907 | 1444.12225 | 1.621062123 | 0.54570069 | 2.97060671 | 0.00297212 | 0.02055442 |
| SETBP1 | 6095.69566 | 0.024453533 | 0.27481902 | 0.0889805 | 0.92909741 | 0.96237725 |
| SLC14A2 | 158.170258 | 0.565848064 | 0.73882545 | 0.76587516 | 0.44375059 | 0.63517875 |
| EPG5 | 1482.5993 | 0.306480267 | 0.23039213 | 1.3302549 | 0.1834343 | 0.36582772 |
| PSTPIP2 | 264.95877 | -0.987604479 | 0.71621582 | -1.3789202 | 0.16791935 | 0.34613516 |
| ATP5F1A | 880.19133 | -0.041361103 | 0.3022209 | -0.1368572 | 0.89114368 | 0.94333086 |
| HAUS1 | 105.94389 | 0.301162329 | 0.36710488 | 0.82037135 | 0.41200444 | 0.60554845 |

| | | | | | | |
|---|---|---|---|---|---|---|
| C18orf25 | 675.932844 | 0.177429377 | 0.2959062 | 0.59961358 | 0.54876379 | 0.71869915 |
| PIAS2 | 1936.18558 | -0.295246855 | 0.34943359 | -0.8449298 | 0.39815003 | 0.59255618 |
| KATNAL2 | 993.479196 | 1.43062871 | 0.51204463 | 2.79395314 | 0.0052068 | 0.03127504 |
| AC012254.1 | 106.499115 | -0.218271891 | 0.67905251 | -0.321436 | 0.74788004 | 0.85769676 |
| HDHD2 | 70.053146 | -0.03527049 | 0.40240313 | -0.0876496 | 0.93015515 | 0.96282888 |
| SMAD2 | 1785.28872 | -0.208703061 | 0.13300659 | -1.5691182 | 0.1166204 | 0.2720878 |
| ZBTB7C | 59.9041089 | -1.211093236 | 0.69384954 | -1.7454695 | 0.08090317 | 0.21362738 |
| CTIF | 806.260477 | -0.439575574 | 0.2540295 | -1.7304116 | 0.08355677 | 0.21827952 |
| SMAD7 | 184.651088 | 0.946108475 | 0.53135067 | 1.78057266 | 0.07498229 | 0.20327972 |
| DYM | 3386.78219 | -0.031254991 | 0.18756819 | -0.1666327 | 0.86765908 | 0.93103202 |
| C18orf32 | 54.5526191 | -0.009136595 | 0.26205234 | -0.0348655 | 0.97218696 | 0.98614421 |
| LIPG | 241.580643 | -1.714627823 | 0.48999152 | -3.499301 | 0.00046648 | 0.00510298 |
| ACAA2 | 748.138182 | 0.074695203 | 0.46208163 | 0.16164936 | 0.87158198 | 0.93353404 |
| SNHG22 | 464.288945 | 0.268099784 | 0.43185006 | 0.62081683 | 0.53472015 | 0.70760852 |
| MYO5B | 3825.07289 | -1.020759073 | 0.31108093 | -3.2813297 | 0.00103319 | 0.00926827 |
| CFAP53 | 139.430997 | 0.658846681 | 0.24642114 | 2.67366137 | 0.00750282 | 0.04097255 |
| MBD1 | 281.809574 | -0.212763587 | 0.24054336 | -0.8845124 | 0.37641967 | 0.57294604 |
| CXXC1 | 202.129565 | 0.450105925 | 0.34039823 | 1.32229221 | 0.18607087 | 0.36923618 |
| MRO | 171.291053 | 5.237872649 | 0.74165815 | 7.06238128 | 1.64E-12 | 4.15E-10 |
| ME2 | 358.998095 | -1.240194126 | 0.4216289 | -2.9414353 | 0.00326695 | 0.02202661 |
| AC091551.1 | 121.585718 | -0.162649629 | 0.43250426 | -0.3760648 | 0.70686872 | 0.83210663 |
| SMAD4 | 539.827774 | -0.205542251 | 0.15717512 | -1.3077277 | 0.19096569 | 0.37527838 |
| MEX3C | 195.308244 | -0.545162798 | 0.18059358 | -3.0187274 | 0.00253839 | 0.0184191 |
| DCC | 1129.26994 | 2.093303309 | 0.7988703 | 2.62032938 | 0.00878449 | 0.04588757 |
| MBD2 | 1567.63555 | -0.767937694 | 0.20992246 | -3.6581969 | 0.000254 | 0.00322065 |
| POLI | 559.746293 | 0.519454889 | 0.31058462 | 1.67250681 | 0.09442443 | 0.2367433 |
| CCDC68 | 171.449209 | -0.02260367 | 0.43826323 | -0.0515756 | 0.9588669 | 0.97893554 |
| TCF4 | 284.40221 | 0.923745896 | 0.32133379 | 2.87472385 | 0.00404381 | 0.02582875 |
| AC009271.1 | 92.0246618 | 0.184572893 | 0.79200952 | 0.23304378 | 0.81572741 | 0.90027629 |
| AC006305.1 | 694.429228 | 3.697853663 | 0.97752934 | 3.78285694 | 0.00015504 | 0.00216848 |
| LINC01905 | 39.8874085 | 1.446701249 | 0.97543843 | 1.48312924 | 0.13804008 | 0.30380786 |
| TXNL1 | 562.169755 | -0.089389151 | 0.16826766 | -0.5312319 | 0.59525808 | 0.75569163 |
| WDR7 | 1556.8137 | 0.665778895 | 0.22503335 | 2.95857873 | 0.00309061 | 0.02111703 |
| FECH | 159.92479 | -0.081608637 | 0.26880682 | -0.3035959 | 0.76143581 | 0.86671654 |
| NARS | 803.457581 | -0.825041896 | 0.24735069 | -3.3355149 | 0.00085142 | 0.00802638 |
| AC027097.2 | 524.037773 | 0.362590366 | 0.28291926 | 1.28160367 | 0.19998171 | 0.38689475 |
| ATP8B1 | 416.715045 | -0.913841346 | 0.26036385 | -3.5098627 | 0.00044834 | 0.00495994 |
| NEDD4L | 12782.6818 | 0.04763782 | 0.37831561 | 0.12592084 | 0.89979458 | 0.94743919 |
| ALPK2 | 1696.19264 | 1.058669137 | 0.40557105 | 2.6103173 | 0.00904583 | 0.04687757 |
| MALT1 | 2745.80443 | 0.809388229 | 0.35988901 | 2.24899403 | 0.02451288 | 0.09496834 |
| AC104365.1 | 303.523023 | -0.247374534 | 0.6344823 | -0.3898841 | 0.69662228 | 0.82536102 |
| ZNF532 | 311.856002 | -0.51389358 | 0.34573889 | -1.4863633 | 0.13718306 | 0.30245588 |
| SEC11C | 194.413799 | -0.754161781 | 0.41578112 | -1.8138432 | 0.06970186 | 0.19344066 |
| LMAN1 | 693.223894 | -0.491360464 | 0.24287339 | -2.0231136 | 0.04306144 | 0.14036697 |
| AC113137.1 | 55.2061104 | 2.137571372 | 0.65258328 | 3.27555337 | 0.00105455 | 0.00941089 |
| CDH20 | 70.9920184 | -0.345343813 | 0.92636263 | -0.3727955 | 0.70930066 | 0.83353773 |
| AC105094.2 | 308.224798 | 1.798248637 | 0.81572887 | 2.20446855 | 0.02749141 | 0.10294526 |
| RNF152 | 7315.30171 | 2.101742719 | 0.72816228 | 2.88636582 | 0.00389719 | 0.02515181 |
| PIGN | 1074.89791 | 0.273335387 | 0.13831893 | 1.97612421 | 0.04814071 | 0.15165818 |
| KIAA1468 | 1222.63693 | 0.067708977 | 0.12992641 | 0.52113328 | 0.60227393 | 0.76094319 |
| ZCCHC2 | 928.791682 | -0.397227169 | 0.17445146 | -2.2770069 | 0.02278581 | 0.09047658 |
| PHLPP1 | 1413.80574 | 0.09853106 | 0.18614636 | 0.52932035 | 0.59658324 | 0.75667312 |
| BCL2 | 3109.04751 | 1.042868135 | 0.43180243 | 2.41515114 | 0.01572868 | 0.06965849 |
| KDSR | 777.662803 | -0.972665609 | 0.31372259 | -3.1004003 | 0.00193259 | 0.0149408 |
| AC036176.1 | 44.2841912 | 0.338403346 | 0.44405297 | 0.76207879 | 0.44601298 | 0.6372179 |
| VPS4B | 741.281388 | -0.333214939 | 0.18119333 | -1.8390022 | 0.06591486 | 0.18652136 |
| SERPINB8 | 70.2395014 | -2.612150752 | 0.61908672 | -4.2193616 | 2.45E-05 | 0.00051521 |
| DSEL | 263.986301 | 0.899249629 | 0.40146538 | 2.23991825 | 0.02509623 | 0.09667393 |
| AC110597.3 | 36.8322949 | -0.124316968 | 0.77507879 | -0.1603927 | 0.87257175 | 0.93410973 |
| AC114689.3 | 337.243556 | 1.583853223 | 0.56446546 | 2.8059347 | 0.00501709 | 0.03037717 |
| TMX3 | 750.259968 | 0.493922678 | 0.24899185 | 1.98369014 | 0.04729039 | 0.15001375 |
| CCDC102B | 243.25968 | 0.202957729 | 0.50865072 | 0.39901197 | 0.68988438 | 0.82071214 |
| DOK6 | 3855.53158 | 0.410832061 | 0.37060068 | 1.10855723 | 0.26762124 | 0.46564381 |
| AC119868.2 | 136.949978 | -0.868027667 | 0.51713973 | -1.6785167 | 0.09324628 | 0.2350794 |
| CD226 | 241.901091 | 2.436200711 | 0.43722049 | 5.57201863 | 2.52E-08 | 1.79E-06 |
| RTTN | 1450.97238 | 1.779630468 | 0.34038394 | 5.22830335 | 1.71E-07 | 8.75E-06 |
| SOCS6 | 641.009996 | -0.063648093 | 0.23211038 | -0.2742148 | 0.78391958 | 0.88133085 |
| LINC01909 | 44.1160292 | 0.347015572 | 0.38311914 | 0.90576413 | 0.36506071 | 0.56248732 |
| AC027458.1 | 111.956777 | 2.592512067 | 1.00352068 | 2.58341668 | 0.00978271 | 0.04964517 |
| FBXO15 | 102.872196 | 1.156178463 | 0.59447879 | 1.94486074 | 0.05179174 | 0.1587111 |
| TIMM21 | 132.654614 | -0.100513314 | 0.19358282 | -0.5192264 | 0.60360287 | 0.76169806 |
| CYB5A | 3552.30109 | 0.776387023 | 0.49286804 | 1.57524319 | 0.11520032 | 0.26983097 |
| CNDP2 | 2577.38162 | -0.300771872 | 0.36023674 | -0.8349284 | 0.40375804 | 0.59790609 |
| CNDP1 | 54.9575382 | 0.607901568 | 0.48697202 | 1.24832957 | 0.21191039 | 0.4015283 |
| LINC00909 | 159.769084 | -0.077967259 | 0.33235005 | -0.2345938 | 0.81452402 | 0.89950699 |
| ZNF407 | 3462.10394 | -0.072903298 | 0.26254257 | -0.2776818 | 0.78125662 | 0.87992477 |
| ZADH2 | 227.023066 | 0.015361489 | 0.31520196 | 0.04873539 | 0.96113018 | 0.97985953 |
| TSHZ1 | 1046.82966 | 0.898561447 | 0.43025761 | 2.08842663 | 0.03675937 | 0.12602818 |
| ZNF516 | 1827.02938 | -0.313292009 | 0.3260566 | -0.9608516 | 0.33662678 | 0.53671552 |
| AC009716.1 | 51.7306381 | -1.328497242 | 0.6306942 | -2.1064047 | 0.03516921 | 0.12210344 |
| ZNF236-DT | 95.3624036 | 0.454765295 | 0.23109998 | 1.96782919 | 0.04908769 | 0.15361759 |
| ZNF236 | 666.194918 | 0.270587909 | 0.2535179 | 1.06733255 | 0.28582169 | 0.48541172 |
| MBP | 2866.38508 | -1.490661029 | 0.37113089 | -4.0165372 | 5.91E-05 | 0.00103388 |
| AC124254.1 | 84.5438773 | -0.981234851 | 1.19269653 | -0.8227029 | 0.41067698 | 0.60443997 |
| ATP9B | 2055.88913 | 0.199825003 | 0.19727935 | 1.01290379 | 0.31110612 | 0.51127879 |
| NFATC1 | 43.4831121 | -1.322679427 | 0.68405064 | -1.9335987 | 0.05316249 | 0.16130504 |
| CTDP1 | 435.302025 | -0.023626388 | 0.34743755 | -0.0680018 | 0.94578418 | 0.97196767 |
| AC021594.2 | 121.499921 | 1.503072549 | 1.12927956 | 1.33100129 | 0.18318859 | 0.36558246 |
| KCNG2 | 69.8851334 | 0.488919732 | 0.96072809 | 0.50890542 | 0.61081852 | 0.76707767 |
| PQLC1 | 1255.3447 | -0.409715186 | 0.3526817 | -1.1617138 | 0.24535175 | 0.4408227 |
| HSBP1L1 | 527.581191 | -0.350514826 | 0.38435193 | -0.9119632 | 0.36178807 | 0.55961134 |
| TXNL4A | 498.614111 | -0.59999914 | 0.25376251 | -2.3644121 | 0.01805871 | 0.0768252 |
| ADNP2 | 882.363937 | -0.477059283 | 0.42165529 | -1.1313964 | 0.25788828 | 0.45523177 |
| PARD6G-AS1 | 232.352603 | -0.441934785 | 0.57322841 | -0.7709576 | 0.44073208 | 0.63268271 |
| PARD6G | 146.264214 | 1.300539413 | 0.31581411 | 4.11805357 | 3.82E-05 | 0.00072639 |
| C20orf96 | 157.583498 | 0.366032229 | 0.38454889 | 0.95184836 | 0.34117389 | 0.54194815 |
| ZCCHC3 | 133.089465 | 0.009253058 | 0.266298 | 0.03474701 | 0.97228148 | 0.98615459 |
| NRSN2-AS1 | 465.456568 | 0.640102973 | 0.24203654 | 2.64465432 | 0.00817744 | 0.04364805 |
| SOX12 | 97.4316856 | 0.065744165 | 0.37616796 | 0.17477343 | 0.86125767 | 0.92699323 |
| NRSN2 | 133.477216 | -0.873112969 | 0.39246412 | -2.224695 | 0.02610172 | 0.09938197 |
| RBCK1 | 525.301616 | -1.009389177 | 0.28589559 | -3.5306218 | 0.00041458 | 0.00470033 |
| TBC1D20 | 390.114637 | -0.598666062 | 0.31494475 | -1.9008606 | 0.05732028 | 0.16976859 |
| CSNK2A1 | 3102.11228 | -0.635047429 | 0.27664943 | -2.2954951 | 0.02170477 | 0.08763099 |
| RSPO4 | 45.625492 | 0.472893061 | 0.62233734 | 0.75986612 | 0.44733461 | 0.63804532 |
| PSMF1 | 705.721217 | -0.604734923 | 0.30516345 | -1.9816755 | 0.04751558 | 0.15042597 |
| AL136531.2 | 304.135525 | -0.819209379 | 0.40604444 | -2.0175362 | 0.04363958 | 0.1415605 |
| SDCBP2-AS1 | 646.613104 | 0.803340348 | 0.34103412 | 2.35560112 | 0.01849276 | 0.078066 |
| NSFL1C | 345.842424 | -0.810611467 | 0.28788428 | -2.8157546 | 0.00486628 | 0.02963223 |
| SIRPA | 307.848883 | -0.322800995 | 0.35448835 | -0.910611 | 0.36250037 | 0.56030588 |
| STK35 | 313.204626 | -0.236063706 | 0.31711928 | -0.7444004 | 0.4566343 | 0.64552736 |
| SNRPB | 276.169729 | -1.333373733 | 0.39507576 | -3.3749824 | 0.0007382 | 0.00722399 |
| AL049650.1 | 69.9597352 | 0.422131172 | 0.3762849 | 1.12183926 | 0.26193079 | 0.45999812 |
| NOP56 | 418.868369 | -0.928070945 | 0.66216946 | -1.4015611 | 0.16104635 | 0.3371137 |

| | | | | | | |
|---|---|---|---|---|---|---|
| IDH3B | 176.567789 | 0.128920296 | 0.30912882 | 0.41704392 | 0.67664629 | 0.81279385 |
| EBF4 | 68.0378594 | -1.103106276 | 0.70277715 | -1.5696388 | 0.11649917 | 0.27203647 |
| PCED1A | 201.805617 | 1.101405642 | 0.26991438 | 4.08057411 | 4.49E-05 | 0.00082878 |
| VPS16 | 95.2390367 | 0.208251872 | 0.42817683 | 0.48636885 | 0.62670567 | 0.77912423 |
| PTPRA | 1490.79724 | -0.203358707 | 0.14968169 | -1.3586077 | 0.17427092 | 0.3540435 |
| MRPS26 | 252.623582 | -0.514810489 | 0.33092661 | -1.5556636 | 0.1197881 | 0.27702309 |
| UBOX5-AS1 | 161.281958 | -0.306485154 | 0.65124896 | -0.4706113 | 0.6379183 | 0.7871047 |
| UBOX5 | 237.158039 | 0.944015084 | 0.48435081 | 1.94903169 | 0.05129164 | 0.15784789 |
| LZTS3 | 285.698292 | 0.190859633 | 0.40512441 | 0.47111364 | 0.63755959 | 0.78694546 |
| DDRGK1 | 308.733496 | -0.842880013 | 0.21560435 | -3.9093832 | 9.25E-05 | 0.00146557 |
| ITPA | 480.36332 | -0.068408345 | 0.28914208 | -0.2365908 | 0.81297429 | 0.8986644 |
| C20orf194 | 4081.55112 | 0.280647377 | 0.27446347 | 1.02253089 | 0.3065297 | 0.50646126 |
| ATRN | 1476.18898 | 0.192493057 | 0.15794196 | 1.21875816 | 0.222936 | 0.4151316 |
| ADAM33 | 122.862914 | -2.703242806 | 1.48716821 | -1.8177115 | 0.06910823 | 0.19255756 |
| C20orf27 | 59.0274025 | 0.060334585 | 0.39551176 | 0.15254814 | 0.87875462 | 0.93706833 |
| CENPB | 197.447156 | -0.966019253 | 0.48238843 | -2.0025755 | 0.04522287 | 0.14543841 |
| CDC25B | 256.173397 | -0.765197221 | 0.569388 | -1.3438942 | 0.1789826 | 0.360146 |
| AP5S1 | 52.8969483 | -0.061116947 | 0.41682121 | -0.1466263 | 0.883427 | 0.93956657 |
| MAVS | 818.166866 | 0.230426006 | 0.24274851 | 0.94923755 | 0.34249981 | 0.54304741 |
| PANK2 | 521.341611 | -0.311895997 | 0.23205873 | -1.344039 | 0.17893576 | 0.36010461 |
| RNF24 | 686.993827 | -2.440042034 | 0.40645977 | -6.0031576 | 1.94E-09 | 2.04E-07 |
| SMOX | 158.030337 | -3.833224118 | 0.44365217 | -8.6401564 | 5.61E-18 | 5.50E-15 |
| PRNP | 575.512977 | 0.149994365 | 0.35861612 | 0.41825884 | 0.67575788 | 0.81215372 |
| SLC23A2 | 243.463681 | 0.477928319 | 0.28081924 | 1.70190731 | 0.08877275 | 0.22722839 |
| TMEM230 | 369.936578 | -0.225640117 | 0.23326973 | -0.9672927 | 0.33339771 | 0.53275938 |
| PCNA | 84.9865657 | -0.019271715 | 0.24340723 | -0.0791748 | 0.9368936 | 0.96611631 |
| CDS2 | 835.593617 | -0.555578443 | 0.20864487 | -2.6627946 | 0.00774947 | 0.04188564 |
| GPCPD1 | 833.369374 | 0.007925768 | 0.49577093 | 0.01598675 | 0.98724496 | 0.99360989 |
| SHLD1 | 325.516051 | 0.893423565 | 0.350274 | 2.55064199 | 0.01075247 | 0.05312686 |
| TRMT6 | 95.941301 | -0.572706671 | 0.45395275 | -1.2615997 | 0.20709285 | 0.39595783 |
| MCM8 | 228.919376 | -0.371013971 | 0.33956516 | -1.092615 | 0.27456289 | 0.47243952 |
| MCM8-AS1 | 36.4532344 | 0.255807602 | 0.57645715 | 0.44375823 | 0.65721739 | 0.80027901 |
| CRLS1 | 1680.45432 | -0.032927145 | 0.30727225 | -0.1071595 | 0.91466243 | 0.95437477 |
| LRRN4 | 58.3785385 | -1.536387305 | 0.50604004 | -3.0360983 | 0.00239661 | 0.01766148 |
| FERMT1 | 209.909987 | 1.041753412 | 0.2884552 | 3.61149116 | 0.00030444 | 0.00371274 |
| BMP2 | 47.4787857 | -0.87316185 | 0.55566677 | -1.5713768 | 0.11609516 | 0.27150936 |
| AL096799.1 | 79.9910292 | -0.910827786 | 0.75064428 | -1.2133947 | 0.22497895 | 0.41717891 |
| LINC01428 | 393.913298 | 3.193591901 | 0.73789315 | 4.32798689 | 1.50E-05 | 0.00035309 |
| TMX4 | 294.242244 | -0.292230215 | 0.31343835 | -0.9323371 | 0.35116232 | 0.55041531 |
| PLCB1 | 1335.08 | -0.436282366 | 0.45502014 | -0.9588199 | 0.33764948 | 0.5378456 |
| PLCB4 | 1544.91229 | -1.800982475 | 0.52972077 | -3.3998714 | 0.00067418 | 0.00672559 |
| PAK5 | 34.533857 | 0.691975673 | 0.37352597 | 1.85255038 | 0.06394683 | 0.18279858 |
| ANKEF1 | 67.9924634 | -0.348169788 | 0.25726487 | -1.3533515 | 0.17594337 | 0.35617425 |
| SNAP25-AS1 | 63.7472993 | -1.446256911 | 0.66774753 | -2.1658739 | 0.03032083 | 0.11009472 |
| SNAP25 | 121.915326 | -2.301078114 | 0.71878224 | -3.2013564 | 0.00136782 | 0.01144612 |
| MKKS | 158.949582 | 0.213993236 | 0.19578719 | 1.09298893 | 0.27439866 | 0.47233234 |
| SLX4IP | 1134.97114 | 0.440685288 | 0.27001877 | 1.63205425 | 0.10266806 | 0.25027051 |
| JAG1 | 268.139003 | -0.196898315 | 0.42669012 | -0.4614551 | 0.64447215 | 0.79162826 |
| AL050403.2 | 247.889048 | -0.875650456 | 0.53805272 | -1.6274436 | 0.10364292 | 0.25157313 |
| BTBD3 | 399.846604 | -0.26092736 | 0.26567958 | -0.982113 | 0.3260442 | 0.52620838 |
| SPTLC3 | 2805.41195 | 0.062237225 | 0.5432858 | 0.11455706 | 0.90879622 | 0.95219603 |
| ISM1 | 127.824332 | 0.507016314 | 0.44260515 | 1.14552737 | 0.25199076 | 0.44840158 |
| TASP1 | 2313.29228 | -0.018362451 | 0.18285953 | -0.1004183 | 0.92001221 | 0.95701657 |
| ESF1 | 886.62099 | -0.138290452 | 0.30186979 | -0.4581129 | 0.64687131 | 0.79280318 |
| NDUFAF5 | 547.234465 | 0.766467018 | 0.27705592 | 2.76647049 | 0.00566667 | 0.03324749 |
| MACROD2 | 4520.42128 | 0.811784052 | 0.42217751 | 1.92285007 | 0.05449888 | 0.16425158 |
| FLRT3 | 772.933407 | 1.47115454 | 0.36927431 | 3.9839071 | 6.78E-05 | 0.0011486 |
| KIF16B | 2184.90545 | 0.494037144 | 0.25409893 | 1.94427083 | 0.0518628 | 0.15885663 |
| SNRPB2 | 312.270365 | -0.319287336 | 0.29808743 | -1.0711198 | 0.28411559 | 0.48341266 |
| DSTN | 1283.1903 | -0.473821321 | 0.19282054 | -2.4573177 | 0.01399788 | 0.0637651 |
| RRBP1 | 1615.72566 | -1.047130082 | 0.33474707 | -3.1281232 | 0.00175926 | 0.01389869 |
| SNX5 | 745.600684 | 0.665122824 | 0.22637031 | 2.93820694 | 0.00330117 | 0.02218092 |
| OVOL2 | 243.764275 | 0.401093963 | 0.41078629 | 0.97640542 | 0.3288636 | 0.52869297 |
| MGME1 | 353.576219 | 0.470056266 | 0.31869325 | 1.47494895 | 0.14022627 | 0.30697512 |
| KAT14 | 168.03013 | 1.157686551 | 0.34500697 | 3.35554539 | 0.00079209 | 0.0076137 |
| ZNF133 | 428.036751 | 0.386788491 | 0.21751647 | 1.77820321 | 0.07537049 | 0.20404963 |
| DZANK1 | 323.088403 | -0.142383708 | 0.26496294 | -0.5373722 | 0.59101057 | 0.75259958 |
| POLR3F | 138.149087 | 0.028626226 | 0.25716106 | 0.11131633 | 0.91136551 | 0.95319043 |
| RBBP9 | 96.9337161 | -0.016932749 | 0.34966738 | -0.0484253 | 0.96137729 | 0.97988733 |
| SEC23B | 697.101961 | -0.533808186 | 0.16530701 | -3.2291927 | 0.0012414 | 0.01064159 |
| DTD1 | 137.23883 | -1.005120792 | 0.373639 | -2.6900853 | 0.00714338 | 0.03954092 |
| SLC24A3 | 64.670033 | -1.845957865 | 0.63617304 | -2.90166 | 0.00371191 | 0.02418938 |
| AL121761.1 | 402.970068 | 0.36583091 | 0.33265329 | 1.09973633 | 0.27144702 | 0.46966982 |
| RIN2 | 1630.82205 | 1.294934923 | 0.35834002 | 3.61370444 | 0.00030185 | 0.0036942 |
| NAA20 | 543.401623 | -0.684922492 | 0.26942226 | -2.5421897 | 0.01101604 | 0.05402391 |
| CRNKL1 | 163.756155 | -0.179576985 | 0.2305623 | -0.7788653 | 0.43605904 | 0.62830455 |
| CFAP61 | 128.431971 | -0.113731608 | 0.3817723 | -0.2979043 | 0.76577621 | 0.86913095 |
| RALGAPA2 | 5153.34247 | -0.40029254 | 0.25850427 | -1.5484949 | 0.12150319 | 0.27953076 |
| KIZ | 545.061023 | 0.433170732 | 0.32954766 | 1.31444031 | 0.18869807 | 0.37263859 |
| KIZ-AS1 | 126.690648 | -0.737079191 | 0.41586264 | -1.7724102 | 0.07632649 | 0.20550131 |
| XRN2 | 1105.11982 | -0.35675691 | 0.23035975 | -1.5486946 | 0.12145515 | 0.27946713 |
| NXT1 | 58.0673988 | -0.812961574 | 0.4874811 | -1.6676781 | 0.09537964 | 0.23817976 |
| GZF1 | 86.4413156 | -0.969670923 | 0.40266574 | -2.4081287 | 0.01603453 | 0.07064779 |
| NAPB | 518.73847 | 0.170491323 | 0.22626629 | 0.75349854 | 0.45115039 | 0.64095339 |
| CST3 | 1442.66319 | -1.309152887 | 0.38536989 | -3.3971333 | 0.00068096 | 0.00676369 |
| SYNDIG1 | 630.283846 | 0.283645585 | 0.84761659 | 0.33463902 | 0.73789741 | 0.85123628 |
| APMAP | 178.735975 | -0.479119712 | 0.28212428 | -1.6982576 | 0.08945915 | 0.22813331 |
| ACSS1 | 627.229074 | 0.057203531 | 0.30442153 | 0.18790895 | 0.85094803 | 0.92167424 |
| ENTPD6 | 324.65689 | -0.367616376 | 0.27899485 | -1.3176457 | 0.18762227 | 0.37126295 |
| PYGB | 354.270525 | -1.451937413 | 0.4364001 | -3.3270786 | 0.00087762 | 0.00821125 |
| ABHD12 | 618.528763 | 0.234898187 | 0.31602246 | 0.74329586 | 0.45730258 | 0.64588182 |
| GINS1 | 57.0806204 | 0.37504622 | 0.35865331 | 1.04570684 | 0.29569641 | 0.4955986 |
| NINL | 3240.50522 | 0.766618415 | 0.32650355 | 2.34796349 | 0.01887637 | 0.07917333 |
| NANP | 94.3677004 | 0.033970804 | 0.30650936 | 0.11083121 | 0.91175019 | 0.95319043 |
| ZNF337-AS1 | 386.555083 | 0.824321405 | 0.4367449 | 1.88742081 | 0.05910375 | 0.17307314 |
| ZNF337 | 290.74405 | 0.337386523 | 0.21869981 | 1.54269234 | 0.12290544 | 0.28169614 |
| FAM182B | 327.368066 | -0.305071904 | 0.36936361 | -0.8259393 | 0.40883855 | 0.60264006 |
| HM13 | 1331.34095 | -1.13283572 | 0.30906591 | -3.6653532 | 0.000247 | 0.00316115 |
| ID1 | 137.675872 | 0.109000604 | 0.59792293 | 0.18229875 | 0.85534828 | 0.92352338 |
| BCL2L1 | 2808.19157 | -0.058521294 | 0.49914591 | -0.1172429 | 0.9066676 | 0.95109493 |
| AL117381.1 | 479.274666 | -0.246438909 | 0.55967835 | -0.4403224 | 0.65970363 | 0.8020214 |
| DUSP15 | 142.147666 | 0.957581986 | 0.66090804 | 1.44888838 | 0.14736876 | 0.31681745 |
| TTLL9 | 78.7294827 | 0.252616555 | 0.46996235 | 0.53752509 | 0.59090496 | 0.75259958 |
| PDRG1 | 75.2327493 | -0.417020305 | 0.33221833 | -1.2552598 | 0.20938448 | 0.39883728 |
| TM9SF4 | 868.310373 | -0.055523186 | 0.16431931 | -0.3378981 | 0.73543996 | 0.85033112 |
| PLAGL2 | 153.54973 | 0.239207253 | 0.27212277 | 0.87904167 | 0.37937868 | 0.57597637 |
| POFUT1 | 257.487808 | 0.078276897 | 0.25248153 | 0.31003019 | 0.756538 | 0.86351 |
| KIF3B | 907.177467 | 0.278456437 | 0.20751008 | 1.34189352 | 0.17963052 | 0.360867 |
| ASXL1 | 2274.0427 | 0.010967903 | 0.21697309 | 0.0505496 | 0.95968443 | 0.97933252 |
| NOL4L | 237.86694 | 0.410858788 | 0.36707138 | 1.11928854 | 0.26301706 | 0.46101981 |
| FO393400.1 | 88.6734401 | -0.357143686 | 0.49371979 | -0.7233732 | 0.4694506 | 0.65640716 |
| DNMT3B | 201.965706 | -0.265739782 | 0.35181567 | -0.7553381 | 0.45004614 | 0.64044668 |
| MAPRE1 | 247.418987 | -0.49054894 | 0.36094123 | -1.3590826 | 0.17412042 | 0.3538951 |

| | | | | | | |
|---|---|---|---|---|---|---|
| AL035071.2 | 66.3659967 | -1.639063132 | 0.38487818 | -4.2586543 | 2.06E-05 | 0.0004532 |
| AL035071.1 | 75.345413 | -0.372317637 | 0.48845674 | -0.7622326 | 0.44592121 | 0.6372179 |
| CDK5RAP1 | 414.263043 | 0.442555271 | 0.2235177 | 1.97995626 | 0.04770844 | 0.15078156 |
| SNTA1 | 166.07968 | 1.219069989 | 0.32857422 | 3.71018143 | 0.00020711 | 0.00275351 |
| CBFA2T2 | 2842.99281 | 0.19489111 | 0.18463388 | 1.05555446 | 0.29117182 | 0.49133228 |
| NECAB3 | 75.4871479 | 0.391089384 | 0.31132685 | 1.25620191 | 0.20904278 | 0.3982971 |
| PXMP4 | 58.3511172 | 1.069716016 | 0.39917027 | 2.6798489 | 0.00736554 | 0.04046471 |
| ZNF341 | 92.8178937 | -0.206519808 | 0.26639852 | -0.7752288 | 0.43820446 | 0.63007117 |
| CHMP4B | 590.312526 | -1.111579331 | 0.37349798 | -2.9761321 | 0.00291909 | 0.02026949 |
| RALY | 944.017308 | -0.595991875 | 0.24332568 | -2.4493587 | 0.01431108 | 0.06482552 |
| EIF2S2 | 671.1355 | -1.176337742 | 0.32572155 | -3.6114827 | 0.00030445 | 0.00371274 |
| ASIP | 200.972337 | 0.790204329 | 0.64861007 | 1.21830413 | 0.22310842 | 0.41522737 |
| AHCY | 518.293084 | 0.872447962 | 0.28614177 | 3.049006 | 0.002296 | 0.0170761 |
| ITCH | 3153.91694 | 0.144932697 | 0.20597401 | 0.70364557 | 0.48165352 | 0.66636403 |
| DYNLRB1 | 803.380773 | -0.295159404 | 0.1865829 | -1.581921 | 0.11366761 | 0.26731977 |
| MAP1LC3A | 193.213755 | -0.646425536 | 0.51360769 | -1.2585979 | 0.20817562 | 0.3973076 |
| PIGU | 429.785451 | 0.275365453 | 0.23367272 | 1.17842361 | 0.23862777 | 0.4338025 |
| NCOA6 | 1739.51105 | 0.033433287 | 0.1672289 | 0.19992529 | 0.84153901 | 0.91634694 |
| TP53INP2 | 126.494003 | 0.381957906 | 0.32135529 | 1.18858445 | 0.23460323 | 0.42901494 |
| GGT7 | 503.930949 | 0.597751857 | 0.23563582 | 2.53676139 | 0.01118832 | 0.0545759 |
| ACSS2 | 2471.99725 | 1.557448163 | 0.49715707 | 3.13270849 | 0.00173201 | 0.01372295 |
| GSS | 427.964742 | 0.583530455 | 0.36597142 | 1.59447004 | 0.1108308 | 0.26301207 |
| TRPC4AP | 1479.41487 | -0.334056999 | 0.17917739 | -1.8643926 | 0.06226659 | 0.17941731 |
| EDEM2 | 316.774009 | 0.148006946 | 0.22473914 | 0.65857219 | 0.51017052 | 0.68954593 |
| PROCR | 94.5605256 | -0.62546173 | 0.38244033 | -1.6354492 | 0.10195492 | 0.24928578 |
| MMP24OS | 250.552734 | 0.211376879 | 0.26365703 | 0.80171151 | 0.42271986 | 0.61620984 |
| MMP24 | 419.502469 | -0.429605063 | 0.44960382 | -0.9555192 | 0.33931522 | 0.53974628 |
| EIF6 | 66.7216324 | -0.44158151 | 0.65486197 | -0.6743123 | 0.50011276 | 0.68141606 |
| UQCC1 | 975.761044 | 0.098504622 | 0.15215896 | 0.64737972 | 0.51738622 | 0.69425481 |
| CEP250 | 387.691086 | -1.863977674 | 0.79769865 | -2.336694 | 0.0194551 | 0.0808339 |
| ERGIC3 | 446.561947 | 0.405649201 | 0.32769418 | 1.23788954 | 0.21575702 | 0.40633004 |
| CPNE1 | 389.132739 | 0.027565796 | 0.22054763 | 0.12498795 | 0.90053309 | 0.94768577 |
| NFS1 | 101.341526 | 1.169741511 | 0.27125725 | 4.31229579 | 1.62E-05 | 0.00037409 |
| ROMO1 | 273.652725 | -0.879543094 | 0.48096707 | -1.8286971 | 0.06744499 | 0.18920764 |
| RBM39 | 3312.74819 | -0.214930424 | 0.19190642 | -1.1199752 | 0.26272434 | 0.46074961 |
| PHF20 | 1424.1101 | 0.04418305 | 0.2915006 | 0.15157104 | 0.87952528 | 0.93716208 |
| SCAND1 | 474.925647 | -0.875253757 | 0.42432356 | -2.0627036 | 0.03914079 | 0.13168946 |
| CNBD2 | 82.8928913 | 0.71928483 | 0.49174098 | 1.4627311 | 0.14354097 | 0.31111893 |
| NORAD | 995.524921 | -0.805682085 | 0.20805361 | -3.8724734 | 0.00010774 | 0.00164679 |
| EPB41L1 | 1820.2493 | 0.126019647 | 0.34413593 | 0.36619148 | 0.7142222 | 0.83652428 |
| AAR2 | 137.854985 | -0.153970339 | 0.28874326 | -0.5332431 | 0.59386533 | 0.75469239 |
| DLGAP4 | 1209.10262 | -0.575165378 | 0.29898736 | -1.9237113 | 0.05439078 | 0.16403396 |
| DLGAP4-AS1 | 415.338053 | 0.599588612 | 0.27265071 | 2.19910889 | 0.02787018 | 0.1038653 |
| MYL9 | 113.91541 | -0.412010756 | 0.38985726 | -1.0568246 | 0.29059163 | 0.49071572 |
| RAB5IF | 353.244081 | -2.089279472 | 0.62138891 | -3.3622735 | 0.00077304 | 0.00749893 |
| NDRG3 | 409.593656 | 0.087797417 | 0.27449284 | 0.31985321 | 0.74907961 | 0.85857831 |
| DSN1 | 77.8089737 | 0.059043415 | 0.29688871 | 0.1988739 | 0.84236139 | 0.91687823 |
| SOGA1 | 273.651875 | -1.186778354 | 0.41601587 | -2.8527238 | 0.00433463 | 0.02723966 |
| SAMHD1 | 143.734717 | -0.862186406 | 0.60255097 | -1.4308937 | 0.15246067 | 0.32411759 |
| RBL1 | 141.6811 | 0.621158411 | 0.35250398 | 1.76213162 | 0.07804706 | 0.20857693 |
| MROH8 | 216.555936 | 1.285205918 | 0.45492684 | 2.82508266 | 0.00472684 | 0.02909332 |
| RPN2 | 680.534372 | -0.079940208 | 0.23183351 | -0.3448173 | 0.73023172 | 0.84648502 |
| MANBAL | 212.029949 | -0.268992678 | 0.24082388 | -1.1169685 | 0.2640078 | 0.46207385 |
| SRC | 362.326829 | -0.685607699 | 0.29830766 | -2.2983241 | 0.02154335 | 0.08731304 |
| BLCAP | 875.617818 | 0.310734139 | 0.30917765 | 1.0050343 | 0.31488033 | 0.51522797 |
| CTNNBL1 | 660.274201 | 0.216346884 | 0.19833956 | 1.09079037 | 0.27536514 | 0.47286769 |
| TTI1 | 341.775182 | 0.121275272 | 0.20938545 | 0.57919628 | 0.56245674 | 0.73014439 |
| RPRD1B | 657.4295 | 0.300073191 | 0.22962832 | 1.3067778 | 0.19128819 | 0.37564287 |
| TGM2 | 1211.49377 | -3.29811805 | 0.70500811 | -4.6781278 | 2.90E-06 | 9.28E-05 |
| KIAA1755 | 350.866117 | 0.487420048 | 1.08293642 | 0.4500911 | 0.65264475 | 0.79674044 |
| LBP | 562.037071 | -0.950455801 | 0.48718709 | -1.9509051 | 0.05106833 | 0.15740806 |
| AL391095.2 | 58.4383292 | -1.3428396 | 0.64918735 | -2.0684932 | 0.03859367 | 0.1306504 |
| RALGAPB | 1159.08823 | 0.006424503 | 0.17791412 | 0.03611014 | 0.97119454 | 0.98569779 |
| ARHGAP40 | 99.5322075 | -1.826062547 | 0.57096298 | -3.1982153 | 0.00138281 | 0.01153632 |
| ACTR5 | 128.450995 | 0.155004057 | 0.33079043 | 0.46858688 | 0.63936496 | 0.78803844 |
| PPP1R16B | 1129.91334 | 1.335123831 | 0.80265697 | 1.66338036 | 0.09623632 | 0.23950313 |
| FAM83D | 55.4417425 | 2.058426553 | 0.61413854 | 3.35172998 | 0.00080308 | 0.00768169 |
| DHX35 | 378.436265 | 0.578153103 | 0.20709705 | 2.79170124 | 0.00524317 | 0.0313972 |
| TOP1 | 1682.40505 | -0.42291167 | 0.17776676 | -2.3790256 | 0.01735847 | 0.07472756 |
| PLCG1-AS1 | 50.8615413 | -0.336356074 | 0.50632287 | -0.6643114 | 0.506491 | 0.68655777 |
| PLCG1 | 371.30291 | -0.461626578 | 0.24017216 | -1.9220653 | 0.05459754 | 0.16434269 |
| ZHX3 | 3251.526 | 0.226005868 | 0.26426742 | 0.85521655 | 0.39243128 | 0.58736138 |
| LPIN3 | 1089.60284 | 0.008976098 | 0.50507333 | 0.01777187 | 0.98582085 | 0.99299282 |
| CHD6 | 1610.12631 | 0.355084127 | 0.27545242 | 1.28909425 | 0.19736532 | 0.38334795 |
| PTPRT | 135.206247 | 1.632308737 | 0.56580352 | 2.88493919 | 0.00391489 | 0.02524056 |
| SRSF6 | 669.564135 | 1.070209658 | 0.23974126 | 4.46401942 | 8.04E-06 | 0.00021203 |
| L3MBTL1 | 301.435552 | -1.065612639 | 0.29115656 | -3.65993 | 0.00025228 | 0.00321082 |
| Z98752.3 | 110.033223 | -0.772204713 | 0.46583603 | -1.6576749 | 0.0973831 | 0.24130272 |
| SGK2 | 719.552145 | 0.992838782 | 0.32664149 | 3.03953669 | 0.00236942 | 0.01749875 |
| IFT52 | 349.856407 | 0.130715585 | 0.24491108 | 0.53372671 | 0.59353063 | 0.75444301 |
| OSER1 | 336.948164 | -0.709858023 | 0.36745135 | -1.9318422 | 0.05337899 | 0.16177838 |
| OSER1-DT | 53.1619505 | -0.625518802 | 0.52869472 | -1.183138 | 0.23675447 | 0.43136226 |
| GDAP1L1 | 43.0254449 | -0.872527158 | 0.809207 | -1.0782496 | 0.28092236 | 0.47950097 |
| FITM2 | 150.416471 | 0.221503478 | 0.24404014 | 0.90765183 | 0.3640622 | 0.56189625 |
| R3HDML | 101.502894 | 2.26900251 | 1.0781518 | 2.10452973 | 0.03533226 | 0.12251438 |
| AL117382.1 | 437.764364 | 3.317734879 | 1.56336491 | 2.12217561 | 0.03382299 | 0.11902742 |
| HNF4A | 1942.5735 | 2.765947672 | 0.77235892 | 3.58116881 | 0.00034206 | 0.0040559 |
| HNF4A-AS1 | 1103.69397 | 3.911287151 | 1.27334315 | 3.0716678 | 0.00212866 | 0.01612914 |
| TTPAL | 129.834887 | -0.392227263 | 0.27067323 | -1.4490804 | 0.14731514 | 0.31679458 |
| SERINC3 | 487.665666 | -0.556826998 | 0.15035035 | -3.7035298 | 0.00021262 | 0.00280769 |
| PKIG | 934.32234 | 0.548543424 | 0.23062807 | 2.37847636 | 0.01738435 | 0.074792 |
| YWHAB | 811.388638 | -0.802661526 | 0.30694118 | -2.6150337 | 0.00892186 | 0.04636532 |
| PABPC1L | 216.061524 | 0.544469279 | 0.24409516 | 2.23056157 | 0.02571018 | 0.09832854 |
| TOMM34 | 138.300884 | -0.623798413 | 0.31963566 | -1.9515921 | 0.05098665 | 0.1573332 |
| STK4 | 633.091901 | -0.219446892 | 0.29339599 | -0.7479547 | 0.45448751 | 0.64389254 |
| SLPI | 910.173191 | -4.438006101 | 0.84726173 | -5.238058 | 1.62E-07 | 8.39E-06 |
| SDC4 | 1832.85554 | -0.224736589 | 0.31659321 | -0.7098592 | 0.47779148 | 0.66346751 |
| SYS1-DBNDD | 133.666464 | -0.377674053 | 0.43519834 | -0.8678205 | 0.38549259 | 0.58174027 |
| TP53TG5 | 42.1435191 | -1.257198701 | 0.45339083 | -2.7728808 | 0.00555625 | 0.03279909 |
| PIGT | 169.984469 | -0.145361771 | 0.3176792 | -0.4575741 | 0.64725846 | 0.79306085 |
| WFDC2 | 2362.90836 | -3.087085965 | 0.63227767 | -4.8824846 | 1.05E-06 | 4.02E-05 |
| DNTTIP1 | 180.85602 | -0.241925931 | 0.22970859 | -1.0531863 | 0.29225563 | 0.49231264 |
| SNX21 | 90.4193469 | 0.640481833 | 0.39715857 | 1.61266022 | 0.10681835 | 0.25646507 |
| ACOT8 | 172.744698 | 0.381369477 | 0.19004415 | 2.00674146 | 0.0447772 | 0.14421078 |
| ZSWIM3 | 87.4511977 | 1.078044705 | 0.35434804 | 3.04233292 | 0.00234752 | 0.01739216 |
| NEURL2 | 95.7491594 | -0.587643648 | 0.68485146 | -0.85806 | 0.39085933 | 0.58629953 |
| CTSA | 1000.17366 | -0.308782197 | 0.3104678 | -0.9945708 | 0.31994507 | 0.520351 |
| PLTP | 304.479478 | -2.546508338 | 0.95881748 | -2.6558843 | 0.00791007 | 0.04251897 |
| PCIF1 | 64.8084312 | 0.106852321 | 0.29377926 | 0.36371635 | 0.71606983 | 0.83776419 |
| ZNF335 | 224.442453 | -0.095475439 | 0.27678076 | -0.3449497 | 0.73013219 | 0.84648502 |
| NCOA5 | 512.572118 | -0.242430203 | 0.315014 | -0.7695855 | 0.44154582 | 0.63336825 |
| CD40 | 62.6693535 | -0.986900915 | 0.55950599 | -1.7638791 | 0.07775234 | 0.20807328 |
| SLC35C2 | 291.284766 | -0.364796799 | 0.25611837 | -1.4243289 | 0.15435128 | 0.32695817 |

| | | | | | | |
|---|---|---|---|---|---|---|
| AL133227.1 | 41.9208672 | -0.079537264 | 0.58974986 | -0.1348661 | 0.89271774 | 0.94395449 |
| ELMO2 | 251.437043 | -0.259058492 | 0.21638542 | -1.1972086 | 0.23122525 | 0.42417083 |
| ZNF334 | 239.303933 | 0.538068871 | 0.50517629 | 1.0651111 | 0.28682566 | 0.48651396 |
| SLC13A3 | 13309.4771 | 0.812430993 | 0.42536242 | 1.90997359 | 0.05613661 | 0.16734766 |
| TP53RK | 82.4199782 | -0.978110405 | 0.31118661 | -3.1431635 | 0.00167132 | 0.01334602 |
| EYA2 | 1274.56323 | 2.026552681 | 0.35410723 | 5.72299156 | 1.05E-08 | 8.40E-07 |
| ZMYND8 | 1692.60367 | 0.438547936 | 0.21974553 | 1.99570811 | 0.0459657 | 0.14700231 |
| NCOA3 | 1355.38903 | 0.154954993 | 0.16355438 | 0.94742184 | 0.34342387 | 0.54383762 |
| SULF2 | 1048.17174 | -1.043112928 | 0.41190682 | -2.5324003 | 0.01132846 | 0.0550845 |
| LINC00494 | 54.4922338 | 0.816310081 | 0.65873077 | 1.23921656 | 0.21526531 | 0.4058096 |
| ARFGEF2 | 1821.62751 | -0.204697038 | 0.20762747 | -0.9858861 | 0.324189 | 0.52469696 |
| CSE1L | 447.911167 | -0.08700772 | 0.21685224 | -0.4012304 | 0.68825047 | 0.81976444 |
| STAU1 | 1515.60221 | -0.195498205 | 0.23000575 | -0.8499709 | 0.39534125 | 0.59017017 |
| DDX27 | 264.525146 | -0.456543969 | 0.2810963 | -1.624155 | 0.10434273 | 0.25237302 |
| ZNFX1 | 463.278604 | -1.117243011 | 0.41677719 | -2.6806722 | 0.00734745 | 0.0404139 |
| ZFAS1 | 440.810464 | -1.395580141 | 0.33627943 | -4.1500609 | 3.32E-05 | 0.0006546 |
| B4GALT5 | 4980.15657 | -0.189319093 | 0.30701729 | -0.6166398 | 0.53747231 | 0.70967393 |
| SLC9A8 | 637.244776 | 0.100405255 | 0.37372003 | 0.26866437 | 0.78818798 | 0.88421655 |
| SPATA2 | 73.0819913 | -0.08393164 | 0.31957392 | -0.2626361 | 0.79283109 | 0.8871958 |
| RNF114 | 860.492954 | -0.269089859 | 0.25513048 | -1.0547147 | 0.29155583 | 0.4916955 |
| CEBPB | 552.597635 | -3.098534664 | 0.33818821 | -9.1621606 | 5.09E-20 | 9.96E-17 |
| PTPN1 | 5140.61042 | -1.922266612 | 0.50146522 | -3.8333 | 0.00012644 | 0.0018711 |
| RIPOR3 | 656.650552 | 1.123279804 | 0.37151143 | 3.02354039 | 0.00249836 | 0.01818639 |
| PARD6B | 2030.78741 | 0.740249395 | 0.30503033 | 2.42680588 | 0.0152324 | 0.06792146 |
| BCAS4 | 87.4872846 | -0.559834703 | 0.44008973 | -1.2720922 | 0.20334037 | 0.39134885 |
| ADNP | 1626.47001 | 0.324358226 | 0.24275359 | 1.33616242 | 0.18149619 | 0.36373182 |
| DPM1 | 480.304712 | -0.057084017 | 0.22060816 | -0.2587575 | 0.79582234 | 0.88843413 |
| NFATC2 | 95.7444451 | -0.335333046 | 0.64598442 | -0.5191039 | 0.60368827 | 0.76169806 |
| ATP9A | 364.003411 | -1.975005058 | 0.53974096 | -3.6591721 | 0.00025303 | 0.00321437 |
| ZFP64 | 552.931543 | 0.325951202 | 0.21684857 | 1.50312822 | 0.13280598 | 0.29622814 |
| TSHZ2 | 149.839175 | -1.265370582 | 0.62089807 | -2.0379683 | 0.0415531 | 0.13691548 |
| AL157838.1 | 42.3628495 | -1.024147813 | 0.55600007 | -1.8419922 | 0.06547629 | 0.18571941 |
| ZNF217 | 696.769593 | -0.479017376 | 0.26696289 | -1.794322 | 0.07276179 | 0.19907103 |
| CYP24A1 | 4399.57887 | -1.703145958 | 0.88952527 | -1.9146684 | 0.0555348 | 0.16620426 |
| PFDN4 | 140.722392 | -0.692426078 | 0.27071772 | -2.557742 | 0.01053542 | 0.05243611 |
| DOK5 | 48.1547825 | -0.945081559 | 0.62776048 | -1.5054811 | 0.13220043 | 0.29551064 |
| FAM210B | 111.476158 | -0.341848201 | 0.31690241 | -1.0787176 | 0.28071365 | 0.47940967 |
| AURKA | 37.8631416 | 0.374674688 | 0.44038141 | 0.85079589 | 0.39488275 | 0.58961414 |
| CSTF1 | 106.085842 | -0.15121811 | 0.19424722 | -0.7784828 | 0.43628447 | 0.62849724 |
| RTF2 | 331.031708 | -0.213178377 | 0.26156203 | -0.8150203 | 0.41506067 | 0.60853959 |
| BMP7 | 74.3065397 | -0.641124933 | 0.7144625 | -0.8973528 | 0.36953069 | 0.56695645 |
| RAE1 | 210.558296 | -0.056256718 | 0.25122147 | -0.2239328 | 0.82280962 | 0.90508398 |
| RBM38 | 98.5521947 | 0.21235429 | 0.37724932 | 0.56290171 | 0.57350182 | 0.73867594 |
| PCK1 | 6748.272 | 1.348849034 | 0.89151963 | 1.51297737 | 0.13028541 | 0.29285373 |
| PMEPA1 | 137.087676 | -1.574700436 | 0.78071513 | -2.0169975 | 0.04369578 | 0.14164143 |
| RAB22A | 811.449333 | -0.538349355 | 0.23364325 | -2.3041511 | 0.02121416 | 0.08628561 |
| VAPB | 963.652915 | -0.25133826 | 0.15087633 | -1.6658561 | 0.09574208 | 0.23882914 |
| STX16-NPEPI | 65.5033411 | 0.441857048 | 0.5088908 | 0.86827478 | 0.38524392 | 0.58152841 |
| NPEPL1 | 182.116987 | 0.423442516 | 0.40765067 | 1.03873866 | 0.29892629 | 0.49896259 |
| GNAS | 1255.15488 | -1.121398874 | 0.23840232 | -4.7038086 | 2.55E-06 | 8.37E-05 |
| NELFCD | 375.786105 | -0.196669951 | 0.20093334 | -0.9787821 | 0.32768767 | 0.52749441 |
| CTSZ | 547.103908 | -1.740413067 | 0.53870122 | -3.2307576 | 0.00123463 | 0.01059012 |
| ATP5F1E | 1135.76264 | -0.988557537 | 0.41462039 | -2.3842473 | 0.0171141 | 0.07395389 |
| PRELID3B | 129.53436 | -0.546590465 | 0.29608865 | -1.8460365 | 0.0648869 | 0.18463873 |
| SYCP2 | 124.617371 | 0.742689798 | 0.57152344 | 1.29949141 | 0.19377534 | 0.37885739 |
| FAM217B | 154.207046 | -0.039305248 | 0.31284074 | -0.1256398 | 0.90001706 | 0.94743919 |
| PPP1R3D | 38.3334864 | -1.047028826 | 0.55093572 | -1.9004555 | 0.05737337 | 0.16988912 |
| CDH26 | 93.2751976 | -0.058933914 | 0.49824317 | -0.1182834 | 0.90584308 | 0.95068694 |
| MIR646HG | 928.438677 | 0.570591176 | 0.79136844 | 0.72101836 | 0.47089822 | 0.65738999 |
| CDH4 | 85.9038376 | -2.203086455 | 0.69887145 | -3.1523486 | 0.00161963 | 0.01308211 |
| TAF4 | 738.198901 | 0.802437263 | 0.27352252 | 2.93371549 | 0.00334931 | 0.0224165 |
| LSM14B | 204.799952 | 0.88874616 | 0.29652814 | 2.99717305 | 0.00272496 | 0.01929288 |
| PSMA7 | 567.411327 | -1.088953234 | 0.29128303 | -3.7384712 | 0.00018514 | 0.00250381 |
| SS18L1 | 705.485758 | 0.057470833 | 0.35088808 | 0.16378679 | 0.86989899 | 0.93283939 |
| MTG2 | 441.205287 | 0.345556888 | 0.21171045 | 1.63221458 | 0.10263429 | 0.25027051 |
| OSBPL2 | 454.765915 | -0.190053479 | 0.20477097 | -0.9281271 | 0.35334166 | 0.55189396 |
| ADRM1 | 136.720635 | -1.014702458 | 0.50751127 | -1.9993693 | 0.04556841 | 0.14614624 |
| AL354836.1 | 116.896656 | 0.982815052 | 0.55419635 | 1.77340586 | 0.07616148 | 0.2054356 |
| LAMA5 | 926.622804 | -0.033651909 | 0.28443466 | -0.1183116 | 0.9058208 | 0.95068694 |
| RPS21 | 134.394704 | -1.133894207 | 0.45737838 | -2.4791163 | 0.01317083 | 0.06121825 |
| CABLES2 | 42.2053352 | -0.503377302 | 0.44893892 | -1.1212601 | 0.26217716 | 0.46007129 |
| SLCO4A1 | 1365.05602 | -4.121110188 | 0.38897448 | -10.594809 | 3.15E-26 | 2.16E-22 |
| MRGBP | 132.077581 | -0.279051295 | 0.38699481 | -0.7210724 | 0.47086494 | 0.65738999 |
| OGFR | 482.474384 | -1.049012463 | 0.29538716 | -3.5513136 | 0.00038331 | 0.00442897 |
| TCFL5 | 99.7174367 | 0.054131575 | 0.3127902 | 0.17306033 | 0.86260401 | 0.92800511 |
| DIDO1 | 1062.51624 | -0.318367997 | 0.17912301 | -1.7773707 | 0.07550726 | 0.20417796 |
| GID8 | 277.399444 | -0.230397949 | 0.2291342 | -1.0055153 | 0.31464878 | 0.51515657 |
| YTHDF1 | 424.891173 | -0.358920839 | 0.32391658 | -1.1080657 | 0.26783345 | 0.46575866 |
| NKAIN4 | 734.8433 | 0.417533282 | 0.79072803 | 0.52803653 | 0.59747398 | 0.75757406 |
| AL121827.2 | 60.8398933 | -0.587594605 | 0.77383272 | -0.7593303 | 0.44765502 | 0.63836947 |
| ARFGAP1 | 589.374212 | -0.306261427 | 0.47529824 | -0.6443563 | 0.5193444 | 0.69597507 |
| CHRNA4 | 105.261686 | 3.966617812 | 0.94523526 | 4.19643444 | 2.71E-05 | 0.00055555 |
| PPDPF | 664.549437 | -0.582533888 | 0.39227834 | -1.4850014 | 0.13754345 | 0.30315293 |
| HELZ2 | 117.243131 | -1.321823755 | 0.90461933 | -1.4611934 | 0.14396239 | 0.3116381 |
| GMEB2 | 228.973617 | -0.108727983 | 0.20689934 | -0.5255115 | 0.59922766 | 0.75897371 |
| MHENCR | 121.129763 | 0.606202086 | 0.43892867 | 1.38109475 | 0.16724983 | 0.34521039 |
| STMN3 | 80.7632985 | -1.222047373 | 0.36673914 | -3.3321978 | 0.00086163 | 0.00810038 |
| ARFRP1 | 195.556655 | 0.418366153 | 0.22870606 | 1.82927441 | 0.06735851 | 0.1890424 |
| SLC2A4RG | 497.643001 | 1.458989741 | 0.49515539 | 2.94652903 | 0.00321362 | 0.02173648 |
| ZBTB46 | 493.282293 | 0.08794953 | 0.36687476 | 0.2397263 | 0.81054244 | 0.89723132 |
| TPD52L2 | 461.820112 | -0.695838977 | 0.3793633 | -1.8342285 | 0.06662008 | 0.1877107 |
| DNAJC5 | 1096.09446 | -0.626655582 | 0.39469872 | -1.5876808 | 0.11235854 | 0.26548122 |
| UCKL1 | 228.215998 | 0.615358049 | 0.27815797 | 2.21226107 | 0.02694863 | 0.101591 |
| ZNF512B | 290.919517 | 1.267144922 | 0.33867252 | 3.74150498 | 0.00018292 | 0.00248036 |
| SAMD10 | 37.6032041 | 0.025931362 | 0.47895217 | 0.05414186 | 0.95682214 | 0.97808644 |
| PRPF6 | 459.263101 | 0.08682974 | 0.21603212 | 0.40192976 | 0.68773572 | 0.81938377 |
| TCEA2 | 112.70485 | 0.879945049 | 0.31789914 | 2.76800071 | 0.00564013 | 0.03313729 |
| PCMTD2 | 499.541425 | 0.424565362 | 0.22238489 | 1.90914663 | 0.05624318 | 0.16761238 |
| LINC00266-1 | 48.2294079 | -0.079136147 | 0.26170073 | -0.3023918 | 0.76235343 | 0.86754491 |
| PLPP2 | 54.4766496 | -3.603374988 | 0.83646351 | -4.3078687 | 1.65E-05 | 0.00037909 |
| MIER2 | 97.3514159 | 0.224716416 | 0.49239126 | 0.45637775 | 0.64811836 | 0.79355195 |
| SHC2 | 130.56683 | -1.503793001 | 0.66232013 | -2.2704927 | 0.02317771 | 0.09147815 |
| MADCAM1 | 52.6763685 | 0.515117977 | 0.37752075 | 1.36447592 | 0.17241783 | 0.35183282 |
| TPGS1 | 219.053179 | -0.774597094 | 0.49800032 | -1.5554148 | 0.1198473 | 0.27702309 |
| CDC34 | 178.700312 | -0.0775846 | 0.29991689 | -0.258687 | 0.79587675 | 0.88843413 |
| BSG | 906.825527 | -0.508190662 | 0.32687801 | -1.5546798 | 0.12002234 | 0.27738091 |
| POLRMT | 86.440799 | 0.950995793 | 0.26791464 | 3.5496223 | 0.00038578 | 0.00444365 |
| RNF126 | 315.007063 | -0.910352643 | 0.2729108 | -3.335715 | 0.0008508 | 0.00802638 |
| FSTL3 | 230.404726 | -1.430950949 | 0.76074784 | -1.8809793 | 0.05997473 | 0.1745825 |
| PALM | 343.846587 | 2.910612187 | 0.56088124 | 5.18935556 | 2.11E-07 | 1.02E-05 |
| MISP | 162.458621 | 0.295814799 | 0.35177972 | 0.84090918 | 0.40039881 | 0.59467619 |
| PTBP1 | 790.006207 | 0.074605205 | 0.29767566 | 0.25062581 | 0.80210342 | 0.89247761 |
| MED16 | 155.339251 | 0.433750014 | 0.36099605 | 1.20153673 | 0.22954307 | 0.42266884 |

| | | | | | | |
|---|---|---|---|---|---|---|
| R3HDM4 | 126.784639 | -0.209460546 | 0.40963795 | -0.5113309 | 0.60911936 | 0.76570058 |
| ARID3A | 387.897694 | 0.558402194 | 0.31889683 | 1.7510434 | 0.07993843 | 0.21173952 |
| WDR18 | 171.31115 | 0.611843981 | 0.41713448 | 1.46677873 | 0.14243626 | 0.30981487 |
| TMEM259 | 649.780518 | -0.106626132 | 0.22832558 | -0.4669916 | 0.64050587 | 0.78888372 |
| CNN2 | 55.5865101 | -0.693599544 | 0.5276489 | -1.3145096 | 0.18867477 | 0.37263859 |
| ABCA7 | 131.35682 | 0.164996845 | 0.51313037 | 0.32154956 | 0.74779396 | 0.85769676 |
| POLR2E | 347.406671 | -0.441685405 | 0.24788052 | -1.781848 | 0.07477403 | 0.20291578 |
| GPX4 | 1318.17806 | -0.02112782 | 0.38151385 | -0.0553789 | 0.9558366 | 0.97756553 |
| SBNO2 | 557.199258 | -1.151530481 | 0.33051975 | -3.483999 | 0.00049398 | 0.0053162 |
| STK11 | 340.041948 | 0.207805368 | 0.32063785 | 0.64809993 | 0.51692032 | 0.6940373 |
| ATP5F1D | 756.665044 | -0.508413692 | 0.3565842 | -1.4257886 | 0.15392936 | 0.32636683 |
| MIDN | 998.480509 | -0.909568013 | 0.44277372 | -2.0542502 | 0.03995148 | 0.13346025 |
| CIRBP | 909.88555 | 0.891700428 | 0.3652012 | 2.44166897 | 0.01461954 | 0.06587446 |
| C19orf24 | 228.590511 | -0.577160521 | 0.46226905 | -1.2485381 | 0.21183406 | 0.40149467 |
| MUM1 | 655.135547 | 0.268244759 | 0.28409869 | 0.94419568 | 0.34506966 | 0.5454262 |
| NDUFS7 | 264.595968 | 0.085506978 | 0.28320929 | 0.30192151 | 0.7627119 | 0.86770535 |
| GAMT | 280.383079 | -0.048354196 | 0.41282176 | -0.1171309 | 0.9067563 | 0.95109493 |
| DAZAP1 | 851.614191 | -0.070886174 | 0.21330841 | -0.3323178 | 0.73964933 | 0.85217898 |
| RPS15 | 1827.271 | -1.02191169 | 0.41234422 | -2.4782976 | 0.0132011 | 0.06129656 |
| C19orf25 | 536.460085 | -0.475696073 | 0.35492438 | -1.3402744 | 0.18015614 | 0.3616579 |
| PCSK4 | 56.6560043 | 1.261923344 | 0.50705042 | 2.48875317 | 0.01281919 | 0.06011381 |
| REEP6 | 56.398414 | 1.015732219 | 0.5293502 | 1.91882845 | 0.05500605 | 0.16516274 |
| MBD3 | 222.946799 | 0.066338616 | 0.44040955 | 0.15062938 | 0.88026809 | 0.937662 |
| UQCR11 | 26.8183948 | -1.140729491 | 0.73847733 | -1.5447048 | 0.12241768 | 0.28088034 |
| TCF3 | 238.302501 | -0.069834629 | 0.43098314 | -0.1620356 | 0.87127779 | 0.93338059 |
| REXO1 | 448.06127 | 0.612398868 | 0.32392755 | 1.89054269 | 0.05868542 | 0.17217487 |
| KLF16 | 79.6213133 | -0.862151759 | 0.55216513 | -1.561402 | 0.11842892 | 0.2747859 |
| AC012615.1 | 32.3522274 | -0.43012807 | 0.41478295 | -1.0369955 | 0.29973794 | 0.50006182 |
| ABHD17A | 234.517949 | -0.22351236 | 0.36105075 | -0.6190608 | 0.53587632 | 0.70860168 |
| SCAMP4 | 282.813891 | -0.208697302 | 0.33903529 | -0.6155622 | 0.53818353 | 0.70990443 |
| CSNK1G2 | 505.514202 | -0.253709753 | 0.35472592 | -0.7152275 | 0.47446845 | 0.6606602 |
| BTBD2 | 462.917857 | 0.706241145 | 0.32374912 | 2.18144576 | 0.02915046 | 0.10692593 |
| MKNK2 | 260.594381 | -0.354654735 | 0.40692572 | -0.8715466 | 0.38345576 | 0.58007152 |
| MOB3A | 300.252245 | -0.174411721 | 0.37380449 | -0.4665854 | 0.64079652 | 0.78905143 |
| AP3D1 | 1150.07553 | -0.225743364 | 0.37040308 | -0.6094533 | 0.54222404 | 0.71320074 |
| DOT1L | 692.173783 | 0.103669554 | 0.45668646 | 0.22700378 | 0.8204208 | 0.90347135 |
| PLEKHJ1 | 425.217278 | -0.287885835 | 0.26447369 | -1.0885235 | 0.27636406 | 0.47404858 |
| SF3A2 | 148.548419 | -0.266950767 | 0.39306926 | -0.6791444 | 0.4970464 | 0.67892526 |
| AC005258.1 | 44.5633708 | -3.253994533 | 0.92896429 | -3.5028198 | 0.00046036 | 0.00505217 |
| LSM7 | 347.776426 | -1.004207989 | 0.39165104 | -2.5640376 | 0.01034623 | 0.05180591 |
| SPPL2B | 513.811443 | 0.38004643 | 0.22885537 | 1.66064018 | 0.09678573 | 0.24042082 |
| TMPRSS9 | 43.5528512 | -0.489742842 | 0.47255449 | -1.0363733 | 0.30002804 | 0.50030223 |
| TIMM13 | 285.591028 | -0.967670377 | 0.38524832 | -2.5118095 | 0.01201139 | 0.05736301 |
| LMNB2 | 84.0739723 | -0.567914484 | 0.42845097 | -1.3255064 | 0.18500326 | 0.36794391 |
| GADD45B | 681.925709 | -1.464593066 | 0.61196238 | -2.3932731 | 0.01669881 | 0.07266389 |
| GNG7 | 575.555035 | 0.500682012 | 0.61981016 | 0.80779897 | 0.41920632 | 0.61258646 |
| SGTA | 273.21329 | -0.356129953 | 0.38152712 | -0.9334329 | 0.35059651 | 0.55003164 |
| THOP1 | 135.960312 | 0.397955552 | 0.27200515 | 1.46304417 | 0.14345529 | 0.3110808 |
| ZNF554 | 82.7455472 | 0.638986016 | 0.25476964 | 2.50809324 | 0.01213846 | 0.05775145 |
| ZNF555 | 110.450288 | -0.310421855 | 0.30506661 | -1.0175544 | 0.30888979 | 0.50906843 |
| ZNF57 | 78.6780684 | -0.087403168 | 0.39152191 | -0.2232395 | 0.82334908 | 0.90531413 |
| AC119403.1 | 33.2697599 | 0.582707251 | 0.4302194 | 1.35444205 | 0.17559539 | 0.35574102 |
| ZNF77 | 49.5450449 | 0.349187345 | 0.35248453 | 0.99064588 | 0.32185853 | 0.52222003 |
| TLE2 | 50.3129959 | -0.680776005 | 0.64039444 | -1.0630573 | 0.28775595 | 0.48754893 |
| AES | 657.466931 | -0.356556019 | 0.37135024 | -0.960161 | 0.33697418 | 0.5370278 |
| GNA11 | 363.346782 | -0.038145264 | 0.32635593 | -0.1168824 | 0.90695325 | 0.95122652 |
| NCLN | 227.138985 | 0.053679916 | 0.36820071 | 0.14578982 | 0.88408731 | 0.9399771 |
| NFIC | 1157.34925 | 0.3052038 | 0.28642327 | 1.06556913 | 0.28661846 | 0.48629422 |
| SMIM24 | 506.245038 | 0.5793179 | 0.57144769 | 1.0137724 | 0.31069137 | 0.51104512 |
| DOHH | 97.0963454 | 0.003534861 | 0.42357444 | 0.00834531 | 0.99334148 | 0.99654041 |
| FZR1 | 175.691685 | -0.007770946 | 0.2825366 | -0.0275042 | 0.97805758 | 0.98906609 |
| MFSD12 | 838.149726 | 0.267080984 | 0.30955376 | 0.86279353 | 0.38825099 | 0.58387159 |
| C19orf71 | 56.2155709 | 0.823211356 | 0.40558089 | 2.02970943 | 0.04238608 | 0.138859 |
| AC005786.3 | 131.332255 | 0.274602011 | 0.35692483 | 0.76935531 | 0.44168241 | 0.63341542 |
| HMG20B | 229.668692 | 0.192085264 | 0.23012536 | 0.83469837 | 0.40388756 | 0.5979787 |
| CACTIN | 70.4457001 | 0.011423511 | 0.39395123 | 0.02899727 | 0.97686677 | 0.9886052 |
| PIP5K1C | 235.180082 | -0.314504076 | 0.33352316 | -0.9429752 | 0.34569358 | 0.54612944 |
| TJP3 | 191.489904 | -0.433921055 | 0.51096071 | -0.8492259 | 0.39575562 | 0.59046721 |
| APBA3 | 91.8131764 | 0.421006826 | 0.26673496 | 1.5783714 | 0.11448031 | 0.2686035 |
| MRPL54 | 190.571129 | -0.758236957 | 0.47044894 | -1.6117306 | 0.10702057 | 0.25681565 |
| DAPK3 | 217.015862 | -0.61648559 | 0.31622649 | -1.9495065 | 0.05123497 | 0.15774433 |
| EEF2 | 1809.87834 | -0.96283116 | 0.42444062 | -2.2684708 | 0.02330053 | 0.09174958 |
| PIAS4 | 201.240504 | 0.226913543 | 0.35002934 | 0.64827007 | 0.51681029 | 0.69403559 |
| ZBTB7A | 569.397117 | -0.41105643 | 0.42468171 | -0.9679165 | 0.33308607 | 0.53261909 |
| MAP2K2 | 444.846089 | -0.420404725 | 0.26407023 | -1.5920186 | 0.11138052 | 0.26403572 |
| SIRT6 | 132.946898 | -0.581501152 | 0.34194351 | -1.7005767 | 0.08902251 | 0.22765514 |
| ANKRD24 | 115.011505 | 2.015045208 | 0.48303025 | 4.17167497 | 3.02E-05 | 0.0006065 |
| YJU2 | 101.464661 | 0.339881742 | 0.36139441 | 0.94047316 | 0.34697491 | 0.54716809 |
| STAP2 | 245.511027 | -0.260697401 | 0.40636618 | -0.6415332 | 0.52117631 | 0.69702056 |
| MPND | 83.4263031 | 0.72122601 | 0.3359196 | 2.14701974 | 0.03179171 | 0.11383724 |
| AC007292.3 | 46.1714289 | -0.007881124 | 0.46514995 | -0.0169432 | 0.98648194 | 0.99351271 |
| SH3GL1 | 890.219282 | -0.533999951 | 0.30750226 | -1.7365725 | 0.08246266 | 0.21657707 |
| CHAF1A | 52.1121778 | 0.005748528 | 0.43909439 | 0.01309178 | 0.98955457 | 0.99493132 |
| UBXN6 | 348.312735 | -0.441191259 | 0.34407326 | -1.2822596 | 0.19975158 | 0.38650408 |
| HDGFL2 | 191.490066 | 0.528434005 | 0.25080456 | 2.10695535 | 0.03512145 | 0.12199942 |
| PLIN4 | 99.9296965 | -0.538118139 | 0.59308718 | -0.9073171 | 0.36423913 | 0.56199352 |
| PLIN5 | 1091.30103 | 0.33564797 | 0.58424985 | 0.57449389 | 0.56563363 | 0.73247225 |
| TNFAIP8L1 | 204.665398 | 1.104823721 | 0.27724421 | 3.98501994 | 6.75E-05 | 0.00114464 |
| MYDGF | 462.409396 | -1.149521534 | 0.30238131 | -3.8015628 | 0.00014379 | 0.00203814 |
| DPP9 | 417.956353 | -0.298508244 | 0.35962335 | -0.830058 | 0.40650599 | 0.60029924 |
| FEM1A | 52.352068 | -0.351957491 | 0.48704527 | -0.7226381 | 0.46990223 | 0.65683764 |
| TICAM1 | 408.42464 | -0.395859998 | 0.33232299 | -1.1911905 | 0.2335788 | 0.42762983 |
| PLIN3 | 267.425918 | -1.731599356 | 0.37472593 | -4.6209756 | 3.82E-06 | 0.00011569 |
| KDM4B | 1271.12916 | -0.538171393 | 0.21253072 | -2.5322052 | 0.01133477 | 0.05509419 |
| PTPRS | 781.687232 | 0.515115938 | 0.355587 | 1.44863544 | 0.14743942 | 0.31691267 |
| TINCR | 94.6112564 | -1.332905228 | 0.3675146 | -3.626809 | 0.00028695 | 0.00353702 |
| SAFB2 | 1499.79201 | 0.903527663 | 0.20112312 | 4.49241075 | 7.04E-06 | 0.0001889 |
| SAFB | 1038.60592 | 0.140572859 | 0.18886872 | 0.74428871 | 0.45670183 | 0.64552736 |
| RPL36 | 1067.29409 | -1.061500468 | 0.43083421 | -2.4638259 | 0.01374629 | 0.06305902 |
| C19orf70 | 346.409523 | -0.382203942 | 0.25799545 | -1.4814368 | 0.13849022 | 0.30456515 |
| LONP1 | 605.14703 | -1.083625625 | 0.45487879 | -2.3822294 | 0.01720817 | 0.07422041 |
| PRR22 | 56.0783635 | 1.072796363 | 0.54027246 | 1.98565804 | 0.04707131 | 0.14956106 |
| DUS3L | 76.1933359 | -0.796077266 | 0.39277826 | -2.0267855 | 0.04268436 | 0.13956918 |
| NRTN | 90.3417286 | 0.9673542 | 0.57721725 | 1.67589275 | 0.0937592 | 0.23571272 |
| FUT6 | 1516.29576 | 2.55991412 | 0.57761665 | 4.43185651 | 9.34E-06 | 0.00024116 |
| FUT3 | 175.314833 | 3.58827124 | 0.52432795 | 6.84356278 | 7.72E-12 | 1.74E-09 |
| AC024592.2 | 37.7405207 | 3.374345549 | 0.60226671 | 5.60274296 | 2.11E-08 | 1.56E-06 |
| AC104532.2 | 62.4937239 | 0.456267622 | 0.39362109 | 1.15915441 | 0.24639325 | 0.44202291 |
| CAPS | 109.659717 | 1.02374061 | 0.48217301 | 2.12318109 | 0.03373868 | 0.11879014 |
| RANBP3 | 254.011503 | -0.049448177 | 0.22161979 | -0.2231217 | 0.82344081 | 0.90534236 |
| RFX2 | 375.948722 | 0.158693293 | 0.41841383 | 0.37927354 | 0.70448475 | 0.83047014 |
| ACSBG2 | 42.6247744 | 1.181257488 | 0.63431323 | 1.86226209 | 0.06256616 | 0.17982686 |
| MLLT1 | 259.296218 | 0.079398414 | 0.34505052 | 0.23010664 | 0.81800891 | 0.90168461 |

| | | | | | | |
|---|---|---|---|---|---|---|
| CLPP | 157.025417 | -0.572843281 | 0.22573399 | -2.5376918 | 0.01115862 | 0.05449168 |
| ALKBH7 | 163.873551 | -0.258587049 | 0.48055627 | -0.5380994 | 0.59050842 | 0.75237952 |
| GTF2F1 | 225.252465 | -0.366478733 | 0.2954816 | -1.240276 | 0.21487333 | 0.40534939 |
| KHSRP | 350.845212 | 0.172968451 | 0.36156531 | 0.47838786 | 0.63237418 | 0.78329594 |
| SLC25A23 | 246.04701 | -0.271257828 | 0.32797314 | -0.8270733 | 0.40819552 | 0.60201592 |
| CRB3 | 76.1012053 | -0.945123883 | 0.4499683 | -2.1004232 | 0.03569163 | 0.12334168 |
| DENND1C | 46.0629385 | 0.061459931 | 0.48339084 | 0.12714335 | 0.89882694 | 0.94743919 |
| AC010503.4 | 113.475575 | -0.164169644 | 0.41184503 | -0.3986199 | 0.69017326 | 0.82084207 |
| C3 | 1550.47717 | -3.249730035 | 0.8100493 | -4.0117682 | 6.03E-05 | 0.00105097 |
| GPR108 | 86.6457693 | -0.670150512 | 0.34594397 | -1.9371649 | 0.05272519 | 0.16051116 |
| TRIP10 | 504.301216 | -0.173867311 | 0.31449962 | -0.5528379 | 0.58037441 | 0.74375393 |
| SH2D3A | 252.715616 | -0.286311039 | 0.24719934 | -1.1582193 | 0.24677457 | 0.44256694 |
| ZNF557 | 129.428795 | 0.201846345 | 0.31344805 | 0.64395471 | 0.5196048 | 0.69597507 |
| INSR | 9557.54471 | 1.453291549 | 0.31823195 | 4.56676825 | 4.95E-06 | 0.00014171 |
| PEX11G | 53.8256537 | 1.190141057 | 0.48258718 | 2.46616797 | 0.01365673 | 0.06275319 |
| ZNF358 | 183.346582 | 0.235748077 | 0.42011137 | 0.56115614 | 0.5746911 | 0.73951285 |
| MCOLN1 | 176.81789 | -0.310085993 | 0.47485712 | -0.653009 | 0.51375045 | 0.69196393 |
| PNPLA6 | 407.025161 | -0.087776639 | 0.33017602 | -0.265848 | 0.79035627 | 0.8860758 |
| CAMSAP3 | 572.847086 | 0.774455849 | 0.27230406 | 2.84408483 | 0.00445392 | 0.02778783 |
| XAB2 | 129.168211 | 0.3120554 | 0.2250164 | 1.38681181 | 0.16549915 | 0.34277681 |
| STXBP2 | 241.971107 | 0.389281006 | 0.37958428 | 1.02554563 | 0.3051058 | 0.50529195 |
| EVI5L | 181.802525 | 0.586215988 | 0.242759 | 2.41480636 | 0.01574358 | 0.06970194 |
| MAP2K7 | 458.251007 | -0.52286445 | 0.46078319 | -1.1347299 | 0.25648852 | 0.45375428 |
| SNAPC2 | 43.0516504 | -0.428048115 | 0.33233311 | -1.2880092 | 0.19774275 | 0.38391783 |
| TIMM44 | 248.668973 | -0.376424975 | 0.30778264 | -1.2230221 | 0.22132134 | 0.4134786 |
| ELAVL1 | 724.086162 | -0.134305989 | 0.2530769 | -0.5306924 | 0.59563195 | 0.75582107 |
| AC008946.1 | 38.3116771 | 1.069327302 | 0.35975415 | 2.97238349 | 0.00295497 | 0.02047716 |
| CERS4 | 1342.17369 | 1.220354862 | 0.28036658 | 4.35271155 | 1.34E-05 | 0.00032222 |
| AC022146.2 | 280.324851 | 0.608683476 | 0.45239803 | 1.34546004 | 0.17847672 | 0.35946591 |
| CD320 | 125.678876 | -0.350645585 | 0.35869215 | -0.9775669 | 0.32828856 | 0.5282489 |
| RPS28 | 492.755486 | -0.570025031 | 0.33871389 | -1.6829101 | 0.09239249 | 0.23365755 |
| KANK3 | 41.1367956 | -0.06166164 | 0.59928392 | -0.1028922 | 0.91804853 | 0.95613488 |
| ANGPTL4 | 1470.25749 | -1.548289531 | 0.9016937 | -1.7170903 | 0.08596267 | 0.22236088 |
| RAB11B-AS1 | 138.945188 | 0.775434225 | 0.33565057 | 2.31024255 | 0.02087473 | 0.08508175 |
| RAB11B | 316.278294 | 0.368423124 | 0.28536778 | 1.29104668 | 0.19668749 | 0.38248 |
| 2-Mar | 194.356352 | 0.187459578 | 0.26518749 | 0.70689449 | 0.47963205 | 0.66481737 |
| HNRNPM | 1396.97666 | -0.017084979 | 0.16921647 | -0.1009652 | 0.91957806 | 0.95690112 |
| ZNF414 | 58.7549707 | 0.078018617 | 0.48332283 | 0.16142134 | 0.87176156 | 0.93353404 |
| ADAMTS10 | 42.8750264 | -1.103664925 | 0.71364849 | -1.5465106 | 0.12198131 | 0.28025441 |
| ZNF558 | 98.1793019 | -0.224286826 | 0.29716082 | -0.7547658 | 0.45038952 | 0.64073577 |
| ZNF317 | 179.213891 | -0.095796657 | 0.20859199 | -0.4592538 | 0.64605194 | 0.79229077 |
| ZNF699 | 57.0972389 | -0.386403763 | 0.33967797 | -1.1375591 | 0.25530463 | 0.45232479 |
| ZNF559-ZNF | 71.0505524 | -0.268575141 | 0.54694498 | -0.491046 | 0.62339392 | 0.77645256 |
| ZNF266 | 190.485972 | -0.10760566 | 0.32499695 | -0.3310974 | 0.74057088 | 0.85259576 |
| ZNF426 | 100.523931 | -0.247841185 | 0.32854633 | -0.7543569 | 0.45063496 | 0.64075861 |
| ZNF121 | 282.742008 | -0.574314727 | 0.30563699 | -1.8790747 | 0.0602343 | 0.17514458 |
| ZNF561 | 192.446724 | -0.025745326 | 0.23233893 | -0.1108093 | 0.91176753 | 0.95319043 |
| ZNF562 | 420.744435 | -0.300304895 | 0.27351078 | -1.0979637 | 0.27222036 | 0.47033143 |
| ZNF846 | 368.318775 | 0.49330157 | 0.33881571 | 1.45595835 | 0.14540413 | 0.31386684 |
| FBXL12 | 245.573778 | 0.453986492 | 0.188052 | 2.41415402 | 0.01577179 | 0.06975927 |
| UBL5 | 218.516874 | -0.724530801 | 0.42057659 | -1.7227083 | 0.08494128 | 0.22059305 |
| PIN1 | 180.934413 | -0.169415997 | 0.27381181 | -0.6187315 | 0.53609324 | 0.70869254 |
| C19orf66 | 254.966408 | -0.655322087 | 0.65612093 | -0.9987825 | 0.31790008 | 0.5180048 |
| EIF3G | 200.584825 | -0.456618078 | 0.44934491 | -1.0161862 | 0.30954074 | 0.50977712 |
| DNMT1 | 166.485091 | -0.343235565 | 0.35125502 | -0.9771691 | 0.32848543 | 0.52840626 |
| MRPL4 | 221.968342 | -0.320172484 | 0.24900212 | -1.2858223 | 0.19850507 | 0.38480603 |
| AC011511.2 | 141.143144 | -2.663268002 | 0.52148728 | -5.1070622 | 3.27E-07 | 1.49E-05 |
| ICAM1 | 531.216988 | -3.213543484 | 0.77863991 | -4.127124 | 3.67E-05 | 0.00070518 |
| RAVER1 | 72.3008413 | 0.362096789 | 0.42402348 | 0.85395458 | 0.39313016 | 0.58802176 |
| TYK2 | 295.747646 | 0.17420196 | 0.24322501 | 0.71621731 | 0.47385718 | 0.66016796 |
| CDC37 | 530.216575 | -0.554713433 | 0.34972713 | -1.5861321 | 0.11270934 | 0.26591198 |
| KEAP1 | 257.037412 | -0.274368836 | 0.42744034 | -0.641888 | 0.52094589 | 0.69678037 |
| ATG4D | 214.125836 | 0.92817672 | 0.25709325 | 3.61027267 | 0.00030588 | 0.00372678 |
| KRI1 | 200.592252 | -0.072412265 | 0.3629519 | -0.1995093 | 0.8418644 | 0.91655563 |
| CDKN2D | 62.0307836 | -0.123126669 | 0.4437951 | -0.2774404 | 0.781442 | 0.88006125 |
| AP1M2 | 286.348547 | 0.385457647 | 0.45879376 | 0.8401545 | 0.40082176 | 0.5949339 |
| SLC44A2 | 128.233855 | -0.033332297 | 0.63959846 | -0.0521144 | 0.95843753 | 0.9788247 |
| ILF3-DT | 224.885435 | 0.177762664 | 0.34414167 | 0.5165392 | 0.60547786 | 0.76301233 |
| ILF3 | 712.139096 | -0.069520355 | 0.3679798 | -0.1889244 | 0.85015209 | 0.92118851 |
| QTRT1 | 354.460212 | 1.280191994 | 0.31490539 | 4.06532256 | 4.80E-05 | 0.00087663 |
| DNM2 | 2414.89571 | 0.677697489 | 0.25994902 | 2.60703996 | 0.00913287 | 0.04722152 |
| TMED1 | 110.384131 | -0.458775346 | 0.31777879 | -1.4436941 | 0.14882505 | 0.31894074 |
| CARM1 | 317.2495 | -0.084930928 | 0.28184081 | -0.3013436 | 0.76315249 | 0.86789421 |
| YIPF2 | 192.467405 | -0.676964987 | 0.37735092 | -1.7939932 | 0.07281425 | 0.19913507 |
| TIMM29 | 132.322247 | 0.413633271 | 0.2276299 | 1.8171307 | 0.0691971 | 0.19266396 |
| SMARCA4 | 1175.13514 | 0.259786659 | 0.34543387 | 0.75205902 | 0.45201557 | 0.64143458 |
| LDLR | 293.602538 | -1.590410447 | 0.60595142 | -2.6246501 | 0.00867381 | 0.04539589 |
| KANK2 | 196.06177 | 0.103381129 | 0.44884546 | 0.23032678 | 0.81783785 | 0.90160299 |
| DOCK6 | 341.063438 | -0.969732018 | 0.4180352 | -2.3197377 | 0.02035507 | 0.08356004 |
| RAB3D | 100.181955 | -0.759159821 | 0.46030562 | -1.6492517 | 0.09909608 | 0.24421252 |
| CCDC159 | 97.2162908 | 0.849614799 | 0.49683762 | 1.71004521 | 0.08725751 | 0.22473483 |
| PLPPR2 | 47.6904029 | -0.185866743 | 0.29298571 | -0.6343884 | 0.52582737 | 0.70084961 |
| SWSAP1 | 67.0739944 | 0.035817218 | 0.44652392 | 0.08021344 | 0.9360675 | 0.96587767 |
| EPOR | 116.605335 | -0.37859736 | 0.45556293 | -0.8310539 | 0.40594317 | 0.60009523 |
| RGL3 | 869.072711 | -0.217653585 | 0.36623211 | -0.594305 | 0.55230811 | 0.72184315 |
| PRKCSH | 355.608403 | 0.142990699 | 0.28877618 | 0.49516099 | 0.62048648 | 0.77455476 |
| ECSIT | 308.252885 | 0.295375904 | 0.25345588 | 1.16539376 | 0.24385964 | 0.43900266 |
| ELOF1 | 105.94057 | -0.707972118 | 0.40397604 | -1.7525102 | 0.07968612 | 0.21135017 |
| ZNF627 | 426.980516 | 0.959684695 | 0.27674973 | 3.46769872 | 0.00052494 | 0.0055221 |
| ACP5 | 206.163379 | 1.155581251 | 0.39483623 | 2.92673562 | 0.0034254 | 0.02281436 |
| AC008543.1 | 68.7066326 | -0.229010009 | 0.44112592 | -0.5191488 | 0.60365695 | 0.76169806 |
| ZNF823 | 385.101515 | -0.298638427 | 0.25108098 | -1.1894108 | 0.23427806 | 0.42856658 |
| ZNF441 | 214.853089 | -0.04097553 | 0.36000845 | -0.1138182 | 0.90938187 | 0.95253371 |
| ZNF491 | 42.2838713 | 0.550348692 | 0.31575337 | 1.7429701 | 0.08133884 | 0.21450053 |
| ZNF440 | 438.31968 | -0.185962568 | 0.40897212 | -0.4547072 | 0.6493199 | 0.79436638 |
| ZNF439 | 176.003678 | 0.891976163 | 0.28404682 | 3.14024349 | 0.00168807 | 0.01344476 |
| ZNF69 | 933.332617 | 0.975070642 | 0.24225407 | 4.02499174 | 5.70E-05 | 0.00100643 |
| AC008770.2 | 88.3504922 | 0.689224552 | 0.43922043 | 1.56919966 | 0.11660142 | 0.2720878 |
| ZNF763 | 31.8410849 | 0.901289094 | 0.65431471 | 1.37745503 | 0.16837161 | 0.34690862 |
| ZNF433 | 67.407261 | 0.316297546 | 0.47267379 | 0.66916667 | 0.50338917 | 0.68404435 |
| ZNF844 | 271.711591 | 0.890006098 | 0.31908104 | 2.78927915 | 0.00528255 | 0.03152282 |
| ZNF788P | 169.035491 | 1.197204673 | 0.40758684 | 2.93729959 | 0.00331084 | 0.02221326 |
| ZNF20 | 70.9137482 | 0.578951646 | 0.50159946 | 1.15421106 | 0.24841365 | 0.4446919 |
| ZNF136 | 153.091231 | 0.192635452 | 0.31294873 | 0.61554956 | 0.53819185 | 0.70990443 |
| ZNF44 | 1243.81245 | 0.623665174 | 0.28897751 | 2.1581789 | 0.03091393 | 0.11148044 |
| ZNF563 | 157.440408 | 0.293344047 | 0.4191499 | 0.69985474 | 0.48401802 | 0.66798581 |
| ZNF442 | 86.4223638 | 1.052474068 | 0.39116701 | 2.69060031 | 0.00713236 | 0.03953225 |
| AC008758.4 | 54.9560245 | -0.18722443 | 0.41301646 | -0.4533099 | 0.65032564 | 0.79482956 |
| AC008758.5 | 182.97356 | -0.502506742 | 0.49069087 | -1.0240801 | 0.30579745 | 0.50583258 |
| AC008758.6 | 73.0693923 | -0.586011589 | 0.37895672 | -1.5463813 | 0.12201251 | 0.28027911 |
| AC008758.1 | 128.854015 | -0.398973367 | 0.54313533 | -0.7345745 | 0.46259867 | 0.65094732 |
| AC010422.6 | 41.3435568 | 0.675012342 | 0.30140609 | 2.2395445 | 0.02512051 | 0.09672102 |
| ZNF490 | 339.436867 | -0.085360547 | 0.27001706 | -0.3161302 | 0.75190369 | 0.8605113 |

| | | | | | | |
|---|---|---|---|---|---|---|
| ZNF791 | 565.093499 | -0.269235823 | 0.24524265 | -1.0978344 | 0.2722768 | 0.47033143 |
| MAN2B1 | 136.954401 | 0.126277151 | 0.27921132 | 0.45226372 | 0.65107901 | 0.79560845 |
| WDR83 | 102.509434 | -0.294459681 | 0.32169324 | -0.9153431 | 0.36001155 | 0.55846014 |
| DHPS | 134.982073 | -0.0528772 | 0.22264831 | -0.237492 | 0.8122751 | 0.89847117 |
| FBXW9 | 74.9560127 | 0.052681596 | 0.43420667 | 0.12132839 | 0.90343093 | 0.94935049 |
| TNPO2 | 486.862933 | -0.084059935 | 0.29031563 | -0.2895467 | 0.77216305 | 0.87334259 |
| TRIR | 536.132846 | -0.436270281 | 0.2315593 | -1.8840542 | 0.05955764 | 0.17383682 |
| ASNA1 | 179.687375 | -0.376341276 | 0.28575912 | -1.3169878 | 0.18784272 | 0.37143107 |
| HOOK2 | 1262.67202 | -0.503999889 | 0.30369911 | -1.6595369 | 0.09700764 | 0.24079749 |
| JUNB | 417.584586 | -0.964434314 | 0.5039259 | -1.9138415 | 0.05564041 | 0.16644764 |
| DNASE2 | 35.0268574 | -0.328343632 | 0.47157235 | -0.6962741 | 0.48625716 | 0.66963057 |
| GCDH | 335.249636 | 1.002441011 | 0.47556221 | 2.10790721 | 0.03503902 | 0.12176973 |
| SYCE2 | 47.4940887 | -0.118641691 | 0.72617477 | -0.163379 | 0.87022005 | 0.93290367 |
| FARSA | 230.068465 | -0.485877706 | 0.36113262 | -1.3454273 | 0.17848727 | 0.35946591 |
| CALR | 801.885965 | -1.045300439 | 0.47690987 | -2.1918197 | 0.02839253 | 0.10506923 |
| AC092069.1 | 147.56887 | -0.97910613 | 0.61206877 | -1.5996669 | 0.1096725 | 0.26107694 |
| RAD23A | 360.277688 | -0.609043597 | 0.35518601 | -1.7147173 | 0.08639707 | 0.22318972 |
| GADD45GIP1 | 406.770771 | -1.345748561 | 0.36185764 | -3.719 | 0.00020001 | 0.0026695 |
| NFIX | 306.01783 | 0.355790143 | 0.41360015 | 0.86022731 | 0.38966375 | 0.5852642 |
| TRMT1 | 177.489953 | -0.063079653 | 0.42162121 | -0.1496121 | 0.88107063 | 0.93815891 |
| NACC1 | 255.112427 | -0.437932614 | 0.47251086 | -0.9268202 | 0.35401989 | 0.55245569 |
| STX10 | 153.346706 | -0.058521474 | 0.34284269 | -0.1706948 | 0.86446373 | 0.92913074 |
| IER2 | 391.631649 | -0.49315277 | 0.24116754 | -2.0448555 | 0.04086911 | 0.13544314 |
| CACNA1A | 64.4097507 | 0.018031486 | 0.39097247 | 0.04611958 | 0.96321494 | 0.98083195 |
| CCDC130 | 578.497268 | -0.185022606 | 0.16716398 | -1.106833 | 0.26836614 | 0.4663406 |
| MRI1 | 385.411533 | 0.669628951 | 0.21756196 | 3.07787706 | 0.00208481 | 0.01585342 |
| AC008686.1 | 40.859331 | 0.992034189 | 0.62760412 | 1.58066869 | 0.11395381 | 0.26768893 |
| C19orf53 | 371.207972 | -1.28213336 | 0.4283194 | -2.9934049 | 0.00275884 | 0.01948035 |
| ZSWIM4 | 257.856142 | -1.415934721 | 0.46135355 | -3.0690881 | 0.00214713 | 0.01623317 |
| CC2D1A | 345.388284 | -0.510597137 | 0.26901248 | -1.8980425 | 0.05769048 | 0.17045895 |
| DCAF15 | 95.0340657 | -0.80648345 | 0.3750795 | -2.1501667 | 0.03154203 | 0.11315013 |
| RFX1 | 358.377348 | 0.632157101 | 0.27205945 | 2.32359914 | 0.02014699 | 0.08282987 |
| IL27RA | 42.3914109 | -0.86157337 | 0.58215962 | -1.4799607 | 0.13888373 | 0.30497906 |
| PALM3 | 319.405899 | 1.77364084 | 0.45660634 | 3.88439821 | 0.00010258 | 0.00158525 |
| MISP3 | 120.165706 | 0.358908451 | 0.42739585 | 0.83975651 | 0.40104492 | 0.59511992 |
| SAMD1 | 96.1755198 | -0.277809772 | 0.45423363 | -0.6116011 | 0.54080173 | 0.71215 |
| PRKACA | 212.453934 | 0.109361583 | 0.2435151 | 0.4490957 | 0.65336262 | 0.79727674 |
| AC022098.1 | 68.2950161 | 0.271618351 | 0.46290192 | 0.586773 | 0.55735617 | 0.7261364 |
| ADGRL1 | 65.1739279 | -0.878281703 | 0.49186252 | -1.7856244 | 0.0741601 | 0.20180911 |
| ADGRE5 | 60.5417175 | -1.793643781 | 0.82907701 | -2.1634224 | 0.0305087 | 0.1105125 |
| DDX39A | 209.761061 | -0.643903083 | 0.57320312 | -1.1233419 | 0.26129231 | 0.45940659 |
| PKN1 | 283.298337 | -0.477257097 | 0.25437666 | -1.8761827 | 0.06063019 | 0.17607162 |
| GIPC1 | 447.813731 | 0.235405873 | 0.29448876 | 0.79937135 | 0.42407512 | 0.61713533 |
| DNAJB1 | 500.286857 | -1.148770923 | 0.6262172 | -1.8344608 | 0.06658562 | 0.1877107 |
| TECR | 375.195984 | 0.802591007 | 0.20400293 | 3.93421317 | 8.35E-05 | 0.00135239 |
| NDUFB7 | 466.143665 | -0.23256087 | 0.31692133 | -0.7338126 | 0.46306296 | 0.65120457 |
| ADGRE3 | 43.1488712 | -0.634260883 | 0.44409371 | -1.4282141 | 0.15323026 | 0.32534099 |
| ZNF333 | 132.651737 | -0.228841148 | 0.5099462 | -0.4487555 | 0.65360806 | 0.79741929 |
| SYDE1 | 43.9436573 | 0.5017861 | 0.45615507 | 1.10003402 | 0.2713173 | 0.46956392 |
| ILVBL | 276.819361 | 0.426507942 | 0.31360922 | 1.3599981 | 0.17383053 | 0.3536616 |
| BRD4 | 1966.51148 | -0.042069773 | 0.22641028 | -0.1858121 | 0.85259211 | 0.92215576 |
| AKAP8 | 527.293274 | -0.03376618 | 0.23939689 | -0.1410469 | 0.88783292 | 0.94209263 |
| AKAP8L | 1235.48024 | 0.612093618 | 0.21691809 | 2.82177309 | 0.0047759 | 0.02929002 |
| WIZ | 117.686866 | -0.047197192 | 0.28828969 | -0.1637145 | 0.86995593 | 0.93283939 |
| CYP4F3 | 2104.8933 | 0.720017467 | 0.61903145 | 1.16313553 | 0.24477452 | 0.44013175 |
| CYP4F12 | 171.1087 | 0.355613921 | 0.84739548 | 0.4196552 | 0.67473736 | 0.81128289 |
| CYP4F2 | 159.415247 | 3.19555198 | 0.93990413 | 3.39987012 | 0.00067418 | 0.00672559 |
| CYP4F11 | 116.801915 | -1.005618752 | 0.84985032 | -1.1832893 | 0.23669453 | 0.43131773 |
| TPM4 | 217.835797 | -1.374079651 | 0.55530097 | -2.4744773 | 0.01334313 | 0.0617798 |
| RAB8A | 171.095393 | -0.227588278 | 0.24178641 | -0.9412782 | 0.3465623 | 0.54670612 |
| FAM32A | 148.902044 | -0.657723777 | 0.30454364 | -2.1597029 | 0.03079568 | 0.11120031 |
| AP1M1 | 490.295203 | 0.471455832 | 0.23187426 | 2.03323915 | 0.04202837 | 0.13799537 |
| KLF2 | 56.2298162 | 0.368395224 | 0.7874847 | 0.46781255 | 0.63991865 | 0.78856044 |
| EPS15L1 | 1280.46483 | -0.037575455 | 0.31697244 | -0.1185449 | 0.90563595 | 0.95066597 |
| AC008764.1 | 51.6894133 | -0.374641935 | 0.40382035 | -0.9277441 | 0.35354034 | 0.55199652 |
| C19orf44 | 193.812692 | 0.654710873 | 0.26685977 | 2.45338919 | 0.01415171 | 0.06428545 |
| AC008764.8 | 58.4705459 | -0.463215265 | 0.32156676 | -1.4404948 | 0.14972745 | 0.32012387 |
| TMEM38A | 147.639312 | 0.35029238 | 0.43937225 | 0.7972565 | 0.42530208 | 0.61832293 |
| SIN3B | 580.030787 | -0.492990065 | 0.30373656 | -1.6230844 | 0.10457137 | 0.25263333 |
| HAUS8 | 58.456667 | 0.162731033 | 0.41669919 | 0.390524 | 0.69614911 | 0.82508568 |
| MYO9B | 709.986192 | -0.867715501 | 0.41703132 | -2.0806962 | 0.03746172 | 0.12780481 |
| USE1 | 130.194005 | 0.072887236 | 0.31316208 | 0.23274605 | 0.81595861 | 0.90043834 |
| OCEL1 | 153.193094 | 0.443586466 | 0.34895961 | 1.27116852 | 0.2036687 | 0.39170576 |
| NR2F6 | 321.525387 | -0.109672152 | 0.33745366 | -0.3249991 | 0.74518172 | 0.85589122 |
| USHBP1 | 96.60135 | -0.625916909 | 0.44961587 | -1.3921148 | 0.16388764 | 0.34080823 |
| ABHD8 | 126.802367 | 0.442730816 | 0.22607728 | 1.95831631 | 0.05019291 | 0.15569002 |
| MRPL34 | 259.553216 | -0.159654728 | 0.36478202 | -0.4376716 | 0.66162436 | 0.80319592 |
| DDA1 | 158.012153 | -0.38747394 | 0.24908136 | -1.555612 | 0.11980039 | 0.27702309 |
| ANO8 | 61.6826191 | 0.242817531 | 0.35490723 | 0.68417183 | 0.49386667 | 0.67606416 |
| GTPBP3 | 85.067291 | 0.340802285 | 0.29110783 | 1.17070806 | 0.24171615 | 0.43681951 |
| PLVAP | 46.9487688 | 1.129756275 | 0.59643984 | 1.89416636 | 0.05820293 | 0.1712176 |
| BST2 | 67.8271724 | -0.506792516 | 0.91171133 | -0.5558695 | 0.57830007 | 0.74211119 |
| MVB12A | 145.025578 | 0.448989933 | 0.33011805 | 1.36008901 | 0.17380176 | 0.3536616 |
| SLC27A1 | 217.224318 | 0.593914921 | 0.28322388 | 2.09698036 | 0.03599531 | 0.12406855 |
| AC010618.3 | 243.367302 | 0.185773356 | 0.32268783 | 0.5757061 | 0.56481385 | 0.73181808 |
| PGLS | 354.763706 | -0.667432949 | 0.29367176 | -2.2727175 | 0.02304321 | 0.0911662 |
| COLGALT1 | 207.53554 | -0.711344435 | 0.41941714 | -1.6960309 | 0.08988002 | 0.22886597 |
| MAP1S | 81.9373988 | -0.595654164 | 0.41848924 | -1.423344 | 0.15463645 | 0.32720003 |
| B3GNT3 | 143.724427 | -1.260322393 | 0.55912224 | -2.2541089 | 0.02418932 | 0.09411383 |
| RPL18A | 100.383429 | -1.702880919 | 0.50542945 | -3.3691763 | 0.00075393 | 0.00735003 |
| CCDC124 | 143.588151 | -0.864938444 | 0.38105617 | -2.269845 | 0.02321699 | 0.09157849 |
| ARRDC2 | 56.6716993 | -1.864180163 | 0.4844832 | -3.8477705 | 0.0001192 | 0.00178953 |
| MAST3 | 483.014724 | 1.41884237 | 0.40239731 | 3.52597383 | 0.00042193 | 0.0047639 |
| AC007192.2 | 76.1283369 | 0.639377228 | 0.71832305 | 0.89009705 | 0.37341378 | 0.57086578 |
| JUND | 1120.27966 | -1.039002383 | 0.50208314 | -2.0693831 | 0.03851015 | 0.13046431 |
| LSM4 | 365.404012 | -0.664220337 | 0.22880792 | -2.9029604 | 0.00369653 | 0.0241393 |
| PGPEP1 | 1887.81322 | 1.180716769 | 0.33497454 | 3.52479551 | 0.00042381 | 0.00476943 |
| GDF15 | 826.769991 | -2.272638528 | 0.32966735 | -6.8937325 | 5.43E-12 | 1.26E-09 |
| SSBP4 | 228.190822 | -0.965892532 | 0.29002479 | -3.3303792 | 0.00086728 | 0.00813674 |
| ISYNA1 | 68.2749617 | -0.743973086 | 0.45237972 | -1.6445766 | 0.10005716 | 0.2460337 |
| ELL | 802.977986 | -0.509850883 | 0.15820093 | -3.2228058 | 0.00126942 | 0.0108343 |
| FKBP8 | 328.237884 | -0.372646697 | 0.2791471 | -1.3349474 | 0.18189356 | 0.36399188 |
| KXD1 | 205.519461 | -0.47669921 | 0.23106996 | -2.0630082 | 0.03911185 | 0.13162864 |
| AC005253.1 | 40.1172907 | -0.992242111 | 0.49589355 | -2.0009176 | 0.04540127 | 0.14577752 |
| UBA52 | 638.042456 | -0.657265348 | 0.51390017 | -1.2789748 | 0.20090594 | 0.38801046 |
| CRLF1 | 57.867095 | -0.732289388 | 0.48527657 | -1.5090145 | 0.13129507 | 0.29439908 |
| REX1BD | 189.518963 | 0.311851661 | 0.42491075 | 0.73392274 | 0.46299583 | 0.65120457 |
| KLHL26 | 43.2533063 | 0.826824368 | 0.40053405 | 2.06430482 | 0.03898882 | 0.13143142 |
| CRTC1 | 536.712624 | 0.561325339 | 0.25716021 | 2.18278458 | 0.02905168 | 0.10673046 |
| UPF1 | 432.06391 | 0.167870128 | 0.29874171 | 0.56192398 | 0.57416782 | 0.73911704 |
| COPE | 305.507921 | -0.499376715 | 0.36762269 | -1.3583947 | 0.17433848 | 0.35411786 |
| DDX49 | 172.487855 | -0.538665078 | 0.38414752 | -1.402235 | 0.1608451 | 0.33687295 |
| HOMER3 | 33.1342887 | -0.041276196 | 0.64298417 | -0.0641947 | 0.94881517 | 0.97382325 |

| | | | | | | |
|---|---|---|---|---|---|---|
| SUGP2 | 1518.8241 | 0.074756417 | 0.20236762 | 0.36940898 | 0.71182291 | 0.83509031 |
| ARMC6 | 123.083867 | -0.039420186 | 0.36808367 | -0.1070957 | 0.91471304 | 0.95437477 |
| SLC25A42 | 1507.84027 | 1.402999357 | 0.42555181 | 3.29689433 | 0.0009776 | 0.00889184 |
| TMEM161A | 224.83497 | 0.671219172 | 0.27004032 | 2.48562579 | 0.01293239 | 0.06043786 |
| BORCS8-MEI | 71.3641858 | 0.7630224 | 0.47947328 | 1.59137628 | 0.11152492 | 0.26429324 |
| RFXANK | 89.5558467 | 0.321979502 | 0.29018587 | 1.10956298 | 0.26718739 | 0.46517688 |
| NR2C2AP | 38.0706972 | -0.741042005 | 0.42401471 | -1.7476799 | 0.08051947 | 0.21296171 |
| TM6SF2 | 119.298208 | 0.002696395 | 0.87559645 | 0.00307949 | 0.99754292 | 0.99870545 |
| SUGP1 | 289.463858 | 0.544355081 | 0.18642801 | 2.9199211 | 0.0035012 | 0.02321768 |
| MAU2 | 637.936614 | 0.21553902 | 0.19831974 | 1.08682583 | 0.27711379 | 0.47500877 |
| GATAD2A | 1925.89872 | -0.241174116 | 0.35129018 | -0.6865382 | 0.49237381 | 0.67462692 |
| YJEFN3 | 37.2948185 | 0.883555271 | 0.568501 | 1.55418419 | 0.12014049 | 0.27756037 |
| ATP13A1 | 198.582159 | 0.029044999 | 0.37194008 | 0.07809053 | 0.93775604 | 0.96652545 |
| ZNF101 | 72.6909658 | 0.254207811 | 0.37255631 | 0.68233393 | 0.49502782 | 0.67697759 |
| ZNF14 | 181.114971 | 0.655640676 | 0.29777499 | 2.20179902 | 0.02767951 | 0.10344639 |
| AC011477.4 | 96.8261666 | -0.428371155 | 0.35011793 | -1.2235053 | 0.2211389 | 0.41341392 |
| ZNF253 | 317.775768 | 0.399327665 | 0.36990817 | 1.0795319 | 0.28035068 | 0.47908824 |
| ZNF93 | 81.1585906 | -0.626694209 | 0.34697258 | -1.8061779 | 0.07089055 | 0.19578819 |
| ZNF682 | 231.633067 | 0.71937013 | 0.3272594 | 2.19816492 | 0.02793736 | 0.10400253 |
| ZNF90 | 649.306032 | -0.870042648 | 0.41124836 | -2.1156137 | 0.03437768 | 0.12031025 |
| AC011447.3 | 189.343959 | 0.381240725 | 0.4098402 | 0.93021798 | 0.35225823 | 0.5513135 |
| ZNF486 | 83.8114657 | 0.487846273 | 0.62289114 | 0.78319668 | 0.43351162 | 0.62582221 |
| AC008554.1 | 1095.77721 | 2.285040643 | 0.70283287 | 3.25118635 | 0.00114925 | 0.01002718 |
| ZNF737 | 282.39331 | 0.771338362 | 0.31813673 | 2.42454988 | 0.01532738 | 0.06823396 |
| AC010636.2 | 77.5892433 | -0.448332951 | 0.65025126 | -0.6894765 | 0.49052347 | 0.67330824 |
| ZNF66 | 53.8312145 | -0.113149034 | 0.40273145 | -0.2809541 | 0.77874564 | 0.87801961 |
| ZNF85 | 58.8302482 | -0.04414425 | 0.55120229 | -0.0800872 | 0.9361679 | 0.96587767 |
| ZNF430 | 220.885289 | -0.338496369 | 0.27889523 | -1.2137044 | 0.22486063 | 0.41705712 |
| ZNF714 | 135.0337 | -0.35913638 | 0.39834413 | -0.9015732 | 0.36728365 | 0.564643 |
| ZNF431 | 457.035904 | -0.141070351 | 0.27586227 | -0.5113796 | 0.60908524 | 0.76570058 |
| ZNF708 | 453.2648 | 0.025669745 | 0.24137042 | 0.10635 | 0.91530466 | 0.95480068 |
| ZNF738 | 40.3427596 | -0.955472022 | 0.42246386 | -2.2616657 | 0.02371806 | 0.09290451 |
| ZNF493 | 203.848418 | 0.744756501 | 0.23121162 | 3.22110323 | 0.00127698 | 0.01087249 |
| AC010615.4 | 311.624599 | -0.30242581 | 0.70277372 | -0.4303317 | 0.66695437 | 0.80652347 |
| ZNF429 | 461.054945 | 0.646048307 | 0.32391807 | 1.99448063 | 0.04609956 | 0.14729291 |
| AC123912.1 | 175.790083 | 0.009520261 | 0.72938339 | 0.01305248 | 0.98958592 | 0.99493132 |
| AC123912.4 | 38.735166 | -0.860460462 | 0.66056699 | -1.3026089 | 0.1927083 | 0.37735039 |
| ZNF100 | 244.922355 | -0.270610686 | 0.4814304 | -0.5620972 | 0.57404978 | 0.73911704 |
| ZNF43 | 400.564742 | -0.07473971 | 0.39447755 | -0.1894651 | 0.84972834 | 0.92090248 |
| ZNF208 | 62.2297718 | -0.873701214 | 0.7327411 | -1.1923737 | 0.23311475 | 0.42706548 |
| ZNF98 | 75.2479362 | 0.510883358 | 0.71707737 | 0.71245221 | 0.47618479 | 0.661907 |
| AC092329.3 | 141.768579 | 1.460178662 | 0.36832923 | 3.96433008 | 7.36E-05 | 0.00122584 |
| ZNF91 | 2173.76844 | 0.557908279 | 0.25993969 | 2.14629889 | 0.03184914 | 0.11401148 |
| ZNF675 | 182.105748 | 0.430179403 | 0.26519721 | 1.62211138 | 0.1047795 | 0.2529434 |
| ZNF254 | 1254.57265 | 0.201587426 | 0.20050238 | 1.00541166 | 0.31469867 | 0.51517671 |
| LINC00662 | 480.529115 | 0.136628164 | 0.24309078 | 0.56204585 | 0.57408478 | 0.73911704 |
| AC006504.5 | 457.968448 | -0.418639729 | 0.53802561 | -0.7781037 | 0.43650787 | 0.62857823 |
| LINC01532 | 30.3555585 | 0.10099255 | 0.85139171 | 0.11862055 | 0.90557599 | 0.95066597 |
| UQCRFS1 | 442.67091 | -0.785750681 | 0.33809216 | -2.3240724 | 0.02012161 | 0.08277943 |
| POP4 | 73.5281882 | -0.36607081 | 0.35896066 | -1.0198076 | 0.30781971 | 0.50791823 |
| PLEKHF1 | 44.5875666 | -1.726469525 | 0.48358698 | -3.5701323 | 0.0003568 | 0.004198 |
| C19orf12 | 741.455374 | 0.54123262 | 0.20596103 | 2.62783999 | 0.00859289 | 0.04506742 |
| URI1 | 1726.53844 | -0.06298237 | 0.1449878 | -0.4343977 | 0.66399965 | 0.80472531 |
| ZNF507 | 538.107022 | 0.328511148 | 0.13284953 | 2.47280631 | 0.01340568 | 0.06195267 |
| DPY19L3 | 448.766004 | -0.788963606 | 0.36508933 | -2.1610152 | 0.03069416 | 0.11095065 |
| PDCD5 | 225.267219 | -0.761441925 | 0.21544113 | -3.5343386 | 0.0004088 | 0.00465701 |
| ANKRD27 | 551.751756 | -0.106383455 | 0.22966191 | -0.4632177 | 0.64320835 | 0.79092642 |
| NUDT19 | 206.062175 | 0.272637973 | 0.23320513 | 1.1690908 | 0.24236705 | 0.43756794 |
| SLC7A9 | 789.549036 | 2.15419202 | 0.4670114 | 4.6127183 | 3.97E-06 | 0.0001192 |
| CEP89 | 518.43599 | 0.082483631 | 0.22008401 | 0.37478248 | 0.70782224 | 0.83258556 |
| RHPN2 | 2426.22247 | -0.043042568 | 0.38575613 | -0.1115797 | 0.91115664 | 0.9531576 |
| GPATCH1 | 191.645634 | 0.213448129 | 0.24707887 | 0.86388661 | 0.38765018 | 0.58371098 |
| WDR88 | 43.8228171 | -0.001707739 | 0.54935652 | -0.0031086 | 0.99751969 | 0.99870545 |
| CEBPG | 176.51804 | -0.802631475 | 0.41997989 | -1.9111188 | 0.05598931 | 0.16710188 |
| PEPD | 2273.33189 | 1.016113555 | 0.26829568 | 3.78728997 | 0.0001523 | 0.00213671 |
| LSM14A | 2439.1976 | 0.383248272 | 0.14064617 | 2.72491086 | 0.00643189 | 0.03651123 |
| KIAA0355 | 1241.02166 | -0.15899514 | 0.22626711 | -0.7026878 | 0.48225034 | 0.66670759 |
| GPI | 1568.34601 | -1.127973501 | 0.27518868 | -4.0989096 | 4.15E-05 | 0.00077836 |
| UBA2 | 830.715623 | -0.418136778 | 0.16519845 | -2.5311181 | 0.01136996 | 0.05522608 |
| WTIP | 42.7951632 | -0.140226983 | 0.4188736 | -0.3347716 | 0.73779739 | 0.85123628 |
| SCGB2B2 | 529.137954 | 0.404951225 | 0.43456426 | 0.9318558 | 0.35141106 | 0.55058411 |
| ZNF302 | 499.665432 | -0.207266611 | 0.36000163 | -0.575738 | 0.5647923 | 0.73181808 |
| ZNF181 | 84.7522795 | 0.342957402 | 0.38980442 | 0.87981918 | 0.37895727 | 0.57574455 |
| ZNF30 | 76.528064 | 1.034301585 | 0.33478633 | 3.08943789 | 0.00200536 | 0.01538188 |
| GRAMD1A | 424.650494 | -0.975717645 | 0.38394617 | -2.5412876 | 0.0110445 | 0.0540972 |
| HPN | 3906.60888 | 0.811996088 | 0.50092134 | 1.6210052 | 0.10501652 | 0.25338171 |
| HPN-AS1 | 51.3397567 | 0.418568213 | 0.75148329 | 0.55698938 | 0.57753469 | 0.74154889 |
| LSR | 317.720487 | -0.931969327 | 0.41014363 | -2.2722999 | 0.0230684 | 0.09120237 |
| USF2 | 273.244623 | 0.152573164 | 0.33473434 | 0.45580375 | 0.64853111 | 0.79376872 |
| DMKN | 121.961589 | -1.271392819 | 0.5593947 | -2.272801 | 0.02303818 | 0.0911662 |
| TMEM147-AS | 38.0087246 | 0.756570973 | 0.50246595 | 1.50571591 | 0.13214012 | 0.2954593 |
| TMEM147 | 175.825058 | -0.192407086 | 0.39463754 | -0.4875539 | 0.62586583 | 0.77848801 |
| HAUS5 | 41.5512123 | 0.077211283 | 0.60516077 | 0.12758805 | 0.89847499 | 0.94743919 |
| RBM42 | 111.528129 | -0.517915649 | 0.34497538 | -1.5013119 | 0.1332749 | 0.29699221 |
| COX6B1 | 593.543616 | -0.488590258 | 0.31228819 | -1.5645493 | 0.11768863 | 0.27360212 |
| KMT2B | 360.086759 | -0.100052411 | 0.26045487 | -0.3841449 | 0.70087105 | 0.82796169 |
| PROSER3 | 162.186777 | 0.181878868 | 0.32160719 | 0.5655311 | 0.57171258 | 0.737619 |
| PRODH2 | 1362.27847 | 2.83706023 | 0.55765814 | 5.08745419 | 3.63E-07 | 1.63E-05 |
| NPHS1 | 43.0711954 | 0.38344799 | 0.54384227 | 0.70507207 | 0.48076539 | 0.66570878 |
| NFKBID | 154.553426 | -1.303478856 | 0.43745003 | -2.9797206 | 0.00288511 | 0.02011501 |
| LRFN3 | 34.370543 | -0.129361553 | 0.41292356 | -0.3132821 | 0.75406636 | 0.86212258 |
| SDHAF1 | 69.2506895 | -0.454242225 | 0.37086996 | -1.2248019 | 0.22064987 | 0.41261225 |
| SYNE4 | 82.5147915 | 0.600023712 | 0.35845048 | 1.67393754 | 0.09414288 | 0.23621021 |
| ALKBH6 | 50.8281481 | 0.644136062 | 0.28484566 | 2.26135114 | 0.02373752 | 0.09290451 |
| THAP8 | 65.028511 | 0.409065155 | 0.33082322 | 1.23650677 | 0.21627025 | 0.40703231 |
| POLR2I | 51.8425951 | -0.437417488 | 0.51131024 | -0.8554835 | 0.39228352 | 0.58726848 |
| TBCB | 167.09708 | -0.411763345 | 0.31686938 | -1.2994734 | 0.1937815 | 0.37885739 |
| CAPNS1 | 563.53894 | -0.75176235 | 0.40099503 | -1.8747423 | 0.06082818 | 0.17645966 |
| COX7A1 | 36.1378617 | 1.255771997 | 0.91620429 | 1.37062444 | 0.17049206 | 0.34957886 |
| ZNF565 | 550.256518 | -0.315203969 | 0.31017585 | -1.0162106 | 0.30952913 | 0.50977712 |
| ZNF146 | 690.285155 | -0.028665651 | 0.2640651 | -0.1085552 | 0.91355526 | 0.95399223 |
| AC012617.1 | 33.898361 | 0.480184627 | 0.44785233 | 1.07219411 | 0.28363285 | 0.48292248 |
| LINC00665 | 337.095715 | -0.030430443 | 0.21607397 | -0.1408335 | 0.88800151 | 0.94209263 |
| ZFP14 | 658.434771 | 0.433313466 | 0.34283291 | 1.26392028 | 0.20625864 | 0.39493857 |
| ZFP82 | 200.602849 | -0.364063768 | 0.26493302 | -1.3741729 | 0.16938802 | 0.34835733 |
| ZNF566 | 351.774512 | 0.250821099 | 0.22301669 | 1.1246741 | 0.26072716 | 0.4589178 |
| ZNF260 | 230.108771 | -0.128036405 | 0.17486079 | -0.7322191 | 0.46403487 | 0.6518268 |
| ZNF529 | 389.954492 | -0.360572251 | 0.20512138 | -1.7578482 | 0.07877334 | 0.20970788 |
| ZNF529-AS1 | 310.205219 | -0.610417042 | 0.24348166 | -2.507035 | 0.01217486 | 0.05782987 |
| ZNF382 | 109.918599 | -0.333425625 | 0.27670648 | -1.2049795 | 0.22821123 | 0.42106492 |
| ZNF461 | 193.851414 | -0.355075382 | 0.3220985 | -1.1023814 | 0.27029591 | 0.46856533 |
| ZNF567 | 202.595768 | -0.154010832 | 0.24093401 | -0.6392241 | 0.52267714 | 0.69848255 |
| ZNF850 | 49.4835029 | -0.18391653 | 0.35684213 | -0.5154003 | 0.60627334 | 0.76359388 |

| | | | | | | |
|---|---|---|---|---|---|---|
| AC020928.1 | 317.301609 | 0.284723777 | 0.26726835 | 1.0653105 | 0.28673544 | 0.48642113 |
| ZNF790 | 351.541582 | 0.249457684 | 0.30325591 | 0.82259793 | 0.41073668 | 0.60446292 |
| ZNF345 | 364.350066 | 0.075087373 | 0.38287217 | 0.19611604 | 0.84451934 | 0.91835198 |
| ZNF829 | 38.2768033 | -0.170570731 | 0.37433687 | -0.455661 | 0.64863374 | 0.79382345 |
| ZNF568 | 186.380819 | -0.145941372 | 0.26480611 | -0.5511254 | 0.58154771 | 0.74455393 |
| ZNF420 | 256.184646 | 0.138070001 | 0.1853938 | 0.74473905 | 0.45642949 | 0.645443 |
| AC012309.1 | 39.6964153 | -0.393195339 | 0.60733324 | -0.6474128 | 0.51736479 | 0.69425481 |
| ZNF383 | 176.10116 | 0.122690228 | 0.20168556 | 0.60832432 | 0.5429724 | 0.71356452 |
| LINC01535 | 29.5295757 | -0.747810631 | 0.64831868 | -1.1534615 | 0.24872101 | 0.44495157 |
| HKR1 | 592.178735 | 0.304692708 | 0.19570011 | 1.55693679 | 0.11948549 | 0.27650132 |
| ZNF527 | 59.8185275 | 0.775403255 | 0.31516226 | 2.46033028 | 0.01388092 | 0.06348541 |
| ZNF569 | 198.396888 | 0.485343075 | 0.25662126 | 1.89128162 | 0.05858676 | 0.17195904 |
| ZNF570 | 64.1026462 | 0.035242739 | 0.23173961 | 0.15207905 | 0.87912459 | 0.93706833 |
| ZNF793 | 176.92681 | 0.6030082 | 0.34984256 | 1.72365591 | 0.08476998 | 0.22023163 |
| ZNF540 | 178.265151 | 0.826167092 | 0.48618042 | 1.69930146 | 0.0892624 | 0.22776623 |
| ZNF571 | 194.63655 | -0.017114676 | 0.28114421 | -0.0608751 | 0.9514587 | 0.97536791 |
| ZFP30 | 331.390759 | 0.108795175 | 0.29229016 | 0.37221634 | 0.70973178 | 0.83388131 |
| AC093227.1 | 38.2893618 | 0.849042638 | 0.30206444 | 2.81079973 | 0.00494185 | 0.0300257 |
| ZNF573 | 297.913713 | 0.484852745 | 0.21270856 | 2.27942279 | 0.02264194 | 0.09011415 |
| SIPA1L3 | 4542.18502 | 0.235482984 | 0.50236327 | 0.4687504 | 0.63924806 | 0.78803844 |
| SPINT2 | 298.280165 | -0.25588464 | 0.25843521 | -0.9901307 | 0.32211023 | 0.52244289 |
| AC011479.1 | 57.4940387 | -0.76434628 | 0.48733618 | -1.5684169 | 0.11678388 | 0.27223752 |
| YIF1B | 63.1397456 | 0.093852117 | 0.31226454 | 0.30055324 | 0.76375519 | 0.86815453 |
| CATSPERG | 233.012996 | 0.934486269 | 0.46733302 | 1.99961531 | 0.04554182 | 0.14614624 |
| PSMD8 | 394.310955 | -0.812715377 | 0.32259605 | -2.5192973 | 0.01175893 | 0.05641571 |
| FAM98C | 58.5347363 | 0.490159189 | 0.25165155 | 1.94776942 | 0.05144256 | 0.15796366 |
| EIF3K | 464.579404 | 0.179312177 | 0.21168722 | 0.8470619 | 0.39696062 | 0.5916211 |
| ACTN4 | 3743.49653 | -0.094146032 | 0.34678546 | -0.2714821 | 0.78602029 | 0.88296838 |
| CAPN12 | 1568.5647 | 1.175325369 | 0.5128963 | 2.29154584 | 0.02193186 | 0.088236 |
| AC022144.1 | 100.582908 | 1.344129201 | 0.41499175 | 3.23892995 | 0.00119979 | 0.01038988 |
| ECH1 | 497.892779 | -0.22401674 | 0.47107002 | -0.4755487 | 0.63439592 | 0.78445331 |
| HNRNPL | 705.850086 | 0.286504345 | 0.25387785 | 1.12851255 | 0.25910352 | 0.45712397 |
| RINL | 34.3069256 | -0.588024277 | 0.39222419 | -1.4992045 | 0.13382058 | 0.29777495 |
| SIRT2 | 350.629612 | 0.33859298 | 0.34166074 | 0.99102103 | 0.32167531 | 0.52210014 |
| NFKBIB | 142.420126 | -0.346903556 | 0.29251688 | -1.1859266 | 0.23565127 | 0.42995285 |
| SARS2 | 33.6524536 | 1.033991984 | 0.48130415 | 2.14831306 | 0.03168889 | 0.11355808 |
| AC011455.2 | 67.8535436 | 0.995810559 | 0.58394074 | 1.70532811 | 0.08813325 | 0.22623195 |
| MRPS12 | 174.763399 | -0.881430687 | 0.34888175 | -2.5264454 | 0.01152233 | 0.05567027 |
| FBXO17 | 1704.60772 | 0.465857924 | 0.31466758 | 1.4804764 | 0.13874615 | 0.30482346 |
| FBXO27 | 115.555858 | -0.560061754 | 0.35650693 | -1.5709702 | 0.11618957 | 0.27158651 |
| PAK4 | 439.649384 | 0.444031449 | 0.26758294 | 1.65941614 | 0.09703197 | 0.24081426 |
| AC011443.1 | 54.11079 | -0.325528202 | 0.50963292 | -0.6387503 | 0.52298536 | 0.69875819 |
| SAMD4B | 626.563938 | -0.370726549 | 0.29171651 | -1.2708453 | 0.2037837 | 0.39181696 |
| PAF1 | 138.18139 | -0.411872682 | 0.29521164 | -1.3951776 | 0.16296228 | 0.33942014 |
| MED29 | 155.663315 | -0.236494183 | 0.37584149 | -0.6292392 | 0.5291925 | 0.70348575 |
| ZFP36 | 733.202269 | -2.1792583 | 0.44371935 | -4.9113438 | 9.05E-07 | 3.55E-05 |
| RPS16 | 806.380792 | -0.648091136 | 0.43813179 | -1.4792151 | 0.13908285 | 0.30514868 |
| SUPT5H | 721.974096 | 0.04736012 | 0.29161431 | 0.16240671 | 0.87098559 | 0.93338059 |
| TIMM50 | 145.584659 | 0.084330234 | 0.2513317 | 0.33553362 | 0.73722259 | 0.85110831 |
| EID2B | 34.1682986 | 0.030908605 | 0.3488822 | 0.08859324 | 0.92940518 | 0.9626233 |
| EID2 | 69.4565131 | 0.224881713 | 0.28857599 | 0.77928074 | 0.43581436 | 0.62808405 |
| DYRK1B | 127.440552 | 0.264429962 | 0.33247275 | 0.79534326 | 0.42641386 | 0.61928954 |
| FBL | 114.767037 | -0.864811961 | 0.30971749 | -2.7922607 | 0.00523412 | 0.03137037 |
| FCGBP | 66.3440554 | 2.035699496 | 0.47805423 | 4.25830244 | 2.06E-05 | 0.0004532 |
| PSMC4 | 276.075436 | -0.281889203 | 0.36807424 | -0.7658488 | 0.4437663 | 0.63517875 |
| ZNF546 | 213.451257 | 0.705610221 | 0.39034323 | 1.80766607 | 0.07065849 | 0.19541679 |
| ZNF780B | 775.127568 | 0.681240824 | 0.27749357 | 2.45497881 | 0.01408929 | 0.06409621 |
| ZNF780A | 164.829598 | 0.317892605 | 0.25384523 | 1.25230874 | 0.21045738 | 0.39989367 |
| MAP3K10 | 224.209088 | 0.081323713 | 0.33551437 | 0.24238519 | 0.80848171 | 0.89601696 |
| AKT2 | 934.466225 | 0.337045168 | 0.31656625 | 1.06469078 | 0.28701589 | 0.48665596 |
| C19orf47 | 61.1764276 | 0.426813256 | 0.50363746 | 0.84746129 | 0.39673805 | 0.59148233 |
| PLD3 | 929.332736 | 0.273012729 | 0.27759192 | 0.98350386 | 0.32535951 | 0.52572237 |
| SERTAD1 | 62.5071187 | -0.615646007 | 0.39430066 | -1.5613618 | 0.1184384 | 0.2747859 |
| SERTAD3 | 68.7187022 | -0.052308295 | 0.32495874 | -0.160969 | 0.87211779 | 0.93380181 |
| BLVRB | 688.965819 | -0.798254036 | 0.33917176 | -2.3535392 | 0.01859564 | 0.07830737 |
| SPTBN4 | 87.1870254 | 0.340623845 | 0.48775303 | 0.69835311 | 0.48495639 | 0.66867491 |
| SHKBP1 | 124.956202 | -0.170447732 | 0.33285083 | -0.5120844 | 0.60859191 | 0.76545874 |
| NUMBL | 77.0747475 | -0.086660163 | 0.31393446 | -0.2760454 | 0.7825132 | 0.88090575 |
| COQ8B | 226.067342 | 0.105732385 | 0.3002276 | 0.35217409 | 0.7247077 | 0.84347189 |
| ITPKC | 450.456172 | -1.045145978 | 0.21298162 | -4.907212 | 9.24E-07 | 3.62E-05 |
| C19orf54 | 65.3750472 | 0.204832867 | 0.25182109 | 0.81340632 | 0.41598513 | 0.60944659 |
| SNRPA | 331.604321 | -0.356080542 | 0.30890093 | -1.1527338 | 0.24901967 | 0.44519533 |
| AC008537.4 | 36.3109061 | -1.118496531 | 0.47799223 | -2.3399889 | 0.01928431 | 0.08031907 |
| AC008537.1 | 361.438762 | -0.674341311 | 0.30423718 | -2.2164987 | 0.02665737 | 0.10088142 |
| CYP2B6 | 180.248469 | 2.101070009 | 0.58929863 | 3.565374 | 0.00036334 | 0.00425664 |
| HNRNPUL1 | 457.147979 | -0.209994963 | 0.27561184 | -0.7619229 | 0.44610604 | 0.63722129 |
| TGFB1 | 227.237083 | -0.816900145 | 0.34062633 | -2.3982296 | 0.01647454 | 0.07198484 |
| CCDC97 | 123.996514 | -0.267356483 | 0.32571467 | -0.8208303 | 0.41174292 | 0.60522897 |
| B9D2 | 39.9851405 | -0.344260144 | 0.39950698 | -0.8617125 | 0.38884576 | 0.58441983 |
| EXOSC5 | 45.7097836 | -0.759732714 | 0.52068529 | -1.4591016 | 0.14453716 | 0.31243823 |
| DMAC2 | 154.781272 | -0.045552089 | 0.19865277 | -0.2293051 | 0.81863181 | 0.90192453 |
| RPS19 | 2284.89428 | -1.403828572 | 0.35111031 | -3.9982551 | 6.38E-05 | 0.00109692 |
| ARHGEF1 | 256.815373 | -0.133024533 | 0.34264074 | -0.3882333 | 0.69784342 | 0.82589622 |
| RABAC1 | 448.509881 | -0.656961224 | 0.45260383 | -1.451515 | 0.14663652 | 0.31578112 |
| ZNF574 | 37.1209027 | -0.071332517 | 0.41684864 | -0.1711233 | 0.86412681 | 0.92885255 |
| POU2F2 | 58.0751823 | -0.447222006 | 0.55681629 | -0.8031769 | 0.42187247 | 0.61556376 |
| DEDD2 | 132.34133 | -0.690557813 | 0.48914124 | -1.4117759 | 0.15801596 | 0.33265624 |
| ZNF526 | 81.0132302 | -0.755507479 | 0.45074377 | -1.6761352 | 0.09371172 | 0.23571272 |
| GSK3A | 133.244478 | -0.074895992 | 0.38484002 | -0.1946159 | 0.84569365 | 0.91865716 |
| CIC | 189.057619 | -0.355954541 | 0.230867 | -1.5418165 | 0.1231182 | 0.28205819 |
| PAFAH1B3 | 80.8690408 | -0.983182464 | 0.46893447 | -2.0966308 | 0.03602627 | 0.12408456 |
| MEGF8 | 251.752451 | 0.645871124 | 0.35035992 | 1.84345037 | 0.06526328 | 0.18536341 |
| LIPE-AS1 | 217.594326 | -0.207896842 | 0.45038368 | -0.4615994 | 0.64436861 | 0.79162826 |
| CEACAM1 | 389.010492 | 0.534830006 | 0.57847257 | 0.92455552 | 0.35519716 | 0.55366752 |
| PHLDB3 | 231.216779 | 0.433710239 | 0.34574146 | 1.25443516 | 0.20968388 | 0.39926448 |
| ETHE1 | 294.147525 | 0.29595239 | 0.41626381 | 0.71097315 | 0.47710087 | 0.66270994 |
| XRCC1 | 80.6699048 | 0.240894156 | 0.2755369 | 0.87427187 | 0.38197022 | 0.57865448 |
| PINLYP | 38.4173149 | -0.342616961 | 0.5068521 | -0.6759703 | 0.49905951 | 0.68080506 |
| ZNF428 | 241.24426 | -0.189923869 | 0.2906799 | -0.6533781 | 0.51351258 | 0.69196393 |
| CADM4 | 124.363521 | 0.324344137 | 0.44250019 | 0.73298078 | 0.46357015 | 0.65157466 |
| SMG9 | 166.928834 | 0.023130062 | 0.21383921 | 0.10816567 | 0.91386427 | 0.95416953 |
| ZNF283 | 139.548395 | 0.019827536 | 0.34491731 | 0.0574849 | 0.95415894 | 0.97682198 |
| ZNF404 | 33.1919759 | -0.563920215 | 0.69150283 | -0.8154995 | 0.41478645 | 0.60833275 |
| ZNF45 | 98.6208782 | 0.038514096 | 0.32574318 | 0.11823454 | 0.90588182 | 0.95068694 |
| ZNF155 | 152.507843 | -0.110006082 | 0.31023544 | -0.354589 | 0.7228975 | 0.84233055 |
| ZNF230 | 44.0497803 | -0.365379997 | 0.33238542 | -1.099266 | 0.27165205 | 0.46990595 |
| ZNF224 | 316.407043 | -0.294159077 | 0.2824943 | -1.0412921 | 0.29774002 | 0.49812308 |
| ZNF225 | 53.597265 | 0.18100612 | 0.26850156 | 0.67413434 | 0.50022591 | 0.68143476 |
| ZNF234 | 132.523594 | 0.040052835 | 0.26310581 | 0.1522309 | 0.87900482 | 0.93706833 |
| ZNF226 | 162.695489 | 0.254340246 | 0.28786162 | 0.88355038 | 0.37693898 | 0.57337727 |
| ZNF227 | 122.399174 | 0.179960828 | 0.31001736 | 0.58048629 | 0.56158673 | 0.72977525 |
| ZNF235 | 646.097204 | 0.119558699 | 0.40788605 | 0.29311789 | 0.76943204 | 0.87154822 |
| ZNF229 | 87.5884078 | -0.634452195 | 0.46945933 | -1.3514529 | 0.17655038 | 0.35692863 |

| | | | | | | |
|---|---|---|---|---|---|---|
| ZNF180 | 87.4467272 | 0.579146962 | 0.2972004 | 1.94867493 | 0.05133426 | 0.15790414 |
| AC243964.2 | 349.684396 | -1.011530337 | 0.32405644 | -3.1214635 | 0.00179955 | 0.01412736 |
| PVR | 181.96772 | -1.729555511 | 0.65475878 | -2.6415156 | 0.0082536 | 0.04389935 |
| CEACAM19 | 408.542701 | -1.416425014 | 0.71821675 | -1.9721414 | 0.04859346 | 0.15252361 |
| BCL3 | 218.523632 | -0.359118844 | 0.64376013 | -0.5578457 | 0.57694973 | 0.74130618 |
| CBLC | 272.274368 | 0.427495335 | 0.45402347 | 0.941571 | 0.34641232 | 0.54665827 |
| BCAM | 598.600398 | 0.031785479 | 0.43730074 | 0.07268563 | 0.94205629 | 0.96969312 |
| NECTIN2 | 551.977437 | -0.215729813 | 0.29382348 | -0.7342157 | 0.46281728 | 0.65111737 |
| TOMM40 | 231.171997 | -0.740608724 | 0.43731612 | -1.6935317 | 0.0903543 | 0.22981749 |
| APOE | 5094.07775 | -0.104972875 | 0.59432315 | -0.1766259 | 0.85980223 | 0.92628466 |
| CLPTM1 | 564.6994 | 0.622463082 | 0.19306944 | 3.22403727 | 0.00126397 | 0.01080127 |
| RELB | 395.647081 | -1.303116538 | 0.5720808 | -2.277854 | 0.02273528 | 0.09035444 |
| CLASRP | 646.864046 | 0.281037152 | 0.27229774 | 1.03209508 | 0.30202758 | 0.50241408 |
| GEMIN7-AS1 | 253.810109 | 0.532841755 | 0.35919985 | 1.48341309 | 0.13796469 | 0.30371689 |
| MARK4 | 443.202181 | 0.10702391 | 0.31353206 | 0.34134917 | 0.73284073 | 0.84847098 |
| TRAPPC6A | 202.23237 | 0.586297195 | 0.29694201 | 1.97445013 | 0.04833058 | 0.15208157 |
| ERCC2 | 136.43624 | 0.753368194 | 0.31873783 | 2.36359834 | 0.01809842 | 0.07695443 |
| PPP1R13L | 398.146579 | -0.869212844 | 0.33318581 | -2.6087931 | 0.00908622 | 0.04701577 |
| CD3EAP | 49.8837354 | -1.946554828 | 0.78810804 | -2.4699086 | 0.01351476 | 0.06226895 |
| ERCC1 | 191.375541 | -1.244925468 | 0.69027807 | -1.803513 | 0.07130769 | 0.19663681 |
| VASP | 315.673902 | -1.166255997 | 0.33026402 | -3.5312838 | 0.00041355 | 0.00469247 |
| OPA3 | 671.236374 | 0.159670817 | 0.35499704 | 0.44978069 | 0.65286858 | 0.79687174 |
| EML2 | 167.204667 | -0.544562191 | 0.33387086 | -1.6310564 | 0.10287843 | 0.25038257 |
| SNRPD2 | 327.017045 | -0.987574373 | 0.3320949 | -2.9737716 | 0.00294164 | 0.02040539 |
| QPCTL | 102.68968 | -0.419837695 | 0.4094556 | -1.0253559 | 0.30519531 | 0.50529195 |
| FBXO46 | 211.275788 | 0.557845155 | 0.26369723 | 2.11547595 | 0.0343894 | 0.12031025 |
| SIX5 | 34.9754703 | 0.41096033 | 0.44770452 | 0.91792758 | 0.35865678 | 0.5571924 |
| DM1-AS | 81.1589985 | 0.117905624 | 0.43458539 | 0.271306 | 0.78615569 | 0.88304811 |
| DMPK | 237.967738 | 0.361795118 | 0.38836079 | 0.93159538 | 0.35154567 | 0.55063839 |
| SYMPK | 797.906256 | 0.022447302 | 0.32712725 | 0.06861948 | 0.94529251 | 0.97196767 |
| IRF2BP1 | 32.0508619 | -0.272876492 | 0.49617846 | -0.5499563 | 0.58234931 | 0.74544846 |
| MYPOP | 229.238588 | 1.051072591 | 0.25223885 | 4.1669734 | 3.09E-05 | 0.00061676 |
| CCDC61 | 64.3257098 | 0.551103249 | 0.26934603 | 2.04607901 | 0.0407486 | 0.13514179 |
| PPP5C | 383.298152 | 0.022470778 | 0.21908062 | 0.10256853 | 0.91830542 | 0.95632976 |
| PPP5D1 | 145.402594 | 0.690920582 | 0.33110973 | 2.08668159 | 0.03691693 | 0.12633733 |
| CALM3 | 321.737679 | -0.296728459 | 0.33429543 | -0.8876234 | 0.37474334 | 0.5715597 |
| PRKD2 | 405.814635 | -0.270157561 | 0.32539545 | -0.8302438 | 0.40640094 | 0.60027346 |
| STRN4 | 193.692338 | -0.013047312 | 0.29384958 | -0.0444013 | 0.9645845 | 0.98155603 |
| FKRP | 135.467436 | 0.352549355 | 0.27721029 | 1.27177585 | 0.20345277 | 0.39145524 |
| SLC1A5 | 228.500685 | -0.290321798 | 0.47648457 | -0.6092995 | 0.54232595 | 0.71322817 |
| AP2S1 | 335.436963 | -0.528903386 | 0.26861763 | -1.9689824 | 0.04895511 | 0.1533524 |
| ARHGAP35 | 1247.54361 | 0.425758954 | 0.1728545 | 2.46310596 | 0.01377392 | 0.06314352 |
| TMEM160 | 218.136305 | -0.595393768 | 0.4287204 | -1.3887694 | 0.16490289 | 0.34222897 |
| ZC3H4 | 254.708067 | 0.268289343 | 0.3107609 | 0.86333042 | 0.38795582 | 0.58387159 |
| SAE1 | 342.260356 | -0.424151399 | 0.50726768 | -0.8361491 | 0.40307104 | 0.59728592 |
| BBC3 | 52.481485 | -2.868895727 | 0.52955574 | -5.417552 | 6.04E-08 | 3.66E-06 |
| CCDC9 | 102.731353 | -0.144170706 | 0.39835438 | -0.3619157 | 0.71741502 | 0.8386157 |
| INAFM1 | 65.494904 | -0.584485714 | 0.4475131 | -1.3060751 | 0.19152703 | 0.37589647 |
| C5AR1 | 513.581448 | -0.096719936 | 0.69139894 | -0.1398902 | 0.88874675 | 0.94260301 |
| DHX34 | 375.616042 | -0.285419326 | 0.39707106 | -0.7188117 | 0.47225694 | 0.65831648 |
| KPTN | 40.1524581 | 0.52866732 | 0.38005895 | 1.39101401 | 0.16422118 | 0.34110921 |
| NAPA-AS1 | 82.4191723 | -0.276931426 | 0.42593152 | -0.6501783 | 0.51557706 | 0.69315508 |
| NAPA | 534.096789 | -0.651123292 | 0.32938908 | -1.9767604 | 0.04806872 | 0.15153587 |
| BICRA-AS1 | 63.4892318 | -0.157472239 | 0.42355098 | -0.3717905 | 0.71004882 | 0.83398794 |
| NOP53 | 152.488231 | -0.529107994 | 0.39363869 | -1.3441463 | 0.17890108 | 0.36010262 |
| SELENOW | 458.06504 | -0.136717252 | 0.26929864 | -0.507679 | 0.6116785 | 0.76786127 |
| PLA2G4C | 650.022794 | -1.901644536 | 0.47182567 | -4.0303965 | 5.57E-05 | 0.00098844 |
| LIG1 | 112.520805 | 0.157768867 | 0.42410714 | 0.37200238 | 0.70989108 | 0.83388131 |
| ZSWIM9 | 161.819553 | 0.156188141 | 0.23380494 | 0.66802756 | 0.504116 | 0.68462479 |
| CARD8 | 389.745471 | 0.398289481 | 0.26652753 | 1.49436526 | 0.13508018 | 0.29948045 |
| EMP3 | 68.0451368 | -1.330899103 | 0.55896198 | -2.381019 | 0.01726482 | 0.07440511 |
| TMEM143 | 210.281832 | 1.416895555 | 0.42730651 | 3.31587633 | 0.00091356 | 0.00845732 |
| KDELR1 | 386.062147 | -0.488387772 | 0.25816623 | -1.891757 | 0.05852337 | 0.17180976 |
| GRWD1 | 67.0565816 | -0.569722756 | 0.52336316 | -1.0885802 | 0.27633906 | 0.47404858 |
| KCNJ14 | 59.4857455 | -0.631147526 | 0.57568647 | -1.096339 | 0.27293045 | 0.47075093 |
| CYTH2 | 236.240085 | -0.604852077 | 0.32798951 | -1.8441202 | 0.06516563 | 0.18520981 |
| LMTK3 | 57.5323728 | -1.147287224 | 0.51828222 | -2.2136341 | 0.02685396 | 0.10132045 |
| SULT2B1 | 51.8887854 | -1.315143815 | 0.70428238 | -1.867353 | 0.06185231 | 0.17848161 |
| RPL18 | 858.839597 | -0.939058956 | 0.45142862 | -2.0801937 | 0.03750777 | 0.12792213 |
| SPHK2 | 263.638743 | 1.155293028 | 0.32770057 | 3.52545326 | 0.00042276 | 0.00476622 |
| DBP | 50.993057 | 0.581204194 | 0.47555891 | 1.22214973 | 0.22165101 | 0.41380692 |
| CA11 | 38.6521656 | 0.3655135 | 0.37019961 | 0.98734166 | 0.32347518 | 0.52401995 |
| BCAT2 | 395.280779 | 0.023726825 | 0.31057121 | 0.07639737 | 0.93910296 | 0.9673318 |
| HSD17B14 | 830.379287 | 1.470341873 | 0.38965695 | 3.77342649 | 0.00016102 | 0.00224072 |
| PLEKHA4 | 139.812238 | -2.158716552 | 0.65355418 | -3.3030415 | 0.00095642 | 0.00875146 |
| PPP1R15A | 258.365976 | -2.455340897 | 0.57674747 | -4.2572201 | 2.07E-05 | 0.00045467 |
| NUCB1 | 1052.57461 | -0.092612453 | 0.4197013 | -0.2206628 | 0.82535502 | 0.90642548 |
| NUCB1-AS1 | 150.20099 | -0.93106744 | 0.73660864 | -1.263992 | 0.20623289 | 0.39493857 |
| DHDH | 239.062126 | 1.01408528 | 0.55616759 | 1.82334478 | 0.06825119 | 0.19083704 |
| BAX | 399.826461 | -0.384313049 | 0.2124312 | -1.8091177 | 0.07043272 | 0.195074 |
| FTL | 24154.4632 | -1.345032288 | 0.61118886 | -2.2006819 | 0.02775855 | 0.1035901 |
| GYS1 | 654.594477 | 0.020463917 | 0.26112181 | 0.07836923 | 0.93753435 | 0.96641041 |
| RUVBL2 | 155.59422 | -0.384239783 | 0.31343745 | -1.2258898 | 0.22024013 | 0.41207091 |
| SNRNP70 | 2322.74154 | 1.053823067 | 0.35670255 | 2.9543469 | 0.00313332 | 0.02132491 |
| LIN7B | 65.3989829 | 0.165178155 | 0.3995911 | 0.41336795 | 0.67933705 | 0.81417094 |
| PPFIA3 | 38.5477815 | 0.484006393 | 0.57427736 | 0.84280946 | 0.39933502 | 0.5934221 |
| TRPM4 | 249.924836 | -0.545140178 | 0.6355262 | -0.8577777 | 0.39101525 | 0.58639453 |
| SLC6A16 | 64.4516717 | -0.653614865 | 0.73907292 | -0.8843713 | 0.37649582 | 0.57294604 |
| PIH1D1 | 232.066682 | -0.074020497 | 0.24454807 | -0.3026828 | 0.7621316 | 0.86743651 |
| RPL13A | 1593.75521 | -0.938772793 | 0.44406723 | -2.114033 | 0.03451245 | 0.12058682 |
| RPS11 | 1051.51665 | -0.732999102 | 0.43757743 | -1.6751301 | 0.09390871 | 0.23588779 |
| FCGRT | 481.732074 | 1.261007778 | 0.37752061 | 3.34023563 | 0.00083707 | 0.00792935 |
| RCN3 | 142.86877 | 2.638729129 | 0.34548179 | 7.63782412 | 2.21E-14 | 9.77E-12 |
| NOSIP | 294.232925 | 0.219594427 | 0.16492497 | 1.33148081 | 0.18303085 | 0.36548738 |
| PRRG2 | 86.6949083 | 0.083137277 | 0.27870818 | 0.29829507 | 0.76547796 | 0.86903543 |
| PRR12 | 228.048789 | 0.947429677 | 0.33136783 | 2.85914803 | 0.00424781 | 0.0267946 |
| RRAS | 93.4145592 | -1.420838699 | 0.51221609 | -2.7739049 | 0.00553879 | 0.03271218 |
| SCAF1 | 189.861275 | -0.105808939 | 0.2721751 | -0.3887532 | 0.69745872 | 0.82563837 |
| IRF3 | 402.409 | -0.011510187 | 0.29944148 | -0.0384389 | 0.96933778 | 0.98471155 |
| PRMT1 | 294.788074 | -0.140406046 | 0.28416581 | -0.494099 | 0.62123625 | 0.77507036 |
| AP2A1 | 547.061352 | 0.015468154 | 0.26150127 | 0.05915135 | 0.95283156 | 0.9761912 |
| FUZ | 58.5847785 | 0.943513425 | 0.47187132 | 1.99951423 | 0.04555274 | 0.14614624 |
| MED25 | 339.684911 | 0.085261646 | 0.28642314 | 0.29767723 | 0.76594952 | 0.86918372 |
| PTOV1-AS1 | 59.8555353 | 0.265009472 | 0.34496475 | 0.7682219 | 0.44235536 | 0.63404423 |
| PTOV1 | 350.393033 | 0.289069059 | 0.21517725 | 1.34339972 | 0.17914258 | 0.36041498 |
| PNKP | 415.620496 | -0.662602844 | 0.41680592 | -1.5897155 | 0.11189894 | 0.26485905 |
| TBC1D17 | 267.190345 | 0.372832192 | 0.21765374 | 1.7129602 | 0.08671986 | 0.22377054 |
| IL4I1 | 55.084936 | -2.885051813 | 0.68108541 | -4.2359619 | 2.28E-05 | 0.00048865 |
| ATF5 | 149.893472 | -1.295670004 | 0.34023157 | -3.8082004 | 0.00013998 | 0.00200494 |
| VRK3 | 881.565074 | -0.624504833 | 0.23881369 | -2.6150294 | 0.00892198 | 0.04636532 |
| ZNF473 | 105.393578 | -0.687794865 | 0.38293554 | -1.7961113 | 0.07247681 | 0.19878777 |
| MYH14 | 484.235899 | 0.006440892 | 0.32785096 | 0.01964579 | 0.98432593 | 0.99236213 |
| KCNC3 | 540.088443 | 0.438149028 | 0.25580632 | 1.71281552 | 0.08674648 | 0.22379711 |

| | | | | | | |
|---|---|---|---|---|---|---|
| NR1H2 | 490.051257 | -0.144246761 | 0.36493293 | -0.3952692 | 0.6926442 | 0.82278136 |
| NAPSA | 1075.41331 | -0.008337665 | 0.52186919 | -0.0159765 | 0.98725311 | 0.99360989 |
| POLD1 | 38.8320044 | -0.504895747 | 0.36801528 | -1.3719423 | 0.17008139 | 0.34910236 |
| EMC10 | 421.502885 | -0.301252254 | 0.36545263 | -0.8243264 | 0.40975412 | 0.60360018 |
| JOSD2 | 100.310152 | 0.436583538 | 0.41056367 | 1.06337597 | 0.28761148 | 0.48746773 |
| ASPDH | 95.99244 | 2.950161322 | 0.52222392 | 5.64922677 | 1.61E-08 | 1.22E-06 |
| C19orf48 | 99.7791625 | -0.988772573 | 0.63381951 | -1.5600223 | 0.11875461 | 0.27533313 |
| CTU1 | 91.1468686 | -0.863950422 | 0.39129957 | -2.2079003 | 0.02725123 | 0.10236573 |
| ETFB | 671.404143 | 0.32752664 | 0.35593119 | 0.92019652 | 0.35747007 | 0.5561453 |
| ZNF175 | 168.177282 | 0.340678943 | 0.21787423 | 1.56364954 | 0.11789989 | 0.27381461 |
| SPACA6 | 277.51297 | 0.406264458 | 0.42376959 | 0.95869186 | 0.337714 | 0.53788587 |
| ZNF577 | 257.596961 | 0.305903697 | 0.46595331 | 0.65651148 | 0.51149509 | 0.69054104 |
| ZNF649 | 50.4718831 | 0.242660442 | 0.44949121 | 0.53985581 | 0.58929648 | 0.75126513 |
| ZNF613 | 55.940235 | 0.017425662 | 0.30081427 | 0.05792831 | 0.95380573 | 0.97675123 |
| ZNF350-AS1 | 44.5322718 | -0.425420908 | 0.39356375 | -1.0809454 | 0.27972142 | 0.4782088 |
| ZNF350 | 89.9509051 | -0.183431105 | 0.33579007 | -0.5462672 | 0.58488227 | 0.74757378 |
| ZNF615 | 106.617678 | 0.567607281 | 0.26018691 | 2.18153661 | 0.02914375 | 0.10692593 |
| ZNF614 | 49.7225903 | 0.02668178 | 0.30417231 | 0.08771929 | 0.93009979 | 0.96282888 |
| ZNF432 | 272.380273 | -0.567014497 | 0.420427 | -1.3486634 | 0.17744511 | 0.35805096 |
| ZNF616 | 97.0966095 | -0.081674954 | 0.24034179 | -0.3398284 | 0.7339858 | 0.84943603 |
| AC011468.3 | 95.5669164 | -0.902905749 | 0.55659926 | -1.6221828 | 0.10476421 | 0.2529434 |
| ZNF836 | 73.0103235 | -0.308609572 | 0.26504951 | -1.1643469 | 0.24428346 | 0.43943757 |
| PPP2R1A | 840.111207 | -0.538067583 | 0.25384428 | -2.1196758 | 0.03403339 | 0.11946114 |
| ZNF766 | 256.422412 | -0.255114722 | 0.27897532 | -0.9144706 | 0.36046963 | 0.5586436 |
| ZNF480 | 260.971243 | -0.472873475 | 0.36092941 | -1.310155 | 0.19014341 | 0.3741919 |
| AC010320.4 | 52.453364 | -0.654555335 | 0.56377706 | -1.161018 | 0.24563457 | 0.44119572 |
| ZNF610 | 42.6345239 | -0.223511959 | 0.5386985 | -0.4149111 | 0.67820702 | 0.81367034 |
| ZNF880 | 45.9908169 | -0.205338868 | 0.40627683 | -0.5054161 | 0.61326658 | 0.76933839 |
| ZNF528 | 98.6812099 | 0.299506323 | 0.42674282 | 0.70184268 | 0.48277728 | 0.66687778 |
| ZNF808 | 241.130586 | 0.499852997 | 0.26855207 | 1.86128892 | 0.06270339 | 0.18014574 |
| ZNF701 | 313.750283 | 0.340821116 | 0.22364459 | 1.52394084 | 0.12752349 | 0.288872 |
| ZNF83 | 2616.51754 | -0.284777762 | 0.44870284 | -0.6346689 | 0.52564435 | 0.70084961 |
| ZNF611 | 1365.25525 | -0.578330903 | 0.43246752 | -1.3372817 | 0.1811307 | 0.36329506 |
| ZNF600 | 144.727279 | -0.30580314 | 0.3300081 | -0.9266534 | 0.35410651 | 0.55250289 |
| ZNF28 | 89.542149 | -0.171276867 | 0.25573581 | -0.6697414 | 0.50302263 | 0.68381743 |
| AC010487.3 | 62.0204976 | -0.185695998 | 0.60468601 | -0.3070949 | 0.75877113 | 0.86482758 |
| ZNF160 | 675.709866 | -0.479155365 | 0.27112436 | -1.76729 | 0.07717968 | 0.20709409 |
| ZNF347 | 116.174693 | -0.717954373 | 0.27449489 | -2.6155474 | 0.00890846 | 0.04635326 |
| ZNF677 | 44.376866 | -0.97712898 | 0.51578768 | -1.8944403 | 0.05816659 | 0.1712176 |
| ZNF845 | 62.9267075 | 0.054682554 | 0.35232054 | 0.15520683 | 0.87665827 | 0.93570743 |
| ZNF525 | 138.799106 | -0.593719392 | 0.28987944 | -2.0481597 | 0.04054435 | 0.13459468 |
| ZNF765 | 272.590364 | -0.484415313 | 0.26321417 | -1.8403846 | 0.06571179 | 0.18621285 |
| ZNF761 | 338.079638 | -0.782674753 | 0.33580634 | -2.3307325 | 0.01976747 | 0.08180939 |
| ZNF813 | 77.0367431 | -0.11548734 | 0.31314762 | -0.3687952 | 0.71228038 | 0.83538853 |
| ZNF331 | 242.87408 | 0.277590115 | 0.2650672 | 1.04724431 | 0.29498691 | 0.49496702 |
| NDUFA3 | 165.848844 | 0.28686505 | 0.41161564 | 0.69692456 | 0.48585001 | 0.66931729 |
| TFPT | 68.3889903 | 0.012134185 | 0.29683812 | 0.04087812 | 0.96739306 | 0.98336311 |
| PRPF31 | 332.137489 | 0.140928235 | 0.23185628 | 0.60782584 | 0.54330299 | 0.71379134 |
| CNOT3 | 186.525839 | 0.094477701 | 0.27935862 | 0.33819504 | 0.73521621 | 0.85021586 |
| LENG1 | 52.407293 | 0.165577243 | 0.33393387 | 0.49583841 | 0.62000842 | 0.77432081 |
| TMC4 | 401.303563 | 0.053502174 | 0.48940073 | 0.10932181 | 0.91294725 | 0.95364791 |
| MBOAT7 | 308.409227 | -0.32840192 | 0.33125626 | -0.9913833 | 0.32149845 | 0.52188291 |
| TSEN34 | 170.759829 | -0.200988098 | 0.30079215 | -0.668196 | 0.50400852 | 0.68461449 |
| RPS9 | 1946.93861 | -0.78665674 | 0.36437379 | -2.1589279 | 0.03085576 | 0.1113407 |
| TTYH1 | 35.7884133 | -1.076605263 | 0.56669103 | -1.8998099 | 0.05745806 | 0.17002973 |
| LENG8-AS1 | 238.583117 | 0.428064887 | 0.30402346 | 1.40799953 | 0.15913121 | 0.33408049 |
| LENG8 | 1217.04883 | 0.771096085 | 0.31685958 | 2.43355772 | 0.01495125 | 0.06690722 |
| LENG9 | 47.0044025 | 0.399960154 | 0.38103663 | 1.04966328 | 0.29387295 | 0.49394984 |
| AC011476.3 | 82.5335058 | -1.40000226 | 0.51998837 | -2.6923723 | 0.00709457 | 0.03937056 |
| RDH13 | 182.245874 | -0.024023832 | 0.3009992 | -0.0798136 | 0.9363855 | 0.96589714 |
| PPP1R12C | 383.698676 | 0.069488538 | 0.32860009 | 0.21146841 | 0.83252178 | 0.91159738 |
| TMEM86B | 43.5102615 | 0.946632842 | 0.5649614 | 1.67557084 | 0.09382229 | 0.23579429 |
| PPP6R1 | 385.646175 | -0.104782065 | 0.4417697 | -0.2371871 | 0.81251166 | 0.89861746 |
| HSPBP1 | 125.702446 | 0.042956376 | 0.40975841 | 0.10483342 | 0.91650799 | 0.95525626 |
| TMEM150B | 213.271751 | 1.139711951 | 0.60415921 | 1.88644306 | 0.05923527 | 0.17320622 |
| KMT5C | 106.30931 | 0.008505051 | 0.42668743 | 0.01993274 | 0.98409703 | 0.99220433 |
| TMEM238 | 112.783731 | -0.604438354 | 0.40559818 | -1.4902393 | 0.13616132 | 0.30097779 |
| RPL28 | 1391.35314 | -1.212008775 | 0.40989197 | -2.956898 | 0.00310751 | 0.02115977 |
| UBE2S | 75.2042447 | -0.806737883 | 0.48543469 | -1.6618876 | 0.09653532 | 0.23992921 |
| ISOC2 | 381.995332 | -0.121365934 | 0.43102249 | -0.2815768 | 0.77826803 | 0.87771268 |
| ZNF628 | 35.9807244 | -0.504596231 | 0.5095485 | -0.9902811 | 0.32203675 | 0.52244289 |
| NAT14 | 57.1943942 | -0.142399599 | 0.49147365 | -0.28974 | 0.77201512 | 0.87329455 |
| ZNF579 | 94.549107 | 0.292780452 | 0.3487831 | 0.83943417 | 0.40122571 | 0.59532376 |
| FIZ1 | 52.7516878 | 0.300274261 | 0.34525879 | 0.8697078 | 0.38446011 | 0.58075763 |
| ZNF524 | 74.5005703 | 0.131700531 | 0.37171824 | 0.35430204 | 0.72311254 | 0.84233055 |
| ZNF865 | 95.5069501 | 0.270827958 | 0.37239467 | 0.72726056 | 0.46706636 | 0.65464135 |
| ZNF784 | 33.9463657 | 0.922878747 | 0.3823501 | 2.41370084 | 0.01579142 | 0.06977854 |
| CCDC106 | 125.870477 | 0.627731777 | 0.33578316 | 1.86945579 | 0.06155943 | 0.17786575 |
| U2AF2 | 294.252451 | 0.126975615 | 0.27480244 | 0.46206145 | 0.64403725 | 0.79148232 |
| AC008735.2 | 120.919086 | 0.561406748 | 0.49503431 | 1.13407644 | 0.25676249 | 0.45400457 |
| EPN1 | 502.044422 | 0.40570834 | 0.32342784 | 1.2544014 | 0.20969614 | 0.39926448 |
| ZNF787 | 308.60365 | -0.4195553 | 0.47807439 | -0.8775942 | 0.38016399 | 0.57657954 |
| ZNF444 | 361.225598 | 0.54205889 | 0.24917502 | 2.17541423 | 0.02959909 | 0.1080623 |
| ZSCAN5A | 748.99957 | 0.412817542 | 0.25332557 | 1.62959288 | 0.10318757 | 0.25087631 |
| EDDM13 | 72.8014849 | -0.479921169 | 0.51443769 | -0.9329044 | 0.35086934 | 0.55033368 |
| ZNF582 | 56.3795075 | 0.376888436 | 0.31709953 | 1.18854933 | 0.23461706 | 0.42901494 |
| ZNF583 | 86.102965 | -0.148204262 | 0.36635057 | -0.4045422 | 0.68581406 | 0.81810131 |
| ZNF667 | 132.593039 | -1.005769046 | 0.46383305 | -2.1683859 | 0.03012934 | 0.10953047 |
| ZNF667-AS1 | 128.799203 | -0.255515166 | 0.27712987 | -0.9220051 | 0.3565259 | 0.55520342 |
| ZNF471 | 166.499818 | -0.190392254 | 0.31751956 | -0.5996237 | 0.54875705 | 0.71869915 |
| ZFP28 | 58.2121559 | -0.190827615 | 0.33913256 | -0.5626933 | 0.57364376 | 0.73871994 |
| AC005498.3 | 89.102812 | -0.302315467 | 0.51468867 | -0.5873754 | 0.55695161 | 0.72581628 |
| ZNF470 | 122.764895 | -1.028648364 | 0.54306637 | -1.8941485 | 0.0582053 | 0.1712176 |
| ZNF71 | 74.1972284 | -0.297516003 | 0.26516621 | -1.1219982 | 0.26186322 | 0.45999812 |
| ZNF264 | 357.144429 | 0.053967057 | 0.16334398 | 0.33038901 | 0.74110604 | 0.85299694 |
| ZNF805 | 75.5858382 | -0.279534883 | 0.30843219 | -0.906309 | 0.3647723 | 0.56229577 |
| AC005261.3 | 38.8375579 | -0.748163467 | 0.39960652 | -1.8722504 | 0.06117197 | 0.17723192 |
| ZNF460-AS1 | 100.126956 | 0.064700635 | 0.26763121 | 0.24175295 | 0.8089716 | 0.8961912 |
| ZNF460 | 161.350126 | -0.477217674 | 0.32959151 | -1.4479065 | 0.14764321 | 0.31730095 |
| AC005261.1 | 494.972997 | -0.666293774 | 0.26674631 | -2.4978556 | 0.01249471 | 0.05897554 |
| ZNF304 | 37.0207349 | 0.383591462 | 0.30798178 | 1.2455005 | 0.21294783 | 0.40288142 |
| AC003002.2 | 49.6088051 | 0.393575332 | 0.31845158 | 1.23590321 | 0.21649454 | 0.40728668 |
| AC004076.1 | 158.289281 | -0.247406824 | 0.37082321 | -0.6671827 | 0.50465545 | 0.6850618 |
| ZNF773 | 34.6911094 | -0.818469234 | 0.53340312 | -1.534429 | 0.12492414 | 0.28485404 |
| ZNF549 | 103.09249 | -0.547889783 | 0.31610695 | -1.7332418 | 0.0830527 | 0.21766795 |
| ZNF550 | 61.6112054 | -0.066050831 | 0.26838316 | -0.2461065 | 0.80559982 | 0.89453456 |
| ZNF134 | 60.2324257 | -0.355307589 | 0.29228075 | -1.215638 | 0.22412285 | 0.41638004 |
| ZNF211 | 45.0597555 | 0.140431541 | 0.28780915 | 0.48793286 | 0.6255974 | 0.77834833 |
| AC003006.1 | 64.9456179 | -0.663869329 | 0.41411675 | -1.603097 | 0.10891323 | 0.25996684 |
| ZNF671 | 39.3571821 | 0.6671907 | 0.32438934 | 2.05675902 | 0.03970941 | 0.13282013 |
| ZNF776 | 80.3526196 | 0.689728542 | 0.25754791 | 2.67805921 | 0.00740501 | 0.04061645 |
| ZNF586 | 1030.97447 | 0.109099099 | 0.25030726 | 0.4358607 | 0.6629378 | 0.80379726 |
| ZNF552 | 266.409119 | 0.582014018 | 0.25524018 | 2.28026019 | 0.02259226 | 0.09002097 |

| | | | | | | |
|---|---|---|---|---|---|---|
| AC010522.1 | 63.8208738 | 0.398852839 | 0.2768303 | 1.44078461 | 0.14964554 | 0.31999867 |
| ZNF814 | 654.689836 | 0.564877261 | 0.25078507 | 2.2524358 | 0.02429475 | 0.0943633 |
| ZNF587 | 343.976309 | 0.498452706 | 0.29088895 | 1.71354983 | 0.08661143 | 0.22357494 |
| AC010326.2 | 37.7040099 | 0.400016041 | 0.43292333 | 0.92398819 | 0.35549247 | 0.55390875 |
| ZNF418 | 78.2041355 | 0.485500823 | 0.44763598 | 1.08458847 | 0.27810397 | 0.47631776 |
| C19orf18 | 74.6825956 | 0.304503859 | 0.31976991 | 0.95225927 | 0.34096551 | 0.5416799 |
| ZNF606 | 128.750262 | 0.736212231 | 0.35235997 | 2.08937533 | 0.03667395 | 0.12582058 |
| ZNF135 | 108.661758 | 0.541897558 | 0.57919748 | 0.93560069 | 0.34947882 | 0.54909516 |
| ZSCAN18 | 262.420193 | 0.110682352 | 0.42137862 | 0.26266722 | 0.79280708 | 0.8871958 |
| ZNF329 | 282.087422 | 0.412629315 | 0.23862619 | 1.72918705 | 0.08377562 | 0.21857428 |
| ZNF274 | 472.829865 | -0.294064336 | 0.2507541 | -1.17272 | 0.24090812 | 0.43618862 |
| AC020915.3 | 138.839059 | 0.182788803 | 0.32046682 | 0.57038293 | 0.568418 | 0.73472607 |
| AC010642.2 | 70.2672987 | -0.141816299 | 0.3446252 | -0.4115088 | 0.6806995 | 0.81442647 |
| ZNF497 | 53.5786315 | 0.694246346 | 0.3774297 | 1.8394057 | 0.06585554 | 0.18643214 |
| ZNF837 | 45.2375402 | 0.719107377 | 0.29736263 | 2.41828433 | 0.01559389 | 0.06917327 |
| RPS5 | 695.704198 | -1.281278836 | 0.43893274 | -2.9190779 | 0.00351069 | 0.02326932 |
| ZNF584 | 140.587601 | -0.122591857 | 0.39383971 | -0.3112735 | 0.75559273 | 0.86293197 |
| ZNF132 | 47.4152445 | 0.36410776 | 0.32475145 | 1.12118903 | 0.26220741 | 0.46007129 |
| ZNF324B | 35.9943394 | -0.116167392 | 0.37147105 | -0.3127226 | 0.75449143 | 0.86224896 |
| ZNF324 | 93.9243966 | -0.737493328 | 0.38018224 | -1.9398416 | 0.05239894 | 0.15967814 |
| ZNF446 | 81.0226757 | 0.241689368 | 0.31213471 | 0.77431109 | 0.43874685 | 0.63058645 |
| SLC27A5 | 105.109513 | 0.91959848 | 0.26490924 | 3.47137183 | 0.00051781 | 0.00546808 |
| ZBTB45 | 46.1888536 | -0.093836443 | 0.42844796 | -0.2190148 | 0.82663852 | 0.90711186 |
| TRIM28 | 331.690391 | -0.621340201 | 0.42039849 | -1.4779791 | 0.13941336 | 0.30565243 |
| CHMP2A | 217.789136 | -0.41923889 | 0.34643633 | -1.210147 | 0.22622249 | 0.41880492 |
| UBE2M | 218.759988 | -0.707790638 | 0.32404842 | -2.1842126 | 0.02894663 | 0.10645867 |
| MZF1-AS1 | 133.211259 | 0.257675825 | 0.32035199 | 0.8043522 | 0.4211936 | 0.61490049 |
| MZF1 | 549.839047 | 0.364477832 | 0.29615779 | 1.23068796 | 0.21843959 | 0.40970874 |
| RPS4Y1 | 832.391021 | 0.533906903 | 1.72032805 | 0.3103518 | 0.75629345 | 0.86345151 |
| ZFY | 361.696548 | 0.700949152 | 1.86358818 | 0.37612878 | 0.70682116 | 0.83210663 |
| ZFY-AS1 | 151.856156 | 2.176953767 | 1.61208942 | 1.35039269 | 0.17689006 | 0.35735135 |
| LINC00278 | 6644.04286 | 1.108413335 | 2.13538674 | 0.51906913 | 0.60371254 | 0.76169806 |
| PCDH11Y | 50.7952807 | -0.24966837 | 0.32606835 | -0.7656934 | 0.44385877 | 0.63517875 |
| TBL1Y | 327.762854 | 1.865713989 | 1.69414931 | 1.10126892 | 0.27077963 | 0.46898433 |
| TTTY15 | 437.403451 | 1.057451966 | 1.71696578 | 0.61588413 | 0.537971 | 0.70986341 |
| USP9Y | 2036.07806 | 1.148308716 | 1.91446866 | 0.59980544 | 0.54863591 | 0.71869915 |
| DDX3Y | 354.795098 | 0.234930154 | 2.09534635 | 0.11211996 | 0.91072829 | 0.95307319 |
| UTY | 2635.23335 | 0.846071198 | 1.97578896 | 0.42821942 | 0.66849138 | 0.80729015 |
| NLGN4Y | 997.43144 | 0.284046097 | 1.64583917 | 0.17258436 | 0.86297815 | 0.92826191 |
| TTTY14 | 4874.60387 | 1.571932477 | 1.40893866 | 1.11568553 | 0.26455676 | 0.46242769 |
| KDM5D | 382.445143 | 1.267817998 | 1.98103098 | 0.63997889 | 0.52218633 | 0.69809889 |
| TTTY10 | 171.750467 | 1.234635577 | 1.351534 | 0.91350686 | 0.36097603 | 0.5590214 |
| EIF1AY | 128.246016 | 0.578754291 | 2.14345323 | 0.27001023 | 0.78715239 | 0.88351603 |
| IL17RA | 249.427543 | -0.815672397 | 0.2707186 | -3.0129898 | 0.00258688 | 0.01867209 |
| HDHD5 | 304.416604 | 0.014616964 | 0.32581602 | 0.04486263 | 0.9642168 | 0.98143402 |
| HDHD5-AS1 | 92.1020516 | 0.376353652 | 0.58979837 | 0.63810561 | 0.52340493 | 0.69911435 |
| ADA2 | 48.0215196 | -0.242910871 | 0.88966522 | -0.2730363 | 0.78482534 | 0.88201288 |
| CECR3 | 46.266017 | -0.044383358 | 1.0871962 | -0.0408237 | 0.96743646 | 0.98336311 |
| CECR2 | 540.287876 | -0.153991569 | 0.84506311 | -0.1822249 | 0.85540621 | 0.92352338 |
| SLC25A18 | 57.9678327 | 0.186582355 | 0.84427267 | 0.22099774 | 0.82509419 | 0.90642548 |
| ATP6V1E1 | 777.445605 | -0.32340324 | 0.25682477 | -1.2592369 | 0.20794476 | 0.39703285 |
| BCL2L13 | 1844.06508 | 0.145997519 | 0.19917605 | 0.7330074 | 0.46355391 | 0.65157466 |
| BID | 138.171882 | -0.727813712 | 0.4270047 | -1.704463 | 0.08829462 | 0.22639124 |
| MICAL3 | 3893.81882 | -0.281984997 | 0.29641974 | -0.951303 | 0.34145057 | 0.54213633 |
| PEX26 | 487.852527 | -0.116124884 | 0.15518724 | -0.7482889 | 0.45428594 | 0.64374004 |
| USP18 | 114.659371 | -0.750387314 | 0.91151735 | -0.8232288 | 0.41037791 | 0.60419441 |
| AC023490.3 | 38.9644206 | -0.698864017 | 0.72786907 | -0.9601507 | 0.33697936 | 0.5370278 |
| AC007326.4 | 63.5599207 | -0.905139455 | 0.75579083 | -1.1976058 | 0.23107053 | 0.42401159 |
| DGCR2 | 845.481855 | 0.238184474 | 0.26771394 | 0.88969767 | 0.37362824 | 0.5710026 |
| ESS2 | 47.6476893 | 0.274093624 | 0.33604094 | 0.81565546 | 0.41469722 | 0.60826697 |
| SLC25A1 | 385.875965 | 0.218553247 | 0.26228293 | 0.83327287 | 0.40469085 | 0.59871533 |
| CLTCL1 | 283.712503 | -0.081234101 | 0.24755759 | -0.3281422 | 0.74280412 | 0.8540591 |
| HIRA | 407.679416 | -0.103754031 | 0.20201638 | -0.5135922 | 0.60753715 | 0.76448285 |
| MRPL40 | 145.348497 | -0.974194974 | 0.34523795 | -2.8218073 | 0.00477539 | 0.02929002 |
| UFD1 | 335.750558 | -0.474036063 | 0.37019678 | -1.2804975 | 0.20037021 | 0.387495 |
| GNB1L | 70.9003609 | -0.501825077 | 0.46532723 | -1.0784348 | 0.28083977 | 0.47944813 |
| TXNRD2 | 278.09072 | 0.196863249 | 0.33398271 | 0.58944145 | 0.55556518 | 0.72476748 |
| COMT | 570.084445 | -0.305567434 | 0.18573284 | -1.6451987 | 0.09992884 | 0.24586692 |
| ARVCF | 213.49031 | 0.009131571 | 0.219789 | 0.04154699 | 0.96685983 | 0.9830686 |
| TANGO2 | 357.913365 | -0.446281128 | 0.3670738 | -1.2157804 | 0.22406859 | 0.41634609 |
| AC006547.1 | 58.1664503 | -1.268052302 | 0.5765416 | -2.1994116 | 0.02784867 | 0.10384158 |
| DGCR8 | 286.037099 | -0.018543982 | 0.33618092 | -0.0551607 | 0.95601042 | 0.97756553 |
| TRMT2A | 108.913416 | 0.050914791 | 0.38791228 | 0.13125336 | 0.89557488 | 0.94595231 |
| RANBP1 | 116.672372 | -0.207449089 | 0.23979492 | -0.8651104 | 0.38697818 | 0.58321165 |
| ZDHHC8 | 212.607821 | 0.510157573 | 0.39115311 | 1.30424011 | 0.19215173 | 0.37658332 |
| DGCR6L | 95.7952855 | 0.089529784 | 0.39664374 | 0.22571838 | 0.82142046 | 0.90404363 |
| ZNF74 | 39.085169 | 0.399552032 | 0.38446008 | 1.03925493 | 0.29868619 | 0.49879283 |
| KLHL22 | 634.630726 | 0.478556802 | 0.27794984 | 1.721738 | 0.08511699 | 0.22086814 |
| MED15 | 1225.27895 | -0.334585308 | 0.24932263 | -1.3419773 | 0.17960335 | 0.36086531 |
| PI4KA | 1509.85107 | -0.156602448 | 0.23073162 | -0.6787212 | 0.49731452 | 0.67922381 |
| SNAP29 | 237.714032 | 0.085039838 | 0.30725974 | 0.27676857 | 0.78195782 | 0.88049748 |
| CRKL | 375.364704 | -0.053470561 | 0.25369213 | -0.2107695 | 0.83306715 | 0.91175754 |
| LZTR1 | 170.06783 | 0.278163097 | 0.20756241 | 1.34014195 | 0.1801992 | 0.36169138 |
| THAP7 | 88.4225567 | 0.331452612 | 0.37152681 | 0.89213646 | 0.3723198 | 0.56984389 |
| THAP7-AS1 | 43.5923761 | 0.335129716 | 0.51547812 | 0.65013374 | 0.51560583 | 0.69315508 |
| GGT2 | 3162.00503 | -0.654822002 | 0.67474095 | -0.9704791 | 0.33180773 | 0.53141196 |
| HIC2 | 291.779291 | 0.293637172 | 0.3764335 | 0.78005059 | 0.4353611 | 0.62776085 |
| UBE2L3 | 850.98737 | -0.222713661 | 0.30085141 | -0.7402779 | 0.45913136 | 0.6475935 |
| YDJC | 85.3559426 | -1.327334904 | 0.46653358 | -2.8451005 | 0.00443974 | 0.0277499 |
| SDF2L1 | 188.019609 | -1.807513706 | 0.58770511 | -3.0755453 | 0.00210118 | 0.01595616 |
| PPIL2 | 603.422606 | 0.783798605 | 0.4388042 | 1.78621493 | 0.07406447 | 0.20162892 |
| YPEL1 | 143.555601 | 0.524337951 | 0.535369 | 0.97939543 | 0.32738464 | 0.52719234 |
| MAPK1 | 1702.2882 | -0.496715729 | 0.22766663 | -2.1817678 | 0.02912667 | 0.10692001 |
| PPM1F | 168.140125 | -0.1915608 | 0.39534769 | -0.4845375 | 0.62800443 | 0.78013927 |
| AC245452.1 | 54.1713707 | -0.960348784 | 0.45281287 | -2.1208514 | 0.03393431 | 0.11926606 |
| TOP3B | 228.936664 | 0.047228545 | 0.27727837 | 0.17032899 | 0.86475141 | 0.92922135 |
| AC245060.6 | 53.8125536 | -0.447228471 | 0.54177492 | -0.8254876 | 0.40909483 | 0.60295299 |
| AC245060.5 | 420.10951 | 0.860587532 | 0.46566276 | 1.8480918 | 0.06458906 | 0.18409697 |
| ZNF280B | 51.9580755 | -0.3276929 | 0.35990538 | -0.9104974 | 0.36256028 | 0.56033529 |
| RSPH14 | 104.497397 | 0.912943257 | 0.58763891 | 1.55357863 | 0.12028496 | 0.27766015 |
| RAB36 | 118.892686 | 0.151658379 | 0.45200449 | 0.33552405 | 0.7372298 | 0.85110831 |
| BCR | 1689.29201 | 0.567568318 | 0.33710719 | 1.68364347 | 0.09225059 | 0.2335141 |
| PCAT14 | 38.0521857 | 1.139428059 | 0.56725394 | 2.00867367 | 0.04457175 | 0.1437856 |
| AP000345.2 | 166.509474 | 2.083227031 | 0.6991713 | 2.97956602 | 0.00288657 | 0.02011501 |
| DRICH1 | 419.445406 | 2.209498843 | 0.49528871 | 4.46103208 | 8.16E-06 | 0.00021459 |
| RGL4 | 62.0995547 | 1.076422906 | 0.66320731 | 1.62305646 | 0.10457734 | 0.25263333 |
| ZNF70 | 57.0296595 | 0.216908435 | 0.49727998 | 0.43618976 | 0.66269906 | 0.80378869 |
| CHCHD10 | 927.100501 | -0.516481557 | 0.4281455 | -1.2063225 | 0.22769316 | 0.42039199 |
| SMARCB1 | 483.308783 | 0.426092941 | 0.20554088 | 2.07303254 | 0.03816925 | 0.12956561 |
| SLC2A11 | 66.5893955 | 0.969441121 | 0.59943784 | 1.61725044 | 0.10582423 | 0.25497148 |
| AP000350.4 | 71.8120376 | 1.288256625 | 0.30347883 | 4.24496368 | 2.19E-05 | 0.00047267 |
| AP000350.6 | 33.3236387 | -0.926942596 | 1.01643418 | -0.9119554 | 0.36179222 | 0.55961134 |
| GSTT2B | 45.9128301 | 0.803836658 | 0.54843938 | 1.46568005 | 0.14273547 | 0.31024713 |

| | | | | | | | |
|---|---|---|---|---|---|---|---|
| DDT | 663.601967 | | 0.011477706 | 0.41456276 | 0.02768629 | 0.97791236 | 0.98902418 |
| CABIN1 | 1215.55154 | | 0.051587939 | 0.26141742 | 0.19733933 | 0.84356201 | 0.91781904 |
| SUSD2 | 106.830368 | | 2.622836695 | 0.92603791 | 2.83232107 | 0.00462114 | 0.02863562 |
| ADORA2A-AS | 1030.90655 | | 0.691167657 | 0.43755831 | 1.57960127 | 0.11419821 | 0.26812519 |
| UPB1 | 1414.04757 | | 2.057727163 | 0.61444556 | 3.34891698 | 0.00081128 | 0.0077385 |
| GUCD1 | 576.054655 | | -0.340347491 | 0.25420379 | -1.3388765 | 0.18061087 | 0.36235848 |
| SNRPD3 | 409.048048 | | -0.021558954 | 0.22093078 | -0.0975824 | 0.92226391 | 0.95826799 |
| GGT1 | 1716.43023 | | 0.708372866 | 0.28040493 | 2.52624968 | 0.01152875 | 0.05567263 |
| LRRC75B | 420.421052 | | 1.462312178 | 0.60022585 | 2.43626992 | 0.01483961 | 0.06664695 |
| SGSM1 | 88.4621999 | | -0.652415347 | 0.49445102 | -1.3194742 | 0.18701064 | 0.370427 |
| KIAA1671 | 3899.49104 | | -0.701453301 | 0.45761929 | -1.5328316 | 0.12531736 | 0.28538379 |
| AL022323.4 | 268.957048 | | -1.206728848 | 0.57807775 | -2.0874854 | 0.03684428 | 0.12619305 |
| AL022323.1 | 60.9134659 | | -0.841989344 | 0.6144025 | -1.3704198 | 0.1705559 | 0.34960516 |
| LRP5L | 337.59475 | | -0.388642938 | 0.37055203 | -1.0488215 | 0.29426028 | 0.49425158 |
| GRK3 | 397.532887 | | -0.588393478 | 0.44474837 | -1.3229806 | 0.18584182 | 0.36901838 |
| HPS4 | 451.058002 | | 0.206746955 | 0.35298702 | 0.58570696 | 0.55807246 | 0.72661361 |
| SRRD | 76.7442757 | | 0.148598119 | 0.25315275 | 0.58698994 | 0.55721046 | 0.72608459 |
| TFIP11 | 167.417392 | | -0.446876568 | 0.31149801 | -1.4346049 | 0.15139972 | 0.32276202 |
| TPST2 | 197.675854 | | -0.358219731 | 0.34094875 | -1.0506556 | 0.29341677 | 0.49335914 |
| MIATNB | 264.898689 | | -0.137997968 | 0.2758248 | -0.5003102 | 0.61685666 | 0.77202832 |
| PITPNB | 1226.73382 | | -0.525161559 | 0.23238318 | -2.2598949 | 0.02382777 | 0.09313011 |
| TTC28 | 7452.62446 | | 0.635296809 | 0.29367994 | 2.16322845 | 0.03052361 | 0.1105125 |
| CHEK2 | 162.020196 | | -0.042309156 | 0.27435259 | -0.1542145 | 0.87744058 | 0.93632371 |
| HSCB | 356.24449 | | -0.155496411 | 0.2467311 | -0.6302262 | 0.52854657 | 0.70289976 |
| CCDC117 | 184.619433 | | 0.020471646 | 0.19397561 | 0.10553722 | 0.91594954 | 0.95496466 |
| XBP1 | 532.982903 | | -1.247226323 | 0.35895587 | -3.4745952 | 0.00051162 | 0.0054279 |
| Z93930.2 | 386.362514 | | 0.478980488 | 0.46167393 | 1.03748653 | 0.29950917 | 0.49980183 |
| ZNRF3 | 4279.03872 | | 0.654181772 | 0.22311899 | 2.93198613 | 0.00336802 | 0.02249309 |
| ZNRF3-AS1 | 64.4741279 | | -0.493292039 | 0.45046288 | -1.0950781 | 0.27348241 | 0.47134709 |
| KREMEN1 | 462.663588 | | 0.702170263 | 0.43220564 | 1.62462079 | 0.10424338 | 0.25231575 |
| EMID1 | 37.2806517 | | 0.198985933 | 0.64643887 | 0.30781864 | 0.75822034 | 0.86448741 |
| RHBDD3 | 215.760487 | | -0.512469214 | 0.4821848 | -1.0628067 | 0.28786964 | 0.48756075 |
| EWSR1 | 1356.80949 | | 0.317182439 | 0.26400539 | 1.20142408 | 0.22958674 | 0.42269247 |
| GAS2L1 | 72.7731232 | | -0.153713544 | 0.4332666 | -0.3547782 | 0.72275576 | 0.84228022 |
| AP1B1 | 559.692554 | | 0.209864119 | 0.19179523 | 1.09420925 | 0.27386322 | 0.47170685 |
| THOC5 | 272.815157 | | 0.404629563 | 0.22609236 | 1.78966494 | 0.07350779 | 0.2005117 |
| NIPSNAP1 | 513.884972 | | 1.444474093 | 0.44776495 | 3.2259651 | 0.00125549 | 0.01073547 |
| NF2 | 612.896707 | | -0.126240562 | 0.37233233 | -0.3390535 | 0.73456946 | 0.84975472 |
| ZMAT5 | 106.137984 | | 0.358389405 | 0.25008117 | 1.43309229 | 0.15183146 | 0.32326093 |
| UQCR10 | 713.835883 | | -1.077317155 | 0.47760585 | -2.2556616 | 0.02409183 | 0.09382649 |
| ASCC2 | 425.82413 | | 0.09185698 | 0.28819271 | 0.31873457 | 0.7499278 | 0.85900284 |
| MTMR3 | 1914.50804 | | 0.231890835 | 0.23671513 | 0.97961984 | 0.32727381 | 0.52707579 |
| LIF | 107.388359 | | -2.763300472 | 0.79669971 | -3.4684341 | 0.0005235 | 0.0055147 |
| TBC1D10A | 332.44505 | | 0.534604418 | 0.31154045 | 1.71600324 | 0.08616145 | 0.22274896 |
| SF3A1 | 164.283433 | | -0.128893295 | 0.19813497 | -0.6505328 | 0.51534813 | 0.69315508 |
| CCDC157 | 71.8727556 | | 0.409906865 | 0.22675635 | 1.80769737 | 0.07065361 | 0.19541679 |
| RNF215 | 497.076784 | | 0.354525831 | 0.4002902 | 0.88567202 | 0.37579429 | 0.57252554 |
| SEC14L2 | 119.672649 | | -0.338440603 | 0.41356019 | -0.8183588 | 0.41315237 | 0.60685942 |
| SEC14L6 | 234.474434 | | -2.227683959 | 0.67964338 | -3.2777248 | 0.00104647 | 0.00936293 |
| GAL3ST1 | 278.486274 | | -0.351219594 | 0.63921363 | -0.5494557 | 0.58269273 | 0.74567914 |
| PES1 | 386.965779 | | -0.208475597 | 0.33371818 | -0.6247055 | 0.53216434 | 0.70560925 |
| TCN2 | 506.963744 | | 0.105627853 | 0.46386215 | 0.22771388 | 0.81986868 | 0.90294063 |
| SLC35E4 | 79.1374761 | | -1.099595319 | 0.39313083 | -2.7970214 | 0.00515761 | 0.03103397 |
| DUSP18 | 176.491721 | | -0.117387845 | 0.27328261 | -0.4295474 | 0.66752488 | 0.80685745 |
| OSBP2 | 470.223778 | | -0.204339678 | 0.24496856 | -0.8341465 | 0.40419842 | 0.59824508 |
| MORC2-AS1 | 39.9987104 | | -0.455098015 | 0.43692051 | -1.0416037 | 0.29759546 | 0.49810722 |
| MORC2 | 346.356958 | | -0.034306012 | 0.28243014 | -0.1214672 | 0.90332096 | 0.9493077 |
| SMTN | 395.027507 | | 0.327171871 | 0.6154274 | 0.53161733 | 0.59499106 | 0.75552751 |
| SELENOM | 256.386398 | | -0.993488944 | 0.44648551 | -2.2251314 | 0.02607242 | 0.09932591 |
| RNF185 | 701.14698 | | 0.322398697 | 0.3403623 | 0.94722212 | 0.34352561 | 0.54391885 |
| LIMK2 | 760.124514 | | -0.504317623 | 0.29795661 | -1.6925874 | 0.09053402 | 0.23023188 |
| PIK3IP1 | 103.170559 | | -0.747272971 | 0.60832071 | -1.2284194 | 0.21928957 | 0.41085321 |
| PIK3IP1-AS1 | 77.6999463 | | -0.617991195 | 0.37669703 | -1.6405523 | 0.10089038 | 0.247308 |
| PATZ1 | 122.03404 | | 0.087826289 | 0.37076642 | 0.23687768 | 0.81275169 | 0.89861746 |
| DRG1 | 531.88991 | | -0.072117245 | 0.21505764 | -0.3353391 | 0.73736927 | 0.85118107 |
| EIF4ENIF1 | 639.257502 | | -0.373181174 | 0.35383087 | -1.054688 | 0.29156804 | 0.4916955 |
| SFI1 | 563.249512 | | -0.196409836 | 0.37657919 | -0.5215632 | 0.60197452 | 0.7609161 |
| PISD | 1272.86237 | | -0.415083778 | 0.37007696 | -1.1216147 | 0.26202628 | 0.46002317 |
| PRR14L | 1147.12721 | | 0.010968005 | 0.30257063 | 0.0362494 | 0.97108349 | 0.98568139 |
| YWHAH | 150.601193 | | -1.579320123 | 0.46279288 | -3.4125852 | 0.0006435 | 0.00649038 |
| SLC5A1 | 375.978286 | | 2.312718241 | 0.85839231 | 2.69424389 | 0.00705485 | 0.03921365 |
| SLC5A4-AS1 | 63.1582758 | | 0.775132604 | 0.73846678 | 1.04965128 | 0.29387847 | 0.49394984 |
| RTCB | 933.973924 | | -0.471993362 | 0.22198851 | -2.1262063 | 0.03348608 | 0.11834101 |
| FBXO7 | 715.902732 | | 0.209704971 | 0.19118514 | 1.09686854 | 0.27269886 | 0.47062144 |
| SYN3 | 66.7096063 | | 0.298526198 | 0.5911015 | 0.50503373 | 0.61353514 | 0.76948725 |
| TIMP3 | 295.2162 | | 0.762523483 | 0.47383907 | 1.60924569 | 0.10756263 | 0.25757529 |
| LARGE1 | 2557.92117 | | -0.361945133 | 0.25047232 | -1.4450504 | 0.14844372 | 0.3183724 |
| HMGXB4 | 347.871691 | | -0.336072744 | 0.2554836 | -1.3154376 | 0.18836286 | 0.37213746 |
| TOM1 | 765.7662 | | -0.503198192 | 0.26369176 | -1.9082818 | 0.0563548 | 0.16770632 |
| HMOX1 | 2980.40684 | | -2.110182964 | 0.59420648 | -3.5512621 | 0.00038339 | 0.00442897 |
| MCM5 | 35.1435331 | | 0.049983055 | 0.44324778 | 0.11276549 | 0.91021648 | 0.9527558 |
| APOL6 | 664.652675 | | -1.065010088 | 0.5170216 | -2.0598948 | 0.0394086 | 0.13226584 |
| RBFOX2 | 2118.06215 | | -0.168314875 | 0.18841533 | -0.8933184 | 0.3716867 | 0.56913135 |
| APOL2 | 397.585998 | | -1.77863486 | 0.5467267 | -3.2532431 | 0.00114096 | 0.00997393 |
| APOL1 | 88.5776332 | | -2.263483973 | 0.59701709 | -3.7913219 | 0.00014985 | 0.00210879 |
| MYH9 | 1984.43665 | | -1.592035544 | 0.54793188 | -2.9055355 | 0.00366625 | 0.02399044 |
| TXN2 | 216.600673 | | 0.022012151 | 0.27541142 | 0.07992461 | 0.93629721 | 0.96589342 |
| FOXRED2 | 60.8858774 | | 0.834571574 | 0.33320385 | 2.50468767 | 0.01225596 | 0.05806858 |
| EIF3D | 415.590316 | | -0.913023271 | 0.35654793 | -2.5607308 | 0.01044523 | 0.05209327 |
| IFT27 | 150.145083 | | 0.421319289 | 0.31654585 | 1.33098977 | 0.18319238 | 0.36558246 |
| TST | 483.50905 | | 0.056244542 | 0.28536745 | 0.19709515 | 0.84375308 | 0.91781904 |
| MPST | 770.850801 | | 0.191064054 | 0.33834261 | 0.56470585 | 0.57227385 | 0.73806372 |
| C1QTNF6 | 85.0157579 | | 0.645132228 | 0.33929419 | 1.90139487 | 0.05725031 | 0.16967136 |
| ELFN2 | 149.256546 | | 2.270059593 | 0.82384213 | 2.7554546 | 0.00586106 | 0.03408468 |
| CARD10 | 291.892423 | | -0.367372672 | 0.27753751 | -1.3236866 | 0.18560716 | 0.36871266 |
| CDC42EP1 | 130.602851 | | -2.476435497 | 0.47530059 | -5.2102513 | 1.89E-07 | 9.27E-06 |
| LGALS2 | 476.710677 | | 1.266133642 | 0.4462654 | 2.83717638 | 0.00455145 | 0.02826764 |
| GGA1 | 542.874547 | | 0.009448203 | 0.21702347 | 0.0435354 | 0.96527475 | 0.98197757 |
| LGALS1 | 84.0734237 | | -1.531977708 | 0.40720542 | -3.7621741 | 0.00016844 | 0.00232278 |
| TRIOBP | 444.412934 | | -0.596477556 | 0.31125249 | -1.9163784 | 0.05531695 | 0.16588908 |
| H1F0 | 465.339598 | | -1.146278447 | 0.59369162 | -1.9307641 | 0.05351224 | 0.16209773 |
| GCAT | 256.120938 | | 0.676268324 | 0.42199627 | 1.60254575 | 0.10903498 | 0.26014664 |
| ANKRD54 | 176.892506 | | 0.320094904 | 0.21406306 | 1.49532996 | 0.13482835 | 0.29928618 |
| EIF3L | 596.155743 | | -0.13473024 | 0.36473257 | -0.3693946 | 0.71183363 | 0.83509031 |
| MICALL1 | 155.46177 | | -0.943872103 | 0.54983905 | -1.7166335 | 0.08604617 | 0.22253487 |
| POLR2F | 165.771358 | | 0.042581273 | 0.26677021 | 0.1596178 | 0.87318216 | 0.93447126 |
| PICK1 | 233.616003 | | 0.081798452 | 0.21083526 | 0.38797331 | 0.69803579 | 0.82596482 |
| AL031587.2 | 84.4366961 | | -0.320095915 | 0.4967358 | -0.6443987 | 0.51931691 | 0.69597507 |
| BAIAP2L2 | 869.119962 | | 0.345763257 | 0.33445608 | 1.03380766 | 0.30122609 | 0.50175064 |
| PLA2G6 | 240.287815 | | 1.253224514 | 0.32112959 | 3.90255075 | 9.52E-05 | 0.00149621 |
| MAFF | 189.39852 | | -3.664422399 | 0.44838947 | -8.1724096 | 3.02E-16 | 2.30E-13 |
| TMEM184B | 382.642739 | | -0.930482445 | 0.40128039 | -2.3187838 | 0.02040676 | 0.08368911 |
| CSNK1E | 104.383445 | | -0.405415803 | 0.43619855 | -0.9294295 | 0.35266654 | 0.55163759 |

| | | | | | | |
|---|---|---|---|---|---|---|
| TPTEP2-CSNI | 940.629743 | -0.422457478 | 0.27590523 | -1.5311688 | 0.12572768 | 0.28603309 |
| DDX17 | 5475.03378 | 0.782067237 | 0.23629422 | 3.309718 | 0.0009339 | 0.00859125 |
| FAM227A | 270.542965 | 0.226178145 | 0.32196917 | 0.70248386 | 0.48237746 | 0.66670759 |
| CBY1 | 291.464942 | 0.390893824 | 0.20210276 | 1.93413401 | 0.05309665 | 0.16120138 |
| AL021707.2 | 44.9736056 | 1.320344225 | 0.35000168 | 3.772394 | 0.00016169 | 0.00224546 |
| TOMM22 | 87.6571588 | -0.618395695 | 0.45512849 | -1.3587277 | 0.17423289 | 0.35401871 |
| JOSD1 | 332.963721 | -1.162149594 | 0.31021347 | -3.7462899 | 0.00017947 | 0.00245507 |
| GTPBP1 | 254.668757 | -0.129795831 | 0.21051538 | -0.6165622 | 0.53752351 | 0.70967393 |
| SUN2 | 442.686829 | 0.621818818 | 0.31394056 | 1.98068967 | 0.04762609 | 0.15062546 |
| AL021707.1 | 154.827671 | -0.252752152 | 0.27886596 | -0.9063571 | 0.36474686 | 0.56229577 |
| NPTXR | 47.7301727 | -2.265387012 | 0.68119503 | -3.3256071 | 0.00088226 | 0.00824347 |
| CBX6 | 665.962875 | -0.728767192 | 0.54931663 | -1.3266796 | 0.18461468 | 0.3674867 |
| CBX7 | 254.176675 | 0.557128878 | 0.42327588 | 1.3162311 | 0.18809649 | 0.37177196 |
| AL031846.2 | 83.0693682 | -0.081454571 | 0.52607031 | -0.1548359 | 0.87695069 | 0.9358738 |
| PDGFB | 88.230935 | 0.121453863 | 0.66336234 | 0.18308827 | 0.85472876 | 0.92322796 |
| AL031590.1 | 62.129355 | 0.721902444 | 0.37827581 | 1.90840235 | 0.05633923 | 0.16770632 |
| RPL3 | 1021.99215 | -0.640351645 | 0.49118134 | -1.303697 | 0.19233691 | 0.3768513 |
| SYNGR1 | 99.2675698 | -1.172944927 | 0.55981779 | -2.0952263 | 0.03615088 | 0.12435953 |
| TAB1 | 259.405199 | 0.268715774 | 0.22066539 | 1.21775223 | 0.22331814 | 0.41556092 |
| MGAT3 | 48.7962277 | 1.135162116 | 0.69894565 | 1.62410642 | 0.1043531 | 0.25237302 |
| MIEF1 | 223.832204 | -0.339428973 | 0.23771892 | -1.4278585 | 0.1533326 | 0.32545768 |
| ATF4 | 722.98424 | -0.639113709 | 0.47321973 | -1.3505644 | 0.17683502 | 0.35735135 |
| RPS19BP1 | 391.812862 | -0.688918312 | 0.3544722 | -1.9435045 | 0.05195523 | 0.15896214 |
| FAM83F | 191.611625 | 0.106991212 | 0.45270604 | 0.23633705 | 0.81317114 | 0.89880951 |
| TNRC6B | 5715.59578 | 0.402135205 | 0.19731816 | 2.03800401 | 0.04154953 | 0.13691548 |
| SGSM3 | 36.3697078 | 0.722306142 | 0.4037492 | 1.78899707 | 0.07361529 | 0.20068511 |
| AL022238.2 | 53.7638551 | -0.19018831 | 0.4991276 | -0.3810415 | 0.70317248 | 0.82953655 |
| MKL1 | 2057.32805 | -0.584227869 | 0.43758201 | -1.3351277 | 0.18183455 | 0.3639885 |
| SLC25A17 | 577.500408 | 0.244018191 | 0.24592919 | 0.99222946 | 0.32108561 | 0.52141674 |
| ST13 | 963.299865 | -0.049276977 | 0.15937504 | -0.3091888 | 0.75717791 | 0.86387001 |
| XPNPEP3 | 784.442027 | 0.312415101 | 0.25232916 | 1.23812524 | 0.21566963 | 0.40631248 |
| RBX1 | 465.94914 | -0.052464309 | 0.2391915 | -0.2193402 | 0.82638505 | 0.90697894 |
| EP300 | 1603.49049 | 0.284804727 | 0.15747512 | 1.80856968 | 0.07051788 | 0.19519156 |
| L3MBTL2 | 157.647752 | 0.142990099 | 0.32496309 | 0.4400195 | 0.65992298 | 0.8020537 |
| AL035681.1 | 48.6086064 | -0.053948748 | 0.39710515 | -0.1358551 | 0.89193585 | 0.94356446 |
| RANGAP1 | 433.966855 | -0.463628124 | 0.46292356 | -1.001522 | 0.31657452 | 0.51664328 |
| ZC3H7B | 566.775597 | 0.388254853 | 0.33546406 | 1.15736648 | 0.24712266 | 0.44301731 |
| TEF | 165.870957 | 2.083237789 | 0.64004285 | 3.25484112 | 0.00113456 | 0.00993064 |
| TOB2 | 577.203203 | 1.190646479 | 0.28907151 | 4.11886487 | 3.81E-05 | 0.00072484 |
| PHF5A | 97.3540586 | -0.02257802 | 0.40072405 | -0.0563431 | 0.95506852 | 0.97731423 |
| ACO2 | 1710.59311 | 0.745883334 | 0.1793047 | 4.15986497 | 3.18E-05 | 0.00063281 |
| POLR3H | 227.288548 | 0.146188381 | 0.41374678 | 0.35332815 | 0.72384245 | 0.84282267 |
| CSDC2 | 543.578376 | 1.040132994 | 0.41155295 | 2.52733701 | 0.01149312 | 0.05558791 |
| PMM1 | 292.671413 | -0.104438887 | 0.36161262 | -0.2888143 | 0.7727235 | 0.87376019 |
| DESI1 | 390.553136 | -0.915737579 | 0.17330752 | -5.2838885 | 1.26E-07 | 6.67E-06 |
| XRCC6 | 841.240947 | -0.271353824 | 0.21099417 | -1.2860726 | 0.19841773 | 0.38479227 |
| SNU13 | 533.36762 | -0.702797617 | 0.29314072 | -2.3974752 | 0.0165085 | 0.07206432 |
| C22orf46 | 61.7315912 | 0.576605334 | 0.48234659 | 1.19541705 | 0.23192413 | 0.42522525 |
| CCDC134 | 101.259853 | 0.331810963 | 0.35443582 | 0.93616658 | 0.34918743 | 0.54887281 |
| SREBF2 | 1348.16239 | -0.552995093 | 0.21966631 | -2.5174324 | 0.01182137 | 0.05665575 |
| WBP2NL | 238.282201 | -0.096559285 | 0.45834938 | -0.2106674 | 0.83314679 | 0.9117719 |
| NAGA | 69.6413912 | 0.167489401 | 0.36789656 | 0.45526221 | 0.64892061 | 0.79404373 |
| SMDT1 | 236.570459 | -0.464504097 | 0.40603708 | -1.1439943 | 0.25262601 | 0.44918209 |
| NDUFA6 | 253.003743 | -0.49360153 | 0.4359451 | -1.1322562 | 0.25752675 | 0.45486445 |
| TCF20 | 2241.93291 | -0.751774432 | 0.37777738 | -1.9899932 | 0.04659168 | 0.14846635 |
| RRP7A | 211.540828 | -0.185047 | 0.46169892 | -0.4007958 | 0.68857045 | 0.81993182 |
| SERHL2 | 58.7609658 | -0.702978368 | 0.34698007 | -2.0259906 | 0.04276575 | 0.13970212 |
| POLDIP3 | 308.72446 | -0.037691844 | 0.18769641 | -0.2008128 | 0.84084494 | 0.9158822 |
| CYB5R3 | 887.421732 | -0.659263093 | 0.28350128 | -2.3254325 | 0.02004884 | 0.08259979 |
| A4GALT | 694.512708 | -0.005047373 | 0.42530706 | -0.0118676 | 0.99053125 | 0.99518064 |
| ARFGAP3 | 1139.6147 | -1.34983012 | 0.27763848 | -4.8618266 | 1.16E-06 | 4.38E-05 |
| PACSIN2 | 2694.22762 | -0.600658374 | 0.31644963 | -1.8981169 | 0.05768069 | 0.17045895 |
| TTLL1 | 93.0457127 | 0.235383286 | 0.44220753 | 0.53229145 | 0.59452415 | 0.75522636 |
| BIK | 57.2110335 | 0.703831463 | 0.42938964 | 1.63914403 | 0.10118327 | 0.24784115 |
| MCAT | 153.263269 | 0.583660254 | 0.29146805 | 2.00248447 | 0.04523265 | 0.14543841 |
| TSPO | 421.979382 | -1.660469627 | 0.31869504 | -5.2102149 | 1.89E-07 | 9.27E-06 |
| TTLL12 | 114.618692 | -0.397546961 | 0.25789096 | -1.5415312 | 0.12318755 | 0.28212727 |
| SCUBE1 | 56.8724412 | -1.26460819 | 1.03700275 | -1.2194839 | 0.22266057 | 0.41477101 |
| EFCAB6 | 1076.12896 | 0.611479907 | 0.36880195 | 1.65801702 | 0.09731403 | 0.24130272 |
| PNPLA3 | 963.453498 | 2.83727017 | 0.70325812 | 4.03446484 | 5.47E-05 | 0.00097724 |
| SAMM50 | 265.560932 | 0.86612586 | 0.20160439 | 4.29616557 | 1.74E-05 | 0.00039766 |
| PARVB | 257.217351 | -0.548941655 | 0.46441445 | -1.1820081 | 0.23720248 | 0.43178411 |
| RTL6 | 54.4790124 | -0.403915313 | 0.40313441 | -1.0019371 | 0.31637399 | 0.51643899 |
| PRR5 | 443.361484 | 1.062954499 | 0.51622644 | 2.05908575 | 0.03948602 | 0.13239602 |
| PRR5-ARHGA | 140.379631 | -0.562387743 | 0.59827158 | -0.9400208 | 0.34720688 | 0.54728204 |
| NUP50-DT | 850.342518 | -0.886195846 | 0.34526156 | -2.5667376 | 0.01026603 | 0.05148789 |
| NUP50 | 307.705603 | -0.413764676 | 0.28181759 | -1.4682003 | 0.14204982 | 0.3093053 |
| KIAA0930 | 410.451853 | -1.631576613 | 0.58148178 | -2.8058947 | 0.00501771 | 0.03037717 |
| FAM118A | 239.276315 | 0.596188099 | 0.53008129 | 1.1247107 | 0.26071164 | 0.4589178 |
| ATXN10 | 1951.66924 | -0.107596692 | 0.14206646 | -0.7573687 | 0.44882899 | 0.639148 |
| MIRLET7BHC | 676.020628 | 0.240413446 | 0.2680431 | 0.89692087 | 0.36976114 | 0.5671795 |
| PPARA | 4877.02453 | 0.985779865 | 0.39563536 | 2.49163744 | 0.01271558 | 0.05974577 |
| TTC38 | 1970.1304 | 1.476167902 | 0.37200312 | 3.96816002 | 7.24E-05 | 0.0012122 |
| TRMU | 450.939688 | 0.257829496 | 0.21283814 | 1.21138765 | 0.22574687 | 0.41815032 |
| CELSR1 | 1905.75459 | 1.140793214 | 0.35310362 | 3.23076042 | 0.00123461 | 0.01059012 |
| AL021392.1 | 54.807228 | 1.516717953 | 0.58250291 | 2.60379464 | 0.0092198 | 0.04753847 |
| GRAMD4 | 489.259422 | 0.0301668 | 0.23163865 | 0.13023215 | 0.89638276 | 0.94620317 |
| CERK | 362.658626 | 1.070367984 | 0.25112146 | 4.26235163 | 2.02E-05 | 0.00044866 |
| AL118516.1 | 71.4082401 | 0.123244323 | 0.61723334 | 0.19967217 | 0.84173698 | 0.91648969 |
| TBC1D22A | 3038.08852 | -0.229745748 | 0.23299301 | -0.9860629 | 0.32410227 | 0.52462178 |
| FAM19A5 | 119.873798 | -0.34216611 | 0.81272319 | -0.4210119 | 0.67374641 | 0.81073146 |
| BRD1 | 668.419984 | -0.06530297 | 0.33445197 | -0.1952537 | 0.84519438 | 0.91854292 |
| ZBED4 | 476.046751 | -0.649205215 | 0.44362078 | -1.4634238 | 0.14335144 | 0.3110032 |
| ALG12 | 121.056783 | -0.028597097 | 0.25251766 | -0.1132479 | 0.90983401 | 0.95264747 |
| CRELD2 | 104.841279 | -0.404135072 | 0.47132258 | -0.857449 | 0.39119679 | 0.58646585 |
| PIM3 | 1001.66833 | -0.907227862 | 0.38383965 | -2.3635595 | 0.01810032 | 0.07695443 |
| PANX2 | 87.8636644 | 0.384781787 | 0.50561792 | 0.76101295 | 0.44664933 | 0.63757779 |
| TRABD | 322.309708 | -0.265726961 | 0.26725463 | -0.9942838 | 0.32008471 | 0.52051264 |
| SELENOO | 923.162665 | 0.195545964 | 0.18604285 | 1.05108023 | 0.29322174 | 0.49326436 |
| TUBGCP6 | 188.92692 | 0.713462431 | 0.28771891 | 2.47972029 | 0.01314855 | 0.0611354 |
| HDAC10 | 82.7317102 | 0.915336971 | 0.34803069 | 2.63004671 | 0.00853731 | 0.04481844 |
| MAPK12 | 172.018601 | -0.548449429 | 0.55956757 | -0.9801308 | 0.32702154 | 0.52691716 |
| MAPK11 | 56.2894494 | 0.185635286 | 0.51719512 | 0.35892699 | 0.71964971 | 0.84036791 |
| PLXNB2 | 1032.17021 | 0.273465706 | 0.30827542 | 0.88708241 | 0.37503453 | 0.5718924 |
| DENND6B | 199.928748 | 0.484800441 | 0.43902549 | 1.1042649 | 0.26947824 | 0.467621 |
| PPP6R2 | 2461.68764 | 0.613223384 | 0.18254758 | 3.35925241 | 0.00078154 | 0.00753872 |
| SBF1 | 303.269947 | 0.278173627 | 0.34014597 | 0.81780662 | 0.41346762 | 0.60717813 |
| ADM2 | 122.528938 | 0.253706998 | 0.37863994 | 0.67004816 | 0.50282709 | 0.68363952 |
| MIOX | 3391.55949 | 2.279184581 | 0.77031289 | 2.95877768 | 0.00308862 | 0.02111703 |
| LMF2 | 230.095518 | -0.442677239 | 0.32339672 | -1.3688365 | 0.17105039 | 0.34999069 |
| NCAPH2 | 67.0771217 | 0.066019829 | 0.46411455 | 0.142249 | 0.88688333 | 0.94159858 |
| TYMP | 166.734419 | -2.201267664 | 0.38229846 | -5.757982 | 8.51E-09 | 7.11E-07 |
| ODF3B | 77.7350953 | -1.608978288 | 0.68419455 | -2.3516386 | 0.01869093 | 0.07858789 |

| | | | | | | |
|---|---|---|---|---|---|---|
| ARSA | 185.900272 | 0.211308605 | 0.26270122 | 0.80436858 | 0.42118414 | 0.61490049 |
| SHANK3 | 439.064918 | 0.54198405 | 0.45781465 | 1.18385038 | 0.2364723 | 0.43108469 |
| RABL2B | 288.895481 | 0.065393715 | 0.23451707 | 0.27884416 | 0.78036443 | 0.87935343 |
| CU633967.1 | 286.274655 | 0.434806993 | 0.59348026 | 0.73263935 | 0.46377842 | 0.65164168 |
| U2AF1L5 | 50.0191038 | -2.032005585 | 0.96627573 | -2.1029252 | 0.03547231 | 0.12281204 |
| FP671120.1 | 5835.36822 | -1.425237848 | 0.67270681 | -2.1186612 | 0.03411911 | 0.11965335 |
| FP236383.1 | 3223.40581 | -0.729738493 | 0.67443527 | -1.0819993 | 0.27925284 | 0.4778675 |
| CR381653.1 | 421.004248 | 0.82284625 | 0.40762666 | 2.01862715 | 0.04352599 | 0.14130997 |
| CR382287.2 | 34.9386066 | 0.468881109 | 0.47044071 | 0.99668481 | 0.31891752 | 0.51910729 |
| HSPA13 | 125.88605 | -0.821677202 | 0.34270624 | -2.3976138 | 0.01650225 | 0.07206002 |
| AF127577.4 | 281.374369 | 0.025281978 | 0.31824621 | 0.07944157 | 0.93668141 | 0.96599895 |
| NRIP1 | 2484.47346 | 0.071589712 | 0.29466132 | 0.24295592 | 0.80803954 | 0.89590037 |
| AF127577.2 | 71.9490381 | -0.554553892 | 0.5433168 | -1.0206824 | 0.30740494 | 0.50760144 |
| AJ009632.2 | 259.363949 | 3.530939987 | 0.66126535 | 5.33967188 | 9.31E-08 | 5.15E-06 |
| USP25 | 2619.04195 | -0.09921896 | 0.2533163 | -0.3916801 | 0.69529458 | 0.82442931 |
| MIR99AHG | 4422.21542 | 2.120547851 | 0.53477529 | 3.96530634 | 7.33E-05 | 0.00122381 |
| CXADR | 386.173947 | -1.767449448 | 0.54284002 | -3.2559306 | 0.00113021 | 0.00991795 |
| BTG3 | 94.0432781 | -2.104199403 | 0.59321041 | -3.5471384 | 0.00038944 | 0.00447073 |
| C21orf91 | 139.146167 | 0.384003637 | 0.25794836 | 1.48868415 | 0.13657056 | 0.30154199 |
| CHODL | 153.111083 | -0.32462374 | 0.62419754 | -0.5200657 | 0.60301777 | 0.7614943 |
| NCAM2 | 152.147074 | 0.405369927 | 0.66083525 | 0.61342056 | 0.5395983 | 0.7111118 |
| LINC01687 | 153.561404 | 1.211826721 | 0.95775612 | 1.26527692 | 0.20577208 | 0.39447802 |
| MRPL39 | 148.174476 | 0.103570407 | 0.27516675 | 0.37639142 | 0.70662592 | 0.83210663 |
| ATP5PF | 609.577852 | -0.491061884 | 0.35583248 | -1.3800367 | 0.16757535 | 0.34566671 |
| GABPA | 278.186585 | 0.16060898 | 0.1899087 | 0.84571682 | 0.39771074 | 0.59222391 |
| APP | 7190.8744 | -1.397099606 | 0.41016952 | -3.4061517 | 0.00065886 | 0.00661122 |
| AP001439.1 | 135.845406 | -2.170172269 | 0.52570538 | -4.128115 | 3.66E-05 | 0.00070313 |
| CYYR1 | 48.3995072 | 0.982000194 | 0.65200761 | 1.50611768 | 0.13203697 | 0.29538611 |
| ADAMTS1 | 1217.40443 | -2.45989824 | 0.47825468 | -5.1434901 | 2.70E-07 | 1.26E-05 |
| ADAMTS5 | 85.0637962 | 1.090525436 | 0.54067964 | 2.01695303 | 0.04370042 | 0.14164143 |
| LINC00113 | 95.3586505 | 2.130315819 | 0.95898567 | 2.22142612 | 0.02632211 | 0.09997152 |
| LINC01695 | 46.7058845 | 0.420422098 | 0.58856854 | 0.7143129 | 0.47503372 | 0.6611115 |
| AF165147.1 | 222.249968 | -0.684302033 | 0.40107138 | -1.7061852 | 0.08797361 | 0.22602705 |
| N6AMT1 | 93.7018329 | 0.614017469 | 0.34857455 | 1.76150977 | 0.07815216 | 0.20877639 |
| LTN1 | 785.888734 | -0.251197991 | 0.20900843 | -1.2018558 | 0.2294194 | 0.42261144 |
| RWDD2B | 103.555716 | 1.064705484 | 0.34851325 | 3.05499286 | 0.00225066 | 0.01678443 |
| USP16 | 437.392807 | -0.169536757 | 0.28127072 | -0.602753 | 0.54667305 | 0.71691997 |
| CCT8 | 363.051323 | -0.495380275 | 0.19060501 | -2.5989887 | 0.00934988 | 0.04798161 |
| MAP3K7CL | 177.119037 | -0.483480607 | 0.46990744 | -1.0288848 | 0.30353383 | 0.5037581 |
| BACH1 | 4197.68086 | -0.839219295 | 0.33931492 | -2.4732755 | 0.01338809 | 0.06189225 |
| GRIK1 | 157.235037 | -1.465664585 | 0.59003576 | -2.4840267 | 0.01299061 | 0.06058599 |
| TIAM1 | 535.574668 | 0.801607327 | 0.56931394 | 1.40802335 | 0.15912416 | 0.33408049 |
| AP000253.1 | 42.7286025 | 0.941154475 | 0.67550615 | 1.39325819 | 0.16354173 | 0.3402651 |
| SOD1 | 983.627116 | -0.568283448 | 0.2836456 | -2.0034982 | 0.04512384 | 0.14519072 |
| SCAF4 | 972.048378 | 0.370574782 | 0.16869151 | 2.19676013 | 0.02803758 | 0.10415702 |
| HUNK | 698.103707 | -1.161374973 | 0.3883264 | -2.9907186 | 0.00278322 | 0.0196041 |
| MIS18A | 48.2155347 | -0.606688765 | 0.49954582 | -1.2144807 | 0.22456421 | 0.41674812 |
| URB1 | 376.462584 | -0.417038781 | 0.40025988 | -1.04192 | 0.29744877 | 0.49796328 |
| URB1-AS1 | 37.5918699 | 0.196722242 | 0.55759162 | 0.35280703 | 0.72423311 | 0.8430627 |
| EVA1C | 719.571838 | -1.360042061 | 0.48850167 | -2.7841094 | 0.00536749 | 0.03194626 |
| SYNJ1 | 670.378772 | 0.573203955 | 0.17403357 | 3.29364023 | 0.00098899 | 0.00895974 |
| PAXBP1-AS1 | 56.8895757 | 0.705165344 | 0.58494015 | 1.20553418 | 0.22799715 | 0.42083988 |
| PAXBP1 | 1121.89013 | 0.029379701 | 0.15572919 | 0.18865892 | 0.85036015 | 0.92134112 |
| C21orf62-AS | 269.317691 | 0.198464503 | 0.56370522 | 0.35207143 | 0.72478469 | 0.84347804 |
| C21orf62 | 74.2356868 | -0.365508429 | 0.70979255 | -0.5149511 | 0.60658722 | 0.76363804 |
| IFNAR1 | 547.815065 | -0.584608126 | 0.23081081 | -2.5328456 | 0.01131408 | 0.05507176 |
| IFNGR2 | 989.108894 | -0.46616304 | 0.23927776 | -1.9482088 | 0.05138998 | 0.15793778 |
| TMEM50B | 611.577802 | 0.344733823 | 0.23224979 | 1.48432349 | 0.13772312 | 0.30335382 |
| DNAJC28 | 45.5343837 | 1.223725842 | 0.2784622 | 4.39458511 | 1.11E-05 | 0.0002771 |
| GART | 445.087511 | -0.092397183 | 0.2468619 | -0.3742869 | 0.70819085 | 0.83287619 |
| SON | 2053.96059 | -0.385502684 | 0.3511381 | -1.0978663 | 0.27226288 | 0.47033143 |
| DONSON | 89.3766054 | -0.350577729 | 0.30867998 | -1.135732 | 0.25606874 | 0.45330418 |
| AP000311.1 | 619.021023 | -0.762025546 | 0.52756766 | -1.4444129 | 0.14862285 | 0.31860705 |
| ITSN1 | 2662.42287 | -0.08157219 | 0.26685147 | -0.3056839 | 0.75984536 | 0.86555309 |
| LINC00649 | 82.8937839 | 0.355780104 | 0.56280467 | 0.63215557 | 0.52728523 | 0.70176703 |
| MRPS6 | 1155.87629 | -0.827356361 | 0.37226817 | -2.2224741 | 0.02625128 | 0.09975778 |
| LINC00310 | 103.89608 | -0.673389424 | 0.48887791 | -1.3774184 | 0.16838293 | 0.34690862 |
| AP000317.1 | 93.3412074 | -0.397360051 | 0.46217528 | -0.8597605 | 0.38992107 | 0.58542916 |
| KCNE1 | 76.6141215 | 0.636115117 | 0.54507279 | 1.16702784 | 0.24319913 | 0.43827642 |
| RCAN1 | 784.864956 | 0.487642317 | 0.28216804 | 1.72819828 | 0.08395268 | 0.21893824 |
| CLIC6 | 61.9578032 | -1.876671209 | 0.7015246 | -2.6751324 | 0.00746997 | 0.04085831 |
| RUNX1 | 1196.8145 | -0.716187302 | 0.52561751 | -1.3625636 | 0.17302008 | 0.35259425 |
| SETD4 | 423.947465 | 0.219995274 | 0.17884328 | 1.23010087 | 0.21865934 | 0.40984049 |
| CBR1 | 499.375599 | -1.516540358 | 0.44122908 | -3.4370816 | 0.00058802 | 0.00603291 |
| AP000688.2 | 47.2493475 | -1.879012646 | 0.56574832 | -3.3212872 | 0.00089603 | 0.00833803 |
| CBR3 | 110.948567 | -1.999386867 | 0.55981086 | -3.57154 | 0.00035489 | 0.00418628 |
| DOPEY2 | 349.648287 | -1.133535293 | 0.46968952 | -2.4133715 | 0.0158057 | 0.06981912 |
| MORC3 | 912.933992 | -0.147931578 | 0.21861437 | -0.6766782 | 0.49861015 | 0.68045095 |
| CLDN14 | 51.4112077 | -1.529441228 | 0.56221161 | -2.7204014 | 0.00652027 | 0.0368252 |
| SIM2 | 64.9388201 | 1.075026516 | 0.85779388 | 1.25324573 | 0.21011629 | 0.3997313 |
| HLCS | 1338.01962 | -0.002675858 | 0.27008384 | -0.0099075 | 0.99209508 | 0.99609195 |
| PIGP | 119.402099 | 0.231177353 | 0.31357931 | 0.73722132 | 0.46098777 | 0.6496103 |
| TTC3 | 2490.98394 | -0.356204257 | 0.23531739 | -1.5137184 | 0.13009728 | 0.2925268 |
| VPS26C | 536.869987 | -0.602028692 | 0.24058069 | -2.5023983 | 0.01233551 | 0.05840484 |
| DYRK1A | 3128.00094 | -0.031389657 | 0.24003526 | -0.130771 | 0.89595645 | 0.9460654 |
| KCNJ6 | 69.1299991 | -2.373448312 | 0.67604316 | -3.5107941 | 0.00044677 | 0.00494659 |
| KCNJ15 | 9530.09046 | 1.596156277 | 0.50379868 | 3.16824226 | 0.00153364 | 0.0125727 |
| ETS2 | 912.887345 | -1.587906827 | 0.38200203 | -4.1568021 | 3.23E-05 | 0.00063834 |
| PSMG1 | 121.705652 | -0.146295717 | 0.27940701 | -0.5235936 | 0.60056125 | 0.76020679 |
| BRWD1 | 2756.66415 | 0.254641831 | 0.14132372 | 1.80183361 | 0.07157159 | 0.19707348 |
| HMGN1 | 638.472341 | -0.462810314 | 0.20237947 | -2.2868441 | 0.02220492 | 0.08891699 |
| WRB | 338.010826 | 0.154137535 | 0.21759873 | 0.70835677 | 0.47872373 | 0.66415649 |
| LCA5L | 146.510805 | 0.671557898 | 0.28119085 | 2.38826371 | 0.01692819 | 0.07347521 |
| SH3BGR | 93.1413608 | 0.69637163 | 0.48606048 | 1.43268515 | 0.15194784 | 0.32340823 |
| B3GALT5 | 57.0192002 | 0.998276726 | 0.84928193 | 1.1754362 | 0.23982025 | 0.43490488 |
| PCP4 | 174.562254 | 0.429264804 | 0.71767707 | 0.59813086 | 0.54975262 | 0.71955844 |
| DSCAM | 50.1330329 | 0.28544677 | 0.334892 | 0.85235469 | 0.39401726 | 0.58879979 |
| BACE2 | 527.860152 | -2.75843668 | 0.42771951 | -6.449172 | 1.12E-10 | 1.73E-08 |
| MX1 | 744.244405 | -2.283537938 | 1.38536514 | -1.6483293 | 0.09928512 | 0.24450253 |
| AP001610.1 | 46.6989646 | -3.339100209 | 1.50728755 | -2.215304 | 0.0267392 | 0.10110738 |
| TMPRSS2 | 345.787199 | 0.130374518 | 0.36352423 | 0.35864052 | 0.71986403 | 0.84047498 |
| RIPK4 | 405.17425 | 0.00578264 | 0.52588711 | 0.01099597 | 0.99122666 | 0.99558466 |
| PRDM15 | 176.480465 | -0.332880974 | 0.46839677 | -0.7106816 | 0.47728155 | 0.66284015 |
| C2CD2 | 320.44051 | -0.776246467 | 0.23553803 | -3.2956311 | 0.00098201 | 0.00891266 |
| ZBTB21 | 291.049581 | -1.22864214 | 0.28416005 | -4.323768 | 1.53E-05 | 0.00035817 |
| ABCG1 | 119.422661 | -1.444274984 | 0.68972439 | -2.0939886 | 0.036261 | 0.12460003 |
| RSPH1 | 90.760539 | -1.0531668 | 0.69987808 | -1.5047861 | 0.13237908 | 0.2956689 |
| SLC37A1 | 309.513335 | -1.815107495 | 0.51489616 | -3.5251914 | 0.00042318 | 0.00476622 |
| LINC01671 | 134.289129 | -0.176537313 | 0.57365479 | -0.3077414 | 0.75827913 | 0.86448741 |
| PDE9A | 703.77746 | -0.275048233 | 0.48140484 | -0.571345 | 0.56776582 | 0.73451775 |
| AP001628.1 | 135.305581 | -1.542718934 | 0.88322398 | -1.7466905 | 0.08069104 | 0.21331381 |
| WDR4 | 264.545861 | -0.4921229 | 0.63253654 | -0.778015 | 0.43656019 | 0.62857823 |
| NDUFV3 | 671.691738 | 0.281574224 | 0.20302199 | 1.38691489 | 0.16546772 | 0.34277681 |

| | | | | | | | |
|---|---|---|---|---|---|---|---|
| PKNOX1 | 617.698988 | | -0.170476198 | 0.29960574 | -0.5690018 | 0.56935494 | 0.73547716 |
| U2AF1 | 63.0963047 | | 1.290917152 | 1.17075654 | 1.10263501 | 0.2701857 | 0.46843352 |
| RRP1B | 249.063706 | | -0.365616925 | 0.43766123 | -0.835388 | 0.4034993 | 0.59761138 |
| PDXK | 1243.64761 | | 0.004414842 | 0.31597631 | 0.01397207 | 0.98885227 | 0.99458453 |
| CSTB | 748.171555 | | -1.704771157 | 0.35133748 | -4.8522325 | 1.22E-06 | 4.51E-05 |
| RRP1 | 148.831489 | | -0.403990522 | 0.43523447 | -0.9282135 | 0.35329682 | 0.55189396 |
| AGPAT3 | 2016.44788 | | 0.780838424 | 0.29889771 | 2.61239342 | 0.00899107 | 0.04664672 |
| TRAPPC10 | 1071.22207 | | 0.128065882 | 0.19514609 | 0.65625647 | 0.51165912 | 0.69069446 |
| GATD3A | 68.8780229 | | -0.837942894 | 1.14392912 | -0.7325129 | 0.46385554 | 0.65164168 |
| PFKL | 1044.1163 | | 0.257284732 | 0.16536455 | 1.55586392 | 0.11974045 | 0.27700969 |
| C21orf2 | 132.383891 | | 0.5359732 | 0.47771414 | 1.12195381 | 0.26188208 | 0.45999812 |
| UBE2G2 | 1246.37839 | | 0.019693791 | 0.18685531 | 0.10539594 | 0.91606164 | 0.95500889 |
| SUMO3 | 231.268425 | | -0.718738223 | 0.43493935 | -1.6525022 | 0.09843222 | 0.24301341 |
| PTTG1IP | 1268.24859 | | -0.811392935 | 0.26905695 | -3.0156922 | 0.00256393 | 0.01855205 |
| FAM207A | 136.137253 | | -0.581816084 | 0.32141015 | -1.8101983 | 0.07026505 | 0.19476534 |
| ADARB1 | 448.736406 | | -1.007779523 | 0.31739886 | -3.1751202 | 0.00149774 | 0.01233008 |
| POFUT2 | 289.296814 | | 0.095279704 | 0.31369793 | 0.30373074 | 0.76133304 | 0.86667154 |
| COL18A1 | 1896.89333 | | 0.281184542 | 0.36974406 | 0.76048427 | 0.44696517 | 0.63774996 |
| COL18A1-AS | 40.4593242 | | 2.151486412 | 0.68079836 | 3.16024028 | 0.00157639 | 0.01283111 |
| SLC19A1 | 112.115212 | | 0.040216985 | 0.44576271 | 0.09022061 | 0.9281119 | 0.96208348 |
| COL6A1 | 219.427525 | | 0.712309919 | 0.29223558 | 2.43745104 | 0.01479122 | 0.06647319 |
| COL6A2 | 69.1028083 | | 0.225112409 | 0.61560101 | 0.36567908 | 0.71460456 | 0.8368246 |
| FTCD | 3225.73346 | | 1.236945433 | 0.40693872 | 3.03963564 | 0.00236865 | 0.01749875 |
| SPATC1L | 37.4312076 | | 0.136726877 | 0.4112799 | 0.3324424 | 0.73955523 | 0.85214219 |
| LSS | 344.240007 | | 0.011183168 | 0.28773914 | 0.03886565 | 0.96899751 | 0.98443884 |
| MCM3AP-AS | 103.692519 | | -0.47770983 | 0.30151562 | -1.5843618 | 0.11311141 | 0.26653225 |
| MCM3AP | 665.949363 | | 0.315254582 | 0.12425803 | 2.53709635 | 0.01117762 | 0.05456254 |
| YBEY | 140.369874 | | 0.615045535 | 0.35335405 | 1.74059287 | 0.08175497 | 0.21527667 |
| C21orf58 | 44.3043918 | | 0.351896665 | 0.41249737 | 0.85308827 | 0.39361036 | 0.58850053 |
| PCNT | 415.882541 | | 0.236065702 | 0.38456456 | 0.61385195 | 0.53931316 | 0.71087273 |
| DIP2A | 938.532113 | | -0.305837396 | 0.19223253 | -1.5909763 | 0.11161492 | 0.26445968 |
| PRMT2 | 923.947498 | | -0.694024808 | 0.28503484 | -2.4348772 | 0.01489685 | 0.06678156 |
| AC011043.1 | 200.662082 | | 0.787937859 | 0.61219171 | 1.28707698 | 0.19806746 | 0.38422171 |
| AL592183.1 | 1308.53242 | | 0.094717849 | 0.49847639 | 0.19001471 | 0.8492976 | 0.92069932 |
| AL354822.1 | 98.6324877 | | 0.384665341 | 0.48661661 | 0.79048954 | 0.42924193 | 0.62214435 |
| AC240274.1 | 63.6942317 | | -0.795636942 | 0.46154901 | -1.7238406 | 0.08473662 | 0.22018669 |

**Thin limbs**

| | baseMean | log2 fold change control/AKI* | lfcSE | stat | pvalue | padj |
|---|---|---|---|---|---|---|
| AL669831.5 | 52.2154787 | 0.340075896 | 0.24264241 | 1.40155176 | 0.16104914 | 0.42721953 |
| NOC2L | 55.91855 | -0.244145408 | 0.29017845 | -0.841363 | 0.40014461 | 0.67469094 |
| HES4 | 57.0154683 | 0.144304531 | 0.65727766 | 0.21954881 | 0.82622256 | 0.93297016 |
| ISG15 | 305.258589 | -1.147474093 | 1.71354113 | -0.6696507 | 0.50308046 | 0.75403031 |
| AGRN | 248.033876 | -0.033915573 | 0.47727619 | -0.0710607 | 0.94334946 | 0.98112027 |
| C1orf159 | 58.7516737 | -0.157478727 | 0.31521239 | -0.4995956 | 0.61735985 | 0.82616513 |
| AL390719.2 | 26.314542 | 0.184552625 | 0.57036289 | 0.32357053 | 0.7462632 | 0.89352127 |
| SDF4 | 94.2080825 | -0.820373022 | 0.28105812 | -2.9188733 | 0.00351299 | 0.04547423 |
| UBE2J2 | 90.0143783 | -0.719439874 | 0.25473828 | -2.8242315 | 0.00473941 | 0.05591589 |
| INTS11 | 71.0480294 | 0.156248668 | 0.2344825 | 0.66635536 | 0.50518399 | 0.75524917 |
| DVL1 | 29.0869628 | -0.317999262 | 0.29902187 | -1.0634649 | 0.28757118 | 0.57450105 |
| AURKAIP1 | 103.585098 | -0.995636505 | 0.4353642 | -2.2869049 | 0.02220138 | 0.14286955 |
| CCNL2 | 182.325605 | 0.094181283 | 0.23253735 | 0.40501572 | 0.68546596 | 0.86578038 |
| MRPL20 | 113.341296 | -0.553003477 | 0.35605293 | -1.5531497 | 0.12038739 | 0.36507762 |
| VWA1 | 41.9851934 | -2.079567388 | 0.6532612 | -3.1833628 | 0.00145575 | 0.02651072 |
| ATAD3B | 41.2815756 | -0.340901758 | 0.27913545 | -1.2212772 | 0.22198108 | 0.50375143 |
| ATAD3A | 25.5240541 | -0.677599995 | 0.39091778 | -1.7333568 | 0.08303227 | 0.29986597 |
| SSU72 | 109.015086 | -0.253980107 | 0.27581803 | -0.9208249 | 0.35714186 | 0.63885377 |
| MIB2 | 30.654244 | 0.8350394 | 0.3366456 | 2.48047026 | 0.01312092 | 0.10500025 |
| CDK11B | 39.7338448 | -0.01284428 | 0.27565333 | -0.0465958 | 0.9628354 | 0.98817751 |
| SLC35E2B | 50.1812219 | 0.226614704 | 0.3022288 | 0.74981175 | 0.45336809 | 0.71320153 |
| SLC35E2A | 50.0978225 | 0.424568429 | 0.35227295 | 1.20522573 | 0.22811617 | 0.51066249 |
| NADK | 47.6562208 | -0.341221888 | 0.28269539 | -1.2070303 | 0.22742049 | 0.51029771 |
| GNB1 | 513.707706 | -0.241320881 | 0.241311 | -1.000041 | 0.31729069 | 0.60364002 |
| PRKCZ | 99.1976964 | 0.14756521 | 0.24015729 | 0.61445234 | 0.53891646 | 0.77708778 |
| FAAP20 | 80.0415192 | -0.087559259 | 0.29610121 | -0.2957072 | 0.7674537 | 0.90412483 |
| SKI | 180.491474 | -0.038926628 | 0.27758865 | -0.1402313 | 0.88847721 | 0.95907089 |
| MORN1 | 60.8365544 | 0.240028865 | 0.29287407 | 0.81956338 | 0.41246506 | 0.68391947 |
| RER1 | 93.0631492 | -0.304517804 | 0.23433103 | -1.2995198 | 0.19376561 | 0.47204496 |
| PEX10 | 27.2832986 | -1.251857101 | 0.32311138 | -3.8743826 | 0.0001069 | 0.00487596 |
| TNFRSF14 | 82.7536551 | -0.628085315 | 0.37233417 | -1.6868861 | 0.09162526 | 0.31554208 |
| ARHGEF16 | 39.2844351 | 0.250248076 | 0.38590149 | 0.64847657 | 0.51667676 | 0.76200872 |
| TPRG1L | 30.4018568 | -0.266950615 | 0.29142483 | -0.9160188 | 0.35965705 | 0.64001011 |
| LRRC47 | 30.8531533 | -0.304512835 | 0.26741743 | -1.1387172 | 0.25482111 | 0.54159654 |
| CEP104 | 78.7240175 | 0.186606396 | 0.21286537 | 0.87664047 | 0.38068195 | 0.65803602 |
| AJAP1 | 49.3217402 | 2.09873186 | 0.69178204 | 3.03380505 | 0.0024149 | 0.03558826 |
| NPHP4 | 52.7220754 | -0.066256172 | 0.26654974 | -0.2485696 | 0.8036937 | 0.92125386 |
| RPL22 | 97.4727796 | 0.126804694 | 0.44424261 | 0.28544019 | 0.77530692 | 0.90712899 |
| RNF207 | 35.4724407 | -0.079740206 | 0.28841171 | -0.2764805 | 0.78217905 | 0.91094142 |
| ICMT | 42.4148785 | -0.503687205 | 0.31771141 | -1.5853608 | 0.11288439 | 0.3518775 |
| NOL9 | 66.3724511 | 0.134789924 | 0.26914699 | 0.50080414 | 0.61650897 | 0.82592881 |
| ZBTB48 | 27.2207303 | -0.032835936 | 0.28070548 | -0.1169765 | 0.9068787 | 0.96640239 |
| KLHL21 | 35.1219449 | -0.281845907 | 0.41139297 | -0.6851014 | 0.49327992 | 0.7455922 |
| DNAJC11 | 139.659154 | 0.338848239 | 0.26644485 | 1.27173876 | 0.20346595 | 0.48230645 |
| CAMTA1 | 528.805391 | 0.110082042 | 0.1666715 | 0.66047311 | 0.50895027 | 0.75796558 |
| VAMP3 | 48.0905403 | -0.461667003 | 0.2463773 | -1.8738212 | 0.06095508 | 0.24952507 |
| PER3 | 136.569854 | 0.124251161 | 0.5403813 | 0.22993238 | 0.81814431 | 0.92999132 |
| PARK7 | 186.269627 | -0.902061705 | 0.38618247 | -2.3358432 | 0.01949941 | 0.13290237 |
| ERRFI1 | 688.8972 | 0.13165967 | 0.71314842 | 0.18461749 | 0.85352907 | 0.9451971 |
| RERE | 1733.27225 | 0.603577132 | 0.15450542 | 3.90651106 | 9.36E-05 | 0.00448767 |
| ENO1 | 810.640303 | -1.48012214 | 0.36926726 | -4.008268 | 6.12E-05 | 0.00332957 |
| MIR34AHG | 65.1425322 | -1.005224667 | 0.56580092 | -1.7766402 | 0.07562746 | 0.28431531 |
| H6PD | 122.004374 | -1.159850556 | 0.36426501 | -3.1840845 | 0.00145213 | 0.02649508 |
| SPSB1 | 147.701827 | -2.824592912 | 0.76560465 | -3.6893623 | 0.00022482 | 0.00818829 |
| SLC25A33 | 61.5537764 | 0.650079811 | 0.5401186 | 1.20358715 | 0.22874919 | 0.51112377 |
| PIK3CD | 33.2398565 | -0.165384463 | 0.28086995 | -0.5888293 | 0.55597577 | 0.78784845 |
| CLSTN1 | 236.701946 | 0.314814603 | 0.29943427 | 1.05136465 | 0.29309114 | 0.58090628 |
| CTNNBIP1 | 45.0622262 | 0.02019188 | 0.32145841 | 0.06281335 | 0.94991513 | 0.98270285 |
| LZIC | 52.2020534 | 0.408051657 | 0.24875642 | 1.64036634 | 0.10092902 | 0.33189121 |
| NMNAT1 | 57.5549054 | 0.060061264 | 0.22395461 | 0.26818499 | 0.78855693 | 0.91447782 |
| UBE4B | 268.72592 | 0.028374809 | 0.19289867 | 0.14709697 | 0.88305548 | 0.95741805 |
| KIF1B | 408.840987 | 0.034691622 | 0.17307933 | 0.20043769 | 0.84113828 | 0.94060981 |
| PGD | 36.5927084 | -0.016505978 | 0.3163835 | -0.0521708 | 0.9583926 | 0.98566059 |
| DFFA | 68.1828516 | -0.579111567 | 0.2130075 | -2.7187379 | 0.00655315 | 0.06867903 |
| PEX14 | 169.071488 | 0.099840209 | 0.21949287 | 0.45486767 | 0.64920444 | 0.84651696 |
| CASZ1 | 135.081393 | -0.17897734 | 0.52468795 | -0.341112 | 0.73301929 | 0.88869659 |
| SRM | 39.0533713 | -1.80918925 | 0.57381008 | -3.1529409 | 0.00161635 | 0.02795985 |
| EXOSC10 | 51.7419569 | 0.158702087 | 0.29359289 | 0.54055153 | 0.58881674 | 0.81143368 |
| MTOR | 106.717495 | -0.221765907 | 0.22406877 | -0.9897225 | 0.32230977 | 0.60955681 |
| AGTRAP | 62.4399309 | -0.804878422 | 0.4792954 | -1.6792951 | 0.09309454 | 0.31780207 |
| MTHFR | 51.100282 | -0.358267745 | 0.46909881 | -0.7637362 | 0.44502446 | 0.70741608 |
| CLCN6 | 47.7383561 | -0.427869194 | 0.24538123 | -1.7436916 | 0.08121288 | 0.29624453 |
| KIAA2013 | 42.0539237 | -0.213030759 | 0.34043065 | -0.6257685 | 0.53146683 | 0.77193643 |
| MFN2 | 53.8389851 | -0.223017944 | 0.23421817 | -0.9521804 | 0.34100552 | 0.62624461 |
| VPS13D | 357.56489 | -0.0472546 | 0.13656636 | -0.3460193 | 0.72932819 | 0.88653304 |
| DHRS3 | 114.57007 | -0.974344663 | 0.63195936 | -1.5417837 | 0.12312615 | 0.36868566 |
| PRDM2 | 211.762686 | 0.396525949 | 0.22532473 | 1.75979771 | 0.07844211 | 0.29033886 |
| KAZN | 4685.04528 | -0.937655986 | 0.35946871 | -2.6084495 | 0.00909534 | 0.08508231 |
| TMEM51-AS | 32.2407885 | -1.264751847 | 0.71948132 | -1.7578661 | 0.07877029 | 0.29095613 |
| TMEM51 | 263.448769 | -0.628277381 | 0.39211304 | -1.6022864 | 0.1090923 | 0.34570701 |
| FHAD1 | 61.7483917 | -0.298282995 | 0.51744389 | -0.5764548 | 0.56430784 | 0.79329392 |
| CASP9 | 42.1720736 | 1.63061978 | 0.44684697 | 3.64916824 | 0.00026309 | 0.00887376 |
| DNAJC16 | 123.54841 | 0.129096426 | 0.1473027 | 0.87640229 | 0.38081138 | 0.65803602 |
| DDI2 | 145.821708 | 0.011515715 | 0.14864758 | 0.07746991 | 0.93824973 | 0.97988379 |
| PLEKHM2 | 92.8596555 | -0.382998132 | 0.21066404 | -1.8180518 | 0.06905622 | 0.26900753 |
| FBLIM1 | 65.6164787 | -0.712316037 | 0.43104039 | -1.6525505 | 0.09842237 | 0.3278122 |
| SPEN | 320.282587 | 0.186466067 | 0.11848955 | 1.57369208 | 0.11555865 | 0.35788437 |
| EPHA2 | 66.5083576 | 0.747561072 | 0.32213022 | 2.32067971 | 0.02030414 | 0.1355161 |
| FBXO42 | 258.89644 | 0.565493238 | 0.23679733 | 2.38808961 | 0.01693621 | 0.12225469 |
| SZRD1 | 83.6736509 | -0.385620064 | 0.27049329 | -1.4256179 | 0.15397868 | 0.41677359 |
| NECAP2 | 44.8584866 | -0.275480745 | 0.29520225 | -0.9331932 | 0.3507202 | 0.6332797 |
| NBPF1 | 63.0307114 | 0.148589829 | 0.21637069 | 0.68673732 | 0.49224827 | 0.74455175 |
| CROCC | 59.9099106 | 0.74074887 | 0.28160543 | 2.63044952 | 0.0085272 | 0.08168208 |
| ATP13A2 | 27.7784801 | -0.782977928 | 0.44488285 | -1.7599643 | 0.07841386 | 0.29033886 |
| SDHB | 80.9296177 | -0.121564503 | 0.24591205 | -0.4943414 | 0.6210651 | 0.82845887 |
| RCC2 | 51.3879274 | -0.477181225 | 0.26320185 | -1.8129858 | 0.06983401 | 0.2709535 |
| ARHGEF10L | 558.977623 | -0.166008582 | 0.44819649 | -0.3703924 | 0.71109012 | 0.87748 |
| UBR4 | 261.720216 | 0.176039678 | 0.15384126 | 1.1442943 | 0.2525016 | 0.5393622 |
| AL035413.1 | 50.0126856 | 0.340623617 | 0.25984497 | 1.31087245 | 0.18990086 | 0.46597228 |
| EMC1 | 39.0455323 | 0.025735272 | 0.28495307 | 0.09031407 | 0.92803764 | 0.97474479 |
| MRTO4 | 38.9048284 | -1.31546376 | 0.45195654 | -2.9105979 | 0.00360738 | 0.04634485 |
| AKR7A2 | 37.1624927 | -0.212208395 | 0.42253763 | -0.5022237 | 0.6155102 | 0.82592881 |
| CAPZB | 283.114069 | -0.413974742 | 0.15685818 | -2.6391657 | 0.00831103 | 0.0803344 |
| NBL1 | 42.0508772 | -0.679419471 | 0.45044804 | -1.5083193 | 0.13147282 | 0.38163088 |
| TMCO4 | 160.359504 | 0.141186829 | 0.39707859 | 0.35556394 | 0.72216715 | 0.88235193 |
| OTUD3 | 41.083136 | 0.361064415 | 0.26745625 | 1.34999432 | 0.17701781 | 0.44872611 |
| CAMK2N1 | 107.855002 | -0.597671157 | 0.37202567 | -1.606532 | 0.10815707 | 0.34396964 |
| PINK1 | 48.0149176 | 0.077148887 | 0.26780801 | 0.28807535 | 0.77328906 | 0.90643453 |
| DDOST | 51.2360372 | -0.445297367 | 0.32071011 | -1.3884731 | 0.16499303 | 0.43324238 |
| HP1BP3 | 329.046103 | 0.392341664 | 0.18958188 | 2.06951034 | 0.03849822 | 0.19501558 |
| EIF4G3 | 1004.29916 | 0.117969524 | 0.17982654 | 0.65601844 | 0.51181226 | 0.75961818 |

*This means that positive fold changes imply higher mean value in controls than in AKI and vice versa.

| | | | | | | |
|---|---|---|---|---|---|---|
| ECE1 | 117.171501 | -0.248305667 | 0.23916691 | -1.0382108 | 0.29917192 | 0.58640635 |
| NBPF3 | 27.6841536 | 0.714112845 | 0.30535518 | 2.33863017 | 0.01935458 | 0.13214365 |
| RAP1GAP | 125.978253 | 0.694181673 | 0.64339808 | 1.07893028 | 0.28061881 | 0.56841775 |
| USP48 | 207.50747 | 0.383702147 | 0.15557728 | 2.46631217 | 0.01365123 | 0.10762663 |
| HSPG2 | 241.633486 | 0.903642622 | 0.40666335 | 2.22209013 | 0.02627722 | 0.15732221 |
| CDC42 | 233.877957 | -0.200375082 | 0.22540767 | -0.8889453 | 0.3740325 | 0.65261518 |
| ZBTB40 | 60.0207708 | 0.336277642 | 0.31799581 | 1.05749079 | 0.29028766 | 0.57714103 |
| EPHB2 | 175.101967 | -1.351041548 | 0.64954899 | -2.0799687 | 0.03752841 | 0.19223776 |
| KDM1A | 147.974466 | 0.267816771 | 0.18417874 | 1.45411337 | 0.14591488 | 0.40399066 |
| AL031428.1 | 38.9406991 | 0.351945314 | 0.33926221 | 1.03738435 | 0.29955677 | 0.58680046 |
| LUZP1 | 138.914955 | -0.138588286 | 0.33513234 | -0.413533 | 0.67921614 | 0.86197819 |
| LINC01355 | 41.8559851 | 0.564472438 | 0.31614014 | 1.78551337 | 0.07417808 | 0.2809616 |
| HNRNPR | 297.485801 | 0.041292463 | 0.16297885 | 0.25336087 | 0.79998936 | 0.92028103 |
| TCEA3 | 83.5415436 | 0.396560814 | 0.54055107 | 0.73362322 | 0.46317841 | 0.72119408 |
| ASAP3 | 40.2871574 | -0.803818915 | 0.55986016 | -1.4357494 | 0.15107365 | 0.41252409 |
| ID3 | 103.071869 | 1.527845375 | 0.83692881 | 1.82553803 | 0.06791988 | 0.26653196 |
| RPL11 | 632.823467 | -0.291072388 | 0.56330893 | -0.5167189 | 0.60535238 | 0.82056119 |
| ELOA | 50.7478309 | -0.862971329 | 0.27273003 | -3.1641963 | 0.00155512 | 0.02753509 |
| ELOA-AS1 | 48.3682283 | 0.100763668 | 0.27449092 | 0.3670929 | 0.71354972 | 0.87865957 |
| PITHD1 | 44.1391474 | -0.105845724 | 0.2452014 | -0.4316685 | 0.66598235 | 0.85763969 |
| LYPLA2 | 48.9175373 | -1.167867886 | 0.41990267 | -2.7812824 | 0.00541446 | 0.06066098 |
| GALE | 35.5043585 | -2.283235161 | 0.41961093 | -5.4413148 | 5.29E-08 | 1.75E-05 |
| HMGCL | 68.098303 | -0.657241859 | 0.2136443 | -3.0763369 | 0.00209561 | 0.03274688 |
| FUCA1 | 47.979151 | -0.528890406 | 0.41047752 | -1.2884759 | 0.19758034 | 0.47630586 |
| PNRC2 | 85.6690606 | -0.130425851 | 0.27418349 | -0.4756882 | 0.63429653 | 0.83690447 |
| SRSF10 | 210.722978 | 0.302390359 | 0.20365327 | 1.48482935 | 0.13758904 | 0.39016145 |
| IFNLR1 | 30.3786586 | 0.419102504 | 0.29101887 | 1.44012142 | 0.14983305 | 0.41050134 |
| STPG1 | 38.0320007 | 0.052962462 | 0.28041706 | 0.18887032 | 0.85019445 | 0.94435157 |
| NIPAL3 | 156.683614 | 0.701942467 | 0.44780448 | 1.56751998 | 0.1169932 | 0.36034658 |
| RCAN3 | 40.1987952 | -0.340230696 | 0.36386949 | -0.935035 | 0.34977028 | 0.63286907 |
| SRRM1 | 192.969848 | 0.231388658 | 0.16910453 | 1.36831733 | 0.17121277 | 0.44163059 |
| CLIC4 | 879.929095 | 0.522246455 | 0.2379996 | 2.19431651 | 0.02821266 | 0.16280067 |
| SYF2 | 69.5679224 | -0.144117753 | 0.37159893 | -0.3878315 | 0.69814076 | 0.87167497 |
| RSRP1 | 383.247482 | 0.082187302 | 0.27006264 | 0.30432682 | 0.76087892 | 0.90125623 |
| TMEM50A | 128.227296 | -0.569344602 | 0.23226233 | -2.4512998 | 0.01423414 | 0.11067271 |
| MACO1 | 115.811048 | 0.2977164 | 0.17401243 | 1.71089154 | 0.08710114 | 0.307082 |
| LDLRAP1 | 28.3535324 | 0.156586005 | 0.40283247 | 0.38871248 | 0.69748885 | 0.8713153 |
| MAN1C1 | 56.0910358 | 0.777078636 | 0.88206773 | 0.88097389 | 0.37833195 | 0.65629152 |
| MTFR1L | 36.9994463 | -0.098093334 | 0.33328789 | -0.2943201 | 0.76851329 | 0.90459435 |
| PAQR7 | 40.2037752 | 0.139247885 | 0.42334372 | 0.32892395 | 0.74221317 | 0.89253455 |
| STMN1 | 58.9064339 | 0.323415144 | 0.66405207 | 0.48703281 | 0.62623508 | 0.83180006 |
| PAFAH2 | 37.9725317 | -0.418640843 | 0.22904722 | -1.8277491 | 0.06758722 | 0.26544403 |
| ZNF593 | 37.4417451 | -1.568507939 | 0.49484588 | -3.1696898 | 0.00152602 | 0.02712008 |
| SH3BGRL3 | 98.2284848 | -1.724505865 | 0.641841 | -2.6868116 | 0.00721376 | 0.0727375 |
| UBXN11 | 41.580528 | 0.132917206 | 0.28232 | 0.47080336 | 0.63778117 | 0.83912075 |
| DHDDS | 53.3789636 | -0.204744337 | 0.32271383 | -0.6344455 | 0.52579013 | 0.76734993 |
| HMGN2 | 138.643183 | -0.286501716 | 0.31206043 | -0.9180969 | 0.35856813 | 0.64001011 |
| RPS6KA1 | 82.6313641 | 0.031485837 | 0.19897694 | 0.15823862 | 0.87426877 | 0.95263572 |
| ARID1A | 238.832712 | 0.082954285 | 0.21350099 | 0.38854286 | 0.69761434 | 0.87135843 |
| GPN2 | 31.7608977 | -0.622613283 | 0.31064144 | -2.0042828 | 0.04503978 | 0.21106218 |
| NUDC | 100.625358 | -0.433842524 | 0.24184361 | -1.793897 | 0.07282961 | 0.27853529 |
| TRNP1 | 38.8680188 | -1.886254834 | 0.65011484 | -2.9014179 | 0.00371478 | 0.04713098 |
| SLC9A1 | 34.94285 | -0.317638056 | 0.26036111 | -1.2199904 | 0.22246851 | 0.50402693 |
| WDTC1 | 96.7161072 | 0.389108469 | 0.21339507 | 1.82341823 | 0.06824007 | 0.26710321 |
| TMEM222 | 37.3974699 | -0.388638494 | 0.27364792 | -1.4202136 | 0.15554551 | 0.41978152 |
| WASF2 | 265.004584 | -0.212438268 | 0.21374175 | -0.9939016 | 0.32027077 | 0.60702226 |
| AHDC1 | 103.912831 | -0.089856239 | 0.39688347 | -0.2264046 | 0.82088676 | 0.93138972 |
| IFI6 | 182.616365 | -1.488315453 | 1.59653873 | -0.9322138 | 0.35122604 | 0.63335467 |
| FAM76A | 49.5930421 | 0.197757954 | 0.27959872 | 0.70729207 | 0.47938499 | 0.73490443 |
| STX12 | 145.801098 | -0.016740548 | 0.21964628 | -0.0762159 | 0.9392473 | 0.97996404 |
| PPP1R8 | 37.8854955 | -0.424066494 | 0.25311379 | -1.6753986 | 0.09385605 | 0.31892412 |
| RPA2 | 47.7312989 | -0.023540914 | 0.32298855 | -0.0728847 | 0.94189789 | 0.98091268 |
| EYA3 | 210.115327 | -0.001620012 | 0.21137524 | -0.0076642 | 0.99388495 | 0.99884271 |
| DNAJC8 | 77.4147257 | -0.431030542 | 0.24545425 | -1.7560524 | 0.07907945 | 0.29145906 |
| ATP5IF1 | 219.74406 | 0.05354894 | 0.35819557 | 0.14949638 | 0.88116197 | 0.95634765 |
| SESN2 | 38.6041552 | -0.672595676 | 0.28566293 | -2.3545081 | 0.01854724 | 0.12902252 |
| PHACTR4 | 365.776054 | 0.110358264 | 0.1822185 | 0.60563698 | 0.54475584 | 0.78068896 |
| RCC1 | 84.8765137 | -1.524583401 | 0.40558227 | -3.7589992 | 0.00017059 | 0.00686607 |
| TRNAU1AP | 80.7563474 | 0.047432429 | 0.22545822 | 0.21038234 | 0.83336927 | 0.93607461 |
| TAF12 | 47.2918339 | 0.354351649 | 0.30920073 | 1.14602463 | 0.25178496 | 0.53906677 |
| GMEB1 | 47.2787702 | -0.121940223 | 0.32707955 | -0.3728152 | 0.709286 | 0.87645132 |
| YTHDF2 | 122.121792 | -0.217937344 | 0.17228339 | -1.2649934 | 0.20587371 | 0.48573011 |
| EPB41 | 265.217138 | -0.459771039 | 0.25402157 | -1.8099685 | 0.07030067 | 0.27219276 |
| SRSF4 | 334.309014 | -0.223189229 | 0.23843923 | -0.9360424 | 0.34925136 | 0.63265484 |
| MECR | 47.27038 | -0.405856677 | 0.24418771 | -1.6620684 | 0.09649906 | 0.32402421 |
| PTPRU | 34.5712101 | -0.219574678 | 0.41366323 | -0.5308054 | 0.59555363 | 0.81525977 |
| PUM1 | 653.674723 | 0.210458734 | 0.11733532 | 1.79365209 | 0.07286872 | 0.27853529 |
| SNRNP40 | 62.9104188 | 0.295994791 | 0.3254702 | 0.90943745 | 0.36311926 | 0.64351459 |
| ZCCHC17 | 71.9321851 | -0.103911211 | 0.26855892 | -0.3869215 | 0.69881435 | 0.8717697 |
| SERINC2 | 148.252558 | -1.495245763 | 0.4595956 | -3.2533944 | 0.00114035 | 0.02270982 |
| TINAGL1 | 120.684174 | 0.432425904 | 0.41647371 | 1.03830301 | 0.29912901 | 0.58640635 |
| PEF1 | 42.5798954 | -0.339724002 | 0.29617076 | -1.1470545 | 0.2513591 | 0.53868382 |
| PTP4A2 | 52.9752532 | 0.129350227 | 0.38500617 | 0.33596923 | 0.73689407 | 0.89057637 |
| KHDRBS1 | 288.166095 | -0.107793994 | 0.2244059 | -0.4803528 | 0.63097658 | 0.83470856 |
| TMEM39B | 26.9837697 | -0.034440128 | 0.35118809 | -0.0980675 | 0.92187872 | 0.97248656 |
| KPNA6 | 214.812582 | -0.203581954 | 0.19818648 | -1.0272242 | 0.30431489 | 0.5922455 |
| TXLNA | 63.4858878 | -0.570576871 | 0.22084544 | -2.5836027 | 0.00977744 | 0.08848555 |
| EIF3I | 65.5454085 | -1.443112591 | 0.44057402 | -3.2755281 | 0.00105465 | 0.02158644 |
| HDAC1 | 89.8106362 | -0.12677987 | 0.26964306 | -0.4701767 | 0.63822881 | 0.83942176 |
| MARCKSL1 | 38.4905563 | -1.605148744 | 0.50767955 | -3.161736 | 0.00156832 | 0.02761563 |
| BSDC1 | 97.1925494 | 0.119377435 | 0.22540952 | 0.52960246 | 0.59638759 | 0.81585918 |
| ZBTB8OS | 132.198746 | 0.051288303 | 0.13668826 | 0.37522099 | 0.70749612 | 0.87544414 |
| RBBP4 | 164.497548 | -0.106151213 | 0.13107372 | -0.8098588 | 0.41802131 | 0.68753883 |
| SYNC | 38.4745747 | -0.128867167 | 0.41549825 | -0.3101509 | 0.75644619 | 0.89917544 |
| KIAA1522 | 151.739252 | -0.10001777 | 0.1885041 | -0.5305867 | 0.59570521 | 0.81525977 |
| YARS | 80.4625303 | -0.912630069 | 0.32582823 | -2.8009546 | 0.00509517 | 0.05887409 |
| S100PBP | 136.253703 | 0.1167906 | 0.18661454 | 0.62583869 | 0.53142075 | 0.77193643 |
| TMEM54 | 48.052993 | -0.668586489 | 0.43826514 | -1.5255297 | 0.12712703 | 0.37451555 |
| RNF19B | 77.7652461 | -0.972331936 | 0.53783048 | -1.8078781 | 0.07062548 | 0.27279822 |
| AK2 | 119.519118 | -0.228569801 | 0.20421363 | -1.1192681 | 0.26302579 | 0.55096925 |
| ZNF362 | 51.0457704 | 0.308818184 | 0.3770203 | 0.81910227 | 0.41272807 | 0.68408231 |
| AL513327.1 | 44.9041214 | -1.044840466 | 0.46486254 | -2.2476332 | 0.02459959 | 0.15217128 |
| PHC2 | 246.58113 | -1.222946349 | 0.38742075 | -3.1566362 | 0.001596 | 0.02784721 |
| SMIM12 | 43.9424992 | -0.352079553 | 0.29784696 | -1.1820821 | 0.23717314 | 0.52000641 |
| ZMYM1 | 79.295495 | 0.345083951 | 0.22360254 | 1.54329172 | 0.12276001 | 0.3680495 |
| SFPQ | 361.384965 | 0.259129139 | 0.21888131 | 1.18387969 | 0.23646069 | 0.51974689 |
| ZMYM4 | 620.965211 | 0.069395566 | 0.12685766 | 0.54703488 | 0.58435477 | 0.80803859 |
| KIAA0319L | 305.428847 | -0.053720256 | 0.1744732 | -0.3078998 | 0.75815861 | 0.89970284 |
| PSMB2 | 86.254854 | -0.442435502 | 0.32579292 | -1.3580268 | 0.17445519 | 0.44586614 |
| AGO4 | 118.83606 | 0.068842312 | 0.24167797 | 0.28485141 | 0.77575798 | 0.90732427 |
| AGO1 | 52.6151596 | 0.502019983 | 0.25815587 | 1.94463907 | 0.05181843 | 0.22800587 |
| AGO3 | 350.542744 | 0.309154052 | 0.12974402 | 2.3828 | 0.01718153 | 0.12346725 |
| ADPRHL2 | 43.5739511 | -0.863119647 | 0.41344223 | -2.0876427 | 0.03683008 | 0.19008369 |

| | | | | | | |
|---|---|---|---|---|---|---|
| TRAPPC3 | 40.7714197 | -0.480093193 | 0.31117624 | -1.5428337 | 0.12287112 | 0.36812373 |
| MAP7D1 | 36.6983021 | -0.913926704 | 0.38620432 | -2.3664331 | 0.01796042 | 0.12664775 |
| THRAP3 | 280.776608 | -0.273548457 | 0.16361111 | -1.6719431 | 0.09453555 | 0.32021075 |
| EVA1B | 56.2370791 | -0.321785511 | 0.66036806 | -0.4872821 | 0.62605846 | 0.83180006 |
| STK40 | 71.5197207 | -1.076714718 | 0.37286044 | -2.8877151 | 0.00388051 | 0.0484634 |
| OSCP1 | 32.3555162 | -0.114903574 | 0.3312189 | -0.3469113 | 0.72865797 | 0.88653304 |
| MRPS15 | 32.4483304 | -1.159193242 | 0.37452843 | -3.0950741 | 0.00196764 | 0.03151251 |
| MEAF6 | 70.0796879 | 0.298559991 | 0.27270744 | 1.09479958 | 0.27360445 | 0.56099915 |
| SNIP1 | 34.3853412 | 0.416383125 | 0.27386049 | 1.52042057 | 0.12840531 | 0.37635066 |
| DNALI1 | 43.5151118 | 0.040628739 | 0.3374539 | 0.12039789 | 0.90416796 | 0.96533938 |
| GNL2 | 27.7594564 | -0.558093363 | 0.43081786 | -1.2954276 | 0.19517275 | 0.47360141 |
| C1orf122 | 49.6950006 | -1.225291564 | 0.454288 | -2.6971691 | 0.00699317 | 0.07109414 |
| MTF1 | 54.5472862 | -0.030981602 | 0.22675846 | -0.1366282 | 0.89132468 | 0.95969529 |
| INPP5B | 80.9674806 | -0.241723069 | 0.24465828 | -0.9880028 | 0.32315127 | 0.61018452 |
| SF3A3 | 53.1899258 | -0.000999794 | 0.30816853 | -0.0032443 | 0.99741142 | 0.99923686 |
| RRAGC | 44.648645 | -0.618083933 | 0.28480219 | -2.1702218 | 0.02999005 | 0.16928383 |
| AL139260.1 | 76.4795603 | -0.177391141 | 0.42592171 | -0.4164877 | 0.67705319 | 0.8613973 |
| AKIRIN1 | 123.512197 | 0.194222343 | 0.335741 | 0.57848861 | 0.56293429 | 0.79260446 |
| NDUFS5 | 201.33101 | -1.095959128 | 0.45859005 | -2.389845 | 0.01685549 | 0.12184079 |
| MACF1 | 1560.1094 | -0.077954317 | 0.27790132 | -0.2805108 | 0.77908565 | 0.90966132 |
| PABPC4 | 68.5163453 | -0.352471859 | 0.28777635 | -1.2248118 | 0.22064616 | 0.50236569 |
| PPIE | 42.315566 | -0.058604705 | 0.28125564 | -0.2083681 | 0.83494156 | 0.93707149 |
| TRIT1 | 104.526439 | 0.279686507 | 0.20752842 | 1.3477022 | 0.17775418 | 0.44938172 |
| CAP1 | 155.76781 | -0.728422778 | 0.39927296 | -1.8243729 | 0.06809571 | 0.26689396 |
| PPT1 | 50.7614585 | -0.255092071 | 0.46651377 | -0.546805 | 0.5845127 | 0.80805991 |
| RLF | 247.609756 | 0.145087719 | 0.26484022 | 0.54783115 | 0.58380785 | 0.80790168 |
| ZMPSTE24 | 69.5766846 | 0.005589081 | 0.2421275 | 0.02308322 | 0.98158389 | 0.9959581 |
| SMAP2 | 46.3374308 | -0.232827289 | 0.26369711 | -0.8829346 | 0.37727163 | 0.65530141 |
| AL603839.3 | 42.0843172 | -0.078105787 | 0.2985129 | -0.2616496 | 0.79359159 | 0.91723627 |
| ZNF684 | 26.6050638 | 0.557275316 | 0.41488118 | 1.34321668 | 0.17920182 | 0.45108076 |
| NFYC | 140.190681 | 0.586099741 | 0.23256784 | 2.52012374 | 0.01173136 | 0.09871458 |
| CITED4 | 84.8032316 | -2.410626376 | 0.51153589 | -4.7125264 | 2.45E-06 | 0.00032459 |
| CTPS1 | 50.0940164 | -0.689437993 | 0.5439442 | -1.2674793 | 0.20498397 | 0.48398853 |
| SLFNL1-AS1 | 36.0845288 | -0.206328582 | 0.3686048 | -0.5597555 | 0.57564619 | 0.80181981 |
| SCMH1 | 840.897399 | -0.319251182 | 0.19748617 | -1.6165749 | 0.10597007 | 0.34040487 |
| AC093151.3 | 35.0011844 | -1.691062013 | 0.76101475 | -2.2221146 | 0.02627556 | 0.15732221 |
| HIVEP3 | 159.566924 | -1.088261542 | 0.32078112 | -3.3925361 | 0.00069249 | 0.01617888 |
| FOXJ3 | 340.60575 | 0.024870149 | 0.1623111 | 0.15322519 | 0.87822069 | 0.95466137 |
| PPCS | 40.5723322 | -0.463171062 | 0.32653288 | -1.4184515 | 0.15605898 | 0.42038498 |
| CCDC30 | 332.827055 | 0.715341131 | 0.35745758 | 2.00119167 | 0.04537174 | 0.21180111 |
| PPIH | 35.37524 | 0.149700075 | 0.29363396 | 0.50981868 | 0.6101785 | 0.822646 |
| YBX1 | 229.702122 | -0.845581367 | 0.5309056 | -1.5927151 | 0.11122412 | 0.34845447 |
| ERMAP | 36.6433142 | 0.169011777 | 0.31144678 | 0.54266663 | 0.58735935 | 0.8106314 |
| SLC2A1 | 52.1225598 | -1.582509427 | 0.75447999 | -2.0974836 | 0.03595079 | 0.18777131 |
| EBNA1BP2 | 59.7831549 | -0.072531093 | 0.25901795 | -0.2800234 | 0.77945954 | 0.90987605 |
| ELOVL1 | 34.8369559 | -0.64331214 | 0.35863162 | -1.7937965 | 0.07284566 | 0.27853529 |
| SZT2 | 68.0968502 | 0.093533123 | 0.23516778 | 0.39772933 | 0.69082972 | 0.86843279 |
| PTPRF | 272.249249 | -0.905507877 | 0.25754273 | -3.515952 | 0.00043818 | 0.01223711 |
| ATP6V0B | 157.652154 | -0.72197237 | 0.29683584 | -2.4322278 | 0.01500627 | 0.11379145 |
| B4GALT2 | 34.2916159 | -1.340135935 | 0.39947812 | -3.3547167 | 0.00079446 | 0.01778667 |
| CCDC24 | 45.828969 | 0.047282107 | 0.37934152 | 0.12464258 | 0.90080652 | 0.96303857 |
| DMAP1 | 27.2130501 | 0.179120435 | 0.40804568 | 0.43897153 | 0.66068217 | 0.85407213 |
| ERI3 | 221.534343 | 0.058387436 | 0.21010734 | 0.27789337 | 0.78109421 | 0.91077326 |
| RNF220 | 50.8315638 | -0.247192495 | 0.26966065 | -0.9166799 | 0.35931041 | 0.64001011 |
| ARMH1 | 38.6277048 | 0.327657271 | 0.49153281 | 0.66660306 | 0.50502572 | 0.75524917 |
| RPS8 | 537.876307 | -0.55447821 | 0.44830879 | -1.2368221 | 0.21615314 | 0.49772378 |
| EIF2B3 | 92.2996744 | 0.053814971 | 0.2839813 | 0.18950181 | 0.84969953 | 0.94414047 |
| UROD | 24.8422876 | -0.346196101 | 0.32164484 | -1.0763303 | 0.28177954 | 0.56931659 |
| ZSWIM5 | 239.017759 | 0.191102519 | 0.30168287 | 0.633455 | 0.52643656 | 0.76812427 |
| TESK2 | 68.2700951 | 0.249126807 | 0.30926723 | 0.80553898 | 0.42050872 | 0.68891139 |
| PRDX1 | 481.051176 | -0.960686324 | 0.50198365 | -1.9137801 | 0.05564826 | 0.23710513 |
| AKR1A1 | 70.486806 | -0.583220472 | 0.29410968 | -1.9830033 | 0.04736706 | 0.21751153 |
| NASP | 114.171192 | 0.235686136 | 0.25296706 | 0.93168705 | 0.35149828 | 0.6334905 |
| GPBP1L1 | 334.882771 | 0.073710035 | 0.15615004 | 0.47204621 | 0.6368938 | 0.83872935 |
| IPP | 70.2513553 | 0.302852932 | 0.27415321 | 1.10468499 | 0.26929611 | 0.55642525 |
| MAST2 | 104.634049 | -0.215349028 | 0.23515648 | -0.915769 | 0.35978804 | 0.64001011 |
| TSPAN1 | 92.2676015 | -1.756213716 | 0.55374349 | -3.1715293 | 0.00151639 | 0.02704927 |
| POMGNT1 | 50.8320882 | -0.745580134 | 0.22557346 | -3.3052653 | 0.00094887 | 0.0201534 |
| LURAP1 | 28.8574237 | 0.053816391 | 0.49899848 | 0.10784881 | 0.91411563 | 0.96861876 |
| LRRC41 | 94.3747275 | -0.041514723 | 0.22528972 | -0.1842726 | 0.8537996 | 0.9452497 |
| UQCRH | 152.366579 | -0.093411805 | 0.38599025 | -0.2420056 | 0.80877582 | 0.92414046 |
| NSUN4 | 61.9846814 | 0.127791306 | 0.18981434 | 0.67324368 | 0.50079228 | 0.75213064 |
| FAAH | 28.4650849 | 0.347064513 | 0.48480953 | 0.71587808 | 0.47406664 | 0.73101808 |
| MKNK1 | 57.1740307 | 0.35778116 | 0.22413702 | 1.59626088 | 0.11043056 | 0.34758554 |
| ATPAF1 | 67.7311263 | 0.254703112 | 0.22918378 | 1.11134877 | 0.26641825 | 0.55430505 |
| EFCAB14 | 126.309826 | -0.093494994 | 0.20782537 | -0.4498729 | 0.65280212 | 0.8484656 |
| PDZK1IP1 | 642.444455 | -2.01981018 | 0.67101102 | -3.0100999 | 0.00261162 | 0.03745013 |
| CMPK1 | 153.986984 | -0.430438822 | 0.24474939 | -1.7586921 | 0.07862981 | 0.29089077 |
| TRABD2B | 1274.79463 | -1.306965833 | 0.40980586 | -3.1892317 | 0.00142651 | 0.02622761 |
| SPATA6 | 286.633888 | 0.184052799 | 0.2431742 | 0.75687635 | 0.44912395 | 0.71037275 |
| AGBL4 | 678.062899 | 0.166944956 | 0.52226257 | 0.31965713 | 0.74922826 | 0.89498161 |
| FAF1 | 786.789592 | -0.109437839 | 0.15205072 | -0.7197456 | 0.47168164 | 0.72945405 |
| RNF11 | 151.126515 | -0.1328843 | 0.171753 | -0.7736942 | 0.43911165 | 0.70285029 |
| TTC39A | 95.0368541 | -2.410722107 | 0.38068905 | -6.3325228 | 2.41E-10 | 1.65E-07 |
| EPS15 | 417.600548 | -0.023596854 | 0.16239112 | -0.1453088 | 0.88446709 | 0.95761929 |
| OSBPL9 | 370.267157 | 0.243971335 | 0.18356526 | 1.32907142 | 0.18382441 | 0.45721925 |
| NRDC | 314.08265 | 0.037593771 | 0.14913865 | 0.25207263 | 0.80098493 | 0.92075299 |
| RAB3B | 35.8329903 | -0.189082467 | 0.50567877 | -0.3739181 | 0.70846522 | 0.87588905 |
| TXNDC12 | 70.5235281 | -0.438442483 | 0.2427806 | -1.8059206 | 0.07093075 | 0.27362662 |
| BTF3L4 | 129.185802 | 0.017281258 | 0.18893748 | 0.09146548 | 0.92712273 | 0.97431801 |
| ZFYVE9 | 224.835795 | 0.183882053 | 0.20696206 | 0.88848195 | 0.37428156 | 0.65269307 |
| CC2D1B | 47.4666625 | -0.336394825 | 0.37700262 | -0.8922878 | 0.37223871 | 0.65197926 |
| PRPF38A | 47.061888 | -0.032361263 | 0.2887817 | -0.1120613 | 0.91077477 | 0.96844354 |
| TUT4 | 576.873357 | 0.401572378 | 0.14850491 | 2.70410174 | 0.00684893 | 0.07024188 |
| ZYG11B | 172.507192 | 0.177421951 | 0.22941691 | 0.7733604 | 0.43930912 | 0.70299733 |
| ECHDC2 | 308.216912 | 0.160251086 | 0.3047432 | 0.52585616 | 0.59898815 | 0.81717937 |
| SCP2 | 201.67368 | 0.063868353 | 0.16175466 | 0.39484706 | 0.69295576 | 0.86938105 |
| CPT2 | 25.7152376 | -0.481846877 | 0.32389918 | -1.4876446 | 0.13684464 | 0.38887471 |
| C1orf123 | 26.2131279 | -0.151676906 | 0.36555108 | -0.4149267 | 0.67819556 | 0.86151115 |
| MAGOH | 44.387942 | 0.036193265 | 0.27709699 | 0.13061587 | 0.89607919 | 0.96163371 |
| GLIS1 | 87.147892 | 0.184330975 | 0.3954092 | 0.46617776 | 0.64108826 | 0.84180733 |
| NDC1 | 59.1370707 | -0.057842856 | 0.34824985 | -0.1660959 | 0.86808152 | 0.95090501 |
| YIPF1 | 39.234008 | -0.62245951 | 0.2393836 | -2.6002596 | 0.00931532 | 0.08613079 |
| HSPB11 | 82.6362223 | 0.006903488 | 0.32046362 | 0.02154219 | 0.98281315 | 0.99616479 |
| LRRC42 | 47.6603854 | -0.050125662 | 0.23681704 | -0.2116641 | 0.83236911 | 0.93582908 |
| TMEM59 | 287.559064 | 0.211704181 | 0.33661847 | 0.62891434 | 0.52940514 | 0.77040597 |
| TCEANC2 | 30.6066357 | 0.154031774 | 0.34681655 | 0.4441304 | 0.65694831 | 0.85164733 |
| MRPL37 | 25.0506842 | -0.033508805 | 0.30672907 | -0.1092456 | 0.91300768 | 0.96844354 |
| SSBP3 | 270.840316 | -0.276161257 | 0.44707517 | -0.6177065 | 0.5367688 | 0.77570079 |
| ACOT11 | 151.233569 | 1.271594538 | 0.4357603 | 2.91810552 | 0.00352165 | 0.04552483 |
| DHCR24 | 108.105885 | 0.266855046 | 0.32404697 | 0.8235073 | 0.41021957 | 0.68208527 |
| USP24 | 381.561784 | 0.279097741 | 0.13139452 | 2.12412013 | 0.0336601 | 0.18043094 |
| PLPP3 | 161.856305 | -1.467158483 | 0.55282636 | -2.6539228 | 0.0079562 | 0.07721628 |

| | | | | | | |
|---|---|---|---|---|---|---|
| PRKAA2 | 383.951175 | 0.715249904 | 0.14140512 | 5.05816122 | 4.23E-07 | 8.82E-05 |
| FYB2 | 73.3070328 | 0.636945021 | 0.77646222 | 0.82031682 | 0.41203552 | 0.68367283 |
| DAB1 | 296.628484 | 0.524382138 | 0.89285099 | 0.58731204 | 0.55699415 | 0.7885674 |
| TACSTD2 | 340.331377 | 0.602656496 | 0.78480372 | 0.76790729 | 0.44254227 | 0.70510851 |
| MYSM1 | 170.414229 | 0.412061668 | 0.21269877 | 1.9373016 | 0.05270849 | 0.23008926 |
| AL136985.3 | 48.5729415 | -0.346333138 | 0.72370228 | -0.4785575 | 0.63225348 | 0.8353707 |
| JUN | 287.830035 | 0.669374238 | 0.32277273 | 2.07382527 | 0.03809554 | 0.19379564 |
| LINC01135 | 61.7535965 | 0.890515722 | 0.46290569 | 1.92375196 | 0.05438568 | 0.23385288 |
| FGGY | 355.068937 | 0.312398969 | 0.33359573 | 0.93645973 | 0.34903654 | 0.63263722 |
| HOOK1 | 231.908085 | 0.487171464 | 0.25209835 | 1.93246593 | 0.05330202 | 0.23167003 |
| NFIA | 1121.83967 | 0.518584751 | 0.2922316 | 1.7745677 | 0.0759693 | 0.28492948 |
| TM2D1 | 151.589241 | 0.43689854 | 0.14800572 | 2.95190307 | 0.00315822 | 0.04214469 |
| PATJ | 1727.61455 | 0.071552057 | 0.21571776 | 0.33169294 | 0.74012114 | 0.89253455 |
| USP1 | 43.5607379 | -0.038069673 | 0.25481296 | -0.1494024 | 0.8812361 | 0.95634765 |
| DOCK7 | 238.719559 | 0.187890129 | 0.21560711 | 0.87144681 | 0.38351024 | 0.66001519 |
| ATG4C | 70.312279 | 0.490230467 | 0.28985276 | 1.69130861 | 0.09077788 | 0.3142614 |
| ALG6 | 37.3408644 | 0.097140027 | 0.22567276 | 0.4304464 | 0.66687095 | 0.85838595 |
| ITGB3BP | 119.88349 | -0.479625359 | 0.42494223 | -1.1286837 | 0.2590313 | 0.54605211 |
| EFCAB7 | 70.6677 | 0.27901168 | 0.35908307 | 0.77701152 | 0.43715197 | 0.70192663 |
| PGM1 | 79.5944965 | -1.11917373 | 0.3705355 | -3.0204224 | 0.00252422 | 0.03652499 |
| ROR1 | 161.366784 | -0.236283625 | 0.5703999 | -0.4142421 | 0.67869685 | 0.86168533 |
| CACHD1 | 84.5697007 | 0.100518566 | 0.6146832 | 0.16352906 | 0.8701019 | 0.95125032 |
| RAVER2 | 139.383949 | 0.584313325 | 0.26737883 | 2.18533877 | 0.02886402 | 0.16526504 |
| JAK1 | 561.086845 | 0.070825404 | 0.26053586 | 0.27184512 | 0.78574111 | 0.91275941 |
| AK4 | 187.46412 | -1.183252492 | 0.42284583 | -2.7983071 | 0.00513712 | 0.05914485 |
| DNAJC6 | 188.125981 | -0.004696928 | 0.34731634 | -0.0135235 | 0.98921015 | 0.99678172 |
| LEPR | 310.318668 | 0.089121745 | 0.32901149 | 0.2708773 | 0.7864854 | 0.91318105 |
| PDE4B | 43.288202 | 1.265671355 | 0.57125069 | 2.21561458 | 0.02671791 | 0.15837305 |
| SGIP1 | 591.888444 | 1.996954818 | 0.67463695 | 2.96004365 | 0.00307595 | 0.04154042 |
| WDR78 | 239.977455 | 0.353242082 | 0.36709244 | 0.9622701 | 0.33591394 | 0.62046272 |
| MIER1 | 224.040024 | 0.054691558 | 0.19970219 | 0.27386559 | 0.78418792 | 0.91212179 |
| SLC35D1 | 62.9448272 | 0.498027082 | 0.20624316 | 2.4147569 | 0.01574571 | 0.11664981 |
| SERBP1 | 243.377981 | -0.573908837 | 0.21146072 | -2.714021 | 0.00664719 | 0.06902242 |
| GADD45A | 90.2397422 | -1.333112616 | 0.46852624 | -2.8453318 | 0.00443652 | 0.05345586 |
| GNG12 | 375.540838 | -0.6395241 | 0.33942029 | -1.8841658 | 0.05954255 | 0.24616233 |
| GNG12-AS1 | 127.451841 | -0.548446855 | 0.36691905 | -1.4947353 | 0.13498353 | 0.38712403 |
| WLS | 174.081886 | -0.138007857 | 0.41776247 | -0.3303501 | 0.74113547 | 0.89253455 |
| LRRC40 | 69.6071607 | 0.418516904 | 0.19599791 | 2.13531311 | 0.03273544 | 0.17715978 |
| SRSF11 | 423.851525 | 0.257979512 | 0.17445855 | 1.47874391 | 0.13920877 | 0.39289358 |
| ANKRD13C | 201.920311 | 0.206974948 | 0.14990771 | 1.3806825 | 0.1673766 | 0.43603495 |
| ZRANB2 | 356.111548 | 0.344577523 | 0.18260422 | 1.88701837 | 0.05915785 | 0.24535287 |
| NEGR1 | 62.1489493 | -0.774509088 | 0.42905236 | -1.8051622 | 0.07104931 | 0.27376561 |
| CRYZ | 176.792058 | 0.201101372 | 0.2570174 | 0.78244262 | 0.43395449 | 0.69898269 |
| TYW3 | 56.9921571 | 0.161245315 | 0.31493485 | 0.51199578 | 0.60865396 | 0.82155842 |
| SLC44A5 | 831.684975 | 1.703711 | 0.62417922 | 2.72952212 | 0.00634262 | 0.06705955 |
| ACADM | 121.431057 | 0.127000352 | 0.20926376 | 0.60689128 | 0.54392306 | 0.78027954 |
| RABGGTB | 57.2005005 | -0.441219284 | 0.42378033 | -1.0411509 | 0.29780551 | 0.58554106 |
| ST6GALNAC3 | 49.7162088 | -0.757520219 | 0.63813282 | -1.1870886 | 0.23519266 | 0.51814869 |
| PIGK | 93.1472474 | 0.361986522 | 0.19832565 | 1.82521285 | 0.06796891 | 0.26661517 |
| AC118549.1 | 507.637521 | 0.319143493 | 0.17796533 | 1.7932903 | 0.07292652 | 0.27864504 |
| USP33 | 312.641294 | 0.249802595 | 0.14932374 | 1.67289265 | 0.09434843 | 0.31991632 |
| MIGA1 | 205.64314 | 0.17691603 | 0.14554706 | 1.21552458 | 0.22416608 | 0.50617113 |
| FUBP1 | 261.878485 | 0.499440924 | 0.19335081 | 2.58308156 | 0.00979222 | 0.08848555 |
| GIPC2 | 102.617484 | -0.312853895 | 0.49658197 | -0.6300146 | 0.52868503 | 0.76999755 |
| PTGFR | 202.646101 | 2.178159417 | 1.04029802 | 2.09378407 | 0.03627922 | 0.18835699 |
| IFI44L | 244.089937 | -1.271182695 | 1.45348176 | -0.8745777 | 0.38180374 | 0.65885391 |
| IFI44 | 91.3519066 | -0.340894497 | 1.20976257 | -0.2817863 | 0.77810739 | 0.90889326 |
| AL136234.1 | 60.8622399 | -0.078377473 | 0.56358821 | -0.1390687 | 0.88939587 | 0.95918866 |
| LINC01781 | 460.558568 | -0.207729781 | 0.68015269 | -0.3054164 | 0.76004904 | 0.90125623 |
| ADGRL2 | 336.999929 | 0.550706793 | 0.38183438 | 1.44226612 | 0.14922731 | 0.40958408 |
| LINC01725 | 207.645142 | 0.097591877 | 0.38202328 | 0.25546055 | 0.79836741 | 0.91949148 |
| TTLL7 | 44.9072568 | -1.183827319 | 0.71303405 | -1.6602676 | 0.09686064 | 0.32454105 |
| PRKACB | 190.381204 | 0.11387854 | 0.20905675 | 0.54472548 | 0.58594233 | 0.80945221 |
| RPF1 | 27.969632 | -0.166050149 | 0.26995351 | -0.6151065 | 0.53848441 | 0.77706269 |
| GNG5 | 114.886757 | -0.623565718 | 0.31138125 | -2.0025796 | 0.04522244 | 0.21161979 |
| SPATA1 | 42.4936269 | 0.306784911 | 0.58175438 | 0.5273444 | 0.59795445 | 0.81658679 |
| CTBS | 65.4001211 | -0.268027658 | 0.2561018 | -1.0465669 | 0.29529939 | 0.5833518 |
| SSX2IP | 58.6363305 | 0.505039141 | 0.23851333 | 2.11744619 | 0.034222 | 0.18232064 |
| SYDE2 | 61.0320524 | 1.304268472 | 0.32895756 | 3.96485336 | 7.34E-05 | 0.00390829 |
| C1orf52 | 55.9035652 | 0.501497091 | 0.21168821 | 2.36903646 | 0.0178345 | 0.12607869 |
| BCL10 | 30.8220022 | -0.030459231 | 0.27382235 | -0.1112372 | 0.91142825 | 0.96844354 |
| AL078459.1 | 95.1907948 | -0.167197732 | 0.38508862 | -0.4341799 | 0.66425782 | 0.85636934 |
| DDAH1 | 487.053884 | 0.310442364 | 0.27441407 | 1.13129173 | 0.25793232 | 0.54515124 |
| CYR61 | 143.10354 | 3.151721128 | 0.56065207 | 5.62152767 | 1.89E-08 | 7.55E-06 |
| ZNHIT6 | 138.01247 | -0.109198276 | 0.20683739 | -0.5279426 | 0.59753916 | 0.81658679 |
| ODF2L | 269.49649 | 0.424283276 | 0.25165402 | 1.68597853 | 0.09179993 | 0.31558127 |
| SH3GLB1 | 183.187612 | -0.170389117 | 0.19551788 | -0.8714759 | 0.38349437 | 0.66001519 |
| SELENOF | 196.006657 | 0.100077496 | 0.16474908 | 0.60745406 | 0.54354962 | 0.78025803 |
| HS2ST1 | 156.177815 | -0.152310013 | 0.22775976 | -0.668731 | 0.50366708 | 0.75479145 |
| LMO4 | 67.0630918 | 0.141841242 | 0.23388794 | 0.60644957 | 0.54421626 | 0.78027954 |
| PKN2-AS1 | 41.965207 | 0.202151978 | 0.44233967 | 0.45700621 | 0.64766658 | 0.84557696 |
| PKN2 | 476.938574 | 0.317385143 | 0.13088915 | 2.42483921 | 0.01531517 | 0.11488175 |
| GTF2B | 70.9863221 | -0.149956938 | 0.21679064 | -0.6917132 | 0.48911749 | 0.74185878 |
| KYAT3 | 97.9257214 | 0.007620981 | 0.21387381 | 0.03563307 | 0.97157494 | 0.99175186 |
| GBP3 | 47.2994934 | -1.465900343 | 0.5352805 | -2.7385648 | 0.0061708 | 0.06585681 |
| GBP2 | 90.3349679 | -2.006979315 | 0.6153133 | -3.2617194 | 0.00110739 | 0.02211903 |
| LRRC8B | 76.2639539 | -0.311220396 | 0.352019 | -0.8841011 | 0.37664162 | 0.65512273 |
| ZNF326 | 142.146336 | 0.125143082 | 0.14913893 | 0.83910404 | 0.40141093 | 0.67562929 |
| ZNF644 | 422.96022 | 0.275865252 | 0.1583767 | 1.74182979 | 0.08153823 | 0.29654614 |
| HFM1 | 57.7653698 | 0.305345166 | 0.33691852 | 0.90628786 | 0.36478351 | 0.64469765 |
| TGFBR3 | 162.543066 | 0.179717941 | 0.4849336 | 0.3706032 | 0.7109331 | 0.87748 |
| BTBD8 | 73.3946296 | 0.33860725 | 0.35733507 | 0.94759032 | 0.34333805 | 0.62811979 |
| GLMN | 80.1591984 | 0.270828476 | 0.17722108 | 1.52819557 | 0.12646399 | 0.37394935 |
| RPAP2 | 245.701606 | -0.054359465 | 0.15137231 | -0.359111 | 0.71951204 | 0.88166633 |
| EVI5 | 409.801073 | 0.307636385 | 0.15924802 | 1.9318067 | 0.05338337 | 0.23183714 |
| RPL5 | 315.503196 | -0.472800857 | 0.39121211 | -1.2085537 | 0.22683434 | 0.50976168 |
| FAM69A | 79.9071348 | -0.1792809 | 0.30704218 | -0.5838966 | 0.55928985 | 0.78981496 |
| MTF2 | 268.497943 | 0.262011954 | 0.2060853 | 1.27137623 | 0.20359484 | 0.48249256 |
| TMED5 | 143.771117 | -0.114280912 | 0.26985145 | -0.4234956 | 0.67193368 | 0.86015189 |
| CCDC18 | 68.7966222 | 0.615858055 | 0.3005417 | 2.04916006 | 0.04044647 | 0.19973796 |
| DR1 | 53.7817126 | -0.109434232 | 0.28304156 | -0.3866366 | 0.69902525 | 0.8717697 |
| FNBP1L | 539.2885 | -0.044030544 | 0.26235473 | -0.1678283 | 0.86671837 | 0.95090501 |
| BCAR3 | 284.931633 | -0.443341543 | 0.46784408 | -0.9476267 | 0.34331953 | 0.62811979 |
| DNTTIP2 | 27.6564744 | -0.652873246 | 0.45376416 | -1.4387942 | 0.15020884 | 0.4109827 |
| GCLM | 89.3269095 | -1.894283432 | 0.31750413 | -5.9661694 | 2.43E-09 | 1.22E-06 |
| ABCA4 | 83.418315 | 3.889459079 | 1.20084206 | 3.23894309 | 0.00119974 | 0.02359807 |
| ARHGAP29 | 2190.23433 | 1.028362599 | 0.29179493 | 3.52426481 | 0.00042466 | 0.01207067 |
| ABCD3 | 258.953052 | 0.280796291 | 0.22988282 | 1.22147575 | 0.22190594 | 0.50375143 |
| SLC44A3 | 187.869211 | 0.842376616 | 0.23447798 | 3.59256177 | 0.00032744 | 0.01021686 |
| CNN3 | 385.333202 | -0.329172148 | 0.3143661 | -1.0470981 | 0.29505433 | 0.58298841 |
| AC105942.1 | 35.4375047 | 0.158268965 | 0.26075607 | 0.60696178 | 0.54387628 | 0.78027954 |
| ALG14 | 68.1697793 | -0.270146491 | 0.35007306 | -0.771686 | 0.44030041 | 0.70364324 |
| PTBP2 | 487.122763 | 0.198230471 | 0.10359111 | 1.91358577 | 0.0556731 | 0.23710513 |

| | | | | | | |
|---|---|---|---|---|---|---|
| DPYD | 432.949683 | -0.571083923 | 0.42582382 | -1.3411272 | 0.17987914 | 0.451844 |
| SNX7 | 84.4753449 | -0.178385814 | 0.27161265 | -0.6567655 | 0.51133173 | 0.75950482 |
| AGL | 101.673619 | 0.18342694 | 0.19960589 | 0.91894552 | 0.35812407 | 0.64001011 |
| TRMT13 | 80.0986668 | 0.069255268 | 0.16570801 | 0.41793556 | 0.67599424 | 0.86116336 |
| DBT | 76.9701072 | 0.138217766 | 0.21280592 | 0.64950151 | 0.51601427 | 0.76140599 |
| RTCA | 52.9546304 | 0.025087436 | 0.21333737 | 0.11759513 | 0.90638846 | 0.96621245 |
| CDC14A | 150.168213 | 0.173371683 | 0.27982338 | 0.61957541 | 0.53553736 | 0.77504588 |
| VCAM1 | 1287.50893 | -0.925531871 | 0.81819792 | -1.1311834 | 0.25797793 | 0.54515124 |
| EXTL2 | 38.1504041 | 0.501878813 | 0.32692629 | 1.53514363 | 0.12474855 | 0.37130091 |
| SLC30A7 | 166.623425 | -0.356150963 | 0.1908506 | -1.8661244 | 0.06202396 | 0.25250472 |
| DPH5 | 44.6765899 | -0.401614065 | 0.24674703 | -1.6276348 | 0.10360234 | 0.33583603 |
| AC093157.1 | 37.5161444 | -0.076276257 | 0.34478651 | -0.2212275 | 0.8249153 | 0.93217972 |
| RNPC3 | 142.309998 | 0.314450432 | 0.18114107 | 1.73594221 | 0.08257405 | 0.29927236 |
| AMY2B | 49.5460648 | 0.419702579 | 0.30832888 | 1.36121721 | 0.17344506 | 0.4445893 |
| VAV3 | 479.119314 | 1.384130891 | 0.66258773 | 2.08897754 | 0.03670974 | 0.18976937 |
| SLC25A24 | 100.453515 | 0.062892816 | 0.23756722 | 0.26473693 | 0.79121213 | 0.91622594 |
| PRPF38B | 110.559686 | -0.090949842 | 0.22693717 | -0.400771 | 0.68858871 | 0.86751885 |
| STXBP3 | 154.093918 | 0.652715161 | 0.17432322 | 3.74428126 | 0.00018091 | 0.00709811 |
| GPSM2 | 33.3237679 | 0.144155871 | 0.28733443 | 0.50170066 | 0.6158781 | 0.82592881 |
| WDR47 | 33.0758161 | 0.515930216 | 0.26005649 | 1.98391594 | 0.04726521 | 0.21714794 |
| TAF13 | 29.1951112 | -0.343447712 | 0.32683862 | -1.0508174 | 0.29334246 | 0.58128412 |
| TMEM167B | 41.2280698 | -0.781897015 | 0.24355599 | -3.2103379 | 0.00132579 | 0.02509831 |
| SARS | 104.408831 | -0.479250422 | 0.20989005 | -2.2833403 | 0.02241033 | 0.14387973 |
| CELSR2 | 62.2270261 | 1.392107327 | 0.34840463 | 3.99566257 | 6.45E-05 | 0.00347177 |
| SORT1 | 226.88837 | 0.610564159 | 0.3017928 | 2.02312367 | 0.0430604 | 0.20605022 |
| PSMA5 | 108.84471 | -0.487423459 | 0.31941601 | -1.5259832 | 0.12701405 | 0.37435927 |
| SYPL2 | 44.3268864 | 0.116026509 | 0.41992814 | 0.27630087 | 0.78231699 | 0.9109913 |
| GNAI3 | 129.697301 | -0.213476151 | 0.175269 | -1.2179915 | 0.22322721 | 0.50502915 |
| AMPD2 | 31.5996147 | -0.00761405 | 0.35918577 | -0.0211981 | 0.98308764 | 0.99619132 |
| GSTM3 | 69.6715534 | 0.581160532 | 0.73453703 | 0.79119296 | 0.4288314 | 0.69479256 |
| CSF1 | 87.3995719 | -5.359529268 | 1.28727257 | -4.1634766 | 3.13E-05 | 0.00212938 |
| AHCYL1 | 304.954046 | -0.722647767 | 0.33912957 | -2.1308899 | 0.03309822 | 0.1783171 |
| STRIP1 | 48.3546841 | 0.089899491 | 0.21590115 | 0.41639191 | 0.67712325 | 0.8613973 |
| RBM15 | 49.9214581 | 0.131870249 | 0.25677892 | 0.51355559 | 0.60756274 | 0.8211189 |
| SLC16A4 | 87.0378387 | -0.227457109 | 0.40856511 | -0.5567218 | 0.57771751 | 0.80353652 |
| LAMTOR5 | 158.831776 | -0.812978158 | 0.36147408 | -2.2490635 | 0.02450846 | 0.15195244 |
| LRIF1 | 33.8088149 | -0.361154932 | 0.40967998 | -0.8815538 | 0.37801816 | 0.65622253 |
| DRAM2 | 89.1269842 | -0.795256907 | 0.35651746 | -2.2306254 | 0.02570595 | 0.15532034 |
| CEPT1 | 123.512763 | -0.268633071 | 0.26615198 | -1.0093221 | 0.31282018 | 0.60019427 |
| DENND2D | 43.5714029 | -0.968874691 | 0.29254313 | -3.3119038 | 0.00092663 | 0.01973385 |
| WDR77 | 28.3812129 | -1.049762459 | 0.33530827 | -3.1307384 | 0.00174367 | 0.0293224 |
| ATP5PB | 110.010513 | -0.476314593 | 0.24760351 | -1.9236989 | 0.05439234 | 0.23385288 |
| RAP1A | 311.046283 | 0.12222512 | 0.21587524 | 0.56618405 | 0.57126867 | 0.79857531 |
| KCND3 | 66.5548702 | -1.803071889 | 0.47148222 | -3.8242627 | 0.00013116 | 0.00565867 |
| CTTNBP2NL | 158.576409 | -0.34851242 | 0.21574879 | -1.6153621 | 0.1062323 | 0.34093088 |
| WNT2B | 36.8037243 | -0.957593196 | 0.52278241 | -1.8317242 | 0.06699252 | 0.26375724 |
| ST7L | 141.202704 | 0.469718903 | 0.21855525 | 2.14919984 | 0.03161856 | 0.17376604 |
| CAPZA1 | 195.295114 | -0.130636302 | 0.13661687 | -0.9562238 | 0.33895919 | 0.62369754 |
| MOV10 | 60.6543683 | -0.587829238 | 0.43053541 | -1.3653447 | 0.17214474 | 0.44309343 |
| RHOC | 115.827277 | -1.674251572 | 0.49129264 | -3.4078499 | 0.00065477 | 0.01564098 |
| SLC16A1-AS1 | 120.224329 | -0.96195395 | 0.79221747 | -1.2142549 | 0.2246504 | 0.50669324 |
| LRIG2 | 117.250042 | 0.446582423 | 0.19899235 | 2.24421902 | 0.02481831 | 0.15268762 |
| MAGI3 | 623.970229 | 0.616312965 | 0.36531761 | 1.68706067 | 0.09159169 | 0.31554208 |
| PHTF1 | 73.8402686 | -0.279761457 | 0.30325233 | -0.9225369 | 0.3562486 | 0.63785147 |
| RSBN1 | 98.04728 | 0.320855418 | 0.19991103 | 1.60499108 | 0.10849577 | 0.3445906 |
| HIPK1 | 118.653949 | 0.11285642 | 0.23610923 | 0.47798393 | 0.63266164 | 0.83561475 |
| TRIM33 | 292.423206 | 0.155369109 | 0.17043048 | 0.91162748 | 0.36196485 | 0.64232332 |
| BCAS2 | 37.5732052 | -0.063244907 | 0.35483889 | -0.1782356 | 0.85853799 | 0.94734879 |
| NRAS | 40.4916067 | -0.463006833 | 0.39356639 | -1.176439 | 0.23941951 | 0.5232488 |
| CSDE1 | 313.898177 | 0.059530994 | 0.16683316 | 0.3568295 | 0.72121945 | 0.88235193 |
| SIKE1 | 39.4917813 | -0.034387117 | 0.35816657 | -0.0960087 | 0.92351364 | 0.97297642 |
| VANGL1 | 25.4532647 | -0.3366248 | 0.35589532 | -0.9458534 | 0.34422337 | 0.62925871 |
| SLC22A15 | 48.6409141 | -0.6707181 | 0.40693879 | -1.6482039 | 0.09931084 | 0.329397 |
| ATP1A1 | 705.836242 | -0.875145743 | 0.2745391 | -3.1876908 | 0.00143414 | 0.02628508 |
| LINC01762 | 50.1118131 | 0.661566882 | 0.66394465 | 0.99641872 | 0.31904674 | 0.60583257 |
| CD58 | 91.6575921 | -0.393876123 | 0.24265833 | -1.6231716 | 0.10455272 | 0.33777911 |
| IGSF3 | 96.7555786 | -0.58730831 | 0.37579962 | -1.562823 | 0.11809422 | 0.36280454 |
| PTGFRN | 54.9170069 | -0.41600442 | 0.29325076 | -1.4185962 | 0.15601677 | 0.42038498 |
| TTF2 | 41.9045358 | -0.567346849 | 0.28430441 | -1.9955612 | 0.0459817 | 0.21391877 |
| VTCN1 | 35.1198223 | 2.718642307 | 0.86405873 | 3.14636287 | 0.00165315 | 0.02837903 |
| MAN1A2 | 388.595544 | 0.32857055 | 0.1601341 | 2.05184627 | 0.0401846 | 0.19892937 |
| GDAP2 | 98.5217183 | 0.137130539 | 0.20397744 | 0.67228288 | 0.50140362 | 0.75245892 |
| WDR3 | 34.5615816 | -0.837707699 | 0.3281026 | -2.5531883 | 0.01067418 | 0.09276247 |
| SPAG17 | 184.918157 | 0.65835872 | 0.50645667 | 1.29993098 | 0.19362463 | 0.47194155 |
| WARS2 | 77.3368962 | 0.116538844 | 0.2174371 | 0.53596578 | 0.5919822 | 0.81275584 |
| AL359915.2 | 232.788662 | 0.221620334 | 0.35297849 | 0.62785791 | 0.53009703 | 0.77088195 |
| ZNF697 | 47.1703428 | 0.10603952 | 0.33715768 | 0.31451017 | 0.75313359 | 0.89718526 |
| PHGDH | 56.7944243 | 0.009246527 | 0.38868259 | 0.0237894 | 0.98102059 | 0.99578216 |
| NOTCH2 | 241.995894 | -0.314611771 | 0.30655787 | -1.0262721 | 0.30476337 | 0.59263668 |
| SEC22B | 129.735938 | -0.200634028 | 0.21765646 | -0.9217922 | 0.35663698 | 0.63842751 |
| NBPF26 | 45.467905 | 0.279578581 | 0.48177983 | 0.58030362 | 0.56170988 | 0.79161674 |
| LINC00623 | 254.287762 | -0.055342782 | 0.32757366 | -0.1689476 | 0.86583786 | 0.95058578 |
| SRGAP2C | 59.0813473 | 0.308259807 | 0.35488989 | 0.8686069 | 0.38506218 | 0.66173495 |
| AC239800.3 | 35.2789346 | 0.220485297 | 0.31905925 | 0.69104813 | 0.4895353 | 0.74220617 |
| NBPF15 | 86.4104161 | -0.314499226 | 0.30074728 | -1.0457259 | 0.29568759 | 0.5839143 |
| FP700111.1 | 58.6484726 | -0.60672586 | 0.28262873 | -2.146724 | 0.03181526 | 0.17444671 |
| SRGAP2B | 71.3925343 | 0.129433383 | 0.31144428 | 0.41559081 | 0.67770945 | 0.86151115 |
| GPR89A | 39.0284331 | 0.093332611 | 0.27256584 | 0.34242226 | 0.73203314 | 0.8885131 |
| PDZK1 | 342.843915 | 0.72756535 | 0.47033316 | 1.54691485 | 0.12188378 | 0.36668381 |
| RNF115 | 298.147076 | 0.202032103 | 0.16498263 | 1.2245659 | 0.22073882 | 0.50236569 |
| POLR3C | 43.4258767 | -0.392905524 | 0.21132082 | -1.8592845 | 0.06298682 | 0.25490103 |
| PIAS3 | 24.8941426 | -0.273579622 | 0.42106156 | -0.6497378 | 0.51586159 | 0.76140599 |
| PEX11B | 31.4026471 | -0.118885145 | 0.31352642 | -0.379187 | 0.704549 | 0.873867 |
| RBM8A | 79.085764 | -0.675288778 | 0.40522048 | -1.6664725 | 0.09561934 | 0.32251327 |
| LIX1L | 30.2903987 | -0.157567851 | 0.38975087 | -0.4042784 | 0.68600802 | 0.8662366 |
| TXNIP | 1439.85098 | -0.494979294 | 0.48411218 | -1.0224475 | 0.30656914 | 0.59421809 |
| NBPF12 | 69.2394617 | 0.032150004 | 0.22677584 | 0.14176997 | 0.8872617 | 0.95896094 |
| PRKAB2 | 30.4477462 | -0.437847109 | 0.300891 | -1.4551685 | 0.14562262 | 0.40362241 |
| AC242426.2 | 92.6884196 | 0.13765694 | 0.2658788 | 0.5177432 | 0.60463746 | 0.82048068 |
| FMO5 | 37.8659737 | 0.064481575 | 0.47255405 | 0.13645333 | 0.89146291 | 0.95969529 |
| CHD1L | 81.1018486 | -0.163085267 | 0.25109226 | -0.6495034 | 0.51601308 | 0.76140599 |
| BCL9 | 88.9036319 | 0.018802524 | 0.38113932 | 0.04933242 | 0.96065439 | 0.98703297 |
| ACP6 | 105.613158 | 0.249967091 | 0.29926564 | 0.83526827 | 0.40356668 | 0.67761003 |
| GPR89B | 42.63211 | 0.362686344 | 0.28129318 | 1.28935351 | 0.19727522 | 0.47630586 |
| NBPF11 | 67.4940936 | 0.28585784 | 0.21773045 | 1.31289783 | 0.18921738 | 0.46534359 |
| AC239809.3 | 66.853158 | 0.027486355 | 0.26746946 | 0.10276446 | 0.91814991 | 0.97057281 |
| NBPF14 | 52.1217715 | 0.042296079 | 0.21457547 | 0.19711516 | 0.84373742 | 0.94193856 |
| PDE4DIP | 315.546667 | 0.4606758 | 0.45309901 | 1.01672215 | 0.30928563 | 0.59586625 |
| NBPF9 | 47.5422155 | 0.068381424 | 0.32760054 | 0.20873416 | 0.83465577 | 0.93697031 |
| AC245297.3 | 465.154514 | -0.215433992 | 0.31338091 | -0.6874509 | 0.49179862 | 0.74434176 |
| NBPF19 | 75.9093062 | -0.083817198 | 0.62630607 | -0.1338279 | 0.8935387 | 0.96030598 |
| SF3B4 | 27.8420085 | -0.844698754 | 0.32334446 | -2.6123805 | 0.00899141 | 0.08477238 |
| OTUD7B | 198.096754 | 0.080902619 | 0.38391169 | 0.21073237 | 0.83309612 | 0.93607461 |

| | | | | | | |
|---|---|---|---|---|---|---|
| VPS45 | 138.186118 | 0.328475131 | 0.13596996 | 2.41579187 | 0.01570103 | 0.11641149 |
| ANP32E | 105.613158 | 0.160491384 | 0.41655506 | 0.38528252 | 0.70002811 | 0.87182648 |
| APH1A | 73.9238959 | -0.39139941 | 0.30245686 | -1.2940669 | 0.19564232 | 0.47420489 |
| MRPS21 | 128.218654 | -0.628000695 | 0.24381372 | -2.5757398 | 0.01000259 | 0.08977568 |
| PRPF3 | 120.267896 | 0.418130239 | 0.22381092 | 1.86822986 | 0.06173004 | 0.25172926 |
| RPRD2 | 382.899337 | 0.299037479 | 0.21661045 | 1.38053116 | 0.16742316 | 0.43603765 |
| TARS2 | 29.4771183 | 0.029018147 | 0.26566138 | 0.10922983 | 0.9130202 | 0.96844354 |
| MCL1 | 193.74444 | -0.288939061 | 0.34288433 | -0.8426721 | 0.39941185 | 0.67417905 |
| ENSA | 334.336638 | 0.161550051 | 0.17094703 | 0.9450299 | 0.34464361 | 0.62954637 |
| GOLPH3L | 95.1943952 | 0.046554937 | 0.2437109 | 0.19102525 | 0.84850581 | 0.94366633 |
| ARNT | 185.548999 | 0.796847484 | 0.16917154 | 4.71029288 | 2.47E-06 | 0.00032459 |
| SETDB1 | 102.059659 | -0.229326912 | 0.21284121 | -1.0774554 | 0.28127687 | 0.56893012 |
| CERS2 | 117.108983 | -0.347755259 | 0.20415331 | -1.7034025 | 0.08849277 | 0.30970853 |
| MINDY1 | 29.8953959 | -0.181781964 | 0.38601841 | -0.4709153 | 0.63770125 | 0.83912075 |
| PRUNE1 | 38.0010817 | 0.614922536 | 0.31721274 | 1.93851777 | 0.05256009 | 0.22987695 |
| C1orf56 | 29.1799089 | 0.533807258 | 0.48765788 | 1.09463474 | 0.27367669 | 0.56099915 |
| CDC42SE1 | 77.5719253 | 0.00519069 | 0.19576287 | 0.02651519 | 0.97884642 | 0.99557142 |
| GABPB2 | 112.102221 | 0.137911719 | 0.4205022 | 0.32796908 | 0.74293504 | 0.89269628 |
| VPS72 | 26.3199992 | -0.030970265 | 0.28353195 | -0.1092302 | 0.91301987 | 0.96844354 |
| PIP5K1A | 328.953739 | 0.287803432 | 0.1883403 | 1.52810329 | 0.1264869 | 0.37394935 |
| PSMD4 | 81.4466445 | -0.224354474 | 0.23049984 | -0.973339 | 0.33038487 | 0.61559165 |
| PI4KB | 77.5739772 | 0.068383652 | 0.17089274 | 0.40015541 | 0.68904206 | 0.86751885 |
| PSMB4 | 58.5246777 | -0.699423319 | 0.37059484 | -1.8872992 | 0.0591201 | 0.24535287 |
| POGZ | 506.261782 | 0.611619874 | 0.20023531 | 3.05450563 | 0.00225432 | 0.0341679 |
| CGN | 68.1511562 | 0.226073628 | 0.37494491 | 0.60295159 | 0.54654089 | 0.78253724 |
| TUFT1 | 86.5624345 | 0.276804282 | 0.36709168 | 0.75404673 | 0.45082116 | 0.71160181 |
| SNX27 | 138.748435 | 0.302179033 | 0.18904488 | 1.59845129 | 0.10994258 | 0.34708436 |
| MRPL9 | 28.5448447 | -0.636460258 | 0.38751122 | -1.6424305 | 0.10050081 | 0.33116522 |
| RORC | 36.6396473 | -0.334882702 | 0.31404625 | -1.0663484 | 0.2862662 | 0.57346076 |
| THEM4 | 58.5782506 | 0.002160634 | 0.23767918 | 0.00909055 | 0.99274689 | 0.99829858 |
| S100A10 | 526.650077 | -0.96991855 | 0.63557924 | -1.5260387 | 0.12700021 | 0.37435927 |
| S100A11 | 867.88199 | -2.057430825 | 0.61478913 | -3.3465634 | 0.0008182 | 0.01810054 |
| FLG-AS1 | 153.021938 | 1.418768753 | 0.31623233 | 4.48647594 | 7.24E-06 | 0.00070063 |
| S100A9 | 88.2615934 | -4.165919806 | 1.13287235 | -3.6773073 | 0.00023571 | 0.0083651 |
| S100A6 | 1265.35628 | -2.069809242 | 0.79542523 | -2.6021418 | 0.00926435 | 0.08582519 |
| S100A4 | 31.0132761 | 0.592354373 | 1.15802404 | 0.51152165 | 0.60898583 | 0.82173197 |
| S100A2 | 64.2981637 | -2.407790283 | 1.32092906 | -1.8228006 | 0.0683336 | 0.26717042 |
| S100A16 | 162.898165 | -1.910832871 | 0.57386674 | -3.3297502 | 0.00086924 | 0.01883601 |
| S100A14 | 116.08328 | -1.293089364 | 0.80361937 | -1.6090819 | 0.10759844 | 0.34308427 |
| S100A13 | 278.682658 | -0.378392024 | 0.53305224 | -0.7098592 | 0.47779147 | 0.73413221 |
| S100A1 | 116.868491 | 0.002975218 | 0.71689145 | 0.00415017 | 0.99668866 | 0.99923686 |
| CHTOP | 90.1086678 | 0.222275018 | 0.26937391 | 0.82515421 | 0.40928405 | 0.68208527 |
| ILF2 | 87.5570504 | 0.065362892 | 0.32381968 | 0.20184966 | 0.84003426 | 0.94021723 |
| NPR1 | 200.382882 | 0.63138194 | 0.54181173 | 1.16531611 | 0.24389106 | 0.52963783 |
| INTS3 | 195.592959 | 0.906165724 | 0.24660813 | 3.67451686 | 0.0002383 | 0.0083651 |
| GATAD2B | 400.878199 | 0.12090746 | 0.17631935 | 0.68572997 | 0.49288341 | 0.74527705 |
| DENND4B | 52.5368406 | -0.089493757 | 0.21876817 | -0.4090803 | 0.6824807 | 0.86417483 |
| CRTC2 | 30.5591919 | 0.020700205 | 0.32210811 | 0.06426477 | 0.9487594 | 0.98262487 |
| SLC39A1 | 72.9419345 | -1.087582072 | 0.41283571 | -2.6344186 | 0.00842815 | 0.08097619 |
| JTB | 67.4858168 | -0.13951233 | 0.38295735 | -0.3643025 | 0.71563211 | 0.88009244 |
| RAB13 | 128.720475 | -1.220829688 | 0.41134333 | -2.9679093 | 0.00299833 | 0.04079685 |
| RPS27 | 140.616161 | -0.28532777 | 0.41050041 | -0.6950731 | 0.48700952 | 0.74072153 |
| TPM3 | 139.103246 | -0.967769336 | 0.32241232 | -3.0016512 | 0.0026852 | 0.03793731 |
| C1orf43 | 109.827998 | -0.437243568 | 0.17708821 | -2.4690721 | 0.01354639 | 0.10741794 |
| UBAP2L | 126.971414 | -0.166833841 | 0.16930788 | -0.9853874 | 0.32443384 | 0.61116061 |
| HAX1 | 30.0915316 | 0.102939816 | 0.37507014 | 0.27445484 | 0.7837351 | 0.91208827 |
| IL6R | 88.8353038 | -1.045602411 | 0.63227177 | -1.6537231 | 0.09818379 | 0.32724515 |
| UBE2Q1 | 40.8782523 | -0.252872024 | 0.33120937 | -0.7634809 | 0.44517665 | 0.70741705 |
| ADAR | 199.757472 | -0.650749552 | 0.28315506 | -2.2982091 | 0.02154989 | 0.14119452 |
| PMVK | 45.3074058 | -0.445470906 | 0.37424345 | -1.1903238 | 0.23391913 | 0.51653098 |
| PBXIP1 | 42.4586482 | 0.044102656 | 0.49075575 | 0.08986682 | 0.92839305 | 0.9749043 |
| ZBTB7B | 23.7026663 | -0.718266185 | 0.3032977 | -2.3681887 | 0.01787542 | 0.12627479 |
| EFNA1 | 180.265876 | -1.551350183 | 0.41404014 | -3.7468593 | 0.00017906 | 0.0070586 |
| DPM3 | 50.5074823 | -0.401532536 | 0.35616524 | -1.1273771 | 0.25958308 | 0.54625358 |
| THBS3 | 35.8557143 | 0.358347632 | 0.32585571 | 1.0997126 | 0.27145736 | 0.55848155 |
| GBA | 29.750251 | -0.407006052 | 0.31825636 | -1.2788623 | 0.20094556 | 0.48001435 |
| SCAMP3 | 48.73878 | -0.50644485 | 0.23553613 | -2.1501791 | 0.03154105 | 0.17373701 |
| CLK2 | 40.5872221 | 0.054058519 | 0.24382828 | 0.22170734 | 0.82454172 | 0.93217972 |
| FDPS | 116.905848 | -0.273342068 | 0.27671351 | -0.9878161 | 0.3232427 | 0.61020115 |
| ASH1L | 805.409959 | 0.022475491 | 0.13066496 | 0.17200856 | 0.8634308 | 0.94929509 |
| YY1AP1 | 159.851588 | 0.035372966 | 0.18217019 | 0.19417538 | 0.84603856 | 0.94271476 |
| DAP3 | 135.164326 | -0.467411806 | 0.26404575 | -1.7701925 | 0.07669508 | 0.28619486 |
| GON4L | 185.627031 | -0.118051287 | 0.18290513 | -0.6454236 | 0.5186527 | 0.7636788 |
| RIT1 | 55.3145895 | 0.312215106 | 0.29383757 | 1.06254319 | 0.28798916 | 0.57459866 |
| KHDC4 | 152.451969 | 0.133106822 | 0.27999779 | 0.47538525 | 0.6345124 | 0.83695873 |
| ARHGEF2 | 67.5164803 | -0.555100259 | 0.23057429 | -2.4074682 | 0.01606356 | 0.11845486 |
| SSR2 | 103.794301 | -0.830629275 | 0.39445613 | -2.1057583 | 0.03522535 | 0.18531364 |
| UBQLN4 | 38.869744 | -0.14462944 | 0.31513071 | -0.4589506 | 0.64626962 | 0.84427583 |
| LAMTOR2 | 29.0283209 | -0.375503976 | 0.39204297 | -0.9578133 | 0.33815691 | 0.62340358 |
| RAB25 | 42.0336426 | -0.6262942 | 0.59045275 | -1.0607016 | 0.28882552 | 0.57571692 |
| LMNA | 218.010214 | -0.178904183 | 0.30906294 | -0.57886 | 0.56268362 | 0.79260446 |
| SEMA4A | 94.605269 | 0.534839103 | 0.52459821 | 1.01952141 | 0.3079555 | 0.59456464 |
| SLC25A44 | 26.9465275 | 0.279594287 | 0.26889285 | 1.03979813 | 0.2984337 | 0.58610017 |
| SMG5 | 102.651489 | -0.383586498 | 0.23622032 | -1.6238506 | 0.1044077 | 0.33754429 |
| CCT3 | 125.00387 | -0.837585278 | 0.34959796 | -2.3958529 | 0.01658174 | 0.12088016 |
| MEF2D | 49.7966076 | 0.356632053 | 0.19957915 | 1.78692038 | 0.07395036 | 0.28056182 |
| NAXE | 32.9501651 | -0.813173229 | 0.55200495 | -1.4731267 | 0.14071688 | 0.39586695 |
| GPATCH4 | 27.2280269 | -1.335829876 | 0.54245389 | -2.4625685 | 0.01379458 | 0.10854146 |
| ISG20L2 | 29.3271994 | -0.847962919 | 0.27400395 | -3.0947106 | 0.00197005 | 0.03151251 |
| RRNAD1 | 31.2764864 | -0.117133991 | 0.25039513 | -0.4677966 | 0.63993006 | 0.8407475 |
| MRPL24 | 28.6082303 | -0.7053363 | 0.41691576 | -1.6917957 | 0.09068493 | 0.3142614 |
| HDGF | 170.690606 | -1.037784796 | 0.36006919 | -2.8821816 | 0.00394932 | 0.04900329 |
| PRCC | 74.3626406 | 0.130346263 | 0.23719773 | 0.54952576 | 0.58264469 | 0.80699154 |
| ARHGEF11 | 174.84091 | 0.172083206 | 0.22052096 | 0.78034853 | 0.43518575 | 0.70008054 |
| KIRREL1 | 381.60514 | 0.356196853 | 0.41360543 | 0.86119964 | 0.3891281 | 0.66513878 |
| IFI16 | 137.527298 | -2.455372735 | 1.31481426 | -1.8674674 | 0.06183635 | 0.25205548 |
| DUSP23 | 197.137491 | -2.626505802 | 0.49961839 | -5.2570238 | 1.46E-07 | 4.01E-05 |
| TAGLN2 | 150.999853 | -1.637704154 | 0.4254609 | -3.8492471 | 0.00011848 | 0.00532833 |
| IGSF8 | 31.0914199 | 0.605796344 | 0.27883348 | 2.1726098 | 0.0298097 | 0.16887967 |
| PEA15 | 50.2105617 | -0.664218332 | 0.46601379 | -1.4253191 | 0.15406499 | 0.41681551 |
| DCAF8 | 53.62542 | 0.399038313 | 0.33340659 | 1.19685191 | 0.23136428 | 0.51432779 |
| COPA | 214.726639 | -0.17304967 | 0.17646367 | -0.9806532 | 0.32676378 | 0.61330512 |
| NCSTN | 57.0059161 | 0.012410729 | 0.37888199 | 0.03275619 | 0.97386902 | 0.99315845 |
| F11R | 79.8713519 | -0.502938195 | 0.39184303 | -1.2835196 | 0.19931011 | 0.4782544 |
| AL591806.3 | 45.3766009 | 0.528288491 | 0.39907205 | 1.32379226 | 0.18557205 | 0.45992099 |
| PFDN2 | 83.8718303 | -0.678402628 | 0.40156489 | -1.6893973 | 0.09114333 | 0.31502156 |
| NIT1 | 60.8379428 | -0.098823132 | 0.34541805 | -0.2860972 | 0.77480369 | 0.90689106 |
| DEDD | 30.1159262 | -0.469798769 | 0.34465342 | -1.3631049 | 0.17284945 | 0.44401526 |
| UFC1 | 61.7138025 | -0.794719508 | 0.41616938 | -1.9096059 | 0.05618398 | 0.23855776 |
| B4GALT3 | 26.1364413 | 0.011541398 | 0.29094093 | 0.03966922 | 0.96835685 | 0.99123492 |
| NDUFS2 | 64.8243761 | -0.050376145 | 0.25105919 | -0.2006545 | 0.84096877 | 0.94053005 |
| SDHC | 152.159669 | -0.026271295 | 0.18316433 | -0.1434302 | 0.88595044 | 0.95827905 |
| AL592295.4 | 28.8212992 | -0.877367876 | 0.39695978 | -2.2102186 | 0.02709 | 0.15919943 |

| | | | | | | |
|---|---|---|---|---|---|---|
| ATF6 | 270.53157 | 0.065496338 | 0.14176083 | 0.46201999 | 0.64406699 | 0.84344334 |
| NOS1AP | 115.255128 | 0.150278142 | 0.34590221 | 0.43445268 | 0.66395975 | 0.85622919 |
| UHMK1 | 118.885388 | 0.25184933 | 0.20323348 | 1.23921182 | 0.21526706 | 0.49731142 |
| UAP1 | 29.9848858 | 0.286220395 | 0.33545154 | 0.85323917 | 0.39352668 | 0.6686322 |
| HSD17B7 | 31.5845433 | 1.205236896 | 0.2999072 | 4.01869949 | 5.85E-05 | 0.00328109 |
| PBX1 | 873.091748 | 0.714034928 | 0.26032133 | 2.74289823 | 0.00608996 | 0.06580904 |
| MGST3 | 326.187885 | -0.505414785 | 0.23488685 | -2.1517373 | 0.03141805 | 0.17346026 |
| ALDH9A1 | 67.9799759 | 0.03249492 | 0.36666822 | 0.08862213 | 0.92938222 | 0.97545894 |
| TMCO1 | 112.139125 | -0.48549841 | 0.20649663 | -2.3511202 | 0.01871698 | 0.1295448 |
| UCK2 | 41.3210431 | -1.624731529 | 0.46723873 | -3.4773049 | 0.00050648 | 0.01358988 |
| POGK | 48.4005897 | -0.181798015 | 0.28034065 | -0.6484897 | 0.51666829 | 0.76200872 |
| ILDR2 | 56.2001577 | 0.736320885 | 0.85402466 | 0.86217755 | 0.38858982 | 0.66481549 |
| POU2F1 | 407.433838 | 0.092151338 | 0.21310515 | 0.43242192 | 0.66543479 | 0.85742752 |
| CREG1 | 66.9781469 | 0.456087469 | 0.41226648 | 1.10629288 | 0.26859977 | 0.55570567 |
| MPZL1 | 205.346422 | -0.849547755 | 0.19979097 | -4.2521829 | 2.12E-05 | 0.00159673 |
| MPC2 | 127.342005 | -0.089153407 | 0.38523432 | -0.2314264 | 0.81698353 | 0.92944005 |
| DCAF6 | 540.303733 | 0.217944962 | 0.15474953 | 1.40837234 | 0.15902085 | 0.42466147 |
| TIPRL | 53.0032259 | -0.210625058 | 0.22201501 | -0.9486974 | 0.34277455 | 0.62779669 |
| SFT2D2 | 125.32898 | -0.361083909 | 0.29353042 | -1.2301414 | 0.21864417 | 0.5006005 |
| LINC00970 | 50.5881898 | 4.040737252 | 0.80134312 | 5.04245581 | 4.60E-07 | 8.90E-05 |
| ATP1B1 | 725.401123 | 1.008639589 | 0.40424882 | 2.49509594 | 0.01259231 | 0.10239535 |
| NME7 | 310.257499 | 0.408220586 | 0.17975917 | 2.27093048 | 0.02315119 | 0.14676719 |
| BLZF1 | 88.4685614 | -0.221097917 | 0.21698618 | -1.0189493 | 0.30822703 | 0.59478378 |
| SLC19A2 | 84.6679862 | 0.257547446 | 0.50630556 | 0.50867987 | 0.61097663 | 0.82279561 |
| F5 | 51.2525461 | 0.319819725 | 0.89566385 | 0.35707562 | 0.7210352 | 0.88235193 |
| SCYL3 | 76.7161929 | -0.100814839 | 0.18777488 | -0.536892 | 0.59134219 | 0.81252197 |
| KIFAP3 | 161.097478 | 0.275676833 | 0.22788613 | 1.20971307 | 0.22638901 | 0.50893695 |
| FMO4 | 39.7955336 | -0.175348288 | 0.50019227 | -0.3505618 | 0.72591714 | 0.88498933 |
| PRRC2C | 495.086348 | -0.144687714 | 0.15429015 | -0.9377638 | 0.34836582 | 0.63229918 |
| VAMP4 | 46.5508853 | 0.702953118 | 0.3022198 | 2.32596645 | 0.02002034 | 0.13480227 |
| DNM3 | 648.695723 | 2.227099811 | 0.60641934 | 3.6725409 | 0.00024015 | 0.0083651 |
| PIGC | 66.6966546 | 0.064174927 | 0.30023867 | 0.21374637 | 0.83074485 | 0.93455137 |
| SUCO | 209.70529 | -0.235124715 | 0.16044462 | -1.4654572 | 0.14279623 | 0.39972096 |
| AL645568.1 | 101.29298 | 1.106521737 | 0.27606252 | 4.00822875 | 6.12E-05 | 0.00332957 |
| PRDX6 | 212.285887 | -0.498943006 | 0.31943119 | -1.5619734 | 0.11829427 | 0.36318615 |
| KLHL20 | 104.923998 | 0.067300191 | 0.16059097 | 0.41907831 | 0.6751589 | 0.86115505 |
| ZBTB37 | 79.1519686 | 0.174360554 | 0.29027239 | 0.6006791 | 0.54805375 | 0.78342416 |
| RC3H1 | 283.164095 | 0.111959621 | 0.1655387 | 0.67633502 | 0.49882796 | 0.75042493 |
| RABGAP1L | 909.599015 | 0.294755404 | 0.18945868 | 1.55577676 | 0.11976119 | 0.36465111 |
| CACYBP | 30.860084 | -0.423541185 | 0.28171114 | -1.5034592 | 0.13272068 | 0.38380408 |
| MRPS14 | 33.7260721 | 0.0920458 | 0.32625852 | 0.28212536 | 0.77784739 | 0.90876938 |
| KIAA0040 | 46.7837586 | -0.691891573 | 0.2508076 | -2.7586547 | 0.00580398 | 0.06375728 |
| COP1 | 454.742845 | 0.080083978 | 0.14560955 | 0.54999125 | 0.58232538 | 0.80678258 |
| PAPPA2 | 115.223073 | -1.609505374 | 1.42400299 | -1.1302683 | 0.25836321 | 0.54540257 |
| SEC16B | 43.2244842 | 1.160018473 | 0.53683211 | 2.16085895 | 0.03070624 | 0.17115672 |
| RASAL2 | 723.535382 | 0.020380779 | 0.32688039 | 0.06234935 | 0.95028463 | 0.98270285 |
| RALGPS2 | 505.73022 | 0.395135633 | 0.32040613 | 1.2332337 | 0.21748859 | 0.49951605 |
| ANGPTL1 | 35.5239783 | 0.130786057 | 0.52208405 | 0.25050767 | 0.80219478 | 0.92119571 |
| FAM20B | 72.2769511 | -0.073901044 | 0.28514081 | -0.2591739 | 0.7955011 | 0.91797597 |
| ABL2 | 280.646559 | 0.239054748 | 0.18364956 | 1.30168973 | 0.19302247 | 0.47104693 |
| SOAT1 | 119.711596 | -0.555214701 | 0.21305653 | -2.6059502 | 0.00916198 | 0.08519229 |
| TOR1AIP2 | 190.946393 | 0.131235507 | 0.27104944 | 0.48417554 | 0.6282613 | 0.8332518 |
| TOR1AIP1 | 77.7788779 | 0.197088526 | 0.23865155 | 0.82584222 | 0.4088936 | 0.68208527 |
| CEP350 | 335.558695 | 0.319355638 | 0.13965253 | 2.28678727 | 0.02220824 | 0.14286955 |
| QSOX1 | 42.4450956 | -0.71154087 | 0.38725498 | -1.8373963 | 0.06615141 | 0.26198834 |
| ACBD6 | 305.566784 | 0.080854198 | 0.17254957 | 0.46858534 | 0.63936606 | 0.84033321 |
| XPR1 | 417.446079 | -0.065928966 | 0.22195022 | -0.2970439 | 0.76643297 | 0.90372312 |
| STX6 | 67.4631146 | -0.083259628 | 0.2622719 | -0.3174554 | 0.75089809 | 0.89599662 |
| MR1 | 29.8063435 | -0.043512584 | 0.37144406 | -0.1171444 | 0.90674564 | 0.9663681 |
| GLUL | 132.639092 | 0.220858406 | 0.4296764 | 0.51401103 | 0.60724428 | 0.8211189 |
| DHX9 | 276.370192 | 0.364985631 | 0.23520968 | 1.55174576 | 0.12072307 | 0.36507762 |
| LAMC1 | 448.751599 | -0.284326885 | 0.24498109 | -1.1606075 | 0.24580157 | 0.53233852 |
| LAMC2 | 89.6910734 | -1.512330639 | 0.58832241 | -2.5705814 | 0.0101528 | 0.09004967 |
| SMG7 | 252.859384 | 0.278907089 | 0.12827574 | 2.1742778 | 0.02968428 | 0.16865107 |
| ARPC5 | 104.633593 | -0.331634997 | 0.33995178 | -0.9755354 | 0.32929474 | 0.61439703 |
| RGL1 | 139.590053 | -0.774919385 | 0.59123335 | -1.3106828 | 0.18996495 | 0.46598572 |
| TSEN15 | 73.0845144 | 0.190496529 | 0.2231955 | 0.85349628 | 0.39338415 | 0.6686322 |
| C1orf21 | 46.0656453 | -0.67812757 | 0.72811204 | -0.9313506 | 0.35167225 | 0.63356563 |
| EDEM3 | 137.747901 | -0.386315718 | 0.22172397 | -1.7423273 | 0.08145119 | 0.29654614 |
| FAM129A | 107.715297 | -2.578928676 | 0.77149668 | -3.3427606 | 0.00082949 | 0.01826604 |
| RNF2 | 61.2747543 | 0.41053393 | 0.21706912 | 1.89125899 | 0.05858978 | 0.24443767 |
| TRMT1L | 68.207907 | -0.01812585 | 0.3688795 | -0.0491376 | 0.96080965 | 0.98708662 |
| SWT1 | 82.6662692 | 0.376426326 | 0.20345125 | 1.85020406 | 0.06428414 | 0.25807955 |
| IVNS1ABP | 412.987478 | 1.37526274 | 0.67771446 | 2.02926575 | 0.04243123 | 0.20434829 |
| HMCN1 | 44.1291415 | 1.471038331 | 1.09274428 | 1.34618717 | 0.17824216 | 0.44977081 |
| TPR | 217.693156 | -0.10433595 | 0.13122174 | -0.7951118 | 0.42654849 | 0.69264418 |
| ODR4 | 118.10008 | 0.18021911 | 0.17284389 | 1.04266982 | 0.29710124 | 0.58520063 |
| PLA2G4A | 40.2640091 | 2.339212269 | 0.90059926 | 2.59739529 | 0.00939338 | 0.08626955 |
| ERVMER61-1 | 54.1164184 | 0.687461226 | 0.28597935 | 2.40388414 | 0.01622191 | 0.11901464 |
| AL136454.1 | 150.000554 | -0.553654549 | 0.49713984 | -1.1136797 | 0.26541662 | 0.55350835 |
| UCHL5 | 76.839715 | 0.088650069 | 0.23371781 | 0.37930387 | 0.70446223 | 0.873867 |
| TROVE2 | 158.689329 | 0.04633521 | 0.17412897 | 0.26609708 | 0.79016445 | 0.9154554 |
| CDC73 | 277.942786 | 0.498986438 | 0.16676769 | 2.99210495 | 0.00277061 | 0.03841511 |
| KCNT2 | 859.244089 | -0.708314623 | 1.04697642 | -0.6765335 | 0.49870198 | 0.75040313 |
| CFH | 816.300827 | -0.927120702 | 1.40340838 | -0.6606208 | 0.50885556 | 0.75796558 |
| ZBTB41 | 49.890668 | 0.010249732 | 0.23204617 | 0.04417109 | 0.96476803 | 0.98965864 |
| CRB1 | 39.8803441 | 0.329122631 | 0.58966205 | 0.55815468 | 0.57673877 | 0.80275223 |
| DENND1B | 331.136321 | 0.698499129 | 0.17061206 | 4.0940783 | 4.24E-05 | 0.00275447 |
| NEK7 | 452.123484 | 0.180584009 | 0.24115713 | 0.74882302 | 0.45396388 | 0.71380828 |
| MIR181A1HG | 73.5065807 | 1.29885496 | 0.46454251 | 2.79598727 | 0.00517414 | 0.05935703 |
| ZNF281 | 31.3078202 | -0.180779566 | 0.30451145 | -0.5936708 | 0.55273229 | 0.78613936 |
| DDX59 | 31.1381874 | -0.112306908 | 0.28029627 | -0.4006721 | 0.68866152 | 0.86751885 |
| CAMSAP2 | 303.010767 | 0.153112168 | 0.22453437 | 0.68190972 | 0.49529604 | 0.74715602 |
| INAVA | 28.370069 | -1.284634204 | 0.36444613 | -3.5248946 | 0.00042365 | 0.01207067 |
| TMEM9 | 60.2517639 | -0.658381776 | 0.35882177 | -1.8348434 | 0.06652889 | 0.26290439 |
| TNNT2 | 62.9413437 | 1.037220219 | 1.21720904 | 0.8521299 | 0.394142 | 0.66881952 |
| LAD1 | 70.0152306 | -0.666705827 | 0.5154205 | -1.2935182 | 0.19583188 | 0.47442426 |
| CSRP1 | 50.1019497 | 0.328073308 | 0.37288569 | 0.87982274 | 0.37895534 | 0.65677822 |
| IPO9 | 151.174973 | -0.257089263 | 0.20028877 | -1.283593 | 0.19928441 | 0.4782544 |
| TIMM17A | 45.8583223 | -0.661139348 | 0.2898132 | -2.2812603 | 0.02253305 | 0.1442808 |
| RNPEP | 82.350276 | -0.247728004 | 0.24272876 | -1.020596 | 0.30744587 | 0.59456464 |
| ELF3 | 352.295957 | -1.227301423 | 0.37021142 | -3.3151366 | 0.00091598 | 0.01958528 |
| ARL8A | 57.6882564 | -0.142134999 | 0.22266011 | -0.6383496 | 0.52324612 | 0.76534368 |
| PPP1R12B | 319.575717 | 0.487974574 | 0.19277795 | 2.53127789 | 0.01136478 | 0.09668135 |
| KDM5B | 314.545716 | -0.084045908 | 0.1696548 | -0.4953936 | 0.62032228 | 0.82828879 |
| KLHL12 | 38.3610763 | -0.023376264 | 0.32830357 | -0.0712032 | 0.94323604 | 0.98112027 |
| ADIPOR1 | 34.4660705 | 0.105757196 | 0.28738145 | 0.36800286 | 0.7128711 | 0.87827554 |
| CYB5R1 | 31.3325024 | -0.543546848 | 0.34391763 | -1.5804565 | 0.11400237 | 0.35432469 |
| BTG2 | 60.538137 | 0.695800359 | 0.70326848 | 0.98938084 | 0.32247685 | 0.60975241 |
| ATP2B4 | 183.461905 | -0.120497835 | 0.18369143 | -0.6559796 | 0.51183724 | 0.75961818 |
| SNRPE | 39.0221695 | -0.210525965 | 0.35619003 | -0.5910496 | 0.55448721 | 0.78734553 |
| SOX13 | 72.1889329 | 0.249599242 | 0.38780535 | 0.6436199 | 0.51982195 | 0.76385303 |
| PLEKHA6 | 261.11317 | -1.079245382 | 0.32904278 | -3.2799546 | 0.00103824 | 0.0213418 |

| | | | | | | |
|---|---|---|---|---|---|---|
| PPP1R15B | 52.7921602 | -0.300487632 | 0.28958972 | -1.0376323 | 0.2994413 | 0.58669425 |
| PIK3C2B | 66.1093343 | 0.217843004 | 0.44009246 | 0.49499372 | 0.62060455 | 0.82839623 |
| MDM4 | 282.310289 | 0.420305141 | 0.19392598 | 2.16734832 | 0.0302083 | 0.16991506 |
| LRRN2 | 122.570222 | -1.280990949 | 0.77350759 | -1.6560806 | 0.09770549 | 0.32610483 |
| RBBP5 | 54.578242 | 0.133126944 | 0.26474813 | 0.50284376 | 0.61507412 | 0.82592881 |
| DSTYK | 98.5422858 | -0.291133656 | 0.19031319 | -1.5297608 | 0.12607594 | 0.37339076 |
| NUAK2 | 190.203881 | 2.036541755 | 0.44745076 | 4.55143213 | 5.33E-06 | 0.00059481 |
| CDK18 | 49.8622142 | -0.557871014 | 0.37647173 | -1.4818404 | 0.13838277 | 0.39113855 |
| ELK4 | 145.965091 | -0.021287472 | 0.2788483 | -0.0763407 | 0.93914805 | 0.97996404 |
| NUCKS1 | 489.037952 | 0.220460427 | 0.20014297 | 1.1015147 | 0.27067271 | 0.5578257 |
| RAB29 | 55.737291 | -0.357159922 | 0.3164151 | -1.1287702 | 0.2589948 | 0.54605211 |
| SLC41A1 | 34.4550382 | -0.203840735 | 0.27257448 | -0.747835 | 0.45455969 | 0.71410256 |
| RHEX | 2007.64222 | 0.256444402 | 0.58113422 | 0.44128257 | 0.65900844 | 0.85262229 |
| SRGAP2 | 141.21943 | -0.016132004 | 0.30089027 | -0.0536142 | 0.95724251 | 0.98521822 |
| EIF2D | 41.1041673 | 0.258559232 | 0.23858551 | 1.08371724 | 0.27849019 | 0.56578102 |
| MAPKAPK2 | 138.504244 | -0.203204451 | 0.27026601 | -0.7518683 | 0.45213024 | 0.71231034 |
| PIGR | 412.0148 | -3.263151791 | 0.7953181 | -4.1029518 | 4.08E-05 | 0.0026763 |
| PFKFB2 | 28.7660671 | -0.63410777 | 0.3159191 | -2.007184 | 0.04473007 | 0.21050788 |
| CD55 | 60.2552532 | 1.025122827 | 0.5239418 | 1.95655858 | 0.05039939 | 0.2253855 |
| CD46 | 331.606316 | 0.426342746 | 0.2261073 | 1.88557712 | 0.05935196 | 0.24597663 |
| MIR29B2CHG | 182.650371 | -0.303560182 | 0.42508325 | -0.7141194 | 0.47515337 | 0.73198684 |
| PLXNA2 | 80.5905119 | 0.396608451 | 0.2877305 | 1.37840255 | 0.16807904 | 0.43712834 |
| IRF6 | 24.4836099 | -0.076486577 | 0.50869156 | -0.1503594 | 0.88048104 | 0.95614192 |
| UTP25 | 28.3427564 | -0.771063926 | 0.33833809 | -2.2789747 | 0.02266857 | 0.14468046 |
| HHAT | 90.6194864 | -0.189214456 | 0.46651228 | -0.4055937 | 0.68504113 | 0.86561619 |
| RCOR3 | 351.833024 | 0.609950498 | 0.2003205 | 3.04487306 | 0.00232779 | 0.03494962 |
| TRAF5 | 75.7761402 | -0.113656766 | 0.31301743 | -0.3631004 | 0.71652986 | 0.88063119 |
| LINC00467 | 61.8884535 | -0.216082534 | 0.29650904 | -0.7287553 | 0.46615136 | 0.72394032 |
| SLC30A1 | 62.8816595 | -0.991325597 | 0.3889848 | -2.5484944 | 0.0108189 | 0.09353273 |
| LPGAT1 | 426.187417 | -1.143062036 | 0.40091385 | -2.8511413 | 0.00435626 | 0.05282104 |
| INTS7 | 46.8314707 | 0.030588229 | 0.48595857 | 0.06294411 | 0.94981101 | 0.98270285 |
| PPP2R5A | 341.085641 | 0.151933572 | 0.17103863 | 0.88829973 | 0.37437955 | 0.65274512 |
| AL360091.3 | 50.1818695 | -0.113983676 | 0.46350463 | -0.245917 | 0.80574647 | 0.92235247 |
| NENF | 71.6490721 | -0.566327532 | 0.31328918 | -1.807683 | 0.07065585 | 0.27279822 |
| ATF3 | 67.2471391 | 1.553938045 | 0.63880327 | 2.43257684 | 0.01499181 | 0.11379145 |
| NSL1 | 102.944165 | -0.011147771 | 0.21269568 | -0.0524118 | 0.95820054 | 0.98566059 |
| TATDN3 | 44.6461044 | 0.013034424 | 0.35055744 | 0.03718199 | 0.9703399 | 0.99175186 |
| FLVCR1 | 69.8357443 | 0.229836234 | 0.24388312 | 0.94240322 | 0.34598624 | 0.63016843 |
| ANGEL2 | 76.1470946 | 0.255594578 | 0.20791201 | 1.22934011 | 0.21894431 | 0.50078023 |
| RPS6KC1 | 257.8316 | 0.026766796 | 0.28382054 | 0.09430888 | 0.9248638 | 0.97343922 |
| SMYD2 | 100.013174 | -0.542303611 | 0.27534154 | -1.969567 | 0.04888802 | 0.22173216 |
| PTPN14 | 970.357347 | 0.652637949 | 0.28787151 | 2.26711546 | 0.02338317 | 0.14774895 |
| KCTD3 | 203.737928 | -0.315025248 | 0.21997639 | -1.4320866 | 0.15211903 | 0.41431567 |
| ESRRG | 1095.46176 | 0.729812503 | 0.34209248 | 2.13337781 | 0.03289375 | 0.17771528 |
| GPATCH2 | 310.463182 | -0.292881184 | 0.24396156 | -1.2005218 | 0.22993674 | 0.51293991 |
| SPATA17 | 182.560915 | 0.088114296 | 0.36126478 | 0.24390503 | 0.80730437 | 0.92342711 |
| RRP15 | 53.900945 | -0.555450019 | 0.31304499 | -1.7743457 | 0.07600599 | 0.28495554 |
| TGFB2 | 80.6394056 | 0.960539086 | 0.44686722 | 2.14949549 | 0.03159514 | 0.17373701 |
| LYPLAL1 | 330.282977 | 0.599181602 | 0.28503676 | 2.10212045 | 0.03554273 | 0.18635127 |
| EPRS | 119.445595 | -0.440558932 | 0.20401626 | -2.1594305 | 0.03081678 | 0.17142506 |
| BPNT1 | 38.0230799 | -0.439654761 | 0.2846633 | -1.544473 | 0.1224738 | 0.36765168 |
| IARS2 | 102.993898 | -0.378292963 | 0.23198695 | -1.6306649 | 0.10296105 | 0.33489437 |
| RAB3GAP2 | 294.87147 | 0.075225561 | 0.13203802 | 0.56972653 | 0.56886319 | 0.79724075 |
| MARK1 | 146.350552 | -0.226153031 | 0.35044027 | -0.6453397 | 0.51870705 | 0.7636788 |
| C1orf115 | 33.5828224 | -0.311307955 | 0.3324355 | -0.9364462 | 0.3490435 | 0.63263722 |
| 2-Mar | 104.868834 | 0.24763476 | 0.28340512 | 0.87378364 | 0.38223609 | 0.65936242 |
| AL513314.2 | 30.678143 | 0.3605515 | 0.43768993 | 0.8237601 | 0.41007588 | 0.68208527 |
| MIA3 | 104.435553 | 0.17426512 | 0.15664464 | 1.11248698 | 0.26592883 | 0.55384374 |
| AIDA | 51.0463669 | 0.178520379 | 0.22382023 | 0.7976061 | 0.42509911 | 0.69181522 |
| BROX | 69.5621269 | 0.205771653 | 0.18838712 | 1.09228088 | 0.27470966 | 0.56187466 |
| FAM177B | 64.0064154 | -1.932057843 | 0.60830974 | -3.1761087 | 0.00149265 | 0.02683835 |
| AC093152.1 | 229.496968 | 1.402333462 | 0.41784296 | 3.35612559 | 0.00079043 | 0.01778667 |
| DISP1 | 29.3439517 | 0.781211688 | 0.49139799 | 1.58977389 | 0.11188578 | 0.3500176 |
| SUSD4 | 43.3312845 | 1.56362908 | 0.81730486 | 1.9131528 | 0.05572849 | 0.23710513 |
| CAPN2 | 268.993043 | -0.429198873 | 0.31768493 | -1.3510206 | 0.17668884 | 0.44872611 |
| TP53BP2 | 81.1166779 | -0.689217305 | 0.24829435 | -2.7758074 | 0.00550648 | 0.06154379 |
| FBXO28 | 145.719141 | -0.014065795 | 0.14872494 | -0.0945759 | 0.92465169 | 0.97332292 |
| DEGS1 | 73.5034181 | -0.932961412 | 0.3025674 | -3.0834829 | 0.00204593 | 0.03226175 |
| NVL | 127.569997 | 0.016463982 | 0.25776991 | 0.06387085 | 0.94907307 | 0.98262487 |
| CNIH4 | 71.9027123 | -0.657603638 | 0.27858041 | -2.3605523 | 0.01824774 | 0.12781151 |
| WDR26 | 269.123116 | 0.449590217 | 0.16876956 | 2.66392958 | 0.00772337 | 0.07595707 |
| CNIH3 | 142.722516 | 0.623771974 | 0.42623692 | 1.46343959 | 0.14334713 | 0.40048158 |
| DNAH14 | 687.733335 | 0.210327508 | 0.19835158 | 1.0603773 | 0.28897298 | 0.57573323 |
| LBR | 267.058823 | 0.565293572 | 0.30380562 | 1.86070807 | 0.06278541 | 0.25462383 |
| ENAH | 179.168179 | -0.044405966 | 0.2549096 | -0.1742028 | 0.86170609 | 0.94853347 |
| SRP9 | 132.426749 | 0.013989681 | 0.26780343 | 0.05223862 | 0.95833856 | 0.98566059 |
| EPHX1 | 43.6231748 | -1.075644951 | 0.4789102 | -2.2460264 | 0.02470232 | 0.15226737 |
| AL591895.1 | 41.2007585 | -0.071207053 | 0.40961901 | -0.1738373 | 0.86199335 | 0.9486379 |
| TMEM63A | 99.8661323 | -0.390675225 | 0.29605609 | -1.3195987 | 0.18696905 | 0.46206921 |
| H3F3A | 33.1537512 | -0.057606372 | 0.30564017 | -0.1884778 | 0.85050215 | 0.94435157 |
| ACBD3 | 169.943243 | -0.321601421 | 0.25698225 | -1.2514538 | 0.21076895 | 0.49146928 |
| PARP1 | 62.9948239 | -0.49471193 | 0.30840828 | -1.6040812 | 0.10869615 | 0.34511119 |
| ITPKB | 61.0207009 | 0.278299224 | 0.30308907 | 0.91820938 | 0.35850926 | 0.64001011 |
| COQ8A | 46.2243626 | 0.049643195 | 0.29886177 | 0.16610755 | 0.86807232 | 0.95090501 |
| CDC42BPA | 789.017001 | 0.415418962 | 0.12163114 | 3.41539962 | 0.00063688 | 0.01548406 |
| ZNF678 | 72.6396772 | 0.440477567 | 0.17906186 | 2.45991835 | 0.01389686 | 0.10898655 |
| SNAP47 | 37.989115 | 0.035111659 | 0.27760351 | 0.12648132 | 0.89935093 | 0.96269027 |
| ARF1 | 188.064808 | -0.661216032 | 0.26681676 | -2.4781653 | 0.01320599 | 0.10541684 |
| MRPL55 | 30.5043363 | -0.335021923 | 0.36732769 | -0.9120519 | 0.36174139 | 0.64204573 |
| GUK1 | 116.227063 | -0.657959217 | 0.35516057 | -1.8525683 | 0.06394426 | 0.25768702 |
| OBSCN | 164.522637 | 0.375213776 | 0.27183555 | 1.38029694 | 0.16749524 | 0.43610679 |
| RNF187 | 56.5791492 | -0.508638106 | 0.36064175 | -1.4103695 | 0.15843062 | 0.42403514 |
| RHOU | 70.3970895 | -1.254551764 | 0.48723535 | -2.5748373 | 0.01002872 | 0.08977568 |
| RAB4A | 61.9590299 | -0.128304134 | 0.20042675 | -0.6401547 | 0.52207201 | 0.7647848 |
| NUP133 | 99.5209109 | 0.333003935 | 0.16024147 | 2.07813827 | 0.03769663 | 0.19289315 |
| ABCB10 | 45.4589583 | 0.327472642 | 0.22606162 | 1.44859903 | 0.14744959 | 0.40633477 |
| GALNT2 | 272.018365 | -0.986858446 | 0.32709359 | -3.0170522 | 0.00255246 | 0.03682229 |
| COG2 | 49.989004 | 0.00367132 | 0.20219946 | 0.01815692 | 0.98551367 | 0.99668665 |
| C1orf198 | 38.2336626 | -0.092585554 | 0.41348134 | -0.2239171 | 0.82282179 | 0.93217972 |
| TTC13 | 68.4056525 | 0.477302418 | 0.26643731 | 1.79142482 | 0.07322515 | 0.27896901 |
| ARV1 | 28.6225796 | 0.403742218 | 0.2830888 | 1.42620342 | 0.15380963 | 0.41655145 |
| C1orf131 | 26.5740598 | -0.073140835 | 0.30332363 | -0.2411313 | 0.80945332 | 0.92449664 |
| GNPAT | 87.3020448 | 0.140771122 | 0.18458909 | 0.76261887 | 0.44569073 | 0.70788783 |
| EGLN1 | 251.340318 | -0.586459633 | 0.31683391 | -1.8510002 | 0.06416952 | 0.25789884 |
| DISC1 | 93.41229 | -0.764371688 | 0.47321235 | -1.6152826 | 0.1062495 | 0.34093088 |
| SIPA1L2 | 262.284594 | 0.086276782 | 0.94048088 | 0.09173688 | 0.92690709 | 0.97419827 |
| NTPCR | 53.3223807 | -0.238769522 | 0.21892451 | -1.0906477 | 0.27542792 | 0.56268552 |
| PCNX2 | 396.929242 | -0.313432659 | 0.24387302 | -1.2852289 | 0.1987123 | 0.47741788 |
| MAP3K21 | 299.277345 | -0.347982147 | 0.2939266 | -1.1839083 | 0.23644936 | 0.51974689 |
| KCNK1 | 52.2086409 | -1.400261967 | 0.56041939 | -2.4985966 | 0.01246862 | 0.1019086 |
| SLC35F3 | 208.325451 | -0.331700036 | 0.6881263 | -0.4820337 | 0.62978204 | 0.83428048 |
| TARBP1 | 82.5483381 | -0.385312224 | 0.28529173 | -1.3505902 | 0.17682674 | 0.44872611 |
| IRF2BP2 | 132.504952 | -0.380395983 | 0.24948115 | -1.5247484 | 0.12732186 | 0.37459861 |

| | | | | | | |
|---|---|---|---|---|---|---|
| TOMM20 | 160.150278 | -0.216626493 | 0.39993624 | -0.5416526 | 0.58805787 | 0.81096757 |
| ARID4B | 629.38089 | 0.246946426 | 0.16062324 | 1.53742649 | 0.1241889 | 0.37041562 |
| B3GALNT2 | 87.3438073 | -0.233239797 | 0.21880136 | -1.0659888 | 0.28642871 | 0.57346076 |
| LYST | 155.455732 | 0.314437359 | 0.18385032 | 1.71028995 | 0.08721227 | 0.30713468 |
| GPR137B | 78.2247005 | -0.168402238 | 0.2696913 | -0.6244259 | 0.53234791 | 0.7728646 |
| ERO1B | 79.3047899 | -0.143141123 | 0.42586069 | -0.3361219 | 0.73677891 | 0.89057637 |
| LGALS8 | 104.417512 | 0.311510749 | 0.22056817 | 1.4123105 | 0.15785856 | 0.42320938 |
| HEATR1 | 85.6574263 | -0.558194516 | 0.25249315 | -2.2107313 | 0.02705445 | 0.15908812 |
| MTR | 421.501502 | -0.043306589 | 0.30072704 | -0.1440063 | 0.88549549 | 0.95800331 |
| LINC01139 | 42.1176774 | -0.022825397 | 0.63146304 | -0.0361468 | 0.97116527 | 0.99175186 |
| FMN2 | 74.9264397 | 0.007516707 | 1.22332853 | 0.00614447 | 0.99509745 | 0.99886097 |
| FH | 28.9019231 | -0.036300817 | 0.29050316 | -0.1249584 | 0.90055647 | 0.96303857 |
| OPN3 | 33.8516802 | -0.878509556 | 0.41292334 | -2.1275367 | 0.03337552 | 0.17930682 |
| CEP170 | 200.700074 | 0.436352772 | 0.24942461 | 1.74943753 | 0.08021542 | 0.2943998 |
| SDCCAG8 | 446.21734 | 0.135581721 | 0.14524958 | 0.93343966 | 0.35059301 | 0.63322593 |
| AKT3 | 787.212304 | -0.041874264 | 0.33386513 | -0.1254227 | 0.90018892 | 0.96303857 |
| ZBTB18 | 45.2703705 | 0.041500995 | 0.30805302 | 0.1347203 | 0.89283303 | 0.95976294 |
| AL645465.1 | 33.7130334 | -0.824600578 | 0.41362075 | -1.9936151 | 0.04619413 | 0.21432487 |
| ADSS | 156.913786 | 0.132255817 | 0.40168451 | 0.32925297 | 0.74196449 | 0.89253455 |
| CATSPERE | 111.687736 | 0.123692565 | 0.3364418 | 0.36764922 | 0.71313481 | 0.8783691 |
| DESI2 | 104.68154 | -0.017151589 | 0.17569465 | -0.0976216 | 0.92223279 | 0.97248656 |
| COX20 | 113.740735 | 0.188887828 | 0.23546083 | 0.80220488 | 0.42243445 | 0.69043214 |
| HNRNPU | 579.774603 | 0.105178575 | 0.16617057 | 0.63295549 | 0.52676271 | 0.76821934 |
| EFCAB2 | 235.773719 | 0.207357347 | 0.35518977 | 0.58379312 | 0.5593595 | 0.78981496 |
| KIF26B | 428.294976 | -1.132833562 | 0.57830924 | -1.9588716 | 0.05012782 | 0.22501594 |
| SMYD3 | 1025.82503 | 0.361781832 | 0.16933459 | 2.13649107 | 0.03263941 | 0.17693995 |
| CNST | 114.725669 | 0.126588135 | 0.16658045 | 0.75992193 | 0.44730125 | 0.70891772 |
| SCCPDH | 270.339851 | 1.317278211 | 0.52058996 | 2.53035656 | 0.01139467 | 0.09684961 |
| AHCTF1 | 198.955778 | 0.326525664 | 0.16332157 | 1.99928071 | 0.04557799 | 0.21266029 |
| ZNF124 | 126.24333 | -0.00494833 | 0.34336603 | -0.0144112 | 0.98850189 | 0.99678172 |
| AL390728.6 | 39.5968142 | 0.083257047 | 0.40560897 | 0.20526431 | 0.83736562 | 0.93836281 |
| ZNF496 | 75.5703652 | 0.007632761 | 0.31367984 | 0.02433297 | 0.98058702 | 0.99578216 |
| ZNF672 | 25.8131499 | -0.318462817 | 0.26931401 | -1.1824963 | 0.23700884 | 0.52000641 |
| ZNF692 | 31.1454975 | 0.443655777 | 0.31727867 | 1.39831578 | 0.16201826 | 0.4283668 |
| FAM110C | 98.512504 | -1.067596378 | 0.45405541 | -2.3512469 | 0.01871061 | 0.1295448 |
| SH3YL1 | 190.780108 | 0.106209872 | 0.19880987 | 0.53422836 | 0.59318355 | 0.81351655 |
| ACP1 | 75.6339456 | -0.271137938 | 0.29628358 | -0.9151298 | 0.36012348 | 0.64036065 |
| SNTG2 | 63.6722942 | 0.817651478 | 0.6222867 | 1.31394658 | 0.18886418 | 0.46479265 |
| PXDN | 108.279446 | 0.257291807 | 0.4632118 | 0.55545175 | 0.5785857 | 0.80404359 |
| EIPR1 | 101.724014 | -0.295625295 | 0.35823469 | -0.825228 | 0.40924216 | 0.68208527 |
| TRAPPC12 | 179.678173 | 0.072045699 | 0.28622509 | 0.25170994 | 0.80126527 | 0.92096473 |
| ADI1 | 58.9435441 | -0.37636613 | 0.28857975 | -1.3042015 | 0.1921649 | 0.46933901 |
| RPS7 | 386.070281 | -0.408461023 | 0.54070046 | -0.7554294 | 0.44999138 | 0.71082906 |
| RNF144A | 63.2565754 | 0.433145307 | 0.40232147 | 1.07661495 | 0.28165232 | 0.56931659 |
| ID2 | 139.516879 | -0.192171772 | 0.40409442 | -0.4755616 | 0.63438677 | 0.83690825 |
| KIDINS220 | 260.875594 | 0.350875892 | 0.15300495 | 2.29323225 | 0.02183464 | 0.14189552 |
| MBOAT2 | 151.898368 | 1.041200643 | 0.50758508 | 2.05128298 | 0.0402394 | 0.19909772 |
| ASAP2 | 370.032646 | 0.699861904 | 0.46373154 | 1.50919625 | 0.13124864 | 0.38163088 |
| ITGB1BP1 | 68.9810522 | 0.439333125 | 0.26628593 | 1.6498548 | 0.09897264 | 0.32904321 |
| CPSF3 | 37.1968102 | 0.030415316 | 0.28105805 | 0.1082172 | 0.9138234 | 0.9684766 |
| IAH1 | 65.1269796 | 0.36009393 | 0.22693148 | 1.58679587 | 0.11255888 | 0.35131051 |
| ADAM17 | 171.01924 | -0.008982982 | 0.1487684 | -0.0603823 | 0.95185113 | 0.9832613 |
| YWHAQ | 204.153561 | -0.254227136 | 0.14839379 | -1.7131926 | 0.08667712 | 0.30671597 |
| TAF1B | 108.20542 | 0.191663009 | 0.29739867 | 0.64446492 | 0.519274 | 0.76385303 |
| GRHL1 | 34.8664857 | 0.002712788 | 0.38059278 | 0.0071278 | 0.99431289 | 0.99885288 |
| KLF11 | 33.1602752 | 0.350723334 | 0.38087227 | 0.92084239 | 0.35713272 | 0.63885377 |
| CYS1 | 172.159239 | 0.600666737 | 0.6418872 | 0.93578239 | 0.34938524 | 0.63273599 |
| HPCAL1 | 196.389853 | -0.019078383 | 0.27932545 | -0.0683016 | 0.94554553 | 0.98119662 |
| ODC1 | 233.109862 | 0.381808436 | 0.56967327 | 0.67022354 | 0.5027153 | 0.75403031 |
| NOL10 | 181.120181 | 0.131799599 | 0.2341771 | 0.56282018 | 0.57355734 | 0.80042333 |
| PDIA6 | 184.011442 | -0.573364413 | 0.32451826 | -1.7668171 | 0.07725886 | 0.28751463 |
| PQLC3 | 27.9598015 | 0.045300199 | 0.4269282 | 0.1061073 | 0.91549722 | 0.9691911 |
| ROCK2 | 549.289465 | 0.882683563 | 0.1890875 | 4.66812217 | 3.04E-06 | 0.00036857 |
| E2F6 | 34.2441228 | 0.026463 | 0.25692375 | 0.10299943 | 0.91796342 | 0.97057281 |
| LPIN1 | 249.086544 | 1.274828322 | 0.54468212 | 2.34049968 | 0.01925796 | 0.13185986 |
| MIR3681HG | 148.745744 | 2.366757588 | 1.13320351 | 2.08855476 | 0.03674782 | 0.18986374 |
| TRIB2 | 52.2104841 | 0.716813769 | 0.51810889 | 1.38351953 | 0.16650562 | 0.43485499 |
| FAM84A | 24.7002863 | -0.118011888 | 0.52960973 | -0.222828 | 0.82366936 | 0.93217972 |
| NBAS | 385.52088 | 0.286813292 | 0.1452153 | 1.97509004 | 0.04825793 | 0.21960224 |
| DDX1 | 119.921732 | -0.003832902 | 0.16745061 | -0.0228897 | 0.98173822 | 0.99598288 |
| SMC6 | 94.4386682 | -0.347630561 | 0.26513725 | -1.3111344 | 0.18981237 | 0.46597228 |
| KCNS3 | 45.1252808 | -0.300363923 | 0.57304058 | -0.5241582 | 0.6001685 | 0.81769468 |
| RDH14 | 32.5682178 | -0.437345529 | 0.36651919 | -1.1932405 | 0.23277521 | 0.51579248 |
| LINC01376 | 186.995295 | -0.147698405 | 0.3031734 | -0.4871747 | 0.62613454 | 0.83180006 |
| TTC32 | 35.2573869 | 0.250281136 | 0.3113635 | 0.80382299 | 0.42149921 | 0.6898802 |
| WDR35 | 171.399867 | 0.303783812 | 0.14634041 | 2.07587103 | 0.03790588 | 0.1933562 |
| LAPTM4A | 466.847748 | -0.005049786 | 0.32890217 | -0.0153535 | 0.98775019 | 0.99678172 |
| SDC1 | 33.5011573 | -0.946186596 | 0.54219396 | -1.7451072 | 0.08096621 | 0.29573375 |
| PUM2 | 576.15312 | 0.402252261 | 0.11049322 | 3.64051516 | 0.00027209 | 0.00898752 |
| RHOB | 138.448793 | -0.134122201 | 0.51714709 | -0.2593502 | 0.79536505 | 0.91797597 |
| HS1BP3 | 30.6959136 | -0.066916038 | 0.35039449 | -0.1909734 | 0.84854641 | 0.94366633 |
| GDF7 | 49.5445456 | 0.649174942 | 0.40150075 | 1.61687104 | 0.10590612 | 0.34040487 |
| LDAH | 183.461687 | -0.155367128 | 0.22737341 | -0.6833127 | 0.49440929 | 0.74652374 |
| KLHL29 | 354.489846 | 0.235581077 | 0.33530485 | 0.70258774 | 0.4823127 | 0.73685381 |
| ATAD2B | 290.752801 | 0.407344338 | 0.1961498 | 2.0767002 | 0.03782924 | 0.19316567 |
| UBXN2A | 118.941182 | 0.071621321 | 0.28595434 | 0.25046419 | 0.8022284 | 0.92119571 |
| SF3B6 | 81.3196679 | -0.444159184 | 0.48116039 | -0.9231001 | 0.35595506 | 0.63744504 |
| FAM228B | 55.718928 | -0.94910795 | 0.45850479 | -2.0700066 | 0.03845172 | 0.1948831 |
| TP53I3 | 28.5725664 | -0.467074191 | 0.34114439 | -1.3691393 | 0.17095573 | 0.44115974 |
| ITSN2 | 374.260344 | -0.228673221 | 0.1771664 | -1.2907257 | 0.19679883 | 0.47592425 |
| NCOA1 | 641.014633 | 0.057563291 | 0.1813396 | 0.31743365 | 0.75091458 | 0.89599662 |
| DNAJC27 | 44.2116593 | 0.523289272 | 0.28807747 | 1.816488 | 0.06929555 | 0.26972046 |
| DNAJC27-AS | 39.6369206 | 1.264451874 | 0.6160083 | 2.05265397 | 0.04010615 | 0.19874641 |
| DNMT3A | 140.838093 | 0.06169264 | 0.30528686 | 0.20208089 | 0.83985349 | 0.94016087 |
| DTNB | 284.153819 | -0.095097183 | 0.30490216 | -0.3118941 | 0.75512101 | 0.89810084 |
| ASXL2 | 194.059017 | 0.086032312 | 0.15926664 | 0.54017785 | 0.58907439 | 0.81148442 |
| RAB10 | 438.140814 | -0.152328761 | 0.17569212 | -0.867021 | 0.3859305 | 0.66275158 |
| HADHA | 147.01358 | -0.508480235 | 0.17992233 | -2.8261096 | 0.00471171 | 0.05572038 |
| HADHB | 203.986987 | 0.054507853 | 0.21916163 | 0.24871075 | 0.80358454 | 0.92125386 |
| SELENOI | 109.566528 | 0.056146895 | 0.29425521 | 0.1908102 | 0.8486743 | 0.94366633 |
| MAPRE3 | 30.5657623 | 0.016461113 | 0.26376347 | 0.06240861 | 0.95023744 | 0.98270285 |
| TMEM214 | 32.248413 | -0.51329584 | 0.35300389 | -1.4540798 | 0.14592418 | 0.40399066 |
| AGBL5 | 48.5888651 | -0.146339567 | 0.3004529 | -0.4870632 | 0.62621351 | 0.83180006 |
| OST4 | 90.9423291 | -0.041892604 | 0.40810726 | -0.102651 | 0.91823999 | 0.97057281 |
| ATRAID | 73.3847418 | -0.292801688 | 0.44892112 | -0.6522342 | 0.51425013 | 0.76112515 |
| CAD | 30.4069318 | -0.317366985 | 0.35935653 | -0.8831535 | 0.37715333 | 0.65530141 |
| MPV17 | 47.2975133 | -0.377690275 | 0.24292606 | -1.554754 | 0.12000466 | 0.36481264 |
| GTF3C2 | 95.2137176 | -0.096550601 | 0.19665498 | -0.4909644 | 0.6234516 | 0.83030289 |
| SNX17 | 41.1268101 | -0.255675689 | 0.40495095 | -0.6313745 | 0.52779571 | 0.76940421 |
| PPM1G | 162.583681 | -1.253333659 | 0.33617007 | -3.7282726 | 0.0001928 | 0.00731604 |
| NRBP1 | 68.1822168 | -0.272121411 | 0.28494165 | -0.9550075 | 0.33957391 | 0.62445354 |
| IFT172 | 71.3787261 | 0.659600624 | 0.35814749 | 1.84170107 | 0.06551889 | 0.26095859 |
| ZNF512 | 101.295632 | -0.047802692 | 0.17934716 | -0.2665372 | 0.78982551 | 0.91528808 |

| | | | | | | |
|---|---|---|---|---|---|---|
| SUPT7L | 34.4922785 | 0.146032641 | 0.25298724 | 0.57723322 | 0.56378192 | 0.79293888 |
| SLC4A1AP | 43.6501037 | 0.142800231 | 0.22708196 | 0.62884886 | 0.52944801 | 0.77040597 |
| MRPL33 | 218.188128 | 0.606984528 | 0.28665558 | 2.11746977 | 0.03422 | 0.18232064 |
| RBKS | 158.147927 | 0.491265886 | 0.38438766 | 1.27804802 | 0.2012325 | 0.48010114 |
| BABAM2 | 836.2662 | -0.142906964 | 0.20683933 | -0.6909081 | 0.48962331 | 0.74222213 |
| FOSL2 | 353.547812 | 0.692395284 | 0.4643053 | 1.49125001 | 0.13589587 | 0.38788633 |
| PPP1CB | 179.006189 | 0.133208915 | 0.25158623 | 0.52947617 | 0.59647517 | 0.81585918 |
| TRMT61B | 61.3884235 | 0.474415886 | 0.22918373 | 2.07002431 | 0.03845007 | 0.1948831 |
| WDR43 | 156.055387 | -0.473955411 | 0.31801662 | -1.4903479 | 0.13613278 | 0.38809998 |
| CLIP4 | 199.587917 | 0.367109327 | 0.49513907 | 0.74142671 | 0.45843476 | 0.71752865 |
| YPEL5 | 87.6478339 | 0.00358945 | 0.26738925 | 0.01342406 | 0.98928947 | 0.99678172 |
| LBH | 70.5624264 | 1.388569803 | 0.60034294 | 2.31296099 | 0.02072478 | 0.13776732 |
| LCLAT1 | 223.07687 | 0.080759863 | 0.17397407 | 0.46420632 | 0.64249994 | 0.84298912 |
| GALNT14 | 1314.28776 | 0.744226655 | 0.23433037 | 3.1759719 | 0.00149335 | 0.02683835 |
| DPY30 | 70.313831 | 0.580817023 | 0.2667301 | 2.17754584 | 0.02943987 | 0.16756059 |
| SPAST | 91.2371521 | 0.209807099 | 0.18530873 | 1.13220301 | 0.25754909 | 0.54484602 |
| SLC30A6 | 100.66942 | -0.048625495 | 0.17498491 | -0.2778839 | 0.78110146 | 0.91077326 |
| YIPF4 | 175.355871 | 0.28547096 | 0.13188589 | 2.16452993 | 0.0304237 | 0.17085095 |
| BIRC6 | 883.209934 | 0.190444443 | 0.11027157 | 1.72704927 | 0.08415882 | 0.30211489 |
| TTC27 | 137.674997 | -0.134857147 | 0.26130518 | -0.5160906 | 0.60579114 | 0.82065808 |
| LINC00486 | 165.167448 | 0.701759441 | 0.1908258 | 3.67748721 | 0.00023554 | 0.0083651 |
| LTBP1 | 124.317699 | 4.820877326 | 1.26940402 | 3.7977486 | 0.00014602 | 0.00613461 |
| RASGRP3 | 79.2244789 | 1.910179827 | 0.61813963 | 3.09020765 | 0.00200017 | 0.03182656 |
| FAM98A | 64.3734776 | 0.583127882 | 0.3047904 | 1.91320945 | 0.05572124 | 0.23710513 |
| LINC01317 | 35.6431598 | 0.724235732 | 0.59751755 | 1.2120744 | 0.2254839 | 0.50767998 |
| AC068599.1 | 64.5653653 | 1.231353676 | 0.51126881 | 2.40842713 | 0.01602142 | 0.11832631 |
| LINC01320 | 2598.7516 | -0.666167899 | 0.38995898 | -1.7083025 | 0.08758023 | 0.30786459 |
| AC073218.2 | 30.4778409 | 0.145675033 | 0.74285092 | 0.19610265 | 0.84452982 | 0.94193856 |
| AC012593.1 | 1743.07318 | -0.395760426 | 0.96058676 | -0.4119986 | 0.68034043 | 0.86244955 |
| CRIM1 | 1211.93078 | 1.044304257 | 0.34890389 | 2.99310005 | 0.00276159 | 0.03841511 |
| FEZ2 | 225.686278 | 0.231942143 | 0.30073403 | 0.77125341 | 0.44055674 | 0.70393545 |
| STRN | 370.09179 | 0.9505722 | 0.30047625 | 3.16355181 | 0.00155857 | 0.02754521 |
| HEATR5B | 136.217821 | 0.093049906 | 0.27104211 | 0.34330424 | 0.73136959 | 0.88793274 |
| EIF2AK2 | 202.799693 | -0.256339302 | 0.61086271 | -0.4196349 | 0.6747522 | 0.86115505 |
| CEBPZOS | 99.6680945 | 0.090742995 | 0.29889669 | 0.30359318 | 0.76143785 | 0.90141135 |
| CEBPZ | 105.21008 | -0.175906759 | 0.28838696 | -0.6099678 | 0.54188313 | 0.77897227 |
| NDUFAF7 | 62.9881528 | 0.040018275 | 0.21873897 | 0.18294991 | 0.85483732 | 0.94566193 |
| PRKD3 | 182.501206 | 0.032420029 | 0.20297181 | 0.15972676 | 0.87309632 | 0.95193455 |
| QPCT | 28.0214944 | -0.740020794 | 0.38682962 | -1.9130407 | 0.05574285 | 0.23710513 |
| CDC42EP3 | 49.2338348 | -0.467608971 | 0.40585535 | -1.1521567 | 0.24925668 | 0.53636804 |
| RMDN2 | 108.404317 | 1.154457409 | 0.3184697 | 3.6250149 | 0.00028894 | 0.00944642 |
| CYP1B1 | 217.879905 | 0.087877489 | 0.59794499 | 0.14696584 | 0.88315898 | 0.9574219 |
| CYP1B1-AS1 | 149.502611 | 0.863604052 | 0.49630524 | 1.74006635 | 0.08184737 | 0.29731359 |
| ATL2 | 202.662106 | 0.501206794 | 0.21368468 | 2.34554385 | 0.01899934 | 0.13055575 |
| HNRNPLL | 134.118085 | 0.976552515 | 0.23931803 | 4.08056392 | 4.49E-05 | 0.00285001 |
| GALM | 95.3734757 | 0.727413548 | 0.28403996 | 2.56095499 | 0.01043849 | 0.0917342 |
| SRSF7 | 136.278417 | -0.180567449 | 0.22501807 | -0.8024576 | 0.42228833 | 0.69043214 |
| DHX57 | 117.615032 | 0.167421372 | 0.18280362 | 0.91585372 | 0.35974362 | 0.64001011 |
| MORN2 | 39.0606734 | 0.123066869 | 0.49276883 | 0.24974565 | 0.80278406 | 0.92125386 |
| SOS1 | 519.939012 | 0.074719866 | 0.17002165 | 0.43947265 | 0.66031909 | 0.85383323 |
| MAP4K3 | 534.897289 | 0.451475954 | 0.2086919 | 2.16336119 | 0.03051341 | 0.17102865 |
| AC007388.1 | 238.050178 | 0.426277565 | 0.30165626 | 1.41312354 | 0.1576194 | 0.42280489 |
| TMEM178A | 40.9580677 | 3.101606988 | 0.70065333 | 4.42673555 | 9.57E-06 | 0.00088118 |
| THUMPD2 | 103.641972 | 0.156870064 | 0.15313923 | 1.02436236 | 0.30566415 | 0.59306399 |
| SLC8A1 | 82.1260764 | 0.47592371 | 1.11504423 | 0.42682047 | 0.66951009 | 0.85924744 |
| EML4 | 486.771635 | 0.132021911 | 0.29060001 | 0.454308 | 0.64960716 | 0.8468409 |
| COX7A2L | 121.006277 | -0.456528853 | 0.25318479 | -1.8031448 | 0.07136547 | 0.27465189 |
| MTA3 | 264.073046 | 0.451120404 | 0.20214157 | 2.2317052 | 0.02563445 | 0.1551184 |
| ZFP36L2 | 296.309125 | -0.501686108 | 0.32172927 | -1.5593424 | 0.11891535 | 0.36415926 |
| THADA | 447.229573 | 0.034350695 | 0.17086894 | 0.20103534 | 0.84067094 | 0.94042285 |
| PLEKHH2 | 573.456228 | 1.279618845 | 0.26522539 | 4.82464695 | 1.40E-06 | 0.00022771 |
| DYNC2LI1 | 358.824121 | 1.299147169 | 0.33022865 | 3.93408378 | 8.35E-05 | 0.00415961 |
| LRPPRC | 260.36409 | -0.125487492 | 0.20723627 | -0.6055286 | 0.54482782 | 0.78068896 |
| PPM1B | 251.478262 | 0.114917749 | 0.29791802 | 0.38573614 | 0.69969209 | 0.87181583 |
| SLC3A1 | 274.830358 | -0.727391104 | 0.64762694 | -1.1231638 | 0.26136795 | 0.54868368 |
| PREPL | 287.064235 | 0.270864732 | 0.24545882 | 1.10350376 | 0.26980846 | 0.55688327 |
| CAMKMT | 944.507601 | 0.346717125 | 0.21017382 | 1.64966849 | 0.09901076 | 0.32904321 |
| SRBD1 | 153.212727 | 0.037475651 | 0.20683386 | 0.18118722 | 0.85622063 | 0.94627061 |
| PRKCE | 583.991281 | -0.175557266 | 0.48891322 | -0.3590765 | 0.71953783 | 0.88166633 |
| EPAS1 | 69.7614512 | -1.315672308 | 0.52807403 | -2.4914543 | 0.01272213 | 0.10301379 |
| RHOQ | 138.080549 | -0.500569004 | 0.27058471 | -1.849953 | 0.06432032 | 0.25811662 |
| PIGF | 67.4063397 | 0.235342412 | 0.28760072 | 0.81829562 | 0.4131884 | 0.68443902 |
| CRIPT | 29.1829976 | 0.283696845 | 0.47886859 | 0.59243151 | 0.55356167 | 0.78684779 |
| SOCS5 | 113.022721 | 0.377057996 | 0.23094848 | 1.63264984 | 0.10254266 | 0.33398714 |
| MCFD2 | 86.0059036 | -0.488084251 | 0.23194602 | -2.104301 | 0.0353522 | 0.18575902 |
| TTC7A | 98.8341514 | -0.519301892 | 0.25252913 | -2.0564039 | 0.0397436 | 0.19750809 |
| AC106869.1 | 93.9145663 | 0.931698394 | 0.48338865 | 1.92743126 | 0.05392591 | 0.2332446 |
| EPCAM | 309.164634 | -0.021900423 | 0.54589058 | -0.0401187 | 0.96799849 | 0.99123492 |
| MSH2 | 114.196505 | 0.243464201 | 0.20035869 | 1.21514169 | 0.22431206 | 0.50628774 |
| MSH6 | 50.5756722 | -0.160294503 | 0.22505715 | -0.7122391 | 0.47631673 | 0.73318945 |
| FBXO11 | 462.300534 | 0.100891454 | 0.13640812 | 0.7396294 | 0.4595249 | 0.71819041 |
| FOXN2 | 61.7134745 | 0.055878822 | 0.31185203 | 0.17918377 | 0.85779341 | 0.94685447 |
| PPP1R21 | 289.43187 | 0.218355763 | 0.25295403 | 0.86322311 | 0.38801481 | 0.66428677 |
| GPR75-ASB3 | 80.9392287 | -0.045866563 | 0.24005803 | -0.1910645 | 0.84847507 | 0.94366633 |
| ERLEC1 | 60.1400994 | 0.102002054 | 0.2050111 | 0.49754405 | 0.61880545 | 0.82733103 |
| PSME4 | 264.206368 | 0.192649422 | 0.18011455 | 1.06959386 | 0.28480216 | 0.57246822 |
| ACYP2 | 407.483791 | 0.105840674 | 0.237146 | 0.44631019 | 0.6553732 | 0.85058543 |
| SPTBN1 | 647.020299 | 0.163845196 | 0.20144638 | 0.81334394 | 0.41602089 | 0.68554346 |
| EML6 | 182.753954 | 2.043268353 | 0.67285082 | 3.03673311 | 0.00239157 | 0.03547399 |
| RTN4 | 812.590984 | 0.295307224 | 0.32475094 | 0.90933448 | 0.3631736 | 0.64351459 |
| CLHC1 | 93.5173658 | 0.409285641 | 0.30804162 | 1.32866994 | 0.18395688 | 0.45721925 |
| RPS27A | 691.819877 | -0.019315507 | 0.49457605 | -0.0390547 | 0.9688468 | 0.99151533 |
| MTIF2 | 42.0571225 | -0.354276969 | 0.27451287 | -1.290566 | 0.19685422 | 0.47593804 |
| AC012358.3 | 48.0162611 | 0.913825358 | 0.35071991 | 2.60557021 | 0.00917215 | 0.08519229 |
| CCDC88A | 112.420028 | 0.402057186 | 0.17475291 | 2.30071809 | 0.02140757 | 0.14045419 |
| CFAP36 | 124.51991 | -0.063309958 | 0.17127173 | -0.3696463 | 0.71164606 | 0.87748 |
| PPP4R3B | 258.749524 | 0.156861206 | 0.16841811 | 0.93137971 | 0.35165718 | 0.63356563 |
| PNPT1 | 109.191329 | -0.441112648 | 0.55389886 | -0.7963776 | 0.42581259 | 0.69215319 |
| EFEMP1 | 86.7913569 | -0.631055184 | 0.55868474 | -1.1295372 | 0.25867129 | 0.54565345 |
| CCDC85A | 172.479608 | 2.322917534 | 0.80409837 | 2.88884746 | 0.00386657 | 0.04842913 |
| VRK2 | 170.180724 | -0.22444368 | 0.22145236 | -1.0135077 | 0.3108177 | 0.59807427 |
| FANCL | 119.871471 | 0.592811765 | 0.28824955 | 2.05659215 | 0.03972547 | 0.19750809 |
| BCL11A | 33.3597731 | -0.338328079 | 0.52451108 | -0.6450351 | 0.51890441 | 0.7636788 |
| PAPOLG | 65.5894205 | 0.19933934 | 0.2502367 | 0.79660313 | 0.42568156 | 0.69215319 |
| REL | 82.6198061 | 0.369285167 | 0.31160539 | 1.18510521 | 0.23597585 | 0.51951567 |
| PUS10 | 42.8651243 | -0.120936504 | 0.41083514 | -0.2943675 | 0.76847712 | 0.90459435 |
| PEX13 | 63.9578224 | -0.200879495 | 0.23027859 | -0.8723325 | 0.38302703 | 0.65965766 |
| KIAA1841 | 46.7702728 | 0.211548212 | 0.26517881 | 0.79775685 | 0.42501161 | 0.69181522 |
| C2orf74 | 32.7764258 | 0.537235757 | 0.54242303 | 0.99043685 | 0.32196064 | 0.60925741 |
| USP34 | 961.066236 | 0.28301522 | 0.12172108 | 2.32511267 | 0.02006594 | 0.13480227 |
| XPO1 | 471.820652 | 0.094671636 | 0.32751138 | 0.28906366 | 0.77253267 | 0.90624314 |
| FAM161A | 29.941112 | 0.40860818 | 0.40542474 | 1.0078521 | 0.31352547 | 0.60065312 |
| CCT4 | 109.313087 | -0.353267283 | 0.26484317 | -1.3338735 | 0.18224532 | 0.4553499 |

| | | | | | | |
|---|---|---|---|---|---|---|
| COMMD1 | 373.076423 | 0.270330666 | 0.12044852 | 2.24436679 | 0.02480881 | 0.15268762 |
| B3GNT2 | 52.4239157 | 0.172214934 | 0.30578714 | 0.56318567 | 0.57330846 | 0.80019259 |
| EHBP1 | 679.127713 | 0.202194013 | 0.22076293 | 0.91588751 | 0.3597259 | 0.64001011 |
| WDPCP | 473.544213 | 0.190197588 | 0.17456297 | 1.08956437 | 0.27590508 | 0.56304683 |
| MDH1 | 110.954693 | -0.427592761 | 0.33384477 | -1.2808131 | 0.20025931 | 0.47897227 |
| UGP2 | 164.475364 | -0.828857321 | 0.27765166 | -2.9852417 | 0.00283354 | 0.03905397 |
| VPS54 | 342.025157 | 0.226574928 | 0.16776744 | 1.35052979 | 0.1768461 | 0.44872611 |
| PELI1 | 188.749923 | 0.306961523 | 0.24982516 | 1.22870539 | 0.21918228 | 0.50096566 |
| AC008074.1 | 47.4838651 | 0.526933065 | 0.43340367 | 1.21580203 | 0.22406035 | 0.50617113 |
| AFTPH | 221.121539 | 0.210207039 | 0.18162891 | 1.15734349 | 0.24713205 | 0.5338317 |
| SERTAD2 | 87.2893335 | 0.073199891 | 0.26360534 | 0.27768744 | 0.7812523 | 0.91077326 |
| CEP68 | 45.9468201 | -0.281573668 | 0.2570957 | -1.0952096 | 0.27342483 | 0.56096305 |
| RAB1A | 394.031905 | 0.043266821 | 0.20418685 | 0.21189818 | 0.83218647 | 0.93582908 |
| ACTR2 | 263.656083 | -0.418212538 | 0.2114269 | -1.9780479 | 0.04792331 | 0.21870289 |
| SPRED2 | 253.832918 | -0.055189719 | 0.14036746 | -0.3931803 | 0.69418633 | 0.86986959 |
| MEIS1 | 88.4027131 | 0.404010954 | 0.78757431 | 0.51298137 | 0.60796435 | 0.8211189 |
| ETAA1 | 43.2075689 | -0.052912451 | 0.22397167 | -0.2362462 | 0.81324165 | 0.92694452 |
| C1D | 72.5343545 | 0.444981271 | 0.25968441 | 1.71354632 | 0.08661208 | 0.30659908 |
| PNO1 | 26.1167936 | -0.84171035 | 0.41459534 | -2.0301973 | 0.04233649 | 0.20409723 |
| APLF | 103.678919 | 0.459273102 | 0.33773122 | 1.35987754 | 0.17386868 | 0.44528422 |
| ANTXR1 | 73.6189424 | -0.588416125 | 0.62112592 | -0.9473379 | 0.34346662 | 0.62823502 |
| GFPT1 | 198.360346 | -0.020262174 | 0.24300224 | -0.0833827 | 0.93354727 | 0.9776374 |
| NFU1 | 59.9212035 | 0.032576397 | 0.2398374 | 0.13582701 | 0.89195804 | 0.95976294 |
| AAK1 | 469.595723 | -0.095416187 | 0.21448151 | -0.4448691 | 0.65641439 | 0.8513124 |
| ANXA4 | 453.504106 | -0.659848602 | 0.29752692 | -2.2177778 | 0.02656999 | 0.15808253 |
| GMCL1 | 98.4288339 | 0.464605597 | 0.21239205 | 2.18749054 | 0.02870673 | 0.1646176 |
| MXD1 | 40.0797048 | 0.12026264 | 0.28792908 | 0.41768146 | 0.67618003 | 0.86116336 |
| PCBP1 | 151.427474 | -0.775421087 | 0.37717541 | -2.0558633 | 0.03979569 | 0.19761687 |
| C2orf42 | 30.7173335 | 0.434983414 | 0.30679265 | 1.41784171 | 0.15623699 | 0.4206279 |
| PCYOX1 | 96.5077885 | -0.141332764 | 0.48506095 | -0.2913711 | 0.77076748 | 0.90547178 |
| SNRPG | 53.9850066 | -0.466610274 | 0.3632558 | -1.2845226 | 0.19895917 | 0.47789115 |
| FAM136A | 43.2537797 | -0.840049043 | 0.30317893 | -2.7708028 | 0.00559183 | 0.06222098 |
| TGFA | 113.562748 | -0.138737175 | 0.47386914 | -0.2927753 | 0.76969392 | 0.90533273 |
| NAGK | 41.5606604 | -0.023392581 | 0.29683222 | -0.0788074 | 0.9371858 | 0.97909629 |
| MCEE | 58.7887365 | 0.156573022 | 0.31041767 | 0.50439467 | 0.61398406 | 0.82514464 |
| MPHOSPH10 | 47.3217818 | -0.429336244 | 0.28940648 | -1.483506 | 0.13794003 | 0.39068755 |
| PAIP2B | 33.3018784 | 0.405910636 | 0.48018626 | 0.84531914 | 0.39793268 | 0.67286798 |
| ZNF638 | 751.189825 | 0.344763822 | 0.16104443 | 2.14079948 | 0.03229021 | 0.17584306 |
| EXOC6B | 746.10235 | 0.266505588 | 0.17445032 | 1.52768762 | 0.12659012 | 0.37394935 |
| SFXN5 | 65.7421842 | 0.041490761 | 0.22940286 | 0.18086419 | 0.85647418 | 0.94627061 |
| RAB11FIP5 | 35.5342209 | -0.352603423 | 0.31118877 | -1.1330853 | 0.25717843 | 0.54454292 |
| CCT7 | 99.8667908 | -0.637249482 | 0.35395132 | -1.8003873 | 0.0717995 | 0.27587942 |
| ALMS1 | 196.280563 | 0.016571814 | 0.17937026 | 0.09238886 | 0.92638909 | 0.97408135 |
| DUSP11 | 33.6507938 | -0.170430606 | 0.26371011 | -0.6462801 | 0.51809796 | 0.76316474 |
| STAMBP | 76.2504373 | 0.000526135 | 0.21052566 | 0.00249915 | 0.99800597 | 0.99923686 |
| DGUOK | 78.4870276 | -0.107214718 | 0.35822706 | -0.2992926 | 0.76471678 | 0.90314131 |
| TET3 | 116.966773 | 0.256754293 | 0.30969781 | 0.82904782 | 0.40707735 | 0.68040675 |
| MOB1A | 145.01518 | 0.027706421 | 0.1984161 | 0.13963797 | 0.88894604 | 0.95908272 |
| AC006030.1 | 34.4870226 | -0.437145811 | 0.32800791 | -1.3327295 | 0.1826206 | 0.45602782 |
| RTKN | 58.7700268 | -0.5188512 | 0.29127764 | -1.7812943 | 0.07486438 | 0.28266689 |
| MOGS | 51.0976372 | -0.230799263 | 0.23820453 | -0.9689122 | 0.33258902 | 0.61719981 |
| AUP1 | 39.3014051 | -0.133303623 | 0.40016892 | -0.3331184 | 0.73904493 | 0.89238766 |
| HK2 | 55.9892884 | -1.158248513 | 0.83035395 | -1.3948853 | 0.16305043 | 0.42986266 |
| POLE4 | 34.6279447 | -0.140520416 | 0.40288023 | -0.3487896 | 0.72724731 | 0.88564265 |
| EVA1A | 89.4516773 | -0.985865171 | 0.4549977 | -2.1667476 | 0.0302541 | 0.17007281 |
| MRPL19 | 119.832542 | 0.145959409 | 0.25398197 | 0.57468413 | 0.56550494 | 0.79444443 |
| GCFC2 | 156.869597 | 0.486011281 | 0.30495627 | 1.59370809 | 0.11100143 | 0.34822126 |
| SUCLG1 | 106.905364 | 0.279142529 | 0.27683581 | 1.00833246 | 0.31329489 | 0.60065312 |
| DNAH6 | 165.883892 | 1.058320492 | 0.49862673 | 2.12247045 | 0.03379825 | 0.18066599 |
| TMSB10 | 920.484419 | -1.413780946 | 0.51373281 | -2.7519771 | 0.00592367 | 0.06458327 |
| KCMF1 | 238.746427 | -0.08077124 | 0.18969961 | -0.425785 | 0.67026455 | 0.85938484 |
| TCF7L1 | 285.138275 | 1.664990371 | 0.54018761 | 3.08224463 | 0.00205446 | 0.03226175 |
| TGOLN2 | 228.861424 | -0.365373033 | 0.26145758 | -1.3974467 | 0.16227929 | 0.42858376 |
| RETSAT | 30.1729431 | -0.59492795 | 0.30667246 | -1.9399458 | 0.05238628 | 0.22955543 |
| ELMOD3 | 78.4718189 | 0.314613089 | 0.18834857 | 1.67037687 | 0.09484482 | 0.3204651 |
| CAPG | 49.861974 | -0.538663416 | 0.30452977 | -1.7688367 | 0.07692114 | 0.28670334 |
| MAT2A | 149.621877 | -0.639160488 | 0.33242601 | -1.922715 | 0.05451585 | 0.23406873 |
| GGCX | 50.8248757 | 0.058308902 | 0.22392662 | 0.2603929 | 0.79456072 | 0.91762696 |
| VAMP8 | 182.820924 | -0.429355219 | 0.38992132 | -1.101133 | 0.27083879 | 0.55804793 |
| VAMP5 | 39.8986487 | -0.290754324 | 0.42925893 | -0.6773402 | 0.49819014 | 0.74986853 |
| USP39 | 100.701666 | 0.288440532 | 0.2551613 | 1.13042429 | 0.25829748 | 0.54540257 |
| C2orf68 | 39.8351073 | 0.024535352 | 0.34516683 | 0.07108259 | 0.94333203 | 0.98112027 |
| ATOH8 | 79.7170297 | 1.580627538 | 1.06568198 | 1.48320753 | 0.13801928 | 0.39068755 |
| ST3GAL5 | 49.4949824 | 0.862834078 | 0.47450937 | 1.81837101 | 0.06900745 | 0.2689269 |
| POLR1A | 64.3175103 | -0.65130549 | 0.28245463 | -2.3058765 | 0.02111754 | 0.13943713 |
| PTCD3 | 128.428 | 0.050187392 | 0.15132914 | 0.33164393 | 0.74015814 | 0.89253455 |
| IMMT | 150.26583 | -0.150947167 | 0.17685543 | -0.853506 | 0.39337878 | 0.6686322 |
| MRPL35 | 26.5881989 | -0.046361043 | 0.35915723 | -0.1290829 | 0.89729208 | 0.96175013 |
| KDM3A | 141.471721 | -0.030047108 | 0.4217005 | -0.0712522 | 0.943197 | 0.98112027 |
| CHMP3 | 54.7534787 | -0.501438141 | 0.43823398 | -1.1442247 | 0.25253047 | 0.5393622 |
| RMND5A | 103.210292 | 0.272872607 | 0.23048785 | 1.18389149 | 0.23645602 | 0.51974689 |
| PLGLB1 | 30.4019689 | 0.909405027 | 0.45613602 | 1.99371458 | 0.04618326 | 0.21432487 |
| CYTOR | 64.8679309 | -1.555336052 | 0.44841151 | -3.4685463 | 0.00052328 | 0.01373303 |
| KRCC1 | 122.958859 | -0.013169716 | 0.18669901 | -0.0705398 | 0.943764 | 0.98112027 |
| THNSL2 | 40.9561663 | 0.300490475 | 0.83066214 | 0.36174813 | 0.71754026 | 0.88106948 |
| EIF2AK3 | 156.577531 | 0.663123931 | 0.60081433 | 1.10370857 | 0.26971957 | 0.55681979 |
| RPIA | 44.989206 | -0.303851841 | 0.30158296 | -1.0075232 | 0.31368339 | 0.60082169 |
| MAL | 274.04545 | 1.21317506 | 0.72277969 | 1.67848528 | 0.0932524 | 0.31811423 |
| MRPS5 | 121.355873 | -0.119489255 | 0.17730322 | -0.673926 | 0.50035834 | 0.75183255 |
| ANKRD36C | 129.672709 | 0.352344078 | 0.27574253 | 1.27780101 | 0.2013196 | 0.48018935 |
| STARD7 | 92.6835283 | -0.065340645 | 0.17055083 | -0.3831154 | 0.70163422 | 0.87200065 |
| TMEM127 | 49.5754917 | -0.668983788 | 0.21430652 | -3.1216212 | 0.00179858 | 0.02983582 |
| CIAO1 | 55.4718052 | -0.41569282 | 0.25976054 | -1.6002924 | 0.10953373 | 0.34627843 |
| SNRNP200 | 131.424616 | -0.642251725 | 0.26210318 | -2.4503775 | 0.01427065 | 0.11086664 |
| NEURL3 | 30.9345517 | -0.462100112 | 0.75439692 | -0.6125424 | 0.54017894 | 0.7777505 |
| KANSL3 | 147.450098 | 0.140885829 | 0.20684723 | 0.68111055 | 0.49580154 | 0.74744774 |
| FER1L5 | 24.3700369 | 0.099630134 | 0.6191143 | 0.16092365 | 0.87215353 | 0.95193455 |
| LMAN2L | 64.9597822 | 0.307562232 | 0.2220871 | 1.38487212 | 0.16609157 | 0.43446007 |
| CNNM4 | 55.7453706 | 0.078514783 | 0.4911487 | 0.15985949 | 0.87299176 | 0.95193455 |
| ANKRD23 | 43.9444281 | 0.153969177 | 0.26087243 | 0.5902087 | 0.55505074 | 0.78784845 |
| ANKRD36 | 198.995664 | 0.830075016 | 0.25407325 | 3.26706968 | 0.00108667 | 0.02186808 |
| AC092683.1 | 110.537101 | -0.251881142 | 0.38942703 | -0.6467993 | 0.51776183 | 0.76300557 |
| ANKRD36B | 59.7381516 | 0.771871422 | 0.38691335 | 1.99494647 | 0.04604872 | 0.21412654 |
| COX5B | 134.988118 | -0.551958835 | 0.4158226 | -1.3273902 | 0.18437965 | 0.45779488 |
| ACTR1B | 48.1470323 | -0.296154489 | 0.25925497 | -1.1423291 | 0.25331726 | 0.54018835 |
| TMEM131 | 355.24955 | 0.590246225 | 0.25074777 | 2.35394404 | 0.01857541 | 0.12912469 |
| INPP4A | 106.309214 | 0.174756136 | 0.18645646 | 0.93724903 | 0.34863047 | 0.63236722 |
| COA5 | 44.8912203 | 0.364123146 | 0.29731595 | 1.22470103 | 0.22068789 | 0.50236569 |
| UNC50 | 52.2025399 | 0.136135784 | 0.33396652 | 0.40763303 | 0.68354311 | 0.86478143 |
| MGAT4A | 412.715302 | 0.223211 | 0.25938041 | 0.86055458 | 0.38948341 | 0.66527489 |
| TSGA10 | 200.878451 | 0.496850741 | 0.32116968 | 1.54700389 | 0.12186231 | 0.36668381 |
| AC079447.1 | 112.227539 | 0.444579712 | 0.27037595 | 1.64430199 | 0.10011384 | 0.33080044 |
| MITD1 | 55.2515974 | -0.292282295 | 0.3110527 | -0.9396552 | 0.34739444 | 0.63096157 |

| | | | | | | |
|---|---|---|---|---|---|---|
| LYG1 | 44.144881 | 0.085035945 | 0.57033209 | 0.149099 | 0.88147551 | 0.95635451 |
| TXNDC9 | 54.7177852 | -0.246640681 | 0.34262856 | -0.7198486 | 0.47161824 | 0.72945405 |
| EIF5B | 248.88661 | -0.564294354 | 0.2286318 | -2.4681359 | 0.01358187 | 0.10755093 |
| REV1 | 245.070107 | 0.605678413 | 0.15072292 | 4.01848908 | 5.86E-05 | 0.00328109 |
| AFF3 | 155.298921 | 0.390064765 | 0.55608478 | 0.70144838 | 0.48302324 | 0.73696727 |
| LONRF2 | 53.8172204 | -0.114231333 | 0.64359684 | -0.177489 | 0.85912434 | 0.94734879 |
| NPAS2 | 362.093257 | 0.495799037 | 0.48814058 | 1.01568903 | 0.30977749 | 0.59657391 |
| RPL31 | 180.799584 | -0.010120095 | 0.35161736 | -0.0287816 | 0.97703881 | 0.99441014 |
| TBC1D8 | 241.088977 | -0.177565808 | 0.31285135 | -0.5675725 | 0.57032532 | 0.79765604 |
| CNOT11 | 71.5790909 | -0.289033183 | 0.18969832 | -1.5236465 | 0.12759703 | 0.37492391 |
| RNF149 | 430.952372 | 0.128410229 | 0.21549695 | 0.59587957 | 0.55125568 | 0.78577398 |
| MAP4K4 | 442.664902 | -1.105217472 | 0.41089811 | -2.6897604 | 0.00715033 | 0.07217391 |
| IL1R1 | 491.290156 | -0.652555074 | 0.4748011 | -1.3743757 | 0.16932509 | 0.43908094 |
| IL18R1 | 31.8033217 | 0.041931001 | 0.5768857 | 0.07268511 | 0.9420567 | 0.98091268 |
| TMEM182 | 42.2801658 | -0.269973629 | 0.29166585 | -0.9256265 | 0.35464013 | 0.63640307 |
| POU3F3 | 267.482428 | 0.000948252 | 0.65657824 | 0.00144423 | 0.99884767 | 0.99936932 |
| MRPS9 | 100.493594 | 0.567703592 | 0.29791872 | 1.90556537 | 0.05670662 | 0.24003212 |
| TGFBRAP1 | 74.4688975 | -0.213402452 | 0.2172461 | -0.9823074 | 0.32594844 | 0.61256819 |
| C2orf49 | 32.4964288 | 0.120903367 | 0.2875839 | 0.42041077 | 0.67418541 | 0.86115505 |
| FHL2 | 170.923971 | -0.482102566 | 0.61542067 | -0.7833708 | 0.43340941 | 0.69833957 |
| NCK2 | 183.575499 | 0.00479867 | 0.2380667 | 0.02015683 | 0.98391826 | 0.99657935 |
| UXS1 | 176.784956 | 0.256521786 | 0.25538698 | 1.00444347 | 0.3151649 | 0.60205159 |
| SULT1C2 | 148.556787 | -1.62835592 | 0.69716227 | -2.3356914 | 0.01950733 | 0.13290237 |
| SULT1C4 | 70.5224103 | 0.111179619 | 0.47814563 | 0.23252251 | 0.81613221 | 0.92869214 |
| GCC2 | 383.170024 | -0.172252814 | 0.22448951 | -0.767309 | 0.44289785 | 0.70544038 |
| GCC2-AS1 | 49.0033654 | 0.060213864 | 0.26432425 | 0.22780302 | 0.81979938 | 0.93087461 |
| LIMS1 | 371.863243 | -0.249199314 | 0.25865496 | -0.963443 | 0.33532524 | 0.61996508 |
| RANBP2 | 219.991938 | 0.164681855 | 0.20827594 | 0.79069073 | 0.42912449 | 0.69498002 |
| CCDC138 | 61.1965461 | 0.267216672 | 0.30065577 | 0.88877947 | 0.37412161 | 0.65265178 |
| EDAR | 31.6470573 | -0.000683413 | 0.35820212 | -0.0019079 | 0.99847772 | 0.99928153 |
| SH3RF3 | 109.422684 | -2.948880785 | 1.00438547 | -2.936005 | 0.00332469 | 0.04368614 |
| 10-Sep | 269.389954 | 0.263191606 | 0.16108398 | 1.6338782 | 0.10228443 | 0.33390378 |
| SOWAHC | 31.1342421 | 0.248509828 | 0.45403315 | 0.54733851 | 0.58414619 | 0.80801969 |
| NPHP1 | 103.202409 | 0.338765659 | 0.29675495 | 1.14156702 | 0.25363404 | 0.54042261 |
| SMIM37 | 52.4408464 | -0.625338482 | 0.41592674 | -1.5034823 | 0.13271472 | 0.38380408 |
| MIR4435-2H | 482.439914 | -1.422695204 | 0.33395578 | -4.2601305 | 2.04E-05 | 0.00156565 |
| BCL2L11 | 48.1097588 | -0.33613491 | 0.35687183 | -0.9418925 | 0.34624767 | 0.6302171 |
| ANAPC1 | 170.699407 | -0.024700399 | 0.21556221 | -0.1145859 | 0.90877332 | 0.96752108 |
| MERTK | 50.4970547 | 0.067653737 | 0.37522335 | 0.18030258 | 0.85691503 | 0.94638973 |
| TMEM87B | 140.152089 | -0.431924082 | 0.19725352 | -2.1896901 | 0.02854672 | 0.16413506 |
| ZC3H8 | 62.2624168 | -0.227719171 | 0.25651763 | -0.887733 | 0.37468439 | 0.65310592 |
| ZC3H6 | 165.498141 | 0.645446607 | 0.34934237 | 1.84760471 | 0.06465955 | 0.25893554 |
| TTL | 39.5060815 | -0.732026729 | 0.36950947 | -1.981077 | 0.04758263 | 0.21810285 |
| POLR1B | 36.9211327 | -0.617993786 | 0.30677118 | -2.0145106 | 0.04395596 | 0.20803069 |
| CHCHD5 | 29.5811209 | -0.71040232 | 0.38645577 | -1.83825 | 0.06602558 | 0.26198834 |
| SLC20A1 | 75.9327416 | -0.304941291 | 0.45642119 | -0.6681138 | 0.50406096 | 0.75508812 |
| PSD4 | 47.095609 | 0.001321776 | 0.41595147 | 0.00317772 | 0.99746455 | 0.99923686 |
| PAX8 | 4716.51084 | 0.473652997 | 0.25835637 | 1.83333202 | 0.06675321 | 0.26332729 |
| CBWD2 | 73.6818128 | 0.18208423 | 0.29055836 | 0.62667009 | 0.53087551 | 0.77154552 |
| RABL2A | 59.1812133 | 0.265527589 | 0.27980264 | 0.94898171 | 0.34262992 | 0.62772374 |
| SLC35F5 | 851.077743 | 0.607636279 | 0.18251984 | 3.32915185 | 0.00087111 | 0.01883601 |
| ACTR3 | 404.846129 | 0.366505684 | 0.19914577 | 1.84038902 | 0.06571114 | 0.26131276 |
| DDX18 | 98.5697671 | -0.485047014 | 0.28151531 | -1.7229863 | 0.084891 | 0.30278126 |
| CCDC93 | 158.593796 | -0.228815543 | 0.17304488 | -1.3222902 | 0.18607154 | 0.46080126 |
| INSIG2 | 101.10922 | -0.325383372 | 0.40360117 | -0.8062003 | 0.42012738 | 0.68863795 |
| C2orf76 | 30.9483216 | 0.141105287 | 0.33608776 | 0.41984655 | 0.67459755 | 0.86115505 |
| DBI | 164.278071 | -1.200594257 | 0.37665807 | -3.1874911 | 0.00143513 | 0.02628508 |
| TMEM37 | 71.6460036 | -1.251529615 | 0.55797705 | -2.2429769 | 0.02489831 | 0.15298324 |
| SCTR | 153.729568 | -0.656963245 | 1.262883 | -0.5202091 | 0.60291783 | 0.81954732 |
| CFAP221 | 428.053139 | -0.036933084 | 0.53419037 | -0.0691384 | 0.94487943 | 0.98119662 |
| PTPN4 | 222.187171 | 0.53968828 | 0.17440755 | 3.09440886 | 0.00197206 | 0.03151251 |
| EPB41L5 | 598.785803 | 0.813997405 | 0.5058542 | 1.60915419 | 0.10758263 | 0.34308427 |
| RALB | 65.1030358 | -0.442903964 | 0.24593912 | -1.8008683 | 0.07172364 | 0.27569852 |
| TFCP2L1 | 187.629924 | 0.433091942 | 0.58738757 | 0.73731888 | 0.46092845 | 0.71918385 |
| CLASP1 | 669.410923 | 0.253354528 | 0.2150012 | 1.17838659 | 0.23864252 | 0.52193243 |
| NIFK-AS1 | 70.7317685 | 0.286381898 | 0.34180217 | 0.83785863 | 0.4021101 | 0.6759938 |
| NIFK | 33.9097206 | -0.6628324 | 0.37709274 | -1.7577437 | 0.07879113 | 0.29095613 |
| TSN | 52.5235649 | -0.866257821 | 0.26239689 | -3.3013265 | 0.00096229 | 0.02030344 |
| BIN1 | 132.785218 | -0.35327632 | 0.38956799 | -0.9068413 | 0.36449075 | 0.644313 |
| ERCC3 | 61.8744904 | 0.104942969 | 0.25026829 | 0.41932187 | 0.67498092 | 0.86115505 |
| MAP3K2 | 404.425099 | 0.188832649 | 0.16154859 | 1.1688907 | 0.24244767 | 0.52725736 |
| IWS1 | 186.055822 | -0.072377354 | 0.23198885 | -0.3119863 | 0.7550509 | 0.89810084 |
| MYO7B | 39.0180153 | -1.269704705 | 0.72749648 | -1.745307 | 0.08093144 | 0.29573375 |
| LIMS2 | 41.0279917 | 0.714334454 | 0.64448004 | 1.10838879 | 0.26769395 | 0.55491026 |
| WDR33 | 303.045762 | 0.268657638 | 0.16515054 | 1.62674396 | 0.10379149 | 0.3359982 |
| POLR2D | 31.2870185 | -0.07814408 | 0.26041447 | -0.3000758 | 0.76411934 | 0.90275027 |
| AMMECR1L | 34.9533571 | -0.123841243 | 0.24303138 | -0.5095689 | 0.61035349 | 0.82270732 |
| SAP130 | 87.4403929 | 0.300196248 | 0.17086044 | 1.75696757 | 0.07892334 | 0.29099563 |
| UGGT1 | 99.300612 | -0.26765587 | 0.24465749 | -1.0940023 | 0.27395395 | 0.5612072 |
| HS6ST1 | 235.858754 | 0.424035504 | 0.4225889 | 1.00342321 | 0.31565671 | 0.60232326 |
| SMPD4 | 82.4214127 | 0.469662088 | 0.27565083 | 1.70382973 | 0.0884129 | 0.30970853 |
| MZT2B | 154.223906 | -0.405735603 | 0.4532939 | -0.8950829 | 0.37074279 | 0.6507871 |
| CCDC115 | 29.37784 | -0.17642594 | 0.3214589 | -0.5488289 | 0.58312287 | 0.80730365 |
| IMP4 | 34.5436943 | -1.064101576 | 0.39874615 | -2.668619 | 0.00761638 | 0.07529132 |
| PTPN18 | 55.418473 | -0.366960518 | 0.28513776 | -1.2869587 | 0.19810869 | 0.47656532 |
| FAM168B | 165.477271 | 0.027261214 | 0.19494341 | 0.13984168 | 0.88878508 | 0.95907089 |
| PLEKHB2 | 144.731053 | -0.3552866 | 0.22152805 | -1.6037996 | 0.10875823 | 0.34519386 |
| MZT2A | 78.997288 | -0.278241486 | 0.27019621 | -1.0297757 | 0.30311531 | 0.59099157 |
| GPR39 | 99.6113783 | -0.129858751 | 0.46757959 | -0.2777254 | 0.78122312 | 0.91077326 |
| LYPD1 | 37.1707268 | -2.262515588 | 0.61591901 | -3.6733979 | 0.00023935 | 0.0083651 |
| NCKAP5 | 177.375182 | -0.357179542 | 0.68240738 | -0.523411 | 0.60068826 | 0.81779318 |
| MGAT5 | 239.553054 | -0.482497848 | 0.31937963 | -1.5107346 | 0.13085609 | 0.38143239 |
| TMEM163 | 476.688517 | 0.132772624 | 0.72251889 | 0.18376353 | 0.85419897 | 0.9452497 |
| CCNT2-AS1 | 77.3279193 | -0.134238213 | 0.32979987 | -0.4070293 | 0.68398647 | 0.86505233 |
| ACMSD | 108.852033 | -0.604609398 | 0.57047274 | -1.0598392 | 0.28921774 | 0.57585049 |
| CCNT2 | 200.419916 | 0.131453016 | 0.18688448 | 0.70339183 | 0.48181159 | 0.73620566 |
| RAB3GAP1 | 348.608602 | 0.188199114 | 0.1209547 | 1.55594715 | 0.11972066 | 0.36465111 |
| ZRANB3 | 164.727721 | -0.162128372 | 0.26088669 | -0.6214513 | 0.53430272 | 0.77394311 |
| R3HDM1 | 303.389929 | 0.148575478 | 0.13090952 | 1.13494788 | 0.25639716 | 0.54336911 |
| UBXN4 | 278.750163 | -0.256886134 | 0.16554753 | -1.5517364 | 0.12072531 | 0.36507762 |
| DARS | 262.498909 | -0.479469695 | 0.21180374 | -2.2637452 | 0.02358979 | 0.14836941 |
| HNMT | 191.760111 | 0.240125082 | 0.47720195 | 0.50319384 | 0.61482799 | 0.82589221 |
| AC114763.1 | 111.789434 | 0.134179661 | 0.41207574 | 0.32561893 | 0.74471269 | 0.89323545 |
| SPOPL | 188.532508 | 0.270351501 | 0.16943959 | 1.5955628 | 0.11058644 | 0.34758554 |
| LRP1B | 46.1447912 | -0.823540682 | 0.34558788 | -2.3830138 | 0.01717155 | 0.12346725 |
| GTDC1 | 293.824956 | -0.004040439 | 0.33932927 | -0.0119071 | 0.99049971 | 0.9976863 |
| ZEB2 | 145.155549 | -0.78363353 | 0.77654932 | -1.0091227 | 0.3129158 | 0.60019427 |
| TEX41 | 128.137206 | 0.828730564 | 0.43744274 | 1.89448923 | 0.0581601 | 0.24328193 |
| ACVR2A | 133.796104 | 0.377455891 | 0.24077176 | 1.5676917 | 0.1169531 | 0.36033893 |
| ORC4 | 263.355607 | 0.698444776 | 0.21894103 | 3.19010458 | 0.00142221 | 0.02622761 |
| MBD5 | 1309.31795 | 0.454212492 | 0.22602314 | 2.00958403 | 0.04447523 | 0.20973445 |
| EPC2 | 272.653311 | 0.433500051 | 0.173442 | 2.4993949 | 0.01244056 | 0.10185309 |
| MMADHC | 58.9222056 | 0.005558353 | 0.38377623 | 0.01448332 | 0.98844439 | 0.99678172 |

| | | | | | | |
|---|---|---|---|---|---|---|
| AC007364.1 | 70.921598 | -0.495981106 | 0.53514057 | -0.926824 | 0.35401794 | 0.63575888 |
| RND3 | 100.284534 | 0.878380199 | 0.5657609 | 1.55256433 | 0.12052725 | 0.36507762 |
| NMI | 48.5445062 | -0.812046592 | 0.54483415 | -1.4904473 | 0.13610666 | 0.38809998 |
| RIF1 | 265.43002 | -0.11956972 | 0.24938647 | -0.4794555 | 0.6316146 | 0.83520655 |
| ARL5A | 70.011349 | 0.420901035 | 0.32982428 | 1.27613721 | 0.20190702 | 0.48041015 |
| STAM2 | 128.213925 | 0.457499748 | 0.16377928 | 2.79339212 | 0.00521584 | 0.05962119 |
| FMNL2 | 967.269152 | -0.908789112 | 0.46250989 | -1.9649074 | 0.04942495 | 0.2235324 |
| PRPF40A | 189.131577 | 0.085441018 | 0.19080691 | 0.44778786 | 0.6543063 | 0.84972886 |
| ARL6IP6 | 91.8332489 | 0.486526743 | 0.2285785 | 2.12848867 | 0.03329659 | 0.17908367 |
| KCNJ3 | 425.09304 | 1.874878072 | 0.93121614 | 2.01336509 | 0.04407624 | 0.20839401 |
| LINC01876 | 157.831886 | -0.83508585 | 0.67384117 | -1.2392918 | 0.21523745 | 0.49731142 |
| GPD2 | 201.197376 | 0.2733213 | 0.20829154 | 1.31220546 | 0.18945082 | 0.46555911 |
| ACVR1 | 253.744354 | -0.245315983 | 0.35059369 | -0.6997159 | 0.48410473 | 0.73727416 |
| UPP2 | 150.335023 | 1.014240096 | 0.53643982 | 1.89068757 | 0.05866606 | 0.24443767 |
| CCDC148 | 424.910679 | 1.103811756 | 0.52362766 | 2.10800887 | 0.03503022 | 0.1847551 |
| PKP4 | 1520.22208 | 0.449894379 | 0.3143577 | 1.43115431 | 0.152386 | 0.41457127 |
| PKP4-AS1 | 50.1972217 | 0.884174783 | 0.5327987 | 1.65949126 | 0.09701684 | 0.32493857 |
| TANC1 | 479.683274 | -0.054284273 | 0.23558627 | -0.2304221 | 0.81776383 | 0.92999132 |
| WDSUB1 | 154.489944 | 0.643809815 | 0.28802569 | 2.23525134 | 0.02540085 | 0.15487891 |
| BAZ2B | 862.154154 | 0.158627193 | 0.17793315 | 0.89149879 | 0.37266164 | 0.65212383 |
| 7-Mar | 325.524664 | 0.165983048 | 0.14527453 | 1.14254752 | 0.25322651 | 0.54018835 |
| PLA2R1 | 171.261625 | -0.746496613 | 0.80355316 | -0.9289947 | 0.35289184 | 0.63441164 |
| ITGB6 | 1080.9893 | -1.81994536 | 0.7451098 | -2.4425197 | 0.01458513 | 0.11197822 |
| LINC02478 | 39.3373202 | -0.057722238 | 0.61755442 | -0.0934691 | 0.92553092 | 0.97349958 |
| RBMS1 | 1047.39058 | -0.036262408 | 0.28788562 | -0.1259612 | 0.89976267 | 0.96299738 |
| TANK | 406.836965 | -0.334310379 | 0.21748061 | -1.5371962 | 0.12424527 | 0.37041562 |
| PSMD14 | 125.98916 | -0.452580605 | 0.28269904 | -1.6009273 | 0.10939302 | 0.34606201 |
| DPP4 | 118.784297 | 0.079344275 | 0.49160146 | 0.16139959 | 0.87177869 | 0.95193455 |
| IFIH1 | 65.7592 | -0.400652776 | 0.86378966 | -0.4638314 | 0.64276854 | 0.84312153 |
| GCA | 84.0954407 | -0.345148934 | 0.31730952 | -1.0877358 | 0.27671174 | 0.56369053 |
| FIGN | 423.289115 | 1.21294301 | 0.24461285 | 4.95862339 | 7.10E-07 | 0.00012831 |
| GRB14 | 232.762354 | 0.3195421 | 0.26762059 | 1.19401166 | 0.23247341 | 0.51579248 |
| COBLL1 | 1803.09891 | 1.149879715 | 0.55269352 | 2.08050153 | 0.03747956 | 0.19219308 |
| AC019197.1 | 811.380298 | 1.422745041 | 0.56298146 | 2.52716144 | 0.01149886 | 0.09721852 |
| SLC38A11 | 248.233916 | 1.673443767 | 1.23443564 | 1.35563468 | 0.17521543 | 0.4470934 |
| SCN3A | 36.9726197 | 0.331967595 | 0.29197853 | 1.13695891 | 0.25555546 | 0.54242539 |
| SCN2A | 111.572759 | 1.375375233 | 0.65009821 | 2.11564224 | 0.03437525 | 0.18283203 |
| CSRNP3 | 132.543297 | 1.299431127 | 0.19499614 | 6.66388115 | 2.67E-11 | 2.84E-08 |
| GALNT3 | 58.6140532 | -0.177924347 | 0.43117437 | -0.4126506 | 0.67986265 | 0.86235436 |
| TTC21B | 152.538341 | 0.799244896 | 0.18815727 | 4.24774921 | 2.16E-05 | 0.00160599 |
| SCN9A | 60.0265784 | 0.013390291 | 0.84639 | 0.01582047 | 0.98737761 | 0.99678172 |
| STK39 | 334.267253 | -1.082648388 | 0.39523774 | -2.7392333 | 0.00615827 | 0.06585681 |
| CERS6 | 195.450269 | -0.331422338 | 0.24083147 | -1.3761588 | 0.16877248 | 0.43835998 |
| NOSTRIN | 190.888495 | -0.513527772 | 0.56821818 | -0.903751 | 0.36612743 | 0.64623818 |
| SPC25 | 37.6285144 | -0.16046905 | 0.54054596 | -0.2968648 | 0.76656976 | 0.90372312 |
| LRP2 | 773.256741 | 0.539793271 | 0.67646333 | 0.79796383 | 0.42489148 | 0.69181522 |
| FASTKD1 | 69.6749352 | 0.306896299 | 0.28742804 | 1.06773264 | 0.28564113 | 0.57312354 |
| PPIG | 223.694271 | 0.251888855 | 0.14601996 | 1.72503033 | 0.08452201 | 0.30235944 |
| CCDC173 | 28.8885213 | 0.284941601 | 0.40814651 | 0.69813558 | 0.48509241 | 0.7383919 |
| KLHL23 | 88.0622627 | 0.2296261 | 0.20305929 | 1.13083279 | 0.25812547 | 0.54534273 |
| SSB | 100.501501 | -0.150456502 | 0.26251819 | -0.5731279 | 0.56655809 | 0.7955233 |
| METTL5 | 38.6361408 | -0.400704933 | 0.25218631 | -1.5889242 | 0.1120775 | 0.35029418 |
| UBR3 | 442.827299 | 0.562652385 | 0.15270138 | 3.68465811 | 0.00022901 | 0.00824694 |
| MYO3B | 162.819358 | 1.62775013 | 0.74733534 | 2.17807195 | 0.02940068 | 0.16751053 |
| GORASP2 | 93.0810338 | -0.295226655 | 0.23954825 | -1.2324309 | 0.21778818 | 0.49980665 |
| TLK1 | 359.364042 | 0.033508506 | 0.17914808 | 0.18704363 | 0.85162643 | 0.94469179 |
| METTL8 | 68.5304389 | -0.299685858 | 0.36491835 | -0.8212409 | 0.41150909 | 0.68316214 |
| DCAF17 | 58.7584717 | 0.025563357 | 0.21176951 | 0.12071312 | 0.90391827 | 0.96520171 |
| CYBRD1 | 48.9172324 | -0.37750674 | 0.80540052 | -0.4687193 | 0.63927032 | 0.84033321 |
| DYNC1I2 | 192.946468 | -0.043733309 | 0.16597806 | -0.2634885 | 0.79217409 | 0.916675 |
| SLC25A12 | 170.577576 | -0.032740285 | 0.1936368 | -0.1690809 | 0.86573301 | 0.95058578 |
| HAT1 | 81.528038 | 0.283672156 | 0.22705482 | 1.24935534 | 0.21153513 | 0.49253647 |
| METAP1D | 43.163764 | 0.123840601 | 0.24474284 | 0.50600295 | 0.61285458 | 0.824514 |
| ITGA6 | 77.4194909 | -0.518082202 | 0.4536901 | -1.1419297 | 0.25348324 | 0.54030172 |
| PDK1 | 128.425521 | -0.642067667 | 0.5923035 | -1.084018 | 0.27835681 | 0.56578102 |
| RAPGEF4 | 33.8096974 | 0.964064321 | 0.41031807 | 2.34955367 | 0.01879593 | 0.12967472 |
| MAP3K20 | 194.58474 | -1.080217927 | 0.40406836 | -2.6733544 | 0.00750969 | 0.07454435 |
| SP3 | 207.6331 | 0.155015339 | 0.17089292 | 0.90709049 | 0.36435895 | 0.644313 |
| OLA1 | 323.939579 | -0.29546864 | 0.2314024 | -1.2768607 | 0.20165142 | 0.48024387 |
| CIR1 | 118.70269 | -0.102404744 | 0.26532705 | -0.3859567 | 0.69952876 | 0.87181583 |
| SCRN3 | 45.4067679 | 0.271840164 | 0.31598987 | 0.86028127 | 0.38963402 | 0.66529505 |
| GPR155 | 130.165063 | 0.723219205 | 0.35721146 | 2.02462489 | 0.0429059 | 0.20590961 |
| AC010894.3 | 227.214373 | -0.828481628 | 0.36855645 | -2.2479097 | 0.02458195 | 0.15217128 |
| WIPF1 | 172.020709 | -0.501098392 | 0.27589324 | -1.8162764 | 0.06932798 | 0.26973709 |
| CHN1 | 109.996945 | -0.351712524 | 0.82298399 | -0.4273625 | 0.66911529 | 0.85924744 |
| ATF2 | 217.52338 | 0.218231022 | 0.18120365 | 1.20434123 | 0.22845772 | 0.51106877 |
| ATP5MC3 | 161.24873 | -0.882636105 | 0.51346946 | -1.7189651 | 0.08562072 | 0.30455287 |
| LNPK | 80.9596711 | 0.172737287 | 0.22224804 | 0.77722749 | 0.43702457 | 0.70192125 |
| HOXD10 | 60.3712017 | 0.475225312 | 0.58994858 | 0.80553683 | 0.42050996 | 0.68891139 |
| HOXD8 | 28.3028196 | 0.097077368 | 0.64537723 | 0.15041957 | 0.8804336 | 0.95614192 |
| HOXD3 | 61.5782337 | 0.759563753 | 0.61339164 | 1.23830145 | 0.21560431 | 0.49731142 |
| MTX2 | 225.555732 | 0.325383244 | 0.15823689 | 2.05630458 | 0.03975317 | 0.19750809 |
| HNRNPA3 | 367.907524 | -0.094333555 | 0.2628417 | -0.3588987 | 0.71967085 | 0.88166633 |
| NFE2L2 | 600.298887 | -0.521460057 | 0.18300607 | -2.8494141 | 0.00437998 | 0.05297457 |
| AGPS | 183.844468 | -0.057648284 | 0.3522133 | -0.1636744 | 0.86998751 | 0.95125032 |
| RBM45 | 38.9842601 | 0.091866093 | 0.27317309 | 0.33629262 | 0.73665022 | 0.89057637 |
| AC009948.1 | 41.7963626 | -0.361981043 | 0.6097559 | -0.5936491 | 0.55274684 | 0.78613936 |
| PRKRA | 110.674219 | -0.077544333 | 0.25594816 | -0.3029689 | 0.76191356 | 0.90141135 |
| PJVK | 27.445105 | -0.225770546 | 0.35139895 | -0.6424907 | 0.52055465 | 0.76418222 |
| PLEKHA3 | 65.8481598 | 0.119720926 | 0.24122119 | 0.49631181 | 0.61967444 | 0.82775923 |
| TTN-AS1 | 267.822384 | -0.124876661 | 0.22355457 | -0.5585959 | 0.57643756 | 0.80245537 |
| SESTD1 | 279.441786 | -0.670285545 | 0.35702109 | -1.8774396 | 0.06045787 | 0.24800908 |
| ZNF385B | 259.681939 | 0.794591761 | 0.58709544 | 1.3534286 | 0.17591875 | 0.44805256 |
| CWC22 | 87.634271 | 0.60571554 | 0.27176509 | 2.22882024 | 0.02582587 | 0.15586278 |
| UBE2E3 | 333.621347 | 0.368671739 | 0.19680047 | 1.87332749 | 0.06102318 | 0.24969714 |
| SSFA2 | 200.855394 | 0.376622052 | 0.25156306 | 1.49712778 | 0.13436001 | 0.38591741 |
| PDE1A | 1432.23757 | -0.547672893 | 1.26582341 | -0.4326614 | 0.6652608 | 0.85742752 |
| DNAJC10 | 135.756178 | -0.58122441 | 0.21127023 | -2.7510947 | 0.00593965 | 0.0646544 |
| NCKAP1 | 523.910478 | 0.074920693 | 0.19852722 | 0.37738246 | 0.7058894 | 0.87473681 |
| NUP35 | 62.2711915 | -0.076331421 | 0.29156562 | -0.2617984 | 0.79347685 | 0.91723627 |
| FSIP2 | 42.1287981 | 0.989650248 | 0.50411081 | 1.96316013 | 0.04962757 | 0.22398317 |
| FSIP2-AS1 | 35.8830447 | 0.877956496 | 0.41374119 | 2.12199442 | 0.03383821 | 0.18077868 |
| LINC01473 | 78.4936881 | 0.55151453 | 0.37358021 | 1.47629482 | 0.13986476 | 0.3940484 |
| ZC3H15 | 125.805932 | -0.317238055 | 0.21759344 | -1.4579394 | 0.14485725 | 0.40282422 |
| ITGAV | 574.245069 | -0.222839337 | 0.33649935 | -0.6622281 | 0.50782503 | 0.75687816 |
| AC017101.1 | 60.2284561 | -0.776295273 | 0.6819707 | -1.1383118 | 0.25499032 | 0.54182615 |
| AC007319.1 | 381.216631 | -0.912947502 | 0.64137971 | -1.4234119 | 0.15461679 | 0.41802467 |
| TFPI | 878.194402 | -2.115635968 | 0.43939434 | -4.8148913 | 1.47E-06 | 0.0002341 |
| LINC01090 | 31.3793956 | -0.816495392 | 0.53310296 | -1.5315904 | 0.12562354 | 0.37252428 |
| GULP1 | 723.277189 | -0.429539181 | 0.52554399 | -0.817323 | 0.41374388 | 0.68448816 |
| WDR75 | 67.1635942 | -0.158147189 | 0.227356 | -0.6955928 | 0.4866839 | 0.74034382 |
| SLC40A1 | 85.276334 | -0.001428566 | 0.46643088 | -0.0030628 | 0.99755628 | 0.99923686 |
| ASNSD1 | 31.5658295 | 0.092667131 | 0.28695516 | 0.32293244 | 0.74674641 | 0.89352127 |

| | | | | | | |
|---|---|---|---|---|---|---|
| ANKAR | 161.930971 | 0.312948547 | 0.29078873 | 1.07620588 | 0.28183519 | 0.56931659 |
| ORMDL1 | 71.4261124 | -0.123013625 | 0.29958208 | -0.4106174 | 0.68135308 | 0.86320343 |
| PMS1 | 102.697556 | 0.103631383 | 0.22401865 | 0.46260158 | 0.64364998 | 0.84343682 |
| C2orf88 | 43.0117827 | -0.210870954 | 0.34381172 | -0.6133327 | 0.53965641 | 0.77746559 |
| HIBCH | 282.009988 | 0.355203691 | 0.24468507 | 1.45167702 | 0.14659143 | 0.40513541 |
| INPP1 | 33.0874481 | -0.85695491 | 0.24472193 | -3.5017496 | 0.00046221 | 0.01267041 |
| MFSD6 | 77.3506531 | -0.273118276 | 0.21700549 | -1.2585777 | 0.2081829 | 0.48842485 |
| NAB1 | 108.788285 | 0.52587679 | 0.23818707 | 2.20783093 | 0.02725606 | 0.15949106 |
| GLS | 1323.9712 | 0.233458 | 0.2970884 | 0.78581999 | 0.43197296 | 0.69731529 |
| STAT1 | 567.634367 | -0.802027991 | 0.72022515 | -1.1135795 | 0.26545961 | 0.55350835 |
| MYO1B | 147.462668 | -0.998537522 | 0.56623063 | -1.763482 | 0.07781923 | 0.28915067 |
| NABP1 | 87.1063691 | -1.575354128 | 0.65406185 | -2.4085706 | 0.01601513 | 0.11832631 |
| AC098617.1 | 136.018712 | 0.204176577 | 0.56773317 | 0.35963475 | 0.71912029 | 0.88155276 |
| SLC39A10 | 122.68021 | 0.207816434 | 0.34920329 | 0.59511591 | 0.55176599 | 0.78577398 |
| DNAH7 | 414.555287 | 0.22384037 | 0.37345178 | 0.59938226 | 0.54891801 | 0.78420366 |
| STK17B | 30.1543289 | -0.063889911 | 0.31334735 | -0.2038948 | 0.83843567 | 0.93912246 |
| GTF3C3 | 58.88712 | 0.19179486 | 0.22049726 | 0.86982876 | 0.38439399 | 0.66091637 |
| PGAP1 | 78.1799927 | 0.659394083 | 0.33657063 | 1.95915516 | 0.05009462 | 0.22501594 |
| ANKRD44 | 314.576153 | 0.320605721 | 0.31399148 | 1.02106503 | 0.30722363 | 0.59456464 |
| SF3B1 | 247.6228 | 0.103377788 | 0.19979013 | 0.51743191 | 0.60485469 | 0.82048068 |
| COQ10B | 45.2255839 | -0.469375589 | 0.27621183 | -1.699332 | 0.08925666 | 0.31101837 |
| HSPD1 | 207.24381 | -0.61514427 | 0.30959735 | -1.9869171 | 0.04693158 | 0.21634149 |
| MOB4 | 32.2733546 | -0.317395706 | 0.42507692 | -0.7466783 | 0.45525778 | 0.71478681 |
| PLCL1 | 443.218092 | 0.506249815 | 0.97439114 | 0.51955502 | 0.60337375 | 0.81981804 |
| SATB2 | 513.670005 | 1.184772197 | 0.58580814 | 2.02245773 | 0.04312909 | 0.20614587 |
| FTCDNL1 | 36.0903049 | -0.306955342 | 0.32845783 | -0.934535 | 0.35002798 | 0.63286907 |
| C2orf69 | 28.4822794 | -0.284735356 | 0.38204371 | -0.7452952 | 0.45609326 | 0.71539502 |
| TYW5 | 59.1467985 | 0.12641235 | 0.17963594 | 0.70371414 | 0.48161082 | 0.73620566 |
| MAIP1 | 29.1001384 | 0.116603951 | 0.26121158 | 0.44639657 | 0.65531081 | 0.85058543 |
| SPATS2L | 577.020476 | -0.767547027 | 0.52547282 | -1.4606788 | 0.1441036 | 0.40162014 |
| AOX1 | 47.691673 | -1.868508429 | 0.83592258 | -2.2352649 | 0.02539996 | 0.15487891 |
| BZW1 | 48.7515292 | -0.686675928 | 0.41446189 | -1.6567891 | 0.09756213 | 0.3259791 |
| CLK1 | 163.947804 | 0.698709137 | 0.19585126 | 3.56754982 | 0.00036033 | 0.0107864 |
| PPIL3 | 37.7118149 | -0.074290491 | 0.34426053 | -0.2157973 | 0.82914576 | 0.93373939 |
| ORC2 | 174.497015 | -0.067032446 | 0.17900489 | -0.3744727 | 0.70805265 | 0.87571806 |
| AC005037.1 | 49.1284114 | -0.016806261 | 0.2893158 | -0.0580897 | 0.9536772 | 0.98368305 |
| FAM126B | 153.642363 | 0.077177364 | 0.19499567 | 0.39579013 | 0.69225986 | 0.86892611 |
| NDUFB3 | 83.3879725 | -0.147441784 | 0.48302628 | -0.3052459 | 0.76017889 | 0.90125623 |
| CFLAR | 537.543121 | -0.334902293 | 0.37094556 | -0.9028341 | 0.36661396 | 0.64650131 |
| CFLAR-AS1 | 46.0053086 | -0.200562568 | 0.58009622 | -0.3457402 | 0.72953799 | 0.88660802 |
| CASP10 | 42.5734582 | 0.061766756 | 0.40428235 | 0.15278123 | 0.8785708 | 0.95482524 |
| CASP8 | 79.2149175 | -0.711500625 | 0.33094463 | -2.1499084 | 0.03156246 | 0.17373701 |
| TRAK2 | 118.443004 | -0.233864756 | 0.23859543 | -0.9801728 | 0.32700082 | 0.61330512 |
| STRADB | 69.7703762 | -0.357809476 | 0.40746979 | -0.8781252 | 0.37987579 | 0.65718444 |
| ALS2 | 118.679867 | 0.45121024 | 0.21481545 | 2.10045526 | 0.03568881 | 0.18676485 |
| SUMO1 | 115.464753 | 0.162911261 | 0.23859079 | 0.68280616 | 0.49472934 | 0.74677156 |
| NOP58 | 143.779291 | -0.134346736 | 0.25612454 | -0.5245368 | 0.59990526 | 0.81763227 |
| BMPR2 | 596.818789 | -0.037342767 | 0.18519538 | -0.2016398 | 0.8401983 | 0.94021723 |
| FAM117B | 165.757212 | 0.042796002 | 0.27613118 | 0.15498432 | 0.87683368 | 0.95377544 |
| ICA1L | 100.831093 | -0.284229958 | 0.33477941 | -0.8490067 | 0.39587757 | 0.67091161 |
| WDR12 | 77.1169009 | -0.006871061 | 0.28745304 | -0.0239032 | 0.98092978 | 0.99578216 |
| CARF | 116.210753 | 0.531890447 | 0.32581784 | 1.63247795 | 0.10257884 | 0.3339914 |
| NBEAL1 | 783.135243 | -0.062222672 | 0.19225596 | -0.323645 | 0.74620684 | 0.89352127 |
| CYP20A1 | 166.641667 | 0.242463494 | 0.18292811 | 1.32545784 | 0.18501934 | 0.45914515 |
| ABI2 | 353.103353 | 0.492139506 | 0.19032235 | 2.58582084 | 0.00971474 | 0.08812264 |
| RAPH1 | 186.487286 | -0.334450173 | 0.24727209 | -1.3525593 | 0.17619646 | 0.44835649 |
| PARD3B | 1832.9897 | 0.83935465 | 0.22775788 | 3.68529353 | 0.00022844 | 0.00824694 |
| NRP2 | 62.8433998 | 0.342746689 | 1.07478367 | 0.3188983 | 0.74980363 | 0.89544558 |
| INO80D | 418.694806 | 0.507081266 | 0.51720254 | 0.98043073 | 0.32687355 | 0.61330512 |
| NDUFS1 | 201.089102 | -0.127819865 | 0.14978439 | -0.853359 | 0.39346023 | 0.6686322 |
| EEF1B2 | 149.572702 | -0.770940978 | 0.58391451 | -1.3202977 | 0.18673565 | 0.4618489 |
| ZDBF2 | 69.6057099 | 0.229662515 | 0.34919509 | 0.65769114 | 0.51073661 | 0.7592095 |
| FASTKD2 | 68.6852952 | -0.006760518 | 0.21495098 | -0.0314514 | 0.97490952 | 0.99357999 |
| KLF7 | 219.697656 | 0.172172924 | 0.43770192 | 0.39335657 | 0.69405614 | 0.86986959 |
| CREB1 | 212.958498 | 0.284071139 | 0.14173175 | 2.00428721 | 0.0450393 | 0.21106218 |
| METTL21A | 68.1911154 | 0.270355292 | 0.19181252 | 1.40947676 | 0.15869424 | 0.42419408 |
| CCNYL1 | 106.364918 | -0.000102656 | 0.28397515 | -0.0003615 | 0.99971157 | 0.9998728 |
| FZD5 | 34.9821996 | -2.148604792 | 0.49558532 | -4.3354892 | 1.45E-05 | 0.00120098 |
| PLEKHM3 | 125.698666 | 0.0310325 | 0.29119351 | 0.10657003 | 0.9151301 | 0.96915768 |
| IDH1 | 31.8211876 | 0.490574872 | 0.33446344 | 1.46675185 | 0.14244357 | 0.39896695 |
| PIKFYVE | 170.139051 | 0.359102616 | 0.14089614 | 2.54870448 | 0.01081239 | 0.09353273 |
| PTH2R | 1061.59442 | 1.532767399 | 0.68174056 | 2.2483148 | 0.02455612 | 0.15214949 |
| MAP2 | 179.287673 | -0.144764561 | 0.69616735 | -0.2079451 | 0.83527187 | 0.93718696 |
| RPE | 73.4197142 | -0.110464796 | 0.18843706 | -0.5862159 | 0.55773045 | 0.78891584 |
| KANSL1L | 765.212911 | 0.431039179 | 0.27169431 | 1.58648583 | 0.11262914 | 0.35131051 |
| AC007038.1 | 95.7918355 | 0.482627658 | 0.22601292 | 2.13539854 | 0.03272847 | 0.17715978 |
| ACADL | 176.199146 | 1.074600607 | 0.48138499 | 2.23231016 | 0.02559447 | 0.1551184 |
| LANCL1 | 99.7899471 | 0.363872176 | 0.17925599 | 2.02990247 | 0.04236645 | 0.20413896 |
| ERBB4 | 957.567994 | -0.252241125 | 0.60796335 | -0.4148953 | 0.67821856 | 0.86151115 |
| IKZF2 | 133.487215 | 0.433639875 | 0.22385199 | 1.9371723 | 0.05272429 | 0.23008926 |
| SPAG16 | 495.071642 | 0.140345186 | 0.31585692 | 0.44433152 | 0.65680292 | 0.8515857 |
| AC068051.1 | 84.3444851 | -0.427566846 | 0.74808785 | -0.5715463 | 0.56762938 | 0.79644381 |
| BARD1 | 44.9333145 | 0.258345187 | 0.23560635 | 1.09651197 | 0.27285479 | 0.56047062 |
| AC016708.1 | 57.9907542 | 0.207745285 | 0.43422934 | 0.47842296 | 0.63234919 | 0.8353707 |
| ATIC | 60.7938186 | -0.537039917 | 0.30847757 | -1.7409367 | 0.08169468 | 0.29698419 |
| MREG | 99.7412997 | 0.488394356 | 0.32417141 | 1.50659295 | 0.13191503 | 0.38233405 |
| PECR | 124.218723 | 0.74580908 | 0.31633103 | 2.35768549 | 0.01838927 | 0.12829627 |
| TMEM169 | 32.2420466 | 0.58025838 | 0.39179185 | 1.48103741 | 0.1385966 | 0.3916274 |
| XRCC5 | 485.167744 | -0.143535781 | 0.2290339 | -0.626701 | 0.53085523 | 0.77154552 |
| SMARCAL1 | 52.8264384 | 0.027280075 | 0.21872639 | 0.12472238 | 0.90074334 | 0.96303857 |
| RPL37A | 354.547205 | -0.549481893 | 0.33436455 | -1.6433617 | 0.10030812 | 0.33096298 |
| IGFBP2 | 238.902756 | 0.454704925 | 0.56481344 | 0.8050533 | 0.42078892 | 0.68925052 |
| IGFBP5 | 440.633421 | 3.299262809 | 1.07714606 | 3.06296699 | 0.00219154 | 0.03385831 |
| AC007563.2 | 43.5126005 | 2.322171653 | 0.91116904 | 2.54856296 | 0.01081678 | 0.09353273 |
| DIRC3 | 138.814422 | 0.703932324 | 0.52129264 | 1.35035922 | 0.17690079 | 0.44872611 |
| TNS1 | 675.717288 | -0.096689661 | 0.46530705 | -0.2077975 | 0.83538705 | 0.93718696 |
| ARPC2 | 187.598438 | -0.472735168 | 0.25105673 | -1.8829814 | 0.05970288 | 0.246506 |
| AAMP | 54.781928 | -0.140749504 | 0.26288027 | -0.535413 | 0.59236434 | 0.81283604 |
| PNKD | 151.379068 | -0.095721547 | 0.30612832 | -0.3126844 | 0.75452045 | 0.89810084 |
| TMBIM1 | 93.5034816 | -0.679221777 | 0.19521806 | -3.4792979 | 0.00050273 | 0.01352709 |
| CTDSP1 | 91.0215465 | -0.186448613 | 0.24652438 | -0.756309 | 0.44946394 | 0.71046454 |
| USP37 | 157.630866 | 0.04735854 | 0.25194108 | 0.18797467 | 0.85089651 | 0.94435157 |
| CNOT9 | 71.6736913 | 0.107272765 | 0.32881526 | 0.32624023 | 0.74424261 | 0.89303519 |
| ZNF142 | 28.3830383 | -0.233751487 | 0.27624638 | -0.8461703 | 0.39745774 | 0.67253979 |
| STK36 | 35.7807511 | 0.598515494 | 0.25748235 | 2.32449134 | 0.02009918 | 0.13491941 |
| TTLL4 | 28.436393 | -0.069807754 | 0.38005837 | -0.1836764 | 0.85426732 | 0.9452497 |
| CYP27A1 | 57.0703058 | -0.512288541 | 0.40206058 | -1.2741576 | 0.20260758 | 0.48158147 |
| CNPPD1 | 36.6894678 | -0.931916642 | 0.26853396 | -3.4703865 | 0.00051971 | 0.01373303 |
| RETREG2 | 59.5175791 | -0.589025747 | 0.36356077 | -1.6201576 | 0.10519842 | 0.33906316 |
| ZFAND2B | 34.3377835 | -0.299287976 | 0.32113795 | -0.9319608 | 0.35135679 | 0.63335467 |
| ANKZF1 | 57.1077984 | 0.253126985 | 0.2691209 | 0.94056978 | 0.34692537 | 0.63046825 |
| TUBA4A | 63.4982485 | -1.607871787 | 0.59404052 | -2.7066702 | 0.00679617 | 0.07001456 |
| DNAJB2 | 47.8960818 | 0.109094835 | 0.341915 | 0.31907005 | 0.74967339 | 0.89540168 |

| | | | | | | |
|---|---|---|---|---|---|---|
| DNPEP | 39.7044681 | -0.532467687 | 0.34990443 | -1.5217518 | 0.12807129 | 0.37588973 |
| OBSL1 | 62.3801799 | -0.038281842 | 0.2756162 | -0.1388955 | 0.88953275 | 0.95918866 |
| EPHA4 | 154.099568 | 1.892597242 | 0.68294544 | 2.77122759 | 0.00558454 | 0.06222098 |
| SGPP2 | 360.692512 | -1.920185588 | 0.59853798 | -3.2081265 | 0.00133603 | 0.02519253 |
| FARSB | 90.1522677 | -0.243283399 | 0.38044614 | -0.6394687 | 0.52251807 | 0.76512314 |
| ACSL3 | 152.181743 | 0.417964753 | 0.23390635 | 1.7868893 | 0.07395538 | 0.28056182 |
| AP1S3 | 231.840451 | 0.350041246 | 0.56574974 | 0.618721 | 0.53610017 | 0.77540842 |
| WDFY1 | 135.676061 | -0.133180789 | 0.27534615 | -0.4836849 | 0.62860948 | 0.83341871 |
| SERPINE2 | 99.1940178 | -3.622910082 | 0.7962987 | -4.5496873 | 5.37E-06 | 0.00059481 |
| CUL3 | 405.35996 | 0.246279168 | 0.11918043 | 2.06643959 | 0.03878699 | 0.19565065 |
| IRS1 | 97.9688471 | -0.332920594 | 0.2817458 | -1.1816346 | 0.23735071 | 0.52003259 |
| RHBDD1 | 187.937781 | 0.218925222 | 0.26528267 | 0.82525263 | 0.40922818 | 0.68208527 |
| COL4A4 | 411.895403 | 1.558505796 | 0.46120073 | 3.3792353 | 0.00072688 | 0.01665732 |
| COL4A3 | 596.418071 | 1.675279252 | 0.50845684 | 3.29483078 | 0.00098481 | 0.02055227 |
| AC097662.1 | 92.6308874 | 1.540004378 | 0.6287088 | 2.44947166 | 0.0143066 | 0.1109659 |
| MFF | 173.207112 | 0.619396224 | 0.22664287 | 2.73291729 | 0.00627761 | 0.06659271 |
| AGFG1 | 315.921293 | 0.01280968 | 0.16968297 | 0.07549184 | 0.93982339 | 0.98003138 |
| TRIP12 | 477.106266 | -0.10099038 | 0.19770375 | -0.5108167 | 0.60947941 | 0.82205058 |
| FBXO36 | 63.2416366 | 0.390400828 | 0.2647091 | 1.47482963 | 0.14025835 | 0.39480892 |
| SP110 | 79.853943 | -0.738155242 | 0.71744696 | -1.0288639 | 0.30354366 | 0.5915859 |
| SP140L | 62.8629344 | -0.13476931 | 0.36427952 | -0.3699613 | 0.71141136 | 0.87748 |
| SP100 | 292.507286 | -0.589793773 | 0.58869727 | -1.0018626 | 0.31640996 | 0.60281198 |
| CAB39 | 293.145593 | 0.270435509 | 0.20553804 | 1.31574433 | 0.18825987 | 0.46394167 |
| ITM2C | 98.7752984 | -0.766974552 | 0.56872463 | -1.3485868 | 0.17746971 | 0.44901805 |
| PSMD1 | 210.378767 | -0.188760566 | 0.26389021 | -0.7152996 | 0.47442393 | 0.73121591 |
| ARMC9 | 325.671452 | -0.283599172 | 0.19483128 | -1.4556142 | 0.14549931 | 0.40339737 |
| NCL | 214.443017 | -1.326831796 | 0.42521179 | -3.1204022 | 0.00180604 | 0.02986263 |
| PTMA | 936.012656 | -1.013269005 | 0.42862339 | -2.3640077 | 0.01807844 | 0.12695993 |
| PDE6D | 58.8716657 | 0.573293306 | 0.2411801 | 2.37703408 | 0.01745248 | 0.12466612 |
| COPS7B | 38.2473958 | 0.441730491 | 0.23295769 | 1.89618333 | 0.05793581 | 0.24313032 |
| DIS3L2 | 338.778456 | -0.066584635 | 0.20597763 | -0.3232615 | 0.74649722 | 0.89352127 |
| EIF4E2 | 78.25627 | -0.349725374 | 0.28466349 | -1.2285572 | 0.21923789 | 0.50097321 |
| EFHD1 | 199.479546 | 1.079068259 | 0.36206596 | 2.98030849 | 0.00287958 | 0.03957434 |
| GIGYF2 | 475.494299 | 0.225730601 | 0.15590183 | 1.44790218 | 0.14764441 | 0.40657927 |
| ATG16L1 | 41.2392028 | 0.057682837 | 0.32680664 | 0.17650448 | 0.85989763 | 0.94745904 |
| DGKD | 99.0053357 | -0.032614948 | 0.31891347 | -0.102269 | 0.91854319 | 0.97062606 |
| USP40 | 263.127424 | 0.34042871 | 0.26829587 | 1.26885556 | 0.20449259 | 0.48318561 |
| ARL4C | 66.3228927 | -2.169248131 | 0.73537407 | -2.9498567 | 0.00317921 | 0.04229679 |
| SH3BP4 | 172.062534 | -0.088485252 | 0.53774769 | -0.1645479 | 0.86929984 | 0.95106604 |
| AGAP1 | 1367.25758 | -0.41838216 | 0.28959896 | -1.444695 | 0.14854359 | 0.40820743 |
| IQCA1 | 112.73161 | 0.491686315 | 0.28972512 | 1.69707863 | 0.0896818 | 0.31211662 |
| COPS8 | 40.7063924 | -0.180000079 | 0.23218931 | -0.7752298 | 0.43820386 | 0.70236452 |
| MLPH | 33.5759705 | 0.447629455 | 0.71247797 | 0.62827129 | 0.52982624 | 0.77060515 |
| RAB17 | 107.275882 | 0.351340338 | 0.37267043 | 0.94276419 | 0.34580153 | 0.63016843 |
| LRRFIP1 | 779.772457 | -0.271948445 | 0.26017983 | -1.0452326 | 0.29591546 | 0.584087 |
| ILKAP | 127.828274 | -0.007412153 | 0.19396886 | -0.0382131 | 0.96951777 | 0.99175186 |
| PER2 | 72.9480868 | 0.019081546 | 0.36337467 | 0.05251204 | 0.9581207 | 0.98566059 |
| TRAF3IP1 | 132.867135 | 0.138517541 | 0.21199463 | 0.65340119 | 0.51349767 | 0.76083448 |
| HDAC4 | 252.038761 | -0.48308442 | 0.25073098 | -1.9267042 | 0.05401651 | 0.23338748 |
| NDUFA10 | 164.739652 | -0.023311555 | 0.16516734 | -0.141139 | 0.88776013 | 0.95904526 |
| COPS9 | 50.617597 | -0.637438072 | 0.40993741 | -1.5549644 | 0.11995455 | 0.36481264 |
| AC124861.1 | 39.7032354 | 1.679658848 | 0.75856206 | 2.21426688 | 0.02681043 | 0.15852905 |
| GPC1 | 57.9798585 | -1.145904786 | 0.55623479 | -2.0601099 | 0.03938804 | 0.19685721 |
| ANKMY1 | 72.9745527 | -0.164265063 | 0.28387504 | -0.5786527 | 0.56282352 | 0.79260446 |
| RNPEPL1 | 50.7835354 | -1.114681645 | 0.32939638 | -3.3840131 | 0.00071435 | 0.01644885 |
| SNED1 | 180.159496 | -0.845603742 | 0.47760541 | -1.770507 | 0.07664271 | 0.2861109 |
| AC005237.1 | 42.8874461 | -0.302958864 | 0.59830848 | -0.506359 | 0.61260467 | 0.82440856 |
| MTERF4 | 100.659088 | 0.265661986 | 0.36097026 | 0.73596641 | 0.46175114 | 0.7200251 |
| PPP1R7 | 61.4772631 | -0.328302526 | 0.20275811 | -1.6191832 | 0.10540785 | 0.33950969 |
| HDLBP | 274.685035 | -0.148377254 | 0.16316163 | -0.9093881 | 0.36314528 | 0.64351459 |
| 2-Sep | 439.262676 | -0.114251123 | 0.20198247 | -0.5656487 | 0.57163261 | 0.79878464 |
| FARP2 | 158.13893 | 0.211054049 | 0.15188409 | 1.38957309 | 0.16465855 | 0.43295753 |
| STK25 | 74.3822565 | -0.155826205 | 0.23950983 | -0.6506046 | 0.51530173 | 0.76135326 |
| BOK | 39.4465596 | -0.163740239 | 0.54080831 | -0.3027695 | 0.76206556 | 0.90141135 |
| THAP4 | 74.9713901 | -0.240494864 | 0.2103686 | -1.143207 | 0.25295264 | 0.53989156 |
| ATG4B | 116.067928 | 0.033830836 | 0.19838785 | 0.17052877 | 0.86459431 | 0.94987371 |
| DTYMK | 25.0467037 | 0.250207402 | 0.32870213 | 0.76119799 | 0.44653882 | 0.70852995 |
| ING5 | 86.571866 | 0.559260843 | 0.21802253 | 2.56515166 | 0.01031307 | 0.09096585 |
| D2HGDH | 129.51922 | 0.610173176 | 0.30160513 | 2.02308623 | 0.04306426 | 0.20605022 |
| CHL1 | 344.733613 | 1.055844211 | 0.65597688 | 1.60957533 | 0.1074906 | 0.34298882 |
| LINC01266 | 184.03426 | 0.620350514 | 0.49926707 | 1.24252238 | 0.21404389 | 0.49620678 |
| CNTN4 | 1181.87309 | 0.246363702 | 0.34340031 | 0.71742423 | 0.47311237 | 0.73010083 |
| TRNT1 | 80.1747575 | -0.243811532 | 0.24485489 | -0.9957389 | 0.31937703 | 0.60604448 |
| CRBN | 112.1734 | 0.378581371 | 0.32646998 | 1.15962077 | 0.24620324 | 0.53308789 |
| AC034195.1 | 142.382727 | 0.046268801 | 0.41186134 | 0.11234073 | 0.91055325 | 0.968393 |
| SUMF1 | 673.429141 | -0.112619867 | 0.31001507 | -0.3632722 | 0.71640153 | 0.88058646 |
| ITPR1 | 270.545493 | 0.941916703 | 0.29498972 | 3.1930492 | 0.00140779 | 0.02608359 |
| EGOT | 87.1189714 | 0.805793184 | 0.7875125 | 1.0232132 | 0.30620705 | 0.59376697 |
| AC018816.1 | 42.2146899 | -0.115415555 | 0.33980563 | -0.3396517 | 0.73411885 | 0.8894668 |
| BHLHE40-AS | 34.7441043 | 0.194748393 | 0.56582248 | 0.34418639 | 0.73070612 | 0.88757722 |
| BHLHE40 | 297.644836 | -0.659103476 | 0.70034085 | -0.9411181 | 0.34664432 | 0.6302171 |
| ARL8B | 205.807164 | -0.13030639 | 0.23699296 | -0.5498323 | 0.58243439 | 0.80681692 |
| AC026202.2 | 36.9548872 | 0.015469466 | 0.42340595 | 0.03653578 | 0.97085515 | 0.99175186 |
| EDEM1 | 52.3568076 | -0.559943819 | 0.27469176 | -2.0384442 | 0.04150553 | 0.20295123 |
| LMCD1-AS1 | 286.351435 | 0.688949487 | 0.27467625 | 2.50822373 | 0.01213398 | 0.10072046 |
| LMCD1 | 39.5110287 | -0.074558328 | 0.52090729 | -0.1431317 | 0.8861862 | 0.95831763 |
| RAD18 | 51.6664083 | 0.384090129 | 0.36036196 | 1.06584537 | 0.28649355 | 0.57346076 |
| SRGAP3 | 300.130371 | 0.601064821 | 0.57311841 | 1.04876202 | 0.29428766 | 0.58223301 |
| THUMPD3-A: | 165.349496 | -0.090794589 | 0.17834766 | -0.5090876 | 0.61069079 | 0.82275767 |
| THUMPD3 | 41.4316241 | -0.631549981 | 0.30186615 | -2.0921524 | 0.03642489 | 0.1889085 |
| SETD5 | 607.435416 | 0.078457388 | 0.15559816 | 0.50423082 | 0.61409919 | 0.82514464 |
| MTMR14 | 64.4498949 | 0.130967722 | 0.21903519 | 0.59793006 | 0.5498866 | 0.78511906 |
| OGG1 | 43.9854212 | -0.29219391 | 0.32212453 | -0.9070837 | 0.36436254 | 0.644313 |
| TADA3 | 42.3128216 | -0.245374512 | 0.22142231 | -1.1081743 | 0.26778654 | 0.55494823 |
| RPUSD3 | 27.6573309 | -0.266875933 | 0.32260407 | -0.8272553 | 0.40809236 | 0.68150571 |
| JAGN1 | 33.5651677 | -0.958578526 | 0.41499114 | -2.3098771 | 0.02089496 | 0.13860998 |
| IL17RC | 30.7700746 | -0.102306679 | 0.29373512 | -0.3482957 | 0.72761812 | 0.88564265 |
| CRELD1 | 48.687805 | 0.160623324 | 0.37609365 | 0.42708332 | 0.66931864 | 0.85924744 |
| EMC3 | 83.3179655 | -0.004889143 | 0.23004052 | -0.0212534 | 0.98304352 | 0.99619132 |
| BRK1 | 88.3432066 | -0.429971306 | 0.40739482 | -1.0554167 | 0.29123477 | 0.5786931 |
| VHL | 59.7091907 | -0.091089264 | 0.25320351 | -0.3597472 | 0.71903617 | 0.88155276 |
| IRAK2 | 88.8645055 | -1.967214369 | 0.51064628 | -3.8524012 | 0.00011697 | 0.00528495 |
| SEC13 | 73.1763894 | -0.440566497 | 0.24762508 | -1.7791675 | 0.07521231 | 0.28328492 |
| HRH1 | 481.020614 | -0.987112454 | 0.61767702 | -1.5981045 | 0.11001971 | 0.34708436 |
| ATG7 | 451.94581 | -0.502915932 | 0.24508532 | -2.0520035 | 0.04016932 | 0.19892937 |
| VGLL4 | 438.727207 | 0.060445332 | 0.27309579 | 0.22133381 | 0.82483252 | 0.93217972 |
| TAMM41 | 114.176662 | -0.232740612 | 0.22447055 | -1.0368425 | 0.29980925 | 0.58681504 |
| PPARG | 280.55073 | 0.065086791 | 0.37385581 | 0.17409597 | 0.86179004 | 0.94853347 |
| TSEN2 | 156.478398 | 0.086955858 | 0.22120339 | 0.39310365 | 0.69424293 | 0.86986959 |
| MKRN2OS | 26.8000085 | 0.156925473 | 0.43387599 | 0.36168277 | 0.7175891 | 0.88106948 |
| MKRN2 | 32.9892453 | 0.191782141 | 0.24437174 | 0.78479674 | 0.43257275 | 0.69746077 |
| RAF1 | 264.973274 | 0.27472737 | 0.10020539 | 2.74164253 | 0.00611328 | 0.06580904 |
| RPL32 | 543.983019 | -0.762403815 | 0.5208997 | -1.4636288 | 0.14329539 | 0.40048158 |

| | | | | | | |
|---|---|---|---|---|---|---|
| IQSEC1 | 47.7621439 | 0.486652956 | 0.32671137 | 1.48955009 | 0.13634257 | 0.38812051 |
| NUP210 | 55.4678014 | 0.22272008 | 0.37825852 | 0.58880387 | 0.55599285 | 0.78784845 |
| HDAC11 | 36.0236832 | 0.182061595 | 0.30314845 | 0.60056911 | 0.54812702 | 0.78342416 |
| TMEM43 | 26.7363642 | -0.379701626 | 0.36820229 | -1.031231 | 0.30243249 | 0.59002054 |
| XPC | 176.300724 | 0.389539153 | 0.28818754 | 1.35168633 | 0.17647568 | 0.44846601 |
| LSM3 | 73.6622234 | -0.041480049 | 0.46593311 | -0.0890258 | 0.92906144 | 0.97539232 |
| SLC6A6 | 523.191462 | -0.439991241 | 0.39350584 | -1.1181314 | 0.26351087 | 0.55111974 |
| CCDC174 | 51.3431524 | -0.045108391 | 0.24256424 | -0.1859647 | 0.85247243 | 0.94470172 |
| FGD5-AS1 | 104.069424 | -0.280472986 | 0.30367024 | -0.9236104 | 0.3556892 | 0.63728305 |
| NR2C2 | 190.2272 | 0.464406273 | 0.16777709 | 2.76799579 | 0.00564022 | 0.062532 |
| MRPS25 | 109.576274 | -0.506984839 | 0.1766973 | -2.8692279 | 0.00411475 | 0.05066222 |
| RBSN | 62.5698402 | 0.198729511 | 0.21039397 | 0.94455896 | 0.34488409 | 0.62973767 |
| CAPN7 | 254.26231 | 0.324047471 | 0.16481046 | 1.96618267 | 0.04927751 | 0.22328727 |
| SH3BP5 | 43.4407262 | 0.393614581 | 0.67501165 | 0.58312265 | 0.55981074 | 0.78994338 |
| METTL6 | 54.9333839 | 0.047183299 | 0.26737091 | 0.17647132 | 0.85992368 | 0.94745904 |
| EAF1 | 28.3459213 | -0.215284758 | 0.27265718 | -0.7895804 | 0.42977287 | 0.69552194 |
| COLQ | 33.8797358 | 0.090512727 | 0.33090521 | 0.27353068 | 0.78444531 | 0.91212179 |
| HACL1 | 84.7302608 | 0.594204567 | 0.18861029 | 3.15043558 | 0.00163027 | 0.02807268 |
| BTD | 27.201576 | 0.281786735 | 0.32722426 | 0.86114256 | 0.38915953 | 0.66513878 |
| ANKRD28 | 434.445064 | 0.093421859 | 0.17332172 | 0.53900837 | 0.58988108 | 0.81183245 |
| OXNAD1 | 106.366955 | 0.35950115 | 0.23931536 | 1.50220678 | 0.13304372 | 0.38380408 |
| RFTN1 | 205.471064 | -1.26586283 | 0.9532501 | -1.3279441 | 0.18419658 | 0.45769625 |
| PLCL2 | 89.9795023 | -0.154739443 | 0.55517919 | -0.2787198 | 0.78045985 | 0.91077326 |
| TBC1D5 | 1189.92114 | 0.591934181 | 0.19087457 | 3.10116844 | 0.00192759 | 0.03108476 |
| KCNH8 | 36.4513574 | -1.432317649 | 0.41901852 | -3.4182681 | 0.00063021 | 0.01539995 |
| RAB5A | 160.635979 | -0.036384752 | 0.16902965 | -0.2152566 | 0.82956725 | 0.93410423 |
| KAT2B | 113.521767 | 0.595066798 | 0.43041548 | 1.38254041 | 0.16680583 | 0.43502125 |
| ZNF385D | 50.0684605 | 0.980911915 | 0.88468144 | 1.10877415 | 0.26752763 | 0.55480561 |
| UBE2E2 | 869.22202 | -0.266320362 | 0.29156886 | -0.9134047 | 0.36102974 | 0.64137683 |
| UBE2E1 | 423.612153 | -0.207050076 | 0.27874308 | -0.742799 | 0.4576034 | 0.71670748 |
| NKIRAS1 | 36.4615458 | 0.107677786 | 0.35754159 | 0.30116157 | 0.76329129 | 0.90232843 |
| RPL15 | 757.142489 | -0.891750681 | 0.50648168 | -1.7606771 | 0.07829308 | 0.29023584 |
| NR1D2 | 105.693667 | 0.439681859 | 0.36778879 | 1.19547378 | 0.23190197 | 0.51504498 |
| THRB | 858.084591 | 1.098176257 | 0.53963742 | 2.03502616 | 0.04184824 | 0.20312121 |
| TOP2B | 246.068902 | 0.41586201 | 0.16068452 | 2.58806516 | 0.00965167 | 0.08782595 |
| NGLY1 | 124.698717 | -0.085358459 | 0.2522955 | -0.3383273 | 0.73511654 | 0.88966284 |
| NEK10 | 91.5496527 | -0.079958496 | 0.46528137 | -0.1718498 | 0.86355564 | 0.94929509 |
| SLC4A7 | 255.622215 | 0.824500951 | 0.50297527 | 1.6392475 | 0.10116173 | 0.33231419 |
| CMC1 | 156.746486 | 0.123922181 | 0.16591436 | 0.74690452 | 0.4551212 | 0.71471019 |
| AZI2 | 96.6434488 | 0.120056754 | 0.29384993 | 0.40856486 | 0.68285902 | 0.8645396 |
| RBMS3 | 226.202057 | -0.74150036 | 0.71159759 | -1.042022 | 0.29740147 | 0.58521131 |
| RBMS3-AS3 | 85.6088847 | -0.923693778 | 0.79819 | -1.1572355 | 0.24717617 | 0.5338317 |
| TGFBR2 | 668.217071 | -0.303295212 | 0.73126059 | -0.4147567 | 0.67832004 | 0.86152581 |
| STT3B | 244.790508 | -0.003296876 | 0.22988289 | -0.0143415 | 0.9885575 | 0.99678172 |
| OSBPL10 | 1165.91587 | 0.719802549 | 0.32575178 | 2.20966575 | 0.02712837 | 0.15922956 |
| CMTM8 | 67.5900083 | -0.617638001 | 0.24385079 | -2.5328521 | 0.01131387 | 0.0963338 |
| CMTM7 | 77.024192 | -1.272981176 | 0.49150957 | -2.5899418 | 0.00959922 | 0.08782595 |
| CMTM6 | 107.039894 | -0.887165204 | 0.29712822 | -2.9857993 | 0.00282838 | 0.03903901 |
| DYNC1LI1 | 74.9775466 | -0.22791703 | 0.25503249 | -0.8936784 | 0.37149398 | 0.6517474 |
| CNOT10 | 148.277656 | 0.033796697 | 0.17220666 | 0.19625662 | 0.84440932 | 0.94193856 |
| GLB1 | 125.811524 | 0.273091011 | 0.18037114 | 1.51405052 | 0.13001303 | 0.37980933 |
| CRTAP | 87.7462306 | -0.134586427 | 0.28098461 | -0.4789815 | 0.6319518 | 0.8353707 |
| FBXL2 | 311.093585 | 0.10925217 | 0.21815844 | 0.50079277 | 0.61651698 | 0.82592881 |
| UBP1 | 107.456787 | 0.174883543 | 0.19382747 | 0.90226399 | 0.36691665 | 0.6466739 |
| CLASP2 | 342.513579 | 0.341138281 | 0.15952642 | 2.13844374 | 0.03248075 | 0.17651588 |
| PDCD6IP | 255.834391 | 0.24450524 | 0.15023528 | 1.62748221 | 0.10363472 | 0.33583603 |
| LINC01811 | 104.391366 | 2.518219445 | 1.30383034 | 1.93140117 | 0.05343346 | 0.23191623 |
| TRANK1 | 72.0512221 | -1.507686937 | 0.73343957 | -2.0556389 | 0.03981733 | 0.19762188 |
| EPM2AIP1 | 97.9449414 | 0.269251328 | 0.21445907 | 1.25549049 | 0.20930076 | 0.48959511 |
| MLH1 | 50.5481193 | 0.578884132 | 0.33812091 | 1.71206249 | 0.08688515 | 0.30691783 |
| LRRFIP2 | 501.039477 | -0.149720211 | 0.24328637 | -0.6154073 | 0.53828576 | 0.77689306 |
| GOLGA4 | 728.87975 | -0.124560135 | 0.19350356 | -0.6437098 | 0.51976365 | 0.76385303 |
| ITGA9 | 65.9105195 | 0.370870556 | 0.49187379 | 0.75399536 | 0.45085201 | 0.71160181 |
| CTDSPL | 588.947294 | 0.085053203 | 0.17363947 | 0.48982644 | 0.62425672 | 0.83072177 |
| ACAA1 | 76.2025078 | -0.566415822 | 0.30090728 | -1.88236 | 0.05978715 | 0.24674758 |
| OXSR1 | 222.902749 | 0.190077025 | 0.18133024 | 1.04823676 | 0.29452954 | 0.58239173 |
| XYLB | 41.0616854 | -1.099686009 | 0.4938186 | -2.2269028 | 0.02595377 | 0.15616281 |
| ACVR2B | 57.3857772 | 0.02056683 | 0.32149101 | 0.06397327 | 0.94899151 | 0.98262487 |
| EXOG | 60.0794854 | 0.317798573 | 0.22863241 | 1.38999793 | 0.16452951 | 0.432737 |
| WDR48 | 109.812142 | 0.397941344 | 0.25645902 | 1.55167615 | 0.12073973 | 0.36507762 |
| GORASP1 | 29.7424167 | -0.147552324 | 0.26115006 | -0.5650097 | 0.57206714 | 0.79915869 |
| TTC21A | 39.1642408 | 0.116953808 | 0.45916 | 0.25471253 | 0.79894513 | 0.91949148 |
| CSRNP1 | 26.2519436 | -0.388113105 | 0.48426698 | -0.8014445 | 0.42287436 | 0.69043214 |
| RPSA | 383.788112 | -0.648322972 | 0.37849074 | -1.7129164 | 0.08672793 | 0.30678244 |
| MYRIP | 180.263639 | 1.63935731 | 0.59845665 | 2.73930836 | 0.00615686 | 0.06585681 |
| EIF1B | 73.3484011 | 0.074968315 | 0.38867223 | 0.19288313 | 0.8470505 | 0.94323402 |
| ENTPD3-AS1 | 49.7714801 | 0.593142174 | 0.58177729 | 1.01953477 | 0.30794915 | 0.59456464 |
| RPL14 | 641.397048 | -0.70108718 | 0.56590899 | -1.2388691 | 0.21539396 | 0.49731142 |
| ZNF621 | 88.7548266 | -0.206533514 | 0.25183998 | -0.8200982 | 0.41216014 | 0.68367283 |
| CTNNB1 | 364.649572 | 0.0686154 | 0.2471456 | 0.27763149 | 0.78129526 | 0.91077326 |
| ULK4 | 383.534292 | 0.008289607 | 0.25927105 | 0.03197274 | 0.97449379 | 0.99326197 |
| TRAK1 | 241.126959 | 0.304113503 | 0.45081725 | 0.67458267 | 0.49994094 | 0.75144113 |
| SEC22C | 50.5127664 | 0.168234489 | 0.26411752 | 0.6369683 | 0.52414549 | 0.76606495 |
| SS18L2 | 30.4594593 | 0.141455908 | 0.36613313 | 0.38635102 | 0.69923672 | 0.87179338 |
| NKTR | 491.068697 | 0.306920093 | 0.11114409 | 2.76146113 | 0.00575434 | 0.06336748 |
| AC006059.1 | 70.2728613 | 0.135992637 | 0.36341023 | 0.37421246 | 0.70824625 | 0.87578784 |
| SNRK | 535.736738 | 1.869944698 | 0.64934381 | 2.87974517 | 0.00397997 | 0.04925595 |
| ANO10 | 355.311121 | -0.153268855 | 0.24392761 | -0.6283375 | 0.5297829 | 0.77060515 |
| ABHD5 | 60.2376345 | -0.351045372 | 0.45707679 | -0.7680227 | 0.44247367 | 0.70510851 |
| TCAIM | 220.019326 | 0.542347085 | 0.25333646 | 2.14081731 | 0.03228877 | 0.17584306 |
| ZNF445 | 52.6067168 | 0.488253725 | 0.2625817 | 1.85943547 | 0.06296544 | 0.25490103 |
| AC099669.1 | 100.134645 | 0.056781545 | 0.33674449 | 0.16861908 | 0.86609627 | 0.95070686 |
| ZNF197 | 78.0538681 | -0.495056653 | 0.21903192 | -2.2602032 | 0.02380864 | 0.14935362 |
| KIAA1143 | 34.2238957 | -0.54386419 | 0.26289867 | -2.0687217 | 0.03857221 | 0.19528708 |
| ZDHHC3 | 85.9374325 | 0.169054757 | 0.21259038 | 0.79521358 | 0.42648928 | 0.69264418 |
| EXOSC7 | 44.2018493 | -0.009540803 | 0.27126664 | -0.0351713 | 0.97194314 | 0.99192876 |
| LARS2 | 101.654011 | -0.247903478 | 0.23951808 | -1.0350095 | 0.3006645 | 0.58788025 |
| LIMD1 | 77.1473104 | -0.514375265 | 0.28778895 | -1.787335 | 0.07388336 | 0.28056182 |
| SACM1L | 129.474191 | 0.43312162 | 0.15302877 | 2.83032807 | 0.00465003 | 0.05512702 |
| LZTFL1 | 48.232001 | 0.289980607 | 0.22819417 | 1.27076253 | 0.20381314 | 0.48277104 |
| FYCO1 | 66.4749032 | -0.249172571 | 0.18127352 | -1.3745669 | 0.16926576 | 0.43904596 |
| LTF | 212.78516 | -5.100819482 | 1.38806833 | -3.6747611 | 0.00023807 | 0.0083651 |
| ALS2CL | 35.0264903 | -0.999997065 | 0.42880025 | -2.3320813 | 0.01969641 | 0.13342561 |
| PTH1R | 57.16233 | 0.09369124 | 0.66993379 | 0.13985149 | 0.88877733 | 0.95907089 |
| CCDC12 | 123.346837 | -0.002224641 | 0.22858896 | -0.0097321 | 0.99223507 | 0.99829858 |
| NBEAL2 | 29.0829853 | 0.236836741 | 0.56535891 | 0.41891396 | 0.67527901 | 0.86115505 |
| SETD2 | 573.369897 | 0.230060659 | 0.14414893 | 1.59599282 | 0.1104904 | 0.34758554 |
| KIF9-AS1 | 144.368305 | 0.107497151 | 0.33543134 | 0.32047438 | 0.74860875 | 0.89457625 |
| KIF9 | 47.5649578 | -0.268274023 | 0.2926145 | -0.9168173 | 0.3592384 | 0.64001011 |
| KLHL18 | 64.8927652 | 0.32546503 | 0.31751095 | 1.02505136 | 0.30533895 | 0.59275349 |
| PTPN23 | 46.8320953 | -0.489442374 | 0.30465352 | -1.6065541 | 0.10815221 | 0.34396964 |
| SCAP | 98.1937212 | 0.081257842 | 0.22169524 | 0.36652949 | 0.71397001 | 0.87895113 |
| ELP6 | 32.8926201 | -0.516013052 | 0.2781452 | -1.8551931 | 0.06356866 | 0.25658128 |

| | | | | | | |
|---|---|---|---|---|---|---|
| SMARCC1 | 432.885873 | -0.163211689 | 0.20135197 | -0.810579 | 0.41760745 | 0.68711498 |
| DHX30 | 85.3003799 | 0.235475196 | 0.1731363 | 1.36005678 | 0.17381196 | 0.44528422 |
| MAP4 | 491.43321 | -0.316443602 | 0.2073322 | -1.5262637 | 0.12694421 | 0.37435927 |
| ZNF589 | 27.9057716 | 0.620074798 | 0.41661257 | 1.48837275 | 0.13665262 | 0.38854124 |
| PLXNB1 | 185.482861 | 0.504898585 | 0.25875866 | 1.95123359 | 0.05102926 | 0.22682567 |
| TMA7 | 118.234014 | 0.129217241 | 0.52329281 | 0.24693105 | 0.80496159 | 0.92227408 |
| SHISA5 | 120.50293 | -1.308208943 | 0.32196652 | -4.0631831 | 4.84E-05 | 0.00294775 |
| PFKFB4 | 60.736574 | -0.785110437 | 0.80331252 | -0.9773412 | 0.32840025 | 0.61394673 |
| UQCRC1 | 56.4606016 | -0.30296773 | 0.2601356 | -1.1646531 | 0.24415944 | 0.5298248 |
| NCKIPSD | 27.7855574 | -0.14819358 | 0.34920109 | -0.4243789 | 0.6712895 | 0.8598933 |
| IP6K2 | 140.732262 | 0.561738084 | 0.24941945 | 2.25218236 | 0.02431075 | 0.15131898 |
| PRKAR2A | 197.567589 | -0.133951554 | 0.18844497 | -0.7108258 | 0.47719218 | 0.73376419 |
| ARIH2 | 180.029381 | 0.084172963 | 0.13896406 | 0.60571748 | 0.54470238 | 0.78068896 |
| P4HTM | 64.8318419 | 0.016557762 | 0.23509656 | 0.07042962 | 0.94385172 | 0.98112027 |
| WDR6 | 118.652715 | 0.35180211 | 0.24082638 | 1.4608122 | 0.14406699 | 0.40162014 |
| DALRD3 | 26.1686095 | 0.346589601 | 0.31251537 | 1.10903219 | 0.2674163 | 0.55480561 |
| NDUFAF3 | 57.0276096 | -0.25559124 | 0.51142959 | -0.4997584 | 0.6172452 | 0.82612711 |
| IMPDH2 | 45.7118595 | -1.17113153 | 0.41098987 | -2.8495387 | 0.00437827 | 0.05297457 |
| QRICH1 | 116.898155 | 0.331625932 | 0.19807119 | 1.67427645 | 0.09407628 | 0.31933263 |
| QARS | 36.1610234 | -0.076463679 | 0.29374176 | -0.2603092 | 0.79462529 | 0.91762696 |
| USP19 | 24.475377 | 0.023055839 | 0.30245573 | 0.07622881 | 0.93923706 | 0.97996404 |
| LAMB2 | 76.0941487 | 0.577599292 | 0.27445587 | 2.10452521 | 0.03533266 | 0.18575825 |
| USP4 | 106.182853 | 0.035659941 | 0.21495589 | 0.16589423 | 0.86824019 | 0.95090501 |
| RHOA | 287.031752 | -0.406601256 | 0.21120723 | -1.9251295 | 0.05421316 | 0.23344987 |
| TCTA | 30.4359581 | -0.402576171 | 0.37841909 | -1.0638368 | 0.28740262 | 0.57443572 |
| DAG1 | 94.3046957 | 0.557932371 | 0.37765187 | 1.47737219 | 0.1395759 | 0.39358184 |
| APEH | 40.6968501 | -0.374182407 | 0.44240698 | -0.8457878 | 0.39767115 | 0.67266325 |
| MST1 | 54.9262888 | 0.879056307 | 0.51253787 | 1.71510508 | 0.08632596 | 0.30615195 |
| RNF123 | 27.1624725 | -0.085030612 | 0.30725964 | -0.2767386 | 0.7819808 | 0.91082127 |
| IP6K1 | 167.613673 | 0.094712284 | 0.21544413 | 0.43961413 | 0.6602166 | 0.85381597 |
| RBM6 | 622.319889 | 0.369740616 | 0.16893717 | 2.18862796 | 0.02862389 | 0.16438145 |
| RBM5 | 240.794284 | 0.606245856 | 0.14173426 | 4.27734179 | 1.89E-05 | 0.00149732 |
| SEMA3F | 53.0853857 | 0.121918622 | 0.5515744 | 0.22103749 | 0.82506324 | 0.93217972 |
| GNAI2 | 122.025325 | -0.517754291 | 0.35590025 | -1.4547736 | 0.14573194 | 0.40369182 |
| SEMA3B | 93.3395475 | 0.362027645 | 0.26039787 | 1.39028649 | 0.1644419 | 0.43262536 |
| IFRD2 | 31.3927004 | -0.679027211 | 0.3502989 | -1.9384223 | 0.05257172 | 0.22987695 |
| HYAL2 | 30.5998851 | -1.279956961 | 0.35757121 | -3.5795862 | 0.00034414 | 0.01043198 |
| TUSC2 | 26.7381918 | -0.615992617 | 0.33177069 | -1.8566819 | 0.06335643 | 0.25585634 |
| HEMK1 | 26.5647757 | -0.160209806 | 0.33453893 | -0.4788974 | 0.63201166 | 0.8353707 |
| MAPKAPK3 | 42.5162587 | -1.136477199 | 0.403717 | -2.8150343 | 0.0048772 | 0.05710298 |
| DOCK3 | 107.464515 | 0.057865358 | 0.618345 | 0.09358102 | 0.92544198 | 0.97349958 |
| MANF | 48.445097 | -1.27784796 | 0.62523938 | -2.043774 | 0.04097588 | 0.2014928 |
| RBM15B | 24.7377599 | -0.800183503 | 0.5102613 | -1.5681838 | 0.11683825 | 0.36010089 |
| DCAF1 | 99.5119368 | -0.088180437 | 0.22235179 | -0.3965807 | 0.69167666 | 0.86887335 |
| RAD54L2 | 110.543894 | 0.490134307 | 0.28063989 | 1.74648835 | 0.08072613 | 0.29548169 |
| TEX264 | 59.3391599 | -0.767969501 | 0.21396745 | -3.5891885 | 0.00033171 | 0.01028298 |
| RPL29 | 710.017589 | -1.23440185 | 0.53169167 | -2.3216498 | 0.0202518 | 0.1355161 |
| WDR82 | 122.047924 | -0.057352702 | 0.1466209 | -0.3911632 | 0.6956766 | 0.87029987 |
| DNAH1 | 27.3528463 | 0.111401906 | 0.38449728 | 0.28973392 | 0.7720198 | 0.90624314 |
| BAP1 | 30.6142642 | -0.637980943 | 0.27067348 | -2.3570131 | 0.0184226 | 0.12843527 |
| NISCH | 110.20664 | 0.301442394 | 0.27207371 | 1.10794385 | 0.26788607 | 0.55494823 |
| SMIM4 | 60.5440625 | -0.343424892 | 0.27530451 | -1.2474365 | 0.21223749 | 0.49369183 |
| PBRM1 | 285.663331 | -0.024696096 | 0.17464538 | -0.1414071 | 0.88754835 | 0.95903278 |
| GNL3 | 71.1900246 | -0.875603217 | 0.42209351 | -2.0744295 | 0.03803944 | 0.19361308 |
| GLT8D1 | 39.8000771 | -0.06165039 | 0.24696165 | -0.2496355 | 0.80286926 | 0.92125386 |
| SPCS1 | 145.015253 | -0.705759262 | 0.40486854 | -1.7431813 | 0.08130195 | 0.29645656 |
| NEK4 | 87.3405305 | 0.490828795 | 0.20202242 | 2.4295759 | 0.0151165 | 0.11411159 |
| SFMBT1 | 71.3901376 | -0.037566014 | 0.29437485 | -0.1276129 | 0.89845536 | 0.9624585 |
| TKT | 168.595065 | -1.8744523 | 0.46079973 | -4.0678242 | 4.75E-05 | 0.00293267 |
| DCP1A | 173.440256 | -0.040678534 | 0.18335626 | -0.2218552 | 0.82442663 | 0.93217972 |
| CACNA1D | 138.166265 | -1.292709138 | 0.5664356 | -2.282182 | 0.0224786 | 0.14412481 |
| CHDH | 78.5050502 | -0.119126056 | 0.30998665 | -0.3842942 | 0.70076044 | 0.87182648 |
| SELENOK | 72.531365 | -0.427352287 | 0.45236863 | -0.9446992 | 0.34481247 | 0.62973472 |
| CACNA2D3 | 794.792034 | -0.122101682 | 0.66465603 | -0.1837066 | 0.85424365 | 0.9452497 |
| ERC2 | 160.74306 | 0.053692495 | 0.24995595 | 0.21480783 | 0.82991715 | 0.93416878 |
| CCDC66 | 190.039086 | 0.372874745 | 0.29900965 | 1.2470325 | 0.21238558 | 0.49391636 |
| FAM208A | 225.801247 | 0.457553067 | 0.16036603 | 2.85317951 | 0.00432842 | 0.05265324 |
| ARHGEF3 | 318.038593 | -2.443708452 | 0.39965785 | -6.1145013 | 9.69E-10 | 5.80E-07 |
| IL17RD | 46.1343795 | 0.504464024 | 0.74636599 | 0.67589364 | 0.49910818 | 0.75042493 |
| APPL1 | 125.698546 | 0.147631081 | 0.17706004 | 0.83379108 | 0.40439872 | 0.67791921 |
| PDE12 | 29.4001968 | -0.461975236 | 0.32994786 | -1.4001462 | 0.16146955 | 0.42791083 |
| ARF4 | 126.595045 | -0.212849168 | 0.32732364 | -0.6502713 | 0.51551699 | 0.76135326 |
| DENND6A | 168.960342 | 0.374274318 | 0.18897718 | 1.98052648 | 0.0476444 | 0.21810285 |
| DENND6A-AS | 37.5635505 | 0.358408147 | 0.3086041 | 1.16138493 | 0.2454854 | 0.53201462 |
| SLMAP | 462.088093 | -0.101790645 | 0.15198241 | -0.6697528 | 0.50301537 | 0.75403031 |
| FLNB | 452.218134 | 0.782595218 | 0.20791773 | 3.76396575 | 0.00016724 | 0.00675945 |
| ABHD6 | 36.0315764 | -0.348557818 | 0.44565463 | -0.7821254 | 0.43414086 | 0.69904779 |
| PXK | 141.312967 | 1.002981337 | 0.32804058 | 3.05749164 | 0.00223198 | 0.03403479 |
| PDHB | 42.812537 | -0.245057782 | 0.34933716 | -0.7014936 | 0.48299503 | 0.73696727 |
| C3orf67 | 42.0426066 | -0.050607127 | 0.49392031 | -0.1024601 | 0.91839148 | 0.97057281 |
| FHIT | 2074.26584 | 0.136239065 | 0.27629982 | 0.49308415 | 0.62195313 | 0.82915174 |
| PTPRG | 2456.90838 | -0.746391056 | 0.33701537 | -2.2147093 | 0.02678003 | 0.15852905 |
| THOC7 | 69.1197649 | -0.19762279 | 0.23075804 | -0.8564069 | 0.39177272 | 0.66731068 |
| ATXN7 | 45.9934821 | 0.02357778 | 0.33936096 | 0.069477 | 0.94460994 | 0.98115679 |
| PSMD6 | 30.8053261 | 0.036223356 | 0.3801247 | 0.09529335 | 0.92408182 | 0.9731617 |
| PRICKLE2 | 299.111637 | -0.12046675 | 0.62081009 | -0.1940477 | 0.84613856 | 0.94271476 |
| ADAMTS9 | 1010.0218 | -1.246862065 | 0.50422307 | -2.4728382 | 0.01340448 | 0.10673443 |
| ADAMTS9-AS | 661.159552 | 1.292652669 | 0.53851592 | 2.40039823 | 0.01637724 | 0.11984539 |
| ADAMTS9-AS | 2256.61488 | 0.913664882 | 0.50955174 | 1.79307579 | 0.07296081 | 0.2786649 |
| MAGI1 | 2380.80088 | 0.954548642 | 0.33070434 | 2.88641097 | 0.00389663 | 0.04847509 |
| MAGI1-AS1 | 36.0099085 | 1.416226298 | 0.48687048 | 2.90883583 | 0.00362777 | 0.04650235 |
| SLC25A26 | 166.833826 | 0.118107792 | 0.26967969 | 0.43795584 | 0.6614183 | 0.85467771 |
| LRIG1 | 98.4028634 | -0.053456191 | 0.33966854 | -0.1573775 | 0.87494733 | 0.95291321 |
| SUCLG2 | 373.057908 | -0.026304685 | 0.17634118 | -0.1491693 | 0.88142007 | 0.95635451 |
| SUCLG2-AS1 | 196.094979 | 0.478411132 | 0.38612291 | 1.23901256 | 0.21534085 | 0.49731142 |
| EOGT | 34.2899908 | -0.39639253 | 0.30501508 | -1.2995834 | 0.19374379 | 0.47204496 |
| TMF1 | 177.172762 | -0.026489394 | 0.14872835 | -0.1781059 | 0.85863982 | 0.94734879 |
| UBA3 | 112.816395 | 0.163219907 | 0.15689601 | 1.04030629 | 0.29819762 | 0.58581522 |
| ARL6IP5 | 116.320887 | -0.676750691 | 0.23963267 | -2.824117 | 0.00474111 | 0.05591589 |
| FRMD4B | 503.52141 | 0.64532898 | 0.30387985 | 2.12363199 | 0.03370093 | 0.18044784 |
| MITF | 483.201337 | 0.714200092 | 0.22603298 | 3.15971625 | 0.00157923 | 0.02770592 |
| FOXP1 | 1139.91691 | 0.106331641 | 0.21610894 | 0.49202797 | 0.62269957 | 0.82971751 |
| EIF4E3 | 50.1779096 | -0.560356086 | 0.36071972 | -1.553439 | 0.12031829 | 0.36507762 |
| RYBP | 216.501413 | 0.319018997 | 0.30292216 | 1.05313854 | 0.2922775 | 0.57977349 |
| SHQ1 | 78.1925775 | -0.045731153 | 0.2874417 | -0.1590971 | 0.87359234 | 0.95210417 |
| GXYLT2 | 67.9145183 | -1.12199276 | 0.61241813 | -1.8320698 | 0.06694102 | 0.26366762 |
| PPP4R2 | 214.946006 | 0.150427125 | 0.1819948 | 0.82654626 | 0.40849429 | 0.68193914 |
| PDZRN3 | 343.166037 | 1.249405671 | 0.68342847 | 1.82814402 | 0.06752794 | 0.26531999 |
| CNTN3 | 150.591029 | 1.108200296 | 0.65734057 | 1.68588454 | 0.09181804 | 0.31558127 |
| ZNF717 | 98.6843673 | 0.045423182 | 0.28902872 | 0.15715802 | 0.87512031 | 0.95291321 |
| ROBO2 | 172.91309 | -0.373791925 | 0.96823343 | -0.3860556 | 0.6994555 | 0.87181583 |
| ROBO1 | 241.988129 | -0.246824432 | 0.97424369 | -0.2533498 | 0.79999793 | 0.92028103 |
| LINC02027 | 87.6977283 | -0.076065912 | 0.78421484 | -0.0969963 | 0.92272936 | 0.97279601 |

| | | | | | | |
|---|---|---|---|---|---|---|
| GBE1 | 614.854457 | 0.042612809 | 0.37937364 | 0.11232412 | 0.91056642 | 0.968393 |
| CADM2 | 199.677188 | 1.567905104 | 1.17532677 | 1.33401633 | 0.1821985 | 0.4553499 |
| CHMP2B | 66.2214552 | 0.00827288 | 0.2285417 | 0.03619856 | 0.97112404 | 0.99175186 |
| CGGBP1 | 176.904925 | 0.173901881 | 0.25194578 | 0.69023534 | 0.49004621 | 0.74227587 |
| ZNF654 | 197.85669 | 0.170674288 | 0.24208179 | 0.70502738 | 0.4807932 | 0.73578929 |
| C3orf38 | 51.2880349 | -0.063039128 | 0.22758699 | -0.2769892 | 0.78178843 | 0.9108187 |
| PROS1 | 167.783748 | -1.601308915 | 0.49483401 | -3.2360526 | 0.00121195 | 0.02363725 |
| ARL13B | 151.373181 | 0.49589565 | 0.26944871 | 1.8404083 | 0.06570832 | 0.26131276 |
| NSUN3 | 72.95416 | 0.158645268 | 0.22072809 | 0.7187362 | 0.47230347 | 0.72983472 |
| MTRNR2L12 | 264.611502 | -0.885056206 | 0.54269951 | -1.6308403 | 0.10292403 | 0.33488767 |
| EPHA6 | 64.0401383 | 0.79678997 | 0.6996848 | 1.13878417 | 0.25479319 | 0.54159654 |
| ARL6 | 56.6355771 | 0.624653835 | 0.2960792 | 2.10975252 | 0.03487968 | 0.18449058 |
| CRYBG3 | 110.290284 | -0.41376046 | 0.28592721 | -1.4470832 | 0.14787363 | 0.40703491 |
| RIOX2 | 48.957168 | -0.515533685 | 0.33811779 | -1.5247162 | 0.1273299 | 0.37459861 |
| CLDND1 | 53.0451924 | 0.350549576 | 0.34591405 | 1.01340081 | 0.31086875 | 0.59807427 |
| ST3GAL6 | 217.446789 | 0.90818679 | 0.50728673 | 1.79028296 | 0.07340843 | 0.27937985 |
| DCBLD2 | 305.255187 | -0.230775718 | 0.46386968 | -0.4975012 | 0.61883566 | 0.82733103 |
| COL8A1 | 190.293826 | -0.440612881 | 0.54791246 | -0.8041666 | 0.42130078 | 0.68985303 |
| CMSS1 | 338.083443 | -0.33518412 | 0.25160635 | -1.3321767 | 0.18280213 | 0.45624325 |
| FILIP1L | 111.209445 | -0.41717369 | 0.34457382 | -1.2106947 | 0.22601245 | 0.50844839 |
| TBC1D23 | 115.851827 | 0.309019672 | 0.18026513 | 1.71425097 | 0.08648264 | 0.30636531 |
| NIT2 | 52.4477127 | 0.267929194 | 0.22382044 | 1.19707203 | 0.23127848 | 0.51432779 |
| TOMM70 | 107.171702 | -0.145060325 | 0.17803 | -0.8148083 | 0.41518205 | 0.68495052 |
| TFG | 215.82046 | -0.097481349 | 0.19904864 | -0.4897363 | 0.62432049 | 0.83072177 |
| ABI3BP | 234.644055 | 0.275473742 | 0.49614414 | 0.55522926 | 0.57873786 | 0.80406395 |
| SENP7 | 225.364133 | 0.374046314 | 0.24138414 | 1.5495894 | 0.1212401 | 0.36572187 |
| TRMT10C | 35.512994 | -1.239049107 | 0.41351917 | -2.9963523 | 0.00273231 | 0.03818317 |
| PCNP | 177.956154 | 0.090843211 | 0.17092593 | 0.53147707 | 0.59508823 | 0.81511365 |
| ZBTB11 | 60.4290818 | 0.105413888 | 0.25297108 | 0.41670331 | 0.67689543 | 0.8613973 |
| RPL24 | 270.848271 | -0.498720777 | 0.48926844 | -1.0193193 | 0.30805139 | 0.59456464 |
| CEP97 | 40.1128968 | 0.238457702 | 0.2784498 | 0.85637592 | 0.39178988 | 0.66731068 |
| NFKBIZ | 556.5116 | -0.931541986 | 0.45748462 | -2.0362258 | 0.04172769 | 0.20312121 |
| ALCAM | 288.588264 | -0.318849107 | 0.24963602 | -1.277256 | 0.20151189 | 0.48024387 |
| CBLB | 475.218245 | -0.320200351 | 0.48058138 | -0.6662771 | 0.50523402 | 0.75524917 |
| LINC00882 | 96.6035987 | 1.444920867 | 0.45541411 | 3.17276259 | 0.00150996 | 0.02700412 |
| BBX | 720.467791 | -0.279034491 | 0.30438071 | -0.9167286 | 0.35928489 | 0.64001011 |
| CD47 | 131.099839 | -0.621988689 | 0.44927524 | -1.3844268 | 0.16622781 | 0.43446007 |
| IFT57 | 107.353943 | 0.138439119 | 0.23975977 | 0.57740761 | 0.56366413 | 0.79293888 |
| DZIP3 | 168.664625 | 0.608136028 | 0.4725187 | 1.28700944 | 0.198091 | 0.47656532 |
| NECTIN3 | 130.042182 | -0.230315083 | 0.31680573 | -0.7269915 | 0.46723114 | 0.72514698 |
| PHLDB2 | 739.123605 | 0.288239937 | 0.1618352 | 1.78107077 | 0.07490089 | 0.28269331 |
| ABHD10 | 37.8126873 | 0.110179901 | 0.34943625 | 0.31530759 | 0.75252813 | 0.89668701 |
| SLC9C1 | 80.0549978 | -0.218732157 | 0.37381637 | -0.5851326 | 0.55845854 | 0.78926435 |
| CD200 | 178.031258 | 0.614026798 | 0.34576799 | 1.77583473 | 0.07576016 | 0.28447926 |
| AC092894.1 | 220.98456 | 1.381910745 | 0.28567083 | 4.83742335 | 1.32E-06 | 0.00021723 |
| ATG3 | 48.1446958 | 0.036938104 | 0.2505146 | 0.14744891 | 0.8827777 | 0.95722522 |
| SLC35A5 | 37.5244908 | 0.10272855 | 0.24218378 | 0.424176 | 0.67143746 | 0.85996783 |
| NEPRO | 61.3359428 | -0.092331674 | 0.19017155 | -0.4855178 | 0.62730909 | 0.83250121 |
| BOC | 128.788691 | 1.048103253 | 0.54698615 | 1.91614222 | 0.055347 | 0.23619173 |
| CFAP44 | 153.296898 | 0.760523958 | 0.38867137 | 1.95672751 | 0.05037951 | 0.2253855 |
| SPICE1 | 95.6752215 | 0.470246702 | 0.17433379 | 2.69739274 | 0.00698848 | 0.07109414 |
| USF3 | 93.5135415 | -0.000954871 | 0.17528339 | -0.0054476 | 0.99565348 | 0.99899312 |
| NAA50 | 194.56855 | 0.105926666 | 0.16917691 | 0.62612958 | 0.53122995 | 0.77193643 |
| ATP6V1A | 245.522254 | 0.071854636 | 0.27186035 | 0.26430716 | 0.79154324 | 0.91648412 |
| GRAMD1C | 125.439248 | 0.374198671 | 0.27173578 | 1.37706809 | 0.1684912 | 0.43774808 |
| ZDHHC23 | 26.8041744 | -0.071972698 | 0.29229562 | -0.2462326 | 0.80550221 | 0.92227408 |
| CCDC191 | 138.896001 | 0.17574034 | 0.39601327 | 0.44377387 | 0.65720608 | 0.85164733 |
| QTRT2 | 58.4643311 | -0.41091643 | 0.29963332 | -1.3713976 | 0.17025103 | 0.44040903 |
| ZBTB20 | 6580.32414 | 0.594807088 | 0.1982157 | 3.00080706 | 0.00269265 | 0.03796996 |
| ZBTB20-AS1 | 30.386651 | 0.63502334 | 0.44490706 | 1.42731683 | 0.15348859 | 0.41627045 |
| ZBTB20-AS5 | 144.568866 | 0.507159725 | 0.5249929 | 0.96603159 | 0.33402837 | 0.61861931 |
| ZBTB20-AS2 | 35.5786702 | 0.426088434 | 0.66661169 | 0.63918537 | 0.52270235 | 0.76512314 |
| LSAMP | 122.161109 | -2.387001089 | 0.8148174 | -2.929492 | 0.00339517 | 0.04424801 |
| IGSF11 | 44.6107406 | 1.119284569 | 0.33302073 | 3.36100572 | 0.00077659 | 0.01758623 |
| B4GALT4 | 53.2383902 | -0.05333023 | 0.36916722 | -0.1444609 | 0.88513653 | 0.95793953 |
| ARHGAP31 | 136.48547 | -0.327296828 | 0.24866339 | -1.3162244 | 0.18809873 | 0.46366385 |
| TMEM39A | 52.8219225 | -0.100518433 | 0.27267394 | -0.3686397 | 0.71239632 | 0.87814237 |
| POGLUT1 | 42.7864708 | 0.219945793 | 0.30643102 | 0.71776608 | 0.47290153 | 0.73007373 |
| TIMMDC1 | 80.7836883 | -0.249978711 | 0.19165893 | -1.3042894 | 0.19213492 | 0.46933901 |
| COX17 | 49.2010946 | -0.014482684 | 0.35834303 | -0.0404157 | 0.96776172 | 0.99123492 |
| MAATS1 | 128.097734 | 0.173297625 | 0.38315829 | 0.45228729 | 0.65106203 | 0.84724399 |
| GSK3B | 554.98759 | 0.163141856 | 0.21397814 | 0.762423 | 0.44580758 | 0.70795604 |
| LRRC58 | 84.582589 | -0.425969191 | 0.27300148 | -1.5603183 | 0.11868469 | 0.3637939 |
| FSTL1 | 163.66664 | -0.256409984 | 0.37040346 | -0.6922451 | 0.48878341 | 0.74185878 |
| NDUFB4 | 83.4566378 | -0.450885625 | 0.36100618 | -1.2489693 | 0.21167632 | 0.49274543 |
| HGD | 86.502557 | -0.732126071 | 0.4868915 | -1.5036739 | 0.13266534 | 0.38380408 |
| RABL3 | 51.8021048 | -0.29761416 | 0.30702776 | -0.9693396 | 0.33237579 | 0.61719981 |
| GOLGB1 | 385.660673 | -0.203036353 | 0.12418732 | -1.6349202 | 0.10206578 | 0.33374822 |
| IQCB1 | 161.464242 | 0.04378153 | 0.15131369 | 0.28934281 | 0.77231905 | 0.90624314 |
| SLC15A2 | 70.3988893 | 1.571054734 | 0.67704119 | 2.32047142 | 0.02031539 | 0.1355161 |
| ILDR1 | 26.8284555 | -0.481293125 | 0.51519158 | -0.9342022 | 0.35019958 | 0.63305563 |
| CASR | 60.049231 | 0.332848741 | 1.29098054 | 0.2578263 | 0.79654096 | 0.91869184 |
| CCDC58 | 28.8273112 | 0.333823428 | 0.28394046 | 1.17568107 | 0.23972235 | 0.52362764 |
| FAM162A | 132.076535 | -0.779329609 | 0.476024 | -1.6371645 | 0.1015961 | 0.33300898 |
| AC083798.2 | 77.9630474 | 0.088718821 | 0.47646127 | 0.18620364 | 0.85228507 | 0.94470172 |
| KPNA1 | 193.527281 | 0.150320253 | 0.23837489 | 0.63060439 | 0.52829923 | 0.76990012 |
| PARP9 | 142.804301 | -0.92550642 | 0.7282338 | -1.2708919 | 0.20376712 | 0.48277104 |
| DTX3L | 65.8228727 | -1.240193994 | 0.57929232 | -2.1408777 | 0.0322839 | 0.17584306 |
| PARP14 | 465.438033 | -0.714682329 | 0.79718022 | -0.8965129 | 0.36997891 | 0.6499226 |
| HSPBAP1 | 66.3066073 | -0.289696219 | 0.30930678 | -0.9365983 | 0.34896521 | 0.63263722 |
| DIRC2 | 104.546183 | -0.322793534 | 0.28397919 | -1.1366802 | 0.25567198 | 0.54255249 |
| SEMA5B | 63.6772922 | 1.204174948 | 0.71177107 | 1.69180092 | 0.09068394 | 0.3142614 |
| PDIA5 | 59.1973622 | -0.67949885 | 0.3049326 | -2.2283575 | 0.02585668 | 0.15587236 |
| SEC22A | 169.328657 | -0.117381943 | 0.14604391 | -0.8037442 | 0.42154473 | 0.6898802 |
| ADCY5 | 182.653543 | 0.474918167 | 0.43148312 | 1.10066453 | 0.27104269 | 0.55822789 |
| HACD2 | 88.9700315 | 0.179017551 | 0.33410357 | 0.53581454 | 0.59208673 | 0.81278286 |
| MYLK-AS1 | 30.2709301 | 0.163077185 | 0.4295147 | 0.37967778 | 0.70418462 | 0.87375446 |
| MYLK | 220.205929 | -0.000288933 | 0.91179344 | -0.0003169 | 0.99974716 | 0.9998728 |
| CCDC14 | 189.299585 | 0.554398399 | 0.15081552 | 3.67600364 | 0.00023692 | 0.0083651 |
| KALRN | 190.082762 | -1.190778893 | 0.39176909 | -3.0394917 | 0.00236978 | 0.0352486 |
| UMPS | 35.087436 | -0.232773396 | 0.32031705 | -0.7266969 | 0.46741168 | 0.72530964 |
| ITGB5 | 111.47045 | -0.46903078 | 0.40711412 | -1.1520867 | 0.24928543 | 0.53636804 |
| HEG1 | 114.430013 | 0.222250971 | 0.47322474 | 0.46965206 | 0.63860363 | 0.83969583 |
| SLC12A8 | 168.472366 | -0.584386209 | 0.34610964 | -1.6884425 | 0.09132633 | 0.31502156 |
| ZNF148 | 827.954937 | 0.41052142 | 0.23007755 | 1.78427415 | 0.07437913 | 0.28138929 |
| SNX4 | 205.056996 | 0.04166891 | 0.17151165 | 0.24295091 | 0.80804343 | 0.92380073 |
| OSBPL11 | 75.2930119 | 0.27688345 | 0.25977539 | 1.06585714 | 0.28648823 | 0.57346076 |
| SLC41A3 | 38.2679155 | -0.360832155 | 0.23681804 | -1.5236683 | 0.12759158 | 0.37492391 |
| ALDH1L1 | 89.5448062 | -0.580314504 | 0.44727191 | -1.2974535 | 0.1944752 | 0.47305178 |
| ZXDC | 192.812784 | 0.189852648 | 0.15906477 | 1.1935556 | 0.23265185 | 0.51579248 |
| TXNRD3NB | 26.6477394 | -0.173944671 | 0.41271107 | -0.4214684 | 0.67341308 | 0.86100159 |
| TXNRD3 | 30.2403193 | 0.510183202 | 0.27556335 | 1.85141896 | 0.0641093 | 0.25781575 |
| CHCHD6 | 71.1590328 | 0.179869218 | 0.25993344 | 0.69198184 | 0.48894873 | 0.74185878 |

| | | | | | | |
|---|---|---|---|---|---|---|
| PLXNA1 | 30.2040939 | 0.29336263 | 0.35254517 | 0.83212778 | 0.40533682 | 0.67879745 |
| TPRA1 | 35.2322071 | -0.278247028 | 0.31160091 | -0.8929596 | 0.3718788 | 0.65192016 |
| MGLL | 225.809391 | -0.858526444 | 0.37420789 | -2.29425 | 0.02177615 | 0.14172056 |
| SEC61A1 | 116.855031 | -0.956077789 | 0.34918383 | -2.7380357 | 0.00618074 | 0.06585681 |
| RUVBL1 | 75.5523778 | -0.496561381 | 0.19010848 | -2.6119897 | 0.0090017 | 0.0847859 |
| EEFSEC | 116.062854 | 0.110253049 | 0.15018297 | 0.73412485 | 0.46287265 | 0.72095238 |
| RPN1 | 92.0869849 | -0.787241592 | 0.31310012 | -2.5143446 | 0.01192538 | 0.10002913 |
| RAB7A | 362.856751 | -0.233538989 | 0.20509153 | -1.1387062 | 0.25482572 | 0.54159654 |
| ACAD9 | 67.7493438 | -0.012163683 | 0.21679051 | -0.056108 | 0.95525577 | 0.98395647 |
| CNBP | 200.808014 | -0.799138353 | 0.35109159 | -2.2761535 | 0.02283682 | 0.14544808 |
| COPG1 | 49.6160305 | -0.71328323 | 0.26621338 | -2.6793665 | 0.00737616 | 0.07378801 |
| HMCES | 44.8511116 | 0.057364707 | 0.30193269 | 0.1899917 | 0.84931563 | 0.94413304 |
| H1FX | 127.474942 | -1.238604001 | 0.63652381 | -1.9458879 | 0.0516682 | 0.22786819 |
| MBD4 | 85.9325735 | -0.429251074 | 0.19837584 | -2.1638274 | 0.0304776 | 0.17102806 |
| IFT122 | 202.309266 | 0.365251408 | 0.2696131 | 1.35472427 | 0.17550542 | 0.4475949 |
| TMCC1 | 581.543959 | 0.035683829 | 0.2002177 | 0.17822515 | 0.85854616 | 0.94734879 |
| PIK3R4 | 73.0474851 | 0.197270053 | 0.22511445 | 0.87631003 | 0.38086152 | 0.65803602 |
| ATP2C1 | 227.748685 | 0.531295667 | 0.2181833 | 2.43508862 | 0.01488814 | 0.11327523 |
| NEK11 | 341.608341 | -0.007876589 | 0.30742061 | -0.0256215 | 0.97955921 | 0.99557142 |
| NUDT16 | 32.8126062 | -0.632775223 | 0.29363193 | -2.1549946 | 0.03116225 | 0.17254522 |
| MRPL3 | 94.9748718 | -0.205460626 | 0.22231268 | -0.9241966 | 0.35538395 | 0.63706583 |
| DNAJC13 | 185.823335 | 0.215692996 | 0.25724528 | 0.83847212 | 0.4017656 | 0.67562929 |
| UBA5 | 97.5507724 | 0.235063094 | 0.26986812 | 0.87102949 | 0.38373805 | 0.66018004 |
| CDV3 | 125.002328 | -0.923915191 | 0.33458116 | -2.7614083 | 0.00575527 | 0.06336748 |
| TOPBP1 | 56.7586271 | 0.304932416 | 0.25644527 | 1.18907403 | 0.23441054 | 0.51702015 |
| SRPRB | 33.9252488 | -0.771635783 | 0.30238659 | -2.5518188 | 0.01071622 | 0.09289657 |
| RAB6B | 32.5877505 | -0.404918375 | 0.49142446 | -0.8239687 | 0.40995734 | 0.68208527 |
| SLCO2A1 | 160.043572 | -0.429627029 | 0.72611969 | -0.5916752 | 0.55406809 | 0.78721719 |
| RYK | 302.233353 | 0.26183175 | 0.12887553 | 2.03166382 | 0.0421877 | 0.20399596 |
| AMOTL2 | 110.13685 | 0.963146774 | 0.52208927 | 1.84479326 | 0.06506762 | 0.26002617 |
| ANAPC13 | 41.9894263 | 0.115486782 | 0.31208093 | 0.37005395 | 0.71134229 | 0.87748 |
| CEP63 | 131.882571 | 0.349239272 | 0.17059777 | 2.04715024 | 0.04064334 | 0.20006295 |
| EPHB1 | 36.8102861 | 0.382180482 | 0.53357865 | 0.71625894 | 0.47383147 | 0.73077309 |
| PPP2R3A | 1084.4156 | -0.805538786 | 0.24662784 | -3.2662118 | 0.00108997 | 0.02188845 |
| MSL2 | 102.535096 | 0.191013352 | 0.19066019 | 1.00185231 | 0.31641493 | 0.60281198 |
| PCCB | 53.8429381 | 0.038899694 | 0.33037594 | 0.11774372 | 0.90627072 | 0.96621245 |
| STAG1 | 1011.51809 | 0.480691291 | 0.13900675 | 3.45804273 | 0.00054412 | 0.01396038 |
| NCK1 | 108.519111 | -0.578526965 | 0.33563595 | -1.723674 | 0.08476671 | 0.30275178 |
| DZIP1L | 82.5371342 | 1.320794813 | 0.27488904 | 4.80482902 | 1.55E-06 | 0.0002355 |
| ARMC8 | 234.348674 | 0.116309322 | 0.17906998 | 0.64951882 | 0.51600309 | 0.76140599 |
| NME9 | 41.9350557 | 0.211002046 | 0.33260574 | 0.63439088 | 0.52582576 | 0.76734993 |
| MRAS | 58.9801686 | -0.861317161 | 0.47909409 | -1.7978038 | 0.07220811 | 0.2768941 |
| CEP70 | 258.690553 | 0.289779613 | 0.51010562 | 0.56807767 | 0.56998223 | 0.79765604 |
| FAIM | 38.9193357 | 0.071525287 | 0.28489284 | 0.25106032 | 0.80176748 | 0.92109986 |
| PIK3CB | 540.265061 | 0.851632496 | 0.18916993 | 4.50194424 | 6.73E-06 | 0.00067187 |
| COPB2 | 113.75368 | -0.246174272 | 0.20817685 | -1.1825247 | 0.23699756 | 0.52000641 |
| AC097103.2 | 226.29732 | 0.582778264 | 0.29455121 | 1.97852954 | 0.047869 | 0.21855917 |
| NMNAT3 | 170.152836 | 2.13657634 | 0.49429252 | 4.32249378 | 1.54E-05 | 0.00126308 |
| CLSTN2 | 274.348188 | 1.99629494 | 0.51473866 | 3.87826893 | 0.0001052 | 0.00484487 |
| SLC25A36 | 291.544508 | 0.259607264 | 0.29414131 | 0.88259369 | 0.37745584 | 0.65536514 |
| PXYLP1 | 29.3578677 | -0.130372204 | 0.32774058 | -0.3977909 | 0.69078436 | 0.86843279 |
| ZBTB38 | 229.116648 | -0.843459487 | 0.270698 | -3.1158689 | 0.00183404 | 0.03020328 |
| RASA2 | 371.911168 | 0.255324856 | 0.21548212 | 1.18490042 | 0.23605682 | 0.51957452 |
| RNF7 | 77.993049 | -0.730318601 | 0.31131403 | -2.3459225 | 0.01898005 | 0.13051681 |
| ATP1B3 | 167.344327 | 0.926664766 | 0.67680398 | 1.36917748 | 0.1709438 | 0.44115974 |
| TFDP2 | 375.111145 | 0.587402961 | 0.2166488 | 2.71131416 | 0.00670171 | 0.069251 |
| GK5 | 211.184296 | -0.00931951 | 0.25979395 | -0.0358727 | 0.97138387 | 0.99175186 |
| XRN1 | 354.297764 | 0.087136541 | 0.34416098 | 0.25318542 | 0.80012493 | 0.92031661 |
| ATR | 280.682185 | -0.218603588 | 0.21188514 | -1.031708 | 0.30220894 | 0.58973509 |
| PLS1 | 120.770876 | -0.342317924 | 0.42151766 | -0.8121081 | 0.41672958 | 0.68612112 |
| TRPC1 | 194.363291 | 0.200303465 | 0.2639179 | 0.75896127 | 0.44787573 | 0.70935873 |
| U2SURP | 247.366859 | 0.360338691 | 0.1881094 | 1.91558045 | 0.05541852 | 0.23625011 |
| SLC9A9 | 125.559989 | 0.003715856 | 0.55051568 | 0.00674977 | 0.9946145 | 0.99886097 |
| C3orf58 | 62.9294291 | -0.190757429 | 0.38135184 | -0.5002137 | 0.61692459 | 0.82592881 |
| AC107021.1 | 86.6984296 | 0.27817836 | 0.72230798 | 0.38512431 | 0.70014532 | 0.87182648 |
| LNCSRLR | 30.72241 | -0.078083433 | 0.48898097 | -0.159686 | 0.8731284 | 0.95193455 |
| PLOD2 | 723.799613 | 0.146137055 | 0.44359921 | 0.32943488 | 0.741827 | 0.89253455 |
| PLSCR4 | 118.359282 | -0.473222932 | 0.38809766 | -1.2193398 | 0.22271524 | 0.50422814 |
| PLSCR2 | 50.4730357 | -0.340608234 | 0.49721692 | -0.6850295 | 0.49332534 | 0.7455922 |
| PLSCR1 | 400.535591 | -1.550451401 | 0.52452299 | -2.9559265 | 0.00311731 | 0.04188043 |
| GYG1 | 34.2077575 | -1.074996422 | 0.30531766 | -3.5209113 | 0.00043007 | 0.01209909 |
| HLTF | 86.111638 | 0.437318574 | 0.24181164 | 1.8085092 | 0.07052729 | 0.27263151 |
| HPS3 | 80.253156 | -0.321902677 | 0.22814956 | -1.4109283 | 0.15826575 | 0.42394509 |
| CP | 82.6972644 | -5.052028847 | 1.18792816 | -4.2528067 | 2.11E-05 | 0.00159673 |
| TM4SF1 | 78.2672612 | -1.290360933 | 0.6995744 | -1.8444942 | 0.06511115 | 0.26007572 |
| WWTR1 | 427.100971 | 0.214812799 | 0.24387329 | 0.88083773 | 0.37840565 | 0.65630052 |
| COMMD2 | 64.7666248 | 0.153491108 | 0.30754622 | 0.49908306 | 0.61772087 | 0.82653278 |
| ANKUB1 | 123.812579 | 0.247763904 | 0.44680723 | 0.5545208 | 0.57922247 | 0.80434504 |
| RNF13 | 276.974538 | -0.004055608 | 0.15049307 | -0.0269488 | 0.97850057 | 0.99538128 |
| TSC22D2 | 291.81042 | 0.342551732 | 0.18126896 | 1.88974295 | 0.05879235 | 0.24454867 |
| SERP1 | 201.132439 | -0.09468589 | 0.33747883 | -0.2805684 | 0.77904147 | 0.90966132 |
| EIF2A | 83.2278201 | -0.098281581 | 0.24781824 | -0.3965874 | 0.69167178 | 0.86887335 |
| SELENOT | 99.9786947 | 0.125566607 | 0.23860668 | 0.52624934 | 0.59871498 | 0.81696449 |
| SIAH2 | 28.0096961 | -0.34744517 | 0.30099189 | -1.154334 | 0.24836327 | 0.53534467 |
| MBNL1 | 701.694437 | 0.179861198 | 0.14778172 | 1.2170734 | 0.22357629 | 0.50557208 |
| DHX36 | 233.426921 | -0.072471132 | 0.14629334 | -0.4953823 | 0.62033028 | 0.82828879 |
| PLCH1 | 203.249582 | -0.494056328 | 0.39889064 | -1.2385759 | 0.21550261 | 0.49731142 |
| GMPS | 100.089683 | -0.120583434 | 0.18726326 | -0.6439247 | 0.51962427 | 0.76385303 |
| SSR3 | 80.3496373 | -0.447627437 | 0.24176651 | -1.8514865 | 0.06409959 | 0.25781575 |
| TIPARP | 40.3319055 | -0.170835453 | 0.57415796 | -0.2975409 | 0.76605362 | 0.90371395 |
| LINC00886 | 89.9557175 | 0.586695789 | 0.40286058 | 1.45632465 | 0.14530289 | 0.40323664 |
| LEKR1 | 125.211172 | 0.93668991 | 0.47009367 | 1.99256013 | 0.04630964 | 0.21459827 |
| CCNL1 | 225.078344 | 0.203384255 | 0.317592 | 0.64039476 | 0.52191599 | 0.76473507 |
| VEPH1 | 154.535939 | 1.842953329 | 0.4839097 | 3.80846532 | 0.00013983 | 0.00595311 |
| RSRC1 | 311.906285 | 0.18096884 | 0.19726041 | 0.91741084 | 0.3589274 | 0.64001011 |
| MLF1 | 86.9656595 | 0.531329759 | 0.30859901 | 1.7217481 | 0.08511516 | 0.30331775 |
| GFM1 | 74.2427621 | -0.257654514 | 0.3086484 | -0.8347832 | 0.40383976 | 0.67765972 |
| RARRES1 | 77.6706735 | -4.125019248 | 1.05640695 | -3.9047635 | 9.43E-05 | 0.00449487 |
| MFSD1 | 39.5712974 | -0.185562847 | 0.29141002 | -0.6367758 | 0.5242709 | 0.76611507 |
| IQCJ-SCHIP1 | 258.417294 | 0.982544472 | 0.19551085 | 5.02552389 | 5.02E-07 | 9.43E-05 |
| SMC4 | 37.6803235 | 0.348256445 | 0.30864986 | 1.12832207 | 0.25918392 | 0.54609557 |
| KPNA4 | 219.61236 | 0.073631073 | 0.20653577 | 0.35650519 | 0.72146227 | 0.88235193 |
| PPM1L | 211.254231 | -0.095718026 | 0.46199851 | -0.2071825 | 0.8358673 | 0.93745146 |
| NMD3 | 170.739055 | 0.011218621 | 0.31393836 | 0.03573511 | 0.97149358 | 0.99175186 |
| LINC01322 | 43.8577679 | -0.768828205 | 0.55087127 | -1.3956586 | 0.16281734 | 0.42941279 |
| PDCD10 | 111.744188 | 0.09533689 | 0.23825699 | 0.4001431 | 0.68905112 | 0.86751885 |
| GOLIM4 | 89.2762332 | 0.338626034 | 0.19919309 | 1.69998884 | 0.08913302 | 0.3108137 |
| MECOM | 1607.41137 | 1.272339342 | 0.76169116 | 1.6704137 | 0.09483754 | 0.3204651 |
| MYNN | 33.8020553 | -0.042732508 | 0.30856701 | -0.138487 | 0.88985557 | 0.95918866 |
| SEC62 | 269.239377 | 0.330946754 | 0.16997825 | 1.94699477 | 0.05153536 | 0.22780675 |
| GPR160 | 107.274547 | 0.192278754 | 0.49990369 | 0.3846316 | 0.70051038 | 0.87182648 |
| PHC3 | 460.819949 | 0.212807603 | 0.19044627 | 1.11741547 | 0.26381672 | 0.55140746 |
| PRKCI | 451.41916 | 1.059269823 | 0.24383911 | 4.34413423 | 1.40E-05 | 0.00116469 |

| | | | | | | |
|---|---|---|---|---|---|---|
| SKIL | 190.852724 | 0.616947358 | 0.25101425 | 2.45781803 | 0.0139784 | 0.10939467 |
| RPL22L1 | 43.033452 | -1.888127571 | 0.52103257 | -3.6238187 | 0.00029029 | 0.00945796 |
| TNIK | 588.531093 | -0.402612548 | 0.37534813 | -1.0726377 | 0.28343371 | 0.57084461 |
| PLD1 | 81.1656203 | -0.221667356 | 0.48933071 | -0.4530011 | 0.65054794 | 0.84724399 |
| FNDC3B | 1777.08811 | -0.211942735 | 0.26275422 | -0.8066197 | 0.41988561 | 0.68835944 |
| TNFSF10 | 1141.95745 | -1.462178628 | 0.55367098 | -2.64088 | 0.0082691 | 0.0800098 |
| NCEH1 | 439.597139 | -0.130089158 | 0.42979139 | -0.3026798 | 0.76213392 | 0.90141135 |
| ECT2 | 43.2256494 | 0.137120787 | 0.44263809 | 0.30978081 | 0.75672764 | 0.89923013 |
| NLGN1 | 1876.85215 | -0.578516729 | 0.1848992 | -3.1288223 | 0.00175508 | 0.029443 |
| NAALADL2 | 955.619163 | 0.058139602 | 0.24666525 | 0.23570244 | 0.81366357 | 0.92720494 |
| TBL1XR1 | 1099.34721 | 0.316059019 | 0.32316747 | 0.97800382 | 0.32807243 | 0.61394673 |
| KCNMB2-AS1 | 281.197082 | 0.021127421 | 0.70384891 | 0.03001698 | 0.97605351 | 0.99411128 |
| ZMAT3 | 89.8296608 | -0.531640305 | 0.29576419 | -1.7975141 | 0.07225404 | 0.27695935 |
| PIK3CA | 211.821033 | 0.140402413 | 0.18076998 | 0.77669097 | 0.43734112 | 0.70195888 |
| KCNMB3 | 28.0397408 | 0.515053768 | 0.37909984 | 1.35862303 | 0.17426607 | 0.44550166 |
| ZNF639 | 38.0864227 | 0.36504879 | 0.28692766 | 1.27226769 | 0.20327802 | 0.48221897 |
| MFN1 | 155.877263 | 0.157246579 | 0.18025417 | 0.87236027 | 0.38301187 | 0.65965766 |
| ACTL6A | 48.1646873 | -0.165333744 | 0.29389862 | -0.5625537 | 0.57373885 | 0.80056001 |
| MRPL47 | 30.4602492 | 0.323519985 | 0.37413967 | 0.86470378 | 0.38720139 | 0.66404483 |
| NDUFB5 | 62.7630008 | 0.184331262 | 0.41576443 | 0.44335505 | 0.65750894 | 0.85176897 |
| USP13 | 38.2258264 | -0.257596638 | 0.34448045 | -0.747783 | 0.45459103 | 0.71410256 |
| TTC14 | 222.790886 | 0.581311955 | 0.19216522 | 3.02506333 | 0.00248581 | 0.03618781 |
| FXR1 | 228.005689 | 0.446585219 | 0.1489624 | 2.99797283 | 0.00271782 | 0.03811711 |
| DNAJC19 | 37.2633076 | 0.252443011 | 0.31528251 | 0.80068827 | 0.42331213 | 0.69062823 |
| ATP11B | 335.738152 | 0.396173248 | 0.20370713 | 1.94481781 | 0.05179691 | 0.22800587 |
| DCUN1D1 | 161.643741 | 0.728331501 | 0.22141668 | 3.2894157 | 0.00100396 | 0.02077084 |
| MCCC1 | 198.537464 | 0.764863979 | 0.23610046 | 3.23956999 | 0.0011971 | 0.02359807 |
| B3GNT5 | 33.7654362 | -0.202609516 | 0.35522591 | -0.5703681 | 0.56842809 | 0.79713855 |
| KLHL24 | 250.510829 | 0.088389904 | 0.29535059 | 0.29927112 | 0.76473319 | 0.90314131 |
| YEATS2 | 169.355757 | -0.514655191 | 0.31064268 | -1.6567433 | 0.09757138 | 0.3259791 |
| ABCC5 | 134.04237 | 0.416423812 | 0.38463233 | 1.0826542 | 0.27896194 | 0.56636829 |
| EIF2B5 | 39.9444615 | -0.289824492 | 0.29343339 | -0.9877011 | 0.32329904 | 0.61020115 |
| DVL3 | 68.8051315 | 0.289238911 | 0.26413153 | 1.09505637 | 0.27349194 | 0.56098058 |
| AP2M1 | 139.023046 | -0.724510885 | 0.38147157 | -1.8992526 | 0.05753127 | 0.24202547 |
| PSMD2 | 69.3752686 | -1.002725835 | 0.45320378 | -2.2125275 | 0.02693024 | 0.15864374 |
| EIF4G1 | 138.143207 | -0.956332561 | 0.25383062 | -3.7676012 | 0.00016482 | 0.00669003 |
| POLR2H | 74.1384482 | -0.460003116 | 0.27990081 | -1.6434504 | 0.10028977 | 0.33096298 |
| MAGEF1 | 37.5219406 | -0.778801196 | 0.32872174 | -2.3691807 | 0.01782754 | 0.12607869 |
| VPS8 | 227.48757 | 0.300559784 | 0.16953018 | 1.77289835 | 0.07624555 | 0.28527949 |
| EHHADH | 100.448472 | 0.379471474 | 0.29703618 | 1.27752609 | 0.20141658 | 0.48024387 |
| MAP3K13 | 329.774859 | -0.205107281 | 0.20279857 | -1.0113842 | 0.31183256 | 0.59932559 |
| TMEM41A | 46.1673426 | -0.416726139 | 0.35613544 | -1.1701339 | 0.2419471 | 0.5268672 |
| LIPH | 93.0901695 | 0.590992492 | 0.98531076 | 0.59980314 | 0.54863744 | 0.78391975 |
| SENP2 | 115.448553 | 0.53396537 | 0.22579482 | 2.36482563 | 0.01803856 | 0.12677284 |
| IGF2BP2 | 467.046501 | -0.008151907 | 0.19971437 | -0.0408178 | 0.96744113 | 0.99123492 |
| TRA2B | 197.038954 | 0.18698824 | 0.1729264 | 1.08131689 | 0.27955618 | 0.56686465 |
| ETV5 | 31.4958433 | -1.47941654 | 0.39526125 | -3.7428828 | 0.00018192 | 0.00709811 |
| EIF4A2 | 235.84026 | 0.095974633 | 0.29985084 | 0.32007458 | 0.74891179 | 0.89482675 |
| ST6GAL1 | 224.699282 | -0.840856463 | 0.99925707 | -0.8414816 | 0.40007817 | 0.67469094 |
| AC072022.1 | 27.9809195 | -0.018219118 | 0.61213691 | -0.0297631 | 0.97625595 | 0.99421176 |
| BCL6 | 367.318436 | -1.258158414 | 0.35883574 | -3.5062238 | 0.00045451 | 0.01258164 |
| LPP | 2354.68376 | -0.468759846 | 0.33312463 | -1.4071606 | 0.15937978 | 0.42483013 |
| TPRG1 | 89.0065664 | -1.689340913 | 0.83328642 | -2.0273232 | 0.04262937 | 0.20489047 |
| P3H2 | 1154.85594 | 1.368830011 | 0.46678018 | 2.93249387 | 0.00336252 | 0.0440021 |
| P3H2-AS1 | 25.4507221 | 0.846480454 | 0.63573744 | 1.33149378 | 0.18302659 | 0.45656555 |
| CLDN1 | 572.144665 | -0.437311484 | 0.67209198 | -0.6506721 | 0.51525819 | 0.76135326 |
| CLDN16 | 47.3053926 | 0.280410862 | 0.43120063 | 0.65030253 | 0.51549683 | 0.76135326 |
| IL1RAP | 68.8409517 | -0.52652743 | 0.82562859 | -0.6377292 | 0.52365 | 0.76569124 |
| CCDC50 | 258.98517 | 0.512903153 | 0.22963364 | 2.23357154 | 0.02551128 | 0.1551184 |
| MB21D2 | 73.0337301 | 0.57730846 | 0.46031479 | 1.25416014 | 0.20978381 | 0.49024618 |
| ATP13A4 | 48.645452 | 1.810858979 | 0.70848984 | 2.55594205 | 0.01059008 | 0.09230421 |
| OPA1 | 205.14901 | -0.036072409 | 0.14189959 | -0.2542108 | 0.7993327 | 0.91973669 |
| HES1 | 188.903297 | 1.024059991 | 0.59797487 | 1.71254688 | 0.08679593 | 0.3067964 |
| ATP13A3 | 637.265001 | 0.104077469 | 0.20935599 | 0.49713155 | 0.61909629 | 0.82744849 |
| LINC00884 | 33.7855516 | 0.044491501 | 0.47348133 | 0.09396675 | 0.92513557 | 0.97344855 |
| LSG1 | 57.7378172 | -0.241508617 | 0.29653909 | -0.8144242 | 0.41540197 | 0.68495052 |
| XXYLT1 | 51.9971003 | 0.027721919 | 0.29124599 | 0.09518386 | 0.92416879 | 0.9731617 |
| ACAP2 | 341.050925 | 0.234111236 | 0.11008717 | 2.1265988 | 0.03345343 | 0.17962468 |
| PPP1R2 | 85.0999051 | 0.430795788 | 0.23257657 | 1.8522751 | 0.06398633 | 0.25774813 |
| MUC20-OT1 | 514.401366 | 0.375373012 | 0.40205135 | 0.93364445 | 0.35048733 | 0.63322593 |
| TFRC | 79.6782754 | 0.262927859 | 0.40373003 | 0.65124673 | 0.51488723 | 0.76135326 |
| PCYT1A | 53.2752373 | -0.164041747 | 0.21023202 | -0.7802891 | 0.43522076 | 0.70008054 |
| UBXN7 | 143.358729 | 0.321142249 | 0.18678314 | 1.7193321 | 0.08555392 | 0.3044283 |
| RNF168 | 100.89399 | 0.095252645 | 0.2136102 | 0.44591805 | 0.65565644 | 0.85068646 |
| PIGX | 40.6428514 | -0.041494568 | 0.24148786 | -0.1718288 | 0.86357212 | 0.94929509 |
| PAK2 | 237.851389 | -0.160825784 | 0.14379409 | -1.118445 | 0.26337699 | 0.55096925 |
| SENP5 | 241.941708 | -0.053292648 | 0.17186286 | -0.3100882 | 0.75649388 | 0.89917544 |
| NCBP2 | 49.256563 | -0.788857548 | 0.28101777 | -2.8071447 | 0.00499828 | 0.05810499 |
| NCBP2-AS2 | 29.4177533 | -0.424218355 | 0.43217181 | -0.9815965 | 0.32629865 | 0.61290706 |
| DLG1 | 731.537584 | -0.022505378 | 0.18032684 | -0.1248033 | 0.9006793 | 0.96303857 |
| BDH1 | 46.3553397 | 0.075005228 | 0.3783211 | 0.19825812 | 0.84284312 | 0.94141724 |
| RUBCN | 51.3905814 | -0.100407144 | 0.31059001 | -0.3232787 | 0.74648416 | 0.89352127 |
| FYTTD1 | 90.1317312 | -0.315820934 | 0.28443233 | -1.1103553 | 0.26684596 | 0.55453954 |
| LRCH3 | 325.911748 | 0.406371824 | 0.17617395 | 2.30665098 | 0.02107429 | 0.13931717 |
| IQCG | 67.0955493 | -0.420767668 | 0.33529183 | -1.2549297 | 0.20950429 | 0.48995155 |
| RPL35A | 312.647342 | -0.324022013 | 0.45157363 | -0.7175397 | 0.47304115 | 0.73010083 |
| LMLN | 81.1439164 | 0.081662781 | 0.30129354 | 0.2710406 | 0.78635981 | 0.91314591 |
| ZNF595 | 50.6118619 | 0.455802782 | 0.22578839 | 2.01871665 | 0.04351668 | 0.2066642 |
| ZNF718 | 151.93688 | 0.349523065 | 0.1716076 | 2.03675755 | 0.04167434 | 0.20312121 |
| ZNF141 | 122.54826 | 0.466978622 | 0.24793062 | 1.88350523 | 0.05963193 | 0.24631922 |
| ZNF721 | 339.025931 | 0.191930478 | 0.17646544 | 1.08763778 | 0.27675504 | 0.56369053 |
| PIGG | 83.7526544 | 0.036696556 | 0.221254 | 0.16585714 | 0.86826938 | 0.95090501 |
| ATP5ME | 49.0714578 | -0.308256504 | 0.36742587 | -0.8389624 | 0.40149038 | 0.67562929 |
| PCGF3 | 113.45616 | 0.149937917 | 0.25513842 | 0.58767282 | 0.55675192 | 0.7883969 |
| AC139887.2 | 190.256853 | -0.373248753 | 0.29234473 | -1.2767419 | 0.2016934 | 0.48024387 |
| GAK | 124.755737 | -0.251083377 | 0.22118821 | -1.1351572 | 0.25630947 | 0.54330348 |
| FGFRL1 | 27.9487812 | -0.274905293 | 0.37540759 | -0.7322849 | 0.46399472 | 0.7217612 |
| RNF212 | 32.7834835 | 0.821470617 | 0.42621462 | 1.92736375 | 0.05393431 | 0.2332446 |
| AC092535.3 | 82.2680709 | 0.429949521 | 0.35041624 | 1.22696803 | 0.21983461 | 0.50155763 |
| SPON2 | 94.4134312 | -1.395896224 | 0.72747814 | -1.9188153 | 0.05500771 | 0.23554932 |
| CTBP1-AS | 33.3951181 | 0.099625575 | 0.49158838 | 0.20266056 | 0.83940035 | 0.93987329 |
| CTBP1 | 91.3262469 | -0.026526981 | 0.23163904 | -0.1145186 | 0.90882669 | 0.96752108 |
| CTBP1-DT | 27.1463707 | -0.149701755 | 0.27103316 | -0.5523374 | 0.5807172 | 0.80525463 |
| MAEA | 86.2305464 | -0.148607401 | 0.15506971 | -0.9583264 | 0.33789819 | 0.62316649 |
| UVSSA | 106.919485 | 0.34533795 | 0.28283186 | 1.2210009 | 0.22208568 | 0.50375143 |
| SLBP | 91.6137515 | -0.197625369 | 0.27150065 | -0.7279002 | 0.4666747 | 0.72440074 |
| TMEM129 | 30.067174 | -0.172985794 | 0.33726615 | -0.5129059 | 0.60801716 | 0.8211189 |
| FGFR3 | 96.1430002 | 0.521510741 | 0.65481043 | 0.79643011 | 0.42578208 | 0.69215319 |
| LETM1 | 53.7755453 | -0.664361054 | 0.31146199 | -2.1330405 | 0.0329214 | 0.17776443 |
| NSD2 | 225.615643 | -0.13345761 | 0.25551357 | -0.5223112 | 0.60145365 | 0.81829073 |
| NELFA | 88.4794121 | 0.465683133 | 0.20205309 | 2.30475626 | 0.02118023 | 0.13968678 |
| MXD4 | 80.9679825 | -0.171876455 | 0.40801686 | -0.4212484 | 0.6735737 | 0.86105131 |
| ZFYVE28 | 73.6041536 | -0.083120063 | 0.48680848 | -0.1707449 | 0.86442436 | 0.94985035 |

| | | | | | | |
|---|---|---|---|---|---|---|
| RNF4 | 99.4876736 | -0.107847301 | 0.17930869 | -0.6014617 | 0.54753254 | 0.78315875 |
| FAM193A | 211.995289 | 0.096599773 | 0.23420133 | 0.41246466 | 0.67999888 | 0.86238758 |
| TNIP2 | 35.2975983 | -1.198477305 | 0.32423384 | -3.6963363 | 0.00021873 | 0.00802776 |
| SH3BP2 | 92.9612107 | -1.171648403 | 0.29214693 | -4.0104765 | 6.06E-05 | 0.00332957 |
| ADD1 | 313.979473 | 0.365680475 | 0.16529405 | 2.21230276 | 0.02694575 | 0.15864374 |
| MFSD10 | 71.9759079 | -0.077410791 | 0.35657436 | -0.2170958 | 0.8281337 | 0.93346403 |
| NOP14 | 32.8570332 | -0.84386857 | 0.36755843 | -2.295876 | 0.02168297 | 0.14167784 |
| GRK4 | 32.0157713 | -0.132732682 | 0.33478082 | -0.3964764 | 0.69175365 | 0.86887335 |
| HTT | 205.782522 | 0.155572515 | 0.17771089 | 0.87542478 | 0.38134282 | 0.65851823 |
| RGS12 | 242.611294 | 0.5474532 | 0.25071642 | 2.18355545 | 0.02899493 | 0.16561863 |
| LRPAP1 | 121.34909 | -0.351212755 | 0.31414182 | -1.118007 | 0.263564 | 0.55111974 |
| TMEM128 | 34.9245843 | 0.174177234 | 0.27768969 | 0.62723696 | 0.53050392 | 0.7712395 |
| STX18 | 216.181759 | 0.008316236 | 0.22794664 | 0.03648326 | 0.97089703 | 0.99175186 |
| STX18-AS1 | 114.189294 | -0.012567545 | 0.31157945 | -0.040335 | 0.96782608 | 0.99123492 |
| STK32B | 399.915799 | 0.777243332 | 0.53125492 | 1.46303272 | 0.14345843 | 0.40048158 |
| EVC2 | 185.984272 | 0.196249206 | 0.36464158 | 0.53819755 | 0.59044067 | 0.81191949 |
| EVC | 716.71641 | 0.936310406 | 0.6258026 | 1.49617533 | 0.13460797 | 0.38651372 |
| CRMP1 | 184.578314 | 1.272886585 | 0.64849423 | 1.9628341 | 0.04966545 | 0.22398559 |
| MRFAP1 | 134.858949 | -0.481653056 | 0.45854035 | -1.050405 | 0.29353197 | 0.58142927 |
| MRFAP1L1 | 34.4155624 | -0.24027446 | 0.29773266 | -0.8070141 | 0.41965835 | 0.68835944 |
| KIAA0232 | 291.254657 | 0.296757312 | 0.21255839 | 1.3961214 | 0.16267794 | 0.42936994 |
| TBC1D14 | 135.242863 | -0.77946162 | 0.25717773 | -3.0308286 | 0.00243884 | 0.03574435 |
| TADA2B | 28.4786053 | -0.671005622 | 0.28973925 | -2.3158948 | 0.02056401 | 0.13698379 |
| GRPEL1 | 37.1346271 | -0.760512349 | 0.47988227 | -1.5847894 | 0.11301418 | 0.35216749 |
| AFAP1 | 170.753926 | 0.525719824 | 0.29804995 | 1.76386483 | 0.07775474 | 0.28902315 |
| ABLIM2 | 156.111096 | -2.096303177 | 1.28245489 | -1.6346019 | 0.10213253 | 0.33378626 |
| SH3TC1 | 74.6288951 | -1.190829207 | 0.42543104 | -2.7991122 | 0.00512433 | 0.05910434 |
| ACOX3 | 60.1326092 | 0.483363723 | 0.2105819 | 2.29537165 | 0.02171183 | 0.14167784 |
| TRMT44 | 60.2738738 | 0.581831869 | 0.2556782 | 2.27564132 | 0.02286749 | 0.14554661 |
| SLC2A9 | 192.637248 | -0.994490685 | 0.50491789 | -1.9696087 | 0.04888323 | 0.22173216 |
| WDR1 | 117.852953 | -0.370540915 | 0.25296545 | -1.4647886 | 0.14297859 | 0.3999438 |
| HS3ST1 | 95.3170362 | -3.878015066 | 0.7000492 | -5.5396322 | 3.03E-08 | 1.16E-05 |
| AC005699.1 | 134.366915 | -2.179885064 | 0.51428511 | -4.2386704 | 2.25E-05 | 0.00164188 |
| RAB28 | 240.065969 | 0.238226884 | 0.15770217 | 1.51061262 | 0.13088718 | 0.38143239 |
| BOD1L1 | 198.495189 | -0.273110959 | 0.14933812 | -1.8288094 | 0.06742817 | 0.26503671 |
| CPEB2 | 258.217852 | -0.14928037 | 0.33129098 | -0.450602 | 0.65227644 | 0.84801249 |
| AC098829.1 | 149.751478 | -0.370280112 | 0.44080466 | -0.8400095 | 0.40090305 | 0.67532043 |
| CC2D2A | 147.144854 | 0.362949787 | 0.20173898 | 1.79910593 | 0.07200193 | 0.27643545 |
| FBXL5 | 288.413056 | 0.277286963 | 0.16160256 | 1.71585744 | 0.08618814 | 0.30586185 |
| FAM200B | 66.7446468 | 0.327964944 | 0.32503863 | 1.00900297 | 0.31297321 | 0.60019427 |
| PROM1 | 876.375033 | 0.874943531 | 0.29966723 | 2.91971707 | 0.00350349 | 0.04544458 |
| TAPT1 | 161.761047 | 0.398620889 | 0.14089827 | 2.82913971 | 0.00466733 | 0.05526375 |
| TAPT1-AS1 | 270.373055 | 0.349866359 | 0.37665494 | 0.92887766 | 0.35295249 | 0.63441164 |
| LDB2 | 91.0252846 | 1.024088722 | 0.81413164 | 1.25789083 | 0.20843124 | 0.48863506 |
| QDPR | 34.0457892 | 0.466377916 | 0.29452602 | 1.5834863 | 0.11331067 | 0.35267249 |
| LAP3 | 74.202511 | -1.553658261 | 0.78110206 | -1.9890592 | 0.04669467 | 0.21587268 |
| MED28 | 79.9241098 | 0.219102386 | 0.22543986 | 0.9718884 | 0.33110607 | 0.61627893 |
| DCAF16 | 82.3745115 | 0.353647562 | 0.26534882 | 1.3327648 | 0.182609 | 0.45602782 |
| LCORL | 225.652704 | 0.251937351 | 0.14208136 | 1.77319072 | 0.07619711 | 0.28527949 |
| AC024230.1 | 79.6712079 | 0.118981462 | 0.56517683 | 0.21052077 | 0.83326124 | 0.93607461 |
| SLIT2 | 79.103815 | 0.291785078 | 0.80478296 | 0.36256369 | 0.71693084 | 0.88101097 |
| PACRGL | 138.116872 | 0.145607147 | 0.17355748 | 0.83895635 | 0.40149381 | 0.67562929 |
| KCNIP4 | 2356.95919 | -2.201609587 | 0.72828983 | -3.0229855 | 0.00250294 | 0.03638192 |
| ADGRA3 | 312.8613 | 0.58537343 | 0.33872415 | 1.72817153 | 0.08395748 | 0.30203879 |
| PPARGC1A | 389.986458 | 0.166488147 | 0.43277701 | 0.3846973 | 0.7004617 | 0.87182648 |
| DHX15 | 279.509172 | 0.182240211 | 0.2049 | 0.88941051 | 0.3737825 | 0.65253555 |
| SOD3 | 185.267771 | -0.804831374 | 1.14686542 | -0.7017662 | 0.48282499 | 0.73696727 |
| CCDC149 | 81.3138599 | 0.387836209 | 0.39305965 | 0.98671083 | 0.32378442 | 0.6106578 |
| LGI2 | 54.0853175 | 2.415356694 | 1.194529 | 2.02201595 | 0.0431747 | 0.20614587 |
| SEPSECS | 74.7117675 | 0.461181014 | 0.32492789 | 1.41933343 | 0.15580183 | 0.4201649 |
| ZCCHC4 | 46.5849439 | 0.259956037 | 0.31587194 | 0.82297921 | 0.41051982 | 0.68234762 |
| ANAPC4 | 102.991085 | 0.054461593 | 0.15636557 | 0.34829657 | 0.72761747 | 0.88564265 |
| SLC34A2 | 364.690055 | -5.322435028 | 0.67588278 | -7.8747901 | 3.41E-15 | 1.09E-11 |
| SEL1L3 | 429.277956 | -0.8344437 | 0.4174678 | -1.9988217 | 0.04562765 | 0.21268479 |
| SMIM20 | 49.1297718 | -0.701993654 | 0.37845316 | -1.8549024 | 0.06361017 | 0.25658128 |
| RBPJ | 295.78378 | -0.053370168 | 0.22062244 | -0.2419073 | 0.80885203 | 0.92414046 |
| TBC1D19 | 224.720851 | 0.261588094 | 0.27040182 | 0.96740508 | 0.33334157 | 0.6179175 |
| STIM2 | 110.788159 | 0.10678039 | 0.1626483 | 0.65651096 | 0.51149542 | 0.75961818 |
| ARAP2 | 331.801663 | 0.289837471 | 0.28039098 | 1.03369043 | 0.30128091 | 0.58849303 |
| C4orf19 | 272.552453 | -0.551962864 | 0.4180041 | -1.3204724 | 0.18667736 | 0.46182398 |
| RELL1 | 82.6559509 | -0.352114976 | 0.23122786 | -1.5228052 | 0.12780746 | 0.37531198 |
| PGM2 | 24.647964 | -0.5858676 | 0.35250153 | -1.6620285 | 0.09650705 | 0.32402421 |
| TBC1D1 | 461.599981 | 0.100775708 | 0.25212693 | 0.39970227 | 0.68937582 | 0.86755279 |
| KLF3 | 169.592661 | -0.001261293 | 0.41049624 | -0.0030726 | 0.99754842 | 0.99923686 |
| FAM114A1 | 78.3756878 | -0.113261816 | 0.31766085 | -0.3565495 | 0.72142909 | 0.88235193 |
| KLHL5 | 101.268378 | -0.4025666 | 0.1697639 | -2.3713321 | 0.0177241 | 0.12575079 |
| WDR19 | 182.078293 | 0.274047854 | 0.18670462 | 1.46781508 | 0.14215446 | 0.39850156 |
| RFC1 | 140.165052 | 0.428853091 | 0.16872086 | 2.5417906 | 0.01102862 | 0.09487502 |
| RPL9 | 35.1400925 | -0.27298086 | 0.34755552 | -0.7854309 | 0.43220095 | 0.69737124 |
| LIAS | 37.9762962 | 0.207701979 | 0.33003596 | 0.62933136 | 0.52913215 | 0.77040597 |
| UGDH | 67.3399333 | 0.028503802 | 0.4972949 | 0.0573177 | 0.95429212 | 0.98390867 |
| UGDH-AS1 | 84.8065799 | 0.155715888 | 0.27135241 | 0.57385113 | 0.56606854 | 0.79495244 |
| SMIM14 | 220.460712 | -0.123142326 | 0.28492795 | -0.4321876 | 0.66560507 | 0.85742752 |
| UBE2K | 493.921494 | 0.15710562 | 0.16048922 | 0.97891695 | 0.32762101 | 0.6137848 |
| PDS5A | 455.880096 | 0.291809495 | 0.13771027 | 2.11901034 | 0.03408959 | 0.18181749 |
| N4BP2 | 112.052817 | 0.422507561 | 0.20618274 | 2.04918981 | 0.04044356 | 0.19973796 |
| RBM47 | 1235.38272 | 0.021949359 | 0.27072173 | 0.08107719 | 0.93538056 | 0.97836695 |
| NSUN7 | 132.229378 | 0.498395857 | 0.26511187 | 1.87994544 | 0.06011551 | 0.24726898 |
| APBB2 | 502.619364 | 0.397994807 | 0.3344677 | 1.18993494 | 0.23407197 | 0.51656729 |
| UCHL1 | 51.4987254 | -1.236244593 | 0.7069202 | -1.7487753 | 0.08032987 | 0.2945893 |
| LIMCH1 | 330.302062 | -0.13488049 | 0.45784915 | -0.2945959 | 0.76830258 | 0.90456864 |
| TMEM33 | 80.2571753 | -0.051166667 | 0.27708162 | -0.1846628 | 0.85349353 | 0.9451971 |
| SLC30A9 | 255.957724 | 0.094033957 | 0.11608599 | 0.81003711 | 0.41791885 | 0.68748834 |
| ATP8A1 | 124.76629 | 0.212326175 | 0.41392103 | 0.51296301 | 0.6079772 | 0.8211189 |
| KCTD8 | 78.3763568 | -0.4732355 | 0.5235204 | -0.9039485 | 0.36602269 | 0.64617238 |
| GUF1 | 45.8276951 | 0.036246152 | 0.28374897 | 0.1277402 | 0.89835457 | 0.96245816 |
| GNPDA2 | 37.3614313 | 0.326119958 | 0.26831967 | 1.21541579 | 0.22420755 | 0.50617113 |
| GABRA2 | 112.559101 | 1.55477486 | 0.75914013 | 2.0480736 | 0.04055279 | 0.19992491 |
| COMMD8 | 29.5590484 | -0.042448214 | 0.40668714 | -0.1043756 | 0.91687129 | 0.9701436 |
| ATP10D | 190.710589 | 0.955520143 | 0.27101951 | 3.52565072 | 0.00042244 | 0.01207067 |
| AC107068.2 | 26.7003075 | 0.475360348 | 0.4719831 | 1.00715545 | 0.31386008 | 0.60093257 |
| NFXL1 | 91.7037798 | 0.262493426 | 0.19565941 | 1.34158346 | 0.17973109 | 0.451844 |
| NIPAL1 | 32.2828701 | 0.046389699 | 0.44007789 | 0.10541247 | 0.91604852 | 0.96948721 |
| TXK | 43.0247864 | -0.193274791 | 0.55516781 | -0.3481376 | 0.72773684 | 0.88565509 |
| TEC | 84.886554 | 0.071037037 | 0.30459919 | 0.2332148 | 0.81559462 | 0.92869214 |
| SLAIN2 | 189.124018 | 0.326040973 | 0.13556145 | 2.4051157 | 0.01616735 | 0.11876305 |
| FRYL | 476.510286 | 0.195740299 | 0.12549384 | 1.55976025 | 0.11881655 | 0.36397304 |
| OCIAD1 | 256.825179 | 0.049195277 | 0.12403618 | 0.39662037 | 0.69164743 | 0.86887335 |
| OCIAD2 | 100.480323 | -0.9896811 | 0.42664784 | -2.3196674 | 0.02035888 | 0.13571167 |
| DCUN1D4 | 246.047118 | 0.174204113 | 0.23795666 | 0.73208338 | 0.46411768 | 0.72183524 |
| SGCB | 46.5690718 | -0.116753786 | 0.31990096 | -0.3649685 | 0.71513489 | 0.87993284 |
| USP46 | 67.3634546 | 0.430147337 | 0.28041613 | 1.53396075 | 0.12503931 | 0.3716263 |
| SCFD2 | 261.987255 | -0.656810542 | 0.42859454 | -1.5324753 | 0.12540519 | 0.37225172 |

| | | | | | | |
|---|---|---|---|---|---|---|
| AC058822.1 | 140.554524 | 0.468829123 | 0.44966914 | 1.04260908 | 0.29712938 | 0.58520063 |
| LNX1 | 101.977545 | 0.71141301 | 0.67407951 | 1.05538441 | 0.29124955 | 0.5786931 |
| CHIC2 | 218.922305 | -0.070222685 | 0.2704328 | -0.2596678 | 0.79512004 | 0.91786634 |
| TMEM165 | 330.00612 | -0.914925597 | 0.32128684 | -2.8476908 | 0.00440377 | 0.05319508 |
| CLOCK | 233.353396 | 0.295100704 | 0.18289159 | 1.61352802 | 0.10662984 | 0.34137943 |
| EXOC1 | 178.348563 | 0.061900565 | 0.21824707 | 0.2836261 | 0.77669693 | 0.90765392 |
| CEP135 | 87.0484831 | 0.027631604 | 0.21074148 | 0.13111611 | 0.89568346 | 0.96142445 |
| AASDH | 83.6012433 | 0.476243066 | 0.19980538 | 2.3835348 | 0.01714726 | 0.12340619 |
| PPAT | 39.1948866 | -0.152481921 | 0.34352594 | -0.4438731 | 0.65713434 | 0.85164733 |
| PAICS | 65.7999163 | -0.889428076 | 0.35543308 | -2.5023784 | 0.0123362 | 0.1012583 |
| SRP72 | 99.7603133 | -0.205401768 | 0.25163664 | -0.8162634 | 0.41434953 | 0.68449528 |
| REST | 135.820437 | -0.084987034 | 0.20185399 | -0.4210322 | 0.67373156 | 0.86105131 |
| NOA1 | 26.1448285 | -0.400319967 | 0.37330653 | -1.0723626 | 0.28355719 | 0.57084461 |
| POLR2B | 131.204947 | -0.121725572 | 0.17724392 | -0.6867687 | 0.49222852 | 0.74455175 |
| IGFBP7 | 2505.91301 | 0.252707072 | 0.49347735 | 0.51209457 | 0.60858482 | 0.82155842 |
| ADGRL3 | 508.464794 | 1.180408352 | 1.03406541 | 1.14152194 | 0.25365279 | 0.54042261 |
| AC116049.2 | 46.0028193 | -0.447273415 | 0.55599325 | -0.8044584 | 0.42113231 | 0.68969506 |
| CENPC | 123.897355 | 0.817522891 | 0.32812417 | 2.4915046 | 0.01272033 | 0.10301379 |
| UBA6 | 221.242625 | 0.000614303 | 0.31552757 | 0.00194691 | 0.99844659 | 0.99928153 |
| UBA6-AS1 | 337.735902 | -0.249837049 | 0.38065762 | -0.6563301 | 0.51161176 | 0.75961818 |
| YTHDC1 | 397.313851 | 0.725364357 | 0.21311919 | 3.40356194 | 0.00066513 | 0.01577057 |
| UGT2A3 | 170.172477 | -0.182062155 | 0.60336935 | -0.3017425 | 0.76284839 | 0.90191616 |
| UGT2B7 | 1651.78485 | 0.817068509 | 0.53620796 | 1.52379034 | 0.12756109 | 0.37492391 |
| AC111000.4 | 124.60864 | 0.889895273 | 0.79137279 | 1.12449567 | 0.26080281 | 0.54773736 |
| UTP3 | 35.1516751 | -1.017140932 | 0.26880512 | -3.7839344 | 0.00015437 | 0.00637369 |
| RUFY3 | 472.749217 | -0.499452271 | 0.25219383 | -1.9804302 | 0.04765521 | 0.21810285 |
| GRSF1 | 88.8241464 | -0.201241 | 0.15532362 | -1.2956239 | 0.19510508 | 0.47360141 |
| MOB1B | 296.415721 | 0.873751331 | 0.25289362 | 3.45501535 | 0.00055026 | 0.01405588 |
| SLC4A4 | 134.143867 | 0.41761552 | 0.44614265 | 0.93605827 | 0.34924319 | 0.63265484 |
| ADAMTS3 | 276.31026 | 0.223199489 | 0.50478786 | 0.44216493 | 0.65836986 | 0.85215492 |
| ANKRD17 | 869.957419 | 0.23772993 | 0.12855822 | 1.8492006 | 0.06442885 | 0.25833569 |
| AC053527.1 | 152.946165 | -0.00631826 | 0.39037577 | -0.0161851 | 0.98708674 | 0.99678172 |
| RASSF6 | 139.332709 | -0.2804542 | 0.42890511 | -0.653884 | 0.51318652 | 0.76076145 |
| CXCL1 | 188.886331 | -4.333253427 | 1.72963172 | -2.5053041 | 0.01223462 | 0.10091982 |
| MTHFD2L | 123.870668 | -0.225484177 | 0.26074016 | -0.864785 | 0.38715681 | 0.66404483 |
| BTC | 31.8836942 | 0.037815952 | 0.60031606 | 0.0629934 | 0.94977176 | 0.98270285 |
| RCHY1 | 106.601788 | 0.761301283 | 0.20795925 | 3.66081959 | 0.00025141 | 0.00863173 |
| THAP6 | 62.715721 | 0.127426273 | 0.18849363 | 0.6760243 | 0.49902522 | 0.75042493 |
| CDKL2 | 205.853941 | 1.729138042 | 0.42564716 | 4.06237418 | 4.86E-05 | 0.00294775 |
| G3BP2 | 156.689018 | -0.053810248 | 0.18212066 | -0.2954648 | 0.76763883 | 0.90423172 |
| USO1 | 273.436957 | -0.078654963 | 0.20869868 | -0.3768829 | 0.70626063 | 0.87473681 |
| NAAA | 43.0104246 | 0.495238644 | 0.4317622 | 1.14701714 | 0.25137454 | 0.53868382 |
| SDAD1 | 95.216319 | -0.014903329 | 0.23017039 | -0.0647491 | 0.94837376 | 0.98262487 |
| NUP54 | 117.703655 | 0.32567835 | 0.22330707 | 1.45843276 | 0.1447213 | 0.40282422 |
| SCARB2 | 302.294104 | -0.225241459 | 0.1911485 | -1.1783585 | 0.23865373 | 0.52193243 |
| SHROOM3 | 534.901335 | 0.862657533 | 0.35388841 | 2.43765411 | 0.01478291 | 0.11283307 |
| 11-Sep | 412.104219 | 0.725176399 | 0.31647718 | 2.29140184 | 0.02194018 | 0.14229081 |
| CCNI | 430.03852 | -0.256478512 | 0.29445805 | -0.8710188 | 0.38374387 | 0.66018004 |
| CCNG2 | 107.561971 | -0.046961784 | 0.34708539 | -0.1353033 | 0.89237211 | 0.95976294 |
| CNOT6L | 202.625905 | 0.268152114 | 0.18312351 | 1.46432381 | 0.14310549 | 0.40011895 |
| MRPL1 | 108.610027 | 0.308404797 | 0.19309988 | 1.59712579 | 0.11023767 | 0.34724322 |
| FRAS1 | 729.600155 | 1.69942835 | 0.40617637 | 4.18396656 | 2.86E-05 | 0.00201769 |
| ANXA3 | 134.630715 | 0.816648841 | 0.49741686 | 1.64177957 | 0.1006357 | 0.33126781 |
| BMP2K | 183.31146 | -0.29830402 | 0.23371532 | -1.2763563 | 0.20182961 | 0.48041015 |
| PAQR3 | 30.5759109 | 0.766891258 | 0.44845962 | 1.71005642 | 0.08725544 | 0.30717378 |
| CFAP299 | 101.381858 | -1.091764123 | 0.52928475 | -2.062716 | 0.03913962 | 0.19618964 |
| RASGEF1B | 869.978313 | 0.882584914 | 0.49358967 | 1.78809439 | 0.07376078 | 0.28049008 |
| HNRNPD | 305.411995 | 0.000994987 | 0.13403338 | 0.00742342 | 0.99407702 | 0.99884271 |
| HNRNPDL | 252.392302 | 0.022999666 | 0.22711891 | 0.10126707 | 0.91933846 | 0.97108563 |
| ENOPH1 | 51.9824731 | -0.935342347 | 0.30184084 | -3.0987932 | 0.00194311 | 0.03122988 |
| TMEM150C | 69.8424134 | -0.603523351 | 0.33933531 | -1.7785457 | 0.07531428 | 0.28328492 |
| SCD5 | 108.535505 | 0.008029597 | 0.5930079 | 0.01354045 | 0.98919661 | 0.99678172 |
| SEC31A | 410.546322 | -0.13537925 | 0.13852297 | -0.9773054 | 0.32841796 | 0.61394673 |
| LIN54 | 69.5564061 | -0.141326276 | 0.19454582 | -0.7264421 | 0.46756777 | 0.72536326 |
| COPS4 | 60.7191314 | 0.518992416 | 0.21239174 | 2.44356214 | 0.01454307 | 0.11197822 |
| HELQ | 65.0651739 | 0.56867981 | 0.20957549 | 2.71348431 | 0.00665797 | 0.06902242 |
| MRPS18C | 38.2789383 | -0.196525082 | 0.3799351 | -0.5172596 | 0.60497495 | 0.82048068 |
| ABRAXAS1 | 44.3717933 | 0.140870257 | 0.23894033 | 0.58956249 | 0.55548401 | 0.78784845 |
| GPAT3 | 223.610697 | 1.53359058 | 0.60900827 | 2.51817695 | 0.0117964 | 0.09912084 |
| CDS1 | 218.366018 | 0.664532691 | 0.2456723 | 2.70495571 | 0.00683135 | 0.07019895 |
| WDFY3 | 1347.84129 | 1.077725747 | 0.19705167 | 5.4692547 | 4.52E-08 | 1.55E-05 |
| ARHGAP24 | 745.378794 | 0.984950736 | 0.32334872 | 3.0460944 | 0.00231835 | 0.0348626 |
| MAPK10 | 1436.04317 | 0.933710803 | 0.30087878 | 3.10327901 | 0.00191389 | 0.03107316 |
| PTPN13 | 967.515051 | 0.87250774 | 0.27782005 | 3.14054998 | 0.00168631 | 0.02869122 |
| AC093827.5 | 37.0842064 | -0.345029984 | 0.40911182 | -0.8433635 | 0.39902517 | 0.67377896 |
| AFF1 | 1545.39591 | 0.40098044 | 0.13992532 | 2.86567473 | 0.00416122 | 0.05110293 |
| KLHL8 | 100.162577 | -0.283613703 | 0.27056114 | -1.0482425 | 0.29452687 | 0.58239173 |
| NUDT9 | 69.28072 | -0.356961392 | 0.23302822 | -1.5318376 | 0.12556251 | 0.37248786 |
| SPP1 | 9555.38623 | -3.470566691 | 0.50505944 | -6.8716005 | 6.35E-12 | 8.69E-09 |
| PKD2 | 521.94612 | 0.489662815 | 0.18394403 | 2.66202075 | 0.00776731 | 0.07623264 |
| PPM1K | 88.201403 | 0.446409066 | 0.57598293 | 0.77503871 | 0.43831679 | 0.70236452 |
| HERC6 | 56.1630034 | -0.685240226 | 1.10735518 | -0.618808 | 0.53604286 | 0.77540842 |
| HERC5 | 55.5827789 | -1.338021704 | 1.05199859 | -1.2718855 | 0.20341382 | 0.48230223 |
| PYURF | 35.4426583 | -0.157172092 | 0.40246491 | -0.3905237 | 0.69614932 | 0.87043654 |
| HERC3 | 108.304197 | 0.415949469 | 0.34479762 | 1.20635829 | 0.22767937 | 0.51041354 |
| FAM13A-AS1 | 40.7765044 | 0.848756801 | 0.41274362 | 2.05637776 | 0.03974612 | 0.19750809 |
| FAM13A | 359.460531 | 0.234514477 | 0.25907664 | 0.90519344 | 0.36536292 | 0.64536444 |
| CCSER1 | 1568.45607 | 1.363422326 | 0.55191457 | 2.47035032 | 0.01349808 | 0.10712353 |
| SMARCAD1 | 172.840642 | 0.402193077 | 0.17846328 | 2.25364608 | 0.02421844 | 0.15123107 |
| PDLIM5 | 478.426067 | -0.367602678 | 0.40125044 | -0.9161427 | 0.35959203 | 0.64001011 |
| BMPR1B | 280.846058 | 0.515733945 | 0.91865311 | 0.56140227 | 0.57452334 | 0.8009242 |
| UNC5C | 192.346286 | -0.151796094 | 1.32112201 | -0.1148994 | 0.90852488 | 0.96752108 |
| STPG2 | 289.551505 | 0.783162043 | 0.41373707 | 1.89289794 | 0.05837144 | 0.243852 |
| RAP1GDS1 | 212.967536 | -0.080400492 | 0.19712011 | -0.4078756 | 0.68336498 | 0.864723 |
| TSPAN5 | 437.954323 | 0.849022281 | 1.18291892 | 0.71773498 | 0.47292071 | 0.73007373 |
| EIF4E | 135.583829 | -0.025050229 | 0.21053151 | -0.1189857 | 0.90528672 | 0.96556525 |
| METAP1 | 69.730342 | -0.221969864 | 0.22783667 | -0.97425 | 0.32993245 | 0.61510761 |
| ADH5 | 96.0551814 | -0.191599401 | 0.27077227 | -0.7076035 | 0.47919153 | 0.73490443 |
| AP002026.1 | 48.1730907 | 0.743230545 | 0.57332222 | 1.29635746 | 0.19485236 | 0.47336819 |
| LAMTOR3 | 46.349778 | -0.138412605 | 0.27656008 | -0.5004793 | 0.6167376 | 0.82592881 |
| DNAJB14 | 224.40297 | 0.094328006 | 0.15306766 | 0.61625039 | 0.53772927 | 0.77644087 |
| H2AFZ | 87.2059115 | 0.1751399 | 0.65064179 | 0.26918022 | 0.787791 | 0.91381085 |
| DDIT4L | 83.3890976 | 2.135663362 | 0.54815956 | 3.8960615 | 9.78E-05 | 0.00459086 |
| PPP3CA | 459.449725 | 0.119638432 | 0.3056995 | 0.3913596 | 0.69553145 | 0.87024545 |
| BANK1 | 75.2346521 | -0.419122014 | 0.42739698 | -0.9806387 | 0.32677095 | 0.61330512 |
| SLC39A8 | 132.466285 | 0.130014748 | 0.50568359 | 0.25710691 | 0.79709624 | 0.9188367 |
| NFKB1 | 278.047989 | -0.153369669 | 0.29537406 | -0.5192388 | 0.60359423 | 0.81988502 |
| MANBA | 112.45627 | 0.264388589 | 0.31663566 | 0.83499309 | 0.4037216 | 0.67765972 |
| UBE2D3 | 538.805465 | 0.001369301 | 0.10451794 | 0.01310111 | 0.98954713 | 0.99683162 |
| CISD2 | 58.2811725 | 0.00157486 | 0.25680658 | 0.00613247 | 0.99510702 | 0.99886097 |
| SLC9B1 | 31.8972835 | 0.807220344 | 0.66547375 | 1.21300102 | 0.22512943 | 0.50741525 |
| SLC9B2 | 28.0166265 | -0.728555782 | 0.38228137 | -1.9058103 | 0.05667482 | 0.24000359 |
| TET2 | 284.886154 | 0.719631969 | 0.2403419 | 2.99420109 | 0.00275165 | 0.03836683 |

| | | | | | | |
|---|---|---|---|---|---|---|
| PPA2 | 328.402718 | 0.066483685 | 0.16740088 | 0.39715255 | 0.69125497 | 0.86884967 |
| ARHGEF38 | 211.052167 | -0.526266208 | 0.53063442 | -0.9917679 | 0.32131074 | 0.60844973 |
| INTS12 | 99.7923471 | -0.31766135 | 0.35870482 | -0.8855787 | 0.37584459 | 0.65434666 |
| GSTCD | 125.565647 | -0.054768165 | 0.29124364 | -0.1880493 | 0.850838 | 0.94435157 |
| NPNT | 318.370214 | 0.469069675 | 0.46537523 | 1.00793864 | 0.31348392 | 0.60065312 |
| TBCK | 559.78509 | 0.468950436 | 0.29748029 | 1.5764084 | 0.11493171 | 0.35651907 |
| AIMP1 | 88.1915664 | 0.0421319 | 0.23201785 | 0.18158905 | 0.85590524 | 0.94621534 |
| PAPSS1 | 112.848404 | -0.027855334 | 0.3200497 | -0.0870344 | 0.93064417 | 0.97602738 |
| SGMS2 | 129.186992 | -0.301667172 | 0.42493391 | -0.7099155 | 0.47775654 | 0.73413221 |
| HADH | 44.6307463 | 0.214772018 | 0.32422663 | 0.66241325 | 0.50770641 | 0.75687816 |
| RPL34-AS1 | 29.8602032 | 0.596772859 | 0.58996936 | 1.01153195 | 0.3117619 | 0.5993111 |
| RPL34 | 304.495871 | -0.242543031 | 0.4504079 | -0.5384964 | 0.59023439 | 0.81183245 |
| OSTC | 77.2039 | -0.900114137 | 0.36742279 | -2.4498049 | 0.01429336 | 0.11095311 |
| SEC24B | 317.455404 | 0.166003303 | 0.16321171 | 1.0171041 | 0.30910391 | 0.59563596 |
| MCUB | 47.2650502 | -1.878020035 | 0.61352276 | -3.0610438 | 0.00220567 | 0.03391348 |
| CASP6 | 43.5052769 | -0.786029578 | 0.31381204 | -2.5047782 | 0.01225282 | 0.10091982 |
| PLA2G12A | 53.8744059 | -0.265677185 | 0.26867954 | -0.9888255 | 0.32274852 | 0.60995011 |
| GAR1 | 30.5961497 | -0.605555048 | 0.33297479 | -1.8186213 | 0.06896922 | 0.26888733 |
| EGF | 49.7878213 | 1.976643876 | 0.62651905 | 3.15496213 | 0.00160519 | 0.02795656 |
| ELOVL6 | 119.806823 | -0.521131445 | 0.31679853 | -1.6449933 | 0.0999712 | 0.33078553 |
| ENPEP | 139.073162 | 0.27527966 | 0.5966371 | 0.46138542 | 0.64452211 | 0.84377167 |
| FAM241A | 33.2555489 | -0.398809206 | 0.46583009 | -0.8561259 | 0.39192815 | 0.6674275 |
| AP1AR | 78.6826212 | -0.178722641 | 0.30367703 | -0.5885287 | 0.55617749 | 0.78799352 |
| ALPK1 | 298.075054 | -1.172360672 | 0.29955991 | -3.91361 | 9.09E-05 | 0.00439891 |
| LARP7 | 104.481515 | 0.262153828 | 0.21806548 | 1.20217943 | 0.22929402 | 0.5117445 |
| ANK2 | 273.932193 | 1.026393822 | 0.50277341 | 2.04146401 | 0.04120473 | 0.20195151 |
| CAMK2D | 554.128702 | 0.448316737 | 0.13839236 | 3.23946159 | 0.00119756 | 0.02359807 |
| ARSJ | 149.906618 | 0.706641559 | 0.25951295 | 2.72295295 | 0.00647013 | 0.06810698 |
| UGT8 | 160.652958 | 0.947623234 | 0.61346957 | 1.54469476 | 0.12242011 | 0.36765168 |
| SNHG8 | 67.7261259 | -0.059926148 | 0.44063909 | -0.1359983 | 0.89182266 | 0.95976294 |
| METTL14 | 63.641508 | 0.321956381 | 0.21339006 | 1.50876936 | 0.13135773 | 0.38163088 |
| SEC24D | 132.970733 | -0.743438085 | 0.35568528 | -2.090157 | 0.0366037 | 0.189528 |
| USP53 | 345.258519 | -0.293878254 | 0.20159569 | -1.4577606 | 0.14490654 | 0.40282422 |
| C4orf3 | 141.701679 | -0.460985882 | 0.39331767 | -1.1720447 | 0.24117912 | 0.52565524 |
| PDE5A | 150.219709 | 0.56744973 | 0.3052729 | 1.85882773 | 0.06305156 | 0.25505528 |
| AC108866.1 | 77.5837153 | -0.603779609 | 0.33906674 | -1.7807102 | 0.0749598 | 0.28280423 |
| PRDM5 | 101.23197 | 0.224650216 | 0.3435518 | 0.65390492 | 0.51317306 | 0.76076145 |
| ANXA5 | 249.25635 | -0.709839459 | 0.37664785 | -1.8846237 | 0.05948066 | 0.24614695 |
| BBS7 | 48.9012672 | 0.112019982 | 0.22565658 | 0.49641797 | 0.61959955 | 0.82775923 |
| KIAA1109 | 467.959661 | 0.556899264 | 0.15297199 | 3.64053089 | 0.00027208 | 0.00898752 |
| FGF2 | 90.5204744 | -0.322085639 | 0.39668875 | -0.8119354 | 0.4168287 | 0.68616638 |
| NUDT6 | 28.9949298 | 0.252105681 | 0.35563558 | 0.70888767 | 0.47839419 | 0.73438108 |
| SPATA5 | 307.514139 | 0.354322558 | 0.17822935 | 1.98801461 | 0.04681007 | 0.21609335 |
| SPRY1 | 37.9909002 | -0.597913999 | 0.54755513 | -1.0919704 | 0.2748461 | 0.56187466 |
| LINC01091 | 51.3050408 | -0.329005049 | 0.50411191 | -0.6526429 | 0.51398653 | 0.76088633 |
| ANKRD50 | 53.2429289 | 0.060550435 | 0.43982526 | 0.1376693 | 0.89050178 | 0.95941026 |
| FAT4 | 160.289753 | 1.401624289 | 0.37651352 | 3.72264004 | 0.00019715 | 0.00743506 |
| INTU | 198.238426 | 0.691805538 | 0.2663879 | 2.59698555 | 0.00940459 | 0.0862898 |
| HSPA4L | 73.8064795 | -0.11149147 | 0.58979432 | -0.1890345 | 0.85006578 | 0.94435157 |
| MFSD8 | 91.1788774 | 0.372664124 | 0.19841772 | 1.87817968 | 0.06035659 | 0.24792272 |
| ABHD18 | 232.249871 | 0.125860938 | 0.15555214 | 0.80912381 | 0.41844393 | 0.68785864 |
| LARP1B | 325.185501 | -0.108948007 | 0.22259567 | -0.4894435 | 0.62452774 | 0.83088211 |
| PGRMC2 | 126.463807 | -0.13247632 | 0.21435984 | -0.6180091 | 0.53656937 | 0.77558443 |
| AC108062.1 | 183.636253 | 0.813959066 | 0.39422465 | 2.06470872 | 0.03895057 | 0.19580915 |
| JADE1 | 184.997474 | 0.570671649 | 0.31046371 | 1.83812675 | 0.06604373 | 0.26198834 |
| SCLT1 | 171.933014 | 0.325276873 | 0.30999507 | 1.04929692 | 0.29404148 | 0.58218755 |
| AC096711.2 | 92.1894046 | 0.412705164 | 0.33031174 | 1.24944139 | 0.21150368 | 0.49253647 |
| AC079380.1 | 45.0458611 | 0.239730292 | 0.42155611 | 0.56867944 | 0.5695737 | 0.79749848 |
| ELF2 | 505.011728 | -0.549137074 | 0.24107306 | -2.2778866 | 0.02273334 | 0.14498178 |
| NDUFC1 | 93.0862294 | -0.341018649 | 0.30242775 | -1.1276037 | 0.25948733 | 0.54618849 |
| NAA15 | 237.17581 | 0.021765353 | 0.19853601 | 0.10962924 | 0.91270342 | 0.96844354 |
| AC097376.2 | 253.231755 | -0.14387802 | 0.22820196 | -0.6304855 | 0.52837701 | 0.76990012 |
| SETD7 | 56.5169262 | -1.032306808 | 0.33236986 | -3.1058977 | 0.00189702 | 0.03090405 |
| MGST2 | 69.8173355 | -0.007799616 | 0.22842573 | -0.0341451 | 0.97276145 | 0.99244669 |
| MAML3 | 1101.30619 | 0.436699408 | 0.46820422 | 0.9327114 | 0.350969 | 0.63331896 |
| SCOC | 148.191662 | 0.172259672 | 0.30275846 | 0.56896734 | 0.56937831 | 0.79749848 |
| ELMOD2 | 63.3511488 | 0.116727818 | 0.18365824 | 0.63557082 | 0.5250562 | 0.76681099 |
| TBC1D9 | 87.7970972 | -0.105668486 | 0.26740208 | -0.395167 | 0.69271962 | 0.86932546 |
| RNF150 | 156.610374 | -0.03526978 | 0.70523666 | -0.0500113 | 0.96011341 | 0.98690056 |
| ZNF330 | 49.1097837 | 0.08746637 | 0.25845148 | 0.33842472 | 0.73504315 | 0.88966284 |
| IL15 | 161.200101 | 0.07095225 | 0.52514835 | 0.13510897 | 0.89252572 | 0.95976294 |
| INPP4B | 70.6289475 | -0.597552869 | 0.47795647 | -1.2502245 | 0.21121756 | 0.49218977 |
| AC104596.1 | 74.5889124 | 0.575511595 | 0.46259319 | 1.24409872 | 0.21346323 | 0.49544024 |
| USP38 | 50.9069962 | 0.085297062 | 0.22551124 | 0.37823863 | 0.70525333 | 0.87462735 |
| GAB1 | 534.860997 | 0.127842267 | 0.35779061 | 0.35731029 | 0.72085953 | 0.88235193 |
| SMARCA5 | 148.737305 | 0.119932371 | 0.21020347 | 0.57055372 | 0.5683022 | 0.79713855 |
| ANAPC10 | 148.590769 | 0.279941658 | 0.20363883 | 1.37469683 | 0.16922545 | 0.43904596 |
| ABCE1 | 83.1358549 | -0.410236355 | 0.26495604 | -1.5483186 | 0.12154561 | 0.3663579 |
| OTUD4 | 155.582568 | -0.118716992 | 0.15620676 | -0.7599991 | 0.44725513 | 0.70891772 |
| SMAD1 | 102.9221 | -0.187129007 | 0.25500197 | -0.7338336 | 0.46305018 | 0.72111164 |
| MMAA | 43.0477557 | -0.219451701 | 0.30052283 | -0.730233 | 0.46524775 | 0.72300587 |
| ZNF827 | 276.132089 | -0.176013736 | 0.41816166 | -0.4209227 | 0.67381152 | 0.86105131 |
| LSM6 | 42.8330365 | 0.257809459 | 0.43687349 | 0.59012384 | 0.55510763 | 0.78784845 |
| SLC10A7 | 125.040379 | -0.218142576 | 0.2104592 | -1.0365077 | 0.29996536 | 0.58700065 |
| TMEM184C | 76.8198966 | 0.292231273 | 0.36893803 | 0.79208769 | 0.42830955 | 0.69444434 |
| PRMT9 | 81.8823928 | 0.779945918 | 0.32957932 | 2.36648927 | 0.01795769 | 0.12664775 |
| ARHGAP10 | 588.935067 | 0.279766833 | 0.32774657 | 0.8536072 | 0.39332267 | 0.6686322 |
| NR3C2 | 630.355017 | 0.641616727 | 0.42949887 | 1.49387291 | 0.13520884 | 0.38731025 |
| DCLK2 | 85.3069582 | 0.442414658 | 0.75028945 | 0.58965864 | 0.55541953 | 0.78784845 |
| LRBA | 1505.10126 | 0.418663357 | 0.19811288 | 2.11325659 | 0.03457881 | 0.18326846 |
| RPS3A | 217.249287 | -0.073725801 | 0.49602894 | -0.1486321 | 0.88184398 | 0.95664592 |
| SH3D19 | 384.032461 | -0.315706621 | 0.29785125 | -1.0599473 | 0.28916858 | 0.57585049 |
| FAM160A1 | 461.242597 | -0.118576556 | 0.28394494 | -0.417604 | 0.67623664 | 0.86116336 |
| GATB | 32.4808224 | -0.325083659 | 0.33960735 | -0.9572339 | 0.33844922 | 0.62345706 |
| FBXW7 | 500.433518 | -0.025335334 | 0.25354427 | -0.0999247 | 0.92040411 | 0.97180033 |
| ARFIP1 | 234.221428 | 0.245371583 | 0.14569282 | 1.68417071 | 0.09214868 | 0.31603732 |
| TRIM2 | 305.911938 | 0.61357015 | 0.38363717 | 1.59935011 | 0.10974283 | 0.34671061 |
| TMEM131L | 92.6045489 | -0.079700604 | 0.47984299 | -0.1660973 | 0.86808042 | 0.95090501 |
| PLRG1 | 49.3501374 | 0.525089998 | 0.25419187 | 2.06572305 | 0.03885464 | 0.19575457 |
| FGB | 300.253085 | -6.980597655 | 1.44075048 | -4.8451122 | 1.27E-06 | 0.00021265 |
| FGA | 115.508155 | -5.888292656 | 1.49625598 | -3.9353511 | 8.31E-05 | 0.00415961 |
| LRAT | 47.9547319 | -0.528531084 | 0.62113539 | -0.8509112 | 0.39481866 | 0.6696972 |
| MAP9 | 75.365823 | 0.348256392 | 0.3196005 | 1.08966159 | 0.27586224 | 0.56304683 |
| GUCY1A1 | 222.528125 | -0.103245972 | 0.32075634 | -0.3218829 | 0.74754143 | 0.89399656 |
| GUCY1B1 | 125.653117 | 0.26924897 | 0.74353646 | 0.36211939 | 0.71726281 | 0.88106948 |
| TDO2 | 27.6831835 | 1.974254454 | 1.14419156 | 1.72545797 | 0.08444497 | 0.3023414 |
| CTSO | 82.0394837 | -0.559911699 | 0.4026801 | -1.3904628 | 0.16438839 | 0.43262536 |
| PDGFC | 385.075405 | 0.229995082 | 0.37161451 | 0.6189077 | 0.53597717 | 0.77540842 |
| TMEM144 | 31.0754398 | -0.1392478 | 0.41244959 | -0.3376117 | 0.73565584 | 0.89020304 |
| ETFDH | 103.496533 | 0.795861714 | 0.2919486 | 2.72603368 | 0.00641005 | 0.06769771 |
| PPID | 45.72793 | -0.042167742 | 0.27575587 | -0.1529169 | 0.87846378 | 0.95481727 |
| FNIP2 | 705.557616 | -0.945045169 | 0.41054204 | -2.3019449 | 0.02133828 | 0.14039576 |
| C4orf45 | 28.1799007 | -0.321048795 | 0.45286607 | -0.7089266 | 0.47837003 | 0.73438108 |

| | | | | | | |
|---|---|---|---|---|---|---|
| RAPGEF2 | 648.397847 | 0.434634594 | 0.22483848 | 1.93309703 | 0.05322425 | 0.23155955 |
| NAF1 | 81.0689076 | 0.17261387 | 0.20187177 | 0.85506689 | 0.39251412 | 0.66806907 |
| TMA16 | 48.9865309 | -0.573724224 | 0.29429476 | -1.9494884 | 0.05123712 | 0.22743299 |
| 1-Mar | 832.584885 | 2.388402661 | 1.10417584 | 2.16306368 | 0.03053628 | 0.17105675 |
| TMEM192 | 101.928594 | -0.618711949 | 0.24529799 | -2.5222871 | 0.01165945 | 0.09840166 |
| KLHL2 | 255.210697 | -0.675086899 | 0.43930912 | -1.5367013 | 0.12436647 | 0.37055989 |
| MSMO1 | 120.298841 | 1.092314706 | 0.52063781 | 2.09803185 | 0.03590233 | 0.18762053 |
| CPE | 602.384037 | 1.606777214 | 0.83121479 | 1.93304696 | 0.05323041 | 0.23155955 |
| DDX60 | 130.934906 | -0.840313068 | 1.01009444 | -0.8319153 | 0.40545673 | 0.67887957 |
| DDX60L | 131.826071 | -0.530189892 | 1.06404102 | -0.4982796 | 0.61828701 | 0.82705925 |
| PALLD | 290.864698 | 1.320932027 | 0.69276799 | 1.90674518 | 0.05655359 | 0.23970215 |
| CBR4 | 294.55259 | 0.465855014 | 0.21009138 | 2.2173923 | 0.0265963 | 0.15808253 |
| SH3RF1 | 433.457396 | 0.535010085 | 0.24775474 | 2.1594343 | 0.03081649 | 0.17142506 |
| NEK1 | 240.87207 | 0.293767674 | 0.19339129 | 1.51903264 | 0.12875428 | 0.37682165 |
| CLCN3 | 233.662857 | 0.033111877 | 0.17158071 | 0.19298135 | 0.84697357 | 0.94323402 |
| HPF1 | 36.2813267 | 0.036327996 | 0.27950642 | 0.12997196 | 0.89658861 | 0.96174942 |
| MFAP3L | 103.949945 | 0.87419733 | 0.53151667 | 1.64472233 | 0.10002709 | 0.33080044 |
| GALNT7 | 158.971736 | -0.560237931 | 0.21892135 | -2.5590831 | 0.01049486 | 0.09189242 |
| HMGB2 | 56.3851959 | -0.756960061 | 0.5390106 | -1.404351 | 0.16021436 | 0.426067 |
| FBXO8 | 40.709199 | 0.453729647 | 0.28706178 | 1.58059931 | 0.11396968 | 0.35432469 |
| CEP44 | 71.4966467 | 0.421689021 | 0.21165091 | 1.99237992 | 0.04632939 | 0.21459827 |
| GPM6A | 134.239703 | 1.460344073 | 0.50244035 | 2.90650237 | 0.00365494 | 0.04674324 |
| WDR17 | 40.3874941 | 0.186495931 | 0.33696369 | 0.55346002 | 0.57994845 | 0.80465327 |
| SPCS3 | 93.4812448 | -0.279823319 | 0.28697912 | -0.9750651 | 0.32952797 | 0.61459277 |
| AC078881.1 | 75.0374536 | 0.097819286 | 0.40326 | 0.24257126 | 0.80833755 | 0.92399348 |
| TENM3 | 677.061415 | -0.573299141 | 0.69538266 | -0.8244369 | 0.40969134 | 0.68208527 |
| DCTD | 71.6903022 | -0.258269997 | 0.20062998 | -1.2872952 | 0.19799143 | 0.47656532 |
| WWC2 | 731.896987 | 0.921725302 | 0.16739546 | 5.50627418 | 3.67E-08 | 1.35E-05 |
| CDKN2AIP | 28.9642482 | 1.113425352 | 0.66549426 | 1.67308033 | 0.09431148 | 0.31990429 |
| RWDD4 | 40.3505307 | 0.133028014 | 0.2752845 | 0.48323829 | 0.62892655 | 0.83372327 |
| TRAPPC11 | 120.997168 | 0.247001332 | 0.22681856 | 1.08898199 | 0.27616183 | 0.56317011 |
| STOX2 | 1617.24577 | 0.01863454 | 0.26804625 | 0.06951987 | 0.94457582 | 0.98115679 |
| IRF2 | 135.498159 | -0.448946362 | 0.26539291 | -1.6916291 | 0.09071672 | 0.3142614 |
| CASP3 | 38.0376419 | -0.348169749 | 0.25784954 | -1.3502826 | 0.17692535 | 0.44872611 |
| PRIMPOL | 59.037448 | 0.564116896 | 0.3874493 | 1.45597603 | 0.14539925 | 0.40323664 |
| ACSL1 | 164.081043 | 0.189694958 | 0.34268648 | 0.5535525 | 0.57988514 | 0.80465327 |
| SLC25A4 | 49.0217984 | -0.069255469 | 0.36820444 | -0.1880897 | 0.85080632 | 0.94435157 |
| CFAP97 | 118.222619 | -0.266397792 | 0.25515391 | -1.0440671 | 0.29645437 | 0.58477205 |
| SNX25 | 267.742936 | 0.926348501 | 0.24120268 | 3.84053979 | 0.00012276 | 0.00547711 |
| LRP2BP | 37.2539985 | 0.735394405 | 0.50188537 | 1.46526368 | 0.14284899 | 0.39975183 |
| AC112722.1 | 27.8678586 | 0.775415743 | 0.35585833 | 2.17900125 | 0.02933158 | 0.16724236 |
| ANKRD37 | 52.5435162 | 0.208546209 | 1.10062338 | 0.18948008 | 0.84971656 | 0.94414047 |
| UFSP2 | 61.9478696 | 0.566760886 | 0.32148388 | 1.76295274 | 0.07790846 | 0.28936997 |
| C4orf47 | 111.306592 | -0.595666487 | 0.65131572 | -0.9145587 | 0.36042333 | 0.64077488 |
| SORBS2 | 289.175438 | 0.141330767 | 0.48424526 | 0.29185782 | 0.77039533 | 0.90533273 |
| FAM149A | 153.25561 | 0.587927713 | 0.36310084 | 1.61918577 | 0.1054073 | 0.33950969 |
| CYP4V2 | 56.5316584 | 0.340947889 | 0.3097159 | 1.10084078 | 0.27096596 | 0.55818988 |
| KLKB1 | 40.387073 | 0.988689971 | 0.26227324 | 3.7696944 | 0.00016345 | 0.0066624 |
| FAT1 | 662.979162 | 1.111025016 | 0.24629206 | 4.51100629 | 6.45E-06 | 0.00066456 |
| FRG1-DT | 67.1605817 | 0.284822258 | 0.48701279 | 0.58483527 | 0.55865849 | 0.78929051 |
| FRG1 | 30.111144 | 0.143735909 | 0.33910062 | 0.42387392 | 0.67165776 | 0.86013159 |
| SDHA | 124.670909 | -0.382279379 | 0.22020827 | -1.7359901 | 0.08256558 | 0.29927236 |
| PDCD6 | 79.8999512 | -0.229525285 | 0.27711517 | -0.8282668 | 0.40751945 | 0.68078632 |
| EXOC3 | 97.8168964 | 0.118592287 | 0.26472542 | 0.44798223 | 0.65416602 | 0.84966187 |
| SLC9A3 | 120.000418 | 1.176168035 | 0.75228744 | 1.56345564 | 0.11794546 | 0.3625801 |
| BRD9 | 45.8373034 | -0.325019315 | 0.26111321 | -1.2447448 | 0.21322556 | 0.49502851 |
| SLC12A7 | 91.3546394 | -1.49091182 | 0.38330559 | -3.8896167 | 0.0001004 | 0.00465146 |
| CLPTM1L | 73.1541737 | -0.512074781 | 0.3115004 | -1.6438977 | 0.10019734 | 0.3309277 |
| LPCAT1 | 59.3962726 | -0.463909945 | 0.45239854 | -1.0254453 | 0.30515313 | 0.59267272 |
| MRPL36 | 30.3265665 | -0.655530244 | 0.39384854 | -1.6644222 | 0.09602809 | 0.32319324 |
| NDUFS6 | 88.4947104 | -0.232698741 | 0.42536803 | -0.5470527 | 0.58434251 | 0.80803859 |
| IRX2 | 25.6007887 | 0.244731196 | 1.10653076 | 0.22116981 | 0.82496022 | 0.93217972 |
| IRX1 | 32.8712976 | 0.356081409 | 1.14533351 | 0.31089757 | 0.75587849 | 0.89874207 |
| ADAMTS16 | 124.029491 | 1.181530805 | 1.02419235 | 1.15362198 | 0.24865518 | 0.53573279 |
| ICE1 | 163.131451 | 0.218675734 | 0.17722881 | 1.2338611 | 0.21725467 | 0.49932392 |
| MED10 | 51.4687296 | -0.133814859 | 0.25621671 | -0.5222722 | 0.60148084 | 0.81829073 |
| NSUN2 | 78.9760723 | -0.171908033 | 0.28992052 | -0.5929488 | 0.5532154 | 0.78647229 |
| TENT4A | 122.289926 | -0.064301524 | 0.24483031 | -0.2626371 | 0.7928303 | 0.91680379 |
| MTRR | 84.5185251 | -0.203283674 | 0.26242446 | -0.7746369 | 0.43855424 | 0.7024935 |
| SEMA5A | 240.057621 | 0.017807531 | 0.70073925 | 0.02541249 | 0.97972595 | 0.99557142 |
| SNHG18 | 30.0542198 | 0.085560504 | 0.50761168 | 0.16855503 | 0.86614665 | 0.95070686 |
| FAM173B | 46.018448 | 0.588392477 | 0.30560474 | 1.92533818 | 0.05418706 | 0.23344987 |
| CCT5 | 101.656861 | -0.906708501 | 0.38153634 | -2.376467 | 0.01747933 | 0.12476488 |
| CMBL | 53.6344569 | -1.142764304 | 0.65344855 | -1.7488206 | 0.08032203 | 0.2945893 |
| 6-Mar | 399.443292 | 0.279234223 | 0.16180389 | 1.72575719 | 0.0843911 | 0.3023414 |
| ANKRD33B | 231.35714 | 0.020368588 | 0.41015344 | 0.0496609 | 0.96039262 | 0.98697574 |
| DAP | 186.914768 | 0.313442132 | 0.26556423 | 1.18028745 | 0.23788591 | 0.52090663 |
| DNAH5 | 220.328425 | -1.573516831 | 0.5147887 | -3.0566266 | 0.00223843 | 0.03403479 |
| TRIO | 1444.86124 | -0.253228667 | 0.17792084 | -1.4232659 | 0.15465909 | 0.41802467 |
| OTULIN | 65.4539475 | -0.146915594 | 0.22170753 | -0.6626549 | 0.50755157 | 0.75682386 |
| ANKH | 117.26069 | 0.525191206 | 0.51209806 | 1.02556766 | 0.30509542 | 0.59267272 |
| LINC02149 | 37.4346317 | 3.357461536 | 1.15001326 | 2.9194981 | 0.00350596 | 0.04544458 |
| FBXL7 | 600.947148 | 1.185678756 | 0.56061472 | 2.11496187 | 0.0344332 | 0.18303862 |
| RETREG1 | 525.485538 | -1.337921381 | 0.43529995 | -3.073562 | 0.0021152 | 0.03294549 |
| MYO10 | 524.656668 | 0.109517824 | 0.33053866 | 0.33133137 | 0.7403942 | 0.89253455 |
| LINC02241 | 81.278346 | 0.843280399 | 0.28641146 | 2.94429695 | 0.00323689 | 0.04294487 |
| C5orf17 | 66.6477688 | 0.034255071 | 0.54521856 | 0.06282814 | 0.94990335 | 0.98270285 |
| CDH9 | 89.5984685 | -0.334118646 | 1.17157791 | -0.2851869 | 0.77550097 | 0.90724522 |
| AC113386.1 | 76.0586411 | -1.485076564 | 0.67992583 | -2.1841744 | 0.02894944 | 0.16552368 |
| CDH6 | 934.624212 | -1.274613389 | 0.73564546 | -1.7326463 | 0.08315855 | 0.30002855 |
| DROSHA | 120.378836 | 0.178733706 | 0.19613192 | 0.91129331 | 0.36214085 | 0.64251662 |
| C5orf22 | 32.1718647 | -0.240656476 | 0.32008136 | -0.7518603 | 0.45213505 | 0.71231034 |
| PDZD2 | 313.463331 | 1.300056011 | 0.43358884 | 2.99836134 | 0.00271436 | 0.03811711 |
| GOLPH3 | 313.877376 | 0.20757952 | 0.20755068 | 1.00013896 | 0.31724326 | 0.60364002 |
| MTMR12 | 165.027209 | 0.046786421 | 0.26584819 | 0.17598924 | 0.86030239 | 0.94776729 |
| ZFR | 430.204743 | 0.310540634 | 0.14821243 | 2.09524011 | 0.03614965 | 0.18804981 |
| SUB1 | 204.44477 | -0.068824569 | 0.33614165 | -0.2047487 | 0.83776846 | 0.93870442 |
| NPR3 | 240.477611 | 1.450109712 | 0.58394353 | 2.48330471 | 0.01301697 | 0.10449615 |
| TARS | 66.4490665 | -0.368320648 | 0.27238964 | -1.352183 | 0.17631677 | 0.44835649 |
| C1QTNF3-AN | 209.474609 | 0.234127844 | 0.18515942 | 1.2644663 | 0.20606271 | 0.48593666 |
| RAI14 | 468.495205 | 0.800814988 | 0.25508518 | 3.13940226 | 0.00169293 | 0.02875277 |
| TTC23L | 31.0681668 | -0.147977955 | 0.57203975 | -0.2586847 | 0.79587849 | 0.91818862 |
| RAD1 | 28.7172351 | 0.128160592 | 0.35585372 | 0.36014965 | 0.71873523 | 0.88155276 |
| BRIX1 | 42.5204 | -0.603652607 | 0.28062247 | -2.15112 | 0.03146673 | 0.17362347 |
| DNAJC21 | 85.0659299 | -0.235298431 | 0.19552 | -1.2034494 | 0.22880246 | 0.51112377 |
| PRLR | 63.1732376 | -0.145827396 | 0.49279864 | -0.2959168 | 0.76729362 | 0.90404743 |
| SPEF2 | 156.465932 | 0.022439262 | 0.43513329 | 0.05156871 | 0.95887235 | 0.98594228 |
| LMBRD2 | 60.5983942 | 0.584982666 | 0.18814989 | 3.10913098 | 0.00187639 | 0.03077721 |
| NADK2 | 94.6147548 | 0.80772011 | 0.39743568 | 2.03232915 | 0.04212035 | 0.20377313 |
| RANBP3L | 112.530602 | 6.008684734 | 1.39372205 | 4.3112504 | 1.62E-05 | 0.00130672 |
| NIPBL | 861.78271 | 0.234922658 | 0.12494803 | 1.88016289 | 0.06008588 | 0.24726898 |
| CPLANE1 | 369.858887 | 0.915268736 | 0.25107234 | 3.64543841 | 0.00026694 | 0.0089405 |
| NUP155 | 65.4500709 | 0.399843602 | 0.28559588 | 1.40003279 | 0.1615035 | 0.42791083 |

| | | | | | | |
|---|---|---|---|---|---|---|
| WDR70 | 419.082559 | 0.295615852 | 0.12894683 | 2.29254062 | 0.02187447 | 0.14205798 |
| LIFR | 514.686228 | 1.246177051 | 0.35084319 | 3.55194876 | 0.00038239 | 0.01120155 |
| LIFR-AS1 | 40.6295976 | 1.020515439 | 0.64456255 | 1.58326828 | 0.11336033 | 0.35267249 |
| OSMR-AS1 | 257.45247 | -1.207810864 | 0.48219708 | -2.5048075 | 0.0122518 | 0.10091982 |
| OSMR | 305.44699 | -1.477686954 | 0.4328274 | -3.4140328 | 0.00064009 | 0.01552255 |
| RICTOR | 443.279435 | 0.218958338 | 0.22434072 | 0.976008 | 0.3290605 | 0.61429707 |
| DAB2 | 46.0076101 | -2.488554163 | 0.48383242 | -5.1434217 | 2.70E-07 | 6.53E-05 |
| TTC33 | 84.7776716 | 0.336054212 | 0.23178271 | 1.44986745 | 0.14709548 | 0.4061769 |
| PRKAA1 | 136.017106 | 0.193336565 | 0.23174369 | 0.83426896 | 0.40412944 | 0.67765972 |
| RPL37 | 241.578301 | -0.682681972 | 0.27791064 | -2.4564801 | 0.01403056 | 0.10948271 |
| OXCT1 | 98.6550788 | 0.749211654 | 0.22748727 | 3.29342239 | 0.00098976 | 0.02061061 |
| C5orf51 | 29.4462532 | 0.423708101 | 0.27441941 | 1.54401651 | 0.12258433 | 0.367753 |
| GHR | 91.4387523 | 0.259879622 | 0.99471963 | 0.26125917 | 0.79389265 | 0.91723627 |
| SELENOP | 96.660551 | 0.562168404 | 0.37259165 | 1.50880569 | 0.13134845 | 0.38163088 |
| ZNF131 | 230.563578 | 0.112755688 | 0.14463836 | 0.77956971 | 0.43564419 | 0.70062219 |
| HMGCS1 | 255.781694 | 1.165054786 | 0.65039426 | 1.79130546 | 0.07324429 | 0.27896901 |
| CCL28 | 101.829091 | -0.825382343 | 0.404333 | -2.041343 | 0.04121675 | 0.20195151 |
| TMEM267 | 154.078147 | -0.161369304 | 0.2107731 | -0.7656067 | 0.44391033 | 0.70661084 |
| PAIP1 | 93.3662284 | -0.014680888 | 0.23818878 | -0.0616355 | 0.95085309 | 0.98286628 |
| NNT-AS1 | 82.9414164 | -0.104070509 | 0.2163741 | -0.4809749 | 0.63053435 | 0.83461713 |
| NNT | 165.802708 | -0.136119356 | 0.23249098 | -0.5854823 | 0.55822347 | 0.78914147 |
| MRPS30-DT | 38.5939905 | 0.337591052 | 0.54191783 | 0.62295616 | 0.53331333 | 0.77348803 |
| MRPS30 | 37.9485015 | -0.343960628 | 0.3090687 | -1.1128938 | 0.26575406 | 0.55384374 |
| PARP8 | 299.219714 | -0.462877053 | 0.35439268 | -1.3061135 | 0.19151397 | 0.46865601 |
| ITGA1 | 374.487528 | 0.669729452 | 0.44730337 | 1.49726002 | 0.13432561 | 0.38591741 |
| ITGA2 | 453.566004 | -2.227788938 | 0.9554651 | -2.3316278 | 0.01972028 | 0.13342561 |
| MOCS2 | 33.1431582 | 0.206594236 | 0.38594368 | 0.53529633 | 0.59244497 | 0.81283604 |
| NDUFS4 | 185.415302 | 0.085707043 | 0.15819096 | 0.54179483 | 0.58795985 | 0.81095283 |
| ARL15 | 1057.37465 | 1.017605555 | 0.20188386 | 5.04054943 | 4.64E-07 | 8.90E-05 |
| SNX18 | 34.780515 | 0.174331916 | 0.30567127 | 0.57032483 | 0.5684574 | 0.79713855 |
| AC026704.1 | 29.2629505 | -0.101967992 | 0.59472329 | -0.1714545 | 0.86386639 | 0.9495096 |
| DHX29 | 118.552193 | 0.065553199 | 0.15389007 | 0.4259742 | 0.67012665 | 0.85932305 |
| MTREX | 241.533275 | 0.267473297 | 0.14452822 | 1.85066489 | 0.06421778 | 0.25792121 |
| PLPP1 | 211.679224 | 0.609334578 | 0.22074828 | 2.76031399 | 0.00577458 | 0.06350716 |
| SLC38A9 | 149.750548 | 0.067104467 | 0.22547163 | 0.29761823 | 0.76599456 | 0.90371395 |
| IL6ST | 478.596591 | -0.681064524 | 0.2828247 | -2.4080801 | 0.01603666 | 0.1183476 |
| LINC01948 | 86.6360893 | 1.374503734 | 0.48382856 | 2.84089001 | 0.00449878 | 0.0540023 |
| MAP3K1 | 868.945136 | 1.001129483 | 0.2917211 | 3.43180341 | 0.00059958 | 0.0149187 |
| AC008937.2 | 38.3564326 | 1.030989775 | 0.51965762 | 1.98397896 | 0.04725819 | 0.21714794 |
| MIER3 | 46.8981829 | 0.467942932 | 0.21768223 | 2.14966072 | 0.03158206 | 0.17373701 |
| GPBP1 | 454.00761 | 0.650788666 | 0.19187014 | 3.39181839 | 0.0006943 | 0.01618185 |
| PDE4D | 3248.84549 | 0.609623154 | 0.3057388 | 1.99393455 | 0.04615921 | 0.21432487 |
| ELOVL7 | 208.03613 | 0.745954338 | 0.37261872 | 2.00192394 | 0.04529291 | 0.21178118 |
| ERCC8 | 92.5228749 | 0.171173865 | 0.1818447 | 0.94131896 | 0.34654143 | 0.6302171 |
| NDUFAF2 | 206.115784 | -0.353354482 | 0.32340038 | -1.0926223 | 0.27455965 | 0.56187466 |
| SMIM15 | 37.4709793 | -0.478325703 | 0.37883176 | -1.2626336 | 0.20672089 | 0.48621411 |
| ZSWIM6 | 775.10996 | -0.083122903 | 0.2324445 | -0.3576032 | 0.72064027 | 0.88235193 |
| KIF2A | 177.527741 | -0.474251832 | 0.20093943 | -2.3601731 | 0.01826641 | 0.12781151 |
| DIMT1 | 31.703595 | -0.212343333 | 0.35952385 | -0.5906238 | 0.55477253 | 0.78763392 |
| IPO11 | 90.0226474 | 0.068412707 | 0.17552538 | 0.38975963 | 0.6967143 | 0.87084339 |
| RNF180 | 378.123359 | 0.734970308 | 0.20745157 | 3.54285244 | 0.00039582 | 0.01152462 |
| SREK1IP1 | 86.7548882 | -0.093540266 | 0.17757307 | -0.5267706 | 0.59835293 | 0.81658679 |
| CWC27 | 206.980464 | 0.313756934 | 0.19640823 | 1.59747347 | 0.11016021 | 0.34724322 |
| PPWD1 | 85.2506264 | 0.628611567 | 0.25978586 | 2.41972971 | 0.01553205 | 0.11578325 |
| TRIM23 | 68.9759095 | 0.341125756 | 0.36247705 | 0.94109615 | 0.34665559 | 0.6302171 |
| TRAPPC13 | 74.4973211 | -0.108231382 | 0.21131026 | -0.5121918 | 0.60851678 | 0.82155842 |
| NLN | 74.3606903 | -0.819883938 | 0.44296724 | -1.8508907 | 0.06418528 | 0.25789884 |
| ERBIN | 730.754019 | 0.306136902 | 0.15051352 | 2.03394957 | 0.04195668 | 0.20339223 |
| SREK1 | 239.445662 | 0.461831686 | 0.11451653 | 4.03288244 | 5.51E-05 | 0.00317935 |
| MAST4 | 429.558482 | -0.503897361 | 0.27311049 | -1.8450311 | 0.06503301 | 0.26002617 |
| PIK3R1 | 374.069863 | -0.519287745 | 0.59887824 | -0.8671007 | 0.38588683 | 0.66275158 |
| SLC30A5 | 97.2404227 | 0.182528219 | 0.26359587 | 0.69245479 | 0.48865177 | 0.74185878 |
| MRPS36 | 63.1571905 | 0.151894436 | 0.33094682 | 0.45896931 | 0.6462562 | 0.84427583 |
| CDK7 | 65.1109256 | 0.002908128 | 0.19780221 | 0.0147022 | 0.98826976 | 0.99678172 |
| CCDC125 | 94.2516392 | 0.160480687 | 0.3629124 | 0.44220226 | 0.65834285 | 0.85215492 |
| AK6 | 45.8670158 | -0.123413475 | 0.34726945 | -0.3553825 | 0.72230308 | 0.88240546 |
| RAD17 | 83.1437107 | 0.321549835 | 0.18841023 | 1.70664747 | 0.0878876 | 0.30849223 |
| MARVELD2 | 66.3766354 | 0.063229916 | 0.4177161 | 0.15137055 | 0.87968342 | 0.95570914 |
| OCLN | 204.868549 | 0.157236206 | 0.30102161 | 0.52234192 | 0.60143229 | 0.81829073 |
| GTF2H2C | 82.0957676 | 0.299997752 | 0.2045553 | 1.46658506 | 0.14248897 | 0.39897744 |
| NAIP | 171.220488 | 0.813485895 | 0.29996961 | 2.71189438 | 0.00668999 | 0.06920456 |
| GTF2H2 | 70.2535039 | 0.313513208 | 0.48859744 | 0.64165954 | 0.52109426 | 0.76440016 |
| BDP1 | 317.106352 | 0.228128464 | 0.19750506 | 1.15505126 | 0.24806943 | 0.53495207 |
| MCCC2 | 133.740742 | -0.094672383 | 0.32444212 | -0.2918005 | 0.77043914 | 0.90533273 |
| MRPS27 | 132.013999 | -0.01085693 | 0.18735879 | -0.0579473 | 0.95379063 | 0.98368305 |
| TNPO1 | 297.69961 | 0.094543224 | 0.15915232 | 0.5940424 | 0.55248376 | 0.78613936 |
| FCHO2 | 377.99333 | 0.01899842 | 0.16339009 | 0.11627645 | 0.90743345 | 0.96677845 |
| TMEM171 | 34.8442855 | 0.030221886 | 0.31591726 | 0.09566393 | 0.92378749 | 0.9730548 |
| BTF3 | 173.872514 | -0.551983692 | 0.4512168 | -1.2233226 | 0.22120788 | 0.50310453 |
| ANKRA2 | 44.0050559 | 0.6839485 | 0.32934612 | 2.07668605 | 0.03783055 | 0.19316567 |
| ARHGEF28 | 868.746507 | 0.171193552 | 0.29324223 | 0.58379569 | 0.55935778 | 0.78981496 |
| HEXB | 71.3754302 | -0.317878608 | 0.30189467 | -1.0529454 | 0.29236601 | 0.57982899 |
| GFM2 | 59.7717179 | 0.542108876 | 0.22046511 | 2.45893272 | 0.01393507 | 0.10914478 |
| NSA2 | 72.976378 | -0.434802427 | 0.28815979 | -1.5088935 | 0.13132601 | 0.38163088 |
| FAM169A | 63.8846368 | 0.209718273 | 0.49586976 | 0.42293015 | 0.67234622 | 0.86015189 |
| HMGCR | 82.2317085 | 0.952233804 | 0.35840608 | 2.65685729 | 0.00788728 | 0.07685888 |
| COL4A3BP | 286.243168 | 0.070608082 | 0.22433397 | 0.31474539 | 0.75295498 | 0.89708405 |
| POLK | 220.43502 | 0.173484263 | 0.25710238 | 0.67476723 | 0.49982367 | 0.75138275 |
| ANKDD1B | 50.8422743 | 0.251805831 | 0.38903782 | 0.64725283 | 0.51746832 | 0.76270643 |
| POC5 | 44.8110133 | -0.16521034 | 0.31083142 | -0.5315111 | 0.59506468 | 0.81511365 |
| IQGAP2 | 68.1697766 | -0.10498637 | 0.74568041 | -0.1407927 | 0.8880337 | 0.95907089 |
| F2RL1 | 42.525241 | 0.236978901 | 0.61279614 | 0.38671735 | 0.69896547 | 0.8717697 |
| ZBED3 | 44.1212749 | -0.425523364 | 0.22442577 | -1.8960539 | 0.05795292 | 0.24313032 |
| WDR41 | 137.046039 | 0.018348405 | 0.18400157 | 0.09971874 | 0.92056762 | 0.97180045 |
| TBCA | 338.269192 | 0.050031013 | 0.21379355 | 0.23401554 | 0.81497291 | 0.92814476 |
| AP3B1 | 446.968779 | 0.077053237 | 0.22343976 | 0.34485016 | 0.73020702 | 0.88730852 |
| SCAMP1 | 166.572638 | -3.44E-05 | 0.21607795 | -0.0001594 | 0.9998728 | 0.9998728 |
| LHFPL2 | 83.2563207 | -0.273171755 | 0.34704737 | -0.787131 | 0.43120516 | 0.69681733 |
| ARSB | 148.808208 | -0.652585405 | 0.4365675 | -1.4948099 | 0.13496407 | 0.38712403 |
| DMGDH | 148.569777 | 0.368929465 | 0.29784034 | 1.23868201 | 0.21546328 | 0.49731142 |
| BHMT2 | 205.261606 | -0.432934717 | 0.42665814 | -1.014711 | 0.3102436 | 0.5972314 |
| BHMT | 76.4332253 | 0.959150978 | 0.70448823 | 1.36148615 | 0.17336011 | 0.44449049 |
| JMY | 156.39633 | 0.677716186 | 0.24616805 | 2.75306311 | 0.00590405 | 0.0645604 |
| HOMER1 | 494.248102 | 2.658943992 | 0.58488201 | 4.54612029 | 5.46E-06 | 0.00059481 |
| TENT2 | 478.074418 | 0.339348573 | 0.15988279 | 2.12248347 | 0.03379716 | 0.18066599 |
| CMYA5 | 40.8267027 | 0.068820174 | 0.57406441 | 0.11988232 | 0.90457637 | 0.9654526 |
| MTX3 | 39.7143566 | 0.370472465 | 0.26486687 | 1.39871197 | 0.16189937 | 0.4283668 |
| THBS4 | 36.9100983 | 0.736383605 | 0.427381 | 1.72301436 | 0.08488593 | 0.30278126 |
| SERINC5 | 265.685949 | 0.879571859 | 0.27427768 | 3.20686632 | 0.00134189 | 0.02525343 |
| ZFYVE16 | 212.070875 | 0.498462168 | 0.18688024 | 2.66728131 | 0.00764676 | 0.07534671 |
| FAM151B | 50.6209919 | 0.710361621 | 0.31280493 | 2.27094127 | 0.02315053 | 0.14676719 |
| DHFR | 382.35303 | -0.768630479 | 0.35205382 | -2.1832755 | 0.02901553 | 0.1656375 |
| MSH3 | 260.377228 | -0.088946989 | 0.19342655 | -0.4598489 | 0.64562467 | 0.84407726 |

| | | | | | | |
|---|---|---|---|---|---|---|
| RASGRF2 | 89.7059179 | 0.43200066 | 0.69322137 | 0.62317851 | 0.53316722 | 0.77346794 |
| CKMT2-AS1 | 118.823258 | 0.506837579 | 0.2089309 | 2.42586227 | 0.01527207 | 0.11471025 |
| ZCCHC9 | 32.3326176 | 0.129287081 | 0.24281681 | 0.532447 | 0.59441644 | 0.81481326 |
| SSBP2 | 556.277144 | 0.178436376 | 0.32883946 | 0.54262459 | 0.58738831 | 0.8106314 |
| ATG10 | 258.525668 | 0.514159818 | 0.26582884 | 1.9341762 | 0.05309147 | 0.23116508 |
| RPS23 | 602.0299 | -0.527972997 | 0.52285192 | -1.0097945 | 0.31259375 | 0.60006724 |
| ATP6AP1L | 103.625963 | 0.361647687 | 0.42385695 | 0.85323053 | 0.39353148 | 0.6686322 |
| TMEM167A | 66.1493917 | 0.089390925 | 0.21446216 | 0.41681444 | 0.67681414 | 0.8613973 |
| XRCC4 | 113.097258 | -0.127291901 | 0.2753771 | -0.4622458 | 0.64390508 | 0.84344334 |
| VCAN | 177.060576 | -1.050048309 | 1.04705323 | -1.0028605 | 0.31592818 | 0.60248379 |
| EDIL3 | 709.504502 | 0.062842823 | 0.23168495 | 0.27124258 | 0.78620447 | 0.91307621 |
| AC113383.1 | 179.502094 | 0.191116565 | 0.28899439 | 0.66131582 | 0.5084098 | 0.75751399 |
| COX7C | 209.729567 | -0.747984721 | 0.36957948 | -2.0238806 | 0.04298244 | 0.20605022 |
| RASA1 | 405.849713 | 0.232082448 | 0.12984384 | 1.78739668 | 0.0738734 | 0.28056182 |
| CCNH | 513.138938 | 0.450977203 | 0.26704782 | 1.68875071 | 0.09126721 | 0.31502156 |
| TMEM161B | 100.368251 | -0.018561872 | 0.24278549 | -0.0764538 | 0.93905808 | 0.97996404 |
| TMEM161B-. | 289.78077 | 0.487342963 | 0.24346727 | 2.00167754 | 0.04531942 | 0.21178118 |
| MEF2C | 50.1909273 | 0.048825194 | 0.81595756 | 0.05983791 | 0.95228473 | 0.98341878 |
| CETN3 | 35.3862203 | 0.613930067 | 0.31994354 | 1.91887003 | 0.05500078 | 0.23554932 |
| LYSMD3 | 38.0902571 | 0.005429524 | 0.28705798 | 0.01891438 | 0.98490941 | 0.99668665 |
| ADGRV1 | 176.447252 | 1.718297838 | 0.64270573 | 2.67353746 | 0.00750559 | 0.07454435 |
| ARRDC3 | 327.87926 | -1.060266973 | 0.4821434 | -2.1990698 | 0.02787296 | 0.16152154 |
| ARRDC3-AS1 | 31.3058505 | -0.112160525 | 0.35948283 | -0.3120052 | 0.75503655 | 0.89810084 |
| AC124854.1 | 194.308132 | 5.832404501 | 1.48410115 | 3.92992385 | 8.50E-05 | 0.00419564 |
| NR2F1-AS1 | 49.3105778 | -0.9039426 | 0.86183293 | -1.0488606 | 0.29424228 | 0.58223301 |
| FAM172A | 628.492184 | 0.101012864 | 0.13453429 | 0.75083361 | 0.4527528 | 0.71261515 |
| KIAA0825 | 359.790835 | 0.245204665 | 0.25894423 | 0.94694006 | 0.34366932 | 0.62848577 |
| SLF1 | 46.6901684 | -0.096559007 | 0.3352623 | -0.2880103 | 0.77333883 | 0.90643453 |
| MCTP1 | 154.962822 | 1.059263892 | 0.54244875 | 1.95274466 | 0.05084986 | 0.22655386 |
| TTC37 | 247.051451 | 0.191101009 | 0.1902326 | 1.004565 | 0.31510636 | 0.60205159 |
| ARSK | 35.1060184 | 0.205160373 | 0.306885 | 0.66852525 | 0.50379837 | 0.7548701 |
| RHOBTB3 | 361.323125 | 1.0450146 | 0.41630046 | 2.51024127 | 0.01206487 | 0.10031173 |
| GLRX | 111.668584 | -0.064870866 | 0.49142454 | -0.1320058 | 0.89497974 | 0.96120764 |
| ELL2 | 212.027257 | -0.3343444 | 0.48225687 | -0.6932911 | 0.48812688 | 0.74136156 |
| AC104123.1 | 101.477166 | -0.337249306 | 0.2698273 | -1.2498709 | 0.21134669 | 0.49233706 |
| CAST | 463.935551 | -0.156877916 | 0.17374062 | -0.9029432 | 0.36655602 | 0.64650131 |
| ERAP1 | 107.516953 | 0.029852821 | 0.25085639 | 0.11900363 | 0.90527248 | 0.96556525 |
| AC009126.1 | 58.2728492 | 0.762144829 | 0.48357656 | 1.57605826 | 0.11501238 | 0.35665379 |
| ERAP2 | 58.6982468 | -0.195612605 | 0.4122934 | -0.47445 | 0.63517904 | 0.83749209 |
| LNPEP | 363.989003 | 0.485504271 | 0.17150681 | 2.83081621 | 0.00464294 | 0.05512702 |
| LIX1-AS1 | 78.8412346 | 0.782631581 | 0.28075993 | 2.78754729 | 0.00531087 | 0.06034736 |
| LIX1 | 40.9182845 | 3.677068341 | 1.08516428 | 3.38849002 | 0.00070279 | 0.01630021 |
| CHD1 | 319.605418 | 0.340245221 | 0.13819325 | 2.46209727 | 0.01381272 | 0.10854146 |
| LINC02062 | 97.4856384 | -0.071506756 | 0.444351 | -0.160924 | 0.87215323 | 0.95193455 |
| FAM174A | 36.3260483 | -0.013987525 | 0.29159509 | -0.047969 | 0.96174095 | 0.9875899 |
| SLCO4C1 | 165.676042 | 1.553447211 | 0.41608449 | 3.73348984 | 0.00018884 | 0.00723578 |
| PAM | 720.668549 | -0.240166713 | 0.27791044 | -0.8641875 | 0.38748493 | 0.66406384 |
| GIN1 | 31.9984679 | 0.157313557 | 0.36872536 | 0.42664154 | 0.66964043 | 0.85924744 |
| PPIP5K2 | 200.073892 | 0.414585004 | 0.1623675 | 2.55337434 | 0.01066848 | 0.09276247 |
| NUDT12 | 29.9766002 | 0.27643115 | 0.29753113 | 0.92908312 | 0.35284601 | 0.63441164 |
| EFNA5 | 995.48974 | -0.830256614 | 0.82415248 | -1.0074066 | 0.31373944 | 0.60082169 |
| FBXL17 | 708.464607 | 0.151912608 | 0.13247418 | 1.14673372 | 0.25149169 | 0.53881433 |
| FER | 786.727351 | 0.661984052 | 0.156161 | 4.23911242 | 2.24E-05 | 0.00164188 |
| PJA2 | 257.877957 | -0.162654024 | 0.18476209 | -0.880343 | 0.3786735 | 0.65652733 |
| MAN2A1 | 298.857381 | -0.581797459 | 0.36341033 | -1.6009381 | 0.10939062 | 0.34606201 |
| TMEM232 | 308.496278 | 0.077716858 | 0.42938722 | 0.18099481 | 0.85637165 | 0.94627061 |
| SLC25A46 | 103.720427 | 0.30045708 | 0.22217891 | 1.35232045 | 0.17627282 | 0.44835649 |
| WDR36 | 75.669084 | 0.162968135 | 0.32509388 | 0.5012956 | 0.6161631 | 0.82592881 |
| STARD4 | 46.1385867 | 0.927017745 | 0.56210574 | 1.64918748 | 0.09910923 | 0.32904321 |
| NREP | 108.493285 | 0.471146902 | 0.55195708 | 0.85359336 | 0.39333034 | 0.6686322 |
| EPB41L4A | 440.221859 | 0.198449015 | 0.19326391 | 1.02682916 | 0.30450092 | 0.59248716 |
| EPB41L4A-A! | 27.828658 | -0.110584793 | 0.55660271 | -0.1986781 | 0.84251452 | 0.94132995 |
| APC | 128.614502 | 0.266856937 | 0.19166753 | 1.3922908 | 0.16383436 | 0.43150104 |
| SRP19 | 62.8904529 | 0.130166144 | 0.25210521 | 0.51631676 | 0.60563319 | 0.820591 |
| REEP5 | 276.652106 | 0.026515143 | 0.21205035 | 0.12504173 | 0.90049051 | 0.96303857 |
| DCP2 | 117.417401 | 0.255370181 | 0.1970262 | 1.29612296 | 0.19493312 | 0.47344432 |
| MCC | 70.3339852 | -0.293685732 | 0.81064876 | -0.3622848 | 0.71713921 | 0.88104097 |
| YTHDC2 | 182.810686 | 0.46340556 | 0.18444066 | 2.51249136 | 0.0119882 | 0.10009484 |
| PGGT1B | 70.1373486 | -0.209400625 | 0.19218944 | -1.0895532 | 0.27590999 | 0.56304683 |
| FEM1C | 50.3641573 | -0.734138323 | 0.339464 | -2.1626397 | 0.0305689 | 0.1711394 |
| TMED7-TICA | 28.4395444 | -0.289329343 | 0.30655997 | -0.9437936 | 0.34527512 | 0.62997912 |
| ATG12 | 64.7790137 | 0.122746191 | 0.27762752 | 0.44212545 | 0.65839843 | 0.85215492 |
| AP3S1 | 101.845549 | -0.017742437 | 0.30330314 | -0.0584974 | 0.95335246 | 0.98368305 |
| COMMD10 | 480.92559 | 0.259102984 | 0.25789945 | 1.00466668 | 0.31505738 | 0.60205159 |
| SEMA6A | 184.978893 | -2.02067722 | 0.47574598 | -4.2473869 | 2.16E-05 | 0.00160599 |
| SEMA6A-AS1 | 78.4903191 | 0.084719949 | 0.31036554 | 0.27296828 | 0.7848776 | 0.91212179 |
| DTWD2 | 111.175387 | 0.420049015 | 0.19232867 | 2.18401661 | 0.02896103 | 0.16552368 |
| AC008629.1 | 60.1075133 | 0.725974166 | 0.52170756 | 1.39153468 | 0.16406335 | 0.43198538 |
| DMXL1 | 404.593726 | 0.455582901 | 0.15829055 | 2.87814341 | 0.00400023 | 0.04944287 |
| TNFAIP8 | 380.8644 | -0.701376922 | 0.31887946 | -2.1995049 | 0.02784204 | 0.16152154 |
| HSD17B4 | 141.785285 | 0.057220741 | 0.2460277 | 0.23257845 | 0.81608876 | 0.92869214 |
| SRFBP1 | 94.9675923 | -0.449287147 | 0.27055214 | -1.6606306 | 0.09678767 | 0.32454105 |
| SNX2 | 192.223231 | 0.725472877 | 0.27264542 | 2.66086585 | 0.007794 | 0.07633817 |
| SNX24 | 226.355322 | 0.15281497 | 0.16365904 | 0.93373985 | 0.35043811 | 0.63322593 |
| PPIC | 46.0536206 | -0.466155554 | 0.36155483 | -1.2893081 | 0.19729101 | 0.47630586 |
| CEP120 | 68.2657963 | 0.530563978 | 0.32552545 | 1.62986941 | 0.1031291 | 0.33528295 |
| CSNK1G3 | 173.726367 | 0.220993588 | 0.16411705 | 1.34656078 | 0.17812173 | 0.44971747 |
| ZNF608 | 329.495154 | 0.366749384 | 0.5964778 | 0.6148584 | 0.53864824 | 0.77706498 |
| GRAMD2B | 485.74216 | 0.007454737 | 0.28968146 | 0.02573426 | 0.9794693 | 0.99557142 |
| ALDH7A1 | 93.1244351 | 0.258039211 | 0.24146052 | 1.06866006 | 0.28522287 | 0.57265771 |
| PHAX | 43.5892902 | -0.332082702 | 0.23077209 | -1.4390072 | 0.15014847 | 0.4109827 |
| C5orf63 | 32.139416 | -0.468311854 | 0.29129607 | -1.6076834 | 0.10790454 | 0.34350867 |
| PRRC1 | 76.1781234 | -0.172366571 | 0.30447308 | -0.5661143 | 0.57131607 | 0.79857531 |
| LINC01184 | 186.369039 | -0.143222383 | 0.25103226 | -0.5705338 | 0.56831572 | 0.79713855 |
| SLC12A2 | 64.1303693 | -0.225149524 | 0.3310509 | -0.6801055 | 0.49643768 | 0.74817127 |
| FBN2 | 94.6640791 | 1.9466412 | 0.40895425 | 4.76004638 | 1.94E-06 | 0.00028091 |
| SLC27A6 | 59.8588172 | 2.202995323 | 0.88994495 | 2.47542877 | 0.01330763 | 0.10613971 |
| HINT1 | 536.086282 | -0.793814417 | 0.4764179 | -1.6662145 | 0.0956707 | 0.3225729 |
| LYRM7 | 48.7888708 | 0.220714875 | 0.25560885 | 0.86348684 | 0.38786985 | 0.66428354 |
| CDC42SE2 | 850.329627 | -0.382914634 | 0.38516187 | -0.9941655 | 0.32014231 | 0.60701567 |
| P4HA2 | 33.9475311 | -0.599531232 | 0.38670548 | -1.5503562 | 0.12105605 | 0.36534213 |
| SLC22A4 | 41.8732414 | -0.584871728 | 0.34335095 | -1.7034225 | 0.08848903 | 0.30970853 |
| SLC22A5 | 101.646185 | -0.660751907 | 0.20049476 | -3.2956069 | 0.00098209 | 0.02054528 |
| AC116366.3 | 182.101624 | -0.302892494 | 0.5767463 | -0.5251746 | 0.59946184 | 0.81763227 |
| IRF1 | 75.75095 | -2.140794548 | 0.60761986 | -3.5232465 | 0.00042629 | 0.01208129 |
| KIF3A | 108.386047 | 0.654626012 | 0.23063811 | 2.83832541 | 0.00453509 | 0.05423427 |
| 8-Sep | 107.561893 | 1.136083271 | 0.40395195 | 2.81242178 | 0.004917 | 0.05736897 |
| UQCRQ | 75.5821885 | -0.667730241 | 0.39644702 | -1.6842862 | 0.09212637 | 0.31603732 |
| AFF4 | 487.523361 | 0.07332375 | 0.15538751 | 0.47187673 | 0.63701478 | 0.83872935 |
| ZCCHC10 | 68.2009293 | -0.036347272 | 0.23044721 | -0.1577249 | 0.87467355 | 0.95285999 |
| HSPA4 | 135.85107 | -0.032672869 | 0.20910111 | -0.1562539 | 0.87583288 | 0.95314737 |
| C5orf15 | 98.0489657 | -0.18926458 | 0.34106233 | -0.5549267 | 0.57894482 | 0.80419264 |
| VDAC1 | 135.930694 | -0.819755903 | 0.3228896 | -2.5388117 | 0.01112297 | 0.09521617 |

| | | | | | | |
|---|---|---|---|---|---|---|
| SKP1 | 269.677213 | 0.172263439 | 0.42075849 | 0.40941168 | 0.68223757 | 0.86398119 |
| CDKL3 | 92.3939757 | 0.897367336 | 0.27992585 | 3.20573226 | 0.00134719 | 0.02530345 |
| UBE2B | 200.674224 | -0.083474319 | 0.20827257 | -0.4007936 | 0.68857207 | 0.86751885 |
| JADE2 | 90.2041167 | -0.165172777 | 0.25021685 | -0.6601185 | 0.50917777 | 0.75806868 |
| SAR1B | 90.9002907 | 0.066910974 | 0.2447518 | 0.27338297 | 0.78455884 | 0.91212179 |
| SEC24A | 100.069245 | 0.014020862 | 0.25012437 | 0.05605556 | 0.95529754 | 0.98395647 |
| CAMLG | 50.5058357 | 0.025378322 | 0.34407833 | 0.0737574 | 0.94120343 | 0.9805098 |
| DDX46 | 190.941249 | -0.010294456 | 0.21619145 | -0.0476173 | 0.96202123 | 0.9875899 |
| C5orf24 | 72.0270578 | 0.07705663 | 0.31019257 | 0.24841546 | 0.80381297 | 0.92125386 |
| TXNDC15 | 51.1725918 | -0.328541797 | 0.2234514 | -1.4703054 | 0.14147906 | 0.39754411 |
| PCBD2 | 140.192858 | -0.169536907 | 0.32177334 | -0.526883 | 0.59827483 | 0.81658679 |
| H2AFY | 177.079391 | -0.358128822 | 0.32001251 | -1.1191088 | 0.26309371 | 0.55096925 |
| CXCL14 | 174.036325 | -1.192552156 | 0.45240528 | -2.6360262 | 0.00838833 | 0.08075639 |
| SLC25A48 | 62.1872102 | 1.593675179 | 0.54707292 | 2.91309461 | 0.00357866 | 0.04613728 |
| SMAD5 | 188.470334 | 0.069201862 | 0.14365625 | 0.48171841 | 0.63000599 | 0.83433967 |
| SPOCK1 | 169.635362 | -0.386782764 | 0.56886601 | -0.6799189 | 0.4965558 | 0.74823156 |
| HNRNPA0 | 87.1809461 | -0.758738484 | 0.39156581 | -1.9377036 | 0.0526594 | 0.23008398 |
| FAM13B | 334.5004 | 0.423384619 | 0.21655842 | 1.95505957 | 0.05057604 | 0.2259645 |
| NME5 | 84.5390582 | 0.651828438 | 0.37730036 | 1.72761148 | 0.0840579 | 0.30203879 |
| BRD8 | 72.9282436 | 0.513736257 | 0.17718818 | 2.89938213 | 0.00373899 | 0.04737536 |
| CDC23 | 37.8399091 | 0.364462209 | 0.24370311 | 1.49551724 | 0.1347795 | 0.38689028 |
| FAM53C | 33.3240479 | 0.226123446 | 0.33069869 | 0.68377484 | 0.49411736 | 0.74643592 |
| KDM3B | 202.65299 | 0.116845847 | 0.16104596 | 0.7255435 | 0.46811867 | 0.72581883 |
| EGR1 | 252.357162 | 2.557731331 | 0.70619235 | 3.62186212 | 0.00029249 | 0.0094975 |
| ETF1 | 119.307538 | -0.231524625 | 0.30808443 | -0.7514973 | 0.45235341 | 0.71244547 |
| HSPA9 | 207.080053 | -0.65075484 | 0.26340974 | -2.4705041 | 0.01349228 | 0.10712353 |
| CTNNA1 | 728.858003 | 0.267106526 | 0.17093579 | 1.56261324 | 0.11814359 | 0.36283985 |
| SIL1 | 302.113131 | -0.297402738 | 0.21459926 | -1.3858516 | 0.16579221 | 0.43415078 |
| PAIP2 | 149.474953 | 0.121507041 | 0.23028606 | 0.52763525 | 0.59775252 | 0.81658679 |
| UBE2D2 | 440.442426 | -0.443842124 | 0.2600083 | -1.7070306 | 0.08781636 | 0.30835519 |
| CXXC5 | 214.860714 | 0.020383649 | 0.39952854 | 0.05101926 | 0.95931018 | 0.98628659 |
| PURA | 195.377642 | -0.321766771 | 0.24721294 | -1.3015774 | 0.19306089 | 0.47104693 |
| IGIP | 32.148248 | 0.341598112 | 0.36882204 | 0.92618683 | 0.3543489 | 0.63611471 |
| CYSTM1 | 524.352624 | 0.382314302 | 0.42919305 | 0.89077468 | 0.37305006 | 0.65217873 |
| PFDN1 | 97.5723224 | 0.211762011 | 0.2480941 | 0.8535552 | 0.39335149 | 0.6686322 |
| SRA1 | 40.0985148 | -0.564485592 | 0.33479367 | -1.6860701 | 0.0917823 | 0.31558127 |
| SLC35A4 | 41.9368217 | -0.774388877 | 0.35572437 | -2.1769352 | 0.02948541 | 0.16772013 |
| NDUFA2 | 71.6378362 | -0.448796574 | 0.40612003 | -1.1050836 | 0.26912338 | 0.55632249 |
| IK | 45.547807 | -0.656529915 | 0.32152879 | -2.0419009 | 0.04116136 | 0.20195151 |
| HARS | 59.8738836 | -0.236492201 | 0.2598415 | -0.9101402 | 0.36274857 | 0.64323742 |
| ZMAT2 | 36.7256328 | -0.467107726 | 0.35251404 | -1.3250755 | 0.1851461 | 0.45922178 |
| TAF7 | 91.7216862 | -0.148107916 | 0.286045 | -0.5177784 | 0.60461291 | 0.82048068 |
| DIAPH1 | 445.248476 | -0.127172872 | 0.33018851 | -0.3851523 | 0.70012456 | 0.87182648 |
| HDAC3 | 61.0738057 | 0.22018771 | 0.29264267 | 0.7524115 | 0.45180363 | 0.71231034 |
| DELE1 | 57.302293 | -0.439799401 | 0.27932623 | -1.5745009 | 0.11537169 | 0.35752108 |
| RNF14 | 57.0290812 | -0.078420659 | 0.28717463 | -0.2730766 | 0.78479437 | 0.91212179 |
| GNPDA1 | 43.783963 | -0.851770417 | 0.47476591 | -1.794085 | 0.0727996 | 0.27853529 |
| NDFIP1 | 284.486906 | -0.011391475 | 0.16390184 | -0.0695018 | 0.94459019 | 0.98115679 |
| SPRY4 | 36.9461429 | -1.285147698 | 0.44487886 | -2.8887588 | 0.00386766 | 0.04842913 |
| SPRY4-AS1 | 160.535549 | -0.017390795 | 0.61117594 | -0.0284546 | 0.97729954 | 0.9945344 |
| FGF1 | 30.378492 | 0.492961354 | 0.35732959 | 1.37957048 | 0.16771893 | 0.43654799 |
| ARHGAP26 | 1250.67734 | -0.559688834 | 0.41229327 | -1.3575017 | 0.17462184 | 0.44605403 |
| NR3C1 | 372.108384 | -0.743473506 | 0.48355275 | -1.5375231 | 0.12416527 | 0.37041562 |
| YIPF5 | 39.4514477 | -0.271975081 | 0.24017442 | -1.1324065 | 0.25746357 | 0.54481971 |
| KCTD16 | 448.53238 | 1.752894637 | 0.49771034 | 3.52191723 | 0.00042844 | 0.01209909 |
| PRELID2 | 240.665679 | 0.26470761 | 0.38483534 | 0.68784641 | 0.49154951 | 0.74421671 |
| SH3RF2 | 46.2392157 | 0.560925541 | 0.73639838 | 0.7617148 | 0.44623024 | 0.70815763 |
| LARS | 181.377089 | 0.157246324 | 0.18598001 | 0.8455012 | 0.39783106 | 0.67281493 |
| TCERG1 | 208.262278 | 0.200108465 | 0.22173703 | 0.90245847 | 0.36681337 | 0.6466739 |
| PPP2R2B | 184.697144 | 1.231557792 | 0.68910596 | 1.78718203 | 0.07390807 | 0.28056182 |
| DPYSL3 | 84.6528832 | -0.963850704 | 1.2303446 | -0.783399 | 0.43339285 | 0.69833957 |
| AC091948.1 | 56.459383 | 1.030315626 | 0.68818208 | 1.49715556 | 0.13435278 | 0.38591741 |
| FBXO38 | 92.8976076 | -0.064085692 | 0.16573326 | -0.3866797 | 0.69899333 | 0.8717697 |
| CSNK1A1 | 669.000982 | -0.076895174 | 0.13048938 | -0.589283 | 0.55567144 | 0.78784845 |
| ARHGEF37 | 48.3657727 | 0.954171668 | 0.49519233 | 1.92687086 | 0.05399573 | 0.23338748 |
| SLC26A2 | 45.9440241 | -0.567264148 | 0.44581532 | -1.2724196 | 0.20322408 | 0.48221041 |
| HMGXB3 | 78.6618025 | -0.228763328 | 0.24530575 | -0.9325641 | 0.35104508 | 0.63331896 |
| PDGFRB | 62.6104188 | 0.251270219 | 1.49958929 | 0.16755936 | 0.86692995 | 0.95090501 |
| TCOF1 | 34.4458409 | -0.418823799 | 0.43065606 | -0.972525 | 0.33078942 | 0.61598598 |
| CD74 | 223.14648 | -2.132953696 | 0.56905946 | -3.7482088 | 0.0001781 | 0.00704975 |
| RPS14 | 485.29131 | -0.332995985 | 0.44672693 | -0.7454128 | 0.45602219 | 0.71539502 |
| NDST1 | 99.0794664 | 0.284431545 | 0.27405569 | 1.03786038 | 0.29933506 | 0.58660608 |
| SYNPO | 37.6440914 | -0.110136045 | 0.61071175 | -0.1803405 | 0.85688528 | 0.94638973 |
| RBM22 | 33.2643796 | -0.013606577 | 0.25311857 | -0.0537557 | 0.95712977 | 0.98520804 |
| DCTN4 | 154.405611 | 0.318795509 | 0.14927532 | 2.135621 | 0.03271032 | 0.17715978 |
| SMIM3 | 34.1612641 | -1.410231117 | 0.64442741 | -2.1883475 | 0.0286443 | 0.16440009 |
| GPX3 | 1413.61785 | -1.49994592 | 0.7657227 | -1.9588631 | 0.05012882 | 0.22501594 |
| TNIP1 | 199.117646 | -1.644957998 | 0.44081573 | -3.7316228 | 0.00019025 | 0.00726059 |
| SLC36A1 | 28.7287273 | -0.441464279 | 0.38483355 | -1.1471564 | 0.25131698 | 0.53868382 |
| SPARC | 134.709833 | 0.830987429 | 1.17147661 | 0.70935042 | 0.47810704 | 0.7341756 |
| ATOX1 | 87.5958171 | -0.085575953 | 0.26102633 | -0.3278441 | 0.74302951 | 0.89269781 |
| G3BP1 | 228.451389 | -0.315759639 | 0.18217757 | -1.733252 | 0.08305088 | 0.29986597 |
| FAM114A2 | 36.0423072 | 0.240325808 | 0.22606695 | 1.0630736 | 0.28774858 | 0.57450105 |
| MFAP3 | 50.2001196 | 0.067182717 | 0.3106596 | 0.2162583 | 0.82878641 | 0.933664 |
| GALNT10 | 198.633443 | 0.27212055 | 0.36898318 | 0.73748769 | 0.46082583 | 0.71918385 |
| SAP30L-AS1 | 44.1847036 | 0.150689193 | 0.30058071 | 0.50132689 | 0.61614109 | 0.82592881 |
| SAP30L | 25.2855158 | 0.0468798 | 0.36109392 | 0.12982717 | 0.89670317 | 0.96175013 |
| LARP1 | 320.616811 | -0.194828103 | 0.25035074 | -0.7782206 | 0.43643897 | 0.70145116 |
| FAXDC2 | 45.2977725 | 0.66819847 | 0.58191362 | 1.14827777 | 0.25085391 | 0.53838647 |
| CNOT8 | 56.9808129 | 0.089525262 | 0.22138668 | 0.40438413 | 0.68593027 | 0.8662366 |
| GEMIN5 | 47.2611763 | -0.330966813 | 0.23305512 | -1.4201225 | 0.15557202 | 0.41978152 |
| MRPL22 | 47.2260203 | -0.174907811 | 0.25577811 | -0.6838264 | 0.49408482 | 0.74643592 |
| SGCD | 100.074248 | 2.045436421 | 1.08146039 | 1.89136508 | 0.05857563 | 0.24443767 |
| TIMD4 | 47.0943776 | -0.971767455 | 0.63890774 | -1.5209824 | 0.12826424 | 0.37610564 |
| HAVCR1 | 157.421324 | -1.050592448 | 0.94563715 | -1.110989 | 0.26657309 | 0.55450677 |
| CYFIP2 | 199.311642 | 1.134404376 | 0.55678769 | 2.03740924 | 0.04160905 | 0.20312121 |
| CLINT1 | 279.101766 | -0.289691705 | 0.26558359 | -1.0907741 | 0.27537229 | 0.56268552 |
| EBF1 | 231.1484 | -3.02000884 | 1.36257319 | -2.2164012 | 0.02666404 | 0.15827108 |
| RNF145 | 135.27202 | 0.042418566 | 0.22610859 | 0.18760262 | 0.85118817 | 0.94435157 |
| UBLCP1 | 45.04094 | 0.309381482 | 0.30188493 | 1.02483248 | 0.30544224 | 0.59275349 |
| TTC1 | 112.151508 | -0.142249127 | 0.22478111 | -0.632834 | 0.52684206 | 0.76821934 |
| PWWP2A | 155.298041 | 0.417924238 | 0.21449271 | 1.94843095 | 0.05136342 | 0.22752396 |
| CCNJL | 69.0769829 | 1.253382779 | 0.71855632 | 1.74430695 | 0.08110558 | 0.29596585 |
| SLU7 | 35.0483533 | 0.245950266 | 0.25220101 | 0.97521524 | 0.3294535 | 0.61457354 |
| AC113414.1 | 92.724953 | -0.082431525 | 0.371796 | -0.2217117 | 0.82453832 | 0.93217972 |
| CCNG1 | 78.1497937 | -0.97748393 | 0.35358454 | -2.7644985 | 0.00570104 | 0.06312976 |
| NUDCD2 | 31.697554 | -0.319272596 | 0.32014015 | -0.9972901 | 0.31862373 | 0.60557475 |
| MAT2B | 66.802831 | 0.144001755 | 0.29973993 | 0.48042233 | 0.63092712 | 0.83470856 |
| TENM2 | 56.1934909 | -0.916760725 | 0.6823645 | -1.3435059 | 0.17910822 | 0.45101935 |
| WWC1 | 1722.61598 | -0.539717082 | 0.39848987 | -1.354406 | 0.17560687 | 0.44761528 |
| RARS | 113.493659 | -0.618207486 | 0.18788178 | -3.2904068 | 0.00100043 | 0.02077084 |
| PANK3 | 106.457504 | 0.085466377 | 0.18582264 | 0.45993523 | 0.64556271 | 0.84407726 |
| SLIT3 | 857.557353 | 1.824002023 | 0.75260016 | 2.42360038 | 0.01536751 | 0.11509409 |

| | | | | | | |
|---|---|---|---|---|---|---|
| RANBP17 | 156.929628 | -0.657724884 | 0.29902429 | -2.1995701 | 0.02783741 | 0.16152154 |
| NPM1 | 177.084114 | -0.503600311 | 0.32220592 | -1.5629766 | 0.1180581 | 0.36280454 |
| FBXW11 | 383.962615 | 0.011398643 | 0.16430872 | 0.06937333 | 0.94469246 | 0.98115679 |
| UBTD2 | 113.170248 | -0.134549882 | 0.23696812 | -0.5677974 | 0.57017255 | 0.79765604 |
| SH3PXD2B | 44.3802729 | -0.874190574 | 0.46900582 | -1.8639226 | 0.06233257 | 0.25332358 |
| DUSP1 | 320.396169 | 0.98744191 | 0.63371227 | 1.55818652 | 0.11918904 | 0.3645149 |
| ERGIC1 | 237.531013 | -0.564657937 | 0.29408271 | -1.920065 | 0.05484969 | 0.23529115 |
| ATP6V0E1 | 272.873726 | -0.631706297 | 0.19864204 | -3.180124 | 0.00147212 | 0.02655639 |
| CREBRF | 231.513615 | 0.257585084 | 0.24169945 | 1.06572473 | 0.2865481 | 0.57346076 |
| BOD1 | 32.7765009 | -0.386948989 | 0.29848003 | -1.2963983 | 0.19483831 | 0.47336819 |
| CPEB4 | 698.54262 | 0.972796381 | 0.3396986 | 2.86370442 | 0.00418719 | 0.05129034 |
| SFXN1 | 129.424687 | -0.275876706 | 0.21518758 | -1.2820289 | 0.1998325 | 0.47866717 |
| THOC3 | 47.8553312 | 0.447627847 | 0.26907302 | 1.66359244 | 0.0961939 | 0.32331278 |
| SIMC1 | 133.389265 | 0.127002546 | 0.32172016 | 0.39476092 | 0.69301934 | 0.86938105 |
| KIAA1191 | 74.3914594 | -0.126652415 | 0.2458039 | -0.515258 | 0.60637277 | 0.82098159 |
| ARL10 | 43.1037572 | 0.257456712 | 0.3515566 | 0.73233361 | 0.46396497 | 0.7217612 |
| CLTB | 62.7581338 | -0.6060283 | 0.41500448 | -1.4602934 | 0.14420946 | 0.40168142 |
| FAF2 | 147.893012 | -0.061141975 | 0.1625415 | -0.3761623 | 0.70679628 | 0.87495497 |
| RNF44 | 38.1440775 | 0.271420623 | 0.23431529 | 1.15835643 | 0.24671861 | 0.53365473 |
| TSPAN17 | 30.0292612 | -0.650621447 | 0.33895593 | -1.9194868 | 0.05492275 | 0.23549912 |
| UIMC1 | 210.326552 | -0.007231641 | 0.18707694 | -0.038656 | 0.96916467 | 0.99152285 |
| ZNF346 | 94.7244087 | 0.215978234 | 0.23892139 | 0.90397193 | 0.36601029 | 0.64617238 |
| NSD1 | 457.691403 | 0.446881373 | 0.17707228 | 2.52372286 | 0.01161194 | 0.09808713 |
| PRELID1 | 64.2223505 | -0.547911539 | 0.45513321 | -1.2038487 | 0.22864805 | 0.51112377 |
| LMAN2 | 87.9210171 | -0.930756288 | 0.3152359 | -2.9525708 | 0.0031514 | 0.04214469 |
| DBN1 | 35.5006766 | -0.010982875 | 0.45734344 | -0.0240145 | 0.98084104 | 0.99578216 |
| PDLIM7 | 31.8342777 | -0.840575921 | 0.59889875 | -1.403536 | 0.16045708 | 0.42647568 |
| FAM193B | 116.477793 | 0.097585818 | 0.2747992 | 0.35511683 | 0.72250207 | 0.88242348 |
| TMED9 | 108.012711 | -1.146358321 | 0.3846888 | -2.9799628 | 0.00288283 | 0.03957434 |
| B4GALT7 | 27.5646957 | -0.598454559 | 0.3364401 | -1.7787849 | 0.07527504 | 0.28328492 |
| RMND5B | 64.1147652 | -0.330576791 | 0.24297702 | -1.360527 | 0.17366321 | 0.44502941 |
| NHP2 | 62.3793727 | -1.38821713 | 0.49861102 | -2.7841686 | 0.00536651 | 0.06041319 |
| HNRNPAB | 82.966649 | -1.604080296 | 0.39521493 | -4.0587543 | 4.93E-05 | 0.00295364 |
| PHYKPL | 222.621338 | -0.551251403 | 0.27021585 | -2.040041 | 0.04134624 | 0.20237899 |
| COL23A1 | 64.9388414 | -1.831955376 | 0.76858307 | -2.3835489 | 0.0171466 | 0.12340619 |
| CLK4 | 96.3770236 | 0.633436131 | 0.34712994 | 1.82478105 | 0.06803408 | 0.26676153 |
| ZNF354A | 48.7085627 | -0.386434828 | 0.25233641 | -1.5314272 | 0.12566385 | 0.37252428 |
| ZNF354B | 36.2962683 | -0.204850471 | 0.28498417 | -0.7188135 | 0.47225583 | 0.72983472 |
| RUFY1 | 173.159282 | 0.03554446 | 0.200788 | 0.17702482 | 0.85948889 | 0.94734879 |
| HNRNPH1 | 418.942379 | 0.784833703 | 0.19514661 | 4.02176439 | 5.78E-05 | 0.00327408 |
| CANX | 375.290057 | -0.691779943 | 0.27669213 | -2.5001793 | 0.01241305 | 0.10171477 |
| MAML1 | 63.9239325 | 0.273714301 | 0.23882492 | 1.14608771 | 0.25175886 | 0.53906677 |
| MGAT4B | 90.3961563 | -0.981344393 | 0.43558073 | -2.2529564 | 0.0242619 | 0.15130515 |
| SQSTM1 | 446.278027 | -1.028401704 | 0.29705298 | -3.4620144 | 0.00053615 | 0.01394437 |
| MRNIP | 75.5876983 | 0.340500639 | 0.25888191 | 1.315274 | 0.18841783 | 0.46421153 |
| TBC1D9B | 95.3896825 | -0.552797579 | 0.25132923 | -2.1994958 | 0.02784269 | 0.16152154 |
| RNF130 | 227.397083 | 0.002336988 | 0.25517481 | 0.00915838 | 0.99269277 | 0.99829858 |
| MAPK9 | 77.6573489 | 0.090021884 | 0.21488593 | 0.4189287 | 0.67526824 | 0.86115505 |
| CNOT6 | 128.360529 | 0.309472089 | 0.14855162 | 2.08326299 | 0.03722726 | 0.19120638 |
| MGAT1 | 56.4912539 | -0.308908971 | 0.26525881 | -1.1645569 | 0.2441984 | 0.5298248 |
| ZFP62 | 25.889056 | 0.217901838 | 0.26918995 | 0.8094724 | 0.41824347 | 0.68778613 |
| TRIM41 | 29.4276641 | -0.274196035 | 0.27529803 | -0.9959971 | 0.31925156 | 0.60592643 |
| RACK1 | 354.095043 | -0.920723221 | 0.41329566 | -2.2277593 | 0.02589657 | 0.15591658 |
| TRIM52 | 43.5725602 | 0.532416317 | 0.25850525 | 2.05959574 | 0.0394372 | 0.19685721 |
| DUSP22 | 58.4916954 | -0.439766908 | 0.26805037 | -1.640613 | 0.10087778 | 0.33183662 |
| EXOC2 | 169.853893 | -0.045927079 | 0.23447859 | -0.195869 | 0.84471273 | 0.94193856 |
| FOXC1 | 264.368437 | 0.415638092 | 0.52467664 | 0.79217952 | 0.42825601 | 0.69444434 |
| GMDS | 1049.15277 | 0.285284457 | 0.23044897 | 1.23795069 | 0.21573435 | 0.49746509 |
| GMDS-DT | 891.589564 | -0.149623894 | 0.21384556 | -0.699682 | 0.4841259 | 0.73727416 |
| WRNIP1 | 70.3830498 | -0.14274716 | 0.20563691 | -0.6941709 | 0.48757503 | 0.74111095 |
| SERPINB1 | 58.9400818 | -0.906665376 | 0.25492513 | -3.5565947 | 0.00037569 | 0.01107312 |
| SERPINB6 | 113.60774 | -0.430654346 | 0.17196911 | -2.504254 | 0.01227099 | 0.10098265 |
| NQO2 | 55.2387446 | -0.27212261 | 0.29696274 | -0.9163527 | 0.35948193 | 0.64001011 |
| RIPK1 | 103.399221 | -0.541348598 | 0.26208167 | -2.0655721 | 0.0388689 | 0.19575457 |
| BPHL | 35.630105 | -0.512281126 | 0.27254634 | -1.8796111 | 0.0601611 | 0.24733183 |
| PSMG4 | 62.2497001 | -0.119684696 | 0.24789647 | -0.4828011 | 0.62923694 | 0.833789 |
| SLC22A23 | 145.005572 | 1.566060266 | 0.53900839 | 2.90544693 | 0.00366729 | 0.04677626 |
| PXDC1 | 55.1231318 | -0.049326096 | 0.45315084 | -0.1088514 | 0.91332037 | 0.96844354 |
| PRPF4B | 226.393325 | 0.121615828 | 0.19024961 | 0.63924353 | 0.52266452 | 0.76512314 |
| ECI2 | 57.4644828 | -0.579808294 | 0.30885727 | -1.8772694 | 0.06048118 | 0.24800908 |
| CDYL | 255.058918 | -0.044392794 | 0.19583132 | -0.2266889 | 0.82066563 | 0.93138972 |
| LYRM4 | 148.233616 | -0.458009629 | 0.29285633 | -1.5639397 | 0.11783173 | 0.36234677 |
| FARS2 | 465.513945 | 0.239284068 | 0.17696218 | 1.35217632 | 0.17631892 | 0.44835649 |
| RREB1 | 380.576666 | 0.304230696 | 0.28661428 | 1.06146386 | 0.28847915 | 0.57533662 |
| SSR1 | 141.413951 | -0.163254219 | 0.27721988 | -0.5888979 | 0.55592974 | 0.78784845 |
| AL139095.4 | 26.4750593 | 0.22644623 | 0.54756947 | 0.41354795 | 0.6792052 | 0.86197819 |
| RIOK1 | 34.0827164 | -0.392882865 | 0.35501861 | -1.1066543 | 0.26844343 | 0.55558531 |
| DSP | 185.730655 | -0.767812958 | 0.22006165 | -3.4890811 | 0.00048468 | 0.01311522 |
| SNRNP48 | 43.1321624 | 0.451055311 | 0.2341575 | 1.92629025 | 0.05406814 | 0.23338941 |
| BMP6 | 149.321697 | 1.250017137 | 0.44891178 | 2.78454963 | 0.00536021 | 0.06041319 |
| BLOC1S5-TXI | 54.2586145 | 0.346994954 | 0.42565552 | 0.81520135 | 0.41495707 | 0.68495052 |
| SLC35B3 | 81.4774682 | 0.568787939 | 0.33587146 | 1.69346909 | 0.09036621 | 0.3135161 |
| GCNT2 | 187.99 | 0.921810478 | 0.53383673 | 1.7267648 | 0.08420991 | 0.30211489 |
| TMEM14C | 71.3927019 | 0.06784481 | 0.36863158 | 0.18404503 | 0.85397813 | 0.9452497 |
| MAK | 107.117136 | 0.421364022 | 0.31303252 | 1.34607107 | 0.1782796 | 0.44977081 |
| AL139807.1 | 178.98264 | -0.611653613 | 0.95740491 | -0.6388662 | 0.52291 | 0.76519323 |
| NEDD9 | 1070.5792 | 0.555185093 | 0.61068994 | 0.90911124 | 0.36329141 | 0.64360151 |
| TMEM170B | 37.2822644 | 0.645097325 | 0.27048695 | 2.38494805 | 0.01708153 | 0.12315032 |
| HIVEP1 | 274.018059 | -0.819670968 | 0.21023286 | -3.8988718 | 9.66E-05 | 0.00456026 |
| PHACTR1 | 313.067713 | -0.565612106 | 0.45745 | -1.2364457 | 0.21629292 | 0.49780633 |
| TBC1D7 | 25.5689398 | 0.292959467 | 0.44001917 | 0.66578797 | 0.50554664 | 0.75528806 |
| GFOD1 | 42.3779095 | -1.073428118 | 0.64294276 | -1.6695547 | 0.0950075 | 0.32056246 |
| SIRT5 | 67.8817371 | 0.468968731 | 0.24675435 | 1.90054897 | 0.05736112 | 0.24173433 |
| NOL7 | 84.5816162 | -0.556605147 | 0.38047238 | -1.4629318 | 0.14348604 | 0.40048158 |
| RANBP9 | 222.231941 | 0.386744755 | 0.14459063 | 2.67475678 | 0.00747835 | 0.07451094 |
| MCUR1 | 75.93539 | 0.697010296 | 0.37758632 | 1.84596279 | 0.06489761 | 0.25978027 |
| JARID2 | 210.536191 | 1.16297231 | 0.26011486 | 4.47099523 | 7.79E-06 | 0.0007384 |
| DTNBP1 | 201.631084 | -0.37805394 | 0.25309937 | -1.4936977 | 0.13525466 | 0.38732567 |
| MYLIP | 64.903854 | -0.009959599 | 0.3252149 | -0.0306247 | 0.97556887 | 0.99407418 |
| GMPR | 81.8220746 | -0.091916131 | 0.69317675 | -0.1326013 | 0.89450871 | 0.96091723 |
| ATXN1 | 699.160936 | 0.405975927 | 0.34343926 | 1.18208945 | 0.23717021 | 0.52000641 |
| CAP2 | 118.238392 | 0.915311426 | 0.22812238 | 4.01237015 | 6.01E-05 | 0.00332957 |
| FAM8A1 | 36.2785351 | -0.13128148 | 0.31042117 | -0.4229141 | 0.67235796 | 0.86015189 |
| NUP153 | 220.608165 | 0.435447502 | 0.18505395 | 2.35308413 | 0.01861842 | 0.12915243 |
| KIF13A | 357.013716 | 0.413866822 | 0.29094545 | 1.42248941 | 0.15488422 | 0.4182791 |
| TPMT | 55.608966 | 0.135190827 | 0.3334435 | 0.40543848 | 0.68515523 | 0.86561619 |
| KDM1B | 63.3982596 | -0.082150305 | 0.32800439 | -0.2504549 | 0.80223558 | 0.92119571 |
| DEK | 185.321407 | -0.14844494 | 0.2203459 | -0.6736905 | 0.50050811 | 0.75193964 |
| RNF144B | 943.326276 | 0.736002414 | 0.34653722 | 2.12387696 | 0.03368043 | 0.18043896 |
| MIR548A1HC | 97.7096682 | -0.753517258 | 0.81093143 | -0.9291997 | 0.35278559 | 0.63441164 |
| AL022068.1 | 297.47134 | 1.192446882 | 0.42936412 | 2.77723927 | 0.00548228 | 0.06134902 |
| ID4 | 260.795666 | 0.691514133 | 0.6868398 | 1.00680556 | 0.31402822 | 0.60099077 |
| MBOAT1 | 50.6968459 | -0.76205154 | 0.63375163 | -1.2024451 | 0.22919113 | 0.5117445 |

| | | | | | | |
|---|---|---|---|---|---|---|
| E2F3 | 116.848873 | -1.030360068 | 0.34318435 | -3.0023516 | 0.00267903 | 0.03790604 |
| CDKAL1 | 1006.14234 | 0.661167325 | 0.22409277 | 2.95041788 | 0.00317344 | 0.04227875 |
| SOX4 | 240.658763 | -0.198237625 | 0.48774153 | -0.4064399 | 0.6844194 | 0.86539363 |
| CASC15 | 1218.88608 | 0.932518428 | 0.31691822 | 2.94245759 | 0.00325618 | 0.04302205 |
| NBAT1 | 47.9261224 | 0.854966648 | 0.36803433 | 2.32306224 | 0.02017581 | 0.13533899 |
| DCDC2 | 1349.33793 | 0.345495848 | 0.38319749 | 0.90161302 | 0.36726247 | 0.64693034 |
| MRS2 | 61.9109058 | -0.076887632 | 0.23484667 | -0.327395 | 0.74336913 | 0.89269973 |
| ALDH5A1 | 33.2451395 | -0.169523264 | 0.38082768 | -0.4451443 | 0.65621549 | 0.85116969 |
| TDP2 | 63.9520187 | -0.321041027 | 0.22483428 | -1.4279007 | 0.15332044 | 0.41605 |
| ACOT13 | 44.9344507 | 0.186997635 | 0.30984559 | 0.60351879 | 0.54616362 | 0.78213504 |
| C6orf62 | 160.60613 | -0.16240899 | 0.2910464 | -0.5580175 | 0.57683242 | 0.80275223 |
| RIPOR2 | 455.64496 | -0.737103199 | 0.50851617 | -1.4495177 | 0.14719306 | 0.40621213 |
| CARMIL1 | 530.646966 | 0.091491373 | 0.20359064 | 0.44938889 | 0.65315114 | 0.84880407 |
| SLC17A1 | 105.152276 | -0.314075934 | 1.05786575 | -0.2968958 | 0.76654604 | 0.90372312 |
| TRIM38 | 208.490777 | -0.720731433 | 0.43981691 | -1.6387079 | 0.10127411 | 0.33245536 |
| HIST1H1C | 82.9933834 | -0.438208126 | 0.61591779 | -0.7114718 | 0.47679193 | 0.73352295 |
| HIST1H4C | 85.1114101 | -2.000754982 | 0.55516815 | -3.603872 | 0.00031351 | 0.00994413 |
| HIST1H2AC | 327.175047 | 0.896819232 | 0.70033461 | 1.2805582 | 0.20034889 | 0.4790669 |
| HIST1H1E | 42.4347813 | -1.432770946 | 0.77694251 | -1.8441145 | 0.06516646 | 0.26009562 |
| HIST1H2BD | 63.2544113 | 0.953077879 | 0.74866868 | 1.27303026 | 0.2030073 | 0.4818154 |
| HIST1H2BG | 38.3174082 | 0.520600852 | 1.10029357 | 0.47314723 | 0.63610814 | 0.83814123 |
| BTN2A1 | 33.8059406 | -0.44942984 | 0.28905143 | -1.5548439 | 0.11998326 | 0.36481264 |
| HCG11 | 35.7087438 | 0.754556592 | 0.50299656 | 1.50012277 | 0.1335826 | 0.38489489 |
| HMGN4 | 29.6442253 | -0.27042055 | 0.35439395 | -0.7630507 | 0.44543317 | 0.70759607 |
| LINC00240 | 102.169262 | 1.091156976 | 0.53725347 | 2.03099103 | 0.0422559 | 0.20401679 |
| ZSCAN16-AS | 34.8340681 | 0.195865134 | 0.3485508 | 0.56194142 | 0.57415593 | 0.80072326 |
| ZKSCAN8 | 42.8268446 | 0.407337408 | 0.26153051 | 1.55751391 | 0.11934852 | 0.3645149 |
| TRIM27 | 112.35491 | -0.355445804 | 0.23582003 | -1.5072757 | 0.13174001 | 0.38194235 |
| GABBR1 | 89.5631506 | 0.601253611 | 0.37242889 | 1.61441185 | 0.10643812 | 0.34105698 |
| HLA-F | 42.3820037 | -1.692831005 | 0.78036282 | -2.1692871 | 0.03006089 | 0.16938429 |
| HLA-A | 837.706712 | -1.049241932 | 0.38831957 | -2.7020063 | 0.00689224 | 0.07061049 |
| PPP1R11 | 33.8750122 | -0.471082808 | 0.32677207 | -1.4416251 | 0.14940818 | 0.40996303 |
| TRIM26 | 97.8977934 | 0.073396719 | 0.28549383 | 0.25708688 | 0.7971117 | 0.9188367 |
| HCG17 | 191.724276 | 0.099538395 | 0.20776721 | 0.47908616 | 0.63187734 | 0.8353707 |
| HLA-E | 230.045158 | -2.095077538 | 0.46064793 | -4.5481101 | 5.41E-06 | 0.00059481 |
| GNL1 | 78.3199892 | -0.083834401 | 0.21828277 | -0.3840633 | 0.70093153 | 0.8718638 |
| PRR3 | 29.932418 | -0.247025841 | 0.31217301 | -0.7913107 | 0.4287627 | 0.69479256 |
| ABCF1 | 73.358229 | -0.320473695 | 0.21977526 | -1.4581882 | 0.14478867 | 0.40282422 |
| PPP1R10 | 88.4502232 | 0.270938575 | 0.26151816 | 1.03602204 | 0.30019186 | 0.58720397 |
| MRPS18B | 62.4784084 | -1.014616646 | 0.23770321 | -4.2684179 | 1.97E-05 | 0.00153789 |
| ATAT1 | 30.8382511 | 0.28098841 | 0.23088419 | 1.21701017 | 0.22360035 | 0.50557208 |
| DHX16 | 31.4142214 | -0.393198915 | 0.27567592 | -1.4263085 | 0.15377931 | 0.41655145 |
| MDC1 | 48.452967 | 0.383063145 | 0.23363101 | 1.63960743 | 0.10108682 | 0.33218203 |
| TUBB | 220.76236 | -1.022811001 | 0.65879455 | -1.5525493 | 0.12053085 | 0.36507762 |
| FLOT1 | 192.426928 | -0.28724457 | 0.25681512 | -1.1184878 | 0.26335873 | 0.55096925 |
| IER3 | 277.218578 | -2.515412939 | 0.98968372 | -2.5416331 | 0.01103359 | 0.09487502 |
| DDR1 | 135.568181 | -0.698599129 | 0.32791612 | -2.1304202 | 0.03313693 | 0.17842535 |
| CCHCR1 | 41.9631413 | 0.696921584 | 0.33837036 | 2.05964132 | 0.03943284 | 0.19685721 |
| POU5F1 | 165.825053 | 1.010605426 | 0.35006301 | 2.88692437 | 0.00389028 | 0.04847509 |
| HLA-C | 336.1179 | -1.185556525 | 0.6000236 | -1.9758498 | 0.04817179 | 0.21931444 |
| HLA-B | 546.436371 | -1.61068225 | 0.52593814 | -3.0624937 | 0.00219501 | 0.03385831 |
| MICA | 49.2245869 | 0.123300999 | 0.32449611 | 0.37997681 | 0.70396263 | 0.87370536 |
| NFKBIL1 | 26.3936454 | -0.431680361 | 0.33566518 | -1.2860445 | 0.19842753 | 0.47709271 |
| PRRC2A | 108.916487 | 0.182303639 | 0.25361294 | 0.71882626 | 0.47224798 | 0.72983472 |
| BAG6 | 54.887353 | -0.361433142 | 0.31185515 | -1.1589776 | 0.2464653 | 0.53341417 |
| DDAH2 | 72.5166033 | -0.316690958 | 0.41440034 | -0.764215 | 0.44473912 | 0.70707984 |
| CLIC1 | 307.539366 | -1.732886734 | 0.39638225 | -4.3717567 | 1.23E-05 | 0.00105027 |
| VARS | 34.7044262 | -1.366006082 | 0.50401591 | -2.710244 | 0.00672337 | 0.06939999 |
| LSM2 | 47.0015543 | -0.675767532 | 0.34876965 | -1.9375755 | 0.05267504 | 0.23008398 |
| HSPA1A | 229.553822 | 0.908295805 | 0.75398096 | 1.20466677 | 0.22833197 | 0.51090678 |
| HSPA1B | 114.53796 | 0.667988217 | 0.75447683 | 0.88536611 | 0.37595921 | 0.65436143 |
| C6orf48 | 70.471127 | 0.052048891 | 0.46617313 | 0.11165142 | 0.9110998 | 0.96844354 |
| NEU1 | 58.477918 | -0.864446016 | 0.47819219 | -1.8077376 | 0.07064734 | 0.27279822 |
| SLC44A4 | 117.689255 | -0.902579305 | 0.46292574 | -1.9497281 | 0.05120854 | 0.2274115 |
| EHMT2 | 40.2160536 | 0.516031466 | 0.35605757 | 1.44929223 | 0.14725599 | 0.40626876 |
| NELFE | 28.3834457 | -0.365962751 | 0.42070623 | -0.8698772 | 0.38436753 | 0.66091637 |
| STK19 | 43.1656853 | -0.67628553 | 0.41209517 | -1.6410907 | 0.1007786 | 0.33162424 |
| ATF6B | 66.3590222 | -0.00651431 | 0.22481299 | -0.0289766 | 0.97688327 | 0.99441014 |
| AGPAT1 | 32.7243008 | -0.177232685 | 0.36087773 | -0.4911156 | 0.62334468 | 0.83030289 |
| RNF5 | 45.8472678 | -0.257471802 | 0.53133373 | -0.4845764 | 0.62797683 | 0.83304114 |
| PBX2 | 52.1282099 | 0.542224409 | 0.20606141 | 2.63137289 | 0.00850407 | 0.081542 |
| AL662796.1 | 65.7684007 | 0.66600283 | 0.5953647 | 1.11864683 | 0.26329084 | 0.55096925 |
| HLA-DRA | 47.7163122 | -1.305875141 | 0.77620891 | -1.6823759 | 0.09249598 | 0.31666153 |
| HLA-DRB1 | 43.4488905 | -1.105532449 | 1.18295649 | -0.9345504 | 0.35002006 | 0.63286907 |
| PSMB8 | 69.2958444 | -1.503317517 | 0.409927 | -3.6672811 | 0.00024514 | 0.00849612 |
| PSMB9 | 42.7819892 | -1.665660087 | 0.80792985 | -2.0616395 | 0.03924207 | 0.19660032 |
| TAP1 | 57.2970886 | -1.774293123 | 0.76080679 | -2.3321205 | 0.01969435 | 0.13342561 |
| BRD2 | 143.768844 | -0.258730686 | 0.25116351 | -1.0301285 | 0.3029497 | 0.59078893 |
| RXRB | 26.4157007 | 0.102968114 | 0.3623649 | 0.28415587 | 0.77629093 | 0.90750438 |
| SLC39A7 | 45.0911371 | -0.625465045 | 0.54005126 | -1.1581587 | 0.2467993 | 0.53365473 |
| RING1 | 53.6651206 | 0.371921659 | 0.27641679 | 1.34551036 | 0.17846048 | 0.45009819 |
| VPS52 | 44.9227431 | 0.29520055 | 0.2174285 | 1.3576902 | 0.17456201 | 0.44602013 |
| RPS18 | 332.947849 | -0.810738658 | 0.49969593 | -1.622464 | 0.10470404 | 0.33803841 |
| PFDN6 | 39.2343338 | -0.642326872 | 0.36463183 | -1.7615765 | 0.07814087 | 0.28987733 |
| RGL2 | 65.0897614 | 0.118003054 | 0.28438567 | 0.41494022 | 0.67818566 | 0.86151115 |
| TAPBP | 223.70391 | -0.717844554 | 0.33595232 | -2.1367453 | 0.03261871 | 0.17693995 |
| PHF1 | 50.2813847 | 0.529634686 | 0.28518059 | 1.85719053 | 0.06328405 | 0.25577972 |
| CUTA | 75.571318 | 0.066787426 | 0.43098081 | 0.15496612 | 0.87684802 | 0.95377544 |
| SYNGAP1 | 27.0604509 | 0.208084457 | 0.29837333 | 0.69739631 | 0.48555481 | 0.73886093 |
| ITPR3 | 149.363511 | -0.328965656 | 0.35856841 | -0.9174418 | 0.35891117 | 0.64001011 |
| UQCC2 | 40.3762109 | 0.020892596 | 0.36061648 | 0.05793578 | 0.95379979 | 0.98368305 |
| LEMD2 | 55.1278239 | -0.489616436 | 0.21742173 | -2.2519204 | 0.0243273 | 0.15131898 |
| HMGA1 | 45.8331926 | -2.480063862 | 0.79598927 | -3.1157001 | 0.00183509 | 0.03020328 |
| NUDT3 | 90.7802989 | -0.34699311 | 0.41590749 | -0.8343036 | 0.40410993 | 0.67765972 |
| C6orf106 | 213.987178 | -0.128760595 | 0.16601346 | -0.7756034 | 0.43798322 | 0.70228343 |
| SNRPC | 45.9219877 | -0.654393329 | 0.44262136 | -1.4784495 | 0.1392875 | 0.39299999 |
| UHRF1BP1 | 98.5837325 | -0.081911323 | 0.19270372 | -0.4250635 | 0.67079037 | 0.8597138 |
| TAF11 | 55.8687578 | 0.178520939 | 0.27722606 | 0.64395439 | 0.51960501 | 0.76385303 |
| ANKS1A | 298.369924 | 0.028519392 | 0.1674142 | 0.17035228 | 0.8647331 | 0.94991724 |
| ZNF76 | 61.2281155 | 0.062847686 | 0.21744803 | 0.28902394 | 0.77256306 | 0.90624314 |
| PPARD | 102.505588 | -0.408986911 | 0.35701852 | -1.1455622 | 0.25197636 | 0.53912923 |
| RPL10A | 366.048712 | -0.460836854 | 0.49911239 | -0.9233128 | 0.35584423 | 0.63736572 |
| TEAD3 | 68.4372596 | -0.616546558 | 0.28484858 | -2.1644713 | 0.0304282 | 0.17085095 |
| FKBP5 | 463.951222 | -3.81570901 | 1.08844258 | -3.5056595 | 0.00045548 | 0.01258164 |
| SRPK1 | 155.098875 | -0.915770495 | 0.32835368 | -2.7889759 | 0.0052875 | 0.06015317 |
| MAPK14 | 248.842741 | -0.800671334 | 0.2928372 | -2.7341858 | 0.00625347 | 0.06648394 |
| MAPK13 | 29.949429 | -0.909124398 | 0.53795361 | -1.689968 | 0.09103407 | 0.31492068 |
| BRPF3 | 35.1082082 | 0.157909534 | 0.33330222 | 0.47377283 | 0.63566191 | 0.83799478 |
| KCTD20 | 140.101772 | -0.052692374 | 0.14254345 | -0.3696583 | 0.71163709 | 0.87748 |
| STK38 | 283.612152 | 0.642823542 | 0.25743811 | 2.49700221 | 0.01252482 | 0.10219355 |
| SRSF3 | 187.723201 | -0.009546412 | 0.29769273 | -0.032068 | 0.97441782 | 0.99326197 |
| CDKN1A | 69.2135544 | -1.71040736 | 0.63565704 | -2.6907707 | 0.00712872 | 0.07203162 |
| C6orf89 | 124.052819 | -0.577445803 | 0.21643976 | -2.6679286 | 0.00763205 | 0.07533214 |

| | | | | | | |
|---|---|---|---|---|---|---|
| MTCH1 | 132.331639 | -0.180207496 | 0.25300926 | -0.7122565 | 0.47630594 | 0.73318945 |
| PIM1 | 35.4549085 | -0.94765697 | 0.46906627 | -2.0203051 | 0.04335175 | 0.20633261 |
| TBC1D22B | 98.0457901 | -0.111916933 | 0.19345873 | -0.5785055 | 0.5629229 | 0.79260446 |
| RNF8 | 63.4141018 | -0.132340278 | 0.33253023 | -0.3979797 | 0.69064516 | 0.86843279 |
| CMTR1 | 51.6817406 | 0.002741189 | 0.37389247 | 0.00733149 | 0.99415037 | 0.99884271 |
| ZFAND3 | 1561.23984 | 0.092739438 | 0.19222843 | 0.48244391 | 0.62949063 | 0.83400979 |
| BTBD9 | 628.304327 | -0.109904077 | 0.24166632 | -0.4547761 | 0.64927029 | 0.84651696 |
| GLO1 | 93.478953 | -0.096895776 | 0.21738132 | -0.445741 | 0.65578432 | 0.85072562 |
| KCNK5 | 75.3266352 | -0.595763239 | 0.19327161 | -3.0825181 | 0.00205257 | 0.03226175 |
| KIF6 | 64.2328012 | 0.195925778 | 0.52810376 | 0.37099864 | 0.71063854 | 0.87748 |
| UNC5CL | 123.009644 | -1.863750902 | 0.57611687 | -3.2350223 | 0.00121633 | 0.02363725 |
| OARD1 | 64.5615538 | -0.264282771 | 0.2918345 | -0.9055913 | 0.36515223 | 0.64511126 |
| NFYA | 55.0510178 | -0.225165263 | 0.23957242 | -0.939863 | 0.34728782 | 0.63088754 |
| FOXP4 | 38.045192 | -0.068348102 | 0.61562694 | -0.1110219 | 0.91159894 | 0.96844354 |
| TFEB | 39.5473831 | -0.41415357 | 0.46109611 | -0.8981936 | 0.36908235 | 0.64880401 |
| USP49 | 66.083892 | 0.448614602 | 0.29242147 | 1.53413702 | 0.12499595 | 0.37161272 |
| CCND3 | 375.870364 | -0.816200835 | 0.55520928 | -1.4700778 | 0.14154068 | 0.39760065 |
| TAF8 | 52.1624227 | 0.125540278 | 0.23233256 | 0.54034732 | 0.58895753 | 0.81148442 |
| C6orf132 | 58.6310719 | 0.042236337 | 0.51204593 | 0.08248545 | 0.93426069 | 0.97768238 |
| MRPS10 | 27.3303282 | 0.074807294 | 0.27142408 | 0.27561038 | 0.78284734 | 0.91149807 |
| TRERF1 | 33.051554 | 0.976246993 | 0.59998645 | 1.62711506 | 0.10371266 | 0.33585652 |
| UBR2 | 392.785497 | 0.258643143 | 0.14048997 | 1.84100791 | 0.0656204 | 0.26125428 |
| BICRAL | 254.404003 | 0.667785474 | 0.26162809 | 2.55242271 | 0.01069766 | 0.09281968 |
| RPL7L1 | 124.100183 | -0.475119466 | 0.23784806 | -1.9975755 | 0.0457627 | 0.21318196 |
| CNPY3 | 32.6119318 | -0.479679173 | 0.35713278 | -1.3431396 | 0.17922679 | 0.45108076 |
| PEX6 | 52.7314216 | -0.52387949 | 0.37256098 | -1.4061577 | 0.1596773 | 0.42532651 |
| PPP2R5D | 27.3581765 | 0.429683922 | 0.25938988 | 1.65651769 | 0.09761703 | 0.3259791 |
| MEA1 | 22.760679 | -0.41790304 | 0.49467724 | -0.8447994 | 0.39822284 | 0.67323978 |
| KLHDC3 | 38.7194796 | 0.074004512 | 0.34096223 | 0.21704607 | 0.82817244 | 0.93346403 |
| RRP36 | 52.3535995 | -0.502912248 | 0.25402313 | -1.9797892 | 0.04772722 | 0.2182239 |
| CUL7 | 32.0258591 | -0.275313938 | 0.29443006 | -0.9350741 | 0.34975009 | 0.63286907 |
| KLC4 | 42.670076 | 0.336316286 | 0.26995012 | 1.24584606 | 0.21282092 | 0.4947372 |
| MRPL2 | 32.1228741 | 0.003676886 | 0.36083785 | 0.01018986 | 0.99186981 | 0.99814428 |
| PTK7 | 89.3760935 | -0.164291806 | 0.3201392 | -0.5131887 | 0.60781936 | 0.8211189 |
| CUL9 | 41.3009962 | 0.146702323 | 0.24651655 | 0.59510132 | 0.55177575 | 0.78577398 |
| DNPH1 | 74.4455185 | -0.821931025 | 0.44042335 | -1.8662294 | 0.06200927 | 0.25250472 |
| ZNF318 | 37.2457171 | 0.574577873 | 0.29108728 | 1.97390239 | 0.04839284 | 0.2201116 |
| ABCC10 | 35.2841992 | -0.091396343 | 0.28809169 | -0.3172474 | 0.75105588 | 0.89599662 |
| TJAP1 | 40.3963099 | 0.072985628 | 0.21563668 | 0.33846574 | 0.73501224 | 0.88966284 |
| POLR1C | 84.7577267 | -0.270371976 | 0.21798465 | -1.2403258 | 0.21485492 | 0.49688442 |
| YIPF3 | 70.2358126 | -0.073702303 | 0.35889178 | -0.2053608 | 0.83729025 | 0.93836281 |
| XPO5 | 261.370461 | -0.34823742 | 0.30481369 | -1.1424599 | 0.25326291 | 0.54018835 |
| POLH | 45.3589713 | 0.204836038 | 0.36270372 | 0.56474756 | 0.57224548 | 0.79929126 |
| GTPBP2 | 50.0363656 | -0.01804917 | 0.24244012 | -0.074448 | 0.94065395 | 0.98016757 |
| MRPS18A | 31.3839565 | -1.022179494 | 0.28451115 | -3.5927573 | 0.0003272 | 0.01021686 |
| VEGFA | 138.767129 | 0.623561979 | 0.44972696 | 1.38653456 | 0.16558373 | 0.43384207 |
| MRPL14 | 71.0518576 | -1.556303542 | 0.44026139 | -3.5349535 | 0.00040785 | 0.01173203 |
| TMEM63B | 46.1045821 | -0.255531697 | 0.23125618 | -1.1049724 | 0.26917154 | 0.55632249 |
| SLC29A1 | 43.1393262 | 0.120900922 | 0.36649684 | 0.32988258 | 0.74148869 | 0.89253455 |
| HSP90AB1 | 488.56661 | -0.946987566 | 0.37769986 | -2.5072489 | 0.0121675 | 0.1007368 |
| SLC35B2 | 31.2413388 | -0.995537782 | 0.46049321 | -2.1618946 | 0.0306263 | 0.17115672 |
| CDC5L | 73.6833114 | 0.036536759 | 0.19804962 | 0.18448286 | 0.85363468 | 0.94520478 |
| SUPT3H | 598.412796 | -0.133146196 | 0.26016044 | -0.5117849 | 0.60880153 | 0.82159902 |
| RUNX2 | 62.5515913 | -2.153027605 | 0.78629211 | -2.7382032 | 0.00617759 | 0.06585681 |
| CLIC5 | 150.264216 | 1.979353412 | 0.77375459 | 2.55811525 | 0.01052412 | 0.09189658 |
| ENPP4 | 45.3018685 | 0.422706846 | 0.31722387 | 1.33251906 | 0.18268968 | 0.45608144 |
| ENPP5 | 46.3209081 | 0.005508504 | 0.32526619 | 0.01693537 | 0.98648817 | 0.99678172 |
| RCAN2 | 143.656609 | 0.182561569 | 0.42702916 | 0.42751546 | 0.66900393 | 0.85924744 |
| CYP39A1 | 33.9590705 | 0.355584165 | 0.5436548 | 0.6540624 | 0.51307161 | 0.76076145 |
| SLC25A27 | 136.480833 | 0.878548492 | 0.52132194 | 1.68523216 | 0.09194379 | 0.31578686 |
| ADGRF1 | 162.535832 | 1.468458337 | 0.57783312 | 2.54131909 | 0.01104351 | 0.09487513 |
| TNFRSF21 | 44.4597439 | -0.7549719 | 0.51153663 | -1.4758902 | 0.13997337 | 0.39412255 |
| CD2AP | 554.027686 | -0.119068837 | 0.18315563 | -0.6500965 | 0.51562988 | 0.7614026 |
| PTCHD4 | 124.237102 | 0.885420618 | 0.60958406 | 1.45249962 | 0.14636274 | 0.40473691 |
| MUT | 54.0029092 | 0.245912054 | 0.3107312 | 0.79139801 | 0.42871177 | 0.69479256 |
| PKHD1 | 4334.9091 | 1.164356754 | 0.57811201 | 2.01406775 | 0.04400242 | 0.20814775 |
| MCM3 | 47.9548138 | -0.374846847 | 0.42085525 | -0.8906788 | 0.37310152 | 0.65217873 |
| EFHC1 | 146.738725 | 0.265711507 | 0.26323793 | 1.00939674 | 0.3127844 | 0.60019427 |
| TRAM2 | 83.2538741 | 0.668979618 | 0.34210568 | 1.9554765 | 0.05052685 | 0.22585009 |
| TMEM14A | 24.5474834 | 0.772236867 | 0.45930101 | 1.68133067 | 0.0926987 | 0.31712889 |
| GSTA4 | 47.4111513 | 0.439283994 | 0.39855749 | 1.10218477 | 0.27038135 | 0.55746512 |
| ICK | 44.3823073 | 0.001125856 | 0.29817955 | 0.00377577 | 0.99698738 | 0.99923686 |
| FBXO9 | 130.552415 | 0.094478755 | 0.21666738 | 0.43605436 | 0.66279729 | 0.85530584 |
| ELOVL5 | 130.835268 | 0.681526718 | 0.33166097 | 2.0548897 | 0.03988966 | 0.19787833 |
| GCLC | 94.5839546 | -1.687634942 | 0.33481569 | -5.0404894 | 4.64E-07 | 8.90E-05 |
| LRRC1 | 169.08026 | 0.176104837 | 0.28415148 | 0.61975688 | 0.53541786 | 0.77504588 |
| TINAG | 113.289505 | -1.260505378 | 0.61071307 | -2.0639895 | 0.03901871 | 0.1958911 |
| FAM83B | 68.1733454 | -0.707726661 | 0.54043604 | -1.3095475 | 0.19034896 | 0.46644991 |
| DST | 607.720553 | -0.583926979 | 0.52613196 | -1.1098489 | 0.26706415 | 0.55453954 |
| ZNF451 | 190.565985 | 0.105274891 | 0.17157678 | 0.61357308 | 0.53949748 | 0.77746559 |
| PRIM2 | 179.727391 | 0.566211442 | 0.28210799 | 2.00707338 | 0.04474185 | 0.21050788 |
| AL021368.2 | 32.2310398 | 0.583808581 | 0.2592456 | 2.25195175 | 0.02432532 | 0.15131898 |
| PTP4A1 | 204.058403 | 0.625678171 | 0.5784551 | 1.08163654 | 0.27941406 | 0.56686465 |
| PHF3 | 363.322882 | 0.023891017 | 0.18893553 | 0.12645063 | 0.89937522 | 0.96269027 |
| LMBRD1 | 283.596832 | 0.415188172 | 0.17602865 | 2.35863974 | 0.01834205 | 0.12806015 |
| COL19A1 | 109.483733 | 1.005584603 | 0.29368652 | 3.42400667 | 0.00061705 | 0.01527321 |
| FAM135A | 408.477036 | -0.865297738 | 0.18668019 | -4.6351878 | 3.57E-06 | 0.000427 |
| SMAP1 | 228.915155 | -0.031124359 | 0.18904319 | -0.1646415 | 0.86922615 | 0.95106604 |
| OGFRL1 | 175.5462 | -0.850857342 | 0.44715169 | -1.9028382 | 0.05706165 | 0.24110877 |
| AL136164.2 | 438.015055 | 1.803160252 | 0.54446901 | 3.31177754 | 0.00092705 | 0.01973385 |
| LINC00472 | 109.189869 | 0.629732689 | 0.50655956 | 1.24315626 | 0.21381026 | 0.49581433 |
| AC019205.1 | 45.0836364 | 0.26808317 | 0.3534164 | 0.75854762 | 0.44812321 | 0.70956008 |
| OOEP | 316.562548 | -0.855459065 | 0.43907928 | -1.9483021 | 0.05137883 | 0.22752396 |
| MTO1 | 52.9290345 | 0.263699669 | 0.18786751 | 1.40364701 | 0.16042399 | 0.42647568 |
| EEF1A1 | 1677.0619 | -0.372120976 | 0.51475289 | -0.7229119 | 0.46973403 | 0.72749915 |
| SLC17A5 | 75.9056175 | 0.315896853 | 0.23938879 | 1.3195975 | 0.18696944 | 0.46206921 |
| COL12A1 | 71.901333 | 2.827606228 | 0.40676694 | 6.95141604 | 3.62E-12 | 5.77E-09 |
| COX7A2 | 98.1141921 | -0.187387758 | 0.42906248 | -0.4367377 | 0.66230159 | 0.85507439 |
| TMEM30A | 127.879651 | -0.097905161 | 0.3085579 | -0.3172991 | 0.75101662 | 0.89599662 |
| SENP6 | 673.830103 | 0.270622665 | 0.14723407 | 1.8380438 | 0.06605595 | 0.26198834 |
| MYO6 | 611.073676 | -0.498826762 | 0.29439336 | -1.6944226 | 0.090185 | 0.31311423 |
| IRAK1BP1 | 145.906289 | 0.279408993 | 0.28919853 | 0.96614942 | 0.33396942 | 0.61861931 |
| PHIP | 537.109448 | 0.198971776 | 0.14968688 | 1.32925328 | 0.18376442 | 0.45721925 |
| HMGN3 | 230.336187 | 0.202108551 | 0.21262172 | 0.9505546 | 0.34183053 | 0.6270562 |
| LCA5 | 57.5400827 | 0.104430948 | 0.36077951 | 0.2894592 | 0.77223 | 0.90624314 |
| SH3BGRL2 | 84.3063908 | 0.890039976 | 0.36716933 | 2.42405859 | 0.01534813 | 0.11503891 |
| BCKDHB | 265.455969 | 0.473999317 | 0.27469176 | 1.72556799 | 0.08442516 | 0.3023414 |
| TENT5A | 58.0138775 | -0.77485792 | 0.60616334 | -1.2782989 | 0.20114406 | 0.48009543 |
| IBTK | 239.984436 | 0.087804845 | 0.26740012 | 0.32836502 | 0.74263569 | 0.89253455 |
| TPBG | 34.8750525 | -0.579072426 | 0.34061008 | -1.7001036 | 0.08911144 | 0.3108137 |
| UBE3D | 186.321895 | -0.002097539 | 0.28322739 | -0.0074058 | 0.99409104 | 0.99884271 |
| DOPEY1 | 237.925493 | 0.371498487 | 0.16857038 | 2.20381828 | 0.02753713 | 0.16064442 |
| PGM3 | 58.3531188 | -0.495663308 | 0.38910107 | -1.2738677 | 0.20271029 | 0.48158147 |

| | | | | | | |
|---|---|---|---|---|---|---|
| ME1 | 54.1242176 | -0.769406424 | 0.26231659 | -2.9331215 | 0.00335573 | 0.04397331 |
| CYB5R4 | 65.7371814 | 0.353354706 | 0.24520658 | 1.44104906 | 0.14957082 | 0.41029179 |
| CEP162 | 136.634913 | 0.385509188 | 0.24143495 | 1.59674143 | 0.11032336 | 0.34739889 |
| SNX14 | 204.238371 | 0.194824316 | 0.14368903 | 1.35587467 | 0.17513905 | 0.44701757 |
| SYNCRIP | 100.512728 | -0.434846733 | 0.29282561 | -1.4850024 | 0.13754319 | 0.39014693 |
| ZNF292 | 770.015828 | -0.170618982 | 0.30123507 | -0.5663981 | 0.57112316 | 0.79853872 |
| SMIM8 | 101.597433 | -0.065545985 | 0.31278611 | -0.2095553 | 0.83401478 | 0.93647023 |
| RARS2 | 159.277214 | 0.267671088 | 0.17029356 | 1.57182156 | 0.11599194 | 0.35899412 |
| ORC3 | 134.449318 | 0.228526863 | 0.22411146 | 1.01970183 | 0.30786989 | 0.59456464 |
| AKIRIN2 | 46.6806732 | -0.604034393 | 0.2910451 | -2.0753979 | 0.03794967 | 0.19346455 |
| RNGTT | 281.776359 | 0.000589656 | 0.17337831 | 0.00340098 | 0.99728641 | 0.99923686 |
| PNRC1 | 146.67081 | -0.50782096 | 0.42581425 | -1.192588 | 0.23303077 | 0.5159967 |
| SRSF12 | 33.4033227 | -0.772812436 | 0.34055016 | -2.2693057 | 0.02324974 | 0.14709991 |
| PM20D2 | 76.9199013 | -0.128634401 | 0.2063725 | -0.6233117 | 0.53307968 | 0.77345809 |
| UBE2J1 | 75.8368834 | -0.151860629 | 0.21707154 | -0.6995879 | 0.48418469 | 0.73727416 |
| RRAGD | 47.3549583 | -1.403985017 | 0.34640187 | -4.0530527 | 5.06E-05 | 0.00298922 |
| ANKRD6 | 250.880142 | -0.015458935 | 0.38835411 | -0.0398063 | 0.96824757 | 0.99123492 |
| LYRM2 | 74.6886564 | -0.356692927 | 0.27280822 | -1.307486 | 0.19104771 | 0.46768362 |
| MDN1 | 202.686918 | -0.034096447 | 0.24265226 | -0.1405157 | 0.88825257 | 0.95907089 |
| CASP8AP2 | 83.648464 | 0.199189441 | 0.2237843 | 0.8900957 | 0.3734145 | 0.65236869 |
| BACH2 | 208.160736 | -0.164775729 | 0.50944559 | -0.3234413 | 0.74636108 | 0.89352127 |
| MAP3K7 | 187.497496 | 0.141195303 | 0.20630357 | 0.68440553 | 0.49371912 | 0.74606965 |
| EPHA7 | 576.041632 | 0.771848665 | 0.3213707 | 2.40173936 | 0.01631733 | 0.11949825 |
| MANEA | 41.3618681 | 0.026231552 | 0.29791989 | 0.08804901 | 0.92983772 | 0.97545894 |
| FUT9 | 96.0936242 | 0.026286299 | 1.01717048 | 0.02584257 | 0.97938291 | 0.99557142 |
| UFL1-AS1 | 66.9634034 | -0.136362412 | 0.41675209 | -0.3272027 | 0.74351457 | 0.89272074 |
| UFL1 | 120.107773 | -0.108228483 | 0.20004669 | -0.5410161 | 0.58849647 | 0.81121355 |
| NDUFAF4 | 28.9045543 | -0.663742279 | 0.4235977 | -1.5669166 | 0.11713418 | 0.36066483 |
| KLHL32 | 69.0169404 | 1.161064953 | 0.48396017 | 2.39909195 | 0.01643579 | 0.12018199 |
| MMS22L | 72.0418978 | -0.032235085 | 0.33601188 | -0.0959344 | 0.9235727 | 0.97297642 |
| FBXL4 | 227.955397 | 0.336685293 | 0.2554882 | 1.31781154 | 0.18756674 | 0.46292038 |
| PNISR | 537.611139 | 0.810556235 | 0.17988363 | 4.50600342 | 6.61E-06 | 0.00066781 |
| AL513550.1 | 28.577104 | 0.271272441 | 0.43595289 | 0.62225173 | 0.53377635 | 0.77365669 |
| USP45 | 108.085863 | 0.114451912 | 0.25225378 | 0.45371734 | 0.65003229 | 0.84716453 |
| CCNC | 178.325293 | 0.06934461 | 0.22752101 | 0.30478333 | 0.76053118 | 0.90125623 |
| SIM1 | 236.827746 | 1.037122872 | 0.71160825 | 1.45743515 | 0.1449963 | 0.40293576 |
| ASCC3 | 416.790711 | -0.217916622 | 0.17959225 | -1.2133966 | 0.22497824 | 0.50731323 |
| HACE1 | 108.240022 | 0.648806366 | 0.25075765 | 2.58738411 | 0.00967077 | 0.08789025 |
| PREP | 69.679379 | -0.176902682 | 0.20681023 | -0.8553865 | 0.39233722 | 0.66799147 |
| ATG5 | 186.857765 | 0.245460232 | 0.16906454 | 1.45187298 | 0.14653693 | 0.40510165 |
| CRYBG1 | 93.7189409 | -0.201433122 | 0.31966987 | -0.6301286 | 0.52861047 | 0.76999755 |
| RTN4IP1 | 43.5771321 | -0.173362541 | 0.46334906 | -0.3741511 | 0.70829193 | 0.87578784 |
| QRSL1 | 64.9436631 | -0.043586671 | 0.30750115 | -0.1417447 | 0.88728164 | 0.95896094 |
| LINC02532 | 942.84189 | 0.924275654 | 0.4551889 | 2.03053204 | 0.04230249 | 0.20409723 |
| CD24 | 1140.62738 | 0.088532949 | 0.3092896 | 0.28624612 | 0.77468962 | 0.90689106 |
| C6orf203 | 32.7790636 | 0.084395938 | 0.25661114 | 0.3288865 | 0.74224148 | 0.89253455 |
| PDSS2 | 337.606018 | 0.239008408 | 0.15776497 | 1.51496497 | 0.12978128 | 0.37924797 |
| SOBP | 91.2658972 | -0.534981322 | 0.81917325 | -0.6530747 | 0.51370812 | 0.76083448 |
| SEC63 | 250.276663 | 0.046457371 | 0.20462645 | 0.22703503 | 0.8203965 | 0.93133185 |
| OSTM1 | 66.6885651 | -0.152774931 | 0.23006444 | -0.6640528 | 0.50665653 | 0.75584222 |
| SNX3 | 151.2504 | -0.202401514 | 0.23512238 | -0.8608348 | 0.38932906 | 0.66513878 |
| AFG1L | 111.721082 | 0.777674526 | 0.27539973 | 2.82380275 | 0.00474576 | 0.05591589 |
| FOXO3 | 615.586533 | -0.365542603 | 0.32037944 | -1.1409677 | 0.25388335 | 0.54067332 |
| ARMC2 | 130.676607 | -0.113412593 | 0.33500918 | -0.3385358 | 0.73495947 | 0.88966284 |
| SESN1 | 158.870611 | -0.096613883 | 0.24147276 | -0.4001026 | 0.68908094 | 0.86751885 |
| CEP57L1 | 55.3978701 | 0.162584169 | 0.30366376 | 0.53540853 | 0.59236739 | 0.81283604 |
| CD164 | 250.392316 | -0.042238309 | 0.19019246 | -0.2220819 | 0.8242501 | 0.93217972 |
| PPIL6 | 103.23514 | 0.450870279 | 0.40075534 | 1.12505121 | 0.26056734 | 0.54748289 |
| AK9 | 97.0745165 | 1.000927153 | 0.39597888 | 2.52772865 | 0.0114803 | 0.09721852 |
| FIG4 | 98.3849926 | 0.046214758 | 0.20197221 | 0.22881741 | 0.81901083 | 0.93042039 |
| WASF1 | 80.7522342 | -0.442896985 | 0.29751843 | -1.4886371 | 0.13658295 | 0.38845845 |
| CDC40 | 66.3151652 | -0.026188784 | 0.27615266 | -0.0948344 | 0.92444633 | 0.97321369 |
| CDK19 | 238.443323 | 0.568430047 | 0.30399754 | 1.86985081 | 0.06150454 | 0.25100302 |
| AMD1 | 119.751539 | -0.451271924 | 0.31519787 | -1.4317099 | 0.15222685 | 0.41449148 |
| GTF3C6 | 80.744456 | -0.912833755 | 0.46501028 | -1.9630399 | 0.04964153 | 0.22398317 |
| RPF2 | 54.9917987 | -0.757597361 | 0.29184883 | -2.5958554 | 0.00943558 | 0.08649129 |
| SLC16A10 | 140.395372 | 0.458651287 | 0.505171 | 0.90791293 | 0.36392422 | 0.644123 |
| MFSD4B | 46.4537259 | 0.221549342 | 0.26871211 | 0.8244859 | 0.40966353 | 0.68208527 |
| REV3L | 430.333091 | -0.044061141 | 0.23231424 | -0.1896618 | 0.84957414 | 0.94414047 |
| TRAF3IP2-AS | 228.670439 | -0.19885812 | 0.28239764 | -0.7041777 | 0.48132211 | 0.73620566 |
| TRAF3IP2 | 49.0768319 | -0.951858182 | 0.39034412 | -2.4385104 | 0.01474794 | 0.11265588 |
| FYN | 160.514734 | 0.323309307 | 0.39226995 | 0.82420105 | 0.40982533 | 0.68208527 |
| TUBE1 | 59.9764848 | 0.422549097 | 0.34649395 | 1.2194992 | 0.22265478 | 0.50421043 |
| FAM229B | 32.3180556 | -0.283863082 | 0.30129221 | -0.9421521 | 0.34611478 | 0.63019075 |
| MARCKS | 133.632527 | -0.566111899 | 0.26257118 | -2.1560321 | 0.03108116 | 0.17239514 |
| HDAC2 | 99.4865115 | -0.243738902 | 0.20927516 | -1.1646815 | 0.24414795 | 0.5298248 |
| FRK | 140.473035 | -0.499624912 | 0.36601942 | -1.365023 | 0.17224583 | 0.44309343 |
| NT5DC1 | 227.576566 | 0.056483573 | 0.22738967 | 0.2483999 | 0.80382501 | 0.92125386 |
| TSPYL4 | 32.2822868 | 0.069246609 | 0.32856431 | 0.21075512 | 0.83307836 | 0.93607461 |
| DSE | 191.784007 | 0.709798474 | 0.342097 | 2.07484568 | 0.03800084 | 0.19361308 |
| TSPYL1 | 88.5692668 | -0.032027406 | 0.29064385 | -0.1101947 | 0.91225498 | 0.96844354 |
| RWDD1 | 57.2751144 | -0.359958713 | 0.37249872 | -0.9663354 | 0.33387637 | 0.61860768 |
| ZUP1 | 38.8389854 | 0.297026051 | 0.34047618 | 0.87238424 | 0.3829988 | 0.65965766 |
| KPNA5 | 141.370304 | 0.298412625 | 0.2659032 | 1.12226037 | 0.26175175 | 0.54924863 |
| DCBLD1 | 87.4535931 | 0.2851026 | 0.35616125 | 0.80048742 | 0.42342845 | 0.69062823 |
| GOPC | 91.7950659 | 0.121198121 | 0.2087233 | 0.58066408 | 0.56146687 | 0.79150717 |
| NUS1 | 94.9020848 | -0.383261434 | 0.24655724 | -1.5544522 | 0.1200766 | 0.36491553 |
| SLC35F1 | 31.4922715 | -0.504992446 | 0.27874266 | -1.8116798 | 0.07003569 | 0.27149813 |
| CEP85L | 129.332603 | -0.516085381 | 0.31592272 | -1.6335811 | 0.10234684 | 0.33390378 |
| MCM9 | 147.752201 | 0.102013368 | 0.20410646 | 0.49980469 | 0.61721261 | 0.82612711 |
| AL365275.1 | 60.9213885 | 0.26289195 | 0.56740011 | 0.46332728 | 0.64312979 | 0.84321657 |
| FAM184A | 51.7787545 | -0.473384231 | 0.45139292 | -1.0487188 | 0.29430757 | 0.58223301 |
| MAN1A1 | 239.167496 | -0.400363092 | 0.35956994 | -1.1134498 | 0.26551528 | 0.55350835 |
| TBC1D32 | 195.414352 | 0.403897698 | 0.2370839 | 1.70360658 | 0.08845461 | 0.30970853 |
| GJA1 | 35.7872941 | -1.821502891 | 0.86664681 | -2.1017823 | 0.03557235 | 0.18640458 |
| HSF2 | 41.4584393 | 0.825676347 | 0.24670552 | 3.34680932 | 0.00081747 | 0.01810054 |
| SERINC1 | 184.849679 | -0.080981282 | 0.23575606 | -0.3434961 | 0.73122529 | 0.88791456 |
| RNF217 | 247.195943 | -0.039272133 | 0.20083972 | -0.1955397 | 0.84497048 | 0.94193856 |
| TPD52L1 | 106.502868 | 0.878644502 | 0.5410747 | 1.6238876 | 0.1043998 | 0.33754429 |
| HDDC2 | 60.1577427 | -0.340013621 | 0.39805222 | -0.8541935 | 0.39299778 | 0.6686322 |
| NCOA7 | 345.108467 | 0.005822507 | 0.32866106 | 0.01771584 | 0.98586554 | 0.99668665 |
| HINT3 | 48.3806751 | -0.114012771 | 0.31716451 | -0.3594752 | 0.71923964 | 0.88158624 |
| TRMT11 | 117.351646 | 0.31310996 | 0.21546861 | 1.4531581 | 0.14617986 | 0.40436474 |
| RNF146 | 67.5559878 | 0.105021085 | 0.27875982 | 0.37674399 | 0.70636387 | 0.87473681 |
| ECHDC1 | 132.677352 | -0.289818319 | 0.16074556 | -1.8029631 | 0.07139401 | 0.27465189 |
| PTPRK | 961.184556 | 0.047286572 | 0.25890436 | 0.18264108 | 0.85507965 | 0.94571158 |
| ARHGAP18 | 134.88607 | 0.618393117 | 0.29391617 | 2.10397787 | 0.03538038 | 0.18580517 |
| L3MBTL3 | 63.8896696 | -0.508347154 | 0.28460208 | -1.7861681 | 0.07407204 | 0.28082432 |
| EPB41L2 | 242.196689 | -0.768355333 | 0.4004883 | -1.9185463 | 0.05504178 | 0.23558767 |
| AKAP7 | 89.1537197 | 0.055176971 | 0.49430605 | 0.11162512 | 0.91112066 | 0.96844354 |
| MED23 | 107.251367 | 0.341529647 | 0.16900026 | 2.02088238 | 0.04329194 | 0.20621259 |
| ENPP1 | 84.7876132 | 2.506141598 | 0.54130404 | 4.62982248 | 3.66E-06 | 0.00042813 |
| CTGF | 555.907138 | 1.933461865 | 1.06646934 | 1.81295588 | 0.06983863 | 0.2709535 |

| | | | | | | |
|---|---|---|---|---|---|---|
| STX7 | 254.088147 | 0.457321059 | 0.25433028 | 1.79813844 | 0.07215507 | 0.27680153 |
| SLC18B1 | 47.6685928 | -0.817084278 | 0.36262354 | -2.2532577 | 0.02424291 | 0.15128521 |
| RPS12 | 301.674313 | -0.375804242 | 0.47217284 | -0.795904 | 0.42608785 | 0.69224822 |
| TBPL1 | 44.674094 | 0.36103615 | 0.23498552 | 1.53641869 | 0.12443572 | 0.37055989 |
| SGK1 | 71.0748797 | -1.448697675 | 0.7549975 | -1.9188112 | 0.05500823 | 0.23554932 |
| ALDH8A1 | 39.1811764 | -0.324331055 | 0.43025518 | -0.7538109 | 0.45096276 | 0.71164003 |
| HBS1L | 116.390705 | 0.294778308 | 0.15177697 | 1.94218075 | 0.05211522 | 0.22878631 |
| AHI1 | 488.65373 | 0.59051879 | 0.20062536 | 2.94339061 | 0.00324639 | 0.04295554 |
| LINC00271 | 285.454416 | 0.699898319 | 0.36778814 | 1.90299319 | 0.05704143 | 0.24110877 |
| PDE7B | 900.119723 | 0.651034448 | 0.30533247 | 2.13221494 | 0.03298918 | 0.17803006 |
| AL138828.1 | 149.895482 | -1.390001396 | 0.51112168 | -2.7195117 | 0.00653784 | 0.06859359 |
| BCLAF1 | 281.337207 | 0.149392108 | 0.12035603 | 1.24125155 | 0.21451284 | 0.49669288 |
| MAP7 | 577.490255 | -0.048996681 | 0.35490378 | -0.1380562 | 0.89019597 | 0.95918866 |
| MAP3K5 | 957.214952 | -1.635417449 | 0.52416256 | -3.1200578 | 0.00180816 | 0.02986263 |
| PEX7 | 66.8228888 | 0.172224877 | 0.21147606 | 0.81439419 | 0.41541916 | 0.68495052 |
| IFNGR1 | 115.43112 | -1.073394537 | 0.46938149 | -2.2868276 | 0.02220589 | 0.14286955 |
| TNFAIP3 | 61.8894273 | -3.638910566 | 1.18635407 | -3.0673057 | 0.00215998 | 0.03353393 |
| PERP | 221.013518 | -0.311568553 | 0.47651579 | -0.6538473 | 0.5132102 | 0.76076145 |
| ARFGEF3 | 92.4251092 | -0.183775596 | 0.48666564 | -0.3776219 | 0.70571151 | 0.87473681 |
| HEBP2 | 84.7232636 | -0.663497164 | 0.28082195 | -2.362697 | 0.0181425 | 0.12731647 |
| NHSL1 | 87.1535639 | -0.940903006 | 0.37823107 | -2.4876407 | 0.01285936 | 0.10368668 |
| CCDC28A | 41.9153394 | 0.359926643 | 0.40854998 | 0.88098559 | 0.37832562 | 0.65629152 |
| REPS1 | 223.998465 | 0.078605429 | 0.15579597 | 0.50454084 | 0.61388137 | 0.82514464 |
| HECA | 33.1500215 | 0.036780161 | 0.33844777 | 0.10867308 | 0.91346179 | 0.96844354 |
| CITED2 | 132.609203 | 0.850685064 | 0.44899195 | 1.89465548 | 0.05813806 | 0.24328193 |
| VTA1 | 98.677858 | -0.208943469 | 0.20769481 | -1.006012 | 0.3144098 | 0.60132156 |
| ADGRG6 | 241.011038 | 0.668289095 | 0.52332909 | 1.27699588 | 0.2016037 | 0.48024387 |
| HIVEP2 | 549.662655 | 0.263364489 | 0.33514188 | 0.78582983 | 0.43196719 | 0.69731529 |
| AIG1 | 344.697143 | -0.017962107 | 0.20110714 | -0.0893161 | 0.92883069 | 0.97525695 |
| ADAT2 | 34.6713642 | -0.598070524 | 0.40631593 | -1.4719347 | 0.14103851 | 0.39653885 |
| PEX3 | 128.95946 | 0.381267959 | 0.26961342 | 1.41412829 | 0.15732424 | 0.42243085 |
| FUCA2 | 44.1307774 | -1.041680574 | 0.27457497 | -3.7937929 | 0.00014836 | 0.00620241 |
| PHACTR2 | 244.737127 | -0.183319582 | 0.24544628 | -0.7468827 | 0.45513437 | 0.71471019 |
| PLAGL1 | 251.506336 | 0.08261963 | 0.26665708 | 0.30983475 | 0.75668663 | 0.89923013 |
| SF3B5 | 94.1840948 | -1.023003647 | 0.39182283 | -2.6108832 | 0.00903087 | 0.0847991 |
| UTRN | 1735.79905 | 0.937446865 | 0.23381133 | 4.00941586 | 6.09E-05 | 0.00332957 |
| EPM2A | 65.8712959 | -0.250457549 | 0.39848603 | -0.6285228 | 0.52966153 | 0.7705996 |
| AL356599.1 | 92.0381086 | 0.262421294 | 0.33340613 | 0.787092 | 0.43122801 | 0.69681733 |
| SHPRH | 336.853583 | 0.521104595 | 0.17089076 | 3.04934335 | 0.00229342 | 0.03454196 |
| STXBP5 | 297.116697 | -0.094374133 | 0.18790283 | -0.5022497 | 0.61549191 | 0.82592881 |
| SAMD5 | 268.920458 | -1.47575076 | 0.60497082 | -2.4393751 | 0.01471269 | 0.11247632 |
| AL033504.1 | 167.607857 | -1.640207686 | 0.70013521 | -2.3427013 | 0.0191447 | 0.13134709 |
| SASH1 | 1546.81349 | 0.251106954 | 0.21631691 | 1.16082907 | 0.24571142 | 0.53226361 |
| UST | 44.8741388 | -1.125194037 | 0.72480412 | -1.5524112 | 0.12056386 | 0.36507762 |
| TAB2 | 249.080533 | 0.340120886 | 0.22019762 | 1.54461654 | 0.12243904 | 0.36765168 |
| PPIL4 | 103.447419 | 0.056396519 | 0.25195741 | 0.22383354 | 0.82288683 | 0.93217972 |
| GINM1 | 125.71796 | -0.214943918 | 0.16533237 | -1.3000716 | 0.19357642 | 0.47194155 |
| KATNA1 | 51.7250901 | 0.541544127 | 0.2607113 | 2.07717934 | 0.03778501 | 0.19316567 |
| LATS1 | 168.961491 | 0.169265124 | 0.14723936 | 1.14959153 | 0.25031215 | 0.53773045 |
| NUP43 | 52.6859958 | 0.394587111 | 0.29286492 | 1.34733486 | 0.17787241 | 0.44944337 |
| PCMT1 | 116.42825 | 0.257870689 | 0.19148611 | 1.34668088 | 0.17808303 | 0.44971747 |
| PPP1R14C | 190.301258 | 0.05621024 | 0.41140108 | 0.13663124 | 0.89132228 | 0.95969529 |
| PLEKHG1 | 66.2314998 | -1.029737584 | 0.44131252 | -2.3333523 | 0.01962966 | 0.13327863 |
| MTHFD1L | 280.823859 | -1.607343362 | 0.38595853 | -4.1645494 | 3.12E-05 | 0.00212938 |
| AKAP12 | 470.936924 | 0.374676263 | 0.49895503 | 0.75092191 | 0.45269965 | 0.71261515 |
| RMND1 | 70.1472569 | -0.027257083 | 0.20915761 | -0.1303184 | 0.89631453 | 0.9616708 |
| ARMT1 | 38.7987749 | -0.236350018 | 0.26412279 | -0.894849 | 0.37086781 | 0.65088728 |
| SYNE1 | 2403.17371 | 0.814216447 | 0.50221787 | 1.6212415 | 0.10496586 | 0.33865541 |
| MTRF1L | 36.2738538 | -0.494628248 | 0.31135121 | -1.5886505 | 0.11213932 | 0.3503531 |
| IPCEF1 | 183.31303 | 0.366326496 | 0.60326218 | 0.6072426 | 0.54368992 | 0.78027954 |
| CNKSR3 | 694.251655 | 0.041239668 | 0.30252831 | 0.13631672 | 0.8915709 | 0.95969858 |
| SCAF8 | 436.085041 | 0.440569993 | 0.1518232 | 2.90186208 | 0.00370952 | 0.04712662 |
| TIAM2 | 240.58573 | 0.00779253 | 0.16295945 | 0.04781883 | 0.96186063 | 0.9875899 |
| TFB1M | 93.9412535 | -0.009979901 | 0.24957924 | -0.0399869 | 0.96810357 | 0.99123492 |
| ARID1B | 1314.02023 | 0.115767946 | 0.18743557 | 0.61764128 | 0.53681183 | 0.77570079 |
| TMEM242 | 53.4140151 | 0.055254671 | 0.23255784 | 0.23759539 | 0.81219493 | 0.92641286 |
| ZDHHC14 | 1212.6484 | 0.583170873 | 0.31752917 | 1.83658991 | 0.06627046 | 0.26209939 |
| SNX9 | 306.78488 | -0.17688063 | 0.2412796 | -0.733094 | 0.4635011 | 0.72133763 |
| SYNJ2 | 133.046009 | 0.242516741 | 0.49967725 | 0.48534677 | 0.62743038 | 0.83254684 |
| SERAC1 | 54.4609757 | 0.737590048 | 0.30439759 | 2.42311393 | 0.0153881 | 0.11515829 |
| GTF2H5 | 73.243301 | -0.130740151 | 0.28365588 | -0.4609111 | 0.64486237 | 0.84398642 |
| TULP4 | 377.998214 | -0.146516375 | 0.21166815 | -0.6921985 | 0.48881268 | 0.74185878 |
| TMEM181 | 83.6520723 | -0.685938137 | 0.26237253 | -2.6143672 | 0.00893929 | 0.0844472 |
| DYNLT1 | 171.672914 | -0.666073718 | 0.38339588 | -1.7373001 | 0.08233421 | 0.29885538 |
| SYTL3 | 56.4315943 | -0.249823434 | 0.54545478 | -0.4580094 | 0.64694566 | 0.84498126 |
| EZR | 484.44969 | 0.133865056 | 0.26615091 | 0.50296675 | 0.61498765 | 0.82592881 |
| RSPH3 | 115.693062 | 0.071391774 | 0.32148015 | 0.2220721 | 0.82425775 | 0.93217972 |
| AL035530.2 | 59.6866534 | 0.588382657 | 0.37791529 | 1.55691678 | 0.11949024 | 0.3645149 |
| SOD2.1 | 46.0356925 | 0.240665429 | 0.37708205 | 0.63823093 | 0.52332337 | 0.76534368 |
| WTAP | 281.528388 | -0.777364749 | 0.31941024 | -2.4337502 | 0.0149433 | 0.11360468 |
| TCP1 | 77.905622 | -0.355412956 | 0.33118966 | -1.0731403 | 0.28320819 | 0.57084461 |
| MRPL18 | 42.9886206 | -0.673444187 | 0.3186979 | -2.1131115 | 0.03459123 | 0.18326846 |
| IGF2R | 421.577459 | -0.006297124 | 0.17360108 | -0.0362735 | 0.97106426 | 0.99175186 |
| SLC22A2 | 153.785782 | -0.339968616 | 0.42507981 | -0.799776 | 0.42384061 | 0.69092389 |
| SLC22A3 | 65.9643206 | -1.082701294 | 0.57424056 | -1.885449 | 0.05936924 | 0.24597663 |
| MAP3K4 | 179.706374 | 0.41955135 | 0.19482409 | 2.15348808 | 0.03128034 | 0.17299906 |
| AGPAT4 | 118.607798 | -0.454576777 | 0.37018942 | -1.2279572 | 0.21946302 | 0.50121948 |
| PRKN | 882.071589 | 0.347490121 | 0.1568844 | 2.21494377 | 0.02676392 | 0.15852905 |
| PACRG | 302.092816 | 0.239010701 | 0.40956187 | 0.58357654 | 0.55950524 | 0.78990431 |
| QKI | 777.24251 | 0.216293188 | 0.18110923 | 1.19426924 | 0.23237266 | 0.51573164 |
| PDE10A | 202.968399 | 0.308105784 | 0.4172487 | 0.7384224 | 0.46025781 | 0.71898395 |
| SFT2D1 | 95.1463699 | 0.049342293 | 0.21777044 | 0.22657939 | 0.82075082 | 0.93138972 |
| MPC1 | 168.357713 | -0.038975611 | 0.33835935 | -0.11519 | 0.90829453 | 0.96748063 |
| RPS6KA2 | 285.528695 | -0.427604038 | 0.30835047 | -1.3867468 | 0.16551898 | 0.43384207 |
| Z94721.1 | 51.3646676 | -0.200450135 | 0.24340237 | -0.823534 | 0.41020437 | 0.68208527 |
| FGFR1OP | 184.073451 | 0.561101053 | 0.19063258 | 2.9433639 | 0.00324667 | 0.04295554 |
| AFDN | 786.241355 | 0.380503063 | 0.30024017 | 1.26732898 | 0.20503768 | 0.48399605 |
| SMOC2 | 111.519673 | 3.346255409 | 1.4230863 | 2.35140722 | 0.01870255 | 0.1295448 |
| WDR27 | 223.392916 | 0.738715177 | 0.28173688 | 2.6220038 | 0.00874145 | 0.08306975 |
| PHF10 | 83.1674268 | 0.542154089 | 0.20944991 | 2.58846661 | 0.00964043 | 0.08782595 |
| ERMARD | 50.8137086 | 0.244616615 | 0.38883669 | 0.6290986 | 0.52928451 | 0.77040597 |
| DLL1 | 35.3828503 | 0.730325914 | 0.68424811 | 1.06734078 | 0.28581798 | 0.57325176 |
| FAM120B | 282.626493 | 0.22744955 | 0.28178624 | 0.80717054 | 0.41956823 | 0.68835944 |
| PSMB1 | 153.185681 | -0.56749667 | 0.31577254 | -1.7971691 | 0.07230877 | 0.27705829 |
| TBP | 40.0767992 | -0.057386286 | 0.27099207 | -0.2117637 | 0.83229138 | 0.93582908 |
| PDCD2 | 48.2856241 | -0.435659478 | 0.30985655 | -1.4060038 | 0.15972301 | 0.42532651 |
| FAM20C | 106.559945 | -1.006747824 | 0.49681945 | -2.0263857 | 0.04272528 | 0.20524846 |
| PDGFA | 70.3388044 | 0.680325285 | 0.50045855 | 1.35940386 | 0.17401865 | 0.44528422 |
| PRKAR1B | 71.6017001 | -0.691355382 | 0.31243659 | -2.2127862 | 0.02691239 | 0.15864374 |
| DNAAF5 | 73.8838614 | -0.344446457 | 0.2812173 | -1.2248409 | 0.22063516 | 0.50236569 |
| SUN1 | 178.786161 | 0.171807321 | 0.16432181 | 1.04555397 | 0.29576701 | 0.5839143 |
| COX19 | 51.6688804 | -0.309642406 | 0.24607025 | -1.2583496 | 0.20826534 | 0.48848523 |
| C7orf50 | 140.039651 | -0.198129235 | 0.21305757 | -0.9299329 | 0.35240584 | 0.63408352 |

| | | | | | | |
|---|---|---|---|---|---|---|
| MICALL2 | 60.7619736 | -0.540040757 | 0.25913953 | -2.0839767 | 0.03716228 | 0.19107758 |
| INTS1 | 74.4423354 | 0.113247552 | 0.31589724 | 0.35849491 | 0.71997298 | 0.88192086 |
| MAFK | 53.0484508 | 1.281114613 | 0.40164563 | 3.18966398 | 0.00142438 | 0.02622761 |
| PSMG3 | 30.0208465 | -0.491571197 | 0.39256326 | -1.2522089 | 0.21049376 | 0.49127431 |
| MAD1L1 | 160.558129 | 0.074907896 | 0.30100313 | 0.24886085 | 0.80346842 | 0.92125386 |
| SNX8 | 78.1555922 | -0.285626898 | 0.33765702 | -0.8459084 | 0.39760387 | 0.67266325 |
| EIF3B | 140.947279 | -0.698263465 | 0.32312619 | -2.1609622 | 0.03069826 | 0.17115672 |
| CHST12 | 36.7334243 | 0.107128837 | 0.32805498 | 0.32655757 | 0.74400254 | 0.89297085 |
| BRAT1 | 35.0927308 | -0.225729235 | 0.33821043 | -0.6674224 | 0.50450233 | 0.7552161 |
| IQCE | 78.5323967 | 0.048744253 | 0.26140421 | 0.1864708 | 0.85207557 | 0.94470172 |
| TTYH3 | 80.0832387 | -0.158811903 | 0.4253144 | -0.3733988 | 0.70885163 | 0.87625367 |
| GNA12 | 133.800478 | 0.325021328 | 0.20015605 | 1.62383966 | 0.10441003 | 0.33754429 |
| SDK1 | 708.717108 | -1.534990108 | 0.58881464 | -2.6069156 | 0.00913619 | 0.08519229 |
| FOXK1 | 212.081556 | -0.246443766 | 0.2577652 | -0.9560785 | 0.33903258 | 0.62369754 |
| AP5Z1 | 27.9981193 | -0.474590843 | 0.33513786 | -1.4161063 | 0.15674437 | 0.4216384 |
| RBAK-RBAKI | 110.635988 | -0.215408168 | 0.22177029 | -0.9713121 | 0.33139289 | 0.61663024 |
| WIPI2 | 92.573928 | -0.078825855 | 0.21012433 | -0.3751391 | 0.707557 | 0.87544414 |
| TNRC18 | 378.646193 | 0.606776722 | 0.30257927 | 2.00534798 | 0.04492586 | 0.21105679 |
| FBXL18 | 33.1110312 | -0.256809523 | 0.29617242 | -0.8670947 | 0.38589014 | 0.66275158 |
| ACTB | 1167.03282 | 0.329903448 | 0.55442478 | 0.59503735 | 0.55181851 | 0.78577398 |
| RNF216 | 250.924454 | -0.060424641 | 0.24835671 | -0.2432978 | 0.80777471 | 0.92368079 |
| CCZ1 | 37.691414 | 0.439633489 | 0.34304772 | 1.28155199 | 0.19999985 | 0.47874144 |
| PMS2 | 57.5110547 | 0.221655269 | 0.30942583 | 0.71634379 | 0.4737791 | 0.73077309 |
| AIMP2 | 37.0955762 | -0.7510125 | 0.37775676 | -1.9880849 | 0.0468023 | 0.21609335 |
| EIF2AK1 | 109.625206 | -0.090502572 | 0.16385418 | -0.5523361 | 0.58071814 | 0.80525463 |
| USP42 | 96.7728463 | 0.106124847 | 0.21129115 | 0.50226831 | 0.6154788 | 0.82592881 |
| CYTH3 | 65.4809736 | 0.212144168 | 0.44536369 | 0.47633917 | 0.63383276 | 0.83652302 |
| RAC1 | 335.799347 | -0.459230115 | 0.20077641 | -2.2872713 | 0.02218 | 0.14286955 |
| DAGLB | 32.4226986 | -0.003440184 | 0.25060242 | -0.0137277 | 0.98904726 | 0.99678172 |
| ZDHHC4 | 25.360159 | -0.207539076 | 0.29299806 | -0.7083292 | 0.47874085 | 0.73456009 |
| ZNF316 | 40.1665498 | -0.00051912 | 0.29241386 | -0.0017753 | 0.99858352 | 0.99928153 |
| AC073343.2 | 35.2625289 | 0.160315485 | 0.3388597 | 0.47310284 | 0.63613981 | 0.83814123 |
| ZNF12 | 56.0985308 | 0.085093887 | 0.2459749 | 0.34594541 | 0.72938374 | 0.88653304 |
| CCZ1B | 42.3253862 | 0.135239744 | 0.24619025 | 0.54933022 | 0.58277885 | 0.80706067 |
| C1GALT1 | 202.733844 | 0.248160698 | 0.32820961 | 0.7561043 | 0.44958666 | 0.71054127 |
| COL28A1 | 96.8347538 | -0.296277461 | 0.65735582 | -0.4507109 | 0.65219789 | 0.84801249 |
| MIOS | 108.957114 | 0.313903236 | 0.36338914 | 0.86382117 | 0.38768614 | 0.66421848 |
| RPA3 | 56.4125578 | 0.391088957 | 0.45884625 | 0.85233116 | 0.39403031 | 0.66874847 |
| UMAD1 | 462.493786 | 0.514020793 | 0.25213749 | 2.03865275 | 0.0414847 | 0.20295123 |
| GLCCI1 | 289.04901 | 0.946688036 | 0.43591088 | 2.17174677 | 0.02987477 | 0.16887967 |
| ICA1 | 153.375858 | -0.545805204 | 0.42804658 | -1.275107 | 0.20227136 | 0.48090279 |
| NDUFA4 | 235.158689 | -0.372995924 | 0.48958981 | -0.7618539 | 0.44614718 | 0.70815763 |
| PHF14 | 323.065721 | 0.171560275 | 0.1748203 | 0.98135214 | 0.32641912 | 0.61297172 |
| THSD7A | 1635.0481 | 0.600632393 | 0.66034777 | 0.9095698 | 0.36304943 | 0.64351459 |
| TMEM106B | 168.25159 | -0.166223203 | 0.33471003 | -0.4966185 | 0.61945808 | 0.82770107 |
| SCIN | 579.227992 | 1.010808372 | 0.50003075 | 2.02149243 | 0.04322882 | 0.20614587 |
| ETV1 | 41.2408239 | -0.659866896 | 0.53044749 | -1.2439816 | 0.21350635 | 0.49544024 |
| ISPD | 128.95551 | 0.616202377 | 0.28792675 | 2.14013591 | 0.03234378 | 0.17593476 |
| ANKMY2 | 71.4706394 | 0.528847684 | 0.27147391 | 1.94806081 | 0.05140769 | 0.22752396 |
| BZW2 | 83.2905141 | -0.688771694 | 0.47282901 | -1.4567035 | 0.14519823 | 0.40323664 |
| AC073332.1 | 53.1694773 | -0.425186743 | 0.37983842 | -1.1193884 | 0.26297446 | 0.55096925 |
| AC019117.1 | 120.408672 | -2.22442506 | 1.26809322 | -1.7541495 | 0.0794049 | 0.29254596 |
| SNX13 | 290.364016 | 0.338082754 | 0.10952328 | 3.08685742 | 0.00202285 | 0.03208086 |
| HDAC9 | 139.647102 | -1.153241965 | 1.17348266 | -0.9827516 | 0.32572971 | 0.61243485 |
| TWISTNB | 30.0363341 | -0.532420056 | 0.41472976 | -1.2837759 | 0.1992204 | 0.4782544 |
| MACC1 | 719.733561 | 0.671888748 | 0.55020548 | 1.22115969 | 0.22202556 | 0.50375143 |
| ITGB8 | 1593.44624 | 0.779748805 | 0.39117951 | 1.99332732 | 0.04622562 | 0.21432487 |
| SP4 | 72.4416813 | 0.950009972 | 0.20945164 | 4.53570085 | 5.74E-06 | 0.00061793 |
| DNAH11 | 175.464294 | 3.178355482 | 0.80413123 | 3.95253334 | 7.73E-05 | 0.0039824 |
| CDCA7L | 30.5019508 | 0.543796362 | 0.35067933 | 1.55069409 | 0.12097501 | 0.36534213 |
| RAPGEF5 | 128.138358 | -1.362182067 | 0.56002143 | -2.4323749 | 0.01500017 | 0.11379145 |
| STEAP1B | 302.254316 | -0.361716584 | 0.42977689 | -0.8416381 | 0.39999058 | 0.67468036 |
| TOMM7 | 126.105534 | 0.097258773 | 0.43401593 | 0.22409033 | 0.82268702 | 0.93217972 |
| FAM126A | 43.6859925 | 0.005596002 | 0.33110906 | 0.01690078 | 0.98651577 | 0.99678172 |
| KLHL7 | 93.1601122 | 0.343820111 | 0.24506181 | 1.40299346 | 0.16061879 | 0.42659999 |
| NUPL2 | 74.369495 | -0.001947247 | 0.21834464 | -0.0089182 | 0.99288438 | 0.99830371 |
| AC005082.1 | 28.7351062 | 1.10259217 | 0.85379879 | 1.2913958 | 0.19656647 | 0.47560247 |
| GPNMB | 62.413217 | -0.111807321 | 1.05585564 | -0.1058926 | 0.91566755 | 0.9691911 |
| MALSU1 | 63.2835478 | 0.069073268 | 0.23999624 | 0.28780979 | 0.77349235 | 0.90643453 |
| TRA2A | 409.733535 | 0.368445163 | 0.22768484 | 1.61822438 | 0.10561425 | 0.33994587 |
| CCDC126 | 38.4057441 | 0.197687804 | 0.24892987 | 0.79415059 | 0.42710777 | 0.69319982 |
| FAM221A | 29.4773046 | 0.681743833 | 0.35998088 | 1.89383346 | 0.05824712 | 0.24353957 |
| MPP6 | 343.715462 | -0.376802164 | 0.29844227 | -1.262563 | 0.20674627 | 0.48621411 |
| OSBPL3 | 234.591139 | 1.129476944 | 0.67105149 | 1.68314497 | 0.09234703 | 0.31660422 |
| CYCS | 76.7859632 | -0.532778721 | 0.48494097 | -1.0986465 | 0.27192227 | 0.55916275 |
| NFE2L3 | 40.6648035 | -0.2654513 | 0.23988845 | -1.1065614 | 0.2684836 | 0.55558531 |
| HNRNPA2B1 | 765.741423 | 0.257195045 | 0.20046757 | 1.28297582 | 0.19950055 | 0.47837581 |
| CBX3 | 144.228204 | -0.861352251 | 0.19248614 | -4.4748794 | 7.65E-06 | 0.00073236 |
| SNX10 | 259.13621 | -2.129005276 | 0.37257406 | -5.7143144 | 1.10E-08 | 5.02E-06 |
| AC004540.1 | 152.262196 | 1.600416629 | 0.46635053 | 3.43178901 | 0.00059961 | 0.0149187 |
| SKAP2 | 669.227858 | 0.235719058 | 0.25745207 | 0.91558422 | 0.35988501 | 0.64005542 |
| HOTAIRM1 | 89.5030723 | 0.022294481 | 0.32350391 | 0.06891565 | 0.94505676 | 0.98119662 |
| HOXA3 | 143.728015 | 1.012956901 | 0.34484637 | 2.93741504 | 0.00330961 | 0.04354772 |
| HOXA-AS2 | 66.1704833 | 0.745768917 | 0.46354804 | 1.60882767 | 0.10765403 | 0.34308427 |
| HOXA7 | 35.55364 | -0.75210303 | 0.50661693 | -1.4845596 | 0.13766052 | 0.39024863 |
| HOXA9 | 77.3921155 | 0.341017237 | 0.2542496 | 1.3412695 | 0.17983297 | 0.451844 |
| HOXA10-AS | 28.2245106 | -0.671076421 | 0.51573383 | -1.3012069 | 0.19318764 | 0.47123616 |
| HIBADH | 343.53762 | 0.282703255 | 0.21276167 | 1.32873209 | 0.18393638 | 0.45721925 |
| TAX1BP1 | 434.411779 | 0.117781473 | 0.20044021 | 0.58761398 | 0.55679142 | 0.7883969 |
| JAZF1 | 363.869958 | -0.117801675 | 0.32326332 | -0.364414 | 0.71554889 | 0.88009244 |
| CREB5 | 224.207354 | -0.330868744 | 0.54218262 | -0.6102533 | 0.54169402 | 0.77887826 |
| AC005162.3 | 121.538966 | 1.238273951 | 0.51815665 | 2.38976757 | 0.01685904 | 0.12184079 |
| CPVL | 406.819571 | 1.02710962 | 0.47901754 | 2.14420042 | 0.03201683 | 0.17525099 |
| WIPF3 | 54.7725272 | 0.464938579 | 0.46395721 | 1.00211522 | 0.31628795 | 0.60281198 |
| SCRN1 | 110.42467 | -0.245791792 | 0.24866587 | -0.988442 | 0.32293622 | 0.61001895 |
| FKBP14 | 25.2823721 | -0.423068667 | 0.33627709 | -1.2580954 | 0.20835724 | 0.48858116 |
| PLEKHA8 | 80.9538553 | 0.067637608 | 0.26986696 | 0.25063316 | 0.80209774 | 0.92119571 |
| MTURN | 110.017218 | 0.978940768 | 0.62701826 | 1.56126357 | 0.11846158 | 0.36346683 |
| ZNRF2 | 110.797982 | -0.64958029 | 0.1809083 | -3.5906605 | 0.00032984 | 0.01025827 |
| AC006978.2 | 89.7474268 | -0.349054317 | 0.26259815 | -1.3292337 | 0.18377087 | 0.45721925 |
| AC005154.6 | 69.7167025 | 0.841253628 | 0.26682637 | 3.15281298 | 0.00161705 | 0.02795985 |
| GARS | 91.0222506 | -0.571901504 | 0.32043543 | -1.7847636 | 0.07429967 | 0.28131087 |
| AQP1 | 603.516193 | -0.252732994 | 0.64566882 | -0.3914282 | 0.69548074 | 0.87024545 |
| PDE1C | 57.808629 | 1.834663742 | 0.72761286 | 2.52148339 | 0.01168612 | 0.09853991 |
| LSM5 | 32.3510718 | -0.096771214 | 0.39687477 | -0.2438331 | 0.80736007 | 0.92342711 |
| AVL9 | 319.901889 | -0.270098326 | 0.16438729 | -1.6430609 | 0.10037034 | 0.33096298 |
| KBTBD2 | 78.0877032 | -0.035911423 | 0.22976954 | -0.1562932 | 0.87580189 | 0.95314737 |
| FKBP9 | 56.6057257 | 0.038959176 | 0.28854489 | 0.13501946 | 0.89259648 | 0.95976294 |
| NT5C3A | 122.357275 | -0.168442936 | 0.38585231 | -0.4365477 | 0.6624394 | 0.85507439 |
| RP9 | 26.8687564 | 0.08738431 | 0.28772964 | 0.30370285 | 0.76135429 | 0.90141135 |
| BBS9 | 345.134458 | 0.498789759 | 0.26460391 | 1.88504307 | 0.05942402 | 0.24609715 |
| DPY19L1 | 89.7576906 | -0.275535639 | 0.29991846 | -0.9187018 | 0.35825155 | 0.64001011 |
| HERPUD2 | 107.650158 | 0.259348937 | 0.15337218 | 1.69097767 | 0.09084107 | 0.31436654 |

| | | | | | | |
|---|---|---|---|---|---|---|
| 7-Sep | 301.691888 | 0.22707003 | 0.12433994 | 1.82620351 | 0.06781961 | 0.26624757 |
| KIAA0895 | 67.2166688 | 0.098687164 | 0.27497452 | 0.35889567 | 0.71967314 | 0.88166633 |
| ELMO1 | 119.98452 | 0.40858507 | 1.19659679 | 0.34145593 | 0.73276038 | 0.88869659 |
| STARD3NL | 76.785246 | 0.013950581 | 0.21364711 | 0.06529731 | 0.94793729 | 0.98239654 |
| VPS41 | 457.82527 | 0.614872101 | 0.24922356 | 2.46715075 | 0.0136193 | 0.10755093 |
| POU6F2 | 96.2550242 | 0.400278927 | 1.12883412 | 0.35459499 | 0.72289303 | 0.88256338 |
| YAE1D1 | 37.0205056 | 0.392662968 | 0.23879173 | 1.6443742 | 0.10009893 | 0.33080044 |
| RALA | 150.51041 | -0.292493097 | 0.19625826 | -1.4903479 | 0.13613278 | 0.38809998 |
| CDK13 | 545.001777 | 0.281019919 | 0.14872448 | 1.8895337 | 0.05882035 | 0.24454867 |
| MPLKIP | 62.5046444 | 0.111779776 | 0.23284357 | 0.48006383 | 0.631182 | 0.83474975 |
| SUGCT | 398.823666 | 0.708761392 | 0.39072318 | 1.81397322 | 0.06968185 | 0.27065319 |
| GLI3 | 152.322248 | 0.001082285 | 0.42776121 | 0.00253011 | 0.99798126 | 0.99923686 |
| MRPL32 | 43.943662 | -0.587815223 | 0.36729703 | -1.6003811 | 0.10951406 | 0.34627843 |
| STK17A | 56.2922959 | -0.280225289 | 0.2893007 | -0.9686298 | 0.33272991 | 0.61719981 |
| COA1 | 357.077488 | 0.231697352 | 0.17123405 | 1.35310325 | 0.17602265 | 0.44807891 |
| BLVRA | 61.3072695 | -0.437558057 | 0.28286591 | -1.5468745 | 0.12189351 | 0.36668381 |
| URGCP | 68.0088529 | 0.158528717 | 0.30058597 | 0.52739892 | 0.59791659 | 0.81658679 |
| UBE2D4 | 27.2463232 | -0.192678694 | 0.31652899 | -0.6087237 | 0.54270761 | 0.77939972 |
| DBNL | 58.0935219 | -0.044023497 | 0.26181041 | -0.1681503 | 0.86646504 | 0.95083843 |
| POLM | 60.5871212 | 0.157029215 | 0.34030618 | 0.4614351 | 0.64448648 | 0.84377167 |
| AEBP1 | 184.934326 | -0.322384414 | 0.51967054 | -0.6203631 | 0.53501878 | 0.77486315 |
| POLD2 | 50.1855142 | -0.064586908 | 0.36598693 | -0.1764733 | 0.85992215 | 0.94745904 |
| YKT6 | 27.7723089 | -0.387324993 | 0.35518048 | -1.0905019 | 0.2754921 | 0.56268552 |
| NUDCD3 | 195.489964 | -0.127813506 | 0.16636377 | -0.7682773 | 0.44232246 | 0.70510851 |
| DDX56 | 40.6775481 | 0.11578787 | 0.28715076 | 0.40323024 | 0.68677885 | 0.86707288 |
| TMED4 | 89.9911332 | -0.255575167 | 0.26654123 | -0.9588579 | 0.33763033 | 0.62279241 |
| OGDH | 226.371222 | 0.008533477 | 0.14989803 | 0.05692855 | 0.95460211 | 0.98390867 |
| ZMIZ2 | 93.675081 | -0.115722674 | 0.30679532 | -0.3771983 | 0.70602624 | 0.87473681 |
| PPIA | 402.782456 | -0.724428884 | 0.44253387 | -1.6370021 | 0.10163002 | 0.33300898 |
| H2AFV | 128.105704 | 0.221972047 | 0.29016147 | 0.76499489 | 0.44427458 | 0.70681054 |
| PURB | 112.922012 | -0.007031541 | 0.19451364 | -0.0361493 | 0.97116328 | 0.99175186 |
| CCM2 | 119.988758 | -0.102552561 | 0.24891576 | -0.4119971 | 0.68034158 | 0.86244955 |
| TBRG4 | 29.9403075 | -0.581144087 | 0.29854907 | -1.9465614 | 0.05158735 | 0.22782627 |
| IGFBP3 | 287.08391 | 0.251568943 | 1.16186577 | 0.21652152 | 0.82858125 | 0.93365249 |
| AC004870.4 | 261.264807 | 2.009683971 | 1.54756683 | 1.29860884 | 0.1940782 | 0.47268627 |
| TNS3 | 1159.50983 | 0.408666668 | 0.22065735 | 1.85204192 | 0.0640198 | 0.25777457 |
| HUS1 | 46.5804498 | 0.154510795 | 0.27806936 | 0.55565559 | 0.57844632 | 0.80396861 |
| UPP1 | 56.5944107 | -2.364621163 | 0.56389081 | -4.1934026 | 2.75E-05 | 0.00194987 |
| ABCA13 | 387.105436 | 1.394707356 | 0.67154736 | 2.07685628 | 0.03781483 | 0.19316567 |
| GRB10 | 176.810755 | -0.844538768 | 0.43072602 | -1.9607331 | 0.04991016 | 0.2246661 |
| COBL | 359.043024 | 0.353017797 | 0.28193937 | 1.25210535 | 0.21053148 | 0.49127431 |
| SEC61G | 61.9743678 | -0.909526644 | 0.28873315 | -3.1500597 | 0.00163237 | 0.02807268 |
| EGFR | 804.903479 | 0.295503639 | 0.26763114 | 1.10414522 | 0.26953015 | 0.55667265 |
| LANCL2 | 64.7054326 | -0.455878664 | 0.2349833 | -1.9400471 | 0.05237397 | 0.22955543 |
| VOPP1 | 175.225071 | -0.476845286 | 0.23534797 | -2.0261288 | 0.04275159 | 0.20527192 |
| NIPSNAP2 | 207.21246 | 0.295731086 | 0.22859411 | 1.29369515 | 0.19577074 | 0.47439614 |
| PSPH | 39.0038141 | 1.168949148 | 0.42415599 | 2.75594164 | 0.00585234 | 0.06406812 |
| CCT6A | 91.7477657 | -0.776245416 | 0.38675711 | -2.0070618 | 0.04474309 | 0.21050788 |
| SUMF2 | 101.892875 | -0.077416914 | 0.18077916 | -0.4282403 | 0.66847621 | 0.85916414 |
| CHCHD2 | 213.120575 | -0.62838147 | 0.5361096 | -1.1721138 | 0.24115136 | 0.52565524 |
| ZNF727 | 36.0346519 | -0.086192239 | 0.45650067 | -0.1888108 | 0.85024114 | 0.94435157 |
| ZNF736 | 119.039879 | 0.118167238 | 0.2762241 | 0.42779481 | 0.66880052 | 0.85924744 |
| ZNF680 | 77.6006463 | 0.279146002 | 0.35990265 | 0.77561529 | 0.43797617 | 0.70228343 |
| ZNF107 | 40.9890282 | 0.670794517 | 0.42690649 | 1.57129144 | 0.11611497 | 0.35925881 |
| ZNF138 | 68.9734401 | 0.264014833 | 0.32719657 | 0.80689977 | 0.41972423 | 0.68835944 |
| ZNF117 | 62.580707 | 0.041485971 | 0.2373135 | 0.17481505 | 0.86122496 | 0.94835188 |
| VKORC1L1 | 179.494317 | 0.012447231 | 0.21863035 | 0.05693277 | 0.95459875 | 0.98390867 |
| GUSB | 38.6876255 | 0.041198507 | 0.26327258 | 0.15648613 | 0.87564985 | 0.95314737 |
| AC068533.4 | 32.9662129 | -0.040429043 | 0.42068728 | -0.0961024 | 0.92343927 | 0.97297642 |
| CRCP | 31.3174483 | -0.618399921 | 0.3305581 | -1.8707753 | 0.06137624 | 0.25071343 |
| AC068533.3 | 28.9475934 | 0.449213987 | 0.4498916 | 0.99849382 | 0.31803996 | 0.60470519 |
| TPST1 | 720.818799 | 0.273173928 | 0.33670791 | 0.81130831 | 0.41718864 | 0.68664089 |
| AC027644.4 | 124.102699 | 0.112889365 | 0.22841843 | 0.49422178 | 0.62114956 | 0.82845887 |
| AC027644.3 | 28.4036838 | -0.030983038 | 0.40955144 | -0.0756512 | 0.93969664 | 0.98003138 |
| TMEM248 | 119.879742 | -0.177205843 | 0.20496801 | -0.8645536 | 0.38728383 | 0.66404483 |
| SBDS | 96.207053 | -0.078216636 | 0.35245732 | -0.221918 | 0.82437769 | 0.93217972 |
| TYW1 | 235.92846 | 0.138764356 | 0.18038636 | 0.76926191 | 0.44173785 | 0.70476463 |
| AUTS2 | 1139.21983 | 0.784329896 | 0.55911062 | 1.40281702 | 0.16067141 | 0.42659999 |
| TYW1B | 166.212507 | 0.260994445 | 0.18489159 | 1.41160796 | 0.15806543 | 0.42352692 |
| POM121 | 175.516795 | 0.068339525 | 0.20729566 | 0.32967176 | 0.74164799 | 0.89253455 |
| NSUN5 | 38.4752908 | -0.134788529 | 0.31230585 | -0.4315914 | 0.66603838 | 0.85763969 |
| BAZ1B | 242.327104 | 0.663988306 | 0.16305555 | 4.07216022 | 4.66E-05 | 0.00292513 |
| BCL7B | 38.0563363 | 0.185055911 | 0.30191389 | 0.61294268 | 0.53991424 | 0.77760314 |
| MLXIPL | 81.3206564 | -0.047271217 | 0.6258139 | -0.0755356 | 0.93978859 | 0.98003138 |
| BUD23 | 69.9711029 | -0.233306932 | 0.34973195 | -0.6671021 | 0.50470689 | 0.75524917 |
| ABHD11 | 41.7648649 | -1.214427559 | 0.22476291 | -5.4031494 | 6.55E-08 | 2.09E-05 |
| CLDN3 | 158.391659 | -0.928912457 | 0.40270588 | -2.3066772 | 0.02107282 | 0.13931717 |
| CLDN4 | 207.126239 | -0.553869517 | 0.4652766 | -1.1904091 | 0.23388562 | 0.51653098 |
| LIMK1 | 32.7724614 | -1.166041118 | 0.42535721 | -2.7413221 | 0.00611925 | 0.06580904 |
| EIF4H | 120.594432 | -0.521318525 | 0.28365816 | -1.8378407 | 0.06608588 | 0.26198834 |
| CLIP2 | 144.960463 | -0.130812357 | 0.3678547 | -0.3556088 | 0.72213357 | 0.88235193 |
| GTF2IRD1 | 495.403035 | 0.385113134 | 0.34153794 | 1.12758523 | 0.25949513 | 0.54618849 |
| GTF2I | 844.069408 | 0.26475732 | 0.20451805 | 1.29454256 | 0.19547808 | 0.47392673 |
| GTF2IRD2 | 65.5043342 | -0.000231849 | 0.38509087 | -0.0006021 | 0.99951962 | 0.9998728 |
| CASTOR2 | 86.8350136 | -0.270974033 | 0.58376491 | -0.4641835 | 0.64251629 | 0.84298912 |
| RCC1L | 76.3492375 | 0.2924241 | 0.35625506 | 0.82082793 | 0.41174429 | 0.68343416 |
| GTF2IRD2B | 102.771227 | 0.128054462 | 0.23193695 | 0.55210894 | 0.58087372 | 0.80535379 |
| TRIM73 | 50.6730221 | 0.549253178 | 0.40114896 | 1.36920004 | 0.17093675 | 0.44115974 |
| POM121C | 151.476886 | 0.214986703 | 0.17257071 | 1.24578907 | 0.21284184 | 0.4947372 |
| SPDYE5 | 41.0553916 | -0.454084326 | 0.45106196 | -1.0067006 | 0.31407869 | 0.60099077 |
| HIP1 | 294.675572 | 0.394516691 | 0.38118097 | 1.03498527 | 0.30067579 | 0.58788025 |
| RHBDD2 | 44.933952 | -0.202200144 | 0.2825759 | -0.7155605 | 0.4742628 | 0.73113966 |
| POR | 152.142074 | -0.082480313 | 0.23802486 | -0.3465197 | 0.72895215 | 0.88653304 |
| TMEM120A | 26.2953323 | 0.342637805 | 0.32483549 | 1.05480411 | 0.29151493 | 0.5787402 |
| STYXL1 | 81.7444526 | 0.051242723 | 0.30655709 | 0.16715557 | 0.86724765 | 0.95090501 |
| MDH2 | 75.3437815 | -0.487415138 | 0.38940272 | -1.2516994 | 0.21067944 | 0.49138016 |
| HSPB1 | 233.880798 | -0.007547202 | 0.39223889 | -0.0192413 | 0.98464858 | 0.99668665 |
| YWHAG | 119.119178 | -0.793771957 | 0.27799624 | -2.8553334 | 0.00429917 | 0.05249369 |
| DTX2 | 61.6103463 | -0.267217122 | 0.24776127 | -1.0785266 | 0.28079881 | 0.56842177 |
| POMZP3 | 40.6921904 | -0.062805255 | 0.38272144 | -0.1641017 | 0.86965105 | 0.95106604 |
| CCDC146 | 205.374305 | 0.070671242 | 0.40762281 | 0.17337411 | 0.86235737 | 0.94877974 |
| GSAP | 137.247058 | -0.32032828 | 0.32860921 | -0.9748001 | 0.32965945 | 0.61471829 |
| PTPN12 | 417.691745 | -0.593215622 | 0.19095317 | -3.1066027 | 0.00189251 | 0.03090405 |
| APTR | 62.0617641 | 0.162559254 | 0.35401859 | 0.45918282 | 0.64610289 | 0.84427583 |
| RSBN1L | 213.231642 | 0.217830276 | 0.17071653 | 1.27597645 | 0.20196384 | 0.48041015 |
| PHTF2 | 148.055979 | 0.792274424 | 0.22224287 | 3.56490373 | 0.00036399 | 0.01082813 |
| MAGI2 | 2154.22563 | 1.011689547 | 0.40349913 | 2.50729053 | 0.01216607 | 0.1007368 |
| GNAI1 | 440.459812 | 0.671800479 | 0.30013086 | 2.23835859 | 0.02519768 | 0.15393404 |
| SEMA3C | 82.3365732 | -0.733017856 | 0.54078902 | -1.35546 | 0.17527106 | 0.44711624 |
| CACNA2D1 | 248.174669 | 0.673524583 | 0.25846415 | 2.6058724 | 0.00916406 | 0.08519229 |
| PCLO | 424.111807 | -0.575405691 | 0.49193423 | -1.1696801 | 0.24212972 | 0.52712741 |
| AC003984.1 | 323.201754 | 2.242824494 | 0.62205082 | 3.60553256 | 0.00031151 | 0.00994413 |
| KIAA1324L | 43.9339827 | 0.460750936 | 0.57423141 | 0.8023785 | 0.42233405 | 0.69043214 |

| | | | | | | |
|---|---|---|---|---|---|---|
| DMTF1 | 283.153305 | 0.607264713 | 0.26447103 | 2.29614833 | 0.0216674 | 0.14167375 |
| TMEM243 | 92.1973579 | -0.306380758 | 0.24252879 | -1.2632758 | 0.20649006 | 0.48621411 |
| CROT | 56.6083727 | 0.317219336 | 0.294099 | 1.07861411 | 0.28075979 | 0.56842177 |
| ABCB1 | 228.128977 | 1.051610358 | 0.28325115 | 3.71264283 | 0.00020511 | 0.00758576 |
| RUNDC3B | 31.1322505 | 0.468779896 | 0.28479292 | 1.64603772 | 0.09975599 | 0.33030164 |
| SLC25A40 | 54.450097 | 0.171251369 | 0.22368915 | 0.76557745 | 0.44392776 | 0.70661084 |
| ADAM22 | 40.4080925 | 1.094613762 | 0.60501837 | 1.80922401 | 0.07041621 | 0.27240724 |
| SRI | 61.9922321 | -0.327276469 | 0.3903722 | -0.8383703 | 0.40182274 | 0.67562929 |
| STEAP2 | 45.9682839 | 0.58301073 | 0.5609455 | 1.03933579 | 0.2986486 | 0.58610017 |
| CFAP69 | 153.49507 | 0.282876913 | 0.36825849 | 0.76814769 | 0.44239944 | 0.70510851 |
| GTPBP10 | 70.1867166 | 0.128827699 | 0.24437324 | 0.52717597 | 0.59807139 | 0.81658679 |
| CLDN12 | 71.1910018 | -0.516767001 | 0.25269681 | -2.045008 | 0.04085407 | 0.20099697 |
| CDK14 | 378.781984 | 0.008990401 | 0.46931024 | 0.01915663 | 0.98471616 | 0.99668665 |
| FZD1 | 33.224457 | -0.181599555 | 0.45639836 | -0.397897 | 0.69070608 | 0.86843279 |
| AC079760.1 | 35.9862943 | -0.438875023 | 0.46933526 | -0.9350992 | 0.34973718 | 0.63286907 |
| MTERF1 | 166.855912 | -0.373716113 | 0.30709393 | -1.216944 | 0.22362552 | 0.50557208 |
| AKAP9 | 608.547471 | 0.316454507 | 0.13509332 | 2.34248818 | 0.01915564 | 0.13134709 |
| KRIT1 | 193.555524 | 0.129839789 | 0.18483737 | 0.70245423 | 0.48239594 | 0.73686345 |
| ANKIB1 | 485.116994 | 0.020357195 | 0.17837992 | 0.11412268 | 0.90914054 | 0.96752108 |
| GATAD1 | 91.0069791 | -0.084961682 | 0.23084555 | -0.3680456 | 0.71283925 | 0.87827554 |
| AC007566.1 | 74.7594863 | -0.012186242 | 0.35472032 | -0.0343545 | 0.97259446 | 0.99238201 |
| PEX1 | 80.3146312 | 0.112266671 | 0.19516641 | 0.57523562 | 0.56513195 | 0.79410283 |
| FAM133B | 103.92547 | 0.118249107 | 0.19860301 | 0.59540441 | 0.55157318 | 0.78577398 |
| CDK6 | 264.278207 | 0.045777187 | 0.44183486 | 0.10360701 | 0.91748123 | 0.97046739 |
| SAMD9L | 79.881083 | -0.499675612 | 1.25051976 | -0.3995743 | 0.68947006 | 0.86755279 |
| VPS50 | 159.743907 | 0.159540592 | 0.17421657 | 0.91576013 | 0.35979272 | 0.64001011 |
| CALCR | 54.2501271 | -0.499961443 | 1.20393453 | -0.4152729 | 0.6779421 | 0.86151115 |
| GNGT1 | 218.438466 | -0.175186479 | 0.42904032 | -0.4083217 | 0.6830375 | 0.86454781 |
| TFPI2 | 72.0789125 | -1.668763798 | 1.02900709 | -1.6217224 | 0.10486281 | 0.33843695 |
| GNG11 | 51.638276 | -0.973084269 | 0.56592756 | -1.7194502 | 0.08553242 | 0.3044283 |
| BET1 | 70.0938168 | -0.079498635 | 0.28654077 | -0.2774427 | 0.78144022 | 0.91077326 |
| CASD1 | 138.919216 | 0.405162576 | 0.17041507 | 2.37750433 | 0.01743024 | 0.12460019 |
| SGCE | 141.863212 | 0.407007391 | 0.37155699 | 1.09541041 | 0.27333688 | 0.56090273 |
| PPP1R9A | 653.583036 | 0.098004297 | 0.18278378 | 0.53617612 | 0.59183683 | 0.81275584 |
| AC002429.2 | 49.8096052 | 0.21529426 | 0.34025647 | 0.63274112 | 0.52690271 | 0.76821934 |
| PON2 | 287.610021 | -0.580733064 | 0.36537025 | -1.5894372 | 0.11196172 | 0.35014082 |
| PDK4 | 140.584531 | -0.621150999 | 1.01889952 | -0.6096293 | 0.5421074 | 0.77900491 |
| SLC25A13 | 296.581643 | -0.336145582 | 0.25165398 | -1.3357451 | 0.18163259 | 0.45460829 |
| SEM1 | 153.434725 | 0.264319943 | 0.22480152 | 1.1757925 | 0.23967781 | 0.52362764 |
| SDHAF3 | 39.6518982 | -0.265623367 | 0.25491084 | -1.0420246 | 0.29740027 | 0.58521131 |
| ASNS | 28.5365651 | 0.025359783 | 0.380159 | 0.06670836 | 0.94681388 | 0.98158791 |
| LMTK2 | 230.913057 | 0.118514382 | 0.21387217 | 0.55413654 | 0.57948541 | 0.80456304 |
| TECPR1 | 29.9881662 | 0.389914393 | 0.36913391 | 1.05629524 | 0.29083335 | 0.57810597 |
| BRI3 | 248.224683 | -0.252612192 | 0.3279495 | -0.7702777 | 0.44113516 | 0.70438969 |
| BAIAP2L1 | 575.097361 | 0.250645181 | 0.49643688 | 0.50488832 | 0.61363728 | 0.82498688 |
| TMEM130 | 60.0386352 | 0.89665269 | 1.23038736 | 0.72875642 | 0.46615067 | 0.72394032 |
| TRRAP | 269.295245 | 0.136665507 | 0.18609891 | 0.73437025 | 0.46272311 | 0.72095238 |
| SMURF1 | 351.973598 | -0.290543126 | 0.25198102 | -1.1530358 | 0.24889571 | 0.53613042 |
| ARPC1B | 43.8906882 | -0.649084666 | 0.49299784 | -1.3166075 | 0.18797022 | 0.46346633 |
| PDAP1 | 60.5582349 | -0.649502321 | 0.31875146 | -2.037645 | 0.04158545 | 0.20312121 |
| BUD31 | 46.4744435 | -0.502158779 | 0.25900789 | -1.9387779 | 0.05252839 | 0.22987695 |
| CPSF4 | 29.1776897 | -0.59483372 | 0.33846364 | -1.7574523 | 0.07884074 | 0.29099563 |
| ZNF789 | 58.8122041 | 0.199894788 | 0.18224952 | 1.09681926 | 0.27272041 | 0.56035796 |
| ZNF394 | 41.8538264 | -0.052401261 | 0.24477619 | -0.2140783 | 0.83048603 | 0.93446017 |
| CYP3A5 | 158.20404 | -2.401247063 | 1.12523555 | -2.133995 | 0.03284319 | 0.17754227 |
| TRIM4 | 77.6166157 | -0.242818511 | 0.2222726 | -1.0924356 | 0.27464166 | 0.56187466 |
| ZKSCAN1 | 226.721916 | -0.003545335 | 0.16400519 | -0.0216172 | 0.9827533 | 0.99616479 |
| ZNF3 | 43.9914929 | -0.284618401 | 0.33038361 | -0.8614786 | 0.38897452 | 0.66513878 |
| COPS6 | 61.0912338 | -0.241295494 | 0.26919649 | -0.8963545 | 0.37006344 | 0.64995191 |
| MCM7 | 32.7025336 | -0.168021781 | 0.37330764 | -0.4500893 | 0.65264604 | 0.84837785 |
| AP4M1 | 34.9477246 | 0.045110902 | 0.27982792 | 0.16120944 | 0.87192845 | 0.95193455 |
| TAF6 | 32.1022244 | -0.304271973 | 0.27719475 | -1.097683 | 0.27234293 | 0.5597024 |
| LAMTOR4 | 125.871243 | -0.169926572 | 0.359292 | -0.4729484 | 0.63625 | 0.83814123 |
| STAG3 | 32.3746219 | -0.538827684 | 0.55141805 | -0.9771673 | 0.32848633 | 0.61394673 |
| CASTOR3 | 200.737832 | -0.540724748 | 0.24827819 | -2.1778987 | 0.02941358 | 0.16751053 |
| SPDYE3 | 110.334086 | 0.249921778 | 0.41179456 | 0.60690888 | 0.54391138 | 0.78027954 |
| PILRB | 159.968567 | -0.473427153 | 0.27806101 | -1.7026017 | 0.08864263 | 0.30992958 |
| PILRA | 36.1381303 | -0.351716219 | 0.2603442 | -1.3509662 | 0.17670625 | 0.44872611 |
| MEPCE | 25.4892665 | -0.531453928 | 0.39605937 | -1.3418542 | 0.17964328 | 0.45177289 |
| PPP1R35 | 40.5671669 | -0.682287234 | 0.35993042 | -1.8956087 | 0.0580118 | 0.24313032 |
| TSC22D4 | 65.3341916 | -0.020482788 | 0.34253524 | -0.0597976 | 0.95231683 | 0.98341878 |
| AGFG2 | 42.580805 | 0.748433739 | 0.55664094 | 1.34455388 | 0.17876934 | 0.45040283 |
| LRCH4 | 26.8468324 | 0.024061073 | 0.35300985 | 0.06815978 | 0.94565845 | 0.98119662 |
| GNB2 | 135.336194 | -1.208518886 | 0.36671355 | -3.2955392 | 0.00098233 | 0.02054528 |
| GIGYF1 | 85.709363 | 0.166209606 | 0.20682121 | 0.80363906 | 0.42160546 | 0.6898802 |
| EPHB4 | 36.8903439 | -0.4871329 | 0.38102005 | -1.2784968 | 0.20107433 | 0.4800825 |
| SLC12A9 | 74.6823994 | 0.186424251 | 0.3996734 | 0.46644147 | 0.64089953 | 0.84179029 |
| TRIP6 | 67.5828564 | -0.133713534 | 0.36352489 | -0.367825 | 0.71300374 | 0.87832598 |
| SRRT | 42.9881558 | -0.215936266 | 0.23993561 | -0.8999759 | 0.36813307 | 0.64798726 |
| TRIM56 | 198.7164 | -1.122083416 | 0.31216662 | -3.5945016 | 0.00032501 | 0.01020754 |
| AP1S1 | 46.6897909 | -1.050084824 | 0.39186435 | -2.6797151 | 0.00736848 | 0.07378801 |
| PLOD3 | 46.3369454 | -1.018782535 | 0.31492745 | -3.2349753 | 0.00121653 | 0.02363725 |
| ZNHIT1 | 79.797594 | -0.564048614 | 0.36222818 | -1.5571638 | 0.1194316 | 0.3645149 |
| FIS1 | 143.555043 | -0.297722506 | 0.3597223 | -0.8276454 | 0.40787135 | 0.68125539 |
| IFT22 | 45.1603884 | -0.490252765 | 0.39824388 | -1.2310365 | 0.2183092 | 0.50052269 |
| COL26A1 | 229.118917 | 2.475976088 | 0.56497222 | 4.382474 | 1.17E-05 | 0.00101506 |
| CUX1 | 1047.37164 | -0.020376081 | 0.26908703 | -0.075723 | 0.93963947 | 0.98003138 |
| PRKRIP1 | 337.640232 | -0.236503367 | 0.24336111 | -0.9718207 | 0.33113975 | 0.61627893 |
| ORAI2 | 122.617425 | -0.309784068 | 0.42134158 | -0.7352326 | 0.46219784 | 0.72048708 |
| POLR2J | 34.624869 | 0.1389676 | 0.38562884 | 0.3603662 | 0.7185733 | 0.88155276 |
| SPDYE2 | 27.4867813 | 0.027737037 | 0.39417895 | 0.07036661 | 0.94390187 | 0.98112027 |
| AC105052.3 | 72.4463726 | 0.547252553 | 0.44194417 | 1.23828437 | 0.21561064 | 0.49731142 |
| FAM185A | 124.461458 | 0.471297504 | 0.27625074 | 1.70604975 | 0.08799882 | 0.30876947 |
| FBXL13 | 158.15764 | -0.24031564 | 0.29788532 | -0.8067388 | 0.419817 | 0.68835944 |
| ARMC10 | 93.7138655 | -0.222749259 | 0.17672283 | -1.2604441 | 0.20750919 | 0.48718884 |
| NAPEPLD | 226.990561 | 1.664390283 | 0.33947162 | 4.90288494 | 9.44E-07 | 0.00016448 |
| PMPCB | 74.8521871 | 0.394180407 | 0.20309662 | 1.94085159 | 0.05227628 | 0.22928319 |
| DNAJC2 | 70.6577316 | -0.080959047 | 0.27129818 | -0.2984135 | 0.76538757 | 0.9036913 |
| PSMC2 | 61.3936167 | -0.338618094 | 0.22442549 | -1.5088219 | 0.1313443 | 0.38163088 |
| ORC5 | 96.839984 | -0.05357845 | 0.21514348 | -0.2490359 | 0.80333301 | 0.92125386 |
| LHFPL3 | 247.689311 | 1.017055062 | 0.52469947 | 1.93835733 | 0.05257964 | 0.22987695 |
| LINC01004 | 205.497771 | 0.382284357 | 0.21176911 | 1.80519417 | 0.0710443 | 0.27376561 |
| KMT2E | 617.70012 | 0.38936495 | 0.21280263 | 1.82969993 | 0.06729482 | 0.26472981 |
| SRPK2 | 688.639292 | 0.071287648 | 0.20100689 | 0.35465276 | 0.72284974 | 0.88256338 |
| PUS7 | 80.3684132 | -0.707367754 | 0.46777294 | -1.5122032 | 0.13048217 | 0.38094749 |
| RINT1 | 41.5371477 | 0.059276965 | 0.27334064 | 0.21686115 | 0.82831656 | 0.93346403 |
| ATXN7L1 | 242.018567 | 0.058965647 | 0.21191158 | 0.2782559 | 0.78081593 | 0.91077326 |
| CDHR3 | 103.618216 | -0.514080795 | 0.61682253 | -0.8334339 | 0.40460007 | 0.67791921 |
| SYPL1 | 170.505573 | -0.148765501 | 0.21551933 | -0.6902652 | 0.49002741 | 0.74227587 |
| NAMPT | 1505.86016 | -3.603874798 | 0.70294362 | -5.1268334 | 2.95E-07 | 6.88E-05 |
| PRKAR2B | 72.6163886 | 1.841355887 | 0.67179226 | 2.74096027 | 0.00612599 | 0.06580904 |
| HBP1 | 143.106991 | 0.263601859 | 0.22116827 | 1.19186109 | 0.23331572 | 0.5160831 |
| COG5 | 388.050372 | 0.175559929 | 0.1342204 | 1.30799741 | 0.19087419 | 0.46742434 |

| | | | | | | |
|---|---|---|---|---|---|---|
| BCAP29 | 74.1572525 | 0.059710113 | 0.31951508 | 0.18687729 | 0.85175684 | 0.94469179 |
| CBLL1 | 29.7625273 | 0.090095512 | 0.23303226 | 0.38662248 | 0.69903571 | 0.8717697 |
| SLC26A3 | 621.333007 | 0.736541015 | 0.56866178 | 1.29521808 | 0.19524501 | 0.47360141 |
| DLD | 102.465339 | 0.165924271 | 0.23463011 | 0.70717383 | 0.47945846 | 0.73490443 |
| LAMB1 | 870.691895 | -0.210685676 | 0.50743802 | -0.4151949 | 0.67799924 | 0.86151115 |
| NRCAM | 107.844023 | -2.846317141 | 0.7318251 | -3.8893407 | 0.00010052 | 0.00465146 |
| PNPLA8 | 130.657693 | 0.09722884 | 0.24157064 | 0.40248616 | 0.68732627 | 0.86733718 |
| DNAJB9 | 36.7707028 | -0.274581972 | 0.52357179 | -0.52444 | 0.59997257 | 0.81763227 |
| IMMP2L | 3479.72174 | 0.726643806 | 0.32727693 | 2.22027201 | 0.02640031 | 0.15746485 |
| DOCK4 | 809.379342 | -0.160890982 | 0.5317362 | -0.3025767 | 0.76221248 | 0.90141135 |
| ZNF277 | 197.633884 | -0.113875866 | 0.21317807 | -0.5341819 | 0.59321569 | 0.81351655 |
| IFRD1 | 131.177979 | 0.425501314 | 0.38342336 | 1.10974281 | 0.26710987 | 0.55453954 |
| TMEM168 | 138.812422 | 0.451970542 | 0.14739722 | 3.06634365 | 0.00216694 | 0.03358759 |
| BMT2 | 163.276597 | 0.434665682 | 0.21384503 | 2.03262002 | 0.04209093 | 0.2037337 |
| SMIM30 | 29.1526231 | -0.475803864 | 0.43952828 | -1.082533 | 0.27901576 | 0.56636829 |
| FOXP2 | 1918.99365 | 0.99882977 | 0.57339944 | 1.74194409 | 0.08151823 | 0.29654614 |
| MDFIC | 187.029561 | 0.518585049 | 0.28298341 | 1.83256345 | 0.06686752 | 0.2635901 |
| TES | 110.859991 | 0.298954574 | 0.38822856 | 0.77004787 | 0.4412715 | 0.70443757 |
| CAV2 | 64.2538148 | 0.351920374 | 0.4316609 | 0.81527044 | 0.41491752 | 0.68495052 |
| MET | 606.48818 | -0.225224317 | 0.38104535 | -0.5910696 | 0.5544738 | 0.78734553 |
| CAPZA2 | 271.776591 | 0.03154895 | 0.21655786 | 0.1456837 | 0.88417109 | 0.95761929 |
| ST7 | 293.705378 | 0.018166419 | 0.24256614 | 0.07489264 | 0.94030013 | 0.98016757 |
| CFTR | 66.2059966 | -1.572569663 | 0.76176461 | -2.0643774 | 0.03898194 | 0.19580915 |
| CTTNBP2 | 696.028709 | 0.551589899 | 0.53978285 | 1.02187369 | 0.30684069 | 0.59449635 |
| LSM8 | 74.4172651 | 0.094603702 | 0.21888717 | 0.43220306 | 0.66559384 | 0.85742752 |
| TSPAN12 | 344.569044 | 0.569711617 | 0.53272091 | 1.06943732 | 0.28487266 | 0.57246822 |
| ING3 | 46.0556506 | 0.36592747 | 0.21904243 | 1.67057806 | 0.09480505 | 0.3204651 |
| FAM3C | 206.250211 | -0.44335915 | 0.30334953 | -1.4615455 | 0.14386579 | 0.40130764 |
| AASS | 99.9008356 | 1.065663528 | 0.32485414 | 3.28043698 | 0.00103646 | 0.0213418 |
| CADPS2 | 429.679765 | 0.459542653 | 0.31997593 | 1.4361788 | 0.15095147 | 0.41230799 |
| AC004594.1 | 27.97976 | 0.943439065 | 0.5850568 | 1.61255977 | 0.10684018 | 0.34182436 |
| SLC13A1 | 105.476656 | -0.673569301 | 1.19889889 | -0.5618233 | 0.57423643 | 0.80072326 |
| IQUB | 90.7058103 | 0.802430841 | 0.41326905 | 1.9416669 | 0.05217744 | 0.2289545 |
| NDUFA5 | 128.252552 | 0.43332451 | 0.37873974 | 1.14412211 | 0.252573 | 0.5393622 |
| WASL | 237.879814 | 0.258843086 | 0.17318718 | 1.49458574 | 0.13502259 | 0.38712403 |
| POT1 | 99.6528281 | 0.336104793 | 0.20194576 | 1.664332 | 0.0960461 | 0.32319324 |
| POT1-AS1 | 39.3706775 | 1.090102612 | 0.39902554 | 2.7319119 | 0.0062968 | 0.06672237 |
| GRM8 | 105.204234 | 0.514674669 | 1.13633341 | 0.45292576 | 0.6506022 | 0.84724399 |
| ZNF800 | 136.915263 | 0.108990084 | 0.24104386 | 0.45215873 | 0.65115463 | 0.84724399 |
| ARF5 | 40.3418045 | -0.332084761 | 0.41120416 | -0.807591 | 0.41932609 | 0.68835944 |
| SND1 | 472.556434 | -0.412095821 | 0.28684554 | -1.4366472 | 0.15081828 | 0.41206169 |
| RBM28 | 64.0983524 | -0.000675415 | 0.26787737 | -0.0025214 | 0.99798825 | 0.99923686 |
| HILPDA | 122.137738 | -4.26967941 | 1.31329717 | -3.2511144 | 0.00114954 | 0.02284523 |
| METTL2B | 70.1539735 | -0.339747092 | 0.23781598 | -1.4286134 | 0.15311538 | 0.41574018 |
| CALU | 115.730743 | 0.125146417 | 0.27621784 | 0.45307144 | 0.6504973 | 0.84724399 |
| KCP | 111.235613 | 0.638261052 | 0.46184098 | 1.38199311 | 0.16697382 | 0.43534082 |
| ATP6V1F | 120.165757 | -1.175040151 | 0.48108471 | -2.4424807 | 0.01458671 | 0.11197822 |
| TNPO3 | 239.000452 | 0.033124473 | 0.15952835 | 0.20764004 | 0.83551004 | 0.93718696 |
| TSPAN33 | 48.9828755 | 0.269440453 | 0.62598748 | 0.43042467 | 0.66688676 | 0.85838595 |
| SMO | 53.4659845 | 0.52164375 | 0.50303082 | 1.03700157 | 0.29973513 | 0.58681504 |
| AHCYL2 | 327.236852 | 0.13735656 | 0.20847295 | 0.65886992 | 0.5099793 | 0.75867242 |
| NRF1 | 271.463827 | 0.221422639 | 0.1598932 | 1.38481584 | 0.16610879 | 0.43446007 |
| UBE2H | 587.485691 | 0.14378329 | 0.17149726 | 0.83839992 | 0.40180614 | 0.67562929 |
| ZC3HC1 | 66.7354413 | 0.278453674 | 0.24324114 | 1.1447639 | 0.25230697 | 0.53930681 |
| KLHDC10 | 269.921577 | -0.140969509 | 0.17143079 | -0.8223115 | 0.41089963 | 0.68247066 |
| TMEM209 | 47.622061 | 0.069815536 | 0.3330141 | 0.20964739 | 0.83394289 | 0.93647023 |
| COPG2 | 88.7411084 | 0.082163541 | 0.3038715 | 0.2703891 | 0.78686092 | 0.91339653 |
| AC016831.7 | 1933.30542 | 0.311340402 | 0.29724342 | 1.0474257 | 0.29490328 | 0.58292995 |
| LINC00513 | 223.474592 | -0.169796379 | 0.48722178 | -0.3484992 | 0.72746535 | 0.88564265 |
| AC058791.1 | 41.0161858 | 0.11678132 | 0.46906034 | 0.24896865 | 0.80338503 | 0.92125386 |
| MKLN1 | 1461.81955 | 0.297398797 | 0.26263442 | 1.13236793 | 0.25747978 | 0.54481971 |
| PODXL | 33.2525347 | -0.795959171 | 0.55655522 | -1.4301531 | 0.15267309 | 0.41499872 |
| PLXNA4 | 46.7855692 | 0.022830088 | 1.25049113 | 0.0182569 | 0.98543391 | 0.99668665 |
| CHCHD3 | 698.744169 | -0.238014589 | 0.2472643 | -0.9625918 | 0.33575242 | 0.62028397 |
| EXOC4 | 1455.14643 | 0.406826743 | 0.14605318 | 2.78546998 | 0.00534502 | 0.06041319 |
| LRGUK | 76.3758678 | 0.087908488 | 0.37946574 | 0.23166383 | 0.81679912 | 0.92934063 |
| SLC35B4 | 42.9444618 | 0.583671385 | 0.25712213 | 2.27001612 | 0.02320661 | 0.14695664 |
| AKR1B1 | 84.6309634 | 0.825041851 | 0.64380199 | 1.28151491 | 0.20001287 | 0.47874144 |
| BPGM | 27.3815781 | -1.325989736 | 0.57779801 | -2.2949019 | 0.02173875 | 0.14167784 |
| CALD1 | 357.174774 | 0.026538269 | 0.27677037 | 0.09588551 | 0.9236115 | 0.97297642 |
| AGBL3 | 132.1447 | 0.390421343 | 0.40298592 | 0.96882131 | 0.33263435 | 0.61719981 |
| CYREN | 79.5604702 | -0.187281489 | 0.30602855 | -0.6119739 | 0.54055502 | 0.77817501 |
| TMEM140 | 65.2699947 | -1.205839901 | 0.7748725 | -1.5561785 | 0.11966566 | 0.36465111 |
| WDR91 | 126.487998 | 0.102600073 | 0.23933208 | 0.42869335 | 0.6681464 | 0.85905308 |
| CNOT4 | 371.740493 | 0.161843762 | 0.15092188 | 1.07236781 | 0.28355486 | 0.57084461 |
| NUP205 | 92.2692163 | 0.487484088 | 0.32447204 | 1.50239164 | 0.13299599 | 0.38380408 |
| STMP1 | 148.702222 | 0.245181484 | 0.33095905 | 0.7408212 | 0.45880186 | 0.71752865 |
| MTPN | 122.28227 | -0.144025329 | 0.23967489 | -0.6009196 | 0.54789357 | 0.7834123 |
| CREB3L2 | 159.80987 | 0.043515862 | 0.36405432 | 0.11953123 | 0.9048545 | 0.96553428 |
| TRIM24 | 177.920731 | 0.301352645 | 0.28442687 | 1.05950834 | 0.28936833 | 0.57591092 |
| ATP6V0A4 | 27.5373332 | 1.891841867 | 0.61248433 | 3.08880045 | 0.00200966 | 0.03192466 |
| KIAA1549 | 87.0881278 | 0.317440659 | 0.42756363 | 0.74244075 | 0.45782034 | 0.71693004 |
| ZC3HAV1 | 156.255918 | -0.555135961 | 0.46286219 | -1.1993547 | 0.23039004 | 0.51368983 |
| TTC26 | 70.854342 | 0.19622079 | 0.27971265 | 0.70150847 | 0.48298575 | 0.73696727 |
| UBN2 | 196.961874 | 0.356330179 | 0.21541387 | 1.65416544 | 0.0980939 | 0.32705935 |
| LUC7L2 | 65.645701 | 0.190558397 | 0.21690582 | 0.87853059 | 0.37965584 | 0.65703312 |
| KLRG2 | 206.728452 | 1.529637076 | 0.37803692 | 4.04626371 | 5.20E-05 | 0.00303967 |
| HIPK2 | 1690.14003 | 0.424158024 | 0.26654835 | 1.59129863 | 0.11154239 | 0.34917141 |
| TBXAS1 | 123.370705 | 0.154527955 | 0.39982433 | 0.38648962 | 0.69913409 | 0.87177889 |
| PARP12 | 94.8538876 | -0.535184596 | 0.72237882 | -0.7408642 | 0.4587758 | 0.71752865 |
| KDM7A | 178.317934 | -0.389667658 | 0.30018044 | -1.2981114 | 0.19424905 | 0.47284316 |
| SLC37A3 | 120.570384 | 0.303787019 | 0.2418567 | 1.25606206 | 0.20909347 | 0.48959511 |
| MKRN1 | 93.9396989 | 0.072476557 | 0.20121627 | 0.36019234 | 0.71870331 | 0.88155276 |
| DENND2A | 121.005742 | 0.353166609 | 0.49095936 | 0.71933981 | 0.47193157 | 0.72972277 |
| NDUFB2 | 128.368862 | -0.11136131 | 0.27181025 | -0.4097024 | 0.68202427 | 0.86382527 |
| BRAF | 763.310506 | 0.698526259 | 0.13805563 | 5.05974499 | 4.20E-07 | 8.82E-05 |
| MRPS33 | 45.1815394 | -0.244694733 | 0.31171553 | -0.7849937 | 0.43245724 | 0.69739191 |
| TMEM178B | 451.55319 | -0.293991411 | 0.71203916 | -0.4128866 | 0.67968972 | 0.86235436 |
| AGK | 94.9121882 | 0.330333295 | 0.18642875 | 1.77190102 | 0.07641099 | 0.28546836 |
| AC004918.1 | 27.3927387 | 0.58286265 | 0.35683257 | 1.63343454 | 0.10237764 | 0.33390378 |
| KIAA1147 | 98.2109546 | 0.469061966 | 0.26882663 | 1.74484931 | 0.0810111 | 0.29573375 |
| SSBP1 | 122.65443 | 0.11993047 | 0.19458943 | 0.61632573 | 0.53767956 | 0.77644087 |
| GSTK1 | 94.3265911 | -0.55984313 | 0.2899299 | -1.9309603 | 0.05348796 | 0.23204765 |
| TMEM139 | 30.85729 | 1.076134565 | 0.46487453 | 2.31489249 | 0.02061882 | 0.1372534 |
| CASP2 | 46.5968628 | 0.905207324 | 0.27100405 | 3.34019845 | 0.00083719 | 0.01839312 |
| ZYX | 56.3816702 | -0.304250918 | 0.33308082 | -0.9134447 | 0.36100869 | 0.64137683 |
| EPHA1 | 23.9639061 | -0.379591044 | 0.45000296 | -0.8435301 | 0.39893204 | 0.67377896 |
| TCAF2 | 70.6029267 | -1.594514577 | 0.66085716 | -2.4127976 | 0.01583061 | 0.11718809 |
| TCAF1 | 133.935748 | -0.263182352 | 0.19315902 | -1.3625165 | 0.17303494 | 0.44428037 |
| AC004889.1 | 189.346663 | 2.113965348 | 0.45810774 | 4.6145593 | 3.94E-06 | 0.00045463 |
| AC074386.1 | 281.28236 | 1.200523885 | 0.37472597 | 3.20373814 | 0.00135656 | 0.02537982 |
| ARHGEF5 | 65.5917981 | 0.580176524 | 0.47521699 | 1.22086655 | 0.22213655 | 0.50375143 |
| TPK1 | 352.710833 | 0.298131419 | 0.27046323 | 1.10229926 | 0.27033159 | 0.55746512 |

| | | | | | | |
|---|---|---|---|---|---|---|
| CNTNAP2 | 32.5524272 | 1.226063584 | 0.41407531 | 2.9609676 | 0.00306674 | 0.04154042 |
| CUL1 | 212.56677 | -0.117846031 | 0.13065385 | -0.9019714 | 0.36707207 | 0.64671387 |
| EZH2 | 32.7489693 | 0.273645818 | 0.31914413 | 0.85743647 | 0.39120371 | 0.66671992 |
| PDIA4 | 71.0897118 | -1.266680198 | 0.54571562 | -2.3211361 | 0.0202795 | 0.1355161 |
| ZNF398 | 65.7590719 | -0.298904011 | 0.2331748 | -1.2818881 | 0.19988189 | 0.47866717 |
| ZNF282 | 42.391429 | -0.350298882 | 0.27899532 | -1.2555726 | 0.20927096 | 0.48959511 |
| ZNF783 | 34.5190485 | 0.039924775 | 0.26035676 | 0.15334642 | 0.8781251 | 0.95466137 |
| ZNF746 | 27.3295798 | -0.240511739 | 0.33652848 | -0.7146847 | 0.47480392 | 0.73162933 |
| ZNF862 | 78.3142777 | 0.175953787 | 0.2517924 | 0.69880501 | 0.4846739 | 0.73787211 |
| ACTR3C | 274.757454 | -0.183468824 | 0.41112594 | -0.4462594 | 0.65540986 | 0.85058543 |
| LRRC61 | 39.4260204 | -0.620762421 | 0.28319365 | -2.1920069 | 0.02837901 | 0.16346515 |
| RARRES2 | 77.4767382 | 0.473757874 | 0.48071827 | 0.98552084 | 0.3243683 | 0.61115735 |
| TMEM176B | 168.685332 | -1.54647794 | 0.58910618 | -2.625126 | 0.00866169 | 0.08247549 |
| TMEM176A | 95.0253895 | -1.490160962 | 0.53160643 | -2.8031282 | 0.00506095 | 0.05854938 |
| ABCB8 | 26.0699857 | -0.38952043 | 0.40156459 | -0.9700069 | 0.33204305 | 0.61700946 |
| SLC4A2 | 48.9754702 | -0.419210711 | 0.32460128 | -1.2914635 | 0.196543 | 0.47560247 |
| FASTK | 34.5544285 | 0.068623115 | 0.34733686 | 0.19756934 | 0.84338203 | 0.94189614 |
| TMUB1 | 41.432882 | -0.303130951 | 0.42648204 | -0.7107707 | 0.47722632 | 0.73376419 |
| AGAP3 | 56.2705624 | -0.371964868 | 0.38535581 | -0.9652504 | 0.33441939 | 0.61889555 |
| SMARCD3 | 78.0830315 | -1.896230472 | 1.18392104 | -1.6016528 | 0.1092324 | 0.34587513 |
| NUB1 | 104.368754 | 0.107443033 | 0.43521911 | 0.24687113 | 0.80500796 | 0.92227408 |
| RHEB | 214.971145 | -0.125291842 | 0.20421509 | -0.6135288 | 0.53952673 | 0.77746559 |
| PRKAG2 | 178.991988 | 0.493435204 | 0.41548612 | 1.18760935 | 0.23498735 | 0.51805188 |
| GALNT11 | 268.572816 | 0.245484534 | 0.21995944 | 1.11604453 | 0.26440307 | 0.55239194 |
| KMT2C | 1057.5349 | 0.411813718 | 0.13638971 | 3.01938985 | 0.00253284 | 0.03659444 |
| ACTR3B | 100.246199 | -0.071027358 | 0.3034884 | -0.2340365 | 0.81495666 | 0.92814476 |
| DPP6 | 453.197273 | -0.530356084 | 0.68071752 | -0.7791133 | 0.43591297 | 0.70084094 |
| PAXIP1-AS2 | 40.2707294 | 0.523641771 | 0.52943301 | 0.98906144 | 0.32263308 | 0.60992743 |
| PAXIP1 | 71.779307 | 0.074567622 | 0.20153307 | 0.37000192 | 0.71138106 | 0.87748 |
| PAXIP1-AS1 | 31.0165627 | 0.509228362 | 0.33738289 | 1.5093485 | 0.13120975 | 0.38163088 |
| INSIG1 | 84.4855418 | 1.744165268 | 0.67525862 | 2.58295891 | 0.0097957 | 0.08848555 |
| AC009403.1 | 62.1363392 | -1.100737259 | 0.21722282 | -5.0673186 | 4.03E-07 | 8.78E-05 |
| RBM33 | 427.474119 | -0.191547768 | 0.17257082 | -1.1099661 | 0.26701361 | 0.55453954 |
| LINC01006 | 96.0231596 | -0.068240231 | 0.36282629 | -0.1880796 | 0.85081424 | 0.94435157 |
| LMBR1 | 320.760729 | 0.137032403 | 0.18366053 | 0.74611787 | 0.45559622 | 0.71496661 |
| NOM1 | 35.3842825 | -0.882650329 | 0.33810324 | -2.6105941 | 0.00903851 | 0.0847991 |
| UBE3C | 357.561755 | -0.114584272 | 0.22303253 | -0.5137559 | 0.60742268 | 0.8211189 |
| DNAJB6 | 186.166299 | 0.006432513 | 0.21413915 | 0.03003894 | 0.976036 | 0.99411128 |
| ESYT2 | 418.897283 | 0.729579937 | 0.37217912 | 1.96029249 | 0.04996161 | 0.22479206 |
| WDR60 | 292.427656 | 0.010386133 | 0.17823116 | 0.05827338 | 0.95353087 | 0.98368305 |
| GTPBP6 | 76.4413195 | 0.046933714 | 0.25120524 | 0.18683413 | 0.85179068 | 0.94469179 |
| IL3RA | 27.0448306 | 0.800735512 | 0.41884458 | 1.91177243 | 0.05590539 | 0.23748015 |
| SLC25A6 | 291.336396 | -0.840151176 | 0.40123004 | -2.0939389 | 0.03626542 | 0.18835699 |
| LINC00106 | 37.0770464 | 0.278324252 | 0.35373691 | 0.78681145 | 0.43139225 | 0.69696301 |
| ASMTL | 52.2770796 | -0.068700439 | 0.22220581 | -0.3091748 | 0.75718856 | 0.89933158 |
| AKAP17A | 50.8943265 | -0.068509313 | 0.27125023 | -0.2525687 | 0.80060153 | 0.9205332 |
| DHRSX | 371.815962 | 0.551873637 | 0.2420313 | 2.2801747 | 0.02259733 | 0.14449921 |
| CD99 | 50.4163315 | -0.610723715 | 0.64344975 | -0.9491397 | 0.34254955 | 0.62772374 |
| ARSD | 127.100657 | -0.047839692 | 0.41829072 | -0.1143695 | 0.90894491 | 0.96752108 |
| ARSE | 37.9216559 | -1.023056824 | 0.70802032 | -1.444954 | 0.1484708 | 0.40820743 |
| PRKX | 241.543178 | -0.276425256 | 0.36015467 | -0.7675182 | 0.44277348 | 0.70535959 |
| BX890604.1 | 37.3609918 | 0.530709988 | 0.40421207 | 1.31294939 | 0.1892 | 0.46534359 |
| PUDP | 75.7294099 | -0.366431427 | 0.33662832 | -1.0885342 | 0.27635936 | 0.56334633 |
| TBL1X | 918.277645 | 1.065753918 | 0.28485754 | 3.74135763 | 0.00018303 | 0.00709811 |
| GPR143 | 32.1379629 | -0.572772729 | 0.41692207 | -1.3738124 | 0.16949992 | 0.43941535 |
| SHROOM2 | 116.614125 | 0.001064193 | 0.47920598 | 0.00222074 | 0.99822811 | 0.9992713 |
| WWC3 | 83.4001313 | 0.359486927 | 0.24091793 | 1.49215515 | 0.13565848 | 0.38788633 |
| MID1 | 399.063682 | -0.378973694 | 0.45470001 | -0.8334587 | 0.40458606 | 0.67791921 |
| AC073529.1 | 43.6742888 | 0.109821541 | 0.32224876 | 0.3407974 | 0.73325611 | 0.88875873 |
| ARHGAP6 | 898.865734 | -0.028058635 | 0.26119016 | -0.1074261 | 0.91445096 | 0.9686812 |
| TMSB4X | 372.152513 | -0.48546435 | 0.52002006 | -0.9335493 | 0.35053645 | 0.63322593 |
| RAB9A | 76.2007075 | 0.172184957 | 0.29267621 | 0.58831209 | 0.55632283 | 0.78804453 |
| TRAPPC2 | 32.3208843 | 0.233434613 | 0.24901011 | 0.93745035 | 0.34852695 | 0.63229918 |
| OFD1 | 158.707951 | 0.190298535 | 0.25148973 | 0.75668511 | 0.44923854 | 0.71037275 |
| GEMIN8 | 51.6322323 | 0.034448962 | 0.24945198 | 0.13809857 | 0.89016252 | 0.95918866 |
| MOSPD2 | 42.661709 | -0.157848366 | 0.22354805 | -0.7061049 | 0.48012293 | 0.73526739 |
| PIR | 60.1118 | 0.27979346 | 0.29923157 | 0.93503991 | 0.34976773 | 0.63286907 |
| CLTRN | 151.713947 | -0.550610889 | 0.51800787 | -1.0629392 | 0.28780951 | 0.57450105 |
| CA5B | 118.536443 | 0.35147082 | 0.3675748 | 0.95618857 | 0.33897698 | 0.62369754 |
| ZRSR2 | 48.4454398 | 0.103114325 | 0.30198102 | 0.34145962 | 0.7327576 | 0.88869659 |
| AP1S2 | 34.633903 | 0.437659278 | 0.30295231 | 1.44464742 | 0.14855695 | 0.40820743 |
| CTPS2 | 108.376859 | 0.212618405 | 0.17182613 | 1.23740441 | 0.21593698 | 0.49746509 |
| SYAP1 | 99.5791177 | -0.104810959 | 0.26782241 | -0.391345 | 0.69554224 | 0.87024545 |
| TXLNG | 73.3733492 | -0.137291575 | 0.28684522 | -0.478626 | 0.63220473 | 0.8353707 |
| RBBP7 | 88.0835452 | -0.069428414 | 0.25046052 | -0.277203 | 0.78162422 | 0.91080813 |
| REPS2 | 223.561161 | -0.956419768 | 0.59388112 | -1.6104566 | 0.10729821 | 0.3426554 |
| NHS | 1064.87641 | -0.090934963 | 0.33894212 | -0.2682905 | 0.78847569 | 0.91447782 |
| SCML1 | 72.6898169 | -0.68098285 | 0.40392224 | -1.6859256 | 0.09181013 | 0.31558127 |
| RAI2 | 67.0615739 | 0.414515451 | 0.31457478 | 1.31770084 | 0.18760381 | 0.46292038 |
| CDKL5 | 160.239936 | 0.392172528 | 0.20445104 | 1.9181733 | 0.05508904 | 0.23568464 |
| PHKA2 | 81.2214146 | 0.052125152 | 0.23144238 | 0.2252187 | 0.82180914 | 0.93192711 |
| PDHA1 | 95.2593038 | 0.079874499 | 0.34776708 | 0.22967815 | 0.81834188 | 0.92999132 |
| MAP3K15 | 66.3021573 | 0.872299932 | 0.52944479 | 1.64757488 | 0.09943994 | 0.32948294 |
| SH3KBP1 | 95.0599434 | 0.387725303 | 0.79031999 | 0.49059281 | 0.62371448 | 0.83030289 |
| BCLAF3 | 91.1066147 | 0.414205704 | 0.1697058 | 2.44072802 | 0.01465769 | 0.11223502 |
| EIF1AX | 168.899894 | -0.217551493 | 0.25766395 | -0.8443226 | 0.39848918 | 0.67357117 |
| RPS6KA3 | 260.765176 | -0.606782039 | 0.17822249 | -3.4046323 | 0.00066253 | 0.01574788 |
| MBTPS2 | 66.133517 | -0.322879916 | 0.18981767 | -1.7010003 | 0.08894294 | 0.31048994 |
| SMS | 172.439828 | -0.076573301 | 0.24532651 | -0.3121281 | 0.75494316 | 0.89810084 |
| PRDX4 | 26.4726697 | -1.031774513 | 0.39262234 | -2.6279058 | 0.00859123 | 0.08196753 |
| ACOT9 | 86.0121148 | 0.808836058 | 0.31835207 | 2.54069675 | 0.01106318 | 0.09494336 |
| SAT1 | 1016.81145 | -1.275523704 | 0.64188745 | -1.9871454 | 0.04690628 | 0.21632898 |
| APOO | 818.561987 | -0.414151284 | 0.50308756 | -0.8232191 | 0.41038341 | 0.68223927 |
| EIF2S3 | 113.424498 | -0.471299446 | 0.21333992 | -2.209148 | 0.02716435 | 0.15924997 |
| ZFX | 157.446114 | 0.490659878 | 0.31556336 | 1.55486963 | 0.11997712 | 0.36481264 |
| PDK3 | 126.006179 | 0.579492445 | 0.45267646 | 1.28014706 | 0.20049342 | 0.47920186 |
| POLA1 | 140.89082 | 0.263155858 | 0.26103285 | 1.0081331 | 0.31339057 | 0.60065312 |
| GK | 52.1689706 | 0.236794993 | 0.30717268 | 0.77088558 | 0.44077475 | 0.70416634 |
| TAB3 | 109.069138 | -0.063063418 | 0.17332485 | -0.3638452 | 0.71597359 | 0.88028636 |
| DMD | 93.2404116 | 0.195803102 | 0.52188733 | 0.37518271 | 0.70752459 | 0.87544414 |
| TMEM47 | 39.749199 | 0.126029194 | 0.42289035 | 0.29801861 | 0.76568896 | 0.90371395 |
| AF241726.2 | 105.449531 | 0.813275279 | 0.3582819 | 2.26993126 | 0.02321175 | 0.14695664 |
| DYNLT3 | 46.8659737 | 0.103517556 | 0.37963898 | 0.27267368 | 0.78510407 | 0.91224065 |
| RPGR | 104.094336 | 0.041419523 | 0.32752184 | 0.12646339 | 0.89936512 | 0.96269027 |
| MID1IP1 | 37.3973659 | -0.603515642 | 0.37894001 | -1.5926416 | 0.11124061 | 0.34845447 |
| BCOR | 172.257037 | 0.244513368 | 0.27752328 | 0.88105534 | 0.37828787 | 0.65629152 |
| ATP6AP2 | 147.587548 | -0.02132 | 0.31872836 | -0.0668908 | 0.94666862 | 0.98158791 |
| CXorf38 | 67.7697684 | -0.097911264 | 0.23875033 | -0.410099 | 0.68173335 | 0.86357099 |
| MED14 | 163.228246 | 0.415312238 | 0.22115054 | 1.8779617 | 0.06038641 | 0.24793887 |
| USP9X | 453.896549 | -0.001476083 | 0.23025712 | -0.0064106 | 0.99488513 | 0.99886097 |
| DDX3X | 200.952937 | 0.32204892 | 0.25926316 | 1.24217001 | 0.21417384 | 0.49638791 |
| CASK | 394.681432 | -0.49021017 | 0.29513417 | -1.6609739 | 0.09671868 | 0.32454105 |
| Z93403.1 | 117.004032 | 1.111637984 | 0.62471004 | 1.77944631 | 0.07516662 | 0.28328492 |

| | | | | | | |
|---|---|---|---|---|---|---|
| MAOA | 311.620084 | -0.971781978 | 0.30073527 | -3.2313535 | 0.00123205 | 0.02374618 |
| MAOB | 126.106973 | -0.234795551 | 0.63169588 | -0.3716908 | 0.71012307 | 0.87725934 |
| EFHC2 | 71.121945 | 0.424051754 | 0.56920893 | 0.74498436 | 0.45628117 | 0.71557259 |
| KDM6A | 467.821367 | 0.475490775 | 0.32893528 | 1.44554509 | 0.14830484 | 0.40798738 |
| MIR222HG | 56.0477308 | 0.01938353 | 0.53377052 | 0.03631435 | 0.97103171 | 0.99175186 |
| KRBOX4 | 56.7993001 | -0.097849633 | 0.21256935 | -0.4603186 | 0.64528752 | 0.84407726 |
| SLC9A7 | 128.703396 | 0.21782099 | 0.40592066 | 0.53660976 | 0.5915372 | 0.81261076 |
| RP2 | 30.5560927 | 0.101669715 | 0.30920301 | 0.32881217 | 0.74229766 | 0.89253455 |
| JADE3 | 95.6012618 | -0.245419026 | 0.24912903 | -0.9851081 | 0.32457098 | 0.61129875 |
| RGN | 37.1134619 | -1.334937103 | 0.77261753 | -1.727811 | 0.08402211 | 0.30203879 |
| NDUFB11 | 84.9042278 | -0.082874507 | 0.45422885 | -0.182451 | 0.85522881 | 0.94576734 |
| RBM10 | 67.0461062 | 0.186154895 | 0.18739909 | 0.99336072 | 0.3205342 | 0.6072779 |
| UBA1 | 127.509335 | 0.139123981 | 0.21063815 | 0.66048805 | 0.50894068 | 0.75796558 |
| CDK16 | 39.6334009 | 0.009766553 | 0.30523757 | 0.03199656 | 0.97447479 | 0.99326197 |
| USP11 | 60.3256647 | 0.439150606 | 0.30705958 | 1.43018044 | 0.15266524 | 0.41499872 |
| ZNF41 | 33.1944868 | -0.004052515 | 0.26897112 | -0.0150667 | 0.98797895 | 0.99678172 |
| ARAF | 40.1307736 | -0.019044158 | 0.30895819 | -0.0616399 | 0.95084959 | 0.98286628 |
| TIMP1 | 212.681379 | -1.520889952 | 0.68383556 | -2.224058 | 0.02614454 | 0.15698891 |
| UXT | 26.0211179 | 0.173657801 | 0.31658181 | 0.54854004 | 0.58332114 | 0.80734478 |
| ZNF81 | 85.2656757 | 0.399805934 | 0.29013194 | 1.37801418 | 0.16819891 | 0.43715387 |
| ZNF630 | 31.5856766 | 0.544365529 | 0.45842296 | 1.18747439 | 0.23504055 | 0.51805188 |
| EBP | 33.6812303 | -0.54580576 | 0.32495967 | -1.6796108 | 0.09303307 | 0.31776118 |
| RBM3 | 102.539807 | -1.068162859 | 0.41720274 | -2.5602969 | 0.01045828 | 0.09182388 |
| WDR13 | 98.575198 | -0.458479356 | 0.26174286 | -1.7516404 | 0.07983566 | 0.29363267 |
| HDAC6 | 54.1124691 | -0.00785166 | 0.23974414 | -0.0327502 | 0.97387382 | 0.99315845 |
| PCSK1N | 68.0831236 | -0.840232473 | 0.57997481 | -1.4487396 | 0.14741032 | 0.40633477 |
| TIMM17B | 32.063132 | -0.405364738 | 0.46234136 | -0.876765 | 0.38061428 | 0.65803602 |
| OTUD5 | 86.2224414 | -0.116041071 | 0.19025962 | -0.6099091 | 0.54192204 | 0.77897227 |
| GRIPAP1 | 75.7498616 | -0.375167316 | 0.16133594 | -2.3253797 | 0.02005167 | 0.13480227 |
| TFE3 | 47.8427324 | -0.537134759 | 0.3982622 | -1.3486963 | 0.17743453 | 0.44901805 |
| CCDC120 | 28.1486282 | -0.089047817 | 0.2704066 | -0.3293108 | 0.74192078 | 0.89253455 |
| WDR45 | 46.2320373 | -0.639956873 | 0.26198299 | -2.442742 | 0.01457615 | 0.11197822 |
| PLP2 | 43.8763135 | -1.305453219 | 0.46889056 | -2.784132 | 0.00536712 | 0.06041319 |
| CLCN5 | 168.873189 | 0.3349334 | 0.40149734 | 0.83421077 | 0.40416223 | 0.67765972 |
| SHROOM4 | 191.039271 | 0.060942544 | 0.27321761 | 0.22305496 | 0.82349273 | 0.93217972 |
| MAGED1 | 226.21774 | -0.029123117 | 0.18190679 | -0.1600991 | 0.87280299 | 0.95193455 |
| FAM156A | 30.776596 | 0.056199297 | 0.40087823 | 0.14019044 | 0.88850952 | 0.95907089 |
| GPR173 | 49.3784949 | -0.076023441 | 0.43930181 | -0.1730552 | 0.86260808 | 0.94877974 |
| KANTR | 165.047146 | 0.436222645 | 0.26204089 | 1.66471213 | 0.0959702 | 0.32319324 |
| KDM5C | 75.8990926 | -0.055720428 | 0.34284907 | -0.1625217 | 0.87089501 | 0.95174591 |
| IQSEC2 | 40.8093317 | 0.187383328 | 0.33355973 | 0.56176843 | 0.5742738 | 0.80072326 |
| SMC1A | 83.4223954 | -0.052668256 | 0.29277858 | -0.1798911 | 0.85723807 | 0.94638973 |
| HSD17B10 | 48.6662423 | -0.732353116 | 0.47380476 | -1.5456854 | 0.12218056 | 0.36721014 |
| HUWE1 | 361.769606 | 0.153687596 | 0.13300864 | 1.15547078 | 0.24789769 | 0.53482252 |
| PHF8 | 95.5770156 | 0.354310288 | 0.20849621 | 1.69936079 | 0.08925122 | 0.31101837 |
| FAM120C | 62.9910729 | 0.027546132 | 0.25281531 | 0.10895753 | 0.91323618 | 0.96844354 |
| WNK3 | 31.1475533 | 0.170570231 | 0.3408379 | 0.50044385 | 0.61676258 | 0.82592881 |
| TSR2 | 31.8745523 | -0.514072718 | 0.3550276 | -1.4479796 | 0.14762276 | 0.40657927 |
| GNL3L | 26.3734163 | -0.127410323 | 0.30418061 | -0.418864 | 0.67531549 | 0.86115505 |
| MAGED2 | 71.1252515 | -0.22030511 | 0.3541763 | -0.622021 | 0.53392803 | 0.77366209 |
| FAM104B | 30.3903275 | 0.108471906 | 0.32643657 | 0.33229091 | 0.7396696 | 0.89253455 |
| RRAGB | 117.205346 | 0.285095103 | 0.19018018 | 1.49907891 | 0.13385316 | 0.38527026 |
| AL445472.1 | 182.774354 | 0.350370757 | 0.17483179 | 2.00404493 | 0.04506525 | 0.21106218 |
| KLF8 | 47.5211161 | 0.029373424 | 0.39027306 | 0.07526378 | 0.94000484 | 0.98003278 |
| UBQLN2 | 42.0308695 | -0.433480438 | 0.32226919 | -1.3450881 | 0.1785968 | 0.45020493 |
| NBDY | 151.845845 | 0.098217297 | 0.16931948 | 0.58007086 | 0.56186683 | 0.79172145 |
| SPIN3 | 40.6543693 | 0.494734885 | 0.24509859 | 2.01851383 | 0.04353778 | 0.2066642 |
| FAAH2 | 220.275654 | -0.042017716 | 0.28131827 | -0.1493601 | 0.88126952 | 0.95634765 |
| ARHGEF9 | 72.0314253 | -0.163900247 | 0.25463745 | -0.6436612 | 0.51979515 | 0.76385303 |
| MTMR8 | 28.8744566 | 0.496916656 | 0.3051622 | 1.62836897 | 0.10344667 | 0.33567605 |
| ZC4H2 | 87.9048593 | 0.40033824 | 0.2309708 | 1.73328506 | 0.08304501 | 0.29986597 |
| ZC3H12B | 57.7839213 | 0.268366011 | 0.27870744 | 0.96289503 | 0.3356002 | 0.62024201 |
| LAS1L | 35.3682807 | -0.43472938 | 0.2825952 | -1.5383467 | 0.12396388 | 0.37032493 |
| MSN | 386.059447 | -0.245831495 | 0.22640254 | -1.085816 | 0.27756042 | 0.56497051 |
| AR | 48.5668177 | 0.243231802 | 0.5997635 | 0.40554619 | 0.68507607 | 0.86561619 |
| OPHN1 | 399.849823 | 1.679867227 | 0.41084496 | 4.08881057 | 4.34E-05 | 0.00278749 |
| YIPF6 | 90.1839903 | -0.223550989 | 0.17774497 | -1.2577064 | 0.20849795 | 0.48867185 |
| EDA | 174.065726 | 0.842822303 | 0.34479542 | 2.44441269 | 0.01450882 | 0.11197822 |
| IGBP1 | 69.707423 | -0.088939455 | 0.245475 | -0.3623157 | 0.7171161 | 0.88104097 |
| DLG3 | 69.0686819 | 0.286004428 | 0.2847591 | 1.00437325 | 0.31519874 | 0.60205159 |
| SNX12 | 33.4959777 | -0.431933337 | 0.26911878 | -1.6049915 | 0.10849569 | 0.3445906 |
| ZMYM3 | 40.5351062 | 0.406616069 | 0.31390928 | 1.2953299 | 0.19520645 | 0.47360141 |
| NONO | 117.716538 | -0.284429322 | 0.22644999 | -1.2560359 | 0.20910295 | 0.48959511 |
| TAF1 | 191.916739 | -0.155583583 | 0.19140036 | -0.8128698 | 0.41629269 | 0.68587335 |
| OGT | 256.629538 | 0.591279389 | 0.20030952 | 2.95182864 | 0.00315898 | 0.04214469 |
| NHSL2 | 123.021992 | 0.399864683 | 0.66920735 | 0.59751986 | 0.55016035 | 0.78536075 |
| PIN4 | 80.6125152 | -0.208594372 | 0.25327279 | -0.8235957 | 0.41016934 | 0.68208527 |
| RPS4X | 494.564723 | -0.422840142 | 0.48420393 | -0.8732687 | 0.38251663 | 0.65946633 |
| HDAC8 | 394.219204 | -0.139004724 | 0.18278639 | -0.7604763 | 0.44696991 | 0.70891772 |
| CHIC1 | 137.156153 | 0.406395456 | 0.15780569 | 2.5752902 | 0.0100156 | 0.08977568 |
| XIST | 687.484043 | 1.03622216 | 3.78442951 | 0.27381199 | 0.78422911 | 0.91212179 |
| JPX | 519.19882 | 0.162635873 | 0.39440698 | 0.41235546 | 0.6800789 | 0.86238758 |
| FTX | 2687.96429 | 0.312075581 | 0.24203215 | 1.2893972 | 0.19726003 | 0.47630586 |
| SLC16A2 | 34.8649663 | -0.271243526 | 0.50374667 | -0.5384522 | 0.59026486 | 0.81183245 |
| RLIM | 71.7815338 | -0.104681924 | 0.24653688 | -0.4246096 | 0.67112129 | 0.85979281 |
| NEXMIF | 86.4814283 | 0.837200196 | 0.72474234 | 1.15516943 | 0.24802105 | 0.53495207 |
| ABCB7 | 129.187801 | 0.391886355 | 0.17327313 | 2.2616684 | 0.0237179 | 0.14897951 |
| ZDHHC15 | 31.6600035 | 0.028706263 | 0.25260858 | 0.1136393 | 0.90952373 | 0.96782135 |
| PBDC1 | 25.1056724 | -0.51339634 | 0.39583058 | -1.2970103 | 0.19462764 | 0.47318227 |
| ATRX | 679.355955 | 0.425023527 | 0.14024252 | 3.03063231 | 0.00244042 | 0.03574435 |
| MAGT1 | 112.814117 | -0.45579043 | 0.23321394 | -1.9543876 | 0.0506554 | 0.22621355 |
| COX7B | 132.458559 | -0.925028291 | 0.38260587 | -2.4177054 | 0.01561872 | 0.11597805 |
| PGK1 | 285.328958 | -1.029456206 | 0.37409908 | -2.7518277 | 0.00592637 | 0.06458327 |
| BRWD3 | 189.55983 | 0.292343582 | 0.24576585 | 1.18952079 | 0.2342348 | 0.51675152 |
| HMGN5 | 35.991606 | 0.251959218 | 0.39414899 | 0.63924867 | 0.52266117 | 0.76512314 |
| SH3BGRL | 134.361688 | 0.135324269 | 0.23445108 | 0.57719618 | 0.56380694 | 0.79293888 |
| RPS6KA6 | 159.281364 | 0.879635022 | 0.39148466 | 2.24692082 | 0.02464509 | 0.15217128 |
| HDX | 71.7539031 | -0.108964276 | 0.41275005 | -0.2639958 | 0.79178316 | 0.91648412 |
| APOOL | 83.3072256 | 0.044541456 | 0.18719305 | 0.23794396 | 0.81192456 | 0.92621476 |
| ZNF711 | 30.2878328 | 1.125335706 | 0.50452923 | 2.2304668 | 0.02571647 | 0.15532034 |
| POF1B | 47.0226492 | 0.01554223 | 0.75517776 | 0.02058089 | 0.98357999 | 0.9964794 |
| CHM | 181.545748 | 0.038682494 | 0.20891328 | 0.18516053 | 0.85310313 | 0.94515269 |
| DIAPH2 | 416.713609 | -0.210130472 | 0.32236206 | -0.6518462 | 0.51450042 | 0.76132352 |
| TSPAN6 | 28.3977985 | -0.12764277 | 0.48358535 | -0.2639509 | 0.79181778 | 0.91648412 |
| SYTL4 | 45.9191547 | 0.567379416 | 0.44711207 | 1.26898702 | 0.20444569 | 0.48318561 |
| TRMT2B | 57.7581948 | 0.148988006 | 0.31820554 | 0.46821311 | 0.6396322 | 0.84047144 |
| HNRNPH2 | 50.722249 | -0.694396587 | 0.40174143 | -1.7284665 | 0.08390463 | 0.30203879 |
| ARMCX4 | 239.608213 | 0.313640053 | 0.35729987 | 0.87780623 | 0.38004889 | 0.65731361 |
| ARMCX3 | 74.1588955 | -0.068550108 | 0.27954203 | -0.2452229 | 0.80628385 | 0.92263409 |
| ARMCX2 | 31.5273515 | 0.019164861 | 0.3521854 | 0.05441697 | 0.95660296 | 0.98498331 |
| ZMAT1 | 156.974349 | 0.576643849 | 0.42323086 | 1.36248063 | 0.17304625 | 0.44428037 |
| GPRASP1 | 28.5402552 | 0.23818776 | 0.56581629 | 0.42096307 | 0.67378205 | 0.86105131 |
| BHLHB9 | 33.1752964 | 0.75444925 | 0.27813652 | 2.71251416 | 0.00667749 | 0.06914995 |

| | | | | | | |
|---|---|---|---|---|---|---|
| LINC00630 | 114.305723 | 0.530265368 | 0.24003278 | 2.20913726 | 0.0271651 | 0.15924997 |
| BEX4 | 90.7552127 | 0.272391701 | 0.36285019 | 0.75070018 | 0.45283311 | 0.71261515 |
| TCEAL8 | 42.5570641 | -0.676969141 | 0.34094259 | -1.985581 | 0.04707987 | 0.2166537 |
| BEX2 | 69.7161073 | -0.373010453 | 0.30493197 | -1.223258 | 0.22123228 | 0.50310453 |
| TCEAL9 | 117.780106 | -0.2951772 | 0.35381352 | -0.8342734 | 0.40412695 | 0.67765972 |
| BEX3 | 219.464519 | 0.519939929 | 0.39617903 | 1.31238629 | 0.18938983 | 0.46552866 |
| TCEAL4 | 114.249036 | -0.631348314 | 0.20699114 | -3.0501225 | 0.00228748 | 0.03450674 |
| TCEAL3 | 43.6986134 | -0.536756284 | 0.35156768 | -1.5267509 | 0.12682296 | 0.37435927 |
| MORF4L2 | 173.758454 | -0.089547537 | 0.27047223 | -0.3310785 | 0.7405852 | 0.89253455 |
| SLC25A53 | 76.510685 | 0.656111982 | 0.21896436 | 2.99643272 | 0.00273159 | 0.03818317 |
| FAM199X | 127.983888 | 0.042910096 | 0.22147497 | 0.19374693 | 0.84637404 | 0.94283253 |
| NRK | 165.647513 | 0.744457865 | 0.43208858 | 1.72292883 | 0.0849014 | 0.30278126 |
| RNF128 | 89.9769426 | 0.557102288 | 0.21648179 | 2.57343718 | 0.01006939 | 0.08977568 |
| TBC1D8B | 60.4235354 | 0.352329558 | 0.33062328 | 1.06565259 | 0.28658072 | 0.57346076 |
| MORC4 | 70.6682077 | -0.537766879 | 0.25841373 | -2.0810306 | 0.03743111 | 0.19204745 |
| CLDN2 | 160.458545 | -2.882155372 | 0.98325321 | -2.9312443 | 0.00337607 | 0.04410166 |
| RBM41 | 106.929986 | 0.079741288 | 0.22801592 | 0.34971807 | 0.72655029 | 0.88522597 |
| TSC22D3 | 74.1079452 | -0.2732119 | 0.58232038 | -0.469178 | 0.63894242 | 0.840026 |
| MID2 | 42.4579078 | -0.035164978 | 0.28083571 | -0.1252155 | 0.90035296 | 0.96303857 |
| COL4A6 | 76.3625524 | 1.664258755 | 0.45876651 | 3.62768145 | 0.00028598 | 0.00938822 |
| COL4A5 | 491.368562 | 1.315090682 | 0.28045745 | 4.68909163 | 2.74E-06 | 0.0003423 |
| ACSL4 | 590.372785 | -0.585468129 | 0.58329854 | -1.0037195 | 0.31551383 | 0.60232326 |
| TMEM164 | 102.084204 | 0.058106072 | 0.28114983 | 0.20667297 | 0.83626527 | 0.93756847 |
| AMMECR1 | 154.78625 | 0.675321055 | 0.28588267 | 2.36223149 | 0.0181653 | 0.12738314 |
| PAK3 | 54.8707177 | 0.360882604 | 0.83488147 | 0.4322561 | 0.66555529 | 0.85742752 |
| ALG13 | 197.36174 | 0.364968639 | 0.26135385 | 1.39645406 | 0.16257781 | 0.4292538 |
| AMOT | 73.0011326 | -0.355602652 | 0.25911026 | -1.372399 | 0.16993927 | 0.44007794 |
| PLS3 | 74.8013555 | -1.128614577 | 0.38137818 | -2.9593056 | 0.00308333 | 0.04154042 |
| DANT2 | 627.84256 | -0.19240445 | 0.23782376 | -0.8090211 | 0.41850299 | 0.68785864 |
| KLHL13 | 400.520151 | 0.333944991 | 0.54648759 | 0.61107516 | 0.54114982 | 0.77844634 |
| WDR44 | 147.532362 | 0.196311194 | 0.17082834 | 1.14917227 | 0.25048495 | 0.53798103 |
| DOCK11 | 556.160133 | 2.207298937 | 0.34017227 | 6.4887679 | 8.65E-11 | 7.54E-08 |
| IL13RA1 | 114.068515 | -0.543596348 | 0.39575866 | -1.3735552 | 0.16957982 | 0.43945897 |
| LONRF3 | 50.0967134 | 0.801702737 | 0.64897751 | 1.23533208 | 0.21670694 | 0.49828031 |
| PGRMC1 | 142.550546 | -0.290237699 | 0.40884626 | -0.7098945 | 0.47776958 | 0.73413221 |
| SLC25A43 | 77.1023927 | -0.615121992 | 0.37672409 | -1.6328183 | 0.10250721 | 0.33398714 |
| SLC25A5 | 100.302435 | -1.107348032 | 0.51379429 | -2.1552362 | 0.03114336 | 0.17254033 |
| UBE2A | 51.0480656 | 0.325679342 | 0.30223199 | 1.07758063 | 0.28122095 | 0.56893012 |
| 6-Sep | 64.3737339 | -0.279936253 | 0.54799141 | -0.5108406 | 0.60946269 | 0.82205058 |
| UPF3B | 88.6110832 | -0.020629198 | 0.25479066 | -0.0809653 | 0.93546956 | 0.97836695 |
| NDUFA1 | 100.685918 | -0.814936038 | 0.30656651 | -2.6582683 | 0.00785433 | 0.0767721 |
| NKAP | 34.7241638 | -0.242073692 | 0.26488173 | -0.9138935 | 0.3607728 | 0.6411582 |
| RHOXF1-AS1 | 59.7918076 | 0.850428354 | 0.3407172 | 2.49599476 | 0.01256045 | 0.10234405 |
| LAMP2 | 167.215391 | -0.534732339 | 0.24314352 | -2.1992457 | 0.02786046 | 0.16152154 |
| CUL4B | 50.6398529 | -0.063257954 | 0.22996025 | -0.2750821 | 0.78325315 | 0.91174892 |
| MCTS1 | 64.4621287 | 0.24366152 | 0.21478631 | 1.13443692 | 0.25661132 | 0.54370269 |
| THOC2 | 332.012555 | 0.078598348 | 0.14473902 | 0.54303496 | 0.58710573 | 0.81058905 |
| XIAP | 240.129102 | -0.129561899 | 0.15286789 | -0.8475416 | 0.39669331 | 0.67195848 |
| STAG2 | 374.989708 | 0.472320299 | 0.17600164 | 2.68361309 | 0.00728313 | 0.0731995 |
| TENM1 | 119.100035 | 0.187942744 | 0.62183895 | 0.30223701 | 0.76247139 | 0.90158172 |
| SMARCA1 | 166.444432 | 0.319134631 | 0.24867869 | 1.28332118 | 0.19937958 | 0.47830126 |
| OCRL | 49.8499563 | -0.032543181 | 0.19783132 | -0.1644996 | 0.86933783 | 0.95106604 |
| ZDHHC9 | 37.7359674 | -1.141409775 | 0.32907456 | -3.4685446 | 0.00052329 | 0.01373303 |
| UTP14A | 29.567941 | -1.160946244 | 0.38148172 | -3.0432552 | 0.00234034 | 0.0349985 |
| BCORL1 | 50.9043365 | -0.271557134 | 0.39114037 | -0.6942703 | 0.48751273 | 0.74111095 |
| AIFM1 | 64.5067705 | -0.266175248 | 0.42776719 | -0.6222433 | 0.53378192 | 0.77365669 |
| ZNF280C | 38.5622244 | 0.062464616 | 0.25863429 | 0.24151715 | 0.80915433 | 0.92426536 |
| ENOX2 | 194.75746 | 0.836785047 | 0.26423101 | 3.16686923 | 0.0015409 | 0.02733378 |
| RAP2C | 35.1362643 | -0.082284852 | 0.3166521 | -0.2598589 | 0.79497265 | 0.9178068 |
| RAP2C-AS1 | 156.81731 | 0.18212103 | 0.31941872 | 0.57016391 | 0.56856653 | 0.79717488 |
| MBNL3 | 77.724248 | -0.07685669 | 0.24834987 | -0.3094694 | 0.75696447 | 0.89930004 |
| GPC4 | 666.9858 | 0.031532956 | 0.52840882 | 0.0596753 | 0.95241424 | 0.98341878 |
| GPC3 | 110.790497 | 0.647799681 | 0.48049523 | 1.34819169 | 0.17759674 | 0.44922079 |
| PHF6 | 98.6958255 | 0.643879001 | 0.28838219 | 2.23272806 | 0.02556688 | 0.1551184 |
| HPRT1 | 92.9161469 | -0.592703723 | 0.2698516 | -2.1964062 | 0.02806289 | 0.16213171 |
| FAM122B | 39.8237118 | -0.006091794 | 0.36189472 | -0.0168331 | 0.9865698 | 0.99678172 |
| RTL8C | 43.8763831 | -0.741700283 | 0.35730031 | -2.0758456 | 0.03790823 | 0.1933562 |
| RTL8A | 36.0900628 | -0.247133276 | 0.40915732 | -0.6040055 | 0.54583998 | 0.78178845 |
| ZNF75D | 88.5921961 | -0.272137516 | 0.25547746 | -1.0652115 | 0.28678025 | 0.57374018 |
| INTS6L | 63.7906462 | 0.296113064 | 0.27307024 | 1.08438424 | 0.27819447 | 0.56578102 |
| MMGT1 | 25.8531123 | 0.149633593 | 0.40942588 | 0.36547175 | 0.71475929 | 0.87969668 |
| SLC9A6 | 38.118671 | 0.526428781 | 0.29708512 | 1.77197963 | 0.07639794 | 0.28546836 |
| FHL1 | 125.958213 | 0.663677001 | 0.55406808 | 1.19782573 | 0.23098487 | 0.51425247 |
| MAP7D3 | 45.5082961 | 0.089328026 | 0.29813967 | 0.29961805 | 0.76446852 | 0.90305142 |
| HTATSF1 | 66.3260655 | 0.090213943 | 0.39276355 | 0.22969021 | 0.8183325 | 0.92999132 |
| AL683813.1 | 69.1043731 | 0.491175937 | 0.32679612 | 1.50300417 | 0.13283797 | 0.38380408 |
| RBMX | 254.371384 | 0.364223791 | 0.11253296 | 3.23659664 | 0.00120964 | 0.02363725 |
| FGF13 | 264.042966 | 0.312506094 | 0.58213655 | 0.5368261 | 0.59138774 | 0.81252197 |
| ATP11C | 132.210521 | 0.260510637 | 0.23810182 | 1.09411445 | 0.27390479 | 0.5612072 |
| SLITRK4 | 35.8163728 | 0.424844011 | 0.63431627 | 0.66976685 | 0.50300643 | 0.75403031 |
| AL589669.1 | 104.751407 | -1.153189069 | 0.57061993 | -2.0209408 | 0.0432859 | 0.20621259 |
| FMR1 | 134.812041 | -0.187985012 | 0.29563083 | -0.6358776 | 0.52485623 | 0.76663584 |
| IDS | 75.9894676 | 0.01462636 | 0.31359885 | 0.04664035 | 0.96279987 | 0.98817751 |
| TMEM185A | 37.7949532 | -0.058389291 | 0.27642492 | -0.2112302 | 0.83270765 | 0.93601742 |
| LINC00894 | 299.386092 | 0.242378956 | 0.38921927 | 0.62273113 | 0.53346122 | 0.773543 |
| MAMLD1 | 82.3807005 | -0.486229543 | 0.42079761 | -1.155495 | 0.24788777 | 0.53482252 |
| MTM1 | 121.842925 | 0.052692788 | 0.26123172 | 0.201709 | 0.84014423 | 0.94021723 |
| MTMR1 | 115.145775 | 0.186352278 | 0.23300528 | 0.79977706 | 0.42383998 | 0.69092389 |
| CD99L2 | 65.5731982 | -0.128049211 | 0.18688105 | -0.685191 | 0.49322341 | 0.7455922 |
| VMA21 | 39.5464784 | -0.343119063 | 0.3355501 | -1.0225569 | 0.3065174 | 0.59421809 |
| CETN2 | 55.7937705 | -0.445006889 | 0.44786244 | -0.993624 | 0.32040593 | 0.60715498 |
| ZNF185 | 39.5236313 | 0.688298517 | 0.45833953 | 1.50172193 | 0.13316894 | 0.38399315 |
| BGN | 190.195305 | 0.376831018 | 1.06595741 | 0.35351414 | 0.72370303 | 0.88310208 |
| SLC6A8 | 67.7668932 | -1.44045711 | 0.34720616 | -4.1487084 | 3.34E-05 | 0.00225549 |
| BCAP31 | 133.979122 | -0.602830425 | 0.39880355 | -1.5115974 | 0.13063631 | 0.38116515 |
| IDH3G | 30.8444558 | 0.008134042 | 0.38990259 | 0.02086173 | 0.98335596 | 0.99635781 |
| SSR4 | 77.4605711 | -0.820995344 | 0.48882535 | -1.679527 | 0.09304939 | 0.31776118 |
| HCFC1 | 36.5133027 | -0.325502422 | 0.28436054 | -1.1446821 | 0.25234086 | 0.53930681 |
| IRAK1 | 43.8928073 | -1.55465019 | 0.64860793 | -2.3969028 | 0.01653431 | 0.12071808 |
| MECP2 | 200.603779 | 0.288797739 | 0.18886428 | 1.52912843 | 0.12623261 | 0.37366569 |
| FLNA | 312.009083 | 0.110581636 | 0.5349597 | 0.20671022 | 0.83623618 | 0.93756847 |
| EMD | 63.4269939 | -0.344540883 | 0.37536827 | -0.9178743 | 0.35868469 | 0.64001011 |
| RPL10 | 740.442882 | -0.396967505 | 0.55549585 | -0.7146183 | 0.47484493 | 0.73162933 |
| TAZ | 52.1661185 | 0.103958277 | 0.22050067 | 0.47146469 | 0.63730893 | 0.8389147 |
| ATP6AP1 | 69.2663238 | -0.124824084 | 0.39423035 | -0.3166273 | 0.75152643 | 0.89611476 |
| GDI1 | 58.9684836 | -0.043320862 | 0.32131276 | -0.1348246 | 0.89275056 | 0.95976294 |
| FAM50A | 47.0311849 | -0.353339712 | 0.3598406 | -0.981934 | 0.32613237 | 0.61279365 |
| PLXNA3 | 41.5078859 | 0.468524799 | 0.37771659 | 1.24041361 | 0.21482245 | 0.49688442 |
| FAM3A | 32.3655968 | -0.224320812 | 0.29024372 | -0.7728705 | 0.43959901 | 0.70322627 |
| DKC1 | 48.0921016 | -0.406964971 | 0.36246933 | -1.122757 | 0.26154072 | 0.54892607 |
| MPP1 | 28.0570957 | -0.219716238 | 0.40867251 | -0.537634 | 0.59082974 | 0.81210477 |
| FUNDC2 | 55.8332595 | -0.082108354 | 0.42844826 | -0.1916412 | 0.84802324 | 0.94366633 |
| BRCC3 | 66.117992 | 0.115515348 | 0.18363531 | 0.62904759 | 0.5293179 | 0.77040597 |

| | | | | | | |
|---|---|---|---|---|---|---|
| VBP1 | 26.2674868 | -0.439151238 | 0.38489304 | -1.1409695 | 0.2538826 | 0.54067332 |
| TMLHE-AS1 | 48.5673474 | -0.002455263 | 0.39199239 | -0.0062635 | 0.99500425 | 0.99886097 |
| TMLHE | 120.736553 | 0.060625757 | 0.27937775 | 0.21700281 | 0.82820616 | 0.93346403 |
| VAMP7 | 49.7777623 | 0.060939781 | 0.2314196 | 0.26333025 | 0.79229605 | 0.91670538 |
| ZNF596 | 27.3645402 | -0.087542798 | 0.27367575 | -0.3198778 | 0.74906096 | 0.89489336 |
| FBXO25 | 89.6917116 | 0.240620757 | 0.26358686 | 0.91287083 | 0.36131048 | 0.64151865 |
| TDRP | 166.80616 | 0.493422355 | 0.33884085 | 1.45620681 | 0.14533546 | 0.40323664 |
| ERICH1 | 182.671418 | -0.372658474 | 0.32824539 | -1.1353045 | 0.25624775 | 0.54329288 |
| KBTBD11 | 75.0543077 | -0.455752168 | 0.49477587 | -0.9211285 | 0.35698333 | 0.63880877 |
| AC246817.1 | 64.4856791 | 0.671267229 | 0.4972478 | 1.34996521 | 0.17702714 | 0.44872611 |
| CSMD1 | 123.677407 | 1.568026712 | 0.81454625 | 1.92503092 | 0.05422549 | 0.23344987 |
| MCPH1 | 284.496135 | -0.109866808 | 0.19942861 | -0.5509079 | 0.58169678 | 0.80614488 |
| ANGPT2 | 67.4378048 | -0.233926335 | 0.32565558 | -0.7183243 | 0.47255732 | 0.72988407 |
| MCPH1-AS1 | 69.111081 | -0.082281802 | 0.25562716 | -0.3218821 | 0.74754203 | 0.89399656 |
| AGPAT5 | 157.180435 | -1.195094213 | 0.32255244 | -3.705116 | 0.00021129 | 0.00778456 |
| AF233439.1 | 63.4183811 | -0.01218762 | 0.4872491 | -0.0250131 | 0.9800445 | 0.99558815 |
| DEFB1 | 520.590174 | -2.101355774 | 0.43158898 | -4.8688819 | 1.12E-06 | 0.00019198 |
| FAM85B | 256.412059 | 0.741606314 | 0.49358496 | 1.5024897 | 0.13297069 | 0.38380408 |
| PRAG1 | 113.387155 | -0.773463381 | 0.42924353 | -1.8019221 | 0.07155767 | 0.27517098 |
| MFHAS1 | 553.088905 | -0.497580471 | 0.2797773 | -1.7784876 | 0.0753238 | 0.28328492 |
| ERI1 | 71.2120876 | -0.742185874 | 0.32783458 | -2.2639036 | 0.02358005 | 0.14836941 |
| PPP1R3B | 37.8790211 | -0.83407163 | 0.52306125 | -1.5945965 | 0.1108025 | 0.34794171 |
| TNKS | 551.015378 | 0.148756339 | 0.16208991 | 0.91773964 | 0.35875519 | 0.64001011 |
| MSRA | 194.831417 | -0.86934714 | 0.35463555 | -2.4513818 | 0.01423089 | 0.11067271 |
| XKR6 | 260.062226 | -0.176678963 | 0.54226211 | -0.3258184 | 0.74456177 | 0.89323545 |
| MTMR9 | 75.180683 | -0.087236471 | 0.20869614 | -0.4180071 | 0.67594191 | 0.86116336 |
| FDFT1 | 153.108816 | 0.246100875 | 0.4548587 | 0.54104906 | 0.58847377 | 0.81121355 |
| CTSB | 248.228602 | -1.221384972 | 0.4014192 | -3.042667 | 0.00234492 | 0.0349985 |
| FAM66D | 82.2693954 | 0.075162715 | 0.67884329 | 0.11072175 | 0.911837 | 0.96844354 |
| AC145124.1 | 32.3093671 | 0.401820121 | 0.66244204 | 0.60657401 | 0.54413365 | 0.78027954 |
| LONRF1 | 123.928226 | 0.011395759 | 0.35255734 | 0.03232314 | 0.97421436 | 0.99326197 |
| TRMT9B | 73.2155191 | -0.667468081 | 0.34667018 | -1.9253692 | 0.05418319 | 0.23344987 |
| DLC1 | 173.750457 | 0.56878255 | 0.61587941 | 0.92352909 | 0.35573155 | 0.63728305 |
| TUSC3 | 295.860768 | -0.0203568 | 0.27374277 | -0.0743647 | 0.9407202 | 0.98016757 |
| MSR1 | 82.7121759 | 0.297095543 | 0.83965741 | 0.35382948 | 0.72346668 | 0.88292615 |
| MICU3 | 123.500918 | 0.076082691 | 0.2344628 | 0.32449792 | 0.7455611 | 0.89349803 |
| ZDHHC2 | 137.854941 | 0.084006591 | 0.30258457 | 0.27763012 | 0.78129631 | 0.91077326 |
| CNOT7 | 66.3352771 | 0.16054513 | 0.1964202 | 0.8173555 | 0.4137253 | 0.68448816 |
| VPS37A | 95.9145144 | 0.020190743 | 0.22885796 | 0.08822391 | 0.92969872 | 0.97545894 |
| SLC7A2 | 54.2505437 | -1.245882567 | 0.60986953 | -2.0428674 | 0.04106557 | 0.20176225 |
| MTUS1 | 152.570283 | -0.771874797 | 0.39420428 | -1.9580579 | 0.05022322 | 0.2251146 |
| PCM1 | 529.302898 | 0.266588415 | 0.16673381 | 1.59888638 | 0.10984585 | 0.34692166 |
| ASAH1 | 246.313005 | 0.396745893 | 0.28637998 | 1.38538278 | 0.16593545 | 0.43440713 |
| PSD3 | 198.21076 | -1.488545094 | 0.56515435 | -2.6338736 | 0.00844169 | 0.08102502 |
| SH2D4A | 76.7013419 | 0.516274542 | 0.36886352 | 1.39963568 | 0.16162244 | 0.42791083 |
| CSGALNACT1 | 139.532127 | -0.512537965 | 1.0674204 | -0.480165 | 0.63111004 | 0.83474975 |
| INTS10 | 215.622756 | 0.057718722 | 0.2306503 | 0.25024344 | 0.8023991 | 0.92125386 |
| ATP6V1B2 | 98.1697627 | -0.25072855 | 0.20371218 | -1.230798 | 0.21839841 | 0.50057436 |
| XPO7 | 156.915981 | 0.095421971 | 0.14242818 | 0.66996555 | 0.50287975 | 0.75403031 |
| DMTN | 64.9643025 | 0.357215044 | 0.34441185 | 1.03717408 | 0.29965474 | 0.58681504 |
| FAM160B2 | 48.0105881 | -0.057161579 | 0.24555794 | -0.2327824 | 0.81593034 | 0.92869214 |
| BMP1 | 91.8514801 | -1.566343246 | 0.7114845 | -2.2015142 | 0.02769964 | 0.16119979 |
| SLC39A14 | 339.562719 | -2.199407013 | 0.96123433 | -2.288107 | 0.02213129 | 0.1428542 |
| PPP3CC | 263.093281 | 0.37533555 | 0.25294656 | 1.48385314 | 0.13784789 | 0.39066417 |
| SORBS3 | 34.5909374 | -0.43577155 | 0.29800295 | -1.4623061 | 0.14365733 | 0.40084288 |
| CCAR2 | 41.2668277 | 0.086850587 | 0.26442267 | 0.32845363 | 0.7425687 | 0.89253455 |
| BIN3 | 73.7500473 | 0.112278265 | 0.24467676 | 0.45888406 | 0.64631743 | 0.84427583 |
| TNFRSF10B | 159.819674 | -0.790826484 | 0.25851203 | -3.0591478 | 0.00221968 | 0.0339653 |
| TNFRSF10D | 65.4909925 | -1.515138379 | 0.47903659 | -3.1628865 | 0.00156213 | 0.02755739 |
| TNFRSF10A | 36.2714488 | -1.909538126 | 0.3609753 | -5.2899413 | 1.22E-07 | 3.55E-05 |
| CHMP7 | 26.4014016 | -0.162004271 | 0.3301442 | -0.4907076 | 0.62363327 | 0.83030289 |
| LOXL2 | 33.4719461 | 0.300222956 | 0.66397502 | 0.45216001 | 0.65115371 | 0.84724399 |
| ENTPD4 | 66.386261 | 0.187229249 | 0.25089373 | 0.74624923 | 0.45551688 | 0.71495923 |
| SLC25A37 | 561.123493 | -2.045151178 | 0.4996539 | -4.0931357 | 4.26E-05 | 0.00275447 |
| DOCK5 | 346.883222 | -1.348935296 | 0.31278846 | -4.3126121 | 1.61E-05 | 0.00130672 |
| KCTD9 | 89.1205949 | 0.363671098 | 0.2361854 | 1.53976959 | 0.12361653 | 0.36947593 |
| PPP2R2A | 298.057738 | 0.343957192 | 0.16795919 | 2.04786173 | 0.04057355 | 0.19992491 |
| BNIP3L | 418.819503 | 0.168150534 | 0.50452521 | 0.3332847 | 0.73891939 | 0.89234856 |
| DPYSL2 | 148.368598 | 0.045754512 | 0.34462031 | 0.13276789 | 0.89437695 | 0.96088344 |
| PTK2B | 125.307016 | -0.566951726 | 0.3492932 | -1.6231399 | 0.1045595 | 0.33777911 |
| EPHX2 | 63.1207312 | 0.033538302 | 0.48084581 | 0.06974856 | 0.94439379 | 0.98115679 |
| CLU | 1218.31833 | -1.068006247 | 0.5445574 | -1.9612372 | 0.04985136 | 0.22450688 |
| SCARA3 | 93.3567015 | 0.270741632 | 0.55956204 | 0.4838456 | 0.62849545 | 0.83338288 |
| CCDC25 | 55.3524324 | -0.017465592 | 0.29748072 | -0.0587117 | 0.95318176 | 0.98368111 |
| ELP3 | 148.016924 | 0.287737313 | 0.24769354 | 1.16166661 | 0.24537091 | 0.53188684 |
| ZNF395 | 57.0420603 | -0.321691438 | 0.37248591 | -0.8636338 | 0.38778907 | 0.66427602 |
| FBXO16 | 66.2764671 | 0.129246444 | 0.38067066 | 0.339523 | 0.73421577 | 0.88947172 |
| FZD3 | 52.0660833 | -0.233308333 | 0.21039509 | -1.1089058 | 0.26747082 | 0.55480561 |
| EXTL3 | 60.0122651 | -0.689381575 | 0.29608402 | -2.3283309 | 0.01989454 | 0.13439337 |
| INTS9 | 78.2381756 | -0.082425748 | 0.27034414 | -0.3048919 | 0.76044846 | 0.90125623 |
| HMBOX1 | 339.279182 | -0.129728722 | 0.1398005 | -0.9279561 | 0.35343036 | 0.63494176 |
| KIF13B | 401.942266 | 0.547578564 | 0.34367063 | 1.5933237 | 0.11108759 | 0.34822126 |
| SARAF | 184.465217 | -0.394900085 | 0.32400011 | -1.218827 | 0.22290986 | 0.50454951 |
| LEPROTL1 | 68.5002299 | -0.575289017 | 0.32682808 | -1.760219 | 0.07837068 | 0.29033886 |
| DCTN6 | 38.2995168 | -0.343906432 | 0.30803864 | -1.1164393 | 0.26423416 | 0.55215947 |
| RBPMS-AS1 | 27.5601426 | -1.02188399 | 0.50715379 | -2.0149391 | 0.04391103 | 0.2079208 |
| RBPMS | 3840.12025 | 0.44583725 | 0.25807744 | 1.72753285 | 0.08407201 | 0.30203879 |
| GTF2E2 | 128.071791 | 0.23140626 | 0.16732365 | 1.38298594 | 0.16666918 | 0.43490657 |
| GSR | 84.8517133 | -0.308521733 | 0.21566253 | -1.4305765 | 0.15255164 | 0.41490407 |
| PPP2CB | 279.048356 | -0.187104118 | 0.186112 | -1.0053308 | 0.31473761 | 0.60177076 |
| PURG | 52.4005924 | 1.756685855 | 0.55900495 | 3.14252288 | 0.00167499 | 0.02854928 |
| WRN | 316.654069 | 0.667556794 | 0.19082243 | 3.49831404 | 0.00046821 | 0.01277772 |
| NRG1 | 435.732555 | 0.355180533 | 0.54546194 | 0.65115548 | 0.51494612 | 0.76135326 |
| FUT10 | 76.3849524 | 0.260295691 | 0.31330775 | 0.83079875 | 0.40608733 | 0.67957905 |
| TTI2 | 29.1566955 | -0.033420622 | 0.40842251 | -0.0818285 | 0.93478305 | 0.97790458 |
| UNC5D | 835.441428 | 0.888696766 | 0.62755532 | 1.41612499 | 0.15673891 | 0.4216384 |
| ERLIN2 | 68.1104908 | -0.119429908 | 0.29625531 | -0.4031317 | 0.68685134 | 0.86707288 |
| PLPBP | 45.6211381 | 0.163984044 | 0.31658836 | 0.51797243 | 0.6044775 | 0.82048068 |
| RAB11FIP1 | 135.02171 | 0.211786545 | 0.35529043 | 0.59609414 | 0.55111234 | 0.78577398 |
| EIF4EBP1 | 45.2946905 | -1.664294384 | 0.41830351 | -3.9786766 | 6.93E-05 | 0.00370851 |
| ASH2L | 66.5499771 | 0.090736639 | 0.19639966 | 0.46199999 | 0.64408132 | 0.84344334 |
| LSM1 | 41.9718297 | -0.118506389 | 0.39088279 | -0.3031763 | 0.76175553 | 0.90141135 |
| BAG4 | 68.8483186 | -0.21887837 | 0.24666563 | -0.8873485 | 0.37489131 | 0.65328067 |
| DDHD2 | 106.669717 | 0.245660929 | 0.20189802 | 1.21675751 | 0.22369649 | 0.50561319 |
| PLPP5 | 43.3255076 | -0.790624899 | 0.37541757 | -2.105988 | 0.03520539 | 0.18531364 |
| NSD3 | 400.921053 | -0.180940855 | 0.17404473 | -1.0396227 | 0.29851523 | 0.58610017 |
| FGFR1 | 263.660429 | -0.924599868 | 0.25380289 | -3.642984 | 0.0002695 | 0.00896353 |
| AC069120.1 | 38.7894484 | 0.848072831 | 0.59004902 | 1.43729215 | 0.150635 | 0.41173164 |
| TACC1 | 270.806189 | -0.344082419 | 0.33988089 | -1.0123618 | 0.31136512 | 0.59878869 |
| PLEKHA2 | 264.910333 | 0.182955468 | 0.29911543 | 0.61165506 | 0.540766 | 0.77836176 |
| ADAM9 | 192.621302 | -0.29331474 | 0.22346131 | -1.3125974 | 0.18931864 | 0.46547311 |
| TCIM | 284.610522 | 0.176076059 | 1.00441757 | 0.17530165 | 0.86084261 | 0.94814435 |
| GOLGA7 | 53.5596178 | 0.014934046 | 0.23699198 | 0.06301499 | 0.94975457 | 0.98270285 |

| | | | | | | |
|---|---|---|---|---|---|---|
| GPAT4 | 75.738166 | -0.237061619 | 0.29783388 | -0.7959525 | 0.42605966 | 0.69224822 |
| KAT6A | 390.339106 | 0.290103017 | 0.14036342 | 2.06679936 | 0.03875306 | 0.19565065 |
| AP3M2 | 40.5754072 | 0.021108106 | 0.25511815 | 0.08273855 | 0.93405943 | 0.97768238 |
| IKBKB | 126.461538 | -0.228121032 | 0.2366792 | -0.9638406 | 0.33512583 | 0.6199188 |
| POLB | 38.4441713 | 0.514016707 | 0.2553552 | 2.0129479 | 0.04412011 | 0.20849854 |
| VDAC3 | 89.5460477 | 0.031165613 | 0.21354142 | 0.14594645 | 0.88396366 | 0.95761929 |
| SLC20A2 | 415.466351 | 0.504492869 | 0.23184254 | 2.17601507 | 0.02955413 | 0.1680113 |
| SMIM19 | 69.5073476 | 0.156538989 | 0.25171635 | 0.62188646 | 0.53401653 | 0.77366209 |
| RNF170 | 108.313963 | -0.052265286 | 0.24330699 | -0.2148121 | 0.82991381 | 0.93416878 |
| HOOK3 | 206.664623 | 0.115061171 | 0.12172535 | 0.9452523 | 0.34453008 | 0.62954637 |
| FNTA | 64.8288303 | -0.44312981 | 0.25341904 | -1.7486051 | 0.08035931 | 0.2945893 |
| HGSNAT | 192.277168 | 0.165067811 | 0.1571485 | 1.05039378 | 0.29353711 | 0.58142927 |
| SPIDR | 1076.78786 | -0.494689861 | 0.29817699 | -1.6590478 | 0.09710617 | 0.32501048 |
| CEBPD | 696.197597 | -1.427291963 | 0.49269551 | -2.8969047 | 0.00376864 | 0.047688 |
| PRKDC | 283.685019 | -0.223568966 | 0.27715641 | -0.8066527 | 0.41986661 | 0.68835944 |
| UBE2V2 | 104.21223 | 0.195397499 | 0.21887591 | 0.89273185 | 0.37200079 | 0.65192016 |
| PCMTD1 | 356.09301 | 0.049773075 | 0.21259113 | 0.23412583 | 0.81488729 | 0.92814476 |
| AC064807.1 | 41.9637494 | 0.032440454 | 0.3204765 | 0.10122569 | 0.91937131 | 0.97108563 |
| RB1CC1 | 283.470939 | 0.193306787 | 0.1413472 | 1.3676025 | 0.17143653 | 0.44204322 |
| ATP6V1H | 195.116035 | -0.02036682 | 0.23989808 | -0.0848978 | 0.93234264 | 0.97701675 |
| TCEA1 | 131.292449 | 0.348755324 | 0.2040083 | 1.70951541 | 0.08735552 | 0.30722583 |
| LYPLA1 | 135.851218 | -0.313412116 | 0.29356286 | -1.067615 | 0.28569421 | 0.57312354 |
| MRPL15 | 35.3356303 | -0.972481223 | 0.40008456 | -2.4306892 | 0.01507013 | 0.11411159 |
| RP1 | 121.126313 | 1.667822649 | 1.03163101 | 1.61668526 | 0.10594624 | 0.34040487 |
| XKR4 | 562.48634 | -2.00096459 | 0.42730906 | -4.6827104 | 2.83E-06 | 0.00034768 |
| TMEM68 | 99.0535063 | 0.532907718 | 0.25712661 | 2.07254987 | 0.03821419 | 0.19425881 |
| TGS1 | 85.8096306 | 0.168519143 | 0.36581816 | 0.46066369 | 0.64503991 | 0.84403456 |
| LYN | 79.7305663 | -1.031304715 | 0.38257545 | -2.6956898 | 0.00702431 | 0.07120193 |
| RPS20 | 246.208998 | -0.424792696 | 0.39190682 | -1.0839125 | 0.2784036 | 0.56578102 |
| PLAG1 | 82.2095539 | -0.145638963 | 0.47066599 | -0.3094317 | 0.75699318 | 0.89930004 |
| CHCHD7 | 32.2858552 | 0.354955188 | 0.3478505 | 1.02042455 | 0.30752716 | 0.59456464 |
| IMPAD1 | 95.3471624 | -0.324365302 | 0.31571359 | -1.0274037 | 0.30423043 | 0.5922455 |
| LINC01606 | 70.4220615 | 1.325328467 | 1.16229443 | 1.14026913 | 0.25417419 | 0.54093192 |
| FAM110B | 370.039087 | 0.138997737 | 0.29449454 | 0.47198748 | 0.63693572 | 0.83872935 |
| UBXN2B | 106.023719 | 0.250461931 | 0.22561676 | 1.11012112 | 0.26694684 | 0.55453954 |
| SDCBP | 129.428691 | -0.83924737 | 0.29523912 | -2.8426022 | 0.00447469 | 0.05380163 |
| NSMAF | 116.488257 | 0.008692326 | 0.24010981 | 0.03620146 | 0.97112172 | 0.99175186 |
| TOX | 67.5546905 | -0.598690847 | 0.66662868 | -0.8980874 | 0.36913893 | 0.64880401 |
| LINC01301 | 40.9107389 | 0.398417819 | 0.56509778 | 0.70504226 | 0.48078393 | 0.73578929 |
| RAB2A | 372.886916 | 0.080327839 | 0.17247295 | 0.46574165 | 0.64140043 | 0.8419864 |
| CHD7 | 341.871672 | -0.098701708 | 0.22605119 | -0.4366343 | 0.66237656 | 0.85507439 |
| ASPH | 336.338137 | 0.185046359 | 0.21706827 | 0.85248001 | 0.39394773 | 0.66874218 |
| YTHDF3 | 149.816492 | 0.184908616 | 0.17914344 | 1.03218191 | 0.30198691 | 0.58963159 |
| CYP7B1 | 141.347487 | -0.94268768 | 0.44343584 | -2.1258716 | 0.03351395 | 0.1798488 |
| ARMC1 | 64.85024 | 0.231911926 | 0.20980979 | 1.10534366 | 0.26901071 | 0.55632249 |
| MTFR1 | 109.411382 | -0.47505662 | 0.26519405 | -1.7913547 | 0.07323639 | 0.27896901 |
| PDE7A | 190.036936 | -0.53603311 | 0.39822429 | -1.3460583 | 0.17828372 | 0.44977081 |
| TRIM55 | 38.9118767 | 0.971940492 | 0.67551153 | 1.43882147 | 0.15020111 | 0.4109827 |
| ADHFE1 | 71.2895204 | 0.520576229 | 0.52080269 | 0.99956516 | 0.31752099 | 0.60395821 |
| VCPIP1 | 58.6601462 | -0.06158543 | 0.22534593 | -0.2732929 | 0.78462811 | 0.91212179 |
| SGK3 | 125.021123 | -0.042962424 | 0.23875136 | -0.1799463 | 0.85719473 | 0.94638973 |
| PPP1R42 | 38.3936521 | 0.361852637 | 0.52655276 | 0.6872106 | 0.49195003 | 0.74445329 |
| CSPP1 | 321.534065 | 0.290699887 | 0.21468389 | 1.35408337 | 0.17570978 | 0.44775565 |
| ARFGEF1 | 559.854992 | 0.038629028 | 0.13047942 | 0.29605455 | 0.76718842 | 0.90403468 |
| PREX2 | 49.6651463 | 3.256809683 | 1.28846177 | 2.52767272 | 0.01148213 | 0.09721852 |
| C8orf34 | 267.044341 | 1.063681977 | 0.61986375 | 1.71599319 | 0.08616329 | 0.30586185 |
| SULF1 | 59.021345 | -0.129169319 | 0.58612276 | -0.2203793 | 0.82557577 | 0.93245965 |
| NCOA2 | 867.71206 | 0.117080213 | 0.21981044 | 0.53264172 | 0.59428161 | 0.81474504 |
| TRAM1 | 224.465724 | -0.37725348 | 0.29879854 | -1.262568 | 0.20674447 | 0.48621411 |
| LACTB2 | 66.0523336 | -1.227484236 | 0.36169829 | -3.3936689 | 0.00068963 | 0.01615151 |
| EYA1 | 114.988554 | 0.517939509 | 0.77132752 | 0.67149103 | 0.50190777 | 0.75309752 |
| MSC-AS1 | 304.411333 | -0.372117238 | 1.15252689 | -0.3228708 | 0.74679312 | 0.89352127 |
| TERF1 | 97.8608481 | 0.29731677 | 0.19548035 | 1.52095479 | 0.12827118 | 0.37610564 |
| RPL7 | 452.629382 | -0.187271873 | 0.58279134 | -0.3213361 | 0.74795573 | 0.89435376 |
| RDH10 | 118.89145 | 0.226030606 | 0.70502041 | 0.32060151 | 0.7485124 | 0.89457271 |
| RDH10-AS1 | 45.9354408 | 0.986471021 | 0.76417306 | 1.29090002 | 0.19673835 | 0.47589814 |
| STAU2 | 329.65487 | 0.106349143 | 0.17851099 | 0.59575684 | 0.55133768 | 0.78577398 |
| AC027018.1 | 49.9187944 | -0.136511099 | 0.57648151 | -0.2368005 | 0.81281158 | 0.92667485 |
| ELOC | 74.9699554 | -0.280277172 | 0.23999609 | -1.1678406 | 0.24287107 | 0.52778176 |
| GDAP1 | 70.9765667 | -0.417886749 | 0.32893216 | -1.2704345 | 0.20392991 | 0.48285556 |
| HNF4G | 64.1573412 | -0.760228155 | 0.47004954 | -1.6173362 | 0.10580572 | 0.34033344 |
| LINC01111 | 44.6709183 | 3.321441934 | 0.7026134 | 4.72726813 | 2.28E-06 | 0.0003114 |
| ZFHX4 | 42.8106171 | 1.196284472 | 0.69944086 | 1.710344 | 0.08720228 | 0.30713468 |
| PEX2 | 69.0846521 | 0.251156619 | 0.21799403 | 1.15212614 | 0.24926924 | 0.53636804 |
| ZC2HC1A | 96.8371308 | 0.313036204 | 0.37964523 | 0.82454928 | 0.40962753 | 0.68208527 |
| IL7 | 59.9555385 | -0.833686515 | 0.41298529 | -2.0186833 | 0.04352014 | 0.2066642 |
| TPD52 | 184.403382 | 0.48470407 | 0.32500008 | 1.49139676 | 0.13585736 | 0.38788633 |
| ZBTB10 | 101.00751 | 0.272042603 | 0.29600024 | 0.9190621 | 0.35806309 | 0.64001011 |
| ZNF704 | 203.388434 | 0.167094684 | 0.51403867 | 0.32506248 | 0.74513378 | 0.89323545 |
| PAG1 | 248.123548 | 0.520804674 | 0.91676666 | 0.56808858 | 0.56997482 | 0.79765604 |
| IMPA1 | 53.6192759 | 0.326606701 | 0.35153228 | 0.92909449 | 0.35284012 | 0.63441164 |
| ZFAND1 | 88.4745076 | 0.319820615 | 0.25769206 | 1.24109611 | 0.21457025 | 0.49670575 |
| CHMP4C | 32.7871041 | -0.856180051 | 0.38353342 | -2.2323479 | 0.02559198 | 0.1551184 |
| SNX16 | 49.4325059 | 0.212385856 | 0.31531725 | 0.67356243 | 0.50058954 | 0.75194406 |
| E2F5 | 49.7297282 | -0.025407578 | 0.26495277 | -0.0958947 | 0.92360417 | 0.97297642 |
| C8orf59 | 53.597773 | 0.123338795 | 0.29311421 | 0.42078749 | 0.67391027 | 0.86106262 |
| CA13 | 66.8626845 | -0.068449728 | 0.24051079 | -0.2846015 | 0.77594948 | 0.90732667 |
| CA3 | 88.2012034 | 0.184637285 | 0.3655967 | 0.50502996 | 0.61353779 | 0.82498688 |
| CA2 | 61.5877995 | 1.059847885 | 0.43617329 | 2.429878 | 0.01510391 | 0.11411159 |
| WWP1 | 394.886433 | -0.248977713 | 0.17687374 | -1.4076579 | 0.15923239 | 0.42480123 |
| RMDN1 | 115.892765 | 0.283310384 | 0.17254551 | 1.64194588 | 0.10060122 | 0.33126781 |
| CPNE3 | 102.287159 | -0.10067361 | 0.26396023 | -0.3813969 | 0.70290877 | 0.87284977 |
| AF117829.1 | 309.633747 | -0.653449455 | 0.42672427 | -1.5313154 | 0.12569146 | 0.37252428 |
| RIPK2 | 41.0023928 | -1.097306238 | 0.46809724 | -2.3441844 | 0.01906874 | 0.13093869 |
| OSGIN2 | 54.7788373 | -0.305150873 | 0.28377233 | -1.075337 | 0.28222389 | 0.56986144 |
| NBN | 105.814874 | -0.364571681 | 0.33841285 | -1.0772986 | 0.2813469 | 0.56893012 |
| DECR1 | 143.558057 | -0.095168397 | 0.24754983 | -0.3844414 | 0.70065134 | 0.87182648 |
| TMEM64 | 130.130696 | -0.288330537 | 0.40538173 | -0.7112569 | 0.47692508 | 0.7335365 |
| NECAB1 | 106.362332 | 0.388496402 | 0.5951215 | 0.65280182 | 0.51388405 | 0.76088633 |
| PIP4P2 | 37.181111 | -0.342364532 | 0.40029749 | -0.8552752 | 0.39239879 | 0.66799147 |
| OTUD6B-AS1 | 41.9088475 | 0.015767587 | 0.33036468 | 0.04772782 | 0.96193316 | 0.9875899 |
| OTUD6B | 45.0712635 | 0.157073392 | 0.29648837 | 0.52977927 | 0.59626498 | 0.81582949 |
| LRRC69 | 65.0643237 | -0.343440358 | 0.26646541 | -1.2888741 | 0.19744186 | 0.47630586 |
| TRIQK | 108.114807 | 0.179024525 | 0.17640393 | 1.01485567 | 0.31017463 | 0.59721866 |
| FAM92A | 41.5033989 | -0.072227547 | 0.3442736 | -0.2097969 | 0.83382616 | 0.93647023 |
| RBM12B | 76.313161 | 0.052940402 | 0.18790543 | 0.2817396 | 0.77814319 | 0.90889326 |
| TMEM67 | 140.343357 | 0.491833652 | 0.26162582 | 1.87991254 | 0.06012 | 0.24726898 |
| VIRMA | 112.029486 | 0.278781399 | 0.16544594 | 1.68503012 | 0.09198276 | 0.31580748 |
| ESRP1 | 97.747885 | 0.360044107 | 0.63534255 | 0.56669289 | 0.57092285 | 0.79837518 |
| DPY19L4 | 102.299596 | 0.28539528 | 0.19158095 | 1.48968503 | 0.13630707 | 0.38812051 |
| INTS8 | 101.916183 | 0.296911912 | 0.21584797 | 1.37556037 | 0.16895778 | 0.43860342 |
| NDUFAF6 | 120.200863 | 0.457812891 | 0.19624684 | 2.33284213 | 0.01965643 | 0.13334913 |
| TP53INP1 | 42.0979278 | -0.152965458 | 0.4476746 | -0.3416889 | 0.732585 | 0.88869659 |

| | | | | | | |
|---|---|---|---|---|---|---|
| PLEKHF2 | 52.4900938 | 0.024362458 | 0.47012904 | 0.05182079 | 0.95867149 | 0.98584157 |
| C8orf37-AS1 | 91.2777194 | 0.314512188 | 0.5089095 | 0.61801202 | 0.53656742 | 0.77558443 |
| UQCRB | 188.287202 | -0.260886217 | 0.42803518 | -0.6094971 | 0.54219496 | 0.77901388 |
| PTDSS1 | 34.9923581 | -0.644384445 | 0.39552272 | -1.629197 | 0.10327132 | 0.33544793 |
| SDC2 | 446.772653 | 0.914355839 | 0.30897544 | 2.95931559 | 0.00308323 | 0.04154042 |
| CPQ | 345.555681 | 0.570493462 | 0.28952146 | 1.97047038 | 0.04878449 | 0.22147232 |
| MTDH | 405.418211 | -0.168182796 | 0.12545405 | -1.3405928 | 0.1800527 | 0.45208652 |
| LAPTM4B | 171.980328 | -1.109110259 | 0.38789257 | -2.8593233 | 0.00424546 | 0.05193774 |
| RPL30 | 324.276424 | -0.188200492 | 0.46106422 | -0.4081872 | 0.68313628 | 0.86454781 |
| RIDA | 58.4880907 | -0.193301081 | 0.40133038 | -0.4816508 | 0.63005406 | 0.83433967 |
| NIPAL2 | 44.3561022 | -0.491156867 | 0.31311129 | -1.5686335 | 0.11673336 | 0.35989342 |
| STK3 | 795.33065 | 0.050997022 | 0.17632545 | 0.289221 | 0.77241226 | 0.90624314 |
| AC104986.2 | 61.6021269 | 0.383675742 | 0.33714401 | 1.13801736 | 0.25511323 | 0.5419671 |
| VPS13B | 799.086454 | 0.180537082 | 0.14046724 | 1.28526113 | 0.19870105 | 0.47741788 |
| COX6C | 130.376338 | -0.31225499 | 0.28876187 | -1.0813581 | 0.27953785 | 0.56686465 |
| POLR2K | 53.9094612 | 0.01878075 | 0.49558854 | 0.03789585 | 0.96977072 | 0.99175186 |
| SPAG1 | 141.934481 | 0.003993794 | 0.38063239 | 0.01049252 | 0.99162833 | 0.99814428 |
| RNF19A | 318.67474 | -0.306439395 | 0.28142095 | -1.0889004 | 0.2761978 | 0.56317011 |
| ANKRD46 | 62.9734836 | 0.264055799 | 0.21271975 | 1.24133186 | 0.21448318 | 0.49669288 |
| PABPC1 | 329.06014 | -0.64471103 | 0.34702874 | -1.857803 | 0.06319699 | 0.25553567 |
| YWHAZ | 419.888328 | 0.067940977 | 0.2387019 | 0.28462687 | 0.77593002 | 0.90732667 |
| ZNF706 | 141.651979 | 0.13095877 | 0.25200152 | 0.51967453 | 0.60329044 | 0.81981804 |
| AP001207.3 | 29.7372225 | 1.258928886 | 1.22628343 | 1.02662146 | 0.30459875 | 0.59255715 |
| GRHL2 | 373.387277 | 0.916852104 | 0.74078802 | 1.23767135 | 0.21583795 | 0.49746509 |
| NCALD | 56.7074294 | 1.320613909 | 0.57835027 | 2.28341537 | 0.02240591 | 0.14387973 |
| RRM2B | 96.2714809 | 0.307158518 | 0.4151567 | 0.73986164 | 0.45938396 | 0.71808729 |
| UBR5 | 557.562105 | 0.286163604 | 0.14789002 | 1.93497577 | 0.05299327 | 0.23089662 |
| KLF10 | 122.752416 | 0.257615384 | 0.47825429 | 0.53865776 | 0.59012302 | 0.81183245 |
| AZIN1 | 118.663941 | 0.232952702 | 0.26359314 | 0.88375858 | 0.37682655 | 0.65512273 |
| AZIN1-AS1 | 255.823303 | 0.153515488 | 0.30087694 | 0.51022682 | 0.60989256 | 0.82246865 |
| AP003354.2 | 70.448432 | 0.276337665 | 0.34659975 | 0.79728179 | 0.42528739 | 0.69200407 |
| ATP6V1C1 | 176.675253 | 0.063786116 | 0.27992767 | 0.22786642 | 0.81975009 | 0.93087461 |
| BAALC-AS1 | 72.9394886 | 0.696985682 | 0.52617398 | 1.32462971 | 0.185294 | 0.45946964 |
| FZD6 | 174.395157 | 0.556875794 | 0.18064726 | 3.08266955 | 0.00205153 | 0.03226175 |
| SLC25A32 | 53.8133284 | 0.15528475 | 0.23033404 | 0.67417194 | 0.50020201 | 0.75171556 |
| DCAF13 | 38.2747228 | -0.602040223 | 0.40519995 | -1.4857855 | 0.13733586 | 0.38978968 |
| RIMS2 | 135.194385 | -2.55619692 | 1.07379274 | -2.380531 | 0.01728771 | 0.1239513 |
| DPYS | 43.736251 | -0.567840316 | 0.82232033 | -0.6905342 | 0.48985831 | 0.74227587 |
| LRP12 | 51.8656477 | 0.117700508 | 0.27445303 | 0.42885484 | 0.66802887 | 0.85905308 |
| ZFPM2 | 685.641815 | 0.005390309 | 1.03082194 | 0.00522914 | 0.99582777 | 0.99899312 |
| OXR1 | 2630.16534 | 0.669653712 | 0.25981679 | 2.57740742 | 0.00995445 | 0.08945002 |
| AC090579.1 | 268.981111 | 0.916051986 | 0.51238981 | 1.78780289 | 0.07380781 | 0.28055756 |
| EIF3E | 237.522677 | -0.036247009 | 0.14236233 | -0.254611 | 0.7990236 | 0.91949148 |
| EMC2 | 135.662303 | 0.111762228 | 0.29185882 | 0.3829325 | 0.70176981 | 0.87200065 |
| NUDCD1 | 88.7962607 | -0.14769465 | 0.40840429 | -0.3616383 | 0.71762231 | 0.88106948 |
| ENY2 | 89.6785177 | -0.391998038 | 0.27503558 | -1.425263 | 0.15408119 | 0.41681551 |
| EBAG9 | 51.2961766 | 0.105474447 | 0.26300986 | 0.4010285 | 0.68839914 | 0.86751885 |
| TRPS1 | 629.099024 | -0.101833944 | 0.23975037 | -0.4247499 | 0.67101901 | 0.85977676 |
| EIF3H | 253.019962 | -0.157038654 | 0.19448619 | -0.807454 | 0.41940496 | 0.68835944 |
| UTP23 | 43.9369197 | -0.555203099 | 0.24176306 | -2.2964761 | 0.02164867 | 0.14164798 |
| RAD21 | 133.418989 | 0.15001695 | 0.19157751 | 0.78306137 | 0.43359107 | 0.69851478 |
| MED30 | 26.0315107 | 0.149711677 | 0.35028984 | 0.42739371 | 0.66909258 | 0.85924744 |
| EXT1 | 1191.32832 | -0.140098062 | 0.4655682 | -0.3009185 | 0.76347668 | 0.90234693 |
| SAMD12 | 883.199804 | -0.243792225 | 0.19209216 | -1.269142 | 0.20439043 | 0.48318561 |
| AC023590.1 | 85.2803121 | -0.256615806 | 0.49269991 | -0.5208359 | 0.60248108 | 0.81930243 |
| SAMD12-AS1 | 58.9113429 | -0.702587339 | 0.31667061 | -2.2186693 | 0.02650923 | 0.15781973 |
| TNFRSF11B | 531.993313 | -0.477875217 | 0.40117597 | -1.191186 | 0.23358056 | 0.51635994 |
| COLEC10 | 116.926336 | 0.431811393 | 0.53572052 | 0.80603855 | 0.42022062 | 0.68867294 |
| MAL2 | 198.480905 | 0.442850924 | 0.4805175 | 0.92161248 | 0.35673074 | 0.63847605 |
| TAF2 | 159.952518 | 0.480955524 | 0.20253533 | 2.37467466 | 0.01756442 | 0.12490692 |
| DEPTOR | 917.474469 | 0.913164577 | 0.59765542 | 1.52791147 | 0.12653452 | 0.37394935 |
| AC091563.1 | 69.0337969 | 0.059182034 | 0.55108376 | 0.10739208 | 0.91447793 | 0.9686812 |
| COL14A1 | 83.7658522 | 1.813648863 | 0.53257247 | 3.40544987 | 0.00066055 | 0.01573986 |
| MRPL13 | 48.3986323 | -0.142917177 | 0.25177161 | -0.5676461 | 0.57027529 | 0.79765604 |
| MTBP | 30.7695402 | 0.873481425 | 0.38972976 | 2.24124899 | 0.02500995 | 0.15333673 |
| SNTB1 | 663.164981 | 0.402470557 | 0.20260564 | 1.98647261 | 0.04698087 | 0.21646453 |
| ZHX2 | 640.931537 | -0.258794783 | 0.2525205 | -1.0248466 | 0.30543556 | 0.59275349 |
| AC104316.1 | 38.4429514 | 0.18559594 | 0.48346803 | 0.38388462 | 0.70106397 | 0.87191532 |
| DERL1 | 126.118686 | 0.115327885 | 0.27048991 | 0.42636667 | 0.66984068 | 0.85924744 |
| WDYHV1 | 28.5063331 | -0.582344403 | 0.41361469 | -1.4079394 | 0.15914902 | 0.42476693 |
| FAM91A1 | 119.527291 | 0.107577578 | 0.23991209 | 0.44840415 | 0.65386155 | 0.84938158 |
| TMEM65 | 184.095744 | 0.332419194 | 0.19424749 | 1.71131785 | 0.08702245 | 0.30691783 |
| RNF139 | 26.6592776 | -0.068118828 | 0.25898485 | -0.2630224 | 0.79253328 | 0.9167584 |
| TATDN1 | 113.149149 | -0.146586989 | 0.19882073 | -0.7372822 | 0.46095074 | 0.71918385 |
| NDUFB9 | 84.4089389 | -0.107298216 | 0.27630191 | -0.3883369 | 0.69776676 | 0.87143518 |
| MTSS1 | 546.617677 | 1.564870173 | 0.4669953 | 3.35093346 | 0.0008054 | 0.01794161 |
| SQLE | 88.4450851 | 0.551656571 | 0.6055875 | 0.91094445 | 0.36232464 | 0.64272366 |
| WASHC5 | 69.7374468 | -0.002138492 | 0.18928592 | -0.0112977 | 0.99098595 | 0.9977564 |
| NSMCE2 | 573.513466 | -0.120236203 | 0.21252988 | -0.5657379 | 0.57157198 | 0.79878464 |
| TRIB1 | 69.4288593 | -0.170895063 | 0.29850048 | -0.5725118 | 0.56697526 | 0.79583763 |
| FAM84B | 121.867898 | 1.288617892 | 0.64796676 | 1.98870988 | 0.04673323 | 0.21594677 |
| PCAT1 | 996.87955 | 1.135127023 | 0.52577535 | 2.15895823 | 0.03085341 | 0.17152919 |
| PVT1 | 240.330931 | -1.425490036 | 0.66662746 | -2.1383608 | 0.03248747 | 0.17651588 |
| FAM49B | 377.420681 | 0.16103386 | 0.2413275 | 0.6672835 | 0.50459106 | 0.7552309 |
| ASAP1 | 400.965952 | -0.554790867 | 0.43671286 | -1.2703791 | 0.20394964 | 0.48285556 |
| EFR3A | 236.851045 | 0.24771316 | 0.19133095 | 1.2946842 | 0.19542919 | 0.47392673 |
| KCNQ3 | 142.50388 | -2.433745133 | 0.94739444 | -2.5688826 | 0.0102027 | 0.09024162 |
| LRRC6 | 172.095743 | 0.421173022 | 0.30998589 | 1.35868451 | 0.17424658 | 0.44550166 |
| PHF20L1 | 135.914461 | 0.34521555 | 0.19728554 | 1.74982689 | 0.08014819 | 0.29426581 |
| NDRG1 | 383.112095 | -1.066524848 | 0.72584857 | -1.469349 | 0.14173816 | 0.397922 |
| ST3GAL1 | 81.7239547 | -1.827774826 | 0.75351467 | -2.4256659 | 0.01528033 | 0.11471025 |
| ZFAT | 62.7414973 | 0.208315532 | 0.26048666 | 0.79971669 | 0.42387496 | 0.69092389 |
| AC083843.3 | 41.534817 | 0.165856015 | 0.3780937 | 0.43866379 | 0.66090517 | 0.85412987 |
| KHDRBS3 | 99.929385 | -0.944113209 | 0.37564104 | -2.5133388 | 0.01195944 | 0.10009484 |
| TRAPPC9 | 549.78407 | 0.005923869 | 0.22976924 | 0.02578182 | 0.97943136 | 0.99557142 |
| CHRAC1 | 28.6229144 | -0.382344086 | 0.35113429 | -1.0888828 | 0.27620557 | 0.56317011 |
| AGO2 | 218.951636 | -0.207867202 | 0.16850016 | -1.2336321 | 0.21734003 | 0.49937639 |
| PTK2 | 1370.16147 | 0.330841792 | 0.19200326 | 1.7231051 | 0.08486952 | 0.30278126 |
| DENND3 | 112.275813 | -0.283292318 | 0.31434072 | -0.9012269 | 0.36746768 | 0.64705385 |
| SLC45A4 | 225.796214 | 1.207986495 | 0.55616686 | 2.17198574 | 0.02985674 | 0.16887967 |
| TSNARE1 | 59.9792485 | 0.031808346 | 0.26355874 | 0.12068788 | 0.90393826 | 0.96520171 |
| JRK | 31.6439404 | 0.173731194 | 0.3776583 | 0.46002218 | 0.6455003 | 0.84407726 |
| LY6E | 239.229268 | -1.47483783 | 1.12426448 | -1.3118246 | 0.18957931 | 0.46563596 |
| RHPN1 | 42.7664723 | 0.007858752 | 0.33385147 | 0.02353967 | 0.9812198 | 0.99585061 |
| ZC3H3 | 52.4653937 | -0.180762055 | 0.28876204 | -0.6259897 | 0.53132171 | 0.77193643 |
| GSDMD | 67.1948772 | -0.47220966 | 0.49470419 | -0.9545293 | 0.33981577 | 0.62462345 |
| EEF1D | 314.778861 | -0.7415388 | 0.40359458 | -1.8373359 | 0.06616033 | 0.26198834 |
| TSTA3 | 37.0728257 | -1.660677539 | 0.48831326 | -3.4008447 | 0.00067178 | 0.01577201 |
| ZNF623 | 36.5954667 | 0.246663836 | 0.35260019 | 0.69955673 | 0.48420417 | 0.73727416 |
| MAPK15 | 31.2562807 | 1.065434521 | 0.55315117 | 1.92611816 | 0.05408962 | 0.23338941 |
| FAM83H | 41.3981341 | -0.794668526 | 0.3560345 | -2.2319987 | 0.02561505 | 0.1551184 |
| IQANK1 | 31.1796069 | 0.052242807 | 0.46879074 | 0.11144163 | 0.91126615 | 0.96844354 |
| SCRIB | 27.6122167 | -0.471198413 | 0.3398164 | -1.3866264 | 0.1655557 | 0.43384207 |

| | | | | | | |
|---|---|---|---|---|---|---|
| PUF60 | 68.557405 | -0.985859069 | 0.35281834 | -2.7942399 | 0.00520219 | 0.05957347 |
| NRBP2 | 79.8308008 | -1.542505489 | 0.59350149 | -2.5989918 | 0.0093498 | 0.08623137 |
| PLEC | 92.3476685 | -0.129203509 | 0.30576508 | -0.4225581 | 0.6726177 | 0.86026394 |
| PARP10 | 49.9989235 | 0.054308195 | 0.69862384 | 0.07773596 | 0.93803809 | 0.97987643 |
| GRINA | 83.3788567 | -0.637723197 | 0.41865028 | -1.5232838 | 0.12768772 | 0.37507533 |
| GPAA1 | 57.1698179 | -0.538047584 | 0.42082381 | -1.2785579 | 0.2010528 | 0.4800825 |
| CYC1 | 54.145749 | -1.090718681 | 0.41833639 | -2.6072766 | 0.00912656 | 0.08519229 |
| SHARPIN | 40.7307194 | 0.161447712 | 0.21568742 | 0.74852632 | 0.45414275 | 0.71384952 |
| MAF1 | 50.9438764 | -0.306086839 | 0.34841309 | -0.878517 | 0.37966319 | 0.65703312 |
| MROH1 | 152.776327 | -0.286434363 | 0.35337969 | -0.8105569 | 0.41762015 | 0.68711498 |
| HSF1 | 89.9968959 | -0.395526552 | 0.22304919 | -1.7732705 | 0.0761839 | 0.28527949 |
| CPSF1 | 25.1049429 | -0.065022144 | 0.48071709 | -0.1352607 | 0.89240573 | 0.95976294 |
| VPS28 | 120.313006 | -0.846338372 | 0.38056863 | -2.2238784 | 0.02615662 | 0.15698891 |
| CYHR1 | 62.3309039 | -0.012241852 | 0.23228553 | -0.0527017 | 0.95796955 | 0.9856488 |
| PPP1R16A | 79.1276542 | 0.120222196 | 0.34370679 | 0.34978126 | 0.72650286 | 0.88522597 |
| ARHGAP39 | 39.4786574 | 0.096182003 | 0.29120863 | 0.33028555 | 0.74118421 | 0.89253455 |
| ZNF251 | 115.660308 | 0.261760105 | 0.20845869 | 1.25569297 | 0.20922731 | 0.48959511 |
| RPL8 | 650.67521 | -0.513497722 | 0.51040939 | -1.0060507 | 0.31439118 | 0.60132156 |
| ZNF7 | 38.5869833 | -0.472129441 | 0.30150238 | -1.5659228 | 0.1173667 | 0.36114862 |
| ZNF250 | 36.8386592 | 0.660805161 | 0.26709081 | 2.4740842 | 0.01335782 | 0.10645139 |
| C8orf33 | 29.5196648 | -0.723795649 | 0.36454225 | -1.9854918 | 0.04708978 | 0.2166537 |
| WASHC1 | 154.500073 | 0.466989017 | 0.24561805 | 1.90128131 | 0.05726518 | 0.24164895 |
| CBWD1 | 49.8739167 | 0.201616387 | 0.3238923 | 0.62247971 | 0.53362648 | 0.77365669 |
| DOCK8 | 163.169727 | 0.651401594 | 0.46206015 | 1.40977663 | 0.15860565 | 0.42414392 |
| KANK1 | 1174.41694 | 1.131751966 | 0.13583561 | 8.33177679 | 7.96E-17 | 3.81E-13 |
| SMARCA2 | 235.184231 | 0.188878543 | 0.31415479 | 0.60122765 | 0.54768837 | 0.7832647 |
| VLDLR-AS1 | 41.9718508 | 0.090129442 | 0.37601979 | 0.23969334 | 0.810568 | 0.92511264 |
| VLDLR | 38.7678349 | -0.599822804 | 0.38409659 | -1.5616458 | 0.11837146 | 0.36330672 |
| PUM3 | 352.39612 | -0.492131611 | 0.3369843 | -1.4603992 | 0.14418038 | 0.40168142 |
| RFX3 | 439.272609 | 0.624582795 | 0.28217827 | 2.21343339 | 0.02686778 | 0.15864374 |
| RFX3-AS1 | 389.11428 | 0.065828993 | 0.33474095 | 0.19665653 | 0.84409633 | 0.94193856 |
| GLIS3 | 2072.41547 | 0.047451395 | 0.30755769 | 0.15428454 | 0.87738539 | 0.95407818 |
| SLC1A1 | 248.431034 | 0.147968827 | 0.3260972 | 0.45375681 | 0.65000387 | 0.84716453 |
| SPATA6L | 280.320884 | -0.09128374 | 0.38502952 | -0.2370824 | 0.81259283 | 0.92664604 |
| CDC37L1 | 80.7821686 | 0.459559682 | 0.22729076 | 2.02190218 | 0.04318646 | 0.20614587 |
| AK3 | 287.497277 | 0.630921754 | 0.24146788 | 2.61285997 | 0.00897881 | 0.08473695 |
| RCL1 | 113.36457 | -0.529857759 | 0.34517592 | -1.5350368 | 0.12477478 | 0.37130091 |
| JAK2 | 130.150801 | -0.230424746 | 0.24906735 | -0.9251504 | 0.3548877 | 0.63648555 |
| PLGRKT | 73.3171745 | -0.099844543 | 0.25469194 | -0.3920208 | 0.69504283 | 0.87024545 |
| RIC1 | 360.635918 | 0.873874698 | 0.23711973 | 3.68537325 | 0.00022837 | 0.00824694 |
| ERMP1 | 531.704329 | 0.925448553 | 0.6788339 | 1.36329159 | 0.17279063 | 0.44401526 |
| KIAA2026 | 344.108507 | 0.233915529 | 0.16560267 | 1.41251064 | 0.15779966 | 0.42316991 |
| RANBP6 | 29.1643685 | -0.520752025 | 0.34279553 | -1.5191331 | 0.128729 | 0.37682165 |
| UHRF2 | 170.770984 | 0.279709635 | 0.1894197 | 1.476666 | 0.13976519 | 0.39399964 |
| GLDC | 92.8396632 | -0.336153404 | 0.63780743 | -0.5270453 | 0.59816213 | 0.81658679 |
| AL162411.1 | 49.6252367 | -0.18419774 | 0.31394278 | -0.5867239 | 0.55738914 | 0.78866035 |
| KDM4C | 559.204465 | -0.246262582 | 0.18640924 | -1.3210857 | 0.18647278 | 0.46155628 |
| DMAC1 | 47.273841 | -0.778889277 | 0.36910193 | -2.1102281 | 0.03483871 | 0.1843757 |
| PTPRD | 470.591558 | 0.968950121 | 0.44659074 | 2.16966014 | 0.0300326 | 0.16932448 |
| MPDZ | 162.104007 | 0.089901687 | 0.34173955 | 0.26307077 | 0.79249603 | 0.9167584 |
| NFIB | 745.772945 | 0.024899536 | 0.32642494 | 0.07627952 | 0.93919672 | 0.97996404 |
| ZDHHC21 | 277.433938 | 0.750393198 | 0.19130978 | 3.92239847 | 8.77E-05 | 0.00428474 |
| TTC39B | 242.213369 | -1.184456667 | 0.48265899 | -2.4540238 | 0.01412676 | 0.11010598 |
| SNAPC3 | 111.422014 | 0.308097001 | 0.18098084 | 1.70237363 | 0.08868535 | 0.30992958 |
| PSIP1 | 103.829921 | 0.694614988 | 0.24687334 | 2.81364923 | 0.00489826 | 0.05722009 |
| CCDC171 | 125.112704 | 0.747485268 | 0.3354019 | 2.22862563 | 0.02583882 | 0.15586278 |
| BNC2 | 833.809455 | -0.01107392 | 0.5015573 | -0.0220791 | 0.98238488 | 0.99616479 |
| CNTLN | 169.220179 | 0.043902763 | 0.26714209 | 0.16434237 | 0.86946163 | 0.95106604 |
| RRAGA | 43.3831898 | -0.967500473 | 0.42832127 | -2.2588196 | 0.02389461 | 0.14976011 |
| HAUS6 | 42.4393176 | -0.098061967 | 0.23006058 | -0.4262441 | 0.66992998 | 0.85924744 |
| PLIN2 | 128.81303 | -2.479106905 | 0.60085152 | -4.1259892 | 3.69E-05 | 0.00247275 |
| DENND4C | 435.473548 | -0.10004757 | 0.18111307 | -0.5524039 | 0.58067164 | 0.80525463 |
| RPS6 | 705.935248 | -0.465913153 | 0.5720166 | -0.8145099 | 0.41535293 | 0.68495052 |
| MLLT3 | 742.492071 | 0.280506565 | 0.28139703 | 0.99683556 | 0.31884433 | 0.60563353 |
| FOCAD | 471.533514 | 0.507648072 | 0.20912975 | 2.42743111 | 0.01520617 | 0.11460263 |
| KLHL9 | 45.5030629 | -0.226630538 | 0.34047773 | -0.6656252 | 0.50565073 | 0.75528806 |
| LINC01239 | 43.8439463 | -0.741763009 | 0.41077647 | -1.8057583 | 0.0709561 | 0.27362662 |
| CAAP1 | 183.383755 | 0.283274076 | 0.2610944 | 1.08494887 | 0.2779443 | 0.56551157 |
| PLAA | 70.7819882 | 0.05074654 | 0.22422241 | 0.22632234 | 0.82095073 | 0.93138972 |
| IFT74 | 227.931964 | 0.721846074 | 0.23719121 | 3.04330871 | 0.00233992 | 0.0349985 |
| LRRC19 | 71.2749357 | 1.114867981 | 0.49717049 | 2.24242591 | 0.02493386 | 0.15310348 |
| MOB3B | 425.937894 | -1.474483053 | 0.58133991 | -2.5363527 | 0.01120138 | 0.09563107 |
| C9orf72 | 75.4119332 | -0.048422751 | 0.37495645 | -0.1291423 | 0.89724503 | 0.96175013 |
| ACO1 | 114.82045 | -0.936726156 | 0.37965346 | -2.4673189 | 0.01361291 | 0.10755093 |
| DDX58 | 167.857414 | -1.019338249 | 0.99363102 | -1.025872 | 0.30495192 | 0.59267272 |
| TOPORS | 44.926508 | -0.161813682 | 0.23108217 | -0.700243 | 0.48377553 | 0.73727416 |
| NDUFB6 | 65.0613391 | -0.019014464 | 0.30454319 | -0.062436 | 0.95021561 | 0.98270285 |
| APTX | 121.272088 | -0.105810422 | 0.32520311 | -0.3253672 | 0.74490318 | 0.89323545 |
| DNAJA1 | 112.198793 | -0.388808099 | 0.29585452 | -1.3141868 | 0.18878335 | 0.46475347 |
| SMU1 | 76.2401567 | 0.245341952 | 0.24521407 | 1.00052152 | 0.31705819 | 0.60343739 |
| B4GALT1 | 314.967426 | -0.637700191 | 0.36916557 | -1.7274097 | 0.08409411 | 0.30203879 |
| BAG1 | 84.9520215 | 0.012085424 | 0.47776693 | 0.02529565 | 0.97981915 | 0.99557142 |
| CHMP5 | 78.7464674 | -0.036699282 | 0.37740924 | -0.09724 | 0.92253579 | 0.97269899 |
| NFX1 | 221.303068 | -0.177256958 | 0.25142454 | -0.7050106 | 0.48080366 | 0.73578929 |
| AQP3 | 73.1309448 | -0.500356549 | 0.77039828 | -0.6494778 | 0.51602962 | 0.76140599 |
| UBE2R2 | 359.887834 | -0.002073714 | 0.24479185 | -0.0084713 | 0.99324093 | 0.9985574 |
| UBAP2 | 421.467932 | 0.584069677 | 0.18281194 | 3.19492083 | 0.00139869 | 0.02596525 |
| DCAF12 | 55.6651988 | -0.410184429 | 0.24873843 | -1.6490594 | 0.09913547 | 0.32904321 |
| UBAP1 | 327.473571 | -0.122573994 | 0.20787764 | -0.5896449 | 0.55542873 | 0.78784845 |
| FAM219A | 72.6177134 | -0.669036593 | 0.33263844 | -2.0113027 | 0.04429349 | 0.20921468 |
| DNAI1 | 44.3721474 | 0.49000864 | 0.58567965 | 0.83664959 | 0.40278957 | 0.67688395 |
| DCTN3 | 39.2779223 | 0.035828478 | 0.46923298 | 0.07635541 | 0.93913634 | 0.97996404 |
| VCP | 147.588216 | -0.886107901 | 0.28965269 | -3.0592083 | 0.00221923 | 0.0339653 |
| STOML2 | 51.6991892 | -0.792366958 | 0.45060658 | -1.7584451 | 0.0786718 | 0.29089077 |
| UNC13B | 185.082611 | 0.088009755 | 0.23722446 | 0.37099781 | 0.71063916 | 0.87748 |
| RUSC2 | 205.138505 | 0.350342771 | 0.40811637 | 0.85843842 | 0.39065042 | 0.66624498 |
| CCDC107 | 40.0895728 | -0.600505876 | 0.33699238 | -1.7819569 | 0.07475627 | 0.28248137 |
| TLN1 | 115.997896 | -0.336377001 | 0.30347943 | -1.1084013 | 0.26768854 | 0.55491026 |
| CREB3 | 28.3502623 | -0.63657624 | 0.46842922 | -1.3589593 | 0.17415949 | 0.44546696 |
| GBA2 | 59.8041127 | -0.107842914 | 0.20017528 | -0.5387424 | 0.59006459 | 0.81183245 |
| RGP1 | 42.6257553 | -0.31804475 | 0.2866306 | -1.109598 | 0.26717227 | 0.55454891 |
| AL133410.1 | 54.0185229 | 0.54943502 | 0.34884437 | 1.57501474 | 0.11525304 | 0.35728443 |
| TMEM8B | 43.662208 | 0.514674827 | 0.38861731 | 1.32437441 | 0.18537873 | 0.45956078 |
| RECK | 98.0070522 | 0.03476629 | 0.38390119 | 0.09056052 | 0.92784181 | 0.97474479 |
| CLTA | 274.581487 | -0.757683672 | 0.27631601 | -2.7420911 | 0.00610494 | 0.06580904 |
| GNE | 176.099227 | 0.631911985 | 0.29676036 | 2.12936788 | 0.03322384 | 0.17879277 |
| RNF38 | 334.915461 | 0.14529782 | 0.14828968 | 0.97982422 | 0.3271729 | 0.61330512 |
| EBLN3P | 93.8700726 | -0.142489055 | 0.21329093 | -0.6680502 | 0.50410153 | 0.75508812 |
| ZCCHC7 | 766.539709 | 0.090555084 | 0.18598612 | 0.48689163 | 0.62633513 | 0.83180006 |
| GRHPR | 62.9433155 | -0.17389116 | 0.32013565 | -0.5431796 | 0.58700613 | 0.81058905 |
| ZBTB5 | 53.1442412 | 0.218464613 | 0.19039005 | 1.14745814 | 0.25119232 | 0.53867677 |
| AL138752.2 | 93.3369309 | 0.538499546 | 0.30524201 | 1.76417246 | 0.07770295 | 0.28902315 |
| TRMT10B | 76.4345226 | 0.194102769 | 0.25894258 | 0.74959772 | 0.45349702 | 0.71320153 |

| | | | | | | |
|---|---|---|---|---|---|---|
| DCAF10 | 246.609871 | -0.002793733 | 0.1719452 | -0.0162478 | 0.98703669 | 0.99678172 |
| FAM95C | 38.7488943 | -0.38100136 | 0.35276245 | -1.0800508 | 0.28011957 | 0.56776669 |
| ANKRD18A | 37.0357987 | -0.817625812 | 0.345572 | -2.3660071 | 0.0179811 | 0.12664775 |
| GLIDR | 50.5473114 | -0.071954859 | 0.31987239 | -0.2249486 | 0.82201923 | 0.93195907 |
| CNTNAP3B | 191.353751 | -2.874168328 | 0.9179806 | -3.1309685 | 0.00174231 | 0.0293224 |
| CBWD5 | 88.553469 | -0.080551562 | 0.37093973 | -0.2171554 | 0.82808725 | 0.93346403 |
| PGM5 | 95.0443582 | -0.363173056 | 0.70794292 | -0.5129976 | 0.60795296 | 0.8211189 |
| AL353616.2 | 108.636035 | -1.176628737 | 0.67959405 | -1.73137 | 0.0833858 | 0.30062195 |
| PIP5K1B | 82.2970502 | 1.157227426 | 0.41591739 | 2.78234921 | 0.00539669 | 0.06060367 |
| APBA1 | 145.763808 | 1.076689567 | 0.67411184 | 1.59719723 | 0.11022175 | 0.34724322 |
| PTAR1 | 259.080853 | 0.491398958 | 0.13740279 | 3.57633898 | 0.00034844 | 0.01052904 |
| MAMDC2-AS | 105.023595 | 0.174986661 | 0.38587423 | 0.45348107 | 0.65020237 | 0.84724399 |
| MAMDC2 | 393.366916 | 0.986281195 | 0.43388041 | 2.27316367 | 0.02301632 | 0.14629948 |
| SMC5-AS1 | 99.0921915 | 0.126858404 | 0.40722257 | 0.31152105 | 0.75540454 | 0.89832652 |
| SMC5 | 310.318584 | 0.146840275 | 0.11028094 | 1.33151088 | 0.18302097 | 0.45656555 |
| KLF9 | 120.260726 | 0.086922727 | 0.44803767 | 0.19400763 | 0.84616991 | 0.94271476 |
| TRPM3 | 307.670026 | -0.50981728 | 0.54461348 | -0.9361084 | 0.34921736 | 0.63265484 |
| CEMIP2 | 134.263707 | -1.329252028 | 0.38284912 | -3.4719997 | 0.0005166 | 0.01373303 |
| ABHD17B | 52.7436881 | -0.141985592 | 0.2758597 | -0.5147022 | 0.60676114 | 0.8211189 |
| C9orf85 | 87.3762688 | -0.122758734 | 0.20807181 | -0.5899825 | 0.55520236 | 0.78784845 |
| ZFAND5 | 295.351669 | -0.790601263 | 0.38838085 | -2.035634 | 0.04178712 | 0.20312121 |
| ALDH1A1 | 213.499664 | -0.2332661 | 0.46886653 | -0.4975107 | 0.61882899 | 0.82733103 |
| ANXA1 | 100.424245 | -0.478570848 | 1.06091095 | -0.4510943 | 0.65192161 | 0.84789641 |
| CARNMT1 | 115.45141 | 0.136861645 | 0.20347009 | 0.67263765 | 0.50117784 | 0.75235583 |
| NMRK1 | 77.9537111 | -0.135470608 | 0.31157012 | -0.4347997 | 0.6637078 | 0.8562037 |
| OSTF1 | 42.9064332 | -0.201303394 | 0.2655505 | -0.7580607 | 0.44841466 | 0.7097429 |
| PCSK5 | 66.5077676 | -0.907265144 | 0.33598225 | -2.7003365 | 0.00692694 | 0.07087956 |
| RFK | 46.3807941 | -0.430489511 | 0.34286845 | -1.255553 | 0.20927809 | 0.48959511 |
| PRUNE2 | 905.32671 | 0.401385267 | 0.37554418 | 1.06880972 | 0.28515542 | 0.57264845 |
| PCA3 | 109.163107 | 0.47319613 | 0.70892047 | 0.66748831 | 0.50446027 | 0.7552161 |
| VPS13A | 444.965456 | 0.241150536 | 0.16169211 | 1.49141811 | 0.13585176 | 0.38788633 |
| GNAQ | 1141.91224 | -0.200564407 | 0.13387852 | -1.4981075 | 0.13410533 | 0.38564845 |
| CEP78 | 41.9506573 | 0.372663831 | 0.34289031 | 1.08683105 | 0.27711148 | 0.56429653 |
| TLE4 | 229.73143 | -0.204109893 | 0.51508686 | -0.3962631 | 0.69191098 | 0.86887982 |
| TLE1 | 158.992845 | -0.750092796 | 0.27557546 | -2.7219143 | 0.0064905 | 0.06817157 |
| RASEF | 83.6283378 | -0.46235549 | 0.52376291 | -0.8827572 | 0.37736746 | 0.65533048 |
| FRMD3 | 481.391915 | 0.923819746 | 0.40940706 | 2.25648219 | 0.02404045 | 0.15036388 |
| IDNK | 26.9104361 | 0.11430406 | 0.37761914 | 0.30269668 | 0.76212103 | 0.90141135 |
| UBQLN1 | 169.421121 | 0.094378052 | 0.15561505 | 0.6064841 | 0.54419334 | 0.78027954 |
| GKAP1 | 105.852707 | 0.865620358 | 0.27202476 | 3.18213812 | 0.00146192 | 0.02655639 |
| KIF27 | 193.172118 | 0.294497693 | 0.28125143 | 1.04709757 | 0.29505458 | 0.58298841 |
| HNRNPK | 377.50979 | -0.08970904 | 0.28238535 | -0.317683 | 0.75072537 | 0.89599662 |
| RMI1 | 39.2978067 | 0.527956891 | 0.27607792 | 1.91234742 | 0.05583165 | 0.23737743 |
| NTRK2 | 173.213172 | 1.885693035 | 1.02521971 | 1.83930626 | 0.06587015 | 0.26170477 |
| AGTPBP1 | 170.125498 | -0.079051862 | 0.19942799 | -0.396393 | 0.69181512 | 0.86887335 |
| NAA35 | 176.004963 | 0.089727931 | 0.1577887 | 0.56865877 | 0.56958773 | 0.79749848 |
| GOLM1 | 84.3549805 | -0.737200553 | 0.39605234 | -1.8613716 | 0.06269172 | 0.25435156 |
| ISCA1 | 41.7645825 | 0.070062794 | 0.22718689 | 0.30839277 | 0.75778348 | 0.89967348 |
| TUT7 | 131.030598 | -0.173634248 | 0.18015081 | -0.9638272 | 0.33513259 | 0.6199188 |
| DAPK1 | 1218.33958 | 1.022311733 | 0.2522245 | 4.05318171 | 5.05E-05 | 0.00298922 |
| CTSL | 173.568797 | -2.392298553 | 0.70819927 | -3.378002 | 0.00073015 | 0.01669227 |
| SPIN1 | 399.208252 | 0.632695802 | 0.23899367 | 2.64733285 | 0.00811295 | 0.07857827 |
| SECISBP2 | 264.806924 | -0.08112953 | 0.15639144 | -0.5187594 | 0.60392854 | 0.82004322 |
| SEMA4D | 673.708241 | 0.118620675 | 0.35744769 | 0.33185464 | 0.73999902 | 0.89253455 |
| LINC01508 | 71.3880151 | 0.82862376 | 0.87354032 | 0.94858101 | 0.34283375 | 0.62779669 |
| SYK | 155.455868 | 0.230048853 | 0.28312782 | 0.81252648 | 0.4164896 | 0.68597057 |
| AL158071.4 | 27.9605041 | 0.639548651 | 0.35530865 | 1.79998053 | 0.07186371 | 0.27601544 |
| AUH | 390.03071 | 0.135605224 | 0.20758773 | 0.65324297 | 0.51359965 | 0.76083448 |
| NFIL3 | 49.9990568 | -2.435282687 | 0.68872885 | -3.5359092 | 0.00040637 | 0.01172488 |
| ROR2 | 42.9420087 | -0.996979611 | 0.84491284 | -1.1799792 | 0.23800847 | 0.52099706 |
| SPTLC1 | 148.948608 | 0.003568815 | 0.16559063 | 0.02155204 | 0.98280529 | 0.99616479 |
| IARS | 154.125651 | -0.198344175 | 0.28597162 | -0.6935799 | 0.48794568 | 0.74125768 |
| NOL8 | 52.3400888 | -0.4463883 | 0.23688129 | -1.8844388 | 0.05950564 | 0.24614695 |
| CENPP | 91.3691997 | -0.311231555 | 0.49884852 | -0.6238999 | 0.53269331 | 0.77315581 |
| IPPK | 35.8867721 | 0.058088808 | 0.23596457 | 0.24617598 | 0.80554601 | 0.92227408 |
| BICD2 | 45.9581116 | -0.170588881 | 0.32587599 | -0.5234779 | 0.60064172 | 0.81779318 |
| NINJ1 | 42.352511 | -0.634120502 | 0.42645012 | -1.4869746 | 0.13702152 | 0.38912812 |
| FAM120AOS | 37.9050649 | -0.128180502 | 0.3650696 | -0.3511125 | 0.72550395 | 0.88462351 |
| FAM120A | 295.013253 | -0.378919344 | 0.20148486 | -1.8806343 | 0.06002168 | 0.24718301 |
| PHF2 | 80.7179099 | 0.060215425 | 0.23370598 | 0.25765462 | 0.79667347 | 0.91869184 |
| PTPDC1 | 77.1439423 | 0.055346432 | 0.28267801 | 0.1957932 | 0.84477203 | 0.94193856 |
| AL158152.2 | 118.122328 | -0.196907014 | 0.31561952 | -0.6238746 | 0.53270992 | 0.77315581 |
| ZNF169 | 37.6823294 | 0.438017974 | 0.22500147 | 1.94673381 | 0.05156666 | 0.22782627 |
| MFSD14B | 89.0860698 | -0.000742988 | 0.16776305 | -0.0044288 | 0.99646635 | 0.99923686 |
| FBP1 | 32.6447594 | -0.250197618 | 0.46676956 | -0.5360196 | 0.59194502 | 0.81275584 |
| C9orf3 | 1089.4998 | 0.256489563 | 0.28867783 | 0.88849761 | 0.37427314 | 0.65269307 |
| FANCC | 561.19125 | 0.378391386 | 0.22742814 | 1.66378434 | 0.09615553 | 0.32329725 |
| PTCH1 | 87.2063357 | 0.023538807 | 0.48283522 | 0.04875122 | 0.96111756 | 0.98729706 |
| LINC00476 | 156.0344 | 0.275634508 | 0.37588443 | 0.73329588 | 0.46337799 | 0.72133763 |
| ERCC6L2 | 215.828724 | 0.042413314 | 0.17500828 | 0.24235033 | 0.80850872 | 0.92407887 |
| SLC35D2 | 41.3124707 | -0.090079191 | 0.27971446 | -0.3220398 | 0.74742254 | 0.89399656 |
| HABP4 | 130.700099 | 0.919178085 | 0.30634242 | 3.00049235 | 0.00269544 | 0.03796996 |
| CDC14B | 182.798009 | 0.250013078 | 0.33788654 | 0.73993205 | 0.45934123 | 0.71808729 |
| ZNF510 | 25.8331071 | 0.219866333 | 0.3109395 | 0.70710325 | 0.47950232 | 0.73490443 |
| ZNF782 | 84.7532495 | -0.104451876 | 0.22543499 | -0.4633348 | 0.64312441 | 0.84321657 |
| MFSD14C | 243.835809 | -0.012940565 | 0.19019526 | -0.0680383 | 0.94575513 | 0.98119662 |
| TDRD7 | 61.1949766 | -0.218729934 | 0.52983766 | -0.4128244 | 0.67973525 | 0.86235436 |
| TSTD2 | 60.9056365 | -0.22456697 | 0.25381459 | -0.8847678 | 0.3762819 | 0.65466091 |
| NCBP1 | 56.0974892 | 0.196259164 | 0.23256445 | 0.84389152 | 0.39873003 | 0.67377896 |
| XPA | 58.7281807 | 0.336003532 | 0.22658945 | 1.4828737 | 0.13810797 | 0.39082312 |
| ANP32B | 119.864088 | -0.730531445 | 0.23187465 | -3.1505447 | 0.00162966 | 0.02807268 |
| TRIM14 | 51.9770599 | -1.780601921 | 0.8743726 | -2.0364338 | 0.04170681 | 0.20312121 |
| ANKS6 | 99.1718588 | -0.148905733 | 0.23131202 | -0.643744 | 0.51974142 | 0.76385303 |
| TGFBR1 | 74.2585431 | -0.530337414 | 0.3201002 | -1.6567856 | 0.09756282 | 0.3259791 |
| ALG2 | 36.6157875 | -0.291879263 | 0.36157735 | -0.8072388 | 0.41952891 | 0.68835944 |
| SEC61B | 127.49887 | -0.704156166 | 0.35253161 | -1.997427 | 0.04577882 | 0.21318196 |
| AL359710.1 | 75.8497302 | 0.748390296 | 0.48777557 | 1.53429228 | 0.12495776 | 0.37161272 |
| STX17 | 123.166948 | 0.336699137 | 0.28467708 | 1.18274058 | 0.23691197 | 0.52000641 |
| ERP44 | 153.077515 | -0.1686049 | 0.18858046 | -0.8940741 | 0.37128226 | 0.65149528 |
| INVS | 341.446701 | 0.069525612 | 0.18147102 | 0.3831224 | 0.70162901 | 0.87200065 |
| TEX10 | 62.0608718 | 0.157217861 | 0.20234652 | 0.77697339 | 0.43717447 | 0.70192663 |
| MRPL50 | 26.5858484 | -0.52267911 | 0.40055018 | -1.3049029 | 0.1919259 | 0.46899444 |
| ZNF189 | 36.9436861 | 0.005024267 | 0.33730198 | 0.01489546 | 0.98811558 | 0.99678172 |
| ALDOB | 96.9613354 | -0.574716038 | 1.0340226 | -0.5558061 | 0.57834344 | 0.80396861 |
| TMEM246 | 30.8142932 | -0.04045569 | 0.29649309 | -0.1364473 | 0.89146766 | 0.95969529 |
| RNF20 | 40.1489145 | 0.097841626 | 0.26217322 | 0.37319459 | 0.70900362 | 0.87632848 |
| NIPSNAP3A | 30.730939 | -0.220976061 | 0.29814376 | -0.7411728 | 0.45858864 | 0.71752865 |
| ABCA1 | 48.4316753 | -0.481334093 | 0.5011097 | -0.9605364 | 0.33678533 | 0.62147307 |
| SLC44A1 | 167.488895 | 0.254359868 | 0.20986765 | 1.21200131 | 0.22551188 | 0.50767998 |
| FSD1L | 114.918444 | -0.032339239 | 0.2339276 | -0.1382447 | 0.89004707 | 0.95918866 |
| FKTN | 112.448742 | 0.115755401 | 0.17936626 | 0.64535771 | 0.5186954 | 0.7636788 |
| TMEM38B | 165.687065 | 0.697697696 | 0.38057111 | 1.83329128 | 0.06675927 | 0.26332729 |
| LINC01505 | 115.906131 | 0.573252236 | 0.39537475 | 1.44989591 | 0.14708755 | 0.4061769 |

| | | | | | | |
|---|---|---|---|---|---|---|
| ZNF462 | 401.723998 | 0.259814503 | 0.20867246 | 1.24508284 | 0.2131013 | 0.49485996 |
| RAD23B | 287.189586 | -0.044107466 | 0.18598519 | -0.2371558 | 0.81253593 | 0.92664604 |
| ELP1 | 73.5100388 | 0.553952704 | 0.17625837 | 3.14284486 | 0.00167314 | 0.02854928 |
| FAM206A | 31.315471 | 0.046183574 | 0.25442427 | 0.18152189 | 0.85595795 | 0.94621534 |
| CTNNAL1 | 131.70741 | -0.693419276 | 0.49176453 | -1.4100636 | 0.1585209 | 0.42411742 |
| TMEM245 | 388.004448 | 0.342055761 | 0.19950826 | 1.71449426 | 0.08643799 | 0.30636531 |
| EPB41L4B | 44.9499466 | 0.220919987 | 0.81704773 | 0.2703881 | 0.78686169 | 0.91339653 |
| PTPN3 | 365.310831 | -0.068157262 | 0.28435824 | -0.239688 | 0.81057213 | 0.92511264 |
| PALM2 | 49.8022538 | 0.092861464 | 0.56850527 | 0.16334319 | 0.87024823 | 0.9512847 |
| TXN | 131.616718 | -1.390286391 | 0.48420981 | -2.8712479 | 0.00408855 | 0.05046933 |
| LPAR1 | 74.4865214 | -1.423878308 | 0.43657612 | -3.2614663 | 0.00110838 | 0.02211903 |
| ECPAS | 307.014861 | -0.211854155 | 0.22898148 | -0.9252021 | 0.35486077 | 0.63648555 |
| ZNF483 | 72.1874984 | 0.756660448 | 0.45571651 | 1.66037531 | 0.09683897 | 0.32454105 |
| PTGR1 | 241.265663 | 0.125688934 | 0.25101044 | 0.5007319 | 0.61655982 | 0.82592881 |
| C9orf84 | 76.7034376 | 0.088511449 | 0.58749353 | 0.15065944 | 0.88024437 | 0.95614192 |
| UGCG | 865.765113 | -0.445929285 | 0.45192249 | -0.9867384 | 0.32377089 | 0.6106578 |
| SUSD1 | 294.245121 | 0.287057053 | 0.40290866 | 0.71246185 | 0.47617883 | 0.73318945 |
| PTBP3 | 922.159092 | 0.279776034 | 0.39587938 | 0.70672039 | 0.47974025 | 0.73503389 |
| HSDL2 | 168.813283 | -0.076440988 | 0.19894768 | -0.3842266 | 0.70081051 | 0.87182648 |
| C9orf147 | 29.6922595 | 0.613964737 | 0.46148857 | 1.33040074 | 0.18338627 | 0.4570849 |
| KIAA1958 | 331.170863 | 0.121590227 | 0.19138669 | 0.63531183 | 0.52522506 | 0.76694068 |
| INIP | 59.7598222 | 0.174860071 | 0.19586912 | 0.89273937 | 0.37199676 | 0.65192016 |
| SNX30 | 152.013115 | 0.252210389 | 0.32059714 | 0.78668945 | 0.43146368 | 0.69696301 |
| FKBP15 | 59.9024964 | 0.066351712 | 0.20960462 | 0.31655653 | 0.75158012 | 0.89611476 |
| SLC31A1 | 42.9586377 | -0.437094672 | 0.34622926 | -1.2624429 | 0.20678947 | 0.48621411 |
| CDC26 | 33.4924868 | -0.010302819 | 0.41378991 | -0.0248987 | 0.98013579 | 0.99558815 |
| PRPF4 | 35.9425518 | -0.208442843 | 0.38436237 | -0.5423081 | 0.58760625 | 0.81081536 |
| WDR31 | 39.8183318 | -0.167725091 | 0.40505367 | -0.4140812 | 0.67881467 | 0.86169702 |
| BSPRY | 34.9086934 | -0.123394206 | 0.42128409 | -0.2929002 | 0.76959841 | 0.90533273 |
| HDHD3 | 62.1787576 | -0.596898846 | 0.22909733 | -2.6054378 | 0.00917569 | 0.08519229 |
| RGS3 | 92.4670353 | -1.318652187 | 0.32075142 | -4.1111343 | 3.94E-05 | 0.002601 |
| ZNF618 | 297.563159 | 0.487342779 | 0.32605563 | 1.49466145 | 0.13500282 | 0.38712403 |
| KIF12 | 235.162958 | 1.048476251 | 0.37385717 | 2.80448345 | 0.00503973 | 0.0583743 |
| COL27A1 | 334.521566 | -0.304768406 | 0.31049651 | -0.9815518 | 0.3263207 | 0.61290706 |
| WHRN | 53.1177175 | 0.124317033 | 0.28338068 | 0.43869268 | 0.66088423 | 0.85412987 |
| ATP6V1G1 | 166.928571 | -0.831972857 | 0.40231409 | -2.0679685 | 0.03864299 | 0.19546419 |
| TNFSF15 | 42.9585067 | -2.183233894 | 0.81610964 | -2.6751723 | 0.00746909 | 0.07451094 |
| TNC | 474.168942 | -1.792065646 | 0.49959564 | -3.5870322 | 0.00033446 | 0.01030168 |
| PAPPA | 177.822022 | -0.991259048 | 0.41616753 | -2.381875 | 0.01722474 | 0.123685 |
| ASTN2 | 802.998073 | 1.226157372 | 0.52540778 | 2.33372519 | 0.01961012 | 0.13327863 |
| CDK5RAP2 | 123.515049 | 0.462567451 | 0.21379791 | 2.16357333 | 0.03049711 | 0.17102865 |
| MEGF9 | 261.550466 | 0.293795573 | 0.32974908 | 0.89096707 | 0.37294684 | 0.65217873 |
| FBXW2 | 83.8458082 | -0.068831168 | 0.23465984 | -0.2933232 | 0.76927515 | 0.90533273 |
| PSMD5 | 85.6242126 | -0.148419701 | 0.27026694 | -0.5491597 | 0.58289589 | 0.80710606 |
| TRAF1 | 68.3997857 | -1.487671346 | 1.04151091 | -1.4283781 | 0.15318307 | 0.41579501 |
| CNTRL | 71.8127412 | 0.304157255 | 0.31131833 | 0.97699757 | 0.32857036 | 0.61394673 |
| RAB14 | 152.296886 | 0.008206548 | 0.18730196 | 0.04381453 | 0.96505224 | 0.98965864 |
| GSN | 111.759355 | -0.329641209 | 0.27282645 | -1.208245 | 0.22695302 | 0.50976168 |
| STOM | 157.272075 | -1.339253795 | 0.29064609 | -4.6078507 | 4.07E-06 | 0.00046396 |
| DAB2IP | 180.589659 | -0.832816396 | 0.50191896 | -1.6592647 | 0.09706247 | 0.32497777 |
| TTLL11 | 48.9940827 | -0.129105865 | 0.30786145 | -0.4193635 | 0.67495047 | 0.86115505 |
| NDUFA8 | 35.5947141 | -0.109994658 | 0.35889319 | -0.306483 | 0.75923693 | 0.9005757 |
| RBM18 | 28.0662202 | -0.298615984 | 0.30566833 | -0.9769281 | 0.32860475 | 0.61394673 |
| MRRF | 112.16129 | -0.326032514 | 0.20265784 | -1.6087831 | 0.10766378 | 0.34308427 |
| PDCL | 31.2348085 | -0.010244637 | 0.27429272 | -0.0373493 | 0.97020651 | 0.99175186 |
| RC3H2 | 153.419221 | -0.17469803 | 0.20217067 | -0.8641117 | 0.38752656 | 0.66406384 |
| RABGAP1 | 406.570148 | 0.388158337 | 0.17514161 | 2.21625421 | 0.0266741 | 0.15827108 |
| STRBP | 256.330076 | 0.180268067 | 0.23903866 | 0.7541377 | 0.45076654 | 0.71160181 |
| CRB2 | 66.2989502 | 1.034186797 | 1.27968991 | 0.80815422 | 0.41900181 | 0.68832418 |
| DENND1A | 439.454873 | 0.137015999 | 0.25020086 | 0.54762401 | 0.5839501 | 0.8079818 |
| NEK6 | 122.392417 | -2.357826556 | 0.75240799 | -3.1337075 | 0.00172613 | 0.02921304 |
| PSMB7 | 136.052974 | -0.354490223 | 0.26907964 | -1.3174175 | 0.18769872 | 0.46303529 |
| NR6A1 | 325.692155 | -0.145870631 | 0.27113273 | -0.5380045 | 0.59057394 | 0.81198619 |
| MIR181A2HC | 91.3824731 | 1.07269752 | 0.29780333 | 3.60203334 | 0.00031574 | 0.00998169 |
| RPL35 | 270.506138 | -0.660540309 | 0.42738023 | -1.5455565 | 0.12221171 | 0.36721014 |
| ARPC5L | 44.7812563 | -1.157398455 | 0.40543184 | -2.85473 | 0.00430734 | 0.05249369 |
| GOLGA1 | 59.1150416 | 0.208375498 | 0.27914267 | 0.74648385 | 0.45537519 | 0.71485397 |
| SCAI | 127.901735 | 0.704739156 | 0.18769998 | 3.75460441 | 0.00017362 | 0.00692943 |
| PPP6C | 117.22908 | 0.225977011 | 0.18365879 | 1.23041761 | 0.21854076 | 0.50057436 |
| RABEPK | 33.2792893 | -0.849686472 | 0.45253445 | -1.8776172 | 0.06043355 | 0.24800908 |
| HSPA5 | 225.228651 | -1.551495856 | 0.59730146 | -2.5975089 | 0.00939027 | 0.08626955 |
| GAPVD1 | 200.727251 | 0.054152869 | 0.17084182 | 0.31697665 | 0.75126132 | 0.89599662 |
| MAPKAP1 | 302.147316 | -0.180061825 | 0.20485241 | -0.8789832 | 0.37941039 | 0.65699285 |
| PBX3 | 299.999942 | -0.145678442 | 0.38612274 | -0.3772853 | 0.70596158 | 0.87473681 |
| MVB12B | 58.9745681 | 0.194563052 | 0.29785325 | 0.65321782 | 0.51361586 | 0.76083448 |
| ZBTB43 | 68.4663097 | -0.384850667 | 0.23841621 | -1.6141967 | 0.10648477 | 0.34105698 |
| RALGPS1 | 190.459238 | 0.556926156 | 0.44800599 | 1.24312211 | 0.21382284 | 0.49581433 |
| GARNL3 | 77.442377 | -0.018800107 | 0.26206975 | -0.071737 | 0.94281118 | 0.98112027 |
| RPL12 | 117.420855 | -0.817666438 | 0.41372055 | -1.9763737 | 0.04811246 | 0.21931444 |
| LRSAM1 | 105.618712 | -0.151415857 | 0.30617932 | -0.4945333 | 0.62092963 | 0.82839623 |
| FAM129B | 235.012415 | -0.998907627 | 0.35477691 | -2.8155937 | 0.00486872 | 0.0570838 |
| STXBP1 | 81.4789038 | -0.458649955 | 0.47314655 | -0.9693613 | 0.33236495 | 0.61719981 |
| CDK9 | 43.277212 | 0.249777879 | 0.36619096 | 0.68209734 | 0.4951774 | 0.74709472 |
| FPGS | 54.6557361 | -0.543221002 | 0.32952258 | -1.6485092 | 0.09924823 | 0.32930335 |
| DPM2 | 28.2564008 | -0.998155606 | 0.44559894 | -2.2400314 | 0.02508888 | 0.1536614 |
| FAM102A | 35.3084414 | -0.921992005 | 0.39654196 | -2.3250806 | 0.02006765 | 0.13480227 |
| SLC25A25 | 30.1002429 | -0.315747406 | 0.39401849 | -0.8013518 | 0.42292804 | 0.69043214 |
| PTGES2 | 53.388387 | -1.4798664 | 0.42632912 | -3.471183 | 0.00051817 | 0.01373303 |
| LCN2 | 461.424183 | -7.869826019 | 1.38921109 | -5.6649605 | 1.47E-08 | 6.12E-06 |
| C9orf16 | 94.1288751 | -1.199888059 | 0.54015448 | -2.2213794 | 0.02632527 | 0.1573452 |
| CIZ1 | 59.002358 | -0.226027906 | 0.24252877 | -0.9319633 | 0.35135551 | 0.63335467 |
| DNM1 | 256.021035 | -0.034503163 | 0.44491169 | -0.0775506 | 0.93818555 | 0.97988379 |
| GOLGA2 | 109.816969 | -0.227612533 | 0.24455422 | -0.9307242 | 0.35199624 | 0.63379173 |
| SWI5 | 31.456303 | -0.092249378 | 0.33507663 | -0.2753083 | 0.78307939 | 0.91165745 |
| TRUB2 | 31.4906289 | -0.932522261 | 0.28756464 | -3.2428266 | 0.0011835 | 0.02342308 |
| COQ4 | 112.604903 | -0.192981507 | 0.18649961 | -1.0347556 | 0.30078308 | 0.58788025 |
| SLC27A4 | 38.1496448 | -0.744402173 | 0.34744372 | -2.1425115 | 0.03215234 | 0.17569156 |
| URM1 | 38.2647877 | -0.732494479 | 0.32976469 | -2.2212642 | 0.02633307 | 0.1573452 |
| CERCAM | 51.3825965 | 0.985273681 | 0.66428124 | 1.48321767 | 0.13801659 | 0.39068755 |
| ODF2 | 50.9076602 | 0.207603416 | 0.22881093 | 0.90731425 | 0.36424064 | 0.644313 |
| GLE1 | 56.0480643 | -0.195838037 | 0.21775131 | -0.8993656 | 0.36845797 | 0.64843999 |
| SPTAN1 | 242.562287 | 0.591901062 | 0.28211262 | 2.09810204 | 0.03589613 | 0.18762053 |
| WDR34 | 59.5596336 | 0.442476944 | 0.39835123 | 1.11077087 | 0.26666699 | 0.55453954 |
| SET | 232.091254 | -0.57566566 | 0.377468 | -1.5250714 | 0.12724128 | 0.37456799 |
| ZDHHC12 | 29.6991877 | -0.614178456 | 0.51471671 | -1.1932359 | 0.232777 | 0.51579248 |
| ZER1 | 150.809426 | -0.051471925 | 0.27700397 | -0.1858166 | 0.85258862 | 0.94470172 |
| TBC1D13 | 37.7405809 | 0.51811139 | 0.32719381 | 1.58349999 | 0.11330755 | 0.35267249 |
| KYAT1 | 46.2484975 | 0.048779599 | 0.44983184 | 0.10843963 | 0.91364696 | 0.96844354 |
| LRRC8A | 31.4193806 | -0.374946803 | 0.3328586 | -1.1264447 | 0.25997734 | 0.54648299 |
| NUP188 | 38.2909164 | -0.315451454 | 0.28718705 | -1.0984181 | 0.27202197 | 0.55916275 |
| SH3GLB2 | 57.0486122 | 0.080858625 | 0.32593963 | 0.24807853 | 0.80407364 | 0.92142857 |
| MIGA2 | 29.0592309 | 0.719159272 | 0.28695344 | 2.50618799 | 0.01220407 | 0.10091982 |
| PTPA | 66.6640955 | -0.636684902 | 0.26486798 | -2.403782 | 0.01622644 | 0.11901464 |

| | | | | | | |
|---|---|---|---|---|---|---|
| IER5L | 97.6758146 | -2.575565759 | 0.37903925 | -6.7949843 | 1.08E-11 | 1.30E-08 |
| LINC01503 | 47.2893635 | -1.691859609 | 0.44666262 | -3.7877797 | 0.000152 | 0.00630305 |
| NTMT1 | 30.9128514 | -1.019634434 | 0.305723 | -3.3351577 | 0.00085251 | 0.01859731 |
| C9orf78 | 52.7387451 | -0.439036687 | 0.36358717 | -1.2075142 | 0.22723417 | 0.50999908 |
| USP20 | 47.1146741 | 0.091819736 | 0.30557155 | 0.30048523 | 0.76380706 | 0.90255018 |
| FNBP1 | 481.80163 | 0.958266218 | 0.27368691 | 3.50132285 | 0.00046295 | 0.01267041 |
| GPR107 | 168.678735 | 0.135436755 | 0.16671348 | 0.81239236 | 0.41656653 | 0.68597057 |
| NCS1 | 74.407543 | -0.198182489 | 0.39617267 | -0.5002427 | 0.61690419 | 0.82592881 |
| ASS1 | 80.1605723 | -0.348232101 | 0.73289471 | -0.4751462 | 0.6346828 | 0.83703065 |
| FUBP3 | 144.109571 | -0.000714757 | 0.1948143 | -0.0036689 | 0.99707263 | 0.99923686 |
| ABL1 | 353.513532 | 0.063528262 | 0.21511708 | 0.29531947 | 0.76774985 | 0.90425129 |
| AIF1L | 545.412759 | 1.320231339 | 0.5004632 | 2.63801883 | 0.0083392 | 0.08052535 |
| NUP214 | 352.35615 | 1.162387564 | 0.29337105 | 3.96217538 | 7.43E-05 | 0.00393056 |
| AL157938.2 | 89.6401825 | 1.326410845 | 0.59821105 | 2.21729579 | 0.02660289 | 0.15808253 |
| AL157938.3 | 27.4367276 | 1.543615637 | 0.48368784 | 3.1913468 | 0.00141611 | 0.02618713 |
| PRRC2B | 420.643102 | -0.035965502 | 0.16776662 | -0.2143782 | 0.83025216 | 0.93443608 |
| POMT1 | 36.1269616 | 0.585766579 | 0.23960151 | 2.44475328 | 0.01449513 | 0.11197822 |
| RAPGEF1 | 233.062547 | -0.027762706 | 0.19069372 | -0.1455879 | 0.88424669 | 0.95761929 |
| MED27 | 190.847773 | -0.246867652 | 0.20956994 | -1.1779726 | 0.23880752 | 0.52214956 |
| SETX | 351.692226 | -0.015049253 | 0.21962067 | -0.0685238 | 0.94536864 | 0.98119662 |
| TTF1 | 66.9664693 | 0.075216394 | 0.23141099 | 0.3250338 | 0.74515549 | 0.89323545 |
| DDX31 | 169.769812 | 0.033251635 | 0.2314568 | 0.14366238 | 0.88576708 | 0.95818892 |
| GTF3C4 | 50.6449645 | -0.183091208 | 0.32744047 | -0.5591588 | 0.57605337 | 0.80227031 |
| AK8 | 46.5567767 | 0.278048107 | 0.42476519 | 0.6545925 | 0.51273015 | 0.7606384 |
| TSC1 | 113.959758 | 0.446707001 | 0.19002043 | 2.35083668 | 0.01873125 | 0.12954995 |
| GTF3C5 | 33.419854 | -0.207864169 | 0.3151448 | -0.6595831 | 0.50952143 | 0.75834459 |
| ABO | 178.911358 | 0.52885262 | 0.3770217 | 1.40271137 | 0.16070292 | 0.42659999 |
| SURF6 | 33.6980257 | -1.060515535 | 0.30672257 | -3.4575726 | 0.00054507 | 0.01396038 |
| RPL7A | 362.809561 | -0.459322879 | 0.47308324 | -0.9709134 | 0.33159138 | 0.61675997 |
| SURF1 | 28.8068535 | -0.416329466 | 0.29058972 | -1.4327054 | 0.15194204 | 0.41406907 |
| SURF4 | 103.300084 | -0.765011907 | 0.39661182 | -1.9288681 | 0.05374723 | 0.23285607 |
| REXO4 | 28.0826785 | -0.508359999 | 0.32787 | -1.5504926 | 0.12102334 | 0.36534213 |
| VAV2 | 234.485092 | 0.268833129 | 0.33841711 | 0.79438398 | 0.42697193 | 0.69309678 |
| BRD3OS | 25.0063528 | -0.290822193 | 0.42130363 | -0.6902912 | 0.49001108 | 0.74227587 |
| BRD3 | 68.3635376 | 0.347601698 | 0.24582021 | 1.41404849 | 0.15734767 | 0.42243085 |
| WDR5 | 42.1836854 | -0.394803363 | 0.32017152 | -1.2330996 | 0.21753862 | 0.49951605 |
| RXRA | 52.2102543 | -0.230574759 | 0.47888714 | -0.4814804 | 0.63017512 | 0.83433967 |
| COL5A1 | 94.1771733 | 0.352225274 | 1.05669403 | 0.33332759 | 0.73888702 | 0.89234856 |
| PPP1R26-AS1 | 32.7339217 | -0.629192852 | 0.36985596 | -1.7011835 | 0.08890854 | 0.31048301 |
| PPP1R26 | 28.500396 | -0.038963708 | 0.29684696 | -0.1312586 | 0.89557076 | 0.96142295 |
| MRPS2 | 29.7686523 | -0.6311019 | 0.32600843 | -1.9358453 | 0.05288665 | 0.2306927 |
| CAMSAP1 | 89.9476918 | -0.127430238 | 0.26383423 | -0.4829936 | 0.6291003 | 0.83372327 |
| UBAC1 | 44.0673667 | -0.211591077 | 0.21529824 | -0.9827812 | 0.32571511 | 0.61243485 |
| NACC2 | 71.0391778 | -0.052361255 | 0.34823468 | -0.150362 | 0.88047904 | 0.95614192 |
| QSOX2 | 26.1348403 | -0.404866685 | 0.28253823 | -1.4329625 | 0.15186856 | 0.41406907 |
| SNAPC4 | 37.3605696 | -0.19011127 | 0.22675416 | -0.8384026 | 0.40180465 | 0.67562929 |
| ENTR1 | 38.0308307 | -0.719315929 | 0.41805081 | -1.7206424 | 0.08531574 | 0.3038065 |
| PMPCA | 36.0162353 | -0.572589438 | 0.25801042 | -2.2192493 | 0.02646977 | 0.15778091 |
| SEC16A | 81.6635972 | -0.19262483 | 0.21890871 | -0.8799322 | 0.37889603 | 0.65677822 |
| NOTCH1 | 67.6961122 | 0.183106153 | 0.35946439 | 0.50938607 | 0.61048164 | 0.82270732 |
| AGPAT2 | 47.4768156 | -1.00305254 | 0.37149671 | -2.7000308 | 0.00693331 | 0.07087956 |
| SNHG7 | 67.9492888 | 0.034859135 | 0.4669047 | 0.07466006 | 0.94048518 | 0.98016757 |
| RABL6 | 121.187566 | -0.041903564 | 0.24550677 | -0.1706819 | 0.86447389 | 0.94985035 |
| PHPT1 | 123.503182 | -0.791112352 | 0.4344191 | -1.8210809 | 0.06859455 | 0.26781626 |
| EDF1 | 174.417008 | -0.653882041 | 0.40506043 | -1.6142827 | 0.10646612 | 0.34105698 |
| TRAF2 | 32.6152045 | 0.122275009 | 0.37080478 | 0.32975575 | 0.74158453 | 0.89253455 |
| FBXW5 | 67.9038146 | -0.729821367 | 0.41101609 | -1.7756516 | 0.07579036 | 0.28448114 |
| PAXX | 37.9098078 | -0.506370134 | 0.50917052 | -0.9945001 | 0.31997946 | 0.60682701 |
| MAN1B1 | 56.9354011 | -0.18358092 | 0.19391057 | -0.9467298 | 0.34377646 | 0.6285617 |
| DPP7 | 49.3162723 | -0.611244527 | 0.32127961 | -1.9025313 | 0.05710173 | 0.24117172 |
| SSNA1 | 53.5956423 | -0.633693274 | 0.38725427 | -1.6363752 | 0.10176109 | 0.33325452 |
| TPRN | 42.5064294 | -0.309074826 | 0.28824702 | -1.0722568 | 0.28360471 | 0.57084461 |
| TMEM203 | 28.2986399 | -0.603907655 | 0.36578594 | -1.6509865 | 0.09874133 | 0.32848864 |
| TUBB4B | 193.828728 | -1.268219393 | 0.52377934 | -2.4212856 | 0.01546572 | 0.11564881 |
| NELFB | 74.6869901 | -0.18982876 | 0.26372509 | -0.7197979 | 0.47164946 | 0.72945405 |
| EXD3 | 102.840975 | 0.029943919 | 0.28247861 | 0.1060042 | 0.91557902 | 0.9691911 |
| NOXA1 | 37.1649993 | 0.119573344 | 0.36383776 | 0.32864468 | 0.74242428 | 0.89253455 |
| NSMF | 69.981553 | 0.522491636 | 0.55633767 | 0.93916278 | 0.34764717 | 0.63118125 |
| PNPLA7 | 69.7947076 | 0.528535639 | 0.69049499 | 0.76544456 | 0.44400686 | 0.70661933 |
| MRPL41 | 100.378889 | -0.893894026 | 0.31472885 | -2.8402036 | 0.00450847 | 0.05405091 |
| DPH7 | 42.9455158 | 0.121161408 | 0.2491939 | 0.48621337 | 0.62681589 | 0.83221501 |
| ARRDC1 | 54.8176845 | -0.051598383 | 0.247618 | -0.208379 | 0.83493307 | 0.93707149 |
| EHMT1 | 400.523319 | 0.113080034 | 0.19678724 | 0.57463092 | 0.56554093 | 0.79444443 |
| BET1L | 31.8601279 | -0.579370391 | 0.30534951 | -1.8974007 | 0.05777507 | 0.24283781 |
| RIC8A | 37.3477943 | -0.978464368 | 0.29925214 | -3.2696988 | 0.00107662 | 0.02175727 |
| SIRT3 | 37.9197546 | -0.431774145 | 0.34221311 | -1.2617113 | 0.20705269 | 0.48648432 |
| PSMD13 | 79.4582963 | -0.400564749 | 0.28847842 | -1.3885432 | 0.1649717 | 0.43324238 |
| AC136475.3 | 27.8259868 | 1.157598928 | 1.29946615 | 0.89082653 | 0.37302224 | 0.65217873 |
| PGGHG | 170.293851 | 0.578870439 | 0.74824288 | 0.77363976 | 0.43914385 | 0.70285029 |
| IFITM2 | 462.453098 | -2.021008493 | 0.62876373 | -3.2142575 | 0.00130782 | 0.02485643 |
| IFITM3 | 851.231518 | -1.626841152 | 0.64919705 | -2.5059281 | 0.01221304 | 0.10091982 |
| SIGIRR | 46.326669 | -0.650540372 | 0.29676078 | -2.1921373 | 0.02836959 | 0.16346515 |
| PTDSS2 | 48.7640499 | -0.571189656 | 0.2251973 | -2.5363966 | 0.01119998 | 0.09563107 |
| RNH1 | 97.8913923 | -0.567366013 | 0.24919446 | -2.2768003 | 0.02279816 | 0.14529842 |
| HRAS | 41.7479284 | -0.961195576 | 0.536544 | -1.7914571 | 0.07321997 | 0.27896901 |
| RASSF7 | 128.059866 | 0.244784043 | 0.28770344 | 0.8508207 | 0.39486896 | 0.6696972 |
| MIR210HG | 29.9300123 | -1.081207443 | 0.65718637 | -1.6452067 | 0.09992718 | 0.33075414 |
| PHRF1 | 62.9654871 | -0.214674089 | 0.23440807 | -0.9158135 | 0.3597647 | 0.64001011 |
| DEAF1 | 144.259592 | 0.442382806 | 0.16628702 | 2.66035677 | 0.00780579 | 0.07637557 |
| EPS8L2 | 287.59054 | 1.013452358 | 0.3242386 | 3.12563762 | 0.0017742 | 0.02957033 |
| AP006621.4 | 52.9473095 | 1.175479968 | 0.65604157 | 1.79177667 | 0.07316875 | 0.27896901 |
| TALDO1 | 151.304538 | -1.075788315 | 0.36194813 | -2.9722168 | 0.00295658 | 0.04028601 |
| GATD1 | 35.9184018 | -0.262501581 | 0.28913582 | -0.9078833 | 0.36393988 | 0.644123 |
| AP006621.3 | 45.3449666 | 0.208773313 | 0.40116568 | 0.52041668 | 0.60277319 | 0.81954732 |
| RPLP2 | 302.735262 | -0.404845736 | 0.44459378 | -0.9105969 | 0.36250778 | 0.64292947 |
| PNPLA2 | 82.2668459 | 0.540456207 | 0.23421958 | 2.30747667 | 0.02102826 | 0.13920506 |
| CD151 | 231.732849 | -0.583264169 | 0.49126036 | -1.1872812 | 0.23511673 | 0.51810057 |
| POLR2L | 154.960036 | -0.8367643 | 0.46745727 | -1.7900338 | 0.07344848 | 0.27941344 |
| TSPAN4 | 89.8433285 | -0.062768782 | 0.32050256 | -0.1958449 | 0.84473159 | 0.94193856 |
| CHID1 | 45.5507092 | 0.147785972 | 0.24046002 | 0.61459686 | 0.53882099 | 0.77708778 |
| AP2A2 | 220.337483 | 0.150275686 | 0.26409461 | 0.56902217 | 0.56934109 | 0.79749848 |
| TOLLIP | 68.0126306 | 0.085100134 | 0.26241736 | 0.32429308 | 0.74571616 | 0.89352127 |
| MOB2 | 40.7640861 | -0.03667121 | 0.22117437 | -0.1658023 | 0.86831257 | 0.95090501 |
| CD81-AS1 | 34.5980759 | 0.254144436 | 0.28929635 | 0.87849169 | 0.37967695 | 0.65703312 |
| CD81 | 434.097178 | -0.755852258 | 0.33078719 | -2.2850107 | 0.0223122 | 0.143442 |
| TSSC4 | 30.3279933 | -0.369157 | 0.34453121 | -1.0714762 | 0.28395535 | 0.57143032 |
| KCNQ1 | 139.253024 | -0.479265096 | 0.46484739 | -1.031016 | 0.30253331 | 0.59009705 |
| KCNQ1OT1 | 541.096251 | -0.748674146 | 0.563951 | -1.3275518 | 0.18432623 | 0.45778091 |
| CDKN1C | 118.903819 | -1.476549367 | 0.81349236 | -1.8150747 | 0.06951244 | 0.27023524 |
| SLC22A18 | 35.4290946 | -0.757106297 | 0.49128794 | -1.5410643 | 0.12330114 | 0.36897896 |
| NAP1L4 | 257.743757 | -0.052494271 | 0.20511126 | -0.2559307 | 0.79800434 | 0.91942309 |
| CARS | 68.7936919 | -0.828365534 | 0.26631181 | -3.11051 | 0.00186765 | 0.03068642 |
| OSBPL5 | 47.7589304 | -0.057570079 | 0.25752634 | -0.2235503 | 0.82310728 | 0.93217972 |

| | | | | | | |
|---|---|---|---|---|---|---|
| ZNF195 | 66.2744073 | -0.089398386 | 0.33427943 | -0.2674361 | 0.78913341 | 0.91492483 |
| AC127526.1 | 46.6906146 | 0.642023177 | 0.42525155 | 1.50974917 | 0.13110744 | 0.38163088 |
| NUP98 | 214.924435 | 0.171151234 | 0.20502689 | 0.83477456 | 0.40384465 | 0.67765972 |
| PGAP2 | 43.9283331 | -0.192095171 | 0.31189097 | -0.6159049 | 0.5379573 | 0.77665304 |
| STIM1 | 222.331896 | -0.757082423 | 0.32999622 | -2.2942155 | 0.02177813 | 0.14172056 |
| RRM1 | 36.996205 | 0.189279085 | 0.33044971 | 0.57279241 | 0.56678526 | 0.79572563 |
| AC104389.4 | 128.639466 | 0.821114635 | 0.60155913 | 1.36497742 | 0.17226015 | 0.44309343 |
| TRIM5 | 39.1357347 | -1.187601303 | 0.73425054 | -1.6174333 | 0.10578477 | 0.34033344 |
| TRIM22 | 133.682246 | -1.346744096 | 0.78019884 | -1.7261549 | 0.08431954 | 0.3023414 |
| FAM160A2 | 52.3147834 | 0.358691019 | 0.2230782 | 1.60791607 | 0.10785356 | 0.34350867 |
| SMPD1 | 43.705857 | -0.581865461 | 0.32757905 | -1.7762597 | 0.07569013 | 0.28432774 |
| APBB1 | 45.8067358 | -0.561783805 | 0.28066825 | -2.0015937 | 0.04532845 | 0.21178118 |
| ARFIP2 | 28.2962663 | -0.753087322 | 0.33873834 | -2.2232125 | 0.02620147 | 0.15715963 |
| AC084337.2 | 45.8858953 | 0.012503413 | 0.31142037 | 0.04014963 | 0.96797383 | 0.99123492 |
| DNHD1 | 65.830875 | 0.115259661 | 0.33552418 | 0.34352118 | 0.73120641 | 0.88791456 |
| ILK | 93.5835784 | -0.022166098 | 0.30177993 | -0.0734512 | 0.94144708 | 0.98065698 |
| TAF10 | 39.2283781 | -0.154991129 | 0.39166745 | -0.3957212 | 0.69231068 | 0.86892611 |
| TPP1 | 46.0214573 | -0.488003531 | 0.28780183 | -1.6956234 | 0.08995721 | 0.31254993 |
| MRPL17 | 46.7418417 | -0.959952569 | 0.41530154 | -2.3114592 | 0.0208075 | 0.13822127 |
| ZNF214 | 26.9460888 | -0.00702202 | 0.39439185 | -0.0178047 | 0.98579467 | 0.99668665 |
| AC107884.1 | 30.178796 | 0.245028066 | 0.40955418 | 0.59827997 | 0.54965314 | 0.78490271 |
| PPFIBP2 | 85.6855236 | -0.027296497 | 0.45913619 | -0.0594519 | 0.95259221 | 0.98349653 |
| EIF3F | 92.6001294 | 0.094449354 | 0.28158014 | 0.3354262 | 0.73730361 | 0.89095891 |
| TUB | 124.333821 | 2.287594114 | 0.7475811 | 3.05999457 | 0.00221341 | 0.0339653 |
| RIC3 | 101.295087 | -0.353694598 | 0.48547692 | -0.7285508 | 0.46627648 | 0.72401725 |
| STK33 | 483.165211 | 0.009193953 | 0.31384047 | 0.02929499 | 0.97662932 | 0.99438056 |
| TRIM66 | 103.891269 | -0.021870467 | 0.36042636 | -0.0606794 | 0.95161451 | 0.98322893 |
| RPL27A | 337.887991 | -0.456141778 | 0.43282412 | -1.0538733 | 0.29194094 | 0.57922582 |
| ST5 | 204.096187 | -1.374539444 | 0.40409898 | -3.401492 | 0.00067019 | 0.01577201 |
| TMEM9B | 51.2011957 | 0.11793389 | 0.3131168 | 0.37664504 | 0.70643741 | 0.87473681 |
| TMEM9B-AS | 48.1210063 | -0.239507767 | 0.45553097 | -0.5257771 | 0.59904307 | 0.81717937 |
| DENND5A | 250.356907 | -0.446925231 | 0.37197675 | -1.201487 | 0.22956234 | 0.51222401 |
| TMEM41B | 148.688266 | 0.359481759 | 0.21366972 | 1.68241789 | 0.09248783 | 0.31666153 |
| IPO7 | 237.124921 | -0.472615198 | 0.29744881 | -1.5888959 | 0.1120839 | 0.35029418 |
| ZNF143 | 63.5214915 | 0.057342755 | 0.2634719 | 0.21764277 | 0.82770746 | 0.93346403 |
| WEE1 | 66.7987832 | 1.389085105 | 0.51996275 | 2.67150891 | 0.00755111 | 0.07480047 |
| SWAP70 | 204.769399 | 0.348825575 | 0.26390025 | 1.32180841 | 0.18623196 | 0.46107933 |
| SBF2-AS1 | 55.1706049 | -0.354150116 | 0.31962426 | -1.1080202 | 0.26785311 | 0.55494823 |
| SBF2 | 1120.02939 | -0.212789171 | 0.20873998 | -1.0193983 | 0.30801393 | 0.59456464 |
| AC011092.2 | 45.358629 | -0.274732283 | 0.33117894 | -0.8295584 | 0.40678849 | 0.68040675 |
| ADM | 96.9284066 | -3.201316256 | 1.10907795 | -2.8864664 | 0.00389594 | 0.04847509 |
| MTRNR2L8 | 101.336151 | -1.590578538 | 0.60726182 | -2.6192632 | 0.00881199 | 0.08357433 |
| RNF141 | 42.3633644 | 0.105066715 | 0.32348169 | 0.32479958 | 0.74533276 | 0.89333615 |
| CTR9 | 70.3654303 | -0.008025604 | 0.23917319 | -0.0335556 | 0.97323151 | 0.99282052 |
| EIF4G2 | 249.364989 | -0.279769941 | 0.25517238 | -1.0963959 | 0.27290557 | 0.56047062 |
| ZBED5 | 98.5179163 | 0.229926064 | 0.22286686 | 1.03167453 | 0.30222461 | 0.58973509 |
| ZBED5-AS1 | 36.1971689 | 0.737740977 | 0.52204101 | 1.41318587 | 0.15760108 | 0.42280489 |
| GALNT18 | 631.548133 | 0.719870108 | 0.39061371 | 1.84292073 | 0.06534059 | 0.26050329 |
| USP47 | 405.813243 | 0.142348902 | 0.11823208 | 1.20397862 | 0.22859784 | 0.51112377 |
| MICAL2 | 77.9753753 | -0.792276015 | 0.71360058 | -1.1102514 | 0.26689072 | 0.55453954 |
| PARVA | 216.159371 | -0.00233944 | 0.29848319 | -0.0078378 | 0.99374644 | 0.99884271 |
| TEAD1 | 924.106068 | 0.302639241 | 0.24455854 | 1.23749202 | 0.21590448 | 0.49746509 |
| LINC00958 | 109.047746 | -0.48741084 | 0.38132686 | -1.278197 | 0.20117998 | 0.48009543 |
| ARNTL | 47.326169 | 0.300015519 | 0.41457324 | 0.72367315 | 0.46926642 | 0.72712764 |
| BTBD10 | 108.514186 | -0.045752912 | 0.25346329 | -0.180511 | 0.85675142 | 0.94638973 |
| FAR1 | 218.75342 | 0.334867976 | 0.15636689 | 2.14155298 | 0.03222947 | 0.17584306 |
| SPON1 | 210.049 | -1.140185331 | 0.66613771 | -1.7116361 | 0.08696376 | 0.30691783 |
| RRAS2 | 290.742646 | -0.026296686 | 0.27644093 | -0.0951259 | 0.92421484 | 0.9731617 |
| COPB1 | 141.739256 | 0.030081668 | 0.20844478 | 0.1443148 | 0.88525189 | 0.95795615 |
| SOX6 | 1311.12093 | 0.725437014 | 0.2574776 | 2.81747626 | 0.00484027 | 0.05681978 |
| C11orf58 | 224.02132 | 0.092077328 | 0.18230534 | 0.50507203 | 0.61350825 | 0.82498688 |
| PLEKHA7 | 1401.39949 | 0.076519186 | 0.21864128 | 0.34997593 | 0.72635676 | 0.88522597 |
| RPS13 | 297.624053 | -0.570528026 | 0.55147595 | -1.0345474 | 0.30088031 | 0.58795033 |
| PIK3C2A | 399.931725 | 0.185121562 | 0.14866154 | 1.24525522 | 0.21303795 | 0.49483281 |
| NUCB2 | 89.0754023 | 0.556401704 | 0.32945514 | 1.68885423 | 0.09124737 | 0.31502156 |
| USH1C | 80.2659914 | -0.832974408 | 0.60189276 | -1.3839249 | 0.16638144 | 0.43474299 |
| SERGEF | 155.833035 | -0.013398919 | 0.16238807 | -0.0825117 | 0.9342398 | 0.97768238 |
| SAA1 | 895.760918 | -6.990404518 | 1.6601011 | -4.2108306 | 2.54E-05 | 0.00181882 |
| HPS5 | 57.7112933 | 0.759296213 | 0.34964261 | 2.17163525 | 0.02988319 | 0.16887967 |
| GTF2H1 | 96.6418253 | 0.079848427 | 0.19411777 | 0.41134013 | 0.68082315 | 0.86276028 |
| LDHA | 396.871454 | -1.492033665 | 0.48537489 | -3.073982 | 0.00211222 | 0.03294549 |
| TSG101 | 136.870716 | 0.053238559 | 0.1824686 | 0.29176833 | 0.77046376 | 0.90533273 |
| UEVLD | 93.6834096 | -0.300941712 | 0.16380948 | -1.8371446 | 0.06618855 | 0.26199177 |
| SPTY2D1OS | 31.513 | 0.375116074 | 0.23887911 | 1.57031761 | 0.11634124 | 0.35938557 |
| SPTY2D1 | 35.7796453 | -0.264482246 | 0.35106998 | -0.7533605 | 0.45123332 | 0.71185178 |
| ZDHHC13 | 85.8162679 | 0.127762912 | 0.20018725 | 0.63821704 | 0.52333241 | 0.76534368 |
| NAV2 | 435.845245 | -0.499634688 | 0.60020608 | -0.8324386 | 0.40516144 | 0.67862239 |
| HTATIP2 | 32.1930538 | -0.98348673 | 0.41238902 | -2.3848519 | 0.017086 | 0.12315032 |
| PRMT3 | 130.536296 | -0.393317901 | 0.31757796 | -1.2384924 | 0.21553352 | 0.49731142 |
| ANO5 | 51.0059033 | 0.480450585 | 0.37854783 | 1.26919387 | 0.20437193 | 0.48318561 |
| GAS2 | 48.9598726 | 0.979594348 | 0.45952386 | 2.13175947 | 0.03302663 | 0.17813178 |
| SVIP | 80.1328293 | 0.200339696 | 0.39932111 | 0.50170074 | 0.61587804 | 0.82592881 |
| MUC15 | 24.1398711 | 1.82832208 | 1.44925615 | 1.26155896 | 0.20710753 | 0.48648432 |
| BBOX1 | 51.2630531 | -1.124726096 | 0.71798043 | -1.5665136 | 0.11722843 | 0.36083905 |
| LGR4 | 979.899968 | 0.730592773 | 0.33670706 | 2.16981722 | 0.0300207 | 0.16932448 |
| LIN7C | 52.4581659 | 0.086436601 | 0.34189328 | 0.25281749 | 0.80040925 | 0.92042259 |
| BDNF-AS | 248.558653 | 0.047798038 | 0.24800834 | 0.19272754 | 0.84717235 | 0.94323402 |
| METTL15 | 471.310027 | 0.18331777 | 0.25313366 | 0.72419356 | 0.46894691 | 0.72686771 |
| ARL14EP | 44.2710436 | -0.325263558 | 0.21564657 | -1.5083178 | 0.13147321 | 0.38163088 |
| MPPED2 | 148.322313 | 0.223327839 | 0.49541074 | 0.4507933 | 0.65213853 | 0.84801249 |
| DCDC1 | 296.992016 | 0.186684756 | 0.37211256 | 0.50168894 | 0.61588635 | 0.82592881 |
| DNAJC24 | 67.0179656 | 0.015312549 | 0.27152468 | 0.05639468 | 0.95502739 | 0.98393042 |
| IMMP1L | 171.270805 | 0.147951876 | 0.23004083 | 0.64315486 | 0.52012362 | 0.76400369 |
| ELP4 | 266.710558 | 0.059384507 | 0.19936255 | 0.29787192 | 0.76580092 | 0.90371395 |
| AC131571.1 | 35.333898 | -0.013020971 | 0.59675044 | -0.0218198 | 0.98259171 | 0.99616479 |
| RCN1 | 88.884528 | -0.961226669 | 0.42876497 | -2.2418498 | 0.02497108 | 0.1532338 |
| WT1 | 148.85842 | 1.454564073 | 1.30781048 | 1.11221319 | 0.2660465 | 0.55384374 |
| EIF3M | 121.922905 | -0.230605584 | 0.43581692 | -0.5291341 | 0.59671244 | 0.81597551 |
| PRRG4 | 86.0096102 | 0.502832398 | 0.29636535 | 1.69666391 | 0.08976022 | 0.31211662 |
| QSER1 | 181.579173 | -0.009921938 | 0.14574562 | -0.0680771 | 0.94572426 | 0.98119662 |
| TCP11L1 | 65.6921678 | 0.055671829 | 0.27047476 | 0.20583004 | 0.83692368 | 0.93808705 |
| CSTF3 | 153.122769 | 0.086657641 | 0.20883131 | 0.41496479 | 0.67816768 | 0.86151115 |
| HIPK3 | 483.231274 | -0.57040171 | 0.27925528 | -2.0425817 | 0.04109387 | 0.20176225 |
| FBXO3 | 95.4481537 | 0.463764831 | 0.2109908 | 2.19803342 | 0.02794672 | 0.16175328 |
| CAPRIN1 | 208.676509 | -0.081263539 | 0.14196259 | -0.5724293 | 0.5670312 | 0.79583763 |
| NAT10 | 69.2940466 | -0.589290328 | 0.30456257 | -1.9348744 | 0.05300571 | 0.23089662 |
| ABTB2 | 171.147585 | -1.035934355 | 0.28407108 | -3.6467435 | 0.00026558 | 0.00892645 |
| CAT | 218.601842 | 1.169622779 | 0.5536651 | 2.1125095 | 0.03464277 | 0.18344008 |
| ELF5 | 35.9166064 | 3.402809268 | 2.23730175 | 1.52094337 | 0.12827405 | 0.37610564 |
| EHF | 201.475349 | 0.522784943 | 0.62719227 | 0.83353219 | 0.40454465 | 0.67791921 |
| APIP | 74.1328131 | 0.190984719 | 0.29323108 | 0.65131131 | 0.51484555 | 0.76135326 |
| PDHX | 127.171523 | 0.274249328 | 0.22345395 | 1.2273192 | 0.21970264 | 0.50143713 |
| TRIM44 | 411.894163 | -0.293096127 | 0.16535134 | -1.7725658 | 0.07630069 | 0.28527949 |

| | | | | | | |
|---|---|---|---|---|---|---|
| LDLRAD3 | 97.6207847 | -0.746073738 | 0.46568875 | -1.6020867 | 0.10913645 | 0.34570701 |
| TRAF6 | 40.9486051 | 0.397556766 | 0.28353067 | 1.40216497 | 0.16086599 | 0.42685189 |
| C11orf74 | 42.1912808 | 0.703597611 | 0.35084026 | 2.00546431 | 0.04491343 | 0.21105679 |
| LRRC4C | 472.753719 | 0.445480567 | 0.48586774 | 0.91687619 | 0.35920752 | 0.64001011 |
| API5 | 111.904239 | 0.075542097 | 0.20091275 | 0.37599455 | 0.70692095 | 0.87499622 |
| TTC17 | 635.291191 | 0.107555997 | 0.15501612 | 0.6938375 | 0.48778412 | 0.74125768 |
| HSD17B12 | 318.975511 | -0.173798345 | 0.20954612 | -0.8294038 | 0.40687596 | 0.68040675 |
| ALKBH3 | 40.4251541 | -0.051434828 | 0.27678368 | -0.1858304 | 0.85257775 | 0.94470172 |
| EXT2 | 124.964985 | -0.083326383 | 0.21804614 | -0.3821502 | 0.70234993 | 0.872382 |
| CD82 | 39.056488 | 0.572101606 | 0.54863885 | 1.0427654 | 0.29705696 | 0.58520063 |
| TSPAN18 | 42.8632949 | -0.070634948 | 0.66660776 | -0.1059618 | 0.91561267 | 0.9691911 |
| TP53I11 | 29.0976642 | 0.525210084 | 0.39217143 | 1.33923596 | 0.18049387 | 0.45284201 |
| PRDM11 | 50.4343442 | 0.290965835 | 0.25307277 | 1.14973189 | 0.25025431 | 0.5377268 |
| CRY2 | 79.7117837 | 0.104661805 | 0.2893038 | 0.36177128 | 0.71752296 | 0.88106948 |
| LARGE2 | 26.5816281 | 1.030615314 | 0.44411952 | 2.32058101 | 0.02030947 | 0.1355161 |
| PHF21A | 585.095435 | 0.183572096 | 0.22138794 | 0.82918743 | 0.40699836 | 0.68040675 |
| DGKZ | 27.2219767 | -0.879407267 | 0.35656819 | -2.4663088 | 0.01365136 | 0.10762663 |
| MDK | 50.7728294 | -2.905883062 | 0.95154688 | -3.0538517 | 0.00225924 | 0.03418836 |
| AMBRA1 | 748.804787 | -0.249670139 | 0.27466857 | -0.9089869 | 0.36335703 | 0.64360151 |
| ATG13 | 78.785728 | 0.076840133 | 0.22676895 | 0.33884769 | 0.73472448 | 0.88966284 |
| ARHGAP1 | 59.6988117 | -0.41900176 | 0.31000364 | -1.3516027 | 0.17650244 | 0.44846601 |
| CKAP5 | 109.877538 | 0.224316877 | 0.17508515 | 1.28118736 | 0.20012786 | 0.47889703 |
| LRP4 | 114.812643 | -0.763354541 | 0.41301489 | -1.8482494 | 0.06456627 | 0.25877835 |
| C11orf49 | 437.003894 | 0.056636957 | 0.1728688 | 0.32762972 | 0.74319165 | 0.89269973 |
| ARFGAP2 | 60.3949904 | -0.130375328 | 0.31933913 | -0.4082661 | 0.68307835 | 0.86454781 |
| PACSIN3 | 28.9670597 | 0.363042512 | 0.3715231 | 0.97717345 | 0.32848329 | 0.61394673 |
| DDB2 | 67.9422753 | -0.262044524 | 0.48245067 | -0.543153 | 0.58702447 | 0.81058905 |
| MADD | 46.159749 | 0.233127245 | 0.29516244 | 0.78982694 | 0.42962884 | 0.69540633 |
| SLC39A13 | 40.944151 | -0.162180074 | 0.30900839 | -0.5248404 | 0.59969417 | 0.81763227 |
| PSMC3 | 28.234122 | -0.760174465 | 0.4298894 | -1.7683024 | 0.07701036 | 0.28692424 |
| CELF1 | 498.596017 | 0.059763255 | 0.17498678 | 0.34153012 | 0.73270453 | 0.88869659 |
| NDUFS3 | 47.2654677 | -0.188785839 | 0.27920359 | -0.6761583 | 0.49894012 | 0.75042493 |
| MTCH2 | 106.68361 | -0.148471451 | 0.24950411 | -0.5950661 | 0.55179926 | 0.78577398 |
| FNBP4 | 305.225742 | -0.31927108 | 0.14820209 | -2.1542954 | 0.03121701 | 0.17274857 |
| NUP160 | 200.730343 | 0.20096844 | 0.1483777 | 1.35443831 | 0.17559658 | 0.44761528 |
| PTPRJ | 887.443975 | -0.021187659 | 0.23279852 | -0.0910129 | 0.92748237 | 0.97458903 |
| TNKS1BP1 | 83.4716332 | -1.299071358 | 0.35948262 | -3.6137251 | 0.00030183 | 0.00973475 |
| SSRP1 | 56.8348224 | -0.462287048 | 0.3240096 | -1.4267696 | 0.15364632 | 0.41634457 |
| TIMM10 | 33.7840962 | -1.440866293 | 0.43209669 | -3.3345923 | 0.00085425 | 0.01859731 |
| UBE2L6 | 50.9606477 | -1.628286213 | 0.61275399 | -2.6573245 | 0.00787636 | 0.07683059 |
| SERPING1 | 353.778172 | -1.870744289 | 0.29096652 | -6.4294142 | 1.28E-10 | 1.02E-07 |
| ZDHHC5 | 109.531633 | -0.766008017 | 0.24825306 | -3.0855934 | 0.00203146 | 0.03216428 |
| CTNND1 | 276.071478 | 0.090797524 | 0.1737676 | 0.52252276 | 0.6013064 | 0.81829073 |
| LPXN | 63.3368036 | 0.404931074 | 0.37756385 | 1.07248369 | 0.28350284 | 0.57084461 |
| GLYAT | 97.632588 | 0.052197251 | 0.76067352 | 0.06861978 | 0.94529227 | 0.98119662 |
| FAM111A | 73.4141055 | 0.566451308 | 0.33916338 | 1.6701429 | 0.09489109 | 0.32049248 |
| OSBP | 140.501355 | 0.118123672 | 0.18081353 | 0.65329 | 0.51356934 | 0.76083448 |
| PATL1 | 98.0677376 | -0.324122535 | 0.32986841 | -0.9825813 | 0.32581355 | 0.61243485 |
| STX3 | 120.093276 | -0.019331225 | 0.30993574 | -0.0623717 | 0.95026682 | 0.98270285 |
| MRPL16 | 33.133513 | -0.292244863 | 0.32856148 | -0.8894678 | 0.37375171 | 0.65253555 |
| PRPF19 | 47.3263462 | -0.714040757 | 0.37580668 | -1.9000215 | 0.0574303 | 0.24181314 |
| TMEM109 | 40.4251756 | -0.11589581 | 0.42682087 | -0.2715327 | 0.78598138 | 0.9129278 |
| VPS37C | 47.8812256 | -0.370282022 | 0.29189334 | -1.2685525 | 0.20460071 | 0.48320272 |
| DDB1 | 129.510291 | -0.188824013 | 0.22243294 | -0.8489031 | 0.3959352 | 0.67091161 |
| CPSF7 | 101.621336 | 0.080856736 | 0.15649174 | 0.51668372 | 0.60537696 | 0.82056119 |
| AP003108.2 | 72.5022202 | -0.068478634 | 0.30732764 | -0.2228196 | 0.82367588 | 0.93217972 |
| MYRF | 68.9700627 | -1.048824835 | 0.64393648 | -1.6287706 | 0.10336158 | 0.33551358 |
| TMEM258 | 73.2689839 | -0.649502512 | 0.33037795 | -1.9659378 | 0.04930579 | 0.22330979 |
| FADS1 | 50.8136849 | -0.367566983 | 0.4960569 | -0.7409775 | 0.45870711 | 0.71752865 |
| FADS3 | 48.3156714 | -1.653756408 | 0.45294375 | -3.6511298 | 0.00026109 | 0.0088687 |
| FTH1 | 2132.14107 | -2.070925275 | 0.55463469 | -3.7338546 | 0.00018857 | 0.00723578 |
| ASRGL1 | 126.744554 | 1.114107595 | 0.37089458 | 3.00383898 | 0.00266596 | 0.03788912 |
| AHNAK | 626.93241 | 0.293064875 | 0.30100076 | 0.97363499 | 0.33023781 | 0.61551418 |
| EML3 | 25.9832086 | -0.722861805 | 0.35958091 | -2.0102897 | 0.04440053 | 0.20957343 |
| B3GAT3 | 44.338629 | -0.493066524 | 0.41887557 | -1.1771193 | 0.23914789 | 0.52277444 |
| GANAB | 149.740191 | -0.376299778 | 0.25641959 | -1.4675157 | 0.14223581 | 0.39850156 |
| UQCC3 | 40.8980113 | -0.558787687 | 0.47053092 | -1.1875685 | 0.23500346 | 0.51805188 |
| UBXN1 | 66.2141545 | -0.472997843 | 0.40903092 | -1.1563865 | 0.24752309 | 0.53437541 |
| TTC9C | 52.6714161 | -0.155781318 | 0.21886675 | -0.7117633 | 0.47661138 | 0.73340729 |
| TMEM223 | 34.2759247 | -0.853809472 | 0.29486138 | -2.89563 | 0.00378398 | 0.047769 |
| TMEM179B | 58.1842153 | -0.581172706 | 0.37780957 | -1.5382689 | 0.12398288 | 0.37032493 |
| NXF1 | 33.1641569 | -0.031127782 | 0.25247741 | -0.1232894 | 0.90187795 | 0.96386131 |
| STX5 | 96.667005 | 0.162817659 | 0.22887106 | 0.71139469 | 0.47683969 | 0.73352295 |
| SLC3A2 | 108.489212 | -1.0570541 | 0.29834924 | -3.5430092 | 0.00039559 | 0.01152462 |
| RARRES3 | 128.394518 | -2.22335582 | 0.9575346 | -2.3219587 | 0.02023516 | 0.1355161 |
| PLA2G16 | 201.692336 | -0.565613882 | 0.40729002 | -1.3887251 | 0.16491635 | 0.43324238 |
| ATL3 | 113.866146 | -0.356552976 | 0.22402346 | -1.5915877 | 0.11147738 | 0.34908199 |
| RTN3 | 236.519593 | 0.185643016 | 0.15588658 | 1.19088514 | 0.23369868 | 0.51635994 |
| MARK2 | 172.05171 | 0.040702806 | 0.15903553 | 0.2559353 | 0.79800079 | 0.91942309 |
| NAA40 | 52.4715733 | 0.171238243 | 0.26222053 | 0.65303141 | 0.51373603 | 0.76083448 |
| OTUB1 | 53.7376721 | -0.148570762 | 0.25310213 | -0.5869992 | 0.55720421 | 0.78863167 |
| MACROD1 | 88.6869356 | -0.054313567 | 0.38742341 | -0.1401918 | 0.88850849 | 0.95907089 |
| STIP1 | 62.2094585 | -0.212826434 | 0.25338648 | -0.8399281 | 0.40094869 | 0.67532043 |
| VEGFB | 67.5602236 | -0.469794564 | 0.22480571 | -2.0897803 | 0.03663754 | 0.18960073 |
| FKBP2 | 83.4965445 | -0.616424332 | 0.39259035 | -1.5701464 | 0.11638105 | 0.35938557 |
| PPP1R14B | 55.0099 | -1.560105588 | 0.59321172 | -2.6299305 | 0.00854023 | 0.08169244 |
| BAD | 46.1665416 | 0.153036763 | 0.29499647 | 0.5187749 | 0.60391772 | 0.82004322 |
| GPR137 | 29.7594339 | 0.07543809 | 0.28859671 | 0.26139622 | 0.79378697 | 0.91723627 |
| ESRRA | 79.1804956 | -1.047424662 | 0.40169594 | -2.6075062 | 0.00912044 | 0.08519229 |
| TRMT112 | 94.743414 | -0.350593089 | 0.4972878 | -0.7050104 | 0.48080374 | 0.73578929 |
| PRDX5 | 95.2308755 | -0.241693291 | 0.47286126 | -0.5111294 | 0.60926045 | 0.82198674 |
| SF1 | 269.553348 | 0.189556213 | 0.18063623 | 1.04938092 | 0.29400284 | 0.58218755 |
| EHD1 | 53.1102879 | -0.748040775 | 0.48245877 | -1.5504761 | 0.12102729 | 0.36534213 |
| ATG2A | 31.0784801 | -0.786131986 | 0.26722776 | -2.9418051 | 0.00326305 | 0.04305342 |
| VPS51 | 93.8580634 | -0.183650454 | 0.22113854 | -0.8304769 | 0.40626919 | 0.67976464 |
| TM7SF2 | 23.4863004 | 0.026104789 | 0.42721482 | 0.0611046 | 0.95127591 | 0.98319213 |
| FAU | 320.487861 | -0.529536726 | 0.47242935 | -1.1208802 | 0.26233884 | 0.54999863 |
| SYVN1 | 31.8631501 | -0.108601462 | 0.32298386 | -0.3362442 | 0.73668669 | 0.89057637 |
| MRPL49 | 29.312858 | -0.526887009 | 0.35146852 | -1.4991016 | 0.13384729 | 0.38527026 |
| CAPN1 | 116.626291 | -0.561524607 | 0.27834363 | -2.0173791 | 0.04365597 | 0.2071226 |
| DPF2 | 60.5853829 | 0.13458223 | 0.21836963 | 0.61630469 | 0.53769344 | 0.77644087 |
| FRMD8 | 38.3388325 | -0.534920889 | 0.25676581 | -2.0833027 | 0.03722365 | 0.19120638 |
| NEAT1 | 7503.38749 | -0.829127661 | 0.34146732 | -2.4281318 | 0.01517683 | 0.11447152 |
| MALAT1 | 87344.7714 | -0.170832818 | 0.3454322 | -0.494548 | 0.62091918 | 0.82839623 |
| SCYL1 | 56.4183066 | -0.378072785 | 0.21388617 | -1.7676355 | 0.07712186 | 0.28711631 |
| LTBP3 | 147.007006 | 0.28051413 | 0.31455967 | 0.89176761 | 0.3725175 | 0.65199071 |
| SSSCA1 | 31.5996363 | -1.375408191 | 0.66721297 | -2.061423 | 0.0392627 | 0.19660083 |
| EHBP1L1 | 43.9726737 | -0.671620697 | 0.5290021 | -1.2695993 | 0.2042274 | 0.48315491 |
| MAP3K11 | 43.3239151 | -0.292637052 | 0.30744344 | -0.9518403 | 0.34117799 | 0.62632205 |
| PCNX3 | 30.7125437 | -0.043683248 | 0.34842112 | -0.1253749 | 0.90022679 | 0.96303857 |
| SIPA1 | 28.8102373 | -0.627608431 | 0.45909363 | -1.3670598 | 0.17160657 | 0.44224358 |
| RELA | 46.7259393 | -0.549159754 | 0.28233971 | -1.9450319 | 0.05177114 | 0.22800587 |
| KAT5 | 45.0906449 | -0.213032053 | 0.2497752 | -0.8528952 | 0.39371745 | 0.6686322 |

| | | | | | | |
|---|---|---|---|---|---|---|
| RNASEH2C | 39.9157116 | 0.13276434 | 0.43098792 | 0.30804654 | 0.75804692 | 0.89970284 |
| CFL1 | 228.339753 | -1.225434738 | 0.50143743 | -2.4438437 | 0.01453172 | 0.11197822 |
| MUS81 | 30.4961348 | -0.216728489 | 0.35582166 | -0.609093 | 0.54246278 | 0.77916494 |
| EFEMP2 | 51.6093308 | -0.291515144 | 0.34710183 | -0.8398548 | 0.40098979 | 0.67532043 |
| DRAP1 | 47.6202462 | -0.179129763 | 0.53109346 | -0.3372848 | 0.7359022 | 0.89027623 |
| SART1 | 31.6644619 | 0.205547101 | 0.30376967 | 0.67665445 | 0.49862522 | 0.75040313 |
| BANF1 | 72.3228159 | -0.52408898 | 0.4601387 | -1.1389804 | 0.25471131 | 0.54159654 |
| SF3B2 | 107.343213 | -0.421244596 | 0.19283285 | -2.1845064 | 0.02892506 | 0.1655156 |
| PACS1 | 493.493063 | -0.009996985 | 0.24126652 | -0.0414354 | 0.96694876 | 0.99123492 |
| KLC2 | 26.7448867 | -0.167079363 | 0.36157087 | -0.462093 | 0.64401464 | 0.84344334 |
| RAB1B | 56.9919613 | -0.735111402 | 0.29810015 | -2.465988 | 0.01366359 | 0.10763448 |
| YIF1A | 25.1973449 | -1.101288259 | 0.33124889 | -3.3246549 | 0.00088528 | 0.01905641 |
| BRMS1 | 27.5196378 | -0.603855789 | 0.34160548 | -1.7676993 | 0.07711118 | 0.28711631 |
| SLC29A2 | 24.2287447 | 0.681083715 | 0.47999531 | 1.41893827 | 0.15591701 | 0.4202389 |
| MRPL11 | 26.9171497 | -0.698635275 | 0.41417805 | -1.6867994 | 0.09164193 | 0.31554208 |
| ZDHHC24 | 30.152718 | -0.312188808 | 0.34577812 | -0.9028588 | 0.36660086 | 0.64650131 |
| CCS | 71.5002844 | 0.199980318 | 0.29846407 | 0.67003147 | 0.50283773 | 0.75403031 |
| RBM14-RBM | 55.1340688 | 0.602123209 | 0.22170133 | 2.71592057 | 0.00660918 | 0.06896439 |
| RBM4 | 48.5169167 | 0.222652808 | 0.27785157 | 0.80133722 | 0.42293645 | 0.69043214 |
| RBM4B | 47.0073585 | 0.2963203 | 0.34453486 | 0.86005898 | 0.38975653 | 0.66529908 |
| C11orf80 | 195.331113 | 0.514709462 | 0.27080077 | 1.90069425 | 0.05734207 | 0.24173433 |
| PC | 38.636502 | 1.151171847 | 0.34407113 | 3.34573799 | 0.00082064 | 0.01811268 |
| RHOD | 37.5096497 | -0.51854308 | 0.43452405 | -1.1933588 | 0.2327289 | 0.51579248 |
| KDM2A | 592.468362 | 0.21301095 | 0.17658341 | 1.20629082 | 0.22770538 | 0.51041354 |
| GRK2 | 57.8111179 | -0.523246873 | 0.360825 | -1.4501403 | 0.1470194 | 0.4061769 |
| ANKRD13D | 63.5380483 | -0.372100224 | 0.24155669 | -1.5404261 | 0.12345653 | 0.36921404 |
| SSH3 | 38.7213038 | -0.435397926 | 0.25727962 | -1.6923141 | 0.0905861 | 0.3141652 |
| RAD9A | 259.348641 | 0.398964449 | 0.25857791 | 1.54291773 | 0.12285074 | 0.36812373 |
| CLCF1 | 23.6813881 | -0.422719062 | 0.63655801 | -0.66407 | 0.50664552 | 0.75584222 |
| PPP1CA | 82.9454779 | -1.020404432 | 0.42164253 | -2.42007 | 0.01551752 | 0.11578325 |
| RPS6KB2 | 29.7515626 | 0.250441593 | 0.34851126 | 0.71860402 | 0.47238493 | 0.72983472 |
| CORO1B | 41.2206647 | -0.74891351 | 0.3157832 | -2.3716066 | 0.01771094 | 0.12575079 |
| TMEM134 | 55.9437865 | -0.230587566 | 0.36182561 | -0.6372892 | 0.52393647 | 0.7659932 |
| AIP | 29.557982 | -0.164500018 | 0.43586425 | -0.3774111 | 0.7058681 | 0.87473681 |
| PITPNM1 | 40.1640904 | -0.39400035 | 0.37678718 | -1.0456841 | 0.29570693 | 0.5839143 |
| CDK2AP2 | 62.7468431 | -0.595372017 | 0.49932676 | -1.1923495 | 0.23312423 | 0.51603997 |
| GSTP1 | 806.958886 | -0.984442165 | 0.59431384 | -1.6564349 | 0.09763379 | 0.3259791 |
| NDUFV1 | 95.9928445 | 0.076725222 | 0.23486329 | 0.32668035 | 0.74390967 | 0.89297085 |
| UNC93B1 | 44.5174732 | -0.84557145 | 0.68431232 | -1.2356514 | 0.21658818 | 0.49824645 |
| NDUFS8 | 86.4673923 | -0.56345226 | 0.29945506 | -1.8815921 | 0.05989143 | 0.24679372 |
| CHKA | 158.040277 | 0.243198821 | 0.35743697 | 0.68039639 | 0.49625351 | 0.74801139 |
| KMT5B | 245.434349 | 0.19719212 | 0.20355313 | 0.96875012 | 0.33266988 | 0.61719981 |
| LRP5 | 116.61524 | -0.368738257 | 0.30801962 | -1.1971259 | 0.2312575 | 0.51432779 |
| PPP6R3 | 689.451698 | 0.481311642 | 0.12176502 | 3.95279066 | 7.72E-05 | 0.0039824 |
| CPT1A | 121.690138 | 0.572891047 | 0.45998769 | 1.24544866 | 0.21296688 | 0.49481974 |
| MRPL21 | 27.7995829 | -0.194925869 | 0.29569459 | -0.6592135 | 0.50975867 | 0.75857982 |
| IGHMBP2 | 31.5949488 | 0.14834714 | 0.25647574 | 0.57840613 | 0.56298995 | 0.79260446 |
| TPCN2 | 42.1825338 | -0.194458687 | 0.37730684 | -0.5153861 | 0.60628327 | 0.82097646 |
| CCND1 | 168.09621 | -0.658195966 | 0.76312175 | -0.8625045 | 0.38840994 | 0.66462645 |
| ORAOV1 | 43.0569469 | 0.484966388 | 0.33573265 | 1.4445017 | 0.1485979 | 0.40820743 |
| PPFIA1 | 280.678483 | -0.08497183 | 0.22807047 | -0.3725683 | 0.70946977 | 0.87656532 |
| AP002336.2 | 103.638812 | 0.006321449 | 0.46226115 | 0.01367506 | 0.98908922 | 0.99678172 |
| CTTN | 301.950275 | -0.767714581 | 0.25132636 | -3.054652 | 0.00225322 | 0.0341679 |
| SHANK2 | 1025.76937 | 0.085898868 | 0.29621433 | 0.2899889 | 0.77182473 | 0.90624314 |
| NADSYN1 | 48.3579685 | -0.446228104 | 0.27057413 | -1.6491898 | 0.09910876 | 0.32904321 |
| AP002495.2 | 121.223957 | 0.640177433 | 0.2786628 | 2.29731931 | 0.02160056 | 0.14142978 |
| RNF121 | 170.740324 | -0.025865484 | 0.17784757 | -0.1454363 | 0.88436644 | 0.95761929 |
| NUMA1 | 175.496983 | 0.337264051 | 0.26930388 | 1.25235498 | 0.21044054 | 0.49127431 |
| LAMTOR1 | 95.6911073 | -0.672082845 | 0.33339552 | -2.0158724 | 0.04381332 | 0.20766342 |
| INPPL1 | 61.2826873 | 0.165210644 | 0.29612137 | 0.5579153 | 0.57690222 | 0.80275223 |
| CLPB | 45.3112122 | -0.210191099 | 0.27058478 | -0.7768031 | 0.43727495 | 0.70195888 |
| ARAP1 | 52.9924725 | -0.266833396 | 0.29921077 | -0.8917907 | 0.3725051 | 0.65199071 |
| ATG16L2 | 30.724751 | 0.658518551 | 0.36182377 | 1.81999802 | 0.06875931 | 0.26828733 |
| FCHSD2 | 531.865935 | -0.387588231 | 0.38801473 | -0.9989008 | 0.31784274 | 0.60445019 |
| P2RY6 | 33.8886624 | -1.607784824 | 0.47271142 | -3.4011973 | 0.00067091 | 0.01577201 |
| ARHGEF17 | 90.9155246 | 0.411512285 | 0.33479431 | 1.22914959 | 0.21901573 | 0.50082398 |
| FAM168A | 213.07255 | 0.557033139 | 0.17787381 | 3.13161979 | 0.00173845 | 0.0293224 |
| RAB6A | 285.089336 | -0.191552504 | 0.15038665 | -1.2737334 | 0.20275791 | 0.48158147 |
| MRPL48 | 44.904387 | 0.573680773 | 0.22442802 | 2.55619051 | 0.01058252 | 0.09230421 |
| COA4 | 61.8612446 | -1.10390838 | 0.40188445 | -2.7468303 | 0.00601743 | 0.06537526 |
| PAAF1 | 81.4243568 | 0.088687854 | 0.24541561 | 0.36137822 | 0.71781673 | 0.88119524 |
| C2CD3 | 108.946766 | 0.3275081 | 0.18951876 | 1.72810388 | 0.0839696 | 0.30203879 |
| PPME1 | 155.887059 | 0.091149191 | 0.2742821 | 0.33231914 | 0.73964829 | 0.89253455 |
| P4HA3 | 29.25487 | 0.006542638 | 0.43598488 | 0.01500657 | 0.98802694 | 0.99678172 |
| PGM2L1 | 73.8756257 | -1.268182082 | 0.46084972 | -2.7518343 | 0.00592625 | 0.06458327 |
| POLD3 | 80.4133577 | -0.244924105 | 0.19460632 | -1.2585619 | 0.20818861 | 0.48842485 |
| RNF169 | 186.91404 | -0.225834678 | 0.22146071 | -1.0197505 | 0.3078468 | 0.59456464 |
| XRRA1 | 167.923139 | 0.689030889 | 0.50680815 | 1.35954975 | 0.17397245 | 0.44528422 |
| SPCS2 | 156.435264 | -0.300205545 | 0.28807427 | -1.0421116 | 0.29735993 | 0.58521131 |
| NEU3 | 32.2031921 | 0.004055651 | 0.26769404 | 0.01515032 | 0.98791225 | 0.99678172 |
| ARRB1 | 37.6796726 | 0.522066698 | 0.52404283 | 0.99622907 | 0.31913886 | 0.60583257 |
| RPS3 | 587.081194 | -0.295019065 | 0.46072989 | -0.6403298 | 0.52195823 | 0.76473507 |
| SERPINH1 | 47.5334622 | -0.396400231 | 0.25533218 | -1.5524883 | 0.12054542 | 0.36507762 |
| UVRAG | 369.915932 | -0.122779209 | 0.19080634 | -0.6434755 | 0.51991559 | 0.76385303 |
| THAP12 | 38.2492816 | 0.341516168 | 0.28575253 | 1.1951466 | 0.23202976 | 0.5151336 |
| EMSY | 183.026028 | 0.286180986 | 0.14095629 | 2.03028182 | 0.0423279 | 0.20409723 |
| ACER3 | 175.097119 | -0.617470072 | 0.2525989 | -2.4444686 | 0.01450657 | 0.11197822 |
| CAPN5 | 25.3048296 | -0.017486588 | 0.50361415 | -0.0347222 | 0.97230126 | 0.99218854 |
| PAK1 | 130.66532 | 0.181433644 | 0.32480089 | 0.55859959 | 0.57643502 | 0.80245537 |
| CLNS1A | 131.076381 | -0.115376817 | 0.15336548 | -0.7522998 | 0.45187079 | 0.71231034 |
| RSF1 | 409.862626 | 0.201351572 | 0.15360465 | 1.31084297 | 0.18991082 | 0.46597228 |
| AAMDC | 110.339196 | 0.433052373 | 0.28677309 | 1.51008718 | 0.13102118 | 0.38159072 |
| AP002812.2 | 46.1282914 | 0.455045326 | 0.32380997 | 1.40528511 | 0.15993651 | 0.42556441 |
| INTS4 | 102.375046 | 0.197559328 | 0.18282305 | 1.08060406 | 0.27987328 | 0.56738754 |
| ALG8 | 59.0521939 | -0.062098338 | 0.31960774 | -0.1942955 | 0.84594453 | 0.94271476 |
| GAB2 | 170.404382 | -1.250615546 | 0.52247441 | -2.3936398 | 0.01668212 | 0.12124283 |
| AP003086.2 | 50.8778039 | -1.44395016 | 0.53199043 | -2.7142408 | 0.00664279 | 0.06902242 |
| NARS2 | 154.873762 | 0.195709526 | 0.2140739 | 0.91421479 | 0.36060399 | 0.64097711 |
| TENM4 | 170.029059 | -1.284980958 | 0.68283246 | -1.8818393 | 0.05985784 | 0.24679372 |
| PRCP | 62.417987 | 0.115907254 | 0.30739696 | 0.37706051 | 0.70612864 | 0.87473681 |
| RAB30 | 33.4784702 | 0.770584358 | 0.38143522 | 2.02022339 | 0.04336022 | 0.20633261 |
| RAB30-AS1 | 174.215233 | 0.25891857 | 0.30589398 | 0.84643238 | 0.39731158 | 0.67251432 |
| PCF11 | 133.139673 | 0.108143954 | 0.19394395 | 0.55760416 | 0.57711472 | 0.80293128 |
| ANKRD42 | 73.2128242 | -0.060486666 | 0.30407247 | -0.1989219 | 0.84232386 | 0.94132995 |
| CCDC90B | 49.4970297 | -0.602869076 | 0.29850335 | -2.0196393 | 0.04342082 | 0.20651839 |
| DLG2 | 1041.87906 | 1.154730924 | 0.38388807 | 3.00798856 | 0.00262983 | 0.03764493 |
| TMEM126B | 42.6526752 | -0.859568242 | 0.28497357 | -3.0163086 | 0.00255873 | 0.03685721 |
| CREBZF | 132.407923 | 0.313948322 | 0.25281634 | 1.2418039 | 0.21430892 | 0.49648737 |
| SYTL2 | 410.024361 | 0.333552203 | 0.27827639 | 1.1986364 | 0.23066936 | 0.51385623 |
| PICALM | 1218.99384 | 0.906999553 | 0.16580842 | 5.47016582 | 4.50E-08 | 1.55E-05 |
| EED | 53.6586639 | 0.438909118 | 0.28279081 | 1.55206288 | 0.12064718 | 0.36507762 |
| HIKESHI | 69.9557584 | 0.167166868 | 0.20158526 | 0.82926139 | 0.40695651 | 0.68040675 |
| AP001831.1 | 46.8965246 | -0.170894444 | 0.47463714 | -0.3600528 | 0.71880763 | 0.88155276 |

| | | | | | | |
|---|---|---|---|---|---|---|
| ME3 | 112.552185 | 0.088601524 | 0.32855507 | 0.26967024 | 0.78741396 | 0.91370543 |
| PRSS23 | 137.011968 | 0.063314435 | 0.4842795 | 0.13073945 | 0.89598142 | 0.96163371 |
| AP001528.2 | 92.6218006 | -0.482797431 | 0.61063297 | -0.7906508 | 0.42914781 | 0.69498002 |
| FZD4 | 49.5467986 | 0.221393706 | 0.41622473 | 0.53190906 | 0.59478898 | 0.81507555 |
| TMEM135 | 260.837389 | 0.371186982 | 0.23831623 | 1.5575397 | 0.1193424 | 0.3645149 |
| CTSC | 310.019092 | 0.156002428 | 0.67469067 | 0.23122067 | 0.81714337 | 0.9295115 |
| GRM5-AS1 | 61.0231764 | 0.829372857 | 0.55385233 | 1.49746207 | 0.13427307 | 0.38591741 |
| GRM5 | 178.771485 | 0.105829443 | 0.59788404 | 0.17700664 | 0.85950318 | 0.94734879 |
| NOX4 | 156.234695 | -0.588920326 | 0.66521401 | -0.8853096 | 0.3759897 | 0.65436143 |
| CHORDC1 | 60.0584233 | 0.735271608 | 0.26401785 | 2.78493142 | 0.0053539 | 0.06041319 |
| FAT3 | 129.380625 | 2.906156838 | 1.07762846 | 2.69680779 | 0.00700077 | 0.07109414 |
| SLC36A4 | 152.257878 | 0.209983431 | 0.35147746 | 0.59743071 | 0.55021985 | 0.78536075 |
| SMCO4 | 44.448281 | 0.545733178 | 0.27074617 | 2.01566349 | 0.04383518 | 0.20766427 |
| CEP295 | 81.2999831 | 0.022393417 | 0.24792627 | 0.09032289 | 0.92803063 | 0.97474479 |
| TAF1D | 296.24599 | -0.392289931 | 0.25531676 | -1.5364833 | 0.1244199 | 0.37055989 |
| MRE11 | 89.7582785 | 0.274705699 | 0.23043175 | 1.19213476 | 0.23320841 | 0.51603997 |
| PIWIL4 | 49.5727429 | -1.677239021 | 0.61194256 | -2.7408439 | 0.00612816 | 0.06580904 |
| AMOTL1 | 98.6044683 | -0.847251193 | 0.4567738 | -1.8548594 | 0.0636163 | 0.25658128 |
| CWC15 | 47.333942 | -0.509219099 | 0.28537716 | -1.7843723 | 0.07436319 | 0.28138929 |
| SRSF8 | 47.5811657 | -0.330704281 | 0.36023279 | -0.9180294 | 0.35860349 | 0.64001011 |
| ENDOD1 | 98.5676437 | -1.140386837 | 0.48768737 | -2.3383563 | 0.01936877 | 0.13214635 |
| AP000787.1 | 64.6129446 | 0.902476822 | 0.58305654 | 1.54783758 | 0.12166141 | 0.36636109 |
| SESN3 | 302.003073 | 1.13000757 | 0.49241117 | 2.29484551 | 0.02174198 | 0.14167784 |
| FAM76B | 50.237196 | 0.473795921 | 0.20476826 | 2.3138153 | 0.02067786 | 0.13755082 |
| CEP57 | 126.38082 | 0.252427005 | 0.23755686 | 1.06259616 | 0.28796513 | 0.57459866 |
| MTMR2 | 167.640912 | -0.024275864 | 0.21483308 | -0.1129987 | 0.91003157 | 0.96825418 |
| MAML2 | 3501.08544 | 0.025399896 | 0.22167076 | 0.11458388 | 0.90877496 | 0.96752108 |
| CCDC82 | 85.6000524 | -0.134967705 | 0.21317894 | -0.6331193 | 0.52665573 | 0.76821934 |
| CNTN5 | 187.663716 | 2.155244789 | 0.60168108 | 3.58203848 | 0.00034092 | 0.01037405 |
| ARHGAP42 | 198.673993 | -0.162796068 | 0.52803991 | -0.3083026 | 0.7578521 | 0.89967348 |
| CEP126 | 119.217622 | -0.140432457 | 0.38649217 | -0.3633514 | 0.71634242 | 0.88058646 |
| YAP1 | 677.989505 | 0.040723393 | 0.20223406 | 0.20136763 | 0.84041112 | 0.94034554 |
| BIRC3 | 263.985746 | -3.144808478 | 0.76449165 | -4.1135943 | 3.90E-05 | 0.00259129 |
| BIRC2 | 117.221271 | 0.06106292 | 0.28015955 | 0.21795766 | 0.8274621 | 0.93346403 |
| TMEM123 | 291.681209 | -0.328360785 | 0.20912156 | -1.570191 | 0.11637069 | 0.35938557 |
| MMP7 | 463.100793 | -3.256085221 | 0.68764655 | -4.7351146 | 2.19E-06 | 0.00030579 |
| DCUN1D5 | 55.0223877 | -0.233619429 | 0.244112 | -0.9570174 | 0.33855848 | 0.62345706 |
| DYNC2H1 | 376.101439 | 0.474166304 | 0.21022466 | 2.25552183 | 0.02410059 | 0.15059334 |
| PDGFD | 150.184866 | 1.183143692 | 0.48157272 | 2.4568329 | 0.01401679 | 0.10948271 |
| CASP4 | 87.9317188 | -1.338431548 | 0.4030767 | -3.3205381 | 0.00089844 | 0.01925316 |
| MSANTD4 | 32.3603273 | -0.066765026 | 0.30007305 | -0.2224959 | 0.82392785 | 0.93217972 |
| KBTBD3 | 29.4736512 | 0.470114333 | 0.29960671 | 1.56910483 | 0.11662352 | 0.35978636 |
| AASDHPPT | 70.2181918 | 0.32093643 | 0.26334776 | 1.21867916 | 0.22296599 | 0.50455734 |
| CWF19L2 | 156.921981 | 0.347066482 | 0.22753987 | 1.52529966 | 0.12718437 | 0.37451555 |
| ALKBH8 | 28.6992363 | -0.005734168 | 0.30358061 | -0.0188885 | 0.98493009 | 0.99668665 |
| SLC35F2 | 58.6298118 | -0.910692555 | 0.36249868 | -2.5122644 | 0.01199592 | 0.10009484 |
| CUL5 | 344.441464 | 0.475725169 | 0.1666281 | 2.85501166 | 0.00430352 | 0.05249369 |
| ACAT1 | 126.095616 | 0.124320068 | 0.24771289 | 0.50187161 | 0.61575784 | 0.82592881 |
| NPAT | 91.7727083 | 0.036140386 | 0.25235068 | 0.14321493 | 0.88612044 | 0.95831763 |
| ATM | 394.568052 | 0.494068421 | 0.30958076 | 1.5959274 | 0.110505 | 0.34758554 |
| C11orf65 | 87.3375569 | 0.837334839 | 0.48260332 | 1.73503746 | 0.08273417 | 0.29973927 |
| EXPH5 | 126.887156 | 1.00921152 | 0.47687107 | 2.11631946 | 0.03431765 | 0.18262708 |
| DDX10 | 199.246088 | -0.161132613 | 0.27704047 | -0.5816212 | 0.56082186 | 0.79071424 |
| RDX | 286.76039 | 0.392242038 | 0.18967994 | 2.0679152 | 0.038648 | 0.19546419 |
| ZC3H12C | 137.533654 | -0.518629524 | 0.339497 | -1.527641 | 0.1266017 | 0.37394935 |
| FDX1 | 70.4345309 | -0.005025691 | 0.27165895 | -0.0185 | 0.98523998 | 0.99668665 |
| SIK2 | 598.711595 | 0.538758011 | 0.35828531 | 1.50371226 | 0.13265547 | 0.38380408 |
| PPP2R1B | 115.955181 | 0.032141958 | 0.24440274 | 0.13151226 | 0.89537009 | 0.96141128 |
| C11orf1 | 26.1361615 | 0.656521818 | 0.54717945 | 1.19982908 | 0.23020573 | 0.51342041 |
| CRYAB | 1938.03333 | -0.101090357 | 0.60501976 | -0.167086 | 0.86730235 | 0.95090501 |
| DIXDC1 | 206.180594 | 0.345070091 | 0.39681291 | 0.869604 | 0.38451685 | 0.66091637 |
| DLAT | 51.8062583 | 0.114811069 | 0.2632906 | 0.43606216 | 0.66279163 | 0.85530584 |
| NKAPD1 | 32.1896983 | 0.069275724 | 0.27877247 | 0.24850275 | 0.80374544 | 0.92125386 |
| TIMM8B | 47.7875354 | -0.977195111 | 0.46621091 | -2.0960365 | 0.03607895 | 0.18804981 |
| IL18 | 55.8175925 | -1.563328303 | 0.89567815 | -1.745413 | 0.080913 | 0.29573375 |
| PTS | 106.444938 | 0.175198902 | 0.31155093 | 0.56234433 | 0.57388144 | 0.80064234 |
| AP002884.1 | 69.2317632 | 0.282364001 | 0.47612027 | 0.59305184 | 0.55314646 | 0.78647229 |
| TTC12 | 94.2642635 | 0.404071959 | 0.23379279 | 1.7283337 | 0.08392841 | 0.30203879 |
| ZW10 | 32.169248 | 0.354353148 | 0.2530822 | 1.40015041 | 0.16146828 | 0.42791083 |
| USP28 | 57.5629151 | 0.431448104 | 0.28424877 | 1.51785392 | 0.12905123 | 0.37757535 |
| ZBTB16 | 773.332373 | -0.532791924 | 0.5621427 | -0.9477877 | 0.34323755 | 0.62811979 |
| AP002518.2 | 74.2614838 | -1.319005388 | 0.7644808 | -1.7253611 | 0.08446243 | 0.3023414 |
| NNMT | 410.68323 | -5.587329648 | 1.1846028 | -4.7166271 | 2.40E-06 | 0.00032351 |
| REXO2 | 67.7536067 | -0.376794891 | 0.31342561 | -1.2021828 | 0.2292927 | 0.5117445 |
| CADM1 | 425.854004 | 0.001227602 | 0.40992378 | 0.00299471 | 0.99761057 | 0.99923686 |
| SIK3 | 1188.11642 | 0.728959725 | 0.21831341 | 3.33905147 | 0.00084065 | 0.01842697 |
| PAFAH1B2 | 144.681969 | 0.157595599 | 0.15393578 | 1.02377499 | 0.30594156 | 0.59348203 |
| SIDT2 | 28.7646326 | 0.262571499 | 0.34707365 | 0.75652963 | 0.44933171 | 0.71037275 |
| TAGLN | 55.4445323 | 0.084818577 | 0.76273061 | 0.11120385 | 0.9114547 | 0.96844354 |
| PCSK7 | 91.0382481 | -0.381467093 | 0.2070136 | -1.8427151 | 0.06537061 | 0.26050329 |
| RNF214 | 75.6571683 | 0.229020873 | 0.24346056 | 0.94068985 | 0.34686382 | 0.63046825 |
| BACE1 | 58.9460453 | -0.214372964 | 0.3346425 | -0.6406029 | 0.5217807 | 0.7647089 |
| CEP164 | 95.9972821 | 0.802678004 | 0.32160164 | 2.49587656 | 0.01256463 | 0.10234405 |
| DSCAML1 | 1160.06851 | 0.10662657 | 0.61611704 | 0.1730622 | 0.86260254 | 0.94877974 |
| FXYD2 | 237.937563 | 0.868620717 | 0.61416732 | 1.41430631 | 0.15727198 | 0.42243085 |
| MPZL3 | 41.6500488 | 0.511016799 | 0.47640232 | 1.07265809 | 0.28342455 | 0.57084461 |
| UBE4A | 103.116086 | 0.283882924 | 0.19343597 | 1.46758084 | 0.14221812 | 0.39850156 |
| ATP5MG | 180.221616 | -0.264937258 | 0.43084456 | -0.6149254 | 0.53860399 | 0.77706498 |
| KMT2A | 468.382629 | -0.070068901 | 0.13185879 | -0.5313935 | 0.59514614 | 0.81511365 |
| IFT46 | 34.8842453 | 0.604836745 | 0.24622714 | 2.45641783 | 0.01403299 | 0.10948271 |
| ARCN1 | 77.7708456 | -0.048292687 | 0.2503363 | -0.1929112 | 0.84702848 | 0.94323402 |
| PHLDB1 | 252.096258 | 0.854086692 | 0.27989157 | 3.05149133 | 0.00227708 | 0.03440396 |
| AP002954.1 | 37.4180241 | 0.562532671 | 0.48123057 | 1.16894625 | 0.24242529 | 0.52725736 |
| DDX6 | 231.563523 | 0.264208479 | 0.18570647 | 1.42272095 | 0.15481707 | 0.41821565 |
| BCL9L | 61.2450985 | -1.205561283 | 0.38623202 | -3.1213396 | 0.0018003 | 0.02983582 |
| CCDC84 | 136.607294 | 0.337779011 | 0.25227286 | 1.3389431 | 0.1805892 | 0.45296253 |
| RPS25 | 387.81766 | -0.519069929 | 0.43124814 | -1.2036456 | 0.22872659 | 0.51112377 |
| TRAPPC4 | 69.0858832 | 0.004810233 | 0.35204429 | 0.01366372 | 0.98909827 | 0.99678172 |
| SLC37A4 | 40.0974971 | -0.012114625 | 0.47478246 | -0.0255162 | 0.97964326 | 0.99557142 |
| HYOU1 | 41.0394194 | -1.333414658 | 0.5166617 | -2.5808274 | 0.00985639 | 0.08873526 |
| VPS11 | 26.2684732 | 0.185686729 | 0.33335764 | 0.55701957 | 0.57751406 | 0.80337019 |
| HINFP | 25.8163327 | -0.228306471 | 0.24720622 | -0.9235466 | 0.35572241 | 0.63728305 |
| NLRX1 | 29.076563 | -0.481420678 | 0.35268199 | -1.3650277 | 0.17224436 | 0.44309343 |
| CBL | 121.738004 | 0.093646872 | 0.23366602 | 0.40077232 | 0.68858776 | 0.86751885 |
| THY1 | 90.3690377 | 0.798245729 | 1.25390769 | 0.63660646 | 0.52438122 | 0.76611507 |
| OAF | 49.4202175 | -0.776110922 | 0.40504108 | -1.9161289 | 0.05534869 | 0.23619173 |
| ARHGEF12 | 807.911766 | -0.003965368 | 0.16660197 | -0.0238014 | 0.98101099 | 0.99578216 |
| SC5D | 37.5912454 | 0.315480911 | 0.42087304 | 0.74958689 | 0.45350355 | 0.71320153 |
| SORL1 | 166.603809 | 1.059935811 | 0.32409513 | 3.27044661 | 0.00107378 | 0.02174571 |
| UBASH3B | 268.793615 | -0.686544986 | 0.49582204 | -1.3846601 | 0.16615643 | 0.43446007 |
| JHY | 137.086526 | 0.194754872 | 0.36954672 | 0.52701015 | 0.59818654 | 0.81658679 |
| HSPA8 | 195.011797 | -0.450177799 | 0.47080117 | -0.9561952 | 0.33897365 | 0.62369754 |
| GRAMD1B | 38.4073202 | 0.518294938 | 0.55067453 | 0.94120013 | 0.34660231 | 0.6302171 |

| | | | | | | |
|---|---|---|---|---|---|---|
| VWA5A | 36.6780984 | -0.374461236 | 0.2873903 | -1.302971 | 0.19258465 | 0.4702443 |
| TBRG1 | 92.8995721 | 0.165443879 | 0.21506175 | 0.76928548 | 0.44172385 | 0.70476463 |
| SIAE | 72.8148037 | 0.213550477 | 0.42053234 | 0.50780987 | 0.61158669 | 0.82338564 |
| ESAM | 43.6769322 | 1.392130069 | 0.45541439 | 3.05684248 | 0.00223682 | 0.03403479 |
| MSANTD2 | 139.580951 | 0.790610818 | 0.20674262 | 3.82413088 | 0.00013123 | 0.00565867 |
| PKNOX2 | 73.4937608 | -0.71163388 | 0.75970987 | -0.936718 | 0.34890366 | 0.63263722 |
| EI24 | 76.7876991 | -0.789755124 | 0.40618072 | -1.9443442 | 0.05185396 | 0.22805742 |
| STT3A | 59.2962429 | 0.044263809 | 0.24033698 | 0.18417394 | 0.853877 | 0.9452497 |
| CDON | 54.1591582 | -0.495100631 | 0.88241462 | -0.5610748 | 0.57474653 | 0.8009242 |
| RPUSD4 | 27.4793696 | 0.005212016 | 0.3194764 | 0.01631424 | 0.98698369 | 0.99678172 |
| FAM118B | 41.554651 | -0.790693536 | 0.21294035 | -3.7132161 | 0.00020464 | 0.00758576 |
| SRPRA | 56.2059427 | -0.628883747 | 0.39470577 | -1.5932976 | 0.11109345 | 0.34822126 |
| FOXRED1 | 38.5999916 | -0.250315825 | 0.23743666 | -1.0542425 | 0.29177189 | 0.57910139 |
| DCPS | 35.470608 | 0.625359254 | 0.48967886 | 1.27708036 | 0.20157388 | 0.48024387 |
| ST3GAL4 | 71.8209724 | 2.182554586 | 0.55490005 | 3.93323915 | 8.38E-05 | 0.00415961 |
| KIRREL3 | 136.511242 | 2.639877559 | 1.0093148 | 2.61551456 | 0.00890931 | 0.08424709 |
| ETS1 | 186.202812 | -0.203080592 | 0.40473918 | -0.5017567 | 0.61583868 | 0.82592881 |
| ARHGAP32 | 296.555982 | 0.223674622 | 0.2847813 | 0.78542595 | 0.43220387 | 0.69737124 |
| TMEM45B | 30.71147 | -0.160539124 | 0.47347846 | -0.3390632 | 0.73456212 | 0.88966284 |
| NFRKB | 57.1262912 | 0.260946078 | 0.2340038 | 1.11513605 | 0.26479213 | 0.55272255 |
| PRDM10 | 67.4705386 | 0.376375134 | 0.28769906 | 1.30822511 | 0.19079697 | 0.46742434 |
| APLP2 | 463.041453 | -0.240231309 | 0.27022582 | -0.8890021 | 0.37400197 | 0.65261518 |
| ST14 | 152.692951 | -0.083411892 | 0.4487715 | -0.1858672 | 0.85254892 | 0.94470172 |
| ZBTB44 | 338.88871 | 1.018216553 | 0.21635954 | 4.706132 | 2.52E-06 | 0.0003268 |
| AP002986.1 | 41.6314385 | 1.431778776 | 0.59771316 | 2.39542788 | 0.01660098 | 0.12092835 |
| ADAMTS15 | 108.908597 | 2.051642578 | 0.65069908 | 3.15298216 | 0.00161612 | 0.02795985 |
| SNX19 | 94.3321814 | 0.826399726 | 0.31998026 | 2.58265847 | 0.00980423 | 0.08848555 |
| NTM | 274.675481 | 1.048085496 | 0.64069555 | 1.63585575 | 0.10186978 | 0.33338252 |
| NCAPD3 | 34.5546547 | -0.187144059 | 0.31871338 | -0.5871861 | 0.55707875 | 0.78857061 |
| VPS26B | 37.5257414 | 0.328515589 | 0.27794648 | 1.18193828 | 0.23723019 | 0.52000641 |
| ZMYND11 | 316.837651 | 0.456476968 | 0.14192506 | 3.21632388 | 0.00129844 | 0.0248259 |
| DIP2C | 689.577121 | 0.152390803 | 0.23152715 | 0.65819842 | 0.51041064 | 0.7589605 |
| LARP4B | 472.454118 | 0.293811218 | 0.18037534 | 1.62888794 | 0.10333674 | 0.33551358 |
| GTPBP4 | 57.12288 | -0.809034801 | 0.36140659 | -2.2385724 | 0.02518375 | 0.15393404 |
| IDI1 | 68.7536094 | 0.688160463 | 0.42040468 | 1.63690009 | 0.10165135 | 0.33300898 |
| WDR37 | 131.739367 | 0.140988414 | 0.14534921 | 0.96999782 | 0.33204758 | 0.61700946 |
| PFKP | 184.562092 | -0.10940082 | 0.50247668 | -0.2177232 | 0.8276448 | 0.93346403 |
| PITRM1 | 73.8378297 | -0.579540241 | 0.27041574 | -2.1431453 | 0.03210143 | 0.17552647 |
| KLF6 | 416.257916 | -0.096903249 | 0.50366031 | -0.192398 | 0.84743044 | 0.94323402 |
| AKR1C1 | 143.979428 | 1.207088222 | 0.80911701 | 1.49185866 | 0.1357362 | 0.38788633 |
| AKR1C2 | 82.9001125 | 2.127069279 | 0.82190861 | 2.58796326 | 0.00965453 | 0.08782595 |
| AKR1C3 | 60.795935 | -0.957968165 | 0.36328512 | -2.6369595 | 0.00836528 | 0.0806959 |
| NET1 | 56.1999578 | -0.268110898 | 0.30366566 | -0.8829148 | 0.37728231 | 0.65530141 |
| ASB13 | 43.7092654 | 0.022470927 | 0.32913325 | 0.06827304 | 0.94556829 | 0.98119662 |
| FAM208B | 314.639327 | -0.035989076 | 0.19448718 | -0.185046 | 0.85319295 | 0.94515269 |
| GDI2 | 196.184258 | -0.665346925 | 0.20144616 | -3.3028523 | 0.00095707 | 0.02023786 |
| ANKRD16 | 31.5401074 | -0.021329733 | 0.40246086 | -0.0529983 | 0.95773328 | 0.98551156 |
| FBH1 | 104.35391 | -0.07063898 | 0.19627705 | -0.3598942 | 0.71892622 | 0.88155276 |
| RBM17 | 124.209329 | -0.129737985 | 0.21349612 | -0.6076831 | 0.54339766 | 0.78025803 |
| PFKFB3 | 854.716961 | 0.428339765 | 0.70176855 | 0.61037185 | 0.54161551 | 0.77887826 |
| AL137145.2 | 146.176806 | 0.120701025 | 0.80433515 | 0.1500631 | 0.88071483 | 0.95628739 |
| PRKCQ | 235.944133 | 0.896871521 | 0.5653821 | 1.58631042 | 0.11266891 | 0.35132015 |
| SFMBT2 | 234.152081 | -0.883586644 | 0.64894153 | -1.3615813 | 0.17333006 | 0.44449049 |
| ITIH5 | 176.308123 | 0.320435284 | 0.69477719 | 0.46120582 | 0.64465095 | 0.84382501 |
| KIN | 44.5712166 | 0.326759054 | 0.21882612 | 1.49323607 | 0.1353754 | 0.38755559 |
| ATP5F1C | 146.69505 | 0.087484442 | 0.16267488 | 0.53778705 | 0.59072407 | 0.81207605 |
| TAF3 | 269.783319 | 0.544429331 | 0.21267888 | 2.55986558 | 0.01047127 | 0.09185371 |
| CELF2 | 97.7268357 | -0.261038997 | 0.38036719 | -0.6862816 | 0.49253556 | 0.74486866 |
| USP6NL | 174.156207 | -0.027275505 | 0.16554934 | -0.1647576 | 0.86913481 | 0.95106604 |
| ECHDC3 | 40.2762718 | -0.772672474 | 0.43675378 | -1.769126 | 0.07687285 | 0.28663488 |
| PROSER2 | 251.844254 | 0.512058389 | 0.50339895 | 1.01720194 | 0.30905737 | 0.59563596 |
| UPF2 | 197.28519 | 0.078499633 | 0.14252375 | 0.55078283 | 0.58178256 | 0.80614713 |
| DHTKD1 | 91.1301664 | -0.050551338 | 0.19176022 | -0.2636174 | 0.79207472 | 0.91667074 |
| SEC61A2 | 35.5832371 | 0.414445165 | 0.27042954 | 1.53254397 | 0.12538826 | 0.37225172 |
| NUDT5 | 73.4732959 | -0.076556096 | 0.20783358 | -0.3683529 | 0.71261013 | 0.87827554 |
| CDC123 | 73.5617096 | -0.177778264 | 0.20997973 | -0.8466449 | 0.39719308 | 0.6724483 |
| CAMK1D | 299.351939 | -0.283236911 | 0.49755011 | -0.5692631 | 0.56917761 | 0.79749848 |
| OPTN | 130.247975 | -0.455687861 | 0.51604188 | -0.8830443 | 0.37721232 | 0.65530141 |
| PHYH | 31.6607092 | -0.16325408 | 0.39207503 | -0.4163848 | 0.67712846 | 0.8613973 |
| SEPHS1 | 53.4897872 | 0.292064322 | 0.33170884 | 0.88048399 | 0.37859717 | 0.65651382 |
| BEND7 | 231.350697 | -0.482575882 | 0.15755716 | -3.0628622 | 0.00219231 | 0.03385831 |
| PRPF18 | 28.6773642 | -0.177287019 | 0.36002517 | -0.4924295 | 0.62241574 | 0.8294547 |
| AL157392.5 | 53.9503891 | 0.006590887 | 0.30404535 | 0.02167732 | 0.98270536 | 0.99616479 |
| FRMD4A | 1275.45524 | 1.289711916 | 0.60261839 | 2.14018015 | 0.03234021 | 0.17593476 |
| FAM107B | 262.029545 | -0.186628565 | 0.22638381 | -0.8243901 | 0.40971795 | 0.68208527 |
| DCLRE1C | 52.2862601 | -0.363937801 | 0.21747341 | -1.6734818 | 0.09423248 | 0.31974955 |
| RPP38 | 86.8348102 | -9.59E-05 | 0.333251 | -0.0002879 | 0.9997703 | 0.9998728 |
| NMT2 | 83.6835973 | 0.119721358 | 0.35590856 | 0.33638235 | 0.73658256 | 0.89057637 |
| FAM171A1 | 461.294579 | 0.340605505 | 0.4743859 | 0.71799247 | 0.47276193 | 0.73006392 |
| MINDY3 | 122.528701 | 0.159347107 | 0.15600042 | 1.02145308 | 0.30703983 | 0.59456464 |
| PTER | 89.6508567 | 0.365812822 | 0.21063976 | 1.73667506 | 0.08244454 | 0.29914253 |
| RSU1 | 162.126019 | -0.093722269 | 0.23871691 | -0.3926084 | 0.69460872 | 0.87010029 |
| CUBN | 167.605507 | 0.480435546 | 0.54845342 | 0.8759824 | 0.3810396 | 0.65812808 |
| TRDMT1 | 44.5916114 | 0.57818375 | 0.29191029 | 1.98068986 | 0.04762607 | 0.21810285 |
| VIM | 320.872052 | -0.828102318 | 0.7517397 | -1.1015812 | 0.27064379 | 0.5578257 |
| HACD1 | 46.9706689 | -0.135356543 | 0.46299883 | -0.2923475 | 0.77002096 | 0.90533273 |
| STAM | 103.958882 | -0.023864431 | 0.27822165 | -0.0857749 | 0.93164537 | 0.97671347 |
| CACNB2 | 155.926122 | 1.217328761 | 0.43536418 | 2.79611605 | 0.00517208 | 0.05935703 |
| AL450384.2 | 55.2804199 | 0.07256027 | 0.32759084 | 0.22149664 | 0.82470575 | 0.93217972 |
| NSUN6 | 132.42512 | 0.247190323 | 0.23563848 | 1.04902358 | 0.29416726 | 0.58223301 |
| ARL5B | 73.3193812 | -0.125973764 | 0.36413473 | -0.3459537 | 0.7293775 | 0.88653304 |
| MALRD1 | 48.7472465 | 0.012335526 | 0.68246967 | 0.01807483 | 0.98557915 | 0.99668665 |
| PLXDC2 | 776.252797 | 0.796247953 | 0.49526651 | 1.60771611 | 0.10789737 | 0.34350867 |
| NEBL | 1374.09315 | 1.361648768 | 0.46169797 | 2.94921975 | 0.00318577 | 0.04232528 |
| MLLT10 | 623.297738 | 0.398905193 | 0.14682806 | 2.71681858 | 0.00659127 | 0.06885256 |
| DNAJC1 | 457.911756 | 0.100844445 | 0.20654032 | 0.4882555 | 0.62536888 | 0.83169854 |
| BMI1 | 29.2737256 | 0.356950573 | 0.30389837 | 1.17457218 | 0.24016592 | 0.52416253 |
| PIP4K2A | 128.857019 | -0.374089853 | 0.46899002 | -0.7976499 | 0.42507367 | 0.69181522 |
| ARMC3 | 70.6885959 | -0.223751362 | 0.73016542 | -0.3064393 | 0.75927021 | 0.9005757 |
| MSRB2 | 69.2199292 | -0.423479052 | 0.35056249 | -1.2079988 | 0.22704774 | 0.50981957 |
| OTUD1 | 33.3442974 | 0.582710715 | 0.46187614 | 1.26161684 | 0.20708669 | 0.48648432 |
| KIAA1217 | 857.060615 | 0.279751379 | 0.32816809 | 0.85246368 | 0.39395679 | 0.66874218 |
| ARHGAP21 | 274.454729 | -0.065389487 | 0.19796348 | -0.3303109 | 0.74116508 | 0.89253455 |
| MYO3A | 71.19025 | -2.402066687 | 0.53894197 | -4.4570043 | 8.31E-06 | 0.00078053 |
| APBB1IP | 75.2399062 | -0.739778212 | 0.77304304 | -0.956969 | 0.33858291 | 0.62345706 |
| ABI1 | 237.135064 | 0.244392171 | 0.16472767 | 1.48361331 | 0.13791154 | 0.39068755 |
| ANKRD26 | 210.237643 | 0.355934238 | 0.24327725 | 1.46308067 | 0.14344531 | 0.40048158 |
| YME1L1 | 195.174079 | -0.173419628 | 0.14351161 | -1.2084014 | 0.2268929 | 0.50976168 |
| MASTL | 28.7055266 | -0.482291326 | 0.33893888 | -1.4229448 | 0.15475215 | 0.41815822 |
| ACBD5 | 95.2491099 | 0.005825451 | 0.24371412 | 0.02390281 | 0.98093014 | 0.99578216 |
| RAB18 | 177.807383 | 0.017254733 | 0.1308591 | 0.13185734 | 0.89509713 | 0.96122594 |
| MPP7 | 163.023507 | 1.029739139 | 0.44743424 | 2.30143123 | 0.02136727 | 0.14045419 |
| AL358234.1 | 102.692442 | 0.655604117 | 1.2206194 | 0.53710773 | 0.5911932 | 0.81248776 |

| | | | | | | |
|---|---|---|---|---|---|---|
| WAC-AS1 | 108.941398 | 0.129433546 | 0.48477466 | 0.26699734 | 0.78947121 | 0.91509495 |
| WAC | 745.340711 | 0.491756109 | 0.13893094 | 3.53957244 | 0.00040078 | 0.01159828 |
| BAMBI | 28.0073323 | 1.386029666 | 0.55549512 | 2.49512482 | 0.01259128 | 0.10239535 |
| SVIL | 631.53911 | -1.160266372 | 0.32893472 | -3.5273454 | 0.00041975 | 0.01203824 |
| MTPAP | 45.0171181 | -0.21020179 | 0.22241064 | -0.9451067 | 0.3446044 | 0.62954637 |
| MAP3K8 | 66.4250261 | -1.865243762 | 0.54011747 | -3.4534039 | 0.00055356 | 0.01407874 |
| ZNF438 | 159.940965 | -0.100473271 | 0.20617193 | -0.4873276 | 0.62602619 | 0.83180006 |
| ZEB1 | 364.521802 | -1.805147571 | 0.59644106 | -3.0265314 | 0.00247377 | 0.03612234 |
| ARHGAP12 | 250.194391 | -0.064099403 | 0.17915549 | -0.3577864 | 0.72050315 | 0.88234463 |
| KIF5B | 223.747489 | 0.116943422 | 0.1634422 | 0.71550325 | 0.47429814 | 0.73113966 |
| EPC1 | 363.821961 | 0.434439917 | 0.14101564 | 3.08079251 | 0.0020645 | 0.03236643 |
| CCDC7 | 197.377445 | 0.099596848 | 0.3347533 | 0.29752312 | 0.76606716 | 0.90371395 |
| ITGB1 | 403.090507 | -0.573982502 | 0.4099731 | -1.4000492 | 0.1614986 | 0.42791083 |
| NRP1 | 1665.43176 | -1.278833793 | 0.50213048 | -2.5468157 | 0.01087108 | 0.09381451 |
| PARD3 | 2684.96717 | 0.300646711 | 0.26866783 | 1.11902759 | 0.26312837 | 0.55096925 |
| CUL2 | 106.613885 | -0.026034805 | 0.21747754 | -0.1197126 | 0.90471081 | 0.9654885 |
| CREM | 183.256487 | 0.446842472 | 0.37149077 | 1.20283601 | 0.22903979 | 0.51153465 |
| CCNY | 654.137722 | 0.155660549 | 0.16963227 | 0.91763523 | 0.35880987 | 0.64001011 |
| AL117336.3 | 74.6883195 | 0.458280816 | 0.37777464 | 1.21310635 | 0.22508916 | 0.50741525 |
| ZNF248 | 148.491081 | 0.25787831 | 0.25086057 | 1.02797466 | 0.30396175 | 0.59203936 |
| ZNF25 | 31.8862512 | -0.080338516 | 0.40915425 | -0.1963526 | 0.84433416 | 0.94193856 |
| ZNF33A | 308.241339 | -0.149708057 | 0.15426696 | -0.970448 | 0.33182325 | 0.61695165 |
| ZNF33B | 249.817277 | 0.059561102 | 0.17710526 | 0.33630341 | 0.73664208 | 0.89057637 |
| BMS1 | 96.4009575 | -0.005395511 | 0.25105164 | -0.0214916 | 0.98285347 | 0.99616479 |
| CSGALNACT2 | 64.186637 | 0.082534898 | 0.25213115 | 0.32734908 | 0.74340388 | 0.89269973 |
| HNRNPF | 203.946425 | -0.985076033 | 0.27155821 | -3.627495 | 0.00028618 | 0.00938822 |
| ZNF487 | 50.278364 | 0.142587588 | 0.26482987 | 0.53841204 | 0.59029262 | 0.81183245 |
| ZNF32 | 41.6038562 | -0.227341658 | 0.28301899 | -0.8032735 | 0.42181666 | 0.69010791 |
| TMEM72-AS | 34.1083708 | -0.302935668 | 0.46580592 | -0.6503474 | 0.51546784 | 0.76135326 |
| RASSF4 | 102.311512 | -0.704885847 | 0.55260087 | -1.2755786 | 0.20210452 | 0.48062542 |
| ZNF22 | 47.6031433 | -0.877745825 | 0.3306231 | -2.654823 | 0.007935 | 0.07708863 |
| 8-Mar | 107.757916 | 0.120471768 | 0.26336202 | 0.45743789 | 0.64735633 | 0.84537847 |
| ZFAND4 | 57.9781159 | 0.424031074 | 0.22452955 | 1.88853126 | 0.05895467 | 0.24500075 |
| WASHC2C | 57.0332765 | -0.276742081 | 0.21480593 | -1.2883354 | 0.19762924 | 0.47630586 |
| AGAP4 | 64.0507918 | 0.13002806 | 0.33947387 | 0.38302819 | 0.70169887 | 0.87200065 |
| TIMM23 | 65.0189332 | 0.082522185 | 0.24201635 | 0.34097773 | 0.73312035 | 0.88870663 |
| NCOA4 | 84.6659716 | -0.38284659 | 0.19894688 | -1.9243659 | 0.05430874 | 0.23370324 |
| AL356056.2 | 49.6948841 | 1.32351094 | 0.60923193 | 2.17242544 | 0.02982359 | 0.16887967 |
| PTPN20 | 38.9480995 | -0.311532538 | 0.48678135 | -0.6399845 | 0.52218265 | 0.76482991 |
| AGAP9 | 52.1795731 | -0.03939379 | 0.31002408 | -0.1270669 | 0.89888747 | 0.96269027 |
| MAPK8 | 511.877765 | 0.287644692 | 0.17751853 | 1.62036435 | 0.10515403 | 0.33906316 |
| ARHGAP22 | 60.3007092 | -2.688022023 | 0.74457186 | -3.6101579 | 0.00030601 | 0.0098365 |
| ERCC6 | 76.8555443 | 0.168324981 | 0.17544315 | 0.95942748 | 0.33734344 | 0.62238305 |
| PARG | 202.858575 | 0.100348048 | 0.16184877 | 0.62001118 | 0.53525042 | 0.77504588 |
| TIMM23B | 155.75631 | 0.163722249 | 0.33893014 | 0.48305603 | 0.62905595 | 0.83372327 |
| LINC00843 | 44.709963 | 0.135535822 | 0.24570962 | 0.55160975 | 0.58121575 | 0.80559481 |
| AGAP6 | 61.4504308 | 0.004407956 | 0.24132206 | 0.01826586 | 0.98542676 | 0.99668665 |
| WASHC2A | 64.0512059 | -0.427684398 | 0.37840652 | -1.1302247 | 0.25838155 | 0.54540257 |
| SGMS1 | 705.69971 | -0.279004432 | 0.16601058 | -1.6806425 | 0.09283237 | 0.31724627 |
| PRKG1 | 551.349902 | 1.098766784 | 0.49212775 | 2.23268611 | 0.02556965 | 0.1551184 |
| IPMK | 64.8823865 | -0.494006187 | 0.32823526 | -1.5050369 | 0.13231458 | 0.38326016 |
| CISD1 | 119.206907 | -0.355217716 | 0.25609276 | -1.3870666 | 0.16542146 | 0.43384207 |
| UBE2D1 | 50.5071196 | -0.149715876 | 0.31331545 | -0.4778439 | 0.63276132 | 0.83561475 |
| TFAM | 40.7385081 | -0.619954877 | 0.27593205 | -2.2467665 | 0.02465496 | 0.15217128 |
| BICC1 | 5875.93154 | 0.744579654 | 0.27910459 | 2.66774423 | 0.00763624 | 0.07533214 |
| PHYHIPL | 530.593774 | 1.292908595 | 0.5441446 | 2.37603866 | 0.01749963 | 0.12481679 |
| SLC16A9 | 67.5406947 | -0.533808558 | 0.80893153 | -0.6598934 | 0.50932225 | 0.75816595 |
| CCDC6 | 168.818899 | -0.43762221 | 0.19609166 | -2.2317227 | 0.02563329 | 0.1551184 |
| ANK3 | 1271.41261 | 0.543766768 | 0.27430737 | 1.98232646 | 0.04744271 | 0.21765027 |
| RHOBTB1 | 43.8644482 | -0.307939006 | 0.42430947 | -0.7257415 | 0.46799723 | 0.72581883 |
| CABCOCO1 | 62.9785238 | 0.780784829 | 0.49774222 | 1.568653 | 0.11672882 | 0.35989342 |
| ARID5B | 564.403237 | 0.209374402 | 0.23539987 | 0.88944143 | 0.37376589 | 0.65253555 |
| JMJD1C | 1180.51466 | 0.170406346 | 0.15453432 | 1.10270874 | 0.27015367 | 0.55735559 |
| REEP3 | 169.495566 | 0.21061526 | 0.19453876 | 1.08263907 | 0.27896866 | 0.56636829 |
| LINC01515 | 142.690958 | -0.08531011 | 0.27257334 | -0.3129804 | 0.75429556 | 0.89810084 |
| SIRT1 | 78.986396 | 0.14964324 | 0.25004358 | 0.59846865 | 0.54952727 | 0.78483998 |
| HERC4 | 602.874033 | -0.020988964 | 0.19503112 | -0.1076185 | 0.91429829 | 0.9686812 |
| PBLD | 51.1112585 | 0.045996609 | 0.39118327 | 0.11758327 | 0.90639786 | 0.96621245 |
| HNRNPH3 | 186.444815 | 0.644968418 | 0.20939382 | 3.08016936 | 0.00206883 | 0.03238123 |
| RUFY2 | 163.987185 | 0.462696143 | 0.16392038 | 2.82268839 | 0.00476228 | 0.05604166 |
| SLC25A16 | 161.84571 | 0.029599528 | 0.24641326 | 0.12012149 | 0.90438691 | 0.96543237 |
| TET1 | 34.2075211 | 0.941180971 | 0.46414627 | 2.02776805 | 0.04258393 | 0.20477482 |
| CCAR1 | 257.819445 | -0.00655703 | 0.20417035 | -0.0321155 | 0.97437995 | 0.99326197 |
| DDX50 | 104.258404 | -0.129155713 | 0.19927957 | -0.6481132 | 0.51691176 | 0.76223795 |
| DDX21 | 143.678799 | -1.458652742 | 0.44218898 | -3.298709 | 0.00097131 | 0.02044865 |
| KIF1BP | 49.4850472 | -0.264039828 | 0.2413976 | -1.0937964 | 0.27404427 | 0.5612722 |
| VPS26A | 123.124308 | 0.219320072 | 0.22638072 | 0.96881075 | 0.33263962 | 0.61719981 |
| SUPV3L1 | 51.2504632 | 0.041307063 | 0.19125172 | 0.21598269 | 0.82900124 | 0.93373939 |
| HKDC1 | 156.222259 | -0.26008589 | 0.35552676 | -0.7315508 | 0.46444278 | 0.7222236 |
| HK1 | 215.915155 | -0.206885653 | 0.38203661 | -0.5415336 | 0.58813986 | 0.81096757 |
| TSPAN15 | 38.6659366 | -1.140232127 | 0.51803107 | -2.2010883 | 0.02772977 | 0.16127718 |
| H2AFY2 | 77.6580385 | 0.037397514 | 0.25787162 | 0.14502377 | 0.8846921 | 0.95767495 |
| AIFM2 | 35.4931573 | -0.208143412 | 0.26414271 | -0.787996 | 0.43069902 | 0.69655005 |
| SAR1A | 82.4156178 | -0.016441502 | 0.20642172 | -0.0796501 | 0.93651558 | 0.97871293 |
| PPA1 | 148.862103 | -1.670065294 | 0.46727886 | -3.5740228 | 0.00035154 | 0.01058926 |
| EIF4EBP2 | 77.5832919 | -0.208815204 | 0.22344397 | -0.9345305 | 0.35003031 | 0.63286907 |
| SGPL1 | 139.797031 | 0.347491362 | 0.1858254 | 1.86998849 | 0.06148542 | 0.25100302 |
| PCBD1 | 79.7809741 | -0.516006148 | 0.43366586 | -1.1898703 | 0.23409736 | 0.51656729 |
| PSAP | 365.233044 | -0.473067593 | 0.28206283 | -1.6771709 | 0.09350906 | 0.31835715 |
| CHST3 | 51.5718688 | -1.778598532 | 0.40757404 | -4.3638661 | 1.28E-05 | 0.00107372 |
| SPOCK2 | 173.161872 | 0.824466215 | 0.64219298 | 1.28382939 | 0.19920166 | 0.4782544 |
| ASCC1 | 136.995175 | 0.411590407 | 0.25111757 | 1.63903471 | 0.10120603 | 0.33234576 |
| ANAPC16 | 338.033416 | 0.384469488 | 0.21993825 | 1.74807923 | 0.08045031 | 0.29481006 |
| DDIT4 | 138.470481 | -2.099629572 | 1.12905029 | -1.8596422 | 0.06293616 | 0.25490103 |
| DNAJB12 | 42.264508 | -0.210802154 | 0.28668184 | -0.7353174 | 0.4621462 | 0.72048708 |
| MICU1 | 328.729588 | 0.157790848 | 0.18990801 | 0.83088044 | 0.40604118 | 0.67957905 |
| MCU | 369.073219 | 0.858748492 | 0.27645366 | 3.10630177 | 0.00189443 | 0.03090405 |
| P4HA1 | 251.820523 | -0.335353099 | 0.51856606 | -0.6466931 | 0.51783059 | 0.76300557 |
| ECD | 65.4304876 | 0.525941971 | 0.25224241 | 2.08506557 | 0.03706335 | 0.19087625 |
| FAM149B1 | 103.012688 | 0.288084837 | 0.26369765 | 1.09248163 | 0.27462146 | 0.56187466 |
| DNAJC9 | 93.7763433 | 0.144359589 | 0.2649849 | 0.5447842 | 0.58590195 | 0.80945221 |
| MRPS16 | 59.5187538 | -0.988462558 | 0.27552959 | -3.5875005 | 0.00033386 | 0.01030168 |
| CFAP70 | 146.318521 | 0.741983468 | 0.58712728 | 1.26375234 | 0.20631893 | 0.48621411 |
| ANXA7 | 210.655343 | -0.15768671 | 0.20191129 | -0.7809702 | 0.43481999 | 0.69978843 |
| PPP3CB | 227.491121 | 0.364149157 | 0.1547393 | 2.35330757 | 0.01860724 | 0.12915243 |
| USP54 | 264.639399 | 0.64686154 | 0.20052728 | 3.22580322 | 0.0012562 | 0.02411445 |
| AGAP5 | 25.7659617 | 0.466048973 | 0.29429033 | 1.58363674 | 0.11327641 | 0.35267249 |
| SEC24C | 58.5647686 | -0.176406315 | 0.20237378 | -0.8716856 | 0.38337989 | 0.66001519 |
| FUT11 | 60.5111226 | -0.225826575 | 0.3833424 | -0.5890989 | 0.55579495 | 0.78784845 |
| CHCHD1 | 45.1354947 | -0.262121873 | 0.41669511 | -0.6290496 | 0.52931662 | 0.77040597 |
| ZSWIM8 | 87.9157475 | -0.523727659 | 0.21910543 | -2.3902998 | 0.01683463 | 0.12184079 |
| CAMK2G | 76.8535259 | 0.135011615 | 0.3520001 | 0.38355562 | 0.70130784 | 0.87200065 |
| VCL | 436.006162 | 0.866566232 | 0.43746489 | 1.9808818 | 0.04760453 | 0.21810285 |

| | | | | | | |
|---|---|---|---|---|---|---|
| AP3M1 | 43.5590954 | 0.02954121 | 0.25078489 | 0.11779501 | 0.90623008 | 0.96621245 |
| ADK | 739.102633 | 0.24875259 | 0.33438783 | 0.74390443 | 0.4569343 | 0.71624507 |
| KAT6B | 263.207579 | 0.506545086 | 0.28541632 | 1.77475866 | 0.07593775 | 0.2849227 |
| SAMD8 | 106.280839 | 0.055019928 | 0.1590376 | 0.34595547 | 0.72937618 | 0.88653304 |
| VDAC2 | 79.7346301 | -0.378860307 | 0.34055889 | -1.1124664 | 0.26593769 | 0.55384374 |
| COMTD1 | 29.2987575 | -0.419245414 | 0.39611715 | -1.0583874 | 0.28987884 | 0.57668732 |
| LRMDA | 1561.24377 | -1.05505504 | 0.75431181 | -1.3986988 | 0.16190332 | 0.4283668 |
| KCNMA1 | 77.5592302 | -0.597033935 | 0.83229351 | -0.7173358 | 0.47316692 | 0.73010083 |
| DLG5 | 93.7782114 | -0.099994127 | 0.22520197 | -0.4440198 | 0.65702829 | 0.85164733 |
| POLR3A | 55.5469885 | -0.299769632 | 0.20002258 | -1.4986789 | 0.13395694 | 0.38545316 |
| RPS24 | 716.842668 | -0.306924351 | 0.46575688 | -0.6589797 | 0.50990878 | 0.75867242 |
| ZMIZ1 | 373.290851 | 0.020137554 | 0.34215902 | 0.05885437 | 0.9530681 | 0.98368111 |
| NUTM2B-AS | 317.32996 | -0.168546819 | 0.22222355 | -0.7584562 | 0.44817795 | 0.70956008 |
| TMEM254 | 33.2717632 | 0.461768378 | 0.26710267 | 1.72880482 | 0.08384403 | 0.30203879 |
| ANXA11 | 237.188808 | -0.219656167 | 0.1799935 | -1.2203561 | 0.22232991 | 0.5039513 |
| FAM213A | 81.6440011 | -0.613180175 | 0.29558926 | -2.0744332 | 0.0380391 | 0.19361308 |
| TSPAN14 | 160.815281 | -1.285142583 | 0.24704014 | -5.2021611 | 1.97E-07 | 5.10E-05 |
| NRG3 | 2081.61391 | -1.405697043 | 1.14937534 | -1.2230096 | 0.22132608 | 0.50310453 |
| GHITM | 94.697044 | -0.699152669 | 0.25348133 | -2.7582018 | 0.00581203 | 0.06377255 |
| CCSER2 | 258.478308 | 0.251017407 | 0.17570877 | 1.4285992 | 0.15311946 | 0.41574018 |
| WAPL | 248.738502 | 0.065830933 | 0.14851031 | 0.44327517 | 0.65756671 | 0.85176897 |
| BMPR1A | 516.216948 | 0.190714294 | 0.16766631 | 1.13746339 | 0.25534462 | 0.54219288 |
| ADIRF | 181.271855 | -0.755171444 | 0.95897998 | -0.7874736 | 0.43100466 | 0.69671579 |
| GLUD1 | 179.014789 | -0.675096007 | 0.27945219 | -2.4157835 | 0.01570139 | 0.11641149 |
| SHLD2 | 292.540104 | 0.27108521 | 0.27726502 | 0.97771154 | 0.32821701 | 0.61394673 |
| NUTM2A-AS | 348.068364 | 0.261884281 | 0.24414589 | 1.07265488 | 0.28342599 | 0.57084461 |
| MINPP1 | 38.3405756 | -0.367137527 | 0.38489698 | -0.9538592 | 0.34015492 | 0.62492213 |
| PAPSS2 | 96.519561 | -0.371661005 | 0.38839086 | -0.9569252 | 0.33860501 | 0.62345706 |
| ATAD1 | 148.944357 | 0.132657615 | 0.17138523 | 0.7740318 | 0.43891198 | 0.70279675 |
| PTEN | 431.994995 | 0.075356252 | 0.19348351 | 0.38947118 | 0.69692763 | 0.87095496 |
| RNLS | 180.28053 | 0.823084711 | 0.43317868 | 1.90010438 | 0.05741942 | 0.24181314 |
| FAS | 42.5507382 | -1.069780732 | 0.32521196 | -3.2894877 | 0.0010037 | 0.02077084 |
| LIPA | 117.809052 | -0.225033083 | 0.38388748 | -0.5861954 | 0.5577442 | 0.78891584 |
| IFIT3 | 129.871263 | -2.44348197 | 1.25226919 | -1.9512434 | 0.0510281 | 0.22682567 |
| IFIT5 | 30.0296944 | -0.992838457 | 0.45290049 | -2.1921779 | 0.02836666 | 0.16346515 |
| SLC16A12 | 251.845601 | -0.278815903 | 0.70722424 | -0.3942397 | 0.69340405 | 0.86938709 |
| PANK1 | 33.469479 | 1.073550526 | 0.41013111 | 2.6175789 | 0.0088556 | 0.08382194 |
| RPP30 | 30.9530457 | 0.290505458 | 0.24498419 | 1.18581306 | 0.23569613 | 0.51901914 |
| PCGF5 | 313.675049 | 0.225304528 | 0.18450543 | 1.22112679 | 0.22203802 | 0.50375143 |
| HECTD2 | 146.578912 | 0.495211776 | 0.24167266 | 2.04910139 | 0.0404522 | 0.19973796 |
| PPP1R3C | 61.4616954 | -0.524452211 | 0.83936012 | -0.6248238 | 0.53208669 | 0.77271958 |
| TNKS2 | 252.309834 | 0.23582205 | 0.17670933 | 1.33451952 | 0.18203365 | 0.45515541 |
| BTAF1 | 233.191215 | 0.364448289 | 0.2370628 | 1.5373491 | 0.12420784 | 0.37041562 |
| CPEB3 | 239.512528 | 0.486865721 | 0.35488478 | 1.37189799 | 0.1700952 | 0.44024369 |
| 5-Mar | 115.155294 | 0.444098852 | 0.15878168 | 2.79691485 | 0.00515931 | 0.059329 |
| IDE | 120.551206 | -0.030590132 | 0.15240091 | -0.2007215 | 0.84091638 | 0.94053005 |
| EXOC6 | 175.208245 | 0.301595377 | 0.23341434 | 1.29210303 | 0.19632147 | 0.4753699 |
| MYOF | 366.632296 | 0.281718331 | 0.31009001 | 0.90850502 | 0.36361146 | 0.64381408 |
| RBP4 | 127.603233 | -1.977532607 | 0.65178571 | -3.0340226 | 0.00241316 | 0.03558826 |
| FRA10AC1 | 104.506432 | 0.650725781 | 0.21031927 | 3.09399025 | 0.00197484 | 0.03151251 |
| PLCE1 | 140.710854 | -0.028328528 | 1.09867303 | -0.0257843 | 0.97942938 | 0.99557142 |
| NOC3L | 51.401684 | -0.194876298 | 0.32434672 | -0.6008271 | 0.54795515 | 0.7834123 |
| TBC1D12 | 103.720584 | 0.184583491 | 0.22077543 | 0.83606897 | 0.4031161 | 0.67708038 |
| HELLS | 48.8940027 | 0.218641055 | 0.39462626 | 0.55404589 | 0.57954744 | 0.80456304 |
| PDLIM1 | 48.1092814 | -2.251207209 | 0.7746809 | -2.90598 | 0.00366105 | 0.04675891 |
| SORBS1 | 151.118788 | -1.037414228 | 0.28243714 | -3.6730801 | 0.00023964 | 0.0083651 |
| ALDH18A1 | 69.888052 | -0.773219498 | 0.28204271 | -2.7414979 | 0.00611597 | 0.06580904 |
| TCTN3 | 75.8336795 | 0.030996096 | 0.22394872 | 0.13840711 | 0.88991868 | 0.95918866 |
| ENTPD1-AS1 | 339.803101 | 0.046035435 | 0.33287111 | 0.13829808 | 0.89000484 | 0.95918866 |
| AL365273.2 | 27.4886157 | 0.317168114 | 0.42939952 | 0.73863173 | 0.46013065 | 0.71894055 |
| ZNF518A | 291.37577 | 0.06778228 | 0.28764473 | 0.23564583 | 0.81370751 | 0.92720494 |
| BLNK | 98.3990016 | -0.002452155 | 0.5132122 | -0.0047781 | 0.99618768 | 0.99921275 |
| TLL2 | 38.7333596 | -1.279490129 | 0.77452041 | -1.6519773 | 0.09853918 | 0.32808718 |
| TM9SF3 | 319.526792 | -0.051310553 | 0.12760277 | -0.4021116 | 0.68760189 | 0.86744877 |
| PIK3AP1 | 217.039976 | -0.870320776 | 0.56523542 | -1.5397492 | 0.1236215 | 0.36947593 |
| LCOR | 603.448219 | 0.320024428 | 0.17875762 | 1.79027017 | 0.07341049 | 0.27937985 |
| RRP12 | 59.9063504 | -0.232525801 | 0.31573788 | -0.736452 | 0.46145565 | 0.71979868 |
| PGAM1 | 127.372286 | -0.80812281 | 0.34030475 | -2.3747033 | 0.01756306 | 0.12490692 |
| EXOSC1 | 47.2692079 | -0.002766247 | 0.21048872 | -0.013142 | 0.98951449 | 0.99683162 |
| ZDHHC16 | 28.4472324 | -0.031887033 | 0.28924271 | -0.1102432 | 0.91221653 | 0.96844354 |
| MMS19 | 122.20761 | 0.058117594 | 0.17573164 | 0.33071787 | 0.7408576 | 0.89253455 |
| UBTD1 | 115.791141 | 0.606066112 | 0.31970922 | 1.89567916 | 0.05800248 | 0.24313032 |
| MORN4 | 42.9269155 | -0.177807859 | 0.30523606 | -0.5825257 | 0.56021261 | 0.78997153 |
| ZFYVE27 | 54.216189 | -0.362581574 | 0.25715733 | -1.4099601 | 0.15855147 | 0.42411742 |
| CRTAC1 | 124.583194 | -4.279199539 | 1.58215757 | -2.7046608 | 0.00683742 | 0.07019895 |
| R3HCC1L | 146.061181 | -0.139388307 | 0.20218609 | -0.689406 | 0.49056779 | 0.74294844 |
| LOXL4 | 48.0798353 | 0.712474059 | 0.4661712 | 1.5283528 | 0.12642497 | 0.37394935 |
| HPS1 | 44.8047886 | -0.636874466 | 0.27386398 | -2.3255138 | 0.0200445 | 0.13480227 |
| CNNM1 | 35.8428781 | 1.399746849 | 0.47935373 | 2.92007083 | 0.00349952 | 0.04544458 |
| GOT1 | 132.900733 | 0.234459836 | 0.31535093 | 0.74348865 | 0.45718589 | 0.71628781 |
| SLC25A28 | 53.5052934 | -0.769199423 | 0.52378335 | -1.468545 | 0.14195624 | 0.39830078 |
| CUTC | 78.4844831 | -0.349712901 | 0.25842707 | -1.3532363 | 0.17598014 | 0.44807891 |
| COX15 | 53.6350415 | 0.133620518 | 0.25849885 | 0.51690953 | 0.60521931 | 0.82056119 |
| DNMBP | 140.04997 | -0.123962919 | 0.45319811 | -0.2735292 | 0.78444644 | 0.91212179 |
| ERLIN1 | 44.1138272 | 0.138398076 | 0.28473046 | 0.48606699 | 0.62691967 | 0.83221501 |
| CHUK | 49.6143597 | 0.473619594 | 0.37464711 | 1.26417524 | 0.20616714 | 0.48606325 |
| CWF19L1 | 62.0240898 | 0.45607897 | 0.24136968 | 1.8895454 | 0.05881879 | 0.24454867 |
| BLOC1S2 | 30.5125412 | -0.730145408 | 0.24553643 | -2.9736745 | 0.00294257 | 0.04015226 |
| HIF1AN | 53.3677171 | -0.063407338 | 0.23817676 | -0.2662197 | 0.79007003 | 0.9154554 |
| PAX2 | 707.891234 | 0.796305703 | 0.32068137 | 2.48316796 | 0.01302197 | 0.10449615 |
| SLF2 | 126.418553 | -0.282234162 | 0.29376352 | -0.9607529 | 0.33667641 | 0.62147307 |
| MRPL43 | 42.1525574 | -0.063457664 | 0.32475784 | -0.1953999 | 0.84507986 | 0.94193856 |
| LZTS2 | 64.0021674 | -0.895739669 | 0.32103772 | -2.7901384 | 0.00526855 | 0.06000886 |
| BTRC | 286.963952 | 0.518612772 | 0.21083987 | 2.45974714 | 0.01390349 | 0.10898655 |
| DPCD | 42.0227835 | -0.205147176 | 0.37170706 | -0.5519055 | 0.58101309 | 0.80543045 |
| POLL | 40.7859889 | -0.213829323 | 0.27209624 | -0.7858592 | 0.43195 | 0.69731529 |
| FBXW4 | 250.926507 | 0.15915151 | 0.18937373 | 0.84040966 | 0.40067873 | 0.67524659 |
| OGA | 438.497607 | 0.03412279 | 0.25285238 | 0.13495143 | 0.89265028 | 0.95976294 |
| ARMH3 | 261.508672 | 0.115242997 | 0.1442808 | 0.79874107 | 0.42444057 | 0.69133076 |
| LDB1 | 32.2562241 | 0.031749022 | 0.39404634 | 0.0805718 | 0.9357825 | 0.97837377 |
| PPRC1 | 39.4109805 | -0.904167151 | 0.49788723 | -1.8160079 | 0.06936916 | 0.26978773 |
| NOLC1 | 45.9800846 | -1.084079399 | 0.41863602 | -2.5895512 | 0.00961011 | 0.08782595 |
| GBF1 | 356.509787 | -0.039698322 | 0.13193458 | -0.300894 | 0.76349534 | 0.90234693 |
| NFKB2 | 42.2157818 | -1.031898729 | 0.57125723 | -1.8063644 | 0.07086144 | 0.27348175 |
| FBXL15 | 30.447218 | -0.086876802 | 0.44363928 | -0.1958276 | 0.84474513 | 0.94193856 |
| CUEDC2 | 45.745118 | -0.229626369 | 0.25602861 | -0.8968778 | 0.36978415 | 0.6497738 |
| ACTR1A | 49.2975527 | -0.275913478 | 0.26568919 | -1.0384821 | 0.29904564 | 0.58639882 |
| SUFU | 150.791846 | 0.090316195 | 0.35963018 | 0.25113631 | 0.80170872 | 0.92109986 |
| TRIM8 | 95.1469455 | -0.534186394 | 0.30399323 | -1.7572312 | 0.07887841 | 0.29099563 |
| ARL3 | 127.982513 | 0.141965909 | 0.22302489 | 0.63654737 | 0.52441972 | 0.76611507 |
| SFXN2 | 38.957545 | 0.474224781 | 0.40944736 | 1.15820696 | 0.24677959 | 0.53365473 |
| WBP1L | 162.491494 | -0.278677874 | 0.31447819 | -0.8861596 | 0.37553152 | 0.65403934 |
| CNNM2 | 187.268626 | 0.743203773 | 0.37959736 | 1.95787393 | 0.05024481 | 0.2251146 |

| | | | | | | |
|---|---|---|---|---|---|---|
| NT5C2 | 443.029618 | -0.271456344 | 0.19810824 | -1.3702425 | 0.1706112 | 0.4409518 |
| PCGF6 | 51.0270131 | 0.390260527 | 0.22598979 | 1.72689449 | 0.08418661 | 0.30211489 |
| ATP5MD | 56.7300152 | -0.149357728 | 0.28859827 | -0.5175282 | 0.60478752 | 0.82048068 |
| PDCD11 | 48.8430023 | -0.686851538 | 0.30440707 | -2.2563587 | 0.02404817 | 0.15036388 |
| SH3PXD2A | 61.6005599 | 0.604574621 | 0.23588877 | 2.56296485 | 0.01037826 | 0.09137253 |
| STN1 | 35.9595075 | -1.074474256 | 0.36371424 | -2.9541716 | 0.0031351 | 0.04206035 |
| SLK | 148.336283 | -0.129458864 | 0.17409768 | -0.743599 | 0.4571191 | 0.71628781 |
| CFAP43 | 173.604131 | 0.340723922 | 0.56169581 | 0.60659865 | 0.54411729 | 0.78027954 |
| GSTO1 | 92.8842925 | -1.378977938 | 0.39690283 | -3.4743465 | 0.0005121 | 0.01366406 |
| GSTO2 | 110.780456 | 0.309870533 | 0.3019924 | 1.0260872 | 0.30485048 | 0.59267272 |
| SORCS1 | 66.0723946 | 0.883176011 | 0.54511551 | 1.62016307 | 0.10519725 | 0.33906316 |
| LINC01435 | 1600.89114 | 1.315894991 | 0.5647935 | 2.32986922 | 0.01981306 | 0.13393743 |
| AL353740.1 | 56.8686587 | 1.797810832 | 0.83364802 | 2.15655862 | 0.03104007 | 0.172267 |
| XPNPEP1 | 83.9659681 | -0.284187059 | 0.27247986 | -1.0429654 | 0.29696433 | 0.58520063 |
| ADD3 | 309.484424 | 0.795784619 | 0.29898342 | 2.66163457 | 0.00777623 | 0.07624203 |
| MXI1 | 170.486619 | -0.37853178 | 0.34611899 | -1.0936464 | 0.2741101 | 0.56128701 |
| SMNDC1 | 50.0770297 | 0.154198704 | 0.25972133 | 0.59370828 | 0.55270724 | 0.78613936 |
| SMC3 | 99.9863314 | 0.148237895 | 0.2460253 | 0.60253111 | 0.54682066 | 0.78253724 |
| PDCD4 | 135.744053 | 0.397686015 | 0.29562902 | 1.34521982 | 0.17855426 | 0.45020493 |
| BBIP1 | 63.9070504 | 0.275311469 | 0.21806479 | 1.26252143 | 0.20676122 | 0.48621411 |
| SHOC2 | 195.343354 | 0.280769457 | 0.18450962 | 1.5217063 | 0.12808269 | 0.37588973 |
| ZDHHC6 | 52.3092631 | -0.007085728 | 0.24101022 | -0.0294001 | 0.97654548 | 0.99438056 |
| VTI1A | 561.193798 | -0.217733589 | 0.15956724 | -1.3645256 | 0.1724022 | 0.44333978 |
| TCF7L2 | 537.799535 | 0.047804406 | 0.28880706 | 0.16552368 | 0.86853182 | 0.95103638 |
| CASP7 | 52.1262294 | -0.478188475 | 0.39331469 | -1.215791 | 0.22406453 | 0.50617113 |
| NHLRC2 | 151.157836 | 0.456147588 | 0.13097844 | 3.4826157 | 0.00049654 | 0.0133982 |
| CCDC186 | 150.717347 | 0.154086778 | 0.20523966 | 0.75076511 | 0.45279403 | 0.71261515 |
| AFAP1L2 | 41.2927173 | 1.708227285 | 0.49562983 | 3.44657882 | 0.00056773 | 0.01434911 |
| ABLIM1 | 602.112696 | -0.146553615 | 0.29726702 | -0.4930033 | 0.62201027 | 0.82915174 |
| FAM160B1 | 90.6529565 | -0.159629717 | 0.30678674 | -0.520328 | 0.60283501 | 0.81954732 |
| TRUB1 | 49.3728211 | -0.239797966 | 0.23326484 | -1.0280073 | 0.30394637 | 0.59203936 |
| ATRNL1 | 242.246522 | -0.579615821 | 0.38926923 | -1.4889844 | 0.13649147 | 0.38831357 |
| HSPA12A | 291.312744 | 0.532546711 | 0.42408105 | 1.25576634 | 0.2092007 | 0.48959511 |
| ENO4 | 37.7061202 | 0.580982387 | 0.42241812 | 1.37537278 | 0.1690159 | 0.43863542 |
| SHTN1 | 299.309736 | -0.043678422 | 0.17820537 | -0.2451016 | 0.80637776 | 0.92263409 |
| PDZD8 | 253.970279 | -0.037079917 | 0.16685736 | -0.2222252 | 0.82413855 | 0.93217972 |
| EMX2OS | 270.814199 | 0.080327154 | 0.42581398 | 0.18864377 | 0.85037202 | 0.94435157 |
| EMX2 | 108.353707 | 0.293258973 | 0.42912311 | 0.68339124 | 0.49435966 | 0.74652374 |
| RAB11FIP2 | 246.22113 | 0.621336546 | 0.28919036 | 2.14853823 | 0.03167102 | 0.17395454 |
| CASC2 | 172.498563 | 0.231057383 | 0.41723715 | 0.55377951 | 0.57972975 | 0.80465327 |
| FAM204A | 117.543626 | 0.47209645 | 0.18927359 | 2.49425419 | 0.01262221 | 0.10255145 |
| CACUL1 | 202.368936 | -0.133720945 | 0.23721957 | -0.5637012 | 0.57295753 | 0.79981931 |
| EIF3A | 271.591706 | -0.473075544 | 0.28799483 | -1.6426529 | 0.10045477 | 0.33112739 |
| FAM45A | 48.0457183 | -0.362293543 | 0.22705787 | -1.5956 | 0.11057812 | 0.34758554 |
| SFXN4 | 52.4150021 | -0.239166747 | 0.27664989 | -0.8645105 | 0.38730749 | 0.66404483 |
| PRDX3 | 104.496236 | 0.094686259 | 0.42521731 | 0.22267734 | 0.82378664 | 0.93217972 |
| TIAL1 | 145.392828 | 0.423859177 | 0.16152844 | 2.6240529 | 0.00868903 | 0.08265363 |
| BAG3 | 43.3029658 | -0.883428698 | 0.28855683 | -3.0615414 | 0.00220201 | 0.03391159 |
| INPP5F | 111.312066 | -0.248804983 | 0.19168479 | -1.2979902 | 0.19429071 | 0.47284316 |
| MCMBP | 84.7587822 | -0.268379898 | 0.208438 | -1.2875766 | 0.19789337 | 0.47656532 |
| SEC23IP | 104.09529 | -0.160609048 | 0.20454141 | -0.7852153 | 0.43232734 | 0.69737124 |
| PLPP4 | 43.4291332 | 0.177299429 | 0.39966852 | 0.4436162 | 0.65732009 | 0.85167986 |
| WDR11-AS1 | 31.0981361 | 0.331398902 | 0.46716905 | 0.70937683 | 0.47809066 | 0.7341756 |
| WDR11 | 166.385933 | 0.169806718 | 0.14185503 | 1.197044 | 0.2312894 | 0.51432779 |
| FGFR2 | 364.636836 | 0.937259987 | 0.35047766 | 2.67423604 | 0.00748997 | 0.07451094 |
| ATE1 | 337.598766 | -0.007399849 | 0.11038565 | -0.0670363 | 0.94655278 | 0.98158791 |
| NSMCE4A | 31.6366387 | 0.357575923 | 0.36019763 | 0.99272147 | 0.32084571 | 0.60774788 |
| TACC2 | 201.181483 | 0.297648514 | 0.37233026 | 0.7994207 | 0.42404651 | 0.69092389 |
| PLEKHA1 | 706.238768 | -1.043515009 | 0.30291273 | -3.4449362 | 0.00057119 | 0.0143986 |
| HTRA1 | 30.4718597 | -1.214555312 | 0.47920386 | -2.5345274 | 0.01125991 | 0.09595969 |
| C10orf88 | 29.8686123 | -0.275332413 | 0.27370403 | -1.0059494 | 0.31443989 | 0.60132156 |
| IKZF5 | 37.517905 | 0.0699073 | 0.27172042 | 0.25727658 | 0.79696527 | 0.9188367 |
| ACADSB | 62.8091545 | 0.074727576 | 0.36214162 | 0.20634904 | 0.83651827 | 0.93774237 |
| BUB3 | 57.4861562 | -0.238759387 | 0.29244832 | -0.8164157 | 0.41426244 | 0.68449528 |
| CHST15 | 82.1368964 | -0.811753803 | 0.5383385 | -1.5078873 | 0.13158337 | 0.381746 |
| OAT | 28.4994086 | 0.076645184 | 0.40384751 | 0.18978744 | 0.8494757 | 0.94414047 |
| LHPP | 33.5894876 | 0.213283398 | 0.40818263 | 0.52251954 | 0.60130865 | 0.81829073 |
| ABRAXAS2 | 44.1025053 | -0.154513129 | 0.31498018 | -0.4905487 | 0.62374565 | 0.83030289 |
| ZRANB1 | 116.76113 | 0.39135786 | 0.15203683 | 2.574099 | 0.01005015 | 0.08977568 |
| CTBP2 | 641.007007 | -0.343860266 | 0.18439245 | -1.8648283 | 0.06220548 | 0.2529698 |
| EDRF1 | 70.1836972 | 0.141378302 | 0.21211488 | 0.66651762 | 0.50508031 | 0.75524917 |
| UROS | 41.8818224 | -0.217328572 | 0.28264121 | -0.7689203 | 0.44194061 | 0.7049707 |
| BCCIP | 77.9029819 | 0.009979016 | 0.25046199 | 0.03984244 | 0.96821874 | 0.99123492 |
| DHX32 | 187.521829 | -0.657761466 | 0.19148701 | -3.4350187 | 0.00059251 | 0.01482276 |
| FANK1 | 155.908132 | -0.053591321 | 0.28090791 | -0.190779 | 0.84869877 | 0.94366633 |
| DOCK1 | 979.79362 | 0.267558093 | 0.2286599 | 1.17011375 | 0.24195519 | 0.5268672 |
| FAM196A | 46.4927463 | 0.255296028 | 0.49320968 | 0.51762169 | 0.60472225 | 0.82048068 |
| PTPRE | 52.4981862 | -2.295071138 | 0.66141219 | -3.4699559 | 0.00052054 | 0.01373303 |
| MGMT | 280.640193 | 0.27655878 | 0.23182096 | 1.19298439 | 0.23287549 | 0.51589137 |
| GLRX3 | 107.211963 | -0.340382619 | 0.23719645 | -1.4350241 | 0.15128023 | 0.41285281 |
| PPP2R2D | 195.884886 | 0.216571391 | 0.19137155 | 1.13168017 | 0.25776892 | 0.54495001 |
| BNIP3 | 176.736499 | -0.873812661 | 0.54036656 | -1.6170739 | 0.10586233 | 0.34040123 |
| STK32C | 37.3244621 | -0.013384459 | 0.29686617 | -0.0450858 | 0.96403889 | 0.98902523 |
| LRRC27 | 38.7308887 | 0.133798147 | 0.37598055 | 0.35586454 | 0.72194201 | 0.88235193 |
| PWWP2B | 22.9469306 | -0.643670763 | 0.68308145 | -0.9423046 | 0.34603674 | 0.63016843 |
| INPP5A | 112.116523 | 0.084201779 | 0.29130534 | 0.2890499 | 0.77254319 | 0.90624314 |
| CFAP46 | 42.4425987 | -0.109439592 | 0.49718109 | -0.2201202 | 0.82577756 | 0.9325776 |
| TUBGCP2 | 120.619144 | 0.010556119 | 0.17976898 | 0.05872047 | 0.95317476 | 0.98368111 |
| ECHS1 | 107.025514 | -0.649853682 | 0.45791926 | -1.4191447 | 0.15585684 | 0.42019495 |
| SLC6A12 | 155.441485 | 1.782594187 | 0.64336085 | 2.7707533 | 0.00559268 | 0.06222098 |
| SLC6A13 | 82.189672 | 1.034348613 | 0.62765369 | 1.64796069 | 0.09936074 | 0.32944843 |
| KDM5A | 257.362027 | 0.050483773 | 0.16989317 | 0.2971501 | 0.76635192 | 0.90372312 |
| B4GALNT3 | 50.9680636 | -0.01829379 | 0.51504911 | -0.0355185 | 0.97166627 | 0.99175186 |
| WNK1 | 389.441583 | -0.179777 | 0.30826217 | -0.5831951 | 0.55976195 | 0.78994338 |
| RAD52 | 67.1557487 | 0.284817645 | 0.27749267 | 1.02639702 | 0.30470448 | 0.59263668 |
| ERC1 | 796.822905 | 0.112312269 | 0.17578966 | 0.63890144 | 0.52288705 | 0.76519323 |
| ADIPOR2 | 139.762209 | 0.185314844 | 0.21977002 | 0.84322167 | 0.39910448 | 0.67377896 |
| DCP1B | 77.2129795 | -0.053490301 | 0.28107047 | -0.1903092 | 0.84906683 | 0.94396601 |
| FKBP4 | 70.0667558 | -0.436643724 | 0.38789064 | -1.1256877 | 0.26029773 | 0.54703642 |
| ITFG2 | 104.465005 | 0.053124203 | 0.2041931 | 0.2601665 | 0.79473535 | 0.91764343 |
| TULP3 | 170.476172 | -0.132510541 | 0.25031293 | -0.5293795 | 0.59654219 | 0.81585918 |
| TEAD4 | 100.598885 | -0.363392147 | 0.36751692 | -0.9887767 | 0.32277243 | 0.60995011 |
| TSPAN9 | 314.693198 | -0.312672351 | 0.36401257 | -0.8589603 | 0.39036241 | 0.66594506 |
| PARP11 | 72.7750472 | 0.202187716 | 0.37926107 | 0.5331096 | 0.59395771 | 0.81441754 |
| CCND2 | 50.8070067 | -0.081067389 | 1.13704313 | -0.0712967 | 0.94316165 | 0.98112027 |
| TIGAR | 28.7141066 | -0.146454801 | 0.39848164 | -0.3675321 | 0.71322214 | 0.8783691 |
| C12orf4 | 54.1214041 | 0.46677506 | 0.20723652 | 2.25237834 | 0.02429837 | 0.15131898 |
| CD9 | 329.49816 | 0.504284392 | 0.41837212 | 1.20534895 | 0.22806862 | 0.51066249 |
| TNFRSF1A | 188.038678 | -1.639271914 | 0.40277623 | -4.0699321 | 4.70E-05 | 0.00292513 |
| SCNN1A | 238.612153 | -0.115748292 | 0.50661204 | -0.2284752 | 0.81927683 | 0.93050186 |
| LTBR | 105.945097 | -1.095777432 | 0.25860452 | -4.2372709 | 2.26E-05 | 0.00164188 |
| MRPL51 | 80.1834659 | -0.52643815 | 0.5575841 | -0.9441412 | 0.34509747 | 0.6298216 |
| NCAPD2 | 40.23528 | 0.213284318 | 0.35004511 | 0.60930524 | 0.54232214 | 0.77907974 |

| | | | | | | |
|---|---|---|---|---|---|---|
| GAPDH | 713.283486 | -1.963949436 | 0.57496148 | -3.415793 | 0.00063597 | 0.01548406 |
| CHD4 | 64.0413169 | -0.538714547 | 0.22645238 | -2.3789308 | 0.01736294 | 0.12439758 |
| AC006064.2 | 27.1784431 | -0.290073542 | 0.31917309 | -0.9088283 | 0.36344076 | 0.64363081 |
| ING4 | 41.7192193 | 0.281953125 | 0.23513414 | 1.19911606 | 0.23048282 | 0.51368983 |
| ZNF384 | 42.6519743 | -0.088448902 | 0.2227744 | -0.3970335 | 0.69134275 | 0.86884967 |
| COPS7A | 39.5526683 | -0.71630557 | 0.31991885 | -2.2390227 | 0.02515444 | 0.15393404 |
| MLF2 | 90.5539715 | -0.464933825 | 0.37682065 | -1.2338332 | 0.21726507 | 0.49932392 |
| PTMS | 153.281827 | -0.346134595 | 0.46503727 | -0.7443158 | 0.45668544 | 0.71603996 |
| USP5 | 34.6767308 | -0.668342479 | 0.32432625 | -2.0607104 | 0.03933067 | 0.19673552 |
| TPI1 | 204.359451 | -1.344113273 | 0.495158 | -2.7145139 | 0.00663731 | 0.06902242 |
| LRRC23 | 65.5107027 | -0.100020315 | 0.27811294 | -0.3596392 | 0.71911697 | 0.88155276 |
| ENO2 | 27.4324187 | -0.213037884 | 0.59880804 | -0.3557699 | 0.72201288 | 0.88235193 |
| ATN1 | 119.92736 | -0.076632046 | 0.17823732 | -0.4299439 | 0.66723647 | 0.85849001 |
| C12orf57 | 51.3346141 | -0.708946086 | 0.46336991 | -1.5299787 | 0.126022 | 0.37338842 |
| PTPN6 | 32.2510706 | -0.141055893 | 0.28962044 | -0.4870371 | 0.62623205 | 0.83180006 |
| PHB2 | 87.5149061 | -0.878667467 | 0.30048635 | -2.924151 | 0.00345397 | 0.04495325 |
| LPCAT3 | 204.732791 | -0.118503613 | 0.23430329 | -0.5057701 | 0.61301802 | 0.82461727 |
| C1S | 155.089437 | -1.01075005 | 0.66214101 | -1.5264876 | 0.12688847 | 0.37435927 |
| C1R | 171.407099 | -2.166728253 | 1.31099978 | -1.6527297 | 0.09838589 | 0.32780467 |
| C1RL | 230.323346 | -1.874062612 | 0.5363405 | -3.4941658 | 0.00047555 | 0.0129044 |
| C1RL-AS1 | 30.0904598 | -1.605227733 | 0.51721207 | -3.1036162 | 0.00191171 | 0.03107316 |
| RBP5 | 41.9169676 | -1.542140736 | 0.61349795 | -2.5136852 | 0.0119477 | 0.10009484 |
| CLSTN3 | 54.6875292 | -0.4320324 | 0.37627925 | -1.1481696 | 0.25089855 | 0.53838647 |
| PEX5 | 37.8917479 | -0.016381492 | 0.24512567 | -0.066829 | 0.94671787 | 0.98158791 |
| NECAP1 | 32.4606869 | -0.040131423 | 0.48438024 | -0.0828511 | 0.93396996 | 0.97768238 |
| LINC00937 | 111.889375 | 0.304140786 | 0.33662114 | 0.90351066 | 0.36625493 | 0.64634414 |
| RIMKLB | 178.073122 | -0.141746144 | 0.33731258 | -0.4202219 | 0.67432334 | 0.86115505 |
| PHC1 | 77.0905514 | 0.441269837 | 0.22009656 | 2.00489203 | 0.04497459 | 0.21106218 |
| M6PR | 106.18117 | -0.786620885 | 0.20506268 | -3.8360022 | 0.00012505 | 0.00549489 |
| CLEC2D | 90.883923 | 0.029741106 | 0.23124322 | 0.12861396 | 0.89766312 | 0.96193255 |
| GABARAPL1 | 233.927079 | 0.907179809 | 0.46961685 | 1.93174461 | 0.05339104 | 0.23183714 |
| KLRD1 | 152.768064 | 0.052663647 | 0.63294223 | 0.08320451 | 0.93368893 | 0.97767886 |
| MAGOHB | 25.3692222 | -0.104959925 | 0.29740984 | -0.3529134 | 0.72415335 | 0.88342651 |
| YBX3 | 365.183611 | -0.428052456 | 0.32030187 | -1.3364032 | 0.18141751 | 0.45456404 |
| SMIM10L1 | 28.5410411 | -0.748198745 | 0.33463917 | -2.2358373 | 0.02536243 | 0.15484175 |
| ETV6 | 665.820205 | -1.303899485 | 0.39057719 | -3.3383913 | 0.00084265 | 0.01842864 |
| BCL2L14 | 76.8932398 | -0.104439787 | 0.25782621 | -0.4050782 | 0.68542 | 0.86578038 |
| LRP6 | 264.659173 | 0.510676449 | 0.19508683 | 2.617688 | 0.00885277 | 0.08382194 |
| BORCS5 | 65.4583942 | -0.179823141 | 0.22015433 | -0.8168049 | 0.41403991 | 0.68449528 |
| DUSP16 | 190.277775 | -0.123130872 | 0.37357702 | -0.3295997 | 0.74170245 | 0.89253455 |
| CREBL2 | 67.7273909 | -0.184663027 | 0.22993423 | -0.8031124 | 0.42190978 | 0.69014238 |
| CDKN1B | 44.7511133 | -0.321557246 | 0.40170529 | -0.8004805 | 0.42343247 | 0.69062823 |
| GPRC5D-AS1 | 49.0657125 | 0.571549709 | 0.41969576 | 1.3618191 | 0.17325498 | 0.44449049 |
| HEBP1 | 74.8701292 | 0.054704567 | 0.34047662 | 0.16067055 | 0.87235288 | 0.95193455 |
| EMP1 | 152.814462 | -0.187344761 | 0.98141606 | -0.1908923 | 0.84860998 | 0.94366633 |
| ATF7IP | 466.770415 | 0.475175424 | 0.2404699 | 1.97602872 | 0.04815152 | 0.21931444 |
| H2AFJ | 120.008732 | -0.954476539 | 0.26765893 | -3.5660179 | 0.00036245 | 0.01081581 |
| WBP11 | 74.3718028 | -0.257571028 | 0.24155871 | -1.0662875 | 0.28629371 | 0.57346076 |
| MGP | 100.868483 | 0.113590162 | 0.76046361 | 0.14936962 | 0.88126198 | 0.95634765 |
| ERP27 | 75.8033854 | 0.170769244 | 1.15750976 | 0.14753158 | 0.88271245 | 0.95722522 |
| RERG | 536.670512 | -0.631051092 | 0.8068056 | -0.78216 | 0.43412053 | 0.69904779 |
| PTPRO | 95.597913 | -0.578784557 | 0.68614273 | -0.8435338 | 0.39892999 | 0.67377896 |
| EPS8 | 423.541465 | -1.262336145 | 0.54782444 | -2.3042713 | 0.02120742 | 0.13971517 |
| STRAP | 96.5214871 | -0.104678359 | 0.26532476 | -0.3945292 | 0.69319039 | 0.86938709 |
| DERA | 171.949277 | -0.502624361 | 0.14464796 | -3.4748114 | 0.00051121 | 0.01366406 |
| MGST1 | 201.365069 | -2.151090729 | 0.62853337 | -3.422397 | 0.00062072 | 0.01532433 |
| PLEKHA5 | 794.751077 | 0.462093909 | 0.396275 | 1.16609401 | 0.24357644 | 0.52907453 |
| AEBP2 | 350.367931 | 0.347817682 | 0.24232118 | 1.43535816 | 0.15118506 | 0.41271067 |
| PDE3A | 161.064282 | 1.135808617 | 1.26823959 | 0.8955789 | 0.37047771 | 0.65044098 |
| SLCO1A2 | 126.702262 | 5.117106921 | 0.84878734 | 6.02872674 | 1.65E-09 | 9.29E-07 |
| PYROXD1 | 55.8891318 | 0.299392082 | 0.35094635 | 0.85309929 | 0.39360424 | 0.6686322 |
| RECQL | 50.408912 | -0.099851332 | 0.25564925 | -0.3905794 | 0.69610816 | 0.87043654 |
| GOLT1B | 44.8026344 | 0.227506149 | 0.32493562 | 0.70015762 | 0.48382887 | 0.73727416 |
| SPX | 120.214195 | 0.854389394 | 0.57433303 | 1.48762015 | 0.1368511 | 0.38887471 |
| LDHB | 401.700592 | -0.444201901 | 0.3424635 | -1.2970781 | 0.19460432 | 0.47318227 |
| CMAS | 51.8973128 | -0.259589646 | 0.2908917 | -0.8923927 | 0.37218246 | 0.65197926 |
| C2CD5 | 184.226307 | 0.555215459 | 0.21957894 | 2.52854608 | 0.01145361 | 0.09717811 |
| ETNK1 | 334.121577 | 0.351157413 | 0.22252328 | 1.57807046 | 0.11454942 | 0.35579408 |
| AC084816.1 | 43.6858421 | -0.965762228 | 0.72606262 | -1.3301363 | 0.18347337 | 0.4570849 |
| CASC1 | 125.118339 | 0.613671512 | 0.51865852 | 1.18318987 | 0.23673391 | 0.52000641 |
| ETFRF1 | 56.429705 | 0.390117991 | 0.32643228 | 1.19509624 | 0.23204943 | 0.5151336 |
| KRAS | 105.838979 | 0.029396914 | 0.22766105 | 0.12912579 | 0.89725811 | 0.96175013 |
| RASSF8-AS1 | 81.5647847 | -0.077655275 | 0.28793301 | -0.2696991 | 0.78739175 | 0.91370543 |
| RASSF8 | 284.308458 | 0.165323879 | 0.27455878 | 0.60214385 | 0.54707839 | 0.78262603 |
| BHLHE41 | 66.1183219 | -0.725704803 | 0.63163278 | -1.1489347 | 0.25058292 | 0.53807079 |
| SSPN | 49.9504873 | -0.800571735 | 0.66344728 | -1.2066848 | 0.22755356 | 0.51041354 |
| ITPR2 | 305.418616 | -0.348165485 | 0.21627309 | -1.6098419 | 0.10743237 | 0.34291724 |
| INTS13 | 32.8475685 | -0.067860642 | 0.27631834 | -0.2455886 | 0.80600069 | 0.92253323 |
| FGFR1OP2 | 110.031021 | 0.57211712 | 0.27139584 | 2.10805415 | 0.03502631 | 0.1847551 |
| TM7SF3 | 129.558497 | -0.312153202 | 0.23333821 | -1.3377715 | 0.18097095 | 0.45368247 |
| MED21 | 52.1499237 | 0.217863783 | 0.3187973 | 0.68339281 | 0.49435866 | 0.74652374 |
| STK38L | 77.6397768 | 0.080821323 | 0.2184972 | 0.36989637 | 0.7114597 | 0.87748 |
| ARNTL2 | 51.2690867 | -1.283747234 | 0.53090279 | -2.4180457 | 0.01560412 | 0.11595952 |
| PPFIBP1 | 1003.73677 | 0.823373192 | 0.23786416 | 3.46152691 | 0.00053712 | 0.01394437 |
| MRPS35 | 80.4779302 | -0.174967651 | 0.22842205 | -0.7659841 | 0.44368579 | 0.70646046 |
| KLHL42 | 70.2001491 | -0.493930591 | 0.30389745 | -1.6253199 | 0.1040944 | 0.33686496 |
| CCDC91 | 500.073779 | 0.014764832 | 0.14222303 | 0.10381463 | 0.91731645 | 0.97040026 |
| FAR2 | 39.0675686 | -0.26388084 | 0.69171688 | -0.3814868 | 0.70284209 | 0.87284977 |
| AC009318.1 | 28.3690184 | -0.305207018 | 0.57119816 | -0.5343277 | 0.59311481 | 0.81351655 |
| ERGIC2 | 96.3208398 | 0.090185171 | 0.16988953 | 0.53084595 | 0.59552553 | 0.81525977 |
| IPO8 | 118.664452 | 0.488694591 | 0.2495729 | 1.95812362 | 0.05021551 | 0.2251146 |
| CAPRIN2 | 62.775318 | 0.571660422 | 0.25703175 | 2.22408482 | 0.02614274 | 0.15698891 |
| SINHCAF | 197.285726 | 0.121582331 | 0.20073517 | 0.60568524 | 0.54472378 | 0.78068896 |
| DENND5B | 231.512268 | -0.015303318 | 0.30748426 | -0.0497694 | 0.96030612 | 0.98697574 |
| AMN1 | 47.5202194 | 0.290479992 | 0.32027935 | 0.90695823 | 0.36442889 | 0.644313 |
| KIAA1551 | 95.6849932 | -0.021179437 | 0.26238359 | -0.0807194 | 0.93566514 | 0.97837377 |
| BICD1 | 764.8229 | 0.249276713 | 0.47517979 | 0.52459452 | 0.5998651 | 0.81763227 |
| FGD4 | 895.369845 | 0.473835072 | 0.34132156 | 1.38823657 | 0.16506502 | 0.43331264 |
| DNM1L | 197.023065 | -0.013832338 | 0.19308672 | -0.071638 | 0.94289003 | 0.98112027 |
| YARS2 | 73.3088888 | 0.059640216 | 0.20880872 | 0.28562129 | 0.7751682 | 0.90707747 |
| PKP2 | 52.3249586 | 0.357817921 | 0.31494815 | 1.13611693 | 0.25590763 | 0.54275365 |
| SYT10 | 38.870982 | 0.620574797 | 1.4672145 | 0.4229612 | 0.67232357 | 0.86015189 |
| ALG10B | 30.1108721 | 0.715486987 | 0.36556736 | 1.95719603 | 0.05032443 | 0.22536591 |
| CPNE8 | 384.105734 | -0.16837658 | 0.16271512 | -1.0347937 | 0.30076526 | 0.58788025 |
| KIF21A | 268.09286 | 0.184278228 | 0.30045223 | 0.61333619 | 0.53965407 | 0.77746559 |
| C12orf40 | 54.7115452 | 1.666688053 | 0.51359201 | 3.2451596 | 0.00117385 | 0.02328011 |
| SLC2A13 | 741.153977 | 1.106620795 | 0.48335822 | 2.28944236 | 0.02205366 | 0.1425647 |
| LINC02471 | 350.987275 | -0.406244175 | 1.00060856 | -0.4059971 | 0.68474473 | 0.86544 |
| AC079630.1 | 141.531303 | 0.31666571 | 0.68774138 | 0.46044301 | 0.64519827 | 0.84407726 |
| LRRK2 | 1039.70673 | 0.559623173 | 0.55715983 | 1.00442126 | 0.31517561 | 0.60205159 |
| GXYLT1 | 103.956349 | 0.422464941 | 0.21620593 | 1.95399332 | 0.05070201 | 0.22628104 |
| YAF2 | 199.899202 | 0.255117246 | 0.16980135 | 1.50244536 | 0.13298213 | 0.38380408 |
| PPHLN1 | 274.550404 | 0.111205598 | 0.12857107 | 0.86493484 | 0.38707455 | 0.66404483 |

| | | | | | | |
|---|---|---|---|---|---|---|
| ZCRB1 | 53.1321661 | 0.361299542 | 0.32060852 | 1.12691809 | 0.25977712 | 0.54640004 |
| PRICKLE1 | 180.240265 | -0.247825638 | 0.63870096 | -0.3880151 | 0.69800483 | 0.87161886 |
| PUS7L | 61.2055765 | 0.094833927 | 0.28858739 | 0.32861424 | 0.74244728 | 0.89253455 |
| IRAK4 | 53.2092642 | -0.034521644 | 0.19871635 | -0.1737232 | 0.86208299 | 0.9486379 |
| TWF1 | 33.4162809 | -0.694194107 | 0.35808603 | -1.9386238 | 0.05254716 | 0.22987695 |
| TMEM117 | 148.297576 | -0.193728985 | 0.39160659 | -0.4947031 | 0.62080973 | 0.82839623 |
| ANO6 | 283.636858 | -0.437056043 | 0.21676727 | -2.0162456 | 0.0437743 | 0.20758121 |
| AC008124.1 | 35.5282374 | 0.442914651 | 0.40175417 | 1.1024519 | 0.27026526 | 0.55746512 |
| ARID2 | 484.852572 | 0.428332965 | 0.13093496 | 3.27134136 | 0.00107039 | 0.02174571 |
| SCAF11 | 550.627366 | 0.152388778 | 0.113515 | 1.342455 | 0.17944851 | 0.45152017 |
| SLC38A1 | 364.561533 | -0.201538151 | 0.43565225 | -0.4626124 | 0.64364219 | 0.84343682 |
| SLC38A2 | 307.524378 | 0.393141814 | 0.39196914 | 1.00299174 | 0.31586485 | 0.60248295 |
| AC008014.1 | 651.871142 | 0.15205424 | 0.33705904 | 0.45112049 | 0.65190271 | 0.84789641 |
| PCED1B | 199.370842 | 0.027928719 | 0.42600994 | 0.06555885 | 0.94772906 | 0.98231681 |
| RPAP3 | 46.144975 | 0.503753363 | 0.33983299 | 1.4823557 | 0.13824567 | 0.39107241 |
| RAPGEF3 | 299.141346 | 0.691879406 | 0.623393 | 1.10986073 | 0.26705904 | 0.55453954 |
| SLC48A1 | 152.071113 | 0.269668712 | 0.62679738 | 0.43023267 | 0.6670264 | 0.85839746 |
| HDAC7 | 182.892461 | 0.266236454 | 0.26079418 | 1.02086809 | 0.30731694 | 0.59456464 |
| VDR | 26.7860154 | -0.395112252 | 0.49843529 | -0.7927052 | 0.4279496 | 0.69421324 |
| TMEM106C | 42.6186123 | -0.085662608 | 0.37488553 | -0.2285034 | 0.81925493 | 0.93050186 |
| SENP1 | 63.9145426 | 0.188740728 | 0.22220623 | 0.84939441 | 0.39566186 | 0.67068572 |
| PFKM | 26.864556 | -0.159602269 | 0.27253583 | -0.5856194 | 0.55813132 | 0.78914147 |
| ASB8 | 33.8771442 | -0.183664248 | 0.22900101 | -0.8020238 | 0.42253922 | 0.69043214 |
| ZNF641 | 28.2261923 | -0.273517934 | 0.30944605 | -0.8838954 | 0.37675269 | 0.65512273 |
| KANSL2 | 39.7420357 | 0.39881477 | 0.26405696 | 1.51033616 | 0.13095767 | 0.38152175 |
| CCNT1 | 76.9340052 | 0.324847968 | 0.1714377 | 1.89484556 | 0.05811287 | 0.24328193 |
| ADCY6 | 75.0491338 | 0.360313144 | 0.21410429 | 1.68288613 | 0.09239714 | 0.31666153 |
| DDX23 | 47.8371912 | -0.478965144 | 0.2115575 | -2.2639951 | 0.02357441 | 0.14836941 |
| CCDC65 | 55.0662616 | -0.326505727 | 0.30692935 | -1.0637814 | 0.28742775 | 0.57443572 |
| PRKAG1 | 61.7622572 | -0.202440543 | 0.23401371 | -0.8650798 | 0.38699497 | 0.66404483 |
| KMT2D | 57.2443977 | -0.10737246 | 0.30829851 | -0.3482743 | 0.72763416 | 0.88564265 |
| LMBR1L | 48.8466021 | -0.459283625 | 0.25716948 | -1.7859181 | 0.07411253 | 0.28082432 |
| TUBA1B | 222.379247 | -1.51542266 | 0.62463045 | -2.4261108 | 0.01526161 | 0.11471025 |
| TUBA1A | 46.1501814 | -0.689775238 | 0.70685539 | -0.9758364 | 0.32914553 | 0.61429707 |
| TUBA1C | 116.178321 | -1.830455334 | 0.58238183 | -3.1430502 | 0.00167197 | 0.02854928 |
| DNAJC22 | 49.1906134 | -0.074288101 | 0.46449906 | -0.1599316 | 0.87293492 | 0.95193455 |
| SPATS2 | 242.099863 | 0.210187411 | 0.24963932 | 0.84196437 | 0.3998079 | 0.67449099 |
| MCRS1 | 28.723857 | -0.338820934 | 0.27715994 | -1.2224744 | 0.22152827 | 0.50332527 |
| PRPF40B | 141.148557 | -0.182015558 | 0.23576877 | -0.7720088 | 0.44010923 | 0.7034987 |
| FMNL3 | 40.2411768 | -0.177637588 | 0.48754961 | -0.3643477 | 0.71559837 | 0.88009244 |
| TMBIM6 | 446.446697 | -0.389677658 | 0.33158251 | -1.1752057 | 0.23991243 | 0.52377013 |
| NCKAP5L | 46.6501566 | -0.016514648 | 0.2568152 | -0.0643056 | 0.94872692 | 0.98262487 |
| AQP2 | 45.4510232 | 1.933109943 | 0.56724529 | 3.4078907 | 0.00065467 | 0.01564098 |
| SMARCD1 | 49.8328962 | -0.28172593 | 0.35410121 | -0.7956085 | 0.42625963 | 0.69240987 |
| AC074032.1 | 142.72007 | 0.047009402 | 0.2613872 | 0.17984585 | 0.85727359 | 0.94638973 |
| CERS5 | 112.427763 | 0.010006834 | 0.2025272 | 0.04940983 | 0.9605927 | 0.98703297 |
| LIMA1 | 219.149655 | -0.137073926 | 0.17200401 | -0.7969229 | 0.42549583 | 0.69215319 |
| LARP4 | 222.355009 | -0.323969176 | 0.20228853 | -1.6015203 | 0.10926173 | 0.34587513 |
| DIP2B | 376.093059 | -0.218112907 | 0.27502527 | -0.793065 | 0.42773998 | 0.69399072 |
| ATF1 | 115.040285 | 0.074208344 | 0.20003952 | 0.37096842 | 0.71066105 | 0.87748 |
| METTL7A | 111.99948 | 0.0579868 | 0.52976144 | 0.10945833 | 0.91283897 | 0.96844354 |
| SLC11A2 | 203.851349 | -0.230987672 | 0.17118464 | -1.3493481 | 0.17722518 | 0.44893092 |
| LETMD1 | 102.397516 | 0.067750618 | 0.15988179 | 0.42375445 | 0.6717449 | 0.86013159 |
| CSRNP2 | 34.8504201 | -0.239714885 | 0.2613925 | -0.9170687 | 0.35910662 | 0.64001011 |
| TFCP2 | 251.331455 | 0.198225334 | 0.21592326 | 0.91803603 | 0.3586 | 0.64001011 |
| POU6F1 | 40.1550596 | 0.599247424 | 0.37158717 | 1.61266986 | 0.10681625 | 0.34182436 |
| DAZAP2 | 192.795131 | -0.300849254 | 0.22173845 | -1.3567753 | 0.17485262 | 0.44640545 |
| SMAGP | 25.7896132 | -0.768930372 | 0.48739542 | -1.5776315 | 0.11465029 | 0.35599194 |
| FIGNL2 | 67.6419236 | -0.184526344 | 0.46796804 | -0.394314 | 0.69334924 | 0.86938709 |
| ACVR1B | 227.537797 | -0.4734728 | 0.31400496 | -1.5078513 | 0.13159258 | 0.381746 |
| NR4A1 | 149.260451 | 5.49806187 | 1.06184088 | 5.17785852 | 2.24E-07 | 5.66E-05 |
| ATG101 | 27.7025529 | -0.772835989 | 0.47491627 | -1.62731 | 0.10367127 | 0.33583603 |
| KRT7 | 251.526259 | -1.249061881 | 0.60849172 | -2.0527179 | 0.04009994 | 0.19874641 |
| KRT8 | 402.665819 | -2.389850551 | 0.49659517 | -4.8124724 | 1.49E-06 | 0.0002341 |
| KRT18 | 475.597721 | -2.863778258 | 0.6044586 | -4.7377574 | 2.16E-06 | 0.00030579 |
| EIF4B | 235.903745 | -0.254203633 | 0.25336862 | -1.0032956 | 0.31571825 | 0.60232326 |
| AC068888.1 | 84.8142592 | -0.294574911 | 0.1941056 | -1.5176013 | 0.12911494 | 0.37764642 |
| TNS2 | 87.4911216 | -0.429130984 | 0.60742707 | -0.7064733 | 0.47989388 | 0.73503413 |
| SPRYD3 | 25.538408 | -1.002732466 | 0.33041361 | -3.0347795 | 0.00240712 | 0.03558826 |
| CSAD | 97.3899423 | 0.4908025 | 0.25442581 | 1.92905942 | 0.05372349 | 0.23285607 |
| ZNF740 | 25.7796037 | 0.178691073 | 0.24711583 | 0.72310655 | 0.46961443 | 0.72743153 |
| RARG | 40.32707 | 0.489254157 | 0.42515303 | 1.15077188 | 0.2498261 | 0.53692712 |
| PFDN5 | 196.618801 | -0.174572339 | 0.41625419 | -0.4193888 | 0.67493202 | 0.86115505 |
| SP1 | 130.749266 | -0.24833482 | 0.26807349 | -0.9263684 | 0.35425455 | 0.63606454 |
| PRR13 | 107.181001 | -0.704614743 | 0.35111448 | -2.0067949 | 0.04477151 | 0.21053819 |
| PCBP2 | 317.351281 | 0.125872747 | 0.12696726 | 0.99137955 | 0.32150028 | 0.60850646 |
| MAP3K12 | 36.0661839 | -0.398087581 | 0.38099545 | -1.0448618 | 0.29608686 | 0.58420198 |
| ATP5MC2 | 237.961883 | -0.183525403 | 0.36873093 | -0.4977218 | 0.61868018 | 0.82733103 |
| CALCOCO1 | 74.507462 | 0.116914866 | 0.41109361 | 0.28439962 | 0.77610415 | 0.90739676 |
| HOXC4 | 51.2939408 | 0.672239113 | 0.32276913 | 2.08272431 | 0.03727636 | 0.19135598 |
| AC023794.1 | 29.4299471 | 0.362176995 | 0.47444891 | 0.76336353 | 0.44524662 | 0.70741705 |
| SMUG1 | 30.3704091 | 0.12605719 | 0.30224151 | 0.41707438 | 0.676624 | 0.8613973 |
| LINC02381 | 63.1653517 | 0.462161753 | 0.52121169 | 0.88670642 | 0.37523698 | 0.65364522 |
| CBX5 | 215.197676 | 0.208091933 | 0.23158936 | 0.89853842 | 0.36889857 | 0.64873865 |
| HNRNPA1 | 179.395139 | -0.275833683 | 0.3509953 | -0.7858615 | 0.43194864 | 0.69731529 |
| COPZ1 | 101.952069 | -0.562656717 | 0.2005537 | -2.8055165 | 0.0050236 | 0.05825796 |
| PPP1R1A | 45.388496 | -1.287505409 | 0.53675266 | -2.3986941 | 0.01645366 | 0.12022087 |
| CD63 | 291.351817 | -1.533320266 | 0.48775204 | -3.1436471 | 0.00166857 | 0.02854928 |
| ORMDL2 | 32.2998208 | -0.938295866 | 0.39587855 | -2.3701609 | 0.01778034 | 0.12595648 |
| RAB5B | 85.1054406 | -0.062882864 | 0.21132068 | -0.2975708 | 0.76603077 | 0.90371395 |
| SUOX | 25.0896927 | -0.232867585 | 0.37930747 | -0.6139283 | 0.53926272 | 0.77736608 |
| RPS26 | 52.6461512 | -1.438054669 | 0.4164481 | -3.4531425 | 0.0005541 | 0.01407874 |
| ERBB3 | 45.0680194 | -0.925793148 | 0.44990629 | -2.0577466 | 0.03961446 | 0.19722814 |
| PA2G4 | 51.2015622 | -1.063952404 | 0.52130541 | -2.0409387 | 0.04125692 | 0.202045 |
| RPL41 | 993.180066 | -0.391566433 | 0.42370666 | -0.9241451 | 0.35541077 | 0.63706583 |
| ESYT1 | 65.4053221 | -0.356132038 | 0.26906792 | -1.3235767 | 0.18564366 | 0.45997946 |
| MYL6B | 24.9231645 | 0.324822507 | 0.32754775 | 0.99167985 | 0.32135373 | 0.60844973 |
| MYL6 | 422.860956 | 0.071257198 | 0.43579914 | 0.16350927 | 0.87011748 | 0.95125032 |
| SMARCC2 | 151.590489 | -0.204068638 | 0.15736119 | -1.2968168 | 0.19469421 | 0.47322402 |
| RNF41 | 48.3705449 | 0.020927639 | 0.31392949 | 0.0666635 | 0.94684959 | 0.98158791 |
| ANKRD52 | 41.5291779 | 0.2439575 | 0.21809906 | 1.1185628 | 0.2633267 | 0.55096925 |
| CS | 58.3829778 | -0.405473925 | 0.30212948 | -1.3420535 | 0.17957866 | 0.45172898 |
| PAN2 | 49.0170401 | 0.617369745 | 0.23847554 | 2.58881793 | 0.0096306 | 0.08782595 |
| STAT2 | 125.310221 | -0.389724595 | 0.51770262 | -0.7527963 | 0.45157235 | 0.71226932 |
| RBMS2 | 224.278652 | 1.145685454 | 0.26136372 | 4.38349076 | 1.17E-05 | 0.00101506 |
| BAZ2A | 152.953763 | -0.237065883 | 0.19256207 | -1.2311141 | 0.21828018 | 0.50052269 |
| ATP5F1B | 198.736726 | -0.636615677 | 0.30269884 | -2.1031322 | 0.03545422 | 0.18598901 |
| PTGES3 | 311.603619 | 0.071165848 | 0.13412151 | 0.53060726 | 0.59569096 | 0.81525977 |
| NACA | 243.590474 | -0.397748915 | 0.48892166 | -0.8135228 | 0.41591838 | 0.68549246 |
| NEMP1 | 36.2666652 | 0.290407223 | 0.28479218 | 1.01971627 | 0.30786304 | 0.59456464 |
| NAB2 | 26.8407627 | 0.090852207 | 0.39615554 | 0.22933469 | 0.8186088 | 0.93007397 |
| STAT6 | 150.846927 | -0.828356178 | 0.20294121 | -4.0817544 | 4.47E-05 | 0.00285001 |
| LRP1 | 40.4850213 | 0.459418145 | 0.48013457 | 0.95685288 | 0.33864151 | 0.62345706 |

| | | | | | | |
|---|---|---|---|---|---|---|
| R3HDM2 | 384.783013 | 0.189556436 | 0.21099927 | 0.89837485 | 0.36898574 | 0.64877284 |
| MARS | 44.556304 | -0.012514089 | 0.35075605 | -0.0356775 | 0.97153953 | 0.99175186 |
| DDIT3 | 47.7304201 | -0.864292785 | 0.61064241 | -1.4153828 | 0.15695628 | 0.42197143 |
| MBD6 | 43.7855977 | -0.033778131 | 0.24438535 | -0.1382167 | 0.89006917 | 0.95918866 |
| DCTN2 | 77.828706 | -0.18367785 | 0.22974348 | -0.799491 | 0.42400574 | 0.69092389 |
| PIP4K2C | 33.2709066 | -0.503320235 | 0.31233567 | -1.6114722 | 0.10707685 | 0.34246715 |
| DTX3 | 35.6388438 | 0.271268881 | 0.32338567 | 0.83884013 | 0.40155903 | 0.67562929 |
| OS9 | 108.456318 | -0.397106969 | 0.2165386 | -1.8338854 | 0.06667101 | 0.26332729 |
| TSPAN31 | 36.7542708 | -0.521760048 | 0.24806802 | -2.1032943 | 0.03544006 | 0.18598901 |
| CDK4 | 38.5436722 | -0.860331699 | 0.40432283 | -2.1278336 | 0.03335088 | 0.17927504 |
| 9-Mar | 35.1427481 | -0.368873721 | 0.29786549 | -1.2383903 | 0.21557139 | 0.49731142 |
| TSFM | 36.8454419 | -0.723205178 | 0.28898596 | -2.5025617 | 0.01232981 | 0.1012583 |
| CTDSP2 | 112.740492 | -0.344077267 | 0.23054209 | -1.4924705 | 0.13557585 | 0.38788633 |
| LRIG3 | 116.528097 | 0.542333687 | 0.39786721 | 1.36310225 | 0.17285029 | 0.44401526 |
| SLC16A7 | 250.954134 | -0.336655524 | 0.37996556 | -0.8860159 | 0.37560896 | 0.65405531 |
| FAM19A2 | 126.673846 | 0.017056885 | 0.43010023 | 0.03965793 | 0.96836584 | 0.99123492 |
| USP15 | 347.042997 | 0.145718576 | 0.21185298 | 0.68782878 | 0.49156061 | 0.74421671 |
| MON2 | 497.627468 | 0.271661309 | 0.12075907 | 2.2496142 | 0.02447345 | 0.15183364 |
| AC048341.1 | 97.3976565 | -0.071738809 | 0.47992914 | -0.1494779 | 0.88117653 | 0.95634765 |
| PPM1H | 542.609431 | 0.737808261 | 0.47615467 | 1.54951387 | 0.12125824 | 0.36572187 |
| DPY19L2 | 184.264938 | 0.655714721 | 0.37801853 | 1.73461 | 0.08280991 | 0.29986597 |
| RXYLT1 | 40.4405551 | -0.188101459 | 0.28269042 | -0.6653974 | 0.50579637 | 0.75528806 |
| SRGAP1 | 604.611057 | -0.43140864 | 0.5285097 | -0.8162738 | 0.41434353 | 0.68449528 |
| XPOT | 62.3122936 | -0.581950561 | 0.30765338 | -1.8915786 | 0.05854714 | 0.24443767 |
| TBK1 | 141.998013 | -0.254788366 | 0.21053237 | -1.2102099 | 0.22619835 | 0.50862769 |
| RASSF3 | 51.6186397 | -0.664849932 | 0.45920561 | -1.4478262 | 0.14766565 | 0.40657927 |
| GNS | 159.861314 | -0.238551867 | 0.24344591 | -0.9798968 | 0.32713706 | 0.61330512 |
| LEMD3 | 176.095626 | 0.36444703 | 0.14459132 | 2.52053192 | 0.01171776 | 0.09871458 |
| MSRB3 | 118.804932 | 0.592064875 | 0.67898978 | 0.87197907 | 0.3832198 | 0.65987101 |
| HMGA2 | 82.4490501 | -1.164041197 | 0.4953178 | -2.3500896 | 0.01876889 | 0.1296231 |
| AC090673.1 | 166.318643 | 1.07291656 | 0.5054938 | 2.12251182 | 0.03379478 | 0.18066599 |
| HELB | 57.3612579 | -0.153958356 | 0.38481477 | -0.4000843 | 0.68909441 | 0.86751885 |
| GRIP1 | 1109.13472 | -0.304134783 | 0.27268977 | -1.1153142 | 0.26471579 | 0.55272255 |
| CAND1 | 199.683837 | 0.329577066 | 0.1966344 | 1.67609061 | 0.09372045 | 0.31857635 |
| DYRK2 | 43.6895522 | -0.123452945 | 0.26081824 | -0.4733294 | 0.63597818 | 0.83814123 |
| MDM1 | 31.6865984 | 1.002832316 | 0.25417007 | 3.94551697 | 7.96E-05 | 0.00405722 |
| RAP1B | 428.404736 | -0.212648633 | 0.28080325 | -0.7572869 | 0.44887797 | 0.71024151 |
| NUP107 | 102.850919 | -0.055332579 | 0.25476617 | -0.2171897 | 0.82806054 | 0.93346403 |
| SLC35E3 | 64.5401746 | -0.343533179 | 0.31459354 | -1.0919906 | 0.27483725 | 0.56187466 |
| MDM2 | 192.090753 | -0.955897432 | 0.32113306 | -2.9766397 | 0.00291426 | 0.0398796 |
| CPM | 342.85712 | -0.870453031 | 0.65996725 | -1.3189337 | 0.18719129 | 0.46249816 |
| CPSF6 | 309.058034 | 0.3030353 | 0.24628092 | 1.23044573 | 0.21853024 | 0.50057436 |
| FRS2 | 208.226144 | 0.363542048 | 0.12057549 | 3.01505761 | 0.0025693 | 0.03695401 |
| CCT2 | 89.5571373 | -0.578438603 | 0.27644995 | -2.0923809 | 0.03640445 | 0.1889048 |
| RAB3IP | 193.75061 | 0.627768675 | 0.29773272 | 2.1084974 | 0.03498799 | 0.1847551 |
| MYRFL | 52.100092 | 1.613615844 | 1.05250125 | 1.53312487 | 0.12524509 | 0.37200705 |
| AC025159.1 | 165.491681 | 0.166921671 | 0.45522422 | 0.36668012 | 0.71385764 | 0.87892575 |
| CNOT2 | 326.931609 | 0.168060515 | 0.15983835 | 1.05144053 | 0.29305631 | 0.58090628 |
| ZFC3H1 | 232.230045 | 0.149708152 | 0.14029203 | 1.06711803 | 0.28591854 | 0.57333341 |
| RAB21 | 232.408571 | 0.027922367 | 0.18027196 | 0.15489024 | 0.87690785 | 0.95377544 |
| TBC1D15 | 190.561114 | 0.2880944 | 0.1830911 | 1.57350299 | 0.1156024 | 0.35790413 |
| TRHDE | 598.267819 | 0.953965977 | 0.37110534 | 2.57060695 | 0.01015205 | 0.09004967 |
| ATXN7L3B | 89.6664228 | -0.254875598 | 0.26026667 | -0.9792864 | 0.3274385 | 0.61368293 |
| CAPS2 | 158.445195 | 0.807015862 | 0.3869631 | 2.08551116 | 0.03702293 | 0.19077063 |
| KRR1 | 67.6914353 | 0.295562565 | 0.24820598 | 1.19079552 | 0.23373387 | 0.51635994 |
| PHLDA1 | 49.0493832 | 0.786585033 | 0.70202153 | 1.12045714 | 0.26251901 | 0.55025592 |
| NAP1L1 | 255.483127 | 0.137474453 | 0.1703859 | 0.80684173 | 0.41975767 | 0.68835944 |
| OSBPL8 | 284.337709 | -0.021946298 | 0.16382533 | -0.1339616 | 0.89343296 | 0.96030008 |
| ZDHHC17 | 189.632951 | 0.49963276 | 0.14881969 | 3.35730289 | 0.00078707 | 0.01778143 |
| NAV3 | 110.206614 | 1.994608461 | 0.83324452 | 2.39378528 | 0.01667551 | 0.12124283 |
| AC027288.3 | 152.32949 | -0.165933208 | 1.03645605 | -0.1600967 | 0.87280489 | 0.95193455 |
| PAWR | 951.408916 | -0.07330299 | 0.56764876 | -0.1291344 | 0.89725129 | 0.96175013 |
| AC073569.2 | 87.6260003 | -0.424066689 | 0.85674208 | -0.4949759 | 0.62061712 | 0.82839623 |
| PPP1R12A | 410.025472 | 0.250215769 | 0.22249143 | 1.12460859 | 0.26075493 | 0.54773736 |
| LIN7A | 66.5621369 | -0.552489242 | 0.84658731 | -0.6526075 | 0.51400934 | 0.76088633 |
| ACSS3 | 175.84158 | -1.313557307 | 0.62309312 | -2.1081236 | 0.0350203 | 0.1847551 |
| PPFIA2 | 70.1772916 | 1.557716871 | 1.35314073 | 1.15118615 | 0.24965566 | 0.5368017 |
| CCDC59 | 47.0702474 | 0.131435434 | 0.30168484 | 0.43567133 | 0.66307521 | 0.85554922 |
| METTL25 | 119.856188 | 0.77801913 | 0.16435136 | 4.73387709 | 2.20E-06 | 0.00030579 |
| TMTC2 | 519.927009 | -0.48252095 | 0.3336113 | -1.4463568 | 0.14807716 | 0.40747807 |
| LRRIQ1 | 218.95416 | 0.403412554 | 0.30795921 | 1.30995449 | 0.19021123 | 0.46635101 |
| MGAT4C | 176.45495 | 1.021791655 | 1.05818757 | 0.96560542 | 0.33424166 | 0.61868616 |
| CEP290 | 208.727797 | 0.299745642 | 0.19087434 | 1.57038206 | 0.11632625 | 0.35938557 |
| TMTC3 | 99.9006695 | 0.271104856 | 0.28211776 | 0.96096344 | 0.33657055 | 0.62147307 |
| KITLG | 193.91501 | -0.057432173 | 0.61154056 | -0.0939139 | 0.92517753 | 0.97344855 |
| DUSP6 | 100.697008 | -0.556853321 | 0.52446733 | -1.0617503 | 0.28834908 | 0.57519697 |
| POC1B | 109.601094 | 0.138858657 | 0.25418178 | 0.54629665 | 0.58486204 | 0.80842619 |
| POC1B-AS1 | 58.840572 | 0.375911121 | 0.24960882 | 1.50600096 | 0.13206693 | 0.38265853 |
| AC025034.1 | 31.8015094 | 0.652505311 | 0.36154416 | 1.80477347 | 0.07111014 | 0.27388983 |
| ATP2B1 | 269.879019 | 0.650016414 | 0.25532044 | 2.54588472 | 0.01090012 | 0.0939804 |
| BTG1 | 235.242324 | -0.518581111 | 0.46516715 | -1.1148275 | 0.26492437 | 0.5528781 |
| EEA1 | 209.499406 | 0.042844012 | 0.18391474 | 0.23295584 | 0.8157957 | 0.92869214 |
| AC138123.1 | 169.105065 | 0.138439179 | 0.53971113 | 0.25650606 | 0.7975601 | 0.91924295 |
| NUDT4 | 108.722947 | -0.548678447 | 0.34141876 | -1.6070542 | 0.10804248 | 0.34383352 |
| UBE2N | 100.480877 | -0.078424669 | 0.247845 | -0.3164263 | 0.75167898 | 0.89612109 |
| MRPL42 | 94.6473388 | 0.087056708 | 0.24514396 | 0.35512483 | 0.72249608 | 0.88242348 |
| CRADD | 392.055056 | -0.075838854 | 0.17334528 | -0.4375017 | 0.66174754 | 0.85475724 |
| AC012085.2 | 34.0869885 | 0.209425584 | 0.58631572 | 0.3571891 | 0.72095024 | 0.88235193 |
| CEP83 | 146.99096 | -0.188780363 | 0.18178251 | -1.0384957 | 0.29903932 | 0.58639882 |
| TMCC3 | 55.672659 | 1.46457712 | 0.37530976 | 3.90231555 | 9.53E-05 | 0.0045181 |
| NDUFA12 | 68.3751382 | -0.265189568 | 0.31700335 | -0.8365513 | 0.40284483 | 0.67688395 |
| NR2C1 | 117.884144 | 0.163451092 | 0.16290174 | 1.00337228 | 0.31568127 | 0.60232326 |
| FGD6 | 115.3003 | -1.273088963 | 0.40128182 | -3.1725558 | 0.00151104 | 0.02700412 |
| VEZT | 343.292583 | 0.076633502 | 0.11773466 | 0.65090008 | 0.51511099 | 0.76135326 |
| METAP2 | 142.86778 | -0.196788343 | 0.15372732 | -1.280113 | 0.20050538 | 0.47920186 |
| NTN4 | 913.045143 | 1.816909743 | 0.34400013 | 5.28171244 | 1.28E-07 | 3.61E-05 |
| SNRPF | 43.4460038 | -1.266004939 | 0.37667302 | -3.3610183 | 0.00077656 | 0.01758623 |
| LTA4H | 66.5645618 | -0.038360647 | 0.23358454 | -0.164226 | 0.86955326 | 0.95106604 |
| ELK3 | 53.9889723 | -2.063990694 | 0.64562043 | -3.1969105 | 0.00138908 | 0.02588716 |
| CDK17 | 199.549593 | -0.069589665 | 0.22183042 | -0.3137066 | 0.75374389 | 0.89777709 |
| CFAP54 | 215.817235 | -0.335444121 | 0.64669725 | -0.5187035 | 0.60396753 | 0.82004322 |
| NEDD1 | 62.763361 | 0.406157988 | 0.2535011 | 1.60219417 | 0.10911268 | 0.34570701 |
| TMPO | 151.214438 | 0.779790091 | 0.30413948 | 2.56392265 | 0.01034966 | 0.0912046 |
| SLC25A3 | 199.526842 | -0.493959606 | 0.42786881 | -1.1544651 | 0.24830954 | 0.53534467 |
| IKBIP | 44.7671715 | -0.266341171 | 0.48774802 | -0.5460631 | 0.5850226 | 0.80853144 |
| APAF1 | 75.3944685 | -0.006173409 | 0.22957933 | -0.0268901 | 0.9785474 | 0.99538128 |
| ANKS1B | 113.751905 | 0.577406337 | 0.68459326 | 0.84342977 | 0.39898813 | 0.67377896 |
| UHRF1BP1L | 118.2296 | 0.234796776 | 0.17602978 | 1.33384687 | 0.18225405 | 0.4553499 |
| ACTR6 | 47.8283865 | 0.173166898 | 0.24704344 | 0.70095729 | 0.48332967 | 0.73723167 |
| SCYL2 | 153.49749 | 0.184198928 | 0.16249313 | 1.13357976 | 0.25697087 | 0.54422373 |
| NR1H4 | 377.091697 | -0.275210659 | 0.32771725 | -0.8397808 | 0.40103129 | 0.67532043 |
| SLC5A8 | 56.9172342 | 2.125467349 | 0.61438894 | 3.45948179 | 0.00054122 | 0.01396038 |
| UTP20 | 85.3090073 | -0.163866425 | 0.34360429 | -0.4769045 | 0.63343014 | 0.83622207 |

| | | | | | | |
|---|---|---|---|---|---|---|
| ARL1 | 87.070883 | -0.446166858 | 0.21581792 | -2.06733 | 0.03870307 | 0.19563942 |
| CHPT1 | 133.285942 | -0.357994253 | 0.37401928 | -0.9571546 | 0.33848925 | 0.62345706 |
| GNPTAB | 99.3823889 | -0.111139758 | 0.1864233 | -0.5961688 | 0.55106246 | 0.78577398 |
| DRAM1 | 175.202205 | -0.002848457 | 0.20937786 | -0.0136044 | 0.9891456 | 0.99678172 |
| WASHC3 | 93.1607399 | 0.323441872 | 0.27671178 | 1.16887641 | 0.24245343 | 0.52725736 |
| NUP37 | 29.932365 | -0.243943224 | 0.27951823 | -0.8727274 | 0.38281168 | 0.65964257 |
| HSP90B1 | 338.406943 | -1.070217384 | 0.46712483 | -2.2910736 | 0.02195915 | 0.14229081 |
| C12orf73 | 27.1372244 | 0.542400783 | 0.30369126 | 1.78602697 | 0.07409489 | 0.28082432 |
| TDG | 70.5377227 | -0.224616821 | 0.21190436 | -1.0599915 | 0.28914846 | 0.57585049 |
| HCFC2 | 54.8833757 | 0.833763817 | 0.23032219 | 3.61998914 | 0.00029462 | 0.00953419 |
| NFYB | 40.9031417 | 0.10345477 | 0.25125326 | 0.41175493 | 0.68051906 | 0.86248903 |
| TXNRD1 | 865.203359 | -0.994881187 | 0.31377381 | -3.1706954 | 0.00152074 | 0.02707661 |
| CHST11 | 79.969549 | -3.182524597 | 0.59572021 | -5.3423143 | 9.18E-08 | 2.84E-05 |
| SLC41A2 | 142.226975 | -0.644680508 | 0.29972554 | -2.1509029 | 0.03148387 | 0.17362347 |
| C12orf45 | 36.7004242 | -0.327350041 | 0.39783193 | -0.822835 | 0.41060182 | 0.68236552 |
| WASHC4 | 312.686106 | 0.014772682 | 0.23132522 | 0.06386109 | 0.94908083 | 0.98262487 |
| APPL2 | 153.214451 | -0.204336388 | 0.22003529 | -0.9286528 | 0.35306905 | 0.63441164 |
| C12orf75 | 133.783921 | -0.738865181 | 0.34183951 | -2.1614388 | 0.03066146 | 0.17115672 |
| NUAK1 | 68.7008784 | 0.572362202 | 0.35633013 | 1.60626946 | 0.10821472 | 0.34403875 |
| CKAP4 | 48.2263594 | -0.377669463 | 0.48900126 | -0.7723282 | 0.43992006 | 0.70338746 |
| TCP11L2 | 65.0152926 | 0.160040017 | 0.482122 | 0.33194921 | 0.73992761 | 0.89253455 |
| POLR3B | 64.9643174 | 0.198945453 | 0.30363416 | 0.65521433 | 0.51232977 | 0.76016215 |
| RIC8B | 262.047265 | 0.122216743 | 0.63838436 | 0.19144696 | 0.84817543 | 0.94366633 |
| TMEM263 | 115.422202 | 0.146172783 | 0.29219892 | 0.50025094 | 0.61689839 | 0.82592881 |
| MTERF2 | 42.7873082 | 0.658668391 | 0.4132855 | 1.59373698 | 0.11099496 | 0.34822126 |
| CRY1 | 103.032137 | 0.152500174 | 0.39190122 | 0.38912912 | 0.69718064 | 0.87104386 |
| BTBD11 | 55.3564404 | -2.155298774 | 1.17244811 | -1.8382893 | 0.06601979 | 0.26198834 |
| PWP1 | 56.3178121 | -0.169031522 | 0.29761801 | -0.5679479 | 0.57007034 | 0.79765604 |
| PRDM4 | 62.3222782 | -0.358057481 | 0.25112058 | -1.4258389 | 0.15391486 | 0.4167186 |
| SART3 | 65.2284721 | -0.200677174 | 0.17622981 | -1.1387243 | 0.25481815 | 0.54159654 |
| ISCU | 131.017114 | -0.03378868 | 0.47318713 | -0.0714066 | 0.94307416 | 0.98112027 |
| CORO1C | 97.9476703 | -0.912868061 | 0.46839964 | -1.9489085 | 0.05130635 | 0.2274958 |
| SSH1 | 104.87423 | -0.013591571 | 0.36489503 | -0.0372479 | 0.97028735 | 0.99175186 |
| USP30 | 35.2434862 | 0.203218768 | 0.31681164 | 0.64144982 | 0.52123047 | 0.76440016 |
| ACACB | 142.270252 | 0.702039635 | 0.45649197 | 1.53790138 | 0.12407273 | 0.37036232 |
| KCTD10 | 39.887408 | -0.198995122 | 0.2749688 | -0.7237007 | 0.46924948 | 0.72712764 |
| UBE3B | 74.26012 | 0.010536418 | 0.1925684 | 0.0547152 | 0.95636536 | 0.98484453 |
| MMAB | 74.4975748 | -0.042661944 | 0.24635879 | -0.17317 | 0.86251783 | 0.94877974 |
| MVK | 45.6576195 | -0.053431427 | 0.37544727 | -0.1423141 | 0.88683194 | 0.95879945 |
| TRPV4 | 33.2816978 | -0.124671066 | 0.59705178 | -0.2088111 | 0.83459567 | 0.93697031 |
| GLTP | 75.3843104 | 0.356370456 | 0.37813676 | 0.94243801 | 0.34596844 | 0.63016843 |
| TCHP | 57.8319082 | 0.324855806 | 0.25906212 | 1.25396878 | 0.20985335 | 0.49028909 |
| GIT2 | 116.701133 | 0.029570351 | 0.21144927 | 0.13984608 | 0.88878161 | 0.95907089 |
| ANKRD13A | 140.476065 | 0.007413075 | 0.31620086 | 0.0234442 | 0.98129595 | 0.99585061 |
| C12orf76 | 35.3606873 | 0.306561566 | 0.25118977 | 1.22043809 | 0.22229884 | 0.5039513 |
| IFT81 | 127.691508 | 0.465645878 | 0.23452007 | 1.98552679 | 0.04708589 | 0.2166537 |
| ATP2A2 | 280.042756 | -0.356179486 | 0.23435173 | -1.51985 | 0.12854867 | 0.37647982 |
| ANAPC7 | 89.2979577 | -0.23254259 | 0.34554818 | -0.6729672 | 0.50096817 | 0.75227686 |
| ARPC3 | 221.845684 | -0.827684536 | 0.33297418 | -2.4857319 | 0.01292853 | 0.10406927 |
| GPN3 | 27.1799425 | -0.130666713 | 0.45488364 | -0.2872531 | 0.77391857 | 0.90650109 |
| VPS29 | 104.635421 | -0.432775193 | 0.1842063 | -2.3494049 | 0.01880344 | 0.12967472 |
| PPTC7 | 74.710656 | -0.460026182 | 0.26766993 | -1.7186323 | 0.08568135 | 0.3045708 |
| TCTN1 | 78.4210194 | 0.208956809 | 0.26535953 | 0.78744792 | 0.43101971 | 0.69671579 |
| PPP1CC | 64.684023 | 0.217894521 | 0.27277658 | 0.79880216 | 0.42440514 | 0.69133076 |
| SH2B3 | 48.232105 | -1.008458445 | 0.58079469 | -1.7363424 | 0.08250331 | 0.29924241 |
| ATXN2 | 506.599524 | 0.053293315 | 0.18801494 | 0.28345256 | 0.77682994 | 0.90769139 |
| BRAP | 63.9814577 | -0.247966284 | 0.2021847 | -1.2264344 | 0.22003523 | 0.50159865 |
| ACAD10 | 63.4361314 | 0.226870884 | 0.29444079 | 0.77051446 | 0.44099478 | 0.70428292 |
| ALDH2 | 88.7639344 | -0.40637721 | 0.38232317 | -1.0629155 | 0.28782029 | 0.57450105 |
| MAPKAPK5 | 85.8545884 | 0.243373921 | 0.2256187 | 1.07869568 | 0.28072341 | 0.56842177 |
| TMEM116 | 87.7573506 | 1.245240375 | 0.50911369 | 2.44589844 | 0.01444917 | 0.11180016 |
| ERP29 | 123.585589 | -0.589886908 | 0.35625509 | -1.6557992 | 0.09776249 | 0.32618142 |
| NAA25 | 162.490332 | -0.073850935 | 0.26482917 | -0.2788625 | 0.78035032 | 0.91077326 |
| TRAFD1 | 60.5150998 | -0.322786398 | 0.26922057 | -1.1989663 | 0.23054103 | 0.51368983 |
| HECTD4 | 406.613585 | 0.326413332 | 0.14579655 | 2.23882762 | 0.02516713 | 0.15393404 |
| RPL6 | 431.790188 | -0.731946763 | 0.56219539 | -1.3019437 | 0.19293563 | 0.47098123 |
| PTPN11 | 159.730582 | -0.412571639 | 0.27264797 | -1.5132027 | 0.13022819 | 0.3803219 |
| RASAL1 | 116.430673 | -0.518959169 | 0.37407092 | -1.3873283 | 0.16534168 | 0.43384207 |
| DDX54 | 50.0595277 | -0.741860961 | 0.43608265 | -1.7011935 | 0.08890666 | 0.31048301 |
| TPCN1 | 268.845716 | -0.303306306 | 0.29583026 | -1.0252714 | 0.30523515 | 0.59271183 |
| SLC8B1 | 39.6948663 | -0.75631758 | 0.2516912 | -3.0049425 | 0.00265631 | 0.03786429 |
| PLBD2 | 34.211816 | -0.886324232 | 0.23243819 | -3.813161 | 0.0001372 | 0.00586717 |
| SDSL | 39.5865182 | -1.553618491 | 0.4847501 | -3.2049885 | 0.00135068 | 0.02531928 |
| RBM19 | 69.9091212 | -0.281984995 | 0.29945553 | -0.941659 | 0.34636726 | 0.6302171 |
| MED13L | 1964.73439 | -0.022837938 | 0.18244015 | -0.1251804 | 0.9003807 | 0.96303857 |
| C12orf49 | 80.8153722 | -0.478892336 | 0.28074642 | -1.7057825 | 0.08804858 | 0.30883096 |
| FBXW8 | 133.053164 | -0.135717493 | 0.1989602 | -0.6821339 | 0.4951543 | 0.74709472 |
| TESC | 45.6739715 | -1.92932862 | 0.49978 | -3.8603558 | 0.00011322 | 0.00514007 |
| FBXO21 | 150.310207 | 0.11338788 | 0.28930676 | 0.39192958 | 0.69511024 | 0.87024545 |
| KSR2 | 34.1618508 | 0.867128009 | 0.7503358 | 1.15565325 | 0.24782302 | 0.53482252 |
| WSB2 | 63.5856128 | -0.084398121 | 0.29022012 | -0.2908073 | 0.77119872 | 0.90586298 |
| VSIG10 | 83.4735904 | 0.688486496 | 0.50251131 | 1.37009154 | 0.17065833 | 0.4409518 |
| PEBP1 | 663.156954 | 0.406940631 | 0.55690406 | 0.73071945 | 0.46495053 | 0.72289582 |
| TAOK3 | 355.592256 | 0.339344973 | 0.32659004 | 1.03905488 | 0.29877921 | 0.58623639 |
| SUDS3 | 88.1577173 | 0.195810051 | 0.20029306 | 0.97761776 | 0.32826341 | 0.61394673 |
| HSPB8 | 94.5451885 | -1.171992571 | 0.53498959 | -2.190683 | 0.02847474 | 0.16381956 |
| AC002070.1 | 40.8859165 | -1.662708255 | 0.4860054 | -3.4211724 | 0.00062352 | 0.01535393 |
| PRKAB1 | 99.4883023 | -0.74448132 | 0.39568105 | -1.8815187 | 0.05990139 | 0.24679372 |
| CIT | 124.046256 | -0.269206334 | 0.54130795 | -0.4973257 | 0.61895942 | 0.827381 |
| BICDL1 | 365.62744 | 1.443596176 | 0.38726131 | 3.72770568 | 0.00019323 | 0.00731604 |
| RAB35 | 29.9191389 | -0.229178964 | 0.33160162 | -0.6911274 | 0.48948549 | 0.74220617 |
| GCN1 | 83.1447447 | -0.560458227 | 0.23866866 | -2.348269 | 0.01886089 | 0.12997734 |
| RPLP0 | 611.77128 | -0.705884908 | 0.51114893 | -1.380977 | 0.16728603 | 0.43591755 |
| PXN | 97.7060667 | -0.290913654 | 0.33896923 | -0.8582303 | 0.39076533 | 0.66627644 |
| GATC | 33.5028757 | -0.41278609 | 0.33380136 | -1.236622 | 0.21622746 | 0.49777526 |
| SRSF9 | 121.757811 | -0.052921183 | 0.40829643 | -0.1296146 | 0.89687134 | 0.96175013 |
| DYNLL1 | 210.868856 | -0.480521047 | 0.44770938 | -1.0732879 | 0.28314198 | 0.57084461 |
| COQ5 | 36.0320529 | 0.066934955 | 0.30250202 | 0.2212711 | 0.82488135 | 0.93217972 |
| RNF10 | 140.216441 | -0.061320228 | 0.13845342 | -0.4428943 | 0.65784219 | 0.85201059 |
| POP5 | 34.9729782 | -0.253538148 | 0.33136064 | -0.7651426 | 0.44418663 | 0.70678799 |
| MLEC | 176.442022 | -0.949073189 | 0.24190633 | -3.9233087 | 8.73E-05 | 0.00428474 |
| UNC119B | 44.367204 | 0.288814094 | 0.28316258 | 1.01995854 | 0.30774813 | 0.59456464 |
| SPPL3 | 477.532329 | 0.302241797 | 0.15681321 | 1.92740009 | 0.05392979 | 0.2332446 |
| P2RX4 | 35.0192943 | -0.468956391 | 0.47723507 | -0.9826528 | 0.32577834 | 0.61243485 |
| CAMKK2 | 52.1672505 | -0.404989068 | 0.31568303 | -1.2828978 | 0.19952788 | 0.47837581 |
| ANAPC5 | 149.985564 | 0.24611697 | 0.18674435 | 1.31793527 | 0.18752532 | 0.46292038 |
| RNF34 | 57.0688518 | -0.11991084 | 0.26515011 | -0.4522376 | 0.65109785 | 0.84724399 |
| KDM2B | 69.1377399 | 0.212882535 | 0.39522997 | 0.53862954 | 0.5901425 | 0.81183245 |
| TMEM120B | 93.0315264 | 0.584880683 | 0.24887394 | 2.35010816 | 0.01876796 | 0.1296231 |
| RHOF | 25.3415703 | 0.347993279 | 0.50405571 | 0.69038655 | 0.48995113 | 0.74227587 |
| SETD1B | 58.3125183 | 0.063569069 | 0.21724573 | 0.29261367 | 0.76981747 | 0.90533273 |
| WDR66 | 143.042956 | -0.107195624 | 0.37282895 | -0.2875196 | 0.77371451 | 0.90643453 |
| BCL7A | 135.764561 | 0.208740488 | 0.31346728 | 0.66590838 | 0.50546966 | 0.75528806 |

| | | | | | | |
|---|---|---|---|---|---|---|
| MLXIP | 279.162104 | -0.685882856 | 0.28343849 | -2.4198649 | 0.01552627 | 0.11578325 |
| CLIP1 | 270.383143 | -0.479533288 | 0.27402297 | -1.7499748 | 0.08012266 | 0.29426581 |
| ZCCHC8 | 92.2927474 | 0.47379473 | 0.18429116 | 2.57090318 | 0.01014337 | 0.09004967 |
| RSRC2 | 152.153931 | 0.283948919 | 0.19156805 | 1.48223523 | 0.13827772 | 0.39107241 |
| DENR | 149.244275 | -0.4662773 | 0.21631622 | -2.1555355 | 0.03111995 | 0.17251043 |
| HIP1R | 162.026287 | 0.152226649 | 0.3365857 | 0.45226713 | 0.65107655 | 0.84724399 |
| VPS37B | 154.231142 | 0.421673289 | 0.38679088 | 1.09018417 | 0.27563203 | 0.5628393 |
| ARL6IP4 | 132.001978 | -0.302055344 | 0.36655325 | -0.8240422 | 0.40991558 | 0.68208527 |
| PITPNM2 | 154.555758 | -0.435848496 | 0.28825396 | -1.5120295 | 0.13052636 | 0.38096039 |
| MPHOSPH9 | 111.295361 | -0.101767661 | 0.25987692 | -0.3915995 | 0.69535419 | 0.87024545 |
| C12orf65 | 56.4438122 | -0.330423391 | 0.25649901 | -1.2882053 | 0.1976745 | 0.47630586 |
| SBNO1 | 168.916894 | 0.008538326 | 0.15759877 | 0.05417762 | 0.95679365 | 0.98507377 |
| KMT5A | 42.4789528 | -0.117930348 | 0.22683425 | -0.5198966 | 0.60313567 | 0.8197271 |
| RILPL2 | 61.1578042 | -0.089016911 | 0.21446885 | -0.4150575 | 0.67809979 | 0.86151115 |
| SNRNP35 | 62.6242016 | 0.265064937 | 0.2210536 | 1.19909802 | 0.23048983 | 0.51368983 |
| RILPL1 | 163.037636 | 0.203591209 | 0.20858085 | 0.97607816 | 0.32902573 | 0.61429707 |
| TMED2 | 82.8989028 | -0.296922897 | 0.26994129 | -1.0999536 | 0.27135234 | 0.55841559 |
| DDX55 | 41.1795798 | -0.252853615 | 0.28691992 | -0.881269 | 0.37817226 | 0.65629152 |
| EIF2B1 | 43.018495 | -0.049663417 | 0.23358234 | -0.2126163 | 0.83162624 | 0.93532321 |
| GTF2H3 | 46.2983982 | -0.055929466 | 0.25677357 | -0.2178163 | 0.82757225 | 0.93346403 |
| TCTN2 | 44.7520723 | 0.566466845 | 0.30967302 | 1.82924183 | 0.06736339 | 0.26489076 |
| ATP6V0A2 | 64.9087031 | -0.00105479 | 0.40364064 | -0.0026132 | 0.99791498 | 0.99923686 |
| CCDC92 | 151.65486 | 0.366456109 | 0.14485817 | 2.52975789 | 0.01141413 | 0.096929 |
| NCOR2 | 289.870051 | 0.330931416 | 0.30597844 | 1.08155143 | 0.27945189 | 0.56686465 |
| SCARB1 | 42.8176702 | -0.036163303 | 0.28582599 | -0.1265221 | 0.89931866 | 0.96269027 |
| UBC | 633.946794 | -0.753640796 | 0.29280743 | -2.5738445 | 0.01005755 | 0.08977568 |
| DHX37 | 27.5089705 | -1.029478548 | 0.38760692 | -2.655986 | 0.00790769 | 0.07690128 |
| BRI3BP | 26.6727951 | 0.208574127 | 0.35620442 | 0.58554615 | 0.55818055 | 0.78914147 |
| AACS | 48.924009 | -0.190132698 | 0.26469634 | -0.718305 | 0.47256927 | 0.72988407 |
| SLC15A4 | 95.5315774 | -0.61699377 | 0.30518873 | -2.0216794 | 0.04320949 | 0.20614587 |
| TMEM132D | 235.357034 | -0.063824878 | 0.54401707 | -0.1173215 | 0.90660532 | 0.96632606 |
| RIMBP2 | 38.4102147 | -0.151606747 | 0.66977531 | -0.2263546 | 0.82092561 | 0.93138972 |
| STX2 | 129.755134 | 1.014801997 | 0.24362895 | 4.16535877 | 3.11E-05 | 0.00212938 |
| RAN | 147.550471 | -1.086278915 | 0.4655537 | -2.3333053 | 0.01963213 | 0.13327863 |
| SFSWAP | 164.903169 | 0.070760937 | 0.16113098 | 0.43915165 | 0.66055166 | 0.85401867 |
| ULK1 | 68.6449813 | 0.581355403 | 0.23868348 | 2.43567504 | 0.01486403 | 0.11321507 |
| PUS1 | 31.3014486 | -0.549929274 | 0.45973512 | -1.1961872 | 0.23162351 | 0.51466519 |
| EP400 | 166.036038 | 0.154041749 | 0.17515981 | 0.87943548 | 0.3791652 | 0.65699285 |
| DDX51 | 25.7734675 | -0.580948053 | 0.27791987 | -2.0903437 | 0.03658694 | 0.189528 |
| FBRSL1 | 75.6015767 | -0.131316328 | 0.23128224 | -0.5677752 | 0.57018759 | 0.79765604 |
| ANKLE2 | 161.702147 | -0.706503991 | 0.32686539 | -2.1614525 | 0.03066041 | 0.17115672 |
| GOLGA3 | 103.504954 | -0.256394457 | 0.28083666 | -0.9129665 | 0.36126017 | 0.64151865 |
| CHFR | 105.488131 | -0.106271236 | 0.20714604 | -0.5130257 | 0.60793337 | 0.8211189 |
| ZNF26 | 68.8765959 | 0.033989311 | 0.20573809 | 0.1652067 | 0.86878129 | 0.95106604 |
| ZNF84 | 104.208635 | 0.37759495 | 0.26799469 | 1.40896428 | 0.15884573 | 0.42441661 |
| ZNF140 | 47.7976323 | 0.407738057 | 0.26713387 | 1.52634353 | 0.12692433 | 0.37435927 |
| ZNF891 | 54.7695043 | 0.903269357 | 0.2097676 | 4.30604803 | 1.66E-05 | 0.00132667 |
| MPHOSPH8 | 211.282511 | -0.029457437 | 0.21527704 | -0.136835 | 0.8911612 | 0.95969529 |
| PSPC1 | 341.784095 | 0.467203601 | 0.19965951 | 2.34000174 | 0.01928365 | 0.13189928 |
| ZMYM5 | 104.955619 | 0.102916047 | 0.23022092 | 0.44703169 | 0.65485218 | 0.85026877 |
| ZMYM2 | 641.95888 | 0.52327782 | 0.17663225 | 2.96252712 | 0.00305125 | 0.04139932 |
| CRYL1 | 171.182496 | -0.309712926 | 0.2626116 | -1.1793574 | 0.2382559 | 0.52130043 |
| IFT88 | 410.099862 | 0.254970418 | 0.22582601 | 1.1290569 | 0.25887383 | 0.54596047 |
| EEF1AKMT1 | 39.6869878 | -0.173435713 | 0.2409569 | -0.719779 | 0.47166108 | 0.72945405 |
| XPO4 | 196.536671 | 0.167449148 | 0.19236439 | 0.87047893 | 0.38403873 | 0.66056868 |
| LATS2 | 196.831584 | 0.955420841 | 0.27551884 | 3.4677151 | 0.0005249 | 0.01373784 |
| SAP18 | 137.208559 | -0.351924061 | 0.44335861 | -0.7937684 | 0.42733027 | 0.69344344 |
| MRPL57 | 82.3709979 | -0.476444951 | 0.42754568 | -1.114372 | 0.26511962 | 0.55316507 |
| ZDHHC20 | 177.86709 | 0.331679474 | 0.21298614 | 1.557282 | 0.11940355 | 0.3645149 |
| MICU2 | 290.365458 | 0.244696522 | 0.13956376 | 1.75329563 | 0.07955127 | 0.29274747 |
| LINC00621 | 122.975155 | -1.879893762 | 0.62201248 | -3.0222766 | 0.00250881 | 0.03641199 |
| AL157931.1 | 50.7495763 | -1.962965766 | 0.61724764 | -3.1801916 | 0.00147178 | 0.02655639 |
| SACS | 66.2849515 | -0.312787042 | 0.44462607 | -0.7034834 | 0.48175457 | 0.73620566 |
| TNFRSF19 | 114.093034 | 1.052521391 | 0.33949168 | 3.10028625 | 0.00193334 | 0.0311251 |
| MIPEP | 192.61824 | -0.131862672 | 0.24795439 | -0.5318021 | 0.59486305 | 0.81507555 |
| PARP4 | 451.803825 | 0.059614893 | 0.36730711 | 0.16230258 | 0.87106758 | 0.95174591 |
| MTMR6 | 69.3684194 | 0.200967211 | 0.25132949 | 0.7996165 | 0.42393302 | 0.69092389 |
| NUP58 | 232.166622 | 0.257327687 | 0.22506273 | 1.14335985 | 0.25288922 | 0.53987649 |
| RNF6 | 30.2392179 | -0.141683813 | 0.33351289 | -0.4248226 | 0.670966 | 0.85977676 |
| CDK8 | 247.654419 | -0.202547594 | 0.21019216 | -0.9636306 | 0.33523117 | 0.6199188 |
| WASF3 | 97.0213189 | -0.600227954 | 0.25743531 | -2.3315681 | 0.01972342 | 0.13342561 |
| USP12 | 153.705089 | -0.097214679 | 0.19612886 | -0.4956674 | 0.62012911 | 0.82825108 |
| LINC02340 | 55.3501906 | 0.156521626 | 0.62078787 | 0.25213384 | 0.80093762 | 0.92075299 |
| RPL21 | 204.114735 | 0.109710405 | 0.47062712 | 0.23311535 | 0.81567184 | 0.92869214 |
| GTF3A | 50.7060052 | -0.177171562 | 0.38350431 | -0.4619806 | 0.64409521 | 0.84344334 |
| MTIF3 | 90.072374 | 0.19048899 | 0.26079336 | 0.73042117 | 0.46513278 | 0.72300587 |
| LNX2 | 119.650449 | 0.750685596 | 0.36456248 | 2.05914111 | 0.03948072 | 0.19697179 |
| POLR1D | 93.6835032 | -0.097704361 | 0.16911902 | -0.5777254 | 0.5634495 | 0.79290184 |
| PAN3 | 718.962733 | 0.181374774 | 0.13240456 | 1.36985297 | 0.1707328 | 0.4409518 |
| POMP | 82.3404897 | -0.296860704 | 0.39557404 | -0.7504555 | 0.45298043 | 0.71272989 |
| SLC7A1 | 91.6299112 | -0.435708626 | 0.44118162 | -0.9875947 | 0.32335119 | 0.61020115 |
| UBL3 | 203.106863 | 0.34735628 | 0.26925155 | 1.29008088 | 0.19702258 | 0.4762249 |
| KATNAL1 | 134.427276 | 0.568448168 | 0.2378604 | 2.38983949 | 0.01685574 | 0.12184079 |
| HMGB1 | 496.728511 | -0.24737939 | 0.23799081 | -1.0394493 | 0.29859581 | 0.58610017 |
| USPL1 | 80.4390806 | 0.095130424 | 0.23872091 | 0.39850059 | 0.69026122 | 0.86817387 |
| HSPH1 | 92.3701532 | 0.319989585 | 0.37721416 | 0.84829686 | 0.39627267 | 0.67136468 |
| B3GLCT | 59.8467964 | 0.451159932 | 0.30985795 | 1.4560218 | 0.14538659 | 0.40323664 |
| FRY | 758.479517 | 1.135135153 | 0.45589688 | 2.48989453 | 0.0127781 | 0.10329235 |
| N4BP2L1 | 135.568283 | -0.07138572 | 0.2860842 | -0.249527 | 0.80295319 | 0.92125386 |
| N4BP2L2 | 1027.4287 | 0.320580095 | 0.18585611 | 1.72488328 | 0.08454851 | 0.30235944 |
| PDS5B | 340.519798 | 0.458721871 | 0.18435008 | 2.48831929 | 0.01283484 | 0.10357622 |
| KL | 48.4290116 | 0.569999776 | 0.43678388 | 1.30499271 | 0.19189533 | 0.46899444 |
| STARD13 | 271.778744 | 0.502190904 | 0.19515876 | 2.57324297 | 0.01007504 | 0.08977568 |
| AL139383.1 | 48.1288019 | 0.167428636 | 0.57980985 | 0.28876473 | 0.77276142 | 0.90636484 |
| RFC3 | 57.8849797 | -0.130397319 | 0.32602436 | -0.3999619 | 0.68918458 | 0.86751885 |
| NBEA | 426.604284 | 0.379016743 | 0.10949474 | 3.46150624 | 0.00053716 | 0.01394437 |
| DCLK1 | 35.5824304 | 0.912278999 | 0.85597011 | 1.0657837 | 0.28652143 | 0.57346076 |
| SPART | 180.492813 | -0.104608278 | 0.19831257 | -0.5274919 | 0.59785203 | 0.81658679 |
| RFXAP | 29.8324441 | 0.613647574 | 0.28399797 | 2.16074635 | 0.03071494 | 0.17115672 |
| SMAD9 | 329.710045 | 1.390220559 | 0.65231494 | 2.13121065 | 0.03307179 | 0.17827501 |
| ALG5 | 69.5984517 | 0.091363932 | 0.19880789 | 0.45955889 | 0.64583287 | 0.84416957 |
| EXOSC8 | 43.7081742 | 0.264143412 | 0.32314568 | 0.81741279 | 0.41369257 | 0.68448816 |
| SUPT20H | 185.245933 | 0.395978084 | 0.22229706 | 1.78130152 | 0.07486321 | 0.28266689 |
| LINC00571 | 53.5261347 | -0.504081037 | 0.41090411 | -1.2267608 | 0.21991252 | 0.50155763 |
| UFM1 | 117.900898 | 0.296310845 | 0.22294867 | 1.32905409 | 0.18383012 | 0.45721925 |
| FREM2 | 293.171516 | 1.463287496 | 0.56095982 | 2.60854243 | 0.00909287 | 0.08508231 |
| PROSER1 | 50.7136679 | 0.254368127 | 0.28750703 | 0.88473706 | 0.37629846 | 0.65466091 |
| NHLRC3 | 93.2481396 | 0.038494632 | 0.3158054 | 0.12189352 | 0.90298335 | 0.96461219 |
| COG6 | 128.491589 | 0.496762526 | 0.18693438 | 2.65741657 | 0.00787421 | 0.07683059 |
| FOXO1 | 409.689086 | -0.387795005 | 0.21639841 | -1.7920418 | 0.07312626 | 0.27896901 |
| MRPS31 | 101.693874 | 0.349104447 | 0.18612038 | 1.87569167 | 0.06069763 | 0.24868375 |
| ELF1 | 464.446936 | 0.304074426 | 0.17768441 | 1.71131742 | 0.08702253 | 0.30691783 |
| WBP4 | 82.875808 | -0.00941593 | 0.20297216 | -0.0463903 | 0.9629992 | 0.98817751 |

| | | | | | | |
|---|---|---|---|---|---|---|
| AL354696.2 | 46.3242346 | 0.896677904 | 0.47520038 | 1.88694693 | 0.05916746 | 0.24535287 |
| MTRF1 | 31.3277353 | -0.1572543 | 0.3015239 | -0.5215318 | 0.60199636 | 0.8187595 |
| NAA16 | 180.666739 | 0.170057728 | 0.16580081 | 1.02567491 | 0.30504484 | 0.59267272 |
| VWA8 | 440.298668 | 0.294549469 | 0.20129296 | 1.46328747 | 0.14338874 | 0.40048158 |
| DGKH | 653.364194 | -0.359382081 | 0.28255683 | -1.2718931 | 0.2034111 | 0.48230223 |
| AKAP11 | 154.040575 | 0.393544531 | 0.20007387 | 1.96699616 | 0.04918365 | 0.22296745 |
| EPSTI1 | 206.911315 | -0.88576912 | 1.14236048 | -0.7753849 | 0.43811223 | 0.70236452 |
| DNAJC15 | 135.0462 | -0.337005642 | 0.26179816 | -1.2872728 | 0.19799923 | 0.47656532 |
| ENOX1 | 210.63142 | -0.749019118 | 0.9900173 | -0.7565717 | 0.44930647 | 0.71037275 |
| CCDC122 | 135.081277 | 0.010646988 | 0.28397063 | 0.03749327 | 0.9700917 | 0.99175186 |
| SMIM2-AS1 | 193.963722 | -2.145993717 | 0.71408148 | -3.0052505 | 0.00265362 | 0.03786429 |
| SERP2 | 33.0055836 | -0.172973682 | 0.28208198 | -0.6132036 | 0.53974174 | 0.7774716 |
| TSC22D1 | 923.200358 | -0.12389086 | 0.31964643 | -0.3875872 | 0.69832155 | 0.8717697 |
| NUFIP1 | 31.4753226 | -0.035658707 | 0.32860977 | -0.1085138 | 0.9135881 | 0.96844354 |
| AL359706.1 | 28.8161236 | -0.123107022 | 0.56527631 | -0.217782 | 0.82759895 | 0.93346403 |
| GPALPP1 | 89.9639633 | 0.295496886 | 0.2223531 | 1.32895327 | 0.18386339 | 0.45721925 |
| GTF2F2 | 161.768515 | -0.084696943 | 0.22509532 | -0.3762715 | 0.7067151 | 0.87495497 |
| TPT1 | 1033.84969 | -0.851371305 | 0.53547994 | -1.5899219 | 0.1118524 | 0.3500176 |
| SLC25A30 | 66.5867028 | 0.437440917 | 0.30597674 | 1.42965415 | 0.15281631 | 0.41527018 |
| COG3 | 123.357507 | 0.349515611 | 0.18967902 | 1.84266882 | 0.06537738 | 0.26050329 |
| ZC3H13 | 193.751833 | 0.115101369 | 0.20591519 | 0.55897464 | 0.57617903 | 0.80232867 |
| LRRC63 | 29.5483969 | 1.89002064 | 1.13106963 | 1.6710029 | 0.0947211 | 0.3204651 |
| AL138686.1 | 40.7401581 | -0.055623206 | 0.56915573 | -0.0977293 | 0.92214723 | 0.97248656 |
| LRCH1 | 455.854095 | -0.472129476 | 0.2697652 | -1.7501497 | 0.08009249 | 0.29426581 |
| ESD | 96.7181953 | -0.160057145 | 0.21703077 | -0.737486 | 0.46082688 | 0.71918385 |
| SUCLA2 | 144.737968 | -0.093462393 | 0.24311508 | -0.3844368 | 0.70065471 | 0.87182648 |
| MED4 | 119.706354 | 0.069634554 | 0.18486174 | 0.37668451 | 0.70640808 | 0.87473681 |
| ITM2B | 1494.62826 | 0.178229132 | 0.37627281 | 0.47366998 | 0.63573526 | 0.83799478 |
| RB1 | 442.660317 | -0.194340615 | 0.2087461 | -0.9309904 | 0.35185853 | 0.63377755 |
| LPAR6 | 70.7489214 | 0.217987624 | 0.48580495 | 0.4487143 | 0.65363777 | 0.84920605 |
| RCBTB2 | 98.4882848 | 0.795006944 | 0.23051998 | 3.4487551 | 0.00056318 | 0.01427163 |
| FNDC3A | 625.133242 | 0.292617351 | 0.20706152 | 1.41319037 | 0.15759976 | 0.42280489 |
| CDADC1 | 43.6383378 | 0.48595313 | 0.30821315 | 1.57667876 | 0.11486946 | 0.35651907 |
| CAB39L | 77.6949564 | 0.923018932 | 0.47384715 | 1.94792546 | 0.05142388 | 0.22752396 |
| SETDB2 | 87.8818628 | 0.463799462 | 0.3112383 | 1.49017477 | 0.13617829 | 0.3881142 |
| PHF11 | 81.6190579 | 0.028792257 | 0.40725519 | 0.07069832 | 0.94363786 | 0.98112027 |
| RCBTB1 | 37.6791241 | 0.31759004 | 0.34196729 | 0.92871467 | 0.35303698 | 0.63441164 |
| EBPL | 50.9123859 | -0.706558662 | 0.29869861 | -2.3654568 | 0.01800784 | 0.12673521 |
| KPNA3 | 151.485724 | -0.160398928 | 0.19463236 | -0.8241123 | 0.40987574 | 0.68208527 |
| SPRYD7 | 33.7089738 | 0.157414937 | 0.2431302 | 0.64745119 | 0.51733997 | 0.76263459 |
| DLEU2 | 284.954018 | -0.154787997 | 0.38670321 | -0.400276 | 0.68895327 | 0.86751885 |
| TRIM13 | 73.5701544 | -0.054540162 | 0.23729644 | -0.2298398 | 0.81821627 | 0.92999132 |
| RNASEH2B | 80.3007597 | 0.116892672 | 0.29304937 | 0.39888388 | 0.68997877 | 0.86793258 |
| INTS6 | 268.36052 | 0.242746385 | 0.305509 | 0.79456376 | 0.4268673 | 0.69304439 |
| INTS6-AS1 | 37.5227197 | -0.114264471 | 0.3699238 | -0.3088865 | 0.75740786 | 0.89936902 |
| WDFY2 | 155.470406 | 0.222430798 | 0.25058064 | 0.88766153 | 0.37472284 | 0.65310592 |
| ATP7B | 37.2161377 | 0.634521621 | 0.31182217 | 2.03488297 | 0.04186265 | 0.20312121 |
| UTP14C | 42.4480086 | -0.045558516 | 0.34712636 | -0.1312448 | 0.89558169 | 0.96142295 |
| NEK5 | 46.3015601 | -0.040884387 | 0.58141901 | -0.0703183 | 0.94394033 | 0.98112027 |
| NEK3 | 49.4919433 | -0.440804003 | 0.37496254 | -1.1755948 | 0.23975684 | 0.52362764 |
| VPS36 | 131.894497 | 0.25512294 | 0.20741231 | 1.23002794 | 0.21868664 | 0.5006005 |
| SUGT1 | 118.641446 | 0.114648416 | 0.18359353 | 0.62446873 | 0.5323198 | 0.7728646 |
| DIAPH3 | 151.788777 | 1.657384501 | 0.34605795 | 4.78932644 | 1.67E-06 | 0.00025046 |
| TDRD3 | 299.890756 | 0.148646631 | 0.17553646 | 0.84681341 | 0.39709913 | 0.67240809 |
| PCDH9 | 268.55162 | -0.559424147 | 1.11030188 | -0.5038487 | 0.61436771 | 0.82538966 |
| DIS3 | 71.3745676 | -0.105800309 | 0.20488887 | -0.516379 | 0.60558974 | 0.820591 |
| PIBF1 | 396.432291 | -0.086514419 | 0.14684956 | -0.5891364 | 0.55576978 | 0.78784845 |
| KLF5 | 59.2682829 | -0.433960469 | 0.52667079 | -0.8239691 | 0.4099571 | 0.68208527 |
| KLF12 | 332.949758 | 0.547785866 | 0.27475621 | 1.99371604 | 0.0461831 | 0.21432487 |
| TBC1D4 | 1080.04018 | 1.049473918 | 0.46430764 | 2.26029862 | 0.02380272 | 0.14935362 |
| COMMD6 | 92.2527746 | -0.161637945 | 0.38660406 | -0.4180969 | 0.6758763 | 0.86116336 |
| UCHL3 | 84.4858531 | -0.176590856 | 0.36905438 | -0.4784955 | 0.63229758 | 0.8353707 |
| LMO7-AS1 | 45.6246776 | 0.216750124 | 0.51242783 | 0.42298664 | 0.67230501 | 0.86015189 |
| LMO7 | 595.398912 | 1.37838796 | 0.31449849 | 4.3828126 | 1.17E-05 | 0.00101506 |
| AC001226.2 | 57.4129377 | 0.526697394 | 0.55961506 | 0.94117803 | 0.34661364 | 0.6302171 |
| FBXL3 | 112.918222 | 0.379987717 | 0.27830887 | 1.36534531 | 0.17214454 | 0.44309343 |
| MYCBP2 | 455.747453 | 0.34246546 | 0.21257411 | 1.61104028 | 0.10717094 | 0.34253936 |
| SLAIN1 | 53.4923111 | 0.317967218 | 0.33032396 | 0.96259206 | 0.33575228 | 0.62028397 |
| RNF219 | 43.8008941 | 0.00200988 | 0.3036158 | 0.00661981 | 0.99471819 | 0.99886097 |
| RBM26 | 407.358589 | 0.261983433 | 0.15139133 | 1.73050491 | 0.0835401 | 0.30106496 |
| NDFIP2 | 146.999186 | 0.041067658 | 0.33569656 | 0.12233565 | 0.9026332 | 0.96445325 |
| SPRY2 | 34.5323718 | 0.393630189 | 0.46510624 | 0.84632317 | 0.39737249 | 0.67251432 |
| GPC5 | 77.9486433 | 1.420253444 | 0.64137985 | 2.21437179 | 0.02680321 | 0.15852905 |
| GPC6 | 343.08103 | -0.492819993 | 0.72657844 | -0.678275 | 0.49759735 | 0.74932951 |
| TGDS | 39.0191755 | 0.024790329 | 0.26340572 | 0.09411462 | 0.92501811 | 0.97344855 |
| GPR180 | 61.2538911 | 0.25949756 | 0.25914966 | 1.00134248 | 0.31666126 | 0.60316131 |
| ABCC4 | 832.051291 | -0.508879651 | 0.47373438 | -1.0741877 | 0.28273856 | 0.57066006 |
| CLDN10 | 164.288719 | -1.341134811 | 0.4968405 | -2.6993267 | 0.00694799 | 0.07095397 |
| DZIP1 | 88.1796119 | -1.053550616 | 0.83635522 | -1.2596928 | 0.20778022 | 0.48770564 |
| DNAJC3 | 196.991257 | -0.362068941 | 0.3308428 | -1.0943836 | 0.27378676 | 0.5611047 |
| UGGT2 | 469.748623 | 0.060207999 | 0.14772546 | 0.40756685 | 0.68359171 | 0.86478143 |
| MBNL2 | 501.628488 | 0.470720955 | 0.26874707 | 1.7515389 | 0.07985313 | 0.29363267 |
| RAP2A | 32.8861155 | -0.030384471 | 0.37683446 | -0.0806308 | 0.93573556 | 0.97837377 |
| IPO5 | 113.009425 | -0.426397993 | 0.24367797 | -1.7498422 | 0.08014554 | 0.29426581 |
| FARP1 | 1941.45582 | 0.429876836 | 0.22735013 | 1.89081414 | 0.05864916 | 0.24443767 |
| STK24 | 314.548517 | -0.275304681 | 0.13330821 | -2.0651742 | 0.03890652 | 0.19580915 |
| SLC15A1 | 39.6125191 | -1.573590138 | 0.70031375 | -2.2469788 | 0.02464138 | 0.15217128 |
| DOCK9 | 342.810127 | -0.091175914 | 0.2574826 | -0.3541051 | 0.72326009 | 0.88289614 |
| UBAC2 | 314.467293 | -0.12994744 | 0.15291456 | -0.8498042 | 0.39543393 | 0.67053666 |
| TM9SF2 | 219.308982 | -0.058854447 | 0.12888243 | -0.4566522 | 0.64792105 | 0.84579391 |
| CLYBL | 224.462927 | -0.050359651 | 0.28451353 | -0.1770027 | 0.8595063 | 0.94734879 |
| PCCA | 822.357618 | -0.424138554 | 0.22781907 | -1.8617342 | 0.06264056 | 0.25425167 |
| GGACT | 66.2750524 | -1.113740483 | 0.3670357 | -3.03442 | 0.00240999 | 0.03558826 |
| TMTC4 | 49.9028736 | -0.705177216 | 0.30309143 | -2.3266155 | 0.01998574 | 0.1347244 |
| FGF14 | 315.757767 | 0.529676475 | 0.6802659 | 0.77863153 | 0.4361968 | 0.70117959 |
| TPP2 | 178.432219 | -0.102796684 | 0.14173925 | -0.7252521 | 0.46829741 | 0.72597847 |
| EFNB2 | 64.26215 | 0.802539382 | 0.51582403 | 1.55583946 | 0.11974627 | 0.36465111 |
| ARGLU1 | 346.970839 | 0.488918681 | 0.14956994 | 3.26882992 | 0.00107993 | 0.02177825 |
| FAM155A | 465.790912 | 0.069640563 | 1.08404313 | 0.06424151 | 0.94877793 | 0.98262487 |
| ABHD13 | 39.633289 | 0.370838574 | 0.22120327 | 1.67646064 | 0.093648 | 0.31857635 |
| TNFSF13B | 40.03946 | 0.01552268 | 0.40137474 | 0.03867378 | 0.96915048 | 0.99152285 |
| IRS2 | 64.4360576 | -0.918952964 | 0.48786488 | -1.8836219 | 0.05961614 | 0.24631922 |
| COL4A1 | 769.346766 | 1.14265766 | 0.21441196 | 5.32926261 | 9.86E-08 | 2.95E-05 |
| COL4A2 | 908.878777 | 0.627192164 | 0.2096174 | 2.99208067 | 0.00277083 | 0.03841511 |
| RAB20 | 66.5011791 | -1.70841879 | 0.46808111 | -3.6498349 | 0.00026241 | 0.00887376 |
| NAXD | 35.1320623 | -0.135764408 | 0.29297275 | -0.4634029 | 0.64307563 | 0.84321657 |
| CARS2 | 123.568313 | -0.02974709 | 0.18538818 | -0.1604584 | 0.87251999 | 0.95193455 |
| ANKRD10 | 419.984268 | -0.087842412 | 0.13701999 | -0.641092 | 0.52146294 | 0.76444589 |
| ARHGEF7 | 254.730865 | 0.344478555 | 0.23096213 | 1.49149365 | 0.13583194 | 0.38788633 |
| TUBGCP3 | 129.865286 | 0.036060203 | 0.28196754 | 0.12788778 | 0.89823778 | 0.96244068 |
| ATP11A | 426.473835 | -0.16807949 | 0.28140612 | -0.5972844 | 0.55031751 | 0.7853831 |
| MCF2L | 216.325035 | 0.831604509 | 0.27916239 | 2.97892747 | 0.00289259 | 0.03963969 |
| PCID2 | 86.5315724 | 0.08086349 | 0.16342329 | 0.49481008 | 0.62073419 | 0.82839623 |

| | | | | | | |
|---|---|---|---|---|---|---|
| CUL4A | 198.888067 | -0.421296916 | 0.21591147 | -1.9512484 | 0.0510275 | 0.22682567 |
| LAMP1 | 189.118467 | -0.511394164 | 0.28257118 | -1.8097888 | 0.07032854 | 0.27219276 |
| GRTP1 | 44.7416194 | -0.196798302 | 0.55343896 | -0.3555917 | 0.72214636 | 0.88235193 |
| DCUN1D2 | 73.2403409 | 0.526930557 | 0.20681219 | 2.54786997 | 0.01083829 | 0.09361582 |
| TMCO3 | 148.237734 | 0.088040548 | 0.29094056 | 0.30260665 | 0.76218965 | 0.90141135 |
| TFDP1 | 62.9505828 | -0.057203716 | 0.31197237 | -0.1833615 | 0.8545144 | 0.94541388 |
| GAS6 | 46.5898397 | -0.830871052 | 0.65265436 | -1.2730644 | 0.20299518 | 0.4818154 |
| RASA3 | 69.378382 | -1.053428222 | 0.38222181 | -2.7560652 | 0.00585013 | 0.06406812 |
| CDC16 | 142.50856 | -0.057990368 | 0.20354641 | -0.2849 | 0.77572077 | 0.90732427 |
| UPF3A | 134.704293 | 0.091172319 | 0.18209617 | 0.50068225 | 0.61659476 | 0.82592881 |
| CHAMP1 | 26.5059016 | -0.408352382 | 0.31834478 | -1.2827362 | 0.1995845 | 0.47837581 |
| TTC5 | 45.2531001 | 0.24024187 | 0.27525624 | 0.87279355 | 0.38277561 | 0.65964257 |
| CCNB1IP1 | 58.8416517 | 0.056019834 | 0.29851434 | 0.18766212 | 0.85114153 | 0.94435157 |
| TEP1 | 37.5486809 | -0.203375656 | 0.27850567 | -0.7302388 | 0.4652442 | 0.72300587 |
| OSGEP | 34.521082 | 0.369605398 | 0.2620638 | 1.41036419 | 0.15843217 | 0.42403514 |
| APEX1 | 59.9426051 | -0.821529505 | 0.48412781 | -1.6969269 | 0.08971048 | 0.31211662 |
| PNP | 30.3043505 | -2.066260726 | 0.45710331 | -4.5203364 | 6.17E-06 | 0.00064828 |
| METTL17 | 33.1268065 | 0.103472292 | 0.26288062 | 0.39360943 | 0.69386941 | 0.86985671 |
| NDRG2 | 33.1487991 | -0.249627169 | 0.5113122 | -0.4882089 | 0.62540187 | 0.83169854 |
| ARHGEF40 | 39.9589021 | -0.354221336 | 0.59908105 | -0.5912745 | 0.55433653 | 0.78734553 |
| ZNF219 | 30.8593906 | 0.06233278 | 0.28211577 | 0.22094753 | 0.82513329 | 0.93217972 |
| HNRNPC | 706.525176 | 0.002952252 | 0.15723689 | 0.01877582 | 0.98501994 | 0.99668665 |
| SUPT16H | 126.82203 | -0.227882189 | 0.20656839 | -1.1031803 | 0.26994886 | 0.55705301 |
| CHD8 | 156.762842 | -0.006953528 | 0.17651265 | -0.0393939 | 0.96857631 | 0.99134443 |
| RAB2B | 61.833398 | 0.351044739 | 0.20796939 | 1.68796346 | 0.09141824 | 0.31522511 |
| TOX4 | 86.0475751 | -0.256871462 | 0.27230636 | -0.9433179 | 0.34551832 | 0.63006282 |
| METTL3 | 38.9973746 | 0.025317128 | 0.31984239 | 0.07915501 | 0.93690933 | 0.97901761 |
| DAD1 | 173.978689 | -0.298403832 | 0.39498684 | -0.7554779 | 0.44996228 | 0.71082906 |
| ABHD4 | 26.4931318 | 0.081282414 | 0.33824089 | 0.24030925 | 0.81009053 | 0.92495996 |
| OXA1L | 41.1559089 | -0.593928622 | 0.27737171 | -2.1412733 | 0.03225201 | 0.17584306 |
| MRPL52 | 44.5554117 | -0.065353551 | 0.42157297 | -0.1550231 | 0.8768031 | 0.95377544 |
| MMP14 | 45.2052731 | -0.356063461 | 0.24733678 | -1.4395896 | 0.14998354 | 0.41071822 |
| LRP10 | 151.301385 | -0.941037087 | 0.20070993 | -4.6885427 | 2.75E-06 | 0.0003423 |
| RBM23 | 99.3268851 | 0.122766893 | 0.17326302 | 0.70855795 | 0.47859884 | 0.73445983 |
| PRMT5 | 29.7808694 | -0.707560072 | 0.26418499 | -2.678275 | 0.00740024 | 0.07391754 |
| PSMB5 | 90.6014039 | -1.032007307 | 0.34491698 | -2.9920455 | 0.00277115 | 0.03841511 |
| ACIN1 | 191.863608 | 0.183658995 | 0.16697488 | 1.09991994 | 0.27136701 | 0.55841559 |
| C14orf119 | 28.2270247 | -1.33689424 | 0.43946373 | -3.0421037 | 0.00234931 | 0.0349985 |
| BCL2L2-PABI | 46.683226 | -0.030480456 | 0.25610612 | -0.1190149 | 0.90526352 | 0.96556525 |
| PABPN1 | 117.791547 | 0.933312484 | 0.29034809 | 3.21446055 | 0.0013069 | 0.02485643 |
| SLC22A17 | 39.1580051 | -0.343079611 | 0.51539641 | -0.6656616 | 0.5056274 | 0.75528806 |
| NGDN | 41.8465249 | -0.297513476 | 0.22356252 | -1.3307842 | 0.18326002 | 0.45690988 |
| ZFHX2 | 45.6518969 | 0.388155553 | 0.50001356 | 0.77629005 | 0.43757775 | 0.70212475 |
| AP1G2 | 34.1471257 | 0.10059714 | 0.34429261 | 0.29218501 | 0.77014517 | 0.90533273 |
| DHRS4-AS1 | 33.5216611 | 0.206399678 | 0.31714239 | 0.65081076 | 0.51516865 | 0.76135326 |
| DHRS4L2 | 38.4351734 | -0.134430781 | 0.42196525 | -0.3185826 | 0.75004306 | 0.89561986 |
| PSME1 | 118.054197 | -0.542838113 | 0.43912669 | -1.2361765 | 0.21639295 | 0.4979169 |
| AL136295.1 | 35.3975802 | -0.245680577 | 0.28087851 | -0.8746863 | 0.38174464 | 0.65885391 |
| GMPR2 | 28.4709012 | -0.573754467 | 0.344515 | -1.6653977 | 0.09583345 | 0.32289433 |
| KHNYN | 37.930923 | -0.623136945 | 0.27434917 | -2.2713279 | 0.02312713 | 0.14676719 |
| SDR39U1 | 34.8627163 | -0.249736027 | 0.30260318 | -0.8252921 | 0.40920575 | 0.68208527 |
| STXBP6 | 81.6414651 | 3.060324382 | 0.33983743 | 9.00525997 | 2.15E-19 | 2.06E-15 |
| NOVA1 | 101.923122 | -0.51496597 | 0.58669887 | -0.8777347 | 0.3800877 | 0.65731361 |
| PRKD1 | 1858.73668 | -0.119627553 | 0.25449074 | -0.4700664 | 0.63830755 | 0.83942176 |
| G2E3 | 82.3888815 | 0.455345832 | 0.17276112 | 2.63569618 | 0.00839649 | 0.08075639 |
| SCFD1 | 289.772221 | 0.190663715 | 0.16844626 | 1.13189642 | 0.25767799 | 0.54487802 |
| COCH | 22.6702848 | -0.769062519 | 0.40592874 | -1.8945752 | 0.05814871 | 0.24328193 |
| STRN3 | 748.043357 | -0.063546825 | 0.25497084 | -0.2492317 | 0.80318153 | 0.92125386 |
| AP4S1 | 68.5457819 | 0.508595189 | 0.31744719 | 1.60214111 | 0.10912441 | 0.34570701 |
| HECTD1 | 374.786454 | 0.05535166 | 0.14381197 | 0.3848891 | 0.70031959 | 0.87182648 |
| HEATR5A | 56.1198077 | 0.317691086 | 0.29952562 | 1.06064746 | 0.28885014 | 0.57571692 |
| AL139353.1 | 32.1514834 | 0.171096778 | 0.35950762 | 0.47591975 | 0.63413155 | 0.83680206 |
| NUBPL | 306.430463 | 0.396598649 | 0.20411498 | 1.94301591 | 0.05201424 | 0.22857606 |
| ARHGAP5 | 489.609555 | 0.017852437 | 0.14246796 | 0.12530844 | 0.90027937 | 0.96303857 |
| AKAP6 | 117.976044 | -0.179133546 | 0.39290662 | -0.4559189 | 0.64844831 | 0.84621977 |
| NPAS3 | 581.704818 | -0.260503481 | 0.62001957 | -0.4201536 | 0.67437322 | 0.86115505 |
| EGLN3 | 168.341954 | -0.538912607 | 0.69138469 | -0.7794685 | 0.43570377 | 0.70062219 |
| EAPP | 89.494837 | 0.218884495 | 0.28817849 | 0.75954488 | 0.44752668 | 0.70904036 |
| SNX6 | 230.523557 | -0.024514144 | 0.20106559 | -0.1219211 | 0.90296148 | 0.96461219 |
| CFL2 | 35.3891579 | -0.011418302 | 0.3973254 | -0.0287379 | 0.97707362 | 0.99441014 |
| BAZ1A | 195.180968 | -0.911819307 | 0.30888947 | -2.9519275 | 0.00315797 | 0.04214469 |
| SRP54 | 189.271773 | -0.105782808 | 0.16981992 | -0.6229117 | 0.53334257 | 0.77348803 |
| FAM177A1 | 115.809161 | -0.109588905 | 0.24224589 | -0.4523871 | 0.65099017 | 0.84724399 |
| PPP2R3C | 61.5420017 | -0.539117917 | 0.27044107 | -1.9934765 | 0.0462093 | 0.21432487 |
| NFKBIA | 218.714206 | -1.698345772 | 0.65048762 | -2.610881 | 0.00903093 | 0.0847991 |
| RALGAPA1 | 540.097074 | 0.485297381 | 0.16183223 | 2.99876851 | 0.00271073 | 0.03811711 |
| BRMS1L | 36.7615258 | -0.043304946 | 0.30340681 | -0.142729 | 0.88650422 | 0.95855333 |
| MBIP | 69.0624399 | 0.075061315 | 0.25896249 | 0.28985401 | 0.77192793 | 0.90624314 |
| SLC25A21 | 79.1722987 | -0.14898401 | 0.27543415 | -0.5409061 | 0.58857231 | 0.81121355 |
| MIPOL1 | 309.522929 | 0.52001586 | 0.17969136 | 2.8939391 | 0.00380442 | 0.0479507 |
| SEC23A | 192.659517 | 0.36228512 | 0.18605327 | 1.94721182 | 0.05150935 | 0.22779687 |
| TRAPPC6B | 82.2560034 | 0.380691988 | 0.18450126 | 2.06335717 | 0.03907871 | 0.19608954 |
| PNN | 202.147423 | 0.105438395 | 0.17764205 | 0.59354411 | 0.55281707 | 0.78613936 |
| MIA2 | 460.546914 | 0.081382572 | 0.22990384 | 0.35398527 | 0.72334992 | 0.88289614 |
| FBXO33 | 90.5326755 | 0.36268531 | 0.21159184 | 1.71407987 | 0.08651406 | 0.30636531 |
| AL049875.1 | 35.9511593 | 0.456761202 | 0.34185193 | 1.33613754 | 0.18150432 | 0.45460445 |
| LRFN5 | 133.943373 | -0.308173912 | 1.28188841 | -0.2404062 | 0.81001538 | 0.92495996 |
| KLHL28 | 103.154958 | 0.12954058 | 0.26934858 | 0.48094027 | 0.63055895 | 0.83461713 |
| TOGARAM1 | 214.341143 | 0.131727433 | 0.17713036 | 0.74367506 | 0.45707308 | 0.71628781 |
| PRPF39 | 78.4839998 | 0.306507302 | 0.18054658 | 1.69766328 | 0.08957133 | 0.31198177 |
| FKBP3 | 60.4188665 | 0.23954631 | 0.29043605 | 0.8247816 | 0.40949559 | 0.68208527 |
| FANCM | 45.7901561 | 0.42969091 | 0.30806742 | 1.39479502 | 0.16307766 | 0.42986266 |
| MIS18BP1 | 71.9887452 | 0.145918554 | 0.26355104 | 0.55366335 | 0.57980926 | 0.80465327 |
| MDGA2 | 124.631879 | 0.325300767 | 1.47494178 | 0.2205516 | 0.82544159 | 0.93241804 |
| RPS29 | 191.628892 | -0.459808658 | 0.2399808 | -1.9160227 | 0.05536221 | 0.23619173 |
| RPL36AL | 233.370061 | -0.950079892 | 0.49695123 | -1.9118172 | 0.05589965 | 0.23748015 |
| DNAAF2 | 27.6780522 | -0.517248937 | 0.30098668 | -1.718511 | 0.08570344 | 0.3045708 |
| POLE2 | 35.4260931 | -0.659553993 | 0.39097337 | -1.6869537 | 0.09161225 | 0.31554208 |
| KLHDC1 | 101.64441 | 0.73374347 | 0.35623114 | 2.05973985 | 0.03942342 | 0.19685721 |
| KLHDC2 | 119.296461 | 0.359890463 | 0.26922588 | 1.33676028 | 0.1813009 | 0.45439071 |
| NEMF | 316.536462 | 0.342922411 | 0.13503949 | 2.53942311 | 0.01110354 | 0.09513493 |
| ARF6 | 78.9722342 | 0.377456496 | 0.37305061 | 1.01181043 | 0.3116287 | 0.5991753 |
| LINC01588 | 29.0463922 | 1.605856143 | 0.43961169 | 3.65289678 | 0.0002593 | 0.00883922 |
| VCPKMT | 39.2620787 | 0.440216493 | 0.21635291 | 2.0347149 | 0.04187957 | 0.20312121 |
| SOS2 | 328.756273 | 0.412200224 | 0.21670339 | 1.90214015 | 0.05715283 | 0.24128118 |
| L2HGDH | 121.051972 | -0.174850267 | 0.30006612 | -0.5827058 | 0.56009138 | 0.78995945 |
| ATP5S | 68.5774545 | 0.200148891 | 0.19627117 | 1.01975694 | 0.30784375 | 0.59456464 |
| CDKL1 | 306.857742 | 1.561743022 | 0.41069359 | 3.80269639 | 0.00014313 | 0.00603982 |
| MAP4K5 | 341.291111 | 0.490896695 | 0.19936958 | 2.46224476 | 0.01380704 | 0.10854146 |
| SAV1 | 86.1144914 | -0.17717203 | 0.23563401 | -0.751895 | 0.45211421 | 0.71231034 |
| NIN | 101.783392 | -0.660506784 | 0.28831877 | -2.2908907 | 0.02196974 | 0.14229081 |
| TMX1 | 54.8325705 | 0.321629578 | 0.30490039 | 1.0548677 | 0.29148584 | 0.5787402 |
| RTRAF | 84.0015567 | -0.436640938 | 0.31498489 | -1.3862282 | 0.16567724 | 0.43396834 |

| | | | | | | |
|---|---|---|---|---|---|---|
| NID2 | 257.605551 | 0.704484753 | 0.55766938 | 1.26326597 | 0.20649361 | 0.48621411 |
| PTGER2 | 34.6710297 | 0.518930182 | 0.46878097 | 1.10697792 | 0.26830348 | 0.55545256 |
| TXNDC16 | 58.0097356 | 0.922958195 | 0.28175781 | 3.27571468 | 0.00105395 | 0.02158644 |
| ERO1A | 154.421159 | -1.223569922 | 0.50724169 | -2.412203 | 0.01585645 | 0.11728876 |
| PSMC6 | 119.686859 | -0.178412834 | 0.18926612 | -0.942656 | 0.3458569 | 0.63016843 |
| STYX | 37.006966 | 0.354347659 | 0.22761449 | 1.55678864 | 0.11952068 | 0.3645149 |
| FERMT2 | 235.415392 | -0.059765342 | 0.24282037 | -0.2461299 | 0.80558171 | 0.92227408 |
| DDHD1 | 85.0275475 | -0.522046996 | 0.21053402 | -2.4796325 | 0.01315178 | 0.10515938 |
| CNIH1 | 47.158157 | -0.376757802 | 0.26731533 | -1.4094134 | 0.15871297 | 0.42419408 |
| GMFB | 56.5221569 | -0.354143971 | 0.24721341 | -1.4325435 | 0.15198834 | 0.41407744 |
| CGRRF1 | 30.1215725 | 0.740684088 | 0.30021 | 2.4672199 | 0.01361667 | 0.10755093 |
| SAMD4A | 645.139683 | -0.345707196 | 0.2856017 | -1.2104522 | 0.22610543 | 0.50853813 |
| GCH1 | 36.7024123 | -1.633440772 | 0.60643566 | -2.6935104 | 0.00707039 | 0.07151774 |
| SOCS4 | 45.2911011 | -0.202886629 | 0.26144538 | -0.7760192 | 0.43773766 | 0.70212475 |
| MAPK1IP1L | 191.695078 | 0.052232563 | 0.21001638 | 0.24870708 | 0.80358737 | 0.92125386 |
| LGALS3 | 123.709287 | -0.664333962 | 0.59373446 | -1.1189075 | 0.26317959 | 0.55096925 |
| FBXO34 | 290.551901 | 0.368187399 | 0.17358853 | 2.12103527 | 0.03391883 | 0.18110843 |
| ATG14 | 65.7124552 | 0.203235274 | 0.30436051 | 0.66774521 | 0.50429624 | 0.75520517 |
| KTN1-AS1 | 36.5011719 | 0.196218024 | 0.45780836 | 0.42860297 | 0.66821219 | 0.85905308 |
| KTN1 | 615.935865 | 0.407651595 | 0.16313588 | 2.49884687 | 0.01245981 | 0.1019086 |
| PELI2 | 417.193593 | 0.606954162 | 0.33292456 | 1.82309817 | 0.06828852 | 0.26710321 |
| TMEM260 | 139.911799 | 0.404294924 | 0.20692111 | 1.95386017 | 0.05071776 | 0.22628104 |
| EXOC5 | 157.312089 | 0.182776316 | 0.17280587 | 1.05769738 | 0.29019343 | 0.57707346 |
| AP5M1 | 78.1969859 | -0.105823892 | 0.20457157 | -0.5172952 | 0.60495011 | 0.82048068 |
| NAA30 | 58.9341762 | 0.015228501 | 0.25218533 | 0.06038615 | 0.95184809 | 0.9832613 |
| CCDC198 | 809.415615 | 1.502464721 | 0.35997917 | 4.17375459 | 3.00E-05 | 0.00209494 |
| SLC35F4 | 94.8117993 | 2.085523932 | 0.80070584 | 2.60460686 | 0.00919797 | 0.08529273 |
| ARMH4 | 232.125184 | 1.033203566 | 0.31550877 | 3.27472217 | 0.00105766 | 0.02160196 |
| ACTR10 | 145.388833 | 0.026206291 | 0.15012096 | 0.17456783 | 0.86141923 | 0.94835188 |
| PSMA3 | 114.959953 | 0.156681893 | 0.22811238 | 0.68686273 | 0.49216923 | 0.74455175 |
| ARID4A | 217.791137 | 0.322237906 | 0.27037889 | 1.19180129 | 0.23333917 | 0.5160831 |
| TIMM9 | 51.4778196 | -0.607923045 | 0.24203012 | -2.5117661 | 0.01201287 | 0.10014906 |
| KIAA0586 | 110.22448 | 0.025824958 | 0.19065232 | 0.13545578 | 0.89225153 | 0.95976294 |
| DAAM1 | 171.951559 | 0.593619283 | 0.3301287 | 1.79814506 | 0.07215402 | 0.27680153 |
| GPR135 | 59.6744096 | -0.682855321 | 0.48829386 | -1.3984516 | 0.1619775 | 0.4283668 |
| L3HYPDH | 54.6178492 | -0.773345788 | 0.52640396 | -1.4691109 | 0.14180272 | 0.39798659 |
| JKAMP | 67.404181 | 0.061584449 | 0.22199678 | 0.27741145 | 0.78146419 | 0.91077326 |
| PCNX4 | 314.319097 | -0.152199687 | 0.29652223 | -0.5132826 | 0.60775368 | 0.8211189 |
| DHRS7 | 76.5062885 | -0.289868424 | 0.43820495 | -0.6614905 | 0.50829778 | 0.75746491 |
| PPM1A | 136.693681 | -0.0964809 | 0.2050871 | -0.4704387 | 0.63804165 | 0.83930252 |
| MNAT1 | 397.712252 | 0.165271555 | 0.20909719 | 0.79040542 | 0.42929104 | 0.69507954 |
| SLC38A6 | 87.9241852 | -0.193276225 | 0.27301098 | -0.707943 | 0.47898063 | 0.73481029 |
| PRKCH | 94.9770566 | 0.305378036 | 0.60351139 | 0.50600211 | 0.61285517 | 0.824514 |
| TMEM30B | 33.8754586 | 0.057639929 | 0.61837008 | 0.09321267 | 0.92573461 | 0.97360692 |
| HIF1A | 2456.24318 | -0.974889934 | 0.43334403 | -2.2496904 | 0.0244686 | 0.15183364 |
| HIF1A-AS2 | 403.222929 | -0.483004806 | 0.54467457 | -0.8867769 | 0.37519905 | 0.65364522 |
| SNAPC1 | 113.694296 | -0.485020403 | 0.23142487 | -2.0958007 | 0.03609988 | 0.18804981 |
| PPP2R5E | 409.934003 | -0.210428408 | 0.17965092 | -1.1713183 | 0.24147087 | 0.52617139 |
| WDR89 | 52.9479826 | 0.02795172 | 0.21681957 | 0.12891696 | 0.89742336 | 0.96178321 |
| SGPP1 | 141.522052 | 0.739326171 | 0.25665789 | 2.88059003 | 0.00396932 | 0.04918768 |
| SYNE2 | 3411.4046 | -0.276783826 | 0.20505217 | -1.3498215 | 0.17707325 | 0.44872611 |
| ESR2 | 268.90352 | 0.604884406 | 0.59460617 | 1.01728578 | 0.3090175 | 0.59563596 |
| MTHFD1 | 73.2783775 | 0.10739355 | 0.33866059 | 0.31711263 | 0.75115814 | 0.89599662 |
| AL122035.1 | 62.5271523 | 0.802849647 | 0.46093068 | 1.74180128 | 0.08154322 | 0.29654614 |
| ZBTB25 | 96.8205989 | -0.159659175 | 0.34192096 | -0.4669476 | 0.64053736 | 0.84142998 |
| AKAP5 | 30.1255385 | 2.686043304 | 0.63520746 | 4.22860791 | 2.35E-05 | 0.00169355 |
| ZBTB1 | 192.843611 | 0.321650842 | 0.225316 | 1.42755437 | 0.15342016 | 0.41620271 |
| HSPA2 | 30.1424228 | 0.978106981 | 0.71214665 | 1.37346286 | 0.1696085 | 0.43945897 |
| PLEKHG3 | 210.856391 | 0.666410103 | 0.3559364 | 1.872273 | 0.06116884 | 0.25014908 |
| CHURC1-FNT | 41.2009999 | -0.937254777 | 0.23082552 | -4.0604469 | 4.90E-05 | 0.00295075 |
| MAX | 160.580106 | -0.740247049 | 0.36446289 | -2.0310629 | 0.04224862 | 0.20401679 |
| FUT8 | 403.497839 | 0.406791199 | 0.18151441 | 2.24109584 | 0.02501987 | 0.15333673 |
| AL359232.1 | 133.249542 | -0.602323026 | 0.51537334 | -1.168712 | 0.24251967 | 0.52725736 |
| GPHN | 1459.0511 | -0.035776128 | 0.33081511 | -0.1081454 | 0.91388037 | 0.9684766 |
| MPP5 | 204.829033 | 0.414658503 | 0.12159622 | 3.41012664 | 0.00064933 | 0.01564098 |
| ATP6V1D | 118.305039 | 0.223617257 | 0.2571281 | 0.86967258 | 0.38447936 | 0.66091637 |
| EIF2S1 | 73.9340328 | -0.564116149 | 0.2552707 | -2.2098743 | 0.02711389 | 0.15922956 |
| PLEKHH1 | 134.799313 | 0.264539953 | 0.39499693 | 0.66972661 | 0.50303208 | 0.75403031 |
| PIGH | 51.3392038 | 1.099473245 | 0.4220072 | 2.60534241 | 0.00917825 | 0.08519229 |
| ARG2 | 240.766972 | 2.372029497 | 1.10681355 | 2.14311571 | 0.03210381 | 0.17552647 |
| VTI1B | 57.5445658 | 0.06070196 | 0.36296533 | 0.167239 | 0.867182 | 0.95090501 |
| ZFYVE26 | 55.6677103 | 0.035438063 | 0.23332665 | 0.15188177 | 0.87928019 | 0.95537942 |
| RAD51B | 765.736197 | 0.459699843 | 0.43180637 | 1.06459717 | 0.28705826 | 0.57406774 |
| ZFP36L1 | 816.174686 | -0.299698725 | 0.26430621 | -1.1339072 | 0.25683346 | 0.54405301 |
| ACTN1 | 215.759104 | -0.376093052 | 0.22265486 | -1.6891302 | 0.09119449 | 0.31502156 |
| DCAF5 | 307.494483 | 0.058316527 | 0.19146294 | 0.30458389 | 0.76068309 | 0.90125623 |
| EXD2 | 61.5179458 | 0.674170222 | 0.20543272 | 3.28170812 | 0.0010318 | 0.02130096 |
| ERH | 111.256112 | -0.289148925 | 0.34589078 | -0.8359544 | 0.40318055 | 0.67708038 |
| SLC39A9 | 100.129734 | -0.13604198 | 0.16926052 | -0.8037431 | 0.42154533 | 0.6898802 |
| SUSD6 | 322.234524 | -1.234139941 | 0.44580975 | -2.7683108 | 0.00563477 | 0.062532 |
| SRSF5 | 341.804547 | 0.804910883 | 0.27656047 | 2.91043359 | 0.00360928 | 0.04634485 |
| AL357153.2 | 63.5176894 | -0.134457068 | 0.38454306 | -0.3496541 | 0.72659828 | 0.88522597 |
| MED6 | 64.0890025 | 0.400081525 | 0.21824258 | 1.83319648 | 0.06677336 | 0.26332729 |
| MAP3K9 | 84.4602411 | -0.452133775 | 0.26822211 | -1.6856693 | 0.09185951 | 0.31561057 |
| PCNX1 | 402.481044 | 0.113927794 | 0.20036662 | 0.56859668 | 0.56962988 | 0.79749848 |
| SIPA1L1 | 765.796608 | -0.004659389 | 0.27089343 | -0.0172001 | 0.986277 | 0.99678172 |
| RGS6 | 235.315961 | 2.283303574 | 1.35221992 | 1.68855934 | 0.09130391 | 0.31502156 |
| ZFYVE1 | 60.2966519 | -0.210999017 | 0.32824185 | -0.6428157 | 0.52034369 | 0.76400554 |
| RBM25 | 282.915485 | 0.219591801 | 0.17827864 | 1.23173366 | 0.21804859 | 0.50016462 |
| PSEN1 | 208.135732 | 0.000270553 | 0.16057133 | 0.00168494 | 0.99865561 | 0.99928153 |
| PAPLN | 184.736834 | -0.442750332 | 0.64826132 | -0.6829813 | 0.49461869 | 0.7467222 |
| NUMB | 706.721991 | -0.059318367 | 0.18914739 | -0.3136092 | 0.75381785 | 0.89777709 |
| DNAL1 | 95.130306 | 0.386676038 | 0.22506016 | 1.71810077 | 0.08577824 | 0.30466063 |
| PNMA1 | 36.2340242 | -0.337534167 | 0.38401471 | -0.8789616 | 0.37942212 | 0.65699285 |
| ELMSAN1 | 115.296009 | -0.076646943 | 0.25855737 | -0.2964408 | 0.7668935 | 0.90390954 |
| LINC02274 | 42.0696873 | -2.467304906 | 0.64394023 | -3.8315744 | 0.00012733 | 0.0055692 |
| AC005480.1 | 53.2576838 | 0.711873711 | 0.37292912 | 1.90887135 | 0.05627869 | 0.23885404 |
| ENTPD5 | 63.4110246 | -0.250855604 | 0.32251725 | -0.7778052 | 0.43668385 | 0.70170049 |
| BBOF1 | 89.464479 | 0.404296582 | 0.35099221 | 1.15186768 | 0.24937545 | 0.53644003 |
| ALDH6A1 | 59.0944279 | 0.275215907 | 0.47230156 | 0.58271225 | 0.56008703 | 0.78995945 |
| LIN52 | 108.439557 | 0.615706488 | 0.25454449 | 2.41885613 | 0.0155694 | 0.11595952 |
| ABCD4 | 117.511428 | -0.043081017 | 0.17942224 | -0.2401097 | 0.81024522 | 0.92495996 |
| NPC2 | 122.506776 | -1.065214203 | 0.42790968 | -2.4893436 | 0.01279792 | 0.10336535 |
| ISCA2 | 31.6343387 | -0.453007927 | 0.48960856 | -0.9252451 | 0.35483843 | 0.63648555 |
| AREL1 | 82.301483 | 0.040046154 | 0.18550157 | 0.2158804 | 0.82908097 | 0.93373939 |
| FCF1 | 46.4407943 | -0.018617806 | 0.24074945 | -0.0773327 | 0.93835887 | 0.97989095 |
| YLPM1 | 222.140899 | 0.07113137 | 0.15041495 | 0.47290095 | 0.63628385 | 0.83814123 |
| DLST | 80.201512 | -0.154954737 | 0.23242564 | -0.6666852 | 0.50497323 | 0.75524917 |
| MLH3 | 148.67549 | 0.13522353 | 0.20418701 | 0.66225335 | 0.50780887 | 0.75687816 |
| ACYP1 | 66.2389705 | 0.95021738 | 0.32420177 | 2.93094448 | 0.00337933 | 0.04410166 |
| NEK9 | 121.652651 | 0.172302089 | 0.14261008 | 1.20820417 | 0.22696874 | 0.50976168 |
| TMED10 | 194.097132 | -0.50895906 | 0.20069273 | -2.5360115 | 0.01121231 | 0.09563908 |
| FOS | 116.055313 | 2.917761891 | 0.77710294 | 3.75466587 | 0.00017357 | 0.00692943 |

| | | | | | | |
|---|---|---|---|---|---|---|
| JDP2 | 30.1472175 | 1.369604209 | 0.50007937 | 2.73877366 | 0.00616688 | 0.06585681 |
| FLVCR2 | 110.928271 | 0.708176314 | 0.52381017 | 1.3519713 | 0.1763845 | 0.44840421 |
| TTLL5 | 299.750187 | 0.233101534 | 0.16653022 | 1.39975517 | 0.16158665 | 0.42791083 |
| ERG28 | 34.2455183 | -0.103273074 | 0.33695378 | -0.3064903 | 0.75923134 | 0.9005757 |
| IFT43 | 213.889909 | -0.089370339 | 0.2936144 | -0.30438 | 0.76083844 | 0.90125623 |
| GPATCH2L | 217.185722 | -0.248736866 | 0.24850922 | -1.000916 | 0.31686741 | 0.60343077 |
| ANGEL1 | 26.9007202 | 0.396536347 | 0.2767713 | 1.43272207 | 0.15193728 | 0.41406907 |
| IRF2BPL | 87.4816284 | 0.035025573 | 0.41482609 | 0.08443435 | 0.93271109 | 0.97718906 |
| POMT2 | 45.9164329 | 0.326037374 | 0.25671589 | 1.27003193 | 0.20407326 | 0.48301501 |
| TMED8 | 115.399229 | -0.316722698 | 0.16599105 | -1.9080709 | 0.05638206 | 0.23918676 |
| VIPAS39 | 32.505102 | -0.033690029 | 0.2311923 | -0.145723 | 0.88414009 | 0.95761929 |
| AHSA1 | 49.2866278 | -0.095106935 | 0.29616916 | -0.3211237 | 0.74811666 | 0.89440069 |
| SPTLC2 | 345.183543 | 0.985930377 | 0.280053 | 3.52051356 | 0.00043071 | 0.01209909 |
| ALKBH1 | 51.9341687 | 0.224428266 | 0.20178414 | 1.11221956 | 0.26604376 | 0.55384374 |
| SLIRP | 81.7472792 | -0.161639562 | 0.23272241 | -0.6945595 | 0.4873314 | 0.74104173 |
| SNW1 | 76.1295587 | -0.127046478 | 0.2149288 | -0.5911096 | 0.55444698 | 0.78734553 |
| NRXN3 | 164.08022 | 0.330359826 | 0.92854104 | 0.35578377 | 0.72200251 | 0.88235193 |
| CEP128 | 103.521477 | 0.600395982 | 0.31233905 | 1.92225721 | 0.0545734 | 0.23421085 |
| AC007262.2 | 53.6998079 | 0.576975892 | 0.57182205 | 1.00901301 | 0.31296839 | 0.60019427 |
| GTF2A1 | 84.4593108 | 0.491708182 | 0.19137174 | 2.5693876 | 0.01018784 | 0.09022316 |
| STON2 | 239.445626 | 0.680259558 | 0.55408707 | 1.22771238 | 0.21955496 | 0.50121948 |
| SEL1L | 168.719234 | 0.107520773 | 0.29035653 | 0.37030603 | 0.71115448 | 0.87748 |
| GALC | 51.9074528 | 0.632045863 | 0.24693995 | 2.55951242 | 0.01048191 | 0.09186297 |
| SPATA7 | 122.981227 | 0.366123658 | 0.28040755 | 1.30568402 | 0.19166004 | 0.46873362 |
| PTPN21 | 134.945426 | 0.727492972 | 0.23041753 | 3.15728134 | 0.00159248 | 0.02783637 |
| ZC3H14 | 198.134611 | 0.250625554 | 0.17050907 | 1.46986638 | 0.14159795 | 0.39764489 |
| EML5 | 101.243057 | -0.183119702 | 0.50824588 | -0.3602975 | 0.7186247 | 0.88155276 |
| TTC8 | 110.429414 | 0.274579711 | 0.23244457 | 1.18126961 | 0.23749564 | 0.52023112 |
| FOXN3 | 190.74953 | 0.290218903 | 0.29884907 | 0.97112199 | 0.33148753 | 0.61668656 |
| EFCAB11 | 80.5450353 | 0.253852145 | 0.36071494 | 0.70374725 | 0.48159019 | 0.73620566 |
| TDP1 | 50.9603014 | -0.477594107 | 0.23056306 | -2.0714251 | 0.03831909 | 0.19441659 |
| PSMC1 | 68.6434967 | -0.602737318 | 0.27431917 | -2.1972118 | 0.02800532 | 0.16199455 |
| NRDE2 | 68.0434047 | 0.265560752 | 0.34368639 | 0.77268334 | 0.43970981 | 0.7032516 |
| CALM1 | 270.313253 | -0.38099723 | 0.33167028 | -1.1487228 | 0.25067029 | 0.53813776 |
| TTC7B | 240.437565 | 0.530730745 | 0.3758333 | 1.41214401 | 0.15790756 | 0.42322231 |
| RPS6KA5 | 104.779172 | 0.023207676 | 0.31235578 | 0.07429885 | 0.94077259 | 0.98016757 |
| DGLUCY | 171.997001 | -0.126350419 | 0.29076545 | -0.4345441 | 0.66389336 | 0.85622919 |
| PPP4R3A | 229.07628 | 0.282620465 | 0.126583 | 2.23268895 | 0.02556946 | 0.1551184 |
| CATSPERB | 83.9875903 | -0.187833492 | 0.37540184 | -0.5003531 | 0.61682644 | 0.82592881 |
| TC2N | 94.2735665 | 0.331968542 | 0.30623176 | 1.08404346 | 0.27834553 | 0.56578102 |
| TRIP11 | 246.738823 | 0.257156966 | 0.17385615 | 1.47913644 | 0.13910385 | 0.39282895 |
| ATXN3 | 155.511016 | 0.191429597 | 0.25222226 | 0.75897187 | 0.44786939 | 0.70935873 |
| NDUFB1 | 65.6197362 | -0.309337389 | 0.32027819 | -0.9658397 | 0.33412441 | 0.61861931 |
| CPSF2 | 88.0753598 | -0.167719676 | 0.21611325 | -0.7760731 | 0.43770583 | 0.70212475 |
| LGMN | 75.9603084 | -0.345294877 | 0.40759788 | -0.8471459 | 0.39691379 | 0.67221309 |
| GOLGA5 | 59.5484905 | -0.107437716 | 0.17932156 | -0.5991344 | 0.54908327 | 0.78432279 |
| ITPK1 | 98.7926733 | 0.351566847 | 0.22000387 | 1.59800304 | 0.1100423 | 0.34708436 |
| MOAP1 | 36.7578512 | -0.506846379 | 0.39240985 | -1.291625 | 0.19648704 | 0.47560247 |
| BTBD7 | 529.022631 | 0.010194498 | 0.22127687 | 0.04607123 | 0.96325347 | 0.9883253 |
| UNC79 | 51.9649716 | 0.155198246 | 0.62917622 | 0.24666896 | 0.80516443 | 0.92227408 |
| DDX24 | 192.627944 | -0.680077535 | 0.20064838 | -3.3893996 | 0.00070046 | 0.01628566 |
| IFI27L2 | 33.9434593 | -0.301960043 | 0.42871009 | -0.7043455 | 0.48121761 | 0.73620566 |
| SERPINA1 | 1320.89685 | -3.089556275 | 0.68573639 | -4.5054576 | 6.62E-06 | 0.00066781 |
| DICER1 | 368.621186 | -0.187409911 | 0.17997414 | -1.0413158 | 0.29772901 | 0.58554106 |
| DICER1-AS1 | 44.5962248 | 0.083859109 | 0.48040863 | 0.17455787 | 0.86142705 | 0.94835188 |
| CLMN | 506.192476 | -0.088294136 | 0.30877164 | -0.2859529 | 0.77491422 | 0.90689106 |
| GLRX5 | 55.7004637 | -0.183286602 | 0.45553472 | -0.4023548 | 0.68742289 | 0.86733718 |
| ATG2B | 195.248402 | 0.585162942 | 0.22727876 | 2.57464863 | 0.0100342 | 0.08977568 |
| GSKIP | 35.4127229 | -0.21106347 | 0.28576475 | -0.7385917 | 0.46015497 | 0.71894055 |
| AK7 | 57.4585792 | -0.193954341 | 0.55683604 | -0.348315 | 0.72760364 | 0.88564265 |
| PAPOLA | 322.328047 | 0.450363067 | 0.12886318 | 3.49489341 | 0.00047425 | 0.0129044 |
| VRK1 | 43.3656498 | 0.270695969 | 0.27715029 | 0.97671183 | 0.32871184 | 0.61402705 |
| LINC01550 | 41.414702 | 0.400764398 | 0.48598892 | 0.8246369 | 0.40957777 | 0.68208527 |
| SETD3 | 256.292668 | 0.139686322 | 0.16889145 | 0.82707751 | 0.40819313 | 0.68155517 |
| CCNK | 62.5124944 | -0.093505891 | 0.20715767 | -0.4513755 | 0.65171896 | 0.84786309 |
| CCDC85C | 38.6168146 | 0.342541848 | 0.34712203 | 0.98680527 | 0.32373811 | 0.6106578 |
| EML1 | 46.1762308 | 0.256056797 | 0.51062379 | 0.50145881 | 0.61604826 | 0.82592881 |
| EVL | 182.757152 | -1.18086723 | 0.38950963 | -3.0316766 | 0.002432 | 0.03574435 |
| YY1 | 224.056423 | -0.122866206 | 0.13205407 | -0.9304235 | 0.35215186 | 0.63395276 |
| SLC25A29 | 141.290042 | -0.041773428 | 0.25724735 | -0.1623862 | 0.87100171 | 0.95174591 |
| WARS | 85.1339942 | -1.493348889 | 0.47653413 | -3.1337711 | 0.00172575 | 0.02921304 |
| WDR25 | 66.9618882 | -0.38387433 | 0.22154385 | -1.7327239 | 0.08314475 | 0.30002855 |
| PPP2R5C | 239.559695 | 0.143834889 | 0.19149333 | 0.75112219 | 0.45257912 | 0.71261515 |
| DYNC1H1 | 251.708862 | -0.4177154 | 0.17654391 | -2.3660708 | 0.017978 | 0.12664775 |
| HSP90AA1 | 858.72466 | 0.328494941 | 0.35036846 | 0.93756996 | 0.34846545 | 0.63229918 |
| WDR20 | 251.392712 | 0.460031182 | 0.18203497 | 2.52715832 | 0.01149897 | 0.09721852 |
| MOK | 201.512812 | 0.597542099 | 0.32037905 | 1.86510977 | 0.06216602 | 0.25296869 |
| ZNF839 | 41.9131094 | 0.093159199 | 0.31368241 | 0.29698573 | 0.76647741 | 0.90372312 |
| CINP | 24.2035772 | -0.545953433 | 0.31337187 | -1.7421903 | 0.08147515 | 0.29654614 |
| TECPR2 | 92.6223558 | 0.037363708 | 0.21389444 | 0.17468293 | 0.86132878 | 0.94835188 |
| RCOR1 | 172.099259 | 0.150563426 | 0.17125965 | 0.87915296 | 0.37931835 | 0.65699285 |
| TRAF3 | 207.132425 | -0.312897716 | 0.30983495 | -1.0098851 | 0.31255032 | 0.60006724 |
| AMN | 45.0849068 | -0.920234213 | 0.61347021 | -1.5000471 | 0.1336022 | 0.38489489 |
| CDC42BPB | 290.180483 | -0.034828849 | 0.20825491 | -0.1672414 | 0.86718008 | 0.95090501 |
| TNFAIP2 | 231.998417 | -2.780700986 | 0.73049432 | -3.8066018 | 0.00014089 | 0.00597159 |
| EIF5 | 138.960747 | -0.807025689 | 0.30698454 | -2.6288805 | 0.00856665 | 0.08181445 |
| MARK3 | 559.277429 | 0.266335004 | 0.19259847 | 1.38285108 | 0.16671053 | 0.43490657 |
| CKB | 107.527072 | -1.119492266 | 0.41883731 | -2.6728571 | 0.00752083 | 0.07457766 |
| BAG5 | 68.6224736 | -0.837144351 | 0.18526816 | -4.5185549 | 6.23E-06 | 0.00064828 |
| KLC1 | 138.938709 | 0.06810073 | 0.20703941 | 0.32892641 | 0.74221131 | 0.89253455 |
| ZFYVE21 | 52.8781963 | 0.065441818 | 0.28338972 | 0.23092516 | 0.81737294 | 0.92966224 |
| PPP1R13B | 458.781531 | 0.15668332 | 0.24594863 | 0.63705709 | 0.52408766 | 0.76606495 |
| ATP5MPL | 86.2900221 | -0.235120378 | 0.31115159 | -0.7556458 | 0.44986161 | 0.71082906 |
| INF2 | 60.4503341 | -0.581408449 | 0.65247616 | -0.89108 | 0.37288626 | 0.65217873 |
| SIVA1 | 47.6048408 | 0.195704012 | 0.36561908 | 0.5352675 | 0.5924649 | 0.81283604 |
| AKT1 | 61.8471611 | -0.903988171 | 0.29304208 | -3.0848408 | 0.00203661 | 0.03219258 |
| CEP170B | 39.2392969 | -0.364172694 | 0.36726836 | -0.9915711 | 0.32140679 | 0.60844973 |
| CLBA1 | 29.4661834 | 0.091269397 | 0.30056276 | 0.3036617 | 0.76138564 | 0.90141135 |
| BRF1 | 60.9869595 | 0.076589439 | 0.24759532 | 0.30933315 | 0.75706812 | 0.89930004 |
| PACS2 | 116.263319 | -0.116061776 | 0.27710112 | -0.4188427 | 0.67533111 | 0.86115505 |
| TEX22 | 27.9088739 | -0.23641312 | 0.35987877 | -0.6569243 | 0.51122957 | 0.75947086 |
| MTA1 | 76.7214636 | 0.386598336 | 0.31439509 | 1.22965767 | 0.21882533 | 0.50062761 |
| CRIP2 | 73.5435509 | 0.08478549 | 0.50806244 | 0.16688006 | 0.86746442 | 0.95090501 |
| NIPA1 | 91.0745987 | 0.196764056 | 0.30682271 | 0.6412956 | 0.52133064 | 0.76440016 |
| NIPA2 | 75.6952502 | -0.312810973 | 0.33183966 | -0.942657 | 0.34585638 | 0.63016843 |
| CYFIP1 | 169.302077 | -0.054510531 | 0.24164016 | -0.2255856 | 0.82152377 | 0.93183851 |
| TUBGCP5 | 53.6100836 | -0.025240193 | 0.34874567 | -0.0723742 | 0.94230411 | 0.98093339 |
| AC124312.1 | 431.470317 | 1.245716254 | 0.31994447 | 3.89353891 | 9.88E-05 | 0.00461625 |
| PWAR6 | 98.0241924 | 1.221191358 | 0.36965044 | 3.30363829 | 0.00095439 | 0.02022586 |
| UBE3A | 341.569539 | 0.189902467 | 0.10803645 | 1.75776296 | 0.07878785 | 0.29095613 |
| ATP10A | 125.846765 | 0.338394055 | 0.66098983 | 0.51195047 | 0.60868568 | 0.82155842 |
| OCA2 | 63.0091073 | 1.223917919 | 0.71682136 | 1.70742389 | 0.08774329 | 0.3082165 |
| HERC2 | 352.77442 | 0.188625881 | 0.18674373 | 1.01007878 | 0.31245755 | 0.60004628 |

| | | | | | | |
|---|---|---|---|---|---|---|
| GOLGA8M | 33.9800262 | 0.598169004 | 0.72950531 | 0.81996525 | 0.41223592 | 0.68367283 |
| FAM189A1 | 367.153361 | -0.294831024 | 1.01883493 | -0.2893806 | 0.77229017 | 0.90624314 |
| TJP1 | 390.450988 | -0.103153096 | 0.23418506 | -0.4404768 | 0.65959178 | 0.85312309 |
| FAN1 | 84.1675898 | 0.452445625 | 0.25197374 | 1.79560626 | 0.07255716 | 0.27778778 |
| MTMR10 | 414.345422 | 1.318368195 | 0.37112321 | 3.5523733 | 0.00038177 | 0.01120155 |
| KLF13 | 117.101502 | -0.148192317 | 0.32710527 | -0.4530417 | 0.65051874 | 0.84724399 |
| OTUD7A | 101.452218 | -0.019344093 | 0.21018803 | -0.0920323 | 0.92667236 | 0.97414308 |
| LINC02256 | 219.591747 | -0.311463098 | 0.2348673 | -1.3261237 | 0.18479871 | 0.45871648 |
| AC123768.3 | 50.2024893 | 0.642220462 | 0.4595608 | 1.39746572 | 0.16227357 | 0.42858376 |
| FMN1 | 76.3460566 | -1.216020433 | 0.48256081 | -2.519932 | 0.01173775 | 0.09871458 |
| RYR3 | 61.9846513 | -1.633934271 | 0.55581324 | -2.9397182 | 0.00328511 | 0.04328482 |
| AVEN | 134.020446 | 0.510280452 | 0.42878001 | 1.19007518 | 0.23401685 | 0.51656729 |
| EMC7 | 57.5325632 | -0.487696148 | 0.32078992 | -1.5202976 | 0.12843619 | 0.37635066 |
| KATNBL1 | 215.570143 | 0.25274126 | 0.23055609 | 1.09622462 | 0.27298049 | 0.56047062 |
| EMC4 | 32.4870891 | -0.695221464 | 0.34736122 | -2.0014366 | 0.04534536 | 0.21178118 |
| SLC12A6 | 126.928964 | 0.55936766 | 0.32192189 | 1.73758813 | 0.08228341 | 0.29878424 |
| NOP10 | 88.4946841 | -0.873416595 | 0.52880843 | -1.6516692 | 0.09860201 | 0.32818229 |
| GOLGA8A | 134.955904 | 0.507115646 | 0.40492057 | 1.25238303 | 0.21043033 | 0.49127431 |
| GOLGA8B | 141.641984 | 0.130287563 | 0.44996108 | 0.28955296 | 0.77215826 | 0.90624314 |
| AQR | 157.352448 | 0.269243102 | 0.16589179 | 1.62300442 | 0.10458846 | 0.33777911 |
| ZNF770 | 94.1204655 | -0.05706735 | 0.21358974 | -0.2671821 | 0.78932898 | 0.91504082 |
| AC021231.1 | 197.160396 | 0.151072911 | 0.18164736 | 0.83168239 | 0.40558824 | 0.67898109 |
| DPH6 | 536.195963 | -0.054327819 | 0.1899015 | -0.2860842 | 0.77481363 | 0.90689106 |
| DPH6-DT | 228.266349 | 1.238043153 | 0.58918881 | 2.10126726 | 0.03561751 | 0.18653917 |
| C15orf41 | 93.8305767 | 0.309476411 | 0.26730281 | 1.15777462 | 0.24695602 | 0.53375265 |
| MEIS2 | 275.673635 | -0.4675714 | 0.906341 | -0.5158891 | 0.6059319 | 0.8207327 |
| SPRED1 | 272.321907 | -0.482155912 | 0.19796444 | -2.4355683 | 0.01486842 | 0.11321507 |
| FAM98B | 57.1055493 | 0.228906186 | 0.23638908 | 0.96834503 | 0.33287208 | 0.61734399 |
| RASGRP1 | 117.807663 | 0.069308122 | 0.43132418 | 0.16068685 | 0.87234005 | 0.95193455 |
| C15orf53 | 77.2545459 | 0.014643713 | 0.71990398 | 0.0203412 | 0.98377119 | 0.9965677 |
| AC013652.1 | 235.404931 | 0.470568507 | 0.37927657 | 1.24070019 | 0.21471653 | 0.49688442 |
| THBS1 | 295.32727 | 2.166237079 | 0.5927981 | 3.6542578 | 0.00025793 | 0.00882387 |
| FSIP1 | 66.5466005 | 1.16775902 | 0.28908738 | 4.0394673 | 5.36E-05 | 0.00311014 |
| GPR176 | 101.681646 | 0.526703378 | 0.81921006 | 0.64294056 | 0.52026267 | 0.76400369 |
| EIF2AK4 | 141.120253 | 0.543324152 | 0.28582054 | 1.90092761 | 0.05731149 | 0.24173433 |
| SRP14 | 374.096081 | -0.595837051 | 0.46643018 | -1.277441 | 0.20144659 | 0.48024387 |
| SRP14-AS1 | 26.4341632 | 0.403467596 | 0.49886822 | 0.80876588 | 0.41864983 | 0.68798193 |
| IVD | 82.9595335 | 0.064063074 | 0.29437585 | 0.2176234 | 0.82772255 | 0.93346403 |
| BAHD1 | 29.192383 | -0.451267417 | 0.41327717 | -1.0919244 | 0.27486634 | 0.56187466 |
| CCDC32 | 86.4882544 | -0.091668268 | 0.35970329 | -0.2548441 | 0.7988435 | 0.91949148 |
| RMDN3 | 47.3446076 | -0.1304533 | 0.30584332 | -0.4265364 | 0.66971705 | 0.85924744 |
| GCHFR | 48.2276844 | -1.167078554 | 0.45445691 | -2.5680731 | 0.01022656 | 0.09036919 |
| DNAJC17 | 69.5193533 | -0.030271123 | 0.20638618 | -0.1466722 | 0.88339073 | 0.95756476 |
| ZFYVE19 | 29.0047018 | -0.423372594 | 0.33685023 | -1.2568571 | 0.20880539 | 0.48927271 |
| SPINT1-AS1 | 35.4227262 | 0.275161477 | 0.46173499 | 0.59592945 | 0.55122236 | 0.78577398 |
| SPINT1 | 97.4887068 | -1.155936424 | 0.30476685 | -3.7928548 | 0.00014893 | 0.00620241 |
| INO80 | 229.920127 | -0.088448763 | 0.30696711 | -0.2881376 | 0.77324142 | 0.90643453 |
| CHP1 | 112.471332 | 0.786669722 | 0.57494129 | 1.36826097 | 0.1712304 | 0.44163059 |
| RTF1 | 140.938235 | -0.367987123 | 0.16679097 | -2.2062772 | 0.0273646 | 0.16002776 |
| RPAP1 | 28.0432824 | -0.297891098 | 0.28867912 | -1.0319108 | 0.30211391 | 0.58973509 |
| MGA | 195.118263 | 0.042119307 | 0.21896985 | 0.19235208 | 0.84746642 | 0.94323402 |
| MAPKBP1 | 75.1943142 | 0.188522424 | 0.25567068 | 0.73736425 | 0.46090087 | 0.71918385 |
| EHD4 | 65.6170042 | -0.55007732 | 0.35392595 | -1.5542159 | 0.12013293 | 0.36497094 |
| VPS39 | 90.145339 | -0.312836825 | 0.15580981 | -2.0078121 | 0.04466326 | 0.21044237 |
| TMEM87A | 199.768273 | -0.229783699 | 0.18525406 | -1.2403707 | 0.21483832 | 0.49688442 |
| GANC | 77.7103259 | 0.222719974 | 0.45744751 | 0.48687548 | 0.62634658 | 0.83180006 |
| ZNF106 | 125.783605 | -0.196812115 | 0.16499769 | -1.1928174 | 0.23294091 | 0.51591698 |
| SNAP23 | 101.414405 | 0.118830345 | 0.21929425 | 0.54187625 | 0.58790376 | 0.81095283 |
| HAUS2 | 31.3931415 | -0.218785992 | 0.29798537 | -0.7342172 | 0.46281636 | 0.72095238 |
| TTBK2 | 221.95941 | 0.324979686 | 0.20381209 | 1.59450645 | 0.11082265 | 0.34794171 |
| UBR1 | 323.658375 | 0.084017801 | 0.16502942 | 0.50910803 | 0.61067651 | 0.82275767 |
| AC068724.3 | 72.0388281 | 0.363702711 | 0.38794775 | 0.93750436 | 0.34849918 | 0.63229918 |
| TMEM62 | 77.7787173 | -0.122536649 | 0.33997166 | -0.360432 | 0.71852413 | 0.88155276 |
| CCNDBP1 | 39.2686272 | 0.151132657 | 0.39004109 | 0.38747881 | 0.69840177 | 0.8717697 |
| ADAL | 31.115425 | 0.068741313 | 0.35179144 | 0.1954036 | 0.845077 | 0.94193856 |
| TUBGCP4 | 47.1254595 | -0.12237528 | 0.29235583 | -0.4185833 | 0.67552067 | 0.86116336 |
| TP53BP1 | 213.592341 | 0.419635225 | 0.19452707 | 2.1572073 | 0.03098952 | 0.17208613 |
| PPIP5K1 | 49.7003736 | -0.112361899 | 0.3183637 | -0.3529356 | 0.72413668 | 0.88342651 |
| CATSPER2 | 66.8267752 | 0.081292395 | 0.46233749 | 0.17582912 | 0.86042819 | 0.94779688 |
| PDIA3 | 210.164014 | -1.05019442 | 0.34335442 | -3.0586309 | 0.00222351 | 0.03396969 |
| SERF2 | 546.145951 | -0.32193106 | 0.42509043 | -0.7573237 | 0.44885595 | 0.71024151 |
| MFAP1 | 41.6835602 | -0.267163317 | 0.29718163 | -0.89899 | 0.36865799 | 0.6485537 |
| FRMD5 | 44.620092 | -0.773989788 | 0.48786164 | -1.5864944 | 0.11262719 | 0.35131051 |
| CASC4 | 474.02782 | 0.163537613 | 0.14887686 | 1.09847571 | 0.27199682 | 0.55916275 |
| CTDSPL2 | 275.912939 | 0.32318056 | 0.14039647 | 2.30191373 | 0.02134004 | 0.14039576 |
| EIF3J-DT | 72.9515082 | 0.080176531 | 0.3573814 | 0.22434444 | 0.8224893 | 0.93217972 |
| EIF3J | 144.813174 | -0.261431419 | 0.29341081 | -0.8910081 | 0.37292481 | 0.65217873 |
| SPG11 | 299.032543 | -0.166747906 | 0.14382211 | -1.1594038 | 0.24629162 | 0.53315875 |
| B2M | 1361.1058 | -0.990412804 | 0.48928442 | -2.0242067 | 0.04294889 | 0.20601272 |
| DUOX1 | 32.9295886 | -0.363359022 | 0.40713528 | -0.8924774 | 0.37213712 | 0.65197926 |
| GATM | 93.8082823 | 0.799493811 | 0.51497518 | 1.55248998 | 0.12054503 | 0.36507762 |
| SPATA5L1 | 35.3834122 | -0.348541682 | 0.28540714 | -1.2212087 | 0.22200701 | 0.50375143 |
| SQOR | 60.196184 | -1.665475522 | 0.53271081 | -3.1264158 | 0.00176951 | 0.02957033 |
| SEMA6D | 65.3104028 | 0.928601904 | 0.71130827 | 1.30548447 | 0.19172793 | 0.46874984 |
| MYEF2 | 312.137214 | 0.745702283 | 0.27000632 | 2.76179569 | 0.00574844 | 0.06336748 |
| SLC12A1 | 305.873592 | 1.621515506 | 1.23816236 | 1.3096146 | 0.19032625 | 0.46644991 |
| DUT | 81.1579537 | 0.373534119 | 0.27152218 | 1.3757039 | 0.16891332 | 0.43860342 |
| CEP152 | 36.6412754 | 0.592993454 | 0.33331447 | 1.77908107 | 0.07522647 | 0.28328492 |
| EID1 | 172.625389 | -0.545457312 | 0.36614676 | -1.4897232 | 0.13629704 | 0.38812051 |
| SECISBP2L | 264.140776 | -0.023461957 | 0.21160185 | -0.1108778 | 0.91171321 | 0.96844354 |
| COPS2 | 85.1392079 | -0.082031846 | 0.24042162 | -0.3412 | 0.73295306 | 0.88869659 |
| GALK2 | 202.331797 | -0.062033068 | 0.24227562 | -0.2560434 | 0.79791735 | 0.91942309 |
| FAM227B | 114.327705 | 0.540336872 | 0.47143071 | 1.14616391 | 0.25172734 | 0.53906677 |
| DTWD1 | 117.394182 | 0.992765011 | 0.35345251 | 2.80876492 | 0.00497319 | 0.05795405 |
| ATP8B4 | 41.6969961 | -0.812450203 | 0.62617466 | -1.2974818 | 0.19446548 | 0.47305178 |
| SLC27A2 | 106.489169 | -1.400072657 | 0.46339212 | -3.0213562 | 0.00251645 | 0.03646761 |
| GABPB1 | 109.889085 | 0.058750159 | 0.21950916 | 0.26764332 | 0.78897388 | 0.9148506 |
| GABPB1-AS1 | 290.668266 | 0.209506195 | 0.35616384 | 0.58822984 | 0.55637803 | 0.78804453 |
| USP8 | 195.837526 | 0.144729976 | 0.1309839 | 1.10494478 | 0.26918352 | 0.55632249 |
| TRPM7 | 381.606113 | 0.229238421 | 0.18140804 | 1.26366183 | 0.20635143 | 0.48621411 |
| SPPL2A | 213.370731 | -0.326661532 | 0.19287969 | -1.6936025 | 0.09034084 | 0.3135161 |
| AC021752.1 | 40.8436485 | 1.453386896 | 0.62975011 | 2.30787875 | 0.02100588 | 0.13915307 |
| AP4E1 | 98.1644869 | 0.376592972 | 0.23372817 | 1.61124338 | 0.10712669 | 0.34251219 |
| DMXL2 | 111.698815 | 0.639364781 | 0.23529295 | 2.71731375 | 0.00658142 | 0.06885256 |
| TMOD3 | 165.775252 | -0.60888969 | 0.23154101 | -2.6297272 | 0.00854534 | 0.08169244 |
| MAPK6 | 124.578122 | 0.072616081 | 0.21368792 | 0.33982306 | 0.73398978 | 0.88942291 |
| GNB5 | 84.2427324 | 0.010819526 | 0.29734259 | 0.03638741 | 0.97097346 | 0.99175186 |
| MYO5C | 163.574743 | 0.20018488 | 0.25293382 | 0.79145161 | 0.42868051 | 0.69479256 |
| MYO5A | 91.0738644 | 0.241829235 | 0.40544651 | 0.59645163 | 0.55087356 | 0.78577398 |
| ARPP19 | 141.502148 | -0.001342652 | 0.25469585 | -0.0052716 | 0.9957939 | 0.99899312 |
| FAM214A | 205.125578 | 0.334021318 | 0.25575342 | 1.30602876 | 0.19154278 | 0.46865601 |
| WDR72 | 224.322414 | 0.132586077 | 0.66757258 | 0.19860923 | 0.84256843 | 0.94132995 |
| UNC13C | 174.580129 | 4.334156248 | 1.21336995 | 3.57199901 | 0.00035427 | 0.010638 |

| | | | | | | |
|---|---|---|---|---|---|---|
| RSL24D1 | 67.7032204 | -0.138262196 | 0.40860954 | -0.3383724 | 0.73508256 | 0.88966284 |
| RAB27A | 129.564815 | -0.098736092 | 0.17597938 | -0.5610663 | 0.57475237 | 0.8009242 |
| PIGB | 60.6859758 | 0.116099627 | 0.26960706 | 0.43062532 | 0.66674083 | 0.85838595 |
| CCPG1 | 220.959534 | 0.350701573 | 0.24945255 | 1.40588491 | 0.1597583 | 0.42532651 |
| DNAAF4 | 125.812841 | 0.442161917 | 0.24264665 | 1.82224616 | 0.06841764 | 0.26728084 |
| PYGO1 | 38.2824949 | 0.702550848 | 0.49892425 | 1.40813129 | 0.1590922 | 0.4247336 |
| PRTG | 100.095088 | -0.065673736 | 0.60539957 | -0.10848 | 0.91361496 | 0.96844354 |
| NEDD4 | 146.17993 | -0.717041965 | 0.46358519 | -1.5467318 | 0.12192793 | 0.36668381 |
| RFX7 | 282.250065 | 0.135370337 | 0.19405287 | 0.69759513 | 0.48543043 | 0.73878902 |
| TEX9 | 246.14483 | 0.364573968 | 0.24478233 | 1.48938023 | 0.13638727 | 0.38813241 |
| MNS1 | 64.483411 | 0.795652941 | 0.43916617 | 1.81173552 | 0.07002707 | 0.27149813 |
| ZNF280D | 145.206975 | 0.584389513 | 0.2392664 | 2.44242198 | 0.01458908 | 0.11197822 |
| AC090517.4 | 104.445569 | 0.508347862 | 0.18507477 | 2.74671619 | 0.00601952 | 0.06537526 |
| TCF12 | 1206.06129 | 0.108833919 | 0.12320176 | 0.8833796 | 0.37703121 | 0.65530141 |
| CGNL1 | 303.431615 | 0.831626204 | 0.49779359 | 1.67062456 | 0.09479586 | 0.3204651 |
| POLR2M | 24.4486571 | -0.218521223 | 0.3031583 | -0.7208156 | 0.47102299 | 0.72902395 |
| ALDH1A2 | 2956.88114 | 0.808893741 | 0.47228522 | 1.71272296 | 0.08676352 | 0.30679504 |
| AQP9 | 50.7099821 | -1.444419118 | 0.74220252 | -1.9461253 | 0.05163969 | 0.22786819 |
| LIPC | 271.715259 | 0.310216804 | 0.25221736 | 1.22995819 | 0.21871276 | 0.5006005 |
| ADAM10 | 1349.05624 | 0.473471808 | 0.19912649 | 2.37774397 | 0.01741892 | 0.12460019 |
| MINDY2 | 528.166628 | 0.164496611 | 0.2123323 | 0.77471309 | 0.43850921 | 0.7024935 |
| RNF111 | 332.752996 | 0.227151329 | 0.18568247 | 1.22333212 | 0.22120428 | 0.50310453 |
| SLTM | 448.754415 | 0.018687272 | 0.14457478 | 0.1292568 | 0.89715446 | 0.96175013 |
| MYO1E | 485.740248 | -0.645477611 | 0.29173095 | -2.2125784 | 0.02692672 | 0.15864374 |
| FAM81A | 41.8775428 | 0.505748903 | 0.47801992 | 1.058008 | 0.29005179 | 0.57691157 |
| GCNT3 | 119.954531 | -1.40477436 | 0.69430773 | -2.0232734 | 0.04304497 | 0.20605022 |
| GTF2A2 | 86.7643676 | -0.302210948 | 0.4253557 | -0.7104899 | 0.47740037 | 0.733914 |
| BNIP2 | 111.797018 | -0.134016478 | 0.14026821 | -0.9554302 | 0.3393602 | 0.62418037 |
| ANXA2 | 420.799381 | -0.718179018 | 0.37980477 | -1.8909163 | 0.05863552 | 0.24443767 |
| ICE2 | 163.302401 | 0.061997549 | 0.22963484 | 0.2699832 | 0.78717318 | 0.91364739 |
| RORA-AS1 | 88.3558698 | 0.706121646 | 0.54817793 | 1.28812491 | 0.19770248 | 0.47630586 |
| RORA | 2093.7286 | 0.682247296 | 0.41753322 | 1.63399523 | 0.10225985 | 0.33390378 |
| AC012404.1 | 58.0487324 | 0.509744118 | 0.5057733 | 1.00785099 | 0.313526 | 0.60065312 |
| AC009554.1 | 109.562724 | 0.603653833 | 0.55430376 | 1.08903073 | 0.27614034 | 0.56317011 |
| VPS13C | 780.288724 | 0.381055579 | 0.17069765 | 2.23234223 | 0.02559235 | 0.1551184 |
| TLN2 | 372.11478 | 0.204347125 | 0.2508959 | 0.81446976 | 0.41537589 | 0.68495052 |
| TPM1 | 1269.28513 | 0.596166739 | 0.25257595 | 2.36034644 | 0.01825788 | 0.12781151 |
| LACTB | 54.2128869 | -1.05528216 | 0.2394456 | -4.4071895 | 1.05E-05 | 0.00095535 |
| RPS27L | 117.730426 | -0.333594211 | 0.29907358 | -1.1154252 | 0.26466826 | 0.55272255 |
| RAB8B | 105.022146 | -0.673433348 | 0.32586921 | -2.0665756 | 0.03877417 | 0.19565065 |
| APH1B | 69.9195475 | -0.094172313 | 0.3607501 | -0.2610458 | 0.79405716 | 0.91723627 |
| CA12 | 141.365154 | -1.868646831 | 0.49453766 | -3.7785734 | 0.00015773 | 0.00645679 |
| USP3 | 308.567573 | -0.129149488 | 0.18446894 | -0.7001151 | 0.48385544 | 0.73727416 |
| USP3-AS1 | 62.9034252 | 0.096554788 | 0.28366188 | 0.34038689 | 0.73356519 | 0.88902087 |
| HERC1 | 312.624662 | 0.197846044 | 0.13690195 | 1.44516597 | 0.14841127 | 0.40816296 |
| FAM96A | 52.1346936 | -0.183839676 | 0.23043725 | -0.7977863 | 0.42499452 | 0.69181522 |
| SNX1 | 165.781414 | 0.014351259 | 0.17082197 | 0.08401296 | 0.93304613 | 0.97732628 |
| PPIB | 247.148993 | -1.273029802 | 0.47051057 | -2.7056349 | 0.0068174 | 0.07014376 |
| CSNK1G1 | 49.2649973 | -0.462629596 | 0.3145028 | -1.4709872 | 0.14129458 | 0.39714225 |
| TRIP4 | 105.572997 | 0.002212983 | 0.23595164 | 0.00937897 | 0.99251678 | 0.99829858 |
| ZNF609 | 575.475746 | -0.03932809 | 0.15049075 | -0.2613323 | 0.79383628 | 0.91723627 |
| OAZ2 | 57.0270977 | -0.343398626 | 0.25025929 | -1.3721714 | 0.1700101 | 0.44014237 |
| SPG21 | 89.3718925 | -0.357086258 | 0.16216387 | -2.2020088 | 0.02766469 | 0.16119224 |
| MTFMT | 49.2811962 | -0.016990151 | 0.22212993 | -0.0764874 | 0.9390313 | 0.97996404 |
| UBAP1L | 32.4036556 | 0.396425005 | 0.280155 | 1.41502026 | 0.15706256 | 0.42213867 |
| PDCD7 | 53.3721181 | -0.08234678 | 0.19880127 | -0.4142166 | 0.67871551 | 0.86168533 |
| CLPX | 103.358914 | -0.17802359 | 0.14409319 | -1.2354754 | 0.21665364 | 0.49827736 |
| PARP16 | 37.4396111 | -0.252779125 | 0.31525101 | -0.8018345 | 0.42264873 | 0.69043214 |
| DPP8 | 164.913905 | 0.071925894 | 0.17461121 | 0.41192025 | 0.68039788 | 0.86244955 |
| HACD3 | 117.201426 | -0.04365649 | 0.22183196 | -0.1967998 | 0.84398419 | 0.94193856 |
| INTS14 | 45.8063765 | -0.413701669 | 0.26561652 | -1.5575148 | 0.1193483 | 0.3645149 |
| SLC24A1 | 37.2075456 | -0.222618817 | 0.3345842 | -0.6653596 | 0.5058205 | 0.75528806 |
| DENND4A | 693.937903 | 0.196812479 | 0.28975945 | 0.67922713 | 0.49699396 | 0.74877401 |
| RAB11A | 397.371821 | 0.141605067 | 0.37960249 | 0.37303513 | 0.7091223 | 0.87636208 |
| MEGF11 | 80.2079697 | -5.26691682 | 0.90259913 | -5.835278 | 5.37E-09 | 2.57E-06 |
| DIS3L | 52.6132974 | 0.270769103 | 0.26334814 | 1.02817927 | 0.30386551 | 0.59203936 |
| TIPIN | 47.5768266 | 1.124513498 | 0.47354384 | 2.37467665 | 0.01756433 | 0.12490692 |
| MAP2K1 | 154.144155 | -0.454291625 | 0.23309807 | -1.9489291 | 0.05130389 | 0.2274958 |
| SNAPC5 | 27.5373578 | -0.170760706 | 0.33818673 | -0.5049302 | 0.61360783 | 0.82498688 |
| RPL4 | 375.319146 | -0.909048499 | 0.52077999 | -1.7455519 | 0.08088885 | 0.29573375 |
| SMAD6 | 142.617512 | 0.848627194 | 0.32653741 | 2.59886667 | 0.00935321 | 0.08623137 |
| SMAD3 | 797.252281 | 0.104393842 | 0.46319356 | 0.22537844 | 0.82168488 | 0.9319109 |
| AAGAB | 148.334323 | -0.031459081 | 0.19700367 | -0.1596878 | 0.87312702 | 0.95193455 |
| IQCH | 205.189752 | 0.283838921 | 0.26346104 | 1.07734686 | 0.28132534 | 0.56893012 |
| IQCH-AS1 | 68.35252 | 0.602756127 | 0.34765663 | 1.73376853 | 0.08295916 | 0.29986597 |
| MAP2K5 | 287.276297 | 0.31456955 | 0.14015624 | 2.24442065 | 0.02480535 | 0.15268762 |
| PIAS1 | 588.849901 | 0.390083211 | 0.26909925 | 1.4495886 | 0.14717328 | 0.40621213 |
| FEM1B | 64.8566494 | -0.019632512 | 0.2264105 | -0.086712 | 0.93090042 | 0.97609039 |
| CORO2B | 60.2832146 | -1.773898556 | 1.08332819 | -1.6374526 | 0.10153594 | 0.33297252 |
| ANP32A | 128.925681 | -0.494451158 | 0.19146596 | -2.5824495 | 0.00981017 | 0.08848555 |
| SPESP1 | 33.0228927 | 0.267676072 | 0.45671699 | 0.58608739 | 0.55781679 | 0.78891584 |
| GLCE | 93.0980673 | 0.271751049 | 0.35063688 | 0.77502129 | 0.43832708 | 0.70236452 |
| PAQR5 | 302.08847 | -0.157621583 | 0.58541283 | -0.2692486 | 0.78773839 | 0.91381085 |
| RPLP1 | 676.060041 | -0.279459618 | 0.45487046 | -0.6143719 | 0.53896961 | 0.77708778 |
| TLE3 | 72.985505 | -0.060145597 | 0.38263978 | -0.157186 | 0.8750983 | 0.95291321 |
| UACA | 394.782442 | 0.080106586 | 0.28006927 | 0.28602419 | 0.77485959 | 0.90689106 |
| LARP6 | 29.4003803 | -0.2363076 | 0.45992895 | -0.5137915 | 0.60739775 | 0.8211189 |
| LRRC49 | 89.745946 | -0.425241019 | 0.46843397 | -0.9077929 | 0.36398766 | 0.644123 |
| THSD4 | 274.346066 | -0.207322418 | 0.46977283 | -0.4413248 | 0.65897785 | 0.85262229 |
| THSD4-AS1 | 140.256307 | -0.55030576 | 0.46627393 | -1.1802199 | 0.23791278 | 0.52090663 |
| MYO9A | 2646.83663 | 0.718921793 | 0.2988428 | 2.40568553 | 0.01614215 | 0.11866898 |
| AC022872.1 | 77.3423331 | 1.156586615 | 0.52240404 | 2.21396949 | 0.02683088 | 0.15855211 |
| SENP8 | 35.1515309 | 0.556048287 | 0.37386588 | 1.48729349 | 0.13693731 | 0.38900431 |
| GRAMD2A | 26.4661265 | 1.238859072 | 0.6000867 | 2.0644668 | 0.03897348 | 0.19580915 |
| PKM | 722.666506 | -1.237740602 | 0.31424091 | -3.9388271 | 8.19E-05 | 0.00414993 |
| PARP6 | 92.9245718 | 0.224457947 | 0.15950034 | 1.40725684 | 0.15935124 | 0.42483013 |
| AC009690.2 | 33.7213165 | -0.146097772 | 0.61003276 | -0.2394917 | 0.81072435 | 0.92512511 |
| AC009690.1 | 124.833855 | -0.347656842 | 0.45354444 | -0.7665331 | 0.44335915 | 0.70605773 |
| ARIH1 | 616.230293 | 0.421853533 | 0.18602399 | 2.26773726 | 0.02334523 | 0.14760655 |
| BBS4 | 135.225807 | 0.298423362 | 0.20536503 | 1.45313622 | 0.14618594 | 0.40436474 |
| ADPGK | 71.1938013 | -0.308368987 | 0.24891264 | -1.2388643 | 0.21539574 | 0.49731142 |
| NEO1 | 290.995028 | -0.112924572 | 0.21519871 | -0.5247456 | 0.59976006 | 0.81763227 |
| NPTN | 271.769441 | 0.465115598 | 0.20171508 | 2.30580475 | 0.02112154 | 0.13943713 |
| CD276 | 31.2152816 | -0.394787197 | 0.45940238 | -0.8593495 | 0.39014773 | 0.66569738 |
| LOXL1 | 50.2166103 | -0.058515938 | 0.59796007 | -0.0978593 | 0.92204404 | 0.97248656 |
| PML | 91.9038958 | -2.214567326 | 0.5952725 | -3.7202581 | 0.00019902 | 0.00745243 |
| ARID3B | 29.0656895 | 1.025611452 | 0.32761456 | 3.130543 | 0.00174483 | 0.0293224 |
| CLK3 | 43.8915333 | 0.396480801 | 0.25960468 | 1.52724827 | 0.12669929 | 0.37412222 |
| EDC3 | 51.8623832 | 0.039039287 | 0.20303261 | 0.19228087 | 0.8475222 | 0.94323402 |
| SCAMP2 | 69.0865247 | -0.030385085 | 0.29801917 | -0.1019568 | 0.91879095 | 0.97078078 |
| FAM219B | 35.000326 | -0.289928973 | 0.26967953 | -1.0750871 | 0.28233575 | 0.56996716 |
| COX5A | 67.0789468 | -0.416819407 | 0.37080117 | -1.1241049 | 0.26096853 | 0.54796526 |
| PPCDC | 94.8876613 | 0.101245991 | 0.28763535 | 0.35199425 | 0.72484257 | 0.88404214 |

| | | | | | | |
|---|---|---|---|---|---|---|
| COMMD4 | 48.171226 | -0.396800523 | 0.35052095 | -1.1320308 | 0.25762148 | 0.54487802 |
| NEIL1 | 25.6693082 | 0.259358541 | 0.31701086 | 0.81813771 | 0.41327856 | 0.68443902 |
| MAN2C1 | 80.5532865 | -0.035719967 | 0.24788548 | -0.1440987 | 0.88542256 | 0.95800331 |
| SIN3A | 159.450257 | 0.146095873 | 0.15341998 | 0.95226106 | 0.3409646 | 0.62624461 |
| PTPN9 | 242.110607 | -0.200837201 | 0.25626008 | -0.7837241 | 0.43320201 | 0.69824029 |
| SNUPN | 26.377725 | -0.371888095 | 0.41211475 | -0.9023897 | 0.3668499 | 0.6466739 |
| IMP3 | 32.4523867 | -1.339466775 | 0.36282842 | -3.6917361 | 0.00022273 | 0.00814319 |
| UBE2Q2 | 96.1259581 | -0.580017889 | 0.30727819 | -1.8875986 | 0.05907985 | 0.24535287 |
| FBXO22 | 109.20093 | -0.289904288 | 0.20474388 | -1.4159363 | 0.15679416 | 0.42165391 |
| ETFA | 190.817891 | 0.06049473 | 0.16430394 | 0.36818794 | 0.7127331 | 0.87827554 |
| SCAPER | 605.894338 | 0.22442435 | 0.19887751 | 1.12845516 | 0.25912774 | 0.54609557 |
| RCN2 | 97.3627728 | 0.459451906 | 0.25910916 | 1.77319821 | 0.07619587 | 0.28527949 |
| TSPAN3 | 164.555603 | -0.370812586 | 0.22370815 | -1.657573 | 0.09740369 | 0.32577861 |
| PEAK1 | 264.826293 | 0.368190908 | 0.26157418 | 1.40759655 | 0.15925057 | 0.42480123 |
| HMG20A | 121.358352 | 0.242072713 | 0.13582625 | 1.78222332 | 0.07471282 | 0.28248137 |
| LINGO1 | 1017.04684 | 0.903990612 | 0.53934142 | 1.67610085 | 0.09371844 | 0.31857635 |
| TBC1D2B | 78.0775497 | -0.31695918 | 0.38238931 | -0.8288913 | 0.4071659 | 0.68040675 |
| IDH3A | 79.1796724 | -0.48543832 | 0.30055134 | -1.6151594 | 0.10627619 | 0.34093088 |
| WDR61 | 70.0244159 | 0.113136401 | 0.182501 | 0.61992208 | 0.53530909 | 0.77504588 |
| IREB2 | 179.872549 | 0.019145392 | 0.13785791 | 0.13887772 | 0.88954677 | 0.95918866 |
| PSMA4 | 83.8856379 | -0.702410232 | 0.32625368 | -2.1529573 | 0.03132204 | 0.17302987 |
| MORF4L1 | 297.081093 | -0.070691701 | 0.18485559 | -0.3824158 | 0.70215296 | 0.87225045 |
| CTSH | 111.548228 | -1.028773986 | 0.28873237 | -3.5630711 | 0.00036654 | 0.01087028 |
| TMED3 | 51.373438 | -1.14506974 | 0.39693567 | -2.8847741 | 0.00391695 | 0.04866463 |
| ZFAND6 | 376.792943 | -0.01819345 | 0.14719449 | -0.1236014 | 0.90163086 | 0.9638123 |
| AC016705.2 | 112.036255 | 0.629630004 | 0.58829353 | 1.07026505 | 0.28450002 | 0.57232018 |
| ARNT2 | 363.648977 | 0.332057932 | 0.53807755 | 0.61711909 | 0.53715618 | 0.7760813 |
| ABHD17C | 40.2836134 | 0.326097358 | 0.42670865 | 0.76421548 | 0.44473884 | 0.70707984 |
| AC027808.1 | 27.0189933 | -0.287531682 | 0.36382741 | -0.7902969 | 0.42935438 | 0.69507954 |
| MESD | 72.9950808 | -0.579676698 | 0.18880969 | -3.0701639 | 0.00213941 | 0.03326857 |
| AC104041.1 | 42.6992074 | 0.731035819 | 0.78321434 | 0.933379 | 0.35062432 | 0.63322593 |
| EFL1 | 84.9459431 | -0.494569606 | 0.30960896 | -1.5974008 | 0.11017639 | 0.34724322 |
| RPS17 | 32.8642658 | -0.354115317 | 0.47310002 | -0.7484999 | 0.45415869 | 0.71384952 |
| WHAMM | 137.527453 | 0.654290364 | 0.28571709 | 2.28999376 | 0.02202168 | 0.14253085 |
| HOMER2 | 75.884685 | 0.226007388 | 0.28656076 | 0.78868924 | 0.43029366 | 0.69612953 |
| C15orf40 | 52.5698019 | 0.300730651 | 0.2445305 | 1.22982881 | 0.21876122 | 0.5006005 |
| BTBD1 | 112.041933 | -0.081373704 | 0.22362018 | -0.3638925 | 0.71593829 | 0.88028636 |
| HDGFL3 | 116.749626 | -0.09807931 | 0.2151075 | -0.4559549 | 0.64842243 | 0.84621977 |
| WDR73 | 110.558773 | -0.066585885 | 0.2471102 | -0.2694583 | 0.78757706 | 0.91378399 |
| SEC11A | 213.657017 | -0.146595602 | 0.18941361 | -0.7739444 | 0.43896366 | 0.70279675 |
| ZNF592 | 125.717746 | -0.060767999 | 0.18468459 | -0.3290367 | 0.74212798 | 0.89253455 |
| ALPK3 | 116.279936 | -0.596470214 | 0.49972576 | -1.1935951 | 0.23263639 | 0.51579248 |
| SLC28A1 | 75.7237237 | -0.506005712 | 0.68735472 | -0.7361639 | 0.46163097 | 0.71995492 |
| PDE8A | 326.348304 | -0.376554324 | 0.29822227 | -1.2626633 | 0.2067102 | 0.48621411 |
| AKAP13 | 823.777805 | -0.169226065 | 0.28465001 | -0.5945057 | 0.55217391 | 0.78604159 |
| MRPS11 | 41.519487 | -0.762573024 | 0.35338595 | -2.1579042 | 0.03093528 | 0.17188461 |
| DET1 | 49.3198534 | -0.074743702 | 0.39752372 | -0.1880233 | 0.85085842 | 0.94435157 |
| ABHD2 | 98.8019685 | -0.280155869 | 0.2617831 | -1.0701832 | 0.28453688 | 0.57232018 |
| POLG | 39.3942544 | -0.262934414 | 0.26703674 | -0.9846376 | 0.32480211 | 0.61161381 |
| ANPEP | 52.596426 | -1.673230096 | 0.73581226 | -2.2739905 | 0.02296656 | 0.14608012 |
| ZNF710 | 63.776976 | -0.488348717 | 0.30903879 | -1.5802182 | 0.11405691 | 0.35437923 |
| IDH2 | 72.0878443 | -0.953767493 | 0.26627648 | -3.5818691 | 0.00034114 | 0.01037405 |
| SEMA4B | 364.33215 | -1.624489387 | 0.41290826 | -3.9342622 | 8.35E-05 | 0.00415961 |
| CIB1 | 192.531523 | -1.763519201 | 0.43580387 | -4.0465892 | 5.20E-05 | 0.00303967 |
| IQGAP1 | 397.038726 | -0.159531951 | 0.18458615 | -0.8642683 | 0.38744053 | 0.66406384 |
| CRTC3 | 221.228452 | -0.395227972 | 0.19406987 | -2.0365241 | 0.04169776 | 0.20312121 |
| FURIN | 50.7207518 | -1.380808045 | 0.43028277 | -3.2090712 | 0.00133165 | 0.02515943 |
| MAN2A2 | 31.430883 | -0.234313492 | 0.26152722 | -0.895943 | 0.37028319 | 0.65021864 |
| UNC45A | 46.2613172 | -0.020930068 | 0.21787001 | -0.0960668 | 0.92346754 | 0.97297642 |
| AC104035.1 | 78.1620098 | -1.475583889 | 0.83917081 | -1.7583832 | 0.07868232 | 0.29089077 |
| SLCO3A1 | 834.042026 | 0.189234836 | 0.21412994 | 0.88373834 | 0.37683748 | 0.65512273 |
| FAM174B | 63.9777181 | 0.380682706 | 0.33926302 | 1.12208724 | 0.26182535 | 0.54928275 |
| AC013394.1 | 257.530125 | 0.168724546 | 0.22416765 | 0.75267124 | 0.4516475 | 0.71227056 |
| CHD2 | 330.219782 | 0.392487279 | 0.23962207 | 1.63794294 | 0.1014336 | 0.33286483 |
| MCTP2 | 454.419106 | -0.517723938 | 0.26822247 | -1.9302034 | 0.05358164 | 0.2323488 |
| LINC01197 | 102.146659 | 0.251875478 | 1.2755851 | 0.19745878 | 0.84346854 | 0.94189614 |
| AC012409.2 | 133.004611 | 0.332725255 | 0.54519724 | 0.61028418 | 0.54167357 | 0.77887826 |
| NR2F2-AS1 | 939.210903 | 0.716609838 | 0.41139983 | 1.74188169 | 0.08152915 | 0.29654614 |
| NR2F2 | 347.303621 | -0.005421108 | 0.46238974 | -0.0117241 | 0.99064573 | 0.99772846 |
| AC087477.2 | 81.376689 | -0.059622897 | 0.64856164 | -0.091931 | 0.92675289 | 0.97414308 |
| ARRDC4 | 77.5800416 | -0.025767981 | 0.4677771 | -0.055086 | 0.95606993 | 0.98464615 |
| IGF1R | 1242.05343 | 0.25705723 | 0.25158507 | 1.02175074 | 0.30689889 | 0.59449635 |
| TTC23 | 103.143019 | 0.203141567 | 0.21078732 | 0.96372763 | 0.3351825 | 0.6199188 |
| LRRC28 | 138.848346 | 0.614212605 | 0.19206202 | 3.19799103 | 0.00138389 | 0.02584063 |
| MEF2A | 716.873666 | 0.188963616 | 0.21163937 | 0.89285662 | 0.37193396 | 0.65192016 |
| AC090825.1 | 102.279424 | 0.831112987 | 0.53453991 | 1.55481934 | 0.1199891 | 0.36481264 |
| LINS1 | 36.2211996 | 0.420810403 | 0.27046828 | 1.55585862 | 0.11974172 | 0.36465111 |
| ASB7 | 57.3721774 | -0.191866744 | 0.2235194 | -0.8583897 | 0.39067732 | 0.66624498 |
| ALDH1A3 | 80.8198215 | 1.71931409 | 1.26447974 | 1.35970078 | 0.17392463 | 0.44528422 |
| LRRK1 | 71.364261 | -0.015398057 | 0.69905178 | -0.0220271 | 0.98242637 | 0.99616479 |
| CHSY1 | 77.9930518 | -0.585639272 | 0.40207533 | -1.4565412 | 0.14524307 | 0.40323664 |
| SELENOS | 85.7833054 | -1.113524318 | 0.34415494 | -3.2355319 | 0.00121416 | 0.02363725 |
| SNRPA1 | 63.983921 | -0.243807291 | 0.21291713 | -1.1450807 | 0.25217574 | 0.53930681 |
| TM2D3 | 67.130042 | 0.060116757 | 0.32004928 | 0.18783594 | 0.85100526 | 0.94435157 |
| TARSL2 | 122.735822 | 0.027141067 | 0.31090195 | 0.08729783 | 0.93043478 | 0.9759784 |
| RHBDF1 | 27.7074983 | -0.947300307 | 0.38951369 | -2.4320077 | 0.01501539 | 0.11379145 |
| MPG | 55.502013 | -1.105460746 | 0.25211427 | -4.3847606 | 1.16E-05 | 0.00101506 |
| NPRL3 | 103.368112 | -0.109883097 | 0.2389635 | -0.4598321 | 0.64563671 | 0.84407726 |
| LUC7L | 188.036749 | 0.565053447 | 0.16084563 | 3.51301704 | 0.00044305 | 0.01233711 |
| AXIN1 | 53.3432326 | -0.347035952 | 0.28617314 | -1.2126783 | 0.22525285 | 0.50741684 |
| MRPL28 | 38.4167609 | -0.807285402 | 0.47586634 | -1.6964541 | 0.08979992 | 0.31211662 |
| TMEM8A | 48.6236849 | -1.135264601 | 0.35940546 | -3.1587294 | 0.00158459 | 0.02774908 |
| NME4 | 28.787467 | -0.298284102 | 0.33523495 | -0.8897763 | 0.37358604 | 0.65253555 |
| RAB11FIP3 | 257.625891 | -0.320065012 | 0.39168395 | -0.8171512 | 0.41384203 | 0.68448816 |
| CAPN15 | 46.6653024 | -0.48746488 | 0.45553443 | -1.0700945 | 0.28457679 | 0.57232018 |
| PIGQ | 39.211038 | -0.204436258 | 0.27316125 | -0.7484087 | 0.45421368 | 0.71384952 |
| RAB40C | 84.1063269 | -0.123292051 | 0.34552349 | -0.3568268 | 0.72122145 | 0.88235193 |
| METTL26 | 31.6573654 | 0.11264235 | 0.4908226 | 0.22949707 | 0.8184826 | 0.9300409 |
| MCRIP2 | 43.5636942 | -0.195846311 | 0.33377629 | -0.5867592 | 0.55736543 | 0.78866035 |
| WDR90 | 30.5211952 | 0.379095334 | 0.26482727 | 1.43148148 | 0.15229227 | 0.41452845 |
| RHOT2 | 53.1372892 | -0.198447668 | 0.26109222 | -0.7600673 | 0.44721434 | 0.70891772 |
| STUB1 | 80.4345581 | -0.265022205 | 0.3119236 | -0.8496382 | 0.39552627 | 0.67057454 |
| JMJD8 | 46.1722233 | -0.276665078 | 0.33584258 | -0.8237939 | 0.41005666 | 0.68208527 |
| METRN | 33.4799938 | -0.485007741 | 0.4580605 | -1.058829 | 0.28967766 | 0.5764068 |
| FAM173A | 48.1624981 | -0.539795308 | 0.47112069 | -1.1457686 | 0.25189089 | 0.53906677 |
| LMF1 | 82.2266601 | -0.208827592 | 0.29774734 | -0.7013584 | 0.48307939 | 0.73696727 |
| CACNA1H | 34.4223868 | 1.622233415 | 0.5929259 | 2.73598001 | 0.00621948 | 0.066196 |
| UBE2I | 136.533948 | -0.463124348 | 0.2326931 | -1.9902797 | 0.04656014 | 0.21545872 |
| TSR3 | 40.2671198 | -0.32973649 | 0.38228783 | -0.8625346 | 0.38839339 | 0.66462645 |
| GNPTG | 52.2705632 | -0.338841928 | 0.26866534 | -1.2612044 | 0.2072352 | 0.48666485 |
| UNKL | 45.0798789 | 0.248305131 | 0.25398988 | 0.97761821 | 0.32826319 | 0.61394673 |
| CLCN7 | 74.9729863 | -0.225330381 | 0.26419956 | -0.8528795 | 0.39372615 | 0.6686322 |
| TELO2 | 26.992513 | -0.177124515 | 0.34482936 | -0.5136584 | 0.60749081 | 0.8211189 |

| | | | | | | |
|---|---|---|---|---|---|---|
| IFT140 | 84.9743238 | 0.383334904 | 0.27284282 | 1.40496607 | 0.16003137 | 0.42569855 |
| JPT2 | 117.498677 | -0.334401265 | 0.22295546 | -1.4998568 | 0.1336515 | 0.38492115 |
| MAPK8IP3 | 139.386285 | 0.192441834 | 0.20488567 | 0.93926451 | 0.34759496 | 0.63118125 |
| NME3 | 107.310633 | 0.163787431 | 0.35281812 | 0.46422625 | 0.64248566 | 0.84298912 |
| MRPS34 | 55.7013565 | -0.715501424 | 0.38891631 | -1.8397311 | 0.06580773 | 0.26156523 |
| SPSB3 | 74.1648525 | 0.69064455 | 0.31774484 | 2.17358227 | 0.02973652 | 0.16866909 |
| NUBP2 | 27.921106 | 0.115549441 | 0.30850724 | 0.3745437 | 0.70799985 | 0.87571806 |
| HAGH | 98.3600265 | 0.305630596 | 0.19107859 | 1.59950201 | 0.1097091 | 0.34671061 |
| FAHD1 | 35.4921047 | -1.452870966 | 0.32721396 | -4.4401253 | 8.99E-06 | 0.00083613 |
| MSRB1 | 32.6484984 | -1.719511032 | 0.443542 | -3.8767716 | 0.00010585 | 0.00485145 |
| NDUFB10 | 76.6088886 | -0.49045911 | 0.49830161 | -0.9842615 | 0.32498694 | 0.61184156 |
| RPS2 | 401.24367 | -0.680289056 | 0.45940418 | -1.4808073 | 0.13865793 | 0.39168515 |
| TSC2 | 86.697245 | 0.242347312 | 0.29396347 | 0.82441302 | 0.40970492 | 0.68208527 |
| PKD1 | 67.007749 | 0.067823063 | 0.25614678 | 0.26478203 | 0.79117738 | 0.91622594 |
| SNHG19 | 34.7121081 | -0.46776845 | 0.32077809 | -1.4582307 | 0.14477697 | 0.40282422 |
| TRAF7 | 32.6221912 | -0.505638951 | 0.39505309 | -1.2799266 | 0.20057096 | 0.47923902 |
| PGP | 46.6191614 | -0.435757105 | 0.2899823 | -1.5027024 | 0.1329158 | 0.38380408 |
| ECI1 | 96.0358969 | -0.332163668 | 0.20160392 | -1.6476052 | 0.09943372 | 0.32948294 |
| RNPS1 | 99.5049289 | -0.212687056 | 0.22368117 | -0.9508492 | 0.34168096 | 0.62700419 |
| ABCA3 | 43.6038911 | -0.448564255 | 0.41795823 | -1.0732275 | 0.28316907 | 0.57084461 |
| PDPK1 | 184.718408 | -0.053119631 | 0.22482501 | -0.236271 | 0.81322238 | 0.92694452 |
| AC093525.4 | 27.3482434 | -0.802496941 | 0.31196084 | -2.5724285 | 0.01009878 | 0.08982009 |
| SRRM2 | 493.79883 | 0.215371819 | 0.14563546 | 1.47884186 | 0.13918259 | 0.39289358 |
| ELOB | 127.121044 | -0.454858745 | 0.41818579 | -1.0876954 | 0.27672961 | 0.56369053 |
| PRSS22 | 31.1477304 | -0.1557365 | 0.85192477 | -0.1828055 | 0.85495066 | 0.94567811 |
| FLYWCH1 | 51.9130514 | 0.553117056 | 0.33308121 | 1.66060719 | 0.09679236 | 0.32454105 |
| TNFRSF12A | 143.361889 | -0.780022902 | 0.72727686 | -1.0725254 | 0.28348411 | 0.57084461 |
| HCFC1R1 | 100.897048 | 0.329021023 | 0.59263329 | 0.55518484 | 0.57876823 | 0.80406395 |
| IL32 | 116.690408 | -1.646050877 | 0.74186451 | -2.2188026 | 0.02650016 | 0.15781973 |
| ZNF213-AS1 | 37.3458161 | 0.420689734 | 0.35600121 | 1.18170873 | 0.23732129 | 0.52003259 |
| ZNF213 | 64.8089451 | 0.190843125 | 0.37073596 | 0.51476832 | 0.60671493 | 0.8211189 |
| ZNF263 | 33.331403 | 0.054298999 | 0.2243526 | 0.24202527 | 0.80876058 | 0.92414046 |
| ZNF75A | 73.7767844 | -0.045932843 | 0.25954217 | -0.1769764 | 0.85952692 | 0.94734879 |
| AC025283.3 | 35.8201024 | 0.04090291 | 0.48301125 | 0.08468314 | 0.9325133 | 0.9770887 |
| NAA60 | 30.4677805 | -0.437636839 | 0.30387479 | -1.4401881 | 0.1498142 | 0.41050134 |
| CLUAP1 | 99.7946199 | 0.02300855 | 0.22273816 | 0.10329865 | 0.91772595 | 0.97057281 |
| DNASE1 | 115.133392 | 0.441871453 | 0.26550358 | 1.66427684 | 0.09605712 | 0.32319324 |
| TRAP1 | 101.211194 | -0.271141369 | 0.22303963 | -1.2156646 | 0.22411273 | 0.50617113 |
| CREBBP | 336.29755 | 0.408611758 | 0.14410675 | 2.83547959 | 0.00457569 | 0.05460848 |
| ADCY9 | 288.964479 | 0.392123534 | 0.27230406 | 1.4400209 | 0.14986149 | 0.41050134 |
| GLIS2 | 56.6542695 | -0.595046533 | 0.24762801 | -2.4029855 | 0.01626183 | 0.11918292 |
| VASN | 72.1819104 | -0.436477558 | 0.58060483 | -0.7517636 | 0.45219326 | 0.71231034 |
| DNAJA3 | 65.2651992 | -0.171431191 | 0.17695144 | -0.9688036 | 0.33264318 | 0.61719981 |
| NMRAL1 | 46.898905 | -0.302010044 | 0.24939493 | -1.210971 | 0.2259065 | 0.50832943 |
| HMOX2 | 47.1394233 | -0.614627179 | 0.26709595 | -2.3011475 | 0.0213833 | 0.14045419 |
| CDIP1 | 39.4341401 | -0.220929584 | 0.38310783 | -0.5766773 | 0.56415748 | 0.7932808 |
| MGRN1 | 115.833529 | -0.213479455 | 0.18939977 | -1.1271368 | 0.25968465 | 0.54634729 |
| NUDT16L1 | 32.3115308 | 0.060608557 | 0.44129206 | 0.13734341 | 0.89075936 | 0.95957984 |
| ANKS3 | 83.3182016 | -0.053738745 | 0.16226122 | -0.3311866 | 0.74050353 | 0.89253455 |
| ZNF500 | 29.2974003 | 0.188385467 | 0.27223514 | 0.69199541 | 0.48894021 | 0.74185878 |
| GLYR1 | 201.341392 | 0.041513076 | 0.22339962 | 0.18582429 | 0.85258255 | 0.94470172 |
| UBN1 | 102.999803 | 0.279707484 | 0.2216984 | 1.26165768 | 0.20707199 | 0.48648432 |
| PPL | 124.407307 | 0.647161147 | 0.47443988 | 1.364053 | 0.17255089 | 0.44360304 |
| RBFOX1 | 2377.46128 | -0.866456647 | 1.34755088 | -0.6429862 | 0.52023304 | 0.76400369 |
| METTL22 | 27.4886687 | -0.041547144 | 0.2612636 | -0.1590239 | 0.87365008 | 0.95210417 |
| ABAT | 55.8806288 | -0.143367728 | 0.45853216 | -0.3126667 | 0.75453391 | 0.89810084 |
| PMM2 | 134.569245 | -0.861121523 | 0.25180712 | -3.4197664 | 0.00062675 | 0.01535456 |
| CARHSP1 | 305.860006 | -1.764482944 | 0.52140311 | -3.3841051 | 0.00071411 | 0.01644885 |
| USP7 | 121.345507 | 0.060088258 | 0.23590797 | 0.25471059 | 0.79894663 | 0.91949148 |
| C16orf72 | 118.193937 | -0.153050651 | 0.17850945 | -0.8573812 | 0.39123423 | 0.66671992 |
| GRIN2A | 48.5799389 | 1.850696229 | 0.90396626 | 2.04730676 | 0.04062797 | 0.20006295 |
| ATF7IP2 | 162.393305 | 0.352076228 | 0.36745922 | 0.95813686 | 0.33799375 | 0.62322274 |
| EMP2 | 45.0829487 | 1.108833835 | 0.43013653 | 2.57786484 | 0.00994129 | 0.08941558 |
| NUBP1 | 29.0584567 | -0.691619674 | 0.24245617 | -2.8525554 | 0.00433693 | 0.05265324 |
| DEXI | 33.5833807 | -0.545266673 | 0.29274145 | -1.862622 | 0.06251546 | 0.25385148 |
| CLEC16A | 373.570644 | -0.024919648 | 0.27104429 | -0.0919394 | 0.92674619 | 0.97414308 |
| LITAF | 214.551428 | -0.074509684 | 0.21383611 | -0.348443 | 0.72750755 | 0.88564265 |
| TXNDC11 | 160.900743 | 0.156045696 | 0.16394121 | 0.95183936 | 0.34117846 | 0.62632205 |
| ZC3H7A | 184.101126 | 0.422716567 | 0.19254382 | 2.19543041 | 0.02813274 | 0.16243731 |
| RSL1D1 | 70.2435891 | -0.796649476 | 0.34318624 | -2.3213328 | 0.02026889 | 0.1355161 |
| AC007216.4 | 101.513779 | 0.515613062 | 0.314056 | 1.64178703 | 0.10063415 | 0.33126781 |
| GSPT1 | 250.963353 | -0.172955531 | 0.25709566 | -0.6727283 | 0.50112015 | 0.75235583 |
| SNX29 | 1299.27815 | 0.118163355 | 0.36076858 | 0.32753228 | 0.74326533 | 0.89269973 |
| CPPED1 | 98.7117059 | -0.832656536 | 0.42548681 | -1.9569503 | 0.05035332 | 0.2253855 |
| SHISA9 | 306.519416 | 0.454194356 | 0.58748845 | 0.77311198 | 0.43945611 | 0.7031151 |
| U91319.1 | 158.047474 | 0.822887412 | 0.59499959 | 1.38300499 | 0.16666333 | 0.43490657 |
| ERCC4 | 39.8141779 | -0.141962251 | 0.2881702 | -0.4926334 | 0.62227168 | 0.8293781 |
| MKL2 | 443.938511 | 0.060104594 | 0.1896441 | 0.31693364 | 0.75129396 | 0.89599662 |
| PARN | 182.458233 | 0.228372853 | 0.21988223 | 1.0386144 | 0.29898411 | 0.58639882 |
| BFAR | 59.2589828 | -0.717379616 | 0.23121806 | -3.1026107 | 0.00191822 | 0.03108476 |
| PDXDC1 | 352.67248 | 0.356502037 | 0.15445389 | 2.30814542 | 0.02099105 | 0.13915105 |
| RRN3 | 62.580876 | -0.143477751 | 0.24234766 | -0.5920328 | 0.55382866 | 0.78699373 |
| C16orf45 | 68.2689204 | -0.250974384 | 0.63378892 | -0.3959905 | 0.69211204 | 0.86892611 |
| MARF1 | 103.488301 | 0.048887653 | 0.18616845 | 0.26259903 | 0.79285965 | 0.91680379 |
| NDE1 | 43.609775 | -0.223686575 | 0.24827582 | -0.90096 | 0.3676096 | 0.64718477 |
| FOPNL | 44.105947 | 0.474163873 | 0.21677493 | 2.18735569 | 0.02871657 | 0.1646176 |
| ABCC1 | 160.890858 | 0.184172421 | 0.22802752 | 0.80767628 | 0.41927696 | 0.68835944 |
| XYLT1 | 102.853628 | 0.038200987 | 0.66479345 | 0.05746294 | 0.95417643 | 0.98390867 |
| RPS15A | 90.5316597 | 0.248385024 | 0.52284591 | 0.47506354 | 0.63474169 | 0.83703065 |
| SMG1 | 453.152985 | 0.177482321 | 0.19353755 | 0.91704336 | 0.35911992 | 0.64001011 |
| COQ7 | 29.8689831 | 0.595566385 | 0.28421783 | 2.09545753 | 0.03613034 | 0.18804981 |
| GDE1 | 30.7707415 | -0.450396849 | 0.33730302 | -1.3352885 | 0.18178194 | 0.45476343 |
| CCP110 | 61.1713302 | 0.474948604 | 0.27783605 | 1.70945638 | 0.08736644 | 0.30722583 |
| VPS35L | 187.192738 | -0.036884648 | 0.20814076 | -0.1772101 | 0.85934335 | 0.94734879 |
| KNOP1 | 44.3862094 | -1.059894718 | 0.3391542 | -3.1251116 | 0.00177738 | 0.02957033 |
| IQCK | 249.300775 | 0.070044247 | 0.2704854 | 0.25895759 | 0.79566796 | 0.9180563 |
| GPRC5B | 78.145986 | -0.085800124 | 0.42851368 | -0.2002273 | 0.84130284 | 0.94068401 |
| UMOD | 148.251134 | 2.0029155 | 0.64582012 | 3.10135197 | 0.00192639 | 0.03108476 |
| ACSM2A | 162.05198 | 0.816562251 | 0.99776785 | 0.81838902 | 0.41313509 | 0.68443902 |
| ACSM2B | 102.696092 | 1.095031094 | 0.99684027 | 1.09850206 | 0.27198532 | 0.55916275 |
| ACSM3 | 238.830624 | -1.914635161 | 1.05555725 | -1.813862 | 0.06969898 | 0.27065319 |
| THUMPD1 | 89.7873454 | -0.185185811 | 0.24765045 | -0.7477709 | 0.45459833 | 0.71410256 |
| ERI2 | 39.576634 | 0.076929076 | 0.48225473 | 0.15951959 | 0.87325953 | 0.95196916 |
| LYRM1 | 47.34765 | 0.126287908 | 0.20661485 | 0.61122378 | 0.54105144 | 0.77844268 |
| TMEM159 | 41.5128613 | -0.046173053 | 0.31777757 | -0.1452999 | 0.88447408 | 0.95761929 |
| CRYM | 32.0384323 | 0.131081084 | 0.4498841 | 0.29136634 | 0.77077116 | 0.90547178 |
| METTL9 | 194.9761 | 0.020705227 | 0.14159968 | 0.14622368 | 0.88374481 | 0.95761929 |
| NPIPB4 | 52.6260604 | 0.46007966 | 0.31593531 | 1.4562464 | 0.14532452 | 0.40323664 |
| UQCRC2 | 135.594694 | -0.068003294 | 0.21775214 | -0.3122968 | 0.75481498 | 0.89810084 |
| MOSMO | 144.708367 | -0.138505509 | 0.13696403 | -1.0112546 | 0.31189457 | 0.59932559 |
| EEF2K | 45.6043499 | -0.634560848 | 0.32046373 | -1.9801331 | 0.04768857 | 0.2181513 |
| POLR3E | 57.6879786 | 0.030858279 | 0.29034222 | 0.10628244 | 0.91535826 | 0.9691911 |
| CDR2 | 677.892375 | 0.955815939 | 0.33081184 | 2.8893039 | 0.00386096 | 0.04842913 |

| | | | | | | |
|---|---|---|---|---|---|---|
| NPIPB5 | 154.949002 | 0.107940064 | 0.25501057 | 0.42327682 | 0.6720933 | 0.86015189 |
| USP31 | 150.226629 | -0.581001956 | 0.35539124 | -1.6348235 | 0.10208605 | 0.33374822 |
| COG7 | 124.87962 | 0.180541832 | 0.19137129 | 0.94341128 | 0.34547057 | 0.63006282 |
| GGA2 | 126.256028 | -0.362239634 | 0.27511485 | -1.3166851 | 0.18794419 | 0.46346633 |
| EARS2 | 25.5257457 | -0.991890989 | 0.35828766 | -2.7684208 | 0.00563287 | 0.062532 |
| UBFD1 | 48.1002536 | -0.431462579 | 0.263133 | -1.6397129 | 0.10106487 | 0.33218203 |
| NDUFAB1 | 55.2173338 | -0.032548262 | 0.31689408 | -0.1027102 | 0.91819296 | 0.97057281 |
| PALB2 | 32.9565367 | -0.110491094 | 0.28351782 | -0.3897148 | 0.69674744 | 0.87084339 |
| DCTN5 | 76.3952854 | -0.258667775 | 0.197155 | -1.3120021 | 0.18951942 | 0.46560824 |
| RBBP6 | 158.129719 | 0.081230113 | 0.19107199 | 0.42512832 | 0.67074313 | 0.8597138 |
| TNRC6A | 704.195928 | 0.179762016 | 0.25547256 | 0.70364511 | 0.48165381 | 0.73620566 |
| ARHGAP17 | 132.848469 | 0.246472981 | 0.20326267 | 1.21258363 | 0.22528905 | 0.50741684 |
| LCMT1 | 124.011079 | 0.284875581 | 0.19066063 | 1.49415001 | 0.13513642 | 0.38731025 |
| NSMCE1 | 64.7994779 | 0.080082623 | 0.23133956 | 0.34616917 | 0.72921558 | 0.88653304 |
| IL4R | 213.731827 | -1.474419279 | 0.43758847 | -3.3694199 | 0.00075327 | 0.01713904 |
| GTF3C1 | 138.507765 | 0.72146588 | 0.25821628 | 2.79403721 | 0.00520545 | 0.05957347 |
| KIAA0556 | 210.707698 | 0.290981429 | 0.23731651 | 1.22613224 | 0.22014891 | 0.50173839 |
| XPO6 | 152.74066 | -0.17287062 | 0.29918414 | -0.5778068 | 0.56339459 | 0.79290184 |
| NUPR1 | 574.584036 | 0.143917191 | 0.62609413 | 0.2298651 | 0.81819659 | 0.92999132 |
| SGF29 | 36.6211605 | -0.147836627 | 0.25939749 | -0.5699231 | 0.56872985 | 0.79717578 |
| SULT1A1 | 28.1481961 | 0.478406831 | 0.38666493 | 1.23726461 | 0.21598886 | 0.49746509 |
| ATXN2L | 85.0126305 | -0.194524778 | 0.23345979 | -0.833226 | 0.40471726 | 0.67799696 |
| TUFM | 113.10652 | -0.690997787 | 0.40154052 | -1.7208669 | 0.08527498 | 0.30377428 |
| SH2B1 | 71.3594231 | 0.50368069 | 0.23225128 | 2.16868853 | 0.03010634 | 0.16944102 |
| RABEP2 | 38.2922613 | -0.061664864 | 0.24767498 | -0.2489749 | 0.80338016 | 0.92125386 |
| NFATC2IP | 147.714331 | -0.03038105 | 0.18057726 | -0.1682441 | 0.86639128 | 0.95083843 |
| SPNS1 | 36.5412015 | -0.253043476 | 0.2642498 | -0.9575919 | 0.33826857 | 0.62345706 |
| MAZ | 57.2227221 | -0.518931993 | 0.34422495 | -1.5075374 | 0.13167296 | 0.38186355 |
| AC009133.1 | 34.7064268 | -1.539813416 | 0.43493161 | -3.5403576 | 0.00039959 | 0.01159828 |
| MVP | 168.048043 | -1.737625198 | 0.45275085 | -3.837928 | 0.00012408 | 0.00547711 |
| CDIPT | 54.3637157 | -0.578266444 | 0.29438411 | -1.9643263 | 0.04949227 | 0.22373121 |
| KCTD13 | 47.4283477 | 0.095703638 | 0.30663201 | 0.31211235 | 0.75495514 | 0.89810084 |
| TMEM219 | 85.5287542 | 0.193480405 | 0.34236745 | 0.56512501 | 0.57198874 | 0.79915869 |
| TAOK2 | 29.8143937 | 0.044086081 | 0.36728156 | 0.12003347 | 0.90445663 | 0.96543237 |
| INO80E | 42.2509507 | 0.311922424 | 0.29561836 | 1.05515239 | 0.29135563 | 0.5787402 |
| PPP4C | 64.4767314 | -0.498419307 | 0.34821373 | -1.4313603 | 0.15232698 | 0.41452845 |
| YPEL3 | 75.2054954 | 0.495314467 | 0.33057547 | 1.49834005 | 0.13404492 | 0.38559049 |
| MAPK3 | 33.0752777 | -0.346984139 | 0.38606195 | -0.8987784 | 0.36877071 | 0.64863288 |
| NPIPB13 | 38.5815193 | 0.162531946 | 0.29688226 | 0.54746265 | 0.58406092 | 0.80801843 |
| CD2BP2 | 41.658138 | -0.597745943 | 0.34839246 | -1.7157258 | 0.08621224 | 0.30586185 |
| TBC1D10B | 25.9655428 | -0.156733537 | 0.28207069 | -0.5556534 | 0.57844782 | 0.80396861 |
| DCTPP1 | 26.8097972 | -1.251025953 | 0.47757248 | -2.619552 | 0.00880453 | 0.08357433 |
| SEPHS2 | 90.5882804 | -1.127098723 | 0.3012368 | -3.7415705 | 0.00018287 | 0.00709811 |
| ZNF768 | 32.5796986 | 0.151300317 | 0.35779301 | 0.42287108 | 0.67238932 | 0.86015189 |
| ZNF785 | 40.1182418 | 0.048928083 | 0.21549345 | 0.22705137 | 0.8203838 | 0.93133185 |
| PRR14 | 24.5209527 | -0.01498486 | 0.38628076 | -0.0387927 | 0.96905569 | 0.99152285 |
| FBRS | 45.7011496 | -0.213831417 | 0.24716973 | -0.8651197 | 0.38697307 | 0.66404483 |
| PHKG2 | 33.3547611 | -0.391942845 | 0.23808067 | -1.6462607 | 0.0997101 | 0.33026383 |
| RNF40 | 51.2842845 | -0.003446757 | 0.30115858 | -0.011445 | 0.99086842 | 0.9977564 |
| BCL7C | 148.972145 | 0.370881924 | 0.28308273 | 1.31015384 | 0.1901438 | 0.46630503 |
| FBXL19 | 41.8325656 | 0.019130467 | 0.26443162 | 0.07234561 | 0.94232687 | 0.98093339 |
| SETD1A | 34.0807627 | -0.029743947 | 0.35522309 | -0.0837331 | 0.9332686 | 0.97745243 |
| STX4 | 33.6408944 | -0.11622249 | 0.35976419 | -0.3230519 | 0.74665597 | 0.89352127 |
| BCKDK | 32.3725054 | -0.564532629 | 0.27251373 | -2.071575 | 0.0383051 | 0.19441659 |
| KAT8 | 63.5101417 | 0.122543989 | 0.30680746 | 0.39941658 | 0.68958628 | 0.86755279 |
| PRSS8 | 58.5607369 | -1.27046798 | 0.66639365 | -1.9064827 | 0.0565876 | 0.23974023 |
| FUS | 317.486808 | 0.363791798 | 0.15877729 | 2.291208 | 0.02195139 | 0.14229081 |
| AC026471.4 | 38.1254736 | -0.23265323 | 0.32220027 | -0.7220765 | 0.47024747 | 0.72817662 |
| C16orf58 | 46.5745243 | -0.581780907 | 0.23350022 | -2.4915647 | 0.01271818 | 0.10301379 |
| ZNF720 | 85.0031066 | -0.222077283 | 0.23538642 | -0.9434584 | 0.3454465 | 0.63006282 |
| ZNF267 | 36.8605536 | -0.892563291 | 0.45544731 | -1.9597509 | 0.05002491 | 0.2248768 |
| VPS35 | 95.1138536 | -0.504074687 | 0.19842014 | -2.5404411 | 0.01107127 | 0.09494336 |
| C16orf87 | 37.0316686 | 0.075456366 | 0.32030375 | 0.23557753 | 0.81376051 | 0.92720494 |
| DNAJA2 | 55.8909821 | -0.322323299 | 0.31574866 | -1.0208224 | 0.30733861 | 0.59456464 |
| ITFG1 | 552.62958 | 0.054801644 | 0.19514208 | 0.28082946 | 0.7788412 | 0.90959765 |
| PHKB | 340.088363 | 0.43522653 | 0.14551043 | 2.99103324 | 0.00278035 | 0.03846189 |
| LONP2 | 272.366544 | 0.181760268 | 0.16521812 | 1.10012307 | 0.27127851 | 0.55841559 |
| SIAH1 | 72.3038906 | 0.08886738 | 0.21628875 | 0.4108738 | 0.68116507 | 0.8630794 |
| N4BP1 | 140.548629 | -0.07788062 | 0.34814027 | -0.2237047 | 0.82298708 | 0.93217972 |
| ZNF423 | 67.9036689 | -0.381594748 | 1.0997887 | -0.346971 | 0.72861313 | 0.88653304 |
| HEATR3 | 26.6573377 | -0.117988795 | 0.41046594 | -0.2874509 | 0.77376711 | 0.90643453 |
| TENT4B | 129.714108 | 0.476354192 | 0.21169103 | 2.25023321 | 0.02443415 | 0.15178643 |
| BRD7 | 106.629774 | 0.179516341 | 0.20538689 | 0.87403991 | 0.38209652 | 0.65924037 |
| CYLD | 94.8924099 | -0.019423785 | 0.27384692 | -0.0709294 | 0.94345398 | 0.98112027 |
| SALL1 | 110.378757 | -0.237167056 | 0.40860684 | -0.5804285 | 0.56162569 | 0.79161455 |
| AC087564.1 | 319.311874 | -0.115417588 | 0.28895487 | -0.3994312 | 0.68957551 | 0.86755279 |
| HNRNPA1P4 | 28.8082032 | -0.645297073 | 0.5574318 | -1.1576251 | 0.24701704 | 0.5337641 |
| TOX3 | 198.371992 | 0.22552869 | 0.52715044 | 0.42782605 | 0.66877777 | 0.85924744 |
| AC007906.2 | 69.8292408 | 1.667472558 | 0.50103866 | 3.32803176 | 0.00087462 | 0.01886931 |
| CHD9 | 826.569645 | 0.426543363 | 0.12641116 | 3.37425413 | 0.00074016 | 0.01688094 |
| RBL2 | 174.427867 | -0.045021251 | 0.2479588 | -0.1815675 | 0.85592218 | 0.94621534 |
| AKTIP | 34.4958489 | 1.032646829 | 0.40421601 | 2.5546906 | 0.01062822 | 0.09255249 |
| RPGRIP1L | 76.2853164 | 0.461947845 | 0.30711842 | 1.5041359 | 0.13254638 | 0.38380408 |
| FTO | 984.326511 | 0.489413237 | 0.17971764 | 2.72323421 | 0.00646462 | 0.06810698 |
| IRX3 | 251.58434 | 0.28348879 | 0.59463133 | 0.47674715 | 0.63354218 | 0.83625473 |
| CRNDE | 218.377842 | 1.525777985 | 0.54798282 | 2.78435368 | 0.00536345 | 0.06041319 |
| IRX5 | 78.5979448 | 0.350605055 | 0.65014185 | 0.5392747 | 0.58969732 | 0.81183245 |
| LPCAT2 | 155.518382 | 1.258312048 | 0.40266111 | 3.1249903 | 0.00177811 | 0.02957033 |
| AMFR | 121.741878 | 0.713301454 | 0.25640333 | 2.78195079 | 0.00540332 | 0.06060706 |
| NUDT21 | 96.7710737 | 0.249157096 | 0.21926542 | 1.13632646 | 0.25581996 | 0.54274627 |
| OGFOD1 | 36.6433878 | 0.078405004 | 0.23119372 | 0.3391312 | 0.7345109 | 0.88966284 |
| BBS2 | 81.7695431 | 0.356200728 | 0.21125848 | 1.68608958 | 0.09177855 | 0.31558127 |
| MT2A | 925.364806 | -3.223035631 | 0.73743274 | -4.3706164 | 1.24E-05 | 0.00105027 |
| MT1E | 361.454742 | -2.783662099 | 0.99147698 | -2.8075913 | 0.00499135 | 0.05809499 |
| MT1F | 68.4223982 | -1.109394739 | 0.83362889 | -1.3308017 | 0.18325427 | 0.45690988 |
| MT1G | 338.025193 | -1.020969494 | 1.04631881 | -0.9757729 | 0.32917704 | 0.61429707 |
| MT1H | 86.2521483 | -0.960578642 | 1.16796302 | -0.8224393 | 0.41082694 | 0.68247066 |
| MT1X | 268.774187 | -2.432025833 | 0.72511661 | -3.353979 | 0.00079658 | 0.01778667 |
| NUP93 | 76.2683626 | 0.133930599 | 0.34995824 | 0.38270451 | 0.70193887 | 0.87209759 |
| SLC12A3 | 37.3914233 | 2.946363254 | 0.57306936 | 5.14137284 | 2.73E-07 | 6.53E-05 |
| HERPUD1 | 127.75763 | 0.094319687 | 0.44079684 | 0.21397542 | 0.83056623 | 0.93446017 |
| NLRC5 | 38.4638119 | -1.484573683 | 0.67474915 | -2.2001861 | 0.0277937 | 0.16152154 |
| FAM192A | 99.6007675 | -0.070811633 | 0.17160761 | -0.4126369 | 0.67987266 | 0.86235436 |
| RSPRY1 | 141.248687 | 0.269959172 | 0.18783381 | 1.43722351 | 0.1506545 | 0.41173164 |
| ARL2BP | 57.7513029 | 0.158327506 | 0.24688155 | 0.64130959 | 0.52132156 | 0.76440016 |
| PLLP | 46.2996922 | 0.120214557 | 0.61358404 | 0.19592191 | 0.84467129 | 0.94193856 |
| CX3CL1 | 60.6482893 | -0.121670105 | 0.39145566 | -0.3108145 | 0.75594163 | 0.89874207 |
| CIAPIN1 | 42.0160878 | -0.341336661 | 0.26986343 | -1.2648496 | 0.20592523 | 0.48573203 |
| COQ9 | 73.269715 | -0.37420856 | 0.21445176 | -1.7449545 | 0.08099279 | 0.29573375 |
| POLR2C | 26.369122 | -0.457273348 | 0.35536107 | -1.2867851 | 0.19816919 | 0.47659117 |
| DOK4 | 28.5800102 | -0.56193879 | 0.45584554 | -1.2327395 | 0.21767298 | 0.49966199 |
| ADGRG1 | 306.195152 | -1.136538043 | 0.5585045 | -2.0349667 | 0.04185422 | 0.20312121 |
| KIFC3 | 424.794021 | 0.771426075 | 0.25497317 | 3.02551858 | 0.00248207 | 0.03618781 |

| | | | | | | |
|---|---|---|---|---|---|---|
| AC092118.1 | 31.4265778 | 0.919196694 | 0.45472514 | 2.02143365 | 0.04323489 | 0.20614587 |
| CSNK2A2 | 86.2957655 | 0.02111477 | 0.26794258 | 0.07880334 | 0.93718905 | 0.97909629 |
| CNOT1 | 234.06669 | 0.146455682 | 0.09850717 | 1.48675145 | 0.13708047 | 0.38918015 |
| GOT2 | 61.3198997 | -0.353083086 | 0.35242009 | -1.0018813 | 0.31640094 | 0.60281198 |
| AC138305.1 | 44.3058095 | -0.163780316 | 0.31901948 | -0.5133866 | 0.60768094 | 0.8211189 |
| TK2 | 28.1301634 | -0.180707703 | 0.4278302 | -0.4223818 | 0.67274634 | 0.86026394 |
| CMTM4 | 185.305542 | 0.26827878 | 0.24791944 | 1.08212077 | 0.27919887 | 0.5666199 |
| DYNC1LI2 | 166.261817 | 0.134935987 | 0.18283496 | 0.73802073 | 0.46050186 | 0.71913063 |
| NAE1 | 67.5060994 | 0.190918593 | 0.18314942 | 1.04241987 | 0.29721706 | 0.58520908 |
| CDH16 | 167.959509 | 0.335877916 | 0.40686592 | 0.82552482 | 0.4090737 | 0.68208527 |
| RRAD | 117.619693 | -2.129530886 | 0.48584247 | -4.3831715 | 1.17E-05 | 0.00101506 |
| FAM96B | 94.9331639 | -0.57098353 | 0.43229339 | -1.3208241 | 0.18656002 | 0.46165292 |
| CES2 | 65.9075163 | -1.188427746 | 0.34358117 | -3.4589432 | 0.0005423 | 0.01396038 |
| CBFB | 253.941874 | 0.02897089 | 0.1866555 | 0.15521048 | 0.87665539 | 0.95377544 |
| C16orf70 | 43.1633874 | -0.061209129 | 0.27685433 | -0.2210879 | 0.82502403 | 0.93217972 |
| NOL3 | 36.5058872 | -1.439763114 | 0.4046138 | -3.5583639 | 0.00037317 | 0.01103277 |
| E2F4 | 25.6431055 | -0.604315232 | 0.38714688 | -1.5609456 | 0.11853658 | 0.36352436 |
| TMEM208 | 51.6567952 | -1.161431932 | 0.40195029 | -2.8894915 | 0.00385865 | 0.04842913 |
| ZDHHC1 | 40.9877132 | -0.092267853 | 0.27929071 | -0.3303649 | 0.74112425 | 0.89253455 |
| ATP6V0D1 | 72.0168136 | -0.168038026 | 0.2345772 | -0.7163442 | 0.47377882 | 0.73077309 |
| AC009061.2 | 34.5885071 | 0.784271841 | 0.34190413 | 2.29383551 | 0.02179995 | 0.14176627 |
| RIPOR1 | 75.9229965 | -0.353575059 | 0.35937797 | -0.9838529 | 0.32518785 | 0.61197925 |
| CTCF | 128.835457 | 0.053669864 | 0.22416168 | 0.23942479 | 0.81077621 | 0.92512511 |
| ENKD1 | 35.1530681 | 0.073752811 | 0.35042894 | 0.21046438 | 0.83330525 | 0.93607461 |
| GFOD2 | 45.9718108 | -0.389638994 | 0.2817689 | -1.3828318 | 0.16671645 | 0.43490657 |
| RANBP10 | 114.888235 | 0.138097134 | 0.15051523 | 0.9174961 | 0.35888274 | 0.64001011 |
| NUTF2 | 33.4788571 | -0.545963903 | 0.35184481 | -1.5517179 | 0.12072973 | 0.36507762 |
| PSKH1 | 35.5693629 | 0.03738933 | 0.23144951 | 0.16154422 | 0.87166479 | 0.95193455 |
| SLC12A4 | 35.2285478 | 0.666788169 | 0.29720295 | 2.24354492 | 0.0248617 | 0.15285635 |
| DUS2 | 29.7415546 | 0.344100997 | 0.35603964 | 0.96646821 | 0.33380995 | 0.61860428 |
| NFATC3 | 304.093583 | 0.341697598 | 0.22908458 | 1.49157835 | 0.13580972 | 0.38788633 |
| ESRP2 | 24.354992 | 0.597795869 | 0.44324495 | 1.3486806 | 0.17743958 | 0.44901805 |
| PLA2G15 | 39.6200913 | 0.35197396 | 0.40887968 | 0.86082526 | 0.38933429 | 0.66513878 |
| SLC7A6 | 94.9050259 | 1.122694373 | 0.31340024 | 3.58230216 | 0.00034058 | 0.01037405 |
| SLC7A6OS | 55.3180862 | 0.156667952 | 0.36038309 | 0.43472615 | 0.66376122 | 0.8562037 |
| PRMT7 | 54.2340714 | 0.374618634 | 0.31054106 | 1.20634169 | 0.22768577 | 0.51041354 |
| ZFP90 | 71.0827254 | 0.046647054 | 0.25100795 | 0.18583895 | 0.85257106 | 0.94470172 |
| CDH1 | 273.300221 | 0.950541183 | 0.32148263 | 2.95674195 | 0.00310908 | 0.04182851 |
| TANGO6 | 134.697364 | -0.205387937 | 0.2723795 | -0.7540506 | 0.45081882 | 0.71160181 |
| CHTF8 | 59.9054757 | -0.043624593 | 0.29522376 | -0.1477679 | 0.88252595 | 0.95722522 |
| UTP4 | 48.7227995 | -0.545712491 | 0.25915729 | -2.1057193 | 0.03522874 | 0.18531364 |
| SNTB2 | 92.1842687 | 0.306317893 | 0.31889123 | 0.96057172 | 0.33676755 | 0.62147307 |
| VPS4A | 60.2735037 | 0.126733627 | 0.24212025 | 0.52343259 | 0.60067325 | 0.81779318 |
| COG8 | 33.1460476 | 0.291907802 | 0.30737465 | 0.94968079 | 0.34227447 | 0.62749229 |
| TERF2 | 69.7711505 | 0.075041554 | 0.26231601 | 0.2860731 | 0.77482213 | 0.90689106 |
| CYB5B | 217.834031 | 0.213409394 | 0.20048748 | 1.0644525 | 0.28712376 | 0.57406774 |
| NFAT5 | 2561.65778 | 0.966489079 | 0.43316459 | 2.23122825 | 0.02566601 | 0.15521133 |
| NQO1 | 37.4232424 | -2.016518745 | 0.60669494 | -3.3237771 | 0.00088807 | 0.01907363 |
| NOB1 | 31.8886656 | -0.718016457 | 0.30153369 | -2.3812147 | 0.01725565 | 0.12381414 |
| WWP2 | 235.4973 | 0.004768479 | 0.26146691 | 0.01823741 | 0.98544946 | 0.99668665 |
| PDPR | 103.527379 | -0.001226258 | 0.22457505 | -0.0054603 | 0.99564329 | 0.99899312 |
| AC009060.1 | 37.9030983 | 0.043608499 | 0.40121432 | 0.10869128 | 0.91344736 | 0.96844354 |
| EXOSC6 | 54.0350696 | -0.139103351 | 0.32178333 | -0.4322889 | 0.66553148 | 0.85742752 |
| AARS | 66.9874083 | -0.030758275 | 0.25404603 | -0.1210736 | 0.90363272 | 0.96509062 |
| ST3GAL2 | 74.7327947 | 0.185738326 | 0.56040404 | 0.33143645 | 0.74031484 | 0.89253455 |
| FUK | 32.3573066 | -0.074508905 | 0.35041317 | -0.2126316 | 0.83161434 | 0.93532321 |
| COG4 | 64.5853518 | 0.168912658 | 0.21079279 | 0.80132084 | 0.42294593 | 0.69043214 |
| SF3B3 | 144.59319 | -0.261069074 | 0.19039582 | -1.3711912 | 0.17031537 | 0.44045651 |
| MTSS1L | 34.3957508 | -0.133074438 | 0.65065373 | -0.2045242 | 0.83794388 | 0.93879116 |
| VAC14 | 98.0863359 | -0.15613561 | 0.37357872 | -0.4179457 | 0.67598684 | 0.86116336 |
| HYDIN | 517.456324 | 0.047031979 | 0.39748576 | 0.11832368 | 0.90581119 | 0.96601708 |
| CMTR2 | 47.2687215 | 0.020107555 | 0.32595361 | 0.06168839 | 0.95081098 | 0.98286628 |
| ZNF19 | 70.2173133 | 0.12515528 | 0.31957355 | 0.39163216 | 0.69533002 | 0.87024545 |
| MARVELD3 | 70.5711883 | 0.313207486 | 0.51244532 | 0.61120177 | 0.54106601 | 0.77844268 |
| PHLPP2 | 87.9160544 | 0.43837945 | 0.24035252 | 1.82390203 | 0.06816689 | 0.26695445 |
| AP1G1 | 254.759063 | 0.417802854 | 0.16774948 | 2.49063578 | 0.01275148 | 0.10316418 |
| IST1 | 178.342664 | 0.051673659 | 0.13605381 | 0.37980309 | 0.70409159 | 0.87375221 |
| ZNF821 | 57.6970022 | 0.352273978 | 0.3538768 | 0.99547068 | 0.3195074 | 0.60617179 |
| TXNL4B | 34.8246121 | -0.338847335 | 0.28886116 | -1.1730457 | 0.24077748 | 0.52525792 |
| DHX38 | 38.8063342 | -0.313360327 | 0.2953068 | -1.0611348 | 0.28862863 | 0.57551492 |
| LINC01572 | 119.698612 | 0.217777134 | 0.34548693 | 0.6303484 | 0.52846666 | 0.76991364 |
| AC004943.2 | 88.5982462 | 0.653576088 | 0.28679026 | 2.27893406 | 0.02267099 | 0.14468046 |
| ZFHX3 | 475.143548 | 0.075730625 | 0.41857613 | 0.18092437 | 0.85642694 | 0.94627061 |
| AC138627.1 | 45.17839 | 0.537284825 | 0.68854014 | 0.78032463 | 0.43519982 | 0.70008054 |
| PSMD7 | 73.6105602 | -0.718049584 | 0.36336203 | -1.9761272 | 0.04814038 | 0.21931444 |
| GLG1 | 509.652468 | 0.145749127 | 0.18928969 | 0.7699792 | 0.44131223 | 0.70443757 |
| RFWD3 | 56.2034783 | 0.263616587 | 0.24664513 | 1.06880922 | 0.28515564 | 0.57264845 |
| WDR59 | 220.001833 | 0.252828835 | 0.15846365 | 1.59550055 | 0.11060035 | 0.34758554 |
| ZNRF1 | 102.064698 | 0.446772403 | 0.2520002 | 1.77290496 | 0.07624446 | 0.28527949 |
| BCAR1 | 148.420994 | -1.132038284 | 0.44471019 | -2.5455641 | 0.01091013 | 0.09398216 |
| CFDP1 | 461.421591 | 0.627639804 | 0.20668914 | 3.03663652 | 0.00239234 | 0.03547399 |
| GABARAPL2 | 108.108307 | 0.259475974 | 0.47573415 | 0.54542221 | 0.58546317 | 0.80902361 |
| ADAT1 | 37.4159305 | -0.63636975 | 0.33671615 | -1.8899294 | 0.05876741 | 0.24454867 |
| KARS | 59.6078904 | -0.750275239 | 0.29328103 | -2.5582126 | 0.01052117 | 0.09189658 |
| TERF2IP | 262.375524 | 0.034394582 | 0.44755391 | 0.07685014 | 0.93874276 | 0.97996404 |
| ADAMTS18 | 44.2526467 | 0.034046305 | 0.83642541 | 0.04070453 | 0.96753145 | 0.99123492 |
| VAT1L | 63.6497237 | 1.323332568 | 1.18999532 | 1.11204855 | 0.26611728 | 0.55384374 |
| WWOX | 2008.1057 | 0.32136143 | 0.39285937 | 0.81800628 | 0.4133536 | 0.68444497 |
| MAF | 81.4041261 | 1.139975398 | 0.85103392 | 1.33951818 | 0.18040204 | 0.4527302 |
| CDYL2 | 37.6295259 | -0.812997977 | 0.61120478 | -1.3301564 | 0.18346673 | 0.4570849 |
| CMC2 | 84.8055757 | -0.090961284 | 0.26613285 | -0.341789 | 0.73250968 | 0.88869659 |
| ATMIN | 44.238345 | -0.450274874 | 0.25965894 | -1.7341012 | 0.08290014 | 0.29986597 |
| GAN | 95.8290697 | -0.052781304 | 0.24103956 | -0.2189736 | 0.8266706 | 0.93336606 |
| CMIP | 424.997343 | -0.46909996 | 0.33510067 | -1.3998777 | 0.16154994 | 0.42791083 |
| PLCG2 | 152.033413 | -0.557846925 | 0.48713931 | -1.1451487 | 0.25214758 | 0.53930681 |
| MPHOSPH6 | 36.329865 | -0.448845393 | 0.31519333 | -1.424032 | 0.1544372 | 0.41766063 |
| HSBP1 | 117.623885 | -0.629419207 | 0.33832511 | -1.8603976 | 0.0628293 | 0.25469397 |
| MBTPS1 | 151.949802 | -0.084002447 | 0.1728006 | -0.4861236 | 0.62687955 | 0.83221501 |
| HSDL1 | 28.4201877 | 0.632868884 | 0.40881651 | 1.5480512 | 0.12160997 | 0.36636109 |
| TAF1C | 30.5781536 | 0.18289199 | 0.25806782 | 0.7086974 | 0.47851228 | 0.73444466 |
| TLDC1 | 79.8937639 | -0.067465333 | 0.37908227 | -0.1779702 | 0.85874641 | 0.94734879 |
| COTL1 | 69.871267 | -0.938650482 | 0.39685758 | -2.3652074 | 0.01801998 | 0.12673521 |
| KLHL36 | 108.265491 | -0.568837865 | 0.30347358 | -1.874423 | 0.06087214 | 0.2492921 |
| USP10 | 186.952922 | -0.323592148 | 0.25228164 | -1.2826623 | 0.19961041 | 0.47837581 |
| ZDHHC7 | 101.240605 | 0.206156473 | 0.2131172 | 0.9673385 | 0.33337485 | 0.6179175 |
| GSE1 | 223.407778 | -0.14031179 | 0.33794085 | -0.4151963 | 0.67799821 | 0.86151115 |
| EMC8 | 31.4205335 | -0.659531364 | 0.31155092 | -2.1169296 | 0.03426582 | 0.18245266 |
| COX4I1 | 294.460331 | -0.692322433 | 0.42117681 | -1.643781 | 0.10022145 | 0.3309277 |
| MTHFSD | 25.3327529 | 0.223564447 | 0.30039121 | 0.7442443 | 0.45672869 | 0.71603996 |
| FBXO31 | 37.7719867 | -0.415331661 | 0.27229211 | -1.5253166 | 0.12718016 | 0.37451555 |
| MAP1LC3B | 94.4877442 | -0.617364117 | 0.31283756 | -1.9734335 | 0.0484462 | 0.22019444 |
| ZCCHC14 | 237.297279 | -0.017588041 | 0.36694569 | -0.0479309 | 0.9617713 | 0.9875899 |
| KLHDC4 | 87.7359814 | 0.330007476 | 0.38223027 | 0.86337348 | 0.38793216 | 0.66428354 |

| | | | | | | |
|---|---|---|---|---|---|---|
| BANP | 103.831474 | 0.050758307 | 0.28494173 | 0.17813575 | 0.85861637 | 0.94734879 |
| ZFPM1 | 42.3865663 | -0.533108478 | 0.27163792 | -1.9625701 | 0.04969614 | 0.22401852 |
| ZC3H18 | 69.416633 | -0.209835138 | 0.23281867 | -0.9012814 | 0.36743872 | 0.64705385 |
| CYBA | 351.964771 | -1.348352661 | 0.52377943 | -2.5742757 | 0.01004502 | 0.08977568 |
| MVD | 40.9370097 | 0.272677551 | 0.42260749 | 0.6452265 | 0.51878041 | 0.7636788 |
| PIEZO1 | 72.785298 | -0.140991125 | 0.45636864 | -0.3089413 | 0.75736617 | 0.89936902 |
| APRT | 100.0527 | -1.312795281 | 0.48350813 | -2.7151462 | 0.00662465 | 0.06902242 |
| GALNS | 45.776609 | -0.014517822 | 0.33249566 | -0.0436632 | 0.96517288 | 0.98965864 |
| TRAPPC2L | 26.6486565 | -0.341131448 | 0.34840297 | -0.979129 | 0.32751625 | 0.61370857 |
| ACSF3 | 51.9983541 | -0.149519926 | 0.25931707 | -0.5765911 | 0.56421569 | 0.7932808 |
| ANKRD11 | 582.89754 | 0.097095554 | 0.20056612 | 0.48410746 | 0.62830961 | 0.8332518 |
| SPG7 | 210.2454 | 0.092657097 | 0.28872214 | 0.32092134 | 0.74827 | 0.89440069 |
| RPL13 | 1093.947 | -0.539916919 | 0.42516264 | -1.2699068 | 0.20411783 | 0.48301501 |
| CHMP1A | 59.8171596 | -0.335974881 | 0.32322234 | -1.0394544 | 0.29859346 | 0.58610017 |
| CDK10 | 89.7437237 | 0.020510379 | 0.27559178 | 0.07442304 | 0.94067377 | 0.98016757 |
| VPS9D1 | 39.0904873 | 0.300468972 | 0.31616189 | 0.95036431 | 0.34192717 | 0.62709561 |
| ZNF276 | 39.9170708 | -0.192446795 | 0.27710508 | -0.6944903 | 0.48737477 | 0.74104173 |
| TCF25 | 305.955192 | -0.323275005 | 0.19137603 | -1.6892137 | 0.09117848 | 0.31502156 |
| DEF8 | 37.9544676 | -0.837214555 | 0.32727679 | -2.5581239 | 0.01052386 | 0.09189658 |
| GAS8 | 84.0192572 | 0.234440805 | 0.26849461 | 0.87316764 | 0.38257171 | 0.65946633 |
| FAM157C | 82.625773 | 1.210220604 | 0.49571345 | 2.44137135 | 0.0146316 | 0.11217436 |
| RPH3AL | 169.422965 | 0.616732047 | 0.46163926 | 1.335961 | 0.18156202 | 0.45460445 |
| VPS53 | 281.392687 | -0.083693555 | 0.20827795 | -0.4018359 | 0.68780481 | 0.86751885 |
| GLOD4 | 50.5777896 | -0.027388789 | 0.26302642 | -0.1041294 | 0.91706664 | 0.97024314 |
| NXN | 167.321337 | 0.708984363 | 0.39030218 | 1.81650118 | 0.06929353 | 0.26972046 |
| ABR | 141.944314 | -0.36480635 | 0.45548221 | -0.8009234 | 0.42317601 | 0.69059047 |
| YWHAE | 447.517139 | -0.105593846 | 0.14940338 | -0.7067701 | 0.47970933 | 0.73503389 |
| CRK | 143.278694 | -0.332254014 | 0.25447446 | -1.3056478 | 0.19167236 | 0.46873362 |
| MYO1C | 168.12425 | -0.002797885 | 0.27359441 | -0.0102264 | 0.99184066 | 0.99814428 |
| INPP5K | 24.2803222 | -0.508322399 | 0.30213609 | -1.6824286 | 0.09248576 | 0.31666153 |
| PITPNA | 173.807787 | -0.084250367 | 0.14168103 | -0.5946482 | 0.55207865 | 0.7860228 |
| SLC43A2 | 75.2412636 | -0.588523958 | 0.58244803 | -1.0104317 | 0.3122885 | 0.5998419 |
| PRPF8 | 157.830863 | -0.033473737 | 0.155669 | -0.2150315 | 0.82974277 | 0.93416878 |
| WDR81 | 31.6624593 | -0.433675827 | 0.33652184 | -1.2887004 | 0.19750227 | 0.47630586 |
| SERPINF2 | 86.2266641 | -0.227920159 | 0.66758857 | -0.3414081 | 0.73279636 | 0.88869659 |
| SMYD4 | 43.4397802 | 0.149433821 | 0.28637062 | 0.52181966 | 0.6017959 | 0.81860309 |
| RPA1 | 95.910607 | 0.144760773 | 0.17739485 | 0.81603706 | 0.41447894 | 0.68453341 |
| SMG6 | 337.224797 | 0.137958197 | 0.20689639 | 0.66679847 | 0.50490087 | 0.75524917 |
| TSR1 | 38.3405251 | -1.047149772 | 0.36948963 | -2.8340437 | 0.0045963 | 0.05471328 |
| SGSM2 | 55.9335662 | 0.088225216 | 0.2562345 | 0.34431435 | 0.73060989 | 0.88757288 |
| MNT | 29.5468957 | -0.653523918 | 0.35671502 | -1.8320617 | 0.06694223 | 0.26366762 |
| METTL16 | 147.35756 | 0.035677494 | 0.21671882 | 0.16462573 | 0.86923858 | 0.95106604 |
| PAFAH1B1 | 498.925984 | 0.352938594 | 0.10952669 | 3.22239808 | 0.00127122 | 0.02435411 |
| CLUH | 65.4066117 | -0.840190459 | 0.37629993 | -2.232768 | 0.02556424 | 0.1551184 |
| RAP1GAP2 | 155.334554 | 0.134206515 | 0.39076297 | 0.34344737 | 0.73126193 | 0.88791456 |
| CTNS | 35.6790046 | -0.109948301 | 0.37642529 | -0.2920853 | 0.77022139 | 0.90533273 |
| TAX1BP3 | 36.1184679 | -0.715398281 | 0.45940159 | -1.5572394 | 0.11941365 | 0.3645149 |
| EMC6 | 38.7593049 | -0.409305344 | 0.50580661 | -0.8092131 | 0.41839257 | 0.68785864 |
| ITGAE | 28.2356987 | 0.56782561 | 0.33213514 | 1.70962221 | 0.08733576 | 0.30722583 |
| NCBP3 | 105.5958 | 0.565781782 | 0.16468858 | 3.43546456 | 0.00059154 | 0.01482276 |
| ZZEF1 | 150.093058 | 0.074984351 | 0.19176729 | 0.39101741 | 0.69578437 | 0.87032103 |
| CYB5D2 | 52.2780616 | -0.198745808 | 0.32897443 | -0.6041376 | 0.54575219 | 0.78177961 |
| ANKFY1 | 145.486795 | -0.132779609 | 0.31154263 | -0.4262005 | 0.66996179 | 0.85924744 |
| UBE2G1 | 183.246909 | -0.099787375 | 0.20579872 | -0.4848785 | 0.62776253 | 0.83287219 |
| SPNS2 | 187.656246 | 0.176035589 | 0.36555931 | 0.48155138 | 0.63012467 | 0.83433967 |
| MYBBP1A | 27.4237937 | -1.023215794 | 0.35238308 | -2.903703 | 0.00368778 | 0.04691265 |
| SMTNL2 | 54.906916 | 0.85400201 | 0.39882508 | 2.14129467 | 0.03225028 | 0.17584306 |
| PELP1 | 46.6786236 | -0.299631315 | 0.30320724 | -0.9882063 | 0.32305161 | 0.61011659 |
| CXCL16 | 41.2319766 | -1.835661015 | 0.50386624 | -3.6431514 | 0.00026932 | 0.00896353 |
| PSMB6 | 66.7078029 | -0.46140157 | 0.49360842 | -0.9347522 | 0.34991601 | 0.63286907 |
| MINK1 | 120.37132 | -0.301495043 | 0.27112896 | -1.1119987 | 0.26613871 | 0.55384374 |
| SLC25A11 | 47.506221 | -0.081239648 | 0.36742166 | -0.2211074 | 0.82500881 | 0.93217972 |
| RNF167 | 43.2269305 | -0.410745763 | 0.36623963 | -1.1215219 | 0.26206578 | 0.54954643 |
| PFN1 | 295.123712 | -1.39796453 | 0.55693581 | -2.5100999 | 0.0120697 | 0.10031173 |
| SPAG7 | 69.9928668 | -0.106388763 | 0.36074879 | -0.2949109 | 0.76806192 | 0.90439645 |
| CAMTA2 | 41.980305 | -0.067421584 | 0.25400739 | -0.2654316 | 0.79067701 | 0.91582769 |
| KIF1C | 96.7307343 | -0.170598507 | 0.28208452 | -0.604778 | 0.54532652 | 0.78128668 |
| ZNF594 | 38.2999573 | 0.280940006 | 0.27456949 | 1.02320183 | 0.30621243 | 0.59376697 |
| AC087500.1 | 78.4674822 | 0.461155188 | 0.30185627 | 1.52773102 | 0.12657934 | 0.37394935 |
| RABEP1 | 231.669964 | 0.001028631 | 0.19845458 | 0.00518321 | 0.99586442 | 0.99899312 |
| NUP88 | 146.020742 | 0.465887738 | 0.25355873 | 1.8373958 | 0.06615149 | 0.26198834 |
| RPAIN | 74.6264329 | 0.188056092 | 0.20595858 | 0.91307725 | 0.36120192 | 0.64151865 |
| C1QBP | 58.1453211 | -1.244557542 | 0.49536396 | -2.5124104 | 0.01199096 | 0.10009484 |
| DERL2 | 53.3721222 | -0.834053122 | 0.36470285 | -2.2869389 | 0.02219939 | 0.14286955 |
| KIAA0753 | 115.082334 | 0.147400041 | 0.21311074 | 0.69165936 | 0.48915127 | 0.74185878 |
| TXNDC17 | 29.5107742 | -1.12067544 | 0.49102919 | -2.282299 | 0.02247169 | 0.14412481 |
| MED31 | 30.0418716 | 0.064063676 | 0.36050826 | 0.17770377 | 0.85895563 | 0.94734879 |
| XAF1 | 126.393915 | -1.050332577 | 1.75055312 | -0.6000004 | 0.54850597 | 0.78384882 |
| ALOX12-AS1 | 41.2419901 | -1.032616802 | 0.5957247 | -1.7333792 | 0.08302829 | 0.29986597 |
| ACADVL | 187.147527 | 0.080664596 | 0.28046812 | 0.287607 | 0.77364759 | 0.90643453 |
| DVL2 | 29.7920653 | -0.204905315 | 0.34017094 | -0.6023598 | 0.54693463 | 0.78253724 |
| PHF23 | 27.7386438 | -1.135920255 | 0.30987022 | -3.6657935 | 0.00024657 | 0.00849612 |
| ELP5 | 31.4626201 | -0.440247474 | 0.27608728 | -1.5945953 | 0.11080276 | 0.34794171 |
| EIF5A | 80.3482079 | -1.600157968 | 0.58344681 | -2.7425944 | 0.00609559 | 0.06580904 |
| GPS2 | 56.8273663 | -1.198846518 | 0.37032568 | -3.2372762 | 0.00120677 | 0.02363725 |
| ZBTB4 | 162.713267 | 0.560702546 | 0.33816327 | 1.65808233 | 0.09730085 | 0.32554832 |
| POLR2A | 134.057082 | -0.199227569 | 0.30612554 | -0.6508035 | 0.51517335 | 0.76135326 |
| EIF4A1 | 96.2680522 | -1.803673682 | 0.47177151 | -3.8231933 | 0.00013173 | 0.00565867 |
| MPDU1 | 32.7382987 | -0.312157081 | 0.5120096 | -0.6096704 | 0.54208019 | 0.77900491 |
| FXR2 | 42.1579485 | -0.009866373 | 0.24182904 | -0.040799 | 0.96745617 | 0.99123492 |
| SAT2 | 88.9732438 | -0.060557632 | 0.27922015 | -0.2168813 | 0.82830084 | 0.93346403 |
| TP53 | 51.053446 | -0.74248846 | 0.32559106 | -2.2804326 | 0.02258205 | 0.14449794 |
| KDM6B | 59.0394651 | -0.404296773 | 0.33542164 | -1.205339 | 0.22807247 | 0.51066249 |
| NAA38 | 142.146422 | 0.205212781 | 0.26243134 | 0.78196752 | 0.43423366 | 0.69907971 |
| CHD3 | 112.914994 | 0.388773278 | 0.29657799 | 1.31086356 | 0.18990386 | 0.46597228 |
| TRAPPC1 | 28.6469039 | -0.899491057 | 0.52823762 | -1.7028152 | 0.08860265 | 0.30992958 |
| PER1 | 86.9514665 | 0.493997304 | 0.56379075 | 0.87620683 | 0.38091761 | 0.65803602 |
| CTC1 | 36.8108858 | 0.601781292 | 0.30861475 | 1.94994341 | 0.05118286 | 0.2274029 |
| NDEL1 | 106.735875 | 0.28690615 | 0.29090413 | 0.98625672 | 0.32400715 | 0.61072469 |
| MYH10 | 193.541068 | 0.276876011 | 0.30873821 | 0.89679865 | 0.36982637 | 0.6497738 |
| STX8 | 302.160603 | 0.188521889 | 0.21876428 | 0.86175809 | 0.38882065 | 0.66509161 |
| USP43 | 78.1046298 | -0.612326041 | 0.32256889 | -1.8982799 | 0.05765922 | 0.24245728 |
| SCO1 | 51.6860003 | -0.300028074 | 0.26536277 | -1.1306336 | 0.25820933 | 0.54539959 |
| TMEM220 | 61.487317 | 0.448577798 | 0.2856719 | 1.57025525 | 0.11635574 | 0.35938557 |
| TMEM220-A: | 38.1028138 | -0.346062695 | 0.46008181 | -0.7521764 | 0.45194497 | 0.71231034 |
| SHISA6 | 37.4186628 | 0.278109933 | 0.59779602 | 0.46522547 | 0.64177 | 0.84235611 |
| DNAH9 | 46.0308862 | -1.06610854 | 0.52365398 | -2.0359027 | 0.04176013 | 0.20312121 |
| MAP2K4 | 372.332117 | 0.088331337 | 0.12578931 | 0.70221656 | 0.48254412 | 0.73696727 |
| ARHGAP44 | 59.3864216 | 0.003241685 | 0.4593303 | 0.00705742 | 0.99436904 | 0.99885288 |
| ELAC2 | 36.0047292 | -0.721721896 | 0.32393314 | -2.2279965 | 0.02588075 | 0.15591658 |
| COX10 | 74.2163223 | -0.00890122 | 0.2923665 | -0.0304454 | 0.97571182 | 0.99408036 |
| TRIM16 | 37.6082142 | 0.086531453 | 0.31828541 | 0.27186748 | 0.78572391 | 0.91275941 |
| ZSWIM7 | 101.271017 | -0.151093466 | 0.21363202 | -0.7072604 | 0.47940468 | 0.73490443 |

| | | | | | | |
|---|---|---|---|---|---|---|
| TTC19 | 228.233405 | 0.381191978 | 0.2056445 | 1.85364541 | 0.06378991 | 0.2571732 |
| NCOR1 | 570.642079 | 0.267640835 | 0.13487791 | 1.98431921 | 0.04722027 | 0.21714794 |
| PIGL | 169.920414 | 0.003724161 | 0.26509217 | 0.01404855 | 0.98879125 | 0.99678172 |
| UBB | 320.962126 | -0.81677323 | 0.40459854 | -2.0187251 | 0.0435158 | 0.2066642 |
| LRRC75A | 30.6047172 | -0.436164054 | 0.44953302 | -0.9702603 | 0.33191675 | 0.61700573 |
| AC098850.3 | 78.8936311 | 0.420584087 | 0.56804751 | 0.74040301 | 0.4590555 | 0.71780813 |
| MPRIP | 320.432496 | 0.463429321 | 0.37891309 | 1.22304912 | 0.22131114 | 0.50310453 |
| COPS3 | 38.5962864 | 0.354332464 | 0.26529373 | 1.3356232 | 0.18167246 | 0.45460829 |
| MED9 | 37.5595167 | -0.209631254 | 0.33709869 | -0.6218691 | 0.53402795 | 0.77366209 |
| RASD1 | 260.443632 | 0.424050816 | 1.01350236 | 0.41840141 | 0.67565366 | 0.86116336 |
| PEMT | 47.5479799 | -0.330129559 | 0.47601786 | -0.6935235 | 0.4879811 | 0.74125768 |
| RAI1 | 224.335418 | -0.622271129 | 0.37084478 | -1.6779827 | 0.09335048 | 0.31833543 |
| SREBF1 | 42.3730374 | -0.811722665 | 0.39278685 | -2.0665729 | 0.03877441 | 0.19565065 |
| TOM1L2 | 218.813332 | 0.632369784 | 0.18586854 | 3.40224221 | 0.00066835 | 0.01577201 |
| DRC3 | 46.3648647 | 0.442435645 | 0.4037799 | 1.0957347 | 0.27319489 | 0.56073149 |
| ATPAF2 | 34.4823165 | 0.167374257 | 0.27071596 | 0.61826519 | 0.53640054 | 0.77557446 |
| DRG2 | 27.0879427 | -0.278737266 | 0.33981274 | -0.8202673 | 0.41206377 | 0.68367283 |
| ALKBH5 | 94.7482548 | -0.450051492 | 0.23142228 | -1.9447198 | 0.05180871 | 0.22800587 |
| FLII | 72.9299483 | -0.606012883 | 0.27272316 | -2.2220807 | 0.02627785 | 0.15732221 |
| TOP3A | 50.3420164 | -0.353976287 | 0.23564942 | -1.5021309 | 0.1330633 | 0.38380408 |
| SHMT1 | 203.413068 | -0.219642977 | 0.28097484 | -0.7817176 | 0.43438056 | 0.69919869 |
| TVP23B | 46.4591767 | -0.222443734 | 0.34682136 | -0.6413784 | 0.52127686 | 0.76440016 |
| PRPSAP2 | 140.730034 | 0.788680163 | 0.26185944 | 3.0118455 | 0.00259665 | 0.03729128 |
| SLC5A10 | 43.2686669 | 0.483400142 | 0.25221539 | 1.91661635 | 0.05528669 | 0.23619173 |
| EPN2 | 310.597027 | 0.224306329 | 0.15691896 | 1.42944056 | 0.15287765 | 0.41531905 |
| B9D1 | 63.8133008 | -0.047932021 | 0.22383901 | -0.2141361 | 0.8304409 | 0.93446017 |
| AC004448.2 | 38.4885301 | -1.386452977 | 0.53697806 | -2.5819546 | 0.00982425 | 0.08852916 |
| SLC47A1 | 73.1840731 | 0.199501991 | 0.34546785 | 0.57748353 | 0.56361286 | 0.79293888 |
| ALDH3A2 | 101.255005 | -0.08399511 | 0.29323401 | -0.286444 | 0.77453811 | 0.90689106 |
| ULK2 | 79.2445676 | 0.066340227 | 0.24229448 | 0.2738 | 0.78423833 | 0.91212179 |
| AKAP10 | 230.43548 | 0.268082522 | 0.17175283 | 1.56086237 | 0.11855623 | 0.36352436 |
| CCDC144NL-/ | 43.2150969 | -0.245973385 | 0.42569377 | -0.5778177 | 0.56338722 | 0.79290184 |
| USP22 | 198.916585 | 0.077585512 | 0.17035807 | 0.45542609 | 0.64880273 | 0.8462529 |
| DHRS7B | 161.377561 | -0.298741839 | 0.23320174 | -1.2810446 | 0.20017799 | 0.47889734 |
| TMEM11 | 25.9348079 | -0.764567789 | 0.34771991 | -2.1988036 | 0.02789189 | 0.16153351 |
| AC087294.1 | 27.585456 | -0.524313699 | 0.30025489 | -1.7462287 | 0.08077122 | 0.29553381 |
| MAP2K3 | 54.7664262 | -0.913198258 | 0.35977329 | -2.5382603 | 0.01114051 | 0.09528119 |
| WSB1 | 774.71289 | 0.255770654 | 0.43852786 | 0.58324836 | 0.55972612 | 0.78994338 |
| KSR1 | 135.314634 | -0.034328513 | 0.30066167 | -0.1141766 | 0.90909784 | 0.96752108 |
| NLK | 505.705681 | 0.746192754 | 0.15626295 | 4.77523782 | 1.79E-06 | 0.00026452 |
| IFT20 | 28.5431549 | 0.223080673 | 0.28897856 | 0.7719627 | 0.44013652 | 0.7034987 |
| TNFAIP1 | 55.0501276 | 0.266542829 | 0.3280954 | 0.8123943 | 0.41656542 | 0.68597057 |
| POLDIP2 | 47.8641633 | -0.674039145 | 0.24215029 | -2.783557 | 0.00537664 | 0.06044935 |
| SARM1 | 35.7432533 | -0.366493913 | 0.5045392 | -0.7263933 | 0.46759768 | 0.72536326 |
| KIAA0100 | 88.8059621 | -0.352653322 | 0.24340222 | -1.44885 | 0.14737947 | 0.40633477 |
| SDF2 | 47.6808537 | -0.175253956 | 0.32942439 | -0.5320005 | 0.59472562 | 0.81507555 |
| SUPT6H | 105.503585 | -0.4987888 | 0.18493678 | -2.6970773 | 0.0069951 | 0.07109414 |
| RAB34 | 65.5568626 | -0.625874246 | 0.38855868 | -1.6107586 | 0.10723234 | 0.34262128 |
| RPL23A | 337.182259 | -0.563348201 | 0.49425915 | -1.1397831 | 0.25437669 | 0.54124263 |
| NEK8 | 39.1292822 | -0.446834327 | 0.54712204 | -0.8166996 | 0.41410015 | 0.68449528 |
| TRAF4 | 81.7168056 | -0.29798691 | 0.31361837 | -0.9501577 | 0.34203213 | 0.62716802 |
| FAM222B | 309.496577 | -0.077519411 | 0.22977194 | -0.3373755 | 0.73583389 | 0.89027623 |
| FLOT2 | 157.763344 | -0.838833555 | 0.26248424 | -3.1957483 | 0.00139469 | 0.02594119 |
| PHF12 | 92.7302677 | 0.191743121 | 0.23433212 | 0.81825368 | 0.41321235 | 0.68443902 |
| MYO18A | 57.8803669 | -0.383639909 | 0.50195622 | -0.7642896 | 0.4446947 | 0.70707984 |
| NUFIP2 | 244.856331 | -0.440287975 | 0.18392178 | -2.3938871 | 0.01667088 | 0.12124283 |
| TAOK1 | 424.092813 | -0.002977204 | 0.20480274 | -0.0145369 | 0.98840161 | 0.99678172 |
| ABHD15-AS1 | 58.9286018 | 0.388100079 | 0.37297946 | 1.0405401 | 0.29808905 | 0.58581522 |
| GIT1 | 37.5775932 | -0.124338613 | 0.38172267 | -0.3257302 | 0.74462847 | 0.89323545 |
| SSH2 | 796.420656 | -0.018597262 | 0.19571014 | -0.0950245 | 0.92429535 | 0.9731617 |
| AC023389.1 | 30.5286526 | -0.157134921 | 0.29743051 | -0.528308 | 0.59728557 | 0.81652612 |
| NSRP1 | 296.287813 | 0.226214579 | 0.16076884 | 1.40707973 | 0.15940375 | 0.42483013 |
| BLMH | 66.5863947 | -0.349836151 | 0.27149148 | -1.2885714 | 0.19754713 | 0.47630586 |
| CPD | 400.255937 | -1.426210769 | 0.40685083 | -3.5054881 | 0.00045577 | 0.01258164 |
| GOSR1 | 150.94124 | 0.08446369 | 0.19141586 | 0.44125753 | 0.65902657 | 0.85262229 |
| CRLF3 | 136.846029 | 0.621870632 | 0.22334641 | 2.78433228 | 0.00536381 | 0.06041319 |
| ATAD5 | 31.7365426 | 0.530564927 | 0.38731909 | 1.36983936 | 0.17073705 | 0.4409518 |
| TEFM | 55.8365319 | 0.076798645 | 0.29423044 | 0.2610153 | 0.79408071 | 0.91723627 |
| RNF135 | 39.5320076 | 0.024146746 | 0.32950317 | 0.07328229 | 0.94158149 | 0.98069034 |
| NF1 | 928.175751 | 0.331795148 | 0.15689956 | 2.11469772 | 0.03445572 | 0.18305675 |
| RAB11FIP4 | 34.015361 | 0.396083245 | 0.34732299 | 1.1403888 | 0.25412436 | 0.54093192 |
| COPRS | 37.0317925 | 0.126751632 | 0.43371563 | 0.29224594 | 0.77009859 | 0.90533273 |
| UTP6 | 58.4870557 | 0.101360639 | 0.25805477 | 0.39278732 | 0.69447657 | 0.87004854 |
| SUZ12 | 243.530037 | 0.333486848 | 0.18485612 | 1.80403464 | 0.07122588 | 0.27422535 |
| LRRC37B | 63.5290217 | 0.048243456 | 0.23770448 | 0.2029556 | 0.83916973 | 0.93972491 |
| RHOT1 | 143.711981 | 0.226103238 | 0.17194736 | 1.31495617 | 0.18852463 | 0.46435523 |
| C17orf75 | 25.5223272 | 0.540688367 | 0.28880612 | 1.87215 | 0.06118585 | 0.25014908 |
| ZNF207 | 366.078824 | 0.159883467 | 0.17668625 | 0.90490043 | 0.36551814 | 0.64551959 |
| PSMD11 | 91.1965618 | -0.267335545 | 0.28392714 | -0.9415639 | 0.34641595 | 0.6302171 |
| MYO1D | 657.444687 | 0.319151141 | 0.24291272 | 1.31385111 | 0.18889632 | 0.46479265 |
| TMEM98 | 51.6614149 | -0.901950134 | 0.55877495 | -1.6141563 | 0.10649352 | 0.34105698 |
| CCL2 | 148.473391 | -3.151187871 | 0.99277367 | -3.1741252 | 0.00150289 | 0.02695912 |
| LIG3 | 32.0232695 | -0.051228884 | 0.25827129 | -0.198353 | 0.84276889 | 0.94141724 |
| RFFL | 34.330975 | -1.519997695 | 0.39745728 | -3.8243046 | 0.00013114 | 0.00565867 |
| SLFN5 | 100.245711 | -2.476167897 | 0.4998196 | -4.9541233 | 7.27E-07 | 0.00012889 |
| SLFN13 | 28.5160906 | 0.562289858 | 0.55005023 | 1.02225183 | 0.30666172 | 0.59427729 |
| LINC02001 | 33.9410204 | -0.232893351 | 0.29712909 | -0.783812 | 0.43315042 | 0.69824029 |
| AP2B1 | 197.895317 | 0.153947176 | 0.18415274 | 0.83597549 | 0.40316869 | 0.67708038 |
| TAF15 | 407.13604 | 0.012054309 | 0.20029768 | 0.06018197 | 0.95201071 | 0.98332009 |
| ZNHIT3 | 30.4191624 | -0.105378891 | 0.42952229 | -0.2453397 | 0.80619338 | 0.92263409 |
| MYO19 | 82.6039197 | -0.638958052 | 0.27771624 | -2.3007587 | 0.02140528 | 0.14045419 |
| GGNBP2 | 177.634385 | -0.177060083 | 0.15669383 | -1.1299748 | 0.25848684 | 0.54550461 |
| AATF | 142.656338 | -0.297568449 | 0.1820776 | -1.6342946 | 0.102197 | 0.33388304 |
| ACACA | 299.608239 | 0.245436721 | 0.22007128 | 1.11526014 | 0.26473896 | 0.55272255 |
| TADA2A | 88.8643572 | 0.088958667 | 0.29131843 | 0.30536574 | 0.76008762 | 0.90125623 |
| SYNRG | 205.203652 | 0.040163949 | 0.17259518 | 0.23270609 | 0.81598964 | 0.92869214 |
| DDX52 | 142.688351 | -0.039040857 | 0.24892739 | -0.1568363 | 0.87537384 | 0.95308093 |
| HNF1B | 564.046406 | 0.922622513 | 0.37678511 | 2.44867032 | 0.01433846 | 0.11107948 |
| MRPL45 | 46.9795135 | 0.162770997 | 0.21835711 | 0.74543485 | 0.45600888 | 0.71539502 |
| SOCS7 | 74.7283926 | 0.582526355 | 0.39456183 | 1.47638802 | 0.13983976 | 0.3940484 |
| ARHGAP23 | 60.5872328 | 0.606747913 | 0.63167031 | 0.96054524 | 0.33678087 | 0.62147307 |
| MLLT6 | 235.488515 | -0.081787103 | 0.20935685 | -0.3906588 | 0.69604943 | 0.87043654 |
| CISD3 | 37.7643532 | -0.319626749 | 0.38560184 | -0.8289036 | 0.40715896 | 0.68040675 |
| PCGF2 | 50.8265995 | -0.319375675 | 0.26999539 | -1.1828931 | 0.23685152 | 0.52000641 |
| PSMB3 | 78.127645 | -0.934072883 | 0.39920593 | -2.3398272 | 0.01929267 | 0.13189928 |
| PIP4K2B | 72.3881329 | 0.404890882 | 0.29634416 | 1.36628602 | 0.17184921 | 0.4426307 |
| CWC25 | 44.0016385 | -0.388856636 | 0.28918854 | -1.3446475 | 0.1787391 | 0.45040283 |
| RPL23 | 262.622266 | -0.554114531 | 0.4299138 | -1.2888968 | 0.19743396 | 0.47630586 |
| LASP1 | 129.433165 | -0.30992301 | 0.30389306 | -1.0198424 | 0.30780323 | 0.59456464 |
| RPL19 | 536.536723 | -0.593636758 | 0.53527339 | -1.1090347 | 0.26741521 | 0.55480561 |
| FBXL20 | 326.498425 | 0.273234505 | 0.27135996 | 1.00690797 | 0.313979 | 0.60099077 |
| MED1 | 71.4036477 | -0.031460337 | 0.18951822 | -0.1660016 | 0.86815565 | 0.95090501 |
| CDK12 | 198.314644 | -0.257088747 | 0.16608221 | -1.5479607 | 0.12163175 | 0.36636109 |

| | | | | | | |
|---|---|---|---|---|---|---|
| STARD3 | 41.8013379 | -0.328476327 | 0.21168628 | -1.5517129 | 0.12073094 | 0.36507762 |
| PGAP3 | 26.2765554 | 0.193376708 | 0.46534788 | 0.415553 | 0.67773712 | 0.86151115 |
| ERBB2 | 171.824765 | 0.593483803 | 0.40707116 | 1.45793624 | 0.14485812 | 0.40282422 |
| MIEN1 | 36.0355071 | -0.635444327 | 0.42052617 | -1.5110696 | 0.13077071 | 0.381325 |
| GRB7 | 111.126989 | 0.567937599 | 0.27532166 | 2.06281481 | 0.03913023 | 0.19618964 |
| PSMD3 | 59.5427339 | -0.772046668 | 0.31078509 | -2.4841818 | 0.01298495 | 0.10443563 |
| MED24 | 57.2763947 | 0.056127244 | 0.23510103 | 0.2387367 | 0.81130976 | 0.92562365 |
| THRA | 68.1425643 | 0.473885875 | 0.3175956 | 1.49210468 | 0.13567171 | 0.38788633 |
| MSL1 | 105.394775 | -0.323448202 | 0.26231564 | -1.2330496 | 0.21755726 | 0.49951605 |
| CASC3 | 104.093974 | 0.200525613 | 0.20036748 | 1.00078922 | 0.31692872 | 0.60343077 |
| WIPF2 | 197.739479 | 0.342373405 | 0.16023972 | 2.13663254 | 0.03262789 | 0.17693995 |
| RARA | 39.4698322 | -1.006640476 | 0.43245453 | -2.3277371 | 0.01992607 | 0.13451147 |
| IGFBP4 | 411.533252 | -1.637556699 | 0.65369148 | -2.5050911 | 0.01224199 | 0.10091982 |
| KRT10 | 89.9937708 | -0.715363137 | 0.45951889 | -1.5567654 | 0.11952618 | 0.3645149 |
| KRT19 | 395.367646 | -1.388737565 | 0.41019488 | -3.3855556 | 0.00071034 | 0.01643569 |
| EIF1 | 408.405884 | -0.777677037 | 0.38072339 | -2.04263 | 0.04108908 | 0.20176225 |
| JUP | 291.361116 | 0.52543141 | 0.23865255 | 2.2016585 | 0.02768944 | 0.16119979 |
| P3H4 | 38.5805985 | 0.097499479 | 0.3960205 | 0.24619806 | 0.80552892 | 0.92227408 |
| FKBP10 | 34.1411769 | -0.024552372 | 0.81682015 | -0.0300585 | 0.97602041 | 0.99411128 |
| NT5C3B | 44.1883447 | -0.091610184 | 0.2322501 | -0.3944463 | 0.6932516 | 0.86938709 |
| ACLY | 161.852556 | -0.163729194 | 0.50870718 | -0.3218535 | 0.74756367 | 0.89399656 |
| TTC25 | 36.4223819 | 0.048188753 | 0.4319489 | 0.11156123 | 0.91117131 | 0.96844354 |
| CNP | 90.1502264 | -0.196928917 | 0.38705816 | -0.5087838 | 0.61090379 | 0.82279561 |
| DNAJC7 | 169.936269 | 0.048155066 | 0.13501119 | 0.35667463 | 0.7213354 | 0.88235193 |
| NKIRAS2 | 26.4102507 | -0.31692084 | 0.30447807 | -1.0408659 | 0.29793778 | 0.5856651 |
| KAT2A | 35.2142665 | 0.372301589 | 0.25610979 | 1.45367964 | 0.14603515 | 0.40418106 |
| GHDC | 23.4080309 | -0.355697543 | 0.35681236 | -0.9968756 | 0.31882489 | 0.60563353 |
| STAT5B | 175.54028 | -0.423405109 | 0.18745876 | -2.2586573 | 0.02390471 | 0.14976011 |
| STAT5A | 27.149819 | -1.386204037 | 0.46165901 | -3.0026579 | 0.00267633 | 0.03790604 |
| STAT3 | 677.881146 | -0.593144129 | 0.27326865 | -2.1705532 | 0.02996497 | 0.16924199 |
| CAVIN1 | 109.949717 | 0.428551615 | 0.54582844 | 0.78513977 | 0.43237162 | 0.69737124 |
| ATP6V0A1 | 192.126096 | -0.298392481 | 0.22453749 | -1.3289206 | 0.18387418 | 0.45721925 |
| NAGLU | 38.3608299 | -0.804506204 | 0.52770348 | -1.5245422 | 0.12737334 | 0.37461135 |
| COASY | 36.2660697 | -0.302732065 | 0.27209512 | -1.1125964 | 0.26588179 | 0.55384374 |
| MLX | 31.5762312 | -0.649241049 | 0.30276408 | -2.1443794 | 0.0320025 | 0.17525099 |
| RETREG3 | 102.732017 | -0.105944953 | 0.16258497 | -0.6516282 | 0.51464105 | 0.76135083 |
| TUBG2 | 31.4215345 | 0.505141072 | 0.23528578 | 2.14692566 | 0.0317992 | 0.17444671 |
| EZH1 | 100.24283 | 0.128701639 | 0.18969729 | 0.67845796 | 0.49748137 | 0.74927265 |
| VPS25 | 49.4968314 | -1.228949879 | 0.33523571 | -3.6659277 | 0.00024644 | 0.00849612 |
| COA3 | 64.8488371 | -0.357458265 | 0.43633549 | -0.819228 | 0.41265632 | 0.68408231 |
| BECN1 | 42.4393183 | -0.130782039 | 0.24253813 | -0.5392226 | 0.58973328 | 0.81183245 |
| PSME3 | 33.9402064 | -0.824587877 | 0.29097013 | -2.8339262 | 0.00459799 | 0.05471328 |
| RUNDC1 | 39.7078739 | -0.165918351 | 0.29568284 | -0.5611362 | 0.57470467 | 0.8009242 |
| RPL27 | 172.65527 | -1.299532058 | 0.46169468 | -2.8147001 | 0.00488228 | 0.05710298 |
| IFI35 | 42.7752329 | -0.793554984 | 0.79304538 | -1.0006426 | 0.31699963 | 0.60343739 |
| VAT1 | 47.6632111 | -0.751239718 | 0.29273713 | -2.5662605 | 0.01028016 | 0.09075913 |
| BRCA1 | 53.9883819 | 0.189469316 | 0.32388139 | 0.584996 | 0.5585504 | 0.78926435 |
| NBR1 | 143.739697 | -0.062697199 | 0.27248827 | -0.2300914 | 0.81802077 | 0.92999132 |
| LINC00854 | 219.45542 | 0.457017909 | 0.54260488 | 0.8422665 | 0.3996388 | 0.67444328 |
| DHX8 | 53.7657979 | -0.296967764 | 0.1978735 | -1.500796 | 0.13340833 | 0.38456768 |
| DUSP3 | 80.9811245 | -0.285888923 | 0.37003689 | -0.7725957 | 0.43976167 | 0.7032516 |
| MPP3 | 48.7124489 | 0.516904042 | 0.44636962 | 1.15801796 | 0.24685671 | 0.5336584 |
| TMEM101 | 31.2650221 | -0.423893212 | 0.2955875 | -1.4340702 | 0.15155224 | 0.41347733 |
| LSM12 | 71.5844347 | -0.201533768 | 0.24660889 | -0.8172202 | 0.41380259 | 0.68448816 |
| G6PC3 | 35.1119569 | -0.881276148 | 0.41813996 | -2.1076104 | 0.0350647 | 0.1847551 |
| HDAC5 | 45.4409491 | 0.572148247 | 0.32435439 | 1.76396025 | 0.07773867 | 0.28902315 |
| TMUB2 | 37.2572072 | -0.404589667 | 0.26053755 | -1.5529035 | 0.12044619 | 0.36507762 |
| ATXN7L3 | 32.3621948 | -0.379219095 | 0.41282486 | -0.9185956 | 0.35830714 | 0.64001011 |
| UBTF | 71.1568148 | 0.035717342 | 0.35290263 | 0.10121019 | 0.91938361 | 0.97108563 |
| SLC25A39 | 51.3962315 | -0.613161575 | 0.39909838 | -1.536367 | 0.1244484 | 0.37055989 |
| GRN | 159.961802 | -0.852695803 | 0.31600302 | -2.6983786 | 0.00696781 | 0.07108061 |
| GPATCH8 | 576.379471 | 0.368302451 | 0.17579141 | 2.09511063 | 0.03616116 | 0.18804981 |
| EFTUD2 | 77.4025589 | -0.103770034 | 0.27290389 | -0.3802439 | 0.70376439 | 0.87357252 |
| DCAKD | 96.4054781 | -0.018983193 | 0.3558548 | -0.0533453 | 0.95745676 | 0.98533286 |
| NMT1 | 123.952629 | -0.190614615 | 0.18584892 | -1.0256428 | 0.30505996 | 0.59267272 |
| PLCD3 | 57.5594297 | 0.310029205 | 0.36352899 | 0.85283214 | 0.3937524 | 0.6686322 |
| ACBD4 | 37.2806575 | 1.182459206 | 0.3361358 | 3.51780203 | 0.00043514 | 0.01218765 |
| HEXIM1 | 51.1374738 | 0.370392347 | 0.30668432 | 1.2077316 | 0.22715052 | 0.50993082 |
| AC138150.1 | 91.2493331 | 0.557192657 | 0.38145368 | 1.46070857 | 0.14409544 | 0.40162014 |
| MAP3K14 | 148.302658 | -0.255078334 | 0.30246653 | -0.8433275 | 0.39904532 | 0.67377896 |
| ARHGAP27 | 35.9165435 | -0.814101685 | 0.27932417 | -2.9145408 | 0.00356212 | 0.04598594 |
| PLEKHM1 | 59.7770632 | -0.478052003 | 0.27021566 | -1.7691499 | 0.07686886 | 0.28663488 |
| LINC02210-C | 52.8196683 | 0.331526772 | 0.37122033 | 0.89307277 | 0.3718182 | 0.65192016 |
| KANSL1 | 1040.52773 | 0.678232556 | 0.30871282 | 2.19696919 | 0.02802265 | 0.16199694 |
| ARL17B | 244.428857 | 2.143018977 | 1.09116188 | 1.96397896 | 0.04953253 | 0.2238076 |
| LRRC37A2 | 61.9658395 | 0.944617985 | 0.32921295 | 2.86932206 | 0.00411353 | 0.05066222 |
| ARL17A | 69.70518 | 0.107380808 | 0.35386335 | 0.30345276 | 0.76154485 | 0.90141135 |
| NSF | 216.213903 | -0.00088933 | 0.37139098 | -0.0023946 | 0.99808939 | 0.99923686 |
| AC005670.2 | 32.0073723 | 0.377311368 | 0.43872151 | 0.86002477 | 0.38977539 | 0.66529908 |
| CDC27 | 270.526361 | 0.071977067 | 0.20902264 | 0.34435058 | 0.73058265 | 0.88757288 |
| AC068234.1 | 763.697634 | -0.311581939 | 0.50554904 | -0.6163239 | 0.53768079 | 0.77644087 |
| AC068234.2 | 142.225806 | -0.519479041 | 0.52195731 | -0.995252 | 0.31961373 | 0.60625346 |
| EFCAB13 | 372.494762 | 0.035741381 | 0.35556914 | 0.10051879 | 0.91993246 | 0.97155822 |
| NPEPPS | 525.344669 | -0.120684576 | 0.19006842 | -0.6349533 | 0.52545886 | 0.7671049 |
| AC025682.1 | 63.0305784 | 0.272725287 | 0.29328466 | 0.92989958 | 0.35242308 | 0.63408352 |
| KPNB1 | 230.977182 | -0.296847692 | 0.21987041 | -1.350103 | 0.17698296 | 0.44872611 |
| SCRN2 | 36.7856587 | 0.025288419 | 0.25886992 | 0.09768775 | 0.92218025 | 0.97248656 |
| SP2 | 24.9138658 | 0.403628377 | 0.2939922 | 1.37292206 | 0.16977658 | 0.43977551 |
| SP2-AS1 | 29.7138379 | -0.381113596 | 0.2743803 | -1.3889977 | 0.16483346 | 0.43317961 |
| PNPO | 29.2570393 | -0.691274853 | 0.33167828 | -2.0841728 | 0.03714445 | 0.19107758 |
| CDK5RAP3 | 58.8195666 | 0.445755152 | 0.25885581 | 1.72202102 | 0.08506571 | 0.30325435 |
| NFE2L1 | 188.544554 | -0.58324852 | 0.19415416 | -3.0040486 | 0.00266413 | 0.03788912 |
| CBX1 | 81.8749149 | 0.01726543 | 0.26350287 | 0.06552274 | 0.9477578 | 0.98231681 |
| SKAP1 | 482.639995 | 0.744284515 | 0.54661199 | 1.36163225 | 0.17331397 | 0.44449049 |
| HOXB2 | 46.3902699 | 0.554395834 | 0.64617578 | 0.85796443 | 0.3909121 | 0.66640808 |
| HOXB3 | 224.441653 | 0.65969268 | 0.57719324 | 1.14293209 | 0.25306679 | 0.54001487 |
| HOXB-AS3 | 53.8575987 | 1.029443433 | 0.69300776 | 1.48547173 | 0.13741891 | 0.38990987 |
| HOXB7 | 62.0945986 | -0.297854989 | 0.51086077 | -0.5830453 | 0.55986278 | 0.78994338 |
| AC103702.2 | 33.8326204 | -0.492279503 | 0.517712 | -0.9508752 | 0.34166773 | 0.62700419 |
| LINC02086 | 66.4364513 | 0.226323758 | 0.52765444 | 0.4289242 | 0.66797839 | 0.85905308 |
| CALCOCO2 | 103.14548 | -0.04309328 | 0.2729029 | -0.157907 | 0.87453008 | 0.95281206 |
| ATP5MC1 | 85.1026838 | -0.830122946 | 0.44837368 | -1.8514087 | 0.06411078 | 0.25781575 |
| UBE2Z | 149.170433 | -0.437352358 | 0.29631999 | -1.4759462 | 0.13995834 | 0.39412255 |
| SNF8 | 61.9481769 | -0.286350594 | 0.21725752 | -1.3180239 | 0.18749566 | 0.46292038 |
| ZNF652 | 221.420987 | 0.29691098 | 0.20708647 | 1.43375366 | 0.15164257 | 0.41360598 |
| PHB | 108.335898 | -0.51817677 | 0.41510799 | -1.2482939 | 0.21192345 | 0.49320085 |
| SPOP | 176.013014 | -0.10657588 | 0.24351801 | -0.4376509 | 0.66163936 | 0.85473405 |
| SLC35B1 | 25.1305365 | -0.547907563 | 0.4582301 | -1.195704 | 0.23181209 | 0.51496475 |
| FAM117A | 27.4231097 | 0.908667259 | 0.35592791 | 2.55295312 | 0.01068139 | 0.09276247 |
| KAT7 | 81.7082595 | 0.230861673 | 0.22686613 | 1.01761189 | 0.30886243 | 0.59563596 |
| ITGA3 | 644.09584 | -1.398238975 | 0.661271 | -2.1144719 | 0.03447498 | 0.18305755 |
| PDK2 | 28.3917456 | 0.305752784 | 0.56607306 | 0.54012954 | 0.5891077 | 0.81148442 |
| PPP1R9B | 24.4323981 | -0.470830782 | 0.41231114 | -1.1419308 | 0.25348279 | 0.54030172 |

| | | | | | | |
|---|---|---|---|---|---|---|
| XYLT2 | 38.2564237 | -0.024642867 | 0.26310238 | -0.0936627 | 0.92537714 | 0.97349958 |
| MRPL27 | 53.5206613 | -0.689766294 | 0.35693904 | -1.9324484 | 0.05330418 | 0.23167003 |
| LRRC59 | 35.1459577 | -1.46168651 | 0.422516 | -3.459482 | 0.00054122 | 0.01396038 |
| ACSF2 | 68.3923555 | 0.286549328 | 0.49060633 | 0.58407181 | 0.559172 | 0.78981496 |
| RSAD1 | 37.402849 | 0.019480635 | 0.24427156 | 0.07974991 | 0.93643616 | 0.97871293 |
| SPATA20 | 35.3452307 | -0.36960779 | 0.32945521 | -1.1218757 | 0.26191529 | 0.54935113 |
| ABCC3 | 556.672426 | -1.575218616 | 0.5998832 | -2.6258755 | 0.00864264 | 0.08237598 |
| ANKRD40 | 41.8144778 | 0.171572338 | 0.28009157 | 0.61255802 | 0.54016862 | 0.7777505 |
| LUC7L3 | 449.72405 | 0.716857666 | 0.09531922 | 7.52059973 | 5.45E-14 | 1.31E-10 |
| TOB1 | 86.6132586 | 0.245549524 | 0.44127712 | 0.55645197 | 0.57790192 | 0.80357578 |
| SPAG9 | 766.9623 | 0.3342324 | 0.25318163 | 1.32012896 | 0.18679197 | 0.46186893 |
| AC005920.2 | 75.0542363 | 0.459445911 | 0.59076845 | 0.77770896 | 0.43674061 | 0.70170049 |
| MBTD1 | 230.203689 | 0.252649921 | 0.18927019 | 1.33486377 | 0.18192094 | 0.45499234 |
| UTP18 | 55.4286665 | -0.145063677 | 0.24074333 | -0.6025657 | 0.54679763 | 0.78253724 |
| TOM1L1 | 318.80774 | 0.403065605 | 0.2185425 | 1.8443351 | 0.06513432 | 0.26007572 |
| COX11 | 56.1017272 | 0.590972116 | 0.21292839 | 2.77545014 | 0.00551254 | 0.06154379 |
| STXBP4 | 210.233607 | 0.041102672 | 0.26104705 | 0.15745312 | 0.87488775 | 0.95291321 |
| HLF | 81.3111192 | -0.480623222 | 0.57990701 | -0.8287936 | 0.4072212 | 0.68040675 |
| DGKE | 29.8978572 | -0.501341997 | 0.25536075 | -1.9632696 | 0.04961486 | 0.22398317 |
| TRIM25 | 48.5053363 | -0.93468599 | 0.40045701 | -2.3340483 | 0.01959319 | 0.13327863 |
| COIL | 39.9021147 | 0.021115961 | 0.23862657 | 0.08848956 | 0.92948758 | 0.97545894 |
| SCPEP1 | 26.3828633 | 0.45170759 | 0.43386132 | 1.04113359 | 0.29781355 | 0.58554106 |
| AKAP1 | 48.2712621 | -0.397726972 | 0.24344979 | -1.6337125 | 0.10231923 | 0.33390378 |
| MSI2 | 1252.50528 | -0.124713946 | 0.20530227 | -0.607465 | 0.54354235 | 0.78025803 |
| MRPS23 | 43.5275374 | -0.829091394 | 0.30265031 | -2.7394368 | 0.00615445 | 0.06585681 |
| CUEDC1 | 152.621336 | -0.277161418 | 0.25284787 | -1.0961588 | 0.27300928 | 0.56047062 |
| VEZF1 | 108.599101 | 0.046440166 | 0.18850365 | 0.24636215 | 0.8054019 | 0.92227408 |
| AC015813.2 | 51.010101 | -0.076763998 | 0.24130817 | -0.318116 | 0.75039693 | 0.89593072 |
| SRSF1 | 25.9925256 | 0.325097335 | 0.32963023 | 0.98624856 | 0.32401116 | 0.61072469 |
| DYNLL2 | 76.6125475 | -0.565284987 | 0.22871885 | -2.4715278 | 0.01345371 | 0.10694861 |
| MKS1 | 27.319259 | 0.173587319 | 0.28060848 | 0.61861038 | 0.53617306 | 0.77540842 |
| TSPOAP1 | 63.5038919 | 0.547388499 | 0.39169693 | 1.39747966 | 0.16226938 | 0.42858376 |
| SUPT4H1 | 36.0514191 | -0.63985851 | 0.32833914 | -1.9487732 | 0.05132252 | 0.2274958 |
| RNF43 | 89.4351484 | 0.536757673 | 0.49441857 | 1.08563413 | 0.27764089 | 0.56501426 |
| MTMR4 | 37.5018791 | 0.13764731 | 0.27057469 | 0.50872204 | 0.61094707 | 0.82279561 |
| RAD51C | 31.2925016 | 0.681489662 | 0.24452427 | 2.78700214 | 0.00531981 | 0.06037735 |
| PPM1E | 59.1071323 | 1.06727002 | 0.51855559 | 2.05815931 | 0.03957484 | 0.19722814 |
| TRIM37 | 108.302126 | 0.435826327 | 0.21402079 | 2.03637379 | 0.04171284 | 0.20312121 |
| SKA2 | 65.9935514 | 0.97639292 | 0.29148124 | 3.34976251 | 0.00080881 | 0.01797582 |
| YPEL2 | 369.023233 | -0.080309752 | 0.33061063 | -0.2429134 | 0.80807249 | 0.92380073 |
| DHX40 | 90.1850714 | 0.002985112 | 0.26200708 | 0.01139325 | 0.9909097 | 0.9977564 |
| CLTC | 409.189573 | 0.129427757 | 0.21680556 | 0.5969762 | 0.55052327 | 0.78544272 |
| PTRH2 | 68.4917332 | -0.268391286 | 0.28428533 | -0.9440912 | 0.34512304 | 0.6298216 |
| VMP1 | 1072.30922 | -0.590255131 | 0.41990327 | -1.4056931 | 0.15981527 | 0.42535996 |
| TUBD1 | 61.6281312 | 0.21400448 | 0.36147175 | 0.59203652 | 0.55382613 | 0.78699373 |
| RPS6KB1 | 97.229731 | 0.264316731 | 0.21479793 | 1.23053667 | 0.21849621 | 0.50057436 |
| HEATR6 | 30.4538672 | 0.169164431 | 0.25625783 | 0.6601337 | 0.50916804 | 0.75806868 |
| USP32 | 380.293262 | -0.018334911 | 0.22420357 | -0.081778 | 0.93482328 | 0.97790458 |
| APPBP2 | 259.074481 | 0.199508842 | 0.17060392 | 1.16942708 | 0.2422316 | 0.52722939 |
| PPM1D | 89.3081402 | 0.724991977 | 0.25102995 | 2.8880696 | 0.00387614 | 0.0484634 |
| BCAS3 | 820.004882 | 0.284758575 | 0.19955621 | 1.42695924 | 0.15359164 | 0.41634457 |
| NACA2 | 36.7670312 | -0.249983796 | 0.66212882 | -0.3775456 | 0.70576821 | 0.87473681 |
| INTS2 | 53.3889638 | 0.237418287 | 0.25107604 | 0.94560313 | 0.34435105 | 0.62937201 |
| MED13 | 503.701585 | 0.092140245 | 0.13023258 | 0.70750536 | 0.47925249 | 0.73490443 |
| METTL2A | 26.2304809 | -0.660481607 | 0.35956412 | -1.8368952 | 0.06622537 | 0.26202924 |
| TLK2 | 301.485344 | 0.287564413 | 0.18566649 | 1.5488224 | 0.12142442 | 0.36610781 |
| MRC2 | 161.449728 | 0.533946341 | 0.61731375 | 0.86495131 | 0.38706551 | 0.66404483 |
| TANC2 | 475.35954 | 0.149681822 | 0.37919276 | 0.39473808 | 0.69303619 | 0.86938105 |
| AC015923.1 | 34.6722384 | 0.135177334 | 0.53784392 | 0.2513319 | 0.80155751 | 0.92108024 |
| CYB561 | 44.7112665 | -0.064389479 | 0.31611602 | -0.2036894 | 0.83859624 | 0.93919249 |
| DCAF7 | 207.514494 | -0.316544975 | 0.19193241 | -1.6492523 | 0.09909596 | 0.32904321 |
| MAP3K3 | 110.767343 | 0.134218453 | 0.2472182 | 0.54291493 | 0.58718837 | 0.81058905 |
| CCDC47 | 52.9602497 | -0.045504954 | 0.32758389 | -0.1389108 | 0.8895206 | 0.95918866 |
| DDX42 | 146.890788 | -0.108129581 | 0.18676779 | -0.578952 | 0.56262159 | 0.79260446 |
| FTSJ3 | 29.5482679 | -0.292008766 | 0.33431234 | -0.8734609 | 0.3824119 | 0.65946633 |
| PSMC5 | 39.2953998 | -0.476546981 | 0.4024392 | -1.1841465 | 0.23635506 | 0.51974689 |
| SMARCD2 | 44.4734996 | -0.497840528 | 0.2256937 | -2.2058238 | 0.02739634 | 0.16011564 |
| ERN1 | 131.105323 | -0.141393234 | 0.57746683 | -0.2448508 | 0.80657193 | 0.92274603 |
| SNHG25 | 41.0655164 | -0.783459217 | 0.50087964 | -1.5641666 | 0.11777844 | 0.36229919 |
| TEX2 | 71.8403135 | -0.444351184 | 0.25764978 | -1.7246325 | 0.08459373 | 0.30235944 |
| POLG2 | 50.2606606 | 0.071917602 | 0.29946588 | 0.2401529 | 0.81021172 | 0.92495996 |
| DDX5 | 468.619596 | -0.069713435 | 0.20742533 | -0.3360893 | 0.73680353 | 0.89057637 |
| CEP95 | 199.337693 | 0.459450021 | 0.27099708 | 1.69540578 | 0.08999847 | 0.31257989 |
| SMURF2 | 295.958996 | 0.253062359 | 0.15529768 | 1.62953086 | 0.10320069 | 0.33533222 |
| LRRC37A3 | 227.648496 | 0.692208817 | 0.28948863 | 2.3911434 | 0.01679599 | 0.12179319 |
| GNA13 | 143.532967 | 0.261155206 | 0.18399716 | 1.41934366 | 0.15579885 | 0.4201649 |
| RGS9 | 45.2912862 | 1.307805221 | 0.55711184 | 2.3474734 | 0.01890122 | 0.13016162 |
| CEP112 | 519.933436 | 0.075489487 | 0.33663163 | 0.22424953 | 0.82256314 | 0.93217972 |
| PRKCA | 517.313623 | 0.054278747 | 0.48875787 | 0.11105447 | 0.91157315 | 0.96844354 |
| HELZ | 461.448555 | 0.259201901 | 0.17656487 | 1.46802646 | 0.14209704 | 0.39848551 |
| PSMD12 | 92.958473 | -0.192628077 | 0.21425545 | -0.899058 | 0.36862177 | 0.6485537 |
| PITPNC1 | 838.565622 | -0.829615701 | 0.7230186 | -1.1474334 | 0.25120254 | 0.53867677 |
| NOL11 | 54.7559575 | 0.246386944 | 0.30509973 | 0.80756199 | 0.41934277 | 0.68835944 |
| BPTF | 581.118462 | 0.321077499 | 0.18758832 | 1.71160706 | 0.08696911 | 0.30691783 |
| ARSG | 62.4429704 | -0.822804551 | 0.4106025 | -2.0038956 | 0.04508124 | 0.21106218 |
| WIPI1 | 78.5018295 | -0.266189224 | 0.36228281 | -0.7347553 | 0.46248851 | 0.72082289 |
| PRKAR1A | 227.491236 | 0.17013967 | 0.21978827 | 0.77410713 | 0.43886744 | 0.70279675 |
| ABCA10 | 34.6426772 | 0.429694151 | 0.38441623 | 1.11778357 | 0.26365944 | 0.55119899 |
| ABCA5 | 141.642661 | 0.249319992 | 0.26898377 | 0.92689606 | 0.35398051 | 0.63575888 |
| AC005208.1 | 188.660775 | 0.468975806 | 0.45356964 | 1.03396649 | 0.30115183 | 0.58836088 |
| KCNJ16 | 684.186324 | 0.438888593 | 0.29039387 | 1.51135626 | 0.13069771 | 0.38122817 |
| SOX9 | 88.0347336 | -1.552109442 | 0.66871081 | -2.3210473 | 0.02028429 | 0.1355161 |
| LINC00511 | 285.116372 | -0.752482776 | 0.92403282 | -0.8143464 | 0.41544655 | 0.68495052 |
| SLC39A11 | 198.086638 | -0.465137334 | 0.22068968 | -2.1076534 | 0.03506099 | 0.1847551 |
| COG1 | 28.0092673 | -0.274127137 | 0.25648053 | -1.0688029 | 0.28515848 | 0.57264845 |
| FAM104A | 45.7625916 | -0.088218523 | 0.21719765 | -0.406167 | 0.68461989 | 0.8653964 |
| C17orf80 | 28.345405 | -0.255651192 | 0.25479515 | -1.0033597 | 0.31568733 | 0.60232326 |
| CDC42EP4 | 55.0784192 | -1.064989397 | 0.39103261 | -2.7235309 | 0.00645882 | 0.06810698 |
| RPL38 | 159.788308 | -0.844395646 | 0.3618879 | -2.3333072 | 0.01963203 | 0.13327863 |
| GPRC5C | 201.52955 | -1.044338599 | 0.23276782 | -4.4866108 | 7.24E-06 | 0.00070063 |
| SLC9A3R1 | 49.6896734 | -1.570794787 | 0.66142422 | -2.3748674 | 0.01755526 | 0.12490692 |
| NAT9 | 29.7470669 | -0.46285861 | 0.27473921 | -1.6847199 | 0.09204263 | 0.31589979 |
| KCTD2 | 39.5196942 | -0.086143845 | 0.25765859 | -0.3343333 | 0.73812807 | 0.89173021 |
| ATP5PD | 63.7035007 | -0.105635197 | 0.37970787 | -0.2782012 | 0.78085789 | 0.91077326 |
| SLC16A5 | 204.05057 | 0.650062207 | 0.77273411 | 0.84124953 | 0.40020816 | 0.67469094 |
| ARMC7 | 72.1991007 | 1.010877674 | 0.55848013 | 1.81005127 | 0.07028784 | 0.27219276 |
| NT5C | 77.2656217 | -0.771573997 | 0.34870462 | -2.2126865 | 0.02691927 | 0.15864374 |
| JPT1 | 80.9005806 | -0.417876717 | 0.6025213 | -0.6935468 | 0.48796647 | 0.74125768 |
| SUMO2 | 61.4389579 | 0.087346424 | 0.2631726 | 0.33189787 | 0.73996638 | 0.89253455 |
| NUP85 | 50.1657891 | 0.278312079 | 0.27323318 | 1.01858816 | 0.30839853 | 0.59499486 |
| GGA3 | 59.0123666 | 0.082730084 | 0.19232132 | 0.43016596 | 0.66707492 | 0.85839746 |
| MRPS7 | 45.5777793 | -0.475830324 | 0.31026432 | -1.5336289 | 0.12512098 | 0.37175369 |
| GRB2 | 180.316865 | -0.11452888 | 0.15167884 | -0.7550749 | 0.45020406 | 0.71104777 |

| | | | | | | |
|---|---|---|---|---|---|---|
| AC011933.4 | 30.5003729 | 0.151360193 | 0.50028243 | 0.30254949 | 0.76223321 | 0.90141135 |
| TMEM94 | 52.1880228 | 0.486230513 | 0.253711 | 1.91647389 | 0.0553048 | 0.23619173 |
| CASKIN2 | 26.9821906 | 0.358115699 | 0.33058793 | 1.08326912 | 0.27868899 | 0.56606485 |
| TSEN54 | 36.6680952 | 0.004082353 | 0.23041406 | 0.01771746 | 0.98586425 | 0.99668665 |
| LLGL2 | 301.90857 | -0.076146969 | 0.24834469 | -0.3066181 | 0.7591341 | 0.9005757 |
| MYO15B | 92.8725754 | 0.558781498 | 0.26863619 | 2.08006786 | 0.03751931 | 0.19223776 |
| RECQL5 | 48.4678619 | 0.475430426 | 0.3447241 | 1.37916212 | 0.16784477 | 0.43666081 |
| SAP30BP | 71.7855944 | -0.449607061 | 0.28873981 | -1.5571357 | 0.11943828 | 0.3645149 |
| H3F3B | 714.631385 | -0.663500848 | 0.42572687 | -1.558513 | 0.1191117 | 0.3645149 |
| UNK | 156.813636 | 0.591386659 | 0.26631359 | 2.22064022 | 0.02637534 | 0.15741394 |
| WBP2 | 100.138986 | -0.894091182 | 0.22594371 | -3.9571413 | 7.59E-05 | 0.00394884 |
| TRIM47 | 35.3671113 | -2.698318547 | 0.47330368 | -5.7010302 | 1.19E-08 | 5.19E-06 |
| TRIM65 | 24.6239062 | -0.487222128 | 0.30699382 | -1.5870747 | 0.11249573 | 0.35123747 |
| ACOX1 | 91.4417945 | 0.082117279 | 0.18372151 | 0.44696606 | 0.65489957 | 0.85026877 |
| EVPL | 44.2042056 | -0.305406341 | 0.41168585 | -0.7418432 | 0.45818236 | 0.71726243 |
| SRP68 | 57.8485214 | -0.273950229 | 0.22330266 | -1.2268113 | 0.21989352 | 0.50155763 |
| EXOC7 | 115.480047 | -0.096799473 | 0.23366848 | -0.4142599 | 0.6786838 | 0.86168533 |
| FOXJ1 | 30.4065855 | -0.872152515 | 0.51861584 | -1.6816928 | 0.09262843 | 0.31700169 |
| RNF157 | 88.8415781 | 0.098900969 | 0.46828353 | 0.21119891 | 0.83273206 | 0.93601742 |
| UBALD2 | 66.6836426 | -0.872479315 | 0.26464516 | -3.2967893 | 0.00097797 | 0.02054377 |
| PRPSAP1 | 76.8358837 | 0.513981079 | 0.18108564 | 2.83833154 | 0.00453501 | 0.05423427 |
| UBE2O | 85.6580324 | -0.131840536 | 0.28692906 | -0.4594883 | 0.64588358 | 0.84416957 |
| RHBDF2 | 46.3359074 | -1.45651014 | 0.39152755 | -3.7200707 | 0.00019917 | 0.00745243 |
| MXRA7 | 97.8212426 | 0.095261323 | 0.45754822 | 0.20819953 | 0.83507318 | 0.93710941 |
| JMJD6 | 28.730883 | -0.333380026 | 0.37926895 | -0.8790069 | 0.37939753 | 0.65699285 |
| SRSF2 | 146.174583 | 0.336606224 | 0.28430234 | 1.18397275 | 0.23642385 | 0.51974689 |
| MFSD11 | 79.7204168 | 0.512157449 | 0.23714973 | 2.15963746 | 0.03080075 | 0.17142506 |
| SEC14L1 | 163.281327 | -0.025117378 | 0.19251717 | -0.1304682 | 0.89619597 | 0.96165131 |
| 9-Sep | 333.938762 | -0.631071962 | 0.37888083 | -1.6656212 | 0.09578888 | 0.32285773 |
| TNRC6C | 146.229815 | 0.318327421 | 0.20984967 | 1.51693079 | 0.12928416 | 0.37802593 |
| TMC6 | 35.4856543 | -0.394177592 | 0.47244011 | -0.8343441 | 0.40408714 | 0.67765972 |
| SYNGR2 | 83.2502866 | -0.357889842 | 0.4398641 | -0.8136373 | 0.41585276 | 0.68549246 |
| AFMID | 88.8720804 | 0.944449448 | 0.39851174 | 2.36994132 | 0.01779091 | 0.12595648 |
| SOCS3 | 86.9753637 | -2.52332506 | 0.53656911 | -4.7027028 | 2.57E-06 | 0.00032791 |
| PGS1 | 76.9549756 | -0.672664951 | 0.30763105 | -2.1865964 | 0.028772 | 0.1648367 |
| DNAH17 | 28.009301 | 2.489396954 | 0.82128903 | 3.03108512 | 0.00243677 | 0.03574435 |
| CYTH1 | 168.295018 | 0.645689893 | 0.31788239 | 2.03122257 | 0.04223242 | 0.20401679 |
| USP36 | 109.991605 | -0.10016137 | 0.33057026 | -0.3029957 | 0.76189313 | 0.90141135 |
| TIMP2 | 82.9785676 | 0.219484388 | 0.32979285 | 0.66552198 | 0.50571669 | 0.75528806 |
| LGALS3BP | 113.140053 | -1.037919145 | 0.70036566 | -1.4819675 | 0.13834894 | 0.39113855 |
| CANT1 | 28.7683613 | -0.732095103 | 0.28153444 | -2.6003749 | 0.0093122 | 0.08613079 |
| TBC1D16 | 400.292769 | 0.187666528 | 0.24672328 | 0.76063567 | 0.44687471 | 0.70891772 |
| CCDC40 | 146.966224 | 0.280953704 | 0.31501106 | 0.8918852 | 0.37245447 | 0.65199071 |
| GAA | 72.7418598 | 0.094170842 | 0.36931944 | 0.2549848 | 0.79873484 | 0.91949148 |
| EIF4A3 | 53.9382364 | -0.674238014 | 0.39718988 | -1.6975206 | 0.08959827 | 0.31198177 |
| AC087741.1 | 29.7135532 | -0.073382754 | 0.42559985 | -0.1724219 | 0.86310582 | 0.94910925 |
| SLC26A11 | 65.7281142 | -0.017702168 | 0.31214263 | -0.0567118 | 0.95477478 | 0.98390867 |
| RNF213 | 680.016433 | -1.339316768 | 0.54883464 | -2.440292 | 0.0146754 | 0.11228085 |
| AC124319.2 | 222.356535 | 0.310998319 | 0.35404574 | 0.87841281 | 0.37971973 | 0.65703312 |
| ENDOV | 37.5175066 | -0.294095901 | 0.27255098 | -1.0790492 | 0.28056582 | 0.56841775 |
| RPTOR | 217.132269 | -0.092056436 | 0.23458953 | -0.3924149 | 0.69475165 | 0.87016554 |
| BAIAP2-DT | 24.0988791 | 0.033119224 | 0.4045219 | 0.08187251 | 0.93474809 | 0.97790458 |
| BAIAP2 | 82.512507 | 1.092757175 | 0.31948588 | 3.42036146 | 0.00062538 | 0.01535456 |
| NDUFAF8 | 47.2526934 | -0.348172421 | 0.28663863 | -1.2146738 | 0.22449054 | 0.50645193 |
| SLC38A10 | 111.472368 | 0.199726148 | 0.28050831 | 0.71201509 | 0.47645543 | 0.73328512 |
| BAHCC1 | 39.468038 | -0.005968954 | 0.37102166 | -0.0160879 | 0.98716428 | 0.99678172 |
| ACTG1 | 1033.51381 | 0.305254141 | 0.56310677 | 0.54208927 | 0.58775701 | 0.81090658 |
| FAAP100 | 25.3568304 | -0.667904604 | 0.38514307 | -1.7341727 | 0.08288744 | 0.29986597 |
| NPLOC4 | 190.127706 | -0.12039237 | 0.2132944 | -0.5644423 | 0.57245319 | 0.79946481 |
| OXLD1 | 26.8094393 | -0.683266521 | 0.38969253 | -1.7533477 | 0.07954233 | 0.29274747 |
| CCDC137 | 33.0405103 | -0.508492673 | 0.45925717 | -1.1072068 | 0.26820452 | 0.55536772 |
| ARL16 | 39.7890821 | -0.052910022 | 0.3168564 | -0.1669842 | 0.86738246 | 0.95090501 |
| HGS | 67.2480718 | -0.337891984 | 0.29534044 | -1.1440763 | 0.25259201 | 0.5393622 |
| MCRIP1 | 62.9109721 | -0.484127109 | 0.47053265 | -1.0288916 | 0.3035306 | 0.5915859 |
| P4HB | 219.647268 | -1.10256032 | 0.34669597 | -3.1801937 | 0.00147177 | 0.02655639 |
| ARHGDIA | 136.078418 | -1.185443259 | 0.46265222 | -2.5622773 | 0.01039883 | 0.09146957 |
| ANAPC11 | 76.9467799 | -0.48387891 | 0.42151384 | -1.147955 | 0.25098715 | 0.53845596 |
| SIRT7 | 26.7224166 | -0.459743017 | 0.27955591 | -1.6445477 | 0.10006313 | 0.33080044 |
| ASPSCR1 | 40.8619819 | -0.065977215 | 0.40029908 | -0.1648198 | 0.86908582 | 0.95106604 |
| CENPX | 29.7406135 | -0.132005615 | 0.51661436 | -0.2555206 | 0.79832103 | 0.91949148 |
| DCXR | 42.102376 | -0.580761297 | 0.36572666 | -1.5879654 | 0.11229416 | 0.35072244 |
| RFNG | 38.3645994 | 0.203058114 | 0.25778307 | 0.78770928 | 0.43086678 | 0.69670372 |
| GPS1 | 38.5515308 | -0.633832851 | 0.45188811 | -1.4026323 | 0.16072652 | 0.42659999 |
| DUS1L | 47.1775919 | -0.777373758 | 0.39321798 | -1.9769538 | 0.04804686 | 0.21916231 |
| CCDC57 | 219.503093 | 0.032521236 | 0.22940244 | 0.141765 | 0.88726563 | 0.95896094 |
| SLC16A3 | 44.8818936 | -0.959857506 | 0.42377188 | -2.2650335 | 0.02351062 | 0.14826087 |
| CSNK1D | 148.288902 | -0.348285994 | 0.15745932 | -2.211911 | 0.02697282 | 0.15870553 |
| OGFOD3 | 43.9296679 | -0.191519857 | 0.26993638 | -0.7095 | 0.47801426 | 0.7341756 |
| HEXDC | 77.946831 | 0.037916292 | 0.26043729 | 0.14558703 | 0.88424741 | 0.95761929 |
| CYBC1 | 40.8734242 | -1.411638272 | 0.3923244 | -3.5981404 | 0.0003205 | 0.01009893 |
| NARF | 58.9585476 | -0.260704914 | 0.21758006 | -1.1982022 | 0.2308383 | 0.51411301 |
| FOXK2 | 202.351018 | 0.134000233 | 0.20559121 | 0.65177999 | 0.51454311 | 0.76132352 |
| WDR45B | 147.967181 | 0.057213771 | 0.31534671 | 0.18143133 | 0.85602903 | 0.94621534 |
| RAB40B | 34.9842926 | 0.406927028 | 0.27237928 | 1.49397203 | 0.13518293 | 0.38731025 |
| FN3KRP | 27.3785321 | 0.111910813 | 0.38232118 | 0.29271413 | 0.76974067 | 0.90533273 |
| FN3K | 27.714537 | 0.550340305 | 0.62890653 | 0.87507488 | 0.38153316 | 0.65862428 |
| TBCD | 104.245573 | 0.200818422 | 0.27239827 | 0.73722356 | 0.46098641 | 0.71918385 |
| METRNL | 46.3366763 | 0.023430218 | 0.36499235 | 0.06419372 | 0.94881598 | 0.98262487 |
| USP14 | 161.444546 | -0.031188805 | 0.21343699 | -0.1461265 | 0.88382151 | 0.95761929 |
| THOC1 | 145.277694 | 0.311071987 | 0.24849696 | 1.25181407 | 0.21063762 | 0.49138016 |
| ENOSF1 | 466.776518 | -0.093669197 | 0.31947936 | -0.2931933 | 0.76937443 | 0.90533273 |
| YES1 | 194.151076 | -0.021658113 | 0.18950405 | -0.1142884 | 0.90900919 | 0.96752108 |
| METTL4 | 44.3576211 | 0.419290464 | 0.25639706 | 1.63531697 | 0.10198262 | 0.33363781 |
| SMCHD1 | 393.602423 | 0.181648377 | 0.23036016 | 0.78854076 | 0.43038047 | 0.6961524 |
| AP001011.1 | 62.4270193 | 0.050372754 | 0.46194808 | 0.10904419 | 0.91316744 | 0.96844354 |
| LPIN2 | 248.510238 | 1.035469825 | 0.41771504 | 2.47889046 | 0.01317918 | 0.10529052 |
| MYOM1 | 41.2750585 | 1.12180171 | 0.49164571 | 2.28172785 | 0.02250541 | 0.14420023 |
| MYL12A | 232.319371 | -0.695621757 | 0.40303243 | -1.7259697 | 0.08435287 | 0.3023414 |
| MYL12B | 447.151909 | -0.62934962 | 0.44375464 | -1.4182378 | 0.15612134 | 0.42043472 |
| TGIF1 | 108.143994 | 0.168497657 | 0.45458932 | 0.37065908 | 0.71089147 | 0.87748 |
| DLGAP1 | 254.072589 | -0.690103499 | 0.72774203 | -0.9482804 | 0.34298673 | 0.62795678 |
| LINC00667 | 47.4364959 | -0.184587137 | 0.2589828 | -0.712739 | 0.47600729 | 0.73306653 |
| EPB41L3 | 127.037464 | -0.142754474 | 0.35668518 | -0.4002254 | 0.68899049 | 0.86751885 |
| L3MBTL4 | 676.278974 | -0.022508646 | 0.26741756 | -0.0841704 | 0.93292094 | 0.97730202 |
| AP001021.2 | 40.8930378 | 0.427580002 | 0.39209947 | 1.0904886 | 0.27549797 | 0.56268552 |
| ARHGAP28 | 62.0517482 | 1.095114123 | 0.57506205 | 1.9043408 | 0.05686581 | 0.24059966 |
| LAMA1 | 216.983432 | 0.040207552 | 0.65933567 | 0.06098192 | 0.95137361 | 0.98319213 |
| PTPRM | 1577.22921 | 0.094199263 | 0.22296858 | 0.42247775 | 0.67267634 | 0.86026394 |
| RAB12 | 116.115174 | -0.016052896 | 0.19474549 | -0.0824301 | 0.93430468 | 0.97768238 |
| MTCL1 | 87.760986 | 0.118460809 | 0.36221004 | 0.32705005 | 0.74363003 | 0.89274747 |
| NDUFV2 | 128.152713 | -0.108040705 | 0.24925418 | -0.4334559 | 0.66468358 | 0.85693189 |
| ANKRD12 | 686.408598 | -0.064751608 | 0.18195506 | -0.3558659 | 0.72194097 | 0.88235193 |
| TWSG1 | 86.728143 | 0.385845275 | 0.22255741 | 1.73368871 | 0.08297333 | 0.29986597 |

| | | | | | | |
|---|---|---|---|---|---|---|
| RALBP1 | 126.584224 | -0.257574948 | 0.21241651 | -1.2125938 | 0.22528514 | 0.50741684 |
| PPP4R1 | 177.899746 | 0.002645133 | 0.26030173 | 0.0101618 | 0.9918922 | 0.99814428 |
| RAB31 | 53.5221218 | -0.98659897 | 0.39257934 | -2.5131199 | 0.01196686 | 0.10009484 |
| VAPA | 315.937619 | 0.129058007 | 0.20056649 | 0.64346743 | 0.51992084 | 0.76385303 |
| NAPG | 92.056208 | -0.155288166 | 0.22169153 | -0.7004695 | 0.48363412 | 0.73727416 |
| MPPE1 | 91.2146895 | 0.277786061 | 0.26677609 | 1.04127044 | 0.29775005 | 0.58554106 |
| IMPA2 | 183.685679 | -0.280763424 | 0.42212043 | -0.6651264 | 0.50596967 | 0.75528806 |
| TUBB6 | 90.4367814 | -0.654612529 | 0.44692413 | -1.4647062 | 0.1430011 | 0.3999438 |
| AFG3L2 | 130.25102 | -0.327907547 | 0.19241011 | -1.7042117 | 0.08834155 | 0.30970853 |
| SPIRE1 | 168.50083 | -0.723776174 | 0.256543 | -2.8212665 | 0.00478344 | 0.05622161 |
| PSMG2 | 80.9836733 | -0.183655857 | 0.17320059 | -1.0603651 | 0.28897853 | 0.57573323 |
| PTPN2 | 236.126923 | -0.34717234 | 0.29005012 | -1.1969391 | 0.23133028 | 0.51432779 |
| SEH1L | 76.8950967 | -0.110107508 | 0.27362395 | -0.4024045 | 0.68738636 | 0.86733718 |
| CEP192 | 176.985298 | 0.204166957 | 0.23521626 | 0.86799679 | 0.38539609 | 0.66218998 |
| LDLRAD4 | 272.576156 | -0.230272147 | 0.69481727 | -0.331414 | 0.74033182 | 0.89253455 |
| FAM210A | 70.494888 | -0.601979442 | 0.36176115 | -1.6640246 | 0.09610752 | 0.32324927 |
| RNMT | 162.293912 | 0.220370541 | 0.30015115 | 0.73419856 | 0.46282773 | 0.72095238 |
| ZNF519 | 29.3518058 | 0.895488972 | 0.35665333 | 2.51081065 | 0.01204543 | 0.10031173 |
| ROCK1 | 398.965874 | 0.0424325 | 0.13936303 | 0.30447459 | 0.76076635 | 0.90125623 |
| GREB1L | 156.47491 | -0.138140094 | 0.3856509 | -0.3581999 | 0.72019376 | 0.88207851 |
| ESCO1 | 131.251262 | 0.133266516 | 0.25788201 | 0.51677322 | 0.60531448 | 0.82056119 |
| SNRPD1 | 56.7062376 | -0.92433092 | 0.38870429 | -2.3779797 | 0.01740778 | 0.12460019 |
| ABHD3 | 70.4640186 | -0.780168746 | 0.33530899 | -2.3267159 | 0.01998039 | 0.1347244 |
| MIB1 | 311.577226 | 0.057980228 | 0.2094932 | 0.27676424 | 0.78196114 | 0.91082127 |
| AC090912.1 | 40.2926541 | -0.736546755 | 0.53761417 | -1.3700285 | 0.17067799 | 0.4409518 |
| RBBP8 | 336.364297 | 0.269468711 | 0.32972321 | 0.81725733 | 0.41378139 | 0.68448816 |
| CABLES1 | 235.917006 | 0.002806061 | 0.34707244 | 0.00808494 | 0.99354922 | 0.99876251 |
| TMEM241 | 155.649355 | -0.495022833 | 0.24320433 | -2.0354195 | 0.04180868 | 0.20312121 |
| RIOK3 | 216.838974 | 0.062694815 | 0.27714178 | 0.22621928 | 0.82103088 | 0.93138972 |
| RMC1 | 61.0745136 | -0.004041438 | 0.22694286 | -0.0178082 | 0.98579189 | 0.99668665 |
| NPC1 | 195.869928 | -0.975005223 | 0.43085249 | -2.2629676 | 0.02363769 | 0.14857309 |
| LAMA3 | 76.1889793 | -0.789734 | 0.72864946 | -1.0838325 | 0.27843905 | 0.56578102 |
| TTC39C | 150.016444 | -0.908224203 | 0.37265898 | -2.4371456 | 0.01480372 | 0.11290192 |
| OSBPL1A | 118.307489 | -0.331426455 | 0.35805806 | -0.9256221 | 0.3546424 | 0.63640307 |
| IMPACT | 34.1907299 | 0.218201351 | 0.28538146 | 0.76459541 | 0.44451251 | 0.70707162 |
| SS18 | 227.582358 | 0.215696021 | 0.18093394 | 1.1921258 | 0.23321192 | 0.51603997 |
| TAF4B | 47.5603787 | -0.063907259 | 0.23259288 | -0.2747602 | 0.7835005 | 0.91192604 |
| KCTD1 | 281.504568 | -0.161298197 | 0.3271794 | -0.4929962 | 0.62201528 | 0.82915174 |
| AQP4-AS1 | 83.0248976 | -0.069754492 | 0.51486361 | -0.1354815 | 0.89223119 | 0.95976294 |
| CHST9 | 246.497491 | 0.409471008 | 0.41787024 | 0.97989991 | 0.32713553 | 0.61330512 |
| CDH2 | 751.402593 | 0.555302307 | 0.27114131 | 2.04801808 | 0.04055823 | 0.19992491 |
| DSC2 | 106.50066 | 0.437391258 | 0.22099745 | 1.97916879 | 0.04779701 | 0.2184387 |
| DSG2 | 145.827282 | -0.209783687 | 0.31572354 | -0.6644537 | 0.50639994 | 0.75581256 |
| TRAPPC8 | 229.484429 | 0.372501986 | 0.17569272 | 2.12019019 | 0.03399001 | 0.18138735 |
| RNF138 | 102.631126 | 0.457119576 | 0.21814524 | 2.09548268 | 0.03612811 | 0.18804981 |
| GAREM1 | 597.857949 | 0.132441028 | 0.28555823 | 0.46379692 | 0.64279325 | 0.84312153 |
| KLHL14 | 92.5104156 | 0.525576189 | 0.51718976 | 1.01621539 | 0.30952683 | 0.59621104 |
| CCDC178 | 104.016658 | 1.417926868 | 0.58924682 | 2.40633776 | 0.01611336 | 0.11854829 |
| ASXL3 | 127.334008 | 0.676771067 | 0.38180332 | 1.77256466 | 0.07630087 | 0.28527949 |
| NOL4 | 81.2829238 | 1.155536118 | 0.60082466 | 1.92325014 | 0.05444864 | 0.23397085 |
| DTNA | 201.819597 | -0.99054926 | 0.93159829 | -1.0632794 | 0.28765527 | 0.57450105 |
| MAPRE2 | 66.3630861 | 0.140739911 | 0.38629503 | 0.3643327 | 0.71560959 | 0.88009244 |
| ZNF397 | 81.8965342 | -0.092268015 | 0.18176566 | -0.5076207 | 0.61171935 | 0.82344852 |
| ZSCAN30 | 71.4042109 | -0.23704629 | 0.24607155 | -0.9633226 | 0.33538564 | 0.61996508 |
| ZNF24 | 109.94428 | -0.080300325 | 0.17629068 | -0.4554996 | 0.64874988 | 0.8462529 |
| GALNT1 | 126.323216 | -0.360627291 | 0.36854492 | -0.9785165 | 0.32781892 | 0.61394673 |
| RPRD1A | 194.639046 | 0.113054242 | 0.16688052 | 0.6774562 | 0.49811655 | 0.74986853 |
| SLC39A6 | 64.4772324 | -0.52909491 | 0.31077041 | -1.7025266 | 0.08865669 | 0.30992958 |
| ELP2 | 156.71758 | 0.134204515 | 0.19160885 | 0.70040875 | 0.48367207 | 0.73727416 |
| FHOD3 | 724.034918 | 0.098410803 | 0.35511365 | 0.27712481 | 0.78168427 | 0.91080813 |
| TPGS2 | 93.7650852 | -0.280724174 | 0.23013319 | -1.2198335 | 0.22252799 | 0.50404247 |
| KIAA1328 | 443.12258 | 0.61999159 | 0.26311378 | 2.35636305 | 0.01845487 | 0.12847326 |
| AC016205.1 | 91.9679919 | -0.368340715 | 0.33697816 | -1.09307 | 0.27436306 | 0.56168493 |
| PIK3C3 | 264.905064 | 0.265191276 | 0.17990691 | 1.47404716 | 0.14046889 | 0.39528541 |
| LINC00907 | 46.069562 | -0.883124378 | 0.42215878 | -2.0919247 | 0.03644525 | 0.18891183 |
| SETBP1 | 1163.81303 | 0.711861551 | 0.24953524 | 2.85274959 | 0.00433428 | 0.05265324 |
| SLC14A2 | 138.442323 | 4.256753042 | 1.08611629 | 3.91924243 | 8.88E-05 | 0.00431919 |
| EPG5 | 197.172354 | 0.541570943 | 0.17132476 | 3.16107806 | 0.00157186 | 0.02762731 |
| PSTPIP2 | 32.6230177 | -0.294727674 | 0.38287129 | -0.7697826 | 0.44142884 | 0.7045063 |
| ATP5F1A | 163.721482 | -0.261869536 | 0.25744271 | -1.0171954 | 0.30906048 | 0.59563596 |
| C18orf25 | 110.505593 | 0.061527449 | 0.23740004 | 0.25917203 | 0.79550251 | 0.91797597 |
| PIAS2 | 224.542895 | 0.087316601 | 0.19642557 | 0.44452767 | 0.65666113 | 0.85151712 |
| KATNAL2 | 66.2664722 | 0.345181739 | 0.47573815 | 0.72557086 | 0.46810189 | 0.72581883 |
| SMAD2 | 445.343808 | 0.151513155 | 0.23170641 | 0.65390145 | 0.5131753 | 0.76076145 |
| ZBTB7C | 195.569855 | 0.277945482 | 1.09053444 | 0.25487089 | 0.79882282 | 0.91949148 |
| CTIF | 208.773518 | 0.385928469 | 0.58666393 | 0.65783569 | 0.51064372 | 0.75918923 |
| SMAD7 | 95.1575392 | 1.098737715 | 0.62377693 | 1.76142731 | 0.07816611 | 0.28987733 |
| DYM | 503.672675 | 0.109100319 | 0.18111797 | 0.60237157 | 0.54692683 | 0.78253724 |
| LIPG | 134.722136 | 0.229707663 | 0.75287131 | 0.3051088 | 0.76028329 | 0.90125623 |
| ACAA2 | 116.392838 | 0.077387091 | 0.38495651 | 0.20102814 | 0.84067657 | 0.94042285 |
| SNHG22 | 97.2804144 | 0.152294192 | 0.36465144 | 0.41764319 | 0.67620801 | 0.86116336 |
| MYO5B | 336.093405 | -0.571969762 | 0.31079408 | -1.8403496 | 0.06571693 | 0.26131276 |
| MBD1 | 45.1794734 | 0.126142886 | 0.2392479 | 0.52724761 | 0.59802165 | 0.81658679 |
| CXXC1 | 34.6580471 | -0.232512734 | 0.27142121 | -0.8566491 | 0.39163884 | 0.66729072 |
| ME2 | 73.5589411 | 0.01411011 | 0.34383444 | 0.04103751 | 0.96726599 | 0.99123492 |
| AC091551.1 | 28.8049115 | -0.247740784 | 0.47021495 | -0.5268671 | 0.59828589 | 0.81658679 |
| SMAD4 | 128.819355 | 0.388504342 | 0.19891669 | 1.95310078 | 0.05080765 | 0.22647116 |
| MEX3C | 31.3504417 | -0.467514674 | 0.25775074 | -1.8138248 | 0.06970471 | 0.27065319 |
| MBD2 | 313.566809 | 0.014251273 | 0.23421177 | 0.06084781 | 0.95148042 | 0.98319643 |
| POLI | 110.40109 | 0.831950017 | 0.22258107 | 3.73773937 | 0.00018568 | 0.00717198 |
| TCF4 | 83.6911944 | 1.045389812 | 0.54359874 | 1.92309094 | 0.05446863 | 0.23397085 |
| TXNL1 | 105.412251 | -0.045723716 | 0.20521679 | -0.2228069 | 0.8236858 | 0.93217972 |
| WDR7 | 239.250309 | 0.367800687 | 0.15720746 | 2.33958805 | 0.01930502 | 0.13189928 |
| FECH | 29.8283958 | 0.12113901 | 0.27477146 | 0.44087188 | 0.65930575 | 0.8528683 |
| NARS | 112.406645 | -0.926752948 | 0.25715252 | -3.6039038 | 0.00031347 | 0.00994413 |
| AC027097.2 | 87.711266 | -0.028007704 | 0.36889952 | -0.0759223 | 0.93948091 | 0.98003138 |
| ATP8B1 | 181.373241 | -0.304867038 | 0.24297573 | -1.2547222 | 0.20957964 | 0.49000814 |
| NEDD4L | 723.278787 | 1.054820081 | 0.20879106 | 5.05203673 | 4.37E-07 | 8.90E-05 |
| ALPK2 | 206.065775 | 0.203505461 | 0.4418132 | 0.46061426 | 0.64507538 | 0.84403456 |
| MALT1 | 305.549245 | 0.386726885 | 0.29567588 | 1.30794196 | 0.190893 | 0.46742434 |
| AC104365.1 | 47.8051541 | 0.333827569 | 0.49639521 | 0.67250361 | 0.50126315 | 0.75236598 |
| ZNF532 | 221.594755 | -0.104955227 | 0.42220526 | -0.2485882 | 0.80367937 | 0.92125386 |
| SEC11C | 38.6370631 | -0.940822978 | 0.47676719 | -1.9733383 | 0.04845703 | 0.22019444 |
| LMAN1 | 187.456023 | 0.174577169 | 0.261468 | 0.66768083 | 0.50433735 | 0.75520517 |
| RNF152 | 54.2560012 | 0.900547045 | 0.51251999 | 1.75709644 | 0.07890137 | 0.29099563 |
| PIGN | 188.205269 | 0.440552208 | 0.15564646 | 2.83046722 | 0.00464801 | 0.05512702 |
| KIAA1468 | 213.147228 | 0.185653266 | 0.13405696 | 1.38488348 | 0.1660881 | 0.43446007 |
| ZCCHC2 | 125.624224 | -0.127396805 | 0.25987165 | -0.4902297 | 0.62397135 | 0.83048793 |
| PHLPP1 | 251.682294 | 0.191001187 | 0.18313491 | 1.04295346 | 0.29696985 | 0.58520063 |
| BCL2 | 1170.98871 | 1.538935047 | 0.24183714 | 6.36351826 | 1.97E-10 | 1.45E-07 |
| KDSR | 139.984287 | -1.060598114 | 0.26779636 | -3.960465 | 7.48E-05 | 0.00393707 |
| VPS4B | 124.609756 | -0.194900256 | 0.2018005 | -0.9658066 | 0.33414096 | 0.61861931 |
| TMX3 | 112.244672 | 0.332866375 | 0.28426986 | 1.17095204 | 0.24161805 | 0.52637238 |

| | | | | | | |
|---|---|---|---|---|---|---|
| CCDC102B | 51.432135 | 0.898945787 | 0.60218748 | 1.49280052 | 0.13548941 | 0.38776608 |
| DOK6 | 50.6661879 | -1.343652162 | 0.69426892 | -1.9353483 | 0.05294757 | 0.23085334 |
| RTTN | 138.648383 | 0.347738647 | 0.2885668 | 1.20505422 | 0.22818237 | 0.51069134 |
| SOCS6 | 54.746111 | 0.353008065 | 0.25286927 | 1.39601015 | 0.16271144 | 0.42936994 |
| CYB5A | 310.885098 | -0.005801347 | 0.37436892 | -0.0154963 | 0.98763621 | 0.99678172 |
| CNDP2 | 406.025748 | -0.383315171 | 0.26618358 | -1.4400407 | 0.1498559 | 0.41050134 |
| LINC00909 | 31.2414985 | -0.484998364 | 0.30259575 | -1.6027931 | 0.10898035 | 0.3456698 |
| ZNF407 | 444.286356 | 0.095496836 | 0.14960835 | 0.63831223 | 0.52327045 | 0.76534368 |
| ZADH2 | 46.729468 | -0.116455327 | 0.27760178 | -0.419505 | 0.67484712 | 0.86115505 |
| TSHZ1 | 89.4661683 | -0.418624509 | 0.24052657 | -1.7404502 | 0.08178 | 0.29718157 |
| ZNF516 | 115.688687 | -0.297172441 | 0.567578 | -0.5235799 | 0.60057076 | 0.81779318 |
| ZNF236 | 116.713084 | 0.241309736 | 0.25601228 | 0.94257093 | 0.34590042 | 0.63016843 |
| MBP | 227.637335 | -0.494892811 | 0.22026395 | -2.2468171 | 0.02465172 | 0.15217128 |
| ATP9B | 358.758855 | 0.061529155 | 0.20732841 | 0.29677146 | 0.76664099 | 0.90372312 |
| CTDP1 | 38.2495069 | -0.623510448 | 0.28683735 | -2.1737422 | 0.0297245 | 0.16866909 |
| PQLC1 | 70.0616319 | -0.403240503 | 0.2874829 | -1.4026591 | 0.16071852 | 0.42659999 |
| HSBP1L1 | 69.4502888 | -0.199701457 | 0.24934818 | -0.800894 | 0.42319303 | 0.69059047 |
| TXNL4A | 91.4318311 | -0.417923634 | 0.30184843 | -1.384548 | 0.16619073 | 0.43446007 |
| ADNP2 | 80.3920809 | -0.353676891 | 0.22913827 | -1.5435086 | 0.12270743 | 0.36800703 |
| NRSN2-AS1 | 66.9121985 | 0.551619785 | 0.33031173 | 1.66999759 | 0.09491984 | 0.32049248 |
| NRSN2 | 32.6557467 | -0.456355276 | 0.43539182 | -1.0481485 | 0.2945702 | 0.58239173 |
| RBCK1 | 99.8668413 | -0.439383077 | 0.25733798 | -1.7074164 | 0.08774469 | 0.3082165 |
| TBC1D20 | 54.4451787 | -0.012555557 | 0.20242015 | -0.0620272 | 0.95054117 | 0.98286203 |
| CSNK2A1 | 298.449085 | -0.259870257 | 0.17702381 | -1.4679961 | 0.14210528 | 0.39848551 |
| PSMF1 | 151.312002 | -0.492056676 | 0.26231755 | -1.8758054 | 0.060682 | 0.24868375 |
| AL136531.2 | 54.3800806 | 0.058290529 | 0.38226589 | 0.15248687 | 0.87880294 | 0.95496919 |
| SDCBP2-AS1 | 86.4291283 | 0.630177217 | 0.26848442 | 2.34716498 | 0.01891688 | 0.13017583 |
| NSFL1C | 60.1428252 | -0.394462195 | 0.29268385 | -1.3477416 | 0.17774151 | 0.44938172 |
| SIRPA | 108.370579 | 0.288737409 | 0.23828765 | 1.21171791 | 0.22562038 | 0.5078049 |
| STK35 | 47.6839654 | -0.065139315 | 0.2477912 | -0.2628799 | 0.79264318 | 0.91677482 |
| SNRPB | 99.7905607 | -1.220930861 | 0.48775032 | -2.5031882 | 0.01230801 | 0.10120034 |
| NOP56 | 54.436641 | -1.232221319 | 0.52630528 | -2.3412673 | 0.0192184 | 0.13168319 |
| IDH3B | 30.603539 | -0.407391797 | 0.31991705 | -1.2734295 | 0.20286569 | 0.48171801 |
| EBF4 | 67.3796763 | 0.313283484 | 0.30931228 | 1.01283881 | 0.31113716 | 0.59847045 |
| PCED1A | 41.2008494 | 0.539214456 | 0.26824321 | 2.01016998 | 0.0444132 | 0.20957343 |
| PTPRA | 337.696875 | -0.0240069 | 0.15524264 | -0.1546411 | 0.87710423 | 0.95388072 |
| MRPS26 | 58.9597342 | -0.262666096 | 0.47484336 | -0.5531637 | 0.58015134 | 0.8048182 |
| UBOX5 | 30.3891488 | 0.337335119 | 0.29693756 | 1.13604732 | 0.25593676 | 0.54275365 |
| LZTS3 | 44.9692954 | 0.814638751 | 0.39451881 | 2.06489205 | 0.03893322 | 0.19580915 |
| DDRGK1 | 61.8351358 | -0.463879257 | 0.32624477 | -1.4218749 | 0.15506256 | 0.4186427 |
| ITPA | 95.0362058 | -0.530021625 | 0.29742887 | -1.7820114 | 0.07474738 | 0.28248137 |
| SLC4A11 | 209.503988 | 2.487592113 | 0.55346679 | 4.49456439 | 6.97E-06 | 0.00068843 |
| C20orf194 | 586.771086 | 0.475869532 | 0.34028115 | 1.39845986 | 0.16197502 | 0.4283668 |
| ATRN | 219.311541 | 0.204331585 | 0.19134253 | 1.06788379 | 0.28557293 | 0.57312029 |
| CENPB | 52.3794765 | -1.325636179 | 0.37852215 | -3.5021363 | 0.00046154 | 0.01267041 |
| CDC25B | 40.4216365 | -0.49678123 | 0.35959854 | -1.3814884 | 0.16712883 | 0.43562642 |
| MAVS | 130.173568 | -0.391540496 | 0.17284307 | -2.2652947 | 0.0234946 | 0.14825743 |
| PANK2 | 143.27432 | 0.759529821 | 0.28639107 | 2.65207228 | 0.00799994 | 0.07756218 |
| RNF24 | 324.446028 | 1.01082409 | 0.56788734 | 1.77997293 | 0.07508039 | 0.28314766 |
| SMOX | 31.5874637 | -3.197642108 | 0.81021774 | -3.9466454 | 7.93E-05 | 0.00405722 |
| PRNP | 98.0907898 | -0.618737899 | 0.33277303 | -1.8593391 | 0.06297909 | 0.25490103 |
| SLC23A2 | 97.7000867 | 0.343417918 | 0.21486089 | 1.59832678 | 0.10997027 | 0.34708436 |
| TMEM230 | 99.3532102 | -0.110041303 | 0.21006636 | -0.5238407 | 0.60038936 | 0.81779318 |
| PCNA | 30.9414441 | -0.675452459 | 0.38323907 | -1.7624833 | 0.07798768 | 0.28954938 |
| CDS2 | 132.932507 | 0.116956738 | 0.16591028 | 0.70493966 | 0.48084779 | 0.73578929 |
| GPCPD1 | 321.538907 | 0.679654821 | 0.40510242 | 1.67773579 | 0.09339869 | 0.31835715 |
| SHLD1 | 91.6255197 | 0.621364231 | 0.29758781 | 2.08800296 | 0.03679757 | 0.19001827 |
| MCM8 | 41.1655 | 1.385472679 | 0.43101941 | 3.21440905 | 0.00130713 | 0.02485643 |
| CRLS1 | 221.466763 | 0.152380469 | 0.28723791 | 0.53050263 | 0.59576348 | 0.81525977 |
| TMX4 | 174.134899 | 0.250579019 | 0.25317725 | 0.98973749 | 0.32230245 | 0.60955681 |
| PLCB1 | 1161.85936 | -0.120215934 | 0.88057073 | -0.1365205 | 0.89140984 | 0.95969529 |
| PLCB4 | 436.394658 | 0.803368081 | 0.72334527 | 1.11062878 | 0.26672817 | 0.55453954 |
| SNAP25 | 44.6095889 | -1.009488817 | 0.69906904 | -1.4440474 | 0.14872565 | 0.40844123 |
| SLX4IP | 177.155205 | 0.325775246 | 0.2345227 | 1.38909898 | 0.16480265 | 0.43317961 |
| JAG1 | 365.469412 | 1.036737138 | 0.51236663 | 2.0234283 | 0.04302901 | 0.20605022 |
| AL050403.2 | 61.3461463 | 0.128449194 | 0.40731011 | 0.3153597 | 0.75248856 | 0.89668701 |
| BTBD3 | 117.680417 | 0.026659665 | 0.2775371 | 0.09605802 | 0.92347449 | 0.97297642 |
| SPTLC3 | 252.012317 | -0.100504972 | 0.41585652 | -0.2416818 | 0.80902669 | 0.92422978 |
| ISM1 | 102.109971 | -0.12234729 | 1.24292615 | -0.0984349 | 0.92158698 | 0.97248656 |
| TASP1 | 362.796639 | 0.050129047 | 0.150917 | 0.33216303 | 0.73976616 | 0.89253455 |
| ESF1 | 121.55405 | -0.236489849 | 0.25772913 | -0.9175907 | 0.3588332 | 0.64001011 |
| NDUFAF5 | 52.4753931 | 0.323557868 | 0.22675583 | 1.42689989 | 0.15360875 | 0.41634457 |
| MACROD2 | 808.342173 | 0.640930755 | 0.32990495 | 1.94277401 | 0.05204347 | 0.22857606 |
| FLRT3 | 161.9254 | 0.458568045 | 0.45073638 | 1.01737526 | 0.30897494 | 0.59563596 |
| KIF16B | 224.255713 | 0.362580169 | 0.22590819 | 1.60498904 | 0.10849622 | 0.3445906 |
| SNRPB2 | 53.1825665 | -0.688786416 | 0.26511832 | -2.5980341 | 0.00937592 | 0.08626955 |
| DSTN | 571.532639 | 0.141921371 | 0.3417886 | 0.41523144 | 0.67797248 | 0.86151115 |
| RRBP1 | 197.743433 | -1.599678289 | 0.39929965 | -4.0062101 | 6.17E-05 | 0.00333916 |
| SNX5 | 123.193648 | 0.20702084 | 0.21181442 | 0.97736895 | 0.32838653 | 0.61394673 |
| OVOL2 | 54.9964541 | 0.570536965 | 0.3969266 | 1.43738658 | 0.15060818 | 0.41173164 |
| MGME1 | 62.151044 | 0.431714277 | 0.30890126 | 1.39758017 | 0.16223918 | 0.42858376 |
| KAT14 | 25.3995293 | 0.003440222 | 0.31465319 | 0.01093338 | 0.9912766 | 0.99794415 |
| ZNF133 | 80.854393 | 0.28755551 | 0.23001042 | 1.25018469 | 0.21123209 | 0.49218977 |
| DZANK1 | 71.0971211 | 0.720648669 | 0.24940692 | 2.88944937 | 0.00385917 | 0.04842913 |
| POLR3F | 27.1443679 | 0.10455717 | 0.34288838 | 0.30493063 | 0.76041899 | 0.90125623 |
| SEC23B | 89.7237459 | -0.219616296 | 0.27575765 | -0.7964105 | 0.42579347 | 0.69215319 |
| DTD1 | 35.1326381 | -1.199023205 | 0.37700606 | -3.1803817 | 0.00147081 | 0.02655639 |
| SLC24A3 | 70.8517576 | -0.358600978 | 0.88234971 | -0.4064159 | 0.68443702 | 0.86539363 |
| AL121761.1 | 96.3864553 | 0.216522013 | 0.32032088 | 0.67595349 | 0.49907018 | 0.75042493 |
| RIN2 | 239.52236 | 0.174423182 | 0.29372402 | 0.59383357 | 0.55262343 | 0.78613936 |
| NAA20 | 97.5994832 | -0.314147559 | 0.23755901 | -1.322398 | 0.18603565 | 0.46080126 |
| CRNKL1 | 30.5359988 | -0.02275186 | 0.33088383 | -0.0687609 | 0.94517997 | 0.98119662 |
| CFAP61 | 44.8133548 | -0.068587476 | 0.58970518 | -0.1163081 | 0.90740838 | 0.96677845 |
| RALGAPA2 | 532.520258 | -0.549442681 | 0.20846321 | -2.6356818 | 0.00839684 | 0.08075639 |
| KIZ | 131.619812 | 0.755791221 | 0.30703358 | 2.46159144 | 0.01383221 | 0.10860555 |
| KIZ-AS1 | 44.903632 | 0.18583724 | 0.47730499 | 0.38934694 | 0.69701951 | 0.87095616 |
| XRN2 | 215.22336 | -0.372056191 | 0.19124504 | -1.9454423 | 0.05172177 | 0.22799945 |
| NAPB | 86.6340681 | 0.41719374 | 0.26771844 | 1.55833024 | 0.11915498 | 0.3645149 |
| CST3 | 311.876216 | -1.445603287 | 0.48064767 | -3.0076153 | 0.00263306 | 0.03764493 |
| APMAP | 55.7203593 | -0.227326181 | 0.34584733 | -0.6573021 | 0.51098668 | 0.75922776 |
| ACSS1 | 119.19664 | -0.983337485 | 0.57016393 | -1.7246575 | 0.08458922 | 0.30235944 |
| ENTPD6 | 63.0059569 | -0.346860372 | 0.23842161 | -1.4548193 | 0.14571929 | 0.40369182 |
| PYGB | 98.380217 | -0.901900672 | 0.29076324 | -3.1018387 | 0.00192323 | 0.03108476 |
| ABHD12 | 166.980349 | 0.556169932 | 0.43899635 | 1.26691243 | 0.2051866 | 0.48422825 |
| NINL | 246.447015 | 0.169639816 | 0.26412286 | 0.64227616 | 0.52069389 | 0.76418222 |
| ZNF337-AS1 | 63.2259858 | 0.297301744 | 0.29408036 | 1.0109541 | 0.3120384 | 0.59948171 |
| ZNF337 | 66.5320326 | 0.296301006 | 0.26835437 | 1.10414078 | 0.26953207 | 0.55667265 |
| FAM182B | 66.612878 | -0.166541958 | 0.39827021 | -0.4181632 | 0.67582777 | 0.86116336 |
| HM13 | 225.751409 | -0.693173156 | 0.28564969 | -2.4266547 | 0.01523875 | 0.11466773 |
| ID1 | 373.339458 | 0.453540638 | 0.7639322 | 0.59369226 | 0.55271796 | 0.78613936 |
| BCL2L1 | 291.420736 | -0.356208748 | 0.433423 | -0.8218501 | 0.4111622 | 0.68270459 |
| AL117381.1 | 51.5595966 | 0.026213556 | 0.4652315 | 0.05634519 | 0.95506682 | 0.98393042 |
| TM9SF4 | 120.127512 | -0.136461531 | 0.17582685 | -0.7761132 | 0.43768218 | 0.70212475 |

| | | | | | | |
|---|---|---|---|---|---|---|
| POFUT1 | 45.6403375 | -0.391348007 | 0.22030131 | -1.7764216 | 0.07566345 | 0.28432774 |
| KIF3B | 131.060051 | -0.359191531 | 0.19701731 | -1.8231471 | 0.06828112 | 0.26710321 |
| ASXL1 | 409.815863 | 0.144928592 | 0.18085977 | 0.80133131 | 0.42293987 | 0.69043214 |
| NOL4L | 55.727939 | 0.107099592 | 0.41572307 | 0.25762244 | 0.79669831 | 0.91869184 |
| MAPRE1 | 53.9943138 | 0.165845342 | 0.37988302 | 0.43656951 | 0.66242358 | 0.85507439 |
| CDK5RAP1 | 58.0947668 | 0.262694325 | 0.19226378 | 1.3663225 | 0.17183776 | 0.4426307 |
| SNTA1 | 33.1144297 | 0.216110971 | 0.40318942 | 0.53600358 | 0.59195608 | 0.81275584 |
| CBFA2T2 | 531.724834 | 0.527961238 | 0.20391814 | 2.58908428 | 0.00962315 | 0.08782595 |
| CHMP4B | 107.318533 | -0.498172635 | 0.26695746 | -1.8661124 | 0.06202564 | 0.25250472 |
| RALY | 150.941682 | -0.217719903 | 0.16843375 | -1.2926145 | 0.19614444 | 0.47506133 |
| EIF2S2 | 122.782629 | -0.777584679 | 0.32154934 | -2.4182437 | 0.01559563 | 0.11595952 |
| AHCY | 51.797557 | -1.034310634 | 0.25695101 | -4.0253224 | 5.69E-05 | 0.00324416 |
| ITCH | 415.53547 | 0.10609994 | 0.13964458 | 0.75978561 | 0.44738275 | 0.70892958 |
| DYNLRB1 | 209.121744 | -0.036664624 | 0.34248141 | -0.1070558 | 0.91474471 | 0.96885665 |
| MAP1LC3A | 36.0912788 | -0.309828639 | 0.42269839 | -0.732978 | 0.46357182 | 0.72133763 |
| PIGU | 95.6359753 | -0.085172434 | 0.20420794 | -0.4170868 | 0.67661492 | 0.8613973 |
| NCOA6 | 250.025992 | 0.077551604 | 0.17531938 | 0.44234473 | 0.65823976 | 0.85215492 |
| GGT7 | 87.7765593 | 0.002630588 | 0.29050419 | 0.00905525 | 0.99277506 | 0.99829858 |
| ACSS2 | 156.210193 | 0.641145868 | 0.31031148 | 2.0661365 | 0.03881559 | 0.19569187 |
| GSS | 56.2497256 | -0.374654599 | 0.24264164 | -1.5440656 | 0.12257245 | 0.367753 |
| TRPC4AP | 242.243788 | -0.019330072 | 0.17136161 | -0.1128028 | 0.91018689 | 0.96831189 |
| EDEM2 | 42.456767 | -0.116974858 | 0.23468777 | -0.4984276 | 0.61818271 | 0.82703521 |
| PROCR | 34.9154484 | 0.00707076 | 0.45397986 | 0.01557505 | 0.98757341 | 0.99678172 |
| MMP24OS | 89.4354705 | 0.432688125 | 0.25772244 | 1.67889194 | 0.09317311 | 0.31795696 |
| MMP24 | 59.669785 | 0.159135736 | 0.24204272 | 0.65746964 | 0.51087899 | 0.75922776 |
| UQCC1 | 142.314862 | 0.182992977 | 0.17924447 | 1.02091279 | 0.30729576 | 0.59456464 |
| CEP250 | 40.6243037 | -0.188022546 | 0.40595558 | -0.4631604 | 0.6432494 | 0.84325798 |
| ERGIC3 | 74.2428433 | -0.130962052 | 0.21000025 | -0.6236281 | 0.53287186 | 0.77327368 |
| CPNE1 | 75.7766151 | 0.114060705 | 0.29103603 | 0.39191265 | 0.69512275 | 0.87024545 |
| ROMO1 | 63.1721979 | -1.15648646 | 0.44675716 | -2.5886243 | 0.00963602 | 0.08782595 |
| RBM39 | 565.79247 | 0.299548442 | 0.19522766 | 1.53435456 | 0.12494245 | 0.37161272 |
| PHF20 | 248.21377 | 0.153712615 | 0.13030136 | 1.17967004 | 0.23813147 | 0.52114722 |
| SCAND1 | 103.573926 | -1.208764839 | 0.37426361 | -3.2297151 | 0.00123914 | 0.02383471 |
| NORAD | 281.941338 | 0.168237483 | 0.18084834 | 0.93026834 | 0.35223217 | 0.63397819 |
| EPB41L1 | 466.197444 | 0.493012098 | 0.3970228 | 1.24177276 | 0.21432042 | 0.49648737 |
| AAR2 | 32.9427893 | 0.204861708 | 0.22936654 | 0.89316302 | 0.37176988 | 0.65192016 |
| DLGAP4 | 191.480595 | -0.138279544 | 0.21481727 | -0.6437078 | 0.51976495 | 0.76385303 |
| DLGAP4-AS1 | 83.7375995 | 0.511747441 | 0.43296527 | 1.18195957 | 0.23722174 | 0.52000641 |
| MYL9 | 123.937338 | 0.877462062 | 0.55416633 | 1.58339114 | 0.11333234 | 0.35267249 |
| RAB5IF | 33.9633089 | -0.943904708 | 0.49865849 | -1.8928881 | 0.05837276 | 0.243852 |
| NDRG3 | 54.1751036 | 0.45187337 | 0.23146016 | 1.95227274 | 0.05090583 | 0.2266978 |
| SOGA1 | 227.015498 | 1.18833392 | 0.59800462 | 1.98716512 | 0.0469041 | 0.21632898 |
| SAMHD1 | 49.4058048 | -0.643473847 | 0.86621626 | -0.7428559 | 0.45756893 | 0.71670748 |
| RBL1 | 41.3970165 | 0.478627237 | 0.39960009 | 1.19776559 | 0.23100829 | 0.51425247 |
| RPN2 | 125.833288 | -0.359303627 | 0.20622608 | -1.7422803 | 0.08145941 | 0.29654614 |
| MANBAL | 39.644787 | -0.654480337 | 0.27196364 | -2.4064994 | 0.01610623 | 0.11854829 |
| SRC | 54.8877642 | -0.052043295 | 0.41150915 | -0.1264694 | 0.89936041 | 0.96269027 |
| BLCAP | 96.1420536 | 0.081263127 | 0.17554132 | 0.46292877 | 0.64341542 | 0.8433602 |
| CTNNBL1 | 82.5968771 | 0.241029736 | 0.17472878 | 1.37945071 | 0.16775583 | 0.43654799 |
| TTI1 | 55.931885 | -0.151503546 | 0.22859813 | -0.6627506 | 0.50749031 | 0.75682386 |
| RPRD1B | 100.106303 | -0.005056209 | 0.22456546 | -0.0225155 | 0.98203673 | 0.99616479 |
| TGM2 | 231.370803 | -3.890242862 | 0.52440287 | -7.4184241 | 1.19E-13 | 2.27E-10 |
| RALGAPB | 180.101279 | 0.112881212 | 0.1488457 | 0.7583774 | 0.44822508 | 0.70956008 |
| DHX35 | 46.661922 | 0.244534968 | 0.23180095 | 1.05493516 | 0.29145498 | 0.5787402 |
| TOP1 | 299.074951 | -0.285034194 | 0.21925949 | -1.2999857 | 0.19360587 | 0.47194155 |
| PLCG1 | 63.7950676 | -0.31655953 | 0.25888857 | -1.2227637 | 0.22141898 | 0.50319629 |
| ZHX3 | 408.245799 | -0.572686022 | 0.28129312 | -2.0359048 | 0.04175991 | 0.20312121 |
| LPIN3 | 63.1127126 | 0.250882862 | 0.26688898 | 0.94002704 | 0.34720369 | 0.63085435 |
| CHD6 | 384.313625 | 0.378803509 | 0.13814763 | 2.74201953 | 0.00610627 | 0.06580904 |
| SRSF6 | 101.978364 | 0.402727492 | 0.34063902 | 1.18227058 | 0.23709835 | 0.52000641 |
| L3MBTL1 | 52.502769 | 0.054444655 | 0.25097616 | 0.21693158 | 0.82826167 | 0.93346403 |
| SGK2 | 26.8215104 | 0.072886178 | 0.36036838 | 0.20225464 | 0.83971766 | 0.94011868 |
| IFT52 | 80.6974039 | 0.268621795 | 0.21879766 | 1.22771786 | 0.2195529 | 0.50121948 |
| OSER1 | 41.1834105 | -0.178595312 | 0.3401912 | -0.5249851 | 0.59959355 | 0.81763227 |
| SERINC3 | 94.9566124 | 0.018437655 | 0.25386436 | 0.07262798 | 0.94210216 | 0.98091268 |
| PKIG | 136.700768 | 0.038075641 | 0.24311846 | 0.15661353 | 0.87554944 | 0.95314737 |
| YWHAB | 265.524535 | -0.968600265 | 0.38959899 | -2.4861468 | 0.01291347 | 0.10403543 |
| PABPC1L | 40.5348733 | 0.687103787 | 0.30935952 | 2.22105264 | 0.0263474 | 0.1573452 |
| STK4 | 107.208118 | 0.198791525 | 0.2053718 | 0.96795921 | 0.33306474 | 0.61758172 |
| SLPI | 1566.61228 | -2.218695583 | 0.60342771 | -3.6768208 | 0.00023616 | 0.0083651 |
| SDC4 | 413.046007 | -1.16476402 | 0.4097708 | -2.8424768 | 0.00447645 | 0.05380163 |
| PIGT | 43.5396866 | -0.054735385 | 0.26804748 | -0.2042003 | 0.83819695 | 0.93896487 |
| WFDC2 | 1191.58307 | -3.469813511 | 0.55892471 | -6.208016 | 5.37E-10 | 3.43E-07 |
| DNTTIP1 | 34.337925 | -0.266934873 | 0.25984968 | -1.0272665 | 0.30429499 | 0.5922455 |
| CTSA | 74.0208003 | -1.101831374 | 0.44374225 | -2.4830437 | 0.01302651 | 0.10449615 |
| ZNF335 | 37.2557435 | -0.140031731 | 0.27567766 | -0.5079546 | 0.61148519 | 0.82336473 |
| NCOA5 | 74.8823364 | 0.08328112 | 0.20811052 | 0.40017736 | 0.68902589 | 0.86751885 |
| CD40 | 51.6650644 | -1.362020411 | 0.55093405 | -2.4722023 | 0.01342835 | 0.1068357 |
| SLC35C2 | 54.0628903 | -0.355542044 | 0.25236653 | -1.408832 | 0.15888485 | 0.42441661 |
| ELMO2 | 41.2911805 | 0.198399781 | 0.25982671 | 0.763585 | 0.44511459 | 0.70741705 |
| ZNF334 | 42.7325033 | 0.36407853 | 0.56446028 | 0.64500293 | 0.51892529 | 0.7636788 |
| SLC13A3 | 61.8917997 | -0.20107034 | 0.53267195 | -0.377475 | 0.70582064 | 0.87473681 |
| EYA2 | 139.980353 | 1.596009713 | 1.14340912 | 1.39583434 | 0.16276439 | 0.42939137 |
| ZMYND8 | 380.54406 | 0.051227053 | 0.36359744 | 0.14088948 | 0.88795725 | 0.95907089 |
| NCOA3 | 269.361728 | 0.129484003 | 0.12074314 | 1.0723922 | 0.28354391 | 0.57084461 |
| SULF2 | 94.4434476 | -2.133079118 | 0.74414059 | -2.8665002 | 0.00415038 | 0.05103528 |
| ARFGEF2 | 239.314946 | -0.143204335 | 0.15791814 | -0.9068264 | 0.36449861 | 0.644313 |
| CSE1L | 91.849808 | -0.007129479 | 0.26240776 | -0.0271695 | 0.97832457 | 0.99536602 |
| STAU1 | 229.751037 | 0.07735286 | 0.16969677 | 0.4558299 | 0.6485123 | 0.84621977 |
| DDX27 | 43.1468913 | -0.623407697 | 0.22943196 | -2.717179 | 0.0065841 | 0.06885256 |
| ZNFX1 | 100.169449 | -0.908852047 | 0.48609559 | -1.8696982 | 0.06152574 | 0.25100302 |
| ZFAS1 | 106.002195 | -0.687044057 | 0.31112734 | -2.2082407 | 0.0272275 | 0.15949106 |
| B4GALT5 | 326.343503 | -0.904481638 | 0.436535 | -2.0719567 | 0.03826948 | 0.1943708 |
| SLC9A8 | 95.3979682 | -0.403954958 | 0.2356607 | -1.714138 | 0.08650339 | 0.30636531 |
| RNF114 | 157.402135 | 0.256177791 | 0.39939864 | 0.64140877 | 0.52125714 | 0.76440016 |
| CEBPB | 123.464313 | -3.068901082 | 0.71909474 | -4.2677285 | 1.97E-05 | 0.00153789 |
| PTPN1 | 389.314981 | -1.396732196 | 0.44297886 | -3.1530448 | 0.00161577 | 0.02795985 |
| RIPOR3 | 155.702926 | 1.329031035 | 0.40631446 | 3.27094201 | 0.0010719 | 0.02174571 |
| PARD6B | 408.780644 | 0.811508539 | 0.32495217 | 2.49731688 | 0.01251371 | 0.10218995 |
| BCAS4 | 48.0260572 | -0.298955995 | 0.5110487 | -0.5849853 | 0.55855758 | 0.78926435 |
| ADNP | 385.686722 | 0.287853377 | 0.17436337 | 1.65088218 | 0.09876263 | 0.32848864 |
| DPM1 | 78.7784998 | 0.023401927 | 0.23437865 | 0.09984667 | 0.92046606 | 0.97180033 |
| ATP9A | 162.146245 | -0.093977156 | 0.2348778 | -0.4001109 | 0.68907487 | 0.86751885 |
| ZFP64 | 67.1334364 | -0.467935164 | 0.23851647 | -1.9618568 | 0.04977916 | 0.22428719 |
| TSHZ2 | 969.116068 | -0.59543734 | 0.61244342 | -0.9722324 | 0.33093494 | 0.61613719 |
| AL109930.1 | 83.5043489 | -1.108617794 | 0.58485926 | -1.8955292 | 0.05802233 | 0.24313032 |
| ZNF217 | 118.873962 | -0.642031421 | 0.20536906 | -3.1262324 | 0.00177062 | 0.02957033 |
| BCAS1 | 98.7063222 | 1.109266754 | 1.00392151 | 1.10493374 | 0.2691883 | 0.55632249 |
| CYP24A1 | 469.357325 | -3.788406776 | 0.74003211 | -5.1192465 | 3.07E-07 | 6.99E-05 |
| PFDN4 | 31.3037746 | -0.317967731 | 0.36717789 | -0.8659773 | 0.3865026 | 0.66361505 |
| DOK5 | 46.2527885 | 1.029707711 | 0.5027043 | 2.04833677 | 0.04052701 | 0.19992491 |
| RTF2 | 51.4175368 | 0.251997288 | 0.26402685 | 0.9544381 | 0.33986193 | 0.62462345 |
| BMP7 | 119.090124 | 1.520991075 | 0.77612291 | 1.95972964 | 0.0500274 | 0.2248768 |

| | | | | | | |
|---|---|---|---|---|---|---|
| RAE1 | 32.9199773 | -0.489368566 | 0.2420659 | -2.0216336 | 0.04321421 | 0.20614587 |
| PCK1 | 120.614003 | 0.371043527 | 1.0587137 | 0.35046635 | 0.72598874 | 0.88498933 |
| PMEPA1 | 66.5522084 | -2.420882089 | 0.86281304 | -2.8058015 | 0.00501916 | 0.05825796 |
| RAB22A | 120.834388 | -0.231713716 | 0.20135078 | -1.1507962 | 0.24981608 | 0.53692712 |
| VAPB | 143.628892 | -0.140177517 | 0.16274663 | -0.8613236 | 0.38905983 | 0.66513878 |
| NPEPL1 | 29.0956025 | 0.040435098 | 0.32945211 | 0.12273437 | 0.90231744 | 0.96422343 |
| GNAS | 533.454197 | -0.330742566 | 0.24331266 | -1.3593315 | 0.17404156 | 0.44528422 |
| NELFCD | 76.429898 | -0.037680405 | 0.19268027 | -0.1955592 | 0.84495518 | 0.94193856 |
| CTSZ | 121.817008 | -0.92241728 | 0.34376339 | -2.6832912 | 0.00729015 | 0.0731995 |
| ATP5F1E | 250.739543 | -0.958052247 | 0.35754928 | -2.679497 | 0.00737329 | 0.07378801 |
| PRELID3B | 25.5496482 | -0.313669359 | 0.34370107 | -0.9126226 | 0.36144107 | 0.64163157 |
| PHACTR3 | 73.7477335 | -0.408015759 | 1.18989729 | -0.3429 | 0.73167371 | 0.88818938 |
| MIR646HG | 67.4736836 | 1.164141124 | 0.77142473 | 1.50907934 | 0.13127851 | 0.38163088 |
| TAF4 | 81.9401993 | 0.336594242 | 0.27402127 | 1.22835077 | 0.21931532 | 0.50103063 |
| LSM14B | 29.4964387 | 0.424461709 | 0.25386883 | 1.67197254 | 0.09452974 | 0.32021075 |
| PSMA7 | 140.503769 | -0.939267806 | 0.3844154 | -2.4433667 | 0.01455094 | 0.11197822 |
| SS18L1 | 59.258377 | 0.140371931 | 0.22178293 | 0.63292485 | 0.52678272 | 0.76821934 |
| MTG2 | 57.0170056 | 0.019923981 | 0.20138001 | 0.09893723 | 0.92118811 | 0.97225548 |
| OSBPL2 | 87.8895799 | 0.020809054 | 0.24014863 | 0.08665073 | 0.93094914 | 0.97609039 |
| ADRM1 | 26.3524358 | -0.668455672 | 0.55865011 | -1.1965552 | 0.23147999 | 0.51446561 |
| LAMA5 | 180.846829 | 0.647562531 | 0.3048594 | 2.12413504 | 0.03365885 | 0.18043094 |
| RPS21 | 35.7065362 | -0.869836682 | 0.42606267 | -2.0415698 | 0.04119422 | 0.20195151 |
| SLCO4A1 | 91.7866442 | 0.106527679 | 0.72225915 | 0.14749232 | 0.88274344 | 0.95722522 |
| OGFR | 75.2974047 | -0.589270786 | 0.36156726 | -1.6297681 | 0.10315052 | 0.33528295 |
| TCFL5 | 28.9176424 | -0.149214733 | 0.35069752 | -0.4254799 | 0.6704869 | 0.85955487 |
| DIDO1 | 180.263495 | 0.086657933 | 0.17734532 | 0.48863952 | 0.62509693 | 0.83152388 |
| GID8 | 63.1396149 | -0.190713226 | 0.29389878 | -0.6489079 | 0.51639793 | 0.7618321 |
| YTHDF1 | 70.0287931 | -0.162385754 | 0.1826509 | -0.8890499 | 0.37397628 | 0.65261518 |
| ARFGAP1 | 53.6608118 | -0.013039328 | 0.39341011 | -0.0331444 | 0.97355946 | 0.99304931 |
| PPDPF | 279.788548 | -0.49329305 | 0.42568635 | -1.1588181 | 0.24653034 | 0.5334344 |
| GMEB2 | 48.8650113 | -0.090323726 | 0.19829743 | -0.4554962 | 0.6487523 | 0.8462529 |
| STMN3 | 29.8853152 | -2.408699983 | 0.54968726 | -4.3819462 | 1.18E-05 | 0.00101506 |
| ARFRP1 | 43.9865508 | -0.184401671 | 0.30977763 | -0.5952711 | 0.55166227 | 0.78577398 |
| SLC2A4RG | 69.6824352 | -0.752754649 | 0.26951727 | -2.7929737 | 0.00522259 | 0.0596272 |
| ZBTB46 | 122.76631 | -0.389540468 | 0.50553937 | -0.7705443 | 0.4409771 | 0.70428292 |
| TPD52L2 | 77.9173221 | -0.456458687 | 0.23925063 | -1.9078682 | 0.05640825 | 0.23919195 |
| DNAJC5 | 147.049901 | -0.444728888 | 0.26213214 | -1.6965829 | 0.08977555 | 0.31211662 |
| UCKL1 | 38.3032898 | 0.805041097 | 0.3030969 | 2.65605189 | 0.00790614 | 0.07690128 |
| ZNF512B | 29.583315 | -0.036469783 | 0.42898246 | -0.0850146 | 0.93224976 | 0.97701675 |
| PRPF6 | 101.466556 | -0.298416021 | 0.19014341 | -1.5694261 | 0.1165487 | 0.35967138 |
| PCMTD2 | 140.330435 | 0.747834804 | 0.26108179 | 2.86436986 | 0.0041784 | 0.05124824 |
| MIER2 | 30.5004418 | 0.374769935 | 0.30717617 | 1.22004886 | 0.22244635 | 0.50402693 |
| SHC2 | 34.8242719 | -0.239620627 | 0.42644039 | -0.5619088 | 0.57417813 | 0.80072326 |
| TPGS1 | 34.624582 | -0.897347198 | 0.41675801 | -2.1531612 | 0.03130601 | 0.17302987 |
| CDC34 | 34.6196108 | -0.180530273 | 0.32003518 | -0.5640951 | 0.57268942 | 0.79959427 |
| BSG | 163.656148 | -0.982471197 | 0.38239259 | -2.5692736 | 0.0101912 | 0.09022316 |
| RNF126 | 42.2116622 | -0.719344043 | 0.3348828 | -2.1480471 | 0.03171001 | 0.17406888 |
| FSTL3 | 70.0641391 | -2.490244834 | 0.73644282 | -3.3814503 | 0.00072104 | 0.01656323 |
| PTBP1 | 146.956465 | -0.205681313 | 0.30189484 | -0.6813012 | 0.49568094 | 0.74738355 |
| MED16 | 42.2066701 | -0.003334366 | 0.34858893 | -0.0095653 | 0.99236809 | 0.99829858 |
| R3HDM4 | 36.1466726 | -0.948505617 | 0.39181746 | -2.4207845 | 0.01548706 | 0.11571803 |
| ARID3A | 94.2119808 | 0.357791303 | 0.35637195 | 1.00398279 | 0.31538691 | 0.60229091 |
| WDR18 | 28.279958 | -0.387229253 | 0.46115037 | -0.8397028 | 0.40107505 | 0.67532043 |
| TMEM259 | 106.701453 | -0.090258474 | 0.3083874 | -0.2926789 | 0.76976763 | 0.90533273 |
| CNN2 | 39.2611381 | 0.051478452 | 0.44679603 | 0.1152169 | 0.90827321 | 0.96748063 |
| POLR2E | 61.2008831 | -0.673358136 | 0.23151101 | -2.9085362 | 0.00363125 | 0.04650235 |
| GPX4 | 195.323036 | -0.416843295 | 0.41122703 | -1.0136573 | 0.31074629 | 0.59807427 |
| SBNO2 | 119.099675 | -1.711249702 | 0.34348749 | -4.9819855 | 6.29E-07 | 0.00011593 |
| STK11 | 66.0751024 | -0.299433141 | 0.24836905 | -1.2055977 | 0.22797266 | 0.51066249 |
| ATP5F1D | 170.019519 | -0.762202446 | 0.36289608 | -2.1003326 | 0.0356996 | 0.18676485 |
| MIDN | 212.581873 | -1.170337754 | 0.35870593 | -3.2626664 | 0.00110369 | 0.02211775 |
| CIRBP | 280.520523 | 0.209750876 | 0.17886396 | 1.17268383 | 0.24092262 | 0.52533525 |
| C19orf24 | 39.8294557 | -1.07244535 | 0.41683481 | -2.5728306 | 0.01008706 | 0.08979917 |
| MUM1 | 111.982362 | 0.158364752 | 0.33194794 | 0.47707707 | 0.63330724 | 0.83617506 |
| NDUFS7 | 51.0926177 | -0.037508568 | 0.37918024 | -0.0989202 | 0.92120167 | 0.97225548 |
| DAZAP1 | 119.66712 | -0.320083132 | 0.23505021 | -1.3617649 | 0.17327208 | 0.44449049 |
| RPS15 | 691.097192 | -0.905004257 | 0.43669154 | -2.0724108 | 0.03822715 | 0.19425881 |
| C19orf25 | 67.0035884 | -0.436544827 | 0.19346291 | -2.2564781 | 0.0240407 | 0.15036388 |
| MBD3 | 42.5723676 | -0.312355413 | 0.42902143 | -0.7280648 | 0.46657391 | 0.72436166 |
| TCF3 | 51.8252077 | -0.110716698 | 0.35953265 | -0.3079462 | 0.7581233 | 0.89970284 |
| REXO1 | 40.0540637 | -0.309012646 | 0.31752275 | -0.9731984 | 0.33045469 | 0.615602 |
| ABHD17A | 47.0167759 | -0.410085997 | 0.27848643 | -1.4725529 | 0.14087163 | 0.39618595 |
| SCAMP4 | 65.6562145 | 0.060697147 | 0.2381581 | 0.25486073 | 0.79883067 | 0.91949148 |
| CSNK1G2 | 91.4089957 | -0.360475081 | 0.22896597 | -1.5743609 | 0.11540403 | 0.35752108 |
| BTBD2 | 106.252703 | -0.014780393 | 0.35927362 | -0.0411397 | 0.96718456 | 0.99123492 |
| MKNK2 | 54.3487432 | -0.712538785 | 0.28592485 | -2.4920492 | 0.01270085 | 0.10301379 |
| MOB3A | 58.7593048 | -0.436248173 | 0.31458834 | -1.3867271 | 0.165525 | 0.43384207 |
| AP3D1 | 123.798426 | -0.389850007 | 0.30551183 | -1.2760553 | 0.20193596 | 0.48041015 |
| DOT1L | 75.7005122 | 0.152000338 | 0.34355597 | 0.44243254 | 0.65817624 | 0.85215492 |
| PLEKHJ1 | 53.6132083 | -0.821890688 | 0.33459741 | -2.456357 | 0.01403537 | 0.10948271 |
| LSM7 | 95.2561804 | -1.004770644 | 0.42638855 | -2.3564672 | 0.0184497 | 0.12847326 |
| SPPL2B | 76.217872 | 0.106616361 | 0.27711713 | 0.38473392 | 0.70043456 | 0.87182648 |
| TIMM13 | 65.2513383 | -0.641443043 | 0.39722943 | -1.6147924 | 0.10635567 | 0.34105698 |
| GADD45B | 113.082402 | -0.471961753 | 0.66120235 | -0.7137932 | 0.47535506 | 0.73217979 |
| GNG7 | 81.1156123 | 1.130001403 | 0.35196803 | 3.21052289 | 0.00132494 | 0.02509831 |
| SGTA | 44.5609075 | -0.441415262 | 0.29610931 | -1.4907173 | 0.13603574 | 0.38809998 |
| THOP1 | 28.0192173 | -0.043865903 | 0.27591412 | -0.1589839 | 0.87368156 | 0.95210417 |
| TLE2 | 73.7988477 | 0.246016679 | 0.43001117 | 0.57211695 | 0.56724274 | 0.79601791 |
| AES | 233.91401 | -0.366429291 | 0.27600888 | -1.3275996 | 0.18431041 | 0.45778091 |
| GNA11 | 85.2001332 | -0.283162104 | 0.22228931 | -1.2738449 | 0.20271838 | 0.48158147 |
| NCLN | 45.9970891 | -0.509159562 | 0.38051861 | -1.3380674 | 0.18087445 | 0.45355926 |
| NFIC | 274.934238 | -0.519079655 | 0.26408823 | -1.9655539 | 0.04935016 | 0.22340509 |
| SMIM24 | 39.9028162 | -1.393676402 | 0.66490058 | -2.0960674 | 0.03607622 | 0.18804981 |
| FZR1 | 36.5936447 | 0.02302651 | 0.2881973 | 0.07989842 | 0.93631804 | 0.97871293 |
| MFSD12 | 66.0706967 | -0.864343203 | 0.31269885 | -2.7641394 | 0.00570732 | 0.06312976 |
| HMG20B | 38.8555455 | -0.084031153 | 0.31527512 | -0.2665328 | 0.78982892 | 0.91528808 |
| PIP5K1C | 40.0255647 | -0.284906566 | 0.31022017 | -0.9184012 | 0.35840888 | 0.64001011 |
| MRPL54 | 45.5262561 | -1.249602382 | 0.46728073 | -2.6742005 | 0.00749076 | 0.07451094 |
| DAPK3 | 38.1794062 | 0.22320617 | 0.27331874 | 0.81665155 | 0.41412759 | 0.68449528 |
| EEF2 | 452.263387 | -0.997043974 | 0.38377518 | -2.5979897 | 0.00937713 | 0.08626955 |
| PIAS4 | 32.8363829 | 0.056034012 | 0.34641781 | 0.16175269 | 0.87150061 | 0.95193455 |
| ZBTB7A | 101.971971 | -0.667978072 | 0.22416337 | -2.9798716 | 0.00288369 | 0.03957434 |
| MAP2K2 | 99.5462486 | -0.51150066 | 0.22591423 | -2.2641365 | 0.02356572 | 0.14836941 |
| STAP2 | 36.519904 | -0.583108284 | 0.2833625 | -2.0578174 | 0.03960766 | 0.19722814 |
| SH3GL1 | 96.5088997 | -0.183246077 | 0.31449865 | -0.5826609 | 0.56012158 | 0.78995945 |
| UBXN6 | 70.4487969 | -0.592975844 | 0.3901646 | -1.5198094 | 0.12855888 | 0.37647982 |
| HDGFL2 | 41.6249551 | 0.060096253 | 0.30145412 | 0.19935456 | 0.8419854 | 0.94122747 |
| PLIN5 | 49.5086408 | -2.1399539 | 0.79688032 | -2.6854144 | 0.00724399 | 0.07296551 |
| MYDGF | 83.703023 | -1.014801492 | 0.33794044 | -3.0029005 | 0.0026742 | 0.03790604 |
| DPP9 | 56.2672932 | -0.556713886 | 0.25763902 | -2.1608291 | 0.03070854 | 0.17115672 |
| PLIN3 | 80.0511136 | -1.191366595 | 0.35511433 | -3.3548818 | 0.00079399 | 0.01778667 |
| KDM4B | 195.584187 | -0.251331128 | 0.2867757 | -0.8764031 | 0.38081091 | 0.65803602 |
| PTPRS | 165.827403 | 0.693319996 | 0.23804132 | 2.91260353 | 0.00358429 | 0.04614778 |

| | | | | | | |
|---|---|---|---|---|---|---|
| TINCR | 30.2995745 | -0.863485153 | 0.4650533 | -1.8567445 | 0.06334751 | 0.25585634 |
| SAFB2 | 131.07381 | 0.238990051 | 0.18588496 | 1.28568794 | 0.19855199 | 0.47727216 |
| SAFB | 158.235913 | -0.139690186 | 0.14640145 | -0.9541585 | 0.34000342 | 0.62476361 |
| RPL36 | 343.613203 | -1.015172575 | 0.41460655 | -2.4485203 | 0.01434443 | 0.11107948 |
| C19orf70 | 73.3715623 | -1.123927559 | 0.36943273 | -3.0423064 | 0.00234773 | 0.0349985 |
| LONP1 | 77.4563221 | -0.563397289 | 0.2847196 | -1.9787794 | 0.04784084 | 0.21853478 |
| RANBP3 | 56.7060503 | -0.018340165 | 0.20797921 | -0.0881827 | 0.92973148 | 0.97545894 |
| RFX2 | 158.233045 | 0.630717093 | 0.53966778 | 1.16871362 | 0.24251903 | 0.52725736 |
| MLLT1 | 74.117279 | -0.02447677 | 0.23878024 | -0.1025075 | 0.91835384 | 0.97057281 |
| CLPP | 38.1872996 | -0.75388942 | 0.37715996 | -1.9988586 | 0.04562366 | 0.21268479 |
| ALKBH7 | 50.3650701 | 0.224500901 | 0.48146776 | 0.46628439 | 0.64101195 | 0.84180733 |
| GTF2F1 | 53.5021388 | -0.484549295 | 0.24900305 | -1.9459573 | 0.05165987 | 0.22786819 |
| KHSRP | 71.6158734 | -0.331406516 | 0.35539155 | -0.932511 | 0.35107252 | 0.63331896 |
| SLC25A23 | 47.9315671 | -0.259053286 | 0.32829201 | -0.7890941 | 0.43005701 | 0.69586421 |
| C3 | 587.657405 | -5.222030216 | 1.12798319 | -4.6295284 | 3.66E-06 | 0.00042813 |
| TRIP10 | 94.460534 | -0.245634018 | 0.32235007 | -0.7620101 | 0.44605395 | 0.70815763 |
| SH2D3A | 58.3124736 | -0.028201594 | 0.64150605 | -0.0439615 | 0.96493506 | 0.98965864 |
| INSR | 380.501648 | 0.053981442 | 0.33371239 | 0.16176037 | 0.87149456 | 0.95193455 |
| ZNF358 | 41.261253 | 0.017738966 | 0.4060084 | 0.04369113 | 0.96515061 | 0.98965864 |
| PNPLA6 | 54.3446612 | 0.461393891 | 0.2751142 | 1.67709952 | 0.09352302 | 0.31835715 |
| CAMSAP3 | 29.1928162 | -0.393528758 | 0.39248763 | -1.0026526 | 0.31602849 | 0.60255512 |
| STXBP2 | 27.6593941 | -0.2493242 | 0.44807425 | -0.556435 | 0.57791351 | 0.80357578 |
| EVI5L | 26.3236341 | -0.036438753 | 0.27991985 | -0.1301757 | 0.89642745 | 0.96168423 |
| MAP2K7 | 70.3091812 | -0.420342853 | 0.27975115 | -1.5025599 | 0.13295257 | 0.38380408 |
| TIMM44 | 35.3933919 | -0.607110585 | 0.25599528 | -2.3715695 | 0.01771272 | 0.12575079 |
| ELAVL1 | 112.484908 | 0.149546594 | 0.19237899 | 0.7773541 | 0.43694989 | 0.70191899 |
| CERS4 | 110.17901 | -0.006863427 | 0.41995481 | -0.0163433 | 0.98696055 | 0.99678172 |
| AC022146.2 | 33.6593585 | -0.149092579 | 0.388786 | -0.3834824 | 0.70136214 | 0.87200065 |
| CD320 | 35.7780386 | -0.494837752 | 0.41115731 | -1.2035242 | 0.22877355 | 0.51112377 |
| RPS28 | 127.29697 | -0.460353808 | 0.39897952 | -1.1538282 | 0.24857063 | 0.5356711 |
| ANGPTL4 | 80.2905984 | -3.183336933 | 0.88798683 | -3.5848921 | 0.00033722 | 0.01035323 |
| RAB11B | 72.09022 | -0.040022743 | 0.36454788 | -0.1097873 | 0.91257803 | 0.96844354 |
| 2-Mar | 35.5086435 | -0.143246184 | 0.34403633 | -0.4163693 | 0.67713975 | 0.8613973 |
| HNRNPM | 294.550583 | 0.148879249 | 0.1809797 | 0.82262954 | 0.4107187 | 0.68244135 |
| ZNF558 | 28.1878212 | -0.161993316 | 0.28139619 | -0.575677 | 0.56483352 | 0.79379993 |
| ZNF317 | 29.6484399 | -0.353901192 | 0.25804971 | -1.3714458 | 0.17023603 | 0.44040903 |
| ZNF559-ZNF | 28.6519132 | 0.141199354 | 0.49731894 | 0.28392113 | 0.77647083 | 0.90760391 |
| ZNF266 | 49.0470644 | -0.013288157 | 0.20870333 | -0.0636701 | 0.94923293 | 0.98267613 |
| ZNF426 | 29.2842186 | -0.593517147 | 0.23925874 | -2.4806498 | 0.01311431 | 0.10500025 |
| ZNF121 | 68.0350434 | -0.343230105 | 0.29978475 | -1.1449218 | 0.25224154 | 0.53930681 |
| ZNF561 | 35.9044737 | 0.015653215 | 0.31118604 | 0.05030179 | 0.9598819 | 0.98676848 |
| ZNF562 | 87.2447095 | -0.230966482 | 0.19855791 | -1.1632198 | 0.24474035 | 0.53076021 |
| ZNF846 | 155.16746 | 1.218537372 | 0.45772175 | 2.6621793 | 0.00776365 | 0.07623264 |
| FBXL12 | 36.1839566 | -0.03253996 | 0.28899839 | -0.1125956 | 0.91035114 | 0.96837907 |
| UBL5 | 68.0706362 | -0.628211472 | 0.45070672 | -1.3938365 | 0.16336699 | 0.43045905 |
| PIN1 | 54.7876105 | -0.318003199 | 0.29489748 | -1.0783517 | 0.28087684 | 0.56845958 |
| C19orf66 | 68.9566836 | -0.844406861 | 0.65037959 | -1.2983293 | 0.19417421 | 0.4727999 |
| EIF3G | 53.7412474 | -0.486921139 | 0.45513616 | -1.0698362 | 0.28469305 | 0.57243381 |
| DNMT1 | 42.0490253 | -0.023089064 | 0.40772099 | -0.0566296 | 0.95484028 | 0.98390867 |
| MRPL4 | 39.6440166 | -0.878039524 | 0.43811491 | -2.0041307 | 0.04505607 | 0.21106218 |
| ICAM1 | 86.9602783 | -3.234553389 | 1.10192065 | -2.9353778 | 0.00333142 | 0.0437146 |
| TYK2 | 42.1858737 | 0.000576725 | 0.23216952 | 0.00248407 | 0.998018 | 0.99923686 |
| CDC37 | 119.014644 | -0.650931386 | 0.344886 | -1.8873813 | 0.05910906 | 0.24535287 |
| KEAP1 | 36.5818454 | -0.526465572 | 0.37170511 | -1.4163528 | 0.15667224 | 0.4216384 |
| KRI1 | 36.8078045 | -0.469543257 | 0.29908099 | -1.5699536 | 0.11642592 | 0.35940827 |
| AP1M2 | 48.4352019 | -0.317202243 | 0.33234378 | -0.9544401 | 0.3398609 | 0.62462345 |
| SLC44A2 | 74.7276043 | 0.128662375 | 0.47044324 | 0.27349182 | 0.78447518 | 0.91212179 |
| ILF3-DT | 60.8113632 | -0.337107505 | 0.51210434 | -0.658279 | 0.5103589 | 0.7589605 |
| ILF3 | 192.554328 | 0.046634941 | 0.23451365 | 0.19885811 | 0.84237374 | 0.94132995 |
| QTRT1 | 49.0643347 | 0.144140513 | 0.27918833 | 0.51628416 | 0.60565595 | 0.820591 |
| DNM2 | 385.685479 | -0.129951548 | 0.23686116 | -0.5486402 | 0.58325241 | 0.80734478 |
| TMED1 | 25.4890952 | -0.801215912 | 0.36604335 | -2.1888553 | 0.02860736 | 0.16438145 |
| CARM1 | 51.1389416 | -0.41825099 | 0.22968516 | -1.8209753 | 0.06861062 | 0.26781626 |
| YIPF2 | 34.8856497 | -0.276113988 | 0.26871597 | -1.027531 | 0.3041705 | 0.5922455 |
| SMARCA4 | 247.267523 | -0.335552897 | 0.28746575 | -1.1672796 | 0.24309749 | 0.52815397 |
| LDLR | 86.4044509 | 0.308423783 | 0.72337913 | 0.42636533 | 0.66984166 | 0.85924744 |
| KANK2 | 88.7644056 | 0.15662761 | 0.40468815 | 0.38703286 | 0.69873188 | 0.8717697 |
| DOCK6 | 102.187967 | -0.107904669 | 0.33624105 | -0.3209146 | 0.74827509 | 0.89440069 |
| RAB3D | 51.8516624 | -0.355779378 | 0.3306283 | -1.0760705 | 0.28189571 | 0.56931879 |
| EPOR | 23.3000782 | 0.658915299 | 0.5572696 | 1.18239949 | 0.23704722 | 0.52000641 |
| RGL3 | 218.996538 | 0.228120879 | 0.52124099 | 0.43764954 | 0.66164035 | 0.85473405 |
| PRKCSH | 77.3992392 | -0.376371123 | 0.383002 | -0.9826871 | 0.32576147 | 0.61243485 |
| ECSIT | 44.6260076 | -0.267841058 | 0.28712892 | -0.9328251 | 0.35091029 | 0.63331896 |
| ELOF1 | 26.3053059 | -1.004567005 | 0.39940841 | -2.5151373 | 0.0118986 | 0.09989195 |
| ZNF627 | 55.4771053 | -0.007701364 | 0.50376541 | -0.0152876 | 0.98780274 | 0.99678172 |
| ZNF823 | 59.1060773 | -0.036459149 | 0.27062031 | -0.1347244 | 0.89282981 | 0.95976294 |
| ZNF441 | 44.7680803 | -0.171115436 | 0.34180328 | -0.5006255 | 0.61663471 | 0.82592881 |
| ZNF440 | 49.905938 | -0.536237342 | 0.23268048 | -2.3046082 | 0.02118853 | 0.13968678 |
| ZNF439 | 68.1872099 | 0.90266607 | 0.29178075 | 3.093645 | 0.00197714 | 0.03151251 |
| ZNF69 | 151.682438 | 1.147402434 | 0.29891527 | 3.83855414 | 0.00012376 | 0.00547711 |
| ZNF844 | 40.3590646 | 0.071288233 | 0.43422163 | 0.16417477 | 0.86959357 | 0.95106604 |
| ZNF136 | 33.6821905 | 0.272792753 | 0.2553613 | 1.06826194 | 0.28540237 | 0.57289801 |
| ZNF44 | 233.072873 | 0.136575837 | 0.22872033 | 0.5971303 | 0.55042039 | 0.78541292 |
| AC008758.5 | 28.037833 | -0.22187199 | 0.3956974 | -0.5607113 | 0.57499439 | 0.80114491 |
| ZNF490 | 40.2955259 | -0.085347295 | 0.28914016 | -0.2951762 | 0.76785928 | 0.904269 |
| ZNF791 | 83.2314973 | -0.093656469 | 0.18382354 | -0.5094912 | 0.610408 | 0.82270732 |
| DHPS | 31.3210222 | 0.555320774 | 0.30259971 | 1.83516622 | 0.06648107 | 0.26282382 |
| TNPO2 | 81.6995621 | -0.419864402 | 0.2510729 | -1.6722808 | 0.09446896 | 0.32021075 |
| TRIR | 132.360163 | -0.149152598 | 0.37269992 | -0.4001949 | 0.68901299 | 0.86751885 |
| ASNA1 | 38.3520444 | -0.755619252 | 0.40217882 | -1.8788141 | 0.06026988 | 0.24767274 |
| HOOK2 | 156.676146 | -0.159995382 | 0.32950898 | -0.485557 | 0.62728131 | 0.83250121 |
| JUNB | 226.726232 | 0.737369817 | 0.62125462 | 1.18690436 | 0.23526535 | 0.51818964 |
| GCDH | 36.9840164 | 0.082145834 | 0.24831247 | 0.33081639 | 0.74078318 | 0.89253455 |
| FARSA | 36.3299371 | -1.338546455 | 0.31391138 | -4.2640902 | 2.01E-05 | 0.00155055 |
| CALR | 220.960231 | -0.93273034 | 0.48684133 | -1.9158816 | 0.05538017 | 0.23619173 |
| AC092069.1 | 41.0322156 | -0.585341989 | 0.60124369 | -0.973552 | 0.33027904 | 0.61551418 |
| RAD23A | 66.3116723 | -0.71554172 | 0.34326789 | -2.0844994 | 0.03711476 | 0.19103831 |
| GADD45GIP1 | 110.888931 | -1.077489563 | 0.51095147 | -2.1087904 | 0.03496268 | 0.1847551 |
| NFIX | 51.8373846 | -0.374187898 | 0.72825536 | -0.5138141 | 0.60738195 | 0.8211189 |
| NACC1 | 25.414581 | -1.086429638 | 0.52259419 | -2.0789164 | 0.03762503 | 0.19262971 |
| IER2 | 182.825816 | 1.317911283 | 0.59235743 | 2.22485819 | 0.02609076 | 0.1568885 |
| CCDC130 | 87.5585528 | -0.181589694 | 0.34964924 | -0.5193482 | 0.60351797 | 0.81988502 |
| MRI1 | 48.6321714 | 0.379398778 | 0.45139603 | 0.84050093 | 0.40062758 | 0.67524659 |
| C19orf53 | 109.98787 | -0.891621591 | 0.40914883 | -2.1792109 | 0.029316 | 0.16724236 |
| ZSWIM4 | 46.757662 | -0.269693554 | 0.24827998 | -1.0862477 | 0.27736943 | 0.56470176 |
| AC020916.1 | 28.7192425 | 1.556766782 | 0.70638967 | 2.20383573 | 0.0275359 | 0.16064442 |
| CC2D1A | 56.2238792 | 0.092776451 | 0.24360245 | 0.38085188 | 0.70331316 | 0.87323872 |
| RFX1 | 46.6845642 | 0.368420972 | 0.29399237 | 1.25316507 | 0.21014564 | 0.49085225 |
| PALM3 | 56.0438879 | 1.251947158 | 0.57723817 | 2.16885718 | 0.03009353 | 0.16944102 |
| PRKACA | 38.7484235 | -0.643876177 | 0.28337616 | -2.2721607 | 0.02307681 | 0.1465867 |
| ADGRL1 | 46.8887895 | 0.001570553 | 0.27670612 | 0.00567589 | 0.99547132 | 0.99899312 |
| PKN1 | 69.369772 | 0.325041459 | 0.30570854 | 1.06323971 | 0.28767326 | 0.57450105 |
| GIPC1 | 126.452737 | -0.277375875 | 0.29223291 | -0.9491603 | 0.3425391 | 0.62772374 |

| | | | | | | |
|---|---|---|---|---|---|---|
| DNAJB1 | 50.7508709 | -0.136259517 | 0.42113628 | -0.3235521 | 0.74627718 | 0.89352127 |
| TECR | 56.5116286 | 0.599308617 | 0.32684871 | 1.83359641 | 0.06671393 | 0.26332729 |
| NDUFB7 | 93.1324745 | -0.459097533 | 0.38061326 | -1.2062048 | 0.22773854 | 0.51041354 |
| ZNF333 | 30.0083516 | -0.026547673 | 0.35287287 | -0.075233 | 0.94002935 | 0.98003278 |
| ILVBL | 40.6398018 | -0.715452762 | 0.32457256 | -2.2042922 | 0.0275038 | 0.16064442 |
| NOTCH3 | 25.7469311 | 0.334624642 | 0.6207176 | 0.5390932 | 0.58982255 | 0.81183245 |
| BRD4 | 297.590944 | -0.238559307 | 0.22851517 | -1.0439539 | 0.29650672 | 0.58477205 |
| AKAP8 | 74.861413 | -0.269169855 | 0.19974785 | -1.3475482 | 0.17780373 | 0.44938837 |
| AKAP8L | 184.032172 | 0.25511772 | 0.1527137 | 1.67056213 | 0.0948082 | 0.3204651 |
| TPM4 | 116.189382 | 0.070355067 | 0.49839564 | 0.14116309 | 0.88774111 | 0.95904526 |
| RAB8A | 35.1530441 | -0.37494273 | 0.32880249 | -1.1403281 | 0.25414963 | 0.54093192 |
| FAM32A | 32.7550208 | -0.723271282 | 0.35563999 | -2.0337175 | 0.04198009 | 0.20340275 |
| AP1M1 | 39.1566397 | -0.146217691 | 0.31385629 | -0.4658747 | 0.64130522 | 0.84197679 |
| EPS15L1 | 109.568355 | 0.164321536 | 0.23151488 | 0.70976662 | 0.47784887 | 0.73413221 |
| C19orf44 | 31.8726412 | 0.499630005 | 0.28386376 | 1.76010494 | 0.07839001 | 0.29033886 |
| SIN3B | 114.015708 | 0.08584966 | 0.28704876 | 0.29907693 | 0.76488135 | 0.90320494 |
| CPAMD8 | 98.7630268 | 0.268520629 | 0.73511388 | 0.36527759 | 0.7149042 | 0.87976199 |
| MYO9B | 107.334204 | -0.049925011 | 0.2946084 | -0.1694623 | 0.86543304 | 0.95046814 |
| NR2F6 | 57.7106926 | -0.40535814 | 0.27807431 | -1.4577332 | 0.14491411 | 0.40282422 |
| ABHD8 | 28.6702712 | -0.0427541 | 0.35925491 | -0.1190077 | 0.90526926 | 0.96556525 |
| MRPL34 | 44.4153096 | -0.632329861 | 0.32406594 | -1.9512383 | 0.05102871 | 0.22682567 |
| DDA1 | 27.1797557 | -0.365534469 | 0.28339742 | -1.2898299 | 0.19710972 | 0.47630586 |
| BST2 | 84.1757023 | 0.003830474 | 1.28482022 | 0.00298133 | 0.99762125 | 0.99923686 |
| MVB12A | 41.9124398 | -0.408521551 | 0.26480697 | -1.5427145 | 0.12290006 | 0.36812373 |
| SLC27A1 | 49.682598 | 0.397722993 | 0.44580308 | 0.8921495 | 0.37231281 | 0.65198983 |
| AC010618.3 | 36.2804645 | -0.075552413 | 0.32261354 | -0.2341886 | 0.81483857 | 0.92814476 |
| PGLS | 73.0125459 | -0.616987943 | 0.30911586 | -1.9959764 | 0.0459365 | 0.21381229 |
| COLGALT1 | 34.8248358 | 0.023640897 | 0.25573406 | 0.09244329 | 0.92634585 | 0.97408135 |
| RPL18A | 49.2182671 | -1.554703564 | 0.48808089 | -3.1853399 | 0.00144584 | 0.02643074 |
| CCDC124 | 33.502765 | -0.755929334 | 0.50737803 | -1.489874 | 0.13625737 | 0.38812051 |
| JUND | 214.177727 | 0.662563007 | 0.31354008 | 2.11316844 | 0.03458635 | 0.18326846 |
| LSM4 | 76.1999375 | -0.64587301 | 0.34919302 | -1.8496161 | 0.06436891 | 0.25820342 |
| PGPEP1 | 79.9262459 | -0.201010446 | 0.3412165 | -0.5890994 | 0.55579458 | 0.78784845 |
| GDF15 | 270.178388 | -0.505978165 | 0.93728918 | -0.5398314 | 0.58931328 | 0.81165089 |
| SSBP4 | 89.4109763 | -0.549007749 | 0.19363074 | -2.8353336 | 0.00457779 | 0.05460848 |
| ISYNA1 | 32.7170383 | 0.320438868 | 0.52400655 | 0.61151691 | 0.54085742 | 0.77837639 |
| ELL | 108.701792 | -0.871470616 | 0.31887482 | -2.7329553 | 0.00627689 | 0.06659271 |
| FKBP8 | 86.0213531 | -0.295798695 | 0.25693643 | -1.1512525 | 0.24962839 | 0.5368017 |
| KXD1 | 34.7957773 | -0.532604123 | 0.26257994 | -2.0283504 | 0.0425245 | 0.20459174 |
| UBA52 | 201.015507 | -0.486192215 | 0.49183887 | -0.9885193 | 0.32289839 | 0.61001895 |
| REX1BD | 33.9049225 | -0.222717039 | 0.36005597 | -0.6185623 | 0.53620476 | 0.77540842 |
| CRTC1 | 108.625189 | 0.813337812 | 0.30178869 | 2.69505732 | 0.00703765 | 0.07126181 |
| UPF1 | 74.047087 | -0.037955236 | 0.21389065 | -0.1774516 | 0.85915369 | 0.94734879 |
| COPE | 50.7720708 | -0.514440424 | 0.41750595 | -1.2321751 | 0.21788369 | 0.49990608 |
| DDX49 | 26.2462158 | -1.048370322 | 0.30566295 | -3.4298246 | 0.00060397 | 0.01498819 |
| SUGP2 | 161.418259 | 0.090475986 | 0.17604368 | 0.51394055 | 0.60729356 | 0.8211189 |
| SLC25A42 | 52.245012 | 0.731901826 | 0.26492662 | 2.76265866 | 0.00573327 | 0.0633437 |
| TMEM161A | 29.3033594 | 0.026065181 | 0.29003247 | 0.08986987 | 0.92839062 | 0.9749043 |
| SUGP1 | 46.0970949 | 0.419553511 | 0.22128383 | 1.89599712 | 0.05796043 | 0.24313032 |
| MAU2 | 120.301825 | 0.132723395 | 0.20189183 | 0.65739853 | 0.5109247 | 0.75922776 |
| GATAD2A | 246.62095 | -0.543150371 | 0.20350979 | -2.6689152 | 0.00760967 | 0.07529132 |
| PBX4 | 39.7445325 | 0.393591158 | 0.517703 | 0.7602644 | 0.44709656 | 0.70891772 |
| ATP13A1 | 36.1651273 | -0.911655189 | 0.37523481 | -2.4295592 | 0.0151172 | 0.11411159 |
| ZNF14 | 34.1846888 | 0.377627968 | 0.3248904 | 1.1623242 | 0.2451038 | 0.53142809 |
| ZNF253 | 63.831228 | 0.323191756 | 0.30932098 | 1.04484267 | 0.29609569 | 0.58420198 |
| ZNF682 | 45.3038396 | -0.283475829 | 0.41585423 | -0.6816711 | 0.49544692 | 0.74726595 |
| ZNF90 | 302.210876 | -0.405358141 | 0.3258948 | -1.2438313 | 0.21356167 | 0.49544858 |
| AC011447.3 | 121.871965 | 1.745280595 | 0.4288105 | 4.07005099 | 4.70E-05 | 0.00292513 |
| ZNF486 | 95.0314146 | 0.841341679 | 0.50820954 | 1.65550153 | 0.09782281 | 0.32626903 |
| AC008554.1 | 149.924566 | 1.329343843 | 0.76108395 | 1.74664548 | 0.08069885 | 0.29548169 |
| ZNF737 | 50.2974185 | 0.215052568 | 0.30639273 | 0.70188536 | 0.48275066 | 0.73696727 |
| ZNF430 | 58.1457527 | -0.215200196 | 0.19103458 | -1.1264986 | 0.25995451 | 0.54648299 |
| ZNF714 | 33.8962896 | 0.314492383 | 0.4356515 | 0.72188981 | 0.47036222 | 0.72823658 |
| ZNF431 | 133.281224 | 0.287783053 | 0.25302803 | 1.13735642 | 0.25538932 | 0.54219288 |
| ZNF708 | 97.3836458 | 0.175784365 | 0.19409508 | 0.90566111 | 0.36511525 | 0.64511126 |
| ZNF493 | 36.8092671 | 0.758766223 | 0.27188302 | 2.7907819 | 0.00525809 | 0.05996099 |
| AC010615.4 | 55.5664675 | -0.353101326 | 0.61501078 | -0.5741384 | 0.56587413 | 0.79479594 |
| ZNF429 | 104.292331 | 0.111569227 | 0.21286613 | 0.52412861 | 0.60018909 | 0.81769468 |
| AC123912.1 | 35.8333503 | 0.029132699 | 0.60606086 | 0.04806893 | 0.96166131 | 0.9875899 |
| ZNF100 | 55.7700739 | -0.307065245 | 0.37992528 | -0.8082254 | 0.41896086 | 0.68832418 |
| ZNF43 | 110.034825 | 0.213840829 | 0.21818846 | 0.98007397 | 0.32704961 | 0.61330512 |
| ZNF208 | 90.0920552 | -0.092933973 | 0.26840647 | -0.3462434 | 0.72915979 | 0.88653304 |
| ZNF98 | 49.7988732 | 0.913082518 | 0.48784732 | 1.8716563 | 0.06125417 | 0.25032154 |
| ZNF91 | 315.757879 | 0.055190495 | 0.18639276 | 0.29609784 | 0.76715536 | 0.90403468 |
| ZNF675 | 33.1600071 | 0.536653526 | 0.24160718 | 2.22118203 | 0.02633863 | 0.1573452 |
| ZNF254 | 256.890707 | 0.261640834 | 0.23219653 | 1.12680769 | 0.2598238 | 0.54640004 |
| LINC00662 | 90.4694364 | 0.125866508 | 0.27521398 | 0.45734054 | 0.6474263 | 0.84537847 |
| AC006504.5 | 99.1138113 | 0.200973241 | 0.3377518 | 0.59503234 | 0.55182185 | 0.78577398 |
| UQCRFS1 | 73.4383395 | -0.683628253 | 0.44085731 | -1.5506792 | 0.12097859 | 0.36534213 |
| C19orf12 | 53.1036358 | -0.483633347 | 0.40842857 | -1.184132 | 0.23636081 | 0.51974689 |
| URI1 | 326.116859 | 0.083894027 | 0.13917178 | 0.6028092 | 0.54663562 | 0.78253724 |
| ZNF536 | 152.704982 | 0.473704154 | 1.40602518 | 0.33691015 | 0.73618463 | 0.89050544 |
| ZNF507 | 101.950489 | 0.076695786 | 0.24305388 | 0.31555056 | 0.75234367 | 0.89668701 |
| DPY19L3 | 156.387093 | -0.112034194 | 0.22956909 | -0.4880195 | 0.62553602 | 0.83176146 |
| PDCD5 | 40.4734529 | -0.705967134 | 0.38265477 | -1.8449192 | 0.06504929 | 0.26002617 |
| ANKRD27 | 81.7219009 | 0.024449293 | 0.22411918 | 0.10909059 | 0.91313064 | 0.96844354 |
| CEP89 | 107.701848 | 0.181584777 | 0.20745097 | 0.87531419 | 0.38140298 | 0.65851823 |
| RHPN2 | 262.879263 | -1.209211797 | 0.31978045 | -3.7813812 | 0.00015596 | 0.00641179 |
| GPATCH1 | 27.6310098 | 0.134925616 | 0.30045066 | 0.44907746 | 0.65337578 | 0.84898082 |
| PEPD | 83.4855962 | -0.337226539 | 0.31990609 | -1.0541423 | 0.29181777 | 0.57910139 |
| LSM14A | 525.827576 | 0.592917097 | 0.13121443 | 4.51868809 | 6.22E-06 | 0.00064828 |
| KIAA0355 | 191.885735 | 0.151624717 | 0.20776292 | 0.72979682 | 0.46551439 | 0.72327407 |
| GPI | 164.163118 | -0.282719242 | 0.43645059 | -0.6477692 | 0.51713425 | 0.76244867 |
| UBA2 | 177.975276 | -0.177212691 | 0.21552878 | -0.8222228 | 0.41095008 | 0.68247066 |
| WTIP | 40.4189364 | 0.442708248 | 0.49716228 | 0.8904703 | 0.37321341 | 0.65225529 |
| SCGB2B2 | 94.3629242 | 0.468370351 | 0.3990253 | 1.17378609 | 0.24048072 | 0.52473002 |
| ZNF302 | 111.295858 | 0.145954313 | 0.28611256 | 0.510129 | 0.60996109 | 0.82246865 |
| GRAMD1A | 94.7064262 | -0.897845874 | 0.49595498 | -1.8103374 | 0.07024347 | 0.27219276 |
| HPN | 256.625642 | -0.439761111 | 0.41122227 | -1.0694 | 0.28488945 | 0.57246822 |
| LSR | 71.7292565 | -0.850271773 | 0.25984314 | -3.2722502 | 0.00106695 | 0.02174537 |
| USF2 | 60.6383614 | 0.008262266 | 0.30210677 | 0.02734883 | 0.97818151 | 0.99532619 |
| DMKN | 161.93928 | -0.009564426 | 0.31273515 | -0.0305831 | 0.97560198 | 0.99407418 |
| TMEM147 | 45.9860459 | -0.649599776 | 0.52159465 | -1.2454111 | 0.21298066 | 0.49481974 |
| RBM42 | 31.1547784 | -0.918529233 | 0.42498817 | -2.1613054 | 0.03067175 | 0.17115672 |
| COX6B1 | 146.661986 | -0.679155392 | 0.37807883 | -1.7963328 | 0.07244161 | 0.27745629 |
| KMT2B | 52.5099986 | -0.42095918 | 0.21966481 | -1.9163706 | 0.05531794 | 0.23619173 |
| PROSER3 | 31.2650099 | 0.106461546 | 0.30228591 | 0.35218825 | 0.72469709 | 0.88397726 |
| NPHS1 | 53.6256209 | 0.936746005 | 1.26237933 | 0.74204796 | 0.45805828 | 0.7171854 |
| NFKBID | 34.015101 | 0.087910015 | 0.30532789 | 0.28792003 | 0.77340795 | 0.90643453 |
| TBCB | 41.0104556 | -0.432303626 | 0.36544507 | -1.182951 | 0.23682856 | 0.52000641 |
| CAPNS1 | 179.958988 | -0.847188516 | 0.39400553 | -2.1501945 | 0.03153983 | 0.17373701 |
| ZNF565 | 104.503544 | -0.017073575 | 0.29044775 | -0.0587836 | 0.95312444 | 0.98368111 |
| ZNF146 | 131.674947 | -0.052056485 | 0.19073158 | -0.2729306 | 0.78490656 | 0.91212179 |

| | | | | | | |
|---|---|---|---|---|---|---|
| LINC00665 | 52.9193766 | -0.736076183 | 0.33865435 | -2.1735323 | 0.02974027 | 0.16866909 |
| ZFP14 | 72.1949104 | 0.474438339 | 0.24603129 | 1.92836583 | 0.05380964 | 0.23302104 |
| ZFP82 | 31.9997621 | -0.539011256 | 0.42472938 | -1.2690699 | 0.20441615 | 0.48318561 |
| ZNF566 | 55.3097134 | -0.072559809 | 0.19354166 | -0.3749054 | 0.70773084 | 0.87554614 |
| ZNF260 | 42.1680519 | -0.132098236 | 0.27082507 | -0.4877622 | 0.62571827 | 0.83180006 |
| ZNF529 | 70.3779759 | 0.38742382 | 0.23938993 | 1.61837979 | 0.10558077 | 0.33994587 |
| ZNF529-AS1 | 62.6594578 | -0.22048747 | 0.30216029 | -0.7297037 | 0.46557135 | 0.72327407 |
| ZNF461 | 29.7710799 | 0.296699019 | 0.26201159 | 1.1323889 | 0.25747097 | 0.54481971 |
| ZNF567 | 33.8302401 | 0.194774282 | 0.31704473 | 0.61434323 | 0.53898854 | 0.77708778 |
| AC020928.1 | 72.9151488 | 0.192454835 | 0.2625272 | 0.73308532 | 0.46350639 | 0.72133763 |
| ZNF790 | 46.2579087 | 0.104554073 | 0.28346033 | 0.36884905 | 0.71224024 | 0.87806297 |
| ZNF345 | 68.2979729 | 0.499311381 | 0.29570225 | 1.68856129 | 0.09130353 | 0.31502156 |
| ZNF568 | 47.3855093 | 0.528070726 | 0.30500141 | 1.73137144 | 0.08338554 | 0.30062195 |
| ZNF420 | 71.3413014 | 0.067287863 | 0.41316734 | 0.16285862 | 0.87062975 | 0.95159315 |
| ZNF383 | 26.8185482 | 0.00461775 | 0.33917301 | 0.01361473 | 0.98913735 | 0.99678172 |
| HKR1 | 107.650562 | 0.329906347 | 0.1969845 | 1.67478326 | 0.09397676 | 0.31910791 |
| ZNF569 | 27.5269209 | 0.416863769 | 0.25886023 | 1.61038166 | 0.10731456 | 0.3426554 |
| ZNF793 | 51.1963131 | 0.422464885 | 0.25188659 | 1.67720274 | 0.09350284 | 0.31835715 |
| ZNF540 | 35.8090599 | 0.266899586 | 0.69327101 | 0.38498593 | 0.70024784 | 0.87182648 |
| ZNF571 | 29.0371194 | -0.108006209 | 0.30010825 | -0.3598908 | 0.71892877 | 0.88155276 |
| ZFP30 | 49.8688158 | -0.132712935 | 0.32670258 | -0.4062194 | 0.68458138 | 0.8653964 |
| ZNF573 | 59.2491161 | -0.160609127 | 0.3271721 | -0.4909011 | 0.62349643 | 0.83030289 |
| SIPA1L3 | 618.316315 | 0.150681331 | 0.31938589 | 0.47178456 | 0.63708057 | 0.83872935 |
| SPINT2 | 41.5844904 | 0.289501627 | 0.35469344 | 0.81620238 | 0.4143844 | 0.68449528 |
| CATSPERG | 34.0776371 | 0.475746473 | 0.48414437 | 0.98265414 | 0.32577769 | 0.61243485 |
| PSMD8 | 89.2637418 | -1.009228849 | 0.37191769 | -2.7135812 | 0.00665603 | 0.06902242 |
| EIF3K | 88.2036474 | -0.181474555 | 0.29478464 | -0.6156174 | 0.53814706 | 0.77680993 |
| ACTN4 | 640.384863 | 0.402384885 | 0.24236411 | 1.66024949 | 0.09686427 | 0.32454105 |
| CAPN12 | 41.8733995 | -0.391186596 | 0.55612581 | -0.7034138 | 0.48179788 | 0.73620566 |
| ECH1 | 75.2289404 | -0.025677285 | 0.45220382 | -0.0567825 | 0.95471842 | 0.98390867 |
| HNRNPL | 92.8912905 | 0.229487104 | 0.20169021 | 1.13781977 | 0.25519575 | 0.54202219 |
| SIRT2 | 40.642831 | 0.142493366 | 0.30415914 | 0.46848293 | 0.63943927 | 0.84033321 |
| MRPS12 | 36.6031482 | -1.053852485 | 0.43780008 | -2.4071546 | 0.01607736 | 0.11846542 |
| FBXO17 | 208.932127 | 0.225764851 | 0.49639412 | 0.45480968 | 0.64924617 | 0.84651696 |
| PAK4 | 73.1160339 | 0.025046792 | 0.31168185 | 0.08036012 | 0.93595084 | 0.97844299 |
| SAMD4B | 162.168398 | -0.017744785 | 0.20400361 | -0.0869827 | 0.93068526 | 0.97602738 |
| PAF1 | 27.4273568 | -0.365653208 | 0.33732906 | -1.0839659 | 0.27837991 | 0.56578102 |
| MED29 | 39.1928211 | -0.494780664 | 0.3516661 | -1.4069615 | 0.15943881 | 0.42483013 |
| ZFP36 | 126.379086 | -0.080586158 | 0.60451346 | -0.1333075 | 0.89395022 | 0.96064047 |
| RPS16 | 306.056714 | -0.434978873 | 0.45882122 | -0.9480356 | 0.34311131 | 0.62806483 |
| SUPT5H | 108.108996 | -0.420578115 | 0.22554358 | -1.8647311 | 0.06221911 | 0.2529698 |
| TIMM50 | 25.7228318 | -0.564318417 | 0.346773 | -1.6273424 | 0.10366439 | 0.33583603 |
| FBL | 42.7571479 | -0.615055537 | 0.34575333 | -1.7788853 | 0.07525856 | 0.28328492 |
| FCGBP | 56.293843 | 3.177351585 | 1.06049267 | 2.99610894 | 0.00273449 | 0.03818317 |
| PSMC4 | 36.3128024 | -0.273649513 | 0.42333323 | -0.6464163 | 0.51800976 | 0.76315219 |
| ZNF546 | 29.1266078 | 0.329289251 | 0.47886975 | 0.68763844 | 0.49168049 | 0.74428057 |
| ZNF780B | 135.346251 | 0.185635123 | 0.29238768 | 0.63489379 | 0.52549769 | 0.7671049 |
| ZNF780A | 36.4432572 | 0.337269508 | 0.34065172 | 0.99007134 | 0.32213925 | 0.60947499 |
| MAP3K10 | 28.8852358 | 0.729222471 | 0.2460818 | 2.96333358 | 0.00304327 | 0.04134956 |
| CNTD2 | 70.0124561 | 2.103597254 | 1.86534053 | 1.12772827 | 0.25943469 | 0.54618849 |
| AKT2 | 269.956343 | 1.13703975 | 0.42633856 | 2.66698781 | 0.00765344 | 0.07534671 |
| PLD3 | 168.688194 | -0.187763989 | 0.24805791 | -0.7569361 | 0.44908814 | 0.71037275 |
| BLVRB | 98.1720755 | -0.661504036 | 0.31663666 | -2.0891581 | 0.0366935 | 0.18976937 |
| COQ8B | 37.3123035 | -0.229621527 | 0.24198274 | -0.948917 | 0.34266285 | 0.62772374 |
| ITPKC | 53.7441251 | -1.005590701 | 0.2618953 | -3.8396668 | 0.0001232 | 0.00547711 |
| SNRPA | 49.9474747 | -0.975725709 | 0.27071983 | -3.60419 | 0.00031313 | 0.00994413 |
| AC008537.1 | 56.0228697 | -0.110546833 | 0.33249286 | -0.3324788 | 0.73952773 | 0.89253455 |
| AXL | 67.3722477 | -0.939064004 | 0.48075444 | -1.9533132 | 0.05078249 | 0.22646438 |
| HNRNPUL1 | 102.084882 | -0.168209273 | 0.27149928 | -0.619557 | 0.5355495 | 0.77504588 |
| TGFB1 | 69.168069 | -0.309139851 | 0.29713419 | -1.0404048 | 0.29815185 | 0.58581522 |
| DMAC2 | 26.7951963 | -0.134426915 | 0.36248426 | -0.370849 | 0.71075002 | 0.87748 |
| RPS19 | 858.50481 | -0.857645246 | 0.35461173 | -2.4185473 | 0.01558262 | 0.11595952 |
| ARHGEF1 | 46.1252776 | 0.238775736 | 0.29991123 | 0.79615469 | 0.42594213 | 0.69224629 |
| RABAC1 | 106.077756 | -0.035918712 | 0.51513876 | -0.0697263 | 0.94441152 | 0.98115679 |
| GRIK5 | 64.5122398 | 0.062389251 | 0.28844665 | 0.2162939 | 0.82875866 | 0.933664 |
| GSK3A | 23.7247958 | -0.435921848 | 0.4065094 | -1.0723537 | 0.2835612 | 0.57084461 |
| CIC | 40.4163902 | -0.211144536 | 0.32063637 | -0.6585171 | 0.5102059 | 0.75889167 |
| MEGF8 | 49.9439662 | 0.096568024 | 0.24390031 | 0.39593235 | 0.69215493 | 0.86892611 |
| LIPE-AS1 | 50.2429758 | 0.084071943 | 0.4670143 | 0.18002006 | 0.85713682 | 0.94638973 |
| PHLDB3 | 39.1284809 | 0.168764058 | 0.33463885 | 0.504317 | 0.61403864 | 0.82514464 |
| ETHE1 | 39.7324109 | -0.488707795 | 0.37965869 | -1.2872293 | 0.19801438 | 0.47656532 |
| ZNF428 | 60.5859204 | -0.231451477 | 0.33964574 | -0.6814497 | 0.49558701 | 0.74735956 |
| SMG9 | 27.1208294 | -0.116711383 | 0.30174501 | -0.3867881 | 0.69891308 | 0.8717697 |
| ZNF283 | 27.973834 | -0.12538311 | 0.35194135 | -0.3562614 | 0.72164483 | 0.88235193 |
| ZNF155 | 40.1129462 | 0.127092387 | 0.40707263 | 0.31221059 | 0.75488048 | 0.89810084 |
| ZNF224 | 49.7995596 | 0.137408958 | 0.29189074 | 0.47075477 | 0.63781587 | 0.83912075 |
| ZNF234 | 27.8612021 | 0.104338295 | 0.31244839 | 0.33393769 | 0.73842657 | 0.89197833 |
| ZNF226 | 36.8587623 | 0.534355384 | 0.31095849 | 1.7184139 | 0.08572115 | 0.3045708 |
| ZNF235 | 99.5306237 | 0.274135923 | 0.37906665 | 0.7231866 | 0.46956525 | 0.72743153 |
| AC243964.2 | 78.2327411 | -0.011229333 | 0.45259791 | -0.0248108 | 0.98020585 | 0.99558815 |
| PVR | 37.2007349 | -0.107197543 | 0.37796449 | -0.283618 | 0.77670312 | 0.90765392 |
| CEACAM19 | 86.0443373 | 0.177446202 | 0.37138512 | 0.47779568 | 0.63279564 | 0.83561475 |
| BCL3 | 42.7154904 | -2.423175092 | 0.47374354 | -5.1149512 | 3.14E-07 | 6.99E-05 |
| CBLC | 45.42175 | -0.371083972 | 0.51502437 | -0.7205173 | 0.47120655 | 0.72919023 |
| BCAM | 570.914137 | 0.562070673 | 0.56983207 | 0.98637951 | 0.32394692 | 0.61072469 |
| NECTIN2 | 210.954386 | 0.215975826 | 0.17776319 | 1.21496368 | 0.22437994 | 0.50632167 |
| TOMM40 | 37.1766011 | -0.957478746 | 0.35676307 | -2.6837944 | 0.00727918 | 0.0731995 |
| APOE | 139.579945 | -0.731422258 | 0.70147971 | -1.0426848 | 0.29709428 | 0.58520063 |
| CLPTM1 | 83.8837969 | -0.337584928 | 0.30313297 | -1.1136529 | 0.2654281 | 0.55350835 |
| RELB | 80.5802843 | -1.007758676 | 0.54092139 | -1.8630409 | 0.06245651 | 0.25371965 |
| CLASRP | 90.8464316 | -0.004585681 | 0.24558984 | -0.0186721 | 0.98510267 | 0.99668665 |
| GEMIN7-AS1 | 29.8333355 | 0.457898872 | 0.30976439 | 1.47821662 | 0.13934981 | 0.39306001 |
| MARK4 | 100.705727 | -0.082055179 | 0.18789176 | -0.4367152 | 0.66231793 | 0.85507439 |
| TRAPPC6A | 49.990674 | -0.069314 | 0.41607096 | -0.1665918 | 0.86769126 | 0.95090501 |
| ERCC2 | 27.4290523 | 0.199882164 | 0.30158648 | 0.66276898 | 0.50747853 | 0.75682386 |
| PPP1R13L | 95.8921375 | -0.031149979 | 0.29744576 | -0.1047249 | 0.9165941 | 0.96995745 |
| ERCC1 | 43.7298124 | -0.454976194 | 0.37115939 | -1.2258243 | 0.2202648 | 0.50176375 |
| VASP | 54.3087159 | -1.014582366 | 0.29767713 | -3.4083316 | 0.00065361 | 0.01564098 |
| OPA3 | 72.3851631 | -0.401741059 | 0.33736651 | -1.1908149 | 0.23372627 | 0.51635994 |
| EML2 | 55.437417 | -0.44660819 | 0.31581528 | -1.4141437 | 0.15731971 | 0.42243085 |
| SNRPD2 | 108.021031 | -0.949263331 | 0.41876149 | -2.2668353 | 0.02340029 | 0.14775964 |
| FBXO46 | 36.9514278 | 0.068094325 | 0.25951709 | 0.2623886 | 0.79302186 | 0.91688068 |
| DMPK | 33.801594 | -0.207008091 | 0.36699614 | -0.5640607 | 0.57271284 | 0.79959427 |
| SYMPK | 154.817618 | -0.045666793 | 0.16680965 | -0.2737659 | 0.78426453 | 0.91212179 |
| HIF3A | 55.8731603 | -0.241350572 | 0.76313921 | -0.3162602 | 0.75180501 | 0.8961598 |
| PPP5C | 87.1308727 | -0.135982586 | 0.16642405 | -0.8170849 | 0.41387989 | 0.68448816 |
| CCDC8 | 23.1744251 | 0.048879385 | 0.54078638 | 0.09038575 | 0.92798068 | 0.97474479 |
| PPP5D1 | 41.533813 | 0.874162288 | 0.43929024 | 1.98994245 | 0.04659728 | 0.21552646 |
| CALM3 | 86.1225353 | -0.716075498 | 0.37169528 | -1.9265122 | 0.05404045 | 0.23338748 |
| PRKD2 | 94.98949 | -0.617497672 | 0.29599518 | -2.0861748 | 0.0369628 | 0.1905633 |
| STRN4 | 34.3021913 | -0.248002933 | 0.30246126 | -0.8199494 | 0.41224494 | 0.68367283 |
| FKRP | 25.8116224 | 0.022661531 | 0.26475782 | 0.08559344 | 0.93178962 | 0.97675779 |
| AP2S1 | 62.972955 | -0.817421768 | 0.40200941 | -2.0333399 | 0.0420182 | 0.20348448 |

| | | | | | | |
|---|---|---|---|---|---|---|
| ARHGAP35 | 172.593143 | 0.216017392 | 0.23914814 | 0.90327859 | 0.36637806 | 0.64644233 |
| TMEM160 | 64.1391956 | -0.435751152 | 0.54693866 | -0.7967094 | 0.42561984 | 0.69215319 |
| ZC3H4 | 51.0184087 | -0.008338136 | 0.32835818 | -0.0253934 | 0.97974116 | 0.99557142 |
| SAE1 | 46.3670907 | -0.023096176 | 0.34049146 | -0.0678319 | 0.94591947 | 0.98126084 |
| C5AR1 | 35.2519481 | -1.030085645 | 0.76812436 | -1.3410402 | 0.1799074 | 0.451844 |
| DHX34 | 50.9967965 | -0.361679837 | 0.28921479 | -1.2505579 | 0.21109583 | 0.49211169 |
| NAPA | 100.472853 | -0.182440998 | 0.28461849 | -0.6410019 | 0.52152144 | 0.76444589 |
| NOP53 | 48.9535903 | -0.609124676 | 0.4922839 | -1.2373443 | 0.21595929 | 0.49746509 |
| SELENOW | 79.5765679 | 0.08410923 | 0.38495019 | 0.2184938 | 0.82704439 | 0.93346403 |
| PLA2G4C | 37.1914985 | -0.96094328 | 0.69736858 | -1.3779561 | 0.16821685 | 0.43715387 |
| LIG1 | 27.9411203 | 0.708974688 | 0.3156175 | 2.24630984 | 0.02468417 | 0.15225348 |
| ZSWIM9 | 31.2637715 | -0.088442926 | 0.25927243 | -0.3411197 | 0.73301349 | 0.88869659 |
| CARD8 | 59.3468041 | 0.327934571 | 0.28461547 | 1.1522022 | 0.249238 | 0.53636804 |
| EMP3 | 37.8505077 | -0.997244764 | 0.81144541 | -1.2289733 | 0.21908181 | 0.50085552 |
| KDELR1 | 101.57788 | -0.905457195 | 0.3467048 | -2.6116085 | 0.00901174 | 0.0847971 |
| CYTH2 | 47.8910649 | -0.269963834 | 0.22972553 | -1.1751582 | 0.23993145 | 0.52377013 |
| LMTK3 | 35.6391936 | 0.088687687 | 0.52304727 | 0.1695596 | 0.8653565 | 0.95046814 |
| SULT2B1 | 45.968213 | 1.915718214 | 1.11936564 | 1.71143203 | 0.08700139 | 0.30691783 |
| RPL18 | 363.497116 | -0.41970587 | 0.43307609 | -0.9691273 | 0.33248167 | 0.61719981 |
| BCAT2 | 45.9520889 | 0.106205183 | 0.29019894 | 0.36597372 | 0.71438469 | 0.87934862 |
| PLEKHA4 | 73.1167419 | -0.593734331 | 0.39186049 | -1.5151676 | 0.12972996 | 0.3792137 |
| PPP1R15A | 37.2681588 | -0.383351424 | 0.41182177 | -0.9308673 | 0.3519222 | 0.63377755 |
| NUCB1 | 112.188311 | -0.949427427 | 0.31362893 | -3.0272316 | 0.00246805 | 0.03609378 |
| BAX | 69.3602041 | -0.650284511 | 0.32656897 | -1.9912624 | 0.04645205 | 0.21506243 |
| FTL | 1610.01798 | -1.464867374 | 0.53646405 | -2.7305974 | 0.00632196 | 0.06691502 |
| GYS1 | 64.33397 | -0.307615972 | 0.3522498 | -0.8732893 | 0.38250542 | 0.65946633 |
| RUVBL2 | 25.9138885 | -0.630159248 | 0.36069796 | -1.7470552 | 0.08062776 | 0.29534735 |
| SNRNP70 | 322.42021 | 0.248784264 | 0.19832678 | 1.25441587 | 0.20969089 | 0.49014861 |
| TRPM4 | 33.3126449 | -0.197467526 | 0.52074885 | -0.3791992 | 0.70453998 | 0.873867 |
| SLC6A16 | 59.4713694 | 0.12239163 | 0.47994272 | 0.25501299 | 0.79871306 | 0.91949148 |
| TEAD2 | 25.6657362 | -1.00993108 | 0.38703811 | -2.6093841 | 0.00907054 | 0.08501632 |
| PIH1D1 | 35.1745485 | 0.128718038 | 0.33592447 | 0.38317553 | 0.70158962 | 0.87200065 |
| RPL13A | 522.336399 | -0.668036657 | 0.39745767 | -1.6807744 | 0.09280675 | 0.31724627 |
| RPS11 | 395.422695 | -0.300928911 | 0.42895708 | -0.7015362 | 0.48296846 | 0.73696727 |
| FCGRT | 96.3611738 | -0.157562976 | 0.40060259 | -0.3933149 | 0.6940869 | 0.86986959 |
| NOSIP | 42.3799113 | 0.153118956 | 0.24281578 | 0.63059721 | 0.52830392 | 0.76990012 |
| PRR12 | 49.6263597 | 0.169230176 | 0.34065612 | 0.49677714 | 0.61934622 | 0.82766705 |
| RRAS | 29.7470891 | -0.737219382 | 0.51186425 | -1.4402635 | 0.14979287 | 0.41050134 |
| SCAF1 | 38.5235577 | -0.202209433 | 0.25751295 | -0.7852399 | 0.43231295 | 0.69737124 |
| IRF3 | 86.5810091 | -0.341971996 | 0.20904264 | -1.6358959 | 0.10186138 | 0.33338252 |
| PRMT1 | 92.8253261 | -0.540139764 | 0.35807877 | -1.5084384 | 0.13144236 | 0.38163088 |
| AP2A1 | 63.1557684 | -0.228944416 | 0.23267494 | -0.9839668 | 0.32513183 | 0.61197925 |
| MED25 | 73.1922357 | -0.271469669 | 0.31555927 | -0.8602811 | 0.38963412 | 0.66529505 |
| PTOV1 | 61.5028898 | 0.355763003 | 0.28516156 | 1.24758401 | 0.21218344 | 0.49368597 |
| PNKP | 52.2742455 | -1.061848911 | 0.31853872 | -3.3335002 | 0.00085761 | 0.01862814 |
| TBC1D17 | 46.2321991 | 0.278057142 | 0.23953253 | 1.16083247 | 0.24571004 | 0.53226361 |
| VRK3 | 52.6246574 | -0.469602474 | 0.2947276 | -1.5933441 | 0.11108302 | 0.34822126 |
| MYH14 | 85.7087894 | 0.556460427 | 0.52860072 | 1.05270463 | 0.29247639 | 0.57992782 |
| KCNC3 | 118.581632 | -0.01827232 | 0.33820462 | -0.0540274 | 0.95691333 | 0.98509111 |
| NR1H2 | 109.053747 | -0.328445182 | 0.3553529 | -0.9242789 | 0.35534114 | 0.63706583 |
| NAPSA | 190.227143 | -0.834706235 | 0.39063392 | -2.1367992 | 0.03261433 | 0.17693995 |
| EMC10 | 86.6499394 | -0.502295102 | 0.32394695 | -1.5505474 | 0.12101019 | 0.36534213 |
| ETFB | 91.8931637 | -0.084041841 | 0.42740334 | -0.1966336 | 0.84411431 | 0.94193856 |
| ZNF175 | 27.3404686 | 0.149438234 | 0.29046648 | 0.5144767 | 0.60691875 | 0.8211189 |
| SPACA6 | 45.4428721 | 0.081904414 | 0.36519607 | 0.22427518 | 0.82254319 | 0.93217972 |
| ZNF577 | 52.62772 | 0.35360361 | 0.31351036 | 1.12788494 | 0.25936851 | 0.54618849 |
| ZNF350-AS1 | 47.9495891 | 1.033314145 | 0.53664633 | 1.92550306 | 0.05416645 | 0.23344987 |
| ZNF432 | 60.9357461 | 0.111383982 | 0.27662029 | 0.4026602 | 0.68719822 | 0.86733718 |
| AC011468.3 | 35.5736693 | -1.053878698 | 0.5547925 | -1.8995908 | 0.05748685 | 0.24194486 |
| PPP2R1A | 211.149038 | -0.459778531 | 0.29847511 | -1.540425 | 0.12345679 | 0.36921404 |
| ZNF766 | 53.1539435 | -0.277863887 | 0.20673008 | -1.3440903 | 0.1789192 | 0.45066186 |
| ZNF480 | 34.8148652 | -0.478247581 | 0.31829365 | -1.5025357 | 0.13295881 | 0.38380408 |
| ZNF528 | 32.4966046 | 0.356197703 | 0.35838456 | 0.99389801 | 0.32027252 | 0.60702226 |
| ZNF701 | 39.8515372 | 0.651241095 | 0.25946209 | 2.50996627 | 0.01207427 | 0.10031173 |
| ZNF83 | 407.787045 | 0.479379193 | 0.2649872 | 1.80906548 | 0.07044083 | 0.27240724 |
| ZNF611 | 196.138116 | 0.245959536 | 0.32009491 | 0.76839565 | 0.44225216 | 0.70510851 |
| ZNF28 | 27.1049922 | 0.389113546 | 0.30347342 | 1.28219975 | 0.19977258 | 0.47864472 |
| AC010487.3 | 57.4573498 | 1.848528831 | 0.54222987 | 3.40912397 | 0.00065172 | 0.01564098 |
| ZNF160 | 127.660014 | 0.078299232 | 0.20582792 | 0.38041114 | 0.70364025 | 0.87353161 |
| ZNF347 | 46.8025726 | -0.088742273 | 0.27928459 | -0.3177486 | 0.75067568 | 0.89599662 |
| ZNF677 | 31.6233174 | -0.173305616 | 0.36072501 | -0.4804369 | 0.63091674 | 0.83470856 |
| ZNF765 | 40.5723972 | -0.067720816 | 0.26944854 | -0.2513312 | 0.80155808 | 0.92108024 |
| ZNF761 | 48.5448966 | -0.010585617 | 0.2913891 | -0.0363281 | 0.97102073 | 0.99175186 |
| ZNF331 | 96.8393708 | 0.185593108 | 0.71109511 | 0.26099618 | 0.79409545 | 0.91723627 |
| MYADM | 40.1960597 | 0.669984624 | 0.50817148 | 1.31842232 | 0.18736231 | 0.46280134 |
| NDUFA3 | 32.1621094 | 0.018065216 | 0.48539042 | 0.03721791 | 0.97031126 | 0.99175186 |
| PRPF31 | 60.6197175 | -0.309194971 | 0.31013787 | -0.9969597 | 0.31878405 | 0.60563353 |
| CNOT3 | 39.4167926 | 0.162979652 | 0.32133977 | 0.50718793 | 0.61202296 | 0.82374146 |
| TMC4 | 81.7849799 | -0.010613831 | 0.3669215 | -0.0289267 | 0.97692305 | 0.99441014 |
| MBOAT7 | 62.5522699 | -0.351577901 | 0.19024946 | -1.8479837 | 0.0646047 | 0.25882408 |
| TSEN34 | 40.8293543 | -0.646017358 | 0.44589978 | -1.448795 | 0.14739485 | 0.40633477 |
| RPS9 | 754.745107 | -0.53838679 | 0.4771961 | -1.1282297 | 0.25922294 | 0.54609557 |
| LENG8-AS1 | 24.3715673 | 0.417539809 | 0.30838864 | 1.35394029 | 0.17575543 | 0.44775565 |
| LENG8 | 182.084608 | 1.006222131 | 0.33644009 | 2.99079141 | 0.00278255 | 0.03846189 |
| RDH13 | 55.9084961 | 0.128563127 | 0.24324423 | 0.52853516 | 0.59712795 | 0.81642715 |
| EPS8L1 | 59.7991052 | 0.985912671 | 0.33948004 | 2.90418448 | 0.00368211 | 0.04690285 |
| PPP1R12C | 60.6242168 | 0.120913799 | 0.29682635 | 0.40735534 | 0.68374702 | 0.86486369 |
| PPP6R1 | 52.0434945 | -0.245343765 | 0.29134931 | -0.8420949 | 0.39973484 | 0.67448652 |
| HSPBP1 | 27.4291273 | -0.726130934 | 0.29023861 | -2.5018412 | 0.01235493 | 0.10132525 |
| TMEM238 | 32.7209259 | -0.650277213 | 0.60404756 | -1.0765331 | 0.28168889 | 0.56931659 |
| RPL28 | 513.680055 | -0.945485044 | 0.44075024 | -2.1451719 | 0.0319391 | 0.17502555 |
| ISOC2 | 42.1823007 | -1.198693401 | 0.51875917 | -2.3106934 | 0.02084979 | 0.13840623 |
| NAT14 | 38.1639562 | 0.181753336 | 0.51045124 | 0.35606405 | 0.7217926 | 0.88235193 |
| ZNF579 | 26.3287856 | -0.223767137 | 0.34734247 | -0.6442262 | 0.51942874 | 0.76385303 |
| U2AF2 | 48.5016545 | -0.04380591 | 0.26251639 | -0.1668692 | 0.86747294 | 0.95090501 |
| EPN1 | 86.5067531 | 0.038973808 | 0.26838596 | 0.14521552 | 0.88454071 | 0.95761929 |
| ZNF787 | 38.045045 | -0.633550388 | 0.35950031 | -1.7623083 | 0.07801722 | 0.28954938 |
| ZNF444 | 42.9442874 | 0.127253549 | 0.29634443 | 0.42941097 | 0.66762418 | 0.85887349 |
| ZSCAN5A | 107.065104 | -0.028582745 | 0.28573977 | -0.1000307 | 0.92031996 | 0.97180033 |
| ZNF667 | 38.4527255 | -0.28881772 | 0.37608329 | -0.7679621 | 0.4425097 | 0.70510851 |
| ZNF667-AS1 | 45.6255004 | 0.409211019 | 0.39487779 | 1.03629787 | 0.3000632 | 0.58707218 |
| ZNF471 | 45.7896522 | 0.505413577 | 0.39840156 | 1.26860341 | 0.20458255 | 0.48320272 |
| ZNF264 | 66.3436747 | 0.090879967 | 0.18516432 | 0.49080711 | 0.62356288 | 0.83030289 |
| AC005261.1 | 92.6405069 | -0.074877024 | 0.26739257 | -0.2800266 | 0.77945712 | 0.90987605 |
| AC004076.1 | 52.0177235 | 0.652821515 | 0.2629396 | 2.4827813 | 0.01303611 | 0.10449615 |
| ZNF549 | 32.2778095 | -0.108537191 | 0.28139421 | -0.3857122 | 0.6997098 | 0.87181583 |
| ZNF586 | 167.47482 | 0.303515031 | 0.17743726 | 1.71054843 | 0.08716451 | 0.30713468 |
| ZNF552 | 39.4163893 | 0.374178316 | 0.29612825 | 1.26356846 | 0.20638496 | 0.48621411 |
| ZNF814 | 119.167248 | -0.201481271 | 0.25428985 | -0.7923292 | 0.42816875 | 0.69444434 |
| ZNF587 | 62.2384828 | 0.608824013 | 0.25084667 | 2.42707635 | 0.01522105 | 0.11462458 |
| ZNF606 | 34.1000564 | 0.515553283 | 0.31593274 | 1.63184504 | 0.10271213 | 0.33431177 |
| ZNF135 | 46.8293282 | 0.293330744 | 0.35815037 | 0.81901561 | 0.41277751 | 0.68408231 |
| ZSCAN18 | 104.410939 | -0.201422046 | 0.35413028 | -0.5687795 | 0.56950578 | 0.79749848 |

| | | | | | | |
|---|---|---|---|---|---|---|
| ZNF329 | 38.6665732 | 0.310509891 | 0.31777187 | 0.97714721 | 0.32849628 | 0.61394673 |
| ZNF274 | 95.8101426 | 0.001724207 | 0.37142975 | 0.00464208 | 0.99629617 | 0.99921694 |
| RPS5 | 255.425514 | -0.943951147 | 0.51796742 | -1.8224141 | 0.06839217 | 0.26728084 |
| TRIM28 | 86.1324718 | -0.551302548 | 0.39999211 | -1.3782836 | 0.16811575 | 0.43712834 |
| CHMP2A | 70.6259986 | -0.64029323 | 0.38345186 | -1.6698139 | 0.0949562 | 0.32050226 |
| UBE2M | 49.4309132 | -0.781045949 | 0.34033198 | -2.2949531 | 0.02173581 | 0.14167784 |
| MZF1-AS1 | 25.918355 | 0.587527248 | 0.43552524 | 1.34900849 | 0.17733424 | 0.44901805 |
| MZF1 | 91.2315333 | -0.03437679 | 0.22416791 | -0.1533529 | 0.87812002 | 0.95466137 |
| RPS4Y1 | 135.85729 | -0.696763804 | 2.42346332 | -0.2875075 | 0.77372378 | 0.90643453 |
| ZFY | 48.0595532 | 0.62798945 | 1.87467798 | 0.33498524 | 0.73763622 | 0.89124841 |
| LINC00278 | 405.930981 | 0.423066617 | 2.62317014 | 0.16128066 | 0.87187235 | 0.95193455 |
| TBL1Y | 68.5713483 | 0.698055642 | 1.56547799 | 0.44590575 | 0.65566533 | 0.85068646 |
| TTTY15 | 73.6199851 | 0.628480959 | 1.93915907 | 0.32409974 | 0.74586252 | 0.89352127 |
| USP9Y | 313.905035 | 0.954259285 | 2.27400249 | 0.41963863 | 0.67474947 | 0.86115505 |
| DDX3Y | 78.2322202 | 0.835982945 | 2.26181928 | 0.36960643 | 0.71167576 | 0.87748 |
| UTY | 499.010239 | 0.648400921 | 1.99348936 | 0.32525928 | 0.74498484 | 0.89323545 |
| NLGN4Y | 216.6342 | -0.64378393 | 1.97289205 | -0.3263148 | 0.74418617 | 0.89303519 |
| TTTY14 | 563.072967 | 0.838277257 | 1.09096411 | 0.76838207 | 0.44226023 | 0.70510851 |
| KDM5D | 54.4140824 | 0.696239345 | 2.15584675 | 0.32295401 | 0.74673007 | 0.89352127 |
| TTTY10 | 34.9879718 | 0.808910178 | 1.47894658 | 0.54695023 | 0.58441292 | 0.80803859 |
| EIF1AY | 27.7465065 | -0.050015944 | 1.98299843 | -0.0252224 | 0.97987758 | 0.99557142 |
| IL17RA | 47.2390268 | -0.576797201 | 0.26320213 | -2.191461 | 0.02841845 | 0.16359395 |
| HDHD5 | 40.1109061 | -0.222649886 | 0.34666542 | -0.6422616 | 0.52070334 | 0.76418222 |
| CECR2 | 54.5606413 | 0.293154377 | 0.84597207 | 0.34652961 | 0.72894473 | 0.88653304 |
| ATP6V1E1 | 95.9050023 | -0.458215665 | 0.24071877 | -1.9035311 | 0.05697128 | 0.24093946 |
| BCL2L13 | 181.274907 | -0.020995809 | 0.13080006 | -0.1605183 | 0.87247277 | 0.95193455 |
| MICAL3 | 582.951424 | -0.205216935 | 0.27696021 | -0.7409618 | 0.4587166 | 0.71752865 |
| PEX26 | 72.1611883 | -0.039400175 | 0.22835871 | -0.1725363 | 0.8630159 | 0.94910925 |
| DGCR2 | 140.758403 | 0.124660359 | 0.2247333 | 0.55470355 | 0.57909745 | 0.80428802 |
| SLC25A1 | 55.0099914 | -0.075756027 | 0.37974576 | -0.1994914 | 0.84187834 | 0.94121763 |
| CLTCL1 | 43.1229191 | 0.071509532 | 0.23803077 | 0.30042138 | 0.76385575 | 0.90255018 |
| HIRA | 56.1018156 | -0.219378852 | 0.32376742 | -0.6775816 | 0.49803701 | 0.74986853 |
| MRPL40 | 47.8751764 | -0.967571249 | 0.59260708 | -1.6327366 | 0.10252441 | 0.33398714 |
| UFD1 | 34.9176825 | -0.296087363 | 0.37427408 | -0.7910977 | 0.42888698 | 0.69479256 |
| TXNRD2 | 46.3393739 | 0.210533268 | 0.24734901 | 0.85115871 | 0.39468119 | 0.66961586 |
| COMT | 162.025012 | -0.083343133 | 0.23685849 | -0.3518689 | 0.7249366 | 0.88404426 |
| TANGO2 | 32.1944919 | -0.211385112 | 0.27749273 | -0.7617681 | 0.44619841 | 0.70815763 |
| DGCR8 | 53.8483746 | -0.116569314 | 0.27333574 | -0.4264693 | 0.66976589 | 0.85924744 |
| RANBP1 | 42.3794302 | -0.004354127 | 0.47742642 | -0.00912 | 0.9927234 | 0.99829858 |
| ZDHHC8 | 39.4762509 | 0.670241893 | 0.43180252 | 1.55219542 | 0.12061547 | 0.36507762 |
| KLHL22 | 103.835516 | 0.471985008 | 0.35330382 | 1.3359182 | 0.18157601 | 0.45460445 |
| MED15 | 199.935305 | -0.001870612 | 0.27359694 | -0.0068371 | 0.99454482 | 0.99886097 |
| PI4KA | 211.631165 | -0.175625307 | 0.20400706 | -0.8608786 | 0.38930492 | 0.66513878 |
| SNAP29 | 47.4489604 | -0.202040843 | 0.26806262 | -0.7537076 | 0.4510248 | 0.71164003 |
| CRKL | 51.0239207 | 0.061293378 | 0.25870158 | 0.23692695 | 0.81271346 | 0.92667328 |
| LZTR1 | 27.3339994 | -0.094352478 | 0.30112366 | -0.3133346 | 0.75402643 | 0.89791387 |
| GGT2 | 228.733539 | -2.535129714 | 0.92378437 | -2.7442873 | 0.00606424 | 0.0657864 |
| HIC2 | 45.546181 | 0.446965309 | 0.22033274 | 2.02859232 | 0.04249983 | 0.20457582 |
| UBE2L3 | 145.539342 | -0.482185112 | 0.20964239 | -2.3000364 | 0.02144616 | 0.14061107 |
| SDF2L1 | 48.2724845 | -2.062939184 | 0.60506588 | -3.4094455 | 0.00065095 | 0.01564098 |
| PPIL2 | 84.166291 | -0.226893309 | 0.22774825 | -0.9962461 | 0.31913057 | 0.60583257 |
| YPEL1 | 36.4372976 | 1.27821915 | 0.42948335 | 2.97617858 | 0.00291865 | 0.03988265 |
| MAPK1 | 308.699625 | 0.013254078 | 0.21084673 | 0.06286119 | 0.94987704 | 0.98270285 |
| PPM1F | 40.9193844 | -0.525606052 | 0.45870673 | -1.1458433 | 0.25185998 | 0.53906677 |
| TOP3B | 31.0515231 | 0.442922163 | 0.28389382 | 1.5601684 | 0.11872009 | 0.3637939 |
| AC245060.5 | 55.3440221 | 0.790386141 | 0.39866224 | 1.98259596 | 0.04741258 | 0.21761624 |
| RAB36 | 40.6911004 | -0.260271506 | 0.31112886 | -0.8365393 | 0.40285159 | 0.67688395 |
| BCR | 197.219269 | -0.268355011 | 0.36360012 | -0.7380498 | 0.46048416 | 0.71913063 |
| PCAT14 | 31.3603221 | 1.674719867 | 0.91817011 | 1.82397558 | 0.06815576 | 0.26695445 |
| CHCHD10 | 160.545899 | -0.760224213 | 0.33210757 | -2.2890903 | 0.02207411 | 0.14258117 |
| SMARCB1 | 78.0671912 | -0.139519329 | 0.20164979 | -0.6918893 | 0.48900687 | 0.74185878 |
| DDT | 107.505 | -0.265724919 | 0.34952398 | -0.760248 | 0.44710634 | 0.70891772 |
| CABIN1 | 188.549017 | -0.264302723 | 0.24826894 | -1.0645823 | 0.28706499 | 0.57406774 |
| GUCD1 | 61.3944693 | -1.285826792 | 0.21359814 | -6.0198409 | 1.75E-09 | 9.29E-07 |
| SNRPD3 | 123.551658 | -0.041407893 | 0.33536745 | -0.1234702 | 0.90173476 | 0.9638158 |
| GGT1 | 92.3374134 | -0.250912042 | 0.38237113 | -0.6562003 | 0.51169526 | 0.75961818 |
| LRRC75B | 34.0176025 | -0.667792801 | 0.39280868 | -1.7000459 | 0.08912229 | 0.3108137 |
| SGSM1 | 70.7538688 | 0.29577022 | 0.59893563 | 0.49382639 | 0.6214288 | 0.82871592 |
| KIAA1671 | 267.498324 | -0.380387478 | 0.22067371 | -1.7237553 | 0.08475202 | 0.30275178 |
| LRP5L | 48.9852121 | -0.243259681 | 0.44734769 | -0.5437821 | 0.58659141 | 0.81023202 |
| GRK3 | 81.5236591 | 0.078587644 | 0.35914789 | 0.21881695 | 0.82679265 | 0.93339384 |
| HPS4 | 83.99794 | -0.051489417 | 0.30937226 | -0.1664319 | 0.86781706 | 0.95090501 |
| TPST2 | 64.9740825 | 0.813135835 | 0.50334842 | 1.61545325 | 0.10621258 | 0.34093088 |
| MIATNB | 67.9992533 | -0.036056309 | 0.3262225 | -0.1105267 | 0.91199164 | 0.96844354 |
| PITPNB | 222.628421 | -0.038172895 | 0.27127117 | -0.1407186 | 0.88809226 | 0.95907089 |
| TTC28 | 845.643982 | -0.398861397 | 0.42627113 | -0.9356988 | 0.34942827 | 0.63273599 |
| HSCB | 62.2669901 | -0.484305757 | 0.21510407 | -2.2514951 | 0.0243542 | 0.15138797 |
| CCDC117 | 33.865159 | 0.491024489 | 0.26638894 | 1.84326152 | 0.06529084 | 0.26048352 |
| XBP1 | 108.442901 | -1.00779407 | 0.41473535 | -2.4299691 | 0.01510011 | 0.11411159 |
| Z93930.2 | 102.355879 | -0.059330194 | 0.48966372 | -0.1211652 | 0.9035602 | 0.96509062 |
| ZNRF3 | 691.754073 | 0.343600047 | 0.42879758 | 0.8013106 | 0.42295186 | 0.69043214 |
| KREMEN1 | 70.2268774 | 0.079446747 | 0.35292422 | 0.22510993 | 0.82189375 | 0.93192711 |
| EMID1 | 40.1947671 | -0.472400804 | 1.10312715 | -0.4282379 | 0.66847796 | 0.85916414 |
| RHBDD3 | 30.3958554 | -0.886362753 | 0.3709232 | -2.3896126 | 0.01686615 | 0.12184079 |
| EWSR1 | 211.838454 | 0.151486195 | 0.16243244 | 0.9326105 | 0.35102111 | 0.63331896 |
| AP1B1 | 72.1002986 | -0.447671351 | 0.31032714 | -1.4425788 | 0.14913916 | 0.40945944 |
| THOC5 | 25.970047 | -0.013908322 | 0.32359534 | -0.0429806 | 0.965717 | 0.99011058 |
| NIPSNAP1 | 41.6863912 | 0.004882806 | 0.26437906 | 0.01846896 | 0.98526474 | 0.99668665 |
| NF2 | 91.5654314 | 0.061172384 | 0.21193771 | 0.28863379 | 0.77286164 | 0.9063714 |
| UQCR10 | 150.049619 | -0.674381942 | 0.4408703 | -1.5296606 | 0.12610075 | 0.37339076 |
| ASCC2 | 57.0745707 | -0.367825197 | 0.22462319 | -1.637521 | 0.10152165 | 0.33297252 |
| MTMR3 | 336.427403 | 0.437627831 | 0.16229193 | 2.69654706 | 0.00700625 | 0.07109414 |
| SF3A1 | 43.5308624 | -0.64115502 | 0.25566056 | -2.507837 | 0.01214727 | 0.1007368 |
| RNF215 | 39.497141 | -0.202762154 | 0.3519736 | -0.576072 | 0.56456648 | 0.79354106 |
| PES1 | 46.7634277 | -0.954827158 | 0.33552635 | -2.8457591 | 0.00443057 | 0.05345142 |
| TCN2 | 36.2621062 | -0.041854001 | 0.50473393 | -0.0829229 | 0.93391285 | 0.97768238 |
| DUSP18 | 40.2946306 | 0.592391752 | 0.32102024 | 1.84534081 | 0.06498798 | 0.26002617 |
| OSBP2 | 68.7914706 | 0.122086027 | 0.31288934 | 0.39018916 | 0.69639668 | 0.87063218 |
| MORC2 | 54.2476844 | -0.038388838 | 0.27504216 | -0.1395744 | 0.88899629 | 0.95908272 |
| SELENOM | 94.5468011 | -0.607141232 | 0.44739545 | -1.3570572 | 0.17476303 | 0.44629567 |
| RNF185 | 81.2284392 | 0.261473578 | 0.1658511 | 1.57655615 | 0.11489769 | 0.35651907 |
| LIMK2 | 201.708025 | 0.025386405 | 0.37584156 | 0.0675455 | 0.94614744 | 0.98139105 |
| PATZ1 | 41.3944397 | -0.105381079 | 0.37891992 | -0.2781091 | 0.78092861 | 0.91077326 |
| DRG1 | 98.0013177 | 0.091854593 | 0.21272618 | 0.4317973 | 0.66588874 | 0.85763969 |
| EIF4ENIF1 | 105.414032 | 0.312357618 | 0.23288062 | 1.34127788 | 0.17983025 | 0.451844 |
| SFI1 | 104.00298 | 0.49184677 | 0.28871134 | 1.70359355 | 0.08845705 | 0.30970853 |
| PISD | 212.293918 | 0.396423896 | 0.19344716 | 2.04926186 | 0.04043652 | 0.19973796 |
| PRR14L | 165.431762 | -0.034915029 | 0.17579135 | -0.1986163 | 0.8425629 | 0.94132995 |
| YWHAH | 36.9121284 | -0.629450242 | 0.35116321 | -1.792472 | 0.07305738 | 0.27892255 |
| RTCB | 90.6015319 | -0.246404762 | 0.18382211 | -1.3404523 | 0.18009836 | 0.45208652 |
| FBXO7 | 116.378388 | 0.313508005 | 0.22608448 | 1.38668519 | 0.16553778 | 0.43384207 |
| TIMP3 | 156.239627 | 1.102942872 | 0.46757242 | 2.35887066 | 0.01833064 | 0.12806015 |
| LARGE1 | 704.235471 | -0.142608004 | 0.27668788 | -0.5154111 | 0.60626578 | 0.82097646 |

| | | | | | | |
|---|---|---|---|---|---|---|
| HMGXB4 | 74.2148177 | -0.102812151 | 0.25706574 | -0.399945 | 0.68919704 | 0.86751885 |
| TOM1 | 67.7143381 | -0.755938181 | 0.23382891 | -3.2328688 | 0.00122554 | 0.02374618 |
| HMOX1 | 80.0687256 | -4.346386043 | 0.6613278 | -6.5722113 | 4.96E-11 | 4.75E-08 |
| APOL6 | 119.09762 | -1.681311639 | 0.7343927 | -2.2893905 | 0.02205668 | 0.1425647 |
| RBFOX2 | 519.309546 | -0.080343938 | 0.18741408 | -0.4286975 | 0.66814341 | 0.85905308 |
| APOL2 | 55.8992278 | -1.234690236 | 0.61444674 | -2.0094341 | 0.04449112 | 0.20973445 |
| APOL1 | 39.4548966 | -1.228849719 | 0.59690316 | -2.0587087 | 0.03952215 | 0.19707585 |
| MYH9 | 325.441279 | -0.127872359 | 0.28261399 | -0.4524629 | 0.65093551 | 0.84724399 |
| TXN2 | 46.833598 | -0.337709582 | 0.35757706 | -0.9444386 | 0.34494556 | 0.62973767 |
| EIF3D | 97.2178435 | -1.070814086 | 0.28567161 | -3.7484092 | 0.00017796 | 0.00704975 |
| IFT27 | 34.7285326 | 0.47582606 | 0.2615217 | 1.81945152 | 0.06884257 | 0.26850285 |
| TST | 49.5162933 | -0.826495818 | 0.3959459 | -2.0873958 | 0.03685237 | 0.19009632 |
| MPST | 75.3940191 | -0.77618901 | 0.32895814 | -2.3595373 | 0.01829774 | 0.12793727 |
| CDC42EP1 | 36.8413635 | -0.859426376 | 0.73769762 | -1.1650117 | 0.24401425 | 0.52978524 |
| GGA1 | 84.440166 | 0.211208285 | 0.18152339 | 1.16353208 | 0.24461369 | 0.53060565 |
| TRIOBP | 142.318347 | -0.410046857 | 0.17134132 | -2.393158 | 0.01670405 | 0.1212996 |
| H1F0 | 172.82134 | -1.020235697 | 0.64514672 | -1.581401 | 0.11378637 | 0.35388299 |
| ANKRD54 | 25.6242212 | 0.041782854 | 0.32853673 | 0.12717864 | 0.89879901 | 0.96269027 |
| EIF3L | 127.740897 | -0.248640135 | 0.24916233 | -0.9979042 | 0.31832581 | 0.60512859 |
| POLR2F | 48.3810137 | 0.342338018 | 0.29313727 | 1.16784202 | 0.2428705 | 0.52778176 |
| PICK1 | 39.5635552 | 0.03275885 | 0.29987006 | 0.10924348 | 0.91300937 | 0.96844354 |
| BAIAP2L2 | 37.8628973 | -0.22736501 | 0.32314748 | -0.7035952 | 0.48168491 | 0.73620566 |
| PLA2G6 | 28.1974084 | 0.869285134 | 0.42713025 | 2.03517579 | 0.04183319 | 0.20312121 |
| MAFF | 30.5000321 | -0.350929862 | 0.43044725 | -0.815268 | 0.41491893 | 0.68495052 |
| TMEM184B | 78.5256527 | 0.239817527 | 0.20815951 | 1.15208537 | 0.24928599 | 0.53636804 |
| CSNK1E | 36.9292904 | 0.67270349 | 0.35698866 | 1.88438338 | 0.05951314 | 0.24614695 |
| TPTEP2-CSNI | 261.473894 | 0.427694826 | 0.26189168 | 1.63309817 | 0.10244835 | 0.33398714 |
| DDX17 | 1246.4215 | 0.646678613 | 0.25027495 | 2.58387268 | 0.00976979 | 0.08848555 |
| FAM227A | 68.5641081 | 0.402930718 | 0.37019774 | 1.08842026 | 0.27640962 | 0.56334633 |
| CBY1 | 50.2921029 | -0.045960858 | 0.21813851 | -0.2106958 | 0.83312469 | 0.93607461 |
| TOMM22 | 24.855222 | -0.32207733 | 0.46004945 | -0.7000928 | 0.48386932 | 0.73727416 |
| JOSD1 | 64.8092312 | -0.036730748 | 0.27624827 | -0.1329628 | 0.89422279 | 0.96082558 |
| GTPBP1 | 51.2195383 | -0.135975881 | 0.41970194 | -0.323982 | 0.74595165 | 0.89352127 |
| SUN2 | 55.8173104 | -0.35050313 | 0.25977095 | -1.3492776 | 0.17724781 | 0.44893092 |
| AL021707.1 | 29.7198025 | -0.309403848 | 0.29838993 | -1.0369112 | 0.29977726 | 0.58681504 |
| CBX6 | 116.950831 | -0.289723956 | 0.43620777 | -0.664188 | 0.50657001 | 0.75584222 |
| CBX7 | 68.2005979 | 0.354473743 | 0.38028943 | 0.93211568 | 0.35127675 | 0.63335467 |
| PDGFB | 29.8579024 | 0.304162957 | 0.64586171 | 0.47094131 | 0.63768265 | 0.83912075 |
| RPL3 | 446.582417 | 0.029676575 | 0.45635077 | 0.06503018 | 0.94814997 | 0.98251066 |
| SYNGR1 | 119.674019 | 0.447901213 | 0.32136462 | 1.39374776 | 0.16339379 | 0.43045905 |
| TAB1 | 52.5696768 | 0.099274688 | 0.27972544 | 0.35490047 | 0.72266416 | 0.88250892 |
| MIEF1 | 30.9219831 | -0.656475885 | 0.31329884 | -2.0953665 | 0.03613843 | 0.18804981 |
| ATF4 | 103.943897 | -0.695876209 | 0.31400842 | -2.2161069 | 0.02668418 | 0.15827108 |
| RPS19BP1 | 86.0455295 | -0.617977064 | 0.43811834 | -1.4105254 | 0.15838459 | 0.42403514 |
| TNRC6B | 1059.58348 | 0.385451538 | 0.18061103 | 2.13415283 | 0.03283027 | 0.17754227 |
| MKL1 | 349.118097 | 0.315750572 | 0.15754311 | 2.00421697 | 0.04504682 | 0.21106218 |
| SLC25A17 | 79.8427659 | 0.07580089 | 0.17746643 | 0.42712806 | 0.66928606 | 0.85924744 |
| ST13 | 189.241825 | -0.144998567 | 0.27303602 | -0.5310602 | 0.59537706 | 0.81525977 |
| XPNPEP3 | 137.870761 | 0.358366866 | 0.18444497 | 1.94294736 | 0.05202252 | 0.22857606 |
| RBX1 | 107.646163 | 0.034959481 | 0.20409934 | 0.17128659 | 0.86399842 | 0.94954576 |
| EP300 | 290.063105 | 0.285618382 | 0.12936615 | 2.20782934 | 0.02725617 | 0.15949106 |
| L3MBTL2 | 29.9487192 | 0.01847224 | 0.31817203 | 0.0580574 | 0.95370291 | 0.98368305 |
| RANGAP1 | 56.8714841 | -0.465140865 | 0.41169727 | -1.1298129 | 0.25855509 | 0.54552846 |
| ZC3H7B | 138.804801 | -0.014079523 | 0.30355978 | -0.0463814 | 0.96300627 | 0.98817751 |
| TEF | 35.7355117 | 0.922149842 | 0.54978988 | 1.67727687 | 0.09348835 | 0.31835715 |
| TOB2 | 77.8619743 | -0.131091674 | 0.19306286 | -0.6790103 | 0.49713133 | 0.74886319 |
| ACO2 | 124.826299 | -0.19883941 | 0.18014437 | -1.1037781 | 0.2696894 | 0.55681979 |
| POLR3H | 30.1408612 | -0.695799373 | 0.29345098 | -2.3710923 | 0.0177356 | 0.12575079 |
| DESI1 | 63.6013597 | -0.680011701 | 0.31303888 | -2.1722915 | 0.02983368 | 0.16887967 |
| XRCC6 | 197.951527 | -0.198073119 | 0.2844513 | -0.696334 | 0.48621967 | 0.73975512 |
| SNU13 | 161.260502 | -0.859239374 | 0.3321845 | -2.586633 | 0.00969188 | 0.08799856 |
| SREBF2 | 353.207461 | 0.583821614 | 0.34516908 | 1.69140762 | 0.09075898 | 0.3142614 |
| WBP2NL | 63.0594311 | 0.043043072 | 0.48449126 | 0.08884179 | 0.92920764 | 0.9754389 |
| SMDT1 | 81.0410894 | -0.010779296 | 0.46763371 | -0.0230507 | 0.98160981 | 0.9959581 |
| NDUFA6 | 53.4898522 | -0.59028835 | 0.33667271 | -1.7533003 | 0.07955048 | 0.29274747 |
| TCF20 | 254.257632 | 0.044862021 | 0.23329072 | 0.19230092 | 0.8475065 | 0.94323402 |
| RRP7A | 48.5108602 | -0.570927086 | 0.37034645 | -1.5416027 | 0.12317017 | 0.3687022 |
| POLDIP3 | 78.1464148 | -0.052620066 | 0.21610576 | -0.2434922 | 0.80762413 | 0.92361885 |
| CYB5R3 | 167.628213 | -1.128343853 | 0.4211477 | -2.6792117 | 0.00737957 | 0.07378801 |
| A4GALT | 156.84557 | -1.68696733 | 0.35109015 | -4.8049406 | 1.55E-06 | 0.0002355 |
| ARFGAP3 | 168.487411 | -0.817069509 | 0.25282502 | -3.2317589 | 0.00123031 | 0.02374618 |
| PACSIN2 | 335.543766 | 0.142069736 | 0.20500344 | 0.69301148 | 0.48830235 | 0.74151049 |
| TSPO | 123.008546 | -1.464785226 | 0.49490335 | -2.95974 | 0.00307899 | 0.04154042 |
| EFCAB6 | 220.954703 | 0.119618436 | 0.34216584 | 0.34959198 | 0.72664493 | 0.88522597 |
| NUP50-DT | 98.4727694 | -0.704746587 | 0.34191388 | -2.0611815 | 0.03928572 | 0.19661335 |
| NUP50 | 61.6607904 | -0.123729783 | 0.22094095 | -0.5600129 | 0.57547064 | 0.80169186 |
| KIAA0930 | 57.5589372 | -1.382407678 | 0.42770811 | -3.2321288 | 0.00122872 | 0.02374618 |
| FAM118A | 40.3132548 | 0.814592136 | 0.42944962 | 1.89682816 | 0.05785062 | 0.24304873 |
| ATXN10 | 564.366464 | 0.437720291 | 0.1773775 | 2.46773289 | 0.01359717 | 0.10755093 |
| WNT7B | 37.7054013 | 1.000633095 | 1.159324 | 0.86311773 | 0.38807274 | 0.66428677 |
| MIRLET7BHG | 182.462787 | -0.043221927 | 0.25574407 | -0.1690046 | 0.86579302 | 0.95058578 |
| PPARA | 160.014128 | -0.210396651 | 0.20612 | -1.0207484 | 0.30737368 | 0.59456464 |
| TTC38 | 72.6579129 | -0.572096476 | 0.4352629 | -1.3143699 | 0.18872174 | 0.46472122 |
| TRMU | 53.3729494 | -0.335707394 | 0.20936956 | -1.6034202 | 0.10884191 | 0.34534502 |
| CELSR1 | 225.575223 | 0.483515915 | 0.39559235 | 1.22225799 | 0.22161008 | 0.50339174 |
| GRAMD4 | 61.5276136 | -0.291616866 | 0.27519821 | -1.0596612 | 0.28929876 | 0.5758921 |
| CERK | 46.1677347 | 0.028229518 | 0.28893233 | 0.09770287 | 0.92216824 | 0.97248656 |
| TBC1D22A | 410.588544 | -0.265885275 | 0.24704241 | -1.0762738 | 0.28180482 | 0.56931659 |
| BRD1 | 99.5757917 | 0.057646271 | 0.27890785 | 0.20668572 | 0.83625531 | 0.93756847 |
| ZBED4 | 66.7248669 | -0.292541145 | 0.41807724 | -0.6997299 | 0.484096 | 0.73727416 |
| PIM3 | 127.623482 | -0.522339215 | 0.39416849 | -1.3251673 | 0.18511565 | 0.45922178 |
| TRABD | 41.5718669 | -0.47700401 | 0.31012595 | -1.5380977 | 0.12402473 | 0.37033446 |
| SELENOO | 80.405314 | -0.324963557 | 0.22608311 | -1.4373633 | 0.1506148 | 0.41173164 |
| PLXNB2 | 336.627382 | -0.240765898 | 0.30041486 | -0.8014447 | 0.42287424 | 0.69043214 |
| DENND6B | 56.6449244 | -0.10879415 | 0.43021903 | -0.2528808 | 0.8003603 | 0.92042259 |
| PPP6R2 | 357.181458 | 0.199970601 | 0.18053349 | 1.10766486 | 0.26800658 | 0.55507785 |
| SBF1 | 27.1895233 | 0.042737069 | 0.29580625 | 0.14447656 | 0.88512416 | 0.95793953 |
| LMF2 | 40.2077186 | -0.748115274 | 0.38209269 | -1.9579419 | 0.05023684 | 0.2251146 |
| RABL2B | 61.1644212 | 0.154683734 | 0.2325563 | 0.6651453 | 0.50595756 | 0.75528806 |
| CU633967.1 | 48.5892436 | 0.702269151 | 0.4470228 | 1.57099181 | 0.11618455 | 0.35935803 |
| FP671120.1 | 853.865247 | -0.779065115 | 0.39641482 | -1.9652775 | 0.04938213 | 0.22344423 |
| FP236383.1 | 489.157491 | -0.741028661 | 0.48372826 | -1.531911 | 0.12554439 | 0.37248786 |
| CR381653.1 | 65.4974238 | 0.131802822 | 0.49924501 | 0.26400429 | 0.79177662 | 0.91648412 |
| HSPA13 | 30.1745696 | 0.115708936 | 0.35079486 | 0.32984786 | 0.74151492 | 0.89253455 |
| NRIP1 | 286.735004 | -0.402065498 | 0.29845889 | -1.3471386 | 0.17793559 | 0.44948443 |
| USP25 | 579.548012 | 0.706010244 | 0.3097852 | 2.27903158 | 0.02266519 | 0.14468046 |
| MIR99AHG | 723.254106 | 1.194123748 | 0.42663782 | 2.79891675 | 0.00512744 | 0.05910434 |
| CXADR | 158.544171 | -0.06920096 | 0.2603875 | -0.2657615 | 0.79042293 | 0.91564412 |
| BTG3 | 54.929253 | -0.816147022 | 0.31394334 | -2.5996634 | 0.00933152 | 0.08619736 |
| C21orf91 | 30.6947864 | -0.102812868 | 0.31625051 | -0.3250995 | 0.7451058 | 0.89323545 |
| CHODL | 68.2196684 | -0.054950735 | 0.62379954 | -0.0880904 | 0.92980485 | 0.97545894 |
| MRPL39 | 34.3927823 | 0.280488218 | 0.28443816 | 0.98611319 | 0.32407757 | 0.6107297 |
| ATP5PF | 114.557259 | -0.409802664 | 0.26497021 | -1.546599 | 0.12195998 | 0.36668381 |

| | | | | | | |
|---|---|---|---|---|---|---|
| GABPA | 56.3098633 | 0.47702203 | 0.19540377 | 2.44121205 | 0.01463806 | 0.11217436 |
| APP | 1451.78866 | 0.272966979 | 0.228681 | 1.19365834 | 0.23261164 | 0.51579248 |
| ADAMTS1 | 654.52106 | -1.27869546 | 0.58029352 | -2.2035322 | 0.02755726 | 0.16066402 |
| LINC00113 | 40.5666297 | 3.290529011 | 0.88626308 | 3.71281292 | 0.00020497 | 0.00758576 |
| AF165147.1 | 59.4094846 | -0.312434425 | 0.44218705 | -0.7065662 | 0.4798361 | 0.73503413 |
| LTN1 | 121.830825 | 0.324919475 | 0.17741073 | 1.83145337 | 0.0670329 | 0.26380779 |
| USP16 | 87.9770826 | -0.041938731 | 0.20200745 | -0.2076098 | 0.83553363 | 0.93718696 |
| CCT8 | 72.5359763 | -0.305858236 | 0.24938782 | -1.2264362 | 0.22003459 | 0.50159865 |
| MAP3K7CL | 36.1540032 | -0.398614319 | 0.46362545 | -0.8597766 | 0.38991219 | 0.66541402 |
| BACH1 | 478.600617 | -0.969985687 | 0.41222386 | -2.3530557 | 0.01861985 | 0.12915243 |
| CLDN8 | 21.5952631 | 1.046288594 | 0.78722223 | 1.32908923 | 0.18381853 | 0.45721925 |
| SOD1 | 166.542402 | -0.411764669 | 0.4331923 | -0.9505355 | 0.34184022 | 0.6270562 |
| SCAF4 | 133.072303 | 0.180373107 | 0.20263041 | 0.89015814 | 0.37338098 | 0.65236869 |
| HUNK | 245.972464 | 1.63552856 | 0.56484323 | 2.89554425 | 0.00378502 | 0.047769 |
| URB1 | 66.2586081 | -0.707647927 | 0.37609858 | -1.881549 | 0.05989727 | 0.24679372 |
| EVA1C | 390.64979 | -0.842217041 | 0.61595794 | -1.3673288 | 0.17152227 | 0.44214526 |
| SYNJ1 | 88.5457211 | 0.435249156 | 0.23024345 | 1.89038668 | 0.05870626 | 0.24449882 |
| PAXBP1 | 217.429122 | 0.372381183 | 0.15561843 | 2.39291178 | 0.01671526 | 0.1212996 |
| C21orf62-AS | 77.9188946 | 0.874695869 | 0.53230654 | 1.64321835 | 0.10033776 | 0.33096298 |
| IFNAR1 | 133.293613 | -0.317033877 | 0.24609011 | -1.2882837 | 0.19764723 | 0.47630586 |
| IFNGR2 | 203.135903 | -0.302534982 | 0.25262953 | -1.197544 | 0.23109458 | 0.51432503 |
| TMEM50B | 119.739953 | 0.570587113 | 0.34318025 | 1.66264555 | 0.09638341 | 0.32383608 |
| GART | 86.9764986 | -0.320914119 | 0.26179438 | -1.2258251 | 0.22026451 | 0.50176375 |
| SON | 416.316948 | -0.010447135 | 0.15003823 | -0.0696298 | 0.9444883 | 0.98115679 |
| AP000311.1 | 77.4877662 | -0.357864211 | 0.54560191 | -0.6559072 | 0.51188384 | 0.75961818 |
| ITSN1 | 262.207727 | 0.285411527 | 0.27376003 | 1.04256098 | 0.29715167 | 0.58520063 |
| LINC00649 | 30.8924642 | 1.136111828 | 0.67583588 | 1.68104693 | 0.0927538 | 0.31720408 |
| MRPS6 | 485.459122 | 2.393839058 | 0.71369512 | 3.35414798 | 0.0007961 | 0.01778667 |
| LINC00310 | 39.8777684 | 2.513014023 | 0.6349157 | 3.95802783 | 7.56E-05 | 0.00394884 |
| AP000317.1 | 56.1244973 | 1.740207129 | 0.50661787 | 3.43495018 | 0.00059266 | 0.01482276 |
| RCAN1 | 71.5153194 | 0.98123083 | 0.46416291 | 2.11397941 | 0.03451703 | 0.18317928 |
| CLIC6 | 51.8148634 | -1.514519836 | 0.72303624 | -2.0946666 | 0.03620064 | 0.18815299 |
| RUNX1 | 244.724896 | -1.495058429 | 0.3827245 | -3.9063567 | 9.37E-05 | 0.00448767 |
| SETD4 | 50.6794634 | 0.24114738 | 0.24708011 | 0.97598864 | 0.3290701 | 0.61429707 |
| CBR1 | 63.6389958 | -1.106675588 | 0.27243549 | -4.0621564 | 4.86E-05 | 0.00294775 |
| DOPEY2 | 63.8331068 | -0.356942868 | 0.36772987 | -0.970666 | 0.33171465 | 0.61686947 |
| MORC3 | 181.091068 | 0.085219055 | 0.17725914 | 0.48075972 | 0.63068728 | 0.83467166 |
| CLDN14 | 111.275014 | 0.178008833 | 0.68955251 | 0.25815124 | 0.79629019 | 0.91855295 |
| SIM2 | 361.675629 | 0.772028316 | 1.230242 | 0.62754183 | 0.53030412 | 0.77106606 |
| HLCS | 314.802015 | 0.031258481 | 0.3036178 | 0.10295339 | 0.91799997 | 0.97057281 |
| PIGP | 25.1995296 | 0.460477882 | 0.38646896 | 1.19150031 | 0.23345723 | 0.51622503 |
| TTC3 | 478.391297 | 0.458532601 | 0.20519101 | 2.23466224 | 0.02543953 | 0.15501607 |
| VPS26C | 81.034811 | -0.171780099 | 0.19041376 | -0.9021412 | 0.36698187 | 0.6466739 |
| DYRK1A | 658.521565 | 0.42569207 | 0.13705044 | 3.1060978 | 0.00189574 | 0.03090405 |
| KCNJ15 | 587.902353 | -0.514507517 | 0.49845985 | -1.0321945 | 0.30198101 | 0.58963159 |
| ETS2 | 222.59247 | -0.487836512 | 0.4520976 | -1.0790513 | 0.28056485 | 0.56841775 |
| BRWD1 | 528.297452 | 0.316015457 | 0.11608939 | 2.72217349 | 0.00648541 | 0.06817157 |
| HMGN1 | 219.935592 | -0.097430567 | 0.22861461 | -0.4261782 | 0.66997799 | 0.85924744 |
| WRB | 56.378624 | 0.4904083 | 0.19996993 | 2.45241027 | 0.01419028 | 0.1105111 |
| PCP4 | 66.1668509 | 0.85601436 | 1.25438066 | 0.68241993 | 0.49497346 | 0.74702234 |
| BACE2 | 281.768737 | -1.011019486 | 0.37353802 | -2.7066039 | 0.00679753 | 0.07001456 |
| MX1 | 191.062352 | -1.762851392 | 1.461816 | -1.2059325 | 0.22784353 | 0.51052939 |
| TMPRSS2 | 96.4455575 | 0.727663436 | 0.62884696 | 1.15713914 | 0.24721552 | 0.5338317 |
| RIPK4 | 53.1179557 | -0.297149935 | 0.3647882 | -0.8145821 | 0.41531156 | 0.68495052 |
| PRDM15 | 24.5398459 | -0.468504061 | 0.43952484 | -1.0659331 | 0.2864539 | 0.57346076 |
| C2CD2 | 68.3941435 | -0.273370555 | 0.23311224 | -1.1726993 | 0.24091642 | 0.52533525 |
| ZBTB21 | 46.2045176 | -0.248223162 | 0.34539012 | -0.7186748 | 0.47234133 | 0.72983472 |
| ABCG1 | 260.280515 | 0.297651396 | 1.00004146 | 0.29763906 | 0.76597866 | 0.90371395 |
| RSPH1 | 44.9243794 | -0.398697085 | 0.40974301 | -0.9730418 | 0.33053252 | 0.61562726 |
| SLC37A1 | 93.9884509 | -0.338910157 | 0.24376907 | -1.3902919 | 0.16444026 | 0.43262536 |
| LINC01671 | 27.7010716 | 0.911195015 | 0.54353545 | 1.67642242 | 0.09365548 | 0.31857635 |
| PDE9A | 330.715702 | 1.509047927 | 0.36191625 | 4.16960536 | 3.05E-05 | 0.00211798 |
| WDR4 | 30.0580344 | -0.8956653 | 0.55017342 | -1.627969 | 0.10353145 | 0.33583603 |
| NDUFV3 | 102.952715 | -0.036390243 | 0.27486679 | -0.1323923 | 0.89467402 | 0.96098704 |
| PKNOX1 | 109.567509 | 0.205860892 | 0.30873096 | 0.66679704 | 0.50490178 | 0.75524917 |
| RRP1B | 64.3520387 | -0.022751022 | 0.26743108 | -0.0850725 | 0.93220378 | 0.97701675 |
| PDXK | 555.886474 | 1.580943276 | 0.39261564 | 4.02669455 | 5.66E-05 | 0.00324416 |
| CSTB | 233.780968 | -0.627551704 | 0.48426462 | -1.2958859 | 0.19501479 | 0.47352261 |
| AGPAT3 | 177.218864 | -0.297614507 | 0.16809346 | -1.77053 | 0.07663889 | 0.2861109 |
| TRAPPC10 | 129.222104 | 0.172163288 | 0.17612269 | 0.97751908 | 0.32831224 | 0.61394673 |
| PFKL | 135.681635 | -0.03765548 | 0.26607485 | -0.1415221 | 0.88745748 | 0.95903278 |
| C21orf2 | 35.6212601 | -0.071767348 | 0.38844885 | -0.1847537 | 0.85342225 | 0.9451971 |
| UBE2G2 | 159.157536 | -0.172927205 | 0.23984665 | -0.7209907 | 0.47091523 | 0.72897495 |
| SUMO3 | 70.1511012 | -0.143502275 | 0.371817 | -0.3859487 | 0.69953469 | 0.87181583 |
| PTTG1IP | 291.702898 | -0.896034344 | 0.37391604 | -2.3963517 | 0.01655919 | 0.1208077 |
| FAM207A | 31.3184829 | -0.414201196 | 0.37653123 | -1.1000447 | 0.27131263 | 0.55841559 |
| ADARB1 | 145.16179 | -1.144814131 | 0.68501597 | -1.6712226 | 0.09467773 | 0.3204651 |
| POFUT2 | 42.293724 | 0.300725687 | 0.21736766 | 1.38348861 | 0.1665151 | 0.43485499 |
| COL18A1 | 397.043151 | 0.271858606 | 0.24833253 | 1.0947362 | 0.27363222 | 0.56099915 |
| COL6A1 | 53.5490166 | 0.011797069 | 0.58791458 | 0.02006596 | 0.98399076 | 0.99657935 |
| COL6A2 | 60.9355472 | 0.072319796 | 1.26903917 | 0.05698784 | 0.95455488 | 0.98390867 |
| LSS | 52.8123196 | 0.4548246 | 0.46408021 | 0.98005601 | 0.32705847 | 0.61330512 |
| MCM3AP-AS | 38.2194706 | -0.028648034 | 0.29143458 | -0.0983001 | 0.92169404 | 0.97248656 |
| MCM3AP | 119.251163 | 0.316381378 | 0.18888099 | 1.67503031 | 0.09392829 | 0.3190564 |
| PCNT | 59.3229666 | 0.474236112 | 0.24063854 | 1.97074045 | 0.04875357 | 0.22143692 |
| DIP2A | 152.328819 | 0.170453357 | 0.15992563 | 1.06582887 | 0.28650101 | 0.57346076 |
| PRMT2 | 198.263055 | 0.085689377 | 0.26103295 | 0.32827035 | 0.74270726 | 0.89253455 |
| AC011043.1 | 55.9380946 | 1.770519423 | 0.33859196 | 5.22906516 | 1.70E-07 | 4.53E-05 |
| AL592183.1 | 146.293065 | -0.251212957 | 0.44078824 | -0.5699176 | 0.56873361 | 0.79717578 |

**Thick ascending limbs**

*This means that positive fold changes imply higher mean value in controls than in AKI and vice versa.

| | baseMean | log2 fold change control/AKI* | lfcSE | stat | pvalue | padj |
|---|---|---|---|---|---|---|
| AL627309.1 | 46.4832515 | -0.166010539 | 0.54028403 | -0.3072653 | 0.75864144 | 0.88316847 |
| AL669831.5 | 439.99619 | 0.213245717 | 0.20945997 | 1.01807382 | 0.30864287 | 0.53250571 |
| SAMD11 | 43.3508878 | -0.668333656 | 0.82592152 | -0.8091975 | 0.41840153 | 0.63753323 |
| NOC2L | 246.032503 | -0.113752371 | 0.32796017 | -0.3468481 | 0.72870542 | 0.86676901 |
| KLHL17 | 42.5399791 | 0.106762229 | 0.38387002 | 0.27812078 | 0.78091965 | 0.89539748 |
| HES4 | 107.829312 | 0.638309001 | 0.48499631 | 1.31611106 | 0.18813677 | 0.39384317 |
| ISG15 | 1233.31999 | 0.182550798 | 1.21438393 | 0.15032379 | 0.88050916 | 0.94639771 |
| AGRN | 623.565741 | -0.477175018 | 0.4561023 | -1.0462017 | 0.2954679 | 0.51851143 |
| C1orf159 | 358.43042 | -0.169125882 | 0.26171044 | -0.6462328 | 0.51812857 | 0.71905445 |
| LINC01342 | 92.9066276 | -0.531589208 | 0.50110631 | -1.0608312 | 0.28876662 | 0.51109718 |
| AL390719.2 | 226.481438 | -0.302422039 | 0.41786546 | -0.7237306 | 0.46923112 | 0.68108245 |
| TTLL10 | 46.6818208 | -0.543174189 | 0.50165772 | -1.0827586 | 0.2789156 | 0.50104668 |
| SDF4 | 373.668797 | -0.408921707 | 0.2124461 | -1.9248257 | 0.05425117 | 0.1756637 |
| UBE2J2 | 376.948358 | -0.321951501 | 0.18189932 | -1.7699434 | 0.07673657 | 0.22097523 |
| ACAP3 | 170.899697 | 0.617274327 | 0.26453566 | 2.33342574 | 0.01962581 | 0.08811603 |
| INTS11 | 425.128513 | 0.39401595 | 0.19994477 | 1.97062391 | 0.04876691 | 0.16328571 |
| CPTP | 67.4225987 | -0.237320882 | 0.36682338 | -0.6469623 | 0.51765637 | 0.71870117 |
| DVL1 | 146.978285 | -0.136633261 | 0.26601528 | -0.5136294 | 0.60751114 | 0.78334778 |
| AURKAIP1 | 378.462361 | -0.867899529 | 0.4341631 | -1.9990173 | 0.04560649 | 0.15600761 |
| CCNL2 | 1074.37096 | -0.255255657 | 0.21547122 | -1.1846392 | 0.23616013 | 0.45391953 |
| MRPL20 | 464.634064 | -0.425621016 | 0.22899134 | -1.8586773 | 0.06307289 | 0.19456557 |
| VWA1 | 143.614962 | -2.268960439 | 0.6733421 | -3.3696994 | 0.0007525 | 0.00807458 |
| ATAD3C | 93.6429029 | -0.717754752 | 0.45163935 | -1.589221 | 0.1120105 | 0.28295022 |
| ATAD3B | 281.766483 | 0.304755593 | 0.23326887 | 1.30645634 | 0.19139742 | 0.39809311 |
| ATAD3A | 140.836864 | 0.179968377 | 0.26439287 | 0.68068543 | 0.49607056 | 0.70141255 |
| SSU72 | 472.675214 | -0.496656498 | 0.1849126 | -2.6858986 | 0.0072335 | 0.04378888 |
| MIB2 | 194.819587 | 0.429805443 | 0.26611783 | 1.61509448 | 0.10629024 | 0.27352187 |
| CDK11B | 263.712491 | -0.343066797 | 0.27453539 | -1.2496269 | 0.21143588 | 0.42405961 |
| FO704657.1 | 41.2857048 | 0.076722084 | 0.53512382 | 0.14337258 | 0.88599593 | 0.94846586 |
| SLC35E2B | 407.982983 | 0.013733991 | 0.3362897 | 0.04083976 | 0.96742364 | 0.98528676 |
| CDK11A | 155.127438 | -0.311707562 | 0.35076912 | -0.8886403 | 0.37419644 | 0.59780424 |
| SLC35E2A | 423.599377 | 0.241161171 | 0.26561998 | 0.90791804 | 0.36392153 | 0.58931463 |
| NADK | 324.730353 | 0.119630299 | 0.17053995 | 0.70147963 | 0.48300374 | 0.69127369 |
| GNB1 | 1981.08381 | -0.659828765 | 0.16936967 | -3.8957905 | 9.79E-05 | 0.00165114 |
| PRKCZ | 664.806574 | -0.224578826 | 0.30531377 | -0.7355673 | 0.46199407 | 0.67527715 |
| FAAP20 | 265.69668 | -0.378151846 | 0.28570754 | -1.3235627 | 0.18564831 | 0.3913547 |
| SKI | 709.808908 | -0.230303529 | 0.28443466 | -0.8096887 | 0.41811912 | 0.63753323 |
| MORN1 | 337.484844 | 0.121974426 | 0.23593204 | 0.51698966 | 0.60516338 | 0.78161758 |
| RER1 | 536.393505 | -0.579774742 | 0.28074637 | -2.0651193 | 0.03891172 | 0.14012673 |
| PEX10 | 137.27454 | -0.954855588 | 0.28201831 | -3.3857929 | 0.00070973 | 0.00772192 |
| PANK4 | 84.762803 | 0.693022108 | 0.24496968 | 2.82901173 | 0.0046692 | 0.03175016 |
| TNFRSF14 | 152.281098 | -0.392499265 | 0.51620007 | -0.7603627 | 0.44703783 | 0.66302748 |
| FAM213B | 66.6313488 | -1.347949143 | 0.43489281 | -3.0994974 | 0.00193849 | 0.01672472 |
| LINC00982 | 1140.99666 | 2.182390215 | 0.73767124 | 2.95848624 | 0.00309154 | 0.02365942 |
| PRDM16 | 3088.24604 | 0.774064845 | 0.37908384 | 2.04193573 | 0.04115791 | 0.14516817 |
| ARHGEF16 | 265.815045 | -0.40059643 | 0.26329903 | -1.5214504 | 0.12814684 | 0.30884182 |
| TPRG1L | 104.351169 | -0.809107769 | 0.32384265 | -2.4984596 | 0.01247344 | 0.06456176 |
| WRAP73 | 125.658482 | 0.099453259 | 0.26426221 | 0.37634309 | 0.70666185 | 0.85221367 |
| LRRC47 | 161.011643 | -0.144758413 | 0.23902377 | -0.6056235 | 0.54476479 | 0.73962362 |
| CEP104 | 549.366667 | 0.060010393 | 0.24777504 | 0.24219708 | 0.80862745 | 0.91010272 |
| DFFB | 105.67844 | 0.797332865 | 0.24843164 | 3.20946581 | 0.00132982 | 0.01251092 |
| C1orf174 | 123.204754 | 0.064547071 | 0.34811964 | 0.18541634 | 0.85290249 | 0.9324606 |
| AJAP1 | 407.943299 | 0.774008286 | 0.59607492 | 1.2985084 | 0.19411269 | 0.40163448 |
| NPHP4 | 362.142763 | 0.155543628 | 0.22877851 | 0.67988741 | 0.49657576 | 0.70182909 |
| RPL22 | 281.28664 | -0.532247375 | 0.21543196 | -2.4706055 | 0.01348845 | 0.06792565 |
| RNF207 | 271.340846 | -0.168998023 | 0.19683335 | -0.8585843 | 0.39056992 | 0.61344091 |
| ICMT | 162.41816 | -0.758002172 | 0.32575376 | -2.3269176 | 0.01996965 | 0.08904071 |
| ACOT7 | 60.7604388 | -0.030847092 | 0.43366918 | -0.0711305 | 0.94329392 | 0.97556917 |
| ESPN | 44.3894082 | 1.482156752 | 0.70177051 | 2.11202484 | 0.03468432 | 0.13011647 |
| PLEKHG5 | 65.4619324 | 0.110359984 | 0.4594341 | 0.24020852 | 0.81016861 | 0.91083524 |
| NOL9 | 406.977548 | 0.601889152 | 0.24669957 | 2.43976572 | 0.01469679 | 0.07224298 |
| ZBTB48 | 194.593165 | -0.210937674 | 0.22820245 | -0.9243445 | 0.35530698 | 0.58081705 |
| KLHL21 | 376.215506 | 0.317466731 | 0.30609361 | 1.03715571 | 0.2996633 | 0.52252876 |
| PHF13 | 103.887466 | 0.04522093 | 0.34219301 | 0.13215036 | 0.89486536 | 0.95172874 |
| THAP3 | 106.412643 | 0.563650526 | 0.19071531 | 2.95545512 | 0.00312208 | 0.02382413 |
| DNAJC11 | 1691.04368 | 1.51708315 | 0.28595984 | 5.30523155 | 1.13E-07 | 6.91E-06 |
| CAMTA1 | 2858.2314 | -0.078484581 | 0.23259951 | -0.3374237 | 0.73579755 | 0.87055499 |
| VAMP3 | 183.397645 | -0.6509247 | 0.21993205 | -2.9596628 | 0.00307976 | 0.0235966 |
| PER3 | 855.527492 | -0.007802808 | 0.78681665 | -0.0099169 | 0.99208756 | 0.99677109 |
| PARK7 | 637.475919 | -0.989933156 | 0.30597439 | -3.2353465 | 0.00121495 | 0.011631 |
| ERRFI1 | 2866.89484 | -1.794920795 | 0.68891735 | -2.6054225 | 0.0091761 | 0.05206631 |
| AL034417.2 | 60.4835317 | -0.75193654 | 0.78988134 | -0.9519614 | 0.34111656 | 0.56653544 |
| RERE | 12891.9394 | 0.374895662 | 0.21332919 | 1.75735754 | 0.07885689 | 0.22453559 |
| AL096855.1 | 46.7644027 | -0.423118929 | 0.41099251 | -1.0295052 | 0.30324233 | 0.52644076 |
| ENO1 | 3209.537 | -1.170814583 | 0.36826297 | -3.1792895 | 0.00147637 | 0.01354258 |
| GPR157 | 213.193704 | 0.386897694 | 0.30084252 | 1.28604724 | 0.19842658 | 0.40699016 |
| MIR34AHG | 229.700908 | -1.28549264 | 0.5576135 | -2.305347 | 0.02114714 | 0.09280154 |
| H6PD | 403.914126 | -0.422862052 | 0.19705037 | -2.1459592 | 0.03187623 | 0.12301462 |
| SPSB1 | 361.481046 | -2.926519941 | 0.54514259 | -5.3683568 | 7.95E-08 | 5.35E-06 |
| SLC25A33 | 682.156208 | 0.73080057 | 0.35801645 | 2.04124857 | 0.04122613 | 0.14525364 |
| TMEM201 | 67.6395888 | 0.848959358 | 0.35721553 | 2.37660259 | 0.0174729 | 0.08143951 |
| PIK3CD | 202.007305 | -0.693730432 | 0.35450235 | -1.9569135 | 0.05035764 | 0.16704521 |
| CLSTN1 | 1079.04305 | -0.112862386 | 0.21663054 | -0.5209902 | 0.6023736 | 0.77961521 |
| CTNNBIP1 | 350.832358 | -0.154706261 | 0.28723734 | -0.5386008 | 0.59016233 | 0.7721373 |
| LZIC | 220.844897 | -0.262643513 | 0.19131813 | -1.3728103 | 0.16981131 | 0.37007716 |
| NMNAT1 | 586.655703 | 0.209200424 | 0.21741643 | 0.96221075 | 0.33594375 | 0.56142219 |
| UBE4B | 1710.44353 | -0.065580287 | 0.19761345 | -0.3318614 | 0.73999388 | 0.87265237 |
| KIF1B | 2279.97157 | 0.112228542 | 0.18472453 | 0.60754542 | 0.54348901 | 0.7391242 |
| PGD | 235.509084 | -0.104922995 | 0.29363776 | -0.3573212 | 0.72085137 | 0.86099338 |
| DFFA | 339.495305 | -0.103124362 | 0.25107057 | -0.4107386 | 0.68126425 | 0.83353275 |
| PEX14 | 1127.06303 | -0.025454316 | 0.11426656 | -0.2227626 | 0.82372028 | 0.91846107 |
| CASZ1 | 1649.75102 | 0.174283699 | 0.50702755 | 0.34373615 | 0.73104472 | 0.86806155 |
| AL139423.1 | 96.2556696 | 0.161410042 | 0.50749723 | 0.31805108 | 0.75044619 | 0.87878048 |
| SRM | 122.461013 | -0.78366339 | 0.37915772 | -2.0668533 | 0.03874798 | 0.13969738 |
| EXOSC10 | 311.850959 | -0.075712253 | 0.2540442 | -0.2980279 | 0.76568189 | 0.88656137 |
| AL109811.1 | 54.2917914 | -0.163042937 | 0.28266818 | -0.5767997 | 0.56407474 | 0.7553629 |
| MTOR | 1082.43368 | 0.621190503 | 0.26648973 | 2.33101104 | 0.01975278 | 0.08847556 |
| UBIAD1 | 106.05204 | 0.511624603 | 0.2762314 | 1.85215946 | 0.06400293 | 0.19624296 |
| FBXO2 | 196.848861 | -1.36297063 | 0.45296993 | -3.0089649 | 0.00262139 | 0.02097274 |
| FBXO44 | 208.428961 | 0.814799328 | 0.26400746 | 3.08627388 | 0.00202682 | 0.01732833 |
| MAD2L2 | 81.7613741 | -1.38876937 | 0.44896667 | -3.0932572 | 0.00197972 | 0.01699743 |
| AGTRAP | 249.053822 | -1.776016202 | 0.33565639 | -5.2911735 | 1.22E-07 | 7.30E-06 |
| MTHFR | 251.91192 | -0.936687707 | 0.30474102 | -3.0737172 | 0.0021141 | 0.0179122 |
| CLCN6 | 346.559879 | -0.521075181 | 0.2280864 | -2.2845518 | 0.02233912 | 0.09605387 |
| KIAA2013 | 156.736317 | -0.044295919 | 0.25941464 | -0.1707533 | 0.86441771 | 0.93833133 |
| PLOD1 | 112.833193 | 0.273093827 | 0.27876619 | 0.97965188 | 0.32725799 | 0.5526381 |
| MFN2 | 387.731519 | 0.170552834 | 0.22806923 | 0.74781169 | 0.45457375 | 0.6698832 |
| MIIP | 53.2930201 | 0.327655008 | 0.32065267 | 1.02183776 | 0.3068577 | 0.53047754 |
| VPS13D | 3632.92704 | -0.062818626 | 0.12550086 | -0.5005434 | 0.61669252 | 0.78978975 |
| DHRS3 | 622.590568 | -1.418801091 | 0.6488304 | -2.1867056 | 0.02876402 | 0.11409728 |
| PRDM2 | 1436.14374 | -0.425833581 | 0.25775555 | -1.652083 | 0.09851762 | 0.26000385 |

| | | | | | | |
|---|---|---|---|---|---|---|
| KAZN | 29880.1677 | -0.926967478 | 0.29932447 | -3.096865 | 0.00195579 | 0.01684095 |
| KAZN-AS1 | 223.742557 | -0.212458619 | 0.76269279 | -0.2785638 | 0.78057958 | 0.89525707 |
| TMEM51-AS | 277.313584 | -1.654760919 | 0.56485209 | -2.9295473 | 0.00339456 | 0.0254188 |
| TMEM51 | 1622.63107 | -1.090363817 | 0.29739595 | -3.6663707 | 0.00024602 | 0.00347184 |
| C1orf195 | 78.1801608 | -1.303354271 | 0.36723098 | -3.5491403 | 0.00038649 | 0.00486206 |
| FHAD1 | 126.001906 | -1.26877207 | 0.35268511 | -3.5974643 | 0.00032133 | 0.0042276 |
| EFHD2 | 79.6567939 | -1.30016626 | 0.33564015 | -3.873691 | 0.0001072 | 0.00176999 |
| CASP9 | 200.064495 | 0.004105729 | 0.20290054 | 0.02023518 | 0.98385576 | 0.99340703 |
| DNAJC16 | 651.515765 | -0.204279806 | 0.20540561 | -0.9945191 | 0.31997021 | 0.54436972 |
| DDI2 | 648.021648 | 0.211751566 | 0.19936649 | 1.06212217 | 0.28818023 | 0.51060665 |
| PLEKHM2 | 479.247076 | -0.381766429 | 0.16640813 | -2.2941573 | 0.02178146 | 0.09452959 |
| FBLIM1 | 56.7718002 | -1.763094688 | 0.36435657 | -4.8389266 | 1.31E-06 | 5.10E-05 |
| UQCRHL | 83.667853 | -0.190101944 | 0.41169009 | -0.4617598 | 0.64425356 | 0.80961167 |
| AL450998.2 | 146.419548 | 0.535643535 | 0.34295343 | 1.561855 | 0.11832215 | 0.29339403 |
| SPEN | 1497.30304 | -0.130756011 | 0.17735431 | -0.7372587 | 0.46096504 | 0.67429537 |
| ZBTB17 | 174.040975 | 0.40025442 | 0.26425354 | 1.5146606 | 0.12985838 | 0.3118726 |
| HSPB7 | 132.849329 | 0.651404374 | 0.73631865 | 0.88467727 | 0.37633072 | 0.59959757 |
| CLCNKA | 1964.20145 | 2.227249243 | 0.84487331 | 2.63619315 | 0.0083842 | 0.04881828 |
| CLCNKB | 1268.41096 | 1.29016642 | 0.81725818 | 1.5786522 | 0.11441585 | 0.28732301 |
| FAM131C | 119.100522 | 0.854655012 | 0.6792933 | 1.25815316 | 0.20833637 | 0.41943532 |
| EPHA2 | 147.419924 | -1.110558031 | 0.39151021 | -2.8366004 | 0.00455966 | 0.03140177 |
| FBXO42 | 1360.37562 | 0.048822995 | 0.25238597 | 0.19344575 | 0.84660989 | 0.92890109 |
| SZRD1 | 397.336621 | -0.111123633 | 0.17185294 | -0.6466205 | 0.51787759 | 0.71887921 |
| SPATA21 | 49.1036907 | -0.073518713 | 0.37699641 | -0.1950117 | 0.84538378 | 0.92854435 |
| NECAP2 | 184.535241 | -0.456155938 | 0.19899709 | -2.2922744 | 0.02188981 | 0.09467668 |
| NBPF1 | 273.557513 | -0.348246142 | 0.30699097 | -1.1343856 | 0.25663284 | 0.47758004 |
| CROCC | 477.125842 | 0.285973541 | 0.36158145 | 0.79089661 | 0.42900433 | 0.64738665 |
| ATP13A2 | 160.601784 | -0.432778344 | 0.2683792 | -1.6125629 | 0.1068395 | 0.27440072 |
| SDHB | 729.477143 | 0.452935436 | 0.16830535 | 2.6911529 | 0.00712056 | 0.04326376 |
| PADI2 | 283.855099 | 1.116055732 | 0.63745446 | 1.75080072 | 0.07998024 | 0.22635474 |
| RCC2 | 272.544087 | -0.438753264 | 0.29674995 | -1.4785285 | 0.13926637 | 0.32604919 |
| ARHGEF10L | 2082.06486 | -0.611567263 | 0.37747724 | -1.6201434 | 0.10520148 | 0.27194033 |
| KLHDC7A | 81.0290699 | 0.975653439 | 0.82561138 | 1.18173449 | 0.23731107 | 0.45502132 |
| UBR4 | 1227.24946 | -0.145999532 | 0.17661294 | -0.8266639 | 0.40842761 | 0.62826602 |
| AL035413.1 | 292.85581 | 0.468850528 | 0.24305511 | 1.92898861 | 0.05373228 | 0.1744714 |
| EMC1 | 164.414003 | 0.261949109 | 0.21267765 | 1.231672 | 0.21807163 | 0.43198983 |
| MRTO4 | 123.322192 | -0.33860332 | 0.31667553 | -1.0692437 | 0.28495986 | 0.50741909 |
| AKR7A2 | 163.357576 | -0.592947564 | 0.3549717 | -1.670408 | 0.09483867 | 0.25363919 |
| PQLC2 | 63.0404902 | 0.019545135 | 0.31230395 | 0.0625837 | 0.95009801 | 0.97831655 |
| CAPZB | 1441.34879 | -0.409060598 | 0.22748294 | -1.7982034 | 0.07214479 | 0.21226201 |
| TMCO4 | 1131.63758 | -0.229303189 | 0.35490057 | -0.6461054 | 0.51821108 | 0.7190934 |
| OTUD3 | 248.217377 | 0.096653424 | 0.22563462 | 0.42836255 | 0.66838718 | 0.82542318 |
| UBXN10 | 50.7198729 | -0.922304085 | 0.41967709 | -2.1976517 | 0.02797393 | 0.11187032 |
| CAMK2N1 | 610.404493 | -1.572926041 | 0.43371058 | -3.6266721 | 0.0002871 | 0.00390553 |
| MUL1 | 110.272123 | 0.120913279 | 0.23301274 | 0.51891273 | 0.6038216 | 0.78087718 |
| CDA | 58.5483082 | 0.640357124 | 0.68003407 | 0.94165448 | 0.34636957 | 0.57154224 |
| PINK1 | 294.621842 | -0.365554962 | 0.24554283 | -1.4887625 | 0.13654992 | 0.32222841 |
| PINK1-AS | 44.1159384 | 0.307558549 | 0.34792874 | 0.88396996 | 0.37671243 | 0.59987878 |
| DDOST | 207.270636 | -0.257204684 | 0.28176586 | -0.9128312 | 0.3613313 | 0.58655834 |
| HP1BP3 | 2076.22625 | 0.275801591 | 0.1463987 | 1.88390742 | 0.0595775 | 0.18714844 |
| EIF4G3 | 7477.46763 | -0.621356251 | 0.3709385 | -1.6750924 | 0.09391611 | 0.25198819 |
| ECE1 | 849.25907 | -0.647826161 | 0.32793503 | -1.9754711 | 0.04821471 | 0.16208414 |
| NBPF3 | 158.56914 | 0.232428785 | 0.28537312 | 0.81447328 | 0.41537387 | 0.63525223 |
| RAP1GAP | 1436.02404 | 1.295991601 | 0.57335872 | 2.26035037 | 0.02379951 | 0.10024483 |
| USP48 | 1317.10882 | 0.189731414 | 0.14419112 | 1.31583288 | 0.18823014 | 0.39384317 |
| HSPG2 | 539.985898 | -0.096998309 | 0.46713665 | -0.2076444 | 0.83550663 | 0.92348178 |
| CDC42 | 1028.48967 | -0.505927908 | 0.18976169 | -2.6661225 | 0.00767317 | 0.04581915 |
| ZBTB40 | 398.858329 | 0.280801408 | 0.28512165 | 0.98484771 | 0.32469888 | 0.54958323 |
| EPHB2 | 584.425577 | -0.984806092 | 0.48776745 | -2.0190074 | 0.04348645 | 0.15092289 |
| KDM1A | 833.547622 | 0.108466759 | 0.11301316 | 0.95977105 | 0.33717045 | 0.56231354 |
| AL031428.1 | 222.771759 | 0.245812741 | 0.41313523 | 0.59499342 | 0.55184787 | 0.74481373 |
| LUZP1 | 772.944826 | 0.429043181 | 0.16216 | 2.64580159 | 0.00814976 | 0.0478658 |
| LINC01355 | 207.502083 | 0.53159711 | 0.23115826 | 2.2997106 | 0.02146462 | 0.09369669 |
| AL109936.6 | 66.2258145 | 0.366407749 | 0.40983606 | 0.89403492 | 0.37130321 | 0.59563048 |
| HNRNPR | 1189.34655 | 0.037849269 | 0.11628676 | 0.3254822 | 0.74481616 | 0.87520849 |
| ZNF436 | 51.4359527 | 0.116073812 | 0.47868409 | 0.24248521 | 0.80840422 | 0.91010272 |
| ZNF436-AS1 | 39.3549025 | 0.180381359 | 0.40933966 | 0.44066426 | 0.65945607 | 0.81980614 |
| TCEA3 | 561.557468 | 1.142453517 | 0.46827202 | 2.43972194 | 0.01469857 | 0.07224298 |
| ASAP3 | 535.433436 | 0.008549362 | 0.34034899 | 0.0251194 | 0.97995973 | 0.9913677 |
| ID3 | 49.7183759 | 0.999683279 | 0.58652209 | 1.70442563 | 0.0883016 | 0.24158571 |
| RPL11 | 1333.02905 | -0.996963219 | 0.39857608 | -2.5013122 | 0.0123734 | 0.06427066 |
| ELOA | 209.376038 | -0.26525062 | 0.19141149 | -1.3857613 | 0.16581982 | 0.36523495 |
| ELOA-AS1 | 286.884546 | 0.421753746 | 0.32719259 | 1.28900764 | 0.19739543 | 0.4057467 |
| PITHD1 | 196.922161 | -0.128667565 | 0.17538371 | -0.7336346 | 0.46317145 | 0.6761003 |
| LYPLA2 | 197.715037 | -0.683185162 | 0.308719 | -2.2129677 | 0.02689988 | 0.10889109 |
| GALE | 263.890284 | -1.213081585 | 0.30064638 | -4.0349117 | 5.46E-05 | 0.00104719 |
| HMGCL | 344.129233 | -0.509362704 | 0.29447563 | -1.7297279 | 0.08367891 | 0.23299214 |
| FUCA1 | 338.172373 | -0.787275292 | 0.32065396 | -2.4552178 | 0.01407993 | 0.07007602 |
| PNRC2 | 360.550642 | -0.249399335 | 0.25917255 | -0.9622907 | 0.3359036 | 0.56142219 |
| SRSF10 | 1431.48887 | 0.437947346 | 0.22474105 | 1.94867535 | 0.05133421 | 0.16901135 |
| IFNLR1 | 317.818634 | 0.454830534 | 0.24355188 | 1.86748932 | 0.06183329 | 0.1919972 |
| STPG1 | 184.276652 | -0.105172254 | 0.21912148 | -0.4799724 | 0.63124705 | 0.80105123 |
| NIPAL3 | 633.34168 | -0.312984399 | 0.26771002 | -1.1691172 | 0.2423564 | 0.46086856 |
| RCAN3 | 173.78702 | -1.199438631 | 0.36113662 | -3.3212877 | 0.00089603 | 0.00916074 |
| NCMAP | 102.325343 | -4.132053992 | 0.56652008 | -7.2937468 | 3.01E-13 | 1.17E-10 |
| SRRM1 | 946.855721 | 0.208402701 | 0.16697178 | 1.2481313 | 0.21198298 | 0.42482955 |
| CLIC4 | 1062.13505 | -1.136798262 | 0.50343096 | -2.2581016 | 0.02393932 | 0.10069367 |
| SYF2 | 272.86708 | -0.584396735 | 0.19393325 | -3.0133912 | 0.00258346 | 0.02075723 |
| RSRP1 | 3062.31768 | 0.023255471 | 0.19929806 | 0.11668689 | 0.90710818 | 0.95771656 |
| RHD | 99.72733 | 0.164072634 | 0.91685745 | 0.17895108 | 0.85797611 | 0.93583868 |
| TMEM50A | 401.776813 | -0.880144095 | 0.23245285 | -3.7863338 | 0.00015289 | 0.00235096 |
| MACO1 | 786.026737 | -0.064482749 | 0.21090004 | -0.3057503 | 0.75979477 | 0.88357016 |
| LDLRAP1 | 156.428727 | -0.090846004 | 0.2550158 | -0.3562368 | 0.72166326 | 0.86143064 |
| MAN1C1 | 490.534412 | 0.35629748 | 0.65573584 | 0.5433552 | 0.58688526 | 0.77047088 |
| AL031280.1 | 43.5316474 | -0.203265695 | 0.60020212 | -0.3386621 | 0.73486432 | 0.86990311 |
| SELENON | 62.0630075 | -0.562460987 | 0.3457328 | -1.6268662 | 0.10376553 | 0.26945789 |
| AL020996.1 | 50.5492431 | -0.390366478 | 0.42461163 | -0.9193495 | 0.35791281 | 0.58337492 |
| MTFR1L | 308.88551 | 1.097027557 | 0.43378264 | 2.52897987 | 0.01143946 | 0.06100275 |
| AL033528.2 | 55.1405465 | -0.151012353 | 0.41993581 | -0.3596082 | 0.71914016 | 0.86003243 |
| PAQR7 | 157.900574 | -0.499508592 | 0.45631378 | -1.0946603 | 0.27366548 | 0.49532027 |
| STMN1 | 178.29508 | -1.163914999 | 0.39448826 | -2.9504427 | 0.00317319 | 0.02407504 |
| PAFAH2 | 220.648269 | -0.263324916 | 0.1812448 | -1.4528688 | 0.14626018 | 0.3368115 |
| PDIK1L | 99.8563535 | 0.152338736 | 0.22433546 | 0.67906668 | 0.49709561 | 0.70224997 |
| ZNF593 | 115.218981 | -1.128221057 | 0.45621126 | -2.4730233 | 0.01339754 | 0.06764837 |
| CNKSR1 | 164.307658 | -0.204734826 | 0.28349545 | -0.7221803 | 0.47018364 | 0.68188686 |
| CEP85 | 165.721319 | 0.616346493 | 0.28203332 | 2.18536768 | 0.0288619 | 0.11432775 |
| SH3BGRL3 | 316.58541 | -2.263315676 | 0.42910912 | -5.2744525 | 1.33E-07 | 7.78E-06 |
| UBXN11 | 128.261238 | 0.2607384 | 0.25673085 | 1.01560993 | 0.30981518 | 0.5337614 |
| DHDDS | 389.333766 | -0.065544045 | 0.2682087 | -0.244377 | 0.80693883 | 0.90969147 |
| HMGN2 | 415.81012 | -0.232280417 | 0.23888444 | -0.9723547 | 0.3308741 | 0.55589409 |
| RPS6KA1 | 247.200303 | -0.809225437 | 0.23427888 | -3.4541117 | 0.00055211 | 0.0063637 |
| ARID1A | 1742.00268 | 0.145566531 | 0.23868796 | 0.60986122 | 0.54195374 | 0.73809463 |
| PIGV | 56.0597952 | 0.182705772 | 0.25910292 | 0.70514747 | 0.48071847 | 0.68948936 |

| | | | | | | |
|---|---|---|---|---|---|---|
| ZDHHC18 | 129.293933 | -0.044886601 | 0.25468441 | -0.176244 | 0.86010225 | 0.93645653 |
| SFN | 101.007161 | -5.685422368 | 0.85861603 | -6.6216122 | 3.55E-11 | 7.22E-09 |
| GPN2 | 203.222827 | -0.152498594 | 0.20295732 | -0.7513826 | 0.45242245 | 0.66816281 |
| GPATCH3 | 69.8203798 | 0.164244295 | 0.20464032 | 0.80259987 | 0.42220604 | 0.64114933 |
| NUDC | 482.633392 | -0.238919129 | 0.10451832 | -2.2859068 | 0.02225971 | 0.0958372 |
| KDF1 | 86.9573896 | -0.128662171 | 0.20499734 | -0.6276285 | 0.53024734 | 0.7282975 |
| TRNP1 | 282.03366 | -2.073942271 | 0.54545116 | -3.8022511 | 0.00014339 | 0.00223349 |
| TENT5B | 43.7563377 | 1.334743961 | 0.76796927 | 1.7380174 | 0.08220775 | 0.23040147 |
| SLC9A1 | 281.201021 | 0.348272331 | 0.20737135 | 1.67946211 | 0.09306202 | 0.25071511 |
| WDTC1 | 638.068313 | -0.089776992 | 0.12794001 | -0.7017116 | 0.48285902 | 0.69127369 |
| TMEM222 | 187.718233 | -0.227069503 | 0.19532351 | -1.1625303 | 0.24502012 | 0.46414323 |
| MAP3K6 | 76.2425555 | -1.262997227 | 0.6921133 | -1.8248417 | 0.06802492 | 0.20444622 |
| WASF2 | 1591.36499 | -0.539013124 | 0.15603596 | -3.4544161 | 0.00055149 | 0.00636207 |
| FO393419.2 | 41.8053953 | -0.224961378 | 0.52610438 | -0.4275984 | 0.66894355 | 0.82587721 |
| AHDC1 | 795.412983 | 0.400465584 | 0.2563169 | 1.5623846 | 0.11819741 | 0.29330634 |
| IFI6 | 971.449025 | -0.962271314 | 1.07176735 | -0.897836 | 0.36927299 | 0.59417383 |
| FAM76A | 231.782771 | 0.253098168 | 0.21783214 | 1.16189542 | 0.24527795 | 0.46435545 |
| STX12 | 855.287486 | -0.234501727 | 0.21652475 | -1.0830251 | 0.2787973 | 0.50097041 |
| PPP1R8 | 212.382771 | -0.029987202 | 0.20463673 | -0.1465387 | 0.88349613 | 0.94755606 |
| RPA2 | 230.326667 | -0.256839164 | 0.17304536 | -1.4842303 | 0.13774783 | 0.32381543 |
| SMPDL3B | 47.2117088 | -2.406313093 | 0.45656218 | -5.2705046 | 1.36E-07 | 7.92E-06 |
| EYA3 | 1309.78674 | -0.074535373 | 0.13134757 | -0.5674667 | 0.57039714 | 0.75945349 |
| PTAFR | 89.3919035 | -2.149331769 | 0.57063876 | -3.7665366 | 0.00016553 | 0.00251317 |
| DNAJC8 | 329.236997 | -0.015597852 | 0.14678886 | -0.1062605 | 0.9153757 | 0.9617561 |
| ATP5IF1 | 668.055922 | -0.116751668 | 0.3536023 | -0.3301779 | 0.74126553 | 0.87368399 |
| AL353622.1 | 73.559887 | 1.309970976 | 0.49478133 | 2.64757562 | 0.00810712 | 0.04772147 |
| SESN2 | 176.421255 | -0.245432963 | 0.31261991 | -0.7850842 | 0.43240419 | 0.65008274 |
| PHACTR4 | 2796.83078 | -0.298782339 | 0.18819575 | -1.5876147 | 0.11237349 | 0.28354182 |
| RCC1 | 329.430386 | -0.706990363 | 0.37895327 | -1.86564 | 0.06209176 | 0.19249428 |
| TRNAU1AP | 493.228782 | -0.72935697 | 0.20025953 | -3.6420587 | 0.00027047 | 0.00372145 |
| SNHG12 | 114.995773 | -0.484841274 | 0.32839443 | -1.4763992 | 0.13983675 | 0.32709468 |
| TAF12 | 300.941201 | -0.055473099 | 0.21625743 | -0.2565142 | 0.79755383 | 0.90505915 |
| GMEB1 | 293.216948 | -0.165171288 | 0.29583129 | -0.5583293 | 0.57661951 | 0.76325956 |
| YTHDF2 | 729.443104 | -0.222380744 | 0.14979191 | -1.4845978 | 0.1376504 | 0.32375786 |
| EPB41 | 1990.75635 | 0.030683087 | 0.17456593 | 0.1757679 | 0.86047629 | 0.93655498 |
| SRSF4 | 2660.87287 | -0.242047169 | 0.15077521 | -1.6053513 | 0.10841652 | 0.27678273 |
| MECR | 301.944065 | 0.113057278 | 0.27232599 | 0.4151542 | 0.67802902 | 0.83118075 |
| PTPRU | 102.575507 | -2.324010414 | 0.46243653 | -5.0255771 | 5.02E-07 | 2.37E-05 |
| PUM1 | 3372.90429 | -0.185941778 | 0.15507133 | -1.1990726 | 0.23049972 | 0.44604027 |
| SNRNP40 | 326.95538 | 0.342562356 | 0.18543768 | 1.84731791 | 0.06470108 | 0.19759391 |
| ZCCHC17 | 665.058167 | -0.433660356 | 0.19216704 | -2.2566843 | 0.02402781 | 0.1008619 |
| AL451070.1 | 99.4419507 | 0.099594851 | 0.36523788 | 0.27268489 | 0.78509545 | 0.89763056 |
| FABP3 | 110.878464 | 0.815495708 | 0.47284862 | 1.72464438 | 0.08459158 | 0.23439694 |
| SERINC2 | 909.479388 | -0.509849608 | 0.26539956 | -1.9210642 | 0.05472361 | 0.17653444 |
| TINAGL1 | 249.176954 | -0.515332043 | 0.41204104 | -1.2506813 | 0.21105076 | 0.42345922 |
| PEF1 | 191.828678 | -0.50905799 | 0.25588257 | -1.9894203 | 0.04665483 | 0.15830044 |
| PTP4A2 | 305.064941 | -0.475020345 | 0.25007842 | -1.8994856 | 0.05750066 | 0.18288617 |
| KHDRBS1 | 1215.58138 | -0.450536457 | 0.23044563 | -1.9550662 | 0.05057526 | 0.1674727 |
| AL445248.1 | 55.8436976 | -0.193925809 | 0.37964274 | -0.5108113 | 0.60948318 | 0.78527735 |
| TMEM39B | 132.423516 | 0.058432328 | 0.26786713 | 0.21813923 | 0.82732063 | 0.91965909 |
| KPNA6 | 1155.21906 | -0.3273986 | 0.15222642 | -2.1507344 | 0.03149717 | 0.12211508 |
| TXLNA | 325.234804 | -0.473284268 | 0.25990092 | -1.8210181 | 0.06860411 | 0.20553225 |
| DCDC2B | 55.0849034 | 0.068919762 | 0.35899218 | 0.19198124 | 0.8477569 | 0.9298365 |
| TMEM234 | 118.533077 | -0.316370425 | 0.19237935 | -1.6445134 | 0.1000702 | 0.26294555 |
| EIF3I | 224.96332 | -1.165450825 | 0.34488957 | -3.3792 | 0.00072697 | 0.00787741 |
| HDAC1 | 441.833448 | -0.018200691 | 0.19090718 | -0.0953379 | 0.92404644 | 0.96654633 |
| TSSK3 | 89.7270849 | 0.708090507 | 0.43256679 | 1.6369507 | 0.10164077 | 0.26554053 |
| FAM229A | 47.3276105 | -0.4780016 | 0.35336039 | -1.3527311 | 0.17614154 | 0.3783991 |
| BSDC1 | 545.767542 | -0.333537574 | 0.17724333 | -1.8818061 | 0.05986235 | 0.1875971 |
| ZBTB8A | 57.7399839 | -0.679188954 | 0.37671972 | -1.8029026 | 0.07140352 | 0.21085827 |
| ZBTB8OS | 894.303585 | 0.10171604 | 0.16806204 | 0.60522913 | 0.54502677 | 0.73990324 |
| RBBP4 | 862.39732 | -0.020435834 | 0.12650321 | -0.161544 | 0.87166496 | 0.94174893 |
| SYNC | 189.737444 | -0.600734028 | 0.37357838 | -1.6080535 | 0.10782346 | 0.2759816 |
| KIAA1522 | 890.479649 | -0.183165247 | 0.26861231 | -0.6818945 | 0.49530568 | 0.70078115 |
| YARS | 408.860007 | -0.351689461 | 0.29240363 | -1.2027534 | 0.22907176 | 0.44457881 |
| S100PBP | 953.954052 | 0.145186042 | 0.12955715 | 1.12063316 | 0.26244404 | 0.4832663 |
| TMEM54 | 92.2089728 | -2.009925097 | 0.46554334 | -4.3173749 | 1.58E-05 | 0.00038841 |
| RNF19B | 398.968543 | -0.140952791 | 0.35373798 | -0.3984667 | 0.69028623 | 0.84106341 |
| AL020995.1 | 379.47875 | -0.580239383 | 0.64315598 | -0.9021752 | 0.3669638 | 0.59249784 |
| AK2 | 1058.26458 | -0.013265237 | 0.30818411 | -0.0430432 | 0.96566708 | 0.98461609 |
| AZIN2 | 169.497989 | -0.236005304 | 0.35647422 | -0.6620543 | 0.5079364 | 0.71103267 |
| TRIM62 | 141.906156 | -0.671887381 | 0.34108104 | -1.9698761 | 0.04885258 | 0.16348966 |
| ZNF362 | 278.570401 | 0.398598378 | 0.26605918 | 1.49815684 | 0.13409251 | 0.31845162 |
| AL513327.1 | 617.718928 | -1.23885529 | 0.55041117 | -2.2507815 | 0.02439938 | 0.10195373 |
| PHC2 | 2352.10526 | -0.663412169 | 0.26615077 | -2.4926178 | 0.01268052 | 0.06540763 |
| AL513327.2 | 54.3917637 | 0.055029198 | 0.41919756 | 0.1312727 | 0.89555958 | 0.95183831 |
| ZSCAN20 | 60.8630913 | -0.163616823 | 0.24880921 | -0.6575995 | 0.5107955 | 0.71294917 |
| CSMD2 | 122.554511 | -1.274027294 | 1.11301697 | -1.1446612 | 0.25234955 | 0.47247132 |
| SMIM12 | 286.029186 | 0.160637538 | 0.15413692 | 1.04217431 | 0.29733087 | 0.52079582 |
| AC114490.3 | 41.7399458 | -0.331189006 | 0.31214613 | -1.0610063 | 0.28868705 | 0.51102482 |
| ZMYM1 | 475.320861 | 0.459589203 | 0.1288026 | 3.56816722 | 0.00035949 | 0.00459133 |
| SFPQ | 2055.68217 | 0.114927155 | 0.19552411 | 0.58779021 | 0.55667312 | 0.74881722 |
| ZMYM4 | 3384.28615 | -0.103347721 | 0.12754619 | -0.8102768 | 0.4177811 | 0.63745762 |
| ZMYM4-AS1 | 80.0287945 | 0.022001747 | 0.60588485 | 0.03631341 | 0.97103245 | 0.98717522 |
| KIAA0319L | 1845.83219 | 0.100817511 | 0.13879428 | 0.7263809 | 0.4676053 | 0.68009231 |
| NCDN | 49.3037794 | -0.221334858 | 0.38596551 | -0.5734576 | 0.56633489 | 0.75654506 |
| PSMB2 | 334.55972 | -0.222595192 | 0.18403735 | -1.209511 | 0.22646659 | 0.44159983 |
| AGO4 | 896.739708 | -0.374766928 | 0.21917508 | -1.7098975 | 0.08728482 | 0.23953446 |
| AGO1 | 280.213784 | 0.133912422 | 0.24652073 | 0.54320959 | 0.58698551 | 0.77047088 |
| AGO3 | 2467.31387 | 0.20908215 | 0.16024417 | 1.30477228 | 0.1919704 | 0.39901448 |
| AL138787.2 | 141.289392 | 0.14642378 | 0.41533113 | 0.35254708 | 0.72442801 | 0.86339165 |
| ADPRHL2 | 171.689987 | -0.824964616 | 0.37294297 | -2.2120396 | 0.02696393 | 0.10908623 |
| TRAPPC3 | 222.795914 | -0.200056567 | 0.17197071 | -1.1633177 | 0.24470064 | 0.46367102 |
| MAP7D1 | 164.830276 | -0.661717284 | 0.35748993 | -1.8510096 | 0.06416818 | 0.196476 |
| THRAP3 | 1563.92863 | -0.231029435 | 0.14405143 | -1.6037983 | 0.10875852 | 0.2771737 |
| SH3D21 | 157.963162 | -0.008156487 | 0.52360926 | -0.0155774 | 0.98757151 | 0.99533283 |
| EVA1B | 170.333146 | -1.000522741 | 0.54610291 | -1.8321139 | 0.06693445 | 0.20199615 |
| AL591845.1 | 123.715231 | 0.146384776 | 0.56511652 | 0.25903468 | 0.79560849 | 0.90424991 |
| STK40 | 427.789151 | -0.354191744 | 0.23563383 | -1.5031447 | 0.13280174 | 0.31658151 |
| LSM10 | 77.8773664 | -0.278614776 | 0.39463532 | -0.7060057 | 0.48018461 | 0.68931285 |
| OSCP1 | 127.395989 | 0.091907271 | 0.24698847 | 0.37211159 | 0.70980977 | 0.85441779 |
| MRPS15 | 168.762305 | -0.468259041 | 0.28175285 | -1.6619496 | 0.09652287 | 0.25648172 |
| LINC01137 | 146.63904 | -0.380973055 | 0.31615037 | -1.2050375 | 0.22818882 | 0.44371354 |
| ZC3H12A | 98.137825 | -1.946608585 | 0.44477012 | -4.3766623 | 1.21E-05 | 0.0003109 |
| MEAF6 | 338.287258 | 0.131528735 | 0.19080874 | 0.68932237 | 0.49062042 | 0.6969611 |
| SNIP1 | 160.226736 | -0.150716411 | 0.22361647 | -0.6739951 | 0.50031441 | 0.70482771 |
| DNALI1 | 116.942243 | -0.283867452 | 0.31458472 | -0.9023561 | 0.36686772 | 0.59241513 |
| GNL2 | 139.178436 | -0.094070189 | 0.21406085 | -0.4394554 | 0.66033162 | 0.82007525 |
| C1orf109 | 97.6596137 | -0.173534903 | 0.31606106 | -0.549055 | 0.5829677 | 0.76842834 |
| YRDC | 50.9347582 | -0.513159163 | 0.31240593 | -1.6426038 | 0.10046494 | 0.26372046 |
| C1orf122 | 290.085457 | -0.421618466 | 0.45790496 | -0.9207554 | 0.35717815 | 0.58271536 |
| MTF1 | 252.29115 | -0.018349888 | 0.17684437 | -0.1037629 | 0.91735749 | 0.96291779 |
| AL929472.2 | 106.118002 | -0.405923911 | 0.37991027 | -1.0684731 | 0.28530715 | 0.50782141 |

| | | | | | | |
|---|---|---|---|---|---|---|
| INPP5B | 815.047291 | -0.185109638 | 0.287553 | -0.6437409 | 0.51974345 | 0.7198387 |
| SF3A3 | 305.693652 | -0.114304824 | 0.20976153 | -0.5449275 | 0.58580338 | 0.77001836 |
| UTP11 | 80.7704052 | 0.086065917 | 0.27148654 | 0.31701726 | 0.75123051 | 0.87930915 |
| RRAGC | 181.882174 | -0.759985016 | 0.25311964 | -3.0024735 | 0.00267795 | 0.02129625 |
| AL139260.1 | 453.509526 | 0.021934973 | 0.3719434 | 0.05897395 | 0.95297285 | 0.97936812 |
| RHBDL2 | 93.6604907 | -0.816499725 | 0.44438799 | -1.8373577 | 0.0661571 | 0.2004287 |
| AKIRIN1 | 1018.93917 | -0.169027964 | 0.2697158 | -0.6266891 | 0.53086302 | 0.72863563 |
| NDUFS5 | 635.398288 | -0.775423067 | 0.37515805 | -2.0669237 | 0.03874135 | 0.13969738 |
| MACF1 | 6891.41 | -0.771439057 | 0.26242717 | -2.939631 | 0.00328603 | 0.02476053 |
| PABPC4 | 256.061004 | -0.241600749 | 0.2048519 | -1.1793923 | 0.23824201 | 0.45614418 |
| HPCAL4 | 113.567035 | -0.738719382 | 0.91278317 | -0.8093043 | 0.41834011 | 0.63753323 |
| PPIE | 206.569681 | 0.143623598 | 0.34500773 | 0.41629096 | 0.67719711 | 0.83063391 |
| TRIT1 | 609.76063 | 0.219050946 | 0.16146886 | 1.35661417 | 0.17490384 | 0.37662539 |
| CAP1 | 523.131065 | -0.848971441 | 0.16365185 | -5.1876678 | 2.13E-07 | 1.16E-05 |
| PPT1 | 150.100041 | 0.017116599 | 0.2663448 | 0.06426481 | 0.94875937 | 0.97784954 |
| RLF | 1413.89895 | -0.325215886 | 0.21749913 | -1.4952514 | 0.13484884 | 0.31968078 |
| AL050341.2 | 40.64586 | 0.875138453 | 0.42813633 | 2.04406492 | 0.04094713 | 0.14473393 |
| ZMPSTE24 | 377.371469 | 0.185798578 | 0.21628925 | 0.85902826 | 0.39032493 | 0.61320085 |
| SMAP2 | 288.091322 | 0.222802594 | 0.18007631 | 1.23726764 | 0.21598774 | 0.42979542 |
| AL603839.3 | 248.153868 | 0.181018058 | 0.25969843 | 0.69703177 | 0.48578291 | 0.69324832 |
| EXO5 | 47.8581551 | -0.141163419 | 0.37556287 | -0.3758716 | 0.70701235 | 0.85248071 |
| ZNF684 | 296.357472 | 0.808238263 | 0.42846898 | 1.88634018 | 0.05924913 | 0.18624982 |
| NFYC | 1541.71995 | 1.245089854 | 0.22544251 | 5.52287069 | 3.34E-08 | 2.56E-06 |
| CITED4 | 113.263234 | -2.738791932 | 0.46860369 | -5.8445804 | 5.08E-09 | 5.39E-07 |
| CTPS1 | 159.910675 | -0.544096337 | 0.53922836 | -1.0090277 | 0.31296136 | 0.53684988 |
| SLFNL1-AS1 | 165.637802 | -0.960192636 | 0.4693531 | -2.0457788 | 0.04077814 | 0.14440321 |
| SLFNL1 | 61.4693503 | -0.874426707 | 0.58872549 | -1.4852877 | 0.13746764 | 0.32350081 |
| SCMH1 | 3249.4724 | -0.66567077 | 0.17220056 | -3.8656713 | 0.00011078 | 0.00181557 |
| AC093151.3 | 129.570802 | -3.943340939 | 0.89673984 | -4.3974191 | 1.10E-05 | 0.00028767 |
| HIVEP3 | 1799.45701 | -1.508545024 | 0.41240631 | -3.6579097 | 0.00025428 | 0.00355428 |
| FOXJ3 | 1965.47437 | -0.124685715 | 0.16740626 | -0.7448092 | 0.4563871 | 0.67145234 |
| RIMKLA | 68.1980323 | -1.400035679 | 0.82912123 | -1.6885778 | 0.09130037 | 0.24700502 |
| ZMYND12 | 61.8580868 | 1.02310367 | 0.5078625 | 2.01452888 | 0.04395404 | 0.15198661 |
| PPCS | 264.733632 | -0.226622571 | 0.2441127 | -0.9283522 | 0.35322489 | 0.57919589 |
| CCDC30 | 1831.66613 | 0.44326211 | 0.19312241 | 2.29523914 | 0.02171942 | 0.09440958 |
| PPIH | 166.349962 | 0.284462952 | 0.24656313 | 1.15371245 | 0.24861808 | 0.46787237 |
| YBX1 | 786.566913 | -0.612941735 | 0.29436506 | -2.0822503 | 0.03731961 | 0.13647696 |
| CLDN19 | 138.57044 | 0.74894637 | 0.80396263 | 0.93156864 | 0.3515595 | 0.57736533 |
| P3H1 | 115.074492 | 0.22678512 | 0.28523973 | 0.79506849 | 0.42657366 | 0.64506142 |
| SVBP | 85.949621 | -0.37424453 | 0.38406633 | -0.9744268 | 0.32984466 | 0.55530437 |
| ERMAP | 259.587132 | 0.348816886 | 0.31193889 | 1.11822188 | 0.26347224 | 0.48439746 |
| ZNF691 | 55.8950109 | 0.367660722 | 0.30700583 | 1.19756919 | 0.23108478 | 0.44684509 |
| SLC2A1 | 595.553961 | -2.482198282 | 0.76256027 | -3.2550847 | 0.00113359 | 0.01102332 |
| SLC2A1-AS1 | 135.347479 | -0.861379261 | 0.78430737 | -1.0982675 | 0.27208772 | 0.49354664 |
| FAM183A | 55.0331405 | -0.959099952 | 0.48247224 | -1.9878863 | 0.04682427 | 0.1587154 |
| EBNA1BP2 | 215.645273 | 0.537024239 | 0.32659744 | 1.6443002 | 0.10011421 | 0.26300887 |
| CFAP57 | 66.0838377 | 0.229494593 | 0.37429392 | 0.61314004 | 0.53978374 | 0.73596237 |
| TMEM125 | 77.670145 | -0.298205798 | 0.43572317 | -0.6843928 | 0.49372717 | 0.69913828 |
| AL139289.2 | 38.3582357 | 0.922462594 | 0.6150223 | 1.4998848 | 0.13364425 | 0.31790147 |
| ELOVL1 | 125.121457 | -0.798072713 | 0.38498941 | -2.0729731 | 0.03817478 | 0.13864467 |
| MED8 | 110.820312 | -0.111544542 | 0.33828424 | -0.3297361 | 0.74159934 | 0.8738293 |
| SZT2 | 479.52192 | -0.002924122 | 0.2162891 | -0.0135195 | 0.98921332 | 0.99594533 |
| HYI | 85.2995125 | 0.044079664 | 0.24778011 | 0.17789832 | 0.85880283 | 0.93618423 |
| PTPRF | 750.834091 | -0.708981487 | 0.3575954 | -1.9826359 | 0.04740811 | 0.15999329 |
| KDM4A-AS1 | 57.9941908 | 0.284792818 | 0.31051594 | 0.91716005 | 0.35905878 | 0.58437745 |
| AL357079.1 | 43.6825209 | -0.353616673 | 0.43548038 | -0.8120152 | 0.41678293 | 0.63667002 |
| IPO13 | 99.5415928 | 0.406158944 | 0.31543791 | 1.28760344 | 0.19788404 | 0.40634007 |
| DPH2 | 43.3632541 | 0.090988664 | 0.44488209 | 0.2045231 | 0.83794474 | 0.92446904 |
| ATP6V0B | 817.33821 | -0.471744617 | 0.38489969 | -1.22563 | 0.22033793 | 0.4341354 |
| B4GALT2 | 104.682629 | -0.038523985 | 0.35660548 | -0.1080297 | 0.91397213 | 0.96119671 |
| CCDC24 | 328.745474 | 0.07772194 | 0.2821889 | 0.27542522 | 0.78298957 | 0.89647912 |
| SLC6A9 | 41.6618301 | -0.319119201 | 0.50044087 | -0.6376761 | 0.52368452 | 0.72325104 |
| DMAP1 | 146.257644 | 0.158404668 | 0.30622866 | 0.51727578 | 0.60496366 | 0.78156518 |
| ERI3 | 1050.78884 | -0.027302575 | 0.21309179 | -0.1281259 | 0.89804935 | 0.9527979 |
| RNF220 | 163.875247 | -0.357809975 | 0.15920982 | -2.2474114 | 0.02461375 | 0.10259482 |
| TMEM53 | 214.871729 | -0.182960042 | 0.30827232 | -0.5935014 | 0.55284568 | 0.74576578 |
| ARMH1 | 206.162581 | 0.752317546 | 0.32332858 | 2.32678954 | 0.01997647 | 0.08904071 |
| RPS8 | 1018.05107 | -1.10758474 | 0.3837077 | -2.8865325 | 0.00389513 | 0.02813051 |
| PLK3 | 64.0714165 | -2.365419586 | 0.48238142 | -4.9036291 | 9.41E-07 | 3.95E-05 |
| EIF2B3 | 612.061328 | 0.344937939 | 0.22535466 | 1.53064478 | 0.12585721 | 0.30481607 |
| HECTD3 | 100.134029 | -0.251101145 | 0.29804754 | -0.8424869 | 0.39951547 | 0.62113914 |
| UROD | 99.5355243 | -0.16411883 | 0.325971 | -0.5034768 | 0.6146291 | 0.78816802 |
| ZSWIM5 | 1053.72968 | 0.108569257 | 0.31694099 | 0.34255353 | 0.73193436 | 0.86826096 |
| MUTYH | 106.247517 | 0.333844647 | 0.22793973 | 1.46461808 | 0.14302515 | 0.33248901 |
| TOE1 | 46.7112277 | -0.262493889 | 0.31033464 | -0.8458414 | 0.39764122 | 0.6192458 |
| TESK2 | 362.260032 | -0.417274996 | 0.24068902 | -1.7336686 | 0.0829769 | 0.23167236 |
| MMACHC | 57.4897024 | 0.663544695 | 0.34657197 | 1.91459425 | 0.05554427 | 0.17839503 |
| PRDX1 | 1283.6066 | -1.874197037 | 0.47947197 | -3.9088772 | 9.27E-05 | 0.0015845 |
| AKR1A1 | 199.466052 | -0.547698273 | 0.27130895 | -2.0187254 | 0.04351577 | 0.15096948 |
| NASP | 489.15187 | 0.130303844 | 0.18187456 | 0.71644898 | 0.47371416 | 0.68438043 |
| GPBP1L1 | 2250.43619 | 0.187704987 | 0.13865632 | 1.35374274 | 0.17581847 | 0.37799253 |
| TMEM69 | 144.477299 | 0.255982235 | 0.19284472 | 1.32740077 | 0.18437615 | 0.38966792 |
| IPP | 469.405829 | 0.236378347 | 0.19112204 | 1.23679274 | 0.21616404 | 0.43001668 |
| MAST2 | 510.587674 | -0.912792096 | 0.26197503 | -3.4842714 | 0.00049348 | 0.00586184 |
| PIK3R3 | 42.4609737 | -0.788491372 | 0.54195131 | -1.4549118 | 0.14569368 | 0.33584931 |
| TSPAN1 | 685.914916 | -2.838601896 | 0.43908055 | -6.4648773 | 1.01E-10 | 1.76E-08 |
| POMGNT1 | 389.994998 | -0.282200757 | 0.19962401 | -1.4136614 | 0.15746135 | 0.35324506 |
| LURAP1 | 413.998005 | 0.340235879 | 0.34712227 | 0.9801615 | 0.32700641 | 0.55240682 |
| LRRC41 | 418.229409 | -0.130265543 | 0.17565124 | -0.7416147 | 0.45832082 | 0.67266311 |
| UQCRH | 921.837596 | 0.413684165 | 0.21628632 | 1.91266916 | 0.05579042 | 0.17899822 |
| NSUN4 | 310.359835 | -0.172379029 | 0.13158039 | -1.3100663 | 0.19017341 | 0.39646473 |
| FAAH | 139.867653 | 0.61062059 | 0.24515773 | 2.49072538 | 0.01274826 | 0.06557701 |
| MKNK1 | 393.014333 | 0.141643193 | 0.13402164 | 1.05686811 | 0.29057179 | 0.51339098 |
| ATPAF1 | 452.447953 | 0.366513885 | 0.16135034 | 2.27154084 | 0.02311426 | 0.09830568 |
| EFCAB14-AS | 44.9008026 | -0.950119499 | 0.58275774 | -1.630385 | 0.10302015 | 0.26790413 |
| EFCAB14 | 599.328906 | -0.285357442 | 0.16012445 | -1.7820979 | 0.07473327 | 0.21714732 |
| CYP4A11 | 52.8046355 | 0.892963713 | 0.88451325 | 1.00955381 | 0.31270911 | 0.5367235 |
| PDZK1IP1 | 480.097845 | -1.834651197 | 0.71562305 | -2.5637117 | 0.01035595 | 0.05687807 |
| CMPK1 | 624.353703 | -0.586240395 | 0.24242439 | -2.4182401 | 0.01559578 | 0.07534919 |
| TRABD2B | 2927.59469 | -1.678148042 | 0.65679537 | -2.5550546 | 0.01061711 | 0.05789965 |
| AL109659.1 | 54.8526834 | -1.73339237 | 0.91158957 | -1.9015053 | 0.05723586 | 0.18228859 |
| SLC5A9 | 54.9100849 | -0.996926738 | 0.42127328 | -2.3664609 | 0.01795907 | 0.08311891 |
| SPATA6 | 1585.63784 | -0.038882422 | 0.12700581 | -0.3061468 | 0.75949289 | 0.88349992 |
| AGBL4 | 4091.24997 | 0.713157052 | 0.46216443 | 1.54308078 | 0.12281118 | 0.300288 |
| AC099788.1 | 68.1739648 | 0.094148617 | 0.70012332 | 0.13447433 | 0.89302751 | 0.95136707 |
| FAF1 | 5534.42809 | 0.069657693 | 0.13838505 | 0.50336141 | 0.6147102 | 0.78816802 |
| RNF11 | 630.458937 | -0.100330272 | 0.1851468 | -0.5418958 | 0.58789031 | 0.7710768 |
| TTC39A | 607.108874 | -2.254542028 | 0.45980761 | -4.903229 | 9.43E-07 | 3.95E-05 |
| TTC39A-AS1 | 73.588066 | -3.216685918 | 0.59092838 | -5.4434446 | 5.23E-08 | 3.77E-06 |
| EPS15 | 2359.58624 | 0.139540141 | 0.14856405 | 0.93925911 | 0.34759772 | 0.57313923 |
| OSBPL9 | 2553.23085 | 0.747573227 | 0.14820282 | 5.04425783 | 4.55E-07 | 2.20E-05 |
| NRDC | 1845.84377 | -0.185044641 | 0.14476843 | -1.2782113 | 0.20117493 | 0.41032344 |
| TXNDC12 | 331.633071 | -0.289422739 | 0.19826865 | -1.4597504 | 0.14435868 | 0.33423906 |

| | | | | | | |
|---|---|---|---|---|---|---|
| BTF3L4 | 556.316134 | -0.065653718 | 0.14422302 | -0.4552236 | 0.64894841 | 0.81204618 |
| ZFYVE9 | 1369.41248 | 0.142824698 | 0.16669141 | 0.85682101 | 0.39154381 | 0.6140943 |
| CC2D1B | 231.882612 | -0.614953901 | 0.25243666 | -2.436072 | 0.01484773 | 0.07277314 |
| PRPF38A | 240.031742 | 0.02099606 | 0.15693541 | 0.13378791 | 0.89357029 | 0.95146032 |
| TUT4 | 2383.51919 | 0.180947751 | 0.15987076 | 1.13183772 | 0.25770267 | 0.47869421 |
| ZYG11B | 890.048259 | 0.3322129 | 0.20459759 | 1.62373809 | 0.10443172 | 0.27047815 |
| ECHDC2 | 1923.17832 | 0.198792258 | 0.29356541 | 0.67716512 | 0.49830119 | 0.70339396 |
| SCP2 | 1134.72779 | 0.143524847 | 0.17515505 | 0.81941598 | 0.41254912 | 0.63186927 |
| CPT2 | 184.16729 | -0.296288141 | 0.27881514 | -1.0626688 | 0.28793218 | 0.5102356 |
| C1orf123 | 110.897143 | 0.30906057 | 0.28203835 | 1.09581044 | 0.27316174 | 0.49481533 |
| MAGOH | 226.31239 | 0.242166367 | 0.18319596 | 1.32189793 | 0.18620214 | 0.39195922 |
| LRP8 | 120.89843 | -2.530022555 | 0.97882729 | -2.5847487 | 0.009745 | 0.05434234 |
| NDC1 | 302.203523 | 0.22324064 | 0.27124579 | 0.82301974 | 0.41049677 | 0.62997812 |
| YIPF1 | 233.043565 | -0.540001308 | 0.224617 | -2.4040981 | 0.01621242 | 0.07725464 |
| HSPB11 | 324.768917 | 0.068324583 | 0.14995441 | 0.4556357 | 0.64865196 | 0.81191151 |
| LRRC42 | 154.412472 | -0.31573212 | 0.21478501 | -1.4699914 | 0.14156408 | 0.33019952 |
| TMEM59 | 1180.45671 | -0.421864776 | 0.27729419 | -1.5213618 | 0.12816907 | 0.30884182 |
| TCEANC2 | 220.156661 | 0.354464749 | 0.21240003 | 1.66885455 | 0.09514621 | 0.25410155 |
| MRPL37 | 125.841839 | 0.100615924 | 0.23799433 | 0.42276606 | 0.67246595 | 0.82807581 |
| SSBP3 | 2874.86718 | -0.400283044 | 0.33854983 | -1.182346 | 0.23706844 | 0.45468811 |
| ACOT11 | 1982.18196 | 1.797429994 | 0.3015568 | 5.96050225 | 2.51E-09 | 2.97E-07 |
| FAM151A | 74.2800167 | 1.708746591 | 0.34792358 | 4.91126979 | 9.05E-07 | 3.86E-05 |
| MROH7-TTC4 | 150.941293 | 2.357062185 | 0.31634395 | 7.45094767 | 9.27E-14 | 4.37E-11 |
| TTC22 | 107.891492 | -1.099440822 | 0.34186045 | -3.2160515 | 0.00129967 | 0.01228877 |
| DHCR24 | 481.846744 | -1.437177517 | 0.43585031 | -3.2974108 | 0.00097581 | 0.00980931 |
| TMEM61 | 196.445422 | 0.270310761 | 0.40888674 | 0.66108958 | 0.50855487 | 0.71123011 |
| BSND | 137.379103 | 0.768741411 | 0.54885938 | 1.40061634 | 0.16132883 | 0.35883595 |
| USP24 | 3618.82586 | 0.637124506 | 0.31718264 | 2.00869915 | 0.04456905 | 0.15372303 |
| AC119674.2 | 91.4601307 | -0.135892725 | 0.52789492 | -0.2574238 | 0.79685162 | 0.9046722 |
| PLPP3 | 1444.46871 | -0.962642959 | 0.43802891 | -2.1976699 | 0.02797264 | 0.11187032 |
| PRKAA2 | 2992.57675 | 0.980852589 | 0.24165784 | 4.05884868 | 4.93E-05 | 0.00096219 |
| DAB1 | 1922.32285 | 1.879787062 | 0.77184909 | 2.43543342 | 0.01487396 | 0.07287469 |
| TACSTD2 | 802.220979 | -1.450759271 | 0.58576873 | -2.4766759 | 0.01326123 | 0.06711401 |
| MYSM1 | 1372.88235 | 0.638372463 | 0.26056381 | 2.44996593 | 0.01428697 | 0.07081299 |
| AL136985.3 | 218.685717 | -1.071162272 | 0.57663576 | -1.8576064 | 0.06322493 | 0.19498906 |
| JUN | 479.891362 | -0.448653948 | 0.60603209 | -0.7403139 | 0.45910958 | 0.67278348 |
| LINC01135 | 440.053757 | 0.48746873 | 0.29503034 | 1.65226644 | 0.09848025 | 0.26000385 |
| LINC01358 | 79.9317675 | -0.07824485 | 0.41471637 | -0.1886708 | 0.85035087 | 0.93136689 |
| FGGY | 1674.81564 | 0.105280617 | 0.30263782 | 0.3478766 | 0.72793286 | 0.86612619 |
| HOOK1 | 953.818972 | 0.275138184 | 0.24886568 | 1.105569 | 0.26891311 | 0.49041465 |
| NFIA | 3380.30261 | 0.495415412 | 0.40337471 | 1.22817667 | 0.21938065 | 0.43356369 |
| TM2D1 | 946.637354 | 0.225149896 | 0.1457772 | 1.54447949 | 0.12247222 | 0.29989549 |
| PATJ | 12520.3222 | -0.084623618 | 0.16313831 | -0.5187232 | 0.60395381 | 0.78089525 |
| USP1 | 207.355981 | -0.308165697 | 0.16118803 | -1.9118399 | 0.05589674 | 0.17920873 |
| DOCK7 | 1581.76343 | -0.19573487 | 0.16841275 | -1.1622331 | 0.2451408 | 0.46420922 |
| ANGPTL3 | 37.4582025 | 3.174997402 | 0.59071576 | 5.37483103 | 7.67E-08 | 5.22E-06 |
| ATG4C | 496.538716 | 0.742247125 | 0.15632658 | 4.74805445 | 2.05E-06 | 7.22E-05 |
| ALG6 | 246.849106 | 0.497486392 | 0.28152923 | 1.76708614 | 0.0772138 | 0.22166902 |
| ITGB3BP | 899.123245 | -0.176947218 | 0.42491131 | -0.4164333 | 0.67709297 | 0.83060787 |
| EFCAB7 | 463.724306 | 0.261401315 | 0.2243449 | 1.16517611 | 0.24394771 | 0.4627096 |
| PGM1 | 1186.6382 | -0.148730059 | 0.22485831 | -0.661439 | 0.50833081 | 0.71121086 |
| ROR1 | 549.263638 | 0.857066299 | 0.587743 | 1.4582331 | 0.1447763 | 0.3347366 |
| UBE2U | 45.3716905 | 2.038098724 | 1.15286478 | 1.76785583 | 0.07708501 | 0.22154393 |
| CACHD1 | 129.277021 | -0.081476908 | 0.54044308 | -0.1507595 | 0.88016547 | 0.94636668 |
| RAVER2 | 785.589875 | 0.478089794 | 0.20123816 | 2.37574126 | 0.01751374 | 0.08160105 |
| JAK1 | 1996.26398 | -0.58816859 | 0.18661672 | -3.1517465 | 0.00162297 | 0.01462334 |
| AK4 | 528.923707 | -2.132558266 | 0.54837826 | -3.8888454 | 0.00010072 | 0.00167664 |
| DNAJC6 | 902.423259 | 1.034402182 | 0.4167727 | 2.48193362 | 0.01306716 | 0.06664086 |
| AL139294.1 | 108.865929 | 0.698263943 | 0.74280742 | 0.94003361 | 0.34720032 | 0.57269843 |
| LEPR | 1214.65749 | -0.909632007 | 0.35944232 | -2.5306759 | 0.0113843 | 0.06078222 |
| SGIP1 | 7160.34328 | 0.081131954 | 0.38313823 | 0.21175636 | 0.83229712 | 0.92160388 |
| AL139147.1 | 119.98758 | -0.473843482 | 0.62987244 | -0.7522848 | 0.45187979 | 0.66765997 |
| WDR78 | 912.905945 | 0.181911336 | 0.21766301 | 0.83574759 | 0.40329692 | 0.62459245 |
| MIER1 | 1366.18999 | -0.076654697 | 0.1644699 | -0.4660713 | 0.64116448 | 0.80747519 |
| SLC35D1 | 549.600983 | 0.647692219 | 0.21921811 | 2.95455622 | 0.00313119 | 0.02387985 |
| IL12RB2 | 43.0492465 | 0.332249159 | 0.6433646 | 0.51642437 | 0.60555805 | 0.78178941 |
| SERBP1 | 849.96493 | -0.31901047 | 0.16504847 | -1.9328291 | 0.05325726 | 0.17348339 |
| GADD45A | 505.743169 | -0.519580736 | 0.2987462 | -1.7392045 | 0.08199879 | 0.2302066 |
| GNG12 | 1709.08539 | -1.049024366 | 0.20399191 | -5.1424803 | 2.71E-07 | 1.40E-05 |
| GNG12-AS1 | 1462.75825 | -0.644152063 | 0.28326846 | -2.2739986 | 0.02296607 | 0.09792087 |
| DIRAS3 | 78.1323009 | 0.097718154 | 0.5587531 | 0.17488611 | 0.86116913 | 0.93697711 |
| WLS | 2788.16622 | 0.531129156 | 0.20533184 | 2.58668673 | 0.00969036 | 0.05412263 |
| LRRC7 | 71.7152312 | 0.38464487 | 0.37885496 | 1.01528267 | 0.3099711 | 0.53392391 |
| LRRC40 | 480.682429 | 0.366575808 | 0.16858036 | 2.17448703 | 0.02966858 | 0.1165999 |
| SRSF11 | 2647.92934 | 0.31516012 | 0.14084084 | 2.23770406 | 0.02524036 | 0.10419747 |
| ANKRD13C | 1220.6955 | -0.0235069 | 0.1739379 | -0.1351454 | 0.89249695 | 0.95118543 |
| HHLA3 | 136.170752 | -0.124079927 | 0.30667085 | -0.4046029 | 0.6857694 | 0.83751486 |
| CTH | 130.568416 | 0.460641361 | 0.34491535 | 1.33552004 | 0.1817062 | 0.38600148 |
| LINC01788 | 63.0728475 | 1.694875827 | 0.59800761 | 2.83420442 | 0.00459399 | 0.03150678 |
| PTGER3 | 9083.73995 | 1.963254996 | 0.47368083 | 4.14467898 | 3.40E-05 | 0.00072733 |
| AL031429.1 | 52.3414179 | 2.018546383 | 0.79457729 | 2.54040281 | 0.01107249 | 0.05964783 |
| ZRANB2-AS1 | 220.143667 | 1.165969874 | 0.48275891 | 2.41522185 | 0.01572563 | 0.07572724 |
| ZRANB2 | 2300.68059 | 0.399398557 | 0.13893951 | 2.87462184 | 0.00404512 | 0.02886653 |
| NEGR1 | 590.580643 | -0.310888655 | 0.36373499 | -0.854712 | 0.39271061 | 0.61497504 |
| LRRIQ3 | 209.277921 | 1.232135762 | 0.45388687 | 2.71463188 | 0.00663495 | 0.04126225 |
| CRYZ | 986.140907 | 0.270154238 | 0.30289626 | 0.8919035 | 0.37244466 | 0.59653468 |
| TYW3 | 287.974277 | 0.406673332 | 0.27444983 | 1.48177658 | 0.13839975 | 0.32460044 |
| SLC44A5 | 134.055136 | -0.755884725 | 0.66072318 | -1.1440263 | 0.25261271 | 0.47267033 |
| ACADM | 878.532269 | 0.785251613 | 0.20765291 | 3.78155839 | 0.00015585 | 0.00238819 |
| RABGGTB | 272.535086 | 0.325958254 | 0.3279577 | 0.99390334 | 0.32026993 | 0.54463101 |
| MSH4 | 68.8646436 | 0.960572958 | 0.56036409 | 1.71419436 | 0.08649304 | 0.23821654 |
| ST6GALNAC3 | 79.9032462 | 0.357162354 | 0.43340338 | 0.82408761 | 0.40988978 | 0.62963532 |
| ST6GALNAC5 | 354.763385 | 2.790539552 | 0.63774214 | 4.37565492 | 1.21E-05 | 0.00031113 |
| PIGK | 567.541696 | -0.080725719 | 0.24161728 | -0.3341057 | 0.73829977 | 0.87200855 |
| AC118549.1 | 2805.2223 | 0.363382443 | 0.12428794 | 2.92371435 | 0.00345882 | 0.02575398 |
| USP33 | 1700.46409 | 0.200723612 | 0.10940768 | 1.83463922 | 0.06655916 | 0.20118535 |
| MIGA1 | 1021.92829 | -0.036045981 | 0.09966692 | -0.3616644 | 0.71760281 | 0.85947463 |
| NEXN | 199.961237 | 0.187915765 | 0.30077391 | 0.62477416 | 0.53211929 | 0.72935185 |
| FUBP1 | 1410.79795 | 0.360490116 | 0.15718387 | 2.29342951 | 0.02182329 | 0.09455259 |
| GIPC2 | 209.601141 | 0.428049425 | 0.40080543 | 1.06797311 | 0.28553264 | 0.50789128 |
| PTGFR | 480.070319 | 0.027017832 | 0.46091278 | 0.0586181 | 0.95325629 | 0.9794308 |
| IFI44L | 998.163173 | -1.149144238 | 1.54984747 | -0.7414563 | 0.45841679 | 0.67266311 |
| IFI44 | 365.190711 | 0.503372728 | 1.14377612 | 0.44009725 | 0.65986667 | 0.81980614 |
| AL136234.1 | 160.178948 | -0.510057768 | 0.91306709 | -0.5586203 | 0.57642092 | 0.76324677 |
| LINC01781 | 792.590153 | -1.836525364 | 1.27339254 | -1.4422303 | 0.1492374 | 0.34127917 |
| ADGRL2 | 2009.68588 | 0.546428545 | 0.3148527 | 1.73550537 | 0.08265133 | 0.23103419 |
| LINC01362 | 218.912976 | 0.181703067 | 0.62021725 | 0.29296681 | 0.76954752 | 0.88901113 |
| LINC01725 | 3016.27557 | 0.621925945 | 0.38242542 | 1.62626729 | 0.10389281 | 0.26963805 |
| TTLL7 | 195.607215 | -0.906764414 | 0.45971781 | -1.972437 | 0.04855974 | 0.16300525 |
| PRKACB | 1110.23941 | 0.319320864 | 0.20298513 | 1.57312439 | 0.11569002 | 0.28931266 |
| SAMD13 | 72.0467889 | 0.34573846 | 0.38351831 | 0.90149141 | 0.3673271 | 0.59270895 |
| RPF1 | 181.902537 | -0.034476044 | 0.19346065 | -0.178207 | 0.85856041 | 0.93614803 |
| GNG5 | 509.236953 | -0.329503632 | 0.27765901 | -1.1867205 | 0.23533789 | 0.45288144 |

| | | | | | | |
|---|---|---|---|---|---|---|
| SPATA1 | 459.812425 | 0.314844203 | 0.46671774 | 0.67459232 | 0.49993481 | 0.70482771 |
| CTBS | 657.022976 | 0.393080337 | 0.1874805 | 2.09664651 | 0.03602488 | 0.133479 |
| SSX2IP | 305.484392 | 0.324561535 | 0.30187961 | 1.07513568 | 0.28231399 | 0.50474898 |
| WDR63 | 65.0766205 | 0.265881361 | 0.54354149 | 0.4891648 | 0.62472503 | 0.79598067 |
| SYDE2 | 230.006008 | 1.600320661 | 0.32620934 | 4.90580884 | 9.30E-07 | 3.94E-05 |
| C1orf52 | 333.502847 | 0.35377367 | 0.21508755 | 1.64478913 | 0.1000133 | 0.26284834 |
| BCL10 | 182.325716 | -0.449748628 | 0.22468691 | -2.0016681 | 0.04532043 | 0.15537776 |
| AL078459.1 | 515.607629 | -0.501534046 | 0.37320699 | -1.3438496 | 0.17899704 | 0.38247715 |
| DDAH1 | 2684.67867 | 0.461165933 | 0.26320249 | 1.75213363 | 0.07975083 | 0.22596069 |
| CYR61 | 168.185072 | 2.43018865 | 0.8871882 | 2.73920307 | 0.00615883 | 0.03901114 |
| ZNHIT6 | 637.480168 | 0.01118599 | 0.18798069 | 0.05950606 | 0.95254904 | 0.97933189 |
| COL24A1 | 219.897517 | 1.542801344 | 0.47076508 | 3.27722129 | 0.00104834 | 0.01036493 |
| AL136382.1 | 154.235645 | 0.065400958 | 0.46013568 | 0.14213407 | 0.88697411 | 0.94913293 |
| ODF2L | 1394.36877 | -0.205645793 | 0.16019578 | -1.2837154 | 0.19924156 | 0.40783848 |
| SH3GLB1 | 844.711263 | -0.403625427 | 0.17365972 | -2.3242317 | 0.02011308 | 0.08933209 |
| SELENOF | 974.636328 | 0.0170089 | 0.13905918 | 0.12231411 | 0.90265026 | 0.95499458 |
| HS2ST1 | 1164.62134 | -0.001732523 | 0.20466295 | -0.0084652 | 0.99324579 | 0.99737436 |
| LMO4 | 517.211174 | -0.338408403 | 0.20386016 | -1.6600027 | 0.09691391 | 0.25726204 |
| PKN2-AS1 | 193.93303 | 0.707482646 | 0.25989213 | 2.72221653 | 0.00648456 | 0.04049863 |
| PKN2 | 2926.84258 | 0.188005744 | 0.12060899 | 1.55880369 | 0.11904285 | 0.29440873 |
| GTF2B | 431.776053 | -0.224529565 | 0.21859984 | -1.0271259 | 0.30436118 | 0.5276817 |
| KYAT3 | 744.12141 | -0.249407971 | 0.18350533 | -1.359132 | 0.17410476 | 0.37589771 |
| GBP3 | 170.326843 | -1.85863222 | 0.49358706 | -3.7655611 | 0.00016618 | 0.00252011 |
| GBP2 | 596.799235 | -5.15314238 | 0.49995874 | -10.307135 | 6.54E-25 | 8.64E-21 |
| LRRC8B | 599.84598 | -0.108387462 | 0.204985 | -0.528758 | 0.59697333 | 0.77654325 |
| LRRC8C-DT | 75.0248616 | -0.053344097 | 0.56293202 | -0.0947612 | 0.92450453 | 0.96673372 |
| LRRC8D | 71.9910534 | 0.338874624 | 0.47237818 | 0.71737993 | 0.4731397 | 0.68383017 |
| ZNF326 | 775.328385 | 0.189508955 | 0.11666147 | 1.62443487 | 0.10428303 | 0.27041118 |
| ZNF644 | 2460.9698 | 0.249237058 | 0.17156294 | 1.45274417 | 0.1462948 | 0.33683249 |
| HFM1 | 403.58366 | 0.837374254 | 0.22683491 | 3.69155805 | 0.00022288 | 0.00320706 |
| CDC7 | 112.218627 | -1.301220429 | 0.6724196 | -1.9351316 | 0.05297415 | 0.17273155 |
| TGFBR3 | 454.819805 | -0.585954823 | 0.3652991 | -1.6040412 | 0.10870496 | 0.27714415 |
| BTBD8 | 371.649503 | -0.535823822 | 0.37577256 | -1.4259259 | 0.15388973 | 0.34807567 |
| GLMN | 469.782863 | 0.36080911 | 0.1847276 | 1.95319542 | 0.05079644 | 0.16799455 |
| RPAP2 | 1558.7208 | 0.102037043 | 0.17294176 | 0.59000813 | 0.5551852 | 0.74754753 |
| EVI5 | 2686.7985 | 0.253579045 | 0.17681202 | 1.43417315 | 0.15152286 | 0.3444863 |
| RPL5 | 817.828164 | -0.837718905 | 0.28844028 | -2.9043062 | 0.00368068 | 0.02694131 |
| FAM69A | 324.696261 | 0.040418095 | 0.31425218 | 0.12861675 | 0.89766091 | 0.95261533 |
| MTF2 | 1402.52987 | 0.213106627 | 0.16013394 | 1.33080239 | 0.18325404 | 0.38804146 |
| TMED5 | 459.759847 | -0.210186873 | 0.27019049 | -0.7779211 | 0.43661556 | 0.65342962 |
| CCDC18 | 215.436677 | 0.339743938 | 0.38005239 | 0.89393976 | 0.37135412 | 0.59563048 |
| DR1 | 257.333035 | 0.099134106 | 0.25209683 | 0.3932382 | 0.69414356 | 0.843734 |
| FNBP1L | 3798.99395 | -0.015742901 | 0.2220893 | -0.0708855 | 0.94348892 | 0.97559625 |
| BCAR3 | 1000.26394 | 0.122639617 | 0.35069804 | 0.34970146 | 0.72656276 | 0.86526092 |
| AL109613.1 | 69.8243779 | 0.309892038 | 0.44277993 | 0.69987823 | 0.48400335 | 0.692203 |
| DNTTIP2 | 116.459892 | -0.577537548 | 0.22405342 | -2.5776779 | 0.00994667 | 0.05520401 |
| GCLM | 537.439555 | -1.36773374 | 0.30987583 | -4.4138123 | 1.02E-05 | 0.00026939 |
| ABCA4 | 189.831078 | 1.146783992 | 0.8141995 | 1.40848034 | 0.15898888 | 0.35534419 |
| ARHGAP29 | 4365.16016 | -0.137190982 | 0.39643092 | -0.3460653 | 0.72929365 | 0.86700908 |
| ABCD3 | 1561.16069 | 0.162760509 | 0.19520463 | 0.83379433 | 0.40439689 | 0.62534284 |
| F3 | 86.4368372 | -2.701495947 | 0.74826671 | -3.6103383 | 0.0003058 | 0.00407103 |
| SLC44A3 | 1293.89064 | 0.294802023 | 0.34399138 | 0.8570041 | 0.39144262 | 0.6140943 |
| CNN3 | 1814.20682 | -0.235799901 | 0.27385034 | -0.8610539 | 0.38920836 | 0.6122488 |
| AC105942.1 | 110.225938 | -0.306745151 | 0.26535402 | -1.1559846 | 0.24768747 | 0.46703404 |
| ALG14 | 397.386055 | 0.102364663 | 0.29794695 | 0.34356675 | 0.73117214 | 0.86806155 |
| AC092802.1 | 99.8587127 | 0.901686665 | 0.39310325 | 2.29376547 | 0.02180397 | 0.09452959 |
| PTBP2 | 2495.41187 | -0.148873617 | 0.13421271 | -1.1092364 | 0.26732823 | 0.48853606 |
| DPYD | 3637.70106 | -1.481260237 | 0.35320824 | -4.1937307 | 2.74E-05 | 0.00061069 |
| DPYD-AS1 | 172.462035 | -1.852094688 | 0.50057291 | -3.6999499 | 0.00021564 | 0.00312326 |
| SNX7 | 431.108292 | -0.696515637 | 0.21711664 | -3.2080251 | 0.0013365 | 0.01255605 |
| FRRS1 | 224.938578 | 0.875550981 | 0.27849172 | 3.14390314 | 0.00166711 | 0.01487892 |
| AGL | 945.290826 | 0.652502541 | 0.11307013 | 5.77077724 | 7.89E-09 | 7.61E-07 |
| AC093019.2 | 51.9559107 | -0.038604276 | 0.55564836 | -0.0694761 | 0.94461066 | 0.9759376 |
| SASS6 | 102.675748 | 0.5344029 | 0.32080238 | 1.66583212 | 0.09574686 | 0.25508678 |
| TRMT13 | 622.839136 | 0.673640588 | 0.16209237 | 4.15590565 | 3.24E-05 | 0.00069703 |
| LRRC39 | 105.672218 | 0.634261844 | 0.30833645 | 2.05704465 | 0.03968193 | 0.14181781 |
| DBT | 679.583413 | 0.512383364 | 0.16642352 | 3.07879178 | 0.00207842 | 0.0176552 |
| RTCA | 405.95189 | 0.328699721 | 0.17543103 | 1.87366923 | 0.06097603 | 0.19001878 |
| CDC14A | 689.206765 | -0.141578563 | 0.27071323 | -0.5229835 | 0.60098571 | 0.7789212 |
| VCAM1 | 107.789119 | -1.152292824 | 0.65644265 | -1.7553595 | 0.07919784 | 0.22487623 |
| EXTL2 | 375.995651 | 0.545174798 | 0.17879706 | 3.04912626 | 0.00229508 | 0.01900672 |
| SLC30A7 | 891.586862 | -0.371285194 | 0.18682964 | -1.9872927 | 0.04688997 | 0.15889419 |
| DPH5 | 277.643663 | 0.739996191 | 0.154749 | 4.78191256 | 1.74E-06 | 6.29E-05 |
| AC093157.1 | 356.429155 | 0.633910126 | 0.1817356 | 3.48808989 | 0.00048648 | 0.00579439 |
| RNPC3 | 1522.64709 | 0.593344905 | 0.18033265 | 3.29028005 | 0.00100088 | 0.01000044 |
| AMY2B | 381.283517 | 0.655365403 | 0.3215752 | 2.03798491 | 0.04155144 | 0.14604923 |
| PRMT6 | 44.5825544 | -0.367635924 | 0.46906751 | -0.7837591 | 0.43318149 | 0.65044973 |
| NTNG1 | 220.342709 | -3.852504066 | 0.67856062 | -5.6774649 | 1.37E-08 | 1.21E-06 |
| VAV3 | 5428.60293 | 0.691123634 | 0.32973098 | 2.09602273 | 0.03608017 | 0.13364639 |
| SLC25A24 | 469.74919 | -0.049411147 | 0.21473714 | -0.2301006 | 0.81801359 | 0.91545722 |
| PRPF38B | 755.767693 | -0.038012442 | 0.22145702 | -0.171647 | 0.86371502 | 0.9382347 |
| STXBP3 | 827.999042 | 0.34428124 | 0.1861167 | 1.84981381 | 0.06434039 | 0.19689524 |
| AL449266.1 | 169.457094 | -0.255975305 | 0.32104085 | -0.7973294 | 0.42525975 | 0.64388538 |
| GPSM2 | 203.247023 | -0.572214317 | 0.53765475 | -1.0642784 | 0.28720261 | 0.50962645 |
| WDR47 | 273.659175 | 0.356470356 | 0.23625287 | 1.50885093 | 0.13133688 | 0.31399617 |
| TAF13 | 160.04235 | -0.117633924 | 0.2819327 | -0.4172412 | 0.67650203 | 0.83016679 |
| TMEM167B | 153.273312 | -0.565993835 | 0.24104228 | -2.3481102 | 0.01886894 | 0.08570832 |
| SARS | 441.803866 | -0.514582716 | 0.21433594 | -2.4008233 | 0.01635823 | 0.07772514 |
| CELSR2 | 141.998855 | -0.630621597 | 0.42255429 | -1.4924037 | 0.13559334 | 0.32074721 |
| SORT1 | 1567.71272 | -0.101411884 | 0.32043162 | -0.3164853 | 0.75163422 | 0.87947193 |
| PSMA5 | 559.378197 | -0.561841628 | 0.28291516 | -1.9859014 | 0.04704427 | 0.15913131 |
| SYPL2 | 351.556585 | 0.666530193 | 0.44704835 | 1.49095771 | 0.1359726 | 0.321247 |
| ATXN7L2 | 41.7794049 | 0.186036552 | 0.40056392 | 0.46443662 | 0.64233496 | 0.80832017 |
| CYB561D1 | 137.235975 | -0.601157482 | 0.28307329 | -2.1236814 | 0.03369679 | 0.1277924 |
| GNAI3 | 786.924849 | -0.058959209 | 0.17605222 | -0.3348961 | 0.73770343 | 0.87166335 |
| AMPD2 | 188.924438 | 0.298573478 | 0.38658603 | 0.77233385 | 0.43991672 | 0.65563127 |
| GSTM4 | 148.764522 | 1.187577358 | 0.37310879 | 3.18292519 | 0.00145795 | 0.01342028 |
| GSTM3 | 413.487641 | -0.049154995 | 0.53507736 | -0.0918652 | 0.92680513 | 0.9678369 |
| AHCYL1 | 1734.48853 | -0.344268799 | 0.31013347 | -1.1100666 | 0.26697034 | 0.48821975 |
| STRIP1 | 262.719709 | 0.170783267 | 0.20975958 | 0.81418577 | 0.41553854 | 0.63535693 |
| KCNC4 | 516.823433 | -1.168650632 | 0.74192863 | -1.5751524 | 0.11522128 | 0.28859915 |
| RBM15-AS1 | 60.3621465 | -0.438660118 | 0.46632654 | -0.9406716 | 0.34687319 | 0.57223029 |
| RBM15 | 266.40979 | 0.434958409 | 0.19835465 | 2.19283192 | 0.02831949 | 0.11284228 |
| SLC16A4 | 277.875756 | -0.48013514 | 0.40982494 | -1.1715615 | 0.24137313 | 0.45974012 |
| AL355488.1 | 119.053144 | 0.953100411 | 0.2462037 | 3.87118636 | 0.00010831 | 0.00178605 |
| LAMTOR5 | 499.921959 | -1.004550615 | 0.35029374 | -2.8677379 | 0.00413418 | 0.02926733 |
| LRIF1 | 221.40039 | -0.27016476 | 0.25585288 | -1.0559379 | 0.29099658 | 0.51373981 |
| DRAM2 | 469.356248 | 0.014572978 | 0.15376164 | 0.09477642 | 0.92449242 | 0.96673372 |
| CEPT1 | 676.776802 | 0.50287284 | 0.17550483 | 2.86529351 | 0.00416623 | 0.02937201 |
| DENND2D | 234.199004 | 0.001251816 | 0.23678493 | 0.00528672 | 0.99578182 | 0.99847083 |
| WDR77 | 114.32306 | 0.212835325 | 0.26767848 | 0.79511557 | 0.42654628 | 0.64506142 |
| ATP5PB | 628.522523 | -0.039012824 | 0.30155269 | -0.1293732 | 0.89706238 | 0.9525128 |
| RAP1A | 1957.42082 | 0.101271028 | 0.20471957 | 0.49468171 | 0.62082482 | 0.7928889 |

| | | | | | | |
|---|---|---|---|---|---|---|
| INKA2 | 131.498425 | -0.321873707 | 0.59420875 | -0.5416846 | 0.58803583 | 0.7710768 |
| DDX20 | 100.653974 | 0.249219259 | 0.2921976 | 0.85291344 | 0.39370731 | 0.61588683 |
| KCND3 | 472.821491 | -1.752357059 | 0.53182096 | -3.2950132 | 0.00098417 | 0.00987479 |
| CTTNBP2NL | 628.811695 | -1.179035303 | 0.17286568 | -6.8205284 | 9.07E-12 | 2.35E-09 |
| WNT2B | 186.930886 | -1.192395827 | 0.45103524 | -2.6436866 | 0.00820085 | 0.04810173 |
| ST7L | 764.480567 | 0.435555296 | 0.15635354 | 2.78570796 | 0.0053411 | 0.03520486 |
| CAPZA1 | 1131.20106 | -0.409468406 | 0.16200017 | -2.52758 | 0.01148516 | 0.0611231 |
| MOV10 | 288.107297 | -0.172732514 | 0.29854859 | -0.5785742 | 0.56287652 | 0.75444301 |
| AL603832.1 | 61.4749675 | -0.973374484 | 0.42998615 | -2.2637345 | 0.02359045 | 0.09974591 |
| RHOC | 319.199721 | -2.187603786 | 0.38726398 | -5.6488698 | 1.62E-08 | 1.39E-06 |
| SLC16A1-AS1 | 74.5441627 | 0.488255601 | 0.43052766 | 1.1340865 | 0.25675827 | 0.47774616 |
| AL357055.3 | 88.5247151 | 1.29527508 | 0.33053826 | 3.91868431 | 8.90E-05 | 0.00154337 |
| LRIG2 | 663.622271 | 0.416555673 | 0.16536735 | 2.51897175 | 0.01176981 | 0.06225259 |
| MAGI3 | 4610.95269 | 0.26539029 | 0.29965979 | 0.88563863 | 0.37581229 | 0.5990954 |
| PHTF1 | 665.369642 | 0.05949995 | 0.1705197 | 0.34893299 | 0.72713962 | 0.8653741 |
| RSBN1 | 620.24851 | 0.050680359 | 0.17056109 | 0.29713905 | 0.76636036 | 0.88690833 |
| AP4B1 | 73.4259059 | 0.287262974 | 0.32729213 | 0.87769594 | 0.38010875 | 0.60281624 |
| HIPK1 | 464.341801 | -0.122964792 | 0.20921007 | -0.5877575 | 0.55669507 | 0.74881722 |
| TRIM33 | 1758.26872 | -0.011488274 | 0.20931471 | -0.0548852 | 0.95622994 | 0.98073153 |
| BCAS2 | 205.236604 | 0.076473521 | 0.27159642 | 0.28157044 | 0.77827291 | 0.89400754 |
| DENND2C | 96.9750427 | -0.400386102 | 0.37098285 | -1.0792577 | 0.28047286 | 0.50294945 |
| NRAS | 136.033689 | -0.446739811 | 0.27582739 | -1.6196355 | 0.10531061 | 0.27208298 |
| CSDE1 | 1246.49651 | -0.196140507 | 0.12407741 | -1.5807914 | 0.11392574 | 0.28641895 |
| SIKE1 | 240.602747 | 0.262003455 | 0.2536978 | 1.03273838 | 0.30172635 | 0.52496248 |
| VANGL1 | 104.200386 | 0.098796359 | 0.33871835 | 0.29167702 | 0.77053358 | 0.88937357 |
| SLC22A15 | 1014.99706 | -0.003748835 | 0.30781847 | -0.0121787 | 0.99028303 | 0.99606252 |
| MAB21L3 | 134.974815 | -3.304007133 | 1.03200881 | -3.20153 | 0.001367 | 0.01276091 |
| ATP1A1 | 15607.88 | -0.240950912 | 0.3225138 | -0.7471026 | 0.4550016 | 0.67027397 |
| LINC01762 | 7477.16215 | 0.618412394 | 0.65352875 | 0.94626654 | 0.34401265 | 0.56886284 |
| AL390066.1 | 69.5046911 | 0.096260449 | 0.53052108 | 0.1814451 | 0.85601822 | 0.93470651 |
| CD58 | 838.138823 | -0.670290125 | 0.16379949 | -4.092138 | 4.27E-05 | 0.00086857 |
| IGSF3 | 1287.91085 | -0.392908065 | 0.30929742 | -1.2703244 | 0.2039691 | 0.41379632 |
| PTGFRN | 322.957413 | -0.442379648 | 0.29628631 | -1.4930816 | 0.13541582 | 0.32061438 |
| AL445231.1 | 41.5537699 | 0.688102594 | 0.39145611 | 1.75780267 | 0.07878109 | 0.22441496 |
| TTF2 | 250.917055 | -0.325953066 | 0.27980242 | -1.1649401 | 0.24404323 | 0.4628237 |
| TRIM45 | 45.7130341 | 1.337731288 | 0.32639729 | 4.09847548 | 4.16E-05 | 0.00085433 |
| VTCN1 | 321.693416 | 0.395369836 | 0.73573961 | 0.5373774 | 0.59100695 | 0.77248385 |
| MAN1A2 | 2073.14117 | 0.198401221 | 0.14776118 | 1.34271545 | 0.17936413 | 0.38281157 |
| TENT5C | 87.7123276 | 0.722684789 | 0.54031158 | 1.3375334 | 0.18104859 | 0.38492165 |
| GDAP2 | 679.54482 | 0.2094347 | 0.18706779 | 1.11956583 | 0.26289882 | 0.48398616 |
| WDR3 | 155.398913 | 0.179472282 | 0.25730854 | 0.69749835 | 0.48549097 | 0.69300247 |
| SPAG17 | 1193.808 | 0.452457121 | 0.30355691 | 1.49051826 | 0.13608802 | 0.32134191 |
| TBX15 | 109.384349 | 0.667119492 | 1.16083428 | 0.57468969 | 0.56550118 | 0.75611956 |
| WARS2 | 576.935982 | 0.416732599 | 0.13648265 | 3.05337424 | 0.00226284 | 0.01876321 |
| AL359915.2 | 1695.12979 | 0.551985392 | 0.32012397 | 1.72428633 | 0.08465617 | 0.2345267 |
| AL359915.1 | 258.888721 | 3.200336109 | 0.7965432 | 4.01778097 | 5.87E-05 | 0.00111177 |
| ZNF697 | 217.819817 | 1.7799625 | 0.35139858 | 5.06536616 | 4.08E-07 | 1.99E-05 |
| PHGDH | 124.391402 | 0.993565889 | 0.49328478 | 2.01418314 | 0.04399031 | 0.15199268 |
| HMGCS2 | 86.5641802 | 2.99076852 | 1.05592607 | 2.83236546 | 0.0046205 | 0.03160652 |
| NOTCH2 | 556.402587 | 0.040554023 | 0.17395546 | 0.23312877 | 0.81566141 | 0.91421906 |
| SEC22B | 674.084533 | 0.285049157 | 0.15826561 | 1.80108083 | 0.07169014 | 0.21127958 |
| NBPF26 | 157.282079 | 0.321594829 | 0.5637402 | 0.57046637 | 0.56836142 | 0.75840853 |
| LINC00623 | 792.811501 | -0.668137663 | 0.58168935 | -1.1486159 | 0.25071438 | 0.47078342 |
| AC244453.3 | 53.0917526 | 0.581180937 | 0.44573048 | 1.3038842 | 0.19227307 | 0.39951784 |
| SRGAP2C | 120.832976 | -0.948890362 | 0.35388841 | -2.6813265 | 0.00733309 | 0.04418924 |
| AC239800.3 | 84.7762755 | -0.989485266 | 0.56046675 | -1.7654665 | 0.0774854 | 0.22216293 |
| NBPF15 | 232.113329 | -0.469700266 | 0.37194322 | -1.2628279 | 0.20665105 | 0.41739901 |
| FP700111.1 | 220.930666 | 0.567430146 | 0.36927154 | 1.53662031 | 0.12438632 | 0.30252603 |
| SRGAP2B | 143.764569 | -1.362286864 | 0.34978741 | -3.8946138 | 9.84E-05 | 0.00165158 |
| GPR89A | 206.982976 | 0.000271578 | 0.23693559 | 0.00114621 | 0.99908546 | 0.99971199 |
| PDZK1 | 288.755536 | 0.901979381 | 0.46421871 | 1.94300523 | 0.05201553 | 0.17074381 |
| RNF115 | 1968.91764 | -0.015233573 | 0.19426242 | -0.0784175 | 0.93749595 | 0.97284815 |
| POLR3C | 270.915421 | -0.37061123 | 0.25603551 | -1.4474993 | 0.14775713 | 0.33919428 |
| PIAS3 | 112.968507 | -0.568745836 | 0.32145459 | -1.7692883 | 0.07684578 | 0.22114508 |
| ITGA10 | 100.864268 | -2.066373914 | 0.71595565 | -2.8861759 | 0.00389954 | 0.02813557 |
| PEX11B | 163.908895 | -0.318222542 | 0.2693161 | -1.1815949 | 0.23736647 | 0.45506149 |
| RBM8A | 268.141513 | -0.752199303 | 0.23213115 | -3.2404066 | 0.00119359 | 0.01146631 |
| LIX1L | 78.3071714 | -0.171319061 | 0.27359245 | -0.6261834 | 0.53119463 | 0.72863563 |
| POLR3GL | 46.236179 | -0.321610348 | 0.52998885 | -0.6068247 | 0.54396722 | 0.73943993 |
| TXNIP | 5191.7281 | 0.522996529 | 0.43602877 | 1.19945418 | 0.23035139 | 0.44592324 |
| NBPF12 | 433.93577 | 0.04740015 | 0.18798695 | 0.25214596 | 0.80092824 | 0.90662963 |
| PRKAB2 | 202.169722 | 0.004021226 | 0.15464679 | 0.02600265 | 0.97925523 | 0.9910179 |
| AC242426.2 | 571.703693 | 0.511276253 | 0.17629562 | 2.90010756 | 0.00373035 | 0.02720105 |
| FMO5 | 307.845487 | 0.396388636 | 0.37032082 | 1.07039251 | 0.28444267 | 0.50683977 |
| CHD1L | 532.258849 | 0.529511498 | 0.21111812 | 2.5081291 | 0.01213723 | 0.063418 |
| LINC00624 | 119.234185 | -0.329758324 | 0.44022466 | -0.7490683 | 0.45381605 | 0.66948056 |
| BCL9 | 566.235133 | 0.351392942 | 0.23813831 | 1.47558343 | 0.14005576 | 0.32747126 |
| ACP6 | 600.31385 | -0.111333526 | 0.17753756 | -0.6270984 | 0.5305947 | 0.72854734 |
| GPR89B | 266.14127 | 0.282347641 | 0.31366354 | 0.90016086 | 0.36803465 | 0.59306694 |
| NBPF11 | 350.914029 | 0.091801787 | 0.32479458 | 0.28264568 | 0.77744846 | 0.89359206 |
| AC239809.3 | 232.702694 | -0.09849026 | 0.36976839 | -0.2663566 | 0.78996457 | 0.90082888 |
| NBPF14 | 149.800103 | -0.094075448 | 0.26045802 | -0.3611924 | 0.71795565 | 0.85963642 |
| NUDT4B | 79.1586284 | 0.713461232 | 0.43345871 | 1.64597278 | 0.09976936 | 0.26241607 |
| PDE4DIP | 3157.38675 | 0.416130735 | 0.33900637 | 1.22750123 | 0.21963426 | 0.43361796 |
| NBPF9 | 158.965232 | -0.293681344 | 0.24202344 | -1.2134417 | 0.22496098 | 0.43942281 |
| AC245297.3 | 2808.98079 | -0.276231023 | 0.21041644 | -1.3127825 | 0.18925624 | 0.39536801 |
| NBPF19 | 283.740692 | 0.095873361 | 0.71513606 | 0.1340631 | 0.89335267 | 0.95146032 |
| BOLA1 | 57.1448981 | -0.153401386 | 0.32938938 | -0.4657144 | 0.64141993 | 0.80757823 |
| SF3B4 | 107.788109 | -0.597524666 | 0.33915573 | -1.7618003 | 0.07810305 | 0.22330371 |
| OTUD7B | 1128.70841 | -0.12329929 | 0.24315958 | -0.5070715 | 0.61210466 | 0.78684784 |
| VPS45 | 1229.76803 | 0.297739867 | 0.11243056 | 2.64821109 | 0.0080919 | 0.04767298 |
| ANP32E | 296.200641 | -0.299637627 | 0.27301913 | -1.0974968 | 0.27242428 | 0.49398089 |
| APH1A | 221.183013 | -0.394571921 | 0.26531558 | -1.4871796 | 0.13696738 | 0.3228412 |
| C1orf54 | 116.493237 | 0.397776094 | 0.65412947 | 0.60809994 | 0.54312119 | 0.73906334 |
| MRPS21 | 523.57681 | -0.290650431 | 0.1471409 | -1.9753205 | 0.04823179 | 0.16208414 |
| PRPF3 | 679.932626 | 0.015275073 | 0.3249211 | 0.04701164 | 0.96250395 | 0.9832997 |
| RPRD2 | 2283.10896 | 0.065431856 | 0.15067888 | 0.43424702 | 0.66410907 | 0.82290964 |
| TARS2 | 166.703103 | -0.11972796 | 0.32470513 | -0.3687283 | 0.71233028 | 0.85574627 |
| AL356356.1 | 42.3357877 | -1.346124702 | 0.55469764 | -2.426772 | 0.01523383 | 0.07418897 |
| ADAMTSL4 | 62.1595237 | -0.800405996 | 0.44333314 | -1.8054279 | 0.07100776 | 0.21020651 |
| MCL1 | 710.910253 | -0.860997771 | 0.36065341 | -2.3873274 | 0.01697137 | 0.07963582 |
| ENSA | 1538.91032 | -0.015283028 | 0.19456703 | -0.0785489 | 0.93739143 | 0.97284815 |
| GOLPH3L | 709.51099 | -0.28615836 | 0.1557279 | -1.8375536 | 0.06612821 | 0.20038714 |
| CTSS | 77.5260272 | -1.459653943 | 0.51402261 | -2.8396688 | 0.00451604 | 0.03115006 |
| CTSK | 49.5476181 | -0.047213929 | 0.33696309 | -0.140116 | 0.88856832 | 0.94975716 |
| ARNT | 825.960966 | 0.152101127 | 0.16888541 | 0.90061732 | 0.36779182 | 0.59294486 |
| SETDB1 | 612.741303 | 0.029508806 | 0.18666245 | 0.15808646 | 0.87438867 | 0.94347633 |
| CERS2 | 333.609571 | -0.307461482 | 0.22734554 | -1.3523972 | 0.17624827 | 0.378495 |
| MINDY1 | 203.208867 | -0.117194495 | 0.27517646 | -0.4258885 | 0.67018908 | 0.82659266 |
| PRUNE1 | 245.700097 | 0.061796921 | 0.20228076 | 0.30550073 | 0.75998481 | 0.88357016 |
| C1orf56 | 173.97059 | 0.414344604 | 0.57226067 | 0.72404872 | 0.46903583 | 0.68108245 |
| CDC42SE1 | 478.859907 | -0.169836784 | 0.24091248 | -0.704973 | 0.48082707 | 0.68948936 |
| GABPB2 | 859.299426 | 0.625352162 | 0.31201901 | 2.00421172 | 0.04504739 | 0.1547717 |

| | | | | | | |
|---|---|---|---|---|---|---|
| SEMA6C | 42.2886733 | 0.360862912 | 0.39720628 | 0.90850254 | 0.36361277 | 0.58901423 |
| SCNM1 | 167.079275 | -0.007448623 | 0.29035512 | -0.0256535 | 0.97953372 | 0.9910179 |
| LYSMD1 | 62.6019042 | 0.385270159 | 0.23546546 | 1.63620667 | 0.10179634 | 0.26584181 |
| VPS72 | 151.973525 | 0.118157766 | 0.22662143 | 0.52138832 | 0.60209629 | 0.77940904 |
| PIP5K1A | 2301.08018 | 0.085420706 | 0.14832223 | 0.57591303 | 0.56467397 | 0.75596786 |
| PSMD4 | 384.343522 | -0.295300496 | 0.13748864 | -2.1478174 | 0.03172827 | 0.12257932 |
| ZNF687 | 66.6560627 | -0.400930311 | 0.33374068 | -1.2013229 | 0.22962598 | 0.44513201 |
| PI4KB | 551.283075 | 0.299073795 | 0.15391396 | 1.94312332 | 0.05200126 | 0.17073941 |
| RFX5 | 65.8077785 | -0.265144311 | 0.26529335 | -0.9994382 | 0.31758245 | 0.54219292 |
| SELENBP1 | 215.055582 | 2.003774734 | 0.70256606 | 2.85208018 | 0.00434341 | 0.03024363 |
| PSMB4 | 205.391502 | -0.515129133 | 0.4149388 | -1.2414581 | 0.21443657 | 0.42793362 |
| POGZ | 3125.11413 | 0.630515323 | 0.15707562 | 4.01408787 | 5.97E-05 | 0.00112029 |
| CGN | 503.343769 | 0.364752804 | 0.25031154 | 1.45719529 | 0.14506248 | 0.33492926 |
| TUFT1 | 506.900618 | 0.135118503 | 0.37328071 | 0.36197559 | 0.71737027 | 0.85947463 |
| SNX27 | 907.183746 | 0.341583261 | 0.16645986 | 2.05204582 | 0.04016521 | 0.14296477 |
| MRPL9 | 92.1029897 | -0.386663626 | 0.29746672 | -1.2998551 | 0.19365064 | 0.40124412 |
| TDRKH | 69.2108793 | -1.13861302 | 0.34944007 | -3.2583928 | 0.00112045 | 0.01091449 |
| RORC | 244.482893 | 0.533152102 | 0.46123526 | 1.15592226 | 0.24771296 | 0.46703404 |
| AL450992.2 | 70.0504036 | -0.307468611 | 0.36001356 | -0.8540473 | 0.39307878 | 0.61533273 |
| THEM4 | 1026.60015 | -0.308208725 | 0.57047204 | -0.5402696 | 0.58901109 | 0.77174702 |
| S100A10 | 1562.53699 | -1.765422822 | 0.48280633 | -3.6565859 | 0.0002556 | 0.00355896 |
| S100A11 | 3061.8223 | -3.247088285 | 0.39815528 | -8.1553316 | 3.48E-16 | 3.07E-13 |
| FLG-AS1 | 1024.82447 | 1.252930412 | 0.36025308 | 3.47791732 | 0.00050533 | 0.00596501 |
| S100A9 | 93.5956255 | -3.751303372 | 0.77898994 | -4.8155993 | 1.47E-06 | 5.51E-05 |
| S100A6 | 5405.33713 | -2.774793428 | 0.56471317 | -4.9136333 | 8.94E-07 | 3.85E-05 |
| S100A4 | 153.178731 | -0.984997097 | 0.62924919 | -1.565353 | 0.11750017 | 0.29215978 |
| S100A2 | 1317.04938 | -2.125322155 | 1.09989382 | -1.9322976 | 0.05332279 | 0.17352568 |
| S100A16 | 318.843976 | -2.184883651 | 0.3696007 | -5.9114705 | 3.39E-09 | 3.86E-07 |
| S100A14 | 152.32635 | -2.429273558 | 0.39436171 | -6.1600138 | 7.27E-10 | 9.80E-08 |
| S100A13 | 675.298388 | -1.772419655 | 0.36362855 | -4.8742587 | 1.09E-06 | 4.44E-05 |
| S100A1 | 68.5407103 | -0.841984424 | 0.60947506 | -1.3814912 | 0.16712799 | 0.36681632 |
| AL162258.2 | 54.761086 | -0.039621166 | 0.5597286 | -0.0707864 | 0.94356777 | 0.97559625 |
| CHTOP | 506.202792 | 0.388400429 | 0.10096629 | 3.84683258 | 0.00011965 | 0.00192746 |
| SNAPIN | 75.636327 | 0.091348066 | 0.27254691 | 0.33516457 | 0.73750095 | 0.87158876 |
| ILF2 | 284.8882 | -0.448068118 | 0.28407147 | -1.5773077 | 0.11472474 | 0.28793447 |
| NPR1 | 46.0459572 | -1.242822903 | 0.66484338 | -1.8693469 | 0.06157457 | 0.19132874 |
| INTS3 | 781.702877 | 0.175974322 | 0.26192508 | 0.67184983 | 0.5016793 | 0.70566164 |
| SLC27A3 | 59.192097 | -0.371244397 | 0.41200397 | -0.90107 | 0.36755113 | 0.59279882 |
| GATAD2B | 2649.39717 | -0.214789009 | 0.21849338 | -0.9830458 | 0.32558488 | 0.55073002 |
| DENND4B | 266.894166 | -0.001011059 | 0.2282374 | -0.0044299 | 0.9964655 | 0.99867719 |
| CRTC2 | 146.565208 | -0.162732314 | 0.2954244 | -0.5508425 | 0.58174165 | 0.7675782 |
| SLC39A1 | 239.424706 | -1.340133026 | 0.32213713 | -4.1601321 | 3.18E-05 | 0.00068537 |
| CREB3L4 | 72.011223 | 0.090808658 | 0.28225629 | 0.32172413 | 0.7476617 | 0.87699701 |
| JTB | 370.553379 | -0.167926984 | 0.38735695 | -0.43352 | 0.66463704 | 0.82323933 |
| RAB13 | 362.315884 | -1.640882694 | 0.35999647 | -4.5580522 | 5.16E-06 | 0.00015642 |
| RPS27 | 347.451812 | -0.81746619 | 0.44533917 | -1.8356036 | 0.0664163 | 0.20089144 |
| NUP210L | 218.492842 | -2.74485874 | 1.29146749 | -2.1253797 | 0.03355495 | 0.1274374 |
| TPM3 | 487.044428 | -0.967071616 | 0.20288685 | -4.7665563 | 1.87E-06 | 6.71E-05 |
| C1orf43 | 580.427573 | -0.26629345 | 0.17310531 | -1.5383321 | 0.12396744 | 0.30190666 |
| UBAP2L | 736.027068 | 0.080965797 | 0.16019628 | 0.50541622 | 0.61326653 | 0.78738701 |
| HAX1 | 157.010241 | -0.036577058 | 0.25423927 | -0.1438686 | 0.8856042 | 0.94843083 |
| ATP8B2 | 109.503336 | 0.34646381 | 0.38339402 | 0.90367558 | 0.36616745 | 0.59142894 |
| IL6R | 227.827696 | -2.741245306 | 0.49271547 | -5.5635463 | 2.64E-08 | 2.09E-06 |
| UBE2Q1 | 174.031494 | -0.173647877 | 0.23988729 | -0.7238728 | 0.46914385 | 0.68108245 |
| ADAR | 973.831901 | -0.229464344 | 0.32538992 | -0.7051981 | 0.48068698 | 0.68948936 |
| KCNN3 | 635.163197 | 1.875461019 | 0.75589952 | 2.4810983 | 0.01309782 | 0.06670975 |
| PMVK | 211.612032 | -0.210581358 | 0.29853864 | -0.7053739 | 0.48057759 | 0.68948936 |
| PBXIP1 | 242.933881 | -0.733870084 | 0.37597818 | -1.9518954 | 0.05095062 | 0.1682517 |
| PYGO2 | 85.5036035 | -0.105571048 | 0.30297723 | -0.3484455 | 0.72750564 | 0.86573172 |
| SHC1 | 51.6837033 | -0.838245266 | 0.37159398 | -2.2558096 | 0.02408256 | 0.10099717 |
| CKS1B | 46.9530346 | -0.014005431 | 0.43108069 | -0.0324891 | 0.97408199 | 0.98832358 |
| FLAD1 | 112.295699 | 0.088727915 | 0.28082033 | 0.31595972 | 0.75203308 | 0.8795471 |
| ZBTB7B | 137.899851 | -0.216696327 | 0.35071093 | -0.6178773 | 0.53665621 | 0.73349204 |
| ADAM15 | 65.7544908 | -0.648856443 | 0.46632747 | -1.391418 | 0.1640987 | 0.36287999 |
| EFNA1 | 540.122766 | -2.429524597 | 0.41322692 | -5.8793957 | 4.12E-09 | 4.61E-07 |
| SLC50A1 | 137.89473 | -0.731405952 | 0.29568263 | -2.4736183 | 0.01337525 | 0.06756165 |
| DPM3 | 180.986197 | -0.508954484 | 0.39395009 | -1.2919263 | 0.19638267 | 0.40443073 |
| MUC1 | 152.173723 | -1.816384825 | 0.63402568 | -2.8648443 | 0.00417214 | 0.02937201 |
| AC234582.1 | 98.3487824 | -0.896746659 | 0.37160844 | -2.4131494 | 0.01581534 | 0.07602133 |
| THBS3 | 131.54972 | 0.113166418 | 0.33028912 | 0.34262836 | 0.73187807 | 0.86826096 |
| MTX1 | 87.1157689 | -0.488944311 | 0.31145626 | -1.5698651 | 0.1164465 | 0.29052291 |
| GBA | 138.115055 | -0.075334575 | 0.23602487 | -0.3191807 | 0.74958952 | 0.87808868 |
| FAM189B | 101.079464 | -0.091443498 | 0.28492367 | -0.3209403 | 0.74825561 | 0.87718567 |
| SCAMP3 | 263.603923 | -0.079874198 | 0.22718288 | -0.3515855 | 0.72514917 | 0.86401269 |
| CLK2 | 280.90995 | 0.223964507 | 0.17931813 | 1.24897858 | 0.21167291 | 0.42447054 |
| HCN3 | 49.5912513 | -0.22261741 | 0.25548904 | -0.8713384 | 0.38356941 | 0.60633896 |
| FDPS | 668.849637 | -0.714612891 | 0.2530929 | -2.8235201 | 0.00474995 | 0.03215891 |
| RUSC1 | 111.309646 | -0.137605418 | 0.32668619 | -0.4212159 | 0.67359744 | 0.82886565 |
| ASH1L | 4612.74488 | -0.032406055 | 0.11096932 | -0.2920272 | 0.77026587 | 0.88929655 |
| ASH1L-AS1 | 46.4519329 | 0.175959374 | 0.3149876 | 0.55862318 | 0.57641892 | 0.76324677 |
| MSTO1 | 78.9873094 | -0.123607564 | 0.30637187 | -0.403456 | 0.68661281 | 0.83821337 |
| YY1AP1 | 819.449875 | 0.159520756 | 0.13756502 | 1.15960258 | 0.24621065 | 0.46546393 |
| DAP3 | 841.118634 | -0.004480443 | 0.18572924 | -0.0241235 | 0.98075408 | 0.99160261 |
| GON4L | 1117.59831 | -0.220676993 | 0.20380081 | -1.0828072 | 0.27889399 | 0.50104668 |
| SYT11 | 63.6516074 | -1.285915556 | 0.47401448 | -2.7128191 | 0.00667135 | 0.04141065 |
| RIT1 | 315.996916 | -0.831852356 | 0.36682285 | -2.2677223 | 0.02334614 | 0.0989334 |
| KHDC4 | 1090.87532 | -0.0056729 | 0.22119184 | -0.025647 | 0.97953892 | 0.9910179 |
| ARHGEF2 | 327.273044 | -0.715346536 | 0.23364953 | -3.061622 | 0.00220141 | 0.01847424 |
| SSR2 | 347.248087 | -0.630725669 | 0.24912377 | -2.5317763 | 0.01134864 | 0.06063912 |
| UBQLN4 | 178.317711 | 0.026593165 | 0.22120373 | 0.12022024 | 0.90430868 | 0.95590696 |
| LAMTOR2 | 127.671542 | -0.6673387 | 0.25940006 | -2.5726235 | 0.01009309 | 0.0558524 |
| RAB25 | 173.287049 | -0.860511724 | 0.34216895 | -2.5148738 | 0.0119075 | 0.06261392 |
| LMNA | 853.80552 | -0.920566199 | 0.3332379 | -2.7624895 | 0.00573624 | 0.03716038 |
| SEMA4A | 289.461709 | 0.09272982 | 0.25339789 | 0.36594551 | 0.71440574 | 0.85740373 |
| SLC25A44 | 174.837959 | 0.396748506 | 0.26224838 | 1.51287304 | 0.13031191 | 0.31253761 |
| SMG5 | 499.332402 | -0.284948652 | 0.17522165 | -1.6262183 | 0.10390322 | 0.26963805 |
| TMEM79 | 47.5865233 | 0.62512642 | 0.35133254 | 1.77930123 | 0.07519039 | 0.21794818 |
| GLMP | 93.5961981 | -0.070974824 | 0.29908674 | -0.2373052 | 0.81242006 | 0.91190984 |
| CCT3 | 507.108301 | -0.297734543 | 0.29986942 | -0.9928806 | 0.32076813 | 0.54511971 |
| MEF2D | 286.122836 | -0.433002883 | 0.26735243 | -1.6195959 | 0.10531912 | 0.27208298 |
| NAXE | 126.074469 | -0.52596567 | 0.48183962 | -1.0915783 | 0.2750185 | 0.49647832 |
| GPATCH4 | 96.9290506 | -0.741290714 | 0.30151589 | -2.4585461 | 0.01395009 | 0.06965503 |
| ISG20L2 | 130.663505 | -0.332405053 | 0.30408584 | -1.093129 | 0.27433716 | 0.49615525 |
| RRNAD1 | 209.162853 | 0.249571371 | 0.20989952 | 1.18900399 | 0.2344381 | 0.45166252 |
| MRPL24 | 102.148522 | -0.06964569 | 0.24381307 | -0.285652 | 0.77514467 | 0.89223749 |
| HDGF | 518.478158 | -0.875186474 | 0.2155711 | -4.0598507 | 4.91E-05 | 0.00096102 |
| PRCC | 403.356623 | 0.167228012 | 0.20792928 | 0.80425427 | 0.42125014 | 0.64039547 |
| ARHGEF11 | 1040.38754 | -0.033157302 | 0.15058043 | -0.2201966 | 0.82571803 | 0.91938879 |
| ETV3 | 96.1599174 | 0.193276072 | 0.23820283 | 0.81139283 | 0.41714011 | 0.63699466 |
| KIRREL1 | 1073.03144 | -0.443716685 | 0.44480708 | -0.9975486 | 0.31849829 | 0.5431247 |
| IFI16 | 209.043311 | -2.443982934 | 0.67457101 | -3.623018 | 0.00029119 | 0.00393682 |
| DUSP23 | 726.411382 | -2.482231523 | 0.54192146 | -4.5804267 | 4.64E-06 | 0.00014188 |
| AL590560.2 | 52.7787673 | 1.029887577 | 0.69419772 | 1.4835652 | 0.13792431 | 0.32399826 |

| | | | | | | |
|---|---|---|---|---|---|---|
| TAGLN2 | 383.456279 | -2.705738882 | 0.35500044 | -7.6217903 | 2.50E-14 | 1.44E-11 |
| KCNJ10 | 247.611949 | 1.233663815 | 0.48992859 | 2.51804823 | 0.01180072 | 0.06230561 |
| IGSF8 | 86.1573787 | 0.296663541 | 0.370329 | 0.80108104 | 0.42308473 | 0.64191663 |
| PEA15 | 116.056347 | -1.364324172 | 0.28823924 | -4.7333048 | 2.21E-06 | 7.66E-05 |
| DCAF8 | 305.738425 | 0.587861634 | 0.26721712 | 2.19993998 | 0.02781115 | 0.11142176 |
| AL139011.2 | 71.4934801 | -0.190512922 | 0.38745724 | -0.4917005 | 0.62293108 | 0.79471114 |
| COPA | 1215.40675 | -0.246640035 | 0.15184966 | -1.6242383 | 0.10432495 | 0.27043463 |
| NCSTN | 190.317295 | -0.17099685 | 0.25441477 | -0.6721184 | 0.50150832 | 0.70555154 |
| VANGL2 | 50.7264729 | -0.166313032 | 0.61817401 | -0.2690392 | 0.78789953 | 0.89956761 |
| F11R | 258.972141 | -0.625101469 | 0.30199005 | -2.0699406 | 0.0384579 | 0.13917687 |
| AL591806.3 | 192.463531 | 0.082990162 | 0.34766534 | 0.23870703 | 0.81133276 | 0.91145556 |
| USF1 | 107.855056 | 0.044141667 | 0.43830357 | 0.10071026 | 0.91978047 | 0.96447683 |
| NECTIN4 | 73.6529267 | -2.984607117 | 0.67149466 | -4.444722 | 8.80E-06 | 0.00024319 |
| KLHDC9 | 57.8990969 | 0.48116602 | 0.39654985 | 1.21338093 | 0.22498421 | 0.43942281 |
| PFDN2 | 312.613195 | -0.25762232 | 0.26463081 | -0.973516 | 0.33029692 | 0.55571163 |
| NIT1 | 344.077914 | -0.342427779 | 0.244478 | -1.4006486 | 0.16131917 | 0.35883595 |
| DEDD | 164.29286 | 0.029967901 | 0.18987988 | 0.15782558 | 0.87459424 | 0.94352461 |
| UFC1 | 252.729262 | -0.897036521 | 0.34970974 | -2.5650887 | 0.01031494 | 0.05676679 |
| USP21 | 50.0938052 | 0.530501984 | 0.31850675 | 1.66559103 | 0.0957949 | 0.2551633 |
| PPOX | 104.820752 | 0.479711143 | 0.21551807 | 2.22585115 | 0.02602415 | 0.10645805 |
| B4GALT3 | 98.38792 | -0.063734936 | 0.32026931 | -0.1990042 | 0.84225947 | 0.92705346 |
| NDUFS2 | 448.270899 | 0.396652448 | 0.24278036 | 1.63379135 | 0.10230267 | 0.26679486 |
| TOMM40L | 44.271264 | -0.139820919 | 0.43521178 | -0.321271 | 0.74800504 | 0.87714848 |
| SDHC | 739.017412 | 0.140185189 | 0.15812153 | 0.88656611 | 0.37531255 | 0.59870627 |
| AL592295.4 | 167.599747 | -0.471947011 | 0.31815403 | -1.4833916 | 0.1379704 | 0.32401004 |
| AL592295.3 | 59.1935588 | -0.590155284 | 0.31520129 | -1.8723124 | 0.0611634 | 0.19027492 |
| DUSP12 | 156.119911 | 0.586437579 | 0.28348784 | 2.0686516 | 0.03857879 | 0.13942196 |
| ATF6 | 1538.86211 | -0.228728469 | 0.20290808 | -1.1272517 | 0.2596361 | 0.48061767 |
| NOS1AP | 5071.18019 | -0.109473698 | 0.47111915 | -0.2323694 | 0.81625108 | 0.91443162 |
| C1orf226 | 113.204485 | -0.965025186 | 0.43040364 | -2.2421399 | 0.02495233 | 0.10351613 |
| UHMK1 | 787.167637 | 0.234984345 | 0.3034434 | 0.77439268 | 0.43869861 | 0.65485026 |
| UAP1 | 341.928731 | 0.48296835 | 0.492112 | 0.98141958 | 0.32638588 | 0.55173164 |
| DDR2 | 123.943396 | -0.582059718 | 0.57807613 | -1.0068911 | 0.31398711 | 0.53787673 |
| HSD17B7 | 319.823452 | -0.307706716 | 0.42782029 | -0.7192429 | 0.47199125 | 0.68308672 |
| RGS5 | 96.1267041 | -0.086937893 | 0.33417808 | -0.2601544 | 0.79474468 | 0.90385363 |
| NUF2 | 87.2677649 | -0.217964966 | 0.54283459 | -0.4015311 | 0.68802915 | 0.83940892 |
| PBX1 | 4883.00623 | 0.968001643 | 0.17885158 | 5.41231795 | 6.22E-08 | 4.42E-06 |
| MGST3 | 1933.95075 | -0.124020531 | 0.23497663 | -0.5277994 | 0.59763855 | 0.77674686 |
| ALDH9A1 | 274.923475 | -0.194323458 | 0.3419271 | -0.5683184 | 0.56981879 | 0.75923656 |
| TMCO1 | 499.268522 | -0.461814229 | 0.19563567 | -2.360583 | 0.01824623 | 0.08397938 |
| AL451074.2 | 40.7398102 | 1.322865467 | 0.45795134 | 2.88865945 | 0.00386888 | 0.0280022 |
| UCK2 | 250.612039 | -0.087992971 | 0.34039705 | -0.258501 | 0.7960203 | 0.90440668 |
| FAM78B | 144.603566 | 0.628533262 | 0.42825648 | 1.46765618 | 0.14219764 | 0.33120942 |
| POGK | 258.137416 | -0.25106206 | 0.30698277 | -0.8178376 | 0.4134499 | 0.63301068 |
| TADA1 | 168.17113 | 0.462857406 | 0.16784521 | 2.7576444 | 0.00582195 | 0.03754366 |
| POU2F1 | 2238.21704 | -0.07554459 | 0.12251101 | -0.6166351 | 0.53747544 | 0.73417922 |
| CREG1 | 208.790052 | -0.309436249 | 0.36448198 | -0.8489754 | 0.39589496 | 0.61804302 |
| MPZL1 | 773.01944 | -0.777983648 | 0.18671103 | -4.1667792 | 3.09E-05 | 0.00066897 |
| ADCY10 | 82.6083262 | 2.03827646 | 0.45735817 | 4.45663067 | 8.33E-06 | 0.00023349 |
| MPC2 | 497.269568 | 0.264890336 | 0.34632156 | 0.76486816 | 0.44435006 | 0.66007871 |
| DCAF6 | 3174.69709 | -0.134251652 | 0.2141658 | -0.6268585 | 0.530752 | 0.72861184 |
| TIPRL | 304.751773 | -0.076034016 | 0.22039228 | -0.344994 | 0.73009887 | 0.86751731 |
| SFT2D2 | 452.702341 | -0.646261999 | 0.13831074 | -4.6725366 | 2.98E-06 | 9.82E-05 |
| TBX19 | 60.6491349 | 0.025394886 | 0.29571856 | 0.08587518 | 0.93156564 | 0.97004734 |
| DPT | 54.7628485 | 2.789560025 | 0.74311017 | 3.75389831 | 0.00017411 | 0.00262098 |
| AL031275.1 | 62.665804 | 2.563023106 | 0.72086136 | 3.55550075 | 0.00037726 | 0.00476864 |
| LINC00626 | 36.5615312 | 2.118693353 | 0.89964045 | 2.35504456 | 0.01852048 | 0.08473747 |
| LINC00970 | 2153.96578 | 2.537540397 | 0.74144034 | 3.42244718 | 0.0006206 | 0.00700643 |
| ATP1B1 | 10315.7315 | 1.190040449 | 0.58264776 | 2.0424698 | 0.04110495 | 0.14502011 |
| NME7 | 1933.31789 | 0.060701776 | 0.15051857 | 0.4032843 | 0.68673909 | 0.83824503 |
| BLZF1 | 485.226448 | -0.035989533 | 0.17580567 | -0.204712 | 0.83779713 | 0.92443926 |
| CCDC181 | 101.637115 | -0.168093337 | 0.28672775 | -0.5862472 | 0.55770942 | 0.74979987 |
| SLC19A2 | 706.736972 | 0.415009199 | 0.43459534 | 0.95493247 | 0.33961185 | 0.56498892 |
| C1orf112 | 120.159636 | 0.245158068 | 0.40065355 | 0.6118954 | 0.54060696 | 0.73655259 |
| SCYL3 | 455.05676 | -0.026702861 | 0.15519595 | -0.172059 | 0.86339114 | 0.93813133 |
| KIFAP3 | 1012.30335 | 0.305949642 | 0.12911527 | 2.36958533 | 0.01780805 | 0.08256457 |
| GORAB | 155.657037 | 0.033013964 | 0.30225693 | 0.10922484 | 0.91302416 | 0.96058432 |
| FMO4 | 597.357708 | 0.93188752 | 0.42142311 | 2.21128716 | 0.02701596 | 0.10918997 |
| PRRC2C | 2427.24826 | -0.172356363 | 0.13312452 | -1.2947004 | 0.19542362 | 0.40302116 |
| MYOCOS | 210.264896 | 0.016144218 | 0.62432517 | 0.02585867 | 0.97937007 | 0.9910179 |
| VAMP4 | 306.90262 | 0.101089327 | 0.24482624 | 0.41290234 | 0.67967816 | 0.83230806 |
| METTL13 | 117.152577 | -0.016216437 | 0.28061701 | -0.0577885 | 0.9539171 | 0.9799573 |
| DNM3 | 3263.42242 | 0.184234281 | 0.22784667 | 0.80858886 | 0.41875168 | 0.63783336 |
| PIGC | 460.351005 | -0.24430951 | 0.26290887 | -0.9292555 | 0.35275671 | 0.57882774 |
| SUCO | 862.693203 | -0.167554754 | 0.2159876 | -0.775761 | 0.43789012 | 0.65419956 |
| AL645568.1 | 885.026638 | 0.915546036 | 0.22740712 | 4.02602183 | 5.67E-05 | 0.00107972 |
| PRDX6 | 998.527253 | -0.696443915 | 0.36998181 | -1.8823734 | 0.05978533 | 0.18751469 |
| SLC9C2 | 55.1637482 | 0.794842147 | 0.53594646 | 1.4830626 | 0.13805778 | 0.3240256 |
| ANKRD45 | 64.021375 | 0.31802699 | 0.47606021 | 0.66803943 | 0.50410843 | 0.70797197 |
| KLHL20 | 758.804281 | 0.027882263 | 0.18682249 | 0.14924468 | 0.88136056 | 0.94645136 |
| DARS2 | 158.444808 | 0.095052397 | 0.27403238 | 0.34686557 | 0.72869231 | 0.86676901 |
| ZBTB37 | 457.345213 | 0.564068563 | 0.29230807 | 1.92970573 | 0.05364331 | 0.17431107 |
| RC3H1 | 1632.87892 | -0.060547723 | 0.19180197 | -0.3156783 | 0.75224669 | 0.87955089 |
| RABGAP1L | 6220.47015 | 0.134624296 | 0.15720127 | 0.85638172 | 0.39178667 | 0.61425577 |
| Z99127.1 | 48.5294556 | 0.373411867 | 0.4334552 | 0.86147743 | 0.38897515 | 0.61210063 |
| CACYBP | 159.537513 | -0.136142542 | 0.24423193 | -0.5574314 | 0.57723274 | 0.76376512 |
| MRPS14 | 183.828735 | 0.147332461 | 0.21773328 | 0.67666488 | 0.4986186 | 0.7035092 |
| KIAA0040 | 357.992019 | -0.997695191 | 0.31983851 | -3.1193717 | 0.00181237 | 0.01580626 |
| AL513329.1 | 68.7773878 | 0.089920254 | 0.61048954 | 0.14729205 | 0.88290151 | 0.94732911 |
| COP1 | 3073.33415 | -0.20684967 | 0.17171671 | -1.2045984 | 0.22835837 | 0.44384722 |
| AL359265.3 | 117.40291 | 0.477883499 | 0.64079573 | 0.74576573 | 0.45580895 | 0.67098931 |
| PAPPA2 | 4100.25713 | -0.447305945 | 1.03774582 | -0.4310361 | 0.6664421 | 0.82409976 |
| SEC16B | 125.44269 | 1.273327947 | 0.53062647 | 2.39966912 | 0.0164099 | 0.07794259 |
| RASAL2 | 3745.6303 | 0.375876177 | 0.26562524 | 1.41506197 | 0.15705033 | 0.35268238 |
| RALGPS2 | 2958.23561 | -0.220735868 | 0.22185576 | -0.9949522 | 0.31975953 | 0.54422157 |
| ANGPTL1 | 285.365744 | 0.366196318 | 0.35327983 | 1.03656163 | 0.2999402 | 0.5227484 |
| FAM20B | 449.094782 | 0.338749336 | 0.13676373 | 2.47689449 | 0.01325311 | 0.06711401 |
| TOR3A | 105.089129 | -0.575837953 | 0.34895564 | -1.6501752 | 0.0989071 | 0.2605632 |
| ABL2 | 1747.99903 | -0.127028207 | 0.14491361 | -0.8765789 | 0.38071541 | 0.60354344 |
| SOAT1 | 447.417534 | -0.909692206 | 0.20442301 | -4.4500481 | 8.59E-06 | 0.00023874 |
| AXDND1 | 140.898082 | -0.026676591 | 0.61399037 | -0.0434479 | 0.9653445 | 0.98449934 |
| TDRD5 | 156.147406 | 0.926931255 | 0.42473799 | 2.18236014 | 0.02908296 | 0.11487944 |
| TOR1AIP2 | 1280.58882 | -0.168584819 | 0.297842 | -0.566021 | 0.57137953 | 0.75974956 |
| TOR1AIP1 | 392.251413 | 0.037800411 | 0.22798261 | 0.16580392 | 0.86831126 | 0.94012487 |
| CEP350 | 1999.29093 | 0.217095187 | 0.1110375 | 1.95515192 | 0.05056514 | 0.1674727 |
| QSOX1 | 198.791186 | -1.070843151 | 0.19360406 | -5.5310986 | 3.18E-08 | 2.47E-06 |
| ACBD6 | 1827.84568 | 0.121118053 | 0.13479355 | 0.89854485 | 0.36889515 | 0.59393582 |
| XPR1 | 2913.5826 | -0.2961613 | 0.22786666 | -1.2997132 | 0.19369929 | 0.40128199 |
| KIAA1614 | 269.305667 | 0.460523526 | 0.40193472 | 1.14576697 | 0.25189158 | 0.47203022 |
| AL162431.2 | 53.050233 | 0.05661297 | 0.31938789 | 0.17725459 | 0.85930842 | 0.93620958 |
| STX6 | 542.30705 | 0.212270442 | 0.23625184 | 0.89849224 | 0.36892318 | 0.59393582 |
| MR1 | 202.774996 | -0.256994487 | 0.17628405 | -1.4578431 | 0.1448838 | 0.33475075 |
| CACNA1E | 190.945904 | -0.327538157 | 0.99381373 | -0.329577 | 0.7417196 | 0.8738293 |

| | | | | | | |
|---|---|---|---|---|---|---|
| GLUL | 711.340882 | -2.129880848 | 0.32890713 | -6.4756299 | 9.44E-11 | 1.66E-08 |
| RNASEL | 111.158645 | -1.029919529 | 0.41726389 | -2.468269 | 0.01357683 | 0.06829257 |
| NPL | 99.0846899 | 1.0422045 | 0.32720967 | 3.18512741 | 0.0014469 | 0.01333715 |
| DHX9 | 1269.90208 | 0.003747468 | 0.24770817 | 0.01512856 | 0.98792962 | 0.99552604 |
| LAMC1 | 1736.93458 | -1.120561438 | 0.22547149 | -4.9698587 | 6.70E-07 | 3.01E-05 |
| LAMC2 | 462.588297 | -2.860939882 | 0.64543926 | -4.432547 | 9.31E-06 | 0.00025207 |
| SMG7 | 1629.95829 | 0.19476102 | 0.16106137 | 1.20923487 | 0.22657263 | 0.44161101 |
| ARPC5 | 439.847736 | -0.844461061 | 0.23446174 | -3.6017009 | 0.00031614 | 0.0041801 |
| RGL1 | 211.154239 | 0.471332287 | 0.3324413 | 1.41779101 | 0.15625179 | 0.35131349 |
| TSEN15 | 494.299889 | 0.67968599 | 0.17429271 | 3.89968113 | 9.63E-05 | 0.00163323 |
| C1orf21 | 142.720677 | -1.157395193 | 0.47815441 | -2.4205469 | 0.01549718 | 0.07503748 |
| EDEM3 | 886.165502 | -0.034482502 | 0.23490141 | -0.1467956 | 0.88329333 | 0.94755606 |
| FAM129A | 86.9895231 | -2.673139937 | 0.53639128 | -4.9835633 | 6.24E-07 | 2.83E-05 |
| RNF2 | 479.458259 | -0.019879319 | 0.17409426 | -0.1141871 | 0.90908948 | 0.95896526 |
| TRMT1L | 420.899641 | 0.291383425 | 0.18559611 | 1.5699867 | 0.1164182 | 0.29052291 |
| SWT1 | 706.080266 | -0.141715709 | 0.21487209 | -0.6595352 | 0.50955214 | 0.71186277 |
| IVNS1ABP | 6950.45927 | 0.521235425 | 0.42044846 | 1.23971301 | 0.21508156 | 0.4288968 |
| LINC01350 | 172.547214 | 0.382940981 | 0.99993094 | 0.38296743 | 0.70174391 | 0.8488446 |
| AL133383.1 | 40.3175489 | 1.470134585 | 0.8550793 | 1.7192962 | 0.08556045 | 0.23643682 |
| HMCN1 | 1691.1738 | 1.137552392 | 0.73264573 | 1.55266365 | 0.12050351 | 0.29668795 |
| TPR | 1347.90001 | -0.161303385 | 0.11994132 | -1.3448525 | 0.17867287 | 0.38189768 |
| ODR4 | 809.518579 | 0.003346099 | 0.13790305 | 0.02426414 | 0.98064191 | 0.99160216 |
| AL596220.1 | 241.333922 | 0.615028165 | 0.39447501 | 1.55910551 | 0.11897141 | 0.29434157 |
| PLA2G4A | 73.8696577 | -0.430862694 | 0.79572877 | -0.5414693 | 0.58818416 | 0.77114884 |
| ERVMER61-1 | 1096.86582 | 0.345746951 | 0.31040005 | 1.11387531 | 0.26533268 | 0.48636961 |
| BRINP3 | 546.210796 | 0.920848617 | 0.72410814 | 1.27170041 | 0.20347959 | 0.41324752 |
| AL954650.1 | 45.9581213 | 5.241216992 | 0.82716693 | 6.33634735 | 2.35E-10 | 3.49E-08 |
| AL390957.1 | 152.965255 | -0.018215282 | 0.35517357 | -0.0512856 | 0.95909796 | 0.98229715 |
| AL136454.1 | 312.343416 | -1.212797133 | 0.43985005 | -2.7572968 | 0.00582814 | 0.03755314 |
| UCHL5 | 431.996835 | 0.13576589 | 0.14316961 | 0.94828707 | 0.34298333 | 0.56815486 |
| TROVE2 | 701.236707 | -0.178207192 | 0.15304805 | -1.1643872 | 0.24426714 | 0.46311535 |
| GLRX2 | 101.539985 | -0.023639313 | 0.35771783 | -0.0660837 | 0.9473112 | 0.97743435 |
| CDC73 | 1560.95917 | 0.091770052 | 0.17108671 | 0.53639499 | 0.59168559 | 0.77313038 |
| LINC01031 | 52.586666 | -1.656610632 | 0.78072264 | -2.1218939 | 0.03384665 | 0.12825025 |
| KCNT2 | 50.2480795 | -0.760162716 | 0.51079699 | -1.4881895 | 0.13670092 | 0.32238573 |
| ZBTB41 | 356.151282 | 0.169570076 | 0.14913517 | 1.13702273 | 0.25552878 | 0.47646523 |
| CRB1 | 226.841998 | 1.098373689 | 0.75454731 | 1.4556724 | 0.1454832 | 0.33560734 |
| DENND1B | 2980.18949 | 1.063037502 | 0.31656963 | 3.35798957 | 0.00078512 | 0.00826862 |
| NEK7 | 2375.21054 | -0.183864713 | 0.13794148 | -1.3329182 | 0.18255865 | 0.38718965 |
| LINC00862 | 167.210534 | 0.475324218 | 0.60153709 | 0.79018273 | 0.42942106 | 0.64764415 |
| AC104461.1 | 167.989857 | 1.467062515 | 0.54374788 | 2.69805652 | 0.00697456 | 0.0427305 |
| ZNF281 | 164.656346 | 0.292234215 | 0.37076443 | 0.78819377 | 0.43058337 | 0.64867424 |
| DDX59 | 310.830633 | 0.062122254 | 0.1912021 | 0.32490362 | 0.74525402 | 0.87533882 |
| CAMSAP2 | 1353.4334 | -0.081434267 | 0.22047005 | -0.3693666 | 0.71185451 | 0.8555078 |
| INAVA | 229.729277 | -1.455322181 | 0.29752399 | -4.8914449 | 1.00E-06 | 4.11E-05 |
| TMEM9 | 234.373344 | -0.586650749 | 0.40592288 | -1.4452271 | 0.14839411 | 0.34019898 |
| LAD1 | 113.192464 | -0.671293257 | 0.50076918 | -1.3405243 | 0.18007495 | 0.38383249 |
| CSRP1 | 155.722194 | -0.397579301 | 0.37870681 | -1.049834 | 0.29379442 | 0.5168794 |
| AC096677.1 | 252.061434 | 0.182538005 | 0.41996042 | 0.43465526 | 0.66381268 | 0.82277392 |
| NAV1 | 147.547689 | -1.030507076 | 0.41265449 | -2.4972637 | 0.01251558 | 0.06470386 |
| IPO9-AS1 | 65.0671427 | -0.591582142 | 0.31326152 | -1.8884609 | 0.0589641 | 0.18561886 |
| IPO9 | 779.854508 | -0.181731562 | 0.2111703 | -0.8605924 | 0.38946256 | 0.61248908 |
| SHISA4 | 43.63191 | -0.581021755 | 0.45125309 | -1.287574 | 0.19789429 | 0.40634007 |
| TIMM17A | 262.236123 | 0.40058451 | 0.27742945 | 1.44391489 | 0.14876292 | 0.34073338 |
| RNPEP | 331.549655 | -0.524663206 | 0.21737352 | -2.4136482 | 0.01579371 | 0.07597198 |
| ELF3 | 1752.72027 | -2.780537805 | 0.36294257 | -7.6610958 | 1.84E-14 | 1.16E-11 |
| ARL8A | 200.42531 | -0.237447874 | 0.21682307 | -1.0951227 | 0.27346286 | 0.49520195 |
| PPP1R12B | 1938.04621 | 0.076183307 | 0.14266801 | 0.53399013 | 0.59334836 | 0.77387448 |
| KDM5B | 1465.17584 | -0.529403355 | 0.19782603 | -2.6761056 | 0.00744832 | 0.04476106 |
| RABIF | 64.776672 | -0.395621797 | 0.40791077 | -0.9698734 | 0.33210962 | 0.55734162 |
| KLHL12 | 244.614449 | 0.127618159 | 0.21510728 | 0.59327679 | 0.55299593 | 0.7458923 |
| ADIPOR1 | 172.064054 | -0.177804177 | 0.17457024 | -1.0185251 | 0.30842847 | 0.53227485 |
| CYB5R1 | 142.751505 | 0.112939703 | 0.35169934 | 0.32112572 | 0.74811512 | 0.87714848 |
| TMEM183A | 138.503723 | 0.132779039 | 0.22357287 | 0.59389603 | 0.55258166 | 0.74556191 |
| PPFIA4 | 89.3048762 | -2.696781465 | 0.95111864 | -2.8353786 | 0.00457714 | 0.03145229 |
| BTG2 | 139.555641 | -0.359572631 | 0.57992321 | -0.6200349 | 0.53523481 | 0.73219616 |
| ATP2B4 | 596.645129 | -1.118560659 | 0.41544458 | -2.6924426 | 0.00709308 | 0.04313648 |
| SNRPE | 147.479358 | 0.084662903 | 0.29853984 | 0.28358997 | 0.77672463 | 0.89337196 |
| SOX13 | 257.837167 | 0.089572849 | 0.24971255 | 0.35870383 | 0.71981667 | 0.86037991 |
| ETNK2 | 104.14743 | -0.438428452 | 0.35749418 | -1.2263933 | 0.22005072 | 0.43389311 |
| GOLT1A | 109.309051 | -0.633639243 | 0.56481753 | -1.1218477 | 0.2619272 | 0.4828072 |
| PLEKHA6 | 1576.45036 | 0.25072854 | 0.2276989 | 1.10114078 | 0.27083539 | 0.49226771 |
| AL606489.1 | 65.1838446 | -0.762993927 | 0.55183654 | -1.3826448 | 0.1667738 | 0.36640505 |
| PPP1R15B | 217.60148 | 0.001002624 | 0.35644791 | 0.00281282 | 0.9977557 | 0.99934448 |
| PIK3C2B | 46.9271851 | -1.067395168 | 0.47065451 | -2.2678953 | 0.02333559 | 0.09892162 |
| MDM4 | 1638.17994 | 0.493832023 | 0.18803871 | 2.62622529 | 0.00863377 | 0.04986594 |
| LRRN2 | 404.409068 | -0.03867414 | 0.6892342 | -0.0561118 | 0.95525278 | 0.98014107 |
| RBBP5 | 322.878633 | 0.094645752 | 0.16712125 | 0.56632983 | 0.57116958 | 0.75965257 |
| DSTYK | 767.430957 | 0.196550195 | 0.17597907 | 1.1168953 | 0.26403909 | 0.48499778 |
| NUAK2 | 638.296671 | 0.99948841 | 0.52265299 | 1.91233653 | 0.05583304 | 0.17904799 |
| LEMD1 | 65.9616045 | -3.071605732 | 0.64242671 | -4.7812547 | 1.74E-06 | 6.29E-05 |
| CDK18 | 296.25464 | -0.572552883 | 0.52047331 | -1.1000619 | 0.27130513 | 0.49275402 |
| MFSD4A | 1505.78628 | 1.592177747 | 0.5840721 | 2.72599522 | 0.00641079 | 0.04013278 |
| ELK4 | 887.92919 | -0.159557379 | 0.33475824 | -0.4766347 | 0.6336223 | 0.80290838 |
| SLC45A3 | 58.8280265 | 0.865688026 | 0.54490082 | 1.58870751 | 0.11212644 | 0.28308069 |
| NUCKS1 | 1761.23255 | 0.119432888 | 0.14161497 | 0.84336341 | 0.39902523 | 0.62081557 |
| RAB29 | 201.468227 | -0.376078809 | 0.29514759 | -1.2742059 | 0.20259044 | 0.41227307 |
| SLC41A1 | 115.223375 | -0.122888397 | 0.19554156 | -0.6284515 | 0.52970818 | 0.7278597 |
| RHEX | 609.308255 | -1.996413481 | 0.62332181 | -3.2028616 | 0.00136069 | 0.01273806 |
| SRGAP2 | 126.934283 | -1.499790164 | 0.52364213 | -2.8641511 | 0.00418129 | 0.02940927 |
| EIF2D | 181.005762 | 0.552802974 | 0.2537166 | 2.17882066 | 0.02934499 | 0.11558224 |
| DYRK3 | 170.492876 | -1.12999714 | 0.32374955 | -3.4903435 | 0.0004824 | 0.00575777 |
| MAPKAPK2 | 1133.42213 | -0.526549879 | 0.23684124 | -2.2232187 | 0.02620105 | 0.10698292 |
| PIGR | 594.338417 | -2.717138971 | 0.72966399 | -3.7238222 | 0.00019623 | 0.0028864 |
| C1orf116 | 86.1619485 | 0.058661217 | 0.56452973 | 0.10391165 | 0.91723946 | 0.96291779 |
| PFKFB2 | 563.036701 | 0.509599036 | 0.27046268 | 1.88417503 | 0.0595413 | 0.1871063 |
| YOD1 | 142.701323 | 0.064809902 | 0.30870851 | 0.20993882 | 0.83371542 | 0.92240112 |
| CD55 | 314.816738 | -1.546839267 | 0.38768266 | -3.9899625 | 6.61E-05 | 0.00121236 |
| CD46 | 1617.42497 | -0.037404704 | 0.19631028 | -0.1905387 | 0.84888703 | 0.93045795 |
| MIR29B2CHG | 2568.69585 | -0.202976447 | 0.32858192 | -0.6177347 | 0.53675025 | 0.73349204 |
| PLXNA2 | 676.758503 | 1.50198397 | 0.47050625 | 3.19227209 | 0.00141158 | 0.01307283 |
| LAMB3 | 239.39009 | -3.470944109 | 0.57659053 | -6.019773 | 1.75E-09 | 2.11E-07 |
| AL031316.1 | 42.1210891 | 1.838405301 | 0.61943931 | 2.9678538 | 0.00299887 | 0.02315141 |
| G0S2 | 606.339449 | -0.210726875 | 0.57563626 | -0.3660764 | 0.71430804 | 0.85736437 |
| IRF6 | 136.786961 | -0.153288747 | 0.27403556 | -0.5593754 | 0.57590555 | 0.76292613 |
| UTP25 | 129.276981 | -0.328411715 | 0.3297243 | -0.9960191 | 0.31924085 | 0.54392231 |
| SYT14 | 87.4678638 | 0.702459226 | 0.53698287 | 1.30815947 | 0.19081923 | 0.39737209 |
| SERTAD4-AS | 40.84922 | -0.296658515 | 0.75802129 | -0.3913591 | 0.69553183 | 0.84441912 |
| SERTAD4 | 40.4105513 | 0.258833687 | 0.79306587 | 0.32637098 | 0.74414369 | 0.87481257 |
| HHAT | 528.900805 | 0.302085051 | 0.4717035 | 0.64041299 | 0.52190414 | 0.7218672 |
| KCNH1 | 85.529346 | 0.576475332 | 0.53862928 | 1.07026364 | 0.28450066 | 0.50687472 |
| RCOR3 | 2113.52099 | 0.499272376 | 0.14821785 | 3.36850359 | 0.00075577 | 0.00808997 |
| TRAF5 | 298.0699 | -0.160653699 | 0.30245957 | -0.5311576 | 0.59530957 | 0.77510539 |

| | | | | | | |
|---|---|---|---|---|---|---|
| LINC00467 | 442.560279 | 0.140740464 | 0.29852395 | 0.47145451 | 0.6373162 | 0.80497022 |
| SLC30A1 | 118.473652 | -0.852151736 | 0.32018317 | -2.6614507 | 0.00778047 | 0.04629381 |
| LPGAT1 | 1525.89858 | -0.874392934 | 0.26798409 | -3.2628538 | 0.00110296 | 0.01077592 |
| INTS7 | 302.58193 | 0.252648942 | 0.34724004 | 0.72759161 | 0.46686362 | 0.67931279 |
| PPP2R5A | 2491.76397 | -0.266431969 | 0.34436205 | -0.7736972 | 0.43910985 | 0.65485132 |
| AL360091.3 | 385.032117 | -0.540728923 | 0.58665143 | -0.921721 | 0.35667414 | 0.58236202 |
| TMEM206 | 107.274698 | -0.206451624 | 0.42150602 | -0.4897952 | 0.62427883 | 0.79564831 |
| NENF | 243.43263 | -0.538085113 | 0.33131187 | -1.6241046 | 0.1043535 | 0.27043463 |
| ATF3 | 130.412374 | 1.230567203 | 0.78301298 | 1.57157956 | 0.11604809 | 0.28978961 |
| NSL1 | 540.583944 | 0.106938888 | 0.15523475 | 0.68888499 | 0.49089565 | 0.69708026 |
| TATDN3 | 410.797948 | 0.241874663 | 0.30135184 | 0.8026321 | 0.42218741 | 0.64114933 |
| FLVCR1-DT | 56.6036944 | 0.204697278 | 0.34473601 | 0.5937798 | 0.5526594 | 0.74559065 |
| FLVCR1 | 320.855865 | -0.062902397 | 0.14749334 | -0.4264762 | 0.66976091 | 0.82642427 |
| ANGEL2 | 443.030364 | 0.303506973 | 0.1554455 | 1.95249764 | 0.05087915 | 0.16818386 |
| RPS6KC1 | 1322.44092 | 0.155135298 | 0.19412853 | 0.79913703 | 0.42421096 | 0.64296071 |
| PROX1 | 1228.5397 | 2.207933327 | 0.55202014 | 3.99973326 | 6.34E-05 | 0.00117152 |
| SMYD2 | 326.963304 | -0.445423022 | 0.31601746 | -1.4094887 | 0.15869072 | 0.35514221 |
| PTPN14 | 5119.94028 | -0.738183945 | 0.20421628 | -3.6147165 | 0.00030068 | 0.00402804 |
| KCTD3 | 1297.0181 | 0.020381139 | 0.2626109 | 0.07760965 | 0.93813857 | 0.97320917 |
| USH2A | 115.180688 | 1.483268502 | 0.37214228 | 3.98575645 | 6.73E-05 | 0.00122892 |
| ESRRG | 51368.916 | 1.476345321 | 0.22968154 | 6.42779259 | 1.29E-10 | 2.19E-08 |
| GPATCH2 | 1802.89408 | -0.35545167 | 0.2468099 | -1.440184 | 0.14981535 | 0.34200906 |
| SPATA17 | 438.167254 | -0.089477482 | 0.31029786 | -0.28836 | 0.77307121 | 0.89082243 |
| RRP15 | 311.320165 | 0.590490008 | 0.31340505 | 1.88411131 | 0.05954992 | 0.1871063 |
| TGFB2 | 106.196315 | 0.01763576 | 0.82364826 | 0.02141176 | 0.98291719 | 0.9928728 |
| LYPLAL1-DT | 107.962876 | 2.219945659 | 0.45767304 | 4.85050559 | 1.23E-06 | 4.86E-05 |
| LYPLAL1 | 3025.35351 | 0.897529998 | 0.30949404 | 2.89999119 | 0.00373173 | 0.02720105 |
| AC096642.1 | 72.0544289 | 0.53541968 | 0.59656676 | 0.8975017 | 0.36945127 | 0.59417383 |
| AL513283.1 | 72.5013315 | -0.094466427 | 0.968644 | -0.0975244 | 0.92230996 | 0.9655855 |
| ZC3H11B | 83.8858386 | 0.436832369 | 0.2840862 | 1.53767544 | 0.12412799 | 0.30213905 |
| SLC30A10 | 48.7271522 | 0.883869659 | 0.84011931 | 1.05207635 | 0.29276452 | 0.51560809 |
| EPRS | 610.476651 | 0.185066891 | 0.15845753 | 1.16792737 | 0.24283606 | 0.46132915 |
| BPNT1 | 195.382957 | 0.023968772 | 0.20326713 | 0.1179176 | 0.90613295 | 0.95706941 |
| IARS2 | 454.629947 | -0.189617915 | 0.22154198 | -0.8559006 | 0.39205276 | 0.614309 |
| RAB3GAP2 | 1812.92064 | -0.10229754 | 0.13729364 | -0.7451004 | 0.45621104 | 0.67128123 |
| MARK1 | 572.200769 | -0.15541772 | 0.47196471 | -0.3292994 | 0.74192936 | 0.87399848 |
| C1orf115 | 75.9897611 | -0.733119938 | 0.43695037 | -1.6778105 | 0.09338409 | 0.25122413 |
| 2-Mar | 679.800086 | 0.830567151 | 0.2907075 | 2.85705439 | 0.00427593 | 0.02994735 |
| 1-Mar | 145.689413 | 0.884942582 | 0.33949353 | 2.60665524 | 0.00914314 | 0.05205672 |
| AL513314.2 | 222.606368 | 0.406735936 | 0.29970684 | 1.3571126 | 0.17474543 | 0.37654099 |
| TAF1A | 92.4775163 | 0.242956639 | 0.29031207 | 0.83688095 | 0.40265949 | 0.62391114 |
| MIA3 | 534.896565 | -0.053314863 | 0.17351101 | -0.3072708 | 0.75863728 | 0.88316847 |
| AIDA | 224.268893 | -0.450262067 | 0.18570344 | -2.4246296 | 0.01532401 | 0.07447199 |
| BROX | 490.981282 | -0.468985513 | 0.15982221 | -2.9344202 | 0.00334171 | 0.02512277 |
| FAM177B | 325.540721 | -3.163578793 | 0.69757484 | -4.5351102 | 5.76E-06 | 0.0001709 |
| AC093152.1 | 413.407421 | -0.069018463 | 0.20153158 | -0.3424697 | 0.73199743 | 0.86826096 |
| TLR5 | 436.393009 | -0.223080423 | 0.2818173 | -0.7915782 | 0.42860668 | 0.6469507 |
| SUSD4 | 1602.60608 | 0.338348931 | 0.43232819 | 0.78262056 | 0.43384996 | 0.65084886 |
| CAPN8 | 96.9068487 | -4.047058515 | 0.51125319 | -7.9159575 | 2.45E-15 | 1.80E-12 |
| CAPN2 | 1514.02259 | -0.508044642 | 0.23815403 | -2.1332608 | 0.03290334 | 0.12575817 |
| TP53BP2 | 491.475302 | -0.303164732 | 0.18212454 | -1.6646013 | 0.09599234 | 0.25543166 |
| FBXO28 | 899.723241 | -0.182830031 | 0.1766985 | -1.0347005 | 0.30080879 | 0.5238475 |
| DEGS1 | 370.881312 | -0.952661688 | 0.36551752 | -2.6063366 | 0.00915165 | 0.05206034 |
| NVL | 718.557041 | 0.198384073 | 0.18944087 | 1.04720839 | 0.29500348 | 0.51817489 |
| CNIH4 | 587.768795 | -0.366650943 | 0.2446139 | -1.4988966 | 0.13390046 | 0.31822439 |
| WDR26 | 1621.60874 | 0.290877052 | 0.20692954 | 1.40568159 | 0.15981869 | 0.35669069 |
| CNIH3 | 690.414945 | 1.12628191 | 0.38155971 | 2.95178416 | 0.00315944 | 0.02401208 |
| DNAH14 | 3853.1588 | 0.174226567 | 0.18615736 | 0.93591017 | 0.34931944 | 0.57533173 |
| LBR | 836.482168 | 0.430542177 | 0.20384408 | 2.11211516 | 0.03467657 | 0.13011647 |
| ENAH | 405.816243 | -0.235043964 | 0.29545276 | -0.7955382 | 0.42630049 | 0.64486981 |
| SRP9 | 511.594737 | -0.364484237 | 0.25282616 | -1.4416397 | 0.14940403 | 0.34155035 |
| EPHX1 | 210.092288 | -0.078348117 | 0.26783922 | -0.2925192 | 0.76988967 | 0.88922989 |
| AL591895.1 | 186.848475 | -1.133345161 | 0.24536544 | -4.6190089 | 3.86E-06 | 0.00012069 |
| TMEM63A | 553.061737 | -1.218837863 | 0.25311654 | -4.815323 | 1.47E-06 | 5.51E-05 |
| AL117348.2 | 49.4972381 | -0.226269996 | 0.51380648 | -0.4403798 | 0.65966205 | 0.81980614 |
| SDE2 | 98.0286482 | 0.337269203 | 0.2066047 | 1.63243726 | 0.1025874 | 0.26722087 |
| H3F3A | 225.848622 | -0.357392285 | 0.16527549 | -2.1624035 | 0.03058709 | 0.11939269 |
| ACBD3 | 881.087302 | -0.236562955 | 0.11296683 | -2.0940921 | 0.03625177 | 0.13396096 |
| LIN9 | 136.657844 | 0.372214616 | 0.26575703 | 1.40058239 | 0.16133899 | 0.35883595 |
| PARP1 | 476.642362 | -0.756875566 | 0.30521478 | -2.479813 | 0.01314513 | 0.06685947 |
| STUM | 571.297783 | 1.420093769 | 0.47325069 | 3.00072204 | 0.0026934 | 0.02134203 |
| ITPKB | 1019.01057 | -0.147235579 | 0.18710405 | -0.7869182 | 0.43132977 | 0.64950239 |
| PSEN2 | 56.0580099 | -0.087729053 | 0.27812602 | -0.3154291 | 0.75243584 | 0.87955089 |
| COQ8A | 431.553906 | 0.390586686 | 0.28649321 | 1.36333661 | 0.17277645 | 0.37425453 |
| CDC42BPA | 5281.57155 | -0.34614073 | 0.28599382 | -1.2103084 | 0.22616057 | 0.44113334 |
| ZNF678 | 516.716094 | 0.050858696 | 0.17880705 | 0.2844334 | 0.77607827 | 0.89296323 |
| SNAP47 | 236.321811 | 0.344469825 | 0.15075053 | 2.28503222 | 0.02231094 | 0.09596392 |
| JMJD4 | 56.3291285 | -0.716465251 | 0.32499513 | -2.2045415 | 0.02748629 | 0.1104666 |
| ARF1 | 670.587393 | -0.84739444 | 0.19737025 | -4.2934253 | 1.76E-05 | 0.00042645 |
| C1orf35 | 108.629533 | -0.415900311 | 0.28195656 | -1.475051 | 0.14019883 | 0.32765152 |
| MRPL55 | 113.612016 | -0.361588873 | 0.35939256 | -1.0061112 | 0.31436209 | 0.53829516 |
| GUK1 | 568.017043 | -0.803678419 | 0.31988285 | -2.5124148 | 0.0119908 | 0.0628518 |
| IBA57 | 60.9306466 | -0.411510335 | 0.28104762 | -1.4642014 | 0.14313892 | 0.33252233 |
| OBSCN-AS1 | 50.4786519 | -0.392539366 | 0.36751411 | -1.0680933 | 0.28547841 | 0.50786322 |
| OBSCN | 796.74138 | 0.039933473 | 0.21502937 | 0.18571172 | 0.85267084 | 0.93236168 |
| TRIM11 | 105.974701 | -0.87385347 | 0.30093622 | -2.903783 | 0.00368684 | 0.02695042 |
| RNF187 | 162.984459 | -0.894406971 | 0.32804777 | -2.7264534 | 0.0064019 | 0.0400961 |
| RAB4A | 340.64648 | -0.126057528 | 0.19857806 | -0.6348009 | 0.52555829 | 0.72472068 |
| CCSAP | 53.8497028 | 0.613949987 | 0.4032997 | 1.522317 | 0.12792967 | 0.30854958 |
| NUP133 | 510.171105 | 0.180469587 | 0.13928088 | 1.29572408 | 0.19507056 | 0.40263651 |
| ABCB10 | 299.291218 | 0.309463265 | 0.18680976 | 1.65656903 | 0.09760664 | 0.2582075 |
| TAF5L | 157.563074 | 0.073236982 | 0.27925707 | 0.2622565 | 0.7931237 | 0.90259076 |
| URB2 | 75.0540392 | -0.059243429 | 0.36077572 | -0.1642112 | 0.86956485 | 0.94056476 |
| GALNT2 | 1158.24557 | -1.003976848 | 0.26437096 | -3.7976063 | 0.0001461 | 0.00226507 |
| PGBD5 | 113.32847 | 0.779250158 | 0.40878063 | 1.90627956 | 0.05661394 | 0.18084972 |
| COG2 | 347.745902 | 0.271223689 | 0.11706412 | 2.31688137 | 0.02051019 | 0.0907906 |
| C1orf198 | 120.103392 | -0.76851705 | 0.21468947 | -3.5796681 | 0.00034403 | 0.00444837 |
| TTC13 | 339.954021 | 0.388350271 | 0.28429095 | 1.36603109 | 0.1719292 | 0.37294355 |
| ARV1 | 213.359781 | 0.529087069 | 0.18806733 | 2.81328545 | 0.00490381 | 0.03294732 |
| FAM89A | 69.494501 | -1.657320414 | 0.33758402 | -4.9093568 | 9.14E-07 | 3.88E-05 |
| C1orf131 | 182.827984 | -0.047182424 | 0.22730824 | -0.2075702 | 0.83556455 | 0.92348178 |
| GNPAT | 543.221695 | 0.14235113 | 0.12570904 | 1.13238579 | 0.25747228 | 0.47845862 |
| EXOC8 | 71.8305045 | -0.372493271 | 0.3319822 | -1.1220278 | 0.26185061 | 0.48277062 |
| SPRTN | 146.320405 | -0.101329557 | 0.29371283 | -0.3449953 | 0.73009788 | 0.86751731 |
| EGLN1 | 1372.77252 | -0.413856455 | 0.285404 | -1.4500724 | 0.14703834 | 0.33786787 |
| TSNAX-DISC1 | 73.0949183 | -0.423136122 | 0.36003832 | -1.175253 | 0.2398935 | 0.4580447 |
| DISC1 | 208.259763 | -1.787635853 | 0.33646385 | -5.3130102 | 1.08E-07 | 6.72E-06 |
| SIPA1L2 | 109.377112 | -1.966907952 | 0.65423232 | -3.0064365 | 0.00264329 | 0.02109683 |
| NTPCR | 321.70168 | 0.266863849 | 0.19605053 | 1.36119931 | 0.17345071 | 0.37516137 |
| PCNX2 | 2078.07472 | -0.33885683 | 0.17255114 | -1.9638052 | 0.04955269 | 0.16520481 |
| MAP3K21 | 1163.59958 | 0.071062302 | 0.25431233 | 0.27942924 | 0.77991543 | 0.8949616 |
| KCNK1 | 984.684799 | -0.292630931 | 0.2639689 | -1.1085811 | 0.26761094 | 0.48878221 |
| SLC35F3 | 120.765822 | -0.762288472 | 0.58296051 | -1.307616 | 0.19100359 | 0.39752425 |

| | | | | | | |
|---|---|---|---|---|---|---|
| AL355472.3 | 46.5194963 | 0.351258915 | 0.62697501 | 0.56024389 | 0.5753131 | 0.76252365 |
| AL355472.1 | 129.598525 | 1.659891368 | 0.43047306 | 3.85597038 | 0.00011527 | 0.00187284 |
| COA6 | 129.943675 | -0.143472131 | 0.30504245 | -0.470335 | 0.63811572 | 0.80525266 |
| TARBP1 | 588.966942 | 0.171785702 | 0.304291 | 0.56454414 | 0.57238387 | 0.76035215 |
| IRF2BP2 | 738.001904 | -0.492514723 | 0.4237522 | -1.1622706 | 0.24512557 | 0.46420922 |
| TOMM20 | 487.834759 | -0.466272263 | 0.28423526 | -1.6404448 | 0.10091272 | 0.26453475 |
| ARID4B | 3707.64261 | -0.178861859 | 0.15573802 | -1.1484791 | 0.25077085 | 0.47078342 |
| TBCE | 67.8224122 | -0.282308225 | 0.38183108 | -0.7393537 | 0.45969224 | 0.67340299 |
| B3GALNT2 | 646.443395 | 0.345444943 | 0.16848712 | 2.05027513 | 0.04033759 | 0.14338515 |
| LYST | 1310.88726 | 0.419428425 | 0.19459271 | 2.15541698 | 0.03112922 | 0.1210082 |
| GPR137B | 216.676833 | -0.729175566 | 0.47304653 | -1.5414458 | 0.12320833 | 0.30093544 |
| ERO1B | 529.054627 | 0.314124132 | 0.20927888 | 1.50098347 | 0.13335984 | 0.31751083 |
| EDARADD | 68.9287105 | -0.332565007 | 0.42335317 | -0.7855498 | 0.43213127 | 0.64979206 |
| LGALS8 | 414.525495 | -0.688888195 | 0.2254304 | -3.0558798 | 0.00224401 | 0.01865397 |
| AL359921.1 | 74.4105698 | 0.088741591 | 0.32282408 | 0.27489148 | 0.78339961 | 0.89665455 |
| HEATR1 | 468.262439 | 0.122096365 | 0.18469461 | 0.66107163 | 0.50856638 | 0.71123011 |
| ACTN2 | 60.2067908 | 1.763677956 | 0.57412011 | 3.07196688 | 0.00212653 | 0.01799448 |
| MTR | 4012.92635 | 0.577772985 | 0.25479447 | 2.26760413 | 0.02335335 | 0.0989334 |
| RYR2 | 119.84253 | 1.799837359 | 0.67567314 | 2.66376929 | 0.00772705 | 0.04604896 |
| LINC01139 | 66.8437152 | 0.799403703 | 0.45515065 | 1.75634969 | 0.07902872 | 0.22471623 |
| CHRM3 | 316.222984 | 0.443161337 | 0.57551898 | 0.77002037 | 0.44128781 | 0.65696085 |
| RGS7 | 510.000329 | 1.64762773 | 0.54906538 | 3.00078605 | 0.00269284 | 0.02134203 |
| FH | 195.133655 | 0.781249043 | 0.26366735 | 2.96301016 | 0.00304647 | 0.02342303 |
| OPN3 | 172.015857 | -0.320093936 | 0.33832624 | -0.9461103 | 0.34409234 | 0.56886284 |
| CEP170 | 744.024761 | 0.050916304 | 0.22957056 | 0.22178935 | 0.82447787 | 0.91879931 |
| SDCCAG8 | 2991.02154 | 0.192454469 | 0.14483827 | 1.32875429 | 0.18392905 | 0.38903423 |
| AKT3 | 4459.82836 | -0.560619269 | 0.27475839 | -2.0404082 | 0.04130969 | 0.14547046 |
| ZBTB18 | 206.839173 | 0.726649571 | 0.24968629 | 2.91025019 | 0.0036114 | 0.02657544 |
| AL645465.1 | 234.736275 | -1.10630097 | 0.41854361 | -2.6432155 | 0.00821227 | 0.04812597 |
| ADSS | 1021.34462 | -0.566223052 | 0.34976559 | -1.6188644 | 0.10547645 | 0.27238287 |
| CATSPERE | 832.452189 | -0.520604513 | 0.3370465 | -1.5446074 | 0.12244126 | 0.29989368 |
| DESI2 | 690.876749 | -0.190454787 | 0.2069574 | -0.9202608 | 0.35743647 | 0.58281427 |
| COX20 | 645.973055 | 0.48682838 | 0.25888149 | 1.88050674 | 0.06003905 | 0.18792791 |
| HNRNPU | 2849.63575 | -0.086187054 | 0.18469691 | -0.4666405 | 0.64075712 | 0.80722564 |
| EFCAB2 | 1628.15843 | 0.241818625 | 0.31516619 | 0.76727336 | 0.442919 | 0.65884202 |
| KIF26B | 708.636771 | -0.364307665 | 0.66524583 | -0.5476286 | 0.58394693 | 0.76885817 |
| SMYD3 | 6033.79455 | 0.173712964 | 0.18330835 | 0.94765441 | 0.34330542 | 0.56833203 |
| TFB2M | 157.631849 | 0.300020382 | 0.22161998 | 1.35376051 | 0.1758128 | 0.37799253 |
| CNST | 701.984822 | -0.000200606 | 0.15946749 | -0.001258 | 0.99899628 | 0.99971199 |
| SCCPDH | 261.712284 | -0.748702053 | 0.31688675 | -2.3626802 | 0.01814332 | 0.08376537 |
| AHCTF1 | 1054.87453 | 0.411731588 | 0.21743363 | 1.89359665 | 0.05827857 | 0.1844278 |
| ZNF124 | 470.35764 | -0.165722977 | 0.23786045 | -0.6967236 | 0.48597582 | 0.6934193 |
| AL390728.6 | 190.053209 | 0.271674531 | 0.27244135 | 0.99718538 | 0.31867454 | 0.5432276 |
| ZNF496 | 224.543939 | -0.087388065 | 0.3608832 | -0.2421506 | 0.80866351 | 0.91010272 |
| LYPD8 | 90.3496398 | 0.589763241 | 0.29864138 | 1.97482091 | 0.04828848 | 0.16221833 |
| SH3BP5L | 126.479749 | 0.128327313 | 0.27991361 | 0.45845328 | 0.64662682 | 0.81106198 |
| ZNF672 | 145.298355 | 0.008142827 | 0.20773326 | 0.03919847 | 0.96873215 | 0.98582596 |
| ZNF692 | 172.072766 | 0.856292097 | 0.22856723 | 3.74634666 | 0.00017943 | 0.00267805 |
| PGBD2 | 70.4498805 | 0.053810691 | 0.38131763 | 0.14111776 | 0.88777691 | 0.94928354 |
| FAM110C | 1478.07195 | -2.248971774 | 0.52341183 | -4.2967538 | 1.73E-05 | 0.00042239 |
| SH3YL1 | 1146.43688 | 0.089116516 | 0.20073351 | 0.44395436 | 0.65707558 | 0.81818546 |
| ACP1 | 375.415263 | -0.046290293 | 0.18128681 | -0.2553429 | 0.7984583 | 0.90515239 |
| ALKAL2 | 61.4618231 | -1.305177929 | 0.83050656 | -1.5715444 | 0.11605625 | 0.28978961 |
| TMEM18 | 85.5846927 | -0.193405308 | 0.26589331 | -0.7273794 | 0.4669936 | 0.67935225 |
| SNTG2 | 131.199138 | 1.62454713 | 0.78437435 | 2.0711375 | 0.03834595 | 0.13904125 |
| EIPR1 | 646.479396 | -0.096136381 | 0.26563587 | -0.3619104 | 0.71741899 | 0.85947463 |
| TRAPPC12 | 1129.54527 | 0.109153009 | 0.19298722 | 0.5655971 | 0.5716677 | 0.75986595 |
| ADI1 | 449.088873 | -0.648820376 | 0.36428088 | -1.7810992 | 0.07489625 | 0.21730765 |
| AC108488.2 | 108.726761 | -0.474588439 | 0.28076976 | -1.6903118 | 0.09096832 | 0.24631864 |
| RNASEH1 | 45.6747662 | -0.022325239 | 0.3753173 | -0.0594836 | 0.9525669 | 0.97933189 |
| RNASEH1-AS | 48.7240375 | 0.592449521 | 0.38386511 | 1.54337944 | 0.12273874 | 0.30023259 |
| RPS7 | 904.976035 | -0.632809722 | 0.41647734 | -1.5194337 | 0.12865336 | 0.30972094 |
| RNF144A | 401.752467 | 1.118189962 | 0.36838597 | 3.03537604 | 0.00240236 | 0.01964877 |
| ID2 | 599.930547 | -1.109425904 | 0.48275899 | -2.2980948 | 0.0215564 | 0.09394208 |
| KIDINS220 | 2009.82646 | 0.765403208 | 0.20949151 | 3.65362391 | 0.00025856 | 0.00358759 |
| MBOAT2 | 1101.43424 | -0.195269197 | 0.40831746 | -0.4782289 | 0.63248732 | 0.80189713 |
| ASAP2 | 3532.92708 | 0.349290104 | 0.37652971 | 0.92765616 | 0.35358596 | 0.57953068 |
| ITGB1BP1 | 949.559526 | 1.062761582 | 0.31370825 | 3.3877387 | 0.00070471 | 0.00769303 |
| CPSF3 | 294.546567 | 0.291611103 | 0.18752144 | 1.55508136 | 0.11992669 | 0.29561832 |
| IAH1 | 354.429358 | 0.177253161 | 0.22310142 | 0.7944959 | 0.42690679 | 0.64526969 |
| ADAM17 | 1099.63808 | -0.175885862 | 0.19599293 | -0.8974092 | 0.36950061 | 0.59417383 |
| AC080162.1 | 56.9508358 | 0.614765657 | 0.40167022 | 1.53052336 | 0.12588724 | 0.30482942 |
| YWHAQ | 1093.83175 | -0.164069601 | 0.14877557 | -1.1027993 | 0.27011431 | 0.49175977 |
| TAF1B | 574.633947 | -0.248578865 | 0.23139437 | -1.0742649 | 0.28270395 | 0.50511067 |
| GRHL1 | 165.210193 | -0.306548821 | 0.53676015 | -0.5711095 | 0.56792542 | 0.75812736 |
| AC104794.2 | 44.5677695 | -0.215754827 | 0.46913567 | -0.4598986 | 0.64558902 | 0.81059872 |
| KLF11 | 261.085976 | 0.92982726 | 0.48486554 | 1.91770127 | 0.0551489 | 0.17750045 |
| AC104794.4 | 78.1150891 | -1.513993315 | 0.67084586 | -2.2568423 | 0.02401793 | 0.1008619 |
| CYS1 | 2692.24884 | -0.177313437 | 0.29383949 | -0.6034364 | 0.54621843 | 0.74053159 |
| HPCAL1 | 858.256092 | -0.056022044 | 0.33487496 | -0.1672924 | 0.86713996 | 0.93961371 |
| ODC1 | 151.565695 | -0.679731707 | 0.37496351 | -1.8127943 | 0.06986356 | 0.20790237 |
| NOL10 | 941.13219 | 0.131870726 | 0.23347172 | 0.56482525 | 0.57219263 | 0.76029498 |
| PDIA6 | 615.844075 | -0.361531531 | 0.31726219 | -1.1395355 | 0.25447986 | 0.47531455 |
| PQLC3 | 96.3598317 | -1.177415788 | 0.35281176 | -3.3372351 | 0.00084616 | 0.00878003 |
| ROCK2 | 1535.67911 | -0.062342647 | 0.15290302 | -0.4077267 | 0.68347432 | 0.83562441 |
| E2F6 | 178.902874 | 0.211575531 | 0.20400951 | 1.03708659 | 0.2996955 | 0.52252876 |
| LPIN1 | 807.861385 | -0.037029431 | 0.22279935 | -0.1662008 | 0.86799893 | 0.93997938 |
| MIR3681HG | 437.15413 | 0.57176325 | 0.71585266 | 0.79871639 | 0.42445488 | 0.64298076 |
| TRIB2 | 639.991228 | -0.099736297 | 0.41633017 | -0.2395606 | 0.81067093 | 0.91117701 |
| LINC00276 | 136.94444 | 1.10602385 | 0.50242732 | 2.20136091 | 0.02771048 | 0.11110055 |
| FAM84A | 78.5811785 | -1.652610922 | 0.54484906 | -3.0331537 | 0.00242012 | 0.01973297 |
| NBAS | 2305.37984 | 0.255120423 | 0.16461778 | 1.54977446 | 0.12119566 | 0.29772614 |
| DDX1 | 640.858011 | 0.261284882 | 0.14426338 | 1.8111657 | 0.07011521 | 0.20845189 |
| SMC6 | 433.744399 | 0.045845075 | 0.30211018 | 0.15174952 | 0.8793845 | 0.94621944 |
| GEN1 | 158.535055 | -0.687718682 | 0.58215937 | -1.1813237 | 0.23747414 | 0.45507241 |
| KCNS3 | 78.5280166 | -0.583919539 | 0.54941731 | -1.0627979 | 0.28787363 | 0.51020029 |
| RDH14 | 118.33937 | 0.09002524 | 0.42336185 | 0.21264372 | 0.83160487 | 0.92158196 |
| LINC01376 | 889.726863 | 0.168590363 | 0.37605859 | 0.44830877 | 0.65393037 | 0.81557608 |
| TTC32 | 194.699911 | 0.632573406 | 0.176936 | 3.57515385 | 0.00035002 | 0.00450053 |
| WDR35 | 774.902504 | 0.384432602 | 0.16201226 | 2.3728612 | 0.0176509 | 0.08197984 |
| LAPTM4A | 916.106331 | -0.880805826 | 0.35176665 | -2.5039492 | 0.01228157 | 0.06394451 |
| AC098828.3 | 70.2724599 | 0.347693266 | 1.29059689 | 0.26940501 | 0.78761803 | 0.89956761 |
| SDC1 | 771.733743 | -1.149311532 | 0.55320806 | -2.0775394 | 0.03775181 | 0.13752443 |
| PUM2 | 3427.74702 | 0.379473929 | 0.14681503 | 2.58470767 | 0.00974616 | 0.05434234 |
| RHOB | 337.934852 | -1.141798759 | 0.51761455 | -2.2058861 | 0.02739197 | 0.11030665 |
| HS1BP3 | 121.444525 | -0.143420997 | 0.32978672 | -0.4348901 | 0.66364216 | 0.82268279 |
| GDF7 | 84.2117636 | -0.916171706 | 0.53479347 | -1.7131318 | 0.08668829 | 0.23858761 |
| LDAH | 1102.99975 | 0.017939939 | 0.20733139 | 0.08652785 | 0.93104682 | 0.96965997 |
| KLHL29 | 410.391288 | -2.272702527 | 0.44010766 | -5.1639695 | 2.42E-07 | 1.29E-05 |
| ATAD2B | 1912.02138 | 0.270863994 | 0.19728196 | 1.37297903 | 0.16975887 | 0.3700239 |
| UBXN2A | 826.127513 | 0.182956253 | 0.18524512 | 0.98764411 | 0.32332698 | 0.54831506 |
| WDCP | 42.8865156 | 0.353770659 | 0.38904279 | 0.90933611 | 0.36317274 | 0.5885693 |
| SF3B6 | 279.463476 | -0.494218889 | 0.33752212 | -1.4642563 | 0.14312394 | 0.33252233 |

| | | | | | | |
|---|---|---|---|---|---|---|
| FAM228B | 207.851402 | 0.170876035 | 0.32974662 | 0.51820405 | 0.60431591 | 0.78119624 |
| TP53I3 | 306.00285 | 0.327727396 | 0.40550361 | 0.80819847 | 0.41897634 | 0.63810198 |
| ITSN2 | 2041.00136 | -0.580771997 | 0.17264442 | -3.3639778 | 0.00076828 | 0.00817742 |
| NCOA1 | 5039.20284 | 0.181828734 | 0.12909137 | 1.40852747 | 0.15897494 | 0.35534419 |
| PTRHD1 | 46.2238805 | -1.288744674 | 0.42808162 | -3.0105115 | 0.00260808 | 0.02090421 |
| CENPO | 54.8910876 | 0.012066944 | 0.35716652 | 0.03378521 | 0.97304843 | 0.98804941 |
| ADCY3 | 89.8917218 | -1.041879819 | 0.46196126 | -2.2553402 | 0.02411198 | 0.10101337 |
| DNAJC27 | 398.980492 | 0.585701548 | 0.22201935 | 2.63806532 | 0.00833805 | 0.04872913 |
| DNAJC27-AS | 265.980969 | 1.649855221 | 0.35139016 | 4.69522312 | 2.66E-06 | 9.02E-05 |
| EFR3B | 98.9188656 | 0.28500953 | 0.47152968 | 0.60443603 | 0.54555379 | 0.74015676 |
| DNMT3A | 1460.33595 | 0.174607997 | 0.2397959 | 0.72815255 | 0.46652021 | 0.67903751 |
| DTNB | 1865.37256 | 0.023918998 | 0.2161113 | 0.11067907 | 0.91187084 | 0.96021221 |
| ASXL2 | 1438.37758 | -0.056964843 | 0.19063536 | -0.2988157 | 0.76508065 | 0.88617593 |
| KIF3C | 72.6840375 | -1.479361205 | 0.40107752 | -3.6884671 | 0.00022561 | 0.00323218 |
| RAB10 | 2419.30399 | -0.315669632 | 0.16469678 | -1.9166716 | 0.05527967 | 0.17774809 |
| HADHA | 1042.29297 | 0.33266895 | 0.26014674 | 1.27877423 | 0.20097657 | 0.41005553 |
| HADHB | 1527.03767 | 0.474056553 | 0.2685783 | 1.76505903 | 0.07755385 | 0.22226269 |
| ADGRF3 | 136.123785 | 1.483595304 | 0.3911771 | 3.7926435 | 0.00014905 | 0.00229735 |
| SELENOI | 742.998979 | 0.378189672 | 0.25785198 | 1.46669294 | 0.1424596 | 0.33152729 |
| AC013472.2 | 49.3307668 | -0.061433425 | 0.57475532 | -0.1068862 | 0.91487924 | 0.96138742 |
| MAPRE3 | 246.62617 | -0.078707235 | 0.22901749 | -0.3436735 | 0.73109186 | 0.86806155 |
| TMEM214 | 138.842855 | -0.287476649 | 0.20598995 | -1.3955858 | 0.16283925 | 0.3609485 |
| AGBL5 | 275.928951 | -0.406989333 | 0.27605248 | -1.4743187 | 0.14039581 | 0.3279958 |
| OST4 | 302.354887 | -0.864529302 | 0.50567528 | -1.7096531 | 0.08733004 | 0.23953446 |
| KHK | 55.0062474 | 1.059600506 | 0.55450062 | 1.91090952 | 0.05601621 | 0.17954819 |
| PREB | 79.8379981 | -0.149028648 | 0.32330644 | -0.4609517 | 0.64483326 | 0.80996601 |
| SLC5A6 | 56.3333888 | -0.063571562 | 0.34502808 | -0.1842504 | 0.85381702 | 0.93291993 |
| ATRAID | 228.219285 | -0.643825998 | 0.35203823 | -1.8288525 | 0.06742171 | 0.20295654 |
| CAD | 125.772569 | 0.173582524 | 0.31561237 | 0.54998643 | 0.58232868 | 0.7679692 |
| MPV17 | 173.494499 | -0.326306443 | 0.27543282 | -1.1847043 | 0.23613438 | 0.45391953 |
| GTF3C2 | 526.557343 | 0.062775556 | 0.19960672 | 0.3144962 | 0.7531442 | 0.88014525 |
| AC074117.1 | 76.084172 | 0.369010601 | 0.23788271 | 1.5512292 | 0.12084677 | 0.2972007 |
| EIF2B4 | 139.396491 | 0.184650046 | 0.19003605 | 0.971658 | 0.33122072 | 0.55622942 |
| SNX17 | 135.745501 | -0.679954338 | 0.35905214 | -1.8937482 | 0.05825844 | 0.18440827 |
| ZNF513 | 77.1271043 | -0.445600727 | 0.35700275 | -1.2481717 | 0.21196818 | 0.42482955 |
| PPM1G | 616.275303 | -0.773023125 | 0.24918061 | -3.1022603 | 0.00192049 | 0.01662369 |
| NRBP1 | 360.876968 | -0.212878116 | 0.24384637 | -0.873001 | 0.38266256 | 0.60541259 |
| KRTCAP3 | 100.808385 | -0.97096622 | 0.29101279 | -3.336507 | 0.00084838 | 0.00878236 |
| IFT172 | 584.685227 | 0.802177304 | 0.24022237 | 3.33931141 | 0.00083986 | 0.00873524 |
| FNDC4 | 97.9634526 | -4.243292251 | 0.63762271 | -6.6548638 | 2.84E-11 | 6.15E-09 |
| ZNF512 | 510.483415 | -0.10773413 | 0.23936349 | -0.4500859 | 0.65264851 | 0.81482365 |
| GPN1 | 116.79109 | 0.052215405 | 0.19872518 | 0.26275183 | 0.79274186 | 0.90238946 |
| SUPT7L | 220.694175 | 0.219248782 | 0.2092312 | 1.04787806 | 0.29469479 | 0.51796358 |
| SLC4A1AP | 280.554261 | 0.261723333 | 0.16482981 | 1.58783977 | 0.11232258 | 0.28352167 |
| MRPL33 | 1213.72749 | 0.539358829 | 0.31230787 | 1.72701004 | 0.08416586 | 0.23370756 |
| RBKS | 998.01072 | 0.063495305 | 0.28484596 | 0.22291102 | 0.82360476 | 0.91846107 |
| BABAM2 | 5696.92351 | -0.178247949 | 0.26449008 | -0.6739306 | 0.50035546 | 0.70482771 |
| FOSL2 | 1207.29391 | -0.67200258 | 0.3787614 | -1.7742108 | 0.07602828 | 0.21975002 |
| AC092164.1 | 80.5401165 | 0.026630969 | 0.31965601 | 0.08331134 | 0.93360399 | 0.97167284 |
| PPP1CB | 1294.39984 | -0.378130611 | 0.26243475 | -1.4408557 | 0.14962545 | 0.341879 |
| SPDYA | 133.422481 | -0.519235436 | 0.48587296 | -1.068665 | 0.28522064 | 0.50777615 |
| TRMT61B | 373.698256 | 0.546752267 | 0.15755174 | 3.47030304 | 0.00051987 | 0.00608236 |
| WDR43 | 587.541973 | -0.015832144 | 0.18659938 | -0.0848456 | 0.93238411 | 0.97074656 |
| CLIP4 | 688.118699 | -0.766697767 | 0.32749172 | -2.3411211 | 0.01922593 | 0.08685203 |
| ALK | 104.676827 | 0.896508708 | 0.50522751 | 1.77446533 | 0.07598621 | 0.21972458 |
| AC016907.2 | 45.1315889 | 1.489069038 | 0.55358457 | 2.68986732 | 0.00714804 | 0.04335439 |
| YPEL5 | 493.518841 | -0.698634674 | 0.26136792 | -2.6729932 | 0.00751778 | 0.04507596 |
| LBH | 135.975413 | -2.546344814 | 0.48974269 | -5.1993524 | 2.00E-07 | 1.09E-05 |
| LINC01936 | 95.9874838 | -0.893084536 | 0.74423469 | -1.2000039 | 0.23013783 | 0.445816 |
| LCLAT1 | 1305.43485 | 0.067697035 | 0.1987841 | 0.34055558 | 0.73343818 | 0.86911141 |
| CAPN13 | 386.330341 | -0.143353899 | 0.47465876 | -0.3020146 | 0.7626409 | 0.88490193 |
| GALNT14 | 4385.16549 | 0.103611882 | 0.31788889 | 0.32593741 | 0.74447171 | 0.8751203 |
| MEMO1 | 102.624539 | 0.356722221 | 0.27932999 | 1.27706379 | 0.20157973 | 0.41084194 |
| DPY30 | 406.360819 | -0.111598075 | 0.21197703 | -0.5264631 | 0.59856651 | 0.77704816 |
| SPAST | 615.803623 | 0.098927466 | 0.21169435 | 0.46731274 | 0.64027615 | 0.80669664 |
| SLC30A6 | 660.994561 | 0.10977793 | 0.17186314 | 0.63875204 | 0.52298425 | 0.7227557 |
| YIPF4 | 938.932856 | -0.010458532 | 0.16293402 | -0.0641888 | 0.94881993 | 0.97784954 |
| BIRC6 | 5815.22295 | 0.125661522 | 0.17094706 | 0.73509027 | 0.46228452 | 0.67537466 |
| TTC27 | 692.696125 | 0.046723359 | 0.20888132 | 0.22368376 | 0.82300338 | 0.9182407 |
| LINC00486 | 1086.26809 | 0.62912512 | 0.21101634 | 2.98140477 | 0.00286929 | 0.02238879 |
| LTBP1 | 420.550889 | 2.051819839 | 0.88555333 | 2.31699183 | 0.02050418 | 0.0907906 |
| RASGRP3 | 336.910264 | 0.205441639 | 0.3835427 | 0.53564215 | 0.59220589 | 0.77343033 |
| FAM98A | 196.717595 | 0.242476555 | 0.24065641 | 1.00756324 | 0.31366419 | 0.53751819 |
| LINC01317 | 89.4604664 | 0.373422581 | 0.41791054 | 0.8935467 | 0.37156448 | 0.59563048 |
| AC068599.1 | 312.934384 | 1.190663251 | 0.53720792 | 2.21639184 | 0.02666468 | 0.10833719 |
| LINC01320 | 15940.276 | -1.378459241 | 0.3295167 | -4.1832758 | 2.87E-05 | 0.00063258 |
| AC073218.2 | 167.841249 | -1.086291444 | 0.6999908 | -1.5518653 | 0.12069445 | 0.29693855 |
| AC073218.1 | 141.458191 | -2.627601476 | 0.80779133 | -3.2528221 | 0.00114265 | 0.01109799 |
| AC012593.1 | 7575.71645 | -1.502537487 | 0.59076417 | -2.5433795 | 0.01097859 | 0.05926285 |
| CRIM1 | 3544.40026 | 0.21311282 | 0.30395989 | 0.70112151 | 0.48322719 | 0.69132684 |
| FEZ2 | 1207.94622 | 0.16847663 | 0.22416413 | 0.7515771 | 0.45230542 | 0.66813937 |
| STRN | 2160.6401 | 0.067352357 | 0.24557927 | 0.27425913 | 0.78388549 | 0.89671287 |
| HEATR5B | 939.017445 | -0.015499569 | 0.22617251 | -0.0685299 | 0.94536386 | 0.97640274 |
| GPATCH11 | 90.0040376 | -0.308435858 | 0.33168195 | -0.9299145 | 0.35241535 | 0.57848321 |
| EIF2AK2 | 904.1361 | 0.569974336 | 0.43150655 | 1.32089383 | 0.18653677 | 0.39228851 |
| CEBPZOS | 588.466495 | -0.113334466 | 0.17602616 | -0.6438501 | 0.51967262 | 0.7198387 |
| CEBPZ | 573.813262 | -0.182357992 | 0.19815858 | -0.9202629 | 0.35743539 | 0.58281427 |
| NDUFAF7 | 386.955189 | 0.029024902 | 0.17909741 | 0.1620621 | 0.87125695 | 0.94147844 |
| PRKD3 | 692.949419 | -0.249292947 | 0.20015001 | -1.2455305 | 0.2129368 | 0.42584137 |
| CDC42EP3 | 241.163342 | 0.37233502 | 0.3585644 | 1.03840486 | 0.29908161 | 0.52207863 |
| RMDN2 | 467.191226 | 0.742804911 | 0.30214148 | 2.45846718 | 0.01395315 | 0.06965503 |
| RMDN2-AS1 | 83.0599031 | 0.559190809 | 0.43520182 | 1.28489998 | 0.19882724 | 0.40743236 |
| CYP1B1 | 235.642701 | 0.99281901 | 0.7168343 | 1.38500488 | 0.16605098 | 0.36543941 |
| CYP1B1-AS1 | 279.150888 | 0.898994691 | 0.50177296 | 1.79163638 | 0.07319123 | 0.21436762 |
| ATL2 | 1077.65559 | 0.673470018 | 0.20816292 | 3.23530248 | 0.00121514 | 0.011631 |
| AC011247.1 | 97.5603963 | 1.257456826 | 0.52152949 | 2.4110944 | 0.01590473 | 0.07631151 |
| HNRNPLL | 608.329909 | 0.002879444 | 0.22509949 | 0.01279187 | 0.98979384 | 0.99606252 |
| GALM | 326.72857 | -0.02591629 | 0.26647662 | -0.0972554 | 0.92252358 | 0.96565607 |
| SRSF7 | 1009.24016 | 0.105911999 | 0.2906017 | 0.3644576 | 0.71551633 | 0.85795708 |
| GEMIN6 | 59.777109 | 0.220390466 | 0.29578383 | 0.74510654 | 0.45620732 | 0.67128123 |
| DHX57 | 569.539738 | -0.098616316 | 0.24690496 | -0.39941 | 0.68959111 | 0.8404511 |
| MORN2 | 119.142662 | -0.006498747 | 0.36641883 | -0.0177358 | 0.98584959 | 0.99426405 |
| ARHGEF33 | 61.3854465 | 0.443860751 | 0.35822049 | 1.23907135 | 0.21531908 | 0.42917605 |
| SOS1 | 3365.04331 | 0.004044219 | 0.10967418 | 0.03687485 | 0.9705848 | 0.98687203 |
| CDKL4 | 51.1470687 | 1.073734337 | 0.52523178 | 2.04430573 | 0.04092335 | 0.14473393 |
| MAP4K3 | 3731.97896 | 0.654487446 | 0.23169626 | 2.82476488 | 0.00473153 | 0.03206712 |
| AC007388.1 | 1891.92522 | 0.690614931 | 0.3593219 | 1.92199512 | 0.05460637 | 0.17622662 |
| TMEM178A | 90.2571533 | 1.029270218 | 0.53207289 | 1.93445343 | 0.0530574 | 0.172875 |
| THUMPD2 | 618.650853 | 0.214798338 | 0.18338778 | 1.17127944 | 0.24148647 | 0.45982339 |
| SLC8A1 | 158.541596 | 1.736978247 | 0.54923367 | 3.16254875 | 0.00156395 | 0.01415459 |
| EML4 | 1774.08154 | -0.213204506 | 0.21695983 | -0.9826912 | 0.32575946 | 0.55095477 |
| COX7A2L | 608.531276 | -0.181894689 | 0.23238797 | -0.7827199 | 0.43379162 | 0.65083524 |
| KCNG3 | 78.2716324 | -1.305778386 | 0.70232148 | -1.8592317 | 0.0629943 | 0.19436853 |

| | | | | | | |
|---|---|---|---|---|---|---|
| MTA3 | 1339.85819 | 0.159791928 | 0.144083 | 1.10902697 | 0.26741855 | 0.48856592 |
| ZFP36L2 | 3244.98737 | -2.165212124 | 0.41948099 | -5.1616454 | 2.45E-07 | 1.29E-05 |
| THADA | 2547.32765 | -0.010499579 | 0.21477209 | -0.0488871 | 0.96100929 | 0.98288592 |
| PLEKHH2 | 2634.09936 | 0.529061446 | 0.28723806 | 1.84189188 | 0.06549097 | 0.19923311 |
| DYNC2LI1 | 1176.81642 | 0.732333214 | 0.25563857 | 2.86472117 | 0.00417377 | 0.02937201 |
| LRPPRC | 1375.25732 | 0.350527671 | 0.2006908 | 1.7466056 | 0.08070577 | 0.22783556 |
| PPM1B | 1547.67527 | 0.213876674 | 0.1498445 | 1.42732417 | 0.15348648 | 0.34739597 |
| SLC3A1 | 1494.62363 | 1.118435145 | 0.66001609 | 1.69455741 | 0.09015941 | 0.24489321 |
| PREPL | 1881.53408 | 0.682254924 | 0.23800713 | 2.86653151 | 0.00414997 | 0.02928255 |
| CAMKMT | 6331.70316 | 0.335105511 | 0.19099099 | 1.75456188 | 0.07933428 | 0.22511846 |
| SRBD1 | 1081.29405 | 0.620245782 | 0.13249541 | 4.68126233 | 2.85E-06 | 9.49E-05 |
| PRKCE | 5662.4027 | 0.010021821 | 0.21269353 | 0.0471186 | 0.96241871 | 0.9832997 |
| EPAS1 | 936.886475 | -2.212321663 | 0.51130503 | -4.3268138 | 1.51E-05 | 0.00037562 |
| ATP6V1E2 | 71.0326568 | -0.248434467 | 0.35328606 | -0.7032105 | 0.48192457 | 0.69035263 |
| RHOQ | 644.289303 | -0.694500919 | 0.27264411 | -2.5472801 | 0.01085662 | 0.05882082 |
| AC018682.1 | 153.333166 | -0.887544985 | 0.54096638 | -1.6406657 | 0.10086683 | 0.26453475 |
| PIGF | 400.984899 | 0.156505304 | 0.18497688 | 0.84608036 | 0.39750791 | 0.6191842 |
| CRIPT | 165.358072 | -0.12573625 | 0.37301013 | -0.3370853 | 0.73605256 | 0.87055499 |
| SOCS5 | 722.81159 | 0.404677092 | 0.23980944 | 1.6874944 | 0.09150833 | 0.24743776 |
| MCFD2 | 420.673868 | -0.158154224 | 0.21960563 | -0.7201738 | 0.47141799 | 0.68263102 |
| TTC7A | 827.270679 | -0.099961181 | 0.40928855 | -0.2442316 | 0.80705148 | 0.90974083 |
| AC106869.1 | 175.746114 | 0.109075868 | 0.46544847 | 0.23434574 | 0.81471658 | 0.91339257 |
| EPCAM | 1036.78267 | -0.484228194 | 0.29976816 | -1.6153423 | 0.10623658 | 0.2735063 |
| MSH2 | 732.742345 | 0.358443185 | 0.23909827 | 1.49914588 | 0.13383579 | 0.31812794 |
| MSH6 | 280.347064 | 0.071065537 | 0.17100977 | 0.4155642 | 0.67772893 | 0.83113187 |
| FBXO11 | 2690.72738 | -0.129072593 | 0.15490607 | -0.8332314 | 0.40471425 | 0.62539431 |
| FOXN2 | 338.141965 | -0.444658458 | 0.1959398 | -2.2693626 | 0.02324628 | 0.09863802 |
| PPP1R21 | 1310.52985 | 0.02462866 | 0.22252014 | 0.11068059 | 0.91186964 | 0.96021221 |
| STON1-GTF2 | 105.214501 | -0.27345363 | 0.73673196 | -0.3711711 | 0.71051008 | 0.85498111 |
| AC009975.1 | 129.333909 | -0.145261759 | 0.85228497 | -0.170438 | 0.86466568 | 0.93833133 |
| NRXN1 | 63.5846463 | 0.256138831 | 0.53493204 | 0.47882499 | 0.63206314 | 0.80154781 |
| AC007402.1 | 54.4413993 | -0.030821177 | 0.31953381 | -0.0964567 | 0.92315786 | 0.96593728 |
| ASB3 | 105.7976 | 0.559131985 | 0.22703156 | 2.46279405 | 0.01378591 | 0.06902883 |
| GPR75-ASB3 | 645.016369 | -0.023847452 | 0.3721588 | -0.0640787 | 0.94890756 | 0.97784954 |
| ERLEC1 | 322.569465 | -0.002687808 | 0.15874686 | -0.0169314 | 0.98649133 | 0.99469954 |
| PSME4 | 2316.89048 | 0.076762722 | 0.24931127 | 0.30789912 | 0.75815909 | 0.88315785 |
| ACYP2 | 2756.16952 | 0.029654384 | 0.2379169 | 0.12464177 | 0.90080716 | 0.95413737 |
| C2orf73 | 64.4248523 | 0.475820064 | 0.56456736 | 0.84280477 | 0.39933764 | 0.62094804 |
| SPTBN1 | 2475.24798 | 0.398230233 | 0.24801812 | 1.60564976 | 0.10835089 | 0.27674263 |
| AC092839.1 | 81.4439405 | 0.555073125 | 0.60452976 | 0.91818991 | 0.35851945 | 0.58385938 |
| AC093110.1 | 78.1541517 | 0.743668407 | 0.5619791 | 1.3233026 | 0.18573477 | 0.39144303 |
| EML6 | 647.19672 | 0.873139516 | 0.27902369 | 3.12926663 | 0.00175243 | 0.01540112 |
| RTN4 | 3331.5258 | -0.113057355 | 0.17892361 | -0.631875 | 0.52746857 | 0.72576379 |
| CLHC1 | 626.228766 | 0.548944315 | 0.24530807 | 2.2377752 | 0.02523572 | 0.10419747 |
| RPS27A | 1444.02193 | -0.736341158 | 0.46697142 | -1.5768442 | 0.11483139 | 0.28809265 |
| MTIF2 | 380.297469 | 0.51868517 | 0.17281955 | 3.00131079 | 0.0026882 | 0.02133921 |
| AC012358.3 | 340.87649 | 0.387524498 | 0.39038135 | 0.99268191 | 0.320865 | 0.54511971 |
| CCDC88A | 398.383743 | -0.046770328 | 0.27498469 | -0.1700834 | 0.86494456 | 0.93847977 |
| CFAP36 | 650.780955 | -0.317541465 | 0.14629271 | -2.1705899 | 0.02996219 | 0.11750907 |
| PPP4R3B | 1400.48472 | -0.007989889 | 0.15859814 | -0.0503782 | 0.95982101 | 0.98229715 |
| PNPT1 | 551.825727 | 0.461864536 | 0.44711026 | 1.03299919 | 0.30160428 | 0.52495597 |
| EFEMP1 | 826.154361 | -1.363247447 | 0.59410046 | -2.2946413 | 0.02175369 | 0.09449014 |
| CCDC85A | 112.787036 | 0.346940527 | 0.44117428 | 0.78640243 | 0.43163176 | 0.64957432 |
| VRK2 | 1136.01588 | -0.220210555 | 0.14057661 | -1.5664808 | 0.1172361 | 0.29185294 |
| FANCL | 816.513922 | 0.303879451 | 0.15951146 | 1.90506343 | 0.05677183 | 0.18126639 |
| LINC01122 | 170.905433 | 0.76712378 | 0.55751272 | 1.37597538 | 0.16882926 | 0.36903287 |
| BCL11A | 145.406763 | 1.200561844 | 0.63966667 | 1.8768554 | 0.06053791 | 0.18908615 |
| PAPOLG | 370.944487 | 0.013007744 | 0.1769995 | 0.07349029 | 0.94141597 | 0.97494048 |
| LINC01185 | 37.4792873 | 0.475077045 | 0.50910084 | 0.93316886 | 0.35073279 | 0.57693534 |
| REL | 450.318907 | -0.467921075 | 0.2132759 | -2.1939707 | 0.02823752 | 0.11265157 |
| PUS10 | 379.617426 | 0.070586806 | 0.27392643 | 0.25768527 | 0.79664981 | 0.90460255 |
| PEX13 | 434.562566 | 0.052464342 | 0.21398445 | 0.24517829 | 0.80631838 | 0.90938008 |
| KIAA1841 | 248.440808 | -0.632517878 | 0.28667355 | -2.2064047 | 0.02735567 | 0.11021426 |
| C2orf74 | 202.576289 | 0.665768544 | 0.31310505 | 2.12634241 | 0.03347476 | 0.12730077 |
| USP34 | 6012.86991 | 0.0657587 | 0.16740396 | 0.39281448 | 0.69445651 | 0.843734 |
| AC016727.1 | 162.973747 | 0.376207769 | 0.28335386 | 1.32769592 | 0.18427858 | 0.38958162 |
| XPO1 | 2591.13533 | 0.259182488 | 0.22059887 | 1.17490397 | 0.24003314 | 0.45806569 |
| FAM161A | 238.14562 | 0.31308105 | 0.21378682 | 1.46445442 | 0.14306983 | 0.33248901 |
| CCT4 | 494.040934 | 0.053233006 | 0.19093325 | 0.27880427 | 0.78039504 | 0.89520087 |
| COMMD1 | 2249.79029 | 0.245202655 | 0.17590557 | 1.39394482 | 0.16333427 | 0.36162962 |
| AC018462.1 | 158.526301 | 0.034965926 | 0.39959107 | 0.08750427 | 0.93027069 | 0.96915731 |
| B3GNT2 | 467.311202 | 0.154715685 | 0.31466317 | 0.49168667 | 0.62294086 | 0.79471114 |
| TMEM17 | 38.5518549 | 0.466259104 | 0.34425692 | 1.35439282 | 0.17561109 | 0.37786639 |
| EHBP1 | 3314.49274 | 0.519779818 | 0.18963539 | 2.74094312 | 0.00612631 | 0.03888633 |
| AC092567.1 | 67.7156707 | 0.343349202 | 0.47075864 | 0.72935294 | 0.4657858 | 0.67871645 |
| WDPCP | 2410.59417 | -0.201876564 | 0.227071 | -0.889046 | 0.37397835 | 0.59778785 |
| MDH1 | 649.605406 | 0.405863158 | 0.2991222 | 1.35684731 | 0.17482973 | 0.37654099 |
| UGP2 | 1383.62938 | -0.432410424 | 0.20736061 | -2.0853064 | 0.03704149 | 0.13567522 |
| VPS54 | 1645.47389 | 0.171277688 | 0.13947996 | 1.22797341 | 0.21945695 | 0.43356369 |
| PELI1 | 1241.60238 | -0.231408443 | 0.27065228 | -0.8550027 | 0.39254963 | 0.61484815 |
| AC012368.1 | 135.713747 | 1.648013249 | 0.73165705 | 2.2524395 | 0.02429451 | 0.10158474 |
| AC008074.1 | 217.087845 | 0.585501263 | 0.40649537 | 1.44036391 | 0.14976447 | 0.34200906 |
| AFTPH | 1758.18199 | 0.302030803 | 0.16845611 | 1.79293466 | 0.07298337 | 0.21394362 |
| SERTAD2 | 320.981431 | -0.224943586 | 0.25067492 | -0.8973518 | 0.36953123 | 0.59417383 |
| CEP68 | 315.623804 | -0.098593008 | 0.25469349 | -0.3871046 | 0.69867881 | 0.84676102 |
| RAB1A | 2128.42628 | 0.099794223 | 0.21188363 | 0.470986 | 0.63765074 | 0.80499195 |
| ACTR2 | 1265.91262 | -0.662814694 | 0.16062291 | -4.1265266 | 3.68E-05 | 0.0007734 |
| SPRED2 | 1609.99334 | -0.103295462 | 0.1784108 | -0.5789754 | 0.56260578 | 0.75438633 |
| MEIS1 | 205.630452 | 0.111004205 | 0.45994846 | 0.24134053 | 0.8092912 | 0.9103008 |
| ETAA1 | 250.660648 | 0.357914693 | 0.17160316 | 2.08571151 | 0.03700476 | 0.13562595 |
| AC007422.2 | 83.9695069 | -3.392622169 | 0.6226529 | -5.4486571 | 5.08E-08 | 3.68E-06 |
| C1D | 325.46038 | 0.128576983 | 0.14632183 | 0.8787273 | 0.37954915 | 0.6023627 |
| WDR92 | 76.886332 | 0.202739644 | 0.27579074 | 0.73512128 | 0.46226564 | 0.67537466 |
| PNO1 | 111.746719 | -0.271245861 | 0.28906776 | -0.938347 | 0.34806613 | 0.57369671 |
| APLF | 512.961058 | 0.078561969 | 0.24481883 | 0.3208984 | 0.74828739 | 0.87718567 |
| ARHGAP25 | 108.327498 | -0.613859018 | 0.48481163 | -1.2661805 | 0.20544848 | 0.41584887 |
| ANTXR1 | 162.669306 | -1.217418952 | 0.57839928 | -2.1048072 | 0.0353081 | 0.13159838 |
| GFPT1 | 1193.18731 | -0.772437572 | 0.23233133 | -3.324724 | 0.00088506 | 0.00908374 |
| NFU1 | 423.871042 | 0.522457212 | 0.16106552 | 3.24375586 | 0.00117965 | 0.0113737 |
| AAK1 | 4076.89243 | 0.109449826 | 0.16413001 | 0.66684834 | 0.50486901 | 0.70861915 |
| ANXA4 | 3951.14543 | -0.804210919 | 0.47503727 | -1.6929428 | 0.09046636 | 0.24558872 |
| GMCL1 | 745.234164 | -0.077397486 | 0.2243867 | -0.344929 | 0.73014773 | 0.86751731 |
| MXD1 | 277.415933 | -0.043764376 | 0.30185337 | -0.1449855 | 0.88472228 | 0.94802439 |
| PCBP1 | 457.884098 | -1.322365168 | 0.35230446 | -3.7534727 | 0.0001744 | 0.00262098 |
| C2orf42 | 243.29956 | 0.21579969 | 0.19643755 | 1.09856641 | 0.27195724 | 0.49344548 |
| PCYOX1 | 313.945552 | -1.270228184 | 0.50661635 | -2.5072783 | 0.01216649 | 0.06354573 |
| SNRPG | 271.039307 | -0.053033823 | 0.27338773 | -0.1939876 | 0.84618561 | 0.92880719 |
| FAM136A | 217.320296 | -0.091153711 | 0.28003383 | -0.3255096 | 0.74479539 | 0.87520849 |
| TGFA | 411.194568 | -1.013626325 | 0.32315922 | -3.1366159 | 0.0017091 | 0.01514117 |
| ATP6V1B1 | 38.0440243 | 1.979776378 | 1.41425085 | 1.39987639 | 0.16155034 | 0.35921571 |
| AC007040.2 | 190.836073 | 0.470032408 | 0.86766466 | 0.54172128 | 0.58801053 | 0.7710768 |
| NAGK | 202.091883 | 0.164128783 | 0.22251153 | 0.73761923 | 0.46074586 | 0.67412407 |
| MCEE | 478.279936 | 0.672640982 | 0.18827281 | 3.57269322 | 0.00035333 | 0.00452859 |

| | | | | | | |
|---|---|---|---|---|---|---|
| MPHOSPH10 | 259.461643 | -0.068127832 | 0.21826679 | -0.312131 | 0.75494096 | 0.88130934 |
| PAIP2B | 185.602773 | 0.210372203 | 0.27712181 | 0.75913261 | 0.44777323 | 0.66352217 |
| ZNF638 | 4853.12927 | -0.051039335 | 0.15593518 | -0.3273112 | 0.7434325 | 0.87467216 |
| DYSF | 47.4679753 | 0.204861261 | 0.65421609 | 0.31314005 | 0.75417426 | 0.88075281 |
| EXOC6B | 4264.3743 | 0.193069237 | 0.21335193 | 0.90493316 | 0.3655008 | 0.59100258 |
| SPR | 60.2881983 | -1.526476998 | 0.53160042 | -2.8714744 | 0.00408562 | 0.02910838 |
| SFXN5 | 671.37183 | -0.209463996 | 0.24372048 | -0.8594436 | 0.39009584 | 0.61313374 |
| RAB11FIP5 | 272.628503 | -0.325988424 | 0.38069491 | -0.8562983 | 0.39183277 | 0.61425577 |
| SMYD5 | 71.1659177 | 0.048108791 | 0.29476701 | 0.16320955 | 0.87035345 | 0.94102469 |
| PRADC1 | 60.2530527 | -0.131719356 | 0.43123818 | -0.3054446 | 0.76002759 | 0.88357016 |
| CCT7 | 352.353086 | -0.513553352 | 0.32984785 | -1.5569401 | 0.1194847 | 0.2951699 |
| ALMS1 | 1059.873 | 0.236420162 | 0.22463045 | 1.05248491 | 0.29257713 | 0.51542428 |
| NAT8 | 113.909642 | 0.246448559 | 1.32323182 | 0.18624746 | 0.85225071 | 0.93200182 |
| TPRKB | 109.127458 | 0.003213626 | 0.20021058 | 0.01605123 | 0.98719352 | 0.99517967 |
| DUSP11 | 204.46656 | -0.139809994 | 0.18971744 | -0.736938 | 0.46116004 | 0.67447531 |
| STAMBP | 581.781482 | -0.205262757 | 0.22182669 | -0.9253294 | 0.35479459 | 0.58044114 |
| DGUOK | 502.500926 | -0.295892708 | 0.1736564 | -1.7038975 | 0.08840024 | 0.24166735 |
| DGUOK-AS1 | 80.0378661 | 0.15112499 | 0.28426133 | 0.53164105 | 0.59497463 | 0.77501911 |
| TET3 | 584.980251 | -0.279026324 | 0.21970417 | -1.2700092 | 0.20408135 | 0.41396047 |
| BOLA3 | 101.723307 | 0.305007323 | 0.39034902 | 0.78137079 | 0.43458445 | 0.65165467 |
| MOB1A | 549.529104 | -0.217408619 | 0.22728873 | -0.9565306 | 0.33880426 | 0.56426858 |
| MTHFD2 | 52.8426511 | -1.940402629 | 0.66667983 | -2.9105465 | 0.00360797 | 0.02657544 |
| AC006030.1 | 190.433249 | 0.043645477 | 0.31044129 | 0.14059172 | 0.88819249 | 0.94943227 |
| DCTN1 | 121.337153 | 0.084654192 | 0.2208461 | 0.38331758 | 0.70148431 | 0.84869711 |
| WDR54 | 67.3214883 | -1.420622594 | 0.5130071 | -2.7692065 | 0.0056193 | 0.03658223 |
| RTKN | 333.717906 | -0.651853289 | 0.26068593 | -2.5005311 | 0.01240072 | 0.06434248 |
| MOGS | 275.790739 | -0.33082146 | 0.22877257 | -1.4460714 | 0.14815718 | 0.33975837 |
| CCDC142 | 122.499672 | 0.046365117 | 0.42230151 | 0.1097915 | 0.91257473 | 0.96042853 |
| TTC31 | 134.43045 | 0.22629887 | 0.32929416 | 0.68722407 | 0.49194154 | 0.69803196 |
| AUP1 | 191.037094 | -0.04427323 | 0.35373101 | -0.1251607 | 0.90039631 | 0.95395051 |
| HTRA2 | 73.9743662 | -0.613832727 | 0.36973097 | -1.6602145 | 0.0968713 | 0.25725232 |
| M1AP | 102.849458 | -0.819163403 | 0.65932846 | -1.2424208 | 0.21408134 | 0.42754769 |
| SEMA4F | 59.6839931 | 0.018884441 | 0.24081006 | 0.07842048 | 0.93749358 | 0.97284815 |
| POLE4 | 131.589063 | -0.700141136 | 0.35102106 | -1.9945844 | 0.04608822 | 0.15694235 |
| TACR1 | 109.314825 | -0.697076611 | 0.49439758 | -1.4099515 | 0.158554 | 0.35503629 |
| AC007681.1 | 140.690904 | -0.300955513 | 0.52384699 | -0.5745103 | 0.5656225 | 0.7561287 |
| EVA1A | 590.954603 | 1.180872112 | 0.36375508 | 3.24633845 | 0.001169 | 0.01129575 |
| AC007099.1 | 157.975998 | 0.978788721 | 0.59914906 | 1.63363142 | 0.10233627 | 0.2668298 |
| MRPL19 | 673.387031 | 0.373859911 | 0.28676657 | 1.30370815 | 0.19233311 | 0.39957975 |
| GCFC2 | 1216.87859 | 0.780721297 | 0.2465023 | 3.16719686 | 0.00153916 | 0.01399423 |
| CTNNA2 | 117.656145 | 0.878758536 | 0.46798746 | 1.87773949 | 0.06041681 | 0.18888655 |
| SUCLG1 | 872.058045 | 0.805799942 | 0.24571399 | 3.27942231 | 0.0010402 | 0.01030756 |
| DNAH6 | 602.187282 | 1.134364061 | 0.40451164 | 2.8042804 | 0.0050429 | 0.03364227 |
| TMSB10 | 1630.20748 | -1.91485132 | 0.36270787 | -5.2793211 | 1.30E-07 | 7.61E-06 |
| KCMF1 | 1283.22853 | -0.162494853 | 0.18792786 | -0.8646661 | 0.38722207 | 0.61065142 |
| TCF7L1 | 184.713135 | -1.416855829 | 0.57118142 | -2.4805706 | 0.01311723 | 0.06674325 |
| TGOLN2 | 855.077544 | -0.550079328 | 0.19857315 | -2.7701596 | 0.00560288 | 0.03651134 |
| RETSAT | 133.556602 | 0.035240808 | 0.24324158 | 0.14487987 | 0.88480572 | 0.94803689 |
| ELMOD3 | 535.072471 | 0.149218374 | 0.17387516 | 0.85819259 | 0.39078613 | 0.61362693 |
| CAPG | 176.061657 | -1.144690961 | 0.33802684 | -3.3863907 | 0.00070818 | 0.00771816 |
| MAT2A | 751.563105 | 0.052491979 | 0.29302865 | 0.179136 | 0.85783092 | 0.93578594 |
| GGCX | 228.158159 | 0.174515646 | 0.18721316 | 0.93217618 | 0.35124549 | 0.57732528 |
| VAMP8 | 641.862229 | -0.592120637 | 0.35235621 | -1.6804603 | 0.09286779 | 0.25024288 |
| VAMP5 | 84.8370568 | -0.045898814 | 0.2478477 | -0.1851896 | 0.85308034 | 0.93250068 |
| RNF181 | 85.118286 | -0.944533247 | 0.42437765 | -2.2256904 | 0.02603493 | 0.10646915 |
| TMEM150A | 75.597784 | 0.017083688 | 0.35081668 | 0.04869691 | 0.96116084 | 0.98296481 |
| USP39 | 594.454498 | 0.126936065 | 0.22120345 | 0.57384307 | 0.566074 | 0.75642605 |
| C2orf68 | 273.030729 | -0.030787129 | 0.23366081 | -0.1317599 | 0.89517419 | 0.95173502 |
| ST3GAL5 | 178.835071 | 1.232829629 | 0.51964369 | 2.37245184 | 0.01767047 | 0.0820419 |
| POLR1A | 383.255611 | 0.227913798 | 0.18760997 | 1.21482776 | 0.22443179 | 0.43912303 |
| PTCD3 | 831.718338 | 0.343526391 | 0.13087231 | 2.62489745 | 0.00866751 | 0.04995161 |
| IMMT | 1239.60036 | 0.556788932 | 0.17387027 | 3.20232394 | 0.00136324 | 0.01274655 |
| MRPL35 | 157.970518 | 0.505926465 | 0.22613764 | 2.23725011 | 0.02527 | 0.10424466 |
| KDM3A | 1217.41742 | -0.362214826 | 0.32820489 | -1.1036241 | 0.26975623 | 0.49134171 |
| CHMP3 | 197.397825 | -1.286887401 | 0.26521945 | -4.8521607 | 1.22E-06 | 4.86E-05 |
| RNF103-CHM | 177.788716 | -3.694595968 | 0.83139022 | -4.4438771 | 8.84E-06 | 0.00024364 |
| AC015971.1 | 67.5668881 | -0.888875109 | 0.64200459 | -1.3845308 | 0.16619599 | 0.36563672 |
| RMND5A | 684.100747 | -0.056099483 | 0.20452009 | -0.2742981 | 0.78385551 | 0.89671287 |
| PLGLB1 | 239.621325 | 0.479882231 | 0.33350361 | 1.43891166 | 0.15017555 | 0.34236605 |
| CYTOR | 454.684932 | -0.852176435 | 0.61770002 | -1.3795959 | 0.16771109 | 0.36756193 |
| KRCC1 | 772.293495 | -0.0561654 | 0.1843144 | -0.304726 | 0.76057481 | 0.88374671 |
| THNSL2 | 170.888441 | 0.517164535 | 0.70637348 | 0.73214036 | 0.4640829 | 0.67705667 |
| EIF2AK3 | 707.092006 | -0.235377732 | 0.24701406 | -0.952892 | 0.34064478 | 0.56605571 |
| RPIA | 262.828193 | 0.300582297 | 0.17552318 | 1.71249349 | 0.08680576 | 0.23862737 |
| MAL | 2516.61421 | 0.305026632 | 0.5382827 | 0.56666624 | 0.57094096 | 0.75965257 |
| MRPS5 | 919.995444 | 0.271744353 | 0.16416681 | 1.65529412 | 0.09786485 | 0.25864282 |
| ZNF514 | 100.467298 | 0.317244815 | 0.22801345 | 1.39134258 | 0.16412157 | 0.36287999 |
| ZNF2 | 83.8455088 | 0.381373485 | 0.31797072 | 1.19939813 | 0.23037318 | 0.44592324 |
| AC092835.1 | 156.079335 | 0.174680352 | 0.24453398 | 0.71433979 | 0.47501709 | 0.68526616 |
| PROM2 | 517.213576 | 0.551453608 | 0.39895658 | 1.38223968 | 0.16689812 | 0.36644902 |
| KCNIP3 | 268.19754 | 1.189823668 | 0.33097698 | 3.59488348 | 0.00032454 | 0.00425703 |
| FAHD2A | 138.453664 | 0.511758194 | 0.2708935 | 1.88914905 | 0.05887186 | 0.18541688 |
| LINC00342 | 41.3791141 | 0.244157409 | 0.4569322 | 0.53434057 | 0.59310593 | 0.77387448 |
| ANKRD36C | 456.409164 | 0.556823465 | 0.35553043 | 1.56617667 | 0.11730726 | 0.291975 |
| STARD7 | 530.06249 | 0.526902385 | 0.16377299 | 3.21727288 | 0.00129415 | 0.01225411 |
| TMEM127 | 332.497147 | -0.267867348 | 0.21459016 | -1.2482741 | 0.21193069 | 0.42482955 |
| CIAO1 | 394.759563 | -0.354896945 | 0.23051036 | -1.5396138 | 0.12365451 | 0.30156806 |
| SNRNP200 | 617.409947 | -0.539887347 | 0.27683126 | -1.9502398 | 0.05114754 | 0.16869111 |
| NEURL3 | 61.0700189 | -0.61784641 | 0.7265484 | -0.8503858 | 0.39511065 | 0.61719685 |
| KANSL3 | 965.505354 | 0.521111928 | 0.23103295 | 2.25557404 | 0.02409732 | 0.10099717 |
| FER1L5 | 170.945029 | -0.185436497 | 0.34033775 | -0.5448602 | 0.58584967 | 0.77001836 |
| LMAN2L | 316.460707 | 0.131116393 | 0.14291053 | 0.91747188 | 0.35889542 | 0.58418356 |
| CNNM4 | 243.491896 | -0.153840982 | 0.3282411 | -0.4686829 | 0.63929633 | 0.80603771 |
| CNNM3 | 103.409858 | 0.126448547 | 0.27634191 | 0.45758006 | 0.64725417 | 0.81146359 |
| ANKRD23 | 236.447241 | 0.135985088 | 0.22610592 | 0.60142207 | 0.5475589 | 0.74151174 |
| SEMA4C | 80.0710736 | -0.523051916 | 0.34536015 | -1.5145115 | 0.12989616 | 0.31190664 |
| FAHD2B | 57.392295 | 0.688033207 | 0.60737261 | 1.13280249 | 0.25729721 | 0.47821004 |
| ANKRD36 | 695.808507 | 0.437145422 | 0.28673931 | 1.52453954 | 0.12737399 | 0.30758373 |
| AC092683.1 | 527.579214 | -0.728769515 | 0.50096129 | -1.4547422 | 0.14574065 | 0.33584931 |
| ANKRD36B | 243.023854 | -0.097206675 | 0.55401732 | -0.1754578 | 0.86071991 | 0.93666578 |
| COX5B | 779.241748 | -0.075128427 | 0.39874959 | -0.18841 | 0.85055523 | 0.93148777 |
| ACTR1B | 239.139635 | -0.42888073 | 0.25516533 | -1.6807955 | 0.09280265 | 0.25016943 |
| C2orf92 | 111.34717 | -0.20656955 | 0.39840834 | -0.518487 | 0.60411852 | 0.78103176 |
| TMEM131 | 2380.15681 | 0.498961444 | 0.2002145 | 2.49213436 | 0.0126978 | 0.06546652 |
| INPP4A | 566.923519 | -0.064320902 | 0.17042122 | -0.3774231 | 0.70585922 | 0.85140119 |
| COA5 | 184.387162 | -0.014720208 | 0.18750045 | -0.0785076 | 0.93742429 | 0.97284815 |
| UNC50 | 236.980562 | -0.048029613 | 0.22659691 | -0.2119606 | 0.83213779 | 0.92158196 |
| MGAT4A | 1883.51738 | -0.068175802 | 0.19115567 | -0.3566507 | 0.72135332 | 0.86135924 |
| KIAA1211L | 406.476584 | -0.115254074 | 0.54370077 | -0.2119807 | 0.83212209 | 0.92158196 |
| TSGA10 | 1684.12321 | 0.471009238 | 0.15900577 | 2.96221469 | 0.00305435 | 0.02346997 |
| AC079447.1 | 817.720827 | 1.079669994 | 0.30314431 | 3.56157099 | 0.00036864 | 0.00467762 |
| MITD1 | 370.085438 | 0.025315245 | 0.16898523 | 0.14980744 | 0.88091654 | 0.94640313 |
| LYG1 | 106.248134 | 0.847813966 | 0.47450016 | 1.78675169 | 0.07397763 | 0.21571093 |

| | | | | | | |
|---|---|---|---|---|---|---|
| TXNDC9 | 302.342382 | 0.110463578 | 0.3596975 | 0.30710132 | 0.75876625 | 0.88316847 |
| EIF5B | 978.033896 | -0.600215094 | 0.16852284 | -3.5616246 | 0.00036857 | 0.00467762 |
| REV1 | 2051.16513 | -0.059449306 | 0.21042596 | -0.2825189 | 0.77754567 | 0.89359206 |
| AFF3 | 94.0679297 | 0.083830462 | 0.47411193 | 0.17681576 | 0.85965311 | 0.93625342 |
| LONRF2 | 548.078577 | 0.199223143 | 0.19189922 | 1.03816546 | 0.29919303 | 0.52213512 |
| PDCL3 | 46.4454675 | -0.878404242 | 0.40853783 | -2.1501173 | 0.03154594 | 0.12226828 |
| NPAS2 | 1166.76594 | -2.877150802 | 0.52493208 | -5.4809964 | 4.23E-08 | 3.12E-06 |
| RPL31 | 630.700245 | 0.087934847 | 0.31799915 | 0.27652541 | 0.78214454 | 0.89588513 |
| TBC1D8 | 2838.66977 | -0.695504178 | 0.3035424 | -2.2912917 | 0.02194655 | 0.09480102 |
| CNOT11 | 569.094242 | -0.739472096 | 0.29237549 | -2.5291863 | 0.01143273 | 0.0609915 |
| RNF149 | 2116.67503 | -0.596464468 | 0.31907397 | -1.8693611 | 0.0615726 | 0.19132874 |
| MAP4K4 | 1746.82472 | -1.884846693 | 0.42480047 | -4.4370165 | 9.12E-06 | 0.00024753 |
| LINC01127 | 48.9107879 | 0.496796884 | 0.73282127 | 0.67792367 | 0.49782009 | 0.70298328 |
| IL1R2 | 207.981754 | -0.008715435 | 0.7306148 | -0.0119289 | 0.99048234 | 0.99606252 |
| IL1R1 | 395.43812 | -0.385812974 | 0.32025619 | -1.2047011 | 0.22831873 | 0.44383548 |
| IL1RL2 | 318.922616 | 1.362187004 | 0.32843084 | 4.14756123 | 3.36E-05 | 0.0007194 |
| SLC9A2 | 44.8678566 | -0.488203505 | 0.51265432 | -0.9523054 | 0.3409421 | 0.56647414 |
| MFSD9 | 117.258775 | -0.1758617 | 0.28698407 | -0.6127925 | 0.54001352 | 0.73609563 |
| TMEM182 | 194.826443 | -0.71500053 | 0.32235113 | -2.2180798 | 0.02654939 | 0.1080046 |
| LINC01965 | 945.643768 | 0.84372995 | 0.648843 | 1.30036073 | 0.19347736 | 0.40113679 |
| POU3F3 | 1177.86096 | 0.047445511 | 0.27946173 | 0.16977462 | 0.86518738 | 0.93858904 |
| AC010884.1 | 51.6864625 | 1.645958846 | 0.38372294 | 4.28944607 | 1.79E-05 | 0.00043236 |
| MRPS9 | 715.934004 | 0.588369596 | 0.14649981 | 4.01617994 | 5.91E-05 | 0.00111455 |
| TGFBRAP1 | 282.141752 | -0.122884863 | 0.22425843 | -0.547961 | 0.58371871 | 0.76885817 |
| AC012360.3 | 93.8221766 | 0.248769522 | 0.288175 | 0.86325853 | 0.38799534 | 0.61116244 |
| C2orf49 | 180.820549 | -0.114003251 | 0.18480136 | -0.6168962 | 0.53730322 | 0.73401988 |
| FHL2 | 786.576348 | -1.55679706 | 1.18646088 | -1.3121352 | 0.18947453 | 0.39556637 |
| AC108058.1 | 48.2470614 | -0.295863253 | 0.91413718 | -0.323653 | 0.74620075 | 0.87598457 |
| NCK2 | 681.612746 | -0.476236984 | 0.19318756 | -2.4651534 | 0.01369546 | 0.06869738 |
| UXS1 | 1256.6741 | 1.359000477 | 0.34733118 | 3.91269353 | 9.13E-05 | 0.00156981 |
| AC018878.1 | 60.5115997 | 1.459393798 | 0.58742526 | 2.48439062 | 0.01297734 | 0.0664152 |
| LINC01594 | 431.261707 | 1.637023841 | 0.45413519 | 3.604706 | 0.00031251 | 0.00414032 |
| SULT1C3 | 50.9560858 | 0.958283598 | 0.76082847 | 1.25952647 | 0.20784024 | 0.41875559 |
| SULT1C2 | 2655.04576 | 0.666429294 | 0.48275806 | 1.38046229 | 0.16744435 | 0.36728204 |
| SULT1C4 | 168.865279 | 0.207930899 | 0.38345598 | 0.54225495 | 0.58764289 | 0.77096904 |
| GCC2 | 2871.95665 | -0.706804726 | 0.14382344 | -4.9143919 | 8.91E-07 | 3.84E-05 |
| GCC2-AS1 | 322.001044 | -0.323030183 | 0.37543318 | -0.8604199 | 0.38955762 | 0.61250643 |
| LIMS1 | 3125.5573 | -0.789424806 | 0.23317277 | -3.3855789 | 0.00071028 | 0.00772192 |
| RANBP2 | 1178.70529 | -0.01685036 | 0.12366991 | -0.1362527 | 0.8916215 | 0.95094295 |
| CCDC138 | 314.258949 | 0.161842876 | 0.25586973 | 0.63252059 | 0.52704676 | 0.72541009 |
| SH3RF3 | 199.623135 | -3.563065089 | 0.96758225 | -3.6824416 | 0.00023101 | 0.00329883 |
| 10-Sep | 1806.61153 | -0.940899329 | 0.33167805 | -2.836785 | 0.00455703 | 0.0314 |
| SOWAHC | 162.764638 | -1.678781797 | 0.46163396 | -3.6366081 | 0.00027625 | 0.00378921 |
| NPHP1 | 492.157153 | 0.278197611 | 0.23670302 | 1.17530232 | 0.23987379 | 0.4580447 |
| SMIM37 | 200.043025 | -0.181610921 | 0.3320967 | -0.5468616 | 0.58447384 | 0.76903227 |
| LINC01106 | 91.3945861 | 1.190061106 | 0.44051041 | 2.70155096 | 0.00690169 | 0.04244154 |
| ACOXL | 90.0867844 | -0.565348817 | 0.55862898 | -1.0120292 | 0.31152411 | 0.53510038 |
| MIR4435-2H( | 2487.72131 | -1.039301671 | 0.53567286 | -1.9401798 | 0.05235784 | 0.17148393 |
| BCL2L11 | 644.033351 | 0.250894827 | 0.34276797 | 0.73196695 | 0.46418874 | 0.67713629 |
| AC017002.5 | 126.345236 | -0.163602029 | 0.51966396 | -0.3148227 | 0.75289625 | 0.8800112 |
| ANAPC1 | 853.822814 | 0.207992885 | 0.16477471 | 1.26228643 | 0.20684574 | 0.4175161 |
| MERTK | 608.682852 | -0.226044054 | 0.33725973 | -0.6702373 | 0.50270653 | 0.70693607 |
| TMEM87B | 772.560521 | -0.8554399 | 0.20221171 | -4.2304172 | 2.33E-05 | 0.00053772 |
| ZC3H8 | 510.312133 | 0.841440471 | 0.25455406 | 3.30554731 | 0.00094791 | 0.00955798 |
| ZC3H6 | 966.351149 | 0.634896966 | 0.36452434 | 1.7417135 | 0.08155859 | 0.22942119 |
| RGPD8 | 82.8378189 | 0.315196057 | 0.33987881 | 0.92737778 | 0.35373043 | 0.57958057 |
| TTL | 366.54129 | 0.306747879 | 0.23675957 | 1.29560918 | 0.19511016 | 0.40263651 |
| POLR1B | 184.725289 | 0.355224873 | 0.27657352 | 1.28437774 | 0.19900982 | 0.40749042 |
| CHCHD5 | 104.158467 | -0.652081622 | 0.29824706 | -2.1863807 | 0.02878776 | 0.11412293 |
| SLC20A1 | 850.911887 | 1.086397421 | 0.46202858 | 2.35136413 | 0.01870472 | 0.08519677 |
| PSD4 | 285.466859 | 0.144469037 | 0.25657816 | 0.56306054 | 0.57339366 | 0.76116346 |
| PAX8 | 14721.0745 | -1.067068979 | 0.30862899 | -3.4574489 | 0.00054532 | 0.00632403 |
| CBWD2 | 382.598068 | 0.396151593 | 0.23646718 | 1.6752921 | 0.09387694 | 0.25193426 |
| RABL2A | 276.793913 | 0.219588167 | 0.28939588 | 0.75878126 | 0.44798342 | 0.66375917 |
| AC017074.1 | 211.791138 | -0.496754117 | 0.41997001 | -1.1828323 | 0.2368756 | 0.45451624 |
| SLC35F5 | 3188.31073 | -0.487421127 | 0.16630534 | -2.9308808 | 0.00338002 | 0.02536747 |
| ACTR3 | 1658.39319 | -0.401022079 | 0.16183672 | -2.4779424 | 0.01321425 | 0.06711401 |
| LINC01191 | 54.4317876 | -3.116813614 | 0.41129742 | -7.5780044 | 3.51E-14 | 1.89E-11 |
| DPP10 | 555.673623 | 1.367945455 | 1.19534564 | 1.14439323 | 0.25246059 | 0.47247832 |
| DDX18 | 403.575792 | -0.702232931 | 0.15873752 | -4.4238623 | 9.70E-06 | 0.00025976 |
| AC009404.1 | 52.9185205 | -0.510680958 | 0.46880484 | -1.0893253 | 0.27601048 | 0.49806317 |
| CCDC93 | 957.586732 | -0.01958378 | 0.20812607 | -0.0940958 | 0.92503309 | 0.9667507 |
| INSIG2 | 518.298763 | -0.251281736 | 0.42554993 | -0.5904871 | 0.55486415 | 0.74754753 |
| STEAP3 | 114.669159 | -1.570730102 | 0.41792565 | -3.758396 | 0.00017101 | 0.00257856 |
| C2orf76 | 195.455236 | 0.094372386 | 0.19992001 | 0.47205072 | 0.63689059 | 0.80480926 |
| DBI | 798.16106 | -1.524927155 | 0.41713367 | -3.6557278 | 0.00025645 | 0.00356578 |
| TMEM37 | 803.52828 | 0.152172939 | 0.41827503 | 0.36381072 | 0.71599936 | 0.85830252 |
| SCTR | 1743.45364 | 0.162094268 | 0.54704915 | 0.29630659 | 0.76699596 | 0.88738281 |
| AC013275.1 | 105.146522 | 1.2646738 | 0.65348064 | 1.93528886 | 0.05295486 | 0.17273155 |
| CFAP221 | 3028.02858 | -0.083539214 | 0.29977372 | -0.2786742 | 0.78049483 | 0.8952376 |
| TMEM177 | 76.1850267 | 0.448208926 | 0.48281983 | 0.92831507 | 0.35324416 | 0.57919589 |
| PTPN4 | 5499.42154 | 1.570872756 | 0.23679343 | 6.63393723 | 3.27E-11 | 6.83E-09 |
| EPB41L5 | 3507.79673 | 0.466814774 | 0.2290757 | 2.03781881 | 0.04156806 | 0.14605549 |
| AC012363.1 | 44.6108512 | 0.389397909 | 0.34569536 | 1.12641926 | 0.2599881 | 0.48104536 |
| RALB | 307.35199 | -0.511742728 | 0.22246186 | -2.3003617 | 0.02142774 | 0.0935691 |
| TFCP2L1 | 5671.24779 | 0.165245939 | 0.45380385 | 0.36413516 | 0.71575708 | 0.85814392 |
| CLASP1 | 6223.78925 | 0.58532056 | 0.24202209 | 2.41845926 | 0.01558639 | 0.07534919 |
| NIFK-AS1 | 272.737162 | 0.691830625 | 0.34114032 | 2.02799431 | 0.04256083 | 0.14841235 |
| NIFK | 111.879332 | -0.256205839 | 0.23459753 | -1.092108 | 0.27478564 | 0.49646334 |
| TSN | 287.169545 | -0.309991423 | 0.20053713 | -1.5458056 | 0.12215152 | 0.2994061 |
| CNTNAP5 | 47.7413664 | 0.16795638 | 0.37964396 | 0.44240499 | 0.65819617 | 0.81888605 |
| LINC01889 | 94.2862993 | 3.089927245 | 0.54426434 | 5.67725458 | 1.37E-08 | 1.21E-06 |
| GYPC | 204.907417 | -0.055550919 | 0.25710505 | -0.2160631 | 0.82893855 | 0.92023714 |
| BIN1 | 148.373099 | -0.921699043 | 0.49202491 | -1.8732772 | 0.06103012 | 0.19003933 |
| ERCC3 | 374.969555 | -0.060499491 | 0.23265323 | -0.2600415 | 0.79483177 | 0.90385363 |
| MAP3K2 | 1861.76567 | -0.019219577 | 0.16920249 | -0.1135892 | 0.90956344 | 0.95919935 |
| IWS1 | 1245.4679 | 0.161059865 | 0.1359109 | 1.18504009 | 0.2360016 | 0.45389346 |
| AC010976.1 | 159.98965 | 0.245821023 | 0.52465614 | 0.4685374 | 0.63940033 | 0.80603771 |
| WDR33 | 1744.02663 | 0.296349758 | 0.20175492 | 1.46886013 | 0.14187073 | 0.33077653 |
| SFT2D3 | 51.1895236 | -0.110671301 | 0.32813937 | -0.3372692 | 0.73591398 | 0.87055499 |
| POLR2D | 144.192775 | 0.061263084 | 0.23112133 | 0.26506893 | 0.79095636 | 0.90144457 |
| AMMECR1L | 210.410426 | -0.012077525 | 0.16904088 | -0.0714474 | 0.94304172 | 0.97556917 |
| SAP130 | 555.197269 | -0.006721287 | 0.20768153 | -0.0323634 | 0.97418222 | 0.98832358 |
| UGGT1 | 612.961741 | 0.108729395 | 0.14370602 | 0.75660987 | 0.44928363 | 0.66516335 |
| HS6ST1 | 546.457025 | -1.051522266 | 0.36408997 | -2.8880836 | 0.00387597 | 0.02803816 |
| SMPD4 | 430.227588 | 0.037778446 | 0.19310189 | 0.19563996 | 0.84489198 | 0.92831294 |
| MZT2B | 389.690781 | -0.499410197 | 0.32570867 | -1.5333033 | 0.12520113 | 0.30383644 |
| CCDC115 | 96.6589412 | -0.322346646 | 0.2908018 | -1.1084754 | 0.26765656 | 0.48879793 |
| IMP4 | 109.360475 | -0.381260245 | 0.29584441 | -1.2887188 | 0.19749588 | 0.40583744 |
| PTPN18 | 351.045228 | -0.380558878 | 0.22024701 | -1.7278731 | 0.08401099 | 0.23347383 |
| FAM168B | 969.840037 | 0.002717314 | 0.14787483 | 0.01837577 | 0.98533908 | 0.99423861 |
| PLEKHB2 | 750.583773 | -0.474857509 | 0.15976854 | -2.972159 | 0.00295713 | 0.02290956 |
| MZT2A | 269.131539 | -0.173661253 | 0.24564029 | -0.7069738 | 0.47958275 | 0.68879075 |

| | | | | | | |
|---|---|---|---|---|---|---|
| CCDC74A | 42.3034563 | -0.142991315 | 0.36981826 | -0.3866529 | 0.69901317 | 0.84693313 |
| ANKRD30BL | 92.3892701 | 0.138809217 | 0.29421386 | 0.471797 | 0.63707169 | 0.80480926 |
| GPR39 | 1099.38008 | 0.58927505 | 0.35145925 | 1.67665254 | 0.09361045 | 0.2517316 |
| LYPD1 | 50.1199544 | -2.517687121 | 0.98765484 | -2.5491569 | 0.01079837 | 0.05855324 |
| NCKAP5 | 912.137965 | 1.56868719 | 0.40657657 | 3.85828235 | 0.00011419 | 0.0018575 |
| AC010890.1 | 39.4840974 | 2.2800944 | 0.68287784 | 3.33894918 | 0.00084096 | 0.00873976 |
| MGAT5 | 5915.0995 | 1.310313117 | 0.31683333 | 4.13565426 | 3.54E-05 | 0.00074565 |
| TMEM163 | 602.076237 | -2.420707077 | 0.67190426 | -3.602756 | 0.00031486 | 0.00416733 |
| CCNT2-AS1 | 452.133989 | -0.692798484 | 0.41128921 | -1.6844558 | 0.09209362 | 0.24861324 |
| ACMSD | 555.332094 | -0.918758621 | 0.69051104 | -1.3305488 | 0.18333752 | 0.38809393 |
| CCNT2 | 1405.48145 | 0.054021456 | 0.1755136 | 0.30779072 | 0.75824159 | 0.88315785 |
| RAB3GAP1 | 2014.33896 | 0.027010922 | 0.1611904 | 0.16757153 | 0.86692037 | 0.93959274 |
| ZRANB3 | 955.286151 | -0.082601226 | 0.20797259 | -0.3971736 | 0.69123943 | 0.84152826 |
| R3HDM1 | 2039.8282 | 0.226551578 | 0.13542632 | 1.67287696 | 0.09435152 | 0.252797 |
| UBXN4 | 1817.19791 | -0.22996974 | 0.16080818 | -1.4300873 | 0.15269196 | 0.34607208 |
| MCM6 | 61.1273252 | -1.014238196 | 0.41670424 | -2.4339522 | 0.01493497 | 0.07314645 |
| DARS | 1543.74385 | -0.429973587 | 0.22650425 | -1.8983025 | 0.05765624 | 0.18310096 |
| DARS-AS1 | 112.948498 | -0.882084777 | 0.64823612 | -1.3607461 | 0.17359394 | 0.37534823 |
| THSD7B | 223.488429 | 1.449020241 | 0.83930656 | 1.72644932 | 0.08426661 | 0.23393814 |
| HNMT | 997.044776 | -0.132898154 | 0.29165806 | -0.4556643 | 0.64863143 | 0.81191151 |
| AC114763.1 | 355.687143 | 0.1869513 | 0.28715986 | 0.65103563 | 0.51502349 | 0.7164072 |
| SPOPL | 763.686012 | -0.047307021 | 0.16573775 | -0.285433 | 0.77531244 | 0.89223749 |
| LRP1B | 8648.92436 | -1.005804546 | 0.48465909 | -2.0752825 | 0.03796036 | 0.13809374 |
| ARHGAP15 | 43.8275822 | -0.212670703 | 0.32253685 | -0.6593687 | 0.50965903 | 0.71186277 |
| GTDC1 | 738.757004 | -0.053877565 | 0.26819155 | -0.2008921 | 0.84078294 | 0.92618646 |
| TEX41 | 412.140227 | 2.929809313 | 0.61701403 | 4.74836741 | 2.05E-06 | 7.22E-05 |
| ACVR2A | 1047.50327 | 0.76780048 | 0.24207245 | 3.17177975 | 0.00151508 | 0.01384002 |
| ORC4 | 1527.35072 | 0.617064255 | 0.17663164 | 3.4935092 | 0.00047672 | 0.0057141 |
| MBD5 | 7239.73393 | 0.049278603 | 0.14798967 | 0.33298677 | 0.73914427 | 0.87227345 |
| EPC2 | 1839.5209 | 0.019782119 | 0.22815976 | 0.08670293 | 0.93090765 | 0.96965997 |
| AC105402.3 | 61.8296273 | 0.242531576 | 1.04183952 | 0.23279168 | 0.81592317 | 0.91427971 |
| LYPD6B | 571.407032 | 0.802910986 | 0.41796218 | 1.9210135 | 0.05473001 | 0.17653444 |
| LYPD6 | 326.748867 | 1.701170464 | 0.40863456 | 4.16306066 | 3.14E-05 | 0.00067774 |
| MMADHC | 328.168573 | 0.213630788 | 0.30194837 | 0.70750767 | 0.47925105 | 0.68864274 |
| AC007364.1 | 185.432468 | -0.354810609 | 0.40080091 | -0.885254 | 0.37601965 | 0.59932418 |
| RND3 | 121.159981 | -0.499779554 | 0.5257337 | -0.9506325 | 0.34179095 | 0.56712537 |
| RBM43 | 84.0178559 | -0.088237271 | 0.42768018 | -0.206316 | 0.83654407 | 0.92398282 |
| NMI | 219.038834 | -0.421054316 | 0.37727658 | -1.1160362 | 0.26440664 | 0.48534566 |
| RIF1 | 1219.15229 | 0.072384898 | 0.19354642 | 0.37399245 | 0.70840994 | 0.85346928 |
| NEB | 327.015322 | 1.778444812 | 0.5254992 | 3.38429591 | 0.00071361 | 0.00775171 |
| ARL5A | 475.181157 | 0.186544018 | 0.33905009 | 0.55019605 | 0.58218491 | 0.76785626 |
| CACNB4 | 1276.21302 | 1.842738373 | 0.58124785 | 3.17031432 | 0.00152274 | 0.01389081 |
| STAM2 | 1297.76083 | 0.552573539 | 0.20251114 | 2.72860817 | 0.00636022 | 0.03994136 |
| FMNL2 | 3668.9885 | -0.837074551 | 0.25091562 | -3.3360799 | 0.00084969 | 0.00878534 |
| PRPF40A | 1042.86686 | 0.042766932 | 0.17242122 | 0.24803752 | 0.80410537 | 0.90804718 |
| ARL6IP6 | 590.363959 | -0.052088987 | 0.24067895 | -0.2164252 | 0.82865634 | 0.92023714 |
| GALNT13 | 536.784055 | 0.335252476 | 0.32504463 | 1.03140444 | 0.3023512 | 0.52556349 |
| AC009227.1 | 58.1811825 | 2.261329526 | 0.60080782 | 3.76381506 | 0.00016734 | 0.00253196 |
| KCNJ3 | 2020.02147 | 1.416080363 | 0.51957284 | 2.72547033 | 0.00642099 | 0.0401637 |
| LINC01876 | 953.991396 | -0.708616613 | 0.65257824 | -1.0858723 | 0.27753551 | 0.49972281 |
| NR4A2 | 97.3102617 | -0.081289671 | 0.37782142 | -0.2151537 | 0.8296475 | 0.92029004 |
| GPD2 | 983.073581 | 0.069321593 | 0.21684543 | 0.31968206 | 0.74920936 | 0.87779905 |
| CYTIP | 190.734917 | 0.029963952 | 0.87241415 | 0.03434602 | 0.97260123 | 0.98801278 |
| ACVR1C | 55.281011 | 0.2925769 | 0.55481634 | 0.52734009 | 0.59795744 | 0.77678206 |
| AC019186.1 | 62.2875716 | 0.272934039 | 0.39901455 | 0.68402027 | 0.49396237 | 0.69925505 |
| ACVR1 | 1522.37916 | -0.135764217 | 0.20763945 | -0.653846 | 0.51321106 | 0.71500948 |
| UPP2 | 1099.11408 | 1.011650083 | 0.47920844 | 2.11108569 | 0.03476495 | 0.13027239 |
| CCDC148-AS1 | 147.704581 | 0.020465059 | 0.56208285 | 0.03640933 | 0.97095598 | 0.98717345 |
| CCDC148 | 5303.09064 | 1.117456396 | 0.30732642 | 3.63605707 | 0.00027684 | 0.00379338 |
| PKP4 | 15092.4726 | 0.806709913 | 0.24537466 | 3.28766591 | 0.00101022 | 0.01006504 |
| PKP4-AS1 | 417.095776 | 0.33133298 | 0.51165361 | 0.64757284 | 0.51726127 | 0.71860582 |
| TANC1 | 1975.50773 | -0.533259534 | 0.22846861 | -2.3340604 | 0.01959256 | 0.08799663 |
| WDSUB1 | 855.305546 | 0.607849269 | 0.25055903 | 2.42597231 | 0.01526743 | 0.07427964 |
| BAZ2B | 5700.21768 | 0.115399102 | 0.17323668 | 0.66613551 | 0.50532449 | 0.70903242 |
| AC009961.1 | 216.2598 | 0.01227521 | 0.18188242 | 0.06748981 | 0.94619177 | 0.9767213 |
| 7-Mar | 1734.70745 | 0.036378128 | 0.12916067 | 0.28165019 | 0.77821175 | 0.89400754 |
| PLA2R1 | 199.082584 | -1.146383986 | 0.36975223 | -3.1004113 | 0.00193252 | 0.0166841 |
| ITGB6 | 14645.9371 | -2.678689368 | 0.70306613 | -3.8100105 | 0.00013896 | 0.00217594 |
| LINC02478 | 80.7009166 | -1.168403671 | 0.56628661 | -2.0632726 | 0.03908673 | 0.14064197 |
| RBMS1 | 5514.03512 | -0.514080094 | 0.30204125 | -1.7020195 | 0.08875171 | 0.24231529 |
| TANK | 1787.51021 | -0.387788486 | 0.23434333 | -1.6547878 | 0.09796755 | 0.25886243 |
| PSMD14 | 638.254431 | -0.072396018 | 0.23863457 | -0.3033761 | 0.76160328 | 0.88431255 |
| DPP4 | 349.147659 | -0.392195476 | 0.52902049 | -0.7413616 | 0.45847422 | 0.67266311 |
| IFIH1 | 247.996332 | 0.29057329 | 0.62676735 | 0.4636063 | 0.64292984 | 0.80857472 |
| GCA | 420.638405 | 0.112771362 | 0.20732098 | 0.54394572 | 0.58647882 | 0.77036771 |
| FIGN | 72.4005533 | -0.120653928 | 0.53085452 | -0.2272825 | 0.8202041 | 0.91658849 |
| GRB14 | 2414.55402 | 0.688558201 | 0.33462949 | 2.05767342 | 0.03962149 | 0.14177464 |
| COBLL1 | 18887.582 | 0.939371633 | 0.3246999 | 2.89304561 | 0.00381526 | 0.02770594 |
| AC019197.1 | 7835.06212 | 0.955095761 | 0.5761322 | 1.65777187 | 0.09736352 | 0.25780996 |
| SLC38A11 | 703.894562 | 1.376716077 | 0.58520322 | 2.35254359 | 0.0186455 | 0.08497037 |
| SCN3A | 693.923574 | -1.112125638 | 0.43051642 | -2.5832363 | 0.00978783 | 0.05452336 |
| SCN2A | 1191.72138 | 1.076714218 | 0.62852973 | 1.71306807 | 0.08670002 | 0.23858761 |
| CSRNP3 | 1322.34234 | 1.475343551 | 0.26807291 | 5.50351593 | 3.72E-08 | 2.79E-06 |
| GALNT3 | 490.091043 | -0.922846812 | 0.22545044 | -4.0933467 | 4.25E-05 | 0.00086633 |
| AC009495.1 | 119.964838 | 0.100579876 | 0.44185698 | 0.22762994 | 0.81993395 | 0.91651491 |
| TTC21B | 950.220452 | 0.44375736 | 0.18161265 | 2.44342763 | 0.01454849 | 0.07170558 |
| AC010127.1 | 95.0234327 | 0.361879468 | 0.71308588 | 0.50748371 | 0.61181546 | 0.78684784 |
| SCN1A | 57.7378051 | 5.08329684 | 0.72080194 | 7.05227965 | 1.76E-12 | 5.41E-10 |
| SCN9A | 156.325254 | -1.951034652 | 0.75119457 | -2.5972428 | 0.00939755 | 0.05295743 |
| SCN7A | 161.922698 | 0.622126948 | 1.01608688 | 0.61227732 | 0.54035429 | 0.73628429 |
| STK39 | 5129.73029 | -1.126924805 | 0.5260802 | -2.142116 | 0.03218415 | 0.1238696 |
| CERS6 | 1192.9435 | -0.184154013 | 0.33636798 | -0.5474778 | 0.58405049 | 0.76885817 |
| NOSTRIN | 155.598758 | -1.007876072 | 0.99370899 | -1.0142568 | 0.31046025 | 0.53438486 |
| LRP2 | 293.978224 | 1.225687511 | 0.82384033 | 1.48777314 | 0.13681073 | 0.3225871 |
| FASTKD1 | 835.624827 | 1.155306505 | 0.20666342 | 5.59028052 | 2.27E-08 | 1.84E-06 |
| PPIG | 1128.50318 | 0.002897158 | 0.11750644 | 0.02465532 | 0.9803299 | 0.99141592 |
| CCDC173 | 84.1209485 | -0.253785091 | 0.39779306 | -0.6379827 | 0.52348494 | 0.723145 |
| KLHL23 | 412.020359 | 0.023124088 | 0.15869249 | 0.14571634 | 0.88414532 | 0.9477505 |
| SSB | 352.512648 | -0.358374836 | 0.27870796 | -1.2858436 | 0.19849766 | 0.40700957 |
| METTL5 | 224.452425 | -0.125573385 | 0.16441177 | -0.7637737 | 0.44500211 | 0.66082438 |
| UBR3 | 3078.08318 | 0.542669515 | 0.17921618 | 3.02801629 | 0.00246165 | 0.01997292 |
| MYO3B | 3892.83594 | 1.265124213 | 0.36452468 | 3.47061333 | 0.00051927 | 0.00608236 |
| AC012594.1 | 81.881344 | 0.855813061 | 0.58308842 | 1.46772434 | 0.14217912 | 0.33120942 |
| AC007277.1 | 84.677229 | 0.964032074 | 0.37559437 | 2.56668401 | 0.01026761 | 0.05655751 |
| AC007405.1 | 244.126472 | 0.538967836 | 0.41078268 | 1.31205103 | 0.18950292 | 0.39556637 |
| AC007405.3 | 73.0430593 | 0.430590316 | 0.36863362 | 1.16807121 | 0.24277804 | 0.46128527 |
| ERICH2 | 48.580392 | 0.652504943 | 0.37719801 | 1.72987377 | 0.08365283 | 0.23299214 |
| GAD1 | 701.898206 | 1.557570425 | 0.40830363 | 3.81473565 | 0.00013633 | 0.00214377 |
| GORASP2 | 540.918914 | -0.190576334 | 0.14202303 | -1.3418692 | 0.1796384 | 0.38325438 |
| TLK1 | 2407.32347 | 0.167101259 | 0.10664474 | 1.56689635 | 0.11713892 | 0.29172097 |
| METTL8 | 428.350992 | 0.346760002 | 0.31467707 | 1.10195508 | 0.2704812 | 0.49198378 |
| DCAF17 | 353.812284 | 0.468374976 | 0.24138872 | 1.94033494 | 0.052339 | 0.17146474 |
| CYBRD1 | 168.200714 | -1.29721815 | 0.57475283 | -2.2570018 | 0.02400796 | 0.1008619 |

| | | | | | | |
|---|---|---|---|---|---|---|
| DYNC1I2 | 1031.14826 | -0.304066582 | 0.08411047 | -3.6150859 | 0.00030025 | 0.00402804 |
| SLC25A12 | 2050.68758 | 0.652179532 | 0.17421552 | 3.74352153 | 0.00018146 | 0.0027053 |
| HAT1 | 462.985393 | -0.222385898 | 0.20080118 | -1.107493 | 0.26808086 | 0.48932276 |
| METAP1D | 185.574352 | 0.49129026 | 0.39014095 | 1.25926351 | 0.20793517 | 0.41888297 |
| ITGA6 | 1103.57229 | 0.772484048 | 0.34499416 | 2.23912208 | 0.02514797 | 0.10407512 |
| AC078883.1 | 153.471899 | -1.243653612 | 0.55761988 | -2.2302892 | 0.02572825 | 0.10557217 |
| PDK1 | 829.945907 | -1.261408055 | 0.59803586 | -2.1092515 | 0.03492288 | 0.13071586 |
| RAPGEF4 | 158.791734 | -0.204023584 | 0.50202998 | -0.4063972 | 0.68445077 | 0.83650168 |
| MAP3K20 | 150.291309 | -2.701230669 | 0.44072086 | -6.1291191 | 8.84E-10 | 1.16E-07 |
| SP3 | 1137.94302 | -0.086530819 | 0.1759002 | -0.4919313 | 0.6227679 | 0.79471114 |
| OLA1 | 1967.69048 | 0.202139985 | 0.22395554 | 0.90258978 | 0.36674365 | 0.59228718 |
| CIR1 | 696.664515 | -0.125184959 | 0.1276019 | -0.9810587 | 0.32656378 | 0.55192222 |
| SCRN3 | 288.543816 | 0.332286009 | 0.26854385 | 1.2373622 | 0.21595265 | 0.42979034 |
| GPR155 | 509.500154 | 0.560168068 | 0.31345379 | 1.78708338 | 0.07392401 | 0.21565688 |
| AC010894.3 | 615.327336 | -1.843393392 | 0.37956849 | -4.8565501 | 1.19E-06 | 4.78E-05 |
| WIPF1 | 559.975977 | -0.401786845 | 0.38062085 | -1.0556091 | 0.29114682 | 0.51373981 |
| CHN1 | 112.627107 | -0.209867062 | 0.29989751 | -0.699796 | 0.48405474 | 0.692203 |
| ATF2 | 1109.17247 | 0.142649702 | 0.14759096 | 0.96652059 | 0.33378375 | 0.5592275 |
| ATP5MC3 | 813.34898 | -0.146850251 | 0.50839082 | -0.2888531 | 0.77269382 | 0.89062065 |
| LNPK | 477.246856 | 0.107181265 | 0.27628034 | 0.38794387 | 0.69805757 | 0.84616339 |
| HOXD10 | 696.521055 | 0.121256947 | 0.48766976 | 0.24864562 | 0.80363492 | 0.90804718 |
| HOXD-AS2 | 180.799842 | 0.973252037 | 0.43881689 | 2.21790015 | 0.02656164 | 0.10802116 |
| HOXD9 | 253.738737 | 0.405903264 | 0.54793323 | 0.74078964 | 0.458821 | 0.67278348 |
| HOXD8 | 255.765079 | 0.304884792 | 0.46856208 | 0.65068174 | 0.51525195 | 0.71656802 |
| HOXD3 | 874.699943 | 0.425381249 | 0.37522819 | 1.13366015 | 0.25693713 | 0.47778941 |
| HAGLR | 235.813876 | -0.749857162 | 0.21890491 | -3.4254926 | 0.00061369 | 0.00694286 |
| HOXD1 | 50.1817132 | -1.118455491 | 0.45511981 | -2.4574968 | 0.0139909 | 0.06971176 |
| MTX2 | 2385.36569 | 0.615969889 | 0.18461191 | 3.33656643 | 0.0008482 | 0.00878236 |
| LINC01116 | 212.179544 | 0.260708976 | 0.45054591 | 0.5786513 | 0.56282448 | 0.75444301 |
| LINC01117 | 222.951076 | -0.565341986 | 0.39735074 | -1.4227782 | 0.15480045 | 0.34929265 |
| AC092162.2 | 49.4739172 | -0.429997639 | 0.48189728 | -0.8923014 | 0.37223139 | 0.5963377 |
| AC079305.3 | 104.725397 | 0.696546104 | 0.33237883 | 2.09563921 | 0.03611421 | 0.13373495 |
| HNRNPA3 | 1639.61865 | 0.154596488 | 0.14212316 | 1.08776425 | 0.27669919 | 0.49896172 |
| NFE2L2 | 3329.41848 | 0.0370523 | 0.27475265 | 0.13485693 | 0.89272499 | 0.95119823 |
| AGPS | 688.372537 | 0.08827369 | 0.13203407 | 0.66856751 | 0.5037714 | 0.7078307 |
| TTC30B | 41.3561907 | -0.15484552 | 0.55151459 | -0.2807641 | 0.77889131 | 0.89433176 |
| TTC30A | 70.7775294 | 0.056657502 | 0.28108754 | 0.20156532 | 0.84025656 | 0.92579761 |
| RBM45 | 268.901483 | 0.704050649 | 0.17009555 | 4.13914803 | 3.49E-05 | 0.00073911 |
| OSBPL6 | 50.2479131 | 0.604979982 | 0.38254581 | 1.58145759 | 0.11377345 | 0.28619948 |
| AC009948.1 | 174.189884 | -0.527048605 | 0.29480275 | -1.7878008 | 0.07380815 | 0.21554455 |
| PRKRA | 654.175904 | -0.03624767 | 0.17737763 | -0.2043531 | 0.83807758 | 0.92446904 |
| PJVK | 164.097707 | 0.142170805 | 0.2474401 | 0.57456656 | 0.56558448 | 0.7561287 |
| PLEKHA3 | 315.070476 | -0.161140499 | 0.18844138 | -0.8551227 | 0.39248322 | 0.61484815 |
| TTN-AS1 | 1617.37887 | -0.063214467 | 0.24356792 | -0.2595353 | 0.79522227 | 0.90396652 |
| TTN | 169.982995 | -0.878322652 | 0.55726991 | -1.5761171 | 0.11499881 | 0.28829365 |
| SESTD1 | 3597.43797 | -1.313447681 | 0.37722121 | -3.481903 | 0.00049786 | 0.00589801 |
| ZNF385B | 1530.96043 | 1.877323661 | 0.36008308 | 5.21358477 | 1.85E-07 | 1.03E-05 |
| CWC22 | 528.49683 | 0.236705904 | 0.20100676 | 1.1776017 | 0.23895543 | 0.45716719 |
| AC009478.1 | 920.3509 | 0.699322313 | 0.26518795 | 2.63708182 | 0.00836227 | 0.04876696 |
| SCHLAP1 | 3269.29313 | -0.21048477 | 0.39295996 | -0.5356392 | 0.59220791 | 0.77343033 |
| UBE2E3 | 2793.65633 | 0.543504914 | 0.21054115 | 2.58146645 | 0.00983816 | 0.05471594 |
| LINC01934 | 53.6519887 | 1.012617495 | 0.4476232 | 2.26220961 | 0.02368446 | 0.10003413 |
| CERKL | 403.874362 | 0.174108427 | 0.36614919 | 0.47551225 | 0.63442191 | 0.80338213 |
| SSFA2 | 1423.41215 | 0.252778683 | 0.39051002 | 0.64730395 | 0.51743524 | 0.71869378 |
| PPP1R1C | 101.35558 | -0.585488408 | 0.4842484 | -1.2090663 | 0.22663739 | 0.44167206 |
| PDE1A | 22859.8803 | 0.864051595 | 0.39611123 | 2.18133577 | 0.02915859 | 0.11507494 |
| DNAJC10 | 592.697125 | -0.4601562 | 0.18848357 | -2.4413597 | 0.01463207 | 0.07206378 |
| NCKAP1 | 2707.09288 | -0.187056697 | 0.19474653 | -0.9605136 | 0.33679678 | 0.56213655 |
| DUSP19 | 42.2807844 | 0.474361018 | 0.35996913 | 1.3177825 | 0.18757647 | 0.39322291 |
| NUP35 | 303.85299 | 0.191273097 | 0.21815886 | 0.87676061 | 0.38061668 | 0.60347686 |
| AC007966.1 | 66.7828097 | 1.211926458 | 0.33263267 | 3.64343785 | 0.00026902 | 0.00371316 |
| FSIP2 | 241.565475 | 0.709826683 | 0.57531118 | 1.23381347 | 0.21727242 | 0.43128814 |
| FSIP2-AS1 | 193.070497 | 1.060912065 | 0.3395276 | 3.12467105 | 0.00178004 | 0.01560427 |
| LINC01473 | 412.655545 | 0.582501506 | 0.26661536 | 2.18480104 | 0.02890344 | 0.11434127 |
| AC017071.1 | 55.1347814 | 0.814083259 | 0.33274533 | 2.44656557 | 0.01442246 | 0.07127057 |
| ZC3H15 | 544.813203 | 0.039423701 | 0.15690414 | 0.25125978 | 0.80161327 | 0.90677969 |
| ITGAV | 2113.06852 | 0.393692523 | 0.37295866 | 1.05559293 | 0.29115423 | 0.51373981 |
| AC017101.1 | 265.792166 | 0.320981776 | 0.76916248 | 0.41731336 | 0.67644922 | 0.83016679 |
| FAM171B | 123.970893 | -2.535823289 | 0.75674553 | -3.3509591 | 0.00080532 | 0.00846935 |
| AC007319.1 | 2512.07907 | -1.99737277 | 0.62002455 | -3.2214414 | 0.00127548 | 0.01212068 |
| CALCRL | 278.920491 | -2.224394392 | 0.75310433 | -2.9536338 | 0.00314056 | 0.02393752 |
| TFPI | 6167.12535 | -2.727116035 | 0.54687393 | -4.9867363 | 6.14E-07 | 2.81E-05 |
| LINC01090 | 175.227099 | -1.613841959 | 0.56256959 | -2.8686975 | 0.00412166 | 0.02922329 |
| GULP1 | 3451.10326 | -1.420155776 | 0.52447228 | -2.7077804 | 0.00677348 | 0.04186754 |
| COL5A2 | 237.806803 | 1.167056399 | 0.44253818 | 2.63718803 | 0.00835965 | 0.04876696 |
| WDR75 | 344.840982 | 0.343249892 | 0.25236263 | 1.36014549 | 0.17378389 | 0.37550853 |
| SLC40A1 | 231.338294 | 0.603504798 | 0.38643083 | 1.5617408 | 0.11834906 | 0.29339403 |
| ASNSD1 | 169.936859 | 0.323511328 | 0.27915207 | 1.15890713 | 0.24649404 | 0.46556654 |
| ANKAR | 883.334277 | 0.18846607 | 0.25434754 | 0.74097855 | 0.45870645 | 0.67278348 |
| OSGEPL1 | 112.522059 | 0.790205621 | 0.23194643 | 3.4068453 | 0.00065718 | 0.00729474 |
| ORMDL1 | 564.041598 | 0.172144185 | 0.24462458 | 0.70370765 | 0.48161486 | 0.69005865 |
| PMS1 | 587.780421 | 0.437813502 | 0.17225057 | 2.54172448 | 0.01103071 | 0.05949556 |
| C2orf88 | 310.14457 | 0.524725487 | 0.32451344 | 1.61696073 | 0.10588676 | 0.27301545 |
| HIBCH | 1938.97887 | 0.495196606 | 0.24002623 | 2.06309368 | 0.03910373 | 0.1406648 |
| INPP1 | 225.681636 | -0.610319592 | 0.27303543 | -2.2353128 | 0.02539682 | 0.10457189 |
| MFSD6 | 490.72969 | -0.972560955 | 0.24069677 | -4.0406066 | 5.33E-05 | 0.00102506 |
| NEMP2 | 95.3060649 | 0.068079273 | 0.36422849 | 0.18691364 | 0.85172834 | 0.93193618 |
| AC108047.1 | 106.593743 | 0.931510789 | 0.29573373 | 3.14982934 | 0.00163366 | 0.0146713 |
| NAB1 | 365.577767 | -0.395056571 | 0.35231653 | -1.1213115 | 0.2621553 | 0.48302544 |
| GLS | 5469.69674 | -0.800422796 | 0.35695979 | -2.2423332 | 0.02493984 | 0.10349682 |
| STAT1 | 1097.79631 | -0.707399616 | 0.62879462 | -1.125009 | 0.26058521 | 0.48174529 |
| MYO1B | 461.227478 | -0.500122716 | 0.41934503 | -1.1926282 | 0.23301501 | 0.44978742 |
| NABP1 | 357.282047 | -1.613922718 | 0.73560503 | -2.1940072 | 0.02823489 | 0.11265157 |
| SLC39A10 | 745.278605 | -0.023072052 | 0.36984003 | -0.0623839 | 0.95025714 | 0.97840413 |
| DNAH7 | 2001.76686 | 0.335767018 | 0.1550322 | 2.16578896 | 0.03032732 | 0.11862408 |
| STK17B | 205.905972 | -0.533180699 | 0.20362662 | -2.6184233 | 0.00883371 | 0.05053466 |
| GTF3C3 | 443.812282 | 0.303847988 | 0.16694962 | 1.81999808 | 0.0687593 | 0.20568756 |
| PGAP1 | 483.808654 | 0.32936028 | 0.18648799 | 1.76612062 | 0.07737561 | 0.22191693 |
| ANKRD44 | 915.640405 | -0.014962007 | 0.29842529 | -0.0501365 | 0.96001359 | 0.98229715 |
| SF3B1 | 1337.38681 | 0.322383677 | 0.17586429 | 1.83313889 | 0.06678192 | 0.20181248 |
| COQ10B | 304.80543 | -0.180992984 | 0.22767584 | -0.7949591 | 0.42663729 | 0.64508083 |
| HSPD1 | 895.716081 | -0.226198417 | 0.26771782 | -0.8449136 | 0.3981591 | 0.61972127 |
| MOB4 | 125.698811 | -0.212651037 | 0.32043332 | -0.6636358 | 0.50692341 | 0.71014438 |
| AC011997.1 | 132.32273 | 0.292403899 | 0.35205689 | 0.83055865 | 0.406223 | 0.62627606 |
| PLCL1 | 18795.9735 | 1.829371156 | 0.38248688 | 4.78283379 | 1.73E-06 | 6.29E-05 |
| SATB2 | 483.020819 | 0.025076146 | 0.46523441 | 0.05390002 | 0.95701482 | 0.98107945 |
| FTCDNL1 | 294.134345 | 0.796427565 | 0.52032359 | 1.53063897 | 0.12585864 | 0.30481607 |
| C2orf69 | 152.977195 | -0.226484662 | 0.30780462 | -0.7358066 | 0.46184843 | 0.67513899 |
| TYW5 | 392.813815 | 0.03024753 | 0.15092408 | 0.20041553 | 0.84115561 | 0.92644247 |
| MAIP1 | 173.827702 | 0.155752337 | 0.24607787 | 0.63293924 | 0.52677332 | 0.72526045 |
| SPATS2L | 1456.52927 | -0.397582287 | 0.55341012 | -0.7184225 | 0.47249682 | 0.68321953 |
| KCTD18 | 179.238021 | 0.021776497 | 0.21005035 | 0.10367275 | 0.91742905 | 0.96291779 |
| SGO2 | 86.5236507 | 0.846934958 | 0.39694723 | 2.13362102 | 0.03287382 | 0.12571808 |

| | | | | | | |
|---|---|---|---|---|---|---|
| AOX1 | 56.9606311 | -0.712160621 | 0.81019835 | -0.8789954 | 0.37940377 | 0.60234909 |
| BZW1 | 238.712243 | -0.892923753 | 0.25188081 | -3.5450249 | 0.00039258 | 0.00492029 |
| CLK1 | 1160.27198 | 0.533937218 | 0.26877142 | 1.98658481 | 0.04696843 | 0.15907844 |
| PPIL3 | 303.453865 | 0.122674718 | 0.32077469 | 0.38243266 | 0.70214047 | 0.8488446 |
| NIF3L1 | 147.157369 | -0.033274864 | 0.24527664 | -0.1356626 | 0.89208802 | 0.95118543 |
| ORC2 | 1364.58223 | 0.105436576 | 0.2366043 | 0.44562409 | 0.65586881 | 0.81726196 |
| AC005037.1 | 316.873549 | 0.16636538 | 0.30437236 | 0.54658504 | 0.58466385 | 0.76905468 |
| FAM126B | 1201.24508 | 0.765347224 | 0.24393107 | 3.13755528 | 0.00170363 | 0.01511301 |
| NDUFB3 | 366.566221 | 0.005386974 | 0.44767409 | 0.01203325 | 0.99039909 | 0.99606252 |
| CFLAR | 3800.1795 | -0.045393579 | 0.18983049 | -0.2391269 | 0.81100718 | 0.91124479 |
| CFLAR-AS1 | 390.958322 | -0.27346196 | 0.55413804 | -0.4934907 | 0.62166592 | 0.79361991 |
| CASP10 | 540.514767 | -0.893784715 | 0.29908512 | -2.9883958 | 0.00280446 | 0.02206666 |
| CASP8 | 505.297177 | -0.393435841 | 0.26184977 | -1.5025251 | 0.13296156 | 0.31682277 |
| ALS2CR12 | 274.830425 | 0.45628721 | 0.40146015 | 1.13656912 | 0.25571846 | 0.47668432 |
| TRAK2 | 1112.46975 | -0.056692579 | 0.22428688 | -0.2527681 | 0.80044739 | 0.90655145 |
| STRADB | 612.249725 | -0.173376428 | 0.35891937 | -0.4830512 | 0.62905939 | 0.79927333 |
| C2CD6 | 100.158652 | 0.728766336 | 0.38944622 | 1.87128879 | 0.06130506 | 0.19062584 |
| TMEM237 | 150.33901 | 0.474676989 | 0.1850645 | 2.56492728 | 0.01031974 | 0.05676679 |
| ALS2 | 731.596882 | 0.287387969 | 0.15646439 | 1.83676283 | 0.06624492 | 0.20051079 |
| SUMO1 | 508.645097 | -0.059563968 | 0.1958292 | -0.3041628 | 0.76100383 | 0.88401192 |
| NOP58 | 753.285045 | 0.025704009 | 0.1636879 | 0.1570306 | 0.87522073 | 0.94384684 |
| BMPR2 | 2851.76435 | -0.662377347 | 0.17157144 | -3.8606503 | 0.00011309 | 0.00184413 |
| FAM117B | 1211.3143 | -0.38126504 | 0.32451435 | -1.1748788 | 0.2400432 | 0.45806569 |
| ICA1L | 432.529796 | -0.313868386 | 0.41771117 | -0.7514005 | 0.45241165 | 0.66816281 |
| WDR12 | 482.216354 | 0.492231893 | 0.235234 | 2.09252022 | 0.036392 | 0.1342368 |
| CARF | 776.320359 | 0.450838691 | 0.24902621 | 1.81040658 | 0.07023276 | 0.20866009 |
| NBEAL1 | 4969.50607 | 0.246725458 | 0.14781308 | 1.66917204 | 0.09508329 | 0.25403624 |
| CYP20A1 | 998.742271 | 0.197624715 | 0.21799592 | 0.90655237 | 0.36464356 | 0.59012212 |
| ABI2 | 1755.27937 | -0.004970295 | 0.14279869 | -0.0348063 | 0.97223419 | 0.98786473 |
| RAPH1 | 895.655897 | -1.131207067 | 0.2228379 | -5.0763675 | 3.85E-07 | 1.90E-05 |
| PARD3B | 7681.73734 | 0.507733424 | 0.15901208 | 3.19304944 | 0.00140779 | 0.01306244 |
| AC007679.1 | 81.3965173 | -0.672712355 | 0.63643892 | -1.0569944 | 0.29051416 | 0.51339098 |
| INO80D | 2260.38872 | 0.552899368 | 0.48938403 | 1.12978629 | 0.25856629 | 0.4795566 |
| NDUFS1 | 1477.85755 | 0.353834248 | 0.13642756 | 2.59356869 | 0.00949856 | 0.0533219 |
| EEF1B2 | 327.472101 | -0.743868087 | 0.3767645 | -1.9743582 | 0.04834103 | 0.16235357 |
| ZDBF2 | 330.53068 | -0.229804596 | 0.24837859 | -0.925219 | 0.354852 | 0.58046316 |
| MDH1B | 86.8558016 | 0.972090765 | 0.38127235 | 2.54959682 | 0.01078476 | 0.05852746 |
| FASTKD2 | 478.527268 | 0.467073183 | 0.20844873 | 2.24071012 | 0.02504486 | 0.10376961 |
| CPO | 152.152822 | 0.59714962 | 0.42659714 | 1.39979752 | 0.16157396 | 0.35921571 |
| KLF7 | 922.671581 | -1.090465499 | 0.45995033 | -2.3708332 | 0.01774804 | 0.08234417 |
| AC009226.1 | 71.047477 | -0.503166809 | 0.64748797 | -0.777106 | 0.43709622 | 0.65356621 |
| CREB1 | 1190.22325 | -0.093207714 | 0.13087317 | -0.7121988 | 0.47634168 | 0.68629987 |
| METTL21A | 304.680074 | 0.485188369 | 0.30373497 | 1.59740702 | 0.11017501 | 0.27970434 |
| CCNYL1 | 433.250353 | 0.044475421 | 0.40662737 | 0.10937636 | 0.91290399 | 0.96054229 |
| FZD5 | 76.0849191 | -1.566397625 | 0.36586828 | -4.2813157 | 1.86E-05 | 0.00044298 |
| PLEKHM3 | 831.604394 | 0.384772449 | 0.17294849 | 2.22478067 | 0.02609597 | 0.10661974 |
| IDH1 | 351.020327 | 0.820844557 | 0.33844268 | 2.42535769 | 0.01529331 | 0.07435014 |
| PIKFYVE | 1168.16883 | 0.014971412 | 0.22268384 | 0.06723169 | 0.94639726 | 0.97676523 |
| PTH2R | 12015.8884 | 1.21875589 | 0.18240319 | 6.68165881 | 2.36E-11 | 5.47E-09 |
| MAP2 | 510.736192 | -1.104082363 | 0.4288272 | -2.5746556 | 0.01003399 | 0.05557192 |
| RPE | 430.13783 | 0.226207707 | 0.2015662 | 1.12225017 | 0.26175609 | 0.48269387 |
| KANSL1L | 5280.92418 | 0.062802185 | 0.22303627 | 0.28157835 | 0.77826684 | 0.89400754 |
| AC007038.2 | 51.2589457 | 0.813942237 | 0.32812172 | 2.4806107 | 0.01311575 | 0.06674325 |
| AC007038.1 | 682.281589 | 0.289759221 | 0.24451123 | 1.18505484 | 0.23599576 | 0.45389346 |
| ACADL | 338.431315 | 0.100117744 | 0.41385923 | 0.24191255 | 0.80884792 | 0.91013478 |
| LANCL1-AS1 | 169.771449 | 0.914160683 | 0.51978166 | 1.75873978 | 0.07862172 | 0.22434959 |
| LANCL1 | 867.890405 | 0.47929685 | 0.15244349 | 3.14409522 | 0.00166601 | 0.01487892 |
| CPS1 | 120.982828 | 0.594504801 | 0.37103585 | 1.60228397 | 0.10909283 | 0.27762728 |
| ERBB4 | 70863.8385 | -0.080972233 | 0.46245066 | -0.1750938 | 0.86100595 | 0.93689987 |
| IKZF2 | 462.8565 | -0.334799279 | 0.21418475 | -1.5631332 | 0.11802128 | 0.29301961 |
| SPAG16 | 3913.94631 | 0.076513161 | 0.27549008 | 0.27773472 | 0.781216 | 0.89545532 |
| AC068051.1 | 535.394202 | -0.646764559 | 0.41897129 | -1.5436966 | 0.12266185 | 0.30015569 |
| VWC2L | 81.6526099 | 0.358675397 | 0.50220799 | 0.71419692 | 0.47510542 | 0.68526616 |
| BARD1 | 258.620994 | -0.257746691 | 0.18970307 | -1.3586849 | 0.17424647 | 0.37589771 |
| AC016708.1 | 270.833151 | -0.068772509 | 0.33683599 | -0.2041721 | 0.83821902 | 0.92451863 |
| ATIC | 253.839047 | -0.086889752 | 0.20588289 | -0.4220348 | 0.67299959 | 0.82864015 |
| FN1 | 94.1941693 | 0.466130016 | 0.29777644 | 1.56536902 | 0.11749641 | 0.29215978 |
| MREG | 231.47851 | -0.216755049 | 0.25795293 | -0.8402892 | 0.40074628 | 0.62224727 |
| PECR | 278.77441 | 0.294560756 | 0.33041353 | 0.89149122 | 0.3726657 | 0.59655016 |
| TMEM169 | 51.8436667 | -0.52464776 | 0.59178076 | -0.8865576 | 0.3753171 | 0.59870627 |
| XRCC5 | 2127.40514 | -0.306226399 | 0.18493291 | -1.6558784 | 0.09774645 | 0.25845947 |
| SMARCAL1 | 335.338955 | 0.017021772 | 0.274989 | 0.06189983 | 0.9506426 | 0.97872472 |
| RPL37A | 827.542498 | -0.656657924 | 0.32853264 | -1.9987601 | 0.04563432 | 0.15600761 |
| IGFBP2 | 287.262076 | -0.729505014 | 0.57276646 | -1.2736518 | 0.20278686 | 0.41232902 |
| IGFBP5 | 226.959411 | -0.522919103 | 0.43370492 | -1.2057025 | 0.22793223 | 0.44334513 |
| DIRC3 | 767.564739 | 0.802722763 | 0.58633757 | 1.36904543 | 0.17098507 | 0.37153181 |
| TNS1 | 2621.12334 | 0.22175689 | 0.39406442 | 0.56274273 | 0.57361008 | 0.761183 |
| ARPC2 | 913.579314 | -0.976033069 | 0.15500763 | -6.2966777 | 3.04E-10 | 4.41E-08 |
| AAMP | 226.803134 | -0.209411796 | 0.29205223 | -0.7170354 | 0.47335224 | 0.68400719 |
| PNKD | 531.697962 | -0.500868943 | 0.29213462 | -1.7145141 | 0.08643434 | 0.2381542 |
| TMBIM1 | 429.586356 | -1.866782637 | 0.20839253 | -8.9580111 | 3.31E-19 | 6.24E-16 |
| CATIP-AS2 | 70.9129144 | -0.601670499 | 0.65336216 | -0.9208836 | 0.3571112 | 0.58271536 |
| CATIP | 69.0006865 | 0.670074711 | 0.66845095 | 1.00242913 | 0.31613638 | 0.54049255 |
| CTDSP1 | 368.790045 | -0.528196865 | 0.16045384 | -3.2918929 | 0.00099516 | 0.00996589 |
| USP37 | 864.289068 | -0.29561579 | 0.26019743 | -1.1361211 | 0.25590589 | 0.47687514 |
| CNOT9 | 422.295899 | 0.081466671 | 0.20104771 | 0.40521063 | 0.68532269 | 0.83718001 |
| PLCD4 | 238.055964 | 0.294771814 | 0.84366694 | 0.34939358 | 0.72679385 | 0.86532561 |
| ZNF142 | 176.520305 | -0.191062357 | 0.22041686 | -0.8668228 | 0.3860391 | 0.60936789 |
| BCS1L | 80.0689388 | -0.083264296 | 0.28332597 | -0.2938816 | 0.76884836 | 0.88857795 |
| RNF25 | 88.1200594 | -0.280863919 | 0.26114787 | -1.0754976 | 0.28215199 | 0.50464019 |
| STK36 | 267.323077 | 0.600078265 | 0.21790479 | 2.75385532 | 0.00588978 | 0.03785796 |
| TTLL4 | 167.026809 | -0.351819487 | 0.24681935 | -1.4254129 | 0.15403787 | 0.3482859 |
| CYP27A1 | 169.818856 | -0.439041367 | 0.27905787 | -1.5732986 | 0.11564969 | 0.28926656 |
| CNPPD1 | 230.48432 | -0.317929942 | 0.21049631 | -1.5103825 | 0.13094586 | 0.31340167 |
| RETREG2 | 238.428334 | -0.649532186 | 0.2930354 | -2.2165656 | 0.02665279 | 0.10832514 |
| ZFAND2B | 147.626673 | -0.134381703 | 0.31097242 | -0.4321338 | 0.66564415 | 0.82357564 |
| ANKZF1 | 402.184134 | -0.05447432 | 0.34495852 | -0.1579156 | 0.87452333 | 0.94352461 |
| GLB1L | 49.4048784 | 0.026405228 | 0.37829271 | 0.06980105 | 0.94435201 | 0.9759376 |
| STK16 | 87.2047778 | -0.412923399 | 0.30348221 | -1.3606182 | 0.17363439 | 0.37537425 |
| TUBA4A | 215.83833 | -1.045982246 | 0.4059984 | -2.5763211 | 0.00998579 | 0.05532813 |
| DNAJB2 | 242.625749 | -0.475820027 | 0.28680385 | -1.6590434 | 0.09710706 | 0.25751598 |
| DNPEP | 180.58741 | -0.46766404 | 0.3022044 | -1.547509 | 0.12174055 | 0.29856496 |
| GMPPA | 104.868626 | 0.359378498 | 0.39050178 | 0.92029925 | 0.3574164 | 0.58281427 |
| CHPF | 107.956243 | -1.423237265 | 0.47972531 | -2.9667754 | 0.00300941 | 0.02320564 |
| OBSL1 | 688.688701 | 0.497708935 | 0.23028191 | 2.16130282 | 0.03067196 | 0.11963711 |
| INHA | 65.9871193 | 0.32544015 | 0.3976947 | 0.81831654 | 0.41317646 | 0.63269742 |
| STK11IP | 152.135387 | 0.312337511 | 0.31907347 | 0.978889 | 0.32763483 | 0.55292302 |
| SLC4A3 | 108.740778 | 0.659209707 | 0.51226262 | 1.2868589 | 0.19814347 | 0.40675481 |
| AC019211.1 | 63.7990333 | 3.917607583 | 0.71105347 | 5.50958227 | 3.60E-08 | 2.73E-06 |
| EPHA4 | 786.668082 | 1.612832666 | 0.51238899 | 3.14767239 | 0.00164576 | 0.0147482 |
| SGPP2 | 2824.45998 | -1.319430418 | 0.31990993 | -4.1243809 | 3.72E-05 | 0.00077858 |
| FARSB | 392.419751 | 0.567286807 | 0.27871303 | 2.03537955 | 0.0418127 | 0.14665859 |
| MOGAT1 | 66.5774613 | 0.898616253 | 0.57297617 | 1.56833094 | 0.11680392 | 0.2910513 |

| | | | | | | |
|---|---|---|---|---|---|---|
| ACSL3 | 1499.29374 | 0.60915754 | 0.27836649 | 2.1883293 | 0.02864562 | 0.11381696 |
| AP1S3 | 1863.97231 | -0.656366698 | 0.43280972 | -1.5165249 | 0.12938669 | 0.31096593 |
| WDFY1 | 802.454347 | -0.204188831 | 0.21848986 | -0.934546 | 0.3500223 | 0.57608659 |
| MRPL44 | 111.172765 | 0.088917188 | 0.21232571 | 0.41877731 | 0.67537889 | 0.82979069 |
| SERPINE2 | 191.892828 | -3.022579516 | 0.83368432 | -3.6255684 | 0.00028833 | 0.00391822 |
| CUL3 | 2379.79198 | -0.037137802 | 0.15949066 | -0.2328525 | 0.81587593 | 0.91427971 |
| DOCK10 | 80.5747278 | -0.785425744 | 0.53905178 | -1.4570506 | 0.1451024 | 0.33496288 |
| AC062015.1 | 49.5534593 | 2.466894072 | 0.74229302 | 3.32334266 | 0.00088946 | 0.00911468 |
| IRS1 | 1285.04518 | 0.313737425 | 0.3677768 | 0.85306474 | 0.3936234 | 0.61588683 |
| AC010735.2 | 38.7732375 | 0.572003799 | 0.31092349 | 1.8396931 | 0.06581331 | 0.19998343 |
| RHBDD1 | 1387.84266 | 0.378512155 | 0.20584113 | 1.83885579 | 0.0659364 | 0.20017327 |
| COL4A4 | 4236.71464 | 0.111607166 | 0.26703967 | 0.41794227 | 0.67598932 | 0.82991711 |
| COL4A3 | 7150.33117 | 0.447891541 | 0.34964533 | 1.2809882 | 0.20019781 | 0.40903525 |
| AC097662.1 | 1038.69225 | 0.132617395 | 0.54664345 | 0.2426031 | 0.80831288 | 0.91010272 |
| MFF | 1269.67558 | 0.682178334 | 0.19196311 | 3.55369501 | 0.00037986 | 0.00479232 |
| AGFG1 | 2199.80587 | -0.3351952 | 0.18449502 | -1.8168251 | 0.06924389 | 0.20674561 |
| SLC19A3 | 335.073878 | -0.017343356 | 0.45400916 | -0.0382005 | 0.96952786 | 0.98632883 |
| DNER | 644.230676 | -1.247266773 | 0.39933364 | -3.1233701 | 0.00178793 | 0.0156503 |
| TRIP12 | 3038.70504 | -0.050717006 | 0.13412744 | -0.3781255 | 0.70533736 | 0.85100486 |
| FBXO36 | 382.828114 | 0.38612309 | 0.18736832 | 2.06077041 | 0.03932495 | 0.14126492 |
| AC009950.1 | 75.0343491 | -0.10248308 | 0.59605022 | -0.171937 | 0.86348707 | 0.93813133 |
| SP110 | 544.639604 | -0.246557928 | 0.49588625 | -0.4972066 | 0.61904335 | 0.79126608 |
| SP140L | 271.613204 | -1.285841806 | 0.40922415 | -3.1421454 | 0.00167715 | 0.01493898 |
| SP100 | 1543.56487 | 0.160011454 | 0.35040506 | 0.45664709 | 0.64792472 | 0.81173905 |
| AC010149.1 | 54.8639922 | 0.146075943 | 0.34202741 | 0.42708841 | 0.66931493 | 0.82610549 |
| CAB39 | 1613.52629 | -0.022509684 | 0.16817994 | -0.1338429 | 0.89352683 | 0.95146032 |
| ITM2C | 955.822516 | -2.201711769 | 0.69551784 | -3.1655719 | 0.00154778 | 0.01406099 |
| PSMD1 | 1082.83162 | -0.123063571 | 0.22493758 | -0.547101 | 0.58430935 | 0.76896903 |
| ARMC9 | 1358.04692 | -0.488087624 | 0.17250546 | -2.829404 | 0.00466348 | 0.03175016 |
| NCL | 600.916786 | -0.914132747 | 0.31409647 | -2.9103566 | 0.00361017 | 0.02657544 |
| PTMA | 2384.64246 | -1.492046834 | 0.32384674 | -4.6072622 | 4.08E-06 | 0.00012651 |
| PDE6D | 459.231805 | 0.21517311 | 0.30471229 | 0.70615172 | 0.48009379 | 0.68929988 |
| COPS7B | 453.239765 | 1.006339641 | 0.29240921 | 3.44154568 | 0.0005784 | 0.00659197 |
| DIS3L2 | 1954.54253 | 0.072208687 | 0.16585582 | 0.43537024 | 0.66329371 | 0.82258481 |
| AC019130.1 | 142.190012 | 0.068064371 | 0.49280979 | 0.13811489 | 0.89014962 | 0.9503707 |
| TIGD1 | 76.4348797 | 0.552845069 | 0.22023554 | 2.51024459 | 0.01206476 | 0.06316874 |
| EIF4E2 | 505.795267 | -0.570132113 | 0.20152669 | -2.8290651 | 0.00466842 | 0.03175016 |
| EFHD1 | 3424.41128 | 0.750506697 | 0.35825011 | 2.09492385 | 0.03617776 | 0.13384823 |
| GIGYF2 | 3230.70026 | 0.401956155 | 0.13577973 | 2.96035466 | 0.00307285 | 0.02355733 |
| KCNJ13 | 42.0571908 | 1.11847025 | 0.62501931 | 1.7894971 | 0.07353479 | 0.21517969 |
| SNORC | 72.0995684 | 0.451579103 | 0.42315617 | 1.0671689 | 0.28589557 | 0.50833149 |
| ATG16L1 | 213.02532 | -0.080059989 | 0.27538144 | -0.290724 | 0.77126243 | 0.8896518 |
| DGKD | 298.465043 | -0.141635602 | 0.34569758 | -0.4097096 | 0.68201902 | 0.8342244 |
| USP40 | 1439.58357 | 0.440198621 | 0.26989363 | 1.63100783 | 0.10288867 | 0.26771179 |
| ARL4C | 151.35314 | -3.583515447 | 0.77649303 | -4.6150002 | 3.93E-06 | 0.00012275 |
| SH3BP4 | 1282.94819 | -0.217213436 | 0.30950471 | -0.7018098 | 0.4827978 | 0.69127369 |
| AGAP1 | 7685.63712 | -0.018736026 | 0.32950174 | -0.0568617 | 0.95465537 | 0.98007893 |
| IQCA1 | 288.841946 | 0.354193242 | 0.50938509 | 0.69533492 | 0.48684544 | 0.69412904 |
| AC105760.2 | 144.98203 | 0.620305968 | 0.3477931 | 1.78354879 | 0.07449701 | 0.21663404 |
| COPS8 | 223.229381 | 0.103407524 | 0.22236576 | 0.46503349 | 0.64190748 | 0.80790432 |
| MLPH | 95.6201178 | 0.439791687 | 0.56153417 | 0.78319667 | 0.43351163 | 0.65072685 |
| RAB17 | 851.771581 | 0.520692122 | 0.41227512 | 1.26297245 | 0.20659908 | 0.41739901 |
| LRRFIP1 | 4314.63659 | -0.948891075 | 0.17171406 | -5.5259954 | 3.28E-08 | 2.53E-06 |
| RBM44 | 52.4872257 | -0.663732355 | 0.75897465 | -0.8745119 | 0.38183955 | 0.60465286 |
| UBE2F-SCLY | 61.9361084 | -0.557910147 | 0.40796113 | -1.3675571 | 0.17145075 | 0.3722987 |
| ERFE | 74.6336826 | -4.318496439 | 0.65796883 | -6.5633754 | 5.26E-11 | 1.04E-08 |
| ILKAP | 773.210934 | -0.164845928 | 0.19793459 | -0.8328303 | 0.40494042 | 0.62552427 |
| PER2 | 312.678466 | -0.604524264 | 0.55218967 | -1.0947765 | 0.27361457 | 0.49529599 |
| TRAF3IP1 | 554.997478 | -0.107001814 | 0.1805889 | -0.592516 | 0.55350511 | 0.74630555 |
| ASB1 | 123.432583 | -0.027149497 | 0.24274651 | -0.111843 | 0.9109479 | 0.95976856 |
| HDAC4 | 1648.30917 | -0.854668897 | 0.25760695 | -3.3177246 | 0.00090754 | 0.00926405 |
| NDUFA10 | 1143.05207 | 0.092515461 | 0.24782645 | 0.37330746 | 0.70891962 | 0.85384511 |
| COPS9 | 250.856224 | -1.025596079 | 0.4906435 | -2.0903081 | 0.03659013 | 0.13464146 |
| AC124861.1 | 134.255902 | 0.617608732 | 0.60915348 | 1.01388033 | 0.31063986 | 0.53450314 |
| GPC1 | 253.142136 | -2.232268123 | 0.44630526 | -5.0016621 | 5.68E-07 | 2.61E-05 |
| ANKMY1 | 536.661745 | -0.14037729 | 0.33336146 | -0.4210963 | 0.67368473 | 0.82886565 |
| RNPEPL1 | 193.714819 | -1.347063533 | 0.32805212 | -4.1062486 | 4.02E-05 | 0.00082997 |
| CAPN10-DT | 48.498142 | -0.066766183 | 0.40423507 | -0.1651667 | 0.86881275 | 0.94020545 |
| CAPN10 | 177.350251 | -0.175976946 | 0.21939309 | -0.802108 | 0.42249051 | 0.64123603 |
| GPR35 | 38.9755581 | 0.665776806 | 0.41305315 | 1.61184294 | 0.10699612 | 0.27469617 |
| SNED1 | 1971.13462 | -0.873076829 | 0.4895342 | -1.7834848 | 0.07450741 | 0.21663404 |
| AC005237.1 | 362.710924 | -0.96265754 | 0.4884303 | -1.970921 | 0.04873291 | 0.1632722 |
| MTERF4 | 603.077189 | 0.156640174 | 0.21909643 | 0.71493715 | 0.47464788 | 0.68512991 |
| PASK | 115.373351 | 0.268642053 | 0.36235101 | 0.74138624 | 0.45845929 | 0.67266311 |
| PPP1R7 | 309.541339 | -0.452443124 | 0.20845584 | -2.1704507 | 0.02997272 | 0.11751548 |
| HDLBP | 1624.12465 | -0.194772088 | 0.13696998 | -1.4220057 | 0.15502459 | 0.34955955 |
| 2-Sep | 2112.5711 | -0.096599751 | 0.15004225 | -0.643817 | 0.51969412 | 0.7198387 |
| FARP2 | 1102.84685 | 0.266763198 | 0.14791137 | 1.80353406 | 0.07130438 | 0.21075399 |
| STK25 | 308.293852 | -0.138090816 | 0.17789206 | -0.7762618 | 0.43759441 | 0.65416304 |
| BOK | 95.9968059 | -0.960655132 | 0.31346753 | -3.0646081 | 0.00217955 | 0.01832573 |
| THAP4 | 541.74802 | 0.207730144 | 0.18173042 | 1.14306752 | 0.25301056 | 0.47317237 |
| ATG4B | 940.489204 | 0.019189537 | 0.23301015 | 0.08235494 | 0.93436447 | 0.97176637 |
| DTYMK | 85.8332277 | -0.174867014 | 0.29637646 | -0.5900165 | 0.55517956 | 0.74754753 |
| ING5 | 480.118914 | 0.249909502 | 0.21291542 | 1.17375013 | 0.24049513 | 0.45857079 |
| D2HGDH | 592.031399 | 0.483146215 | 0.31047804 | 1.55613654 | 0.11967563 | 0.29536727 |
| CHL1 | 1505.47555 | 0.345449015 | 0.43950427 | 0.78599694 | 0.43186928 | 0.64957432 |
| LINC01266 | 1502.99165 | 0.078172838 | 0.23550765 | 0.33193333 | 0.7399396 | 0.87265237 |
| CNTN6 | 106.563361 | 0.736141716 | 0.34912736 | 2.10851912 | 0.03498611 | 0.13084552 |
| CNTN4 | 2013.52582 | -0.699807853 | 0.62738152 | -1.1154423 | 0.26466095 | 0.48574496 |
| TRNT1 | 603.49844 | 0.279334871 | 0.21095492 | 1.32414486 | 0.18545494 | 0.39112862 |
| CRBN | 647.643181 | 0.233707536 | 0.1672587 | 1.3972818 | 0.16232885 | 0.36024895 |
| AC024060.1 | 68.6227281 | 0.313730187 | 0.26645905 | 1.17740487 | 0.23903395 | 0.45716719 |
| AC034195.1 | 1069.09432 | 0.713561391 | 0.39588765 | 1.80243406 | 0.07147714 | 0.21093422 |
| SUMF1 | 4434.82446 | 0.159811947 | 0.32068527 | 0.49834514 | 0.6182408 | 0.79053759 |
| LRRN1 | 104.583855 | 0.10424561 | 0.3459315 | 0.30134755 | 0.76314948 | 0.88525876 |
| SETMAR | 176.087828 | 0.860744375 | 0.27989847 | 3.07520218 | 0.0021036 | 0.01785761 |
| ITPR1 | 4777.55664 | 0.963102211 | 0.28516288 | 3.37737577 | 0.00073181 | 0.00791038 |
| EGOT | 194.99765 | -0.957600823 | 0.60400365 | -1.5854223 | 0.11287043 | 0.28452395 |
| AC018816.1 | 230.142419 | 0.108474146 | 0.20090916 | 0.53991637 | 0.58925471 | 0.77174702 |
| BHLHE40-AS | 126.589413 | -0.479867502 | 0.48301494 | -0.9934838 | 0.32047426 | 0.54486506 |
| BHLHE40 | 1530.55252 | -1.913923076 | 0.6102909 | -3.1360833 | 0.00171221 | 0.01515651 |
| ARL8B | 1168.07586 | -0.193253964 | 0.1929786 | -1.0014269 | 0.31662045 | 0.54097006 |
| AC026202.2 | 287.266777 | 0.027858045 | 0.37041704 | 0.07520725 | 0.94004982 | 0.97411372 |
| EDEM1 | 274.548002 | -0.247495387 | 0.25709193 | -0.9626727 | 0.3357118 | 0.56131863 |
| AC087857.1 | 109.825898 | 0.816701612 | 0.78398325 | 1.0417335 | 0.29753525 | 0.52089373 |
| AC026167.1 | 168.44185 | 1.52712514 | 0.78559223 | 1.9439158 | 0.05190561 | 0.17046771 |
| AC069277.1 | 279.634703 | 0.480805959 | 1.34367845 | 0.35782814 | 0.72047193 | 0.86069582 |
| GRM7 | 216.009943 | 1.050842096 | 0.86647582 | 1.21277717 | 0.22521503 | 0.43967858 |
| LMCD1-AS1 | 1114.82175 | 0.789935198 | 0.28954834 | 2.72816347 | 0.0063688 | 0.03996462 |
| LMCD1 | 110.394492 | 0.960191679 | 0.74316447 | 1.2920312 | 0.19634634 | 0.40441896 |
| RAD18 | 251.599599 | -0.248047894 | 0.32051189 | -0.7739117 | 0.43898302 | 0.65485132 |
| SRGAP3 | 1846.31703 | -0.191251447 | 0.4468958 | -0.4279553 | 0.66868363 | 0.82563489 |
| THUMPD3-A: | 1087.24096 | 0.154784336 | 0.20882399 | 0.74121914 | 0.45856058 | 0.67271509 |

| | | | | | | |
|---|---|---|---|---|---|---|
| THUMPD3 | 250.836004 | 0.043935737 | 0.22622299 | 0.19421429 | 0.84600809 | 0.9287667 |
| SETD5 | 3039.23771 | -0.015989975 | 0.10713936 | -0.1492446 | 0.8813606 | 0.94645136 |
| LHFPL4 | 46.7683147 | -0.277896832 | 0.78670884 | -0.3532397 | 0.72390871 | 0.86292845 |
| MTMR14 | 420.389758 | -0.181945833 | 0.2011596 | -0.904485 | 0.3657383 | 0.59121933 |
| BRPF1 | 61.5960754 | -0.094695492 | 0.31153425 | -0.303965 | 0.76115459 | 0.8841093 |
| OGG1 | 222.894712 | -0.144944848 | 0.21908927 | -0.6615789 | 0.50824111 | 0.71121086 |
| CAMK1 | 116.32076 | -0.297154563 | 0.32675789 | -0.9094029 | 0.3631375 | 0.5885693 |
| TADA3 | 233.350911 | -0.421210778 | 0.17236258 | -2.4437484 | 0.01453556 | 0.07166861 |
| TTLL3 | 84.9729226 | -0.14770335 | 0.29879249 | -0.4943342 | 0.62107017 | 0.79308932 |
| RPUSD3 | 127.766492 | 0.034163894 | 0.26856153 | 0.12721067 | 0.89877366 | 0.9532601 |
| JAGN1 | 100.040337 | -0.761209302 | 0.44116537 | -1.7254512 | 0.08444619 | 0.23419059 |
| IL17RE | 164.660496 | 0.953964485 | 0.44541838 | 2.14172682 | 0.03221547 | 0.12393503 |
| IL17RC | 146.115072 | 0.422505885 | 0.29497491 | 1.43234514 | 0.15204507 | 0.34519823 |
| CRELD1 | 260.154178 | 0.233223899 | 0.27128744 | 0.85969294 | 0.38995833 | 0.61299055 |
| EMC3 | 487.86416 | -0.139930506 | 0.16695288 | -0.8381437 | 0.40194999 | 0.62338352 |
| FANCD2 | 75.2501999 | -0.49873871 | 0.356472 | -1.3990964 | 0.16178406 | 0.35922099 |
| BRK1 | 305.076072 | -0.722851923 | 0.23808344 | -3.0361286 | 0.00239637 | 0.0196241 |
| VHL | 308.413469 | -0.256262723 | 0.22287736 | -1.1497925 | 0.25022934 | 0.47016775 |
| IRAK2 | 241.663338 | -3.448286738 | 0.52587459 | -6.5572416 | 5.48E-11 | 1.05E-08 |
| GHRL | 45.4248998 | -0.051690823 | 0.36214133 | -0.1427366 | 0.88649821 | 0.94892665 |
| SEC13 | 407.642999 | -0.104507674 | 0.22608266 | -0.4622543 | 0.64389899 | 0.80933122 |
| HRH1 | 229.578237 | -3.804244853 | 0.71590843 | -5.3138708 | 1.07E-07 | 6.72E-06 |
| ATG7 | 2689.29042 | -0.516223883 | 0.25472304 | -2.0266086 | 0.04270246 | 0.14879075 |
| VGLL4 | 2384.43209 | -0.15208581 | 0.16788457 | -0.9058951 | 0.36499138 | 0.59048436 |
| TAMM41 | 707.116835 | 0.194205459 | 0.31569718 | 0.61516374 | 0.53844659 | 0.73492731 |
| PPARG | 307.90322 | -0.921412816 | 0.47153594 | -1.954067 | 0.05069329 | 0.16769539 |
| TSEN2 | 679.18596 | 0.253029358 | 0.30254497 | 0.83633637 | 0.4029657 | 0.62429908 |
| MKRN2OS | 116.290261 | 0.150837374 | 0.40221774 | 0.37501422 | 0.70764989 | 0.85293799 |
| MKRN2 | 188.861992 | 0.262922891 | 0.19056249 | 1.3797201 | 0.16767284 | 0.36753909 |
| RAF1 | 1484.23826 | 0.229221301 | 0.14102365 | 1.62541036 | 0.10407515 | 0.27003116 |
| CAND2 | 54.7070695 | 0.139921749 | 0.4282198 | 0.32675217 | 0.74385534 | 0.87470713 |
| RPL32 | 1066.19788 | -1.052839454 | 0.41298792 | -2.5493227 | 0.01079324 | 0.05854944 |
| IQSEC1 | 282.657867 | 0.312634835 | 0.25962654 | 1.20417133 | 0.22852336 | 0.4440498 |
| NUP210 | 517.467026 | 0.37004687 | 0.21009002 | 1.76137287 | 0.07817531 | 0.22341361 |
| HDAC11 | 181.085804 | -0.100050928 | 0.3041789 | -0.3289213 | 0.74221515 | 0.87402695 |
| CHCHD4 | 46.1410444 | 0.353585057 | 0.35044209 | 1.00896858 | 0.3129897 | 0.53684988 |
| TMEM43 | 128.730259 | -0.765166982 | 0.32082379 | -2.385007 | 0.0170788 | 0.07996945 |
| XPC | 544.509025 | -0.356219825 | 0.15777436 | -2.2577801 | 0.02395937 | 0.10072544 |
| LSM3 | 222.729881 | -0.416713504 | 0.30677576 | -1.3583652 | 0.17434784 | 0.3759322 |
| SLC6A6 | 320.383351 | -3.013768437 | 0.70960255 | -4.2471218 | 2.17E-05 | 0.00050356 |
| CCDC174 | 364.898083 | 0.048636928 | 0.17556205 | 0.27703554 | 0.78175282 | 0.89582484 |
| C3orf20 | 66.7670914 | 0.444000424 | 0.94105137 | 0.47181316 | 0.63706016 | 0.80480926 |
| FGD5-AS1 | 505.821279 | -0.07190979 | 0.28742567 | -0.2501857 | 0.80244375 | 0.90725666 |
| NR2C2 | 1100.76047 | -0.077348987 | 0.2254319 | -0.3431146 | 0.73151221 | 0.86826096 |
| MRPS25 | 843.819052 | 0.210935112 | 0.20107613 | 1.04903108 | 0.29416382 | 0.51725371 |
| RBSN | 343.16306 | 0.152149415 | 0.18958425 | 0.80254247 | 0.42223923 | 0.64114933 |
| CAPN7 | 1378.40458 | 0.083239327 | 0.17783591 | 0.46806815 | 0.63973585 | 0.80624663 |
| SH3BP5-AS1 | 93.7563685 | 0.346138896 | 0.27463181 | 1.26037435 | 0.20753435 | 0.41837037 |
| SH3BP5 | 91.2110502 | -1.155134037 | 0.39275612 | -2.9410975 | 0.00327052 | 0.02466685 |
| METTL6 | 349.281655 | 0.054353123 | 0.25402124 | 0.21397078 | 0.83056984 | 0.92100378 |
| EAF1 | 132.467326 | 0.067302319 | 0.20131168 | 0.334319 | 0.73813885 | 0.87200855 |
| COLQ | 219.529337 | -0.418464413 | 0.41953058 | -0.9974587 | 0.31854193 | 0.54312899 |
| HACL1 | 511.099883 | -0.109917771 | 0.15229065 | -0.7217631 | 0.47044014 | 0.68204755 |
| BTD | 151.172738 | 0.414270104 | 0.23314663 | 1.77686507 | 0.07559044 | 0.21881967 |
| ANKRD28 | 3553.76879 | -0.297455208 | 0.20046944 | -1.4837933 | 0.13786378 | 0.32397129 |
| GALNT15 | 47.1002864 | -1.483709102 | 0.5175941 | -2.8665495 | 0.00414973 | 0.02928255 |
| DPH3 | 80.8624994 | -0.621523955 | 0.46878189 | -1.3258276 | 0.18489682 | 0.39045597 |
| OXNAD1 | 1172.28835 | 0.411235084 | 0.21493642 | 1.91328711 | 0.05571131 | 0.17878782 |
| RFTN1 | 111.010641 | -0.282618363 | 0.45990759 | -0.6145112 | 0.53887758 | 0.73506352 |
| PLCL2 | 704.061267 | 0.863329181 | 0.29848183 | 2.89240109 | 0.0038231 | 0.02773161 |
| TBC1D5 | 7098.9762 | 0.319358256 | 0.177209 | 1.80215593 | 0.07152088 | 0.21101614 |
| KCNH8 | 105.510087 | -2.66493598 | 0.55176426 | -4.8298452 | 1.37E-06 | 5.28E-05 |
| EFHB | 42.9782734 | 0.668308058 | 0.66909949 | 0.99881716 | 0.31788327 | 0.54242606 |
| RAB5A | 1109.9975 | -0.223012608 | 0.12745365 | -1.7497546 | 0.08016067 | 0.22663577 |
| KAT2B | 1017.1274 | 1.095567126 | 0.24692142 | 4.43690605 | 9.13E-06 | 0.00024753 |
| ZNF385D | 2049.72553 | 1.360949634 | 0.68404176 | 1.98957096 | 0.04663822 | 0.15828474 |
| UBE2E2 | 3258.33315 | -0.426683678 | 0.3395802 | -1.2565034 | 0.20893351 | 0.42044445 |
| UBE2E1 | 2391.1681 | -0.168297814 | 0.16714892 | -1.0068735 | 0.31399557 | 0.53787673 |
| NKIRAS1 | 185.146426 | 0.156528924 | 0.23468942 | 0.66696198 | 0.50479642 | 0.70859746 |
| RPL15 | 1708.79708 | -1.765539812 | 0.48653071 | -3.6288353 | 0.0002847 | 0.00388496 |
| NR1D2 | 481.299774 | -0.21264594 | 0.42546936 | -0.4997914 | 0.61722194 | 0.7902755 |
| THRB | 5496.76239 | 1.279038517 | 0.22755735 | 5.62073034 | 1.90E-08 | 1.59E-06 |
| AC092422.1 | 69.2928822 | -1.109485425 | 0.64204575 | -1.7280473 | 0.08397975 | 0.23347383 |
| RARB | 778.47997 | 0.314027929 | 0.28013718 | 1.12097912 | 0.26229675 | 0.48321866 |
| TOP2B | 1640.30707 | 0.324323342 | 0.19826601 | 1.63579898 | 0.10188166 | 0.26601204 |
| NGLY1 | 1171.52654 | 0.164526337 | 0.26200129 | 0.62796004 | 0.53003012 | 0.72815058 |
| OXSM | 135.104741 | 0.143427658 | 0.32124115 | 0.44647972 | 0.65525076 | 0.81660603 |
| NEK10 | 2862.13875 | -0.416857238 | 1.3668421 | -0.3049783 | 0.76038265 | 0.88360116 |
| SLC4A7 | 7179.52871 | -0.828723321 | 0.80233478 | -1.0328897 | 0.30165552 | 0.52496248 |
| CMC1 | 1757.05001 | 0.647280045 | 0.18960614 | 3.41381366 | 0.0006406 | 0.00717096 |
| AZI2 | 610.935704 | 0.275025143 | 0.25028281 | 1.09885749 | 0.27183023 | 0.49335058 |
| AC098650.1 | 63.2053576 | 1.08617938 | 0.32226568 | 3.37044698 | 0.00075046 | 0.00807237 |
| RBMS3 | 71.3414783 | -0.5802704 | 0.5525407 | -1.0501858 | 0.29363269 | 0.51666368 |
| TGFBR2 | 1728.7962 | 0.397621495 | 0.29857464 | 1.33173232 | 0.18294816 | 0.38776674 |
| GADL1 | 127.966945 | 7.919764129 | 0.81379424 | 9.73189998 | 2.20E-22 | 9.71E-19 |
| STT3B | 1363.29734 | 0.375208096 | 0.22911137 | 1.63766688 | 0.1014912 | 0.26525471 |
| OSBPL10 | 3172.15694 | -0.189604375 | 0.33367545 | -0.5682299 | 0.56987886 | 0.75923656 |
| OSBPL10-AS | 66.3445226 | -1.037776123 | 0.32234975 | -3.2194103 | 0.00128455 | 0.01218059 |
| ZNF860 | 54.3146341 | -0.83593683 | 0.47462341 | -1.7612634 | 0.07819384 | 0.22341821 |
| GPD1L | 307.141049 | 0.973243854 | 0.29797279 | 3.26621716 | 0.00108995 | 0.01068034 |
| CMTM8 | 361.631905 | -1.141326899 | 0.39854632 | -2.8637246 | 0.00418692 | 0.02943321 |
| CMTM7 | 341.930332 | -3.811698682 | 0.47786742 | -7.9764774 | 1.51E-15 | 1.17E-12 |
| CMTM6 | 439.307655 | -1.402301124 | 0.28302785 | -4.95464 | 7.25E-07 | 3.20E-05 |
| DYNC1LI1 | 461.863239 | -0.088129856 | 0.17153811 | -0.5137625 | 0.60741802 | 0.78330417 |
| CNOT10 | 866.92194 | -0.162338899 | 0.14976288 | -1.0839729 | 0.27837684 | 0.50055858 |
| GLB1 | 771.485191 | 0.110751581 | 0.12393448 | 0.89363008 | 0.37151985 | 0.59563048 |
| CRTAP | 273.631964 | -0.354021912 | 0.26232073 | -1.3495766 | 0.17715185 | 0.37959992 |
| FBXL2 | 1271.22834 | -0.47517256 | 0.23355996 | -2.0344778 | 0.04190344 | 0.14677872 |
| UBP1 | 615.706071 | 0.135522813 | 0.22342742 | 0.60656304 | 0.54414094 | 0.73944091 |
| CLASP2 | 2895.99225 | 0.537183099 | 0.22582085 | 2.37880203 | 0.017369 | 0.08104102 |
| PDCD6IP | 1634.15325 | -0.165448319 | 0.11575998 | -1.4292359 | 0.15293643 | 0.34644783 |
| LINC01811 | 1031.6298 | -3.411860707 | 1.26183521 | -2.7038877 | 0.00685334 | 0.04218352 |
| STAC | 314.882496 | 0.30483331 | 0.59711495 | 0.51051026 | 0.60969403 | 0.7853192 |
| TRANK1 | 193.006867 | -1.56113781 | 0.80018338 | -1.950975 | 0.05106001 | 0.16848657 |
| EPM2AIP1 | 485.828377 | 0.42866162 | 0.25862179 | 1.65748458 | 0.09742154 | 0.257884 |
| MLH1 | 328.165049 | 0.374343976 | 0.21147749 | 1.77013631 | 0.07670444 | 0.22093085 |
| LRRFIP2 | 2524.73524 | -0.305341151 | 0.25099545 | -1.2165207 | 0.22378665 | 0.4383227 |
| GOLGA4 | 4444.56376 | 0.255954516 | 0.17517796 | 1.46111138 | 0.14398488 | 0.33360748 |
| ITGA9 | 179.862873 | 0.294779547 | 0.51746853 | 0.56965695 | 0.56891039 | 0.75867193 |
| CTDSPL | 4533.05479 | 0.223448787 | 0.20919634 | 1.06812954 | 0.28546208 | 0.50786322 |
| PLCD1 | 165.974866 | -0.087227745 | 0.41873001 | -0.208315 | 0.83498302 | 0.92318496 |
| ACAA1 | 393.580145 | -0.011791594 | 0.35547558 | -0.0331713 | 0.97353797 | 0.98804941 |
| MYD88 | 50.6347442 | -1.474755557 | 0.38624353 | -3.8182013 | 0.00013443 | 0.00212209 |

| | | | | | | |
|---|---|---|---|---|---|---|
| OXSR1 | 1367.73238 | 0.06058233 | 0.18139294 | 0.33398394 | 0.73839168 | 0.87200855 |
| XYLB | 148.198546 | 1.155690096 | 0.46496627 | 2.48553537 | 0.01293567 | 0.06630474 |
| ACVR2B | 319.784505 | -0.215255282 | 0.24713788 | -0.8709927 | 0.38375816 | 0.60643561 |
| EXOG | 342.078593 | -0.010367851 | 0.2248868 | -0.0461025 | 0.96322853 | 0.98370849 |
| SCN11A | 91.162745 | 0.747096787 | 0.32082476 | 2.32867561 | 0.01987626 | 0.08881782 |
| WDR48 | 718.524477 | 0.212803783 | 0.1682542 | 1.26477547 | 0.20595183 | 0.41660301 |
| GORASP1 | 204.175516 | 0.042953085 | 0.17716276 | 0.24244986 | 0.8084316 | 0.91010272 |
| TTC21A | 189.201776 | -0.054222431 | 0.32542057 | -0.1666226 | 0.86766699 | 0.93988956 |
| CSRNP1 | 114.9114 | -1.237494938 | 0.49007277 | -2.5251249 | 0.01156572 | 0.06140933 |
| SLC25A38 | 144.026508 | 0.077510517 | 0.26703298 | 0.29026571 | 0.77161297 | 0.88991843 |
| RPSA | 836.109144 | -0.98592147 | 0.3600121 | -2.7385787 | 0.00617054 | 0.03903576 |
| MYRIP | 324.625608 | 0.664113405 | 0.43324933 | 1.53286655 | 0.12530874 | 0.30392546 |
| EIF1B | 288.196585 | -0.286030141 | 0.4373372 | -0.6540266 | 0.5130947 | 0.71499819 |
| ENTPD3-AS1 | 411.019334 | 0.51017436 | 0.42569946 | 1.19843787 | 0.2307466 | 0.44643252 |
| RPL14 | 1368.26679 | -1.192735658 | 0.41948496 | -2.8433335 | 0.00446443 | 0.0308747 |
| ZNF619 | 42.4119288 | 0.586684167 | 0.35169285 | 1.668172 | 0.09528159 | 0.25429643 |
| ZNF620 | 81.937948 | 0.264071566 | 0.28478222 | 0.92727547 | 0.35378353 | 0.57958057 |
| ZNF621 | 531.232772 | -0.06397034 | 0.24973275 | -0.2561552 | 0.79783101 | 0.90506918 |
| CTNNB1 | 1397.23693 | 0.09483266 | 0.13432971 | 0.70596935 | 0.4802072 | 0.68931285 |
| ULK4 | 1621.92943 | -0.450553567 | 0.28401817 | -1.5863548 | 0.11265886 | 0.28420757 |
| TRAK1 | 1360.28527 | 0.320286665 | 0.19909898 | 1.60868057 | 0.10768621 | 0.27583544 |
| SEC22C | 252.584004 | -0.013973949 | 0.23660261 | -0.0590608 | 0.95290365 | 0.97936812 |
| SS18L2 | 137.953341 | -0.185544569 | 0.32182002 | -0.5765476 | 0.56424509 | 0.75551073 |
| NKTR | 2901.90552 | 0.159548213 | 0.12390731 | 1.2876416 | 0.19787075 | 0.40634007 |
| AC006059.1 | 409.589983 | -0.308978537 | 0.44979961 | -0.6869249 | 0.49213008 | 0.69803196 |
| CCDC13 | 97.9916945 | -0.36662044 | 0.46673632 | -0.785498 | 0.43216165 | 0.64979206 |
| AC099329.3 | 131.536791 | -0.696142325 | 0.38916512 | -1.7888097 | 0.07364548 | 0.21536044 |
| AC092042.3 | 72.9901724 | 0.321990092 | 0.45383475 | 0.70948752 | 0.47802199 | 0.68767071 |
| POMGNT2 | 79.2803815 | 0.121867684 | 0.22618612 | 0.53879382 | 0.59002913 | 0.7721373 |
| SNRK | 1525.57703 | 0.657733208 | 0.36573932 | 1.79836614 | 0.072119 | 0.21225934 |
| AC104184.1 | 35.1684728 | 1.765326997 | 0.50808875 | 3.47444617 | 0.00051191 | 0.00603194 |
| ANO10 | 1963.90399 | 0.330928789 | 0.24857725 | 1.33129156 | 0.1830931 | 0.3878872 |
| ABHD5 | 362.80456 | -0.241220829 | 0.36573301 | -0.6595544 | 0.5095398 | 0.71186277 |
| TCAIM | 2910.52816 | 0.503671103 | 0.24294694 | 2.07317326 | 0.03815616 | 0.13861515 |
| ZNF445 | 422.953137 | 0.441491032 | 0.19090428 | 2.31263036 | 0.02074297 | 0.09136176 |
| ZNF852 | 113.686014 | 0.412357445 | 0.23142786 | 1.78179694 | 0.07478235 | 0.21724216 |
| ZKSCAN7 | 114.857615 | 0.431639176 | 0.2163169 | 1.99540196 | 0.04599906 | 0.15675994 |
| AC099669.1 | 769.088941 | 0.582707993 | 0.25745049 | 2.26337886 | 0.02361234 | 0.09980655 |
| ZNF197 | 409.880274 | 0.052571712 | 0.23972701 | 0.21929824 | 0.82641773 | 0.91938879 |
| ZNF35 | 91.7757603 | 0.144420614 | 0.26358866 | 0.54790146 | 0.58375957 | 0.76885817 |
| ZNF502 | 70.0237116 | 0.166366645 | 0.36834953 | 0.45165429 | 0.65151805 | 0.81384903 |
| KIAA1143 | 149.722501 | -0.446964664 | 0.26734725 | -1.6718506 | 0.09455379 | 0.25303098 |
| ZDHHC3 | 483.045575 | -0.174825148 | 0.15124062 | -1.1559404 | 0.24770554 | 0.46703404 |
| EXOSC7 | 296.02437 | 0.033933758 | 0.24371886 | 0.13923321 | 0.88926587 | 0.95012225 |
| CDCP1 | 429.511771 | -3.47636723 | 0.51172381 | -6.7934443 | 1.09E-11 | 2.73E-09 |
| LARS2 | 753.29581 | 0.183166701 | 0.19654405 | 0.93193715 | 0.35136901 | 0.57733962 |
| LIMD1 | 549.00398 | -0.022437033 | 0.22357092 | -0.1003576 | 0.92006046 | 0.96453005 |
| SACM1L | 947.055111 | 0.441663873 | 0.1680743 | 2.62778944 | 0.00859417 | 0.04970244 |
| SLC6A20 | 224.957137 | -5.210833664 | 0.5557652 | -9.3759626 | 6.85E-21 | 1.81E-17 |
| LZTFL1 | 271.343363 | 0.241006563 | 0.18152966 | 1.32764286 | 0.18429612 | 0.38958162 |
| FYCO1 | 370.651104 | -0.084823835 | 0.1585622 | -0.5349562 | 0.59268013 | 0.77381752 |
| LTF | 1199.36616 | -5.295139762 | 0.81395649 | -6.5054334 | 7.75E-11 | 1.40E-08 |
| LRRC2 | 689.689149 | 0.716428152 | 0.44427173 | 1.61258999 | 0.10683361 | 0.27440072 |
| TDGF1 | 80.5088823 | 1.167111621 | 0.73136751 | 1.59579365 | 0.11053487 | 0.28040237 |
| AC104304.1 | 95.4937774 | 0.550683258 | 0.5212512 | 1.05646425 | 0.29075617 | 0.51351761 |
| ALS2CL | 398.355755 | -1.574505628 | 0.24765379 | -6.3576885 | 2.05E-10 | 3.11E-08 |
| MYL3 | 86.4547131 | 0.197743624 | 0.80622368 | 0.24527142 | 0.80624628 | 0.90938008 |
| PTH1R | 2119.66941 | 0.381500518 | 0.85342913 | 0.44702074 | 0.65486009 | 0.81619616 |
| CCDC12 | 791.509075 | 0.234834239 | 0.24329711 | 0.96521589 | 0.33443669 | 0.55954075 |
| NBEAL2 | 274.338103 | 0.359953881 | 0.34943035 | 1.03011625 | 0.30295544 | 0.5262179 |
| SETD2 | 3389.04519 | 0.171591573 | 0.12323036 | 1.39244565 | 0.16378749 | 0.36245082 |
| KIF9-AS1 | 1190.87388 | 0.174549611 | 0.26248404 | 0.66499133 | 0.50605604 | 0.70953129 |
| KIF9 | 163.36553 | -0.200090881 | 0.2287623 | -0.8746672 | 0.38175501 | 0.60462853 |
| KLHL18 | 413.946021 | 0.107267076 | 0.17300711 | 0.62001541 | 0.53524764 | 0.73219616 |
| AC099778.1 | 38.057616 | 0.356545877 | 0.41907779 | 0.85078685 | 0.39488777 | 0.61706761 |
| PTPN23 | 223.062416 | -0.377554942 | 0.22832205 | -1.653607 | 0.09820739 | 0.25939241 |
| SCAP | 646.095684 | 0.249548783 | 0.1711659 | 1.45793519 | 0.14485841 | 0.33475075 |
| ELP6 | 270.922122 | 0.134459263 | 0.30054994 | 0.44737744 | 0.65460256 | 0.81602918 |
| SMARCC1 | 2291.03778 | -0.446373034 | 0.21846722 | -2.0432037 | 0.04103228 | 0.1448411 |
| DHX30 | 513.177822 | 0.452339732 | 0.14357418 | 3.15056472 | 0.00162955 | 0.01465265 |
| MAP4 | 3099.81676 | -0.371106084 | 0.20765432 | -1.7871339 | 0.07391585 | 0.21565688 |
| ZNF589 | 184.467856 | 0.579877042 | 0.2407238 | 2.40888958 | 0.01600114 | 0.07653081 |
| NME6 | 144.776569 | -0.173314076 | 0.23902655 | -0.725083 | 0.46840114 | 0.68050051 |
| PLXNB1 | 1085.05553 | 0.310055413 | 0.22833741 | 1.3578827 | 0.17450091 | 0.37613944 |
| TMA7 | 433.014015 | -0.570480125 | 0.38260635 | -1.4910367 | 0.13595185 | 0.321247 |
| ATRIP | 52.4467692 | 0.348942239 | 0.32132275 | 1.08595559 | 0.27749864 | 0.49972281 |
| SHISA5 | 508.330888 | -1.915091359 | 0.4460542 | -4.2934051 | 1.76E-05 | 0.00042645 |
| PFKFB4 | 349.04908 | -1.97232725 | 0.7576816 | -2.6031083 | 0.00923828 | 0.05228295 |
| COL7A1 | 393.202615 | -3.707594871 | 1.16111795 | -3.1931251 | 0.00140742 | 0.01306244 |
| UQCRC1 | 426.909073 | 0.271508058 | 0.25292195 | 1.07348557 | 0.28305331 | 0.50545507 |
| SLC26A6 | 56.6420225 | 0.052290577 | 0.32711969 | 0.15985151 | 0.87299804 | 0.94280363 |
| NCKIPSD | 113.155884 | -0.984771094 | 0.33739638 | -2.9187364 | 0.00351453 | 0.02606596 |
| IP6K2 | 809.600403 | 0.331305376 | 0.13977436 | 2.37028726 | 0.01777427 | 0.08243691 |
| PRKAR2A | 980.317068 | -0.149359234 | 0.18452605 | -0.8094208 | 0.41827312 | 0.63753323 |
| SLC25A20 | 181.813471 | 1.774051003 | 0.57731917 | 3.072912 | 0.00211981 | 0.01794908 |
| ARIH2 | 1194.58634 | 0.123581713 | 0.16022437 | 0.77130408 | 0.44052671 | 0.65623647 |
| P4HTM | 309.030825 | -0.055619805 | 0.28046481 | -0.198313 | 0.84280022 | 0.92739954 |
| WDR6 | 566.717396 | 0.338523755 | 0.31988173 | 1.05827788 | 0.28992878 | 0.51274189 |
| DALRD3 | 155.655154 | 0.247752619 | 0.27272374 | 0.90843805 | 0.36364683 | 0.58901423 |
| NDUFAF3 | 138.705218 | -1.455620174 | 0.49309556 | -2.9520042 | 0.00315719 | 0.02401208 |
| IMPDH2 | 94.0689173 | -0.378755172 | 0.37693872 | -1.004819 | 0.31498403 | 0.53915046 |
| QRICH1 | 795.03484 | 0.08875443 | 0.16800695 | 0.52827833 | 0.59730617 | 0.77659469 |
| QARS | 213.395635 | 0.483017994 | 0.25908864 | 1.8642963 | 0.06228011 | 0.1928215 |
| USP19 | 131.603709 | 0.190215663 | 0.18772051 | 1.01329186 | 0.31092077 | 0.53451627 |
| LAMB2 | 323.621352 | 0.370251938 | 0.3050646 | 1.21368374 | 0.22486852 | 0.43942281 |
| KLHDC8B | 59.9747357 | 0.264441494 | 0.39249953 | 0.67373709 | 0.50047848 | 0.70482771 |
| C3orf62 | 95.8584593 | 0.167319887 | 0.2421728 | 0.69091115 | 0.48962138 | 0.69631877 |
| USP4 | 744.230469 | -0.28499975 | 0.17300795 | -1.6473217 | 0.09949194 | 0.26173852 |
| RHOA | 1171.78441 | -0.916093325 | 0.19654082 | -4.6610842 | 3.15E-06 | 0.00010267 |
| TCTA | 160.74896 | -0.45734477 | 0.33704851 | -1.3569108 | 0.17480955 | 0.37654099 |
| DAG1 | 610.639618 | 0.449483501 | 0.24706991 | 1.81925634 | 0.06887233 | 0.20591549 |
| BSN | 40.7949794 | 0.341402797 | 0.47014318 | 0.72616771 | 0.46773596 | 0.68013258 |
| APEH | 171.182568 | -0.249947806 | 0.43233222 | -0.5781383 | 0.56317077 | 0.75468425 |
| MST1 | 249.052582 | 0.48678042 | 0.3890432 | 1.25122459 | 0.21085255 | 0.42321096 |
| RNF123 | 181.581761 | 0.093868446 | 0.1841395 | 0.50976811 | 0.61021393 | 0.78568241 |
| GMPPB | 50.6268659 | -0.580255927 | 0.37187187 | -1.5603652 | 0.1186736 | 0.29393579 |
| IP6K1 | 1323.76812 | 0.06647819 | 0.22313725 | 0.29792512 | 0.76576032 | 0.88657446 |
| MON1A | 68.1256057 | 0.408474272 | 0.30161611 | 1.35428532 | 0.17564537 | 0.37786639 |
| RBM6 | 4210.07524 | 0.224228962 | 0.16655467 | 1.34627847 | 0.17821273 | 0.3814636 |
| RBM5 | 1385.48209 | 0.464650628 | 0.12075088 | 3.84801041 | 0.00011908 | 0.001921 |
| SEMA3F | 87.5045522 | -0.22072838 | 0.55814182 | -0.3954701 | 0.69249601 | 0.84221582 |
| GNAI2 | 334.497415 | -1.292059703 | 0.26206114 | -4.9303751 | 8.21E-07 | 3.57E-05 |
| SEMA3B | 510.556502 | -1.586937839 | 0.39506486 | -4.0169045 | 5.90E-05 | 0.00111431 |

| | | | | | | |
|---|---|---|---|---|---|---|
| IFRD2 | 109.221437 | -0.787593151 | 0.2825286 | -2.7876582 | 0.00530905 | 0.03501112 |
| HYAL1 | 147.206547 | 0.48028055 | 0.49726583 | 0.96584265 | 0.33412292 | 0.5593519 |
| HYAL2 | 74.2312808 | -0.771637584 | 0.28645151 | -2.6937808 | 0.00706466 | 0.04308269 |
| TUSC2 | 111.888533 | -0.744260121 | 0.3673002 | -2.0262992 | 0.04273413 | 0.14882023 |
| RASSF1 | 112.545547 | -0.806859461 | 0.31810825 | -2.5364305 | 0.0111989 | 0.06010818 |
| NPRL2 | 59.7317876 | 0.630577599 | 0.31654199 | 1.99208202 | 0.04636206 | 0.1575499 |
| TMEM115 | 75.5671185 | -0.708825863 | 0.37686526 | -1.8808469 | 0.05999274 | 0.18787201 |
| C3orf18 | 45.2864203 | 0.438725327 | 0.45646144 | 0.96114434 | 0.3364796 | 0.56189115 |
| HEMK1 | 165.905152 | 0.515087582 | 0.25538146 | 2.01693414 | 0.04370239 | 0.15147332 |
| CISH | 216.108995 | -0.694772104 | 0.45238723 | -1.5357907 | 0.12458972 | 0.30285345 |
| MAPKAPK3 | 293.574899 | -0.573952606 | 0.3306137 | -1.7360218 | 0.08255997 | 0.23089873 |
| DOCK3 | 1030.37159 | -0.627831339 | 0.47988453 | -1.3082967 | 0.1907727 | 0.39735942 |
| MANF | 223.342701 | -0.000182221 | 0.45220711 | -0.000403 | 0.99967848 | 0.99975417 |
| RBM15B | 75.2236207 | -0.506753369 | 0.26026473 | -1.9470689 | 0.05152647 | 0.1694925 |
| DCAF1 | 620.679608 | -0.154450203 | 0.23948226 | -0.6449338 | 0.51897009 | 0.71946641 |
| RAD54L2 | 713.348904 | 0.154512769 | 0.19931011 | 0.77523798 | 0.43819905 | 0.65440037 |
| TEX264 | 295.129496 | -0.349599847 | 0.28717412 | -1.2173794 | 0.22345991 | 0.43819507 |
| RRP9 | 39.1564446 | 0.079753581 | 0.51148515 | 0.15592551 | 0.87609174 | 0.94421473 |
| PARP3 | 58.8001493 | 0.122709759 | 0.40303869 | 0.30446149 | 0.76077633 | 0.88388009 |
| PCBP4 | 56.4920281 | 0.132712966 | 0.40472417 | 0.32790966 | 0.74297997 | 0.87445523 |
| ABHD14A | 42.4433359 | -0.21197681 | 0.4052144 | -0.5231226 | 0.60088894 | 0.7789212 |
| RPL29 | 1411.7906 | -1.721303195 | 0.462063 | -3.7252565 | 0.00019512 | 0.00287324 |
| POC1A | 45.7254135 | -0.476538007 | 0.48575699 | -0.9810214 | 0.32658219 | 0.55192222 |
| ALAS1 | 108.840292 | 0.260184533 | 0.24136994 | 1.07794921 | 0.28105642 | 0.50360245 |
| WDR82 | 471.010035 | 0.020690577 | 0.1551451 | 0.13336275 | 0.89390651 | 0.95146032 |
| GLYCTK | 47.1347532 | -0.208785321 | 0.33843918 | -0.6169065 | 0.53729642 | 0.73401988 |
| DNAH1 | 117.251553 | -0.034227849 | 0.33117221 | -0.1033536 | 0.91768232 | 0.96310708 |
| BAP1 | 150.78645 | -0.469900205 | 0.30279765 | -1.5518621 | 0.12069521 | 0.29693855 |
| PHF7 | 152.956721 | 0.66424304 | 0.37316947 | 1.78000372 | 0.07507535 | 0.21771028 |
| NISCH | 628.780895 | 0.194860139 | 0.20552147 | 0.94812546 | 0.34306559 | 0.56819785 |
| NT5DC2 | 61.7047093 | -0.461255801 | 0.49116597 | -0.9391037 | 0.34767748 | 0.57319918 |
| SMIM4 | 402.875097 | 0.556755786 | 0.27679809 | 2.01141486 | 0.04428166 | 0.15279947 |
| PBRM1 | 1711.99504 | 0.084499028 | 0.21948373 | 0.38498994 | 0.70024487 | 0.84787759 |
| GNL3 | 254.029486 | -0.150406915 | 0.26157785 | -0.5749987 | 0.5652922 | 0.75611701 |
| GLT8D1 | 180.063332 | 0.282662809 | 0.19027327 | 1.48556235 | 0.13739492 | 0.32338729 |
| SPCS1 | 464.574888 | -0.994277597 | 0.42232521 | -2.3542937 | 0.01855794 | 0.08485161 |
| NEK4 | 445.790474 | 0.286889324 | 0.13404008 | 2.14032497 | 0.03232851 | 0.12413585 |
| SFMBT1 | 375.432253 | 0.044427766 | 0.15113329 | 0.29396413 | 0.76878531 | 0.88857795 |
| AC096887.1 | 76.2422774 | 0.526820779 | 0.44931579 | 1.17249557 | 0.24099815 | 0.4592908 |
| RFT1 | 112.624406 | 0.01534482 | 0.25708732 | 0.05968719 | 0.95240477 | 0.97933189 |
| PRKCD | 129.675669 | -0.256405835 | 0.26985681 | -0.9501551 | 0.34203343 | 0.56736401 |
| TKT | 629.192145 | -1.664685343 | 0.56912387 | -2.9249965 | 0.0034446 | 0.02569155 |
| DCP1A | 1065.16234 | -0.026346691 | 0.1484364 | -0.1774948 | 0.85911975 | 0.93618423 |
| CACNA1D | 2577.2151 | -0.032975608 | 0.39363746 | -0.0837715 | 0.93323809 | 0.97148254 |
| CHDH | 269.503945 | -0.316959172 | 0.57747699 | -0.5488689 | 0.58309541 | 0.76851998 |
| IL17RB | 102.76796 | -0.279025247 | 0.63998302 | -0.4359885 | 0.66284506 | 0.8221918 |
| ACTR8 | 128.344241 | 0.270744578 | 0.23647754 | 1.14490609 | 0.25224806 | 0.47234826 |
| SELENOK | 210.481939 | -0.989717241 | 0.43142784 | -2.2940505 | 0.0217876 | 0.09452959 |
| CACNA2D3 | 17909.8562 | 0.351289378 | 0.30135397 | 1.16570349 | 0.24373435 | 0.46243709 |
| CACNA2D3-A | 128.221047 | 0.450275303 | 0.45326106 | 0.99341273 | 0.32050886 | 0.54486506 |
| LRTM1 | 293.133274 | -0.141072737 | 0.46246234 | -0.305047 | 0.76033038 | 0.88360116 |
| ERC2 | 1308.68353 | 0.213908323 | 0.34164153 | 0.62611921 | 0.53123675 | 0.72863563 |
| CCDC66 | 1417.85679 | 0.421131963 | 0.20367203 | 2.06769662 | 0.03866856 | 0.13955547 |
| FAM208A | 1452.40932 | 0.283479796 | 0.20622398 | 1.37462092 | 0.169249 | 0.36964452 |
| ARHGEF3 | 5010.76077 | -1.210151904 | 0.33043905 | -3.6622545 | 0.00025001 | 0.00350937 |
| ARHGEF3-AS | 52.6634199 | -1.638483793 | 0.71630912 | -2.2873976 | 0.02217262 | 0.09555569 |
| SPATA12 | 132.327211 | -1.757883861 | 0.54573307 | -3.2211423 | 0.00127681 | 0.01212462 |
| IL17RD | 253.810984 | -0.250468304 | 0.42862627 | -0.5843513 | 0.55898401 | 0.75089633 |
| HESX1 | 70.2356529 | 0.792593747 | 0.4370202 | 1.81363182 | 0.06973443 | 0.20769381 |
| APPL1 | 806.621492 | 0.075231041 | 0.11824039 | 0.63625499 | 0.52461024 | 0.72386677 |
| ASB14 | 134.013579 | 0.212110652 | 0.43333271 | 0.48948682 | 0.62449708 | 0.79577291 |
| DNAH12 | 79.1845606 | 0.520850837 | 0.38016528 | 1.37006421 | 0.17066686 | 0.37096241 |
| PDE12 | 135.987357 | 0.050925999 | 0.22730458 | 0.22404299 | 0.82272385 | 0.91813562 |
| ARF4 | 660.951304 | -0.270820498 | 0.27746678 | -0.9760466 | 0.32904138 | 0.55451743 |
| DENND6A | 955.352556 | -0.207186058 | 0.14239404 | -1.4550192 | 0.14566395 | 0.33584931 |
| DENND6A-AS | 217.186159 | -0.282623127 | 0.37453216 | -0.7546031 | 0.45048716 | 0.66649697 |
| SLMAP | 3213.83866 | -0.345544871 | 0.16988614 | -2.0339791 | 0.0419537 | 0.14683946 |
| FLNB | 2938.43033 | 0.265719823 | 0.33039076 | 0.80425925 | 0.42124727 | 0.64039547 |
| ABHD6 | 196.259233 | 0.5877629 | 0.44990792 | 1.3064071 | 0.19141415 | 0.39809311 |
| RPP14 | 107.358836 | 0.482848218 | 0.30807216 | 1.56732179 | 0.11703949 | 0.29152832 |
| PXK | 908.800548 | 0.638624981 | 0.28231041 | 2.26213757 | 0.02368891 | 0.10003413 |
| PDHB | 250.407664 | -0.022315377 | 0.28155059 | -0.0792589 | 0.93682673 | 0.97284815 |
| KCTD6 | 211.466021 | -0.318728578 | 0.26251927 | -1.214115 | 0.22470382 | 0.43932988 |
| ACOX2 | 57.4314044 | 0.51523143 | 0.40839039 | 1.26161498 | 0.20708736 | 0.41787611 |
| FAM107A | 164.864523 | -1.598219076 | 0.88984453 | -1.7960655 | 0.0724841 | 0.21290693 |
| C3orf67 | 200.383076 | -0.530385294 | 0.46602453 | -1.138106 | 0.25507624 | 0.47589013 |
| C3orf67-AS1 | 52.4018098 | -0.716468689 | 0.53329657 | -1.3434714 | 0.17911938 | 0.3825983 |
| FHIT | 17971.034 | 0.53191626 | 0.17020276 | 3.12519171 | 0.00177689 | 0.01559533 |
| PTPRG | 20125.5208 | -0.574357982 | 0.31699915 | -1.8118597 | 0.07000788 | 0.20825264 |
| C3orf14 | 43.2677855 | 0.15630745 | 0.38544998 | 0.40551942 | 0.68509575 | 0.83713504 |
| THOC7 | 339.388571 | -0.152703188 | 0.17872746 | -0.8543913 | 0.39288822 | 0.61518024 |
| ATXN7 | 288.61505 | 0.169530222 | 0.16459388 | 1.02999101 | 0.30301423 | 0.5262179 |
| PSMD6-AS2 | 134.709936 | 0.020772308 | 0.24828582 | 0.08366289 | 0.93332447 | 0.97149589 |
| PSMD6 | 197.627551 | 0.08441322 | 0.26253785 | 0.32152781 | 0.74781044 | 0.8770936 |
| PSMD6-AS1 | 49.1943944 | -0.099887037 | 0.37633793 | -0.2654185 | 0.79068711 | 0.90121546 |
| PRICKLE2 | 3223.32092 | 0.277251604 | 0.39358318 | 0.7044295 | 0.48116534 | 0.68948936 |
| ADAMTS9 | 3723.90908 | -1.682684511 | 0.53717676 | -3.1324596 | 0.00173348 | 0.01526505 |
| ADAMTS9-AS | 2222.14455 | 1.590920554 | 0.46444958 | 3.42538916 | 0.00061392 | 0.00694286 |
| ADAMTS9-AS | 13061.9409 | -0.032356788 | 0.47556007 | -0.0680393 | 0.94575433 | 0.9765845 |
| MAGI1 | 20414.0492 | -0.087194426 | 0.2559106 | -0.3407222 | 0.73331271 | 0.86911141 |
| MAGI1-AS1 | 385.04892 | -0.698867668 | 0.67693834 | -1.0323949 | 0.30188718 | 0.52503327 |
| SLC25A26 | 1041.67046 | 0.302297578 | 0.21702239 | 1.39293265 | 0.16364016 | 0.36224616 |
| LRIG1 | 256.284031 | 0.010024616 | 0.30114488 | 0.03328835 | 0.97344464 | 0.98804941 |
| SUCLG2 | 2867.05222 | 0.300074435 | 0.17611682 | 1.70383748 | 0.08841145 | 0.24166735 |
| SUCLG2-AS1 | 1899.29113 | 0.901179696 | 0.27845591 | 3.23634615 | 0.0012107 | 0.01160537 |
| FAM19A1 | 754.84331 | 0.667236996 | 0.51869149 | 1.28638507 | 0.19830871 | 0.40681157 |
| EOGT | 135.720687 | -0.752876003 | 0.30573297 | -2.462528 | 0.01379614 | 0.06905388 |
| AC109587.1 | 130.960696 | -0.274324925 | 0.67612668 | -0.4057301 | 0.68494095 | 0.83702331 |
| TMF1 | 973.059337 | -0.270834063 | 0.11761609 | -2.3026957 | 0.02129597 | 0.09326873 |
| UBA3 | 690.117772 | 0.065479308 | 0.14214344 | 0.46065657 | 0.64504502 | 0.81007404 |
| ARL6IP5 | 397.889035 | -0.916913528 | 0.22894626 | -4.0049291 | 6.20E-05 | 0.00115576 |
| FRMD4B | 1513.71724 | 0.541240979 | 0.23825444 | 2.27169312 | 0.02310505 | 0.09830568 |
| MITF | 4097.23022 | 0.935138444 | 0.1887447 | 4.95451489 | 7.25E-07 | 3.20E-05 |
| FOXP1 | 8997.18307 | -0.385572158 | 0.2782286 | -1.3858106 | 0.16580473 | 0.36523495 |
| AC097634.3 | 48.7435631 | -0.926207602 | 0.5284341 | -1.75274 | 0.07964664 | 0.22581079 |
| EIF4E3 | 340.82623 | -0.878949718 | 0.34696697 | -2.5332374 | 0.01130144 | 0.060462 |
| RYBP | 1205.94785 | -0.151346989 | 0.25985291 | -0.5824333 | 0.56027485 | 0.75183103 |
| SHQ1 | 472.582841 | -0.016370177 | 0.26712869 | -0.061282 | 0.95113464 | 0.97892609 |
| GXYLT2 | 153.788252 | -1.557266514 | 0.36776642 | -4.2343901 | 2.29E-05 | 0.00053015 |
| PPP4R2 | 1078.01442 | 0.014874332 | 0.18642903 | 0.07978549 | 0.93640786 | 0.97280247 |
| PDZRN3 | 1383.59338 | 0.706629917 | 0.6045308 | 1.16888986 | 0.24244801 | 0.4609234 |
| CNTN3 | 505.704276 | 0.850171559 | 0.29631726 | 2.869126 | 0.00411608 | 0.02922329 |
| LINC02018 | 69.9704538 | 0.30882846 | 0.54491567 | 0.56674542 | 0.57088716 | 0.75965257 |

| | | | | | | |
|---|---|---|---|---|---|---|
| ZNF717 | 669.465843 | 0.401713647 | 0.25640437 | 1.56671918 | 0.11718035 | 0.29176912 |
| ROBO2 | 492.077228 | 1.700740228 | 0.6900298 | 2.46473446 | 0.01371148 | 0.06872334 |
| ROBO1 | 185.005291 | -0.795996558 | 0.42579846 | -1.8694209 | 0.06156427 | 0.19132874 |
| LINC02027 | 128.115046 | -0.298557179 | 0.83701485 | -0.3566928 | 0.72132179 | 0.86135924 |
| GBE1 | 4875.501 | 0.442057988 | 0.38582488 | 1.14574775 | 0.25189954 | 0.47203022 |
| CADM2 | 111.648809 | -0.561697975 | 0.63706878 | -0.8816913 | 0.37794379 | 0.60077175 |
| CHMP2B | 410.768897 | -0.005612955 | 0.17047173 | -0.032926 | 0.97373358 | 0.98817201 |
| AC117513.1 | 72.5209484 | 2.981195309 | 1.23946608 | 2.4052254 | 0.01616249 | 0.07713473 |
| CGGBP1 | 992.836697 | -0.010161686 | 0.2581523 | -0.0393631 | 0.96860087 | 0.98582596 |
| ZNF654 | 1320.84957 | -0.168966627 | 0.18470616 | -0.9147861 | 0.36030393 | 0.58548052 |
| C3orf38 | 241.352195 | 0.250606809 | 0.18536897 | 1.35193507 | 0.17639609 | 0.37867358 |
| PROS1 | 212.679813 | -3.200634921 | 0.32343636 | -9.8957178 | 4.34E-23 | 2.87E-19 |
| ARL13B | 573.796486 | -0.185898248 | 0.21235125 | -0.8754281 | 0.38134103 | 0.60411773 |
| STX19 | 47.4373782 | 0.053436103 | 0.56710777 | 0.09422566 | 0.9249299 | 0.9667507 |
| DHFR2 | 91.4749955 | 0.590480192 | 0.2371476 | 2.48992694 | 0.01277694 | 0.06569503 |
| NSUN3 | 607.004602 | 0.333589975 | 0.22927325 | 1.45498868 | 0.1456724 | 0.33584931 |
| MTRNR2L12 | 888.673909 | -1.112246895 | 0.52952576 | -2.1004585 | 0.03568852 | 0.13247052 |
| EPHA6 | 263.52881 | 0.611676148 | 0.66654459 | 0.91768226 | 0.35878524 | 0.5841433 |
| ARL6 | 381.709276 | 0.37434561 | 0.20248704 | 1.84873859 | 0.06449557 | 0.1971129 |
| CRYBG3 | 1602.25414 | -0.649579731 | 0.28337098 | -2.2923298 | 0.02188662 | 0.09467668 |
| AC110491.2 | 49.9527044 | -0.56733693 | 0.74009338 | -0.7665748 | 0.44333437 | 0.65901459 |
| RIOX2 | 520.841738 | -0.275679451 | 0.26987335 | -1.0215142 | 0.3070109 | 0.53066042 |
| CLDND1 | 280.945502 | 0.031737136 | 0.25356958 | 0.12516145 | 0.90039574 | 0.95395051 |
| CPOX | 106.845461 | 0.319777665 | 0.31559634 | 1.01324898 | 0.31094125 | 0.53451627 |
| ST3GAL6-AS | 92.7408013 | 0.709187879 | 0.31908843 | 2.22254341 | 0.0262466 | 0.10713578 |
| ST3GAL6 | 1588.7113 | 0.566645533 | 0.26595616 | 2.13059749 | 0.03312232 | 0.12633902 |
| DCBLD2 | 846.14391 | -0.047841507 | 0.2148473 | -0.2226768 | 0.82378707 | 0.91846107 |
| AC091212.1 | 45.9612224 | -0.012587208 | 0.56084095 | -0.0224435 | 0.98209422 | 0.99261478 |
| COL8A1 | 749.443338 | -0.729740633 | 0.53262117 | -1.3700932 | 0.17065782 | 0.37096241 |
| CMSS1 | 1465.8166 | -0.096194226 | 0.18950744 | -0.5076013 | 0.61173298 | 0.78684784 |
| FILIP1L | 457.173438 | -0.155550827 | 0.43426132 | -0.3581964 | 0.72019637 | 0.86060013 |
| TBC1D23 | 1003.83326 | -0.108622427 | 0.14775875 | -0.7351336 | 0.46225811 | 0.67537466 |
| NIT2 | 196.098407 | 0.183590019 | 0.21008091 | 0.87390148 | 0.38217191 | 0.60492616 |
| TOMM70 | 463.219057 | 0.508427103 | 0.16427202 | 3.09503157 | 0.00196792 | 0.01693436 |
| LNP1 | 69.8505722 | 1.366189421 | 0.39537607 | 3.45541757 | 0.00054944 | 0.00635349 |
| TFG | 1190.01932 | 0.034716397 | 0.18598953 | 0.1866578 | 0.85192894 | 0.93193618 |
| ABI3BP | 314.385677 | -1.138588982 | 0.46814488 | -2.4321295 | 0.01501034 | 0.07335241 |
| IMPG2 | 65.5307891 | 0.251357171 | 0.45361106 | 0.55412487 | 0.57949339 | 0.76560594 |
| SENP7 | 1619.0823 | 0.319816291 | 0.17493828 | 1.82816646 | 0.06752457 | 0.20312733 |
| TRMT10C | 171.726429 | -0.678161667 | 0.35897274 | -1.8891732 | 0.05886863 | 0.18541688 |
| PCNP | 930.116715 | -0.156978454 | 0.12495635 | -1.2562664 | 0.20901941 | 0.42044445 |
| ZBTB11 | 398.975165 | 0.219235628 | 0.20419102 | 1.07367909 | 0.28296653 | 0.50543677 |
| RPL24 | 618.257488 | -0.639815691 | 0.33054714 | -1.9356262 | 0.0529135 | 0.17266168 |
| CEP97 | 265.016119 | 0.570421179 | 0.16700103 | 3.41567461 | 0.00063624 | 0.0071434 |
| NXPE3 | 99.7642338 | -0.097720172 | 0.24905329 | -0.3923665 | 0.69478742 | 0.84398078 |
| NFKBIZ | 1219.43317 | -1.96620153 | 0.40796203 | -4.8195699 | 1.44E-06 | 5.47E-05 |
| ZPLD1 | 256.31923 | -3.279409113 | 0.83749171 | -3.9157512 | 9.01E-05 | 0.0015541 |
| ALCAM | 1377.08878 | -1.566812144 | 0.42358046 | -3.6989717 | 0.00021647 | 0.00312846 |
| CBLB | 2452.82777 | 0.242372633 | 0.11430571 | 2.12038954 | 0.03397321 | 0.12854543 |
| LINC00882 | 452.512685 | 1.164720163 | 0.36618253 | 3.18070923 | 0.00146915 | 0.01349513 |
| DUBR | 98.6995175 | -0.426885091 | 0.33960816 | -1.256993 | 0.20875615 | 0.42021641 |
| AC063944.1 | 105.874813 | 0.002953781 | 0.43440466 | 0.00679961 | 0.99457474 | 0.99786448 |
| BBX | 4284.96289 | -0.632576064 | 0.21953725 | -2.8814065 | 0.00395905 | 0.02846763 |
| CD47 | 1091.39873 | -0.920427276 | 0.29058495 | -3.1674981 | 0.00153757 | 0.01399423 |
| IFT57 | 526.573974 | -0.204311373 | 0.21292894 | -0.9595284 | 0.3372926 | 0.56232461 |
| CIP2A | 67.529113 | 0.142500473 | 0.45746824 | 0.31149807 | 0.75542201 | 0.88163716 |
| DZIP3 | 730.081699 | 0.837105908 | 0.42474748 | 1.97083195 | 0.0487431 | 0.1632722 |
| AC078980.1 | 2068.1277 | 4.073025753 | 1.37754384 | 2.95673041 | 0.0031092 | 0.02375327 |
| NECTIN3-AS: | 91.5551479 | 0.681074497 | 0.58985272 | 1.15465179 | 0.24823305 | 0.46734754 |
| NECTIN3 | 484.887399 | 0.47423073 | 0.25550324 | 1.8560654 | 0.06344424 | 0.19518591 |
| PHLDB2 | 4217.09368 | 0.710815874 | 0.14266155 | 4.98253301 | 6.28E-07 | 2.84E-05 |
| ABHD10 | 186.579507 | 0.620434481 | 0.26791499 | 2.31578859 | 0.02056981 | 0.09092934 |
| C3orf52 | 166.149866 | -1.422003429 | 0.52966116 | -2.6847418 | 0.00725859 | 0.04390049 |
| SLC9C1 | 114.717251 | -1.477102742 | 0.46790154 | -3.1568666 | 0.00159474 | 0.01440832 |
| ATG3 | 316.892915 | 0.249495128 | 0.192135 | 1.29854078 | 0.19410157 | 0.40163448 |
| SLC35A5 | 195.481507 | -0.099366422 | 0.21400609 | -0.4643159 | 0.64242147 | 0.80832017 |
| GTPBP8 | 200.656148 | 0.211741341 | 0.28851972 | 0.73388861 | 0.46301663 | 0.67594901 |
| NEPRO | 535.985355 | 0.202417157 | 0.17446554 | 1.16021284 | 0.24596216 | 0.46524008 |
| AC078785.1 | 141.519497 | 0.416204174 | 0.23205128 | 1.79358708 | 0.0728791 | 0.21378193 |
| CFAP44 | 794.41609 | 0.225809335 | 0.20487146 | 1.10220008 | 0.2703747 | 0.49185779 |
| CFAP44-AS1 | 51.3749774 | -0.402043635 | 0.5395372 | -0.7451639 | 0.45617266 | 0.67128123 |
| SPICE1 | 586.903433 | 0.078097232 | 0.22044944 | 0.35426369 | 0.72314128 | 0.86248065 |
| SIDT1 | 187.46228 | -0.024375752 | 0.98979463 | -0.0246271 | 0.98035242 | 0.99141592 |
| USF3 | 559.685156 | 0.021874772 | 0.15663538 | 0.13965409 | 0.8889333 | 0.94999353 |
| NAA50 | 1346.03666 | -0.164028232 | 0.22936052 | -0.7151546 | 0.47451351 | 0.6850551 |
| ATP6V1A | 2658.40095 | -0.175250466 | 0.34733614 | -0.5045558 | 0.61387085 | 0.78755051 |
| GRAMD1C | 2781.60777 | -0.084530036 | 0.386749 | -0.2185656 | 0.82698844 | 0.91955386 |
| ZDHHC23 | 213.522443 | -0.11892039 | 0.22516335 | -0.5281516 | 0.5973941 | 0.77659469 |
| CCDC191 | 705.93737 | -0.023834645 | 0.31735971 | -0.0751029 | 0.94013281 | 0.97411372 |
| QTRT2 | 356.666993 | -0.025746368 | 0.32564327 | -0.0790631 | 0.93698243 | 0.97284815 |
| ZBTB20 | 26385.9683 | 0.080481412 | 0.12627226 | 0.63736413 | 0.52388769 | 0.72325104 |
| ZBTB20-AS1 | 130.135247 | -0.608459945 | 0.61702325 | -0.9861216 | 0.32407345 | 0.54894668 |
| ZBTB20-AS5 | 624.911204 | -0.484643522 | 0.62476424 | -0.7757223 | 0.437913 | 0.65419956 |
| ZBTB20-AS2 | 113.643474 | 0.030748981 | 0.55878192 | 0.05502859 | 0.95611569 | 0.98070674 |
| LSAMP | 676.370468 | 1.671103988 | 0.55261925 | 3.02396991 | 0.00249481 | 0.02020026 |
| IGSF11 | 72.6541452 | 0.683144686 | 0.44967674 | 1.51919061 | 0.12871453 | 0.30980142 |
| B4GALT4 | 178.017192 | 0.029309646 | 0.21143321 | 0.13862366 | 0.88974755 | 0.9503707 |
| ARHGAP31 | 48.1253815 | -1.342084578 | 0.47648452 | -2.8166384 | 0.00485291 | 0.03268849 |
| TMEM39A | 358.184995 | 0.175644057 | 0.25043006 | 0.70136971 | 0.48307232 | 0.69127369 |
| POGLUT1 | 207.112445 | 0.017025728 | 0.26945844 | 0.06318499 | 0.9496192 | 0.97821544 |
| TIMMDC1 | 485.812087 | 0.102221232 | 0.14035073 | 0.72832702 | 0.46641343 | 0.67903174 |
| PLA1A | 48.7985195 | 2.18498196 | 0.6221878 | 3.51177243 | 0.00044513 | 0.00541766 |
| COX17 | 227.726144 | -0.135202089 | 0.283738 | -0.4765033 | 0.63371587 | 0.80294992 |
| MAATS1 | 534.118243 | -0.132103066 | 0.28460684 | -0.4641598 | 0.64253323 | 0.80838379 |
| GSK3B | 2742.32587 | -0.048511243 | 0.17100372 | -0.2836853 | 0.77665156 | 0.89337196 |
| AC092910.3 | 144.521441 | 0.652271664 | 0.23341594 | 2.79446071 | 0.00519864 | 0.03448959 |
| LRRC58 | 336.047243 | -0.084965236 | 0.30695486 | -0.2768004 | 0.78193335 | 0.89588513 |
| FSTL1 | 164.696499 | -1.103350664 | 0.41559087 | -2.6548963 | 0.00793328 | 0.04707578 |
| NDUFB4 | 386.579031 | -0.272050435 | 0.25671613 | -1.0597325 | 0.2892663 | 0.51170731 |
| HGD | 712.150891 | -0.221131341 | 0.57280353 | -0.3860509 | 0.69945894 | 0.84724009 |
| RABL3 | 317.731667 | -0.321254683 | 0.1945017 | -1.6516806 | 0.09859968 | 0.2600774 |
| GTF2E1 | 118.92288 | 0.227222124 | 0.20672067 | 1.09917467 | 0.27169188 | 0.49316725 |
| GOLGB1 | 2402.98654 | -0.342697444 | 0.16746382 | -2.0463969 | 0.04071733 | 0.1443853 |
| IQCB1 | 1135.66065 | -0.206695598 | 0.18150929 | -1.1387604 | 0.25480309 | 0.47558202 |
| EAF2 | 248.341023 | 0.963119135 | 0.2912752 | 3.30656076 | 0.00094449 | 0.00953075 |
| SLC15A2 | 503.594205 | 1.32615399 | 0.47927791 | 2.76698333 | 0.00565776 | 0.03674209 |
| ILDR1 | 328.112274 | 0.111822081 | 0.24491277 | 0.45657921 | 0.64797352 | 0.81173905 |
| CASR | 8411.38639 | 1.118523727 | 0.3172661 | 3.52550664 | 0.00042267 | 0.00520811 |
| CCDC58 | 204.513035 | 0.802809623 | 0.32837653 | 2.44478379 | 0.0144939 | 0.07154332 |
| FAM162A | 948.53431 | 0.183625286 | 0.42057701 | 0.43660324 | 0.66239911 | 0.82190799 |
| WDR5B | 40.596762 | 0.079117916 | 0.3704608 | 0.21356623 | 0.83088534 | 0.92119896 |
| AC083798.2 | 509.10807 | 0.547525352 | 0.28254553 | 1.93783051 | 0.0526439 | 0.17207952 |
| KPNA1 | 1087.85596 | -0.087369021 | 0.25481775 | -0.3428687 | 0.73169727 | 0.86826096 |

| | | | | | | |
|---|---|---|---|---|---|---|
| PARP9 | 667.913822 | -0.986887431 | 0.52994993 | -1.8622277 | 0.062571 | 0.19355981 |
| DTX3L | 251.112187 | -1.401144527 | 0.491393 | -2.8513726 | 0.00435309 | 0.03027909 |
| PARP14 | 1688.29393 | -0.329269197 | 0.57544527 | -0.572199 | 0.56718718 | 0.75730066 |
| HSPBAP1 | 929.92343 | -0.507501353 | 0.35612585 | -1.4250618 | 0.15413933 | 0.34845566 |
| DIRC2 | 1369.66938 | -0.929986781 | 0.36603238 | -2.5407227 | 0.01106236 | 0.0596176 |
| LINC02035 | 112.481326 | -0.617167045 | 0.51946517 | -1.1880817 | 0.23480124 | 0.45204629 |
| SEMA5B | 90.2189851 | -2.288321266 | 1.06463231 | -2.1494006 | 0.03160266 | 0.12230869 |
| PDIA5 | 515.161285 | 0.475745798 | 0.35456979 | 1.34175505 | 0.17967543 | 0.38325438 |
| SEC22A | 1251.71224 | 0.259610475 | 0.19208537 | 1.35153696 | 0.17652349 | 0.37870696 |
| ADCY5 | 1320.96721 | 0.350944673 | 0.49762628 | 0.70523742 | 0.4806625 | 0.68948936 |
| HACD2 | 775.652521 | 0.392182355 | 0.32984431 | 1.18899231 | 0.23444269 | 0.45166252 |
| MYLK-AS1 | 229.70661 | 0.640977719 | 0.28861067 | 2.2209079 | 0.0263572 | 0.1074544 |
| MYLK | 1695.81302 | 0.346802162 | 0.45100609 | 0.76895229 | 0.44192164 | 0.65772878 |
| CCDC14 | 1204.50981 | 0.507005399 | 0.15904915 | 3.1877279 | 0.00143395 | 0.01322702 |
| KALRN | 413.591776 | -3.280636645 | 0.76641673 | -4.2804867 | 1.86E-05 | 0.00044383 |
| UMPS | 235.347024 | 0.370161122 | 0.18361315 | 2.01598368 | 0.04380168 | 0.15161856 |
| ITGB5 | 1560.06482 | 0.838187146 | 0.29149573 | 2.87546971 | 0.00403427 | 0.02880468 |
| ITGB5-AS1 | 97.4772688 | 0.640388807 | 0.3435127 | 1.86423621 | 0.06228854 | 0.1928215 |
| HEG1 | 63.0885995 | -1.120422754 | 0.43151178 | -2.5965056 | 0.00941774 | 0.05304857 |
| SLC12A8 | 666.033429 | 0.150439373 | 0.31749597 | 0.47383081 | 0.63562057 | 0.80389813 |
| ZNF148 | 3518.67657 | 0.021057694 | 0.11919769 | 0.17666193 | 0.85977394 | 0.93625342 |
| SNX4 | 1109.9452 | 0.008983119 | 0.13316874 | 0.06745667 | 0.94621816 | 0.9767213 |
| OSBPL11 | 416.012934 | 0.262945345 | 0.26652121 | 0.98658318 | 0.32384702 | 0.54888141 |
| AC092902.3 | 42.293619 | -0.36999507 | 0.66517545 | -0.5562368 | 0.57804898 | 0.76438572 |
| AC092903.2 | 39.7143794 | 0.646370391 | 0.36754698 | 1.75860619 | 0.07864442 | 0.22436591 |
| SLC41A3 | 185.660829 | -0.264832761 | 0.24626329 | -1.075405 | 0.28219346 | 0.50464019 |
| AC079848.1 | 49.3401322 | 0.001914217 | 0.37592574 | 0.00509201 | 0.99593718 | 0.99850734 |
| ALDH1L1 | 976.627631 | 0.703261213 | 0.48435552 | 1.45195251 | 0.14651481 | 0.33711064 |
| ALDH1L1-AS2 | 57.6910791 | 0.809298116 | 0.48283452 | 1.67613971 | 0.09371083 | 0.25184394 |
| KLF15 | 138.329335 | 0.501330411 | 0.45757431 | 1.09562623 | 0.27324238 | 0.49489354 |
| ZXDC | 1678.46745 | -0.18070496 | 0.21592772 | -0.8368771 | 0.40266166 | 0.62391114 |
| TXNRD3NB | 124.790833 | 0.485086342 | 0.29005874 | 1.67237278 | 0.09445084 | 0.25284294 |
| TXNRD3 | 211.407424 | 0.758065076 | 0.22470966 | 3.37353138 | 0.00074211 | 0.00798898 |
| CHCHD6 | 412.831733 | 0.153909674 | 0.23148508 | 0.66487945 | 0.5061276 | 0.7095563 |
| PLXNA1 | 110.45147 | -0.61900218 | 0.29683699 | -2.085327 | 0.03703963 | 0.13567522 |
| TPRA1 | 147.374782 | 0.207561083 | 0.2227694 | 0.93173069 | 0.35147572 | 0.57736509 |
| PODXL2 | 46.8433562 | 0.608576799 | 0.50312559 | 1.20959223 | 0.2264354 | 0.44159983 |
| ABTB1 | 155.393782 | -0.423910933 | 0.32118262 | -1.3198439 | 0.18688713 | 0.39252537 |
| MGLL | 4473.65281 | -1.090292595 | 0.34035906 | -3.20336 | 0.00135834 | 0.01272505 |
| KBTBD12 | 362.937252 | -0.95646476 | 0.32903022 | -2.9069207 | 0.00365006 | 0.02677046 |
| SEC61A1 | 563.03973 | -0.337155517 | 0.22339953 | -1.5092042 | 0.1312466 | 0.31392317 |
| RUVBL1 | 393.745612 | -0.027605714 | 0.18147916 | -0.1521151 | 0.87909618 | 0.94598627 |
| EEFSEC | 675.201809 | 0.026849892 | 0.1568928 | 0.17113527 | 0.8641174 | 0.93827593 |
| RPN1 | 374.705269 | -0.327498851 | 0.23764271 | -1.3781144 | 0.16816796 | 0.36813567 |
| RAB7A | 1813.16776 | -0.199504821 | 0.14399709 | -1.3854781 | 0.16590631 | 0.36530363 |
| ACAD9 | 378.246051 | 0.017450838 | 0.1519405 | 0.1148531 | 0.90856156 | 0.95871462 |
| KIAA1257 | 64.3872157 | -0.300733521 | 0.40103948 | -0.7498851 | 0.45332392 | 0.66912009 |
| CNBP | 822.101772 | -1.151919961 | 0.4316653 | -2.6685489 | 0.00761797 | 0.04559391 |
| COPG1 | 331.830511 | -0.12173376 | 0.19905794 | -0.6115494 | 0.54083593 | 0.73678855 |
| HMCES | 211.603286 | -0.155138522 | 0.20949456 | -0.7405372 | 0.45897408 | 0.67278348 |
| H1FX | 264.569845 | -1.834033356 | 0.53760025 | -3.4115188 | 0.00064602 | 0.00720717 |
| MBD4 | 467.582572 | -0.159047838 | 0.11572384 | -1.3743741 | 0.16932558 | 0.36972673 |
| IFT122 | 636.036257 | 0.236249675 | 0.17191435 | 1.37422895 | 0.16937061 | 0.36972673 |
| TMCC1 | 4514.70741 | 0.074680618 | 0.1669795 | 0.44724423 | 0.65469873 | 0.81607205 |
| LINC02021 | 49.7559358 | 0.486874638 | 0.48891732 | 0.99582202 | 0.31933663 | 0.54392231 |
| COL6A6 | 45.9096049 | 0.517315037 | 0.66682939 | 0.77578319 | 0.43787701 | 0.65419956 |
| PIK3R4 | 511.878815 | 0.283902234 | 0.20783782 | 1.36597963 | 0.17194536 | 0.37294355 |
| ATP2C1 | 1885.19592 | 0.168997342 | 0.24021156 | 0.70353542 | 0.48172214 | 0.6901375 |
| AC055733.2 | 92.6052879 | -0.148664477 | 0.67900188 | -0.2189456 | 0.82669242 | 0.91944889 |
| ASTE1 | 80.3102378 | 0.23123352 | 0.30808768 | 0.75054453 | 0.45292681 | 0.66868339 |
| NEK11 | 1085.05308 | -0.535767471 | 0.30786436 | -1.7402712 | 0.08181142 | 0.22992489 |
| NUDT16 | 130.278773 | -0.318511207 | 0.39270121 | -0.8110777 | 0.41732104 | 0.63704999 |
| MRPL3 | 461.359164 | 0.119957921 | 0.16784806 | 0.7146816 | 0.47480581 | 0.68513327 |
| CPNE4 | 121.334077 | 0.94554094 | 0.41918536 | 2.25566309 | 0.02409174 | 0.10099717 |
| ACPP | 1808.87395 | 0.601463505 | 0.39601041 | 1.51880731 | 0.12881101 | 0.30997715 |
| DNAJC13 | 985.911858 | 0.143415461 | 0.18140085 | 0.79059972 | 0.42917761 | 0.64747247 |
| NPHP3-ACAD | 75.6010595 | 0.050162538 | 0.4117343 | 0.1218323 | 0.90303183 | 0.95509228 |
| ACKR4 | 44.3037664 | -0.062754284 | 0.64329539 | -0.0975513 | 0.92228862 | 0.9655855 |
| UBA5 | 681.538352 | 0.201947679 | 0.2120984 | 0.95214147 | 0.34102524 | 0.56647414 |
| TMEM108 | 264.172009 | 0.466296874 | 0.67246 | 0.6934195 | 0.48804633 | 0.69502858 |
| CDV3 | 426.840994 | -0.989356148 | 0.20478573 | -4.8311771 | 1.36E-06 | 5.26E-05 |
| TOPBP1 | 370.708318 | -0.071409831 | 0.24808004 | -0.28785 | 0.77346159 | 0.89119454 |
| TF | 58.2116722 | 0.725212578 | 0.44258211 | 1.63859444 | 0.10129775 | 0.26501129 |
| SRPRB | 167.076087 | 0.182750369 | 0.30491372 | 0.59935108 | 0.54893879 | 0.74226986 |
| RAB6B | 114.349841 | 0.297166567 | 0.56794391 | 0.52323225 | 0.60081264 | 0.7789212 |
| SLCO2A1 | 121.98715 | -1.615213663 | 0.61631477 | -2.6207609 | 0.00877338 | 0.05041939 |
| RYK | 1663.11335 | -0.064757047 | 0.18355471 | -0.3527942 | 0.7242427 | 0.86324867 |
| AMOTL2 | 287.439112 | 1.059852416 | 0.44833229 | 2.36398857 | 0.01807937 | 0.08352935 |
| ANAPC13 | 126.026641 | -0.419928224 | 0.28606452 | -1.4679493 | 0.14211799 | 0.33114068 |
| CEP63 | 693.398757 | 0.277455785 | 0.18201633 | 1.52434554 | 0.12742242 | 0.30764444 |
| EPHB1 | 319.869443 | -0.833939447 | 0.57776591 | -1.4433864 | 0.14891166 | 0.34089673 |
| PPP2R3A | 9811.2073 | -0.452075256 | 0.19160107 | -2.3594611 | 0.0183015 | 0.08414359 |
| MSL2 | 722.093693 | 0.56890566 | 0.21037551 | 2.70423899 | 0.0068461 | 0.04217825 |
| PCCB | 317.961354 | 0.30226821 | 0.21147879 | 1.42930744 | 0.15291589 | 0.34644783 |
| STAG1 | 5213.7948 | -0.105182593 | 0.21521167 | -0.4887402 | 0.62502565 | 0.79598067 |
| SLC35G2 | 239.937498 | -0.846886418 | 0.37464502 | -2.2605036 | 0.02379001 | 0.10024483 |
| NCK1-DT | 113.346585 | 0.094518329 | 0.26975178 | 0.35039001 | 0.72604602 | 0.86492983 |
| NCK1 | 602.681295 | -0.111335908 | 0.21064657 | -0.5285437 | 0.59712205 | 0.77654325 |
| IL20RB | 57.98475 | -1.161100634 | 0.55256151 | -2.1013057 | 0.03561414 | 0.13239418 |
| CLDN18 | 54.4518466 | -0.551199369 | 0.73695821 | -0.7479384 | 0.45449729 | 0.6698832 |
| DZIP1L | 440.810362 | 0.374787797 | 0.26865537 | 1.39505045 | 0.16300063 | 0.36109189 |
| DBR1 | 118.043951 | 0.309606285 | 0.21515568 | 1.43898728 | 0.15015413 | 0.34236605 |
| ARMC8 | 1513.1402 | 0.087330877 | 0.2122396 | 0.41147306 | 0.68072569 | 0.83310531 |
| NME9 | 190.988814 | -0.405992843 | 0.34807828 | -1.1663837 | 0.24345934 | 0.46211444 |
| MRAS | 549.832502 | -0.795314138 | 0.22912702 | -3.4710622 | 0.0005184 | 0.00608135 |
| ESYT3 | 87.2935494 | -0.580650224 | 0.33450053 | -1.7358723 | 0.08258641 | 0.23092377 |
| CEP70 | 2090.55988 | 0.251316295 | 0.2946304 | 0.85298832 | 0.39366578 | 0.61588683 |
| FAIM | 230.36797 | -0.223045858 | 0.25449333 | -0.876431 | 0.38079574 | 0.60354344 |
| PIK3CB | 2078.16181 | 0.377794561 | 0.1601814 | 2.35854195 | 0.01834689 | 0.08423497 |
| MRPS22 | 127.480746 | 0.282471953 | 0.22368714 | 1.26279925 | 0.20666133 | 0.41739901 |
| COPB2 | 700.97136 | -0.134563632 | 0.15694104 | -0.8574152 | 0.39121547 | 0.6140943 |
| AC097103.2 | 964.226481 | 0.47805747 | 0.40197841 | 1.18926157 | 0.23433675 | 0.45161281 |
| NMNAT3 | 428.832929 | 0.557343932 | 0.27641328 | 2.0163428 | 0.04376414 | 0.15152832 |
| CLSTN2 | 80.9376164 | 0.239178515 | 0.40514559 | 0.59035201 | 0.55495468 | 0.74754753 |
| SLC25A36 | 1962.21929 | 0.202558966 | 0.2827785 | 0.71631671 | 0.47379581 | 0.68442354 |
| PXYLP1 | 118.743868 | -0.272428253 | 0.25834489 | -1.0545138 | 0.29164775 | 0.51433579 |
| AC117383.1 | 102.282978 | -2.490884046 | 0.63791601 | -3.904721 | 9.43E-05 | 0.00160162 |
| ZBTB38 | 1259.08356 | -1.175971756 | 0.276562 | -4.252109 | 2.12E-05 | 0.00049421 |
| RASA2 | 2030.983 | -0.74552807 | 0.2118978 | -3.518338 | 0.00043426 | 0.0053409 |
| RNF7 | 320.446039 | -0.72365882 | 0.27355592 | -2.645378 | 0.00815997 | 0.04789524 |
| ATP1B3 | 1130.83193 | 0.65370837 | 0.26130053 | 2.50174911 | 0.01235815 | 0.06426616 |
| TFDP2 | 3045.62092 | 0.427485424 | 0.20513966 | 2.08387507 | 0.03717153 | 0.13601074 |
| GK5 | 861.959793 | 0.215777407 | 0.22544098 | 0.9571348 | 0.33849922 | 0.56390922 |

| | | | | | | |
|---|---|---|---|---|---|---|
| XRN1 | 2256.48774 | 0.199063944 | 0.19398674 | 1.02617292 | 0.30481008 | 0.52816954 |
| ATR | 1787.13843 | 0.095267411 | 0.22420385 | 0.42491424 | 0.67089919 | 0.82705622 |
| PLS1 | 404.034246 | -0.467565833 | 0.17952473 | -2.6044648 | 0.00920179 | 0.05214346 |
| TRPC1 | 597.905124 | -0.880170558 | 0.39011977 | -2.2561547 | 0.02406094 | 0.10095966 |
| PCOLCE2 | 55.7557442 | -1.279829675 | 0.35747437 | -3.5801998 | 0.00034333 | 0.00444614 |
| U2SURP | 1490.40674 | 0.407909776 | 0.15434199 | 2.64289558 | 0.00822004 | 0.0481501 |
| SLC9A9 | 166.577537 | -1.570436902 | 0.58509424 | -2.6840751 | 0.00727308 | 0.04394788 |
| C3orf58 | 285.685478 | -0.353885861 | 0.32270067 | -1.0966381 | 0.27279961 | 0.49436274 |
| AC107021.1 | 463.007263 | 0.03829993 | 0.7374393 | 0.05193638 | 0.95857938 | 0.9819501 |
| LNCSRLR | 125.81761 | -0.147202358 | 0.52575092 | -0.279985 | 0.77948903 | 0.894861 |
| PLOD2 | 3239.64106 | -0.297348394 | 0.43457073 | -0.6842347 | 0.49382695 | 0.69913828 |
| PLSCR4 | 327.159122 | -0.684794482 | 0.48591694 | -1.409283 | 0.1587515 | 0.35514221 |
| PLSCR2 | 446.077345 | -0.184161848 | 0.33141177 | -0.5556889 | 0.57842353 | 0.76458087 |
| PLSCR1 | 2251.61474 | -1.162730015 | 0.35766984 | -3.2508472 | 0.00115062 | 0.01115895 |
| GYG1 | 140.864065 | -0.914423555 | 0.23311859 | -3.9225681 | 8.76E-05 | 0.00152069 |
| HLTF | 499.311337 | 0.853673759 | 0.25168689 | 3.39180859 | 0.00069433 | 0.00762377 |
| HPS3 | 460.401792 | -0.176141359 | 0.18859878 | -0.9339475 | 0.35033098 | 0.57642276 |
| CP | 602.910048 | -4.471307689 | 1.1004556 | -4.0631423 | 4.84E-05 | 0.00095463 |
| TM4SF1 | 169.800538 | -1.262550607 | 0.66697767 | -1.8929428 | 0.05836547 | 0.18452596 |
| WWTR1 | 2716.64912 | -0.525524809 | 0.21469189 | -2.4478093 | 0.01437277 | 0.07110485 |
| COMMD2 | 283.24511 | 0.001133228 | 0.28644662 | 0.00395616 | 0.99684345 | 0.99880946 |
| ANKUB1 | 687.700728 | -0.271265643 | 0.50032704 | -0.5421767 | 0.58769681 | 0.77096904 |
| RNF13 | 1613.07377 | 0.140994135 | 0.12617914 | 1.11741236 | 0.26381805 | 0.48477502 |
| PFN2 | 41.9253733 | -0.248457034 | 0.34211071 | -0.7262475 | 0.46768708 | 0.68013258 |
| TSC22D2 | 2716.53473 | 0.424241077 | 0.35596738 | 1.19179761 | 0.23334061 | 0.45021855 |
| SERP1 | 944.177693 | 0.128894462 | 0.26514665 | 0.48612518 | 0.62687841 | 0.79757628 |
| EIF2A | 477.803091 | 0.124757666 | 0.17427219 | 0.71587822 | 0.47406655 | 0.68459432 |
| SELENOT | 508.550785 | 0.284405174 | 0.22698307 | 1.25297967 | 0.2102131 | 0.42231253 |
| ERICH6 | 60.2755061 | -0.069125426 | 0.6312018 | -0.109514 | 0.91279483 | 0.96054229 |
| ERICH6-AS1 | 91.3632368 | 0.023125598 | 0.25983882 | 0.08899978 | 0.92908209 | 0.96900671 |
| SIAH2 | 196.467273 | -0.259791439 | 0.25114589 | -1.0344244 | 0.30093781 | 0.52396873 |
| MED12L | 241.23055 | -1.383205719 | 0.57416942 | -2.409055 | 0.01599389 | 0.07653081 |
| SUCNR1 | 468.272619 | 1.135510278 | 0.9251701 | 1.22735298 | 0.21968995 | 0.43361796 |
| MBNL1 | 5162.56033 | 0.112531476 | 0.18735185 | 0.60064244 | 0.54807816 | 0.74186733 |
| RAP2B | 92.4412262 | -2.209328688 | 0.41261577 | -5.3544456 | 8.58E-08 | 5.67E-06 |
| ARHGEF26 | 146.512815 | -0.551713585 | 0.39761777 | -1.3875476 | 0.16527485 | 0.36452087 |
| DHX36 | 1336.43899 | 0.002675383 | 0.14553253 | 0.0183834 | 0.98533299 | 0.99423861 |
| MME | 110.156936 | 0.865872576 | 0.49798719 | 1.73874466 | 0.08207969 | 0.23028687 |
| PLCH1 | 566.712137 | -1.298699707 | 0.73151574 | -1.7753544 | 0.07583938 | 0.21949221 |
| GMPS | 683.988888 | 0.354948102 | 0.18405436 | 1.92849605 | 0.05379346 | 0.17451057 |
| KCNAB1 | 116.151182 | 0.068538584 | 0.56729352 | 0.12081679 | 0.90383615 | 0.95548393 |
| SSR3 | 410.475793 | 0.35013766 | 0.22619521 | 1.54794463 | 0.12163563 | 0.29836662 |
| TIPARP | 233.110073 | 0.566054522 | 0.32461935 | 1.74374856 | 0.08120294 | 0.22874648 |
| LINC00886 | 719.320059 | 0.626818268 | 0.41176121 | 1.52228586 | 0.12793747 | 0.30854958 |
| LEKR1 | 495.977899 | 1.306316106 | 0.39646596 | 3.294901 | 0.00098456 | 0.00987479 |
| CCNL1 | 1648.542 | -0.437319991 | 0.3935128 | -1.1113234 | 0.26642916 | 0.48750606 |
| VEPH1 | 984.762549 | 2.28356552 | 0.41569409 | 5.49337986 | 3.94E-08 | 2.94E-06 |
| RSRC1 | 1769.6878 | -0.141067021 | 0.21310067 | -0.6619736 | 0.50798814 | 0.71103267 |
| MLF1 | 229.793881 | -0.030155249 | 0.31107691 | -0.0969382 | 0.92277544 | 0.96580501 |
| GFM1 | 387.709785 | 0.122004448 | 0.28970459 | 0.42113398 | 0.67365725 | 0.82886565 |
| LXN | 125.048478 | 0.290398097 | 0.39041149 | 0.7438257 | 0.45698193 | 0.67196642 |
| RARRES1 | 118.931692 | -0.685821284 | 0.51239871 | -1.3384524 | 0.180749 | 0.38452466 |
| MFSD1 | 241.732224 | 0.237505156 | 0.16782322 | 1.41521031 | 0.15700684 | 0.35264468 |
| IQCJ-SCHIP1 | 941.652797 | -0.623793292 | 0.31726491 | -1.9661591 | 0.04928024 | 0.16458726 |
| AC079594.2 | 60.6185101 | -0.020105117 | 0.78279412 | -0.0256838 | 0.97950956 | 0.9910179 |
| SMC4 | 235.164553 | -0.209232115 | 0.27980981 | -0.7477655 | 0.45460163 | 0.6698832 |
| KPNA4 | 1242.47238 | -0.041490501 | 0.24724324 | -0.1678125 | 0.8667308 | 0.93959274 |
| PPM1L | 3422.4354 | 1.468355419 | 0.38408898 | 3.82295639 | 0.00013186 | 0.00209597 |
| B3GALNT1 | 219.640797 | 0.57561129 | 0.26774901 | 2.1498167 | 0.03156972 | 0.12228868 |
| NMD3 | 1525.00732 | 0.758724968 | 0.26484629 | 2.86477479 | 0.00417306 | 0.02937201 |
| LINC01322 | 140.056125 | 0.366157131 | 0.36542716 | 1.00199757 | 0.31634477 | 0.54070885 |
| PDCD10 | 663.590612 | -0.100194782 | 0.17944553 | -0.5583576 | 0.57660021 | 0.76325956 |
| SERPINI1 | 124.374553 | 0.424274189 | 0.33532441 | 1.26526485 | 0.2057764 | 0.41631192 |
| GOLIM4 | 592.149679 | 0.712023242 | 0.28574363 | 2.49182546 | 0.01270885 | 0.06549792 |
| MECOM | 47927.8807 | 0.385620851 | 0.27076616 | 1.42418405 | 0.1543932 | 0.34881127 |
| AC074033.1 | 231.814313 | 0.04227777 | 0.54288461 | 0.07787616 | 0.93792656 | 0.97306566 |
| MYNN | 219.401486 | 0.319492049 | 0.24604263 | 1.29852317 | 0.19410762 | 0.40163448 |
| LRRC34 | 80.1492532 | -0.362131703 | 0.38125783 | -0.9498341 | 0.34219653 | 0.56742669 |
| SEC62 | 965.74377 | 0.08603986 | 0.18139956 | 0.47431129 | 0.63527794 | 0.80389813 |
| GPR160 | 236.583128 | -0.666349618 | 0.39708574 | -1.6781001 | 0.09332756 | 0.25117608 |
| PHC3 | 2399.76965 | 0.149622413 | 0.15545533 | 0.96247852 | 0.33580929 | 0.56141057 |
| PRKCI | 1819.77902 | -0.017453531 | 0.24488298 | -0.0712729 | 0.94318053 | 0.97556917 |
| SKIL | 769.089733 | -0.609435201 | 0.31935664 | -1.9083217 | 0.05634965 | 0.18026702 |
| RPL22L1 | 100.608054 | -0.229383655 | 0.39331161 | -0.583211 | 0.55975127 | 0.75155057 |
| TNIK | 437.276736 | -2.186603871 | 0.60451478 | -3.6171222 | 0.0002979 | 0.00400704 |
| PLD1 | 453.008445 | -0.216065112 | 0.25857026 | -0.8356147 | 0.4033717 | 0.62463502 |
| FNDC3B | 10286.0786 | -0.941255703 | 0.21814581 | -4.3148008 | 1.60E-05 | 0.00039148 |
| TNFSF10 | 2119.57228 | -1.934851736 | 0.42835736 | -4.5169102 | 6.27E-06 | 0.00018216 |
| LINC02068 | 75.4005717 | -0.904605018 | 0.57170026 | -1.5823065 | 0.11357962 | 0.28587524 |
| NCEH1 | 1895.17795 | -1.12370463 | 0.42986178 | -2.6141069 | 0.00894611 | 0.05104497 |
| ECT2 | 269.454141 | -0.96915601 | 0.37330066 | -2.5961808 | 0.00942664 | 0.0530761 |
| NLGN1 | 3491.42944 | -3.036618916 | 0.76900754 | -3.9487505 | 7.86E-05 | 0.00139852 |
| NAALADL2 | 10571.8593 | 0.607628675 | 0.37860173 | 1.60492843 | 0.10850956 | 0.27683877 |
| AC092920.1 | 76.3836785 | 0.622228947 | 0.60869799 | 1.02222935 | 0.30667236 | 0.53032741 |
| TBL1XR1 | 5512.40486 | -0.80938015 | 0.33266975 | -2.4329839 | 0.01497497 | 0.0732348 |
| LINC00501 | 55.8303759 | -0.905765583 | 0.61026037 | -1.4842281 | 0.13774842 | 0.32381543 |
| LINC00578 | 128.909519 | -0.76474026 | 0.67368326 | -1.1351629 | 0.25630706 | 0.47722602 |
| KCNMB2-AS1 | 487.476713 | -0.010266478 | 0.52838996 | -0.0194297 | 0.98449829 | 0.99382789 |
| ZMAT3 | 414.889613 | -0.65757389 | 0.37291884 | -1.7633163 | 0.07784715 | 0.22291846 |
| AC076966.1 | 39.6413806 | 1.267298229 | 0.6839991 | 1.85277764 | 0.06391424 | 0.19615315 |
| PIK3CA | 1288.58188 | -0.120868314 | 0.22598107 | -0.5348603 | 0.59274643 | 0.77382759 |
| KCNMB3 | 237.878429 | -0.315507685 | 0.30876957 | -1.0218225 | 0.30686493 | 0.53047754 |
| ZNF639 | 173.472519 | 0.0160053 | 0.24222563 | 0.06607599 | 0.94731732 | 0.97743435 |
| MFN1 | 1135.42644 | 0.084515798 | 0.18416529 | 0.45891273 | 0.64629684 | 0.81080206 |
| GNB4 | 215.95424 | -0.946540421 | 0.44392556 | -2.1322053 | 0.03298997 | 0.12597993 |
| ACTL6A | 236.773411 | -0.281689719 | 0.27557859 | -1.0221756 | 0.30669779 | 0.53032741 |
| MRPL47 | 182.684884 | 0.059222242 | 0.24067373 | 0.24606857 | 0.80562915 | 0.90906846 |
| NDUFB5 | 461.347275 | 0.476026288 | 0.25785299 | 1.84611506 | 0.0648755 | 0.19777071 |
| USP13 | 727.658605 | 1.138513849 | 0.2770785 | 4.10899379 | 3.97E-05 | 0.00082145 |
| PEX5L | 125.621467 | 1.052034565 | 0.37040777 | 2.8402065 | 0.00450843 | 0.03113011 |
| AC125618.1 | 57.6881289 | 1.414868214 | 0.61158223 | 2.3134554 | 0.02069761 | 0.09130981 |
| TTC14 | 1693.34224 | 0.796102188 | 0.14292258 | 5.57016372 | 2.55E-08 | 2.04E-06 |
| CCDC39 | 51.3214943 | 0.644224853 | 0.67853174 | 0.94943952 | 0.34239712 | 0.56753966 |
| FXR1 | 1402.58505 | 0.562326215 | 0.20873143 | 2.69401791 | 0.00705964 | 0.04307194 |
| DNAJC19 | 383.806888 | 0.798228124 | 0.29154469 | 2.7379272 | 0.00618278 | 0.03909444 |
| ATP11B | 1357.30114 | -0.144076857 | 0.150172 | -0.9594122 | 0.33735112 | 0.5623512 |
| DCUN1D1 | 855.443669 | 0.523491 | 0.12905192 | 4.05643725 | 4.98E-05 | 0.00096931 |
| MCCC1 | 1511.0446 | 0.423181161 | 0.20797536 | 2.03476588 | 0.04187444 | 0.14672183 |
| MCF2L2 | 156.965185 | -0.429866556 | 0.47853126 | -0.898304 | 0.36902349 | 0.59393582 |
| B3GNT5 | 184.333534 | -1.005680072 | 0.38374043 | -2.62073 | 0.00877417 | 0.05041939 |
| KLHL24 | 2202.67719 | 0.157000678 | 0.1930996 | 0.81305545 | 0.41618627 | 0.63597923 |
| YEATS2 | 1135.91291 | -0.112319349 | 0.29162527 | -0.3851496 | 0.7001266 | 0.84785721 |
| ABCC5 | 2261.73657 | 0.202644374 | 0.16492393 | 1.22871421 | 0.21917898 | 0.43353326 |

| | | | | | | |
|---|---|---|---|---|---|---|
| EIF2B5 | 212.75918 | 0.328253498 | 0.23416947 | 1.40177752 | 0.16098169 | 0.35824533 |
| DVL3 | 442.340285 | 0.728086762 | 0.39111207 | 1.86158093 | 0.06266218 | 0.19370578 |
| AP2M1 | 518.53897 | -0.985656053 | 0.36742445 | -2.6826088 | 0.00730504 | 0.04412083 |
| ABCF3 | 104.57326 | 0.077235018 | 0.2662584 | 0.29007542 | 0.77175854 | 0.88993091 |
| ALG3 | 84.8514524 | -0.652032267 | 0.37045088 | -1.7601045 | 0.07839008 | 0.22378531 |
| PSMD2 | 277.722303 | -0.638678218 | 0.30554857 | -2.0902674 | 0.03659379 | 0.13464146 |
| EIF4G1 | 550.85567 | -0.628384346 | 0.22777433 | -2.7588023 | 0.00580136 | 0.0374903 |
| FAM131A | 129.669498 | -0.404277647 | 0.27074402 | -1.4932099 | 0.13538226 | 0.32059237 |
| CLCN2 | 76.4219381 | -0.27096393 | 0.33900747 | -0.799286 | 0.42412459 | 0.64296071 |
| POLR2H | 414.312502 | -0.283144981 | 0.24450455 | -1.1580356 | 0.2468495 | 0.46607134 |
| EPHB3 | 69.6842434 | 2.26977573 | 0.58670874 | 3.86865846 | 0.00010944 | 0.00180017 |
| MAGEF1 | 179.563504 | -0.419638203 | 0.32943214 | -1.2738229 | 0.20272619 | 0.41232902 |
| VPS8 | 1453.7469 | 0.163848315 | 0.23050564 | 0.71082128 | 0.477195 | 0.68692989 |
| EHHADH | 384.965703 | 0.596635099 | 0.2497493 | 2.38893606 | 0.01689724 | 0.07956857 |
| MAP3K13 | 1931.69266 | 0.132487964 | 0.24187011 | 0.54776492 | 0.58385333 | 0.76885817 |
| TMEM41A | 256.571024 | -0.248309456 | 0.21288658 | -1.1663932 | 0.24345551 | 0.46211444 |
| LIPH | 890.657927 | -0.068505547 | 0.46796277 | -0.146391 | 0.88361272 | 0.94760416 |
| SENP2 | 684.545523 | 0.196253612 | 0.17318683 | 1.13319019 | 0.25713439 | 0.4779747 |
| IGF2BP2 | 1720.17966 | -0.452109342 | 0.47269341 | -0.9564537 | 0.33884309 | 0.56426858 |
| TRA2B | 959.566795 | 0.387561841 | 0.16408779 | 2.36191764 | 0.01818068 | 0.08384041 |
| ETV5 | 332.455719 | -1.841053698 | 0.55513586 | -3.3164021 | 0.00091185 | 0.00928648 |
| CRYGS | 76.1246195 | 0.540647478 | 0.25522107 | 2.11834972 | 0.03414546 | 0.12893865 |
| TBCCD1 | 141.321951 | 0.390847465 | 0.24185381 | 1.61604842 | 0.10608385 | 0.27325692 |
| DNAJB11 | 133.62322 | 0.660289957 | 0.29860046 | 2.21128244 | 0.02701628 | 0.10918997 |
| AC068631.1 | 1187.76516 | 0.464162488 | 0.39200369 | 1.18407682 | 0.23638266 | 0.45392878 |
| AC068631.2 | 72.9564662 | -1.426659689 | 0.54796839 | -2.6035438 | 0.00922655 | 0.05223895 |
| HRG | 61.0662896 | 0.603387064 | 0.60953052 | 0.989921 | 0.32221274 | 0.54705759 |
| KNG1 | 2181.03806 | 2.08664149 | 0.58897303 | 3.54284726 | 0.00039583 | 0.00493259 |
| EIF4A2 | 1052.88371 | -0.161311473 | 0.28629462 | -0.5634457 | 0.57313142 | 0.76101836 |
| RFC4 | 47.3446639 | -0.045225492 | 0.30428848 | -0.148627 | 0.88184795 | 0.94672712 |
| ST6GAL1 | 956.389544 | -1.812480301 | 0.57803097 | -3.1356111 | 0.00171496 | 0.01515651 |
| AC072022.1 | 152.413446 | -2.181234463 | 0.86625023 | -2.5180189 | 0.0118017 | 0.06230561 |
| BCL6 | 1803.98613 | -2.618077533 | 0.47673782 | -5.4916506 | 3.98E-08 | 2.95E-06 |
| LPP | 9219.96667 | -0.484036252 | 0.21488163 | -2.2525716 | 0.02428617 | 0.10158203 |
| TPRG1 | 370.681771 | -1.438303287 | 0.53634887 | -2.6816562 | 0.00732587 | 0.04418924 |
| P3H2 | 4752.31337 | 1.015113988 | 0.41728804 | 2.43264577 | 0.01498896 | 0.07327503 |
| P3H2-AS1 | 133.883505 | 1.270470233 | 0.57329665 | 2.21607826 | 0.02668614 | 0.10833817 |
| CLDN1 | 207.334861 | -2.752192683 | 0.50648728 | -5.4338831 | 5.51E-08 | 3.94E-06 |
| CLDN16 | 1251.80332 | 1.228528734 | 0.63673947 | 1.92940565 | 0.05368052 | 0.17434621 |
| TMEM207 | 336.974143 | 3.536254979 | 0.78625828 | 4.49757423 | 6.87E-06 | 0.00019598 |
| IL1RAP | 170.595489 | -0.727207362 | 0.5534927 | -1.3138518 | 0.18889609 | 0.39486629 |
| CCDC50 | 1807.21159 | 0.144526485 | 0.2653886 | 0.54458437 | 0.58603941 | 0.77001836 |
| FGF12 | 56.124146 | 0.161606726 | 0.36367564 | 0.4443705 | 0.65677474 | 0.81801439 |
| MB21D2 | 112.589625 | -0.080218483 | 0.52896176 | -0.1516527 | 0.87946086 | 0.94622453 |
| OPA1 | 1374.15672 | 0.315464887 | 0.15888946 | 1.98543624 | 0.04709596 | 0.15923345 |
| OPA1-AS1 | 72.6423358 | -0.235177537 | 0.44016144 | -0.5342984 | 0.59313513 | 0.77387448 |
| HES1 | 245.039948 | -0.396581483 | 0.58219247 | -0.6811862 | 0.49575367 | 0.70114155 |
| ATP13A3 | 1044.61499 | -0.301984528 | 0.24617151 | -1.2267241 | 0.2199263 | 0.43371253 |
| LINC00884 | 74.0034125 | 0.561875187 | 0.41494198 | 1.35410542 | 0.17570275 | 0.37792828 |
| TMEM44-AS | 118.331399 | 0.470558038 | 0.46274786 | 1.01687783 | 0.30921155 | 0.53306909 |
| TMEM44 | 53.5568475 | -1.679569654 | 0.4423802 | -3.7966655 | 0.00014666 | 0.00227101 |
| LSG1 | 366.916386 | -0.011112797 | 0.2663966 | -0.0417152 | 0.96672571 | 0.98510234 |
| FAM43A | 238.378286 | 1.801506135 | 0.99688794 | 1.80713004 | 0.07074201 | 0.20970179 |
| LINC01968 | 38.1649636 | 2.416613056 | 0.93321995 | 2.58954285 | 0.00961035 | 0.05378943 |
| XXYLT1 | 336.897386 | 0.337462252 | 0.29111197 | 1.15921807 | 0.24636731 | 0.46556019 |
| ACAP2 | 2031.2278 | 0.170310477 | 0.16937945 | 1.00549667 | 0.31465775 | 0.53873159 |
| PPP1R2 | 472.812119 | 0.055786214 | 0.14187255 | 0.39321357 | 0.69416174 | 0.843734 |
| MUC20-OT1 | 5542.76703 | 0.676953747 | 0.42637782 | 1.58768518 | 0.11235755 | 0.28354182 |
| TNK2 | 131.225244 | 0.094670196 | 0.24595592 | 0.38490717 | 0.7003062 | 0.84787759 |
| TFRC | 667.305641 | 0.600248176 | 0.30305728 | 1.98064264 | 0.04763136 | 0.16041884 |
| LINC00885 | 89.8963877 | 1.905600097 | 1.11526177 | 1.70865724 | 0.08751446 | 0.23987933 |
| SLC51A | 46.3227013 | -0.358081185 | 0.4220198 | -0.8484938 | 0.39616302 | 0.61805306 |
| PCYT1A | 267.97326 | -0.319390238 | 0.19339067 | -1.6515287 | 0.09863066 | 0.2600774 |
| UBXN7 | 770.1342 | -0.022336663 | 0.19385553 | -0.1152232 | 0.90826817 | 0.95863478 |
| RNF168 | 598.154746 | -0.467952384 | 0.26179633 | -1.7874673 | 0.07386199 | 0.21561173 |
| WDR53 | 110.016362 | 0.035669059 | 0.27442476 | 0.12997755 | 0.89658419 | 0.95230008 |
| FBXO45 | 58.3595167 | 0.494866538 | 0.25391211 | 1.94896787 | 0.05129926 | 0.16898054 |
| PIGX | 173.447549 | -0.048946607 | 0.18771056 | -0.2607557 | 0.79428087 | 0.90358458 |
| PAK2 | 952.245536 | -0.683873808 | 0.14575336 | -4.6919936 | 2.71E-06 | 9.09E-05 |
| SENP5 | 1562.19967 | -0.264991894 | 0.24864239 | -1.0657551 | 0.28653436 | 0.50879744 |
| NCBP2 | 248.628894 | -0.281243419 | 0.24924917 | -1.1283625 | 0.25916686 | 0.48023578 |
| NCBP2-AS2 | 107.68335 | -0.284287417 | 0.51664746 | -0.5502542 | 0.58214505 | 0.76785626 |
| MELTF | 272.376863 | 2.330247259 | 1.32058334 | 1.76455903 | 0.07763791 | 0.22245534 |
| DLG1 | 3504.04681 | -0.2328862 | 0.16117377 | -1.4449386 | 0.14847514 | 0.34031027 |
| BDH1 | 658.674856 | 0.968981759 | 0.51158644 | 1.89407239 | 0.0582154 | 0.18431621 |
| AC024560.1 | 54.4265126 | 1.672808939 | 1.89557478 | 0.88248111 | 0.37751669 | 0.60053769 |
| RUBCN | 413.728135 | -0.263627247 | 0.16286981 | -1.6186379 | 0.10552519 | 0.27245547 |
| FYTTD1 | 508.183644 | -0.140579883 | 0.25421471 | -0.5529966 | 0.58026571 | 0.76631971 |
| LRCH3 | 1672.63361 | -0.088871759 | 0.13448657 | -0.6608225 | 0.50872613 | 0.71131189 |
| IQCG | 296.628441 | -0.157413145 | 0.20519035 | -0.7671567 | 0.44298838 | 0.65884991 |
| RPL35A | 686.128525 | -0.898137045 | 0.41012203 | -2.1899263 | 0.02852958 | 0.11348594 |
| LMLN | 474.340934 | 0.288483272 | 0.2277882 | 1.26645395 | 0.20535061 | 0.41584887 |
| ZNF595 | 395.947088 | 0.420945144 | 0.16728199 | 2.51638058 | 0.01185671 | 0.06247118 |
| ZNF718 | 943.210186 | 0.168573543 | 0.17643734 | 0.95543006 | 0.33936026 | 0.56485001 |
| ZNF141 | 763.04768 | 0.271187555 | 0.13250165 | 2.04667298 | 0.0406902 | 0.14432785 |
| AC092574.2 | 140.932043 | 0.439877006 | 0.49502992 | 0.8885867 | 0.37422524 | 0.59780424 |
| ZNF721 | 2139.92785 | 0.27103278 | 0.17618811 | 1.5383148 | 0.12397166 | 0.30190666 |
| PIGG | 440.791409 | -0.221310229 | 0.20091889 | -1.1014904 | 0.27068328 | 0.49221578 |
| ATP5ME | 258.775558 | 0.315722806 | 0.26456961 | 1.19334496 | 0.2327343 | 0.44950832 |
| SLC49A3 | 55.7437155 | -0.583954857 | 0.35654586 | -1.6378113 | 0.10146105 | 0.26522839 |
| PCGF3 | 676.324 | 0.114633714 | 0.23917968 | 0.47927866 | 0.6317404 | 0.80129239 |
| AC139887.2 | 1394.32134 | 0.373893083 | 0.36842738 | 1.01483521 | 0.31018438 | 0.53418847 |
| GAK | 840.996315 | -0.481346379 | 0.26779159 | -1.7974664 | 0.07226161 | 0.21251609 |
| TMEM175 | 155.474038 | 0.677336154 | 0.1825339 | 3.71074166 | 0.00020665 | 0.00301622 |
| DGKQ | 78.0968674 | -0.475505322 | 0.45045434 | -1.0556127 | 0.29114519 | 0.51373981 |
| IDUA | 135.625291 | 0.251144765 | 0.30496726 | 0.82351387 | 0.41021583 | 0.62984319 |
| FGFRL1 | 124.115639 | -0.333226802 | 0.30289105 | -1.100154 | 0.27126502 | 0.49275402 |
| RNF212 | 525.8986 | 1.71593046 | 0.67705755 | 2.5343938 | 0.01126421 | 0.06034355 |
| AC092535.3 | 1619.64124 | 0.90549627 | 0.51506907 | 1.75800941 | 0.07874591 | 0.22441496 |
| AC092535.1 | 48.9055662 | 1.153955421 | 0.71724246 | 1.60887773 | 0.10764308 | 0.27582104 |
| SPON2 | 134.185465 | 0.312565204 | 0.56495754 | 0.55325433 | 0.58008927 | 0.7661633 |
| CTBP1-AS | 179.959062 | 0.564067746 | 0.46503909 | 1.21294696 | 0.2251501 | 0.4396723 |
| CTBP1 | 445.144339 | -0.010348722 | 0.20237076 | -0.0511374 | 0.95921601 | 0.98229715 |
| CTBP1-DT | 141.182582 | 0.290377641 | 0.25581722 | 1.13509811 | 0.25633421 | 0.47722602 |
| MAEA | 665.770897 | 0.138598485 | 0.15036091 | 0.92177205 | 0.35664749 | 0.58236202 |
| UVSSA | 500.359479 | 0.247578298 | 0.27449016 | 0.90195692 | 0.36707975 | 0.5925402 |
| FAM53A | 63.461393 | 0.206324097 | 0.34912292 | 0.59097838 | 0.55453491 | 0.74735757 |
| SLBP | 418.637221 | -0.209860075 | 0.28820749 | -0.7281562 | 0.46651797 | 0.67903751 |
| TMEM129 | 164.767748 | 0.384289006 | 0.3197134 | 1.20197968 | 0.2293714 | 0.44481325 |
| FGFR3 | 227.498416 | 2.862828439 | 0.65318344 | 4.38288585 | 1.17E-05 | 0.0003032 |
| LETM1 | 356.523761 | 0.107751503 | 0.19130053 | 0.56325772 | 0.57325941 | 0.76109996 |
| NSD2 | 1379.45194 | -0.22669508 | 0.16589852 | -1.3664684 | 0.17179199 | 0.37285581 |
| NELFA | 636.488778 | 0.139123311 | 0.20380788 | 0.68261989 | 0.49484706 | 0.70028229 |

| | | | | | | |
|---|---|---|---|---|---|---|
| C4orf48 | 108.112842 | -1.201514851 | 0.65798447 | -1.8260535 | 0.0678422 | 0.20403635 |
| NAT8L | 187.548347 | 1.833761105 | 0.65276113 | 2.80923757 | 0.0049659 | 0.03322926 |
| POLN | 262.178645 | 0.769690059 | 0.37383109 | 2.05892467 | 0.03950146 | 0.14151742 |
| MXD4 | 526.78885 | -0.328184313 | 0.25768446 | -1.2735898 | 0.20280882 | 0.41232902 |
| ZFYVE28 | 933.830007 | -0.103698461 | 0.26456655 | -0.3919561 | 0.69509068 | 0.84411644 |
| AL158068.2 | 66.0967673 | 1.071237268 | 0.49661341 | 2.15708487 | 0.03099905 | 0.12067978 |
| RNF4 | 582.654893 | -0.058067383 | 0.23474085 | -0.247368 | 0.80462341 | 0.90847684 |
| FAM193A | 1408.18683 | -0.033272468 | 0.17437955 | -0.1908049 | 0.84867847 | 0.9303692 |
| TNIP2 | 162.652349 | -1.159709488 | 0.23154798 | -5.0085061 | 5.49E-07 | 2.55E-05 |
| SH3BP2 | 244.412833 | -2.370734583 | 0.37203371 | -6.3723649 | 1.86E-10 | 2.86E-08 |
| ADD1 | 1331.56456 | -0.067889301 | 0.16147021 | -0.4204447 | 0.67416058 | 0.82909435 |
| MFSD10 | 249.084557 | -0.620536335 | 0.37298687 | -1.6636948 | 0.09617343 | 0.25581048 |
| NOP14-AS1 | 92.4284116 | -0.398387704 | 0.28204435 | -1.4125002 | 0.15780273 | 0.35383118 |
| NOP14 | 167.993304 | -0.124831455 | 0.20578894 | -0.6065994 | 0.54411678 | 0.73944091 |
| GRK4 | 275.773914 | 0.531654238 | 0.3119957 | 1.70404347 | 0.08837297 | 0.24166735 |
| HTT | 1091.30679 | 0.1058365 | 0.17933643 | 0.59015616 | 0.55508596 | 0.74754753 |
| RGS12 | 503.384243 | -0.712842622 | 0.34587801 | -2.0609654 | 0.03930634 | 0.14126492 |
| LRPAP1 | 422.334125 | -0.478981559 | 0.34159663 | -1.4021847 | 0.16086011 | 0.35821031 |
| TMEM128 | 164.623859 | 0.091964213 | 0.20506687 | 0.44845963 | 0.65382151 | 0.81551732 |
| ZBTB49 | 119.471386 | -0.066615395 | 0.26276848 | -0.2535136 | 0.79987132 | 0.90620981 |
| STX18 | 1785.88796 | 0.470330063 | 0.17446151 | 2.69589579 | 0.00701996 | 0.04290916 |
| STX18-AS1 | 841.369159 | 0.760709844 | 0.38019959 | 2.00081712 | 0.0454121 | 0.15560366 |
| STK32B | 3157.75803 | 1.045549441 | 0.44324589 | 2.35884745 | 0.01833179 | 0.08422421 |
| EVC2 | 755.492725 | -0.647780493 | 0.29371377 | -2.2054822 | 0.02742028 | 0.11035784 |
| EVC | 1574.5666 | -0.62662512 | 0.4651563 | -1.3471281 | 0.17793898 | 0.38100113 |
| CRMP1 | 496.66327 | -0.611704592 | 0.69283156 | -0.8829052 | 0.37728749 | 0.60028794 |
| WFS1 | 71.6939994 | 0.287174072 | 0.25475489 | 1.12725639 | 0.2596341 | 0.48061767 |
| MAN2B2 | 160.225707 | -0.253561199 | 0.39338255 | -0.6445665 | 0.51920818 | 0.71956991 |
| MRFAP1 | 327.779801 | -1.219196442 | 0.423652 | -2.8778253 | 0.00400427 | 0.02868001 |
| LINC02482 | 94.5493448 | 0.774971772 | 0.42632261 | 1.81780594 | 0.0690938 | 0.20643745 |
| AC093323.1 | 64.1575086 | 0.151828349 | 0.2792697 | 0.5436621 | 0.58667402 | 0.77046899 |
| MRFAP1L1 | 143.032018 | -0.113858042 | 0.29916846 | -0.3805817 | 0.70351366 | 0.84973591 |
| BLOC1S4 | 88.2132456 | -0.83440673 | 0.38921642 | -2.1438117 | 0.03204798 | 0.12352545 |
| KIAA0232 | 1935.6934 | 0.273992895 | 0.16020481 | 1.71026631 | 0.08721664 | 0.23941076 |
| TBC1D14 | 1732.88525 | -0.226618812 | 0.28609346 | -0.7921146 | 0.42829385 | 0.64670021 |
| TADA2B | 118.927953 | -0.464958848 | 0.30369167 | -1.5310227 | 0.12576377 | 0.30469803 |
| GRPEL1 | 196.953991 | -0.505245566 | 0.3275888 | -1.5423164 | 0.12299674 | 0.30052976 |
| SORCS2 | 880.512007 | 1.038316232 | 0.66943928 | 1.55102377 | 0.12089599 | 0.29724838 |
| AFAP1 | 257.289322 | -0.721399411 | 0.28897758 | -2.4963854 | 0.01254662 | 0.06478823 |
| ABLIM2 | 81.0090124 | -0.197872605 | 0.36015717 | -0.5494063 | 0.58272668 | 0.76826397 |
| SH3TC1 | 244.997399 | -0.834243632 | 0.66567151 | -1.2532362 | 0.21011976 | 0.42219501 |
| ACOX3 | 308.237751 | 0.130473808 | 0.20519868 | 0.63584137 | 0.52487982 | 0.72408747 |
| TRMT44 | 262.600955 | 0.02576076 | 0.19921618 | 0.12931058 | 0.8971119 | 0.9525128 |
| SLC2A9 | 269.267505 | 1.685083504 | 0.5511895 | 3.05717638 | 0.00223433 | 0.01862729 |
| WDR1 | 718.821001 | -0.60557108 | 0.26870673 | -2.2536506 | 0.02421816 | 0.10136174 |
| ZNF518B | 98.1221138 | -0.079828313 | 0.33615288 | -0.2374762 | 0.81228737 | 0.91190752 |
| CLNK | 41.5891405 | 0.256447234 | 0.35423364 | 0.72394941 | 0.4690968 | 0.68108245 |
| HS3ST1 | 562.219729 | -3.099856702 | 0.54696507 | -5.667376 | 1.45E-08 | 1.27E-06 |
| AC005699.1 | 1003.74798 | -0.111558793 | 0.53469921 | -0.2086384 | 0.83473052 | 0.92298305 |
| RAB28 | 1993.6472 | 0.27636199 | 0.16794142 | 1.64558561 | 0.0998491 | 0.26257352 |
| BOD1L1 | 1358.11589 | -0.395943913 | 0.29578592 | -1.3386165 | 0.18069556 | 0.38451679 |
| LINC01182 | 1718.45095 | 0.644442297 | 0.52842402 | 1.21955526 | 0.22263351 | 0.43689884 |
| AC073848.1 | 40.3251354 | 1.144074613 | 0.52306319 | 2.1872589 | 0.02872363 | 0.11400554 |
| AC092546.1 | 252.802757 | 1.744141228 | 0.70190961 | 2.4848516 | 0.01296055 | 0.06636926 |
| LINC00504 | 147.020394 | 0.837702499 | 0.6463166 | 1.29611787 | 0.19493488 | 0.40245308 |
| CPEB2-DT | 234.393469 | 0.363376743 | 0.47459142 | 0.76566226 | 0.44387729 | 0.65959895 |
| CPEB2 | 4404.05896 | 0.514379412 | 0.26528169 | 1.93899325 | 0.05250216 | 0.1718288 |
| AC098829.1 | 1142.60755 | 0.133208382 | 0.42579977 | 0.31284277 | 0.75440012 | 0.88091152 |
| C1QTNF7 | 213.850312 | 2.076908171 | 0.80405015 | 2.58305799 | 0.00979289 | 0.05452336 |
| CC2D2A | 717.065715 | 0.49115082 | 0.29424238 | 1.6692049 | 0.09507678 | 0.25403624 |
| FBXL5 | 1310.64315 | 0.208761779 | 0.15106813 | 1.38190481 | 0.16700093 | 0.36661381 |
| FAM200B | 364.154138 | 0.387542838 | 0.20206558 | 1.91790627 | 0.0551229 | 0.17745999 |
| PROM1 | 2052.06394 | -1.470336355 | 0.44433651 | -3.3090604 | 0.0009361 | 0.00946052 |
| TAPT1 | 1139.42918 | 0.477057678 | 0.18603235 | 2.56438028 | 0.01033602 | 0.05679223 |
| TAPT1-AS1 | 2924.43854 | 1.276933762 | 0.19532362 | 6.53752874 | 6.25E-11 | 1.18E-08 |
| AC097515.1 | 663.944843 | 1.691421103 | 0.47914671 | 3.53006936 | 0.00041545 | 0.00513829 |
| LDB2 | 217.212702 | 1.687100044 | 0.47761816 | 3.53231971 | 0.00041193 | 0.00510431 |
| QDPR | 103.300234 | 0.737918563 | 0.34089302 | 2.16466315 | 0.03041349 | 0.11885556 |
| LAP3 | 320.070048 | 0.079107087 | 0.55982563 | 0.14130665 | 0.88762769 | 0.94928354 |
| AC006160.1 | 80.1922238 | 0.83711598 | 0.51158898 | 1.63630572 | 0.10177562 | 0.26584025 |
| MED28 | 388.607449 | -0.094270808 | 0.13563565 | -0.6950297 | 0.48703667 | 0.69421251 |
| FAM184B | 81.7541166 | -0.363642274 | 0.44122234 | -0.8241701 | 0.4098429 | 0.62963532 |
| DCAF16 | 414.439888 | 0.464970909 | 0.18772433 | 2.47688139 | 0.01325359 | 0.06711401 |
| LCORL | 1648.32599 | 0.324797596 | 0.221131 | 1.46880171 | 0.14188658 | 0.33077653 |
| AC024230.1 | 142.643356 | -0.266937885 | 0.53919074 | -0.4950713 | 0.62054976 | 0.79280799 |
| SLIT2 | 813.633017 | 0.950761884 | 0.59418764 | 1.60010376 | 0.10957557 | 0.27845011 |
| PACRGL | 674.398225 | 0.022895492 | 0.21897362 | 0.10455822 | 0.91672637 | 0.96263921 |
| KCNIP4 | 84786.6426 | -0.745363299 | 0.51226706 | -1.4550287 | 0.14566131 | 0.33584931 |
| AC110296.1 | 97.2564257 | -1.046531984 | 0.76299108 | -1.3716176 | 0.17018252 | 0.37051935 |
| ADGRA3 | 1640.06756 | 0.704911053 | 0.27888581 | 2.52759739 | 0.0114846 | 0.0611231 |
| AC093607.1 | 78.3725903 | 0.161618814 | 0.35241958 | 0.45859771 | 0.64652308 | 0.81100887 |
| PPARGC1A | 7234.8801 | 1.414325405 | 0.38326765 | 3.6901768 | 0.0002241 | 0.00322102 |
| DHX15 | 1849.41665 | 0.089781649 | 0.23275484 | 0.38573483 | 0.69969306 | 0.84744596 |
| SOD3 | 760.357734 | -4.019035991 | 0.88418407 | -4.5454743 | 5.48E-06 | 0.00016416 |
| CCDC149 | 494.364225 | 0.10553574 | 0.28517422 | 0.37007462 | 0.71132689 | 0.8551462 |
| SEPSECS | 420.000228 | 0.311620291 | 0.21595563 | 1.44298293 | 0.14902528 | 0.34109772 |
| AC104662.2 | 137.302129 | 0.8891316 | 0.31103986 | 2.85857771 | 0.00425545 | 0.02983557 |
| PI4K2B | 60.9774377 | 0.060195003 | 0.33959596 | 0.17725477 | 0.85930828 | 0.93620958 |
| ZCCHC4 | 237.319795 | 0.406496798 | 0.26178591 | 1.55278331 | 0.12047491 | 0.29667284 |
| ANAPC4 | 1165.40272 | -0.223790256 | 0.38088159 | -0.5875586 | 0.5568286 | 0.74892058 |
| SLC34A2 | 867.013449 | -4.505300961 | 0.66532062 | -6.7716238 | 1.27E-11 | 3.06E-09 |
| SEL1L3 | 424.622915 | -3.065019021 | 0.60897375 | -5.0330889 | 4.83E-07 | 2.31E-05 |
| SMIM20 | 169.323191 | -0.531754681 | 0.3562705 | -1.4925588 | 0.1355527 | 0.32070852 |
| RBPJ | 1733.21993 | 0.056940402 | 0.17320742 | 0.32874113 | 0.74235137 | 0.87402792 |
| TBC1D19 | 1214.65808 | 0.210770418 | 0.20086813 | 1.04929744 | 0.29404125 | 0.51714864 |
| STIM2 | 836.244873 | -0.466077148 | 0.33466888 | -1.3926516 | 0.16372517 | 0.36237362 |
| PCDH7 | 305.135545 | 0.71479638 | 0.77014408 | 0.92813332 | 0.35333842 | 0.57927854 |
| AC016687.3 | 108.288137 | 4.305269246 | 0.80993893 | 5.31554796 | 1.06E-07 | 6.72E-06 |
| LINC02484 | 364.30876 | 4.451255324 | 0.77196044 | 5.76617026 | 8.11E-09 | 7.71E-07 |
| ARAP2 | 1572.03062 | -0.015225208 | 0.21386161 | -0.0711919 | 0.94324505 | 0.97556917 |
| C4orf19 | 4196.1255 | -0.781160655 | 0.2763219 | -2.8269951 | 0.00469871 | 0.03189373 |
| RELL1 | 611.814784 | -0.280832578 | 0.21341571 | -1.3158946 | 0.18820942 | 0.39384317 |
| PGM2 | 203.817665 | -1.093928295 | 0.32575871 | -3.3580937 | 0.00078482 | 0.00826862 |
| TBC1D1 | 4084.50813 | -0.988159083 | 0.28535734 | -3.462883 | 0.00053442 | 0.00621953 |
| KLF3 | 886.23673 | -1.493184454 | 0.28132291 | -5.3077244 | 1.11E-07 | 6.88E-06 |
| TLR1 | 66.8337184 | -0.31962063 | 0.43063086 | -0.7422149 | 0.45795716 | 0.67250337 |
| FAM114A1 | 336.924415 | -0.343877879 | 0.38183154 | -0.900601 | 0.36780048 | 0.59294486 |
| KLHL5 | 1177.55155 | -0.485219556 | 0.22916972 | -2.1172935 | 0.03423495 | 0.12911907 |
| WDR19 | 921.992469 | -0.373247547 | 0.20319014 | -1.8369373 | 0.06621916 | 0.20049878 |
| RFC1 | 732.165861 | 0.230270929 | 0.17323871 | 1.32921175 | 0.18377812 | 0.38883953 |
| RPL9 | 129.441633 | -0.082272793 | 0.30648346 | -0.2684412 | 0.78835972 | 0.89964955 |
| LIAS | 344.944435 | 0.806176238 | 0.21348266 | 3.77630789 | 0.00015917 | 0.00243625 |
| UGDH | 358.450329 | 0.029288066 | 0.3273289 | 0.08947596 | 0.92870366 | 0.96900671 |

| | | | | | | |
|---|---|---|---|---|---|---|
| UGDH-AS1 | 607.831294 | 0.693786738 | 0.27445041 | 2.52791291 | 0.01147428 | 0.06111443 |
| SMIM14 | 1217.49915 | -0.066547046 | 0.10953894 | -0.6075195 | 0.54350619 | 0.7391242 |
| AC108471.2 | 50.3413154 | 0.874383761 | 0.38679412 | 2.26059216 | 0.02378452 | 0.10024483 |
| UBE2K | 2774.08612 | 0.043595678 | 0.17239875 | 0.25287699 | 0.80036327 | 0.90653391 |
| PDS5A | 2243.66177 | -0.059309537 | 0.16841529 | -0.3521624 | 0.72471645 | 0.86357957 |
| N4BP2 | 644.275038 | 0.286698719 | 0.13618229 | 2.10525701 | 0.03526894 | 0.13148954 |
| RBM47 | 9608.24714 | -0.301955447 | 0.26711277 | -1.1304418 | 0.2582901 | 0.47942539 |
| NSUN7 | 763.624172 | 0.124149259 | 0.21574563 | 0.57544276 | 0.5649919 | 0.75605085 |
| APBB2 | 5283.88109 | 0.555050958 | 0.23993706 | 2.31331898 | 0.02070511 | 0.09130981 |
| AC131953.1 | 41.2019148 | -0.064531973 | 0.41798169 | -0.1543895 | 0.87730265 | 0.94505715 |
| UCHL1 | 769.442112 | -1.130211209 | 0.7217241 | -1.5659879 | 0.11735145 | 0.291975 |
| LIMCH1 | 3208.03834 | 0.283561141 | 0.33437868 | 0.84802398 | 0.39642462 | 0.61829883 |
| TMEM33 | 415.95255 | -0.335414164 | 0.29203343 | -1.1485472 | 0.25074276 | 0.47078342 |
| SLC30A9 | 1617.42375 | -0.011013372 | 0.10791951 | -0.1020517 | 0.91871562 | 0.9638852 |
| AC024022.1 | 824.242917 | 1.71535232 | 0.76713574 | 2.23604799 | 0.02534863 | 0.10446452 |
| SHISA3 | 142.663278 | -0.201205489 | 0.5918871 | -0.339939 | 0.73390249 | 0.86935939 |
| ATP8A1 | 1224.74778 | -0.2472033 | 0.30948431 | -0.7987588 | 0.42443031 | 0.64298076 |
| KCTD8 | 895.066074 | -0.235399089 | 0.59212938 | -0.3975467 | 0.69096435 | 0.84134845 |
| YIPF7 | 47.5558133 | 0.70818333 | 0.34706395 | 2.04049813 | 0.04130074 | 0.14547046 |
| GUF1 | 364.61025 | 0.609544109 | 0.21914644 | 2.78144661 | 0.00541172 | 0.03556391 |
| GNPDA2 | 297.545949 | 0.551562366 | 0.26926996 | 2.04836208 | 0.04052453 | 0.14385611 |
| GABRA2 | 174.340408 | -1.145595345 | 0.76051373 | -1.5063441 | 0.13197887 | 0.31507481 |
| GABRB1 | 489.547506 | -0.023777636 | 0.61035143 | -0.0389573 | 0.96892444 | 0.98586681 |
| COMMD8 | 94.4197505 | -0.141592081 | 0.33670486 | -0.4205228 | 0.67410356 | 0.82909435 |
| ATP10D | 933.689157 | 0.421800203 | 0.23579633 | 1.78883274 | 0.07364176 | 0.21536044 |
| CORIN | 85.8177676 | 0.652489493 | 0.4767081 | 1.36874011 | 0.17108053 | 0.37167807 |
| AC107068.2 | 191.047857 | -0.312834828 | 0.5669157 | -0.551819 | 0.5810724 | 0.76707829 |
| NFXL1 | 604.208781 | 0.247521752 | 0.18790622 | 1.31726217 | 0.18775076 | 0.3934634 |
| NIPAL1 | 330.103496 | -0.066973081 | 0.28263506 | -0.2369596 | 0.81268816 | 0.91196992 |
| CNGA1 | 117.700456 | -1.3477187 | 0.56361886 | -2.3911881 | 0.01679395 | 0.07930914 |
| TXK | 442.721409 | -0.391280574 | 0.40321881 | -0.9703927 | 0.3318508 | 0.55704883 |
| TEC | 511.658721 | -0.587719487 | 0.23371615 | -2.5146721 | 0.01191431 | 0.06262482 |
| SLAIN2 | 947.01826 | 0.247298207 | 0.14811599 | 1.66962535 | 0.09499351 | 0.25400188 |
| FRYL | 2920.51012 | -0.088133155 | 0.13656741 | -0.6453455 | 0.51870334 | 0.71939843 |
| OCIAD1 | 1484.78148 | 0.300312403 | 0.12409285 | 2.42006217 | 0.01551785 | 0.07511005 |
| OCIAD2 | 260.615999 | -0.739030797 | 0.22815876 | -3.2391077 | 0.00119904 | 0.01151029 |
| DCUN1D4 | 1209.5704 | 0.076872438 | 0.15065659 | 0.51024941 | 0.60987674 | 0.78542196 |
| SGCB | 158.236475 | 0.152607087 | 0.24482727 | 0.62332552 | 0.53307063 | 0.73027691 |
| SPATA18 | 44.3282912 | -1.56432656 | 0.58800575 | -2.6603933 | 0.00780495 | 0.04641852 |
| USP46 | 320.768291 | 0.298873496 | 0.21741342 | 1.37467826 | 0.16923121 | 0.36964452 |
| SCFD2 | 1529.96559 | 0.369125928 | 0.34014609 | 1.08519822 | 0.27783387 | 0.49998741 |
| AC058822.1 | 5971.79728 | 0.9110098 | 0.6527813 | 1.39558196 | 0.16284042 | 0.3609485 |
| LNX1 | 4596.82291 | -0.080752856 | 0.37446471 | -0.2156488 | 0.82926153 | 0.92023714 |
| CHIC2 | 1137.10353 | -0.165453333 | 0.18209095 | -0.9086302 | 0.36354537 | 0.58899433 |
| SRD5A3 | 69.0996195 | 0.474386262 | 0.51058663 | 0.92910044 | 0.35283704 | 0.57885329 |
| TMEM165 | 1682.81097 | -0.606598062 | 0.16559929 | -3.6630474 | 0.00024923 | 0.00350282 |
| CLOCK | 1148.28397 | 0.144013314 | 0.19769751 | 0.72845284 | 0.46633643 | 0.67903174 |
| EXOC1 | 1384.90085 | 0.140700053 | 0.20346233 | 0.69152877 | 0.48923331 | 0.69606665 |
| CEP135 | 412.55957 | 0.04276299 | 0.24317789 | 0.17585065 | 0.86041128 | 0.93655498 |
| AASDH | 719.958839 | 0.187288748 | 0.22243856 | 0.84197969 | 0.39979932 | 0.62134844 |
| PPAT | 254.105352 | -0.270950304 | 0.3895017 | -0.6956332 | 0.48665858 | 0.69400145 |
| PAICS | 366.818921 | -0.349206474 | 0.27225061 | -1.2826655 | 0.19960928 | 0.40831558 |
| SRP72 | 541.658766 | -0.074467367 | 0.16106131 | -0.4623542 | 0.64382737 | 0.80931821 |
| REST | 959.536439 | -0.041607791 | 0.19286836 | -0.2157316 | 0.829197 | 0.92023714 |
| NOA1 | 139.080093 | 0.060685491 | 0.23449506 | 0.2587922 | 0.79579558 | 0.90430693 |
| POLR2B | 731.914832 | -0.063666208 | 0.19290618 | -0.3300372 | 0.74137188 | 0.87369479 |
| IGFBP7 | 9523.13073 | -1.159612133 | 0.40529015 | -2.86119 | 0.00422054 | 0.02963999 |
| ADGRL3 | 423.675526 | 0.128250113 | 0.73640281 | 0.17415756 | 0.86174164 | 0.93723716 |
| LINC02232 | 52.7586048 | 1.00085866 | 0.83231928 | 1.20249367 | 0.22917232 | 0.44457881 |
| AC116049.2 | 73.8055687 | 0.161915779 | 0.39109721 | 0.41400392 | 0.67887124 | 0.83168338 |
| AC104806.2 | 217.461801 | 0.207241799 | 0.3841712 | 0.53945168 | 0.58957523 | 0.77174702 |
| CENPC | 883.755397 | 0.466281458 | 0.16986047 | 2.74508512 | 0.00604952 | 0.03860295 |
| UBA6 | 1281.50363 | -0.341646013 | 0.24332822 | -1.4040542 | 0.1603027 | 0.35743389 |
| UBA6-AS1 | 3176.6753 | 0.266932744 | 0.40036303 | 0.66672676 | 0.50494669 | 0.70865287 |
| TMPRSS11D | 64.29443 | 0.121806848 | 0.73274287 | 0.16623409 | 0.86797273 | 0.93997938 |
| YTHDC1 | 1977.81536 | 0.374150357 | 0.19317457 | 1.93685096 | 0.05276357 | 0.17234271 |
| UGT2B7 | 4183.51499 | -0.832155608 | 0.61056259 | -1.3629325 | 0.17290378 | 0.37444497 |
| AC111000.4 | 135.305084 | 0.94152721 | 0.76232226 | 1.23507769 | 0.21680159 | 0.43096046 |
| ENAM | 171.143529 | 0.249957005 | 0.46880017 | 0.53318455 | 0.59390584 | 0.77421753 |
| UTP3 | 125.37083 | -0.622425869 | 0.32041553 | -1.9425584 | 0.05206954 | 0.1708362 |
| RUFY3 | 3139.09083 | -0.80035288 | 0.30587201 | -2.6166267 | 0.00888034 | 0.05073547 |
| GRSF1 | 477.399503 | -0.147453112 | 0.16595957 | -0.8884882 | 0.37427823 | 0.59780424 |
| MOB1B | 1196.14346 | -0.133846091 | 0.24780985 | -0.5401161 | 0.58911696 | 0.77174702 |
| SLC4A4 | 438.240646 | 1.153514354 | 0.45148534 | 2.55493203 | 0.01062085 | 0.05789965 |
| ADAMTS3 | 156.11889 | -1.955747017 | 0.66399939 | -2.9454048 | 0.00322532 | 0.02437541 |
| COX18 | 144.743155 | 0.18324627 | 0.22962923 | 0.79800934 | 0.42486507 | 0.64350908 |
| ANKRD17 | 6561.16528 | 0.370727843 | 0.21054682 | 1.76078575 | 0.07827467 | 0.22355247 |
| AC053527.1 | 1202.23314 | 0.60619753 | 0.44078193 | 1.37527764 | 0.16904538 | 0.36932195 |
| RASSF6 | 490.467698 | -0.024993975 | 0.4815693 | -0.0519011 | 0.9586075 | 0.9819501 |
| MTHFD2L | 765.830752 | -0.325943977 | 0.37599377 | -0.8668866 | 0.38600413 | 0.60936789 |
| BTC | 224.599534 | 0.631652296 | 0.43739173 | 1.44413407 | 0.14870127 | 0.34065124 |
| RCHY1 | 578.193146 | 0.721259239 | 0.25685139 | 2.80807997 | 0.00498378 | 0.03331519 |
| THAP6 | 423.720553 | 0.295875718 | 0.18116709 | 1.63316484 | 0.10243433 | 0.26697087 |
| CDKL2 | 431.307263 | 0.188800438 | 0.41294635 | 0.45720332 | 0.64752491 | 0.81157193 |
| G3BP2 | 624.382 | -0.225040269 | 0.17501089 | -1.2858644 | 0.19849039 | 0.40700957 |
| USO1 | 1641.03517 | 0.352769122 | 0.17584922 | 2.00608866 | 0.04484679 | 0.15433459 |
| NAAA | 411.419623 | 1.095108726 | 0.45818658 | 2.39009344 | 0.01684409 | 0.07944774 |
| SDAD1 | 418.028228 | -0.040387942 | 0.19595042 | -0.2061131 | 0.83670259 | 0.92408065 |
| NUP54 | 745.365326 | 0.398220982 | 0.20651328 | 1.92830687 | 0.05381697 | 0.17453187 |
| SCARB2 | 827.847381 | -0.440073985 | 0.23034313 | -1.9105149 | 0.05606695 | 0.17962366 |
| FAM47E-STB | 59.246863 | -0.044432729 | 0.34237191 | -0.1297791 | 0.89674118 | 0.95232788 |
| SHROOM3 | 5477.98426 | -0.247355744 | 0.33341911 | -0.7418763 | 0.45816227 | 0.67265372 |
| AC107072.1 | 79.5365681 | -1.140679261 | 0.67484372 | -1.6902866 | 0.09097315 | 0.24631864 |
| 11-Sep | 1249.90663 | 0.209178615 | 0.48402315 | 0.43216655 | 0.66562037 | 0.82357564 |
| CCNI | 1732.11227 | -0.430607809 | 0.27805394 | -1.5486485 | 0.12146625 | 0.29822447 |
| CCNG2 | 656.627662 | -0.757849311 | 0.27708667 | -2.7350623 | 0.00623685 | 0.03936097 |
| CNOT6L | 1678.39224 | 0.540955367 | 0.30387896 | 1.78016723 | 0.0750486 | 0.21768048 |
| MRPL1 | 822.6463 | 0.423146692 | 0.18607324 | 2.27408672 | 0.02296077 | 0.09792087 |
| FRAS1 | 4070.37287 | 1.504546975 | 0.52963865 | 2.84070464 | 0.0045014 | 0.03109779 |
| ANXA3 | 272.746474 | -1.173646853 | 0.65621307 | -1.7885149 | 0.07369297 | 0.21540763 |
| BMP2K | 887.939831 | 0.069561857 | 0.27700153 | 0.25112446 | 0.80171789 | 0.90677969 |
| PAQR3 | 166.157245 | 0.237506928 | 0.25883393 | 0.91760354 | 0.35882646 | 0.5841433 |
| LINC01088 | 109.615636 | 0.375259472 | 0.41721142 | 0.8994468 | 0.36841472 | 0.5935133 |
| ANTXR2 | 54.1260916 | -0.341384766 | 0.74966129 | -0.4553853 | 0.64883204 | 0.81203546 |
| CFAP299 | 283.859151 | -0.234371479 | 0.71577781 | -0.3274361 | 0.74333808 | 0.87465031 |
| PRKG2 | 61.9175253 | 2.04474072 | 0.64560777 | 3.16715631 | 0.00153938 | 0.01399423 |
| RASGEF1B | 3721.22569 | 0.798059711 | 0.56433198 | 1.41416707 | 0.15731285 | 0.35297188 |
| HNRNPD | 1756.81356 | -0.043178499 | 0.10247432 | -0.4213592 | 0.6734928 | 0.82886112 |
| AC124016.1 | 78.1867545 | 1.599226698 | 0.35694998 | 4.48025434 | 7.46E-06 | 0.00021087 |
| HNRNPDL | 1363.73688 | 0.303992657 | 0.20724327 | 1.46683971 | 0.14241966 | 0.33149275 |
| ENOPH1 | 264.912893 | -0.290244368 | 0.22364671 | -1.2977807 | 0.19434272 | 0.40190425 |
| TMEM150C | 369.017747 | 0.6396353 | 0.26454442 | 2.41787488 | 0.01561144 | 0.07539728 |
| SCD5 | 1672.04611 | 0.718854813 | 0.27645545 | 2.60025553 | 0.00931544 | 0.05262943 |

| | | | | | | |
|---|---|---|---|---|---|---|
| SEC31A | 2349.95022 | -0.056731482 | 0.13624081 | -0.4164059 | 0.67711299 | 0.83060787 |
| THAP9 | 112.41173 | 0.204057634 | 0.2853534 | 0.71510497 | 0.47454418 | 0.6850551 |
| LIN54 | 372.631814 | -0.225128322 | 0.16801285 | -1.3399471 | 0.18026256 | 0.38404648 |
| COPS4 | 412.360181 | 0.454185883 | 0.16619517 | 2.73284646 | 0.00627896 | 0.03951347 |
| AC114781.2 | 61.7937105 | 0.270995524 | 0.44009832 | 0.61576133 | 0.53805205 | 0.73484533 |
| COQ2 | 94.8903914 | 0.363512902 | 0.35265705 | 1.03078304 | 0.30264257 | 0.52586237 |
| HPSE | 85.244545 | -1.265482488 | 0.6308746 | -2.0059176 | 0.04486503 | 0.15433459 |
| HELQ | 562.170432 | 0.377334507 | 0.18245018 | 2.06815091 | 0.03862583 | 0.13947749 |
| MRPS18C | 207.476237 | 0.209880507 | 0.21871069 | 0.95962619 | 0.33724338 | 0.56231354 |
| ABRAXAS1 | 225.407105 | 0.612860624 | 0.25227071 | 2.42937682 | 0.0151248 | 0.07377531 |
| GPAT3 | 2349.33552 | 2.226495058 | 0.48058605 | 4.63287493 | 3.61E-06 | 0.00011478 |
| CDS1 | 658.934755 | 0.021381638 | 0.21299032 | 0.10038784 | 0.92003642 | 0.96453005 |
| WDFY3 | 5034.78904 | 0.232852983 | 0.21451046 | 1.08550876 | 0.27769638 | 0.4998762 |
| WDFY3-AS1 | 61.502138 | 0.088797695 | 0.65527179 | 0.13551277 | 0.89220647 | 0.95118543 |
| ARHGAP24 | 12617.9106 | 0.657571951 | 0.26483249 | 2.48297311 | 0.01302909 | 0.0665769 |
| MAPK10 | 7040.81042 | 0.575887937 | 0.19286023 | 2.98603784 | 0.00282618 | 0.0221812 |
| AC104827.1 | 150.455922 | -0.20201976 | 0.50313689 | -0.4015205 | 0.68803697 | 0.83940892 |
| PTPN13 | 5101.62269 | 1.046159209 | 0.21009558 | 4.97944409 | 6.38E-07 | 2.87E-05 |
| AC093827.5 | 261.554839 | -0.149175015 | 0.34241071 | -0.4356611 | 0.66308266 | 0.82240928 |
| AFF1 | 7420.79631 | -0.740604135 | 0.19517693 | -3.7945269 | 0.00014793 | 0.00228531 |
| KLHL8 | 713.140033 | -0.163475326 | 0.32248541 | -0.5069232 | 0.61220872 | 0.78684784 |
| HSD17B11 | 172.507697 | -0.775563783 | 0.38426808 | -2.0182883 | 0.04356124 | 0.15106339 |
| NUDT9 | 397.733727 | 0.304525326 | 0.1801672 | 1.69023732 | 0.09098256 | 0.24631864 |
| SPARCL1 | 107.15341 | 0.138517714 | 0.4686995 | 0.29553629 | 0.76758423 | 0.88759696 |
| AC093895.1 | 275.484106 | -0.465713507 | 0.81830004 | -0.5691232 | 0.56927256 | 0.75882548 |
| SPP1 | 54676.147 | -4.267181822 | 0.60262187 | -7.0810272 | 1.43E-12 | 4.50E-10 |
| PKD2 | 3592.76213 | 0.017990117 | 0.22998673 | 0.07822242 | 0.93765113 | 0.97293274 |
| PPM1K | 1167.41799 | 1.561171577 | 0.38618471 | 4.04255149 | 5.29E-05 | 0.00101956 |
| AC107067.1 | 55.2499898 | 2.796597804 | 0.62029907 | 4.50846685 | 6.53E-06 | 0.00018873 |
| HERC6 | 371.903951 | 0.169088718 | 0.82580619 | 0.20475594 | 0.83776282 | 0.92443926 |
| HERC5 | 258.197091 | -0.062141473 | 0.59298123 | -0.104795 | 0.91653847 | 0.96251842 |
| PYURF | 136.075894 | -0.659323554 | 0.34140982 | -1.9311792 | 0.05346089 | 0.17384662 |
| AC098582.1 | 108.095474 | 0.661962863 | 0.47310955 | 1.39917459 | 0.16176063 | 0.35922099 |
| HERC3 | 1237.053 | -0.499936341 | 0.21454467 | -2.3302203 | 0.01979451 | 0.0885424 |
| FAM13A-AS1 | 460.310998 | -0.532053704 | 0.26516088 | -2.0065317 | 0.04479955 | 0.15429683 |
| FAM13A | 3773.62403 | 0.945727304 | 0.34653247 | 2.72911598 | 0.00635044 | 0.03990624 |
| GPRIN3 | 756.949425 | -0.296387595 | 0.52277033 | -0.5669557 | 0.57074431 | 0.75965155 |
| CCSER1 | 31289.809 | 1.036536967 | 0.34695324 | 2.98754084 | 0.00281232 | 0.02208816 |
| AC004054.1 | 71.5973106 | 0.485819063 | 0.5035406 | 0.96480615 | 0.33464192 | 0.55980428 |
| GRID2 | 46.967277 | 0.47946927 | 0.4001385 | 1.19825827 | 0.23081649 | 0.44645701 |
| SMARCAD1 | 1280.83055 | 0.486468179 | 0.14259111 | 3.41163047 | 0.00064576 | 0.00720717 |
| PDLIM5 | 2261.26467 | -0.34652418 | 0.16855232 | -2.055885 | 0.0397936 | 0.14210156 |
| BMPR1B | 99.605337 | -0.142921018 | 0.40373188 | -0.3539998 | 0.72333901 | 0.86257046 |
| UNC5C | 111.087227 | 0.746613058 | 0.78903521 | 0.94623541 | 0.34402853 | 0.56886284 |
| STPG2 | 5690.76324 | 1.583011846 | 0.30913807 | 5.12072766 | 3.04E-07 | 1.54E-05 |
| STPG2-AS1 | 160.379436 | 1.169484556 | 0.54236105 | 2.1562842 | 0.03106148 | 0.12085157 |
| RAP1GDS1 | 1412.92796 | 0.112037782 | 0.16765751 | 0.66825388 | 0.50397155 | 0.70796131 |
| TSPAN5 | 102.855626 | -0.586822254 | 0.42079859 | -1.3945443 | 0.16315331 | 0.36135012 |
| EIF4E | 671.891028 | 0.230481015 | 0.23287709 | 0.98971097 | 0.32231541 | 0.54716158 |
| METAP1 | 353.039473 | 0.055589077 | 0.20158912 | 0.27575435 | 0.78273675 | 0.89633028 |
| ADH5 | 442.860021 | 0.352879223 | 0.2102202 | 1.6786171 | 0.09322669 | 0.25105634 |
| AP002026.1 | 213.453931 | 0.827748164 | 0.57393221 | 1.4422403 | 0.1492346 | 0.34127917 |
| TRMT10A | 118.478439 | 0.64283983 | 0.36091003 | 1.78116367 | 0.07488572 | 0.21730765 |
| MTTP | 40.0766263 | 1.171681856 | 0.55083076 | 2.1271177 | 0.0334103 | 0.12714397 |
| LAMTOR3 | 205.607004 | -0.126770719 | 0.22418679 | -0.5654692 | 0.57175469 | 0.75986595 |
| DNAJB14 | 1203.99389 | 0.009151703 | 0.13388823 | 0.06835331 | 0.94550439 | 0.97640274 |
| H2AFZ | 315.673904 | -1.33707725 | 0.55392118 | -2.4138403 | 0.01578538 | 0.0759596 |
| AC097460.1 | 90.5426062 | 0.651331283 | 0.29023286 | 2.24416795 | 0.0248216 | 0.10320066 |
| DDIT4L | 187.181202 | -0.683533395 | 0.81402267 | -0.8396982 | 0.40107761 | 0.62261536 |
| EMCN | 81.2554905 | 0.651449596 | 0.52817903 | 1.23338785 | 0.21743109 | 0.43128814 |
| PPP3CA | 3225.56813 | -0.105897767 | 0.2710698 | -0.390666 | 0.6960441 | 0.84465287 |
| BANK1 | 164.455467 | 0.367469777 | 0.55317196 | 0.66429574 | 0.50650105 | 0.70992915 |
| AP002075.1 | 148.474417 | 0.231707989 | 0.61194528 | 0.37864168 | 0.70495396 | 0.85073346 |
| SLC39A8 | 163.834499 | 1.250276574 | 0.45632246 | 2.73989709 | 0.00614584 | 0.03899156 |
| NFKB1 | 1293.67622 | -0.137054266 | 0.161899 | -0.8465418 | 0.39725059 | 0.61900236 |
| MANBA | 652.687779 | 0.177333269 | 0.18806807 | 0.94292065 | 0.34572149 | 0.57104354 |
| UBE2D3 | 2825.75949 | -0.122991219 | 0.14059629 | -0.8747828 | 0.3816921 | 0.60460139 |
| AC018797.2 | 114.425289 | 0.014056065 | 0.30612895 | 0.0459155 | 0.9633776 | 0.98378467 |
| CISD2 | 259.595356 | -0.276387957 | 0.27029733 | -1.022533 | 0.30652868 | 0.53024323 |
| SLC9B1 | 178.427539 | 0.694677997 | 0.46579109 | 1.49139391 | 0.13585811 | 0.32114347 |
| SLC9B2 | 89.6850209 | -0.805061707 | 0.31414838 | -2.5626798 | 0.01038678 | 0.05702367 |
| BDH2 | 137.655137 | 0.467617668 | 0.22853319 | 2.04616958 | 0.04073969 | 0.14438667 |
| AC004053.1 | 41.8205427 | 0.822530057 | 0.51363089 | 1.60140302 | 0.10928768 | 0.27793243 |
| AC096577.1 | 44.0296466 | 1.404018337 | 0.77347685 | 1.81520407 | 0.06949255 | 0.20725381 |
| TET2 | 1939.3741 | 0.264329663 | 0.143947 | 1.83629851 | 0.06631352 | 0.20067246 |
| PPA2 | 1984.24714 | 0.126266041 | 0.1944738 | 0.64927018 | 0.51616376 | 0.71746821 |
| ARHGEF38 | 5184.5471 | -1.04868556 | 0.29822065 | -3.5164754 | 0.00043732 | 0.00536353 |
| INTS12 | 764.010026 | -0.450986223 | 0.27994923 | -1.6109572 | 0.10718906 | 0.27497771 |
| GSTCD | 781.525183 | -0.199636676 | 0.2596027 | -0.7690085 | 0.44188828 | 0.65772878 |
| NPNT | 1296.61935 | -0.288755472 | 0.68824643 | -0.4195524 | 0.67481243 | 0.82939913 |
| AC109361.1 | 81.1654925 | -0.038173829 | 0.39421343 | -0.0968354 | 0.92285708 | 0.96580501 |
| TBCK | 3518.53992 | 0.301926631 | 0.12461098 | 2.42295373 | 0.01539489 | 0.07470649 |
| AIMP1 | 428.952896 | -0.180493106 | 0.1589623 | -1.135446 | 0.25618851 | 0.47715651 |
| PAPSS1 | 619.451181 | -0.078536372 | 0.12931823 | -0.6073109 | 0.5436446 | 0.7391242 |
| SGMS2 | 992.174188 | -0.062927723 | 0.29655621 | -0.2121949 | 0.83195497 | 0.92158196 |
| AC096564.1 | 213.672124 | -0.628469599 | 0.46208884 | -1.3600623 | 0.17381022 | 0.37550853 |
| CYP2U1 | 44.6114121 | 0.415416385 | 0.47889857 | 0.86744127 | 0.38570028 | 0.60912422 |
| HADH | 334.127765 | 0.211076661 | 0.26916468 | 0.78419153 | 0.43292774 | 0.65020381 |
| LEF1 | 81.9101901 | -1.077234889 | 0.88648982 | -1.2151689 | 0.22430167 | 0.43906353 |
| RPL34-AS1 | 556.097533 | 0.811131915 | 0.36250959 | 2.2375461 | 0.02525067 | 0.10419747 |
| RPL34 | 697.778797 | -0.751142445 | 0.40546023 | -1.8525675 | 0.06394437 | 0.19619792 |
| OSTC | 317.220984 | -0.023023902 | 0.28092728 | -0.0819568 | 0.93468106 | 0.97183581 |
| COL25A1 | 134.079666 | -0.441061971 | 0.38574556 | -1.1434013 | 0.25287202 | 0.47304724 |
| SEC24B-AS1 | 137.77534 | 0.59400648 | 0.48591609 | 1.22244661 | 0.22153878 | 0.43559181 |
| SEC24B | 1821.60803 | -0.039553224 | 0.1441071 | -0.274471 | 0.78372267 | 0.89671287 |
| MCUB | 286.407989 | -3.389652179 | 0.44745021 | -7.5754847 | 3.58E-14 | 1.89E-11 |
| CASP6 | 325.278556 | -0.544362818 | 0.18021299 | -3.0206636 | 0.00252221 | 0.02035182 |
| PLA2G12A | 479.782615 | -0.042934486 | 0.28414957 | -0.1510982 | 0.87989826 | 0.94636668 |
| AC126283.2 | 164.989414 | -0.61780826 | 0.48912065 | -1.2630999 | 0.20655327 | 0.41739901 |
| CFI | 45.9874647 | -0.471183331 | 0.6030892 | -0.781283 | 0.43463607 | 0.6516581 |
| GAR1 | 158.316636 | 0.278205596 | 0.29144739 | 0.9545654 | 0.33979752 | 0.56521666 |
| RRH | 44.2083311 | 0.224227671 | 0.54032316 | 0.41498808 | 0.67815063 | 0.83118075 |
| LRIT3 | 37.1232975 | 2.288991381 | 0.71606334 | 3.19663256 | 0.00139042 | 0.01295208 |
| EGF | 16888.5984 | 4.012902525 | 0.52434963 | 7.65310458 | 1.96E-14 | 1.18E-11 |
| ELOVL6 | 1227.77996 | -0.061839241 | 0.20035894 | -0.3086423 | 0.75759366 | 0.88292347 |
| ENPEP | 175.521701 | 0.403867793 | 0.5237589 | 0.77109485 | 0.44065071 | 0.6562809 |
| FAM241A | 195.8995 | -0.286113624 | 0.33859858 | -0.8449936 | 0.39811443 | 0.61972127 |
| AP1AR | 311.879289 | -0.400966073 | 0.2925076 | -1.3707885 | 0.17044088 | 0.37070833 |
| TIFA | 62.4384409 | -0.24979439 | 0.40613886 | -0.6150468 | 0.53852384 | 0.73492731 |
| ALPK1 | 1849.16775 | -0.90887434 | 0.25104065 | -3.6204269 | 0.00029412 | 0.00396428 |
| ZGRF1 | 100.038384 | 0.017933084 | 0.53484401 | 0.03352956 | 0.97325229 | 0.98804941 |
| LARP7 | 708.663518 | 0.114997436 | 0.14264847 | 0.80615963 | 0.42015081 | 0.63958123 |
| ANK2 | 4566.99999 | 1.513760697 | 0.60182164 | 2.51529787 | 0.01189319 | 0.06260025 |

| | | | | | | |
|---|---|---|---|---|---|---|
| CAMK2D | 1960.17573 | -0.200453411 | 0.22540049 | -0.8893211 | 0.37383054 | 0.59773969 |
| ARSJ | 464.569654 | -0.38928617 | 0.54042232 | -0.720337 | 0.47131755 | 0.68256042 |
| UGT8 | 3380.05827 | 1.301371303 | 0.25029861 | 5.19927493 | 2.00E-07 | 1.09E-05 |
| TRAM1L1 | 40.8126464 | 0.75214553 | 0.54638378 | 1.37658833 | 0.16863956 | 0.36880132 |
| LINC01378 | 61.1669907 | 3.311404397 | 0.58796601 | 5.63196574 | 1.78E-08 | 1.52E-06 |
| NDST3 | 129.850682 | 1.620578538 | 0.53711694 | 3.01718006 | 0.00255138 | 0.02053699 |
| SNHG8 | 199.852788 | 0.491155676 | 0.48497727 | 1.01273957 | 0.31118457 | 0.53472577 |
| METTL14 | 407.192712 | 0.261234183 | 0.19308657 | 1.35293816 | 0.17607539 | 0.3783991 |
| SEC24D | 609.471941 | -1.086258153 | 0.29617192 | -3.6676609 | 0.00024478 | 0.00345807 |
| USP53 | 4454.39558 | -0.435536865 | 0.3447876 | -1.2632034 | 0.20651609 | 0.41739901 |
| C4orf3 | 408.667928 | -1.471840965 | 0.42389463 | -3.4721859 | 0.00051624 | 0.00606672 |
| PDE5A | 893.364747 | -0.369074736 | 0.30064881 | -1.2275942 | 0.21959934 | 0.43361796 |
| AC108866.1 | 739.681898 | 0.033724153 | 0.53241639 | 0.06334169 | 0.94949442 | 0.97821544 |
| PRDM5 | 576.637975 | 0.359046204 | 0.25599939 | 1.40252756 | 0.16075777 | 0.35808589 |
| ANXA5 | 875.571969 | -0.988885406 | 0.32915914 | -3.0042775 | 0.00266212 | 0.02120868 |
| EXOSC9 | 130.438328 | 0.551647314 | 0.24131451 | 2.2860097 | 0.02225369 | 0.0958372 |
| BBS7 | 231.894702 | -0.07424421 | 0.28388834 | -0.2615261 | 0.79368683 | 0.9031538 |
| KIAA1109 | 3245.55412 | 0.183457793 | 0.1590399 | 1.15353317 | 0.24869161 | 0.46792613 |
| BBS12 | 74.1725618 | -0.262619878 | 0.37271722 | -0.7046089 | 0.48105364 | 0.68948936 |
| FGF2 | 450.180298 | 0.078352404 | 0.32894003 | 0.23819662 | 0.8117286 | 0.91159012 |
| NUDT6 | 160.963145 | 0.574119732 | 0.25758722 | 2.22883621 | 0.0258248 | 0.1058064 |
| SPATA5 | 1611.33006 | 0.167731649 | 0.21377262 | 0.78462642 | 0.43267263 | 0.65015341 |
| SPRY1 | 234.277914 | -1.240294097 | 0.45980304 | -2.6974465 | 0.00698735 | 0.0427692 |
| LINC01091 | 607.745133 | 0.044583862 | 0.75819133 | 0.05880292 | 0.95310909 | 0.97941614 |
| ANKRD50 | 171.608127 | -0.016620229 | 0.40097751 | -0.0414493 | 0.96693773 | 0.9851354 |
| FAT4 | 454.499295 | 2.182993071 | 0.53314009 | 4.09459562 | 4.23E-05 | 0.00086473 |
| INTU | 719.423037 | 0.388774492 | 0.28479932 | 1.36508224 | 0.1722272 | 0.37349354 |
| HSPA4L | 686.721292 | 0.109405993 | 0.447162 | 0.24466746 | 0.80671392 | 0.90951555 |
| MFSD8 | 580.713054 | 0.152195384 | 0.21922775 | 0.69423412 | 0.4875354 | 0.69477345 |
| ABHD18 | 1500.28259 | 0.062641482 | 0.21334341 | 0.29361808 | 0.76904976 | 0.88859152 |
| LARP1B | 1586.90081 | 0.087674341 | 0.21860592 | 0.40106115 | 0.68837511 | 0.83951129 |
| PGRMC2 | 498.147084 | -0.060493705 | 0.20711571 | -0.2920769 | 0.77022786 | 0.88929655 |
| AC108062.1 | 632.428256 | 0.570170147 | 0.25012462 | 2.27954425 | 0.02263473 | 0.0968398 |
| AC078850.2 | 132.742435 | 1.477465011 | 0.76994497 | 1.91892288 | 0.0549941 | 0.17713168 |
| JADE1 | 1156.41112 | -0.500366054 | 0.36053141 | -1.387857 | 0.16518059 | 0.36449555 |
| SCLT1 | 1372.15099 | 0.356577373 | 0.25722965 | 1.38622192 | 0.16567915 | 0.36504686 |
| C4orf33 | 189.065963 | 0.76618793 | 0.3108873 | 2.46451989 | 0.01371969 | 0.06872334 |
| AC096711.2 | 624.467886 | 0.256345999 | 0.30448134 | 0.84191038 | 0.39983812 | 0.62134844 |
| AC018680.1 | 68.1049491 | 1.081309873 | 0.59323723 | 1.8227276 | 0.06834466 | 0.205 |
| AC109927.1 | 173.250699 | -0.635626571 | 0.40155308 | -1.5829204 | 0.1134396 | 0.28563167 |
| NOCT | 112.980182 | -0.390187892 | 0.34729759 | -1.1234973 | 0.26122636 | 0.48232304 |
| ELF2 | 3912.3093 | -0.60156099 | 0.21459071 | -2.8032946 | 0.00505834 | 0.03369423 |
| NDUFC1 | 409.447917 | 0.092629508 | 0.30251123 | 0.30620188 | 0.75945094 | 0.88349992 |
| NAA15 | 1213.38567 | 0.1560682 | 0.2231293 | 0.69945183 | 0.4842697 | 0.69235581 |
| AC097376.2 | 1494.64513 | -0.590713973 | 0.25665901 | -2.3015517 | 0.02136047 | 0.0934273 |
| RAB33B | 92.0199424 | 0.34789843 | 0.24919346 | 1.39609777 | 0.16268505 | 0.36088961 |
| SETD7 | 287.763742 | -0.695157962 | 0.31570666 | -2.201911 | 0.0276716 | 0.1109973 |
| MGST2 | 376.57506 | 0.147032218 | 0.21075607 | 0.69764168 | 0.48540131 | 0.69300247 |
| MAML3 | 2767.24972 | -0.715255748 | 0.34034021 | -2.1015905 | 0.03558916 | 0.13239418 |
| SCOC | 701.431733 | -0.34355665 | 0.17811872 | -1.9288071 | 0.05375481 | 0.17448302 |
| SCOC-AS1 | 146.640919 | 0.261773908 | 0.43077574 | 0.60768025 | 0.54339956 | 0.7391242 |
| ELMOD2 | 344.817982 | 0.237386033 | 0.15463873 | 1.53510073 | 0.12475909 | 0.30309781 |
| TBC1D9 | 1064.85322 | 0.500748814 | 0.35396661 | 1.41467809 | 0.1571629 | 0.35273406 |
| RNF150 | 4828.75911 | 1.114135929 | 0.32006825 | 3.48093233 | 0.00049967 | 0.00591413 |
| ZNF330 | 272.930841 | 0.353941694 | 0.19466519 | 1.8182074 | 0.06903244 | 0.20630079 |
| LINC02432 | 478.783481 | 1.856982463 | 0.47165195 | 3.93718818 | 8.24E-05 | 0.00145209 |
| IL15 | 689.766791 | 1.101691587 | 0.285076 | 3.86455392 | 0.00011129 | 0.00181938 |
| INPP4B | 3555.78298 | 1.32811345 | 0.60547885 | 2.19349273 | 0.0282719 | 0.11272064 |
| AC139720.1 | 99.4635453 | 2.579124382 | 0.67603726 | 3.81506247 | 0.00013615 | 0.00214349 |
| AC104596.1 | 626.778894 | 1.566449464 | 0.64516493 | 2.42798296 | 0.01518306 | 0.0740048 |
| USP38 | 331.481723 | -0.00598861 | 0.22993682 | -0.0260446 | 0.97922177 | 0.9910179 |
| GAB1 | 2633.50267 | -0.323727908 | 0.17198406 | -1.8823135 | 0.05979347 | 0.18751469 |
| SMARCA5 | 781.232984 | -0.14841178 | 0.16296004 | -0.910725 | 0.36244031 | 0.58778073 |
| HHIP | 48.0945856 | -0.05284928 | 0.63510819 | -0.083213 | 0.93368215 | 0.97167284 |
| ANAPC10 | 1125.85607 | 0.300908515 | 0.14549487 | 2.06817264 | 0.03862379 | 0.13947749 |
| ABCE1 | 345.877819 | 0.214589145 | 0.20654195 | 1.03896157 | 0.29882261 | 0.5219024 |
| OTUD4 | 969.911514 | -0.113613342 | 0.10344273 | -1.0983212 | 0.27206425 | 0.49354664 |
| SMAD1 | 821.602907 | -0.56953779 | 0.13457768 | -4.2320376 | 2.32E-05 | 0.00053479 |
| MMAA | 224.725072 | -0.249588247 | 0.20506853 | -1.2170968 | 0.22356739 | 0.43821184 |
| ZNF827 | 1751.96608 | 0.311462477 | 0.35214875 | 0.88446283 | 0.37644642 | 0.59959975 |
| LSM6 | 202.005267 | 0.31480695 | 0.2186982 | 1.43945836 | 0.1500207 | 0.34219019 |
| SLC10A7 | 1115.28435 | 0.080649643 | 0.22764424 | 0.35427931 | 0.72312957 | 0.86248065 |
| TMEM184C | 492.672677 | 0.478890212 | 0.28583721 | 1.67539494 | 0.09385677 | 0.25193426 |
| PRMT9 | 521.634706 | 0.394236837 | 0.20217252 | 1.95000214 | 0.05117586 | 0.16874238 |
| ARHGAP10 | 542.728143 | -1.169468263 | 0.51738155 | -2.2603594 | 0.02379895 | 0.10024483 |
| NR3C2 | 10353.0557 | 0.450871506 | 0.34258175 | 1.31609902 | 0.18814081 | 0.39384317 |
| AC069272.1 | 108.986285 | -0.218370374 | 0.36264742 | -0.6021561 | 0.5470702 | 0.74124838 |
| DCLK2 | 764.326769 | 1.151800518 | 0.40883637 | 2.81726532 | 0.00484345 | 0.03264139 |
| LRBA | 14033.9613 | 0.719479454 | 0.16409507 | 4.38452813 | 1.16E-05 | 0.00030165 |
| RPS3A | 493.28327 | -0.650030065 | 0.35931465 | -1.8090831 | 0.07043809 | 0.20912191 |
| SH3D19 | 2253.20596 | -0.609005106 | 0.25733958 | -2.3665427 | 0.0179551 | 0.08311891 |
| PRSS48 | 93.9418493 | -1.080783656 | 0.60590225 | -1.7837591 | 0.07446282 | 0.21663404 |
| FAM160A1 | 3652.73151 | 0.093163007 | 0.23839939 | 0.39078543 | 0.69595585 | 0.84465287 |
| AC092611.2 | 53.4562136 | 0.353747643 | 0.29426681 | 1.2021323 | 0.22931227 | 0.44478499 |
| GATB | 274.442974 | 0.290608076 | 0.26895344 | 1.08051444 | 0.27991316 | 0.50229221 |
| FBXW7 | 2946.06449 | -0.057325242 | 0.2178577 | -0.2631316 | 0.79244916 | 0.90221177 |
| TMEM154 | 65.2745507 | -4.073964379 | 0.49547726 | -8.2223035 | 2.00E-16 | 2.02E-13 |
| TIGD4 | 75.802465 | 0.581565382 | 0.28628907 | 2.03139221 | 0.04221522 | 0.14747974 |
| ARFIP1 | 1740.6441 | 0.045254405 | 0.15684331 | 0.28853258 | 0.7729391 | 0.89074912 |
| FHDC1 | 146.139395 | -0.889643421 | 0.35712669 | -2.4911143 | 0.01273431 | 0.06557701 |
| TRIM2 | 5660.60203 | 0.207923632 | 0.19341986 | 1.07498596 | 0.28238101 | 0.50480049 |
| MND1 | 135.26363 | 0.213366357 | 0.74314513 | 0.28711264 | 0.77402608 | 0.89176716 |
| TMEM131L | 446.085307 | 0.625688087 | 0.22257719 | 2.81110603 | 0.00493715 | 0.03310204 |
| TLR2 | 1073.2145 | -2.052039633 | 0.68938799 | -2.9766107 | 0.00291454 | 0.02262834 |
| PLRG1 | 294.292486 | 0.550320586 | 0.22043099 | 2.49656629 | 0.01254022 | 0.06478562 |
| FGB | 91.5020639 | -5.205410916 | 1.23625 | -4.2106458 | 2.55E-05 | 0.00057643 |
| MAP9 | 404.62446 | 0.224858304 | 0.18989219 | 1.18413668 | 0.23635896 | 0.45392878 |
| GUCY1A1 | 2108.58759 | 0.498140885 | 0.26157936 | 1.90435853 | 0.05686351 | 0.18138824 |
| GUCY1B1 | 167.464472 | -1.078289339 | 0.52458442 | -2.0555116 | 0.03982962 | 0.14211692 |
| TDO2 | 62.5642475 | 2.464448166 | 0.61362205 | 4.01623144 | 5.91E-05 | 0.00111455 |
| CTSO | 323.064317 | 0.057719315 | 0.30678825 | 0.18814057 | 0.85076646 | 0.93159033 |
| PDGFC | 1606.40375 | 0.07391983 | 0.1858205 | 0.39780234 | 0.69077589 | 0.84134845 |
| GLRB | 257.135865 | 1.088517941 | 0.28444697 | 3.82678694 | 0.00012983 | 0.00207111 |
| TMEM144 | 105.534099 | -0.820338534 | 0.47549751 | -1.7252215 | 0.08448756 | 0.23425612 |
| ETFDH | 1049.82165 | 1.182226747 | 0.28463327 | 4.15350868 | 3.27E-05 | 0.00070323 |
| PPID | 303.137804 | -0.054480998 | 0.24785807 | -0.2198072 | 0.82602129 | 0.91938879 |
| FNIP2 | 4998.10169 | -0.707842191 | 0.47918001 | -1.4771947 | 0.13962345 | 0.32671144 |
| C4orf45 | 236.004936 | -0.782039541 | 0.70749702 | -1.1053609 | 0.26900324 | 0.49051129 |
| RAPGEF2 | 3183.03222 | -0.189248087 | 0.19241056 | -0.9835639 | 0.32532997 | 0.55051032 |
| FSTL5 | 43.6178804 | 0.840434824 | 0.46138112 | 1.82156311 | 0.0685213 | 0.20537733 |
| NAF1 | 376.027205 | 0.041328331 | 0.20607679 | 0.20054821 | 0.84105186 | 0.92640544 |
| NPY1R | 435.717738 | 0.578092419 | 0.65475208 | 0.88291804 | 0.37728056 | 0.60028794 |
| TMA16 | 245.690991 | 0.031696303 | 0.23924506 | 0.13248467 | 0.89460095 | 0.95172874 |

| | | | | | | |
|---|---|---|---|---|---|---|
| 1-Mar | 92.2475214 | 0.270157119 | 0.30170931 | 0.8954219 | 0.3705616 | 0.59532317 |
| TMEM192 | 477.489987 | -0.197864285 | 0.34452692 | -0.5743072 | 0.56575995 | 0.75623591 |
| KLHL2 | 2295.68659 | -1.914000138 | 0.46412673 | -4.123874 | 3.73E-05 | 0.00077865 |
| MSMO1 | 444.559365 | -1.007083182 | 0.53223459 | -1.8921791 | 0.05846712 | 0.18461291 |
| CPE | 103.949873 | 0.434068813 | 0.94202297 | 0.46078368 | 0.64495381 | 0.8100404 |
| DDX60 | 577.606263 | -0.051836584 | 0.91089325 | -0.0569074 | 0.95461894 | 0.98007893 |
| DDX60L | 452.416461 | -0.292203135 | 0.89044632 | -0.3281536 | 0.74279556 | 0.87441041 |
| PALLD | 267.934559 | -0.327343099 | 0.31467376 | -1.0402618 | 0.29821828 | 0.52146747 |
| CBR4 | 2677.53076 | 0.465440758 | 0.17373141 | 2.67908245 | 0.00738242 | 0.04444595 |
| SH3RF1 | 3197.13442 | -0.000537476 | 0.23861431 | -0.0022525 | 0.99820278 | 0.99948912 |
| AC096741.1 | 85.4255029 | -0.152286231 | 0.43632538 | -0.3490199 | 0.72707439 | 0.8653741 |
| NEK1 | 1795.55156 | 0.601860265 | 0.12515868 | 4.80877762 | 1.52E-06 | 5.68E-05 |
| CLCN3 | 1564.92255 | 0.00183684 | 0.21769997 | 0.00843748 | 0.99326794 | 0.99737436 |
| HPF1 | 243.405593 | 0.265682463 | 0.21695264 | 1.22461043 | 0.22072204 | 0.43450333 |
| LINC02275 | 128.245721 | 2.428570136 | 0.50391675 | 4.81938764 | 1.44E-06 | 5.47E-05 |
| MFAP3L | 805.96929 | 1.057018414 | 0.36299948 | 2.91190065 | 0.00359237 | 0.02650927 |
| LINC01612 | 71.610653 | 0.259021907 | 0.38007194 | 0.68150757 | 0.49555038 | 0.70097718 |
| LINC02512 | 69.3505531 | 0.004338176 | 0.48530631 | 0.00893905 | 0.99286777 | 0.99724662 |
| GALNTL6 | 49.7084456 | 0.103351616 | 0.40280274 | 0.25658121 | 0.79750208 | 0.90505915 |
| AC105285.1 | 59.8382571 | -0.438108054 | 0.3806472 | -1.1509557 | 0.24975047 | 0.46940982 |
| GALNT7 | 1138.15189 | -0.77342654 | 0.28291375 | -2.7337892 | 0.00626101 | 0.03944087 |
| HMGB2 | 231.646216 | -0.301790204 | 0.29024515 | -1.0397769 | 0.29844356 | 0.52159011 |
| SAP30 | 74.9002006 | -1.148574925 | 0.46929973 | -2.447423 | 0.01438819 | 0.07115446 |
| FBXO8 | 227.680372 | 0.28188654 | 0.21982378 | 1.28232959 | 0.19972705 | 0.40838926 |
| CEP44 | 506.342516 | 0.428639606 | 0.17314087 | 2.47566965 | 0.01329866 | 0.06727765 |
| GPM6A | 51.1562253 | 0.674556054 | 0.37438628 | 1.80176488 | 0.07158241 | 0.21105625 |
| WDR17 | 400.116094 | 1.178961759 | 0.3527635 | 3.3420741 | 0.00083155 | 0.00867226 |
| AC019163.1 | 50.8870231 | 1.494355813 | 0.39666487 | 3.76730066 | 0.00016502 | 0.00251063 |
| SPCS3 | 553.058631 | 0.286090061 | 0.24581074 | 1.16386315 | 0.24447948 | 0.46332521 |
| AGA | 87.3924594 | 0.606753881 | 0.3130082 | 1.93846001 | 0.05256712 | 0.1719562 |
| AC078881.1 | 602.660641 | 0.491954425 | 0.42892651 | 1.14694339 | 0.25140502 | 0.47183985 |
| TENM3 | 332.671186 | 0.311850984 | 0.38537752 | 0.80920907 | 0.4183949 | 0.63753323 |
| DCTD | 232.304315 | -0.528802065 | 0.19651407 | -2.6909119 | 0.0071257 | 0.04327512 |
| WWC2 | 4767.2367 | 0.570045207 | 0.15857132 | 3.59488213 | 0.00032454 | 0.00425703 |
| CDKN2AIP | 117.069409 | 0.003050913 | 0.28061897 | 0.01087208 | 0.9913255 | 0.99653109 |
| AC107214.1 | 96.8436203 | -0.110371322 | 0.35909208 | -0.3073622 | 0.75856772 | 0.88316847 |
| ING2 | 77.0158195 | -0.815389766 | 0.39580345 | -2.0600876 | 0.03939017 | 0.14131036 |
| RWDD4 | 211.989076 | 0.081647871 | 0.15652568 | 0.52162604 | 0.60193073 | 0.77940904 |
| TRAPPC11 | 727.156881 | 0.148109031 | 0.20736387 | 0.71424704 | 0.47507444 | 0.68526616 |
| STOX2 | 3636.40305 | -0.457398602 | 0.39902499 | -1.1462906 | 0.25167492 | 0.47199621 |
| IRF2 | 719.108951 | -0.593022782 | 0.19611501 | -3.0238521 | 0.00249578 | 0.02020026 |
| LINC02427 | 141.588431 | 1.556702068 | 0.56841223 | 2.73868504 | 0.00616854 | 0.03903576 |
| CASP3 | 276.370486 | -0.556687005 | 0.3795799 | -1.4665872 | 0.14248839 | 0.33153588 |
| PRIMPOL | 355.033569 | 0.571014317 | 0.28446407 | 2.00733371 | 0.04471414 | 0.15409056 |
| ACSL1 | 2316.24252 | 1.129577603 | 0.37943976 | 2.97696162 | 0.0029112 | 0.02262006 |
| AC084871.2 | 44.1698781 | 0.722775277 | 0.6016132 | 1.2013953 | 0.2295979 | 0.44513201 |
| LINC02437 | 210.357683 | 1.738341092 | 0.6832499 | 2.54422444 | 0.01095207 | 0.0591753 |
| LINC02436 | 46.9697997 | 2.057848618 | 0.65310789 | 3.15085557 | 0.00162793 | 0.01464804 |
| SLC25A4 | 581.393086 | 0.476548305 | 0.43828948 | 1.08729122 | 0.27690813 | 0.49900129 |
| CFAP97 | 479.288086 | -0.558539461 | 0.22121859 | -2.5248305 | 0.01157541 | 0.06143013 |
| SNX25 | 1616.4384 | 0.034247288 | 0.34041181 | 0.10060547 | 0.91986366 | 0.96447683 |
| LRP2BP | 241.443312 | 0.134870143 | 0.40448857 | 0.33343375 | 0.73880689 | 0.87211464 |
| AC112722.1 | 244.882419 | 0.153367712 | 0.41943284 | 0.365655 | 0.71462253 | 0.85752192 |
| ANKRD37 | 294.346764 | -1.068729919 | 0.7832233 | -1.3645277 | 0.17240153 | 0.37368753 |
| UFSP2 | 501.182366 | 0.206247471 | 0.18965313 | 1.0874984 | 0.27681661 | 0.49897002 |
| C4orf47 | 751.169204 | -0.621131032 | 0.60158711 | -1.0324873 | 0.30184391 | 0.52503327 |
| CCDC110 | 57.7437938 | -0.006874513 | 0.38923722 | -0.0176615 | 0.98590889 | 0.99426405 |
| SORBS2 | 2604.54304 | 0.245325289 | 0.46540676 | 0.52712017 | 0.59811014 | 0.77678206 |
| TLR3 | 97.1195307 | -0.198698804 | 0.30282378 | -0.6561532 | 0.51172554 | 0.71384334 |
| FAM149A | 481.494316 | -0.208339516 | 0.30125322 | -0.6915761 | 0.4892036 | 0.69606665 |
| CYP4V2 | 448.922394 | 0.476806914 | 0.25826307 | 1.84620635 | 0.06486225 | 0.19777071 |
| KLKB1 | 477.542282 | 0.923181176 | 0.38190092 | 2.41733161 | 0.01563477 | 0.0754676 |
| F11 | 582.498466 | 1.882195449 | 0.53589034 | 3.51227727 | 0.00044428 | 0.00541766 |
| F11-AS1 | 105.280306 | -0.398594419 | 0.7275587 | -0.5478519 | 0.5837936 | 0.76885817 |
| AC018709.1 | 56.2874988 | -2.878321892 | 0.81653026 | -3.5250646 | 0.00042338 | 0.00521195 |
| FAT1 | 1129.08764 | -0.109379405 | 0.21745623 | -0.502995 | 0.61496776 | 0.78834522 |
| LINC01060 | 97.6145182 | -0.883313819 | 0.69575931 | -1.2695681 | 0.20423852 | 0.41414152 |
| FRG1-DT | 652.523408 | 0.874858576 | 0.55435367 | 1.57815961 | 0.11452895 | 0.28749769 |
| FRG1 | 192.51464 | 0.060895376 | 0.23429192 | 0.2599124 | 0.79493134 | 0.90385363 |
| PLEKHG4B | 203.197946 | 2.25521575 | 1.40119774 | 1.60949143 | 0.10750893 | 0.27558421 |
| CCDC127 | 89.5408453 | -0.103918128 | 0.30116184 | -0.3450574 | 0.7300512 | 0.86751731 |
| SDHA | 909.745351 | 0.216471259 | 0.13042882 | 1.65968888 | 0.09697706 | 0.25732623 |
| PDCD6 | 493.347308 | -0.265791907 | 0.2515699 | -1.056533 | 0.29072476 | 0.51351761 |
| AHRR | 48.5634292 | -0.123706329 | 0.685457 | -0.1804728 | 0.85678142 | 0.93522187 |
| EXOC3-AS1 | 47.0865768 | 0.22516518 | 0.31954678 | 0.70463917 | 0.48103481 | 0.68948936 |
| EXOC3 | 723.905421 | -0.228307881 | 0.19654936 | -1.1615804 | 0.24540595 | 0.46447446 |
| SLC9A3 | 1542.485 | 0.894092933 | 0.46889521 | 1.90680756 | 0.05654551 | 0.18067481 |
| CEP72 | 65.8480937 | -0.124807057 | 0.31418247 | -0.3972439 | 0.69118764 | 0.84152826 |
| TPPP | 81.0603319 | 0.192552857 | 0.7869388 | 0.24468594 | 0.80669962 | 0.90951555 |
| ZDHHC11B | 212.149211 | 0.059271316 | 0.78984681 | 0.07504153 | 0.94018167 | 0.97411372 |
| ZDHHC11 | 64.5346311 | -0.971706056 | 0.56288883 | -1.7262841 | 0.08429631 | 0.23395814 |
| BRD9 | 243.431667 | -0.756378186 | 0.22500639 | -3.3615853 | 0.00077496 | 0.00822209 |
| SLC12A7 | 254.041763 | -0.30881684 | 0.29596789 | -1.0434133 | 0.2967569 | 0.51998178 |
| CLPTM1L | 258.680209 | -0.499678908 | 0.26157273 | -1.9102867 | 0.05609631 | 0.17967388 |
| LPCAT1 | 417.446582 | -0.557108325 | 0.3580596 | -1.5559095 | 0.11972962 | 0.29536727 |
| MRPL36 | 296.861032 | -1.051961884 | 0.6561972 | -1.6031185 | 0.10890849 | 0.27734186 |
| NDUFS6 | 664.035067 | -0.408646271 | 0.30365282 | -1.3457681 | 0.17837732 | 0.38168976 |
| AC126768.2 | 71.1817803 | -0.022119811 | 0.46855962 | -0.0472081 | 0.96234737 | 0.9832997 |
| IRX2 | 1256.87285 | -0.136296222 | 0.41708568 | -0.3267823 | 0.74383254 | 0.87470713 |
| C5orf38 | 214.573373 | -0.752646207 | 0.42287032 | -1.7798511 | 0.07510033 | 0.21773492 |
| IRX1 | 652.128319 | -0.454445282 | 0.38837745 | -1.1701124 | 0.24195573 | 0.46023859 |
| AC106799.2 | 47.3751845 | -0.22869326 | 0.41983944 | -0.544716 | 0.58594886 | 0.77001836 |
| AC022424.1 | 89.6472261 | 1.447753848 | 0.63399154 | 2.28355388 | 0.02239776 | 0.09618077 |
| ADAMTS16 | 679.181735 | 1.911062423 | 0.56182872 | 3.40150363 | 0.00067016 | 0.00741388 |
| AC022424.2 | 48.6157403 | 1.823634426 | 0.63197534 | 2.88561012 | 0.00390656 | 0.02813616 |
| ICE1 | 1119.19596 | 0.361018822 | 0.16257855 | 2.22058093 | 0.02637936 | 0.10751155 |
| LINC02145 | 66.7726909 | 0.181492974 | 0.4764033 | 0.38096498 | 0.70322924 | 0.84970316 |
| MED10 | 232.124361 | -0.123403551 | 0.17791005 | -0.6936289 | 0.48791497 | 0.69502858 |
| NSUN2 | 455.133921 | 0.311753859 | 0.22274816 | 1.39957996 | 0.16163914 | 0.35921571 |
| SRD5A1 | 177.210595 | 0.544758286 | 0.33487794 | 1.62673683 | 0.103793 | 0.26945789 |
| TENT4A | 726.420487 | 0.034045377 | 0.19608593 | 0.17362478 | 0.86216035 | 0.93753817 |
| ADCY2 | 37.8271402 | 0.109380403 | 0.3858701 | 0.28346431 | 0.77682093 | 0.89337196 |
| MTRR | 423.10318 | -0.054414443 | 0.21636043 | -0.2514991 | 0.80142829 | 0.90672945 |
| FASTKD3 | 80.1302942 | 0.585365262 | 0.23582926 | 2.48215711 | 0.01305897 | 0.06664086 |
| MIR4458HG | 86.2548718 | 0.368398779 | 0.43085421 | 0.85504277 | 0.39252747 | 0.61484815 |
| SEMA5A | 338.708177 | -0.164900825 | 0.63769949 | -0.258587 | 0.79595388 | 0.90440668 |
| SNHG18 | 74.3472684 | -0.71877457 | 0.42187148 | -1.7037761 | 0.08842292 | 0.24166735 |
| FAM173B | 354.501118 | 0.17362979 | 0.21112212 | 0.82241402 | 0.4108413 | 0.63014283 |
| CCT5 | 386.770589 | -0.668531798 | 0.29381969 | -2.2753131 | 0.02288715 | 0.09777374 |
| CMBL | 388.44469 | -0.102816752 | 0.4535045 | -0.2267161 | 0.82064454 | 0.91677045 |
| 6-Mar | 2780.92443 | 0.152124284 | 0.16492834 | 0.92236597 | 0.35633772 | 0.58195659 |
| ANKRD33B | 261.371839 | -1.301712316 | 0.59105755 | -2.2023444 | 0.02764099 | 0.1109082 |
| DAP | 1316.25011 | -0.672072299 | 0.29549371 | -2.2744047 | 0.02294166 | 0.09791161 |

| | | | | | | |
|---|---|---|---|---|---|---|
| DNAH5 | 3219.98365 | -3.209585206 | 0.73554489 | -4.3635477 | 1.28E-05 | 0.00032382 |
| AC016576.1 | 183.014467 | -2.803491298 | 0.70272927 | -3.9894329 | 6.62E-05 | 0.00121339 |
| TRIO | 7392.8602 | -1.160381941 | 0.25759724 | -4.5046365 | 6.65E-06 | 0.0001905 |
| OTULINL | 98.0026585 | 1.315214101 | 0.51801416 | 2.53895398 | 0.01111844 | 0.05982365 |
| OTULIN | 407.423978 | -0.087976022 | 0.17474937 | -0.5034411 | 0.61465416 | 0.78816802 |
| ANKH | 778.783702 | 0.616473971 | 0.3435653 | 1.79434295 | 0.07275845 | 0.21355982 |
| FBXL7 | 524.010402 | 1.348205116 | 0.72235753 | 1.86639587 | 0.061986 | 0.1923357 |
| ZNF622 | 82.0278519 | -0.436108397 | 0.27097899 | -1.6093809 | 0.10753309 | 0.27559266 |
| RETREG1 | 2130.2054 | -3.187994596 | 0.62422198 | -5.1071489 | 3.27E-07 | 1.64E-05 |
| MYO10 | 4223.14953 | -0.080189479 | 0.29735738 | -0.2696737 | 0.78741126 | 0.89956761 |
| CDH18 | 181.872394 | 3.796184645 | 0.76997436 | 4.93027413 | 8.21E-07 | 3.57E-05 |
| LINC02241 | 504.405144 | 0.457603475 | 0.25097122 | 1.82333051 | 0.06825335 | 0.2048531 |
| AC093274.1 | 73.6773215 | 0.881140684 | 0.39215476 | 2.24692078 | 0.02464509 | 0.10269306 |
| CDH12 | 235.74593 | 1.033934621 | 0.43558938 | 2.37364514 | 0.01761347 | 0.0818924 |
| C5orf17 | 536.201623 | -0.118380866 | 0.57656587 | -0.2053206 | 0.83732163 | 0.9242234 |
| CDH9 | 104.286562 | 0.116899591 | 0.68644151 | 0.17029796 | 0.86477581 | 0.93837375 |
| CDH6 | 159.603781 | -0.980167968 | 0.81187906 | -1.2072832 | 0.22732309 | 0.44274708 |
| DROSHA | 800.282072 | 0.12737067 | 0.18905755 | 0.67371374 | 0.50049332 | 0.70482771 |
| C5orf22 | 227.355398 | 0.171581853 | 0.18563516 | 0.92429611 | 0.35533217 | 0.58081705 |
| PDZD2 | 800.487013 | -1.197093569 | 0.280558 | -4.266831 | 1.98E-05 | 0.00046683 |
| AC025178.1 | 88.981059 | 0.120184077 | 0.55728332 | 0.21566064 | 0.82925228 | 0.92023714 |
| GOLPH3 | 1936.13114 | -0.420962314 | 0.31415535 | -1.3399814 | 0.18025141 | 0.38404648 |
| MTMR12 | 1011.01713 | -0.141861311 | 0.12625926 | -1.1235715 | 0.26111485 | 0.48232304 |
| ZFR | 2400.41535 | 0.142722085 | 0.12748903 | 1.1194852 | 0.2629332 | 0.48398616 |
| SUB1 | 824.950622 | 0.260597972 | 0.29026517 | 0.89779278 | 0.36929603 | 0.59417383 |
| NPR3 | 372.874658 | 1.835077559 | 0.80194261 | 2.28829038 | 0.02212062 | 0.09542496 |
| AC010343.3 | 69.8240855 | 1.821503729 | 0.50798377 | 3.58575184 | 0.00033611 | 0.00437405 |
| TARS | 310.846642 | 0.164659003 | 0.24326263 | 0.67687751 | 0.49848368 | 0.70339396 |
| SLC45A2 | 59.6695576 | 0.554070165 | 0.53946552 | 1.02707244 | 0.30438636 | 0.5276817 |
| C1QTNF3-AN | 1866.12237 | 0.029882334 | 0.26653849 | 0.11211264 | 0.91073409 | 0.95976856 |
| RAI14 | 1466.13858 | -0.078479131 | 0.24117709 | -0.3254004 | 0.74487803 | 0.87520849 |
| TTC23L | 139.583885 | 0.834435648 | 0.48341869 | 1.72611374 | 0.08432695 | 0.23395814 |
| RAD1 | 192.033258 | 0.050245539 | 0.30093165 | 0.16696661 | 0.86739632 | 0.93967342 |
| BRIX1 | 164.360191 | -0.17146745 | 0.21271445 | -0.8060922 | 0.42018971 | 0.63958123 |
| DNAJC21 | 508.005225 | -0.065958861 | 0.15469138 | -0.42639 | 0.66982368 | 0.82642453 |
| PRLR | 333.33774 | 0.339579416 | 0.63622679 | 0.53373957 | 0.59352173 | 0.77392186 |
| SPEF2 | 789.339448 | 0.843037099 | 0.4379652 | 1.92489517 | 0.05424247 | 0.1756637 |
| AC137810.1 | 44.4090452 | 0.544635796 | 0.65298303 | 0.83407344 | 0.4042396 | 0.62531923 |
| LMBRD2 | 427.479226 | 0.203039898 | 0.10678407 | 1.90140624 | 0.05724882 | 0.18228859 |
| SKP2 | 132.403366 | 0.906914536 | 0.20388196 | 4.44823346 | 8.66E-06 | 0.00024026 |
| NADK2 | 808.3382 | 0.650206011 | 0.40775911 | 1.59458364 | 0.11080537 | 0.28084062 |
| RANBP3L | 4491.23661 | 1.897334083 | 0.63169906 | 3.00354108 | 0.00266858 | 0.02123446 |
| NIPBL | 4999.10051 | 0.010695498 | 0.13028831 | 0.082091 | 0.93457435 | 0.97183581 |
| CPLANE1 | 1609.55776 | 0.063725389 | 0.16288002 | 0.39124129 | 0.69561889 | 0.84444719 |
| NUP155 | 331.093393 | 0.371591912 | 0.24185433 | 1.53642859 | 0.1244333 | 0.30252889 |
| WDR70 | 2743.86444 | 0.216901006 | 0.16707567 | 1.29822019 | 0.19421168 | 0.40177637 |
| LIFR | 2788.00855 | 1.085489477 | 0.24576661 | 4.41674913 | 1.00E-05 | 0.00026683 |
| LIFR-AS1 | 142.600018 | 1.095056619 | 0.3460854 | 3.16412259 | 0.00155551 | 0.01409243 |
| OSMR-AS1 | 1946.73568 | -1.662442509 | 0.3542685 | -4.6926061 | 2.70E-06 | 9.09E-05 |
| OSMR | 2251.08 | -2.588851811 | 0.40599998 | -6.3764826 | 1.81E-10 | 2.82E-08 |
| RICTOR | 2955.27398 | 0.042482059 | 0.17339323 | 0.24500414 | 0.80645323 | 0.90945451 |
| FYB1 | 409.82635 | -2.343639758 | 0.65825154 | -3.5604015 | 0.00037029 | 0.00469399 |
| C9 | 69.3055786 | -1.272947933 | 0.47259704 | -2.6935165 | 0.00707026 | 0.0430878 |
| DAB2 | 129.16265 | -2.850048033 | 0.70145438 | -4.0630554 | 4.84E-05 | 0.00095463 |
| TTC33 | 522.779072 | 0.176047464 | 0.21749025 | 0.80944993 | 0.41825639 | 0.63753323 |
| PRKAA1 | 834.32389 | 0.009663563 | 0.19287056 | 0.05010388 | 0.96003961 | 0.98229715 |
| RPL37 | 622.047799 | -0.339440041 | 0.26108612 | -1.3001076 | 0.19356411 | 0.40119134 |
| CARD6 | 64.4418307 | -1.410091928 | 0.87035598 | -1.6201324 | 0.10520384 | 0.27194033 |
| OXCT1 | 1182.11115 | 1.49223095 | 0.34563499 | 4.31736073 | 1.58E-05 | 0.00038841 |
| C5orf51 | 184.771215 | 0.282120418 | 0.21016832 | 1.3423546 | 0.17948105 | 0.38299922 |
| FBXO4 | 146.178056 | -0.649984677 | 0.31542426 | -2.060668 | 0.03933472 | 0.14126492 |
| GHR | 797.334846 | 1.845736753 | 0.40032653 | 4.61057816 | 4.02E-06 | 0.0001248 |
| SELENOP | 369.942459 | 0.067910184 | 0.37987348 | 0.17877053 | 0.85811789 | 0.93591604 |
| ZNF131 | 1184.91867 | 0.058655627 | 0.18430681 | 0.31824991 | 0.75029538 | 0.87868177 |
| NIM1K | 69.9354946 | -0.254972281 | 0.52296015 | -0.4875558 | 0.62586447 | 0.79661594 |
| HMGCS1 | 712.653759 | -0.440872776 | 0.40035029 | -1.1012176 | 0.27080197 | 0.49226771 |
| CCL28 | 215.894177 | -1.216208034 | 0.72312545 | -1.681877 | 0.0925927 | 0.24975637 |
| TMEM267 | 583.283257 | -0.261180014 | 0.19296368 | -1.353519 | 0.17588988 | 0.37808453 |
| C5orf34 | 76.5181528 | -0.490719581 | 0.30035244 | -1.6338125 | 0.10229822 | 0.26679486 |
| PAIP1 | 435.39747 | -0.050251449 | 0.18294619 | -0.2746788 | 0.78356299 | 0.89665455 |
| NNT-AS1 | 798.611682 | 0.401522302 | 0.20460758 | 1.9624019 | 0.04971571 | 0.16558115 |
| NNT | 2218.20414 | 0.881273938 | 0.21820665 | 4.03871265 | 5.37E-05 | 0.00103186 |
| MRPS30-DT | 380.565651 | 0.478158308 | 0.27472605 | 1.74049134 | 0.08177278 | 0.22991414 |
| MRPS30 | 216.102801 | 0.026256174 | 0.20192721 | 0.13002792 | 0.89654434 | 0.95230008 |
| AC093297.2 | 69.3821593 | 0.477101448 | 0.25207434 | 1.89270134 | 0.0583976 | 0.18453921 |
| PARP8 | 1201.99955 | -0.912302193 | 0.32055713 | -2.8459894 | 0.00442737 | 0.03066322 |
| ITGA1 | 604.566037 | 0.731751565 | 0.20031331 | 3.65303514 | 0.00025916 | 0.00358879 |
| ITGA2 | 3129.93654 | -0.348201961 | 0.55427883 | -0.6282072 | 0.5298682 | 0.72800386 |
| MOCS2 | 242.333219 | 0.540715437 | 0.34951994 | 1.54702313 | 0.12185767 | 0.29879672 |
| NDUFS4 | 1442.90868 | 0.660579145 | 0.1653007 | 3.99622707 | 6.44E-05 | 0.00118733 |
| ARL15 | 7280.76812 | 0.002368255 | 0.3509672 | 0.0067478 | 0.99461608 | 0.99786448 |
| SNX18 | 284.314184 | 0.554595348 | 0.316804 | 1.7505945 | 0.08001578 | 0.22635818 |
| AC026704.1 | 151.921249 | -0.787075429 | 0.47233429 | -1.6663525 | 0.09564322 | 0.2549135 |
| DHX29 | 664.416621 | 0.121567255 | 0.16355037 | 0.74330163 | 0.45729908 | 0.67216888 |
| MTREX | 1238.94343 | 0.163350505 | 0.11433055 | 1.42875641 | 0.15307426 | 0.3467006 |
| PLPP1 | 824.14631 | 0.34161594 | 0.25963949 | 1.31573183 | 0.18826407 | 0.3938514 |
| SLC38A9 | 808.046379 | -0.278986696 | 0.22675264 | -1.230357 | 0.21856346 | 0.4326703 |
| IL6ST | 1415.41217 | -0.249727474 | 0.27284593 | -0.9152692 | 0.36005032 | 0.58534212 |
| AC008892.1 | 238.704955 | 1.191912196 | 0.42860184 | 2.78093114 | 0.00542032 | 0.03560271 |
| LINC01948 | 390.829311 | 0.559606893 | 0.50530763 | 1.10745783 | 0.26809604 | 0.48932276 |
| MAP3K1 | 2559.68458 | 0.254636556 | 0.21786955 | 1.16875699 | 0.24250155 | 0.46095884 |
| AC008937.2 | 116.814784 | 0.168644889 | 0.55637482 | 0.30311381 | 0.76180313 | 0.88431827 |
| SETD9 | 64.4359281 | -0.073127488 | 0.37253249 | -0.1962983 | 0.8443767 | 0.92790115 |
| MIER3 | 257.361832 | 0.324385972 | 0.24200526 | 1.34040878 | 0.18011248 | 0.38385056 |
| GPBP1 | 2578.58769 | 0.412482557 | 0.19570688 | 2.10765485 | 0.03506086 | 0.13104664 |
| PDE4D | 16278.1017 | 0.079088277 | 0.46205652 | 0.17116581 | 0.86409339 | 0.93827593 |
| ELOVL7 | 1130.79774 | 0.356753572 | 0.32165312 | 1.10912519 | 0.26737618 | 0.48855609 |
| ERCC8 | 602.586087 | 0.575941049 | 0.18829883 | 3.05865445 | 0.00222333 | 0.01857529 |
| NDUFAF2 | 946.714592 | -0.200005077 | 0.23364541 | -0.8560197 | 0.39198688 | 0.61430622 |
| SMIM15 | 135.109143 | -0.799749909 | 0.2378643 | -3.3622107 | 0.00077321 | 0.00821156 |
| SMIM15-AS1 | 138.203735 | 1.060416128 | 0.32787691 | 3.23418972 | 0.00121988 | 0.01166796 |
| ZSWIM6 | 4441.06647 | -0.418785185 | 0.26857603 | -1.5592798 | 0.11893017 | 0.29433982 |
| KIF2A | 775.506419 | -0.403215533 | 0.16863444 | -2.3910628 | 0.01679968 | 0.07930914 |
| DIMT1 | 126.389679 | 0.235996151 | 0.26941356 | 0.87596243 | 0.38105046 | 0.60380225 |
| IPO11 | 507.33608 | 0.223726602 | 0.17924317 | 1.24817366 | 0.21196747 | 0.42482955 |
| RNF180 | 1220.21924 | 0.170283773 | 0.15988056 | 1.06506867 | 0.28684486 | 0.50912842 |
| SREK1IP1 | 448.997106 | -0.090893548 | 0.12996721 | -0.6993575 | 0.48432862 | 0.69235581 |
| CWC27 | 1217.23133 | 0.349883445 | 0.12588411 | 2.77940908 | 0.00544579 | 0.03573444 |
| PPWD1 | 567.813805 | 0.390831103 | 0.1852244 | 2.11004116 | 0.03485481 | 0.13053507 |
| TRIM23 | 423.627403 | 0.465013805 | 0.21198829 | 2.1935825 | 0.02826544 | 0.11272064 |
| TRAPPC13 | 551.678091 | 0.272065786 | 0.18968971 | 1.43426746 | 0.15149595 | 0.34448442 |
| SGTB | 70.2629214 | -0.421938654 | 0.37858125 | -1.114526 | 0.26505359 | 0.4859929 |
| NLN | 311.44988 | -0.664456854 | 0.34340844 | -1.934888 | 0.05300404 | 0.17278638 |

| | | | | | | |
|---|---|---|---|---|---|---|
| ERBIN | 4431.5847 | -0.051999544 | 0.15199042 | -0.3421238 | 0.73225771 | 0.8684917 |
| SREK1 | 1675.54235 | 0.458458181 | 0.21089237 | 2.17389648 | 0.02971291 | 0.11672734 |
| MAST4 | 16002.1959 | 0.525702185 | 0.35914326 | 1.46376737 | 0.14325752 | 0.3325639 |
| AC112206.2 | 125.63018 | 0.48587892 | 0.5845358 | 0.83122184 | 0.40584832 | 0.62597507 |
| AC106798.1 | 118.307675 | 0.043237268 | 0.66894483 | 0.06463503 | 0.94846459 | 0.97784954 |
| PIK3R1 | 1355.98694 | 0.802815188 | 0.28759528 | 2.79147553 | 0.00524683 | 0.03475301 |
| AC093523.1 | 179.518567 | 2.328195656 | 0.57157975 | 4.07326476 | 4.64E-05 | 0.00092503 |
| LINC02198 | 153.475196 | 1.835096377 | 0.58598702 | 3.13163314 | 0.00173837 | 0.01529788 |
| SLC30A5 | 667.955803 | 0.020061843 | 0.26522107 | 0.07564197 | 0.93970395 | 0.97411372 |
| AC010273.1 | 44.9741493 | 0.501698225 | 0.43494689 | 1.15347008 | 0.24871749 | 0.46792613 |
| MRPS36 | 358.921466 | 0.215879952 | 0.21299796 | 1.01353063 | 0.31080677 | 0.53450314 |
| CDK7 | 362.230781 | -0.184080874 | 0.17181646 | -1.0713809 | 0.28399821 | 0.50652611 |
| CCDC125 | 766.951586 | 0.397462923 | 0.35638693 | 1.11525674 | 0.26474042 | 0.48582331 |
| AK6 | 245.380092 | 0.10800186 | 0.23392889 | 0.46168672 | 0.644306 | 0.80961167 |
| RAD17 | 586.27131 | 0.305815924 | 0.13983607 | 2.18696022 | 0.02874543 | 0.11405778 |
| MARVELD2 | 613.716671 | 0.866687803 | 0.20207473 | 4.28894699 | 1.80E-05 | 0.00043236 |
| AC145146.1 | 90.1288728 | 0.97156721 | 0.49942448 | 1.94537362 | 0.05173003 | 0.17010255 |
| OCLN | 1717.23203 | 0.63336254 | 0.27220812 | 2.32675843 | 0.01997813 | 0.08904071 |
| GTF2H2C | 696.728168 | 0.715460769 | 0.32142163 | 2.22592604 | 0.02601914 | 0.10645805 |
| AC146944.4 | 37.2677992 | 1.06331339 | 0.60618317 | 1.75411237 | 0.07941125 | 0.22520518 |
| NAIP | 2080.32957 | 1.20971419 | 0.28217314 | 4.28713441 | 1.81E-05 | 0.0004331 |
| GTF2H2 | 657.85909 | 0.727785882 | 0.51164084 | 1.42245463 | 0.15489431 | 0.34938507 |
| LINC02197 | 96.4271479 | 1.470646613 | 1.50045839 | 0.98013155 | 0.32702119 | 0.55240682 |
| BDP1 | 1703.40809 | -0.076826317 | 0.14994445 | -0.5123652 | 0.60839545 | 0.78418184 |
| MCCC2 | 798.638106 | -0.231527997 | 0.09540575 | -2.4267719 | 0.01523383 | 0.07418897 |
| MRPS27 | 705.911032 | 0.122213505 | 0.13604235 | 0.89834895 | 0.36899954 | 0.59393582 |
| PTCD2 | 163.954058 | 0.348102661 | 0.25094981 | 1.38714056 | 0.16539891 | 0.36461184 |
| AC063979.2 | 78.3692571 | 1.537009034 | 0.67817799 | 2.26638 | 0.02342813 | 0.0991548 |
| AC035140.1 | 41.1919218 | 1.078187371 | 0.39390224 | 2.73719533 | 0.00619655 | 0.03915169 |
| TNPO1 | 2021.45798 | -0.055867965 | 0.14243369 | -0.3922384 | 0.69488207 | 0.84401813 |
| FCHO2 | 2469.10528 | -1.016057449 | 0.19351084 | -5.2506486 | 1.52E-07 | 8.70E-06 |
| TMEM171 | 459.441864 | -0.20735137 | 0.54447243 | -0.3808299 | 0.70332949 | 0.84973591 |
| AC116345.1 | 293.297774 | -2.980591936 | 0.82482725 | -3.6135954 | 0.00030198 | 0.0040373 |
| BTF3 | 453.606709 | -0.8676193 | 0.31062428 | -2.7931471 | 0.0052198 | 0.03459522 |
| ANKRA2 | 245.531281 | 0.105009323 | 0.20793243 | 0.50501658 | 0.61354719 | 0.78755051 |
| UTP15 | 67.2412374 | 0.405516813 | 0.3061076 | 1.32475251 | 0.18525325 | 0.39092552 |
| ARHGEF28 | 5554.77275 | -0.011612648 | 0.20017623 | -0.0580121 | 0.95373897 | 0.97985052 |
| HEXB | 176.474818 | -0.195003719 | 0.2199402 | -0.8866215 | 0.3752827 | 0.59870627 |
| GFM2 | 396.243485 | 0.568624798 | 0.19305723 | 2.94536908 | 0.0032257 | 0.02437541 |
| NSA2 | 242.536735 | -0.416038949 | 0.26068753 | -1.5959296 | 0.11050451 | 0.28037919 |
| FAM169A | 383.114611 | 0.663849306 | 0.46129209 | 1.43910837 | 0.15011982 | 0.34235717 |
| AC116337.3 | 47.8252303 | 0.114722816 | 0.45733979 | 0.2508481 | 0.80193155 | 0.90683279 |
| ANKRD31 | 88.4563019 | 0.85651303 | 0.55883081 | 1.53268755 | 0.12535286 | 0.30392546 |
| HMGCR | 437.267552 | -0.053010761 | 0.43706615 | -0.1212877 | 0.90346314 | 0.95532916 |
| COL4A3BP | 1602.99541 | 0.036670352 | 0.18925619 | 0.19376038 | 0.84636351 | 0.92885801 |
| POLK | 1387.41073 | 0.197160166 | 0.1544474 | 1.27655219 | 0.20176039 | 0.41114671 |
| ANKDD1B | 374.993341 | 0.406403825 | 0.48336781 | 0.84077552 | 0.4004737 | 0.62204341 |
| POC5 | 287.860273 | -0.323579902 | 0.21393337 | -1.5125266 | 0.13039996 | 0.31260492 |
| IQGAP2 | 73.6087852 | 0.516828483 | 0.45906533 | 1.12582775 | 0.26023844 | 0.48117155 |
| F2RL1 | 129.75889 | -0.807587677 | 0.34460131 | -2.3435421 | 0.01910161 | 0.08651251 |
| AC008581.2 | 121.518109 | -0.148742309 | 0.42599824 | -0.3491618 | 0.72696785 | 0.86532561 |
| ZBED3 | 249.487666 | 0.055045861 | 0.2880943 | 0.1910689 | 0.84847161 | 0.9303692 |
| AC022414.1 | 152.200001 | 1.588889224 | 0.40153476 | 3.95704025 | 7.59E-05 | 0.00135453 |
| PDE8B | 42.558914 | 1.218163517 | 0.65609996 | 1.85667365 | 0.0633576 | 0.1951062 |
| WDR41 | 851.477584 | 0.276766339 | 0.17503346 | 1.5812196 | 0.11382783 | 0.28628177 |
| TBCA | 1682.68724 | 0.06597367 | 0.20295463 | 0.3250661 | 0.74513104 | 0.87531397 |
| AP3B1 | 3560.0006 | 0.226576521 | 0.25214536 | 0.89859483 | 0.36886851 | 0.59393582 |
| SCAMP1-AS1 | 56.5993234 | -0.049516051 | 0.43945933 | -0.1126749 | 0.91028828 | 0.95961675 |
| SCAMP1 | 769.305788 | -0.165979476 | 0.15195236 | -1.0923126 | 0.27469572 | 0.49636878 |
| LHFPL2 | 119.207709 | -0.486843029 | 0.35674745 | -1.3646714 | 0.17235636 | 0.37368753 |
| ARSB | 228.511399 | -0.10050668 | 0.3291841 | -0.3053206 | 0.76012201 | 0.88357016 |
| DMGDH | 1797.64484 | 1.388459528 | 0.288269 | 4.81654122 | 1.46E-06 | 5.51E-05 |
| BHMT2 | 1132.65197 | 0.565314491 | 0.42266089 | 1.33751313 | 0.1810552 | 0.38492165 |
| BHMT | 184.266503 | 1.597554093 | 0.40668548 | 3.92822993 | 8.56E-05 | 0.00149318 |
| JMY | 999.809024 | 0.089786803 | 0.22381486 | 0.40116552 | 0.68829827 | 0.83951129 |
| HOMER1 | 1562.94046 | 2.031389619 | 0.46867321 | 4.33434127 | 1.46E-05 | 0.00036574 |
| TENT2 | 3510.92515 | 0.519610599 | 0.17602383 | 2.95193323 | 0.00315791 | 0.02401208 |
| CMYA5 | 915.003831 | 1.036415663 | 0.37615966 | 2.75525467 | 0.00586465 | 0.03776364 |
| MTX3 | 223.392059 | 0.386343488 | 0.18489817 | 2.08949327 | 0.03666334 | 0.13482352 |
| THBS4 | 366.016579 | 1.083089988 | 0.44067524 | 2.45779634 | 0.01397924 | 0.06967993 |
| SERINC5 | 930.380403 | -0.736967341 | 0.32210224 | -2.287992 | 0.02213799 | 0.09546307 |
| AC010260.1 | 84.4428489 | -0.937343536 | 0.57784926 | -1.6221247 | 0.10477666 | 0.27110575 |
| ZFYVE16 | 1495.77873 | 0.319677356 | 0.22548673 | 1.41772138 | 0.15627213 | 0.35131349 |
| AC008771.1 | 102.342903 | -0.054081729 | 0.301959 | -0.1791029 | 0.85785691 | 0.93578594 |
| FAM151B | 370.142498 | -0.162303279 | 0.41799409 | -0.3882908 | 0.69780081 | 0.84608508 |
| DHFR | 1506.82598 | -0.768936001 | 0.31833118 | -2.4155221 | 0.01571267 | 0.07569243 |
| MSH3 | 1488.53611 | -0.081043959 | 0.2426011 | -0.3340626 | 0.73833231 | 0.87200855 |
| RASGRF2 | 363.176346 | -0.964664981 | 0.54837356 | -1.7591384 | 0.07855401 | 0.22420482 |
| CKMT2-AS1 | 700.539864 | 0.392582605 | 0.28485767 | 1.3781711 | 0.16815047 | 0.36813567 |
| CKMT2 | 241.544706 | 0.922334111 | 0.45005681 | 2.04937264 | 0.04042569 | 0.14362102 |
| ZCCHC9 | 190.378796 | -0.009193216 | 0.20799544 | -0.0441991 | 0.96474568 | 0.98435763 |
| SSBP2 | 5010.22956 | 0.335467071 | 0.19771531 | 1.69671771 | 0.08975004 | 0.24423327 |
| ATG10 | 1756.76338 | 0.721281998 | 0.22283853 | 3.23679215 | 0.00120881 | 0.01159566 |
| RPS23 | 1309.15406 | -1.207476735 | 0.49775465 | -2.4258472 | 0.0152727 | 0.07427964 |
| ATP6AP1L | 746.749074 | 1.00809836 | 0.38021314 | 2.65140324 | 0.00801581 | 0.04737396 |
| TMEM167A | 444.682457 | -0.147959297 | 0.20583098 | -0.7188388 | 0.47224025 | 0.68319499 |
| XRCC4 | 1036.04053 | -0.211138475 | 0.26597029 | -0.7938423 | 0.42728723 | 0.64571091 |
| VCAN | 173.162978 | -3.844198162 | 0.74455266 | -5.1630977 | 2.43E-07 | 1.29E-05 |
| EDIL3 | 162.331182 | -1.193136854 | 0.68616298 | -1.7388534 | 0.08206055 | 0.23028208 |
| COX7C | 861.499113 | -0.612138829 | 0.42754825 | -1.4317421 | 0.15221764 | 0.34548783 |
| RASA1 | 2117.47841 | -0.020738125 | 0.15554404 | -0.1333264 | 0.89393525 | 0.95146032 |
| CCNH | 3193.91136 | 0.435955198 | 0.23902089 | 1.82392088 | 0.06816403 | 0.20463153 |
| TMEM161B | 936.166996 | 0.53351729 | 0.20992411 | 2.54147692 | 0.01103852 | 0.05951341 |
| TMEM161B-. | 3414.51685 | 0.861035336 | 0.26226057 | 3.28312924 | 0.00102662 | 0.01019592 |
| MEF2C | 1249.47182 | 0.477852781 | 0.47942762 | 0.99671517 | 0.31890278 | 0.54353379 |
| MEF2C-AS1 | 529.624211 | 0.834268963 | 0.63675043 | 1.31019773 | 0.19012895 | 0.39643462 |
| CETN3 | 181.509046 | 1.046427678 | 0.26540652 | 3.94273531 | 8.06E-05 | 0.00142639 |
| MBLAC2 | 75.3671437 | 0.264160745 | 0.32158007 | 0.82144626 | 0.41139212 | 0.63062302 |
| POLR3G | 67.1065632 | 0.507835343 | 0.24830863 | 2.04517801 | 0.04083731 | 0.14450041 |
| LYSMD3 | 295.108505 | 0.173216278 | 0.25893535 | 0.66895571 | 0.50352373 | 0.70763326 |
| ADGRV1 | 1984.99139 | 0.588695595 | 0.2894628 | 2.03375215 | 0.04197659 | 0.14687915 |
| LUCAT1 | 103.115713 | -6.942080642 | 1.12931812 | -6.1471436 | 7.89E-10 | 1.05E-07 |
| ARRDC3 | 2896.41996 | -0.551829855 | 0.36527004 | -1.5107449 | 0.13085345 | 0.31323728 |
| ARRDC3-AS1 | 159.720783 | -0.277380763 | 0.34625502 | -0.8010881 | 0.42308066 | 0.64191663 |
| NR2F1-AS1 | 972.322219 | -0.005733041 | 0.57773873 | -0.0099232 | 0.99208253 | 0.99677109 |
| NR2F1 | 98.80792 | -0.080208083 | 0.61477136 | -0.1304682 | 0.89619605 | 0.95213172 |
| FAM172A | 4355.26984 | 0.193939219 | 0.14715159 | 1.31795533 | 0.1875186 | 0.393164 |
| KIAA0825 | 1479.68279 | 0.203853507 | 0.3091381 | 0.65942537 | 0.50962266 | 0.71186277 |
| SLF1 | 240.302556 | 0.054612746 | 0.40312158 | 0.13547463 | 0.89223662 | 0.95118543 |
| MCTP1 | 94.4062715 | -0.278512947 | 0.30450324 | -0.9146469 | 0.36037703 | 0.58548052 |
| TTC37 | 1321.30947 | 0.322194891 | 0.17674955 | 1.82288947 | 0.06832013 | 0.205 |
| ARSK | 220.333109 | 0.606136588 | 0.27928335 | 2.17032837 | 0.02998198 | 0.1175169 |
| RFESD | 41.8834828 | 0.725315448 | 0.38992647 | 1.86013389 | 0.06286659 | 0.1941105 |

| | | | | | | |
|---|---|---|---|---|---|---|
| RHOBTB3 | 4662.50544 | 0.157105171 | 0.29090324 | 0.54005989 | 0.58915573 | 0.77174702 |
| GLRX | 953.113095 | -0.119940977 | 0.4603706 | -0.2605314 | 0.79445393 | 0.90363747 |
| ELL2 | 1681.07504 | -0.983930058 | 0.39818569 | -2.4710332 | 0.01347233 | 0.06789623 |
| AC104123.1 | 1021.50372 | -0.098771765 | 0.36663567 | -0.2694003 | 0.78762164 | 0.89956761 |
| CAST | 2407.35403 | -0.213784472 | 0.15609533 | -1.3695763 | 0.17081919 | 0.37123242 |
| ERAP1 | 511.548514 | -0.183255955 | 0.30230695 | -0.6061917 | 0.54438748 | 0.73944091 |
| AC009126.1 | 276.614375 | 0.893720001 | 0.48315759 | 1.84974843 | 0.06434982 | 0.19689524 |
| ERAP2 | 223.51256 | -1.437983198 | 0.58235503 | -2.4692552 | 0.01353946 | 0.06813057 |
| LNPEP | 2689.42807 | 0.059895612 | 0.27300219 | 0.21939609 | 0.82634152 | 0.91938879 |
| LIX1-AS1 | 940.639415 | 0.348232777 | 0.2412227 | 1.44361527 | 0.14884723 | 0.34086738 |
| LIX1 | 63.9687634 | -1.290123571 | 0.85708971 | -1.5052375 | 0.13226303 | 0.31566403 |
| RIOK2 | 179.970835 | 0.650998525 | 0.23025881 | 2.82724702 | 0.00469501 | 0.03188503 |
| AC008883.1 | 55.383284 | 0.71129348 | 0.50400178 | 1.4112916 | 0.15815865 | 0.35438806 |
| LINC01340 | 92.2520225 | -0.50867809 | 0.52688683 | -0.9654409 | 0.33432402 | 0.55944639 |
| RGMB | 82.2691389 | -0.039509495 | 0.25288778 | -0.1562333 | 0.87584912 | 0.94418437 |
| CHD1 | 1792.27906 | -0.075284685 | 0.22534535 | -0.3340858 | 0.7383148 | 0.87200855 |
| LINC02062 | 629.065259 | 0.329009809 | 0.30523102 | 1.07790424 | 0.28107649 | 0.50360245 |
| AC021086.1 | 162.43419 | 0.786304043 | 0.48007403 | 1.63788082 | 0.10144656 | 0.26522839 |
| FAM174A | 275.308087 | -0.193374291 | 0.26172201 | -0.7388538 | 0.4599958 | 0.67354889 |
| SLCO4C1 | 1434.18991 | 1.713427196 | 0.3634837 | 4.71390385 | 2.43E-06 | 8.34E-05 |
| PAM | 4442.26924 | -0.583576336 | 0.23317984 | -2.5026878 | 0.01232542 | 0.06414756 |
| GIN1 | 261.071701 | 0.471893926 | 0.19625658 | 2.40447445 | 0.01619574 | 0.07721365 |
| PPIP5K2 | 1327.37493 | 0.508097334 | 0.15507882 | 3.27638128 | 0.00105147 | 0.01038027 |
| C5orf30 | 85.5366615 | 1.445650896 | 0.34414614 | 4.20068893 | 2.66E-05 | 0.00059732 |
| NUDT12 | 220.71173 | 0.461036589 | 0.18203731 | 2.53264886 | 0.01132043 | 0.06051459 |
| LINC01950 | 351.980005 | -0.091301592 | 0.50612658 | -0.1803928 | 0.8568442 | 0.93522187 |
| EFNA5 | 23426.0176 | -0.554381771 | 0.3190397 | -1.7376576 | 0.08227116 | 0.23053027 |
| FBXL17 | 6468.15408 | 0.21977269 | 0.15006004 | 1.46456508 | 0.14303962 | 0.33248901 |
| LINC01023 | 44.4431269 | 0.978148684 | 0.56143841 | 1.74221902 | 0.08147013 | 0.22935612 |
| FER | 3547.16876 | -0.285478356 | 0.17763313 | -1.6071234 | 0.1080273 | 0.27626962 |
| AC008467.1 | 171.519265 | -0.964750896 | 0.48531696 | -1.987878 | 0.04682519 | 0.1587154 |
| PJA2 | 1346.92893 | -0.209173352 | 0.16767009 | -1.2475293 | 0.21220348 | 0.42488177 |
| MAN2A1 | 2252.3812 | 0.088964484 | 0.2342766 | 0.37974123 | 0.70413751 | 0.85017199 |
| TMEM232 | 2094.27355 | 0.65217848 | 0.39840316 | 1.63698118 | 0.1016344 | 0.26554053 |
| SLC25A46 | 705.619394 | 0.438763168 | 0.20988499 | 2.09049335 | 0.03657351 | 0.13464146 |
| WDR36 | 394.578495 | 0.280602168 | 0.18782218 | 1.49397776 | 0.13518143 | 0.32034652 |
| STARD4 | 159.230269 | 0.104795368 | 0.46054659 | 0.22754564 | 0.81999948 | 0.91651491 |
| STARD4-AS1 | 137.844875 | 0.209039715 | 0.4154928 | 0.50311273 | 0.61488502 | 0.78831566 |
| NREP | 560.987794 | 0.040172005 | 0.55936354 | 0.07181735 | 0.94274727 | 0.97554394 |
| EPB41L4A | 3532.35329 | 0.270142598 | 0.27698878 | 0.97528356 | 0.32941962 | 0.55488483 |
| EPB41L4A-AS | 81.0538979 | -0.059502325 | 0.31591357 | -0.18835 | 0.85060228 | 0.93148777 |
| AC010261.2 | 75.6628045 | 0.135718233 | 0.59790673 | 0.22698897 | 0.82043232 | 0.91661089 |
| AC104126.1 | 83.5837154 | -0.066679429 | 0.55166332 | -0.1208698 | 0.90379417 | 0.95548393 |
| APC | 919.471066 | 0.118528997 | 0.22511571 | 0.52652478 | 0.59852364 | 0.77704816 |
| SRP19 | 399.543817 | 0.466213125 | 0.25068554 | 1.85975277 | 0.06292051 | 0.19418623 |
| REEP5 | 1114.53053 | -0.522537269 | 0.21018207 | -2.4861173 | 0.01291454 | 0.06622211 |
| DCP2 | 1050.26057 | -0.079649575 | 0.25680223 | -0.3101592 | 0.75643989 | 0.88220153 |
| MCC | 724.279498 | 0.505284215 | 0.28496978 | 1.77311511 | 0.07620964 | 0.22012391 |
| YTHDC2 | 1236.52342 | 0.469704152 | 0.17699564 | 2.65376112 | 0.00796002 | 0.04718482 |
| KCNN2 | 44.86399 | 1.055910799 | 0.44461584 | 2.37488346 | 0.01755449 | 0.08173327 |
| PGGT1B | 438.611836 | 0.161633967 | 0.12307083 | 1.31334101 | 0.18906807 | 0.39509573 |
| CCDC112 | 161.470476 | -0.038333387 | 0.34299683 | -0.1117602 | 0.91101356 | 0.95976856 |
| FEM1C | 263.59251 | -0.131498496 | 0.24216454 | -0.543013 | 0.58712085 | 0.77051955 |
| TMED7-TICAI | 115.03023 | -0.331711474 | 0.30969882 | -1.0710776 | 0.28413454 | 0.50670083 |
| AC010226.1 | 61.80929 | -0.39989794 | 0.48634893 | -0.8222449 | 0.41093752 | 0.63014283 |
| ATG12 | 438.412572 | 0.112576765 | 0.17398342 | 0.64705456 | 0.51759663 | 0.71869378 |
| AP3S1 | 593.083436 | -0.11139587 | 0.17956438 | -0.6203673 | 0.535016 | 0.73210674 |
| COMMD10 | 2867.42952 | 0.093489362 | 0.16048543 | 0.58254112 | 0.56020226 | 0.75183103 |
| SEMA6A | 259.353367 | -1.130789096 | 0.38376687 | -2.9465521 | 0.00321338 | 0.02433805 |
| SEMA6A-AS1 | 377.656923 | 0.310134302 | 0.2406114 | 1.28894266 | 0.19741802 | 0.4057467 |
| LINC02147 | 51.2375021 | 0.416687424 | 0.62960341 | 0.66182523 | 0.50808324 | 0.71109043 |
| DTWD2 | 914.376154 | 0.836315699 | 0.20589294 | 4.06189601 | 4.87E-05 | 0.00095678 |
| AC008629.1 | 564.313082 | 1.132755912 | 0.49597145 | 2.28391353 | 0.02237661 | 0.09615247 |
| DMXL1 | 2475.79533 | 0.159853485 | 0.12663259 | 1.2623408 | 0.20682618 | 0.4175161 |
| TNFAIP8 | 1395.75637 | -1.579797722 | 0.38958665 | -4.0550612 | 5.01E-05 | 0.0009736 |
| HSD17B4 | 1166.5311 | 0.438011531 | 0.23138112 | 1.89303054 | 0.05835381 | 0.18452596 |
| SRFBP1 | 438.296328 | -0.316539587 | 0.25561178 | -1.2383607 | 0.21558234 | 0.42945552 |
| AC119150.1 | 81.686067 | -0.206476679 | 0.4073392 | -0.5068913 | 0.61223112 | 0.78684784 |
| AC022101.1 | 181.240631 | -0.723232986 | 0.74015463 | -0.9771377 | 0.32850099 | 0.55367737 |
| SNX2 | 981.435363 | 0.395509461 | 0.23161382 | 1.70762464 | 0.08770601 | 0.24030466 |
| SNX24 | 1998.39435 | 0.19806517 | 0.23821598 | 0.83145205 | 0.4057183 | 0.6259207 |
| PPIC | 183.430355 | 0.228816384 | 0.29080989 | 0.78682464 | 0.43138453 | 0.6495108 |
| AC106786.2 | 45.2216951 | 0.594007259 | 0.37384708 | 1.58890439 | 0.11208198 | 0.28302254 |
| PRDM6 | 135.460381 | 0.241584485 | 0.35208997 | 0.68614419 | 0.49262219 | 0.69855572 |
| CEP120 | 636.767453 | 0.565896187 | 0.19487996 | 2.90381924 | 0.00368641 | 0.02695042 |
| CSNK1G3 | 934.767243 | -0.04907489 | 0.24912702 | -0.1969874 | 0.84383738 | 0.92769172 |
| ZNF608 | 1892.34743 | -0.159269989 | 0.27697101 | -0.5750421 | 0.56526283 | 0.75611701 |
| AC112196.1 | 71.8347307 | -0.389848182 | 0.37295856 | -1.0452855 | 0.29589103 | 0.5190471 |
| LINC02240 | 71.5184599 | 0.634129764 | 0.437316 | 1.45004932 | 0.14704477 | 0.33786787 |
| AC010235.1 | 116.478727 | -0.008219114 | 0.46208869 | -0.0177869 | 0.98580887 | 0.99426405 |
| GRAMD2B | 5226.09763 | 0.066050093 | 0.21446967 | 0.30796938 | 0.75810563 | 0.88315785 |
| AC093535.1 | 274.569319 | -0.628198429 | 0.66525763 | -0.9442935 | 0.34501971 | 0.57016956 |
| ALDH7A1 | 481.96942 | 0.44844449 | 0.32365801 | 1.3855504 | 0.16588423 | 0.36530363 |
| PHAX | 197.038705 | 0.087752549 | 0.17325312 | 0.50649911 | 0.61250632 | 0.78687125 |
| 3-Mar | 177.1312 | -0.341610077 | 0.43537767 | -0.7846293 | 0.43267095 | 0.65015341 |
| C5orf63 | 273.669325 | 0.24080972 | 0.22037887 | 1.09270785 | 0.27452209 | 0.49616155 |
| PRRC1 | 362.397483 | 0.04460515 | 0.16996399 | 0.26243882 | 0.79298315 | 0.90258633 |
| CCDC192 | 49.7398328 | 0.253339728 | 0.50017955 | 0.50649757 | 0.6125074 | 0.78687125 |
| LINC01184 | 1213.88228 | 0.382670709 | 0.20562273 | 1.86103312 | 0.0627395 | 0.19385405 |
| SLC12A2 | 423.294973 | 0.291498711 | 0.35693124 | 0.81668029 | 0.41411116 | 0.6338348 |
| SLC27A6 | 142.966992 | 1.772548355 | 1.08097457 | 1.63976879 | 0.10105325 | 0.26468617 |
| ISOC1 | 192.261813 | 0.731212809 | 0.224098 | 3.26291537 | 0.00110272 | 0.01077592 |
| CHSY3 | 146.050038 | -0.901679783 | 0.44510743 | -2.0257577 | 0.04278962 | 0.14893495 |
| HINT1 | 1340.06924 | -1.511124053 | 0.4003251 | -3.7747422 | 0.00016017 | 0.00244593 |
| LYRM7 | 304.007983 | 0.474494085 | 0.20200844 | 2.34888243 | 0.01882985 | 0.08556925 |
| CDC42SE2 | 4532.12266 | -2.08502797 | 0.39328256 | -5.3016029 | 1.15E-07 | 6.99E-06 |
| RAPGEF6 | 45.4825629 | -0.045484713 | 0.34478623 | -0.1319215 | 0.89504639 | 0.95172874 |
| P4HA2 | 146.709335 | -0.524608595 | 0.47992964 | -1.0930948 | 0.27435218 | 0.49615525 |
| PDLIM4 | 97.4633856 | -1.937469746 | 0.53943366 | -3.5916738 | 0.00032856 | 0.00429699 |
| SLC22A4 | 158.432243 | 0.304442271 | 0.36589594 | 0.83204605 | 0.40538295 | 0.62573634 |
| SLC22A5 | 363.462442 | 0.44888817 | 0.23385859 | 1.91948548 | 0.05492292 | 0.17694558 |
| AC116366.3 | 814.453102 | 0.624845699 | 0.66482202 | 0.93986913 | 0.34728469 | 0.57276608 |
| IRF1 | 154.352692 | -1.913370299 | 0.43575396 | -4.3909418 | 1.13E-05 | 0.00029404 |
| TH2LCRR | 110.19507 | -0.435735836 | 0.52234101 | -0.834198 | 0.4041694 | 0.62528387 |
| AC004039.1 | 90.3400448 | 0.431902595 | 0.62585858 | 0.69009614 | 0.49013373 | 0.69682235 |
| KIF3A | 568.849688 | 0.34614037 | 0.17330328 | 1.99731003 | 0.04579151 | 0.1562946 |
| 8-Sep | 256.596843 | 0.468188061 | 0.28944567 | 1.6175335 | 0.10576317 | 0.27294329 |
| UQCRQ | 329.481501 | -0.302388677 | 0.43958362 | -0.687898 | 0.49151704 | 0.69766264 |
| AFF4 | 2547.31444 | -0.226478733 | 0.14896948 | -1.5203029 | 0.12843487 | 0.30935378 |
| ZCCHC10 | 372.036889 | -0.194241127 | 0.23292992 | -0.8339037 | 0.40433524 | 0.62532611 |
| HSPA4 | 569.387641 | -0.030869323 | 0.18413665 | -0.1676436 | 0.8668637 | 0.93959274 |
| FSTL4 | 213.016053 | 0.236822528 | 0.63831449 | 0.3710123 | 0.71062837 | 0.85498857 |
| C5orf15 | 383.395607 | -0.027795722 | 0.19720302 | -0.1409498 | 0.88790962 | 0.94928354 |

| | | | | | | |
|---|---|---|---|---|---|---|
| VDAC1 | 721.529168 | -0.257015124 | 0.29744436 | -0.864078 | 0.38754505 | 0.61086904 |
| TCF7 | 142.252371 | -0.999026472 | 0.27045544 | -3.6938672 | 0.00022087 | 0.00318153 |
| SKP1 | 939.365494 | -0.780203936 | 0.37321812 | -2.0904771 | 0.03657496 | 0.13464146 |
| CDKL3 | 539.532355 | 0.266393852 | 0.20203221 | 1.31857117 | 0.18731252 | 0.39298287 |
| UBE2B | 1242.85932 | -0.380296614 | 0.15517012 | -2.4508366 | 0.01425246 | 0.07066846 |
| CDKN2AIPNL | 73.2414928 | 0.412757585 | 0.28831113 | 1.43163943 | 0.15224704 | 0.34548783 |
| JADE2 | 406.257039 | -0.413375334 | 0.30720881 | -1.3455843 | 0.17843663 | 0.38169546 |
| SAR1B | 522.996193 | 0.125721624 | 0.24307311 | 0.51721732 | 0.60500447 | 0.78156518 |
| SEC24A | 674.780534 | -0.296524906 | 0.22870635 | -1.2965311 | 0.19479256 | 0.40241084 |
| CAMLG | 230.344759 | -0.219663093 | 0.18475614 | -1.1889353 | 0.23446513 | 0.45166252 |
| DDX46 | 1087.41449 | 0.019036642 | 0.17778288 | 0.10707804 | 0.91472707 | 0.96131774 |
| C5orf24 | 294.289128 | 0.090525341 | 0.30714985 | 0.29472695 | 0.76820247 | 0.88823411 |
| TXNDC15 | 320.006729 | -0.04995814 | 0.30240507 | -0.1652027 | 0.86878442 | 0.94020545 |
| PCBD2 | 1332.84579 | 0.348695359 | 0.28368426 | 1.22916709 | 0.21900916 | 0.43326376 |
| C5orf66 | 85.0183996 | -0.269631453 | 0.44794222 | -0.6019336 | 0.54721837 | 0.74135461 |
| H2AFY | 800.959316 | -0.37037677 | 0.15918393 | -2.326722 | 0.01998006 | 0.08904071 |
| CXCL14 | 508.616415 | -0.253281913 | 0.58899718 | -0.4300223 | 0.66717943 | 0.82469498 |
| SLC25A48 | 248.450726 | 2.165171322 | 0.44870795 | 4.82534648 | 1.40E-06 | 5.37E-05 |
| SMAD5 | 770.557283 | 0.011017959 | 0.193022 | 0.05708136 | 0.95448038 | 0.98007893 |
| SPOCK1 | 256.821825 | -1.394258417 | 0.86615978 | -1.6097012 | 0.10746311 | 0.27552022 |
| KLHL3 | 248.279682 | 0.623662844 | 0.60242449 | 1.03525479 | 0.30054994 | 0.52367288 |
| HNRNPA0 | 254.394671 | -0.738793759 | 0.28141119 | -2.6253177 | 0.00865682 | 0.04991178 |
| AC106791.1 | 234.13711 | 0.64693322 | 0.40072336 | 1.61441354 | 0.10643776 | 0.27378589 |
| PKD2L2 | 88.9800835 | -0.138464864 | 0.58810639 | -0.2354419 | 0.8138658 | 0.91290365 |
| FAM13B | 1824.05973 | -0.008015158 | 0.14110808 | -0.0568016 | 0.95470328 | 0.98007893 |
| NME5 | 553.04523 | 0.675142549 | 0.39817117 | 1.69560884 | 0.08995998 | 0.24458483 |
| BRD8 | 605.90334 | 0.093829025 | 0.22329936 | 0.42019388 | 0.67434383 | 0.82921315 |
| CDC23 | 205.622469 | 0.149131017 | 0.17809014 | 0.83739065 | 0.40237302 | 0.62374665 |
| FAM53C | 218.671048 | 0.390297762 | 0.35093423 | 1.11216782 | 0.266066 | 0.48710545 |
| KDM3B | 1192.73696 | -0.126056862 | 0.13589343 | -0.9276156 | 0.35360701 | 0.57953068 |
| EGR1 | 320.353726 | 0.799247691 | 0.74243081 | 1.07652819 | 0.2816911 | 0.50418127 |
| ETF1 | 681.053705 | -0.208753268 | 0.36732482 | -0.568307 | 0.56982652 | 0.75923656 |
| HSPA9 | 1074.94175 | -0.063131143 | 0.16515989 | -0.3822426 | 0.70228143 | 0.8488446 |
| CTNNA1 | 4264.85979 | -0.327660349 | 0.17608918 | -1.8607637 | 0.06277755 | 0.19392626 |
| SIL1 | 2360.14791 | 0.61324797 | 0.18086049 | 3.39072382 | 0.00069708 | 0.00764393 |
| AC011405.1 | 95.7186862 | 0.414543487 | 0.57451973 | 0.72154787 | 0.4705725 | 0.68213352 |
| MATR3 | 52.2288104 | -0.194358129 | 0.44203227 | -0.4396922 | 0.66016007 | 0.81993929 |
| PAIP2 | 884.678569 | -0.267520369 | 0.25598734 | -1.0450531 | 0.29599842 | 0.51906534 |
| SPATA24 | 44.5041427 | 0.232573072 | 0.37141266 | 0.62618509 | 0.53119355 | 0.72863563 |
| DNAJC18 | 90.0249737 | 0.005180627 | 0.2558286 | 0.02025038 | 0.98384364 | 0.99340703 |
| UBE2D2 | 4218.35491 | -1.412194838 | 0.5035145 | -2.8046756 | 0.00503672 | 0.03361803 |
| CXXC5 | 747.114111 | -0.52254692 | 0.44553281 | -1.1728584 | 0.24085258 | 0.45907961 |
| AC008667.1 | 71.1566198 | 0.325793642 | 0.24747109 | 1.31649174 | 0.18800905 | 0.39381724 |
| NRG2 | 302.188799 | -0.981620208 | 0.63889384 | -1.5364371 | 0.12443121 | 0.30252889 |
| PURA | 957.323796 | 0.00867746 | 0.15927586 | 0.05448069 | 0.95655219 | 0.9809097 |
| IGIP | 135.010697 | 0.165782011 | 0.28184481 | 0.58820317 | 0.55639593 | 0.74856731 |
| CYSTM1 | 2303.84694 | 0.23918337 | 0.30006161 | 0.79711419 | 0.42538471 | 0.64392696 |
| PFDN1 | 532.621864 | 0.451984106 | 0.1979877 | 2.28288987 | 0.02243686 | 0.09631733 |
| SLC4A9 | 39.1132995 | 1.197744114 | 0.6841405 | 1.75072827 | 0.07999273 | 0.22635474 |
| AC008438.1 | 176.635883 | 1.381293567 | 0.3715249 | 3.71790308 | 0.00020088 | 0.00294176 |
| ANKHD1-EIF4 | 178.19086 | -0.045119799 | 0.26780414 | -0.1684806 | 0.86620521 | 0.93926847 |
| SRA1 | 231.280581 | -0.358976355 | 0.28214067 | -1.2723311 | 0.2032555 | 0.4129194 |
| APBB3 | 109.524766 | 0.586597925 | 0.3613801 | 1.62321592 | 0.10454326 | 0.27066089 |
| SLC35A4 | 132.027619 | -1.113278779 | 0.38238196 | -2.9114312 | 0.00359777 | 0.02651951 |
| NDUFA2 | 277.763454 | -0.651134509 | 0.41397381 | -1.5728882 | 0.11574472 | 0.28933988 |
| IK | 190.346475 | -0.645817442 | 0.22790765 | -2.8336804 | 0.00460153 | 0.03154211 |
| WDR55 | 147.832376 | -0.347143476 | 0.20754583 | -1.6726112 | 0.09440386 | 0.25284294 |
| HARS | 344.466623 | 0.294970605 | 0.13774072 | 2.14149163 | 0.03223441 | 0.12393503 |
| HARS2 | 92.2842795 | -0.092905726 | 0.32542728 | -0.2854884 | 0.77526996 | 0.89223749 |
| ZMAT2 | 148.979776 | -0.54515837 | 0.27511859 | -1.9815396 | 0.0475308 | 0.16021958 |
| AC005609.5 | 106.453638 | 0.179597548 | 0.50429572 | 0.35613538 | 0.72173918 | 0.86143064 |
| AC244517.11 | 46.9020016 | -0.08524946 | 0.52384767 | -0.1627371 | 0.87072542 | 0.94134831 |
| AC244517.10 | 40.1240946 | 0.233078557 | 0.51522567 | 0.45238149 | 0.65099418 | 0.81352716 |
| TAF7 | 365.447952 | -0.127495258 | 0.27351936 | -0.4661288 | 0.64112329 | 0.80747519 |
| DIAPH1 | 2979.91395 | -0.637140337 | 0.25121359 | -2.5362495 | 0.01120469 | 0.06011483 |
| AC008781.2 | 115.485073 | -0.832947716 | 0.51533765 | -1.6163145 | 0.10602633 | 0.2732349 |
| AC008781.1 | 64.7743215 | -0.236225318 | 0.38840548 | -0.6081925 | 0.54305978 | 0.73905591 |
| HDAC3 | 371.35391 | 0.398322074 | 0.20702295 | 1.9240479 | 0.05434858 | 0.1757642 |
| FCHSD1 | 63.3686195 | -0.733345101 | 0.32305802 | -2.2700105 | 0.02320694 | 0.09856609 |
| PCDH1 | 305.636369 | -0.751355321 | 0.57817435 | -1.2995307 | 0.19376186 | 0.40134866 |
| DELE1 | 349.732075 | -0.090820642 | 0.22068561 | -0.4115386 | 0.68067766 | 0.83310531 |
| RNF14 | 342.925013 | -0.040605602 | 0.18223659 | -0.2228181 | 0.82367712 | 0.91846107 |
| GNPDA1 | 199.521451 | -0.459649437 | 0.27972179 | -1.6432378 | 0.10033373 | 0.2634284 |
| NDFIP1 | 1515.42111 | -0.428858687 | 0.16560094 | -2.5897117 | 0.00960563 | 0.05378585 |
| SPRY4 | 105.146554 | -1.902053409 | 0.41959765 | -4.5330412 | 5.81E-06 | 0.00017145 |
| SPRY4-AS1 | 236.578948 | -1.651885788 | 0.40719978 | -4.0566962 | 4.98E-05 | 0.00096931 |
| ARHGAP26 | 1863.86746 | -1.606055746 | 0.39230825 | -4.0938617 | 4.24E-05 | 0.00086613 |
| NR3C1 | 1894.97688 | 0.243556416 | 0.18132334 | 1.34321601 | 0.17920204 | 0.38271298 |
| YIPF5 | 229.408218 | 0.062534042 | 0.19822449 | 0.31547081 | 0.75240421 | 0.87955089 |
| KCTD16 | 2734.75325 | 2.384276637 | 0.49503127 | 4.81641619 | 1.46E-06 | 5.51E-05 |
| PRELID2 | 1221.26426 | -0.336746203 | 0.42612855 | -0.7902456 | 0.42938436 | 0.64764415 |
| PLAC8L1 | 42.0463624 | 0.839571135 | 0.39286837 | 2.13702907 | 0.03259563 | 0.12490735 |
| AC091887.1 | 48.9679579 | 0.189512242 | 0.58231256 | 0.32544763 | 0.74484231 | 0.87520849 |
| LARS | 890.069105 | 0.414668894 | 0.14333341 | 2.89303715 | 0.00381536 | 0.02770594 |
| AC091959.3 | 54.8602678 | -0.451096575 | 0.29320079 | -1.5385244 | 0.12392044 | 0.30189323 |
| TCERG1 | 1173.32504 | 0.364083095 | 0.19333429 | 1.88317915 | 0.05967609 | 0.18736903 |
| PPP2R2B | 1454.26601 | 1.343741191 | 0.50456726 | 2.66315574 | 0.00774116 | 0.04608065 |
| STK32A-AS1 | 58.1557258 | 0.211871395 | 0.73937545 | 0.28655454 | 0.77445342 | 0.8920261 |
| STK32A | 787.593046 | 0.445371482 | 0.76758166 | 0.58022684 | 0.56176166 | 0.75348393 |
| AC091948.1 | 551.278569 | 1.357357748 | 0.29338752 | 4.62650127 | 3.72E-06 | 0.0001178 |
| FBXO38 | 676.074493 | 0.143707075 | 0.22205603 | 0.64716584 | 0.51752462 | 0.71869378 |
| GRPEL2 | 44.5739566 | 0.045555455 | 0.3154359 | 0.14442064 | 0.88516832 | 0.94815256 |
| PCYOX1L | 36.3478233 | 0.818373522 | 0.53502543 | 1.52959744 | 0.1261164 | 0.30521649 |
| CSNK1A1 | 3793.73313 | -0.282454346 | 0.13593829 | -2.077813 | 0.03772659 | 0.13750842 |
| ARHGEF37 | 284.970542 | 0.255968105 | 0.43383065 | 0.59001849 | 0.55517825 | 0.74754753 |
| SLC26A2 | 347.98549 | -2.434489667 | 0.75983061 | -3.20399 | 0.00135537 | 0.01271528 |
| HMGXB3 | 469.396773 | -0.261818602 | 0.28633607 | -0.9143752 | 0.36051973 | 0.58556839 |
| TCOF1 | 148.700308 | -0.240165002 | 0.31019367 | -0.7742421 | 0.43878762 | 0.65485132 |
| CD74 | 1454.7124 | -1.739527746 | 0.45980004 | -3.7832266 | 0.00015481 | 0.00237499 |
| RPS14 | 1115.18195 | -0.898787093 | 0.35433978 | -2.536512 | 0.01119629 | 0.06010818 |
| NDST1 | 382.343702 | 0.747497819 | 0.21388259 | 3.49489786 | 0.00047424 | 0.00569998 |
| SYNPO | 105.482684 | -0.506302762 | 0.37187374 | -1.361491 | 0.17335857 | 0.37508606 |
| RBM22 | 165.039712 | -0.000290318 | 0.23213096 | -0.0012507 | 0.99900211 | 0.99971199 |
| DCTN4 | 764.800333 | 0.010952715 | 0.1233777 | 0.08877387 | 0.92926163 | 0.96900671 |
| SMIM3 | 98.2383191 | -0.906295324 | 0.48371251 | -1.8736239 | 0.06098228 | 0.19001878 |
| ZNF300 | 92.2050127 | -0.071734219 | 0.43172936 | -0.1661555 | 0.86803456 | 0.93997938 |
| GPX3 | 4458.56018 | -1.324685486 | 0.66858711 | -1.9813207 | 0.04755532 | 0.16024444 |
| TNIP1 | 1027.66743 | -0.696648871 | 0.32789112 | -2.1246348 | 0.0336171 | 0.12763676 |
| ANXA6 | 196.726275 | 0.584792743 | 0.52073764 | 1.1230084 | 0.26143393 | 0.48237074 |
| GM2A | 41.1449678 | -0.14682645 | 0.47690497 | -0.3078736 | 0.75817851 | 0.88315785 |
| SLC36A1 | 105.31331 | -1.086397886 | 0.4520386 | -2.4033299 | 0.01624652 | 0.07738922 |
| SPARC | 72.4556977 | 0.262166839 | 0.38003894 | 0.68984204 | 0.49029353 | 0.69684897 |
| ATOX1 | 319.264948 | -0.307334961 | 0.20244986 | -1.5180794 | 0.12899438 | 0.31029263 |

| | | | | | | |
|---|---|---|---|---|---|---|
| G3BP1 | 1248.91997 | 0.0605076 | 0.28902646 | 0.20934969 | 0.83417526 | 0.92260077 |
| LINC01933 | 349.103244 | -0.256642515 | 0.60557436 | -0.4238002 | 0.67171156 | 0.82759425 |
| LINC01470 | 133.779246 | -1.015135715 | 0.68572911 | -1.4803742 | 0.1387734 | 0.32507075 |
| FAM114A2 | 222.049422 | 0.124338872 | 0.16597026 | 0.74916355 | 0.45375864 | 0.66948056 |
| MFAP3 | 328.826439 | -0.142980915 | 0.24172423 | -0.5915043 | 0.55418259 | 0.74695896 |
| GALNT10 | 1206.46813 | -0.180449397 | 0.52144654 | -0.3460554 | 0.72930107 | 0.86700908 |
| SAP30L-AS1 | 329.467599 | 0.107918541 | 0.27672721 | 0.38998168 | 0.6965501 | 0.84496018 |
| SAP30L | 151.918074 | -0.306934922 | 0.20917156 | -1.4673836 | 0.14227172 | 0.33124766 |
| LARP1 | 1770.89633 | 0.27263459 | 0.23288372 | 1.17068975 | 0.2417235 | 0.46003068 |
| FAXDC2 | 714.574148 | 1.032828098 | 0.42785031 | 2.41399401 | 0.01577872 | 0.07595522 |
| CNOT8 | 362.404753 | -0.093199157 | 0.23426953 | -0.3978288 | 0.69075641 | 0.84134845 |
| GEMIN5 | 227.101982 | 0.450779595 | 0.2379731 | 1.89424602 | 0.05819236 | 0.18428744 |
| MRPL22 | 229.103555 | 0.321133197 | 0.20187005 | 1.5907917 | 0.11165646 | 0.28237377 |
| SGCD | 169.88218 | 3.256453155 | 0.7049307 | 4.61953655 | 3.85E-06 | 0.00012067 |
| TIMD4 | 114.009378 | -3.904597809 | 0.82247426 | -4.7473799 | 2.06E-06 | 7.22E-05 |
| HAVCR1 | 580.684223 | -4.250432489 | 0.8466967 | -5.0200178 | 5.17E-07 | 2.42E-05 |
| HAVCR2 | 204.100842 | -0.86145935 | 0.36387833 | -2.3674379 | 0.01791172 | 0.08295791 |
| AC009185.1 | 42.1954434 | 2.335498589 | 0.63933722 | 3.65299958 | 0.00025919 | 0.00358879 |
| CYFIP2 | 4251.82817 | 1.684243741 | 0.34675916 | 4.85709955 | 1.19E-06 | 4.78E-05 |
| AC008676.1 | 72.2784591 | 1.797905458 | 0.6625517 | 2.7136078 | 0.00665549 | 0.04135107 |
| THG1L | 45.7244323 | -0.332014512 | 0.40054589 | -0.8289051 | 0.40715813 | 0.62711657 |
| LSM11 | 43.4308915 | -0.385278022 | 0.34056117 | -1.1313034 | 0.2579274 | 0.47897694 |
| CLINT1 | 3445.3167 | -0.417491886 | 0.26764387 | -1.5598784 | 0.11878862 | 0.29411039 |
| RNF145 | 1090.99519 | -0.839168615 | 0.2491732 | -3.3678125 | 0.00075767 | 0.00810371 |
| UBLCP1 | 295.503558 | 0.195265954 | 0.24640037 | 0.79247426 | 0.42808419 | 0.64645754 |
| AC008691.1 | 84.6935364 | -0.436597756 | 0.55920501 | -0.7807472 | 0.43495118 | 0.65190856 |
| TTC1 | 694.448849 | -0.165544992 | 0.21215459 | -0.7803036 | 0.43521219 | 0.65215177 |
| PWWP2A | 832.682871 | 0.299334926 | 0.1561083 | 1.91748241 | 0.05517667 | 0.17751599 |
| CCNJL | 471.704591 | 0.80998104 | 0.28064587 | 2.88613203 | 0.00390009 | 0.02813557 |
| ZBED8 | 70.4566744 | 0.693970755 | 0.25165052 | 2.75767664 | 0.00582138 | 0.03754366 |
| SLU7 | 215.362285 | -0.139200108 | 0.16667292 | -0.8351693 | 0.40362239 | 0.62487671 |
| GABRB2 | 177.829054 | -0.036646347 | 0.97782388 | -0.0374775 | 0.97010432 | 0.98653537 |
| AC113414.1 | 1326.39122 | -0.977075609 | 0.51565653 | -1.8948186 | 0.05811643 | 0.18409112 |
| CCNG1 | 281.84525 | -0.794442747 | 0.3603558 | -2.2046065 | 0.02748172 | 0.1104666 |
| NUDCD2 | 137.510908 | -0.247329448 | 0.32223514 | -0.7675434 | 0.44275851 | 0.65867746 |
| MAT2B | 332.030756 | -0.093870255 | 0.24783629 | -0.3787591 | 0.70486675 | 0.85073346 |
| AC008415.1 | 205.489367 | 3.647045541 | 0.55839144 | 6.53134217 | 6.52E-11 | 1.21E-08 |
| TENM2 | 81.2958792 | 0.714202986 | 0.3701473 | 1.9295102 | 0.05366755 | 0.17434621 |
| WWC1 | 6635.38695 | -0.381967181 | 0.34645151 | -1.1025127 | 0.27023886 | 0.49175977 |
| RARS | 606.565034 | -0.174005226 | 0.23446751 | -0.7421294 | 0.45800895 | 0.67250337 |
| PANK3 | 540.120724 | 0.065311923 | 0.27151108 | 0.24054975 | 0.8099041 | 0.91064395 |
| SLIT3 | 173.395729 | 2.837888141 | 0.72850663 | 3.89548707 | 9.80E-05 | 0.00165114 |
| SPDL1 | 65.1091659 | 0.098256764 | 0.29775616 | 0.3299907 | 0.74140699 | 0.87369479 |
| KCNIP1 | 84.5699013 | -0.42938261 | 0.77910903 | -0.55112 | 0.58155138 | 0.767524 |
| GABRP | 71.5390562 | -3.090915624 | 0.76135035 | -4.0597809 | 4.91E-05 | 0.00096102 |
| RANBP17 | 641.076604 | -0.156681495 | 0.39155087 | -0.4001562 | 0.6890415 | 0.84010064 |
| NPM1 | 612.859806 | -0.163823088 | 0.22059754 | -0.7426333 | 0.4577037 | 0.67227935 |
| FBXW11 | 3510.70147 | -0.105669536 | 0.31869689 | -0.3315675 | 0.74021585 | 0.87268348 |
| STK10 | 215.708702 | -0.965807311 | 0.35097612 | -2.751775 | 0.00592732 | 0.03798836 |
| UBTD2 | 599.488311 | -0.293284369 | 0.20568118 | -1.4259174 | 0.15389219 | 0.34807567 |
| SH3PXD2B | 121.506208 | -0.405180298 | 0.48061106 | -0.8430524 | 0.39919915 | 0.6208833 |
| DUSP1 | 865.593433 | -1.132350485 | 0.62465177 | -1.8127708 | 0.06986718 | 0.20790237 |
| ERGIC1 | 1261.81293 | -1.11678484 | 0.29232652 | -3.8203337 | 0.00013327 | 0.00210824 |
| RPL26L1 | 72.7759324 | -0.40505438 | 0.31883983 | -1.2704008 | 0.20394189 | 0.41379632 |
| ATP6V0E1 | 2313.91974 | -0.852952051 | 0.36424672 | -2.3416877 | 0.01919677 | 0.08674997 |
| CREBRF | 2092.19037 | -0.610670409 | 0.26883846 | -2.2715143 | 0.02311586 | 0.09830568 |
| BNIP1 | 110.558848 | 0.477111894 | 0.25479053 | 1.87256525 | 0.06112844 | 0.1902558 |
| BOD1 | 164.373555 | 0.112385365 | 0.23882589 | 0.47057446 | 0.63794465 | 0.80514149 |
| CPEB4 | 6988.76159 | 0.791503313 | 0.34483729 | 2.295295 | 0.02171622 | 0.09440958 |
| SFXN1 | 793.987863 | 0.045989664 | 0.21657012 | 0.21235461 | 0.83183039 | 0.92158196 |
| THOC3 | 276.858567 | -0.185335247 | 0.3368353 | -0.5502251 | 0.58216496 | 0.76785626 |
| SIMC1 | 704.836429 | -0.191037561 | 0.15730959 | -1.2144051 | 0.22459308 | 0.43924342 |
| KIAA1191 | 346.6725 | -0.303088219 | 0.1926781 | -1.5730289 | 0.11571213 | 0.28931317 |
| ARL10 | 206.933805 | -0.169841255 | 0.29375181 | -0.5781794 | 0.56314299 | 0.75468425 |
| CLTB | 271.537742 | -0.567060512 | 0.28375554 | -1.9984121 | 0.045672 | 0.15600761 |
| FAF2 | 846.527934 | -0.047997867 | 0.17612673 | -0.2725189 | 0.78522304 | 0.89769873 |
| RNF44 | 210.661556 | 0.116824866 | 0.27905918 | 0.41863832 | 0.67548048 | 0.82983832 |
| CDHR2 | 74.756913 | -0.50152726 | 0.59588149 | -0.841656 | 0.39998051 | 0.6214966 |
| TSPAN17 | 142.33039 | -0.867102696 | 0.35880144 | -2.4166645 | 0.01566345 | 0.07556555 |
| UIMC1 | 1121.8719 | -0.008682598 | 0.15220401 | -0.0570458 | 0.95450872 | 0.98007893 |
| ZNF346 | 616.635309 | 0.367769433 | 0.1825008 | 2.01516614 | 0.04388724 | 0.15187493 |
| FGFR4 | 68.9247924 | 0.049019469 | 0.52792482 | 0.09285313 | 0.92602025 | 0.96724667 |
| NSD1 | 2824.54663 | 0.139125634 | 0.17644131 | 0.78850942 | 0.43039879 | 0.64854411 |
| PRELID1 | 211.417383 | -0.967265844 | 0.41527649 | -2.3292093 | 0.01984798 | 0.08872149 |
| LMAN2 | 361.608401 | -0.46323862 | 0.26287842 | -1.7621782 | 0.0780392 | 0.22316946 |
| RGS14 | 74.1308847 | -0.824402485 | 0.33247967 | -2.4795576 | 0.01315455 | 0.06688162 |
| GRK6 | 59.4732732 | -0.69951285 | 0.30491907 | -2.2940934 | 0.02178513 | 0.09452959 |
| DBN1 | 238.846201 | -0.224139559 | 0.37936672 | -0.5908256 | 0.5546373 | 0.74738138 |
| PDLIM7 | 158.790166 | -1.756817288 | 0.78510469 | -2.2376854 | 0.02524158 | 0.10419747 |
| DDX41 | 96.6618441 | -0.690925324 | 0.37706875 | -1.832359 | 0.06689795 | 0.20193399 |
| FAM193B | 687.313644 | 0.096915783 | 0.18580744 | 0.52159259 | 0.60195402 | 0.77940904 |
| TMED9 | 292.296818 | -1.143414231 | 0.33196642 | -3.4443672 | 0.0005724 | 0.00653483 |
| B4GALT7 | 97.0960419 | -0.236092254 | 0.43820132 | -0.5387758 | 0.59004158 | 0.7721373 |
| AC106795.3 | 97.4763487 | 0.446840079 | 0.31240605 | 1.43031826 | 0.1526257 | 0.34604065 |
| N4BP3 | 51.4441321 | 0.879087309 | 0.48877044 | 1.79856889 | 0.0720869 | 0.21221706 |
| RMND5B | 492.114242 | 0.043361163 | 0.29409316 | 0.14744023 | 0.88278455 | 0.94732911 |
| NHP2 | 156.079928 | -1.376245619 | 0.41818805 | -3.2909731 | 0.00099841 | 0.00998339 |
| HNRNPAB | 219.944829 | -1.23770767 | 0.2643431 | -4.6822015 | 2.84E-06 | 9.49E-05 |
| PHYKPL | 1193.81077 | -0.360441065 | 0.30053911 | -1.199315 | 0.23040549 | 0.44592324 |
| COL23A1 | 181.414172 | -0.603699605 | 0.55934966 | -1.0792884 | 0.28045918 | 0.50294945 |
| CLK4 | 656.271332 | 0.957159156 | 0.19850924 | 4.82173609 | 1.42E-06 | 5.45E-05 |
| ZNF354A | 251.871378 | -0.344485065 | 0.33290065 | -1.0347984 | 0.30076306 | 0.52383693 |
| ZNF354B | 222.651753 | -0.060783046 | 0.24377802 | -0.2493377 | 0.80309959 | 0.90758619 |
| ZFP2 | 179.806125 | 0.721828585 | 0.2463248 | 2.93039347 | 0.00338533 | 0.02538379 |
| ZNF454 | 86.0408296 | -0.36026196 | 0.26639044 | -1.3523832 | 0.17625276 | 0.378495 |
| AC104117.3 | 57.74046 | -0.173971576 | 0.33066292 | -0.5261297 | 0.59879811 | 0.77721524 |
| ZNF879 | 40.0800292 | 0.264368826 | 0.32060225 | 0.82460066 | 0.40959835 | 0.62940723 |
| ZNF354C | 156.957603 | 0.363277634 | 0.22147094 | 1.6402948 | 0.10094389 | 0.26453475 |
| RUFY1 | 862.176229 | 0.113080828 | 0.1692436 | 0.66815422 | 0.50403515 | 0.70797197 |
| AC136604.2 | 102.167419 | -0.325273612 | 0.32196177 | -1.0102864 | 0.31235807 | 0.53625394 |
| HNRNPH1 | 2851.14635 | 0.742237285 | 0.25397418 | 2.9224911 | 0.00347244 | 0.02584079 |
| CANX | 1371.07481 | -0.38875197 | 0.31369281 | -1.239276 | 0.21524331 | 0.42913181 |
| MAML1 | 426.566182 | 0.165972096 | 0.20538449 | 0.80810435 | 0.41903051 | 0.63811091 |
| MGAT4B | 316.203982 | -1.275456629 | 0.29364976 | -4.3434622 | 1.40E-05 | 0.00035154 |
| SQSTM1 | 2005.35318 | -1.261892442 | 0.33121572 | -3.8098809 | 0.00013903 | 0.00217594 |
| MRNIP | 433.291246 | 0.057856522 | 0.1879344 | 0.30785488 | 0.75819276 | 0.88315785 |
| TBC1D9B | 491.108502 | -0.551519793 | 0.29434912 | -1.8736927 | 0.0609728 | 0.19001878 |
| RNF130 | 1610.51655 | -0.290479935 | 0.26582341 | -1.0927553 | 0.27450126 | 0.49616155 |
| MAPK9 | 506.890503 | 0.225663739 | 0.14087466 | 1.60187597 | 0.10918304 | 0.27771977 |
| AC008610.1 | 52.6223934 | 0.491420268 | 0.41524247 | 1.18345378 | 0.23662935 | 0.4542417 |
| GFPT2 | 340.366021 | -1.791706164 | 0.30614244 | -5.8525246 | 4.84E-09 | 5.24E-07 |
| CNOT6 | 700.488978 | -0.095839517 | 0.13976697 | -0.6857093 | 0.49289641 | 0.69868807 |
| MGAT1 | 304.941581 | 0.000270601 | 0.1737693 | 0.00155724 | 0.9987575 | 0.99966567 |

| | | | | | | |
|---|---|---|---|---|---|---|
| LINC00847 | 44.1876423 | 0.364576412 | 0.35748231 | 1.01984463 | 0.30780215 | 0.53140225 |
| ZFP62 | 148.433941 | 0.198949968 | 0.23101634 | 0.86119437 | 0.389131 | 0.6122488 |
| BTNL9 | 95.0298605 | 2.326280424 | 0.4943399 | 4.70583184 | 2.53E-06 | 8.59E-05 |
| TRIM41 | 187.575359 | -0.032834472 | 0.25564425 | -0.1284381 | 0.89780225 | 0.95268878 |
| RACK1 | 858.0182 | -1.034317322 | 0.36303116 | -2.8491145 | 0.00438411 | 0.0303988 |
| TRIM52 | 267.078854 | 0.21176857 | 0.19303804 | 1.09703023 | 0.27262818 | 0.49418767 |
| AC138035.1 | 112.720613 | 0.203804419 | 0.30264718 | 0.67340598 | 0.50068905 | 0.70491515 |
| DUSP22 | 367.647286 | -1.25597459 | 0.15130527 | -8.3009309 | 1.03E-16 | 1.14E-13 |
| EXOC2 | 1085.82527 | -0.259199421 | 0.21854409 | -1.186028 | 0.23561123 | 0.45334141 |
| FOXQ1 | 178.028967 | -1.016348283 | 0.53835721 | -1.8878697 | 0.05904344 | 0.18569162 |
| FOXC1 | 294.198995 | -0.2445323 | 0.48871503 | -0.5003576 | 0.61682327 | 0.78988062 |
| GMDS | 2941.48296 | -0.47571677 | 0.17574496 | -2.7068586 | 0.00679232 | 0.0419447 |
| GMDS-DT | 5167.46369 | -0.081532855 | 0.33301951 | -0.2448291 | 0.80658879 | 0.90951555 |
| LINC01600 | 83.1603824 | -0.27684585 | 0.53051208 | -0.5218465 | 0.60177724 | 0.77940904 |
| MYLK4 | 72.4730802 | 0.033041204 | 0.42365386 | 0.07799104 | 0.93783518 | 0.97304728 |
| WRNIP1 | 420.187209 | 0.128769222 | 0.17674784 | 0.72854766 | 0.46627841 | 0.67903174 |
| SERPINB1 | 318.136361 | -2.012643597 | 0.30687422 | -6.5585294 | 5.43E-11 | 1.05E-08 |
| SERPINB6 | 585.645952 | -0.783198937 | 0.23626811 | -3.3148736 | 0.00091685 | 0.00932303 |
| NQO2 | 366.405272 | 0.256179904 | 0.35427017 | 0.72312016 | 0.46960607 | 0.68135177 |
| RIPK1 | 530.145696 | -0.510082778 | 0.2547839 | -2.0020212 | 0.04528245 | 0.15535466 |
| BPHL | 147.764976 | -0.263243075 | 0.27604064 | -0.9536388 | 0.3402665 | 0.56556941 |
| TUBB2A | 79.8520433 | -0.153215261 | 0.48565704 | -0.3154804 | 0.75239696 | 0.87955089 |
| LINC02525 | 81.9472391 | -1.396395888 | 0.9099229 | -1.534631 | 0.12487449 | 0.30326663 |
| TUBB2B | 119.329139 | -0.424023592 | 0.59338404 | -0.7145854 | 0.47486525 | 0.68514419 |
| PSMG4 | 456.027378 | -0.179616044 | 0.4041806 | -0.4443955 | 0.65675666 | 0.81801439 |
| SLC22A23 | 190.697411 | -0.453199346 | 0.43436218 | -1.0433674 | 0.29677815 | 0.51998178 |
| PXDC1 | 67.8115154 | -1.332007347 | 0.41058881 | -3.2441394 | 0.00117806 | 0.01136669 |
| PRPF4B | 1424.19601 | 0.047341822 | 0.18547295 | 0.2552492 | 0.79853063 | 0.90515671 |
| FAM217A | 105.493893 | 0.761045863 | 0.49404194 | 1.5404479 | 0.12345121 | 0.30130582 |
| ECI2 | 630.550483 | -0.388018032 | 0.2183026 | -1.7774321 | 0.07549718 | 0.21863598 |
| CDYL | 1581.75826 | -0.703065928 | 0.22540719 | -3.1190928 | 0.00181409 | 0.01580626 |
| LYRM4 | 677.204436 | -0.194297137 | 0.26367664 | -0.7368766 | 0.46119742 | 0.67447531 |
| FARS2 | 2852.28347 | 0.233565293 | 0.15479515 | 1.50886698 | 0.13133278 | 0.31399617 |
| AL022097.1 | 62.5499335 | 0.087101732 | 0.51663021 | 0.16859589 | 0.86611451 | 0.93926847 |
| LY86-AS1 | 168.771756 | 0.858322846 | 0.4742361 | 1.80990617 | 0.07031034 | 0.20879705 |
| RREB1 | 1842.67542 | -0.430032159 | 0.27765754 | -1.5487862 | 0.12143314 | 0.2981986 |
| SSR1 | 491.818836 | -0.233848087 | 0.27485137 | -0.8508165 | 0.39487127 | 0.61706761 |
| AL139095.4 | 99.7383962 | -0.171535361 | 0.52852236 | -0.3245565 | 0.74551676 | 0.87556957 |
| RIOK1 | 148.709964 | 0.114674384 | 0.27337339 | 0.41947896 | 0.67486613 | 0.82939913 |
| DSP | 610.537604 | -1.085842891 | 0.26242094 | -4.1377906 | 3.51E-05 | 0.00074111 |
| SNRNP48 | 254.356735 | 0.230624284 | 0.18383332 | 1.25452928 | 0.20964969 | 0.42143703 |
| BMP6 | 2663.44946 | 1.073290167 | 0.67208621 | 1.59695312 | 0.11027616 | 0.27985356 |
| BLOC1S5-TXI | 246.970069 | 0.129276731 | 0.40089952 | 0.32246666 | 0.74709919 | 0.87660707 |
| SLC35B3 | 384.122164 | 0.192895699 | 0.20258034 | 0.95219358 | 0.34099881 | 0.56647414 |
| TFAP2A | 143.316729 | 0.269575325 | 0.30206992 | 0.89242692 | 0.37216415 | 0.59630231 |
| TFAP2A-AS1 | 44.9704888 | -0.273506044 | 0.44003677 | -0.6215527 | 0.53423603 | 0.73149412 |
| GCNT2 | 3601.64116 | 0.850607686 | 0.40342192 | 2.1084816 | 0.03498935 | 0.13084552 |
| C6orf52 | 50.7506241 | 0.509344886 | 0.43750505 | 1.16420345 | 0.24434157 | 0.46318998 |
| PAK1IP1 | 85.0897253 | 0.155618522 | 0.27854901 | 0.55867555 | 0.57638317 | 0.76324677 |
| TMEM14C | 296.728963 | -0.430850701 | 0.22772737 | -1.8919584 | 0.05849653 | 0.18461291 |
| AL024498.2 | 70.4724902 | 0.28056595 | 0.37867386 | 0.74091713 | 0.45874369 | 0.67278348 |
| MAK | 568.683445 | 0.003197581 | 0.43392024 | 0.00736905 | 0.9941204 | 0.99783229 |
| SMIM13 | 132.858735 | 0.315870247 | 0.27856524 | 1.13391837 | 0.25682879 | 0.47778145 |
| AL139807.1 | 761.396971 | 0.448855877 | 0.70762658 | 0.63431178 | 0.52587737 | 0.72494586 |
| NEDD9 | 2216.11085 | 0.637615976 | 0.42150015 | 1.51273012 | 0.13034823 | 0.31253761 |
| TMEM170B | 234.185125 | 0.638097714 | 0.26225087 | 2.43315769 | 0.01496778 | 0.0732348 |
| AL157373.2 | 104.629802 | -0.131819455 | 0.70777805 | -0.1862441 | 0.85225338 | 0.93200182 |
| HIVEP1 | 1790.49909 | -0.920213346 | 0.33213621 | -2.7705903 | 0.00559548 | 0.03648415 |
| EDN1 | 71.7064967 | -1.438089645 | 0.56988709 | -2.5234641 | 0.01162049 | 0.06161986 |
| PHACTR1 | 6461.09862 | -0.364947397 | 0.43575324 | -0.8375093 | 0.40230634 | 0.62371648 |
| TBC1D7 | 197.821071 | 0.099752931 | 0.21855542 | 0.45641939 | 0.64808842 | 0.81173905 |
| GFOD1 | 411.950352 | -2.914593651 | 0.90772258 | -3.2108859 | 0.00132326 | 0.01246719 |
| SIRT5 | 512.369347 | 0.94970648 | 0.27238706 | 3.4866064 | 0.00048919 | 0.00582137 |
| NOL7 | 333.540026 | -0.496552613 | 0.27649386 | -1.7958902 | 0.07251198 | 0.21294147 |
| RANBP9 | 1415.82109 | 0.038615193 | 0.25315349 | 0.15253668 | 0.87876366 | 0.94585961 |
| MCUR1 | 303.119347 | 0.258818338 | 0.2100366 | 1.2322535 | 0.2178544 | 0.43175376 |
| CD83 | 253.381117 | -0.110676288 | 0.66529095 | -0.1663577 | 0.86787544 | 0.93997938 |
| JARID2 | 1861.29665 | 0.51178471 | 0.28215199 | 1.81386177 | 0.06969901 | 0.20763514 |
| DTNBP1 | 1153.93214 | -0.689014019 | 0.25903259 | -2.6599511 | 0.0078152 | 0.04643769 |
| MYLIP | 406.85398 | -0.033507863 | 0.24329374 | -0.137726 | 0.890457 | 0.95041347 |
| GMPR | 1008.8466 | 0.028044076 | 0.34009256 | 0.08246013 | 0.93428082 | 0.97176637 |
| ATXN1 | 5170.95447 | 0.154431693 | 0.32075844 | 0.48145792 | 0.63019108 | 0.79994176 |
| CAP2 | 430.025245 | 0.498842524 | 0.5054892 | 0.986851 | 0.32371569 | 0.54883334 |
| FAM8A1 | 181.474685 | -0.02587747 | 0.41884948 | -0.0617823 | 0.95073623 | 0.97874482 |
| NUP153 | 1109.52624 | 0.325725249 | 0.21156606 | 1.5395912 | 0.12366003 | 0.30156806 |
| KIF13A | 1291.54205 | 0.280040009 | 0.28646732 | 0.97756353 | 0.32829024 | 0.55346341 |
| TPMT | 261.237923 | 0.150852678 | 0.20546298 | 0.73420856 | 0.46282164 | 0.67581373 |
| KDM1B | 228.843233 | -0.350653721 | 0.24140054 | -1.4525805 | 0.14634025 | 0.33687843 |
| DEK | 727.864158 | -0.41949336 | 0.2006928 | -2.0902262 | 0.03659748 | 0.13464146 |
| RNF144B | 2474.49112 | -1.083866281 | 0.31775909 | -3.4109686 | 0.00064733 | 0.00721563 |
| MIR548A1HC | 86.3620463 | -2.960057952 | 0.67696514 | -4.3725412 | 1.23E-05 | 0.00031376 |
| AL022068.1 | 804.724136 | 1.157531341 | 0.74214318 | 1.55971431 | 0.11882741 | 0.29415128 |
| ID4 | 145.036328 | -0.524382756 | 0.59081489 | -0.8875585 | 0.37477831 | 0.59824129 |
| MBOAT1 | 249.463122 | -1.683212559 | 0.73745101 | -2.2824737 | 0.02246139 | 0.09637486 |
| E2F3 | 424.434408 | -1.425977442 | 0.30723574 | -4.6413137 | 3.46E-06 | 0.00011073 |
| CDKAL1 | 4070.42403 | -0.043928991 | 0.19755553 | -0.2223627 | 0.82403151 | 0.91855769 |
| AL513188.1 | 95.047178 | -0.594495356 | 0.57238257 | -1.0386329 | 0.29897551 | 0.52196239 |
| AL512380.2 | 98.0501942 | -0.278729977 | 0.36177263 | -0.7704562 | 0.4410293 | 0.65677069 |
| SOX4 | 644.048636 | -0.287424996 | 0.45678941 | -0.6292287 | 0.52919935 | 0.72746324 |
| CASC15 | 1028.82826 | 0.724524779 | 0.35750719 | 2.02660196 | 0.04270314 | 0.14879075 |
| NBAT1 | 83.5519434 | 1.340675701 | 0.5541899 | 2.419163 | 0.01555627 | 0.0752684 |
| DCDC2 | 7343.70497 | -1.154430892 | 0.39126285 | -2.9505252 | 0.00317234 | 0.02407504 |
| MRS2 | 544.29884 | 0.554829292 | 0.20894244 | 2.65541696 | 0.00792104 | 0.0470243 |
| GPLD1 | 65.616051 | 0.631757605 | 0.29848138 | 2.11657294 | 0.03429611 | 0.12913878 |
| ALDH5A1 | 142.035246 | 0.344295163 | 0.32133213 | 1.07146199 | 0.28396175 | 0.50652611 |
| TDP2 | 305.594069 | -0.24220384 | 0.22767501 | -1.0638139 | 0.28741301 | 0.50981103 |
| ACOT13 | 211.575519 | 0.21207671 | 0.24750477 | 0.85685909 | 0.39152276 | 0.6140943 |
| C6orf62 | 879.710998 | 0.389182283 | 0.21542592 | 1.80657129 | 0.07082915 | 0.20986591 |
| GMNN | 101.071496 | 0.130543526 | 0.32630487 | 0.40006612 | 0.68910781 | 0.84010064 |
| RIPOR2 | 77.8554695 | -1.340640719 | 0.58459798 | -2.2932695 | 0.02183249 | 0.09455259 |
| CARMIL1 | 4008.76271 | 0.284897378 | 0.22386092 | 1.27265349 | 0.20314103 | 0.41281382 |
| SLC17A1 | 98.2894198 | 1.05847044 | 0.85812509 | 1.23346871 | 0.21740095 | 0.43128814 |
| TRIM38 | 689.848871 | -0.445042689 | 0.32726656 | -1.3598783 | 0.17386845 | 0.3755115 |
| U91328.2 | 76.7099417 | -0.144205076 | 0.46952905 | -0.3071271 | 0.75874666 | 0.88316847 |
| HIST1H1C | 160.463309 | -1.301138169 | 0.48321934 | -2.6926451 | 0.00708877 | 0.04313648 |
| HFE | 66.2832447 | -0.066792393 | 0.44180284 | -0.1511814 | 0.87983259 | 0.94636668 |
| HIST1H4C | 299.99102 | -3.108912766 | 0.72526151 | -4.2866093 | 1.81E-05 | 0.00043334 |
| HIST1H2AC | 303.294089 | 0.082188472 | 0.63268946 | 0.12990334 | 0.8966429 | 0.95230008 |
| HIST1H1E | 86.1126358 | -2.182327371 | 0.70415532 | -3.0992131 | 0.00194035 | 0.01672986 |
| HIST1H2BD | 73.1545286 | 0.644654067 | 0.74721089 | 0.86274715 | 0.3882765 | 0.61143829 |
| BTN3A2 | 83.6654082 | -1.490656475 | 0.80813598 | -1.8445615 | 0.06510136 | 0.19827619 |
| BTN2A2 | 48.8179749 | -1.520038669 | 0.4950158 | -3.0706872 | 0.00213567 | 0.01803711 |
| BTN3A1 | 90.1876312 | -0.359436543 | 0.7647917 | -0.4699797 | 0.63836955 | 0.80536995 |

| | | | | | | |
|---|---|---|---|---|---|---|
| BTN2A1 | 175.925498 | -0.260690994 | 0.2520259 | -1.0343817 | 0.30095774 | 0.52396873 |
| HCG11 | 52.5767508 | -0.49205054 | 0.49793676 | -0.9881788 | 0.3230651 | 0.54801228 |
| HMGN4 | 125.582317 | -0.7059941 | 0.32411721 | -2.1782062 | 0.02939069 | 0.11571435 |
| ABT1 | 64.6982289 | -0.044317492 | 0.36225747 | -0.122337 | 0.90263213 | 0.95499458 |
| ZNF322 | 163.457113 | 0.216908452 | 0.24709444 | 0.87783624 | 0.3800326 | 0.60276784 |
| AL513548.1 | 44.5745814 | 1.31366326 | 0.6219258 | 2.1122508 | 0.03466494 | 0.13011647 |
| LINC00240 | 377.87305 | 0.79345977 | 0.31981137 | 2.48102429 | 0.01310054 | 0.06670975 |
| ZNF391 | 58.5129673 | 0.184455161 | 0.25001881 | 0.73776515 | 0.46065718 | 0.67406897 |
| ZNF184 | 52.835649 | 0.052937467 | 0.24462772 | 0.21640012 | 0.82867588 | 0.92023714 |
| HIST1H2BN | 45.6386326 | -0.257711656 | 0.48841947 | -0.5276441 | 0.59774637 | 0.77674686 |
| AL121944.1 | 102.099576 | -0.284574605 | 0.37891083 | -0.7510332 | 0.45263269 | 0.66839857 |
| ZNF165 | 75.1260973 | -1.030106419 | 0.51029258 | -2.0186584 | 0.04352273 | 0.15096948 |
| ZSCAN16-AS | 188.807875 | 0.188924118 | 0.26242189 | 0.71992514 | 0.47157109 | 0.68270303 |
| ZSCAN16 | 44.6556491 | -0.300958462 | 0.38358967 | -0.7845844 | 0.4326973 | 0.65015341 |
| ZKSCAN8 | 291.963725 | 0.225584695 | 0.17516263 | 1.28785859 | 0.19779519 | 0.40634007 |
| ZSCAN9 | 143.26355 | -0.144023863 | 0.20410037 | -0.7056522 | 0.48040448 | 0.68948936 |
| PGBD1 | 96.8588985 | 0.472572791 | 0.28239023 | 1.67347431 | 0.09423396 | 0.25263576 |
| ZSCAN31 | 50.4064216 | -0.236498349 | 0.6671565 | -0.3544871 | 0.7229739 | 0.86248065 |
| ZKSCAN3 | 70.1455119 | 0.478786293 | 0.31326545 | 1.52837249 | 0.12642008 | 0.30578335 |
| ZSCAN12 | 97.3937017 | -0.01185253 | 0.21820292 | -0.0543188 | 0.95668114 | 0.98096505 |
| ZSCAN23 | 119.256268 | 0.338009896 | 0.25499653 | 1.32554707 | 0.18498977 | 0.39058981 |
| ZBED9 | 42.6208681 | -0.089634534 | 0.37447768 | -0.2393588 | 0.81082736 | 0.9112091 |
| TRIM27 | 441.252951 | -0.449932254 | 0.17317865 | -2.5980815 | 0.00937462 | 0.05285079 |
| ZNF311 | 36.9617634 | 0.042644427 | 0.48731673 | 0.08750865 | 0.93026721 | 0.96915731 |
| GABBR1 | 246.700201 | -0.194406679 | 0.54546731 | -0.3564039 | 0.7215381 | 0.86142415 |
| HLA-F | 144.31607 | -1.474689533 | 0.68872723 | -2.1411808 | 0.03225946 | 0.12394275 |
| HLA-A | 2344.20027 | -1.389817154 | 0.49710489 | -2.7958227 | 0.00517678 | 0.03436184 |
| ZNRD1 | 75.2802305 | 0.2328498 | 0.40516002 | 0.5747107 | 0.56548697 | 0.75611956 |
| PPP1R11 | 132.470102 | -1.032531123 | 0.27532096 | -3.7502815 | 0.00017664 | 0.00264534 |
| TRIM26 | 662.120874 | 0.320604842 | 0.22943385 | 1.39737373 | 0.16230121 | 0.36024815 |
| HCG17 | 1415.52183 | 0.276689952 | 0.29740866 | 0.93033589 | 0.3521972 | 0.57826884 |
| HLA-E | 916.826265 | -1.92727635 | 0.44131825 | -4.3670896 | 1.26E-05 | 0.00031923 |
| GNL1 | 393.869448 | -0.480695724 | 0.23257634 | -2.0668299 | 0.03875019 | 0.13969738 |
| PRR3 | 164.524962 | 0.058079976 | 0.27089798 | 0.21439797 | 0.83023673 | 0.92071169 |
| ABCF1 | 327.902422 | -0.577899439 | 0.1765311 | -3.2736409 | 0.00106171 | 0.01045801 |
| PPP1R10 | 508.78375 | 0.522708918 | 0.28720886 | 1.81996095 | 0.06876495 | 0.20568756 |
| MRPS18B | 286.237047 | -0.311501276 | 0.19274729 | -1.6161124 | 0.10607002 | 0.27325692 |
| ATAT1 | 195.953697 | -0.209703252 | 0.19930061 | -1.0521957 | 0.29270976 | 0.51558917 |
| C6orf136 | 236.707435 | 0.755885946 | 0.22749893 | 3.32259118 | 0.00089186 | 0.00913218 |
| DHX16 | 230.218739 | -0.249947014 | 0.22144486 | -1.1287099 | 0.25902022 | 0.48019623 |
| PPP1R18 | 61.6217543 | -1.642033648 | 0.31737296 | -5.1738297 | 2.29E-07 | 1.23E-05 |
| MDC1 | 283.501998 | 0.120946848 | 0.24356875 | 0.49656144 | 0.61949835 | 0.79177104 |
| MDC1-AS1 | 51.9350807 | 0.260961208 | 0.38383176 | 0.67988436 | 0.49657769 | 0.70182909 |
| TUBB | 412.516839 | -1.062301354 | 0.48042052 | -2.2111906 | 0.02702264 | 0.10918997 |
| FLOT1 | 718.087528 | -0.64145438 | 0.23140623 | -2.7719841 | 0.00557158 | 0.03637911 |
| IER3 | 1939.72672 | -3.346390068 | 0.48775972 | -6.8607347 | 6.85E-12 | 1.83E-09 |
| DDR1 | 739.726412 | -1.317252224 | 0.33285223 | -3.9574686 | 7.57E-05 | 0.00135394 |
| VARS2 | 114.177204 | 0.509758078 | 0.37116024 | 1.37341781 | 0.1696225 | 0.3699733 |
| PSORS1C1 | 106.071133 | -3.118201883 | 0.98028428 | -3.1809159 | 0.0014681 | 0.01349489 |
| CCHCR1 | 161.199148 | -0.034501434 | 0.29416373 | -0.1172865 | 0.90663302 | 0.95744448 |
| POU5F1 | 898.520508 | -0.98905278 | 0.42954503 | -2.302559 | 0.02130367 | 0.09327151 |
| HCG27 | 64.4064534 | -0.503630298 | 0.42591416 | -1.182469 | 0.23701965 | 0.45466055 |
| HLA-C | 1233.35966 | -1.536952891 | 0.66201986 | -2.3216115 | 0.02025387 | 0.08980642 |
| HLA-B | 1751.92813 | -2.126228354 | 0.62551915 | -3.3991419 | 0.00067598 | 0.00745323 |
| MICA | 270.677404 | 0.292044989 | 0.35819034 | 0.81533464 | 0.41488079 | 0.63486565 |
| NFKBIL1 | 121.837098 | 0.081886903 | 0.17755823 | 0.46118337 | 0.64466705 | 0.80987296 |
| PRRC2A | 407.492146 | 0.046527254 | 0.22771848 | 0.20431918 | 0.83810408 | 0.92446904 |
| BAG6 | 253.199153 | -0.298562347 | 0.21592215 | -1.3827314 | 0.16674723 | 0.36640505 |
| C6orf47 | 74.6034143 | -0.766080344 | 0.30636669 | -2.500534 | 0.01240062 | 0.06434248 |
| GPANK1 | 90.1455113 | -0.581198586 | 0.26471308 | -2.1955794 | 0.02812206 | 0.11232668 |
| DDAH2 | 189.973719 | -0.911955834 | 0.44836141 | -2.0339748 | 0.04195413 | 0.14683946 |
| CLIC1 | 994.533192 | -2.217091473 | 0.43960407 | -5.0433824 | 4.57E-07 | 2.20E-05 |
| VARS | 128.349462 | -0.906285803 | 0.38417036 | -2.3590727 | 0.01832067 | 0.08420239 |
| LSM2 | 158.97854 | -0.67465339 | 0.22323059 | -3.0222264 | 0.00250923 | 0.0202718 |
| HSPA1A | 392.356573 | -1.182087584 | 0.45136322 | -2.6189276 | 0.00882067 | 0.05053466 |
| HSPA1B | 184.853451 | -0.919451815 | 0.56665019 | -1.6226092 | 0.10467297 | 0.27094361 |
| C6orf48 | 160.093941 | -0.04066015 | 0.32389943 | -0.1255333 | 0.9001014 | 0.95395051 |
| NEU1 | 143.610409 | -1.014284634 | 0.40472864 | -2.5060856 | 0.01220761 | 0.0637101 |
| SLC44A4 | 72.1305355 | -2.309821822 | 0.47835644 | -4.8286625 | 1.37E-06 | 5.29E-05 |
| EHMT2 | 189.030829 | 0.335082786 | 0.28606752 | 1.1713416 | 0.2414615 | 0.45982339 |
| C2 | 92.7256048 | 0.533941544 | 0.37458457 | 1.42542321 | 0.1540349 | 0.3482859 |
| NELFE | 120.6069 | -0.33858812 | 0.21476903 | -1.5765221 | 0.11490553 | 0.28822392 |
| SKIV2L | 101.965102 | 0.07891301 | 0.23989361 | 0.32895003 | 0.74219346 | 0.87402695 |
| DXO | 54.6512608 | 0.596108103 | 0.36332304 | 1.64071098 | 0.10085742 | 0.26453475 |
| STK19 | 175.377298 | -0.251544528 | 0.18108626 | -1.3890868 | 0.16480637 | 0.36385214 |
| ATF6B | 394.335743 | -0.175367126 | 0.25478794 | -0.6882866 | 0.49127232 | 0.69746518 |
| AGPAT1 | 118.608374 | -0.38667807 | 0.27235633 | -1.4197506 | 0.1556803 | 0.35065265 |
| RNF5 | 141.478568 | -0.740786487 | 0.3478383 | -2.1296864 | 0.03319752 | 0.12651644 |
| AGER | 41.011567 | 0.324676077 | 0.44002276 | 0.73786201 | 0.46059831 | 0.67405751 |
| PBX2 | 224.871307 | 0.127407489 | 0.24347244 | 0.52329328 | 0.60077018 | 0.7789212 |
| AL662796.1 | 50.0904332 | 0.954582829 | 0.69308415 | 1.37729716 | 0.16842039 | 0.36849761 |
| HLA-DRA | 405.917 | -1.300306356 | 0.64218456 | -2.0248172 | 0.04288614 | 0.14919225 |
| HLA-DRB5 | 57.7847007 | -0.614340403 | 0.98489626 | -0.6237615 | 0.53278421 | 0.73003595 |
| HLA-DRB1 | 231.969391 | -0.267637512 | 0.8892948 | -0.3009548 | 0.76344898 | 0.88541384 |
| HLA-DQB1 | 72.2907463 | -0.526817551 | 0.87108871 | -0.6047806 | 0.54532479 | 0.7400858 |
| PSMB8 | 223.870075 | -1.83812391 | 0.47838386 | -3.8423619 | 0.00012186 | 0.00196053 |
| PSMB9 | 105.817217 | -1.767475248 | 0.71160608 | -2.4837832 | 0.01299949 | 0.06650282 |
| TAP1 | 151.378097 | -1.699076639 | 0.61145821 | -2.778729 | 0.0054572 | 0.0357738 |
| HLA-DMB | 59.0277281 | -1.411856898 | 0.66704897 | -2.1165716 | 0.03429622 | 0.12913878 |
| HLA-DMA | 100.303069 | 1.560866558 | 0.55157532 | 2.82983396 | 0.00465722 | 0.03175016 |
| BRD2 | 607.006071 | -0.198696202 | 0.20331488 | -0.9772831 | 0.32842901 | 0.55362669 |
| HLA-DPA1 | 66.5946219 | -0.846755011 | 0.78651182 | -1.0765954 | 0.28166106 | 0.50418127 |
| HLA-DPB1 | 47.5209389 | -0.46863432 | 0.71871928 | -0.6520408 | 0.51437483 | 0.71598841 |
| RXRB | 149.90174 | -0.088269451 | 0.23689647 | -0.3726077 | 0.70944043 | 0.85423871 |
| SLC39A7 | 181.756953 | -0.416769445 | 0.31476537 | -1.3240638 | 0.18548184 | 0.39112862 |
| HSD17B8 | 41.7114339 | 0.517386484 | 0.39988123 | 1.29385038 | 0.19571711 | 0.4034387 |
| RING1 | 242.941883 | -0.020743223 | 0.2394916 | -0.0866136 | 0.93097868 | 0.96965997 |
| HCG25 | 76.4229434 | 0.140498026 | 0.32998296 | 0.42577358 | 0.67027284 | 0.82659266 |
| VPS52 | 301.49093 | 0.371823006 | 0.15668377 | 2.37307926 | 0.01764048 | 0.0819669 |
| RPS18 | 650.442136 | -0.864951644 | 0.35782176 | -2.4172696 | 0.01563743 | 0.0754676 |
| B3GALT4 | 54.2954575 | -1.277404635 | 0.34032781 | -3.7534536 | 0.00017441 | 0.00262098 |
| WDR46 | 118.264578 | -0.20971473 | 0.33441057 | -0.6271175 | 0.5305822 | 0.72854734 |
| PFDN6 | 114.460493 | -0.519259515 | 0.32430728 | -1.6011343 | 0.10934717 | 0.27803017 |
| RGL2 | 283.31348 | 0.378698227 | 0.26378801 | 1.43561576 | 0.15111171 | 0.34386696 |
| TAPBP | 663.577241 | -1.379953022 | 0.33326968 | -4.1406498 | 3.46E-05 | 0.00073665 |
| DAXX | 96.9028239 | -0.575882079 | 0.26761051 | -2.1519412 | 0.03140199 | 0.12185336 |
| PHF1 | 270.549479 | -0.152948693 | 0.20384129 | -0.7503322 | 0.45305464 | 0.66879735 |
| CUTA | 249.74824 | -0.411955188 | 0.44738642 | -0.920804 | 0.35715279 | 0.58271536 |
| SYNGAP1 | 192.777083 | 0.062487392 | 0.20390999 | 0.30644595 | 0.75926512 | 0.88339056 |
| SYNGAP1-AS | 84.0574335 | 0.351784547 | 0.32572784 | 1.07999534 | 0.28014425 | 0.50263861 |
| ITPR3 | 840.116702 | -1.9205711 | 0.33122512 | -5.798386 | 6.70E-09 | 6.60E-07 |
| UQCC2 | 237.212007 | 0.723033607 | 0.17561757 | 4.11709147 | 3.84E-05 | 0.00079562 |
| LEMD2 | 272.539761 | -0.387533279 | 0.22967316 | -1.687325 | 0.09154087 | 0.2474751 |

| | | | | | | |
|---|---|---|---|---|---|---|
| HMGA1 | 161.569238 | -2.38974784 | 0.6068922 | -3.9376809 | 8.23E-05 | 0.00145209 |
| SMIM29 | 59.9634966 | -1.145803772 | 0.46243142 | -2.477781 | 0.01322023 | 0.06711401 |
| NUDT3 | 265.109518 | -0.864870923 | 0.29237384 | -2.9580995 | 0.00309542 | 0.02367541 |
| C6orf106 | 924.703498 | -0.125903181 | 0.12555912 | -1.0027403 | 0.3159862 | 0.54030573 |
| SNRPC | 186.812609 | -0.294680507 | 0.2443612 | -1.2059218 | 0.22784763 | 0.44334483 |
| UHRF1BP1 | 626.002958 | 0.18276228 | 0.14483286 | 1.26188407 | 0.20699051 | 0.41774448 |
| TAF11 | 374.329165 | 0.081113475 | 0.22472783 | 0.36094094 | 0.7181436 | 0.85963642 |
| ANKS1A | 1946.90893 | -0.005582734 | 0.14243533 | -0.0391949 | 0.96873502 | 0.98582596 |
| SCUBE3 | 54.8852481 | -0.043399825 | 0.53788895 | -0.0806855 | 0.93569209 | 0.97236337 |
| ZNF76 | 495.06533 | 0.079238939 | 0.22080361 | 0.35886614 | 0.71969523 | 0.8603126 |
| PPARD | 461.387174 | 0.098531183 | 0.16851347 | 0.58470804 | 0.55874405 | 0.75065603 |
| RPL10A | 781.095913 | -1.035913107 | 0.43385226 | -2.3877094 | 0.01695374 | 0.079621 |
| TEAD3 | 328.322863 | -0.929159632 | 0.30494771 | -3.0469474 | 0.00231178 | 0.01911947 |
| FKBP5 | 5601.25984 | -1.49486981 | 0.7488617 | -1.9961894 | 0.04591331 | 0.1565997 |
| SRPK1 | 927.934983 | -0.548548946 | 0.31203186 | -1.7579902 | 0.07874918 | 0.22441496 |
| SLC26A8 | 70.5281198 | -2.039421289 | 0.46153086 | -4.4188189 | 9.92E-06 | 0.00026483 |
| MAPK14 | 1978.26574 | -0.890602704 | 0.35822653 | -2.4861439 | 0.01291357 | 0.06622211 |
| MAPK13 | 127.569133 | -1.22079438 | 0.22727681 | -5.3713989 | 7.81E-08 | 5.29E-06 |
| Z84485.1 | 67.3054082 | 1.141675106 | 0.30950643 | 3.68869596 | 0.00022541 | 0.00323218 |
| BRPF3 | 194.143463 | 0.003550528 | 0.28572535 | 0.01242637 | 0.99008545 | 0.99606252 |
| KCTD20 | 637.30458 | -0.154943913 | 0.15061132 | -1.0287667 | 0.30358932 | 0.52681441 |
| STK38 | 1112.31626 | -0.55431799 | 0.15904233 | -3.4853488 | 0.0004915 | 0.00584354 |
| SRSF3 | 839.257493 | 0.20175206 | 0.18974484 | 1.06328086 | 0.2876546 | 0.51002814 |
| CDKN1A | 221.47363 | -2.858003196 | 0.53146678 | -5.3775764 | 7.55E-08 | 5.17E-06 |
| PPIL1 | 87.144088 | 0.310598422 | 0.37442713 | 0.8295297 | 0.40680474 | 0.62679153 |
| C6orf89 | 715.225296 | -0.291350912 | 0.12684697 | -2.2968693 | 0.02162623 | 0.0941532 |
| MTCH1 | 662.22275 | -0.689879613 | 0.2355727 | -2.928521 | 0.00340579 | 0.02547398 |
| PIM1 | 118.322124 | -0.088054205 | 0.26138746 | -0.3368723 | 0.73621314 | 0.87060335 |
| TMEM217 | 108.377147 | -0.385482356 | 0.3436514 | -1.121725 | 0.2619794 | 0.48283603 |
| TBC1D22B | 591.496306 | -0.237352283 | 0.16879828 | -1.4061297 | 0.1596856 | 0.35649376 |
| RNF8 | 729.972878 | -0.695828574 | 0.34048716 | -2.0436265 | 0.04099046 | 0.14480959 |
| CMTR1 | 252.853659 | 0.321111051 | 0.28069806 | 1.14397319 | 0.25263476 | 0.47267033 |
| ZFAND3 | 9941.2268 | -0.004336422 | 0.18153801 | -0.0238871 | 0.98094264 | 0.99160261 |
| BTBD9 | 5982.38957 | -0.308873408 | 0.21130942 | -1.4617115 | 0.14382029 | 0.33340158 |
| GLO1 | 464.303669 | -0.344435877 | 0.23261464 | -1.4807145 | 0.13868266 | 0.32505424 |
| SAYSD1 | 107.455858 | -0.587872179 | 0.27426033 | -2.1434823 | 0.03207439 | 0.12359118 |
| KCNK5 | 129.787981 | -2.215459406 | 0.73841003 | -3.0003106 | 0.00269704 | 0.02135807 |
| KIF6 | 216.361542 | 0.34453458 | 0.67134564 | 0.5132 | 0.60781143 | 0.783582 |
| DAAM2 | 45.7888442 | 0.395377203 | 0.60034837 | 0.65857962 | 0.51016575 | 0.71241986 |
| MOCS1 | 132.43409 | -0.548338703 | 0.42683739 | -1.2846548 | 0.19891294 | 0.40748588 |
| UNC5CL | 159.655113 | -2.139434143 | 0.66819123 | -3.2018291 | 0.00136558 | 0.01275668 |
| OARD1 | 452.5919 | 0.071336329 | 0.27041364 | 0.26380448 | 0.79193058 | 0.90193232 |
| NFYA | 225.328098 | -0.209024029 | 0.18101017 | -1.154764 | 0.24818709 | 0.46734754 |
| FOXP4 | 128.608597 | -0.730757741 | 0.25323197 | -2.8857246 | 0.00390514 | 0.02813616 |
| TFEB | 701.398312 | 0.155139152 | 0.58952205 | 0.2631609 | 0.79242657 | 0.90221177 |
| FRS3 | 49.2478905 | 0.376598132 | 0.28899278 | 1.30314026 | 0.19252688 | 0.39989594 |
| USP49 | 462.934527 | 0.150508137 | 0.16805513 | 0.89558787 | 0.37047292 | 0.59525323 |
| MED20 | 83.8807573 | 0.218164923 | 0.21526328 | 1.0134795 | 0.31083118 | 0.53450314 |
| CCND3 | 2846.82741 | -1.089544931 | 0.34164321 | -3.189131 | 0.00142701 | 0.01317219 |
| TAF8 | 390.418181 | 0.137872281 | 0.18807087 | 0.73308683 | 0.46350547 | 0.67636365 |
| C6orf132 | 160.478224 | -1.78373729 | 0.42672723 | -4.1800409 | 2.91E-05 | 0.00064057 |
| MRPS10 | 140.585045 | -0.121721469 | 0.17963831 | -0.6775919 | 0.49803045 | 0.70320516 |
| TRERF1 | 327.003274 | -0.167248573 | 0.52852805 | -0.3164422 | 0.7516669 | 0.87947193 |
| UBR2 | 2830.58055 | 0.195068201 | 0.15747569 | 1.23871946 | 0.21544941 | 0.42930627 |
| TBCC | 52.3324047 | -0.064715097 | 0.48457336 | -0.1335507 | 0.89375789 | 0.95146032 |
| BICRAL | 1463.88031 | 0.260100128 | 0.26749293 | 0.97236264 | 0.33087017 | 0.55589409 |
| RPL7L1 | 503.479719 | -0.324781776 | 0.21477684 | -1.5121825 | 0.13048744 | 0.31270112 |
| C6orf226 | 62.9491508 | 0.056004132 | 0.40153346 | 0.13947563 | 0.88907431 | 0.95006736 |
| CNPY3 | 128.033014 | -0.941976254 | 0.33054384 | -2.8497771 | 0.00437499 | 0.0303988 |
| AL035587.1 | 66.9798008 | -0.519453493 | 0.49886413 | -1.0412725 | 0.2977491 | 0.52106092 |
| PEX6 | 278.83182 | -0.152177823 | 0.27009055 | -0.5634326 | 0.57314033 | 0.76101836 |
| PPP2R5D | 163.191748 | 0.317173926 | 0.23297639 | 1.36139942 | 0.1733875 | 0.37508606 |
| MEA1 | 74.1095141 | -0.912974707 | 0.37708015 | -2.4211688 | 0.01547069 | 0.07499168 |
| KLHDC3 | 256.501715 | -0.398806142 | 0.19444916 | -2.0509533 | 0.0402715 | 0.14322731 |
| RRP36 | 285.338482 | -0.243217937 | 0.16689873 | -1.4572785 | 0.14503951 | 0.33492926 |
| CUL7 | 170.471621 | 0.019875833 | 0.2236266 | 0.08887956 | 0.92917763 | 0.96900671 |
| KLC4 | 314.57059 | 0.431282057 | 0.1973542 | 2.18531993 | 0.0288654 | 0.11432775 |
| MRPL2 | 177.004148 | 0.169825288 | 0.21973764 | 0.77285481 | 0.43960831 | 0.65531951 |
| PTK7 | 839.089358 | 0.081015416 | 0.32806526 | 0.24694909 | 0.80494763 | 0.90860991 |
| SRF | 82.2676452 | 0.44766912 | 0.46636984 | 0.95990151 | 0.33710478 | 0.56229538 |
| CUL9 | 269.977784 | 0.117824989 | 0.18886498 | 0.62385832 | 0.53272063 | 0.73002457 |
| DNPH1 | 194.371286 | -0.801546816 | 0.47825103 | -1.675996 | 0.09373897 | 0.25184394 |
| ZNF318 | 215.488813 | 0.709771266 | 0.1850894 | 3.8347483 | 0.00012569 | 0.00201246 |
| ABCC10 | 220.426137 | 0.21418653 | 0.23967996 | 0.89363555 | 0.37151692 | 0.59563048 |
| TJAP1 | 261.633676 | -0.10732507 | 0.1733205 | -0.6192289 | 0.53576555 | 0.73267701 |
| POLR1C | 390.645365 | 0.130376145 | 0.24106697 | 0.54082957 | 0.58862506 | 0.77157373 |
| YIPF3 | 283.656399 | -0.878168148 | 0.31877845 | -2.7547914 | 0.00587296 | 0.03778659 |
| XPO5 | 772.528324 | -0.456862759 | 0.24853199 | -1.8382453 | 0.06602627 | 0.20026199 |
| POLH | 428.405484 | 0.820375655 | 0.25693223 | 3.19296509 | 0.0014082 | 0.01306244 |
| GTPBP2 | 339.031622 | 0.164819184 | 0.29362151 | 0.56133211 | 0.57457116 | 0.76207555 |
| MAD2L1BP | 54.7127773 | -0.287658048 | 0.32848439 | -0.875713 | 0.38118608 | 0.6039447 |
| MRPS18A | 123.402538 | -0.714403276 | 0.31815015 | -2.245491 | 0.02473663 | 0.1029446 |
| VEGFA | 1601.80441 | -0.020866994 | 0.47158747 | -0.0442484 | 0.9647064 | 0.98435763 |
| AL109615.3 | 58.5536568 | -1.273436114 | 0.64428356 | -1.976515 | 0.04809648 | 0.16178954 |
| MRPL14 | 208.478359 | -0.787651837 | 0.3228335 | -2.4398083 | 0.01469506 | 0.07224298 |
| TMEM63B | 269.690723 | 0.083988166 | 0.23391183 | 0.35905907 | 0.7195509 | 0.86029578 |
| SLC29A1 | 284.61892 | -0.810496739 | 0.44890941 | -1.8054795 | 0.07099968 | 0.21020651 |
| HSP90AB1 | 1396.69838 | -1.191708056 | 0.42005605 | -2.8370215 | 0.00455365 | 0.03139312 |
| SLC35B2 | 88.8702751 | -0.668450235 | 0.38691963 | -1.7276204 | 0.08405631 | 0.23354285 |
| NFKBIE | 50.4506746 | -1.427300604 | 0.46657745 | -3.0590861 | 0.00222013 | 0.01856273 |
| AARS2 | 88.0166441 | 0.308592832 | 0.34820506 | 0.88623878 | 0.37548887 | 0.59876913 |
| CDC5L | 429.951957 | 0.222015425 | 0.17995389 | 1.23373508 | 0.21730164 | 0.43128814 |
| SUPT3H | 3418.1856 | 0.577835118 | 0.19777355 | 2.92170068 | 0.00348126 | 0.02586274 |
| RUNX2 | 157.605568 | -0.019171596 | 0.45003575 | -0.0426002 | 0.96602027 | 0.98480834 |
| CLIC5 | 90.3077343 | -1.344593481 | 0.49987315 | -2.6898694 | 0.007148 | 0.04335439 |
| AL035701.1 | 63.5105623 | 0.178268044 | 0.49758647 | 0.35826545 | 0.72014467 | 0.86060013 |
| ENPP4 | 279.460305 | -0.783055518 | 0.33467286 | -2.339764 | 0.01929593 | 0.08707889 |
| ENPP5 | 204.60112 | -0.159412258 | 0.38147946 | -0.417879 | 0.67603558 | 0.82991711 |
| RCAN2 | 526.359637 | 1.045508967 | 0.37826246 | 2.76397762 | 0.00571015 | 0.03702764 |
| CYP39A1 | 230.297506 | 0.572904205 | 0.29880071 | 1.91734553 | 0.05519405 | 0.17751599 |
| SLC25A27 | 1127.4652 | 0.5765563 | 0.3515366 | 1.6401032 | 0.10098371 | 0.26455649 |
| AL591242.1 | 65.5540238 | -0.043104887 | 0.86324269 | -0.0499337 | 0.96017523 | 0.98233713 |
| TDRD6 | 115.039346 | 0.676631584 | 0.53115085 | 1.27389721 | 0.20269985 | 0.41232902 |
| ADGRF5 | 49.9245831 | 0.276121152 | 0.55278208 | 0.49951177 | 0.6174189 | 0.7902755 |
| ADGRF1 | 1504.38355 | -0.947461107 | 0.34133166 | -2.7757786 | 0.00550697 | 0.03602851 |
| TNFRSF21 | 475.862089 | 0.427142913 | 0.34960911 | 1.22177282 | 0.22179355 | 0.43589808 |
| AL355353.1 | 58.1699076 | 0.482355547 | 0.46496287 | 1.03740659 | 0.29954641 | 0.52246938 |
| CD2AP | 3157.39084 | -0.568499949 | 0.22435885 | -2.5338869 | 0.01128051 | 0.06037451 |
| PTCHD4 | 186.049225 | -1.056276423 | 0.65066052 | -1.623391 | 0.10450585 | 0.2706171 |
| MUT | 363.889597 | 0.145807072 | 0.18603901 | 0.78374463 | 0.43318997 | 0.65044973 |
| CENPQ | 59.727941 | -0.414769044 | 0.35075865 | -1.1824913 | 0.23701083 | 0.45466055 |
| TFAP2B | 1428.61284 | 0.307300399 | 0.44638746 | 0.68841629 | 0.49119067 | 0.69742422 |
| AL355997.1 | 114.617985 | 0.388926908 | 0.63903608 | 0.60861494 | 0.5427797 | 0.73882698 |

| | | | | | | |
|---|---|---|---|---|---|---|
| PKHD1 | 21867.5805 | 0.776916743 | 0.2499364 | 3.10845782 | 0.00188066 | 0.01632175 |
| MCM3 | 191.156494 | -0.06755138 | 0.28338196 | -0.2383757 | 0.81158969 | 0.91158913 |
| EFHC1 | 717.171257 | -0.092553353 | 0.14206332 | -0.6514936 | 0.51472788 | 0.71629167 |
| TRAM2 | 212.435954 | 0.192959008 | 0.34424362 | 0.56053038 | 0.57511772 | 0.76252365 |
| TRAM2-AS1 | 45.5938659 | 0.040448389 | 0.29445253 | 0.13736812 | 0.89073983 | 0.95046308 |
| TMEM14A | 107.63683 | 0.5600374 | 0.25630361 | 2.18505464 | 0.02888484 | 0.11433041 |
| GSTA4 | 300.9713 | 0.842379613 | 0.34399471 | 2.44881562 | 0.01433268 | 0.07098626 |
| ICK | 273.530096 | -0.118474819 | 0.19095932 | -0.6204192 | 0.53498186 | 0.73210674 |
| FBXO9 | 770.395007 | 0.181847306 | 0.1389259 | 1.30895176 | 0.19055069 | 0.39706328 |
| ELOVL5 | 298.999396 | -0.603811918 | 0.46611197 | -1.2954225 | 0.19517453 | 0.40269608 |
| GCLC | 506.361566 | -0.947418382 | 0.468544 | -2.0220478 | 0.04317141 | 0.14994771 |
| AL513211.1 | 39.9603005 | 1.048429376 | 0.33930623 | 3.08992082 | 0.0020021 | 0.01713916 |
| LRRC1 | 521.93911 | -0.005158949 | 0.15367515 | -0.0335705 | 0.97321966 | 0.98804941 |
| TINAG | 975.424887 | -1.22418547 | 0.78234143 | -1.5647714 | 0.11763653 | 0.29235389 |
| FAM83B | 1803.78616 | -0.430567243 | 0.23200248 | -1.8558734 | 0.0634716 | 0.19518591 |
| HCRTR2 | 103.77357 | 3.439572043 | 0.527327 | 6.52265488 | 6.91E-11 | 1.27E-08 |
| DST | 4208.89732 | -1.279905037 | 0.3284039 | -3.8973503 | 9.73E-05 | 0.0016448 |
| KIAA1586 | 103.649972 | 0.124769374 | 0.27383951 | 0.45562955 | 0.64865639 | 0.81191151 |
| ZNF451 | 1268.66062 | 0.227417467 | 0.11666685 | 1.94928957 | 0.05126085 | 0.16889614 |
| ZNF451-AS1 | 188.987121 | 0.191932478 | 0.56680821 | 0.33861979 | 0.73489617 | 0.86990311 |
| BAG2 | 62.2488825 | 0.349103441 | 0.39560861 | 0.88244652 | 0.37753539 | 0.60053769 |
| RAB23 | 99.812207 | 0.337532849 | 0.40266916 | 0.83823863 | 0.4018967 | 0.62337405 |
| PRIM2 | 991.979672 | -0.071962774 | 0.30486589 | -0.2360473 | 0.81339596 | 0.91245411 |
| AL021368.2 | 165.628754 | 0.633248784 | 0.25779924 | 2.45636406 | 0.01403509 | 0.06987919 |
| AL445250.1 | 76.9263899 | 0.750219628 | 0.57204672 | 1.31146566 | 0.18970049 | 0.39572864 |
| PTP4A1 | 839.740491 | -1.260367379 | 0.64376515 | -1.9578062 | 0.05025276 | 0.16685843 |
| PHF3 | 2236.94699 | 0.03834431 | 0.1712609 | 0.22389413 | 0.82283969 | 0.91813562 |
| EYS | 246.190066 | -0.361811526 | 0.53726609 | -0.6734308 | 0.50067327 | 0.70491515 |
| LMBRD1 | 1726.57321 | 0.257266295 | 0.1435887 | 1.791689 | 0.0731828 | 0.21436762 |
| COL19A1 | 447.945161 | 0.239991422 | 0.3074564 | 0.78057059 | 0.4350551 | 0.65199033 |
| FAM135A | 4544.6158 | -1.207489837 | 0.25910969 | -4.6601493 | 3.16E-06 | 0.0001028 |
| SDHAF4 | 160.817203 | 0.703809862 | 0.17886844 | 3.93479066 | 8.33E-05 | 0.00146276 |
| SMAP1 | 1974.97926 | 0.078460055 | 0.17695325 | 0.44339427 | 0.65748058 | 0.81859324 |
| OGFRL1 | 272.190896 | -0.912013753 | 0.42755004 | -2.1331158 | 0.03291523 | 0.12576721 |
| AL136164.2 | 2060.20589 | 0.349323808 | 0.38801364 | 0.90028745 | 0.3679673 | 0.59306694 |
| LINC00472 | 917.114702 | -0.212202998 | 0.32791219 | -0.6471336 | 0.51754548 | 0.71869378 |
| RIMS1 | 126.604468 | -1.231211613 | 0.87276733 | -1.4106986 | 0.15833352 | 0.35465956 |
| KHDC1 | 71.0283248 | -0.198612701 | 0.34106128 | -0.5823373 | 0.56033952 | 0.75183103 |
| AC019205.1 | 189.960598 | 0.438874703 | 0.19721508 | 2.22536078 | 0.02605703 | 0.10649359 |
| OOEP | 666.556024 | -1.131823737 | 0.39471126 | -2.8674727 | 0.00413765 | 0.02926733 |
| MTO1 | 414.442066 | 0.644917562 | 0.240516 | 2.68139153 | 0.00733167 | 0.04418924 |
| EEF1A1 | 3472.3364 | -1.352687401 | 0.47475772 | -2.8492162 | 0.00438271 | 0.0303988 |
| AL121972.1 | 48.4534306 | -0.022934017 | 0.45774458 | -0.0501022 | 0.96004093 | 0.98229715 |
| SLC17A5 | 283.693229 | -0.498329167 | 0.29732826 | -1.6760236 | 0.09373358 | 0.25184394 |
| AL357507.1 | 55.7176335 | 0.713570822 | 0.33679279 | 2.11872357 | 0.03411384 | 0.12889292 |
| COX7A2 | 491.18205 | 0.020438056 | 0.35711526 | 0.05723098 | 0.9543612 | 0.98007893 |
| TMEM30A | 543.699784 | -0.437260078 | 0.26200432 | -1.6689041 | 0.09513639 | 0.25410155 |
| AL080250.1 | 101.044491 | 1.026437876 | 0.33536544 | 3.06065493 | 0.00220853 | 0.01852224 |
| FILIP1 | 86.787868 | 0.845545545 | 0.42910471 | 1.97048767 | 0.04878251 | 0.16329654 |
| SENP6 | 4050.453 | 0.176079724 | 0.12959238 | 1.35871972 | 0.17423542 | 0.37589771 |
| MYO6 | 7032.49763 | -1.085056974 | 0.33335606 | -3.254949 | 0.00113413 | 0.01102332 |
| MEI4 | 252.944934 | 1.166012462 | 0.36523795 | 3.19247343 | 0.0014106 | 0.01307283 |
| IRAK1BP1 | 1011.63046 | 0.568404784 | 0.2425476 | 2.34347725 | 0.01910493 | 0.08651251 |
| PHIP | 2693.93393 | -0.291825982 | 0.1996348 | -1.4617992 | 0.14379625 | 0.33340158 |
| HMGN3 | 990.036025 | 0.029778762 | 0.20046766 | 0.14854646 | 0.88191152 | 0.94672712 |
| LCA5 | 139.289704 | -0.2660704 | 0.30275453 | -0.8788321 | 0.37949232 | 0.60235468 |
| SH3BGRL2 | 149.269979 | 0.518903718 | 0.28313356 | 1.8327171 | 0.06684465 | 0.20190968 |
| BCKDHB | 2443.20254 | 0.586800269 | 0.31268956 | 1.87662252 | 0.06056985 | 0.18914117 |
| AL359715.1 | 115.598024 | 0.826302331 | 0.47684158 | 1.7328655 | 0.08311958 | 0.23192364 |
| TENT5A | 260.344589 | -1.290605149 | 0.65004377 | -1.9854127 | 0.04709857 | 0.15923345 |
| IBTK | 1556.46054 | 0.011273871 | 0.24304816 | 0.04638534 | 0.96300312 | 0.98360994 |
| TPBG | 183.658145 | -2.63682719 | 0.44945236 | -5.8667557 | 4.44E-09 | 4.85E-07 |
| UBE3D | 1228.32072 | 0.238010558 | 0.19528459 | 1.2187882 | 0.22292459 | 0.43734011 |
| DOPEY1 | 1733.76661 | 0.068590315 | 0.15178794 | 0.45188252 | 0.65135362 | 0.81378687 |
| PGM3 | 410.384146 | 0.101559178 | 0.25615518 | 0.39647521 | 0.69175449 | 0.84169408 |
| RWDD2A | 72.4378977 | 0.172465266 | 0.35204428 | 0.48989652 | 0.62420713 | 0.79563369 |
| ME1 | 280.957488 | -0.03622058 | 0.48039711 | -0.0753972 | 0.93989872 | 0.97411372 |
| CYB5R4 | 519.055403 | -0.24572235 | 0.27417485 | -0.896225 | 0.37013262 | 0.59485116 |
| MRAP2 | 205.414197 | -0.889373152 | 0.542824 | -1.638419 | 0.10133432 | 0.26505446 |
| CEP162 | 951.766273 | 0.218544912 | 0.14582721 | 1.49865663 | 0.13396273 | 0.31830389 |
| NT5E | 55.6719273 | -3.602436216 | 0.66771199 | -5.3951947 | 6.84E-08 | 4.78E-06 |
| SNX14 | 1534.00185 | 0.097885738 | 0.10946375 | 0.89422968 | 0.37119902 | 0.59563048 |
| AL589666.1 | 84.5397058 | 0.061390496 | 0.34701813 | 0.17690861 | 0.85958017 | 0.93625342 |
| SYNCRIP | 535.587485 | -0.013755469 | 0.18310045 | -0.0751253 | 0.94011505 | 0.97411372 |
| ZNF292 | 5250.59933 | -0.439696497 | 0.27521425 | -1.5976516 | 0.11012053 | 0.27961977 |
| SMIM8 | 956.047829 | 0.851667729 | 0.24272276 | 3.50880871 | 0.00045012 | 0.00546472 |
| SLC35A1 | 68.8973998 | 0.512941922 | 0.40686116 | 1.26072965 | 0.20740627 | 0.41826403 |
| RARS2 | 1015.65454 | 0.076980098 | 0.16883012 | 0.45596187 | 0.64841739 | 0.81184316 |
| ORC3 | 742.9493 | 0.257051487 | 0.34861572 | 0.73734909 | 0.46091009 | 0.67428966 |
| AKIRIN2 | 331.284348 | 0.209558434 | 0.19036266 | 1.1008379 | 0.27096721 | 0.49232544 |
| RNGTT | 1733.75005 | 0.27414322 | 0.2197351 | 1.24760777 | 0.21217473 | 0.42488177 |
| PNRC1 | 630.221363 | -0.666094042 | 0.38946504 | -1.7102794 | 0.08721422 | 0.23941076 |
| SRSF12 | 97.3328414 | -0.407144237 | 0.40753575 | -0.9990393 | 0.31777564 | 0.54237242 |
| PM20D2 | 427.928075 | -0.278811529 | 0.22557581 | -1.2359992 | 0.21645885 | 0.43040869 |
| UBE2J1 | 544.860089 | 0.119590209 | 0.19922566 | 0.60027513 | 0.54832289 | 0.74193783 |
| RRAGD | 686.692341 | 0.581092529 | 0.40021741 | 1.45194216 | 0.14651769 | 0.33711064 |
| ANKRD6 | 1734.42014 | 0.109481193 | 0.17792081 | 0.61533663 | 0.53833243 | 0.73492731 |
| LYRM2 | 413.879072 | 0.130843462 | 0.28093447 | 0.4657437 | 0.64139897 | 0.80757823 |
| MDN1 | 989.896388 | 0.129576379 | 0.13053122 | 0.99268499 | 0.32086349 | 0.54511971 |
| CASP8AP2 | 568.12191 | 0.27282208 | 0.1738095 | 1.56966147 | 0.11649389 | 0.29055283 |
| BACH2 | 1847.76413 | -0.99377763 | 0.44971596 | -2.2097895 | 0.02711977 | 0.10948199 |
| MAP3K7 | 1070.97371 | -0.083059903 | 0.19220575 | -0.4321406 | 0.66563925 | 0.82357564 |
| EPHA7 | 50.9050683 | 1.055287551 | 0.57344095 | 1.8402724 | 0.06572826 | 0.19977094 |
| MANEA | 175.471497 | 0.696324402 | 0.25717427 | 2.7075975 | 0.00677722 | 0.04187102 |
| UFL1-AS1 | 372.147498 | -0.322956982 | 0.53287223 | -0.6060683 | 0.54446938 | 0.73944091 |
| UFL1 | 691.35388 | 0.131348991 | 0.17895447 | 0.73397992 | 0.46296098 | 0.67594247 |
| NDUFAF4 | 174.028821 | 0.459140922 | 0.3395372 | 1.35225515 | 0.17629371 | 0.37852139 |
| KLHL32 | 614.01196 | 0.976604962 | 0.3763484 | 2.59494913 | 0.00946049 | 0.05324399 |
| MMS22L | 996.239343 | -1.248229546 | 0.48763975 | -2.5597371 | 0.01047514 | 0.05738951 |
| AL589740.1 | 309.072756 | 0.539649338 | 0.50764561 | 1.06304345 | 0.28776225 | 0.51007132 |
| FBXL4 | 2166.82424 | 0.47598191 | 0.20614962 | 2.30891484 | 0.02094831 | 0.09214325 |
| FAXC | 167.666599 | 0.499252367 | 0.43921156 | 1.13670133 | 0.25566317 | 0.47664852 |
| COQ3 | 122.804873 | 1.304897081 | 0.24600667 | 5.30431596 | 1.13E-07 | 6.92E-06 |
| PNISR | 3905.20646 | 0.694095618 | 0.14577195 | 4.76151687 | 1.92E-06 | 6.84E-05 |
| AL513550.1 | 282.861869 | 0.258078639 | 0.31491113 | 0.81952847 | 0.41248496 | 0.63185827 |
| USP45 | 770.883923 | 0.095129521 | 0.26900602 | 0.35363343 | 0.72361361 | 0.86271256 |
| TSTD3 | 131.054916 | -0.314307267 | 0.23930001 | -1.3134444 | 0.18903323 | 0.39508544 |
| CCNC | 1096.54887 | -0.292889335 | 0.20635323 | -1.4193591 | 0.15579434 | 0.35075635 |
| SIM1 | 3599.36576 | 1.001936071 | 0.47121674 | 2.12627437 | 0.03348042 | 0.12730077 |
| ASCC3 | 2428.00062 | -0.261507521 | 0.19008651 | -1.375729 | 0.16890555 | 0.36913855 |
| GRIK2 | 551.258183 | 1.366114163 | 0.57844412 | 2.36170465 | 0.01819113 | 0.08384041 |
| HACE1 | 941.137569 | 0.703236197 | 0.10536136 | 6.67451713 | 2.48E-11 | 5.65E-09 |
| PREP | 481.531892 | 0.104998635 | 0.14291364 | 0.7346999 | 0.46252228 | 0.67560066 |

| | | | | | | |
|---|---|---|---|---|---|---|
| ATG5 | 1262.26178 | 0.133591966 | 0.19812788 | 0.6742714 | 0.50013878 | 0.70482771 |
| CRYBG1 | 942.35229 | -0.051596606 | 0.45249185 | -0.1140277 | 0.90921583 | 0.95902196 |
| RTN4IP1 | 364.386332 | 0.356689384 | 0.34304614 | 1.03977087 | 0.29844637 | 0.52159011 |
| QRSL1 | 619.428312 | 0.583455156 | 0.29767246 | 1.96005757 | 0.04998906 | 0.16632381 |
| LINC02532 | 11016.7901 | -0.205523668 | 0.33563672 | -0.6123396 | 0.54031307 | 0.73628429 |
| CD24 | 5013.94885 | -1.190593184 | 0.26243261 | -4.5367578 | 5.71E-06 | 0.00016995 |
| C6orf203 | 252.505756 | 0.30316294 | 0.18701497 | 1.62106238 | 0.10500426 | 0.27158827 |
| BEND3 | 56.8623142 | -0.078457685 | 0.38518656 | -0.2036875 | 0.83859772 | 0.92470467 |
| PDSS2 | 2205.35012 | 0.11728979 | 0.20451563 | 0.57350038 | 0.56630594 | 0.75654506 |
| SOBP | 151.990838 | 0.40417397 | 0.71799564 | 0.56291981 | 0.57348949 | 0.76117591 |
| SEC63 | 1092.45754 | -0.017268615 | 0.18333379 | -0.0941922 | 0.92495647 | 0.9667507 |
| OSTM1 | 388.634158 | -0.138277482 | 0.27392479 | -0.5048009 | 0.61369868 | 0.78755051 |
| SNX3 | 765.581752 | -0.168442421 | 0.11541043 | -1.4595078 | 0.14442541 | 0.33427636 |
| AFG1L | 1022.78427 | 0.987497364 | 0.22586206 | 4.37212596 | 1.23E-05 | 0.00031376 |
| FOXO3 | 5197.82915 | -0.451430506 | 0.27878234 | -1.6192938 | 0.10538407 | 0.27219754 |
| LINC00222 | 45.7219749 | -0.287773765 | 0.50884728 | -0.5655405 | 0.57170616 | 0.75986595 |
| ARMC2 | 691.968622 | 0.16843607 | 0.25712422 | 0.65507664 | 0.51241841 | 0.71435723 |
| SESN1 | 1233.71445 | -0.325476905 | 0.21835503 | -1.4905858 | 0.13607028 | 0.32134191 |
| CEP57L1 | 288.961152 | 0.150276151 | 0.20297942 | 0.74035167 | 0.45908664 | 0.67278348 |
| CD164 | 692.542893 | -0.393875367 | 0.19118389 | -2.0601912 | 0.03938027 | 0.14131036 |
| PPIL6 | 280.321985 | 0.150988667 | 0.29635269 | 0.50948979 | 0.61040895 | 0.78585691 |
| SMPD2 | 58.6562694 | -0.445203448 | 0.2960456 | -1.503834 | 0.1326241 | 0.31632932 |
| MICAL1 | 115.207165 | -0.459458125 | 0.34506705 | -1.3315039 | 0.18302326 | 0.38786366 |
| ZBTB24 | 165.693558 | -0.161684266 | 0.1823727 | -0.8865596 | 0.37531606 | 0.59870627 |
| AK9 | 780.302355 | 0.220064947 | 0.22162629 | 0.99295505 | 0.32073186 | 0.54511971 |
| FIG4 | 675.830703 | -0.098856583 | 0.18632995 | -0.5305459 | 0.59573352 | 0.77558092 |
| WASF1 | 444.872336 | -0.780450346 | 0.31931947 | -2.4441051 | 0.01452119 | 0.07162452 |
| CDC40 | 401.176877 | 0.30667377 | 0.1502208 | 2.04148671 | 0.04120247 | 0.14520904 |
| METTL24 | 80.3547867 | 1.181286056 | 0.65468042 | 1.80437053 | 0.07117324 | 0.21050266 |
| CDK19 | 1609.5822 | -0.192050937 | 0.23270934 | -0.8252825 | 0.40921124 | 0.62915416 |
| AMD1 | 573.074609 | -0.178771452 | 0.26446024 | -0.6759861 | 0.49904946 | 0.70390435 |
| GTF3C6 | 280.336637 | -0.913394391 | 0.39440283 | -2.3158921 | 0.02056416 | 0.09092934 |
| RPF2 | 238.259705 | -0.123352071 | 0.27879144 | -0.4424529 | 0.65816153 | 0.81888605 |
| SLC16A10 | 411.055848 | 1.112437788 | 0.44325505 | 2.50970132 | 0.01208333 | 0.06318635 |
| MFSD4B | 268.554519 | 0.495921045 | 0.22761462 | 2.17877501 | 0.02934839 | 0.11558224 |
| AL080317.2 | 57.3409546 | 0.366165929 | 0.2965353 | 1.23481396 | 0.21689975 | 0.4310907 |
| REV3L | 2699.25829 | 0.300047521 | 0.16811428 | 1.78478312 | 0.0742965 | 0.2163675 |
| AL080317.3 | 105.009103 | 0.486669571 | 0.25246014 | 1.92770856 | 0.05389139 | 0.17473032 |
| TRAF3IP2-AS | 1246.94917 | 0.11495529 | 0.2157185 | 0.53289491 | 0.59410633 | 0.77437838 |
| TRAF3IP2 | 236.402721 | -0.300447431 | 0.21809728 | -1.3775845 | 0.16833162 | 0.36843286 |
| FYN | 82.881749 | -1.365896012 | 0.49278373 | -2.7717961 | 0.00557479 | 0.03638215 |
| TUBE1 | 417.484819 | 0.635345763 | 0.20246509 | 3.1380509 | 0.00170075 | 0.01509762 |
| FAM229B | 161.142557 | 0.179049353 | 0.26651084 | 0.67182765 | 0.50169342 | 0.70566164 |
| LAMA4 | 286.960945 | -1.561950115 | 0.94292566 | -1.6564934 | 0.09762194 | 0.2582075 |
| MARCKS | 156.9321 | -0.880177253 | 0.34197078 | -2.5738376 | 0.01005775 | 0.05568012 |
| HDAC2 | 480.072146 | -0.091494576 | 0.20324352 | -0.4501722 | 0.6525863 | 0.814823 |
| HDAC2-AS2 | 73.6511766 | -0.120868816 | 0.39287674 | -0.3076507 | 0.75834812 | 0.88316847 |
| FRK | 1946.97877 | 0.471143104 | 0.24355048 | 1.9344782 | 0.05305436 | 0.172875 |
| NT5DC1 | 1989.11633 | 0.723760211 | 0.23502244 | 3.07953657 | 0.00207323 | 0.01762245 |
| COL10A1 | 99.0469709 | 0.014220653 | 0.67486922 | 0.02107172 | 0.98318845 | 0.99288503 |
| TSPYL4 | 138.830843 | 0.276914038 | 0.28085 | 0.98598552 | 0.32414022 | 0.54898937 |
| DSE | 1299.36187 | 1.001617141 | 0.19520689 | 5.13105412 | 2.88E-07 | 1.48E-05 |
| TSPYL1 | 291.49272 | -0.902465413 | 0.45225254 | -1.99549 | 0.04598947 | 0.15675994 |
| RWDD1 | 237.380879 | 0.214894417 | 0.23634813 | 0.90922835 | 0.3632296 | 0.5885693 |
| ZUP1 | 250.885336 | 0.289148786 | 0.18344829 | 1.57618691 | 0.11498273 | 0.28829365 |
| KPNA5 | 761.199018 | 0.366819598 | 0.14806689 | 2.47739114 | 0.01323468 | 0.06711401 |
| DCBLD1 | 300.929851 | 0.259049474 | 0.30034019 | 0.86252017 | 0.38840134 | 0.61156196 |
| GOPC | 501.790875 | 0.210093213 | 0.19920542 | 1.05465608 | 0.29158264 | 0.51428964 |
| NUS1 | 475.856446 | -0.020594275 | 0.19079273 | -0.1079406 | 0.91404284 | 0.96119671 |
| SLC35F1 | 114.191484 | 0.505281681 | 0.39166323 | 1.2900922 | 0.19701865 | 0.40536126 |
| CEP85L | 752.159411 | -0.265735172 | 0.25208074 | -1.0541669 | 0.2918065 | 0.51440973 |
| MCM9 | 1246.09308 | -0.139482668 | 0.21996884 | -0.6341019 | 0.5260143 | 0.72497874 |
| ASF1A | 90.0983956 | 0.41563938 | 0.3142661 | 1.32257147 | 0.18597793 | 0.39174123 |
| AL365275.1 | 491.124205 | 0.090266911 | 0.46462958 | 0.19427715 | 0.84595887 | 0.9287667 |
| FAM184A | 1732.42012 | 1.117148922 | 0.33081499 | 3.37695977 | 0.00073292 | 0.00791587 |
| MAN1A1 | 2922.55492 | 0.793709919 | 0.24239381 | 3.27446449 | 0.00105862 | 0.01044314 |
| TBC1D32 | 1482.23764 | 0.329357282 | 0.26081442 | 1.26280319 | 0.20665992 | 0.41739901 |
| GJA1 | 70.1001106 | -2.418638659 | 0.67787509 | -3.567971 | 0.00035976 | 0.00459133 |
| HSF2 | 218.149342 | 0.363656366 | 0.22848184 | 1.59162043 | 0.11147002 | 0.28207039 |
| SERINC1 | 808.76686 | -0.607644183 | 0.208153 | -2.9192189 | 0.0035091 | 0.02604026 |
| PKIB | 260.820957 | 0.837656926 | 0.38374729 | 2.18283475 | 0.02904798 | 0.11477777 |
| SMPDL3A | 108.072447 | 0.433888449 | 0.29737692 | 1.4590522 | 0.14455074 | 0.33439067 |
| NKAIN2 | 526.830502 | 1.869955246 | 0.56456715 | 3.31219281 | 0.00092568 | 0.00939836 |
| RNF217-AS1 | 98.8729657 | -1.305862447 | 0.51066951 | -2.5571577 | 0.01055314 | 0.05767331 |
| RNF217 | 1288.19169 | -0.991167795 | 0.38787798 | -2.5553598 | 0.01060781 | 0.05789597 |
| TPD52L1 | 933.027657 | 0.358909497 | 0.31811768 | 1.12822871 | 0.25922334 | 0.48023578 |
| HDDC2 | 354.604008 | 0.150377126 | 0.21201239 | 0.70928461 | 0.47814788 | 0.6877769 |
| AL365259.1 | 74.3696162 | 1.90594284 | 0.68544407 | 2.7805957 | 0.00542593 | 0.0356218 |
| NCOA7 | 4171.98304 | -1.34338971 | 0.41732707 | -3.2190332 | 0.00128624 | 0.01218787 |
| NCOA7-AS1 | 63.3131173 | -1.816065029 | 0.7894728 | -2.3003516 | 0.02142831 | 0.0935691 |
| HINT3 | 260.58486 | -0.342435975 | 0.16804644 | -2.037746 | 0.04157534 | 0.14605549 |
| TRMT11 | 745.456637 | 0.221419615 | 0.24906246 | 0.88901238 | 0.37399642 | 0.59778785 |
| AC020559.1 | 265.722441 | 0.326727487 | 0.40363635 | 0.80946002 | 0.41825059 | 0.63753323 |
| RNF146 | 413.596017 | 0.298435307 | 0.25222407 | 1.183215 | 0.23672395 | 0.45435726 |
| ECHDC1 | 1057.48004 | 0.233468248 | 0.19221325 | 1.21463139 | 0.22450671 | 0.43913952 |
| PTPRK | 4596.12016 | -0.550802432 | 0.23518755 | -2.3419711 | 0.0191822 | 0.08671379 |
| LAMA2 | 269.773635 | 1.765819143 | 0.4538535 | 3.89072497 | 9.99E-05 | 0.00167111 |
| AL356124.1 | 337.402523 | 1.083177501 | 0.40185404 | 2.69545007 | 0.00702936 | 0.04292004 |
| ARHGAP18 | 2326.69946 | 0.983140955 | 0.21265834 | 4.62310084 | 3.78E-06 | 0.0001189 |
| L3MBTL3 | 428.366819 | -0.872014289 | 0.30754279 | -2.8354242 | 0.00457649 | 0.03145229 |
| EPB41L2 | 620.584871 | -1.291314219 | 0.30563136 | -4.225071 | 2.39E-05 | 0.00054873 |
| AKAP7 | 516.672902 | 0.653100933 | 0.33313543 | 1.96046672 | 0.04994126 | 0.16624853 |
| MED23 | 724.902118 | 0.513368039 | 0.21458622 | 2.39236252 | 0.0167403 | 0.07911363 |
| ENPP3 | 47.3834399 | 0.71030334 | 0.38360046 | 1.8516749 | 0.06407252 | 0.19636518 |
| ENPP1 | 81.0577903 | 1.545494281 | 0.45862405 | 3.36985008 | 0.00075209 | 0.00807458 |
| CTGF | 128.143721 | 0.26927697 | 0.67365488 | 0.39972541 | 0.68935878 | 0.84032301 |
| AL133346.1 | 94.3602183 | 1.635961012 | 0.68714152 | 2.38082107 | 0.0172741 | 0.08065521 |
| STX7 | 1103.30245 | 0.058696942 | 0.19336757 | 0.30355112 | 0.7614699 | 0.88431255 |
| SLC18B1 | 578.256757 | 0.770347375 | 0.25467651 | 3.02480729 | 0.00248792 | 0.02016727 |
| RPS12 | 747.843324 | -0.528753763 | 0.44122465 | -1.1983777 | 0.23077002 | 0.44643252 |
| TBPL1 | 305.011148 | 0.40739655 | 0.26112577 | 1.56015453 | 0.11872337 | 0.29400393 |
| SLC2A12 | 213.198625 | 1.428356764 | 0.36774406 | 3.88410563 | 0.00010271 | 0.00170435 |
| SGK1 | 454.906778 | -2.406402392 | 0.51491699 | -4.6733792 | 2.96E-06 | 9.81E-05 |
| ALDH8A1 | 85.6803372 | 0.201985951 | 0.400262 | 0.50463434 | 0.61381569 | 0.78755051 |
| HBS1L | 662.100582 | 0.224835796 | 0.19062769 | 1.17944985 | 0.2382191 | 0.45614418 |
| AHI1 | 3289.2781 | -0.522697452 | 0.24255294 | -2.154983 | 0.03116317 | 0.1210689 |
| LINC00271 | 1587.54212 | -0.104733034 | 0.39926609 | -0.2623139 | 0.79307947 | 0.90259076 |
| PDE7B | 7114.95948 | -0.406587392 | 0.31737515 | -1.281094 | 0.20016066 | 0.40902261 |
| AL138828.1 | 927.749098 | -1.370052416 | 0.59188027 | -2.3147459 | 0.02062684 | 0.09107013 |
| BCLAF1 | 1421.10996 | 0.195666029 | 0.1735818 | 1.12722666 | 0.25964666 | 0.48061767 |
| MAP7 | 2923.26935 | -0.254758358 | 0.17034711 | -1.495525 | 0.13477748 | 0.31961862 |
| MAP3K5 | 1644.92635 | -1.442595546 | 0.64685236 | -2.2301775 | 0.02573566 | 0.10557217 |
| PEX7 | 409.470345 | 0.162465121 | 0.19142126 | 0.84873082 | 0.39603109 | 0.61804994 |

| | | | | | | |
|---|---|---|---|---|---|---|
| IFNGR1 | 293.579474 | -0.372748451 | 0.2146012 | -1.7369355 | 0.08239855 | 0.23069744 |
| TNFAIP3 | 84.6470503 | -2.504550457 | 0.56222446 | -4.4547163 | 8.40E-06 | 0.00023459 |
| PERP | 353.663266 | -1.708603295 | 0.3190377 | -5.3554903 | 8.53E-08 | 5.66E-06 |
| ARFGEF3 | 1314.50297 | 0.296580928 | 0.38286911 | 0.77462746 | 0.43855982 | 0.65476683 |
| HEBP2 | 517.782956 | -0.895652207 | 0.2209823 | -4.0530496 | 5.06E-05 | 0.00098058 |
| NHSL1 | 1316.79501 | 0.421046386 | 0.38710395 | 1.08768301 | 0.27673507 | 0.49896172 |
| CCDC28A | 234.68135 | -0.060153637 | 0.18985618 | -0.3168379 | 0.75136659 | 0.87939051 |
| REPS1 | 1329.9678 | 0.032180303 | 0.1951904 | 0.16486622 | 0.86904928 | 0.94023032 |
| ABRACL | 59.5737722 | -1.94618015 | 0.54631413 | -3.562383 | 0.0003675 | 0.00467214 |
| HECA | 227.46019 | -0.640146684 | 0.32974169 | -1.9413581 | 0.05221486 | 0.1711902 |
| AL031772.1 | 86.9656721 | -0.365532185 | 0.42393219 | -0.8622421 | 0.3885543 | 0.61165699 |
| CITED2 | 262.546287 | -0.205903492 | 0.28605232 | -0.7198106 | 0.47164161 | 0.68273031 |
| VTA1 | 558.776711 | 0.103171723 | 0.19815475 | 0.52066238 | 0.60260198 | 0.77979671 |
| HIVEP2 | 2876.85993 | -0.953938961 | 0.39593551 | -2.4093292 | 0.01598188 | 0.07651593 |
| LINC01277 | 91.146014 | -0.354738858 | 0.54104808 | -0.6556513 | 0.51204853 | 0.71407594 |
| AIG1 | 1955.77166 | 0.154313914 | 0.14186032 | 1.0877877 | 0.27668884 | 0.49896172 |
| ADAT2 | 186.10141 | 0.34939554 | 0.27586023 | 1.26656728 | 0.20531006 | 0.41584887 |
| PEX3 | 695.860685 | 0.276629088 | 0.21242806 | 1.30222481 | 0.19283955 | 0.40019129 |
| FUCA2 | 226.206415 | -0.274319051 | 0.18724159 | -1.4650541 | 0.14290617 | 0.33233231 |
| PHACTR2 | 1195.19465 | -0.752982318 | 0.25809299 | -2.9174846 | 0.00352867 | 0.0261268 |
| PLAGL1 | 887.793155 | -0.242401886 | 0.23554059 | -1.02913 | 0.30341859 | 0.52658733 |
| SF3B5 | 292.93057 | -1.110883815 | 0.4797233 | -2.3156762 | 0.02057596 | 0.09092934 |
| UTRN | 4862.12056 | 0.362117138 | 0.22519462 | 1.60801861 | 0.1078311 | 0.2759816 |
| AL024474.2 | 107.336593 | -0.463770285 | 0.43992727 | -1.0541976 | 0.29179245 | 0.51440973 |
| EPM2A | 793.512311 | 0.933191729 | 0.35393273 | 2.63663583 | 0.00837327 | 0.04880957 |
| FBXO30 | 189.557528 | 0.25658378 | 0.34760063 | 0.73815683 | 0.46041915 | 0.67395297 |
| AL356599.1 | 598.790628 | 0.650851869 | 0.30447884 | 2.1375931 | 0.03254978 | 0.12476787 |
| SHPRH | 2400.0171 | 0.375257035 | 0.1058704 | 3.54449437 | 0.00039337 | 0.00492029 |
| GRM1 | 364.048104 | 2.980264526 | 0.43934835 | 6.78337475 | 1.17E-11 | 2.87E-09 |
| RAB32 | 143.31008 | 0.435806246 | 0.38778096 | 1.12384641 | 0.26107819 | 0.48225168 |
| ADGB | 74.4884942 | 2.558454644 | 0.55094804 | 4.64373126 | 3.42E-06 | 0.0001097 |
| STXBP5-AS1 | 205.890043 | -0.972815596 | 0.41907679 | -2.3213302 | 0.02026903 | 0.0898435 |
| STXBP5 | 1574.08698 | -0.393993715 | 0.1671081 | -2.3577177 | 0.01838767 | 0.0843248 |
| SAMD5 | 1115.79415 | -0.197783679 | 0.30330504 | -0.6520949 | 0.51433994 | 0.71598841 |
| AL033504.1 | 701.280042 | -0.439713101 | 0.44327953 | -0.9919544 | 0.32121975 | 0.54565222 |
| AL513164.1 | 53.0444104 | -0.880354998 | 0.69696327 | -1.2631297 | 0.20654258 | 0.41739901 |
| SASH1 | 7176.80114 | -1.177031167 | 0.34422991 | -3.419317 | 0.00062779 | 0.00706339 |
| UST | 993.678985 | 0.588019035 | 0.28309731 | 2.07709155 | 0.03779311 | 0.13759387 |
| TAB2 | 1664.71772 | 0.155717545 | 0.14673717 | 1.06120041 | 0.28859884 | 0.51093714 |
| PPIL4 | 642.214521 | 0.359956777 | 0.16773893 | 2.14593467 | 0.03187819 | 0.12301462 |
| GINM1 | 716.496347 | 0.041954131 | 0.16689739 | 0.25137679 | 0.80152281 | 0.90675872 |
| AL078581.3 | 61.6749054 | 0.310153239 | 0.50148779 | 0.61846618 | 0.53626808 | 0.73306066 |
| KATNA1 | 359.344028 | 0.019282675 | 0.24173106 | 0.07976912 | 0.93642088 | 0.97280247 |
| LATS1 | 1049.50596 | 0.379703959 | 0.13424722 | 2.82839351 | 0.00467823 | 0.03178739 |
| NUP43 | 358.068631 | 0.235859063 | 0.21828302 | 1.08051952 | 0.2799109 | 0.50229221 |
| PCMT1 | 587.570022 | -0.053572837 | 0.20173565 | -0.2655596 | 0.79057841 | 0.90116933 |
| AL355312.2 | 45.7449525 | 0.33374255 | 0.34753451 | 0.96031484 | 0.33689678 | 0.56223242 |
| LRP11 | 244.732111 | -0.431413409 | 0.24140845 | -1.7870684 | 0.07392644 | 0.21565688 |
| PPP1R14C | 174.319397 | -3.002998506 | 0.56301614 | -5.3337699 | 9.62E-08 | 6.20E-06 |
| PLEKHG1 | 1557.7673 | -0.825327402 | 0.46878438 | -1.7605693 | 0.07831132 | 0.22360879 |
| MTHFD1L | 1112.886 | -2.382178621 | 0.40069559 | -5.9451082 | 2.76E-09 | 3.23E-07 |
| AL138733.1 | 56.7414174 | -0.668506202 | 0.29855802 | -2.2391165 | 0.02514834 | 0.10407512 |
| AKAP12 | 253.581148 | -1.746678543 | 0.58597338 | -2.9808155 | 0.00287482 | 0.02241646 |
| ZBTB2 | 136.34512 | 0.264128078 | 0.29351356 | 0.89988373 | 0.36818213 | 0.59323222 |
| RMND1 | 431.428269 | 0.147881209 | 0.26894774 | 0.5498511 | 0.58242151 | 0.76801495 |
| ARMT1 | 224.060533 | 0.254029629 | 0.27429237 | 0.9261272 | 0.35437988 | 0.58012193 |
| ESR1 | 226.817537 | 0.345052744 | 0.71274256 | 0.48411974 | 0.62830089 | 0.79869372 |
| SYNE1 | 6847.69627 | -0.525772216 | 0.38644815 | -1.3605246 | 0.17366397 | 0.37537677 |
| AL080276.2 | 82.2394206 | 0.571451659 | 0.31564504 | 1.81042497 | 0.07022991 | 0.20866009 |
| MTRF1L | 209.965765 | 0.01749108 | 0.2229933 | 0.07843769 | 0.93747989 | 0.97284815 |
| RGS17 | 49.9684191 | -0.086027981 | 0.6450805 | -0.1333601 | 0.89390862 | 0.95146032 |
| OPRM1 | 173.788183 | -0.087227889 | 0.4960117 | -0.1758585 | 0.86040508 | 0.93655498 |
| IPCEF1 | 1665.57359 | 0.10391496 | 0.60404548 | 0.17203168 | 0.86341262 | 0.93813133 |
| CNKSR3 | 2878.90827 | 0.508520103 | 0.24344573 | 2.0888438 | 0.03672179 | 0.13485073 |
| SCAF8 | 2667.43076 | 0.385419904 | 0.19683444 | 1.95809182 | 0.05021924 | 0.16683751 |
| TIAM2 | 1548.04839 | 0.284324516 | 0.15616243 | 1.82069736 | 0.06865287 | 0.20553849 |
| TFB1M | 837.665318 | 0.302987552 | 0.23701371 | 1.27835454 | 0.20112444 | 0.41029387 |
| CLDN20 | 57.9612537 | 0.254419779 | 0.39120754 | 0.65034478 | 0.51546954 | 0.71679515 |
| ARID1B | 9049.34524 | 0.164793618 | 0.15436267 | 1.0675743 | 0.28571258 | 0.50814292 |
| TMEM242 | 310.776978 | 0.119607717 | 0.16561373 | 0.72220896 | 0.47016602 | 0.68188686 |
| ZDHHC14 | 4810.67987 | 0.481680486 | 0.29036605 | 1.65887329 | 0.09714133 | 0.25753652 |
| SNX9 | 2291.91608 | -0.361121136 | 0.26852276 | -1.3448437 | 0.17867572 | 0.38189768 |
| SYNJ2 | 429.427073 | -1.915806504 | 0.4682485 | -4.0914312 | 4.29E-05 | 0.00086989 |
| SERAC1 | 222.306107 | -0.15312604 | 0.25375626 | -0.6034375 | 0.54621769 | 0.74053159 |
| GTF2H5 | 238.126888 | -0.005710762 | 0.32175451 | -0.0177488 | 0.98583924 | 0.99426405 |
| TULP4 | 1839.91724 | -0.165449753 | 0.19885803 | -0.8319993 | 0.40540931 | 0.62573634 |
| TMEM181 | 554.987337 | -0.622950771 | 0.20378819 | -3.0568542 | 0.00223673 | 0.01862861 |
| DYNLT1 | 604.385458 | -0.945588301 | 0.23248996 | -4.067222 | 4.76E-05 | 0.00094503 |
| SYTL3 | 497.885521 | -0.639873086 | 0.64925263 | -0.9855533 | 0.32435235 | 0.54927823 |
| EZR | 1682.8765 | -0.812210102 | 0.29416386 | -2.7610805 | 0.00576105 | 0.03726625 |
| C6orf99 | 44.7640122 | 0.518317576 | 0.47665068 | 1.087416 | 0.276853 | 0.49897002 |
| RSPH3 | 548.997878 | -0.199806331 | 0.19516414 | -1.0237861 | 0.30593633 | 0.52956532 |
| AL035530.2 | 533.559101 | 1.496902067 | 0.43396332 | 3.44937461 | 0.00056189 | 0.00643739 |
| SOD2 | 131.243612 | -2.204516353 | 0.58605659 | -3.76161 | 0.00016882 | 0.00254855 |
| SOD2.1 | 253.550839 | -0.5334341 | 0.50563131 | -1.0549863 | 0.29143159 | 0.51409186 |
| WTAP | 1598.00067 | -0.489755403 | 0.27954155 | -1.751995 | 0.07977466 | 0.22597974 |
| ACAT2 | 143.664835 | -0.976964701 | 0.39054869 | -2.5015183 | 0.0123662 | 0.06426616 |
| TCP1 | 349.172045 | -0.277468685 | 0.24198646 | -1.146629 | 0.251535 | 0.47188266 |
| MRPL18 | 194.944588 | -0.127885723 | 0.26514593 | -0.482322 | 0.6295772 | 0.79962358 |
| IGF2R | 2550.15724 | 0.361706961 | 0.20892006 | 1.73131752 | 0.08339515 | 0.23244704 |
| SLC22A1 | 149.682129 | 1.310919351 | 0.48887531 | 2.68150042 | 0.00732928 | 0.04418924 |
| SLC22A2 | 464.240151 | 1.156612277 | 0.60945598 | 1.8977782 | 0.05772531 | 0.18320364 |
| SLC22A3 | 679.99071 | 0.715032346 | 0.53337142 | 1.34058993 | 0.18005363 | 0.38383249 |
| MAP3K4 | 1073.29369 | 0.417696725 | 0.15475539 | 2.69907709 | 0.00695321 | 0.04269 |
| AGPAT4 | 1366.04118 | 1.322726697 | 0.32501964 | 4.06968232 | 4.71E-05 | 0.00093651 |
| PRKN | 8073.70409 | 0.871359401 | 0.26568895 | 3.2796223 | 0.00103946 | 0.01030756 |
| PACRG | 1697.26104 | 0.110035555 | 0.24820517 | 0.443325 | 0.65753068 | 0.81859324 |
| CAHM | 47.8448759 | 1.103321536 | 0.31419136 | 3.51162279 | 0.00044538 | 0.00541766 |
| QKI | 5162.54236 | 0.457985126 | 0.17489441 | 2.61863784 | 0.00882816 | 0.05053466 |
| PDE10A | 2232.66195 | 0.487778632 | 0.50498803 | 0.96592118 | 0.33408362 | 0.5593519 |
| SFT2D1 | 466.523265 | -0.038706505 | 0.15319035 | -0.2526694 | 0.80052373 | 0.90656019 |
| MPC1 | 2022.3764 | 0.979320687 | 0.26138219 | 3.74670021 | 0.00017918 | 0.00267805 |
| AL022069.1 | 89.5547913 | 0.982810719 | 0.4399591 | 2.23386838 | 0.02549173 | 0.10486463 |
| RPS6KA2 | 1287.82737 | -0.692973493 | 0.39693942 | -1.7457916 | 0.08084718 | 0.22811304 |
| AL159163.1 | 169.098856 | -0.598041228 | 0.56354854 | -1.0612062 | 0.28859619 | 0.51093714 |
| Z94721.1 | 233.91486 | -0.155282261 | 0.45805808 | -0.3390012 | 0.73460879 | 0.8697963 |
| FGFR1OP | 999.098263 | 0.35589914 | 0.20246889 | 1.75779665 | 0.07878211 | 0.22441496 |
| LINC01558 | 47.647918 | -1.023191526 | 0.65761425 | -1.5559145 | 0.11972843 | 0.29536727 |
| AFDN | 3777.70255 | -0.019102251 | 0.19126999 | -0.0998706 | 0.92044705 | 0.96454603 |
| WDR27 | 1211.82694 | 0.464062644 | 0.20693243 | 2.24258053 | 0.02492388 | 0.1034956 |
| C6orf120 | 69.2052196 | -0.439953484 | 0.34926371 | -1.2596599 | 0.20779208 | 0.41872245 |
| PHF10 | 510.328832 | 0.141210792 | 0.23058564 | 0.61240062 | 0.54027273 | 0.73628429 |
| TCTE3 | 83.8773704 | 0.623581408 | 0.28477418 | 2.18973995 | 0.0285431 | 0.11349363 |

| | | | | | | |
|---|---|---|---|---|---|---|
| ERMARD | 328.403905 | 0.316331241 | 0.35389085 | 0.89386669 | 0.37139322 | 0.59563048 |
| DLL1 | 61.90343 | 0.253722443 | 0.44810432 | 0.56621289 | 0.57124907 | 0.75965257 |
| FAM120B | 1679.03019 | -0.086138483 | 0.22294775 | -0.3863618 | 0.69922877 | 0.8470493 |
| PSMB1 | 553.041684 | -0.689302996 | 0.30796592 | -2.2382444 | 0.02520512 | 0.10419747 |
| TBP | 255.921724 | 0.148858399 | 0.21712247 | 0.68559648 | 0.4929676 | 0.69868807 |
| PDCD2 | 208.72061 | -0.474193807 | 0.27405688 | -1.7302751 | 0.08358113 | 0.23286714 |
| AC093627.5 | 55.5760518 | 0.265671474 | 0.41928311 | 0.63363266 | 0.52632058 | 0.72497874 |
| FAM20C | 654.383997 | -2.854886425 | 0.56507191 | -5.0522533 | 4.37E-07 | 2.12E-05 |
| PDGFA | 316.256808 | -0.598097449 | 0.31819957 | -1.87963 | 0.06015853 | 0.18821269 |
| PRKAR1B | 624.974508 | 0.688091007 | 0.28637754 | 2.40274081 | 0.01627272 | 0.07745813 |
| DNAAF5 | 419.390631 | 0.086054868 | 0.20196052 | 0.42609749 | 0.67003681 | 0.82659266 |
| SUN1 | 1180.17665 | 0.020675625 | 0.17340296 | 0.11923456 | 0.90508953 | 0.95650273 |
| AC073957.3 | 61.398798 | -0.054342674 | 0.5701469 | -0.0953135 | 0.92406585 | 0.96654633 |
| ADAP1 | 83.509212 | 0.611151358 | 0.66536961 | 0.91851408 | 0.35834979 | 0.58372701 |
| COX19 | 320.456014 | -0.144124458 | 0.19959071 | -0.7221 | 0.47023298 | 0.68188686 |
| C7orf50 | 665.590239 | -0.038506138 | 0.23870444 | -0.161313 | 0.87184685 | 0.94186841 |
| ZFAND2A | 47.1071981 | -0.6490506 | 0.34640292 | -1.8736869 | 0.0609736 | 0.19001878 |
| MICALL2 | 156.576986 | -1.033949003 | 0.43109421 | -2.3984293 | 0.01646555 | 0.07817882 |
| INTS1 | 324.750138 | 0.215435545 | 0.30582879 | 0.70443186 | 0.48116387 | 0.68948936 |
| MAFK | 215.179519 | -0.006002922 | 0.21631114 | -0.0277513 | 0.97786048 | 0.99038178 |
| TMEM184A | 83.288927 | -0.140710023 | 0.41424212 | -0.3396806 | 0.73409705 | 0.86945001 |
| PSMG3 | 78.6333143 | -0.212283684 | 0.34620838 | -0.6131674 | 0.53976568 | 0.73596237 |
| PSMG3-AS1 | 125.465747 | 0.314939499 | 0.24013128 | 1.3115305 | 0.1896786 | 0.39572864 |
| ELFN1 | 127.566915 | 1.468894297 | 0.38163633 | 3.84893723 | 0.00011863 | 0.00192035 |
| MAD1L1 | 698.528276 | -0.265703015 | 0.21233623 | -1.2513315 | 0.21081355 | 0.423197 |
| MRM2 | 87.0811155 | -0.509364316 | 0.24979603 | -2.0391209 | 0.04143796 | 0.1458444 |
| SNX8 | 179.596739 | -0.465917102 | 0.42256812 | -1.1025846 | 0.27020761 | 0.49175977 |
| EIF3B | 528.883097 | -0.403463752 | 0.27370322 | -1.4740921 | 0.1404568 | 0.32804946 |
| CHST12 | 217.856471 | 0.138390629 | 0.25295243 | 0.54710139 | 0.58430908 | 0.76896903 |
| BRAT1 | 184.766121 | 0.227927361 | 0.30076047 | 0.75783683 | 0.44854867 | 0.66444762 |
| IQCE | 374.212115 | 0.034864502 | 0.21628706 | 0.16119551 | 0.87193942 | 0.94189138 |
| TTYH3 | 192.408666 | -0.977663311 | 0.26975659 | -3.6242426 | 0.00028981 | 0.00393028 |
| AMZ1 | 70.3846155 | -0.972565686 | 0.46099629 | -2.109704 | 0.03488386 | 0.13060685 |
| GNA12 | 460.108404 | -0.661758711 | 0.18472165 | -3.5824643 | 0.00034037 | 0.00441643 |
| SDK1 | 2424.2462 | -1.223028536 | 0.29980952 | -4.079352 | 4.52E-05 | 0.00091074 |
| FOXK1 | 1134.57882 | -0.376521846 | 0.23325552 | -1.6142033 | 0.10648333 | 0.27380538 |
| AP5Z1 | 199.681648 | -0.298985305 | 0.30829944 | -0.9697887 | 0.33215184 | 0.55734167 |
| RBAK-RBAKD | 635.028894 | -0.38240918 | 0.25018976 | -1.5284765 | 0.12639427 | 0.30577691 |
| WIPI2 | 611.798665 | -0.335390222 | 0.22381778 | -1.4984967 | 0.13400424 | 0.31830389 |
| SLC29A4 | 43.8626948 | 0.328651872 | 0.4797159 | 0.6850969 | 0.49328278 | 0.69881955 |
| TNRC18 | 2145.97745 | 0.090460265 | 0.29971852 | 0.3018174 | 0.76279127 | 0.88492094 |
| AC092171.4 | 44.3101347 | 0.140228686 | 0.52045021 | 0.26943727 | 0.78759321 | 0.89956761 |
| FBXL18 | 247.551968 | -0.280833633 | 0.28210985 | -0.9954762 | 0.31950473 | 0.54413848 |
| ACTB | 2797.50851 | -0.914525941 | 0.48628865 | -1.8806237 | 0.06002313 | 0.18792261 |
| RNF216 | 1418.79929 | 0.013171772 | 0.20081228 | 0.06559246 | 0.94770229 | 0.97760554 |
| CCZ1 | 212.992408 | -0.013954531 | 0.33590715 | -0.0415428 | 0.96686315 | 0.9851354 |
| PMS2 | 367.529886 | 0.14372123 | 0.23069742 | 0.62298587 | 0.53329381 | 0.73050689 |
| AIMP2 | 229.431413 | -0.007529291 | 0.22539665 | -0.0334046 | 0.97335192 | 0.98804941 |
| EIF2AK1 | 548.70747 | -0.119281891 | 0.11855472 | -1.0061336 | 0.3143513 | 0.53829516 |
| USP42 | 548.598868 | -0.105969126 | 0.17719261 | -0.5980448 | 0.54981001 | 0.74303667 |
| CYTH3 | 638.269082 | 0.045093654 | 0.16216194 | 0.27807791 | 0.78095255 | 0.89539748 |
| FAM220A | 61.6090877 | 0.385727094 | 0.33866263 | 1.13897152 | 0.25471503 | 0.47549755 |
| RAC1 | 1339.62425 | -0.768977528 | 0.16015146 | -4.8015642 | 1.57E-06 | 5.83E-05 |
| DAGLB | 184.451786 | -0.554998271 | 0.2656318 | -2.0893517 | 0.03667607 | 0.1348328 |
| GRID2IP | 79.9888703 | -0.591254201 | 0.37775023 | -1.5651988 | 0.11753631 | 0.29215978 |
| ZDHHC4 | 125.786209 | -0.235946814 | 0.17822277 | -1.323887 | 0.18554057 | 0.39119 |
| C7orf26 | 164.90223 | 0.316064316 | 0.31639121 | 0.9989668 | 0.31781077 | 0.54237242 |
| ZNF316 | 181.120751 | 0.072472845 | 0.23548276 | 0.30776285 | 0.75826279 | 0.88315785 |
| AC073343.2 | 204.950957 | 0.024577847 | 0.41160211 | 0.05971264 | 0.95238451 | 0.97933189 |
| ZNF12 | 307.52434 | 0.111448439 | 0.18444831 | 0.60422585 | 0.5456935 | 0.74015676 |
| CCZ1B | 299.904525 | -0.064667415 | 0.25263641 | -0.2559703 | 0.79797378 | 0.90506918 |
| C1GALT1 | 762.76025 | -0.353508281 | 0.29987334 | -1.1788586 | 0.23845447 | 0.45641864 |
| COL28A1 | 1612.1055 | -0.191519966 | 0.28083962 | -0.6819549 | 0.49526745 | 0.70078115 |
| AC004982.1 | 106.841545 | 0.292021779 | 0.30839076 | 0.9469213 | 0.34367889 | 0.56845159 |
| AC004982.2 | 300.795148 | 0.161897487 | 0.3287534 | 0.49245875 | 0.62239508 | 0.79439719 |
| MIOS | 811.357876 | 0.25512645 | 0.35736507 | 0.71390988 | 0.47528291 | 0.68532684 |
| RPA3 | 358.268047 | -0.545972792 | 0.46389123 | -1.1769414 | 0.2392189 | 0.45728544 |
| UMAD1 | 3178.8905 | -0.104336347 | 0.16168052 | -0.6453242 | 0.51871713 | 0.71939843 |
| GLCCI1 | 2024.79616 | 0.326937218 | 0.16355075 | 1.99899549 | 0.04560884 | 0.15600761 |
| ICA1 | 1794.31511 | -0.640876741 | 0.34656794 | -1.8492095 | 0.06442757 | 0.19704184 |
| AC004949.1 | 105.991303 | 3.243346108 | 0.93554462 | 3.46680004 | 0.00052669 | 0.00614584 |
| NDUFA4 | 978.662951 | -0.256806054 | 0.47279892 | -0.5431613 | 0.58701878 | 0.77047088 |
| PHF14 | 1994.44041 | 0.067073578 | 0.20225068 | 0.33163587 | 0.74016423 | 0.87268348 |
| THSD7A | 7583.58765 | -0.290629893 | 0.5773013 | -0.5034284 | 0.61466308 | 0.78816802 |
| TMEM106B | 960.426121 | 0.305979229 | 0.2749687 | 1.11277839 | 0.26580362 | 0.48682751 |
| SCIN | 2160.90017 | -0.70396233 | 0.21222528 | -3.3170523 | 0.00090973 | 0.00927921 |
| AC011287.1 | 340.515571 | -1.041446235 | 0.44182125 | -2.3571664 | 0.01841499 | 0.0843248 |
| ETV1 | 361.057793 | -1.25694355 | 0.56348417 | -2.2306635 | 0.02570343 | 0.10552301 |
| ISPD | 1236.43117 | 0.982099212 | 0.21605871 | 4.54552017 | 5.48E-06 | 0.00016416 |
| SOSTDC1 | 41.2108536 | 1.282784367 | 0.8983544 | 1.42792685 | 0.15331291 | 0.34712209 |
| ANKMY2 | 460.126837 | 0.810162462 | 0.2344728 | 3.45525137 | 0.00054978 | 0.00635349 |
| BZW2 | 957.673104 | -0.882469793 | 0.25552077 | -3.4536128 | 0.00055313 | 0.00636394 |
| AGR2 | 117.70722 | -3.737685723 | 1.06366833 | -3.5139579 | 0.00044148 | 0.00539958 |
| AHR | 135.591891 | 0.124597828 | 0.57266921 | 0.21757382 | 0.82776118 | 0.91981743 |
| AC073332.1 | 218.896509 | -0.900702213 | 0.48065981 | -1.8738871 | 0.06094599 | 0.19001878 |
| AC019117.1 | 226.290789 | -3.4989578 | 0.86842893 | -4.0290664 | 5.60E-05 | 0.00106891 |
| SNX13 | 2039.48372 | -0.084134877 | 0.17434598 | -0.4825742 | 0.62939809 | 0.79954985 |
| HDAC9 | 1948.29378 | 0.29781037 | 0.64225627 | 0.46369399 | 0.642867 | 0.80857472 |
| TWISTNB | 111.442561 | -0.301141698 | 0.26868451 | -1.1208004 | 0.26237284 | 0.4832663 |
| MACC1 | 1188.38257 | -0.443776502 | 0.46432787 | -0.9557395 | 0.33920385 | 0.56472695 |
| ITGB8 | 2490.68917 | -0.124759155 | 0.38165343 | -0.3268912 | 0.74375016 | 0.87470713 |
| SP4 | 519.489231 | 0.148394409 | 0.21404402 | 0.69328922 | 0.48812807 | 0.69502858 |
| DNAH11 | 87.7885205 | -0.544188956 | 0.68835382 | -0.7905658 | 0.42919744 | 0.64747247 |
| CDCA7L | 105.636173 | -0.216590141 | 0.33815343 | -0.6405085 | 0.52184207 | 0.72185694 |
| RAPGEF5 | 679.100098 | -3.560060674 | 0.60653336 | -5.8695216 | 4.37E-09 | 4.81E-07 |
| STEAP1B | 644.588376 | -1.345619678 | 0.43552135 | -3.0896755 | 0.00200375 | 0.01714221 |
| TOMM7 | 373.975523 | -0.462069831 | 0.3813514 | -1.2116642 | 0.22564095 | 0.44031487 |
| FAM126A | 89.7391309 | 0.217682367 | 0.35385029 | 0.61518211 | 0.53843446 | 0.73492731 |
| KLHL7 | 513.216255 | 0.030723932 | 0.22221951 | 0.13825938 | 0.89003542 | 0.9503707 |
| NUPL2 | 582.750113 | 0.090369122 | 0.24327565 | 0.37146801 | 0.71028898 | 0.85479293 |
| AC005082.1 | 135.409876 | -0.474809842 | 0.42534977 | -1.1162809 | 0.2643019 | 0.48528287 |
| GPNMB | 201.730332 | -0.64803118 | 0.62808168 | -1.0317626 | 0.30218335 | 0.52534086 |
| MALSU1 | 308.870903 | -0.027062517 | 0.17051 | -0.1587151 | 0.87389331 | 0.94333543 |
| TRA2A | 3074.43393 | 0.557906617 | 0.16681558 | 3.34445152 | 0.00082445 | 0.00860887 |
| CCDC126 | 260.668743 | 0.062752476 | 0.27244831 | 0.230328 | 0.8178369 | 0.91533703 |
| FAM221A | 260.782099 | 0.760514318 | 0.23419256 | 3.24738886 | 0.00116469 | 0.01126238 |
| STK31 | 59.7513894 | 0.531750664 | 0.58118282 | 0.9149456 | 0.36022019 | 0.58547416 |
| MPP6 | 1623.08143 | 0.116654835 | 0.27664705 | 0.42167388 | 0.67326307 | 0.82881006 |
| GSDME | 270.156389 | 0.598284425 | 0.28979419 | 2.06451489 | 0.03896892 | 0.1402945 |
| OSBPL3 | 6782.15885 | 0.535010361 | 0.31952717 | 1.67438144 | 0.09405566 | 0.25227697 |
| CYCS | 415.140273 | 0.193312565 | 0.47791357 | 0.40449273 | 0.68585042 | 0.83751486 |
| C7orf31 | 102.298597 | 0.100529142 | 0.31288415 | 0.32129829 | 0.74798435 | 0.87714848 |
| NFE2L3 | 250.744799 | -0.472137221 | 0.41740622 | -1.1311217 | 0.25800388 | 0.47905162 |

| | | | | | | |
|---|---|---|---|---|---|---|
| AC004520.1 | 45.5886602 | 0.681608821 | 0.58208472 | 1.17097873 | 0.24160733 | 0.45992091 |
| HNRNPA2B1 | 4353.82998 | 0.219459626 | 0.1248557 | 1.75770609 | 0.07879753 | 0.22441496 |
| CBX3 | 774.011008 | -0.919151674 | 0.12393511 | -7.4163945 | 1.20E-13 | 5.30E-11 |
| SNX10 | 2818.0173 | -0.956200447 | 0.33722661 | -2.8354834 | 0.00457564 | 0.03145229 |
| AC004540.2 | 47.3535304 | 0.165665256 | 0.5180162 | 0.31980709 | 0.74911457 | 0.87779905 |
| AC004540.1 | 464.999279 | 0.60833866 | 0.52672748 | 1.15494006 | 0.24811497 | 0.46734754 |
| SKAP2 | 2699.07487 | 0.005155436 | 0.21525275 | 0.02395061 | 0.980892 | 0.99160261 |
| HOTAIRM1 | 517.706023 | -0.09208641 | 0.3847421 | -0.2393458 | 0.81083744 | 0.9112091 |
| HOXA3 | 737.207125 | 0.803848385 | 0.30495275 | 2.63597686 | 0.00838955 | 0.04881828 |
| HOXA-AS2 | 320.345083 | 0.587198912 | 0.37183689 | 1.57918412 | 0.11429383 | 0.28723501 |
| HOXA-AS3 | 113.892201 | 0.42856968 | 0.43583007 | 0.98334124 | 0.32543952 | 0.55057386 |
| HOXA7 | 108.657955 | -0.657988336 | 0.53324808 | -1.2339254 | 0.21723072 | 0.43128814 |
| HOXA9 | 233.595694 | 0.515039107 | 0.49559051 | 1.03924329 | 0.2986916 | 0.52181158 |
| HOXA10-AS | 73.892168 | -0.258347707 | 0.54735976 | -0.4719889 | 0.63693474 | 0.80480926 |
| HIBADH | 3562.85164 | 0.896966279 | 0.26351464 | 3.40385752 | 0.00066441 | 0.00736262 |
| TAX1BP1 | 2880.07943 | -0.025349272 | 0.14280268 | -0.1775126 | 0.85910578 | 0.93618423 |
| JAZF1 | 1327.68296 | -0.14146091 | 0.41109328 | -0.344109 | 0.73076429 | 0.86804546 |
| CREB5 | 474.767239 | -0.381230308 | 0.64344694 | -0.5924813 | 0.55352831 | 0.74630555 |
| AC005162.3 | 373.804399 | 0.841785717 | 0.4536586 | 1.85554889 | 0.06351788 | 0.19518591 |
| CPVL | 1130.32392 | 0.381927511 | 0.43817341 | 0.87163553 | 0.38340724 | 0.60615515 |
| AC007255.1 | 187.220959 | -0.487533808 | 0.34383364 | -1.4179352 | 0.1562097 | 0.35131349 |
| PRR15 | 78.1037485 | -1.822601888 | 0.34895977 | -5.2229571 | 1.76E-07 | 9.86E-06 |
| WIPF3 | 127.905692 | -0.185123574 | 0.60325918 | -0.3068724 | 0.75894052 | 0.88316847 |
| SCRN1 | 733.228702 | -0.381627252 | 0.16720327 | -2.282415 | 0.02246485 | 0.09637486 |
| AC007285.1 | 112.2504 | -0.142997861 | 0.31351539 | -0.4561111 | 0.64831006 | 0.81178572 |
| FKBP14 | 194.796887 | -0.036397528 | 0.2535677 | -0.1435417 | 0.88586241 | 0.94846586 |
| PLEKHA8 | 590.238273 | -0.539070176 | 0.18203339 | -2.9613808 | 0.00306263 | 0.02351993 |
| MTURN | 1388.3835 | 0.252519775 | 0.25390497 | 0.99454445 | 0.31995788 | 0.54436972 |
| ZNRF2 | 907.235413 | -0.464509418 | 0.21774446 | -2.1332777 | 0.03290195 | 0.12575817 |
| AC006978.2 | 816.134882 | -0.494611905 | 0.23315586 | -2.1213788 | 0.03388993 | 0.12834064 |
| NOD1 | 144.08592 | -0.14412167 | 0.47210152 | -0.3052769 | 0.7601553 | 0.88357016 |
| GGCT | 58.816901 | -0.317069215 | 0.46149745 | -0.6870443 | 0.49205478 | 0.69803196 |
| AC005154.6 | 426.500276 | 0.647392672 | 0.23502479 | 2.75457183 | 0.0058769 | 0.03779355 |
| GARS | 366.524503 | -0.190002939 | 0.19657668 | -0.9665589 | 0.3337646 | 0.5592275 |
| AQP1 | 178.894346 | 0.051718687 | 0.65507868 | 0.07895034 | 0.93707212 | 0.97284815 |
| PDE1C | 53.2448092 | -0.539658306 | 0.48211131 | -1.1193645 | 0.26298465 | 0.48401342 |
| LSM5 | 171.926538 | 0.047005351 | 0.27749522 | 0.16939157 | 0.86548865 | 0.93876167 |
| AVL9 | 2004.26777 | -0.292012915 | 0.14838264 | -1.9679722 | 0.04907123 | 0.16405515 |
| KBTBD2 | 568.916946 | -0.279360615 | 0.21174305 | -1.3193378 | 0.18705621 | 0.39270408 |
| FKBP9 | 349.970977 | 0.393958764 | 0.24247811 | 1.62471886 | 0.10422248 | 0.27030723 |
| NT5C3A | 745.852509 | 0.543695205 | 0.26835321 | 2.02604321 | 0.04276036 | 0.14887231 |
| RP9 | 146.258599 | 0.492873002 | 0.21916104 | 2.24890789 | 0.02451836 | 0.10232637 |
| BBS9 | 2382.86616 | 0.502824761 | 0.13735385 | 3.66079849 | 0.00025143 | 0.00351851 |
| BMPER | 133.963386 | 0.021509039 | 0.49901636 | 0.04310287 | 0.96561953 | 0.98461609 |
| NPSR1 | 109.129265 | -0.378522357 | 0.86051433 | -0.4398792 | 0.6600246 | 0.81992523 |
| DPY19L1 | 1149.51274 | -0.822203588 | 0.27667567 | -2.9717235 | 0.00296133 | 0.02292863 |
| HERPUD2 | 753.14215 | 0.020258533 | 0.16911902 | 0.11978861 | 0.9046506 | 0.95619187 |
| AC018647.2 | 48.4600867 | -0.345544631 | 0.40717268 | -0.8486439 | 0.39607945 | 0.61805238 |
| SEPT7-AS1 | 229.014209 | 0.307849039 | 0.33749022 | 0.91217175 | 0.36167831 | 0.58697738 |
| 7-Sep | 1602.57766 | -0.126826946 | 0.16139575 | -0.7858134 | 0.43197679 | 0.64966201 |
| EEPD1 | 379.391722 | -0.757628255 | 0.5799622 | -1.3063407 | 0.19143671 | 0.39809311 |
| KIAA0895 | 451.305441 | -0.118219537 | 0.30831482 | -0.3834377 | 0.70139523 | 0.84869711 |
| ELMO1 | 335.660377 | -0.231486738 | 0.94408015 | -0.2451982 | 0.80630298 | 0.90938008 |
| EPDR1 | 266.379556 | 0.664580204 | 0.35797311 | 1.85650872 | 0.06338108 | 0.1951062 |
| SFRP4 | 100.577569 | 0.912200791 | 0.65986971 | 1.38239531 | 0.16685036 | 0.36640505 |
| STARD3NL | 336.521115 | 0.021463127 | 0.13940936 | 0.15395758 | 0.87764319 | 0.9452698 |
| AMPH | 279.873104 | 4.150315855 | 0.76718619 | 5.40978956 | 6.31E-08 | 4.43E-06 |
| VPS41 | 1952.50646 | 0.277012511 | 0.17205701 | 1.61000426 | 0.10739693 | 0.275404 |
| YAE1D1 | 218.362245 | 0.579175469 | 0.23915 | 2.42180839 | 0.01544349 | 0.07488733 |
| RALA | 793.535906 | -0.302392372 | 0.20223682 | -1.4952389 | 0.13485209 | 0.31968078 |
| LINC00265 | 175.804636 | 0.218807298 | 0.30327248 | 0.72148748 | 0.47060964 | 0.68213352 |
| CDK13 | 3423.52709 | 0.078574823 | 0.16581196 | 0.4738791 | 0.63558612 | 0.80389813 |
| MPLKIP | 309.830629 | 0.56122958 | 0.21287964 | 2.63637041 | 0.00837982 | 0.04881828 |
| SUGCT | 1242.24818 | 0.669730066 | 0.17149317 | 3.90528713 | 9.41E-05 | 0.00159993 |
| MRPL32 | 226.475163 | -0.298327646 | 0.269646 | -1.1063678 | 0.26856737 | 0.48998707 |
| AC005537.1 | 83.5506327 | -0.695799397 | 0.78918371 | -0.8816697 | 0.37795543 | 0.60077175 |
| STK17A | 836.559988 | 0.396097702 | 0.25802188 | 1.53513223 | 0.12475135 | 0.30309781 |
| COA1 | 2151.0298 | 0.348014244 | 0.15476929 | 2.24860007 | 0.02453795 | 0.10237581 |
| BLVRA | 231.858395 | -0.484015658 | 0.22874827 | -2.1159314 | 0.03435064 | 0.12926998 |
| URGCP | 429.286743 | 0.428345219 | 0.21046914 | 2.03519256 | 0.0418315 | 0.14665859 |
| UBE2D4 | 124.830816 | 0.234687154 | 0.26896393 | 0.87255995 | 0.38290298 | 0.6054789 |
| SPDYE1 | 129.59931 | 0.067336105 | 0.28222537 | 0.23858984 | 0.81142365 | 0.91148014 |
| AC004951.1 | 120.753649 | 0.274208832 | 0.38164181 | 0.71849789 | 0.47245035 | 0.68321953 |
| DBNL | 298.810187 | -0.248095747 | 0.1759611 | -1.4099466 | 0.15855546 | 0.35503629 |
| POLM | 293.01239 | 0.187902599 | 0.27400138 | 0.68577246 | 0.4928566 | 0.69868807 |
| POLD2 | 142.992737 | -0.391553988 | 0.23387088 | -1.6742315 | 0.09408512 | 0.25228793 |
| YKT6 | 134.982743 | -0.233714655 | 0.24469818 | -0.955114 | 0.33952006 | 0.56496857 |
| CAMK2B | 51.8768924 | 2.26253667 | 0.69118444 | 3.27341958 | 0.00106255 | 0.0104584 |
| NUDCD3 | 1096.04092 | -0.099162313 | 0.25032818 | -0.3961293 | 0.69200968 | 0.8419228 |
| DDX56 | 189.71093 | 0.18169857 | 0.20240864 | 0.89768189 | 0.36935517 | 0.59417383 |
| TMED4 | 371.01355 | -0.29602113 | 0.22302676 | -1.3272897 | 0.18441286 | 0.38968316 |
| OGDH | 2223.6061 | 0.536007936 | 0.11625376 | 4.61067164 | 4.01E-06 | 0.0001248 |
| ZMIZ2 | 557.007782 | -0.071415756 | 0.25436364 | -0.2807624 | 0.77889262 | 0.89433176 |
| PPIA | 1351.51426 | -1.094068808 | 0.36853311 | -2.9687124 | 0.0029905 | 0.02311384 |
| H2AFV | 556.991293 | -0.118710544 | 0.22994653 | -0.5162528 | 0.60567785 | 0.78178941 |
| PURB | 462.884683 | -0.085469731 | 0.19239971 | -0.44423 | 0.65687627 | 0.81801439 |
| SNHG15 | 59.4298117 | 0.327180165 | 0.42736722 | 0.7655715 | 0.44393131 | 0.65960502 |
| CCM2 | 582.175183 | -0.358457818 | 0.15227641 | -2.3539945 | 0.01857289 | 0.08485965 |
| TBRG4 | 137.872807 | 0.15965208 | 0.24420566 | 0.65376076 | 0.51326595 | 0.71501054 |
| ADCY1 | 48.0075482 | 1.832051431 | 0.68697779 | 2.66682775 | 0.00765709 | 0.04576584 |
| IGFBP3 | 328.391599 | -0.994247329 | 0.53342526 | -1.8638925 | 0.06233681 | 0.1928805 |
| AC011294.1 | 84.2281257 | 1.532257244 | 1.52450322 | 1.00508626 | 0.31485531 | 0.53899998 |
| AC004870.4 | 755.042551 | 1.036441662 | 0.88818313 | 1.16692338 | 0.24324131 | 0.46195412 |
| AC004870.3 | 47.4851753 | 0.91497591 | 0.66621233 | 1.37339984 | 0.16962808 | 0.3699733 |
| TNS3 | 6356.96111 | -1.175784663 | 0.37062696 | -3.1724208 | 0.00151174 | 0.01382188 |
| HUS1 | 285.372412 | 0.317344974 | 0.21389375 | 1.48365704 | 0.13789993 | 0.32399826 |
| UPP1 | 1101.76682 | -2.161043563 | 0.74301844 | -2.9084656 | 0.00363207 | 0.02666817 |
| ZPBP | 144.575934 | 1.11476686 | 0.48927614 | 2.27840019 | 0.02270275 | 0.09704873 |
| FIGNL1 | 51.6136834 | 0.470386518 | 0.39101151 | 1.20299917 | 0.22897664 | 0.44451188 |
| GRB10 | 681.040825 | -0.398673593 | 0.48143552 | -0.8280934 | 0.40761759 | 0.62748459 |
| COBL | 2388.73922 | 0.346846343 | 0.4342666 | 0.7986945 | 0.42446758 | 0.64298076 |
| SEC61G | 229.064498 | -0.37866945 | 0.39858723 | -0.950029 | 0.34209749 | 0.56739902 |
| EGFR | 1390.07039 | -1.098359749 | 0.27977616 | -3.9258519 | 8.64E-05 | 0.00150404 |
| LANCL2 | 387.073102 | 0.206342071 | 0.33398947 | 0.61781011 | 0.53670052 | 0.73349204 |
| VOPP1 | 2158.5191 | -1.351365725 | 0.26852706 | -5.0325122 | 4.84E-07 | 2.31E-05 |
| AC092647.5 | 42.4422705 | 0.593874788 | 0.29713093 | 1.99869729 | 0.04564112 | 0.15600761 |
| NIPSNAP2 | 1135.32165 | 0.324128663 | 0.18019384 | 1.79877775 | 0.07205384 | 0.21221706 |
| PSPH | 235.418543 | 0.638590563 | 0.42633922 | 1.49784617 | 0.13417322 | 0.31858603 |
| CCT6A | 476.863449 | -0.279150366 | 0.253887 | -1.0995064 | 0.27154726 | 0.49297248 |
| SUMF2 | 652.124717 | -0.579120279 | 0.16195178 | -3.5758808 | 0.00034905 | 0.00449816 |
| PHKG1 | 163.774563 | -0.53938063 | 0.34908289 | -1.5451362 | 0.12231331 | 0.29963586 |
| CHCHD2 | 801.900849 | -0.950283384 | 0.38460535 | -2.4708013 | 0.01348107 | 0.06791437 |
| NUPR2 | 56.9702397 | -0.588269757 | 0.53666431 | -1.0961596 | 0.27300892 | 0.49467419 |

| | | | | | | |
|---|---|---|---|---|---|---|
| ZNF727 | 145.040234 | 0.205689721 | 0.35743806 | 0.57545556 | 0.56498324 | 0.75605085 |
| ZNF736 | 865.073053 | 0.082547189 | 0.21582336 | 0.38247568 | 0.70210856 | 0.8488446 |
| ZNF680 | 905.027826 | 0.5475381 | 0.21232147 | 2.57881643 | 0.00991395 | 0.05506867 |
| ZNF107 | 308.381175 | 0.26805771 | 0.28920565 | 0.92687577 | 0.35399104 | 0.57979331 |
| ZNF138 | 680.851897 | 0.564581632 | 0.19679637 | 2.86886203 | 0.00411951 | 0.02922329 |
| ZNF273 | 101.386306 | -0.057178025 | 0.23566177 | -0.2426275 | 0.80829398 | 0.91010272 |
| ZNF117 | 325.38803 | 0.024420671 | 0.18047644 | 0.13531224 | 0.89236501 | 0.95118543 |
| ZNF92 | 114.400957 | 0.180206741 | 0.2203581 | 0.8177904 | 0.41347688 | 0.63301068 |
| VKORC1L1 | 881.54906 | -0.087649316 | 0.18641052 | -0.4701951 | 0.63821562 | 0.80525266 |
| GUSB | 222.90169 | 0.176960329 | 0.1755326 | 1.00813368 | 0.31339029 | 0.53711851 |
| ASL | 159.70188 | -0.643112338 | 0.3563867 | -1.8045352 | 0.07114745 | 0.21047854 |
| AC068533.4 | 186.837028 | -0.774896393 | 0.37666029 | -2.0572819 | 0.03965912 | 0.14177464 |
| CRCP | 139.85714 | -0.521304697 | 0.20228279 | -2.5771084 | 0.00996307 | 0.0552691 |
| AC068533.3 | 183.250851 | 0.087199811 | 0.45843085 | 0.19021366 | 0.8491417 | 0.93065987 |
| TPST1 | 7403.12251 | -0.576481162 | 0.30602447 | -1.8837747 | 0.05959545 | 0.18716032 |
| AC027644.4 | 863.440789 | 0.013896196 | 0.28981961 | 0.04794774 | 0.96175789 | 0.9832997 |
| AC027644.3 | 128.909468 | -0.21704319 | 0.41276716 | -0.5258248 | 0.59900997 | 0.77729304 |
| AC073335.2 | 52.8668335 | 0.693171748 | 0.35362596 | 1.96018344 | 0.04997435 | 0.16631676 |
| TMEM248 | 533.121913 | -0.367714418 | 0.16587471 | -2.2168203 | 0.02663538 | 0.10828768 |
| SBDS | 297.46211 | -0.889737414 | 0.3317409 | -2.6820251 | 0.0073178 | 0.04417769 |
| TYW1 | 1358.22943 | 0.241266207 | 0.16741023 | 1.44116764 | 0.14953733 | 0.34173678 |
| AC006480.2 | 186.042086 | 0.604259454 | 0.29242662 | 2.06636265 | 0.03879425 | 0.13977994 |
| LINC01372 | 112.08989 | 1.228298971 | 0.39454788 | 3.11318105 | 0.00185082 | 0.0160945 |
| AUTS2 | 3677.03197 | 0.353450464 | 0.4897119 | 0.72175184 | 0.47044706 | 0.68204755 |
| GALNT17 | 289.92556 | 2.349901054 | 0.97788179 | 2.40305226 | 0.01625886 | 0.07742009 |
| CALN1 | 136.219616 | 0.479072411 | 0.33006233 | 1.45146044 | 0.14665169 | 0.33725691 |
| TYW1B | 1002.7919 | 0.29738382 | 0.17136206 | 1.73541222 | 0.08266782 | 0.23103419 |
| POM121 | 1015.09495 | 0.167831226 | 0.18517784 | 0.90632455 | 0.36476409 | 0.59024488 |
| NSUN5 | 206.842771 | 0.347960597 | 0.27308251 | 1.27419583 | 0.20259402 | 0.41227307 |
| BAZ1B | 922.044274 | 0.0342544 | 0.17523277 | 0.19547941 | 0.84501765 | 0.9283738 |
| BCL7B | 174.173672 | -0.285668105 | 0.18241405 | -1.5660422 | 0.11733874 | 0.291975 |
| TBL2 | 114.436088 | 0.111534534 | 0.2112276 | 0.52803012 | 0.59747843 | 0.77662789 |
| MLXIPL | 249.138077 | -0.068326584 | 0.60562199 | -0.1128205 | 0.91017285 | 0.95961675 |
| VPS37D | 56.6955195 | -0.44141566 | 0.5808781 | -0.759911 | 0.44730779 | 0.66320446 |
| DNAJC30 | 70.3531724 | -0.422022762 | 0.30237946 | -1.3956727 | 0.16281308 | 0.3609485 |
| BUD23 | 303.128632 | -0.111608481 | 0.22027958 | -0.5066674 | 0.61238822 | 0.78687125 |
| ABHD11 | 243.699623 | -1.038421374 | 0.3581989 | -2.8990078 | 0.00374346 | 0.02725872 |
| CLDN3 | 393.726873 | -3.005904433 | 0.52407991 | -5.7355842 | 9.72E-09 | 8.98E-07 |
| CLDN4 | 622.540721 | -2.042980194 | 0.4506196 | -4.5337136 | 5.80E-06 | 0.00017145 |
| METTL27 | 77.3692139 | -0.331163803 | 0.3594542 | -0.9212962 | 0.35689579 | 0.58255058 |
| LIMK1 | 121.198385 | -0.943338204 | 0.39233438 | -2.404424 | 0.01619797 | 0.07721365 |
| EIF4H | 497.500875 | -0.389469913 | 0.15490759 | -2.514208 | 0.01193 | 0.06268234 |
| RFC2 | 101.037628 | -0.732116504 | 0.3221221 | -2.2727919 | 0.02303872 | 0.09813559 |
| CLIP2 | 768.019195 | -0.067304125 | 0.339712 | -0.1981211 | 0.84295029 | 0.92739954 |
| GTF2IRD1 | 3697.1005 | -0.372734846 | 0.32163416 | -1.1588783 | 0.24650579 | 0.46556654 |
| GTF2I | 4260.80318 | 0.05823497 | 0.18672476 | 0.311876 | 0.75513476 | 0.88145768 |
| GTF2IRD2 | 500.157274 | -0.220892816 | 0.36778318 | -0.6006061 | 0.54810239 | 0.74186733 |
| CASTOR2 | 556.606893 | -0.764329055 | 0.38811203 | -1.9693516 | 0.04891273 | 0.16364949 |
| RCC1L | 370.807305 | -0.230726459 | 0.23994195 | -0.9615928 | 0.33625418 | 0.5617555 |
| GTF2IRD2B | 738.678568 | 0.151129358 | 0.35202345 | 0.42931616 | 0.66769315 | 0.82486496 |
| AC211486.5 | 171.428464 | -1.124837382 | 0.54986019 | -2.0456789 | 0.04078797 | 0.14440321 |
| TRIM73 | 348.121856 | 0.535768696 | 0.39692839 | 1.34978679 | 0.17708438 | 0.37954042 |
| POM121C | 895.289516 | 0.221493358 | 0.20270541 | 1.09268596 | 0.27453171 | 0.49616155 |
| SPDYE5 | 244.517577 | -0.280544191 | 0.33149305 | -0.8463049 | 0.39738268 | 0.61906213 |
| HIP1 | 8619.42008 | 1.052638668 | 0.33688399 | 3.12463249 | 0.00178027 | 0.01560427 |
| RHBDD2 | 195.389941 | -0.618644405 | 0.24991196 | -2.4754494 | 0.01330686 | 0.06729339 |
| POR | 412.748687 | -0.562633801 | 0.25941359 | -2.1688679 | 0.03009272 | 0.11784604 |
| TMEM120A | 118.918857 | 0.488248985 | 0.23283725 | 2.09695395 | 0.03599765 | 0.13341554 |
| STYXL1 | 558.851072 | 0.141377103 | 0.27375225 | 0.51644179 | 0.60554589 | 0.78178941 |
| MDH2 | 289.135998 | -0.245376189 | 0.32223616 | -0.7614794 | 0.44637079 | 0.66218686 |
| HSPB1 | 717.175781 | -0.87747692 | 0.46469432 | -1.8882885 | 0.05898723 | 0.18564745 |
| YWHAG | 470.025117 | -0.327877835 | 0.1808183 | -1.8133 | 0.06978556 | 0.20779925 |
| SSC4D | 73.03033 | -0.250574343 | 0.65569064 | -0.3821533 | 0.70234766 | 0.8488446 |
| ZP3 | 51.9140707 | -0.030234015 | 0.42888424 | -0.0704946 | 0.94380001 | 0.97568955 |
| DTX2 | 336.082275 | -0.014160279 | 0.21858461 | -0.0647817 | 0.94834782 | 0.97784954 |
| SPDYE16 | 76.2568784 | 0.269879089 | 0.36516632 | 0.73905799 | 0.45987179 | 0.67351662 |
| POMZP3 | 238.449219 | 0.076645738 | 0.36705957 | 0.20881008 | 0.8345965 | 0.92298305 |
| CCDC146 | 1229.70418 | 0.678747024 | 0.27607755 | 2.45853758 | 0.01395042 | 0.06965503 |
| FGL2 | 232.256271 | -0.997724891 | 0.84016547 | -1.1875338 | 0.23501713 | 0.45233007 |
| GSAP | 1447.47545 | 0.622153064 | 0.26917804 | 2.31130695 | 0.02081591 | 0.09162189 |
| PTPN12 | 2202.72022 | -1.175401924 | 0.28226649 | -4.1641568 | 3.13E-05 | 0.00067559 |
| APTR | 428.061581 | 0.518537211 | 0.22137295 | 2.34236936 | 0.01916174 | 0.08666407 |
| RSBN1L | 1118.69764 | 0.010299353 | 0.14060476 | 0.07325039 | 0.94160687 | 0.97495762 |
| TMEM60 | 79.5054642 | 0.309392861 | 0.34976359 | 0.88457711 | 0.37638476 | 0.59959757 |
| PHTF2 | 813.014657 | 0.349963943 | 0.17027055 | 2.05534036 | 0.03984614 | 0.14213548 |
| MAGI2 | 4687.75902 | 0.040426993 | 0.40872494 | 0.09891002 | 0.92120971 | 0.96504145 |
| GNAI1 | 4065.80511 | 0.308015766 | 0.33940813 | 0.90750852 | 0.36413795 | 0.58952055 |
| CD36 | 161.728334 | 1.049397507 | 0.52831457 | 1.98631189 | 0.0469987 | 0.1590994 |
| SEMA3C | 583.179285 | -0.456981101 | 0.70041808 | -0.6524405 | 0.51411706 | 0.71596592 |
| CACNA2D1 | 891.862426 | -0.826030156 | 0.56249876 | -1.4685013 | 0.14196811 | 0.33090819 |
| PCLO | 3686.35036 | 0.193475621 | 0.4812583 | 0.40202033 | 0.68766906 | 0.8391926 |
| KIAA1324L | 247.120476 | 0.188975209 | 0.34035316 | 0.55523271 | 0.5787355 | 0.76468112 |
| DMTF1 | 1882.26237 | 0.578939926 | 0.23423168 | 2.47165507 | 0.01344892 | 0.06780998 |
| TMEM243 | 530.495032 | -0.781880693 | 0.19708906 | -3.9671441 | 7.27E-05 | 0.00130762 |
| TP53TG1 | 90.3902906 | 0.226880452 | 0.26252995 | 0.86420788 | 0.3874737 | 0.61086904 |
| CROT | 404.787088 | 0.060972987 | 0.21314106 | 0.2860687 | 0.7748255 | 0.89222126 |
| ABCB1 | 135.282498 | 0.214438473 | 0.59646525 | 0.35951545 | 0.71920952 | 0.86003243 |
| RUNDC3B | 86.3359995 | -0.083006445 | 0.45556001 | -0.1822075 | 0.8554199 | 0.93436215 |
| SLC25A40 | 496.337309 | 0.839400631 | 0.1539592 | 5.45209792 | 4.98E-08 | 3.63E-06 |
| DBF4 | 142.909188 | -0.171496934 | 0.28221988 | -0.6076714 | 0.54340547 | 0.7391242 |
| ADAM22 | 462.625026 | 1.124217655 | 0.4076481 | 2.75781403 | 0.00581893 | 0.03754366 |
| SRI | 409.513019 | -0.457054978 | 0.25544752 | -1.7892324 | 0.07357739 | 0.21525665 |
| AC003991.1 | 80.7424843 | -0.036009914 | 0.44619252 | -0.0807049 | 0.93567666 | 0.97236337 |
| ZNF804B | 64.4928367 | 2.763079846 | 0.69689147 | 3.96486392 | 7.34E-05 | 0.0013162 |
| STEAP2 | 347.254998 | 0.663485433 | 0.2842409 | 2.33423634 | 0.01958335 | 0.08799176 |
| CFAP69 | 816.963776 | 0.792469265 | 0.22448621 | 3.5301467 | 0.00041533 | 0.00513829 |
| AC002064.1 | 52.9991356 | 0.951118763 | 0.46088187 | 2.06369315 | 0.03904682 | 0.14053663 |
| GTPBP10 | 465.960413 | -0.001663722 | 0.24083714 | -0.0069081 | 0.99448819 | 0.99786448 |
| CLDN12 | 474.741852 | -0.440702518 | 0.18858569 | -2.3368821 | 0.01944532 | 0.08757353 |
| CDK14 | 2469.04946 | 0.385157293 | 0.26452227 | 1.45604865 | 0.14537917 | 0.33542594 |
| AC079760.1 | 341.034946 | -0.016976924 | 0.64914205 | -0.0261529 | 0.97913541 | 0.9910179 |
| MTERF1 | 1437.74823 | -0.214974291 | 0.40124874 | -0.5357632 | 0.59212225 | 0.77343033 |
| AKAP9 | 4502.88906 | 0.59909169 | 0.1554743 | 3.85331646 | 0.00011653 | 0.00189094 |
| KRIT1 | 1515.4212 | 0.144435489 | 0.17427768 | 0.82876642 | 0.40723659 | 0.6271643 |
| ANKIB1 | 2800.78366 | 0.060956963 | 0.15327737 | 0.39769055 | 0.6908583 | 0.84134845 |
| GATAD1 | 447.665383 | -0.106186273 | 0.30367522 | -0.3496705 | 0.72658598 | 0.86526092 |
| AC007566.1 | 407.864012 | 0.045615242 | 0.34538993 | 0.13206882 | 0.89492985 | 0.95172874 |
| PEX1 | 599.865831 | 0.235799438 | 0.17908099 | 1.31671951 | 0.18793266 | 0.39374474 |
| RBM48 | 163.996411 | 0.261863613 | 0.26737835 | 0.97937477 | 0.32739484 | 0.55270302 |
| FAM133B | 699.020992 | 0.272084276 | 0.14164272 | 1.92091957 | 0.05474185 | 0.17653444 |
| CDK6 | 1619.17002 | 1.184160723 | 0.33428576 | 3.54236069 | 0.00039656 | 0.00493346 |
| SAMD9 | 140.793899 | -0.563495715 | 0.39682473 | -1.4200116 | 0.15560431 | 0.35056751 |
| SAMD9L | 307.509086 | -0.048064809 | 0.76380923 | -0.0629278 | 0.94982402 | 0.97825441 |

| | | | | | | |
|---|---|---|---|---|---|---|
| VPS50 | 1297.01672 | 0.31596396 | 0.14324346 | 2.20578286 | 0.02739921 | 0.11030665 |
| CALCR | 2479.44112 | -0.903341153 | 0.56173216 | -1.608135 | 0.10780561 | 0.2759816 |
| GNGT1 | 1872.03806 | -0.220947861 | 0.36229969 | -0.6098483 | 0.54196228 | 0.73809463 |
| TFPI2 | 76.3021157 | -1.099082967 | 0.73286292 | -1.4997115 | 0.13368914 | 0.31792719 |
| GNG11 | 65.4015811 | -0.163917114 | 0.42052015 | -0.3897961 | 0.69668733 | 0.84496018 |
| BET1 | 414.024 | -0.044067665 | 0.31099015 | -0.1417012 | 0.88731606 | 0.94926361 |
| CASD1 | 833.943587 | 0.33323605 | 0.14535646 | 2.29254384 | 0.02187428 | 0.09467149 |
| SGCE | 1459.37692 | 0.231704253 | 0.24681658 | 0.93877101 | 0.34784833 | 0.57340928 |
| PEG10 | 90.0582861 | -0.414798323 | 0.58611899 | -0.7077033 | 0.47912955 | 0.68864274 |
| PPP1R9A | 3857.41847 | -0.177469973 | 0.22408663 | -0.7919704 | 0.42837795 | 0.64675327 |
| AC002429.2 | 326.743205 | -0.398973168 | 0.48162451 | -0.8283905 | 0.40744938 | 0.62741885 |
| PON2 | 1629.2063 | -1.315536104 | 0.41212531 | -3.1920779 | 0.00141253 | 0.01307283 |
| PDK4 | 3190.92568 | 1.015045108 | 1.19213398 | 0.85145221 | 0.3945182 | 0.61682844 |
| AC002451.1 | 66.8501048 | 2.311344773 | 0.64570688 | 3.57955728 | 0.00034418 | 0.00444837 |
| SLC25A13 | 2088.96205 | -0.229812927 | 0.16321642 | -1.4080257 | 0.15912345 | 0.35555175 |
| SEM1 | 968.295143 | 0.353331782 | 0.20162297 | 1.7524381 | 0.07969851 | 0.22590935 |
| SDHAF3 | 372.796151 | -0.051058698 | 0.46897623 | -0.1088727 | 0.91330348 | 0.96075537 |
| ASNS | 154.405505 | 0.020955838 | 0.34739689 | 0.06032247 | 0.95189881 | 0.97925478 |
| LMTK2 | 910.08115 | -0.36119037 | 0.23887258 | -1.5120629 | 0.13051786 | 0.31271729 |
| TECPR1 | 166.507573 | -0.039495722 | 0.41754199 | -0.094591 | 0.92463968 | 0.9667507 |
| BRI3 | 1015.73837 | -0.436813361 | 0.25506032 | -1.7125884 | 0.08678828 | 0.23862737 |
| BAIAP2L1 | 1811.32942 | -0.630203666 | 0.31529109 | -1.9987995 | 0.04563005 | 0.15600761 |
| TRRAP | 1466.93779 | 0.245800696 | 0.13644496 | 1.80146411 | 0.07162976 | 0.21114874 |
| SMURF1 | 2078.05595 | -0.60035819 | 0.20150318 | -2.9793982 | 0.00288815 | 0.02248061 |
| AC004834.1 | 68.6535111 | 0.846447355 | 0.3640692 | 2.32496286 | 0.02007395 | 0.08927837 |
| ARPC1B | 183.347993 | -2.149594063 | 0.43389118 | -4.9542239 | 7.26E-07 | 3.20E-05 |
| PDAP1 | 206.850732 | -0.769642424 | 0.25800925 | -2.9830032 | 0.00285435 | 0.02229634 |
| BUD31 | 168.288298 | -0.253599193 | 0.20935206 | -1.2113528 | 0.22576023 | 0.44048256 |
| CPSF4 | 147.01832 | -0.091751633 | 0.23825209 | -0.3851032 | 0.70016098 | 0.84785721 |
| ZNF789 | 378.732846 | 0.289228058 | 0.18680086 | 1.54832292 | 0.12154457 | 0.29836131 |
| ZNF394 | 247.357284 | -0.336763606 | 0.19383053 | -1.7374126 | 0.08231436 | 0.23060242 |
| ZKSCAN5 | 199.233246 | 0.094030367 | 0.2369949 | 0.39676113 | 0.69154361 | 0.84169408 |
| FAM200A | 169.480133 | 0.289404703 | 0.24333476 | 1.18932743 | 0.23431084 | 0.45161281 |
| ZSCAN25 | 151.003886 | 0.272699391 | 0.20183652 | 1.35109044 | 0.17666646 | 0.37882911 |
| CYP3A5 | 77.5099432 | -1.008537738 | 0.67945036 | -1.4843435 | 0.13771781 | 0.32381543 |
| TRIM4 | 454.277328 | -0.201604501 | 0.18906679 | -1.0663137 | 0.28628187 | 0.50853916 |
| ZKSCAN1 | 1316.43175 | -0.044133239 | 0.11962112 | -0.3689419 | 0.71217107 | 0.85565469 |
| ZSCAN21 | 123.767651 | 0.124488803 | 0.26065256 | 0.47760438 | 0.63293181 | 0.80230473 |
| ZNF3 | 266.514183 | 0.091845751 | 0.33425278 | 0.27477931 | 0.78348579 | 0.89665455 |
| COPS6 | 274.499077 | -0.35588887 | 0.22790385 | -1.5615746 | 0.11838824 | 0.29339403 |
| MCM7 | 143.872188 | -0.237998459 | 0.26730411 | -0.8903659 | 0.37326946 | 0.59720401 |
| AP4M1 | 228.283911 | -0.370572906 | 0.22686794 | -1.63343 | 0.10237858 | 0.26688745 |
| TAF6 | 117.578046 | -0.617800203 | 0.2998883 | -2.060101 | 0.03938888 | 0.14131036 |
| AC073842.2 | 35.219415 | 0.763695691 | 0.50924681 | 1.4996573 | 0.1337032 | 0.31792719 |
| LAMTOR4 | 433.879217 | -0.722456737 | 0.30160712 | -2.395357 | 0.01660419 | 0.07866741 |
| C7orf43 | 63.3048231 | 0.427203387 | 0.43879177 | 0.97359024 | 0.33026003 | 0.55571163 |
| STAG3 | 161.165276 | -0.838180015 | 0.44537063 | -1.8819831 | 0.05983831 | 0.18757511 |
| CASTOR3 | 974.673502 | -1.389898619 | 0.3163513 | -4.3935291 | 1.12E-05 | 0.00029171 |
| SPDYE3 | 380.435205 | 0.224494747 | 0.30324736 | 0.74030239 | 0.45911653 | 0.67278348 |
| PILRB | 980.76789 | -0.296383801 | 0.32445339 | -0.9134865 | 0.36098671 | 0.58607098 |
| PILRA | 263.105025 | 0.856550577 | 0.38843848 | 2.20511257 | 0.0274462 | 0.11042851 |
| ZCWPW1 | 132.167764 | 0.662211735 | 0.52164454 | 1.26946931 | 0.20427373 | 0.41414152 |
| MEPCE | 118.170037 | -0.551858956 | 0.26824351 | -2.0573059 | 0.03965681 | 0.14177464 |
| PPP1R35 | 178.15005 | -0.591833794 | 0.32199134 | -1.8380426 | 0.06605613 | 0.20028899 |
| TSC22D4 | 505.674765 | -0.276189127 | 0.27378715 | -1.0087732 | 0.31308343 | 0.53694093 |
| AGFG2 | 358.575765 | -0.695032062 | 0.39576191 | -1.7561873 | 0.07905642 | 0.22471623 |
| SAP25 | 76.3764383 | 0.271699151 | 0.44779077 | 0.60675469 | 0.54401372 | 0.73943993 |
| LRCH4 | 159.235434 | -0.300889055 | 0.30505602 | -0.9863403 | 0.32396614 | 0.54890568 |
| PCOLCE-AS1 | 49.9692374 | 1.126434353 | 0.59836552 | 1.88251882 | 0.05976561 | 0.18751469 |
| PCOLCE | 107.708401 | 0.099535861 | 0.30754001 | 0.32365175 | 0.74620171 | 0.87598457 |
| MOSPD3 | 53.8182418 | -0.930216598 | 0.35355189 | -2.631061 | 0.00851188 | 0.04937785 |
| GNB2 | 548.18476 | -1.38287004 | 0.35355515 | -3.9113277 | 9.18E-05 | 0.00157367 |
| GIGYF1 | 602.493135 | -0.109465364 | 0.23231626 | -0.4711911 | 0.63750424 | 0.80497022 |
| POP7 | 62.8399525 | -1.108912262 | 0.50453891 | -2.1978726 | 0.02795818 | 0.11187032 |
| EPHB4 | 178.008397 | -0.102597505 | 0.32345515 | -0.3171924 | 0.75109764 | 0.87923155 |
| SLC12A9 | 434.385097 | 0.725896627 | 0.38923279 | 1.86494212 | 0.06218952 | 0.19265041 |
| TRIP6 | 216.471881 | 0.13825716 | 0.25279105 | 0.54692269 | 0.58443185 | 0.76903227 |
| SRRT | 275.588119 | -0.113861075 | 0.22655677 | -0.5025719 | 0.61526527 | 0.78857354 |
| TRIM56 | 856.354045 | -1.183818019 | 0.45004687 | -2.6304328 | 0.00852762 | 0.04944749 |
| AP1S1 | 205.977598 | -0.612797238 | 0.45227902 | -1.3549097 | 0.17544632 | 0.37768423 |
| PLOD3 | 165.387218 | -0.53892261 | 0.22897189 | -2.3536628 | 0.01858947 | 0.08488512 |
| ZNHIT1 | 292.361443 | -0.66496368 | 0.22733659 | -2.9250183 | 0.00344436 | 0.02569155 |
| CLDN15 | 73.7403896 | 0.051431261 | 0.28710197 | 0.17913935 | 0.85782828 | 0.93578594 |
| FIS1 | 475.235459 | -0.67829763 | 0.28771369 | -2.3575438 | 0.01839628 | 0.0843248 |
| IFT22 | 132.377364 | -0.584495796 | 0.25316681 | -2.3087379 | 0.02095813 | 0.09215577 |
| COL26A1 | 126.623505 | -0.067206591 | 0.54469373 | -0.1233842 | 0.90180288 | 0.95457905 |
| CUX1 | 5220.11396 | -0.418225587 | 0.20581861 | -2.0320105 | 0.04215259 | 0.14729989 |
| SH2B2 | 84.5698568 | -1.948257013 | 0.4107101 | -4.7436306 | 2.10E-06 | 7.32E-05 |
| PRKRIP1 | 1584.69618 | -0.15227146 | 0.21856961 | -0.6966726 | 0.48600771 | 0.6934193 |
| ORAI2 | 234.548506 | -0.837583373 | 0.39332601 | -2.1294889 | 0.03321383 | 0.12653816 |
| ALKBH4 | 92.5180953 | 0.072501648 | 0.28743213 | 0.25223919 | 0.80085619 | 0.90662963 |
| LRWD1 | 41.4273172 | 0.635941366 | 0.41303292 | 1.53968687 | 0.1236367 | 0.30156806 |
| POLR2J | 137.825657 | -0.804930685 | 0.33704618 | -2.3881911 | 0.01693154 | 0.0796186 |
| AC093668.2 | 78.2413277 | 0.798180978 | 0.52939431 | 1.50772489 | 0.13162495 | 0.31440036 |
| SPDYE2 | 208.20004 | 0.23779072 | 0.54627559 | 0.43529442 | 0.66334873 | 0.82258481 |
| AC105052.1 | 83.631027 | 0.690068135 | 0.49839338 | 1.38458528 | 0.16617931 | 0.36563672 |
| AC105052.3 | 492.972939 | 0.585046756 | 0.48876465 | 1.19699073 | 0.23131017 | 0.44721546 |
| FAM185A | 932.923031 | 0.487697389 | 0.16081797 | 3.03260505 | 0.00242453 | 0.0197445 |
| FBXL13 | 690.14367 | -0.527870073 | 0.26336305 | -2.0043437 | 0.04503325 | 0.1547717 |
| LRRC17 | 114.963283 | 0.971481873 | 0.25489762 | 3.811263 | 0.00013826 | 0.00216895 |
| ARMC10 | 452.57596 | -0.369683327 | 0.21336425 | -1.7326395 | 0.08315977 | 0.23195161 |
| NAPEPLD | 1056.01835 | 1.271057837 | 0.36692418 | 3.46408852 | 0.00053203 | 0.00620265 |
| PMPCB | 465.107425 | 0.413629519 | 0.2051294 | 2.01643217 | 0.0437548 | 0.15152832 |
| DNAJC2 | 413.142836 | -0.040646235 | 0.19976437 | -0.2034709 | 0.83876699 | 0.92481412 |
| PSMC2 | 316.666256 | 0.191393719 | 0.16108807 | 1.18813092 | 0.23478183 | 0.45204629 |
| SLC26A5 | 121.031568 | -0.946712163 | 0.44148495 | -2.1443815 | 0.03200233 | 0.12338552 |
| ORC5 | 566.066305 | 0.144664972 | 0.18290593 | 0.79092553 | 0.42898745 | 0.64738665 |
| LHFPL3 | 572.211638 | -0.122324554 | 0.34163457 | -0.3580567 | 0.72030087 | 0.86064266 |
| LINC01004 | 1761.27676 | 0.622152876 | 0.21442076 | 2.90155153 | 0.0037132 | 0.02712811 |
| KMT2E-AS1 | 117.933192 | -0.044736937 | 0.27562319 | -0.162312 | 0.8710602 | 0.94147844 |
| KMT2E | 3314.09994 | 0.175502444 | 0.16957174 | 1.03497459 | 0.30068078 | 0.52376268 |
| SRPK2 | 4810.373 | 0.197457146 | 0.16290452 | 1.21210354 | 0.22547275 | 0.44005165 |
| PUS7 | 271.256576 | -0.451443732 | 0.29903354 | -1.5096759 | 0.13112614 | 0.31371948 |
| RINT1 | 263.557492 | 0.324568529 | 0.25728383 | 1.26151934 | 0.2071218 | 0.41788175 |
| EFCAB10 | 77.679984 | -1.586881834 | 0.44994967 | -3.5267985 | 0.00042062 | 0.00518761 |
| ATXN7L1 | 1592.06456 | -0.084716131 | 0.24669674 | -0.3434019 | 0.73129613 | 0.86813072 |
| CDHR3 | 398.44581 | 0.465808031 | 0.59133145 | 0.78772748 | 0.43085613 | 0.64893712 |
| SYPL1 | 539.902436 | -0.001018303 | 0.28302368 | -0.0035979 | 0.99712926 | 0.99894429 |
| NAMPT | 2947.96909 | -1.54521833 | 0.56005767 | -2.7590343 | 0.00579724 | 0.03748204 |
| AC005050.1 | 63.0272016 | 2.831828826 | 0.55676146 | 5.08625149 | 3.65E-07 | 1.81E-05 |
| CCDC71L | 167.493069 | 0.859189431 | 0.69804147 | 1.23085729 | 0.21837624 | 0.43246354 |
| HBP1 | 987.202306 | -0.368561094 | 0.19932908 | -1.8490081 | 0.06445664 | 0.19708512 |
| COG5 | 2616.2276 | 0.002506833 | 0.21765919 | 0.01151724 | 0.99081078 | 0.99624112 |

| | | | | | | |
|---|---|---|---|---|---|---|
| DUS4L | 128.914395 | 0.006292276 | 0.21189153 | 0.02969574 | 0.97630971 | 0.98947549 |
| BCAP29 | 489.649924 | -0.061025051 | 0.24849145 | -0.2455821 | 0.80600575 | 0.90934469 |
| CBLL1 | 199.736484 | 0.162143329 | 0.26987765 | 0.6008031 | 0.54797114 | 0.74186733 |
| SLC26A3 | 2881.76756 | 0.838715142 | 0.63017535 | 1.33092344 | 0.1832142 | 0.3880193 |
| DLD | 674.249533 | 0.402918324 | 0.24160824 | 1.66765141 | 0.09538495 | 0.25441407 |
| LAMB1 | 5123.07176 | 0.460706257 | 0.48274873 | 0.95433966 | 0.33991174 | 0.56526428 |
| LAMB4 | 141.92334 | 0.982488623 | 0.4209087 | 2.33420842 | 0.01958481 | 0.08799176 |
| NRCAM | 2125.12146 | 0.901375663 | 0.48290643 | 1.86656381 | 0.06196253 | 0.19230804 |
| PNPLA8 | 1298.49443 | -0.325739791 | 0.34553398 | -0.9427142 | 0.34582711 | 0.57105512 |
| DNAJB9 | 158.422638 | -0.586851543 | 0.46173091 | -1.2709817 | 0.20373515 | 0.41350055 |
| IMMP2L | 25545.2378 | 0.486313936 | 0.18746319 | 2.59418368 | 0.00948158 | 0.05329491 |
| LRRN3 | 170.86298 | -0.198012344 | 0.65832609 | -0.3007816 | 0.76358108 | 0.88541384 |
| DOCK4 | 2728.12895 | 0.491780065 | 0.36053459 | 1.36403019 | 0.17255806 | 0.37384278 |
| ZNF277 | 1545.90283 | -0.029535134 | 0.2419241 | -0.1220843 | 0.90283225 | 0.95509228 |
| AC004112.1 | 87.7288428 | -0.092986704 | 0.42164079 | -0.2205354 | 0.82545424 | 0.91938879 |
| IFRD1 | 1336.24176 | 1.24729467 | 0.4536512 | 2.74945742 | 0.0059694 | 0.03818394 |
| LSMEM1 | 156.390906 | 0.887862718 | 0.70546552 | 1.2585487 | 0.20819339 | 0.41927526 |
| TMEM168 | 753.304585 | 0.32335789 | 0.20684066 | 1.5633188 | 0.11797763 | 0.2929811 |
| BMT2 | 1036.5416 | -0.234072944 | 0.15989204 | -1.4639437 | 0.14320933 | 0.33255675 |
| SMIM30 | 119.299085 | -0.150642751 | 0.39789235 | -0.3786018 | 0.7049836 | 0.85073346 |
| FOXP2 | 6965.77193 | 0.125870935 | 0.63588563 | 0.19794587 | 0.84308741 | 0.9274102 |
| MDFIC | 814.541483 | 0.055806867 | 0.35812996 | 0.15582853 | 0.87616818 | 0.94422008 |
| TES | 523.306777 | -0.329049593 | 0.2465888 | -1.3344061 | 0.18207081 | 0.38654023 |
| AC002066.1 | 59.7102881 | -0.445467087 | 0.54683662 | -0.8146256 | 0.41528667 | 0.63525223 |
| CAV2 | 108.1447 | -1.483356595 | 0.46147394 | -3.2143887 | 0.00130723 | 0.01235132 |
| MET | 2327.30067 | -1.597637357 | 0.51361725 | -3.1105602 | 0.00186733 | 0.01622733 |
| CAPZA2 | 1349.03644 | -0.351679028 | 0.19531005 | -1.8006192 | 0.07176291 | 0.21144687 |
| ST7 | 1955.22332 | 0.126577024 | 0.19625396 | 0.64496546 | 0.51894957 | 0.71946641 |
| ST7-AS2 | 126.950548 | -0.242641171 | 0.5841903 | -0.4153461 | 0.67788855 | 0.83117329 |
| CFTR | 57.3689886 | -1.42544907 | 0.43916479 | -3.2458182 | 0.00117114 | 0.01130814 |
| CTTNBP2 | 828.151989 | -0.228537689 | 0.34734442 | -0.657957 | 0.51056577 | 0.7127524 |
| LSM8 | 493.540697 | -0.096149286 | 0.16037685 | -0.599521 | 0.54882552 | 0.74226986 |
| ANKRD7 | 401.654168 | 0.768532314 | 0.29115888 | 2.63956339 | 0.00830129 | 0.04856144 |
| TSPAN12 | 1135.30676 | 0.436160149 | 0.37595059 | 1.16015286 | 0.24598658 | 0.46524008 |
| ING3 | 328.388984 | 0.579138895 | 0.23947858 | 2.41833281 | 0.01559181 | 0.07534919 |
| FAM3C | 1017.72738 | -0.705927701 | 0.25503006 | -2.7680177 | 0.00563984 | 0.03666173 |
| AASS | 642.449167 | 0.156518044 | 0.34803893 | 0.44971418 | 0.65291655 | 0.81499693 |
| CADPS2 | 6101.65263 | 0.939596406 | 0.35900964 | 2.61718988 | 0.0088657 | 0.05067375 |
| AC004594.1 | 377.877944 | 0.618617871 | 0.64764233 | 0.95518443 | 0.33948444 | 0.56496857 |
| SLC13A1 | 68.3572108 | 1.173398811 | 0.95346272 | 1.23067089 | 0.21844598 | 0.43253679 |
| IQUB | 307.944251 | -0.220621687 | 0.30015009 | -0.7350379 | 0.46231642 | 0.67537466 |
| NDUFA5 | 704.484782 | 0.415914491 | 0.25863066 | 1.60814067 | 0.10780437 | 0.2759816 |
| WASL | 1168.71297 | 0.195333466 | 0.19467326 | 1.00339135 | 0.31567208 | 0.53997831 |
| AC006333.2 | 52.9191271 | 0.276035851 | 0.41041023 | 0.67258521 | 0.50121121 | 0.70528379 |
| POT1 | 635.808821 | 0.043049306 | 0.18805795 | 0.22891511 | 0.8189349 | 0.9158675 |
| POT1-AS1 | 342.560906 | 0.745362988 | 0.3388638 | 2.19959462 | 0.02783567 | 0.11148616 |
| GRM8 | 639.626703 | 1.338109922 | 0.56595984 | 2.36431955 | 0.01806322 | 0.08348395 |
| ZNF800 | 818.30068 | -0.169438681 | 0.25180776 | -0.672889 | 0.50101789 | 0.70523715 |
| GCC1 | 51.0763242 | -0.284602381 | 0.30300707 | -0.9392599 | 0.34759733 | 0.57313923 |
| ARF5 | 159.195199 | -0.937599658 | 0.43219845 | -2.169373 | 0.03005438 | 0.1177308 |
| SND1 | 2765.15998 | -0.055615936 | 0.21411109 | -0.2597527 | 0.79505453 | 0.90385363 |
| RBM28 | 443.719341 | 0.492750245 | 0.20627395 | 2.38881471 | 0.01690282 | 0.07956857 |
| HILPDA | 472.522694 | -3.135625792 | 1.15050756 | -2.7254282 | 0.00642181 | 0.0401637 |
| METTL2B | 368.067407 | -0.287472919 | 0.20840997 | -1.3793626 | 0.16778298 | 0.3676585 |
| AC090114.2 | 111.350873 | -0.203553635 | 0.32375507 | -0.6287272 | 0.52952764 | 0.72768733 |
| AC018638.7 | 60.9357213 | 0.704140417 | 0.26883925 | 2.61918752 | 0.00881395 | 0.05053466 |
| FAM71F2 | 164.808487 | 0.689683506 | 0.29572717 | 2.33216145 | 0.0196922 | 0.08832403 |
| FAM71F1 | 44.0750314 | 0.807382286 | 0.58003374 | 1.39195746 | 0.16393528 | 0.36265636 |
| CALU | 578.264552 | -0.0151178 | 0.19882951 | -0.076034 | 0.93939206 | 0.97411372 |
| KCP | 1269.95072 | 0.698858038 | 0.6072037 | 1.15094495 | 0.24975488 | 0.46940982 |
| ATP6V1F | 368.761903 | -1.460614032 | 0.42170357 | -3.4636036 | 0.00053299 | 0.00620836 |
| IRF5 | 55.7685745 | 0.444674499 | 0.46449451 | 0.95732994 | 0.33840075 | 0.56381629 |
| TNPO3 | 1291.78874 | -0.085594456 | 0.18176354 | -0.470911 | 0.63770428 | 0.80499195 |
| TSPAN33 | 509.091508 | 1.454803413 | 0.4964853 | 2.9302044 | 0.00338739 | 0.02538379 |
| SMO | 411.639839 | 1.708108017 | 0.27856098 | 6.13189985 | 8.68E-10 | 1.15E-07 |
| AHCYL2 | 2114.70614 | 0.21467422 | 0.13932031 | 1.54086806 | 0.1233489 | 0.3011674 |
| AC078846.1 | 93.9111772 | 0.555264992 | 0.24907601 | 2.22929938 | 0.02579399 | 0.10574576 |
| NRF1 | 1697.71199 | 0.114234377 | 0.15575662 | 0.73341589 | 0.46330481 | 0.67622024 |
| UBE2H | 6443.81367 | -0.272758935 | 0.27895629 | -0.9777838 | 0.32818128 | 0.55346341 |
| AC073320.1 | 48.7183062 | 0.03183877 | 0.32597253 | 0.09767317 | 0.92219182 | 0.9655855 |
| ZC3HC1 | 359.873242 | 0.337298553 | 0.22123368 | 1.5246257 | 0.12735248 | 0.30758373 |
| KLHDC10 | 1942.28206 | 0.202653364 | 0.16512981 | 1.22723667 | 0.21973365 | 0.43361796 |
| TMEM209 | 388.719311 | 0.095948443 | 0.19852268 | 0.48331224 | 0.62887405 | 0.79919158 |
| AC087071.2 | 103.986895 | -0.052560533 | 0.46206014 | -0.1137526 | 0.90943393 | 0.95917541 |
| CEP41 | 159.094178 | 0.431017661 | 0.24451693 | 1.76273134 | 0.07794581 | 0.22304728 |
| MEST | 267.50429 | 0.321224574 | 0.72678089 | 0.44198269 | 0.65850173 | 0.81895766 |
| COPG2 | 599.010693 | 0.096368718 | 0.22246099 | 0.43319377 | 0.664874 | 0.82323966 |
| AC016831.7 | 15047.5189 | -1.34153116 | 0.38810908 | -3.4565828 | 0.00054707 | 0.00633883 |
| LINC00513 | 1983.64782 | -1.403228732 | 0.66775296 | -2.101419 | 0.0356042 | 0.13239418 |
| AC058791.1 | 406.448947 | -1.541020699 | 0.71962943 | -2.1414087 | 0.0322411 | 0.12393503 |
| MKLN1 | 10904.0717 | -0.554767969 | 0.35699817 | -1.5539799 | 0.12018922 | 0.29613493 |
| PODXL | 142.166059 | -0.589174065 | 0.42105234 | -1.3992894 | 0.16172622 | 0.35922099 |
| PLXNA4 | 418.770925 | -0.244097412 | 0.58488506 | -0.4173425 | 0.67642788 | 0.83016679 |
| CHCHD3 | 3553.2331 | 0.328534493 | 0.16141828 | 2.03529917 | 0.04182078 | 0.14665859 |
| EXOC4 | 7612.95075 | 0.04824618 | 0.12650837 | 0.38136749 | 0.70293058 | 0.84942 |
| LRGUK | 457.042638 | 0.23779345 | 0.27742522 | 0.8571443 | 0.39136514 | 0.6140943 |
| SLC35B4 | 537.101924 | 0.992258587 | 0.19759938 | 5.02156727 | 5.13E-07 | 2.41E-05 |
| AKR1B1 | 621.124786 | 0.548422133 | 0.82423181 | 0.66537366 | 0.50581153 | 0.70948881 |
| BPGM | 149.615158 | -0.341721317 | 0.29957417 | -1.1406902 | 0.25399886 | 0.47461749 |
| CALD1 | 241.916679 | -0.604472404 | 0.41985314 | -1.4397234 | 0.14994566 | 0.34213137 |
| AGBL3 | 404.262357 | 0.215546977 | 0.2709287 | 0.79558561 | 0.42627293 | 0.64486981 |
| CYREN | 411.00815 | -0.684377799 | 0.22908007 | -2.9875048 | 0.00281265 | 0.02208816 |
| TMEM140 | 343.22637 | -1.312127662 | 0.44603691 | -2.9417468 | 0.00326367 | 0.02463416 |
| WDR91 | 1285.23064 | -0.610376595 | 0.18657899 | -3.2714112 | 0.00107012 | 0.01051729 |
| AC009542.1 | 222.863727 | -1.484153706 | 0.60671437 | -2.4462148 | 0.01443649 | 0.07131326 |
| CNOT4 | 2042.59052 | 0.165579621 | 0.13076082 | 1.26627855 | 0.20541338 | 0.41584887 |
| NUP205 | 436.727495 | 0.159444349 | 0.22005287 | 0.7245729 | 0.46871409 | 0.68088028 |
| STMP1 | 804.223689 | -0.650873379 | 0.27419398 | -2.3737698 | 0.01760752 | 0.0818924 |
| SLC13A4 | 164.069556 | -1.176326488 | 0.52840852 | -2.2261687 | 0.02600288 | 0.10643697 |
| MTPN | 494.55935 | -0.367508847 | 0.13885136 | -2.6467789 | 0.00812625 | 0.04779144 |
| DGKI | 137.368779 | 1.224239369 | 0.48722726 | 2.512666 | 0.01198227 | 0.0628518 |
| CREB3L2 | 1846.70991 | 0.970278446 | 0.32184822 | 3.01470814 | 0.00257227 | 0.02069249 |
| TRIM24 | 1080.562 | 0.40088306 | 0.23807543 | 1.68384893 | 0.09221087 | 0.24887891 |
| SVOPL | 677.253948 | 0.714785044 | 0.64725974 | 1.10432489 | 0.26945223 | 0.49092338 |
| ATP6V0A4 | 2639.2944 | 0.520494426 | 0.38035587 | 1.36844062 | 0.1711742 | 0.37175928 |
| TMEM213 | 1518.6696 | 0.561796299 | 0.32359001 | 1.7361361 | 0.08253977 | 0.23089112 |
| KIAA1549 | 336.179959 | 0.361525031 | 0.32948137 | 1.09725486 | 0.27253 | 0.49407751 |
| ZC3HAV1L | 74.3313151 | 0.435465206 | 0.44751721 | 0.97306918 | 0.33051892 | 0.55580197 |
| ZC3HAV1 | 467.148751 | -0.838658961 | 0.33546848 | -2.4999635 | 0.01242061 | 0.06441054 |
| TTC26 | 304.310018 | -0.348455798 | 0.19056211 | -1.8285681 | 0.06746433 | 0.20303858 |
| UBN2 | 1338.77777 | 0.270047603 | 0.15243733 | 1.77153192 | 0.07647229 | 0.22055076 |
| LUC7L2 | 323.266107 | 0.590616362 | 0.2551572 | 2.31471568 | 0.0206285 | 0.09107013 |
| HIPK2 | 4938.28114 | -0.316003211 | 0.15739666 | -2.0076869 | 0.04467657 | 0.15400127 |

| | | | | | | |
|---|---|---|---|---|---|---|
| TBXAS1 | 359.590134 | -1.116001762 | 0.36670269 | -3.0433422 | 0.00233966 | 0.01924321 |
| PARP12 | 455.860384 | -0.178125218 | 0.59955844 | -0.297094 | 0.76639474 | 0.88690833 |
| KDM7A | 2268.31907 | -0.987538475 | 0.4132482 | -2.3896982 | 0.01686222 | 0.07944774 |
| SLC37A3 | 676.80277 | -0.160554365 | 0.1981375 | -0.8103179 | 0.41775749 | 0.63745762 |
| RAB19 | 54.0672626 | 0.520851512 | 0.2978214 | 1.74887199 | 0.08031315 | 0.22694435 |
| MKRN1 | 801.092307 | 0.043632359 | 0.19171219 | 0.22759304 | 0.81996263 | 0.91651491 |
| DENND2A | 3348.51712 | 0.373825849 | 0.31976945 | 1.16904805 | 0.24238427 | 0.46086856 |
| ADCK2 | 47.3899106 | -0.020391052 | 0.40461942 | -0.0503956 | 0.95980712 | 0.98229715 |
| NDUFB2 | 625.039624 | -0.086531533 | 0.34060135 | -0.2540552 | 0.79945294 | 0.90589122 |
| BRAF | 5047.99884 | 0.198265639 | 0.17404345 | 1.13917321 | 0.25463092 | 0.47546222 |
| MRPS33 | 194.9018 | 0.264026639 | 0.26882568 | 0.98214814 | 0.32602687 | 0.55126587 |
| TMEM178B | 884.951416 | -0.901389314 | 0.52812282 | -1.7067797 | 0.087863 | 0.24068486 |
| AC073878.1 | 57.6404416 | 1.210080909 | 0.52629187 | 2.29925821 | 0.02149028 | 0.09377267 |
| AGK | 539.408657 | 0.283364027 | 0.18825809 | 1.50518911 | 0.13227546 | 0.31566403 |
| AC004918.1 | 179.91052 | 0.339632751 | 0.39442621 | 0.86108058 | 0.38919366 | 0.6122488 |
| KIAA1147 | 523.71931 | 0.002446224 | 0.2991244 | 0.00817795 | 0.99347501 | 0.99743207 |
| WEE2-AS1 | 126.847596 | 0.502798983 | 0.47985525 | 1.04781386 | 0.29472437 | 0.51796358 |
| SSBP1 | 729.477811 | 0.304098215 | 0.19763605 | 1.53867785 | 0.12388296 | 0.30188476 |
| MGAM | 234.878174 | 0.977451073 | 0.79212286 | 1.23396398 | 0.21721633 | 0.43128814 |
| GSTK1 | 544.070949 | 0.237241255 | 0.24605626 | 0.96417483 | 0.33495828 | 0.56020055 |
| AC073342.1 | 127.03043 | 0.718520416 | 0.45391662 | 1.58293479 | 0.11343633 | 0.28563167 |
| TMEM139 | 88.0690862 | 0.520812027 | 0.58582106 | 0.8890292 | 0.37398739 | 0.59778785 |
| CASP2 | 157.427562 | -0.054755999 | 0.23958539 | -0.2285448 | 0.81922272 | 0.91603427 |
| ZYX | 138.319867 | -1.386120021 | 0.44672397 | -3.1028557 | 0.00191663 | 0.01660116 |
| EPHA1 | 70.4925629 | -0.354924549 | 0.34151364 | -1.039269 | 0.29867964 | 0.52181158 |
| TCAF2 | 230.68152 | -2.19243629 | 0.80632428 | -2.7190503 | 0.00654696 | 0.04083043 |
| TCAF1 | 703.500522 | 0.002578966 | 0.17428535 | 0.01479738 | 0.98819383 | 0.99552604 |
| AC004889.1 | 455.014462 | 0.779012486 | 0.34303039 | 2.27097223 | 0.02314866 | 0.09841346 |
| AC074386.1 | 939.007572 | -0.066582805 | 0.31207685 | -0.2133539 | 0.83105095 | 0.92124425 |
| ARHGEF5 | 189.747545 | -0.461394931 | 0.32403194 | -1.4239181 | 0.15447019 | 0.34881127 |
| TPK1 | 543.721591 | -0.330056143 | 0.37567342 | -0.878572 | 0.37963339 | 0.60242401 |
| CNTNAP2 | 156.907886 | 0.288717699 | 0.33603246 | 0.85919586 | 0.39023247 | 0.61320085 |
| CUL1 | 1516.16819 | 0.066625106 | 0.13141307 | 0.50698996 | 0.61216186 | 0.78684784 |
| EZH2 | 246.288933 | 0.065154193 | 0.4213198 | 0.15464308 | 0.8771027 | 0.94491882 |
| PDIA4 | 208.547841 | -0.336544617 | 0.37054499 | -0.9082423 | 0.36375024 | 0.58910948 |
| ZNF786 | 180.185608 | 0.014822511 | 0.35067668 | 0.04226831 | 0.96628481 | 0.984926 |
| ZNF425 | 151.330279 | 0.25641361 | 0.34429956 | 0.74473987 | 0.45642899 | 0.67145234 |
| ZNF398 | 424.060496 | 0.077089775 | 0.23088812 | 0.33388369 | 0.73846733 | 0.87201993 |
| ZNF282 | 249.286207 | -0.075669322 | 0.1956389 | -0.3867806 | 0.69891867 | 0.84693191 |
| ZNF212 | 172.79853 | 0.505939049 | 0.25114329 | 2.0145434 | 0.04395251 | 0.15198661 |
| ZNF783 | 226.864574 | 0.265891061 | 0.20769929 | 1.28017321 | 0.20048422 | 0.40930388 |
| ZNF777 | 126.423318 | 0.014503216 | 0.24695109 | 0.0587291 | 0.95316788 | 0.97941614 |
| ZNF746 | 184.882853 | 0.111232953 | 0.18697947 | 0.59489395 | 0.55191436 | 0.74481373 |
| KRBA1 | 77.4617202 | 0.025366926 | 0.27704517 | 0.09156242 | 0.92704571 | 0.96785859 |
| ZNF467 | 120.605914 | 0.840538948 | 0.48202546 | 1.74376462 | 0.08120014 | 0.22874648 |
| SSPO | 43.5566515 | 0.35076565 | 0.61820285 | 0.56739572 | 0.57044536 | 0.75945349 |
| ZNF862 | 651.14448 | 0.320579312 | 0.23768177 | 1.34877537 | 0.17740913 | 0.38004614 |
| ATP6V0E2-A: | 49.7939125 | 0.882437305 | 0.46344077 | 1.90409945 | 0.05689723 | 0.1814475 |
| ATP6V0E2 | 79.3646572 | -0.284892955 | 0.57653006 | -0.4941511 | 0.62119948 | 0.79317776 |
| AC092666.1 | 135.311956 | -0.883852738 | 0.72761006 | -1.2147341 | 0.22446753 | 0.4391279 |
| ACTR3C | 2648.35978 | -0.301516681 | 0.24162123 | -1.2478899 | 0.21207139 | 0.42488177 |
| LRRC61 | 132.971818 | -0.782411022 | 0.37779455 | -2.0709961 | 0.03835916 | 0.13904125 |
| RARRES2 | 162.171838 | 1.120591047 | 0.72349245 | 1.5488635 | 0.12141454 | 0.2981986 |
| AC073111.5 | 55.9204807 | 1.647138811 | 0.8354636 | 1.97152673 | 0.04866366 | 0.16318819 |
| TMEM176B | 581.935331 | -1.236298223 | 0.46339038 | -2.667941 | 0.00763176 | 0.04565579 |
| TMEM176A | 264.092476 | -1.432303685 | 0.50630522 | -2.8289333 | 0.00467034 | 0.03175016 |
| AOC1 | 110.568798 | -0.633556739 | 0.76020605 | -0.8334013 | 0.40461842 | 0.62539431 |
| ABCB8 | 188.795903 | -0.060741128 | 0.24149788 | -0.2515183 | 0.80141344 | 0.90672945 |
| SLC4A2 | 291.474351 | 0.529374999 | 0.27230803 | 1.94403005 | 0.05189183 | 0.17046484 |
| FASTK | 218.772577 | 0.752898463 | 0.33697515 | 2.23428483 | 0.02546434 | 0.10481721 |
| TMUB1 | 177.690928 | -0.415835327 | 0.3322277 | -1.2516576 | 0.21069465 | 0.42302259 |
| AGAP3 | 263.538408 | -0.374125229 | 0.24522124 | -1.525664 | 0.12709356 | 0.30701881 |
| CHPF2 | 96.0974553 | -0.889796233 | 0.31521406 | -2.8228317 | 0.00476016 | 0.03219503 |
| SMARCD3 | 329.387368 | -1.346426559 | 0.44954527 | -2.9950856 | 0.00274368 | 0.02166243 |
| AC021097.1 | 53.6374731 | 0.051556001 | 0.31265492 | 0.16489745 | 0.8690247 | 0.94023032 |
| NUB1 | 483.424425 | 0.153679728 | 0.29015657 | 0.52964415 | 0.59635868 | 0.77616532 |
| RHEB | 1165.13472 | -0.293711583 | 0.16833637 | -1.7447898 | 0.08102146 | 0.22843384 |
| PRKAG2 | 1694.57084 | 0.429450963 | 0.36589577 | 1.17369753 | 0.2405162 | 0.45857079 |
| GALNT11 | 1220.04156 | 0.029098521 | 0.12779815 | 0.22769125 | 0.81988627 | 0.91651491 |
| KMT2C | 5385.95873 | 0.008287649 | 0.13974157 | 0.05930697 | 0.95270761 | 0.97936812 |
| ACTR3B | 744.660868 | 0.29115812 | 0.15923898 | 1.82843501 | 0.06748429 | 0.20305239 |
| DPP6 | 1994.61632 | 0.427961127 | 0.40533427 | 1.05582271 | 0.29104922 | 0.51373981 |
| PAXIP1-AS2 | 227.631617 | 0.454346522 | 0.36694299 | 1.23819377 | 0.21564422 | 0.42946242 |
| PAXIP1 | 453.754495 | 0.332317328 | 0.18811013 | 1.7666105 | 0.07729348 | 0.22180526 |
| PAXIP1-AS1 | 138.583289 | 0.167891249 | 0.30113775 | 0.55752308 | 0.57717009 | 0.76375874 |
| INSIG1 | 228.068052 | -1.453811628 | 0.6994694 | -2.0784492 | 0.037668 | 0.13733278 |
| AC009403.1 | 315.215296 | -0.103810554 | 0.29178993 | -0.3557715 | 0.72201166 | 0.86152231 |
| RBM33 | 2313.52325 | -0.260476492 | 0.17778815 | -1.4650948 | 0.14289507 | 0.33233231 |
| SHH | 77.865437 | -1.746850118 | 0.41643274 | -4.1947953 | 2.73E-05 | 0.00060939 |
| LINC01006 | 640.600355 | -0.061156441 | 0.44515679 | -0.1373818 | 0.89072902 | 0.95046308 |
| RNF32 | 64.6004688 | 0.073459453 | 0.42655318 | 0.1722164 | 0.8632674 | 0.93813133 |
| LMBR1 | 2056.72862 | 0.028273528 | 0.21321065 | 0.13260842 | 0.89450307 | 0.95172874 |
| NOM1 | 198.593366 | -0.041582184 | 0.30157689 | -0.1378825 | 0.89033326 | 0.95041347 |
| UBE3C | 2404.08182 | 0.148641328 | 0.20621463 | 0.72080882 | 0.47102715 | 0.68235944 |
| DNAJB6 | 1224.05692 | -0.289423683 | 0.20100741 | -1.4398658 | 0.14990538 | 0.34210439 |
| PTPRN2 | 53.413058 | -0.47583164 | 0.42137399 | -1.1292383 | 0.25879733 | 0.47985036 |
| NCAPG2 | 98.0015228 | 0.316283997 | 0.34223371 | 0.92417546 | 0.35539497 | 0.58084783 |
| ESYT2 | 2048.7388 | -0.528093712 | 0.25425858 | -2.0769947 | 0.03780205 | 0.13759387 |
| WDR60 | 1998.06428 | -0.115870911 | 0.16438445 | -0.7048776 | 0.48088642 | 0.68948936 |
| PLCXD1 | 59.5263419 | 0.709684087 | 0.34760314 | 2.04165038 | 0.04118622 | 0.14519362 |
| GTPBP6 | 308.759529 | 0.17978801 | 0.26247819 | 0.68496363 | 0.49336687 | 0.69886145 |
| LINC00685 | 188.302824 | -0.249703238 | 0.5141402 | -0.4856715 | 0.6272001 | 0.79775504 |
| PPP2R3B | 134.969888 | 0.004170694 | 0.42987386 | 0.00970213 | 0.99225894 | 0.9967867 |
| IL3RA | 218.844066 | 0.686296316 | 0.45244657 | 1.51685606 | 0.12930303 | 0.31082142 |
| SLC25A6 | 953.280806 | -0.800898948 | 0.42148739 | -1.900173 | 0.05741042 | 0.18268713 |
| LINC00106 | 414.749037 | 0.390952807 | 0.37187094 | 1.05131316 | 0.29311479 | 0.51602735 |
| ASMTL-AS1 | 202.279013 | 0.384395023 | 0.30050994 | 1.27914245 | 0.2008469 | 0.40991759 |
| ASMTL | 334.523879 | -0.166050715 | 0.23195208 | -0.7158837 | 0.47406315 | 0.68459432 |
| AKAP17A | 325.489241 | 0.383279756 | 0.21614965 | 1.77321482 | 0.07619312 | 0.22012391 |
| DHRSX | 1605.15013 | 0.159178171 | 0.2496144 | 0.63769626 | 0.52367142 | 0.72325104 |
| CD99 | 306.834929 | -0.683035622 | 0.35606021 | -1.918315 | 0.05507109 | 0.17733642 |
| GYG2 | 49.7113346 | -0.059847479 | 0.44374286 | -0.1348697 | 0.89271487 | 0.95119823 |
| ARSD | 652.685655 | -0.214525363 | 0.39741833 | -0.5397974 | 0.58933679 | 0.77174702 |
| ARSE | 60.1586635 | -0.055353037 | 0.5592419 | -0.0989787 | 0.92115518 | 0.96504145 |
| MXRA5 | 47.2011905 | 0.691516938 | 0.71030269 | 0.97355247 | 0.33027879 | 0.55571163 |
| PRKX | 2688.40377 | -0.527205143 | 0.38927068 | -1.3543407 | 0.1756277 | 0.37786639 |
| PRKX-AS1 | 204.820647 | -0.995931038 | 0.64260242 | -1.5498402 | 0.12117989 | 0.29772614 |
| BX890604.1 | 203.268172 | 0.202653908 | 0.38787931 | 0.5224664 | 0.60134563 | 0.77912452 |
| FAM239A | 58.0904187 | -4.308094175 | 1.6320246 | -2.6397238 | 0.00829736 | 0.04855997 |
| NLGN4X | 224.363667 | -1.544086749 | 1.00468535 | -1.5368859 | 0.12432125 | 0.30242347 |
| AC097347.1 | 58.8974533 | -0.034588887 | 0.55026958 | -0.0628581 | 0.94987952 | 0.97825441 |
| PUDP | 404.559377 | -0.367640432 | 0.3029846 | -1.2133964 | 0.22497829 | 0.43942281 |
| STS | 126.265953 | -1.050134346 | 0.48288862 | -2.1746927 | 0.02965315 | 0.11658408 |

| | | | | | | |
|---|---|---|---|---|---|---|
| PNPLA4 | 104.100305 | 0.228794687 | 0.24301558 | 0.94148154 | 0.34645814 | 0.57161699 |
| AC078960.1 | 58.394651 | 0.287333504 | 0.45442265 | 0.63230454 | 0.5271879 | 0.72545317 |
| TBL1X | 3631.0499 | -0.046386743 | 0.33465088 | -0.1386123 | 0.88975649 | 0.9503707 |
| GPR143 | 93.9668973 | -0.056348995 | 0.54621305 | -0.103163 | 0.91783358 | 0.9631893 |
| SHROOM2 | 382.983138 | -0.039780925 | 0.29308665 | -0.1357309 | 0.89203399 | 0.95118543 |
| WWC3 | 356.155492 | 0.22747074 | 0.28669805 | 0.79341573 | 0.42753565 | 0.64592455 |
| CLCN4 | 71.3089607 | 1.276741724 | 0.43412085 | 2.94098225 | 0.00327173 | 0.02466685 |
| MID1 | 683.089457 | -1.577895037 | 0.48687318 | -3.2408749 | 0.00119163 | 0.01145997 |
| AC073529.1 | 236.834141 | -0.008095193 | 0.42531733 | -0.0190333 | 0.98481454 | 0.99405563 |
| HCCS | 59.4174964 | -0.078284885 | 0.31717389 | -0.2468201 | 0.80504747 | 0.90864497 |
| ARHGAP6 | 19651.9323 | 0.72785539 | 0.28092445 | 2.59092929 | 0.00957172 | 0.05368696 |
| MSL3 | 98.394113 | -0.26898773 | 0.32706664 | -0.8224248 | 0.41083514 | 0.63014283 |
| PRPS2 | 150.914347 | 0.762219285 | 0.34711788 | 2.19585143 | 0.02810258 | 0.11228281 |
| TMSB4X | 857.377309 | -1.364302648 | 0.48038803 | -2.8400014 | 0.00451133 | 0.03113386 |
| TCEANC | 38.3389062 | 0.747920567 | 0.41508421 | 1.8018526 | 0.0715686 | 0.21105625 |
| RAB9A | 429.970217 | -0.2729624 | 0.28078167 | -0.9721518 | 0.33097504 | 0.55593201 |
| TRAPPC2 | 219.109694 | -0.022666951 | 0.23684086 | -0.0957054 | 0.92375454 | 0.96642544 |
| OFD1 | 1110.03162 | -0.12927101 | 0.21071237 | -0.6134951 | 0.53954901 | 0.73579422 |
| GPM6B | 183.611191 | -1.079737847 | 0.42424758 | -2.5450655 | 0.01092573 | 0.05909824 |
| GEMIN8 | 311.40054 | 0.094251454 | 0.19877643 | 0.47415809 | 0.63538718 | 0.80389813 |
| MOSPD2 | 231.649567 | 0.093888522 | 0.22737873 | 0.4129169 | 0.6796675 | 0.83230806 |
| PIGA | 123.082595 | -0.357159822 | 0.30408302 | -1.1745471 | 0.24017596 | 0.45822612 |
| PIR | 175.299648 | 0.324431762 | 0.51389956 | 0.63131356 | 0.52783552 | 0.72611741 |
| BMX | 45.0163287 | -0.560654669 | 0.34729783 | -1.6143339 | 0.10645502 | 0.27378589 |
| CLTRN | 154.58266 | -0.643748389 | 0.65253215 | -0.986539 | 0.32386871 | 0.54888141 |
| CA5B | 608.205725 | -0.095901647 | 0.23272887 | -0.4120746 | 0.68028478 | 0.8328016 |
| ZRSR2 | 253.43186 | 0.172782811 | 0.28724735 | 0.60151229 | 0.54749883 | 0.74150641 |
| AP1S2 | 386.17475 | -0.799049943 | 0.36630582 | -2.1813738 | 0.02915578 | 0.11507494 |
| CTPS2 | 571.548817 | 0.045576975 | 0.1775531 | 0.2566949 | 0.79741431 | 0.90505915 |
| SYAP1 | 519.2376 | -0.534965161 | 0.13129106 | -4.0746503 | 4.61E-05 | 0.0009237 |
| TXLNG | 362.424644 | 0.016775904 | 0.23111172 | 0.07258786 | 0.94213409 | 0.97528599 |
| RBBP7 | 418.364895 | -0.208412202 | 0.17124755 | -1.2170229 | 0.22359549 | 0.43821184 |
| REPS2 | 305.603554 | -0.731751986 | 0.3343828 | -2.1883661 | 0.02864294 | 0.11381696 |
| NHS | 8926.37083 | -0.52291492 | 0.42930263 | -1.2180566 | 0.22320245 | 0.43782019 |
| NHS-AS1 | 47.7209454 | -1.331407493 | 0.47212633 | -2.8200238 | 0.00480201 | 0.03239516 |
| SCML1 | 572.416724 | -0.330485718 | 0.33956196 | -0.9732707 | 0.33041876 | 0.55577504 |
| RAI2 | 230.325335 | -1.06746717 | 0.2626166 | -4.0647361 | 4.81E-05 | 0.00095282 |
| SCML2 | 66.208424 | 0.831315252 | 0.48881433 | 1.70067692 | 0.08900367 | 0.24280246 |
| CDKL5 | 1017.1716 | 0.691459857 | 0.2477152 | 2.79135017 | 0.00524887 | 0.03475301 |
| PHKA2 | 361.324899 | 0.109593176 | 0.1780288 | 0.6155924 | 0.53816357 | 0.7348912 |
| ADGRG2 | 61.0814945 | -0.449119338 | 0.52117593 | -0.8617423 | 0.38882935 | 0.6119441 |
| PDHA1 | 740.325722 | 0.037543675 | 0.34328192 | 0.10936689 | 0.9129115 | 0.96054229 |
| MAP3K15 | 735.471667 | 0.70826518 | 0.34472903 | 2.05455625 | 0.03992189 | 0.14232879 |
| SH3KBP1 | 163.129529 | -0.595117221 | 0.28644715 | -2.0775812 | 0.03774795 | 0.13752443 |
| BCLAF3 | 780.44802 | 0.122980106 | 0.23684131 | 0.51925108 | 0.60358566 | 0.78064849 |
| EIF1AX | 605.557426 | -0.338183593 | 0.2632813 | -1.2844953 | 0.19896871 | 0.40748588 |
| RPS6KA3 | 1040.16225 | -0.822601726 | 0.2129792 | -3.8623571 | 0.0001123 | 0.00183356 |
| MBTPS2 | 452.909104 | 0.475147838 | 0.21529586 | 2.20695294 | 0.02731735 | 0.11017858 |
| SMS | 1400.25108 | -0.162946509 | 0.25902335 | -0.6290804 | 0.52929642 | 0.72752096 |
| PHEX | 116.160481 | -2.077286496 | 0.9816553 | -2.1161058 | 0.03433581 | 0.12925098 |
| PTCHD1-AS | 396.302828 | -0.363700278 | 0.63165609 | -0.5757884 | 0.56475818 | 0.75596786 |
| PRDX4 | 116.376748 | -0.840214533 | 0.3485719 | -2.4104482 | 0.01593293 | 0.07637249 |
| ACOT9 | 293.177088 | -0.246159113 | 0.34694533 | -0.709504 | 0.47801175 | 0.68767071 |
| SAT1 | 7840.22703 | -1.836545766 | 0.38317953 | -4.792912 | 1.64E-06 | 6.03E-05 |
| APOO | 2057.46872 | -0.454267902 | 0.33073091 | -1.3735272 | 0.16958851 | 0.3699733 |
| KLHL15 | 120.945569 | 0.573667732 | 0.39970419 | 1.4352307 | 0.15122137 | 0.34403773 |
| EIF2S3 | 530.983404 | 0.077672474 | 0.19590984 | 0.39647051 | 0.69175796 | 0.84169408 |
| ZFX | 890.347139 | 0.460797168 | 0.30125124 | 1.52961087 | 0.12611308 | 0.30521649 |
| PDK3 | 586.264566 | 0.343716259 | 0.41389168 | 0.8304498 | 0.40628452 | 0.6262822 |
| POLA1 | 748.713466 | 0.077922953 | 0.28121675 | 0.27709215 | 0.78170935 | 0.89582484 |
| IL1RAPL1 | 78.2552679 | 0.740915361 | 0.36101535 | 2.05230984 | 0.04013956 | 0.14295051 |
| GK | 452.87037 | 0.677390726 | 0.20173553 | 3.35781571 | 0.00078561 | 0.00826862 |
| GK-AS1 | 44.735881 | 0.596044704 | 0.45679329 | 1.30484557 | 0.19194544 | 0.39901448 |
| TAB3 | 723.809938 | -0.079163822 | 0.17649265 | -0.4485389 | 0.65376432 | 0.81551732 |
| DMD | 727.619062 | 0.73247479 | 0.24763538 | 2.95787613 | 0.00309767 | 0.02367886 |
| TMEM47 | 422.920925 | -1.402865809 | 0.49484731 | -2.8349468 | 0.00458333 | 0.03146633 |
| CFAP47 | 197.299555 | -0.973239868 | 0.53150967 | -1.8310859 | 0.06708772 | 0.20232002 |
| AF241726.2 | 661.893924 | -0.312744043 | 0.37568999 | -0.8324524 | 0.40515362 | 0.6256341 |
| DYNLT3 | 247.600909 | -1.191375671 | 0.49265225 | -2.4182893 | 0.01559367 | 0.07534919 |
| RPGR | 469.502927 | 0.059504469 | 0.27281011 | 0.2181168 | 0.8273381 | 0.91965909 |
| MID1IP1 | 125.323303 | -0.799815957 | 0.34971622 | -2.2870428 | 0.02219333 | 0.09561372 |
| BCOR | 793.51138 | -0.270038043 | 0.30880707 | -0.8744555 | 0.38187024 | 0.60465286 |
| ATP6AP2 | 403.840227 | -0.486980428 | 0.27475676 | -1.7724056 | 0.07632725 | 0.22032488 |
| CXorf38 | 255.137898 | -0.306630826 | 0.17641862 | -1.7380865 | 0.08219557 | 0.23040147 |
| MED14 | 1204.19317 | -0.416455466 | 0.23713282 | -1.7562119 | 0.07905224 | 0.22471623 |
| USP9X | 2072.12209 | -0.183855462 | 0.19790396 | -0.9290135 | 0.35288207 | 0.57885329 |
| DDX3X | 955.930203 | -0.02761027 | 0.30637472 | -0.0901193 | 0.92819242 | 0.96874951 |
| CASK | 1477.46947 | 0.176728834 | 0.17127507 | 1.03184214 | 0.30214608 | 0.52534086 |
| Z93403.1 | 147.200906 | 2.178592815 | 0.58679576 | 3.71269349 | 0.00020507 | 0.00299636 |
| MAOA | 2222.82356 | -1.435312863 | 0.40799633 | -3.5179553 | 0.00043489 | 0.00534152 |
| MAOB | 611.295844 | 0.384574966 | 0.33925103 | 1.13360001 | 0.25696237 | 0.47778941 |
| EFHC2 | 408.491221 | -0.287344712 | 0.45935242 | -0.625543 | 0.53161471 | 0.7289628 |
| FUNDC1 | 94.5532053 | -0.118327971 | 0.27538141 | -0.4296876 | 0.66742292 | 0.82469498 |
| KDM6A | 2955.42299 | 0.128084975 | 0.39232099 | 0.32648005 | 0.74406118 | 0.87481257 |
| MIR222HG | 724.947341 | 0.149161479 | 0.50783307 | 0.29372148 | 0.76897074 | 0.88857795 |
| KRBOX4 | 449.471331 | 0.467934144 | 0.14882055 | 3.14428455 | 0.00166493 | 0.01487892 |
| ZNF674 | 132.455139 | 0.404634901 | 0.17195766 | 2.35310768 | 0.01861724 | 0.08488614 |
| SLC9A7 | 552.796221 | -0.510778988 | 0.29219773 | -1.7480594 | 0.08045374 | 0.22726977 |
| RP2 | 138.498987 | -0.452653413 | 0.26559808 | -1.7042797 | 0.08832885 | 0.24161022 |
| JADE3 | 547.174044 | -0.703915853 | 0.24224895 | -2.905754 | 0.00366369 | 0.02684067 |
| RGN | 383.735114 | -0.652816902 | 0.72174844 | -0.9044937 | 0.36573368 | 0.59121933 |
| NDUFB11 | 349.961497 | -0.631813602 | 0.51700552 | -1.2220635 | 0.2216836 | 0.43577101 |
| RBM10 | 465.875936 | 0.080773843 | 0.16758237 | 0.48199487 | 0.62980959 | 0.79976494 |
| UBA1 | 832.018398 | 0.251452734 | 0.20368505 | 1.23451735 | 0.21701018 | 0.43120432 |
| CDK16 | 255.37914 | 0.09008435 | 0.22831891 | 0.39455493 | 0.69317138 | 0.84287037 |
| USP11 | 303.350608 | 0.160879952 | 0.21210221 | 0.75850203 | 0.4481505 | 0.66393225 |
| ZNF41 | 255.002776 | 0.247658377 | 0.15750239 | 1.57241031 | 0.11585543 | 0.28950706 |
| ARAF | 201.304803 | 0.226203983 | 0.2836182 | 0.79756512 | 0.4251229 | 0.64375196 |
| TIMP1 | 540.086293 | -4.244029775 | 0.44294889 | -9.5813081 | 9.58E-22 | 3.16E-18 |
| ELK1 | 105.796903 | -0.065936425 | 0.1929889 | -0.3416592 | 0.73260741 | 0.8685945 |
| UXT | 141.789219 | -0.19895277 | 0.21630952 | -0.9197596 | 0.35769838 | 0.58316933 |
| ZNF81 | 553.617058 | 0.023235271 | 0.18520148 | 0.12545942 | 0.90015985 | 0.95395051 |
| ZNF182 | 113.021245 | -0.143321734 | 0.18685846 | -0.7670069 | 0.44307739 | 0.65885504 |
| ZNF630 | 206.914691 | 0.329845229 | 0.30122761 | 1.0950033 | 0.27351519 | 0.49520195 |
| FTSJ1 | 58.6742354 | -0.080898194 | 0.27795769 | -0.291045 | 0.77101691 | 0.88961935 |
| PORCN | 37.8248813 | 0.463560358 | 0.36352135 | 1.27519431 | 0.20224047 | 0.4117439 |
| EBP | 169.620477 | 0.280518221 | 0.3600845 | 0.77903443 | 0.43595943 | 0.65305582 |
| TBC1D25 | 172.613869 | 0.132090385 | 0.22862552 | 0.5777587 | 0.56342704 | 0.75487451 |
| RBM3 | 368.445186 | -0.524533914 | 0.25672069 | -2.0432085 | 0.0410318 | 0.1448411 |
| WDR13 | 327.537295 | 0.09644362 | 0.24317473 | 0.39660215 | 0.69166086 | 0.84169408 |
| AC231533.1 | 47.5623124 | 0.080181749 | 0.41403566 | 0.19365904 | 0.84644287 | 0.92885801 |
| HDAC6 | 245.587094 | 0.214114397 | 0.23755173 | 0.90133797 | 0.36740865 | 0.59270895 |
| PCSK1N | 772.519498 | -1.300375049 | 0.53812777 | -2.4164801 | 0.01567138 | 0.07557623 |

| | | | | | | |
|---|---|---|---|---|---|---|
| TIMM17B | 104.333269 | -0.878810989 | 0.31340538 | -2.8040711 | 0.00504618 | 0.03364564 |
| PQBP1 | 74.3426484 | -0.058024442 | 0.42141372 | -0.13769 | 0.89048545 | 0.95041347 |
| SLC35A2 | 62.1438878 | -0.094950193 | 0.28665932 | -0.3312301 | 0.74047069 | 0.87282504 |
| OTUD5 | 491.039942 | -0.34912641 | 0.19168072 | -1.8213955 | 0.06854676 | 0.20540701 |
| GRIPAP1 | 432.782379 | -0.424517272 | 0.2299286 | -1.8463004 | 0.0648486 | 0.19777071 |
| TFE3 | 238.958323 | -0.742556321 | 0.26190102 | -2.8352556 | 0.0045789 | 0.03145229 |
| CCDC120 | 147.597883 | -0.533938563 | 0.30170778 | -1.7697209 | 0.07677365 | 0.22103381 |
| WDR45 | 326.570395 | -0.337622646 | 0.20696899 | -1.6312717 | 0.10283301 | 0.2676495 |
| GPKOW | 63.9514137 | -0.344365054 | 0.22492395 | -1.5310289 | 0.12576226 | 0.30469803 |
| MAGIX | 78.851389 | 0.308460337 | 0.30233439 | 1.02026216 | 0.30760415 | 0.53112982 |
| PLP2 | 141.616677 | -2.558070975 | 0.35379303 | -7.2304165 | 4.82E-13 | 1.79E-10 |
| PRICKLE3 | 53.0266643 | -0.361446519 | 0.3059258 | -1.1814843 | 0.2374104 | 0.45507241 |
| SYP | 68.2964092 | 0.524914297 | 0.46436775 | 1.13038492 | 0.25831407 | 0.47942539 |
| CCDC22 | 78.7736389 | -0.260395667 | 0.3126298 | -0.8329202 | 0.40488975 | 0.62551915 |
| PPP1R3F | 84.3998901 | 0.526383962 | 0.23636568 | 2.22698986 | 0.02594795 | 0.10624504 |
| CLCN5 | 5924.59299 | 1.066339888 | 0.41716926 | 2.55613244 | 0.01058428 | 0.0578196 |
| CCNB3 | 55.3548498 | 0.429909789 | 0.53372683 | 0.80548655 | 0.42053896 | 0.63996534 |
| SHROOM4 | 1387.69809 | -0.754987814 | 0.30412566 | -2.4824864 | 0.0130469 | 0.06661637 |
| AL158055.1 | 74.6122823 | 0.429166591 | 0.6922909 | 0.61992233 | 0.53530892 | 0.73220416 |
| GSPT2 | 49.3844258 | -0.736288605 | 0.45692652 | -1.6113939 | 0.1070939 | 0.27484035 |
| MAGED1 | 2100.01626 | 0.483650171 | 0.20530688 | 2.35574262 | 0.01848572 | 0.08460772 |
| FAM156A | 151.258678 | 0.363433732 | 0.20606574 | 1.76367859 | 0.07778611 | 0.22283164 |
| GPR173 | 67.5473102 | -1.669280261 | 0.57953015 | -2.8804028 | 0.00397167 | 0.0285183 |
| TSPYL2 | 175.224294 | -0.816327867 | 0.43112808 | -1.8934695 | 0.05829546 | 0.18443706 |
| KANTR | 1052.15956 | 0.435442257 | 0.25443243 | 1.71142589 | 0.08700252 | 0.23902169 |
| KDM5C | 419.965581 | -0.049096918 | 0.28235644 | -0.1738828 | 0.8619576 | 0.93739486 |
| IQSEC2 | 155.947626 | -0.543810877 | 0.38927634 | -1.396979 | 0.16241988 | 0.36039043 |
| SMC1A | 421.724597 | -0.084643034 | 0.25318669 | -0.3343108 | 0.73814507 | 0.87200855 |
| HSD17B10 | 153.682375 | -0.652192619 | 0.49123114 | -1.3276695 | 0.1842873 | 0.38958162 |
| HUWE1 | 1907.63918 | 0.066379772 | 0.13011566 | 0.51015974 | 0.60993955 | 0.78542196 |
| PHF8 | 1112.99299 | 0.091722968 | 0.39904278 | 0.22985748 | 0.81820252 | 0.91549538 |
| FAM120C | 414.407507 | 0.139020735 | 0.14465724 | 0.96103542 | 0.33653436 | 0.56191156 |
| WNK3 | 369.103622 | 0.289013478 | 0.23592707 | 1.22501193 | 0.22057073 | 0.43435181 |
| TSR2 | 93.4391895 | -0.929290266 | 0.31026542 | -2.9951461 | 0.00274313 | 0.02166243 |
| FGD1 | 131.863373 | -0.346319201 | 0.2330482 | -1.4860411 | 0.13726826 | 0.32320435 |
| GNL3L | 157.432274 | -0.068957651 | 0.27473427 | -0.2509976 | 0.80181594 | 0.90677969 |
| MAGED2 | 289.292349 | -0.848630303 | 0.38294911 | -2.2160394 | 0.0266888 | 0.10833817 |
| TRO | 40.3329378 | 0.217814751 | 0.60994674 | 0.35710454 | 0.72101354 | 0.86110921 |
| APEX2 | 55.1889937 | -0.198795929 | 0.38124618 | -0.5214372 | 0.60206226 | 0.77940904 |
| FAM104B | 96.7056751 | 0.816103366 | 0.34599498 | 2.35871446 | 0.01833836 | 0.0842251 |
| MAGEH1 | 46.8866375 | -0.093060561 | 0.53967551 | -0.172438 | 0.86309321 | 0.93813133 |
| USP51 | 46.282563 | 0.162193348 | 0.54858961 | 0.29565516 | 0.76749344 | 0.88756968 |
| RRAGB | 730.241737 | 0.011664406 | 0.14475872 | 0.08057826 | 0.93577736 | 0.97236337 |
| AL445472.1 | 750.244137 | -0.473768966 | 0.19844627 | -2.3873916 | 0.0169684 | 0.07963582 |
| KLF8 | 106.305381 | -0.882204465 | 0.42780358 | -2.0621718 | 0.03919138 | 0.14094173 |
| UBQLN2 | 207.064492 | -0.312603313 | 0.25207568 | -1.2401169 | 0.21493216 | 0.42879308 |
| AL354793.1 | 45.4095951 | -0.357481141 | 0.41363447 | -0.8642441 | 0.38745383 | 0.61086904 |
| NBDY | 946.960942 | 0.006855319 | 0.22674626 | 0.03023344 | 0.97588088 | 0.9893257 |
| SPIN3 | 276.614797 | 0.148467947 | 0.21285668 | 0.69750195 | 0.48548872 | 0.69300247 |
| FAAH2 | 2384.26541 | -0.832086405 | 0.31444175 | -2.6462338 | 0.00813935 | 0.04782594 |
| ZXDB | 47.9026787 | 0.179243205 | 0.32370648 | 0.5537214 | 0.57976953 | 0.76589416 |
| ARHGEF9 | 643.226955 | 0.194284367 | 0.26664108 | 0.72863629 | 0.46622417 | 0.67903174 |
| AMER1 | 40.4740629 | 0.494473332 | 0.41449602 | 1.19295074 | 0.23288867 | 0.44960924 |
| MTMR8 | 331.80803 | 0.116164433 | 0.18547657 | 0.62630246 | 0.53111657 | 0.72863563 |
| ZC4H2 | 761.825031 | 0.62390011 | 0.19909167 | 3.13373285 | 0.00172598 | 0.01521926 |
| ZC3H12B | 306.926426 | 0.244594769 | 0.19653452 | 1.24453843 | 0.21330146 | 0.42637697 |
| LAS1L | 132.855396 | -0.003412662 | 0.28602065 | -0.0119315 | 0.99048025 | 0.99606252 |
| MSN | 1361.95631 | -1.334049561 | 0.29139614 | -4.5781304 | 4.69E-06 | 0.00014279 |
| OPHN1 | 1309.74823 | 0.270685484 | 0.19557705 | 1.38403498 | 0.16634774 | 0.36590963 |
| YIPF6 | 450.543442 | -0.136932343 | 0.13864012 | -0.987682 | 0.32330844 | 0.54831506 |
| STARD8 | 236.295795 | 1.237185099 | 0.32257321 | 3.83536221 | 0.00012538 | 0.00200987 |
| PJA1 | 42.6477988 | -0.159196938 | 0.40830458 | -0.3898975 | 0.69661234 | 0.84496018 |
| EDA | 1094.1319 | 0.093631519 | 0.29100614 | 0.32175102 | 0.74764133 | 0.87699701 |
| IGBP1 | 416.756236 | -0.0516401 | 0.1905614 | -0.2709893 | 0.78639926 | 0.89881006 |
| PDZD11 | 48.3708223 | -1.079503017 | 0.3531038 | -3.0571832 | 0.00223428 | 0.01862729 |
| DLG3 | 831.610488 | 0.169185864 | 0.21607445 | 0.78299799 | 0.43362829 | 0.65072685 |
| SNX12 | 114.636085 | -0.359949812 | 0.24748024 | -1.4544588 | 0.14581915 | 0.3359716 |
| MED12 | 122.332763 | 0.019547251 | 0.21862101 | 0.08941158 | 0.92875482 | 0.96900671 |
| ZMYM3 | 139.613798 | 0.277098832 | 0.32863002 | 0.84319391 | 0.39912 | 0.6208833 |
| NONO | 461.352417 | -0.390091253 | 0.23139421 | -1.6858298 | 0.09182858 | 0.24804985 |
| TAF1 | 1066.97791 | -0.103908186 | 0.24715795 | -0.4204121 | 0.67418445 | 0.82909435 |
| OGT | 1680.11529 | 0.21151667 | 0.33134927 | 0.63834959 | 0.52324614 | 0.72294254 |
| GCNA | 230.494125 | 0.348692224 | 0.3526593 | 0.98875098 | 0.322785 | 0.54781794 |
| NHSL2 | 122.452665 | -0.574027421 | 0.39548869 | -1.4514383 | 0.14665786 | 0.33725691 |
| PIN4 | 367.68746 | -0.151069518 | 0.30071679 | -0.5023648 | 0.61541097 | 0.78868375 |
| RPS4X | 1160.40735 | -1.476569017 | 0.46943285 | -3.1454318 | 0.00165842 | 0.0148415 |
| HDAC8 | 2952.92779 | -0.181026527 | 0.20306645 | -0.8914645 | 0.37268004 | 0.59655016 |
| PHKA1 | 212.921149 | 0.467638992 | 0.27171249 | 1.72108021 | 0.08523627 | 0.235788 |
| NAP1L2 | 45.2536729 | -0.076290845 | 0.59218239 | -0.12883 | 0.89749218 | 0.9525128 |
| CHIC1 | 994.595162 | 0.360307906 | 0.21759963 | 1.65582956 | 0.09775634 | 0.25845947 |
| TSIX | 100.418882 | -1.031728815 | 2.50377297 | -0.4120696 | 0.68028838 | 0.8328016 |
| XIST | 3707.09931 | 0.842321746 | 3.8866352 | 0.21672262 | 0.82842452 | 0.92023714 |
| JPX | 2612.09954 | 0.27674873 | 0.33288998 | 0.83135193 | 0.40577484 | 0.62593483 |
| FTX | 17205.9332 | -0.139090148 | 0.21786402 | -0.6384264 | 0.52319613 | 0.72294254 |
| SLC16A2 | 316.322947 | -0.452670622 | 0.29398613 | -1.5397686 | 0.12361676 | 0.30156806 |
| RLIM | 372.346629 | -0.328868166 | 0.2016401 | -1.6309661 | 0.10289748 | 0.26771179 |
| NEXMIF | 1202.85644 | -0.914730781 | 0.26983525 | -3.3899603 | 0.00069903 | 0.00765332 |
| ABCB7 | 1004.53343 | 0.239086016 | 0.17975685 | 1.33005229 | 0.18350104 | 0.38837771 |
| UPRT | 217.489297 | 0.018191127 | 0.16021674 | 0.11354074 | 0.90960187 | 0.95919935 |
| ZDHHC15 | 262.999776 | -0.216857717 | 0.25149111 | -0.8622878 | 0.38852915 | 0.61165699 |
| PBDC1 | 90.7942957 | -0.900241742 | 0.36013884 | -2.4997074 | 0.01242959 | 0.06441054 |
| ATRX | 3764.42049 | 0.162298079 | 0.16977071 | 0.95598399 | 0.33908033 | 0.56459246 |
| MAGT1 | 580.676459 | -0.022241018 | 0.24556621 | -0.0905704 | 0.92783399 | 0.96852846 |
| COX7B | 783.226211 | 0.342522334 | 0.41087072 | 0.8336499 | 0.4044783 | 0.62539431 |
| PGK1 | 1603.04953 | -1.086180857 | 0.46399086 | -2.3409531 | 0.01923458 | 0.0868614 |
| TAF9B | 61.3013359 | -0.385709448 | 0.37072586 | -1.0404169 | 0.29814626 | 0.52143889 |
| TENT5D | 48.1068536 | 0.535086175 | 0.40215455 | 1.33054861 | 0.18333758 | 0.38809393 |
| BRWD3 | 1273.78031 | -0.088664548 | 0.15823203 | -0.5603451 | 0.57524404 | 0.76252365 |
| HMGN5 | 177.88125 | 0.224337484 | 0.1803811 | 1.24368622 | 0.21361507 | 0.42681008 |
| SH3BGRL | 876.522256 | -0.142471291 | 0.22440667 | -0.6348799 | 0.52550673 | 0.72472068 |
| RPS6KA6 | 1850.06498 | 0.980678215 | 0.28285033 | 3.46712767 | 0.00052605 | 0.00614378 |
| HDX | 444.053255 | -0.381331321 | 0.27174593 | -1.4032641 | 0.16053809 | 0.35771722 |
| APOOL | 646.607143 | 0.13553216 | 0.12228953 | 1.10828915 | 0.26773696 | 0.48887718 |
| SATL1 | 110.643307 | 1.658275747 | 0.34187998 | 4.85046173 | 1.23E-06 | 4.86E-05 |
| ZNF711 | 338.499855 | 1.47763435 | 0.23706889 | 6.23293231 | 4.58E-10 | 6.39E-08 |
| POF1B | 170.600976 | -1.037805669 | 0.53191412 | -1.9510775 | 0.05104783 | 0.16848657 |
| CHM | 1109.96749 | 0.053196172 | 0.18842923 | 0.28231381 | 0.7777029 | 0.89359206 |
| PCDH11X | 82.8706711 | -0.293882371 | 0.40602636 | -0.7238012 | 0.46918779 | 0.68108245 |
| DIAPH2 | 1173.74656 | 0.330398727 | 0.42696441 | 0.77383201 | 0.43903014 | 0.65485132 |
| DIAPH2-AS1 | 117.120463 | 0.677977025 | 0.66837099 | 1.01437231 | 0.31040513 | 0.53435962 |
| PCDH19 | 43.8586654 | 0.589526781 | 0.45650375 | 1.29139528 | 0.19656665 | 0.40474651 |
| TSPAN6 | 180.01124 | 0.597898424 | 0.2634141 | 2.26980421 | 0.02321946 | 0.09858756 |
| SRPX2 | 164.894615 | 0.436719298 | 0.46329087 | 0.94264603 | 0.34586199 | 0.57105512 |

| | | | | | | |
|---|---|---|---|---|---|---|
| SYTL4 | 980.408138 | 1.236096846 | 0.26403481 | 4.68156767 | 2.85E-06 | 9.49E-05 |
| CSTF2 | 46.8117967 | 0.040246333 | 0.4028488 | 0.09990431 | 0.92042029 | 0.96454603 |
| TRMT2B | 440.866422 | -0.037790137 | 0.2474043 | -0.1527465 | 0.87859821 | 0.94585961 |
| GLA | 59.8887649 | -0.348903817 | 0.4039209 | -0.8637924 | 0.38770193 | 0.61089763 |
| HNRNPH2 | 173.295676 | -1.208918715 | 0.39383669 | -3.0695939 | 0.0021435 | 0.01809169 |
| ARMCX4 | 1636.27533 | 0.528468669 | 0.37114716 | 1.42387905 | 0.15448149 | 0.34881127 |
| ARMCX1 | 69.5181143 | -0.148227199 | 0.29318893 | -0.5055689 | 0.61315933 | 0.7873259 |
| ARMCX6 | 95.4197732 | -0.31780424 | 0.25744178 | -1.2344703 | 0.2170277 | 0.43120432 |
| ARMCX3 | 341.834206 | -0.345770637 | 0.18822946 | -1.8369635 | 0.06621529 | 0.20049878 |
| ARMCX2 | 165.662317 | -0.445285302 | 0.28270876 | -1.5750672 | 0.11524093 | 0.28859915 |
| ZMAT1 | 1418.09302 | 0.520525763 | 0.33167395 | 1.56938997 | 0.1165571 | 0.2906556 |
| TCEAL2 | 53.6257854 | -0.948632524 | 0.75282998 | -1.2600887 | 0.20763738 | 0.41847454 |
| BEX5 | 46.709883 | -0.8872591 | 0.53479404 | -1.6590669 | 0.09710231 | 0.25751598 |
| ARMCX5 | 85.9270178 | -0.237467164 | 0.2743836 | -0.8654568 | 0.3867881 | 0.6104043 |
| AL035427.1 | 112.049558 | -0.494357465 | 0.71465085 | -0.6917468 | 0.48909633 | 0.6960367 |
| GPRASP1 | 82.6413437 | 0.109887908 | 0.41593303 | 0.26419615 | 0.79162877 | 0.90190943 |
| BHLHB9 | 160.809138 | 0.291543734 | 0.18185044 | 1.60320607 | 0.10888916 | 0.27734186 |
| LINC00630 | 612.302787 | 0.322061512 | 0.2071072 | 1.55504738 | 0.11993479 | 0.29561832 |
| Z68871.1 | 72.6675728 | 0.529475664 | 0.2851007 | 1.85715317 | 0.06328936 | 0.19505368 |
| BEX4 | 307.346366 | -0.502961907 | 0.24447162 | -2.0573427 | 0.03965327 | 0.14177464 |
| TCEAL8 | 122.695843 | -0.820188917 | 0.28745673 | -2.8532604 | 0.00432731 | 0.03017925 |
| BEX2 | 254.613763 | -0.652625043 | 0.39930863 | -1.6343875 | 0.10217751 | 0.26662637 |
| TCEAL9 | 331.839346 | -0.676980259 | 0.40002517 | -1.6923442 | 0.09058037 | 0.24578392 |
| BEX3 | 638.977535 | 0.083302889 | 0.31598689 | 0.26362768 | 0.79206683 | 0.90200972 |
| RAB40A | 51.0901855 | 0.534519484 | 0.37755001 | 1.41575809 | 0.15684634 | 0.35240404 |
| TCEAL4 | 385.961139 | -0.909608343 | 0.21638446 | -4.2036676 | 2.63E-05 | 0.00059299 |
| TCEAL3 | 204.700419 | -0.768393544 | 0.31913439 | -2.4077428 | 0.01605148 | 0.07668139 |
| TCEAL1 | 58.7564796 | -0.181009057 | 0.3295969 | -0.5491831 | 0.58287978 | 0.76838912 |
| MORF4L2 | 936.374398 | -0.030984221 | 0.22025139 | -0.1406766 | 0.88812541 | 0.94943227 |
| MORF4L2-AS | 44.5979394 | 0.051994577 | 0.37601517 | 0.13827787 | 0.89002082 | 0.9503707 |
| SLC25A53 | 508.192834 | 0.344116563 | 0.19218756 | 1.79052469 | 0.0733696 | 0.21483908 |
| FAM199X | 675.915064 | 0.065656144 | 0.13878545 | 0.47307656 | 0.63615856 | 0.80446022 |
| IL1RAPL2 | 110.719988 | 0.694757091 | 0.47099524 | 1.47508305 | 0.14019022 | 0.32765152 |
| NRK | 2243.64241 | 1.184851347 | 0.29809423 | 3.97475436 | 7.05E-05 | 0.00127479 |
| CXorf57 | 212.726627 | 0.074375877 | 0.49638871 | 0.14983394 | 0.88089563 | 0.94640313 |
| RNF128 | 253.203103 | -0.161473663 | 0.72106386 | -0.2239381 | 0.82280548 | 0.91813562 |
| TBC1D8B | 448.027809 | 0.065952035 | 0.33550436 | 0.19657579 | 0.84415952 | 0.92788526 |
| MORC4 | 395.352668 | -0.227608354 | 0.32995579 | -0.6898147 | 0.49031072 | 0.69684897 |
| RBM41 | 861.825183 | 0.027988702 | 0.16388602 | 0.17078151 | 0.86439557 | 0.93833133 |
| NUP62CL | 194.037169 | -0.286765227 | 0.43838246 | -0.6541439 | 0.51301908 | 0.71496824 |
| PRPS1 | 71.5621844 | -0.241247001 | 0.3410986 | -0.7072647 | 0.479402 | 0.68864274 |
| TSC22D3 | 346.391236 | 0.051485468 | 0.5076671 | 0.1014158 | 0.91922039 | 0.96418504 |
| MID2 | 180.796551 | -0.084479063 | 0.28344449 | -0.2980445 | 0.76566922 | 0.88656137 |
| PSMD10 | 57.278707 | -0.409477696 | 0.38144579 | -1.0734886 | 0.28305197 | 0.50545507 |
| ATG4A | 113.203554 | 0.200968196 | 0.21922833 | 0.91670721 | 0.35929608 | 0.58469163 |
| COL4A6 | 159.533489 | 0.984256329 | 0.67521349 | 1.45769648 | 0.14492422 | 0.3347856 |
| COL4A5 | 1913.22092 | 0.68765766 | 0.45580353 | 1.50867122 | 0.13138282 | 0.31404917 |
| ACSL4 | 9408.76758 | -1.443458034 | 0.65422923 | -2.206349 | 0.02735957 | 0.11021426 |
| TMEM164 | 1650.18257 | 0.32166612 | 0.27387602 | 1.17449539 | 0.24019666 | 0.45822612 |
| AMMECR1 | 1124.13766 | 0.581007091 | 0.20245805 | 2.86976535 | 0.00410777 | 0.02920608 |
| CAPN6 | 43.6947583 | -0.01042201 | 0.72673651 | -0.0143408 | 0.98855806 | 0.99571934 |
| ALG13 | 759.742383 | -0.004018889 | 0.24940248 | -0.0161141 | 0.98714339 | 0.99517967 |
| ALG13-AS1 | 68.0309573 | -0.706260144 | 0.55372874 | -1.2754623 | 0.20214565 | 0.41161429 |
| AMOT | 81.4600865 | -1.769149087 | 0.51541409 | -3.432481 | 0.00059809 | 0.00678704 |
| LRCH2 | 128.013857 | -0.597291209 | 0.53746309 | -1.1113158 | 0.26643245 | 0.48750606 |
| PLS3-AS1 | 196.906872 | 0.933508591 | 0.57756636 | 1.61627937 | 0.10603392 | 0.2732349 |
| PLS3 | 1228.58646 | 0.193464099 | 0.22256255 | 0.86925719 | 0.38470647 | 0.60777273 |
| DANT2 | 5262.89091 | 0.084937339 | 0.23529911 | 0.36097603 | 0.71811736 | 0.85963642 |
| KLHL13 | 6035.14239 | 0.00091872 | 0.27459639 | 0.00334571 | 0.99733051 | 0.99907013 |
| WDR44 | 706.937299 | -0.230382709 | 0.24431508 | -0.9429738 | 0.34569432 | 0.57104354 |
| DOCK11 | 469.956547 | 2.461495355 | 1.40001784 | 1.75818856 | 0.07871543 | 0.22441496 |
| IL13RA1 | 605.816597 | -1.092998518 | 0.40019429 | -2.7311697 | 0.006311 | 0.03967728 |
| LONRF3 | 247.902676 | -0.525502194 | 0.53999612 | -0.9731592 | 0.33047419 | 0.55579749 |
| PGRMC1 | 430.476827 | -1.084560796 | 0.39170275 | -2.7688363 | 0.00562569 | 0.03658775 |
| SLC25A43 | 471.100637 | -0.10224021 | 0.39911959 | -0.2561643 | 0.79782394 | 0.90506918 |
| SLC25A5 | 494.596833 | 0.036238339 | 0.52119115 | 0.06952984 | 0.94456788 | 0.9759376 |
| CXorf56 | 120.246723 | 0.086975215 | 0.28105318 | 0.30946177 | 0.75697029 | 0.88250844 |
| UBE2A | 301.039219 | -0.257485844 | 0.27049686 | -0.9518996 | 0.34114792 | 0.56653544 |
| NKRF | 170.698214 | -0.031339879 | 0.21737872 | -0.1441718 | 0.88536482 | 0.94825135 |
| 6-Sep | 1041.87631 | 0.044038019 | 0.32615953 | 0.13501988 | 0.89259616 | 0.95119823 |
| RPL39 | 47.5968822 | 0.305311316 | 0.46189468 | 0.66099769 | 0.5086138 | 0.71123011 |
| UPF3B | 675.477928 | 0.1395555 | 0.37812684 | 0.3690706 | 0.7120751 | 0.85561721 |
| NDUFA1 | 452.603341 | -0.396739163 | 0.36110836 | -1.0986707 | 0.27191174 | 0.49343071 |
| NKAP | 174.98007 | -0.167275151 | 0.19912291 | -0.8400598 | 0.40087486 | 0.62237376 |
| RHOXF1-AS1 | 250.607441 | 0.798828223 | 0.42815066 | 1.86576432 | 0.06207435 | 0.19249428 |
| ZBTB33 | 58.0251099 | -0.25656734 | 0.34120529 | -0.7519442 | 0.45208461 | 0.66788789 |
| LAMP2 | 517.837338 | -0.334985115 | 0.15135764 | -2.2132025 | 0.02688368 | 0.10888294 |
| CUL4B | 307.380375 | 0.300052105 | 0.18806169 | 1.59549827 | 0.11060086 | 0.28051588 |
| MCTS1 | 368.077264 | 0.58610341 | 0.19377129 | 3.02471744 | 0.00248866 | 0.02016727 |
| C1GALT1C1 | 88.3748447 | -0.508153256 | 0.2827142 | -1.7974098 | 0.07227059 | 0.21251609 |
| GRIA3 | 68.8021832 | -0.675459016 | 0.58429017 | -1.1560335 | 0.24766745 | 0.46703404 |
| THOC2 | 1686.08901 | -0.035697186 | 0.17296873 | -0.2063794 | 0.83649454 | 0.92398282 |
| XIAP | 1230.83464 | -0.319702767 | 0.15435551 | -2.0712106 | 0.03833912 | 0.13904125 |
| STAG2 | 2062.36342 | 0.021342565 | 0.19678183 | 0.10845801 | 0.91363239 | 0.96091809 |
| TENM1 | 637.351103 | -0.470925832 | 0.55483891 | -0.8487614 | 0.39601409 | 0.61804994 |
| AL442647.1 | 107.278387 | -0.415391962 | 0.52687414 | -0.7884083 | 0.4304579 | 0.64855918 |
| SMARCA1 | 879.987878 | 0.072163561 | 0.19700903 | 0.36629571 | 0.71414443 | 0.85724589 |
| OCRL | 261.716084 | 0.202652316 | 0.21673183 | 0.93503717 | 0.34976914 | 0.57578522 |
| ZDHHC9 | 174.88478 | -1.783863245 | 0.30687171 | -5.8130586 | 6.13E-09 | 6.28E-07 |
| AL034405.1 | 52.9937009 | 0.471661123 | 0.38639614 | 1.22066729 | 0.22221202 | 0.43639586 |
| UTP14A | 124.997153 | 0.004816673 | 0.30897702 | 0.0155891 | 0.9875622 | 0.99533283 |
| BCORL1 | 151.697999 | -0.241162958 | 0.33690659 | -0.7158155 | 0.47410529 | 0.68459432 |
| ELF4 | 121.354826 | -0.793280876 | 0.27823239 | -2.851145 | 0.00435621 | 0.03028483 |
| AIFM1 | 287.500262 | 0.30712141 | 0.23569269 | 1.30305869 | 0.19255472 | 0.39989594 |
| ZNF280C | 185.709938 | 0.302419823 | 0.17697798 | 1.70879913 | 0.08748816 | 0.23985703 |
| SLC25A14 | 157.967821 | 0.46344887 | 0.17888773 | 2.59072477 | 0.00957741 | 0.05369607 |
| RBMX2 | 97.5983937 | 0.278747349 | 0.22464416 | 1.24083949 | 0.21466505 | 0.42832487 |
| ENOX2 | 718.362648 | -0.330336371 | 0.28059799 | -1.1772585 | 0.23909235 | 0.45716719 |
| STK26 | 111.395757 | 0.275501144 | 0.45979762 | 0.59917915 | 0.54905343 | 0.74226986 |
| RAP2C | 748.198449 | 2.145612808 | 0.39993878 | 5.36485305 | 8.10E-08 | 5.43E-06 |
| RAP2C-AS1 | 1396.25168 | 0.963452056 | 0.29429013 | 3.27381705 | 0.00106105 | 0.01045801 |
| MBNL3 | 710.930531 | 0.8127037 | 0.28293683 | 2.87238569 | 0.00407385 | 0.02904023 |
| HS6ST2 | 4185.52505 | 0.743598917 | 0.20664534 | 3.59843057 | 0.00032014 | 0.00422033 |
| GPC4 | 950.169463 | -1.194027919 | 0.32288041 | -3.69805 | 0.00021726 | 0.00313641 |
| GPC3 | 1522.50275 | 1.09054406 | 0.47320065 | 2.30461233 | 0.02118829 | 0.09294933 |
| CCDC160 | 143.246338 | 0.69403448 | 0.36197373 | 1.91736147 | 0.05519203 | 0.17751599 |
| PHF6 | 434.551107 | 0.2356373 | 0.19969893 | 1.17996275 | 0.23801503 | 0.45603216 |
| HPRT1 | 1005.39196 | -0.249564831 | 0.28948052 | -0.8621127 | 0.3886255 | 0.61169616 |
| MIR503HG | 369.251479 | 0.969284928 | 0.53998945 | 1.79500717 | 0.07265256 | 0.21330688 |
| FAM122B | 199.450482 | -0.182090426 | 0.22355709 | -0.8145142 | 0.41535044 | 0.63525223 |
| FAM122C | 164.317287 | 0.041841934 | 0.26395804 | 0.15851737 | 0.87404913 | 0.94334544 |
| MOSPD1 | 103.870027 | 0.122537539 | 0.22947619 | 0.53398803 | 0.59334981 | 0.77387448 |
| RTL8C | 257.992366 | -0.484206006 | 0.42274171 | -1.1453945 | 0.25204579 | 0.47214352 |

| | | | | | | |
|---|---|---|---|---|---|---|
| RTL8A | 149.67991 | -1.405179259 | 0.33014986 | -4.2561862 | 2.08E-05 | 0.00048701 |
| ZNF75D | 711.044971 | -0.206999192 | 0.30179821 | -0.6858861 | 0.49278495 | 0.69868807 |
| ZNF449 | 124.615104 | -0.209131446 | 0.22862415 | -0.9147391 | 0.36032861 | 0.58548052 |
| INTS6L | 245.67964 | -0.2239959 | 0.23880322 | -0.9379937 | 0.34824767 | 0.57385273 |
| MMGT1 | 102.314288 | -0.665226811 | 0.29247032 | -2.2745105 | 0.02293531 | 0.09791161 |
| SLC9A6 | 205.921139 | 0.161848762 | 0.22233484 | 0.72795053 | 0.46664387 | 0.67914267 |
| MAP7D3 | 51.9968144 | -1.008673234 | 0.3943384 | -2.5578874 | 0.01053102 | 0.0576005 |
| HTATSF1 | 186.418709 | -0.441025472 | 0.25793886 | -1.7098063 | 0.08730171 | 0.23953446 |
| ARHGEF6 | 48.4339527 | -0.03338929 | 0.32992686 | -0.1012021 | 0.91939003 | 0.9642864 |
| AL683813.1 | 268.429664 | 0.440082088 | 0.42234852 | 1.04198798 | 0.29741725 | 0.52079582 |
| RBMX | 1376.6609 | 0.519160431 | 0.11798963 | 4.40005134 | 1.08E-05 | 0.00028477 |
| FGF13 | 3505.65567 | -0.410358085 | 0.70757945 | -0.5799463 | 0.56195083 | 0.75366113 |
| ATP11C | 538.681337 | -0.300102225 | 0.19438044 | -1.5438911 | 0.12261473 | 0.30009596 |
| LDOC1 | 58.8166202 | -1.52342821 | 0.50685626 | -3.0056415 | 0.00265021 | 0.02112653 |
| SLITRK4 | 179.551057 | 0.669288358 | 0.45286059 | 1.47791258 | 0.13943118 | 0.32631935 |
| AL589669.1 | 61.3071363 | 0.481084173 | 0.64278161 | 0.7484411 | 0.45419414 | 0.66965626 |
| FMR1 | 934.347707 | -0.042382014 | 0.22413937 | -0.1890878 | 0.85002403 | 0.93124056 |
| IDS | 319.886374 | -0.557410766 | 0.23319113 | -2.3903601 | 0.01683186 | 0.07943267 |
| LINC00893 | 66.6922756 | -0.149343934 | 0.34577921 | -0.4319055 | 0.66581011 | 0.82362669 |
| TMEM185A | 244.834538 | -0.293854104 | 0.22206858 | -1.3232583 | 0.18574948 | 0.39144303 |
| LINC00894 | 1295.46923 | 0.035864915 | 0.36320474 | 0.09874572 | 0.92134016 | 0.96504145 |
| MAMLD1 | 79.981322 | -1.193686933 | 0.52001818 | -2.2954715 | 0.02170612 | 0.09440958 |
| MTM1 | 630.289889 | -0.001319475 | 0.16210218 | -0.0081398 | 0.99350547 | 0.99743207 |
| MTMR1 | 579.954938 | 0.126694223 | 0.1344361 | 0.94241222 | 0.34598163 | 0.57105512 |
| CD99L2 | 407.901613 | 0.007244708 | 0.19223014 | 0.03768768 | 0.9699367 | 0.98653537 |
| VMA21 | 152.577788 | -0.597178472 | 0.26887302 | -2.2210428 | 0.02634806 | 0.1074544 |
| CETN2 | 88.6391476 | -1.501095231 | 0.50572632 | -2.9681968 | 0.00299553 | 0.02313912 |
| NSDHL | 72.1261765 | -0.279540168 | 0.32348698 | -0.8641466 | 0.38750738 | 0.61086904 |
| ZNF185 | 103.669902 | -0.323337157 | 0.48619245 | -0.6650395 | 0.50602526 | 0.70953129 |
| ZNF275 | 79.3398308 | 0.752579081 | 0.35343101 | 2.12935212 | 0.03322514 | 0.12653816 |
| BGN | 46.12554 | 0.56149655 | 0.82942241 | 0.67697296 | 0.49842311 | 0.70339396 |
| CCNQ | 70.5446769 | -0.452953373 | 0.36844214 | -1.2293745 | 0.21893143 | 0.43323824 |
| DUSP9 | 529.544623 | 1.144293626 | 0.66238426 | 1.72753747 | 0.08407118 | 0.23354285 |
| SLC6A8 | 705.540557 | -0.831038907 | 0.40495236 | -2.0521893 | 0.04015127 | 0.14295366 |
| BCAP31 | 371.810612 | -1.257194164 | 0.35770556 | -3.5146062 | 0.00044041 | 0.00539298 |
| ABCD1 | 42.8193506 | -0.297924721 | 0.40041918 | -0.7440321 | 0.45685706 | 0.67185759 |
| IDH3G | 150.691182 | -0.095158271 | 0.31710653 | -0.300083 | 0.76411386 | 0.88555042 |
| SSR4 | 219.914842 | -0.914752325 | 0.47427284 | -1.928747 | 0.05376228 | 0.17448302 |
| ARHGAP4 | 106.103488 | 0.120459497 | 0.45249285 | 0.26621304 | 0.79007514 | 0.90082888 |
| HCFC1 | 149.419552 | -0.149337177 | 0.24799962 | -0.602167 | 0.54706301 | 0.74124838 |
| TMEM187 | 54.3548931 | 0.151227355 | 0.36617789 | 0.41298876 | 0.67961485 | 0.83230806 |
| IRAK1 | 139.907935 | -1.603194954 | 0.44765724 | -3.5813001 | 0.00034189 | 0.0044318 |
| MECP2 | 1108.39932 | -0.054839965 | 0.17330151 | -0.3164425 | 0.75166666 | 0.87947193 |
| FLNA | 450.593728 | -0.986635658 | 0.64335515 | -1.5335786 | 0.12513337 | 0.30372779 |
| EMD | 222.635277 | -0.546970421 | 0.36881219 | -1.4830595 | 0.1380586 | 0.3240256 |
| RPL10 | 1660.12011 | -1.024778471 | 0.49013275 | -2.0908182 | 0.03654436 | 0.13464146 |
| DNASE1L1 | 114.447264 | -0.299060107 | 0.32342005 | -0.9246802 | 0.35513229 | 0.58081705 |
| TAZ | 296.79571 | 0.249875674 | 0.2625747 | 0.95163654 | 0.34128134 | 0.56661454 |
| AC244090.1 | 42.1715153 | 0.686641078 | 0.33346296 | 2.0591225 | 0.03948251 | 0.1414879 |
| ATP6AP1 | 259.5389 | -0.463726095 | 0.40435127 | -1.1468397 | 0.25144787 | 0.47185323 |
| GDI1 | 231.927837 | -0.228327051 | 0.2941262 | -0.7762894 | 0.43757813 | 0.65416304 |
| FAM50A | 137.512276 | -0.409949029 | 0.31349381 | -1.3076782 | 0.19098247 | 0.39752425 |
| PLXNA3 | 111.690914 | -1.61948714 | 0.39023138 | -4.1500689 | 3.32E-05 | 0.00071272 |
| UBL4A | 61.7222794 | -0.774908589 | 0.33819911 | -2.2912792 | 0.02194727 | 0.09480102 |
| SLC10A3 | 45.7848288 | -0.837747789 | 0.44645491 | -1.8764443 | 0.06059429 | 0.18917277 |
| FAM3A | 169.472896 | -0.239190238 | 0.19933198 | -1.1999592 | 0.23015519 | 0.445816 |
| G6PD | 96.3054115 | -1.661597827 | 0.56534145 | -2.9391049 | 0.00329162 | 0.02478846 |
| IKBKG | 134.339964 | -0.37268517 | 0.25960497 | -1.4355857 | 0.15112027 | 0.34386696 |
| DKC1 | 140.264111 | -0.265163393 | 0.23915273 | -1.1087617 | 0.26753298 | 0.48870739 |
| MPP1 | 83.8728813 | 0.898606629 | 0.34174234 | 2.62948581 | 0.00855141 | 0.04950124 |
| F8 | 219.806426 | 0.39919213 | 0.24745824 | 1.61316971 | 0.10670764 | 0.27422203 |
| FUNDC2 | 290.680662 | -0.094020271 | 0.2760465 | -0.3405958 | 0.73340791 | 0.86911141 |
| MTCP1 | 119.871205 | -0.264497825 | 0.64704443 | -0.4087785 | 0.68270225 | 0.83498278 |
| BRCC3 | 431.509922 | -0.213349774 | 0.18909922 | -1.1282425 | 0.25921753 | 0.48023578 |
| VBP1 | 140.314334 | 0.113376309 | 0.22400931 | 0.50612321 | 0.61277016 | 0.78690269 |
| CLIC2 | 53.3973388 | 0.490608555 | 0.47729723 | 1.02788896 | 0.30400206 | 0.52732281 |
| TMLHE-AS1 | 368.129351 | -0.353869454 | 0.53506939 | -0.6613525 | 0.50838631 | 0.71121317 |
| TMLHE | 797.026241 | -0.263285862 | 0.20422806 | -1.2891758 | 0.19733699 | 0.4057467 |
| VAMP7 | 348.009274 | -0.165441044 | 0.20943357 | -0.7899452 | 0.42955978 | 0.64772318 |
| ZNF596 | 211.149854 | 0.123207893 | 0.21757845 | 0.56626882 | 0.57121105 | 0.75965257 |
| FBXO25 | 667.027234 | 0.118204641 | 0.22745353 | 0.519687 | 0.60328175 | 0.78033183 |
| TDRP | 1030.51597 | 0.511266818 | 0.2884902 | 1.77221557 | 0.07635878 | 0.22033505 |
| ERICH1 | 2433.58611 | 0.808021669 | 0.47038775 | 1.71777789 | 0.08583714 | 0.23695355 |
| DLGAP2 | 108.910792 | 2.002518183 | 0.77913103 | 2.57019437 | 0.01016415 | 0.0561985 |
| AC100810.1 | 58.5136362 | -0.040215312 | 0.49311821 | -0.0815531 | 0.93500211 | 0.9719401 |
| AC019257.8 | 49.2015391 | 0.794508157 | 0.5049542 | 1.57342617 | 0.11562017 | 0.28924751 |
| KBTBD11 | 336.412518 | -0.741789462 | 0.35767072 | -2.0739452 | 0.0380844 | 0.13846873 |
| MYOM2 | 49.2950852 | 0.339384423 | 0.48367121 | 0.70168416 | 0.48287616 | 0.69127369 |
| AC246817.2 | 276.818407 | 0.660442937 | 0.44598507 | 1.48086335 | 0.13864299 | 0.32505064 |
| AC246817.1 | 611.227362 | 0.414374182 | 0.31096794 | 1.33253022 | 0.18268602 | 0.38733541 |
| CSMD1 | 467.39957 | 1.216294238 | 0.84845794 | 1.43353511 | 0.15170497 | 0.34478166 |
| MCPH1 | 1718.7373 | 0.140928433 | 0.19366624 | 0.72768714 | 0.46680512 | 0.67930251 |
| ANGPT2 | 301.722583 | -0.153405648 | 0.48931372 | -0.3135118 | 0.75389182 | 0.88075281 |
| MCPH1-AS1 | 416.875239 | 0.273563424 | 0.20848605 | 1.31214261 | 0.18947202 | 0.39556637 |
| AGPAT5 | 711.427617 | -0.182357765 | 0.44988872 | -0.4053397 | 0.68522782 | 0.83714154 |
| AF233439.1 | 1280.82535 | -0.375178029 | 0.81919576 | -0.4579834 | 0.64696439 | 0.81129186 |
| DEFB1 | 2895.09626 | -0.227554537 | 0.52945863 | -0.4297872 | 0.66735045 | 0.82469498 |
| FAM66B | 59.5971812 | 0.215254223 | 0.45679468 | 0.47122752 | 0.63747827 | 0.80497022 |
| FAM85B | 1022.96736 | 0.34242099 | 0.385377 | 0.88853509 | 0.37425299 | 0.59780424 |
| PRAG1 | 629.97825 | -2.823258502 | 0.81152378 | -3.4789597 | 0.00050336 | 0.00594717 |
| MFHAS1 | 5403.10283 | -0.658803846 | 0.28735471 | -2.2926503 | 0.02186815 | 0.09467149 |
| ERI1 | 480.565207 | -0.97332274 | 0.32562609 | -2.989081 | 0.00279818 | 0.02204004 |
| PPP1R3B | 210.143768 | -0.745562453 | 0.36970733 | -2.0166288 | 0.04373427 | 0.15150431 |
| AC022784.3 | 105.632134 | 1.913510641 | 0.5467433 | 3.49983372 | 0.00046555 | 0.00561592 |
| AC022784.1 | 38.0053268 | 2.754410482 | 0.56206236 | 4.90054249 | 9.56E-07 | 3.97E-05 |
| TNKS | 3113.57406 | -0.132379776 | 0.19205078 | -0.6892957 | 0.49063721 | 0.6969611 |
| MSRA | 1514.39342 | -0.445162964 | 0.25083769 | -1.7747052 | 0.07594657 | 0.21965804 |
| PINX1 | 46.1447649 | 0.458074604 | 0.4279438 | 1.07040832 | 0.28443555 | 0.50683977 |
| XKR6 | 2863.285 | -1.063609796 | 0.30447339 | -3.4932767 | 0.00047713 | 0.0057141 |
| MTMR9 | 469.572583 | -0.120587803 | 0.17551926 | -0.6870346 | 0.49206094 | 0.69803196 |
| FAM167A | 119.647276 | -0.711498727 | 0.66701426 | -1.0666919 | 0.28611097 | 0.50853442 |
| NEIL2 | 76.8912512 | -0.429857331 | 0.34477843 | -1.246764 | 0.21248404 | 0.42519341 |
| FDFT1 | 866.218002 | -0.566029854 | 0.33992291 | -1.6651712 | 0.0958786 | 0.25518041 |
| CTSB | 564.974882 | -1.342934509 | 0.31806692 | -4.222176 | 2.42E-05 | 0.0005539 |
| FAM66D | 437.416194 | 0.982428077 | 0.61934756 | 1.58623064 | 0.112687 | 0.28422429 |
| AC145124.1 | 120.112717 | 0.002466422 | 0.54993051 | 0.00448497 | 0.99642152 | 0.99867719 |
| FAM66A | 51.913566 | 1.767732924 | 0.8872651 | 1.99233907 | 0.04633387 | 0.15749461 |
| LONRF1 | 598.273237 | -0.51405507 | 0.36588199 | -1.4049751 | 0.16002869 | 0.35694343 |
| TRMT9B | 2849.59551 | -0.657484313 | 0.2579617 | -2.5487672 | 0.01081044 | 0.05859465 |
| DLC1 | 508.241775 | 0.823707472 | 0.68337642 | 1.20534956 | 0.22806838 | 0.44354465 |
| SGCZ | 41.3803128 | 0.521223395 | 0.41167025 | 1.26611869 | 0.20547059 | 0.41584887 |
| TUSC3 | 1482.18994 | -0.252081318 | 0.2829327 | -0.8909586 | 0.37295138 | 0.59676739 |
| MSR1 | 1264.90501 | -0.221825649 | 0.74953495 | -0.295951 | 0.76726747 | 0.88754147 |

| | | | | | | |
|---|---|---|---|---|---|---|
| AC011586.1 | 91.1757377 | -0.554164467 | 0.61535787 | -0.9005564 | 0.36782422 | 0.59294486 |
| AC068992.1 | 617.272899 | -0.19469423 | 0.75867641 | -0.2566235 | 0.7974694 | 0.90505915 |
| MICU3 | 1121.08837 | 0.528101402 | 0.21536824 | 2.45208584 | 0.01420308 | 0.07045004 |
| AC079193.2 | 59.4292641 | 0.395015451 | 0.36975528 | 1.06831593 | 0.28537803 | 0.50782141 |
| ZDHHC2 | 1290.61446 | -0.005097052 | 0.23673491 | -0.0215306 | 0.98282237 | 0.9928728 |
| CNOT7 | 385.38491 | 0.203315715 | 0.18169473 | 1.11899624 | 0.26314174 | 0.48419683 |
| VPS37A | 586.874595 | 0.065995448 | 0.22612149 | 0.29185837 | 0.77039492 | 0.88936781 |
| MTMR7 | 256.343947 | 0.652289092 | 0.52703095 | 1.23766753 | 0.21583936 | 0.42962962 |
| SLC7A2 | 379.228208 | -0.958513844 | 0.59386658 | -1.6140222 | 0.10652261 | 0.27385309 |
| MTUS1 | 2032.87809 | -0.054584599 | 0.26656589 | -0.2047696 | 0.83775212 | 0.92443926 |
| AC027117.2 | 349.379307 | -0.215709857 | 0.42716706 | -0.5049777 | 0.61357447 | 0.78755051 |
| PCM1 | 3337.84653 | 0.358829228 | 0.10912699 | 3.28818053 | 0.00100837 | 0.01006504 |
| ASAH1 | 479.171716 | -0.524614929 | 0.29453329 | -1.7811736 | 0.07488409 | 0.21730765 |
| NAT1 | 39.8876893 | 0.712046073 | 0.34399416 | 2.06993651 | 0.03845829 | 0.13917687 |
| PSD3 | 594.18778 | -1.496296143 | 0.36734626 | -4.0732581 | 4.64E-05 | 0.00092503 |
| SH2D4A | 740.642601 | -0.512401427 | 0.38612565 | -1.3270329 | 0.18449781 | 0.38980031 |
| CSGALNACT1 | 4171.09214 | -2.257100727 | 0.70275987 | -3.2117667 | 0.00131921 | 0.01243791 |
| INTS10 | 1533.84438 | -0.118639583 | 0.22561934 | -0.5258396 | 0.59899965 | 0.77729304 |
| LPL | 242.185239 | 3.090696059 | 0.6873144 | 4.49677188 | 6.90E-06 | 0.00019598 |
| ATP6V1B2 | 381.716454 | 0.074361775 | 0.20048896 | 0.37090209 | 0.71071046 | 0.85498857 |
| GFRA2 | 46.5248023 | 0.81149829 | 0.81260317 | 0.99864032 | 0.31796896 | 0.54243213 |
| XPO7 | 931.636816 | 0.190469005 | 0.1447725 | 1.31564356 | 0.18829371 | 0.3938514 |
| NPM2 | 39.9272504 | -0.082857096 | 0.52097516 | -0.1590423 | 0.87363553 | 0.94326073 |
| DMTN | 445.392106 | 0.686753338 | 0.32236399 | 2.13036614 | 0.0331414 | 0.12633902 |
| FAM160B2 | 268.205518 | -0.588230615 | 0.21089868 | -2.7891622 | 0.00528446 | 0.03488293 |
| BMP1 | 81.8630821 | -1.1963807 | 0.40092898 | -2.9840215 | 0.00284487 | 0.02227907 |
| POLR3D | 68.0462579 | -0.340451267 | 0.29383162 | -1.1586611 | 0.24659435 | 0.46565614 |
| SLC39A14 | 627.33079 | -1.690125923 | 0.48681742 | -3.471786 | 0.00051701 | 0.00607037 |
| PPP3CC | 1229.32836 | 0.283471135 | 0.20367446 | 1.39178541 | 0.16398739 | 0.36271089 |
| SORBS3 | 98.6172515 | -0.448935975 | 0.37234158 | -1.20571 | 0.22792932 | 0.44334513 |
| CCAR2 | 190.598748 | 0.17956012 | 0.19604495 | 0.91591299 | 0.35971253 | 0.58502113 |
| AC037459.3 | 45.6116727 | -0.539674691 | 0.3880319 | -1.3907998 | 0.16428614 | 0.36306425 |
| BIN3 | 614.761816 | -0.744252309 | 0.2366742 | -3.1446279 | 0.00166298 | 0.01487226 |
| PEBP4 | 71.8150046 | 0.898795246 | 0.4808938 | 1.86900983 | 0.06162145 | 0.19142937 |
| RHOBTB2 | 80.8651706 | -0.440394631 | 0.46548636 | -0.9460957 | 0.34409979 | 0.56886284 |
| TNFRSF10B | 724.71483 | -0.899716116 | 0.26169255 | -3.4380654 | 0.00058589 | 0.00667153 |
| TNFRSF10D | 457.919462 | -2.507452037 | 0.34864855 | -7.1919187 | 6.39E-13 | 2.28E-10 |
| TNFRSF10A | 74.9150973 | -2.87994393 | 0.40450351 | -7.1197008 | 1.08E-12 | 3.66E-10 |
| CHMP7 | 131.709268 | 0.019862045 | 0.21032034 | 0.09443711 | 0.92476194 | 0.9667507 |
| R3HCC1 | 80.1831939 | -0.111592597 | 0.30086512 | -0.3709057 | 0.71070775 | 0.85498857 |
| LOXL2 | 71.1510224 | -0.510474779 | 0.42506823 | -1.2009243 | 0.22978055 | 0.4453663 |
| ENTPD4 | 420.920812 | -0.399878324 | 0.20965738 | -1.9072942 | 0.0564825 | 0.18057989 |
| SLC25A37 | 678.453488 | -0.744291223 | 0.35806898 | -2.0786253 | 0.03765181 | 0.13733278 |
| AC120193.1 | 68.5750584 | 1.322781308 | 0.60924483 | 2.17118183 | 0.02991743 | 0.11736838 |
| DOCK5 | 1343.87327 | -1.557796102 | 0.38025479 | -4.0967166 | 4.19E-05 | 0.00085951 |
| KCTD9 | 546.141018 | 0.002715531 | 0.1907233 | 0.01423807 | 0.98864005 | 0.99572599 |
| PPP2R2A | 1701.02303 | 0.203229033 | 0.18231657 | 1.1147041 | 0.26497724 | 0.48592037 |
| BNIP3L | 1627.48975 | -1.795905752 | 0.54242661 | -3.3108732 | 0.00093005 | 0.0094283 |
| DPYSL2 | 275.567564 | 0.131616299 | 0.55465134 | 0.23729556 | 0.81242751 | 0.91190984 |
| TRIM35 | 127.709309 | -0.17610629 | 0.30431191 | -0.5787032 | 0.56278943 | 0.75444301 |
| PTK2B | 387.313748 | -0.64890094 | 0.42364165 | -1.5317213 | 0.12559122 | 0.30439164 |
| EPHX2 | 710.169287 | 1.332846746 | 0.42483433 | 3.13733294 | 0.00170492 | 0.01511432 |
| CLU | 1188.07582 | -3.618984537 | 0.56052711 | -6.4563952 | 1.07E-10 | 1.84E-08 |
| CCDC25 | 241.633126 | -0.026723607 | 0.21340707 | -0.1252236 | 0.90034651 | 0.95395051 |
| ELP3 | 759.913049 | -0.093934977 | 0.15001477 | -0.6261715 | 0.53120244 | 0.72863563 |
| ZNF395 | 272.305072 | -1.087074393 | 0.45540371 | -2.3870565 | 0.01698388 | 0.07966622 |
| FBXO16 | 431.890032 | 0.271507332 | 0.2909976 | 0.93302258 | 0.35080831 | 0.57693534 |
| FZD3 | 258.597424 | -0.183713194 | 0.32115691 | -0.5720356 | 0.56729783 | 0.75737185 |
| EXTL3 | 295.942091 | -0.64737732 | 0.24445429 | -2.6482551 | 0.00809084 | 0.04767298 |
| INTS9 | 642.709774 | -0.019458838 | 0.16342412 | -0.1190696 | 0.90522025 | 0.95656434 |
| HMBOX1 | 2061.74462 | -0.604009319 | 0.20726133 | -2.9142403 | 0.00356555 | 0.02634083 |
| KIF13B | 3177.09275 | -0.596386849 | 0.19980377 | -2.9848629 | 0.00283705 | 0.02225336 |
| AC108449.1 | 165.272504 | -0.844095417 | 0.47380629 | -1.7815201 | 0.07482753 | 0.21730765 |
| SARAF | 745.813575 | -0.685084514 | 0.29586981 | -2.3154931 | 0.02058596 | 0.09094313 |
| LEPROTL1 | 264.926051 | -0.177100929 | 0.24898668 | -0.7112868 | 0.47690656 | 0.6867392 |
| DCTN6 | 152.729304 | -0.447259488 | 0.24935567 | -1.7936608 | 0.07286732 | 0.21378193 |
| RBPMS-AS1 | 135.658094 | -1.862584862 | 0.49480291 | -3.7642965 | 0.00016702 | 0.00252999 |
| RBPMS | 13308.2277 | -1.171875244 | 0.22750864 | -5.1509043 | 2.59E-07 | 1.36E-05 |
| GTF2E2 | 583.596024 | -0.117007326 | 0.1958732 | -0.5973626 | 0.5502653 | 0.74342379 |
| GSR | 427.455655 | 0.100767852 | 0.23570651 | 0.42751407 | 0.66900494 | 0.82587721 |
| UBXN8 | 77.9010464 | 0.187157532 | 0.23734869 | 0.78853407 | 0.43038438 | 0.64854411 |
| PPP2CB | 1097.78935 | -0.154637564 | 0.18523271 | -0.8348286 | 0.40381422 | 0.62502719 |
| WRN | 1135.3906 | 0.179186596 | 0.23368159 | 0.76679809 | 0.44320157 | 0.65889134 |
| NRG1 | 1237.11717 | 0.875833736 | 0.27106453 | 3.23108942 | 0.00123319 | 0.0117782 |
| FUT10 | 583.748698 | 0.585664039 | 0.22920104 | 2.55524158 | 0.01061141 | 0.05789597 |
| TTI2 | 227.132591 | -0.043220822 | 0.32617384 | -0.1325085 | 0.89458206 | 0.95172874 |
| MAK16 | 90.0165552 | 0.36927673 | 0.37677412 | 0.98010111 | 0.32703621 | 0.55240682 |
| RNF122 | 167.085412 | 0.497180236 | 0.36597872 | 1.35849492 | 0.17430669 | 0.37592646 |
| UNC5D | 106.175664 | -0.188392853 | 0.31407794 | -0.5998283 | 0.54862064 | 0.74226468 |
| ERLIN2 | 264.66435 | -0.026807537 | 0.20445317 | -0.1311182 | 0.89568179 | 0.9518916 |
| PLPBP | 239.589719 | -0.142460476 | 0.16164662 | -0.8813081 | 0.37815107 | 0.60093811 |
| RAB11FIP1 | 611.539678 | -0.740818414 | 0.26356156 | -2.8107984 | 0.00494187 | 0.03310204 |
| EIF4EBP1 | 96.3709677 | -0.602764958 | 0.34029263 | -1.7713136 | 0.07650857 | 0.22060722 |
| ASH2L | 429.276609 | 0.281433134 | 0.23490686 | 1.19806267 | 0.23089262 | 0.44653889 |
| LSM1 | 222.846242 | -0.149221387 | 0.25539665 | -0.5842731 | 0.5590366 | 0.75089633 |
| BAG4 | 516.3164 | 0.342425729 | 0.18042195 | 1.89791609 | 0.05770714 | 0.18319 |
| DDHD2 | 676.851523 | 0.249296671 | 0.15531684 | 1.60508457 | 0.10847519 | 0.27682551 |
| PLPP5 | 414.38673 | 0.015894275 | 0.36461919 | 0.04359144 | 0.96523008 | 0.98449934 |
| NSD3 | 2411.53423 | -0.242585054 | 0.12173852 | -1.9926729 | 0.04629728 | 0.15746395 |
| LETM2 | 145.14087 | -0.147439982 | 0.31250764 | -0.4717964 | 0.63707212 | 0.80480926 |
| FGFR1 | 610.651936 | -1.369138751 | 0.39686536 | -3.4498822 | 0.00056083 | 0.00643616 |
| AC069120.1 | 98.2888076 | 0.587580151 | 0.75578537 | 0.77744314 | 0.43689737 | 0.65349081 |
| TACC1 | 5376.78773 | 1.300075296 | 0.48540632 | 2.67832382 | 0.00739916 | 0.04452645 |
| PLEKHA2 | 763.423308 | 0.061240285 | 0.35928292 | 0.17045142 | 0.86465514 | 0.93833133 |
| TM2D2 | 78.043784 | -0.575029888 | 0.40398264 | -1.4234025 | 0.15461952 | 0.34900362 |
| ADAM9 | 823.760052 | -0.625920148 | 0.24232493 | -2.5829788 | 0.00979514 | 0.05452336 |
| ADAM32 | 93.9413191 | -1.383872194 | 0.33114841 | -4.1790091 | 2.93E-05 | 0.00064242 |
| TCIM | 4788.39262 | -1.316301104 | 0.53353343 | -2.4671389 | 0.01361975 | 0.06843031 |
| AC105999.2 | 233.506323 | 4.100588468 | 1.48281091 | 2.76541563 | 0.00568503 | 0.03688289 |
| ZMAT4 | 218.485633 | 0.742242344 | 1.19415108 | 0.62156486 | 0.53422803 | 0.73149412 |
| SFRP1 | 1737.71023 | -0.383235301 | 0.56861178 | -0.6739841 | 0.50032142 | 0.70482771 |
| GOLGA7 | 229.007002 | 0.083379776 | 0.20711058 | 0.40258579 | 0.68725297 | 0.8387623 |
| AC009630.1 | 68.8189723 | -0.027620398 | 0.42431617 | -0.0650939 | 0.94809923 | 0.97778285 |
| GPAT4 | 280.373937 | -0.495417535 | 0.22091359 | -2.2425852 | 0.02492358 | 0.1034956 |
| KAT6A | 2268.58326 | -0.029549634 | 0.13980192 | -0.2113679 | 0.83260023 | 0.92170771 |
| AP3M2 | 188.998921 | 0.554448829 | 0.29857829 | 1.85696295 | 0.06331642 | 0.19508903 |
| IKBKB | 839.385472 | -0.504589279 | 0.20072103 | -2.5138834 | 0.01194099 | 0.06271512 |
| POLB | 222.979138 | 0.158405612 | 0.219214 | 0.7226072 | 0.46992124 | 0.68173418 |
| VDAC3 | 582.293006 | 0.369054459 | 0.24909635 | 1.48157312 | 0.13845391 | 0.32466497 |
| SLC20A2 | 4015.07362 | -0.457170287 | 0.32947353 | -1.3875782 | 0.16526553 | 0.36452087 |
| SMIM19 | 490.157569 | -0.419847374 | 0.25422744 | -1.6514637 | 0.09864394 | 0.2600774 |
| THAP1 | 44.4658813 | -0.193538817 | 0.38183649 | -0.5068631 | 0.6122509 | 0.78684784 |
| RNF170 | 730.988589 | 0.029568946 | 0.24159705 | 0.12238951 | 0.90259054 | 0.95499458 |

| | | | | | | |
|---|---|---|---|---|---|---|
| HOOK3 | 1151.61587 | -0.218137303 | 0.11524352 | -1.8928378 | 0.05837944 | 0.18452596 |
| FNTA | 314.513374 | -0.380176714 | 0.24337519 | -1.5621014 | 0.11826411 | 0.29330654 |
| POMK | 80.199291 | 0.453112571 | 0.22815265 | 1.98600621 | 0.04703263 | 0.15913131 |
| HGSNAT | 1169.70517 | -0.366411995 | 0.19779658 | -1.8524688 | 0.06395853 | 0.19619792 |
| SPIDR | 7075.00129 | -0.876549213 | 0.31602684 | -2.7736543 | 0.00554305 | 0.0362287 |
| CEBPD | 1944.34012 | -2.418624752 | 0.47562404 | -5.0851609 | 3.67E-07 | 1.82E-05 |
| PRKDC | 1361.38088 | -0.03697103 | 0.19215853 | -0.1923986 | 0.84743 | 0.92955513 |
| MCM4 | 159.884589 | -0.350098959 | 0.33502087 | -1.0450064 | 0.29602001 | 0.51906534 |
| UBE2V2 | 534.427517 | 0.094215366 | 0.22208709 | 0.42422711 | 0.67140019 | 0.82736497 |
| SNTG1 | 48.7567846 | 0.353602647 | 0.26502075 | 1.33424513 | 0.18212353 | 0.38654023 |
| PXDNL | 1581.57656 | 2.304327056 | 0.51927597 | 4.43757695 | 9.10E-06 | 0.00024753 |
| PCMTD1 | 3199.50387 | 0.179206464 | 0.14154438 | 1.26607967 | 0.20548456 | 0.41584887 |
| AC064807.1 | 250.154769 | 0.994665291 | 0.32408058 | 3.0691913 | 0.00214639 | 0.01809665 |
| RB1CC1 | 2066.48276 | -0.072023423 | 0.21364702 | -0.3371141 | 0.73603088 | 0.87055499 |
| ATP6V1H | 2047.51895 | 0.500570759 | 0.21554461 | 2.32235337 | 0.02021392 | 0.08968949 |
| TCEA1 | 614.411649 | 0.219060659 | 0.17782622 | 1.23188051 | 0.21799372 | 0.43190027 |
| LYPLA1 | 1123.54733 | -0.120607345 | 0.16286171 | -0.7405506 | 0.45896595 | 0.67278348 |
| MRPL15 | 299.59954 | 0.382117858 | 0.24929741 | 1.53277909 | 0.1253303 | 0.30392546 |
| RP1 | 2645.11131 | 0.461968206 | 0.50174953 | 0.92071478 | 0.35719936 | 0.58271536 |
| XKR4 | 243.326469 | -3.320381744 | 0.65509287 | -5.0685664 | 4.01E-07 | 1.97E-05 |
| TMEM68 | 402.910095 | 0.407208633 | 0.18809342 | 2.16492756 | 0.03039323 | 0.11881155 |
| TGS1 | 461.970385 | 0.033621994 | 0.1561695 | 0.21529167 | 0.82953992 | 0.92024799 |
| LYN | 610.844988 | -1.86701921 | 0.30083728 | -6.2060766 | 5.43E-10 | 7.47E-08 |
| RPS20 | 650.5988 | -0.81476236 | 0.3870085 | -2.1052828 | 0.03526669 | 0.13148954 |
| PLAG1 | 434.771581 | -0.600818929 | 0.44195119 | -1.359469 | 0.17399802 | 0.37572992 |
| CHCHD7 | 149.710169 | -0.178539414 | 0.26252489 | -0.6800857 | 0.49645022 | 0.70179911 |
| IMPAD1 | 378.56715 | 0.02177693 | 0.1959268 | 0.11114829 | 0.91149875 | 0.96012656 |
| LINC01606 | 5043.68312 | 0.816481709 | 0.54429513 | 1.50007169 | 0.13359583 | 0.31784355 |
| FAM110B | 1022.56408 | -0.087671774 | 0.23682383 | -0.3701983 | 0.71123476 | 0.8551462 |
| AC027698.1 | 83.4293554 | -0.290911229 | 0.54962722 | -0.5292882 | 0.59660551 | 0.77627853 |
| UBXN2B | 559.673911 | 0.153218835 | 0.15384785 | 0.99591142 | 0.31929318 | 0.54392231 |
| SDCBP | 744.08208 | -1.093137996 | 0.35044594 | -3.1192771 | 0.00181295 | 0.01580626 |
| NSMAF | 736.838898 | 0.062538091 | 0.16472054 | 0.37966176 | 0.70419651 | 0.85017199 |
| TOX | 2015.58418 | 0.962369611 | 0.38342609 | 2.50992208 | 0.01207578 | 0.06317188 |
| CA8 | 939.870708 | -3.723997523 | 0.63994674 | -5.8192304 | 5.91E-09 | 6.10E-07 |
| LINC01301 | 356.516534 | 0.578809346 | 0.45444739 | 1.27365533 | 0.20278559 | 0.41232902 |
| RAB2A | 2074.38152 | -0.370981139 | 0.16526192 | -2.2448071 | 0.02478052 | 0.10309477 |
| CHD7 | 1856.09174 | -0.062799662 | 0.1863414 | -0.337014 | 0.73610634 | 0.87055499 |
| CLVS1 | 68.0129455 | -0.486665778 | 0.23168343 | -2.1005636 | 0.03567929 | 0.13247052 |
| ASPH | 1917.10999 | -0.061561952 | 0.15667142 | -0.3929367 | 0.69436624 | 0.843734 |
| NKAIN3 | 63.5624675 | -0.466398229 | 0.40718772 | -1.1454133 | 0.252038 | 0.47214352 |
| YTHDF3 | 1056.94696 | 0.236694219 | 0.17149201 | 1.38020553 | 0.16752337 | 0.36733337 |
| AC022639.1 | 40.6454369 | -0.004732856 | 0.34723233 | -0.0136302 | 0.98912499 | 0.99594533 |
| CYP7B1 | 1549.90405 | 0.80707041 | 0.24096605 | 3.34931163 | 0.00081013 | 0.00849957 |
| ARMC1 | 334.286764 | 0.446837151 | 0.16495197 | 2.70889256 | 0.00675082 | 0.04178612 |
| MTFR1 | 613.350341 | -0.110183534 | 0.31497005 | -0.3498223 | 0.72647209 | 0.86526092 |
| PDE7A | 1079.0252 | 0.498818114 | 0.24514965 | 2.03474943 | 0.04187609 | 0.14672183 |
| TRIM55 | 240.794984 | 0.216201742 | 0.4527525 | 0.47752743 | 0.63298659 | 0.80230473 |
| RRS1-AS1 | 97.4610238 | 0.667891474 | 0.39937269 | 1.67235139 | 0.09445505 | 0.25284294 |
| ADHFE1 | 652.048841 | 0.96597491 | 0.50736821 | 1.90389326 | 0.05692409 | 0.18148912 |
| VXN | 167.174428 | 1.957170472 | 0.37469732 | 5.22333727 | 1.76E-07 | 9.86E-06 |
| VCPIP1 | 320.18143 | -0.173249306 | 0.15218317 | -1.1384262 | 0.25494258 | 0.47570794 |
| SGK3 | 788.882423 | -0.151926906 | 0.23543667 | -0.6452984 | 0.51873383 | 0.71939843 |
| MCMDC2 | 113.066201 | 1.418064287 | 0.3811413 | 3.72057368 | 0.00019877 | 0.00292054 |
| PPP1R42 | 191.124733 | 0.407805916 | 0.35315999 | 1.1547342 | 0.24819929 | 0.46734754 |
| COPS5 | 101.023011 | 0.164462604 | 0.3430851 | 0.47936388 | 0.63167979 | 0.80129239 |
| CSPP1 | 1658.09409 | 0.19331836 | 0.18770185 | 1.02992253 | 0.30304637 | 0.5262179 |
| ARFGEF1 | 3963.24998 | -0.415262564 | 0.16325089 | -2.5437079 | 0.01096828 | 0.05923139 |
| C8orf34 | 84.8705317 | 0.075519321 | 0.29253531 | 0.25815455 | 0.79628763 | 0.90454795 |
| SULF1 | 72.9089168 | 0.264365291 | 0.54672829 | 0.48354054 | 0.62871198 | 0.7991394 |
| NCOA2 | 8331.62049 | -0.101545326 | 0.16857684 | -0.6023682 | 0.54692909 | 0.74124838 |
| TRAM1 | 898.100846 | -0.716316284 | 0.27899946 | -2.5674468 | 0.01024505 | 0.05650278 |
| LACTB2-AS1 | 122.162831 | -1.300212179 | 0.71334407 | -1.8226999 | 0.06834885 | 0.205 |
| LACTB2 | 674.616438 | -1.49916567 | 0.41272639 | -3.6323475 | 0.00028085 | 0.00383641 |
| XKR9 | 199.135821 | -0.148980107 | 0.53160998 | -0.2802432 | 0.77929089 | 0.89471129 |
| TERF1 | 491.769202 | 0.15998944 | 0.14307919 | 1.11818803 | 0.26348669 | 0.48439746 |
| AC022893.1 | 77.6912602 | -0.157937584 | 0.35292633 | -0.4475086 | 0.65450785 | 0.81598813 |
| SBSPON | 126.465523 | -1.638737365 | 0.42587229 | -3.8479549 | 0.00011911 | 0.001921 |
| RPL7 | 1016.12977 | -0.916101404 | 0.49926584 | -1.834897 | 0.06652095 | 0.20111587 |
| RDH10 | 368.444143 | -0.282476675 | 0.46395097 | -0.6088503 | 0.54262369 | 0.73869074 |
| RDH10-AS1 | 145.009614 | -0.274261415 | 0.63226619 | -0.4337752 | 0.66445167 | 0.82323933 |
| STAU2 | 1962.70019 | -0.029445191 | 0.1974665 | -0.1491149 | 0.88146299 | 0.94645136 |
| AC027018.1 | 323.5202 | 0.002789066 | 0.5330379 | 0.0052324 | 0.99582517 | 0.99847083 |
| ELOC | 345.422236 | -0.106621113 | 0.19501567 | -0.546731 | 0.58456356 | 0.76905468 |
| TMEM70 | 94.2396458 | -0.217555978 | 0.26904591 | -0.8086203 | 0.4187336 | 0.63783336 |
| GDAP1 | 1022.58147 | 0.644593174 | 0.46474765 | 1.38697456 | 0.16544952 | 0.36466256 |
| MIR2052HG | 274.60997 | -0.277534147 | 0.65649898 | -0.4227488 | 0.67247856 | 0.82807581 |
| CRISPLD1 | 47.3285161 | 0.038155874 | 0.50916733 | 0.07493779 | 0.94026421 | 0.97411372 |
| CASC9 | 77.8936919 | 2.273306178 | 0.80017584 | 2.84100827 | 0.00449711 | 0.03108445 |
| HNF4G | 126.269176 | -1.016918507 | 0.60946193 | -1.6685513 | 0.09520633 | 0.25421072 |
| PEX2 | 388.047129 | 0.08711842 | 0.19322488 | 0.45086543 | 0.65208654 | 0.81435295 |
| PKIA | 73.2879634 | 0.142210877 | 0.47471657 | 0.29957007 | 0.76450512 | 0.88572451 |
| ZC2HC1A | 536.571086 | 0.01204644 | 0.29733728 | 0.0405144 | 0.96768303 | 0.98528676 |
| IL7 | 528.034851 | -0.89451066 | 0.37533655 | -2.3832229 | 0.0171618 | 0.08024433 |
| AC083837.1 | 59.4953843 | -1.142454847 | 0.786569 | -1.4524534 | 0.14637557 | 0.33690101 |
| LINC01607 | 37.7817409 | 1.223317523 | 0.69439972 | 1.7616907 | 0.07812157 | 0.22330833 |
| TPD52 | 1343.93162 | 0.213116933 | 0.33124853 | 0.64337472 | 0.51998098 | 0.71990992 |
| ZBTB10 | 737.587093 | -0.303579138 | 0.3820852 | -0.7945326 | 0.42688546 | 0.64526969 |
| ZNF704 | 2781.79156 | 0.284712157 | 0.25531821 | 1.11512669 | 0.26479613 | 0.48585805 |
| PAG1 | 171.41973 | 0.08638448 | 0.69321379 | 0.12461448 | 0.90082876 | 0.95413737 |
| IMPA1 | 393.988445 | 0.604575008 | 0.16983376 | 3.55980455 | 0.00037113 | 0.00470016 |
| ZFAND1 | 477.780292 | 0.346340165 | 0.19862862 | 1.74365687 | 0.08121894 | 0.22874648 |
| CHMP4C | 212.222489 | -0.556656194 | 0.27672119 | -2.0116139 | 0.04426066 | 0.15276692 |
| SNX16 | 354.481372 | -0.114533663 | 0.23124039 | -0.4953013 | 0.62038747 | 0.79267731 |
| RALYL | 2442.94828 | 2.561820551 | 0.67254282 | 3.80915605 | 0.00013944 | 0.00217736 |
| LRRCC1 | 174.235519 | -0.08680235 | 0.35919793 | -0.241656 | 0.80904669 | 0.91023512 |
| E2F5 | 279.589653 | -0.323608916 | 0.2773349 | -1.1668525 | 0.24326996 | 0.46195412 |
| C8orf59 | 282.806904 | -0.044620341 | 0.24680053 | -0.1807952 | 0.85652836 | 0.93503166 |
| CA13 | 467.584655 | -0.143816816 | 0.16218321 | -0.8867553 | 0.37521067 | 0.59870627 |
| CA1 | 151.358614 | -0.356810495 | 0.44465862 | -0.8024369 | 0.42230026 | 0.64115803 |
| CA3 | 588.290622 | -0.212036347 | 0.24734077 | -0.857264 | 0.39129898 | 0.6140943 |
| CA2 | 383.359158 | 0.596438815 | 0.43905405 | 1.3584633 | 0.17431672 | 0.37592646 |
| ATP6V0D2 | 239.916548 | 2.242926716 | 0.78650252 | 2.85177308 | 0.00434761 | 0.03025691 |
| WWP1 | 2825.577 | -0.232503567 | 0.14617234 | -1.5906126 | 0.11169679 | 0.28237377 |
| RMDN1 | 1013.62697 | 0.675834154 | 0.13905418 | 4.86022171 | 1.17E-06 | 4.72E-05 |
| CPNE3 | 782.047071 | 0.329633056 | 0.1801793 | 1.82947239 | 0.06732887 | 0.20284564 |
| CNGB3 | 115.484204 | -0.629576883 | 0.41603679 | -1.5132721 | 0.13021055 | 0.31253761 |
| CNBD1 | 47.1206024 | 0.14712754 | 0.33859322 | 0.43452595 | 0.66390656 | 0.82281276 |
| AF117829.1 | 2419.71853 | 0.081462476 | 0.27133662 | 0.30022662 | 0.7640043 | 0.88555042 |
| RIPK2 | 186.830585 | -0.532401152 | 0.31923823 | -1.6677236 | 0.0953706 | 0.25441407 |
| OSGIN2 | 278.277247 | -0.402238705 | 0.18899469 | -2.1283069 | 0.03331164 | 0.12680504 |
| NBN | 524.455048 | -0.055570315 | 0.22848475 | -0.2432124 | 0.8078409 | 0.91010272 |
| DECR1 | 1441.7702 | 0.523556192 | 0.17783526 | 2.94405174 | 0.00323946 | 0.02446541 |

| | | | | | | |
|---|---|---|---|---|---|---|
| CALB1 | 50.3849536 | 2.162591626 | 0.64094866 | 3.37404814 | 0.00074071 | 0.0079805 |
| LINC00534 | 381.986243 | 1.05321486 | 0.64030489 | 1.64486462 | 0.09999773 | 0.26284834 |
| TMEM64 | 158.322674 | 0.192718698 | 0.54826985 | 0.35150337 | 0.72521075 | 0.86401269 |
| NECAB1 | 64.0801347 | -0.650766607 | 0.75053081 | -0.8670751 | 0.38590084 | 0.60936789 |
| C8orf88 | 66.3825359 | 0.232480341 | 0.65674469 | 0.35398892 | 0.72334719 | 0.86257046 |
| PIP4P2 | 425.991423 | 0.408829433 | 0.3281785 | 1.24575324 | 0.212855 | 0.42574224 |
| OTUD6B-AS1 | 230.708132 | 0.200749246 | 0.34107394 | 0.58857985 | 0.55614316 | 0.74830345 |
| OTUD6B | 256.128343 | 0.161133489 | 0.16865128 | 0.95542407 | 0.33936329 | 0.56485001 |
| LRRC69 | 445.502167 | -0.473669972 | 0.36453439 | -1.2993835 | 0.19381235 | 0.40136505 |
| SLC26A7 | 216.891441 | 1.333417157 | 0.57861588 | 2.30449459 | 0.0211949 | 0.09294933 |
| AC117834.1 | 132.1194 | -0.439604345 | 0.55148532 | -0.7971279 | 0.42537674 | 0.64392696 |
| TRIQK | 591.071189 | -0.022829221 | 0.14559806 | -0.1567962 | 0.87540548 | 0.94393722 |
| LINC00535 | 78.4737727 | 0.940509383 | 0.3693988 | 2.54605423 | 0.01089483 | 0.05897941 |
| FAM92A | 466.5592 | 0.619470379 | 0.32434942 | 1.90988591 | 0.0561479 | 0.17975222 |
| AC120053.1 | 63.6985639 | 0.61034276 | 0.28709764 | 2.12590658 | 0.03351104 | 0.12730725 |
| RBM12B | 520.219462 | 0.356408308 | 0.16837292 | 2.11677925 | 0.03427859 | 0.12913878 |
| TMEM67 | 803.973894 | 0.163759911 | 0.14171469 | 1.15556057 | 0.24786094 | 0.46717968 |
| PDP1 | 107.716503 | -0.494141653 | 0.23297905 | -2.1209704 | 0.0339243 | 0.12843394 |
| GEM | 69.2496929 | -0.425262501 | 0.40898578 | -1.0397978 | 0.29843386 | 0.52159011 |
| RAD54B | 50.4298126 | 0.83120536 | 0.38757366 | 2.14463839 | 0.03198177 | 0.12334225 |
| VIRMA | 606.822775 | 0.097573615 | 0.13743332 | 0.70997059 | 0.47772238 | 0.68746521 |
| AC023632.2 | 97.0445041 | 0.592694641 | 0.33312788 | 1.77918054 | 0.07521017 | 0.21795768 |
| AC108860.2 | 56.161689 | 0.612409263 | 0.3300181 | 1.85568386 | 0.06349863 | 0.19518591 |
| ESRP1 | 1040.13389 | -0.043468196 | 0.26825111 | -0.1620429 | 0.87127205 | 0.94147844 |
| DPY19L4 | 660.46626 | 0.251053769 | 0.19403643 | 1.29384864 | 0.19571771 | 0.4034387 |
| INTS8 | 608.528943 | 0.198804438 | 0.14365963 | 1.38385735 | 0.16640214 | 0.3659452 |
| NDUFAF6 | 671.005104 | 0.425764605 | 0.1781714 | 2.38963494 | 0.01686513 | 0.07944774 |
| TP53INP1 | 317.859271 | 0.417218083 | 0.34510919 | 1.20894514 | 0.22668392 | 0.44169759 |
| PLEKHF2 | 182.69912 | 0.579512916 | 0.24170968 | 2.39755778 | 0.01650478 | 0.07830877 |
| C8orf37-AS1 | 466.647525 | -0.271811161 | 0.40753956 | -0.6669565 | 0.50479992 | 0.70859746 |
| C8orf37 | 60.4161863 | 0.445899642 | 0.29482413 | 1.51242585 | 0.13042556 | 0.31260956 |
| UQCRB | 805.016884 | -0.192560093 | 0.53209833 | -0.3618882 | 0.7174356 | 0.85947463 |
| MTERF3 | 202.927776 | 0.394774177 | 0.19312223 | 2.04416745 | 0.04093701 | 0.14473393 |
| PTDSS1 | 176.258011 | -0.202637129 | 0.31981569 | -0.633606 | 0.52633801 | 0.72497874 |
| SDC2 | 1217.4133 | -0.392948823 | 0.37647289 | -1.0437639 | 0.29659463 | 0.51979812 |
| CPQ | 1765.8757 | 0.717326958 | 0.34865526 | 2.05741041 | 0.03964676 | 0.14177464 |
| TSPYL5 | 100.782155 | -0.183714795 | 0.34110228 | -0.5385915 | 0.59016874 | 0.7721373 |
| MTDH | 2235.45848 | -0.023193767 | 0.10664432 | -0.2174871 | 0.82782873 | 0.91981743 |
| LAPTM4B | 925.448573 | -1.164488874 | 0.27803146 | -4.188335 | 2.81E-05 | 0.00062175 |
| MATN2 | 45.5611811 | -0.809247766 | 0.54686169 | -1.4798034 | 0.13892573 | 0.32536701 |
| RPL30 | 761.833406 | -0.643321163 | 0.34072739 | -1.8880817 | 0.05901498 | 0.18569053 |
| RIDA | 199.372168 | 0.175577529 | 0.43136797 | 0.40702495 | 0.68398966 | 0.83609286 |
| POP1 | 138.272107 | -0.054764007 | 0.27363274 | -0.2001369 | 0.84137352 | 0.92645072 |
| NIPAL2 | 310.933762 | -0.182720817 | 0.3028751 | -0.6032877 | 0.54631732 | 0.74058965 |
| AP003438.1 | 45.4751093 | -0.004472983 | 0.3731366 | -0.0119875 | 0.99043557 | 0.99606252 |
| STK3 | 4106.79434 | -0.355116733 | 0.19664522 | -1.8058752 | 0.07093783 | 0.21009369 |
| AC104986.2 | 394.226707 | 0.538586782 | 0.24587738 | 2.19046898 | 0.02849024 | 0.11338584 |
| VPS13B | 4855.52424 | -0.003861024 | 0.13224871 | -0.0291952 | 0.97670893 | 0.98948597 |
| COX6C | 705.455934 | 0.338323245 | 0.28889922 | 1.17107702 | 0.24156782 | 0.45991198 |
| POLR2K | 159.950375 | -0.448911464 | 0.41866292 | -1.0722504 | 0.2836076 | 0.50610278 |
| SPAG1 | 530.999084 | -0.682150541 | 0.27862622 | -2.448264 | 0.01435464 | 0.0710684 |
| RNF19A | 2400.65009 | -1.478904965 | 0.34044996 | -4.3439717 | 1.40E-05 | 0.00035139 |
| AP000424.1 | 127.980485 | -5.836569205 | 0.9696911 | -6.0189985 | 1.75E-09 | 2.11E-07 |
| ANKRD46 | 339.68232 | 0.435053557 | 0.16936944 | 2.56866626 | 0.01020907 | 0.05632901 |
| PABPC1 | 1152.72843 | -0.580378737 | 0.25092425 | -2.3129639 | 0.02072462 | 0.09134186 |
| YWHAZ | 1816.43926 | -0.346918406 | 0.18267956 | -1.8990543 | 0.05755733 | 0.18298338 |
| AP003469.4 | 43.1383958 | 0.584756803 | 0.34343322 | 1.70267981 | 0.08862801 | 0.24205091 |
| ZNF706 | 563.548632 | 0.10935378 | 0.19516573 | 0.56031242 | 0.57526636 | 0.76252365 |
| AP001207.3 | 58.4687709 | 1.537950501 | 0.44076505 | 3.48927504 | 0.00048433 | 0.00577396 |
| GRHL2 | 1995.45421 | 0.479031321 | 0.2478702 | 1.93258941 | 0.0532868 | 0.17349404 |
| NCALD | 86.1423059 | -0.668520284 | 0.46210463 | -1.446686 | 0.1479849 | 0.33942221 |
| RRM2B | 399.12668 | -0.050158037 | 0.43635385 | -0.1149481 | 0.90848628 | 0.95871259 |
| UBR5-AS1 | 106.885766 | 0.369749666 | 0.54009805 | 0.6845973 | 0.49359807 | 0.69903901 |
| UBR5 | 3356.41035 | 0.185537046 | 0.19493196 | 0.95180413 | 0.34119633 | 0.5665446 |
| KLF10 | 180.811675 | -0.26588234 | 0.55368765 | -0.4802028 | 0.63108322 | 0.80092028 |
| AZIN1 | 615.382026 | -0.277416076 | 0.25611872 | -1.0831542 | 0.27873998 | 0.50097041 |
| AZIN1-AS1 | 2078.06139 | -0.179606405 | 0.29556888 | -0.6076634 | 0.54341071 | 0.7391242 |
| AP003354.2 | 593.599425 | -0.412006793 | 0.42583977 | -0.967516 | 0.33328614 | 0.55874814 |
| ATP6V1C1 | 843.832028 | 0.168434556 | 0.20481092 | 0.82239051 | 0.41085468 | 0.63014283 |
| BAALC-AS1 | 550.183957 | 0.710739787 | 0.30045442 | 2.36554945 | 0.01800334 | 0.0832363 |
| FZD6 | 1023.66184 | 0.461609258 | 0.22562166 | 2.04594391 | 0.04076189 | 0.14438825 |
| SLC25A32 | 336.731418 | 0.108333215 | 0.19879276 | 0.54495555 | 0.58578409 | 0.77001836 |
| DCAF13 | 197.072032 | 0.154598033 | 0.27223803 | 0.56787817 | 0.5701177 | 0.75944783 |
| RIMS2 | 1184.62679 | 0.60470275 | 0.7744306 | 0.7808353 | 0.43489937 | 0.65190487 |
| DPYS | 76.7094895 | 0.943460337 | 0.73317403 | 1.28681637 | 0.1981583 | 0.40675481 |
| LRP12 | 290.44804 | 0.167483565 | 0.25543727 | 0.65567395 | 0.51203394 | 0.71407594 |
| ZFPM2 | 85.7913253 | -0.870322034 | 0.46037815 | -1.8904504 | 0.05869775 | 0.18495674 |
| AC027031.2 | 380.383274 | 0.116084847 | 0.46732125 | 0.24840481 | 0.80382121 | 0.90804718 |
| OXR1 | 18792.1481 | 0.213783604 | 0.41537988 | 0.5146701 | 0.60678357 | 0.78286816 |
| AC090579.1 | 1597.24432 | -0.013492172 | 0.45688543 | -0.0295308 | 0.97644129 | 0.98947549 |
| ANGPT1 | 47.3998209 | 1.627755746 | 0.58751806 | 2.77056293 | 0.00559595 | 0.03648415 |
| EIF3E | 938.018536 | 0.261926707 | 0.14911408 | 1.75655243 | 0.07899413 | 0.2246842 |
| EMC2 | 781.996197 | 0.400690212 | 0.22594975 | 1.77335987 | 0.07616909 | 0.22010885 |
| NUDCD1 | 363.723412 | -0.064141476 | 0.25429887 | -0.2522287 | 0.80086429 | 0.90662963 |
| ENY2 | 456.68916 | 0.060330304 | 0.26298727 | 0.22940389 | 0.81855501 | 0.91559451 |
| EBAG9 | 282.701015 | 0.478625047 | 0.20136919 | 2.37685342 | 0.01746103 | 0.08141288 |
| SYBU | 203.87946 | 0.464915115 | 0.81017414 | 0.57384591 | 0.56607208 | 0.75642605 |
| TRPS1 | 7059.12187 | 0.367872412 | 0.36314468 | 1.01301886 | 0.31105115 | 0.53456605 |
| EIF3H | 1166.13463 | -0.186090006 | 0.15716543 | -1.184039 | 0.23639761 | 0.45392878 |
| UTP23 | 210.848768 | 0.255257572 | 0.27225793 | 0.93755789 | 0.34847166 | 0.57409675 |
| RAD21 | 680.626396 | -0.144593795 | 0.22209876 | -0.6510338 | 0.51502468 | 0.7164072 |
| MED30 | 160.930379 | 0.111208268 | 0.28535578 | 0.38971795 | 0.69674512 | 0.84496018 |
| EXT1 | 6098.38909 | 0.059064271 | 0.22883939 | 0.2581036 | 0.79632695 | 0.90454795 |
| SAMD12 | 7765.15821 | -0.818955087 | 0.11764046 | -6.9615089 | 3.37E-12 | 9.46E-10 |
| AC023590.1 | 756.793608 | -1.128978369 | 0.49345824 | -2.2878904 | 0.0221439 | 0.09546307 |
| SAMD12-AS1 | 826.903446 | -2.048413752 | 0.46265324 | -4.4275357 | 9.53E-06 | 0.00025642 |
| TNFRSF11B | 596.912685 | -1.219689671 | 0.67474376 | -1.8076339 | 0.0706635 | 0.20951609 |
| COLEC10 | 311.690676 | -0.953395396 | 0.59591104 | -1.5998955 | 0.10962177 | 0.27851393 |
| MAL2 | 953.205486 | 0.013809164 | 0.28911032 | 0.04776434 | 0.96190406 | 0.9832997 |
| MAL2-AS1 | 113.888572 | -0.133030517 | 0.72449977 | -0.1836171 | 0.85431388 | 0.93338561 |
| ENPP2 | 93.2283369 | 0.700965651 | 0.53199168 | 1.31762523 | 0.18762914 | 0.39327091 |
| TAF2 | 1027.37074 | 0.097119387 | 0.19866436 | 0.48886165 | 0.62493965 | 0.79598067 |
| DEPTOR | 11464.5596 | -1.1428276 | 0.37306885 | -3.0633155 | 0.00218899 | 0.01838168 |
| AC091563.1 | 556.582515 | -1.564048387 | 0.43774205 | -3.5729909 | 0.00035293 | 0.00452859 |
| COL14A1 | 853.535314 | -0.621617809 | 0.55079601 | -1.1285808 | 0.25907471 | 0.48022983 |
| MRPL13 | 303.859524 | 0.209192829 | 0.23692082 | 0.88296516 | 0.3772551 | 0.60028794 |
| MTBP | 178.938061 | 0.014285139 | 0.38051584 | 0.03754151 | 0.97005324 | 0.98653537 |
| SNTB1 | 1195.51087 | -1.781002214 | 0.40190167 | -4.4314378 | 9.36E-06 | 0.00025285 |
| ZHX2 | 3106.39215 | -0.74066322 | 0.22376244 | -3.3100426 | 0.00093282 | 0.00944183 |
| AC104316.1 | 175.41104 | -0.355069045 | 0.38409069 | -0.9244406 | 0.35525694 | 0.58081705 |
| DERL1 | 393.849546 | -0.245082454 | 0.1987075 | -1.233383 | 0.2174329 | 0.43128814 |
| TBC1D31 | 215.378622 | 0.605271495 | 0.22128483 | 2.7352598 | 0.00623311 | 0.03935616 |
| C8orf76 | 82.4610276 | 0.464340428 | 0.31366519 | 1.48036966 | 0.13877462 | 0.32507075 |

| | | | | | | |
|---|---|---|---|---|---|---|
| ATAD2 | 127.686121 | -0.054283314 | 0.26494798 | -0.2048829 | 0.83766361 | 0.92443926 |
| WDYHV1 | 219.865725 | 0.060024862 | 0.30308242 | 0.19804799 | 0.84300752 | 0.92739954 |
| FBXO32 | 414.043798 | -1.08013394 | 0.36378163 | -2.9691822 | 0.00298593 | 0.02309204 |
| FAM91A1 | 776.460239 | 0.17437955 | 0.20575436 | 0.84751328 | 0.39670909 | 0.61845042 |
| TMEM65 | 1422.51223 | 0.391950324 | 0.18363304 | 2.13442162 | 0.03280828 | 0.12557653 |
| TRMT12 | 104.21839 | 0.583057654 | 0.2041468 | 2.85607056 | 0.0042892 | 0.03002439 |
| RNF139-AS1 | 58.753482 | 0.63372105 | 0.34281412 | 1.84858502 | 0.06451776 | 0.1971351 |
| RNF139 | 131.086351 | 0.737964793 | 0.2760509 | 2.67329245 | 0.00751107 | 0.04505621 |
| TATDN1 | 681.824399 | 0.017741546 | 0.15464277 | 0.114726 | 0.9086623 | 0.95874434 |
| NDUFB9 | 435.743707 | -0.066860835 | 0.28299558 | -0.2362611 | 0.81323011 | 0.91242304 |
| MTSS1 | 1895.83245 | 1.240999549 | 0.36290187 | 3.41965601 | 0.000627 | 0.00706061 |
| SQLE | 289.76934 | -1.340516346 | 0.45281282 | -2.9604204 | 0.00307219 | 0.02355733 |
| WASHC5 | 525.435679 | 0.074653485 | 0.18739336 | 0.3983785 | 0.6903512 | 0.84106341 |
| NSMCE2 | 3904.11933 | -0.019123093 | 0.19180444 | -0.099701 | 0.92058171 | 0.96454603 |
| TRIB1 | 189.857525 | -1.169666424 | 0.29291819 | -3.9931505 | 6.52E-05 | 0.00120117 |
| FAM84B | 260.394804 | -0.547846548 | 0.34630266 | -1.5819877 | 0.11365238 | 0.28600387 |
| PCAT1 | 3077.10487 | 0.037418833 | 0.49038448 | 0.07630509 | 0.93917637 | 0.97405628 |
| CASC8 | 46.2748919 | 0.106451802 | 0.59302821 | 0.17950546 | 0.85754083 | 0.93574122 |
| PVT1 | 126.709885 | -2.132127866 | 0.52949238 | -4.0267394 | 5.66E-05 | 0.00107798 |
| CCDC26 | 56.5136458 | 0.45265684 | 0.41609703 | 1.08786366 | 0.2766553 | 0.49896172 |
| FAM49B | 3168.37382 | -0.087259296 | 0.16403944 | -0.5319409 | 0.5947669 | 0.77500996 |
| ASAP1 | 2161.8776 | -0.320119216 | 0.28026305 | -1.1422099 | 0.2533668 | 0.47377152 |
| EFR3A | 1582.16531 | 0.032711681 | 0.27988653 | 0.1168748 | 0.90695927 | 0.95763589 |
| KCNQ3 | 490.662145 | -0.966081067 | 0.64187004 | -1.5051038 | 0.13229738 | 0.31566403 |
| LRRC6 | 736.672749 | 0.395371639 | 0.1621458 | 2.43837111 | 0.01475362 | 0.07241938 |
| PHF20L1 | 1003.81036 | 0.369229846 | 0.17957341 | 2.0561499 | 0.03976807 | 0.14204879 |
| TG | 109.527407 | 1.016161436 | 0.68368579 | 1.48629889 | 0.13720008 | 0.32320435 |
| NDRG1 | 2860.45813 | -1.285390074 | 0.7581852 | -1.695351 | 0.09000885 | 0.24458483 |
| ST3GAL1 | 358.856164 | -1.449337006 | 0.37371816 | -3.8781552 | 0.00010525 | 0.00174001 |
| ZFAT | 517.831739 | 0.055683938 | 0.23754122 | 0.234418 | 0.81466048 | 0.91339257 |
| AC083843.3 | 531.280921 | 0.203383293 | 0.35112293 | 0.57923671 | 0.56242946 | 0.75422648 |
| KHDRBS3 | 1363.76692 | 0.014064891 | 0.34584841 | 0.04066779 | 0.96756074 | 0.98528676 |
| LINC02055 | 147.489354 | 1.859074436 | 0.90756331 | 2.04842397 | 0.04051847 | 0.14385611 |
| TRAPPC9 | 4696.33401 | -0.184132083 | 0.19232191 | -0.957416 | 0.33835731 | 0.56381503 |
| CHRAC1 | 86.674866 | -0.4532361 | 0.29912252 | -1.5152189 | 0.12971698 | 0.31164633 |
| AGO2 | 1183.30675 | -0.573652839 | 0.17148533 | -3.3452007 | 0.00082223 | 0.00860562 |
| PTK2 | 8080.56831 | -0.019912032 | 0.20607001 | -0.0966275 | 0.92302221 | 0.96587185 |
| DENND3 | 392.838426 | -1.296792961 | 0.50672429 | -2.5591687 | 0.01049228 | 0.05745959 |
| SLC45A4 | 919.3537 | -0.43653168 | 0.36176857 | -1.20666 | 0.22756309 | 0.44308386 |
| TSNARE1 | 483.240584 | 0.190403775 | 0.22019405 | 0.86470901 | 0.38719852 | 0.61065142 |
| JRK | 159.197057 | 0.084489732 | 0.28760515 | 0.29376989 | 0.76893375 | 0.88857795 |
| THEM6 | 58.3356398 | 0.373253464 | 0.40857849 | 0.91354165 | 0.36095774 | 0.58607098 |
| LY6E | 699.088255 | -2.971710104 | 0.68852295 | -4.3160654 | 1.59E-05 | 0.00038997 |
| GLI4 | 96.3512514 | 0.204900422 | 0.29317295 | 0.69890631 | 0.48461058 | 0.69239889 |
| TOP1MT | 120.66793 | 0.33953707 | 0.24147302 | 1.4061077 | 0.15969215 | 0.35649376 |
| RHPN1 | 261.458401 | 0.08429227 | 0.36654394 | 0.22996498 | 0.81811898 | 0.91549538 |
| ZC3H3 | 330.412099 | -0.026975839 | 0.30980045 | -0.0870749 | 0.93061199 | 0.96943642 |
| GSDMD | 204.078611 | -0.225763814 | 0.41605484 | -0.5426299 | 0.58738462 | 0.77078914 |
| NAPRT | 112.687425 | -0.218731326 | 0.42519688 | -0.5144236 | 0.60695584 | 0.78294756 |
| EEF1D | 915.941355 | -0.584706937 | 0.25453247 | -2.2971802 | 0.02160849 | 0.09410702 |
| TIGD5 | 40.100707 | -0.240804157 | 0.47721487 | -0.5046032 | 0.61383755 | 0.78755051 |
| TSTA3 | 106.88974 | -1.278360852 | 0.51124227 | -2.5004991 | 0.01240184 | 0.06434248 |
| ZNF623 | 165.309164 | -0.254506297 | 0.1854917 | -1.372063 | 0.17004382 | 0.37027842 |
| ZNF707 | 69.8249144 | -0.033960264 | 0.2757072 | -0.1231751 | 0.90196844 | 0.95457905 |
| FAM83H | 163.000228 | -0.883233191 | 0.29915759 | -2.952401 | 0.00315313 | 0.0240056 |
| IQANK1 | 186.479607 | -0.456638696 | 0.39730035 | -1.1493539 | 0.25041009 | 0.47037357 |
| SCRIB | 158.694541 | -0.150538094 | 0.3382893 | -0.4449981 | 0.65632113 | 0.81763141 |
| PUF60 | 191.888727 | -0.800738073 | 0.37023128 | -2.162805 | 0.03055618 | 0.11934258 |
| NRBP2 | 218.619071 | -0.578865017 | 0.36861691 | -1.5703702 | 0.11632901 | 0.29036089 |
| PLEC | 438.260873 | -0.859364951 | 0.30456961 | -2.8215716 | 0.0047789 | 0.03227222 |
| PARP10 | 226.297149 | 1.132788197 | 0.46430955 | 2.43972624 | 0.0146984 | 0.07224298 |
| GRINA | 274.357687 | 0.258852672 | 0.41556443 | 0.6228942 | 0.53335405 | 0.73051365 |
| SPATC1 | 332.227549 | 1.573379109 | 0.69679481 | 2.25802357 | 0.02394419 | 0.10069367 |
| OPLAH | 146.242784 | 1.356368108 | 0.46988192 | 2.88661482 | 0.00389411 | 0.02813051 |
| EXOSC4 | 64.7808882 | -0.591763648 | 0.43775059 | -1.3518283 | 0.17643024 | 0.37867358 |
| GPAA1 | 249.169922 | -0.162791516 | 0.45290244 | -0.3594406 | 0.71926552 | 0.86003243 |
| CYC1 | 356.670874 | 0.077539292 | 0.43527492 | 0.17813866 | 0.85861408 | 0.93614803 |
| SHARPIN | 270.145861 | -0.128157926 | 0.22702581 | -0.5645082 | 0.57240834 | 0.76035215 |
| MAF1 | 212.808972 | -0.99085619 | 0.36848939 | -2.688968 | 0.00716733 | 0.04343104 |
| MROH1 | 1056.95729 | -0.268016193 | 0.27565691 | -0.9722818 | 0.33091038 | 0.55589409 |
| SCX | 85.3642146 | 0.496387398 | 0.36553426 | 1.35797776 | 0.17447074 | 0.3761358 |
| HSF1 | 525.723022 | -0.374981685 | 0.1587649 | -2.3618677 | 0.01818313 | 0.08384041 |
| DGAT1 | 123.75316 | 0.25831031 | 0.26128262 | 0.98862417 | 0.32284705 | 0.54785287 |
| SLC52A2 | 76.2619903 | -1.246767186 | 0.44625613 | -2.7938377 | 0.00520866 | 0.03453876 |
| ADCK5 | 60.6110608 | 0.323956709 | 0.33066907 | 0.97970067 | 0.3272339 | 0.5526381 |
| CPSF1 | 141.638641 | 0.305807596 | 0.2889528 | 1.05833062 | 0.28990474 | 0.51274189 |
| SLC39A4 | 53.2219704 | -1.785478923 | 0.43664399 | -4.0890954 | 4.33E-05 | 0.00087734 |
| VPS28 | 433.376841 | -1.008203568 | 0.34656407 | -2.9091404 | 0.00362424 | 0.02665512 |
| CYHR1 | 335.767143 | 0.038384534 | 0.34965518 | 0.10977825 | 0.91258524 | 0.96042853 |
| KIFC2 | 166.181857 | 0.584520223 | 0.39914295 | 1.46443829 | 0.14307423 | 0.33248901 |
| PPP1R16A | 369.866805 | 0.014545587 | 0.2394772 | 0.06073892 | 0.95156713 | 0.97914234 |
| GPT | 39.336184 | 1.717658075 | 0.58249489 | 2.94879512 | 0.00319015 | 0.02418986 |
| MFSD3 | 61.636849 | 0.353971927 | 0.3971975 | 0.89117361 | 0.37283604 | 0.5967274 |
| LRRC14 | 66.828552 | -0.066781709 | 0.38613631 | -0.1729485 | 0.86269188 | 0.93788453 |
| ARHGAP39 | 200.737205 | 0.245893126 | 0.24039791 | 1.02285882 | 0.3063746 | 0.53011734 |
| ZNF251 | 798.731387 | 0.241429799 | 0.16218107 | 1.4886435 | 0.13658127 | 0.32222841 |
| ZNF34 | 57.1929075 | -0.07324091 | 0.39173082 | -0.1869674 | 0.85168616 | 0.93193618 |
| RPL8 | 1387.20047 | -0.997376237 | 0.48640637 | -2.0504999 | 0.04031567 | 0.14334581 |
| ZNF517 | 41.3067828 | -0.233317782 | 0.36204558 | -0.6444431 | 0.51928813 | 0.71960522 |
| ZNF7 | 219.766982 | -0.016740097 | 0.22194948 | -0.075423 | 0.93987815 | 0.97411372 |
| COMMD5 | 76.6026171 | -0.041280117 | 0.2651449 | -0.1556889 | 0.87627825 | 0.94426166 |
| ZNF250 | 178.539457 | 0.669091116 | 0.22174847 | 3.01734269 | 0.00255001 | 0.02053699 |
| ZNF16 | 158.49174 | -0.118372528 | 0.32435363 | -0.364949 | 0.71514949 | 0.85767293 |
| C8orf33 | 116.531651 | -1.056233843 | 0.2848444 | -3.7081082 | 0.00020881 | 0.00304103 |
| WASHC1 | 1049.12929 | 0.426146704 | 0.2010672 | 2.11942428 | 0.03405463 | 0.12874286 |
| CBWD1 | 241.833161 | 0.156298483 | 0.24459959 | 0.63899731 | 0.52282468 | 0.7226864 |
| DOCK8 | 61.9920242 | 0.491201924 | 0.32948321 | 1.49082536 | 0.13600735 | 0.32126629 |
| AL161725.1 | 64.2126711 | 2.302468264 | 0.68558857 | 3.35838191 | 0.000784 | 0.00826862 |
| KANK1 | 5742.86387 | 1.277283715 | 0.32199765 | 3.96674857 | 7.29E-05 | 0.00130762 |
| SMARCA2 | 2196.26789 | 0.027806381 | 0.19870618 | 0.13993717 | 0.88870963 | 0.94983134 |
| VLDLR-AS1 | 1151.04873 | 0.65788828 | 0.5552422 | 1.18486721 | 0.23606996 | 0.45389346 |
| VLDLR | 505.128541 | 0.018400573 | 0.39168193 | 0.04697836 | 0.96253048 | 0.9832997 |
| PUM3 | 1786.67477 | -0.291465301 | 0.17622825 | -1.6539079 | 0.09814622 | 0.2592827 |
| RFX3 | 2822.82669 | 0.288954847 | 0.26851264 | 1.07613128 | 0.28186855 | 0.50430463 |
| RFX3-AS1 | 2325.39135 | 0.015431996 | 0.33955469 | 0.04544775 | 0.96375042 | 0.98401324 |
| GLIS3 | 10436.9176 | -0.302349287 | 0.18853311 | -1.6036933 | 0.10878168 | 0.27717924 |
| SLC1A1 | 647.125301 | -0.775939732 | 0.40937389 | -1.8954304 | 0.0580354 | 0.18392265 |
| SPATA6L | 1938.67391 | -1.35109448 | 0.26111951 | -5.1742379 | 2.29E-07 | 1.23E-05 |
| CDC37L1 | 687.678099 | -0.118353493 | 0.31136319 | -0.380114 | 0.70386082 | 0.84992208 |
| AK3 | 5374.98805 | 0.890225892 | 0.38174754 | 2.33197544 | 0.01970198 | 0.08833791 |
| RCL1 | 1023.90955 | -0.424681973 | 0.25341244 | -1.6758529 | 0.09376702 | 0.25184394 |
| JAK2 | 1179.17374 | -0.458322944 | 0.23944704 | -1.914089 | 0.05560879 | 0.17854558 |
| PLGRKT | 581.500877 | -0.082930811 | 0.20340182 | -0.4077191 | 0.68347991 | 0.83562441 |

| | | | | | | |
|---|---|---|---|---|---|---|
| CD274 | 82.484131 | -1.408036762 | 0.558698 | -2.5202109 | 0.01172845 | 0.06209265 |
| RIC1 | 2074.53064 | 0.183651931 | 0.27931316 | 0.65751264 | 0.51085135 | 0.71294917 |
| ERMP1 | 2136.50204 | -0.172751269 | 0.38012844 | -0.454455 | 0.64950138 | 0.81243027 |
| KIAA2026 | 2899.35559 | -0.163195714 | 0.30005647 | -0.5438833 | 0.58652176 | 0.77036771 |
| MLANA | 43.7616708 | -0.042714278 | 0.70545731 | -0.0605484 | 0.95171891 | 0.97922223 |
| RANBP6 | 216.72961 | -0.246242627 | 0.37968228 | -0.6485492 | 0.51662982 | 0.71787958 |
| UHRF2 | 1147.60167 | 0.159677316 | 0.17031585 | 0.93753645 | 0.34848268 | 0.57409675 |
| AL133480.1 | 131.873695 | 0.326118618 | 0.47715459 | 0.68346532 | 0.49431285 | 0.69960125 |
| GLDC | 735.87666 | -0.301870633 | 0.57261359 | -0.5271804 | 0.59806835 | 0.77678206 |
| AL162411.1 | 250.926821 | -0.614523683 | 0.61748363 | -0.9952064 | 0.31963587 | 0.54422157 |
| KDM4C | 4424.37949 | -0.241030974 | 0.19639598 | -1.2272704 | 0.21972096 | 0.43361796 |
| DMAC1 | 197.725531 | 0.150698409 | 0.30937607 | 0.48710429 | 0.62618443 | 0.7969236 |
| PTPRD | 285.31955 | 0.36440313 | 0.42424482 | 0.85894538 | 0.39037065 | 0.61320085 |
| TYRP1 | 114.376839 | 1.689877214 | 0.43798412 | 3.85830705 | 0.00011418 | 0.0018575 |
| LURAP1L-AS | 74.890405 | -1.120842162 | 0.64417938 | -1.7399535 | 0.08186719 | 0.22998377 |
| LURAP1L | 327.037928 | -2.205799576 | 0.56739205 | -3.887611 | 0.00010124 | 0.00168204 |
| MPDZ | 1702.43031 | 0.711554315 | 0.20068911 | 3.54555524 | 0.00039179 | 0.00491931 |
| NFIB | 3244.67992 | 0.246784596 | 0.25536908 | 0.966384 | 0.33385207 | 0.55927102 |
| ZDHHC21 | 2408.03322 | 0.558392858 | 0.25326478 | 2.20477893 | 0.02746961 | 0.1104666 |
| FREM1 | 1986.35412 | 2.966993812 | 0.48619961 | 6.1024192 | 1.04E-09 | 1.33E-07 |
| TTC39B | 1060.85908 | -0.168706695 | 0.30794003 | -0.5478557 | 0.58379101 | 0.76885817 |
| SNAPC3 | 721.023343 | 0.034108678 | 0.18511802 | 0.18425369 | 0.85381444 | 0.93291993 |
| PSIP1 | 641.950089 | 0.02750798 | 0.20672457 | 0.13306585 | 0.89414131 | 0.95155595 |
| CCDC171 | 1578.23224 | 0.417872515 | 0.28624789 | 1.45982741 | 0.14433751 | 0.33423906 |
| BNC2 | 2211.00232 | -1.230793837 | 0.69454646 | -1.7720828 | 0.07638081 | 0.22033505 |
| CNTLN | 1018.06645 | 0.040503328 | 0.14584209 | 0.27772043 | 0.78122698 | 0.89545532 |
| SH3GL2 | 457.113114 | 0.722371418 | 0.35305493 | 2.04605957 | 0.04075051 | 0.14438667 |
| ADAMTSL1 | 89.1848451 | 0.695105176 | 0.31531177 | 2.20450122 | 0.02748911 | 0.1104666 |
| SAXO1 | 67.4829467 | 0.360817443 | 0.54673563 | 0.65994866 | 0.50928678 | 0.71164382 |
| RRAGA | 162.28843 | -1.097848891 | 0.46654296 | -2.3531572 | 0.01861477 | 0.08488614 |
| HAUS6 | 192.333193 | -0.097396762 | 0.2109499 | -0.4617057 | 0.64429241 | 0.80961167 |
| PLIN2 | 381.502542 | -1.355595441 | 0.58065641 | -2.3345914 | 0.01956478 | 0.08796159 |
| DENND4C | 2937.16621 | -0.254528223 | 0.23553916 | -1.0806196 | 0.27986637 | 0.50229221 |
| RPS6 | 1425.47088 | -1.021495194 | 0.46685637 | -2.1880288 | 0.0286675 | 0.11381696 |
| ACER2 | 104.738992 | -1.620459994 | 0.32220789 | -5.0292375 | 4.92E-07 | 2.34E-05 |
| MLLT3 | 3119.26048 | 0.366627123 | 0.23895562 | 1.53428962 | 0.12495842 | 0.30341466 |
| FOCAD | 2233.62144 | 0.34691534 | 0.14626041 | 2.37190195 | 0.01769679 | 0.08213523 |
| HACD4 | 40.2841798 | 0.04538139 | 0.5253879 | 0.08637692 | 0.93116679 | 0.96970846 |
| KLHL9 | 200.407055 | 0.009155347 | 0.29532779 | 0.03100063 | 0.97526904 | 0.98899114 |
| LINC01239 | 249.696249 | -0.755127516 | 0.43935319 | -1.7187255 | 0.08566437 | 0.23657552 |
| AL391117.1 | 109.272028 | 1.366156793 | 0.50874853 | 2.68532823 | 0.00724586 | 0.0438436 |
| TUSC1 | 62.1123463 | -0.916427744 | 0.45251501 | -2.0251875 | 0.04284811 | 0.14909925 |
| CAAP1 | 1449.66161 | 0.445579582 | 0.21345425 | 2.08747112 | 0.03684557 | 0.13519253 |
| PLAA | 459.295056 | 0.008127704 | 0.22033773 | 0.03688748 | 0.97057472 | 0.98687203 |
| IFT74 | 1412.40759 | 0.302295351 | 0.15203353 | 1.98834664 | 0.04677337 | 0.15866189 |
| LRRC19 | 154.322376 | 0.221763429 | 0.44448882 | 0.4989179 | 0.61783722 | 0.79042527 |
| TEK | 43.2664663 | 0.416434195 | 0.67735228 | 0.61479707 | 0.53868875 | 0.73496797 |
| MOB3B | 1975.52902 | -0.998456848 | 0.34983974 | -2.8540406 | 0.0043167 | 0.03016353 |
| C9orf72 | 428.615376 | -0.142166989 | 0.2635373 | -0.5394568 | 0.58957169 | 0.77174702 |
| LINGO2 | 464.835593 | 1.763201284 | 0.60303356 | 2.92388581 | 0.00345692 | 0.02575398 |
| ACO1 | 388.090465 | 0.109761852 | 0.18168829 | 0.60412178 | 0.54576268 | 0.74015676 |
| DDX58 | 796.691531 | 0.387749949 | 0.67468769 | 0.57471028 | 0.56548726 | 0.75611956 |
| TOPORS | 271.095465 | -0.123773344 | 0.22118792 | -0.5595846 | 0.57576283 | 0.76289009 |
| SMIM27 | 71.8296749 | 0.251115766 | 0.35289843 | 0.71158085 | 0.47672437 | 0.68662656 |
| NDUFB6 | 407.585951 | 0.32356241 | 0.23076019 | 1.4021587 | 0.16086786 | 0.35821031 |
| APTX | 699.869912 | -0.014167053 | 0.22576085 | -0.0627525 | 0.94996361 | 0.97825441 |
| DNAJA1 | 486.24619 | -0.497050541 | 0.31254152 | -1.5903505 | 0.11175583 | 0.28246896 |
| SMU1 | 410.909175 | 0.222042525 | 0.17819859 | 1.24603973 | 0.21274981 | 0.42566075 |
| B4GALT1 | 1068.32251 | -0.432743343 | 0.20278116 | -2.1340412 | 0.03283941 | 0.12562288 |
| BAG1 | 334.952368 | -0.437529266 | 0.33093953 | -1.3220822 | 0.18614078 | 0.3918925 |
| CHMP5 | 311.216577 | -0.082472908 | 0.19051456 | -0.4328956 | 0.66509065 | 0.8233644 |
| NFX1 | 1361.96836 | -0.308615102 | 0.22361665 | -1.3801079 | 0.16755344 | 0.36733831 |
| AQP7 | 51.5434645 | 0.546938018 | 0.70709419 | 0.77350093 | 0.43922598 | 0.65489738 |
| AQP3 | 137.242759 | -0.788703237 | 0.6307802 | -1.2503614 | 0.21116755 | 0.42358574 |
| NOL6 | 45.1965907 | -0.303329782 | 0.36859288 | -0.8229399 | 0.41054218 | 0.62997812 |
| UBE2R2 | 2025.70436 | -0.235223754 | 0.26008047 | -0.9044268 | 0.36576913 | 0.59121933 |
| UBAP2 | 1450.82425 | -0.304180783 | 0.16963123 | -1.7931885 | 0.07294279 | 0.21387375 |
| DCAF12 | 283.369888 | -0.288571976 | 0.25702881 | -1.1227223 | 0.26155546 | 0.48239198 |
| UBAP1 | 2163.70218 | -0.605885894 | 0.21029104 | -2.8811779 | 0.00396192 | 0.02847279 |
| KIF24 | 118.305254 | 0.002644202 | 0.37389466 | 0.00707205 | 0.99435737 | 0.99783229 |
| NUDT2 | 155.640518 | -0.097390666 | 0.39004991 | -0.2496877 | 0.80282887 | 0.9075253 |
| C9orf24 | 58.9628519 | 0.625549084 | 0.40157994 | 1.55771994 | 0.11929965 | 0.2949886 |
| FAM219A | 403.402786 | -0.712481155 | 0.23660422 | -3.0112784 | 0.0026015 | 0.02087682 |
| DNAI1 | 189.406938 | 0.907419964 | 0.44827047 | 2.02426887 | 0.04294249 | 0.14934898 |
| RPP25L | 68.1289483 | -0.090512218 | 0.42735741 | -0.2117951 | 0.83226687 | 0.92160388 |
| DCTN3 | 132.865055 | -0.627846864 | 0.46198882 | -1.3590088 | 0.17414381 | 0.37589771 |
| SIGMAR1 | 50.1143072 | -0.159772949 | 0.36875224 | -0.4332799 | 0.66481141 | 0.82323933 |
| IL11RA | 56.315191 | 0.891820818 | 0.48300869 | 1.84638667 | 0.06483608 | 0.19777071 |
| AL162231.2 | 61.4263006 | 0.309200123 | 0.3368577 | 0.91789538 | 0.35867364 | 0.58403848 |
| VCP | 610.713988 | -0.772664829 | 0.21313627 | -3.6252151 | 0.00028872 | 0.00391954 |
| PIGO | 88.0674096 | 0.021988148 | 0.25013255 | 0.08790598 | 0.9299514 | 0.96915731 |
| STOML2 | 206.115365 | -0.606188351 | 0.39279225 | -1.5432798 | 0.1227629 | 0.30023608 |
| FAM214B | 101.740718 | -0.551538131 | 0.3201819 | -1.7225775 | 0.08496496 | 0.23528348 |
| UNC13B | 1542.38746 | 0.420504492 | 0.20224134 | 2.07922129 | 0.03759702 | 0.1372634 |
| RUSC2 | 982.942346 | 0.421795212 | 0.30873362 | 1.3662108 | 0.17187281 | 0.37294355 |
| TESK1 | 94.533498 | -0.568089911 | 0.32089132 | -1.7703499 | 0.07666887 | 0.22087658 |
| CD72 | 129.245842 | 0.401200652 | 0.37260165 | 1.07675489 | 0.28158978 | 0.50413655 |
| CCDC107 | 199.280554 | -1.622619165 | 0.40663235 | -3.9903839 | 6.60E-05 | 0.00121189 |
| TLN1 | 443.321519 | -0.6790271 | 0.19251312 | -3.527173 | 0.00042002 | 0.00518512 |
| CREB3 | 103.530214 | -1.170040145 | 0.39203549 | -2.9845261 | 0.00284018 | 0.02226465 |
| GBA2 | 514.232579 | 0.48232011 | 0.27891469 | 1.72927465 | 0.08375995 | 0.23315857 |
| RGP1 | 204.926254 | -0.55418915 | 0.17524087 | -3.1624424 | 0.00156452 | 0.01415459 |
| AL133410.1 | 244.001247 | -0.284786742 | 0.23834868 | -1.1948324 | 0.2321525 | 0.44864702 |
| HINT2 | 101.025035 | 0.069280379 | 0.3900322 | 0.17762733 | 0.85901566 | 0.93618423 |
| TMEM8B | 207.339376 | 0.227949648 | 0.30733325 | 0.74170187 | 0.45826799 | 0.67266311 |
| RECK | 389.90321 | 0.031274637 | 0.23798157 | 0.13141621 | 0.89544606 | 0.95183831 |
| CLTA | 1206.25573 | -0.71775324 | 0.20775317 | -3.4548365 | 0.00055063 | 0.00635772 |
| GNE | 907.226882 | -0.476589989 | 0.26376501 | -1.8068734 | 0.07078202 | 0.20977332 |
| RNF38 | 1993.3065 | -0.207373186 | 0.13874245 | -1.4946629 | 0.13500244 | 0.31997977 |
| EBLN3P | 498.690627 | -0.127846392 | 0.19504416 | -0.6554741 | 0.51216257 | 0.71407594 |
| ZCCHC7 | 4631.29125 | 0.105235929 | 0.16349695 | 0.64365683 | 0.51979799 | 0.7198387 |
| GRHPR | 242.58258 | -0.045467042 | 0.25620333 | -0.1774647 | 0.85914341 | 0.93618423 |
| ZBTB5 | 278.28553 | 0.114589204 | 0.15891189 | 0.7210864 | 0.47085636 | 0.68226652 |
| AL138752.2 | 563.052547 | 0.135511068 | 0.20125431 | 0.67333251 | 0.50073578 | 0.70491515 |
| TRMT10B | 545.093302 | 0.391386683 | 0.21103869 | 1.85457313 | 0.06365721 | 0.1955006 |
| DCAF10 | 1416.31515 | 0.007708776 | 0.14215877 | 0.05422652 | 0.95675469 | 0.98096505 |
| FAM95C | 253.435986 | -0.181597636 | 0.30309299 | -0.5991483 | 0.54907402 | 0.74226986 |
| ANKRD18A | 240.701672 | -0.368488401 | 0.3036702 | -1.2134493 | 0.22495808 | 0.43942281 |
| CNTNAP3 | 201.741462 | 0.363093665 | 0.63409445 | 0.57261764 | 0.56690361 | 0.75714258 |
| GLIDR | 374.310314 | 0.798603071 | 0.41368286 | 1.93047173 | 0.05354841 | 0.17404552 |
| FAM95B1 | 48.5246577 | 0.110921795 | 0.81556308 | 0.1360064 | 0.89181622 | 0.95107383 |
| BMS1P14 | 307.135094 | -0.719458815 | 0.53223671 | -1.3517647 | 0.17645059 | 0.37867358 |
| CBWD6 | 138.786555 | 0.555714014 | 0.40461666 | 1.37343335 | 0.16961767 | 0.3699733 |

| | | | | | | |
|---|---|---|---|---|---|---|
| AL591926.2 | 70.4507945 | -0.79274004 | 0.56253001 | -1.4092404 | 0.15876408 | 0.35514221 |
| CNTNAP3B | 281.055604 | -0.413810214 | 0.71058752 | -0.5823494 | 0.56033136 | 0.75183103 |
| FAM27C | 107.579003 | -0.561186 | 0.41609601 | -1.3486935 | 0.17743543 | 0.38004614 |
| AL391987.2 | 100.604597 | -0.366126947 | 0.29363862 | -1.2468624 | 0.21244797 | 0.42518563 |
| AL512625.3 | 139.924688 | 0.875507664 | 0.44747209 | 1.95656376 | 0.05039878 | 0.16713971 |
| AL512625.1 | 40.1814196 | 0.32641512 | 0.60061645 | 0.54346684 | 0.58680842 | 0.77047088 |
| CBWD5 | 456.306006 | 0.127262004 | 0.32512437 | 0.39142561 | 0.69548267 | 0.84441912 |
| ZNF658 | 198.441933 | 0.618360742 | 0.34111386 | 1.81276931 | 0.06986741 | 0.20790237 |
| CBWD3 | 95.0629521 | 0.298534122 | 0.3202832 | 0.93209421 | 0.35128784 | 0.57732528 |
| PGM5 | 385.903575 | -0.295733628 | 1.18637135 | -0.2492758 | 0.80314748 | 0.90758619 |
| AL161457.2 | 142.370325 | 0.315841134 | 0.27352332 | 1.15471373 | 0.24820768 | 0.46734754 |
| AL353616.2 | 493.774316 | -2.912220323 | 0.84547655 | -3.444472 | 0.00057218 | 0.00653483 |
| PIP5K1B | 728.062524 | 0.514362025 | 0.4656582 | 1.10459137 | 0.26933669 | 0.49084828 |
| FAM189A2 | 184.756652 | -1.054350019 | 0.67878353 | -1.5532935 | 0.12035304 | 0.29648328 |
| APBA1 | 83.2293377 | 0.588329292 | 0.3663678 | 1.60584332 | 0.10830834 | 0.27674263 |
| PTAR1 | 1441.71472 | 0.633229902 | 0.15962958 | 3.96687063 | 7.28E-05 | 0.00130762 |
| C9orf135 | 83.7422287 | -0.090245006 | 0.84205351 | -0.1071725 | 0.91465211 | 0.96131774 |
| MAMDC2-AS | 950.794016 | 0.751294299 | 0.25750826 | 2.91755417 | 0.00352788 | 0.0261268 |
| MAMDC2 | 848.070309 | -0.307079772 | 0.46160137 | -0.6652488 | 0.50589135 | 0.70948881 |
| SMC5-AS1 | 1175.13733 | 0.602124151 | 0.20997372 | 2.8676167 | 0.00413576 | 0.02926733 |
| SMC5 | 1912.74832 | -0.199782315 | 0.17948652 | -1.113077 | 0.26567536 | 0.48676739 |
| KLF9 | 806.430764 | 0.309535802 | 0.42412795 | 0.72981703 | 0.46550203 | 0.67845265 |
| TRPM3 | 291.121161 | 0.614320953 | 0.40431368 | 1.5194167 | 0.12865764 | 0.30972094 |
| CEMIP2 | 1484.91121 | -1.619719207 | 0.3798554 | -4.2640416 | 2.01E-05 | 0.00047186 |
| ABHD17B | 358.890666 | 0.221586096 | 0.2847227 | 0.77825231 | 0.43642028 | 0.65341766 |
| C9orf85 | 659.476442 | 0.182012139 | 0.1711918 | 1.06320594 | 0.28768857 | 0.51002814 |
| GDA | 138.704273 | -1.315676324 | 0.63095219 | -2.0852235 | 0.03704902 | 0.13567522 |
| LINC01504 | 135.1045 | -0.18951811 | 1.3858864 | -0.1367487 | 0.89122946 | 0.9506034 |
| ZFAND5 | 1839.00497 | -0.310724291 | 0.41835475 | -0.7427292 | 0.45764565 | 0.67227935 |
| TMC1 | 49.3025596 | -0.928422312 | 0.46093457 | -2.0142171 | 0.04398674 | 0.15199268 |
| ALDH1A1 | 1061.34527 | 0.171804589 | 0.29760803 | 0.5772848 | 0.56374708 | 0.75507353 |
| ANXA1 | 109.89004 | -1.970479503 | 0.84768691 | -2.3245369 | 0.02009674 | 0.08931958 |
| C9orf41-AS1 | 86.4594195 | 0.067856565 | 0.44309092 | 0.15314366 | 0.87828499 | 0.94565262 |
| CARNMT1 | 692.810533 | 0.555905809 | 0.13065265 | 4.25483777 | 2.09E-05 | 0.00048909 |
| NMRK1 | 471.624766 | -0.654525265 | 0.24474708 | -2.6742925 | 0.00748871 | 0.04497657 |
| OSTF1 | 306.845491 | -0.524577709 | 0.2746972 | -1.909658 | 0.05617726 | 0.17980263 |
| PCSK5 | 421.494318 | -2.622717395 | 0.51374503 | -5.1050954 | 3.31E-07 | 1.65E-05 |
| RFK | 155.694782 | -1.460905693 | 0.37567998 | -3.8886972 | 0.00010078 | 0.00167664 |
| GCNT1 | 127.690049 | -1.858329817 | 0.45113538 | -4.1192287 | 3.80E-05 | 0.00078951 |
| PRUNE2 | 630.837776 | -1.781695794 | 0.49113845 | -3.6276854 | 0.00028597 | 0.00389425 |
| PCA3 | 83.1872729 | -2.047739194 | 0.66607043 | -3.0743584 | 0.00210956 | 0.01788521 |
| VPS13A | 1869.44185 | 0.251579893 | 0.16361389 | 1.53764388 | 0.12413571 | 0.30213905 |
| GNA14 | 60.263281 | -0.000950401 | 0.44908414 | -0.0021163 | 0.99831143 | 0.99952215 |
| GNAQ | 4907.12012 | -0.705819121 | 0.17365007 | -4.064606 | 4.81E-05 | 0.00095282 |
| CEP78 | 201.47884 | 0.780226321 | 0.37035825 | 2.10668002 | 0.03514533 | 0.13121385 |
| PSAT1 | 75.0615589 | 2.285521721 | 0.39393321 | 5.80180003 | 6.56E-09 | 6.52E-07 |
| TLE4 | 800.652032 | -0.339517133 | 0.264301 | -1.2845851 | 0.1989373 | 0.40748588 |
| AL161782.1 | 59.2974565 | 0.862602345 | 0.53401537 | 1.6153137 | 0.10624278 | 0.2735063 |
| TLE1 | 958.165211 | -0.286732893 | 0.34405936 | -0.8333821 | 0.40462929 | 0.62539431 |
| AL591368.1 | 94.898428 | 2.277873756 | 0.71780773 | 3.17337593 | 0.00150677 | 0.01379277 |
| RASEF | 348.030971 | -0.592743122 | 0.35971056 | -1.6478336 | 0.09938683 | 0.26151408 |
| AL137847.1 | 81.7745074 | -0.067184168 | 0.49680326 | -0.1352329 | 0.8924277 | 0.95118543 |
| FRMD3 | 719.640702 | 0.268581506 | 0.20995027 | 1.2792625 | 0.20080463 | 0.40989467 |
| AL137847.2 | 57.997891 | 0.023238131 | 0.52178363 | 0.04453595 | 0.96447719 | 0.98434112 |
| IDNK | 176.757679 | -0.04283239 | 0.24538458 | -0.1745521 | 0.8614316 | 0.93697711 |
| UBQLN1 | 1029.65073 | 0.10717924 | 0.18906498 | 0.56689102 | 0.57078823 | 0.75965155 |
| GKAP1 | 662.932589 | 0.559632051 | 0.24811622 | 2.25552384 | 0.02410047 | 0.10099717 |
| AL354733.2 | 48.9871648 | 0.576695482 | 0.42406559 | 1.35992047 | 0.17385509 | 0.3755115 |
| KIF27 | 826.751351 | 0.366841133 | 0.18599609 | 1.97230559 | 0.04857473 | 0.16301413 |
| C9orf64 | 105.10441 | 0.115837349 | 0.24224535 | 0.47818193 | 0.63252072 | 0.80189713 |
| HNRNPK | 1304.9811 | -0.202647417 | 0.26692326 | -0.7591973 | 0.44773455 | 0.66352217 |
| AL354733.3 | 113.190238 | 0.068696368 | 0.53889308 | 0.12747681 | 0.89856303 | 0.95318977 |
| RMI1 | 212.991357 | 0.513996132 | 0.23524319 | 2.18495649 | 0.02889204 | 0.11433041 |
| NTRK2 | 1742.79458 | 0.819260068 | 0.27248373 | 3.00663851 | 0.00264154 | 0.02109556 |
| AGTPBP1 | 1101.9606 | -0.193138868 | 0.23746906 | -0.8133222 | 0.41603334 | 0.63589266 |
| NAA35 | 1220.13333 | 0.121114701 | 0.14233314 | 0.85092414 | 0.3948115 | 0.61706761 |
| GOLM1 | 1493.93579 | -0.579352705 | 0.37156957 | -1.5592039 | 0.11894814 | 0.29433982 |
| C9orf153 | 160.669858 | 1.52080082 | 0.390212 | 3.89737072 | 9.72E-05 | 0.0016448 |
| ISCA1 | 338.451693 | 0.245761048 | 0.28868813 | 0.85130292 | 0.3946011 | 0.61683857 |
| TUT7 | 1029.91175 | -0.375328457 | 0.21790732 | -1.7224224 | 0.08499303 | 0.23531186 |
| AL513318.2 | 192.642533 | -0.025605048 | 0.33689782 | -0.0760024 | 0.93941718 | 0.97411372 |
| DAPK1 | 7442.71946 | -0.783929914 | 0.33349534 | -2.3506473 | 0.01874079 | 0.08530223 |
| CTSL | 407.164323 | 0.108203771 | 0.38754464 | 0.27920337 | 0.78008876 | 0.89508273 |
| CDK20 | 58.7925362 | -0.033600466 | 0.31522796 | -0.106591 | 0.91511345 | 0.96155704 |
| SPIN1 | 2962.32465 | 0.21826126 | 0.17941583 | 1.21651062 | 0.22379047 | 0.4383227 |
| SECISBP2 | 1427.49355 | -0.185891998 | 0.24209197 | -0.7678569 | 0.44257219 | 0.65847444 |
| SEMA4D | 2095.79782 | -0.565042324 | 0.27425199 | -2.0603034 | 0.03936954 | 0.14131036 |
| GADD45G | 45.5295189 | 0.376485256 | 0.61207887 | 0.61509272 | 0.53849349 | 0.73492731 |
| LINC01508 | 62.9540252 | -1.014399163 | 0.82965936 | -1.2226695 | 0.22145456 | 0.43555589 |
| SYK | 417.968751 | -0.231917471 | 0.28836822 | -0.8042408 | 0.42125795 | 0.64039547 |
| AL158071.1 | 265.481274 | 0.773986595 | 0.79039272 | 0.97924307 | 0.32745989 | 0.55270302 |
| AL158071.4 | 798.433841 | 1.002410286 | 0.51118577 | 1.96095106 | 0.04988473 | 0.1661022 |
| AL158071.5 | 66.9195218 | 1.035621349 | 0.43767711 | 2.36617665 | 0.01797287 | 0.08312451 |
| AUH | 3764.0865 | 0.376312633 | 0.25529184 | 1.47404882 | 0.14046845 | 0.32804946 |
| NFIL3 | 368.89816 | -0.385429095 | 0.56072629 | -0.6873748 | 0.4918466 | 0.69798042 |
| SPTLC1 | 758.353957 | -0.119829261 | 0.1489251 | -0.8046277 | 0.42103455 | 0.64038167 |
| IARS | 618.260288 | 0.246308933 | 0.1957239 | 1.25845099 | 0.2082287 | 0.41928245 |
| NOL8 | 293.725531 | -0.034166367 | 0.14339855 | -0.2382616 | 0.81167821 | 0.91159012 |
| CENPP | 617.536236 | 0.584800551 | 0.57987788 | 1.00848915 | 0.31321969 | 0.53703517 |
| IPPK | 240.256913 | 0.422518553 | 0.18516022 | 2.28190778 | 0.02249479 | 0.09644338 |
| BICD2 | 244.977614 | 0.148139695 | 0.26103337 | 0.56751248 | 0.57036605 | 0.75945349 |
| ZNF484 | 156.401735 | 0.058925017 | 0.24297678 | 0.24251296 | 0.80838272 | 0.91010272 |
| CARD19 | 137.481063 | -0.71971426 | 0.21914766 | -3.2841521 | 0.0010229 | 0.01016663 |
| NINJ1 | 246.664068 | -0.197277391 | 0.35507244 | -0.5555976 | 0.57848598 | 0.76458087 |
| WNK2 | 74.9854757 | 0.076494553 | 0.35989011 | 0.21254975 | 0.83167817 | 0.92158196 |
| FAM120AOS | 165.307288 | -0.45432595 | 0.19695584 | -2.3067402 | 0.02106931 | 0.09252144 |
| FAM120A | 1386.58472 | -0.146127199 | 0.13841078 | -1.0557501 | 0.29108239 | 0.51373981 |
| PHF2 | 372.13701 | -0.174707672 | 0.16684469 | -1.0471275 | 0.29504076 | 0.51817489 |
| PTPDC1 | 594.495731 | 0.184259338 | 0.2426748 | 0.759285 | 0.44768209 | 0.66352217 |
| AL158152.2 | 739.472972 | 0.584786929 | 0.19967731 | 2.9286599 | 0.00340427 | 0.02547398 |
| MIRLET7DHG | 57.5572738 | 0.506728384 | 0.74857519 | 0.67692383 | 0.49845428 | 0.70339396 |
| ZNF169 | 292.508049 | 0.321532395 | 0.19943452 | 1.61222033 | 0.106914 | 0.2745387 |
| MFSD14B | 658.851177 | 0.193762143 | 0.22610905 | 0.85694113 | 0.39147742 | 0.6140943 |
| FBP1 | 83.004497 | 0.743956246 | 0.49350922 | 1.50748195 | 0.13168717 | 0.3144921 |
| C9orf3 | 7143.50054 | -0.032631177 | 0.23826334 | -0.1369543 | 0.89106696 | 0.95058176 |
| AL353768.1 | 87.8753264 | -0.763214365 | 0.45952635 | -1.6608718 | 0.09673919 | 0.25695314 |
| FANCC | 1617.2909 | 0.117917569 | 0.21505369 | 0.54831687 | 0.58347435 | 0.76878929 |
| PTCH1 | 403.503364 | -0.235725231 | 0.27284646 | -0.8639483 | 0.3876163 | 0.61088911 |
| LINC00476 | 1320.63137 | 0.552782363 | 0.34428114 | 1.60561324 | 0.10835891 | 0.27674263 |
| ERCC6L2 | 1314.92535 | 0.241568033 | 0.15818423 | 1.5271309 | 0.12672846 | 0.30636096 |
| AL160269.1 | 50.0156766 | 0.131308389 | 0.28509639 | 0.46057542 | 0.64510325 | 0.81007404 |
| SLC35D2 | 284.676943 | -0.135567975 | 0.19327281 | -0.7014332 | 0.48303268 | 0.69127369 |
| ZNF367 | 69.7206355 | 1.575201662 | 0.44468285 | 3.54230362 | 0.00039665 | 0.00493346 |

| | | | | | | |
|---|---|---|---|---|---|---|
| HABP4 | 523.026186 | 0.252285943 | 0.22174129 | 1.13774906 | 0.25522528 | 0.47610094 |
| CDC14B | 1179.30337 | 0.121524009 | 0.36540917 | 0.33256968 | 0.73945914 | 0.87241119 |
| AAED1 | 82.5016073 | -0.743948434 | 0.30620635 | -2.4295657 | 0.01511693 | 0.07377531 |
| AL589843.1 | 218.933053 | -0.385212481 | 0.60071772 | -0.6412537 | 0.52135785 | 0.7213382 |
| ZNF510 | 176.226382 | -0.006844018 | 0.22673076 | -0.0301857 | 0.97591899 | 0.9893257 |
| ZNF782 | 679.370211 | -0.192178695 | 0.24177115 | -0.7948785 | 0.42668417 | 0.64508083 |
| MFSD14C | 1556.8323 | -0.374613839 | 0.20676302 | -1.8118029 | 0.07001665 | 0.20825264 |
| AL590705.1 | 103.439621 | -0.579451461 | 0.38990389 | -1.4861392 | 0.1372423 | 0.32320435 |
| CCDC180 | 185.488125 | 0.18436295 | 0.39263079 | 0.46955805 | 0.6386708 | 0.80559613 |
| TDRD7 | 369.45315 | -0.029782174 | 0.26577733 | -0.1120569 | 0.91077832 | 0.95976856 |
| TMOD1 | 80.6270208 | -4.183791273 | 1.06328725 | -3.9347705 | 8.33E-05 | 0.00146276 |
| TSTD2 | 455.310636 | -0.108620165 | 0.20213471 | -0.5373652 | 0.59101535 | 0.77248385 |
| NCBP1 | 431.157915 | 0.345568417 | 0.27505739 | 1.25635024 | 0.20898901 | 0.42044445 |
| XPA | 789.012209 | 1.483553426 | 0.22518841 | 6.58805406 | 4.46E-11 | 8.92E-09 |
| TRMO | 128.237284 | 0.067891099 | 0.25317922 | 0.26815431 | 0.78858055 | 0.89971274 |
| ANP32B | 463.439151 | -0.408741216 | 0.17838501 | -2.2913429 | 0.02194359 | 0.09480102 |
| NANS | 136.268358 | -0.575392746 | 0.29927182 | -1.9226426 | 0.05452495 | 0.17600687 |
| TRIM14 | 156.599644 | -0.502496066 | 0.61108429 | -0.8223024 | 0.41090483 | 0.63014283 |
| TBC1D2 | 126.517289 | -0.908113735 | 0.35674195 | -2.5455759 | 0.01090976 | 0.05903607 |
| ANKS6 | 515.146693 | 0.343397943 | 0.32820558 | 1.04628915 | 0.29542755 | 0.51851143 |
| GALNT12 | 90.7132056 | -0.908193287 | 0.60455337 | -1.5022549 | 0.13303128 | 0.31689995 |
| TGFBR1 | 610.609923 | -1.063750323 | 0.4061734 | -2.6189562 | 0.00881993 | 0.05053466 |
| ALG2 | 166.898085 | -0.169442862 | 0.30956552 | -0.547357 | 0.58413347 | 0.76889078 |
| SEC61B | 426.551134 | -0.494757404 | 0.26521297 | -1.8655099 | 0.06210997 | 0.19249428 |
| AL359710.1 | 689.990606 | 0.86711258 | 0.3617616 | 2.39691713 | 0.01653366 | 0.0783895 |
| STX17 | 668.389795 | 0.246341616 | 0.1809421 | 1.3614389 | 0.17337503 | 0.37508606 |
| ERP44 | 966.225071 | 0.054867033 | 0.16563246 | 0.33125773 | 0.74044982 | 0.87282504 |
| INVS | 2207.71579 | 0.130812008 | 0.15035904 | 0.86999762 | 0.3843017 | 0.60720588 |
| TEX10 | 383.783094 | 0.281759935 | 0.26418816 | 1.06651233 | 0.28619211 | 0.50853442 |
| MSANTD3 | 72.8315459 | 0.156260555 | 0.39651003 | 0.39408979 | 0.69351475 | 0.84313266 |
| MSANTD3-TI | 50.9955011 | 0.184348191 | 0.4206148 | 0.43828271 | 0.66118136 | 0.82074472 |
| PLPPR1 | 1510.83788 | 2.565545398 | 0.38621687 | 6.6427585 | 3.08E-11 | 6.56E-09 |
| MRPL50 | 136.504982 | 0.125412667 | 0.34832054 | 0.36004959 | 0.71881005 | 0.85990372 |
| ZNF189 | 271.153584 | 0.224411183 | 0.18356465 | 1.22251851 | 0.22151161 | 0.43559181 |
| ALDOB | 359.404896 | -0.150222472 | 0.94668541 | -0.1586826 | 0.87391897 | 0.94333543 |
| TMEM246 | 63.235942 | -0.540491289 | 0.53113877 | -1.0176084 | 0.30886408 | 0.53272133 |
| RNF20 | 230.848022 | 0.38460749 | 0.17951496 | 2.14248157 | 0.03215475 | 0.12379251 |
| SMC2 | 161.237216 | 0.31065564 | 0.29009811 | 1.07086407 | 0.28423056 | 0.50680365 |
| NIPSNAP3A | 186.914926 | -0.014112581 | 0.25167991 | -0.0560735 | 0.95528323 | 0.98014107 |
| NIPSNAP3B | 105.673124 | 0.32348332 | 0.31124547 | 1.03931897 | 0.29865642 | 0.52181158 |
| ABCA1 | 586.481808 | -0.430549088 | 0.35855812 | -1.2007791 | 0.22983691 | 0.44541017 |
| SLC44A1 | 1094.67528 | -0.335992423 | 0.21601915 | -1.5553826 | 0.11985498 | 0.29558709 |
| FSD1L | 625.850358 | -0.041423361 | 0.17479582 | -0.2369814 | 0.8126712 | 0.91196992 |
| FKTN | 646.685377 | 0.313263617 | 0.21109727 | 1.48397757 | 0.13781487 | 0.32391399 |
| TMEM38B | 643.890141 | 0.932654167 | 0.2169397 | 4.29914006 | 1.71E-05 | 0.00041942 |
| LINC01505 | 1056.37684 | 0.365657602 | 0.39232023 | 0.93203862 | 0.35131657 | 0.57732528 |
| ZNF462 | 1207.1416 | 0.505122846 | 0.29720307 | 1.69958824 | 0.0892084 | 0.24300965 |
| AL807761.4 | 47.9805868 | 0.161811456 | 0.62072747 | 0.26068035 | 0.79433902 | 0.90358458 |
| RAD23B | 1238.61168 | -0.034036266 | 0.15706723 | -0.2166987 | 0.82844315 | 0.92023714 |
| ELP1 | 460.799836 | 0.404034262 | 0.13206752 | 3.05930084 | 0.00221854 | 0.01856273 |
| FAM206A | 145.338781 | -0.005014662 | 0.20229717 | -0.0247886 | 0.98022359 | 0.99141592 |
| CTNNAL1 | 872.453265 | 0.416993437 | 0.34672876 | 1.20265026 | 0.22911169 | 0.44457881 |
| TMEM245 | 1951.47853 | 0.203815392 | 0.19199733 | 1.06155327 | 0.28843854 | 0.51092727 |
| EPB41L4B | 494.463496 | 0.538308681 | 0.28346326 | 1.89904215 | 0.05755893 | 0.18298338 |
| PTPN3 | 3275.75017 | 0.178243005 | 0.23901092 | 0.74575256 | 0.4558169 | 0.67098931 |
| PALM2 | 718.994354 | 0.221595714 | 0.37601254 | 0.58933064 | 0.55563949 | 0.7478543 |
| PALM2-AKAF | 201.246502 | 0.202712914 | 0.39223778 | 0.51681129 | 0.6052879 | 0.78170198 |
| TXN | 407.916919 | -0.499364226 | 0.24495999 | -2.0385542 | 0.04149454 | 0.14589537 |
| LPAR1 | 596.79281 | -0.091652257 | 0.25449224 | -0.3601377 | 0.71874414 | 0.85990372 |
| ECPAS | 1683.40421 | 0.058265804 | 0.1692988 | 0.34415957 | 0.73072628 | 0.86804546 |
| ZNF483 | 679.019509 | -0.239673802 | 0.23384336 | -1.0249331 | 0.30539474 | 0.52869714 |
| PTGR1 | 975.466385 | -0.240007956 | 0.18900481 | -1.2698511 | 0.20413768 | 0.41401116 |
| C9orf84 | 349.122706 | -0.547786511 | 0.56175194 | -0.9751395 | 0.32949106 | 0.55488483 |
| UGCG | 2554.47838 | -1.556504904 | 0.37632809 | -4.1360317 | 3.53E-05 | 0.00074562 |
| SUSD1 | 488.922791 | -0.759641164 | 0.34065526 | -2.2299411 | 0.02575135 | 0.10560374 |
| PTBP3 | 1612.48156 | -1.130594942 | 0.28937679 | -3.9069995 | 9.34E-05 | 0.00159069 |
| HSDL2 | 1343.44945 | 0.624620641 | 0.28719963 | 2.17486575 | 0.02964018 | 0.11658408 |
| C9orf147 | 309.249262 | 0.486178575 | 0.22272884 | 2.18282726 | 0.02904853 | 0.11477777 |
| KIAA1958 | 4008.12715 | 0.267869754 | 0.15457344 | 1.73296112 | 0.08310258 | 0.23192364 |
| INIP | 405.542858 | 0.446079054 | 0.17444093 | 2.55719262 | 0.01055208 | 0.05767331 |
| SNX30 | 498.958905 | 0.075910647 | 0.18247111 | 0.4160146 | 0.67739932 | 0.83080479 |
| ZNF883 | 50.6753866 | 0.085773186 | 0.44970544 | 0.19073193 | 0.84873562 | 0.9303692 |
| ZFP37 | 65.1082134 | 0.161051744 | 0.38409334 | 0.41930367 | 0.67499422 | 0.82939913 |
| FKBP15 | 382.553787 | 0.113254587 | 0.15344829 | 0.73806351 | 0.46047585 | 0.67395297 |
| SLC31A1 | 265.978973 | 0.123291039 | 0.20411724 | 0.60402069 | 0.54582989 | 0.74015676 |
| CDC26 | 174.653441 | -0.07390229 | 0.34943698 | -0.2114896 | 0.83250524 | 0.92169608 |
| PRPF4 | 189.00501 | -0.423409933 | 0.39723074 | -1.0659043 | 0.28646692 | 0.50879744 |
| WDR31 | 189.227914 | 0.16130261 | 0.31356669 | 0.51441245 | 0.60696366 | 0.78294756 |
| BSPRY | 250.776893 | -0.31620162 | 0.25745472 | -1.2281834 | 0.21937811 | 0.43356369 |
| HDHD3 | 333.134021 | -0.303324988 | 0.36349172 | -0.8344756 | 0.40401301 | 0.62518836 |
| ALAD | 132.436389 | 0.611670525 | 0.30238161 | 2.02284297 | 0.04308934 | 0.14970202 |
| POLE3 | 60.1020314 | -0.772403474 | 0.3291579 | -2.3466047 | 0.01894534 | 0.08596669 |
| RGS3 | 1130.02403 | -0.405306337 | 0.3894883 | -1.0406124 | 0.29805549 | 0.52143889 |
| ZNF618 | 2562.09376 | 0.625166441 | 0.26183259 | 2.38765706 | 0.01695616 | 0.079621 |
| KIF12 | 1078.67997 | 0.763174183 | 0.41796082 | 1.82594671 | 0.06785829 | 0.20403828 |
| COL27A1 | 1067.06002 | -0.701240585 | 0.41252662 | -1.6998675 | 0.08915585 | 0.24298145 |
| AKNA | 79.6629161 | -1.102890517 | 0.32971598 | -3.3449714 | 0.00082291 | 0.00860562 |
| WHRN | 501.965017 | -0.593935371 | 0.26840196 | -2.2128578 | 0.02690745 | 0.10889109 |
| ATP6V1G1 | 523.985672 | -0.915961469 | 0.29867621 | -3.0667372 | 0.00216409 | 0.01821891 |
| TMEM268 | 118.243478 | 0.150342372 | 0.46059656 | 0.32640794 | 0.74411573 | 0.87481257 |
| TNFSF15 | 52.3008217 | -2.243582282 | 0.57417515 | -3.9074876 | 9.33E-05 | 0.00158953 |
| TNC | 128.77585 | -1.628190812 | 0.9054773 | -1.7981575 | 0.07215205 | 0.21226201 |
| PAPPA | 638.484414 | 0.06566513 | 0.34756186 | 0.18893077 | 0.85014707 | 0.93129812 |
| AL137024.1 | 136.176557 | 0.58502474 | 0.47260446 | 1.23787394 | 0.21576281 | 0.42958827 |
| ASTN2 | 4916.2306 | 0.569966628 | 0.40444115 | 1.40926962 | 0.15875546 | 0.35514221 |
| ASTN2-AS1 | 143.324282 | -0.056510925 | 0.72050948 | -0.0784319 | 0.9374845 | 0.97284815 |
| AL133284.1 | 40.3442013 | -0.03931411 | 0.83301725 | -0.0471948 | 0.96235795 | 0.9832997 |
| TRIM32 | 75.499051 | -0.166297496 | 0.36724129 | -0.452829 | 0.6506719 | 0.8132783 |
| CDK5RAP2 | 676.540614 | 0.254547297 | 0.20590647 | 1.23622776 | 0.21637391 | 0.43031868 |
| MEGF9 | 781.790366 | 0.001921262 | 0.22869187 | 0.00840109 | 0.99329698 | 0.99737436 |
| FBXW2 | 501.243091 | 0.075033666 | 0.16721522 | 0.4487251 | 0.65362998 | 0.81543241 |
| PSMD5 | 440.16694 | -0.08056588 | 0.22595492 | -0.3565573 | 0.72142324 | 0.86136487 |
| TRAF1 | 57.6091925 | 0.161616648 | 0.39628824 | 0.40782601 | 0.68340142 | 0.83562441 |
| C5 | 119.234723 | 0.91705563 | 0.32481752 | 2.82329488 | 0.00475328 | 0.03216502 |
| CNTRL | 359.350451 | 0.043924115 | 0.27599384 | 0.15914889 | 0.87355157 | 0.94324717 |
| RAB14 | 706.835336 | -0.058811135 | 0.18618694 | -0.3158714 | 0.75210009 | 0.8795476 |
| GSN | 1359.23147 | -1.119096176 | 0.39555276 | -2.8291957 | 0.00466652 | 0.03175016 |
| STOM | 521.360858 | -2.20905347 | 0.4450039 | -4.9641216 | 6.90E-07 | 3.09E-05 |
| AL359644.1 | 97.9171103 | -0.215078898 | 0.57632674 | -0.3731892 | 0.70900766 | 0.85387328 |
| DAB2IP | 1692.1447 | -0.844756526 | 0.43635392 | -1.9359435 | 0.05287462 | 0.17257744 |
| TTLL11 | 313.613537 | 0.455288959 | 0.20309294 | 2.24177645 | 0.02497582 | 0.10358106 |
| NDUFA8 | 266.740704 | 0.565298645 | 0.20458293 | 2.76317596 | 0.00572419 | 0.0371005 |
| RBM18 | 157.289452 | -0.063992939 | 0.27758377 | -0.2305356 | 0.81767561 | 0.91531162 |

| | | | | | | |
|---|---|---|---|---|---|---|
| MRRF | 865.493004 | 0.378189543 | 0.17558213 | 2.15391824 | 0.03124658 | 0.12132162 |
| PDCL | 156.416963 | -0.444070884 | 0.20759762 | -2.1390943 | 0.03242803 | 0.12440948 |
| RC3H2 | 906.383462 | -0.033776264 | 0.11785772 | -0.2865851 | 0.77443005 | 0.8920261 |
| ZBTB6 | 53.835585 | 0.181804935 | 0.25653371 | 0.70869803 | 0.47851189 | 0.68816298 |
| ZBTB26 | 144.813019 | 0.209973635 | 0.24690686 | 0.85041637 | 0.39509364 | 0.61719685 |
| RABGAP1 | 2339.03031 | 0.20453681 | 0.1470154 | 1.39126112 | 0.16414626 | 0.36287999 |
| STRBP | 2252.74676 | 0.494254658 | 0.18268206 | 2.70554574 | 0.00681923 | 0.04208894 |
| DENND1A | 3268.60788 | 0.584709296 | 0.16519395 | 3.53953218 | 0.00040084 | 0.00498086 |
| AL390774.2 | 58.8352796 | 0.139456818 | 0.68361402 | 0.20399935 | 0.838354 | 0.92456015 |
| NEK6 | 273.374495 | -2.731640555 | 0.70111491 | -3.8961382 | 9.77E-05 | 0.00165093 |
| PSMB7 | 647.963572 | -0.232882479 | 0.17907348 | -1.3004856 | 0.19343459 | 0.40113679 |
| NR6A1 | 4191.84742 | -0.022998129 | 0.26016306 | -0.0883989 | 0.92955963 | 0.96910444 |
| MIR181A2HC | 564.308471 | -0.008238533 | 0.41291131 | -0.0199523 | 0.98408142 | 0.99355893 |
| RPL35 | 594.014849 | -0.843911553 | 0.42288187 | -1.9956201 | 0.04597529 | 0.15675984 |
| ARPC5L | 266.164306 | -0.626801326 | 0.31231216 | -2.0069706 | 0.04475279 | 0.15418352 |
| GOLGA1 | 501.690332 | 0.47439458 | 0.29962511 | 1.58329378 | 0.11335452 | 0.28558074 |
| SCAI | 859.9686 | 0.498539956 | 0.27853779 | 1.78984674 | 0.07347855 | 0.21506275 |
| PPP6C | 689.225332 | 0.16700304 | 0.17635687 | 0.94696077 | 0.34365877 | 0.56845159 |
| RABEPK | 209.894981 | 0.015707596 | 0.41806272 | 0.03757234 | 0.97002866 | 0.98653537 |
| HSPA5 | 677.852273 | -0.569121991 | 0.52574104 | -1.0825139 | 0.27902424 | 0.50110553 |
| GAPVD1 | 1191.53523 | 0.017479774 | 0.12752143 | 0.13707323 | 0.89097291 | 0.95055821 |
| MAPKAP1 | 2069.31496 | -0.135106272 | 0.17373498 | -0.7776573 | 0.4367711 | 0.65344993 |
| PBX3 | 1656.26902 | -0.614338529 | 0.44430836 | -1.3826851 | 0.16676146 | 0.36640505 |
| MVB12B | 376.003062 | 0.324597656 | 0.18497184 | 1.75484908 | 0.07928513 | 0.22502734 |
| ZBTB43 | 649.404382 | -0.01883397 | 0.21159024 | -0.0890115 | 0.92907275 | 0.96900671 |
| ZBTB34 | 151.385731 | 0.120609743 | 0.23743021 | 0.50797976 | 0.61146754 | 0.78684784 |
| RALGPS1 | 2456.8131 | 0.31875699 | 0.22141423 | 1.43964094 | 0.14996901 | 0.34213137 |
| ANGPTL2 | 58.725755 | -0.90600439 | 0.54377717 | -1.6661317 | 0.09568718 | 0.25497922 |
| GARNL3 | 333.895076 | -0.21982366 | 0.32378879 | -0.6789107 | 0.49719447 | 0.70224997 |
| SLC2A8 | 58.154175 | -0.190865353 | 0.31422606 | -0.6074142 | 0.5435761 | 0.7391242 |
| ZNF79 | 68.824888 | 0.09406209 | 0.30743991 | 0.30595276 | 0.75964061 | 0.88355676 |
| RPL12 | 287.294268 | -0.556773479 | 0.34629166 | -1.6078166 | 0.10787535 | 0.27604136 |
| LRSAM1 | 591.361365 | -0.179673087 | 0.24279648 | -0.7400152 | 0.45929076 | 0.67296414 |
| FAM129B | 791.979627 | -2.185134633 | 0.35789568 | -6.1055071 | 1.02E-09 | 1.31E-07 |
| STXBP1 | 448.302614 | -1.023819793 | 0.32652175 | -3.1355333 | 0.00171542 | 0.01515651 |
| PTRH1 | 109.432763 | -0.871878036 | 0.26520058 | -3.2876174 | 0.00101039 | 0.01006504 |
| CDK9 | 215.325733 | 0.128995729 | 0.22660802 | 0.5692461 | 0.56918914 | 0.75882311 |
| FPGS | 316.040908 | -0.903197028 | 0.33547561 | -2.6922882 | 0.00709636 | 0.04313659 |
| ENG | 109.961057 | -0.319530884 | 0.21285783 | -1.501147 | 0.13331755 | 0.3174823 |
| AL162586.1 | 90.1903189 | 0.571705985 | 0.45935616 | 1.24458107 | 0.21328578 | 0.42637697 |
| ST6GALNAC4 | 47.4885633 | -1.701850902 | 0.5085021 | -3.3467923 | 0.00081752 | 0.00857038 |
| DPM2 | 143.527247 | -0.94619628 | 0.34883903 | -2.7124152 | 0.00667949 | 0.04144169 |
| FAM102A | 338.608902 | -0.73375333 | 0.51036264 | -1.4377097 | 0.15051643 | 0.34284731 |
| SLC25A25 | 369.194883 | -0.213051152 | 0.40603651 | -0.5247093 | 0.59978526 | 0.77794221 |
| SLC25A25-AS | 202.651106 | 0.516545753 | 0.33965071 | 1.52081459 | 0.12830637 | 0.30910065 |
| PTGES2 | 266.74749 | -0.386640683 | 0.25468182 | -1.5181323 | 0.12898106 | 0.31029263 |
| LCN2 | 200.557956 | -7.113717453 | 0.84217272 | -8.4468628 | 2.99E-17 | 3.59E-14 |
| C9orf16 | 259.497901 | -1.42259852 | 0.38130526 | -3.7308652 | 0.00019082 | 0.00282249 |
| CIZ1 | 247.79584 | -0.103126285 | 0.3049899 | -0.3381302 | 0.7352651 | 0.87026136 |
| DNM1 | 240.844049 | -0.097720059 | 0.38170364 | -0.2560103 | 0.79794289 | 0.90506918 |
| GOLGA2 | 591.768909 | -0.271997309 | 0.19665887 | -1.383092 | 0.16663666 | 0.36635414 |
| SWI5 | 111.149426 | -0.475085335 | 0.25152754 | -1.8888004 | 0.05891857 | 0.18551977 |
| TRUB2 | 138.315414 | -0.194457075 | 0.21983741 | -0.8845495 | 0.37639965 | 0.59959757 |
| COQ4 | 706.577494 | 0.394045247 | 0.22431692 | 1.75664526 | 0.07897829 | 0.2246842 |
| SLC27A4 | 286.990117 | -0.041061331 | 0.27441797 | -0.1496306 | 0.88105605 | 0.94645136 |
| URM1 | 210.124736 | -0.495671048 | 0.2295572 | -2.1592485 | 0.03083089 | 0.12013133 |
| CERCAM | 53.9156486 | -0.291514752 | 0.42575163 | -0.6847061 | 0.49352938 | 0.69901668 |
| ODF2 | 336.459982 | -0.408117286 | 0.24078204 | -1.6949656 | 0.09008194 | 0.2447331 |
| GLE1 | 362.550163 | 0.279725381 | 0.25060463 | 1.11620195 | 0.2643357 | 0.48528287 |
| AL356481.1 | 49.6021543 | -0.002152167 | 0.49921152 | -0.0043111 | 0.99656023 | 0.99867719 |
| SPTAN1 | 1142.05893 | 0.153273557 | 0.36425101 | 0.42079103 | 0.67390769 | 0.82906274 |
| WDR34 | 312.441273 | 0.184237803 | 0.28995609 | 0.63539898 | 0.52516824 | 0.72440969 |
| SET | 602.089102 | -0.910008384 | 0.27300249 | -3.3333336 | 0.00085812 | 0.00884848 |
| PKN3 | 48.9158795 | -1.483237943 | 0.326344 | -4.5450137 | 5.49E-06 | 0.00016416 |
| ZDHHC12 | 98.7753541 | -1.795129297 | 0.44798076 | -4.0071571 | 6.15E-05 | 0.00115076 |
| ZER1 | 1126.11216 | -0.271866796 | 0.25725679 | -1.0567915 | 0.29060673 | 0.51339098 |
| TBC1D13 | 200.956318 | 0.11287556 | 0.20611883 | 0.54762373 | 0.58395029 | 0.76885817 |
| ENDOG | 121.39909 | 0.070958774 | 0.37359781 | 0.18993359 | 0.84936117 | 0.93082317 |
| SPOUT1 | 77.5418377 | -0.193286394 | 0.32190227 | -0.6004505 | 0.54820601 | 0.74186733 |
| KYAT1 | 268.690841 | 1.053092288 | 0.19815984 | 5.31435769 | 1.07E-07 | 6.72E-06 |
| LRRC8A | 124.322884 | -0.352107297 | 0.19325782 | -1.8219562 | 0.06846163 | 0.20524503 |
| PHYHD1 | 242.968285 | 0.046169486 | 0.2596824 | 0.17779213 | 0.85888623 | 0.93618423 |
| NUP188 | 186.841342 | 0.004743517 | 0.32321911 | 0.01467586 | 0.98829078 | 0.99552604 |
| SH3GLB2 | 350.675105 | 0.295055309 | 0.26544432 | 1.11155253 | 0.26633059 | 0.48748675 |
| MIGA2 | 298.156565 | 0.926986227 | 0.16503583 | 5.61687853 | 1.94E-08 | 1.61E-06 |
| CRAT | 84.7949004 | -0.180583622 | 0.27064202 | -0.6672416 | 0.50461781 | 0.70856773 |
| PTPA | 440.530846 | -0.816069452 | 0.30828383 | -2.6471368 | 0.00811765 | 0.04776216 |
| IER5L | 393.36938 | -2.073389954 | 0.38804907 | -5.3431128 | 9.14E-08 | 5.97E-06 |
| LINC01503 | 979.803931 | -2.253045557 | 0.63573455 | -3.5440036 | 0.0003941 | 0.00492029 |
| NTMT1 | 212.836671 | -0.649223014 | 0.21046669 | -3.0846829 | 0.00203769 | 0.01741002 |
| ASB6 | 123.415693 | -0.102592272 | 0.34309522 | -0.2990198 | 0.76492492 | 0.88607325 |
| TOR1B | 178.358015 | -0.005506538 | 0.2815955 | -0.0195548 | 0.98439854 | 0.99380314 |
| TOR1A | 82.9140256 | -0.721527948 | 0.29767121 | -2.4239091 | 0.01535445 | 0.07456505 |
| C9orf78 | 200.713361 | -0.509431297 | 0.26028509 | -1.9572051 | 0.05032337 | 0.16698912 |
| USP20 | 305.193895 | -0.094269576 | 0.20730216 | -0.4547448 | 0.64929286 | 0.81224637 |
| FNBP1 | 1397.83473 | -0.246194499 | 0.22548361 | -1.091851 | 0.27489862 | 0.49647832 |
| GPR107 | 778.287535 | -0.004499007 | 0.18296715 | -0.0245891 | 0.98038267 | 0.99141592 |
| NCS1 | 307.302454 | 0.425197905 | 0.35376531 | 1.20192086 | 0.22939419 | 0.44481325 |
| ASS1 | 208.191515 | -0.60731728 | 0.47534968 | -1.2776221 | 0.20138269 | 0.41056705 |
| FUBP3 | 791.372067 | 0.046850562 | 0.18608308 | 0.25177228 | 0.80121708 | 0.90672348 |
| EXOSC2 | 93.1305638 | 0.254929337 | 0.26076482 | 0.97762166 | 0.32826148 | 0.55346341 |
| ABL1 | 2065.34979 | 0.363605756 | 0.12488028 | 2.91163461 | 0.00359543 | 0.02651705 |
| AIF1L | 1200.98659 | 0.038016324 | 0.3810047 | 0.09977915 | 0.92051966 | 0.96454603 |
| NUP214 | 1804.78082 | 0.296044327 | 0.16416543 | 1.80332926 | 0.07133652 | 0.21075466 |
| AL157938.2 | 498.858573 | 0.295473502 | 0.52214317 | 0.56588599 | 0.57147129 | 0.75979509 |
| AL157938.3 | 133.473063 | 0.405643306 | 0.37822641 | 1.07248805 | 0.28350088 | 0.5059807 |
| PLPP7 | 79.2801277 | 0.398851253 | 0.30453178 | 1.30971964 | 0.1902907 | 0.39664665 |
| PRRC2B | 2062.41301 | -0.354906401 | 0.18485203 | -1.9199487 | 0.05486438 | 0.17680011 |
| POMT1 | 197.533425 | 0.466865226 | 0.2208907 | 2.11355761 | 0.03455307 | 0.12988375 |
| UCK1 | 95.8625202 | -0.404811255 | 0.25203753 | -1.6061547 | 0.10823993 | 0.27665271 |
| RAPGEF1 | 1691.37415 | 0.259411075 | 0.28816113 | 0.90022925 | 0.36799826 | 0.59306694 |
| MED27 | 1156.0967 | -0.033444689 | 0.1477878 | -0.2263021 | 0.82096648 | 0.91698603 |
| SETX | 1724.1758 | -0.104589682 | 0.17516383 | -0.5970964 | 0.55044305 | 0.74358788 |
| TTF1 | 345.108544 | -0.164034519 | 0.22977975 | -0.7138772 | 0.47530314 | 0.68532684 |
| CFAP77 | 45.3506784 | 0.06059527 | 0.49192621 | 0.12317959 | 0.90196489 | 0.95457905 |
| DDX31 | 443.217685 | -0.104088456 | 0.2267414 | -0.4590624 | 0.64618934 | 0.81074421 |
| GTF3C4 | 163.215418 | -0.045009918 | 0.20824062 | -0.2161438 | 0.82887566 | 0.92023714 |
| AK8 | 212.0951 | 0.214786722 | 0.36698167 | 0.58527916 | 0.55836002 | 0.75039889 |
| SPACA9 | 82.8081878 | 0.121673846 | 0.37773627 | 0.32211322 | 0.74736692 | 0.87680697 |
| TSC1 | 911.247036 | 0.611053967 | 0.19898648 | 3.07083151 | 0.00213464 | 0.01803711 |
| GTF3C5 | 211.506497 | -0.220295049 | 0.27649661 | -0.7967369 | 0.42560385 | 0.64411105 |
| RALGDS | 105.32469 | -0.514356628 | 0.29092035 | -1.7680325 | 0.07705547 | 0.22150724 |
| ABO | 587.155274 | 0.029878864 | 0.46693045 | 0.06398997 | 0.94897822 | 0.97784954 |

| | | | | | | |
|---|---|---|---|---|---|---|
| SURF6 | 121.018159 | -0.477345267 | 0.1932936 | -2.4695347 | 0.01352889 | 0.06810331 |
| MED22 | 93.3271035 | -0.484984373 | 0.25901175 | -1.8724416 | 0.06114554 | 0.19026418 |
| RPL7A | 706.78801 | -0.963053587 | 0.45709379 | -2.1069059 | 0.03512574 | 0.13117781 |
| SURF1 | 141.874078 | 0.029046977 | 0.26453126 | 0.10980546 | 0.91256366 | 0.96042853 |
| SURF2 | 70.6662192 | -0.695373734 | 0.37855984 | -1.8368925 | 0.06622577 | 0.20049878 |
| SURF4 | 510.534962 | -0.316131234 | 0.251138 | -1.2587949 | 0.20810442 | 0.41916001 |
| REXO4 | 160.347606 | -0.128678666 | 0.22872875 | -0.562582 | 0.57371958 | 0.76125182 |
| ADAMTS13 | 80.4097663 | 0.656428236 | 0.38499018 | 1.7050519 | 0.08818475 | 0.24141602 |
| CACFD1 | 115.586834 | -0.452854422 | 0.35587372 | -1.2725144 | 0.20319042 | 0.41285069 |
| ADAMTSL2 | 93.1632259 | 2.425626719 | 0.76343767 | 3.17724263 | 0.00148683 | 0.01362907 |
| VAV2 | 1000.55643 | -0.764256713 | 0.25676451 | -2.9764889 | 0.0029157 | 0.02262834 |
| BRD3OS | 89.4192295 | -0.530431173 | 0.32738883 | -1.6201871 | 0.10519209 | 0.27194033 |
| BRD3 | 308.786092 | -0.243208493 | 0.15982435 | -1.5217236 | 0.12807834 | 0.30877657 |
| WDR5 | 215.990668 | -0.080325404 | 0.20915697 | -0.3840436 | 0.70094611 | 0.84841905 |
| RXRA | 323.997046 | 0.261007389 | 0.34168614 | 0.76388053 | 0.44493844 | 0.66082438 |
| COL5A1 | 74.4913034 | 0.039188167 | 0.84608547 | 0.04631703 | 0.96305757 | 0.98360994 |
| PPP1R26-AS1 | 334.687843 | 0.221243925 | 0.45113797 | 0.490413 | 0.62384168 | 0.79539814 |
| PPP1R26 | 135.973609 | 0.106199739 | 0.23263766 | 0.45650278 | 0.64802847 | 0.81173905 |
| MRPS2 | 100.346502 | -0.302218332 | 0.39666939 | -0.7618897 | 0.44612583 | 0.66204652 |
| CAMSAP1 | 431.858286 | -0.620865769 | 0.21634961 | -2.8697338 | 0.00410818 | 0.02920608 |
| AL355574.1 | 43.9884616 | -0.666088027 | 0.42108937 | -1.5818211 | 0.11369043 | 0.28604511 |
| UBAC1 | 421.949588 | 0.136992747 | 0.2125287 | 0.6445847 | 0.51919635 | 0.71956991 |
| NACC2 | 421.646397 | -0.55315958 | 0.2373651 | -2.3304167 | 0.01978414 | 0.0885424 |
| TMEM250 | 64.1605887 | -0.629956856 | 0.39981545 | -1.5756191 | 0.11511361 | 0.28847195 |
| QSOX2 | 119.812377 | -0.017593425 | 0.28565491 | -0.0615898 | 0.95088951 | 0.97879278 |
| SNAPC4 | 188.765492 | -0.414845725 | 0.21112589 | -1.9649211 | 0.04942336 | 0.16493559 |
| ENTR1 | 189.976121 | -0.132218009 | 0.31631305 | -0.4179973 | 0.67594907 | 0.82991711 |
| PMPCA | 192.872152 | -0.320124515 | 0.24264316 | -1.3193222 | 0.1870614 | 0.39270408 |
| INPP5E | 115.041726 | 0.173156183 | 0.32850921 | 0.52709689 | 0.5981263 | 0.77678206 |
| SEC16A | 482.833545 | -0.456210589 | 0.21861716 | -2.0868014 | 0.03690609 | 0.13533942 |
| NOTCH1 | 350.804951 | -0.905904719 | 0.29874616 | -3.032356 | 0.00242653 | 0.01974862 |
| EGFL7 | 69.6233357 | -1.023810472 | 0.41540488 | -2.4646086 | 0.0137163 | 0.06872334 |
| AGPAT2 | 362.813265 | -0.908483779 | 0.44890511 | -2.0237768 | 0.04299312 | 0.14944634 |
| FAM69B | 90.301619 | 1.311386214 | 0.57750715 | 2.27077051 | 0.02316087 | 0.09843371 |
| SNHG7 | 289.495545 | -0.208713141 | 0.40435258 | -0.5161662 | 0.60573833 | 0.78178941 |
| RABL6 | 686.195671 | 0.043794228 | 0.23517297 | 0.18622136 | 0.85227118 | 0.93200182 |
| AJM1 | 82.3534623 | 0.942085224 | 0.61361189 | 1.53531124 | 0.12470739 | 0.30308371 |
| PHPT1 | 419.535158 | -0.007682703 | 0.43490757 | -0.0176651 | 0.98590599 | 0.99426405 |
| MAMDC4 | 45.3173922 | 1.73422278 | 0.53684326 | 3.23040801 | 0.00123614 | 0.01179778 |
| EDF1 | 632.295925 | -0.215390065 | 0.33152423 | -0.6496963 | 0.51588841 | 0.71730211 |
| TRAF2 | 111.716533 | -0.315054754 | 0.26515166 | -1.1882059 | 0.23475231 | 0.45204629 |
| FBXW5 | 324.958703 | -0.544218129 | 0.33910408 | -1.6048705 | 0.10852231 | 0.27683877 |
| LCN12 | 111.795543 | 1.541089251 | 0.61675754 | 2.49869544 | 0.01246514 | 0.06456176 |
| PAXX | 149.118402 | 0.348072226 | 0.45962121 | 0.75730235 | 0.44886874 | 0.66462361 |
| ABCA2 | 171.584657 | 0.859730938 | 0.49748168 | 1.72816602 | 0.08395846 | 0.23347383 |
| UAP1L1 | 87.4455719 | 1.24792485 | 0.55573553 | 2.24553726 | 0.02473366 | 0.1029446 |
| MAN1B1 | 249.73428 | -0.404276495 | 0.28741357 | -1.406602 | 0.15954545 | 0.3562867 |
| DPP7 | 219.391113 | -0.552652579 | 0.34391103 | -1.6069638 | 0.10806231 | 0.27630566 |
| ANAPC2 | 45.5760375 | 0.106379023 | 0.34148134 | 0.31152222 | 0.75540366 | 0.88163716 |
| SSNA1 | 203.808639 | -1.044057381 | 0.34316922 | -3.0423981 | 0.00234701 | 0.01929165 |
| TPRN | 193.349802 | -0.166187735 | 0.27814616 | -0.5974835 | 0.55018464 | 0.74339084 |
| TMEM203 | 87.8639214 | -0.718571551 | 0.41585844 | -1.7279235 | 0.08400195 | 0.23347383 |
| NDOR1 | 93.772072 | -0.356869828 | 0.34621152 | -1.0307855 | 0.30264141 | 0.52586237 |
| BX255925.3 | 75.1616761 | -0.170371227 | 0.450523 | -0.3781632 | 0.70530935 | 0.85100486 |
| RNF208 | 44.5806961 | 0.004739266 | 0.46837196 | 0.01011859 | 0.99192667 | 0.99677109 |
| TUBB4B | 498.959208 | -1.409014782 | 0.45552649 | -3.0931566 | 0.0019804 | 0.01699743 |
| NELFB | 323.761511 | -0.157354807 | 0.22428172 | -0.7015944 | 0.48293212 | 0.69127369 |
| EXD3 | 597.799223 | -0.231450114 | 0.21767518 | -1.0632821 | 0.28765405 | 0.51002814 |
| NOXA1 | 179.484699 | -0.492114652 | 0.38049099 | -1.2933674 | 0.19588401 | 0.40352961 |
| NSMF | 109.898667 | -0.463788814 | 0.2955411 | -1.569287 | 0.11658108 | 0.29066052 |
| PNPLA7 | 381.473993 | 0.179035355 | 0.41752057 | 0.42880607 | 0.66806436 | 0.82510165 |
| MRPL41 | 458.512583 | -0.579772016 | 0.39090225 | -1.4831637 | 0.13803093 | 0.3240256 |
| DPH7 | 208.20713 | 0.533249488 | 0.18944482 | 2.81480114 | 0.00488074 | 0.03284245 |
| ZMYND19 | 56.6265509 | -0.521489904 | 0.43164002 | -1.2081593 | 0.226986 | 0.44215574 |
| ARRDC1 | 296.298395 | -0.689948923 | 0.27019544 | -2.553518 | 0.01066408 | 0.05802348 |
| ARRDC1-AS1 | 90.8082235 | 0.007252397 | 0.22134526 | 0.03276509 | 0.97386192 | 0.98822632 |
| EHMT1 | 2322.28735 | -0.228023721 | 0.18615672 | -1.2249019 | 0.22061218 | 0.43435181 |
| LINC01001 | 87.8067737 | 0.303955246 | 0.47740341 | 0.63668428 | 0.52433051 | 0.72362308 |
| AC069287.3 | 103.46682 | 0.59377082 | 0.53690703 | 1.10590994 | 0.2687655 | 0.49028083 |
| BET1L | 141.657266 | -0.338885687 | 0.19507177 | -1.7372359 | 0.08234554 | 0.23064084 |
| RIC8A | 127.988102 | -0.918034895 | 0.21217143 | -4.3268545 | 1.51E-05 | 0.00037562 |
| SIRT3 | 241.47198 | -0.145036199 | 0.22878029 | -0.6339541 | 0.52611079 | 0.72497874 |
| PSMD13 | 365.116117 | -0.239571232 | 0.26962969 | -0.8885195 | 0.37426137 | 0.59780424 |
| PGGHG | 158.88241 | -1.240223086 | 0.49056969 | -2.5281282 | 0.01146725 | 0.0611016 |
| IFITM2 | 1607.78534 | -3.143336257 | 0.51598712 | -6.091889 | 1.12E-09 | 1.40E-07 |
| IFITM3 | 2461.40073 | -2.973025687 | 0.40112792 | -7.4116648 | 1.25E-13 | 5.31E-11 |
| SIGIRR | 243.919499 | 0.003338292 | 0.31093222 | 0.0107364 | 0.99143376 | 0.99656408 |
| ANO9 | 87.7729712 | -0.990748605 | 0.34416855 | -2.8786727 | 0.00399353 | 0.02862207 |
| PTDSS2 | 258.634783 | -0.049422687 | 0.14700226 | -0.3362036 | 0.73671735 | 0.87092675 |
| RNH1 | 384.781103 | -0.928972036 | 0.27590352 | -3.3670177 | 0.00075986 | 0.00812052 |
| HRAS | 170.081957 | -0.608623315 | 0.33002703 | -1.844162 | 0.06515953 | 0.19836189 |
| LMNTD2 | 54.3922797 | 0.389867866 | 0.33117665 | 1.17722027 | 0.2391076 | 0.45716719 |
| AP006284.1 | 91.00676 | 0.293589441 | 0.51670416 | 0.5681964 | 0.56990162 | 0.75923656 |
| RASSF7 | 624.420171 | -0.228642988 | 0.32204618 | -0.7099696 | 0.47772301 | 0.68746521 |
| MIR210HG | 200.17505 | -1.885662205 | 0.58619721 | -3.216771 | 0.00129642 | 0.01226677 |
| PHRF1 | 314.621 | -0.030757727 | 0.25235115 | -0.1218846 | 0.90299039 | 0.95509228 |
| IRF7 | 131.931566 | -0.4224883 | 0.53955978 | -0.7830241 | 0.43361297 | 0.65072685 |
| DEAF1 | 783.363796 | -0.225670306 | 0.19869759 | -1.1357476 | 0.25606221 | 0.47705582 |
| AC131934.1 | 169.234683 | 0.082535324 | 0.4465293 | 0.18483742 | 0.85335656 | 0.93272542 |
| EPS8L2 | 1297.68465 | -0.044055792 | 0.30468394 | -0.1445951 | 0.8850306 | 0.94815256 |
| AP006621.4 | 250.897568 | -0.664414392 | 0.56097964 | -1.1843824 | 0.23626173 | 0.45391953 |
| TALDO1 | 583.790981 | -1.316037075 | 0.40697463 | -3.2337079 | 0.00122194 | 0.0116792 |
| GATD1 | 176.956485 | -0.257859933 | 0.20421284 | -1.2627019 | 0.20669635 | 0.41740591 |
| AP006621.3 | 229.321563 | 0.156389446 | 0.27902758 | 0.56048024 | 0.57515191 | 0.76252365 |
| PIDD1 | 114.298114 | 0.740527653 | 0.29105036 | 2.54432822 | 0.01094882 | 0.0591753 |
| RPLP2 | 690.402731 | -0.80847554 | 0.33042863 | -2.4467479 | 0.01441516 | 0.07126119 |
| PNPLA2 | 460.370752 | -0.661091936 | 0.2345872 | -2.8181075 | 0.00483076 | 0.03257251 |
| CRACR2B | 96.4864934 | 1.010326605 | 0.47422604 | 2.13047477 | 0.03313244 | 0.12633902 |
| CD151 | 568.566045 | -1.524342648 | 0.40517463 | -3.7621868 | 0.00016843 | 0.00254559 |
| POLR2L | 459.406372 | -1.341525726 | 0.51489135 | -2.605454 | 0.00917526 | 0.05206631 |
| TSPAN4 | 147.739225 | -0.738720716 | 0.34523833 | -2.1397413 | 0.03237568 | 0.12428083 |
| CHID1 | 232.604209 | 0.239401612 | 0.24447757 | 0.97923753 | 0.32746263 | 0.55270302 |
| AP2A2 | 1599.02139 | 0.051742047 | 0.26278236 | 0.19690077 | 0.84390519 | 0.92769172 |
| MUC6 | 59.7768063 | 0.805794396 | 0.33029024 | 2.43965551 | 0.01470127 | 0.07224298 |
| TOLLIP | 272.65764 | -0.04295942 | 0.1872306 | -0.2294466 | 0.81852185 | 0.91559451 |
| MOB2 | 270.334486 | -0.022434344 | 0.19279256 | -0.1163652 | 0.90736312 | 0.95790916 |
| MRPL23 | 81.5022156 | -0.633430165 | 0.44201557 | -1.4330494 | 0.15184371 | 0.34497826 |
| CD81-AS1 | 98.5541512 | -0.145585244 | 0.39294116 | -0.3705014 | 0.71100895 | 0.85503616 |
| CD81 | 1466.00317 | -1.628266051 | 0.40453968 | -4.0249848 | 5.70E-05 | 0.00108293 |
| TSSC4 | 104.831049 | -0.69784524 | 0.331366 | -2.1059651 | 0.03520738 | 0.13140839 |
| KCNQ1 | 2918.38457 | -0.459434688 | 0.30467329 | -1.5079585 | 0.13156515 | 0.3143712 |
| KCNQ1OT1 | 2828.9102 | -0.661322419 | 0.21713152 | -3.0457228 | 0.00232122 | 0.01913918 |
| CDKN1C | 255.271757 | -1.917997549 | 0.62970392 | -3.045872 | 0.00232007 | 0.01913918 |

| | | | | | | |
|---|---|---|---|---|---|---|
| SLC22A18 | 146.881833 | -1.256407847 | 0.66816119 | -1.8803963 | 0.06005408 | 0.18793044 |
| NAP1L4 | 1859.3702 | -0.169924694 | 0.16236693 | -1.0465474 | 0.29530837 | 0.51843809 |
| CARS | 496.670458 | -1.012429028 | 0.276854 | -3.6569059 | 0.00025528 | 0.00355896 |
| OSBPL5 | 405.82813 | -0.785372023 | 0.28833626 | -2.723806 | 0.00645344 | 0.0403424 |
| ZNF195 | 377.166313 | 0.553710747 | 0.27986379 | 1.97850087 | 0.04787223 | 0.16114788 |
| AC127526.1 | 188.742072 | 0.373935562 | 0.48080828 | 0.7777228 | 0.43673245 | 0.65344993 |
| NUP98 | 1194.58668 | -0.14771863 | 0.2747163 | -0.5377134 | 0.59077494 | 0.77232247 |
| PGAP2 | 198.515975 | 0.237007997 | 0.49862787 | 0.4753204 | 0.63455862 | 0.80347822 |
| RHOG | 70.4977895 | -0.983060345 | 0.27165698 | -3.618756 | 0.00029602 | 0.00398589 |
| STIM1 | 542.376057 | -2.043843966 | 0.35187968 | -5.8083602 | 6.31E-09 | 6.31E-07 |
| RRM1 | 191.725397 | -0.080215245 | 0.27974438 | -0.2867448 | 0.77430773 | 0.89201385 |
| TRIM21 | 52.8497043 | -0.485632088 | 0.51531419 | -0.9424 | 0.34598789 | 0.57105512 |
| TRIM68 | 58.5963154 | -0.223852847 | 0.34983197 | -0.6398868 | 0.52224623 | 0.72221624 |
| HBB | 63.4470885 | -0.834051171 | 1.05078306 | -0.7937425 | 0.42734535 | 0.64571091 |
| AC104389.4 | 476.074085 | 1.297916448 | 0.4802005 | 2.70286361 | 0.00687449 | 0.04229399 |
| TRIM5 | 153.81823 | 0.410656168 | 0.45539442 | 0.90175933 | 0.36718473 | 0.59256482 |
| TRIM22 | 407.828396 | -0.884281183 | 0.78376363 | -1.1282498 | 0.25921443 | 0.48023578 |
| FAM160A2 | 308.056687 | 0.290905421 | 0.19944295 | 1.45858962 | 0.14467809 | 0.33456809 |
| CAVIN3 | 75.4404535 | -2.328676796 | 0.62302279 | -3.7377073 | 0.00018571 | 0.00275927 |
| SMPD1 | 118.106204 | -0.164457559 | 0.445556 | -0.3691064 | 0.71204844 | 0.85561721 |
| APBB1 | 219.849216 | -0.470417009 | 0.22323533 | -2.1072695 | 0.03509422 | 0.13109716 |
| TRIM3 | 44.7159299 | -0.417696474 | 0.44709197 | -0.9342518 | 0.35017403 | 0.5762363 |
| ARFIP2 | 149.758888 | -0.190387943 | 0.22361785 | -0.8513987 | 0.39454792 | 0.61682844 |
| AC084337.2 | 259.676494 | -0.012028802 | 0.37150345 | -0.0323787 | 0.97417004 | 0.98832358 |
| DNHD1 | 469.961625 | 0.099611026 | 0.34081268 | 0.292275 | 0.77007637 | 0.88929655 |
| RRP8 | 103.665051 | 0.056332206 | 0.24253886 | 0.23226054 | 0.81633566 | 0.91443162 |
| ILK | 390.070755 | -0.739829211 | 0.1806652 | -4.0950289 | 4.22E-05 | 0.00086445 |
| TAF10 | 170.09997 | -0.571036932 | 0.33344027 | -1.7125614 | 0.08679326 | 0.23862737 |
| TPP1 | 226.121187 | -0.415832007 | 0.26336782 | -1.5789021 | 0.11435851 | 0.28726204 |
| MRPL17 | 141.046391 | -1.05152239 | 0.40457144 | -2.5991019 | 0.0093468 | 0.05271645 |
| AC087280.2 | 217.126597 | 0.374089229 | 0.55340951 | 0.67597181 | 0.49905854 | 0.70390435 |
| ZNF215 | 95.7446259 | 0.121409281 | 0.38758208 | 0.31324792 | 0.75409231 | 0.88075281 |
| ZNF214 | 139.544404 | 0.523324909 | 0.19349403 | 2.70460495 | 0.00683857 | 0.04215148 |
| NLRP14 | 37.4365818 | 0.124550169 | 0.51554973 | 0.24158711 | 0.80910011 | 0.91023512 |
| SYT9 | 73.0672255 | -0.68218112 | 0.49157467 | -1.3877467 | 0.1652142 | 0.36450883 |
| AC107884.1 | 229.749906 | 0.173865011 | 0.29198053 | 0.59546783 | 0.5515308 | 0.74467698 |
| PPFIBP2 | 250.078892 | -0.052527516 | 0.27290387 | -0.1924763 | 0.84736916 | 0.92955513 |
| OVCH2 | 71.8742186 | -3.363992816 | 1.02318715 | -3.2877591 | 0.00100988 | 0.01006504 |
| AC044810.3 | 542.802373 | -3.180893109 | 1.11525753 | -2.8521602 | 0.00434232 | 0.03024363 |
| EIF3F | 381.169188 | -0.39920746 | 0.25865295 | -1.5434096 | 0.12273141 | 0.30023259 |
| TUB | 104.910931 | 0.801285511 | 0.47399952 | 1.69047746 | 0.09093665 | 0.24631864 |
| RIC3 | 1063.29925 | -0.080980436 | 0.24984032 | -0.3241288 | 0.74584054 | 0.87571624 |
| LMO1 | 139.32373 | 1.293849519 | 0.40877909 | 3.1651558 | 0.00155 | 0.01407144 |
| STK33 | 1695.33204 | 0.094847408 | 0.31901688 | 0.29731156 | 0.76622865 | 0.88688348 |
| TRIM66 | 927.460402 | 0.314611052 | 0.45026219 | 0.69872857 | 0.48472168 | 0.69239889 |
| RPL27A | 844.040259 | -1.015255121 | 0.33773767 | -3.0060464 | 0.00264669 | 0.02111115 |
| ST5 | 2516.57113 | -2.123919712 | 0.40215513 | -5.2813443 | 1.28E-07 | 7.56E-06 |
| AC026894.1 | 248.776649 | -1.972734953 | 0.57427562 | -3.435171 | 0.00059218 | 0.0067258 |
| AKIP1 | 123.05058 | -0.450292783 | 0.26446791 | -1.7026367 | 0.08863607 | 0.24205091 |
| TMEM9B | 238.750596 | -0.02815932 | 0.19611538 | -0.1435855 | 0.88582781 | 0.94846586 |
| TMEM9B-AS | 253.97087 | 0.711540365 | 0.50181369 | 1.41793734 | 0.15620906 | 0.35131349 |
| SCUBE2 | 59.283484 | 0.293418565 | 0.44198653 | 0.66386313 | 0.50677791 | 0.71009117 |
| DENND5A | 1214.18103 | 0.133619955 | 0.27730758 | 0.48184747 | 0.6299143 | 0.79979406 |
| TMEM41B | 995.47386 | -0.182206456 | 0.24159934 | -0.7541679 | 0.45074844 | 0.66654333 |
| IPO7 | 911.082252 | -0.31266079 | 0.20797435 | -1.5033622 | 0.13274566 | 0.31650494 |
| AC132192.2 | 48.5296776 | 1.225179458 | 0.51439578 | 2.38178363 | 0.01722902 | 0.08047316 |
| ZNF143 | 321.559172 | -0.100976225 | 0.32532925 | -0.3103816 | 0.75627076 | 0.88220153 |
| WEE1 | 440.695709 | 0.118579135 | 0.41861721 | 0.28326388 | 0.77697457 | 0.89337196 |
| SWAP70 | 1087.00993 | -0.932955999 | 0.23086896 | -4.041063 | 5.32E-05 | 0.00102455 |
| SBF2-AS1 | 401.816349 | 0.160408441 | 0.30041203 | 0.53396143 | 0.59336822 | 0.77387448 |
| SBF2 | 5992.97782 | 0.04236657 | 0.18112163 | 0.23391227 | 0.81505309 | 0.91369229 |
| AC011092.2 | 265.174361 | -0.042944264 | 0.46149514 | -0.0930546 | 0.92586015 | 0.96724667 |
| AC100763.1 | 77.9323171 | 0.122836209 | 0.59773728 | 0.205502 | 0.83717993 | 0.92422146 |
| ADM | 339.97079 | -3.589150531 | 1.18872853 | -3.0193189 | 0.00253344 | 0.0204299 |
| AMPD3 | 495.522973 | -0.036721266 | 0.52297776 | -0.0702157 | 0.94402195 | 0.97578731 |
| MTRNR2L8 | 367.049984 | -1.53143279 | 0.68929178 | -2.2217482 | 0.02630033 | 0.1073219 |
| RNF141 | 189.233141 | 0.065644072 | 0.17504291 | 0.37501703 | 0.7076478 | 0.85293799 |
| MRVI1-AS1 | 56.527658 | 1.633110038 | 0.52457412 | 3.11321125 | 0.00185064 | 0.0160945 |
| CTR9 | 453.490101 | -0.045951454 | 0.19946494 | -0.2303736 | 0.81780148 | 0.91533703 |
| EIF4G2 | 1079.15885 | -0.288054637 | 0.20360009 | -1.4148061 | 0.15712537 | 0.35273096 |
| ZBED5 | 692.518491 | 0.206922327 | 0.15749375 | 1.31384466 | 0.18889848 | 0.39486629 |
| ZBED5-AS1 | 242.468157 | 1.024183224 | 0.31319095 | 3.27015586 | 0.00107488 | 0.01055622 |
| GALNT18 | 3618.057 | 0.91603314 | 0.45461982 | 2.01494325 | 0.0439106 | 0.15191594 |
| USP47 | 2413.57006 | 0.207628083 | 0.15726061 | 1.32028028 | 0.18674146 | 0.39251209 |
| MICAL2 | 87.7992498 | -1.668789636 | 0.3364205 | -4.9604279 | 7.03E-07 | 3.14E-05 |
| PARVA | 1214.0404 | -0.480458093 | 0.23863688 | -2.0133438 | 0.04407847 | 0.15221766 |
| TEAD1 | 4330.68053 | -0.189678749 | 0.27667031 | -0.6855768 | 0.49298 | 0.69868807 |
| LINC00958 | 628.520025 | 0.026332535 | 0.27967217 | 0.09415501 | 0.92498602 | 0.9667507 |
| AC084859.1 | 63.0595108 | 0.988644918 | 0.51756592 | 1.91018165 | 0.05610983 | 0.17967388 |
| RASSF10 | 49.2890049 | 0.966865312 | 0.55816208 | 1.73223039 | 0.08323255 | 0.23209178 |
| ARNTL | 333.365894 | 0.772023118 | 0.29480001 | 2.61880286 | 0.00882389 | 0.05053466 |
| BTBD10 | 717.82153 | 0.030722687 | 0.20965107 | 0.146542 | 0.88349353 | 0.94755606 |
| FAR1 | 1157.7042 | 0.304280336 | 0.17149655 | 1.77426509 | 0.07601931 | 0.21975002 |
| SPON1 | 197.744541 | -1.56506149 | 0.51570761 | -3.0347845 | 0.00240708 | 0.019663 |
| SPON1-AS1 | 44.087768 | -0.221538411 | 0.55376379 | -0.4000594 | 0.68911276 | 0.84010064 |
| RRAS2 | 957.367297 | -0.244021514 | 0.26999377 | -0.9038042 | 0.36609922 | 0.59142894 |
| COPB1 | 812.167475 | -0.091165866 | 0.16880783 | -0.5400571 | 0.58915765 | 0.77174702 |
| PSMA1 | 46.5670608 | 0.654039254 | 0.45940025 | 1.42368067 | 0.15453893 | 0.34888134 |
| PDE3B | 69.2810153 | 0.354202053 | 0.3793449 | 0.93372034 | 0.35044817 | 0.57654376 |
| CYP2R1 | 176.228701 | 0.334700188 | 0.24957724 | 1.34106855 | 0.1798982 | 0.38357954 |
| SOX6 | 7661.43115 | 0.754354584 | 0.21789171 | 3.46206183 | 0.00053605 | 0.00623304 |
| C11orf58 | 1094.57956 | -0.144314373 | 0.13205104 | -1.0928682 | 0.27445169 | 0.49616155 |
| PLEKHA7 | 6310.98247 | -0.084267294 | 0.25566118 | -0.3296054 | 0.74169817 | 0.8738293 |
| RPS13 | 652.593024 | -1.097448607 | 0.43788464 | -2.5062505 | 0.01220191 | 0.06370556 |
| PIK3C2A | 2429.0507 | 0.119376306 | 0.0783421 | 1.52378237 | 0.12756309 | 0.30792778 |
| NUCB2 | 391.947904 | 0.513242583 | 0.30188009 | 1.70015383 | 0.08910199 | 0.24292017 |
| NCR3LG1 | 64.4415566 | -0.771105698 | 0.43514036 | -1.772085 | 0.07638045 | 0.22033505 |
| KCNC1 | 134.697007 | -0.634932551 | 0.46297662 | -1.3714138 | 0.17024598 | 0.37059498 |
| SERGEF | 1245.78757 | -0.318493263 | 0.25887117 | -1.2303157 | 0.21857892 | 0.4326703 |
| SAAL1 | 143.756196 | 0.77996695 | 0.27081354 | 2.88008842 | 0.00397564 | 0.02852481 |
| SAA1 | 210.404974 | -8.456566465 | 1.56309716 | -5.4101349 | 6.30E-08 | 4.43E-06 |
| HPS5 | 289.922181 | 0.210490486 | 0.1944208 | 1.08265413 | 0.27896197 | 0.50106183 |
| GTF2H1 | 497.469762 | 0.208404695 | 0.19448521 | 1.07157094 | 0.28391279 | 0.50651054 |
| LDHA | 2345.53088 | -2.782716951 | 0.39284467 | -7.0835044 | 1.41E-12 | 4.50E-10 |
| TSG101 | 852.852524 | -0.236843507 | 0.18245245 | -1.2981109 | 0.19424923 | 0.40179112 |
| UEVLD | 617.771099 | -0.161474247 | 0.16390751 | -0.9851547 | 0.3245481 | 0.54946883 |
| SPTY2D1OS | 267.271132 | 0.459343576 | 0.17482901 | 2.62738762 | 0.00860433 | 0.04972678 |
| SPTY2D1 | 210.124276 | -0.090541485 | 0.20339415 | -0.4451528 | 0.6562093 | 0.8175692 |
| TMEM86A | 48.1978563 | 0.254314892 | 0.42051244 | 0.60477376 | 0.54532933 | 0.7400858 |
| ZDHHC13 | 715.944408 | -0.620509229 | 0.18186212 | -3.4119762 | 0.00064494 | 0.00720717 |
| NAV2 | 3687.32446 | 0.313350766 | 0.37364668 | 0.83862853 | 0.40167779 | 0.62318088 |
| HTATIP2 | 152.939504 | -1.884026133 | 0.38710068 | -4.8670185 | 1.13E-06 | 4.58E-05 |
| PRMT3 | 776.494107 | 0.246085524 | 0.2201784 | 1.11766423 | 0.26371043 | 0.48467386 |

| | | | | | | |
|---|---|---|---|---|---|---|
| NELL1 | 2920.727 | 2.703724981 | 0.58735193 | 4.60324522 | 4.16E-06 | 0.00012867 |
| ANO5 | 595.294289 | 0.70386786 | 0.19168782 | 3.67194883 | 0.00024071 | 0.00341208 |
| FANCF | 59.2691475 | 0.137051356 | 0.3925101 | 0.34916645 | 0.72696435 | 0.86532561 |
| GAS2 | 350.852552 | 1.437879705 | 0.3109236 | 4.62454352 | 3.75E-06 | 0.00011844 |
| SVIP | 436.925627 | -0.274816924 | 0.2319984 | -1.1845639 | 0.23618993 | 0.45391953 |
| AC006299.1 | 162.708781 | 0.315186975 | 0.45663184 | 0.69024309 | 0.49004133 | 0.69682235 |
| MUC15 | 228.875653 | 1.157802581 | 0.47047609 | 2.46091692 | 0.01385825 | 0.06931222 |
| SLC5A12 | 94.0633557 | 2.647782363 | 0.91082007 | 2.90703119 | 0.00364877 | 0.02677046 |
| FIBIN | 115.629745 | -1.164820241 | 0.86559839 | -1.3456821 | 0.17840507 | 0.38168976 |
| BBOX1 | 2166.26685 | -0.400983246 | 0.79101604 | -0.5069218 | 0.61220972 | 0.78684784 |
| BBOX1-AS1 | 406.53675 | -0.702269934 | 0.64170279 | -1.094385 | 0.27378614 | 0.49547077 |
| CCDC34 | 134.248833 | 0.055726305 | 0.41641697 | 0.13382333 | 0.89354228 | 0.95146032 |
| LGR4 | 9093.09289 | -0.128813436 | 0.26575445 | -0.4847085 | 0.62788314 | 0.7982736 |
| LIN7C | 262.573653 | 0.07671396 | 0.30322068 | 0.25299712 | 0.80027044 | 0.90650649 |
| BDNF-AS | 1196.47845 | 0.245641431 | 0.18987423 | 1.29370601 | 0.19576699 | 0.40345184 |
| METTL15 | 2795.11005 | 0.138765603 | 0.21140186 | 0.65640674 | 0.51156246 | 0.71369122 |
| AC090833.1 | 212.505637 | 0.103560331 | 0.42747417 | 0.24226102 | 0.80857791 | 0.91010272 |
| ARL14EP | 304.344596 | 0.155309844 | 0.21402721 | 0.72565468 | 0.46805049 | 0.6801407 |
| MPPED2 | 1956.17556 | 1.401331796 | 0.55108327 | 2.54286761 | 0.01099469 | 0.0593255 |
| AL136088.1 | 51.0268061 | 2.576978082 | 0.62385505 | 4.13073209 | 3.62E-05 | 0.00076059 |
| DCDC1 | 1122.24822 | -0.391488785 | 0.27621706 | -1.417323 | 0.15638852 | 0.35149497 |
| DNAJC24 | 466.466609 | 0.056983239 | 0.26464155 | 0.21532235 | 0.82951601 | 0.92024799 |
| IMMP1L | 1292.44143 | 0.566576408 | 0.18927672 | 2.99337602 | 0.0027591 | 0.02177114 |
| ELP4 | 1899.7824 | 0.336446039 | 0.12508026 | 2.68984131 | 0.0071486 | 0.04335439 |
| AC131571.1 | 248.048846 | -0.250262737 | 0.55387906 | -0.4518364 | 0.65138682 | 0.81378687 |
| RCN1 | 193.603663 | -1.364078526 | 0.35548037 | -3.8372823 | 0.0001244 | 0.00199665 |
| EIF3M | 432.323575 | 0.047779581 | 0.21875392 | 0.21841703 | 0.8271042 | 0.91955386 |
| CCDC73 | 81.2649126 | -0.852324694 | 0.47081815 | -1.8103055 | 0.07024842 | 0.20866009 |
| PRRG4 | 575.887707 | -0.555346832 | 0.37176102 | -1.4938275 | 0.13522071 | 0.32038214 |
| QSER1 | 812.941372 | -0.062714296 | 0.1190551 | -0.526767 | 0.59835541 | 0.77692683 |
| TCP11L1 | 381.029325 | -0.12341192 | 0.17898829 | -0.6894972 | 0.49051044 | 0.6969611 |
| CSTF3 | 778.482648 | 0.350041651 | 0.18863614 | 1.85564467 | 0.06350422 | 0.19518591 |
| HIPK3 | 2360.51826 | -0.793980441 | 0.23285017 | -3.4098341 | 0.00065002 | 0.0072348 |
| KIAA1549L | 78.8434304 | 0.457972528 | 0.43075633 | 1.06318234 | 0.28769927 | 0.51002814 |
| AL049629.1 | 238.877552 | -0.363190348 | 0.31531223 | -1.1518435 | 0.24938538 | 0.46884878 |
| CD59 | 185.740704 | -0.234042151 | 0.24668513 | -0.9487485 | 0.34274853 | 0.56790834 |
| FBXO3 | 553.521086 | 0.238315529 | 0.18403327 | 1.29495891 | 0.1953344 | 0.40290008 |
| LMO2 | 51.1772651 | 0.078240664 | 0.47356356 | 0.16521682 | 0.86887333 | 0.94020545 |
| CAPRIN1 | 1040.15681 | 0.154889084 | 0.14249269 | 1.08699667 | 0.27703828 | 0.49909966 |
| NAT10 | 340.50875 | -0.053532669 | 0.25113532 | -0.2131626 | 0.8312001 | 0.92131595 |
| ABTB2 | 2684.73431 | 0.292671157 | 0.30894556 | 0.94732275 | 0.34347434 | 0.56837065 |
| CAT | 1027.14979 | 1.118282069 | 0.50507045 | 2.21411106 | 0.02682114 | 0.10871927 |
| EHF | 1162.09971 | -0.264583345 | 0.20579415 | -1.2856699 | 0.1985583 | 0.4070707 |
| APIP | 394.860755 | 0.44681674 | 0.41635912 | 1.07315228 | 0.2832028 | 0.50565366 |
| PDHX | 962.341728 | 0.28217028 | 0.18547532 | 1.52133596 | 0.12817555 | 0.30884182 |
| CD44 | 1112.54527 | -4.306858658 | 0.87874697 | -4.9011363 | 9.53E-07 | 3.97E-05 |
| TRIM44 | 2108.14618 | -0.136138891 | 0.14604885 | -0.9321463 | 0.35126093 | 0.57732528 |
| LDLRAD3 | 106.251381 | -0.297390681 | 0.55748994 | -0.5334458 | 0.593725 | 0.77411051 |
| COMMD9 | 176.947266 | 0.42726014 | 0.24491167 | 1.74454787 | 0.0810636 | 0.22850386 |
| PRR5L | 215.448122 | -0.153122008 | 0.46810042 | -0.3271136 | 0.74358198 | 0.87467216 |
| TRAF6 | 330.382003 | 0.27809512 | 0.14891391 | 1.86748922 | 0.0618333 | 0.1919972 |
| RAG1 | 86.3335979 | 0.73945465 | 0.2723808 | 2.71478262 | 0.00663193 | 0.04126225 |
| C11orf74 | 327.998881 | 1.477329215 | 0.23303045 | 6.33964027 | 2.30E-10 | 3.46E-08 |
| LRRC4C | 73.0805596 | 0.151723146 | 0.33486015 | 0.45309407 | 0.65048101 | 0.81311665 |
| API5 | 688.90654 | 0.125807872 | 0.1624092 | 0.77463511 | 0.4385553 | 0.65476683 |
| TTC17 | 3510.17387 | 0.202390809 | 0.15917116 | 1.27152939 | 0.20354038 | 0.41330744 |
| AC068205.1 | 96.1101845 | 1.129818075 | 0.21203685 | 5.32840436 | 9.91E-08 | 6.35E-06 |
| HSD17B12 | 2691.06421 | 0.018512557 | 0.26080269 | 0.070983 | 0.94341129 | 0.97559625 |
| AC087521.2 | 67.4931772 | -0.040974419 | 0.49585157 | -0.0826344 | 0.93414221 | 0.97173449 |
| ALKBH3 | 274.799996 | 0.281908538 | 0.22468392 | 1.25468944 | 0.20959152 | 0.42138423 |
| ACCS | 138.008848 | -0.150919455 | 0.5097537 | -0.2960635 | 0.7671816 | 0.88751986 |
| EXT2 | 832.400147 | -0.722721613 | 0.34230511 | -2.1113375 | 0.03474331 | 0.13022828 |
| CD82 | 196.658723 | -1.104966656 | 0.41433651 | -2.6668339 | 0.00765695 | 0.04576584 |
| PRDM11 | 221.365918 | 0.651141977 | 0.24850491 | 2.62023789 | 0.00878684 | 0.05041939 |
| SYT13 | 136.299951 | 0.470430327 | 0.38654661 | 1.21700801 | 0.22360117 | 0.43821184 |
| CRY2 | 368.289789 | 0.0851144 | 0.36918003 | 0.23054985 | 0.81766453 | 0.91531162 |
| MAPK8IP1 | 79.7147982 | 0.42769099 | 0.50459865 | 0.84758648 | 0.39666831 | 0.61845042 |
| PEX16 | 103.137496 | -0.204929019 | 0.25548733 | -0.8021103 | 0.42248916 | 0.64123603 |
| LARGE2 | 258.033828 | 1.083010895 | 0.52929866 | 2.04612439 | 0.04074413 | 0.14438667 |
| PHF21A | 3246.21578 | -0.224092263 | 0.16860782 | -1.329074 | 0.18382357 | 0.38887341 |
| DGKZ | 189.349844 | -0.015083314 | 0.31050762 | -0.0485763 | 0.96125695 | 0.982987 |
| MDK | 75.4057476 | -2.177744104 | 0.75814899 | -2.8724487 | 0.00407304 | 0.02904023 |
| AMBRA1 | 2759.10419 | -0.167151513 | 0.18197401 | -0.9185461 | 0.35833305 | 0.58372701 |
| HARBI1 | 47.914902 | 0.159501821 | 0.39763074 | 0.40113051 | 0.68832404 | 0.83951129 |
| ATG13 | 457.910934 | -0.231354083 | 0.14911594 | -1.5515047 | 0.12078078 | 0.29709373 |
| ARHGAP1 | 376.83495 | -0.84321527 | 0.24045642 | -3.506728 | 0.00045365 | 0.00549752 |
| CKAP5 | 657.679965 | 0.260364649 | 0.14864306 | 1.75160981 | 0.07984093 | 0.22607048 |
| LRP4 | 521.342139 | -0.512052791 | 0.4323632 | -1.1843117 | 0.23628971 | 0.45391953 |
| C11orf49 | 2579.84768 | -0.193317995 | 0.13906039 | -1.390173 | 0.16447636 | 0.36336649 |
| ARFGAP2 | 312.068559 | 0.035599374 | 0.23332222 | 0.15257602 | 0.87873264 | 0.94585961 |
| PACSIN3 | 49.6383076 | -0.066536688 | 0.39018458 | -0.1705262 | 0.86459634 | 0.93833133 |
| DDB2 | 172.612066 | -0.654561331 | 0.38655822 | -1.6933059 | 0.09039725 | 0.24548875 |
| ACP2 | 93.4250499 | 0.37027951 | 0.31476739 | 1.17635917 | 0.23945138 | 0.45753121 |
| NR1H3 | 170.690051 | 0.475895635 | 0.27477555 | 1.73194314 | 0.08328369 | 0.23218536 |
| MADD | 399.304401 | 0.69185665 | 0.24312439 | 2.84569003 | 0.00443153 | 0.03066322 |
| SPI1 | 51.4498453 | 2.286284206 | 0.68082449 | 3.35811097 | 0.00078477 | 0.00826862 |
| AC090559.1 | 40.5365162 | 0.776172828 | 0.4984869 | 1.55705762 | 0.11945681 | 0.29515618 |
| SLC39A13 | 270.416214 | 0.211699937 | 0.34743331 | 0.60932538 | 0.54230879 | 0.73841426 |
| PSMC3 | 117.235555 | -0.542220027 | 0.35186806 | -1.5409754 | 0.12332277 | 0.30115928 |
| CELF1 | 2816.93074 | 0.080385362 | 0.16251237 | 0.49464151 | 0.6208532 | 0.7928889 |
| NDUFS3 | 241.775397 | 0.217663509 | 0.35405253 | 0.61477744 | 0.53870171 | 0.73496797 |
| KBTBD4 | 66.7200079 | -0.391063851 | 0.25511898 | -1.5328685 | 0.12530825 | 0.30392546 |
| MTCH2 | 616.765538 | 0.082393267 | 0.23571126 | 0.34955168 | 0.72667518 | 0.86528914 |
| AGBL2 | 82.6734091 | 0.53429578 | 0.41816184 | 1.27772486 | 0.20134646 | 0.41055655 |
| FNBP4 | 1902.31581 | -0.175258313 | 0.1497346 | -1.1704597 | 0.24181604 | 0.46005301 |
| NUP160 | 1055.14402 | 0.116921213 | 0.21852869 | 0.53503827 | 0.5926234 | 0.77381752 |
| PTPRJ | 4221.20595 | -0.396652852 | 0.3510696 | -1.1298411 | 0.2585432 | 0.4795566 |
| FOLH1 | 55.8792807 | 4.369339355 | 0.50046391 | 8.73057836 | 2.53E-18 | 3.35E-15 |
| TNKS1BP1 | 438.917263 | -1.026164271 | 0.14960199 | -6.8592955 | 6.92E-12 | 1.83E-09 |
| SSRP1 | 185.682007 | -0.571405902 | 0.27910874 | -2.0472519 | 0.04063336 | 0.14416494 |
| SLC43A1 | 99.7271575 | 2.554343332 | 0.60174997 | 4.24485825 | 2.19E-05 | 0.00050777 |
| TIMM10 | 106.433959 | -1.152523289 | 0.42480043 | -2.7130935 | 0.00666583 | 0.04139584 |
| UBE2L6 | 130.240929 | -1.050952906 | 0.64116539 | -1.6391292 | 0.10118636 | 0.26492976 |
| SERPING1 | 1728.2134 | -2.792573183 | 0.38631713 | -7.2287066 | 4.88E-13 | 1.79E-10 |
| ZDHHC5 | 557.345659 | -0.481418911 | 0.20048948 | -2.4012178 | 0.01634061 | 0.07766933 |
| MED19 | 88.799546 | -0.533951388 | 0.27758511 | -1.9235592 | 0.05440986 | 0.1757642 |
| CTNND1 | 1608.25276 | -0.181430327 | 0.16569273 | -1.0949806 | 0.27352513 | 0.49520195 |
| LPXN | 509.232959 | 0.988471681 | 0.21778361 | 4.5387791 | 5.66E-06 | 0.00016871 |
| GLYAT | 61.2460331 | 1.698588343 | 0.84929125 | 2.00000687 | 0.04549952 | 0.15579927 |
| GLYATL1 | 99.1586944 | 1.183993142 | 0.49975234 | 2.36915978 | 0.01782855 | 0.08263064 |
| FAM111A-D1 | 67.1942888 | 0.334400454 | 0.3405186 | 0.98203286 | 0.32608366 | 0.55129132 |
| FAM111A | 431.223669 | 0.663149473 | 0.17975571 | 3.68917054 | 0.00022499 | 0.00323027 |
| DTX4 | 82.0526621 | -2.396475861 | 0.81924176 | -2.9252364 | 0.00344195 | 0.02569155 |

| | | | | | | |
|---|---|---|---|---|---|---|
| OSBP | 1030.63401 | 0.380222374 | 0.27388043 | 1.38827871 | 0.16505219 | 0.36427308 |
| AP000442.2 | 57.013272 | 0.224700531 | 0.50043745 | 0.44900823 | 0.65342572 | 0.81539269 |
| PATL1 | 576.517897 | -0.002900579 | 0.22855572 | -0.0126909 | 0.9898744 | 0.99606252 |
| STX3 | 778.879825 | -0.620554085 | 0.27575684 | -2.250367 | 0.02442566 | 0.10200397 |
| MRPL16 | 169.94894 | 0.037243873 | 0.27033492 | 0.13776938 | 0.89042269 | 0.95041347 |
| PRPF19 | 189.368804 | -0.45057712 | 0.34073927 | -1.3223516 | 0.1860511 | 0.39176614 |
| TMEM109 | 122.971972 | -0.665623706 | 0.42401613 | -1.5698075 | 0.1164599 | 0.29052291 |
| TMEM132A | 46.8255455 | -0.829234264 | 0.43637457 | -1.9002809 | 0.05739627 | 0.18268609 |
| VPS37C | 285.167476 | -0.62710397 | 0.28408822 | -2.2074268 | 0.02728425 | 0.11007871 |
| DDB1 | 731.769447 | -0.071141699 | 0.23024002 | -0.3089893 | 0.75732968 | 0.88284951 |
| TKFC | 128.112425 | -0.490967347 | 0.31222689 | -1.5724698 | 0.11584164 | 0.28950706 |
| CYB561A3 | 98.9699331 | 0.024717566 | 0.23027593 | 0.1073389 | 0.91452012 | 0.96131774 |
| TMEM138 | 119.843716 | 0.129647888 | 0.26455318 | 0.49006361 | 0.62408889 | 0.79555975 |
| TMEM216 | 74.6524018 | 0.092231771 | 0.34500705 | 0.267333 | 0.78921278 | 0.90023416 |
| CPSF7 | 658.217662 | 0.179941195 | 0.09463918 | 1.90133934 | 0.05725758 | 0.18228859 |
| AP003108.2 | 590.744327 | 0.05604161 | 0.37241488 | 0.15048166 | 0.88038462 | 0.94639771 |
| SYT7 | 307.385823 | 1.056122828 | 0.50474421 | 2.09239215 | 0.03640345 | 0.13424153 |
| DAGLA | 65.299267 | 1.437906198 | 0.47189394 | 3.04709615 | 0.00231064 | 0.01911947 |
| MYRF | 99.7945365 | -1.745046095 | 0.55431973 | -3.1480859 | 0.00164343 | 0.01473735 |
| TMEM258 | 190.531413 | -0.92209944 | 0.32363582 | -2.8491884 | 0.00438309 | 0.0303988 |
| FADS2 | 102.006538 | -0.590626643 | 0.34905788 | -1.6920593 | 0.09063466 | 0.24583023 |
| FADS1 | 109.740935 | -1.187971639 | 0.4680259 | -2.5382604 | 0.0111405 | 0.05990255 |
| FADS3 | 136.637924 | -1.541336484 | 0.34047238 | -4.5270529 | 5.98E-06 | 0.00017557 |
| FTH1 | 5963.00085 | -2.165541361 | 0.45219061 | -4.7890011 | 1.68E-06 | 6.12E-05 |
| INCENP | 68.7644822 | -0.125573992 | 0.37251682 | -0.3370962 | 0.7360444 | 0.87055499 |
| ASRGL1 | 756.240186 | 0.2720451 | 0.24633852 | 1.10435468 | 0.26943931 | 0.49092338 |
| AHNAK | 1103.71727 | -1.404735813 | 0.2655259 | -5.290391 | 1.22E-07 | 7.30E-06 |
| MTA2 | 54.8074979 | -0.126576905 | 0.29108454 | -0.4348459 | 0.66367432 | 0.82268279 |
| EML3 | 148.812567 | -0.701816589 | 0.4021566 | -1.7451326 | 0.0809618 | 0.22831434 |
| B3GAT3 | 231.928455 | -0.20230354 | 0.26326187 | -0.7684498 | 0.44221999 | 0.65802453 |
| GANAB | 552.604122 | -0.066227871 | 0.17886 | -0.3702777 | 0.71117558 | 0.8551462 |
| INTS5 | 53.0175106 | -0.664783369 | 0.49705917 | -1.3374331 | 0.18108132 | 0.38492165 |
| CSKMT | 58.7015274 | 0.150655177 | 0.2314109 | 0.65102888 | 0.51502785 | 0.7164072 |
| UQCC3 | 153.842852 | -0.16425387 | 0.31751698 | -0.5173074 | 0.60494162 | 0.78156518 |
| UBXN1 | 207.298538 | -0.795904941 | 0.30535297 | -2.606508 | 0.00914707 | 0.05205672 |
| TTC9C | 302.060829 | -0.131085519 | 0.23034875 | -0.5690742 | 0.56930582 | 0.75882548 |
| ZBTB3 | 38.7140394 | 1.001534906 | 0.29809266 | 3.35981067 | 0.00077996 | 0.00825243 |
| POLR2G | 88.2594691 | -0.162794826 | 0.30630573 | -0.5314782 | 0.59508743 | 0.77501911 |
| TAF6L | 154.042725 | 0.179914955 | 0.2995101 | 0.60069745 | 0.54804152 | 0.74186733 |
| TMEM223 | 169.503865 | -0.336182638 | 0.22252731 | -1.5107478 | 0.13085272 | 0.31323728 |
| TMEM179B | 297.973749 | -0.749025313 | 0.26253958 | -2.8529996 | 0.00433087 | 0.03018809 |
| NXF1 | 312.250869 | 0.474607926 | 0.38710715 | 1.2260376 | 0.22018453 | 0.43409215 |
| STX5 | 616.802189 | -0.231817727 | 0.20215976 | -1.1467056 | 0.25150332 | 0.47188266 |
| WDR74 | 73.6881809 | -0.163228323 | 0.2458731 | -0.6638722 | 0.50677209 | 0.71009117 |
| SLC3A2 | 354.853371 | -0.356788843 | 0.31338156 | -1.1385126 | 0.25490652 | 0.47570785 |
| SLC22A6 | 81.7987307 | 0.354213015 | 0.92540691 | 0.38276461 | 0.70189431 | 0.8488446 |
| RARRES3 | 257.054116 | -2.046781219 | 0.64340316 | -3.1811799 | 0.00146677 | 0.01349199 |
| PLA2G16 | 134.813123 | -1.731371849 | 0.49263273 | -3.5145287 | 0.00044054 | 0.00539298 |
| ATL3 | 573.586474 | -0.644090473 | 0.16170427 | -3.9831383 | 6.80E-05 | 0.00123626 |
| RTN3 | 1233.43059 | -0.222397546 | 0.28836771 | -0.771229 | 0.4405712 | 0.65623647 |
| C11orf95 | 48.7697243 | -0.301169595 | 0.56903418 | -0.5292645 | 0.59662197 | 0.77627853 |
| SPINDOC | 89.0950689 | -0.113157084 | 0.28603901 | -0.3956002 | 0.6924 | 0.84221582 |
| MARK2 | 933.673086 | -0.068229197 | 0.18867077 | -0.361631 | 0.7176278 | 0.85947463 |
| NAA40 | 358.182655 | 0.120859439 | 0.22129534 | 0.54614543 | 0.58496598 | 0.76929665 |
| OTUB1 | 289.885273 | 0.090345382 | 0.19049246 | 0.47427275 | 0.63530542 | 0.80389813 |
| MACROD1 | 518.828532 | 0.914021839 | 0.40098018 | 2.27946887 | 0.02263921 | 0.0968398 |
| AP000721.2 | 178.012857 | 1.891673481 | 0.61635487 | 3.06913041 | 0.00214683 | 0.01809665 |
| FLRT1 | 55.9730058 | 2.056848748 | 0.57285338 | 3.59053262 | 0.00033 | 0.0043099 |
| STIP1 | 277.507616 | -0.217855269 | 0.20216607 | -1.0776055 | 0.28120984 | 0.50372942 |
| TRPT1 | 105.18451 | 0.166412023 | 0.30933618 | 0.53796496 | 0.59060124 | 0.77217181 |
| NUDT22 | 212.338656 | 0.186961242 | 0.17127013 | 1.09161616 | 0.27500185 | 0.49647832 |
| AP001453.1 | 100.828565 | 0.556379838 | 0.33153391 | 1.67819889 | 0.09330828 | 0.25117608 |
| DNAJC4 | 154.006853 | 0.924559573 | 0.3874155 | 2.38648058 | 0.01701051 | 0.0797062 |
| VEGFB | 421.612263 | 0.366076976 | 0.27260596 | 1.34287956 | 0.17931098 | 0.38275997 |
| FKBP2 | 365.262304 | -0.086875941 | 0.37523022 | -0.231527 | 0.81690538 | 0.91483008 |
| PPP1R14B | 165.270495 | -1.006312332 | 0.39341579 | -2.557885 | 0.01053109 | 0.0576005 |
| PLCB3 | 61.5288988 | -0.336669501 | 0.32475742 | -1.03668 | 0.29988504 | 0.52272123 |
| BAD | 174.392544 | -0.299661886 | 0.2895112 | -1.0350615 | 0.30064021 | 0.52376109 |
| GPR137 | 208.081621 | 0.049869443 | 0.24964892 | 0.19975829 | 0.84166962 | 0.92654505 |
| ESRRA | 616.919387 | 0.222928955 | 0.26463454 | 0.84240308 | 0.39956237 | 0.62113914 |
| TRMT112 | 226.662286 | -1.39372771 | 0.4462527 | -3.1231805 | 0.00178908 | 0.0156503 |
| PRDX5 | 367.978128 | -0.749958278 | 0.41694599 | -1.7986941 | 0.07206709 | 0.21221706 |
| RPS6KA4 | 76.6526135 | -0.876433058 | 0.40803688 | -2.1479261 | 0.03171963 | 0.12257932 |
| SLC22A12 | 49.6141485 | 0.173769682 | 0.76333256 | 0.2276461 | 0.81992138 | 0.91651491 |
| SF1 | 1851.51854 | -0.080255471 | 0.1932778 | -0.4152338 | 0.67797078 | 0.83118075 |
| MAP4K2 | 110.660397 | 0.638065538 | 0.26103727 | 2.44434655 | 0.01451148 | 0.07160334 |
| MEN1 | 98.6118766 | -0.135442197 | 0.30748275 | -0.4404871 | 0.65958432 | 0.81980614 |
| CDC42BPG | 71.1297988 | 0.238211816 | 0.2600289 | 0.91609746 | 0.35961578 | 0.58499567 |
| EHD1 | 148.333514 | -0.187428078 | 0.31782209 | -0.5897264 | 0.55537409 | 0.74764945 |
| ATG2A | 225.431014 | -0.589131338 | 0.240154 | -2.4531399 | 0.01416153 | 0.07032316 |
| PPP2R5B | 112.569527 | 0.538709065 | 0.23430757 | 2.29915349 | 0.02149622 | 0.09377267 |
| ZFPL1 | 94.1237927 | 0.028882354 | 0.32958544 | 0.08763237 | 0.93016887 | 0.96915731 |
| VPS51 | 555.164938 | -0.241824398 | 0.20005 | -1.2088198 | 0.2267321 | 0.44172629 |
| TM7SF2 | 104.459588 | -0.101097488 | 0.41754763 | -0.242122 | 0.8086856 | 0.91010272 |
| FAU | 772.820219 | -0.959337185 | 0.40859018 | -2.3479203 | 0.01887856 | 0.08572255 |
| SYVN1 | 150.557353 | 0.083350408 | 0.30083027 | 0.27706789 | 0.78172798 | 0.89582484 |
| MRPL49 | 96.4987993 | -0.821982665 | 0.39348708 | -2.0889699 | 0.03671043 | 0.13485073 |
| AP003068.2 | 49.7052194 | 0.0030962 | 0.43162393 | 0.00717337 | 0.99427653 | 0.99783229 |
| CAPN1 | 504.829983 | -0.791856698 | 0.26577129 | -2.9794666 | 0.00288751 | 0.02248061 |
| POLA2 | 87.7429678 | -0.236907712 | 0.47417972 | -0.4996159 | 0.61734558 | 0.7902755 |
| DPF2 | 298.76189 | -0.02374689 | 0.18155417 | -0.1307978 | 0.89593525 | 0.95200843 |
| SLC25A45 | 97.928657 | -1.281453828 | 0.63329619 | -2.0234668 | 0.04302504 | 0.14951796 |
| FRMD8 | 277.978553 | -0.986949055 | 0.28971888 | -3.4065749 | 0.00065784 | 0.00729584 |
| NEAT1 | 78404.0362 | -2.166767822 | 0.41022241 | -5.2819344 | 1.28E-07 | 7.56E-06 |
| MALAT1 | 1039802.28 | -1.15800011 | 0.40362074 | -2.8690303 | 0.00411732 | 0.02922329 |
| SCYL1 | 250.390332 | -0.497466773 | 0.27464851 | -1.8112852 | 0.07009672 | 0.20844384 |
| LTBP3 | 623.845876 | 0.236554029 | 0.36974711 | 0.63977249 | 0.52232052 | 0.72221624 |
| SSSCA1 | 87.6425503 | -1.173841836 | 0.49027147 | -2.3942691 | 0.01665353 | 0.0788446 |
| FAM89B | 81.6510387 | -1.158036736 | 0.41613327 | -2.7828507 | 0.00538836 | 0.035428 |
| EHBP1L1 | 247.027044 | -2.555325924 | 0.47797048 | -5.3462003 | 8.98E-08 | 5.90E-06 |
| MAP3K11 | 148.410162 | -0.616619232 | 0.30872297 | -1.9973221 | 0.0457902 | 0.1562946 |
| PCNX3 | 143.426179 | -0.106473063 | 0.32456479 | -0.3280487 | 0.74287483 | 0.87441041 |
| SIPA1 | 94.0187647 | -1.724481676 | 0.32907945 | -5.2403201 | 1.60E-07 | 9.13E-06 |
| RELA | 204.201016 | -0.89174165 | 0.27199346 | -3.2785408 | 0.00104345 | 0.01033206 |
| KAT5 | 242.960323 | -0.154566943 | 0.25039979 | -0.6172806 | 0.53704963 | 0.73382524 |
| RNASEH2C | 158.693685 | -0.284294273 | 0.29611737 | -0.9600729 | 0.3370185 | 0.56227228 |
| AP5B1 | 71.4827027 | -0.651116422 | 0.39453523 | -1.6503378 | 0.09887386 | 0.26052758 |
| CFL1 | 675.615446 | -1.566831289 | 0.34401104 | -4.5545959 | 5.25E-06 | 0.00015865 |
| MUS81 | 152.429035 | -0.278191249 | 0.22144735 | -1.256241 | 0.2090286 | 0.42044445 |
| EFEMP2 | 74.3918142 | -0.553713973 | 0.35223124 | -1.5720183 | 0.1159463 | 0.28962455 |
| FIBP | 61.5396273 | -0.518447508 | 0.44067441 | -1.1764865 | 0.23940051 | 0.45750019 |
| CCDC85B | 68.9982851 | -1.120796022 | 0.41087756 | -2.7278103 | 0.00637563 | 0.03998844 |
| C11orf68 | 56.2082127 | -0.214253878 | 0.44582334 | -0.4805802 | 0.63081487 | 0.80065664 |

| | | | | | | |
|---|---|---|---|---|---|---|
| DRAP1 | 131.808212 | -0.365247809 | 0.37509027 | -0.9737597 | 0.33017585 | 0.55571163 |
| SART1 | 158.371149 | -0.09419469 | 0.28935025 | -0.3255387 | 0.74477343 | 0.87520849 |
| EIF1AD | 70.110196 | 0.124558397 | 0.24974766 | 0.49873699 | 0.61796468 | 0.79042527 |
| BANF1 | 192.42414 | -0.81851857 | 0.39856613 | -2.0536581 | 0.04000879 | 0.14260014 |
| SF3B2 | 426.521205 | -0.387317502 | 0.18523032 | -2.0910048 | 0.03652763 | 0.13462429 |
| PACS1 | 3684.92721 | -0.826310951 | 0.18058546 | -4.5757336 | 4.75E-06 | 0.0001441 |
| KLC2 | 131.659069 | -0.559346712 | 0.30651202 | -1.8248769 | 0.06801961 | 0.20444622 |
| RAB1B | 241.265509 | -0.868764827 | 0.33864313 | -2.5654287 | 0.01030484 | 0.0567389 |
| YIF1A | 100.343021 | -0.669518744 | 0.3969192 | -1.6867885 | 0.09164403 | 0.24770329 |
| BRMS1 | 102.626988 | -0.454779026 | 0.33115201 | -1.3733241 | 0.16965162 | 0.3699733 |
| B4GAT1 | 54.3933571 | -0.060167674 | 0.50652041 | -0.1187863 | 0.90544468 | 0.95668833 |
| AP001107.9 | 46.9421468 | 0.291842414 | 0.35046829 | 0.83272132 | 0.40500192 | 0.62554611 |
| SLC29A2 | 238.595042 | 0.735163959 | 0.2925222 | 2.51319029 | 0.01196448 | 0.06280275 |
| MRPL11 | 89.6622149 | -0.344349299 | 0.36551112 | -0.9421035 | 0.34613966 | 0.57123423 |
| DPP3 | 56.142303 | -0.60945337 | 0.46006528 | -1.3247106 | 0.18526715 | 0.39092552 |
| ZDHHC24 | 143.069204 | -0.606026129 | 0.26901415 | -2.2527667 | 0.02427386 | 0.1015627 |
| CTSF | 162.093809 | -0.335761831 | 0.45721981 | -0.7343554 | 0.46273217 | 0.67575779 |
| CCS | 292.984865 | 0.132761175 | 0.2111037 | 0.6288908 | 0.52942055 | 0.72761586 |
| RBM14-RBM | 368.636432 | 0.19246708 | 0.26041818 | 0.73906928 | 0.45986493 | 0.67351662 |
| RBM4 | 220.694188 | 0.383982191 | 0.34223612 | 1.12198032 | 0.26187081 | 0.48277062 |
| RBM4B | 293.238862 | 0.435442374 | 0.20143943 | 2.1616541 | 0.03064485 | 0.11958281 |
| AP001157.1 | 67.8768717 | 0.292287217 | 0.41482775 | 0.70459899 | 0.48105983 | 0.68948936 |
| SPTBN2 | 142.636343 | -0.001198792 | 0.5014443 | -0.0023907 | 0.99809252 | 0.99945448 |
| C11orf80 | 1032.51635 | -1.020848129 | 0.24392339 | -4.1851178 | 2.85E-05 | 0.00062852 |
| RCE1 | 63.9722085 | -0.205937643 | 0.33991468 | -0.605851 | 0.54461371 | 0.73949449 |
| PC | 118.855238 | 0.308542801 | 0.25221868 | 1.22331464 | 0.22121088 | 0.43520621 |
| RHOD | 130.173861 | -0.362576238 | 0.32197187 | -1.1261115 | 0.26011831 | 0.48104792 |
| KDM2A | 3584.28063 | -0.205700058 | 0.16599169 | -1.239219 | 0.21526439 | 0.42913181 |
| GRK2 | 262.08251 | -0.836383994 | 0.27841684 | -3.0040711 | 0.00266393 | 0.02121028 |
| ANKRD13D | 380.633723 | -0.787462853 | 0.28744541 | -2.7395214 | 0.00615287 | 0.03900524 |
| SSH3 | 140.188829 | -1.10852054 | 0.28502006 | -3.8892721 | 0.00010055 | 0.00167664 |
| RAD9A | 1368.27165 | 0.304327849 | 0.17207584 | 1.76856816 | 0.07696597 | 0.2212981 |
| CLCF1 | 67.5325737 | -2.395279638 | 0.5586507 | -4.287616 | 1.81E-05 | 0.00043295 |
| PPP1CA | 305.640494 | -1.435640391 | 0.42611875 | -3.3691087 | 0.00075412 | 0.00807877 |
| RPS6KB2 | 142.652485 | 0.014029132 | 0.3691509 | 0.03800379 | 0.96968466 | 0.98641238 |
| CORO1B | 171.430058 | -1.331309961 | 0.3503703 | -3.7997227 | 0.00014486 | 0.00224845 |
| TMEM134 | 230.19089 | -0.34579914 | 0.22884517 | -1.5110616 | 0.13077277 | 0.31315763 |
| AIP | 98.1959397 | -0.52950651 | 0.38396832 | -1.379037 | 0.16788335 | 0.36781741 |
| PITPNM1 | 157.403234 | -1.351566872 | 0.41196203 | -3.2808045 | 0.00103511 | 0.0102726 |
| CDK2AP2 | 214.069847 | -1.042837785 | 0.46595316 | -2.2380743 | 0.02521621 | 0.10419747 |
| GSTP1 | 1980.77843 | -1.859599491 | 0.5213021 | -3.5672204 | 0.00036079 | 0.00460005 |
| NDUFV1 | 656.264749 | 0.563811621 | 0.28211477 | 1.99851864 | 0.04566046 | 0.15600761 |
| NUDT8 | 71.8284934 | -0.429889453 | 0.40774278 | -1.0543153 | 0.29173859 | 0.51440973 |
| AP003716.1 | 44.5713072 | 0.397022846 | 0.57349426 | 0.69228739 | 0.48875687 | 0.69568852 |
| UNC93B1 | 144.932859 | -1.406893482 | 0.49703485 | -2.8305731 | 0.00464647 | 0.03173485 |
| NDUFS8 | 473.705864 | 0.018207818 | 0.32335513 | 0.05630904 | 0.95509561 | 0.98014107 |
| TCIRG1 | 60.1177543 | 0.214241024 | 0.58210597 | 0.36804471 | 0.71283989 | 0.85606893 |
| CHKA | 1420.10125 | 0.875284519 | 0.1645878 | 5.31804006 | 1.05E-07 | 6.66E-06 |
| KMT5B | 1441.93716 | 0.207226692 | 0.15102612 | 1.37212487 | 0.17002457 | 0.37027842 |
| C11orf24 | 44.5294574 | -0.418058456 | 0.34412118 | -1.2148583 | 0.22442015 | 0.43912303 |
| LRP5 | 539.509788 | 0.065900562 | 0.31580929 | 0.20867202 | 0.83470429 | 0.92298305 |
| PPP6R3 | 3835.78081 | 0.269528487 | 0.1551758 | 1.7369235 | 0.08240068 | 0.23069744 |
| AP003096.1 | 57.6431939 | 0.008152217 | 0.65255623 | 0.01249274 | 0.99003249 | 0.99606252 |
| TESMIN | 99.1102762 | -0.064005788 | 0.45039859 | -0.1421092 | 0.88699374 | 0.94913293 |
| CPT1A | 1319.5998 | 1.391880933 | 0.51693857 | 2.69254613 | 0.00709087 | 0.04313648 |
| MRPL21 | 116.738354 | -0.033146089 | 0.235139 | -0.1409638 | 0.88789854 | 0.94928354 |
| IGHMBP2 | 183.560827 | -0.040738151 | 0.30895359 | -0.1318585 | 0.89509623 | 0.95172874 |
| TPCN2 | 171.827321 | 0.210321869 | 0.21785686 | 0.96541311 | 0.33433794 | 0.55944639 |
| CCND1 | 275.260106 | -1.150671118 | 0.62890748 | -1.829635 | 0.06730454 | 0.20284564 |
| ORAOV1 | 234.370065 | 0.056775481 | 0.28781568 | 0.19726334 | 0.84362147 | 0.92769172 |
| FADD | 47.7717824 | -1.4379416 | 0.3220646 | -4.4647614 | 8.02E-06 | 0.00022528 |
| PPFIA1 | 1419.84185 | -0.022609462 | 0.2267719 | -0.0997013 | 0.92058144 | 0.96454603 |
| AP002336.2 | 562.912959 | -0.184705108 | 0.43305552 | -0.426516 | 0.66973189 | 0.82642427 |
| AP000487.1 | 136.182799 | 0.515629477 | 0.24900589 | 2.07075211 | 0.03838197 | 0.13904125 |
| CTTN | 1489.36459 | -1.188924629 | 0.20608465 | -5.7691081 | 7.97E-09 | 7.63E-07 |
| SHANK2 | 4619.56821 | 0.235441087 | 0.32659602 | 0.72089393 | 0.47097478 | 0.68235944 |
| SHANK2-AS1 | 49.2893989 | 0.018147836 | 0.66940817 | 0.02711027 | 0.97837178 | 0.99074769 |
| DHCR7 | 54.4982852 | -0.676616952 | 0.55046875 | -1.2291651 | 0.21900993 | 0.43326376 |
| NADSYN1 | 309.931802 | -0.410852951 | 0.27861324 | -1.4746354 | 0.14031058 | 0.32785467 |
| FAM86C1 | 41.1800113 | -0.540359127 | 0.37227015 | -1.4515242 | 0.14663395 | 0.33725691 |
| AP002495.2 | 915.969631 | 0.487557967 | 0.14429386 | 3.37892395 | 0.0007277 | 0.00787855 |
| RNF121 | 857.407008 | -0.236198724 | 0.19231712 | -1.2281732 | 0.21938197 | 0.43356369 |
| IL18BP | 68.9505531 | 0.073955823 | 0.36187042 | 0.20437101 | 0.83806358 | 0.92446904 |
| NUMA1 | 1012.37534 | 0.093546224 | 0.21770509 | 0.4296924 | 0.66741941 | 0.82469498 |
| LAMTOR1 | 362.782263 | -0.742078367 | 0.20968873 | -3.538952 | 0.00040172 | 0.00498713 |
| ANAPC15 | 75.6706464 | -0.288822977 | 0.33890536 | -0.8522231 | 0.39409027 | 0.61640479 |
| FOLR3 | 67.8302012 | -0.216571227 | 0.62023615 | -0.3491754 | 0.72695761 | 0.86532561 |
| FOLR1 | 85.8740651 | -1.196160968 | 0.68072138 | -1.7571961 | 0.07888439 | 0.22456549 |
| INPPL1 | 281.394743 | 0.16022938 | 0.24164792 | 0.66306957 | 0.507286 | 0.71048657 |
| CLPB | 197.128327 | -0.090623626 | 0.20449475 | -0.4431587 | 0.65765096 | 0.81859324 |
| ARAP1 | 307.060344 | 0.122086504 | 0.256962 | 0.47511502 | 0.63470499 | 0.80358653 |
| ATG16L2 | 217.690194 | 0.044514461 | 0.28745446 | 0.15485744 | 0.87693371 | 0.94481381 |
| FCHSD2 | 2546.22362 | -0.418236848 | 0.26893769 | -1.5551441 | 0.11991176 | 0.29561832 |
| P2RY6 | 350.704962 | -2.400362784 | 0.89119063 | -2.6934336 | 0.00707202 | 0.0430878 |
| ARHGEF17 | 501.324474 | 0.106475481 | 0.25391136 | 0.41934115 | 0.67496683 | 0.82939913 |
| FAM168A | 1525.27391 | 0.344412198 | 0.11876488 | 2.89994986 | 0.00373222 | 0.02720105 |
| RAB6A | 1860.90397 | -0.449122473 | 0.1908568 | -2.3531909 | 0.01861308 | 0.08488614 |
| MRPL48 | 379.999545 | 0.655995606 | 0.17965621 | 3.65139403 | 0.00026082 | 0.00360752 |
| COA4 | 221.381223 | -0.724123563 | 0.24825564 | -2.9168463 | 0.0035359 | 0.02615323 |
| PAAF1 | 376.900641 | 0.312987215 | 0.32602373 | 0.9600136 | 0.33704837 | 0.56227228 |
| UCP2 | 46.7731839 | -0.72441946 | 0.5981911 | -1.2110168 | 0.22588898 | 0.44066866 |
| C2CD3 | 844.840772 | 0.322053216 | 0.13957665 | 2.30735732 | 0.02103491 | 0.09241548 |
| PPME1 | 997.098161 | -0.327456577 | 0.27816962 | -1.1771831 | 0.23912243 | 0.45716719 |
| P4HA3 | 182.197514 | -0.446427827 | 0.47296349 | -0.9438949 | 0.34522335 | 0.57043473 |
| PGM2L1 | 478.982507 | -1.915390593 | 0.46993034 | -4.0759033 | 4.58E-05 | 0.00092154 |
| KCNE3 | 54.6894492 | 1.495754279 | 0.54433611 | 2.74785054 | 0.00599873 | 0.03829738 |
| POLD3 | 471.811818 | -0.086734029 | 0.29837309 | -0.2906898 | 0.77128854 | 0.8896518 |
| RNF169 | 1128.43015 | -0.086654858 | 0.22271846 | -0.389078 | 0.69721843 | 0.84545655 |
| XRRA1 | 1149.31049 | 0.594723696 | 0.45047887 | 1.32020331 | 0.18676715 | 0.39251209 |
| SPCS2 | 749.916835 | -0.082774028 | 0.23958487 | -0.3454894 | 0.72972649 | 0.8673587 |
| NEU3 | 199.247171 | 0.358378238 | 0.21418234 | 1.67323898 | 0.09428026 | 0.25270859 |
| ARRB1 | 183.297602 | -0.819826314 | 0.35043974 | -2.3394216 | 0.01931362 | 0.08709923 |
| RPS3 | 1241.69737 | -1.0300745 | 0.38838355 | -2.6522094 | 0.00799669 | 0.04732452 |
| SERPINH1 | 120.534624 | -0.80357427 | 0.25574178 | -3.1421313 | 0.00167723 | 0.01493898 |
| MAP6 | 102.751729 | -0.345703649 | 0.44562659 | -0.7757698 | 0.43788492 | 0.65419956 |
| UVRAG | 1638.61237 | 0.13862035 | 0.16840886 | 0.8231179 | 0.41044095 | 0.62997812 |
| THAP12 | 206.033637 | 0.282956357 | 0.23780728 | 1.18985574 | 0.2341031 | 0.45136007 |
| GVQW3 | 104.189487 | 1.076858891 | 0.39565388 | 2.72171952 | 0.00649432 | 0.04054042 |
| AP002360.1 | 104.953667 | 1.112514484 | 0.39246985 | 2.83464953 | 0.0045876 | 0.03147927 |
| EMSY | 1120.11697 | 0.027261038 | 0.23178798 | 0.11761196 | 0.90637513 | 0.95724867 |
| ACER3 | 1491.07535 | -0.81872755 | 0.44018014 | -1.8599829 | 0.06288794 | 0.19413106 |
| CAPN5 | 147.450386 | -1.497708743 | 0.64035828 | -2.3388606 | 0.01934265 | 0.08717061 |
| PAK1 | 881.35009 | 0.134152484 | 0.29651136 | 0.45243623 | 0.65095475 | 0.81352716 |

| | | | | | | |
|---|---|---|---|---|---|---|
| CLNS1A | 642.867893 | 0.235124364 | 0.21107497 | 1.11393768 | 0.26530592 | 0.48636961 |
| AQP11 | 50.6916784 | 0.237236253 | 0.34566855 | 0.68631135 | 0.49251679 | 0.69848124 |
| RSF1 | 2265.66716 | 0.018903274 | 0.13381217 | 0.14126722 | 0.88765885 | 0.94928354 |
| AAMDC | 694.188145 | 0.187534068 | 0.16254186 | 1.15375857 | 0.24859916 | 0.46787237 |
| AP002812.2 | 297.473409 | 0.423514179 | 0.29324653 | 1.44422572 | 0.14867549 | 0.34065124 |
| INTS4 | 597.158571 | 0.044927259 | 0.22675128 | 0.19813453 | 0.84293981 | 0.92739954 |
| ALG8 | 305.729875 | -0.036857361 | 0.26784119 | -0.137609 | 0.89054944 | 0.95041347 |
| KCTD21-AS1 | 59.0264566 | 0.271070476 | 0.74196074 | 0.36534342 | 0.71485507 | 0.85755341 |
| KCTD21 | 52.4384004 | 0.011381069 | 0.33782656 | 0.03368909 | 0.97312508 | 0.98804941 |
| GAB2 | 1293.04935 | -1.424021798 | 0.31550706 | -4.5134388 | 6.38E-06 | 0.00018477 |
| AP003086.2 | 496.267725 | -1.627320094 | 0.62742298 | -2.5936571 | 0.00949611 | 0.0533219 |
| NARS2 | 801.431336 | 0.415034191 | 0.24695216 | 1.68062589 | 0.0928356 | 0.2502072 |
| TENM4 | 3488.10339 | -0.21661633 | 0.39776045 | -0.5445899 | 0.58603559 | 0.77001836 |
| MIR4300HG | 47.0450843 | 0.02045628 | 0.60859771 | 0.03361215 | 0.97318643 | 0.98804941 |
| PRCP | 505.116459 | 0.494799395 | 0.28614416 | 1.7291962 | 0.08377399 | 0.23315857 |
| RAB30 | 289.802804 | 0.721787135 | 0.26651769 | 2.70821477 | 0.00676462 | 0.04185195 |
| RAB30-AS1 | 994.946058 | 0.313545362 | 0.23077096 | 1.35868638 | 0.17424599 | 0.37589771 |
| AP001767.3 | 97.9768755 | 0.251357949 | 0.57600192 | 0.43638387 | 0.66255824 | 0.8219904 |
| PCF11 | 708.801101 | 0.184067656 | 0.2186166 | 0.84196559 | 0.39980722 | 0.62134844 |
| PCF11-AS1 | 75.4902579 | 0.336220711 | 0.34480695 | 0.97509843 | 0.32951144 | 0.55488483 |
| ANKRD42 | 414.190441 | -0.490496231 | 0.24150543 | -2.0309947 | 0.04225554 | 0.14754253 |
| CCDC90B | 317.650056 | -0.287602502 | 0.19074099 | -1.507817 | 0.13160137 | 0.31440036 |
| AP000446.1 | 369.631216 | 0.483093617 | 0.31160634 | 1.55033309 | 0.12106159 | 0.29750745 |
| DLG2 | 2499.69121 | 0.36763457 | 0.24094058 | 1.52583084 | 0.127052 | 0.3070091 |
| TMEM126B | 205.671657 | -0.138698397 | 0.20588114 | -0.6736819 | 0.50051358 | 0.70482771 |
| TMEM126A | 87.8749809 | 0.169532231 | 0.32568862 | 0.52053471 | 0.60269094 | 0.77979671 |
| CREBZF | 835.963618 | 0.215988166 | 0.15530735 | 1.39071434 | 0.16431207 | 0.36306425 |
| SYTL2 | 1851.54727 | -0.229296217 | 0.4366628 | -0.5251105 | 0.59950638 | 0.77773323 |
| PICALM | 4336.15958 | 0.265470312 | 0.14720533 | 1.80340153 | 0.07132518 | 0.21075466 |
| EED | 479.695975 | 0.696427903 | 0.22787322 | 3.05620781 | 0.00224156 | 0.01865397 |
| HIKESHI | 463.452899 | 0.460585123 | 0.16257901 | 2.83299262 | 0.00461144 | 0.03156092 |
| CCDC81 | 51.5691152 | 0.317554103 | 0.33453433 | 0.94924219 | 0.34249745 | 0.56763059 |
| AP001831.1 | 379.726593 | 0.324725675 | 0.38386741 | 0.84593187 | 0.39759075 | 0.61924021 |
| ME3 | 1326.42618 | 0.986450323 | 0.29371527 | 3.35852589 | 0.00078359 | 0.00826862 |
| AP003059.1 | 148.672544 | 0.8365951 | 0.46562018 | 1.7967329 | 0.07237803 | 0.21270422 |
| AP003059.2 | 153.240621 | 1.205447736 | 0.55146958 | 2.18588257 | 0.0288242 | 0.11423307 |
| PRSS23 | 387.937387 | 0.58050727 | 0.40876605 | 1.42014551 | 0.15556532 | 0.35053947 |
| AP001528.2 | 319.258514 | 0.682296259 | 0.66330897 | 1.0286251 | 0.30365588 | 0.5268607 |
| FZD4 | 168.060379 | 0.48420361 | 0.30154482 | 1.60574339 | 0.1083303 | 0.27674263 |
| AP001528.1 | 172.890557 | 0.385625292 | 0.38050761 | 1.01344962 | 0.31084545 | 0.53450314 |
| TMEM135 | 1551.60048 | 0.333992617 | 0.20897359 | 1.59825278 | 0.10998673 | 0.27933373 |
| RAB38 | 146.695901 | 0.658538158 | 0.41522634 | 1.58597396 | 0.11274522 | 0.2842713 |
| CTSC | 506.024933 | 0.325531183 | 0.36864261 | 0.88305361 | 0.37720731 | 0.60028794 |
| GRM5 | 128.375008 | 0.861081888 | 0.43827266 | 1.96471733 | 0.04944696 | 0.16493559 |
| NOX4 | 906.68353 | 0.120550241 | 0.46398203 | 0.25981662 | 0.79500523 | 0.90385363 |
| CHORDC1 | 452.455619 | 0.433612262 | 0.37416672 | 1.15887449 | 0.24650735 | 0.46556654 |
| SLC36A4 | 1168.51693 | 0.056812157 | 0.22900413 | 0.24808355 | 0.80406976 | 0.90804718 |
| DEUP1 | 72.146123 | 0.252009343 | 0.42552899 | 0.59222603 | 0.55369924 | 0.74638364 |
| SMCO4 | 344.223752 | -0.329708601 | 0.33973978 | -0.9704739 | 0.33181032 | 0.55704883 |
| CEP295 | 459.318738 | 0.140654833 | 0.25218537 | 0.55774382 | 0.57701933 | 0.76363574 |
| TAF1D | 1491.44067 | -0.181060319 | 0.18519946 | -0.9776504 | 0.32824728 | 0.55346341 |
| AP001273.2 | 56.1970161 | 0.239326634 | 0.29576327 | 0.80918309 | 0.41840984 | 0.63753323 |
| PANX1 | 119.08214 | -0.682431393 | 0.25341146 | -2.6929776 | 0.0070817 | 0.04312688 |
| MRE11 | 552.775307 | -0.027863617 | 0.24713863 | -0.1127449 | 0.91023281 | 0.95961675 |
| ANKRD49 | 131.255441 | 0.362794344 | 0.32439909 | 1.1183581 | 0.26341408 | 0.48439746 |
| PIWIL4 | 101.663421 | -1.027022485 | 0.41360224 | -2.4831163 | 0.01302385 | 0.06657589 |
| AMOTL1 | 287.828386 | -1.147006961 | 0.64154904 | -1.7878711 | 0.07379681 | 0.21554455 |
| CWC15 | 249.649173 | -0.11273051 | 0.15476475 | -0.7283991 | 0.4663693 | 0.67903174 |
| KDM4D | 46.9383625 | 0.634504621 | 0.49052284 | 1.29352716 | 0.1958288 | 0.4034788 |
| SRSF8 | 232.890483 | -0.692762655 | 0.31705978 | -2.1849591 | 0.02889184 | 0.11433041 |
| ENDOD1 | 478.975697 | -1.264805391 | 0.40913532 | -3.0914109 | 0.00199208 | 0.01707551 |
| AP000787.1 | 345.511826 | -0.234230656 | 0.37085698 | -0.631593 | 0.52765289 | 0.72594178 |
| SESN3 | 594.87022 | -0.554746426 | 0.32628436 | -1.7001931 | 0.08909461 | 0.24292017 |
| FAM76B | 290.069543 | 0.118806423 | 0.22436226 | 0.52952944 | 0.59643823 | 0.77619237 |
| CEP57 | 795.408186 | -0.102301359 | 0.16325151 | -0.6266488 | 0.53088949 | 0.72863563 |
| MTMR2 | 621.730323 | 0.082694243 | 0.15858593 | 0.52144753 | 0.60205505 | 0.77940904 |
| MAML2 | 14811.2758 | 0.509726178 | 0.21784547 | 2.33985205 | 0.01929138 | 0.08707889 |
| CCDC82 | 482.340639 | -0.188553 | 0.21428727 | -0.8799076 | 0.37890937 | 0.6017088 |
| JRKL | 136.488336 | -0.251361313 | 0.28623294 | -0.8781705 | 0.37985122 | 0.60262489 |
| CNTN5 | 168.781632 | 1.022988785 | 0.50432058 | 2.0284494 | 0.0425144 | 0.14828961 |
| ARHGAP42 | 109.402566 | -0.188240505 | 0.40672319 | -0.4628222 | 0.64349185 | 0.80905043 |
| TRPC6 | 135.949211 | 1.465216728 | 0.4284415 | 3.41987585 | 0.0006265 | 0.00706061 |
| CEP126 | 435.108674 | -0.327834569 | 0.26854783 | -1.2207679 | 0.22217391 | 0.43639586 |
| CFAP300 | 61.5311987 | -1.721296088 | 0.79372549 | -2.168629 | 0.03011086 | 0.11788216 |
| YAP1 | 4108.63794 | -0.610889396 | 0.19542858 | -3.1258959 | 0.00177264 | 0.01556838 |
| BIRC3 | 554.191611 | -3.199011127 | 0.58123953 | -5.5037742 | 3.72E-08 | 2.79E-06 |
| BIRC2 | 543.499138 | -0.08551569 | 0.22380443 | -0.3821001 | 0.70238713 | 0.8488446 |
| TMEM123 | 1140.50365 | -0.234374274 | 0.25300097 | -0.926377 | 0.35425011 | 0.58005327 |
| MMP7 | 621.028705 | -3.564781805 | 0.72960419 | -4.8859119 | 1.03E-06 | 4.21E-05 |
| DCUN1D5 | 346.396108 | 0.118850599 | 0.20898053 | 0.56871614 | 0.56954879 | 0.75907275 |
| DYNC2H1 | 2202.18679 | 0.143371704 | 0.1852621 | 0.77388578 | 0.43899834 | 0.65485132 |
| PDGFD | 462.190349 | 0.013713135 | 0.39930888 | 0.03434217 | 0.97260429 | 0.98801278 |
| CASP4 | 623.990657 | -1.790279472 | 0.3562369 | -5.0255307 | 5.02E-07 | 2.37E-05 |
| MSANTD4 | 166.017363 | 0.00911653 | 0.18644021 | 0.04889788 | 0.96100068 | 0.98288592 |
| KBTBD3 | 209.983763 | 0.127592134 | 0.25972552 | 0.49125759 | 0.62324427 | 0.79494337 |
| AASDHPPT | 322.022104 | 0.353746457 | 0.26202105 | 1.35006886 | 0.1769939 | 0.37941536 |
| GUCY1A2 | 56.6373169 | -0.884687261 | 0.39974552 | -2.2131261 | 0.02688895 | 0.10888294 |
| AP000766.1 | 74.6973884 | 0.246930431 | 0.35613582 | 0.69336027 | 0.48808349 | 0.69502858 |
| CWF19L2 | 925.320656 | 0.059322714 | 0.20488254 | 0.28954499 | 0.77216436 | 0.89032114 |
| ALKBH8 | 224.652908 | 0.472985153 | 0.20607451 | 2.2952142 | 0.02172085 | 0.09440958 |
| SLC35F2 | 299.187422 | -0.781099751 | 0.39253893 | -1.9898657 | 0.04660573 | 0.15821514 |
| CUL5 | 1731.86264 | 0.305913453 | 0.12838693 | 2.38274613 | 0.01718404 | 0.08031988 |
| AP002433.1 | 42.2075869 | 0.417440425 | 0.55443664 | 0.75290916 | 0.4515045 | 0.66729785 |
| ACAT1 | 743.551512 | 0.655032244 | 0.27112241 | 2.41600181 | 0.01569198 | 0.07562036 |
| NPAT | 522.087439 | -0.169780942 | 0.15913753 | -1.0668818 | 0.28602519 | 0.50849351 |
| ATM | 1624.31426 | 0.116969937 | 0.23417644 | 0.4994949 | 0.61743078 | 0.7902755 |
| C11orf65 | 646.428474 | 0.583432186 | 0.36209585 | 1.61126449 | 0.10712209 | 0.27485929 |
| KDELC2 | 111.302816 | 1.15063759 | 0.33656964 | 3.41872073 | 0.00062916 | 0.00707286 |
| EXPH5 | 852.31908 | 1.030184165 | 0.40100885 | 2.56898112 | 0.0101998 | 0.05631792 |
| DDX10 | 1041.78861 | 0.304051108 | 0.24841359 | 1.22397129 | 0.22096306 | 0.43484819 |
| AP003049.2 | 38.1055624 | 0.462283215 | 0.51961728 | 0.88966098 | 0.37364795 | 0.59763858 |
| RDX | 1492.0998 | 0.333309388 | 0.20896489 | 1.59504971 | 0.11070112 | 0.28071628 |
| ZC3H12C | 718.457125 | -0.658635756 | 0.29546538 | -2.2291469 | 0.02580413 | 0.1057545 |
| FDX1 | 463.888334 | 0.619377543 | 0.18164709 | 3.40978508 | 0.00065014 | 0.0072348 |
| ARHGAP20 | 42.8138592 | 0.448366612 | 0.39353234 | 1.13933866 | 0.25456193 | 0.47540061 |
| LINC02550 | 399.374672 | 6.7889769 | 1.58208105 | 4.29116884 | 1.78E-05 | 0.00042998 |
| COLCA1 | 192.074174 | 1.699131723 | 0.56695031 | 2.99696762 | 0.0027268 | 0.02155491 |
| COLCA2 | 100.534341 | 0.733255411 | 0.58509325 | 1.25322828 | 0.21012264 | 0.42219501 |
| POU2AF1 | 60.040102 | 2.514737287 | 0.84601821 | 2.97243874 | 0.00295444 | 0.02290212 |
| SIK2 | 2996.68787 | -0.723996127 | 0.29034917 | -2.493536 | 0.01264777 | 0.06528505 |
| PPP2R1B | 724.836653 | -0.602482388 | 0.31133668 | -1.9351475 | 0.0529722 | 0.17273155 |
| C11orf1 | 84.3768411 | 0.376575059 | 0.33163847 | 1.13549871 | 0.25616642 | 0.47715651 |
| CRYAB | 2394.05325 | -2.441854604 | 0.61228087 | -3.9881282 | 6.66E-05 | 0.00121839 |

| | | | | | | |
|---|---|---|---|---|---|---|
| DIXDC1 | 587.830249 | 0.140294005 | 0.30927337 | 0.45362459 | 0.65009905 | 0.81297534 |
| DLAT | 382.922579 | 0.59207534 | 0.21612799 | 2.73946623 | 0.0061539 | 0.03900524 |
| PIH1D2 | 86.6147809 | 0.623139659 | 0.28085629 | 2.21871357 | 0.02650622 | 0.10792868 |
| NKAPD1 | 156.718904 | -0.01007313 | 0.23412948 | -0.0430238 | 0.9656826 | 0.98461609 |
| TIMM8B | 191.769576 | -0.634518614 | 0.35924561 | -1.7662529 | 0.07735343 | 0.22191693 |
| AP002884.3 | 279.418864 | 0.283012775 | 0.55803417 | 0.5071603 | 0.61204235 | 0.78684784 |
| IL18 | 70.212472 | -0.155501075 | 0.53500043 | -0.290656 | 0.77131444 | 0.8896518 |
| BCO2 | 106.391827 | 0.086364583 | 0.62030798 | 0.13922855 | 0.88926955 | 0.95012225 |
| PTS | 690.502088 | -0.230941082 | 0.23012229 | -1.0035581 | 0.31559168 | 0.5399107 |
| AP002884.1 | 412.441584 | -0.433548366 | 0.39318318 | -1.1026626 | 0.27017373 | 0.49175977 |
| AP003063.1 | 60.4890603 | -0.147012514 | 0.57645639 | -0.255028 | 0.79870148 | 0.90525349 |
| TTC12 | 488.140773 | 0.230056348 | 0.20021187 | 1.14906448 | 0.25052939 | 0.47053075 |
| TMPRSS5 | 41.0320475 | -0.828235308 | 0.67612576 | -1.2249723 | 0.22058567 | 0.43435181 |
| ZW10 | 264.330816 | 0.353311264 | 0.2320868 | 1.52232381 | 0.12792796 | 0.30854958 |
| USP28 | 418.290201 | -0.013916397 | 0.21247781 | -0.0654958 | 0.94777928 | 0.97760554 |
| ZBTB16 | 2710.17512 | -2.120366062 | 0.48542482 | -4.3680627 | 1.25E-05 | 0.00031842 |
| AP002518.2 | 607.196582 | -5.130112139 | 1.15596103 | -4.4379629 | 9.08E-06 | 0.00024753 |
| NNMT | 2624.16997 | -6.471535216 | 1.07322499 | -6.0299893 | 1.64E-09 | 2.01E-07 |
| C11orf71 | 69.8164698 | 0.017854717 | 0.33100172 | 0.05394146 | 0.9569818 | 0.98107945 |
| AP002373.1 | 53.7931008 | -1.805880301 | 0.63459818 | -2.8457067 | 0.0044313 | 0.03066322 |
| REXO2 | 542.979208 | -0.10024847 | 0.41762932 | -0.2400417 | 0.8102979 | 0.91083524 |
| CADM1 | 6608.9211 | 1.148757368 | 0.40069586 | 2.86690599 | 0.00414506 | 0.02927921 |
| AP000462.1 | 299.609896 | 1.31521574 | 0.58194375 | 2.26003928 | 0.02381881 | 0.1002791 |
| BUD13 | 125.987436 | 0.184963581 | 0.26474278 | 0.69865393 | 0.48476833 | 0.69239889 |
| ZPR1 | 74.5116819 | -0.671789145 | 0.32863412 | -2.0441856 | 0.04093521 | 0.14473393 |
| APOA1-AS | 60.3841872 | 0.07327337 | 0.59273395 | 0.12361933 | 0.90161669 | 0.95457905 |
| SIK3 | 8061.50504 | 0.213990221 | 0.25928025 | 0.82532403 | 0.40918765 | 0.62915416 |
| PAFAH1B2 | 800.030937 | -0.309128048 | 0.2136563 | -1.4468473 | 0.14793969 | 0.33937745 |
| SIDT2 | 180.745764 | 0.172983493 | 0.2136551 | 0.80963896 | 0.41814771 | 0.63753323 |
| TAGLN | 86.774375 | 0.484318244 | 0.33621941 | 1.4404827 | 0.14973088 | 0.34200906 |
| PCSK7 | 688.683506 | 0.132145196 | 0.24313972 | 0.5434949 | 0.5867891 | 0.77047088 |
| RNF214 | 484.295934 | 0.110265972 | 0.21614773 | 0.5101417 | 0.60995219 | 0.78542196 |
| BACE1 | 340.844839 | 0.159348188 | 0.24563081 | 0.64873045 | 0.51651262 | 0.71779223 |
| BACE1-AS | 39.3907773 | 0.703873934 | 0.36838522 | 1.91070079 | 0.05604304 | 0.17959063 |
| CEP164 | 492.481006 | 0.424864801 | 0.16402689 | 2.59021439 | 0.00959162 | 0.05375294 |
| AP000892.1 | 71.8820333 | 1.020268947 | 0.48288971 | 2.11284052 | 0.03461442 | 0.13000338 |
| DSCAML1 | 3597.57653 | -0.239606698 | 0.63164266 | -0.379339 | 0.70443614 | 0.85038357 |
| FXYD2 | 971.078912 | -0.121732923 | 0.5420913 | -0.2245617 | 0.82232029 | 0.91802136 |
| AP000757.2 | 89.7311406 | -0.059140101 | 0.43247979 | -0.1367465 | 0.89123117 | 0.9506034 |
| FXYD6-FXYD: | 241.048463 | 1.280354298 | 0.89465057 | 1.4311222 | 0.1523952 | 0.3456367 |
| AP000757.1 | 281.490951 | 0.073142844 | 0.58462367 | 0.12511099 | 0.90043568 | 0.95395051 |
| SMIM35 | 43.4213389 | -0.507750943 | 0.42585736 | -1.1923029 | 0.23314252 | 0.44996778 |
| TMPRSS4 | 844.664922 | -2.87031048 | 0.76612168 | -3.7465465 | 0.00017929 | 0.00267805 |
| MPZL3 | 246.447141 | -0.41975207 | 0.23545075 | -1.7827596 | 0.07462545 | 0.21688176 |
| MPZL2 | 83.3531525 | -1.560639589 | 0.40651672 | -3.8390539 | 0.00012351 | 0.00198471 |
| UBE4A | 616.687824 | 0.05335784 | 0.14200043 | 0.37575831 | 0.70709658 | 0.85250445 |
| ATP5MG | 924.066057 | 0.092988562 | 0.39226148 | 0.23705759 | 0.81261211 | 0.91196992 |
| KMT2A | 2275.12157 | -0.51171152 | 0.08084287 | -6.3297052 | 2.46E-10 | 3.61E-08 |
| AP001267.3 | 139.508592 | 0.420617507 | 0.35208158 | 1.19465922 | 0.2322202 | 0.44871221 |
| TMEM25 | 99.4659444 | -0.024477815 | 0.22798583 | -0.1073655 | 0.91449901 | 0.96131774 |
| IFT46 | 196.798167 | 0.455933583 | 0.20877045 | 2.18389901 | 0.02896967 | 0.11456897 |
| ARCN1 | 383.824821 | 0.021904141 | 0.18559647 | 0.11802025 | 0.90605162 | 0.95706004 |
| PHLDB1 | 648.605161 | 0.089120478 | 0.43901403 | 0.20300144 | 0.83913391 | 0.92514146 |
| AP002954.1 | 123.060868 | -0.379708341 | 0.33805443 | -1.1232166 | 0.26134552 | 0.48237074 |
| DDX6 | 1170.19369 | 0.008306527 | 0.11567252 | 0.07181072 | 0.94275254 | 0.97554394 |
| BCL9L | 179.2146 | -0.984225448 | 0.26083343 | -3.7733868 | 0.00016105 | 0.00245642 |
| CCDC84 | 892.432921 | -0.109008443 | 0.2041027 | -0.5340862 | 0.59328187 | 0.77387448 |
| AP003392.1 | 58.9645851 | -0.419773097 | 0.43621792 | -0.9623014 | 0.33589825 | 0.56142219 |
| RPS25 | 853.018954 | -0.858010897 | 0.40470593 | -2.1200848 | 0.03399889 | 0.12860578 |
| TRAPPC4 | 216.515451 | -0.408774769 | 0.34226003 | -1.1943398 | 0.23234509 | 0.44888786 |
| SLC37A4 | 106.083022 | -0.308060228 | 0.43703784 | -0.7048823 | 0.4808835 | 0.68948936 |
| HYOU1 | 131.949982 | -0.093892746 | 0.38759894 | -0.242242 | 0.80859265 | 0.91010272 |
| VPS11 | 147.437247 | -0.055527398 | 0.21572458 | -0.2573995 | 0.7968704 | 0.9046722 |
| HMBS | 54.8349053 | 0.162839297 | 0.40956387 | 0.39759195 | 0.690931 | 0.84134845 |
| H2AFX | 51.4332457 | -0.913353679 | 0.42496054 | -2.149267 | 0.03161324 | 0.12231379 |
| DPAGT1 | 68.0585375 | 0.083444704 | 0.37224671 | 0.22416505 | 0.82262888 | 0.91813562 |
| C2CD2L | 74.5835585 | 0.177287316 | 0.30311065 | 0.58489307 | 0.55861962 | 0.75064156 |
| HINFP | 177.099743 | -0.17153733 | 0.22728774 | -0.7547144 | 0.45042036 | 0.66647279 |
| NLRX1 | 141.921524 | 0.032710122 | 0.35525541 | 0.09207494 | 0.9266385 | 0.9677394 |
| CBL | 611.010031 | -0.398670654 | 0.28485468 | -1.399558 | 0.16164571 | 0.35921571 |
| USP2 | 405.934953 | 1.894924106 | 0.69096197 | 2.74244342 | 0.0060984 | 0.03878772 |
| USP2-AS1 | 393.315748 | 1.365506113 | 0.51442878 | 2.65441236 | 0.00794467 | 0.04712219 |
| THY1 | 155.835197 | 3.088124503 | 0.92504143 | 3.33836345 | 0.00084273 | 0.00875132 |
| NECTIN1 | 66.4886886 | 1.18984366 | 0.43804312 | 2.71627063 | 0.00660219 | 0.04113602 |
| OAF | 276.120605 | 1.110186914 | 0.40320818 | 2.75338392 | 0.00589827 | 0.03787566 |
| TMEM136 | 84.0948529 | 0.070423158 | 0.37202642 | 0.18929612 | 0.84986074 | 0.93113889 |
| ARHGEF12 | 4672.68222 | -0.193374082 | 0.20652745 | -0.9363118 | 0.34911268 | 0.57506291 |
| TECTA | 55.4066972 | 0.422726607 | 0.39340194 | 1.07454124 | 0.28258017 | 0.50501982 |
| SC5D | 148.917799 | -0.257548398 | 0.37258251 | -0.691252 | 0.4894072 | 0.69623906 |
| SORL1 | 1617.07168 | 1.175934075 | 0.49052741 | 2.39728513 | 0.01651707 | 0.07833893 |
| UBASH3B | 363.974248 | -2.54712154 | 0.43538966 | -5.8502114 | 4.91E-09 | 5.27E-07 |
| JHY | 312.97139 | -0.290744916 | 0.38777686 | -0.7497738 | 0.45339097 | 0.66914428 |
| HSPA8 | 573.888994 | -0.627234912 | 0.52014836 | -1.2058769 | 0.22786494 | 0.44334483 |
| CLMP | 46.0427737 | -0.188123039 | 0.4354845 | -0.4319856 | 0.66575187 | 0.82362669 |
| GRAMD1B | 346.903558 | 2.808870048 | 0.75510697 | 3.71983065 | 0.00019936 | 0.00292448 |
| ZNF202 | 100.539491 | 0.057615591 | 0.27711546 | 0.20791186 | 0.83529779 | 0.9233784 |
| VWA5A | 306.275154 | -1.066416013 | 0.26766965 | -3.9840751 | 6.77E-05 | 0.00123424 |
| TBRG1 | 681.61638 | 0.062829082 | 0.18855839 | 0.33320756 | 0.73897761 | 0.87215469 |
| SIAE | 498.996983 | 0.514654692 | 0.30226772 | 1.70264522 | 0.08863448 | 0.24205091 |
| SPA17 | 92.0828779 | -0.251987223 | 0.2599185 | -0.9694855 | 0.33230301 | 0.55752451 |
| ESAM | 173.652484 | 1.543400947 | 0.47361749 | 3.25874987 | 0.00111904 | 0.01091218 |
| AP000866.2 | 69.2264894 | 1.059981239 | 0.43695509 | 2.42583565 | 0.01527318 | 0.07427964 |
| MSANTD2 | 1070.36417 | 0.918745868 | 0.22009749 | 4.1742679 | 2.99E-05 | 0.00065168 |
| AP000866.1 | 67.2867813 | 1.670142579 | 0.29789211 | 5.60653511 | 2.06E-08 | 1.69E-06 |
| ROBO3 | 46.3163553 | 0.447036913 | 0.41365676 | 1.0806953 | 0.27983268 | 0.50229221 |
| CCDC15 | 55.947144 | -0.538517922 | 0.62253054 | -0.8650466 | 0.38701322 | 0.61054074 |
| TMEM218 | 105.711668 | 0.411188228 | 0.3371315 | 1.21966718 | 0.22259107 | 0.43688045 |
| PKNOX2 | 391.97594 | 0.686082366 | 0.35633995 | 1.92535911 | 0.05418445 | 0.17555123 |
| EI24 | 273.522638 | -0.893890535 | 0.30865432 | -2.8960895 | 0.00377845 | 0.02749835 |
| STT3A-AS1 | 80.5686712 | 0.402097579 | 0.35804797 | 1.12302713 | 0.26142597 | 0.48237074 |
| STT3A | 271.066529 | 0.041987067 | 0.20053481 | 0.20937546 | 0.83415515 | 0.92260077 |
| HYLS1 | 59.7318683 | 0.638921674 | 0.40570805 | 1.57483116 | 0.11529542 | 0.2886277 |
| PUS3 | 61.4516486 | -0.283615697 | 0.37599367 | -0.7543098 | 0.45066324 | 0.66654333 |
| DDX25 | 59.1485698 | 1.178187514 | 0.47303387 | 2.49070433 | 0.01274902 | 0.06557701 |
| CDON | 63.7350072 | -0.220245975 | 0.42593734 | -0.5170854 | 0.60509655 | 0.7816077 |
| RPUSD4 | 138.9017 | 0.420132904 | 0.2931558 | 1.43313862 | 0.15181823 | 0.34497826 |
| FAM118B | 258.046134 | -0.448616105 | 0.20906738 | -2.1457967 | 0.0318892 | 0.12302116 |
| SRPRA | 224.282932 | -0.629818426 | 0.19734025 | -3.1915356 | 0.00141519 | 0.01308132 |
| FOXRED1 | 256.74048 | 0.320080031 | 0.28635888 | 1.11775834 | 0.26367022 | 0.48466739 |
| TIRAP | 118.351698 | 0.850935074 | 0.25177742 | 3.37971161 | 0.00072562 | 0.00786922 |
| AP001318.2 | 80.8286052 | 0.26376872 | 0.40291567 | 0.65464993 | 0.51269317 | 0.71466484 |
| DCPS | 431.907285 | 0.677670012 | 0.3796992 | 1.78475491 | 0.07430108 | 0.2163675 |
| GSEC | 46.3102666 | 1.16249213 | 0.50254241 | 2.31322194 | 0.02071044 | 0.09130981 |

| | | | | | | |
|---|---|---|---|---|---|---|
| ST3GAL4 | 703.205432 | 0.799187147 | 0.60422309 | 1.322669 | 0.18594548 | 0.39174123 |
| ETS1 | 290.984382 | -2.420081802 | 0.54486262 | -4.4416367 | 8.93E-06 | 0.00024568 |
| KCNJ1 | 1964.89306 | 1.270166469 | 0.59924711 | 2.11960383 | 0.03403947 | 0.1287224 |
| KCNJ5 | 122.874369 | 1.953220332 | 0.71467158 | 2.73303206 | 0.00627542 | 0.03951004 |
| ARHGAP32 | 2076.71195 | -0.15414196 | 0.21716487 | -0.7097923 | 0.47783293 | 0.68754849 |
| BARX2 | 141.078935 | 0.420747548 | 1.01407434 | 0.41490799 | 0.67820926 | 0.83118075 |
| TMEM45B | 979.808789 | 0.357971455 | 0.2807589 | 1.27501376 | 0.20230437 | 0.41181052 |
| NFRKB | 342.400685 | 0.584488527 | 0.16702943 | 3.49931466 | 0.00046646 | 0.00562173 |
| PRDM10 | 499.823641 | 0.186617357 | 0.32553858 | 0.57325726 | 0.56647052 | 0.75664972 |
| APLP2 | 1578.14772 | -0.605603345 | 0.28091573 | -2.1558185 | 0.03109784 | 0.1209574 |
| ST14 | 615.602158 | -0.39891218 | 0.2348078 | -1.6988881 | 0.08934026 | 0.24331867 |
| ZBTB44 | 1730.22185 | 0.125644558 | 0.28611903 | 0.43913388 | 0.66056453 | 0.82026629 |
| AP002986.1 | 297.96575 | 1.846338169 | 0.50160477 | 3.68086245 | 0.00023245 | 0.00331575 |
| ADAMTS15 | 288.241269 | 1.722916197 | 0.41086146 | 4.19342373 | 2.75E-05 | 0.00061069 |
| AP003486.1 | 45.7649608 | 0.18562331 | 0.32358184 | 0.57365182 | 0.56620344 | 0.75652248 |
| SNX19 | 727.938642 | 0.204567346 | 0.50212791 | 0.40740087 | 0.68371359 | 0.83583275 |
| NTM | 199.42978 | -1.371504728 | 0.74072976 | -1.8515588 | 0.0640892 | 0.19637073 |
| OPCML | 353.224097 | 2.307523354 | 0.76106124 | 3.03198116 | 0.00242954 | 0.01976099 |
| IGSF9B | 54.3972112 | -0.106131629 | 0.70440828 | -0.1506678 | 0.88023779 | 0.94636668 |
| JAM3 | 168.081277 | 0.32237207 | 0.40867928 | 0.78881432 | 0.43022054 | 0.64842345 |
| NCAPD3 | 206.70807 | 0.090700869 | 0.35159697 | 0.25796829 | 0.79643138 | 0.90456252 |
| VPS26B | 232.601418 | 0.143218557 | 0.19023249 | 0.75286064 | 0.45153366 | 0.66729785 |
| THYN1 | 66.86654 | 0.223351549 | 0.31683506 | 0.70494581 | 0.48084396 | 0.68948936 |
| ACAD8 | 143.571946 | 0.432605092 | 0.28353167 | 1.52577343 | 0.1270663 | 0.3070091 |
| TUBB8 | 67.0680637 | 1.163529042 | 0.8291194 | 1.4033311 | 0.16051813 | 0.35771722 |
| ZMYND11 | 1954.79902 | 0.439988752 | 0.09609345 | 4.57875925 | 4.68E-06 | 0.00014269 |
| DIP2C | 4876.10868 | -0.85580773 | 0.22400275 | -3.8205234 | 0.00013317 | 0.00210824 |
| PRR26 | 101.460486 | -0.373903596 | 0.45966492 | -0.8134264 | 0.41597361 | 0.63587494 |
| LARP4B | 2805.62516 | -0.125956341 | 0.15528845 | -0.8111121 | 0.41730128 | 0.63704999 |
| GTPBP4 | 250.941664 | -0.361986797 | 0.20821889 | -1.7384916 | 0.08212422 | 0.23032164 |
| IDI1 | 296.892943 | -0.856604683 | 0.46954982 | -1.8243105 | 0.06810514 | 0.2045477 |
| WDR37 | 1068.78131 | -0.020110905 | 0.18160022 | -0.1107427 | 0.91182036 | 0.96021221 |
| ADARB2 | 236.69507 | 2.883268883 | 0.98320905 | 2.9325085 | 0.00336236 | 0.02526358 |
| PFKP | 1413.3648 | -1.190563667 | 0.5662728 | -2.1024561 | 0.03551335 | 0.13225142 |
| PITRM1 | 561.940723 | -0.764394779 | 0.27203362 | -2.8099276 | 0.00495526 | 0.0331749 |
| PITRM1-AS1 | 88.3037198 | -0.446285434 | 0.47987014 | -0.9300129 | 0.35236439 | 0.57847145 |
| KLF6 | 1261.04754 | -0.728905439 | 0.56228773 | -1.2963211 | 0.19486489 | 0.40243438 |
| AKR1C1 | 100.735191 | -1.963702239 | 0.64451638 | -3.0467841 | 0.00231304 | 0.01911947 |
| AKR1C3 | 65.9688177 | -2.09241805 | 0.55792029 | -3.7503889 | 0.00017656 | 0.00264534 |
| NET1 | 137.224216 | 0.089915379 | 0.3255085 | 0.27623051 | 0.78237103 | 0.89606684 |
| ASB13 | 197.198555 | 0.291235194 | 0.41833201 | 0.69618195 | 0.48631489 | 0.6936328 |
| FAM208B | 1639.94631 | 0.181531993 | 0.17315511 | 1.04837791 | 0.29446453 | 0.51771355 |
| AL365356.1 | 112.80676 | 0.08703684 | 0.46918307 | 0.18550721 | 0.85283123 | 0.93245987 |
| GDI2 | 956.718099 | -0.316982337 | 0.15250389 | -2.0785197 | 0.03766152 | 0.13733278 |
| ANKRD16 | 158.788727 | 0.533627081 | 0.26921843 | 1.98213429 | 0.04746421 | 0.16014171 |
| FBH1 | 741.696469 | 0.080243663 | 0.18665613 | 0.42990103 | 0.66726764 | 0.82469498 |
| RBM17 | 678.047537 | 0.279520091 | 0.245749 | 1.13742108 | 0.2553623 | 0.47622669 |
| PFKFB3 | 1748.7283 | -1.215110817 | 0.65882677 | -1.8443556 | 0.06513134 | 0.19832178 |
| AL137145.2 | 324.15738 | -0.878935188 | 0.96824131 | -0.9077646 | 0.3640026 | 0.58937367 |
| PRKCQ | 2931.81556 | -0.146389921 | 0.31624354 | -0.4629025 | 0.64343427 | 0.80905043 |
| SFMBT2 | 1119.76265 | 0.202104756 | 0.27495153 | 0.73505594 | 0.46230543 | 0.67537466 |
| KIN | 301.427239 | 0.10146741 | 0.23750571 | 0.42722093 | 0.66921842 | 0.82606355 |
| ATP5F1C | 1214.8335 | 0.682029185 | 0.22510891 | 3.0297743 | 0.00244737 | 0.01986924 |
| TAF3 | 1184.24189 | 0.061003173 | 0.20626711 | 0.29574843 | 0.76742221 | 0.88756968 |
| CELF2 | 218.710722 | -1.157214303 | 0.56765362 | -2.0385923 | 0.04149073 | 0.14589537 |
| USP6NL | 1064.26077 | -0.387649082 | 0.17609935 | -2.201309 | 0.02771416 | 0.11110055 |
| ECHDC3 | 188.540408 | -0.184154198 | 0.40049232 | -0.4598195 | 0.64564575 | 0.81059872 |
| PROSER2 | 696.382253 | -0.423653251 | 0.43016796 | -0.9848554 | 0.32469508 | 0.54958323 |
| PROSER2-AS | 51.6741833 | 0.449720497 | 0.63017715 | 0.71364139 | 0.47544896 | 0.68538649 |
| UPF2 | 1119.29251 | 0.042833353 | 0.18398354 | 0.23281078 | 0.81590834 | 0.91427971 |
| DHTKD1 | 486.263307 | 0.011390156 | 0.26638149 | 0.04275881 | 0.9658938 | 0.98475542 |
| SEC61A2 | 259.397765 | 0.453028091 | 0.2521442 | 1.79670237 | 0.07238288 | 0.21270422 |
| NUDT5 | 544.224063 | 0.144354345 | 0.18967064 | 0.76107901 | 0.44660988 | 0.66246714 |
| CDC123 | 459.968069 | 0.008446079 | 0.17203361 | 0.04909551 | 0.96084318 | 0.98286824 |
| CAMK1D | 214.502372 | -2.048858444 | 0.58830842 | -3.4826264 | 0.00049652 | 0.00589267 |
| OPTN | 367.649821 | -0.873007472 | 0.41615808 | -2.0977785 | 0.03592472 | 0.13318259 |
| PHYH | 182.348144 | 0.152480678 | 0.3041745 | 0.50129343 | 0.61616463 | 0.78919021 |
| SEPHS1 | 296.008151 | 0.122591076 | 0.25754336 | 0.47600169 | 0.63407317 | 0.8030945 |
| BEND7 | 884.541786 | -0.723080612 | 0.24489389 | -2.9526283 | 0.00315081 | 0.02400177 |
| PRPF18 | 155.519841 | 0.065433429 | 0.29533634 | 0.22155563 | 0.82465982 | 0.91884704 |
| AL157392.5 | 317.164011 | 0.049746854 | 0.18900087 | 0.26320965 | 0.79238899 | 0.90221177 |
| FRMD4A | 2279.76238 | 0.32297617 | 0.35737983 | 0.90373362 | 0.36613667 | 0.59142894 |
| FAM107B | 1191.3333 | -0.168206494 | 0.30066798 | -0.5594427 | 0.57585966 | 0.76292613 |
| CDNF | 62.6159841 | 1.068072661 | 0.47232998 | 2.26128494 | 0.02374162 | 0.10019266 |
| HSPA14.1 | 133.647709 | -0.110224097 | 0.28699111 | -0.384068 | 0.70092806 | 0.84841905 |
| SUV39H2 | 63.3832967 | 0.361137893 | 0.41371832 | 0.87290767 | 0.3827134 | 0.60542052 |
| DCLRE1C | 353.205121 | -0.399825995 | 0.25697064 | -1.5559209 | 0.1197269 | 0.29536727 |
| RPP38 | 499.978483 | 0.199541927 | 0.28876499 | 0.69101841 | 0.48955397 | 0.69629788 |
| NMT2 | 291.718585 | 0.07026692 | 0.24670814 | 0.284818 | 0.77578358 | 0.89270192 |
| FAM171A1 | 2668.62907 | 0.929292329 | 0.26040613 | 3.56862692 | 0.00035886 | 0.00458871 |
| MINDY3 | 793.679097 | 0.064963055 | 0.1869845 | 0.34742481 | 0.7282722 | 0.86640975 |
| PTER | 713.151092 | 0.031639121 | 0.18363924 | 0.17228954 | 0.86320991 | 0.93813133 |
| RSU1 | 1014.05859 | -0.353154275 | 0.31801734 | -1.1104875 | 0.26678901 | 0.4879557 |
| CUBN | 378.555184 | 1.027053049 | 0.53418543 | 1.92265269 | 0.05452368 | 0.17600687 |
| TRDMT1 | 206.865201 | 0.149480267 | 0.26560323 | 0.56279537 | 0.57357423 | 0.761183 |
| VIM | 249.787939 | -1.678632571 | 0.38831886 | -4.3228201 | 1.54E-05 | 0.00038105 |
| ST8SIA6 | 164.198898 | 3.650042933 | 1.05618742 | 3.45586669 | 0.00054853 | 0.00635012 |
| HACD1 | 109.306554 | 0.282000692 | 0.36199093 | 0.77902696 | 0.43596383 | 0.65305582 |
| STAM-AS1 | 40.0597025 | 0.771096312 | 0.42653248 | 1.80782555 | 0.07063366 | 0.20947462 |
| STAM | 433.204983 | -0.079950967 | 0.21005037 | -0.3806276 | 0.70347961 | 0.84973591 |
| CACNB2 | 2833.09278 | 2.036000582 | 0.45795571 | 4.445846 | 8.75E-06 | 0.00024243 |
| AL450384.2 | 370.06572 | 0.695294831 | 0.32255155 | 2.15560839 | 0.03111426 | 0.12098564 |
| NSUN6 | 859.524792 | 0.532650205 | 0.2133555 | 2.49653838 | 0.01254121 | 0.06478562 |
| ARL5B | 360.659648 | 0.011653526 | 0.38509128 | 0.03026172 | 0.97585832 | 0.9893257 |
| MALRD1 | 1313.27754 | 1.503799675 | 0.35796476 | 4.20097133 | 2.66E-05 | 0.00059732 |
| PLXDC2 | 3491.13115 | 0.364937843 | 0.19247955 | 1.89598243 | 0.05796237 | 0.18377939 |
| NEBL | 5453.88131 | 0.704135437 | 0.34223649 | 2.05745281 | 0.03964269 | 0.14177464 |
| MLLT10 | 3223.86407 | 0.254350546 | 0.14519343 | 1.75180478 | 0.07980738 | 0.22602395 |
| DNAJC1 | 2213.15091 | -0.198700764 | 0.19155029 | -1.0373295 | 0.29958234 | 0.52246938 |
| BMI1 | 129.574371 | -0.153099326 | 0.32292599 | -0.4741004 | 0.63542835 | 0.80389813 |
| PIP4K2A | 1134.55506 | -0.456656426 | 0.29675168 | -1.5388504 | 0.12384083 | 0.30188476 |
| MSRB2 | 218.125296 | -0.401321323 | 0.22000418 | -1.8241532 | 0.06812891 | 0.20457259 |
| C10orf67 | 79.4163648 | 0.611974204 | 0.5505782 | 1.11151186 | 0.26634809 | 0.48748675 |
| OTUD1 | 90.0617972 | 0.498796363 | 0.28103463 | 1.77485732 | 0.07592145 | 0.21963347 |
| KIAA1217 | 8556.73959 | 0.815233276 | 0.3575199 | 2.2802459 | 0.02259311 | 0.09676795 |
| ARHGAP21 | 1487.9273 | -0.253263548 | 0.19121816 | -1.3244743 | 0.18534557 | 0.39096608 |
| PRTFDC1 | 95.1241414 | 1.009448525 | 0.3491892 | 2.89083546 | 0.00384219 | 0.02785484 |
| ENKUR | 52.4872955 | -1.806382587 | 0.62100334 | -2.908813 | 0.00362804 | 0.02666817 |
| THNSL1 | 73.3292575 | 0.671050066 | 0.41499247 | 1.61701744 | 0.10587452 | 0.27301545 |
| PDSS1 | 215.7753 | 0.632523007 | 0.33430894 | 1.89203138 | 0.0584868 | 0.18461291 |
| ABI1 | 1827.72234 | 0.090127031 | 0.14157336 | 0.63661011 | 0.52437884 | 0.72362308 |
| ANKRD26 | 1572.43314 | 0.614282324 | 0.15077057 | 4.07428538 | 4.62E-05 | 0.00092375 |
| YME1L1 | 1013.65213 | -0.070345178 | 0.141914 | -0.4956888 | 0.62011399 | 0.79240454 |

| | | | | | | |
|---|---|---|---|---|---|---|
| MASTL | 225.954479 | -0.123633917 | 0.34709828 | -0.3561928 | 0.72169617 | 0.86143064 |
| ACBD5 | 570.547364 | -0.160853369 | 0.14791816 | -1.0874484 | 0.27683868 | 0.49897002 |
| RAB18 | 888.802243 | -0.253313298 | 0.09633592 | -2.6294793 | 0.00855157 | 0.04950124 |
| MPP7 | 1666.40065 | 1.083061014 | 0.38197407 | 2.8354307 | 0.00457639 | 0.03145229 |
| AL358234.1 | 220.452121 | 1.332049593 | 0.80354332 | 1.6577197 | 0.09737405 | 0.25780996 |
| WAC-AS1 | 422.482614 | -0.139383448 | 0.28833099 | -0.4834147 | 0.62880129 | 0.79917601 |
| WAC | 3625.64164 | 0.075660748 | 0.1718246 | 0.4403371 | 0.65969297 | 0.81980614 |
| BAMBI | 42.4665239 | 0.907262273 | 0.52714723 | 1.72107945 | 0.08523641 | 0.235788 |
| SVIL | 3590.60075 | -0.986907226 | 0.34580436 | -2.8539467 | 0.00431798 | 0.03016353 |
| MTPAP | 265.739331 | 0.630511699 | 0.163804 | 3.84918389 | 0.00011851 | 0.00192035 |
| MAP3K8 | 231.968102 | -1.465504259 | 0.39955345 | -3.6678553 | 0.00024459 | 0.00345807 |
| ZNF438 | 897.455491 | -0.479956424 | 0.22470705 | -2.1359206 | 0.03268588 | 0.12521689 |
| ZEB1-AS1 | 76.9498486 | -1.032560314 | 0.45413044 | -2.2737087 | 0.02298351 | 0.09796359 |
| ZEB1 | 1376.03321 | -2.916683661 | 0.70483596 | -4.1381028 | 3.50E-05 | 0.00074111 |
| ARHGAP12 | 1828.52228 | 0.174376441 | 0.1752456 | 0.99504036 | 0.31971664 | 0.54422157 |
| KIF5B | 966.950668 | -0.083374049 | 0.17710461 | -0.4707616 | 0.63781099 | 0.80504973 |
| EPC1 | 1950.92755 | 0.306862281 | 0.15561243 | 1.9719651 | 0.04861359 | 0.1631031 |
| AL391839.1 | 64.6444785 | 0.205456093 | 0.47601759 | 0.4316145 | 0.66602162 | 0.82381118 |
| CCDC7 | 1797.96673 | 0.404686968 | 0.33377152 | 1.21246704 | 0.22533365 | 0.43984516 |
| ITGB1 | 1570.07404 | -0.684174926 | 0.24852517 | -2.7529402 | 0.00590627 | 0.0379086 |
| NRP1 | 4698.62108 | -3.25794736 | 0.50730709 | -6.4220418 | 1.34E-10 | 2.23E-08 |
| PARD3 | 9733.78973 | -0.378423639 | 0.15631833 | -2.4208526 | 0.01548416 | 0.07500191 |
| CUL2 | 745.749985 | -0.240350379 | 0.16675228 | -1.4413619 | 0.14948248 | 0.34167054 |
| CREM | 1462.62915 | -0.466373 | 0.27255759 | -1.7110989 | 0.08706285 | 0.2391377 |
| CCNY | 3461.87766 | 0.110885747 | 0.23544905 | 0.47095433 | 0.63767335 | 0.80499195 |
| AL117336.3 | 460.419689 | 0.413785174 | 0.25365089 | 1.63131767 | 0.10282331 | 0.2676495 |
| FZD8 | 91.5787439 | 0.080312394 | 0.45175475 | 0.17777875 | 0.85889674 | 0.93618423 |
| ZNF248 | 1121.34797 | 0.19667956 | 0.23157417 | 0.8493156 | 0.3957057 | 0.61783411 |
| ZNF25 | 212.435509 | 0.817553686 | 0.21021966 | 3.88904481 | 0.00010064 | 0.00167664 |
| ZNF33A | 2284.84073 | -0.24414999 | 0.19881727 | -1.228012 | 0.21944248 | 0.43356369 |
| AL117339.5 | 155.35304 | -0.08940541 | 0.30757114 | -0.2906821 | 0.7712945 | 0.8896518 |
| ZNF33B | 1390.57892 | -0.097557586 | 0.15210309 | -0.6413912 | 0.52126855 | 0.7212902 |
| BMS1 | 518.950004 | 0.188775287 | 0.2055479 | 0.91840049 | 0.35840924 | 0.58375186 |
| CSGALNACT2 | 363.53747 | 0.006466575 | 0.18211121 | 0.03550893 | 0.97167392 | 0.98744736 |
| FXYD4 | 61.1868011 | 0.457213707 | 0.72881528 | 0.62733825 | 0.53043753 | 0.72848299 |
| HNRNPF | 668.515594 | -0.999483276 | 0.18643608 | -5.3609972 | 8.28E-08 | 5.52E-06 |
| ZNF487 | 186.500059 | -0.280062591 | 0.22952846 | -1.220165 | 0.22240233 | 0.43657488 |
| ZNF239 | 72.4480763 | 0.724723221 | 0.28562069 | 2.53736249 | 0.01116912 | 0.05999714 |
| ZNF32 | 232.665197 | -0.028748931 | 0.23309075 | -0.1233379 | 0.90183951 | 0.95457905 |
| CXCL12 | 459.335747 | 3.929123992 | 0.60537694 | 6.49037607 | 8.56E-11 | 1.53E-08 |
| TMEM72-AS | 1116.67032 | 2.064864461 | 0.52729002 | 3.91599383 | 9.00E-05 | 0.0015541 |
| TMEM72 | 1609.94746 | 1.319673475 | 0.62491842 | 2.1117532 | 0.03470762 | 0.13013142 |
| AL353801.1 | 89.6307145 | 1.313411218 | 0.60771215 | 2.16123904 | 0.03067688 | 0.11963711 |
| RASSF4 | 250.01186 | 1.055580048 | 0.26747078 | 3.9465248 | 7.93E-05 | 0.00140887 |
| DEPP1 | 585.36729 | -2.784813315 | 0.7200497 | -3.8675293 | 0.00010994 | 0.00180403 |
| ZNF22 | 158.723195 | -0.279953913 | 0.32017174 | -0.8743867 | 0.3819077 | 0.60465286 |
| 8-Mar | 912.703407 | -0.34027567 | 0.17266153 | -1.9707671 | 0.04875052 | 0.1632722 |
| ZFAND4 | 348.871827 | 0.24945594 | 0.18720448 | 1.33253186 | 0.18268548 | 0.38733541 |
| WASHC2C | 350.195847 | -0.1007691 | 0.22773864 | -0.442477 | 0.6581441 | 0.81888605 |
| AGAP4 | 411.989678 | 0.447948522 | 0.22481393 | 1.99253008 | 0.04631293 | 0.15746395 |
| TIMM23 | 403.856178 | 0.405999126 | 0.20744248 | 1.95716487 | 0.05032809 | 0.16698912 |
| NCOA4 | 414.777307 | 0.162639078 | 0.29277403 | 0.5555106 | 0.57854546 | 0.76458299 |
| AL356056.2 | 71.6099826 | 0.643716579 | 0.3274599 | 1.96578748 | 0.04932316 | 0.16464738 |
| PTPN20 | 398.611289 | 0.126678258 | 0.53946865 | 0.23482043 | 0.81434811 | 0.91328954 |
| AGAP9 | 397.48674 | 0.32667138 | 0.31347185 | 1.04210755 | 0.29736182 | 0.52079582 |
| FRMPD2 | 56.9902419 | 0.381757501 | 0.77166157 | 0.49472141 | 0.62079678 | 0.7928889 |
| MAPK8 | 3560.83889 | -0.04750933 | 0.23482886 | -0.2023147 | 0.83967071 | 0.92557877 |
| ARHGAP22 | 244.153953 | -4.474764454 | 0.61069886 | -7.3272848 | 2.35E-13 | 9.40E-11 |
| AL138760.1 | 88.0823999 | 0.236420257 | 0.63503472 | 0.37229501 | 0.70967322 | 0.85441779 |
| ERCC6 | 643.857567 | 0.397523185 | 0.13403922 | 2.9657228 | 0.00301973 | 0.02325771 |
| OGDHL | 599.044636 | 0.572963521 | 0.66752295 | 0.85834281 | 0.3907032 | 0.61357729 |
| PARG | 1285.52624 | 0.142964678 | 0.16379862 | 0.87280757 | 0.38276797 | 0.60543434 |
| TIMM23B | 1024.82973 | 0.092942887 | 0.33426552 | 0.27805108 | 0.78097315 | 0.89539748 |
| LINC00843 | 277.782481 | 0.103320659 | 0.20413936 | 0.50612807 | 0.61276675 | 0.78690269 |
| AGAP6 | 428.831343 | 0.084076592 | 0.16416267 | 0.51215415 | 0.60854313 | 0.78429566 |
| WASHC2A | 345.224453 | 0.160059639 | 0.30623996 | 0.52266085 | 0.60121029 | 0.77902557 |
| SGMS1 | 5566.60956 | -0.83269635 | 0.3196005 | -2.6054288 | 0.00917593 | 0.05206631 |
| SGMS1-AS1 | 140.315428 | -0.628778496 | 0.61984929 | -1.0144055 | 0.31038932 | 0.53435962 |
| ASAH2B | 100.228673 | -0.601774387 | 0.4087575 | -1.4722039 | 0.14096582 | 0.32909465 |
| PRKG1 | 4735.33513 | 0.451867164 | 0.45612134 | 0.99067315 | 0.32184521 | 0.5465741 |
| CSTF2T | 70.5936861 | -0.021887117 | 0.39640142 | -0.0552145 | 0.95596756 | 0.98069086 |
| PCDH15 | 73.8971022 | 0.949052309 | 0.45366802 | 2.09195328 | 0.03644269 | 0.13434875 |
| IPMK | 541.953692 | -1.056925694 | 0.3756621 | -2.813501 | 0.00490052 | 0.03294199 |
| CISD1 | 849.708389 | 0.250477812 | 0.23107474 | 1.0839688 | 0.27837863 | 0.50055858 |
| UBE2D1 | 343.282148 | -0.426474016 | 0.32793128 | -1.3004981 | 0.1934303 | 0.40113679 |
| TFAM | 195.35968 | 0.096429615 | 0.17080906 | 0.56454624 | 0.57238244 | 0.76035215 |
| BICC1 | 22779.7227 | -0.132890879 | 0.25185697 | -0.5276442 | 0.59774628 | 0.77674686 |
| PHYHIPL | 937.618177 | 0.60006229 | 0.53069648 | 1.13070712 | 0.25817838 | 0.47930826 |
| SLC16A9 | 101.471737 | 1.649939606 | 0.71227689 | 2.31643007 | 0.0205348 | 0.09084009 |
| CCDC6 | 977.875022 | -0.311961416 | 0.1820471 | -1.7136303 | 0.08659665 | 0.23845219 |
| ANK3 | 13636.3987 | 0.221594076 | 0.23692389 | 0.93529645 | 0.34963554 | 0.57570879 |
| AL592430.1 | 95.7811503 | -0.928550121 | 0.67429999 | -1.3770579 | 0.16849435 | 0.36854477 |
| RHOBTB1 | 151.562017 | -0.183135872 | 0.25623895 | -0.7147074 | 0.47478987 | 0.68513327 |
| ARID5B | 5109.54879 | -0.482640478 | 0.36300043 | -1.3295865 | 0.18365455 | 0.38864033 |
| ADO | 49.8715206 | -0.510785121 | 0.36614459 | -1.3950366 | 0.16300479 | 0.36109189 |
| NRBF2 | 165.387535 | -0.089721231 | 0.30207176 | -0.2970196 | 0.76645155 | 0.88690833 |
| JMJD1C | 7762.96213 | 0.311508247 | 0.13381694 | 2.32786852 | 0.01991909 | 0.08894902 |
| REEP3 | 1381.74876 | -0.043374958 | 0.26226863 | -0.1653837 | 0.86864198 | 0.94020545 |
| AC022387.1 | 425.743644 | 1.802471679 | 0.64768629 | 2.78293937 | 0.00538689 | 0.035428 |
| LINC01515 | 858.039334 | 0.732434453 | 0.35322658 | 2.07355419 | 0.03812073 | 0.13856268 |
| CTNNA3 | 262.470845 | 1.777146473 | 0.39601122 | 4.48761644 | 7.20E-06 | 0.00020416 |
| SIRT1 | 493.160141 | 0.385878871 | 0.21816232 | 1.76876953 | 0.07693234 | 0.2212981 |
| HERC4 | 3389.4308 | -0.029443104 | 0.13863353 | -0.2123808 | 0.83180995 | 0.92158196 |
| PBLD | 682.283236 | 0.081115342 | 0.35995392 | 0.22534924 | 0.8217076 | 0.91749245 |
| HNRNPH3 | 1132.10667 | 0.348698852 | 0.17168284 | 2.0310641 | 0.04224849 | 0.14754253 |
| RUFY2 | 932.190621 | 0.091683221 | 0.15016635 | 0.61054437 | 0.54150126 | 0.73761887 |
| DNA2 | 173.538024 | 0.098533923 | 0.38180544 | 0.25807365 | 0.79635007 | 0.90454795 |
| SLC25A16 | 1294.27738 | 0.259256499 | 0.18716608 | 1.38516822 | 0.16600104 | 0.36540834 |
| TET1 | 292.960866 | 1.590970821 | 0.37572374 | 4.23441658 | 2.29E-05 | 0.00053015 |
| CCAR1 | 1839.14478 | 0.055248601 | 0.20540272 | 0.26897697 | 0.78794741 | 0.89956761 |
| STOX1 | 226.541542 | 0.50286068 | 0.3831799 | 1.31233576 | 0.18940687 | 0.39556637 |
| DDX50 | 654.740148 | -0.01231989 | 0.14648372 | -0.0841042 | 0.93297362 | 0.97128377 |
| DDX21 | 439.658361 | -1.028724628 | 0.25528597 | -4.0296951 | 5.58E-05 | 0.0010676 |
| KIF1BP | 376.021564 | -0.207503682 | 0.19042063 | -1.0897122 | 0.27583995 | 0.49782347 |
| SRGN | 42.9029771 | 0.305380997 | 0.60195188 | 0.50731796 | 0.61193174 | 0.78684784 |
| VPS26A | 598.928755 | -0.069699017 | 0.15372328 | -0.4534057 | 0.65025661 | 0.81301431 |
| SUPV3L1 | 339.361116 | 0.466046553 | 0.19512375 | 2.38846654 | 0.01691885 | 0.07961563 |
| HKDC1 | 974.512349 | -2.244724945 | 0.63989253 | -3.5079718 | 0.00045154 | 0.00547691 |
| HK1 | 1477.39119 | -0.694273219 | 0.20435299 | -3.3974213 | 0.00068024 | 0.00748776 |
| TSPAN15 | 302.687676 | -0.912229931 | 0.34062798 | -2.6780828 | 0.00740449 | 0.04453822 |
| H2AFY2 | 495.773423 | -0.071478109 | 0.19170156 | -0.3728614 | 0.70925162 | 0.85408922 |
| AIFM2 | 141.62442 | -0.154519021 | 0.28161855 | -0.548682 | 0.58322373 | 0.76857514 |
| TYSND1 | 114.853814 | 0.102088862 | 0.3799911 | 0.26866119 | 0.78819042 | 0.89961465 |

| | | | | | | |
|---|---|---|---|---|---|---|
| SAR1A | 420.288287 | 0.117603752 | 0.17760927 | 0.66214871 | 0.50787592 | 0.71102629 |
| PPA1 | 576.180189 | -1.478404528 | 0.32714204 | -4.5191518 | 6.21E-06 | 0.00018064 |
| EIF4EBP2 | 415.215689 | -0.29069363 | 0.21644125 | -1.3430602 | 0.17925247 | 0.3827588 |
| SGPL1 | 777.581617 | -0.400663018 | 0.16739901 | -2.393461 | 0.01669025 | 0.07896188 |
| PCBD1 | 259.642513 | -0.892670334 | 0.42733805 | -2.0889091 | 0.03671591 | 0.13485073 |
| UNC5B | 85.7232425 | -1.715807355 | 0.83597746 | -2.0524565 | 0.04012532 | 0.14293833 |
| SLC29A3 | 92.5184874 | 0.600082745 | 0.56259028 | 1.06664257 | 0.28613327 | 0.50853442 |
| PSAP | 1416.9469 | -0.588728615 | 0.24440168 | -2.4088567 | 0.01600258 | 0.07653081 |
| CHST3 | 162.657012 | -0.974578082 | 0.69257944 | -1.4071715 | 0.15937653 | 0.35596966 |
| ASCC1 | 769.480657 | -0.158239906 | 0.23370606 | -0.6770894 | 0.49834921 | 0.70339396 |
| ANAPC16 | 1892.56546 | -0.101810245 | 0.21846659 | -0.466022 | 0.64119973 | 0.80747519 |
| DDIT4 | 358.352098 | -2.615211921 | 0.70554306 | -3.7066652 | 0.00021001 | 0.00305504 |
| DNAJB12 | 250.713199 | -0.350462524 | 0.25469966 | -1.3759835 | 0.16882676 | 0.36903287 |
| MICU1 | 1614.71317 | -0.202089389 | 0.10927854 | -1.8493053 | 0.06441374 | 0.19704184 |
| MCU | 1521.45003 | -0.324388742 | 0.25362543 | -1.2790072 | 0.20089453 | 0.40995146 |
| P4HA1 | 2399.64733 | -1.430805442 | 0.56071452 | -2.5517538 | 0.01071822 | 0.05821422 |
| NUDT13 | 137.637421 | 0.274404285 | 0.33429346 | 0.8208485 | 0.41173257 | 0.63092523 |
| ECD | 416.663425 | 0.413535447 | 0.20298755 | 2.03724542 | 0.04162545 | 0.14619267 |
| FAM149B1 | 708.87219 | 0.23709512 | 0.15340166 | 1.54558377 | 0.12220512 | 0.29942636 |
| DNAJC9 | 517.673313 | 0.01040297 | 0.22768905 | 0.04568937 | 0.96355784 | 0.98389266 |
| DNAJC9-AS1 | 51.5217759 | 0.286302178 | 0.3682175 | 0.77753549 | 0.4368429 | 0.65348334 |
| MRPS16 | 260.423791 | -0.140264376 | 0.22666899 | -0.6188071 | 0.53604345 | 0.73290528 |
| CFAP70 | 1178.7507 | 1.140152785 | 0.40676881 | 2.80295036 | 0.00506375 | 0.03371322 |
| ANXA7 | 1051.07092 | -0.286711984 | 0.10825654 | -2.6484494 | 0.00808619 | 0.04767298 |
| MSS51 | 62.081433 | 0.366388794 | 0.29192245 | 1.25508949 | 0.20944627 | 0.42115631 |
| PPP3CB | 1306.53791 | 0.447755398 | 0.17369359 | 2.57784637 | 0.00994182 | 0.05520029 |
| USP54 | 1054.98398 | 0.004681893 | 0.12959984 | 0.03612576 | 0.97118208 | 0.98725135 |
| AC073389.2 | 86.7275374 | 0.073562187 | 0.40588036 | 0.18124106 | 0.85617837 | 0.9348041 |
| AGAP5 | 198.813435 | 0.553602948 | 0.15139787 | 3.65660997 | 0.00025557 | 0.00355896 |
| AC022400.6 | 75.6615926 | 0.421661874 | 0.21755604 | 1.93817586 | 0.05260177 | 0.17202693 |
| SEC24C | 354.911484 | -0.069346362 | 0.16004507 | -0.4332927 | 0.66480213 | 0.82323933 |
| FUT11 | 229.711548 | -1.602242884 | 0.48245905 | -3.3209925 | 0.00089698 | 0.00916334 |
| CHCHD1 | 177.564827 | -0.772963079 | 0.40708377 | -1.8987814 | 0.05759323 | 0.18300431 |
| ZSWIM8 | 641.539441 | 0.09972396 | 0.19981426 | 0.49908331 | 0.6177207 | 0.79041681 |
| CAMK2G | 468.170046 | 0.314856756 | 0.3277774 | 0.96058105 | 0.33676286 | 0.56213655 |
| PLAU | 589.36597 | 1.162077698 | 0.67341307 | 1.72565362 | 0.08440975 | 0.23413867 |
| VCL | 2242.74862 | 0.65129426 | 0.46660621 | 1.39581139 | 0.1627713 | 0.3609485 |
| AP3M1 | 269.036857 | -0.104330072 | 0.16381822 | -0.6368649 | 0.52421286 | 0.72354522 |
| ADK | 6583.12695 | 0.142032344 | 0.20315831 | 0.69912151 | 0.4844761 | 0.69235581 |
| AC022540.1 | 67.8477493 | -0.060984882 | 0.82214706 | -0.0741776 | 0.94086908 | 0.97451107 |
| KAT6B | 2410.31587 | 0.563475292 | 0.24603735 | 2.29020226 | 0.02200959 | 0.09498231 |
| SAMD8 | 520.393433 | -0.344696889 | 0.16960252 | -2.0323808 | 0.04211512 | 0.14722087 |
| VDAC2 | 515.224949 | 0.126836263 | 0.2544908 | 0.49839233 | 0.61820754 | 0.79053759 |
| COMTD1 | 204.059158 | 0.396185132 | 0.4495953 | 0.88120391 | 0.37820746 | 0.60095542 |
| AC010997.3 | 52.9024201 | 0.651513112 | 0.51375344 | 1.26814354 | 0.20474669 | 0.4148641 |
| LRMDA | 3084.00398 | -0.044027841 | 0.38974292 | -0.1129664 | 0.91005722 | 0.95960292 |
| KCNMA1 | 535.11553 | 1.237239125 | 0.76041016 | 1.62706811 | 0.10372263 | 0.26943407 |
| DLG5 | 926.230526 | -0.254950706 | 0.2869223 | -0.8885706 | 0.37423392 | 0.59780424 |
| POLR3A | 450.772674 | -0.049872085 | 0.20258569 | -0.2461777 | 0.80554465 | 0.90905077 |
| RPS24 | 1547.14354 | -0.620197408 | 0.4263129 | -1.4547939 | 0.14572633 | 0.33584931 |
| ZMIZ1-AS1 | 102.19648 | -1.775238172 | 0.54334222 | -3.2672561 | 0.00108595 | 0.01064912 |
| ZMIZ1 | 2312.38781 | -0.445134658 | 0.43990537 | -1.0118873 | 0.31159194 | 0.53514731 |
| PPIF | 186.657945 | 0.446162368 | 0.28539207 | 1.56333133 | 0.11797468 | 0.2929811 |
| ZCCHC24 | 50.2733412 | -0.5971011 | 0.29465897 | -2.0264141 | 0.04272237 | 0.1488185 |
| NUTM2B-AS | 2009.42445 | -0.370247361 | 0.2862029 | -1.2936534 | 0.19578517 | 0.40345184 |
| AL135925.1 | 64.1340202 | 0.464793511 | 0.32738866 | 1.41969947 | 0.15569519 | 0.35065265 |
| SFTPD | 49.1420126 | -1.155458996 | 0.41397599 | -2.7911256 | 0.00525251 | 0.03475971 |
| TMEM254-A: | 97.411712 | 0.539942205 | 0.66588895 | 0.81085924 | 0.41744651 | 0.63711196 |
| TMEM254 | 180.631419 | 0.148559524 | 0.24837524 | 0.59812533 | 0.54975631 | 0.74303667 |
| ANXA11 | 2055.82321 | -0.712567489 | 0.30958389 | -2.3016944 | 0.02135241 | 0.093423 |
| FAM213A | 692.075378 | -0.164293391 | 0.14576128 | -1.1271402 | 0.25968323 | 0.48061767 |
| TSPAN14 | 1106.88765 | -0.807675403 | 0.19504605 | -4.1409472 | 3.46E-05 | 0.00073665 |
| NRG3 | 1998.65598 | -0.613410994 | 0.56987246 | -1.0764005 | 0.28174819 | 0.50421512 |
| GHITM | 488.705588 | -0.34945737 | 0.31690171 | -1.1027311 | 0.27014395 | 0.49175977 |
| CCSER2 | 1867.32903 | 0.332012228 | 0.19902232 | 1.66821608 | 0.09527284 | 0.25429643 |
| WAPL | 1481.44973 | 0.014012796 | 0.16342061 | 0.08574681 | 0.93166769 | 0.97007713 |
| AL731569.1 | 40.11894 | 1.300526744 | 0.38252411 | 3.39985566 | 0.00067421 | 0.00744001 |
| BMPR1A | 2624.1828 | -0.131421241 | 0.15277298 | -0.8602388 | 0.38965743 | 0.61259045 |
| MMRN2 | 78.3668401 | 0.862863684 | 0.62930134 | 1.37114548 | 0.17032961 | 0.37064919 |
| ADIRF | 302.451028 | -0.958992072 | 0.72397484 | -1.3246207 | 0.18529698 | 0.39092602 |
| GLUD1 | 1386.89074 | 0.021608074 | 0.21979084 | 0.09831199 | 0.92168456 | 0.96523677 |
| SHLD2 | 2487.05892 | 0.707621406 | 0.20514561 | 3.44936172 | 0.00056191 | 0.00643739 |
| NUTM2A-AS | 2237.1203 | 0.134230246 | 0.22587033 | 0.59428011 | 0.55232478 | 0.74529145 |
| LINC00864 | 100.005021 | 3.532764229 | 0.93090372 | 3.79498347 | 0.00014765 | 0.00228378 |
| MINPP1 | 196.819305 | 0.280273357 | 0.27448317 | 1.02109488 | 0.30720949 | 0.53079531 |
| PAPSS2 | 747.17648 | 0.696384456 | 0.56816728 | 1.22566801 | 0.22032363 | 0.4341354 |
| ATAD1 | 1022.58795 | 0.41213534 | 0.2285847 | 1.8029874 | 0.07139019 | 0.21085827 |
| PTEN | 2744.49588 | 0.156722094 | 0.18772846 | 0.83483398 | 0.40381119 | 0.62502719 |
| RNLS | 941.486034 | 0.350853958 | 0.24918875 | 1.40798476 | 0.15913558 | 0.35555175 |
| STAMBPL1 | 67.1437103 | -0.395369583 | 0.739837 | -0.5344009 | 0.59306417 | 0.77387448 |
| ACTA2 | 79.5614586 | -1.478205501 | 0.51555651 | -2.8672036 | 0.00414116 | 0.02926733 |
| FAS | 142.867214 | -2.041794185 | 0.51652167 | -3.952969 | 7.72E-05 | 0.00137594 |
| LIPA | 1190.95702 | 0.711144699 | 0.45233827 | 1.5721524 | 0.11591521 | 0.28960167 |
| IFIT2 | 252.688547 | 0.118692136 | 0.89061035 | 0.13327056 | 0.89397941 | 0.95146032 |
| IFIT3 | 559.549116 | -0.330185643 | 0.77888339 | -0.4239218 | 0.67162285 | 0.82756215 |
| IFIT1 | 555.915683 | 1.480957159 | 1.1418081 | 1.29702807 | 0.19462153 | 0.40218332 |
| IFIT5 | 113.174608 | -0.420784093 | 0.46176171 | -0.9112581 | 0.36215941 | 0.58754159 |
| SLC16A12 | 4018.03959 | 1.099352645 | 0.52311796 | 2.1015387 | 0.0355937 | 0.13239418 |
| SLC16A12-AS | 167.148054 | 0.823551865 | 0.63320972 | 1.30059891 | 0.19339578 | 0.40113679 |
| PANK1 | 648.580965 | 1.693728338 | 0.36456212 | 4.64592523 | 3.39E-06 | 0.00010907 |
| AL157400.3 | 80.1528392 | 1.277638652 | 0.39048495 | 3.27192803 | 0.00106817 | 0.0105059 |
| KIF20B | 134.847238 | 0.013738398 | 0.3377633 | 0.04067463 | 0.96755529 | 0.98528676 |
| LINC00865 | 40.1851422 | 0.60113882 | 0.70158015 | 0.85683556 | 0.39153577 | 0.6140943 |
| RPP30 | 208.351688 | 0.582995405 | 0.21870275 | 2.6656977 | 0.00768287 | 0.04583699 |
| PCGF5 | 2203.93777 | -0.06400616 | 0.23823579 | -0.2686673 | 0.78818574 | 0.89961465 |
| HECTD2 | 952.646233 | 0.406363652 | 0.22076237 | 1.84072879 | 0.06566131 | 0.19961341 |
| PPP1R3C | 58.7055616 | -0.487969754 | 1.02040671 | -0.478211 | 0.63250001 | 0.80189713 |
| TNKS2 | 1556.6984 | 0.028315059 | 0.16707193 | 0.16947826 | 0.86542047 | 0.93876167 |
| BTAF1 | 1387.95628 | 0.023678116 | 0.2650996 | 0.08931781 | 0.92882934 | 0.96900671 |
| CPEB3 | 4932.68465 | 1.097145169 | 0.37614432 | 2.91681973 | 0.0035362 | 0.02615323 |
| 5-Mar | 822.988722 | -0.011481715 | 0.22851748 | -0.0502444 | 0.95992766 | 0.98229715 |
| IDE | 670.600879 | -0.042607923 | 0.17491726 | -0.243589 | 0.80754914 | 0.91010272 |
| EXOC6 | 947.974912 | 0.134162417 | 0.20989178 | 0.63919804 | 0.52269411 | 0.72258153 |
| MYOF | 2844.15441 | -1.72244494 | 0.32961833 | -5.2255738 | 1.74E-07 | 9.80E-06 |
| FRA10AC1 | 604.492402 | 0.412173281 | 0.18536757 | 2.22354584 | 0.02617902 | 0.10692598 |
| SLC35G1 | 73.3818737 | 0.425755603 | 0.50875021 | 0.8368657 | 0.40266806 | 0.62391114 |
| PLCE1 | 74.328958 | -1.235232398 | 0.46019106 | -2.6841729 | 0.00727095 | 0.04394788 |
| NOC3L | 201.812668 | 0.26391035 | 0.2286316 | 1.15430393 | 0.24837559 | 0.46754925 |
| TBC1D12 | 494.403968 | -0.05358774 | 0.20624408 | -0.2598268 | 0.79499737 | 0.90385363 |
| HELLS | 168.172085 | -0.371298274 | 0.3061313 | -1.2128726 | 0.22517852 | 0.4396723 |
| AL157834.1 | 57.2261499 | 0.848156456 | 0.51338583 | 1.65208388 | 0.09851745 | 0.26000385 |
| PDLIM1 | 89.2779538 | -2.26230722 | 0.7383281 | -3.0640947 | 0.0021833 | 0.01834553 |
| SORBS1 | 2664.34268 | -0.681833219 | 0.34126451 | -1.9979611 | 0.04572088 | 0.15613419 |

| | | | | | | |
|---|---|---|---|---|---|---|
| ALDH18A1 | 590.360159 | -0.788101402 | 0.27320929 | -2.8846069 | 0.00391903 | 0.02821059 |
| TCTN3 | 480.5932 | 0.104480908 | 0.18248569 | 0.57254303 | 0.56695414 | 0.75714258 |
| ENTPD1 | 65.7263564 | -0.695726958 | 0.59675839 | -1.1658436 | 0.24367768 | 0.46243709 |
| ENTPD1-AS1 | 1493.75718 | 0.422497534 | 0.32353531 | 1.30587766 | 0.19159416 | 0.39835783 |
| AL365273.2 | 97.6045204 | -0.098650743 | 0.46007419 | -0.2144236 | 0.83021678 | 0.92071169 |
| CC2D2B | 113.640606 | -0.047707016 | 0.52850469 | -0.0902679 | 0.92807432 | 0.96870278 |
| CCNJ | 79.3631248 | 0.120945085 | 0.27941556 | 0.43285022 | 0.66512358 | 0.8233644 |
| ZNF518A | 2123.60612 | 0.371939969 | 0.19191355 | 1.93806001 | 0.0526159 | 0.17203055 |
| BLNK | 2614.89664 | -0.251632607 | 0.37791549 | -0.6658436 | 0.50551108 | 0.70921889 |
| TM9SF3 | 1763.18226 | 0.005623208 | 0.1340554 | 0.0419469 | 0.96654103 | 0.98510234 |
| PIK3AP1 | 84.4074429 | -0.309871944 | 0.46079911 | -0.6724665 | 0.50128679 | 0.705315 |
| LCOR | 3410.92659 | 0.17714709 | 0.21507141 | 0.82366638 | 0.41012915 | 0.6297833 |
| FRAT2 | 52.1114302 | -0.760309508 | 0.43131668 | -1.762764 | 0.0779403 | 0.22304728 |
| RRP12 | 210.540833 | -0.327421596 | 0.24443868 | -1.3394836 | 0.1804133 | 0.38418173 |
| PGAM1 | 634.879891 | -0.960884685 | 0.33062534 | -2.9062645 | 0.00365772 | 0.02681179 |
| EXOSC1 | 307.908374 | -0.015997741 | 0.17445841 | -0.0916995 | 0.92693683 | 0.96785223 |
| ZDHHC16 | 230.978612 | 0.015302192 | 0.22781549 | 0.06716923 | 0.94644698 | 0.97676523 |
| MMS19 | 762.057035 | 0.326875514 | 0.18631515 | 1.75442266 | 0.07935811 | 0.22513773 |
| UBTD1 | 683.335955 | 0.20679231 | 0.34111242 | 0.60622921 | 0.54436256 | 0.73944091 |
| ANKRD2 | 105.406308 | 3.472036078 | 0.94955462 | 3.65648905 | 0.00025569 | 0.00355896 |
| AL355315.1 | 174.413996 | 0.984116602 | 0.41470548 | 2.37304941 | 0.0176419 | 0.0819669 |
| MORN4 | 328.569412 | 0.878945029 | 0.25576859 | 3.43648541 | 0.00058931 | 0.00669901 |
| PI4K2A | 75.5309858 | -0.431043646 | 0.32691563 | -1.3185165 | 0.18733081 | 0.39298287 |
| AVPI1 | 46.9949757 | 0.147766693 | 0.37617398 | 0.39281476 | 0.6944563 | 0.843734 |
| ZFYVE27 | 313.202271 | -0.446331278 | 0.19475173 | -2.2917963 | 0.0219174 | 0.09476497 |
| R3HCC1L | 1189.76321 | -0.272375587 | 0.17081708 | -1.5945454 | 0.11081394 | 0.28084062 |
| PYROXD2 | 66.0746084 | 0.394864722 | 0.37706614 | 1.04720282 | 0.29500605 | 0.51817489 |
| HPS1 | 236.824451 | -0.338205741 | 0.2253666 | -1.5006915 | 0.13343538 | 0.31763343 |
| HPSE2 | 44.6657914 | 0.6437383 | 0.38109473 | 1.68918186 | 0.09118458 | 0.24676443 |
| GOT1 | 1689.44266 | 0.474381493 | 0.31250279 | 1.51800723 | 0.12901258 | 0.31029263 |
| AL391684.1 | 112.808142 | 1.702250307 | 0.43292883 | 3.93194024 | 8.43E-05 | 0.00147429 |
| SLC25A28 | 310.593511 | -0.169511419 | 0.29388469 | -0.5767957 | 0.56407749 | 0.7553629 |
| ENTPD7 | 140.043673 | -0.168410142 | 0.4078935 | -0.4128777 | 0.6796962 | 0.83230806 |
| CUTC | 365.809662 | -0.007740509 | 0.21623096 | -0.0357974 | 0.97144389 | 0.98736553 |
| COX15 | 312.89931 | -0.031874181 | 0.24127423 | -0.1321077 | 0.89489911 | 0.95172874 |
| ABCC2 | 117.388609 | -0.39908878 | 0.56075114 | -0.7117039 | 0.47664814 | 0.68659164 |
| DNMBP | 924.395416 | -0.257203463 | 0.28714166 | -0.8957372 | 0.37039315 | 0.59519746 |
| ERLIN1 | 300.017214 | 0.15751691 | 0.15464593 | 1.01856487 | 0.30840959 | 0.53227485 |
| CHUK | 325.844162 | 0.492139845 | 0.31966706 | 1.53953881 | 0.12367281 | 0.30156806 |
| CWF19L1 | 355.792898 | 0.177796765 | 0.22933366 | 0.7752755 | 0.43817688 | 0.65440037 |
| BLOC1S2 | 156.761885 | -0.309290938 | 0.20939569 | -1.4770645 | 0.13965836 | 0.32673526 |
| SCD | 62.4467376 | -2.803866834 | 0.58456556 | -4.7964968 | 1.61E-06 | 5.96E-05 |
| OLMALINC | 124.736963 | 0.081889748 | 0.57110334 | 0.14338867 | 0.88598323 | 0.94846586 |
| SEC31B | 173.956297 | 0.771074705 | 0.29067999 | 2.65265834 | 0.00798607 | 0.04728281 |
| HIF1AN | 306.547305 | -0.013273139 | 0.14065017 | -0.0943699 | 0.92481535 | 0.9667507 |
| PAX2 | 3186.64625 | 0.2863168 | 0.32528157 | 0.88021217 | 0.37874438 | 0.60159146 |
| SLF2 | 754.215908 | 0.056869092 | 0.24069003 | 0.23627523 | 0.81321911 | 0.91242304 |
| MRPL43 | 184.554581 | -0.458292397 | 0.32983346 | -1.3894661 | 0.16469108 | 0.36365838 |
| SEMA4G | 68.0706012 | 0.211176905 | 0.40568972 | 0.52053798 | 0.60268866 | 0.77979671 |
| LZTS2 | 255.253651 | -0.334248006 | 0.22491414 | -1.4861138 | 0.13724902 | 0.32320435 |
| SFXN3 | 58.5149263 | -1.605422774 | 0.44741715 | -3.5882012 | 0.00033297 | 0.00433744 |
| BTRC | 1817.11797 | 0.251884901 | 0.14673569 | 1.71658923 | 0.08605425 | 0.23735448 |
| DPCD | 176.515647 | -0.236995036 | 0.30420847 | -0.7790547 | 0.4359475 | 0.65305582 |
| POLL | 234.513166 | -0.176954046 | 0.27342909 | -0.6471661 | 0.51752443 | 0.71869378 |
| FBXW4 | 1337.01905 | 0.000813141 | 0.15037282 | 0.0054075 | 0.99568546 | 0.99847083 |
| OGA | 2646.89401 | -0.167096731 | 0.2000213 | -0.8353947 | 0.40349551 | 0.62475352 |
| KCNIP2-AS1 | 41.3157338 | 0.336626228 | 0.64025614 | 0.52576806 | 0.59904937 | 0.77729304 |
| KCNIP2 | 42.2316951 | -0.062580385 | 0.38011197 | -0.1646367 | 0.86922993 | 0.94034875 |
| ARMH3 | 1653.33748 | -0.055646576 | 0.16194062 | -0.3436233 | 0.73112958 | 0.86806155 |
| LDB1 | 239.060448 | 0.099391152 | 0.32177962 | 0.30887957 | 0.75741314 | 0.88285773 |
| PPRC1 | 163.482819 | -0.586082121 | 0.32586217 | -1.7985583 | 0.07208857 | 0.21221706 |
| NOLC1 | 146.176066 | -0.551629807 | 0.20339741 | -2.7120788 | 0.00668627 | 0.04146429 |
| GBF1 | 2446.76624 | -0.152631279 | 0.19317508 | -0.7901189 | 0.42945833 | 0.64764415 |
| NFKB2 | 166.235328 | -1.867314753 | 0.3214457 | -5.8091142 | 6.28E-09 | 6.31E-07 |
| FBXL15 | 178.912356 | -0.558303863 | 0.37533341 | -1.4874878 | 0.13688603 | 0.32270705 |
| CUEDC2 | 179.42548 | -0.426535256 | 0.22021326 | -1.9369191 | 0.05275524 | 0.17234271 |
| MFSD13A | 39.7647284 | 0.13424015 | 0.31575604 | 0.42513881 | 0.67073548 | 0.82700878 |
| ACTR1A | 236.549659 | -0.251620769 | 0.24958128 | -1.0081716 | 0.31337208 | 0.53711851 |
| SUFU | 746.692818 | 0.288487425 | 0.25442751 | 1.13386882 | 0.25684958 | 0.47778145 |
| TRIM8 | 491.685114 | -0.641470292 | 0.33192092 | -1.9325998 | 0.05328552 | 0.17349404 |
| ARL3 | 494.28708 | -0.108881013 | 0.2060286 | -0.5284752 | 0.59716954 | 0.77654325 |
| SFXN2 | 269.683906 | 0.755020431 | 0.27812026 | 2.71472648 | 0.00663305 | 0.04126225 |
| WBP1L | 893.292103 | -0.438330523 | 0.24806565 | -1.7669941 | 0.07722922 | 0.22166902 |
| BORCS7-ASN | 53.4895181 | -0.06132726 | 0.41553778 | -0.1475853 | 0.88267007 | 0.94732911 |
| CNNM2 | 1705.72486 | 1.039513941 | 0.18297048 | 5.68132047 | 1.34E-08 | 1.20E-06 |
| NT5C2 | 2841.3357 | -0.556127589 | 0.20681997 | -2.6889454 | 0.00716782 | 0.04343104 |
| PCGF6 | 299.356013 | 0.282157626 | 0.26128892 | 1.07986833 | 0.28020082 | 0.50267182 |
| TAF5 | 63.1982737 | 0.353677606 | 0.44997329 | 0.78599689 | 0.43186931 | 0.64957432 |
| ATP5MD | 385.759513 | 0.436776607 | 0.30222505 | 1.44520317 | 0.14840082 | 0.34019898 |
| PDCD11 | 225.040167 | -0.109134909 | 0.2467959 | -0.4422071 | 0.65833933 | 0.81895766 |
| SH3PXD2A | 356.764334 | 0.344781682 | 0.25518067 | 1.35112772 | 0.17665452 | 0.37882911 |
| STN1 | 324.757637 | -0.236401411 | 0.36703828 | -0.6440784 | 0.51952463 | 0.71978191 |
| SLK | 1004.08727 | -0.077252446 | 0.19478811 | -0.3965973 | 0.69166441 | 0.84169408 |
| CFAP43 | 452.004092 | 0.041198499 | 0.30802874 | 0.13374888 | 0.89360115 | 0.95146032 |
| GSTO1 | 371.457264 | -0.68751609 | 0.23751807 | -2.8945843 | 0.00379661 | 0.02760012 |
| GSTO2 | 977.743855 | 0.596524948 | 0.2661197 | 2.24156629 | 0.02498942 | 0.1036049 |
| ITPRIP | 67.4717078 | -0.587403698 | 0.47674083 | -1.2321238 | 0.21790286 | 0.43178501 |
| SORCS3 | 804.908425 | -0.983506715 | 0.814889 | -1.2069211 | 0.22746254 | 0.44295336 |
| SORCS1 | 61.1943648 | 1.011263749 | 0.47736798 | 2.11841555 | 0.03413989 | 0.12893865 |
| LINC01435 | 137.067182 | 0.660645856 | 0.72594229 | 0.91005286 | 0.36279463 | 0.58828312 |
| XPNPEP1 | 469.912225 | -0.326699853 | 0.25216209 | -1.2955947 | 0.19511517 | 0.40263651 |
| ADD3 | 2122.2009 | 0.531250131 | 0.15995073 | 3.32133609 | 0.00089588 | 0.00916074 |
| MXI1 | 1081.98807 | -0.773184934 | 0.29345496 | -2.6347653 | 0.00841955 | 0.04894974 |
| SMNDC1 | 240.750738 | 0.119093674 | 0.21672791 | 0.5495078 | 0.58265702 | 0.7682488 |
| DUSP5 | 176.350617 | -2.053680785 | 0.73290474 | -2.8021115 | 0.00507693 | 0.03378398 |
| SMC3 | 550.226228 | 0.163187103 | 0.19246899 | 0.84786181 | 0.39651493 | 0.61836668 |
| RBM20 | 553.274585 | 0.823565598 | 0.82701226 | 0.9958324 | 0.31933159 | 0.54392231 |
| PDCD4 | 752.723339 | 0.173828231 | 0.21725721 | 0.80010341 | 0.42365089 | 0.64248043 |
| BBIP1 | 441.072283 | -0.019364573 | 0.11905788 | -0.1626484 | 0.87079529 | 0.94134831 |
| SHOC2 | 1310.93049 | -0.144489169 | 0.19455441 | -0.7426672 | 0.45768321 | 0.67227935 |
| GPAM | 120.736536 | 0.550632276 | 0.22924714 | 2.40191552 | 0.01630947 | 0.07754926 |
| ACSL5 | 52.1485826 | -0.774114072 | 0.47520376 | -1.629015 | 0.10330984 | 0.26857304 |
| ZDHHC6 | 302.198725 | 0.093131899 | 0.21406305 | 0.43506761 | 0.66351335 | 0.82268279 |
| VTI1A | 4113.02425 | -0.122513794 | 0.18285912 | -0.6699901 | 0.50286411 | 0.70694289 |
| TCF7L2 | 4116.86221 | -0.334787863 | 0.19735694 | -1.6963572 | 0.08981826 | 0.24431824 |
| AL158212.2 | 47.1706524 | -0.189120661 | 0.45065861 | -0.4196539 | 0.67473828 | 0.82939913 |
| CASP7 | 278.832069 | -0.414271084 | 0.25022533 | -1.6555921 | 0.09780445 | 0.25853493 |
| DCLRE1A | 52.1451186 | 0.739345694 | 0.31013975 | 2.38391141 | 0.01712973 | 0.08012271 |
| NHLRC2 | 821.875221 | 0.158568301 | 0.14951741 | 1.06053402 | 0.28890172 | 0.5112678 |
| CCDC186 | 1071.92839 | 0.267545036 | 0.22804767 | 1.17319785 | 0.24071647 | 0.4588864 |
| TDRD1 | 52.4682068 | 1.933085877 | 0.73775029 | 2.62024414 | 0.00878668 | 0.05041939 |
| VWA2 | 59.319986 | -0.520971317 | 0.84083971 | -0.6195846 | 0.53553132 | 0.73243252 |
| AFAP1L2 | 282.479882 | 0.219471304 | 0.85179829 | 0.25765643 | 0.79667208 | 0.90460255 |

| | | | | | | |
|---|---|---|---|---|---|---|
| ABLIM1 | 6744.33254 | -0.09056558 | 0.38833209 | -0.2332168 | 0.81559304 | 0.91421906 |
| FAM160B1 | 598.285125 | 0.249998825 | 0.18429089 | 1.35654464 | 0.17492594 | 0.37662539 |
| TRUB1 | 269.956735 | 0.54371305 | 0.1533832 | 3.54480181 | 0.00039291 | 0.00492029 |
| ATRNL1 | 705.045437 | 0.217530022 | 0.66928377 | 0.32501912 | 0.7451666 | 0.87531397 |
| HSPA12A | 1204.35343 | 0.288575421 | 0.30237049 | 0.95437693 | 0.33989288 | 0.56526428 |
| ENO4 | 160.603132 | -0.117895795 | 0.37364089 | -0.3155324 | 0.75235748 | 0.87955089 |
| SHTN1 | 1318.11331 | -0.402632973 | 0.1893855 | -2.1259968 | 0.03350353 | 0.12730725 |
| PDZD8 | 984.67552 | -0.259288046 | 0.23512188 | -1.1027815 | 0.27012208 | 0.49175977 |
| AC005871.2 | 78.5483087 | -1.049209551 | 0.56875336 | -1.8447531 | 0.06507346 | 0.19827619 |
| EMX2OS | 1455.24414 | -0.451831984 | 0.24583087 | -1.837979 | 0.0660655 | 0.20028899 |
| EMX2 | 497.549375 | -0.266545121 | 0.25980754 | -1.025933 | 0.30492318 | 0.52822225 |
| RAB11FIP2 | 982.046616 | 0.208072398 | 0.13715794 | 1.51702774 | 0.12925968 | 0.31077378 |
| AC022395.1 | 84.8729498 | -0.105002604 | 0.52535796 | -0.1998687 | 0.84158328 | 0.92654505 |
| CASC2 | 1188.0661 | 0.239668671 | 0.40691667 | 0.58898711 | 0.55586992 | 0.748078 |
| FAM204A | 673.530107 | 0.239800131 | 0.13396801 | 1.78998048 | 0.07345705 | 0.21504746 |
| CACUL1 | 1183.39343 | -0.238695337 | 0.18198935 | -1.3115896 | 0.18965864 | 0.39572864 |
| EIF3A | 1302.04083 | -0.297232891 | 0.15225568 | -1.9521958 | 0.05091497 | 0.16821806 |
| FAM45A | 291.707303 | 0.188068502 | 0.22759124 | 0.82634333 | 0.40860936 | 0.62839924 |
| SFXN4 | 376.410749 | 0.612949663 | 0.33779973 | 1.8145357 | 0.06959529 | 0.20746202 |
| PRDX3 | 435.612331 | -0.265886898 | 0.44839211 | -0.5929785 | 0.55319552 | 0.74600915 |
| GRK5 | 114.260771 | -0.054950077 | 0.4265296 | -0.1288306 | 0.89749167 | 0.9525128 |
| RGS10 | 61.5846429 | 0.475240031 | 0.57081921 | 0.83255788 | 0.40509412 | 0.62561537 |
| TIAL1 | 796.108157 | 0.537009803 | 0.15338195 | 3.5011278 | 0.00046329 | 0.00559383 |
| BAG3 | 283.604434 | 0.033749113 | 0.2307871 | 0.14623483 | 0.88373601 | 0.94765944 |
| INPP5F | 669.405073 | -0.255013479 | 0.15557349 | -1.6391833 | 0.10117508 | 0.26492976 |
| MCMBP | 476.442056 | 0.108669753 | 0.18434053 | 0.58950548 | 0.55552224 | 0.74777267 |
| SEC23IP | 664.12071 | 0.096103338 | 0.19020233 | 0.50526897 | 0.61336993 | 0.78744323 |
| WDR11-AS1 | 137.102549 | 0.177840104 | 0.37858744 | 0.46974644 | 0.63853618 | 0.80550324 |
| WDR11 | 1055.78713 | 0.292558167 | 0.13575602 | 2.15502912 | 0.03115956 | 0.1210689 |
| FGFR2 | 1932.95385 | 0.946731546 | 0.36084079 | 2.6236822 | 0.00869849 | 0.05008647 |
| ATE1 | 1676.23239 | -0.010253711 | 0.22741295 | -0.0450885 | 0.96403676 | 0.98420487 |
| NSMCE4A | 169.570749 | 0.450372156 | 0.24499233 | 1.83831123 | 0.06601656 | 0.20026199 |
| TACC2 | 1011.14765 | 0.030404619 | 0.45105045 | 0.06740847 | 0.94625653 | 0.9767213 |
| PLEKHA1 | 5035.29153 | -2.055091894 | 0.25020428 | -8.213656 | 2.15E-16 | 2.02E-13 |
| HTRA1 | 93.0824893 | -1.20046446 | 0.48796948 | -2.460122 | 0.01388898 | 0.06943963 |
| FAM24B | 55.8934201 | -0.438712928 | 0.27343071 | -1.6044757 | 0.10860923 | 0.27695355 |
| C10orf88 | 204.916618 | 0.238167912 | 0.20776151 | 1.14635244 | 0.25164935 | 0.47199621 |
| PSTK | 199.949865 | 0.321081132 | 0.20468287 | 1.56867615 | 0.11672342 | 0.29096049 |
| IKZF5 | 251.088446 | 0.271860109 | 0.21076342 | 1.28988279 | 0.19709136 | 0.40544771 |
| ACADSB | 475.035763 | 0.203859997 | 0.1939973 | 1.05083938 | 0.29333237 | 0.51627278 |
| BUB3 | 312.762645 | 0.030920667 | 0.23731347 | 0.13029461 | 0.89633334 | 0.95220099 |
| CPXM2 | 541.343206 | -0.670201751 | 0.70757126 | -0.9471862 | 0.34354389 | 0.56837065 |
| CHST15 | 56.1674119 | -2.922726584 | 0.61977688 | -4.7157722 | 2.41E-06 | 8.28E-05 |
| OAT | 212.90627 | 0.000120829 | 0.27557237 | 0.00043847 | 0.99965015 | 0.99975417 |
| LHPP | 277.857855 | 0.383065776 | 0.38956443 | 0.98331814 | 0.32545088 | 0.55057386 |
| ABRAXAS2 | 280.795439 | 0.005277244 | 0.18108669 | 0.02914208 | 0.97675127 | 0.98948597 |
| ZRANB1 | 620.931754 | -0.044924913 | 0.23142601 | -0.1941221 | 0.84608024 | 0.92876871 |
| CTBP2 | 4495.33617 | -0.365182031 | 0.25861476 | -1.4120696 | 0.15792948 | 0.35399465 |
| EDRF1 | 460.914686 | 0.253492956 | 0.20863687 | 1.21499599 | 0.22436762 | 0.43912303 |
| UROS | 231.579478 | 0.065786659 | 0.2612228 | 0.25184119 | 0.80116382 | 0.90672348 |
| BCCIP | 472.481915 | 0.021243236 | 0.14034048 | 0.15136928 | 0.87968442 | 0.9463109 |
| DHX32 | 1233.0692 | -0.445714039 | 0.23561866 | -1.8916754 | 0.05853424 | 0.18461766 |
| FANK1 | 660.590758 | -0.478487652 | 0.25910011 | -1.8467289 | 0.06478644 | 0.19768171 |
| ADAM12 | 161.329566 | -2.749286276 | 0.58176052 | -4.7258042 | 2.29E-06 | 7.93E-05 |
| DOCK1 | 8170.94607 | 0.063481279 | 0.17733456 | 0.35797466 | 0.72036227 | 0.86064266 |
| AL359094.1 | 119.506841 | -0.147455477 | 0.59848659 | -0.2463806 | 0.80538763 | 0.90895123 |
| FAM196A | 350.24224 | 0.223667735 | 0.58747476 | 0.38072739 | 0.70340554 | 0.84973591 |
| MGMT | 1204.89379 | -0.118032902 | 0.21934332 | -0.5381194 | 0.59049459 | 0.77217181 |
| AL157832.3 | 165.507792 | -0.166116124 | 0.44716622 | -0.3714863 | 0.71027536 | 0.85479293 |
| C10orf143 | 167.956073 | 0.600913528 | 0.41650009 | 1.44276927 | 0.14908548 | 0.34111729 |
| GLRX3 | 592.567038 | -0.065230228 | 0.19230047 | -0.3392099 | 0.73445159 | 0.86968813 |
| PPP2R2D | 1132.91693 | 0.018206572 | 0.17240949 | 0.10560076 | 0.91589912 | 0.96214624 |
| BNIP3 | 1052.95805 | -1.611937487 | 0.59489915 | -2.7095979 | 0.00673648 | 0.04173649 |
| STK32C | 252.207576 | -0.199769603 | 0.3006875 | -0.6643762 | 0.50644959 | 0.70992915 |
| LRRC27 | 204.247361 | -0.084755202 | 0.38421216 | -0.2205948 | 0.82540796 | 0.91938879 |
| PWWP2B | 90.4384895 | -0.526217022 | 0.25762866 | -2.0425407 | 0.04109793 | 0.14502011 |
| INPP5A | 843.081329 | -0.371845485 | 0.28906142 | -1.2863892 | 0.19830727 | 0.40681157 |
| CFAP46 | 75.4987259 | -0.30372052 | 0.45843861 | -0.6625108 | 0.50764393 | 0.71077684 |
| KNDC1 | 94.6172429 | 0.358404187 | 0.59690495 | 0.60043761 | 0.54821463 | 0.74186733 |
| ADAM8 | 43.7289009 | 1.126474264 | 0.43290542 | 2.60212555 | 0.00926479 | 0.05241056 |
| TUBGCP2 | 747.595486 | -0.031627917 | 0.20263649 | -0.156082 | 0.87596835 | 0.94420235 |
| ECHS1 | 359.56551 | -0.813769085 | 0.433554 | -1.8769729 | 0.06052181 | 0.18908057 |
| PAOX | 61.5131344 | -0.870302514 | 0.21915283 | -3.9712127 | 7.15E-05 | 0.00129036 |
| SCART1 | 145.246506 | 0.891865767 | 0.43056316 | 2.0713936 | 0.03832203 | 0.13904125 |
| SLC6A12 | 489.234256 | -0.898898895 | 0.54073513 | -1.6623645 | 0.09643971 | 0.25636387 |
| SLC6A13 | 61.3434858 | 1.472069715 | 0.59688811 | 2.46624066 | 0.01365396 | 0.06855002 |
| KDM5A | 1574.39567 | -0.003004218 | 0.12018601 | -0.0249964 | 0.98005784 | 0.99139102 |
| CCDC77 | 126.285054 | -0.295447826 | 0.31862084 | -0.9272709 | 0.35378591 | 0.57958057 |
| B4GALNT3 | 730.31664 | -0.187828149 | 0.31828616 | -0.5901235 | 0.55510784 | 0.74754753 |
| WNK1 | 4596.51604 | 0.523101406 | 0.30941507 | 1.69061388 | 0.09091057 | 0.24631864 |
| RAD52 | 498.379901 | 0.31943702 | 0.18395933 | 1.73645462 | 0.08248348 | 0.23078253 |
| ERC1 | 4005.72562 | -0.286777768 | 0.11883978 | -2.4131462 | 0.01581548 | 0.07602133 |
| WNT5B | 139.731024 | 0.852810709 | 0.38972173 | 2.1882555 | 0.028651 | 0.11381696 |
| FBXL14 | 145.050368 | 0.466549614 | 0.19354931 | 2.41049487 | 0.0159309 | 0.07637249 |
| ADIPOR2 | 1236.5347 | 0.136812454 | 0.29884962 | 0.45779698 | 0.6470983 | 0.81134518 |
| DCP1B | 535.375946 | 0.284611798 | 0.1715489 | 1.65907095 | 0.0971015 | 0.25751598 |
| CACNA1C | 717.826539 | -0.129418739 | 0.78976024 | -0.1638709 | 0.86983276 | 0.94069272 |
| FKBP4 | 237.904104 | -0.38461093 | 0.45635393 | -0.8427909 | 0.39934541 | 0.62094804 |
| ITFG2-AS1 | 83.7070997 | 0.424690493 | 0.33901401 | 1.25272253 | 0.2103067 | 0.42243632 |
| ITFG2 | 910.013997 | 0.389550571 | 0.23073394 | 1.68831066 | 0.09135161 | 0.24706458 |
| TULP3 | 1429.20691 | -0.089443999 | 0.19661997 | -0.454908 | 0.64917543 | 0.81217638 |
| TEAD4 | 584.036594 | -1.549699586 | 0.40456749 | -3.8305095 | 0.00012788 | 0.0020425 |
| TSPAN9 | 4068.44352 | -0.195669891 | 0.37899448 | -0.5162869 | 0.60565404 | 0.78178941 |
| LINC02417 | 61.8148951 | 2.759983757 | 0.97865391 | 2.82018366 | 0.00479962 | 0.03239516 |
| PRMT8 | 146.065339 | 3.016995174 | 0.89674359 | 3.36439002 | 0.00076713 | 0.0081718 |
| PARP11 | 292.269871 | -0.010037991 | 0.29012562 | -0.0345988 | 0.97239968 | 0.98795688 |
| CCND2 | 48.72158 | -2.403724968 | 0.95762577 | -2.510088 | 0.01207011 | 0.06316874 |
| TIGAR | 180.16532 | -0.492296242 | 0.40422019 | -1.2178913 | 0.2232653 | 0.43787844 |
| C12orf4 | 349.814209 | 0.409256353 | 0.16335048 | 2.50538817 | 0.01223171 | 0.06375249 |
| AKAP3 | 44.6026082 | 0.537871789 | 0.52686204 | 1.02089682 | 0.30730332 | 0.53081857 |
| AC005833.2 | 65.2971636 | 0.845244125 | 0.28368939 | 2.97947035 | 0.00288747 | 0.02248061 |
| GALNT8 | 50.1958888 | 0.670364616 | 0.39750541 | 1.68642891 | 0.09171322 | 0.24783958 |
| ANO2 | 53.4987558 | -0.983559477 | 0.52394629 | -1.8772143 | 0.06048872 | 0.18906663 |
| VWF | 44.8872081 | 0.277134697 | 0.41950521 | 0.66062278 | 0.50885426 | 0.71134045 |
| CD9 | 994.993848 | -0.904168759 | 0.41467609 | -2.1804218 | 0.02922621 | 0.11527293 |
| PLEKHG6 | 42.2282007 | -1.237308319 | 0.44334371 | -2.7908557 | 0.00525689 | 0.03477129 |
| TNFRSF1A | 501.635658 | -1.322458624 | 0.39003336 | -3.3906295 | 0.00069732 | 0.00764393 |
| SCNN1A | 2735.57698 | -0.176644896 | 0.31409011 | -0.562402 | 0.57384218 | 0.76133803 |
| LTBR | 282.260566 | -0.814706845 | 0.22858441 | -3.56414 | 0.00036505 | 0.00464543 |
| TAPBPL | 139.436606 | 0.111969665 | 0.37316897 | 0.30005085 | 0.76413837 | 0.88555042 |
| MRPL51 | 253.780192 | -1.011682566 | 0.44721109 | -2.2622037 | 0.02368482 | 0.10003413 |
| NCAPD2 | 216.02849 | -0.261938429 | 0.29989101 | -0.8734454 | 0.38242034 | 0.60517434 |
| GAPDH | 2272.90492 | -1.915874644 | 0.50241816 | -3.8133069 | 0.00013712 | 0.00215364 |

| | | | | | | |
|---|---|---|---|---|---|---|
| CHD4 | 313.446609 | -0.282804864 | 0.19496491 | -1.4505423 | 0.14690734 | 0.33777181 |
| AC006064.2 | 140.976316 | -0.360152678 | 0.42937167 | -0.83879 | 0.40158714 | 0.62311342 |
| ING4 | 274.881844 | -0.067519917 | 0.18508743 | -0.3648001 | 0.71526061 | 0.85772831 |
| ZNF384 | 262.439445 | 0.009931298 | 0.21319718 | 0.04658269 | 0.96284583 | 0.98354579 |
| COPS7A | 200.479013 | -0.076028561 | 0.21364151 | -0.3558698 | 0.72193808 | 0.86151234 |
| MLF2 | 345.552249 | -0.850327101 | 0.38084645 | -2.2327295 | 0.02556679 | 0.1050752 |
| PTMS | 511.922283 | -0.693104131 | 0.52413865 | -1.3223679 | 0.18604567 | 0.39176614 |
| P3H3 | 50.7170868 | -0.215214991 | 0.43892112 | -0.4903273 | 0.62390234 | 0.79539871 |
| USP5 | 159.377478 | -0.44525771 | 0.36629504 | -1.2155712 | 0.2241483 | 0.4388934 |
| TPI1 | 828.351336 | -1.119243038 | 0.42700204 | -2.6211655 | 0.00876297 | 0.0504138 |
| SPSB2 | 56.668704 | -0.565403735 | 0.32085175 | -1.7621962 | 0.07803615 | 0.22316946 |
| LRRC23 | 280.908117 | -0.652176069 | 0.26922215 | -2.4224458 | 0.01541642 | 0.07478353 |
| ENO2 | 137.727865 | -2.627697053 | 0.59528646 | -4.4141724 | 1.01E-05 | 0.00026939 |
| ATN1 | 606.99151 | -0.47561934 | 0.33023223 | -1.4402572 | 0.14979465 | 0.34200906 |
| C12orf57 | 197.166055 | -0.922785734 | 0.46228036 | -1.9961604 | 0.04591647 | 0.1565997 |
| PTPN6 | 231.962095 | -0.250827796 | 0.37228363 | -0.6737546 | 0.50046737 | 0.70482771 |
| PHB2 | 322.09736 | -0.463886433 | 0.32907812 | -1.4096544 | 0.15864175 | 0.35514221 |
| EMG1.1 | 107.132232 | 0.022321316 | 0.31694096 | 0.07042736 | 0.94385351 | 0.97568955 |
| LPCAT3 | 988.929714 | -0.082791951 | 0.24006362 | -0.344875 | 0.73018831 | 0.86751731 |
| C1S | 463.994264 | -0.353908559 | 0.34006009 | -1.0407236 | 0.29800385 | 0.52143766 |
| C1R | 252.372735 | -3.328109797 | 0.54863826 | -6.0661278 | 1.31E-09 | 1.62E-07 |
| C1RL | 884.923836 | -1.551888523 | 0.29486438 | -5.2630586 | 1.42E-07 | 8.21E-06 |
| C1RL-AS1 | 97.7125155 | -1.668993513 | 0.43914023 | -3.8005935 | 0.00014435 | 0.00224584 |
| RBP5 | 139.442483 | -0.056277093 | 0.30142106 | -0.1867059 | 0.85189122 | 0.93193618 |
| CLSTN3 | 308.834224 | -0.213116222 | 0.30110947 | -0.7077699 | 0.47908816 | 0.68864274 |
| PEX5 | 156.396882 | 0.064429383 | 0.21275061 | 0.30283994 | 0.76201184 | 0.88448281 |
| CD163L1 | 65.9805737 | -0.616931818 | 0.3544817 | -1.740377 | 0.08179283 | 0.22992159 |
| FOXJ2 | 141.484351 | 0.066201964 | 0.25903108 | 0.25557537 | 0.79827874 | 0.90515239 |
| NECAP1 | 182.224815 | 0.264601063 | 0.20657779 | 1.28087854 | 0.20023633 | 0.40905068 |
| FAM66C | 198.803613 | 0.396737809 | 0.4068312 | 0.97519022 | 0.32946591 | 0.55488483 |
| LINC00937 | 212.660707 | -0.965136533 | 0.26271782 | -3.6736622 | 0.0002391 | 0.00339597 |
| RIMKLB | 1602.19931 | -0.30644234 | 0.53021326 | -0.5779605 | 0.56329076 | 0.75476848 |
| A2ML1-AS1 | 253.150596 | -0.098013937 | 0.65250843 | -0.150211 | 0.88059816 | 0.94639771 |
| AC006581.2 | 103.687649 | -0.078005687 | 0.3501742 | -0.2227625 | 0.82372034 | 0.91846107 |
| PHC1 | 357.390998 | 0.314440563 | 0.18611814 | 1.68946758 | 0.09112986 | 0.24666687 |
| M6PR | 361.589219 | -0.43395292 | 0.22991854 | -1.8874203 | 0.05910382 | 0.18583726 |
| KLRG1 | 98.6489584 | 0.708336222 | 0.39622527 | 1.78771089 | 0.07382266 | 0.21554455 |
| A2M | 170.46574 | -0.866062714 | 0.57631161 | -1.5027681 | 0.13289885 | 0.31675585 |
| CLEC2D | 449.043793 | -0.028994119 | 0.31049905 | -0.0933791 | 0.9256024 | 0.96719264 |
| KLRF1 | 75.6508255 | 0.444834766 | 0.67656946 | 0.65748573 | 0.51086864 | 0.71294917 |
| TMEM52B | 684.610192 | 1.19514288 | 1.14937962 | 1.03981562 | 0.29842557 | 0.52159011 |
| GABARAPL1 | 1166.08907 | -0.298049981 | 0.38702818 | -0.7700989 | 0.44124121 | 0.65696085 |
| KLRD1 | 579.617698 | -0.326224644 | 0.57660364 | -0.5657693 | 0.57155061 | 0.75982407 |
| AC022075.1 | 54.5345406 | -0.770769049 | 0.28822518 | -2.6741905 | 0.00749099 | 0.04497657 |
| EIF2S3B | 70.8451214 | 0.155078774 | 0.31721261 | 0.48887959 | 0.62492694 | 0.79598067 |
| MAGOHB | 152.319993 | 0.399979441 | 0.28466891 | 1.40506893 | 0.16000078 | 0.35694145 |
| YBX3 | 2018.90374 | -1.411227168 | 0.32905514 | -4.2887256 | 1.80E-05 | 0.00043236 |
| PRH1 | 139.300245 | 0.184873734 | 0.50960036 | 0.3627818 | 0.71676789 | 0.85906788 |
| SMIM10L1 | 120.885363 | -0.195055988 | 0.2764534 | -0.7055655 | 0.48045837 | 0.68948936 |
| AC244131.2 | 337.070627 | -0.38577613 | 0.55091801 | -0.7002424 | 0.48377596 | 0.69195416 |
| PRB2 | 57.9146849 | -0.153115305 | 0.7560721 | -0.2025142 | 0.83951479 | 0.92548413 |
| LINC01252 | 146.19877 | 0.024919564 | 0.70007737 | 0.03559544 | 0.97160494 | 0.98744736 |
| ETV6 | 3606.60611 | -1.440538273 | 0.27416384 | -5.2542972 | 1.49E-07 | 8.57E-06 |
| BCL2L14 | 703.350756 | 0.544377844 | 0.44412369 | 1.22573477 | 0.2202985 | 0.4341354 |
| LRP6 | 2304.30104 | 1.005164837 | 0.16511523 | 6.08765675 | 1.15E-09 | 1.43E-07 |
| MANSC1 | 124.530955 | -0.726915921 | 0.21732876 | -3.3447755 | 0.00082349 | 0.00860562 |
| BORCS5 | 588.935846 | -0.106529523 | 0.23502943 | -0.4532604 | 0.65036128 | 0.81304393 |
| DUSP16 | 833.319118 | -0.291857529 | 0.35245794 | -0.8280634 | 0.40763459 | 0.62748459 |
| AC007619.1 | 52.0066792 | 0.251487186 | 0.42227108 | 0.59555862 | 0.55147012 | 0.74467698 |
| CREBL2 | 345.303084 | -0.374593812 | 0.16975933 | -2.2066169 | 0.02734084 | 0.11021426 |
| CDKN1B | 189.731644 | -0.345400809 | 0.333402 | -1.035989 | 0.30020727 | 0.52314483 |
| APOLD1 | 305.651201 | 0.648962262 | 0.52630315 | 1.23305791 | 0.21755416 | 0.43128814 |
| GPRC5A | 132.992315 | -2.330671595 | 0.8731273 | -2.6693377 | 0.0076001 | 0.04550758 |
| GPRC5D-AS1 | 414.994207 | 0.508262006 | 0.37363772 | 1.360307 | 0.1737328 | 0.37546409 |
| HEBP1 | 159.141224 | 0.191911382 | 0.32820097 | 0.58473741 | 0.5587243 | 0.75065603 |
| AC023790.2 | 91.142559 | 0.79999123 | 0.38865192 | 2.05837457 | 0.03955419 | 0.14166792 |
| FAM234B | 114.801665 | 0.402988331 | 0.32448091 | 1.24194772 | 0.21425585 | 0.42776686 |
| EMP1 | 1053.77714 | -1.088951404 | 0.35655554 | -3.0540863 | 0.00225747 | 0.01873049 |
| ATF7IP | 2449.35244 | 0.285925699 | 0.25689031 | 1.1130264 | 0.26569709 | 0.48676739 |
| PLBD1 | 133.468481 | -0.194440659 | 0.26065455 | -0.7459707 | 0.4556851 | 0.67098025 |
| PLBD1-AS1 | 119.692505 | 0.379307078 | 0.45819677 | 0.82782573 | 0.4077692 | 0.62761866 |
| AC010168.1 | 230.27998 | 0.434902342 | 0.47810147 | 0.90964444 | 0.36301005 | 0.58856018 |
| H2AFJ | 298.558907 | -1.286952404 | 0.41723454 | -3.084482 | 0.00203907 | 0.01741052 |
| WBP11 | 321.917346 | -0.057500592 | 0.20448176 | -0.2812016 | 0.7785558 | 0.8941782 |
| C12orf60 | 306.336645 | 0.580874367 | 0.36905716 | 1.57394146 | 0.11550098 | 0.28905883 |
| MGP | 127.523759 | -0.466915407 | 0.5050446 | -0.9245033 | 0.35522433 | 0.58081705 |
| ERP27 | 55.0747579 | -0.209322101 | 0.701392 | -0.2984381 | 0.76536881 | 0.88643197 |
| RERG | 792.526143 | 0.260549365 | 0.50346725 | 0.51751006 | 0.60480015 | 0.78156518 |
| PTPRO | 51.8172095 | -0.368025496 | 0.48797294 | -0.7541924 | 0.45073368 | 0.66654333 |
| EPS8 | 710.211692 | -0.532165058 | 0.39066137 | -1.3622157 | 0.17312982 | 0.37472092 |
| STRAP | 489.982972 | -0.163954699 | 0.25444949 | -0.6443507 | 0.51934806 | 0.71961278 |
| DERA | 1289.64619 | -0.345598945 | 0.20552949 | -1.6815054 | 0.09266479 | 0.24984879 |
| MGST1 | 583.309863 | -0.185471839 | 0.63079934 | -0.2940267 | 0.76873751 | 0.88857795 |
| LMO3 | 177.060789 | 1.979270465 | 0.55083239 | 3.59323546 | 0.0003266 | 0.00427554 |
| PIK3C2G | 64.5649344 | 1.054432356 | 0.42769069 | 2.46540872 | 0.0136857 | 0.06868331 |
| PLEKHA5 | 9481.82787 | 0.44269081 | 0.21574882 | 2.05188055 | 0.04018127 | 0.14298087 |
| AEBP2 | 3269.6958 | 0.615945102 | 0.29572029 | 2.08286389 | 0.03726363 | 0.13630998 |
| AC024901.1 | 541.569731 | 0.146372869 | 0.60136064 | 0.24340281 | 0.80769337 | 0.91010272 |
| LINC02398 | 270.54487 | -0.250538502 | 0.73502393 | -0.3408576 | 0.73321078 | 0.86911141 |
| PDE3A | 451.049749 | 0.232600463 | 0.84803421 | 0.27428194 | 0.78386797 | 0.89671287 |
| PYROXD1 | 376.363423 | -0.190766253 | 0.13273738 | -1.4371706 | 0.15066953 | 0.34308559 |
| RECQL | 288.23495 | -0.485963685 | 0.2268106 | -2.1425969 | 0.03214548 | 0.12379251 |
| GOLT1B | 277.298461 | 0.236240902 | 0.17261688 | 1.3685852 | 0.17112897 | 0.37172219 |
| SPX | 93.3631681 | 0.099285423 | 0.66127926 | 0.15014144 | 0.88065302 | 0.94639771 |
| LDHB | 3187.10101 | -0.621824837 | 0.47060741 | -1.3213239 | 0.18639337 | 0.39217427 |
| AC010197.1 | 453.215327 | -0.969698531 | 0.41284092 | -2.3488431 | 0.01883184 | 0.08556925 |
| AC010185.1 | 60.4447803 | -1.137956483 | 0.50320249 | -2.2614286 | 0.02373273 | 0.10018717 |
| ABCC9 | 228.476408 | -1.858897676 | 0.61792566 | -3.0082869 | 0.00262725 | 0.02100687 |
| AC008250.1 | 419.647345 | -0.954712018 | 0.41074866 | -2.3243217 | 0.02010826 | 0.08933209 |
| CMAS | 265.589096 | -0.284906227 | 0.19384759 | -1.4697434 | 0.14163125 | 0.33029789 |
| ST8SIA1 | 50.184627 | 1.196966936 | 0.41834892 | 2.86116891 | 0.00422082 | 0.02963999 |
| C2CD5 | 1843.13499 | 0.651051682 | 0.1967363 | 3.3092605 | 0.00093543 | 0.00946052 |
| ETNK1 | 3577.63361 | 0.281485149 | 0.28920602 | 0.9733032 | 0.33040263 | 0.55577504 |
| AC084816.1 | 213.937016 | -0.618901679 | 0.48102127 | -1.286641 | 0.19821945 | 0.40675481 |
| SOX5 | 260.606304 | -0.16750282 | 0.55492415 | -0.3018481 | 0.76276783 | 0.88492094 |
| CASC1 | 541.77525 | 0.72258476 | 0.38813712 | 1.86167393 | 0.06264906 | 0.19370578 |
| ETFRF1 | 428.380303 | 0.461020921 | 0.21213244 | 2.1732693 | 0.02976005 | 0.1168551 |
| KRAS | 810.087553 | -0.256457146 | 0.21011713 | -1.2205437 | 0.22225883 | 0.43642291 |
| LMNTD1 | 58.0243092 | 2.467716729 | 0.8947437 | 2.7580152 | 0.00581535 | 0.03754366 |
| RASSF8-AS1 | 988.408846 | 0.086565885 | 0.42955867 | 0.20152284 | 0.84028977 | 0.92579761 |
| RASSF8 | 3625.66123 | 0.630632368 | 0.49012877 | 1.28666671 | 0.19821048 | 0.40675481 |
| BHLHE41 | 68.3039343 | -2.621857299 | 0.71314363 | -3.6764786 | 0.00023648 | 0.00336233 |
| SSPN | 94.6589534 | -0.213676079 | 0.40180401 | -0.5317918 | 0.59487021 | 0.77501911 |

| | | | | | | |
|---|---|---|---|---|---|---|
| AC022509.2 | 59.9991429 | 0.238202333 | 0.43258113 | 0.55065355 | 0.58187119 | 0.76767245 |
| AC024145.1 | 177.501514 | 0.141794072 | 0.60274171 | 0.23524848 | 0.81401588 | 0.91299446 |
| ITPR2 | 7494.05068 | 0.455873884 | 0.23923127 | 1.90557818 | 0.05670495 | 0.18109665 |
| AC023051.1 | 193.928995 | 0.246204118 | 0.58160858 | 0.42331583 | 0.67206485 | 0.82787509 |
| INTS13 | 185.822575 | 0.159953355 | 0.25006929 | 0.63963614 | 0.52240918 | 0.72226322 |
| FGFR1OP2 | 648.220675 | 0.655053844 | 0.19347283 | 3.38576665 | 0.0007098 | 0.00772192 |
| TM7SF3 | 541.102801 | 0.032854896 | 0.25456887 | 0.12906094 | 0.89730943 | 0.9525128 |
| MED21 | 220.465283 | 0.166869429 | 0.26307726 | 0.6342982 | 0.52588623 | 0.72494586 |
| STK38L | 707.917724 | 0.32342961 | 0.31083773 | 1.0405095 | 0.29810325 | 0.52143889 |
| ARNTL2 | 75.8293059 | -2.648086757 | 0.6594415 | -4.0156508 | 5.93E-05 | 0.00111546 |
| PPFIBP1 | 1724.35893 | 0.015673426 | 0.29139011 | 0.05378846 | 0.9571037 | 0.98109443 |
| MRPS35 | 429.845509 | 0.066107551 | 0.15017526 | 0.44020268 | 0.65979032 | 0.81980614 |
| KLHL42 | 491.436552 | -0.162372745 | 0.18279908 | -0.888258 | 0.37440197 | 0.59792959 |
| CCDC91 | 3294.46417 | -0.097206557 | 0.16665865 | -0.5832674 | 0.5597133 | 0.75155057 |
| AC022364.1 | 84.239593 | 0.443744518 | 0.2545123 | 1.74350909 | 0.08124473 | 0.22877033 |
| FAR2 | 138.560043 | -0.24921447 | 0.42797635 | -0.582309 | 0.56035858 | 0.75183103 |
| AC009318.1 | 205.280709 | -0.225507199 | 0.57024439 | -0.3954571 | 0.69250557 | 0.84221582 |
| ERGIC2 | 602.239387 | 0.068552415 | 0.16947486 | 0.40449903 | 0.68584579 | 0.83751486 |
| TMTC1 | 86.6400381 | 0.142941873 | 0.69448582 | 0.20582403 | 0.83692837 | 0.9240982 |
| IPO8 | 757.730125 | 0.374675911 | 0.14412242 | 2.59970591 | 0.00933037 | 0.05269125 |
| CAPRIN2 | 503.426736 | 0.211700708 | 0.16301701 | 1.29864185 | 0.19406687 | 0.40163448 |
| AC010198.1 | 160.332587 | 1.066691264 | 0.4582517 | 2.32774098 | 0.01992586 | 0.08894921 |
| DDX11-AS1 | 50.7344581 | 0.398905553 | 0.4008639 | 0.99511469 | 0.31968049 | 0.54422157 |
| DDX11 | 145.396316 | 0.547947034 | 0.53672654 | 1.02090542 | 0.30729925 | 0.53081857 |
| SINHCAF | 1241.06546 | -0.187988377 | 0.17457569 | -1.0768302 | 0.28155614 | 0.50413655 |
| DENND5B | 1445.11832 | -0.356946077 | 0.19185563 | -1.8604931 | 0.06281579 | 0.19399901 |
| DENND5B-A! | 69.5619271 | -1.186019475 | 0.43656509 | -2.7167071 | 0.00659349 | 0.0411012 |
| ETFBKMT | 103.818622 | 0.160929602 | 0.21375327 | 0.75287553 | 0.45152472 | 0.66729785 |
| AMN1 | 451.522638 | 0.452917988 | 0.21640385 | 2.09292946 | 0.03635545 | 0.13417691 |
| KIAA1551 | 516.447604 | -0.522283031 | 0.15707163 | -3.3251265 | 0.00088378 | 0.00907769 |
| BICD1 | 2085.86793 | -2.76815617 | 0.509029 | -5.4381109 | 5.38E-08 | 3.87E-06 |
| FGD4 | 11626.6108 | -0.064602343 | 0.17903281 | -0.3608408 | 0.71821845 | 0.85963642 |
| DNM1L | 1559.17378 | 0.503265444 | 0.15090521 | 3.33497727 | 0.00085306 | 0.00880322 |
| YARS2 | 454.90734 | 0.123643543 | 0.15365738 | 0.80467037 | 0.42100992 | 0.64038167 |
| PKP2 | 142.257885 | -0.443371115 | 0.26151229 | -1.6954122 | 0.08999726 | 0.24458483 |
| SYT10 | 79.0346718 | 1.54769527 | 0.58365981 | 2.6517078 | 0.00800858 | 0.04737365 |
| ALG10 | 42.6076518 | 0.428223836 | 0.36447831 | 1.17489524 | 0.24003663 | 0.45806569 |
| ALG10B | 155.119753 | 0.560108723 | 0.28222263 | 1.98463435 | 0.04718517 | 0.1594038 |
| CPNE8 | 2050.62365 | -0.463726163 | 0.28597561 | -1.6215584 | 0.10489793 | 0.27136637 |
| KIF21A | 702.351186 | 0.337994171 | 0.35282051 | 0.95797768 | 0.33807401 | 0.56341403 |
| C12orf40 | 205.591604 | -0.154710996 | 0.49774728 | -0.3108224 | 0.75593565 | 0.88214393 |
| SLC2A13 | 2491.52819 | 0.063183552 | 0.24400809 | 0.2589404 | 0.79568123 | 0.90425478 |
| LINC02471 | 106.095068 | -2.662248925 | 0.92930389 | -2.8647775 | 0.00417302 | 0.02937201 |
| AC079630.1 | 125.6449 | -0.407626533 | 0.49003595 | -0.8318298 | 0.40550499 | 0.62581089 |
| LRRK2 | 3576.64906 | -0.68828928 | 0.3326155 | -2.0693241 | 0.03851568 | 0.1392701 |
| CNTN1 | 634.207962 | 0.768503612 | 0.6851923 | 1.12158822 | 0.26203756 | 0.48287586 |
| PDZRN4 | 87.0845857 | 3.067268544 | 0.6256821 | 4.9022795 | 9.47E-07 | 3.96E-05 |
| AC090630.1 | 79.2033288 | 0.158283863 | 0.69725025 | 0.22701155 | 0.82041476 | 0.91661089 |
| GXYLT1 | 537.461515 | 0.216246678 | 0.19081994 | 1.13324988 | 0.25710933 | 0.4779747 |
| YAF2 | 1132.4728 | -0.062920371 | 0.2280662 | -0.2758864 | 0.78263529 | 0.89629179 |
| PPHLN1 | 1597.67487 | -0.03906951 | 0.11584131 | -0.3372675 | 0.73591525 | 0.87055499 |
| ZCRB1 | 249.980653 | 0.091455109 | 0.20024789 | 0.45670947 | 0.64787988 | 0.81173905 |
| PRICKLE1 | 365.330097 | -3.537090535 | 0.63263248 | -5.5910669 | 2.26E-08 | 1.84E-06 |
| PUS7L | 384.390455 | 0.593847296 | 0.19918574 | 2.98137459 | 0.00286958 | 0.02238879 |
| IRAK4 | 478.66463 | -0.244613251 | 0.20334778 | -1.2029305 | 0.2290032 | 0.44451188 |
| TWF1 | 194.16418 | -0.306125737 | 0.28169167 | -1.0867405 | 0.27715152 | 0.49923558 |
| TMEM117 | 2419.30097 | 0.330987569 | 0.20140726 | 1.64337453 | 0.10030547 | 0.26340654 |
| ANO6 | 2197.56001 | -0.278627664 | 0.20881296 | -1.3343409 | 0.18209216 | 0.38654023 |
| AC008124.1 | 209.344631 | 0.699192167 | 0.29452274 | 2.37398368 | 0.01759733 | 0.081875 |
| ARID2 | 3092.48551 | 0.164024836 | 0.16037286 | 1.02277177 | 0.30641577 | 0.53011734 |
| SCAF11 | 2763.64145 | 0.120165886 | 0.11809261 | 1.01755635 | 0.30888884 | 0.53272133 |
| SLC38A1 | 1224.88881 | -1.51061914 | 0.30700587 | -4.9204895 | 8.63E-07 | 3.74E-05 |
| SLC38A2 | 1324.0007 | -0.480761156 | 0.44355837 | -1.0838735 | 0.27842089 | 0.50056644 |
| AC008014.1 | 4034.56389 | 0.290124579 | 0.28611597 | 1.01401044 | 0.31057777 | 0.53450314 |
| SLC38A4 | 96.0721815 | 0.434227022 | 0.62962782 | 0.68965667 | 0.49041012 | 0.69691526 |
| PCED1B | 321.594419 | -2.121189505 | 0.61416694 | -3.453767 | 0.00055281 | 0.00636394 |
| RPAP3 | 304.130388 | 0.631976768 | 0.20760702 | 3.04410108 | 0.00233377 | 0.01921866 |
| AC004241.1 | 61.3300144 | 0.630661261 | 0.31438224 | 2.00603335 | 0.04485269 | 0.15433459 |
| RAPGEF3 | 1186.95181 | -1.856418181 | 0.45526218 | -4.0776903 | 4.55E-05 | 0.00091588 |
| SLC48A1 | 421.422593 | -1.36992877 | 0.24230784 | -5.6536709 | 1.57E-08 | 1.36E-06 |
| HDAC7 | 1189.61056 | -1.802467005 | 0.38657416 | -4.6626681 | 3.12E-06 | 0.00010231 |
| VDR | 290.761991 | -0.651089805 | 0.66530595 | -0.9786322 | 0.32776176 | 0.55299353 |
| TMEM106C | 162.321138 | -0.307453306 | 0.32324775 | -0.9511383 | 0.34153418 | 0.56696305 |
| SENP1 | 400.797442 | -0.041959766 | 0.16418554 | -0.2555631 | 0.79828821 | 0.90515239 |
| PFKM | 202.529593 | 0.841613842 | 0.26271648 | 3.20350607 | 0.00135765 | 0.01272505 |
| ASB8 | 198.711628 | -0.063056962 | 0.19180084 | -0.3287627 | 0.74233506 | 0.87402792 |
| ZNF641 | 189.354508 | 0.014503192 | 0.25625733 | 0.0565962 | 0.95486686 | 0.98009452 |
| AC090115.1 | 152.361905 | 0.596182347 | 0.36859562 | 1.61744284 | 0.10578272 | 0.27294329 |
| KANSL2 | 289.639566 | 0.631700568 | 0.24869579 | 2.54005331 | 0.01108356 | 0.05968313 |
| CCNT1 | 460.373735 | -0.149843684 | 0.14420374 | -1.0391109 | 0.29875314 | 0.52185008 |
| ADCY6 | 533.493076 | 0.434640309 | 0.24949178 | 1.74210273 | 0.08149047 | 0.2293645 |
| DDX23 | 237.049442 | -0.573920079 | 0.22943255 | -2.5014763 | 0.01236767 | 0.06426616 |
| AC073610.3 | 67.5182293 | -0.580872834 | 0.39795147 | -1.4596575 | 0.14438424 | 0.33423964 |
| CCDC65 | 229.795369 | -0.586247118 | 0.38994225 | -1.5034204 | 0.13273068 | 0.31650494 |
| PRKAG1 | 328.076841 | -0.251039365 | 0.2783711 | -0.9018155 | 0.3671549 | 0.59256482 |
| KMT2D | 344.760832 | -0.289164925 | 0.24999358 | -1.1566894 | 0.24739928 | 0.46677577 |
| LMBR1L | 329.747355 | -0.823931333 | 0.24891474 | -3.3100946 | 0.00093264 | 0.00944183 |
| AC011603.2 | 161.73724 | 0.54585546 | 0.29841197 | 1.82920097 | 0.06736951 | 0.20284564 |
| TUBA1B | 447.421675 | -1.324704609 | 0.5631093 | -2.3524822 | 0.01864858 | 0.08497037 |
| TUBA1A | 187.683335 | -1.829019358 | 0.48450505 | -3.7750264 | 0.00015999 | 0.00244593 |
| TUBA1C | 377.3182 | -0.845928485 | 0.32067713 | -2.6379446 | 0.00834102 | 0.04872913 |
| AC010173.1 | 68.7128603 | 0.629311989 | 0.37644522 | 1.6717226 | 0.09457903 | 0.25303098 |
| SPATS2 | 1000.9256 | -0.341957002 | 0.17709202 | -1.9309566 | 0.05348842 | 0.17389332 |
| KCNH3 | 41.9386099 | 0.808400242 | 0.7097735 | 1.13895523 | 0.25472183 | 0.47549755 |
| MCRS1 | 145.816189 | -0.086346714 | 0.25255817 | -0.3418884 | 0.73243487 | 0.86854584 |
| PRPF40B | 893.851107 | 0.103441917 | 0.17821649 | 0.58042843 | 0.56162574 | 0.75337812 |
| FMNL3 | 55.370702 | -1.352931777 | 0.51922415 | -2.6056796 | 0.00916922 | 0.05206631 |
| TMBIM6 | 1604.8832 | -0.81718012 | 0.31994853 | -2.5540987 | 0.01064631 | 0.05800659 |
| NCKAP5L | 239.109794 | -0.096082513 | 0.28584698 | -0.3361327 | 0.73677081 | 0.87092675 |
| BCDIN3D | 87.9150925 | -0.229076837 | 0.352828 | -0.6492592 | 0.51617083 | 0.71746821 |
| AQP2 | 239.840419 | 0.368424435 | 0.65600843 | 0.5616154 | 0.57437809 | 0.76189598 |
| RACGAP1 | 43.2232607 | 0.385279201 | 0.4759856 | 0.80943457 | 0.41826522 | 0.63753323 |
| SMARCD1 | 232.06023 | -0.264453079 | 0.23871973 | -1.1077973 | 0.26794935 | 0.48919736 |
| AC074032.1 | 1081.09068 | -0.016850415 | 0.22462036 | -0.0750173 | 0.94020094 | 0.97411372 |
| CERS5 | 709.502541 | -0.312447478 | 0.1691526 | -1.8471338 | 0.06472775 | 0.19759391 |
| LIMA1 | 976.890118 | -0.439629506 | 0.20879149 | -2.1055911 | 0.03523989 | 0.13145542 |
| LARP4 | 1497.14239 | 0.380177588 | 0.17139534 | 2.21813253 | 0.0265458 | 0.1080046 |
| DIP2B | 2418.32527 | -0.050110734 | 0.22341495 | -0.2242945 | 0.82252819 | 0.91813562 |
| ATF1 | 693.296584 | 0.258632556 | 0.23280414 | 1.11094482 | 0.2665921 | 0.48773062 |
| METTL7A | 446.085891 | 0.825826553 | 0.52650075 | 1.56851923 | 0.11676001 | 0.29099678 |
| SLC11A2 | 919.84309 | -0.263259682 | 0.18728961 | -1.4056288 | 0.15983437 | 0.35669069 |
| LETMD1 | 932.682179 | 0.560158511 | 0.21737012 | 2.57698025 | 0.00996676 | 0.0552691 |
| CSRNP2 | 308.177056 | 0.178980624 | 0.24417502 | 0.73300138 | 0.46355759 | 0.67636498 |

| | | | | | | |
|---|---|---|---|---|---|---|
| TFCP2 | 1900.04074 | 0.383336481 | 0.20845321 | 1.83895695 | 0.06592152 | 0.20017327 |
| POU6F1 | 363.506673 | 0.366907926 | 0.27303237 | 1.34382572 | 0.17900475 | 0.38247715 |
| DAZAP2 | 910.963162 | -0.270782696 | 0.24287242 | -1.1149174 | 0.2648858 | 0.48588759 |
| SMAGP | 255.726057 | 0.669400415 | 0.52733326 | 1.26940678 | 0.20429602 | 0.41414152 |
| GALNT6 | 148.34049 | 0.125377825 | 0.46299166 | 0.27079932 | 0.78654539 | 0.89889929 |
| SLC4A8 | 743.885984 | 0.71690174 | 0.59557807 | 1.20370742 | 0.22870268 | 0.44425496 |
| SCN8A | 46.1664769 | 0.077497762 | 0.52613765 | 0.1472956 | 0.8828987 | 0.94732911 |
| AC068987.3 | 46.3512585 | -0.144101127 | 0.56324127 | -0.2558426 | 0.79807236 | 0.90510327 |
| FIGNL2 | 239.139999 | -0.492250937 | 0.4702878 | -1.0467015 | 0.29523728 | 0.51838219 |
| ACVR1B | 2513.74072 | -0.642027197 | 0.25870799 | -2.4816674 | 0.01307693 | 0.06664086 |
| GRASP | 48.0386192 | -1.175816979 | 0.6112681 | -1.92357 | 0.05440851 | 0.1757642 |
| NR4A1 | 182.923269 | 0.885918805 | 0.79851717 | 1.10945492 | 0.26723398 | 0.48843139 |
| ATG101 | 123.461998 | -0.521837738 | 0.29432031 | -1.7730266 | 0.0762243 | 0.22012391 |
| AC078864.2 | 70.7720275 | -0.615934578 | 0.91933578 | -0.6699778 | 0.50287193 | 0.70694289 |
| AC078864.1 | 131.954254 | 0.084965748 | 0.88797324 | 0.09568503 | 0.92377073 | 0.96642544 |
| KRT7 | 1174.66954 | -2.209241018 | 0.65660228 | -3.3646563 | 0.00076639 | 0.00817103 |
| KRT8 | 705.396598 | -1.285208432 | 0.33015319 | -3.8927639 | 9.91E-05 | 0.00166133 |
| KRT18 | 768.287777 | -2.381150944 | 0.45689349 | -5.2116106 | 1.87E-07 | 1.04E-05 |
| EIF4B | 913.831035 | -0.271543649 | 0.11503292 | -2.3605733 | 0.01824671 | 0.08397938 |
| AC068888.1 | 472.792678 | -0.370454581 | 0.24513444 | -1.5112302 | 0.1307298 | 0.31311151 |
| TNS2 | 504.017586 | -0.827443923 | 0.32933981 | -2.5124322 | 0.01199022 | 0.0628518 |
| SPRYD3 | 78.0776244 | -0.913530188 | 0.38679907 | -2.3617694 | 0.01818795 | 0.08384041 |
| IGFBP6 | 79.5813602 | -3.164269805 | 0.73791257 | -4.2881365 | 1.80E-05 | 0.00043272 |
| CSAD | 887.486653 | 0.292359541 | 0.17100745 | 1.70963041 | 0.08733424 | 0.23953446 |
| ZNF740 | 144.620525 | 0.356477564 | 0.18191772 | 1.95955384 | 0.05004796 | 0.16639404 |
| RARG | 156.183212 | -1.122943916 | 0.42509652 | -2.6416211 | 0.00825103 | 0.04831023 |
| MFSD5 | 52.4183565 | -1.23234862 | 0.47654526 | -2.5860054 | 0.00970954 | 0.05418392 |
| PFDN5 | 624.285029 | -0.280646366 | 0.33811258 | -0.8300382 | 0.40651717 | 0.62649938 |
| C12orf10 | 87.5284178 | -0.792475733 | 0.34077167 | -2.3255329 | 0.02004348 | 0.08923302 |
| AAAS | 141.990888 | 0.806381776 | 0.23732897 | 3.39773849 | 0.00067945 | 0.00748532 |
| SP1 | 833.847032 | -0.733600403 | 0.18381843 | -3.9908969 | 6.58E-05 | 0.00121096 |
| PRR13 | 543.094672 | 0.06002531 | 0.30293587 | 0.19814527 | 0.8429314 | 0.92739954 |
| PCBP2 | 1764.49055 | 0.292947724 | 0.22482627 | 1.30299599 | 0.19257613 | 0.39989594 |
| MAP3K12 | 165.028136 | -0.464338383 | 0.36842902 | -1.2603198 | 0.20755402 | 0.41837037 |
| TARBP2 | 55.7940574 | 0.299441314 | 0.36317873 | 0.82450123 | 0.40965482 | 0.62942078 |
| ATP5MC2 | 871.060475 | -0.427369186 | 0.20791437 | -2.0555058 | 0.03983018 | 0.14211692 |
| CALCOCO1 | 489.69665 | -0.146973564 | 0.23114044 | -0.6358626 | 0.52486597 | 0.72408747 |
| HOXC4 | 395.998186 | 1.003393919 | 0.32014497 | 3.13418608 | 0.00172332 | 0.01520593 |
| AC023794.1 | 213.122366 | 0.47379822 | 0.33290723 | 1.42321397 | 0.15467414 | 0.34906726 |
| SMUG1 | 225.394689 | 0.173782492 | 0.2030198 | 0.85598789 | 0.39200448 | 0.61430622 |
| LINC02381 | 730.118922 | 0.012708904 | 0.21033268 | 0.06042287 | 0.95181885 | 0.97924878 |
| CBX5 | 804.049898 | 0.009851061 | 0.16201419 | 0.06080369 | 0.95151555 | 0.97914234 |
| AC078778.1 | 80.2402295 | 0.744673269 | 0.35060361 | 2.12397489 | 0.03367224 | 0.12774861 |
| HNRNPA1 | 552.820864 | -0.169955383 | 0.27146172 | -0.6260749 | 0.53126579 | 0.72863563 |
| COPZ1 | 441.411585 | -0.211453 | 0.14885616 | -1.4205189 | 0.15545666 | 0.35035438 |
| PPP1R1A | 1004.36797 | 0.472183363 | 0.50904693 | 0.92758316 | 0.35362384 | 0.57953068 |
| AC079310.1 | 173.779492 | 1.519664246 | 1.53667703 | 0.98892885 | 0.32269796 | 0.54774059 |
| CD63 | 812.286479 | -1.510888724 | 0.34555404 | -4.372366 | 1.23E-05 | 0.00031376 |
| GDF11 | 55.8218866 | -0.118328941 | 0.49787822 | -0.2376664 | 0.81213982 | 0.91189688 |
| ORMDL2 | 118.852261 | -0.754139033 | 0.38480647 | -1.9597878 | 0.0500206 | 0.16638315 |
| PYM1 | 127.748014 | 0.343499628 | 0.20503386 | 1.67533126 | 0.09386926 | 0.25193426 |
| DGKA | 65.3604719 | -0.30636103 | 0.36116125 | -0.8482666 | 0.39628951 | 0.6181611 |
| CDK2 | 61.4796089 | -0.967766062 | 0.25902256 | -3.7362231 | 0.0001868 | 0.00277248 |
| RAB5B | 474.306454 | -0.382855699 | 0.17726901 | -2.1597441 | 0.03079249 | 0.12001711 |
| SUOX | 161.19456 | 0.33484811 | 0.32267 | 1.03774168 | 0.29939034 | 0.52234143 |
| IKZF4 | 176.034171 | -0.032452142 | 0.23454857 | -0.13836 | 0.8899559 | 0.9503707 |
| RPS26 | 184.277742 | -0.554302967 | 0.35925786 | -1.5429112 | 0.12285232 | 0.300288 |
| ERBB3 | 289.669514 | -0.857928432 | 0.29359713 | -2.9221281 | 0.00347648 | 0.02584181 |
| PA2G4 | 184.705334 | -0.567813229 | 0.33838703 | -1.6779994 | 0.09334722 | 0.25117608 |
| RPL41 | 2352.01135 | -0.985806531 | 0.40466589 | -2.4360999 | 0.01484658 | 0.07277314 |
| ESYT1 | 185.458256 | -0.875899941 | 0.25167153 | -3.4803298 | 0.0005008 | 0.00592214 |
| AC034102.5 | 45.5920353 | 0.822934167 | 0.47918198 | 1.71737293 | 0.08591105 | 0.23705852 |
| MYL6B | 143.269724 | 0.195891832 | 0.29120053 | 0.67270423 | 0.50113547 | 0.70528379 |
| MYL6 | 1321.4919 | -0.363418144 | 0.35734638 | -1.0169912 | 0.3091576 | 0.53306909 |
| SMARCC2 | 957.629878 | -0.339236342 | 0.19557483 | -1.7345603 | 0.08281872 | 0.23127959 |
| RNF41 | 207.053417 | -0.205384072 | 0.24117217 | -0.8516077 | 0.39443188 | 0.61679303 |
| NABP2 | 68.7478802 | -0.342801669 | 0.26221845 | -1.3073133 | 0.19110633 | 0.39765651 |
| ANKRD52 | 272.622887 | 0.306567037 | 0.13448628 | 2.27954135 | 0.0226349 | 0.0968398 |
| COQ10A | 123.472306 | 0.762780265 | 0.31720878 | 2.40466313 | 0.01618738 | 0.07721365 |
| CS | 406.063147 | 0.327345122 | 0.21434412 | 1.52719432 | 0.1267127 | 0.30636096 |
| AC073896.1 | 105.17047 | 0.390434998 | 0.30057943 | 1.29894116 | 0.19396412 | 0.4015787 |
| AC073896.2 | 146.904806 | 0.658616159 | 0.28990658 | 2.27182202 | 0.02309726 | 0.09830568 |
| PAN2 | 397.428728 | 0.752873193 | 0.18254752 | 4.12425871 | 3.72E-05 | 0.00077858 |
| STAT2 | 602.961562 | 0.088339773 | 0.31278547 | 0.28242928 | 0.77761437 | 0.89359206 |
| TIMELESS | 83.5479954 | -0.386944138 | 0.33276316 | -1.1628214 | 0.24490196 | 0.46398595 |
| SPRYD4 | 82.9395293 | -0.182163358 | 0.27489664 | -0.6626613 | 0.50754752 | 0.71071718 |
| GLS2 | 69.7048283 | -0.062585928 | 0.28453025 | -0.2199623 | 0.82590051 | 0.91938879 |
| RBMS2 | 824.774027 | -0.420116222 | 0.16819136 | -2.4978466 | 0.01249503 | 0.06464817 |
| BAZ2A | 686.265051 | -0.251665634 | 0.17635674 | -1.4270259 | 0.15357242 | 0.34747142 |
| ATP5F1B | 1212.8345 | -0.081885416 | 0.43778091 | -0.1870466 | 0.85162412 | 0.93193618 |
| PTGES3 | 1387.3772 | -0.18759443 | 0.13868126 | -1.3527021 | 0.17615081 | 0.3783991 |
| NACA | 589.307388 | -0.719618154 | 0.35215676 | -2.0434597 | 0.04100695 | 0.14482909 |
| PRIM1 | 52.0080213 | 0.499640645 | 0.28798032 | 1.73498188 | 0.08274401 | 0.23116738 |
| NEMP1 | 145.251263 | -0.497775873 | 0.27404816 | -1.816381 | 0.06931195 | 0.20685529 |
| NAB2 | 95.2636522 | -0.991830836 | 0.62972575 | -1.5750203 | 0.11525176 | 0.28859915 |
| STAT6 | 791.016568 | -1.117085551 | 0.21098364 | -5.2946547 | 1.19E-07 | 7.19E-06 |
| LRP1 | 96.9161886 | 0.049719121 | 0.32829285 | 0.15144747 | 0.87962275 | 0.9463109 |
| SHMT2 | 65.0284477 | -0.78186238 | 0.54441239 | -1.4361583 | 0.1509573 | 0.34361451 |
| R3HDM2 | 2715.19171 | 0.125985793 | 0.1418605 | 0.88809637 | 0.37448891 | 0.59799614 |
| MARS | 270.325171 | -0.032491289 | 0.26326462 | -0.1234168 | 0.90177702 | 0.95457905 |
| DDIT3 | 349.633122 | 0.641974634 | 0.63331197 | 1.01367836 | 0.31073625 | 0.53450314 |
| MBD6 | 271.692758 | 0.278420807 | 0.23210647 | 1.19953918 | 0.23031836 | 0.44592324 |
| DCTN2 | 493.782684 | -0.61524722 | 0.18024742 | -3.4133483 | 0.0006417 | 0.00717714 |
| PIP4K2C | 312.714338 | -0.364854885 | 0.36049479 | -1.0120947 | 0.31149275 | 0.53510038 |
| DTX3 | 155.591487 | 0.201971601 | 0.33197539 | 0.60839329 | 0.54292666 | 0.73895088 |
| OS9 | 482.682293 | -0.427388784 | 0.21982864 | -1.9441907 | 0.05187246 | 0.17045649 |
| AC025165.2 | 62.6258679 | -0.290737313 | 0.49022533 | -0.5930687 | 0.55313516 | 0.74600391 |
| TSPAN31 | 163.755125 | -0.729874782 | 0.24956709 | -2.9245634 | 0.0034494 | 0.02571281 |
| CDK4 | 127.89779 | -0.275946455 | 0.20381997 | -1.3538735 | 0.17577675 | 0.37799253 |
| 9-Mar | 185.981281 | -0.671088521 | 0.23304438 | -2.8796597 | 0.00398105 | 0.02854812 |
| TSFM | 242.692937 | -0.734017469 | 0.33681318 | -2.1793015 | 0.02930928 | 0.11551991 |
| AVIL | 723.514706 | -0.452719811 | 0.39600412 | -1.14322 | 0.25294728 | 0.47312102 |
| CTDSP2 | 668.745749 | -0.707569046 | 0.17167492 | -4.1215633 | 3.76E-05 | 0.00078278 |
| AC084033.3 | 43.9689028 | 0.703951579 | 0.33182358 | 2.12146341 | 0.03388282 | 0.12834064 |
| ATP23 | 54.0229352 | 0.839471528 | 0.33826216 | 2.48171872 | 0.01307504 | 0.06664086 |
| LRIG3 | 461.426991 | 0.703766978 | 0.30590767 | 2.30058628 | 0.02141503 | 0.0935691 |
| AC068305.2 | 92.755536 | 1.160409746 | 0.49609144 | 2.33910457 | 0.01933002 | 0.08714343 |
| SLC16A7 | 4725.03118 | 0.409182364 | 0.44073708 | 0.92840467 | 0.3531977 | 0.57919589 |
| FAM19A2 | 815.96968 | 0.834471267 | 0.43819066 | 1.9043566 | 0.05686376 | 0.18138824 |
| USP15 | 2070.48242 | 0.161119993 | 0.20575874 | 0.78305298 | 0.433596 | 0.65072685 |
| MON2 | 3219.92224 | 0.311938718 | 0.15720665 | 1.98425911 | 0.04722696 | 0.1595042 |
| AC048341.1 | 1120.59062 | -1.214980188 | 0.46090807 | -2.6360575 | 0.00838755 | 0.04881828 |
| PPM1H | 2629.81194 | -0.775559329 | 0.2473452 | -3.1355342 | 0.00171541 | 0.01515651 |

| | | | | | | |
|---|---|---|---|---|---|---|
| DPY19L2 | 671.904921 | 0.966645373 | 0.45524567 | 2.12334885 | 0.03372463 | 0.12786126 |
| RXYLT1 | 218.75083 | 0.116875377 | 0.31768821 | 0.36789334 | 0.71295277 | 0.85612664 |
| SRGAP1 | 2138.98595 | -0.16010942 | 0.42498062 | -0.3767452 | 0.70636295 | 0.85193099 |
| AC020611.2 | 106.37042 | -0.207792752 | 0.51776733 | -0.4013246 | 0.68818118 | 0.83950731 |
| C12orf66 | 87.4866834 | 0.521953907 | 0.23788385 | 2.19415442 | 0.02822431 | 0.11265157 |
| C12orf56 | 44.6171269 | -0.040703202 | 0.37744302 | -0.1078393 | 0.91412314 | 0.96120463 |
| XPOT | 251.533433 | -0.0319149 | 0.23095072 | -0.1381892 | 0.89009087 | 0.9503707 |
| TBK1 | 916.80224 | -0.414834696 | 0.21868835 | -1.8969218 | 0.05783827 | 0.18348602 |
| RASSF3 | 174.216873 | -0.795547771 | 0.22112812 | -3.5976779 | 0.00032107 | 0.0042276 |
| GNS | 849.045618 | -0.487955667 | 0.20832216 | -2.3423129 | 0.01916464 | 0.08666407 |
| AC025262.1 | 40.6205564 | 0.221357186 | 0.48393598 | 0.45741006 | 0.64737633 | 0.81148066 |
| TBC1D30 | 582.476712 | 0.184065582 | 0.22362028 | 0.82311666 | 0.41044165 | 0.62997812 |
| LINC02389 | 309.206997 | 0.734207669 | 0.32335623 | 2.27058456 | 0.02317214 | 0.09844992 |
| LINC02231 | 41.5597454 | 1.179173559 | 0.46920879 | 2.51311055 | 0.01196718 | 0.06280275 |
| LEMD3 | 1043.52476 | 0.079784249 | 0.20355415 | 0.39195589 | 0.69509081 | 0.84411644 |
| MSRB3 | 71.7675468 | 0.005213714 | 0.50590814 | 0.01030565 | 0.99177742 | 0.99675782 |
| HMGA2 | 394.811637 | -3.443912003 | 0.91815752 | -3.7508945 | 0.0001762 | 0.00264487 |
| AC090673.1 | 143.499826 | -0.029061004 | 0.54739189 | -0.0530899 | 0.95766025 | 0.98151258 |
| LLPH | 90.8982263 | -0.315407432 | 0.32569162 | -0.9684235 | 0.33283289 | 0.55812995 |
| HELB | 491.484233 | -0.189659593 | 0.27516841 | -0.6892491 | 0.4906665 | 0.6969611 |
| GRIP1 | 4984.28122 | -1.831454847 | 0.46432545 | -3.9443344 | 8.00E-05 | 0.0014188 |
| CAND1 | 1036.63494 | 0.39575362 | 0.14550164 | 2.71992545 | 0.00652966 | 0.04074177 |
| DYRK2 | 138.227337 | -0.751650925 | 0.21168378 | -3.5508197 | 0.00038403 | 0.00483575 |
| MDM1 | 167.041867 | 0.884115731 | 0.28505544 | 3.10155711 | 0.00192506 | 0.01663053 |
| RAP1B | 1700.90194 | -0.638115751 | 0.17081116 | -3.7357965 | 0.00018712 | 0.00277407 |
| NUP107 | 528.099527 | -0.300606435 | 0.25564786 | -1.1758614 | 0.23965028 | 0.45777883 |
| SLC35E3 | 505.030945 | -0.177364988 | 0.37232621 | -0.4763699 | 0.6338109 | 0.8029933 |
| MDM2 | 1244.22657 | -0.767392924 | 0.28456924 | -2.6966827 | 0.0070034 | 0.04282772 |
| CPM | 3705.29496 | 0.403558796 | 0.37625294 | 1.07257313 | 0.28346269 | 0.5059807 |
| CPSF6 | 1511.69618 | 0.576063208 | 0.1771313 | 3.25218191 | 0.00114523 | 0.01111484 |
| YEATS4 | 127.586337 | 0.130199678 | 0.1811166 | 0.71887212 | 0.47221972 | 0.68319499 |
| FRS2 | 1420.7087 | 0.34456317 | 0.15562421 | 2.21407179 | 0.02682384 | 0.10871927 |
| CCT2 | 396.40864 | -0.297991961 | 0.28176193 | -1.0576019 | 0.29023697 | 0.51314953 |
| RAB3IP | 1378.09638 | 0.365625352 | 0.20600092 | 1.77487241 | 0.07591896 | 0.21963347 |
| AC025263.2 | 54.1948847 | 0.739568946 | 0.65798111 | 1.12399722 | 0.2610142 | 0.48220092 |
| MYRFL | 618.191064 | 0.752959908 | 0.58593076 | 1.28506635 | 0.1987691 | 0.40737642 |
| AC025159.1 | 967.711793 | 0.172358595 | 0.55633472 | 0.30981096 | 0.75670472 | 0.88227669 |
| CNOT2 | 1992.67407 | 0.275426886 | 0.19373835 | 1.42164361 | 0.15512974 | 0.34973694 |
| TSPAN8 | 883.459666 | -0.229052377 | 0.30639787 | -0.7475652 | 0.45472248 | 0.66998653 |
| LGR5 | 190.279988 | 3.068868867 | 1.70788488 | 1.79688274 | 0.07235424 | 0.21270422 |
| ZFC3H1 | 1416.1555 | -0.245216611 | 0.16272814 | -1.5069097 | 0.13183381 | 0.3147854 |
| THAP2 | 77.3346538 | 0.531274961 | 0.42362041 | 1.25412974 | 0.20979485 | 0.42166467 |
| TMEM19 | 83.1940973 | -0.216556016 | 0.33997898 | -0.6369688 | 0.52414514 | 0.72352734 |
| RAB21 | 1174.39604 | -0.37831225 | 0.22231235 | -1.7017149 | 0.08880882 | 0.2424211 |
| TBC1D15 | 1357.17144 | -0.465654778 | 0.26377419 | -1.7653538 | 0.07750431 | 0.22216894 |
| TRHDE | 3268.00988 | 0.416293199 | 0.28476519 | 1.46188235 | 0.14377345 | 0.33340158 |
| ATXN7L3B | 314.23231 | -0.423553662 | 0.27997738 | -1.5128138 | 0.13032695 | 0.31253761 |
| CAPS2 | 1316.13548 | 1.109189587 | 0.2420278 | 4.58290165 | 4.59E-06 | 0.00014054 |
| GLIPR1L1 | 124.398638 | 0.913922031 | 0.58470493 | 1.56304826 | 0.11804124 | 0.29301961 |
| GLIPR1L2 | 122.158773 | 1.507147735 | 0.66475115 | 2.267236 | 0.02337581 | 0.09899683 |
| KRR1 | 343.901496 | 0.198135102 | 0.23390771 | 0.84706529 | 0.39695873 | 0.61869356 |
| NAP1L1 | 1163.46466 | -0.053806635 | 0.18594172 | -0.2893737 | 0.77229546 | 0.89039458 |
| BBS10 | 59.3796015 | 0.506456728 | 0.25886896 | 1.95642122 | 0.05041555 | 0.16715338 |
| OSBPL8 | 1683.29927 | 0.062603923 | 0.20503976 | 0.30532578 | 0.76011804 | 0.88357016 |
| AC107032.2 | 182.500157 | 0.610975766 | 0.58493647 | 1.04451646 | 0.29624651 | 0.51932583 |
| ZDHHC17 | 1321.3381 | 0.562438679 | 0.22032204 | 2.55280263 | 0.010686 | 0.05811092 |
| CSRP2 | 351.434418 | 0.549861474 | 0.52994354 | 1.03758502 | 0.2994633 | 0.52239972 |
| NAV3 | 57.8179449 | 0.064804324 | 0.38686078 | 0.1675133 | 0.86696619 | 0.93959274 |
| SYT1 | 442.0408 | 1.208731683 | 0.49276072 | 2.45297895 | 0.01416786 | 0.07032818 |
| AC090709.1 | 682.118026 | 2.415073662 | 0.50298362 | 4.80149566 | 1.57E-06 | 5.83E-05 |
| AC027288.3 | 451.310694 | 0.426134972 | 0.5515427 | 0.77262372 | 0.4397451 | 0.65544945 |
| PAWR | 3463.23383 | 0.479859693 | 0.24573323 | 1.95276676 | 0.05084724 | 0.16812044 |
| AC073569.2 | 283.919197 | 0.559555055 | 0.38661156 | 1.44733143 | 0.14780412 | 0.3392432 |
| PPP1R12A | 2368.62141 | 0.111577744 | 0.12472166 | 0.89461399 | 0.37099347 | 0.59561104 |
| LIN7A | 375.352874 | 0.467344175 | 0.50500141 | 0.92543142 | 0.35474154 | 0.58042624 |
| ACSS3 | 2276.67006 | 0.560839779 | 0.26537712 | 2.113369 | 0.0345692 | 0.1299074 |
| AC078955.1 | 50.99925 | -0.771312541 | 0.85570051 | -0.9013814 | 0.36738556 | 0.59270895 |
| PPFIA2 | 1146.89966 | 0.453061984 | 0.44602918 | 1.01576758 | 0.30974008 | 0.5337614 |
| CCDC59 | 191.572484 | 0.128971696 | 0.20351708 | 0.63371436 | 0.52626725 | 0.72497874 |
| METTL25 | 1070.04639 | 0.501341026 | 0.19358151 | 2.5898188 | 0.00960265 | 0.05378585 |
| TMTC2 | 3130.68096 | -0.24842056 | 0.17982937 | -1.3814237 | 0.16714872 | 0.36681632 |
| LRRIQ1 | 527.735199 | 0.047113278 | 0.38248705 | 0.12317614 | 0.90196762 | 0.95457905 |
| RASSF9 | 123.652183 | 0.218063835 | 0.55667663 | 0.39172443 | 0.69526184 | 0.84424652 |
| MGAT4C | 89.038883 | 1.064187896 | 0.64932978 | 1.63890204 | 0.10123366 | 0.26494858 |
| C12orf29 | 117.191479 | -0.175805248 | 0.36856684 | -0.4769969 | 0.63336432 | 0.80265847 |
| CEP290 | 1121.3905 | 0.302833867 | 0.1797101 | 1.68512432 | 0.09196459 | 0.24831567 |
| TMTC3 | 778.012996 | 0.099256161 | 0.15873974 | 0.62527606 | 0.53178989 | 0.72906475 |
| KITLG | 946.768412 | 0.271008111 | 0.42244377 | 0.64152469 | 0.52118184 | 0.72124578 |
| AC024941.2 | 58.5496545 | 1.429163002 | 0.60492281 | 2.36255432 | 0.01814948 | 0.08376537 |
| AC024909.1 | 69.5091016 | -0.364857485 | 0.57226908 | -0.6375628 | 0.52375831 | 0.72325104 |
| DUSP6 | 235.069336 | -1.202018681 | 0.39167263 | -3.0689371 | 0.00214822 | 0.01809682 |
| POC1B | 741.096417 | -0.29298259 | 0.14027863 | -2.0885761 | 0.0367459 | 0.13490177 |
| POC1B-AS1 | 337.959209 | 0.05744362 | 0.27718634 | 0.20723828 | 0.83582377 | 0.92365065 |
| AC025034.1 | 166.963857 | 0.750488931 | 0.34242854 | 2.19166583 | 0.02840364 | 0.11309997 |
| ATP2B1 | 1274.77644 | 0.099677681 | 0.25994532 | 0.38345634 | 0.70138144 | 0.84869711 |
| ATP2B1-AS1 | 36.9497496 | -0.00545517 | 0.41914728 | -0.0130149 | 0.98961589 | 0.99606252 |
| BTG1 | 1059.88392 | -1.3409926 | 0.31487164 | -4.2588548 | 2.05E-05 | 0.00048209 |
| AC025164.1 | 165.877717 | 0.263250928 | 0.52783816 | 0.49873417 | 0.61796667 | 0.79042527 |
| LINC02391 | 118.558671 | 0.298494584 | 0.52375551 | 0.56991207 | 0.56873734 | 0.75864147 |
| PLEKHG7 | 212.844507 | 0.40693565 | 0.66124441 | 0.61540883 | 0.53828476 | 0.73492731 |
| EEA1 | 2350.74997 | 0.700838517 | 0.18958203 | 3.69675607 | 0.00021837 | 0.00314899 |
| LINC02413 | 72.9434008 | 2.479209402 | 0.49512211 | 5.00726859 | 5.52E-07 | 2.56E-05 |
| AC138123.1 | 3384.23489 | 2.155357856 | 0.37638854 | 5.72641728 | 1.03E-08 | 9.41E-07 |
| AC138123.2 | 129.79482 | 2.245500236 | 0.48129697 | 4.6655192 | 3.08E-06 | 0.00010115 |
| AC124947.1 | 756.305342 | 1.437910766 | 0.43352787 | 3.31676665 | 0.00091066 | 0.00928153 |
| NUDT4 | 2991.86162 | 1.616460551 | 0.44476238 | 3.63443632 | 0.00027859 | 0.00381335 |
| UBE2N | 631.673952 | 0.293791487 | 0.21735473 | 1.35166827 | 0.17648146 | 0.3786783 |
| MRPL42 | 579.086661 | 0.334302317 | 0.14797756 | 2.2591419 | 0.02387456 | 0.10046482 |
| SOCS2 | 272.729445 | -0.296262042 | 0.65855679 | -0.4498656 | 0.65280736 | 0.81494494 |
| CRADD | 5188.0218 | -0.246700962 | 0.31785057 | -0.7761539 | 0.43765812 | 0.65418424 |
| AC012085.2 | 493.807075 | -0.740517506 | 0.73414926 | -1.0086743 | 0.31313085 | 0.53695254 |
| AC012464.1 | 47.342055 | -0.18081205 | 0.67384423 | -0.2683292 | 0.78844597 | 0.89967025 |
| PLXNC1 | 72.6225486 | -0.463982547 | 0.25973138 | -1.7863939 | 0.07403551 | 0.21578443 |
| CEP83 | 701.915443 | 0.281515328 | 0.16034309 | 1.75570601 | 0.07913862 | 0.22478731 |
| TMCC3 | 118.065004 | 0.809694426 | 0.58989222 | 1.37261418 | 0.16987232 | 0.37014905 |
| NDUFA12 | 432.104924 | 0.859172163 | 0.29628214 | 2.89984459 | 0.00373348 | 0.02720105 |
| NR2C1 | 850.86253 | 0.244001393 | 0.19681043 | 1.23977874 | 0.21505724 | 0.4288968 |
| FGD6 | 498.397923 | -0.545707822 | 0.28503012 | -1.9145619 | 0.05554839 | 0.17839503 |
| VEZT | 1677.39294 | 0.058663185 | 0.14018731 | 0.41846288 | 0.67560872 | 0.82986765 |
| METAP2 | 733.98003 | 0.132622813 | 0.10077682 | 1.31600514 | 0.18817232 | 0.39384317 |
| USP44 | 221.806514 | 0.187520003 | 0.37591781 | 0.49883246 | 0.61789742 | 0.79042527 |
| NTN4 | 3046.46605 | 1.153321222 | 0.40513887 | 2.84673063 | 0.00441707 | 0.03061128 |
| AC090001.1 | 75.7380032 | 0.248418763 | 0.69203276 | 0.35896966 | 0.71961779 | 0.86029789 |

| | | | | | | |
|---|---|---|---|---|---|---|
| SNRPF | 166.988974 | -0.163937144 | 0.25721269 | -0.6373602 | 0.52389022 | 0.72325104 |
| LTA4H | 371.09778 | 0.332324697 | 0.18315517 | 1.81444344 | 0.06960948 | 0.20746202 |
| ELK3 | 393.880735 | -2.607497807 | 0.46373364 | -5.6228352 | 1.88E-08 | 1.58E-06 |
| CDK17 | 922.902442 | -0.244860389 | 0.15777875 | -1.5519225 | 0.12068078 | 0.29693855 |
| CFAP54 | 220.831119 | -1.464502656 | 0.64559306 | -2.268461 | 0.02330113 | 0.09880724 |
| NEDD1 | 336.193997 | 0.131432087 | 0.17619583 | 0.74594324 | 0.45570171 | 0.67098025 |
| TMPO | 647.663135 | 0.255541203 | 0.30179779 | 0.84672987 | 0.39714571 | 0.61891194 |
| SLC25A3 | 1058.61616 | -0.397706263 | 0.40305452 | -0.9867307 | 0.32377468 | 0.54886291 |
| IKBIP | 126.470748 | -1.032661335 | 0.39157765 | -2.6371815 | 0.00835981 | 0.04876696 |
| APAF1 | 375.612339 | -0.412246203 | 0.19117658 | -2.1563635 | 0.03105529 | 0.12085157 |
| ANKS1B | 322.841421 | -0.102008378 | 0.50515118 | -0.2019363 | 0.8399665 | 0.92566636 |
| UHRF1BP1L | 625.628704 | -0.094555162 | 0.19572498 | -0.4831022 | 0.6290232 | 0.79927333 |
| ACTR6 | 312.915199 | 0.243632491 | 0.18996381 | 1.28252054 | 0.1996601 | 0.40831558 |
| DEPDC4 | 124.411148 | 0.608813191 | 0.26771046 | 2.27414792 | 0.02295709 | 0.09792087 |
| SCYL2 | 1121.96508 | 0.091271761 | 0.214342 | 0.42582303 | 0.67023681 | 0.82659266 |
| NR1H4 | 2069.9682 | -0.26399995 | 0.35403881 | -0.7456808 | 0.45586025 | 0.67098931 |
| GAS2L3 | 156.28882 | -2.013630262 | 0.50852489 | -3.9597477 | 7.50E-05 | 0.0013429 |
| SLC5A8 | 92.4478315 | 3.683169124 | 0.94557111 | 3.8951794 | 9.81E-05 | 0.00165114 |
| UTP20 | 407.491478 | 0.149118097 | 0.23912386 | 0.62360192 | 0.53288905 | 0.73010387 |
| ARL1 | 310.133179 | -0.532573716 | 0.21265681 | -2.5043812 | 0.01226658 | 0.06389167 |
| CHPT1 | 704.422847 | 0.720158479 | 0.36851857 | 1.95419862 | 0.05067773 | 0.16768591 |
| SYCP3 | 43.95146 | 0.501145472 | 0.63131225 | 0.79381553 | 0.42730283 | 0.64571091 |
| GNPTAB | 734.213467 | 0.027310632 | 0.27394318 | 0.09969451 | 0.92058686 | 0.96454603 |
| DRAM1 | 303.073554 | -2.638279805 | 0.37569891 | -7.0223248 | 2.18E-12 | 6.41E-10 |
| WASHC3 | 612.435536 | 0.044569705 | 0.20173743 | 0.22092928 | 0.8251475 | 0.91931286 |
| NUP37 | 200.065639 | 0.109678132 | 0.20379474 | 0.53817942 | 0.59045319 | 0.77217181 |
| PARPBP | 58.9239828 | 0.131208946 | 0.53448445 | 0.24548693 | 0.80607943 | 0.90934469 |
| NT5DC3 | 131.945676 | 0.590940731 | 0.35921821 | 1.64507455 | 0.09995444 | 0.26279821 |
| HSP90B1 | 1223.70442 | -0.120041357 | 0.38658288 | -0.3105191 | 0.75616626 | 0.88219397 |
| C12orf73 | 141.753131 | 0.534346202 | 0.26206689 | 2.03896875 | 0.04145315 | 0.14585898 |
| TDG | 472.22714 | 0.036853647 | 0.15293225 | 0.24098022 | 0.80957045 | 0.91048242 |
| HCFC2 | 336.706951 | 0.663294486 | 0.18702306 | 3.54659202 | 0.00039025 | 0.00490465 |
| NFYB | 263.856134 | 0.639451514 | 0.2332311 | 2.7417077 | 0.00611207 | 0.03882844 |
| TXNRD1 | 2190.84698 | -0.84040969 | 0.23072364 | -3.642495 | 0.00027001 | 0.00372145 |
| CHST11 | 273.405935 | -4.110303552 | 0.54541074 | -7.5361617 | 4.84E-14 | 2.37E-11 |
| SLC41A2 | 814.278643 | -0.701344203 | 0.29798798 | -2.353599 | 0.01859266 | 0.08488512 |
| C12orf45 | 173.921008 | 0.372117942 | 0.24521571 | 1.51751265 | 0.1291373 | 0.31053607 |
| WASHC4 | 1647.39364 | 0.199580079 | 0.19452428 | 1.0259906 | 0.30489601 | 0.52822225 |
| APPL2 | 851.910325 | -0.164364915 | 0.19695081 | -0.834548 | 0.40397223 | 0.62518836 |
| C12orf75 | 446.987671 | -1.825598998 | 0.47168666 | -3.8703638 | 0.00010867 | 0.00178985 |
| NUAK1 | 383.724836 | 0.533010443 | 0.34637977 | 1.53880362 | 0.12385225 | 0.30188476 |
| CKAP4 | 146.926609 | -0.024927291 | 0.32856692 | -0.0758667 | 0.93952514 | 0.97411372 |
| TCP11L2 | 647.844679 | 0.155491189 | 0.23066264 | 0.67410652 | 0.50024359 | 0.70482771 |
| POLR3B | 459.107823 | 0.818106297 | 0.1843453 | 4.43790165 | 9.08E-06 | 0.00024753 |
| AC079385.1 | 49.8260646 | 0.726319922 | 0.30466243 | 2.38401543 | 0.01712488 | 0.08012271 |
| RIC8B | 1134.90946 | 0.445634586 | 0.16970078 | 2.62600206 | 0.00863943 | 0.04987079 |
| AC007541.1 | 46.2672131 | 1.544710787 | 0.50875681 | 3.03624593 | 0.00239544 | 0.0196241 |
| TMEM263 | 771.161005 | 0.446591226 | 0.19250639 | 2.31987738 | 0.02034751 | 0.09016111 |
| MTERF2 | 214.185673 | 0.833517638 | 0.29506102 | 2.82489915 | 0.00472955 | 0.03206712 |
| CRY1 | 862.21031 | 0.072325018 | 0.5750339 | 0.12577522 | 0.89990985 | 0.95395051 |
| BTBD11 | 1556.43044 | 1.401490979 | 0.49490036 | 2.83186495 | 0.00462774 | 0.03163965 |
| PWP1 | 309.939364 | -0.147622446 | 0.15523597 | -0.9509552 | 0.34162713 | 0.56704608 |
| PRDM4 | 380.356514 | -0.398677132 | 0.21302545 | -1.8715 | 0.06127581 | 0.19057975 |
| FICD | 41.6707862 | -0.797143239 | 0.42999447 | -1.8538453 | 0.06376129 | 0.19577473 |
| SART3 | 350.596307 | 0.043706722 | 0.18269859 | 0.23922857 | 0.81092835 | 0.91123374 |
| ISCU | 607.307938 | -1.247245906 | 0.4102255 | -3.040391 | 0.00236271 | 0.01939656 |
| CORO1C | 368.041541 | -1.571854962 | 0.30528441 | -5.1488216 | 2.62E-07 | 1.37E-05 |
| SSH1 | 1230.79924 | -0.016056963 | 0.28990331 | -0.0553873 | 0.95582991 | 0.98062581 |
| USP30 | 210.394531 | 0.453923103 | 0.23251856 | 1.9522016 | 0.05091427 | 0.16821806 |
| ALKBH2 | 51.3939906 | -0.080845473 | 0.37458982 | -0.215824 | 0.82912494 | 0.92023714 |
| UNG | 154.259022 | 0.332928261 | 0.30336188 | 1.09746241 | 0.27243931 | 0.49398089 |
| ACACB | 3947.00808 | 1.600871122 | 0.45618087 | 3.50929034 | 0.0004493 | 0.00545985 |
| KCTD10 | 187.880445 | -0.859029984 | 0.23309527 | -3.6853171 | 0.00022842 | 0.00326888 |
| UBE3B | 440.119898 | 0.137831338 | 0.24480796 | 0.56301821 | 0.57342248 | 0.76116346 |
| MMAB | 418.641479 | -0.025019286 | 0.23745621 | -0.1053638 | 0.91608715 | 0.96219745 |
| MVK | 388.322529 | -0.09573461 | 0.31640909 | -0.3025659 | 0.76222068 | 0.88460822 |
| FAM222A | 252.06765 | 0.617173599 | 0.33739604 | 1.829226 | 0.06736576 | 0.20284564 |
| FAM222A-AS | 104.544592 | 0.429958112 | 0.38479694 | 1.11736365 | 0.26383887 | 0.48477502 |
| TRPV4 | 152.695358 | 0.822995749 | 0.46676772 | 1.76318053 | 0.07787005 | 0.22292706 |
| GLTP | 621.113672 | 0.851275462 | 0.26669655 | 3.19192532 | 0.00141328 | 0.01307283 |
| TCHP | 464.213007 | -0.0274247 | 0.28194777 | -0.0972687 | 0.92251299 | 0.96565607 |
| GIT2 | 757.291494 | -0.141883222 | 0.18092446 | -0.7842125 | 0.43291544 | 0.65020381 |
| ANKRD13A | 1093.11411 | -0.279782963 | 0.28023804 | -0.9983761 | 0.31809701 | 0.5425805 |
| C12orf76 | 203.368637 | 0.017541332 | 0.2357026 | 0.07442146 | 0.94067503 | 0.97438649 |
| IFT81 | 764.956638 | 0.067174823 | 0.18612907 | 0.36090453 | 0.71817082 | 0.85963642 |
| ATP2A2 | 2426.13029 | -0.09600642 | 0.35663382 | -0.2692017 | 0.7877745 | 0.89956761 |
| ANAPC7 | 464.546313 | -0.02728412 | 0.24246744 | -0.1125269 | 0.91040561 | 0.95966385 |
| ARPC3 | 954.920416 | -1.092557802 | 0.23003384 | -4.7495526 | 2.04E-06 | 7.22E-05 |
| GPN3 | 116.005601 | -0.037913765 | 0.25841965 | -0.1467139 | 0.88335782 | 0.94755606 |
| FAM216A | 68.0027341 | 0.208361479 | 0.43483088 | 0.4791782 | 0.63181186 | 0.80130609 |
| VPS29 | 597.721235 | -0.567340801 | 0.18572156 | -3.0547924 | 0.00225216 | 0.01869819 |
| PPTC7 | 787.423049 | 0.320460468 | 0.218948 | 1.46363736 | 0.14329306 | 0.33258795 |
| TCTN1 | 417.521524 | 0.236055808 | 0.23917562 | 0.98695597 | 0.32366422 | 0.54881652 |
| HVCN1 | 125.990931 | 0.312111842 | 0.62165526 | 0.5020658 | 0.61562124 | 0.78887669 |
| PPP1CC | 302.212862 | 0.308659 | 0.24192775 | 1.27583132 | 0.20201515 | 0.41147543 |
| SH2B3 | 141.297444 | -1.528025697 | 0.36606032 | -4.1742456 | 2.99E-05 | 0.00065168 |
| ATXN2 | 3392.91125 | 0.047603755 | 0.17991534 | 0.26458975 | 0.79132552 | 0.90178749 |
| BRAP | 400.935971 | -0.032846315 | 0.17616282 | -0.1864543 | 0.85208851 | 0.93200182 |
| ACAD10 | 509.961843 | 0.457693035 | 0.2595664 | 1.76329844 | 0.07785017 | 0.22291846 |
| ALDH2 | 564.588812 | -0.568111011 | 0.29926818 | -1.8983341 | 0.05765208 | 0.18310096 |
| MAPKAPK5-A | 112.531006 | 0.183496883 | 0.34529371 | 0.53142261 | 0.59512596 | 0.77501911 |
| MAPKAPK5 | 447.595082 | 0.188898957 | 0.16013621 | 1.17961429 | 0.23815366 | 0.45614418 |
| TMEM116 | 1267.24957 | 1.215336145 | 0.336219 | 3.61471588 | 0.00030068 | 0.00402804 |
| ERP29 | 441.882182 | -0.99921711 | 0.37001127 | -2.7005045 | 0.00692344 | 0.04253568 |
| NAA25 | 816.571119 | 0.093500893 | 0.23143376 | 0.40400715 | 0.68620746 | 0.83787339 |
| TRAFD1 | 253.185632 | -0.178736027 | 0.13374414 | -1.3364027 | 0.1814177 | 0.38551261 |
| HECTD4 | 2815.35181 | -0.258232298 | 0.13308515 | -1.940354 | 0.05233668 | 0.17146474 |
| RPL6 | 1039.86086 | -1.064635338 | 0.38832701 | -2.7415949 | 0.00611417 | 0.03882844 |
| PTPN11 | 792.693524 | -0.254609678 | 0.19810168 | -1.2852475 | 0.19870582 | 0.40730993 |
| DTX1 | 246.96701 | 1.557251881 | 0.36896951 | 4.22054358 | 2.44E-05 | 0.00055696 |
| RASAL1 | 601.349254 | -2.230107547 | 0.48998508 | -4.5513785 | 5.33E-06 | 0.00016072 |
| DDX54 | 184.695653 | -0.614442539 | 0.31165587 | -1.9715417 | 0.04866195 | 0.16318819 |
| RITA1 | 58.3789133 | -0.638155156 | 0.30454811 | -2.0954166 | 0.03613397 | 0.13377065 |
| IQCD | 99.6132477 | -0.44617542 | 0.48269659 | -0.9243393 | 0.35530969 | 0.58081705 |
| TPCN1 | 1707.28093 | -0.721228951 | 0.28262836 | -2.5518634 | 0.01071485 | 0.05821422 |
| SLC8B1 | 531.275206 | -0.808638443 | 0.32062162 | -2.5220958 | 0.01166579 | 0.06178567 |
| PLBD2 | 159.392389 | -0.529169706 | 0.18239979 | -2.901153 | 0.00371792 | 0.02714762 |
| SDSL | 130.359374 | -0.090361709 | 0.43951847 | -0.2055925 | 0.83710922 | 0.92422064 |
| RBM19 | 395.333246 | -0.046935342 | 0.26542368 | -0.1768318 | 0.85964052 | 0.93625342 |
| MED13L | 7553.72104 | -0.385851746 | 0.19790928 | -1.9496395 | 0.0512191 | 0.16880068 |
| MAP1LC3B2 | 47.5598679 | -1.399716374 | 0.41600795 | -3.3646385 | 0.00076644 | 0.00817103 |
| C12orf49 | 386.791267 | -1.096846957 | 0.33124529 | -3.3112832 | 0.00092869 | 0.00942173 |
| RNFT2 | 86.4503126 | -0.076233057 | 0.35342585 | -0.2156975 | 0.82922358 | 0.92023714 |

| | | | | | | |
|---|---|---|---|---|---|---|
| FBXW8 | 915.008105 | -0.202548512 | 0.20497168 | -0.988178 | 0.32306546 | 0.54801228 |
| AC127164.1 | 71.8796562 | -0.007025408 | 0.55157468 | -0.012737 | 0.98983762 | 0.99606252 |
| TESC | 813.210035 | -1.307478191 | 0.43095631 | -3.0338996 | 0.00241415 | 0.0196964 |
| FBXO21 | 1074.95835 | 0.298012774 | 0.26762953 | 1.11352726 | 0.26548205 | 0.48657588 |
| NOS1 | 752.485918 | 0.618312616 | 0.89174698 | 0.69337226 | 0.48807597 | 0.69502858 |
| KSR2 | 585.42299 | 0.588455585 | 0.28014455 | 2.1005427 | 0.03568113 | 0.13247052 |
| RFC5 | 86.5895769 | 0.348306324 | 0.34792434 | 1.00109788 | 0.31677949 | 0.54117179 |
| WSB2 | 330.860854 | -0.16906527 | 0.21499981 | -0.7863508 | 0.431662 | 0.64957432 |
| VSIG10 | 740.224038 | 0.694799118 | 0.36729053 | 1.89168807 | 0.05853255 | 0.18461766 |
| PEBP1 | 2362.93858 | -0.598630384 | 0.48152133 | -1.2432064 | 0.2137918 | 0.42709858 |
| TAOK3 | 3892.02729 | 0.401871085 | 0.17308287 | 2.32184203 | 0.02024144 | 0.08978147 |
| SUDS3 | 543.982525 | 0.011049116 | 0.1704639 | 0.06481792 | 0.94831897 | 0.97784954 |
| HSPB8 | 60.5456686 | -2.724602495 | 0.56348421 | -4.8352774 | 1.33E-06 | 5.18E-05 |
| CCDC60 | 237.237657 | -1.850162211 | 0.50504474 | -3.663363 | 0.00024893 | 0.00350282 |
| AC002070.1 | 1467.64447 | -1.350265991 | 0.3615508 | -3.7346508 | 0.00018798 | 0.0027836 |
| TMEM233 | 97.2152602 | -1.824670039 | 0.41446704 | -4.4024491 | 1.07E-05 | 0.0002822 |
| PRKAB1 | 952.841393 | -0.627767657 | 0.19068878 | -3.2921059 | 0.0009944 | 0.00996589 |
| CIT | 1979.98217 | -1.051022249 | 0.21917265 | -4.7954079 | 1.62E-06 | 5.97E-05 |
| BICDL1 | 7748.22127 | 1.444685008 | 0.39050094 | 3.69956855 | 0.00021597 | 0.00312453 |
| RAB35 | 240.703407 | -0.509497784 | 0.21086501 | -2.4162273 | 0.01568227 | 0.07560113 |
| GCN1 | 388.72835 | -0.039788634 | 0.17191332 | -0.2314459 | 0.8169684 | 0.91483008 |
| RPLP0 | 1429.54557 | -1.489706915 | 0.47235097 | -3.1538136 | 0.00161152 | 0.01454 |
| PXN-AS1 | 85.2779205 | 0.175544987 | 0.31500296 | 0.55728043 | 0.57733584 | 0.7637681 |
| PXN | 864.422279 | -0.455131956 | 0.17746214 | -2.5646708 | 0.01032737 | 0.05676832 |
| SIRT4 | 79.176607 | 1.321684993 | 0.37571207 | 3.51781354 | 0.00043512 | 0.00534152 |
| GATC | 156.219137 | -0.433665881 | 0.22861905 | -1.896893 | 0.05784206 | 0.18348602 |
| SRSF9 | 468.267765 | -0.254418599 | 0.26672956 | -0.9538448 | 0.34016222 | 0.56546725 |
| DYNLL1 | 683.774872 | -1.079538522 | 0.51065527 | -2.114026 | 0.03451305 | 0.12977024 |
| COQ5 | 248.600866 | 0.665867253 | 0.2704231 | 2.46231651 | 0.01380428 | 0.06906845 |
| RNF10 | 1696.98093 | 0.529616908 | 0.17862835 | 2.96490962 | 0.00302772 | 0.02329245 |
| POP5 | 162.593669 | -0.052713976 | 0.34638521 | -0.1521831 | 0.87904252 | 0.94598627 |
| AC125616.1 | 51.0498034 | 1.721304159 | 0.66730002 | 2.57950562 | 0.00989418 | 0.05498203 |
| CABP1 | 425.913988 | 1.08390252 | 0.61705436 | 1.75657542 | 0.07899021 | 0.2246842 |
| MLEC | 672.398941 | -0.278347453 | 0.35405141 | -0.7861781 | 0.43176317 | 0.64957432 |
| UNC119B | 170.284291 | -0.170307372 | 0.26681735 | -0.638292 | 0.52328364 | 0.72294254 |
| ACADS | 146.969929 | 0.217552002 | 0.2811146 | 0.7738908 | 0.43899537 | 0.65485132 |
| SPPL3 | 2212.83527 | -0.119760457 | 0.16755813 | -0.7147398 | 0.47476986 | 0.68513327 |
| C12orf43 | 78.1471019 | -0.573088782 | 0.29010258 | -1.9754694 | 0.0482149 | 0.16208414 |
| P2RX4 | 150.076443 | -0.510708064 | 0.52779236 | -0.9676306 | 0.33322886 | 0.55872304 |
| CAMKK2 | 191.59975 | -0.361776248 | 0.20795856 | -1.7396555 | 0.08191953 | 0.23006759 |
| ANAPC5 | 981.080514 | 0.437459976 | 0.16509036 | 2.6498215 | 0.00805343 | 0.04757503 |
| RNF34 | 468.118698 | -0.162179204 | 0.27236033 | -0.5954582 | 0.5515372 | 0.74467698 |
| KDM2B | 797.101574 | 0.734837643 | 0.3077487 | 2.38778472 | 0.01695027 | 0.079621 |
| ORAI1 | 81.7450406 | -1.971404873 | 0.50213317 | -3.9260598 | 8.63E-05 | 0.00150404 |
| MORN3 | 59.3333664 | -0.609008985 | 0.36335276 | -1.6760819 | 0.09372215 | 0.25184394 |
| TMEM120B | 833.404645 | 0.325170126 | 0.45934808 | 0.70789482 | 0.47901058 | 0.68864274 |
| RHOF | 141.641798 | -0.729967398 | 0.29995504 | -2.4335893 | 0.01494995 | 0.07319267 |
| SETD1B | 310.027006 | -0.620991213 | 0.22634463 | -2.7435651 | 0.0060776 | 0.03870155 |
| HPD | 48.1381422 | 0.400494646 | 0.56687835 | 0.70649135 | 0.47988263 | 0.68907161 |
| WDR66 | 424.744551 | 0.058270578 | 0.33972169 | 0.17152446 | 0.8638114 | 0.93825218 |
| AC069503.1 | 39.1115884 | 2.03241317 | 0.60494217 | 3.35968173 | 0.00078032 | 0.00825243 |
| BCL7A | 806.170846 | 0.342181966 | 0.32472642 | 1.05375463 | 0.29199527 | 0.51467381 |
| MLXIP | 1295.57628 | -1.315797216 | 0.24795691 | -5.306556 | 1.12E-07 | 6.90E-06 |
| CLIP1 | 1870.44584 | -0.727605499 | 0.26423084 | -2.7536737 | 0.00589305 | 0.03786055 |
| ZCCHC8 | 675.425151 | 0.266716225 | 0.16155673 | 1.65091376 | 0.09875619 | 0.26026944 |
| RSRC2 | 893.868094 | 0.181460098 | 0.15900654 | 1.14121154 | 0.2537819 | 0.47441341 |
| KNTC1 | 147.488602 | 0.048368231 | 0.33460318 | 0.14455401 | 0.88506301 | 0.94815256 |
| DENR | 936.589197 | -0.331184248 | 0.19309739 | -1.7151151 | 0.08632413 | 0.23790016 |
| CCDC62 | 52.9872905 | 0.112649382 | 0.38651106 | 0.2914519 | 0.77070572 | 0.88937357 |
| HIP1R | 750.40745 | 0.228304273 | 0.30927563 | 0.73819031 | 0.46039881 | 0.67395297 |
| VPS37B | 1377.45496 | -0.402188905 | 0.22608013 | -1.7789662 | 0.0752453 | 0.21801167 |
| ABCB9 | 184.476093 | 0.751178187 | 0.31570339 | 2.37937953 | 0.01734181 | 0.08094275 |
| OGFOD2 | 55.0993025 | 0.504817666 | 0.323493 | 1.56052115 | 0.11863678 | 0.2938997 |
| ARL6IP4 | 524.183578 | -0.442190125 | 0.27456251 | -1.6105263 | 0.10728302 | 0.27516532 |
| PITPNM2 | 2175.01868 | -0.485505579 | 0.33021781 | -1.4702586 | 0.14149173 | 0.33008905 |
| MPHOSPH9 | 867.67821 | 0.525796039 | 0.21094343 | 2.49259261 | 0.01268143 | 0.06540763 |
| C12orf65 | 309.333182 | -0.520139135 | 0.23210029 | -2.2410103 | 0.02502541 | 0.10372157 |
| AC068768.1 | 61.8790096 | 0.424429813 | 0.41404688 | 1.0250767 | 0.305327 | 0.52868668 |
| SBNO1 | 949.638897 | -0.108255692 | 0.15569841 | -0.6952909 | 0.48687301 | 0.69412904 |
| KMT5A | 223.4143 | -0.217142832 | 0.21434947 | -1.0130318 | 0.31104497 | 0.53456605 |
| RILPL2 | 255.812823 | -0.497770002 | 0.1999553 | -2.4894064 | 0.01279566 | 0.06576571 |
| SNRNP35 | 364.984127 | 0.056866037 | 0.17978734 | 0.31629611 | 0.75177777 | 0.87948211 |
| RILPL1 | 978.695077 | 0.405386425 | 0.20790492 | 1.94986456 | 0.05119226 | 0.16875434 |
| AC055713.1 | 90.9005696 | 0.531347919 | 0.28805529 | 1.84460389 | 0.06509518 | 0.19827619 |
| TMED2 | 310.97637 | -0.478779768 | 0.24453423 | -1.9579253 | 0.05023878 | 0.16685843 |
| DDX55 | 214.657139 | 0.012657556 | 0.29006351 | 0.04363719 | 0.96519361 | 0.98449934 |
| EIF2B1 | 267.619736 | 0.239976984 | 0.19492356 | 1.23113383 | 0.21827281 | 0.43232353 |
| GTF2H3 | 258.996486 | 0.228722191 | 0.17482849 | 1.30826613 | 0.19078306 | 0.39735942 |
| TCTN2 | 228.736843 | 0.370060453 | 0.19087025 | 1.93880638 | 0.05252492 | 0.1718607 |
| ATP6V0A2 | 274.4175 | -0.042906355 | 0.33468652 | -0.1281986 | 0.89799178 | 0.9527979 |
| CCDC92 | 790.346895 | -0.09341471 | 0.12489065 | -0.747972 | 0.45447704 | 0.6698832 |
| RFLNA | 167.386882 | 0.608458959 | 0.5206954 | 1.16855068 | 0.24258471 | 0.46105057 |
| NCOR2 | 1216.86211 | 0.042076284 | 0.33241611 | 0.12657715 | 0.89927508 | 0.95371534 |
| SCARB1 | 63.5993352 | -0.280794881 | 0.38121656 | -0.7365758 | 0.46138037 | 0.67460408 |
| UBC | 2190.87201 | -1.779151478 | 0.3459042 | -5.1434804 | 2.70E-07 | 1.40E-05 |
| DHX37 | 123.143621 | -0.55329229 | 0.32584337 | -1.6980315 | 0.08950183 | 0.24365821 |
| BRI3BP | 217.415597 | 0.543062458 | 0.31083269 | 1.74712142 | 0.08061628 | 0.22763155 |
| AACS | 530.737764 | 0.085016052 | 0.40109029 | 0.21196238 | 0.83213638 | 0.92158196 |
| LINC02405 | 42.9169371 | 0.126752541 | 0.59705347 | 0.2122968 | 0.83187549 | 0.92158196 |
| TMEM132C | 150.419689 | 2.240636216 | 0.62966958 | 3.55843173 | 0.00037308 | 0.00472027 |
| SLC15A4 | 379.670297 | -0.32189032 | 0.24822625 | -1.2967618 | 0.19471315 | 0.40230972 |
| TMEM132D | 45.2650628 | -0.083538953 | 0.29196509 | -0.2861265 | 0.77478123 | 0.89222126 |
| RIMBP2 | 1404.23931 | -1.399321491 | 0.61088087 | -2.2906618 | 0.02198298 | 0.09492422 |
| STX2 | 344.420108 | -0.062790403 | 0.41634949 | -0.1508118 | 0.8801242 | 0.94636668 |
| RAN | 352.484089 | -1.195817716 | 0.47663974 | -2.5088502 | 0.01211248 | 0.06331373 |
| SFSWAP | 1043.04436 | 0.248071665 | 0.20663208 | 1.20054767 | 0.22992671 | 0.44551883 |
| ULK1 | 744.642103 | -0.321048825 | 0.24466192 | -1.3122141 | 0.1894479 | 0.39556637 |
| PUS1 | 155.360807 | 0.10804836 | 0.20605167 | 0.52437507 | 0.60001769 | 0.77816728 |
| EP400 | 1008.68591 | 0.144096706 | 0.19238741 | 0.74899238 | 0.4538618 | 0.66948056 |
| DDX51 | 112.371881 | -0.241413246 | 0.31694725 | -0.7616827 | 0.44624939 | 0.66215549 |
| NOC4L | 41.2335667 | -0.022008157 | 0.49964724 | -0.0440474 | 0.96486663 | 0.98439201 |
| GALNT9 | 209.179081 | 0.492329649 | 0.51796061 | 0.95051562 | 0.34185032 | 0.56713148 |
| AC148477.2 | 45.6481679 | 1.385289014 | 0.67466211 | 2.05330787 | 0.04004273 | 0.1426826 |
| FBRSL1 | 546.95735 | 0.448891864 | 0.23580874 | 1.9036269 | 0.05695879 | 0.18151235 |
| POLE | 136.067671 | 0.250338135 | 0.39557328 | 0.63284895 | 0.52683229 | 0.72526604 |
| AC135586.2 | 37.3334693 | 0.520279486 | 0.33637631 | 1.54671859 | 0.12193112 | 0.29892133 |
| PGAM5 | 54.1218091 | -0.652029111 | 0.32847501 | -1.9850189 | 0.04714237 | 0.15929996 |
| ANKLE2 | 731.085784 | -0.949177721 | 0.28456804 | -3.3355035 | 0.00085145 | 0.00879344 |
| GOLGA3 | 587.582662 | -0.383834105 | 0.22271532 | -1.7234293 | 0.08481092 | 0.23490616 |
| CHFR | 738.837824 | -0.159806365 | 0.20922184 | -0.763813 | 0.44497868 | 0.66082438 |
| ZNF26 | 548.239153 | 0.142408933 | 0.22590171 | 0.6304022 | 0.52843147 | 0.72663454 |
| ZNF84 | 611.650754 | 0.201902312 | 0.16322856 | 1.23693003 | 0.21611306 | 0.42998003 |
| ZNF140 | 333.755726 | -0.010659643 | 0.18556757 | -0.0574435 | 0.95419195 | 0.98007893 |

| | | | | | | |
|---|---|---|---|---|---|---|
| ZNF891 | 283.303928 | 0.507675114 | 0.1666504 | 3.04634804 | 0.0023164 | 0.01913267 |
| ZNF268 | 106.01996 | -0.379757565 | 0.32146876 | -1.1813203 | 0.23747552 | 0.45507241 |
| TPTE2 | 87.957782 | 0.151377133 | 0.40246048 | 0.37612918 | 0.70682086 | 0.85232762 |
| MPHOSPH8 | 1383.95057 | -0.051041925 | 0.2590923 | -0.1970029 | 0.8438253 | 0.92769172 |
| PSPC1 | 2009.74688 | 0.133305703 | 0.247698 | 0.53817835 | 0.59045392 | 0.77217181 |
| ZMYM5 | 818.377416 | 0.148711422 | 0.13088463 | 1.13620234 | 0.25587189 | 0.47687514 |
| AL355001.1 | 67.7904418 | 0.000894923 | 0.45487554 | 0.0019674 | 0.99843024 | 0.99953295 |
| ZMYM2 | 4292.591 | 0.249759897 | 0.20160397 | 1.23886397 | 0.21539588 | 0.42926436 |
| CRYL1 | 1943.5245 | 0.725804802 | 0.27698509 | 2.62037497 | 0.00878331 | 0.05041939 |
| IFT88 | 2325.61779 | 0.284574208 | 0.19519268 | 1.45791431 | 0.14486416 | 0.33475075 |
| EEF1AKMT1 | 217.057948 | 0.092954554 | 0.22432485 | 0.41437475 | 0.67859968 | 0.8314278 |
| XPO4 | 1189.21494 | 0.359765907 | 0.13840533 | 2.59936444 | 0.00933966 | 0.05271645 |
| LATS2 | 808.008319 | 0.599016736 | 0.25145188 | 2.38223208 | 0.01720805 | 0.08040365 |
| LATS2-AS1 | 51.6570372 | 0.59742379 | 0.53515826 | 1.11634975 | 0.26427245 | 0.48528287 |
| SAP18 | 452.366069 | -0.874339053 | 0.34035817 | -2.5688792 | 0.0102028 | 0.05631792 |
| MRPL57 | 305.754198 | -0.619114225 | 0.29475833 | -2.100413 | 0.03569253 | 0.13247052 |
| ZDHHC20 | 1244.45729 | -0.179695871 | 0.14932329 | -1.2034015 | 0.22882099 | 0.44441942 |
| MICU2 | 2277.22152 | 0.103076424 | 0.17740316 | 0.58102923 | 0.56122075 | 0.75291133 |
| FGF9 | 1205.74082 | 1.910248676 | 0.44165568 | 4.32519897 | 1.52E-05 | 0.00037767 |
| AL136962.1 | 464.233568 | 0.074299394 | 0.92788832 | 0.08007364 | 0.93617869 | 0.97270387 |
| LINC00621 | 437.232914 | -1.651834396 | 0.4452813 | -3.7096424 | 0.00020755 | 0.003026 |
| AL157931.1 | 189.900169 | -3.229353915 | 0.51562359 | -6.2630066 | 3.78E-10 | 5.36E-08 |
| SACS | 126.90408 | -1.479477624 | 0.4513663 | -3.277776 | 0.00104628 | 0.01035233 |
| TNFRSF19 | 215.652817 | 0.739739677 | 0.5004043 | 1.47828401 | 0.13933178 | 0.3261445 |
| MIPEP | 1127.68409 | 0.218477616 | 0.24254924 | 0.90075573 | 0.36771821 | 0.59294486 |
| PARP4 | 1031.92953 | -1.633943776 | 0.43310772 | -3.7726037 | 0.00016155 | 0.0024613 |
| CENPJ | 85.808644 | 0.163619208 | 0.3955532 | 0.41364653 | 0.67913299 | 0.8319269 |
| MTMR6 | 338.382213 | -0.220462019 | 0.13220433 | -1.6675854 | 0.09539805 | 0.25441407 |
| NUP58 | 954.789446 | 0.183649751 | 0.26549338 | 0.69173005 | 0.48910687 | 0.6960367 |
| RNF6 | 144.586076 | -0.059562091 | 0.22424615 | -0.2656103 | 0.79053934 | 0.90116933 |
| CDK8 | 1640.81693 | 0.010022966 | 0.17480288 | 0.05733868 | 0.95427541 | 0.98007893 |
| WASF3 | 370.329536 | 0.174963691 | 0.17466379 | 1.001717 | 0.31648029 | 0.54080054 |
| USP12 | 1212.7421 | -0.076535026 | 0.23374547 | -0.3274289 | 0.7433435 | 0.87465031 |
| LINC02340 | 420.298062 | 0.538647476 | 0.43683833 | 1.2330591 | 0.21755371 | 0.43128814 |
| RPL21 | 508.026803 | -0.426177587 | 0.39050136 | -1.0913601 | 0.27511448 | 0.49658201 |
| GTF3A | 199.611675 | -0.441077871 | 0.20743704 | -2.1263217 | 0.03347648 | 0.12730077 |
| MTIF3 | 492.912212 | -0.090312285 | 0.14331407 | -0.6301704 | 0.5285831 | 0.72676737 |
| LNX2 | 637.568916 | -0.368479836 | 0.23881446 | -1.5429545 | 0.12284182 | 0.300288 |
| POLR1D | 477.528151 | 0.080222517 | 0.14739948 | 0.54425237 | 0.58626782 | 0.77024186 |
| PAN3 | 4766.29292 | -0.072942251 | 0.15988417 | -0.4562194 | 0.64823225 | 0.81176524 |
| FLT1 | 67.4176943 | -0.01032949 | 0.37172666 | -0.0277879 | 0.97783135 | 0.99038178 |
| POMP | 320.009727 | -0.197954225 | 0.20867858 | -0.9486083 | 0.34281988 | 0.56795532 |
| SLC46A3 | 194.67833 | -0.635019675 | 0.39275201 | -1.6168464 | 0.10591144 | 0.27302579 |
| MTUS2 | 51.4051174 | -1.059563161 | 0.62230818 | -1.7026341 | 0.08863656 | 0.24205091 |
| SLC7A1 | 381.354705 | -1.433916452 | 0.30147313 | -4.7563658 | 1.97E-06 | 7.00E-05 |
| UBL3 | 1157.59999 | 0.049182842 | 0.15559602 | 0.31609319 | 0.75193178 | 0.8795471 |
| KATNAL1 | 399.113961 | 0.054708138 | 0.3011378 | 0.18167144 | 0.85584058 | 0.93464491 |
| HMGB1 | 2185.78762 | 0.239289884 | 0.24602329 | 0.97263103 | 0.33073672 | 0.55588514 |
| USPL1 | 524.819555 | 0.133163679 | 0.26647049 | 0.49973143 | 0.6172642 | 0.7902755 |
| HSPH1 | 534.105647 | -0.125992361 | 0.39235838 | -0.3211155 | 0.74812286 | 0.87714848 |
| B3GLCT | 662.996534 | 0.659356778 | 0.27650365 | 2.38462232 | 0.01709666 | 0.08002474 |
| FRY | 2092.3387 | 2.069051752 | 0.73552118 | 2.81304169 | 0.00490753 | 0.03295554 |
| FRY-AS1 | 39.4491821 | 2.638606643 | 1.13371134 | 2.32740606 | 0.01994366 | 0.0889986 |
| N4BP2L1 | 771.982651 | -0.110979089 | 0.33829964 | -0.3280497 | 0.7428741 | 0.87441041 |
| N4BP2L2 | 8650.78926 | 0.071216616 | 0.17452266 | 0.40806516 | 0.68322584 | 0.83554579 |
| PDS5B | 2106.31216 | 0.210194805 | 0.19710713 | 1.06639877 | 0.28624342 | 0.50853916 |
| KL | 184.64491 | 1.483540384 | 0.26637755 | 5.56931456 | 2.56E-08 | 2.04E-06 |
| STARD13 | 3704.51943 | 0.908952273 | 0.22699426 | 4.00429633 | 6.22E-05 | 0.00115648 |
| AL139383.1 | 1487.20299 | 1.17577018 | 0.33621691 | 3.49705846 | 0.00047042 | 0.00565916 |
| RFC3 | 301.999632 | 0.068217829 | 0.36306679 | 0.18789333 | 0.85096027 | 0.93170782 |
| LINC02343 | 2342.72494 | 0.654100488 | 0.42979293 | 1.52189681 | 0.12803494 | 0.30872828 |
| LINC00457 | 136.357497 | 0.299569415 | 0.72622427 | 0.41250262 | 0.67997106 | 0.83256746 |
| NBEA | 2744.78501 | -0.548938557 | 0.30321729 | -1.8103801 | 0.07023686 | 0.20866009 |
| DCLK1 | 65.3353945 | 1.389113561 | 0.68630404 | 2.02404981 | 0.04296503 | 0.14938801 |
| SPART | 899.552111 | 0.31287447 | 0.25766048 | 1.21428971 | 0.22463711 | 0.43926449 |
| SPART-AS1 | 77.1062308 | -0.120286421 | 0.51300978 | -0.234472 | 0.81461858 | 0.91339257 |
| RFXAP | 168.857358 | 0.690362709 | 0.23593712 | 2.92604533 | 0.00343301 | 0.02564854 |
| SMAD9 | 172.690029 | -2.856763303 | 0.60161348 | -4.7485028 | 2.05E-06 | 7.22E-05 |
| ALG5 | 432.754433 | 0.256292674 | 0.12610693 | 2.0323441 | 0.04211883 | 0.14722087 |
| EXOSC8 | 236.927587 | 0.590251542 | 0.25173923 | 2.34469431 | 0.01904269 | 0.08631943 |
| SUPT20H | 1135.67424 | 0.203307984 | 0.14801071 | 1.37360315 | 0.16956492 | 0.3699733 |
| LINC02334 | 125.274933 | -0.728315276 | 0.55915138 | -1.3025368 | 0.19273293 | 0.40009576 |
| LINC00571 | 422.746043 | -0.155223587 | 0.3065578 | -0.5063436 | 0.61261544 | 0.78690269 |
| UFM1 | 553.544876 | 0.08405553 | 0.18723095 | 0.44894036 | 0.65347468 | 0.81539269 |
| FREM2 | 1397.28842 | 1.171665144 | 0.34564042 | 3.38983834 | 0.00069934 | 0.00765332 |
| PROSER1 | 400.472213 | 0.552976961 | 0.26531598 | 2.08422032 | 0.03714013 | 0.1359335 |
| NHLRC3 | 833.370491 | 0.652725248 | 0.29511813 | 2.21174225 | 0.02698448 | 0.10913594 |
| AL354809.1 | 57.978497 | 0.402966396 | 0.32644786 | 1.23439742 | 0.21705485 | 0.43120432 |
| LHFPL6 | 58.5410372 | 0.459979619 | 0.58019019 | 0.79280834 | 0.4278895 | 0.64638522 |
| COG6 | 1069.45484 | 0.536581455 | 0.21213055 | 2.52948689 | 0.01142294 | 0.06096391 |
| LINC00598 | 302.170852 | 0.058868941 | 0.32415544 | 0.18160713 | 0.85589105 | 0.93464491 |
| FOXO1 | 4280.27885 | -0.674884528 | 0.29249987 | -2.3072985 | 0.02103818 | 0.09241548 |
| MRPS31 | 915.774197 | 0.468640392 | 0.19225134 | 2.43764432 | 0.01478331 | 0.07253817 |
| SLC25A15 | 87.1822504 | 0.023592171 | 0.31341888 | 0.07527361 | 0.93999702 | 0.97411372 |
| ELF1 | 2985.75905 | -0.182533215 | 0.23081187 | -0.7908312 | 0.42904252 | 0.64738665 |
| WBP4 | 508.582428 | 0.346259388 | 0.15388273 | 2.25015113 | 0.02443935 | 0.10202889 |
| KBTBD6 | 52.8659317 | 0.264952457 | 0.50136444 | 0.5284628 | 0.59717816 | 0.77654325 |
| AL354696.2 | 212.098528 | 0.502814089 | 0.25095414 | 2.00360944 | 0.04511191 | 0.15493585 |
| KBTBD7 | 58.3738093 | 0.598892291 | 0.45194401 | 1.32514709 | 0.18512237 | 0.39074487 |
| MTRF1 | 290.126141 | 0.440661303 | 0.23243916 | 1.89581352 | 0.05798471 | 0.1838061 |
| NAA16 | 1185.20342 | 0.214222414 | 0.16967858 | 1.26251887 | 0.20676214 | 0.41747495 |
| VWA8 | 4894.70441 | 0.593226027 | 0.20689452 | 2.86728733 | 0.00414007 | 0.02926733 |
| DGKH | 4028.65492 | -2.071985104 | 0.42670724 | -4.8557533 | 1.20E-06 | 4.79E-05 |
| AKAP11 | 891.881496 | 0.2353216 | 0.09539287 | 2.46686777 | 0.01363007 | 0.06845611 |
| LINC02341 | 72.4634516 | 0.579247427 | 0.49769754 | 1.1638543 | 0.24448306 | 0.46332521 |
| EPSTI1 | 625.33324 | 0.42888074 | 0.75275658 | 0.56974691 | 0.56884937 | 0.75867193 |
| DNAJC15 | 833.157291 | -0.450565589 | 0.26295362 | -1.7134793 | 0.0866244 | 0.23847888 |
| ENOX1 | 4013.33868 | 0.849043221 | 0.75597643 | 1.12310807 | 0.2613916 | 0.48237074 |
| CCDC122 | 693.615415 | -0.076368839 | 0.27485956 | -0.2778468 | 0.78112999 | 0.89545532 |
| LINC00284 | 66.977256 | -0.932527159 | 0.8976467 | -1.0388577 | 0.29887094 | 0.5219178 |
| SMIM2-AS1 | 188.423956 | 0.542139467 | 0.48137479 | 1.12623152 | 0.26006753 | 0.48104792 |
| SERP2 | 178.747381 | -0.513964413 | 0.31270872 | -1.6435884 | 0.10026126 | 0.2633428 |
| TSC22D1 | 4836.7496 | -0.770169322 | 0.29544879 | -2.6067777 | 0.00913987 | 0.05205672 |
| TSC22D1-AS1 | 47.7166911 | 0.339693865 | 0.46955051 | 0.72344478 | 0.46940667 | 0.68113728 |
| NUFIP1 | 158.588282 | 0.083169642 | 0.24265155 | 0.34275339 | 0.73178399 | 0.86826096 |
| AL359706.1 | 183.94988 | 0.423142344 | 0.46398682 | 0.91197061 | 0.36178419 | 0.58707707 |
| GPALPP1 | 481.065575 | 0.026274531 | 0.1229698 | 0.21366653 | 0.83080712 | 0.92118956 |
| GTF2F2 | 985.974595 | -0.33314801 | 0.21818475 | -1.5269079 | 0.12678393 | 0.30643895 |
| TPT1 | 2831.54807 | -1.639815525 | 0.46159502 | -3.5524983 | 0.00038159 | 0.00480958 |
| AL138963.1 | 57.5850399 | -0.360816081 | 0.40165233 | -0.8983294 | 0.36900998 | 0.59393582 |
| SLC25A30 | 548.337122 | 0.676436171 | 0.40615663 | 1.66545642 | 0.09582173 | 0.25517074 |
| COG3 | 707.717291 | 0.325315324 | 0.16417002 | 1.98157575 | 0.04752674 | 0.16021958 |
| LINC01055 | 115.639239 | 0.042405444 | 1.07662041 | 0.03938755 | 0.9685814 | 0.98582596 |

| | | | | | | |
|---|---|---|---|---|---|---|
| SIAH3 | 198.345277 | 0.58735549 | 0.90035403 | 0.65236059 | 0.51416858 | 0.71596592 |
| ZC3H13 | 1298.53428 | 0.193155845 | 0.10447046 | 1.84890399 | 0.06447168 | 0.19708549 |
| CPB2-AS1 | 74.7320667 | 1.144846621 | 0.28039387 | 4.08299444 | 4.45E-05 | 0.00089933 |
| AL138686.1 | 139.356465 | -0.991061381 | 0.58872694 | -1.6833974 | 0.09229819 | 0.24906371 |
| LRCH1 | 3039.66663 | 0.457277495 | 0.21035668 | 2.17381968 | 0.02971868 | 0.11672734 |
| ESD | 447.905874 | 0.096537175 | 0.19684062 | 0.49043321 | 0.62382738 | 0.79539814 |
| SUCLA2 | 1320.65168 | 0.362504663 | 0.2787587 | 1.30042455 | 0.1934555 | 0.40113679 |
| NUDT15 | 58.5751227 | -0.643582185 | 0.28393128 | -2.2666829 | 0.0234096 | 0.09910816 |
| MED4 | 658.403477 | 0.158343644 | 0.19076795 | 0.83003276 | 0.40652026 | 0.62649938 |
| ITM2B | 3922.82138 | -0.606215161 | 0.31292106 | -1.9372782 | 0.05271134 | 0.17225733 |
| RB1 | 2455.39434 | -0.523462836 | 0.25306734 | -2.0684725 | 0.03859561 | 0.13944459 |
| LPAR6 | 160.73702 | -0.426071216 | 0.5123354 | -0.8316256 | 0.40562032 | 0.62584264 |
| RCBTB2 | 424.169484 | 0.159826608 | 0.19582546 | 0.8161687 | 0.41440366 | 0.634209 |
| FNDC3A | 3365.08671 | -0.520557109 | 0.30792104 | -1.6905539 | 0.09092204 | 0.24631864 |
| CDADC1 | 295.857124 | 0.564199216 | 0.21601285 | 2.61187808 | 0.00900464 | 0.05133459 |
| CAB39L | 521.68625 | 0.4389124 | 0.39546741 | 1.10985733 | 0.26706051 | 0.48822301 |
| SETDB2 | 1049.52568 | 1.103017166 | 0.28662859 | 3.84824548 | 0.00011897 | 0.001921 |
| PHF11 | 564.405969 | 0.080986319 | 0.30693054 | 0.26385879 | 0.79188874 | 0.90193232 |
| RCBTB1 | 207.673831 | 0.130511398 | 0.26707916 | 0.48866186 | 0.62508111 | 0.79598067 |
| EBPL | 214.4556 | -0.226949171 | 0.3387378 | -0.6699848 | 0.50286749 | 0.70694289 |
| KPNA3 | 930.893835 | -0.022538209 | 0.19261343 | -0.1170127 | 0.90685003 | 0.95759709 |
| SPRYD7 | 178.921191 | 0.054822488 | 0.2518035 | 0.21771932 | 0.82764781 | 0.91981743 |
| DLEU2 | 1232.09777 | -1.095432061 | 0.43415953 | -2.5231096 | 0.01163221 | 0.06165727 |
| TRIM13 | 407.605486 | -0.249011122 | 0.18888037 | -1.3183536 | 0.18738531 | 0.39298287 |
| RNASEH2B | 450.73286 | 0.091017286 | 0.23820528 | 0.38209601 | 0.70239015 | 0.8488446 |
| FAM124A | 152.07743 | 1.279215512 | 0.57199114 | 2.23642538 | 0.02532392 | 0.10440189 |
| INTS6 | 2184.03882 | 0.658000805 | 0.37435142 | 1.75770885 | 0.07879706 | 0.22441496 |
| INTS6-AS1 | 303.596302 | 0.778706362 | 0.30166912 | 2.58132602 | 0.00984216 | 0.05471594 |
| WDFY2 | 2502.91299 | 1.252310943 | 0.26509734 | 4.72396638 | 2.31E-06 | 7.98E-05 |
| ATP7B | 145.378991 | 0.320972354 | 0.23844054 | 1.34613163 | 0.17826007 | 0.38150312 |
| ALG11 | 141.025232 | 0.424176349 | 0.21179566 | 2.00276222 | 0.04520282 | 0.15516736 |
| UTP14C | 162.910555 | -0.223798086 | 0.29537087 | -0.757685 | 0.44863957 | 0.66445974 |
| NEK5 | 118.949875 | 0.495280807 | 0.48822144 | 1.01445936 | 0.31036361 | 0.53435962 |
| NEK3 | 437.475785 | 0.065479938 | 0.37851257 | 0.17299277 | 0.86265711 | 0.93788453 |
| THSD1 | 59.8700564 | 0.00045043 | 0.40223824 | 0.00111981 | 0.99910652 | 0.99971199 |
| VPS36 | 872.455225 | 0.26358012 | 0.12733427 | 2.06998574 | 0.03845368 | 0.13917687 |
| CKAP2 | 93.9138195 | 0.386198988 | 0.26378036 | 1.46409304 | 0.14316853 | 0.33253263 |
| HNRNPA1L2 | 38.7018358 | 1.062640112 | 0.37947841 | 2.80026498 | 0.00510607 | 0.03396075 |
| SUGT1 | 616.651786 | 0.271698933 | 0.20377449 | 1.33333146 | 0.18242305 | 0.38696421 |
| OLFM4 | 336.745939 | -2.972659419 | 1.30074298 | -2.285355 | 0.02229202 | 0.09594504 |
| DIAPH3 | 772.66819 | -0.631217005 | 0.57997 | -1.0883615 | 0.27643556 | 0.49876211 |
| TDRD3 | 2488.05587 | 0.572226929 | 0.17191359 | 3.32857291 | 0.00087292 | 0.00898708 |
| PCDH9 | 9224.21489 | 1.511142701 | 0.4324204 | 3.4946147 | 0.00047475 | 0.00570084 |
| PCDH9-AS2 | 201.888277 | 0.93735899 | 0.68015278 | 1.37815946 | 0.16815406 | 0.36813567 |
| DACH1 | 2248.78114 | 0.542548309 | 0.34142895 | 1.58905184 | 0.11204869 | 0.28299257 |
| MZT1 | 95.07591 | -0.305334282 | 0.28666452 | -1.0651275 | 0.28681821 | 0.50912842 |
| BORA | 97.2240209 | -0.210104222 | 0.32657959 | -0.6433477 | 0.51999853 | 0.71990992 |
| DIS3 | 421.080224 | -0.169508803 | 0.13064294 | -1.2974969 | 0.19446028 | 0.40198581 |
| PIBF1 | 2600.37141 | -0.064167293 | 0.14593546 | -0.4396964 | 0.66015702 | 0.81993929 |
| KLF5 | 475.697168 | -1.768099722 | 0.30119224 | -5.8703363 | 4.35E-09 | 4.81E-07 |
| KLF12 | 4413.82942 | 0.789284895 | 0.24702853 | 3.19511636 | 0.00139774 | 0.01300198 |
| TBC1D4 | 6519.06486 | 0.791772664 | 0.20258462 | 3.90835533 | 9.29E-05 | 0.00158587 |
| COMMD6 | 401.564589 | 0.186334827 | 0.41590661 | 0.44802084 | 0.65413816 | 0.81575821 |
| UCHL3 | 504.242609 | -0.389634512 | 0.28053518 | -1.3888971 | 0.16486403 | 0.36391861 |
| LMO7-AS1 | 289.979661 | 0.116971763 | 0.4483307 | 0.26090509 | 0.7941657 | 0.90354304 |
| LMO7 | 5624.82234 | 1.057646448 | 0.36271035 | 2.91595335 | 0.00354604 | 0.02621129 |
| KCTD12 | 92.8766434 | -0.856595818 | 0.58615545 | -1.4613799 | 0.1439112 | 0.33355379 |
| AC001226.2 | 323.563842 | 0.129646987 | 0.4125162 | 0.31428338 | 0.75330582 | 0.88025624 |
| FBXL3 | 785.131444 | 0.223602739 | 0.24733944 | 0.90403189 | 0.36597849 | 0.59141301 |
| MYCBP2 | 2457.70697 | 0.237659557 | 0.13686024 | 1.73651276 | 0.08247321 | 0.23078253 |
| SLAIN1 | 333.643806 | 0.346414403 | 0.18397528 | 1.88293998 | 0.0597085 | 0.18742624 |
| RNF219 | 231.939885 | 0.135748859 | 0.18310428 | 0.74137458 | 0.45846635 | 0.67266311 |
| RBM26 | 2516.78113 | 0.554578622 | 0.07169419 | 7.73533599 | 1.03E-14 | 6.81E-12 |
| RBM26-AS1 | 90.9572187 | 0.99229143 | 0.40456277 | 2.45275024 | 0.01417687 | 0.07034648 |
| NDFIP2 | 1046.51964 | -0.189899665 | 0.29239984 | -0.649452 | 0.51604626 | 0.71744606 |
| SPRY2 | 89.6081044 | 0.040970133 | 0.28101239 | 0.14579476 | 0.88408341 | 0.9477505 |
| LINC00379 | 142.44302 | 3.510201047 | 0.76393328 | 4.59490529 | 4.33E-06 | 0.00013299 |
| GPC5 | 19699.5831 | 0.219002095 | 1.24399121 | 0.17604794 | 0.86025628 | 0.93654703 |
| GPC5-AS1 | 74.5760128 | 0.870913048 | 0.62676522 | 1.38953634 | 0.16466972 | 0.36365838 |
| GPC6 | 4177.1307 | 0.287722595 | 1.1494887 | 0.25030485 | 0.80235161 | 0.90723013 |
| TGDS | 274.804988 | 0.528569095 | 0.18252519 | 2.89586924 | 0.0037811 | 0.0275025 |
| GPR180 | 296.818265 | 0.092125151 | 0.29476772 | 0.31253473 | 0.75463417 | 0.88110694 |
| ABCC4 | 591.245015 | -1.253584786 | 0.42402156 | -2.9564176 | 0.00311235 | 0.02376363 |
| CLDN10 | 4339.36755 | -0.320728183 | 0.41861513 | -0.7661648 | 0.44357824 | 0.65930291 |
| CLDN10-AS1 | 6061.76382 | -0.388259044 | 0.46592917 | -0.8333006 | 0.40467523 | 0.62539431 |
| DZIP1 | 2100.27829 | 0.523947152 | 0.3226467 | 1.62390364 | 0.10439638 | 0.27047815 |
| DNAJC3-DT | 224.366974 | -0.06338438 | 0.33746735 | -0.1878237 | 0.85101482 | 0.93170782 |
| DNAJC3 | 1266.72096 | 0.127015555 | 0.24661059 | 0.51504501 | 0.60652157 | 0.78260657 |
| UGGT2 | 2698.85835 | 0.075518369 | 0.24808953 | 0.30439966 | 0.76082343 | 0.88388009 |
| MBNL2 | 3807.20714 | 0.445627187 | 0.22698267 | 1.96326527 | 0.04961536 | 0.16537201 |
| RAP2A | 141.683349 | -0.255243073 | 0.28679775 | -0.8899759 | 0.37347885 | 0.59746664 |
| IPO5 | 519.76623 | -0.05894499 | 0.1899448 | -0.3103269 | 0.75631234 | 0.88220153 |
| FARP1 | 10572.457 | 0.381437477 | 0.25552894 | 1.49273692 | 0.13550606 | 0.32065561 |
| STK24 | 2571.52171 | -0.987880513 | 0.1926704 | -5.1273082 | 2.94E-07 | 1.50E-05 |
| SLC15A1 | 164.13156 | -3.368690394 | 0.71476507 | -4.7130037 | 2.44E-06 | 8.35E-05 |
| DOCK9 | 1621.67068 | -0.11132257 | 0.2279837 | -0.4882918 | 0.62534317 | 0.7961598 |
| UBAC2 | 2270.03947 | 0.051248658 | 0.15456639 | 0.33156405 | 0.74021847 | 0.87268348 |
| TM9SF2 | 1120.44848 | -0.139488468 | 0.13687558 | -1.0190895 | 0.30816047 | 0.53195133 |
| CLYBL | 1947.97992 | 0.40301138 | 0.28793755 | 1.39964858 | 0.16161858 | 0.35921571 |
| AL355338.1 | 181.514658 | 0.70332023 | 0.33131428 | 2.12281897 | 0.03376902 | 0.12799284 |
| PCCA | 9502.18095 | 0.031757487 | 0.22358485 | 0.14203774 | 0.8870502 | 0.94913293 |
| PCCA-AS1 | 115.551996 | -0.408929975 | 0.49082559 | -0.8331472 | 0.40476171 | 0.62539448 |
| GGACT | 268.04119 | -0.476635162 | 0.31807715 | -1.4984892 | 0.1340062 | 0.31830389 |
| TMTC4 | 332.67897 | -0.186186189 | 0.34785216 | -0.5352452 | 0.59248033 | 0.77370961 |
| FGF14 | 4288.37788 | 1.594595864 | 0.51113898 | 3.11969134 | 0.00181041 | 0.01580546 |
| TPP2 | 1129.58088 | 0.245217015 | 0.16995429 | 1.44284098 | 0.14906527 | 0.34111729 |
| TEX30 | 143.286233 | -0.269822281 | 0.31407376 | -0.8591048 | 0.39028269 | 0.61320085 |
| KDELC1 | 57.103656 | 0.481300068 | 0.33840054 | 1.42227926 | 0.1549452 | 0.34944017 |
| ARGLU1 | 1694.08175 | 0.021160322 | 0.1073217 | 0.19716723 | 0.84369668 | 0.92769172 |
| FAM155A | 3886.78623 | -1.934281681 | 0.7677426 | -2.5194404 | 0.01175416 | 0.06220378 |
| LIG4 | 213.667536 | 0.68805465 | 0.35018696 | 1.96482087 | 0.04943497 | 0.16493559 |
| ABHD13 | 279.422588 | 0.034397573 | 0.299247 | 0.11494709 | 0.90848706 | 0.95871259 |
| TNFSF13B | 368.797148 | -0.063257291 | 0.29343885 | -0.2155723 | 0.82932114 | 0.92023714 |
| MYO16 | 73.1928401 | 0.594860858 | 0.50062537 | 1.18823555 | 0.23474062 | 0.45204629 |
| IRS2 | 665.106675 | -0.347332276 | 0.36669141 | -0.9472059 | 0.34353386 | 0.56837065 |
| COL4A1 | 1690.22541 | 0.994328897 | 0.3564626 | 2.789434 | 0.00528003 | 0.03487193 |
| COL4A2 | 2650.09257 | 0.891484154 | 0.38546766 | 2.31273397 | 0.02073727 | 0.09136176 |
| RAB20 | 412.404564 | -0.087505128 | 0.40462832 | -0.2162605 | 0.82878469 | 0.92023714 |
| NAXD | 321.873693 | 0.077623303 | 0.1779163 | 0.43629114 | 0.66262551 | 0.82199665 |
| CARS2 | 645.982747 | 0.067101143 | 0.2011767 | 0.33354332 | 0.7387242 | 0.87211464 |
| ING1 | 73.8512999 | 0.010588661 | 0.34802507 | 0.03042499 | 0.97572811 | 0.9893257 |
| ANKRD10 | 3521.93076 | -0.385490359 | 0.21235293 | -1.8153286 | 0.06947342 | 0.20724355 |
| ARHGEF7 | 2217.81906 | 0.381654966 | 0.24711916 | 1.54441674 | 0.12248741 | 0.29989549 |

| | | | | | | | |
|---|---|---|---|---|---|---|---|
| TUBGCP3 | 734.552916 | | 0.333365611 | 0.18859074 | 1.76766688 | 0.07711661 | 0.22158653 |
| ATP11A | 2663.89115 | | -0.995798128 | 0.3337575 | -2.9835977 | 0.00284881 | 0.02227942 |
| MCF2L | 1249.32589 | | 0.819772458 | 0.28560138 | 2.87033786 | 0.00410033 | 0.02918174 |
| F10 | 48.9140346 | | 1.162476764 | 0.58023248 | 2.00346724 | 0.04512716 | 0.15494792 |
| PCID2 | 439.098524 | | -0.007082275 | 0.17441155 | -0.0406067 | 0.96760945 | 0.98528676 |
| CUL4A | 1270.19627 | | -0.496186137 | 0.2021238 | -2.4548625 | 0.01409385 | 0.07009249 |
| LAMP1 | 550.898313 | | -0.541325343 | 0.25301881 | -2.1394668 | 0.03239788 | 0.12432994 |
| GRTP1 | 359.825725 | | 0.139075069 | 0.23022256 | 0.60408966 | 0.54578404 | 0.74015676 |
| ADPRHL1 | 63.7943894 | | 0.473851881 | 0.33190311 | 1.42768133 | 0.1533836 | 0.34722262 |
| DCUN1D2 | 480.610941 | | 0.028439026 | 0.18680576 | 0.15223849 | 0.87899884 | 0.94598627 |
| TMCO3 | 687.03551 | | -0.895753696 | 0.24803753 | -3.6113636 | 0.00030459 | 0.00405989 |
| TFDP1 | 334.558783 | | -0.325087689 | 0.24418227 | -1.3313321 | 0.18307977 | 0.3878872 |
| GAS6-AS1 | 60.0186958 | | 0.821483982 | 0.63223638 | 1.29933046 | 0.19383055 | 0.40136505 |
| GAS6 | 570.845369 | | 1.018737526 | 0.37756644 | 2.69816756 | 0.00697223 | 0.0427305 |
| RASA3 | 374.858524 | | -2.069741171 | 0.60969123 | -3.3947367 | 0.00068695 | 0.00755527 |
| CDC16 | 737.205641 | | 0.225505524 | 0.2006275 | 1.12400106 | 0.26101258 | 0.48220092 |
| UPF3A | 940.684368 | | 0.395744853 | 0.15975173 | 2.4772492 | 0.01323994 | 0.06711401 |
| CHAMP1 | 151.062333 | | 0.270694853 | 0.24445817 | 1.10732586 | 0.26815307 | 0.48932276 |
| TTC5 | 209.908712 | | 0.266756192 | 0.21798017 | 1.22376356 | 0.22104144 | 0.43493763 |
| CCNB1IP1 | 492.951645 | | 1.384194666 | 0.26800256 | 5.1648562 | 2.41E-07 | 1.29E-05 |
| PARP2 | 98.8707967 | | 0.478907395 | 0.25870538 | 1.85116905 | 0.06414524 | 0.196476 |
| TEP1 | 299.982266 | | -1.166529296 | 0.2516014 | -4.6364182 | 3.54E-06 | 0.00011311 |
| OSGEP | 223.455515 | | 0.235259611 | 0.32184056 | 0.73098187 | 0.46479023 | 0.67778915 |
| APEX1 | 195.878731 | | -0.398273485 | 0.40695766 | -0.9786607 | 0.32774764 | 0.55299353 |
| PIP4P1 | 101.916148 | | -0.15760717 | 0.25988786 | -0.606443 | 0.54422063 | 0.73944091 |
| PNP | 207.706819 | | -1.372715002 | 0.37909957 | -3.620988 | 0.00029348 | 0.00395973 |
| METTL17 | 232.45905 | | 0.616709663 | 0.21994532 | 2.80392261 | 0.0050485 | 0.03364564 |
| NDRG2 | 385.071226 | | 1.145231803 | 0.43162137 | 2.65332509 | 0.00797031 | 0.04721066 |
| ARHGEF40 | 167.071174 | | 0.31934109 | 0.32576199 | 0.9802896 | 0.32694319 | 0.55240682 |
| ZNF219 | 99.3289263 | | 0.268531341 | 0.33556508 | 0.80023625 | 0.42357393 | 0.64243748 |
| HNRNPC | 3955.55116 | | 0.153212898 | 0.15532337 | 0.98641241 | 0.32393078 | 0.54890568 |
| SUPT16H | 553.684365 | | -0.071865237 | 0.16327049 | -0.4401606 | 0.6598208 | 0.81980614 |
| CHD8 | 981.597913 | | 0.009418311 | 0.14457664 | 0.06514407 | 0.94805929 | 0.97778285 |
| RAB2B | 382.494469 | | 0.23780028 | 0.2233612 | 1.06464452 | 0.28703682 | 0.5094007 |
| TOX4 | 458.215134 | | -0.2542689 | 0.11602082 | -2.19158 | 0.02840985 | 0.11309997 |
| METTL3 | 214.101417 | | 0.270091214 | 0.17151027 | 1.5747816 | 0.11530687 | 0.2886277 |
| DAD1 | 522.451339 | | -0.512490995 | 0.25624225 | -2.0000253 | 0.04549753 | 0.15579927 |
| ABHD4 | 219.033347 | | -0.140103324 | 0.25213679 | -0.5556639 | 0.57844061 | 0.76458087 |
| OXA1L | 220.675496 | | -0.154288779 | 0.20321356 | -0.7592445 | 0.4477063 | 0.66352217 |
| MRPL52 | 179.445413 | | -0.042701575 | 0.26655323 | -0.1601991 | 0.87272428 | 0.94258503 |
| MMP14 | 149.378118 | | -0.982803605 | 0.27231873 | -3.6090195 | 0.00030736 | 0.00408438 |
| LRP10 | 663.164924 | | -1.567547025 | 0.27241894 | -5.7541778 | 8.71E-09 | 8.10E-07 |
| RBM23 | 714.466629 | | 0.030746847 | 0.15523897 | 0.19806139 | 0.84299703 | 0.92739954 |
| PRMT5 | 127.611777 | | 0.050129325 | 0.22445228 | 0.22334068 | 0.82327037 | 0.91844587 |
| AL132780.1 | 345.417922 | | 0.2675016 | 0.53674138 | 0.49838081 | 0.61821566 | 0.79053759 |
| AL132780.3 | 42.3442842 | | 0.690372673 | 0.48273686 | 1.43012213 | 0.15268197 | 0.34607208 |
| AL132780.2 | 69.2306232 | | 1.25531316 | 0.53052231 | 2.36618356 | 0.01797253 | 0.08312451 |
| C14orf93 | 217.661444 | | 0.823769002 | 0.2349599 | 3.50599819 | 0.0004549 | 0.00550756 |
| PSMB5 | 326.462134 | | -0.780366949 | 0.29126129 | -2.6792676 | 0.00737834 | 0.04444162 |
| ACIN1 | 1259.71965 | | 0.184705803 | 0.15178921 | 1.21685728 | 0.22365852 | 0.43825922 |
| C14orf119 | 153.085611 | | -0.933060578 | 0.31463344 | -2.965548 | 0.00302144 | 0.02325771 |
| SLC7A8 | 320.409475 | | 0.587559602 | 0.31320656 | 1.87594922 | 0.06066225 | 0.18934019 |
| RNF212B | 77.4737185 | | 0.478574016 | 0.45470335 | 1.05249722 | 0.29257149 | 0.51542428 |
| HOMEZ | 123.189229 | | 0.082300322 | 0.2799446 | 0.29398789 | 0.76876716 | 0.88857795 |
| PPP1R3E | 106.627177 | | 0.381057828 | 0.52142642 | 0.73079885 | 0.46490203 | 0.67787735 |
| BCL2L2 | 42.1097689 | | 0.304200314 | 0.51923937 | 0.58585756 | 0.55797124 | 0.74999919 |
| BCL2L2-PABI | 268.322858 | | 0.052762179 | 0.18854604 | 0.27983712 | 0.77960247 | 0.8948704 |
| PABPN1 | 569.073113 | | 1.180354662 | 0.33017114 | 3.57497831 | 0.00035026 | 0.00450053 |
| NGDN | 209.825107 | | -0.0394891 | 0.23894438 | -0.1652648 | 0.86873555 | 0.94020545 |
| ZFHX2 | 326.023584 | | 0.967430812 | 0.35197105 | 2.74860907 | 0.00598487 | 0.03822736 |
| THTPA | 51.9912267 | | -0.054347268 | 0.29898627 | -0.1817718 | 0.85576183 | 0.93464491 |
| AP1G2 | 220.831237 | | -0.26696971 | 0.39059682 | -0.6834918 | 0.49429615 | 0.69960125 |
| AL135999.1 | 79.3545055 | | 0.450922754 | 0.33008063 | 1.36609879 | 0.17190796 | 0.37294355 |
| DHRS4-AS1 | 251.902907 | | 0.127364356 | 0.19867575 | 0.64106644 | 0.52147952 | 0.72143097 |
| DHRS4 | 92.8377811 | | 0.206929848 | 0.26732339 | 0.77408059 | 0.43888314 | 0.65485132 |
| DHRS4L2 | 173.649554 | | 0.533629458 | 0.29764529 | 1.7928369 | 0.07299901 | 0.21394362 |
| NRL | 127.388963 | | -0.086968621 | 0.39008257 | -0.2229493 | 0.823575 | 0.91846107 |
| PCK2 | 64.1777289 | | -0.283336412 | 0.31975579 | -0.8861025 | 0.37556229 | 0.59876913 |
| DCAF11 | 72.9070143 | | -0.320764602 | 0.34427901 | -0.9316995 | 0.35149182 | 0.57736509 |
| PSME1 | 283.772357 | | -1.54476421 | 0.44479911 | -3.4729481 | 0.00051477 | 0.00605491 |
| EMC9 | 88.3723954 | | -0.140022658 | 0.37424002 | -0.374152 | 0.70829122 | 0.85346928 |
| PSME2 | 81.3805608 | | -0.131075851 | 0.38479754 | -0.3406359 | 0.73337771 | 0.86911141 |
| REC8 | 68.1383085 | | -0.798385594 | 0.34778394 | -2.2956367 | 0.02169666 | 0.09440958 |
| IPO4 | 45.4076698 | | -0.869292952 | 0.45145616 | -1.9255313 | 0.05416293 | 0.17552456 |
| AL136295.1 | 236.329337 | | -0.222190828 | 0.25719912 | -0.8638864 | 0.38765028 | 0.61088911 |
| GMPR2 | 136.024315 | | -0.444560136 | 0.32391544 | -1.3724574 | 0.1699211 | 0.37016988 |
| TINF2 | 79.8868059 | | -0.857346152 | 0.28731777 | -2.983965 | 0.00284539 | 0.02227907 |
| RABGGTA | 47.9662638 | | 0.61485215 | 0.36748637 | 1.67312913 | 0.09430188 | 0.25271526 |
| DHRS1 | 137.508814 | | -0.402018643 | 0.32356514 | -1.2424659 | 0.21406472 | 0.42754769 |
| LTB4R | 59.662209 | | -0.757481418 | 0.6438334 | -1.1765177 | 0.23938805 | 0.45750019 |
| NFATC4 | 189.058814 | | 0.714468672 | 0.49984182 | 1.42938955 | 0.1528923 | 0.34644783 |
| NYNRIN | 97.3742596 | | 1.206027152 | 0.5834786 | 2.06696038 | 0.03873789 | 0.13969738 |
| KHNYN | 191.025642 | | -0.591750794 | 0.24118905 | -2.4534729 | 0.01414842 | 0.07031095 |
| SDR39U1 | 223.710837 | | 0.081760756 | 0.2654492 | 0.30800905 | 0.75807545 | 0.88315785 |
| NOVA1 | 1100.023 | | -0.047813882 | 0.34525544 | -0.1384884 | 0.88985442 | 0.9503707 |
| AL110292.1 | 159.60113 | | 0.597466156 | 0.70870995 | 0.84303339 | 0.39920977 | 0.6208833 |
| LINC00645 | 1055.02258 | | 1.505924992 | 0.27618035 | 5.45268687 | 4.96E-08 | 3.63E-06 |
| PRKD1 | 22141.5827 | | -0.148142809 | 0.38009995 | -0.389747 | 0.69672365 | 0.84496018 |
| AL133372.3 | 41.3575574 | | 1.51040889 | 0.56647336 | 2.666337 | 0.00766828 | 0.04581198 |
| G2E3-AS1 | 70.4254058 | | 0.217841327 | 0.57532845 | 0.3786382 | 0.70495655 | 0.85073346 |
| G2E3 | 458.658855 | | 0.010014841 | 0.17778078 | 0.05633253 | 0.95507691 | 0.98014107 |
| SCFD1 | 1743.41715 | | 0.193262834 | 0.17184945 | 1.12460548 | 0.26075625 | 0.48199403 |
| COCH | 129.156714 | | -0.731970344 | 0.39551907 | -1.8506575 | 0.06421884 | 0.19658555 |
| STRN3 | 3609.71048 | | -0.491970473 | 0.26154631 | -1.8810071 | 0.05997095 | 0.18784831 |
| AP4S1 | 267.289632 | | 0.16390341 | 0.22651299 | 0.72359386 | 0.46931511 | 0.68108245 |
| HECTD1 | 3005.46106 | | 0.200879658 | 0.22146272 | 0.90705856 | 0.36437583 | 0.5897611 |
| AL136418.1 | 62.0123343 | | -0.639642979 | 0.40649457 | -1.5735585 | 0.11558956 | 0.28922571 |
| HEATR5A | 488.242355 | | -0.003385318 | 0.21342895 | -0.0158616 | 0.98734483 | 0.99525625 |
| AL139353.1 | 152.018859 | | 0.360359875 | 0.24206571 | 1.48868618 | 0.13657003 | 0.32222841 |
| AL139353.2 | 52.1226321 | | 0.394133296 | 0.44710193 | 0.88152896 | 0.37803159 | 0.60082051 |
| NUBPL | 1602.26541 | | 0.334895251 | 0.24960194 | 1.34171733 | 0.17968767 | 0.38325438 |
| ARHGAP5 | 2515.64517 | | 0.077194572 | 0.14565194 | 0.52999345 | 0.59611647 | 0.776003 |
| AKAP6 | 1381.54121 | | 0.935579723 | 0.3315468 | 2.82186325 | 0.00477455 | 0.03225937 |
| NPAS3 | 2109.01319 | | 0.140125382 | 0.42032007 | 0.3333778 | 0.73884912 | 0.87211464 |
| EGLN3 | 389.21125 | | 0.023614122 | 0.60602424 | 0.03896564 | 0.96891778 | 0.98586681 |
| EAPP | 524.189022 | | -0.231261746 | 0.1764019 | -1.3109935 | 0.18985995 | 0.39599874 |
| SNX6 | 1167.6765 | | -0.041517282 | 0.17586095 | -0.2360802 | 0.81337046 | 0.91245411 |
| CFL2 | 112.207405 | | 0.105668685 | 0.37289622 | 0.28337291 | 0.77689099 | 0.89337196 |
| BAZ1A | 651.930596 | | -1.074733245 | 0.27330128 | -3.932412 | 8.41E-05 | 0.00147429 |
| SRP54 | 975.800243 | | -0.200388996 | 0.16820556 | -1.1913339 | 0.23352252 | 0.45043793 |
| FAM177A1 | 579.059426 | | -0.142079634 | 0.18072793 | -0.7861521 | 0.43177841 | 0.64957432 |
| PPP2R3C | 398.206929 | | -0.445666963 | 0.21626207 | -2.0607726 | 0.03932474 | 0.14126492 |
| NFKBIA | 475.068328 | | -1.158997509 | 0.51166252 | -2.2651601 | 0.02350286 | 0.09943922 |

| | | | | | | |
|---|---|---|---|---|---|---|
| RALGAPA1 | 3538.96534 | 0.227567021 | 0.15310205 | 1.4863748 | 0.13718001 | 0.32320435 |
| BRMS1L | 252.800028 | 0.442370362 | 0.325089 | 1.36076693 | 0.17358735 | 0.37534823 |
| PTCSC3 | 41.732299 | 3.681772343 | 0.54996688 | 6.6945347 | 2.16E-11 | 5.10E-09 |
| MBIP | 436.819247 | 0.500098567 | 0.19809952 | 2.52448144 | 0.01158691 | 0.06146648 |
| SLC25A21 | 216.524907 | 0.249181258 | 0.38287612 | 0.6508143 | 0.51516637 | 0.71652443 |
| MIPOL1 | 1518.54564 | 0.32832138 | 0.21701773 | 1.51287816 | 0.13031061 | 0.31253761 |
| SEC23A | 1023.13573 | -0.03802417 | 0.199232 | -0.1908537 | 0.8486402 | 0.9303692 |
| GEMIN2 | 128.565884 | 0.403100216 | 0.1925549 | 2.09343001 | 0.03631079 | 0.13408699 |
| TRAPPC6B | 694.687618 | 0.453336347 | 0.18907397 | 2.39766659 | 0.01649988 | 0.07830877 |
| PNN | 1137.89834 | 0.2519887 | 0.17603462 | 1.43147237 | 0.15229488 | 0.34552784 |
| MIA2 | 2248.10404 | 0.065454229 | 0.20004656 | 0.32719497 | 0.74352043 | 0.87467216 |
| FBXO33 | 804.111577 | -0.186551271 | 0.18754564 | -0.994698 | 0.31988318 | 0.54436188 |
| AL049875.1 | 380.086724 | 0.222430694 | 0.31302831 | 0.71057693 | 0.47734644 | 0.68707302 |
| LRFN5 | 259.282165 | 1.047408651 | 0.78182356 | 1.33969953 | 0.18034305 | 0.38409406 |
| C14orf28 | 71.343524 | 0.293700671 | 0.22758083 | 1.29053345 | 0.19686551 | 0.40510928 |
| KLHL28 | 746.076394 | 0.204809523 | 0.24126263 | 0.84890694 | 0.39593308 | 0.61804302 |
| TOGARAM1 | 1234.70351 | 0.216937175 | 0.15332247 | 1.41490787 | 0.15709551 | 0.35272388 |
| PRPF39 | 519.355219 | 0.446671681 | 0.22642716 | 1.97269481 | 0.04853034 | 0.16294797 |
| FKBP3 | 397.954018 | 0.509840853 | 0.21826708 | 2.33585774 | 0.01949866 | 0.08778383 |
| FANCM | 338.337678 | 0.43421731 | 0.38422294 | 1.13011813 | 0.25842645 | 0.47954387 |
| MIS18BP1 | 428.056043 | -0.253013323 | 0.27629689 | -0.9157299 | 0.35980859 | 0.58502113 |
| LINC00871 | 173.218787 | 0.771698719 | 0.42100658 | 1.83298494 | 0.06680481 | 0.20183549 |
| AL358335.2 | 112.43914 | 3.101233418 | 0.78893107 | 3.93093079 | 8.46E-05 | 0.00147846 |
| AL110505.1 | 35.3210152 | 2.957239024 | 1.0337036 | 2.86081911 | 0.00422548 | 0.02965695 |
| RPS29 | 757.359446 | 0.080272802 | 0.24847759 | 0.32305852 | 0.74665093 | 0.87635381 |
| LRR1 | 66.3868382 | -0.059566577 | 0.3411925 | -0.1745835 | 0.86140692 | 0.93697711 |
| RPL36AL | 702.112525 | -1.516924859 | 0.39655344 | -3.8252722 | 0.00013063 | 0.00207887 |
| MGAT2 | 48.8302026 | -0.185923376 | 0.42978027 | -0.432601 | 0.66530466 | 0.82338698 |
| DNAAF2 | 172.001562 | 0.458878126 | 0.24901469 | 1.84277531 | 0.06536182 | 0.19888605 |
| POLE2 | 155.947792 | -0.459827413 | 0.3353795 | -1.371066 | 0.17035439 | 0.37064919 |
| KLHDC1 | 597.995477 | 0.474524748 | 0.20324112 | 2.33478716 | 0.01955454 | 0.08796159 |
| KLHDC2 | 719.618266 | 0.442481002 | 0.22850812 | 1.93639073 | 0.05281986 | 0.17244132 |
| NEMF | 2096.29551 | 0.01621218 | 0.16102417 | 0.10068165 | 0.91980318 | 0.96447683 |
| ARF6 | 189.936572 | -0.854568467 | 0.31791925 | -2.6880048 | 0.00718804 | 0.0435336 |
| LINC01588 | 103.552918 | 0.261486885 | 0.35536954 | 0.73581681 | 0.46184219 | 0.67513899 |
| VCPKMT | 252.792522 | 0.630810211 | 0.1933039 | 3.26330821 | 0.0011012 | 0.0107746 |
| SOS2 | 2185.80017 | 0.183932323 | 0.13818133 | 1.3310939 | 0.18315812 | 0.38796273 |
| L2HGDH | 775.782498 | 0.500315745 | 0.18983791 | 2.63548915 | 0.00840161 | 0.04886697 |
| ATP5S | 443.247945 | 0.385570765 | 0.17506999 | 2.20238068 | 0.02763843 | 0.1109082 |
| AL359397.2 | 73.3652937 | 1.179543229 | 0.34230226 | 3.44591133 | 0.00056914 | 0.00650886 |
| CDKL1 | 1836.00425 | 1.503001166 | 0.3418485 | 4.39668787 | 1.10E-05 | 0.00028807 |
| MAP4K5 | 1713.78793 | -0.116533872 | 0.09498972 | -1.2268051 | 0.21989587 | 0.43371253 |
| ATL1 | 178.437658 | 0.515406869 | 0.45400403 | 1.13524736 | 0.25627169 | 0.47722602 |
| SAV1 | 580.608444 | -0.08225299 | 0.28225712 | -0.2914116 | 0.77073657 | 0.88937357 |
| NIN | 602.248487 | 0.041527497 | 0.16287773 | 0.25496118 | 0.79875308 | 0.90525349 |
| PYGL | 241.960066 | -0.895739033 | 0.42303684 | -2.1174019 | 0.03422575 | 0.12911907 |
| AL358334.2 | 38.8902777 | 2.947567619 | 0.70226965 | 4.19720203 | 2.70E-05 | 0.00060398 |
| TRIM9 | 111.310922 | -1.290685102 | 0.61639461 | -2.0939266 | 0.03626652 | 0.13396096 |
| TMX1 | 285.695787 | 0.189369387 | 0.25353858 | 0.7469056 | 0.45512054 | 0.67027397 |
| FRMD6 | 67.7037001 | -1.993025323 | 0.65145541 | -3.0593427 | 0.00221823 | 0.01856273 |
| GNG2 | 111.788461 | -1.121422496 | 0.80635306 | -1.3907339 | 0.16430615 | 0.36306425 |
| RTRAF | 311.247425 | -0.354589393 | 0.31484253 | -1.1262436 | 0.2600624 | 0.48104792 |
| TXNDC16 | 338.468583 | 0.428274922 | 0.22602162 | 1.8948405 | 0.05811354 | 0.18409112 |
| GPR137C | 92.3218454 | 0.084936406 | 0.3530463 | 0.24058149 | 0.80987949 | 0.91064395 |
| ERO1A | 893.710524 | -2.025277359 | 0.61617963 | -3.2868294 | 0.00101322 | 0.01008564 |
| AL133453.1 | 83.494887 | -2.550696783 | 0.78706463 | -3.2407717 | 0.00119207 | 0.01145997 |
| PSMC6 | 586.087504 | -0.060580315 | 0.12954679 | -0.4676327 | 0.64004727 | 0.80648521 |
| STYX | 293.72253 | 0.316746188 | 0.18037897 | 1.75600398 | 0.07908773 | 0.22475685 |
| GNPNAT1 | 207.558632 | -0.053080491 | 0.23378987 | -0.2270436 | 0.82038985 | 0.91661089 |
| FERMT2 | 1585.50755 | 0.52304469 | 0.30401322 | 1.72046692 | 0.0853476 | 0.23604616 |
| DDHD1 | 484.84909 | 0.103424221 | 0.18971535 | 0.54515472 | 0.58564711 | 0.77001836 |
| LINC02331 | 39.3492839 | 2.504136106 | 0.5762686 | 4.34543217 | 1.39E-05 | 0.00034973 |
| CNIH1 | 235.605726 | -0.006471163 | 0.24358645 | -0.0265662 | 0.97880574 | 0.99083348 |
| GMFB | 241.621972 | -0.471624649 | 0.18775155 | -2.5119614 | 0.01200622 | 0.06290766 |
| CGRRF1 | 192.79728 | 0.40771193 | 0.32680075 | 1.24758567 | 0.21218283 | 0.42488177 |
| SAMD4A | 4725.51627 | -1.271425906 | 0.39407038 | -3.226393 | 0.00125361 | 0.01193868 |
| GCH1 | 130.361688 | -0.820528645 | 0.39180081 | -2.0942495 | 0.03623776 | 0.13396096 |
| WDHD1 | 103.961685 | -0.063576014 | 0.49199413 | -0.1292211 | 0.89718271 | 0.9525128 |
| SOCS4 | 259.245818 | -0.106623804 | 0.23929229 | -0.4455798 | 0.65590083 | 0.81726196 |
| MAPK1IP1L | 1077.32016 | -0.322089744 | 0.15386757 | -2.093292 | 0.0363231 | 0.13409497 |
| LGALS3 | 634.34682 | -2.416549934 | 0.49898526 | -4.8429285 | 1.28E-06 | 5.01E-05 |
| FBXO34 | 2292.57404 | -0.074064009 | 0.22500318 | -0.3291687 | 0.74202817 | 0.87402695 |
| AL158801.3 | 145.030525 | -0.005622884 | 0.45239859 | -0.012429 | 0.99008331 | 0.99606252 |
| ATG14 | 556.549746 | 0.303413642 | 0.32148907 | 0.94377593 | 0.34528416 | 0.57046385 |
| KTN1-AS1 | 107.729647 | 0.522869017 | 0.34556554 | 1.51308205 | 0.13025882 | 0.31253761 |
| KTN1 | 2416.05654 | -0.002431248 | 0.16305702 | -0.0149104 | 0.98810365 | 0.99552604 |
| PELI2 | 3222.59911 | 0.051723811 | 0.17211111 | 0.30052569 | 0.7637762 | 0.88551917 |
| LINC02284 | 71.4004865 | -0.372206029 | 0.51016792 | -0.7295755 | 0.46564968 | 0.67859297 |
| TMEM260 | 1145.00246 | 0.29606463 | 0.11625461 | 2.54669153 | 0.01087495 | 0.05889596 |
| EXOC5 | 748.931151 | -0.07083284 | 0.11654831 | -0.6077552 | 0.54334985 | 0.7391242 |
| AP5M1 | 730.502523 | 0.666835199 | 0.24652651 | 2.70492284 | 0.00683203 | 0.04213082 |
| NAA30 | 272.876174 | 0.082486743 | 0.18116176 | 0.45532094 | 0.64887836 | 0.81203546 |
| CCDC198 | 2689.4997 | 0.58551133 | 0.36799972 | 1.59106463 | 0.11159503 | 0.28227859 |
| ARMH4 | 1046.8436 | 0.821371986 | 0.23443964 | 3.50355422 | 0.00045909 | 0.00555326 |
| ACTR10 | 771.088393 | -0.080016211 | 0.15148159 | -0.528224 | 0.59734387 | 0.77659469 |
| PSMA3 | 654.884188 | 0.049249724 | 0.23798893 | 0.20694124 | 0.83605575 | 0.92369276 |
| ARID4A | 1255.90778 | -0.08192376 | 0.15665815 | -0.522946 | 0.6010118 | 0.7789212 |
| AL139021.2 | 56.8710823 | -0.194806882 | 0.46194978 | -0.4217058 | 0.6732398 | 0.82881006 |
| AL139021.1 | 167.416005 | 0.323248394 | 0.36171718 | 0.89364955 | 0.37150942 | 0.59563048 |
| TIMM9 | 256.69421 | 0.126933337 | 0.18419868 | 0.68911099 | 0.49075342 | 0.6969611 |
| KIAA0586 | 821.859645 | 0.046356748 | 0.13527018 | 0.34269746 | 0.73182608 | 0.86826096 |
| DAAM1 | 1731.81469 | 0.218698798 | 0.22669379 | 0.9647322 | 0.33467896 | 0.55980428 |
| GPR135 | 231.316761 | -0.027792694 | 0.50524181 | -0.0550087 | 0.95613153 | 0.98070674 |
| L3HYPDH | 201.318218 | -0.004258942 | 0.3248806 | -0.0131093 | 0.98954063 | 0.99606252 |
| JKAMP | 429.605625 | 0.442919611 | 0.18939445 | 2.33860924 | 0.01935567 | 0.08719952 |
| CCDC175 | 43.2875696 | -0.164210597 | 0.79750428 | -0.2059056 | 0.83686465 | 0.9240982 |
| RTN1 | 85.2551062 | 0.536375803 | 0.41790674 | 1.28348205 | 0.19932324 | 0.40794248 |
| LRRC9 | 73.5156391 | 1.263112522 | 0.50406435 | 2.50585572 | 0.01221555 | 0.06372636 |
| AL157911.1 | 406.201399 | 1.815307965 | 0.4293093 | 4.22843853 | 2.35E-05 | 0.00054152 |
| PCNX4 | 1660.1409 | 0.056560363 | 0.2725492 | 0.2075235 | 0.83560104 | 0.92348178 |
| DHRS7 | 193.653377 | -0.683136609 | 0.31901537 | -2.1413909 | 0.03224253 | 0.12393503 |
| PPM1A | 898.103682 | 0.186552614 | 0.26752371 | 0.69733115 | 0.48559558 | 0.69305586 |
| SIX4 | 121.771666 | 0.046215385 | 0.36862051 | 0.12537388 | 0.90022756 | 0.95395051 |
| MNAT1 | 3310.93093 | 0.486504302 | 0.1767855 | 2.75194689 | 0.00592421 | 0.03798686 |
| TRMT5 | 103.140673 | 0.577970361 | 0.28476582 | 2.02963388 | 0.04239377 | 0.14794697 |
| SLC38A6 | 605.784069 | 0.338234563 | 0.13817494 | 2.44787199 | 0.01437027 | 0.07110485 |
| PRKCH | 447.168509 | -0.503577193 | 0.29602802 | -1.7011132 | 0.08892172 | 0.24262901 |
| TMEM30B | 151.299523 | -0.998327878 | 0.42406998 | -2.3541583 | 0.0185647 | 0.08485161 |
| AL359220.1 | 54.2547304 | -1.374575081 | 0.69588835 | -1.9752811 | 0.04823626 | 0.16208414 |
| HIF1A-AS1 | 46.4843392 | -0.996494241 | 0.42385593 | -2.3510211 | 0.01872197 | 0.08524595 |
| HIF1A | 13074.5635 | -1.75177389 | 0.36707529 | -4.7722469 | 1.82E-06 | 6.54E-05 |
| HIF1A-AS2 | 2515.76921 | -2.011873225 | 0.595991 | -3.3756772 | 0.00073634 | 0.00793988 |
| SNAPC1 | 896.476226 | -0.938813271 | 0.40252358 | -2.3323187 | 0.01968393 | 0.08831693 |

| | | | | | | |
|---|---|---|---|---|---|---|
| SYT16 | 300.675476 | -2.11974571 | 0.68217211 | -3.1073474 | 0.00188774 | 0.01636169 |
| PPP2R5E | 2222.33762 | -0.269695398 | 0.23397231 | -1.1526808 | 0.24904141 | 0.46836355 |
| WDR89 | 288.223515 | 0.296504532 | 0.15412273 | 1.92382097 | 0.05437703 | 0.1757642 |
| SGPP1 | 551.347334 | 0.345632498 | 0.21793285 | 1.58595868 | 0.11274868 | 0.2842713 |
| SYNE2 | 16886.4949 | -0.714522639 | 0.23037183 | -3.101606 | 0.00192474 | 0.01663053 |
| ESR2 | 1367.69822 | -0.536668814 | 0.56243254 | -0.9541923 | 0.3399863 | 0.56531709 |
| MTHFD1 | 286.198257 | 0.063198488 | 0.20551607 | 0.30751118 | 0.75845432 | 0.88316847 |
| AL122035.1 | 198.138381 | -0.730477148 | 0.38624095 | -1.8912473 | 0.05859134 | 0.18475365 |
| ZBTB25 | 439.527743 | -0.236845758 | 0.28654676 | -0.8265519 | 0.40849111 | 0.62829053 |
| ZBTB1 | 1204.40067 | 0.033157782 | 0.20816797 | 0.15928379 | 0.8734453 | 0.94320952 |
| AL049869.3 | 176.776193 | 0.262057054 | 0.59495634 | 0.44046434 | 0.65960083 | 0.81980614 |
| HSPA2 | 286.386077 | 0.645691025 | 0.72256002 | 0.89361576 | 0.37152751 | 0.59563048 |
| PPP1R36 | 80.9693064 | 0.970871761 | 0.56705165 | 1.71213992 | 0.08687089 | 0.2387097 |
| PLEKHG3 | 759.538175 | 0.153499163 | 0.41989579 | 0.3655649 | 0.71468977 | 0.85752192 |
| SPTB | 55.7248553 | 0.106511689 | 0.35266861 | 0.30201636 | 0.76263959 | 0.88490193 |
| CHURC1-FNT | 307.048414 | -0.772480339 | 0.32720638 | -2.3608352 | 0.01823383 | 0.08397938 |
| RAB15 | 131.964844 | -1.083982285 | 0.30312439 | -3.5760312 | 0.00034885 | 0.00449816 |
| FNTB | 44.2697438 | 0.232623669 | 0.34286311 | 0.6784739 | 0.49747127 | 0.70256581 |
| MAX | 736.420618 | -0.40985111 | 0.22593619 | -1.8140127 | 0.06967578 | 0.20761275 |
| AL139022.1 | 47.8497897 | -0.760516072 | 0.63214648 | -1.2030694 | 0.22894948 | 0.44451188 |
| FUT8 | 758.059536 | -0.479725169 | 0.16147728 | -2.9708523 | 0.00296975 | 0.0229803 |
| AL359232.1 | 600.837779 | 0.09301638 | 0.49026622 | 0.18972627 | 0.84952364 | 0.93092398 |
| GPHN | 6339.78913 | 0.262942015 | 0.15302277 | 1.71831956 | 0.08573834 | 0.23673031 |
| MPP5 | 960.855412 | -0.355416763 | 0.15744562 | -2.2573938 | 0.02398348 | 0.10079472 |
| ATP6V1D | 630.802249 | 0.268562751 | 0.31027399 | 0.86556644 | 0.38672797 | 0.61038233 |
| EIF2S1 | 369.774678 | -0.104126752 | 0.17489038 | -0.5953829 | 0.55158752 | 0.74467698 |
| PLEKHH1 | 364.394794 | -0.458182067 | 0.26957829 | -1.6996252 | 0.08920144 | 0.24300965 |
| PIGH | 286.575344 | -0.100734925 | 0.35282391 | -0.2855105 | 0.77525307 | 0.89223749 |
| ARG2 | 895.180813 | -0.516362905 | 0.87972127 | -0.5869619 | 0.55722927 | 0.7492306 |
| VTI1B | 212.035025 | -0.283063327 | 0.3164201 | -0.8945807 | 0.37101125 | 0.59561104 |
| RDH11 | 81.6747918 | -0.590086111 | 0.31640219 | -1.8649874 | 0.06218316 | 0.19265041 |
| ZFYVE26 | 354.738156 | 0.137648463 | 0.29566448 | 0.46555631 | 0.64153312 | 0.80758706 |
| RAD51B | 4178.96407 | 0.383176144 | 0.37162714 | 1.03107686 | 0.30250478 | 0.52576127 |
| ZFP36L1 | 2297.18055 | -0.894638028 | 0.28403049 | -3.1497958 | 0.00163385 | 0.0146713 |
| ACTN1 | 531.039331 | -1.489015102 | 0.26969103 | -5.5211888 | 3.37E-08 | 2.57E-06 |
| DCAF5 | 1910.81442 | -0.061689905 | 0.147923 | -0.4170407 | 0.67664866 | 0.83026959 |
| AL359317.2 | 176.007854 | 0.287872411 | 0.47848775 | 0.60162964 | 0.5474207 | 0.74147662 |
| EXD2 | 408.349776 | 0.431101564 | 0.20708221 | 2.08178943 | 0.03736171 | 0.13659308 |
| AL359317.1 | 46.0212774 | 0.166924305 | 0.57775436 | 0.28891916 | 0.77264324 | 0.89062065 |
| GALNT16 | 471.32013 | 0.073360213 | 0.78970801 | 0.09289536 | 0.92598669 | 0.96724667 |
| ERH | 389.027224 | -0.445804383 | 0.25566714 | -1.7436906 | 0.08121305 | 0.22874648 |
| SLC39A9 | 787.870116 | 0.092276586 | 0.14657244 | 0.62956301 | 0.52898053 | 0.72723811 |
| PLEKHD1 | 44.0754096 | 0.374025356 | 0.8567535 | 0.43656122 | 0.66242959 | 0.82190799 |
| SUSD6 | 1516.73459 | -0.540825131 | 0.20251753 | -2.6705102 | 0.00757361 | 0.04536951 |
| SRSF5 | 2029.17999 | 0.693667682 | 0.23998642 | 2.89044558 | 0.00384696 | 0.02787412 |
| AL357153.2 | 419.588986 | -0.322834371 | 0.40235367 | -0.8023647 | 0.42234204 | 0.64115803 |
| MED6 | 351.523066 | 0.071639476 | 0.25088256 | 0.28554984 | 0.77522292 | 0.89223749 |
| MAP3K9 | 481.785785 | -0.025887131 | 0.19122623 | -0.1353744 | 0.89231588 | 0.95118543 |
| AC004816.1 | 41.0047753 | 0.638046336 | 0.45000917 | 1.41785186 | 0.15623402 | 0.35131349 |
| PCNX1 | 2629.19877 | 0.428175351 | 0.22023689 | 1.94415818 | 0.05187638 | 0.17045649 |
| AC005476.2 | 40.2016091 | 0.677597 | 0.48339611 | 1.40174277 | 0.16099207 | 0.35824533 |
| SIPA1L1 | 3729.28396 | -0.353355671 | 0.26418245 | -1.3375441 | 0.18104511 | 0.38492165 |
| RGS6 | 3070.80805 | -2.258897974 | 0.76668393 | -2.9463223 | 0.00321577 | 0.02434219 |
| DPF3 | 98.3522997 | -0.011084609 | 0.37460037 | -0.0295905 | 0.97639365 | 0.98947549 |
| DCAF4 | 115.379405 | -0.50606535 | 0.23036265 | -2.1968203 | 0.02803328 | 0.11203983 |
| ZFYVE1 | 436.044434 | -0.220831297 | 0.20055842 | -1.1010821 | 0.27086091 | 0.49226771 |
| RBM25 | 1481.1116 | 0.299471731 | 0.1853476 | 1.61573032 | 0.10615263 | 0.2733808 |
| PSEN1 | 1120.88573 | -0.489679656 | 0.19907611 | -2.4597611 | 0.01390295 | 0.06948321 |
| PAPLN | 430.392929 | -1.305515264 | 0.749772 | -1.7412164 | 0.08164566 | 0.22960561 |
| NUMB | 3341.12781 | -0.289810383 | 0.1506634 | -1.9235619 | 0.05440952 | 0.1757642 |
| HEATR4 | 153.140725 | -0.680785935 | 0.41483113 | -1.6411158 | 0.10077337 | 0.26447755 |
| AC005225.3 | 49.0189513 | 0.014080131 | 0.47597758 | 0.0295815 | 0.97640082 | 0.98947549 |
| ACOT2 | 50.1095288 | 0.320819855 | 0.55745739 | 0.57550561 | 0.5649494 | 0.75605085 |
| AC005225.2 | 60.228348 | 1.007913752 | 0.53763094 | 1.87473166 | 0.06082965 | 0.18977299 |
| DNAL1 | 385.548144 | -0.045536138 | 0.1985717 | -0.2293184 | 0.81862148 | 0.91559451 |
| PNMA1 | 83.634893 | -0.869667298 | 0.51825738 | -1.6780606 | 0.09333527 | 0.25117608 |
| ELMSAN1 | 521.254654 | -0.503626132 | 0.27052649 | -1.8616518 | 0.06265219 | 0.19370578 |
| AC005520.2 | 72.5292414 | 0.539434689 | 0.86907056 | 0.62070299 | 0.53479506 | 0.73203192 |
| AC005480.1 | 313.314211 | 0.150740109 | 0.36376466 | 0.4143891 | 0.67858917 | 0.8314278 |
| COQ6 | 160.989011 | 0.523217378 | 0.19078749 | 2.7424092 | 0.00609903 | 0.03878772 |
| ENTPD5 | 508.0951 | 0.312890694 | 0.25375158 | 1.23305907 | 0.21755373 | 0.43128814 |
| BBOF1 | 428.125309 | -0.036811127 | 0.17470916 | -0.2106995 | 0.83312179 | 0.92205326 |
| ALDH6A1 | 371.066823 | 0.062201099 | 0.30769665 | 0.20215072 | 0.8397989 | 0.92558662 |
| LIN52 | 746.431295 | 0.490852088 | 0.21839752 | 2.24751674 | 0.02460702 | 0.10259482 |
| ABCD4 | 643.022049 | -0.172097129 | 0.17519905 | -0.9822949 | 0.32595461 | 0.55121424 |
| NPC2 | 341.193624 | -1.078365964 | 0.37713406 | -2.8593704 | 0.00424483 | 0.02977692 |
| ISCA2 | 123.363938 | -0.439730014 | 0.43697943 | -1.0062945 | 0.31427391 | 0.53828372 |
| LTBP2 | 287.986632 | 1.510751228 | 0.64709029 | 2.33468381 | 0.01955994 | 0.08796159 |
| AREL1 | 543.497873 | 0.474391348 | 0.19138319 | 2.47875145 | 0.01318431 | 0.06700716 |
| FCF1 | 244.790164 | 0.044154838 | 0.19657818 | 0.22461719 | 0.82227709 | 0.91802136 |
| YLPM1 | 1282.80555 | 0.30742719 | 0.14313351 | 2.14783514 | 0.03172685 | 0.12257932 |
| DLST | 545.981795 | 0.255182544 | 0.20321981 | 1.25569717 | 0.20922578 | 0.42077701 |
| RPS6KL1 | 45.6967642 | -0.479314065 | 0.66785376 | -0.7176931 | 0.47294651 | 0.68364527 |
| EIF2B2 | 133.909404 | -0.210854202 | 0.25375309 | -0.8309424 | 0.40600617 | 0.62607231 |
| AL049780.1 | 40.9033503 | 0.798173794 | 0.36210944 | 2.20423361 | 0.02750792 | 0.11047495 |
| MLH3 | 897.923902 | 0.176438622 | 0.16877724 | 1.04539346 | 0.29584116 | 0.51903949 |
| ACYP1 | 579.772158 | 1.015526858 | 0.34477109 | 2.94551046 | 0.00322422 | 0.02437541 |
| ZC2HC1C | 205.394295 | 0.907274458 | 0.54694249 | 1.65881144 | 0.09715379 | 0.25753652 |
| NEK9 | 724.943439 | 0.160017317 | 0.11286904 | 1.41772547 | 0.15627093 | 0.35131349 |
| AL049780.2 | 119.204251 | 0.30927731 | 0.39162484 | 0.78972853 | 0.42968633 | 0.64784005 |
| TMED10 | 817.562723 | -0.459092942 | 0.22945302 | -2.0008145 | 0.04541238 | 0.15560366 |
| FOS | 233.436859 | -0.834024096 | 0.73994858 | -1.1271379 | 0.25968418 | 0.48061767 |
| JDP2 | 184.0059 | 1.351839689 | 0.35338074 | 3.82544814 | 0.00013053 | 0.00207887 |
| FLVCR2 | 58.013952 | -0.576925799 | 0.73524926 | -0.784667 | 0.43264886 | 0.65015341 |
| TTLL5 | 1592.6864 | -0.056572992 | 0.212484 | -0.2662459 | 0.79004984 | 0.90082888 |
| ERG28 | 123.358628 | -0.218449403 | 0.30086705 | -0.7260662 | 0.46779817 | 0.6801328 |
| AF107885.2 | 78.7320535 | -0.079359779 | 0.4850339 | -0.163617 | 0.87003268 | 0.94075489 |
| IFT43 | 1017.04502 | -0.450113019 | 0.14776211 | -3.0462006 | 0.00231753 | 0.01913267 |
| GPATCH2L | 2276.22393 | 0.011191468 | 0.221332 | 0.05056416 | 0.95967282 | 0.98229715 |
| AC016526.1 | 305.853362 | 1.778611657 | 0.69411926 | 2.56240065 | 0.01039513 | 0.05704583 |
| ESRRB | 6085.30994 | 1.381742814 | 0.37290593 | 3.70533882 | 0.00021111 | 0.00306769 |
| AC008050.1 | 43.5093016 | 0.667316916 | 0.61132524 | 1.09159065 | 0.27501307 | 0.49647832 |
| VASH1-AS1 | 54.9312823 | 0.166291253 | 0.33783088 | 0.49223224 | 0.62255517 | 0.79452476 |
| ANGEL1 | 198.409887 | 0.498864852 | 0.18486328 | 2.69856107 | 0.006964 | 0.04272523 |
| IRF2BPL | 346.64851 | -0.853940655 | 0.45774896 | -1.8655218 | 0.0621083 | 0.19249428 |
| AC007686.2 | 64.9516229 | -2.598076923 | 0.97537702 | -2.6636643 | 0.00772947 | 0.04604896 |
| POMT2 | 195.951478 | -0.09542443 | 0.26135634 | -0.3651124 | 0.71502753 | 0.85760497 |
| GSTZ1 | 87.029198 | -0.624998062 | 0.2611661 | -2.3931056 | 0.01670643 | 0.07898184 |
| TMED8 | 556.473567 | 0.021058946 | 0.13441066 | 0.15667615 | 0.87550008 | 0.94396217 |
| SAMD15 | 64.9365156 | -0.873867358 | 0.39937765 | -2.1880728 | 0.0286643 | 0.11381696 |
| NOXRED1 | 126.189665 | 0.190656345 | 0.35868937 | 0.53153609 | 0.59504734 | 0.77501911 |
| VIPAS39 | 265.399853 | 0.028449264 | 0.2702988 | 0.10525117 | 0.91617651 | 0.9622148 |
| AHSA1 | 204.674784 | -0.068109737 | 0.28316751 | -0.2405281 | 0.8099209 | 0.91064395 |

| | | | | | | |
|---|---|---|---|---|---|---|
| SPTLC2 | 1206.40488 | 0.259804943 | 0.20481711 | 1.26847284 | 0.20462914 | 0.41468953 |
| ALKBH1 | 365.789904 | -0.013263321 | 0.23640513 | -0.0561042 | 0.9552588 | 0.98014107 |
| SLIRP | 412.76478 | 0.153426322 | 0.27797852 | 0.55193589 | 0.58099228 | 0.76704917 |
| SNW1 | 438.593529 | -0.19144766 | 0.15898927 | -1.2041546 | 0.22852983 | 0.4440498 |
| ADCK1 | 240.222159 | 0.821076862 | 0.36179049 | 2.2694816 | 0.02323906 | 0.09863802 |
| NRXN3 | 996.208854 | -1.87065708 | 0.74871427 | -2.4984926 | 0.01247227 | 0.06456176 |
| CEP128 | 528.571893 | -0.307147081 | 0.38232586 | -0.8033647 | 0.421764 | 0.64072233 |
| TSHR | 62.7970367 | 1.202633197 | 0.52720078 | 2.28116733 | 0.02253855 | 0.09659691 |
| AC007262.2 | 254.61471 | 1.570477881 | 0.49505008 | 3.17236165 | 0.00151205 | 0.01382188 |
| GTF2A1 | 472.812003 | 0.104190421 | 0.15759992 | 0.66110707 | 0.50854365 | 0.71123011 |
| STON2 | 1476.49051 | 0.217822286 | 0.2157874 | 1.00943006 | 0.31276843 | 0.5367235 |
| SEL1L | 747.390986 | 0.135552587 | 0.13125574 | 1.03273646 | 0.30172724 | 0.52496248 |
| AL355838.1 | 60.5603153 | 1.447110298 | 0.68358369 | 2.11694678 | 0.03426436 | 0.12913878 |
| LINC02301 | 52.3918722 | 1.237510092 | 0.55888745 | 2.21423847 | 0.02681238 | 0.10871927 |
| GALC | 245.526806 | 0.393162614 | 0.25218695 | 1.55901254 | 0.11899341 | 0.29434157 |
| SPATA7 | 651.719922 | 0.369663807 | 0.19830567 | 1.86411116 | 0.0623061 | 0.19283065 |
| PTPN21 | 570.934283 | 0.114556974 | 0.27067434 | 0.42322806 | 0.67212888 | 0.82787676 |
| AL162171.2 | 46.5140847 | 0.791954435 | 0.30274875 | 2.61588012 | 0.00889978 | 0.05081356 |
| ZC3H14 | 1069.641 | 0.330601655 | 0.2225075 | 1.48580005 | 0.13733202 | 0.32329685 |
| EML5 | 459.373773 | -0.143739438 | 0.4590577 | -0.3131185 | 0.75419067 | 0.88075281 |
| TTC8 | 586.714917 | 0.273757901 | 0.17529912 | 1.56166161 | 0.11836773 | 0.29339403 |
| FOXN3 | 2172.96693 | 0.54157188 | 0.16564066 | 3.26955886 | 0.00107715 | 0.01057067 |
| AL356805.1 | 66.3542623 | 0.127372944 | 0.48755239 | 0.26124976 | 0.79389991 | 0.90331845 |
| EFCAB11 | 539.769615 | -0.243079632 | 0.48859806 | -0.4975043 | 0.61883348 | 0.79107437 |
| TDP1 | 246.820993 | -0.466251653 | 0.24226526 | -1.9245502 | 0.05428565 | 0.1756637 |
| KCNK13 | 569.538788 | 2.453969314 | 0.46005387 | 5.33409118 | 9.60E-08 | 6.20E-06 |
| PSMC1 | 316.767855 | -0.376455153 | 0.23017026 | -1.6355508 | 0.10193364 | 0.26604257 |
| NRDE2 | 420.213644 | -0.045743193 | 0.23839608 | -0.191879 | 0.84783702 | 0.92984716 |
| CALM1 | 669.980969 | -0.389119274 | 0.27620474 | -1.4088074 | 0.15889214 | 0.35524819 |
| TTC7B | 1355.91247 | 0.38550517 | 0.24416632 | 1.578863 | 0.11436748 | 0.28726204 |
| RPS6KA5 | 740.120489 | -0.77461031 | 0.18713641 | -4.1392818 | 3.48E-05 | 0.00073911 |
| DGLUCY | 1312.11286 | 0.493434561 | 0.36563676 | 1.34952119 | 0.17716962 | 0.37959992 |
| CCDC88C | 63.8060983 | -0.002459537 | 0.33500986 | -0.0073417 | 0.99414223 | 0.99783229 |
| PPP4R3A | 1246.14138 | 0.110344872 | 0.16711141 | 0.66030722 | 0.5090567 | 0.71154814 |
| CATSPERB | 217.356383 | -0.417420299 | 0.38860201 | -1.0741589 | 0.28275148 | 0.50512096 |
| TC2N | 293.724982 | -0.267206108 | 0.24004659 | -1.1131427 | 0.26564715 | 0.48676739 |
| FBLN5 | 50.4492182 | 0.848271171 | 0.49110681 | 1.72726413 | 0.08412024 | 0.23362999 |
| TRIP11 | 1481.23501 | 0.218707816 | 0.13731933 | 1.592695 | 0.11122863 | 0.28156745 |
| ATXN3 | 1068.87343 | -0.104487617 | 0.21799522 | -0.4793115 | 0.63171703 | 0.80129239 |
| NDUFB1 | 448.015182 | 0.366895997 | 0.23701871 | 1.54796214 | 0.12163141 | 0.29836662 |
| CPSF2 | 490.058022 | 0.251466565 | 0.17551416 | 1.43274235 | 0.15193148 | 0.34499965 |
| RIN3 | 55.5869773 | -1.670880744 | 0.46258354 | -3.6120627 | 0.00030377 | 0.00405714 |
| LGMN | 255.47554 | 0.036171945 | 0.40821905 | 0.08860916 | 0.92939253 | 0.96900671 |
| GOLGA5 | 360.399716 | 0.120960249 | 0.15124127 | 0.79978337 | 0.42383632 | 0.64268786 |
| ITPK1 | 781.537416 | -0.630743656 | 0.38771307 | -1.626831 | 0.10377299 | 0.26945789 |
| MOAP1 | 176.938607 | -1.641897872 | 0.45506285 | -3.6080684 | 0.00030849 | 0.00409526 |
| AL110118.2 | 65.4110469 | 0.003370846 | 0.3401195 | 0.00991077 | 0.99209248 | 0.99677109 |
| GON7 | 103.962525 | -0.125100186 | 0.22701243 | -0.551072 | 0.58158435 | 0.767524 |
| UBR7 | 122.861333 | -0.23780621 | 0.20549342 | -1.1572449 | 0.24717231 | 0.46641415 |
| BTBD7 | 3267.62636 | -0.505758405 | 0.1325224 | -3.8163993 | 0.00013541 | 0.00213446 |
| UNC79 | 175.891703 | -0.505778546 | 0.50760273 | -0.9964063 | 0.31905278 | 0.54371929 |
| DDX24 | 1100.82034 | -0.762065958 | 0.15653342 | -4.8683916 | 1.13E-06 | 4.56E-05 |
| IFI27L1 | 106.846285 | 0.212852849 | 0.26973971 | 0.78910462 | 0.43005086 | 0.64824168 |
| IFI27 | 219.266661 | 1.294386662 | 0.81174201 | 1.59457887 | 0.11080644 | 0.28084062 |
| IFI27L2 | 74.0881746 | -1.188329715 | 0.50978312 | -2.3310496 | 0.01975075 | 0.08847556 |
| PPP4R4 | 91.0507926 | -0.293370547 | 1.07309012 | -0.2733885 | 0.78455456 | 0.89708978 |
| SERPINA1 | 1214.05281 | -2.869505387 | 1.05217196 | -2.7272209 | 0.00638703 | 0.04004092 |
| DICER1 | 2553.26145 | -0.044350312 | 0.13560542 | -0.3270541 | 0.74362695 | 0.87467216 |
| DICER1-AS1 | 237.036368 | 0.685055252 | 0.32120613 | 2.13275897 | 0.0329445 | 0.12584267 |
| CLMN | 5653.37622 | -0.425268885 | 0.40592383 | -1.0476569 | 0.29479673 | 0.51799872 |
| SNHG10 | 55.487455 | 0.40120699 | 0.28600762 | 1.40278426 | 0.16068118 | 0.35797567 |
| GLRX5 | 255.679655 | -0.296868317 | 0.35364157 | -0.839461 | 0.40121065 | 0.62271905 |
| C14orf132 | 133.798295 | -1.033392696 | 0.34208854 | -3.0208341 | 0.00252079 | 0.02035182 |
| AL355102.2 | 111.539652 | 0.418329136 | 0.27117836 | 1.54263466 | 0.12291944 | 0.30039647 |
| AL355102.4 | 81.0552881 | 1.777068667 | 0.51865743 | 3.42628597 | 0.0006119 | 0.0069378 |
| ATG2B | 1532.68602 | 0.319703229 | 0.18585818 | 1.72014612 | 0.08540588 | 0.23615791 |
| GSKIP | 258.600416 | -0.1023212 | 0.23238819 | -0.4403029 | 0.65971773 | 0.81980614 |
| AK7 | 311.814868 | -0.559145498 | 0.24645563 | -2.2687471 | 0.02328371 | 0.09876509 |
| PAPOLA | 1867.30649 | 0.344696319 | 0.20869919 | 1.65164186 | 0.09860758 | 0.2600774 |
| VRK1 | 200.932062 | 0.260216494 | 0.26753452 | 0.97264641 | 0.33072907 | 0.55588514 |
| LINC01550 | 418.18147 | 1.043644222 | 0.33140218 | 3.14917734 | 0.00163731 | 0.01469239 |
| BCL11B | 236.649877 | 1.196317023 | 0.48489079 | 2.4671886 | 0.01361786 | 0.06843031 |
| SETD3 | 1312.83243 | 0.308110379 | 0.16110603 | 1.91246962 | 0.05581599 | 0.17903675 |
| CCNK | 377.101895 | -0.037551883 | 0.17602307 | -0.213335 | 0.83106568 | 0.92124425 |
| CCDC85C | 180.831481 | -0.502459892 | 0.25098821 | -2.0019262 | 0.04529266 | 0.15535466 |
| CYP46A1 | 109.85311 | 0.814136168 | 0.4383781 | 1.8571552 | 0.06328908 | 0.19505368 |
| EML1 | 570.390074 | -0.570965538 | 0.30156226 | -1.8933587 | 0.05831018 | 0.18443945 |
| EVL | 910.229459 | -0.641792349 | 0.32893976 | -1.9510939 | 0.05104588 | 0.16848657 |
| DEGS2 | 58.2289664 | -0.026185354 | 0.59265482 | -0.0441831 | 0.96475842 | 0.98435763 |
| AL133523.1 | 91.6972718 | 0.762067743 | 0.22614067 | 3.36988361 | 0.000752 | 0.00807458 |
| YY1 | 1131.29338 | -0.404549997 | 0.1530397 | -2.6434316 | 0.00820703 | 0.0481166 |
| SLC25A29 | 1172.49628 | -0.390567859 | 0.30497317 | -1.280663 | 0.20031205 | 0.40914209 |
| WARS | 447.030089 | -0.577792966 | 0.40571258 | -1.4241436 | 0.15440491 | 0.34881127 |
| AL157871.4 | 84.6607953 | -0.400682734 | 0.39269408 | -1.0203432 | 0.30756572 | 0.53112982 |
| WDR25 | 424.050522 | -0.021464574 | 0.25829858 | -0.0830999 | 0.93377214 | 0.97167284 |
| PPP2R5C | 1706.93983 | -0.223665873 | 0.16744556 | -1.3357528 | 0.18163009 | 0.38590187 |
| DYNC1H1 | 1051.5289 | -0.386751636 | 0.25916978 | -1.4922714 | 0.13562802 | 0.3207718 |
| HSP90AA1 | 2781.39242 | -0.913039427 | 0.30078099 | -3.0355622 | 0.00240088 | 0.01964877 |
| WDR20 | 1511.14846 | -0.042644457 | 0.17173615 | -0.2483138 | 0.80389162 | 0.90804718 |
| MOK | 1348.8393 | 0.707765157 | 0.25735974 | 2.75010055 | 0.0059577 | 0.03812754 |
| ZNF839 | 259.584178 | 0.28098505 | 0.27390358 | 1.02585387 | 0.30496047 | 0.52822225 |
| CINP | 135.384247 | -0.052244204 | 0.20449192 | -0.255483 | 0.79835009 | 0.90515239 |
| TECPR2 | 710.908017 | -0.144536746 | 0.17856759 | -0.8094232 | 0.41827179 | 0.63753323 |
| ANKRD9 | 255.385989 | -0.011039277 | 0.35314261 | -0.0312601 | 0.9750621 | 0.98891242 |
| RCOR1 | 1477.42099 | -0.057967247 | 0.21357265 | -0.271417 | 0.78607033 | 0.89851185 |
| TRAF3 | 996.82927 | -1.432991183 | 0.26623391 | -5.3824517 | 7.35E-08 | 5.08E-06 |
| AMN | 100.365044 | -0.516012543 | 0.46371299 | -1.1127843 | 0.26580107 | 0.48682751 |
| CDC42BPB | 1458.20899 | -0.642064174 | 0.24246495 | -2.64807 | 0.00809528 | 0.04767298 |
| AL117209.1 | 59.9792797 | -0.663833253 | 0.45347804 | -1.4638708 | 0.14322926 | 0.33255675 |
| TNFAIP2 | 323.503325 | -2.503243266 | 0.41738649 | -5.9974229 | 2.00E-09 | 2.39E-07 |
| AL161669.3 | 68.9682414 | -0.354062948 | 0.57628356 | -0.6143902 | 0.53895754 | 0.73506352 |
| EIF5 | 795.110593 | -0.641298927 | 0.2753054 | -2.3294092 | 0.0198374 | 0.0887042 |
| MARK3 | 3266.71628 | -0.148103489 | 0.18013772 | -0.8221681 | 0.41098121 | 0.63014283 |
| CKB | 1931.98168 | -0.658923742 | 0.52687681 | -1.250622 | 0.21107241 | 0.42345922 |
| TRMT61A | 57.4677925 | -0.565011362 | 0.39430061 | -1.4329457 | 0.15187335 | 0.34498626 |
| AL139300.2 | 51.4535905 | -1.718534422 | 0.46933573 | -3.6616314 | 0.00025061 | 0.00351419 |
| BAG5 | 410.299334 | -1.294128392 | 0.33483244 | -3.8650031 | 0.00011109 | 0.00181829 |
| KLC1 | 765.398953 | -0.12922435 | 0.16177942 | -0.7987688 | 0.4244245 | 0.64298076 |
| AL049840.1 | 82.3218057 | -0.841438185 | 0.52024842 | -1.6173777 | 0.10579678 | 0.27294329 |
| ZFYVE21 | 229.061856 | -0.925546185 | 0.21684329 | -4.2682721 | 1.97E-05 | 0.00046549 |
| PPP1R13B | 2245.42526 | 0.036001069 | 0.1481009 | 0.24308474 | 0.80793976 | 0.91010272 |
| ATP5MPL | 432.433954 | 0.047065012 | 0.27360753 | 0.1720165 | 0.86342455 | 0.93813133 |
| TDRD9 | 187.840063 | -0.993104303 | 0.70375361 | -1.4111534 | 0.15819939 | 0.35441922 |

| | | | | | | |
|---|---|---|---|---|---|---|
| AL590079.1 | 111.925669 | 1.394729598 | 0.76736345 | 1.81756064 | 0.06913131 | 0.20650283 |
| INF2 | 115.706951 | -1.76030298 | 0.37657644 | -4.6744904 | 2.95E-06 | 9.78E-05 |
| SIVA1 | 178.826196 | -0.596871696 | 0.24737377 | -2.4128334 | 0.01582905 | 0.07604413 |
| AKT1 | 288.945087 | -0.3305961 | 0.23911896 | -1.3825591 | 0.16680009 | 0.36640505 |
| AL583810.1 | 62.5465617 | -0.166826139 | 0.31399095 | -0.5313088 | 0.59520484 | 0.77504542 |
| CEP170B | 275.067929 | 0.079236309 | 0.25569337 | 0.30988801 | 0.75664612 | 0.88227669 |
| CLBA1 | 127.253241 | 0.070497035 | 0.23250246 | 0.30320985 | 0.76172995 | 0.88431255 |
| CDCA4 | 46.3277531 | -0.101469368 | 0.37059211 | -0.2738034 | 0.78423573 | 0.89695816 |
| GPR132 | 505.930353 | 0.299776346 | 0.64988313 | 0.46127732 | 0.64459966 | 0.80987296 |
| JAG2 | 58.979112 | -0.80835487 | 0.464121 | -1.7416899 | 0.08156271 | 0.22942119 |
| NUDT14 | 122.544344 | -0.106790949 | 0.47693561 | -0.2239106 | 0.82282685 | 0.91813562 |
| BRF1 | 364.438024 | 0.043525402 | 0.22150752 | 0.19649628 | 0.84422175 | 0.92788526 |
| BTBD6 | 91.5639183 | -1.029226749 | 0.33798419 | -3.045192 | 0.00232532 | 0.01916103 |
| PACS2 | 684.061267 | 0.001027425 | 0.23553525 | 0.00436208 | 0.99651957 | 0.99867719 |
| TEX22 | 141.334435 | -0.061604148 | 0.32929823 | -0.1870771 | 0.85160021 | 0.93193618 |
| MTA1 | 401.502982 | 0.261464467 | 0.29937931 | 0.87335518 | 0.38246951 | 0.60517966 |
| CRIP2 | 328.540928 | -0.443460337 | 0.46688531 | -0.9498271 | 0.34220009 | 0.56742669 |
| AC087386.1 | 102.784795 | 0.912866314 | 0.42748242 | 2.13544763 | 0.03272446 | 0.12530982 |
| NIPA1 | 549.51209 | 0.041511615 | 0.15346592 | 0.27049403 | 0.78678021 | 0.89905515 |
| NIPA2 | 433.034707 | -0.268101345 | 0.16691384 | -1.606226 | 0.10822427 | 0.27665271 |
| CYFIP1 | 931.760962 | -0.540529927 | 0.16365518 | -3.3028586 | 0.00095705 | 0.00963481 |
| TUBGCP5 | 428.708393 | 0.247902169 | 0.19395584 | 1.27813715 | 0.20120107 | 0.41032344 |
| SNRPN | 77.8203624 | 1.400595591 | 0.50580611 | 2.76903654 | 0.00562223 | 0.03658329 |
| AC124312.1 | 2557.36613 | 0.551237968 | 0.24983726 | 2.20638817 | 0.02735683 | 0.11021426 |
| AC124312.2 | 81.87391 | 0.612108116 | 0.92647738 | 0.66068328 | 0.50881545 | 0.71134045 |
| PWAR6 | 634.479322 | 0.562548639 | 0.30699325 | 1.83244626 | 0.06688496 | 0.20193399 |
| UBE3A | 1894.11035 | 0.25168729 | 0.14643021 | 1.71882078 | 0.08564701 | 0.23657552 |
| LINC02250 | 59.3218341 | -0.413362838 | 0.59265631 | -0.6974748 | 0.48550571 | 0.69300247 |
| GABRB3 | 1084.60389 | 0.2079539 | 0.38752156 | 0.53662536 | 0.59152642 | 0.77302885 |
| OCA2 | 59.0709274 | 0.923142007 | 0.58954155 | 1.5658642 | 0.11738041 | 0.29199207 |
| HERC2 | 2233.97012 | 0.138358615 | 0.16732276 | 0.82689656 | 0.40829569 | 0.62813625 |
| GOLGA8M | 88.3308701 | 1.492020115 | 0.49697511 | 3.00220292 | 0.00268033 | 0.02130078 |
| FAM189A1 | 124.090169 | -1.210776261 | 0.43246706 | -2.7996959 | 0.00511508 | 0.03400354 |
| TJP1 | 1836.15768 | -0.325360293 | 0.18639909 | -1.7455037 | 0.08089724 | 0.228181 |
| AC091057.2 | 50.9076893 | 1.269087915 | 0.37886605 | 3.34970085 | 0.00080899 | 0.00849438 |
| FAN1 | 548.414226 | 0.496535184 | 0.23353639 | 2.12615765 | 0.03349014 | 0.12730106 |
| MTMR10 | 1737.53477 | 0.368730994 | 0.24925936 | 1.47930653 | 0.13905841 | 0.32562002 |
| KLF13 | 361.163892 | -0.378959533 | 0.3441609 | -1.1011115 | 0.27084812 | 0.49226771 |
| OTUD7A | 1005.19451 | 0.157948335 | 0.3093264 | 0.51062029 | 0.60961696 | 0.78529652 |
| CHRNA7 | 55.7431976 | 0.374629171 | 0.60260654 | 0.62168123 | 0.5341515 | 0.73149412 |
| LINC02256 | 1645.40566 | 0.315409734 | 0.32563629 | 0.96859515 | 0.33274722 | 0.55805715 |
| AC123768.3 | 264.246402 | 0.68630414 | 0.27811714 | 2.46768014 | 0.01359918 | 0.06837896 |
| GOLGA8N | 199.455519 | 0.937401998 | 0.267624 | 3.50268283 | 0.0004606 | 0.00556636 |
| FMN1 | 2698.15342 | 0.39501536 | 0.57995012 | 0.68111954 | 0.49579585 | 0.70114155 |
| RYR3 | 129.706194 | -0.678045777 | 0.52652469 | -1.2877758 | 0.197824 | 0.40634007 |
| AC010809.1 | 55.4322451 | 0.041864676 | 0.38117374 | 0.10983096 | 0.91254344 | 0.96042853 |
| AVEN | 619.737511 | 0.283668889 | 0.26729354 | 1.06126352 | 0.28857016 | 0.51093714 |
| CHRM5 | 108.792823 | 0.498224861 | 0.44344726 | 1.12352675 | 0.26121385 | 0.48232304 |
| EMC7 | 243.726571 | -0.215668283 | 0.24940493 | -0.8647314 | 0.38718621 | 0.61065142 |
| PGBD4 | 53.1854081 | 1.172573357 | 0.2760034 | 4.24840185 | 2.15E-05 | 0.00050157 |
| KATNBL1 | 1188.36174 | -0.011176246 | 0.15516673 | -0.0720273 | 0.94258015 | 0.97554394 |
| EMC4 | 139.464699 | -0.284623427 | 0.27039318 | -1.052628 | 0.29251153 | 0.51542428 |
| SLC12A6 | 742.271857 | -0.131489329 | 0.28067782 | -0.4684707 | 0.63944804 | 0.80603771 |
| NOP10 | 257.345475 | -1.358766648 | 0.39454332 | -3.4438972 | 0.00057339 | 0.00654055 |
| LPCAT4 | 93.6396442 | -0.508484614 | 0.42655967 | -1.1920598 | 0.23323782 | 0.45008595 |
| GOLGA8A | 903.614568 | 0.09428768 | 0.234741 | 0.40166686 | 0.68792922 | 0.83940892 |
| GOLGA8B | 970.246829 | 0.351091752 | 0.34783033 | 1.00937647 | 0.31279412 | 0.5367235 |
| AQR | 991.451556 | 0.105555112 | 0.19687995 | 0.53613947 | 0.59186217 | 0.77328146 |
| ZNF770 | 457.965297 | -0.286470316 | 0.26545205 | -1.0791791 | 0.28050788 | 0.50294945 |
| AC021231.1 | 1317.86685 | 0.522725706 | 0.42601018 | 1.22702633 | 0.21981269 | 0.43361796 |
| DPH6 | 3869.0526 | 0.203845984 | 0.29389048 | 0.69361207 | 0.48792552 | 0.69502858 |
| DPH6-DT | 2224.11943 | 1.895701386 | 0.52801094 | 3.59026915 | 0.00033034 | 0.0043099 |
| AC021351.1 | 47.3920039 | 0.713149703 | 0.77240716 | 0.9232821 | 0.35586021 | 0.58124861 |
| C15orf41 | 783.365825 | 0.754181882 | 0.25027019 | 3.01347072 | 0.00258278 | 0.02075723 |
| MEIS2 | 67.5108898 | 0.296788717 | 0.56671933 | 0.52369612 | 0.60048991 | 0.77870324 |
| SPRED1 | 1336.73347 | -0.798003624 | 0.29575174 | -2.6982212 | 0.00697111 | 0.0427305 |
| FAM98B | 240.666641 | 0.371148573 | 0.18862636 | 1.96763898 | 0.04910959 | 0.16410433 |
| RASGRP1 | 114.719463 | -1.170653752 | 0.3252949 | -3.5987461 | 0.00031976 | 0.00421943 |
| C15orf53 | 231.800428 | 0.510803946 | 0.64986141 | 0.78601981 | 0.43185588 | 0.64957432 |
| AC013652.1 | 506.173325 | 1.332894716 | 0.52937936 | 2.51784413 | 0.01180756 | 0.06231163 |
| THBS1 | 208.146469 | 1.278160834 | 0.53479617 | 2.38999624 | 0.01684855 | 0.07944774 |
| FSIP1 | 234.620267 | 0.417427281 | 0.1624611 | 2.5693983 | 0.01018753 | 0.05630421 |
| EIF2AK4 | 542.951024 | 0.117795395 | 0.15617372 | 0.75425877 | 0.45069385 | 0.66654333 |
| SRP14 | 1287.85953 | -1.639733914 | 0.44145736 | -3.7143653 | 0.00020371 | 0.00297991 |
| SRP14-AS1 | 207.682007 | 0.600560241 | 0.31550053 | 1.9035158 | 0.05697327 | 0.18151471 |
| BMF | 130.100194 | -1.600395527 | 0.48457751 | -3.3026616 | 0.00095772 | 0.00963481 |
| PAK6 | 41.6427893 | -0.416620494 | 0.60812134 | -0.6850943 | 0.49328439 | 0.69881955 |
| CCDC9B | 77.8156412 | 0.448649649 | 0.4155491 | 1.07965495 | 0.28029586 | 0.50277404 |
| KNSTRN | 45.9711976 | 0.417883093 | 0.40665174 | 1.02761911 | 0.30412903 | 0.52747379 |
| IVD | 455.984132 | -0.133355406 | 0.22255376 | -0.5992054 | 0.54903594 | 0.74226986 |
| BAHD1 | 231.597991 | -0.106876021 | 0.25489195 | -0.4192993 | 0.6749974 | 0.82939913 |
| AC091045.1 | 105.243762 | -0.725111209 | 0.63352283 | -1.14457 | 0.25238733 | 0.4724751 |
| CCDC32 | 476.890505 | -0.488091363 | 0.23147819 | -2.1085847 | 0.03498045 | 0.13084552 |
| RPUSD2 | 44.1675868 | 0.070614306 | 0.32321255 | 0.21847638 | 0.82705796 | 0.91955386 |
| KNL1 | 45.9296567 | 1.044448716 | 0.55245954 | 1.89054337 | 0.05868533 | 0.18495674 |
| RMDN3 | 231.590024 | -0.437620088 | 0.22222897 | -1.9692306 | 0.04892661 | 0.16365451 |
| GCHFR | 107.821078 | -2.348937596 | 0.5293455 | -4.4374375 | 9.10E-06 | 0.00024753 |
| DNAJC17 | 457.470698 | -0.190799853 | 0.22931754 | -0.8320334 | 0.40539008 | 0.62573634 |
| ZFYVE19 | 142.147107 | 0.010285074 | 0.30934285 | 0.03324814 | 0.97347671 | 0.98804941 |
| SPINT1-AS1 | 183.744214 | -0.352077676 | 0.26949751 | -1.3064227 | 0.19140883 | 0.39809311 |
| SPINT1 | 470.960304 | -1.149996817 | 0.24506644 | -4.692592 | 2.70E-06 | 9.09E-05 |
| VPS18 | 79.1406342 | -0.625534599 | 0.29572622 | -2.115249 | 0.03440873 | 0.12945169 |
| INO80 | 1079.21194 | 0.164845622 | 0.2155176 | 0.76488242 | 0.44434156 | 0.66007871 |
| AC020661.3 | 43.1374708 | -0.131933288 | 0.33816088 | -0.3901495 | 0.69642602 | 0.84496018 |
| INO80-AS1 | 46.4663324 | 0.151991549 | 0.51048657 | 0.29773859 | 0.76590269 | 0.88666158 |
| CHP1 | 879.149567 | -0.40977126 | 0.28628646 | -1.431333 | 0.1523348 | 0.34555905 |
| OIP5 | 57.4558135 | 0.664288105 | 0.43599154 | 1.52362614 | 0.12760213 | 0.30794192 |
| NDUFAF1 | 158.285056 | 0.165927307 | 0.20637819 | 0.80399632 | 0.4213991 | 0.64046063 |
| RTF1 | 609.908858 | -0.503318874 | 0.1971857 | -2.5525121 | 0.01069492 | 0.05813548 |
| RPAP1 | 141.428634 | 0.304406572 | 0.23832944 | 1.27725122 | 0.20151357 | 0.41077048 |
| MGA | 1070.15984 | 0.120942196 | 0.18549433 | 0.65199942 | 0.51440154 | 0.71598841 |
| MAPKBP1 | 727.333992 | 0.209427202 | 0.29441195 | 0.71134071 | 0.47687313 | 0.6867392 |
| EHD4 | 236.075133 | -0.831549975 | 0.28815385 | -2.8857847 | 0.00390439 | 0.02813616 |
| PLA2G4F | 97.7409113 | 0.708554486 | 0.88907164 | 0.79695995 | 0.42547429 | 0.64398876 |
| VPS39 | 762.530268 | -0.083614976 | 0.15374388 | -0.5438589 | 0.58653857 | 0.77036771 |
| TMEM87A | 1062.36809 | -0.263561218 | 0.26028916 | -1.0125709 | 0.31126519 | 0.53479473 |
| GANC | 660.900751 | 0.393520554 | 0.31852282 | 1.23545483 | 0.21666128 | 0.43074636 |
| ZNF106 | 896.462191 | 0.000694994 | 0.17018018 | 0.00408387 | 0.99674155 | 0.99878313 |
| SNAP23 | 606.622498 | -0.293407913 | 0.18772461 | -1.5629699 | 0.11805968 | 0.29301961 |
| LRRC57 | 101.036402 | -0.269453486 | 0.19576366 | -1.3764224 | 0.1686909 | 0.36885253 |
| HAUS2 | 166.185448 | -0.147489999 | 0.22377633 | -0.6590956 | 0.50983437 | 0.71203237 |
| STARD9 | 45.9094929 | -1.286526285 | 0.31378765 | -4.0999902 | 4.13E-05 | 0.00085008 |
| CDAN1 | 82.6752686 | 0.145915587 | 0.28963987 | 0.5037828 | 0.61441401 | 0.78809426 |

| | | | | | | |
|---|---|---|---|---|---|---|
| TTBK2 | 1258.63323 | -0.02522067 | 0.19043822 | -0.1324349 | 0.89464029 | 0.95172874 |
| UBR1 | 2064.69452 | -0.251509633 | 0.13289893 | -1.892488 | 0.05842599 | 0.18458477 |
| AC068724.3 | 289.365363 | -0.194815417 | 0.33868589 | -0.5752097 | 0.56514947 | 0.75610852 |
| TMEM62 | 566.122678 | -0.099537344 | 0.25459647 | -0.3909612 | 0.69582591 | 0.84462089 |
| CCNDBP1 | 272.636998 | -0.485893372 | 0.22555195 | -2.1542415 | 0.03122124 | 0.12125885 |
| ADAL | 411.248852 | 0.979377373 | 0.28491643 | 3.43741974 | 0.00058728 | 0.00668169 |
| ZSCAN29 | 123.80451 | 0.161397904 | 0.26254281 | 0.6147489 | 0.53872056 | 0.73496797 |
| TUBGCP4 | 297.412961 | 0.036747505 | 0.1924851 | 0.19091091 | 0.84859539 | 0.9303692 |
| TP53BP1 | 1428.72597 | 0.150579394 | 0.16345186 | 0.92124614 | 0.35692194 | 0.58255058 |
| PPIP5K1 | 514.079594 | -0.106053245 | 0.29337907 | -0.3614888 | 0.71773408 | 0.85952398 |
| AC011330.3 | 71.0301313 | 0.001081485 | 0.58181321 | 0.00185882 | 0.99851688 | 0.99953295 |
| CATSPER2 | 579.113085 | 0.496772347 | 0.44247164 | 1.12272134 | 0.26155586 | 0.48239198 |
| PDIA3 | 812.633982 | -0.273696121 | 0.29780177 | -0.9190547 | 0.35806695 | 0.58348526 |
| AC018512.1 | 45.7967386 | -0.828314551 | 0.40260976 | -2.0573633 | 0.03965129 | 0.14177464 |
| SERF2 | 2177.88609 | -1.06676424 | 0.45473343 | -2.3459112 | 0.01898063 | 0.08609723 |
| MFAP1 | 218.372336 | -0.496205936 | 0.21563511 | -2.3011371 | 0.02138388 | 0.09346781 |
| WDR76 | 53.999489 | 0.15844648 | 0.34947929 | 0.45337874 | 0.65027604 | 0.81301431 |
| FRMD5 | 82.9258467 | -0.852342171 | 0.26394423 | -3.229251 | 0.00124115 | 0.01183707 |
| CASC4 | 2385.35377 | -0.636154116 | 0.18330357 | -3.470495 | 0.0005195 | 0.00608236 |
| AC025043.1 | 115.744894 | -0.680085566 | 0.42980004 | -1.5823302 | 0.11357422 | 0.28587524 |
| CTDSPL2 | 1458.94243 | -0.099340735 | 0.19247567 | -0.516121 | 0.6057699 | 0.78178941 |
| EIF3J-DT | 424.448967 | -0.278217331 | 0.27368904 | -1.0165454 | 0.30936974 | 0.53327221 |
| EIF3J | 561.254691 | -0.146827744 | 0.17803308 | -0.8247217 | 0.4095296 | 0.62937481 |
| SPG11 | 1862.15419 | -0.128953388 | 0.2077392 | -0.6207465 | 0.53476641 | 0.73203192 |
| B2M | 4055.12428 | -1.414377838 | 0.47403307 | -2.9837114 | 0.00284775 | 0.02227942 |
| TRIM69 | 130.104227 | 0.223359124 | 0.43321577 | 0.51558401 | 0.60614498 | 0.78219705 |
| SORD | 59.9941384 | -0.717605164 | 0.61875874 | -1.1597495 | 0.2461508 | 0.46541739 |
| DUOX1 | 168.992394 | -0.213935678 | 0.30955483 | -0.6911075 | 0.48949796 | 0.6962932 |
| SHF | 58.9501272 | -0.122169471 | 0.39381115 | -0.3102235 | 0.75639101 | 0.88220153 |
| AC051619.5 | 68.42027 | 1.5214238 | 0.56184821 | 2.70789114 | 0.00677122 | 0.04186754 |
| GATM | 208.180842 | 1.152463051 | 0.39255033 | 2.93583512 | 0.00332651 | 0.02503235 |
| SPATA5L1 | 174.03513 | -0.089408878 | 0.23273518 | -0.3841657 | 0.70085562 | 0.84841905 |
| AC025580.2 | 79.2367119 | -1.411624773 | 0.67412658 | -2.0940055 | 0.03625949 | 0.13396096 |
| SLC30A4 | 111.494208 | 0.518598649 | 0.34825944 | 1.48911586 | 0.13645686 | 0.32215526 |
| SQOR | 133.34889 | -2.524506878 | 0.33421144 | -7.5536221 | 4.23E-14 | 2.15E-11 |
| SEMA6D | 1930.90753 | 2.292304461 | 0.3443459 | 6.65698203 | 2.80E-11 | 6.15E-09 |
| AC092078.2 | 1190.2981 | 3.486929166 | 0.88682434 | 3.93192769 | 8.43E-05 | 0.00147429 |
| SLC24A5 | 91.349018 | -0.181274849 | 0.63967124 | -0.2833875 | 0.77687979 | 0.89337196 |
| MYEF2 | 1848.24336 | 0.224329746 | 0.23404895 | 0.95847361 | 0.33782399 | 0.5630684 |
| SLC12A1 | 33457.3288 | 2.748719497 | 0.5202829 | 5.28312481 | 1.27E-07 | 7.56E-06 |
| AC023355.1 | 100.422528 | 0.683251336 | 0.63360575 | 1.07835406 | 0.28087578 | 0.50347241 |
| DUT | 353.664548 | 0.221566146 | 0.24906064 | 0.88960724 | 0.37367682 | 0.59763858 |
| FBN1 | 139.800206 | 0.00752829 | 0.34250195 | 0.02198028 | 0.98246368 | 0.99281397 |
| CEP152 | 619.40339 | 0.630053641 | 0.30399914 | 2.07255073 | 0.03821411 | 0.13874936 |
| EID1 | 470.52377 | -1.187538839 | 0.4315857 | -2.7515714 | 0.00593101 | 0.03799355 |
| SECISBP2L | 2100.99131 | 0.022418983 | 0.1022479 | 0.21926107 | 0.82644668 | 0.91938879 |
| COPS2 | 412.308101 | 0.196842133 | 0.16985785 | 1.15886394 | 0.24651165 | 0.46556654 |
| GALK2 | 1184.54563 | 0.321861292 | 0.19841632 | 1.62215131 | 0.10477095 | 0.27110575 |
| FAM227B | 1070.20165 | 0.615331068 | 0.31272725 | 1.96762857 | 0.04911079 | 0.16410433 |
| FGF7 | 48.005022 | -0.356548992 | 0.54176258 | -0.6581278 | 0.51045604 | 0.71269123 |
| DTWD1 | 743.080921 | 0.892352917 | 0.21192801 | 4.21064161 | 2.55E-05 | 0.00057643 |
| SLC27A2 | 75.9175694 | -1.448530477 | 0.75260811 | -1.9246809 | 0.05426929 | 0.1756637 |
| GABPB1 | 525.585347 | -0.078393966 | 0.19643275 | -0.3990881 | 0.68982832 | 0.84058508 |
| GABPB1-AS1 | 1165.18995 | 0.970991869 | 0.40370632 | 2.40519362 | 0.0161639 | 0.07713473 |
| USP8 | 1433.84137 | 0.235174099 | 0.16413988 | 1.43276638 | 0.15192461 | 0.34499965 |
| USP50 | 263.030534 | 0.280466592 | 0.51841159 | 0.54101143 | 0.58849971 | 0.77148597 |
| TRPM7 | 3518.87428 | 0.492360037 | 0.27230897 | 1.80809332 | 0.07059198 | 0.20944507 |
| SPPL2A | 1919.24654 | -0.198710982 | 0.17114028 | -1.1611 | 0.24560123 | 0.4647176 |
| AC021752.1 | 88.7251104 | -0.574674008 | 0.44520052 | -1.2908206 | 0.19676589 | 0.40509362 |
| AP4E1 | 627.143911 | -0.222848113 | 0.241285 | -0.9235888 | 0.35570046 | 0.58105953 |
| DMXL2 | 710.030892 | 0.714310607 | 0.19147371 | 3.73059356 | 0.00019103 | 0.00282249 |
| LYSMD2 | 52.4445062 | -0.439249851 | 0.43247996 | -1.0156537 | 0.30979434 | 0.5337614 |
| TMOD2 | 117.706411 | 0.043693939 | 0.29469382 | 0.14826893 | 0.88213053 | 0.94686025 |
| TMOD3 | 1205.86726 | -0.410286959 | 0.1496529 | -2.7415905 | 0.00611425 | 0.03882844 |
| AC090971.2 | 76.2380314 | -0.29801624 | 0.42665772 | -0.6984902 | 0.48487067 | 0.69247018 |
| LEO1 | 154.446165 | 0.108679914 | 0.23053938 | 0.47141583 | 0.63734381 | 0.80497022 |
| MAPK6 | 665.362523 | 0.066193276 | 0.24917526 | 0.26564947 | 0.79050918 | 0.90116933 |
| GNB5 | 592.413729 | -1.011302697 | 0.2092797 | -4.8323018 | 1.35E-06 | 5.24E-05 |
| CERNA1 | 119.718973 | -0.007843101 | 0.51263438 | -0.0152996 | 0.98779316 | 0.99548027 |
| MYO5C | 1383.16027 | -0.111179294 | 0.23415265 | -0.4748154 | 0.63491852 | 0.80377983 |
| MYO5A | 955.085387 | -0.012681465 | 0.21440464 | -0.0591473 | 0.95283475 | 0.97936812 |
| ARPP19 | 647.14753 | -0.472503154 | 0.175287 | -2.6955972 | 0.00702626 | 0.04292004 |
| AC025917.1 | 67.9579236 | -0.832129892 | 0.67903261 | -1.2254638 | 0.22040051 | 0.43419394 |
| FAM214A | 1916.77405 | 0.029879538 | 0.21475829 | 0.13913101 | 0.88934662 | 0.95012776 |
| WDR72 | 3390.6106 | 0.401995389 | 0.54287108 | 0.74049882 | 0.45899738 | 0.67278348 |
| UNC13C | 691.537079 | 4.933380959 | 1.24212321 | 3.97173237 | 7.14E-05 | 0.00128931 |
| RSL24D1 | 229.926554 | -0.141309508 | 0.28307643 | -0.4991921 | 0.61764408 | 0.79039534 |
| RAB27A | 1022.24437 | -1.091331693 | 0.27188389 | -4.0139623 | 5.97E-05 | 0.00112029 |
| PIGB | 408.440045 | -0.043057101 | 0.18360382 | -0.2345109 | 0.81458836 | 0.91339257 |
| CCPG1 | 1177.7879 | 0.018511993 | 0.14642567 | 0.12642587 | 0.89939483 | 0.95376576 |
| C15orf65 | 76.01349 | 0.048805669 | 0.40197918 | 0.12141343 | 0.90336358 | 0.95532916 |
| DNAAF4 | 500.437209 | 0.117193986 | 0.2725289 | 0.43002407 | 0.66717813 | 0.82469498 |
| PYGO1 | 67.6461885 | 0.322552691 | 0.43065879 | 0.74897506 | 0.45387224 | 0.66948056 |
| PRTG | 205.460627 | -0.994866409 | 0.91581956 | -1.0863127 | 0.27734068 | 0.49950818 |
| NEDD4 | 273.372641 | -2.085591854 | 0.35834593 | -5.8200517 | 5.88E-09 | 6.10E-07 |
| RFX7 | 1450.40422 | -0.271804967 | 0.1375192 | -1.9764875 | 0.04809959 | 0.16178954 |
| TEX9 | 992.194908 | 0.015775122 | 0.19026513 | 0.08291126 | 0.9339221 | 0.97167284 |
| MNS1 | 176.644387 | -0.499759839 | 0.36459959 | -1.3707087 | 0.17046577 | 0.37070833 |
| AC090518.1 | 93.4590516 | 0.743557888 | 0.50180425 | 1.4817688 | 0.13840182 | 0.32460044 |
| ZNF280D | 803.046732 | 0.617388495 | 0.20357958 | 3.03266413 | 0.00242405 | 0.0197445 |
| AC090517.4 | 484.203946 | 0.722323508 | 0.30997676 | 2.33025054 | 0.01979291 | 0.0885424 |
| TCF12 | 6997.18492 | 0.091928814 | 0.19764644 | 0.46511747 | 0.64184733 | 0.80790432 |
| CGNL1 | 11725.1648 | 0.277387474 | 0.37094122 | 0.74779361 | 0.45458466 | 0.6698832 |
| POLR2M | 125.221748 | -0.233189332 | 0.32969451 | -0.7072891 | 0.47938683 | 0.68864274 |
| ALDH1A2 | 4184.30232 | -0.265133924 | 0.57279789 | -0.4628752 | 0.64345384 | 0.80905043 |
| LIPC | 662.948973 | -0.466465224 | 0.56244923 | -0.8293464 | 0.40690844 | 0.62687819 |
| ADAM10 | 3487.55584 | -0.829754804 | 0.29286753 | -2.8332086 | 0.00460833 | 0.03155596 |
| MINDY2 | 1772.86279 | -0.460373727 | 0.21976571 | -2.0948387 | 0.03618533 | 0.13384823 |
| RNF111 | 2012.36768 | 0.011086841 | 0.18474401 | 0.06001191 | 0.95214615 | 0.97933189 |
| SLTM | 2386.43273 | -0.239694615 | 0.16444986 | -1.4575544 | 0.1449634 | 0.33481755 |
| MYO1E | 4319.23873 | -0.981903812 | 0.38355247 | -2.5600247 | 0.01046647 | 0.05736582 |
| FAM81A | 222.867346 | 0.222659224 | 0.41251692 | 0.5397578 | 0.58936407 | 0.77174702 |
| GCNT3 | 87.4437492 | -4.469895734 | 0.76532412 | -5.8405264 | 5.20E-09 | 5.46E-07 |
| GTF2A2 | 351.368551 | -0.204358154 | 0.2543034 | -0.8035998 | 0.42162815 | 0.64058963 |
| BNIP2 | 628.009675 | -0.269889726 | 0.11581578 | -2.3303364 | 0.01978838 | 0.0885424 |
| ANXA2 | 971.545496 | -3.113469307 | 0.50898669 | -6.1169956 | 9.54E-10 | 1.23E-07 |
| ICE2 | 994.444126 | 0.40152836 | 0.16641788 | 2.41277171 | 0.01583173 | 0.07604413 |
| RORA-AS1 | 685.176397 | 0.871003131 | 0.33662966 | 2.58742241 | 0.0096697 | 0.05403005 |
| RORA | 11124.5279 | 0.777110422 | 0.29677846 | 2.61848663 | 0.00883208 | 0.05053466 |
| AC012404.1 | 328.401398 | 0.536217371 | 0.38409616 | 1.39604981 | 0.1626995 | 0.36088961 |
| AC009554.1 | 529.219699 | -0.448168762 | 0.68506263 | -0.6542012 | 0.51298222 | 0.71496824 |
| VPS13C | 3882.84046 | -0.616603353 | 0.33668363 | -1.8314028 | 0.06704044 | 0.2022236 |
| C2CD4A | 280.818788 | -8.39943321 | 1.04564296 | -8.0327928 | 9.53E-16 | 7.87E-13 |

| | | | | | | |
|---|---|---|---|---|---|---|
| TLN2 | 2051.6531 | 0.540446171 | 0.23212301 | 2.32827485 | 0.01989751 | 0.08888274 |
| TPM1 | 1064.58516 | -1.004378532 | 0.41452167 | -2.422982 | 0.01539369 | 0.07470649 |
| LACTB | 275.231509 | -0.406402573 | 0.20108699 | -2.0210287 | 0.0432768 | 0.15027424 |
| RPS27L | 505.525909 | -0.43986593 | 0.33362632 | -1.3184389 | 0.18735676 | 0.39298287 |
| RAB8B | 463.739951 | -0.540880947 | 0.18230642 | -2.9668782 | 0.0030084 | 0.02320564 |
| APH1B | 332.537419 | -0.420789989 | 0.37616129 | -1.1186425 | 0.2632927 | 0.48437789 |
| CA12 | 7892.31211 | 0.511161046 | 0.6512746 | 0.78486255 | 0.43253416 | 0.65015341 |
| LINC02568 | 381.529714 | 2.507959275 | 0.71414842 | 3.51181799 | 0.00044505 | 0.00541766 |
| USP3 | 2136.9024 | -0.055235398 | 0.16620377 | -0.3323354 | 0.739636 | 0.87254192 |
| USP3-AS1 | 448.262577 | 0.139514001 | 0.33096955 | 0.42153123 | 0.67336721 | 0.82886101 |
| HERC1 | 2436.88098 | 0.409298663 | 0.19267111 | 2.12433852 | 0.03364185 | 0.12769401 |
| AC073167.1 | 108.962901 | 0.187555492 | 0.54274489 | 0.34556842 | 0.72966708 | 0.8673587 |
| DAPK2 | 55.3761907 | -2.935025388 | 0.81919117 | -3.5828333 | 0.00033989 | 0.00441453 |
| FAM96A | 390.895264 | 0.221896995 | 0.20446911 | 1.08523483 | 0.27781766 | 0.49998741 |
| SNX1 | 977.452948 | -0.086088551 | 0.10646024 | -0.8086451 | 0.41871933 | 0.63783336 |
| PPIB | 678.857363 | -0.942087564 | 0.37434769 | -2.5166112 | 0.01184895 | 0.06245523 |
| CSNK1G1 | 254.17175 | -0.485186572 | 0.26901717 | -1.8035525 | 0.0713015 | 0.21075399 |
| TRIP4 | 629.838098 | -0.105898975 | 0.20904452 | -0.5065857 | 0.61244551 | 0.78687125 |
| ZNF609 | 3199.21902 | -0.222289574 | 0.15639727 | -1.4213137 | 0.15522559 | 0.34989331 |
| OAZ2 | 274.97543 | -0.264712695 | 0.28377178 | -0.9328366 | 0.35090435 | 0.57700679 |
| AC100830.1 | 62.8179816 | 0.132062844 | 0.55295821 | 0.2388297 | 0.81123764 | 0.91142621 |
| RBPMS2 | 74.3049511 | 0.103666169 | 0.4460286 | 0.23242045 | 0.81621146 | 0.91443162 |
| ANKDD1A | 64.9864981 | -0.771823694 | 0.43639717 | -1.7686267 | 0.07695619 | 0.2212981 |
| AC103691.1 | 41.4346449 | 0.628499321 | 0.46949363 | 1.33867487 | 0.18067654 | 0.38451679 |
| SPG21 | 415.694845 | 0.000111719 | 0.27695327 | 0.00040338 | 0.99967815 | 0.99975417 |
| MTFMT | 365.829578 | 0.213977588 | 0.19092782 | 1.12072504 | 0.26240492 | 0.4832663 |
| UBAP1L | 186.692217 | 0.119066806 | 0.3671909 | 0.32426404 | 0.74573814 | 0.87570052 |
| PDCD7 | 279.639193 | -0.332659638 | 0.24939426 | -1.3338705 | 0.18224631 | 0.38671349 |
| CLPX | 634.944346 | -0.00174391 | 0.15701523 | -0.0111066 | 0.99113837 | 0.99641881 |
| PARP16 | 238.862408 | 0.118682172 | 0.33486667 | 0.3544162 | 0.723027 | 0.86248065 |
| DPP8 | 912.028568 | -0.097094587 | 0.17348394 | -0.5596748 | 0.57570128 | 0.76288505 |
| HACD3 | 597.418411 | 0.144459062 | 0.19294934 | 0.74868908 | 0.45404463 | 0.66958468 |
| INTS14 | 255.430268 | 0.19197657 | 0.16922907 | 1.1344184 | 0.25661909 | 0.47758004 |
| SLC24A1 | 232.032276 | -0.022508715 | 0.31305282 | -0.0719007 | 0.94268094 | 0.97554394 |
| DENND4A | 3339.2538 | -0.451637911 | 0.29884683 | -1.5112689 | 0.13071996 | 0.31311151 |
| RAB11A | 1869.2386 | -0.721571378 | 0.42307336 | -1.7055467 | 0.08809251 | 0.24126352 |
| MEGF11 | 1356.21021 | -5.167788371 | 0.71905468 | -7.1869199 | 6.63E-13 | 2.30E-10 |
| DIS3L | 300.845331 | 0.416961458 | 0.21225686 | 1.96441925 | 0.04948149 | 0.1650091 |
| TIPIN | 305.352127 | 0.944760589 | 0.32530438 | 2.90423566 | 0.00368151 | 0.02694131 |
| MAP2K1 | 1120.2646 | -0.683944237 | 0.27451862 | -2.4914311 | 0.01272296 | 0.06554509 |
| SNAPC5 | 119.982411 | -0.183957141 | 0.26567179 | -0.6924226 | 0.48867201 | 0.69564269 |
| RPL4 | 773.477105 | -1.423677643 | 0.44822843 | -3.1762323 | 0.00149201 | 0.01366714 |
| ZWILCH | 108.930849 | 0.056676891 | 0.33797333 | 0.16769634 | 0.86682217 | 0.93959274 |
| SMAD6 | 305.008387 | -0.069326121 | 0.53074362 | -0.1306207 | 0.89607534 | 0.95208005 |
| AC087482.1 | 67.0058853 | 0.695220843 | 0.84275289 | 0.82494033 | 0.40940547 | 0.62933048 |
| SMAD3 | 3344.31541 | -1.373475565 | 0.4922519 | -2.7901885 | 0.00526774 | 0.03481445 |
| AAGAB | 873.068276 | -0.257050373 | 0.19729391 | -1.3028804 | 0.19261559 | 0.39991502 |
| IQCH | 1043.3635 | 0.095415876 | 0.25493783 | 0.37427115 | 0.70820259 | 0.85344841 |
| IQCH-AS1 | 691.244397 | 0.646536128 | 0.23741595 | 2.72322112 | 0.00646488 | 0.04039478 |
| C15orf61 | 76.0119784 | 0.505660076 | 0.33762626 | 1.49769178 | 0.13421335 | 0.31862403 |
| MAP2K5 | 1873.18244 | 0.059556508 | 0.1766832 | 0.33708075 | 0.73605603 | 0.87055499 |
| PIAS1 | 2445.71569 | 0.095314992 | 0.14844911 | 0.64207182 | 0.52082655 | 0.72082962 |
| FEM1B | 247.055054 | 0.186308666 | 0.27558125 | 0.67605711 | 0.49900439 | 0.70390435 |
| CORO2B | 53.0851575 | -2.520303264 | 0.75448945 | -3.3404089 | 0.00083655 | 0.00870788 |
| ANP32A | 520.154336 | -0.396395725 | 0.23900411 | -1.658531 | 0.09721034 | 0.25760256 |
| SPESP1 | 192.67815 | 0.56467929 | 0.26279362 | 2.14875574 | 0.03165377 | 0.12243473 |
| NOX5 | 58.8282059 | 0.360164586 | 0.43548618 | 0.82704023 | 0.40821426 | 0.62808412 |
| GLCE | 782.643812 | 0.503186526 | 0.2297776 | 2.18988503 | 0.02853258 | 0.11348594 |
| AC026992.2 | 159.265677 | 1.483929675 | 0.44402743 | 3.34197751 | 0.00083184 | 0.00867226 |
| AC026992.1 | 119.856199 | 1.23799382 | 0.65760539 | 1.88257858 | 0.0597575 | 0.18751469 |
| PAQR5 | 3535.86869 | 0.034604727 | 0.3124221 | 0.11076274 | 0.91180449 | 0.96021221 |
| AC027237.3 | 116.216579 | -0.129357414 | 0.61170352 | -0.2114708 | 0.83251994 | 0.92169608 |
| KIF23 | 518.382818 | 0.27598237 | 0.71126624 | 0.38801556 | 0.69800452 | 0.84616339 |
| RPLP1 | 1686.09696 | -0.858186216 | 0.4306844 | -1.9926104 | 0.04630413 | 0.15746395 |
| DRAIC | 60.4259364 | 2.085895343 | 0.44752819 | 4.66092506 | 3.15E-06 | 0.00010267 |
| TLE3 | 325.572396 | -0.730824499 | 0.27887492 | -2.6206175 | 0.00877707 | 0.05041939 |
| UACA | 1679.10257 | -0.532479473 | 0.17518995 | -3.0394407 | 0.00237018 | 0.01944577 |
| AC009269.2 | 47.261654 | -0.7804922 | 0.58348392 | -1.3376413 | 0.18101339 | 0.38492165 |
| LARP6 | 141.27679 | 0.059871282 | 0.41150059 | 0.14549501 | 0.88432006 | 0.9477505 |
| LRRC49 | 506.863722 | -0.188679602 | 0.24999081 | -0.7547462 | 0.45040131 | 0.66647279 |
| THSD4 | 2816.29811 | 0.618436194 | 0.54239295 | 1.14019955 | 0.25420317 | 0.47493207 |
| THSD4-AS1 | 423.507706 | -0.735683034 | 0.21539651 | -3.4154826 | 0.00063669 | 0.0071434 |
| MYO9A | 10171.5166 | 0.237581166 | 0.16855597 | 1.40950905 | 0.1586847 | 0.35514221 |
| AC022872.1 | 330.175568 | -0.022280004 | 0.50770952 | -0.0438834 | 0.96499737 | 0.98444936 |
| SENP8 | 259.200889 | 0.484884943 | 0.25812497 | 1.87848909 | 0.06031429 | 0.18865532 |
| AC020779.2 | 43.3511737 | -0.049099561 | 0.4734135 | -0.1037139 | 0.9173964 | 0.96291779 |
| GRAMD2A | 176.740332 | -0.120898608 | 0.32334435 | -0.3739005 | 0.70847836 | 0.85346928 |
| PKM | 3701.03428 | -1.296081971 | 0.3202664 | -4.0468871 | 5.19E-05 | 0.00100526 |
| PARP6 | 534.849646 | 0.218187774 | 0.22593857 | 0.9656951 | 0.33419677 | 0.5593519 |
| AC009690.3 | 45.5586221 | 1.136585172 | 0.70532378 | 1.61143747 | 0.10708441 | 0.27484035 |
| AC009690.2 | 610.6491 | 1.186990537 | 0.69135195 | 1.7169121 | 0.08599523 | 0.23724124 |
| AC009690.1 | 2004.9591 | 1.46947856 | 0.46784095 | 3.14097889 | 0.00168384 | 0.01497769 |
| TMEM202-A! | 117.415191 | 0.716432108 | 0.44941842 | 1.59413162 | 0.11090656 | 0.28096754 |
| ARIH1 | 3613.49603 | 0.07067726 | 0.19412835 | 0.3640749 | 0.71580208 | 0.85814392 |
| BBS4 | 757.216588 | 0.030521453 | 0.24361049 | 0.12528792 | 0.90029561 | 0.95395051 |
| ADPGK | 432.345058 | -0.241211385 | 0.25088994 | -0.9614231 | 0.33633947 | 0.5617992 |
| NEO1 | 1211.10903 | -0.083776363 | 0.20021416 | -0.4184338 | 0.67563001 | 0.82986765 |
| NPTN | 1257.75134 | 0.203983877 | 0.14808437 | 1.37748418 | 0.1683626 | 0.36843963 |
| CD276 | 92.6080545 | -0.526128464 | 0.25875763 | -2.0332868 | 0.04202356 | 0.14700455 |
| LOXL1-AS1 | 40.7972721 | -0.062485136 | 0.39801174 | -0.1569932 | 0.87525021 | 0.94384684 |
| LOXL1 | 78.1547666 | 0.527082347 | 0.30608585 | 1.72200823 | 0.08506803 | 0.23547016 |
| PML | 219.957827 | -0.721792566 | 0.36090395 | -1.9999575 | 0.04550485 | 0.15579927 |
| UBL7 | 64.7090067 | -0.026901851 | 0.23266952 | -0.1156226 | 0.90795166 | 0.9583773 |
| UBL7-AS1 | 63.0423219 | -0.330020134 | 0.43309178 | -0.7620097 | 0.44605422 | 0.66204652 |
| AC012435.2 | 89.3799412 | 0.25502838 | 0.31063741 | 0.82098411 | 0.41165532 | 0.63088004 |
| ARID3B | 189.097156 | 0.179042392 | 0.23208829 | 0.77144087 | 0.44044566 | 0.65619747 |
| CLK3 | 281.128356 | 0.584297164 | 0.16301185 | 3.58438456 | 0.00033787 | 0.00439269 |
| EDC3 | 376.809837 | 0.401410353 | 0.17499704 | 2.29381227 | 0.02180128 | 0.09452959 |
| CSK | 93.6291913 | -0.434349993 | 0.2780292 | -1.562246 | 0.11823005 | 0.29330654 |
| ULK3 | 187.143403 | 1.026029538 | 0.29951379 | 3.4256504 | 0.00061333 | 0.00694286 |
| SCAMP2 | 315.51804 | -0.254340616 | 0.17029578 | -1.4935227 | 0.13530042 | 0.32045602 |
| MPI | 76.3703717 | 0.06979926 | 0.33752684 | 0.20679618 | 0.83616904 | 0.92372308 |
| FAM219B | 162.463801 | -0.286821228 | 0.20553176 | -1.3955081 | 0.16286268 | 0.3609485 |
| COX5A | 504.6355 | 0.436246817 | 0.24115741 | 1.80897125 | 0.07045547 | 0.20912191 |
| SCAMP5 | 172.620665 | 0.181978064 | 0.34891315 | 0.52155691 | 0.60197887 | 0.77940904 |
| PPCDC | 478.45582 | -0.180746708 | 0.35342072 | -0.5114208 | 0.60905643 | 0.78488062 |
| COMMD4 | 207.380634 | -0.088998469 | 0.26142943 | -0.3404302 | 0.73353259 | 0.86914532 |
| NEIL1 | 258.941003 | -0.074239263 | 0.29777211 | -0.2493157 | 0.80311658 | 0.90758619 |
| MAN2C1 | 424.970023 | 0.148664713 | 0.28332381 | 0.52471663 | 0.5997802 | 0.77794221 |
| SIN3A | 788.347334 | -0.323703252 | 0.17823912 | -1.8161179 | 0.06935229 | 0.20692895 |
| PTPN9 | 897.102209 | -0.784330917 | 0.18595655 | -4.2178181 | 2.47E-05 | 0.00056082 |
| SNUPN | 157.837218 | -0.228870392 | 0.27602991 | -0.8291507 | 0.40701913 | 0.6269756 |
| IMP3 | 107.572812 | -0.93761632 | 0.39309788 | -2.3851981 | 0.01706993 | 0.07995627 |

| | | | | | | |
|---|---|---|---|---|---|---|
| SNX33 | 77.1770859 | -0.884044883 | 0.26500326 | -3.3359774 | 0.00085 | 0.00878534 |
| UBE2Q2 | 404.680888 | -1.153006828 | 0.20159161 | -5.719518 | 1.07E-08 | 9.73E-07 |
| FBXO22 | 615.381388 | 0.037157452 | 0.14974499 | 0.2481382 | 0.80402748 | 0.90804718 |
| NRG4 | 101.733492 | 1.119484997 | 0.5359287 | 2.08886927 | 0.03671949 | 0.13485073 |
| TMEM266 | 209.155504 | 2.166833832 | 0.74497257 | 2.90860887 | 0.00363041 | 0.02666817 |
| ETFA | 1229.36879 | 0.59180078 | 0.21033951 | 2.81355018 | 0.00489977 | 0.03294199 |
| SCAPER | 2786.45164 | 0.054167849 | 0.20514394 | 0.26404801 | 0.79174293 | 0.9019242 |
| RCN2 | 568.515181 | 0.573747186 | 0.19378144 | 2.96079531 | 0.00306846 | 0.02355099 |
| TSPAN3 | 599.261175 | -0.330093222 | 0.17832769 | -1.8510486 | 0.06416256 | 0.196476 |
| PEAK1 | 1365.71673 | 0.028471175 | 0.20761599 | 0.13713382 | 0.89092502 | 0.95055821 |
| HMG20A | 913.998053 | 0.407045597 | 0.10805346 | 3.76707616 | 0.00016517 | 0.00251063 |
| LINGO1 | 4882.46187 | 1.077243779 | 0.55327528 | 1.94703036 | 0.05153109 | 0.1694925 |
| TBC1D2B | 201.081367 | 0.139595136 | 0.22897688 | 0.60964731 | 0.54209547 | 0.7381999 |
| IDH3A | 540.372831 | 0.07835517 | 0.24676708 | 0.31752683 | 0.75084388 | 0.87901239 |
| DNAJA4 | 85.3560532 | -0.242753364 | 0.27230856 | -0.8914643 | 0.37268014 | 0.59655016 |
| WDR61 | 358.080848 | -0.041692428 | 0.17157014 | -0.2430052 | 0.8080014 | 0.91010272 |
| IREB2 | 1002.74455 | 0.037629982 | 0.13667722 | 0.27532008 | 0.78307034 | 0.89647912 |
| HYKK | 167.454064 | 0.841193642 | 0.32797316 | 2.5648246 | 0.0103228 | 0.05676679 |
| PSMA4 | 332.952264 | -0.32668285 | 0.27819638 | -1.1742886 | 0.24027943 | 0.45831784 |
| MORF4L1 | 1121.65877 | -0.312534981 | 0.13910556 | -2.2467468 | 0.02465621 | 0.10270701 |
| CTSH | 372.704169 | -0.24027072 | 0.33966863 | -0.707368 | 0.47933781 | 0.68864274 |
| TMED3 | 292.863945 | -0.926253558 | 0.31696261 | -2.9222802 | 0.00347479 | 0.02584181 |
| MTHFS | 71.4728221 | 0.343426951 | 0.26038072 | 1.31894153 | 0.18718866 | 0.39289501 |
| ZFAND6 | 2319.11432 | -0.286106898 | 0.19498616 | -1.467319 | 0.14228929 | 0.33124766 |
| FAH | 89.9807952 | 0.344235942 | 0.30616082 | 1.12436315 | 0.260859 | 0.48204904 |
| CTXND1 | 108.78154 | 0.773728969 | 0.77330633 | 1.00054653 | 0.31704609 | 0.5415572 |
| LINC00927 | 41.3623807 | 1.452079968 | 0.58688474 | 2.47421662 | 0.01335287 | 0.0674744 |
| AC016705.2 | 1445.42948 | 1.144135369 | 0.55089113 | 2.07688111 | 0.03781253 | 0.13759387 |
| ARNT2 | 3162.6338 | 0.167426347 | 0.29337491 | 0.57069076 | 0.56820929 | 0.75835874 |
| ABHD17C | 993.157384 | 1.155856113 | 0.29883019 | 3.86793627 | 0.00010976 | 0.00180326 |
| CEMIP | 144.25806 | 0.330640245 | 0.57902312 | 0.57103116 | 0.56797852 | 0.75812736 |
| AC027808.1 | 242.488725 | 0.601164949 | 0.30707476 | 1.95771526 | 0.05026343 | 0.16685843 |
| MESD | 426.021804 | -0.020254219 | 0.20514569 | -0.0987309 | 0.92135193 | 0.96504145 |
| TLNRD1 | 60.2388978 | -1.583534783 | 0.43646575 | -3.6280848 | 0.00028553 | 0.00389224 |
| CFAP161 | 70.4367497 | -0.038858266 | 0.56812183 | -0.0683978 | 0.945469 | 0.97640274 |
| AC104041.1 | 441.851097 | 0.569440391 | 0.73207149 | 0.77784806 | 0.43665859 | 0.65342962 |
| EFL1 | 509.059462 | 0.115308073 | 0.23737027 | 0.48577302 | 0.62712811 | 0.79774029 |
| SAXO2 | 123.228485 | 1.351703109 | 0.49569403 | 2.72689002 | 0.00639343 | 0.04006209 |
| GOLGA6L10 | 39.7318262 | 0.85061735 | 0.72662784 | 1.17063688 | 0.24174477 | 0.46003068 |
| GOLGA6L9 | 70.389284 | 0.271163535 | 0.26245247 | 1.03319104 | 0.30151451 | 0.52486889 |
| RPS17 | 77.5903755 | -0.275847092 | 0.5329217 | -0.5176128 | 0.60472845 | 0.78156518 |
| AP3B2 | 108.275239 | -1.188816167 | 0.5995838 | -1.9827356 | 0.04739697 | 0.15999329 |
| AC105339.3 | 80.4348403 | 0.493926623 | 0.34344149 | 1.43816817 | 0.15038634 | 0.3427283 |
| SNHG21 | 73.6852946 | 0.44315899 | 0.28906374 | 1.53308399 | 0.12525516 | 0.30391173 |
| WHAMM | 570.726605 | -0.283310385 | 0.21154074 | -1.339271 | 0.18048246 | 0.38426706 |
| HOMER2 | 438.585568 | -0.027226519 | 0.27201044 | -0.1000937 | 0.92026997 | 0.96454603 |
| AC022558.2 | 141.572498 | 0.095817937 | 0.49078256 | 0.19523501 | 0.84520897 | 0.92842955 |
| C15orf40 | 285.455816 | 0.197458289 | 0.2170949 | 0.90954826 | 0.36306079 | 0.5885693 |
| BTBD1 | 643.255004 | -0.117765772 | 0.17342452 | -0.6790607 | 0.49709942 | 0.70224997 |
| HDGFL3 | 617.627076 | -0.286286565 | 0.17958478 | -1.5941583 | 0.11090059 | 0.28096754 |
| AC103876.1 | 94.8708716 | -0.269982223 | 0.41070696 | -0.6573597 | 0.51094963 | 0.71298687 |
| ADAMTSL3 | 644.745388 | 0.887401752 | 0.73567101 | 1.2062481 | 0.22772184 | 0.44332761 |
| UBE2Q2L | 49.4712712 | 0.08842511 | 0.66831001 | 0.13231152 | 0.8947379 | 0.95172874 |
| GOLGA6L4 | 58.0614171 | 0.219820668 | 0.39352253 | 0.55859741 | 0.57643651 | 0.76324677 |
| AC048382.5 | 116.654897 | -0.193463942 | 0.34591167 | -0.5592871 | 0.57596578 | 0.7629294 |
| ZSCAN2 | 98.196604 | 0.020231657 | 0.27156058 | 0.07450145 | 0.94061139 | 0.97438649 |
| WDR73 | 672.172603 | -0.139821437 | 0.20573626 | -0.6796149 | 0.49674831 | 0.70199513 |
| SEC11A | 1095.66846 | -0.131853798 | 0.15043367 | -0.8764913 | 0.38076303 | 0.60354344 |
| ZNF592 | 756.83261 | -0.03549974 | 0.16836743 | -0.2108468 | 0.83300679 | 0.92200324 |
| ALPK3 | 1039.91483 | -1.150622108 | 0.38265631 | -3.0069336 | 0.00263897 | 0.02108785 |
| SLC28A1 | 53.9192858 | 1.086457756 | 0.61355996 | 1.77074424 | 0.07660324 | 0.22078382 |
| PDE8A | 2521.62692 | 0.031135091 | 0.24975534 | 0.12466236 | 0.90079086 | 0.95413737 |
| AKAP13 | 5269.38703 | -0.724370723 | 0.22971893 | -3.1532914 | 0.00161441 | 0.0145561 |
| AC087286.2 | 180.97693 | -0.951008567 | 0.51462681 | -1.8479577 | 0.06460846 | 0.1973666 |
| KLHL25 | 91.9945457 | -0.174151363 | 0.25570177 | -0.6810722 | 0.49582582 | 0.70114155 |
| MRPL46 | 149.475652 | -0.070730398 | 0.2786266 | -0.2538537 | 0.79960857 | 0.90598984 |
| MRPS11 | 242.158706 | -0.257389264 | 0.25081467 | -1.0262129 | 0.30479122 | 0.52816954 |
| DET1 | 286.843323 | 0.397862749 | 0.44730334 | 0.88946966 | 0.37375072 | 0.59768442 |
| ABHD2 | 688.484106 | -0.656378454 | 0.31062019 | -2.1131223 | 0.0345903 | 0.12994973 |
| FANCI | 47.1504342 | -0.3448618 | 0.53627321 | -0.6430711 | 0.52017797 | 0.72000742 |
| POLG | 203.73429 | -0.158633644 | 0.21910087 | -0.7240211 | 0.46905281 | 0.68108245 |
| PEX11A | 106.563952 | -0.038755818 | 0.27377373 | -0.1415615 | 0.88742638 | 0.94928354 |
| WDR93 | 63.1405996 | 0.101009667 | 0.54066262 | 0.18682569 | 0.8517973 | 0.93193618 |
| ANPEP | 111.017306 | -0.664985747 | 0.66834418 | -0.994975 | 0.31974843 | 0.54422157 |
| AP3S2 | 50.4326254 | -0.351161399 | 0.38519627 | -0.9116428 | 0.36195679 | 0.58728501 |
| ZNF710 | 562.173996 | -0.070012681 | 0.27170586 | -0.2576782 | 0.79665526 | 0.90460255 |
| ZNF710-AS1 | 347.403821 | 0.486117634 | 0.31814777 | 1.52796176 | 0.12652203 | 0.30597392 |
| IDH2 | 2753.49942 | 0.106577496 | 0.28750897 | 0.37069277 | 0.71086638 | 0.85502039 |
| IDH2-DT | 525.408033 | 0.742624137 | 0.5759253 | 1.28944525 | 0.19724334 | 0.40569718 |
| SEMA4B | 2089.51362 | -2.150540776 | 0.42985619 | -5.0029308 | 5.65E-07 | 2.60E-05 |
| CIB1 | 772.169221 | -1.986503415 | 0.45467101 | -4.3691006 | 1.25E-05 | 0.00031752 |
| AC091167.6 | 106.618013 | 0.087975167 | 0.35441104 | 0.24822919 | 0.80395708 | 0.90804718 |
| ZNF774 | 69.3525115 | 0.749172621 | 0.25070872 | 2.98821919 | 0.00280608 | 0.02206666 |
| IQGAP1 | 1844.18164 | -0.939713244 | 0.20114348 | -4.6718553 | 2.98E-06 | 9.83E-05 |
| CRTC3 | 1241.12016 | -0.163176922 | 0.21741066 | -0.750547 | 0.45292531 | 0.66868339 |
| CRTC3-AS1 | 215.82379 | -0.054905384 | 0.56720434 | -0.0968 | 0.92288522 | 0.96580501 |
| FURIN | 193.268183 | -1.319998426 | 0.39809668 | -3.3157735 | 0.0009139 | 0.00930022 |
| MAN2A2 | 185.572028 | 0.110531375 | 0.23216167 | 0.47609657 | 0.63400558 | 0.80308589 |
| HDDC3 | 79.6671671 | -0.220843921 | 0.27985982 | -0.7891234 | 0.43003991 | 0.64824168 |
| UNC45A | 277.933468 | -0.411333778 | 0.19881759 | -2.0689003 | 0.03855544 | 0.1393757 |
| RCCD1 | 94.3199471 | 0.29742894 | 0.22603721 | 1.31584062 | 0.18822754 | 0.39384317 |
| AC104035.1 | 700.511957 | 1.854448059 | 0.9206154 | 2.01435698 | 0.04397207 | 0.15199268 |
| SLCO3A1 | 2796.17444 | -0.168177001 | 0.26086751 | -0.6446836 | 0.51913225 | 0.71956991 |
| AC116903.2 | 60.210779 | -0.403278678 | 0.51774301 | -0.7789167 | 0.43602879 | 0.65307906 |
| FAM174B | 568.834307 | -0.028502116 | 0.42965583 | -0.0663371 | 0.94710946 | 0.97737256 |
| AC013394.1 | 1643.21469 | 0.070309909 | 0.16826065 | 0.41786306 | 0.67604724 | 0.82991711 |
| CHD2 | 2345.71202 | -0.245759873 | 0.28997293 | -0.8475269 | 0.39670148 | 0.61845042 |
| MCTP2 | 255.344738 | -1.942255969 | 0.73011132 | -2.660219 | 0.00780899 | 0.04642165 |
| LINC01197 | 276.000891 | 2.480115468 | 0.84160806 | 2.94687705 | 0.00321001 | 0.02432645 |
| AC012409.2 | 986.635611 | 1.053147151 | 0.4233184 | 2.48783692 | 0.01285227 | 0.06595399 |
| AC024337.2 | 62.8927247 | 1.345343936 | 0.34344642 | 3.9171873 | 8.96E-05 | 0.00155094 |
| NR2F2-AS1 | 6802.41482 | 1.180989338 | 0.26523758 | 4.45257164 | 8.48E-06 | 0.00023645 |
| NR2F2 | 1174.50733 | 0.148781514 | 0.25877458 | 0.57494639 | 0.56532755 | 0.75611701 |
| AC087477.2 | 262.72894 | -0.075351075 | 0.49353454 | -0.1526764 | 0.87865347 | 0.94585961 |
| LINC00923 | 84.2189171 | 1.269930498 | 0.76209679 | 1.66636378 | 0.09564098 | 0.2549135 |
| ARRDC4 | 472.165188 | 0.42523053 | 0.34295889 | 1.23988777 | 0.21501691 | 0.4288968 |
| FAM169B | 208.02445 | -3.8453139 | 0.88767645 | -4.3318868 | 1.48E-05 | 0.00036873 |
| IGF1R | 9809.4182 | 0.438208375 | 0.24969862 | 1.75494913 | 0.07926801 | 0.22502711 |
| AC036108.1 | 138.415627 | 0.150700934 | 0.54735221 | 0.27532717 | 0.78306489 | 0.89647912 |
| TTC23 | 636.680406 | 0.189514482 | 0.17293015 | 1.09590191 | 0.2731217 | 0.49481067 |
| LRRC28 | 1050.75993 | 0.743896978 | 0.15165642 | 4.90514653 | 9.34E-07 | 3.94E-05 |
| MEF2A | 4462.14988 | 0.247482812 | 0.23037578 | 1.07425706 | 0.28270749 | 0.50511067 |
| LYSMD4 | 119.661061 | 0.283359888 | 0.25366286 | 1.11707281 | 0.2639632 | 0.484936 |

| | | | | | | |
|---|---|---|---|---|---|---|
| AC090825.1 | 204.620203 | 0.010343364 | 0.54558638 | 0.01895825 | 0.98487441 | 0.99405563 |
| ADAMTS17 | 1298.00015 | 2.00332664 | 0.5461381 | 3.66816862 | 0.00024429 | 0.00345807 |
| AC027020.2 | 73.377735 | 1.411325065 | 0.47066047 | 2.9986055 | 0.00271218 | 0.02146508 |
| LINS1 | 319.862436 | 0.892781454 | 0.22623028 | 3.94633937 | 7.94E-05 | 0.00140887 |
| ASB7 | 413.26083 | 0.190507076 | 0.17245797 | 1.104658 | 0.26930781 | 0.49084828 |
| AC087762.1 | 48.806775 | 1.249981204 | 0.45075782 | 2.77306604 | 0.00555308 | 0.03627631 |
| CHSY1 | 806.08526 | -0.526758009 | 0.2736987 | -1.9245908 | 0.05428057 | 0.1756637 |
| SELENOS | 370.525576 | -0.907540376 | 0.31464814 | -2.8843024 | 0.00392282 | 0.0282225 |
| SNRPA1 | 339.272691 | -0.138800045 | 0.17134069 | -0.8100822 | 0.41789293 | 0.63753323 |
| AC023024.1 | 49.224642 | 0.668371593 | 0.30893617 | 2.16346175 | 0.03050568 | 0.11918059 |
| PCSK6 | 1195.8407 | -1.04341606 | 0.41120826 | -2.5374395 | 0.01116667 | 0.05999714 |
| TM2D3 | 373.022891 | -0.025420318 | 0.19968967 | -0.1272991 | 0.89870366 | 0.9532601 |
| TARSL2 | 628.242003 | 0.070150833 | 0.2193997 | 0.31973988 | 0.74916552 | 0.87779905 |
| POLR3K | 41.1045766 | -0.389533026 | 0.37125043 | -1.049246 | 0.29406491 | 0.51714864 |
| SNRNP25 | 71.3101916 | -0.130582929 | 0.48759658 | -0.2678094 | 0.78884606 | 0.89989357 |
| RHBDF1 | 56.9919095 | -1.515026974 | 0.45092513 | -3.3598194 | 0.00077993 | 0.00825243 |
| MPG | 208.660535 | -1.208800447 | 0.21648115 | -5.5838601 | 2.35E-08 | 1.89E-06 |
| NPRL3 | 651.108065 | -0.212827031 | 0.24823683 | -0.8573548 | 0.39124884 | 0.6140943 |
| LUC7L | 1110.51391 | 0.358409168 | 0.16446646 | 2.17922351 | 0.02931507 | 0.11551991 |
| FAM234A | 269.047181 | 0.603296152 | 0.33806841 | 1.78453865 | 0.07433618 | 0.216422 |
| AXIN1 | 285.581159 | -0.612126711 | 0.22679841 | -2.6989903 | 0.00695502 | 0.04269 |
| MRPL28 | 170.527574 | -0.221407871 | 0.46719266 | -0.4739113 | 0.63556318 | 0.80389813 |
| TMEM8A | 596.757372 | -0.536149998 | 0.31708485 | -1.6908723 | 0.09086119 | 0.24629294 |
| NME4 | 136.040961 | 0.119631253 | 0.50358906 | 0.23755729 | 0.81222448 | 0.91190752 |
| DECR2 | 79.9331498 | 0.033867669 | 0.32096871 | 0.10551704 | 0.91596555 | 0.96214624 |
| RAB11FIP3 | 1845.06024 | 0.393826669 | 0.3070715 | 1.28252431 | 0.19965878 | 0.40831558 |
| CAPN15 | 275.534141 | -0.361945979 | 0.30173831 | -1.199536 | 0.23031958 | 0.44592324 |
| PIGQ | 279.486957 | 0.23767229 | 0.3305637 | 0.7189909 | 0.47214653 | 0.68319499 |
| RAB40C | 670.533921 | -0.197413248 | 0.30461873 | -0.6480667 | 0.51694183 | 0.71823757 |
| WFIKKN1 | 39.5783757 | 1.977773715 | 0.67017506 | 2.95112996 | 0.00316614 | 0.02404917 |
| METTL26 | 101.483274 | -0.041040866 | 0.44469358 | -0.0922902 | 0.92646746 | 0.96763728 |
| MCRIP2 | 347.152495 | 1.115578518 | 0.27874355 | 4.00216798 | 6.28E-05 | 0.00116278 |
| WDR90 | 149.622973 | -0.315835282 | 0.27175993 | -1.1621849 | 0.24516038 | 0.46420922 |
| RHOT2 | 336.418425 | 0.413126553 | 0.25612623 | 1.61298025 | 0.1067488 | 0.27427444 |
| STUB1 | 293.678466 | -0.571594874 | 0.32980322 | -1.7331392 | 0.08307094 | 0.23188589 |
| JMJD8 | 148.332657 | -0.551429396 | 0.4294789 | -1.2839499 | 0.19915948 | 0.40773366 |
| WDR24 | 46.3627079 | -0.22257321 | 0.36536412 | -0.6091819 | 0.54240388 | 0.73846762 |
| METRN | 224.151211 | -0.145278648 | 0.28656338 | -0.5069686 | 0.61217682 | 0.78684784 |
| FAM173A | 180.958129 | -0.271105038 | 0.40823206 | -0.6640954 | 0.50662924 | 0.71003349 |
| HAGHL | 84.7352903 | 1.064175806 | 0.53763381 | 1.9793692 | 0.04777445 | 0.16085974 |
| NARFL | 181.65487 | -0.004552935 | 0.26400387 | -0.0172457 | 0.9862406 | 0.99452264 |
| PRR25 | 45.8578202 | 0.039569207 | 0.40247714 | 0.09831417 | 0.92168282 | 0.96523677 |
| LMF1 | 861.104778 | 0.556236702 | 0.5338476 | 1.04193913 | 0.2974399 | 0.52079582 |
| CACNA1H | 482.21616 | 0.000354331 | 0.4553085 | 0.00077822 | 0.99937907 | 0.99975417 |
| UBE2I | 697.931808 | -0.59904321 | 0.2441878 | -2.4532069 | 0.01415889 | 0.07032316 |
| BAIAP3 | 119.94984 | -0.222288374 | 0.34466672 | -0.6449372 | 0.51896791 | 0.71946641 |
| TSR3 | 125.377361 | -0.642542194 | 0.37119439 | -1.7310127 | 0.0834495 | 0.23254946 |
| GNPTG | 367.513 | 0.019757527 | 0.19379787 | 0.10194915 | 0.91879704 | 0.96389406 |
| UNKL | 377.831733 | -0.545450831 | 0.27710256 | -1.9684078 | 0.04902114 | 0.16392918 |
| C16orf91 | 45.5719543 | -0.024866528 | 0.41777527 | -0.0595213 | 0.9525369 | 0.97933189 |
| CCDC154 | 61.6811252 | 0.524604892 | 0.36568211 | 1.43459272 | 0.15140319 | 0.34433277 |
| CLCN7 | 477.876489 | -0.032685571 | 0.2818056 | -0.1159862 | 0.90766345 | 0.95814964 |
| TELO2 | 129.488423 | -0.507866147 | 0.26631945 | -1.9069811 | 0.05652304 | 0.1806467 |
| IFT140 | 479.071193 | 0.041213675 | 0.27495981 | 0.14988982 | 0.88085155 | 0.94640313 |
| AL133297.2 | 55.7622751 | -0.181689898 | 0.38502283 | -0.4718938 | 0.63700258 | 0.80480926 |
| CRAMP1 | 107.792902 | 0.102542333 | 0.32604823 | 0.31450051 | 0.75314093 | 0.88014525 |
| JPT2 | 773.474933 | -1.288186389 | 0.28577591 | -4.5076801 | 6.55E-06 | 0.00018901 |
| MAPK8IP3 | 1278.39712 | -0.223075576 | 0.25963335 | -0.8591946 | 0.39023315 | 0.61320085 |
| NME3 | 398.037593 | 0.108472533 | 0.33974024 | 0.31928079 | 0.74951359 | 0.87807761 |
| MRPS34 | 189.656682 | -0.58185188 | 0.3189176 | -1.8244584 | 0.0680828 | 0.20452712 |
| EME2 | 116.178856 | 0.348852026 | 0.32586876 | 1.07052922 | 0.28438116 | 0.50683977 |
| SPSB3 | 504.95678 | 0.001984376 | 0.21202361 | 0.00935922 | 0.99253253 | 0.99698572 |
| NUBP2 | 114.722352 | -0.239624371 | 0.26002948 | -0.9215277 | 0.35677498 | 0.58245467 |
| HAGH | 627.482414 | -0.170301637 | 0.18069613 | -0.9424753 | 0.34594935 | 0.57105512 |
| FAHD1 | 151.207215 | -0.578020504 | 0.44788738 | -1.2905488 | 0.19686019 | 0.40510928 |
| MSRB1 | 138.52177 | -2.099974236 | 0.45632709 | -4.6019057 | 4.19E-06 | 0.0001292 |
| NDUFB10 | 305.954442 | -0.463936104 | 0.42427456 | -1.0934809 | 0.27418273 | 0.49598462 |
| RPS2 | 867.329272 | -0.995242808 | 0.39250166 | -2.5356397 | 0.01122421 | 0.06019514 |
| SNHG9 | 88.6373778 | -0.294580242 | 0.4379816 | -0.6725859 | 0.50121079 | 0.70528379 |
| TBL3 | 48.478632 | -0.403885465 | 0.35509065 | -1.1374151 | 0.2553648 | 0.47622669 |
| GFER | 48.3627866 | -0.122068398 | 0.41790059 | -0.2920991 | 0.77021084 | 0.88929655 |
| ZNF598 | 95.1707936 | -0.102137362 | 0.2857202 | -0.3574734 | 0.72073746 | 0.86093517 |
| SLC9A3R2 | 51.7093589 | -0.074128858 | 0.40439133 | -0.1833097 | 0.85455501 | 0.93357185 |
| TSC2 | 677.824764 | 0.225653042 | 0.3190689 | 0.70722356 | 0.47942756 | 0.68864274 |
| PKD1 | 428.873176 | 0.148258664 | 0.20668474 | 0.7173179 | 0.47317797 | 0.68383017 |
| SNHG19 | 155.687598 | 0.256300196 | 0.3081846 | 0.83164505 | 0.40560933 | 0.62584264 |
| TRAF7 | 124.437117 | -1.04041479 | 0.32419098 | -3.2092651 | 0.00133075 | 0.01251092 |
| MLST8 | 71.3398836 | -0.193746876 | 0.3990811 | -0.4854825 | 0.62733415 | 0.79784872 |
| BRICD5 | 59.7326861 | 0.775290633 | 0.45274106 | 1.71243719 | 0.08681613 | 0.23862737 |
| PGP | 409.05456 | -0.008762917 | 0.21889933 | -0.0400317 | 0.96806783 | 0.98560259 |
| E4F1 | 188.210151 | 0.015458589 | 0.31368363 | 0.04928083 | 0.9606955 | 0.98279328 |
| ECI1 | 669.696615 | -0.102809709 | 0.24688125 | -0.4164339 | 0.67709256 | 0.83060787 |
| RNPS1 | 473.469488 | -0.080090244 | 0.15966636 | -0.50161 | 0.61594187 | 0.78898859 |
| ABCA3 | 217.411975 | -0.59222512 | 0.18262369 | -3.2428713 | 0.00118332 | 0.01140075 |
| CCNF | 54.5879648 | -0.75900201 | 0.39141515 | -1.9391227 | 0.0524864 | 0.17181979 |
| TBC1D24 | 103.723245 | 0.744331734 | 0.60334589 | 1.23367333 | 0.21732466 | 0.43128814 |
| AMDHD2 | 75.6535185 | 0.671397592 | 0.40531207 | 1.65649545 | 0.09762153 | 0.2582075 |
| PDPK1 | 1034.43317 | -0.100155323 | 0.14452786 | -0.6929828 | 0.48832033 | 0.69521699 |
| AC093525.4 | 101.482409 | -0.395639102 | 0.34703446 | -1.1400571 | 0.25426251 | 0.47497574 |
| KCTD5 | 107.785927 | 0.008241363 | 0.30704054 | 0.02684129 | 0.97858632 | 0.99081303 |
| SRRM2-AS1 | 38.1883626 | -0.031312233 | 0.52043998 | -0.0601649 | 0.95202428 | 0.97930759 |
| SRRM2 | 3256.56835 | 0.128356802 | 0.19404364 | 0.6614842 | 0.50830185 | 0.71121086 |
| ELOB | 440.291019 | -1.130041312 | 0.47867182 | -2.3607851 | 0.01823629 | 0.08397938 |
| PRSS22 | 152.912823 | -1.192332432 | 0.51420302 | -2.318797 | 0.02040604 | 0.09039015 |
| FLYWCH2 | 127.771414 | 0.013055986 | 0.31326539 | 0.04167708 | 0.96675613 | 0.98510234 |
| FLYWCH1 | 339.325734 | 0.36883636 | 0.30247442 | 1.21939688 | 0.22269359 | 0.43695182 |
| TNFRSF12A | 505.745518 | -1.534877306 | 0.60471107 | -2.5381994 | 0.01114245 | 0.05990255 |
| HCFC1R1 | 165.936058 | -0.513887648 | 0.44870971 | -1.1452564 | 0.25210298 | 0.47214352 |
| THOC6 | 51.8335218 | 0.28824818 | 0.50871102 | 0.56662461 | 0.57096925 | 0.75965257 |
| BICDL2 | 190.280323 | -0.754180431 | 0.45492159 | -1.6578251 | 0.09735277 | 0.25780996 |
| MMP25-AS1 | 96.6201906 | -0.894140358 | 0.37413831 | -2.3898658 | 0.01685453 | 0.07944774 |
| IL32 | 113.071696 | -1.345029904 | 0.48871298 | -2.7521878 | 0.00591986 | 0.03797736 |
| ZNF213-AS1 | 218.821788 | 0.169287545 | 0.2137803 | 0.79187627 | 0.42843282 | 0.64676219 |
| ZNF205 | 82.0470988 | 0.151441982 | 0.31218249 | 0.48510723 | 0.62760028 | 0.79811035 |
| ZNF213 | 384.684409 | 0.082239583 | 0.29467967 | 0.2790813 | 0.78018244 | 0.89511246 |
| ZNF200 | 73.3468372 | 0.328792697 | 0.30714744 | 1.07047189 | 0.28440695 | 0.50683977 |
| ZNF263 | 192.382028 | -0.289817016 | 0.26496841 | -1.0937795 | 0.27405169 | 0.49588339 |
| TIGD7 | 154.565783 | -0.056092314 | 0.31195818 | -0.1798072 | 0.85730396 | 0.93564637 |
| ZNF75A | 554.538115 | -0.230479281 | 0.30195564 | -0.7632885 | 0.44529134 | 0.66110524 |
| AC025283.3 | 196.93246 | 0.214694469 | 0.45780229 | 0.46896766 | 0.63909275 | 0.80589749 |
| ZSCAN32 | 120.622008 | 0.222400154 | 0.23461113 | 0.94795229 | 0.34315374 | 0.56819785 |
| NAA60 | 156.075546 | -0.26185501 | 0.24144656 | -1.0845257 | 0.27813177 | 0.50031901 |
| CLUAP1 | 484.899405 | -0.004086202 | 0.22692156 | -0.0180071 | 0.98563318 | 0.99426405 |

| | | | | | | |
|---|---|---|---|---|---|---|
| SLX4 | 75.3185673 | 0.298714658 | 0.35345512 | 0.84512753 | 0.39803964 | 0.61972013 |
| DNASE1 | 1733.12347 | 1.155296012 | 0.3436171 | 3.36216097 | 0.00077335 | 0.00821156 |
| TRAP1 | 519.177814 | 0.424639562 | 0.14492067 | 2.93015186 | 0.00338796 | 0.02538379 |
| CREBBP | 2180.67551 | 0.169209187 | 0.17395957 | 0.97269259 | 0.33070611 | 0.55588514 |
| ADCY9 | 698.186221 | -0.695393443 | 0.31106572 | -2.2355194 | 0.02538327 | 0.10454869 |
| TFAP4 | 47.4905659 | 0.227408193 | 0.53302212 | 0.42663932 | 0.66964206 | 0.82642427 |
| GLIS2 | 289.041913 | -0.521658458 | 0.48475961 | -1.0761178 | 0.28187456 | 0.50430463 |
| GLIS2-AS1 | 40.2421224 | -0.32372583 | 0.46616256 | -0.6944484 | 0.48740107 | 0.69465696 |
| PAM16 | 51.3242197 | -0.701380045 | 0.46755958 | -1.500087 | 0.13359187 | 0.31784355 |
| VASN | 133.49764 | -1.334724659 | 0.62684622 | -2.1292697 | 0.03323195 | 0.12653816 |
| DNAJA3 | 375.206802 | 0.128027678 | 0.16528848 | 0.77457109 | 0.43859315 | 0.65476683 |
| NMRAL1 | 288.982515 | -0.166709635 | 0.20966003 | -0.7951427 | 0.42653052 | 0.64506142 |
| HMOX2 | 313.514064 | -0.234125523 | 0.24078897 | -0.9723266 | 0.33088809 | 0.55589409 |
| CDIP1 | 242.988377 | 0.067075262 | 0.26614685 | 0.25202351 | 0.80102289 | 0.90665907 |
| UBALD1 | 122.055207 | -0.368361767 | 0.27000595 | -1.3642728 | 0.17248172 | 0.3737929 |
| MGRN1 | 707.616389 | 0.082998457 | 0.18110227 | 0.45829606 | 0.64673976 | 0.81112661 |
| NUDT16L1 | 138.989077 | 0.155072013 | 0.36219562 | 0.42814436 | 0.66854602 | 0.82554215 |
| ANKS3 | 422.809397 | 0.071201471 | 0.17773011 | 0.40061568 | 0.6887031 | 0.83983375 |
| ZNF500 | 201.504236 | 0.417299918 | 0.26813074 | 1.55632999 | 0.11962964 | 0.29536727 |
| ROGDI | 88.5082276 | 0.5534699 | 0.5411362 | 1.02279224 | 0.30640609 | 0.53011734 |
| GLYR1 | 1151.29794 | -0.167365022 | 0.14194913 | -1.1790493 | 0.23837855 | 0.45633945 |
| UBN1 | 507.306051 | 0.298814706 | 0.21273566 | 1.40462915 | 0.16013158 | 0.35711263 |
| PPL | 402.089519 | -0.775931182 | 0.29456231 | -2.6341836 | 0.00843399 | 0.04901212 |
| NAGPA | 107.581473 | 0.091684931 | 0.31722954 | 0.28901763 | 0.77256789 | 0.89062065 |
| ALG1 | 166.92408 | 1.088020024 | 0.25455977 | 4.27412405 | 1.92E-05 | 0.00045454 |
| C16orf89 | 157.185208 | 1.405048616 | 0.90280165 | 1.55632039 | 0.11963193 | 0.29536727 |
| EEF2KMT | 43.0382419 | -0.3194285 | 0.4966157 | -0.6432106 | 0.52008743 | 0.71995755 |
| AC026458.2 | 73.9981157 | 0.899303356 | 0.3550512 | 2.53288359 | 0.01131285 | 0.06049857 |
| RBFOX1 | 8820.73171 | 0.137615221 | 0.53701838 | 0.25625794 | 0.79775168 | 0.90506918 |
| METTL22 | 191.340564 | -0.054900787 | 0.31147326 | -0.1762616 | 0.8600884 | 0.93645653 |
| ABAT | 233.288539 | 0.250893305 | 0.28568673 | 0.87821128 | 0.37982907 | 0.60262489 |
| TMEM186 | 55.7111304 | 0.140376648 | 0.31356695 | 0.4476768 | 0.65438646 | 0.8159138 |
| PMM2 | 788.211999 | -0.314846198 | 0.19434417 | -1.6200445 | 0.10522273 | 0.27194033 |
| AC022167.2 | 67.6263212 | 0.396507281 | 0.37081756 | 1.06927859 | 0.28494416 | 0.50741909 |
| CARHSP1 | 496.764535 | -0.944626793 | 0.30976542 | -3.0494908 | 0.0022923 | 0.01899558 |
| USP7 | 592.53472 | 0.044024383 | 0.16363182 | 0.26904536 | 0.78789477 | 0.89956761 |
| C16orf72 | 607.013792 | -0.034710481 | 0.16119209 | -0.2153361 | 0.82950526 | 0.92024799 |
| AC087190.1 | 92.5931768 | -0.270161086 | 0.56988171 | -0.4740652 | 0.63545341 | 0.80389813 |
| ATF7IP2 | 1067.49118 | 0.786132023 | 0.29674524 | 2.64918158 | 0.0080687 | 0.04762262 |
| NUBP1 | 171.323445 | -0.383726662 | 0.22503403 | -1.7051939 | 0.08815826 | 0.24139355 |
| CIITA | 290.296024 | -1.482451885 | 0.5356863 | -2.7673881 | 0.00565074 | 0.03671455 |
| DEXI | 152.634929 | -0.626679714 | 0.26089623 | -2.4020267 | 0.01630452 | 0.07754926 |
| CLEC16A | 3266.20681 | -0.715595278 | 0.20894363 | -3.4248246 | 0.0006152 | 0.00695135 |
| AC007014.1 | 210.34245 | -1.044934745 | 0.5209302 | -2.0059016 | 0.04486674 | 0.15433459 |
| AC007220.1 | 225.441804 | -0.202093316 | 0.38637683 | -0.5230472 | 0.60094141 | 0.7789212 |
| RMI2 | 58.7693638 | 0.638127087 | 0.52296593 | 1.22020775 | 0.22238613 | 0.43657488 |
| SOCS1 | 322.079868 | -2.125377183 | 0.49178651 | -4.3217476 | 1.55E-05 | 0.00038219 |
| AC099489.1 | 237.155348 | 0.742150093 | 0.37271461 | 1.99120202 | 0.04645868 | 0.15779705 |
| LITAF | 1512.96648 | -0.352636134 | 0.2512795 | -1.4033621 | 0.16050888 | 0.35771722 |
| SNN | 53.1732257 | -1.037128496 | 0.39307294 | -2.6385141 | 0.00832702 | 0.04869042 |
| TXNDC11 | 1121.9402 | 0.308011132 | 0.16836193 | 1.82945831 | 0.06733098 | 0.20284564 |
| ZC3H7A | 1091.25192 | -0.010177076 | 0.13333208 | -0.0763288 | 0.93915752 | 0.97405628 |
| RSL1D1 | 197.447065 | -0.324978049 | 0.21132616 | -1.5378033 | 0.12409673 | 0.30213905 |
| AC007216.4 | 579.881028 | 0.396948336 | 0.26760053 | 1.48336152 | 0.13797838 | 0.32401004 |
| GSPT1 | 1310.87104 | -0.141767055 | 0.1730242 | -0.8193481 | 0.41258782 | 0.63186927 |
| NPIPB2 | 128.726701 | 0.694471644 | 0.9937521 | 0.69883791 | 0.48465333 | 0.69239889 |
| SNX29 | 7638.09221 | 0.218020408 | 0.25937859 | 0.84054897 | 0.40060066 | 0.62209429 |
| AC010333.2 | 86.0514585 | 0.458305783 | 0.41670809 | 1.09982454 | 0.27140858 | 0.49278844 |
| CPPED1 | 1413.42484 | -0.41786452 | 0.33597128 | -1.2437507 | 0.21359132 | 0.42681008 |
| AC109597.1 | 95.4445987 | -0.469871797 | 0.52276147 | -0.8988264 | 0.36874515 | 0.59392204 |
| SHISA9 | 1233.39979 | 1.370272702 | 0.61402496 | 2.23162379 | 0.02563984 | 0.1053278 |
| U91319.1 | 720.981678 | 1.351506771 | 0.51508666 | 2.6238435 | 0.00869437 | 0.05008459 |
| ERCC4 | 224.380147 | 0.251804153 | 0.1760108 | 1.43061767 | 0.15253982 | 0.34590532 |
| MKL2 | 2744.8196 | 0.475006629 | 0.2165034 | 2.19399154 | 0.02823602 | 0.11265157 |
| AC130650.1 | 60.4509211 | 0.254777222 | 0.56120285 | 0.45398419 | 0.6498402 | 0.81277713 |
| MIR193BHG | 72.561095 | 0.049981734 | 0.38730501 | 0.12905006 | 0.89731804 | 0.9525128 |
| PARN | 1179.79844 | 0.014018858 | 0.19821263 | 0.07072636 | 0.94361554 | 0.97559625 |
| BFAR | 304.599042 | -0.317773853 | 0.21783745 | -1.458766 | 0.14462954 | 0.33451437 |
| NOMO1 | 228.076426 | 0.678037408 | 0.29994435 | 2.26054402 | 0.02378751 | 0.10024483 |
| NPIPA1 | 99.887537 | 0.53763007 | 0.38928452 | 1.3810723 | 0.16725673 | 0.36693143 |
| PDXDC1 | 2386.32897 | 0.090415473 | 0.17829635 | 0.50710783 | 0.61207916 | 0.78684784 |
| NTAN1 | 153.692496 | -0.118648559 | 0.29979717 | -0.3957628 | 0.69228004 | 0.84217416 |
| RRN3 | 343.430397 | 0.516006127 | 0.26502278 | 1.94702553 | 0.05153167 | 0.1694925 |
| C16orf45 | 833.789296 | -0.710918877 | 0.25177261 | -2.8236545 | 0.00474795 | 0.03215891 |
| MARF1 | 650.333628 | 0.116246061 | 0.24093095 | 0.48248704 | 0.62946001 | 0.79955161 |
| NDE1 | 232.617783 | -0.294724024 | 0.18978285 | -1.5529539 | 0.12043415 | 0.29662775 |
| MYH11 | 59.5485382 | 0.038958194 | 0.36388596 | 0.10706155 | 0.91474015 | 0.96131774 |
| FOPNL | 268.258949 | 0.255599547 | 0.17532465 | 1.45786429 | 0.14487795 | 0.33475075 |
| ABCC1 | 721.939366 | -0.178496326 | 0.24011927 | -0.7433653 | 0.45726057 | 0.67216888 |
| ABCC6 | 159.375691 | 1.059138081 | 0.65582056 | 1.61498151 | 0.1063147 | 0.27353153 |
| NOMO3 | 124.226198 | 0.982498811 | 0.40438625 | 2.42960491 | 0.01511529 | 0.07377531 |
| NOMO2 | 142.410841 | 0.274158526 | 0.22434662 | 1.222031 | 0.2216959 | 0.43577101 |
| RPS15A | 203.739139 | -0.197444414 | 0.44665891 | -0.4420474 | 0.6584549 | 0.81895766 |
| AC138811.1 | 53.869723 | 1.236644664 | 0.62258305 | 1.98631276 | 0.04699861 | 0.1590994 |
| SMG1 | 2802.12616 | 0.277014272 | 0.16804282 | 1.6484743 | 0.09925538 | 0.26127228 |
| COQ7 | 184.221745 | 0.106479809 | 0.1972184 | 0.53990808 | 0.58926043 | 0.77174702 |
| SYT17 | 300.410364 | 0.676022354 | 0.58394921 | 1.1576732 | 0.24699743 | 0.46618627 |
| GDE1 | 158.718788 | -0.56936388 | 0.25017821 | -2.2758332 | 0.02285599 | 0.09767221 |
| CCP110 | 332.515794 | 0.113478087 | 0.18429918 | 0.61572756 | 0.53807435 | 0.73484533 |
| VPS35L | 1077.67854 | 0.159298229 | 0.25142208 | 0.63358887 | 0.52634917 | 0.72497874 |
| KNOP1 | 165.819179 | -0.536669084 | 0.41281712 | -1.3000165 | 0.1935953 | 0.40119239 |
| IQCK | 991.275788 | -0.414118517 | 0.27524543 | -1.5045427 | 0.13244168 | 0.31595125 |
| GPRC5B | 215.775552 | -1.919996614 | 0.46908201 | -4.0930937 | 4.26E-05 | 0.00086633 |
| GP2 | 1668.92653 | 2.445329279 | 1.09626574 | 2.230599 | 0.0257077 | 0.10552301 |
| UMOD | 16978.8089 | 2.271125616 | 0.6650065 | 3.41519312 | 0.00063737 | 0.0071434 |
| ACSM2A | 117.110458 | 1.633168718 | 0.92888437 | 1.75820454 | 0.07871271 | 0.22441496 |
| ACSM2B | 130.030619 | 1.558398566 | 0.90627013 | 1.71957401 | 0.0855099 | 0.23639605 |
| ACSM3 | 468.50894 | 0.241090049 | 0.38818286 | 0.6210734 | 0.53455133 | 0.73184997 |
| THUMPD1 | 481.520663 | 0.153472067 | 0.17922197 | 0.85632398 | 0.3918186 | 0.61425577 |
| ERI2 | 305.892264 | 0.638173043 | 0.32651194 | 1.95451672 | 0.05064014 | 0.16764552 |
| REXO5 | 51.0979486 | 0.507870683 | 0.33949564 | 1.49595642 | 0.13466501 | 0.31940926 |
| LYRM1 | 210.296112 | -0.215620788 | 0.27739797 | -0.7772976 | 0.4369832 | 0.65354518 |
| TMEM159 | 140.555062 | -0.690537322 | 0.37739707 | -1.8297369 | 0.0672893 | 0.20284564 |
| CRYM | 54.5136578 | -0.032786836 | 0.6119311 | -0.0535793 | 0.95727035 | 0.98118911 |
| NPIPB3 | 78.320936 | 1.041462902 | 0.47966026 | 2.17125119 | 0.02991219 | 0.11736838 |
| METTL9 | 944.877848 | -0.166562657 | 0.25233527 | -0.6600847 | 0.50919946 | 0.71159709 |
| IGSF6 | 56.1146004 | -0.312947835 | 0.62389666 | -0.501602 | 0.61594749 | 0.78898859 |
| NPIPB4 | 274.016431 | 0.615805151 | 0.37101615 | 1.65977991 | 0.09695874 | 0.25732623 |
| UQCRC2 | 1271.66323 | 0.71968744 | 0.17660077 | 4.0752226 | 4.60E-05 | 0.00092283 |
| MOSMO | 948.268747 | 0.058774259 | 0.20819182 | 0.28230821 | 0.77770719 | 0.89359206 |
| EEF2K | 200.107966 | -0.365347378 | 0.27875763 | -1.3106274 | 0.18998369 | 0.39619428 |
| POLR3E | 293.123141 | 0.046439433 | 0.18684877 | 0.24854021 | 0.80371647 | 0.90804718 |
| CDR2 | 2128.02317 | 0.03366145 | 0.27125685 | 0.12409438 | 0.90124055 | 0.95449699 |

| | | | | | | |
|---|---|---|---|---|---|---|
| AC092338.2 | 57.0626173 | 0.099113109 | 0.70096218 | 0.1413958 | 0.88755727 | 0.94928354 |
| NPIPB5 | 742.572946 | 0.049326713 | 0.19018738 | 0.2593585 | 0.79535865 | 0.90404375 |
| USP31 | 540.781285 | -0.350847475 | 0.25401276 | -1.3812199 | 0.16721136 | 0.36689282 |
| COG7 | 846.662678 | 0.22715263 | 0.182536 | 1.24442648 | 0.21334264 | 0.42639476 |
| GGA2 | 1014.58874 | 0.352172688 | 0.29897047 | 1.17795143 | 0.23881597 | 0.45697814 |
| EARS2 | 159.480022 | -0.168237502 | 0.23776491 | -0.7075792 | 0.47920663 | 0.68864274 |
| UBFD1 | 206.099477 | -0.022698491 | 0.17967853 | -0.1263283 | 0.89947202 | 0.95377105 |
| NDUFAB1 | 299.95422 | 0.343174205 | 0.2141461 | 1.60252374 | 0.10903984 | 0.27762284 |
| PALB2 | 172.034659 | -0.080352133 | 0.33159362 | -0.2423211 | 0.80853136 | 0.91010272 |
| DCTN5 | 405.284786 | -0.13601631 | 0.21439726 | -0.6344125 | 0.52581163 | 0.72494586 |
| RBBP6 | 861.681334 | 0.232641634 | 0.19026926 | 1.22269688 | 0.22144421 | 0.43555589 |
| TNRC6A | 5747.5994 | 0.591467577 | 0.19646121 | 3.01060749 | 0.00260726 | 0.02090421 |
| AC008731.1 | 182.711884 | 0.310337372 | 0.44754677 | 0.69341887 | 0.48804672 | 0.69502858 |
| SLC5A11 | 89.009356 | 1.272638308 | 0.60773657 | 2.09406241 | 0.03625442 | 0.13396096 |
| ARHGAP17 | 757.178623 | 0.05985856 | 0.16923566 | 0.35369945 | 0.72356413 | 0.86271256 |
| LCMT1 | 536.976674 | 0.030654743 | 0.17305922 | 0.17713441 | 0.85940281 | 0.93623519 |
| AC008741.2 | 101.101577 | 0.207116248 | 0.34597551 | 0.59864425 | 0.54941014 | 0.74264824 |
| HS3ST4 | 52.3971539 | 0.162606864 | 0.37060504 | 0.43876054 | 0.66083506 | 0.82039194 |
| AC009035.1 | 35.1006855 | 3.440179039 | 0.66665026 | 5.16039556 | 2.46E-07 | 1.30E-05 |
| NSMCE1 | 278.965872 | -0.195549651 | 0.22657676 | -0.8630614 | 0.38810372 | 0.61123907 |
| AC106739.1 | 87.9436822 | 0.373250845 | 0.51752451 | 0.72122351 | 0.47077201 | 0.68221914 |
| IL4R | 281.543055 | -3.196295552 | 0.55227775 | -5.7874785 | 7.15E-09 | 6.99E-07 |
| GTF3C1 | 455.757628 | 0.0347838 | 0.12469017 | 0.27896185 | 0.7802741 | 0.89513988 |
| KIAA0556 | 871.578387 | -0.442045263 | 0.18352459 | -2.4086433 | 0.01601194 | 0.07654785 |
| XPO6 | 794.893675 | -0.135607605 | 0.19724038 | -0.6875246 | 0.49175223 | 0.69792148 |
| SBK1 | 52.0921022 | 0.117643081 | 0.48585044 | 0.24213847 | 0.80867287 | 0.91010272 |
| NUPR1 | 207.941904 | -1.332717717 | 0.6206466 | -2.1473053 | 0.03176899 | 0.12266488 |
| SGF29 | 177.222506 | 0.260051482 | 0.31342155 | 0.82971794 | 0.40669828 | 0.6267006 |
| SULT1A2 | 49.5027466 | 0.065204647 | 0.70069969 | 0.09305648 | 0.92585869 | 0.96724667 |
| SULT1A1 | 385.477784 | 1.303920843 | 0.41532495 | 3.13951963 | 0.00169225 | 0.01504236 |
| ATXN2L | 447.181333 | -0.219750351 | 0.21878614 | -1.0044071 | 0.31518242 | 0.53935026 |
| TUFM | 486.911464 | -0.526952917 | 0.45037018 | -1.170044 | 0.24198327 | 0.46023859 |
| SH2B1 | 724.106251 | 0.427176823 | 0.18116478 | 2.35794625 | 0.01837635 | 0.08431166 |
| AC133550.2 | 119.511259 | 0.393246511 | 0.37537839 | 1.0476003 | 0.2948228 | 0.51799872 |
| ATP2A1 | 103.574283 | 0.247502347 | 0.3416817 | 0.72436524 | 0.46884153 | 0.68099052 |
| RABEP2 | 493.200812 | 0.06219942 | 0.32002412 | 0.19435854 | 0.84589515 | 0.9287667 |
| NFATC2IP | 913.608985 | -0.287479703 | 0.30241685 | -0.9506074 | 0.3418037 | 0.56712537 |
| AC109460.2 | 75.0103915 | 0.164955072 | 0.37569058 | 0.43907162 | 0.66060964 | 0.82026629 |
| SPNS1 | 188.832457 | -0.138670083 | 0.25842734 | -0.5365922 | 0.59154937 | 0.77302885 |
| LAT | 77.6276889 | 0.647909803 | 0.29958612 | 2.16268298 | 0.03056557 | 0.11934396 |
| NPIPB11 | 63.3191476 | 0.591025891 | 0.38452914 | 1.53701196 | 0.12429038 | 0.30242347 |
| QPRT | 85.3359867 | -0.281225762 | 0.56053203 | -0.5017122 | 0.61586998 | 0.78898859 |
| KIF22 | 158.744333 | -0.218047046 | 0.28463785 | -0.7660508 | 0.44364609 | 0.65932956 |
| MAZ | 165.027354 | -0.926739953 | 0.37702666 | -2.4580224 | 0.01397045 | 0.06966238 |
| AC009133.1 | 211.218484 | -2.121194549 | 0.36855793 | -5.7553898 | 8.64E-09 | 8.10E-07 |
| MVP | 863.713417 | -2.056383065 | 0.43684654 | -4.7073352 | 2.51E-06 | 8.54E-05 |
| CDIPT | 221.677962 | -0.395495993 | 0.26630718 | -1.485112 | 0.13751416 | 0.32355263 |
| ASPHD1 | 89.3971429 | -0.734562291 | 0.34139879 | -2.1516253 | 0.03142688 | 0.12191413 |
| KCTD13 | 332.263586 | -0.083712349 | 0.24876712 | -0.3365089 | 0.73648715 | 0.87077146 |
| TMEM219 | 341.443064 | -0.489446217 | 0.32639817 | -1.4995373 | 0.13373431 | 0.31794394 |
| TAOK2 | 152.821186 | -0.040208649 | 0.30095973 | -0.1336014 | 0.89371776 | 0.95146032 |
| HIRIP3 | 41.9881465 | -0.268801582 | 0.41194698 | -0.652515 | 0.51406901 | 0.71596592 |
| INO80E | 244.899098 | 0.324091634 | 0.19062314 | 1.70016941 | 0.08909906 | 0.24292017 |
| DOC2A | 64.8317724 | 1.640620377 | 0.76829988 | 2.13539064 | 0.03272911 | 0.12530982 |
| PPP4C | 263.39549 | -0.609513062 | 0.19944638 | -3.0560248 | 0.00224293 | 0.01865397 |
| YPEL3 | 508.051448 | -0.761240793 | 0.34890414 | -2.181805 | 0.02912393 | 0.11500687 |
| MAPK3 | 106.051645 | -0.742291712 | 0.37037458 | -2.0041649 | 0.0450524 | 0.1547717 |
| NPIPB13 | 219.769758 | 0.315303997 | 0.46410596 | 0.67937933 | 0.49689754 | 0.7021309 |
| CD2BP2 | 169.618138 | -0.250327636 | 0.32174358 | -0.7780346 | 0.43654862 | 0.65342962 |
| TBC1D10B | 147.224666 | -0.15098912 | 0.26493478 | -0.5699105 | 0.56873838 | 0.75864147 |
| ZNF48 | 69.860575 | 0.01458082 | 0.37000199 | 0.03940741 | 0.96856557 | 0.98582596 |
| 1-Sep | 58.3838071 | -0.213522907 | 0.32094679 | -0.6652907 | 0.50586459 | 0.70948881 |
| ZNF771 | 132.222267 | -0.176352561 | 0.36218778 | -0.4869092 | 0.62632268 | 0.79697262 |
| DCTPP1 | 80.7757761 | -1.212245144 | 0.405682 | -2.988166 | 0.00280657 | 0.02206666 |
| SEPHS2 | 427.922609 | -1.172882121 | 0.4142121 | -2.8315979 | 0.0046316 | 0.0316497 |
| ZNF768 | 114.55019 | 0.291927785 | 0.27961741 | 1.04402578 | 0.29647346 | 0.51965472 |
| AC002310.4 | 50.6171327 | 0.600682615 | 0.32273178 | 1.86124405 | 0.06270972 | 0.19380737 |
| ZNF688 | 81.1094301 | -0.357056142 | 0.29159152 | -1.224508 | 0.22076065 | 0.4345146 |
| AC002310.2 | 41.4266537 | 0.625874125 | 0.48328209 | 1.29504928 | 0.19530323 | 0.4028987 |
| ZNF785 | 236.949146 | 0.117191961 | 0.17374394 | 0.67450963 | 0.49998736 | 0.70482771 |
| PRR14 | 138.70225 | -0.061685356 | 0.29792291 | -0.2070514 | 0.83596971 | 0.92369276 |
| FBRS | 241.803575 | -0.293013039 | 0.20249 | -1.4470495 | 0.14788307 | 0.33936545 |
| SRCAP | 51.7859764 | 0.326746199 | 0.30918411 | 1.05680138 | 0.29060224 | 0.51339098 |
| PHKG2 | 192.572335 | -0.163074441 | 0.24473835 | -0.6663216 | 0.50520558 | 0.70894088 |
| CCDC189 | 56.3209737 | 0.187474405 | 0.27292912 | 0.68689778 | 0.49214714 | 0.69803196 |
| RNF40 | 297.798239 | -0.077376458 | 0.25056796 | -0.3088043 | 0.75747041 | 0.88285773 |
| ZNF629 | 87.1157358 | -0.490128268 | 0.3139167 | -1.5613322 | 0.11844538 | 0.29348059 |
| BCL7C | 798.533698 | 0.659711864 | 0.28884155 | 2.28399226 | 0.02237199 | 0.09615247 |
| MIR762HG | 172.417871 | 0.611928326 | 0.3396094 | 1.80185921 | 0.07156756 | 0.21105625 |
| CTF1 | 42.4893948 | 0.21971651 | 0.28999541 | 0.75765514 | 0.44865746 | 0.66445974 |
| FBXL19 | 222.659601 | -0.37817487 | 0.38050939 | -0.9938648 | 0.32028872 | 0.54463101 |
| ORAI3 | 74.0332472 | -1.05936334 | 0.30026778 | -3.528062 | 0.00041861 | 0.00517257 |
| SETD1A | 173.400056 | -0.091640505 | 0.26903405 | -0.3406279 | 0.73338372 | 0.86911141 |
| HSD3B7 | 52.8494694 | -0.885732778 | 0.53230968 | -1.6639426 | 0.09612389 | 0.25573021 |
| STX4 | 153.581351 | -0.235368302 | 0.1844676 | -1.275933 | 0.20197921 | 0.41146566 |
| AC135050.3 | 67.4927224 | -0.02932408 | 0.33396505 | -0.0878058 | 0.930031 | 0.96915731 |
| ZNF668 | 49.9037428 | -0.3228491 | 0.36195694 | -0.8919544 | 0.37241736 | 0.59653468 |
| ZNF646 | 47.6577786 | -0.331970815 | 0.37002648 | -0.8971542 | 0.36963664 | 0.59419866 |
| BCKDK | 154.307735 | -1.011130343 | 0.35262906 | -2.8674051 | 0.00413853 | 0.02926733 |
| KAT8 | 365.936769 | 0.183467553 | 0.25503149 | 0.71939175 | 0.47189958 | 0.68302887 |
| PRSS8 | 368.68579 | -0.219332372 | 0.42170013 | -0.5201146 | 0.60298374 | 0.78002274 |
| FUS | 1781.1325 | 0.276116455 | 0.13792378 | 2.00194965 | 0.04529015 | 0.15535466 |
| AC026471.4 | 165.867985 | 0.101850174 | 0.34577745 | 0.29455412 | 0.76833451 | 0.88830902 |
| AC026471.1 | 46.6377263 | -0.203062777 | 0.41554966 | -0.4886607 | 0.62508195 | 0.79598067 |
| ARMC5 | 42.6227377 | -0.034768545 | 0.37226149 | -0.0933982 | 0.92558724 | 0.96719264 |
| C16orf58 | 243.554436 | -0.108419689 | 0.19961221 | -0.5431516 | 0.58702543 | 0.77047088 |
| ZNF720 | 528.094216 | 0.13184755 | 0.12799651 | 1.03008709 | 0.30296912 | 0.5262179 |
| ZNF267 | 145.594314 | -0.430790357 | 0.21993891 | -1.9586819 | 0.05015005 | 0.16664956 |
| VPS35 | 444.743769 | -0.642989545 | 0.17997859 | -3.572589 | 0.00035347 | 0.00452859 |
| C16orf87 | 171.528903 | 0.348763155 | 0.23753197 | 1.46827879 | 0.14202851 | 0.33099057 |
| GPT2 | 104.702465 | 0.233548529 | 0.39429595 | 0.59231785 | 0.55363776 | 0.74637694 |
| DNAJA2 | 269.376689 | -0.030363717 | 0.20929463 | -0.1450764 | 0.88465053 | 0.94802439 |
| NETO2 | 80.0163421 | -0.101735766 | 0.39927919 | -0.2547986 | 0.79887868 | 0.90531816 |
| ITFG1 | 4123.71241 | -0.16683779 | 0.18061173 | -0.9237373 | 0.35562311 | 0.58105953 |
| AC007533.1 | 49.7405621 | -0.213415619 | 0.57599919 | -0.3705137 | 0.71099974 | 0.85503616 |
| PHKB | 2593.40995 | -0.150309603 | 0.14208729 | -1.057868 | 0.29011562 | 0.51300364 |
| LINC02133 | 120.615832 | -1.00054521 | 0.63476119 | -1.5762545 | 0.11496715 | 0.28829365 |
| LONP2 | 1861.89617 | 0.291555899 | 0.13406975 | 2.17465836 | 0.02965573 | 0.11658408 |
| SIAH1 | 395.300105 | -0.038999276 | 0.24673145 | -0.1580637 | 0.87440664 | 0.94347633 |
| N4BP1 | 817.392523 | 0.167440478 | 0.19313767 | 0.86694884 | 0.38597004 | 0.60936789 |
| ZNF423 | 87.8198853 | 0.450599945 | 0.6440489 | 0.69963623 | 0.48415451 | 0.69227072 |
| CNEP1R1 | 120.922581 | 0.116123821 | 0.25357776 | 0.45794166 | 0.64699435 | 0.81129186 |
| HEATR3 | 172.115622 | 0.551186271 | 0.30683854 | 1.79633977 | 0.0724405 | 0.21282618 |

| | | | | | | |
|---|---|---|---|---|---|---|
| TENT4B | 699.996276 | 0.536715124 | 0.22805002 | 2.35349736 | 0.01859774 | 0.08488512 |
| BRD7 | 559.49568 | -0.268029568 | 0.18820216 | -1.4241578 | 0.1544008 | 0.34881127 |
| CYLD | 488.180825 | 0.038365184 | 0.14108111 | 0.27193708 | 0.7856704 | 0.89813244 |
| SALL1 | 851.559498 | 0.572122833 | 0.4184672 | 1.3671868 | 0.17156676 | 0.37248937 |
| AC087564.1 | 2909.80008 | 0.670447339 | 0.37941385 | 1.76706078 | 0.07721805 | 0.22166902 |
| AC007344.1 | 406.707978 | -1.743346261 | 0.89800901 | -1.9413461 | 0.05221632 | 0.1711902 |
| HNRNPA1P4 | 61.4129295 | -0.725037407 | 0.36599451 | -1.9810063 | 0.04759057 | 0.16032233 |
| LINC01571 | 44.202502 | 0.880137327 | 0.64765815 | 1.35895352 | 0.17416132 | 0.37589771 |
| TOX3 | 942.579763 | 0.249552229 | 0.28347631 | 0.88032833 | 0.37868147 | 0.60156387 |
| AC007906.2 | 225.204453 | -0.682277232 | 0.57825672 | -1.1798864 | 0.2380454 | 0.45603216 |
| CHD9 | 6269.59058 | 0.159544432 | 0.16141982 | 0.98838194 | 0.32296563 | 0.54798369 |
| RBL2 | 1423.26213 | -0.061441246 | 0.1426766 | -0.4306329 | 0.66673528 | 0.82438513 |
| AKTIP | 221.777874 | 0.550247314 | 0.20056722 | 2.74345587 | 0.00607962 | 0.03870155 |
| RPGRIP1L | 272.744341 | 0.378816361 | 0.27260767 | 1.38960274 | 0.16464954 | 0.36365838 |
| FTO | 3280.87681 | -0.224963515 | 0.14656192 | -1.5349384 | 0.12479897 | 0.30313894 |
| IRX3 | 70.3037707 | -1.869341047 | 0.46685496 | -4.0041153 | 6.23E-05 | 0.00115648 |
| CRNDE | 289.719476 | 0.568811756 | 0.39856383 | 1.42715349 | 0.15353566 | 0.34744774 |
| IRX5 | 60.7868498 | -0.381199492 | 0.35145868 | -1.0846211 | 0.27808952 | 0.50031115 |
| AMFR | 1513.76415 | 0.337206917 | 0.4543421 | 0.74218725 | 0.45797389 | 0.67250337 |
| AC092140.1 | 96.568423 | 0.978016197 | 0.75778926 | 1.29061765 | 0.19683629 | 0.40510928 |
| NUDT21 | 611.482187 | 0.207152386 | 0.24199024 | 0.85603612 | 0.3919778 | 0.61430622 |
| OGFOD1 | 273.319159 | 0.104443531 | 0.25025236 | 0.41735283 | 0.67642035 | 0.83016679 |
| BBS2 | 616.756658 | 0.348289886 | 0.18918205 | 1.84103033 | 0.06561711 | 0.19952497 |
| MT2A | 3555.52351 | -2.590269213 | 0.52343851 | -4.9485645 | 7.48E-07 | 3.28E-05 |
| MT1E | 907.322968 | -1.756816968 | 0.50663 | -3.4676528 | 0.00052503 | 0.00613722 |
| MT1M | 95.4149411 | -2.356971099 | 0.74481253 | -3.1645159 | 0.00155341 | 0.01408306 |
| MT1F | 134.984883 | -1.253137762 | 0.52912068 | -2.3683402 | 0.0178681 | 0.08278489 |
| MT1G | 1271.11317 | -0.338665875 | 0.67214954 | -0.503855 | 0.61436331 | 0.78809426 |
| MT1H | 403.585295 | 0.080086562 | 0.7793049 | 0.10276666 | 0.91814817 | 0.96339767 |
| MT1X | 504.490651 | -1.266106093 | 0.57905641 | -2.1864987 | 0.02877914 | 0.11412293 |
| NUP93 | 443.223207 | 0.133075115 | 0.26521421 | 0.50176464 | 0.61583309 | 0.78898859 |
| SLC12A3 | 91.0073739 | 2.402138306 | 0.60047555 | 4.00039318 | 6.32E-05 | 0.00116989 |
| HERPUD1 | 907.706634 | 0.425583132 | 0.35645656 | 1.19392705 | 0.23250651 | 0.44913402 |
| NLRC5 | 224.302087 | -1.169299167 | 0.61252274 | -1.908989 | 0.05626351 | 0.18003507 |
| FAM192A | 642.012674 | 0.210224333 | 0.14404974 | 1.45938709 | 0.1444586 | 0.33429018 |
| RSPRY1 | 820.148893 | 0.15573918 | 0.13082012 | 1.19048337 | 0.23385646 | 0.45095037 |
| ARL2BP | 287.02372 | -0.088805046 | 0.27516337 | -0.3227357 | 0.74689541 | 0.87648733 |
| PLLP | 94.7843975 | -1.557962876 | 0.74747434 | -2.0843028 | 0.03713263 | 0.1359335 |
| CX3CL1 | 63.900228 | 0.087523436 | 0.56804149 | 0.1540793 | 0.87754721 | 0.9452435 |
| CIAPIN1 | 332.106409 | 0.063360314 | 0.23065508 | 0.27469724 | 0.78354885 | 0.89665455 |
| COQ9 | 649.555684 | 0.537316014 | 0.17159918 | 3.13122711 | 0.00174077 | 0.01530885 |
| POLR2C | 164.639105 | -0.161774002 | 0.22061169 | -0.7332975 | 0.463377 | 0.67625089 |
| DOK4 | 92.6809767 | -1.088022996 | 0.50978323 | -2.1342856 | 0.03281941 | 0.12558274 |
| CCDC102A | 45.6788199 | -0.42280704 | 0.40707059 | -1.0386578 | 0.29896392 | 0.52196239 |
| ADGRG1 | 1430.04544 | -0.627705936 | 0.63697146 | -0.9854538 | 0.32440122 | 0.54929057 |
| KATNB1 | 49.1718347 | -0.12227873 | 0.33896131 | -0.3607454 | 0.71828979 | 0.85964391 |
| KIFC3 | 3063.52007 | 1.255040612 | 0.31042034 | 4.04303607 | 5.28E-05 | 0.00101956 |
| AC092118.1 | 76.5675539 | 1.548065482 | 0.45708447 | 3.38682579 | 0.00070706 | 0.0077123 |
| USB1 | 144.71379 | -0.569958809 | 0.31360437 | -1.8174453 | 0.06914895 | 0.20650881 |
| MMP15 | 63.7813152 | -0.670875261 | 0.51299494 | -1.3077619 | 0.19095407 | 0.39752425 |
| CFAP20 | 97.4885935 | 0.116360547 | 0.22944646 | 0.50713595 | 0.61205943 | 0.78684784 |
| CSNK2A2 | 428.250298 | -0.135064906 | 0.16396477 | -0.8237435 | 0.41008534 | 0.6297833 |
| CCDC113 | 82.2173378 | -0.732661695 | 0.32145547 | -2.2792012 | 0.02265511 | 0.09687644 |
| SETD6 | 147.466933 | -0.001135339 | 0.29405659 | -0.003861 | 0.99691941 | 0.99880981 |
| CNOT1 | 1504.32567 | 0.027113331 | 0.1877921 | 0.1443795 | 0.8852008 | 0.94815256 |
| AC009118.3 | 42.1143356 | 0.448077306 | 0.41007966 | 1.09265917 | 0.27454347 | 0.49616155 |
| SLC38A7 | 157.975282 | -0.266035871 | 0.27667117 | -0.9615598 | 0.33627078 | 0.5617555 |
| GOT2 | 424.001584 | 0.220640325 | 0.1672185 | 1.31947316 | 0.18701098 | 0.39270408 |
| AC092378.1 | 301.749015 | 0.6154507 | 0.37019925 | 1.662485 | 0.09641557 | 0.2563513 |
| AC138305.1 | 5287.932 | 0.399068655 | 0.34205138 | 1.16669213 | 0.24333472 | 0.46201068 |
| BEAN1 | 72.9552336 | -0.839433149 | 0.30400723 | -2.7612276 | 0.00575845 | 0.03726625 |
| AC010542.3 | 156.302403 | -0.573606672 | 0.56159804 | -1.021383 | 0.30707303 | 0.53069838 |
| TK2 | 200.576713 | -0.37157567 | 0.29128651 | -1.2756364 | 0.20208409 | 0.41155238 |
| CMTM2 | 71.8683614 | 0.35459685 | 0.62159061 | 0.57046688 | 0.56836108 | 0.75840853 |
| CMTM4 | 2088.6512 | 0.382256149 | 0.18152277 | 2.10583032 | 0.03521909 | 0.13141498 |
| DYNC1LI2 | 1338.18771 | 0.191517008 | 0.1658343 | 1.15486967 | 0.2481438 | 0.46734754 |
| AC018557.1 | 113.147188 | 0.105868826 | 0.39488002 | 0.26810378 | 0.78861944 | 0.89971274 |
| NAE1 | 413.879659 | 0.033142793 | 0.151309 | 0.21904046 | 0.82661853 | 0.91944889 |
| PDP2 | 322.359389 | -0.317462728 | 0.3138595 | -1.0114804 | 0.31178657 | 0.53534237 |
| CDH16 | 1988.44177 | -0.219042629 | 0.41634505 | -0.5261084 | 0.5988129 | 0.77721524 |
| RRAD | 826.673724 | -3.897232798 | 0.56672669 | -6.8767412 | 6.12E-12 | 1.69E-09 |
| FAM96B | 337.964298 | -0.883045162 | 0.30709162 | -2.8755105 | 0.00403375 | 0.02880468 |
| CES2 | 216.116277 | -1.135088117 | 0.3132839 | -3.6231933 | 0.00029099 | 0.00393682 |
| CBFB | 1519.19401 | -0.426349087 | 0.19067902 | -2.2359518 | 0.02535493 | 0.10446452 |
| C16orf70 | 264.487053 | 0.326795336 | 0.28060335 | 1.16461667 | 0.24417419 | 0.46300558 |
| TRADD | 131.972867 | -0.386368867 | 0.33846439 | -1.1415348 | 0.25364746 | 0.47422919 |
| FBXL8 | 42.213452 | -0.855185087 | 0.40691733 | -2.1016188 | 0.03558669 | 0.13239418 |
| HSF4 | 94.9420129 | 0.782175965 | 0.43781689 | 1.78653677 | 0.07401239 | 0.21576466 |
| NOL3 | 246.408855 | -0.570595425 | 0.47888476 | -1.1915088 | 0.23345389 | 0.45037132 |
| KIAA0895L | 228.372568 | 0.321798463 | 0.28210556 | 1.1407023 | 0.25399383 | 0.47461749 |
| E2F4 | 220.083743 | 0.444541865 | 0.33674101 | 1.32012988 | 0.18679166 | 0.39251209 |
| ELMO3 | 157.20639 | 0.688909172 | 0.25059343 | 2.74911109 | 0.00597571 | 0.03820581 |
| LRRC29 | 167.892028 | 0.336960585 | 0.41346731 | 0.81496305 | 0.41509346 | 0.63511751 |
| TMEM208 | 236.503999 | -1.001263438 | 0.40225072 | -2.4891526 | 0.0128048 | 0.06578707 |
| FHOD1 | 107.296446 | 0.730250466 | 0.28919771 | 2.52509074 | 0.01156684 | 0.06140933 |
| SLC9A5 | 39.7023977 | 0.405314063 | 0.34625198 | 1.17057544 | 0.24176947 | 0.46003068 |
| LRRC36 | 118.612646 | 0.778199365 | 0.62353221 | 1.24804999 | 0.21201275 | 0.42482955 |
| ZDHHC1 | 300.91336 | -0.039050375 | 0.25451409 | -0.1534311 | 0.87805833 | 0.94548565 |
| HSD11B2 | 615.349596 | 0.852264122 | 1.10163315 | 0.77363696 | 0.43914551 | 0.65485132 |
| ATP6V0D1 | 446.918234 | -0.242299243 | 0.27400518 | -0.8842871 | 0.37654125 | 0.59967849 |
| AC009061.2 | 446.430067 | 0.723714306 | 0.40030199 | 1.80792081 | 0.07061883 | 0.20947462 |
| AC027682.6 | 66.6864184 | 0.39658153 | 0.66168722 | 0.59934893 | 0.54894022 | 0.74226986 |
| RIPOR1 | 403.71634 | -0.324461412 | 0.31487664 | -1.0304398 | 0.30280362 | 0.526073 |
| AC027682.4 | 161.304863 | -0.399706053 | 0.36607418 | -1.0918717 | 0.27488948 | 0.49647832 |
| CTCF | 792.589861 | -0.10019211 | 0.20308797 | -0.4933434 | 0.62176996 | 0.79367602 |
| ENKD1 | 177.550636 | -0.008573119 | 0.32310162 | -0.0265338 | 0.97883156 | 0.99083348 |
| C16orf86 | 39.9914723 | -0.054366151 | 0.52099508 | -0.1043506 | 0.91689112 | 0.96273568 |
| GFOD2 | 273.235113 | -0.416238444 | 0.25365185 | -1.6409833 | 0.10080089 | 0.26449721 |
| RANBP10 | 918.834304 | 0.036000731 | 0.20183513 | 0.17836702 | 0.85843475 | 0.93614803 |
| TSNAXIP1 | 72.1391995 | 0.545668735 | 0.23366233 | 2.33528754 | 0.0195284 | 0.0878878 |
| CENPT | 104.574734 | -0.092906558 | 0.24587481 | -0.3778612 | 0.70553369 | 0.85116398 |
| NUTF2 | 147.175137 | -0.713023696 | 0.23944855 | -2.9777742 | 0.0029035 | 0.02257346 |
| PSKH1 | 250.12684 | 0.084496859 | 0.19431608 | 0.43484233 | 0.66367688 | 0.82268279 |
| LCAT | 88.8449643 | 0.55260408 | 0.39933775 | 1.38380124 | 0.16641932 | 0.3659452 |
| SLC12A4 | 89.160959 | -0.163908118 | 0.39532742 | -0.4146136 | 0.6784248 | 0.83136777 |
| DUS2 | 154.223524 | 0.224783656 | 0.22271711 | 1.00927879 | 0.31284095 | 0.53673413 |
| NFATC3 | 1460.03223 | 0.279654198 | 0.12842425 | 2.17758091 | 0.02943725 | 0.11586313 |
| AC020978.5 | 86.2229216 | 0.743054008 | 0.38452149 | 1.93241216 | 0.05330865 | 0.17352243 |
| ESRP2 | 127.737889 | 0.281275988 | 0.38518385 | 0.73023828 | 0.46524455 | 0.67822705 |
| PLA2G15 | 181.723002 | 0.014964754 | 0.29763231 | 0.05027933 | 0.95989979 | 0.98229715 |
| SLC7A6 | 613.344867 | 0.872018399 | 0.2806285 | 3.10737646 | 0.00188756 | 0.01636169 |
| SLC7A6OS | 330.285695 | -0.399825028 | 0.29615392 | -1.3500582 | 0.17699731 | 0.37941536 |
| PRMT7 | 371.618564 | 0.910561722 | 0.36689632 | 2.4817957 | 0.01307222 | 0.06664086 |

| | | | | | | |
|---|---|---|---|---|---|---|
| ZFP90 | 435.702307 | 0.214987101 | 0.21140592 | 1.01693983 | 0.30918206 | 0.53306909 |
| CDH3 | 593.688225 | -0.949618259 | 0.66180939 | -1.4348818 | 0.15132078 | 0.34420461 |
| CDH1 | 979.081345 | 0.150935609 | 0.37798696 | 0.39931433 | 0.68966161 | 0.84045947 |
| TANGO6 | 859.760495 | 0.160066515 | 0.17837131 | 0.89737815 | 0.36951717 | 0.59417383 |
| CHTF8 | 335.151585 | -0.441663962 | 0.22065049 | -2.001645 | 0.04532292 | 0.15537776 |
| UTP4 | 203.389588 | 0.117400726 | 0.22991565 | 0.51062521 | 0.60961352 | 0.78529652 |
| SNTB2 | 519.62333 | -0.355304697 | 0.45204324 | -0.7859971 | 0.43186916 | 0.64957432 |
| VPS4A | 212.125068 | -0.250106919 | 0.18863247 | -1.3258954 | 0.18487435 | 0.39045597 |
| COG8 | 114.815526 | -0.001272604 | 0.25539674 | -0.0049829 | 0.99602428 | 0.99851887 |
| NIP7 | 53.4775208 | -0.190912449 | 0.395939 | -0.4821764 | 0.62968062 | 0.79967805 |
| TERF2 | 414.813761 | 0.284848283 | 0.22942754 | 1.241561 | 0.21439858 | 0.42792245 |
| CYB5B | 865.951335 | -0.04655513 | 0.16939155 | -0.2748374 | 0.78344118 | 0.89665455 |
| NFAT5 | 6935.33582 | -0.041448361 | 0.1416167 | -0.2926799 | 0.76976683 | 0.88918672 |
| NQO1 | 417.29665 | -1.591372456 | 0.78194378 | -2.0351495 | 0.04183584 | 0.14665859 |
| AC092115.3 | 64.9255422 | 0.139952823 | 0.32622541 | 0.4290065 | 0.66791849 | 0.82499863 |
| NOB1 | 92.8562083 | 0.039490229 | 0.24128457 | 0.16366662 | 0.8699936 | 0.94075489 |
| WWP2 | 2030.3039 | -0.082129751 | 0.21986828 | -0.3735407 | 0.70874606 | 0.85371391 |
| PDPR | 689.216041 | 0.231751489 | 0.32242083 | 0.71878572 | 0.47227296 | 0.68319499 |
| AC009060.1 | 306.758151 | -0.005191723 | 0.40934771 | -0.0126829 | 0.98988077 | 0.99606252 |
| EXOSC6 | 176.988062 | -0.11190307 | 0.25247724 | -0.4432204 | 0.65760631 | 0.81859324 |
| AARS | 404.785389 | 0.179215831 | 0.19236319 | 0.93165347 | 0.35151564 | 0.57736509 |
| AC012184.2 | 73.1930526 | 0.393473701 | 0.31123814 | 1.26422069 | 0.20615083 | 0.41687787 |
| DDX19A | 87.7202849 | 0.024410257 | 0.27462156 | 0.08888689 | 0.9291718 | 0.96900671 |
| ST3GAL2 | 364.323532 | 0.094649198 | 0.17317361 | 0.54655672 | 0.58468331 | 0.76905468 |
| FUK | 227.720355 | -0.125477001 | 0.19641895 | -0.6388233 | 0.52293788 | 0.7227557 |
| COG4 | 508.208034 | 0.080770227 | 0.26633952 | 0.30326039 | 0.76169143 | 0.88431255 |
| SF3B3 | 639.861979 | -0.107561155 | 0.10721155 | -1.0032609 | 0.31573501 | 0.54001603 |
| MTSS1L | 212.140433 | 1.128977854 | 0.31762813 | 3.55440132 | 0.00037884 | 0.00478404 |
| VAC14 | 426.553639 | -0.298073552 | 0.33636731 | -0.8861549 | 0.37553404 | 0.59876913 |
| HYDIN | 1198.29201 | -0.188062789 | 0.29741724 | -0.6323197 | 0.52717797 | 0.72545317 |
| CMTR2 | 355.17522 | 0.006190523 | 0.29226596 | 0.02118113 | 0.98310117 | 0.9928728 |
| ZNF19 | 435.506266 | -0.316135629 | 0.27590913 | -1.1457962 | 0.25187949 | 0.47203022 |
| MARVELD3 | 166.039721 | -0.353556512 | 0.47190621 | -0.7492093 | 0.45373108 | 0.66948056 |
| PHLPP2 | 343.277391 | -0.052324463 | 0.29162177 | -0.1794258 | 0.85760339 | 0.93574122 |
| AP1G1 | 1512.38779 | 0.114595952 | 0.20191983 | 0.56753195 | 0.57035283 | 0.75945349 |
| IST1 | 1012.44987 | -0.045088979 | 0.1326351 | -0.3399476 | 0.73389602 | 0.86935939 |
| ZNF821 | 389.976568 | 0.041694106 | 0.27611682 | 0.15100169 | 0.87997438 | 0.94636668 |
| DHODH | 116.045546 | -0.486502523 | 0.53814153 | -0.904042 | 0.36597315 | 0.59141301 |
| TXNL4B | 207.585727 | -0.066902277 | 0.1546086 | -0.4327203 | 0.66521799 | 0.8233644 |
| DHX38 | 205.803199 | -0.266307803 | 0.20752646 | -1.2832475 | 0.19940539 | 0.40801923 |
| LINC01572 | 665.239227 | 0.146893852 | 0.37182156 | 0.39506545 | 0.69279459 | 0.84248976 |
| AC004158.1 | 46.361624 | -0.491092399 | 0.55117689 | -0.8909887 | 0.37293521 | 0.59676739 |
| AC004943.2 | 436.685452 | 0.215678622 | 0.17727353 | 1.2166431 | 0.22374004 | 0.4383227 |
| ZFHX3 | 3190.01979 | -0.977134833 | 0.29402086 | -3.323352 | 0.00088943 | 0.00911468 |
| AC138627.1 | 515.71923 | 1.419615589 | 0.54841905 | 2.58855993 | 0.00963782 | 0.05389752 |
| PSMD7 | 342.479359 | -0.402500324 | 0.21103058 | -1.9073081 | 0.05648071 | 0.18057989 |
| NPIPB15 | 165.870934 | -1.099504293 | 0.90404883 | -1.2162001 | 0.2239087 | 0.43848926 |
| AC009053.3 | 208.215814 | 0.793050142 | 0.59321074 | 1.3368776 | 0.18126259 | 0.38524499 |
| GLG1 | 3040.66577 | 0.128276282 | 0.14288346 | 0.89776857 | 0.36930894 | 0.59417383 |
| RFWD3 | 451.504853 | -0.163296497 | 0.1839228 | -0.8878535 | 0.37461956 | 0.59813246 |
| MLKL | 74.667336 | -0.251146375 | 0.59199677 | -0.4242361 | 0.67139367 | 0.82736497 |
| FA2H | 570.906714 | 0.705391184 | 0.70693204 | 0.99782036 | 0.31836647 | 0.54297001 |
| WDR59 | 1267.2463 | 0.060007876 | 0.18002003 | 0.33334 | 0.73887765 | 0.87211464 |
| ZNRF1 | 660.426869 | -0.392793561 | 0.31848656 | -1.2333128 | 0.21745907 | 0.43128814 |
| LDHD | 43.421992 | 0.953964625 | 0.45979173 | 2.07477551 | 0.03800735 | 0.1382266 |
| ZFP1 | 95.5275349 | 0.324947805 | 0.28028399 | 1.15935199 | 0.24631274 | 0.46552367 |
| BCAR1 | 737.996839 | -1.22148265 | 0.28043252 | -4.3557097 | 1.33E-05 | 0.00033435 |
| CFDP1 | 2357.86919 | 0.000806275 | 0.30703743 | 0.00262598 | 0.99790477 | 0.99934986 |
| AC009163.4 | 45.6939588 | 0.254445187 | 0.29411976 | 0.86510741 | 0.38697984 | 0.61054074 |
| TMEM231 | 78.109525 | -0.006329373 | 0.43066371 | -0.0146968 | 0.98827408 | 0.99552604 |
| GABARAPL2 | 417.809424 | -0.834570964 | 0.37080862 | -2.2506784 | 0.02440591 | 0.10195373 |
| ADAT1 | 199.083281 | -0.515708935 | 0.25353671 | -2.0340602 | 0.04194552 | 0.14683946 |
| KARS | 266.441992 | -0.437487028 | 0.1950934 | -2.2424491 | 0.02493236 | 0.10349682 |
| TERF2IP | 1069.99121 | -0.165733296 | 0.28871413 | -0.5740394 | 0.56594111 | 0.75640152 |
| MON1B | 84.9328481 | -0.444404812 | 0.37715571 | -1.1783059 | 0.23867467 | 0.45677393 |
| NUDT7 | 164.043784 | 1.48674331 | 0.36997115 | 4.01853847 | 5.86E-05 | 0.00110979 |
| VAT1L | 808.758706 | 2.958796856 | 1.20542073 | 2.45457605 | 0.01410508 | 0.07012196 |
| WWOX | 18088.1508 | 0.519388973 | 0.25514036 | 2.03569897 | 0.04178059 | 0.14662364 |
| AC046158.1 | 128.30765 | 0.561695982 | 0.58529915 | 0.95967333 | 0.33721965 | 0.56231354 |
| MAF | 95.5282406 | 0.718359216 | 0.5758612 | 1.24745202 | 0.2122318 | 0.42488177 |
| AC105411.1 | 106.47572 | 0.115347262 | 0.49907727 | 0.23112105 | 0.81722076 | 0.9150351 |
| DYNLRB2 | 95.2729087 | -0.041991959 | 0.52886438 | -0.0794002 | 0.93671428 | 0.97284815 |
| CDYL2 | 1013.55467 | -0.011726814 | 0.54166578 | -0.0216495 | 0.98272752 | 0.9928728 |
| CMC2 | 706.454617 | 0.750788556 | 0.28570178 | 2.627875 | 0.00859201 | 0.04970244 |
| ATMIN | 178.42705 | -0.446543445 | 0.25522799 | -1.7495865 | 0.08018969 | 0.22666931 |
| GAN | 473.265818 | -0.219517762 | 0.16453027 | -1.3342089 | 0.18213539 | 0.38654023 |
| CMIP | 1873.56519 | -0.524484324 | 0.24988346 | -2.0989157 | 0.03582433 | 0.13288502 |
| PLCG2 | 1315.08117 | -1.006806593 | 0.48622159 | -2.0706744 | 0.03838924 | 0.13904125 |
| SDR42E1 | 59.1953562 | 0.001279105 | 0.41642973 | 0.0030716 | 0.99754922 | 0.99921344 |
| AC092142.1 | 135.141122 | 0.41368224 | 0.34400409 | 1.20255036 | 0.22915037 | 0.44457881 |
| MPHOSPH6 | 228.386273 | 0.253765269 | 0.27571426 | 0.92039226 | 0.35736781 | 0.58281427 |
| CDH13 | 137.774979 | 2.299214614 | 0.55904103 | 4.11278333 | 3.91E-05 | 0.00080935 |
| HSBP1 | 470.367831 | -0.409726888 | 0.34869162 | -1.1750408 | 0.23997839 | 0.45806569 |
| MLYCD | 100.303948 | -0.078261297 | 0.35504537 | -0.2204262 | 0.82553925 | 0.91938879 |
| AC009119.2 | 108.693506 | -0.180441179 | 0.5407257 | -0.3337019 | 0.73860453 | 0.87210399 |
| MBTPS1 | 885.924605 | 0.066526233 | 0.12968326 | 0.51299014 | 0.60795822 | 0.78369475 |
| HSDL1 | 179.479309 | 0.436943827 | 0.31040345 | 1.4076642 | 0.15923053 | 0.35570371 |
| DNAAF1 | 57.1330172 | 0.293138652 | 0.67000812 | 0.43751507 | 0.66173785 | 0.82128115 |
| TAF1C | 285.218232 | 0.587268188 | 0.24172868 | 2.42945184 | 0.01512167 | 0.07377531 |
| ATP2C2 | 51.5044093 | 0.855156044 | 0.47947065 | 1.783542 | 0.07449812 | 0.21663404 |
| ATP2C2-AS1 | 39.3329607 | 0.157875405 | 0.4754776 | 0.33203542 | 0.73986251 | 0.87265237 |
| TLDC1 | 408.701236 | -0.521339721 | 0.37854556 | -1.3772179 | 0.16844489 | 0.36849761 |
| COTL1 | 172.892941 | -1.263336389 | 0.5087582 | -2.4831765 | 0.01302166 | 0.06657589 |
| KLHL36 | 445.651238 | -0.324585196 | 0.19897588 | -1.6312791 | 0.10283144 | 0.2676495 |
| USP10 | 996.378378 | 0.027296776 | 0.161338 | 0.16919 | 0.8656472 | 0.93885654 |
| ZDHHC7 | 976.242313 | 0.107017234 | 0.26078735 | 0.41036206 | 0.68154037 | 0.83379335 |
| KIAA0513 | 322.36792 | -0.501589683 | 0.36527847 | -1.3731707 | 0.1696993 | 0.36997702 |
| GSE1 | 1492.61069 | 0.05340167 | 0.35430275 | 0.15072327 | 0.88019401 | 0.94636668 |
| EMC8 | 171.927794 | -0.156039716 | 0.25465918 | -0.6127394 | 0.54004866 | 0.73609563 |
| COX4I1 | 1245.56282 | -0.512192733 | 0.45132123 | -1.134874 | 0.25642812 | 0.47733357 |
| IRF8 | 52.5816468 | 0.438855551 | 0.45403902 | 0.9665591 | 0.33376449 | 0.5592275 |
| MTHFSD | 175.66265 | 0.133634041 | 0.21178168 | 0.63099907 | 0.52804114 | 0.72617338 |
| C16orf95 | 62.0795881 | 1.045637395 | 0.28854134 | 3.62387383 | 0.00029022 | 0.00393185 |
| FBXO31 | 213.884007 | 0.11697484 | 0.23594172 | 0.49577853 | 0.62005068 | 0.79240029 |
| MAP1LC3B | 464.868584 | -0.544251954 | 0.37363245 | -1.4566507 | 0.14521284 | 0.33510069 |
| ZCCHC14 | 1391.98732 | -0.788798607 | 0.33464573 | -2.3571154 | 0.01841752 | 0.0843248 |
| KLHDC4 | 517.482089 | -0.13968619 | 0.30406564 | -0.4593948 | 0.64595065 | 0.81059872 |
| SLC7A5 | 425.478491 | -0.193686 | 0.68925794 | -0.2810066 | 0.77870537 | 0.89427224 |
| BANP | 729.719083 | 0.024060862 | 0.29827996 | 0.08066537 | 0.93570808 | 0.97236337 |
| AC138512.1 | 72.5586644 | 2.199979754 | 0.68777238 | 3.19870325 | 0.00138047 | 0.01286849 |
| ZFPM1 | 290.753227 | -0.425123878 | 0.22482816 | -1.8908836 | 0.05863989 | 0.18486259 |
| AC116552.1 | 58.6260796 | 0.58474634 | 0.34540755 | 1.69291709 | 0.09047125 | 0.24558872 |
| ZC3H18 | 398.012859 | -0.407241447 | 0.22148213 | -1.8387102 | 0.06595783 | 0.20019231 |

| | | | | | | |
|---|---|---|---|---|---|---|
| CYBA | 649.778902 | -2.304526893 | 0.64066777 | -3.5970701 | 0.00032182 | 0.00422979 |
| MVD | 212.262051 | -0.379400639 | 0.35457401 | -1.0700182 | 0.28461112 | 0.50700314 |
| SNAI3-AS1 | 78.7799626 | 0.253199332 | 0.30682412 | 0.82522629 | 0.40924313 | 0.62915416 |
| RNF166 | 63.0564679 | 0.922936539 | 0.32618682 | 2.82947221 | 0.00466248 | 0.03175016 |
| PIEZO1 | 141.300027 | -1.182184364 | 0.4300687 | -2.7488268 | 0.0059809 | 0.03822046 |
| APRT | 257.313085 | -1.674855969 | 0.4495377 | -3.7257297 | 0.00019475 | 0.00287106 |
| GALNS | 271.616789 | -0.059396479 | 0.2854877 | -0.2080527 | 0.83518784 | 0.92333413 |
| TRAPPC2L | 87.5146498 | -0.328468335 | 0.36793197 | -0.892742 | 0.37199533 | 0.59617645 |
| ACSF3 | 324.369128 | -0.053379081 | 0.25345738 | -0.2106038 | 0.83319646 | 0.92205865 |
| AC009113.1 | 43.6960125 | 0.636628266 | 0.38922432 | 1.63563331 | 0.10191635 | 0.26604257 |
| ZNF778 | 114.685676 | -0.020782343 | 0.23286956 | -0.0892446 | 0.92888755 | 0.96900671 |
| ANKRD11 | 3229.32112 | -0.375365451 | 0.18225975 | -2.0595082 | 0.03944558 | 0.14144318 |
| SPG7 | 1448.37689 | -0.221221349 | 0.31552895 | -0.7011127 | 0.4832327 | 0.69132684 |
| AC092123.1 | 45.2039542 | 0.183154215 | 0.39857923 | 0.45951771 | 0.64586244 | 0.81059872 |
| RPL13 | 2155.47629 | -1.034309735 | 0.40903846 | -2.5286369 | 0.01145064 | 0.06103775 |
| CHMP1A | 250.106955 | -0.555383243 | 0.25819056 | -2.1510594 | 0.03147151 | 0.12205142 |
| SPATA33 | 110.447673 | -0.767254383 | 0.27293971 | -2.8110764 | 0.00493761 | 0.03310204 |
| CDK10 | 391.771004 | -0.153569831 | 0.21033525 | -0.7301193 | 0.46531724 | 0.67825816 |
| SPATA2L | 73.4905952 | -0.641361805 | 0.26361177 | -2.4329787 | 0.01497518 | 0.0732348 |
| VPS9D1 | 418.404823 | -0.20646708 | 0.27640267 | -0.7469793 | 0.45507604 | 0.67027397 |
| ZNF276 | 241.785783 | 0.18589843 | 0.27773329 | 0.66934156 | 0.50327762 | 0.70736264 |
| FANCA | 62.261132 | 0.08779981 | 0.50006474 | 0.17557689 | 0.86062636 | 0.93664115 |
| SPIRE2 | 104.911041 | 0.354230928 | 0.31717748 | 1.11682243 | 0.26407026 | 0.48499778 |
| TCF25 | 2141.69331 | -0.837108243 | 0.33958896 | -2.4650632 | 0.01369891 | 0.06869738 |
| MC1R | 88.8200278 | -1.111315909 | 0.46542136 | -2.387763 | 0.01695127 | 0.079621 |
| DEF8 | 178.541557 | -0.131966171 | 0.22514664 | -0.5861343 | 0.55778526 | 0.74982551 |
| DBNDD1 | 56.0469467 | -0.778290865 | 0.46448339 | -1.6756054 | 0.09381552 | 0.25192298 |
| GAS8 | 458.029747 | -0.016465811 | 0.29071578 | -0.0566389 | 0.95483288 | 0.98009452 |
| FAM157C | 461.35776 | 0.661683674 | 0.33393157 | 1.98149481 | 0.04753581 | 0.16021958 |
| AC133919.2 | 69.9933056 | 0.716863685 | 0.66487817 | 1.07818803 | 0.28094985 | 0.50353686 |
| LINC02193 | 123.604843 | -0.175361633 | 0.29777121 | -0.588914 | 0.55591897 | 0.748078 |
| AC240565.1 | 47.9446334 | 0.592013248 | 0.43186232 | 1.37083794 | 0.17042548 | 0.37070833 |
| RPH3AL | 701.508936 | -0.562249762 | 0.37421612 | -1.5024734 | 0.1329749 | 0.31682277 |
| AC129507.2 | 151.62977 | 0.245634683 | 0.53807119 | 0.45650963 | 0.64802354 | 0.81173905 |
| C17orf97 | 79.2631884 | 1.114181676 | 0.4457022 | 2.49983439 | 0.01242514 | 0.06441054 |
| VPS53 | 1728.16166 | -0.20152308 | 0.20062709 | -1.004466 | 0.31515407 | 0.53935026 |
| AC015853.3 | 81.5604078 | -0.12579577 | 0.44456474 | -0.2829639 | 0.77720451 | 0.89355857 |
| FAM57A | 96.0430339 | -2.380341244 | 0.46005005 | -5.174092 | 2.29E-07 | 1.23E-05 |
| GLOD4 | 223.711395 | -0.470181743 | 0.20097714 | -2.3394787 | 0.01931067 | 0.08709923 |
| MRM3 | 57.5976065 | -0.249463598 | 0.28386177 | -0.8788207 | 0.3794985 | 0.60235468 |
| NXN | 318.89521 | 0.027872394 | 0.43998883 | 0.06334796 | 0.94948943 | 0.97821544 |
| TIMM22 | 84.601068 | -0.103663245 | 0.28637627 | -0.3619827 | 0.71736499 | 0.85947463 |
| ABR | 399.84804 | -1.190845295 | 0.38489692 | -3.0939331 | 0.00197522 | 0.01698612 |
| YWHAE | 1858.3813 | -0.055957281 | 0.09532614 | -0.5870088 | 0.55719779 | 0.7492306 |
| CRK | 820.488738 | -0.335120681 | 0.16163304 | -2.0733426 | 0.0381404 | 0.13859604 |
| MYO1C | 617.137492 | -0.657883285 | 0.250551 | -2.625746 | 0.00864593 | 0.04987079 |
| INPP5K | 158.361219 | -0.667433826 | 0.28606561 | -2.3331494 | 0.0196403 | 0.08815113 |
| PITPNA-AS1 | 44.3426944 | 0.209964771 | 0.37683868 | 0.55717415 | 0.57740845 | 0.7637681 |
| PITPNA | 1038.07534 | -0.507384322 | 0.16669873 | -3.0437203 | 0.00233672 | 0.01923101 |
| SLC43A2 | 1347.10124 | 0.328922742 | 0.42043816 | 0.78233323 | 0.43401876 | 0.65102814 |
| SCARF1 | 122.712622 | 1.328557013 | 0.3803193 | 3.4932674 | 0.00047715 | 0.0057141 |
| RILP | 63.2808825 | 0.523306953 | 0.397951 | 1.3150035 | 0.18850873 | 0.39417631 |
| PRPF8 | 830.811991 | -0.126976589 | 0.18162241 | -0.699124 | 0.48447453 | 0.69235581 |
| MIR22HG | 222.295155 | -0.46620976 | 0.52861897 | -0.8819391 | 0.37780972 | 0.60068471 |
| WDR81 | 131.068343 | -0.504896606 | 0.47309098 | -1.0672294 | 0.28586826 | 0.50833149 |
| SERPINF2 | 74.5383325 | -0.16014645 | 0.70449185 | -0.2273219 | 0.82017342 | 0.91658849 |
| SMYD4 | 209.333991 | -0.066657434 | 0.29487355 | -0.2260543 | 0.82115918 | 0.91711268 |
| RPA1 | 332.896426 | -0.514283379 | 0.22248774 | -2.3115134 | 0.02080452 | 0.09160228 |
| RTN4RL1 | 185.048013 | -1.101426927 | 0.74793858 | -1.4726168 | 0.14085439 | 0.32889263 |
| DPH1 | 104.654089 | 0.975591448 | 0.31568862 | 3.09035988 | 0.00199914 | 0.01712494 |
| SMG6 | 2606.23809 | -0.174509521 | 0.19749886 | -0.8835976 | 0.37691347 | 0.6000254 |
| AC090617.4 | 70.3568663 | -0.443738629 | 0.40414131 | -1.0979789 | 0.2722137 | 0.49370737 |
| SRR | 144.694525 | 0.739374562 | 0.27012756 | 2.73713111 | 0.00619776 | 0.03915169 |
| TSR1 | 180.938219 | -0.418092293 | 0.24660102 | -1.6954199 | 0.08999578 | 0.24458483 |
| SGSM2 | 394.053382 | 0.084616907 | 0.18300443 | 0.46237629 | 0.6438115 | 0.80931821 |
| MNT | 185.629467 | -1.123402395 | 0.26681235 | -4.2104588 | 2.55E-05 | 0.00057643 |
| METTL16 | 839.220266 | 0.070489774 | 0.24947867 | 0.2825483 | 0.77752312 | 0.89359206 |
| PAFAH1B1 | 2738.96918 | -0.081669297 | 0.12894324 | -0.633374 | 0.52648942 | 0.72502073 |
| CLUH | 583.58213 | 0.29407538 | 0.28054835 | 1.04821638 | 0.29453892 | 0.51777544 |
| RAP1GAP2 | 1723.58339 | 0.168965061 | 0.20779684 | 0.81312622 | 0.4161457 | 0.63597923 |
| TRPV1 | 87.0932739 | 1.082678264 | 0.34948442 | 3.09792999 | 0.00194877 | 0.0167915 |
| CTNS | 209.22497 | -0.147123045 | 0.38080227 | -0.3863502 | 0.69923731 | 0.8470493 |
| TAX1BP3 | 119.512403 | -1.597227539 | 0.38288739 | -4.1715334 | 3.03E-05 | 0.00065829 |
| EMC6 | 146.964152 | -0.732900542 | 0.54109706 | -1.3544715 | 0.175586 | 0.37786639 |
| ITGAE | 171.347594 | 0.217520372 | 0.27216941 | 0.79920948 | 0.42416896 | 0.64296071 |
| NCBP3 | 719.628762 | 0.461579733 | 0.17875404 | 2.58220591 | 0.0098171 | 0.05462261 |
| CAMKK1 | 67.7105593 | -0.757365875 | 0.43133507 | -1.7558644 | 0.07911157 | 0.22477624 |
| ZZEF1 | 1113.37539 | 0.21489791 | 0.24003182 | 0.89528927 | 0.37063247 | 0.59532317 |
| CYB5D2 | 404.258137 | -0.151184014 | 0.2969489 | -0.5091247 | 0.61066483 | 0.78610971 |
| ANKFY1 | 811.624906 | 0.174569408 | 0.19728166 | 0.88487398 | 0.3762246 | 0.59953562 |
| UBE2G1 | 1152.8202 | 0.040484347 | 0.18775674 | 0.21562127 | 0.82928297 | 0.92023714 |
| SPNS2 | 668.820744 | -0.881454023 | 0.64091226 | -1.3753115 | 0.16903488 | 0.36932195 |
| MYBBP1A | 113.299409 | -0.338164144 | 0.28854597 | -1.1719593 | 0.24121339 | 0.45956838 |
| GGT6 | 129.648902 | 0.742773265 | 0.60237329 | 1.23307803 | 0.21754665 | 0.43128814 |
| SMTNL2 | 618.771416 | 1.796466175 | 0.29142674 | 6.16438339 | 7.08E-10 | 9.64E-08 |
| PELP1 | 197.699181 | -0.089493456 | 0.31016907 | -0.2885312 | 0.77294015 | 0.89074912 |
| AC091153.2 | 58.549444 | -0.058381703 | 0.37062962 | -0.1575203 | 0.87483478 | 0.94367599 |
| ARRB2 | 46.5520185 | 0.113350486 | 0.41965137 | 0.27010632 | 0.78707846 | 0.8992751 |
| MED11 | 93.6266461 | -0.427689819 | 0.3021294 | -1.4155849 | 0.15689707 | 0.35245806 |
| CXCL16 | 72.6953844 | -2.040801677 | 0.522016 | -3.9094619 | 9.25E-05 | 0.00158272 |
| PSMB6 | 211.449275 | -0.88362242 | 0.48537617 | -1.8204899 | 0.06868443 | 0.20558165 |
| PLD2 | 67.9934863 | -1.403542648 | 0.40400959 | -3.474033 | 0.0005127 | 0.00603585 |
| MINK1 | 679.453448 | -0.673153055 | 0.23100677 | -2.9139971 | 0.00356833 | 0.02634662 |
| SLC25A11 | 160.588414 | 0.106817577 | 0.40070984 | 0.26657089 | 0.78979957 | 0.90082571 |
| RNF167 | 167.8895 | -0.171768667 | 0.23113357 | -0.7431576 | 0.45738628 | 0.67216888 |
| PFN1 | 657.471165 | -1.616143096 | 0.34776505 | -4.6472268 | 3.36E-06 | 0.00010892 |
| SPAG7 | 253.999842 | 0.073517138 | 0.29280717 | 0.25107697 | 0.8017546 | 0.90677969 |
| CAMTA2 | 226.097456 | 0.089039249 | 0.18870069 | 0.47185439 | 0.63703073 | 0.80480926 |
| KIF1C | 285.526687 | -0.341338936 | 0.30505044 | -1.118959 | 0.26315762 | 0.48419683 |
| ZFP3 | 109.777036 | 0.504230026 | 0.21462715 | 2.34933015 | 0.01880722 | 0.08554566 |
| ZNF232 | 87.9910482 | -1.254293646 | 0.76096111 | -1.6483019 | 0.09929074 | 0.26131328 |
| USP6 | 61.4627421 | -1.633744423 | 0.68010848 | -2.4021821 | 0.01629759 | 0.0775486 |
| ZNF594 | 261.451246 | 0.24359916 | 0.1870486 | 1.3023308 | 0.19280333 | 0.400179 |
| AC087500.1 | 551.982357 | 0.575747648 | 0.27821415 | 2.06944054 | 0.03850477 | 0.13926874 |
| SCIMP | 107.177031 | 0.32892127 | 0.32936363 | 0.99865693 | 0.31796091 | 0.54243213 |
| RABEP1 | 1463.64104 | 0.152392309 | 0.22239085 | 0.68524542 | 0.49318907 | 0.69881955 |
| NUP88 | 1333.18964 | -0.093596745 | 0.22022163 | -0.4250116 | 0.67082823 | 0.82704593 |
| RPAIN | 593.967964 | -0.278536924 | 0.22492733 | -1.2383418 | 0.21558934 | 0.42945552 |
| AC004148.2 | 67.0112409 | -0.510091402 | 0.55918886 | -0.9121988 | 0.36166408 | 0.58697738 |
| C1QBP | 183.406293 | -0.997658155 | 0.39070675 | -2.5534705 | 0.01066553 | 0.05802348 |
| DHX33 | 97.1625058 | -0.150933574 | 0.35639073 | -0.4235059 | 0.67192621 | 0.8277815 |
| DERL2 | 251.797324 | -0.044219139 | 0.20291935 | -0.2179148 | 0.82749546 | 0.91975661 |
| MIS12 | 173.730712 | 0.104628929 | 0.22996156 | 0.45498443 | 0.64912044 | 0.81217638 |

| | | | | | | |
|---|---|---|---|---|---|---|
| AC007846.2 | 43.7536743 | 2.217933037 | 0.60207031 | 3.68384389 | 0.00022974 | 0.00328428 |
| WSCD1 | 218.69755 | 1.154691488 | 0.41910301 | 2.75514962 | 0.00586653 | 0.03776364 |
| PITPNM3 | 95.8368523 | -0.158039056 | 0.53153427 | -0.2973262 | 0.7662175 | 0.88688348 |
| KIAA0753 | 613.060339 | -0.2262096 | 0.18067477 | -1.2520265 | 0.21056021 | 0.42281694 |
| TXNDC17 | 125.466462 | -0.813337883 | 0.34981455 | -2.3250544 | 0.02006905 | 0.08927837 |
| MED31 | 154.571092 | -0.024083187 | 0.26273821 | -0.0916623 | 0.92696635 | 0.96785223 |
| XAF1 | 461.208151 | -0.732529835 | 1.44721731 | -0.5061644 | 0.61274127 | 0.78690269 |
| ALOX12-AS1 | 298.427425 | -0.027540167 | 0.40203161 | -0.0685025 | 0.94538564 | 0.97640274 |
| SLC16A13 | 39.6762157 | -0.257665875 | 0.33922776 | -0.759566 | 0.44751405 | 0.66343581 |
| DLG4 | 56.1122044 | 0.184089038 | 0.73160118 | 0.25162485 | 0.80133105 | 0.90672945 |
| ACADVL | 1586.75515 | 0.570737583 | 0.47704621 | 1.19639896 | 0.23154091 | 0.44759606 |
| DVL2 | 137.992834 | 0.368499415 | 0.18152762 | 2.02999092 | 0.04235746 | 0.14785933 |
| PHF23 | 87.1063896 | -0.608693126 | 0.3376788 | -1.8025802 | 0.07145417 | 0.21091354 |
| AC003688.1 | 45.2820363 | -1.16089298 | 0.36890109 | -3.1468949 | 0.00165014 | 0.01477744 |
| ELP5 | 154.630235 | -0.135978956 | 0.28478803 | -0.4774743 | 0.63302444 | 0.80230473 |
| CLDN7 | 41.4181151 | -1.974831142 | 0.42704754 | -4.6243825 | 3.76E-06 | 0.00011844 |
| EIF5A | 204.754054 | -1.369833523 | 0.41170347 | -3.3272333 | 0.00087713 | 0.00901634 |
| GPS2 | 171.368925 | -0.990145454 | 0.33929723 | -2.9182244 | 0.00352031 | 0.02609414 |
| TNK1 | 41.6500372 | -0.233610858 | 0.277796 | -0.8409439 | 0.40037935 | 0.62197 |
| NLGN2 | 77.1060022 | -1.025730626 | 0.441165 | -2.3250499 | 0.02006929 | 0.08927837 |
| FGF11 | 90.197922 | 0.37917583 | 0.68504739 | 0.55350307 | 0.57991898 | 0.76601498 |
| CHRNB1 | 59.7519887 | 0.540015038 | 0.42083514 | 1.28319854 | 0.19942253 | 0.40801923 |
| ZBTB4 | 573.193865 | -0.065717894 | 0.33930308 | -0.1936849 | 0.84642259 | 0.92885801 |
| POLR2A | 953.969517 | -0.455792048 | 0.22865366 | -1.993373 | 0.04622061 | 0.15731205 |
| EIF4A1 | 275.062291 | -1.364943682 | 0.42333553 | -3.2242597 | 0.00126299 | 0.01201932 |
| CD68 | 67.8259405 | -2.753600236 | 0.58164552 | -4.7341553 | 2.20E-06 | 7.65E-05 |
| MPDU1 | 207.968269 | 0.102443195 | 0.23385042 | 0.43807147 | 0.66133448 | 0.82085765 |
| FXR2 | 192.307354 | 0.280932552 | 0.26139449 | 1.0747455 | 0.28248869 | 0.50492463 |
| SAT2 | 446.312009 | 0.755818227 | 0.35863448 | 2.10748901 | 0.03507521 | 0.13106322 |
| TP53 | 265.921482 | -0.24699375 | 0.34269927 | -0.7207303 | 0.47107546 | 0.68235944 |
| WRAP53 | 52.4658475 | -0.173402445 | 0.42773682 | -0.4053952 | 0.68518705 | 0.83714154 |
| KDM6B | 230.333341 | -0.637433023 | 0.28761361 | -2.2162825 | 0.02667216 | 0.10833719 |
| NAA38 | 652.009117 | -0.213878112 | 0.22012376 | -0.9716266 | 0.33123632 | 0.55622942 |
| CYB5D1 | 89.5813991 | 0.012722363 | 0.30814321 | 0.04128718 | 0.96706696 | 0.98519107 |
| CHD3 | 295.18443 | -0.219549543 | 0.25839187 | -0.8496767 | 0.39550488 | 0.61766658 |
| KCNAB3 | 42.3122475 | -0.269436842 | 0.42919888 | -0.6277669 | 0.53015667 | 0.7282487 |
| TRAPPC1 | 84.144675 | -1.667273739 | 0.56792753 | -2.9357156 | 0.00332779 | 0.02503235 |
| CNTROB | 147.649412 | 0.077350596 | 0.35123184 | 0.22022661 | 0.82569468 | 0.91938879 |
| PER1 | 323.699696 | -0.422346887 | 0.41889115 | -1.0082497 | 0.31333459 | 0.53711851 |
| TMEM107 | 108.017572 | -0.771946782 | 0.25649917 | -3.0095489 | 0.00261636 | 0.02094515 |
| CTC1 | 243.194977 | -0.128667691 | 0.17224456 | -0.7470058 | 0.45506003 | 0.67027397 |
| PFAS | 73.9108409 | 0.297300091 | 0.36047648 | 0.824742 | 0.40951808 | 0.62937481 |
| SLC25A35 | 75.3837043 | 0.270811515 | 0.39887578 | 0.67893697 | 0.4971778 | 0.70224997 |
| RANGRF | 80.9826122 | -0.06466539 | 0.4017265 | -0.1609687 | 0.87211806 | 0.94200731 |
| AC135178.3 | 64.1880937 | 0.738627414 | 0.39985817 | 1.84722354 | 0.06471475 | 0.19759391 |
| NDEL1 | 738.495287 | -0.064711222 | 0.29457413 | -0.2196772 | 0.82612256 | 0.91938879 |
| MYH10 | 1314.13298 | 0.078382934 | 0.28650751 | 0.27358073 | 0.78440684 | 0.89699853 |
| STX8 | 2063.57062 | 0.061994985 | 0.21641845 | 0.28645888 | 0.77452668 | 0.89203269 |
| AC005695.2 | 121.858008 | -0.144969046 | 0.42424626 | -0.3417097 | 0.7325694 | 0.8685945 |
| AC118755.2 | 53.322034 | -0.03540947 | 0.40325875 | -0.0878083 | 0.93002903 | 0.96915731 |
| USP43 | 436.508027 | -1.551574548 | 0.35320252 | -4.3928751 | 1.12E-05 | 0.00029201 |
| GAS7 | 82.434237 | 0.915484032 | 0.52769124 | 1.73488578 | 0.08276104 | 0.23116738 |
| MYH3 | 55.0168661 | -0.298087362 | 0.47037706 | -0.63372 | 0.52626355 | 0.72497874 |
| SCO1 | 246.638739 | -0.275978744 | 0.20898246 | -1.3205833 | 0.18664036 | 0.39244388 |
| ADPRM | 72.7224584 | 0.612868416 | 0.26760695 | 2.29018121 | 0.02201081 | 0.09498231 |
| TMEM220 | 301.20143 | 0.139744728 | 0.28427128 | 0.49158934 | 0.62300968 | 0.7947209 |
| TMEM220-A! | 203.985312 | -0.15883551 | 0.49526409 | -0.3207087 | 0.74843114 | 0.87725423 |
| SHISA6 | 573.753228 | 0.996745506 | 0.60168913 | 1.65657888 | 0.09760465 | 0.2582075 |
| DNAH9 | 97.0731235 | -0.72280656 | 0.2774249 | -2.6054134 | 0.00917635 | 0.05206631 |
| ZNF18 | 88.2994039 | 0.205094782 | 0.2769527 | 0.74054084 | 0.45897189 | 0.67278348 |
| MAP2K4 | 2282.24453 | -0.12205956 | 0.13986471 | -0.8726973 | 0.38282807 | 0.6054569 |
| ARHGAP44 | 880.910134 | -0.362941009 | 0.15475932 | -2.3451965 | 0.01901705 | 0.08623283 |
| ELAC2 | 206.573862 | -0.256969442 | 0.22352135 | -1.1496416 | 0.25029151 | 0.4702177 |
| COX10 | 777.466715 | 0.800089051 | 0.18966735 | 4.21838043 | 2.46E-05 | 0.00056042 |
| AC005224.1 | 36.2023027 | 1.791011586 | 0.47210426 | 3.79367807 | 0.00014843 | 0.00229047 |
| PMP22 | 43.8388021 | -1.026872358 | 0.58605494 | -1.7521776 | 0.07974327 | 0.22596069 |
| TRIM16 | 206.538055 | 0.478283996 | 0.27927343 | 1.71260115 | 0.08678594 | 0.23862737 |
| ZSWIM7 | 505.634739 | -0.161039677 | 0.1672412 | -0.9629187 | 0.33558833 | 0.56118321 |
| TTC19 | 1315.34124 | 0.212581444 | 0.23981375 | 0.88644393 | 0.37537836 | 0.59870627 |
| NCOR1 | 3669.62156 | 0.342991911 | 0.13481424 | 2.54418166 | 0.01095341 | 0.0591753 |
| PIGL | 1078.93353 | 0.229440218 | 0.26579144 | 0.86323404 | 0.3880088 | 0.61116244 |
| CENPV | 88.8673563 | 1.12913476 | 0.44766679 | 2.52226609 | 0.01166015 | 0.06178054 |
| UBB | 1041.39444 | -1.673072365 | 0.47931767 | -3.4905293 | 0.00048206 | 0.00575777 |
| LRRC75A | 126.162517 | 0.174305745 | 0.33955456 | 0.51333649 | 0.60771596 | 0.7835354 |
| ZNF287 | 92.7844984 | 0.28922169 | 0.23824853 | 1.21394952 | 0.224767 | 0.43938839 |
| ZNF624 | 77.5498632 | 0.543640566 | 0.32004574 | 1.69863398 | 0.08938817 | 0.24339896 |
| AC098850.3 | 479.408826 | 0.379295099 | 0.61731129 | 0.61443085 | 0.53893066 | 0.73506352 |
| MPRIP | 1803.0888 | 0.037248314 | 0.22114231 | 0.16843595 | 0.86624033 | 0.93926847 |
| MPRIP-AS1 | 45.7413049 | -0.036700233 | 0.50156131 | -0.073172 | 0.94166927 | 0.97495762 |
| AC055811.1 | 47.5057102 | 0.12817609 | 0.53068238 | 0.24153071 | 0.80914382 | 0.91023512 |
| COPS3 | 218.006694 | 0.114487557 | 0.1898206 | 0.60313557 | 0.54641851 | 0.7406508 |
| NT5M | 101.802433 | 0.939975432 | 0.35714344 | 2.63192694 | 0.00849021 | 0.04927382 |
| MED9 | 481.349857 | -0.752777909 | 0.28262322 | -2.6635388 | 0.00773235 | 0.04604896 |
| RASD1 | 2067.38401 | -3.62636645 | 0.61151495 | -5.9301354 | 3.03E-09 | 3.51E-07 |
| PEMT | 176.874643 | -0.156195024 | 0.43402136 | -0.3598787 | 0.71893788 | 0.85995204 |
| RAI1 | 932.33271 | -0.946725068 | 0.35096659 | -2.6974791 | 0.00698667 | 0.0427692 |
| SMCR5 | 107.613709 | -0.794005075 | 0.26767541 | -2.9662982 | 0.00301408 | 0.02322812 |
| SREBF1 | 541.845291 | -0.397319697 | 0.48326077 | -0.8221642 | 0.41098346 | 0.63014283 |
| TOM1L2 | 2486.83922 | 1.046939811 | 0.1860717 | 5.62653976 | 1.84E-08 | 1.56E-06 |
| DRC3 | 175.513031 | 0.158722612 | 0.42447462 | 0.37392721 | 0.70845847 | 0.85346928 |
| ATPAF2 | 192.02053 | 0.420090034 | 0.18305975 | 2.29482466 | 0.02174317 | 0.09447552 |
| GID4 | 146.076766 | -0.172054635 | 0.23960597 | -0.7180732 | 0.47271212 | 0.68345605 |
| DRG2 | 136.58573 | -0.229069839 | 0.25877629 | -0.8852041 | 0.37604655 | 0.59932418 |
| ALKBH5 | 436.465667 | -0.654823535 | 0.21001956 | -3.1179168 | 0.00182134 | 0.015859 |
| FLII | 321.254694 | -0.56566379 | 0.23632472 | -2.3935871 | 0.01668452 | 0.07896188 |
| MIEF2 | 109.851107 | -0.117887578 | 0.24754394 | -0.4762289 | 0.63391131 | 0.80304349 |
| TOP3A | 309.992641 | -0.192421059 | 0.20789072 | -0.9255875 | 0.35466038 | 0.58039381 |
| SMCR8 | 116.088631 | -0.166098312 | 0.21800337 | -0.7619071 | 0.44611547 | 0.66204652 |
| SHMT1 | 552.066163 | -0.118097095 | 0.27292314 | -0.4327119 | 0.66522408 | 0.8233644 |
| ZNF286B | 87.9047313 | 0.680217796 | 0.29809312 | 2.28189699 | 0.02249542 | 0.09644338 |
| TRIM16L | 86.7250569 | 0.974325808 | 0.40457253 | 2.40828462 | 0.01602768 | 0.07659538 |
| TVP23B | 292.193634 | 0.190108608 | 0.19675094 | 0.9662399 | 0.33392415 | 0.55932084 |
| PRPSAP2 | 761.658038 | 0.477746389 | 0.14629331 | 3.26567484 | 0.00109204 | 0.01069288 |
| SLC5A10 | 94.1359406 | -0.059886528 | 0.32067501 | -0.1867515 | 0.8518555 | 0.93193618 |
| FAM83G | 63.4095304 | -1.840607565 | 0.34472904 | -5.3392879 | 9.33E-08 | 6.07E-06 |
| EPN2 | 1583.11101 | -0.366022025 | 0.25624548 | -1.4284038 | 0.15317566 | 0.34687078 |
| EPN2-AS1 | 82.3651685 | -0.87366688 | 0.45348726 | -1.9265522 | 0.05403546 | 0.17515446 |
| B9D1 | 333.307331 | -0.053672373 | 0.22575478 | -0.2377463 | 0.81207784 | 0.91189688 |
| MAPK7 | 55.9761494 | -0.594694956 | 0.37629488 | -1.5803961 | 0.11401619 | 0.28659179 |
| AC004448.2 | 88.8492097 | -1.247541999 | 0.36821289 | -3.3880997 | 0.00070379 | 0.00768926 |
| SLC47A1 | 332.100744 | 1.516966585 | 0.38649903 | 3.92489104 | 8.68E-05 | 0.00150807 |
| ALDH3A2 | 439.50035 | 0.34408931 | 0.3175626 | 1.08353221 | 0.27857226 | 0.50077042 |
| SLC47A2 | 64.2968092 | -0.191942417 | 0.564656 | -0.3399281 | 0.73391071 | 0.86935939 |

| | | | | | | |
|---|---|---|---|---|---|---|
| ULK2 | 583.150921 | 0.113800964 | 0.24508179 | 0.46433872 | 0.64240509 | 0.80832017 |
| AKAP10 | 1495.86678 | 0.205907776 | 0.1657874 | 1.24199892 | 0.21423696 | 0.42776686 |
| SPECC1 | 155.602066 | -1.113490842 | 0.28249311 | -3.9416567 | 8.09E-05 | 0.0014309 |
| CCDC144NL-/ | 186.423287 | -0.857660267 | 0.41699219 | -2.0567778 | 0.03970761 | 0.14187118 |
| USP22 | 1061.21733 | -0.0667525 | 0.117671 | -0.5672808 | 0.57052343 | 0.75945349 |
| DHRS7B | 932.037754 | -0.342085183 | 0.19127051 | -1.7884889 | 0.07369717 | 0.21540763 |
| TMEM11 | 111.416341 | -0.118976232 | 0.26075955 | -0.456268 | 0.64819729 | 0.81176524 |
| AC087294.1 | 165.948036 | 0.271425225 | 0.25200059 | 1.0770817 | 0.28144377 | 0.5040801 |
| NATD1 | 61.4644046 | -1.284234465 | 0.41118883 | -3.1232231 | 0.00178882 | 0.0156503 |
| MAP2K3 | 320.515048 | 0.462118428 | 0.3191444 | 1.44799164 | 0.14761939 | 0.33893699 |
| KCNJ12 | 66.8186596 | 1.166671583 | 0.89990238 | 1.29644238 | 0.19482312 | 0.40241103 |
| C17orf51 | 54.456754 | 0.077203462 | 0.46194431 | 0.16712721 | 0.86726996 | 0.93961371 |
| WSB1 | 5360.54835 | 0.138768903 | 0.50145972 | 0.27672991 | 0.78198751 | 0.89588513 |
| KSR1 | 908.698797 | -0.061323672 | 0.29250462 | -0.2096503 | 0.83394064 | 0.92257303 |
| AC005697.1 | 104.916437 | 1.835353605 | 0.71437649 | 2.56916854 | 0.01019429 | 0.05631792 |
| NLK | 2788.63633 | 0.489129091 | 0.25436136 | 1.92296935 | 0.0544839 | 0.17596035 |
| IFT20 | 170.912517 | 0.332252335 | 0.20848276 | 1.59366816 | 0.11101038 | 0.28112272 |
| TNFAIP1 | 275.692204 | -0.52155021 | 0.18278902 | -2.8532907 | 0.0043269 | 0.03017925 |
| POLDIP2 | 229.163588 | -0.273799448 | 0.25734601 | -1.0639351 | 0.28735811 | 0.50981103 |
| TMEM199 | 93.9495736 | 0.482414827 | 0.24053654 | 2.00557818 | 0.04490127 | 0.15441315 |
| SARM1 | 239.281374 | 0.000958544 | 0.36635442 | 0.00261644 | 0.99791239 | 0.99934986 |
| AC005726.1 | 187.867448 | 0.041057136 | 0.27451869 | 0.14956044 | 0.88111142 | 0.94645136 |
| KIAA0100 | 499.324676 | -0.258745001 | 0.21397225 | -1.2092456 | 0.22656851 | 0.44161101 |
| SDF2 | 221.341947 | -0.505039972 | 0.22101535 | -2.28509 | 0.02230755 | 0.09596392 |
| SUPT6H | 723.340089 | -0.310590153 | 0.21973876 | -1.4134518 | 0.15752291 | 0.35332317 |
| PROCA1 | 60.4756169 | 0.230864976 | 0.36936297 | 0.62503552 | 0.53194775 | 0.72919238 |
| RAB34 | 258.372329 | -1.318164683 | 0.43610312 | -3.0225986 | 0.00250614 | 0.02025928 |
| RPL23A | 821.404391 | -1.080000731 | 0.3502038 | -3.0839207 | 0.00204292 | 0.01742088 |
| TLCD1 | 67.8601587 | -0.761551267 | 0.52961588 | -1.4379313 | 0.15045354 | 0.34276316 |
| NEK8 | 314.15623 | -0.595301274 | 0.5934502 | -1.0031192 | 0.31580336 | 0.540063 |
| TRAF4 | 493.04879 | -0.465248209 | 0.40426658 | -1.1508451 | 0.24979597 | 0.46942026 |
| FAM222B | 1826.68123 | -0.441420011 | 0.21348179 | -2.0677174 | 0.0386666 | 0.13955547 |
| ERAL1 | 94.8474419 | 0.333116608 | 0.23595841 | 1.41175981 | 0.15802069 | 0.35413901 |
| FLOT2 | 802.862988 | -1.256881683 | 0.29407767 | -4.2739786 | 1.92E-05 | 0.00045454 |
| DHRS13 | 144.655565 | -1.721550761 | 0.46812071 | -3.6775787 | 0.00023546 | 0.00335148 |
| PHF12 | 606.8947 | -0.144651923 | 0.20960361 | -0.6901213 | 0.4901179 | 0.69682235 |
| PIPOX | 63.5670084 | -0.190886708 | 0.40368013 | -0.4728662 | 0.63630861 | 0.80446022 |
| MYO18A | 420.768915 | -0.599880122 | 0.28019303 | -2.1409531 | 0.03227782 | 0.12397725 |
| AC024619.3 | 75.7817036 | -0.892286481 | 0.48096103 | -1.8552157 | 0.06356542 | 0.19526411 |
| NUFIP2 | 1368.14273 | -0.332131046 | 0.16039309 | -2.0707316 | 0.03838389 | 0.13904125 |
| TAOK1 | 2467.87069 | -0.133410033 | 0.14805828 | -0.9010643 | 0.36755412 | 0.59279882 |
| ABHD15-AS1 | 248.767518 | -0.534762838 | 0.34453646 | -1.5521226 | 0.12063288 | 0.29693855 |
| ABHD15 | 43.0921386 | -0.507841608 | 0.29438303 | -1.7251049 | 0.08450856 | 0.23426519 |
| GIT1 | 160.285899 | -0.51139502 | 0.28092484 | -1.820398 | 0.06869842 | 0.20558165 |
| SSH2 | 4010.93117 | -0.576149108 | 0.20163692 | -2.8573592 | 0.00427182 | 0.02993447 |
| AC023389.1 | 164.365842 | -0.67398541 | 0.30031189 | -2.2442848 | 0.02481408 | 0.10320066 |
| EFCAB5 | 146.584126 | -1.813216789 | 0.56298939 | -3.2206944 | 0.0012788 | 0.01213486 |
| NSRP1 | 1741.41118 | 0.093612871 | 0.15478564 | 0.60479041 | 0.54531827 | 0.7400858 |
| AC104984.3 | 85.2606573 | -0.51981411 | 0.50346769 | -1.0324677 | 0.30185308 | 0.52503327 |
| BLMH | 317.106197 | -0.007459325 | 0.220092 | -0.0338918 | 0.97296339 | 0.98804941 |
| CPD | 3076.45342 | -1.924063521 | 0.45556061 | -4.2235072 | 2.41E-05 | 0.00055159 |
| GOSR1 | 939.26293 | 0.228627645 | 0.1773773 | 1.28893408 | 0.197421 | 0.4057467 |
| AC127024.2 | 56.055201 | -0.758792609 | 0.49803516 | -1.5235724 | 0.12761557 | 0.30794192 |
| CRLF3 | 761.807347 | 0.312481589 | 0.24785754 | 1.26073059 | 0.20740593 | 0.41826403 |
| ATAD5 | 184.813549 | 0.142888318 | 0.35874892 | 0.39829617 | 0.69041189 | 0.84106341 |
| AC130324.2 | 85.7873791 | 0.243080332 | 0.4685197 | 0.51882628 | 0.60388189 | 0.7808787 |
| TEFM | 385.972778 | 0.371987852 | 0.2388742 | 1.55725418 | 0.11941015 | 0.2950961 |
| ADAP2 | 84.9051489 | 0.539740516 | 0.48881469 | 1.10418227 | 0.26951408 | 0.49096835 |
| RNF135 | 175.565885 | -0.158490766 | 0.2254456 | -0.7030111 | 0.48204882 | 0.69045574 |
| NF1 | 5387.73222 | 0.023028294 | 0.15415436 | 0.14938464 | 0.88125013 | 0.94645136 |
| OMG | 142.51393 | -0.282192425 | 0.58449395 | -0.4827979 | 0.62923928 | 0.79942499 |
| RAB11FIP4 | 292.91984 | -0.216403489 | 0.31815144 | -0.6801902 | 0.49638404 | 0.70178067 |
| COPRS | 126.849123 | -0.102581589 | 0.4467962 | -0.2295937 | 0.81840751 | 0.91558777 |
| UTP6 | 336.767478 | 0.392784932 | 0.2322505 | 1.69121243 | 0.09079624 | 0.24620512 |
| AC004253.1 | 122.949654 | 0.381719558 | 0.40295375 | 0.94730365 | 0.34348407 | 0.56837065 |
| SUZ12 | 1136.29819 | 0.319712095 | 0.15727164 | 2.03286552 | 0.04206611 | 0.14711445 |
| LRRC37B | 391.120139 | -0.011268377 | 0.22058907 | -0.0510831 | 0.95925929 | 0.98229715 |
| RHOT1 | 1061.18422 | 0.548412778 | 0.14131649 | 3.88074147 | 0.00010414 | 0.00172593 |
| C17orf75 | 180.006014 | 0.797092368 | 0.23392037 | 3.40753721 | 0.00065552 | 0.00728852 |
| ZNF207 | 1919.69422 | 0.325129848 | 0.15537481 | 2.09255192 | 0.03638917 | 0.1342368 |
| PSMD11 | 542.479598 | -0.039733923 | 0.27906971 | -0.1423799 | 0.88677992 | 0.94907438 |
| MYO1D | 2301.49413 | -1.112513671 | 0.23736711 | -4.6868906 | 2.77E-06 | 9.30E-05 |
| AC079336.4 | 50.3315721 | -1.553952209 | 0.43591629 | -3.564795 | 0.00036414 | 0.00463832 |
| ASIC2 | 818.810002 | -0.961330882 | 0.58825935 | -1.6341957 | 0.10221777 | 0.26667876 |
| CCL2 | 165.4027 | -2.102587297 | 0.97092494 | -2.1655508 | 0.03034553 | 0.11866018 |
| CCT6B | 140.722111 | 0.128153438 | 0.58359427 | 0.21959338 | 0.82618785 | 0.91938879 |
| ZNF830 | 80.6482299 | 0.074476801 | 0.32276473 | 0.23074641 | 0.81751182 | 0.9152834 |
| LIG3 | 162.748798 | -0.086311894 | 0.19559256 | -0.4412841 | 0.65900731 | 0.81950928 |
| RFFL | 92.5542185 | -0.418340872 | 0.28136769 | -1.486812 | 0.13706446 | 0.32301239 |
| SLFN5 | 242.595565 | -2.409967482 | 0.49710034 | -4.8480503 | 1.25E-06 | 4.90E-05 |
| SLFN12 | 54.3460318 | 0.421551612 | 0.32929019 | 1.28018274 | 0.20048087 | 0.40930388 |
| SLFN13 | 76.6340515 | 0.349630211 | 0.7790283 | 0.44880297 | 0.6535738 | 0.81543241 |
| LINC02001 | 164.387589 | -0.026116744 | 0.2435169 | -0.1072482 | 0.9145921 | 0.96131774 |
| AP2B1 | 1422.80804 | 0.029257412 | 0.17082304 | 0.17127322 | 0.86400893 | 0.93827593 |
| TAF15 | 2175.22522 | -0.015293215 | 0.18843939 | -0.0811572 | 0.93531693 | 0.97219085 |
| AC015849.3 | 154.909748 | 0.315550383 | 0.48390664 | 0.65208938 | 0.51434351 | 0.71598841 |
| ZNHIT3 | 158.634065 | -0.193809573 | 0.24299904 | -0.7975734 | 0.42511809 | 0.64375196 |
| MYO19 | 902.303238 | 0.071144948 | 0.31493416 | 0.22590419 | 0.82127594 | 0.91716553 |
| GGNBP2 | 998.056633 | -0.241101124 | 0.17290727 | -1.3943955 | 0.1631982 | 0.36138895 |
| DHRS11 | 80.3720652 | 0.555796383 | 0.50291105 | 1.10515842 | 0.26909095 | 0.4906035 |
| MRM1 | 58.699701 | -0.033384149 | 0.37956058 | -0.0879547 | 0.92991266 | 0.96915731 |
| AATF | 796.315482 | -0.319542788 | 0.19983345 | -1.5990456 | 0.10981048 | 0.27893974 |
| ACACA | 2042.63596 | 0.33519199 | 0.31703372 | 1.05727551 | 0.29038587 | 0.51334407 |
| TADA2A | 390.666336 | -0.204506005 | 0.20999513 | -0.9738607 | 0.33012572 | 0.55570671 |
| DUSP14 | 43.0337507 | 0.240765558 | 0.37409482 | 0.64359501 | 0.51983809 | 0.7198387 |
| SYNRG | 1254.46474 | 0.005624326 | 0.14228733 | 0.03952794 | 0.96846948 | 0.98582596 |
| DDX52 | 1025.76501 | -0.651223883 | 0.59907025 | -1.0870576 | 0.27701135 | 0.49909966 |
| AC243585.2 | 61.8812331 | -1.527153073 | 0.82463977 | -1.8519033 | 0.06403972 | 0.19631019 |
| AC243571.2 | 100.80852 | -2.21696467 | 1.00731547 | -2.2008643 | 0.02774563 | 0.11119297 |
| HNF1B | 2664.38254 | -0.139574664 | 0.27519168 | -0.5071907 | 0.61202101 | 0.78684784 |
| MRPL45 | 317.173118 | 0.494466446 | 0.15164006 | 3.26079028 | 0.00111102 | 0.01084663 |
| SOCS7 | 321.243782 | 0.215319526 | 0.23091706 | 0.93245397 | 0.35110197 | 0.57725988 |
| ARHGAP23 | 45.0914318 | -0.892999105 | 0.3980732 | -2.2433038 | 0.02487723 | 0.1033669 |
| MLLT6 | 1650.31069 | -0.185280047 | 0.22041195 | -0.840608 | 0.40056757 | 0.62209429 |
| CISD3 | 138.20438 | -0.459870627 | 0.35435322 | -1.2977747 | 0.19436478 | 0.40190425 |
| PCGF2 | 246.124032 | -0.224432236 | 0.23165151 | -0.9688356 | 0.33262722 | 0.55792672 |
| PSMB3 | 293.350216 | -0.718582788 | 0.30672066 | -2.3427923 | 0.01914004 | 0.08661211 |
| PIP4K2B | 382.302195 | 0.595568212 | 0.22797614 | 2.61241466 | 0.00899051 | 0.05127621 |
| CWC25 | 276.693437 | -0.501169407 | 0.18581969 | -2.6970737 | 0.00699518 | 0.04279728 |
| RPL23 | 621.988302 | -0.887209241 | 0.39461026 | -2.2483177 | 0.02455594 | 0.1024185 |
| LASP1 | 383.267403 | -1.352114039 | 0.21084398 | -6.4128653 | 1.43E-10 | 2.32E-08 |
| RPL19 | 1157.27046 | -1.277452444 | 0.4597135 | -2.7788013 | 0.00545599 | 0.0357738 |
| STAC2 | 116.430715 | -4.001557314 | 0.88984969 | -4.4968913 | 6.90E-06 | 0.00019598 |

| | | | | | | |
|---|---|---|---|---|---|---|
| FBXL20 | 2278.40182 | -0.206284862 | 0.19308139 | -1.068383 | 0.2853478 | 0.50782141 |
| MED1 | 384.931839 | 0.132267955 | 0.13630088 | 0.97041157 | 0.33184139 | 0.55704883 |
| CDK12 | 1274.24794 | 0.06278909 | 0.16235442 | 0.38674088 | 0.69894805 | 0.84693191 |
| PPP1R1B | 166.935224 | -2.266950892 | 1.01782212 | -2.2272565 | 0.02593014 | 0.10620502 |
| STARD3 | 268.1807 | -0.256135565 | 0.24347438 | -1.0520021 | 0.29279857 | 0.51560809 |
| PGAP3 | 180.079413 | 0.383092335 | 0.33720971 | 1.13606556 | 0.25592913 | 0.47687514 |
| ERBB2 | 1140.47945 | 0.070463182 | 0.33625929 | 0.20955015 | 0.83401879 | 0.92258222 |
| MIEN1 | 165.518544 | -1.379246781 | 0.42786741 | -3.2235378 | 0.00126618 | 0.01204097 |
| GRB7 | 1136.90895 | -1.117337332 | 0.21789561 | -5.1278561 | 2.93E-07 | 1.50E-05 |
| GSDMB | 54.2548826 | 0.281297322 | 0.36909427 | 0.76212867 | 0.44598321 | 0.66204652 |
| ORMDL3 | 71.5834064 | -1.148168278 | 0.44098235 | -2.6036604 | 0.00922341 | 0.05223895 |
| AC090844.2 | 82.7171715 | 0.979051159 | 0.44415475 | 2.20430191 | 0.02750312 | 0.11047495 |
| PSMD3 | 227.815984 | -0.43108293 | 0.25290074 | -1.7045539 | 0.08827766 | 0.24157026 |
| MED24 | 337.717143 | 0.204990913 | 0.18453527 | 1.11084948 | 0.26663314 | 0.48773815 |
| THRA | 238.484815 | 0.001877499 | 0.2902357 | 0.00646888 | 0.99483862 | 0.997919 |
| NR1D1 | 57.9124834 | -2.07469062 | 0.5957926 | -3.4822363 | 0.00049724 | 0.00589597 |
| MSL1 | 496.661037 | -0.263212293 | 0.24797973 | -1.0614267 | 0.28849605 | 0.51093714 |
| CASC3 | 489.225317 | 0.093946831 | 0.11453236 | 0.82026451 | 0.41206533 | 0.6313619 |
| RAPGEFL1 | 69.8689284 | 0.33611386 | 0.3618128 | 0.92897171 | 0.35290375 | 0.57885329 |
| WIPF2 | 1214.41823 | -0.201996537 | 0.20481329 | -0.9862472 | 0.3240118 | 0.54891265 |
| RARA | 165.175276 | -0.853644182 | 0.50428848 | -1.6927695 | 0.09049934 | 0.24561451 |
| IGFBP4 | 194.583988 | -0.599023196 | 0.3293436 | -1.8188396 | 0.0689359 | 0.20605891 |
| KRT10 | 247.196136 | -0.99146539 | 0.47880346 | -2.0707148 | 0.03838546 | 0.13904125 |
| TMEM99 | 81.9422615 | 0.404992923 | 0.27924085 | 1.45033551 | 0.14696498 | 0.33784554 |
| KRT19 | 393.098847 | -3.49734907 | 0.6046798 | -5.7838034 | 7.30E-09 | 7.09E-07 |
| EIF1 | 1535.70173 | -1.418581868 | 0.3626553 | -3.9116535 | 9.17E-05 | 0.00157367 |
| HAP1 | 139.320211 | -3.02702769 | 0.5258149 | -5.7568313 | 8.57E-09 | 8.09E-07 |
| JUP | 1506.70108 | -0.167093566 | 0.27750153 | -0.6021357 | 0.54708384 | 0.74124838 |
| P3H4 | 133.612043 | 0.256730704 | 0.28681965 | 0.89509454 | 0.37073655 | 0.59532317 |
| FKBP10 | 49.6644255 | -0.627965259 | 0.7355618 | -0.853722 | 0.39325907 | 0.61554201 |
| NT5C3B | 231.563482 | -0.049744417 | 0.30728812 | -0.161882 | 0.87139877 | 0.94153836 |
| ACLY | 453.378026 | -0.560092466 | 0.25424916 | -2.2029275 | 0.02759986 | 0.11081049 |
| TTC25 | 147.145257 | 0.235922457 | 0.36342584 | 0.64916258 | 0.51623329 | 0.71747954 |
| CNP | 292.071901 | 0.092593461 | 0.45401243 | 0.20394477 | 0.83839666 | 0.92456015 |
| DNAJC7 | 912.260872 | 0.02788393 | 0.11046472 | 0.25242385 | 0.80071346 | 0.90662963 |
| NKIRAS2 | 113.699485 | -0.482993565 | 0.20979692 | -2.3021956 | 0.02132414 | 0.09333021 |
| DHX58 | 69.4875207 | -0.376082023 | 0.6065492 | -0.6200355 | 0.53523443 | 0.73219616 |
| KAT2A | 207.909628 | 1.27461347 | 0.41350899 | 3.08243227 | 0.00205316 | 0.01749694 |
| GHDC | 57.8810941 | -0.823257316 | 0.44088147 | -1.8672985 | 0.06185991 | 0.19203468 |
| STAT5B | 1304.13736 | -0.513830912 | 0.19395462 | -2.6492326 | 0.00806748 | 0.04762262 |
| STAT5A | 202.310167 | -1.161245916 | 0.32113167 | -3.6161052 | 0.00029907 | 0.00401872 |
| STAT3 | 5065.11645 | -1.057648268 | 0.25190953 | -4.1985243 | 2.69E-05 | 0.00060148 |
| CAVIN1 | 103.751683 | -1.470746062 | 0.647108 | -2.2727984 | 0.02303833 | 0.09813559 |
| ATP6V0A1 | 676.240861 | -0.089519946 | 0.24301509 | -0.368372 | 0.7125959 | 0.8559282 |
| NAGLU | 242.010075 | -0.306780159 | 0.37042006 | -0.8281953 | 0.4075599 | 0.62748459 |
| COASY | 224.304636 | -0.022622571 | 0.351476 | -0.0643645 | 0.94868001 | 0.97784954 |
| MLX | 262.701663 | -0.352997433 | 0.26765098 | -1.3188722 | 0.18721184 | 0.39289501 |
| RETREG3 | 611.885496 | -0.161747684 | 0.18142734 | -0.8915287 | 0.37264558 | 0.59655016 |
| TUBG1 | 57.6260413 | -0.696064015 | 0.49524341 | -1.4054988 | 0.15987301 | 0.35671664 |
| TUBG2 | 218.090684 | 0.067187309 | 0.22973564 | 0.29245488 | 0.76993885 | 0.88922989 |
| PLEKHH3 | 93.7542988 | 0.139554955 | 0.36444609 | 0.38292345 | 0.70177652 | 0.8488446 |
| AC100793.2 | 65.212676 | -0.727615755 | 0.62854072 | -1.1576271 | 0.24701625 | 0.46618627 |
| EZH1 | 723.222397 | -0.046201733 | 0.21046669 | -0.2195204 | 0.82624469 | 0.91938879 |
| VPS25 | 222.232931 | -1.009151507 | 0.24361449 | -4.1424117 | 3.44E-05 | 0.00073337 |
| WNK4 | 1462.51123 | 1.139676451 | 0.51459918 | 2.21468766 | 0.02678151 | 0.10864772 |
| COA3 | 401.521951 | -0.078713721 | 0.41571256 | -0.1893465 | 0.84982125 | 0.93113889 |
| CNTD1 | 79.1849846 | 1.52645916 | 0.44115765 | 3.46012171 | 0.00053993 | 0.00627261 |
| BECN1 | 477.24622 | 0.345915898 | 0.19171967 | 1.80427966 | 0.07118748 | 0.21050266 |
| PSME3 | 214.85665 | -0.192294158 | 0.18723416 | -1.027025 | 0.3044087 | 0.5276817 |
| LINC00671 | 179.356499 | 3.29106935 | 0.72795421 | 4.52098401 | 6.16E-06 | 0.00017948 |
| RUNDC1 | 230.927322 | -0.54684162 | 0.20813626 | -2.6273251 | 0.00860591 | 0.04972678 |
| RPL27 | 420.476013 | -1.156805558 | 0.3853395 | -3.0020425 | 0.00268175 | 0.02130078 |
| IFI35 | 149.915954 | -0.749406817 | 0.60093045 | -1.2470775 | 0.2123691 | 0.4250922 |
| VAT1 | 92.8169591 | -1.758969829 | 0.34281421 | -5.1309711 | 2.88E-07 | 1.48E-05 |
| BRCA1 | 268.075754 | -0.479581461 | 0.38999707 | -1.2297053 | 0.21880748 | 0.43305784 |
| NBR1 | 838.509836 | -0.231250074 | 0.14646715 | -1.5788528 | 0.11436983 | 0.28726204 |
| TMEM106A | 76.6849028 | -0.543981058 | 0.39863658 | -1.364604 | 0.17237756 | 0.37368753 |
| LINC00854 | 611.892569 | -0.157605408 | 0.77973718 | -0.2021263 | 0.83981797 | 0.92558662 |
| ARL4D | 145.319371 | -1.901044185 | 0.60331168 | -3.1510151 | 0.00162704 | 0.01464804 |
| DHX8 | 363.31763 | -0.328534162 | 0.15074192 | -2.1794479 | 0.02929841 | 0.11551991 |
| ETV4 | 136.026067 | -1.636670768 | 0.79622221 | -2.0555452 | 0.03982637 | 0.14211692 |
| AC068675.1 | 64.7813815 | -1.587056316 | 0.52222695 | -3.0390165 | 0.00237352 | 0.01946108 |
| DUSP3 | 287.444507 | -0.630057407 | 0.26026087 | -2.4208687 | 0.01548347 | 0.07500191 |
| MPP3 | 55.4247076 | -1.02122356 | 0.36616012 | -2.7890082 | 0.00528697 | 0.03488293 |
| TMEM101 | 155.449645 | -1.222704904 | 0.50137863 | -2.4386857 | 0.01474078 | 0.07238327 |
| LSM12 | 344.316914 | -0.107123992 | 0.19771659 | -0.5418058 | 0.5879523 | 0.7710768 |
| G6PC3 | 115.906722 | -0.889449175 | 0.4004687 | -2.2210205 | 0.02634958 | 0.1074544 |
| HDAC5 | 469.579281 | -0.290689577 | 0.28037118 | -1.0368026 | 0.29982785 | 0.52269052 |
| ASB16-AS1 | 106.849797 | 0.064249266 | 0.29139026 | 0.22049215 | 0.82548788 | 0.91938879 |
| TMUB2 | 241.708601 | -0.087142735 | 0.22920632 | -0.3801934 | 0.70380184 | 0.84992208 |
| ATXN7L3 | 122.065897 | -0.79284793 | 0.31284464 | -2.5343184 | 0.01126663 | 0.06034355 |
| AC004596.1 | 55.8132618 | -0.754100184 | 0.29270371 | -2.576326 | 0.00998565 | 0.05532813 |
| UBTF | 362.07191 | -0.248479698 | 0.278662 | -0.8916885 | 0.37255993 | 0.59655016 |
| AC003102.1 | 54.0158435 | -0.853648581 | 0.4688522 | -1.82072 | 0.06864943 | 0.20553849 |
| SLC25A39 | 300.698866 | -0.183962751 | 0.30571127 | -0.6017533 | 0.54733839 | 0.74144117 |
| GRN | 388.904952 | -1.685260629 | 0.34451673 | -4.8916656 | 1.00E-06 | 4.11E-05 |
| GPATCH8 | 3057.13705 | 0.030182545 | 0.14952626 | 0.20185448 | 0.84003049 | 0.92566636 |
| CCDC43 | 91.4323197 | -0.081000219 | 0.26130768 | -0.3099802 | 0.75657598 | 0.88227669 |
| DBF4B | 151.49533 | 0.125518743 | 0.4179908 | 0.30029068 | 0.76395544 | 0.88555042 |
| EFTUD2 | 393.273191 | -0.304869738 | 0.25897094 | -1.1772353 | 0.2391016 | 0.45716719 |
| DCAKD | 596.287585 | 0.019395156 | 0.25341899 | 0.07653395 | 0.93899431 | 0.9740204 |
| NMT1 | 626.993266 | -0.246481196 | 0.12757231 | -1.9320901 | 0.05334839 | 0.17356623 |
| PLCD3 | 67.0416128 | -1.760925895 | 0.46072437 | -3.822081 | 0.00013233 | 0.0021009 |
| ACBD4 | 142.663079 | 0.216640925 | 0.24161132 | 0.89665056 | 0.36990541 | 0.59455835 |
| AC142472.1 | 44.5009327 | 0.650952903 | 0.43728387 | 1.48862773 | 0.13658542 | 0.32222841 |
| HEXIM1 | 155.923409 | -0.27419971 | 0.4698604 | -0.583577 | 0.55950496 | 0.75137261 |
| AC138150.1 | 623.132701 | 0.624924256 | 0.37153856 | 1.6819903 | 0.09257072 | 0.24974809 |
| HEXIM2 | 49.3861328 | 0.328054358 | 0.31217824 | 1.05085594 | 0.29332476 | 0.51627278 |
| MAP3K14 | 663.85862 | -0.765111766 | 0.29375006 | -2.6046353 | 0.00919721 | 0.05213989 |
| ARHGAP27 | 128.736423 | 0.091052897 | 0.27044493 | 0.33667814 | 0.73635954 | 0.87069852 |
| PLEKHM1 | 400.343758 | -0.414136901 | 0.241127 | -1.7175053 | 0.08588688 | 0.23704134 |
| LINC02210-C | 297.937984 | 0.518779256 | 0.35469773 | 1.46259537 | 0.14357813 | 0.33319107 |
| KANSL1 | 6688.35147 | 0.341652325 | 0.26665171 | 1.2812681 | 0.20009951 | 0.40902261 |
| KANSL1-AS1 | 47.8523278 | 0.391798176 | 0.48305086 | 0.81109095 | 0.41731344 | 0.63704999 |
| ARL17B | 1419.17039 | 1.442785338 | 1.09442676 | 1.31830233 | 0.18740246 | 0.39298287 |
| LRRC37A | 76.932987 | 0.586896941 | 0.30827847 | 1.90378829 | 0.05693776 | 0.18148912 |
| LRRC37A2 | 388.331887 | 0.809557976 | 0.29085203 | 2.78340149 | 0.00537922 | 0.03540314 |
| ARL17A | 466.570779 | 0.050237017 | 0.45454478 | 0.1105216 | 0.91199572 | 0.96026715 |
| NSF | 1689.7212 | 0.159245416 | 0.30610039 | 0.52023918 | 0.60289688 | 0.77998677 |
| AC005670.2 | 288.25159 | -0.002381616 | 0.33577147 | -0.007093 | 0.99434068 | 0.99783229 |
| CDC27 | 1599.52816 | -0.422579233 | 0.19941391 | -2.1191061 | 0.0340815 | 0.12880758 |
| AC068234.1 | 708.781355 | -4.908326207 | 0.76723746 | -6.3974016 | 1.58E-10 | 2.52E-08 |
| AC068234.2 | 129.849942 | -3.267666433 | 0.56226068 | -5.8116574 | 6.19E-09 | 6.29E-07 |

| | | | | | | |
|---|---|---|---|---|---|---|
| EFCAB13 | 781.490988 | -1.902824275 | 0.39797679 | -4.7812444 | 1.74E-06 | 6.29E-05 |
| AC040934.1 | 134.298705 | 0.753653366 | 0.34117519 | 2.2089923 | 0.02717518 | 0.10967214 |
| NPEPPS | 3142.8486 | -0.518102221 | 0.19222082 | -2.6953491 | 0.00703149 | 0.04292004 |
| AC025682.1 | 415.19643 | 0.293135109 | 0.27019873 | 1.08488707 | 0.27797167 | 0.50016726 |
| KPNB1 | 1188.54932 | -0.447297645 | 0.17147417 | -2.6085425 | 0.00909287 | 0.05181525 |
| TBKBP1 | 85.1366076 | -0.628359672 | 0.36770823 | -1.708854 | 0.087478 | 0.23985703 |
| OSBPL7 | 70.4196486 | -0.100764089 | 0.45261623 | -0.2226259 | 0.82382669 | 0.91846107 |
| MRPL10 | 73.7987927 | -0.593628334 | 0.32891554 | -1.8048048 | 0.07110524 | 0.21040078 |
| SCRN2 | 139.924013 | 0.180333022 | 0.29758184 | 0.60599471 | 0.54451827 | 0.73944091 |
| SP2 | 180.218756 | 0.137120375 | 0.24459735 | 0.56059633 | 0.57507275 | 0.76252365 |
| SP2-AS1 | 128.798158 | -0.062761104 | 0.27044909 | -0.2320625 | 0.81648944 | 0.91452633 |
| PNPO | 136.659699 | -0.578671446 | 0.33222729 | -1.7417938 | 0.08154452 | 0.22942119 |
| PRR15L | 177.220594 | -0.425186851 | 0.67379378 | -0.6310341 | 0.52801823 | 0.72617338 |
| CDK5RAP3 | 397.508692 | 0.264612281 | 0.20785929 | 1.2730356 | 0.20300541 | 0.41262777 |
| NFE2L1 | 662.221643 | -0.532998506 | 0.25644046 | -2.0784493 | 0.03766799 | 0.13733278 |
| CBX1 | 409.562946 | -0.118302679 | 0.24570379 | -0.481485 | 0.63017187 | 0.79994176 |
| SNX11 | 119.828119 | -0.241246393 | 0.19008537 | -1.2691476 | 0.20438843 | 0.41426527 |
| SKAP1 | 4952.90701 | 0.231039854 | 0.20658203 | 1.11839279 | 0.26339927 | 0.48439746 |
| SKAP1-AS1 | 103.053322 | -0.103521355 | 0.47560038 | -0.2176646 | 0.82769047 | 0.91981743 |
| HOXB2 | 240.939742 | -0.904860679 | 0.40142758 | -2.2541069 | 0.02418944 | 0.10127365 |
| HOXB-AS1 | 216.380948 | 0.597768098 | 0.32273805 | 1.85217731 | 0.06400037 | 0.19624296 |
| HOXB3 | 2656.15095 | -0.123473698 | 0.25174979 | -0.490462 | 0.62380703 | 0.79539814 |
| HOXB-AS3 | 599.801943 | -0.113446575 | 0.35430781 | -0.3201921 | 0.74882268 | 0.87757952 |
| HOXB7 | 396.514278 | -1.332024699 | 0.2653806 | -5.0192994 | 5.19E-07 | 2.42E-05 |
| AC103702.2 | 60.8758681 | -1.489229985 | 0.63178404 | -2.357182 | 0.01841422 | 0.0843248 |
| LINC02086 | 128.473745 | -0.861035815 | 0.67748203 | -1.2709353 | 0.20375167 | 0.41350055 |
| TTLL6 | 42.4536363 | 0.664143509 | 0.8529261 | 0.77866478 | 0.43617721 | 0.6532273 |
| CALCOCO2 | 459.920983 | -0.020350085 | 0.17770343 | -0.1145171 | 0.90882787 | 0.95876587 |
| ATP5MC1 | 399.578563 | 0.328661729 | 0.43623261 | 0.75340936 | 0.45120396 | 0.66703448 |
| UBE2Z | 745.102531 | -0.627438884 | 0.18486381 | -3.3940602 | 0.00068865 | 0.00756765 |
| SNF8 | 325.583061 | -0.159819417 | 0.14897044 | -1.0728263 | 0.28334904 | 0.50584639 |
| AC091133.1 | 64.8927211 | -0.466119365 | 0.28571347 | -1.6314224 | 0.10280121 | 0.2676495 |
| B4GALNT2 | 144.26396 | 0.698031929 | 0.63947159 | 1.09157614 | 0.27501945 | 0.49647832 |
| ZNF652 | 1258.68395 | -0.154336732 | 0.13970669 | -1.1047197 | 0.26928106 | 0.49084828 |
| AC091180.5 | 68.4105593 | -0.591327634 | 0.53755138 | -1.1000393 | 0.27131501 | 0.49275402 |
| PHB | 381.950457 | -0.898323215 | 0.4522969 | -1.9861361 | 0.04701821 | 0.15912467 |
| SPOP | 982.116413 | 0.116206497 | 0.17396441 | 0.66799007 | 0.50413993 | 0.70797197 |
| SLC35B1 | 117.208297 | 0.240150599 | 0.28438403 | 0.8444588 | 0.39841308 | 0.62000923 |
| FAM117A | 180.669411 | 0.695613327 | 0.3140452 | 2.21501024 | 0.02675936 | 0.10859122 |
| KAT7 | 398.982996 | 0.079488318 | 0.16396644 | 0.48478406 | 0.62782953 | 0.7982736 |
| TAC4 | 79.620686 | -0.989289356 | 0.44331781 | -2.2315579 | 0.0256442 | 0.1053278 |
| ITGA3 | 2580.37267 | -2.445447128 | 0.49790709 | -4.9114527 | 9.04E-07 | 3.86E-05 |
| PDK2 | 189.735912 | 0.805102822 | 0.33663781 | 2.39159954 | 0.01677513 | 0.07924991 |
| PPP1R9B | 96.0225322 | -1.059952818 | 0.35587347 | -2.9784541 | 0.00289706 | 0.0225367 |
| COL1A1 | 55.8124936 | -2.174069727 | 0.53512015 | -4.0627694 | 4.85E-05 | 0.00095463 |
| XYLT2 | 171.846775 | 0.045038299 | 0.30008272 | 0.15008628 | 0.88069654 | 0.94639771 |
| MRPL27 | 265.883545 | -0.188813445 | 0.26893782 | -0.702071 | 0.48263488 | 0.69122023 |
| LRRC59 | 117.437694 | -0.92833345 | 0.32129892 | -2.889314 | 0.00386083 | 0.02795929 |
| ACSF2 | 251.073585 | 0.592302074 | 0.4269449 | 1.38730332 | 0.16534929 | 0.36456331 |
| RSAD1 | 164.188009 | 0.417594829 | 0.21353144 | 1.95565971 | 0.05050525 | 0.16736676 |
| SPATA20 | 120.601911 | 0.081837753 | 0.41126898 | 0.19898839 | 0.84227183 | 0.92705346 |
| ABCC3 | 916.859695 | -3.677305122 | 0.68262663 | -5.3869933 | 7.16E-08 | 4.98E-06 |
| ANKRD40 | 214.263905 | -0.329554058 | 0.22845911 | -1.4425078 | 0.14915916 | 0.34122677 |
| AC005921.2 | 59.546552 | -0.820841784 | 0.45451522 | -1.8059721 | 0.0709227 | 0.21009369 |
| LUC7L3 | 2008.88312 | 0.606914338 | 0.13768703 | 4.40792656 | 1.04E-05 | 0.00027626 |
| TOB1 | 672.592574 | 0.045112081 | 0.40346273 | 0.11181226 | 0.91097226 | 0.95976856 |
| SPAG9 | 2757.78914 | -0.025326219 | 0.2304032 | -0.1099213 | 0.91247179 | 0.96042853 |
| AC005920.2 | 261.099934 | -0.332702423 | 0.55068865 | -0.604157 | 0.54573924 | 0.74015676 |
| MBTD1 | 1547.469 | 0.056499568 | 0.18787304 | 0.30073271 | 0.76361833 | 0.88541384 |
| UTP18 | 293.516951 | -0.086063889 | 0.15569153 | -0.5527846 | 0.58041088 | 0.76635819 |
| CA10 | 3011.07034 | 0.160268563 | 0.32093066 | 0.49938689 | 0.61750686 | 0.7902963 |
| TOM1L1 | 2118.38615 | 0.436343846 | 0.18993793 | 2.29729711 | 0.02160183 | 0.09410702 |
| COX11 | 493.997354 | 0.527262104 | 0.2283292 | 2.30921886 | 0.02093144 | 0.09209973 |
| STXBP4 | 3266.48907 | 1.116146322 | 0.34937118 | 3.19472926 | 0.00139962 | 0.01301027 |
| HLF | 531.257251 | -0.047623162 | 0.54279877 | -0.0877363 | 0.93008626 | 0.96915731 |
| MMD | 106.88957 | 0.529828433 | 0.33718752 | 1.57131685 | 0.11610907 | 0.2898667 |
| PCTP | 131.103801 | -0.39628873 | 0.32560418 | -1.2170874 | 0.22357098 | 0.43821184 |
| C17orf67 | 77.2201668 | -0.62461145 | 0.39105503 | -1.5972469 | 0.11021067 | 0.27974112 |
| DGKE | 244.297442 | -0.111032177 | 0.21137465 | -0.5252862 | 0.59938428 | 0.77765121 |
| TRIM25 | 207.625439 | -0.014083649 | 0.44611093 | -0.0315698 | 0.9748151 | 0.98880477 |
| COIL | 189.472051 | 0.072636446 | 0.17376046 | 0.41802631 | 0.67592788 | 0.82991711 |
| SCPEP1 | 254.536502 | 0.458367525 | 0.23389803 | 1.95968952 | 0.05003209 | 0.16638315 |
| AC007114.1 | 50.5565363 | 0.513236624 | 0.41232064 | 1.24475122 | 0.21322321 | 0.4263496 |
| AKAP1 | 547.945128 | 0.569998399 | 0.24958946 | 2.28374382 | 0.02238659 | 0.09616406 |
| AC007114.2 | 83.1731226 | 0.469833081 | 0.48897504 | 0.96085289 | 0.33662614 | 0.56199377 |
| MSI2 | 14816.6134 | -0.208757096 | 0.3332462 | -0.626435 | 0.53102964 | 0.72863563 |
| AC015845.2 | 63.1349244 | 1.136381145 | 0.41559467 | 2.73434963 | 0.00625036 | 0.03942744 |
| MRPS23 | 195.253281 | -0.190615218 | 0.23713553 | -0.8038239 | 0.42149866 | 0.64046656 |
| CUEDC1 | 1177.87305 | 0.003991933 | 0.26753086 | 0.01492139 | 0.98809489 | 0.99552604 |
| VEZF1 | 607.569323 | -0.025591626 | 0.19401412 | -0.131906 | 0.89505865 | 0.95172874 |
| AC015813.2 | 372.112929 | 0.882132934 | 0.21709266 | 4.06339361 | 4.84E-05 | 0.00095463 |
| SRSF1 | 175.558285 | 0.443186364 | 0.25223394 | 1.75704493 | 0.07891015 | 0.22459043 |
| DYNLL2 | 264.417668 | -0.093268327 | 0.31046864 | -0.3004114 | 0.76386335 | 0.88554248 |
| MKS1 | 155.994799 | 0.308077947 | 0.21812243 | 1.41240835 | 0.15782976 | 0.35383118 |
| TSPOAP1 | 370.074352 | 0.482383351 | 0.32213407 | 1.49746146 | 0.13427323 | 0.31870891 |
| TSPOAP1-AS | 151.501733 | 0.761089985 | 0.43857945 | 1.73535259 | 0.08267837 | 0.23103419 |
| SUPT4H1 | 121.746906 | -0.421032697 | 0.2247386 | -1.873433 | 0.06100862 | 0.19001878 |
| RNF43 | 813.227476 | 1.066011334 | 0.29304996 | 3.63764365 | 0.00027514 | 0.00377794 |
| MTMR4 | 295.58628 | 0.567547263 | 0.33750633 | 1.68158999 | 0.09264837 | 0.24984879 |
| SEPT4-AS1 | 166.931166 | 0.620364202 | 0.46302883 | 1.33979606 | 0.18031166 | 0.38408913 |
| TEX14 | 89.4096636 | 0.81168611 | 0.39012005 | 2.08060598 | 0.03746999 | 0.13691315 |
| RAD51C | 219.350055 | 0.579612555 | 0.20820677 | 2.78383147 | 0.00537209 | 0.03537387 |
| PPM1E | 1769.82217 | 2.098563653 | 0.53581054 | 3.91661511 | 8.98E-05 | 0.00155259 |
| TRIM37 | 589.613862 | 0.370559261 | 0.26620303 | 1.39201744 | 0.16391712 | 0.36265636 |
| AC099850.1 | 46.6714291 | 0.408076709 | 0.41058571 | 0.99388922 | 0.3202768 | 0.54463101 |
| SKA2 | 306.468807 | 0.67479032 | 0.20200874 | 3.34040158 | 0.00083657 | 0.00870788 |
| GDPD1 | 158.207832 | 0.861995583 | 0.27515205 | 3.13279722 | 0.00173149 | 0.01525767 |
| YPEL2 | 1334.75144 | -1.227517431 | 0.24627706 | -4.9842947 | 6.22E-07 | 2.83E-05 |
| DHX40 | 577.810048 | -0.357322833 | 0.17876737 | -1.9988146 | 0.04562842 | 0.15600761 |
| CLTC | 1980.76933 | 0.025762355 | 0.17139697 | 0.15030811 | 0.88052154 | 0.94639771 |
| PTRH2 | 334.646066 | 0.01867558 | 0.26859195 | 0.06953142 | 0.94456662 | 0.9759376 |
| VMP1 | 5593.71542 | -1.204316175 | 0.381565 | -3.1562543 | 0.00159809 | 0.01442873 |
| TUBD1 | 384.75419 | -0.137287786 | 0.27805793 | -0.4937381 | 0.62149117 | 0.79347351 |
| RPS6KB1 | 579.952734 | 0.307046088 | 0.20924585 | 1.46739391 | 0.14226893 | 0.33124766 |
| RNFT1 | 144.711203 | -0.098453282 | 0.37294188 | -0.2639909 | 0.79178689 | 0.9019242 |
| RNFT1-DT | 62.5282277 | -0.208493328 | 0.37417761 | -0.5572042 | 0.57738794 | 0.7637681 |
| HEATR6 | 153.023745 | 0.438466212 | 0.21083187 | 2.07969606 | 0.03755342 | 0.13714214 |
| AC025048.4 | 50.9204398 | 0.314867569 | 0.56441871 | 0.55786167 | 0.57693884 | 0.76360573 |
| CA4 | 354.364208 | -0.648939554 | 0.83699557 | -0.7753202 | 0.43815049 | 0.65440037 |
| USP32 | 2139.49311 | -0.517250953 | 0.19774054 | -2.6158064 | 0.0089017 | 0.05081356 |
| APPBP2 | 1413.32967 | -0.104104876 | 0.14009656 | -0.7430937 | 0.45742494 | 0.67216888 |
| AC011921.1 | 191.692924 | 0.185948257 | 0.32990404 | 0.56364347 | 0.57299679 | 0.76098076 |
| PPM1D | 562.694933 | 0.285689668 | 0.29549344 | 0.96682236 | 0.33363285 | 0.5592275 |

| | | | | | | |
|---|---|---|---|---|---|---|
| BCAS3 | 5899.96147 | -0.106009732 | 0.17491511 | -0.6060639 | 0.54447233 | 0.73944091 |
| AC005884.1 | 44.5036693 | -0.725574459 | 0.46592655 | -1.5572722 | 0.11940587 | 0.2950961 |
| AC005746.2 | 70.8241401 | -0.260215612 | 0.49353946 | -0.5272438 | 0.59802431 | 0.77678206 |
| NACA2 | 71.0178653 | -1.368649545 | 0.56744917 | -2.4119333 | 0.01586819 | 0.07616383 |
| INTS2 | 317.223355 | 0.408055449 | 0.25215629 | 1.61826398 | 0.10560572 | 0.27261011 |
| MED13 | 3168.57485 | 0.033248491 | 0.22287132 | 0.14918245 | 0.88140967 | 0.94645136 |
| EFCAB3 | 62.8724757 | -0.817801438 | 0.47667512 | -1.7156369 | 0.08622852 | 0.23768624 |
| METTL2A | 159.209399 | -0.193923223 | 0.24622224 | -0.7875943 | 0.43093408 | 0.64898053 |
| TLK2 | 1648.63156 | 0.055407487 | 0.206306 | 0.26856944 | 0.78826103 | 0.89961465 |
| 10-Mar | 57.6675658 | -0.356021582 | 0.41742277 | -0.8529041 | 0.3937125 | 0.61588683 |
| TANC2 | 5970.80781 | -1.136075243 | 0.24404961 | -4.6550996 | 3.24E-06 | 0.0001051 |
| AC006270.1 | 58.6835451 | -1.204620282 | 0.57744316 | -2.0861279 | 0.03696704 | 0.13552529 |
| AC015923.1 | 468.298917 | -1.659453616 | 0.55458432 | -2.9922476 | 0.00276932 | 0.02183874 |
| CYB561 | 175.090239 | -0.338429137 | 0.32329027 | -1.0468275 | 0.29517916 | 0.51834904 |
| DCAF7 | 1000.39627 | -0.303864954 | 0.17255473 | -1.7609772 | 0.07824226 | 0.22350822 |
| TACO1 | 76.4802654 | 0.104162074 | 0.34148847 | 0.3050237 | 0.7603481 | 0.88360116 |
| MAP3K3 | 1567.82027 | -0.407994067 | 0.34431709 | -1.184937 | 0.23604234 | 0.45389346 |
| CCDC47 | 297.256671 | 0.128783655 | 0.22307044 | 0.57732283 | 0.56372139 | 0.75507353 |
| DDX42 | 786.647054 | 0.047574073 | 0.14200976 | 0.33500565 | 0.73762082 | 0.87164372 |
| FTSJ3 | 148.587354 | 0.062664808 | 0.24834336 | 0.25233132 | 0.80078498 | 0.90662963 |
| PSMC5 | 168.282572 | -0.259754693 | 0.36007841 | -0.7213837 | 0.47067347 | 0.68215118 |
| SMARCD2 | 206.580514 | -0.228727955 | 0.25993438 | -0.879945 | 0.37888912 | 0.6017088 |
| ERN1 | 572.843617 | -0.967734676 | 0.38622336 | -2.5056347 | 0.01222318 | 0.06374102 |
| SNHG25 | 226.832611 | 0.196890868 | 0.35150968 | 0.5601293 | 0.57539125 | 0.76255072 |
| TEX2 | 1016.80504 | -0.278639615 | 0.33329384 | -0.8360179 | 0.40314484 | 0.62443013 |
| PECAM1 | 53.3800779 | -0.297460687 | 0.31381063 | -0.9478987 | 0.34318103 | 0.56819785 |
| POLG2 | 277.233317 | 0.455882124 | 0.23939017 | 1.90434769 | 0.05686492 | 0.18138824 |
| DDX5 | 3673.84447 | -0.20075389 | 0.23561549 | -0.8520403 | 0.39419174 | 0.61649049 |
| CEP95 | 1273.8714 | 0.420932985 | 0.20439624 | 2.05939695 | 0.03945623 | 0.14144318 |
| SMURF2 | 1977.70051 | 0.296055907 | 0.11912601 | 2.48523307 | 0.01294666 | 0.06633533 |
| LRRC37A3 | 1665.14246 | 0.56347021 | 0.30347107 | 1.85675097 | 0.06334659 | 0.1951062 |
| AC103810.2 | 57.3524491 | -0.412157538 | 0.61576076 | -0.6693469 | 0.50327424 | 0.70736264 |
| GNA13 | 801.089783 | 0.173349159 | 0.32337747 | 0.53605824 | 0.5919183 | 0.77328146 |
| AC004805.1 | 62.7046121 | -0.095632092 | 0.47769435 | -0.2001952 | 0.84132796 | 0.92645072 |
| CEP112 | 1962.93759 | -0.321729973 | 0.19702521 | -1.6329381 | 0.10248202 | 0.26699901 |
| PRKCA | 2252.55417 | -0.707209669 | 0.42237876 | -1.6743495 | 0.09406193 | 0.25227697 |
| PRKCA-AS1 | 90.6018087 | -1.001303605 | 0.39758433 | -2.5184685 | 0.01178664 | 0.06230082 |
| HELZ | 2664.98926 | 0.094208532 | 0.15185856 | 0.62037025 | 0.53501406 | 0.73210674 |
| PSMD12 | 481.015487 | -0.019020966 | 0.1572932 | -0.1209268 | 0.903749 | 0.95548393 |
| PITPNC1 | 4123.10325 | -1.429000777 | 0.53893343 | -2.6515349 | 0.00801268 | 0.04737396 |
| NOL11 | 301.098148 | 0.433653357 | 0.22135067 | 1.95912379 | 0.05009829 | 0.16651946 |
| BPTF | 3480.30159 | -0.079314739 | 0.2073243 | -0.3825636 | 0.70204334 | 0.8488446 |
| KPNA2 | 67.1369487 | -0.450945165 | 0.4819451 | -0.9356775 | 0.34943928 | 0.57545736 |
| AC005332.7 | 123.814994 | 0.191192465 | 0.16895924 | 1.13158931 | 0.25780713 | 0.47882093 |
| AC005332.6 | 43.5846687 | -0.128374165 | 0.35834106 | -0.3582458 | 0.72015941 | 0.86060013 |
| AMZ2 | 97.6284968 | 0.362732862 | 0.26146384 | 1.38731561 | 0.16534554 | 0.36456331 |
| ARSG | 322.530521 | -0.021047526 | 0.48409094 | -0.0434785 | 0.96532014 | 0.98449934 |
| WIPI1 | 372.14792 | -0.371196306 | 0.34302988 | -1.0821107 | 0.27920333 | 0.501359 |
| PRKAR1A | 1124.91586 | -0.302753413 | 0.28149176 | -1.075532 | 0.28213659 | 0.50464019 |
| FAM20A | 239.228095 | -2.613439575 | 0.29804389 | -8.76864 | 1.81E-18 | 2.65E-15 |
| LINC01482 | 277.033214 | -2.22572686 | 0.71344128 | -3.1197057 | 0.00181032 | 0.01580546 |
| ABCA9 | 139.552531 | 0.604262608 | 0.51077265 | 1.18303634 | 0.23679475 | 0.45442711 |
| ABCA6 | 160.805637 | 0.635647886 | 0.3708113 | 1.71420849 | 0.08649044 | 0.23821654 |
| ABCA10 | 592.544845 | 0.482475039 | 0.30137866 | 1.60089319 | 0.10940058 | 0.27811244 |
| ABCA5 | 1748.09049 | 0.277374822 | 0.27182437 | 1.02041926 | 0.30752966 | 0.53112982 |
| MAP2K6 | 396.246071 | 2.305647162 | 0.66903961 | 3.4462043 | 0.00056852 | 0.00650744 |
| AC005208.1 | 2549.53565 | 0.398402657 | 0.37384752 | 1.06568223 | 0.28656731 | 0.50879744 |
| KCNJ16 | 8368.17125 | 0.532507179 | 0.2895145 | 1.83931091 | 0.06586947 | 0.20010806 |
| SOX9 | 277.590223 | -2.991894633 | 0.64395215 | -4.6461443 | 3.38E-06 | 0.00010907 |
| LINC00511 | 615.306919 | -4.76322417 | 0.69873168 | -6.8169575 | 9.30E-12 | 2.36E-09 |
| SLC39A11 | 1322.21137 | -0.245380357 | 0.34505391 | -0.7111363 | 0.47699978 | 0.68679857 |
| COG1 | 193.551787 | 0.000220336 | 0.22873413 | 0.00096328 | 0.99923141 | 0.99975417 |
| FAM104A | 258.67984 | 0.075006189 | 0.15359017 | 0.4883528 | 0.62529997 | 0.7961598 |
| C17orf80 | 164.615337 | 0.110555639 | 0.24170972 | 0.45739012 | 0.64739066 | 0.81148066 |
| CDC42EP4 | 130.619421 | -1.279804492 | 0.22716248 | -5.6338726 | 1.76E-08 | 1.51E-06 |
| RPL38 | 340.791877 | -0.852808881 | 0.34881114 | -2.4449015 | 0.01448917 | 0.07154332 |
| GPRC5C | 852.432902 | -1.526774432 | 0.34919226 | -4.3723032 | 1.23E-05 | 0.00031376 |
| SLC9A3R1 | 135.080974 | -1.13353717 | 0.40369607 | -2.8078975 | 0.00498661 | 0.03331646 |
| NAT9 | 157.0348 | -0.323738711 | 0.28370782 | -1.141099 | 0.25382873 | 0.4744338 |
| TMEM104 | 114.020926 | -0.40749562 | 0.34155829 | -1.1930485 | 0.23285037 | 0.44960101 |
| HID1 | 84.0719958 | -0.226533265 | 0.24648605 | -0.9190511 | 0.35806886 | 0.58348526 |
| CDR2L | 126.777742 | -0.55648351 | 0.36373762 | -1.5299036 | 0.12604059 | 0.30514481 |
| MRPL58 | 100.29249 | 0.042792782 | 0.23839863 | 0.17950096 | 0.85754436 | 0.93574122 |
| KCTD2 | 251.372822 | -0.091887111 | 0.25036048 | -0.3670192 | 0.71360467 | 0.85667582 |
| ATP5PD | 310.740367 | 0.226634884 | 0.33619485 | 0.67411766 | 0.50023652 | 0.70482771 |
| SLC16A5 | 1160.96731 | -1.60607853 | 0.47573857 | -3.3759687 | 0.00073556 | 0.00793795 |
| ARMC7 | 284.156602 | -0.840201189 | 0.47106226 | -1.7836309 | 0.07448366 | 0.21663404 |
| NT5C | 339.640567 | -0.659879362 | 0.28653926 | -2.3029283 | 0.02128288 | 0.0932423 |
| JPT1 | 234.731474 | -0.981953974 | 0.38425009 | -2.5555074 | 0.01060331 | 0.05789597 |
| SUMO2 | 237.341234 | 0.011315141 | 0.22123843 | 0.05114456 | 0.95921033 | 0.98229715 |
| NUP85 | 282.21864 | 0.042115468 | 0.21313938 | 0.1975959 | 0.84336124 | 0.92755692 |
| GGA3 | 348.132967 | 0.014253386 | 0.17157774 | 0.08307247 | 0.93379392 | 0.97167284 |
| MRPS7 | 196.650444 | -0.168383808 | 0.31169316 | -0.540223 | 0.58904326 | 0.77174702 |
| MIF4GD | 47.3381514 | -0.944134303 | 0.36397047 | -2.593986 | 0.00948703 | 0.05330253 |
| SLC25A19 | 57.5285569 | 1.193568657 | 0.29245712 | 4.08117493 | 4.48E-05 | 0.00090501 |
| GRB2 | 935.641624 | -0.349731306 | 0.18264651 | -1.9147988 | 0.05551817 | 0.17839503 |
| AC011933.4 | 156.247037 | -0.669729738 | 0.46192621 | -1.4498631 | 0.14709671 | 0.33791313 |
| TMEM94 | 311.986073 | 0.289431742 | 0.17114598 | 1.69113955 | 0.09081015 | 0.24620512 |
| CASKIN2 | 147.353403 | 0.257995952 | 0.35549555 | 0.72573609 | 0.46800057 | 0.6801407 |
| TSEN54 | 245.252313 | 0.00226802 | 0.26238074 | 0.008644 | 0.99310317 | 0.99737436 |
| LLGL2 | 3073.35177 | -0.358412847 | 0.36772112 | -0.9746866 | 0.32971574 | 0.55515809 |
| MYO15B | 464.743574 | 0.356847406 | 0.40460337 | 0.88196845 | 0.37779387 | 0.60068471 |
| RECQL5 | 262.165108 | 0.360445169 | 0.26927969 | 1.33855312 | 0.1807162 | 0.38451679 |
| SMIM5 | 87.7322003 | 0.476065962 | 0.51399516 | 0.92620709 | 0.35433837 | 0.58012193 |
| SAP30BP | 441.820252 | -0.179430054 | 0.22637584 | -0.7926201 | 0.42799917 | 0.64642414 |
| ITGB4 | 91.7337382 | -2.016491721 | 0.51983751 | -3.8790809 | 0.00010485 | 0.00173557 |
| GALK1 | 58.9321205 | -0.220736754 | 0.46674394 | -0.472929 | 0.63626382 | 0.80446022 |
| H3F3B | 1624.2767 | -1.301703381 | 0.32291092 | -4.0311532 | 5.55E-05 | 0.00106254 |
| UNK | 873.458519 | 0.406663398 | 0.24659383 | 1.64912236 | 0.09912257 | 0.26099533 |
| UNC13D | 52.2546328 | -0.191167214 | 0.40275104 | -0.4746536 | 0.63503393 | 0.80384889 |
| WBP2 | 506.111771 | -1.277394177 | 0.26912759 | -4.7464259 | 2.07E-06 | 7.23E-05 |
| TRIM47 | 137.450874 | -2.843351956 | 0.51373745 | -5.5346402 | 3.12E-08 | 2.44E-06 |
| TRIM65 | 95.323189 | -0.626232402 | 0.28334614 | -2.2101321 | 0.02709599 | 0.1094529 |
| FBF1 | 103.43038 | -0.324591211 | 0.25336099 | -1.2811412 | 0.20014406 | 0.40902261 |
| ACOX1 | 572.275044 | 0.054728065 | 0.17478893 | 0.31310944 | 0.75419752 | 0.88075281 |
| TEN1 | 106.287124 | -0.376212865 | 0.20998365 | -1.7916293 | 0.07319237 | 0.21436762 |
| CDK3 | 94.1775003 | 0.714273795 | 0.43546921 | 1.64023947 | 0.10095539 | 0.26453475 |
| EVPL | 247.614239 | 0.389748169 | 0.48639718 | 0.80129611 | 0.42296024 | 0.64187521 |
| SRP68 | 294.499673 | 0.033683046 | 0.16401933 | 0.20536022 | 0.8372907 | 0.9242234 |
| EXOC7 | 797.851106 | 0.05474037 | 0.25745122 | 0.21262424 | 0.83162006 | 0.92158196 |
| RNF157 | 111.67072 | -1.152733173 | 0.36380423 | -3.1685536 | 0.001532 | 0.01395595 |
| UBALD2 | 306.70734 | -1.095216267 | 0.2611907 | -4.1931672 | 2.75E-05 | 0.00061069 |
| QRICH2 | 158.075433 | 1.385529544 | 0.44196668 | 3.13491852 | 0.00171902 | 0.01517815 |

| | | | | | | |
|---|---|---|---|---|---|---|
| PRPSAP1 | 578.487271 | 0.848210183 | 0.29477108 | 2.87752176 | 0.00400812 | 0.02868001 |
| UBE2O | 677.090949 | 0.173682879 | 0.20699021 | 0.83908743 | 0.40142025 | 0.62292764 |
| RHBDF2 | 191.617936 | -1.082801876 | 0.31831176 | -3.4017024 | 0.00066968 | 0.00741388 |
| AC015802.4 | 75.0768559 | 0.847230974 | 0.42515563 | 1.9927549 | 0.04628829 | 0.15746395 |
| ST6GALNAC2 | 119.609982 | -0.190025667 | 0.54818433 | -0.3466456 | 0.72885761 | 0.86681233 |
| AC005837.1 | 35.2356279 | 1.656426266 | 0.67966118 | 2.4371353 | 0.01480414 | 0.07261339 |
| MXRA7 | 339.914248 | -0.02720704 | 0.43068194 | -0.063172 | 0.94962954 | 0.97821544 |
| JMJD6 | 214.037177 | -0.181934803 | 0.30580501 | -0.5949373 | 0.5518854 | 0.74481373 |
| METTL23 | 117.889537 | 0.070212457 | 0.3338599 | 0.21030515 | 0.83342951 | 0.92223929 |
| AC005837.2 | 139.779578 | 1.375792909 | 0.66806668 | 2.05936466 | 0.03945932 | 0.14144318 |
| SRSF2 | 959.260489 | 0.857667239 | 0.28491047 | 3.01030436 | 0.00260986 | 0.02090579 |
| MFSD11 | 470.964957 | 0.181839762 | 0.2012059 | 0.90374967 | 0.36612815 | 0.59142894 |
| SEC14L1 | 2651.09858 | -1.140829368 | 0.38130044 | -2.9919435 | 0.00277208 | 0.02184746 |
| 9-Sep | 2228.96195 | -1.195575833 | 0.37209813 | -3.2130659 | 0.00131326 | 0.01239062 |
| TNRC6C | 1117.56711 | 0.171068758 | 0.22656952 | 0.75503872 | 0.45022575 | 0.66633411 |
| TMC6 | 100.281404 | -2.030707988 | 0.41862079 | -4.8509487 | 1.23E-06 | 4.86E-05 |
| SYNGR2 | 240.666767 | -0.893388658 | 0.42132038 | -2.1204497 | 0.03396814 | 0.12854543 |
| AFMID | 736.610429 | 1.371821147 | 0.39065441 | 3.51159776 | 0.00044542 | 0.00541766 |
| SOCS3 | 246.992854 | -3.829674564 | 0.57549354 | -6.6545918 | 2.84E-11 | 6.15E-09 |
| AC061992.1 | 50.7971284 | -3.844631484 | 0.79755471 | -4.8205238 | 1.43E-06 | 5.47E-05 |
| PGS1 | 429.505799 | -0.66660035 | 0.24018074 | -2.7754113 | 0.00551319 | 0.03605137 |
| CYTH1 | 1038.19891 | 0.142314485 | 0.24089124 | 0.59078314 | 0.55466573 | 0.74738138 |
| USP36 | 528.541104 | -0.438274764 | 0.22784429 | -1.9235714 | 0.05440833 | 0.1757642 |
| TIMP2 | 315.472849 | -1.032485126 | 0.30868191 | -3.344819 | 0.00082336 | 0.00860562 |
| LGALS3BP | 405.13426 | -0.678416407 | 0.58837077 | -1.1530423 | 0.24889301 | 0.46818965 |
| CANT1 | 122.791812 | -0.384334876 | 0.26722098 | -1.4382661 | 0.15035857 | 0.34272413 |
| ENGASE | 164.073658 | 0.688235863 | 0.22316563 | 3.08396889 | 0.00204259 | 0.01742088 |
| CBX8 | 45.0911669 | 0.482537721 | 0.62205369 | 0.77571716 | 0.43791601 | 0.65419956 |
| CBX4 | 167.510739 | -0.497978387 | 0.47960502 | -1.0383094 | 0.29912605 | 0.5220872 |
| TBC1D16 | 1687.90423 | 0.134750187 | 0.39794012 | 0.33861926 | 0.73489657 | 0.86990311 |
| CCDC40 | 441.456562 | 0.249605351 | 0.33984991 | 0.73445761 | 0.46266989 | 0.67574154 |
| GAA | 244.910735 | 0.362175018 | 0.30902029 | 1.17201047 | 0.24119285 | 0.45956838 |
| EIF4A3 | 163.502718 | -0.298149926 | 0.26745998 | -1.1147459 | 0.26495932 | 0.48592037 |
| AC087741.1 | 157.234853 | 0.315262092 | 0.41395219 | 0.76159058 | 0.4463044 | 0.66216273 |
| SGSH | 96.6675841 | -0.479774322 | 0.28934826 | -1.6581206 | 0.09729312 | 0.25775067 |
| SLC26A11 | 309.329242 | -0.571529639 | 0.18381275 | -3.1093036 | 0.00187529 | 0.01628579 |
| RNF213 | 3809.50191 | -0.939118582 | 0.45947292 | -2.0439041 | 0.04096302 | 0.14475136 |
| AC124319.2 | 1980.11681 | 0.496398808 | 0.32405718 | 1.53182473 | 0.12556569 | 0.30438562 |
| ENDOV | 243.036802 | -0.123032561 | 0.27362659 | -0.4496367 | 0.65297241 | 0.81499693 |
| RPTOR | 1279.90006 | 0.023276624 | 0.23022215 | 0.10110506 | 0.91946706 | 0.96429063 |
| CHMP6 | 85.3649991 | -0.35293762 | 0.33027169 | -1.0686281 | 0.28523727 | 0.50777615 |
| BAIAP2-DT | 59.0451244 | -0.841637067 | 0.58562582 | -1.4371584 | 0.15067299 | 0.34308559 |
| BAIAP2 | 145.550525 | -0.355898244 | 0.25785034 | -1.3802512 | 0.16750931 | 0.36733337 |
| CEP131 | 71.9493221 | 0.25453493 | 0.24359207 | 1.04492287 | 0.29605862 | 0.51906534 |
| TEPSIN | 162.413305 | 0.047120363 | 0.32014816 | 0.14718299 | 0.88298759 | 0.94732911 |
| NDUFAF8 | 204.901567 | -0.455722637 | 0.37550613 | -1.2136224 | 0.22489196 | 0.43942281 |
| SLC38A10 | 596.631595 | -0.057747076 | 0.20383038 | -0.2833095 | 0.77693962 | 0.89337196 |
| AC110285.6 | 68.1940651 | 0.900336294 | 0.45356173 | 1.98503584 | 0.04714048 | 0.15929996 |
| AC110285.2 | 89.4538389 | 0.320446224 | 0.55984737 | 0.57238141 | 0.56706361 | 0.75721221 |
| BAHCC1 | 176.288498 | 0.424421416 | 0.37875336 | 1.12057466 | 0.26246895 | 0.4832663 |
| ACTG1 | 2192.28282 | -0.7579853 | 0.51191946 | -1.4806729 | 0.13869375 | 0.32505424 |
| AC139149.1 | 45.5030741 | -0.040344964 | 0.42363043 | -0.0952362 | 0.9241272 | 0.96654633 |
| FAAP100 | 102.533303 | -0.226961054 | 0.32040285 | -0.7083615 | 0.47872077 | 0.68837608 |
| NPLOC4 | 961.575432 | -0.25633832 | 0.19087643 | -1.3429543 | 0.17928677 | 0.38275997 |
| TSPAN10 | 49.6284123 | 1.042408703 | 0.36246866 | 2.87585888 | 0.0040293 | 0.02880032 |
| OXLD1 | 108.811718 | -0.753076618 | 0.27128592 | -2.7759517 | 0.00550404 | 0.03602718 |
| CCDC137 | 192.21501 | -0.423983557 | 0.26642298 | -1.5913926 | 0.11152126 | 0.282146 |
| ARL16 | 222.563262 | 0.322835755 | 0.26287874 | 1.22807862 | 0.21941745 | 0.43356369 |
| HGS | 425.69348 | -0.58497576 | 0.25878968 | -2.2604292 | 0.02379462 | 0.10024483 |
| AC139530.1 | 38.2950303 | -0.133098769 | 0.33771913 | -0.3941108 | 0.69349921 | 0.84313266 |
| SLC25A10 | 41.6814542 | -0.063650425 | 0.43720675 | -0.1455843 | 0.88424959 | 0.9477505 |
| GCGR | 404.823385 | 1.878177718 | 0.47172228 | 3.98153277 | 6.85E-05 | 0.00124067 |
| MCRIP1 | 233.677222 | -0.39576254 | 0.28628531 | -1.3824061 | 0.16684706 | 0.36640505 |
| P4HB | 835.441259 | -0.577469258 | 0.2686635 | -2.1494146 | 0.03160155 | 0.12230869 |
| ARHGDIA | 412.675783 | -1.387528066 | 0.41227144 | -3.3655692 | 0.00076386 | 0.00815668 |
| ALYREF | 45.2912018 | -0.270026841 | 0.34967487 | -0.7722226 | 0.43998259 | 0.65564599 |
| ANAPC11 | 282.646232 | -0.647969999 | 0.35587786 | -1.8207651 | 0.06864257 | 0.20553849 |
| PCYT2 | 93.1716959 | 0.389797983 | 0.51787777 | 0.75268337 | 0.45164021 | 0.66738064 |
| SIRT7 | 210.60637 | -0.640456528 | 0.25592816 | -2.5024856 | 0.01233246 | 0.06415893 |
| MAFG | 132.013281 | -1.468771565 | 0.35088285 | -4.1859315 | 2.84E-05 | 0.00062731 |
| ASPSCR1 | 316.750447 | 0.010576459 | 0.33589846 | 0.03148707 | 0.9748811 | 0.98880477 |
| CENPX | 102.829378 | -0.395044874 | 0.34888865 | -1.132295 | 0.25751044 | 0.47845862 |
| LRRC45 | 38.9797384 | 0.621443506 | 0.45553919 | 1.36419329 | 0.17250674 | 0.3737929 |
| DCXR | 170.485798 | 0.104974293 | 0.31235621 | 0.33607237 | 0.73681629 | 0.87092675 |
| RFNG | 171.723807 | 0.359248271 | 0.23792484 | 1.50992333 | 0.13106299 | 0.31362519 |
| GPS1 | 184.086108 | -0.194663094 | 0.38404239 | -0.5068792 | 0.61223959 | 0.78684784 |
| DUS1L | 199.487173 | -0.131762012 | 0.29742014 | -0.4430164 | 0.65775385 | 0.81863236 |
| FASN | 92.0084915 | -1.23581219 | 0.45035839 | -2.7440639 | 0.00606837 | 0.03868587 |
| CCDC57 | 1236.74661 | -0.094041427 | 0.20282387 | -0.4636606 | 0.64289096 | 0.80857472 |
| SLC16A3 | 268.604944 | -1.431440865 | 0.39102149 | -3.6607729 | 0.00025146 | 0.00351851 |
| CSNK1D | 695.886206 | -0.778747098 | 0.22857617 | -3.4069478 | 0.00065694 | 0.00729474 |
| AC132872.1 | 63.5985592 | 0.230530086 | 0.3228968 | 0.71394355 | 0.47526209 | 0.68532684 |
| SECTM1 | 239.45143 | -3.275097974 | 0.88048046 | -3.7196714 | 0.00019948 | 0.00292448 |
| OGFOD3 | 230.029362 | -0.099264166 | 0.2961741 | -0.3351548 | 0.73750833 | 0.87158876 |
| HEXDC | 495.38968 | 0.270618742 | 0.25305989 | 1.06938616 | 0.2848957 | 0.50741909 |
| CYBC1 | 231.472489 | -1.01049456 | 0.28881374 | -3.4987759 | 0.0004674 | 0.00562796 |
| NARF | 439.082986 | -0.531231669 | 0.18460676 | -2.8776393 | 0.00400663 | 0.02868001 |
| FOXK2 | 1316.76274 | 0.174979833 | 0.22307645 | 0.784394 | 0.43280896 | 0.65020381 |
| WDR45B | 823.982964 | -0.27024844 | 0.36573795 | -0.7389128 | 0.45995997 | 0.67354889 |
| RAB40B | 617.812742 | -0.222477282 | 0.38536432 | -0.5773168 | 0.5637255 | 0.75507353 |
| FN3KRP | 152.726633 | -0.146081387 | 0.22620841 | -0.6457823 | 0.51842035 | 0.71930578 |
| FN3K | 386.39203 | 0.800103121 | 0.59403314 | 1.34689981 | 0.1780125 | 0.38109678 |
| TBCD | 571.657016 | 0.084470833 | 0.27200633 | 0.3105473 | 0.75614479 | 0.88219397 |
| B3GNTL1 | 79.9889928 | -0.002574864 | 0.43892331 | -0.0058663 | 0.99531938 | 0.998191 |
| METRNL | 167.196182 | -0.018232427 | 0.25598013 | -0.0712259 | 0.94321793 | 0.97556917 |
| USP14 | 873.608623 | -0.040583508 | 0.1511022 | -0.2685832 | 0.78825047 | 0.89961465 |
| THOC1 | 852.018622 | 0.14044917 | 0.21183812 | 0.66300236 | 0.50732904 | 0.71048657 |
| CLUL1 | 43.5046551 | 1.124307734 | 0.40031239 | 2.80857593 | 0.00497611 | 0.03328076 |
| TYMS | 56.1483394 | -0.453134679 | 0.52163726 | -0.8686777 | 0.38502342 | 0.60812798 |
| ENOSF1 | 2871.62286 | -0.150052579 | 0.40701521 | -0.3686658 | 0.71237686 | 0.85574627 |
| YES1 | 1059.85039 | -0.217367957 | 0.21441713 | -1.0137621 | 0.31069629 | 0.53450314 |
| LINC00470 | 107.912308 | 0.557357576 | 0.97882564 | 0.56941456 | 0.56907484 | 0.7587473 |
| AP005230.1 | 50.3183338 | -0.103920243 | 0.56873812 | -0.1827207 | 0.85501715 | 0.93399947 |
| METTL4 | 342.470134 | -0.082815395 | 0.25262209 | -0.3278233 | 0.74304531 | 0.87445523 |
| SMCHD1 | 2202.02491 | 0.097247552 | 0.20184546 | 0.48179213 | 0.62995362 | 0.79979406 |
| AP001011.1 | 376.31982 | -0.240461017 | 0.54526119 | -0.4410015 | 0.65921189 | 0.81968652 |
| LPIN2 | 927.651312 | -0.039571826 | 0.38082502 | -0.1039108 | 0.91724015 | 0.96291779 |
| AP000919.2 | 87.2054443 | -0.284544012 | 0.60751421 | -0.4683743 | 0.63951698 | 0.80604769 |
| MYOM1 | 193.414983 | 1.419591226 | 0.47022492 | 3.01896213 | 0.00253642 | 0.02044149 |
| AP005329.2 | 47.218921 | 0.873135164 | 0.61312821 | 1.4240662 | 0.15442731 | 0.34881127 |
| MYL12A | 646.134641 | -1.231486539 | 0.37273264 | -3.3039407 | 0.00095336 | 0.00960559 |
| MYL12B | 1310.08155 | -1.402049483 | 0.37504451 | -3.7383549 | 0.00018523 | 0.00275527 |
| TGIF1 | 357.369665 | -1.138995904 | 0.34984966 | -3.2556725 | 0.00113124 | 0.01101147 |

| | | | | | | |
|---|---|---|---|---|---|---|
| DLGAP1 | 348.87187 | -1.558154511 | 0.55685916 | -2.7981124 | 0.00514022 | 0.03415352 |
| DLGAP1-AS1 | 132.124283 | -0.770138714 | 0.31673897 | -2.4314618 | 0.01503803 | 0.07346057 |
| DLGAP1-AS2 | 60.0017235 | -1.017915403 | 0.41946834 | -2.4266799 | 0.01523769 | 0.07418897 |
| LINC00667 | 210.726671 | 0.558693692 | 0.28756893 | 1.9428166 | 0.05203832 | 0.1707762 |
| ZBTB14 | 42.2953374 | -0.151073926 | 0.39376377 | -0.3836664 | 0.70122572 | 0.84867973 |
| EPB41L3 | 1102.21706 | 0.734919828 | 0.45914818 | 1.60061581 | 0.10946204 | 0.27821514 |
| L3MBTL4 | 578.832343 | -0.096971191 | 0.43638737 | -0.2222136 | 0.82414764 | 0.91858629 |
| LAMA1 | 2507.34339 | 1.320412822 | 0.41467174 | 3.18423635 | 0.00145136 | 0.01336895 |
| PTPRM | 8444.64438 | -0.212768581 | 0.20617224 | -1.0319943 | 0.30207477 | 0.52529036 |
| RAB12 | 727.228163 | 0.107715885 | 0.17814126 | 0.60466557 | 0.54540123 | 0.74010734 |
| MTCL1 | 370.977776 | 1.085446282 | 0.51706198 | 2.09925759 | 0.0357942 | 0.13281056 |
| NDUFV2 | 786.926981 | 0.159052276 | 0.19784722 | 0.80391465 | 0.42144627 | 0.64046063 |
| AP005899.1 | 62.0059729 | -0.048665963 | 0.38025676 | -0.1279818 | 0.89816334 | 0.9528423 |
| NDUFV2-AS1 | 124.337677 | -0.319324834 | 0.32917223 | -0.9700844 | 0.33200444 | 0.55723592 |
| ANKRD12 | 4075.63798 | -0.568814197 | 0.18882828 | -3.0123358 | 0.00259246 | 0.02081687 |
| TWSG1 | 372.225992 | -0.220015275 | 0.17928363 | -1.2271911 | 0.21975076 | 0.43361796 |
| RALBP1 | 674.466177 | -0.531786382 | 0.14482576 | -3.6719047 | 0.00024075 | 0.00341208 |
| PPP4R1 | 1089.52513 | -0.652696182 | 0.24498623 | -2.6642158 | 0.0077168 | 0.04601863 |
| RAB31 | 330.788208 | -1.703002565 | 0.51416111 | -3.3121963 | 0.00092567 | 0.00939836 |
| VAPA | 2236.22851 | 0.67417986 | 0.31027072 | 2.17287619 | 0.02978964 | 0.1169365 |
| NAPG | 774.800092 | -0.465173382 | 0.26599203 | -1.7488245 | 0.08032136 | 0.22694435 |
| GNAL | 230.020253 | -1.530141414 | 0.27600952 | -5.5437993 | 2.96E-08 | 2.33E-06 |
| MPPE1 | 677.895007 | 0.519303299 | 0.25617468 | 2.02714533 | 0.04264755 | 0.14867551 |
| IMPA2 | 1606.21106 | 0.455723617 | 0.36704088 | 1.24161543 | 0.21437849 | 0.42792245 |
| TUBB6 | 218.257806 | -1.901784983 | 0.45083455 | -4.2183657 | 2.46E-05 | 0.00056042 |
| AFG3L2 | 1083.83187 | 0.34102986 | 0.15228062 | 2.23948299 | 0.02512451 | 0.10406698 |
| SPIRE1 | 2734.12494 | -0.058364375 | 0.30049434 | -0.1942279 | 0.84599746 | 0.9287667 |
| PSMG2 | 617.950177 | 0.034563004 | 0.26312183 | 0.13135742 | 0.89549257 | 0.95183831 |
| CEP76 | 185.250386 | -0.012214569 | 0.27297536 | -0.044746 | 0.96430973 | 0.98427998 |
| PTPN2 | 1515.10797 | -0.627379787 | 0.2583581 | -2.4283341 | 0.01516836 | 0.07396047 |
| SEH1L | 298.480267 | -0.206787223 | 0.22618891 | -0.9142235 | 0.36059941 | 0.58558614 |
| CEP192 | 1623.92784 | 0.460679669 | 0.24526145 | 1.87832076 | 0.0603373 | 0.18868262 |
| LDLRAD4 | 3791.23316 | -0.132710214 | 0.38802187 | -0.3420174 | 0.73233783 | 0.86850875 |
| AP005131.1 | 67.1211858 | -0.251980466 | 0.86842619 | -0.2901576 | 0.77169567 | 0.88993091 |
| FAM210A | 504.274307 | 0.03274509 | 0.36803922 | 0.08897174 | 0.92910437 | 0.96900671 |
| RNMT | 911.635913 | -0.080816275 | 0.31643464 | -0.2553964 | 0.79841693 | 0.90515239 |
| ZNF519 | 88.8476307 | 0.510440047 | 0.50227273 | 1.01626072 | 0.30950525 | 0.53343618 |
| ANKRD30B | 59.3315475 | 0.01894723 | 0.56827063 | 0.03334191 | 0.97340193 | 0.98804941 |
| ROCK1 | 2030.76975 | -0.233620944 | 0.12942279 | -1.805099 | 0.07105919 | 0.21031165 |
| GREB1L | 137.604427 | 0.114854354 | 0.38348403 | 0.29950231 | 0.76455681 | 0.88572451 |
| ESCO1 | 939.524926 | -0.077536603 | 0.18663193 | -0.415452 | 0.67781107 | 0.83115544 |
| SNRPD1 | 203.450376 | -0.051051055 | 0.25809403 | -0.1978002 | 0.84320138 | 0.92745832 |
| ABHD3 | 701.644977 | -0.631701596 | 0.21961977 | -2.8763421 | 0.00402313 | 0.02877183 |
| MIB1 | 2090.21813 | 0.233666138 | 0.21524286 | 1.08559297 | 0.27765911 | 0.4998762 |
| AC090912.1 | 226.206426 | -1.002083572 | 0.38456968 | -2.6057269 | 0.00916795 | 0.05206631 |
| RBBP8 | 3213.18196 | -0.508481259 | 0.27018674 | -1.8819623 | 0.05984113 | 0.18757511 |
| CABLES1 | 798.055701 | -1.815767813 | 0.40285648 | -4.5072325 | 6.57E-06 | 0.00018901 |
| TMEM241 | 430.728831 | -0.949012915 | 0.22751587 | -4.1711944 | 3.03E-05 | 0.00065829 |
| RIOK3 | 1145.51647 | -0.577321358 | 0.25085834 | -2.3013839 | 0.02136994 | 0.09343778 |
| RMC1 | 324.422455 | -0.421994091 | 0.22039058 | -1.9147555 | 0.05552369 | 0.17839503 |
| NPC1 | 1243.94798 | -1.161682945 | 0.34892309 | -3.3293381 | 0.00087053 | 0.00896941 |
| LAMA3 | 142.704375 | -1.763521471 | 0.71738065 | -2.4582786 | 0.01396048 | 0.06965579 |
| TTC39C | 549.815484 | -0.584479499 | 0.33959279 | -1.7211187 | 0.08522928 | 0.235788 |
| OSBPL1A | 810.583792 | 0.744463825 | 0.29154393 | 2.55352196 | 0.01066396 | 0.05802348 |
| IMPACT | 172.671391 | 0.270900831 | 0.22461869 | 1.20604761 | 0.22779913 | 0.44334483 |
| SS18 | 1693.82943 | 0.235435844 | 0.23118128 | 1.01840357 | 0.30848621 | 0.53230494 |
| TAF4B | 347.584518 | 0.596148787 | 0.12071772 | 4.93837034 | 7.88E-07 | 3.45E-05 |
| KCTD1 | 4740.78716 | 0.421564916 | 0.24371086 | 1.72977482 | 0.08367052 | 0.23299214 |
| AQP4-AS1 | 540.483365 | 0.300425498 | 0.29697 | 1.01163583 | 0.31171221 | 0.53528426 |
| AQP4 | 66.2334911 | -0.767296832 | 0.90940704 | -0.8437331 | 0.39881856 | 0.62056713 |
| CHST9 | 887.165174 | -0.168556071 | 0.43143167 | -0.3906901 | 0.69602634 | 0.84465287 |
| CDH2 | 61.3545031 | 0.434234832 | 0.49709855 | 0.87353872 | 0.38236951 | 0.60516641 |
| DSC2 | 180.515234 | -0.557151445 | 0.31500719 | -1.7686944 | 0.07694489 | 0.2212981 |
| DSG2 | 530.131344 | 0.224885574 | 0.37744464 | 0.59581076 | 0.55130166 | 0.7446716 |
| B4GALT6 | 565.982561 | 0.868609286 | 0.40064485 | 2.16802809 | 0.03015655 | 0.11802603 |
| TRAPPC8 | 1464.38105 | 0.206904451 | 0.21524057 | 0.9612707 | 0.33641607 | 0.5618561 |
| RNF138 | 529.151428 | 0.280075897 | 0.19154793 | 1.46217134 | 0.14369426 | 0.333319 |
| GAREM1 | 4032.47848 | -0.612953346 | 0.23830942 | -2.5720902 | 0.01010865 | 0.05591508 |
| AC025887.2 | 151.176414 | 0.877715973 | 0.41992812 | 2.09015765 | 0.03660364 | 0.13464146 |
| KLHL14 | 628.618421 | 0.846281902 | 0.48457868 | 1.74642826 | 0.08073656 | 0.22787378 |
| CCDC178 | 2880.5269 | 0.8442726 | 0.73105312 | 1.15487176 | 0.24814295 | 0.46734754 |
| ASXL3 | 1222.63467 | 0.802667483 | 0.27125705 | 2.95906591 | 0.00308573 | 0.02362865 |
| DTNA | 444.125084 | -0.001110231 | 0.61059611 | -0.0018183 | 0.99854923 | 0.99953295 |
| MAPRE2 | 378.74909 | 0.273798412 | 0.32316439 | 0.8472419 | 0.3968603 | 0.61861314 |
| ZNF397 | 555.50555 | 0.058709048 | 0.18106997 | 0.32423404 | 0.74576085 | 0.87570052 |
| ZSCAN30 | 432.884829 | -0.045017977 | 0.14333455 | -0.3140762 | 0.75346314 | 0.88036219 |
| AC011815.2 | 47.9219054 | 0.188706734 | 0.52175247 | 0.36167866 | 0.71759217 | 0.85947463 |
| ZNF24 | 455.127954 | -0.063446517 | 0.17863352 | -0.355177 | 0.722457 | 0.86197584 |
| ZNF396 | 69.8001524 | -0.306116865 | 0.25829496 | -1.1851445 | 0.2359603 | 0.45389346 |
| GALNT1 | 632.885245 | 0.013993185 | 0.19966368 | 0.07008378 | 0.94412698 | 0.9758195 |
| C18orf21 | 58.5855885 | 0.025861644 | 0.32704319 | 0.07907715 | 0.93697126 | 0.97284815 |
| RPRD1A | 1386.62481 | 0.023673183 | 0.16771334 | 0.14115265 | 0.88774935 | 0.94928354 |
| SLC39A6 | 212.63357 | -0.841852786 | 0.22152463 | -3.8002673 | 0.00014454 | 0.00224615 |
| ELP2 | 839.936817 | 0.091288234 | 0.12956492 | 0.70457523 | 0.48107462 | 0.68948936 |
| MOCOS | 162.910263 | -2.792483329 | 0.52439178 | -5.3251852 | 1.01E-07 | 6.44E-06 |
| FHOD3 | 4349.48612 | 0.822865004 | 0.32160725 | 2.55860217 | 0.01050939 | 0.05752945 |
| TPGS2 | 500.815014 | -0.699062524 | 0.17873539 | -3.911159 | 9.19E-05 | 0.00157367 |
| KIAA1328 | 2474.04741 | 0.327911601 | 0.24978476 | 1.31277663 | 0.18925823 | 0.39536801 |
| AC016493.1 | 95.1980603 | 0.370531269 | 0.69696721 | 0.53163372 | 0.59497971 | 0.77501911 |
| CELF4 | 60.7230313 | 0.011521844 | 0.54220082 | 0.02125014 | 0.98304612 | 0.9928728 |
| AC015961.2 | 44.8001385 | 0.984977593 | 0.321913 | 3.05976329 | 0.00221512 | 0.01856273 |
| AC016205.1 | 195.581374 | -0.132071619 | 0.44663108 | -0.2957063 | 0.7674544 | 0.88756968 |
| AC021504.1 | 94.7384771 | 0.98909469 | 0.85279395 | 1.15982845 | 0.24611866 | 0.46541739 |
| PIK3C3 | 1847.52618 | 0.114026405 | 0.15835204 | 0.72008168 | 0.47147471 | 0.68263832 |
| LINC00907 | 376.738598 | -0.274602777 | 0.52651913 | -0.5215438 | 0.60198801 | 0.77940904 |
| SETBP1 | 7432.25649 | 0.11699578 | 0.21681521 | 0.53961058 | 0.58946562 | 0.77174702 |
| SLC14A2 | 187.821958 | 0.416142756 | 0.4892511 | 0.85057092 | 0.39500775 | 0.61718208 |
| EPG5 | 935.44854 | -0.045729321 | 0.12693907 | -0.3602462 | 0.718663 | 0.85990372 |
| PSTPIP2 | 311.437514 | -0.637689356 | 0.39667068 | -1.607604 | 0.10792194 | 0.27605362 |
| ATP5F1A | 1024.40522 | 0.372797164 | 0.37204702 | 1.00201626 | 0.31633574 | 0.54070885 |
| HAUS1 | 151.031458 | -0.280222999 | 0.363931 | -0.7699894 | 0.4413062 | 0.65696085 |
| C18orf25 | 691.809516 | 0.359657339 | 0.19660371 | 1.82935174 | 0.06734693 | 0.20284564 |
| RNF165 | 246.699653 | 1.492683225 | 0.34459391 | 4.33171684 | 1.48E-05 | 0.00036873 |
| ST8SIA5 | 396.533033 | -0.661535296 | 0.38546336 | -1.716208 | 0.08612398 | 0.23749723 |
| PIAS2 | 2759.30838 | 0.255234465 | 0.28187759 | 0.90547982 | 0.36521125 | 0.590679 |
| KATNAL2 | 1639.98011 | 1.115682702 | 0.3196554 | 3.49026699 | 0.00048254 | 0.00575777 |
| AC012254.1 | 129.253662 | 0.010548453 | 0.57272935 | 0.01841787 | 0.9853055 | 0.99423861 |
| HDHD2 | 72.2557816 | 0.091437603 | 0.36993468 | 0.24717229 | 0.80477489 | 0.90857022 |
| SMAD2 | 2261.51951 | -0.26370047 | 0.17686752 | -1.4909491 | 0.13597487 | 0.321247 |
| ZBTB7C | 152.466615 | -0.029274289 | 0.81545565 | -0.0358993 | 0.97136265 | 0.98735893 |
| CTIF | 1261.99798 | 0.625993148 | 0.24127677 | 2.59450236 | 0.0094728 | 0.05329053 |
| SMAD7 | 307.579493 | 0.525086213 | 0.41276282 | 1.27212576 | 0.20332844 | 0.41300406 |
| DYM | 3444.43045 | 0.201627095 | 0.12467968 | 1.61716086 | 0.10584356 | 0.27301066 |

| | | | | | | |
|---|---|---|---|---|---|---|
| C18orf32 | 49.0552897 | 0.398960138 | 0.2456924 | 1.62381962 | 0.10441431 | 0.27047815 |
| LIPG | 472.787542 | -2.501345772 | 0.59543853 | -4.2008463 | 2.66E-05 | 0.00059732 |
| ACAA2 | 423.624469 | -0.409049689 | 0.37110355 | -1.1022521 | 0.27035207 | 0.49185779 |
| SNHG22 | 362.981024 | -0.439651366 | 0.27858181 | -1.5781768 | 0.11452499 | 0.28749769 |
| MYO5B | 1249.73833 | -0.691410004 | 0.25556586 | -2.7054083 | 0.00682205 | 0.04208894 |
| CFAP53 | 118.989454 | 0.323686596 | 0.29428234 | 1.0999185 | 0.27136763 | 0.49278183 |
| MBD1 | 281.434729 | -0.085635024 | 0.15608646 | -0.5486384 | 0.58325363 | 0.76857514 |
| CXXC1 | 178.558595 | 0.281180313 | 0.25432182 | 1.10560829 | 0.2688961 | 0.49041465 |
| MAPK4 | 337.62477 | 0.998162214 | 0.71205771 | 1.40179961 | 0.1609751 | 0.35824533 |
| MRO | 41.9240117 | 1.989407922 | 0.98644419 | 2.01674655 | 0.04372197 | 0.15150144 |
| ME2 | 487.767294 | -0.087315671 | 0.20557189 | -0.4247452 | 0.67102244 | 0.82713096 |
| AC091551.1 | 230.202602 | -0.17950415 | 0.36601151 | -0.4904331 | 0.62382748 | 0.79539814 |
| SMAD4 | 670.240165 | 0.094819359 | 0.24316938 | 0.38993132 | 0.69658733 | 0.84496018 |
| MEX3C | 157.551352 | -0.483794856 | 0.16952882 | -2.8537617 | 0.00432049 | 0.03016353 |
| DCC | 78.6961588 | 0.56803493 | 0.36054958 | 1.57546966 | 0.11514808 | 0.28850359 |
| AC103949.1 | 92.950734 | 0.789935645 | 0.71649791 | 1.10249539 | 0.27024636 | 0.49175977 |
| MBD2 | 1525.98783 | -0.835474299 | 0.17546381 | -4.7615192 | 1.92E-06 | 6.84E-05 |
| POLI | 572.276781 | 0.175014551 | 0.19838377 | 0.88220195 | 0.37766761 | 0.60060335 |
| TCF4 | 366.884848 | 0.916235389 | 0.59185591 | 1.54807171 | 0.12160503 | 0.29836662 |
| AC009271.1 | 65.6734038 | -0.180835268 | 0.7143358 | -0.2531516 | 0.80015105 | 0.90644899 |
| TXNL1 | 562.045762 | 0.22474278 | 0.16031032 | 1.40192334 | 0.16093814 | 0.35824533 |
| WDR7 | 1895.32349 | 0.357943845 | 0.15471232 | 2.31360911 | 0.02068917 | 0.09130748 |
| FECH | 540.065372 | 1.564470158 | 0.32744803 | 4.77776624 | 1.77E-06 | 6.38E-05 |
| NARS | 698.697779 | -0.378999013 | 0.24035198 | -1.57685 | 0.11483004 | 0.28809265 |
| AC027097.2 | 527.444805 | 0.165320126 | 0.31906762 | 0.51813508 | 0.60436402 | 0.78119624 |
| ATP8B1 | 1490.30356 | -0.586197331 | 0.23707132 | -2.4726624 | 0.01341108 | 0.06766498 |
| NEDD4L | 7701.71628 | 1.08508028 | 0.24546813 | 4.4204528 | 9.85E-06 | 0.00026336 |
| MALT1 | 694.708053 | -0.158269157 | 0.19639469 | -0.8058729 | 0.42031614 | 0.63969995 |
| AC104365.1 | 120.60686 | 0.007166258 | 0.6190451 | 0.01157631 | 0.99076365 | 0.99624112 |
| ZNF532 | 2013.54117 | -0.324731262 | 0.32236552 | -1.0073387 | 0.31377204 | 0.53763326 |
| SEC11C | 203.442167 | -0.35562594 | 0.32807074 | -1.0839916 | 0.2783685 | 0.50055858 |
| LMAN1 | 702.181475 | 0.193346432 | 0.21112937 | 0.91577233 | 0.35978632 | 0.58502113 |
| CDH20 | 99.3094568 | -0.574743165 | 0.64400494 | -0.8924515 | 0.37215099 | 0.59630231 |
| AC105094.2 | 116.480708 | 0.66060263 | 0.59380179 | 1.11249687 | 0.26592458 | 0.48691402 |
| RNF152 | 2058.78316 | -0.082715188 | 0.58038754 | -0.1425172 | 0.88667152 | 0.94903526 |
| PIGN | 1442.66438 | 0.284050521 | 0.12218094 | 2.32483488 | 0.02008079 | 0.08927875 |
| KIAA1468 | 1517.76908 | -0.079234339 | 0.14871061 | -0.5328089 | 0.59416586 | 0.77437956 |
| ZCCHC2 | 750.522493 | 0.300229329 | 0.18446036 | 1.62760892 | 0.10360784 | 0.2692845 |
| PHLPP1 | 1501.69716 | -0.11835284 | 0.18027383 | -0.656517 | 0.51149153 | 0.71366764 |
| BCL2 | 3254.3508 | 0.568880815 | 0.25742051 | 2.20992808 | 0.02711015 | 0.10947662 |
| KDSR | 616.01125 | -1.010064608 | 0.19726993 | -5.1202157 | 3.05E-07 | 1.54E-05 |
| AC036176.1 | 57.4139762 | 0.861048555 | 0.34652979 | 2.48477499 | 0.01296333 | 0.06636926 |
| VPS4B | 675.88967 | -0.30710958 | 0.1338906 | -2.2937352 | 0.02180571 | 0.09452959 |
| SERPINB8 | 82.1081443 | -2.977863795 | 0.52489601 | -5.6732452 | 1.40E-08 | 1.23E-06 |
| TMX3 | 726.240092 | 0.379400171 | 0.26307923 | 1.44215172 | 0.14925958 | 0.34127917 |
| CCDC102B | 277.643011 | 1.404115375 | 0.49555937 | 2.83339487 | 0.00460565 | 0.03155393 |
| DOK6 | 85.2766772 | 0.0191365 | 0.29897164 | 0.06400774 | 0.94896406 | 0.97784954 |
| CD226 | 183.540107 | 1.755554361 | 0.70880295 | 2.47678761 | 0.01325708 | 0.06711401 |
| RTTN | 914.510869 | 0.989236428 | 0.34958731 | 2.82972635 | 0.00465878 | 0.03175016 |
| SOCS6 | 377.195414 | 0.394760096 | 0.24831053 | 1.58978398 | 0.11188351 | 0.28273756 |
| FBXO15 | 133.88376 | 0.830393291 | 0.39791807 | 2.08684487 | 0.03690216 | 0.13533942 |
| TIMM21 | 149.484083 | 0.757962013 | 0.21015613 | 3.60666142 | 0.00031016 | 0.00411338 |
| CYB5A | 1329.56114 | 0.54651895 | 0.33465421 | 1.63308554 | 0.10245101 | 0.26697087 |
| CNDP2 | 1543.54137 | -1.194132296 | 0.30503453 | -3.9147446 | 9.05E-05 | 0.00155856 |
| LINC00909 | 159.522313 | 0.230186795 | 0.29400808 | 0.78292678 | 0.43367011 | 0.65072685 |
| ZNF407 | 2698.7301 | 0.028014487 | 0.19747929 | 0.14186038 | 0.88719029 | 0.94920594 |
| ZADH2 | 389.296599 | 0.080635966 | 0.21671681 | 0.37207989 | 0.70983337 | 0.85441779 |
| TSHZ1 | 879.363628 | -0.233384413 | 0.25215644 | -0.925554 | 0.35467779 | 0.58039381 |
| ZNF516 | 997.771935 | 0.291354511 | 0.2587353 | 1.12607173 | 0.26013516 | 0.48104792 |
| ZNF236-DT | 99.3266298 | 0.494358257 | 0.26399628 | 1.87259554 | 0.06112426 | 0.1902558 |
| ZNF236 | 659.930437 | 0.35775296 | 0.26969141 | 1.32652711 | 0.18466516 | 0.3900915 |
| MBP | 1338.98066 | -0.659739557 | 0.30686813 | -2.1499122 | 0.03156216 | 0.12228868 |
| ATP9B | 2284.95186 | -0.038640027 | 0.2312015 | -0.1671271 | 0.86727007 | 0.93961371 |
| NFATC1 | 191.507799 | -0.247123969 | 0.38272201 | -0.645701 | 0.51847304 | 0.71930578 |
| AC068473.1 | 42.832726 | 1.195744819 | 0.49898097 | 2.39637361 | 0.0165582 | 0.07847768 |
| CTDP1 | 236.11813 | -0.536919621 | 0.2637531 | -2.0356903 | 0.04178147 | 0.14662364 |
| PQLC1 | 306.979662 | 0.00948668 | 0.18207319 | 0.05210366 | 0.9584461 | 0.98194195 |
| HSBP1L1 | 392.818803 | -0.435279138 | 0.24259482 | -1.794264 | 0.07277105 | 0.21355982 |
| TXNL4A | 480.120652 | -0.265246846 | 0.15377752 | -1.724874 | 0.08455019 | 0.23433139 |
| ADNP2 | 567.656206 | -0.003686163 | 0.21940129 | -0.016801 | 0.98659537 | 0.99472851 |
| PARD6G-AS1 | 106.905187 | 0.142031036 | 0.34541472 | 0.41118988 | 0.68093331 | 0.8332822 |
| PARD6G | 58.8126893 | -0.126723809 | 0.36872303 | -0.3436829 | 0.73108476 | 0.86806155 |
| C20orf96 | 143.233683 | 0.365530388 | 0.29930203 | 1.221276 | 0.22198154 | 0.43613776 |
| ZCCHC3 | 104.985969 | 0.479124381 | 0.28889116 | 1.65849441 | 0.09721771 | 0.25760256 |
| NRSN2-AS1 | 750.83376 | 0.662335456 | 0.33010436 | 2.00644265 | 0.04480904 | 0.15429683 |
| SOX12 | 128.590211 | 0.768048743 | 0.32780113 | 2.34303263 | 0.01912771 | 0.08658599 |
| NRSN2 | 137.727307 | -0.813056565 | 0.31267912 | -2.6002906 | 0.00931449 | 0.05262943 |
| RBCK1 | 459.134858 | -0.180119054 | 0.21769709 | -0.8273838 | 0.40801954 | 0.6279308 |
| TBC1D20 | 277.299173 | 0.053451281 | 0.19217957 | 0.27813196 | 0.78091106 | 0.89539748 |
| CSNK2A1 | 1563.73373 | 0.292016146 | 0.12354027 | 2.36373251 | 0.01809187 | 0.08355787 |
| RSPO4 | 101.028042 | 0.059069847 | 0.63540211 | 0.09296451 | 0.92593175 | 0.96724667 |
| PSMF1 | 651.159979 | -0.407284533 | 0.20973371 | -1.9419126 | 0.05214768 | 0.17105009 |
| AL136531.2 | 341.571063 | -0.441933201 | 0.30869335 | -1.4316253 | 0.15225109 | 0.34548783 |
| SDCBP2-AS1 | 517.178964 | 0.572485048 | 0.18887585 | 3.03101249 | 0.00243735 | 0.01981229 |
| NSFL1C | 260.430654 | -0.277877755 | 0.26474451 | -1.0496072 | 0.29389873 | 0.51699405 |
| SIRPA | 161.031591 | -0.019321082 | 0.36962215 | -0.0522725 | 0.95831155 | 0.98194195 |
| STK35 | 332.73591 | -0.242171604 | 0.20389248 | -1.1877417 | 0.23493519 | 0.45223825 |
| SNRPB | 251.182304 | -1.208156602 | 0.26098231 | -4.6292662 | 3.67E-06 | 0.00011652 |
| AL049650.1 | 75.2639601 | -0.048317034 | 0.25593957 | -0.188783 | 0.85026291 | 0.93134777 |
| NOP56 | 186.189556 | -0.3430888 | 0.40435933 | -0.8484751 | 0.39617346 | 0.61805306 |
| IDH3B | 165.754476 | 0.175493985 | 0.32915952 | 0.53315786 | 0.59392431 | 0.77421753 |
| EBF4 | 222.312473 | -0.339206892 | 0.39895823 | -0.8502316 | 0.39519633 | 0.6172577 |
| PCED1A | 192.310773 | 0.400287729 | 0.2167307 | 1.84693596 | 0.06475642 | 0.19763576 |
| VPS16 | 76.190027 | -0.063522436 | 0.35944243 | -0.1767249 | 0.85972446 | 0.93625342 |
| PTPRA | 1794.87321 | -0.159097028 | 0.16417065 | -0.9690954 | 0.33249758 | 0.5577801 |
| MRPS26 | 230.343499 | -0.203280084 | 0.30402423 | -0.6686312 | 0.50373077 | 0.7078307 |
| UBOX5-AS1 | 179.300496 | 0.474881747 | 0.36970961 | 1.2844723 | 0.19897675 | 0.40748588 |
| UBOX5 | 199.437473 | 0.573626325 | 0.27714625 | 2.06976039 | 0.03847479 | 0.13919843 |
| LZTS3 | 230.677856 | 0.483684047 | 0.35790605 | 1.35142743 | 0.17655855 | 0.37872068 |
| DDRGK1 | 235.909633 | -0.926731457 | 0.18932485 | -4.8949277 | 9.83E-07 | 4.06E-05 |
| ITPA | 415.850788 | 0.020993782 | 0.25713895 | 0.08164373 | 0.93493003 | 0.9719401 |
| C20orf194 | 4610.63663 | 0.663315949 | 0.24263815 | 2.73376611 | 0.00626145 | 0.03944087 |
| ATRN | 1971.0916 | 0.649002188 | 0.15539961 | 4.17634364 | 2.96E-05 | 0.00064784 |
| CENPB | 143.639317 | -1.525821839 | 0.34650074 | -4.4035168 | 1.07E-05 | 0.00028138 |
| CDC25B | 166.939468 | -0.569145199 | 0.35449636 | -1.6055036 | 0.10838301 | 0.27675067 |
| AP5S1 | 63.4231137 | -0.300647708 | 0.37781367 | -0.7957566 | 0.42617355 | 0.64484366 |
| MAVS | 840.448713 | -0.310452836 | 0.1814621 | -1.7108412 | 0.08711043 | 0.23921863 |
| AL353194.1 | 53.4011007 | 0.1571371 | 0.46551954 | 0.3375521 | 0.73570074 | 0.87055499 |
| PANK2 | 719.156923 | 0.139665448 | 0.28527714 | 0.48957812 | 0.62443246 | 0.79576733 |
| RNF24 | 2883.32381 | -0.229603426 | 0.4084192 | -0.5621759 | 0.57399619 | 0.76146587 |
| SMOX | 73.986843 | -3.077083063 | 0.47992335 | -6.4116136 | 1.44E-10 | 2.32E-08 |
| PRNP | 183.264152 | -0.6439228 | 0.34535658 | -1.8645158 | 0.06224931 | 0.19279041 |
| SLC23A2 | 539.771327 | 0.179968372 | 0.22195326 | 0.81083907 | 0.4174581 | 0.63711196 |

| | | | | | | |
|---|---|---|---|---|---|---|
| TMEM230 | 443.574464 | -0.226850801 | 0.16644986 | -1.3628777 | 0.17292106 | 0.37444497 |
| PCNA | 121.418037 | -0.231982355 | 0.42104643 | -0.5509662 | 0.58165684 | 0.76754298 |
| CDS2 | 575.150966 | -0.230106888 | 0.11914123 | -1.9313791 | 0.05343619 | 0.17380906 |
| GPCPD1 | 1455.6764 | 0.057228666 | 0.28648626 | 0.19976059 | 0.84166782 | 0.92654505 |
| SHLD1 | 591.293409 | 0.359735344 | 0.25017209 | 1.43795157 | 0.1504478 | 0.34276316 |
| TRMT6 | 97.1221972 | 0.041315193 | 0.36909061 | 0.11193781 | 0.91087272 | 0.95976856 |
| MCM8 | 225.501268 | 0.363725968 | 0.40705068 | 0.89356433 | 0.37155504 | 0.59563048 |
| CRLS1 | 945.92075 | -0.053636161 | 0.33032442 | -0.1623742 | 0.87101119 | 0.94147844 |
| TMX4 | 691.868465 | -0.364542141 | 0.36453671 | -1.0000149 | 0.31730329 | 0.5418564 |
| PLCB1 | 33273.0819 | 0.17029741 | 0.37728997 | 0.4513701 | 0.65172283 | 0.81397569 |
| PLCB4 | 445.98564 | -0.481036918 | 0.5443411 | -0.8837049 | 0.37685551 | 0.6000254 |
| ANKEF1 | 81.2963185 | 0.058475058 | 0.26385485 | 0.22161828 | 0.82461105 | 0.91884704 |
| SNAP25-AS1 | 49.4240955 | -0.463048216 | 0.53521976 | -0.8651553 | 0.38695355 | 0.61054074 |
| SNAP25 | 65.5007758 | -2.323692732 | 0.71308179 | -3.2586623 | 0.00111939 | 0.01091218 |
| MKKS | 171.831018 | 0.53180137 | 0.2240086 | 2.37402216 | 0.01759549 | 0.081875 |
| SLX4IP | 1699.32113 | 0.464540132 | 0.17918749 | 2.59248086 | 0.00952865 | 0.0534681 |
| JAG1 | 442.418134 | 1.10238538 | 0.36736038 | 3.00082821 | 0.00269246 | 0.02134203 |
| AL050403.2 | 381.416004 | 1.443458284 | 0.57391887 | 2.51509117 | 0.01190016 | 0.06260025 |
| BTBD3 | 1259.24901 | -0.335507475 | 0.21476964 | -1.5621737 | 0.11824708 | 0.29330654 |
| SPTLC3 | 2395.35274 | -0.39204157 | 0.34716254 | -1.1292738 | 0.25878234 | 0.47985036 |
| ISM1 | 112.45129 | -0.555607059 | 0.50176948 | -1.1072955 | 0.26816621 | 0.48932276 |
| TASP1 | 2008.43888 | -0.051794802 | 0.16931557 | -0.3059069 | 0.75967553 | 0.88355676 |
| ESF1 | 592.377565 | 0.169764041 | 0.18962735 | 0.8952508 | 0.37065303 | 0.59532317 |
| NDUFAF5 | 530.534401 | 0.857984587 | 0.28210247 | 3.04139341 | 0.00235486 | 0.01934411 |
| MACROD2 | 7480.92954 | 0.427204965 | 0.26065267 | 1.63898173 | 0.10121706 | 0.26494858 |
| FLRT3 | 728.390509 | -0.34921927 | 0.53064034 | -0.6581092 | 0.51046799 | 0.71269123 |
| KIF16B | 2946.18291 | 0.003524927 | 0.36096669 | 0.00976524 | 0.99220859 | 0.9967867 |
| SNRPB2 | 233.369484 | 0.098961486 | 0.18108302 | 0.54649786 | 0.58472376 | 0.76905468 |
| BFSP1 | 52.9703916 | -0.820005896 | 0.60410722 | -1.3573847 | 0.17465901 | 0.3764188 |
| DSTN | 2126.01835 | -0.78639071 | 0.34099809 | -2.306144 | 0.02110258 | 0.09263677 |
| RRBP1 | 1513.337 | -0.543816047 | 0.24505237 | -2.2191829 | 0.02647428 | 0.10786514 |
| SNX5 | 464.782173 | 0.083121428 | 0.21846577 | 0.38047803 | 0.7035906 | 0.84975114 |
| OVOL2 | 682.687152 | 0.37987848 | 0.25462581 | 1.4919088 | 0.13572306 | 0.32093911 |
| MGME1 | 705.093969 | 0.583223866 | 0.17755751 | 3.28470393 | 0.0010209 | 0.01015438 |
| KAT14 | 138.804099 | 0.496611628 | 0.25441429 | 1.95198007 | 0.05094057 | 0.1682517 |
| ZNF133 | 535.536724 | 0.37193955 | 0.16914475 | 2.19894232 | 0.02788202 | 0.11160414 |
| DZANK1 | 349.268423 | -0.052567335 | 0.31356896 | -0.167642 | 0.86686491 | 0.93959274 |
| POLR3F | 140.126736 | 0.224291968 | 0.3475985 | 0.6452616 | 0.51875767 | 0.71939843 |
| RBBP9 | 86.6028592 | -0.547789021 | 0.28523423 | -1.9204884 | 0.05479624 | 0.1766273 |
| SEC23B | 605.893421 | -0.16500354 | 0.17309523 | -0.953253 | 0.34046192 | 0.56582304 |
| DTD1 | 103.928458 | -1.204913219 | 0.32894114 | -3.6630056 | 0.00024927 | 0.00350282 |
| AL035563.1 | 38.823512 | 0.824448174 | 0.37155563 | 2.21890912 | 0.02649291 | 0.10790775 |
| SLC24A3 | 109.520154 | -1.977610978 | 0.581654 | -3.3999783 | 0.00067391 | 0.00744001 |
| AL121761.1 | 402.080166 | -0.7579984 | 0.30130375 | -2.5157284 | 0.01187867 | 0.06256193 |
| RIN2 | 1048.91992 | -0.345857444 | 0.32455032 | -1.0656512 | 0.28658134 | 0.50879744 |
| NAA20 | 481.599586 | -0.297259101 | 0.22096037 | -1.3453051 | 0.17852674 | 0.38170277 |
| CRNKL1 | 169.513778 | -0.05138848 | 0.17620649 | -0.2916378 | 0.77056355 | 0.88937357 |
| CFAP61 | 78.6336084 | 0.064481131 | 0.2775633 | 0.23231144 | 0.81629613 | 0.91443162 |
| RALGAPA2 | 6178.40774 | -1.084522301 | 0.25349867 | -4.278217 | 1.88E-05 | 0.00044758 |
| KIZ | 1527.85504 | 0.578928638 | 0.20793449 | 2.78418759 | 0.0053662 | 0.03535268 |
| KIZ-AS1 | 327.875031 | 0.108840054 | 0.55731027 | 0.19529526 | 0.84516181 | 0.92842955 |
| XRN2 | 1070.81648 | -0.402370892 | 0.21312817 | -1.8879292 | 0.05903545 | 0.18569162 |
| NXT1 | 44.4166461 | -0.655688914 | 0.36096799 | -1.8164738 | 0.06929772 | 0.20685529 |
| GZF1 | 94.81612 | 0.24727019 | 0.3498785 | 0.70673159 | 0.47973329 | 0.68893206 |
| NAPB | 738.65964 | 0.914343674 | 0.22818913 | 4.00695539 | 6.15E-05 | 0.00115076 |
| CST3 | 1267.42423 | -1.687777302 | 0.47208398 | -3.5751633 | 0.00035001 | 0.00450053 |
| SYNDIG1 | 762.381418 | 0.759220057 | 0.3899172 | 1.94713149 | 0.05151897 | 0.1694925 |
| AL035661.1 | 49.0896774 | -0.074234047 | 0.62078256 | -0.1195814 | 0.90481476 | 0.95628885 |
| APMAP | 284.684454 | -0.447881415 | 0.22453021 | -1.9947491 | 0.04607026 | 0.15692161 |
| ACSS1 | 1354.19199 | 0.650199245 | 0.3636986 | 1.78774194 | 0.07381765 | 0.21554455 |
| ENTPD6 | 454.316482 | -0.73321362 | 0.27717673 | -2.6452928 | 0.00816202 | 0.04789524 |
| PYGB | 808.895852 | -1.50324574 | 0.54653306 | -2.750512 | 0.00595022 | 0.03809815 |
| ABHD12 | 804.588244 | -0.036482629 | 0.21979439 | -0.1659853 | 0.86816855 | 0.94004741 |
| GINS1 | 91.3387306 | 1.162224206 | 0.36993027 | 3.14173863 | 0.00167948 | 0.01494895 |
| NINL | 2757.89729 | 0.569391733 | 0.24239706 | 2.34900431 | 0.01882369 | 0.08556925 |
| NANP | 58.8790154 | 0.553944347 | 0.27864692 | 1.98797941 | 0.04681397 | 0.1587154 |
| ZNF337-AS1 | 404.024796 | 0.602382049 | 0.26723448 | 2.25413294 | 0.02418781 | 0.10127365 |
| ZNF337 | 401.511857 | 0.049900052 | 0.20497786 | 0.24344118 | 0.80766365 | 0.91010272 |
| FAM182B | 313.048285 | -0.026932543 | 0.2511992 | -0.1072159 | 0.91461772 | 0.96131774 |
| HM13 | 832.016391 | -0.4726038 | 0.22880986 | -2.0654871 | 0.03887694 | 0.14003967 |
| ID1 | 221.843907 | -1.317255968 | 0.68305895 | -1.928466 | 0.05379719 | 0.17451057 |
| BCL2L1 | 1622.69118 | -0.323609646 | 0.40667972 | -0.7957359 | 0.42618557 | 0.64484366 |
| AL117381.1 | 352.616675 | -0.494620065 | 0.28426327 | -1.7400069 | 0.0818578 | 0.22998377 |
| DUSP15 | 184.896047 | 0.762697294 | 0.62479882 | 1.22070861 | 0.22219637 | 0.43639586 |
| TTLL9 | 178.411159 | 0.015568013 | 0.37307533 | 0.04172887 | 0.96671484 | 0.98510234 |
| PDRG1 | 77.9224426 | -0.272712986 | 0.26276672 | -1.0378521 | 0.29933892 | 0.52232071 |
| TM9SF4 | 717.997986 | 0.092565628 | 0.16651019 | 0.55591569 | 0.57826849 | 0.76458087 |
| PLAGL2 | 115.231502 | -0.158798565 | 0.20434498 | -0.7771102 | 0.43709377 | 0.65356621 |
| POFUT1 | 224.796056 | -0.024037008 | 0.22695055 | -0.105913 | 0.9156514 | 0.96196925 |
| KIF3B | 742.590714 | -0.170797406 | 0.21381358 | -0.7988146 | 0.42439794 | 0.64298076 |
| ASXL1 | 2299.59863 | -0.008263415 | 0.21401473 | -0.0386114 | 0.96920019 | 0.98607143 |
| NOL4L | 183.386493 | -0.476969669 | 0.26178925 | -1.8219605 | 0.06846098 | 0.20524503 |
| FO393400.1 | 63.8699881 | -0.232211568 | 0.32314218 | -0.7186049 | 0.47238437 | 0.68320652 |
| DNMT3B | 73.7105233 | 0.305497977 | 0.306365 | 0.99716997 | 0.31868201 | 0.5432276 |
| MAPRE1 | 223.934756 | 0.094924244 | 0.24466971 | 0.38796892 | 0.69803904 | 0.84616339 |
| AL035071.2 | 80.1204828 | -0.130443462 | 0.53893739 | -0.2420382 | 0.80875053 | 0.91010272 |
| AL035071.1 | 60.376151 | 0.274211586 | 0.38199888 | 0.71783348 | 0.47285997 | 0.68359498 |
| CDK5RAP1 | 390.669873 | 0.341315867 | 0.16417418 | 2.0789863 | 0.03761861 | 0.13730429 |
| SNTA1 | 128.436911 | 0.550088106 | 0.36957283 | 1.48844305 | 0.13663409 | 0.32228566 |
| CBFA2T2 | 2595.91261 | 0.286567286 | 0.15279302 | 1.87552607 | 0.06072039 | 0.18947687 |
| NECAB3 | 90.6558531 | 0.886407249 | 0.34706595 | 2.55400236 | 0.01064925 | 0.05800659 |
| PXMP4 | 77.5886648 | 0.028513291 | 0.46334688 | 0.06153768 | 0.95093101 | 0.97879278 |
| ZNF341 | 110.646055 | 0.341090504 | 0.3593594 | 0.9491626 | 0.34253792 | 0.56763059 |
| CHMP4B | 492.819304 | -0.794135286 | 0.16570839 | -4.7923663 | 1.65E-06 | 6.03E-05 |
| RALY | 903.618452 | -0.504162985 | 0.22945854 | -2.1971855 | 0.0280072 | 0.11196945 |
| EIF2S2 | 600.398577 | -0.878955344 | 0.24482624 | -3.5901191 | 0.00033053 | 0.0043099 |
| ASIP | 89.7647752 | 0.115414336 | 0.62572911 | 0.18444776 | 0.8536622 | 0.93290511 |
| AHCY | 144.0282 | -0.488612253 | 0.25618704 | -1.9072481 | 0.05648846 | 0.18057989 |
| ITCH | 2448.96043 | -0.000726731 | 0.1495794 | -0.0048585 | 0.9961235 | 0.99854256 |
| DYNLRB1 | 642.572843 | -0.583895015 | 0.23107311 | -2.5268843 | 0.01150794 | 0.061195 |
| MAP1LC3A | 195.644074 | -1.375287096 | 0.61450788 | -2.23803 | 0.0252191 | 0.10419747 |
| PIGU | 557.162887 | 0.362285347 | 0.29498132 | 1.22816368 | 0.21938553 | 0.43356369 |
| NCOA6 | 1467.68349 | 0.057250885 | 0.13609598 | 0.4206655 | 0.67399936 | 0.82909435 |
| TP53INP2 | 187.237371 | 0.111990121 | 0.40038026 | 0.2797094 | 0.77970047 | 0.8948704 |
| GGT7 | 596.907108 | 0.014372166 | 0.32308676 | 0.04448392 | 0.96451867 | 0.98434112 |
| ACSS2 | 668.948652 | -0.294160318 | 0.21450795 | -1.371326 | 0.17027337 | 0.37059498 |
| GSS | 182.645644 | -1.148150445 | 0.35731184 | -3.2133008 | 0.00131219 | 0.01238934 |
| MYH7B | 45.7860861 | 0.816653193 | 0.3793459 | 2.152793 | 0.03133495 | 0.12162896 |
| TRPC4AP | 1458.47706 | -0.170093606 | 0.17021324 | -0.9992972 | 0.31765076 | 0.54223944 |
| EDEM2 | 294.649891 | -0.077696731 | 0.24884257 | -0.3122325 | 0.75486385 | 0.88129721 |
| PROCR | 187.918312 | 0.252253641 | 0.54931887 | 0.45921168 | 0.64608217 | 0.81068674 |
| MMP24OS | 514.627571 | 0.235902756 | 0.20978219 | 1.12451281 | 0.26079554 | 0.4819992 |
| MMP24 | 438.530089 | -0.193042492 | 0.32422167 | -0.5954028 | 0.55157425 | 0.74467698 |

| | | | | | | |
|---|---|---|---|---|---|---|
| UQCC1 | 897.686273 | 0.376503527 | 0.16514476 | 2.27983943 | 0.02261721 | 0.0968398 |
| CEP250 | 201.990363 | -0.365248732 | 0.31699764 | -1.1522128 | 0.24923364 | 0.4686302 |
| ERGIC3 | 365.844156 | -0.034447086 | 0.27831882 | -0.1237684 | 0.90149862 | 0.95457905 |
| CPNE1 | 359.480747 | -0.003007443 | 0.26747385 | -0.0112439 | 0.99102887 | 0.99638456 |
| NFS1 | 111.975493 | 0.356031543 | 0.24027082 | 1.48179266 | 0.13839547 | 0.32460044 |
| ROMO1 | 208.925614 | -0.890141362 | 0.41431284 | -2.1484764 | 0.03167593 | 0.12248458 |
| RBM39 | 3538.87384 | 0.22994332 | 0.14337043 | 1.60384058 | 0.1087492 | 0.2771737 |
| PHF20 | 1322.21312 | -0.244775205 | 0.13823044 | -1.7707764 | 0.07659789 | 0.22078382 |
| SCAND1 | 378.783899 | -1.534909295 | 0.3654129 | -4.2004792 | 2.66E-05 | 0.00059732 |
| CNBD2 | 93.7434755 | 1.056469214 | 0.2988278 | 3.53537794 | 0.00040719 | 0.00505033 |
| NORAD | 875.706297 | -0.941721636 | 0.28109625 | -3.350175 | 0.00080761 | 0.0084866 |
| EPB41L1 | 3652.25085 | -0.11930375 | 0.54418685 | -0.2192331 | 0.8264685 | 0.91938879 |
| AAR2 | 181.46143 | 0.103053973 | 0.16660857 | 0.61853946 | 0.53621979 | 0.73306066 |
| DLGAP4 | 847.128485 | -0.48405229 | 0.19537105 | -2.477605 | 0.01322675 | 0.06711401 |
| DLGAP4-AS1 | 423.344697 | 0.46338815 | 0.3140502 | 1.47552252 | 0.14007212 | 0.32747126 |
| MYL9 | 101.124018 | 0.097439371 | 0.39446682 | 0.24701538 | 0.80489633 | 0.90860991 |
| RAB5IF | 122.662666 | -0.706128999 | 0.47338252 | -1.4916668 | 0.13578651 | 0.32103168 |
| NDRG3 | 343.645795 | 0.501919637 | 0.15838883 | 3.16890802 | 0.00153013 | 0.01394856 |
| DSN1 | 95.7713611 | 0.158813594 | 0.30139992 | 0.52691983 | 0.59824926 | 0.77686537 |
| SOGA1 | 639.46228 | 0.348859441 | 0.32243437 | 1.08195488 | 0.27927258 | 0.50141518 |
| SAMHD1 | 133.509613 | 0.651739842 | 0.60868406 | 1.07073585 | 0.28428822 | 0.50683804 |
| RBL1 | 298.292017 | 0.494619268 | 0.23361854 | 2.11720899 | 0.03424212 | 0.12911907 |
| MROH8 | 146.689118 | 0.868909747 | 0.25394968 | 3.42158241 | 0.00062258 | 0.00702275 |
| RPN2 | 552.9283 | -0.159859328 | 0.16957114 | -0.9427272 | 0.34582045 | 0.57105512 |
| MANBAL | 159.07575 | -0.161866438 | 0.19489283 | -0.8305407 | 0.40623312 | 0.62627606 |
| SRC | 235.869966 | -0.246714646 | 0.21248514 | -1.1610913 | 0.24560478 | 0.4647176 |
| BLCAP | 619.682051 | 0.163680395 | 0.21984667 | 0.74452069 | 0.45656153 | 0.67157252 |
| CTNNBL1 | 555.120133 | 0.132557852 | 0.1566205 | 0.84636337 | 0.39735007 | 0.61906213 |
| TTI1 | 357.868316 | 0.102885223 | 0.17660695 | 0.58256609 | 0.56018544 | 0.75183103 |
| RPRD1B | 648.795628 | 0.034853716 | 0.20426499 | 0.17062991 | 0.86451478 | 0.93833133 |
| TGM2 | 607.989359 | -5.513527413 | 0.5937621 | -9.2857516 | 1.61E-20 | 3.53E-17 |
| RALGAPB | 1281.95638 | -0.014343045 | 0.16264223 | -0.0881877 | 0.92972749 | 0.96915731 |
| ARHGAP40 | 68.6664279 | -4.239774947 | 0.9674209 | -4.3825546 | 1.17E-05 | 0.0003032 |
| ACTR5 | 90.7983906 | 0.342315228 | 0.30346222 | 1.12803242 | 0.25930623 | 0.48032197 |
| DHX35 | 317.110301 | 0.659159668 | 0.24839631 | 2.65366129 | 0.00796237 | 0.04718482 |
| MAFB | 52.8667765 | -0.385358588 | 0.5420856 | -0.7108814 | 0.47715771 | 0.68692989 |
| TOP1 | 1773.43653 | -0.377275837 | 0.22343542 | -1.6885229 | 0.09131089 | 0.24700502 |
| PLCG1-AS1 | 73.193033 | -0.263751721 | 0.37606829 | -0.70134 | 0.48309088 | 0.69127369 |
| PLCG1 | 480.415002 | -0.680168836 | 0.25864845 | -2.6297039 | 0.00854593 | 0.04950124 |
| ZHX3 | 4936.64678 | -0.344348506 | 0.35295651 | -0.9756117 | 0.32925692 | 0.55480987 |
| LPIN3 | 517.163087 | -0.66220037 | 0.25418021 | -2.6052397 | 0.009181 | 0.05207035 |
| CHD6 | 2578.37668 | 0.185702228 | 0.12700063 | 1.46221496 | 0.14368231 | 0.333319 |
| PTPRT | 48.6197802 | 0.033692268 | 0.48350681 | 0.06968313 | 0.94444587 | 0.9759376 |
| SRSF6 | 699.118703 | 1.080401417 | 0.26860303 | 4.02229795 | 5.76E-05 | 0.00109379 |
| L3MBTL1 | 279.188203 | -0.360845022 | 0.22637539 | -1.5940117 | 0.11093341 | 0.28098167 |
| Z98752.3 | 90.2697152 | -0.431771642 | 0.31219629 | -1.3830134 | 0.16666077 | 0.36635414 |
| SGK2 | 1045.25786 | 1.070411687 | 0.23942783 | 4.47070697 | 7.80E-06 | 0.00021957 |
| IFT52 | 389.208124 | -0.104872422 | 0.15326727 | -0.6842454 | 0.49382024 | 0.69913828 |
| OSER1 | 245.451569 | -0.636969902 | 0.26688708 | -2.3866644 | 0.017002 | 0.0797062 |
| OSER1-DT | 66.7684328 | 0.369890067 | 0.33824922 | 1.09354301 | 0.27415546 | 0.49598462 |
| FITM2 | 77.9238417 | 0.358919868 | 0.30532829 | 1.17552115 | 0.23978628 | 0.45797238 |
| TTPAL | 123.376322 | -0.152428184 | 0.28332903 | -0.53799 | 0.59058395 | 0.77217181 |
| SERINC3 | 511.273979 | -0.424742787 | 0.17998868 | -2.3598305 | 0.01828329 | 0.08408911 |
| PKIG | 717.222996 | -0.194046459 | 0.1662792 | -1.1669918 | 0.24321369 | 0.46195412 |
| YWHAB | 885.503068 | -1.327582354 | 0.33141085 | -4.0058507 | 6.18E-05 | 0.00115289 |
| PABPC1L | 289.500557 | 0.48650537 | 0.21165117 | 2.2986189 | 0.02152659 | 0.09387412 |
| TOMM34 | 110.144619 | 0.041085752 | 0.25923324 | 0.15848952 | 0.87407108 | 0.94334544 |
| STK4 | 587.978276 | -0.178329332 | 0.22638256 | -0.7877344 | 0.43085208 | 0.64893712 |
| SLPI | 5909.96203 | -5.687769257 | 0.639281 | -8.8971348 | 5.73E-19 | 9.46E-16 |
| SDC4 | 1787.06434 | -0.920196089 | 0.37140469 | -2.4776103 | 0.01322655 | 0.06711401 |
| SYS1-DBNDD | 104.308793 | -0.357313358 | 0.42825745 | -0.8343424 | 0.40408805 | 0.62523124 |
| TP53TG5 | 59.3464561 | -0.486399879 | 0.62269793 | -0.7811169 | 0.43473374 | 0.65173056 |
| PIGT | 177.30668 | 0.054125487 | 0.28351412 | 0.19090931 | 0.84859664 | 0.9303692 |
| WFDC2 | 9006.98496 | -4.209287932 | 0.74090661 | -5.6812666 | 1.34E-08 | 1.20E-06 |
| DNTTIP1 | 198.166545 | -0.305182779 | 0.24011769 | -1.2709717 | 0.20373873 | 0.41350055 |
| SNX21 | 200.776866 | 0.669451005 | 0.33262262 | 2.01264425 | 0.04415207 | 0.15243196 |
| ACOT8 | 187.52416 | 0.24337257 | 0.21238839 | 1.14588452 | 0.25184294 | 0.47203022 |
| ZSWIM3 | 82.7256827 | 0.620317526 | 0.33853569 | 1.83235489 | 0.06689856 | 0.20193399 |
| CTSA | 304.232028 | 0.037825552 | 0.31980213 | 0.11827798 | 0.9058474 | 0.95693796 |
| PCIF1 | 72.9841044 | 0.065411354 | 0.24872614 | 0.26298544 | 0.7925618 | 0.90226224 |
| ZNF335 | 224.281919 | -0.101504208 | 0.22476756 | -0.4515963 | 0.6515598 | 0.81384903 |
| NCOA5 | 515.248154 | -0.054165436 | 0.26176885 | -0.2069209 | 0.83607167 | 0.92369276 |
| CD40 | 58.0588208 | -1.832288649 | 0.51227978 | -3.5767343 | 0.00034791 | 0.00449227 |
| SLC35C2 | 272.389344 | -0.520549825 | 0.2306738 | -2.2566491 | 0.02403001 | 0.1008619 |
| AL133227.1 | 58.1033207 | 0.116905636 | 0.52360945 | 0.22326877 | 0.82332633 | 0.91844587 |
| ELMO2 | 242.120552 | -0.009682139 | 0.20500586 | -0.0472286 | 0.96233104 | 0.9832997 |
| ZNF334 | 183.341252 | 0.606864054 | 0.58953491 | 1.02939461 | 0.30329428 | 0.52644076 |
| SLC13A3 | 206.541704 | -0.057789091 | 0.51633641 | -0.1119214 | 0.91088573 | 0.95976856 |
| EYA2 | 1017.2734 | 1.518895693 | 1.70049606 | 0.89320742 | 0.37174611 | 0.59584933 |
| ZMYND8 | 4737.98141 | -0.27137851 | 0.30361937 | -0.8938116 | 0.3714227 | 0.59563048 |
| NCOA3 | 2265.25458 | 0.04029307 | 0.16131356 | 0.24978105 | 0.80275667 | 0.9075253 |
| SULF2 | 674.964104 | -2.43035731 | 0.55363275 | -4.3898366 | 1.13E-05 | 0.00029496 |
| PREX1 | 137.054411 | -0.697969426 | 0.352154 | -1.9820006 | 0.04747918 | 0.16015129 |
| ARFGEF2 | 2358.2825 | 0.067682125 | 0.13331658 | 0.50767971 | 0.61167798 | 0.78684784 |
| CSE1L | 501.821691 | 0.265027177 | 0.23121773 | 1.14622341 | 0.25170272 | 0.47199621 |
| STAU1 | 1350.99493 | -0.184083606 | 0.18335514 | -1.003973 | 0.31539164 | 0.53963837 |
| DDX27 | 225.136307 | -0.476941686 | 0.167382 | -2.8494205 | 0.0043799 | 0.0303988 |
| ZNFX1 | 415.601512 | -0.985222103 | 0.33647165 | -2.9280984 | 0.00341042 | 0.0254942 |
| ZFAS1 | 375.954375 | -0.447722451 | 0.13947888 | -3.209966 | 0.00132751 | 0.01249825 |
| B4GALT5 | 2110.27879 | -0.392615849 | 0.20916363 | -1.8770751 | 0.0605078 | 0.18908057 |
| SLC9A8 | 792.23794 | -0.081213796 | 0.26460171 | -0.3069285 | 0.75889782 | 0.88316847 |
| SPATA2 | 52.3220149 | 0.040264566 | 0.30585868 | 0.13164435 | 0.89526561 | 0.9517556 |
| RNF114 | 624.209012 | -0.433743208 | 0.18174983 | -2.3864849 | 0.01701031 | 0.0797062 |
| CEBPB | 320.432274 | -3.224780596 | 0.59961323 | -5.3781011 | 7.53E-08 | 5.17E-06 |
| PTPN1 | 1632.4205 | -0.883298822 | 0.28880414 | -3.05847 | 0.0022247 | 0.01857529 |
| RIPOR3 | 966.404631 | 0.679277678 | 1.05098746 | 0.64632329 | 0.51807 | 0.71904872 |
| AL353653.1 | 51.3296125 | 0.35713228 | 1.123816 | 0.31778537 | 0.75064775 | 0.87890283 |
| PARD6B | 1704.94396 | 0.161925295 | 0.28414416 | 0.56987021 | 0.56876573 | 0.75864147 |
| BCAS4 | 315.07123 | -0.907926314 | 0.2794082 | -3.249462 | 0.00115624 | 0.01120522 |
| ADNP | 2198.79929 | 0.093067155 | 0.14901738 | 0.62453892 | 0.53227372 | 0.72948782 |
| ADNP-AS1 | 58.4774565 | 0.100941288 | 0.31642809 | 0.3190023 | 0.74972477 | 0.87816924 |
| DPM1 | 547.239706 | 0.244250007 | 0.14547752 | 1.67895358 | 0.09316109 | 0.25093084 |
| NFATC2 | 496.724956 | -0.563031144 | 0.38032646 | -1.4803891 | 0.13876943 | 0.32507075 |
| ATP9A | 885.661895 | -0.599844449 | 0.24591417 | -2.4392431 | 0.01471806 | 0.07229858 |
| ZFP64 | 443.643207 | -0.373170389 | 0.25551437 | -1.4604673 | 0.14416168 | 0.33389998 |
| TSHZ2 | 85.6804322 | -0.803401917 | 0.48978354 | -1.6403204 | 0.10093857 | 0.26453475 |
| AL157838.1 | 62.5849369 | -0.908064087 | 0.70913869 | -1.2805169 | 0.2003634 | 0.40918369 |
| ZNF217 | 679.8475 | -0.982690959 | 0.2357967 | -4.1675348 | 3.08E-05 | 0.00066785 |
| BCAS1 | 109.825523 | -1.135330771 | 0.72895732 | -1.5574722 | 0.11935841 | 0.29507866 |
| CYP24A1 | 1108.31711 | -6.370169799 | 0.90868808 | -7.0102931 | 2.38E-12 | 6.83E-10 |
| PFDN4 | 153.242004 | -0.342832464 | 0.29611553 | -1.1577659 | 0.24695959 | 0.46618627 |
| DOK5 | 617.504162 | 0.319060626 | 0.45062459 | 0.70804087 | 0.47891988 | 0.68858742 |
| FAM210B | 113.221747 | -0.936909289 | 0.34138458 | -2.7444394 | 0.00606143 | 0.0386603 |

| | | | | | | |
|---|---|---|---|---|---|---|
| CSTF1 | 114.793824 | 0.047047373 | 0.16646514 | 0.28262597 | 0.77746357 | 0.89359206 |
| RTF2 | 338.773823 | -0.174056579 | 0.18786346 | -0.9265057 | 0.35418321 | 0.58001563 |
| RAE1 | 210.036967 | 0.218640641 | 0.21260724 | 1.02837816 | 0.30377198 | 0.52699292 |
| RBM38 | 76.3666514 | -0.202955017 | 0.46799432 | -0.4336698 | 0.66452822 | 0.82323933 |
| PCK1 | 1785.25718 | -0.873160493 | 0.66725108 | -1.3085936 | 0.19067205 | 0.39725348 |
| PMEPA1 | 299.140137 | -1.500182124 | 0.84230012 | -1.7810542 | 0.0749036 | 0.21730765 |
| RAB22A | 627.098227 | -0.371734679 | 0.17713275 | -2.0986219 | 0.03585024 | 0.1329438 |
| VAPB | 827.228636 | -0.150561441 | 0.13354507 | -1.1274204 | 0.25956477 | 0.48061767 |
| STX16-NPEPI | 99.97429 | -0.456190482 | 0.62578018 | -0.7289948 | 0.46600486 | 0.67888587 |
| NPEPL1 | 256.853518 | 0.094282097 | 0.3103087 | 0.30383323 | 0.76125495 | 0.88414812 |
| GNAS-AS1 | 49.235506 | 0.259451696 | 0.51420268 | 0.50457088 | 0.61386027 | 0.78755051 |
| GNAS | 3331.24294 | -0.207184715 | 0.24891185 | -0.8323618 | 0.40520476 | 0.62563994 |
| AL121917.1 | 64.7665035 | 0.512513676 | 0.36378595 | 1.40883306 | 0.15888454 | 0.35524819 |
| NELFCD | 407.446932 | 0.278403356 | 0.15933341 | 1.74730055 | 0.08058522 | 0.22759251 |
| CTSZ | 243.173759 | -1.241806534 | 0.37731859 | -3.2911353 | 0.00099784 | 0.00998339 |
| ATP5F1E | 996.591923 | -0.973846744 | 0.47395993 | -2.0547027 | 0.03990773 | 0.14231674 |
| PRELID3B | 122.124877 | 0.229673463 | 0.3218705 | 0.71355859 | 0.47550017 | 0.68538649 |
| PHACTR3 | 490.384003 | -3.13004648 | 1.09084946 | -2.8693661 | 0.00411295 | 0.02922329 |
| SYCP2 | 202.749065 | 0.286505079 | 0.3247405 | 0.88225854 | 0.37763701 | 0.60060335 |
| FAM217B | 132.218792 | -0.068819862 | 0.30953108 | -0.2223359 | 0.82405243 | 0.91855769 |
| CDH26 | 56.9885364 | 0.088647519 | 0.51228664 | 0.17304281 | 0.86261778 | 0.93788453 |
| MIR646HG | 1665.61008 | 0.596695165 | 0.82025332 | 0.7274523 | 0.46694892 | 0.67935225 |
| CDH4 | 101.298222 | 0.217562427 | 0.50185368 | 0.43351765 | 0.66463875 | 0.82323933 |
| TAF4 | 534.756734 | 0.018808416 | 0.22848389 | 0.08231834 | 0.93439357 | 0.97176637 |
| LSM14B | 129.730167 | -0.073820272 | 0.24517023 | -0.301098 | 0.76333975 | 0.88540171 |
| PSMA7 | 469.638206 | -1.142947188 | 0.29934755 | -3.8181278 | 0.00013447 | 0.00212209 |
| SS18L1 | 439.725541 | -0.207842092 | 0.18507657 | -1.123006 | 0.26143495 | 0.48237074 |
| MTG2 | 505.691202 | 0.376539405 | 0.19259053 | 1.9551294 | 0.0505678 | 0.1674727 |
| OSBPL2 | 725.71633 | -0.234437937 | 0.23970088 | -0.9780437 | 0.3280527 | 0.55334544 |
| ADRM1 | 90.7779992 | -0.920437757 | 0.4111868 | -2.2384905 | 0.02518908 | 0.10419747 |
| AL354836.1 | 74.1724675 | -0.01436574 | 0.48382898 | -0.0296918 | 0.97631287 | 0.98947549 |
| LAMA5 | 516.534637 | -0.428573929 | 0.26332759 | -1.6275314 | 0.10362428 | 0.2692845 |
| RPS21 | 102.375314 | -0.208154048 | 0.37648371 | -0.5528899 | 0.58033879 | 0.76633961 |
| CABLES2 | 50.1434642 | -0.146087577 | 0.34426223 | -0.4243497 | 0.67131079 | 0.82736497 |
| SLCO4A1 | 468.082153 | -2.440566715 | 0.4176386 | -5.8437288 | 5.10E-09 | 5.39E-07 |
| NTSR1 | 46.974436 | 1.365261881 | 0.64500767 | 2.11665993 | 0.03428872 | 0.12913878 |
| LINC00659 | 46.0565915 | 1.144411648 | 0.50181596 | 2.28054057 | 0.02257565 | 0.09672453 |
| MRGBP | 83.6892407 | -0.12745013 | 0.28455989 | -0.4478851 | 0.65423613 | 0.81580337 |
| OGFR | 335.038639 | -1.375704361 | 0.32013605 | -4.2972491 | 1.73E-05 | 0.00042223 |
| TCFL5 | 164.876766 | -0.217275161 | 0.20466094 | -1.0616347 | 0.28840155 | 0.51092727 |
| DIDO1 | 1025.52978 | 0.06956335 | 0.14719029 | 0.47260829 | 0.63649267 | 0.80461591 |
| GID8 | 270.184021 | -0.168116583 | 0.15148298 | -1.1098051 | 0.26708301 | 0.48822301 |
| YTHDF1 | 343.421755 | -0.150244817 | 0.15822171 | -0.9495841 | 0.34232362 | 0.56753966 |
| NKAIN4 | 44.5303461 | 0.006150037 | 0.59640989 | 0.01031176 | 0.99177255 | 0.99675782 |
| ARFGAP1 | 275.135317 | -0.061827651 | 0.31727216 | -0.1948726 | 0.84549268 | 0.92858675 |
| PPDPF | 867.610691 | -1.645148345 | 0.45519381 | -3.6141711 | 0.00030131 | 0.00403243 |
| HELZ2 | 97.5501768 | -1.285995677 | 0.87109701 | -1.4762945 | 0.13986486 | 0.32710251 |
| GMEB2 | 196.334912 | -0.391132323 | 0.18415456 | -2.1239351 | 0.03367557 | 0.12774861 |
| MHENCR | 84.8021538 | 0.243629879 | 0.37405171 | 0.65132674 | 0.51483559 | 0.7163661 |
| STMN3 | 138.475327 | -2.529986492 | 0.70266636 | -3.6005516 | 0.00031754 | 0.00419442 |
| ARFRP1 | 177.276362 | -0.24705708 | 0.28048376 | -0.8808249 | 0.37841261 | 0.60120906 |
| SLC2A4RG | 250.251618 | -0.355869239 | 0.42314824 | -0.8410037 | 0.40034585 | 0.62197 |
| ZBTB46 | 266.716935 | -1.107227557 | 0.47665044 | -2.3229341 | 0.02018269 | 0.08958104 |
| TPD52L2 | 428.342727 | -0.247497603 | 0.27011673 | -0.9162617 | 0.35952967 | 0.58492762 |
| DNAJC5 | 917.127411 | -0.593101701 | 0.19572346 | -3.0303045 | 0.00244307 | 0.01984658 |
| UCKL1 | 240.755685 | 0.691454174 | 0.27361818 | 2.52707683 | 0.01150163 | 0.0611861 |
| ZNF512B | 165.091599 | 0.08210229 | 0.39071214 | 0.21013499 | 0.83356232 | 0.92230899 |
| SAMD10 | 41.3225133 | -0.081707907 | 0.47395654 | -0.1723954 | 0.86312672 | 0.93813133 |
| PRPF6 | 532.962446 | -0.209048865 | 0.227505 | -0.9188759 | 0.35816049 | 0.58356258 |
| TCEA2 | 107.241982 | -0.050291047 | 0.32133424 | -0.156507 | 0.87563343 | 0.94402889 |
| PCMTD2 | 923.370122 | -0.155022769 | 0.30012932 | -0.5165199 | 0.60549134 | 0.78178941 |
| LINC00266-1 | 48.7673364 | 0.470454235 | 0.30609134 | 1.53697335 | 0.12429984 | 0.30242347 |
| PLPP2 | 143.969395 | -2.791602768 | 0.57242587 | -4.8767935 | 1.08E-06 | 4.40E-05 |
| MIER2 | 154.515596 | -0.133906232 | 0.26193053 | -0.511228 | 0.60919139 | 0.78497795 |
| SHC2 | 88.142538 | -1.137460705 | 0.63755849 | -1.7840884 | 0.0744093 | 0.21658714 |
| MADCAM1 | 53.6312463 | 0.285594752 | 0.36699531 | 0.77819728 | 0.43645272 | 0.65341766 |
| TPGS1 | 176.064434 | -0.965123493 | 0.34851561 | -2.7692404 | 0.00561872 | 0.03658223 |
| CDC34 | 125.743023 | -0.091267694 | 0.24971385 | -0.3654891 | 0.71474633 | 0.85752192 |
| BSG | 781.879418 | -0.37700933 | 0.38527331 | -0.9785503 | 0.32780221 | 0.55299353 |
| POLRMT | 69.9808587 | 0.660127532 | 0.2607505 | 2.53164436 | 0.01135291 | 0.06063912 |
| RNF126 | 183.755622 | -0.438837919 | 0.205171 | -2.1388886 | 0.03244469 | 0.12443725 |
| FSTL3 | 131.800481 | -0.965315173 | 0.63511323 | -1.5199104 | 0.12853351 | 0.30953493 |
| PALM | 521.115271 | 1.201783784 | 0.42519667 | 2.82641861 | 0.00470717 | 0.03193478 |
| LINC01836 | 84.8163409 | 1.205917773 | 0.61188324 | 1.97082987 | 0.04874334 | 0.1632722 |
| PTBP1 | 592.203649 | -0.245585839 | 0.24006195 | -1.0230103 | 0.30630299 | 0.53011734 |
| MED16 | 178.93478 | -0.255404495 | 0.3583382 | -0.712747 | 0.4760023 | 0.68588572 |
| R3HDM4 | 193.071415 | -1.061112176 | 0.33988695 | -3.1219562 | 0.00179654 | 0.01570513 |
| ARID3A | 733.900813 | -0.609910014 | 0.42539115 | -1.4337628 | 0.15163996 | 0.3446932 |
| WDR18 | 127.342783 | 0.022045092 | 0.49104441 | 0.04489429 | 0.96419157 | 0.98423542 |
| TMEM259 | 548.079566 | -0.053201477 | 0.30432074 | -0.1748204 | 0.86122075 | 0.93697711 |
| CNN2 | 173.669233 | -0.284062907 | 0.33158647 | -0.8566782 | 0.39162274 | 0.61414517 |
| ABCA7 | 134.56848 | 0.245450191 | 0.41247961 | 0.59506018 | 0.55180324 | 0.74481373 |
| POLR2E | 215.428662 | -0.596281521 | 0.27882516 | -2.1385499 | 0.03247214 | 0.12450638 |
| GPX4 | 691.418205 | -0.943135118 | 0.38141438 | -2.4727309 | 0.01340851 | 0.06766498 |
| SBNO2 | 1091.57107 | -2.293141707 | 0.35712588 | -6.421102 | 1.35E-10 | 2.23E-08 |
| STK11 | 412.320637 | -0.158014752 | 0.21837601 | -0.7235903 | 0.46931732 | 0.68108245 |
| ATP5F1D | 597.875606 | -0.198076212 | 0.42197724 | -0.4694002 | 0.63878359 | 0.80566146 |
| MIDN | 696.209946 | -1.910972795 | 0.37952397 | -5.0351834 | 4.77E-07 | 2.29E-05 |
| CIRBP | 1192.75233 | 0.485071664 | 0.26383349 | 1.8385523 | 0.06598106 | 0.20021683 |
| C19orf24 | 134.664358 | -0.918419311 | 0.36913627 | -2.4880224 | 0.01284556 | 0.06595294 |
| MUM1 | 564.50179 | -0.111541518 | 0.18553513 | -0.6011881 | 0.54771469 | 0.74164667 |
| NDUFS7 | 230.988536 | 0.217492171 | 0.32604065 | 0.66707072 | 0.50472696 | 0.70859746 |
| GAMT | 57.3376622 | -0.154241206 | 0.56259821 | -0.2741587 | 0.78396265 | 0.89672347 |
| DAZAP1 | 741.303633 | 0.203243906 | 0.19902404 | 1.02120282 | 0.30715836 | 0.53077639 |
| RPS15 | 1426.97057 | -1.359343759 | 0.42126598 | -3.2268064 | 0.0012518 | 0.01193004 |
| C19orf25 | 438.058912 | -0.440060525 | 0.24626319 | -1.7869521 | 0.07394523 | 0.21566406 |
| PCSK4 | 71.5074596 | 0.681564326 | 0.38006218 | 1.79329691 | 0.07292546 | 0.21387043 |
| MBD3 | 163.403868 | -0.337049383 | 0.36045629 | -0.9350631 | 0.34975577 | 0.57578522 |
| TCF3 | 257.578774 | -0.304230251 | 0.31502288 | -0.9657402 | 0.33417421 | 0.5593519 |
| REXO1 | 220.882526 | 0.020640971 | 0.25166778 | 0.08201674 | 0.9346334 | 0.97183581 |
| KLF16 | 47.0902523 | -1.680898605 | 0.51838431 | -3.2425723 | 0.00118456 | 0.0114044 |
| AC012615.1 | 47.9718542 | 0.106093569 | 0.33087449 | 0.32064596 | 0.7484787 | 0.87725423 |
| ABHD17A | 224.676249 | -0.429917477 | 0.29587459 | -1.4530395 | 0.14621278 | 0.33676104 |
| SCAMP4 | 353.508609 | -0.191434158 | 0.28778148 | -0.6652067 | 0.50591832 | 0.70948881 |
| CSNK1G2 | 414.218239 | -0.823044095 | 0.20057921 | -4.103337 | 4.07E-05 | 0.00083918 |
| BTBD2 | 463.287013 | -0.237966526 | 0.31314142 | -0.7599331 | 0.44729458 | 0.66320446 |
| MKNK2 | 184.099622 | -0.711482785 | 0.349017 | -2.0385333 | 0.04149663 | 0.14589537 |
| MOB3A | 255.777611 | -0.61039226 | 0.34543488 | -1.7670256 | 0.07722395 | 0.22166902 |
| AP3D1 | 614.167874 | -0.364365685 | 0.21029853 | -1.7326116 | 0.08316472 | 0.23195161 |
| DOT1L | 481.558813 | 0.08315499 | 0.32363778 | 0.25693845 | 0.79722629 | 0.90499845 |
| PLEKHJ1 | 269.197651 | 0.01076993 | 0.30469387 | 0.03534672 | 0.97180327 | 0.98750283 |
| SF3A2 | 120.113107 | -0.182268449 | 0.33422374 | -0.5453486 | 0.58551378 | 0.76994042 |
| LSM7 | 306.167549 | -0.636750685 | 0.34310271 | -1.8558603 | 0.06347347 | 0.19518591 |

| | | | | | | |
|---|---|---|---|---|---|---|
| SPPL2B | 492.242197 | 0.173819363 | 0.18968526 | 0.91635675 | 0.35947981 | 0.58491856 |
| TMPRSS9 | 41.672413 | -0.419634131 | 0.44269964 | -0.9478981 | 0.34318135 | 0.56819785 |
| TIMM13 | 212.90643 | -0.654788342 | 0.41417146 | -1.5809596 | 0.11388728 | 0.28637676 |
| LMNB2 | 66.2726146 | -0.131644569 | 0.2700232 | -0.4875306 | 0.62588237 | 0.79661594 |
| GADD45B | 701.177508 | -1.736606033 | 0.38585109 | -4.5007156 | 6.77E-06 | 0.00019363 |
| GNG7 | 1310.5726 | 1.077052686 | 0.33766334 | 3.18972347 | 0.00142409 | 0.01315441 |
| SGTA | 230.712602 | -0.260335716 | 0.2464745 | -1.0562379 | 0.29085953 | 0.51363149 |
| THOP1 | 141.824899 | 0.60587442 | 0.29675848 | 2.04164148 | 0.04118711 | 0.14519362 |
| ZNF554 | 104.148402 | 0.002051955 | 0.33574702 | 0.00611161 | 0.99512367 | 0.99807051 |
| ZNF555 | 107.413822 | -0.232324926 | 0.31034352 | -0.7486057 | 0.45409491 | 0.66958468 |
| ZNF57 | 71.6178268 | 0.198162914 | 0.23635429 | 0.83841472 | 0.40179782 | 0.62329389 |
| AC119403.1 | 48.7305819 | 0.480396229 | 0.25220762 | 1.9047649 | 0.05681064 | 0.18134648 |
| ZNF77 | 65.7781799 | 0.166654535 | 0.23973312 | 0.69516693 | 0.48695069 | 0.69416487 |
| TLE2 | 236.370871 | -0.913324253 | 0.62088039 | -1.4710148 | 0.14128711 | 0.32966993 |
| AES | 1001.74033 | -0.419065357 | 0.40053434 | -1.0462657 | 0.29543835 | 0.51851143 |
| GNA11 | 296.756771 | -0.519979348 | 0.31222805 | -1.6653832 | 0.09583633 | 0.25517074 |
| NCLN | 208.201058 | 0.298668015 | 0.2507255 | 1.19121518 | 0.23356912 | 0.45046205 |
| NFIC | 2037.58185 | -0.305227847 | 0.2307847 | -1.3225654 | 0.18597996 | 0.39174123 |
| SMIM24 | 78.3133899 | -0.127892282 | 0.58203237 | -0.219734 | 0.82607835 | 0.91938879 |
| DOHH | 46.1671298 | -0.20666943 | 0.28344728 | -0.7291283 | 0.46592319 | 0.67884176 |
| FZR1 | 182.261347 | -0.164239941 | 0.22097323 | -0.7432572 | 0.45732599 | 0.67216888 |
| MFSD12 | 281.335884 | -0.336507682 | 0.20190515 | -1.6666622 | 0.0955816 | 0.25485211 |
| C19orf71 | 70.3877927 | 0.434124014 | 0.42284496 | 1.0266742 | 0.3045739 | 0.52789879 |
| AC005786.3 | 115.49429 | 0.425977077 | 0.30897267 | 1.37868852 | 0.16799081 | 0.36799181 |
| HMG20B | 216.872138 | -0.225320628 | 0.2541894 | -0.8864281 | 0.37538688 | 0.59870627 |
| CACTIN | 66.2033013 | 0.34253498 | 0.32776143 | 1.04507408 | 0.29598873 | 0.51906534 |
| PIP5K1C | 211.236591 | -0.343182406 | 0.26000206 | -1.3199219 | 0.18686111 | 0.39252537 |
| TJP3 | 235.882823 | 0.127708567 | 0.29251745 | 0.43658444 | 0.66241275 | 0.82190799 |
| APBA3 | 81.8121964 | 0.062640469 | 0.29657475 | 0.21121309 | 0.832721 | 0.92176416 |
| MRPL54 | 168.501203 | -0.930407187 | 0.36969273 | -2.516704 | 0.01184583 | 0.06245523 |
| DAPK3 | 236.031666 | -0.591244831 | 0.21441034 | -2.7575389 | 0.00582383 | 0.03754366 |
| EEF2 | 1097.37519 | -1.127542539 | 0.4346429 | -2.5941814 | 0.00948164 | 0.05329491 |
| PIAS4 | 218.296698 | 0.365241644 | 0.28495718 | 1.28174219 | 0.1999331 | 0.4087473 |
| ZBTB7A | 548.686907 | -0.50122259 | 0.3034738 | -1.6516174 | 0.09861258 | 0.2600774 |
| MAP2K2 | 460.942635 | -0.324883533 | 0.21179211 | -1.5339737 | 0.12503612 | 0.30354754 |
| SIRT6 | 128.392033 | -0.607149181 | 0.25188671 | -2.4104058 | 0.01593478 | 0.07637249 |
| ANKRD24 | 128.815267 | 1.110719849 | 0.63872587 | 1.73896173 | 0.0820415 | 0.23027754 |
| YJU2 | 119.585139 | 0.216354063 | 0.35695248 | 0.60611447 | 0.54443874 | 0.73944091 |
| STAP2 | 82.9760162 | -0.844297772 | 0.30943522 | -2.7285122 | 0.00636207 | 0.03994136 |
| MPND | 41.5358611 | -0.044364683 | 0.2795507 | -0.1586999 | 0.87390528 | 0.94333543 |
| SH3GL1 | 366.983648 | -0.586548436 | 0.28587344 | -2.0517766 | 0.04019138 | 0.14298087 |
| CHAF1A | 55.434177 | -0.378652287 | 0.50210439 | -0.7541306 | 0.4507708 | 0.66654333 |
| UBXN6 | 259.208094 | -0.724580739 | 0.33303312 | -2.1757017 | 0.02957757 | 0.11638074 |
| HDGFL2 | 195.6566 | 0.012360907 | 0.26091372 | 0.04737546 | 0.96221399 | 0.9832997 |
| PLIN4 | 111.162512 | 0.316706754 | 0.30152853 | 1.05033761 | 0.29356292 | 0.51660973 |
| PLIN5 | 912.751432 | -0.006695573 | 0.29548511 | -0.0226596 | 0.98192181 | 0.99251646 |
| MYDGF | 282.310441 | -0.882836719 | 0.31393512 | -2.812163 | 0.00492096 | 0.03302892 |
| DPP9 | 507.839501 | -0.268836822 | 0.21242922 | -1.265536 | 0.20567926 | 0.41617913 |
| FEM1A | 52.4853316 | 0.248102429 | 0.28822399 | 0.86079729 | 0.3893497 | 0.61239821 |
| TICAM1 | 156.89975 | -0.804987835 | 0.29105255 | -2.7657817 | 0.00567865 | 0.03685962 |
| PLIN3 | 429.49948 | -1.756381096 | 0.26483129 | -6.6320756 | 3.31E-11 | 6.83E-09 |
| KDM4B | 1157.12801 | -0.581482815 | 0.25986317 | -2.23765 | 0.02524389 | 0.10419747 |
| PTPRS | 667.670848 | 0.708423619 | 0.3157416 | 2.24368161 | 0.02485289 | 0.10329826 |
| TINCR | 88.0878207 | -0.463247615 | 0.32797925 | -1.4124296 | 0.15782349 | 0.35383118 |
| SAFB2 | 783.131798 | 0.268050551 | 0.1503275 | 1.78311052 | 0.07456831 | 0.21676339 |
| SAFB | 933.96109 | 0.092601455 | 0.15110133 | 0.6128434 | 0.53997989 | 0.73609563 |
| RPL36 | 830.245596 | -1.187610467 | 0.45278558 | -2.6228981 | 0.00871853 | 0.05018 |
| C19orf70 | 311.7152 | -0.27167685 | 0.30604311 | -0.8877078 | 0.37469796 | 0.59818533 |
| LONP1 | 431.603367 | -0.497461887 | 0.22272015 | -2.2335738 | 0.02551113 | 0.10491174 |
| DUS3L | 53.4591227 | -0.516237221 | 0.32980722 | -1.5652696 | 0.1175197 | 0.29215978 |
| AC104532.2 | 122.483928 | 0.271647949 | 0.43329116 | 0.62694089 | 0.53069798 | 0.72861184 |
| CAPS | 172.162774 | -0.350963664 | 0.46451498 | -0.7555486 | 0.44991986 | 0.66603064 |
| RANBP3 | 325.840629 | 0.082803808 | 0.22560496 | 0.36703008 | 0.71359658 | 0.85667582 |
| AC011444.1 | 51.3427033 | 0.353604817 | 0.31230952 | 1.13222556 | 0.25753961 | 0.47845862 |
| RFX2 | 617.685049 | -0.277766143 | 0.41204432 | -0.6741172 | 0.50023683 | 0.70482771 |
| MLLT1 | 308.272261 | -0.203950053 | 0.22785558 | -0.8950847 | 0.3707418 | 0.59532317 |
| CLPP | 150.786722 | -1.028034062 | 0.31090429 | -3.3065933 | 0.00094438 | 0.00953075 |
| ALKBH7 | 132.214332 | -0.787115949 | 0.49666065 | -1.5848164 | 0.11300805 | 0.28481651 |
| GTF2F1 | 189.009023 | -0.716423838 | 0.25803619 | -2.7764472 | 0.00549566 | 0.03599015 |
| KHSRP | 287.837537 | -0.042277363 | 0.26848533 | -0.1574662 | 0.87487745 | 0.94367599 |
| SLC25A23 | 273.548905 | -0.651117837 | 0.33289302 | -1.9559372 | 0.05047256 | 0.16730038 |
| CRB3 | 69.3971843 | -1.048662889 | 0.33910182 | -3.092472 | 0.00198497 | 0.01702562 |
| AC010503.4 | 91.4557457 | -0.675961064 | 0.31336387 | -2.1571123 | 0.03099691 | 0.12067978 |
| C3 | 382.579571 | -4.96153294 | 0.83880409 | -5.915008 | 3.32E-09 | 3.81E-07 |
| GPR108 | 84.2961553 | -0.497043525 | 0.3389845 | -1.4662721 | 0.14257417 | 0.33167704 |
| TRIP10 | 334.542203 | -1.125555115 | 0.26940776 | -4.1778867 | 2.94E-05 | 0.00064452 |
| SH2D3A | 393.806052 | -1.263232899 | 0.28446497 | -4.4407327 | 8.97E-06 | 0.0002462 |
| ZNF557 | 125.992022 | 0.092803662 | 0.23928992 | 0.38782939 | 0.69814229 | 0.84618843 |
| INSR | 2481.32984 | 0.385945132 | 0.32721702 | 1.17947756 | 0.23820807 | 0.45614418 |
| AC119396.1 | 52.7014145 | -0.413699505 | 0.43322921 | -0.9549206 | 0.33961784 | 0.56498892 |
| PEX11G | 46.0522486 | -0.140201247 | 0.38161989 | -0.3673845 | 0.71333221 | 0.85650442 |
| ZNF358 | 129.25693 | -0.324728277 | 0.40000489 | -0.8118108 | 0.41690023 | 0.63677557 |
| MCOLN1 | 110.451089 | 0.009939163 | 0.36886483 | 0.02694527 | 0.97850338 | 0.990805 |
| PNPLA6 | 253.289503 | -0.284967174 | 0.30596037 | -0.9313859 | 0.35165398 | 0.57744872 |
| CAMSAP3 | 223.709006 | 0.068855455 | 0.24635335 | 0.27949876 | 0.77986209 | 0.8949616 |
| XAB2 | 103.915201 | -0.167366247 | 0.33629441 | -0.4976778 | 0.61871119 | 0.79099459 |
| PCP2 | 73.0893778 | 1.475479123 | 0.49217075 | 2.99790088 | 0.00271846 | 0.02150189 |
| AC008763.1 | 53.421624 | 0.529164264 | 0.43086389 | 1.22814715 | 0.21939173 | 0.43356369 |
| STXBP2 | 129.268249 | -0.265922213 | 0.37284837 | -0.7132181 | 0.4757108 | 0.68561527 |
| EVI5L | 186.904955 | -0.182824086 | 0.29214346 | -0.6258024 | 0.53144455 | 0.72880513 |
| MAP2K7 | 395.238232 | -0.633181569 | 0.30031499 | -2.1083915 | 0.03499714 | 0.13084552 |
| TIMM44 | 223.731033 | 0.295662834 | 0.20344879 | 1.45325431 | 0.14615316 | 0.33668243 |
| ELAVL1 | 607.646281 | -0.080640945 | 0.17108413 | -0.4713526 | 0.63738897 | 0.80497022 |
| AC008946.1 | 51.8447398 | 0.689199451 | 0.41785313 | 1.64938204 | 0.09906939 | 0.2609387 |
| FBN3 | 42.5707487 | 0.571119762 | 0.75413826 | 0.75731439 | 0.44886153 | 0.66462361 |
| CERS4 | 534.346053 | 0.040847806 | 0.33485606 | 0.12198616 | 0.90290998 | 0.95509228 |
| AC022146.2 | 170.6657 | -0.214666667 | 0.50733733 | -0.4231241 | 0.6722047 | 0.82789295 |
| CD320 | 95.3654408 | -0.30410468 | 0.39340033 | -0.7730158 | 0.43951301 | 0.6552514 |
| RPS28 | 365.283089 | -0.461780522 | 0.39005698 | -1.1838796 | 0.23646071 | 0.45398395 |
| KANK3 | 45.030563 | 0.555424952 | 0.5332711 | 1.04154331 | 0.29762346 | 0.52095615 |
| ANGPTL4 | 455.642664 | -1.579828255 | 1.17592578 | -1.3434762 | 0.17911784 | 0.3825983 |
| RAB11B-AS1 | 127.248806 | -0.253264782 | 0.46258863 | -0.5474946 | 0.58403897 | 0.76885817 |
| RAB11B | 320.935626 | -0.260409222 | 0.33947382 | -0.7670966 | 0.44302406 | 0.65884991 |
| 2-Mar | 181.600332 | -0.288114138 | 0.24495457 | -1.1761942 | 0.2395173 | 0.45759098 |
| HNRNPM | 1473.90627 | 0.131171019 | 0.13485734 | 0.97266507 | 0.33071979 | 0.55588514 |
| ZNF414 | 56.273374 | -0.00801404 | 0.37754459 | -0.0212267 | 0.98306478 | 0.9928728 |
| ZNF558 | 134.759729 | 0.166502813 | 0.21902334 | 0.76020579 | 0.44713159 | 0.66309208 |
| ZNF317 | 165.383885 | 0.172988871 | 0.15971888 | 1.08308343 | 0.27877139 | 0.50097041 |
| ZNF699 | 105.611452 | -0.029943132 | 0.30590001 | -0.0978854 | 0.92202333 | 0.96551499 |
| ZNF559-ZNF | 281.060516 | 0.578203565 | 0.29647683 | 1.95024874 | 0.05114648 | 0.16869111 |
| ZNF266 | 316.44181 | -0.346305901 | 0.20860491 | -1.6601043 | 0.09689348 | 0.2572595 |
| ZNF426 | 152.384098 | -0.291743479 | 0.22157444 | -1.3166838 | 0.18794463 | 0.39374474 |
| ZNF121 | 298.736387 | 0.013137529 | 0.18392474 | 0.07142883 | 0.94305647 | 0.97556917 |

| | | | | | | |
|---|---|---|---|---|---|---|
| ZNF561 | 231.359018 | 0.343456202 | 0.24991246 | 1.37430603 | 0.1693467 | 0.36972673 |
| ZNF562 | 495.950433 | 0.041259837 | 0.19186285 | 0.2150486 | 0.82972942 | 0.92030363 |
| ZNF846 | 588.814454 | 0.162656864 | 0.19803585 | 0.82135062 | 0.41144658 | 0.63063331 |
| FBXL12 | 177.224957 | 0.137087433 | 0.17754001 | 0.77214953 | 0.44002587 | 0.65564599 |
| UBL5 | 223.44407 | -1.215324615 | 0.44992057 | -2.7011982 | 0.00690902 | 0.04246682 |
| PIN1 | 200.67079 | -0.481516902 | 0.26237738 | -1.8352074 | 0.06647497 | 0.20102287 |
| OLFM2 | 161.611365 | 0.248820477 | 0.71894457 | 0.34609132 | 0.72927408 | 0.86700908 |
| RDH8 | 292.414297 | 2.152680006 | 1.26009959 | 1.70834117 | 0.08757306 | 0.23999015 |
| C19orf66 | 422.983079 | -0.262900826 | 0.37921438 | -0.6932776 | 0.48813537 | 0.69502858 |
| EIF3G | 162.745262 | -0.545301241 | 0.30680553 | -1.7773514 | 0.07551044 | 0.21863598 |
| DNMT1 | 215.829337 | -0.064539925 | 0.32759877 | -0.1970091 | 0.84382045 | 0.92769172 |
| MRPL4 | 202.549658 | -0.201470283 | 0.27750748 | -0.7259995 | 0.46783909 | 0.6801328 |
| AC011511.2 | 67.4095271 | 0.004275342 | 0.56356839 | 0.0075862 | 0.99394715 | 0.99779966 |
| ICAM1 | 104.520654 | -3.230847984 | 0.51839332 | -6.232426 | 4.59E-10 | 6.39E-08 |
| ZGLP1 | 37.9818508 | 1.033417044 | 0.60106566 | 1.71930807 | 0.08555829 | 0.23643682 |
| RAVER1 | 75.9110829 | 0.004683348 | 0.34700062 | 0.01349666 | 0.98923155 | 0.99594533 |
| TYK2 | 365.437819 | 0.117162827 | 0.3418713 | 0.34271033 | 0.73181639 | 0.86826096 |
| CDC37 | 484.143503 | -0.599367424 | 0.23043839 | -2.6009878 | 0.00929558 | 0.05256219 |
| PDE4A | 148.411323 | 0.177137925 | 0.54939136 | 0.32242576 | 0.74713017 | 0.87660707 |
| KEAP1 | 152.042388 | -0.563069536 | 0.29696153 | -1.8961026 | 0.05794648 | 0.18377311 |
| ATG4D | 168.340666 | -0.008247882 | 0.19496991 | -0.0423034 | 0.96625687 | 0.984926 |
| KRI1 | 181.559986 | -0.513212658 | 0.246754 | -2.0798555 | 0.03753879 | 0.13712662 |
| CDKN2D | 44.0053053 | -0.882413192 | 0.39499641 | -2.2339777 | 0.02548454 | 0.10486463 |
| AP1M2 | 275.078796 | -0.670571159 | 0.32229284 | -2.0806269 | 0.03746807 | 0.13691315 |
| SLC44A2 | 416.140864 | -0.225593904 | 0.39174806 | -0.5758648 | 0.56470659 | 0.75596786 |
| ILF3-DT | 263.588857 | -0.136446147 | 0.32030452 | -0.4259888 | 0.67011598 | 0.82659266 |
| ILF3 | 797.116073 | 0.137343038 | 0.23284135 | 0.58985673 | 0.5552867 | 0.747608 |
| QTRT1 | 358.296035 | 1.042955959 | 0.28889709 | 3.6101297 | 0.00030604 | 0.00407103 |
| DNM2 | 2811.7189 | -0.285355592 | 0.31503451 | -0.9057915 | 0.3650462 | 0.59048436 |
| TMED1 | 95.7313976 | -0.897717517 | 0.40942481 | -2.192631 | 0.02833398 | 0.11286596 |
| CARM1 | 373.99099 | 0.095197369 | 0.30389458 | 0.31325787 | 0.75408476 | 0.88075281 |
| YIPF2 | 216.548734 | 0.009573027 | 0.21261425 | 0.04502533 | 0.96408712 | 0.98420487 |
| TIMM29 | 137.335181 | 0.223753322 | 0.27871865 | 0.80279279 | 0.42209451 | 0.64114933 |
| SMARCA4 | 1343.01484 | -0.019735281 | 0.37341822 | -0.0528503 | 0.95785115 | 0.98155592 |
| LDLR | 538.704832 | -1.496443452 | 0.54771754 | -2.7321445 | 0.00629235 | 0.03957891 |
| KANK2 | 378.717689 | 0.121259665 | 0.26392118 | 0.45945408 | 0.64590812 | 0.81059872 |
| DOCK6 | 640.333756 | -0.431989281 | 0.38826173 | -1.1126239 | 0.26586999 | 0.48688155 |
| AC011472.1 | 41.762712 | 0.566206645 | 0.39938667 | 1.41769039 | 0.15628118 | 0.35131349 |
| RAB3D | 202.463224 | -1.167427804 | 0.28756743 | -4.0596663 | 4.91E-05 | 0.00096102 |
| CCDC159 | 64.6503954 | 0.496653548 | 0.38318341 | 1.29612485 | 0.19493247 | 0.40245308 |
| PLPPR2 | 53.9574545 | -0.141839298 | 0.33034545 | -0.4293666 | 0.66765646 | 0.82486496 |
| SWSAP1 | 63.8349983 | 0.380468314 | 0.2565886 | 1.48279506 | 0.13812887 | 0.32413293 |
| EPOR | 252.264418 | 0.592209194 | 0.47829892 | 1.23815707 | 0.21565783 | 0.42946242 |
| RGL3 | 2354.5512 | -0.05153518 | 0.30025985 | -0.1716353 | 0.86372428 | 0.9382347 |
| CCDC151 | 45.5779376 | 1.025338394 | 0.39448739 | 2.59916645 | 0.00934504 | 0.05271645 |
| PRKCSH | 251.215923 | -0.426641061 | 0.25098914 | -1.6998388 | 0.08916126 | 0.24298145 |
| ECSIT | 310.810106 | 0.394940776 | 0.2645574 | 1.49283584 | 0.13548016 | 0.32065176 |
| ELOF1 | 96.7905947 | -0.692735552 | 0.38075176 | -1.819389 | 0.06885211 | 0.20590164 |
| ZNF627 | 378.198237 | 0.463396561 | 0.26471901 | 1.75052239 | 0.08002821 | 0.22635818 |
| AC008543.1 | 182.027254 | 0.325089695 | 0.46388881 | 0.70079227 | 0.48343266 | 0.69153801 |
| ZNF823 | 345.14218 | -0.32948794 | 0.19420971 | -1.6965575 | 0.08978035 | 0.24426542 |
| ZNF441 | 265.987499 | -0.249299112 | 0.2698795 | -0.9237423 | 0.35562049 | 0.58105953 |
| ZNF491 | 73.2897757 | 0.628379244 | 0.2389759 | 2.62946699 | 0.00855188 | 0.04950124 |
| ZNF440 | 325.640552 | -0.470257914 | 0.2119688 | -2.2185242 | 0.02651911 | 0.10794791 |
| ZNF439 | 547.980821 | 0.268401517 | 0.18849233 | 1.4239387 | 0.15446422 | 0.34881127 |
| ZNF69 | 819.818732 | 0.044509724 | 0.24112982 | 0.18458822 | 0.85355203 | 0.93286188 |
| AC008770.2 | 121.19395 | 0.369659351 | 0.22226989 | 1.6631103 | 0.09629036 | 0.25606993 |
| ZNF763 | 57.482723 | 0.581307108 | 0.48446623 | 1.19989191 | 0.23018132 | 0.445816 |
| ZNF433 | 106.211308 | 0.399919462 | 0.35277127 | 1.13365089 | 0.25694102 | 0.47778941 |
| AC008770.1 | 39.7340958 | 0.172482897 | 0.35985563 | 0.47931138 | 0.63171713 | 0.80129239 |
| ZNF844 | 348.788496 | 0.720319079 | 0.25453438 | 2.82994806 | 0.00465556 | 0.03175016 |
| ZNF788P | 207.865481 | 1.041349857 | 0.28133013 | 3.70152266 | 0.00021431 | 0.00311078 |
| ZNF20 | 96.8950511 | 0.607062467 | 0.40225285 | 1.50915642 | 0.13125881 | 0.31392317 |
| ZNF136 | 185.491941 | 0.21211539 | 0.17001931 | 1.24759587 | 0.21217909 | 0.42488177 |
| ZNF44 | 2087.98228 | 0.242105783 | 0.19559185 | 1.23781122 | 0.21578607 | 0.42958827 |
| ZNF563 | 115.505564 | -0.180652819 | 0.36504582 | -0.4948771 | 0.62068687 | 0.7928889 |
| ZNF442 | 90.0041138 | 0.154239196 | 0.34253595 | 0.45028616 | 0.65250412 | 0.8147974 |
| AC008758.5 | 156.072099 | -0.744294825 | 0.55818514 | -1.3334193 | 0.18239425 | 0.38696421 |
| AC008758.6 | 81.3665552 | -0.552803693 | 0.4187862 | -1.3200141 | 0.1868303 | 0.39252537 |
| AC008758.1 | 147.521944 | -0.122167971 | 0.44645824 | -0.2736381 | 0.78436277 | 0.89699853 |
| AC010422.6 | 43.529012 | -0.162277363 | 0.30146951 | -0.5382878 | 0.59037837 | 0.77217181 |
| ZNF490 | 391.78003 | -0.206500385 | 0.24064355 | -0.8581173 | 0.39082772 | 0.61362693 |
| ZNF791 | 503.381653 | -0.335142826 | 0.12951033 | -2.587769 | 0.00965997 | 0.05399856 |
| MAN2B1 | 124.219086 | -0.098252376 | 0.27025061 | -0.3635602 | 0.71618642 | 0.85844886 |
| WDR83 | 102.09712 | -0.167837215 | 0.24493222 | -0.6852394 | 0.49319285 | 0.69881955 |
| DHPS | 156.120383 | -0.004095154 | 0.22312547 | -0.0183536 | 0.98535678 | 0.99423861 |
| FBXW9 | 56.8593935 | -0.318663325 | 0.38684319 | -0.8237532 | 0.41007979 | 0.6297833 |
| TNPO2 | 412.818254 | -0.038916659 | 0.21001062 | -0.1853081 | 0.85298742 | 0.93247628 |
| TRIR | 423.152057 | -0.684241602 | 0.2151531 | -3.1802545 | 0.00147146 | 0.01350694 |
| ASNA1 | 132.401251 | -0.839165948 | 0.36941509 | -2.2716071 | 0.02311025 | 0.09830568 |
| HOOK2 | 1239.79671 | -0.490685674 | 0.24553729 | -1.9984161 | 0.04567157 | 0.15600761 |
| JUNB | 441.241433 | -1.703042643 | 0.50374254 | -3.3807799 | 0.0007228 | 0.00784513 |
| DNASE2 | 62.1509931 | -1.108932084 | 0.43904911 | -2.5257586 | 0.01154488 | 0.06136671 |
| GCDH | 147.189056 | 0.042764919 | 0.31735599 | 0.13475378 | 0.89280655 | 0.9512084 |
| FARSA | 180.490145 | -0.191759979 | 0.26432941 | -0.7254584 | 0.46817088 | 0.68024081 |
| CALR | 650.734289 | -0.471035246 | 0.51472099 | -0.9151273 | 0.36012479 | 0.58539113 |
| AC092069.1 | 123.183178 | -0.088392669 | 0.59516902 | -0.1485169 | 0.88193483 | 0.94672712 |
| RAD23A | 286.279351 | -0.499364599 | 0.1949597 | -2.5613735 | 0.01042592 | 0.05716728 |
| GADD45GIP1 | 310.800177 | -1.407812237 | 0.45697165 | -3.080743 | 0.00206485 | 0.01757382 |
| NFIX | 111.638848 | -0.227331855 | 0.3129404 | -0.7264382 | 0.4675702 | 0.68009231 |
| TRMT1 | 100.503115 | 0.054466305 | 0.35445422 | 0.15366245 | 0.87787589 | 0.94536627 |
| NACC1 | 151.877571 | -0.5129342 | 0.28733582 | -1.7851384 | 0.07423888 | 0.21628172 |
| STX10 | 119.075625 | -0.012723381 | 0.26924778 | -0.0472553 | 0.96230977 | 0.9832997 |
| IER2 | 357.536702 | -0.445001319 | 0.43832031 | -1.0152423 | 0.30999034 | 0.53392391 |
| CACNA1A | 63.4679673 | -0.064534189 | 0.32866372 | -0.1963532 | 0.84433369 | 0.92790115 |
| CCDC130 | 680.635971 | -0.265688851 | 0.23008701 | -1.1547321 | 0.24820016 | 0.46734754 |
| MRI1 | 370.0698 | 0.564780406 | 0.2916307 | 1.93662875 | 0.05279074 | 0.17238885 |
| AC008686.1 | 44.979373 | 0.658880202 | 0.68519204 | 0.96159933 | 0.33625091 | 0.5617555 |
| C19orf53 | 361.097141 | -1.44024983 | 0.45510913 | -3.1646253 | 0.00155283 | 0.01408306 |
| ZSWIM4 | 291.706396 | -1.487858555 | 0.32870123 | -4.526477 | 6.00E-06 | 0.00017566 |
| AC020916.1 | 107.576097 | -0.276596938 | 0.51750251 | -0.5344842 | 0.59300655 | 0.77387448 |
| CC2D1A | 406.113962 | -0.323131339 | 0.22434166 | -1.4403537 | 0.14976735 | 0.34200906 |
| DCAF15 | 131.021546 | -0.709346222 | 0.37029083 | -1.9156462 | 0.05541014 | 0.17812425 |
| RFX1 | 335.151821 | -0.029701523 | 0.25928514 | -0.1145516 | 0.90880056 | 0.95876587 |
| PALM3 | 153.498534 | 2.076512392 | 0.56970042 | 3.6449199 | 0.00026748 | 0.00369569 |
| MISP3 | 83.5641175 | 0.276152768 | 0.51648257 | 0.53467974 | 0.59287133 | 0.77387448 |
| SAMD1 | 99.876217 | -0.871058431 | 0.48980762 | -1.7783685 | 0.07534336 | 0.2182479 |
| PRKACA | 265.205777 | -0.33305369 | 0.26342615 | -1.2643152 | 0.20611691 | 0.41687311 |
| AC022098.1 | 103.331378 | 0.105513684 | 0.2450778 | 0.43053138 | 0.66680914 | 0.82439929 |
| ADGRL1 | 241.03816 | -0.11294002 | 0.36344199 | -0.3107512 | 0.7559898 | 0.88214393 |
| DDX39A | 89.9914767 | -0.085425416 | 0.30888018 | -0.2765649 | 0.78211422 | 0.89588513 |
| PKN1 | 191.112995 | -0.601147942 | 0.35947286 | -1.6723041 | 0.09446437 | 0.25284294 |
| GIPC1 | 762.512861 | -0.886461946 | 0.33369694 | -2.6564881 | 0.00789592 | 0.04689625 |

| | | | | | | |
|---|---|---|---|---|---|---|
| DNAJB1 | 183.756387 | -1.186427084 | 0.31146891 | -3.8091349 | 0.00013945 | 0.00217736 |
| TECR | 277.631428 | -0.139031357 | 0.25668887 | -0.5416337 | 0.58807084 | 0.7710768 |
| NDUFB7 | 414.089535 | -0.275813458 | 0.36578781 | -0.7540258 | 0.45083372 | 0.66656174 |
| ADGRE3 | 53.6386167 | -0.674432546 | 0.31221427 | -2.1601592 | 0.03076034 | 0.11992721 |
| ZNF333 | 138.208933 | 0.000263789 | 0.4122513 | 0.00063987 | 0.99948945 | 0.99975417 |
| SYDE1 | 42.4634893 | -0.083452804 | 0.36317122 | -0.2297891 | 0.81825562 | 0.91549538 |
| ILVBL | 188.519325 | 0.06652483 | 0.29398124 | 0.22628937 | 0.82097636 | 0.91698603 |
| NOTCH3 | 234.093445 | 0.650743027 | 0.49608052 | 1.31176896 | 0.1895981 | 0.39570253 |
| BRD4 | 1987.99152 | -0.699160993 | 0.20561467 | -3.4003459 | 0.00067301 | 0.00743912 |
| AKAP8 | 623.858715 | -0.387618249 | 0.21050365 | -1.8413849 | 0.06556518 | 0.19941294 |
| AKAP8L | 1209.84192 | 0.057205051 | 0.14568349 | 0.39266668 | 0.69456568 | 0.84378903 |
| WIZ | 118.901149 | -0.175533648 | 0.2936403 | -0.5977846 | 0.54998366 | 0.74319532 |
| TPM4 | 146.619194 | -1.096019216 | 0.49080696 | -2.2330963 | 0.02554259 | 0.10500844 |
| RAB8A | 135.808103 | -0.189492153 | 0.24247844 | -0.7814804 | 0.43451999 | 0.65163199 |
| FAM32A | 117.876508 | -0.731934956 | 0.29775338 | -2.4581919 | 0.01396385 | 0.06965579 |
| AP1M1 | 195.25158 | -0.264892179 | 0.17112577 | -1.5479386 | 0.12163709 | 0.29836662 |
| KLF2 | 60.3857416 | -1.167378859 | 0.88325196 | -1.3216827 | 0.18627382 | 0.39204763 |
| EPS15L1 | 802.837163 | 0.517187876 | 0.19608102 | 2.63762341 | 0.00834892 | 0.04875373 |
| AC008764.1 | 55.2329087 | 0.606067678 | 0.33400409 | 1.81455168 | 0.06959283 | 0.20746202 |
| C19orf44 | 173.528185 | 0.345656413 | 0.23655863 | 1.4611871 | 0.1439641 | 0.33360748 |
| AC008764.8 | 47.5629608 | -0.073549187 | 0.33600118 | -0.2188956 | 0.82673136 | 0.91944889 |
| AC024075.3 | 41.6581957 | 0.401838645 | 0.43508292 | 0.92359095 | 0.35569933 | 0.58105953 |
| TMEM38A | 236.72314 | 0.476176256 | 0.28522507 | 1.66947546 | 0.09502319 | 0.25402981 |
| NWD1 | 93.6608077 | 1.740558553 | 0.4276234 | 4.07030707 | 4.70E-05 | 0.00093541 |
| SIN3B | 550.093922 | 0.111185822 | 0.14222109 | 0.78178155 | 0.43434297 | 0.65144047 |
| CPAMD8 | 68.3729989 | -2.060621181 | 0.75229833 | -2.7391011 | 0.00616074 | 0.03901114 |
| HAUS8 | 72.839034 | -0.113855885 | 0.31610421 | -0.3601847 | 0.71870905 | 0.85990372 |
| MYO9B | 704.704187 | -0.727024905 | 0.28501826 | -2.5508012 | 0.01074756 | 0.05834958 |
| USE1 | 102.632452 | -0.427123532 | 0.28541201 | -1.4965156 | 0.13451934 | 0.31917836 |
| OCEL1 | 83.5575951 | -0.343976353 | 0.30761156 | -1.1182166 | 0.2634745 | 0.48439746 |
| NR2F6 | 208.102617 | -0.96827251 | 0.37715464 | -2.567309 | 0.01024912 | 0.05650278 |
| USHBP1 | 113.849357 | -0.024349604 | 0.26719371 | -0.0911309 | 0.92738858 | 0.96814003 |
| ABHD8 | 148.279661 | 0.095748964 | 0.22306573 | 0.42924103 | 0.66774782 | 0.82486496 |
| MRPL34 | 215.529009 | -0.200077286 | 0.43807337 | -0.456721 | 0.64787163 | 0.81173905 |
| DDA1 | 145.289977 | -0.678210738 | 0.21179116 | -3.2022618 | 0.00136353 | 0.01274655 |
| ANO8 | 64.473951 | 0.390891336 | 0.26971266 | 1.44928805 | 0.14725716 | 0.33822288 |
| GTPBP3 | 92.4426899 | 0.315503863 | 0.28656282 | 1.1009937 | 0.2708994 | 0.49226994 |
| BST2 | 307.78168 | -0.310723849 | 0.75422265 | -0.4119789 | 0.68035486 | 0.8328058 |
| MVB12A | 212.847542 | -0.45953092 | 0.26588937 | -1.7282787 | 0.08393828 | 0.23347383 |
| SLC27A1 | 337.615975 | 0.696279915 | 0.30809194 | 2.25997448 | 0.02382283 | 0.1002791 |
| AC010618.3 | 262.037344 | 0.245946881 | 0.25465455 | 0.96580595 | 0.33414129 | 0.5593519 |
| PGLS | 276.468397 | -0.847679632 | 0.2732754 | -3.1019244 | 0.00192267 | 0.01663053 |
| COLGALT1 | 158.332296 | -0.300654492 | 0.28056885 | -1.071589 | 0.28390467 | 0.50651054 |
| MAP1S | 71.3906036 | -0.611840601 | 0.25619026 | -2.3882274 | 0.01692986 | 0.0796186 |
| RPL18A | 93.4160866 | -1.712723827 | 0.48333544 | -3.5435511 | 0.00039478 | 0.00492409 |
| CCDC124 | 116.493225 | -0.93279071 | 0.30368116 | -3.0716121 | 0.00212906 | 0.01800434 |
| ARRDC2 | 136.708257 | -2.334012851 | 0.48579686 | -4.8045038 | 1.55E-06 | 5.79E-05 |
| MAST3 | 83.8032279 | -0.642225639 | 0.31330867 | -2.0498176 | 0.04038223 | 0.14350521 |
| IQCN | 91.4946402 | 0.278647348 | 0.51365462 | 0.54247998 | 0.5874879 | 0.77084808 |
| JUND | 663.555445 | -0.781251552 | 0.45529458 | -1.7159255 | 0.08617569 | 0.2375902 |
| LSM4 | 250.182434 | -1.02054332 | 0.33190946 | -3.0747642 | 0.00210669 | 0.01787237 |
| PGPEP1 | 377.36737 | -0.650737112 | 0.24573602 | -2.6481144 | 0.00809421 | 0.04767298 |
| GDF15 | 1038.76319 | -2.608087331 | 0.49811333 | -5.2359316 | 1.64E-07 | 9.31E-06 |
| SSBP4 | 295.338865 | -0.445838789 | 0.22986191 | -1.939594 | 0.05242905 | 0.17167459 |
| ISYNA1 | 189.533533 | 1.199465231 | 0.51780901 | 2.31642404 | 0.02053513 | 0.09084009 |
| ELL | 713.433595 | -0.144394978 | 0.21318329 | -0.6773279 | 0.49819796 | 0.70336649 |
| FKBP8 | 302.76186 | -0.672234346 | 0.31941972 | -2.1045486 | 0.03533062 | 0.13164516 |
| KXD1 | 183.052198 | -0.594305513 | 0.18936003 | -3.138495 | 0.00169818 | 0.01508489 |
| AC005253.1 | 46.3462034 | -0.015782211 | 0.54909075 | -0.0287424 | 0.97707 | 0.98973295 |
| UBA52 | 488.23207 | -0.791696728 | 0.45094421 | -1.7556423 | 0.0791495 | 0.22478731 |
| CRLF1 | 54.1112068 | 0.14915479 | 0.33991369 | 0.43880195 | 0.66080505 | 0.82039194 |
| REX1BD | 170.08394 | 0.356684619 | 0.40805981 | 0.87409886 | 0.38206442 | 0.60482849 |
| KLHL26 | 79.0162381 | -0.027409269 | 0.45094398 | -0.060782 | 0.95153284 | 0.97914234 |
| CRTC1 | 970.526413 | 0.174718704 | 0.24653958 | 0.70868419 | 0.47852048 | 0.68816298 |
| UPF1 | 415.801648 | -0.328310699 | 0.1988315 | -1.6512006 | 0.09869762 | 0.26016701 |
| COPE | 241.014496 | -0.630238854 | 0.25020577 | -2.5188822 | 0.0117728 | 0.06225259 |
| DDX49 | 128.099084 | -0.343727842 | 0.37593465 | -0.9143287 | 0.36054415 | 0.58556839 |
| SUGP2 | 1060.54859 | 0.345278344 | 0.16708 | 2.06654497 | 0.03877705 | 0.13975609 |
| ARMC6 | 93.1400299 | 0.447380766 | 0.26683007 | 1.67665051 | 0.09361084 | 0.2517316 |
| SLC25A42 | 221.67288 | 0.306333351 | 0.38323556 | 0.79933435 | 0.42409656 | 0.64296071 |
| TMEM161A | 152.970061 | 0.353435083 | 0.23435993 | 1.50808663 | 0.13153236 | 0.31434972 |
| BORCS8-MEI | 85.7683967 | 0.136732419 | 0.37889385 | 0.36087263 | 0.71819467 | 0.85963642 |
| RFXANK | 96.2613306 | -0.070677679 | 0.23596903 | -0.299521 | 0.76454255 | 0.88572451 |
| SUGP1 | 255.367746 | 0.137016781 | 0.20101233 | 0.6816337 | 0.4954706 | 0.70093939 |
| MAU2 | 747.58842 | -0.074544651 | 0.20942279 | -0.3559529 | 0.72187583 | 0.86151234 |
| GATAD2A | 1326.63148 | -0.546233679 | 0.20163672 | -2.708999 | 0.00674865 | 0.04178612 |
| YJEFN3 | 58.3121445 | 1.184182938 | 0.44975478 | 2.63295241 | 0.00846462 | 0.0491685 |
| PBX4 | 412.022252 | 0.752284458 | 0.36890809 | 2.03921919 | 0.04142816 | 0.1458444 |
| ATP13A1 | 179.816515 | -0.405928059 | 0.36149318 | -1.1229204 | 0.26147131 | 0.48237074 |
| ZNF101 | 129.2743 | -0.323815478 | 0.26508456 | -1.2215554 | 0.22187579 | 0.43599484 |
| ZNF14 | 245.016799 | 0.381332439 | 0.27669985 | 1.37814472 | 0.16815861 | 0.36813567 |
| AC011477.4 | 166.997244 | -0.362255989 | 0.30649674 | -1.1819244 | 0.23723568 | 0.45494281 |
| ZNF253 | 432.296212 | 0.466891677 | 0.16173151 | 2.88683179 | 0.00389142 | 0.02813051 |
| ZNF93 | 137.381814 | -0.015581289 | 0.38191698 | -0.0407976 | 0.96745727 | 0.98528676 |
| ZNF682 | 338.385954 | 0.577509294 | 0.40593584 | 1.42266149 | 0.15483431 | 0.34930937 |
| ZNF90 | 712.995929 | -0.306024128 | 0.36185714 | -0.8457043 | 0.39771774 | 0.61929195 |
| AC011447.3 | 487.935559 | 0.556530043 | 0.29579119 | 1.88149633 | 0.05990443 | 0.18768446 |
| ZNF486 | 429.871656 | -0.709935442 | 0.53195231 | -1.3345848 | 0.18201229 | 0.38652738 |
| AC008554.1 | 534.771359 | -0.292198898 | 0.6725867 | -0.4344405 | 0.6639686 | 0.82281276 |
| ZNF737 | 416.326822 | 0.583457007 | 0.23023242 | 2.53420868 | 0.01127016 | 0.06034355 |
| AC010636.2 | 85.5235845 | -0.890063264 | 0.37760243 | -2.3571439 | 0.01841611 | 0.0843248 |
| ZNF66 | 138.818831 | 0.128221608 | 0.34242742 | 0.37444901 | 0.70807029 | 0.85336683 |
| ZNF85 | 169.504143 | 0.484696771 | 0.44003472 | 1.10149664 | 0.27068056 | 0.49221578 |
| ZNF430 | 428.320717 | -0.1668782 | 0.17789049 | -0.9380951 | 0.34819554 | 0.57383841 |
| ZNF714 | 244.219611 | 0.222093864 | 0.28702144 | 0.77378841 | 0.43905593 | 0.65485132 |
| ZNF431 | 906.349541 | 0.157817542 | 0.15153104 | 1.04148656 | 0.29764979 | 0.52095615 |
| ZNF708 | 642.300497 | -0.113466691 | 0.19480017 | -0.5824774 | 0.56024517 | 0.75183103 |
| ZNF738 | 115.477064 | 0.272761393 | 0.32586442 | 0.83703949 | 0.40257037 | 0.62391114 |
| ZNF493 | 382.323377 | 0.825005982 | 0.20713015 | 3.9830318 | 6.80E-05 | 0.00123626 |
| AC010615.4 | 537.567027 | -0.455879112 | 0.68213076 | -0.6683163 | 0.5039317 | 0.70796131 |
| ZNF429 | 678.939658 | 0.016054106 | 0.2210984 | 0.07261068 | 0.94211593 | 0.97528599 |
| AC123912.1 | 242.021582 | -0.312886226 | 0.60893599 | -0.5138245 | 0.6073747 | 0.78330417 |
| AC123912.4 | 62.2408196 | 0.988155667 | 0.4505663 | 2.19314155 | 0.02829718 | 0.11278741 |
| ZNF100 | 428.813238 | -0.488773244 | 0.44037117 | -1.109912 | 0.26703695 | 0.48822301 |
| ZNF43 | 631.93004 | -0.346141102 | 0.28085144 | -1.2324705 | 0.21777339 | 0.43165797 |
| ZNF208 | 661.758602 | 0.480597073 | 0.44627298 | 1.07691277 | 0.28151924 | 0.50413655 |
| ZNF257 | 134.268211 | -0.067914371 | 0.30850494 | -0.2201403 | 0.82576189 | 0.91938879 |
| ZNF676 | 157.612577 | 1.030333093 | 0.58302034 | 1.76723353 | 0.07718913 | 0.22166902 |
| ZNF98 | 1343.79733 | 1.15866592 | 0.36870431 | 3.14253426 | 0.00167492 | 0.01493858 |
| ZNF99 | 63.5359701 | 0.624584313 | 0.68950349 | 0.90584648 | 0.36501711 | 0.59048436 |
| AC092329.3 | 138.487853 | 0.198344234 | 0.37157035 | 0.53379995 | 0.59347995 | 0.77392186 |
| ZNF91 | 1959.75554 | 0.162729023 | 0.15132558 | 1.07535698 | 0.28221493 | 0.50464019 |
| ZNF675 | 220.940897 | 0.150700129 | 0.20613385 | 0.73107901 | 0.46473089 | 0.67777745 |

| | | | | | | |
|---|---|---|---|---|---|---|
| ZNF681 | 53.0775334 | 0.072738377 | 0.31422151 | 0.23148758 | 0.81693603 | 0.91483008 |
| AC139769.2 | 98.5941979 | 0.165462101 | 0.55166089 | 0.29993444 | 0.76422716 | 0.88557563 |
| ZNF726 | 138.064424 | 0.734293599 | 0.31319335 | 2.34453766 | 0.01905069 | 0.08632608 |
| AC011503.1 | 78.5933664 | 0.262227618 | 0.59201094 | 0.44294387 | 0.65780633 | 0.81863236 |
| ZNF254 | 1784.87747 | -0.229037299 | 0.17652433 | -1.2974829 | 0.19446509 | 0.40198581 |
| LINC00662 | 536.912121 | 0.339555018 | 0.20823559 | 1.63062911 | 0.1029686 | 0.26784409 |
| AC006504.5 | 403.038001 | 0.301017038 | 0.43274352 | 0.69560149 | 0.48667843 | 0.69400145 |
| UQCRFS1 | 500.723859 | 0.379338189 | 0.43286575 | 0.87634143 | 0.38084445 | 0.60354821 |
| POP4 | 75.4704283 | -0.434284522 | 0.29089855 | -1.4929071 | 0.1354615 | 0.32065176 |
| C19orf12 | 346.449115 | -0.414700549 | 0.20870088 | -1.987057 | 0.04691608 | 0.1589419 |
| URI1 | 2045.97879 | 0.30114542 | 0.11731967 | 2.56687923 | 0.01026183 | 0.05654926 |
| TSHZ3 | 80.11672 | -1.039102536 | 0.48751579 | -2.1314234 | 0.03305428 | 0.12618902 |
| ZNF507 | 505.971364 | 0.507410773 | 0.14006514 | 3.62267699 | 0.00029157 | 0.00393798 |
| DPY19L3 | 797.038928 | -0.419520508 | 0.21044077 | -1.9935324 | 0.04620318 | 0.15729324 |
| PDCD5 | 177.474225 | -0.164464495 | 0.19630488 | -0.8378014 | 0.40214227 | 0.6235353 |
| ANKRD27 | 410.349352 | -0.233309565 | 0.18021608 | -1.2946102 | 0.19545475 | 0.40302244 |
| NUDT19 | 144.251242 | 0.706107681 | 0.17637319 | 4.00348644 | 6.24E-05 | 0.00115794 |
| CEP89 | 432.599732 | 0.011100569 | 0.20951363 | 0.05298256 | 0.9577458 | 0.98152411 |
| RHPN2 | 1081.53523 | -1.845759229 | 0.26211444 | -7.0418068 | 1.90E-12 | 5.70E-10 |
| GPATCH1 | 170.625586 | 0.185592393 | 0.22807982 | 0.81371685 | 0.41580718 | 0.6356941 |
| WDR88 | 60.1864328 | 0.550317402 | 0.401591 | 1.37034297 | 0.17057986 | 0.37089538 |
| LRP3 | 81.7021174 | -0.239929163 | 0.40851118 | -0.5873258 | 0.5569849 | 0.74905453 |
| CEBPG | 80.1730352 | -0.076333325 | 0.39463166 | -0.1934293 | 0.84662277 | 0.92890109 |
| PEPD | 350.706797 | -0.29635108 | 0.24574132 | -1.2059473 | 0.22783781 | 0.44334483 |
| KCTD15 | 157.48288 | 0.915386264 | 0.34965619 | 2.617961 | 0.00884569 | 0.05058127 |
| LSM14A | 2487.22217 | 0.139778442 | 0.11402136 | 1.225897 | 0.22023744 | 0.43413167 |
| KIAA0355 | 1295.15615 | -0.433873427 | 0.15000063 | -2.8924774 | 0.00382217 | 0.02773161 |
| AC010504.1 | 43.395341 | 0.618890858 | 0.43597538 | 1.41955462 | 0.15573739 | 0.35068788 |
| GPI | 2012.43435 | -0.065565234 | 0.42020708 | -0.1560308 | 0.87600877 | 0.94420235 |
| UBA2 | 875.567108 | -0.020957085 | 0.17649762 | -0.1187386 | 0.90548243 | 0.95668833 |
| WTIP | 135.723142 | -0.086276203 | 0.34564136 | -0.2496119 | 0.80288748 | 0.9075253 |
| SCGB2B2 | 894.46819 | 0.494481628 | 0.32637718 | 1.5150619 | 0.12975673 | 0.31168515 |
| ZNF302 | 809.677892 | 0.039872242 | 0.19845067 | 0.20091765 | 0.84076297 | 0.92618646 |
| ZNF181 | 126.71304 | 0.342291305 | 0.19421689 | 1.76241784 | 0.07799873 | 0.22315035 |
| ZNF599 | 56.4523462 | 0.163293569 | 0.30869768 | 0.5289757 | 0.59682231 | 0.77646271 |
| ZNF30 | 94.6945959 | 1.328209468 | 0.39803798 | 3.33689129 | 0.00084721 | 0.00878236 |
| GRAMD1A | 422.233021 | -1.155450113 | 0.25825751 | -4.4740233 | 7.68E-06 | 0.00021665 |
| SCN1B | 49.0093034 | 0.298151573 | 0.41599477 | 0.71671953 | 0.47354717 | 0.68421403 |
| HPN | 1964.80456 | 0.546288745 | 0.38617369 | 1.41461927 | 0.15718015 | 0.35273406 |
| LSR | 337.273683 | -1.011526875 | 0.1976129 | -5.1187291 | 3.08E-07 | 1.55E-05 |
| USF2 | 293.032012 | -0.105399978 | 0.27530147 | -0.3828529 | 0.7018288 | 0.8488446 |
| DMKN | 451.051441 | -0.807920625 | 0.26626304 | -3.0342951 | 0.00241099 | 0.01968276 |
| TMEM147-A! | 55.1610749 | 0.475271214 | 0.27138121 | 1.75130481 | 0.07989342 | 0.22617063 |
| TMEM147 | 153.668355 | -0.534778593 | 0.463958 | -1.1526444 | 0.24905637 | 0.46836355 |
| HAUS5 | 83.3732456 | -0.146506715 | 0.41859216 | -0.3499987 | 0.72633967 | 0.86520161 |
| RBM42 | 106.955875 | -1.027487269 | 0.2899092 | -3.5441692 | 0.00039385 | 0.00492029 |
| COX6B1 | 663.743709 | 0.08211756 | 0.34207579 | 0.24005662 | 0.81028636 | 0.91083524 |
| KMT2B | 297.28194 | -0.503763201 | 0.23870125 | -2.1104338 | 0.034821 | 0.13044545 |
| PROSER3 | 179.410789 | -0.473480962 | 0.2832406 | -1.6716564 | 0.09459209 | 0.25303098 |
| ARHGAP33 | 45.4523654 | 0.842889297 | 0.52574031 | 1.60324266 | 0.10888109 | 0.27734186 |
| NFKBID | 73.4675516 | -1.052489554 | 0.29286013 | -3.5938301 | 0.00032585 | 0.00427002 |
| SDHAF1 | 56.4511334 | -0.687419684 | 0.43853792 | -1.5675262 | 0.11699175 | 0.29146435 |
| SYNE4 | 89.6516337 | -0.011321711 | 0.33222583 | -0.0340784 | 0.97281467 | 0.98804941 |
| ALKBH6 | 60.2375244 | -0.219974368 | 0.2659258 | -0.827202 | 0.40812255 | 0.62801617 |
| AC002116.2 | 53.8845981 | -0.302474904 | 0.42086451 | -0.718699 | 0.4723264 | 0.68319748 |
| THAP8 | 51.6751634 | 0.092780144 | 0.38453706 | 0.24127751 | 0.80934004 | 0.9103008 |
| TBCB | 171.926025 | -1.095326668 | 0.29895634 | -3.6638349 | 0.00024847 | 0.00350267 |
| CAPNS1 | 525.442816 | -1.471274371 | 0.32329281 | -4.5509035 | 5.34E-06 | 0.00016072 |
| COX7A1 | 55.299423 | 1.202189903 | 0.63282299 | 1.8997254 | 0.05746917 | 0.18283001 |
| ZNF565 | 632.613831 | 0.101038916 | 0.26917397 | 0.37536659 | 0.70738785 | 0.85277778 |
| ZNF146 | 661.051829 | 0.246382091 | 0.12803761 | 1.92429461 | 0.05431767 | 0.17572423 |
| LINC00665 | 310.57484 | 0.021574367 | 0.32905432 | 0.06556476 | 0.94772435 | 0.97760554 |
| ZFP14 | 486.510908 | -0.043638942 | 0.22408205 | -0.1947454 | 0.84559229 | 0.92861894 |
| ZFP82 | 231.630968 | 0.086487158 | 0.28756726 | 0.30075454 | 0.76360168 | 0.88541384 |
| ZNF566 | 293.224532 | 0.077986486 | 0.21359614 | 0.36511187 | 0.7150279 | 0.85760497 |
| ZNF260 | 241.738825 | -0.02014445 | 0.15060403 | -0.1337577 | 0.89359417 | 0.95146032 |
| ZNF529 | 447.229221 | 0.427576309 | 0.21289574 | 2.00838358 | 0.04460255 | 0.15378623 |
| ZNF529-AS1 | 419.000363 | -0.20416913 | 0.19871687 | -1.0274373 | 0.30421457 | 0.52755288 |
| ZNF382 | 122.373997 | 0.327942634 | 0.23140946 | 1.41715307 | 0.15643818 | 0.35154677 |
| ZNF461 | 213.547501 | -0.0079207 | 0.25571255 | -0.030975 | 0.97528946 | 0.98899114 |
| ZNF567 | 246.244241 | 0.111789531 | 0.21102475 | 0.52974607 | 0.596288 | 0.77614981 |
| ZNF850 | 78.4401243 | 0.1240741 | 0.32847693 | 0.37772546 | 0.70563455 | 0.85120792 |
| AC020928.1 | 498.439045 | 0.285845731 | 0.20427992 | 1.39928454 | 0.16172767 | 0.35922099 |
| ZNF790 | 416.670135 | 0.344428759 | 0.23399848 | 1.47192731 | 0.14104051 | 0.32915266 |
| ZNF345 | 658.658279 | 0.217872905 | 0.22558561 | 0.96581031 | 0.33413911 | 0.5593519 |
| ZNF829 | 62.0818187 | 0.202135892 | 0.37416017 | 0.54023894 | 0.58903226 | 0.77174702 |
| ZNF568 | 371.529133 | 0.237426306 | 0.14083132 | 1.6858914 | 0.09181672 | 0.24804985 |
| ZNF420 | 721.239127 | 0.355521836 | 0.33420204 | 1.06379314 | 0.28742242 | 0.50981103 |
| AC010632.1 | 40.4461133 | 0.634620392 | 0.4195198 | 1.5127305 | 0.13034813 | 0.31253761 |
| AC012309.1 | 60.2157694 | -0.064718505 | 0.58643389 | -0.1103594 | 0.91212433 | 0.96032603 |
| ZNF383 | 240.009652 | 0.322043861 | 0.237331 | 1.35693975 | 0.17480036 | 0.37654099 |
| LINC01535 | 126.944302 | 0.469219146 | 0.30495861 | 1.53863222 | 0.12389411 | 0.30188476 |
| HKR1 | 787.785539 | 0.315805193 | 0.14120559 | 2.23649209 | 0.02531955 | 0.10440189 |
| ZNF527 | 92.9173978 | 0.884482287 | 0.22730721 | 3.89113162 | 9.98E-05 | 0.00167042 |
| ZNF569 | 226.822477 | 0.057285456 | 0.26503216 | 0.2161453 | 0.82887449 | 0.92023714 |
| ZNF570 | 83.1878087 | -0.046192451 | 0.2940301 | -0.1571011 | 0.87516517 | 0.94384684 |
| ZNF793 | 386.045236 | 0.533501164 | 0.21254353 | 2.51007949 | 0.0120704 | 0.06316874 |
| ZNF540 | 282.715528 | 0.324482152 | 0.35688301 | 0.90921155 | 0.36323847 | 0.5885693 |
| ZNF571 | 241.93316 | 0.166639414 | 0.16635098 | 1.00173391 | 0.31647213 | 0.54080054 |
| ZFP30 | 371.343977 | 0.122330114 | 0.20940541 | 0.58417837 | 0.55910031 | 0.75090554 |
| AC093227.1 | 52.6456326 | 0.771211851 | 0.32704541 | 2.35811859 | 0.01836782 | 0.0843018 |
| ZNF573 | 392.694559 | 0.417058164 | 0.25002012 | 1.66809843 | 0.09529619 | 0.25429643 |
| SIPA1L3 | 2812.49983 | 0.222765034 | 0.23971064 | 0.92930808 | 0.35272945 | 0.57882774 |
| SPINT2 | 231.311519 | -0.429908599 | 0.21115364 | -2.035999 | 0.04175046 | 0.14659271 |
| AC011479.1 | 48.7825008 | -0.396106528 | 0.48623823 | -0.8146347 | 0.41528145 | 0.63525223 |
| YIF1B | 48.7084941 | 0.100036164 | 0.3500592 | 0.28576928 | 0.77505484 | 0.89223749 |
| CATSPERG | 203.06888 | -0.121289178 | 0.42461924 | -0.2856422 | 0.77515217 | 0.89223749 |
| PSMD8 | 363.280721 | -0.725856856 | 0.29566311 | -2.4550133 | 0.01408794 | 0.07008949 |
| SPRED3 | 143.308701 | -3.483903168 | 0.47528096 | -7.3301972 | 2.30E-13 | 9.40E-11 |
| FAM98C | 81.3166985 | -0.22345743 | 0.31013634 | -0.7205135 | 0.47120887 | 0.68247785 |
| EIF3K | 624.57408 | 0.543766561 | 0.23710428 | 2.29336456 | 0.02182702 | 0.09455259 |
| ACTN4 | 2480.96174 | -0.162852371 | 0.29238321 | -0.5569826 | 0.57753929 | 0.76386467 |
| CAPN12 | 105.341652 | -0.49775018 | 0.49115182 | -1.0134345 | 0.31085269 | 0.53450314 |
| ECH1 | 296.529004 | -0.592531144 | 0.54926986 | -1.0787614 | 0.28069409 | 0.50321502 |
| HNRNPL | 567.782688 | 0.145065997 | 0.20266573 | 0.71578949 | 0.47412134 | 0.68459432 |
| SIRT2 | 324.527713 | 0.311496014 | 0.28899763 | 1.07784972 | 0.28110083 | 0.50360245 |
| NFKBIB | 103.126076 | -0.617414023 | 0.22232702 | -2.7770535 | 0.00548542 | 0.0359409 |
| AC011455.2 | 40.9596163 | 0.911371975 | 0.45117137 | 2.02001288 | 0.04338205 | 0.15060014 |
| MRPS12 | 166.618148 | -0.328571308 | 0.40417764 | -0.8129379 | 0.41625368 | 0.63600866 |
| FBXO17 | 1221.12404 | -0.054542005 | 0.53775364 | -0.1014256 | 0.91921259 | 0.96418504 |
| FBXO27 | 78.0500834 | 0.266379243 | 0.26879087 | 0.99102788 | 0.32167197 | 0.54635014 |
| PAK4 | 684.805931 | 0.538495178 | 0.30491023 | 1.76607779 | 0.0773828 | 0.22191693 |
| AC011443.1 | 109.345543 | 0.48149139 | 0.33356956 | 1.44345122 | 0.1488934 | 0.34089673 |

| | | | | | | |
|---|---|---|---|---|---|---|
| SAMD4B | 843.697497 | -0.308111378 | 0.21149864 | -1.4568008 | 0.14517138 | 0.33506356 |
| PAF1 | 128.680771 | -0.76651481 | 0.38881469 | -1.9714142 | 0.04867652 | 0.16318988 |
| MED29 | 175.154609 | -0.606169599 | 0.17259524 | -3.5120877 | 0.0004446 | 0.00541766 |
| ZFP36 | 366.542641 | -1.637916405 | 0.53543273 | -3.0590517 | 0.00222039 | 0.01856273 |
| PLEKHG2 | 51.4299931 | -1.41392776 | 0.43154045 | -3.2764663 | 0.00105115 | 0.01038027 |
| RPS16 | 648.870663 | -0.728151172 | 0.42903148 | -1.6971975 | 0.08965932 | 0.24403667 |
| SUPT5H | 607.060419 | -0.197136481 | 0.15640368 | -1.2604338 | 0.20751293 | 0.41837037 |
| TIMM50 | 119.331423 | -0.01281825 | 0.18017634 | -0.0711428 | 0.9432841 | 0.97556917 |
| EID2 | 53.7929515 | -0.237554727 | 0.42781403 | -0.5552757 | 0.57870611 | 0.76468112 |
| DYRK1B | 113.953861 | -0.341495532 | 0.24479591 | -1.3950214 | 0.16300938 | 0.36109189 |
| FBL | 141.678375 | -0.492923263 | 0.26048537 | -1.8923261 | 0.05844754 | 0.1846087 |
| FCGBP | 439.082338 | 1.870775114 | 0.89006068 | 2.1018512 | 0.03556631 | 0.13239418 |
| PSMC4 | 204.513243 | -0.331624323 | 0.28954425 | -1.1453321 | 0.25207162 | 0.47214352 |
| ZNF546 | 187.884752 | 0.443196993 | 0.33894207 | 1.30758923 | 0.19101267 | 0.39752425 |
| ZNF780B | 1011.20877 | 0.362587206 | 0.20302369 | 1.78593546 | 0.07410971 | 0.21595304 |
| ZNF780A | 178.7449 | 0.078217902 | 0.21402262 | 0.36546558 | 0.7147639 | 0.85752192 |
| MAP3K10 | 182.857359 | -0.028146172 | 0.27398451 | -0.1027291 | 0.91817801 | 0.96339767 |
| CNTD2 | 52.2284771 | -0.144744672 | 0.99079825 | -0.1460889 | 0.88385117 | 0.94770599 |
| AKT2 | 717.788687 | 0.005104647 | 0.17412965 | 0.02931521 | 0.9766132 | 0.98948597 |
| C19orf47 | 54.6536683 | 0.31235685 | 0.31425387 | 0.99396341 | 0.32024068 | 0.54463101 |
| PLD3 | 941.667524 | -0.219016231 | 0.22265665 | -0.9836501 | 0.3252876 | 0.55050915 |
| SERTAD1 | 96.4449787 | -0.784701954 | 0.46823957 | -1.6758557 | 0.09376646 | 0.25184394 |
| SERTAD3 | 54.5203146 | -0.212461596 | 0.29800258 | -0.7129522 | 0.47587534 | 0.68577759 |
| BLVRB | 417.305576 | -0.967005327 | 0.31378215 | -3.0817729 | 0.00205772 | 0.01752443 |
| SPTBN4 | 150.769883 | 0.964001163 | 0.78513115 | 1.22782182 | 0.21951386 | 0.43361128 |
| SHKBP1 | 108.412339 | -0.71584816 | 0.28194703 | -2.5389456 | 0.01111871 | 0.05982365 |
| NUMBL | 48.910985 | -0.61927615 | 0.33187984 | -1.8659649 | 0.06204627 | 0.1924775 |
| COQ8B | 196.392101 | -0.404578611 | 0.26085905 | -1.5509472 | 0.12091434 | 0.29724838 |
| ITPKC | 267.202084 | -1.249760192 | 0.24515687 | -5.097798 | 3.44E-07 | 1.71E-05 |
| C19orf54 | 50.8991262 | -0.142450003 | 0.30460719 | -0.4676515 | 0.64003385 | 0.80648521 |
| SNRPA | 195.215908 | -0.49752312 | 0.22625092 | -2.1989883 | 0.02787875 | 0.11160414 |
| AC008537.4 | 49.0531364 | -0.711425503 | 0.44411674 | -1.6018885 | 0.10918026 | 0.27771977 |
| AC008537.1 | 231.199794 | -0.199774194 | 0.49496082 | -0.4036162 | 0.68649499 | 0.83814699 |
| AXL | 93.8882226 | -1.562277201 | 0.75779214 | -2.0616171 | 0.03924421 | 0.14109329 |
| HNRNPUL1 | 504.542841 | -0.362443291 | 0.16876673 | -2.1475992 | 0.03174561 | 0.12261045 |
| TGFB1 | 186.905792 | -0.691905917 | 0.35648645 | -1.9409039 | 0.05226994 | 0.17132348 |
| CCDC97 | 105.017363 | -0.343636301 | 0.23861894 | -1.4401049 | 0.14983772 | 0.34200906 |
| B9D2 | 46.4798637 | -1.129377092 | 0.34759542 | -3.2491138 | 0.00115765 | 0.01121072 |
| EXOSC5 | 40.8679111 | -0.759242249 | 0.49249279 | -1.5416312 | 0.12316324 | 0.30088094 |
| DMAC2 | 120.132406 | 0.069362119 | 0.23790819 | 0.29154994 | 0.77063076 | 0.88937357 |
| RPS19 | 1910.65501 | -1.346495707 | 0.36592648 | -3.6796892 | 0.00023352 | 0.00332745 |
| ARHGEF1 | 264.189215 | -0.413979306 | 0.25631458 | -1.615122 | 0.10628429 | 0.27352187 |
| RABAC1 | 369.825642 | -0.833294295 | 0.47942938 | -1.738096 | 0.08219389 | 0.23040147 |
| GRIK5 | 486.526267 | 0.079622646 | 0.24652994 | 0.32297354 | 0.74671528 | 0.87635381 |
| POU2F2 | 48.9503145 | -1.122971242 | 0.55339285 | -2.0292478 | 0.04243305 | 0.14804496 |
| DEDD2 | 102.414572 | -0.359825869 | 0.24868939 | -1.4468887 | 0.14792809 | 0.33937745 |
| ZNF526 | 50.191189 | -0.653342428 | 0.42028426 | -1.5545251 | 0.12005922 | 0.29586982 |
| GSK3A | 96.1985776 | -0.647962786 | 0.33157428 | -1.954201 | 0.05067745 | 0.16768591 |
| CIC | 211.603905 | -0.845252275 | 0.23263725 | -3.6333488 | 0.00027977 | 0.0038255 |
| PAFAH1B3 | 86.5690515 | -0.762474258 | 0.44814809 | -1.7013891 | 0.08886996 | 0.24253786 |
| MEGF8 | 252.388251 | 0.07951196 | 0.26223558 | 0.30320813 | 0.76173126 | 0.88431255 |
| LIPE-AS1 | 160.402284 | -0.941463372 | 0.43459615 | -2.1662948 | 0.03028867 | 0.11850801 |
| PHLDB3 | 107.70865 | -1.012500044 | 0.43961607 | -2.3031461 | 0.02127062 | 0.09321953 |
| ETHE1 | 243.758787 | -1.222255871 | 0.26963465 | -4.5330074 | 5.81E-06 | 0.00017145 |
| XRCC1 | 82.6416226 | -0.274474584 | 0.28397592 | -0.9665418 | 0.33377317 | 0.5592275 |
| PINLYP | 94.7058191 | -0.269878396 | 0.33526761 | -0.8049641 | 0.42084039 | 0.64027651 |
| ZNF428 | 213.944636 | -0.604175406 | 0.30078442 | -2.0086659 | 0.04457258 | 0.15372303 |
| CADM4 | 76.5512289 | -0.1727517 | 0.3533229 | -0.4889343 | 0.62488818 | 0.79598067 |
| PLAUR | 86.3907988 | -2.690915554 | 0.57944065 | -4.6439882 | 3.42E-06 | 0.0001097 |
| SMG9 | 133.934254 | -0.086275716 | 0.30650355 | -0.2814836 | 0.77833952 | 0.89400754 |
| ZNF283 | 214.390556 | 0.51241738 | 0.17231487 | 2.97372702 | 0.00294207 | 0.02281959 |
| ZNF404 | 61.8540934 | 1.018745111 | 0.41217638 | 2.47162419 | 0.01345008 | 0.06780998 |
| AC006213.3 | 78.8085087 | 0.433226421 | 0.45285623 | 0.95665334 | 0.33874226 | 0.56424293 |
| ZNF45 | 132.876324 | -0.072866646 | 0.29682658 | -0.2454856 | 0.80608047 | 0.90934469 |
| ZNF155 | 248.812027 | -0.006851438 | 0.28496635 | -0.024043 | 0.98081833 | 0.99160261 |
| ZNF230 | 57.2325334 | -0.054094832 | 0.26259566 | -0.2060005 | 0.83679053 | 0.9240982 |
| ZNF224 | 368.217666 | -0.202621334 | 0.16954604 | -1.1950815 | 0.2320552 | 0.4485246 |
| ZNF225 | 66.0202961 | 0.166213323 | 0.23864665 | 0.69648295 | 0.48612643 | 0.69351377 |
| ZNF234 | 162.536028 | 0.185179205 | 0.25030245 | 0.73982177 | 0.45940815 | 0.67306147 |
| ZNF226 | 267.76016 | 0.385132917 | 0.27148701 | 1.41860531 | 0.15601412 | 0.35119129 |
| ZNF227 | 179.038992 | 0.337639765 | 0.21190026 | 1.59339003 | 0.11107272 | 0.28122667 |
| ZNF235 | 633.928263 | 0.129633559 | 0.28208451 | 0.45955575 | 0.64583512 | 0.81059872 |
| ZNF233 | 49.5104449 | 0.639859 | 0.49642064 | 1.28894519 | 0.19741714 | 0.4057467 |
| ZNF229 | 154.910367 | -0.44495366 | 0.3081016 | -1.4441783 | 0.14868882 | 0.34065124 |
| ZNF180 | 128.366766 | 0.358723085 | 0.28550843 | 1.25643607 | 0.20895791 | 0.42044445 |
| AC243964.2 | 361.699873 | -0.313842967 | 0.34794606 | -0.9019874 | 0.36706356 | 0.5925402 |
| PVR | 133.314946 | -0.98463236 | 0.26012603 | -3.7852128 | 0.00015358 | 0.00235884 |
| CEACAM19 | 227.184618 | -1.095201285 | 0.44255555 | -2.4747205 | 0.01333405 | 0.06740507 |
| BCL3 | 248.552106 | -2.441908548 | 0.56036842 | -4.3576841 | 1.31E-05 | 0.00033198 |
| CBLC | 144.142686 | -1.208652749 | 0.39417164 | -3.0663107 | 0.00216718 | 0.0182333 |
| BCAM | 1487.72286 | -1.270150472 | 0.47642903 | -2.6659804 | 0.00767642 | 0.04581915 |
| NECTIN2 | 637.791091 | -0.362975451 | 0.22263956 | -1.6303277 | 0.10303225 | 0.26790413 |
| TOMM40 | 180.638227 | -0.383049859 | 0.30580176 | -1.2526084 | 0.21034825 | 0.42245553 |
| APOE | 504.385931 | 0.108314271 | 0.62052613 | 0.17455231 | 0.86143142 | 0.93697711 |
| CLPTM1 | 394.946671 | -0.1554152 | 0.19816566 | -0.7842691 | 0.43288224 | 0.65020381 |
| RELB | 339.414845 | -2.619429361 | 0.44352456 | -5.9059399 | 3.51E-09 | 3.96E-07 |
| CLASRP | 663.200646 | -0.01479156 | 0.2160776 | -0.0684549 | 0.94542356 | 0.97640274 |
| GEMIN7-AS1 | 224.195024 | 0.03246932 | 0.27456447 | 0.11825755 | 0.90586359 | 0.95693796 |
| MARK4 | 502.822057 | -0.289185924 | 0.20604163 | -1.4035316 | 0.16045839 | 0.35771722 |
| TRAPPC6A | 192.132796 | -0.241911527 | 0.27054603 | -0.8941603 | 0.37123612 | 0.59563048 |
| KLC3 | 46.132574 | 0.11571953 | 0.40831878 | 0.28340487 | 0.7768665 | 0.89337196 |
| ERCC2 | 111.084037 | 0.117282495 | 0.32606162 | 0.35969427 | 0.71907578 | 0.86003243 |
| PPP1R13L | 387.646167 | -0.715991072 | 0.30465437 | -2.3501749 | 0.01876459 | 0.08538114 |
| ERCC1 | 203.785418 | 0.208265623 | 0.22771213 | 0.91460049 | 0.3604014 | 0.58548052 |
| VASP | 285.662637 | -0.904622952 | 0.23856699 | -3.7919034 | 0.0001495 | 0.00230152 |
| OPA3 | 711.767718 | 0.227945836 | 0.29850461 | 0.76362586 | 0.44509023 | 0.66088095 |
| EML2 | 414.171609 | -0.430087502 | 0.40895807 | -1.0516665 | 0.29295259 | 0.51581055 |
| SNRPD2 | 306.428182 | -1.166223547 | 0.3355067 | -3.4760067 | 0.00050894 | 0.00600231 |
| QPCTL | 101.706227 | -0.360577054 | 0.24866947 | -1.4500254 | 0.14705143 | 0.33786787 |
| FBXO46 | 269.724309 | 0.016743163 | 0.20220851 | 0.08280148 | 0.9340094 | 0.97167284 |
| SIX5 | 55.3464925 | -0.005342354 | 0.39130489 | -0.0136527 | 0.98910709 | 0.99594533 |
| DM1-AS | 69.3392814 | 0.142640318 | 0.43850128 | 0.32529054 | 0.74496119 | 0.87522835 |
| DMPK | 185.715669 | -0.260581817 | 0.3328131 | -0.7829674 | 0.43364624 | 0.65072685 |
| SYMPK | 740.117227 | -0.237366802 | 0.21006379 | -1.1299749 | 0.25848682 | 0.47954387 |
| MYPOP | 240.055845 | 0.539909942 | 0.2925712 | 1.84539674 | 0.06497985 | 0.1980431 |
| CCDC61 | 67.4886624 | 0.274770235 | 0.34153806 | 0.80450839 | 0.42110343 | 0.64038167 |
| HIF3A | 120.665894 | 0.798826971 | 0.77932001 | 1.02503075 | 0.30534868 | 0.52868668 |
| PPP5C | 513.846781 | 0.190733238 | 0.11626292 | 1.64053373 | 0.10089424 | 0.26453475 |
| AC007193.1 | 47.3447593 | 0.072963588 | 0.57820788 | 0.1261892 | 0.89958216 | 0.95381127 |
| CCDC8 | 122.924119 | 0.662149903 | 0.38667931 | 1.71240065 | 0.08682286 | 0.23862737 |
| PNMA8A | 51.3150756 | -0.449087604 | 0.6699473 | -0.6703327 | 0.5026457 | 0.7069258 |
| PPP5D1 | 294.578654 | 0.615185428 | 0.34744553 | 1.77059529 | 0.07662803 | 0.22080707 |
| CALM3 | 197.42787 | -1.041209085 | 0.38901572 | -2.6765219 | 0.00743907 | 0.04472585 |

| | | | | | | |
|---|---|---|---|---|---|---|
| PRKD2 | 457.358082 | -1.040033294 | 0.32919203 | -3.1593514 | 0.00158121 | 0.0142958 |
| STRN4 | 164.015335 | 0.038043257 | 0.18995234 | 0.20027791 | 0.84126324 | 0.92645072 |
| FKRP | 183.96142 | 0.088229433 | 0.18925308 | 0.46619813 | 0.64107369 | 0.80747519 |
| SLC1A5 | 156.744252 | -0.952654225 | 0.5189028 | -1.8359011 | 0.06637228 | 0.20080429 |
| AP2S1 | 274.333872 | -0.527604919 | 0.28708867 | -1.8377769 | 0.06609529 | 0.20033333 |
| ARHGAP35 | 1108.133 | -0.076022806 | 0.2100566 | -0.3619158 | 0.71741494 | 0.85947463 |
| TMEM160 | 291.94962 | -1.260752652 | 0.50323581 | -2.505292 | 0.01223503 | 0.06375249 |
| ZC3H4 | 321.697963 | -0.037955343 | 0.20313125 | -0.1868513 | 0.8517772 | 0.93193618 |
| SAE1 | 290.375397 | -0.006490417 | 0.29748027 | -0.021818 | 0.98259316 | 0.99281519 |
| BBC3 | 50.8438882 | -2.090272042 | 0.4018384 | -5.2017728 | 1.97E-07 | 1.09E-05 |
| CCDC9 | 119.94049 | -0.6198447 | 0.26851833 | -2.3083888 | 0.02097752 | 0.09221035 |
| INAFM1 | 92.6387054 | -0.998950548 | 0.50786791 | -1.9669495 | 0.04918903 | 0.16432421 |
| C5AR1 | 424.536096 | 0.457981019 | 0.49412548 | 0.92685165 | 0.35400356 | 0.57979331 |
| DHX34 | 287.741746 | -0.108891336 | 0.26495718 | -0.4109771 | 0.68108932 | 0.83339591 |
| KPTN | 41.6577885 | 0.311753484 | 0.38748287 | 0.80456069 | 0.42107324 | 0.64038167 |
| NAPA-AS1 | 112.896264 | 0.174561802 | 0.32403897 | 0.5387062 | 0.5900896 | 0.7721373 |
| NAPA | 535.996066 | -0.372651424 | 0.28467818 | -1.309027 | 0.1905252 | 0.39706328 |
| AC010331.1 | 39.5710702 | 0.900234748 | 0.44711203 | 2.01344337 | 0.04406801 | 0.15221766 |
| BICRA-AS1 | 100.934888 | 0.319662582 | 0.48199776 | 0.66320345 | 0.50720026 | 0.71045687 |
| NOP53 | 133.028898 | -0.714351691 | 0.40820232 | -1.7499942 | 0.08011931 | 0.22656734 |
| SELENOW | 613.59498 | -0.712083036 | 0.30539681 | -2.331665 | 0.01971832 | 0.08838118 |
| PLA2G4C | 85.9941502 | -1.341939859 | 0.62614697 | -2.1431707 | 0.03209939 | 0.12365146 |
| LIG1 | 122.266676 | -0.271247054 | 0.4762208 | -0.5695825 | 0.56896087 | 0.75867193 |
| ZSWIM9 | 158.252663 | 0.04192066 | 0.25529913 | 0.16420213 | 0.86957203 | 0.94056476 |
| CARD8 | 457.178062 | 0.550506388 | 0.15120356 | 3.64082957 | 0.00027176 | 0.00373537 |
| EMP3 | 60.9962591 | -0.475777822 | 0.40814268 | -1.1657145 | 0.24372991 | 0.46243709 |
| TMEM143 | 204.578144 | 0.765694431 | 0.38446162 | 1.99160171 | 0.04641478 | 0.15768847 |
| KDELR1 | 317.610401 | -0.853076854 | 0.22797467 | -3.7419808 | 0.00018258 | 0.00271887 |
| GRIN2D | 38.2884338 | 0.702762036 | 0.60148092 | 1.16838625 | 0.242651 | 0.46111021 |
| GRWD1 | 59.1220418 | -0.671990024 | 0.43943779 | -1.529204 | 0.12621388 | 0.30539643 |
| KCNJ14 | 71.0046246 | -0.104555842 | 0.37380121 | -0.2797097 | 0.7797002 | 0.8948704 |
| CYTH2 | 321.587028 | -0.392449558 | 0.29723452 | -1.3203364 | 0.18672273 | 0.39251209 |
| LMTK3 | 301.657603 | 0.316240332 | 0.4065212 | 0.77791843 | 0.4366171 | 0.65342962 |
| SULT2B1 | 103.12882 | 1.128661987 | 0.89123491 | 1.26640236 | 0.20536907 | 0.41584887 |
| RPL18 | 802.441215 | -0.988939102 | 0.4367236 | -2.2644508 | 0.0235464 | 0.09959156 |
| SPHK2 | 160.105554 | 0.195827255 | 0.31027478 | 0.63114139 | 0.52794808 | 0.72617338 |
| CA11 | 55.0546119 | -0.28414984 | 0.42811692 | -0.6637202 | 0.50686942 | 0.71014406 |
| MAMSTR | 51.8092745 | 1.310027349 | 0.40988539 | 3.1960821 | 0.00139307 | 0.01296767 |
| BCAT2 | 239.492226 | 0.079916358 | 0.36563442 | 0.21856902 | 0.82698578 | 0.91955386 |
| HSD17B14 | 125.761926 | -0.678795293 | 0.52324233 | -1.2972867 | 0.19453257 | 0.4020624 |
| PLEKHA4 | 204.300645 | -1.550782336 | 0.3702349 | -4.1886444 | 2.81E-05 | 0.00062175 |
| PPP1R15A | 98.4495297 | -1.567046633 | 0.38763409 | -4.0425924 | 5.29E-05 | 0.00101956 |
| NUCB1 | 441.744173 | -0.453515667 | 0.27501064 | -1.6490841 | 0.09913041 | 0.26099533 |
| NUCB1-AS1 | 60.8421689 | -0.098269629 | 0.44259597 | -0.2220301 | 0.82429045 | 0.91866795 |
| BAX | 190.105551 | -0.60306008 | 0.32493526 | -1.8559392 | 0.06346223 | 0.19518591 |
| FTL | 6121.14695 | -1.424253203 | 0.58756239 | -2.4240034 | 0.01535046 | 0.07456505 |
| GYS1 | 508.409119 | -0.265096376 | 0.38005378 | -0.6975233 | 0.48547534 | 0.69300247 |
| RUVBL2 | 109.789232 | -0.243190689 | 0.23787286 | -1.0223558 | 0.30661255 | 0.53031886 |
| SNRNP70 | 1763.06072 | 0.48827516 | 0.2806848 | 1.73958531 | 0.08193186 | 0.23006759 |
| LIN7B | 52.056142 | 0.270035475 | 0.34344008 | 0.78626663 | 0.4317113 | 0.64957432 |
| PPFIA3 | 45.1897459 | 0.657778383 | 0.41528317 | 1.58392739 | 0.11321024 | 0.28527167 |
| TRPM4 | 464.419516 | 0.571662972 | 0.45704368 | 1.2507841 | 0.21101326 | 0.42345922 |
| SLC6A16 | 641.030324 | 1.017373284 | 0.374763 | 2.71471112 | 0.00663336 | 0.04126225 |
| AC011450.1 | 83.6919594 | 0.670205425 | 0.47898863 | 1.39920946 | 0.16175018 | 0.35922099 |
| TEAD2 | 141.726316 | -0.975252508 | 0.50135968 | -1.9452153 | 0.05174907 | 0.17012283 |
| PIH1D1 | 236.122427 | -0.024719093 | 0.19604082 | -0.1260916 | 0.89965945 | 0.95381665 |
| RPL13A | 1062.54901 | -1.131421028 | 0.42753295 | -2.646395 | 0.00813548 | 0.04782444 |
| RPS11 | 846.831224 | -0.9751552 | 0.41962732 | -2.3238601 | 0.02013299 | 0.08939048 |
| FCGRT | 280.370068 | -0.15336568 | 0.33926741 | -0.4520495 | 0.65123329 | 0.81374899 |
| NOSIP | 329.505619 | 0.252537461 | 0.20268839 | 1.24593943 | 0.21278663 | 0.42566994 |
| PRRG2 | 119.062857 | -0.683031717 | 0.28383786 | -2.4064151 | 0.01610995 | 0.07693286 |
| PRR12 | 268.626808 | 0.095527829 | 0.31263428 | 0.30555776 | 0.75994139 | 0.88357016 |
| RRAS | 80.3008421 | -2.253417734 | 0.58792493 | -3.8328324 | 0.00012668 | 0.00202575 |
| SCAF1 | 169.09276 | -0.409533159 | 0.26186323 | -1.56392 | 0.11783635 | 0.29274033 |
| IRF3 | 469.830712 | -0.7151183 | 0.28318571 | -2.5252627 | 0.01156118 | 0.06140933 |
| BCL2L12 | 43.8034131 | 0.387383638 | 0.33246812 | 1.1651753 | 0.24394804 | 0.4627096 |
| PRMT1 | 316.966582 | 0.048006145 | 0.29257763 | 0.16408002 | 0.86966815 | 0.94059171 |
| AP2A1 | 358.793522 | -0.265132473 | 0.25081876 | -1.0570679 | 0.29048058 | 0.51339098 |
| FUZ | 86.0578673 | 0.465774297 | 0.34072907 | 1.36699311 | 0.17162747 | 0.37255993 |
| MED25 | 460.941994 | -0.147428642 | 0.26491865 | -0.5565053 | 0.57786544 | 0.76421953 |
| PTOV1-AS1 | 122.864274 | 0.33496774 | 0.29270009 | 1.14440598 | 0.25245531 | 0.47247832 |
| PTOV1 | 400.155632 | -0.015172723 | 0.22410461 | -0.0677038 | 0.94602146 | 0.97670764 |
| PNKP | 453.158656 | -0.867048093 | 0.31996767 | -2.709799 | 0.0067324 | 0.04173076 |
| TBC1D17 | 315.794645 | -0.34612667 | 0.29834145 | -1.1601696 | 0.24597978 | 0.46524008 |
| AC011452.1 | 36.4182733 | 2.579769702 | 0.48710613 | 5.29611423 | 1.18E-07 | 7.17E-06 |
| ATF5 | 68.3832156 | -0.863037883 | 0.41022524 | -2.1038147 | 0.03539462 | 0.13184644 |
| VRK3 | 368.465506 | -0.57219414 | 0.24597769 | -2.3262034 | 0.0200077 | 0.08910376 |
| ZNF473 | 81.6098968 | -0.424792032 | 0.29957393 | -1.4179873 | 0.15619447 | 0.35131349 |
| MYH14 | 273.04036 | -1.144229323 | 0.55884335 | -2.0474957 | 0.04060944 | 0.14411879 |
| KCNC3 | 215.963226 | -1.449808164 | 0.61030453 | -2.3755487 | 0.01752288 | 0.08161484 |
| NR1H2 | 324.276136 | -0.812124772 | 0.28448548 | -2.8547143 | 0.00430756 | 0.03013692 |
| NAPSA | 171.833055 | -1.635263154 | 0.63158812 | -2.589129 | 0.0096219 | 0.05383131 |
| EMC10 | 396.086443 | -0.817501087 | 0.31133072 | -2.6258285 | 0.00864384 | 0.04987079 |
| JOSD2 | 94.2155983 | -0.355974879 | 0.34052201 | -1.0453799 | 0.2958474 | 0.51903949 |
| CLEC11A | 113.430774 | -0.301371875 | 0.32301196 | -0.9330053 | 0.35081722 | 0.57693534 |
| C19orf48 | 73.2448704 | 0.069233608 | 0.437225 | 0.15834778 | 0.87418276 | 0.9433889 |
| CTU1 | 64.6740084 | -0.381787928 | 0.30040518 | -1.2709099 | 0.2037607 | 0.41350055 |
| ETFB | 366.682163 | -0.158093294 | 0.46115384 | -0.3428212 | 0.73173301 | 0.86826096 |
| ZNF175 | 194.837685 | 0.318695009 | 0.16354254 | 1.94869794 | 0.05133151 | 0.16901135 |
| SPACA6 | 244.379671 | -0.196690323 | 0.26695049 | -0.7368045 | 0.46124124 | 0.67447531 |
| ZNF577 | 481.081671 | 0.598940979 | 0.27054732 | 2.21381227 | 0.0268417 | 0.10875827 |
| ZNF649 | 65.3153118 | -0.120378583 | 0.29734161 | -0.4048494 | 0.68558819 | 0.83742689 |
| ZNF613 | 95.6413751 | 0.305728892 | 0.22686658 | 1.34761537 | 0.17778212 | 0.38072699 |
| ZNF350-AS1 | 305.734667 | 1.221785832 | 0.66965476 | 1.82450109 | 0.06807635 | 0.20452712 |
| ZNF350 | 201.943502 | 0.312080073 | 0.3626628 | 0.86052408 | 0.38950021 | 0.61248908 |
| ZNF615 | 159.83709 | 0.523088328 | 0.22165412 | 2.35993057 | 0.01827836 | 0.08408911 |
| ZNF614 | 84.9441582 | 0.497757992 | 0.31062894 | 1.60241989 | 0.10906279 | 0.27762728 |
| ZNF432 | 359.92722 | 0.074264796 | 0.29002353 | 0.25606473 | 0.79790086 | 0.90506918 |
| ZNF616 | 105.892566 | 0.185212848 | 0.22152644 | 0.83607557 | 0.40311239 | 0.62443013 |
| AC011468.3 | 83.8916834 | -1.316593988 | 0.55923848 | -2.3542622 | 0.01855952 | 0.08485161 |
| ZNF836 | 90.2462561 | -0.336613697 | 0.27285793 | -1.2336592 | 0.21732993 | 0.43128814 |
| PPP2R1A | 739.330907 | -0.576645836 | 0.29156744 | -1.9777443 | 0.04795756 | 0.161394 |
| ZNF766 | 324.617765 | 0.014299259 | 0.24192482 | 0.05910621 | 0.95286752 | 0.97936812 |
| AC010320.3 | 47.9013251 | -0.236929437 | 0.48884623 | -0.4846707 | 0.62790996 | 0.7982736 |
| ZNF480 | 266.334931 | -0.074413981 | 0.24597838 | -0.3025224 | 0.76225382 | 0.88460822 |
| AC010320.4 | 75.1083126 | 0.421120543 | 0.51837777 | 0.81238156 | 0.41657272 | 0.63642252 |
| ZNF610 | 112.30859 | 0.211952098 | 0.33461675 | 0.63341747 | 0.52646106 | 0.72502073 |
| ZNF880 | 94.4851367 | 0.085636834 | 0.29176953 | 0.29350849 | 0.76913351 | 0.88861056 |
| ZNF528 | 232.407078 | 0.254152692 | 0.33654098 | 0.75519091 | 0.45013444 | 0.66627362 |
| ZNF578 | 58.6420114 | -0.021306124 | 0.44710039 | -0.047654 | 0.96199199 | 0.9832997 |
| ZNF808 | 301.665461 | 0.735640213 | 0.24079355 | 3.05506605 | 0.00225011 | 0.01869289 |
| ZNF701 | 414.93005 | 0.78085989 | 0.17259707 | 4.52417821 | 6.06E-06 | 0.00017718 |
| ZNF83 | 2381.39559 | 0.243444063 | 0.21608911 | 1.12659106 | 0.25991542 | 0.48097825 |

| | | | | | | |
|---|---|---|---|---|---|---|
| ZNF611 | 1136.14096 | 0.090192051 | 0.30250297 | 0.29815261 | 0.76558669 | 0.88656137 |
| ZNF600 | 164.991938 | -0.022373794 | 0.2743148 | -0.0815625 | 0.93499464 | 0.9719401 |
| ZNF28 | 177.068329 | 0.087128165 | 0.17927945 | 0.48599081 | 0.62697368 | 0.79762067 |
| ZNF320 | 88.5287572 | 1.093186441 | 0.54789739 | 1.99523936 | 0.04601678 | 0.15677989 |
| AC010487.3 | 326.740568 | 0.318785703 | 0.49302146 | 0.646596 | 0.51789345 | 0.71887921 |
| ZNF160 | 813.134816 | -0.458291062 | 0.28562722 | -1.6045077 | 0.1086022 | 0.27695355 |
| ZNF415 | 90.964292 | 0.348578326 | 0.23680389 | 1.47201264 | 0.14101746 | 0.32915266 |
| ZNF347 | 260.967727 | -0.504529361 | 0.20602535 | -2.4488703 | 0.0143305 | 0.07098626 |
| ZNF665 | 97.744299 | -0.340348793 | 0.40546142 | -0.839411 | 0.40123869 | 0.62271905 |
| ZNF677 | 155.158487 | -0.165521778 | 0.31104958 | -0.5321395 | 0.59462936 | 0.77490718 |
| ZNF845 | 73.5917541 | 0.073328091 | 0.29604528 | 0.24769215 | 0.80437261 | 0.90827131 |
| ZNF525 | 197.988006 | -0.509899951 | 0.22703321 | -2.2459267 | 0.0247087 | 0.1028932 |
| ZNF765 | 276.403442 | -0.225289535 | 0.24732773 | -0.9108948 | 0.36235083 | 0.58770777 |
| ZNF761 | 321.249696 | -0.277794654 | 0.33334473 | -0.8333555 | 0.40464428 | 0.62539431 |
| ZNF813 | 99.2280539 | -0.237535089 | 0.37259947 | -0.6375079 | 0.5237941 | 0.72325104 |
| ZNF331 | 352.597546 | -0.191705964 | 0.27786683 | -0.6899203 | 0.49024431 | 0.69684897 |
| MYADM | 57.3802863 | -0.316436339 | 0.75369365 | -0.4198474 | 0.67459692 | 0.82939913 |
| NDUFA3 | 142.076608 | -0.152014672 | 0.45702617 | -0.332617 | 0.73942341 | 0.87241119 |
| TFPT | 69.4604225 | -0.124310161 | 0.30561211 | -0.406758 | 0.68418576 | 0.83625517 |
| PRPF31 | 291.770418 | -0.272390011 | 0.207059 | -1.3155188 | 0.1883356 | 0.39387665 |
| CNOT3 | 186.080663 | 0.222864876 | 0.26119309 | 0.85325717 | 0.39351671 | 0.6158723 |
| LENG1 | 59.113576 | 0.062435906 | 0.28386995 | 0.21994546 | 0.82591363 | 0.91938879 |
| TMC4 | 672.12296 | 0.060990811 | 0.15124882 | 0.40324817 | 0.68676567 | 0.83824503 |
| MBOAT7 | 356.211925 | -0.588962734 | 0.2697788 | -2.183132 | 0.02902609 | 0.11475774 |
| TSEN34 | 170.037602 | -1.002258232 | 0.35655565 | -2.8109447 | 0.00493963 | 0.03310204 |
| RPS9 | 1680.57774 | -1.048061558 | 0.37136215 | -2.8222089 | 0.00476941 | 0.03224111 |
| LENG8-AS1 | 209.146522 | -0.026060934 | 0.24008401 | -0.1085492 | 0.91356003 | 0.96091809 |
| LENG8 | 916.604469 | 0.672412783 | 0.30836756 | 2.18055615 | 0.02921626 | 0.1152681 |
| LENG9 | 38.9412411 | 1.145266963 | 0.61132117 | 1.87342925 | 0.06100913 | 0.19001878 |
| NLRP2 | 56.2759961 | -0.054542289 | 0.61174008 | -0.0891593 | 0.92895535 | 0.96900671 |
| AC011476.3 | 168.227953 | -0.175255917 | 0.50560673 | -0.346625 | 0.72887309 | 0.86681233 |
| GP6 | 52.2678281 | 0.142932591 | 0.78912312 | 0.18112838 | 0.85626681 | 0.9348234 |
| RDH13 | 384.218731 | 0.326490419 | 0.25647334 | 1.27299945 | 0.20301824 | 0.41262777 |
| EPS8L1 | 240.237297 | 0.435503839 | 0.51981251 | 0.83780946 | 0.40213773 | 0.6235353 |
| PPP1R12C | 382.283024 | -0.176671059 | 0.21186183 | -0.8338975 | 0.40433873 | 0.62532611 |
| PPP6R1 | 285.111622 | -0.320867714 | 0.19581043 | -1.6386651 | 0.10128302 | 0.26501129 |
| HSPBP1 | 148.104004 | -0.087965085 | 0.27684794 | -0.3177379 | 0.75068376 | 0.87890283 |
| KMT5C | 84.1099215 | -0.301989014 | 0.30960658 | -0.975396 | 0.32936387 | 0.55488483 |
| TMEM238 | 148.614807 | -0.914689637 | 0.54928813 | -1.6652274 | 0.0958674 | 0.25518041 |
| RPL28 | 1260.05757 | -1.289050699 | 0.4021316 | -3.2055444 | 0.00134807 | 0.01265579 |
| UBE2S | 71.395261 | -0.141554166 | 0.3825468 | -0.370031 | 0.71135941 | 0.8551462 |
| ISOC2 | 124.32576 | -0.905582841 | 0.42824205 | -2.1146518 | 0.03445963 | 0.12960629 |
| NAT14 | 97.6733641 | 0.207373967 | 0.46674285 | 0.44430025 | 0.65682552 | 0.81801439 |
| ZNF579 | 94.274459 | -0.298748447 | 0.4014763 | -0.7441247 | 0.45680102 | 0.67184997 |
| FIZ1 | 55.4514856 | 0.114553503 | 0.30788955 | 0.37206038 | 0.70984789 | 0.85441779 |
| ZNF524 | 76.4380383 | -0.706090597 | 0.39642512 | -1.7811449 | 0.07488878 | 0.21730765 |
| ZNF865 | 97.8110269 | -0.16110616 | 0.28987885 | -0.5557707 | 0.57836764 | 0.76458087 |
| CCDC106 | 89.9730213 | 0.014875356 | 0.36672706 | 0.04056247 | 0.9676447 | 0.98528676 |
| U2AF2 | 265.625809 | 0.130424886 | 0.20383974 | 0.63984033 | 0.52227641 | 0.72221624 |
| AC008735.2 | 120.727947 | 0.43636606 | 0.51648648 | 0.84487412 | 0.39818112 | 0.61972127 |
| EPN1 | 767.198973 | 0.760056235 | 0.38664201 | 1.96578802 | 0.0493231 | 0.16464738 |
| ZNF787 | 172.807598 | -0.216653422 | 0.26068667 | -0.8310875 | 0.40592422 | 0.62601903 |
| ZNF444 | 311.945274 | 0.267002599 | 0.21760048 | 1.22703132 | 0.21981082 | 0.43361796 |
| ZSCAN5A | 972.668394 | 0.229587597 | 0.2808505 | 0.81747264 | 0.41365838 | 0.63321515 |
| EDDM13 | 140.951666 | -0.32774741 | 0.49643825 | -0.6601977 | 0.50912695 | 0.71157103 |
| ZNF582 | 97.346538 | 0.410546671 | 0.30923659 | 1.32761351 | 0.18430582 | 0.38958162 |
| ZNF583 | 100.779728 | 0.126718617 | 0.41274167 | 0.30701678 | 0.7588306 | 0.88316847 |
| ZNF667 | 249.75896 | -0.057844099 | 0.37638971 | -0.1536814 | 0.87786095 | 0.94536627 |
| ZNF667-AS1 | 216.412073 | -0.072844121 | 0.19779076 | -0.3682888 | 0.71265791 | 0.8559282 |
| ZNF471 | 228.459918 | -0.003116202 | 0.2570706 | -0.012122 | 0.9903283 | 0.99606252 |
| ZFP28 | 79.0623975 | 0.121100807 | 0.29180297 | 0.41500883 | 0.67813544 | 0.83118075 |
| AC005498.3 | 155.907182 | -0.088748659 | 0.39324737 | -0.2256815 | 0.82144914 | 0.9172814 |
| ZNF470 | 95.0189294 | -0.481739836 | 0.4630413 | -1.040382 | 0.29816246 | 0.52143889 |
| ZNF71 | 65.2535487 | 0.329721717 | 0.28088606 | 1.17386286 | 0.24044996 | 0.45857079 |
| AC006115.3 | 71.55336 | 0.888691754 | 0.25754892 | 3.45057461 | 0.00055939 | 0.00642525 |
| ZIM2 | 40.3409476 | 0.56515438 | 0.46732941 | 1.20932766 | 0.22653699 | 0.44161101 |
| ZNF264 | 352.19307 | 0.093764424 | 0.16202501 | 0.57870339 | 0.56278933 | 0.75444301 |
| ZNF805 | 76.7052546 | -0.140949563 | 0.25157977 | -0.5602579 | 0.57530351 | 0.76252365 |
| ZNF460-AS1 | 182.854845 | 0.395887234 | 0.32024305 | 1.23620868 | 0.21638101 | 0.43031868 |
| ZNF460 | 132.121238 | 0.250109128 | 0.26396934 | 0.94749308 | 0.34338758 | 0.56837065 |
| AC005261.1 | 422.036879 | 0.001290439 | 0.2017688 | 0.00639563 | 0.99489706 | 0.997919 |
| ZNF304 | 70.9724085 | 0.320439234 | 0.33888938 | 0.94555703 | 0.34437457 | 0.56924587 |
| AC003002.2 | 112.461932 | 0.877462474 | 0.28365855 | 3.09337573 | 0.00197893 | 0.01699743 |
| ZNF749 | 45.1348376 | -0.298752243 | 0.34296268 | -0.8710926 | 0.38370362 | 0.60643561 |
| AC004076.1 | 249.306226 | -0.14015462 | 0.32487769 | -0.4314073 | 0.66617221 | 0.82384316 |
| ZNF773 | 47.8327172 | -0.077131851 | 0.46680476 | -0.1652336 | 0.86876009 | 0.94020545 |
| ZNF549 | 168.572046 | -0.208425325 | 0.23990782 | -0.8687725 | 0.38497156 | 0.6081188 |
| ZNF550 | 102.514614 | 0.749472739 | 0.26861569 | 2.79013012 | 0.00526869 | 0.03481445 |
| ZNF134 | 79.2394297 | 0.056940283 | 0.3404893 | 0.16723075 | 0.86718849 | 0.93961371 |
| ZNF211 | 86.8991947 | 0.376044752 | 0.23641074 | 1.59064156 | 0.11169027 | 0.28237377 |
| AC003006.1 | 144.321235 | 0.211441903 | 0.40009997 | 0.52847267 | 0.59717131 | 0.77654325 |
| ZNF154 | 97.0393499 | 0.651163144 | 0.5358338 | 1.21523342 | 0.22427708 | 0.43906353 |
| ZNF671 | 56.9416653 | 0.524767353 | 0.49845295 | 1.05279215 | 0.29243626 | 0.51538234 |
| ZNF776 | 92.5792362 | 0.832185926 | 0.25254927 | 3.2951429 | 0.00098372 | 0.00987479 |
| ZNF586 | 1329.01217 | 0.016350417 | 0.207502 | 0.07879643 | 0.93719454 | 0.97284815 |
| ZNF552 | 475.801261 | 0.4349848 | 0.26964361 | 1.61318418 | 0.1067045 | 0.27422203 |
| AC010522.1 | 71.8409597 | 0.557139032 | 0.25035122 | 2.22542969 | 0.02605241 | 0.10649359 |
| ZNF814 | 764.283053 | 0.106115179 | 0.2230774 | 0.47568772 | 0.63429687 | 0.80330081 |
| ZNF587 | 393.703445 | 0.425217438 | 0.16599231 | 2.56166956 | 0.01041704 | 0.0571423 |
| ZNF418 | 118.071218 | 0.04591176 | 0.35101527 | 0.13079704 | 0.89593586 | 0.95200843 |
| C19orf18 | 160.285818 | 0.636593349 | 0.47054874 | 1.3528744 | 0.17609576 | 0.3783991 |
| ZNF606 | 235.96824 | 0.152496617 | 0.20877474 | 0.73043614 | 0.46512364 | 0.67812562 |
| ZSCAN1 | 50.4209741 | 0.230332961 | 0.58589922 | 0.39312727 | 0.69422548 | 0.843734 |
| ZNF135 | 462.123751 | 0.486282518 | 0.35691414 | 1.36246359 | 0.17305163 | 0.37466627 |
| ZSCAN18 | 543.927072 | -0.599619316 | 0.31222378 | -1.9204793 | 0.05479738 | 0.1766273 |
| ZNF329 | 294.154451 | -0.154613019 | 0.27402464 | -0.5642304 | 0.57259737 | 0.76052676 |
| ZNF274 | 377.858849 | -0.502094178 | 0.23359676 | -2.1494055 | 0.03160227 | 0.12230869 |
| AC020915.3 | 186.705638 | 0.279265455 | 0.25668594 | 1.08796553 | 0.27661032 | 0.49896172 |
| ERVK3-1 | 44.3603265 | -0.052119358 | 0.42452713 | -0.1227704 | 0.90228893 | 0.95484173 |
| AC010642.2 | 64.8599718 | -0.341900467 | 0.35608327 | -0.96017 | 0.33696968 | 0.56227228 |
| ZNF497 | 72.3345158 | 0.478428655 | 0.27405469 | 1.74574153 | 0.08085587 | 0.22811304 |
| ZNF837 | 44.275689 | -0.129161314 | 0.3088624 | -0.418184 | 0.67581259 | 0.82991711 |
| RPS5 | 523.417452 | -1.076679706 | 0.43275242 | -2.4879808 | 0.01284707 | 0.06595294 |
| ZNF584 | 112.620373 | -0.163056066 | 0.30279043 | -0.5385113 | 0.59022411 | 0.7721373 |
| ZNF132 | 43.440324 | 0.086078459 | 0.37644589 | 0.22866091 | 0.81913248 | 0.91601091 |
| ZNF324 | 94.9404741 | -0.371162281 | 0.24725645 | -1.5011227 | 0.13332382 | 0.3174823 |
| ZNF446 | 92.1417826 | -0.449323446 | 0.27991613 | -1.6052074 | 0.10844817 | 0.27681002 |
| SLC27A5 | 113.056861 | 0.315927666 | 0.19877687 | 1.58935829 | 0.11197953 | 0.28292608 |
| ZBTB45 | 54.5000239 | -0.271836793 | 0.42884546 | -0.6338805 | 0.52615878 | 0.72497874 |
| TRIM28 | 282.976432 | -0.527467119 | 0.40526787 | -1.3015271 | 0.19307809 | 0.40062339 |
| CHMP2A | 222.809475 | -1.024938284 | 0.30491834 | -3.3613534 | 0.00077562 | 0.00822239 |
| UBE2M | 187.898457 | -0.64338445 | 0.3019681 | -2.1306371 | 0.03311905 | 0.12633902 |
| MZF1-AS1 | 211.882354 | 0.070352757 | 0.48250415 | 0.14580757 | 0.8840733 | 0.9477505 |

| | | | | | | |
|---|---|---|---|---|---|---|
| MZF1 | 637.895282 | -0.314974164 | 0.23122398 | -1.3622037 | 0.1731336 | 0.37472092 |
| RPS4Y1 | 483.867644 | -0.033997448 | 1.91701271 | -0.0177346 | 0.98585058 | 0.99426405 |
| ZFY | 331.24576 | 0.71862614 | 1.79879377 | 0.39950446 | 0.68952153 | 0.84044385 |
| ZFY-AS1 | 46.297898 | 0.039741666 | 1.81355256 | 0.02191371 | 0.98251679 | 0.99281397 |
| LINC00278 | 1415.34061 | 0.059836331 | 2.24614786 | 0.02663953 | 0.97874724 | 0.99083348 |
| PCDH11Y | 47.9341188 | -0.005802044 | 0.4300178 | -0.0134926 | 0.98923482 | 0.99594533 |
| TBL1Y | 633.909327 | 1.440075234 | 1.87775943 | 0.76691147 | 0.44313416 | 0.65886527 |
| TTTY15 | 329.050073 | 0.099094667 | 1.97888355 | 0.05007605 | 0.96006179 | 0.98229715 |
| USP9Y | 1359.75984 | 0.365725599 | 2.13458594 | 0.17133327 | 0.86396172 | 0.93827593 |
| DDX3Y | 315.535266 | -0.000378859 | 2.05194114 | -0.0001846 | 0.99985268 | 0.99985268 |
| UTY | 3264.18608 | -0.137905582 | 2.15091217 | -0.0641149 | 0.94887871 | 0.97784954 |
| NLGN4Y | 2311.08104 | -1.51444642 | 1.9107415 | -0.7925962 | 0.42801313 | 0.64642414 |
| TTTY14 | 2702.44213 | 0.402186209 | 1.18411369 | 0.33965168 | 0.73411885 | 0.86945001 |
| KDM5D | 374.914957 | 0.439772406 | 2.4662243 | 0.17831809 | 0.85847317 | 0.93614803 |
| TTTY10 | 199.272312 | 0.570221954 | 1.61873393 | 0.35226416 | 0.72464016 | 0.86356657 |
| EIF1AY | 126.561631 | -0.001170875 | 2.26003718 | -0.0005181 | 0.99958663 | 0.99975417 |
| AP000547.3 | 126.489981 | -0.296870272 | 0.25066422 | -1.1843344 | 0.2362807 | 0.45391953 |
| IL17RA | 308.147959 | -0.662120604 | 0.20384112 | -3.248219 | 0.0011613 | 0.0112378 |
| HDHD5 | 269.979335 | 0.115661243 | 0.35164234 | 0.32891728 | 0.74221821 | 0.87402695 |
| HDHD5-AS1 | 107.998102 | 0.26636551 | 0.54708695 | 0.48687966 | 0.62634361 | 0.79697262 |
| ADA2 | 119.318294 | 0.528860705 | 0.64354901 | 0.82178777 | 0.4111977 | 0.63039814 |
| CECR3 | 64.0327134 | 0.663784482 | 0.66393706 | 0.99977019 | 0.31742173 | 0.54198858 |
| CECR2 | 1774.69206 | 0.30626456 | 0.34663074 | 0.88354704 | 0.37694078 | 0.6000254 |
| ATP6V1E1 | 486.680515 | -0.649994128 | 0.1576981 | -4.1217626 | 3.76E-05 | 0.00078278 |
| BCL2L13 | 1028.48167 | 0.068960779 | 0.15203227 | 0.45359303 | 0.65012177 | 0.81297534 |
| MICAL3 | 2371.32027 | 0.352139477 | 0.33217722 | 1.06009521 | 0.28910129 | 0.51148392 |
| PEX26 | 371.122029 | -0.30573019 | 0.15356437 | -1.9908928 | 0.04649268 | 0.15787192 |
| USP18 | 201.942629 | -0.684150953 | 0.78550089 | -0.8709741 | 0.38376828 | 0.60643561 |
| AC023490.3 | 103.334616 | -1.519317867 | 0.94506339 | -1.6076359 | 0.10791494 | 0.27605362 |
| AC007326.4 | 38.1508644 | 1.829101474 | 0.5428567 | 3.36940019 | 0.00075332 | 0.00807679 |
| DGCR2 | 648.300978 | -0.105754325 | 0.21224309 | -0.4982698 | 0.61829388 | 0.79053759 |
| ESS2 | 53.4938072 | -0.313088662 | 0.34506183 | -0.9073408 | 0.36422662 | 0.58959184 |
| SLC25A1 | 210.513749 | -0.676494824 | 0.37528919 | -1.8025961 | 0.07145167 | 0.21091354 |
| CLTCL1 | 244.526237 | -0.221484774 | 0.30388005 | -0.7288559 | 0.4660898 | 0.67893474 |
| HIRA | 257.651482 | 0.319369207 | 0.25322756 | 1.26119448 | 0.20723879 | 0.41805393 |
| MRPL40 | 173.155475 | -1.292064306 | 0.47607743 | -2.7139794 | 0.00664803 | 0.04132416 |
| UFD1 | 237.549417 | -0.212455452 | 0.24199748 | -0.8779242 | 0.37998484 | 0.60276447 |
| GNB1L | 45.3591292 | -0.002166556 | 0.33774899 | -0.0064147 | 0.99488185 | 0.997919 |
| TXNRD2 | 298.673571 | -0.114603494 | 0.26446558 | -0.4333399 | 0.66476784 | 0.82323933 |
| COMT | 781.571143 | -0.276766115 | 0.25951122 | -1.06649 | 0.2862022 | 0.50853442 |
| ARVCF | 70.1992482 | -0.313071356 | 0.36619 | -0.8549424 | 0.39258304 | 0.61484815 |
| TANGO2 | 338.291804 | 0.208796833 | 0.21989749 | 0.94951895 | 0.34235674 | 0.56753966 |
| AC006547.1 | 73.6241095 | 0.18675075 | 0.44521832 | 0.41945881 | 0.67488085 | 0.82939913 |
| DGCR8 | 309.325212 | -0.02589011 | 0.32922235 | -0.0786402 | 0.93731881 | 0.97284815 |
| TRMT2A | 77.017455 | 0.213453177 | 0.29980217 | 0.7119801 | 0.4764771 | 0.68642011 |
| RANBP1 | 125.954774 | -0.158776046 | 0.30371651 | -0.5227771 | 0.60112936 | 0.77899712 |
| ZDHHC8 | 166.937722 | 0.87267606 | 0.46127052 | 1.89189644 | 0.05850478 | 0.18461291 |
| DGCR6L | 87.1374512 | -0.261450215 | 0.48448068 | -0.5396504 | 0.58943812 | 0.77174702 |
| ZNF74 | 46.6026512 | 0.387106395 | 0.42097573 | 0.91954563 | 0.35781025 | 0.58327972 |
| KLHL22 | 635.46121 | 0.362105155 | 0.19515258 | 1.85549769 | 0.06352519 | 0.19518591 |
| MED15 | 941.429145 | -0.39607428 | 0.24720181 | -1.6022305 | 0.10910464 | 0.27762728 |
| PI4KA | 1371.81878 | 0.168017478 | 0.14338558 | 1.17178784 | 0.24128225 | 0.45963328 |
| SNAP29 | 217.449036 | -0.394861812 | 0.26593438 | -1.4848092 | 0.13759437 | 0.32368372 |
| CRKL | 256.707013 | 0.245379116 | 0.23934993 | 1.02518984 | 0.30527362 | 0.52868668 |
| LZTR1 | 138.645013 | -0.130300104 | 0.24717016 | -0.5271676 | 0.59807719 | 0.77678206 |
| THAP7 | 77.7082727 | -0.335801414 | 0.27892874 | -1.2038968 | 0.22862948 | 0.44417808 |
| GGT2 | 3305.37979 | -3.392903573 | 0.85201549 | -3.9822088 | 6.83E-05 | 0.00123885 |
| HIC2 | 197.824231 | 0.889754711 | 0.27799107 | 3.2006593 | 0.00137114 | 0.01279049 |
| UBE2L3 | 634.971122 | -0.193600643 | 0.16067507 | -1.2049203 | 0.22823409 | 0.44373624 |
| YDJC | 68.5321893 | -1.378411768 | 0.36989547 | -3.72649 | 0.00019416 | 0.00286561 |
| SDF2L1 | 122.298727 | -1.306578913 | 0.54965259 | -2.3770995 | 0.01744938 | 0.08138731 |
| PPIL2 | 716.103622 | 0.60092145 | 0.31647017 | 1.89882493 | 0.0575875 | 0.18300431 |
| YPEL1 | 255.121624 | 0.323122491 | 0.33988637 | 0.950678 | 0.34176786 | 0.56712537 |
| MAPK1 | 1586.207 | -0.118052109 | 0.20810225 | -0.5672793 | 0.57052441 | 0.75945349 |
| PPM1F | 169.953163 | -0.294707447 | 0.3665273 | -0.8040532 | 0.42136625 | 0.64046063 |
| AC245452.1 | 71.8201885 | 0.232637984 | 0.30273268 | 0.76846009 | 0.44221389 | 0.65802453 |
| TOP3B | 188.663366 | 0.212637198 | 0.28400121 | 0.74871935 | 0.45402638 | 0.66958468 |
| AC245060.5 | 328.062167 | 1.055441924 | 0.39033402 | 2.7039455 | 0.00685215 | 0.04218352 |
| RSPH14 | 91.8135287 | 0.336371662 | 0.52983851 | 0.63485695 | 0.52552172 | 0.72472068 |
| RAB36 | 109.384503 | 0.071845846 | 0.49381626 | 0.14549105 | 0.88432318 | 0.9477505 |
| BCR | 2646.89386 | -1.281699536 | 0.33697145 | -3.8035849 | 0.00014262 | 0.00222412 |
| PCAT14 | 96.522281 | 1.029727829 | 0.42175755 | 2.44151604 | 0.01462574 | 0.07205945 |
| AP000345.2 | 35.9927282 | 1.187878358 | 0.71031445 | 1.67232746 | 0.09445977 | 0.25284294 |
| DRICH1 | 86.167955 | 1.342581064 | 0.59961033 | 2.23908928 | 0.02515011 | 0.10407512 |
| CHCHD10 | 854.52342 | -0.01834764 | 0.33606806 | -0.054595 | 0.95646111 | 0.98089246 |
| SMARCB1 | 404.561205 | 0.04706053 | 0.25145039 | 0.18715632 | 0.85153807 | 0.93193618 |
| AP000350.4 | 44.0917619 | 0.952483568 | 0.38761005 | 2.45732426 | 0.01399763 | 0.06971895 |
| GSTT2B | 99.4936935 | 0.241819064 | 0.65227122 | 0.37073391 | 0.71083573 | 0.85502039 |
| DDT | 437.412247 | -0.336239786 | 0.40855845 | -0.8229907 | 0.41051331 | 0.62997812 |
| CABIN1 | 1526.96357 | 0.017196571 | 0.25330138 | 0.06788976 | 0.94587339 | 0.97663109 |
| GUCD1 | 249.151184 | -0.792408912 | 0.18108022 | -4.3760104 | 1.21E-05 | 0.00031113 |
| SNRPD3 | 418.70745 | -0.105221563 | 0.20185354 | -0.5212768 | 0.60217397 | 0.77943322 |
| GGT1 | 281.609598 | -1.713586759 | 0.72987666 | -2.3477758 | 0.01888589 | 0.08572635 |
| LRRC75B | 90.17143 | -0.721014375 | 0.62813724 | -1.1478612 | 0.25102586 | 0.4711952 |
| SGSM1 | 706.199274 | 0.264021389 | 0.54082194 | 0.48818542 | 0.62541851 | 0.79617898 |
| KIAA1671 | 1873.03091 | -0.260490299 | 0.19716513 | -1.3211784 | 0.1864419 | 0.3922139 |
| AL022323.4 | 145.561929 | -0.042417656 | 0.3497626 | -0.1212756 | 0.90347278 | 0.95532916 |
| AL022323.1 | 43.683098 | -0.196520615 | 0.42203368 | -0.4656515 | 0.64146497 | 0.80757823 |
| LRP5L | 621.000347 | -0.119744221 | 0.49051709 | -0.2441183 | 0.80713916 | 0.90976203 |
| GRK3 | 437.473658 | -0.324995829 | 0.28031315 | -1.1594027 | 0.24629207 | 0.46552367 |
| ASPHD2 | 76.0341874 | -0.526325083 | 0.49868032 | -1.0554358 | 0.29122604 | 0.51379788 |
| HPS4 | 445.410751 | -0.079379446 | 0.26778867 | -0.2964257 | 0.766905 | 0.8873553 |
| SRRD | 60.1164869 | 0.028748927 | 0.30207644 | 0.09517104 | 0.92417898 | 0.96654633 |
| TFIP11 | 174.04062 | -0.375142361 | 0.2511591 | -1.4936443 | 0.1352686 | 0.32043812 |
| TPST2 | 832.884024 | 0.249991858 | 0.48310408 | 0.51746998 | 0.60482812 | 0.78156518 |
| MIATNB | 433.074164 | -0.157605894 | 0.39739122 | -0.3966013 | 0.69166146 | 0.84169408 |
| PITPNB | 1223.81412 | -0.256865657 | 0.22224014 | -1.1558023 | 0.24776204 | 0.4670599 |
| TTC28 | 2925.93334 | -0.52468853 | 0.21134306 | -2.4826391 | 0.01304131 | 0.06661358 |
| CHEK2 | 120.602836 | -0.824903712 | 0.40328646 | -2.0454536 | 0.04081016 | 0.14444305 |
| HSCB | 415.220432 | -0.39867731 | 0.23510105 | -1.69577 | 0.08992945 | 0.24457033 |
| CCDC117 | 171.690759 | 0.026985461 | 0.30451064 | 0.08861911 | 0.92938462 | 0.96900671 |
| XBP1 | 327.903262 | -0.889742607 | 0.36927599 | -2.4094245 | 0.0159777 | 0.07651593 |
| Z93930.2 | 749.32726 | 0.148430627 | 0.42847256 | 0.34641804 | 0.72902857 | 0.86691919 |
| ZNRF3 | 2752.86521 | 0.102150039 | 0.32082776 | 0.31839527 | 0.75018513 | 0.87863055 |
| ZNRF3-AS1 | 61.9965027 | -0.163586316 | 0.51703143 | -0.3163953 | 0.75170248 | 0.87947193 |
| KREMEN1 | 465.837806 | -0.176483826 | 0.34328439 | -0.5141039 | 0.60717938 | 0.78314935 |
| RHBDD3 | 140.479203 | 0.01207145 | 0.25413251 | 0.04750061 | 0.96211424 | 0.9832997 |
| EWSR1 | 1233.53132 | 0.452081909 | 0.15153972 | 2.98325691 | 0.00285198 | 0.02229105 |
| GAS2L1 | 74.2742634 | 0.250407116 | 0.35833132 | 0.69881449 | 0.48466797 | 0.69239889 |
| AP1B1 | 416.078557 | -0.013583645 | 0.1639403 | -0.0828573 | 0.93396503 | 0.97167284 |
| THOC5 | 170.843137 | 0.181971666 | 0.22566291 | 0.80638803 | 0.42001915 | 0.6395427 |
| NIPSNAP1 | 220.34969 | 0.220491146 | 0.30380191 | 0.72577274 | 0.4679781 | 0.6801407 |
| NF2 | 511.360996 | 0.13221159 | 0.19614866 | 0.6740377 | 0.50028734 | 0.70482771 |

| | | | | | | |
|---|---|---|---|---|---|---|
| ZMAT5 | 99.2975674 | 0.182512307 | 0.20918228 | 0.87250368 | 0.38293366 | 0.6054789 |
| UQCR10 | 608.63943 | -1.031394221 | 0.47782308 | -2.1585274 | 0.03088685 | 0.12031389 |
| ASCC2 | 361.925523 | 0.016597187 | 0.2196146 | 0.07557415 | 0.9397579 | 0.97411372 |
| MTMR3 | 2070.77245 | -0.085258221 | 0.13776348 | -0.6188739 | 0.53599945 | 0.73290528 |
| LIF | 93.3232969 | -3.016914833 | 0.89697755 | -3.3634229 | 0.00076982 | 0.00818728 |
| TBC1D10A | 219.917506 | -0.071175106 | 0.26317771 | -0.270445 | 0.78681789 | 0.89905515 |
| SF3A1 | 176.181851 | -0.273663549 | 0.20400276 | -1.3414698 | 0.17976796 | 0.38336373 |
| CCDC157 | 66.3160085 | -0.34059582 | 0.26112105 | -1.3043599 | 0.19211092 | 0.39924372 |
| RNF215 | 256.755023 | 0.121304965 | 0.35756145 | 0.33925628 | 0.73441668 | 0.86968813 |
| SEC14L2 | 69.6487333 | -0.848885373 | 0.40075167 | -2.1182329 | 0.03415535 | 0.12893912 |
| SEC14L6 | 226.76367 | 0.226980613 | 0.49390148 | 0.45956658 | 0.64582735 | 0.81059872 |
| GAL3ST1 | 169.100907 | 1.645333811 | 0.32510957 | 5.06085935 | 4.17E-07 | 2.03E-05 |
| PES1 | 189.780145 | -0.299064689 | 0.2585736 | -1.156594 | 0.24743826 | 0.46678263 |
| TCN2 | 128.389028 | 0.457218915 | 0.48360822 | 0.94543248 | 0.34443813 | 0.56927968 |
| SLC35E4 | 92.3325508 | 0.025100179 | 0.31595739 | 0.07944166 | 0.93668134 | 0.97284815 |
| DUSP18 | 235.434433 | -0.10377667 | 0.27074506 | -0.3833003 | 0.70149709 | 0.84869711 |
| OSBP2 | 790.963771 | 0.744366352 | 0.20111459 | 3.7012051 | 0.00021458 | 0.00311126 |
| MORC2-AS1 | 48.2766384 | -0.152304483 | 0.33915228 | -0.449074 | 0.65337825 | 0.81539269 |
| MORC2 | 342.249206 | -0.355876772 | 0.22873446 | -1.5558512 | 0.11974349 | 0.29536727 |
| SMTN | 99.863834 | 0.56852722 | 0.54542515 | 1.04235608 | 0.29724662 | 0.52073351 |
| SELENOM | 236.845346 | -0.739714637 | 0.54980121 | -1.345422 | 0.178489 | 0.38170277 |
| INPP5J | 78.4072273 | 1.700972587 | 0.58247279 | 2.92026102 | 0.00349738 | 0.02596792 |
| RNF185 | 452.94792 | -0.091319705 | 0.16767917 | -0.5446097 | 0.58602196 | 0.77001836 |
| LIMK2 | 495.907592 | -1.191445554 | 0.22898902 | -5.2030684 | 1.96E-07 | 1.08E-05 |
| PIK3IP1 | 168.131972 | -1.377758602 | 0.38576534 | -3.571494 | 0.00035495 | 0.00454316 |
| PIK3IP1-AS1 | 107.033872 | -1.561602789 | 0.31789381 | -4.9123409 | 9.00E-07 | 3.86E-05 |
| PATZ1 | 271.662418 | 0.165103969 | 0.335792 | 0.49168524 | 0.62294187 | 0.79471114 |
| DRG1 | 643.536922 | 0.268014054 | 0.19173484 | 1.39783698 | 0.16216203 | 0.3599997 |
| EIF4ENIF1 | 563.008435 | 0.051296932 | 0.13412643 | 0.38245208 | 0.70212606 | 0.8488446 |
| SFI1 | 585.121772 | -0.173006221 | 0.26393787 | -0.6554808 | 0.51215821 | 0.71407594 |
| PISD | 1134.41375 | -0.005116789 | 0.17467479 | -0.0292932 | 0.97663072 | 0.98948597 |
| PRR14L | 1042.15946 | -0.146997657 | 0.17027545 | -0.8632933 | 0.38797622 | 0.61116244 |
| YWHAH | 160.999203 | -2.10555378 | 0.47251446 | -4.4560621 | 8.35E-06 | 0.00023362 |
| RTCB | 478.093601 | -0.278478061 | 0.17138284 | -1.6248888 | 0.10418626 | 0.27026636 |
| FBXO7 | 534.812231 | 0.008663152 | 0.17040458 | 0.05083873 | 0.95945403 | 0.98229715 |
| SYN3 | 897.718364 | 3.608597729 | 0.84047673 | 4.29351293 | 1.76E-05 | 0.00042645 |
| TIMP3 | 689.494374 | -0.758600598 | 0.46707692 | -1.6241449 | 0.10434489 | 0.27043463 |
| LARGE1 | 1991.04067 | -0.672423052 | 0.2459657 | -2.7338083 | 0.00626065 | 0.03944087 |
| HMGXB4 | 351.027748 | -0.097985873 | 0.29516724 | -0.3319673 | 0.73991395 | 0.87265237 |
| TOM1 | 620.171151 | -0.196944856 | 0.24464913 | -0.8050094 | 0.42081424 | 0.64027651 |
| HMOX1 | 185.925782 | -4.209610444 | 0.92979733 | -4.5274495 | 5.97E-06 | 0.00017557 |
| MCM5 | 61.4892464 | -0.552758405 | 0.45043044 | -1.2271782 | 0.21975563 | 0.43361796 |
| APOL6 | 213.116344 | -1.797078575 | 0.64004407 | -2.8077419 | 0.00498902 | 0.03331646 |
| RBFOX2 | 3231.39361 | -0.077860813 | 0.14566334 | -0.5345258 | 0.5929778 | 0.77387448 |
| APOL4 | 154.452176 | -1.738776962 | 1.00610935 | -1.7282187 | 0.08394903 | 0.23347383 |
| APOL2 | 487.650705 | -1.157143555 | 0.55420116 | -2.0879486 | 0.03680247 | 0.13507191 |
| APOL1 | 227.500087 | -3.248441609 | 0.76130476 | -4.2669398 | 1.98E-05 | 0.00046683 |
| MYH9 | 1740.10039 | 0.355757913 | 0.29572412 | 1.20300606 | 0.22897398 | 0.44451188 |
| TXN2 | 257.292643 | -0.238914972 | 0.3070098 | -0.7781998 | 0.43645122 | 0.65341766 |
| EIF3D | 366.506422 | -0.571106222 | 0.25367465 | -2.2513335 | 0.02436442 | 0.10184483 |
| IFT27 | 197.575654 | 0.335679949 | 0.28222987 | 1.18938491 | 0.23428824 | 0.45161281 |
| PVALB | 55.3888435 | 1.681618962 | 0.85683842 | 1.96258586 | 0.04969432 | 0.16555163 |
| TST | 211.824432 | -1.002658412 | 0.35853504 | -2.7965424 | 0.00516526 | 0.03430265 |
| MPST | 255.498156 | -0.230653421 | 0.2860761 | -0.806266 | 0.42008951 | 0.63957611 |
| C1QTNF6 | 91.7184953 | -0.556135463 | 0.40501052 | -1.3731383 | 0.16970935 | 0.36997702 |
| CARD10 | 87.1085077 | -0.594834705 | 0.66137776 | -0.8993872 | 0.36844644 | 0.5935133 |
| CDC42EP1 | 52.385687 | -2.759389573 | 0.56361058 | -4.8959151 | 9.78E-07 | 4.05E-05 |
| GGA1 | 557.36506 | 0.048057801 | 0.17371962 | 0.27664002 | 0.78205653 | 0.89588513 |
| LGALS1 | 56.0374501 | -1.960402413 | 0.42644293 | -4.5971038 | 4.28E-06 | 0.00013191 |
| TRIOBP | 661.409708 | -0.479524878 | 0.31126603 | -1.5405628 | 0.12342322 | 0.30129317 |
| H1F0 | 440.419022 | -1.155680446 | 0.6655119 | -1.7365286 | 0.08247041 | 0.23078253 |
| GCAT | 77.0122943 | 0.301345167 | 0.5778981 | 0.52145035 | 0.60205309 | 0.77940904 |
| ANKRD54 | 156.799597 | 0.183093954 | 0.17987928 | 1.0178713 | 0.30873912 | 0.5326022 |
| EIF3L | 558.589647 | -0.141746952 | 0.21624856 | -0.6554816 | 0.51215773 | 0.71407594 |
| MICALL1 | 115.203423 | -0.826084626 | 0.33163971 | -2.4909099 | 0.01274164 | 0.06557701 |
| POLR2F | 157.599017 | 0.13824117 | 0.29813815 | 0.46368159 | 0.64287589 | 0.80857472 |
| PICK1 | 219.65799 | 0.014409542 | 0.27658709 | 0.05209766 | 0.95845087 | 0.98194195 |
| AL031587.2 | 115.176788 | 0.233372149 | 0.38931686 | 0.59944014 | 0.54887942 | 0.74226986 |
| BAIAP2L2 | 91.5059327 | 0.83772474 | 0.34770112 | 2.40932423 | 0.01598209 | 0.07651593 |
| PLA2G6 | 260.149099 | 0.762075161 | 0.22555214 | 3.37870945 | 0.00072827 | 0.00787855 |
| MAFF | 155.644004 | -1.598325474 | 0.59779365 | -2.6737077 | 0.00750178 | 0.04502091 |
| TMEM184B | 389.498314 | -0.272158782 | 0.21322581 | -1.2763876 | 0.20181854 | 0.41120177 |
| CSNK1E | 143.8392 | 0.313020257 | 0.43326136 | 0.72247444 | 0.47000283 | 0.68177766 |
| TPTEP2-CSNI | 1807.46029 | 0.315477482 | 0.23975432 | 1.31583648 | 0.18822893 | 0.39384317 |
| DDX17 | 7845.14199 | 0.613792296 | 0.2093314 | 2.93215584 | 0.00336618 | 0.02527791 |
| FAM227A | 300.51802 | 0.135961416 | 0.24059916 | 0.56509515 | 0.57200905 | 0.76012752 |
| CBY1 | 292.719148 | 0.257608359 | 0.177855 | 1.44841787 | 0.14750022 | 0.33872225 |
| AL021707.2 | 69.6981694 | 0.930095271 | 0.25752224 | 3.61170855 | 0.00030419 | 0.00405858 |
| TOMM22 | 71.2653128 | -0.404578572 | 0.44414987 | -0.9109055 | 0.36234515 | 0.58770777 |
| JOSD1 | 292.37141 | -0.556891867 | 0.20295957 | -2.7438562 | 0.00607221 | 0.03869169 |
| GTPBP1 | 250.243791 | -0.301162037 | 0.19418952 | -1.5508666 | 0.12093367 | 0.29724838 |
| SUN2 | 237.214412 | -0.804924954 | 0.26520495 | -3.0351054 | 0.00240452 | 0.01965425 |
| AL021707.1 | 182.751335 | -0.358266771 | 0.2835846 | -1.2633506 | 0.20646321 | 0.41739901 |
| CBX6 | 402.067336 | -1.035213978 | 0.49985571 | -2.0710256 | 0.0383564 | 0.13904125 |
| CBX7 | 310.227043 | -0.419013237 | 0.26589913 | -1.5758353 | 0.11506377 | 0.28840177 |
| AL031846.2 | 111.371851 | -0.630935208 | 0.37962321 | -1.6620038 | 0.09651202 | 0.25648172 |
| PDGFB | 63.1915453 | -0.717354072 | 0.64335244 | -1.115025 | 0.2648397 | 0.4858705 |
| AL031590.1 | 43.457381 | -0.569384539 | 0.64768082 | -0.8791129 | 0.37934009 | 0.60232038 |
| RPL3 | 814.185163 | -0.850268659 | 0.49871723 | -1.7049113 | 0.08821096 | 0.24143775 |
| SYNGR1 | 233.621045 | 0.192486768 | 0.41020406 | 0.46924637 | 0.63889354 | 0.8057232 |
| TAB1 | 240.526581 | 0.31471041 | 0.19880011 | 1.58304944 | 0.1134102 | 0.28563167 |
| MIEF1 | 125.046692 | -0.311138686 | 0.23931782 | -1.3001066 | 0.19356442 | 0.40119134 |
| ATF4 | 426.383676 | -0.11557335 | 0.24089437 | -0.4797678 | 0.63139255 | 0.8011589 |
| RPS19BP1 | 281.30407 | -0.706487621 | 0.38315324 | -1.8438775 | 0.065201 | 0.1984424 |
| FAM83F | 229.585749 | -0.193000605 | 0.4579776 | -0.4214193 | 0.67344892 | 0.82886112 |
| TNRC6B | 7556.61751 | 0.285170622 | 0.16495519 | 1.72877634 | 0.08384913 | 0.23331856 |
| AL022238.2 | 74.098154 | 0.658607406 | 0.62120705 | 1.06020595 | 0.28905091 | 0.5114633 |
| MKL1 | 2550.30403 | -0.086439276 | 0.22004135 | -0.392832 | 0.69444359 | 0.843734 |
| SLC25A17 | 588.504802 | 0.114361336 | 0.18831937 | 0.60727335 | 0.54366952 | 0.7391242 |
| ST13 | 760.244933 | -0.277718124 | 0.20846578 | -1.3322001 | 0.18279445 | 0.38750311 |
| XPNPEP3 | 831.340455 | 0.231125961 | 0.1668609 | 1.38514154 | 0.16600919 | 0.36540834 |
| RBX1 | 511.651233 | 0.086354386 | 0.19530582 | 0.44214958 | 0.65838097 | 0.81895766 |
| EP300 | 1684.01138 | 0.055330446 | 0.180365 | 0.30676931 | 0.75901897 | 0.88318196 |
| L3MBTL2 | 154.338311 | -0.097547852 | 0.26373541 | -0.3698701 | 0.71147925 | 0.85521244 |
| AL035681.1 | 60.0187324 | -0.073915701 | 0.37770368 | -0.1956976 | 0.84484687 | 0.92831294 |
| RANGAP1 | 266.866624 | -0.464691447 | 0.26920027 | -1.7261924 | 0.08431281 | 0.23395814 |
| ZC3H7B | 486.150745 | -0.032068234 | 0.22355565 | -0.1434463 | 0.88593771 | 0.94846586 |
| TEF | 177.673502 | 0.754555335 | 0.56508604 | 1.33529282 | 0.18178053 | 0.38609728 |
| TOB2 | 337.198609 | 0.372288491 | 0.24882882 | 1.49616305 | 0.13461117 | 0.31933889 |
| PHF5A | 89.2624831 | -0.178526476 | 0.25950657 | -0.6879459 | 0.49148687 | 0.69766264 |
| ACO2 | 1015.68088 | 0.418814567 | 0.16769008 | 2.49755124 | 0.01250544 | 0.06467672 |
| POLR3H | 140.463749 | -0.417780182 | 0.25028339 | -1.6692285 | 0.09507209 | 0.25403624 |
| CSDC2 | 206.127967 | -0.086925005 | 0.56015544 | -0.1551801 | 0.8766793 | 0.94461676 |

| | | | | | | |
|---|---|---|---|---|---|---|
| PMM1 | 212.077621 | 0.705321115 | 0.33397238 | 2.11191449 | 0.03469378 | 0.13011647 |
| DESI1 | 331.605135 | -0.079335422 | 0.21458895 | -0.3697088 | 0.71159948 | 0.85527912 |
| XRCC6 | 828.677386 | -0.417773818 | 0.19510087 | -2.1413222 | 0.03224806 | 0.12393503 |
| SNU13 | 597.642879 | -0.555078174 | 0.24095676 | -2.3036422 | 0.02124273 | 0.0931282 |
| C22orf46 | 96.6834927 | -0.165591393 | 0.40392896 | -0.4099518 | 0.68184132 | 0.83408428 |
| CCDC134 | 172.107061 | -0.162940045 | 0.46084435 | -0.3535685 | 0.72366229 | 0.86271256 |
| SREBF2 | 2919.72269 | -0.177051333 | 0.31048435 | -0.5702424 | 0.56851332 | 0.75853459 |
| WBP2NL | 403.620318 | 0.437476821 | 0.4875868 | 0.89722859 | 0.36959695 | 0.59419866 |
| NAGA | 50.6707148 | -0.545432274 | 0.31730723 | -1.7189406 | 0.08562519 | 0.23656624 |
| SMDT1 | 271.256645 | -0.645158618 | 0.38279253 | -1.6854002 | 0.09191138 | 0.24822274 |
| NDUFA6 | 264.286911 | -0.183668015 | 0.32885333 | -0.5585104 | 0.57649588 | 0.76324888 |
| TCF20 | 1306.92202 | -0.559401581 | 0.19601252 | -2.8539074 | 0.00431851 | 0.03016353 |
| AL022316.1 | 51.7309093 | -2.969068589 | 0.74801045 | -3.9692876 | 7.21E-05 | 0.00129906 |
| RRP7A | 196.791606 | -0.319690994 | 0.39944358 | -0.8003408 | 0.42351337 | 0.6424194 |
| SERHL2 | 79.4704546 | -0.274769762 | 0.29562139 | -0.9294651 | 0.35264811 | 0.57879335 |
| POLDIP3 | 354.417848 | -0.460828949 | 0.17255205 | -2.6706663 | 0.00757009 | 0.045369 |
| CYB5R3 | 641.44004 | -1.664784488 | 0.22382255 | -7.4379657 | 1.02E-13 | 4.66E-11 |
| A4GALT | 423.97491 | -3.965315307 | 0.62164203 | -6.3787761 | 1.79E-10 | 2.81E-08 |
| ARFGAP3 | 2058.15646 | -1.403699282 | 0.31727177 | -4.4242804 | 9.68E-06 | 0.00025976 |
| PACSIN2 | 2049.33193 | -0.431404089 | 0.20593321 | -2.0948738 | 0.03618221 | 0.13384823 |
| TTLL1 | 104.497615 | -0.220145018 | 0.34803561 | -0.6325359 | 0.52703675 | 0.72541009 |
| MCAT | 106.894386 | 0.130175612 | 0.3615791 | 0.36001974 | 0.71883237 | 0.85990372 |
| TSPO | 408.461824 | -1.863322708 | 0.39574241 | -4.7084231 | 2.50E-06 | 8.52E-05 |
| TTLL12 | 84.119372 | -0.469291719 | 0.35513304 | -1.3214533 | 0.18635028 | 0.39214606 |
| EFCAB6 | 1289.37139 | -0.104085029 | 0.24473522 | -0.4252965 | 0.67062055 | 0.82694426 |
| SAMM50 | 168.585408 | 0.47007742 | 0.23694416 | 1.98391645 | 0.04726515 | 0.15959239 |
| PARVB | 65.2213306 | 0.230421252 | 0.7292419 | 0.31597369 | 0.75202248 | 0.8795471 |
| RTL6 | 53.3805942 | -0.703598092 | 0.42572549 | -1.6527037 | 0.09839118 | 0.2598259 |
| PHF21B | 1200.06176 | 0.487106474 | 0.38440038 | 1.26718521 | 0.20508907 | 0.41549411 |
| NUP50-DT | 690.594548 | -0.15938852 | 0.28143709 | -0.566338 | 0.57116403 | 0.75965257 |
| NUP50 | 294.469966 | -0.298804094 | 0.16518593 | -1.8088956 | 0.07046723 | 0.20912191 |
| KIAA0930 | 164.752012 | -1.511213763 | 0.30195343 | -5.0047909 | 5.59E-07 | 2.58E-05 |
| FAM118A | 261.206799 | 0.195580285 | 0.38901698 | 0.50275514 | 0.61513644 | 0.78848493 |
| FBLN1 | 283.7204 | -4.815570183 | 1.16831043 | -4.1218242 | 3.76E-05 | 0.00078278 |
| ATXN10 | 2542.40882 | 0.175623699 | 0.21644616 | 0.81139669 | 0.4171379 | 0.63699466 |
| MIRLET7BHG | 1515.31292 | -0.46744941 | 0.3122442 | -1.4970635 | 0.13437673 | 0.31889727 |
| PPARA | 2187.4203 | 0.709082585 | 0.23194885 | 3.05706447 | 0.00223516 | 0.01862729 |
| TTC38 | 385.280016 | 0.107335393 | 0.23276076 | 0.46114041 | 0.64469787 | 0.80987296 |
| TRMU | 402.916954 | 0.273617851 | 0.21482836 | 1.27365794 | 0.20278467 | 0.41232902 |
| CELSR1 | 726.561681 | -0.027706766 | 0.37821647 | -0.0732564 | 0.94160211 | 0.97495762 |
| GRAMD4 | 369.288901 | 0.814312828 | 0.22973073 | 3.5446404 | 0.00039315 | 0.00492029 |
| CERK | 268.543783 | 0.025020989 | 0.36276316 | 0.06897335 | 0.94501084 | 0.97627469 |
| AL118516.1 | 87.1992642 | 0.514383936 | 0.49969452 | 1.02939679 | 0.30329326 | 0.52644076 |
| TBC1D22A | 3017.11744 | -0.419366345 | 0.31641571 | -1.3253651 | 0.18505008 | 0.3906547 |
| BRD1 | 642.918118 | -0.060146126 | 0.22766669 | -0.264185 | 0.79163736 | 0.90190943 |
| ZBED4 | 365.87593 | -0.037160414 | 0.33290179 | -0.1116258 | 0.91112015 | 0.95980429 |
| ALG12 | 123.949646 | 0.248133692 | 0.26808847 | 0.92556644 | 0.35467134 | 0.58039381 |
| CRELD2 | 97.8955457 | 0.6341746 | 0.40536088 | 1.56446917 | 0.11770743 | 0.29247506 |
| PIM3 | 502.269609 | -0.646689418 | 0.36544267 | -1.7696057 | 0.07679286 | 0.22104093 |
| TRABD | 303.399867 | 0.195548178 | 0.27964597 | 0.6992705 | 0.484383 | 0.69235581 |
| SELENOO | 552.442261 | -0.10960112 | 0.25400056 | -0.4314995 | 0.6661052 | 0.82383742 |
| TUBGCP6 | 180.480581 | 0.376853991 | 0.28011909 | 1.34533492 | 0.1785171 | 0.38170277 |
| HDAC10 | 43.9589174 | 0.22262481 | 0.47065642 | 0.47300919 | 0.63620662 | 0.80446022 |
| PLXNB2 | 819.748075 | -0.581025602 | 0.27651961 | -2.1012094 | 0.03562259 | 0.13239418 |
| DENND6B | 173.259639 | 0.106586352 | 0.40303011 | 0.2644625 | 0.79142356 | 0.90182141 |
| PPP6R2 | 2761.66171 | 0.387097605 | 0.1701875 | 2.27453611 | 0.02293377 | 0.09791161 |
| SBF1 | 126.515902 | 0.169192232 | 0.19296308 | 0.87681141 | 0.38058908 | 0.60347686 |
| MIOX | 108.406552 | 1.143688748 | 0.78221388 | 1.46211769 | 0.14370896 | 0.333319 |
| LMF2 | 132.200362 | -0.707555202 | 0.38110972 | -1.8565656 | 0.06337298 | 0.1951062 |
| NCAPH2 | 72.0978214 | 0.242143114 | 0.32590234 | 0.74299287 | 0.457486 | 0.67218382 |
| TYMP | 82.4399368 | -1.086230058 | 0.43157259 | -2.5169116 | 0.01183885 | 0.06245184 |
| ARSA | 106.952718 | -0.367009807 | 0.24459834 | -1.5004591 | 0.13349552 | 0.31771933 |
| RABL2B | 310.604085 | 0.147586108 | 0.24431967 | 0.6040697 | 0.54579731 | 0.74015676 |
| CU633967.1 | 210.643805 | 0.23708323 | 0.68160772 | 0.34782944 | 0.72796828 | 0.86612619 |
| U2AF1L5 | 50.5895253 | -2.19712055 | 1.04011395 | -2.1123845 | 0.03465349 | 0.1301131 |
| FP475955.1 | 47.3819069 | -2.081505127 | 0.84035949 | -2.4769223 | 0.01325208 | 0.06711401 |
| FP671120.1 | 3531.63061 | -0.070673499 | 0.41359385 | -0.1708766 | 0.86432082 | 0.93833133 |
| FP236383.1 | 2071.25225 | -0.307769807 | 0.49222575 | -0.6252615 | 0.53179945 | 0.72906475 |
| CR381653.1 | 399.172673 | 0.506736366 | 0.53122669 | 0.95389855 | 0.340135 | 0.56546725 |
| AP001347.1 | 100.206076 | -0.062860331 | 0.48752017 | -0.1289389 | 0.89740597 | 0.9525128 |
| RBM11 | 109.420129 | -0.584480078 | 0.34542161 | -1.6920774 | 0.09063123 | 0.24583023 |
| HSPA13 | 115.218574 | -0.009214287 | 0.33449466 | -0.0275469 | 0.97802354 | 0.99047098 |
| AF127577.4 | 127.127558 | -0.057890438 | 0.35097302 | -0.1649427 | 0.86898908 | 0.94023032 |
| NRIP1 | 1386.37426 | -0.415440724 | 0.25529709 | -1.6272834 | 0.10367692 | 0.2693683 |
| AF127577.2 | 54.7229935 | -0.713976109 | 0.54051429 | -1.32092 | 0.18652806 | 0.39228851 |
| USP25 | 3680.76022 | 0.703464776 | 0.20755726 | 3.38925637 | 0.00070082 | 0.00766324 |
| MIR99AHG | 4895.10665 | 1.30532782 | 0.3258411 | 4.00602566 | 6.17E-05 | 0.00115289 |
| CXADR | 511.662633 | -0.692053554 | 0.20002979 | -3.4597525 | 0.00054067 | 0.00627569 |
| BTG3 | 230.268005 | -1.014720614 | 0.26554021 | -3.8213445 | 0.00013273 | 0.00210466 |
| C21orf91 | 162.382489 | 0.101722924 | 0.29599129 | 0.34366863 | 0.7310955 | 0.86806155 |
| CHODL | 508.990441 | 0.971310648 | 0.54311531 | 1.78840594 | 0.07371054 | 0.21540763 |
| MRPL39 | 201.729276 | 0.343655866 | 0.2136285 | 1.60866111 | 0.10769046 | 0.27583544 |
| JAM2 | 112.059954 | 1.623732279 | 0.64064278 | 2.53453614 | 0.01125963 | 0.06034355 |
| ATP5PF | 714.133769 | 0.242524089 | 0.37132159 | 0.65313759 | 0.51366758 | 0.71549458 |
| GABPA | 338.754302 | 0.200237689 | 0.17234804 | 1.16182164 | 0.24530792 | 0.46435545 |
| APP | 8889.13847 | -0.210511567 | 0.21048544 | -1.0001241 | 0.31725045 | 0.5418362 |
| AP001439.1 | 206.714325 | -0.009217175 | 0.41800754 | -0.0220503 | 0.98240786 | 0.99281397 |
| AP000229.1 | 47.5490811 | 1.223613923 | 0.56394157 | 2.16975302 | 0.03002556 | 0.11765281 |
| CYYR1 | 282.505045 | -0.684434912 | 0.46901525 | -1.459302 | 0.14448199 | 0.33429018 |
| ADAMTS1 | 5363.47415 | -4.736921145 | 0.59936961 | -7.903172 | 2.72E-15 | 1.89E-12 |
| ADAMTS5 | 121.488937 | -1.580141081 | 1.01558111 | -1.5558985 | 0.11973224 | 0.29536727 |
| LINC00113 | 196.743752 | 3.161925985 | 0.81189559 | 3.89449827 | 9.84E-05 | 0.00165158 |
| LINC01695 | 84.8819798 | 0.060931416 | 0.4925042 | 0.12371756 | 0.90153891 | 0.95457905 |
| AF165147.1 | 298.459267 | -0.411841384 | 0.56751854 | -0.725688 | 0.46803007 | 0.6801407 |
| N6AMT1 | 114.902436 | 0.439751677 | 0.19133608 | 2.2983207 | 0.02154354 | 0.09391705 |
| LTN1 | 920.325796 | 0.335319201 | 0.16363011 | 2.04925119 | 0.04043756 | 0.14362455 |
| RWDD2B | 97.8174432 | 0.559554945 | 0.28024193 | 1.99668531 | 0.04585938 | 0.1564858 |
| USP16 | 486.993065 | 0.100232709 | 0.1672545 | 0.59928259 | 0.54898445 | 0.74226986 |
| CCT8 | 335.807577 | -0.168533253 | 0.23613799 | -0.7137067 | 0.47540859 | 0.68538649 |
| MAP3K7CL | 397.805079 | 0.045255949 | 0.30750749 | 0.14717023 | 0.88299766 | 0.94732911 |
| BACH1 | 3551.26314 | -0.380211565 | 0.27423562 | -1.3864412 | 0.16561223 | 0.36496028 |
| GRIK1 | 210.072485 | -1.180435729 | 0.55404004 | -2.1305964 | 0.03312241 | 0.12633902 |
| CLDN8 | 37.1133839 | 0.958440043 | 0.63291612 | 1.51432396 | 0.1299437 | 0.31196406 |
| TIAM1 | 126.966046 | -0.054632788 | 0.50197141 | -0.1088365 | 0.91333221 | 0.96075537 |
| SOD1 | 671.618395 | -0.586456193 | 0.38694112 | -1.5156213 | 0.12961515 | 0.31145834 |
| SCAF4 | 731.482481 | 0.307731181 | 0.16606864 | 1.85303603 | 0.06387719 | 0.19608502 |
| HUNK | 598.970523 | -1.904977076 | 0.6897918 | -2.7616697 | 0.00575066 | 0.03723554 |
| MIS18A | 44.1653132 | -0.18683363 | 0.28668501 | -0.6517035 | 0.51459245 | 0.71617867 |
| URB1 | 278.254909 | 0.058656549 | 0.27116273 | 0.21631494 | 0.82874227 | 0.92023714 |
| EVA1C | 498.124584 | -1.473842307 | 0.36989678 | -3.9844692 | 6.76E-05 | 0.00123389 |
| SYNJ1 | 854.008754 | 0.122748975 | 0.18772508 | 0.65387626 | 0.51319153 | 0.71500948 |
| PAXBP1-AS1 | 97.8537645 | 0.265264648 | 0.39307889 | 0.6748382 | 0.49977857 | 0.70482771 |
| PAXBP1 | 1410.48093 | 0.36377416 | 0.19006703 | 1.9139256 | 0.05562966 | 0.17856919 |

| | | | | | | |
|---|---|---|---|---|---|---|
| C21orf62-AS | 346.336988 | 0.872328428 | 0.36160306 | 2.41239229 | 0.01584822 | 0.07609565 |
| C21orf62 | 281.215134 | 0.73933368 | 0.61961513 | 1.19321438 | 0.23278542 | 0.44954133 |
| AP000282.1 | 521.928675 | 1.15560981 | 0.74768163 | 1.54559076 | 0.12220344 | 0.29942636 |
| IFNAR1 | 636.69712 | -0.573270231 | 0.11557856 | -4.9600048 | 7.05E-07 | 3.14E-05 |
| IFNGR2 | 811.965246 | -0.203633779 | 0.27379666 | -0.7437409 | 0.45703323 | 0.67196705 |
| TMEM50B | 702.641874 | 0.452377924 | 0.15757235 | 2.87092202 | 0.00409276 | 0.02914357 |
| DNAJC28 | 41.945149 | 0.99810157 | 0.38591153 | 2.58634811 | 0.00969989 | 0.05415294 |
| GART | 493.000494 | 0.163382485 | 0.20064916 | 0.81426949 | 0.41549059 | 0.63535693 |
| SON | 2207.21604 | -0.235105512 | 0.25158084 | -0.9345128 | 0.35003944 | 0.57608659 |
| DONSON | 92.5618532 | 0.197753381 | 0.41687558 | 0.47437027 | 0.63523589 | 0.80389813 |
| AP000311.1 | 470.804431 | 0.030013657 | 0.41568541 | 0.07220282 | 0.9424405 | 0.97552673 |
| ITSN1 | 1609.87952 | 0.148275904 | 0.1493537 | 0.99278358 | 0.32081543 | 0.54511971 |
| LINC00649 | 136.851206 | 1.26581514 | 0.60946845 | 2.07691661 | 0.03780926 | 0.13759387 |
| MRPS6 | 5740.90708 | 1.541184866 | 0.54736379 | 2.81565 | 0.00486787 | 0.0327725 |
| LINC00310 | 881.564984 | 2.129438902 | 0.58253795 | 3.65545096 | 0.00025673 | 0.00356588 |
| AP000317.1 | 938.636934 | 1.557485161 | 0.44604178 | 3.49179208 | 0.00047979 | 0.00574055 |
| KCNE2 | 48.4715642 | 0.622060049 | 0.61627506 | 1.00938703 | 0.31278906 | 0.5367235 |
| RCAN1 | 444.571748 | -0.324949423 | 0.57242746 | -0.5676692 | 0.57025963 | 0.75945349 |
| CLIC6 | 192.366066 | -1.842322846 | 0.58091801 | -3.1713991 | 0.00151707 | 0.0138486 |
| LINC01426 | 49.1366249 | -2.905674545 | 0.84139618 | -3.4533964 | 0.00055357 | 0.00636394 |
| RUNX1 | 6036.74426 | -0.267843735 | 0.5695523 | -0.4702707 | 0.63816166 | 0.80525266 |
| SETD4 | 436.944391 | 0.463777348 | 0.29134457 | 1.59185167 | 0.11141804 | 0.28199288 |
| CBR1 | 656.99856 | -0.537308759 | 0.34792615 | -1.5443184 | 0.12251122 | 0.2998982 |
| AP000688.2 | 171.351492 | -0.031957906 | 0.43497279 | -0.073471 | 0.94143128 | 0.97494048 |
| CBR3 | 290.841819 | -0.013686255 | 0.4320143 | -0.0316801 | 0.97472717 | 0.98880049 |
| DOPEY2 | 271.758793 | -0.97009685 | 0.36851333 | -2.6324607 | 0.00847688 | 0.04921809 |
| MORC3 | 1020.98387 | -0.008168582 | 0.12753746 | -0.0640485 | 0.94893162 | 0.97784954 |
| CHAF1B | 48.183944 | -0.24309756 | 0.33232116 | -0.7315139 | 0.4644653 | 0.6774649 |
| AP000695.1 | 45.7823836 | -1.442406713 | 0.7843576 | -1.8389657 | 0.06592023 | 0.20017327 |
| AP000695.2 | 356.431044 | -0.598554417 | 0.43613596 | -1.3724033 | 0.16993793 | 0.37016988 |
| CLDN14 | 1700.88282 | 0.464267331 | 0.41084212 | 1.13003831 | 0.25846008 | 0.47954387 |
| AP000696.1 | 148.581442 | 0.937747456 | 0.31015587 | 3.02347159 | 0.00249892 | 0.02021329 |
| SIM2 | 4962.1858 | 0.37665619 | 0.41130508 | 0.91575866 | 0.35979349 | 0.58502113 |
| HLCS | 2198.92894 | -0.13782229 | 0.16132659 | -0.8543061 | 0.39293541 | 0.6151812 |
| PIGP | 154.650673 | 0.343638653 | 0.28047437 | 1.22520518 | 0.22049793 | 0.43432107 |
| TTC3 | 2863.29249 | 0.464514769 | 0.15082891 | 3.07974627 | 0.00207177 | 0.01762139 |
| VPS26C | 564.855776 | -0.347374015 | 0.14505972 | -2.3946965 | 0.01663413 | 0.07878099 |
| DYRK1A | 4048.04541 | 0.059310877 | 0.17825766 | 0.33272554 | 0.73934147 | 0.87241119 |
| KCNJ6 | 55.2760935 | -1.844723456 | 0.45442376 | -4.0594785 | 4.92E-05 | 0.00096102 |
| KCNJ15 | 178.860042 | 1.096098693 | 0.47118635 | 2.32625307 | 0.02000505 | 0.08910376 |
| ETS2 | 575.994841 | -2.477790767 | 0.39382137 | -6.2916615 | 3.14E-10 | 4.51E-08 |
| PSMG1 | 90.5126046 | 0.162001698 | 0.28606484 | 0.56631111 | 0.57118231 | 0.75965257 |
| BRWD1 | 3781.74776 | 0.230290711 | 0.17198421 | 1.3390224 | 0.18056338 | 0.38437739 |
| HMGN1 | 747.28889 | -0.295597326 | 0.1779509 | -1.6611173 | 0.09668989 | 0.25687384 |
| WRB | 254.779463 | -0.013463845 | 0.26839355 | -0.0501646 | 0.95999125 | 0.98229715 |
| LCA5L | 165.65142 | 0.473411019 | 0.22350575 | 2.1181156 | 0.03416528 | 0.12893976 |
| SH3BGR | 136.484569 | 1.596175364 | 0.28412235 | 5.61791554 | 1.93E-08 | 1.61E-06 |
| B3GALT5 | 43.5990218 | -0.832822146 | 0.59111555 | -1.4088991 | 0.15886501 | 0.35524819 |
| PCP4 | 466.503567 | 2.246339159 | 0.89210316 | 2.51802624 | 0.01180145 | 0.06230561 |
| DSCAM | 48.4163845 | 0.738892208 | 0.30277014 | 2.44043946 | 0.0146694 | 0.07222071 |
| BACE2 | 4682.13474 | 0.033536836 | 0.53429479 | 0.06276841 | 0.94995092 | 0.97825441 |
| PLAC4 | 51.4715497 | -0.057871194 | 0.96536552 | -0.0599474 | 0.95219749 | 0.97933189 |
| FAM3B | 181.24265 | -0.604659035 | 0.33015341 | -1.8314487 | 0.06703359 | 0.2022236 |
| MX1 | 856.321375 | -0.60138291 | 1.09124643 | -0.5510973 | 0.58156701 | 0.767524 |
| TMPRSS2 | 1195.63427 | 0.018329355 | 0.38531579 | 0.0475697 | 0.96205918 | 0.9832997 |
| RIPK4 | 296.557098 | -0.786522784 | 0.31270436 | -2.5152281 | 0.01189554 | 0.06260025 |
| PRDM15 | 174.673423 | -0.383898831 | 0.26283576 | -1.4606035 | 0.14412429 | 0.33387193 |
| C2CD2 | 212.092298 | -1.310977781 | 0.24995043 | -5.2449511 | 1.56E-07 | 8.94E-06 |
| ZBTB21 | 219.514783 | -0.384082802 | 0.35020827 | -1.0967268 | 0.27276083 | 0.49436029 |
| ABCG1 | 302.535203 | -1.119018719 | 0.53258615 | -2.1011037 | 0.03563187 | 0.13239418 |
| TMPRSS3 | 109.334267 | -0.51547827 | 0.56954882 | -0.9050642 | 0.36543136 | 0.59096263 |
| RSPH1 | 121.067515 | -0.962868766 | 0.47710511 | -2.0181481 | 0.04357584 | 0.15107436 |
| SLC37A1 | 415.130882 | -1.223273762 | 0.17234021 | -7.0980171 | 1.27E-12 | 4.18E-10 |
| LINC01671 | 205.389891 | -0.030369861 | 0.46094115 | -0.0658866 | 0.94746808 | 0.97751354 |
| PDE9A | 2267.74045 | 0.481187821 | 0.24001194 | 2.00484951 | 0.04497914 | 0.15464066 |
| AP001628.1 | 77.1398283 | -1.498171789 | 0.62601115 | -2.393203 | 0.016702 | 0.07898184 |
| WDR4 | 185.563936 | -0.291655277 | 0.36412331 | -0.8009794 | 0.42314356 | 0.64193216 |
| NDUFV3 | 670.296538 | 0.606852684 | 0.13701862 | 4.42897958 | 9.47E-06 | 0.00025523 |
| PKNOX1 | 619.049368 | -0.449133312 | 0.23660584 | -1.8982343 | 0.05766523 | 0.18310096 |
| U2AF1 | 62.7419433 | 1.489594899 | 1.0370959 | 1.43631355 | 0.15091314 | 0.34357319 |
| HSF2BP | 46.2477568 | 0.088734051 | 0.54705517 | 0.16220311 | 0.87114591 | 0.94147844 |
| RRP1B | 270.892402 | 0.059639517 | 0.22151645 | 0.2692329 | 0.78775047 | 0.89956761 |
| PDXK | 1562.92828 | 0.660580055 | 0.22934166 | 2.88033174 | 0.00397257 | 0.0285183 |
| CSTB | 774.76256 | -2.055386753 | 0.42809272 | -4.8012653 | 1.58E-06 | 5.83E-05 |
| RRP1 | 98.4439281 | -0.267720798 | 0.32650362 | -0.8199627 | 0.41223736 | 0.63155223 |
| AGPAT3 | 1104.10865 | 0.127193214 | 0.20094179 | 0.63298536 | 0.5267432 | 0.72526045 |
| TRAPPC10 | 1067.49792 | 0.049830753 | 0.18701467 | 0.26645371 | 0.7898898 | 0.90082888 |
| PFKL | 1237.66492 | -0.270178645 | 0.27581365 | -0.9795695 | 0.32729865 | 0.5526381 |
| C21orf2 | 130.542328 | 0.335084308 | 0.33844605 | 0.99006714 | 0.32214131 | 0.54700662 |
| AP001062.1 | 47.886367 | -0.216271463 | 0.37591475 | -0.5753205 | 0.56507455 | 0.75608487 |
| UBE2G2 | 937.519713 | 0.387673105 | 0.15006215 | 2.5834169 | 0.0097827 | 0.05452309 |
| SUMO3 | 170.949406 | -1.550383631 | 0.34403396 | -4.5064843 | 6.59E-06 | 0.00018926 |
| PTTG1IP | 1660.02842 | -1.162913509 | 0.34052172 | -3.4150935 | 0.0006376 | 0.0071434 |
| FAM207A | 132.92206 | -0.254408894 | 0.22199168 | -1.146029 | 0.25178316 | 0.47203022 |
| ADARB1 | 471.291029 | 0.597547379 | 0.34066049 | 1.75408476 | 0.07941599 | 0.22520518 |
| POFUT2 | 265.200872 | -0.039712462 | 0.20540731 | -0.1933352 | 0.84669647 | 0.92890479 |
| BX322562.1 | 48.7020859 | -1.234491069 | 0.75371698 | -1.6378708 | 0.10144864 | 0.26522839 |
| COL18A1 | 1906.46476 | -0.324121087 | 0.38157979 | -0.8494189 | 0.39564824 | 0.61781743 |
| SLC19A1 | 56.2229573 | 0.275069448 | 0.39917077 | 0.68910218 | 0.49075897 | 0.6969611 |
| PCBP3 | 533.642412 | -0.833191919 | 1.42366066 | -0.5852461 | 0.55838222 | 0.75039889 |
| COL6A1 | 467.606951 | 1.24772175 | 0.63560228 | 1.96305425 | 0.04963987 | 0.16541197 |
| FTCD | 71.8018202 | 1.782508494 | 0.45273872 | 3.93716822 | 8.24E-05 | 0.00145209 |
| LSS | 367.216702 | 0.063200638 | 0.37217479 | 0.1698144 | 0.8651561 | 0.93858904 |
| MCM3AP-AS | 151.289254 | 0.205041615 | 0.29446153 | 0.69632735 | 0.48622385 | 0.69357784 |
| MCM3AP | 709.402048 | 0.244579985 | 0.13796845 | 1.77272404 | 0.07627445 | 0.22022058 |
| YBEY | 160.980861 | 0.664832928 | 0.35798609 | 1.85714739 | 0.06329019 | 0.19505368 |
| C21orf58 | 50.6526293 | 0.691158858 | 0.39756592 | 1.73847612 | 0.08212695 | 0.23032164 |
| PCNT | 372.396488 | -0.089652029 | 0.27712383 | -0.3235089 | 0.74630986 | 0.87603367 |
| DIP2A | 1164.14807 | 0.092313895 | 0.19701455 | 0.46856385 | 0.63938142 | 0.80603771 |
| PRMT2 | 908.46061 | -0.7345085 | 0.20167075 | -3.6421172 | 0.0002704 | 0.00372145 |
| AC011043.1 | 180.500103 | 1.248537171 | 0.37513267 | 3.32825496 | 0.00087392 | 0.00899033 |
| AL592183.1 | 898.697904 | -0.027815528 | 0.48900577 | -0.0568818 | 0.95463935 | 0.98007893 |
| AL354822.1 | 81.7576291 | 0.460243424 | 0.62355531 | 0.73809559 | 0.46045636 | 0.67395297 |

**Distal convoluted tubules**

*This means that positive fold changes imply higher mean value in controls than in AKI and vice versa.

| | baseMean | log2 fold change control/AKI* | lfcSE | stat | pvalue | padj |
|---|---|---|---|---|---|---|
| AL669831.5 | 37.7512535 | 0.188654176 | 0.27900594 | 0.67616545 | 0.4989356 | 0.77558432 |
| NOC2L | 22.2146633 | -0.246245197 | 0.5293676 | -0.4651686 | 0.64181071 | 0.85468927 |
| ISG15 | 83.7123448 | -0.097399883 | 1.48155867 | -0.0657415 | 0.94758363 | 0.98633762 |
| AGRN | 81.1115278 | -0.277615264 | 0.47128077 | -0.5890655 | 0.5558173 | 0.81104802 |
| C1orf159 | 32.4346201 | -0.568723351 | 0.32271986 | -1.7622818 | 0.07802169 | 0.29875739 |
| AL390719.2 | 23.2684139 | 0.114913117 | 0.61984095 | 0.1853913 | 0.85292213 | 0.95137449 |
| SDF4 | 41.711187 | -0.752451846 | 0.44111476 | -1.7057961 | 0.08804605 | 0.32206409 |
| UBE2J2 | 41.0119848 | -0.689109324 | 0.42731331 | -1.6126559 | 0.10681929 | 0.35932624 |
| ACAP3 | 17.6617193 | 0.283813094 | 0.48732972 | 0.58238413 | 0.56030797 | 0.81330344 |
| INTS11 | 38.9613273 | 0.394796817 | 0.26142742 | 1.51015843 | 0.131003 | 0.40219691 |
| AURKAIP1 | 31.3272465 | -0.145408238 | 0.46091082 | -0.3154802 | 0.75239708 | 0.90954886 |
| CCNL2 | 119.184147 | 0.139435992 | 0.25561372 | 0.54549495 | 0.58541315 | 0.82884893 |
| MRPL20 | 37.9688801 | -0.188693099 | 0.36007618 | -0.5240366 | 0.60025308 | 0.83602169 |
| ATAD3B | 34.263048 | 0.012140591 | 0.32117565 | 0.03780047 | 0.96984677 | 0.99156233 |
| ATAD3A | 16.2446781 | -0.22044014 | 0.55395228 | -0.3979407 | 0.69067392 | 0.8771239 |
| SSU72 | 42.685554 | -0.161682746 | 0.40101696 | -0.4031818 | 0.68681448 | 0.87523044 |
| MIB2 | 20.3861767 | 0.428868896 | 0.3850609 | 1.11376902 | 0.26537829 | 0.58079175 |
| CDK11B | 27.4942725 | -0.21166192 | 0.49732619 | -0.4255998 | 0.67039949 | 0.86777458 |
| SLC35E2B | 26.8833805 | 0.096529129 | 0.37086962 | 0.2602778 | 0.79464949 | 0.9268423 |
| CDK11A | 18.457347 | 0.012914015 | 0.4023415 | 0.03209715 | 0.97439458 | 0.99357756 |
| SLC35E2A | 32.2079983 | 0.322135407 | 0.29455548 | 1.09363238 | 0.27411624 | 0.58894058 |
| NADK | 29.9025058 | 0.335161677 | 0.40606679 | 0.8253856 | 0.40915271 | 0.70777268 |
| GNB1 | 220.314499 | -0.817752056 | 0.36238338 | -2.2565937 | 0.02403347 | 0.14714173 |
| PRKCZ | 65.2804535 | -0.483912441 | 0.38522246 | -1.2561896 | 0.20904725 | 0.5174675 |
| FAAP20 | 26.0485945 | -0.192744853 | 0.36224426 | -0.5320853 | 0.59466691 | 0.83416944 |
| SKI | 85.7238378 | -0.118661683 | 0.5929668 | -0.2001152 | 0.84139047 | 0.94776435 |
| MORN1 | 28.8976537 | 0.289870755 | 0.31325655 | 0.92534619 | 0.35478586 | 0.66121742 |
| RER1 | 55.8600929 | -0.262419487 | 0.3538828 | -0.7415435 | 0.45836397 | 0.747377 |
| PEX10 | 12.3028773 | -0.416279481 | 0.47734224 | -0.8720776 | 0.38316604 | 0.68720476 |
| TNFRSF14 | 25.8165348 | -0.304858362 | 0.80993699 | -0.3763976 | 0.70662131 | 0.88603006 |
| LINC00982 | 59.6924952 | 1.434161617 | 0.58182257 | 2.4649467 | 0.01370336 | 0.10221613 |
| PRDM16 | 280.54792 | -0.297339456 | 0.50945654 | -0.5836405 | 0.55946222 | 0.81314937 |
| ARHGEF16 | 31.1733735 | -0.242165154 | 0.42239665 | -0.5733122 | 0.56643333 | 0.81595078 |
| LRRC47 | 17.0226089 | 0.059775956 | 0.40332199 | 0.14820902 | 0.88217782 | 0.96161232 |
| CEP104 | 60.3455956 | -0.37273399 | 0.34654662 | -1.0755667 | 0.2821211 | 0.59642935 |
| DFFB | 11.6469611 | 0.494307251 | 0.63451617 | 0.77903019 | 0.43596193 | 0.72891111 |
| AJAP1 | 52.8823468 | 0.709013089 | 0.97377546 | 0.72810737 | 0.46654786 | 0.75300195 |
| NPHP4 | 42.7468475 | 0.360112015 | 0.38969467 | 0.92408762 | 0.3554407 | 0.66163487 |
| RPL22 | 26.3328227 | -0.296609816 | 0.47990767 | -0.618056 | 0.53653842 | 0.80030143 |
| RNF207 | 22.2928014 | -0.54705815 | 0.43865878 | -1.2471155 | 0.21235516 | 0.52134789 |
| ICMT | 15.3380692 | -0.503294672 | 0.53465559 | -0.9413437 | 0.34652875 | 0.65471653 |
| NOL9 | 39.4444687 | 0.41571941 | 0.27492983 | 1.51209275 | 0.13051027 | 0.40125903 |
| ZBTB48 | 17.096483 | -0.026408518 | 0.429141 | -0.0615381 | 0.95093068 | 0.98684281 |
| KLHL21 | 33.4805719 | -0.579363348 | 0.3784918 | -1.5307157 | 0.12583967 | 0.39241775 |
| DNAJC11 | 126.268844 | 1.09075978 | 0.23931662 | 4.55781047 | 5.17E-06 | 0.0002221 |
| CAMTA1 | 426.101238 | -1.381810745 | 0.56538127 | -2.4440335 | 0.01452408 | 0.1059672 |
| VAMP3 | 17.7993446 | -0.510332273 | 0.35752642 | -1.4273974 | 0.15346538 | 0.43892689 |
| PER3 | 99.2047566 | -0.111550012 | 0.55116621 | -0.2023891 | 0.83961258 | 0.9474978 |
| PARK7 | 52.9308574 | -0.695544979 | 0.38925966 | -1.7868406 | 0.07396325 | 0.29120952 |
| ERRFI1 | 121.731756 | -1.799460047 | 0.55754365 | -3.2274783 | 0.00124886 | 0.01783219 |
| RERE | 1351.91417 | -0.010661181 | 0.216544 | -0.0492333 | 0.96073336 | 0.98983463 |
| ENO1 | 235.601653 | -1.367988813 | 0.38366783 | -3.5655551 | 0.00036309 | 0.00683186 |
| GPR157 | 33.3378966 | 0.875989526 | 0.3516339 | 2.49119762 | 0.01273133 | 0.0975406 |
| H6PD | 34.6219096 | -0.4247076 | 0.3148874 | -1.3487602 | 0.17741399 | 0.47467647 |
| SPSB1 | 45.8485804 | -3.365034456 | 0.72665034 | -4.6308854 | 3.64E-06 | 0.00016642 |
| SLC25A33 | 80.9286143 | 0.543199038 | 0.35319053 | 1.53797734 | 0.12405416 | 0.38919278 |
| PIK3CD | 21.4933439 | -0.674951642 | 0.65848738 | -1.0250031 | 0.3053617 | 0.61882037 |
| CLSTN1 | 104.001204 | -0.022075041 | 0.30290158 | -0.0728786 | 0.94190273 | 0.98467329 |
| CTNNBIP1 | 34.2452033 | 0.579751726 | 0.28847992 | 2.00967793 | 0.04446529 | 0.21534941 |
| LZIC | 23.0201226 | -0.004828775 | 0.39600288 | -0.0121938 | 0.99027101 | 0.99857194 |
| NMNAT1 | 63.9459435 | 0.276114803 | 0.27590544 | 1.00075882 | 0.31694342 | 0.63093651 |
| UBE4B | 177.993772 | -0.038525559 | 0.22530486 | -0.170993 | 0.86422925 | 0.95486142 |
| KIF1B | 203.631003 | 0.748176247 | 0.21600971 | 3.46362315 | 0.00053295 | 0.0090186 |
| PGD | 27.2746927 | 0.01370814 | 0.35892483 | 0.03819223 | 0.96953442 | 0.99156233 |
| DFFA | 36.8410371 | -0.155155471 | 0.34275917 | -0.4526661 | 0.65078916 | 0.86113455 |
| PEX14 | 128.786202 | -0.102895887 | 0.25777341 | -0.3991718 | 0.68976659 | 0.87680759 |
| CASZ1 | 218.424067 | -0.057124676 | 0.46375109 | -0.1231796 | 0.90196488 | 0.96803829 |
| EXOSC10 | 31.5505061 | -0.020128165 | 0.38038643 | -0.052915 | 0.9577996 | 0.98865531 |
| MTOR | 86.0107608 | 0.237990178 | 0.2835583 | 0.83929894 | 0.40130158 | 0.70114138 |
| FBXO44 | 22.4467998 | 1.162919604 | 0.46546101 | 2.49842537 | 0.01247464 | 0.09619433 |
| MAD2L2 | 14.4256247 | -0.553421526 | 0.58111246 | -0.9523484 | 0.34092032 | 0.65083559 |
| AGTRAP | 17.1388518 | -1.548789479 | 0.51376028 | -3.0146151 | 0.00257306 | 0.03067925 |
| MTHFR | 22.1808484 | -1.25794992 | 0.38356693 | -3.27961 | 0.00103951 | 0.0153847 |
| CLCN6 | 31.6619692 | -1.40373734 | 0.52152094 | -2.6916222 | 0.00711054 | 0.06436769 |
| KIAA2013 | 16.3176484 | -0.261980763 | 0.46599889 | -0.5621918 | 0.57398533 | 0.82117002 |
| PLOD1 | 13.7166917 | 0.123044652 | 0.48519072 | 0.25360059 | 0.79980414 | 0.92926776 |
| MFN2 | 45.7761056 | 0.294570711 | 0.34951934 | 0.84278801 | 0.39934701 | 0.69959441 |
| VPS13D | 408.235957 | -0.453070339 | 0.20506158 | -2.2094355 | 0.02714437 | 0.1597816 |
| DHRS3 | 94.1616146 | -0.436652495 | 0.5548008 | -0.7870437 | 0.43125627 | 0.72490237 |
| PRDM2 | 105.661076 | -0.620580559 | 0.21201768 | -2.9270227 | 0.00342224 | 0.03844452 |
| KAZN | 1471.97281 | -1.022498111 | 0.40742802 | -2.5096411 | 0.01208539 | 0.09413884 |
| TMEM51-AS | 33.3392523 | -2.169524182 | 0.70940535 | -3.0582292 | 0.00222649 | 0.02735153 |
| TMEM51 | 251.611706 | -1.399588228 | 0.58349161 | -2.3986433 | 0.01645594 | 0.11476078 |
| CASP9 | 24.5750924 | 0.027771522 | 0.36092651 | 0.07694509 | 0.93866723 | 0.98331142 |
| DNAJC16 | 62.4305497 | -0.310138065 | 0.23821644 | -1.3019172 | 0.19294471 | 0.49423529 |
| DDI2 | 67.1823243 | 0.319864868 | 0.2363244 | 1.35349915 | 0.17589622 | 0.47294748 |
| PLEKHM2 | 60.0012419 | -1.010794087 | 0.53113381 | -1.9030875 | 0.05702913 | 0.24905839 |
| AL450998.2 | 11.7912633 | -0.200318496 | 0.46735827 | -0.4286187 | 0.66820073 | 0.86777458 |
| SPEN | 148.052062 | -0.05617322 | 0.39557258 | -0.1420048 | 0.88707618 | 0.96253908 |
| ZBTB17 | 21.4068263 | 0.176528548 | 0.33114374 | 0.53308739 | 0.59397309 | 0.83416944 |
| CLCNKA | 123.072369 | 2.73631422 | 0.6608999 | 4.14028541 | 3.47E-05 | 0.0010745 |
| CLCNKB | 184.920379 | 1.592126703 | 0.87347387 | 1.82275253 | 0.06834088 | 0.27855451 |
| FBXO42 | 151.238878 | 0.039124124 | 0.35170838 | 0.11124024 | 0.91142584 | 0.97128978 |
| SZRD1 | 39.6984238 | -0.313535783 | 0.29758312 | -1.0536074 | 0.29206269 | 0.60690719 |
| NECAP2 | 22.029023 | -0.335899401 | 0.30782216 | -1.0912125 | 0.27517937 | 0.58956168 |
| NBPF1 | 34.4114376 | -0.319614807 | 0.35796656 | -0.8928622 | 0.37193097 | 0.67534262 |
| CROCC | 37.7465235 | 0.440361819 | 0.42041043 | 1.04745694 | 0.29488888 | 0.61006078 |
| ATP13A2 | 14.5549482 | -0.455614276 | 0.47369968 | -0.961821 | 0.33613955 | 0.64728732 |
| SDHB | 76.7735569 | 0.444002716 | 0.21253112 | 2.08911864 | 0.03669704 | 0.19211861 |
| PADI2 | 12.9277227 | 1.61015402 | 1.00134702 | 1.60798803 | 0.1078378 | 0.36142331 |
| RCC2 | 30.9798382 | -1.065700135 | 0.44891186 | -2.373963 | 0.01759832 | 0.12052024 |
| ARHGEF10L | 234.870454 | -1.008426896 | 0.35744585 | -2.8212019 | 0.00478441 | 0.04859458 |
| UBR4 | 141.651676 | -0.47259091 | 0.32885456 | -1.4370818 | 0.15069474 | 0.43488337 |
| AL035413.1 | 27.0527767 | 0.09350788 | 0.37760818 | 0.24763203 | 0.80441912 | 0.93242217 |
| EMC1 | 16.2127884 | 0.678813644 | 0.39912662 | 1.70074762 | 0.08899039 | 0.32396747 |
| AKR7A2 | 14.038456 | 0.168900393 | 0.55033739 | 0.30690336 | 0.75891693 | 0.9118739 |
| CAPZB | 150.217445 | -0.386574358 | 0.35416867 | -1.0914979 | 0.27505386 | 0.58956168 |
| TMCO4 | 99.910813 | -0.243690735 | 0.5171801 | -0.4711912 | 0.63750417 | 0.8538689 |
| OTUD3 | 39.1230079 | -1.179980489 | 0.40866161 | -2.8874268 | 0.00388407 | 0.04191558 |
| CAMK2N1 | 68.4821145 | -1.463472186 | 0.51166734 | -2.8602025 | 0.00423371 | 0.04430424 |
| MUL1 | 14.6938437 | -0.250737896 | 0.43729337 | -0.573386 | 0.56638336 | 0.81595078 |

| | | | | | | |
|---|---|---|---|---|---|---|
| PINK1 | 26.8085725 | -0.025382362 | 0.35484302 | -0.0715312 | 0.94297496 | 0.98507485 |
| DDOST | 18.5390663 | -1.027577699 | 0.40099894 | -2.5625447 | 0.01039082 | 0.08461487 |
| HP1BP3 | 229.421898 | 0.233189895 | 0.16820424 | 1.38634968 | 0.16564015 | 0.45785473 |
| EIF4G3 | 901.01262 | -0.936263619 | 0.51582776 | -1.8150702 | 0.06951312 | 0.28106692 |
| ECE1 | 79.5324165 | 0.594622063 | 0.48766802 | 1.21931731 | 0.22272378 | 0.53302786 |
| NBPF3 | 16.7439629 | 0.169347363 | 0.59979174 | 0.28234361 | 0.77768005 | 0.91971851 |
| RAP1GAP | 50.6853698 | 0.054989883 | 0.6239402 | 0.08813326 | 0.92977076 | 0.97876742 |
| USP48 | 136.156381 | 0.22954891 | 0.23040257 | 0.99629492 | 0.31910687 | 0.63100017 |
| HSPG2 | 64.5897302 | 0.120202727 | 0.38296528 | 0.31387369 | 0.75361697 | 0.91019749 |
| CDC42 | 107.717484 | -0.227400218 | 0.25384145 | -0.8958356 | 0.37034055 | 0.67442486 |
| ZBTB40 | 50.3977142 | 0.042269277 | 0.34796905 | 0.12147424 | 0.90331541 | 0.96843447 |
| EPHB2 | 43.7277707 | -1.755965485 | 0.82951658 | -2.116854 | 0.03427224 | 0.18418162 |
| KDM1A | 81.1732075 | 0.401989721 | 0.34175399 | 1.17625466 | 0.23949313 | 0.5513664 |
| AL031428.1 | 21.6652599 | -0.118017419 | 0.48241759 | -0.2446375 | 0.80673715 | 0.93325451 |
| LUZP1 | 147.586827 | 1.768761906 | 0.25178288 | 7.02494894 | 2.14E-12 | 7.40E-10 |
| LINC01355 | 20.903928 | 0.642478883 | 0.37031292 | 1.73496211 | 0.08274752 | 0.30935759 |
| HNRNPR | 114.461203 | 0.086856948 | 0.23840353 | 0.36432744 | 0.71561351 | 0.88904248 |
| TCEA3 | 56.467264 | 1.416642621 | 0.65863466 | 2.15087774 | 0.03148585 | 0.17567711 |
| ASAP3 | 63.3049494 | 1.284437911 | 0.32605116 | 3.93937536 | 8.17E-05 | 0.00209839 |
| RPL11 | 88.7359097 | -0.743964093 | 0.37802476 | -1.9680301 | 0.04906458 | 0.22737481 |
| ELOA | 24.8713467 | -0.458492919 | 0.38571796 | -1.188674 | 0.23456796 | 0.54657 |
| ELOA-AS1 | 24.8993049 | 0.476575483 | 0.60486974 | 0.78789771 | 0.43075654 | 0.7245261 |
| PITHD1 | 19.5640683 | -0.321639504 | 0.36266164 | -0.8868859 | 0.37514032 | 0.67888822 |
| LYPLA2 | 18.3897874 | -0.316489064 | 0.56436656 | -0.5607864 | 0.57494319 | 0.82146824 |
| GALE | 23.0275384 | -2.231077029 | 0.47716164 | -4.6757259 | 2.93E-06 | 0.00013794 |
| HMGCL | 32.9843768 | -0.491408302 | 0.32607407 | -1.507045 | 0.13179912 | 0.40351308 |
| FUCA1 | 24.2563681 | -0.065460063 | 0.42002736 | -0.1558471 | 0.87615351 | 0.95866717 |
| PNRC2 | 34.9888794 | -0.329153065 | 0.48988834 | -0.6718941 | 0.50165114 | 0.776256 |
| SRSF10 | 141.095368 | -0.062090988 | 0.30517526 | -0.2034601 | 0.83877543 | 0.94741614 |
| IFNLR1 | 19.0011207 | 0.498890824 | 0.46488208 | 1.07315564 | 0.28320129 | 0.59695715 |
| STPG1 | 14.5221871 | -0.333483105 | 0.51903532 | -0.6425056 | 0.52054495 | 0.78804369 |
| NIPAL3 | 49.9663732 | -0.575396137 | 0.45221857 | -1.272385 | 0.20323635 | 0.50790023 |
| RCAN3 | 26.2240626 | -1.70423666 | 0.45127938 | -3.7764559 | 0.00015908 | 0.00349817 |
| NCMAP | 22.5721907 | -4.407232156 | 0.70109106 | -6.2862478 | 3.25E-10 | 6.74E-08 |
| SRRM1 | 92.1078946 | 0.148612657 | 0.27680683 | 0.53688219 | 0.591349 | 0.83234048 |
| CLIC4 | 61.9799372 | -0.319561891 | 0.73560798 | -0.4344187 | 0.66398439 | 0.86599391 |
| SYF2 | 24.1632212 | -0.085292621 | 0.39815593 | -0.2142191 | 0.83037617 | 0.94393022 |
| RSRP1 | 279.556805 | -0.101704756 | 0.36181102 | -0.2810991 | 0.77863439 | 0.92013725 |
| TMEM50A | 33.6930384 | -0.630655901 | 0.38691889 | -1.6299434 | 0.10311345 | 0.35088341 |
| MACO1 | 69.9311308 | -0.205204437 | 0.22823089 | -0.899109 | 0.36859463 | 0.67276052 |
| LDLRAP1 | 15.7306427 | 0.411810319 | 0.43416802 | 0.9485045 | 0.34287268 | 0.65243773 |
| MAN1C1 | 27.2694307 | 0.525877354 | 0.59376316 | 0.88566855 | 0.37579616 | 0.67918655 |
| MTFR1L | 26.3457302 | 1.188985253 | 0.68594639 | 1.73335011 | 0.08303346 | 0.31017787 |
| STMN1 | 15.7159396 | -0.660794471 | 0.4798061 | -1.3772115 | 0.16844688 | 0.46207669 |
| PAFAH2 | 23.9734828 | 0.285344349 | 0.3971232 | 0.71852853 | 0.47243147 | 0.75855881 |
| CNKSR1 | 19.0479271 | -0.080645282 | 0.37367337 | -0.2158176 | 0.82912994 | 0.94324153 |
| CEP85 | 22.5257108 | 0.023892024 | 0.5364435 | 0.04453782 | 0.96447571 | 0.99082982 |
| SH3BGRL3 | 22.5682654 | -1.218214385 | 0.68383562 | -1.7814433 | 0.07484006 | 0.29289497 |
| DHDDS | 39.5185136 | -0.071242138 | 0.32136916 | -0.2216832 | 0.82456052 | 0.94153354 |
| HMGN2 | 33.8601836 | 0.168054282 | 0.40997122 | 0.40991727 | 0.68186664 | 0.87271442 |
| RPS6KA1 | 31.0707828 | -0.857866485 | 0.29609412 | -2.8972764 | 0.00376418 | 0.04122633 |
| ARID1A | 186.581077 | 0.296902793 | 0.25387226 | 1.1694968 | 0.24220353 | 0.55446835 |
| ZDHHC18 | 14.1473709 | -0.28379587 | 0.50736169 | -0.5593561 | 0.5759187 | 0.82147339 |
| GPN2 | 18.8076684 | -1.260215139 | 0.53726852 | -2.3455965 | 0.01899666 | 0.12678946 |
| NUDC | 47.3816428 | -0.201021327 | 0.34927228 | -0.5755433 | 0.56492391 | 0.81537933 |
| TRNP1 | 58.1375021 | -2.930261958 | 0.50788639 | -5.7695225 | 7.95E-09 | 8.72E-07 |
| SLC9A1 | 36.9971572 | 0.081075169 | 0.50411523 | 0.16082666 | 0.87222993 | 0.95690422 |
| WDTC1 | 60.0110137 | -0.175085057 | 0.23837917 | -0.7344813 | 0.46265542 | 0.75091259 |
| TMEM222 | 19.4470846 | -0.297037222 | 0.43062496 | -0.6897817 | 0.49033147 | 0.77196791 |
| WASF2 | 129.346353 | -0.26937662 | 0.20574191 | -1.3092939 | 0.19043483 | 0.49126283 |
| AHDC1 | 58.572462 | 0.029639284 | 0.34081395 | 0.08696617 | 0.9306984 | 0.97888844 |
| IFI6 | 79.6207149 | -0.654149613 | 1.59157889 | -0.4110067 | 0.68106761 | 0.87265005 |
| FAM76A | 20.767325 | 0.587243626 | 0.5007651 | 1.17269279 | 0.24091902 | 0.55327012 |
| STX12 | 84.0024666 | -0.5478645 | 0.40658369 | -1.3474827 | 0.17782482 | 0.47521885 |
| PPP1R8 | 20.5568664 | -0.234076032 | 0.35965609 | -0.650833 | 0.51515433 | 0.78381327 |
| RPA2 | 20.0112751 | 0.223974376 | 0.36113348 | 0.62019831 | 0.53512723 | 0.79931057 |
| EYA3 | 254.445494 | 0.041137349 | 0.29840124 | 0.13785917 | 0.89035172 | 0.96323092 |
| DNAJC8 | 38.371278 | 0.02347774 | 0.37558094 | 0.06251047 | 0.95015633 | 0.98678754 |
| ATP5IF1 | 58.0276126 | -0.245889434 | 0.48228487 | -0.5098427 | 0.61016165 | 0.84109213 |
| PHACTR4 | 247.936219 | -0.584806031 | 0.20835892 | -2.8067243 | 0.00500481 | 0.05012332 |
| RCC1 | 27.1530613 | -1.131086409 | 0.32139885 | -3.5192609 | 0.00043275 | 0.00775956 |
| TRNAU1AP | 48.8421078 | -0.47586546 | 0.39952389 | -1.1910814 | 0.23362165 | 0.54598215 |
| TAF12 | 33.8489906 | -0.076662116 | 0.39331252 | -0.194914 | 0.84546028 | 0.94907846 |
| GMEB1 | 32.590517 | -0.396916103 | 0.42694423 | -0.9296673 | 0.35254335 | 0.65972795 |
| YTHDF2 | 75.5971025 | -0.583737772 | 0.29130545 | -2.0038683 | 0.04508416 | 0.2175801 |
| EPB41 | 195.187213 | -0.219385849 | 0.19503558 | -1.1248504 | 0.26065243 | 0.57426715 |
| SRSF4 | 272.243983 | -0.414600154 | 0.30419589 | -1.3629381 | 0.17290203 | 0.46823658 |
| MECR | 26.4402998 | 0.22213807 | 0.44334369 | 0.50105162 | 0.6163348 | 0.84223226 |
| PUM1 | 358.505001 | -0.396956592 | 0.25081234 | -1.5826837 | 0.11349358 | 0.37119783 |
| SNRNP40 | 37.0275602 | 0.284742737 | 0.35654537 | 0.79861572 | 0.42451327 | 0.72151771 |
| ZCCHC17 | 82.2601577 | -0.939177671 | 0.28223723 | -3.327618 | 0.00087592 | 0.01352164 |
| FABP3 | 18.4426252 | 0.488244187 | 0.57979138 | 0.84210323 | 0.39973018 | 0.69959441 |
| SERINC2 | 92.7426203 | -0.576141982 | 0.29593153 | -1.946876 | 0.0515496 | 0.23455111 |
| TINAGL1 | 44.9480644 | 0.172521966 | 0.60678517 | 0.28432133 | 0.77616414 | 0.91956219 |
| PEF1 | 21.7312176 | -0.148013106 | 0.38659637 | -0.3828621 | 0.701822 | 0.88424653 |
| PTP4A2 | 34.4069963 | -0.625881861 | 0.45999695 | -1.3606218 | 0.17363325 | 0.46960235 |
| KHDRBS1 | 128.929642 | -0.724812995 | 0.33676161 | -2.1523029 | 0.03137351 | 0.17537564 |
| TMEM39B | 15.0949307 | 0.934648558 | 0.41684369 | 2.24220395 | 0.02494819 | 0.15046375 |
| KPNA6 | 117.833799 | -0.339582135 | 0.35915305 | -0.9455082 | 0.34439951 | 0.65359197 |
| TXLNA | 36.2072299 | -0.379952904 | 0.37544998 | -1.0119934 | 0.3115412 | 0.62642275 |
| EIF3I | 16.8414226 | -0.67183458 | 0.41927568 | -1.6023696 | 0.10907391 | 0.36312941 |
| HDAC1 | 48.0880916 | 0.18107374 | 0.31237542 | 0.57966706 | 0.56213916 | 0.81388399 |
| BSDC1 | 63.4424215 | -0.449121781 | 0.26421088 | -1.699861 | 0.08915707 | 0.32396747 |
| ZBTB8OS | 113.911972 | -0.168067222 | 0.31543383 | -0.5328129 | 0.59416308 | 0.83416944 |
| RBBP4 | 92.7599476 | -0.364863171 | 0.33800289 | -1.0794676 | 0.28037933 | 0.59468991 |
| SYNC | 15.8788712 | -1.025318773 | 0.47958152 | -2.1379447 | 0.03252123 | 0.17931872 |
| KIAA1522 | 91.6533006 | 0.064204097 | 0.27566782 | 0.23290385 | 0.81583607 | 0.93717111 |
| YARS | 50.3447139 | -1.075130295 | 0.59921099 | -1.7942433 | 0.07277435 | 0.28837569 |
| S100PBP | 101.779862 | 0.356955007 | 0.24322812 | 1.46757296 | 0.14222026 | 0.4225818 |
| RNF19B | 41.2857587 | -0.108706416 | 0.57317516 | -0.1896565 | 0.84957828 | 0.95003258 |
| AL020995.1 | 32.5567013 | -0.173646634 | 0.55397466 | -0.3134559 | 0.75393431 | 0.91026551 |
| AK2 | 112.457236 | 0.592545741 | 0.27431401 | 2.16010019 | 0.03076491 | 0.17290661 |
| ZNF362 | 22.826344 | 0.405171326 | 0.66131689 | 0.61267349 | 0.54009226 | 0.80166126 |
| AL513327.1 | 43.3456108 | -1.301435463 | 0.7058257 | -1.8438482 | 0.06520526 | 0.27058097 |
| PHC2 | 206.127224 | -0.905546468 | 0.44138494 | -2.0516025 | 0.04020831 | 0.20290676 |
| SMIM12 | 33.9251853 | -0.301147921 | 0.42691003 | -0.7054131 | 0.4805532 | 0.76551071 |
| ZMYM1 | 42.1784265 | 0.508540422 | 0.42181197 | 1.20560928 | 0.22796818 | 0.54017161 |
| SFPQ | 221.051841 | 0.166586942 | 0.28588564 | 0.58270483 | 0.56009202 | 0.81330344 |
| ZMYM4 | 324.997492 | 0.03423555 | 0.1963151 | 0.17439082 | 0.86155833 | 0.95417909 |
| KIAA0319L | 162.469001 | 0.048925074 | 0.24261992 | 0.20165316 | 0.84018788 | 0.9474978 |
| PSMB2 | 25.7651951 | 0.24898317 | 0.3594887 | 0.69260361 | 0.48855835 | 0.77039033 |
| AGO4 | 78.6596924 | 0.097525027 | 0.29566109 | 0.32985411 | 0.7415102 | 0.90383185 |
| AGO1 | 23.6530401 | 0.639809984 | 0.35469707 | 1.80382088 | 0.07125939 | 0.28479321 |

| | | | | | | |
|---|---|---|---|---|---|---|
| AGO3 | 241.764855 | 0.04376636 | 0.24527076 | 0.178441 | 0.85837665 | 0.95271398 |
| ADPRHL2 | 18.6947661 | -0.44888939 | 0.46488588 | -0.9655905 | 0.33424913 | 0.64554754 |
| TRAPPC3 | 32.4700073 | -0.754135776 | 0.51115339 | -1.475361 | 0.14011551 | 0.41923311 |
| MAP7D1 | 19.9639947 | 0.001800179 | 0.60847521 | 0.00295851 | 0.99763946 | 0.99922221 |
| THRAP3 | 174.573584 | -0.300826586 | 0.29216144 | -1.0296588 | 0.30317022 | 0.61760086 |
| SH3D21 | 24.5424977 | 0.069728785 | 0.52416712 | 0.13302777 | 0.89417142 | 0.96482804 |
| EVA1B | 16.3134184 | -1.339218647 | 0.54609799 | -2.4523413 | 0.014193 | 0.10453042 |
| AL591845.1 | 23.6028297 | -0.695439979 | 0.499342 | -1.3927128 | 0.16370667 | 0.45523441 |
| STK40 | 50.8067846 | -1.046572779 | 0.50798286 | -2.0602521 | 0.03937444 | 0.20050645 |
| LINC01137 | 17.2249122 | -0.371679045 | 0.43175057 | -0.8608652 | 0.38931229 | 0.69247382 |
| MEAF6 | 27.8321551 | 0.545139917 | 0.3358338 | 1.62324317 | 0.10453743 | 0.35443892 |
| SNIP1 | 14.1151572 | 0.059981815 | 0.41186804 | 0.14563357 | 0.88421066 | 0.96178024 |
| GNL2 | 13.2480086 | 0.015199054 | 0.53445829 | 0.02843824 | 0.97731262 | 0.99497547 |
| C1orf122 | 24.0102175 | -0.454700896 | 0.52311089 | -0.8692247 | 0.38472425 | 0.68851611 |
| MTF1 | 28.0474046 | -0.019736993 | 0.36346589 | -0.0543022 | 0.95669441 | 0.98859719 |
| INPP5B | 103.003987 | 0.135874294 | 0.37733321 | 0.360091 | 0.71877908 | 0.8907358 |
| SF3A3 | 30.9061999 | 0.138900729 | 0.33834412 | 0.41053094 | 0.68141651 | 0.87270068 |
| RRAGC | 22.8907961 | -1.55093653 | 0.67114491 | -2.3108817 | 0.02083939 | 0.13454463 |
| AL139260.1 | 76.3005875 | -2.559784601 | 0.84694932 | -3.0223587 | 0.00250813 | 0.03021382 |
| AKIRIN1 | 105.522215 | -0.551153101 | 0.2902725 | -1.8987438 | 0.05759817 | 0.25013758 |
| NDUFS5 | 53.7495369 | -0.066509762 | 0.39638747 | -0.1677898 | 0.86674867 | 0.95578708 |
| MACF1 | 899.953329 | -0.742969645 | 0.37930714 | -1.9587547 | 0.05014152 | 0.23113573 |
| PABPC4 | 34.2764283 | -0.360379288 | 0.43808879 | -0.822617 | 0.41072583 | 0.70931795 |
| PPIE | 22.7212737 | 0.617552105 | 0.36962743 | 1.67074207 | 0.09477263 | 0.3357371 |
| TRIT1 | 58.6961508 | 0.272089923 | 0.37355192 | 0.72838583 | 0.46637743 | 0.75300195 |
| CAP1 | 45.9651933 | -0.801173332 | 0.33128801 | -2.418359 | 0.01559069 | 0.11096761 |
| PPT1 | 17.810095 | 0.313399431 | 0.44615876 | 0.70243926 | 0.48240527 | 0.76628579 |
| RLF | 173.200669 | -0.538152529 | 0.23127974 | -2.3268469 | 0.01997342 | 0.13082568 |
| ZMPSTE24 | 37.0754174 | -0.179582543 | 0.33080856 | -0.5428594 | 0.5872266 | 0.8298425 |
| SMAP2 | 29.0996384 | 0.019281177 | 0.34540503 | 0.05582193 | 0.95548366 | 0.98853497 |
| AL603839.3 | 26.0444723 | 0.188766679 | 0.44818859 | 0.4211769 | 0.67362591 | 0.86846218 |
| ZNF684 | 32.4484966 | 0.444133987 | 0.67264887 | 0.66027612 | 0.50907665 | 0.78082426 |
| NFYC | 172.958259 | 1.089052398 | 0.26232887 | 4.15147756 | 3.30E-05 | 0.00104056 |
| SCMH1 | 305.269698 | -0.190123689 | 0.1828584 | -1.0397318 | 0.29846453 | 0.61229556 |
| HIVEP3 | 160.75948 | -1.495828275 | 0.36678821 | -4.0781798 | 4.54E-05 | 0.00130621 |
| FOXJ3 | 212.074421 | -0.392993006 | 0.37174299 | -1.0571632 | 0.29043712 | 0.60501244 |
| PPCS | 21.6311736 | -0.458661295 | 0.48835433 | -0.9391978 | 0.34762922 | 0.6553345 |
| CCDC30 | 138.848626 | 0.258997379 | 0.49880274 | 0.51923808 | 0.60359473 | 0.83748768 |
| PPIH | 15.8786324 | 0.572751889 | 0.5072692 | 1.12908864 | 0.25886044 | 0.57194663 |
| YBX1 | 63.8764799 | -0.729982078 | 0.30573638 | -2.3876193 | 0.0169579 | 0.11753644 |
| ERMAP | 24.5834951 | -0.06485809 | 0.41269983 | -0.1571556 | 0.87512222 | 0.95805533 |
| SLC2A1 | 62.8819231 | -2.303190457 | 0.60955601 | -3.7784722 | 0.00015779 | 0.00347986 |
| SLC2A1-AS1 | 19.6843178 | -1.252796619 | 0.7287359 | -1.7191367 | 0.08558949 | 0.31592889 |
| EBNA1BP2 | 17.3045671 | 1.026950552 | 0.44721467 | 2.29632571 | 0.02165727 | 0.13787238 |
| SZT2 | 49.2863376 | -0.094030514 | 0.33616557 | -0.2797149 | 0.77969626 | 0.920472 |
| PTPRF | 95.6618066 | -0.30766886 | 0.46089023 | -0.6675534 | 0.50441868 | 0.77792612 |
| ATP6V0B | 84.3810146 | -0.101882001 | 0.42652718 | -0.238864 | 0.81121102 | 0.93552094 |
| CCDC24 | 29.0877672 | -0.506913041 | 0.47920995 | -1.0578099 | 0.29014211 | 0.60493852 |
| DMAP1 | 13.7403295 | 0.38565908 | 0.41671368 | 0.92547738 | 0.35471765 | 0.66121742 |
| ERI3 | 91.1584706 | 0.166438804 | 0.44781001 | 0.37167281 | 0.71013647 | 0.88614996 |
| RNF220 | 16.0747664 | -0.529422722 | 0.44120606 | -1.1999443 | 0.23016096 | 0.54292511 |
| TMEM53 | 22.5171112 | -0.215356571 | 0.37174603 | -0.579311 | 0.56237933 | 0.81397468 |
| ARMH1 | 15.7150239 | 1.001218485 | 0.58803839 | 1.70264137 | 0.0886352 | 0.32323227 |
| RPS8 | 78.709234 | -0.922741698 | 0.39554081 | -2.3328609 | 0.01965545 | 0.12988475 |
| EIF2B3 | 53.6830756 | 0.272869581 | 0.23767561 | 1.14807563 | 0.25093734 | 0.56353121 |
| ZSWIM5 | 94.5670927 | 0.400779341 | 0.43440605 | 0.92259154 | 0.35622011 | 0.66255661 |
| TESK2 | 24.442997 | -0.831782616 | 0.34308685 | -2.4244083 | 0.01533336 | 0.110145 |
| PRDX1 | 94.4405285 | -1.312498332 | 0.43934152 | -2.9874216 | 0.00281341 | 0.03299658 |
| AKR1A1 | 18.9964463 | -0.805663133 | 0.47658669 | -1.690486 | 0.09093501 | 0.32812618 |
| NASP | 51.5646612 | 0.219967847 | 0.28015565 | 0.78516298 | 0.43235802 | 0.72599466 |
| GPBP1L1 | 226.549744 | 0.388040348 | 0.19153334 | 2.02596768 | 0.0427681 | 0.21080021 |
| TMEM69 | 14.5962113 | -0.08142473 | 0.36936191 | -0.220447 | 0.82552304 | 0.94182399 |
| IPP | 40.8761234 | 0.202076177 | 0.26081521 | 0.77478678 | 0.43846566 | 0.73069774 |
| MAST2 | 69.3379514 | -0.799027456 | 0.50076619 | -1.5956098 | 0.11057593 | 0.36599573 |
| TSPAN1 | 66.8925765 | -3.020435452 | 0.38743965 | -7.7958863 | 6.40E-15 | 3.51E-12 |
| POMGNT1 | 39.4090349 | -0.304185839 | 0.38187523 | -0.7965582 | 0.42570767 | 0.72208446 |
| LURAP1 | 42.2662934 | 0.374704158 | 0.47084851 | 0.7958062 | 0.42614469 | 0.72211376 |
| LRRC41 | 42.1540822 | -0.528742042 | 0.36818269 | -1.4360861 | 0.15097785 | 0.43488337 |
| UQCRH | 81.769519 | 0.572829628 | 0.35969806 | 1.5925291 | 0.11126587 | 0.36720011 |
| NSUN4 | 32.4272862 | 0.175618588 | 0.31895925 | 0.55059882 | 0.58190872 | 0.82595782 |
| FAAH | 14.7748871 | 1.028762254 | 0.42509254 | 2.42009013 | 0.01551666 | 0.11077898 |
| MKNK1 | 51.4029195 | 0.614104669 | 0.30023559 | 2.04540934 | 0.04081452 | 0.20490426 |
| ATPAF1 | 55.9727505 | 0.65225307 | 0.27146712 | 2.4026964 | 0.01627469 | 0.11409416 |
| EFCAB14 | 79.4475307 | -0.090594037 | 0.39700208 | -0.2281954 | 0.81949436 | 0.9384641 |
| PDZK1IP1 | 43.5141714 | -1.494498461 | 0.64556541 | -2.3150225 | 0.0206117 | 0.13383178 |
| CMPK1 | 55.8641546 | 0.032443743 | 0.33089787 | 0.0980476 | 0.92189449 | 0.97590194 |
| TRABD2B | 774.049077 | -0.283173449 | 0.32912378 | -0.8603859 | 0.38957638 | 0.69267928 |
| SPATA6 | 152.074991 | 0.07785248 | 0.46200301 | 0.16851076 | 0.86618148 | 0.95565922 |
| AGBL4 | 456.512184 | 0.303930397 | 0.74533181 | 0.40777865 | 0.68343619 | 0.87340448 |
| FAF1 | 625.60612 | 0.176643551 | 0.18238024 | 0.96854547 | 0.33277202 | 0.64492217 |
| RNF11 | 68.3627481 | -0.177530454 | 0.31074335 | -0.5713089 | 0.56779024 | 0.81673499 |
| TTC39A | 76.0285556 | -3.282517004 | 0.50946217 | -6.4431025 | 1.17E-10 | 2.73E-08 |
| EPS15 | 239.051589 | 0.491364854 | 0.22515396 | 2.18235047 | 0.02908368 | 0.16626377 |
| OSBPL9 | 215.78403 | 0.751827436 | 0.2451912 | 3.0662904 | 0.00216733 | 0.02673639 |
| NRDC | 185.35788 | -0.123648447 | 0.21014198 | -0.5884043 | 0.55626094 | 0.81141693 |
| TXNDC12 | 35.7175787 | -0.03638632 | 0.30865484 | -0.1178868 | 0.90615738 | 0.96959048 |
| BTF3L4 | 56.9323199 | 0.018309236 | 0.28271496 | 0.06476218 | 0.94836336 | 0.98641632 |
| ZFYVE9 | 162.434378 | 0.555254736 | 0.23718169 | 2.34105223 | 0.01922948 | 0.12791313 |
| CC2D1B | 25.6625455 | -1.231608108 | 0.47505798 | -2.5925427 | 0.00952693 | 0.07938261 |
| PRPF38A | 26.012217 | -0.024218246 | 0.3589673 | -0.0674664 | 0.94621038 | 0.98574808 |
| TUT4 | 300.599298 | 0.48304048 | 0.23782201 | 2.03110084 | 0.04224477 | 0.20951607 |
| ZYG11B | 89.4066198 | 0.564433193 | 0.22524557 | 2.50585698 | 0.0122155 | 0.0949813 |
| ECHDC2 | 219.099153 | 0.269744656 | 0.46637948 | 0.5783802 | 0.56300746 | 0.81448184 |
| SCP2 | 124.514349 | 0.572552819 | 0.22103454 | 2.59033193 | 0.00958834 | 0.07960971 |
| CPT2 | 15.6374669 | -0.211904276 | 0.55233929 | -0.3836487 | 0.70123881 | 0.88421489 |
| MAGOH | 20.9700687 | 0.346617026 | 0.43003591 | 0.80601879 | 0.42023202 | 0.71864146 |
| NDC1 | 33.2595432 | 0.23735506 | 0.34544739 | 0.68709468 | 0.49202307 | 0.77284018 |
| YIPF1 | 27.6790789 | -0.517619207 | 0.31359378 | -1.6506042 | 0.09881942 | 0.34341868 |
| HSPB11 | 33.1194213 | -0.007293683 | 0.35266303 | -0.0206817 | 0.98349954 | 0.99742652 |
| LRRC42 | 15.5760827 | -0.129600958 | 0.4080649 | -0.3175989 | 0.75078922 | 0.9089517 |
| TMEM59 | 111.502826 | -0.28166546 | 0.29726448 | -0.9475248 | 0.34337143 | 0.65298699 |
| TCEANC2 | 23.4569451 | 0.194827261 | 0.56567059 | 0.34441823 | 0.73053178 | 0.89731942 |
| MRPL37 | 13.309099 | 0.072557225 | 0.40010471 | 0.18134559 | 0.85609633 | 0.95219406 |
| SSBP3 | 360.067094 | 0.135876929 | 0.23391158 | 0.58089014 | 0.5613145 | 0.81384822 |
| ACOT11 | 241.741919 | 1.871194588 | 0.29295649 | 6.38727824 | 1.69E-10 | 3.75E-08 |
| MROH7-TTC | 20.6486472 | 1.752854729 | 0.49590887 | 3.53463069 | 0.00040835 | 0.00746552 |
| DHCR24 | 34.9753101 | -1.745997402 | 0.68126885 | -2.562861 | 0.01038136 | 0.08461171 |
| BSND | 11.5751771 | 1.350893605 | 0.71786899 | 1.88181079 | 0.05986171 | 0.25568052 |
| USP24 | 640.252642 | 1.030830159 | 0.32223028 | 3.19904807 | 0.00137882 | 0.01933253 |
| PLPP3 | 288.89634 | -1.696226217 | 0.76938909 | -2.2046403 | 0.02747934 | 0.16065021 |
| PRKAA2 | 272.233442 | 0.121732692 | 0.35996353 | 0.33818063 | 0.73522708 | 0.90007689 |
| DAB1 | 216.574928 | 1.751827167 | 0.91415883 | 1.91632692 | 0.0553235 | 0.24479327 |
| TACSTD2 | 47.2706421 | -1.583152891 | 0.56605129 | -2.7968365 | 0.00516056 | 0.0513523 |

| | | | | | | |
|---|---|---|---|---|---|---|
| MYSM1 | 134.561282 | 0.61582099 | 0.27227659 | 2.26174787 | 0.02371299 | 0.14623008 |
| AL136985.3 | 21.4826754 | -2.14921137 | 0.70311354 | -3.056706 | 0.00223784 | 0.02738266 |
| JUN | 38.9548985 | -1.794135377 | 0.6972931 | -2.5730003 | 0.01008211 | 0.08275141 |
| LINC01135 | 34.3320659 | 0.086360521 | 0.64954196 | 0.13295603 | 0.89422815 | 0.96482804 |
| FGGY | 138.703501 | -0.406956494 | 0.42159038 | -0.9652888 | 0.33440016 | 0.64554754 |
| HOOK1 | 105.161139 | 0.665045206 | 0.26871825 | 2.47487916 | 0.01332813 | 0.1002998 |
| NFIA | 20.6767898 | 1.007953924 | 0.63110313 | 1.59713028 | 0.11023667 | 0.365422 |
| TM2D1 | 80.4136527 | 0.21138517 | 0.26483233 | 0.79818491 | 0.4247632 | 0.72151771 |
| PATJ | 1296.62041 | -0.219003208 | 0.17535641 | -1.2489034 | 0.21170042 | 0.52063276 |
| USP1 | 18.4852948 | -0.217685788 | 0.42446271 | -0.5128502 | 0.60805611 | 0.83943073 |
| DOCK7 | 310.2562 | -0.604510104 | 0.36707507 | -1.6468296 | 0.09959307 | 0.34443834 |
| AC103923.1 | 16.8063484 | -2.975037184 | 0.62455376 | -4.7634605 | 1.90E-06 | 9.70E-05 |
| ATG4C | 55.6090137 | 0.459725394 | 0.41986862 | 1.09492678 | 0.27354872 | 0.58894058 |
| ALG6 | 19.1407255 | 0.581125142 | 0.39720907 | 1.46302082 | 0.14346168 | 0.42447522 |
| ITGB3BP | 67.1696184 | -0.172222015 | 0.72075621 | -0.2389463 | 0.81114724 | 0.93552094 |
| EFCAB7 | 37.0944637 | 0.315680262 | 0.623041 | 0.50667655 | 0.61238179 | 0.84174925 |
| PGM1 | 98.5219581 | 0.03713811 | 0.39263825 | 0.09458607 | 0.92464361 | 0.97735117 |
| CACHD1 | 308.261427 | 1.334515285 | 0.27001855 | 4.94230961 | 7.72E-07 | 4.67E-05 |
| RAVER2 | 88.7773546 | 0.714129909 | 0.33746286 | 2.11617336 | 0.03433007 | 0.18432213 |
| JAK1 | 165.106164 | -0.740323875 | 0.31049293 | -2.3843502 | 0.01710931 | 0.11799351 |
| AK4 | 31.7665314 | -1.411330595 | 0.82127182 | -1.7184695 | 0.08571101 | 0.31615828 |
| DNAJC6 | 23.7983621 | -1.560508384 | 0.61288165 | -2.5461823 | 0.01089083 | 0.08716404 |
| LEPR | 99.189432 | -1.5122596 | 0.4831366 | -3.130087 | 0.00174755 | 0.02285289 |
| SGIP1 | 313.764654 | 0.504307696 | 0.58527952 | 0.86165273 | 0.38887864 | 0.69247382 |
| WDR78 | 60.1718359 | -0.410086956 | 0.56306408 | -0.7283131 | 0.46642193 | 0.75300195 |
| MIER1 | 133.514937 | -0.170442766 | 0.2504426 | -0.6805662 | 0.49614603 | 0.77475164 |
| SLC35D1 | 60.6983186 | 0.59138421 | 0.33221725 | 1.78011287 | 0.07505749 | 0.29306367 |
| SERBP1 | 83.5575657 | -0.56728337 | 0.32097586 | -1.7673708 | 0.07716615 | 0.29743577 |
| GADD45A | 22.2173585 | -1.61845689 | 0.35659568 | -4.538633 | 5.66E-06 | 0.00023997 |
| GNG12 | 178.288916 | -0.600523066 | 0.28462222 | -2.1098952 | 0.03486738 | 0.18591921 |
| GNG12-AS1 | 94.4474938 | -0.398154231 | 0.42644577 | -0.9336574 | 0.35048067 | 0.65765381 |
| WLS | 203.520435 | 0.819983829 | 0.27045595 | 3.03185723 | 0.00243054 | 0.02946992 |
| LRRC40 | 41.3598275 | 1.061581979 | 0.32303077 | 3.28631845 | 0.00101506 | 0.01518241 |
| SRSF11 | 230.341021 | 0.119743425 | 0.21368329 | 0.56037806 | 0.57522159 | 0.82146824 |
| ANKRD13C | 142.517999 | 0.258944154 | 0.22756691 | 1.1378814 | 0.25517001 | 0.56782939 |
| PTGER3 | 928.528996 | 0.911127584 | 0.61928978 | 1.47124595 | 0.14122462 | 0.42136906 |
| ZRANB2-AS1 | 16.9300226 | 0.25106153 | 0.58712852 | 0.42760915 | 0.6689357 | 0.86777458 |
| ZRANB2 | 250.25174 | 0.32435854 | 0.19149481 | 1.69382421 | 0.09029869 | 0.326842 |
| NEGR1 | 18.3033385 | -0.682225102 | 0.56718732 | -1.2028215 | 0.22904541 | 0.54176037 |
| LRRIQ3 | 18.124793 | -0.061750004 | 1.0318468 | -0.0598442 | 0.95227976 | 0.98738095 |
| CRYZ | 93.051584 | 0.803478184 | 0.45759388 | 1.75587616 | 0.07910956 | 0.30106837 |
| TYW3 | 28.1890896 | 0.3895453 | 0.38205482 | 1.01960578 | 0.30791546 | 0.62178394 |
| ACADM | 69.1529173 | 0.610522363 | 0.26107695 | 2.33847672 | 0.01936253 | 0.12858707 |
| RABGGTB | 24.8226168 | 0.170542617 | 0.46088698 | 0.37003132 | 0.71135916 | 0.88696346 |
| PIGK | 45.7031387 | 0.027864419 | 0.36731541 | 0.07585965 | 0.93953076 | 0.98358617 |
| AC118549.1 | 255.869004 | 0.295708757 | 0.15052637 | 1.96449809 | 0.04947236 | 0.22858288 |
| USP33 | 196.734073 | 0.501803357 | 0.16017454 | 3.13285344 | 0.00173116 | 0.02273426 |
| MIGA1 | 116.612142 | 0.60360454 | 0.21223548 | 2.84403224 | 0.00445465 | 0.04609899 |
| NEXN | 51.4213964 | 1.193035313 | 0.43592868 | 2.73676722 | 0.00620462 | 0.05790977 |
| FUBP1 | 158.634635 | 0.295504866 | 0.20995133 | 1.40749222 | 0.15928148 | 0.44760113 |
| GIPC2 | 27.6179942 | 0.740263117 | 0.75325823 | 0.98274814 | 0.32573141 | 0.63751463 |
| PTGFR | 63.6773293 | 1.847994153 | 0.46485326 | 3.97543549 | 7.03E-05 | 0.00188835 |
| IFI44L | 90.2922598 | -0.556312969 | 2.01305064 | -0.2763532 | 0.78227681 | 0.92141851 |
| IFI44 | 28.3670274 | 0.581139766 | 1.56858113 | 0.37048767 | 0.71101916 | 0.88677671 |
| LINC01781 | 93.4603819 | -3.22527679 | 1.39038146 | -2.3197064 | 0.02035677 | 0.13268199 |
| ADGRL2 | 701.549089 | 0.070847237 | 0.63752714 | 0.11112819 | 0.9115147 | 0.97128978 |
| LINC01362 | 17.3264278 | -0.757095857 | 0.91974709 | -0.8231566 | 0.41041896 | 0.70916447 |
| LINC01725 | 198.306171 | 0.492490874 | 0.56775593 | 0.86743414 | 0.38570418 | 0.68921713 |
| TTLL7 | 19.005611 | -0.529146796 | 0.43203116 | -1.2247885 | 0.22065492 | 0.53073441 |
| PRKACB | 124.818769 | 0.722059972 | 0.4557755 | 1.58424481 | 0.11313802 | 0.37079048 |
| RPF1 | 15.8519345 | -0.525791785 | 0.39097677 | -1.3448159 | 0.17868469 | 0.47656064 |
| GNG5 | 37.0690804 | -0.187574217 | 0.35350853 | -0.5306073 | 0.59569091 | 0.83416944 |
| SPATA1 | 21.8100468 | -0.686345851 | 0.65207138 | -1.0525624 | 0.29254158 | 0.60745829 |
| CTBS | 41.30191 | -0.145417652 | 0.33126332 | -0.438979 | 0.66067673 | 0.86492722 |
| SSX2IP | 16.2095454 | -0.798439938 | 0.41847398 | -1.9079799 | 0.05639381 | 0.2473063 |
| SYDE2 | 20.1996731 | 1.056155603 | 0.40637345 | 2.59897788 | 0.00935018 | 0.07861233 |
| C1orf52 | 32.5350852 | 0.199621229 | 0.29381027 | 0.67942222 | 0.49687037 | 0.77475164 |
| BCL10 | 21.1480926 | -0.170200398 | 0.36169617 | -0.4705618 | 0.63795369 | 0.8538689 |
| AL078459.1 | 54.2524004 | -0.172371913 | 0.41933188 | -0.4110632 | 0.68102618 | 0.87265005 |
| DDAH1 | 321.551971 | 0.844020793 | 0.45146592 | 1.86951165 | 0.06155166 | 0.2599337 |
| CYR61 | 25.011754 | 0.919750493 | 0.96365136 | 0.95444321 | 0.33985934 | 0.65027011 |
| ZNHIT6 | 62.6633087 | -0.122588822 | 0.22565279 | -0.543263 | 0.58694871 | 0.8297013 |
| ODF2L | 126.633107 | -0.508148616 | 0.26621566 | -1.9087856 | 0.05628976 | 0.24721889 |
| SH3GLB1 | 71.8756243 | -0.246748013 | 0.24612241 | -1.0025418 | 0.31608198 | 0.63009595 |
| SELENOF | 93.9581402 | 0.117256214 | 0.21610564 | 0.54258748 | 0.58741386 | 0.82985559 |
| HS2ST1 | 118.856426 | 0.217908473 | 0.21139823 | 1.03079612 | 0.30263644 | 0.61691783 |
| PKN2-AS1 | 26.5893967 | 1.39675289 | 0.65357346 | 2.1371016 | 0.03258973 | 0.17956997 |
| PKN2 | 319.697363 | -0.146153034 | 0.3612699 | -0.4045536 | 0.68580568 | 0.87448517 |
| GTF2B | 41.0753907 | -0.518317257 | 0.41752363 | -1.2414082 | 0.21445499 | 0.52290228 |
| KYAT3 | 67.2571134 | -0.246685489 | 0.40922589 | -0.6028101 | 0.54663504 | 0.80581909 |
| GBP3 | 18.3873542 | -1.847464751 | 0.46709307 | -3.9552391 | 7.65E-05 | 0.00199691 |
| GBP2 | 42.0721574 | -5.494284681 | 0.92367332 | -5.9482985 | 2.71E-09 | 3.77E-07 |
| LRRC8B | 81.3951445 | -0.136641505 | 0.28385613 | -0.4813759 | 0.63024935 | 0.84981126 |
| LRRC8C-DT | 11.7339324 | 0.426978385 | 0.81535905 | 0.52366916 | 0.60050867 | 0.83619218 |
| ZNF326 | 64.9878728 | 0.030141655 | 0.29038464 | 0.10379907 | 0.91732881 | 0.97460959 |
| ZNF644 | 288.242643 | -0.031968925 | 0.32086744 | -0.0996328 | 0.92063585 | 0.9754826 |
| HFM1 | 18.490416 | 0.793647664 | 0.62771907 | 1.26433575 | 0.20610954 | 0.51308606 |
| TGFBR3 | 18.1703106 | -1.769103831 | 0.65684743 | -2.6933254 | 0.00707432 | 0.06416435 |
| BTBD8 | 27.6064078 | -0.567964741 | 0.63174788 | -0.899037 | 0.36863294 | 0.67276052 |
| GLMN | 44.3668177 | 0.611180002 | 0.34231368 | 1.78543842 | 0.07419023 | 0.29129403 |
| RPAP2 | 161.893115 | 0.037218741 | 0.17802912 | 0.20905985 | 0.83440151 | 0.94646712 |
| EVI5 | 296.014352 | 0.123813235 | 0.24079922 | 0.51417623 | 0.60712878 | 0.83902013 |
| RPL5 | 76.7615627 | -0.538174853 | 0.28323229 | -1.9001183 | 0.0574176 | 0.24970229 |
| FAM69A | 32.5437548 | -0.043338726 | 0.31292254 | -0.1384967 | 0.88984791 | 0.9631927 |
| MTF2 | 150.004753 | 0.17161416 | 0.22281375 | 0.77021351 | 0.44117325 | 0.73330634 |
| TMED5 | 44.2303444 | -0.319679206 | 0.38303983 | -0.8345848 | 0.40395154 | 0.70295711 |
| CCDC18 | 21.158835 | -0.112909896 | 0.5516058 | -0.2046931 | 0.83781192 | 0.94722424 |
| DR1 | 22.7234163 | 0.073769633 | 0.34499995 | 0.21382506 | 0.83068349 | 0.94408737 |
| FNBP1L | 373.3459 | 0.172046753 | 0.20515032 | 0.8386375 | 0.40167276 | 0.70114138 |
| BCAR3 | 177.721566 | -0.914659908 | 0.46722754 | -1.9576327 | 0.05027313 | 0.23147983 |
| GCLM | 46.9301781 | -1.711152728 | 0.38786658 | -4.4117044 | 1.03E-05 | 0.00039679 |
| ARHGAP29 | 424.11885 | -0.513532619 | 0.6770313 | -0.7585065 | 0.44814784 | 0.7399821 |
| ABCD3 | 166.616761 | 0.382569489 | 0.2633832 | 1.45252049 | 0.14635694 | 0.42885986 |
| SLC44A3 | 150.245493 | 0.299074939 | 0.50519633 | 0.59199745 | 0.5538523 | 0.81056645 |
| CNN3 | 154.587482 | -0.452069208 | 0.26333796 | -1.7166883 | 0.08603613 | 0.3168629 |
| ALG14 | 39.4431572 | 0.075763102 | 0.28170236 | 0.26894735 | 0.7879702 | 0.92377626 |
| PTBP2 | 229.933226 | 0.062344144 | 0.19687429 | 0.31666981 | 0.75149416 | 0.90896158 |
| DPYD | 363.540689 | -1.018353753 | 0.38607297 | -2.6377235 | 0.00834646 | 0.07252786 |
| SNX7 | 38.6858327 | -0.263960718 | 0.33172192 | -0.7957289 | 0.42618962 | 0.72211376 |
| PALMD | 12.109348 | 2.540807183 | 0.62702104 | 4.0521881 | 5.07E-05 | 0.00143366 |
| FRRS1 | 16.6030406 | 0.39049058 | 0.46458028 | 0.84052337 | 0.40061501 | 0.70092619 |
| AGL | 98.0018243 | 0.503892669 | 0.21261129 | 2.37001844 | 0.0177872 | 0.12114524 |
| TRMT13 | 66.4353835 | 1.01569957 | 0.27845711 | 3.64759794 | 0.0002647 | 0.00526246 |

| | | | | | | |
|---|---|---|---|---|---|---|
| LRRC39 | 12.2469573 | 0.495258784 | 0.53989402 | 0.91732594 | 0.35897187 | 0.66541823 |
| DBT | 75.7205573 | 0.883934503 | 0.26882806 | 3.28810356 | 0.00100865 | 0.01514433 |
| RTCA | 44.173198 | 0.757962094 | 0.29719632 | 2.55037511 | 0.01076071 | 0.08656845 |
| CDC14A | 68.780213 | -0.417579455 | 0.24193643 | -1.7259883 | 0.08434952 | 0.31297129 |
| EXTL2 | 29.3906204 | -0.086332842 | 0.33021014 | -0.2614482 | 0.7937469 | 0.92658341 |
| SLC30A7 | 101.241713 | -0.597978327 | 0.28568297 | -2.0931536 | 0.03633545 | 0.19065374 |
| DPH5 | 24.9515473 | 0.124027712 | 0.31103499 | 0.39875807 | 0.69007147 | 0.87691291 |
| AC093157.1 | 33.1666297 | -0.072056099 | 0.31334354 | -0.2299588 | 0.8181238 | 0.9380844 |
| RNPC3 | 428.153059 | 2.265981334 | 0.46987785 | 4.82249025 | 1.42E-06 | 7.61E-05 |
| AMY2B | 140.0477 | 2.701939267 | 0.57338191 | 4.71228553 | 2.45E-06 | 0.00011896 |
| NTNG1 | 59.8087709 | -5.567165399 | 1.06469557 | -5.22888 | 1.71E-07 | 1.30E-05 |
| VAV3 | 652.343686 | 1.125385387 | 0.50984015 | 2.20732984 | 0.02729102 | 0.16044231 |
| SLC25A24 | 51.5538611 | 0.058479818 | 0.30158013 | 0.19391137 | 0.84624528 | 0.94909204 |
| PRPF38B | 80.8430994 | -0.188703866 | 0.43413873 | -0.4346626 | 0.66380735 | 0.86588413 |
| STXBP3 | 77.729593 | 0.360477691 | 0.24780461 | 1.45468519 | 0.14575644 | 0.42828171 |
| AL449266.1 | 18.8567857 | -0.365706103 | 0.35839743 | -1.0203926 | 0.30754228 | 0.62161058 |
| GPSM2 | 17.0846476 | -1.664575328 | 0.59205465 | -2.8115231 | 0.00493076 | 0.04959478 |
| WDR47 | 23.2882667 | 0.644027213 | 0.35939364 | 1.79198275 | 0.07313573 | 0.28899744 |
| TAF13 | 14.7304028 | 0.306583387 | 0.56586027 | 0.54180052 | 0.58795593 | 0.830244 |
| TMEM167B | 13.7965773 | -0.203196879 | 0.37685349 | -0.5391933 | 0.58975348 | 0.83091183 |
| SARS | 40.4238405 | -0.609871856 | 0.43850273 | -1.3908052 | 0.16428452 | 0.45563826 |
| SORT1 | 79.9963937 | 0.13535713 | 0.34232186 | 0.39540895 | 0.6925411 | 0.87849579 |
| PSMA5 | 51.2482255 | -0.577920917 | 0.50210829 | -1.1509886 | 0.24973692 | 0.5624056 |
| GNAI3 | 79.6640313 | -0.393176362 | 0.2493002 | -1.5771201 | 0.11476789 | 0.37337606 |
| AMPD2 | 17.5919861 | -0.366444746 | 0.46221089 | -0.7928085 | 0.42788938 | 0.72328509 |
| GSTM4 | 14.3132533 | 0.428720549 | 0.62955374 | 0.68099119 | 0.49587706 | 0.77475164 |
| GSTM3 | 60.720757 | -0.089981991 | 0.62118446 | -0.1448555 | 0.88482496 | 0.96211171 |
| AHCYL1 | 199.55695 | 0.1706479 | 0.36271778 | 0.47047018 | 0.63801913 | 0.8538689 |
| STRIP1 | 26.3241896 | 0.380303527 | 0.38117596 | 0.99771121 | 0.31841942 | 0.63093651 |
| SLC6A17 | 21.2253452 | 0.727577079 | 0.71427528 | 1.01862279 | 0.30838208 | 0.62223642 |
| KCNC4 | 69.0737876 | -0.580551465 | 0.5550486 | -1.0459471 | 0.29558547 | 0.61055359 |
| RBM15 | 29.4884844 | 0.30398111 | 0.36296892 | 0.83748524 | 0.40231987 | 0.70146019 |
| SLC16A4 | 32.5640898 | -1.137682386 | 0.55185316 | -2.0615672 | 0.03924896 | 0.20036945 |
| AL355488.1 | 14.0606863 | 0.702891965 | 0.43537787 | 1.61444118 | 0.10643177 | 0.35877432 |
| LAMTOR5 | 41.9672692 | -0.555069291 | 0.36895022 | -1.5044558 | 0.13246405 | 0.40428636 |
| LRIF1 | 17.8784358 | -0.238355263 | 0.45712373 | -0.521424 | 0.60207147 | 0.83691476 |
| DRAM2 | 48.7829558 | 0.163951272 | 0.32155768 | 0.50986583 | 0.61014546 | 0.84109213 |
| CEPT1 | 66.527549 | 0.667158451 | 0.24796454 | 2.69053979 | 0.00713365 | 0.06445172 |
| DENND2D | 26.9771693 | -0.630577705 | 0.45501401 | -1.3858424 | 0.16579503 | 0.45790073 |
| ATP5PB | 63.1153393 | 0.319936721 | 0.39878968 | 0.80226931 | 0.42239719 | 0.72013823 |
| RAP1A | 145.695764 | -0.192356923 | 0.27358746 | -0.7030911 | 0.48199895 | 0.76628579 |
| KCND3 | 21.923194 | -1.720394167 | 0.6747623 | -2.5496299 | 0.01078373 | 0.08667889 |
| CTTNBP2NL | 63.9797164 | -0.649252547 | 0.32555639 | -1.994286 | 0.04612081 | 0.22049887 |
| WNT2B | 17.959338 | -1.505658047 | 0.62717192 | -2.4007102 | 0.01636329 | 0.11437129 |
| ST7L | 76.2000197 | 0.381346639 | 0.25269567 | 1.50911424 | 0.13126959 | 0.40268134 |
| CAPZA1 | 127.219054 | -0.451758161 | 0.33232707 | -1.3593782 | 0.17402679 | 0.47019002 |
| MOV10 | 31.6087368 | 0.387873085 | 0.35114494 | 1.1045954 | 0.26933494 | 0.58564558 |
| RHOC | 32.9295933 | -1.687969705 | 0.56802263 | -2.9716593 | 0.00296195 | 0.03430713 |
| LRIG2 | 60.670102 | 0.433700032 | 0.25533666 | 1.69854193 | 0.08940553 | 0.32467397 |
| MAGI3 | 286.883152 | 0.260274011 | 0.26225899 | 0.99243122 | 0.32098722 | 0.63269097 |
| PHTF1 | 64.736194 | 0.236434686 | 0.24819121 | 0.95263117 | 0.34077698 | 0.65083559 |
| RSBN1 | 63.2940421 | -0.013427812 | 0.28889011 | -0.0464807 | 0.96292712 | 0.9905486 |
| HIPK1 | 45.0647719 | 0.005588947 | 0.37750974 | 0.01480477 | 0.98818793 | 0.9977555 |
| TRIM33 | 222.361339 | -0.629905925 | 0.53720623 | -1.1725589 | 0.24097276 | 0.55327012 |
| BCAS2 | 20.3209238 | -0.387749378 | 0.57479284 | -0.6745898 | 0.49993642 | 0.77558432 |
| CSDE1 | 130.112775 | -0.028044084 | 0.24470276 | -0.1146047 | 0.90875846 | 0.97059151 |
| SIKE1 | 25.3831678 | -0.216386707 | 0.33141531 | -0.652917 | 0.51380976 | 0.78323295 |
| SLC22A15 | 149.960278 | 0.525425236 | 0.27364677 | 1.92008568 | 0.05484708 | 0.24363704 |
| ATP1A1 | 1298.34454 | -0.347967475 | 0.35702689 | -0.9746254 | 0.32974611 | 0.64177654 |
| LINC01762 | 852.294932 | 0.254301565 | 0.59249671 | 0.42920334 | 0.66777525 | 0.86777458 |
| CD58 | 79.4141141 | -1.024202666 | 0.23592205 | -4.3412758 | 1.42E-05 | 0.00052833 |
| IGSF3 | 144.707155 | -0.167207406 | 0.27737012 | -0.6028314 | 0.54662088 | 0.80581909 |
| PTGFRN | 44.1686804 | -0.540572082 | 0.45857082 | -1.1788192 | 0.23847018 | 0.55037028 |
| TTF2 | 32.0701747 | -0.110890046 | 0.41677029 | -0.2660699 | 0.79018535 | 0.92448611 |
| VTCN1 | 33.9698522 | -3.904163427 | 0.98382072 | -3.9683688 | 7.24E-05 | 0.00192235 |
| MAN1A2 | 199.295018 | 0.256223023 | 0.19177562 | 1.33605627 | 0.18153088 | 0.47998877 |
| GDAP2 | 73.362802 | 0.068366355 | 0.21464265 | 0.31851245 | 0.75009625 | 0.90865239 |
| WDR3 | 12.0956125 | 0.476699977 | 0.52102748 | 0.91492291 | 0.3602321 | 0.66607353 |
| SPAG17 | 68.7176676 | -0.669160317 | 0.57865594 | -1.1564045 | 0.24751576 | 0.56027856 |
| WARS2 | 44.2990373 | -0.042572994 | 0.32535808 | -0.1308497 | 0.89589424 | 0.96519206 |
| AL359915.2 | 131.716554 | 0.161980945 | 0.4355508 | 0.37189908 | 0.70996799 | 0.88614996 |
| ZNF697 | 27.072743 | 1.009020898 | 0.47035039 | 2.14525365 | 0.03193257 | 0.17712034 |
| PHGDH | 22.1324514 | -0.983367597 | 0.63576099 | -1.5467567 | 0.12192191 | 0.38601016 |
| NOTCH2 | 52.1389875 | 0.297599164 | 0.28334309 | 1.05031382 | 0.29357385 | 0.60876592 |
| SEC22B | 68.1322032 | 0.190152734 | 0.25376649 | 0.74932168 | 0.45366334 | 0.74470138 |
| NBPF26 | 18.6395227 | 0.296493042 | 0.60442339 | 0.49053867 | 0.62375278 | 0.84597807 |
| LINC00623 | 123.547925 | -0.85608528 | 0.56318166 | -1.5200873 | 0.12848904 | 0.39735716 |
| SRGAP2C | 22.2849223 | 0.170865029 | 0.36312113 | 0.47054554 | 0.63796531 | 0.8538689 |
| NBPF15 | 21.2478899 | -0.590550489 | 0.40225291 | -1.4681075 | 0.14207504 | 0.42241953 |
| FP700111.1 | 24.6775805 | 0.862962259 | 0.36630258 | 2.3558727 | 0.01847925 | 0.12417062 |
| SRGAP2B | 27.5598843 | -0.170253178 | 0.3622223 | -0.470024 | 0.63833787 | 0.8538689 |
| GPR89A | 19.7206397 | 0.04942026 | 0.40105014 | 0.12322714 | 0.90192724 | 0.96803829 |
| PDZK1 | 22.0099438 | 0.893354398 | 0.53127605 | 1.68152582 | 0.09266082 | 0.33202083 |
| RNF115 | 217.123942 | 0.256396648 | 0.18791366 | 1.36443862 | 0.17242956 | 0.46814182 |
| POLR3C | 26.2335028 | -0.598245772 | 0.34590415 | -1.7295131 | 0.0837173 | 0.31173327 |
| PEX11B | 12.9413919 | 0.214465437 | 0.45960296 | 0.46663197 | 0.64076321 | 0.85468927 |
| RBM8A | 20.0036257 | -0.693543315 | 0.42472805 | -1.6329115 | 0.10248761 | 0.34990644 |
| TXNIP | 453.287708 | 1.454917521 | 0.5481279 | 2.65433945 | 0.00794638 | 0.06963542 |
| NBPF12 | 50.0263996 | -0.651049953 | 0.29800909 | -2.1846647 | 0.02891344 | 0.16590086 |
| PRKAB2 | 23.1845511 | 0.917323305 | 0.48884499 | 1.87651164 | 0.06058506 | 0.2573479 |
| AC242426.2 | 43.457836 | 0.278943604 | 0.50621121 | 0.55104193 | 0.58160493 | 0.82565231 |
| FMO5 | 23.2031836 | -0.495516384 | 0.7346078 | -0.6745319 | 0.49997322 | 0.77558432 |
| CHD1L | 54.7777393 | 0.419985774 | 0.35988572 | 1.1669976 | 0.24321134 | 0.55555126 |
| BCL9 | 87.2898982 | 0.574210594 | 0.3746368 | 1.53271272 | 0.12534665 | 0.39153507 |
| ACP6 | 66.6061607 | 0.440674273 | 0.36574278 | 1.20487483 | 0.22825163 | 0.54043123 |
| GPR89B | 25.2257615 | 0.355609809 | 0.35148613 | 1.01173214 | 0.31166615 | 0.62642275 |
| NBPF11 | 39.5068263 | -0.126751128 | 0.32163259 | -0.3940867 | 0.69351703 | 0.87889881 |
| NBPF14 | 17.1377677 | 0.061500612 | 0.41148858 | 0.14945885 | 0.88119158 | 0.9612999 |
| PDE4DIP | 215.280355 | 0.798520017 | 0.28478519 | 2.80393805 | 0.00504826 | 0.05050427 |
| NBPF9 | 18.2100706 | -0.280223651 | 0.36749223 | -0.7625295 | 0.44574407 | 0.7388166 |
| AC245297.3 | 334.496742 | 0.026851697 | 0.28057071 | 0.09570385 | 0.92375578 | 0.97676332 |
| NBPF19 | 27.1468337 | -0.207173021 | 0.899385 | -0.2303497 | 0.81782008 | 0.93805075 |
| OTUD7B | 142.623103 | -0.312998251 | 0.36084565 | -0.867402 | 0.38572176 | 0.68921713 |
| VPS45 | 121.24982 | 0.276159181 | 0.29421891 | 0.93861806 | 0.34792688 | 0.65536772 |
| ANP32E | 25.404478 | -0.199797497 | 0.38658775 | -0.5168231 | 0.60527962 | 0.83802715 |
| APH1A | 25.071575 | -0.133342296 | 0.48942397 | -0.2724474 | 0.78527802 | 0.92276669 |
| C1orf54 | 32.2317152 | -0.252176358 | 0.56873475 | -0.4433989 | 0.65747723 | 0.86357172 |
| MRPS21 | 54.9645868 | -0.210304782 | 0.22569112 | -0.9318257 | 0.35142662 | 0.65871783 |
| PRPF3 | 68.5309829 | 0.337463514 | 0.37284821 | 0.90509624 | 0.36541441 | 0.67012584 |
| RPRD2 | 238.138585 | 0.141750848 | 0.16926462 | 0.83745114 | 0.40233903 | 0.70146019 |
| TARS2 | 14.6724253 | 0.383885375 | 0.41419811 | 0.92681586 | 0.35402215 | 0.6607207 |
| MCL1 | 51.927109 | -1.602539917 | 0.32680439 | -4.903667 | 9.41E-07 | 5.59E-05 |
| ENSA | 187.46707 | -0.870791982 | 0.51498804 | -1.6908975 | 0.09085638 | 0.32805373 |

| | | | | | | |
|---|---|---|---|---|---|---|
| GOLPH3L | 66.315524 | -0.012006916 | 0.26281239 | -0.0456863 | 0.96356031 | 0.99076272 |
| ARNT | 84.1505647 | 0.31160872 | 0.2727604 | 1.14242652 | 0.25327677 | 0.56559484 |
| SETDB1 | 63.3532821 | -0.024755061 | 0.24079803 | -0.1028043 | 0.91811833 | 0.97482633 |
| CERS2 | 29.2867935 | -0.727422689 | 0.39939572 | -1.8213082 | 0.06856002 | 0.27887855 |
| MINDY1 | 19.1603228 | -0.192027747 | 0.55154945 | -0.3481605 | 0.72771963 | 0.89598017 |
| PRUNE1 | 25.5270863 | 0.1247932 | 0.40574307 | 0.30756705 | 0.75841179 | 0.91161939 |
| CDC42SE1 | 40.7873743 | -0.336169655 | 0.39043999 | -0.8610021 | 0.38923688 | 0.69247382 |
| GABPB2 | 75.0366046 | 1.093881878 | 0.43692702 | 2.50358028 | 0.01229438 | 0.09519681 |
| SCNM1 | 13.543945 | -0.799080461 | 0.53063881 | -1.5058839 | 0.13209697 | 0.40375585 |
| VPS72 | 14.0222728 | 0.141654174 | 0.41044867 | 0.34512031 | 0.73000392 | 0.89725238 |
| PIP5K1A | 251.552121 | 0.211444124 | 0.20419396 | 1.03550627 | 0.30043254 | 0.61414654 |
| PSMD4 | 41.3843764 | 0.004792163 | 0.26731146 | 0.01792727 | 0.98569688 | 0.99746215 |
| PI4KB | 51.3906621 | 0.226208334 | 0.33948459 | 0.66632872 | 0.50520101 | 0.77885156 |
| PSMB4 | 18.4841159 | -0.297284951 | 0.47552581 | -0.625171 | 0.53185883 | 0.79676281 |
| POGZ | 323.66991 | 0.762203104 | 0.20146769 | 3.78325238 | 0.00015479 | 0.00344459 |
| CGN | 52.9338179 | 0.501719887 | 0.41707382 | 1.20295224 | 0.2289948 | 0.54176037 |
| TUFT1 | 43.6945759 | 0.03660836 | 0.33853478 | 0.10813766 | 0.91388649 | 0.97217087 |
| SNX27 | 78.5807829 | 0.75603553 | 0.26322411 | 2.87221237 | 0.00407609 | 0.04333576 |
| RORC | 37.4963299 | 0.487712684 | 0.4771775 | 1.02207811 | 0.30674393 | 0.62067717 |
| THEM4 | 108.328526 | -0.430096485 | 0.62754205 | -0.6853668 | 0.49311249 | 0.77312609 |
| S100A10 | 103.16598 | -1.400370245 | 0.47434903 | -2.9521937 | 0.00315525 | 0.0361864 |
| S100A11 | 211.570238 | -2.967957409 | 0.49976121 | -5.9387511 | 2.87E-09 | 3.94E-07 |
| FLG-AS1 | 83.8065696 | 1.578224273 | 0.44885576 | 3.51610562 | 0.00043793 | 0.00779242 |
| S100A6 | 302.943367 | -2.397146473 | 0.72118116 | -3.3239172 | 0.00088763 | 0.0136122 |
| S100A2 | 54.0410976 | -1.824153079 | 1.42605706 | -1.2791585 | 0.20084123 | 0.50467291 |
| S100A16 | 26.9682245 | -2.379323837 | 0.51775501 | -4.5954627 | 4.32E-06 | 0.00019263 |
| S100A13 | 33.1134929 | -1.404502483 | 0.43893144 | -3.199822 | 0.00137512 | 0.01930973 |
| CHTOP | 42.7689189 | 0.257745105 | 0.34007068 | 0.75791628 | 0.44850111 | 0.74006308 |
| ILF2 | 23.179734 | -0.690451904 | 0.3361153 | -2.0542114 | 0.03995523 | 0.20257889 |
| INTS3 | 56.9557886 | 0.104509401 | 0.28633936 | 0.3649844 | 0.71512305 | 0.88904248 |
| GATAD2B | 300.297691 | -0.04997617 | 0.36521363 | -0.1368409 | 0.89115654 | 0.96328012 |
| DENND4B | 30.6064564 | -0.097097194 | 0.43333216 | -0.2240711 | 0.82270202 | 0.94066053 |
| CRTC2 | 16.6943596 | -0.271724085 | 0.53946676 | -0.5036901 | 0.61447914 | 0.84174925 |
| SLC39A1 | 28.8007929 | -1.953870512 | 0.62028845 | -3.1499386 | 0.00163305 | 0.02180518 |
| JTB | 31.4033548 | -0.189428009 | 0.41386361 | -0.4577064 | 0.64716342 | 0.85917726 |
| RAB13 | 28.256187 | -1.308332308 | 0.33500277 | -3.9054373 | 9.41E-05 | 0.00230782 |
| RPS27 | 27.0399203 | -0.147330795 | 0.50173673 | -0.2936416 | 0.76903175 | 0.91600052 |
| TPM3 | 55.1357501 | -1.276752993 | 0.44257161 | -2.8848506 | 0.00391599 | 0.04212483 |
| C1orf43 | 65.3186915 | -0.631636333 | 0.31346433 | -2.0150182 | 0.04390274 | 0.21387104 |
| UBAP2L | 82.2435295 | 0.150263308 | 0.41528951 | 0.36182784 | 0.71748068 | 0.89028919 |
| HAX1 | 17.8572458 | 0.079447075 | 0.45736531 | 0.17370595 | 0.86209656 | 0.95458251 |
| ATP8B2 | 14.3601226 | 0.811039051 | 0.49595241 | 1.63531629 | 0.10198276 | 0.34893691 |
| IL6R | 26.574408 | -3.391212955 | 0.91304035 | -3.7141983 | 0.00020385 | 0.0043002 |
| UBE2Q1 | 27.415611 | -1.66614815 | 0.68616041 | -2.4282196 | 0.01517315 | 0.10916241 |
| ADAR | 104.832623 | -0.014362632 | 0.34219571 | -0.041972 | 0.96652104 | 0.99156233 |
| KCNN3 | 81.9430389 | 2.228301693 | 0.54200014 | 4.11125671 | 3.94E-05 | 0.00117224 |
| PMVK | 19.7739269 | 0.079386288 | 0.40873473 | 0.19422448 | 0.84600012 | 0.94909204 |
| PBXIP1 | 28.9249775 | 0.933445877 | 0.55363465 | 1.68603226 | 0.09178959 | 0.33043047 |
| ZBTB7B | 12.0701818 | 0.550548555 | 0.48591647 | 1.1330107 | 0.25720976 | 0.5697847 |
| EFNA1 | 41.0834853 | -2.406765198 | 0.40271455 | -5.9763551 | 2.28E-09 | 3.38E-07 |
| SLC50A1 | 15.4102118 | -1.667773335 | 0.45110216 | -3.6971078 | 0.00021807 | 0.00452847 |
| DPM3 | 15.464502 | -0.340297392 | 0.50348077 | -0.6758896 | 0.49911077 | 0.77558432 |
| SCAMP3 | 26.0687252 | 0.08043399 | 0.50880958 | 0.1580827 | 0.87439164 | 0.95761805 |
| CLK2 | 26.4155754 | -0.179716667 | 0.32138367 | -0.5591966 | 0.57602753 | 0.82147339 |
| FDPS | 60.1462172 | -0.822615593 | 0.32452827 | -2.5348041 | 0.01125102 | 0.08920454 |
| RUSC1 | 13.3206307 | 0.883524953 | 0.46700275 | 1.89190526 | 0.0585036 | 0.25277461 |
| ASH1L | 537.183166 | -0.069875572 | 0.22430529 | -0.3115199 | 0.75540539 | 0.91060567 |
| YY1AP1 | 89.2003682 | 0.264210589 | 0.28820631 | 0.91674116 | 0.35927829 | 0.66545706 |
| DAP3 | 95.3282581 | -0.007477466 | 0.23208363 | -0.0322188 | 0.97429753 | 0.99357756 |
| GON4L | 116.58701 | -0.434298262 | 0.319949 | -1.3573984 | 0.17465466 | 0.47119666 |
| RIT1 | 36.9876677 | -1.16498337 | 0.4845535 | -2.404241 | 0.01620609 | 0.11387005 |
| KHDC4 | 103.981975 | -0.492133424 | 0.29838374 | -1.6493305 | 0.09907994 | 0.34368353 |
| ARHGEF2 | 31.411436 | -1.464477113 | 0.47111768 | -3.1085166 | 0.00188029 | 0.02404915 |
| SSR2 | 32.5615431 | -0.89615997 | 0.42786005 | -2.0945166 | 0.03621399 | 0.19023055 |
| UBQLN4 | 15.250341 | -0.288463946 | 0.39371103 | -0.7326794 | 0.46375402 | 0.75139985 |
| RAB25 | 14.0612749 | -0.164752636 | 0.59979827 | -0.2746801 | 0.78356204 | 0.92211693 |
| LMNA | 84.8846923 | -0.281083143 | 0.39845961 | -0.7054244 | 0.48054614 | 0.76551071 |
| SLC25A44 | 17.8695893 | 0.930951892 | 0.35341573 | 2.63415525 | 0.00843469 | 0.072922 |
| SMG5 | 53.1591246 | -0.471611809 | 0.25813196 | -1.8270183 | 0.06769702 | 0.27769777 |
| CCT3 | 48.3883758 | 0.125522891 | 0.42490551 | 0.29541366 | 0.7676779 | 0.91600052 |
| RHBG | 58.0153479 | 2.450523462 | 0.76972039 | 3.18365408 | 0.00145429 | 0.02017823 |
| MEF2D | 43.2791159 | 0.165129935 | 0.57149966 | 0.28894144 | 0.77262619 | 0.91791018 |
| ISG20L2 | 12.8915062 | -1.023803334 | 0.53402105 | -1.9171591 | 0.05521772 | 0.24446818 |
| RRNAD1 | 18.5076641 | 0.020446386 | 0.36253748 | 0.05639799 | 0.95502476 | 0.98852696 |
| HDGF | 42.5417415 | -0.850592676 | 0.28661724 | -2.9676954 | 0.00300041 | 0.0346236 |
| PRCC | 47.4488261 | 0.146765246 | 0.32401012 | 0.452965 | 0.65057394 | 0.86113455 |
| ARHGEF11 | 107.504895 | 0.133151453 | 0.24995508 | 0.53270153 | 0.59424021 | 0.83416944 |
| KIRREL1 | 72.1151215 | -0.926622684 | 0.42683151 | -2.1709332 | 0.02993622 | 0.16937218 |
| IFI16 | 30.7146053 | -2.136493554 | 0.75713364 | -2.8218183 | 0.00477522 | 0.04855417 |
| DUSP23 | 39.6714806 | -1.41961941 | 0.5211344 | -2.7240946 | 0.0064478 | 0.05976075 |
| TAGLN2 | 39.0719832 | -2.290737999 | 0.54123878 | -4.2323981 | 2.31E-05 | 0.0007727 |
| KCNJ10 | 48.3117258 | 0.660998856 | 0.77567006 | 0.85216498 | 0.39412253 | 0.69624829 |
| IGSF8 | 14.6139774 | 0.806579328 | 0.54567031 | 1.47814405 | 0.13936923 | 0.41770449 |
| DCAF8 | 29.8195568 | 1.010072107 | 0.38392801 | 2.63088935 | 0.00851618 | 0.07325168 |
| COPA | 125.442676 | -0.194856621 | 0.27983933 | -0.6963161 | 0.48623092 | 0.76833754 |
| NCSTN | 21.2387333 | -0.232279539 | 0.50648034 | -0.4586151 | 0.64651058 | 0.8589047 |
| F11R | 37.289245 | -1.175152231 | 0.6099299 | -1.9267005 | 0.05401697 | 0.24144497 |
| AL591806.3 | 21.3078062 | -0.065452882 | 0.42601502 | -0.1536398 | 0.87789372 | 0.95961091 |
| PFDN2 | 39.7869036 | -0.678796535 | 0.35697479 | -1.9015251 | 0.05723326 | 0.24962036 |
| NIT1 | 30.5006825 | -0.813988023 | 0.31621776 | -2.5741376 | 0.01004903 | 0.08269829 |
| DEDD | 15.8552199 | -0.098729597 | 0.38904676 | -0.2537731 | 0.79967087 | 0.92922871 |
| UFC1 | 18.943181 | -1.288601479 | 0.43848393 | -2.9387656 | 0.00329522 | 0.03746908 |
| NDUFS2 | 47.9833627 | 0.770657376 | 0.29585779 | 2.60482371 | 0.00919215 | 0.07756348 |
| SDHC | 79.2540425 | 0.080557859 | 0.2578792 | 0.31238602 | 0.75474717 | 0.91026551 |
| AL592295.4 | 17.3679961 | -0.171741571 | 0.54250257 | -0.3165728 | 0.75156776 | 0.90896158 |
| DUSP12 | 15.867992 | -0.095195343 | 0.39198596 | -0.242854 | 0.80811853 | 0.9342712 |
| ATF6 | 176.291314 | -0.329879347 | 0.41501079 | -0.7948693 | 0.42668952 | 0.72230448 |
| NOS1AP | 823.521198 | 0.077396173 | 0.44980008 | 0.17206794 | 0.86338412 | 0.95458251 |
| C1orf226 | 15.4540475 | -0.794246247 | 0.4782459 | -1.6607487 | 0.09676393 | 0.33969491 |
| UHMK1 | 70.3386403 | 0.138748929 | 0.2455119 | 0.56514137 | 0.57197761 | 0.81947131 |
| UAP1 | 35.4632392 | 0.36945032 | 0.50329446 | 0.73406395 | 0.46290977 | 0.75091259 |
| HSD17B7 | 38.4108443 | -0.814318936 | 0.46179288 | -1.7633856 | 0.07783547 | 0.29864393 |
| PBX1 | 228.232244 | 0.605389798 | 0.39541609 | 1.53101964 | 0.12576454 | 0.39231467 |
| MGST3 | 160.230978 | -0.100436745 | 0.3138757 | -0.3199889 | 0.74897674 | 0.90776798 |
| ALDH9A1 | 21.6332824 | 0.136338048 | 0.4543975 | 0.30004137 | 0.7641456 | 0.91485536 |
| TMCO1 | 45.4585565 | -0.367201848 | 0.30661699 | -1.1975913 | 0.23107616 | 0.54380466 |
| UCK2 | 31.1145556 | 0.250574165 | 0.34251426 | 0.73157293 | 0.46442927 | 0.75205602 |
| FAM78B | 11.9073003 | 0.633423646 | 0.47317046 | 1.33867962 | 0.18067499 | 0.4792642 |
| POGK | 34.0423388 | 0.347467981 | 0.31956569 | 1.08731315 | 0.27689844 | 0.59188469 |
| TADA1 | 17.7015834 | 0.96988156 | 0.3802709 | 2.55050169 | 0.0107568 | 0.08656845 |
| POU2F1 | 229.188417 | 0.188875585 | 0.29981626 | 0.62997112 | 0.52871348 | 0.79434813 |
| CREG1 | 22.8344125 | -0.261138824 | 0.40667534 | -0.642131 | 0.52078815 | 0.78815594 |
| MPZL1 | 69.3682253 | -0.559548468 | 0.25531248 | -2.1916221 | 0.02840681 | 0.16420667 |

| | | | | | | |
|---|---|---|---|---|---|---|
| MPC2 | 42.3497932 | 0.192718229 | 0.44726923 | 0.43087746 | 0.66655747 | 0.86777458 |
| DCAF6 | 294.800038 | 0.131233286 | 0.29950211 | 0.43817149 | 0.66126198 | 0.86492722 |
| TIPRL | 27.1442783 | 0.01662942 | 0.35940759 | 0.04626897 | 0.96309587 | 0.99061289 |
| SFT2D2 | 40.1211627 | -0.772046794 | 0.30027159 | -2.5711616 | 0.0101358 | 0.08297056 |
| LINC00970 | 257.120814 | 2.291874724 | 0.69056733 | 3.31882875 | 0.00090396 | 0.01379461 |
| ATP1B1 | 1530.61446 | 0.905876347 | 0.58655356 | 1.54440517 | 0.12249021 | 0.38683625 |
| NME7 | 194.279901 | 0.015223353 | 0.20115185 | 0.0756809 | 0.93967297 | 0.98358617 |
| BLZF1 | 54.10012 | 0.050340096 | 0.35169292 | 0.14313651 | 0.88618237 | 0.96249798 |
| SLC19A2 | 89.0544055 | 0.097149704 | 0.5657961 | 0.17170444 | 0.86366989 | 0.95458251 |
| SCYL3 | 38.357797 | -0.218106468 | 0.38419679 | -0.5676947 | 0.57024231 | 0.81874643 |
| KIFAP3 | 90.0482271 | 0.65768784 | 0.28842711 | 2.28025667 | 0.02259247 | 0.14147226 |
| GORAB | 15.0356812 | -0.262884874 | 0.5560575 | -0.4727656 | 0.6363804 | 0.85375696 |
| FMO4 | 57.0420853 | 1.810000856 | 0.48856389 | 3.70473724 | 0.00021161 | 0.00443382 |
| PRRC2C | 288.245146 | -0.395012916 | 0.37703464 | -1.0476834 | 0.29478452 | 0.60998021 |
| VAMP4 | 25.1481497 | -0.202817596 | 0.35557679 | -0.5703904 | 0.56841292 | 0.81712643 |
| DNM3 | 264.489926 | 0.595647337 | 0.4940031 | 1.20575627 | 0.22791148 | 0.54017161 |
| PIGC | 34.161491 | -0.271626002 | 0.41224375 | -0.6588966 | 0.50996217 | 0.78153966 |
| SUCO | 95.0998568 | -0.348358296 | 0.46844177 | -0.7436534 | 0.45708622 | 0.7472269 |
| AL645568.1 | 68.6679647 | 0.901980535 | 0.50874361 | 1.772957 | 0.07623583 | 0.29596141 |
| PRDX6 | 82.0624341 | 0.047711872 | 0.41710656 | 0.11438773 | 0.90893044 | 0.97066401 |
| KLHL20 | 77.9941195 | -0.042055563 | 0.30251563 | -0.1390195 | 0.88943476 | 0.96308489 |
| DARS2 | 16.7791371 | -0.274247129 | 0.5119653 | -0.5356752 | 0.59218303 | 0.83293326 |
| ZBTB37 | 41.3754015 | -0.067702569 | 0.29463353 | -0.2297857 | 0.8182583 | 0.9380844 |
| RC3H1 | 160.342884 | -0.247467179 | 0.24937528 | -0.9923485 | 0.32102756 | 0.63269097 |
| RABGAP1L | 781.166627 | 0.737045324 | 0.19441145 | 3.79116219 | 0.00014994 | 0.0033527 |
| CACYBP | 13.4511336 | 0.156711582 | 0.61584627 | 0.25446542 | 0.79913601 | 0.92922871 |
| MRPS14 | 14.548537 | -0.184346258 | 0.45128262 | -0.408494 | 0.68291101 | 0.87314216 |
| KIAA0040 | 23.2601056 | -0.778453236 | 0.3995471 | -1.9483391 | 0.0513744 | 0.23455111 |
| COP1 | 320.691801 | -0.325477941 | 0.22793217 | -1.4279596 | 0.15330348 | 0.43873592 |
| PAPPA2 | 396.318428 | -3.728489414 | 0.61867577 | -6.0265645 | 1.67E-09 | 2.56E-07 |
| RASAL2 | 326.6233 | -2.225865734 | 0.76932552 | -2.8932691 | 0.00381255 | 0.04152772 |
| RALGPS2 | 217.410139 | -0.20339645 | 0.34251439 | -0.5938333 | 0.55262362 | 0.80962553 |
| FAM20B | 43.2010865 | -0.045008043 | 0.27111659 | -0.1660099 | 0.86814914 | 0.95588054 |
| ABL2 | 174.936833 | -0.124959476 | 0.24765281 | -0.5045753 | 0.6138572 | 0.84174925 |
| SOAT1 | 53.230779 | -1.907340389 | 0.43389605 | -4.3958464 | 1.10E-05 | 0.00042339 |
| TOR1AIP2 | 101.077798 | -0.1297397 | 0.19131529 | -0.678146 | 0.49767913 | 0.77553656 |
| TOR1AIP1 | 46.0126206 | -0.315409423 | 0.49022988 | -0.6433909 | 0.51997051 | 0.7877576 |
| CEP350 | 220.854437 | 0.183120187 | 0.25012932 | 0.73210206 | 0.46410628 | 0.75166353 |
| QSOX1 | 31.556147 | -0.939906404 | 0.35698138 | -2.6329284 | 0.00846522 | 0.0729557 |
| ACBD6 | 178.62684 | 0.216731661 | 0.29581553 | 0.73265815 | 0.46376696 | 0.75139985 |
| XPR1 | 194.374367 | -0.540352295 | 0.33979901 | -1.5902115 | 0.11178714 | 0.36830503 |
| KIAA1614 | 29.426976 | -0.365039907 | 0.53708245 | -0.679672 | 0.49671217 | 0.77475164 |
| STX6 | 55.6152544 | -0.252170972 | 0.23480882 | -1.0739417 | 0.28284882 | 0.59668285 |
| MR1 | 18.4215301 | -0.65278352 | 0.41477828 | -1.5738132 | 0.11553065 | 0.3743642 |
| GLUL | 54.0702166 | -2.332029559 | 0.33697921 | -6.9203959 | 4.50E-12 | 1.35E-09 |
| DHX9 | 127.050132 | 0.006399264 | 0.36552111 | 0.01750724 | 0.98603196 | 0.99746215 |
| LAMC1 | 160.181474 | -1.074490796 | 0.56504742 | -1.901594 | 0.05722425 | 0.24962036 |
| LAMC2 | 21.6551915 | -5.128017347 | 1.0740399 | -4.7745129 | 1.80E-06 | 9.33E-05 |
| SMG7 | 154.189973 | 0.214765191 | 0.23814577 | 0.90182239 | 0.36715122 | 0.67131445 |
| ARPC5 | 34.2628045 | -0.977906016 | 0.34135791 | -2.8647527 | 0.00417335 | 0.04411827 |
| RGL1 | 15.688218 | 0.578978955 | 0.49068515 | 1.17993983 | 0.23802415 | 0.54997949 |
| TSEN15 | 48.8618962 | 0.52870707 | 0.37926975 | 1.39401328 | 0.16331359 | 0.45454804 |
| EDEM3 | 86.1480551 | 0.230657822 | 0.34702115 | 0.66467944 | 0.50625555 | 0.77934214 |
| RNF2 | 48.3457604 | 0.306894299 | 0.27671173 | 1.10907584 | 0.26739747 | 0.58364201 |
| TRMT1L | 37.8928915 | 0.306842008 | 0.37429744 | 0.8197812 | 0.41234085 | 0.71110141 |
| SWT1 | 72.3240736 | -0.589264173 | 0.39485246 | -1.4923654 | 0.13560337 | 0.41064171 |
| IVNS1ABP | 699.180668 | 0.720288622 | 0.48655211 | 1.48039358 | 0.13876824 | 0.41617082 |
| HMCN1 | 210.959261 | 0.588400711 | 0.2971162 | 1.98037239 | 0.0476617 | 0.22360104 |
| TPR | 135.560969 | -0.157164418 | 0.24410716 | -0.6438337 | 0.51968327 | 0.7877576 |
| ODR4 | 83.1795258 | 0.174041779 | 0.25270021 | 0.68872826 | 0.49099429 | 0.77215443 |
| AL596220.1 | 17.8200964 | -0.161230478 | 0.7412668 | -0.2175067 | 0.8278135 | 0.94254892 |
| ERVMER61-1 | 126.157236 | 1.272972939 | 0.40428228 | 3.14872309 | 0.00163986 | 0.02181171 |
| BRINP3 | 54.0848082 | 1.0931532 | 0.49677292 | 2.20050882 | 0.02777081 | 0.16155622 |
| AL390957.1 | 22.4581093 | -0.681475323 | 0.54220167 | -1.2568669 | 0.20880184 | 0.51711493 |
| AL136454.1 | 19.0871578 | -0.815248774 | 0.5000299 | -1.6304001 | 0.10301697 | 0.35071178 |
| UCHL5 | 42.6932814 | 0.179689617 | 0.43231795 | 0.41564227 | 0.67767179 | 0.87114221 |
| TROVE2 | 70.2604475 | -0.356893681 | 0.25673267 | -1.3901374 | 0.16448717 | 0.45563826 |
| CDC73 | 169.060196 | 0.03898105 | 0.23098348 | 0.1687612 | 0.86598448 | 0.95565922 |
| ZBTB41 | 36.7601316 | 0.169460063 | 0.32994507 | 0.51360084 | 0.60753109 | 0.83917773 |
| CRB1 | 18.3478104 | 0.014220729 | 1.09775181 | 0.01295441 | 0.98966416 | 0.99842088 |
| DENND1B | 229.654862 | 1.01696062 | 0.25542385 | 3.98146311 | 6.85E-05 | 0.00185108 |
| NEK7 | 202.558409 | 0.26071978 | 0.17937352 | 1.45350207 | 0.1460844 | 0.42860392 |
| AC104461.1 | 24.1258647 | 1.174919182 | 0.56580045 | 2.07656106 | 0.03784209 | 0.19628857 |
| ZNF281 | 15.9362497 | -0.322367549 | 0.49108484 | -0.6564396 | 0.5115413 | 0.78254489 |
| DDX59 | 32.017981 | 0.014728759 | 0.35756673 | 0.04119164 | 0.96714312 | 0.99156233 |
| CAMSAP2 | 154.903181 | -0.126575987 | 0.26574565 | -0.476305 | 0.63385709 | 0.85221104 |
| INAVA | 26.806218 | -0.645090257 | 0.4575123 | -1.4099954 | 0.15854103 | 0.44659715 |
| TMEM9 | 24.0203367 | -0.096414635 | 0.46970956 | -0.2052644 | 0.83736558 | 0.94706413 |
| CSRP1 | 25.259063 | -0.403900274 | 0.66359541 | -0.6086544 | 0.54275353 | 0.80340275 |
| AC096677.1 | 38.9278991 | 0.427608565 | 0.34121376 | 1.25319849 | 0.21013348 | 0.51886901 |
| NAV1 | 40.2578257 | -1.194691987 | 0.40617953 | -2.9412905 | 0.00326848 | 0.03725587 |
| IPO9 | 81.2627569 | -0.418570537 | 0.34138813 | -1.226084 | 0.22016706 | 0.5299013 |
| TIMM17A | 27.5433453 | 0.085229378 | 0.40275549 | 0.21161569 | 0.83240687 | 0.94520892 |
| RNPEP | 27.8211298 | -0.427159405 | 0.3128809 | -1.365246 | 0.17217574 | 0.46801268 |
| ELF3 | 111.001087 | -3.268668883 | 0.4298038 | -7.6050256 | 2.85E-14 | 1.33E-11 |
| ARL8A | 23.9472521 | -0.190099489 | 0.40285448 | -0.4718813 | 0.63701153 | 0.8538689 |
| PPP1R12B | 218.429884 | 0.425022022 | 0.23552149 | 1.80459977 | 0.07113733 | 0.28467146 |
| KDM5B | 144.812846 | -0.534554097 | 0.22908162 | -2.3334657 | 0.01962372 | 0.12983501 |
| KLHL12 | 27.8713302 | 0.281643158 | 0.3355451 | 0.83936006 | 0.40126729 | 0.70114138 |
| ADIPOR1 | 15.8715559 | 0.277218877 | 0.40981557 | 0.67644789 | 0.49875632 | 0.77558432 |
| CYB5R1 | 16.1566009 | 0.60153722 | 0.48295756 | 1.24552811 | 0.21293769 | 0.52201944 |
| TMEM183A | 14.9470901 | 0.338772214 | 0.47900111 | 0.70724724 | 0.47941284 | 0.76437164 |
| BTG2 | 25.4070726 | -0.113875833 | 0.6263937 | -0.1817959 | 0.85574287 | 0.9521686 |
| ATP2B4 | 171.562905 | 0.130606527 | 0.27792133 | 0.46994064 | 0.63839743 | 0.8538689 |
| SNRPE | 12.5487514 | -0.069180285 | 0.41624378 | -0.1662014 | 0.86799848 | 0.95588054 |
| SOX13 | 36.6962526 | -0.101927847 | 0.34838639 | -0.2925713 | 0.76984988 | 0.91655198 |
| GOLT1A | 18.7080876 | -0.672592268 | 0.50995339 | -1.3189289 | 0.18719288 | 0.48725107 |
| PLEKHA6 | 126.40223 | -0.534960011 | 0.34436718 | -1.5534582 | 0.12031371 | 0.38339202 |
| PPP1R15B | 24.8520981 | -0.26484056 | 0.51003951 | -0.519255 | 0.60358294 | 0.83748768 |
| MDM4 | 170.739358 | 0.405223858 | 0.21094095 | 1.92102988 | 0.05472794 | 0.24343712 |
| LRRN2 | 52.1278076 | 0.525007491 | 0.5237764 | 1.00235042 | 0.31617438 | 0.63009595 |
| RBBP5 | 41.7318742 | 0.146792844 | 0.3641127 | 0.40315222 | 0.68683624 | 0.87523044 |
| DSTYK | 77.744107 | 0.3082607 | 0.2974647 | 1.03629339 | 0.30006529 | 0.61374288 |
| NUAK2 | 55.1380385 | 0.409666597 | 0.60683679 | 0.6750853 | 0.49962157 | 0.77558432 |
| CDK18 | 27.9294698 | -1.469639216 | 0.51597325 | -2.8482857 | 0.00439554 | 0.04563926 |
| MFSD4A | 92.9214632 | 1.313413028 | 0.63624395 | 2.064323 | 0.0389871 | 0.19962424 |
| ELK4 | 60.8096879 | -0.384939896 | 0.35799126 | -1.0752774 | 0.28225057 | 0.59642935 |
| NUCKS1 | 170.767728 | 0.004260634 | 0.27567662 | 0.01545519 | 0.98766904 | 0.9977555 |
| RAB29 | 14.9345029 | -0.892671463 | 0.46376319 | -1.9248433 | 0.05424897 | 0.2422497 |
| SRGAP2 | 44.5206266 | -0.087740341 | 0.29551544 | -0.2969061 | 0.76653818 | 0.91564296 |
| EIF2D | 21.2897551 | 0.252863702 | 0.37827338 | 0.66846814 | 0.50383481 | 0.77790293 |
| DYRK3 | 22.8239888 | -2.022701612 | 0.81122946 | -2.4933779 | 0.01265341 | 0.0971032 |
| MAPKAPK2 | 127.51982 | -0.56679964 | 0.42375966 | -1.3375498 | 0.18104325 | 0.47966815 |

| | | | | | | |
|---|---|---|---|---|---|---|
| PIGR | 44.3014489 | -2.699487876 | 0.92763671 | -2.9100701 | 0.00361348 | 0.03987226 |
| PFKFB2 | 97.8683008 | -0.713549408 | 0.57400424 | -1.2431083 | 0.21382794 | 0.52235998 |
| CD55 | 36.843581 | -1.06466852 | 0.71083761 | -1.4977662 | 0.13419402 | 0.40823002 |
| CD46 | 146.351317 | -0.142273191 | 0.29620657 | -0.4803175 | 0.63100166 | 0.85057966 |
| MIR29B2CHC | 246.269776 | 0.229176865 | 0.39878793 | 0.57468355 | 0.56550533 | 0.81595078 |
| PLXNA2 | 14.4824694 | -0.079374572 | 0.53454274 | -0.1484906 | 0.8819556 | 0.96157087 |
| LAMB3 | 57.4873008 | -6.949919481 | 1.63622338 | -4.2475371 | 2.16E-05 | 0.00074363 |
| G0S2 | 93.0450224 | -1.31952907 | 0.74800894 | -1.7640552 | 0.07772268 | 0.29864393 |
| IRF6 | 18.0909935 | 0.748687578 | 0.45701019 | 1.63822952 | 0.10137382 | 0.34801529 |
| UTP25 | 13.0521574 | -0.438103108 | 0.48409871 | -0.9049871 | 0.3654722 | 0.67012584 |
| SYT14 | 20.3030299 | -0.226062863 | 0.82944026 | -0.2725487 | 0.78520015 | 0.92276669 |
| HHAT | 81.9112777 | 1.363592976 | 0.67327452 | 2.02531498 | 0.04283503 | 0.21080021 |
| KCNH1 | 16.0692997 | 2.758739253 | 0.6722046 | 4.10401721 | 4.06E-05 | 0.00119807 |
| RCOR3 | 174.471839 | 0.768819643 | 0.25204506 | 3.05032623 | 0.00228593 | 0.02789791 |
| TRAF5 | 33.5992945 | 0.079605937 | 0.46629991 | 0.17071832 | 0.86444526 | 0.95487238 |
| LINC00467 | 29.9735374 | -0.015901388 | 0.48334089 | -0.0328989 | 0.9737552 | 0.99340049 |
| LPGAT1 | 129.679233 | -0.73910658 | 0.38234095 | -1.9331086 | 0.05322282 | 0.23892614 |
| INTS7 | 32.6103507 | 0.23493259 | 0.37923677 | 0.6194879 | 0.53559499 | 0.79972923 |
| PPP2R5A | 188.917381 | -0.033839116 | 0.29727742 | -0.1138301 | 0.90937249 | 0.97090822 |
| AL360091.3 | 25.6535627 | -1.385281961 | 0.71077862 | -1.948964 | 0.05129972 | 0.23455111 |
| NENF | 29.5133846 | 0.937000083 | 0.64503919 | 1.45262505 | 0.14632789 | 0.42885986 |
| NSL1 | 61.7562788 | 0.281022389 | 0.25578314 | 1.09867442 | 0.2719101 | 0.58786226 |
| TATDN3 | 45.7870445 | 0.232519569 | 0.27290061 | 0.85203023 | 0.39419732 | 0.69624849 |
| FLVCR1 | 37.4836292 | -0.009940927 | 0.35472976 | -0.0280239 | 0.97764305 | 0.99503808 |
| ANGEL2 | 51.0742012 | 0.38146263 | 0.31245328 | 1.22086293 | 0.22213792 | 0.53271965 |
| RPS6KC1 | 158.089036 | 0.163876161 | 0.26408726 | 0.62053793 | 0.53490369 | 0.79921039 |
| PROX1 | 57.3184084 | 1.068948944 | 0.52205119 | 2.04759413 | 0.04059979 | 0.20429163 |
| SMYD2 | 38.6046405 | -0.706149641 | 0.38586461 | -1.8300451 | 0.0672432 | 0.27620069 |
| PTPN14 | 529.810489 | -0.714493155 | 0.45721904 | -1.5626934 | 0.11812474 | 0.37883307 |
| KCTD3 | 95.4517717 | 0.244666324 | 0.25404926 | 0.96306647 | 0.33551417 | 0.64662829 |
| ESRRG | 5317.15071 | 1.102522335 | 0.28513026 | 3.86673209 | 0.0001103 | 0.00261697 |
| GPATCH2 | 176.263884 | -0.264593537 | 0.19989137 | -1.3236867 | 0.18560713 | 0.4858669 |
| SPATA17 | 28.3687775 | 0.284304423 | 0.83351188 | 0.34109223 | 0.73303415 | 0.89840827 |
| RRP15 | 24.3416274 | 0.026540061 | 0.36751404 | 0.07221509 | 0.94243073 | 0.98488741 |
| LYPLAL1 | 221.713407 | 0.420880975 | 0.3511797 | 1.1984775 | 0.23073118 | 0.54354156 |
| EPRS | 58.4520277 | -0.132176719 | 0.30163222 | -0.4382049 | 0.66123775 | 0.86492722 |
| BPNT1 | 16.3660979 | 0.480811014 | 0.42963332 | 1.11911948 | 0.26308917 | 0.57713562 |
| IARS2 | 50.9004773 | -0.362907197 | 0.29311894 | -1.2380885 | 0.21568323 | 0.52507323 |
| RAB3GAP2 | 203.068353 | 0.157083357 | 0.22967315 | 0.68394307 | 0.49401111 | 0.7738663 |
| MARK1 | 161.708683 | 0.001471629 | 0.64719841 | 0.00227385 | 0.99818574 | 0.99922221 |
| 2-Mar | 71.8236333 | 1.065294762 | 0.40635742 | 2.62157082 | 0.00875256 | 0.074665 |
| 1-Mar | 13.1371093 | -0.22341604 | 0.49196168 | -0.454133 | 0.64973309 | 0.86077172 |
| AL513314.2 | 19.2034698 | 0.828581972 | 0.38656713 | 2.14343621 | 0.03207809 | 0.17765195 |
| MIA3 | 45.1708839 | 0.262790587 | 0.27268696 | 0.96370794 | 0.33519237 | 0.64662829 |
| AIDA | 18.7414452 | -0.258032898 | 0.33930397 | -0.7604771 | 0.44696946 | 0.73972874 |
| BROX | 43.8287312 | -0.031591711 | 0.26223103 | -0.1204728 | 0.90410861 | 0.9686272 |
| FAM177B | 32.1579362 | -2.338436179 | 1.04863437 | -2.2299824 | 0.02574861 | 0.15406587 |
| AC093152.1 | 35.8156783 | 0.004525788 | 0.35363257 | 0.01279799 | 0.98978896 | 0.99842088 |
| TLR5 | 26.547939 | 0.486650476 | 0.36408878 | 1.33662585 | 0.1813448 | 0.47994859 |
| SUSD4 | 109.581939 | 0.851303983 | 0.34769345 | 2.44843263 | 0.01434793 | 0.10525575 |
| CAPN2 | 149.860457 | -0.252738879 | 0.32234139 | -0.7840721 | 0.43299782 | 0.72648177 |
| TP53BP2 | 69.7010667 | -1.289487958 | 0.62719448 | -2.0559619 | 0.03978619 | 0.20205143 |
| FBXO28 | 110.156777 | -0.31124281 | 0.36023191 | -0.8640068 | 0.38758417 | 0.69098179 |
| DEGS1 | 46.4045904 | -1.22197491 | 0.50635862 | -2.4132598 | 0.01581055 | 0.11184941 |
| NVL | 80.5166083 | 0.129398653 | 0.26756338 | 0.4836187 | 0.6286565 | 0.84817317 |
| CNIH4 | 65.9108348 | -0.472922613 | 0.26362279 | -1.7939368 | 0.07282326 | 0.28844693 |
| WDR26 | 164.481996 | 0.115561309 | 0.25340545 | 0.45603324 | 0.64836608 | 0.85981586 |
| CNIH3 | 49.4903327 | 0.337163275 | 0.66106995 | 0.51002662 | 0.61003281 | 0.84109213 |
| DNAH14 | 374.791787 | 0.09297839 | 0.24093761 | 0.38590235 | 0.69956899 | 0.88336691 |
| LBR | 50.2700249 | 0.612634124 | 0.31152799 | 1.966546 | 0.04923557 | 0.22771453 |
| ENAH | 75.6507191 | -0.325307831 | 0.3614993 | -0.8998851 | 0.3681814 | 0.67246295 |
| SRP9 | 48.0023295 | -0.200875284 | 0.29855498 | -0.6728251 | 0.50105857 | 0.7760974 |
| EPHX1 | 19.186864 | -0.685286991 | 0.44177789 | -1.5512026 | 0.12085315 | 0.3843229 |
| AL591895.1 | 18.9046352 | -1.119481546 | 0.41622811 | -2.6895866 | 0.00715406 | 0.06457351 |
| TMEM63A | 61.698654 | -1.1126813 | 0.29301901 | -3.7973008 | 0.00014628 | 0.00330246 |
| H3F3A | 22.27977 | -0.582234323 | 0.3379546 | -1.7228182 | 0.08492141 | 0.31408459 |
| ACBD3 | 87.2327726 | -0.26581782 | 0.29064984 | -0.9145638 | 0.36042068 | 0.66624949 |
| LIN9 | 15.5384638 | 0.197789486 | 0.40816126 | 0.48458662 | 0.62796961 | 0.84784081 |
| PARP1 | 46.8671505 | -0.397195746 | 0.32564741 | -1.2197111 | 0.22257442 | 0.53302786 |
| STUM | 45.2770436 | 0.654083937 | 0.610603 | 1.07120983 | 0.28407509 | 0.59746288 |
| ITPKB | 292.795765 | 1.683780852 | 0.49511735 | 3.40077122 | 0.00067196 | 0.0109921 |
| COQ8A | 37.8880812 | 0.43855189 | 0.32061228 | 1.36785744 | 0.1713567 | 0.46718638 |
| CDC42BPA | 374.012199 | -0.726203369 | 0.27543452 | -2.6365735 | 0.00837481 | 0.07270641 |
| ZNF678 | 53.0525023 | -0.386977556 | 0.24968756 | -1.5498471 | 0.12117821 | 0.38456965 |
| SNAP47 | 26.1512346 | 0.388216438 | 0.35904409 | 1.08125003 | 0.27958591 | 0.59383074 |
| ARF1 | 72.9900005 | -0.983169072 | 0.44266051 | -2.2210453 | 0.02634789 | 0.15607861 |
| GUK1 | 57.7865469 | -0.328154433 | 0.4036323 | -0.8130034 | 0.4162161 | 0.71390709 |
| OBSCN | 133.510115 | 0.009609489 | 0.30891551 | 0.03110717 | 0.97518407 | 0.99383717 |
| TRIM11 | 12.6088143 | -1.194583973 | 0.44360558 | -2.6928966 | 0.00708342 | 0.06418448 |
| RNF187 | 15.8462815 | -0.623633582 | 0.60645094 | -1.0283331 | 0.30379316 | 0.61793 |
| RAB4A | 26.9637735 | -0.044203867 | 0.38920825 | -0.1135738 | 0.90957564 | 0.97090822 |
| NUP133 | 48.5106777 | 0.348375886 | 0.24648837 | 1.41335627 | 0.157551 | 0.44434528 |
| ABCB10 | 23.5709533 | 0.662959882 | 0.484383 | 1.36866877 | 0.17110284 | 0.46688992 |
| TAF5L | 17.1759512 | -0.326943443 | 0.56286609 | -0.5808548 | 0.56133835 | 0.81384822 |
| GALNT2 | 105.677671 | -0.717363949 | 0.430844 | -1.6650202 | 0.09590873 | 0.33852036 |
| PGBD5 | 24.5001905 | 0.350741294 | 0.41032666 | 0.85478553 | 0.39266989 | 0.69579134 |
| COG2 | 36.1319413 | 0.38168743 | 0.27388873 | 1.39358575 | 0.16344273 | 0.45477172 |
| TTC13 | 25.0783532 | 0.391158402 | 0.35942446 | 1.08829098 | 0.27646667 | 0.59163995 |
| ARV1 | 17.7883574 | 0.889878265 | 0.45711816 | 1.94671389 | 0.05156905 | 0.23455111 |
| C1orf131 | 19.3943063 | -0.259177969 | 0.47423589 | -0.546517 | 0.58471061 | 0.82845288 |
| GNPAT | 54.5766201 | 0.10832588 | 0.24345798 | 0.44494692 | 0.65635811 | 0.86294007 |
| SPRTN | 13.532094 | 0.351975963 | 0.4150944 | 0.84794197 | 0.39647029 | 0.6969028 |
| EGLN1 | 142.438201 | 0.120867902 | 0.27324154 | 0.44234819 | 0.65823726 | 0.86357172 |
| DISC1 | 20.9823098 | -1.51985434 | 0.65183221 | -2.331665 | 0.01971832 | 0.13002379 |
| NTPCR | 28.7398464 | 0.323115429 | 0.38329093 | 0.84300306 | 0.39922673 | 0.69959441 |
| PCNX2 | 184.434658 | -0.383688006 | 0.24108593 | -1.591499 | 0.11149732 | 0.36760999 |
| MAP3K21 | 89.8505785 | 0.125623911 | 0.3378907 | 0.3717886 | 0.71005025 | 0.88614996 |
| KCNK1 | 40.4549629 | -1.917698219 | 0.41836008 | -4.583846 | 4.57E-06 | 0.00020077 |
| AL355472.1 | 11.4112237 | 0.999099147 | 0.75463535 | 1.32394957 | 0.18551979 | 0.4858669 |
| TARBP1 | 63.6755579 | 0.266898844 | 0.4614896 | 0.57834206 | 0.56303321 | 0.81448184 |
| IRF2BP2 | 75.7398002 | -0.724996587 | 0.44489449 | -1.6295922 | 0.10318772 | 0.35088341 |
| TOMM20 | 48.3109607 | -0.5282562 | 0.32649297 | -1.6179711 | 0.10566882 | 0.35687176 |
| ARID4B | 397.274835 | -0.209002702 | 0.26447968 | -0.7902411 | 0.42938701 | 0.7236272 |
| B3GALNT2 | 71.7296518 | -0.005199226 | 0.27508279 | -0.0189006 | 0.98492041 | 0.99742652 |
| LYST | 144.531971 | 0.130261938 | 0.24006327 | 0.54261503 | 0.58739489 | 0.82985559 |
| GPR137B | 19.8124903 | -1.485890963 | 0.42998934 | -3.455646 | 0.00054898 | 0.00927292 |
| ERO1B | 44.9352426 | -0.24472645 | 0.35648948 | -0.68649 | 0.49240419 | 0.77284018 |
| LGALS8 | 40.2300885 | -0.830337549 | 0.36432911 | -2.2790865 | 0.02266193 | 0.14162184 |
| HEATR1 | 46.4361412 | 0.412670968 | 0.33843848 | 1.2193382 | 0.22271585 | 0.53302786 |
| MTR | 388.694462 | 1.046109647 | 0.29541687 | 3.54113034 | 0.00039842 | 0.00734158 |
| CHRM3 | 158.917575 | -0.099088819 | 0.36910427 | -0.2684575 | 0.78834717 | 0.92377626 |
| RGS7 | 31.3984904 | 1.408119815 | 0.5133615 | 2.74294003 | 0.00608918 | 0.05734901 |
| FH | 17.7804858 | 0.730445537 | 0.49810309 | 1.46645455 | 0.14252449 | 0.42308208 |

| | | | | | | |
|---|---|---|---|---|---|---|
| CEP170 | 79.4800167 | 0.077311875 | 0.241071 | 0.32070168 | 0.74843647 | 0.90723111 |
| SDCCAG8 | 285.111223 | 0.217303664 | 0.32241163 | 0.6739945 | 0.50031481 | 0.77558432 |
| AKT3 | 316.845597 | 0.284154097 | 0.29983552 | 0.94769991 | 0.34328225 | 0.65295057 |
| ZBTB18 | 15.0625272 | 1.033917428 | 0.4597283 | 2.24897493 | 0.02451409 | 0.14871138 |
| ADSS | 60.2953563 | -0.189746255 | 0.29962618 | -0.6332766 | 0.52655302 | 0.79346524 |
| CATSPERE | 45.3034965 | 0.071571165 | 0.5036058 | 0.14211744 | 0.88698725 | 0.96253908 |
| DESI2 | 85.4908816 | -0.702318682 | 0.3477207 | -2.0197782 | 0.0434064 | 0.21196707 |
| COX20 | 65.8641727 | 0.607444415 | 0.29762409 | 2.04097863 | 0.04125295 | 0.20635328 |
| HNRNPU | 308.853218 | -0.151208002 | 0.20352665 | -0.7429396 | 0.45751827 | 0.747377 |
| EFCAB2 | 182.437877 | -0.086512845 | 0.51540587 | -0.1678538 | 0.86669828 | 0.95578708 |
| KIF26B | 49.4590927 | -1.620200177 | 0.51453106 | -3.148887 | 0.00163894 | 0.02181171 |
| SMYD3 | 502.893829 | 0.279005622 | 0.4117492 | 0.67761061 | 0.49801862 | 0.77558432 |
| TFB2M | 15.1489658 | 0.159784368 | 0.45061902 | 0.3545886 | 0.72289782 | 0.89240027 |
| CNST | 90.6016762 | -0.286616134 | 0.39784406 | -0.7204233 | 0.47126441 | 0.7577288 |
| SCCPDH | 20.5160811 | -0.238893182 | 0.44187652 | -0.5406333 | 0.58876033 | 0.83025227 |
| AHCTF1 | 108.557571 | 0.248253053 | 0.36640651 | 0.6775345 | 0.49806689 | 0.77558432 |
| ZNF124 | 52.9267208 | -1.016392007 | 0.33052052 | -3.0751252 | 0.00210414 | 0.02629895 |
| AL390728.6 | 15.413382 | -0.242919946 | 0.44786876 | -0.5423909 | 0.58754924 | 0.82992109 |
| ZNF496 | 26.9876047 | 0.174873053 | 0.38955498 | 0.44890468 | 0.65350042 | 0.86199934 |
| SH3BP5L | 10.4617998 | -0.040552145 | 0.46419065 | -0.087361 | 0.93038459 | 0.97888806 |
| ZNF672 | 14.2420436 | 0.27192876 | 0.47090407 | 0.57746106 | 0.56362804 | 0.81450672 |
| ZNF692 | 18.846058 | 0.59600446 | 0.399271 | 1.49273165 | 0.13550745 | 0.41064171 |
| FAM110C | 110.956442 | -3.632880129 | 0.71702862 | -5.0665762 | 4.05E-07 | 2.66E-05 |
| SH3YL1 | 102.646344 | 0.113636226 | 0.36854123 | 0.3083406 | 0.75782317 | 0.91161939 |
| ACP1 | 31.7326943 | -0.052675669 | 0.29922799 | -0.1760386 | 0.86026364 | 0.95363316 |
| EIPR1 | 80.703525 | -0.541785173 | 0.33103806 | -1.636625 | 0.10170884 | 0.34880722 |
| TRAPPC12 | 113.799008 | -0.254190112 | 0.2455484 | -1.0351935 | 0.30057854 | 0.61419993 |
| ADI1 | 36.7148848 | -0.777856144 | 0.40506505 | -1.920324 | 0.05481699 | 0.24361944 |
| RPS7 | 71.2785662 | -0.805009523 | 0.39648888 | -2.0303458 | 0.04232141 | 0.20978458 |
| RNF144A | 18.5904901 | 0.361601416 | 0.38971705 | 0.92785628 | 0.35348212 | 0.66036212 |
| ID2 | 46.4960316 | -1.434261926 | 0.42907858 | -3.3426556 | 0.00082981 | 0.01291675 |
| KIDINS220 | 162.500018 | 0.605185672 | 0.20066862 | 3.0158461 | 0.00256263 | 0.0305941 |
| MBOAT2 | 83.6849269 | -0.39498136 | 0.46348216 | -0.852204 | 0.39410086 | 0.69624829 |
| ASAP2 | 497.858576 | 0.24327944 | 0.21262689 | 1.14416119 | 0.25255679 | 0.56511628 |
| ITGB1BP1 | 67.5115624 | -0.341166907 | 0.26079122 | -1.3081994 | 0.19080569 | 0.49135864 |
| CPSF3 | 28.3410563 | -0.208111174 | 0.39937989 | -0.5210858 | 0.60230703 | 0.83691476 |
| IAH1 | 26.9736634 | -0.313779934 | 0.31570296 | -0.9939087 | 0.3202673 | 0.63212442 |
| ADAM17 | 118.568004 | -0.47208648 | 0.38877427 | -1.2142946 | 0.22463526 | 0.53581458 |
| YWHAQ | 80.1242535 | -0.4559266 | 0.30928168 | -1.4741468 | 0.14044207 | 0.41997495 |
| TAF1B | 99.7109872 | -0.489414486 | 0.31570906 | -1.5502073 | 0.12109177 | 0.38450126 |
| GRHL1 | 15.4373904 | 0.416116958 | 0.63360932 | 0.65674059 | 0.51134773 | 0.78246322 |
| KLF11 | 23.4745558 | 0.352199426 | 0.50128239 | 0.70259685 | 0.48230703 | 0.76628579 |
| CYS1 | 261.241435 | 0.140600082 | 0.71879568 | 0.19560507 | 0.84491929 | 0.94907846 |
| HPCAL1 | 109.225604 | -0.272283336 | 0.30064922 | -0.9056512 | 0.36512049 | 0.67012584 |
| NOL10 | 117.565082 | 0.315889237 | 0.27652209 | 1.14236527 | 0.25330222 | 0.56559484 |
| PDIA6 | 59.1723857 | -0.519189507 | 0.54898327 | -0.9457292 | 0.34428672 | 0.65359197 |
| ROCK2 | 183.346374 | -0.015622195 | 0.38025704 | -0.0410832 | 0.96722953 | 0.99156233 |
| E2F6 | 20.9177405 | -0.020071794 | 0.44308386 | -0.0453002 | 0.96386802 | 0.99082982 |
| LPIN1 | 70.5469307 | -0.013364181 | 0.47160401 | -0.0283377 | 0.9773928 | 0.99497547 |
| TRIB2 | 58.2625597 | -2.526859946 | 0.45862864 | -5.5095991 | 3.60E-08 | 3.13E-06 |
| NBAS | 287.61242 | 0.263310734 | 0.24616536 | 1.06964984 | 0.28477695 | 0.59819624 |
| DDX1 | 67.0587646 | 0.460475516 | 0.26410753 | 1.74351527 | 0.08124365 | 0.30643842 |
| SMC6 | 53.6372313 | -0.008070382 | 0.34377427 | -0.0234758 | 0.98127073 | 0.99655465 |
| GEN1 | 17.508111 | -0.749821101 | 0.60223298 | -1.2450682 | 0.2131067 | 0.52207222 |
| LINC01376 | 65.5331033 | 0.488532272 | 0.52181945 | 0.9362094 | 0.34916538 | 0.65632016 |
| TTC32 | 16.8573681 | 0.24838401 | 0.40099495 | 0.61941929 | 0.53564018 | 0.79972923 |
| WDR35 | 59.925694 | 0.106147363 | 0.34458299 | 0.30804586 | 0.75804744 | 0.91161939 |
| LAPTM4A | 70.5218852 | -0.640926498 | 0.28078419 | -2.2826303 | 0.02245216 | 0.14099969 |
| SDC1 | 25.3529966 | -1.464766183 | 0.73612664 | -1.9898291 | 0.04660977 | 0.22172933 |
| PUM2 | 330.975842 | 0.148199888 | 0.24858698 | 0.59616914 | 0.55106224 | 0.80853147 |
| RHOB | 39.7139504 | -1.045252644 | 0.6947236 | -1.504559 | 0.13243749 | 0.40428636 |
| LDAH | 97.7248176 | 0.224852255 | 0.46468821 | 0.48387769 | 0.62847268 | 0.84817317 |
| KLHL29 | 115.176011 | -5.968792837 | 1.35901933 | -4.3919852 | 1.12E-05 | 0.00042921 |
| ATAD2B | 197.921588 | 0.458088132 | 0.24137564 | 1.89782254 | 0.05771946 | 0.25043102 |
| UBXN2A | 87.0061875 | -0.061566624 | 0.27934203 | -0.2203987 | 0.82556064 | 0.94182399 |
| SF3B6 | 21.3751057 | -0.233429898 | 0.39621946 | -0.5891429 | 0.55576539 | 0.81104802 |
| FAM228B | 13.9243003 | -0.026910969 | 0.49014925 | -0.0549036 | 0.95621525 | 0.98853497 |
| TP53I3 | 18.2998328 | -0.225177094 | 0.59805861 | -0.3765134 | 0.70653524 | 0.88603006 |
| ITSN2 | 214.283233 | -0.550140873 | 0.29626786 | -1.8569037 | 0.06332486 | 0.26477174 |
| NCOA1 | 510.845546 | 0.191920068 | 0.1807083 | 1.06204345 | 0.28821596 | 0.60280969 |
| DNAJC27 | 40.8973 | 0.269837525 | 0.25906292 | 1.04159067 | 0.29760149 | 0.61141484 |
| DNAJC27-AS | 18.8923076 | 1.057406626 | 0.6750923 | 1.56631415 | 0.11727509 | 0.3774501 |
| DNMT3A | 122.026564 | 0.184270475 | 0.27262102 | 0.67592175 | 0.49909033 | 0.77558432 |
| DTNB | 168.905733 | -0.117081692 | 0.27587274 | -0.4244047 | 0.67127068 | 0.86777458 |
| ASXL2 | 171.771616 | -0.289393603 | 0.30745981 | -0.9412404 | 0.34658167 | 0.65471653 |
| RAB10 | 294.752945 | -0.327102863 | 0.35120539 | -0.931372 | 0.35166118 | 0.65871783 |
| HADHA | 88.2680613 | 0.245249471 | 0.25600154 | 0.95799999 | 0.33806276 | 0.64897646 |
| HADHB | 122.259577 | 0.378379452 | 0.30289032 | 1.24922925 | 0.21158124 | 0.52063276 |
| SELENOI | 72.9658728 | 0.363891328 | 0.33128897 | 1.09841065 | 0.27202521 | 0.58786226 |
| MAPRE3 | 32.2576232 | 0.037864547 | 0.49734922 | 0.07613272 | 0.93931351 | 0.98358617 |
| TMEM214 | 13.5865995 | -0.214689429 | 0.40498864 | -0.5301122 | 0.59603412 | 0.83416944 |
| AGBL5 | 25.2785392 | -0.908006204 | 0.55693436 | -1.6303649 | 0.10302441 | 0.35071178 |
| OST4 | 25.9172644 | -0.70017564 | 0.52613221 | -1.3307979 | 0.18325552 | 0.48210665 |
| ATRAID | 20.8790262 | -0.496181418 | 0.41899695 | -1.1842125 | 0.23632896 | 0.54909824 |
| MPV17 | 14.5683551 | -0.310768473 | 0.55627117 | -0.5586636 | 0.5763913 | 0.82175421 |
| GTF3C2 | 53.881943 | 0.259868149 | 0.25435903 | 1.02165883 | 0.3069424 | 0.62080932 |
| EIF2B4 | 16.22778 | 0.020139534 | 0.38973185 | 0.05167536 | 0.95878737 | 0.9891274 |
| PPM1G | 50.4445052 | -0.464801311 | 0.34473961 | -1.3482678 | 0.17757226 | 0.47495232 |
| NRBP1 | 47.8133629 | -0.654071239 | 0.49959666 | -1.3091986 | 0.19046709 | 0.49126283 |
| IFT172 | 54.0167285 | 0.916162535 | 0.52081181 | 1.75910476 | 0.07855972 | 0.29970982 |
| ZNF512 | 50.5906205 | -0.129637395 | 0.3639575 | -0.3561883 | 0.72169957 | 0.89217596 |
| SUPT7L | 20.8116477 | -0.013255634 | 0.33378145 | -0.0397135 | 0.96832153 | 0.99156233 |
| SLC4A1AP | 31.2943301 | 0.224393862 | 0.32552299 | 0.68933338 | 0.49061349 | 0.77203035 |
| MRPL33 | 94.5955897 | -0.238002303 | 0.31117835 | -0.7648421 | 0.44436558 | 0.73698311 |
| RBKS | 110.044526 | -0.360399189 | 0.30317598 | -1.1887459 | 0.23453969 | 0.54657 |
| BABAM2 | 635.164353 | -0.365954395 | 0.2695523 | -1.3576378 | 0.17457864 | 0.47119666 |
| FOSL2 | 182.407858 | -0.782206806 | 0.58515201 | -1.3367583 | 0.18130155 | 0.47994859 |
| PPP1CB | 125.153728 | -0.467531955 | 0.44789998 | -1.0438312 | 0.29656351 | 0.61093809 |
| TRMT61B | 41.9863963 | 0.330610747 | 0.29936073 | 1.10438917 | 0.26942436 | 0.58564558 |
| WDR43 | 68.6556414 | -0.34065029 | 0.39307967 | -0.866619 | 0.38615082 | 0.68921713 |
| CLIP4 | 117.174801 | -1.149876889 | 0.3442439 | -3.3402971 | 0.00083689 | 0.01300524 |
| YPEL5 | 62.6411902 | 0.014438793 | 0.29044302 | 0.049713 | 0.9603511 | 0.98975502 |
| LCLAT1 | 103.364446 | 0.515767053 | 0.28585701 | 1.80428336 | 0.0711869 | 0.28474759 |
| CAPN13 | 47.0780169 | -4.136111797 | 1.51056891 | -2.7381153 | 0.00617924 | 0.05784663 |
| GALNT14 | 224.822612 | -0.299004682 | 0.414648 | -0.7211049 | 0.47084501 | 0.75744631 |
| DPY30 | 35.8036893 | 0.086413633 | 0.30456962 | 0.28372375 | 0.7766221 | 0.9196152 |
| SPAST | 64.1777552 | 0.180111825 | 0.23176573 | 0.7771288 | 0.43708278 | 0.72995878 |
| SLC30A6 | 74.9702746 | 0.230372065 | 0.33150651 | 0.69492472 | 0.48710248 | 0.76866947 |
| YIPF4 | 94.674674 | -0.280780864 | 0.27998738 | -1.002834 | 0.31594095 | 0.63009595 |
| BIRC6 | 575.593722 | 0.096239819 | 0.22560745 | 0.42658085 | 0.66968464 | 0.86777458 |
| TTC27 | 67.0312761 | 0.048912969 | 0.30363397 | 0.16109189 | 0.87202103 | 0.9568935 |
| LINC00486 | 106.361877 | 0.607089951 | 0.29718709 | 2.04278708 | 0.04107352 | 0.20567643 |
| RASGRP3 | 42.0911511 | 0.799229107 | 0.60831425 | 1.3138425 | 0.18889921 | 0.48924896 |

| | | | | | | |
|---|---|---|---|---|---|---|
| FAM98A | 15.3612184 | 0.669763559 | 0.54254369 | 1.23448778 | 0.2170212 | 0.52654323 |
| AC068599.1 | 23.56259 | 2.239418183 | 0.63424814 | 3.5308234 | 0.00041427 | 0.00751486 |
| LINC01320 | 360.856037 | -2.616479798 | 0.65134282 | -4.0170548 | 5.89E-05 | 0.00162563 |
| AC012593.1 | 106.950232 | -3.721826587 | 1.01312167 | -3.6736225 | 0.00023914 | 0.0048897 |
| CRIM1 | 227.958937 | -0.335835741 | 0.32752885 | -1.0253623 | 0.30519226 | 0.61874596 |
| FEZ2 | 108.700522 | 0.175712399 | 0.25768417 | 0.68189054 | 0.49530817 | 0.77475164 |
| STRN | 157.471121 | 0.125459262 | 0.26721802 | 0.4695015 | 0.63871121 | 0.8538689 |
| HEATR5B | 91.8151745 | 0.150323995 | 0.25820453 | 0.5821896 | 0.56043898 | 0.81331253 |
| EIF2AK2 | 88.9555167 | 0.722543381 | 0.55769017 | 1.29559998 | 0.19511333 | 0.49766572 |
| CEBPZOS | 54.2626253 | -0.19327099 | 0.29461182 | -0.6560191 | 0.51181181 | 0.78271634 |
| CEBPZ | 56.8692144 | -0.189083748 | 0.37525204 | -0.5038847 | 0.61434243 | 0.84174925 |
| NDUFAF7 | 43.1692403 | 0.28984749 | 0.26798033 | 1.08159987 | 0.27943036 | 0.59383074 |
| PRKD3 | 52.2089981 | -0.641683308 | 0.27981713 | -2.2932238 | 0.02183512 | 0.13851354 |
| RMDN2 | 33.2775593 | 0.505841024 | 0.44025979 | 1.14896032 | 0.25057234 | 0.56339619 |
| ATL2 | 186.530603 | 0.732500425 | 0.32844618 | 2.23019922 | 0.02573422 | 0.15406587 |
| AC011247.1 | 16.457146 | -1.22601775 | 0.84405147 | -1.4525391 | 0.14635177 | 0.42885986 |
| HNRNPLL | 62.235253 | -0.321497997 | 0.27246509 | -1.1799603 | 0.23801599 | 0.54997949 |
| GALM | 31.5465575 | -0.453541013 | 0.4596416 | -0.9867275 | 0.32377624 | 0.63622543 |
| SRSF7 | 94.2416827 | 0.107961391 | 0.37714277 | 0.28626133 | 0.77467797 | 0.91864687 |
| DHX57 | 58.3282552 | -0.107420505 | 0.23105751 | -0.4649081 | 0.64199728 | 0.85468927 |
| SOS1 | 339.939219 | 0.351973353 | 0.22354863 | 1.57448226 | 0.115376 | 0.37430961 |
| MAP4K3 | 568.630913 | 0.709812013 | 0.2355361 | 3.01360181 | 0.00258166 | 0.03074257 |
| AC007388.1 | 306.40082 | 0.420004652 | 0.2008751 | 2.09087464 | 0.0365393 | 0.19150785 |
| TMEM178A | 47.9990164 | 2.296189424 | 0.55589756 | 4.13059813 | 3.62E-05 | 0.00109533 |
| THUMPD2 | 77.7635883 | -0.039612489 | 0.19847925 | -0.19958 | 0.84180907 | 0.94776435 |
| SLC8A1-AS1 | 79.8350602 | 2.159056571 | 0.59765262 | 3.61256105 | 0.00030319 | 0.00587718 |
| SLC8A1 | 2520.77743 | 2.537862618 | 0.60530444 | 4.19270448 | 2.76E-05 | 0.00089814 |
| EML4 | 217.552826 | -0.522694158 | 0.33477858 | -1.561313 | 0.11844991 | 0.37913731 |
| COX7A2L | 64.0535784 | -0.375076265 | 0.52191194 | -0.7186581 | 0.47235158 | 0.75855881 |
| MTA3 | 137.530117 | 0.383186577 | 0.30831812 | 1.24282861 | 0.213931 | 0.52235998 |
| ZFP36L2 | 178.274468 | -1.785764074 | 0.40122511 | -4.4507784 | 8.56E-06 | 0.00033803 |
| THADA | 237.230293 | -0.026812724 | 0.22655309 | -0.1183507 | 0.90578976 | 0.96930836 |
| PLEKHH2 | 132.561462 | -0.069202016 | 0.43102493 | -0.1605522 | 0.87244608 | 0.95690422 |
| DYNC2LI1 | 95.5430541 | 0.508621141 | 0.53621886 | 0.94853274 | 0.34285832 | 0.65243773 |
| LRPPRC | 184.37996 | 0.372667981 | 0.27336259 | 1.36327355 | 0.17279632 | 0.468183 |
| PPM1B | 220.101046 | 0.64191336 | 0.18151155 | 3.53648761 | 0.00040549 | 0.00742779 |
| SLC3A1 | 190.41764 | 0.790954156 | 0.44866752 | 1.76289597 | 0.07791804 | 0.29864393 |
| PREPL | 208.959967 | 0.621281381 | 0.21582039 | 2.87869634 | 0.00399323 | 0.04274723 |
| CAMKMT | 619.383899 | 0.101047294 | 0.24522406 | 0.4120611 | 0.68029464 | 0.87237893 |
| SRBD1 | 114.678085 | 0.574970629 | 0.31399591 | 1.83114052 | 0.06707957 | 0.27565003 |
| PRKCE | 545.191565 | -0.496552658 | 0.34731848 | -1.4296753 | 0.15281023 | 0.43799648 |
| EPAS1 | 129.106628 | -2.826743498 | 0.55474237 | -5.0955969 | 3.48E-07 | 2.37E-05 |
| RHOQ | 67.5495474 | -0.921361464 | 0.58991424 | -1.5618566 | 0.11832177 | 0.37909258 |
| PIGF | 32.0325373 | 0.226943305 | 0.35790999 | 0.63407926 | 0.52602909 | 0.79338325 |
| CRIPT | 14.3190632 | 0.100419561 | 0.47246508 | 0.21254388 | 0.83168275 | 0.94487754 |
| SOCS5 | 75.0572803 | 0.338465421 | 0.32752926 | 1.03338989 | 0.30142147 | 0.61540442 |
| MCFD2 | 45.3831474 | -0.346018775 | 0.33115447 | -1.0448863 | 0.29607551 | 0.61093809 |
| TTC7A | 87.9033775 | 0.461228821 | 0.30890783 | 1.49309527 | 0.13541225 | 0.41064171 |
| AC106869.1 | 13.4202433 | 0.071106182 | 0.67588375 | 0.10520475 | 0.91621334 | 0.97396301 |
| EPCAM | 108.909279 | -0.723966533 | 0.31283144 | -2.3142384 | 0.02065465 | 0.13383178 |
| MSH2 | 98.4724786 | 0.576312588 | 0.46992678 | 1.22638804 | 0.22005268 | 0.52989521 |
| MSH6 | 27.5043812 | -0.255439029 | 0.35573608 | -0.7180577 | 0.47272171 | 0.75889416 |
| FBXO11 | 279.622546 | -0.468557435 | 0.33572588 | -1.3956548 | 0.16281848 | 0.45357618 |
| FOXN2 | 41.0020926 | -0.96990731 | 0.68545629 | -1.4149805 | 0.15707421 | 0.44367158 |
| PPP1R21 | 113.703241 | 0.117707846 | 0.22604362 | 0.52073068 | 0.6025544 | 0.83691476 |
| GPR75-ASB3 | 53.1672352 | -0.045300481 | 0.42629179 | -0.1062664 | 0.91537101 | 0.97341689 |
| ERLEC1 | 31.558542 | -0.163901982 | 0.28201886 | -0.5811738 | 0.5611233 | 0.81384822 |
| PSME4 | 193.933214 | 0.243355307 | 0.3119268 | 0.780168 | 0.435292 | 0.72827249 |
| ACYP2 | 266.84996 | -0.316636214 | 0.42470416 | -0.7455454 | 0.4559421 | 0.74621803 |
| SPTBN1 | 686.964855 | -1.717747583 | 0.46823252 | -3.6685782 | 0.0002439 | 0.0049654 |
| EML6 | 52.34588 | 1.071001678 | 0.43629865 | 2.45474443 | 0.01409848 | 0.10421524 |
| RTN4 | 357.418533 | -0.544084469 | 0.31579198 | -1.7229205 | 0.08490291 | 0.31408459 |
| CLHC1 | 46.1893544 | 0.37005323 | 0.34124349 | 1.08442577 | 0.27817606 | 0.59352714 |
| RPS27A | 109.319878 | -0.501857399 | 0.52587802 | -0.9543228 | 0.33992026 | 0.65027011 |
| MTIF2 | 37.101935 | 1.027714116 | 0.30714144 | 3.34606139 | 0.00081968 | 0.01282335 |
| AC012358.3 | 27.6132348 | 0.149788975 | 0.69351079 | 0.21598651 | 0.82899826 | 0.94320681 |
| CCDC88A | 49.6087815 | 0.199359782 | 0.29317956 | 0.67999208 | 0.49650948 | 0.77475164 |
| CFAP36 | 67.2100094 | -0.629817421 | 0.34133073 | -1.8451823 | 0.06501103 | 0.27000571 |
| PPP4R3B | 150.293053 | -0.02598269 | 0.17627471 | -0.1473989 | 0.88281721 | 0.96161232 |
| PNPT1 | 66.9894428 | 0.671625698 | 0.48567843 | 1.38286087 | 0.16670753 | 0.4589256 |
| EFEMP1 | 109.78772 | -1.79092756 | 0.56728698 | -3.1570045 | 0.00159399 | 0.02150847 |
| VRK2 | 119.292227 | 0.12588656 | 0.27737537 | 0.45384909 | 0.64993745 | 0.86079784 |
| FANCL | 83.7342943 | 1.083547986 | 0.44284533 | 2.44678653 | 0.01441362 | 0.10557154 |
| PAPOLG | 40.0146214 | 0.017657422 | 0.26932205 | 0.06556248 | 0.94772616 | 0.98633762 |
| REL | 51.2646584 | -0.47673961 | 0.5811477 | -0.8203416 | 0.41202142 | 0.71102864 |
| PUS10 | 29.4104374 | -0.097664867 | 0.302423 | -0.3229413 | 0.74673973 | 0.90661036 |
| PEX13 | 45.2105617 | -0.335755724 | 0.3099732 | -1.0831766 | 0.27873004 | 0.59369058 |
| KIAA1841 | 20.5048375 | -0.995427393 | 0.4985727 | -1.9965542 | 0.04587364 | 0.21973316 |
| C2orf74 | 13.7813864 | 0.685905611 | 0.47245406 | 1.45179325 | 0.1465591 | 0.42918249 |
| USP34 | 658.099202 | -0.071169035 | 0.24796882 | -0.287008 | 0.77410619 | 0.91862688 |
| AC016727.1 | 14.4216415 | 0.517568107 | 0.48138587 | 1.07516265 | 0.28230191 | 0.59642935 |
| XPO1 | 257.297356 | 0.618666654 | 0.22713868 | 2.72373976 | 0.00645473 | 0.05976559 |
| FAM161A | 19.5590676 | -0.508711335 | 0.5296912 | -0.9603923 | 0.33685783 | 0.64773405 |
| CCT4 | 50.2453929 | 0.050547433 | 0.34263237 | 0.14752673 | 0.88271628 | 0.96161232 |
| COMMD1 | 260.194108 | 0.091974863 | 0.22666518 | 0.40577412 | 0.68490858 | 0.87408051 |
| AC018462.1 | 17.755405 | -0.574732773 | 0.56093537 | -1.0245971 | 0.30555332 | 0.61893963 |
| B3GNT2 | 60.4629844 | -0.030439083 | 0.40947954 | -0.074336 | 0.94074301 | 0.98412294 |
| EHBP1 | 305.394179 | 0.557287388 | 0.2586381 | 2.15469951 | 0.03118536 | 0.17474295 |
| WDPCP | 195.619361 | 0.159439137 | 0.40005746 | 0.3985406 | 0.69023174 | 0.87692066 |
| MDH1 | 67.5448194 | 0.753924518 | 0.32452118 | 2.32319046 | 0.02016892 | 0.13178349 |
| UGP2 | 109.193564 | -0.625459052 | 0.24141389 | -2.5908163 | 0.00957486 | 0.0795686 |
| VPS54 | 155.683092 | 0.351006128 | 0.2322205 | 1.51152084 | 0.1306558 | 0.40139529 |
| PELI1 | 54.7895794 | -0.201604009 | 0.47272589 | -0.4264713 | 0.66976448 | 0.86777458 |
| AC008074.1 | 29.7410336 | -0.553463897 | 0.65451819 | -0.8456051 | 0.3977731 | 0.69815955 |
| AFTPH | 158.162259 | -0.168595101 | 0.25892543 | -0.6511338 | 0.51496012 | 0.78381327 |
| SERTAD2 | 33.4512709 | -0.722615498 | 0.38382549 | -1.8826668 | 0.05974554 | 0.2554971 |
| CEP68 | 23.5744481 | -0.114955555 | 0.32020218 | -0.3590093 | 0.71958815 | 0.89114993 |
| RAB1A | 222.729481 | -0.210742928 | 0.42671073 | -0.4938777 | 0.62139253 | 0.84470972 |
| ACTR2 | 122.085465 | -0.601177638 | 0.25987273 | -2.3133541 | 0.02070318 | 0.13396005 |
| SPRED2 | 189.915697 | -0.258173719 | 0.23627701 | -1.0926739 | 0.27453701 | 0.5891499 |
| LINC01828 | 85.6055555 | 7.204146087 | 1.17795453 | 6.11580998 | 9.61E-10 | 1.64E-07 |
| ETAA1 | 27.4542193 | 0.016285444 | 0.34370706 | 0.04738176 | 0.96220897 | 0.99029318 |
| C1D | 34.7319982 | -0.190298654 | 0.33973104 | -0.560145 | 0.57538051 | 0.82147339 |
| APLF | 59.5421833 | 0.233433953 | 0.32441232 | 0.71955946 | 0.47179628 | 0.75793048 |
| ANTXR1 | 23.455802 | -0.220777438 | 0.36229171 | -0.6093914 | 0.54226506 | 0.80340275 |
| GFPT1 | 134.559694 | -1.733560574 | 0.27890648 | -6.2155622 | 5.11E-10 | 9.93E-08 |
| NFU1 | 48.2522135 | 0.293780178 | 0.30639484 | 0.95882874 | 0.33764503 | 0.64844536 |
| AAK1 | 335.206277 | -0.022182047 | 0.19084661 | -0.1162297 | 0.90747049 | 0.9700616 |
| ANXA4 | 523.946852 | -1.056378759 | 0.62117014 | -1.7006271 | 0.08901304 | 0.32396747 |
| GMCL1 | 97.3815696 | -0.443401517 | 0.53733242 | -0.8251903 | 0.40926354 | 0.70777268 |
| MXD1 | 48.807262 | -0.008645606 | 0.37433708 | -0.0230958 | 0.98157387 | 0.99656336 |
| PCBP1 | 37.8389715 | -0.954576403 | 0.45146048 | -2.1144185 | 0.03447953 | 0.18465662 |
| C2orf42 | 20.7847329 | -0.201658806 | 0.47101146 | -0.4281399 | 0.66854926 | 0.86777458 |

| | | | | | | |
|---|---|---|---|---|---|---|
| PCYOX1 | 38.5384037 | -0.840328005 | 0.76753621 | -1.0948383 | 0.2735875 | 0.58894058 |
| SNRPG | 25.7702644 | -0.388631405 | 0.33364781 | -1.1647953 | 0.24410186 | 0.55607274 |
| FAM136A | 18.8952339 | 0.154654376 | 0.40799922 | 0.37905557 | 0.7046466 | 0.88570031 |
| TGFA | 60.3092789 | -1.028071614 | 0.52772766 | -1.9481101 | 0.05140179 | 0.23455111 |
| AC007040.2 | 46.020087 | 0.079424005 | 0.97048365 | 0.08183961 | 0.93477426 | 0.98085023 |
| NAGK | 18.2043713 | 0.017224383 | 0.46004047 | 0.03744102 | 0.97013337 | 0.99156233 |
| MCEE | 41.1656065 | 0.904943433 | 0.38875765 | 2.32778295 | 0.01992363 | 0.13082568 |
| MPHOSPH10 | 24.911671 | -0.36860899 | 0.39404625 | -0.935446 | 0.3495585 | 0.6567164 |
| PAIP2B | 27.4225263 | -0.418686426 | 0.59227724 | -0.7069095 | 0.4796227 | 0.76457549 |
| ZNF638 | 554.807594 | 0.023148687 | 0.17054124 | 0.13573659 | 0.89202952 | 0.96387568 |
| EXOC6B | 612.19228 | -0.448870062 | 0.4963599 | -0.9043238 | 0.36582374 | 0.67012584 |
| EMX1 | 175.722901 | 1.492476394 | 0.4962588 | 3.00745579 | 0.00263444 | 0.03113252 |
| SFXN5 | 198.339587 | 0.061305498 | 0.27400601 | 0.22373778 | 0.82296134 | 0.9407002 |
| RAB11FIP5 | 49.3147979 | -0.077916586 | 0.40238244 | -0.1936381 | 0.84645923 | 0.94917449 |
| CCT7 | 27.9087949 | -0.287168984 | 0.42187327 | -0.6806996 | 0.49606156 | 0.77475164 |
| ALMS1 | 101.624111 | 0.314343049 | 0.19896054 | 1.57992661 | 0.11412367 | 0.372426 |
| DUSP11 | 20.6844694 | 0.356775299 | 0.3982237 | 0.89591679 | 0.37029721 | 0.67442486 |
| STAMBP | 58.5065002 | -0.173015622 | 0.26460673 | -0.6538595 | 0.51320234 | 0.7830488 |
| DGUOK | 50.9009049 | -0.608188024 | 0.30804791 | -1.9743293 | 0.04834431 | 0.22526853 |
| TET3 | 75.2178982 | -1.190281627 | 0.61490964 | -1.9357017 | 0.05290425 | 0.23818408 |
| MOB1A | 53.0680137 | -0.289712001 | 0.34067748 | -0.8503996 | 0.39510295 | 0.69642725 |
| AC006030.1 | 22.7211738 | -0.48230506 | 0.44093929 | -1.0938128 | 0.27403707 | 0.58894058 |
| DCTN1 | 13.58214 | 0.49200467 | 0.42291288 | 1.16337122 | 0.24467892 | 0.55684312 |
| RTKN | 34.1812674 | -0.702549464 | 0.43543502 | -1.6134427 | 0.10664836 | 0.35932624 |
| MOGS | 24.0635326 | -0.103589456 | 0.43969299 | -0.235595 | 0.81374698 | 0.93633777 |
| CCDC142 | 10.3674596 | -0.104497217 | 0.63392556 | -0.1648415 | 0.86906877 | 0.95612718 |
| TTC31 | 12.4390195 | -0.039483412 | 0.41469221 | -0.0952114 | 0.92414694 | 0.97706612 |
| AUP1 | 18.7955676 | -0.096413741 | 0.40863195 | -0.2359427 | 0.81347711 | 0.93633777 |
| POLE4 | 17.5916605 | -0.430810919 | 0.45738576 | -0.9418984 | 0.34624466 | 0.65471206 |
| TACR1 | 17.6064129 | -1.165626838 | 0.94200298 | -1.2373919 | 0.21594163 | 0.52525486 |
| AC007681.1 | 26.4530146 | -0.282361097 | 0.71531395 | -0.3947373 | 0.69303677 | 0.87856898 |
| EVA1A | 65.1944139 | 0.312351292 | 0.42091935 | 0.74206921 | 0.45804541 | 0.747377 |
| MRPL19 | 67.8437012 | 0.301609134 | 0.43823778 | 0.6882317 | 0.49130689 | 0.77224299 |
| GCFC2 | 122.663824 | 0.478236126 | 0.28198022 | 1.69599175 | 0.08988744 | 0.32574025 |
| SUCLG1 | 98.3687327 | 0.941288048 | 0.3792755 | 2.48180557 | 0.01307186 | 0.09891352 |
| DNAH6 | 33.5792308 | 0.638065778 | 0.97546531 | 0.65411427 | 0.51303819 | 0.7830488 |
| TMSB10 | 124.912461 | -1.449608875 | 0.45574401 | -3.1807525 | 0.00146893 | 0.02032094 |
| KCMF1 | 121.608931 | -0.031084712 | 0.28484146 | -0.1091299 | 0.91309948 | 0.97166624 |
| TGOLN2 | 89.9522237 | -0.536929887 | 0.3927657 | -1.3670488 | 0.17161001 | 0.4673165 |
| RETSAT | 13.0474722 | -0.51953208 | 0.42265573 | -1.2292087 | 0.21899358 | 0.52913753 |
| ELMOD3 | 45.9645843 | 0.126312658 | 0.26708089 | 0.47293783 | 0.63625754 | 0.85375696 |
| CAPG | 17.5346603 | -1.583399934 | 0.60725193 | -2.6074844 | 0.00912102 | 0.07710283 |
| MAT2A | 116.549758 | -0.923362917 | 0.70632737 | -1.3072733 | 0.1911199 | 0.49188042 |
| GGCX | 25.547051 | 0.260847464 | 0.46056628 | 0.56636249 | 0.57114739 | 0.81895612 |
| VAMP8 | 44.1259599 | -0.566635678 | 0.34172221 | -1.6581763 | 0.09728188 | 0.34011144 |
| USP39 | 55.0859367 | 0.049246495 | 0.37049534 | 0.13292069 | 0.8942561 | 0.96482804 |
| C2orf68 | 27.7967533 | -0.002375195 | 0.32456482 | -0.0073181 | 0.99416106 | 0.9992177 |
| POLR1A | 37.8406144 | 0.138642945 | 0.27487813 | 0.50437969 | 0.61399459 | 0.84174925 |
| PTCD3 | 90.8512629 | 0.023899843 | 0.21574633 | 0.11077752 | 0.91179277 | 0.97128978 |
| IMMT | 110.774468 | 0.405654374 | 0.25153891 | 1.61269034 | 0.1068118 | 0.35932624 |
| MRPL35 | 15.0860518 | 1.05134592 | 0.41434108 | 2.53739241 | 0.01116817 | 0.08869847 |
| KDM3A | 92.3581886 | -0.483784155 | 0.33996573 | -1.423038 | 0.15472513 | 0.44032584 |
| CHMP3 | 15.6203002 | -0.942300871 | 0.48178572 | -1.9558506 | 0.05048276 | 0.23198682 |
| RMND5A | 54.1840969 | 0.060736049 | 0.36411122 | 0.16680631 | 0.86752246 | 0.95588054 |
| PLGLB1 | 19.5376351 | 0.398241436 | 0.46773027 | 0.85143395 | 0.39452834 | 0.69642725 |
| CYTOR | 60.2756752 | -2.713135729 | 0.77806862 | -3.4870134 | 0.00048845 | 0.00848097 |
| KRCC1 | 62.9011044 | -0.000290585 | 0.36415352 | -0.000798 | 0.99936331 | 0.99969609 |
| THNSL2 | 12.641161 | 1.045267254 | 0.79666765 | 1.31204933 | 0.18950349 | 0.48972576 |
| EIF2AK3 | 97.0291306 | -0.488664769 | 0.49170501 | -0.9938169 | 0.320312 | 0.63212442 |
| RPIA | 33.5683722 | 0.643958243 | 0.36213331 | 1.77823532 | 0.07536521 | 0.29365034 |
| MAL | 211.477447 | 0.212102148 | 0.7087142 | 0.29927741 | 0.76472839 | 0.91531804 |
| MRPS5 | 93.8587938 | 0.346376104 | 0.27305891 | 1.26850322 | 0.2046183 | 0.51080617 |
| AC092835.1 | 16.0770937 | 0.744574075 | 0.39439288 | 1.88789936 | 0.05903946 | 0.25401736 |
| PROM2 | 54.1186737 | 1.385255602 | 0.59524528 | 2.32720133 | 0.01995455 | 0.13082568 |
| KCNIP3 | 23.2055788 | 0.921006904 | 0.55170995 | 1.66936795 | 0.09504448 | 0.33657225 |
| FAHD2A | 13.7286988 | 0.394896481 | 0.51191393 | 0.77141187 | 0.44046284 | 0.73297797 |
| ANKRD36C | 57.8500629 | 0.932552071 | 0.49879885 | 1.86959547 | 0.06154001 | 0.2599337 |
| STARD7 | 54.0990179 | 0.782622761 | 0.31053073 | 2.52027478 | 0.01172633 | 0.09234481 |
| TMEM127 | 30.7684809 | 0.049632626 | 0.38277884 | 0.12966398 | 0.89683228 | 0.96570784 |
| CIAO1 | 38.868185 | -0.77281051 | 0.30955376 | -2.4965309 | 0.01254148 | 0.09648244 |
| SNRNP200 | 63.2205412 | -0.859959547 | 0.43082376 | -1.996082 | 0.045925 | 0.2198176 |
| KANSL3 | 111.609732 | 0.456064572 | 0.30096326 | 1.51534966 | 0.12968388 | 0.39972644 |
| FER1L5 | 12.9392506 | 0.653477677 | 0.43364069 | 1.50695654 | 0.1318218 | 0.40351308 |
| LMAN2L | 29.2826033 | 0.200201722 | 0.31616767 | 0.63321378 | 0.52659405 | 0.79346524 |
| CNNM4 | 24.0666954 | 0.0681972 | 0.32539573 | 0.20958235 | 0.83399366 | 0.94640649 |
| ANKRD23 | 16.8194493 | -0.031328247 | 0.49190629 | -0.0636874 | 0.94921912 | 0.98641632 |
| FAHD2B | 11.105899 | 1.249393552 | 0.48151793 | 2.5946979 | 0.00946741 | 0.07895718 |
| ANKRD36 | 77.9785738 | 1.099350757 | 0.40123423 | 2.73992263 | 0.00614536 | 0.05758732 |
| AC092683.1 | 112.899741 | -0.074048832 | 0.43199359 | -0.1714119 | 0.86389992 | 0.95472358 |
| ANKRD36B | 33.3363549 | 0.748450324 | 0.34678852 | 2.15823273 | 0.03090974 | 0.173407 |
| COX5B | 75.4501664 | 0.251831561 | 0.34901688 | 0.7215455 | 0.47057396 | 0.75727158 |
| ACTR1B | 29.5238345 | -0.664749434 | 0.42079684 | -1.5797396 | 0.11416651 | 0.372426 |
| C2orf92 | 46.6655527 | -2.918553263 | 1.01778476 | -2.8675545 | 0.00413658 | 0.0438289 |
| TMEM131 | 242.633801 | 0.284509194 | 0.25785419 | 1.10337237 | 0.26986549 | 0.58625952 |
| INPP4A | 60.154591 | 0.341682617 | 0.26862995 | 1.27194536 | 0.20339253 | 0.50815434 |
| COA5 | 18.6825609 | 0.069041418 | 0.39834286 | 0.17332159 | 0.86239865 | 0.95458251 |
| UNC50 | 25.3325275 | 0.408759032 | 0.42765025 | 0.95582553 | 0.33916039 | 0.64971064 |
| MGAT4A | 65.4915426 | -0.197695737 | 0.51896435 | -0.3809428 | 0.70324568 | 0.88513266 |
| KIAA1211L | 19.6222064 | 0.176685143 | 0.72102619 | 0.24504678 | 0.80642021 | 0.93325451 |
| TSGA10 | 139.899515 | -0.020306229 | 0.37767828 | -0.0537659 | 0.95712164 | 0.98859719 |
| AC079447.1 | 60.3701263 | 0.258035496 | 0.35284401 | 0.73130191 | 0.46459476 | 0.75219336 |
| MITD1 | 38.3972627 | -0.395416367 | 0.27905232 | -1.4169972 | 0.15648373 | 0.44294303 |
| TXNDC9 | 26.647117 | -0.494789493 | 0.43331647 | -1.1418663 | 0.25350958 | 0.56559484 |
| EIF5B | 94.8596536 | -0.784715523 | 0.33903373 | -2.3145648 | 0.02063676 | 0.13383178 |
| REV1 | 238.53983 | -0.022444605 | 0.18259697 | -0.1229188 | 0.90217138 | 0.96803829 |
| LONRF2 | 30.8354838 | -0.099878978 | 0.60623604 | -0.1647526 | 0.8691387 | 0.95612718 |
| NPAS2 | 308.798996 | -2.102643408 | 0.65592219 | -3.2056293 | 0.00134767 | 0.01903897 |
| RPL31 | 58.1485589 | -0.511599224 | 0.36447433 | -1.4036632 | 0.16041915 | 0.44957865 |
| TBC1D8 | 223.512074 | -1.035207131 | 0.32582816 | -3.1771567 | 0.00148727 | 0.02045321 |
| CNOT11 | 73.2545972 | -0.892209901 | 0.55833703 | -1.5979773 | 0.11004803 | 0.36528581 |
| RNF149 | 255.957172 | -0.568995213 | 0.34012028 | -1.6729235 | 0.09434236 | 0.33484499 |
| MAP4K4 | 133.854426 | -0.8610066 | 0.47886877 | -1.7980012 | 0.07217682 | 0.28698366 |
| IL1R2 | 19.3147076 | -1.43473577 | 1.09941793 | -1.3049958 | 0.19189428 | 0.49284602 |
| IL1R1 | 88.9919611 | -1.044529085 | 0.60481721 | -1.7270161 | 0.08416477 | 0.31289963 |
| IL1RL2 | 48.0183767 | -0.609751293 | 0.8645203 | -0.7053059 | 0.48061987 | 0.76551071 |
| SLC9A2 | 27.3325089 | 0.858521824 | 0.58881871 | 1.45804101 | 0.14482924 | 0.42666282 |
| MFSD9 | 13.7507671 | -0.096695858 | 0.36927444 | -0.2618536 | 0.79343428 | 0.92649882 |
| TMEM182 | 31.0801568 | -0.329926409 | 0.39488512 | -0.8354997 | 0.4034364 | 0.70245397 |
| POU3F3 | 82.8774869 | 0.045383412 | 0.32354983 | 0.14026715 | 0.88844892 | 0.96307703 |
| MRPS9 | 80.1909614 | 0.449351875 | 0.38162634 | 1.17746556 | 0.23900973 | 0.55079752 |
| TGFBRAP1 | 25.4796018 | 0.109838347 | 0.34213668 | 0.32103645 | 0.74818277 | 0.90723111 |
| C2orf49 | 18.2606268 | -0.421832623 | 0.42696756 | -0.9879735 | 0.32316565 | 0.63569546 |
| NCK2 | 93.4110242 | -0.465750934 | 0.27390619 | -1.7004031 | 0.08905512 | 0.32396747 |

| | | | | | | |
|---|---|---|---|---|---|---|
| UXS1 | 127.830278 | 1.203292012 | 0.27774605 | 4.33234608 | 1.48E-05 | 0.00054803 |
| LINC01594 | 36.810031 | 0.667045291 | 0.50991267 | 1.30815593 | 0.19082042 | 0.49135864 |
| SULT1C2 | 348.042013 | 0.286404165 | 0.45074268 | 0.63540503 | 0.52516429 | 0.79274974 |
| GCC2 | 257.865562 | -0.829671014 | 0.23752453 | -3.4929908 | 0.00047764 | 0.00835562 |
| GCC2-AS1 | 25.2641834 | -1.09234083 | 0.74946797 | -1.4574883 | 0.14498163 | 0.42697684 |
| LIMS1 | 364.594987 | -1.044853058 | 0.28701123 | -3.6404606 | 0.00027215 | 0.00537613 |
| RANBP2 | 139.170435 | 0.23685725 | 0.25061244 | 0.94511371 | 0.34460083 | 0.65359197 |
| CCDC138 | 37.3067358 | -0.153728725 | 0.44984149 | -0.3417398 | 0.73254675 | 0.89812832 |
| SH3RF3 | 30.7393783 | -3.131148202 | 0.94677869 | -3.3071596 | 0.00094247 | 0.01424248 |
| 10-Sep | 242.248636 | -3.492221241 | 0.6698866 | -5.2131528 | 1.86E-07 | 1.40E-05 |
| NPHP1 | 44.2925663 | -0.469194077 | 0.67199661 | -0.698209 | 0.48504651 | 0.76770898 |
| SMIM37 | 21.4607343 | -0.65744929 | 0.65305827 | -1.0067238 | 0.31406753 | 0.62887712 |
| ACOXL | 25.1984755 | -0.609645406 | 0.68819779 | -0.8858578 | 0.37569414 | 0.67913381 |
| MIR4435-2H | 338.066897 | -2.488695385 | 0.42556663 | -5.8479571 | 4.98E-09 | 6.03E-07 |
| BCL2L11 | 54.0165895 | 0.147555697 | 0.42560858 | 0.34669343 | 0.72882165 | 0.8966266 |
| ANAPC1 | 87.2150914 | 0.402980995 | 0.25860417 | 1.55829271 | 0.11916388 | 0.38057334 |
| MERTK | 72.7003925 | -0.029105618 | 0.28934188 | -0.1005925 | 0.91987396 | 0.97520237 |
| TMEM87B | 76.0525188 | -0.472486599 | 0.22518296 | -2.0982342 | 0.03588446 | 0.18935297 |
| ZC3H8 | 63.6323561 | 1.043255991 | 0.27547481 | 3.78711942 | 0.0001524 | 0.00339955 |
| ZC3H6 | 75.5189545 | 0.413619093 | 0.58033289 | 0.7127273 | 0.47601452 | 0.76162498 |
| TTL | 36.1897253 | 0.618730066 | 0.34914889 | 1.77210948 | 0.07637639 | 0.29596141 |
| POLR1B | 17.0587562 | 0.532571861 | 0.42897965 | 1.24148513 | 0.21442659 | 0.52290228 |
| SLC20A1 | 191.503379 | -0.356919931 | 0.59520183 | -0.599662 | 0.5487315 | 0.80661714 |
| PSD4 | 36.74901 | 0.04726162 | 0.34607545 | 0.1365645 | 0.89137504 | 0.96328012 |
| PAX8 | 801.145814 | -1.408544557 | 0.4369412 | -3.2236478 | 0.00126569 | 0.01801723 |
| CBWD2 | 39.7761452 | 0.151758844 | 0.30934158 | 0.49058663 | 0.62371884 | 0.84597807 |
| RABL2A | 27.5185619 | 0.129320073 | 0.35766067 | 0.36157197 | 0.71767191 | 0.8903104 |
| AC017074.1 | 27.3321485 | -0.944343593 | 0.50747663 | -1.8608612 | 0.06276377 | 0.26390468 |
| SLC35F5 | 418.221912 | -0.568004477 | 0.31427608 | -1.8073424 | 0.07070891 | 0.28380968 |
| ACTR3 | 194.271036 | -0.47685261 | 0.24975687 | -1.9092672 | 0.05622763 | 0.24714736 |
| DDX18 | 40.0130037 | -0.854625694 | 0.34463401 | -2.4798066 | 0.01314537 | 0.09932528 |
| CCDC93 | 128.418606 | 0.007810289 | 0.388356 | 0.02011116 | 0.9839547 | 0.99742652 |
| INSIG2 | 65.5003079 | -0.286475648 | 0.54945382 | -0.5213826 | 0.60210029 | 0.83691476 |
| C2orf76 | 22.9428425 | 0.491582698 | 0.35733329 | 1.37569803 | 0.16891514 | 0.46258659 |
| DBI | 67.4426915 | -1.830575025 | 0.54618814 | -3.3515467 | 0.00080361 | 0.01261432 |
| TMEM37 | 63.8394924 | -0.24955191 | 0.66089895 | -0.3775947 | 0.70573174 | 0.88599027 |
| SCTR | 41.1427231 | -0.674768289 | 0.9606428 | -0.7024133 | 0.48242145 | 0.76628579 |
| AC013275.1 | 17.3078698 | 2.286402177 | 0.73173958 | 3.12461187 | 0.0017804 | 0.02317532 |
| CFAP221 | 192.595356 | 0.155282378 | 0.59815902 | 0.2596005 | 0.79517195 | 0.9268423 |
| PTPN4 | 437.419848 | 0.540215125 | 0.41282153 | 1.30859242 | 0.19067245 | 0.49135864 |
| EPB41L5 | 340.830706 | -0.159623253 | 0.33907897 | -0.4707554 | 0.63781542 | 0.8538689 |
| RALB | 29.3505231 | -0.524267793 | 0.30017782 | -1.7465241 | 0.08071993 | 0.30557556 |
| TFCP2L1 | 602.053013 | 0.764707824 | 0.47757435 | 1.60123302 | 0.10932532 | 0.36375272 |
| CLASP1 | 868.346513 | 1.248737588 | 0.3234928 | 3.86017115 | 0.00011331 | 0.00265447 |
| NIFK-AS1 | 31.7866234 | 0.209069443 | 0.46790627 | 0.44681907 | 0.6550057 | 0.8623656 |
| TSN | 26.4934635 | -0.245446573 | 0.33741847 | -0.7274248 | 0.46696575 | 0.75315494 |
| ERCC3 | 39.6814787 | -0.006747811 | 0.36879643 | -0.0182968 | 0.98540204 | 0.99746215 |
| MAP3K2 | 182.232842 | 0.120288132 | 0.24462557 | 0.49172346 | 0.62291485 | 0.84581917 |
| IWS1 | 173.942634 | -0.437852642 | 0.29886598 | -1.4650468 | 0.14290815 | 0.42395023 |
| AC010976.1 | 17.540281 | -1.210051879 | 0.6843495 | -1.7681782 | 0.07703111 | 0.2971468 |
| WDR33 | 213.268339 | 0.042819776 | 0.29030866 | 0.14749741 | 0.88273942 | 0.96161232 |
| POLR2D | 13.5538381 | -0.051556903 | 0.36744425 | -0.1403122 | 0.88841334 | 0.96307703 |
| AMMECR1L | 27.131059 | -0.216264567 | 0.36774972 | -0.5880754 | 0.55648167 | 0.81161194 |
| SAP130 | 68.2959393 | 0.151989028 | 0.42180239 | 0.3603323 | 0.71859864 | 0.8907358 |
| UGGT1 | 62.6940589 | 0.028211188 | 0.32049618 | 0.08802348 | 0.92985802 | 0.97876742 |
| HS6ST1 | 81.6152106 | -2.320047733 | 0.42007524 | -5.5229337 | 3.33E-08 | 2.93E-06 |
| SMPD4 | 42.7543663 | 0.35134224 | 0.32143458 | 1.09304432 | 0.27437434 | 0.5891499 |
| MZT2B | 33.6972984 | -0.217010698 | 0.4548096 | -0.4771463 | 0.63325798 | 0.8520131 |
| PTPN18 | 23.1700443 | -0.217928893 | 0.49581841 | -0.4395337 | 0.66027488 | 0.86490629 |
| FAM168B | 108.478588 | -0.223459923 | 0.38373522 | -0.5823284 | 0.56034549 | 0.81330344 |
| PLEKHB2 | 98.4482557 | -0.580641304 | 0.34261554 | -1.6947313 | 0.0901264 | 0.32647184 |
| MZT2A | 24.6673422 | 0.101342526 | 0.48858482 | 0.20742054 | 0.83568144 | 0.94654164 |
| ANKRD30BL | 14.2256884 | 0.078712709 | 0.48471294 | 0.16239036 | 0.87099846 | 0.95629149 |
| GPR39 | 132.398188 | 0.400755938 | 0.33823587 | 1.18484161 | 0.23608008 | 0.54879348 |
| NCKAP5 | 210.053006 | 1.213974043 | 0.6669378 | 1.82022079 | 0.06872539 | 0.27897062 |
| MGAT5 | 181.349278 | -0.124510812 | 0.23225172 | -0.5361029 | 0.59188746 | 0.83276878 |
| TMEM163 | 39.5769283 | -2.489539214 | 1.06834053 | -2.3302862 | 0.01979103 | 0.13033374 |
| CCNT2-AS1 | 44.2368052 | -1.723082472 | 0.69986236 | -2.4620305 | 0.01381529 | 0.10272232 |
| ACMSD | 28.6552482 | -0.897773011 | 0.90953587 | -0.9870672 | 0.3236097 | 0.63611921 |
| CCNT2 | 137.581072 | 0.024837899 | 0.2811282 | 0.08835079 | 0.92959787 | 0.97876742 |
| RAB3GAP1 | 192.189411 | 0.104242808 | 0.20731511 | 0.50282301 | 0.61508871 | 0.84190945 |
| ZRANB3 | 85.8593507 | -0.141294293 | 0.31477798 | -0.4488697 | 0.65352567 | 0.86199934 |
| R3HDM1 | 197.80239 | 0.289745367 | 0.19811877 | 1.46248316 | 0.14360885 | 0.42454311 |
| UBXN4 | 180.762578 | -0.449336611 | 0.34687793 | -1.2953739 | 0.19519128 | 0.49766572 |
| DARS | 167.095878 | -0.550403732 | 0.21533925 | -2.5559843 | 0.01058879 | 0.08585207 |
| THSD7B | 18.9248008 | -0.848129474 | 0.66392502 | -1.2774477 | 0.20144424 | 0.50571062 |
| HNMT | 82.3525356 | -0.172505389 | 0.53606071 | -0.321802 | 0.74760273 | 0.90704591 |
| AC114763.1 | 27.0336946 | 0.623267212 | 0.55531746 | 1.122362 | 0.26170856 | 0.57564769 |
| SPOPL | 81.9732324 | 0.220578325 | 0.22002864 | 1.00249825 | 0.31610301 | 0.63009595 |
| LRP1B | 738.258554 | -1.210620525 | 0.64341424 | -1.8815569 | 0.0598962 | 0.2557107 |
| GTDC1 | 88.1795164 | -0.140026802 | 0.30034608 | -0.4662182 | 0.64105934 | 0.85468927 |
| TEX41 | 530.448126 | 3.714174228 | 0.48364044 | 7.6796188 | 1.60E-14 | 7.83E-12 |
| ACVR2A | 102.113981 | 0.638768835 | 0.39911184 | 1.60047577 | 0.10949308 | 0.36409182 |
| ORC4 | 150.329214 | 0.82326479 | 0.20268034 | 4.0618878 | 4.87E-05 | 0.00138798 |
| MBD5 | 740.029229 | 0.098424101 | 0.30029947 | 0.32775317 | 0.7430983 | 0.90463949 |
| EPC2 | 179.913317 | 0.094215191 | 0.31767122 | 0.29658082 | 0.76678656 | 0.91566571 |
| LYPD6 | 23.4392392 | 1.785131229 | 0.75355625 | 2.36894225 | 0.01783904 | 0.12140963 |
| MMADHC | 28.6848264 | -0.000960394 | 0.37698753 | -0.0025475 | 0.99796735 | 0.99922221 |
| AC007364.1 | 30.5583775 | -1.498182902 | 0.84340083 | -1.7763593 | 0.07567372 | 0.29457168 |
| RND3 | 34.6366426 | -3.169268701 | 1.08198814 | -2.9291159 | 0.00339928 | 0.03837149 |
| NMI | 16.1534194 | -0.340663768 | 0.6565724 | -0.5188518 | 0.6038641 | 0.83754753 |
| RIF1 | 129.014007 | -0.003035169 | 0.3304894 | -0.0091839 | 0.99267244 | 0.9992177 |
| NEB | 19.2424563 | 0.022741724 | 0.53440342 | 0.04255535 | 0.96605599 | 0.99156233 |
| ARL5A | 49.4578341 | 0.120200657 | 0.31349005 | 0.38342734 | 0.70140293 | 0.88421489 |
| CACNB4 | 544.840118 | 1.638109161 | 0.31596102 | 5.18452922 | 2.17E-07 | 1.58E-05 |
| STAM2 | 142.506585 | 0.732663668 | 0.26202191 | 2.79619242 | 0.00517086 | 0.0513999 |
| FMNL2 | 492.903384 | -2.683672122 | 0.43789452 | -6.1285811 | 8.87E-10 | 1.59E-07 |
| PRPF40A | 111.687851 | -0.04493389 | 0.21560396 | -0.2084094 | 0.83490932 | 0.94654164 |
| ARL6IP6 | 50.4384559 | -0.375005212 | 0.42460888 | -0.883178 | 0.37714012 | 0.68029685 |
| GALNT13 | 31.6910432 | 1.30047851 | 0.78280592 | 1.66130387 | 0.09665243 | 0.33969491 |
| KCNJ3 | 128.098378 | 1.743395963 | 0.45725027 | 3.8127828 | 0.00013741 | 0.00312492 |
| LINC01876 | 126.744187 | -0.047684263 | 1.02519481 | -0.0465124 | 0.96290186 | 0.9905486 |
| GPD2 | 93.4565747 | 0.074487 | 0.27840859 | 0.26754562 | 0.78904909 | 0.92402584 |
| CYTIP | 19.8007943 | 1.890666368 | 0.84993704 | 2.22447813 | 0.02611629 | 0.15539777 |
| ACVR1C | 13.6470038 | 1.285951636 | 0.54747055 | 2.34889647 | 0.01882914 | 0.12585155 |
| ACVR1 | 164.720786 | 0.230310667 | 0.21923866 | 1.05050209 | 0.29348733 | 0.60876592 |
| UPP2 | 52.581794 | 0.004206769 | 0.7621101 | 0.0055199 | 0.99559578 | 0.99922221 |
| CCDC148 | 356.230189 | 1.809295664 | 0.44278214 | 4.08619837 | 4.38E-05 | 0.00126974 |
| PKP4 | 1126.89727 | 1.443226652 | 0.24757474 | 5.82945847 | 5.56E-09 | 6.56E-07 |
| PKP4-AS1 | 27.0548377 | 0.954724879 | 0.68209537 | 1.39969412 | 0.16160493 | 0.45167998 |
| TANC1 | 375.13787 | -0.080780479 | 0.20698671 | -0.3902689 | 0.6963377 | 0.88095695 |
| WDSUB1 | 80.9043536 | 0.48344348 | 0.24404453 | 1.98096425 | 0.04759528 | 0.22360104 |
| BAZ2B | 599.612295 | -0.029411485 | 0.1483675 | -0.198234 | 0.84286197 | 0.94799096 |

| | | | | | | |
|---|---|---|---|---|---|---|
| AC009961.1 | 29.6777454 | -1.255836179 | 0.5562007 | -2.2578831 | 0.02395295 | 0.14714173 |
| 7-Mar | 211.060316 | -0.003091967 | 0.34201438 | -0.0090405 | 0.99278686 | 0.9992177 |
| PLA2R1 | 44.5702926 | -1.602830802 | 0.33866682 | -4.732766 | 2.21E-06 | 0.00010985 |
| ITGB6 | 361.338407 | -3.178159763 | 0.57797873 | -5.4987486 | 3.82E-08 | 3.30E-06 |
| RBMS1 | 505.858948 | -0.965583468 | 0.44449383 | -2.1723214 | 0.02983143 | 0.16898435 |
| TANK | 149.616955 | -0.422140869 | 0.2870703 | -1.4705139 | 0.14142262 | 0.42163989 |
| PSMD14 | 66.7899174 | 0.099779128 | 0.35429387 | 0.28162815 | 0.77822865 | 0.91990415 |
| IFIH1 | 29.4184633 | 0.61657642 | 0.63700624 | 0.96792838 | 0.33308014 | 0.64492217 |
| GCA | 48.4484624 | 0.092321213 | 0.34655742 | 0.26639514 | 0.7899349 | 0.92448262 |
| KCNH7 | 137.974986 | 1.397113925 | 0.81477669 | 1.71472005 | 0.08639657 | 0.31765047 |
| FIGN | 19.5318788 | 0.006693589 | 0.64752593 | 0.01033717 | 0.99175227 | 0.9988777 |
| GRB14 | 488.895987 | 1.421027972 | 0.27377229 | 5.19054713 | 2.10E-07 | 1.54E-05 |
| COBLL1 | 2465.0763 | 0.945115653 | 0.24498304 | 3.85788191 | 0.00011437 | 0.00267274 |
| AC019197.1 | 5568.98734 | 0.388528273 | 0.55312183 | 0.70242802 | 0.48241228 | 0.76628579 |
| SLC38A11 | 56.3053887 | 0.653184847 | 0.73381203 | 0.89012557 | 0.37339847 | 0.67707283 |
| SCN3A | 186.697013 | -0.621088989 | 0.81672672 | -0.7604612 | 0.44697895 | 0.73972874 |
| SCN2A | 903.324381 | 0.525975563 | 0.59319945 | 0.88667574 | 0.3752535 | 0.67888822 |
| CSRNP3 | 73.7154769 | 1.04150915 | 0.66189379 | 1.57352911 | 0.11559635 | 0.3743642 |
| GALNT3 | 36.7560261 | -0.756258583 | 0.31149914 | -2.4278031 | 0.01519059 | 0.10920357 |
| TTC21B | 65.7658702 | 0.085547576 | 0.34811288 | 0.24574665 | 0.80587836 | 0.93308942 |
| SCN7A | 58.4179221 | -0.098966459 | 0.59798427 | -0.1655001 | 0.86855037 | 0.95601034 |
| STK39 | 1008.88829 | -1.556082888 | 0.36300414 | -4.2866808 | 1.81E-05 | 0.00064543 |
| CERS6 | 104.038574 | -0.839730056 | 0.34521293 | -2.4324988 | 0.01499504 | 0.1082988 |
| LRP2 | 43.8727376 | -0.187989828 | 0.75512254 | -0.2489527 | 0.80339733 | 0.93158522 |
| FASTKD1 | 100.850115 | 1.269393031 | 0.26118831 | 4.86006829 | 1.17E-06 | 6.63E-05 |
| PPIG | 134.986821 | 0.269412169 | 0.23367423 | 1.15293915 | 0.24893536 | 0.56145944 |
| KLHL23 | 38.5528939 | 0.153152297 | 0.29425524 | 0.52047433 | 0.60273301 | 0.83691476 |
| SSB | 33.9824561 | -0.15196729 | 0.36904984 | -0.4117799 | 0.68050078 | 0.87252328 |
| METTL5 | 23.6780447 | 0.205491936 | 0.39457794 | 0.52078921 | 0.60251362 | 0.83691476 |
| UBR3 | 357.211125 | 0.49643708 | 0.23151223 | 2.14432332 | 0.03200699 | 0.17742757 |
| MYO3B | 260.241087 | 2.85353314 | 0.55161522 | 5.17305001 | 2.30E-07 | 1.66E-05 |
| AC007405.1 | 19.8145319 | 0.537819249 | 0.67657066 | 0.79491955 | 0.4266603 | 0.72230448 |
| GAD1 | 126.809207 | 2.842178296 | 0.55467106 | 5.12407891 | 2.99E-07 | 2.10E-05 |
| GORASP2 | 53.7824997 | -0.005176076 | 0.2654764 | -0.0194973 | 0.98444438 | 0.99742652 |
| TLK1 | 311.755697 | 0.123448796 | 0.29143288 | 0.42359254 | 0.671863 | 0.86777458 |
| METTL8 | 54.0244084 | 0.17115325 | 0.27488534 | 0.62263505 | 0.53352437 | 0.79784784 |
| DCAF17 | 30.7255858 | 0.514422251 | 0.30720275 | 1.67453663 | 0.09402518 | 0.3344198 |
| CYBRD1 | 14.4646465 | 0.213647694 | 0.54467024 | 0.39225146 | 0.69487243 | 0.87978644 |
| DYNC1I2 | 110.847978 | -0.295416933 | 0.30235696 | -0.9770469 | 0.32854594 | 0.64006734 |
| SLC25A12 | 227.68763 | 0.886159395 | 0.40471906 | 2.18956677 | 0.02855567 | 0.16455691 |
| HAT1 | 50.5073866 | -0.411463567 | 0.3695828 | -1.113319 | 0.26557143 | 0.58092849 |
| METAP1D | 16.7276599 | -0.264947943 | 0.67060772 | -0.3950863 | 0.69277918 | 0.87849579 |
| ITGA6 | 176.760772 | -0.432652975 | 0.29567779 | -1.4632583 | 0.14339672 | 0.42447522 |
| PDK1 | 69.2567931 | -1.503791381 | 0.6558716 | -2.2928137 | 0.02185873 | 0.13855257 |
| RAPGEF4 | 18.1080027 | -0.706568509 | 0.62475571 | -1.1309517 | 0.25807544 | 0.57102406 |
| MAP3K20 | 25.0057674 | -2.420256453 | 0.57105805 | -4.2381969 | 2.25E-05 | 0.00075834 |
| SP3 | 106.510905 | -0.211963731 | 0.2689797 | -0.7880287 | 0.4306799 | 0.7245261 |
| OLA1 | 199.863048 | -0.071326786 | 0.30737382 | -0.2320523 | 0.81649743 | 0.93735334 |
| CIR1 | 58.8367166 | 0.072749962 | 0.27916219 | 0.26060105 | 0.79440018 | 0.9267672 |
| SCRN3 | 22.9382466 | 0.58307313 | 0.60333467 | 0.96641741 | 0.33383536 | 0.64554754 |
| GPR155 | 35.6513896 | -0.390915144 | 0.48894925 | -0.7995004 | 0.42400029 | 0.72151771 |
| AC010894.3 | 35.880725 | -1.976771006 | 0.53269509 | -3.7108865 | 0.00020653 | 0.00433723 |
| ATF2 | 112.914559 | 0.175896846 | 0.27275771 | 0.64488314 | 0.51900292 | 0.78749931 |
| ATP5MC3 | 71.6776585 | 0.142606973 | 0.64921623 | 0.21966021 | 0.8261358 | 0.94213432 |
| LNPK | 44.5442267 | 0.431948361 | 0.44103659 | 0.97939348 | 0.3273856 | 0.63895602 |
| HOXD10 | 79.3100555 | 0.342365803 | 0.53194187 | 0.64361507 | 0.51982508 | 0.7877576 |
| HOXD-AS2 | 27.5170611 | 1.490316052 | 0.61598699 | 2.41939533 | 0.01554633 | 0.11082452 |
| HOXD9 | 38.9894542 | 1.160316922 | 0.5967707 | 1.94432622 | 0.05185612 | 0.235032 |
| HOXD8 | 73.7893078 | 0.929447685 | 0.72136708 | 1.28845315 | 0.19758827 | 0.49994926 |
| HOXD3 | 148.650867 | 0.448695623 | 0.45257513 | 0.99142792 | 0.32147667 | 0.63312885 |
| HAGLR | 40.0337841 | -0.400546875 | 0.40318896 | -0.993447 | 0.32049215 | 0.63217026 |
| MTX2 | 293.278012 | 0.582162413 | 0.22623124 | 2.57330693 | 0.01007318 | 0.08275141 |
| LINC01116 | 34.6781958 | 0.337219034 | 0.62793228 | 0.53703089 | 0.59124628 | 0.83234048 |
| LINC01117 | 28.3163397 | -0.825472007 | 0.52991481 | -1.5577447 | 0.11929377 | 0.38066225 |
| AC079305.3 | 16.0472595 | 0.407481843 | 0.64516708 | 0.63159118 | 0.52765405 | 0.79403853 |
| HNRNPA3 | 187.740266 | 0.094375339 | 0.24677823 | 0.38242976 | 0.70214262 | 0.88441219 |
| NFE2L2 | 855.563434 | 0.920404864 | 0.50700629 | 1.81537169 | 0.06946681 | 0.28100153 |
| AGPS | 77.7314271 | -0.045307732 | 0.28384616 | -0.1596207 | 0.87317985 | 0.95703878 |
| RBM45 | 27.2200894 | 0.370708842 | 0.3499153 | 1.0594245 | 0.2894065 | 0.60448615 |
| AC009948.1 | 15.3587874 | -0.257885824 | 0.62373215 | -0.413456 | 0.67927253 | 0.87202768 |
| PRKRA | 67.9213807 | -0.309616423 | 0.2344572 | -1.3205669 | 0.18664581 | 0.48675708 |
| PJVK | 13.3683673 | -0.017465711 | 0.53249826 | -0.0327996 | 0.97383442 | 0.99340049 |
| PLEKHA3 | 31.5448141 | 0.063838694 | 0.31960411 | 0.19974303 | 0.84168156 | 0.94776435 |
| TTN-AS1 | 196.823397 | 0.211337767 | 0.3645486 | 0.57972453 | 0.5621004 | 0.81388399 |
| TTN | 22.357554 | -0.25008341 | 0.66889606 | -0.3738748 | 0.70849743 | 0.88614996 |
| SESTD1 | 462.821482 | -2.270284021 | 0.47027461 | -4.8275709 | 1.38E-06 | 7.49E-05 |
| ZNF385B | 65.4685528 | 0.310868204 | 0.54905574 | 0.56618697 | 0.57126669 | 0.81895612 |
| CWC22 | 64.4578299 | 0.244043036 | 0.33865611 | 0.72062198 | 0.47114212 | 0.75766284 |
| SCHLAP1 | 63.525505 | -2.157661538 | 0.37362629 | -5.7749189 | 7.70E-09 | 8.55E-07 |
| UBE2E3 | 210.498346 | 0.590543341 | 0.22991929 | 2.56848106 | 0.01021453 | 0.08339779 |
| CERKL | 35.5305264 | -0.175194325 | 0.64784031 | -0.2704283 | 0.78683081 | 0.92339968 |
| SSFA2 | 58.8680268 | -0.248074929 | 0.2705915 | -0.9167876 | 0.35925396 | 0.66545706 |
| PPP1R1C | 41.0960437 | -0.742456799 | 0.89441234 | -0.8301057 | 0.40647901 | 0.70556143 |
| PDE1A | 68.0275687 | -0.897112788 | 0.51832357 | -1.7307968 | 0.08348801 | 0.31112797 |
| DNAJC10 | 61.0565652 | 0.072875012 | 0.27974101 | 0.26050886 | 0.79447128 | 0.9267672 |
| NCKAP1 | 264.787436 | -0.04866639 | 0.22327136 | -0.2179697 | 0.82745273 | 0.9425007 |
| NUP35 | 30.941209 | -0.052603181 | 0.4205854 | -0.1250713 | 0.90046707 | 0.967235 |
| FSIP2 | 19.3964613 | 0.184186134 | 0.64074951 | 0.28745419 | 0.77376457 | 0.91835296 |
| FSIP2-AS1 | 17.2543741 | 0.945559479 | 0.52206799 | 1.81118073 | 0.07011288 | 0.28228674 |
| LINC01473 | 37.465212 | 0.794960345 | 0.66866034 | 1.18888515 | 0.23448487 | 0.54657 |
| ZC3H15 | 54.0146449 | -0.112833923 | 0.27944071 | -0.4037848 | 0.68637095 | 0.87487665 |
| ITGAV | 278.764888 | 0.355348414 | 0.43759729 | 0.81204436 | 0.41676618 | 0.71471912 |
| AC017101.1 | 24.8004122 | -0.07850597 | 0.66054497 | -0.1188503 | 0.90539396 | 0.96921851 |
| AC007319.1 | 66.8037041 | -0.096510574 | 0.81162673 | -0.11891 | 0.90534662 | 0.96921851 |
| TFPI | 59.7220507 | -2.481583852 | 0.91611241 | -2.7088202 | 0.00675229 | 0.06172389 |
| GULP1 | 473.775747 | -2.355011545 | 0.78877705 | -2.9856492 | 0.00282977 | 0.03302226 |
| COL5A2 | 23.928691 | 2.112509762 | 0.67270629 | 3.14031516 | 0.00168766 | 0.02232022 |
| WDR75 | 35.1060361 | 0.446470544 | 0.39813256 | 1.12141178 | 0.26211262 | 0.57612873 |
| SLC40A1 | 15.4484055 | 0.929668013 | 0.62380679 | 1.49031404 | 0.13614168 | 0.41123731 |
| ASNSD1 | 16.3189836 | 0.147783357 | 0.49462489 | 0.29877865 | 0.76510894 | 0.91549415 |
| ANKAR | 75.9220455 | 0.144310099 | 0.43976492 | 0.32815282 | 0.74279612 | 0.90463949 |
| ORMDL1 | 58.0269244 | -0.309238457 | 0.37472619 | -0.8252384 | 0.40923624 | 0.70777268 |
| PMS1 | 56.1354753 | 0.588214237 | 0.295675 | 1.98939456 | 0.04665767 | 0.22184402 |
| C2orf88 | 29.3566176 | -0.451254872 | 0.50186372 | -0.8991582 | 0.36856841 | 0.67276052 |
| HIBCH | 200.668312 | 0.572217751 | 0.42267242 | 1.35380906 | 0.1757973 | 0.47294748 |
| INPP1 | 24.4522979 | -0.658343774 | 0.44893598 | -1.4664536 | 0.14252476 | 0.42308208 |
| MFSD6 | 45.9933566 | -0.819450799 | 0.25486773 | -3.2152003 | 0.00130354 | 0.01847137 |
| NAB1 | 35.7652188 | -0.093969272 | 0.52671182 | -0.1784074 | 0.85840306 | 0.95271398 |
| GLS | 907.19636 | -0.019718404 | 0.5849223 | -0.0337112 | 0.97310749 | 0.99302333 |
| STAT1 | 127.270431 | -0.423354688 | 0.80040386 | -0.5289263 | 0.59685654 | 0.83416944 |
| MYO1B | 292.332629 | 0.279595834 | 0.37742396 | 0.74080043 | 0.45881445 | 0.747377 |
| NABP1 | 127.426579 | -2.1604078 | 0.53182365 | -4.0622635 | 4.86E-05 | 0.00138798 |
| SLC39A10 | 107.007052 | -0.050207426 | 0.7206482 | -0.0696698 | 0.94445647 | 0.98552865 |

| | | | | | | |
|---|---|---|---|---|---|---|
| DNAH7 | 159.792849 | -0.110246348 | 0.30615366 | -0.3601014 | 0.71877134 | 0.8907358 |
| STK17B | 16.3350305 | -0.662289372 | 0.50272268 | -1.317405 | 0.18770291 | 0.48778065 |
| GTF3C3 | 44.3412433 | 0.456542587 | 0.30607777 | 1.49159014 | 0.13580663 | 0.41099025 |
| PGAP1 | 28.8719122 | -0.484033597 | 0.32579939 | -1.4856799 | 0.13736382 | 0.41315491 |
| ANKRD44 | 47.3242883 | 0.03807002 | 0.34042447 | 0.11183103 | 0.91095738 | 0.97128978 |
| SF3B1 | 157.495222 | 0.672971558 | 0.21226186 | 3.17047797 | 0.00152188 | 0.02077605 |
| COQ10B | 41.6486961 | -0.73329882 | 0.53910134 | -1.3602244 | 0.17375891 | 0.46960235 |
| HSPD1 | 87.3418601 | -0.001100737 | 0.35854947 | -0.00307 | 0.99755052 | 0.99922221 |
| PLCL1 | 1482.40687 | 1.361957834 | 0.23912665 | 5.6955501 | 1.23E-08 | 1.25E-06 |
| FTCDNL1 | 38.8149348 | 1.289841686 | 0.73183078 | 1.76248624 | 0.07798718 | 0.29874793 |
| C2orf69 | 15.6384765 | -0.263454962 | 0.50443674 | -0.5222755 | 0.60147851 | 0.83679851 |
| TYW5 | 43.9274766 | -0.288856852 | 0.28584102 | -1.0105507 | 0.3122315 | 0.6271535 |
| MAIP1 | 20.4108015 | 0.01635181 | 0.38144075 | 0.04286854 | 0.96580632 | 0.99156233 |
| SPATS2L | 161.729066 | -0.480849612 | 0.56049796 | -0.8578972 | 0.39094925 | 0.69372012 |
| KCTD18 | 15.361941 | 0.113871005 | 0.43106689 | 0.26416087 | 0.79165596 | 0.92533536 |
| BZW1 | 30.0205229 | -1.571343257 | 0.63252371 | -2.484244 | 0.01298268 | 0.09865568 |
| CLK1 | 115.506853 | 0.156525064 | 0.32685734 | 0.47887883 | 0.63202483 | 0.85146176 |
| PPIL3 | 27.4925872 | -0.289028191 | 0.500744 | -0.5771975 | 0.56380604 | 0.81450672 |
| NIF3L1 | 13.3359263 | 0.257722591 | 0.42085506 | 0.6123785 | 0.54028736 | 0.80166126 |
| ORC2 | 127.940482 | -0.291468996 | 0.2644965 | -1.1019768 | 0.27047177 | 0.58716619 |
| AC005037.1 | 26.5767814 | -0.746290779 | 0.45553764 | -1.6382637 | 0.1013667 | 0.34801529 |
| FAM126B | 106.552665 | 0.289209012 | 0.27626809 | 1.04684189 | 0.29517251 | 0.61025281 |
| NDUFB3 | 30.2079807 | 0.092626778 | 0.48755874 | 0.18998075 | 0.84932421 | 0.95003258 |
| CFLAR | 252.812797 | -0.403133098 | 0.20267819 | -1.9890305 | 0.04669783 | 0.22187604 |
| CFLAR-AS1 | 19.2110909 | -0.769824475 | 0.62665505 | -1.2284661 | 0.21927205 | 0.52935683 |
| CASP10 | 18.0088068 | -1.745778377 | 0.44208192 | -3.9489929 | 7.85E-05 | 0.00203265 |
| CASP8 | 28.8874741 | -1.476661262 | 0.32949906 | -4.481534 | 7.41E-06 | 0.00029913 |
| ALS2CR12 | 21.2416971 | -0.555290102 | 0.4714957 | -1.1777204 | 0.2389081 | 0.55079752 |
| TRAK2 | 139.607674 | -0.922964735 | 0.49073375 | -1.8807851 | 0.06000115 | 0.25589532 |
| STRADB | 58.9611364 | -0.725660266 | 0.35218626 | -2.0604446 | 0.03935606 | 0.20050645 |
| TMEM237 | 14.3863312 | 0.449062028 | 0.54032024 | 0.83110348 | 0.40591518 | 0.70505833 |
| ALS2 | 78.627669 | -0.149123197 | 0.31538703 | -0.472826 | 0.63633731 | 0.85375696 |
| SUMO1 | 47.4856293 | -0.084967933 | 0.25449315 | -0.3338712 | 0.73847676 | 0.90241447 |
| NOP58 | 86.4305367 | -0.196856954 | 0.38668155 | -0.5090932 | 0.61068689 | 0.84144297 |
| BMPR2 | 271.437811 | -0.300199369 | 0.16860618 | -1.7804766 | 0.07499799 | 0.29304439 |
| FAM117B | 79.7333466 | -0.318895929 | 0.30463014 | -1.0468299 | 0.29517806 | 0.61025281 |
| ICA1L | 21.8343348 | -0.560611845 | 0.60362554 | -0.9287411 | 0.35302328 | 0.6597693 |
| WDR12 | 43.3629492 | 0.003706723 | 0.35925443 | 0.01031782 | 0.99176771 | 0.9988777 |
| CARF | 64.3865364 | 0.303841221 | 0.58711239 | 0.51751798 | 0.60479462 | 0.83778698 |
| NBEAL1 | 433.401004 | 0.52121431 | 0.30528309 | 1.70731469 | 0.08776358 | 0.32115683 |
| CYP20A1 | 101.812478 | -0.286973313 | 0.35832171 | -0.8008817 | 0.42320012 | 0.72111072 |
| ABI2 | 179.371392 | 0.076002192 | 0.21373715 | 0.35558718 | 0.72214974 | 0.89217596 |
| RAPH1 | 101.403212 | -1.29879034 | 0.32161262 | -4.0383687 | 5.38E-05 | 0.00151439 |
| PARD3B | 892.408906 | 0.281869482 | 0.27954094 | 1.00832989 | 0.31329612 | 0.62820925 |
| INO80D | 250.112453 | 0.243592603 | 0.57345472 | 0.42478089 | 0.67099641 | 0.86777458 |
| NDUFS1 | 162.689432 | 0.349139082 | 0.2450609 | 1.42470333 | 0.15424298 | 0.44013998 |
| EEF1B2 | 23.2638206 | -0.900572226 | 0.40873827 | -2.203298 | 0.02757375 | 0.16098788 |
| ZDBF2 | 36.9079702 | -0.060506578 | 0.37141552 | -0.1629081 | 0.87059083 | 0.95629149 |
| FASTKD2 | 48.8133316 | -0.022901376 | 0.28454299 | -0.0804848 | 0.93585171 | 0.98164938 |
| CPO | 11.9936406 | -0.220022619 | 0.42203011 | -0.5213434 | 0.60212756 | 0.83691476 |
| KLF7 | 41.2021548 | -0.96155276 | 0.52193961 | -1.8422682 | 0.06543593 | 0.27116648 |
| CREB1 | 108.832411 | 0.075167511 | 0.18279884 | 0.41120344 | 0.68092337 | 0.87265005 |
| METTL21A | 22.6358237 | -0.244495 | 0.43506271 | -0.5619765 | 0.57413206 | 0.82117002 |
| CCNYL1 | 40.8659445 | -0.581443312 | 0.55455007 | -1.0484956 | 0.29441032 | 0.60947643 |
| PLEKHM3 | 84.551507 | 0.306864804 | 0.27171167 | 1.1293766 | 0.258739 | 0.57194663 |
| IDH1 | 28.4286066 | 0.846926113 | 0.41906824 | 2.02097423 | 0.04328243 | 0.21173421 |
| PIKFYVE | 118.04745 | 0.006444782 | 0.2488423 | 0.02589906 | 0.97933785 | 0.99567616 |
| PTH2R | 1270.49578 | 0.976015305 | 0.52805994 | 1.84830403 | 0.06455837 | 0.268828 |
| RPE | 43.8695991 | 0.39558779 | 0.27460148 | 1.44058872 | 0.14970091 | 0.433886 |
| KANSL1L | 450.219119 | 0.102562165 | 0.3688991 | 0.27802227 | 0.78099527 | 0.92083964 |
| AC007038.1 | 49.4613486 | 0.12849294 | 0.30566276 | 0.42037487 | 0.67421163 | 0.86876026 |
| LANCL1 | 64.7905502 | 0.095988009 | 0.34404748 | 0.27899641 | 0.78024758 | 0.9206566 |
| ERBB4 | 6360.59089 | 0.027731601 | 0.38538838 | 0.07195754 | 0.9426357 | 0.98488741 |
| IKZF2 | 38.2833835 | 0.049012755 | 0.58037978 | 0.08444945 | 0.93269909 | 0.97959629 |
| SPAG16 | 278.373437 | -0.447098242 | 0.58910555 | -0.7589442 | 0.44788593 | 0.7399821 |
| AC068051.1 | 73.6609159 | 0.055666993 | 0.8071109 | 0.06897069 | 0.94501295 | 0.98552865 |
| BARD1 | 26.1186955 | -0.785958565 | 0.41588045 | -1.8898666 | 0.05877581 | 0.25348085 |
| AC016708.1 | 17.6672158 | 0.156481214 | 0.65840946 | 0.2376655 | 0.81214054 | 0.93566416 |
| ATIC | 23.2129928 | -0.04778751 | 0.34910632 | -0.1368853 | 0.89112149 | 0.96328012 |
| MREG | 16.0475106 | 0.045885427 | 0.41116046 | 0.1115998 | 0.91114073 | 0.97128978 |
| PECR | 24.7906555 | 0.276194154 | 0.64690573 | 0.42694653 | 0.66941828 | 0.86777458 |
| XRCC5 | 203.833805 | -0.371749936 | 0.31772951 | -1.1700202 | 0.24199285 | 0.55444462 |
| SMARCAL1 | 32.4157517 | -0.045956976 | 0.44726278 | -0.1027516 | 0.9181601 | 0.97482633 |
| RPL37A | 76.7586595 | -0.67572934 | 0.44644274 | -1.5135857 | 0.13013095 | 0.40070379 |
| IGFBP5 | 21.9936077 | -0.140762638 | 0.79601051 | -0.1768351 | 0.85963788 | 0.95342671 |
| DIRC3 | 66.5558095 | 0.749082476 | 0.80632112 | 0.9290126 | 0.35288256 | 0.65972795 |
| TNS1 | 201.178877 | 0.791031749 | 0.43820387 | 1.80516831 | 0.07104835 | 0.28455962 |
| ARPC2 | 116.484866 | -0.932566485 | 0.44157524 | -2.1119084 | 0.0346943 | 0.18538091 |
| AAMP | 19.3386484 | -0.225445144 | 0.49312656 | -0.457175 | 0.64754525 | 0.85921615 |
| PNKD | 56.2240462 | -0.254406876 | 0.4472796 | -0.5687871 | 0.56950062 | 0.81831157 |
| TMBIM1 | 54.6690583 | -1.760242072 | 0.29560152 | -5.9547802 | 2.60E-09 | 3.68E-07 |
| CTDSP1 | 38.6767757 | -0.532349436 | 0.32698837 | -1.6280378 | 0.10351687 | 0.3517461 |
| USP37 | 99.565933 | -0.572436821 | 0.32227788 | -1.7762213 | 0.07569646 | 0.29457168 |
| CNOT9 | 47.9190363 | 0.09208605 | 0.29640124 | 0.31068038 | 0.75604361 | 0.9108865 |
| PLCD4 | 57.6491411 | 0.382912466 | 0.40757517 | 0.93948919 | 0.34747964 | 0.6553345 |
| ZNF142 | 20.2106873 | -0.067639627 | 0.34512028 | -0.1959886 | 0.84461912 | 0.94905142 |
| STK36 | 18.1628447 | 0.415062787 | 0.63342142 | 0.65527116 | 0.51229319 | 0.7830488 |
| TTLL4 | 24.1897637 | -0.980305883 | 0.52143407 | -1.8800189 | 0.06010551 | 0.25601816 |
| CNPPD1 | 22.4261974 | 0.123622136 | 0.46233954 | 0.26738387 | 0.78917362 | 0.92405561 |
| RETREG2 | 22.5052439 | -0.407248748 | 0.48574177 | -0.8384059 | 0.4018028 | 0.70114138 |
| ZFAND2B | 12.5626206 | 0.390156842 | 0.49890602 | 0.78202472 | 0.43420005 | 0.72762064 |
| ANKZF1 | 33.6636584 | -0.028365181 | 0.51199665 | -0.0554011 | 0.95581891 | 0.98853497 |
| TUBA4A | 21.9125943 | -0.374781445 | 0.53176558 | -0.704787 | 0.48094282 | 0.76563273 |
| DNAJB2 | 26.0196148 | -0.19685618 | 0.38776255 | -0.507672 | 0.6116834 | 0.84169124 |
| DNPEP | 18.553168 | -0.305555778 | 0.38930683 | -0.7848714 | 0.432529 | 0.72599466 |
| OBSL1 | 36.1611971 | 0.469169267 | 0.65161473 | 0.72001022 | 0.4715187 | 0.75776258 |
| SGPP2 | 249.816357 | -0.543628772 | 0.42610822 | -1.2757998 | 0.20202631 | 0.50618429 |
| FARSB | 46.1715505 | 0.581097363 | 0.30659526 | 1.89532402 | 0.05804949 | 0.25127828 |
| MOGAT1 | 24.0326638 | -3.097197991 | 1.49394321 | -2.0731698 | 0.03815648 | 0.19688489 |
| ACSL3 | 208.143997 | 0.375054002 | 0.53311005 | 0.70352078 | 0.48173126 | 0.76623375 |
| AP1S3 | 49.5152517 | -1.176977296 | 0.58549103 | -2.0102397 | 0.04440583 | 0.21534941 |
| WDFY1 | 90.4812663 | -0.756051428 | 0.42043819 | -1.7982463 | 0.07213798 | 0.28695159 |
| CUL3 | 237.931096 | -0.122377412 | 0.26867249 | -0.4554892 | 0.64875735 | 0.8599651 |
| IRS1 | 47.1272478 | -0.955856146 | 0.48379387 | -1.9757508 | 0.048183 | 0.22485402 |
| RHBDD1 | 124.455705 | 0.21417133 | 0.33938976 | 0.6310483 | 0.52800895 | 0.7943404 |
| COL4A4 | 581.197847 | -0.122958942 | 0.37748075 | -0.3257357 | 0.74462436 | 0.90519915 |
| COL4A3 | 1070.71572 | 0.325610455 | 0.5569607 | 0.58462016 | 0.55880315 | 0.81279041 |
| AC097662.1 | 124.4742 | -0.687728952 | 0.6489478 | -1.0597601 | 0.28925378 | 0.60443796 |
| MFF | 149.378788 | -0.030621185 | 0.29851081 | -0.1025798 | 0.91829646 | 0.97486009 |
| AGFG1 | 302.965499 | -0.913008209 | 0.48340523 | -1.8887016 | 0.05893183 | 0.25380155 |
| SLC19A3 | 222.274787 | -0.321269326 | 0.73439077 | -0.4374637 | 0.66177507 | 0.86492722 |
| PID1 | 98.1185331 | 2.764719059 | 0.7045953 | 3.92383976 | 8.71E-05 | 0.00219615 |
| DNER | 254.942589 | -0.764385468 | 0.63287128 | -1.2078056 | 0.22712205 | 0.53871433 |

| | | | | | | |
|---|---|---|---|---|---|---|
| TRIP12 | 310.833833 | -0.161388308 | 0.23018708 | -0.701118 | 0.48322937 | 0.76664595 |
| FBXO36 | 33.3633775 | 0.220629796 | 0.36457463 | 0.60517046 | 0.54506574 | 0.80452635 |
| SP110 | 40.5656955 | -0.144182098 | 0.66061464 | -0.2182545 | 0.82723085 | 0.9425007 |
| SP140L | 16.2287004 | -1.83000017 | 0.69178221 | -2.6453415 | 0.00816085 | 0.07111381 |
| SP100 | 152.281009 | -0.018386996 | 0.50645731 | -0.0363051 | 0.97103906 | 0.99199827 |
| CAB39 | 223.165142 | 0.172369104 | 0.38740387 | 0.44493388 | 0.65636754 | 0.86294007 |
| ITM2C | 55.7211301 | -1.701454207 | 0.74599072 | -2.2807981 | 0.0225604 | 0.14136636 |
| PSMD1 | 118.801702 | -0.042082202 | 0.301229 | -0.1397017 | 0.88889568 | 0.96307703 |
| ARMC9 | 103.497871 | -0.525961375 | 0.21899846 | -2.401667 | 0.01632056 | 0.11432974 |
| NCL | 53.3576628 | -0.749054221 | 0.40086786 | -1.8685814 | 0.06168108 | 0.26023274 |
| PTMA | 175.34039 | -0.908878197 | 0.40271582 | -2.2568724 | 0.02401605 | 0.14714173 |
| PDE6D | 31.3223637 | -0.080343323 | 0.33962143 | -0.2365673 | 0.81299249 | 0.93607582 |
| COPS7B | 37.183521 | 0.564768071 | 0.32325319 | 1.74713844 | 0.08061333 | 0.30529596 |
| DIS3L2 | 202.819602 | 0.221584571 | 0.18168711 | 1.21959435 | 0.22261869 | 0.53302786 |
| EIF4E2 | 48.7773313 | -0.464811649 | 0.27765077 | -1.6740874 | 0.09411343 | 0.33444204 |
| EFHD1 | 182.724545 | 1.229377678 | 0.52172162 | 2.35638628 | 0.01845372 | 0.12417062 |
| GIGYF2 | 305.193499 | -0.045994557 | 0.29666572 | -0.1550383 | 0.8767911 | 0.95879545 |
| ATG16L1 | 25.187954 | -0.230250532 | 0.37503689 | -0.613941 | 0.5392543 | 0.80165929 |
| DGKD | 36.4889687 | -0.32724141 | 0.52741074 | -0.6204679 | 0.53494981 | 0.79921039 |
| USP40 | 151.982993 | 0.39949446 | 0.30796478 | 1.29720828 | 0.19455953 | 0.49702866 |
| AC114812.2 | 27.1248739 | -3.399522733 | 0.9595338 | -3.54289 | 0.00039577 | 0.0073217 |
| SH3BP4 | 183.061393 | -0.114806963 | 0.59152448 | -0.1940866 | 0.84610809 | 0.94909204 |
| AGAP1 | 965.425204 | -0.221120868 | 0.48918616 | -0.4520178 | 0.65125613 | 0.86120024 |
| COPS8 | 19.8664527 | 0.203995385 | 0.37189653 | 0.54852726 | 0.58332992 | 0.82696794 |
| RAB17 | 101.820407 | -0.396651426 | 0.49621006 | -0.7993619 | 0.42408058 | 0.72151771 |
| LRRFIP1 | 544.056348 | -0.652452016 | 0.33466167 | -1.9495869 | 0.05122537 | 0.23447491 |
| ILKAP | 75.3761515 | -0.058992933 | 0.3433361 | -0.1718227 | 0.86357692 | 0.95458251 |
| PER2 | 23.3293878 | -0.4740379 | 0.5761059 | -0.8228312 | 0.410604 | 0.70923892 |
| TRAF3IP1 | 48.8594309 | 0.199599246 | 0.30791078 | 0.64823727 | 0.5168315 | 0.78561084 |
| ASB1 | 14.3221453 | 0.202847716 | 0.4535183 | 0.4472757 | 0.65467601 | 0.86217501 |
| HDAC4 | 193.891908 | -1.348915079 | 0.43926354 | -3.070856 | 0.00213446 | 0.02650028 |
| NDUFA10 | 126.694906 | 0.460030318 | 0.24765208 | 1.85756694 | 0.06323054 | 0.26477174 |
| COPS9 | 17.323694 | -0.348798463 | 0.66332666 | -0.5258321 | 0.59900486 | 0.83572068 |
| GPC1 | 27.1471111 | -2.925704265 | 0.74081959 | -3.9492804 | 7.84E-05 | 0.00203265 |
| ANKMY1 | 48.7892357 | -0.464890349 | 0.38110869 | -1.2198366 | 0.22252681 | 0.53302786 |
| RNPEPL1 | 17.2012704 | -1.229183794 | 0.54087571 | -2.2725809 | 0.02305145 | 0.14328781 |
| CAPN10 | 17.6645614 | -0.393360818 | 0.55085354 | -0.7140933 | 0.47516949 | 0.76125092 |
| SNED1 | 78.9197648 | -3.290683048 | 0.84347598 | -3.9013358 | 9.57E-05 | 0.00234112 |
| MTERF4 | 55.087631 | -0.123393531 | 0.41219272 | -0.2993588 | 0.76466627 | 0.91531804 |
| PPP1R7 | 28.6811966 | -0.345109817 | 0.31098921 | -1.1097164 | 0.26712127 | 0.58335197 |
| HDLBP | 173.006029 | -0.424095327 | 0.34556515 | -1.2272514 | 0.2197281 | 0.52962456 |
| 2-Sep | 212.62788 | 0.045519914 | 0.18361816 | 0.2479053 | 0.80420768 | 0.93229297 |
| FARP2 | 106.651232 | -0.379157099 | 0.32130108 | -1.1800679 | 0.2379732 | 0.54997949 |
| STK25 | 27.3165364 | -0.127454195 | 0.33249006 | -0.3833324 | 0.70147335 | 0.88421489 |
| THAP4 | 46.0153702 | 0.076643534 | 0.32561018 | 0.23538433 | 0.81391045 | 0.93633777 |
| ATG4B | 106.573868 | -0.154878559 | 0.3066729 | -0.5050285 | 0.61353881 | 0.84174925 |
| ING5 | 49.649783 | 0.190207849 | 0.26215369 | 0.72555853 | 0.46810945 | 0.75473846 |
| D2HGDH | 51.7020525 | 0.536463737 | 0.49498743 | 1.08379265 | 0.27845675 | 0.59359815 |
| LINC01266 | 164.836125 | -0.206867649 | 0.48298624 | -0.4283096 | 0.66842572 | 0.86777458 |
| CNTN4 | 43.2663459 | -0.834314557 | 0.44230792 | -1.8862754 | 0.05925785 | 0.25450031 |
| TRNT1 | 49.700002 | -0.080069069 | 0.28753796 | -0.2784643 | 0.78065594 | 0.92083964 |
| CRBN | 59.8474579 | 0.179621938 | 0.28911366 | 0.62128485 | 0.53441222 | 0.79873873 |
| AC034195.1 | 108.326433 | 0.554879751 | 0.65723302 | 0.84426639 | 0.39852056 | 0.69875231 |
| SUMF1 | 461.414311 | 0.228526377 | 0.33259339 | 0.68710439 | 0.49201695 | 0.77284018 |
| SETMAR | 15.7412263 | 0.755543507 | 0.50822534 | 1.48663093 | 0.13711231 | 0.41253153 |
| ITPR1 | 961.800612 | 0.486539417 | 0.297419 | 1.63587203 | 0.10186637 | 0.34880722 |
| EGOT | 18.2162894 | -0.036298321 | 0.44213143 | -0.0820985 | 0.9345684 | 0.98085023 |
| AC018816.1 | 24.5259123 | -0.80009674 | 0.44647308 | -1.792038 | 0.07312688 | 0.28899744 |
| BHLHE40 | 143.108847 | -1.612894228 | 0.56997609 | -2.8297577 | 0.00465833 | 0.04771644 |
| ARL8B | 150.911575 | -1.16475573 | 0.62096802 | -1.8757097 | 0.06069515 | 0.25747117 |
| AC026202.2 | 21.7359478 | -0.848405578 | 0.53149532 | -1.5962616 | 0.1104304 | 0.36577373 |
| EDEM1 | 30.6652754 | -0.36550224 | 0.40644406 | -0.8992683 | 0.36850979 | 0.67276052 |
| AC069277.1 | 53.9847139 | 0.567522996 | 1.5119726 | 0.3753527 | 0.70739818 | 0.88614996 |
| LMCD1-AS1 | 76.6286542 | 0.045360983 | 0.71558117 | 0.06339041 | 0.94945562 | 0.98641632 |
| RAD18 | 24.137243 | 0.034043313 | 0.49366174 | 0.06896081 | 0.94502082 | 0.98552865 |
| SRGAP3 | 110.630798 | 0.004090982 | 0.47034046 | 0.00869792 | 0.99306015 | 0.9992177 |
| THUMPD3-A: | 88.6335218 | 0.010665113 | 0.27259394 | 0.03912454 | 0.96879109 | 0.99156233 |
| THUMPD3 | 22.2145721 | 0.60413624 | 0.35931298 | 1.6813649 | 0.09269206 | 0.33202083 |
| SETD5 | 291.437307 | 0.007541346 | 0.1587075 | 0.04751726 | 0.96210097 | 0.99029318 |
| MTMR14 | 39.5010605 | -0.059821497 | 0.294298 | -0.2032684 | 0.83892522 | 0.94741614 |
| OGG1 | 25.8970885 | -0.426857745 | 0.45515706 | -0.9378252 | 0.34833427 | 0.65547624 |
| TADA3 | 24.0082601 | -0.574801599 | 0.37449803 | -1.5348588 | 0.12481852 | 0.39027762 |
| IL17RE | 17.7699395 | 0.821038673 | 0.63673094 | 1.28945936 | 0.19723844 | 0.49934704 |
| IL17RC | 11.1957266 | 0.718447555 | 0.75366256 | 0.95327483 | 0.34045084 | 0.65083559 |
| CRELD1 | 24.118379 | -0.201227711 | 0.49395364 | -0.4073818 | 0.68372761 | 0.87343121 |
| EMC3 | 47.1021296 | -0.067262214 | 0.35198623 | -0.1910933 | 0.84845249 | 0.95003258 |
| BRK1 | 28.0268638 | -0.354108789 | 0.34964978 | -1.0127528 | 0.31117826 | 0.62611697 |
| VHL | 29.2081332 | -0.176384162 | 0.32405518 | -0.5443029 | 0.58623308 | 0.8294442 |
| SEC13 | 37.5294287 | -0.445724097 | 0.32206662 | -1.38395 | 0.16637377 | 0.45864087 |
| ATG7 | 292.70553 | -0.25256956 | 0.27703521 | -0.9116876 | 0.36193321 | 0.66719361 |
| VGLL4 | 267.232102 | 0.306463623 | 0.26631772 | 1.15074439 | 0.24983741 | 0.5624056 |
| TAMM41 | 78.2993242 | 0.496237398 | 0.3217436 | 1.54233807 | 0.12299146 | 0.38755404 |
| SYN2 | 20.5547847 | 0.038014322 | 0.77659502 | 0.04894999 | 0.96095915 | 0.98983463 |
| PPARG | 23.1387743 | -1.836656118 | 0.74156259 | -2.4767378 | 0.01325893 | 0.09993884 |
| TSEN2 | 62.4484078 | 0.377997497 | 0.39468151 | 0.95772791 | 0.33819998 | 0.64897646 |
| MKRN2 | 15.3323601 | 0.495988508 | 0.41978598 | 1.18152709 | 0.2373934 | 0.54992696 |
| RAF1 | 159.815038 | 0.174443372 | 0.29379449 | 0.59375985 | 0.55267274 | 0.80962553 |
| RPL32 | 75.3156747 | -0.43613778 | 0.42895139 | -1.0167534 | 0.30927076 | 0.62335507 |
| IQSEC1 | 62.7365657 | 0.775565554 | 0.46840372 | 1.65576302 | 0.09776982 | 0.34066808 |
| NUP210 | 46.7161777 | 1.161661727 | 0.33000992 | 3.52008124 | 0.00043141 | 0.00775956 |
| HDAC11 | 12.7172981 | 0.633589682 | 0.46325833 | 1.36768114 | 0.17141191 | 0.46718638 |
| XPC | 52.304227 | -0.499277031 | 0.33813975 | -1.4765405 | 0.13979885 | 0.4184541 |
| LSM3 | 17.2716517 | -0.228583034 | 0.46805054 | -0.4883725 | 0.62528598 | 0.8471532 |
| SLC6A6 | 28.552309 | -3.223942255 | 0.69912104 | -4.6114222 | 4.00E-06 | 0.0001819 |
| CCDC174 | 40.6658553 | 0.072329256 | 0.38784721 | 0.18648904 | 0.85206127 | 0.95090866 |
| FGD5-AS1 | 42.9594583 | 0.233531859 | 0.35878876 | 0.65088956 | 0.51511778 | 0.78381327 |
| NR2C2 | 115.866954 | -0.136333734 | 0.32544208 | -0.4189186 | 0.67527564 | 0.86960234 |
| MRPS25 | 89.3134351 | 0.186884794 | 0.26257555 | 0.71173723 | 0.47662751 | 0.76227699 |
| RBSN | 35.758004 | 0.233957293 | 0.36431676 | 0.64218098 | 0.52075568 | 0.78815594 |
| CAPN7 | 149.536424 | -0.13504959 | 0.26378673 | -0.5119651 | 0.60867546 | 0.84008076 |
| METTL6 | 39.8170254 | -0.218025099 | 0.41292573 | -0.5280008 | 0.59749881 | 0.83436856 |
| COLQ | 17.8688454 | -0.965166727 | 0.68571169 | -1.4075401 | 0.15926729 | 0.44760113 |
| HACL1 | 50.7474681 | -0.248174983 | 0.37971626 | -0.6535801 | 0.51338238 | 0.7830488 |
| ANKRD28 | 400.122083 | 0.028445101 | 0.16520628 | 0.1721793 | 0.86329657 | 0.95458251 |
| OXNAD1 | 145.382073 | 0.235863469 | 0.35773005 | 0.65933368 | 0.50968152 | 0.78123796 |
| PLCL2 | 68.2182987 | 2.078419695 | 0.4866631 | 4.27075674 | 1.95E-05 | 0.00068544 |
| TBC1D5 | 568.364702 | 0.39096223 | 0.37694131 | 1.03719656 | 0.29964426 | 0.61356712 |
| RAB5A | 122.291855 | -0.624204811 | 0.3003617 | -2.0781771 | 0.03769305 | 0.19601227 |
| KAT2B | 87.1843231 | 1.253691705 | 0.46062503 | 2.72171859 | 0.00649434 | 0.0598944 |
| ZNF385D | 961.81339 | -0.621243868 | 0.59542651 | -1.0433594 | 0.29678185 | 0.61093809 |
| UBE2E2 | 529.914577 | 0.228949672 | 0.38372163 | 0.59665563 | 0.55073732 | 0.80829133 |
| UBE2E1 | 227.23672 | 0.093442989 | 0.21959061 | 0.42553271 | 0.67044838 | 0.86777458 |
| NKIRAS1 | 17.2502838 | -0.025300867 | 0.4274598 | -0.0591889 | 0.95280167 | 0.98743167 |

| | | | | | | |
|---|---|---|---|---|---|---|
| RPL15 | 120.012011 | -1.519549821 | 0.51725812 | -2.9377012 | 0.00330655 | 0.03755215 |
| NR1D2 | 62.6208826 | -0.64584718 | 0.57911362 | -1.115234 | 0.26475017 | 0.57977023 |
| THRB | 479.520451 | 1.305310783 | 0.47625315 | 2.74079189 | 0.00612913 | 0.05756707 |
| RARB | 25.7111743 | 0.245679005 | 0.44716893 | 0.54940983 | 0.58272423 | 0.82657506 |
| TOP2B | 158.363337 | 0.635808279 | 0.22915237 | 2.77460925 | 0.00552681 | 0.05345806 |
| NGLY1 | 128.841389 | -0.023542968 | 0.25480622 | -0.0923956 | 0.92638375 | 0.97735117 |
| SLC4A7 | 45.9750554 | -3.343832912 | 1.04703853 | -3.1936102 | 0.00140506 | 0.01961191 |
| CMC1 | 181.684564 | 0.671928781 | 0.26831812 | 2.50422443 | 0.01227202 | 0.09511578 |
| AZI2 | 55.9141324 | 0.095947261 | 0.31519682 | 0.30440428 | 0.76081991 | 0.91310141 |
| TGFBR2 | 86.9927086 | 0.595362429 | 0.35194788 | 1.69162102 | 0.09071826 | 0.32772453 |
| STT3B | 137.767952 | 0.15379499 | 0.35024421 | 0.43910787 | 0.66058337 | 0.86492722 |
| OSBPL10 | 173.773732 | -0.621427611 | 0.27849262 | -2.231397 | 0.02565484 | 0.15373119 |
| GPD1L | 30.3621085 | 1.500205557 | 0.44852178 | 3.34477751 | 0.00082349 | 0.01285117 |
| CMTM8 | 122.208597 | -0.714223121 | 0.40911455 | -1.7457779 | 0.08084956 | 0.30571131 |
| CMTM7 | 73.8743533 | -4.562362047 | 0.48852772 | -9.3390035 | 9.72E-21 | 2.27E-17 |
| CMTM6 | 51.3779428 | -1.164898335 | 0.38592041 | -3.0184938 | 0.00254035 | 0.03036691 |
| DYNC1LI1 | 54.4049411 | -0.058865551 | 0.34099471 | -0.1726289 | 0.86294311 | 0.95458251 |
| CNOT10 | 96.1607169 | -0.032285039 | 0.24519643 | -0.1316701 | 0.89524523 | 0.96511349 |
| TRIM71 | 40.6717894 | 0.381986395 | 0.67959499 | 0.56207947 | 0.57406187 | 0.82117002 |
| GLB1 | 77.7947463 | 0.388136951 | 0.32113061 | 1.20865759 | 0.22679442 | 0.53826211 |
| CRTAP | 36.1957423 | 0.22279322 | 0.33293907 | 0.66917114 | 0.50338631 | 0.77790293 |
| FBXL2 | 122.737796 | -0.114008563 | 0.3781328 | -0.301504 | 0.76303018 | 0.91422428 |
| UBP1 | 65.3280919 | 0.345323526 | 0.2991459 | 1.1543649 | 0.2483506 | 0.56107171 |
| CLASP2 | 325.652949 | 0.673158744 | 0.28514405 | 2.36076725 | 0.01823717 | 0.12340471 |
| PDCD6IP | 166.108409 | -0.059459313 | 0.18462073 | -0.322062 | 0.74740576 | 0.90692529 |
| STAC | 142.276923 | 0.702938034 | 0.44679519 | 1.57328918 | 0.11565187 | 0.3743642 |
| TRANK1 | 18.2538531 | -0.363892899 | 1.04883001 | -0.3469513 | 0.72862793 | 0.89661959 |
| EPM2AIP1 | 48.3968599 | 0.203005872 | 0.28437502 | 0.71386676 | 0.47530958 | 0.76134453 |
| MLH1 | 31.1856351 | 0.345388785 | 0.31452385 | 1.09813226 | 0.27214674 | 0.58786226 |
| LRRFIP2 | 296.221593 | -0.005245305 | 0.28132873 | -0.0186448 | 0.9851245 | 0.99742652 |
| GOLGA4 | 523.963912 | -0.312228724 | 0.40497884 | -0.7709754 | 0.44072151 | 0.73302253 |
| ITGA9 | 56.903164 | -0.765595413 | 0.68594712 | -1.1161143 | 0.2643732 | 0.57918602 |
| CTDSPL | 525.221493 | 0.395947703 | 0.35553176 | 1.11367743 | 0.26541759 | 0.58079175 |
| PLCD1 | 18.0796763 | 0.346643744 | 0.56237473 | 0.61639282 | 0.53763529 | 0.80078458 |
| ACAA1 | 37.1676948 | -0.074559957 | 0.41665406 | -0.1789493 | 0.8579775 | 0.95266335 |
| OXSR1 | 155.287177 | 0.417833112 | 0.28035531 | 1.4903699 | 0.136127 | 0.41123731 |
| XYLB | 16.4026073 | 2.085855198 | 0.50686748 | 4.11518841 | 3.87E-05 | 0.00115613 |
| ACVR2B | 29.627672 | -0.016122797 | 0.37480349 | -0.0430167 | 0.96568825 | 0.99156233 |
| EXOG | 24.0460268 | 0.182829636 | 0.62401949 | 0.29298706 | 0.76953204 | 0.91636229 |
| WDR48 | 81.8574124 | 0.290467916 | 0.33272583 | 0.8729948 | 0.38266591 | 0.68680981 |
| GORASP1 | 17.7383667 | -0.02490364 | 0.36170706 | -0.0688503 | 0.94510878 | 0.98552865 |
| TTC21A | 13.4444848 | -0.208690966 | 0.69940009 | -0.2983857 | 0.76540882 | 0.91549415 |
| RPSA | 62.0063669 | -0.650695373 | 0.44313297 | -1.4683976 | 0.14199626 | 0.42241953 |
| MYRIP | 23.9298777 | 0.364099339 | 0.74810665 | 0.48669443 | 0.6264749 | 0.8472036 |
| EIF1B | 20.3199465 | -0.669061869 | 0.47245291 | -1.4161451 | 0.15673303 | 0.44337948 |
| ENTPD3-AS1 | 28.723764 | 0.720062667 | 0.61958895 | 1.16216189 | 0.24516971 | 0.55755248 |
| RPL14 | 93.4730163 | -0.766813062 | 0.49904248 | -1.5365687 | 0.12439896 | 0.38965272 |
| ZNF621 | 49.2732165 | -0.045545381 | 0.30927223 | -0.1472663 | 0.88292182 | 0.96161232 |
| CTNNB1 | 122.671738 | -0.209553061 | 0.21022198 | -0.996818 | 0.31885285 | 0.63100017 |
| ULK4 | 168.741872 | -0.616858968 | 0.32214154 | -1.9148694 | 0.05550916 | 0.24529263 |
| TRAK1 | 240.037084 | 0.272081364 | 0.23235948 | 1.17095015 | 0.24161882 | 0.55407128 |
| SEC22C | 23.7866013 | 0.167960433 | 0.41711756 | 0.40266929 | 0.68719153 | 0.87556351 |
| NKTR | 329.481762 | 0.038934519 | 0.17899747 | 0.21751436 | 0.82780752 | 0.94254892 |
| AC006059.1 | 31.1625437 | -0.853318059 | 0.5426741 | -1.5724319 | 0.11585042 | 0.37454555 |
| SNRK | 206.887498 | -0.151100985 | 0.48330384 | -0.3126418 | 0.75455281 | 0.91026551 |
| ANO10 | 254.279199 | 0.472690879 | 0.34280518 | 1.37889073 | 0.16792845 | 0.4611973 |
| ABHD5 | 34.8791694 | -1.290621599 | 0.71446123 | -1.8064264 | 0.07085177 | 0.28413844 |
| TCAIM | 253.674626 | 0.47968992 | 0.43388201 | 1.10557687 | 0.2689097 | 0.58532868 |
| ZNF445 | 35.8652173 | 0.589385362 | 0.36548272 | 1.6126217 | 0.10682672 | 0.35932624 |
| AC099669.1 | 90.8987426 | 0.510180585 | 0.28348535 | 1.79967178 | 0.07191248 | 0.28629887 |
| ZNF197 | 41.3044761 | -0.098279037 | 0.26320408 | -0.3733948 | 0.70885462 | 0.88614996 |
| KIAA1143 | 11.32944 | -0.120677787 | 0.42559631 | -0.2835499 | 0.77675534 | 0.9196152 |
| ZDHHC3 | 77.3411511 | 0.428839866 | 0.22486913 | 1.90706417 | 0.05651228 | 0.2474967 |
| EXOSC7 | 38.2336162 | 0.336172437 | 0.33544652 | 1.00216403 | 0.31626437 | 0.63009595 |
| CDCP1 | 131.195727 | -1.433269563 | 0.43684424 | -3.2809625 | 0.00103454 | 0.01533546 |
| LARS2 | 79.327246 | 0.114682009 | 0.26175401 | 0.43812895 | 0.66129281 | 0.86492722 |
| LIMD1 | 45.7973469 | -0.294286233 | 0.38318607 | -0.7679983 | 0.44248822 | 0.73477475 |
| SACM1L | 92.1238153 | 0.719261621 | 0.25322794 | 2.84037223 | 0.00450609 | 0.04652802 |
| LZTFL1 | 23.2842038 | -0.291374475 | 0.38570148 | -0.7554404 | 0.44998479 | 0.74152951 |
| FYCO1 | 36.4994912 | 0.214891437 | 0.33439049 | 0.64263621 | 0.52046018 | 0.78804331 |
| LTF | 46.1363324 | -4.326231351 | 1.21663813 | -3.55589 | 0.0003767 | 0.00702473 |
| ALS2CL | 22.4889966 | -2.859722033 | 0.67365688 | -4.2450721 | 2.19E-05 | 0.00074908 |
| PTH1R | 236.844389 | 0.759706091 | 0.5109813 | 1.48675908 | 0.13707845 | 0.41253153 |
| CCDC12 | 84.5191874 | 0.167225353 | 0.26669106 | 0.62703772 | 0.53063451 | 0.79595177 |
| NBEAL2 | 42.8359298 | -0.744283382 | 0.46937895 | -1.585677 | 0.11281261 | 0.37037491 |
| SETD2 | 361.229794 | 0.078354961 | 0.21047994 | 0.37226808 | 0.70969327 | 0.88614996 |
| KIF9-AS1 | 132.053787 | 0.500023397 | 0.30305701 | 1.64993178 | 0.09895689 | 0.34359002 |
| KLHL18 | 47.8044753 | -0.31492899 | 0.38651578 | -0.8147895 | 0.41519283 | 0.71267635 |
| PTPN23 | 23.8697223 | -0.454695072 | 0.53741354 | -0.8460804 | 0.39750789 | 0.6979969 |
| SCAP | 76.0545079 | 0.587448887 | 0.27670443 | 2.12301944 | 0.03375222 | 0.18286213 |
| ELP6 | 23.697795 | 0.019776042 | 0.32865382 | 0.06017286 | 0.95201796 | 0.98738095 |
| SMARCC1 | 276.973798 | -0.740644254 | 0.35042684 | -2.1135489 | 0.03455382 | 0.1848421 |
| DHX30 | 54.5231672 | 0.81639715 | 0.30713013 | 2.65814735 | 0.00785715 | 0.06924395 |
| MAP4 | 334.578499 | -0.046778915 | 0.25355077 | -0.1844953 | 0.85362495 | 0.95137449 |
| ZNF589 | 28.2162269 | -0.255693201 | 0.45860877 | -0.557541 | 0.57715784 | 0.82233385 |
| NME6 | 15.3747928 | -0.33687948 | 0.49094798 | -0.6861816 | 0.49259858 | 0.77284018 |
| PLXNB1 | 130.851674 | 0.253762077 | 0.41151096 | 0.61665935 | 0.53745944 | 0.80078458 |
| TMA7 | 42.8037608 | 0.011900471 | 0.52673436 | 0.02259293 | 0.98197499 | 0.99659802 |
| SHISA5 | 68.138324 | -2.241833871 | 0.42155083 | -5.3180629 | 1.05E-07 | 8.36E-06 |
| PFKFB4 | 51.889432 | -2.910143224 | 0.80345148 | -3.6220522 | 0.00029228 | 0.00567744 |
| COL7A1 | 27.8624963 | -3.473463151 | 0.86837948 | -3.9999369 | 6.34E-05 | 0.00172738 |
| UQCRC1 | 44.4602978 | 0.231360763 | 0.38603559 | 0.59932496 | 0.54895621 | 0.80682025 |
| NCKIPSD | 14.447431 | -0.326744076 | 0.58070724 | -0.5626657 | 0.57366251 | 0.82110314 |
| IP6K2 | 83.1184864 | -0.010373997 | 0.31711895 | -0.0327133 | 0.97390324 | 0.99340049 |
| PRKAR2A | 120.508842 | -0.287581985 | 0.29815017 | -0.9645541 | 0.33476818 | 0.64611437 |
| SLC25A20 | 18.3556951 | 0.940416461 | 0.57547207 | 1.63416525 | 0.10222416 | 0.34939078 |
| ARIH2 | 123.829213 | 0.128348591 | 0.29105534 | 0.44097659 | 0.65922995 | 0.86450914 |
| P4HTM | 27.7452052 | 0.305287176 | 0.4591001 | 0.66496865 | 0.50607054 | 0.77934214 |
| WDR6 | 55.4827171 | 0.090015989 | 0.38286848 | 0.23510943 | 0.8141238 | 0.93633777 |
| DALRD3 | 14.668395 | -0.029018041 | 0.45325972 | -0.0640208 | 0.94895368 | 0.98641632 |
| QRICH1 | 92.4187624 | -0.146515205 | 0.3146822 | -0.4655974 | 0.64150372 | 0.85468927 |
| QARS | 19.7505873 | 0.991816106 | 0.40458985 | 2.45141126 | 0.01422973 | 0.10471821 |
| LAMB2 | 29.7049669 | 0.905691618 | 0.43065231 | 2.10306923 | 0.03545972 | 0.18774924 |
| USP4 | 79.8380873 | 0.030767829 | 0.28437669 | 0.10819392 | 0.91384187 | 0.97217087 |
| RHOA | 108.345043 | -0.331105965 | 0.34023614 | -0.9731652 | 0.33047122 | 0.64255236 |
| TCTA | 15.2689043 | -0.676070637 | 0.51520461 | -1.3122372 | 0.18944012 | 0.48969773 |
| DAG1 | 76.1983536 | 0.81808938 | 0.38551541 | 2.12206661 | 0.03383215 | 0.18318869 |
| APEH | 14.3360563 | 0.219503782 | 0.46827374 | 0.46875099 | 0.63924763 | 0.85391761 |
| MST1 | 20.1137053 | 0.821667107 | 0.47935765 | 1.71410033 | 0.0865103 | 0.31781798 |
| RNF123 | 16.2712881 | 0.109285834 | 0.45132678 | 0.24214347 | 0.80866899 | 0.93433178 |
| IP6K1 | 133.773653 | 0.268191717 | 0.27518627 | 0.97458246 | 0.32976742 | 0.64177654 |
| RBM6 | 453.849863 | 0.110745183 | 0.22729852 | 0.48722352 | 0.62609994 | 0.84715656 |
| RBM5 | 141.047907 | 0.451280014 | 0.23943596 | 1.88476292 | 0.05946185 | 0.25478967 |

| | | | | | | |
|---|---|---|---|---|---|---|
| GNAI2 | 29.4720762 | -1.339628717 | 0.39770101 | -3.3684318 | 0.00075597 | 0.01209035 |
| SEMA3B | 63.6693204 | -1.662811596 | 0.46568152 | -3.5707055 | 0.00035602 | 0.00671972 |
| HYAL1 | 16.3141593 | 1.056248954 | 0.57080011 | 1.85047083 | 0.06424572 | 0.26796228 |
| HEMK1 | 18.7041384 | 0.803293742 | 0.46871706 | 1.71381375 | 0.08656294 | 0.31788611 |
| CISH | 23.1254851 | -0.873171739 | 0.48136627 | -1.8139446 | 0.06968626 | 0.28152282 |
| MAPKAPK3 | 26.8657388 | -0.820382148 | 0.43705589 | -1.8770646 | 0.06050923 | 0.25726772 |
| DOCK3 | 103.286423 | -0.767737774 | 0.55717319 | -1.3779159 | 0.16822927 | 0.46175145 |
| MANF | 20.1262945 | -0.557450379 | 0.67299708 | -0.8283102 | 0.40749482 | 0.70619047 |
| DCAF1 | 71.1275367 | -0.047108753 | 0.38486193 | -0.1224043 | 0.90257884 | 0.96803829 |
| RAD54L2 | 104.963975 | 0.627073457 | 0.28442161 | 2.20473212 | 0.0274729 | 0.16065021 |
| TEX264 | 27.4052851 | -0.35354657 | 0.38605716 | -0.9157882 | 0.35977802 | 0.66572082 |
| RPL29 | 101.54116 | -1.518423292 | 0.47102197 | -3.2236782 | 0.00126556 | 0.01801723 |
| ALAS1 | 16.0426421 | 0.011319983 | 0.36786421 | 0.03077218 | 0.97545123 | 0.99397457 |
| WDR82 | 53.0269827 | -0.059461746 | 0.36857723 | -0.1613278 | 0.87183525 | 0.9568935 |
| DNAH1 | 12.8182411 | -0.645488561 | 0.54584293 | -1.1825537 | 0.23698608 | 0.54992696 |
| BAP1 | 13.4380991 | -0.555539987 | 0.49054951 | -1.1324851 | 0.25743057 | 0.57013838 |
| PHF7 | 16.789385 | 1.48528541 | 0.43825601 | 3.38908169 | 0.00070127 | 0.01137157 |
| NISCH | 75.1387159 | 0.418836691 | 0.2215073 | 1.89084825 | 0.0586446 | 0.25314921 |
| SMIM4 | 38.5390397 | 0.131615372 | 0.44237291 | 0.29752132 | 0.76606853 | 0.91564296 |
| PBRM1 | 188.894848 | -0.431038021 | 0.34568405 | -1.2469132 | 0.21242931 | 0.52134789 |
| GNL3 | 22.6838716 | -0.994654001 | 0.39604013 | -2.511498 | 0.012022 | 0.09375173 |
| GLT8D1 | 17.0243701 | 0.404185359 | 0.40519915 | 0.99749805 | 0.31852282 | 0.63093651 |
| SPCS1 | 32.674874 | -0.845774124 | 0.44194605 | -1.9137497 | 0.05565214 | 0.24569157 |
| NEK4 | 47.9678308 | 0.512322608 | 0.28959137 | 1.76912249 | 0.07687344 | 0.29667546 |
| SFMBT1 | 39.879568 | 0.621125049 | 0.37418219 | 1.65995353 | 0.0969238 | 0.33969491 |
| RFT1 | 11.3979607 | -0.528178556 | 0.49977839 | -1.0568255 | 0.29059122 | 0.60519825 |
| TKT | 55.5639279 | -1.605458951 | 0.46624469 | -3.4433828 | 0.00057449 | 0.0095823 |
| DCP1A | 138.362708 | 0.328694663 | 0.25942914 | 1.26699209 | 0.20515812 | 0.51160585 |
| CACNA1D | 894.428844 | -0.327872529 | 0.41533297 | -0.7894209 | 0.42986604 | 0.72387049 |
| CHDH | 35.5750252 | -1.178695873 | 0.44758663 | -2.6334475 | 0.00845229 | 0.0729557 |
| SELENOK | 16.4615133 | -0.737917212 | 0.45935732 | -1.6064122 | 0.10818337 | 0.36175274 |
| CACNA2D3 | 1265.81226 | 0.007892857 | 0.38027905 | 0.02075544 | 0.98344075 | 0.99742652 |
| ERC2 | 96.7676789 | -0.228950167 | 0.41249887 | -0.5550322 | 0.57887262 | 0.82340325 |
| CCDC66 | 142.456529 | 0.629678097 | 0.25354319 | 2.48351416 | 0.01300932 | 0.09877758 |
| FAM208A | 153.437138 | 0.220671842 | 0.24693729 | 0.89363515 | 0.37151713 | 0.67534262 |
| ARHGEF3 | 964.730308 | -1.394484673 | 0.35680452 | -3.9082595 | 9.30E-05 | 0.00228705 |
| SPATA12 | 17.863779 | -2.765058717 | 0.55699012 | -4.9642868 | 6.90E-07 | 4.26E-05 |
| IL17RD | 15.9330749 | -1.563156412 | 0.53391598 | -2.9277198 | 0.00341458 | 0.03844452 |
| APPL1 | 71.5229676 | 0.196808736 | 0.27716786 | 0.71007055 | 0.47766039 | 0.7631764 |
| ARF4 | 66.4741195 | -0.426738796 | 0.43271018 | -0.9862 | 0.32403496 | 0.63646555 |
| DENND6A | 91.6449552 | 0.038452095 | 0.24790341 | 0.15510918 | 0.87673525 | 0.95879545 |
| DENND6A-AS | 17.5210228 | -0.464005063 | 0.55900499 | -0.8300553 | 0.40650751 | 0.70556143 |
| SLMAP | 341.1461 | -0.171348027 | 0.29491981 | -0.5809987 | 0.56124133 | 0.81384822 |
| FLNB | 301.088495 | -0.135519943 | 0.47752461 | -0.2837968 | 0.77656613 | 0.9196152 |
| ABHD6 | 13.4091996 | 0.434876996 | 0.43097585 | 1.00905188 | 0.31294975 | 0.62764971 |
| PXK | 83.9797997 | 0.938250781 | 0.33355368 | 2.81289292 | 0.0049098 | 0.04949078 |
| PDHB | 23.3869591 | 0.006597488 | 0.37340149 | 0.01766862 | 0.98590322 | 0.99746215 |
| KCTD6 | 26.6871258 | -0.217216654 | 0.47141977 | -0.4607712 | 0.64496276 | 0.85774252 |
| FAM107A | 19.6924593 | -0.112640361 | 0.74175094 | -0.1518574 | 0.87929942 | 0.96035935 |
| FAM3D-AS1 | 13.4577058 | 2.149229118 | 0.58303525 | 3.68627644 | 0.00022756 | 0.00469416 |
| FHIT | 2499.2198 | 0.422704389 | 0.17755237 | 2.3807308 | 0.01727833 | 0.11889534 |
| PTPRG | 2422.37625 | -0.520501004 | 0.28317376 | -1.8380976 | 0.06604803 | 0.27225103 |
| THOC7 | 28.0694464 | 0.187546196 | 0.3197336 | 0.58657017 | 0.55749242 | 0.81230403 |
| ATXN7 | 32.2139951 | 0.148033112 | 0.29261901 | 0.50589028 | 0.61293368 | 0.84174925 |
| PSMD6-AS2 | 16.4574137 | -0.219303136 | 0.39081495 | -0.5611432 | 0.57469995 | 0.82146824 |
| PSMD6 | 22.0695468 | 0.371139838 | 0.45929058 | 0.80807196 | 0.41904916 | 0.71744663 |
| PRICKLE2 | 393.895157 | 0.437927675 | 0.51585213 | 0.84894033 | 0.3959145 | 0.69664216 |
| ADAMTS9 | 677.358883 | -1.056510466 | 0.52422226 | -2.0153865 | 0.04386417 | 0.21382372 |
| ADAMTS9-AS | 254.014042 | 1.636700075 | 0.60653997 | 2.69842081 | 0.00696693 | 0.06337528 |
| ADAMTS9-AS | 1572.95676 | -0.158915648 | 0.38532411 | -0.4124207 | 0.68003108 | 0.87237893 |
| MAGI1 | 1946.76498 | 0.045672797 | 0.2401925 | 0.19015081 | 0.84919096 | 0.95003258 |
| MAGI1-AS1 | 26.68443 | -0.440867997 | 0.77682508 | -0.5675254 | 0.57035725 | 0.81878537 |
| SLC25A26 | 112.35712 | -0.3270928 | 0.48524184 | -0.674082 | 0.50025917 | 0.77558432 |
| LRIG1 | 26.6305814 | -0.256412616 | 0.46772599 | -0.5482112 | 0.58354691 | 0.8271498 |
| SUCLG2 | 292.547443 | 0.461483061 | 0.22602288 | 2.0417537 | 0.04117597 | 0.20607876 |
| SUCLG2-AS1 | 186.819331 | 0.86309119 | 0.69498592 | 1.24188299 | 0.21427974 | 0.52288517 |
| FAM19A1 | 72.8553284 | 1.184994897 | 0.70389374 | 1.68348548 | 0.09228115 | 0.33118916 |
| TMF1 | 95.5699852 | -0.381250284 | 0.21670008 | -1.7593453 | 0.07851887 | 0.29970982 |
| UBA3 | 65.0414076 | -0.007698082 | 0.24013499 | -0.0320573 | 0.97442635 | 0.99357756 |
| ARL6IP5 | 38.817216 | -0.463896902 | 0.29382397 | -1.5788259 | 0.11437599 | 0.37262114 |
| FRMD4B | 130.336263 | 0.311688531 | 0.25743844 | 1.21073034 | 0.22599877 | 0.53714314 |
| MITF | 701.64415 | 1.911669938 | 0.41146465 | 4.64601258 | 3.38E-06 | 0.00015621 |
| FOXP1 | 820.096987 | -0.156604429 | 0.3796776 | -0.4124669 | 0.67999727 | 0.87237893 |
| EIF4E3 | 39.7965843 | -0.617258406 | 0.34185043 | -1.8056388 | 0.07097477 | 0.28450937 |
| RYBP | 184.251586 | -0.093690825 | 0.33571903 | -0.2790751 | 0.78018719 | 0.9206566 |
| SHQ1 | 54.3321866 | -0.301468363 | 0.28364374 | -1.0628416 | 0.2878538 | 0.6024252 |
| PPP4R2 | 119.175085 | 0.106864611 | 0.26164031 | 0.40844092 | 0.68294999 | 0.87314216 |
| PDZRN3 | 311.026322 | 1.361674419 | 0.54215507 | 2.51159585 | 0.01201866 | 0.09375173 |
| CNTN3 | 20.5201748 | -1.285447275 | 0.85812026 | -1.4979803 | 0.13413836 | 0.40819388 |
| ZNF717 | 68.3389754 | 0.450679755 | 0.35951276 | 1.25358487 | 0.20999293 | 0.51870552 |
| ROBO1 | 55.8172752 | -1.188462014 | 0.66299007 | -1.7925789 | 0.07304028 | 0.28899744 |
| GBE1 | 779.741066 | -0.685436205 | 0.50501931 | -1.3572475 | 0.17470258 | 0.47119666 |
| CHMP2B | 37.4826572 | -0.380332523 | 0.33524476 | -1.1344921 | 0.25658819 | 0.56935259 |
| CGGBP1 | 82.1638449 | -0.41015904 | 0.34421871 | -1.1915652 | 0.23343177 | 0.54590364 |
| ZNF654 | 135.298389 | -0.150467883 | 0.21375262 | -0.7039347 | 0.48147345 | 0.76608505 |
| C3orf38 | 26.7419369 | -0.108309168 | 0.36457748 | -0.2970813 | 0.76640441 | 0.91564296 |
| ARL13B | 45.4500212 | -0.073954726 | 0.30036374 | -0.2462172 | 0.80551408 | 0.93308942 |
| NSUN3 | 48.706733 | 0.579981979 | 0.28150262 | 2.06030759 | 0.03936914 | 0.20050645 |
| MTRNR2L12 | 97.3161765 | -0.873669674 | 0.58964588 | -1.4816854 | 0.13842403 | 0.41527892 |
| ARL6 | 30.3615466 | 0.281631143 | 0.56352917 | 0.4997632 | 0.61724182 | 0.84238331 |
| CRYBG3 | 232.067441 | -1.334337451 | 0.34097558 | -3.9132933 | 9.10E-05 | 0.00225159 |
| RIOX2 | 58.8259212 | -0.762323996 | 0.47576526 | -1.6023112 | 0.10908682 | 0.36312941 |
| CLDND1 | 31.9622872 | -0.230817374 | 0.33753625 | -0.6838299 | 0.49408259 | 0.7738663 |
| ST3GAL6 | 125.961614 | -1.397969204 | 0.78243956 | -1.7866801 | 0.0739892 | 0.29120952 |
| DCBLD2 | 73.8784008 | 0.13848151 | 0.31790174 | 0.43561105 | 0.66311895 | 0.86556259 |
| COL8A1 | 45.325338 | 2.629840859 | 0.83463931 | 3.15087107 | 0.00162784 | 0.02180146 |
| CMSS1 | 153.988208 | -0.041615586 | 0.29421476 | -0.1414463 | 0.88751739 | 0.96279373 |
| FILIP1L | 43.5076164 | -0.908392385 | 0.68536142 | -1.325421 | 0.18503157 | 0.48500871 |
| TBC1D23 | 109.111738 | -0.107359337 | 0.19880635 | -0.5400196 | 0.58918348 | 0.83046815 |
| NIT2 | 13.2370996 | 0.497130985 | 0.5763927 | 0.8624866 | 0.3884198 | 0.69192328 |
| TOMM70 | 48.7891369 | 0.787600622 | 0.29950111 | 2.62970857 | 0.00854581 | 0.0733037 |
| TFG | 103.438893 | 0.106378082 | 0.27048138 | 0.39329171 | 0.69410404 | 0.8793242 |
| SENP7 | 146.692687 | 0.541122719 | 0.26895251 | 2.01196379 | 0.04422375 | 0.21487352 |
| TRMT10C | 14.0423553 | -0.729378645 | 0.46117347 | -1.5815711 | 0.11374751 | 0.37174264 |
| PCNP | 100.558022 | -0.152150263 | 0.29864125 | -0.509475 | 0.61041929 | 0.84119857 |
| ZBTB11 | 42.9166905 | -0.025619127 | 0.37260413 | -0.068757 | 0.94518308 | 0.98552865 |
| RPL24 | 47.6172973 | -0.492367617 | 0.37724327 | -1.3051727 | 0.19183405 | 0.49284602 |
| CEP97 | 25.1754101 | 0.991272732 | 0.34880802 | 2.84188626 | 0.00448475 | 0.04635898 |
| NFKBIZ | 151.692356 | -3.06702341 | 0.61134065 | -5.0168812 | 5.25E-07 | 3.33E-05 |
| ALCAM | 61.2396944 | -2.182258444 | 0.57541612 | -3.7924875 | 0.00014915 | 0.00334287 |
| CBLB | 471.759775 | 1.590651587 | 0.24676262 | 6.44607995 | 1.15E-10 | 2.73E-08 |
| LINC00882 | 56.5062812 | 0.77875469 | 0.69560592 | 1.1195343 | 0.26291226 | 0.57700771 |
| BBX | 451.64842 | -0.241587003 | 0.29136844 | -0.8291461 | 0.40702174 | 0.70602034 |

| | | | | | | |
|---|---|---|---|---|---|---|
| CD47 | 123.194428 | -1.045715812 | 0.43201115 | -2.420576 | 0.01549594 | 0.11071582 |
| IFT57 | 52.6077997 | 0.427869389 | 0.37678057 | 1.13559304 | 0.25612692 | 0.56895381 |
| DZIP3 | 62.375567 | 0.900760985 | 0.57176962 | 1.57539148 | 0.11516611 | 0.37388886 |
| AC078980.1 | 474.372592 | 4.367195404 | 0.55617942 | 7.85213419 | 4.09E-15 | 2.38E-12 |
| NECTIN3 | 46.0050955 | -0.307210887 | 0.29411257 | -1.0445351 | 0.29623791 | 0.61093809 |
| PHLDB2 | 468.382576 | 0.16998635 | 0.22147649 | 0.76751419 | 0.44277586 | 0.73499059 |
| ABHD10 | 18.5909184 | 0.674192288 | 0.52693338 | 1.27946401 | 0.20073371 | 0.50462148 |
| C3orf52 | 131.50721 | 1.091500949 | 0.77941893 | 1.40040343 | 0.16139254 | 0.45135695 |
| AC128688.2 | 23.3803269 | 1.661027639 | 0.73112805 | 2.27186968 | 0.02309438 | 0.14345905 |
| SLC9C1 | 68.6538985 | 0.955017583 | 0.81855136 | 1.16671676 | 0.24332477 | 0.55555126 |
| AC112487.1 | 27.5093077 | 1.184930894 | 0.59519909 | 1.99081436 | 0.0465013 | 0.22155244 |
| ATG3 | 29.8731884 | 0.33934257 | 0.33966462 | 0.99905185 | 0.31776957 | 0.63093651 |
| SLC35A5 | 18.6220875 | -0.251810869 | 0.45540431 | -0.5529391 | 0.58030508 | 0.82443464 |
| GTPBP8 | 22.3491299 | -0.37501744 | 0.4654986 | -0.8056253 | 0.42045895 | 0.71867263 |
| NEPRO | 49.5754332 | -0.06268962 | 0.28633578 | -0.2189374 | 0.82669879 | 0.9422064 |
| CFAP44 | 60.8306677 | -0.285941494 | 0.50069126 | -0.5710934 | 0.5679363 | 0.81681908 |
| SPICE1 | 54.6007693 | -0.151574558 | 0.35260614 | -0.4298693 | 0.66729071 | 0.86777458 |
| SIDT1 | 37.7297027 | 1.275841271 | 0.73631276 | 1.73274366 | 0.08314124 | 0.31045612 |
| USF3 | 60.3886158 | -0.192135203 | 0.2350277 | -0.8175002 | 0.41364261 | 0.71143975 |
| NAA50 | 126.288533 | -0.247069088 | 0.28285025 | -0.8734979 | 0.38239177 | 0.68644992 |
| ATP6V1A | 299.996399 | -0.060216328 | 0.24627725 | -0.2445063 | 0.80683876 | 0.93325451 |
| GRAMD1C | 221.708182 | -0.048680248 | 0.57678293 | -0.0843996 | 0.93273873 | 0.97959629 |
| ZDHHC23 | 23.7012413 | 0.167359168 | 0.37391414 | 0.44758717 | 0.65445117 | 0.86217501 |
| CCDC191 | 61.8670873 | -0.035189508 | 0.67512729 | -0.0521228 | 0.95843087 | 0.98911021 |
| QTRT2 | 37.9648685 | -0.652538893 | 0.43495973 | -1.5002283 | 0.13355528 | 0.40681783 |
| ZBTB20 | 1832.68118 | 0.353603981 | 0.20494783 | 1.72533655 | 0.08446684 | 0.31327319 |
| ZBTB20-AS5 | 34.0118187 | -0.679612767 | 0.69280834 | -0.9809535 | 0.32661567 | 0.63830738 |
| LSAMP | 119.226161 | -0.629805445 | 1.40617951 | -0.4478841 | 0.65423684 | 0.86217501 |
| B4GALT4 | 20.6295813 | 0.219287507 | 0.33635148 | 0.6519594 | 0.51442737 | 0.7835482 |
| TMEM39A | 32.695499 | 0.083300217 | 0.3308447 | 0.25178042 | 0.80121079 | 0.93020663 |
| POGLUT1 | 21.7371437 | 0.661499768 | 0.37348561 | 1.77115195 | 0.07653544 | 0.29596141 |
| TIMMDC1 | 46.3743864 | 0.300256982 | 0.30912001 | 0.9713282 | 0.33138487 | 0.64324815 |
| COX17 | 20.2941898 | -0.028198225 | 0.41016686 | -0.0687482 | 0.94519007 | 0.98552865 |
| MAATS1 | 46.1308363 | 0.47722076 | 0.32895342 | 1.45072444 | 0.14685661 | 0.4297124 |
| GSK3B | 345.337948 | -0.142954902 | 0.25612664 | -0.5581415 | 0.57674778 | 0.82212008 |
| AC092910.3 | 11.521932 | -0.03868958 | 0.4609224 | -0.0839395 | 0.93310456 | 0.97987014 |
| LRRC58 | 32.6412834 | -0.242528917 | 0.38101064 | -0.6365411 | 0.52442383 | 0.79185875 |
| FSTL1 | 20.528956 | -1.76467913 | 0.6550021 | -2.694158 | 0.00705667 | 0.0640666 |
| NDUFB4 | 36.9477993 | 0.0872632 | 0.37683799 | 0.23156688 | 0.81687443 | 0.93753535 |
| HGD | 66.233213 | -1.22596734 | 0.75906239 | -1.6151075 | 0.10628742 | 0.35841733 |
| RABL3 | 30.2600673 | 0.012904995 | 0.34997328 | 0.03687423 | 0.97058529 | 0.99175202 |
| GOLGB1 | 232.077925 | -0.512199818 | 0.26330813 | -1.9452488 | 0.05174504 | 0.23500769 |
| IQCB1 | 105.600734 | -0.171148352 | 0.27761111 | -0.616504 | 0.53756196 | 0.80078458 |
| EAF2 | 14.593732 | -0.435509364 | 0.51096636 | -0.8523249 | 0.39403378 | 0.69624829 |
| SLC15A2 | 16.4671696 | 0.815879889 | 0.5930874 | 1.37564867 | 0.16893043 | 0.46258659 |
| ILDR1 | 23.1881296 | 0.483630305 | 0.50877452 | 0.95057886 | 0.3418182 | 0.65136173 |
| CASR | 290.791906 | 1.929285808 | 0.34638096 | 5.56983786 | 2.55E-08 | 2.33E-06 |
| CCDC58 | 18.1564852 | 0.799763619 | 0.45420087 | 1.76081482 | 0.07826975 | 0.29933846 |
| FAM162A | 82.8952241 | 0.61188994 | 0.45542886 | 1.34354668 | 0.17909502 | 0.47750284 |
| AC083798.2 | 35.4624274 | 0.496645802 | 0.53464846 | 0.92892029 | 0.35293039 | 0.65972795 |
| KPNA1 | 116.195953 | -0.218647296 | 0.39074457 | -0.5595658 | 0.57577564 | 0.82147339 |
| PARP9 | 53.4561485 | -0.983853588 | 0.75441962 | -1.3041198 | 0.19219273 | 0.49325764 |
| DTX3L | 19.7639558 | -1.916824084 | 0.60837162 | -3.1507454 | 0.00162854 | 0.02180146 |
| PARP14 | 171.222444 | 0.033526359 | 0.70031568 | 0.04787321 | 0.96181729 | 0.99029318 |
| HSPBAP1 | 81.7385387 | -1.39701097 | 0.5649736 | -2.4727013 | 0.01340962 | 0.10067724 |
| DIRC2 | 145.704983 | -1.107210661 | 0.4785164 | -2.3138406 | 0.02067647 | 0.13388014 |
| PDIA5 | 26.7089749 | 0.361787667 | 0.38525142 | 0.93909497 | 0.34768199 | 0.6553345 |
| SEC22A | 118.675699 | 0.25765257 | 0.22231773 | 1.15893848 | 0.24648126 | 0.55930671 |
| ADCY5 | 85.5264704 | 0.397304467 | 0.49756032 | 0.79850512 | 0.42457742 | 0.72151771 |
| HACD2 | 66.6587843 | 0.071775121 | 0.35894598 | 0.19996079 | 0.84151125 | 0.94776435 |
| MYLK-AS1 | 17.8321589 | 0.291289554 | 0.4314942 | 0.67507178 | 0.49963016 | 0.77558432 |
| MYLK | 49.9839096 | -0.638599852 | 0.45535368 | -1.402426 | 0.16078806 | 0.45020658 |
| CCDC14 | 83.2118985 | -0.082941989 | 0.32076177 | -0.2585782 | 0.79596074 | 0.92698936 |
| KALRN | 74.3679783 | -1.746217599 | 0.82220939 | -2.1238113 | 0.03368592 | 0.18264262 |
| UMPS | 24.5025714 | 0.21570216 | 0.29773753 | 0.72447085 | 0.46877672 | 0.75516139 |
| ITGB5 | 165.034017 | 1.103680405 | 0.2890923 | 3.81774406 | 0.00013468 | 0.00307025 |
| SLC12A8 | 32.6498802 | 1.013355352 | 0.53329991 | 1.90016035 | 0.05741208 | 0.24970229 |
| ZNF148 | 315.908544 | 0.120377333 | 0.18990931 | 0.63386746 | 0.52616732 | 0.79346338 |
| SNX4 | 134.663694 | 0.493542344 | 0.23395315 | 2.10957769 | 0.03489475 | 0.18591921 |
| OSBPL11 | 34.1711433 | 0.505120013 | 0.33221374 | 1.52046694 | 0.12839366 | 0.39719393 |
| SLC41A3 | 15.2389213 | -0.386189602 | 0.50144702 | -0.7701504 | 0.4412107 | 0.73330634 |
| ALDH1L1 | 133.769438 | -0.350344488 | 0.68916854 | -0.5083582 | 0.61120215 | 0.84169124 |
| ZXDC | 162.274449 | -0.232639755 | 0.28773131 | -0.8085312 | 0.41878483 | 0.71712575 |
| TXNRD3NB | 14.4724811 | 0.415141054 | 0.52885204 | 0.78498526 | 0.43246221 | 0.72599466 |
| TXNRD3 | 23.9861129 | 1.128157199 | 0.37119329 | 3.03927151 | 0.00237151 | 0.02882915 |
| CHCHD6 | 46.0959137 | 0.429978841 | 0.36189335 | 1.18813689 | 0.23477948 | 0.54672425 |
| TPRA1 | 20.4733813 | -0.59316581 | 0.67960432 | -0.8728105 | 0.38276635 | 0.68685787 |
| MGLL | 330.157328 | -5.013841696 | 0.58835507 | -8.5217957 | 1.57E-17 | 1.46E-14 |
| SEC61A1 | 53.4611867 | -0.418097627 | 0.38762554 | -1.0786122 | 0.28076066 | 0.5950926 |
| RUVBL1 | 32.5510947 | -0.408178421 | 0.32530829 | -1.2547434 | 0.20957193 | 0.51817785 |
| EEFSEC | 69.3291835 | 0.080966148 | 0.21102169 | 0.38368639 | 0.7012109 | 0.88421489 |
| RPN1 | 40.6768553 | -0.194079939 | 0.32286925 | -0.60111 | 0.54776672 | 0.80618229 |
| RAB7A | 219.424925 | -0.506074768 | 0.40116554 | -1.2615111 | 0.20712478 | 0.51472054 |
| ACAD9 | 38.3546557 | 0.340047698 | 0.303542 | 1.12026572 | 0.26260054 | 0.57679328 |
| CNBP | 76.2021175 | -0.9800914 | 0.39399075 | -2.4876 | 0.01286083 | 0.09812959 |
| COPG1 | 33.9191864 | -0.161723271 | 0.31399008 | -0.5150585 | 0.60651212 | 0.83829217 |
| HMCES | 24.2175334 | 0.025079126 | 0.37372203 | 0.06710636 | 0.94649703 | 0.98574808 |
| MBD4 | 30.3234291 | -0.175069924 | 0.29963451 | -0.5842782 | 0.55903313 | 0.81279041 |
| IFT122 | 31.3698473 | -0.172237635 | 0.54161003 | -0.3180104 | 0.75047703 | 0.9088775 |
| TMCC1 | 493.807251 | 0.050280029 | 0.24480924 | 0.20538452 | 0.83727171 | 0.94706413 |
| PIK3R4 | 54.5578007 | 0.20316825 | 0.32763496 | 0.62010552 | 0.53518832 | 0.79931057 |
| ATP2C1 | 183.752022 | 0.502920023 | 0.21731317 | 2.31426392 | 0.02065325 | 0.13383178 |
| NEK11 | 112.614487 | -0.55866357 | 0.45445468 | -1.2293053 | 0.21895735 | 0.52913753 |
| NUDT16 | 15.3631401 | 0.108045555 | 0.50886342 | 0.21232722 | 0.83185176 | 0.94495441 |
| MRPL3 | 47.5505076 | 0.131800234 | 0.27163091 | 0.48521811 | 0.62752164 | 0.84783129 |
| ACPP | 51.1240663 | -2.621958475 | 0.75638904 | -3.4664152 | 0.00052745 | 0.00895797 |
| DNAJC13 | 112.343296 | 0.096243938 | 0.26096431 | 0.36880115 | 0.71227594 | 0.88727601 |
| UBA5 | 79.7962464 | 0.491786679 | 0.32286821 | 1.52318088 | 0.12771346 | 0.39587776 |
| TMEM108 | 62.8644922 | 1.434311616 | 0.5156958 | 2.78131336 | 0.00541395 | 0.05285825 |
| CDV3 | 64.0222781 | -0.490408955 | 0.46654572 | -1.0511488 | 0.29319027 | 0.6084367 |
| TOPBP1 | 35.2712535 | -0.050080309 | 0.33243068 | -0.1506489 | 0.8802527 | 0.96097414 |
| SRPRB | 12.8085692 | 0.445626957 | 0.45443984 | 0.98060716 | 0.3267865 | 0.6384266 |
| RYK | 165.035168 | -0.064462303 | 0.22498779 | -0.2865147 | 0.77448396 | 0.91864687 |
| AMOTL2 | 33.9537217 | 1.060152474 | 0.53654765 | 1.97587759 | 0.04816864 | 0.22485402 |
| CEP63 | 56.5169771 | 0.376128114 | 0.315275 | 1.19301598 | 0.23286311 | 0.54525758 |
| EPHB1 | 59.8594935 | -0.78606246 | 0.74272168 | -1.058354 | 0.28989409 | 0.60488981 |
| PPP2R3A | 934.099257 | -0.303075839 | 0.22489854 | -1.3476114 | 0.17778339 | 0.47521885 |
| MSL2 | 72.1958641 | 0.462211894 | 0.28245831 | 1.6363898 | 0.10175803 | 0.34880722 |
| PCCB | 38.7354509 | 0.54382732 | 0.3795632 | 1.43277147 | 0.15192316 | 0.43628364 |
| STAG1 | 575.904219 | -0.439632468 | 0.34159254 | -1.2870084 | 0.19809136 | 0.50081448 |
| SLC35G2 | 24.7000844 | -0.844141701 | 0.61632134 | -1.3696454 | 0.17079762 | 0.46658085 |
| NCK1-DT | 15.5522665 | -0.61391763 | 0.54010218 | -1.1366694 | 0.25567651 | 0.56843403 |
| NCK1 | 73.5314774 | -0.117128389 | 0.26768735 | -0.4375567 | 0.66170768 | 0.86492722 |

| | | | | | | |
|---|---|---|---|---|---|---|
| DZIP1L | 28.3324386 | 0.980564938 | 0.43275994 | 2.26584038 | 0.02346116 | 0.14515716 |
| ARMC8 | 133.606636 | 0.413591819 | 0.24678459 | 1.67592237 | 0.0937534 | 0.33428555 |
| MRAS | 57.6108819 | -1.952324019 | 0.74849828 | -2.6083213 | 0.00909875 | 0.07698434 |
| CEP70 | 161.120148 | 0.1744414 | 0.44166093 | 0.39496679 | 0.6928674 | 0.87849579 |
| FAIM | 13.7407983 | 0.258047729 | 0.53950635 | 0.47830341 | 0.63243427 | 0.85152594 |
| PIK3CB | 171.648509 | 0.543570185 | 0.27737261 | 1.95971108 | 0.05002957 | 0.23092856 |
| COPB2 | 68.8200371 | -0.032419448 | 0.25368776 | -0.1277927 | 0.89831301 | 0.96607503 |
| AC097103.2 | 86.415223 | 0.289117212 | 0.55991147 | 0.51636237 | 0.60560134 | 0.83802715 |
| NMNAT3 | 74.6487782 | 1.056829644 | 0.33811242 | 3.12567532 | 0.00177397 | 0.0231336 |
| SLC25A36 | 233.43992 | 0.757467461 | 0.33672161 | 2.24953626 | 0.0244784 | 0.14869351 |
| SPSB4 | 14.5503211 | 2.636508987 | 0.62219667 | 4.23742058 | 2.26E-05 | 0.00075834 |
| PXYLP1 | 13.2776836 | 0.490207967 | 0.51181622 | 0.95778122 | 0.33817309 | 0.64897646 |
| ZBTB38 | 190.130116 | -1.570940963 | 0.4596779 | -3.417482 | 0.00063203 | 0.01043022 |
| RASA2 | 237.524423 | -0.858185071 | 0.32880556 | -2.6100078 | 0.00905402 | 0.07674514 |
| RNF7 | 27.903556 | -0.715690685 | 0.40430895 | -1.7701579 | 0.07670084 | 0.29625461 |
| ATP1B3 | 189.134596 | 0.161709737 | 0.51964627 | 0.31119195 | 0.7556547 | 0.91065328 |
| TFDP2 | 361.880395 | 0.650871161 | 0.34413771 | 1.89131021 | 0.05858295 | 0.25300018 |
| GK5 | 82.1616419 | 0.80946388 | 0.30120688 | 2.68740167 | 0.00720103 | 0.06480927 |
| XRN1 | 258.706389 | 0.527793709 | 0.34982651 | 1.50872989 | 0.13136782 | 0.40278645 |
| ATR | 181.935536 | -0.275528468 | 0.37325472 | -0.7381781 | 0.46040621 | 0.74872989 |
| PLS1 | 41.9168053 | -0.135270485 | 0.43187626 | -0.3132158 | 0.75411669 | 0.91026551 |
| TRPC1 | 83.1336728 | 0.110471847 | 0.40595835 | 0.27212607 | 0.78552509 | 0.92291279 |
| U2SURP | 147.726211 | 0.233862337 | 0.17441154 | 1.34086504 | 0.17996428 | 0.47860436 |
| AC107021.1 | 21.0425131 | -1.282440918 | 0.64776904 | -1.9797811 | 0.04772813 | 0.2237391 |
| PLOD2 | 217.535092 | -0.750204055 | 0.44801844 | -1.6744937 | 0.09403361 | 0.3344198 |
| PLSCR4 | 27.0271885 | 0.070946311 | 0.37476654 | 0.18930802 | 0.84985141 | 0.95003258 |
| PLSCR2 | 46.2795457 | -0.253060671 | 0.36106983 | -0.7008635 | 0.4833882 | 0.76664595 |
| PLSCR1 | 170.342316 | -0.645485218 | 0.42838347 | -1.506793 | 0.13186373 | 0.40351308 |
| HLTF | 43.3656623 | 1.349741106 | 0.38907227 | 3.46912701 | 0.00052215 | 0.00888422 |
| HPS3 | 64.3835781 | -0.334539463 | 0.33550465 | -0.9971232 | 0.31870472 | 0.63093651 |
| WWTR1 | 667.117544 | -1.062045426 | 0.41800699 | -2.5407361 | 0.01106194 | 0.08807986 |
| COMMD2 | 26.3715921 | -0.460653423 | 0.42065906 | -1.0950755 | 0.27348357 | 0.58894058 |
| ANKUB1 | 55.3544446 | -0.90078373 | 0.56177428 | -1.603462 | 0.10883269 | 0.36293132 |
| RNF13 | 140.25467 | -0.003106987 | 0.24433785 | -0.0127159 | 0.98985442 | 0.99842088 |
| AC117386.2 | 93.1916642 | 1.520896103 | 0.66729638 | 2.27919131 | 0.0226557 | 0.14162184 |
| TSC22D2 | 686.675393 | 1.389003186 | 0.19853429 | 6.99628861 | 2.63E-12 | 8.75E-10 |
| SERP1 | 89.0192506 | 0.083816691 | 0.3683329 | 0.2275569 | 0.81999073 | 0.93868552 |
| EIF2A | 45.4387331 | 0.026646143 | 0.27394103 | 0.09726963 | 0.92251227 | 0.97611757 |
| SELENOT | 48.9608155 | 0.495681367 | 0.38806623 | 1.27731125 | 0.20149238 | 0.50571062 |
| SIAH2 | 23.9481306 | 0.226019361 | 0.59442171 | 0.38023403 | 0.7037717 | 0.88543615 |
| MED12L | 95.6823181 | -1.099039486 | 0.4263021 | -2.5780767 | 0.0099352 | 0.08190607 |
| MBNL1 | 602.359702 | 0.621253909 | 0.25637721 | 2.42320254 | 0.01538435 | 0.11034127 |
| DHX36 | 138.82119 | 0.156556453 | 0.29853229 | 0.52442049 | 0.59998611 | 0.83601151 |
| PLCH1 | 34.853532 | -1.314983194 | 0.78484977 | -1.6754585 | 0.09384431 | 0.3344198 |
| GMPS | 59.3747941 | 0.319244786 | 0.27584435 | 1.15733667 | 0.24713484 | 0.55983606 |
| SSR3 | 39.0647096 | -0.068457105 | 0.37074264 | -0.1846486 | 0.85350467 | 0.95137449 |
| TIPARP | 28.2970883 | -0.720116265 | 0.47399805 | -1.5192389 | 0.12870238 | 0.39737957 |
| LINC00886 | 59.9392434 | 0.714645752 | 0.7673166 | 0.93135708 | 0.35166888 | 0.65871783 |
| LEKR1 | 32.6411028 | 1.05183617 | 0.49628207 | 2.11943215 | 0.03405396 | 0.18374951 |
| CCNL1 | 212.235656 | -0.700635334 | 0.43065462 | -1.6269077 | 0.1037567 | 0.35243258 |
| VEPH1 | 62.3301223 | 1.808136173 | 0.62671823 | 2.88508628 | 0.00391306 | 0.04212483 |
| RSRC1 | 148.243907 | 0.338839363 | 0.23206501 | 1.46010536 | 0.14426112 | 0.42566161 |
| MLF1 | 31.2930707 | 0.484351526 | 0.53374584 | 0.90745724 | 0.36416505 | 0.6696613 |
| GFM1 | 39.8644732 | 0.070984453 | 0.37779617 | 0.18789088 | 0.8509622 | 0.95042076 |
| MFSD1 | 24.3339219 | 0.033711777 | 0.46769566 | 0.07208058 | 0.94253778 | 0.98488741 |
| IQCJ-SCHIP1 | 173.568286 | -1.194895735 | 0.79025753 | -1.5120333 | 0.13052538 | 0.40125903 |
| SMC4 | 19.1065008 | -0.478280617 | 0.53361244 | -0.8963071 | 0.37008877 | 0.67436147 |
| KPNA4 | 153.757108 | -0.658136445 | 0.47564752 | -1.3836642 | 0.1664613 | 0.45864087 |
| PPM1L | 223.056284 | 0.330992555 | 0.47354479 | 0.69896779 | 0.48457216 | 0.76759625 |
| B3GALNT1 | 15.7670773 | 0.117275539 | 0.56426517 | 0.20783764 | 0.83535574 | 0.94654164 |
| NMD3 | 109.292459 | 0.602231155 | 0.44492586 | 1.35355394 | 0.17587873 | 0.47294748 |
| LINC01322 | 14.6415771 | 1.11379342 | 0.53670458 | 2.07524486 | 0.03796385 | 0.19654159 |
| PDCD10 | 69.2684231 | -0.137470528 | 0.23018747 | -0.5972112 | 0.5503664 | 0.80796749 |
| SERPINI1 | 13.98431 | 0.087872197 | 0.50665457 | 0.17343611 | 0.86230864 | 0.95458251 |
| GOLIM4 | 55.8010519 | 0.665353018 | 0.4197651 | 1.58506035 | 0.11295262 | 0.37070407 |
| MECOM | 6209.8852 | 0.127039399 | 0.29907452 | 0.42477507 | 0.67100065 | 0.86777458 |
| AC074033.1 | 25.4218703 | -0.808004444 | 0.8491987 | -0.9514904 | 0.34135547 | 0.65087901 |
| MYNN | 22.6217743 | 0.428139711 | 0.45595707 | 0.93899127 | 0.34773522 | 0.6553345 |
| SEC62 | 97.6948389 | 0.085555162 | 0.26978239 | 0.31712656 | 0.75114757 | 0.9089517 |
| GPR160 | 61.5679913 | 0.129718235 | 0.35328163 | 0.36718081 | 0.71348416 | 0.88813689 |
| PHC3 | 261.151918 | 0.158399134 | 0.18067308 | 0.87671685 | 0.38064045 | 0.68449211 |
| PRKCI | 136.997272 | -0.362188045 | 0.28148597 | -1.2867002 | 0.19819882 | 0.50095033 |
| SKIL | 60.6592064 | -1.000769329 | 0.4143275 | -2.4154065 | 0.01571766 | 0.11161571 |
| PLD1 | 79.8796027 | 0.102774624 | 0.32716275 | 0.31413914 | 0.75341537 | 0.91007188 |
| FNDC3B | 1804.69679 | -0.565871791 | 0.29640494 | -1.9091173 | 0.05624697 | 0.24714736 |
| TNFSF10 | 155.201674 | -0.956476328 | 0.66566459 | -1.4368743 | 0.15075372 | 0.43488337 |
| NCEH1 | 154.118336 | -1.225871066 | 0.56029585 | -2.1878996 | 0.02867692 | 0.16498593 |
| ECT2 | 35.1499791 | -1.229757041 | 0.53948215 | -2.2795138 | 0.02263654 | 0.14162184 |
| NLGN1 | 385.233277 | -2.861968824 | 0.76892721 | -3.7220283 | 0.00019763 | 0.00417844 |
| NAALADL2 | 1338.57864 | -0.279837586 | 0.46535248 | -0.6013454 | 0.54760994 | 0.80618229 |
| TBL1XR1 | 615.787598 | -1.299027356 | 0.44905315 | -2.8928142 | 0.00381807 | 0.04152772 |
| ZMAT3 | 32.6011965 | -0.247946163 | 0.5400278 | -0.4591359 | 0.64613656 | 0.85869117 |
| PIK3CA | 116.62417 | -0.072971799 | 0.20047815 | -0.3639888 | 0.71586638 | 0.88904248 |
| ZNF639 | 17.6577951 | -0.114092873 | 0.41595568 | -0.2742909 | 0.78386105 | 0.92232893 |
| MFN1 | 100.714691 | -0.038089921 | 0.31955196 | -0.1191979 | 0.90511858 | 0.96914626 |
| GNB4 | 43.6518356 | 0.759013252 | 0.35273212 | 2.15181213 | 0.03141215 | 0.17538139 |
| ACTL6A | 22.5740353 | -0.616076532 | 0.38658118 | -1.5936537 | 0.11101362 | 0.36676814 |
| MRPL47 | 18.5102926 | -0.166391126 | 0.52191512 | -0.3188088 | 0.74987152 | 0.90857724 |
| NDUFB5 | 39.8551661 | 0.545511111 | 0.4142479 | 1.31687117 | 0.18788181 | 0.48810533 |
| TTC14 | 155.7793 | 0.605394889 | 0.21732174 | 2.78570793 | 0.0053411 | 0.0524767 |
| FXR1 | 159.967837 | 0.484817034 | 0.19007073 | 2.55071909 | 0.01075009 | 0.08656845 |
| DNAJC19 | 23.8548647 | 0.621253372 | 0.43183913 | 1.43862223 | 0.15025758 | 0.4348261 |
| ATP11B | 144.477351 | -0.157177907 | 0.2544719 | -0.6176631 | 0.53679743 | 0.80030369 |
| DCUN1D1 | 113.757556 | 1.312083696 | 0.22515382 | 5.82749919 | 5.63E-09 | 6.56E-07 |
| MCCC1 | 137.350225 | 0.288114006 | 0.26632527 | 1.08181249 | 0.27933585 | 0.59383074 |
| KLHL24 | 186.200471 | -0.167577212 | 0.31723946 | -0.5282357 | 0.59733574 | 0.8342658 |
| YEATS2 | 91.2835089 | -0.198352156 | 0.46603433 | -0.425617 | 0.67038692 | 0.86777458 |
| ABCC5 | 313.323897 | 0.087324911 | 0.50805536 | 0.1718807 | 0.86353132 | 0.95458251 |
| EIF2B5 | 22.6585278 | 0.100814128 | 0.40255522 | 0.25043552 | 0.80225056 | 0.93060267 |
| DVL3 | 46.2147204 | 0.665956798 | 0.48046829 | 1.38605775 | 0.16572927 | 0.45785473 |
| AP2M1 | 45.121933 | -0.102164588 | 0.50308726 | -0.2030753 | 0.83907619 | 0.94741614 |
| PSMD2 | 25.4371935 | -0.471183861 | 0.45461897 | -1.0364369 | 0.29999838 | 0.61374288 |
| EIF4G1 | 52.1722766 | -0.923979481 | 0.36573532 | -2.5263611 | 0.0115251 | 0.0909137 |
| FAM131A | 15.9048569 | 0.095365928 | 0.50572137 | 0.18857405 | 0.85042667 | 0.950309 |
| POLR2H | 47.0720672 | 0.116586046 | 0.35627562 | 0.32723554 | 0.74348974 | 0.90463949 |
| VPS8 | 177.951341 | 0.08684382 | 0.22735488 | 0.38197473 | 0.70248011 | 0.88464675 |
| EHHADH | 27.9419659 | 0.396858014 | 0.51398028 | 0.77212693 | 0.44003925 | 0.73240378 |
| MAP3K13 | 241.250193 | 0.366836067 | 0.22249435 | 1.64874327 | 0.09920024 | 0.343767 |
| TMEM41A | 34.7788851 | -0.681392476 | 0.42879879 | -1.5890727 | 0.11204397 | 0.3686302 |
| LIPH | 15.4641373 | 0.3078316 | 0.65570205 | 0.46946872 | 0.63873464 | 0.8538689 |
| SENP2 | 84.3477566 | -0.038649515 | 0.33500992 | -0.1153683 | 0.90815323 | 0.97050094 |
| IGF2BP2 | 396.599662 | -0.295500509 | 0.30526033 | -0.9680279 | 0.33303044 | 0.64492217 |
| TRA2B | 100.684958 | 0.275033286 | 0.2227549 | 1.23469017 | 0.21694583 | 0.52649739 |

| | | | | | | |
|---|---|---|---|---|---|---|
| ETV5 | 33.3226154 | -3.528901765 | 0.71828578 | -4.9129495 | 8.97E-07 | 5.36E-05 |
| TBCCD1 | 14.6050051 | 0.355794871 | 0.42617644 | 0.83485346 | 0.40380022 | 0.70290417 |
| AC068631.1 | 104.894237 | -0.022871726 | 0.6066123 | -0.037704 | 0.96992367 | 0.99156233 |
| KNG1 | 347.082851 | 2.265567346 | 0.62893171 | 3.60224699 | 0.00031548 | 0.00607752 |
| EIF4A2 | 105.926741 | 0.032094497 | 0.36566857 | 0.08776936 | 0.93005999 | 0.97876742 |
| ST6GAL1 | 239.954184 | -1.889287284 | 0.45662702 | -4.1374847 | 3.51E-05 | 0.00107696 |
| BCL6 | 167.635758 | -3.901583234 | 0.77164336 | -5.0562001 | 4.28E-07 | 2.79E-05 |
| LPP | 1455.52481 | -0.536216117 | 0.36895647 | -1.4533317 | 0.14613169 | 0.42860392 |
| TPRG1 | 26.7347309 | -2.948119301 | 0.81361965 | -3.6234613 | 0.00029069 | 0.00567744 |
| P3H2 | 130.742294 | 1.157800945 | 0.54263706 | 2.13365622 | 0.03287093 | 0.1802874 |
| CCDC50 | 176.824597 | 0.15980271 | 0.3156876 | 0.50620522 | 0.61271259 | 0.84174925 |
| MB21D2 | 51.1740949 | 0.881677515 | 0.54448934 | 1.61927414 | 0.10538829 | 0.35680481 |
| OPA1 | 153.308409 | 0.522345423 | 0.20114441 | 2.59686765 | 0.00940782 | 0.07888124 |
| HES1 | 27.3140677 | -0.729300271 | 0.52438166 | -1.3907814 | 0.16429172 | 0.45563826 |
| ATP13A3 | 145.461857 | -0.98679339 | 0.51466832 | -1.9173385 | 0.05519494 | 0.24446818 |
| LSG1 | 33.0095653 | -0.13449803 | 0.3874193 | -0.347164 | 0.72846812 | 0.89654656 |
| XXYLT1 | 26.2717753 | 0.155807413 | 0.38408884 | 0.40565462 | 0.68499639 | 0.87408051 |
| ACAP2 | 237.781628 | 0.245956403 | 0.22606559 | 1.08798692 | 0.27660088 | 0.59165557 |
| PPP1R2 | 45.2506968 | -0.175854426 | 0.29618262 | -0.5937365 | 0.55268838 | 0.80962553 |
| MUC20-OT1 | 576.669295 | 0.77928292 | 0.42979325 | 1.81315764 | 0.06980751 | 0.28168095 |
| TNK2 | 17.8560371 | 0.428052244 | 0.47924037 | 0.89318904 | 0.37175595 | 0.67534262 |
| TFRC | 103.166828 | 0.212925655 | 0.42744508 | 0.4981357 | 0.6183884 | 0.84283824 |
| PCYT1A | 26.9584708 | 0.090676381 | 0.31008576 | 0.29242355 | 0.7699628 | 0.91655198 |
| UBXN7 | 73.6393249 | -0.169008897 | 0.24788559 | -0.681802 | 0.49536414 | 0.77475164 |
| RNF168 | 58.8798194 | -0.482316424 | 0.34689164 | -1.3903951 | 0.16440894 | 0.45563826 |
| PIGX | 20.154882 | -0.473889685 | 0.39031908 | -1.2141084 | 0.22470632 | 0.53584699 |
| PAK2 | 111.583405 | -0.877057418 | 0.36757594 | -2.3860578 | 0.01703008 | 0.11780307 |
| SENP5 | 177.7802 | -0.476469519 | 0.38638323 | -1.2331527 | 0.21751881 | 0.52693657 |
| NCBP2 | 24.7690927 | -0.437230912 | 0.39973654 | -1.0937977 | 0.27404371 | 0.58894058 |
| DLG1 | 375.024608 | -0.266658076 | 0.22358898 | -1.1926262 | 0.23301581 | 0.54547813 |
| BDH1 | 66.9018568 | 0.857173625 | 0.42241597 | 2.02921691 | 0.0424362 | 0.21013019 |
| RUBCN | 41.8326517 | -0.650277301 | 0.35017392 | -1.8570124 | 0.06330939 | 0.26477174 |
| FYTTD1 | 41.9655931 | -0.118877511 | 0.35148308 | -0.3382169 | 0.73519975 | 0.90007689 |
| LRCH3 | 184.639392 | -0.277846792 | 0.32932036 | -0.8436976 | 0.39883841 | 0.69914822 |
| IQCG | 25.4175644 | -0.210948342 | 0.43623013 | -0.4835712 | 0.62869019 | 0.84817317 |
| RPL35A | 55.5344132 | -0.469780814 | 0.39054032 | -1.2028996 | 0.22901516 | 0.54176037 |
| LMLN | 42.1625066 | 0.567361536 | 0.43872443 | 1.29320708 | 0.19593945 | 0.49845696 |
| ZNF595 | 45.7594705 | 0.430909618 | 0.2569708 | 1.67688163 | 0.09356563 | 0.33374367 |
| ZNF718 | 112.514717 | 0.114098649 | 0.22210729 | 0.51370961 | 0.60745503 | 0.83917773 |
| ZNF141 | 94.0705612 | 0.278048563 | 0.26382034 | 1.05393147 | 0.29191429 | 0.60677688 |
| ZNF721 | 251.084183 | 0.203937043 | 0.22320245 | 0.9136864 | 0.36088165 | 0.66644099 |
| PIGG | 44.3014366 | -0.208809418 | 0.3040776 | -0.6866978 | 0.49227319 | 0.77284018 |
| ATP5ME | 25.581679 | 0.509441709 | 0.47060801 | 1.08251815 | 0.27902236 | 0.59380484 |
| PCGF3 | 69.7170963 | 0.116654141 | 0.23322574 | 0.50017695 | 0.61695049 | 0.84223226 |
| AC139887.2 | 124.035481 | 0.19844684 | 0.36383458 | 0.5454315 | 0.58545678 | 0.82884893 |
| GAK | 106.612442 | -0.780245614 | 0.50386696 | -1.5485151 | 0.12149832 | 0.38493045 |
| TMEM175 | 12.12438 | 0.520533156 | 0.52126087 | 0.99860393 | 0.31798659 | 0.63093651 |
| IDUA | 14.1456227 | 0.502763396 | 0.45564314 | 1.10341483 | 0.26984706 | 0.58625952 |
| RNF212 | 24.2539525 | 1.538229312 | 0.49371791 | 3.11560361 | 0.00183569 | 0.0236504 |
| AC092535.3 | 87.9955971 | 0.80583473 | 0.39176603 | 2.05692856 | 0.0396931 | 0.20168853 |
| CTBP1-AS | 14.6544328 | 0.362240286 | 0.5255313 | 0.68928394 | 0.49064461 | 0.77203035 |
| CTBP1 | 47.4310659 | -0.029408761 | 0.36188692 | -0.0812651 | 0.93523116 | 0.98121924 |
| CTBP1-DT | 12.1894585 | 0.212427696 | 0.47696639 | 0.44537247 | 0.65605061 | 0.8628884 |
| MAEA | 76.1911993 | 0.012215009 | 0.35892417 | 0.03403228 | 0.97285141 | 0.99287068 |
| UVSSA | 42.4032348 | 0.57656282 | 0.28783102 | 2.00312954 | 0.04516339 | 0.21784967 |
| SLBP | 44.7307561 | -0.336635925 | 0.49948946 | -0.67396 | 0.50033673 | 0.77558432 |
| TMEM129 | 16.5529456 | 0.730212422 | 0.47618338 | 1.53346893 | 0.12516036 | 0.39121528 |
| LETM1 | 33.541305 | 0.202096723 | 0.42772763 | 0.47248929 | 0.63657759 | 0.8538689 |
| NSD2 | 155.128583 | -0.464413827 | 0.313087 | -1.483338 | 0.13798464 | 0.41448738 |
| NELFA | 56.7831015 | 0.633368346 | 0.45507265 | 1.39179613 | 0.16398414 | 0.45563826 |
| NAT8L | 19.6266428 | 2.093983577 | 0.90186561 | 2.32183548 | 0.0202418 | 0.13216702 |
| POLN | 16.361608 | 0.222971399 | 0.35296558 | 0.63170861 | 0.5275773 | 0.79403853 |
| MXD4 | 25.7112459 | 0.210487434 | 0.50820809 | 0.41417569 | 0.67874545 | 0.87147102 |
| ZFYVE28 | 56.9584266 | -0.41050133 | 0.24732582 | -1.6597593 | 0.09696289 | 0.33969491 |
| RNF4 | 61.3593442 | -0.372451627 | 0.40608131 | -0.9171849 | 0.35904578 | 0.66542294 |
| FAM193A | 154.891262 | -0.112789344 | 0.3351289 | -0.3365551 | 0.7364523 | 0.90136552 |
| TNIP2 | 17.8435044 | -1.120590962 | 0.5050628 | -2.2187161 | 0.02650604 | 0.15651828 |
| ADD1 | 123.044754 | -0.01446748 | 0.26404709 | -0.0547913 | 0.95630474 | 0.98853497 |
| MFSD10 | 28.0546681 | -0.993799934 | 0.54268832 | -1.8312536 | 0.0670627 | 0.27565003 |
| NOP14 | 19.7581963 | -0.478020251 | 0.42508221 | -1.124536 | 0.26078571 | 0.57429522 |
| GRK4 | 28.3426798 | -0.005653686 | 0.54709341 | -0.010334 | 0.99175477 | 0.9988777 |
| HTT | 133.555544 | 0.243187645 | 0.20267333 | 1.19989956 | 0.23017835 | 0.54292511 |
| RGS12 | 66.6913195 | -0.999512208 | 0.55375573 | -1.8049695 | 0.07107946 | 0.28456198 |
| LRPAP1 | 42.4286858 | -0.576174652 | 0.46352649 | -1.2430242 | 0.21385891 | 0.52235998 |
| TMEM128 | 18.3131191 | 0.13593296 | 0.3641245 | 0.37331451 | 0.70891438 | 0.88614996 |
| ZBTB49 | 13.3906623 | 0.17447067 | 0.45573403 | 0.38283441 | 0.70184255 | 0.88424653 |
| STX18 | 213.891102 | 0.804406434 | 0.25440557 | 3.16190578 | 0.0015674 | 0.02124195 |
| STX18-AS1 | 92.6693569 | 1.052311967 | 0.36761196 | 2.86256186 | 0.00420231 | 0.04426547 |
| STK32B | 239.28782 | 1.072301554 | 0.45097344 | 2.37774881 | 0.01741869 | 0.11959635 |
| EVC2 | 58.7222614 | -0.908912727 | 0.32785216 | -2.772325 | 0.00556574 | 0.05366598 |
| EVC | 75.7344234 | -0.645080015 | 0.37433691 | -1.7232605 | 0.08484143 | 0.31408459 |
| CRMP1 | 19.5261743 | -1.231673307 | 0.69839547 | -1.7635757 | 0.07780344 | 0.29864393 |
| MAN2B2 | 17.9879761 | -0.419993896 | 0.50089594 | -0.8384853 | 0.40175818 | 0.70114138 |
| MRFAP1 | 23.8978264 | -0.697621725 | 0.43189502 | -1.6152576 | 0.10625492 | 0.35841733 |
| KIAA0232 | 198.29514 | 0.465060826 | 0.25518177 | 1.82246884 | 0.06838388 | 0.27855451 |
| TBC1D14 | 178.330326 | -0.145639685 | 0.38841852 | -0.3749556 | 0.70769351 | 0.88614996 |
| TADA2B | 12.7128954 | -0.473973199 | 0.63241161 | -0.7494695 | 0.45357429 | 0.74469566 |
| GRPEL1 | 18.6054404 | -0.735693252 | 0.5549583 | -1.325673 | 0.18494802 | 0.48494245 |
| SORCS2 | 26.2233204 | -2.343740933 | 1.02652048 | -2.2831896 | 0.0224192 | 0.14095524 |
| SH3TC1 | 43.5748981 | -0.349256731 | 0.31936494 | -1.0935976 | 0.2741315 | 0.58894058 |
| ACOX3 | 34.0978533 | -0.300356898 | 0.36312744 | -0.8271391 | 0.40815823 | 0.70684758 |
| TRMT44 | 26.0251785 | -0.215433542 | 0.31285761 | -0.6885993 | 0.49107544 | 0.77215443 |
| WDR1 | 71.727846 | -0.685389843 | 0.33664537 | -2.0359402 | 0.04175636 | 0.20827251 |
| HS3ST1 | 66.9700656 | -2.998582957 | 0.50609531 | -5.9249372 | 3.12E-09 | 4.22E-07 |
| AC005699.1 | 54.5273886 | -1.853590813 | 0.74652363 | -2.4829634 | 0.01302945 | 0.09881314 |
| RAB28 | 231.031308 | 0.316075984 | 0.3072111 | 1.02885601 | 0.30354734 | 0.61784922 |
| BOD1L1 | 147.227209 | -0.552359186 | 0.44967643 | -1.2283481 | 0.21931633 | 0.52935683 |
| LINC01182 | 211.846352 | 0.774924105 | 0.44803927 | 1.72958968 | 0.08370361 | 0.31173327 |
| AC092546.1 | 34.0963599 | 1.788418454 | 0.51440305 | 3.47668712 | 0.00050765 | 0.00871699 |
| CPEB2 | 450.022757 | 0.828732051 | 0.37517926 | 2.2088962 | 0.02718186 | 0.15990137 |
| AC098829.1 | 139.705908 | 0.508628126 | 0.57534075 | 0.88404676 | 0.37667097 | 0.68015091 |
| CC2D2A | 67.2313135 | 0.015626193 | 0.58092295 | 0.02689891 | 0.97854037 | 0.9951912 |
| FBXL5 | 163.118649 | 0.145477298 | 0.18799088 | 0.77385296 | 0.43901775 | 0.73148704 |
| FAM200B | 38.0859179 | 0.169923166 | 0.37737333 | 0.45027868 | 0.65250951 | 0.86179129 |
| PROM1 | 49.1218785 | -2.705216857 | 0.97644861 | -2.7704652 | 0.00559763 | 0.05377622 |
| TAPT1 | 94.5658937 | 0.273047798 | 0.25527595 | 1.0696182 | 0.2847912 | 0.59819624 |
| TAPT1-AS1 | 242.988447 | 1.171547483 | 0.59086899 | 1.98275336 | 0.04739499 | 0.22326436 |
| AC097515.1 | 52.6428974 | 1.613171336 | 0.6676294 | 2.41626766 | 0.01568053 | 0.11143694 |
| LDB2 | 16.0917473 | 0.770418917 | 0.5813241 | 1.32528294 | 0.18507733 | 0.48500871 |
| QDPR | 13.7182892 | -0.037068157 | 0.7145668 | -0.051875 | 0.95862829 | 0.9891274 |
| LAP3 | 31.9097378 | 0.141158931 | 0.59809917 | 0.23601258 | 0.8134229 | 0.93633777 |
| MED28 | 50.7310541 | -0.113081661 | 0.30632572 | -0.369155 | 0.71201223 | 0.88721964 |
| DCAF16 | 42.9717214 | 0.149378967 | 0.28873343 | 0.51735944 | 0.60490527 | 0.83778698 |

| | | | | | | |
|---|---|---|---|---|---|---|
| LCORL | 204.030216 | -0.08951159 | 0.25298473 | -0.3538221 | 0.72347221 | 0.89263661 |
| SLIT2 | 61.7748262 | 0.740199974 | 0.58985976 | 1.2548745 | 0.20952432 | 0.51817785 |
| PACRGL | 60.7823387 | 0.16719614 | 0.34064893 | 0.49081657 | 0.62355619 | 0.84597807 |
| KCNIP4 | 6748.14251 | -1.753491345 | 0.48755137 | -3.5965264 | 0.00032249 | 0.00618712 |
| ADGRA3 | 115.458815 | 0.190271867 | 0.41373449 | 0.45988883 | 0.64559602 | 0.85821747 |
| PPARGC1A | 802.494464 | 0.621494579 | 0.34655757 | 1.79333721 | 0.07291902 | 0.28870359 |
| DHX15 | 195.128794 | 0.107572197 | 0.30400379 | 0.3538515 | 0.72345018 | 0.89263661 |
| CCDC149 | 26.9785273 | -0.0930633 | 0.35814122 | -0.2598508 | 0.79497883 | 0.9268423 |
| SEPSECS | 26.7687318 | 0.152595636 | 0.33919471 | 0.44987622 | 0.65279969 | 0.86183588 |
| ZCCHC4 | 21.2073322 | 0.240759049 | 0.42042224 | 0.57266011 | 0.56687485 | 0.81606004 |
| ANAPC4 | 129.376653 | -0.374082127 | 0.4499334 | -0.8314167 | 0.40573829 | 0.70501376 |
| SEL1L3 | 39.590668 | -4.42473881 | 0.88763409 | -4.9848681 | 6.20E-07 | 3.91E-05 |
| SMIM20 | 15.5017811 | -1.338858828 | 0.78759416 | -1.6999349 | 0.08914317 | 0.32396747 |
| RBPJ | 180.649319 | 0.136815954 | 0.20700706 | 0.66092408 | 0.508661 | 0.7804435 |
| TBC1D19 | 100.545759 | 0.39063549 | 0.49424985 | 0.79036036 | 0.42931735 | 0.7236272 |
| STIM2 | 70.3914629 | -0.940541661 | 0.49084497 | -1.9161685 | 0.05534365 | 0.24479327 |
| ARAP2 | 82.0054939 | -0.646096703 | 0.49699254 | -1.3000129 | 0.19359656 | 0.49536068 |
| C4orf19 | 572.996788 | -0.386470296 | 0.23963635 | -1.6127365 | 0.10680176 | 0.35932624 |
| RELL1 | 163.594331 | 0.000870048 | 0.45432975 | 0.00191501 | 0.99847204 | 0.99922221 |
| PGM2 | 22.6834698 | -0.24206468 | 0.3658934 | -0.6615716 | 0.50824582 | 0.78013108 |
| TBC1D1 | 406.776196 | -0.086218226 | 0.26418847 | -0.3263512 | 0.74415865 | 0.90498699 |
| KLF3 | 72.4221673 | -1.232858827 | 0.3676761 | -3.3531111 | 0.00079909 | 0.01260025 |
| FAM114A1 | 15.9393766 | -0.135296015 | 0.53984343 | -0.2506208 | 0.80210727 | 0.93060267 |
| KLHL5 | 154.386008 | -0.147232718 | 0.398033 | -0.3699008 | 0.71145642 | 0.88696612 |
| WDR19 | 81.9868845 | -0.642920878 | 0.22596723 | -2.8451952 | 0.00443842 | 0.04598205 |
| RFC1 | 65.8702033 | 0.330544417 | 0.25154665 | 1.31404819 | 0.18882999 | 0.48924896 |
| LIAS | 27.8875252 | 0.858566986 | 0.47358591 | 1.81290651 | 0.06984624 | 0.28168095 |
| UGDH | 43.1717425 | -0.773484057 | 0.42603502 | -1.815541 | 0.06944081 | 0.28100153 |
| UGDH-AS1 | 44.5351534 | 0.311955715 | 0.43634853 | 0.71492326 | 0.47465646 | 0.76075482 |
| SMIM14 | 104.518468 | -0.270368542 | 0.34164831 | -0.7913651 | 0.42873097 | 0.7236272 |
| UBE2K | 373.594389 | -0.265909632 | 0.24739347 | -1.074845 | 0.28244413 | 0.59642935 |
| PDS5A | 235.600117 | -0.114074212 | 0.20971685 | -0.5439439 | 0.58648005 | 0.82948466 |
| N4BP2 | 59.7260903 | 0.428180106 | 0.27778834 | 1.54138976 | 0.12322196 | 0.38814916 |
| RBM47 | 882.966167 | -0.724397104 | 0.49821718 | -1.4539786 | 0.14595225 | 0.42860392 |
| NSUN7 | 51.4940885 | -0.464329732 | 0.44523338 | -1.0428906 | 0.29699894 | 0.61093809 |
| APBB2 | 1433.83321 | 1.558815012 | 0.27612774 | 5.64526771 | 1.65E-08 | 1.62E-06 |
| UCHL1 | 45.5654963 | -2.045963197 | 0.61713212 | -3.3152758 | 0.00091553 | 0.01392041 |
| LIMCH1 | 971.391199 | 0.120057081 | 0.44852473 | 0.26767104 | 0.78895255 | 0.92402584 |
| TMEM33 | 49.9539933 | -0.503817712 | 0.47602338 | -1.0583886 | 0.28987832 | 0.60488981 |
| SLC30A9 | 214.362146 | 0.252725949 | 0.21998198 | 1.14884841 | 0.25061849 | 0.56339619 |
| AC024022.1 | 93.7889908 | 0.920436476 | 0.69810573 | 1.31847718 | 0.18734396 | 0.48725107 |
| SHISA3 | 40.3835262 | -0.103203918 | 0.72067731 | -0.1432041 | 0.88612902 | 0.96249798 |
| ATP8A1 | 189.865746 | 0.841127126 | 0.37768676 | 2.22704952 | 0.02594396 | 0.15456754 |
| KCTD8 | 262.681032 | 0.705066126 | 0.44502958 | 1.58431293 | 0.11312252 | 0.37079048 |
| GUF1 | 36.0862188 | 0.890355069 | 0.40496491 | 2.19859804 | 0.02790652 | 0.16193939 |
| GNPDA2 | 30.5191031 | 0.385457989 | 0.42716165 | 0.90237031 | 0.36686019 | 0.67113782 |
| GABRA2 | 51.6945199 | 0.713660929 | 0.67541758 | 1.05662179 | 0.29068423 | 0.60525676 |
| ATP10D | 79.7727522 | 0.659103507 | 0.32506389 | 2.02761218 | 0.04259985 | 0.21039469 |
| AC107068.2 | 15.3171809 | -0.815399474 | 0.78068583 | -1.0444656 | 0.29627004 | 0.61093809 |
| NFXL1 | 57.8609141 | 0.323343241 | 0.3648361 | 0.88626987 | 0.37547212 | 0.67888822 |
| NIPAL1 | 28.3022429 | 0.116892523 | 0.45217341 | 0.2585126 | 0.79601133 | 0.92698936 |
| TXK | 64.73365 | -1.432986959 | 0.48997611 | -2.9246058 | 0.00344893 | 0.03860482 |
| TEC | 74.0992811 | -1.205245455 | 0.36205326 | -3.3289175 | 0.00087184 | 0.01348102 |
| SLAIN2 | 97.6399804 | 0.524652636 | 0.2252195 | 2.32951695 | 0.0198317 | 0.13040249 |
| FRYL | 330.027057 | -0.131731513 | 0.18305265 | -0.7196373 | 0.47174834 | 0.75793048 |
| OCIAD1 | 164.071481 | 0.178128028 | 0.19273294 | 0.924222 | 0.35537074 | 0.66163487 |
| OCIAD2 | 23.9575629 | -1.180035549 | 0.52190685 | -2.261008 | 0.02375876 | 0.14631883 |
| CWH43 | 28.355788 | 0.817524147 | 0.65464657 | 1.24880231 | 0.21173739 | 0.52063276 |
| DCUN1D4 | 123.613614 | 0.061171506 | 0.24345123 | 0.25126801 | 0.8016069 | 0.9304348 |
| SGCB | 13.1219279 | -0.21491122 | 0.41335322 | -0.5199215 | 0.6031183 | 0.8372477 |
| USP46 | 30.2944708 | 0.750560056 | 0.34928947 | 2.14881957 | 0.03164871 | 0.17592332 |
| SCFD2 | 127.838115 | -0.140328043 | 0.32948703 | -0.4258985 | 0.67018178 | 0.86777458 |
| AC058822.1 | 80.3724698 | -0.134932364 | 0.57850983 | -0.2332413 | 0.81557406 | 0.93717111 |
| LNX1 | 227.408167 | 0.492118123 | 0.210767 | 2.3348917 | 0.01954908 | 0.12964126 |
| CHIC2 | 108.312114 | 0.023811105 | 0.23062304 | 0.10324686 | 0.91776705 | 0.97481433 |
| TMEM165 | 176.12529 | 0.01442433 | 0.30260718 | 0.04766685 | 0.96198176 | 0.99029318 |
| CLOCK | 99.8378676 | 0.059653139 | 0.27349618 | 0.21811324 | 0.82734088 | 0.9425007 |
| EXOC1 | 134.981436 | 0.287659922 | 0.23531225 | 1.22246046 | 0.22153355 | 0.53195437 |
| CEP135 | 31.3017906 | 0.140761855 | 0.40205106 | 0.3501094 | 0.72625659 | 0.8946514 |
| AASDH | 54.865546 | 0.225987193 | 0.37273884 | 0.60628828 | 0.54432334 | 0.80393962 |
| PPAT | 17.1767006 | -0.524289725 | 0.42510804 | -1.2333093 | 0.21746037 | 0.52693657 |
| PAICS | 31.2463457 | -0.457424124 | 0.29468529 | -1.5522462 | 0.12060332 | 0.38405237 |
| SRP72 | 52.3576967 | -0.012619751 | 0.22867362 | -0.0551867 | 0.9559897 | 0.98853497 |
| REST | 100.973256 | 0.342870832 | 0.29415894 | 1.16559717 | 0.24377735 | 0.55580983 |
| NOA1 | 12.6750892 | -0.432049012 | 0.45401312 | -0.9516223 | 0.34128856 | 0.65087901 |
| POLR2B | 69.7044095 | -0.033966813 | 0.25822805 | -0.131538 | 0.89534969 | 0.96511451 |
| IGFBP7 | 1511.7823 | -0.498333402 | 0.38091707 | -1.3082465 | 0.19078973 | 0.49135864 |
| ADGRL3 | 52.9958814 | 1.387155758 | 0.87288858 | 1.58915558 | 0.11202527 | 0.3686302 |
| AC104806.2 | 18.8315846 | -0.654892154 | 0.59593684 | -1.0989288 | 0.27179912 | 0.58786226 |
| CENPC | 88.0677739 | 0.657486704 | 0.29350229 | 2.24014167 | 0.02508173 | 0.15107366 |
| UBA6 | 140.198736 | -0.395085126 | 0.29636213 | -1.3331161 | 0.1824937 | 0.48134973 |
| UBA6-AS1 | 353.577307 | 0.179220943 | 0.4993196 | 0.35893032 | 0.71964722 | 0.89114993 |
| YTHDC1 | 196.906302 | 0.172817542 | 0.21883499 | 0.78971623 | 0.42969351 | 0.72374389 |
| UGT2B7 | 114.719141 | -1.748993671 | 0.68657796 | -2.5474072 | 0.01085267 | 0.08700799 |
| RUFY3 | 312.316921 | -1.496012042 | 0.50598996 | -2.9566042 | 0.00311047 | 0.03576083 |
| GRSF1 | 47.0931119 | 0.226128302 | 0.30809938 | 0.73394598 | 0.46298166 | 0.75091259 |
| MOB1B | 135.431313 | -0.085744359 | 0.43957548 | -0.1950617 | 0.84534461 | 0.94907846 |
| SLC4A4 | 22.1639343 | 0.173952935 | 0.45959751 | 0.37848973 | 0.70506682 | 0.88596489 |
| ADAMTS3 | 23.3254025 | -2.382485972 | 0.84230426 | -2.8285337 | 0.00467618 | 0.04780777 |
| COX18 | 12.009547 | -0.01849251 | 0.49487831 | -0.0373678 | 0.97019175 | 0.99156233 |
| ANKRD17 | 739.279953 | 0.3058937 | 0.24939891 | 1.22652378 | 0.22000163 | 0.52989521 |
| AC053527.1 | 123.541618 | 0.62328118 | 0.57004989 | 1.09338005 | 0.27422697 | 0.58900997 |
| RASSF6 | 88.2506505 | -1.967462206 | 0.4080139 | -4.8220471 | 1.42E-06 | 7.61E-05 |
| MTHFD2L | 63.3942461 | 0.071533176 | 0.28138845 | 0.25421504 | 0.79932943 | 0.92922871 |
| BTC | 13.1606713 | 0.340404737 | 0.82756815 | 0.41133137 | 0.68082957 | 0.87265005 |
| RCHY1 | 61.0410851 | 0.370914873 | 0.34142681 | 1.08636714 | 0.27731659 | 0.59237111 |
| THAP6 | 36.7580261 | 0.433911362 | 0.3530572 | 1.2290115 | 0.2190675 | 0.52916718 |
| G3BP2 | 55.7989423 | -0.244435701 | 0.34676762 | -0.7048977 | 0.48087389 | 0.76563273 |
| USO1 | 143.5293 | 0.671113446 | 0.25576503 | 2.62394533 | 0.00869177 | 0.0742824 |
| NAAA | 29.3107231 | 0.631736605 | 0.59556316 | 1.06073821 | 0.28880889 | 0.6036436 |
| SDAD1 | 47.7495475 | -0.356326406 | 0.29156142 | -1.2221316 | 0.22165788 | 0.53197892 |
| NUP54 | 77.5965618 | 0.103545504 | 0.30776986 | 0.33643809 | 0.73654053 | 0.90136552 |
| SCARB2 | 89.0347655 | -0.297479636 | 0.38298258 | -0.7767446 | 0.43730949 | 0.73004974 |
| SHROOM3 | 816.59023 | 0.549511157 | 0.52711408 | 1.04249001 | 0.29718456 | 0.61114883 |
| 11-Sep | 186.638842 | -0.157482201 | 0.45406294 | -0.346829 | 0.72871978 | 0.89661959 |
| CCNI | 180.0398 | -0.61564795 | 0.31942345 | -1.9273724 | 0.05393323 | 0.24118632 |
| CCNG2 | 63.1123239 | -1.022463553 | 0.33284803 | -3.071863 | 0.00212727 | 0.02650028 |
| CNOT6L | 309.38086 | 1.31279008 | 0.32640877 | 4.02192031 | 5.77E-05 | 0.00159713 |
| MRPL1 | 111.092745 | 0.780702537 | 0.40980364 | 1.90506492 | 0.05677163 | 0.24816629 |
| FRAS1 | 501.215672 | 0.306635767 | 0.70376524 | 0.43570746 | 0.66304899 | 0.86556259 |
| ANXA3 | 45.1079074 | -0.501886768 | 0.30585697 | -1.6409199 | 0.10081405 | 0.34685987 |
| BMP2K | 91.8791201 | -0.108504443 | 0.27872841 | -0.3892838 | 0.69706625 | 0.88175902 |
| PAQR3 | 14.9613982 | 0.151963563 | 0.40452121 | 0.37566278 | 0.70716761 | 0.88614996 |

| | | | | | | |
|---|---|---|---|---|---|---|
| RASGEF1B | 573.311084 | 1.397627195 | 0.7401257 | 1.88836464 | 0.05897701 | 0.25387889 |
| HNRNPD | 165.934535 | -0.133603301 | 0.21293092 | -0.627449 | 0.53036492 | 0.7958814 |
| HNRNPDL | 135.710039 | 0.144846192 | 0.22976684 | 0.63040511 | 0.52842957 | 0.7943404 |
| ENOPH1 | 31.4499298 | -0.557160602 | 0.31601066 | -1.763107 | 0.07788244 | 0.29864393 |
| TMEM150C | 39.9752422 | 0.533537209 | 0.51450766 | 1.03698594 | 0.29974241 | 0.61356712 |
| SCD5 | 117.591787 | 0.568801438 | 0.39538345 | 1.43860709 | 0.15026188 | 0.4348261 |
| SEC31A | 207.867393 | -0.022365684 | 0.22045814 | -0.1014509 | 0.91919251 | 0.97513464 |
| LIN54 | 36.6135763 | 0.308718104 | 0.33766956 | 0.91426097 | 0.36057973 | 0.66638631 |
| COPS4 | 47.907262 | 0.316316674 | 0.27594857 | 1.14628852 | 0.25167579 | 0.56401021 |
| HELQ | 51.2259041 | -0.64738776 | 0.32269883 | -2.006167 | 0.04483843 | 0.2166184 |
| MRPS18C | 20.3600418 | 0.021273822 | 0.32691739 | 0.065074 | 0.94811508 | 0.98641208 |
| ABRAXAS1 | 18.3607669 | 0.843945178 | 0.63527122 | 1.3284801 | 0.18401955 | 0.48318736 |
| GPAT3 | 254.729792 | 2.987471964 | 0.61577035 | 4.85160087 | 1.22E-06 | 6.88E-05 |
| CDS1 | 60.2447378 | 0.442013869 | 0.31852837 | 1.38767505 | 0.16523602 | 0.45716932 |
| WDFY3 | 405.305833 | -1.40E-05 | 0.14706361 | -9.54E-05 | 0.99992389 | 0.99992389 |
| ARHGAP24 | 973.22839 | 0.379188145 | 0.32058874 | 1.18278685 | 0.23689363 | 0.54986214 |
| MAPK10 | 590.544998 | 0.463202698 | 0.60796427 | 0.76189132 | 0.44612487 | 0.73900835 |
| PTPN13 | 387.390133 | 0.56905552 | 0.3226584 | 1.76364698 | 0.07779143 | 0.29864393 |
| AC093827.5 | 20.4079594 | 0.150148785 | 0.45258647 | 0.33175712 | 0.74007266 | 0.90304252 |
| AFF1 | 697.987571 | -0.064233569 | 0.272376 | -0.2358268 | 0.81356705 | 0.93633777 |
| KLHL8 | 53.501468 | -0.333899543 | 0.47086337 | -0.7091219 | 0.47824882 | 0.76374064 |
| HSD17B11 | 18.6720974 | -0.29274088 | 0.39884666 | -0.7339685 | 0.46296794 | 0.75091259 |
| NUDT9 | 52.812601 | 0.405078306 | 0.33400267 | 1.21279961 | 0.22520644 | 0.53616259 |
| SPARCL1 | 23.2672015 | 0.447199555 | 0.69385515 | 0.64451428 | 0.519242 | 0.78759816 |
| AC093895.1 | 74.979436 | 1.569130912 | 0.79766293 | 1.96716037 | 0.04916472 | 0.22760049 |
| SPP1 | 6572.93451 | -2.126553117 | 0.59519994 | -3.5728383 | 0.00035313 | 0.00667872 |
| PKD2 | 283.487177 | -0.059000153 | 0.22285792 | -0.2647434 | 0.79120718 | 0.92515873 |
| PPM1K | 82.5915255 | 0.095865797 | 0.4771823 | 0.20089973 | 0.84077698 | 0.94770364 |
| HERC6 | 33.2656291 | 0.723238073 | 0.97471293 | 0.74200111 | 0.45808667 | 0.747377 |
| HERC5 | 43.2207239 | 0.110378081 | 0.60866184 | 0.18134549 | 0.8560964 | 0.95219406 |
| PYURF | 14.5236331 | -0.812998507 | 0.48478156 | -1.6770409 | 0.09353448 | 0.33374367 |
| HERC3 | 139.325464 | -0.258428105 | 0.34147378 | -0.7568022 | 0.44916838 | 0.74080155 |
| FAM13A-AS1 | 55.2730549 | -0.50048885 | 0.29068844 | -1.7217363 | 0.08511729 | 0.31443489 |
| FAM13A | 211.89028 | 0.022931995 | 0.58019718 | 0.03952449 | 0.96847223 | 0.99156233 |
| GPRIN3 | 328.651038 | 0.110122479 | 0.52233012 | 0.21082927 | 0.8330205 | 0.94570597 |
| CCSER1 | 3464.41782 | 0.869110499 | 0.37753485 | 2.30206694 | 0.0213314 | 0.1369229 |
| SMARCAD1 | 114.700051 | 0.309643705 | 0.24842883 | 1.2464081 | 0.21261461 | 0.52155186 |
| PDLIM5 | 281.952347 | -0.848923195 | 0.25338069 | -3.3503863 | 0.00080699 | 0.012646 |
| BMPR1B | 37.1037445 | -1.821744569 | 0.39149733 | -4.6532746 | 3.27E-06 | 0.00015155 |
| UNC5C | 298.09196 | 1.117656461 | 0.5612692 | 1.99130197 | 0.0464477 | 0.2214102 |
| STPG2 | 318.01196 | 1.468184772 | 0.66026712 | 2.22362242 | 0.02617386 | 0.15554178 |
| RAP1GDS1 | 158.032981 | 0.478271145 | 0.25314104 | 1.88934656 | 0.05884541 | 0.25354648 |
| TSPAN5 | 53.8008623 | 2.450201508 | 0.57818366 | 4.23775643 | 2.26E-05 | 0.00075834 |
| EIF4E | 89.0780396 | -0.155677828 | 0.38023202 | -0.4094285 | 0.68222522 | 0.87274146 |
| METAP1 | 56.5605304 | 1.133580037 | 0.29080933 | 3.89801816 | 9.70E-05 | 0.00235523 |
| ADH5 | 36.7966645 | 0.628320127 | 0.37549746 | 1.67330059 | 0.09426814 | 0.33471291 |
| LAMTOR3 | 21.7962232 | -0.436464735 | 0.35003055 | -1.2469333 | 0.21242197 | 0.52134789 |
| DNAJB14 | 121.488259 | -0.119276531 | 0.19633626 | -0.6075115 | 0.54351153 | 0.8036317 |
| H2AFZ | 25.5810725 | -1.222831449 | 0.48852362 | -2.5031163 | 0.01231051 | 0.09519681 |
| PPP3CA | 463.105103 | -0.259576682 | 0.3338683 | -0.7774823 | 0.4368743 | 0.72974131 |
| BANK1 | 30.3258389 | 1.312778596 | 0.81525198 | 1.61027342 | 0.10733818 | 0.36052636 |
| AP002075.1 | 16.0600645 | -0.296610714 | 0.81019823 | -0.3660965 | 0.71429309 | 0.88883875 |
| NFKB1 | 152.540958 | 0.042942988 | 0.28562803 | 0.15034585 | 0.88049176 | 0.96097414 |
| MANBA | 81.0589081 | -0.096586491 | 0.2910465 | -0.3318593 | 0.73999549 | 0.90304252 |
| UBE2D3 | 292.695718 | -0.198735306 | 0.24914584 | -0.7976666 | 0.425064 | 0.72151771 |
| CISD2 | 25.7615473 | -0.301432047 | 0.40475247 | -0.7447318 | 0.45643385 | 0.74649873 |
| SLC9B1 | 13.1837827 | 0.933021454 | 0.65646889 | 1.42127291 | 0.15523744 | 0.4407533 |
| BDH2 | 14.183372 | 0.086432628 | 0.43868023 | 0.19702877 | 0.84380502 | 0.94858974 |
| TET2 | 190.315001 | 0.216952972 | 0.20544084 | 1.05603625 | 0.29095165 | 0.60554313 |
| PPA2 | 205.05693 | 0.103128293 | 0.2860052 | 0.36058188 | 0.71841203 | 0.8907358 |
| ARHGEF38 | 392.815995 | -1.535969061 | 0.40939403 | -3.7518111 | 0.00017556 | 0.00378921 |
| INTS12 | 73.124678 | -0.062425002 | 0.3632332 | -0.1718593 | 0.86354815 | 0.95458251 |
| GSTCD | 70.405825 | -0.308059878 | 0.34367345 | -0.8963738 | 0.37005315 | 0.67436147 |
| NPNT | 183.568076 | -0.260317166 | 0.55265668 | -0.4710287 | 0.63762023 | 0.8538689 |
| TBCK | 370.992367 | 0.174253637 | 0.27634425 | 0.63056726 | 0.52832351 | 0.7943404 |
| AIMP1 | 37.8500751 | -0.248710804 | 0.31438227 | -0.7911095 | 0.4288801 | 0.7236272 |
| PAPSS1 | 71.8663622 | 0.342283033 | 0.24447871 | 1.40005253 | 0.16149759 | 0.45151529 |
| SGMS2 | 142.135334 | 0.066588056 | 0.47762082 | 0.13941615 | 0.88912131 | 0.96307703 |
| AC096564.1 | 21.3651447 | -0.46423306 | 0.70778059 | -0.6558997 | 0.51188868 | 0.78271634 |
| HADH | 27.0652501 | 1.141034732 | 0.39824182 | 2.8651806 | 0.00416771 | 0.04410871 |
| RPL34-AS1 | 56.9910684 | 0.651497566 | 0.60590134 | 1.07525355 | 0.28226123 | 0.59642935 |
| RPL34 | 53.2543178 | -0.509041629 | 0.36704275 | -1.3868729 | 0.16548054 | 0.4575743 |
| OSTC | 31.9534753 | -0.437924134 | 0.33846943 | -1.2938366 | 0.19572187 | 0.49824525 |
| SEC24B-AS1 | 13.1354985 | 0.027678024 | 0.70278051 | 0.0393836 | 0.96858456 | 0.99156233 |
| SEC24B | 195.535491 | 0.143812831 | 0.19595201 | 0.73391863 | 0.46299833 | 0.75091259 |
| MCUB | 27.9318265 | -0.614998182 | 0.71610676 | -0.858808 | 0.39044648 | 0.69343295 |
| CASP6 | 30.3191764 | -0.860785213 | 0.4223407 | -2.0381299 | 0.04153694 | 0.20743999 |
| PLA2G12A | 52.4025298 | -0.150457331 | 0.49607953 | -0.3032928 | 0.76166676 | 0.91342791 |
| GAR1 | 18.9549867 | -0.117538948 | 0.37028988 | -0.3174241 | 0.75092179 | 0.9089517 |
| EGF | 2416.16125 | 5.047307806 | 0.53983958 | 9.34964384 | 8.79E-21 | 2.27E-17 |
| ELOVL6 | 69.8933381 | -0.89862612 | 0.45539345 | -1.9732961 | 0.04846184 | 0.22559069 |
| ENPEP | 18.5514044 | 0.509753866 | 0.6067464 | 0.8401432 | 0.4008281 | 0.70092619 |
| FAM241A | 22.6068097 | -0.618459687 | 0.48314026 | -1.2800831 | 0.2005159 | 0.50442784 |
| AP1AR | 40.5613955 | -0.667972178 | 0.66765528 | -1.0004746 | 0.31708086 | 0.63093651 |
| ALPK1 | 170.987692 | -1.0905185 | 0.30004344 | -3.6345354 | 0.00027848 | 0.00547799 |
| LARP7 | 61.7278837 | -0.313126906 | 0.25096182 | -1.2477074 | 0.21213825 | 0.52120606 |
| ANK2 | 171.279084 | 1.480023828 | 0.46708246 | 3.16865643 | 0.00153145 | 0.02087612 |
| CAMK2D | 155.111831 | -0.295706347 | 0.39826491 | -0.7424866 | 0.45779258 | 0.747377 |
| ARSJ | 63.9607352 | -0.096260013 | 0.37394295 | -0.257419 | 0.79685535 | 0.92769126 |
| UGT8 | 118.141447 | -0.189883654 | 0.33144457 | -0.5728972 | 0.56671432 | 0.81606004 |
| SNHG8 | 16.3417093 | 0.085625414 | 0.59565443 | 0.14375015 | 0.88569777 | 0.96249798 |
| METTL14 | 46.1089557 | 0.056210007 | 0.27589689 | 0.20373556 | 0.83856015 | 0.94741614 |
| SEC24D | 47.0588094 | -1.240515826 | 0.37403472 | -3.3165793 | 0.00091127 | 0.01388342 |
| USP53 | 457.611271 | -1.070315509 | 0.46453159 | -2.3040747 | 0.02121845 | 0.13661886 |
| C4orf3 | 31.7226122 | -1.517492009 | 0.48997668 | -3.0970699 | 0.00195444 | 0.02479344 |
| PDE5A | 90.3937151 | -1.23182758 | 0.28155426 | -4.3750983 | 1.21E-05 | 0.00045819 |
| AC108866.1 | 51.1970463 | 0.798514713 | 0.39345679 | 2.02948516 | 0.0424089 | 0.21010658 |
| PRDM5 | 43.4469911 | 0.876292789 | 0.50198204 | 1.74566561 | 0.08086907 | 0.30571131 |
| ANXA5 | 80.8230322 | -1.190465035 | 0.46636374 | -2.5526535 | 0.01069058 | 0.08645183 |
| EXOSC9 | 13.04722 | 0.701338492 | 0.41765928 | 1.67921204 | 0.09311073 | 0.3326491 |
| BBS7 | 18.3170457 | -0.800381611 | 0.3899511 | -2.0525179 | 0.04011936 | 0.20285949 |
| KIAA1109 | 298.613762 | 0.586813888 | 0.23542521 | 2.49257029 | 0.01268222 | 0.09724428 |
| FGF2 | 36.0502815 | -3.930993723 | 1.0279497 | -3.824111 | 0.00013124 | 0.0030141 |
| NUDT6 | 11.0175327 | 0.212106592 | 0.50081317 | 0.42352439 | 0.67191271 | 0.86777458 |
| SPATA5 | 157.542337 | -0.211311683 | 0.26612311 | -0.7940373 | 0.4271737 | 0.72259933 |
| ANKRD50 | 32.4899543 | -0.399320741 | 0.4544774 | -0.8786372 | 0.37959803 | 0.68343046 |
| FAT4 | 24.6982184 | 0.500634398 | 0.53048996 | 0.94372077 | 0.34531236 | 0.65406518 |
| INTU | 65.2560342 | 0.211758479 | 0.58009068 | 0.36504375 | 0.71507875 | 0.88904248 |
| HSPA4L | 106.15236 | 0.613607894 | 0.42161886 | 1.45536158 | 0.14556919 | 0.4281546 |
| MFSD8 | 53.4564252 | 0.422014073 | 0.24299627 | 1.73671006 | 0.08243836 | 0.30885595 |
| ABHD18 | 155.601616 | 0.250953062 | 0.26503725 | 0.94685957 | 0.34371034 | 0.65306073 |
| LARP1B | 141.648098 | -0.18700759 | 0.30856327 | -0.6060591 | 0.54447548 | 0.80403696 |
| PGRMC2 | 58.5112169 | -0.184209814 | 0.43118132 | -0.4272212 | 0.6692182 | 0.86777458 |

| | | | | | | |
|---|---|---|---|---|---|---|
| AC108062.1 | 64.4710981 | 0.859615151 | 0.41543681 | 2.06918386 | 0.03852884 | 0.19803907 |
| AC078850.2 | 13.9421302 | 1.131607216 | 1.09506767 | 1.03336739 | 0.301432 | 0.61540442 |
| JADE1 | 121.92143 | -0.241155062 | 0.49457149 | -0.4876041 | 0.62583033 | 0.8471532 |
| SCLT1 | 113.36584 | 0.351355484 | 0.38799053 | 0.90557748 | 0.36515953 | 0.67012584 |
| C4orf33 | 15.9774807 | 0.414071513 | 0.64801741 | 0.63898208 | 0.52283459 | 0.79024931 |
| AC096711.2 | 73.9947162 | 0.181708365 | 0.42875068 | 0.42380893 | 0.67170517 | 0.86777458 |
| AC109927.1 | 23.3868121 | -0.535285907 | 0.49701567 | -1.0770001 | 0.28148024 | 0.59579794 |
| NOCT | 20.2141531 | -1.487830523 | 0.42006965 | -3.5418663 | 0.00039731 | 0.00733562 |
| ELF2 | 430.436733 | -0.353639232 | 0.19436441 | -1.819465 | 0.06884052 | 0.27931638 |
| NDUFC1 | 32.7285223 | 0.532376982 | 0.43040184 | 1.23693007 | 0.21611305 | 0.52528481 |
| NAA15 | 125.456193 | 0.138836547 | 0.29171069 | 0.47593918 | 0.63411771 | 0.85231563 |
| AC097376.2 | 183.140797 | -0.720584852 | 0.42592104 | -1.6918273 | 0.09067891 | 0.32770935 |
| SETD7 | 18.9830975 | 0.288597165 | 0.46266805 | 0.62376723 | 0.53278047 | 0.79724684 |
| MGST2 | 29.3156412 | 0.22053799 | 0.59104311 | 0.37313351 | 0.70904908 | 0.88614996 |
| MAML3 | 285.159296 | -0.33682434 | 0.35883438 | -0.9386624 | 0.3479041 | 0.65536772 |
| SCOC | 61.4570356 | -0.379803439 | 0.26442316 | -1.4363471 | 0.15090359 | 0.43488337 |
| ELMOD2 | 41.126201 | 0.339496235 | 0.32530236 | 1.04363287 | 0.29665528 | 0.61093809 |
| TBC1D9 | 524.545935 | 0.125678553 | 0.42792526 | 0.29369277 | 0.76899268 | 0.91600052 |
| RNF150 | 421.220855 | 0.580328631 | 0.39431106 | 1.47175337 | 0.14108749 | 0.42122951 |
| ZNF330 | 27.6002587 | -0.044121246 | 0.4177156 | -0.1056251 | 0.91587982 | 0.97373586 |
| IL15 | 23.9849514 | 0.048681567 | 0.57286438 | 0.08497922 | 0.93227791 | 0.9794317 |
| INPP4B | 300.105636 | 0.2334137 | 0.32978019 | 0.70778569 | 0.47907836 | 0.76419703 |
| AC104596.1 | 37.6023281 | 1.249169369 | 1.05532637 | 1.18368062 | 0.23653951 | 0.54945053 |
| USP38 | 33.8546627 | -0.614154353 | 0.42000491 | -1.4622552 | 0.14367129 | 0.42455984 |
| GAB1 | 405.523452 | -0.692422986 | 0.39505003 | -1.7527476 | 0.07964534 | 0.30261336 |
| SMARCA5 | 79.808525 | 0.096924469 | 0.27731567 | 0.34950953 | 0.72670681 | 0.89508776 |
| ANAPC10 | 118.153432 | 0.367801801 | 0.27575853 | 1.33378212 | 0.18227527 | 0.48131823 |
| ABCE1 | 37.7732101 | -0.477324794 | 0.39024535 | -1.2231402 | 0.22127673 | 0.5316115 |
| OTUD4 | 96.7350925 | -0.085334979 | 0.22922159 | -0.3722816 | 0.7096832 | 0.88614996 |
| SMAD1 | 86.1101185 | -0.091729927 | 0.31193977 | -0.2940629 | 0.76870981 | 0.91600052 |
| MMAA | 19.7819343 | -0.258344382 | 0.39442728 | -0.6549861 | 0.51247671 | 0.7830488 |
| ZNF827 | 158.932274 | 0.256846932 | 0.29935981 | 0.85798735 | 0.39089945 | 0.69372012 |
| LSM6 | 17.3208013 | 0.287087978 | 0.436019 | 0.65842998 | 0.51026188 | 0.78158783 |
| SLC10A7 | 96.9754153 | 0.258287451 | 0.28060647 | 0.9204615 | 0.35733164 | 0.66396179 |
| TMEM184C | 43.5749818 | -0.02321039 | 0.33764962 | -0.0687411 | 0.94519574 | 0.98552865 |
| PRMT9 | 44.351475 | -0.031279861 | 0.2584885 | -0.1210106 | 0.9036826 | 0.96849846 |
| ARHGAP10 | 26.2200971 | -2.492725418 | 0.7078828 | -3.5213815 | 0.0004293 | 0.00774243 |
| NR3C2 | 1325.04746 | 0.107909599 | 0.35281973 | 0.30584911 | 0.75971954 | 0.91236458 |
| DCLK2 | 85.6270487 | 0.891773132 | 0.59562529 | 1.49720495 | 0.13433994 | 0.40854063 |
| LRBA | 1591.25331 | 0.722476134 | 0.1275892 | 5.66251815 | 1.49E-08 | 1.48E-06 |
| RPS3A | 38.5347259 | -0.431598239 | 0.41607084 | -1.0373191 | 0.29958716 | 0.61356712 |
| SH3D19 | 296.645635 | -0.253671677 | 0.51289553 | -0.4945874 | 0.62089138 | 0.84452097 |
| FAM160A1 | 415.984527 | -0.143650325 | 0.40349412 | -0.3560159 | 0.72182865 | 0.89217596 |
| GATB | 31.2334005 | 0.538484382 | 0.401817 | 1.34012344 | 0.18020522 | 0.47864023 |
| FBXW7 | 308.022935 | 0.101036895 | 0.31190571 | 0.3239341 | 0.74598793 | 0.90626599 |
| ARFIP1 | 176.855299 | 0.257247713 | 0.19916498 | 1.29163123 | 0.19648488 | 0.49845696 |
| FHDC1 | 18.9938831 | -1.345194104 | 0.71471762 | -1.8821337 | 0.05981787 | 0.25561036 |
| TRIM2 | 420.956641 | 0.075958431 | 0.19807565 | 0.38348192 | 0.70136248 | 0.88421489 |
| TMEM131L | 30.0730233 | 0.654811405 | 0.34569767 | 1.89417362 | 0.05820197 | 0.25182142 |
| PLRG1 | 26.514761 | 0.620570435 | 0.41112118 | 1.50945869 | 0.1311816 | 0.40261267 |
| MAP9 | 27.220781 | -0.214939295 | 0.40457764 | -0.5312684 | 0.59523283 | 0.83416944 |
| GUCY1A1 | 48.5862963 | 0.469110849 | 0.52447608 | 0.89443708 | 0.37108808 | 0.67522591 |
| CTSO | 13.858839 | -0.475021237 | 0.55910037 | -0.8496171 | 0.395538 | 0.69642725 |
| PDGFC | 88.3846463 | -1.73920718 | 0.654478 | -2.657396 | 0.00787469 | 0.06926755 |
| ETFDH | 87.1166011 | 1.310580801 | 0.33284487 | 3.93751243 | 8.23E-05 | 0.00210733 |
| PPID | 33.7726638 | -0.193982074 | 0.30182416 | -0.642699 | 0.52041945 | 0.78804331 |
| FNIP2 | 888.893715 | -1.951830777 | 0.77701966 | -2.5119452 | 0.01200677 | 0.09375173 |
| C4orf45 | 30.1117908 | -1.699070601 | 0.70361091 | -2.4147872 | 0.01574441 | 0.11172058 |
| RAPGEF2 | 378.975308 | 0.062126187 | 0.27062972 | 0.22956158 | 0.81843246 | 0.93816073 |
| NAF1 | 48.1725923 | -0.000760086 | 0.37005748 | -0.002054 | 0.99836117 | 0.99922221 |
| NPY1R | 69.054956 | 2.068948141 | 0.63186089 | 3.27437284 | 0.00105897 | 0.01562313 |
| TMA16 | 23.9581413 | -0.249901972 | 0.33800226 | -0.73935 | 0.4596945 | 0.74802978 |
| TMEM192 | 42.1296133 | -0.226690545 | 0.45811126 | -0.4948373 | 0.62071496 | 0.84449549 |
| KLHL2 | 136.541507 | -1.960930677 | 0.4753712 | -4.1250515 | 3.71E-05 | 0.00111483 |
| MSMO1 | 36.4326905 | -0.661121981 | 0.35993482 | -1.8367825 | 0.06624201 | 0.27268896 |
| DDX60 | 52.5818969 | 0.848604513 | 1.03022299 | 0.82370954 | 0.41010461 | 0.70890164 |
| DDX60L | 43.3480366 | 0.889422982 | 1.46356504 | 0.60770991 | 0.54337989 | 0.80361674 |
| PALLD | 163.717519 | 0.982189935 | 0.41301399 | 2.37810329 | 0.01740195 | 0.11956948 |
| CBR4 | 392.656185 | -0.065288914 | 0.40190297 | -0.1624494 | 0.87095194 | 0.95629149 |
| SH3RF1 | 616.26788 | -0.574140539 | 0.36269466 | -1.5829859 | 0.11342467 | 0.37119783 |
| NEK1 | 183.681511 | 0.503101566 | 0.18771657 | 2.68011278 | 0.00735974 | 0.06598286 |
| CLCN3 | 128.319624 | 0.186820766 | 0.28211889 | 0.6622058 | 0.50783934 | 0.78008138 |
| HPF1 | 20.4402172 | 0.347601979 | 0.36174937 | 0.96089174 | 0.3366066 | 0.64764721 |
| MFAP3L | 36.3403799 | -0.351572097 | 0.28985177 | -1.2129376 | 0.22515369 | 0.53616259 |
| LINC02512 | 12.5061403 | -0.471371475 | 0.96720304 | -0.4873552 | 0.6260066 | 0.8471532 |
| GALNT7 | 91.3871799 | -0.338697745 | 0.34946387 | -0.9691925 | 0.33244917 | 0.644708 |
| HMGB2 | 18.6379921 | 0.766146173 | 0.46625027 | 1.643208 | 0.10033991 | 0.34581665 |
| FBXO8 | 23.6241144 | 0.257719201 | 0.37815466 | 0.68151798 | 0.4955438 | 0.77475164 |
| CEP44 | 48.4703348 | 0.403297499 | 0.36736813 | 1.09780208 | 0.27229092 | 0.58796679 |
| WDR17 | 28.7668248 | 0.658047716 | 0.48911669 | 1.34537979 | 0.17850262 | 0.47621127 |
| SPCS3 | 51.5833883 | -0.007237636 | 0.37830077 | -0.019132 | 0.98473583 | 0.99742652 |
| AC078881.1 | 51.0075955 | 0.544927884 | 0.31611782 | 1.72381262 | 0.08474168 | 0.31408459 |
| LINC01099 | 50.3241111 | 1.447928057 | 0.92909059 | 1.55843582 | 0.11912997 | 0.38057334 |
| DCTD | 14.8743066 | -1.544796992 | 0.52340115 | -2.9514589 | 0.00316277 | 0.03622707 |
| WWC2 | 241.179483 | -0.202021668 | 0.24962926 | -0.8092868 | 0.41835019 | 0.7167764 |
| RWDD4 | 27.0825469 | 0.765166002 | 0.3322671 | 2.3028642 | 0.02128648 | 0.13687944 |
| TRAPPC11 | 78.7779102 | 0.354966532 | 0.21378395 | 1.66039845 | 0.09683432 | 0.33969491 |
| STOX2 | 192.144996 | -1.401616315 | 0.70887261 | -1.9772471 | 0.04801371 | 0.22451344 |
| IRF2 | 69.6965586 | -0.029640965 | 0.22802469 | -0.1299902 | 0.89657422 | 0.96554146 |
| LINC02427 | 12.0745946 | 0.985966781 | 0.76802884 | 1.2837627 | 0.199225 | 0.50272636 |
| CASP3 | 29.4428482 | -0.707173591 | 0.43658167 | -1.6197968 | 0.10527594 | 0.35668344 |
| PRIMPOL | 30.5860172 | 0.499358259 | 0.56993426 | 0.87616818 | 0.38093861 | 0.68476415 |
| ACSL1 | 210.956029 | 0.001397721 | 0.45940128 | 0.00304248 | 0.99757245 | 0.99922221 |
| LINC02437 | 19.6979232 | 1.030894344 | 0.8814942 | 1.16948511 | 0.24220824 | 0.55446835 |
| SLC25A4 | 47.2725762 | 0.160070258 | 0.59516136 | 0.26895271 | 0.78796608 | 0.92377626 |
| CFAP97 | 65.1842209 | -1.024030793 | 0.39327052 | -2.603884 | 0.0092174 | 0.07756589 |
| SNX25 | 189.629536 | -0.431259947 | 0.32853974 | -1.3126569 | 0.1892986 | 0.48954068 |
| LRP2BP | 22.8125874 | 0.35002228 | 0.49261893 | 0.71053355 | 0.47737333 | 0.76307714 |
| AC112722.1 | 25.6184554 | -0.801038626 | 0.34513404 | -2.3209493 | 0.02028958 | 0.1323863 |
| ANKRD37 | 25.5663585 | -1.719701661 | 0.93459097 | -1.8400581 | 0.06575971 | 0.27167064 |
| UFSP2 | 51.126939 | 0.414067874 | 0.34144049 | 1.21270876 | 0.22524119 | 0.53616259 |
| C4orf47 | 72.7863994 | -1.923989795 | 0.57905786 | -3.322621 | 0.00089176 | 0.01365315 |
| SORBS2 | 265.611162 | 0.426321651 | 0.69480547 | 0.61358419 | 0.53949013 | 0.80166126 |
| FAM149A | 15.7477535 | -1.170634741 | 0.54314737 | -2.1552802 | 0.03113992 | 0.17459326 |
| CYP4V2 | 25.6317388 | 1.00056303 | 0.36288716 | 2.75722905 | 0.00582935 | 0.0554621 |
| KLKB1 | 14.2805854 | 0.008464116 | 0.44886745 | 0.0188566 | 0.9849555 | 0.99742652 |
| F11 | 30.0607927 | 2.12741037 | 0.45598404 | 4.66553696 | 3.08E-06 | 0.0001438 |
| FAT1 | 146.814777 | -0.41060832 | 0.31745576 | -1.2934348 | 0.19586073 | 0.49841852 |
| FRG1-DT | 39.7914018 | 1.129126417 | 0.65419619 | 1.72597522 | 0.08435187 | 0.31297129 |
| FRG1 | 17.696095 | -0.138370475 | 0.38782661 | -0.3567844 | 0.72125323 | 0.89190519 |
| SDHA | 99.625257 | -0.132803785 | 0.3093884 | -0.4292462 | 0.66774409 | 0.86777458 |
| PDCD6 | 61.9090135 | -0.264582717 | 0.39848519 | -0.6639713 | 0.5067087 | 0.77942663 |
| EXOC3 | 56.2718539 | -0.461850358 | 0.3992705 | -1.1567355 | 0.24738044 | 0.56012026 |

| | | | | | | |
|---|---|---|---|---|---|---|
| ZDHHC11B | 38.4740899 | 0.890795628 | 0.4919529 | 1.81073357 | 0.0701821 | 0.28242466 |
| BRD9 | 30.5727385 | -0.736557321 | 0.36511861 | -2.0173097 | 0.0436632 | 0.21303805 |
| SLC12A7 | 51.211024 | -1.979588925 | 0.63871487 | -3.0993312 | 0.00193958 | 0.02463849 |
| CLPTM1L | 27.6078348 | -0.888409407 | 0.43608363 | -2.0372455 | 0.04162545 | 0.2077707 |
| LPCAT1 | 45.619478 | -1.329879876 | 0.65749883 | -2.0226346 | 0.04311084 | 0.21134556 |
| NDUFS6 | 39.8680037 | -0.470017983 | 0.4932992 | -0.9528051 | 0.34068884 | 0.65083559 |
| AC022424.1 | 39.5309026 | -0.009006735 | 0.72315662 | -0.0124548 | 0.9900628 | 0.99852305 |
| ADAMTS16 | 301.72658 | 0.236262427 | 0.48579575 | 0.48634108 | 0.62672536 | 0.84726508 |
| ICE1 | 104.814421 | 0.17998556 | 0.20803313 | 0.86517739 | 0.38694144 | 0.69039914 |
| MED10 | 23.8148526 | -0.061446402 | 0.36450702 | -0.168574 | 0.86613174 | 0.95565922 |
| NSUN2 | 50.7991666 | 0.39599432 | 0.38540088 | 1.02748682 | 0.30419129 | 0.61818174 |
| SRD5A1 | 33.5420106 | 1.595293543 | 0.5044444 | 3.16247647 | 0.00156433 | 0.02123122 |
| TENT4A | 75.36676 | -0.124317317 | 0.27179233 | -0.4573982 | 0.64738487 | 0.85921615 |
| MTRR | 37.7564208 | -0.099329779 | 0.32846985 | -0.3024015 | 0.762346 | 0.91387428 |
| SEMA5A | 80.7009897 | -0.464972487 | 0.84700311 | -0.548962 | 0.58303155 | 0.82679638 |
| FAM173B | 40.8853684 | 0.088452825 | 0.4005105 | 0.22085021 | 0.82520907 | 0.94180992 |
| CCT5 | 32.8269227 | -0.40369079 | 0.41953767 | -0.9622278 | 0.33593521 | 0.6470947 |
| CMBL | 39.9632828 | -0.516089615 | 0.59813032 | -0.8628381 | 0.3882265 | 0.69173015 |
| 6-Mar | 311.461068 | 0.042491338 | 0.25250712 | 0.16827778 | 0.86636475 | 0.95574834 |
| ANKRD33B | 50.8799878 | 1.362301759 | 0.65630696 | 2.07570823 | 0.03792094 | 0.19653968 |
| DAP | 78.4203769 | -1.028807821 | 0.27117216 | -3.7939287 | 0.00014828 | 0.00333153 |
| TRIO | 630.90415 | -0.445907787 | 0.33508137 | -1.3307448 | 0.183273 | 0.48210665 |
| OTULIN | 39.6506672 | -0.053205822 | 0.26447689 | -0.2011738 | 0.84056267 | 0.94769122 |
| ANKH | 42.0793675 | 0.131939217 | 0.27214626 | 0.48480996 | 0.62781116 | 0.84784081 |
| RETREG1 | 320.995686 | -0.671603976 | 0.63466659 | -1.0581997 | 0.28996442 | 0.60488981 |
| MYO10 | 430.434672 | 0.181439771 | 0.41657603 | 0.4355502 | 0.66316311 | 0.86556259 |
| LINC02241 | 45.3512051 | 0.538664855 | 0.33278322 | 1.6186659 | 0.10551917 | 0.35685917 |
| CDH12 | 17.5126851 | -0.097216481 | 0.72348231 | -0.134373 | 0.89310764 | 0.96459349 |
| C5orf17 | 81.8169563 | -0.252129471 | 0.71357456 | -0.353333 | 0.72383877 | 0.89280599 |
| CDH6 | 12.7987287 | -0.813102543 | 0.89091333 | -0.9126618 | 0.36142045 | 0.66677568 |
| DROSHA | 79.3631243 | 0.415530743 | 0.2584843 | 1.60756666 | 0.10793012 | 0.36147286 |
| C5orf22 | 26.209926 | -0.152152952 | 0.29115144 | -0.5225904 | 0.60125932 | 0.83679851 |
| PDZD2 | 48.2184837 | -0.775627899 | 0.50472121 | -1.5367452 | 0.12435572 | 0.38965272 |
| GOLPH3 | 133.931463 | 0.058896588 | 0.24696816 | 0.23847846 | 0.81151002 | 0.93552094 |
| MTMR12 | 90.5694512 | 0.461641072 | 0.32805774 | 1.40719457 | 0.1593697 | 0.4477141 |
| ZFR | 248.821404 | 0.038606728 | 0.20117118 | 0.19190984 | 0.84781283 | 0.94989268 |
| SUB1 | 77.6652085 | 0.115469617 | 0.24339182 | 0.47441865 | 0.6352014 | 0.85291155 |
| NPR3 | 33.7844629 | 2.030297095 | 0.53721917 | 3.77927147 | 0.00015729 | 0.00347986 |
| TARS | 37.3541095 | -0.101178392 | 0.36918412 | -0.2740594 | 0.78403894 | 0.92232893 |
| C1QTNF3-AN | 161.810543 | 0.061367435 | 0.29549234 | 0.2076786 | 0.83547993 | 0.94654164 |
| RAI14 | 155.034178 | -0.422706285 | 0.43527789 | -0.9711182 | 0.33148942 | 0.64324815 |
| RAD1 | 15.5370911 | 0.010327148 | 0.42994678 | 0.0240196 | 0.98083698 | 0.99646326 |
| BRIX1 | 17.0936966 | -0.156537398 | 0.44364916 | -0.3528405 | 0.72420801 | 0.89307175 |
| DNAJC21 | 55.5451006 | 0.15115814 | 0.23496657 | 0.64331763 | 0.52001802 | 0.7877576 |
| PRLR | 46.6447772 | 1.584071167 | 0.33691236 | 4.70173067 | 2.58E-06 | 0.00012463 |
| SPEF2 | 64.4851059 | 0.728336781 | 0.77624054 | 0.93828748 | 0.34809669 | 0.65547624 |
| LMBRD2 | 39.9599044 | 0.6186806 | 0.32016173 | 1.93240025 | 0.05331012 | 0.23903633 |
| SKP2 | 14.0326264 | 1.386591482 | 0.51619875 | 2.68615815 | 0.00722789 | 0.06498825 |
| NADK2 | 55.3042857 | 0.277243344 | 0.37114887 | 0.74698689 | 0.45507147 | 0.74558223 |
| RANBP3L | 159.746119 | 0.095124333 | 0.62406681 | 0.15242652 | 0.87885054 | 0.96026761 |
| NIPBL | 539.966587 | -0.230252427 | 0.32851476 | -0.7008891 | 0.4833722 | 0.76664595 |
| CPLANE1 | 165.278049 | 0.466404227 | 0.25092529 | 1.85873746 | 0.06306436 | 0.26451287 |
| NUP155 | 35.8134314 | 0.343990412 | 0.29943553 | 1.14879625 | 0.25064 | 0.56339619 |
| WDR70 | 288.068571 | 0.164426899 | 0.16276069 | 1.01023716 | 0.31238168 | 0.62732 |
| LIFR | 228.44605 | 1.186395451 | 0.19426257 | 6.10717459 | 1.01E-09 | 1.68E-07 |
| OSMR-AS1 | 188.834406 | -1.634591989 | 0.38658365 | -4.2283009 | 2.35E-05 | 0.0007813 |
| OSMR | 222.852402 | -2.904614862 | 0.32489651 | -8.9401232 | 3.89E-19 | 7.25E-16 |
| RICTOR | 274.461466 | 0.171233697 | 0.22974003 | 0.74533681 | 0.45606814 | 0.74627681 |
| TTC33 | 43.5749531 | -0.034792283 | 0.39463581 | -0.088163 | 0.92974711 | 0.97876742 |
| PRKAA1 | 66.8250721 | 0.038251409 | 0.27091259 | 0.14119465 | 0.88771617 | 0.96289734 |
| RPL37 | 52.8941833 | -0.481387751 | 0.38944793 | -1.2360773 | 0.21642983 | 0.52579252 |
| OXCT1 | 71.2301957 | 1.38356332 | 0.58334228 | 2.37178645 | 0.01770232 | 0.12083193 |
| C5orf51 | 15.6865337 | -0.214441133 | 0.44039746 | -0.4869264 | 0.62631052 | 0.8471956 |
| GHR | 198.355804 | 2.064545891 | 0.49711792 | 4.15303054 | 3.28E-05 | 0.00103702 |
| SELENOP | 42.9582425 | 0.375661206 | 0.42004515 | 0.89433531 | 0.37114252 | 0.67522591 |
| ZNF131 | 121.963963 | -0.025330749 | 0.29127769 | -0.0869643 | 0.93069992 | 0.97888844 |
| HMGCS1 | 63.4352998 | -0.020990808 | 0.36082692 | -0.0581742 | 0.95360989 | 0.98771968 |
| TMEM267 | 44.3468799 | -0.260517875 | 0.30438481 | -0.8558833 | 0.39206234 | 0.6951429 |
| PAIP1 | 51.6431643 | 0.01189725 | 0.2963541 | 0.04014539 | 0.96797722 | 0.99156233 |
| NNT-AS1 | 89.9479689 | 0.386653324 | 0.25943991 | 1.49033865 | 0.13613521 | 0.41123731 |
| NNT | 278.635232 | 1.139805096 | 0.22403545 | 5.08761035 | 3.63E-07 | 2.43E-05 |
| MRPS30-DT | 39.861086 | 0.318672074 | 0.4747378 | 0.67125911 | 0.50205548 | 0.77657022 |
| MRPS30 | 24.0691271 | -0.350549659 | 0.35131424 | -0.9978236 | 0.31836488 | 0.63093651 |
| PARP8 | 113.976088 | -0.70498335 | 0.31266819 | -2.2547332 | 0.02415008 | 0.14746257 |
| ITGA1 | 56.7218675 | -0.657300284 | 0.55091213 | -1.1931127 | 0.23282522 | 0.54525758 |
| ITGA2 | 798.783465 | -1.998969409 | 0.88548452 | -2.2574866 | 0.02397769 | 0.14714173 |
| MOCS2 | 23.3312026 | 0.915065899 | 0.372629 | 2.45570231 | 0.01406096 | 0.10405109 |
| NDUFS4 | 151.60266 | 0.865812572 | 0.22543118 | 3.84069579 | 0.00012269 | 0.00283852 |
| ARL15 | 439.048143 | 1.06481505 | 0.59691201 | 1.78387272 | 0.07444435 | 0.29200098 |
| SNX18 | 24.5830587 | -0.029344064 | 0.49705032 | -0.0590364 | 0.95292311 | 0.98744778 |
| DHX29 | 62.0148953 | 0.121164105 | 0.25420465 | 0.47664 | 0.6336185 | 0.8520131 |
| MTREX | 126.454414 | 0.297817866 | 0.19444138 | 1.53165885 | 0.12560664 | 0.39208447 |
| PLPP1 | 102.088353 | 0.123293044 | 0.28530448 | 0.43214549 | 0.66563568 | 0.86729836 |
| SLC38A9 | 66.0378339 | 0.050755781 | 0.35381597 | 0.14345249 | 0.88593283 | 0.96249798 |
| IL6ST | 145.202742 | 0.252794942 | 0.32012226 | 0.78968247 | 0.42971323 | 0.72374389 |
| AC008892.1 | 46.6486915 | 1.670869614 | 0.42507672 | 3.93074833 | 8.47E-05 | 0.0021424 |
| LINC01948 | 15.0304605 | 0.007776825 | 0.5268343 | 0.01476142 | 0.98822251 | 0.9977555 |
| MAP3K1 | 197.47001 | 0.079158662 | 0.27922835 | 0.28349077 | 0.77680066 | 0.9196152 |
| MIER3 | 28.5538899 | 0.271023619 | 0.35912341 | 0.75468101 | 0.4504404 | 0.74156119 |
| GPBP1 | 251.186796 | 0.430946323 | 0.22876422 | 1.88380129 | 0.05959186 | 0.25511226 |
| PDE4D | 4568.64577 | 0.208577936 | 0.25948837 | 0.80380458 | 0.42150985 | 0.71947943 |
| DEPDC1B | 159.219719 | 1.09442644 | 0.71828762 | 1.52366046 | 0.12759355 | 0.39563761 |
| ELOVL7 | 294.995628 | 0.578973567 | 0.36804062 | 1.57312408 | 0.11569009 | 0.3743642 |
| ERCC8 | 61.6831459 | 0.54181748 | 0.30394215 | 1.78263356 | 0.07464597 | 0.2925595 |
| NDUFAF2 | 117.883166 | -0.270484132 | 0.3380921 | -0.8000309 | 0.42369288 | 0.72151771 |
| SMIM15-AS1 | 11.5465515 | 0.430050441 | 0.55451127 | 0.7755486 | 0.43801556 | 0.7305377 |
| ZSWIM6 | 627.933031 | -0.125648275 | 0.42989749 | -0.292275 | 0.7700764 | 0.91655198 |
| KIF2A | 71.8725649 | -0.489023289 | 0.31569356 | -1.5490442 | 0.12137109 | 0.38478887 |
| IPO11 | 53.3900378 | 0.195507395 | 0.32160345 | 0.60791448 | 0.5432442 | 0.80361674 |
| RNF180 | 138.447653 | 0.292810906 | 0.23703113 | 1.23532679 | 0.21670891 | 0.52616463 |
| SREK1IP1 | 52.0559714 | 0.0260604 | 0.23393428 | 0.11140052 | 0.91129875 | 0.97128978 |
| CWC27 | 117.217859 | 0.654142499 | 0.2941196 | 2.22406975 | 0.02614375 | 0.15546195 |
| PPWD1 | 56.7526841 | 0.254101684 | 0.26155335 | 0.97150997 | 0.33129439 | 0.64324815 |
| TRIM23 | 39.3641583 | 0.411701554 | 0.33287871 | 1.23679149 | 0.2161645 | 0.52528481 |
| TRAPPC13 | 59.3261709 | 0.248029352 | 0.24510935 | 1.01191305 | 0.31157963 | 0.62642275 |
| NLN | 32.3237433 | -0.836759354 | 0.34596646 | -2.4186141 | 0.01557976 | 0.11096761 |
| ERBIN | 467.72294 | -0.137673877 | 0.18350196 | -0.7502583 | 0.45309913 | 0.74451091 |
| SREK1 | 150.085099 | 0.308200933 | 0.29789049 | 1.03461153 | 0.30085037 | 0.61462069 |
| MAST4 | 1325.86291 | 0.242030791 | 0.56088171 | 0.43151842 | 0.66609146 | 0.86745649 |
| PIK3R1 | 141.702336 | 0.786838174 | 0.43461253 | 1.81043602 | 0.0702282 | 0.28248823 |
| AC093523.1 | 16.7958793 | 3.069021631 | 0.59758674 | 5.13569231 | 2.81E-07 | 1.99E-05 |
| SLC30A5 | 64.6427388 | -0.3409885 | 0.37652883 | -0.9056106 | 0.36514199 | 0.67012584 |
| MRPS36 | 35.6273122 | 0.183353616 | 0.3126444 | 0.58646058 | 0.55756604 | 0.81230403 |

| | | | | | | |
|---|---|---|---|---|---|---|
| CDK7 | 36.6774765 | -0.122188862 | 0.35990053 | -0.3395073 | 0.73422759 | 0.89939324 |
| CCDC125 | 72.4130939 | 0.217603135 | 0.40897913 | 0.53206416 | 0.59468156 | 0.83416944 |
| AK6 | 23.125924 | 0.176068373 | 0.35199882 | 0.50019592 | 0.61693713 | 0.84223226 |
| RAD17 | 49.3040656 | 0.061232642 | 0.25492695 | 0.24019682 | 0.81017768 | 0.93552094 |
| MARVELD2 | 45.1328318 | 0.565758342 | 0.33034403 | 1.71263378 | 0.08677993 | 0.31843215 |
| OCLN | 118.020243 | 0.298995444 | 0.24315406 | 1.22965434 | 0.21882657 | 0.52913355 |
| GTF2H2C | 34.3067603 | 0.172818926 | 0.41476351 | 0.41666859 | 0.67692083 | 0.87044681 |
| NAIP | 149.072342 | 1.107788876 | 0.44857843 | 2.46955451 | 0.01352814 | 0.10123305 |
| GTF2H2 | 31.5225295 | 0.376916425 | 0.55869074 | 0.67464233 | 0.49990303 | 0.77558432 |
| BDP1 | 182.052724 | 0.108008937 | 0.23074059 | 0.46809682 | 0.63971535 | 0.85413611 |
| MCCC2 | 70.3298924 | -0.044821033 | 0.25034587 | -0.1790364 | 0.85790909 | 0.95266335 |
| MRPS27 | 63.7483578 | 0.138864649 | 0.30805655 | 0.45077649 | 0.65215065 | 0.8617705 |
| PTCD2 | 18.991911 | 0.53544582 | 0.54969545 | 0.97407723 | 0.33001819 | 0.64199659 |
| TNPO1 | 204.443898 | 0.177143286 | 0.19619947 | 0.90287343 | 0.36659307 | 0.67113782 |
| FCHO2 | 186.633564 | -0.052319887 | 0.25220357 | -0.207451 | 0.83565765 | 0.94654164 |
| BTF3 | 33.6465487 | -0.904499348 | 0.32599281 | -2.774599 | 0.00552698 | 0.05345806 |
| ANKRA2 | 20.5824936 | -0.126503559 | 0.32644799 | -0.3875152 | 0.69837483 | 0.88293518 |
| ARHGEF28 | 471.85136 | -0.323332162 | 0.32535269 | -0.9937897 | 0.32032524 | 0.63212442 |
| HEXB | 20.0426972 | -0.855088251 | 0.44166148 | -1.9360716 | 0.05285893 | 0.23809501 |
| GFM2 | 40.9691899 | 0.466274421 | 0.27577553 | 1.69077518 | 0.09087975 | 0.32805373 |
| NSA2 | 24.487941 | -0.593975994 | 0.32863023 | -1.8074296 | 0.07069533 | 0.28380968 |
| FAM169A | 58.0369766 | 0.826778302 | 0.43243908 | 1.91189543 | 0.05588961 | 0.24615716 |
| HMGCR | 56.9288433 | 0.401623322 | 0.40060873 | 1.00253263 | 0.31608642 | 0.63009595 |
| COL4A3BP | 172.29221 | -0.096861203 | 0.33840325 | -0.2862301 | 0.77470188 | 0.91864687 |
| POLK | 130.724119 | 0.055682568 | 0.31015419 | 0.17953189 | 0.85752008 | 0.95266335 |
| ANKDD1B | 20.5154246 | -1.180798239 | 0.8961073 | -1.3176974 | 0.18760497 | 0.48778065 |
| POC5 | 23.4914034 | -0.333676361 | 0.47842462 | -0.6974481 | 0.48552238 | 0.76820138 |
| IQGAP2 | 15.4224472 | 1.104200209 | 0.57815823 | 1.90985815 | 0.05615148 | 0.24696056 |
| ZBED3 | 24.0518995 | -0.026446019 | 0.37035597 | -0.071407 | 0.94307382 | 0.98507485 |
| WDR41 | 84.9138965 | 0.205757678 | 0.22144599 | 0.92915513 | 0.3528087 | 0.65972795 |
| TBCA | 163.605642 | 0.138407717 | 0.27682347 | 0.49998548 | 0.6170853 | 0.84229298 |
| AP3B1 | 340.682829 | 0.478879652 | 0.21303966 | 2.24784273 | 0.02458622 | 0.14891004 |
| SCAMP1 | 82.0637926 | 0.387684595 | 0.26757473 | 1.44888344 | 0.14737014 | 0.43061083 |
| LHFPL2 | 20.3440501 | -0.649584093 | 0.39190536 | -1.6575024 | 0.09741794 | 0.34032404 |
| ARSB | 20.8125637 | -0.153367237 | 0.62043304 | -0.2471939 | 0.8047582 | 0.93258332 |
| DMGDH | 105.002199 | 1.075795781 | 0.54244388 | 1.98323887 | 0.04734075 | 0.22326436 |
| BHMT2 | 40.693288 | -0.263697881 | 0.56152603 | -0.4696094 | 0.63863413 | 0.8538689 |
| BHMT | 12.1159062 | 0.86850781 | 0.84718019 | 1.02517484 | 0.3052807 | 0.6187907 |
| JMY | 82.7856182 | 0.275979577 | 0.27110212 | 1.0179912 | 0.30868214 | 0.62245953 |
| HOMER1 | 98.9118188 | -1.849149432 | 0.9244663 | -2.0002345 | 0.04547494 | 0.21867374 |
| TENT2 | 369.953034 | 0.457988511 | 0.25591034 | 1.78964439 | 0.0735111 | 0.29006241 |
| CMYA5 | 100.599374 | 0.845188704 | 0.3114404 | 2.7138056 | 0.00665152 | 0.06110223 |
| MTX3 | 21.7191346 | 0.631919113 | 0.47794384 | 1.32216185 | 0.18611426 | 0.48628295 |
| THBS4 | 33.6104539 | 1.102135365 | 0.5324571 | 2.06990451 | 0.03846129 | 0.19801935 |
| SERINC5 | 70.2121384 | -0.470772545 | 0.32773582 | -1.4364391 | 0.15087743 | 0.43488337 |
| ZFYVE16 | 151.523873 | 0.108680545 | 0.24826428 | 0.4377615 | 0.66155918 | 0.86492722 |
| FAM151B | 29.6926577 | -0.546874829 | 0.53393095 | -1.0242426 | 0.3057207 | 0.61914418 |
| DHFR | 174.071921 | -1.036368326 | 0.49424921 | -2.0968538 | 0.03600652 | 0.18961413 |
| MSH3 | 130.106632 | 0.208924113 | 0.29784859 | 0.70144401 | 0.48302596 | 0.7665951 |
| RASGRF2 | 155.076907 | -1.900515068 | 0.59720778 | -3.1823348 | 0.00146093 | 0.02024026 |
| CKMT2-AS1 | 67.6802369 | 0.174539918 | 0.39972396 | 0.43665113 | 0.66236438 | 0.8654548 |
| CKMT2 | 69.7361189 | -0.127857255 | 0.33794311 | -0.3783396 | 0.70517834 | 0.88596489 |
| ZCCHC9 | 23.7697085 | -0.213838573 | 0.43045717 | -0.4967708 | 0.61935065 | 0.84360228 |
| SSBP2 | 198.215696 | -0.338295752 | 0.40017116 | -0.8453776 | 0.39790003 | 0.69815955 |
| ATG10 | 119.0637 | 0.592710886 | 0.44502486 | 1.33186016 | 0.18290614 | 0.48189231 |
| RPS23 | 82.8211069 | -0.638501907 | 0.41443814 | -1.5406447 | 0.12340329 | 0.38858908 |
| ATP6AP1L | 78.1882586 | 0.819277644 | 0.6213424 | 1.31856067 | 0.18731603 | 0.48725107 |
| TMEM167A | 37.8676986 | -0.104015264 | 0.33391643 | -0.3115009 | 0.75541986 | 0.91060567 |
| XRCC4 | 88.8946796 | -0.190112847 | 0.35415835 | -0.5368018 | 0.59140452 | 0.83234048 |
| COX7C | 70.7402121 | -0.45720032 | 0.46237894 | -0.9888001 | 0.32276098 | 0.635042 |
| RASA1 | 221.849326 | 0.056052903 | 0.21414391 | 0.26175343 | 0.79351154 | 0.92649882 |
| CCNH | 297.030131 | 0.346339853 | 0.3442568 | 1.00605087 | 0.3143911 | 0.62905207 |
| TMEM161B | 102.2279 | 0.434508936 | 0.29162073 | 1.48997961 | 0.1362296 | 0.41133574 |
| TMEM161B-, | 326.830374 | 0.580102488 | 0.24231746 | 2.39397721 | 0.01666678 | 0.11605758 |
| MEF2C | 72.8041257 | 1.334579786 | 0.62266849 | 2.14332315 | 0.03208716 | 0.17765195 |
| MEF2C-AS1 | 24.1463012 | 0.42896675 | 0.92188052 | 0.46531708 | 0.64170441 | 0.85468927 |
| LYSMD3 | 20.83187 | 0.000868647 | 0.33876886 | 0.00256413 | 0.99795412 | 0.99922221 |
| ADGRV1 | 82.059493 | 0.207415453 | 0.39829468 | 0.52075878 | 0.60253482 | 0.83691476 |
| ARRDC3 | 262.786719 | -0.306981683 | 0.49872522 | -0.6155327 | 0.53820298 | 0.80104055 |
| ARRDC3-AS1 | 18.6521379 | -1.092108178 | 0.57364958 | -1.9037897 | 0.05693757 | 0.24877504 |
| NR2F1-AS1 | 111.332744 | 0.054660732 | 0.603836 | 0.09052248 | 0.92787203 | 0.97812694 |
| FAM172A | 391.444758 | 0.467695493 | 0.30628905 | 1.52697423 | 0.12676742 | 0.39465089 |
| KIAA0825 | 106.194959 | 0.360103195 | 0.6948115 | 0.51827466 | 0.60426665 | 0.83754753 |
| SLF1 | 22.7984553 | 0.013952423 | 0.35145195 | 0.03969938 | 0.9683328 | 0.99156233 |
| TTC37 | 139.273923 | 0.137883646 | 0.27231925 | 0.50633088 | 0.61262439 | 0.84174925 |
| ARSK | 19.0930023 | 0.142878637 | 0.54245271 | 0.26339372 | 0.79224713 | 0.92567823 |
| RHOBTB3 | 501.508691 | -0.037008181 | 0.42120173 | -0.0878633 | 0.92998531 | 0.97876742 |
| GLRX | 63.8310198 | -0.701646661 | 0.68637382 | -1.0222515 | 0.30666189 | 0.62067717 |
| ELL2 | 105.948373 | -2.141864352 | 0.75111781 | -2.8515691 | 0.0043504 | 0.04532196 |
| AC104123.1 | 38.0129065 | -1.033184049 | 0.58006708 | -1.7811458 | 0.07488863 | 0.29289497 |
| CAST | 209.784447 | -0.784633958 | 0.31173509 | -2.5169896 | 0.01183623 | 0.0928965 |
| ERAP1 | 40.4191794 | -0.538702601 | 0.46595591 | -1.1561236 | 0.24763062 | 0.56027856 |
| AC009126.1 | 17.1836986 | 0.369267238 | 0.74396773 | 0.49634846 | 0.61964858 | 0.84360228 |
| ERAP2 | 18.0404784 | -1.029347944 | 0.62084294 | -1.6579845 | 0.0973206 | 0.34011144 |
| LNPEP | 238.329304 | 0.228979187 | 0.37609166 | 0.60883878 | 0.54263131 | 0.80340275 |
| LIX1-AS1 | 54.1129645 | 0.07965266 | 0.35975794 | 0.22140626 | 0.82477612 | 0.94162024 |
| RIOK2 | 12.8265034 | 0.340656845 | 0.42582591 | 0.79999088 | 0.42371608 | 0.72151771 |
| CHD1 | 177.647843 | -0.320290353 | 0.32378022 | -0.9892215 | 0.32255478 | 0.63502972 |
| LINC02062 | 63.4289708 | -0.445247825 | 0.59596442 | -0.7471047 | 0.45500036 | 0.74558223 |
| AC021086.1 | 12.2432786 | 0.748866046 | 0.64316866 | 1.16433851 | 0.24428685 | 0.55622238 |
| FAM174A | 27.0715776 | 0.549859214 | 0.34179591 | 1.60873549 | 0.10767419 | 0.36113459 |
| SLCO4C1 | 29.9853731 | 1.106722624 | 0.36885865 | 3.0003976 | 0.00269627 | 0.03178263 |
| PAM | 402.483766 | -0.925418638 | 0.32354903 | -2.8602114 | 0.00423359 | 0.04430424 |
| GIN1 | 24.4339358 | 0.165098739 | 0.43661615 | 0.37813246 | 0.70533219 | 0.88596489 |
| PPIP5K2 | 124.361653 | 0.763512625 | 0.24311457 | 3.14054661 | 0.00168633 | 0.02232022 |
| NUDT12 | 19.2413617 | 0.584958299 | 0.51192345 | 1.14266753 | 0.25317666 | 0.56559484 |
| LINC01950 | 27.5842341 | -0.493966902 | 0.83541066 | -0.5912863 | 0.55432859 | 0.81072612 |
| EFNA5 | 2474.06132 | -0.266054887 | 0.36422489 | -0.7304687 | 0.46510374 | 0.75251779 |
| FBXL17 | 723.769517 | 0.405701968 | 0.28381014 | 1.42948368 | 0.15286526 | 0.43801957 |
| FER | 333.545508 | -0.436968869 | 0.19120496 | -2.2853428 | 0.02229274 | 0.1402547 |
| AC008467.1 | 21.7475524 | -2.178355594 | 0.81545362 | -2.6713421 | 0.00755486 | 0.06715111 |
| PJA2 | 135.588821 | -0.106361044 | 0.2055039 | -0.5175622 | 0.60476378 | 0.83778698 |
| MAN2A1 | 246.612418 | -0.272189912 | 0.41425734 | -0.6570551 | 0.51114546 | 0.78232441 |
| TMEM232 | 144.349445 | 0.537812145 | 0.79023867 | 0.68056926 | 0.49614408 | 0.77475164 |
| SLC25A46 | 63.7570755 | 0.074595485 | 0.24976536 | 0.29866225 | 0.76519776 | 0.91549415 |
| WDR36 | 39.5831605 | 0.731888096 | 0.29142737 | 2.51139105 | 0.01202564 | 0.09375173 |
| STARD4 | 15.4729021 | 0.009207695 | 0.60858426 | 0.0151297 | 0.98792871 | 0.9977555 |
| NREP | 53.9467432 | -0.469468583 | 0.52273907 | -0.8980935 | 0.36913568 | 0.67341441 |
| EPB41L4A | 250.313608 | 0.047022815 | 0.26744965 | 0.17581932 | 0.86043589 | 0.95363316 |
| APC | 106.226795 | 0.554162451 | 0.2667798 | 2.07722797 | 0.03778052 | 0.19628857 |
| SRP19 | 31.8936315 | -0.075612604 | 0.41467431 | -0.1823421 | 0.85531423 | 0.95200547 |
| REEP5 | 100.6197 | -0.325937091 | 0.25395595 | -1.2834395 | 0.19933815 | 0.50273976 |
| DCP2 | 116.853294 | -0.524002415 | 0.42152356 | -1.2431154 | 0.21382533 | 0.52235998 |

| | | | | | | |
|---|---|---|---|---|---|---|
| MCC | 138.458763 | 0.160616062 | 0.32217538 | 0.49853611 | 0.61810622 | 0.84283824 |
| YTHDC2 | 130.253348 | 0.67392596 | 0.291852 | 2.309136 | 0.02093604 | 0.13499834 |
| PGGT1B | 43.0168608 | -0.036268927 | 0.29187514 | -0.1242618 | 0.901108 | 0.96762997 |
| CCDC112 | 15.5084316 | 0.006614115 | 0.46425302 | 0.01424679 | 0.98863309 | 0.99783665 |
| FEM1C | 34.1771645 | -1.208046359 | 0.58900922 | -2.0509804 | 0.04026886 | 0.20306481 |
| ATG12 | 49.0239302 | -0.327946957 | 0.27428994 | -1.1956215 | 0.23184428 | 0.54477728 |
| AP3S1 | 97.032533 | 0.279453064 | 0.25449771 | 1.0980573 | 0.27217947 | 0.58786226 |
| COMMD10 | 312.032356 | -0.061262765 | 0.23684432 | -0.2586626 | 0.79589558 | 0.92698936 |
| SEMA6A | 118.01747 | -3.903979715 | 0.74459527 | -5.2430896 | 1.58E-07 | 1.22E-05 |
| SEMA6A-AS1 | 37.9090126 | 0.060905832 | 0.31573192 | 0.19290363 | 0.84703445 | 0.94947694 |
| DTWD2 | 89.4279365 | 1.164396927 | 0.39086354 | 2.97903698 | 0.00289156 | 0.03365904 |
| AC008629.1 | 35.4337262 | 0.66082197 | 0.73260685 | 0.90201446 | 0.36704919 | 0.67131445 |
| DMXL1 | 252.928535 | 0.235203658 | 0.21639858 | 1.08690018 | 0.27708093 | 0.59213903 |
| TNFAIP8 | 73.4265749 | -4.449350639 | 0.73451887 | -6.0575035 | 1.38E-09 | 2.18E-07 |
| HSD17B4 | 94.2860765 | 0.784705327 | 0.40879216 | 1.91957037 | 0.05491219 | 0.24363853 |
| SRFBP1 | 44.8185724 | -0.454256443 | 0.37585292 | -1.2086016 | 0.22681594 | 0.53826211 |
| SNX2 | 95.1169574 | 0.234549369 | 0.26737149 | 0.87724151 | 0.38035547 | 0.68437561 |
| SNX24 | 189.003616 | 0.768789375 | 0.31975672 | 2.40429468 | 0.01620371 | 0.11387005 |
| PPIC | 18.2536397 | -0.247396316 | 0.4215683 | -0.5868475 | 0.55730612 | 0.81230403 |
| CEP120 | 48.9290732 | 0.562240968 | 0.27169326 | 2.06939609 | 0.03850893 | 0.19803907 |
| CSNK1G3 | 101.770369 | 0.393235257 | 0.39484493 | 0.99592327 | 0.31928743 | 0.6311291 |
| ZNF608 | 199.060193 | -1.372296577 | 0.42869532 | -3.2010999 | 0.00136904 | 0.01928238 |
| GRAMD2B | 294.189174 | 0.132705552 | 0.28369858 | 0.46776952 | 0.63994942 | 0.85413987 |
| ALDH7A1 | 37.1343774 | 0.555953221 | 0.48958216 | 1.13556676 | 0.25613793 | 0.56895381 |
| PHAX | 18.8748237 | 0.007594163 | 0.37208963 | 0.0204095 | 0.98371671 | 0.99742652 |
| C5orf63 | 21.5270307 | 0.277546883 | 0.5503621 | 0.50429869 | 0.6140515 | 0.84174925 |
| PRRC1 | 37.3373934 | 0.195693681 | 0.30108015 | 0.64997204 | 0.51571029 | 0.78429012 |
| LINC01184 | 138.704184 | 0.642955067 | 0.33748116 | 1.90515842 | 0.05675948 | 0.24816629 |
| SLC12A2 | 124.165179 | 1.872903988 | 0.36024819 | 5.19892687 | 2.00E-07 | 1.48E-05 |
| SLC27A6 | 17.0991845 | 1.813162649 | 1.80036708 | 1.0071072 | 0.31388326 | 0.62884777 |
| ISOC1 | 20.8690485 | 0.876713849 | 0.49492073 | 1.77142276 | 0.07649043 | 0.29596141 |
| HINT1 | 93.2527851 | -1.103600727 | 0.32862491 | -3.3582382 | 0.00078441 | 0.0124385 |
| LYRM7 | 28.0351374 | 0.751975132 | 0.39562093 | 1.90074657 | 0.05733522 | 0.24969339 |
| CDC42SE2 | 334.514846 | -1.729098916 | 0.4418764 | -3.9130827 | 9.11E-05 | 0.00225159 |
| SLC22A5 | 32.2976877 | 0.389940205 | 0.43108088 | 0.9045639 | 0.36569647 | 0.67012584 |
| AC116366.3 | 103.753918 | 1.034432888 | 0.56298478 | 1.83740827 | 0.06614965 | 0.27250233 |
| IRF1 | 23.4109332 | -3.348268307 | 0.89142998 | -3.7560643 | 0.00017261 | 0.00374275 |
| KIF3A | 61.8033198 | 0.548437465 | 0.31929106 | 1.71767247 | 0.08585637 | 0.31653809 |
| 8-Sep | 20.9090081 | 0.286970981 | 0.48573603 | 0.59079616 | 0.55465701 | 0.81072612 |
| UQCRQ | 35.4477673 | -0.211002389 | 0.457244 | -0.4614656 | 0.64446458 | 0.85720224 |
| AFF4 | 319.690317 | -0.590304082 | 0.3872542 | -1.5243323 | 0.12742572 | 0.3953479 |
| ZCCHC10 | 40.3016861 | -0.108216801 | 0.35147111 | -0.3078967 | 0.75816092 | 0.91161939 |
| HSPA4 | 61.2140445 | 0.003340317 | 0.31587255 | 0.01057489 | 0.99156262 | 0.9988777 |
| FSTL4 | 130.556976 | 1.159393756 | 0.81792095 | 1.41748876 | 0.15634008 | 0.44267078 |
| C5orf15 | 54.7704987 | 0.522800001 | 0.28122246 | 1.85902648 | 0.06302339 | 0.26445997 |
| VDAC1 | 78.4923322 | 0.31219937 | 0.2952718 | 1.05732878 | 0.29036156 | 0.60499022 |
| TCF7 | 20.936284 | -1.842926357 | 0.42038105 | -4.3839425 | 1.17E-05 | 0.00044176 |
| SKP1 | 78.757094 | -0.436220248 | 0.40677616 | -1.072384 | 0.28354759 | 0.59716825 |
| CDKL3 | 42.9898745 | -0.180252257 | 0.27566573 | -0.6538798 | 0.51318924 | 0.7830488 |
| UBE2B | 149.62844 | -0.416778763 | 0.41007222 | -1.0163545 | 0.30946059 | 0.62360288 |
| JADE2 | 31.0443557 | -0.235301161 | 0.48146651 | -0.4887176 | 0.62504163 | 0.8471532 |
| SAR1B | 50.7751289 | -0.192300451 | 0.42600387 | -0.4514054 | 0.65169739 | 0.86141572 |
| SEC24A | 105.142075 | -0.973882546 | 0.61975111 | -1.5714091 | 0.11608765 | 0.37466295 |
| CAMLG | 22.6649686 | 0.080076548 | 0.36086743 | 0.22190018 | 0.82439159 | 0.94153354 |
| DDX46 | 118.893308 | -0.002253173 | 0.26493588 | -0.0085046 | 0.99321439 | 0.9992177 |
| C5orf24 | 23.6718014 | -0.503892993 | 0.39501147 | -1.2756414 | 0.2020823 | 0.50618429 |
| TXNDC15 | 32.9899801 | 0.303358942 | 0.53169734 | 0.57054816 | 0.56830597 | 0.81709866 |
| PCBD2 | 114.076262 | 0.404389971 | 0.28980758 | 1.39537403 | 0.16290308 | 0.45367631 |
| H2AFY | 88.8209323 | -0.354867738 | 0.353655 | -1.0034292 | 0.31565384 | 0.63009595 |
| CXCL14 | 35.2378175 | -0.484495176 | 0.74335404 | -0.6517691 | 0.51455015 | 0.7835482 |
| SMAD5 | 82.3149352 | -0.124526487 | 0.22421505 | -0.5553886 | 0.57862888 | 0.82318213 |
| SPOCK1 | 180.773744 | -1.012079253 | 1.10372356 | -0.9169681 | 0.35915938 | 0.66545706 |
| KLHL3 | 386.294043 | 3.623285856 | 0.48839425 | 7.41877257 | 1.18E-13 | 5.19E-11 |
| HNRNPA0 | 28.9998617 | -0.84998041 | 0.47979619 | -1.7715447 | 0.07647016 | 0.29596141 |
| AC106791.1 | 15.7386529 | -0.025291833 | 0.75451821 | -0.0335205 | 0.97325951 | 0.99306979 |
| FAM13B | 227.222954 | 0.003724044 | 0.25064361 | 0.01485793 | 0.98814553 | 0.9977555 |
| NME5 | 44.9902019 | 0.328247979 | 0.6194749 | 0.529881 | 0.59619444 | 0.83416944 |
| BRD8 | 53.4189222 | -0.044285103 | 0.3185639 | -0.1390148 | 0.88943844 | 0.96308489 |
| CDC23 | 21.683299 | 0.47249166 | 0.35905823 | 1.31591931 | 0.18820113 | 0.4882768 |
| FAM53C | 19.2071517 | 0.075837797 | 0.37672172 | 0.20130986 | 0.84045629 | 0.94768587 |
| KDM3B | 124.070847 | -0.098376074 | 0.27909629 | -0.3524808 | 0.72447774 | 0.89328623 |
| ETF1 | 69.4920746 | -0.426200559 | 0.52953976 | -0.8048509 | 0.42090572 | 0.71917261 |
| HSPA9 | 145.057695 | -0.677344121 | 0.50923257 | -1.3301273 | 0.18347634 | 0.48210665 |
| CTNNA1 | 371.426148 | -0.378690377 | 0.26187106 | -1.4460948 | 0.14815062 | 0.43194383 |
| SIL1 | 195.839937 | 0.482152649 | 0.24186515 | 1.99347716 | 0.04620922 | 0.22049887 |
| PAIP2 | 77.6625701 | -0.131845525 | 0.26391214 | -0.4995811 | 0.61737004 | 0.84243499 |
| UBE2D2 | 330.770314 | -1.060969846 | 0.47645347 | -2.2268069 | 0.02596018 | 0.15456754 |
| CXXC5 | 74.8804916 | 0.000656011 | 0.40589125 | 0.00161622 | 0.99871044 | 0.99924629 |
| NRG2 | 73.9384033 | 1.291785255 | 0.41099018 | 3.1431049 | 0.00167166 | 0.02217148 |
| PURA | 87.3850511 | -0.111711889 | 0.33569891 | -0.3327741 | 0.73930484 | 0.90291434 |
| CYSTM1 | 217.850831 | 0.297073053 | 0.33102323 | 0.89743869 | 0.36948488 | 0.67364366 |
| PFDN1 | 56.5593786 | 0.424509616 | 0.26877682 | 1.57941306 | 0.11424134 | 0.372426 |
| AC008438.1 | 14.7394923 | 0.95076767 | 0.56838709 | 1.67274677 | 0.09437716 | 0.33484499 |
| ANKHD1-EIF4 | 15.8479676 | -0.647367163 | 0.45963464 | -1.4084386 | 0.15900124 | 0.44748795 |
| SRA1 | 17.170158 | -0.247967904 | 0.37902885 | -0.6542191 | 0.51297068 | 0.7830488 |
| NDUFA2 | 22.9138741 | 0.387459872 | 0.48558043 | 0.7979314 | 0.4249103 | 0.72151771 |
| IK | 18.3244546 | -0.345540327 | 0.39698596 | -0.8704094 | 0.38407668 | 0.68808461 |
| WDR55 | 14.6465728 | -0.208964424 | 0.43816875 | -0.476904 | 0.63343049 | 0.8520131 |
| HARS | 35.2230157 | 0.355821399 | 0.3042328 | 1.1695695 | 0.24217426 | 0.55446835 |
| TAF7 | 25.5837692 | -0.695840729 | 0.43327597 | -1.6059989 | 0.10827415 | 0.36184524 |
| DIAPH1 | 321.926844 | -0.562216893 | 0.28046448 | -2.0045922 | 0.04500666 | 0.21731856 |
| HDAC3 | 31.6219511 | 0.058463865 | 0.44673338 | 0.1308697 | 0.89587838 | 0.96519206 |
| PCDH1 | 39.5171656 | -0.247641851 | 0.37179479 | -0.6660713 | 0.50536551 | 0.77892236 |
| DELE1 | 33.6346864 | -0.240947034 | 0.34644582 | -0.6954826 | 0.48675292 | 0.76847808 |
| RNF14 | 39.4528629 | -0.005342048 | 0.27890198 | -0.0191539 | 0.98471837 | 0.99742652 |
| GNPDA1 | 22.1936552 | -0.946073948 | 0.62281179 | -1.5190367 | 0.12875326 | 0.39738346 |
| NDFIP1 | 187.687847 | -0.162442973 | 0.27552935 | -0.5895668 | 0.55548108 | 0.81104802 |
| SPRY4-AS1 | 20.6368205 | -3.098711372 | 0.71657604 | -4.3243302 | 1.53E-05 | 0.00055943 |
| ARHGAP26 | 77.3875617 | -1.715535794 | 0.45716922 | -3.7525182 | 0.00017507 | 0.0037873 |
| YIPF5 | 15.082396 | -0.280338196 | 0.49724869 | -0.5637787 | 0.57290478 | 0.82067356 |
| PRELID2 | 78.0254076 | -1.322636023 | 0.85643711 | -1.5443469 | 0.12250431 | 0.38683625 |
| LARS | 87.8813367 | 0.597034288 | 0.28474208 | 2.09675469 | 0.0360153 | 0.18961413 |
| TCERG1 | 123.29571 | 0.279381466 | 0.19965787 | 1.39930107 | 0.16172272 | 0.45171558 |
| STK32A | 72.5504082 | -0.291113731 | 0.71563192 | -0.4067925 | 0.68416036 | 0.87360405 |
| AC091948.1 | 51.8057109 | 1.039127223 | 0.58256783 | 1.78370169 | 0.07447215 | 0.29200098 |
| FBXO38 | 65.6227073 | 0.439190527 | 0.28138623 | 1.56081028 | 0.11856852 | 0.37938672 |
| CSNK1A1 | 436.902575 | -0.461972814 | 0.27169271 | -1.7003504 | 0.08906503 | 0.32396747 |
| ARHGEF37 | 20.2669003 | -0.4469249 | 0.60593308 | -0.7375813 | 0.46076893 | 0.74896364 |
| SLC26A2 | 33.566518 | -2.89699336 | 0.79197017 | -3.6579577 | 0.00025423 | 0.00513088 |
| HMGXB3 | 46.1294518 | -0.464575496 | 0.31311319 | -1.4837302 | 0.13788051 | 0.41430805 |
| TCOF1 | 13.507042 | -0.124512427 | 0.49666283 | -0.2506981 | 0.80204753 | 0.93060267 |
| CD74 | 74.3838706 | -0.490460139 | 0.49093607 | -0.9990306 | 0.31777989 | 0.63093651 |
| RPS14 | 92.0663943 | -0.78342127 | 0.42488747 | -1.8438323 | 0.06520758 | 0.27058097 |

| | | | | | | |
|---|---|---|---|---|---|---|
| NDST1 | 30.1585219 | 0.400540703 | 0.31725786 | 1.26250836 | 0.20676592 | 0.51396573 |
| RBM22 | 13.5768916 | -0.224634498 | 0.43428236 | -0.5172545 | 0.60497852 | 0.83778698 |
| DCTN4 | 71.6687215 | -0.004336036 | 0.24644163 | -0.0175946 | 0.98596228 | 0.99746215 |
| GPX3 | 302.562688 | -1.052228201 | 0.69890514 | -1.5055379 | 0.13218583 | 0.40383377 |
| TNIP1 | 92.6725902 | -0.652110197 | 0.40110361 | -1.6257899 | 0.10399436 | 0.35311123 |
| ANXA6 | 27.8225436 | 0.922725482 | 0.40786274 | 2.26234317 | 0.02367621 | 0.14609992 |
| ATOX1 | 26.1274314 | 0.310040795 | 0.33531418 | 0.92462775 | 0.35515957 | 0.66163487 |
| G3BP1 | 137.390245 | 0.339342608 | 0.29553546 | 1.14822975 | 0.25087373 | 0.56353121 |
| FAM114A2 | 22.0733479 | 0.759419437 | 0.33790138 | 2.24745879 | 0.02461072 | 0.14891004 |
| MFAP3 | 35.9393661 | 0.366196244 | 0.43688869 | 0.83819117 | 0.40192335 | 0.70114138 |
| GALNT10 | 109.640157 | 0.890608222 | 0.42039198 | 2.11851857 | 0.03413117 | 0.18384695 |
| SAP30L-AS1 | 29.8563983 | 0.056871022 | 0.37910904 | 0.15001231 | 0.88075491 | 0.96104842 |
| SAP30L | 12.9141287 | 0.345062353 | 0.3980005 | 0.86698976 | 0.38594762 | 0.68921713 |
| LARP1 | 166.866521 | 0.361725336 | 0.1857905 | 1.9469528 | 0.05154039 | 0.23455111 |
| FAXDC2 | 40.1433449 | 0.606364501 | 0.63122195 | 0.96062012 | 0.33674321 | 0.64764721 |
| CNOT8 | 30.0228388 | 0.063671575 | 0.30408361 | 0.20938839 | 0.83414506 | 0.94640649 |
| GEMIN5 | 21.8154069 | 0.573504004 | 0.31721448 | 1.80793766 | 0.0706162 | 0.28368181 |
| MRPL22 | 21.8293968 | 0.267570941 | 0.35645464 | 0.75064513 | 0.45286625 | 0.74451091 |
| SGCD | 61.3635203 | 2.585230769 | 0.98477991 | 2.62518635 | 0.00866016 | 0.07408008 |
| HAVCR1 | 54.9238555 | -4.371326236 | 1.65061681 | -2.6482986 | 0.0080898 | 0.07056452 |
| HAVCR2 | 26.5031884 | -1.162862778 | 0.6285469 | -1.8500812 | 0.06430185 | 0.26801541 |
| CYFIP2 | 322.822221 | 1.339614295 | 0.31956131 | 4.19204155 | 2.76E-05 | 0.00089814 |
| CLINT1 | 345.173777 | -0.238069011 | 0.3455133 | -0.68903 | 0.49080442 | 0.772104 |
| RNF145 | 106.947003 | -1.444850102 | 0.30596404 | -4.7222873 | 2.33E-06 | 0.00011444 |
| UBLCP1 | 28.2430613 | -0.046752425 | 0.40251928 | -0.1161495 | 0.90753404 | 0.9700616 |
| TTC1 | 75.9589009 | -0.240139871 | 0.2614951 | -0.9183341 | 0.35844397 | 0.6648362 |
| PWWP2A | 116.07616 | 0.272894287 | 0.32196338 | 0.84759419 | 0.39666401 | 0.69690885 |
| CCNJL | 21.9405624 | 0.705283265 | 0.52260819 | 1.349545 | 0.17716198 | 0.47439927 |
| SLU7 | 17.2906116 | -0.477218523 | 0.47726285 | -0.9999071 | 0.31735546 | 0.63093651 |
| AC113414.1 | 102.059173 | -1.199885717 | 0.71658481 | -1.6744504 | 0.09404212 | 0.3344198 |
| CCNG1 | 24.8348093 | -1.08674726 | 0.33572299 | -3.2370356 | 0.00120778 | 0.01732519 |
| MAT2B | 22.914712 | 0.008248854 | 0.46666978 | 0.017676 | 0.98589733 | 0.99746215 |
| WWC1 | 553.503257 | -1.305924974 | 0.49652845 | -2.6301111 | 0.0085357 | 0.0733037 |
| RARS | 54.0223256 | -0.232746921 | 0.27295486 | -0.8526938 | 0.39382912 | 0.69624829 |
| PANK3 | 65.3154523 | -0.527724583 | 0.33548484 | -1.5730207 | 0.11571403 | 0.3743642 |
| KCNIP1 | 30.7470077 | 0.328423911 | 1.15770933 | 0.28368426 | 0.77665236 | 0.9196152 |
| RANBP17 | 31.3794924 | 0.182716164 | 0.63335886 | 0.28848758 | 0.77297354 | 0.91791018 |
| NPM1 | 63.9740486 | -0.357267768 | 0.30162513 | -1.1844761 | 0.23622464 | 0.54899267 |
| FBXW11 | 296.44922 | -0.176327822 | 0.36863507 | -0.4783262 | 0.63241803 | 0.85152594 |
| STK10 | 19.5818622 | -1.147093107 | 0.53543002 | -2.1423773 | 0.03216314 | 0.17776472 |
| UBTD2 | 62.0029124 | -0.225974854 | 0.29236872 | -0.7729105 | 0.43957535 | 0.73189296 |
| SH3PXD2B | 15.2985659 | 0.074284212 | 0.45044232 | 0.16491393 | 0.86901173 | 0.95612718 |
| DUSP1 | 60.1835052 | 0.710834853 | 0.66223632 | 1.07338548 | 0.2830982 | 0.59692618 |
| ERGIC1 | 113.563558 | -0.634059298 | 0.28941132 | -2.1908587 | 0.02846202 | 0.16422022 |
| ATP6V0E1 | 256.530922 | -0.648280502 | 0.24113037 | -2.6885062 | 0.00717725 | 0.06472019 |
| CREBRF | 216.517575 | -0.581285894 | 0.35166283 | -1.6529637 | 0.09833826 | 0.34225677 |
| BOD1 | 18.9102458 | -0.080390187 | 0.4928256 | -0.163121 | 0.87042319 | 0.95629149 |
| LINC01485 | 18.3753254 | 5.762500738 | 0.90047131 | 6.39942735 | 1.56E-10 | 3.55E-08 |
| CPEB4 | 1238.20832 | 1.91455809 | 0.2349938 | 8.14727077 | 3.72E-16 | 2.67E-13 |
| SFXN1 | 127.353145 | 0.386195843 | 0.24589545 | 1.57056929 | 0.11628273 | 0.37503292 |
| THOC3 | 36.6318049 | -0.395321544 | 0.33214488 | -1.1902082 | 0.23396459 | 0.54619074 |
| SIMC1 | 52.2282608 | -0.238460615 | 0.24533509 | -0.9719792 | 0.33106088 | 0.64324815 |
| KIAA1191 | 31.1078051 | -0.31466775 | 0.29339331 | -1.0725117 | 0.28349028 | 0.59716825 |
| ARL10 | 16.4905991 | 0.176785136 | 0.47543527 | 0.3718385 | 0.7100131 | 0.88614996 |
| CLTB | 27.301204 | -0.90547959 | 0.45497042 | -1.9901944 | 0.04656952 | 0.22165096 |
| FAF2 | 92.6681925 | 0.199183382 | 0.36839081 | 0.540685 | 0.58872472 | 0.83025227 |
| RNF44 | 24.6651184 | 0.249513854 | 0.40583634 | 0.61481398 | 0.53867758 | 0.80128358 |
| UIMC1 | 123.912266 | 0.058431415 | 0.25497513 | 0.22916515 | 0.81874056 | 0.93817586 |
| ZNF346 | 62.6918669 | 0.373325275 | 0.24573628 | 1.51921103 | 0.12870939 | 0.39737957 |
| FGFR4 | 16.0143883 | -1.192111493 | 0.67105615 | -1.7764705 | 0.07565539 | 0.29457168 |
| NSD1 | 321.006263 | 0.160215879 | 0.19334523 | 0.82865184 | 0.40730144 | 0.70615074 |
| PRELID1 | 19.3419408 | -0.607144152 | 0.53796076 | -1.128603 | 0.25906532 | 0.57214086 |
| LMAN2 | 46.1601782 | -0.070143797 | 0.39542853 | -0.1773868 | 0.85920459 | 0.95328983 |
| DBN1 | 27.4112817 | 0.232971187 | 0.46919127 | 0.49653777 | 0.61951505 | 0.84360228 |
| PDLIM7 | 14.9320734 | -1.226642729 | 0.77839645 | -1.5758586 | 0.1150584 | 0.3738073 |
| FAM193B | 71.7427789 | 0.17674462 | 0.25325242 | 0.69789902 | 0.48524034 | 0.76788543 |
| TMED9 | 25.5216387 | -1.008617888 | 0.51247876 | -1.9681165 | 0.04905464 | 0.22737481 |
| RMND5B | 42.7860814 | 0.064111002 | 0.45946416 | 0.13953428 | 0.88902797 | 0.96307703 |
| HNRNPAB | 20.410858 | -1.011466097 | 0.51940895 | -1.9473405 | 0.05149392 | 0.23455111 |
| PHYKPL | 104.657849 | -0.571274012 | 0.27939628 | -2.0446729 | 0.04088711 | 0.20496314 |
| CLK4 | 57.9835729 | 0.983543375 | 0.25125873 | 3.91446442 | 9.06E-05 | 0.00225159 |
| ZNF354A | 17.0761277 | -0.327936257 | 0.42898849 | -0.7644407 | 0.44460467 | 0.73723884 |
| ZNF354B | 20.311291 | 0.263432691 | 0.43011619 | 0.61246867 | 0.54022772 | 0.80166126 |
| ZFP2 | 14.5498413 | 0.48804141 | 0.52801144 | 0.92430083 | 0.35532971 | 0.66163487 |
| ZNF354C | 13.4453044 | 0.435696473 | 0.49789502 | 0.87507699 | 0.38153202 | 0.6854344 |
| RUFY1 | 94.1169295 | 0.037307306 | 0.2448699 | 0.15235562 | 0.87890646 | 0.96026761 |
| HNRNPH1 | 259.457583 | 0.72712019 | 0.26533663 | 2.74036869 | 0.00613703 | 0.05756707 |
| CANX | 143.30932 | -0.668582989 | 0.30936134 | -2.1611718 | 0.03068207 | 0.17278271 |
| MAML1 | 47.5749987 | 0.016175644 | 0.32043616 | 0.05048008 | 0.95973982 | 0.9895734 |
| MGAT4B | 33.7037749 | -1.135375155 | 0.50401095 | -2.2526796 | 0.02427936 | 0.1479791 |
| SQSTM1 | 246.716962 | -1.965513146 | 0.60649553 | -3.240771 | 0.00119207 | 0.01712612 |
| MRNIP | 41.3975185 | 0.040200058 | 0.36831477 | 0.10914593 | 0.91308675 | 0.97166624 |
| TBC1D9B | 42.8376655 | -0.523597553 | 0.4438437 | -1.179689 | 0.23812391 | 0.54997949 |
| RNF130 | 293.685328 | -1.344747973 | 0.63778358 | -2.1084707 | 0.0349903 | 0.18621549 |
| MAPK9 | 53.5795421 | 0.393210887 | 0.30386598 | 1.29402734 | 0.19565598 | 0.49824525 |
| GFPT2 | 37.9306271 | -3.164433328 | 0.74660864 | -4.2384097 | 2.25E-05 | 0.00075834 |
| CNOT6 | 85.3307278 | -0.393226799 | 0.28052922 | -1.4017321 | 0.16099527 | 0.45065144 |
| MGAT1 | 29.5930119 | 0.036483833 | 0.32969553 | 0.11065917 | 0.91188663 | 0.97128978 |
| ZFP62 | 15.5042269 | 0.075406572 | 0.39658339 | 0.19014052 | 0.84919902 | 0.95003258 |
| TRIM41 | 18.3674366 | 0.447448788 | 0.37477324 | 1.19391871 | 0.23250977 | 0.54506955 |
| RACK1 | 73.670098 | -0.76880825 | 0.36152063 | -2.1265958 | 0.03345368 | 0.18219751 |
| TRIM52 | 27.9751598 | 0.47268283 | 0.30338029 | 1.55805386 | 0.11922048 | 0.38057334 |
| AC138035.1 | 10.8890443 | 0.230599613 | 0.45371866 | 0.50824361 | 0.61128251 | 0.84169124 |
| DUSP22 | 41.8507179 | -1.050434433 | 0.27686986 | -3.7939645 | 0.00014826 | 0.00333153 |
| EXOC2 | 117.597705 | 0.193045497 | 0.22788518 | 0.84711738 | 0.3969297 | 0.69724426 |
| FOXQ1 | 28.7491632 | -2.723696137 | 0.88142192 | -3.0901162 | 0.00200078 | 0.02524397 |
| GMDS | 255.145656 | -1.126134649 | 0.29066534 | -3.8743342 | 0.00010692 | 0.0025627 |
| GMDS-DT | 471.650872 | -0.285128332 | 0.37143734 | -0.7676351 | 0.44270401 | 0.73499059 |
| WRNIP1 | 35.5070219 | 0.448091216 | 0.3471798 | 1.2906604 | 0.19682146 | 0.49895685 |
| SERPINB1 | 27.3351155 | -1.680613821 | 0.47369251 | -3.5479003 | 0.00038832 | 0.00721016 |
| SERPINB6 | 57.0169121 | -0.420737558 | 0.32282123 | -1.3033144 | 0.19246745 | 0.49370242 |
| NQO2 | 49.2752453 | 0.425729572 | 0.33714835 | 1.26273663 | 0.20668385 | 0.51389871 |
| RIPK1 | 61.5610808 | -0.384731144 | 0.35975741 | -1.0694183 | 0.28488124 | 0.5982506 |
| BPHL | 17.8440515 | -0.226944883 | 0.49175004 | -0.4615046 | 0.64443666 | 0.85720224 |
| PSMG4 | 73.6462552 | -0.137890945 | 0.43494005 | -0.3170344 | 0.75121752 | 0.9089517 |
| SLC22A23 | 186.51645 | -0.360467281 | 0.66347202 | -0.5433044 | 0.58692022 | 0.8297013 |
| PXDC1 | 18.7375095 | -1.938895821 | 0.61394685 | -3.1580842 | 0.0015881 | 0.02147789 |
| PRPF4B | 146.916545 | -0.392742083 | 0.3298733 | -1.1905846 | 0.23381668 | 0.54598215 |
| ECI2 | 50.5063845 | -0.127210888 | 0.33116505 | -0.3841314 | 0.70088107 | 0.88418551 |
| CDYL | 310.129068 | -1.427382316 | 0.68448475 | -2.0853384 | 0.0370386 | 0.1934722 |
| LYRM4 | 71.4396212 | -0.373108262 | 0.32027061 | -1.1649782 | 0.24402784 | 0.55603997 |
| FARS2 | 233.943809 | 0.298478068 | 0.42240819 | 0.70661051 | 0.47980856 | 0.76474103 |
| RREB1 | 186.464878 | -0.324983005 | 0.25727851 | -1.2631564 | 0.20653296 | 0.51366054 |

| | | | | | | |
|---|---|---|---|---|---|---|
| SSR1 | 49.9147335 | -0.148614154 | 0.39589004 | -0.3753925 | 0.70736858 | 0.88614996 |
| RIOK1 | 16.5658914 | 0.131652842 | 0.47838451 | 0.27520298 | 0.7831603 | 0.92183302 |
| DSP | 86.9438424 | -0.670902224 | 0.30727175 | -2.1834165 | 0.02900515 | 0.16620962 |
| SNRNP48 | 23.3038382 | 0.200360789 | 0.32361477 | 0.61913364 | 0.53582832 | 0.79974391 |
| BLOC1S5-TXI | 19.0579367 | 0.374872555 | 0.49064343 | 0.76404275 | 0.44484177 | 0.73750082 |
| SLC35B3 | 33.4027479 | 0.097959544 | 0.33937169 | 0.28864972 | 0.77284944 | 0.91791018 |
| TFAP2A | 87.4850459 | 1.433360935 | 0.38922231 | 3.68262791 | 0.00023084 | 0.00475137 |
| GCNT2 | 469.149333 | 0.546183999 | 0.41312078 | 1.32209277 | 0.18613726 | 0.48628295 |
| TMEM14C | 22.0541546 | -0.250082505 | 0.42077769 | -0.594334 | 0.55228871 | 0.80943179 |
| MAK | 45.9606763 | -0.318490985 | 0.60011726 | -0.5307146 | 0.59561657 | 0.83416944 |
| SMIM13 | 14.1539115 | -0.167089095 | 0.44589889 | -0.3747242 | 0.70786561 | 0.88614996 |
| AL139807.1 | 35.6150856 | -0.512996403 | 0.61060831 | -0.8401399 | 0.40082995 | 0.70092619 |
| NEDD9 | 143.796908 | -0.743134282 | 0.40908591 | -1.8165727 | 0.06928257 | 0.28060883 |
| TMEM170B | 18.9033266 | 0.848339368 | 0.51918891 | 1.6339705 | 0.10226504 | 0.34940245 |
| HIVEP1 | 145.618329 | -0.791253845 | 0.43932618 | -1.8010624 | 0.07169305 | 0.28591361 |
| PHACTR1 | 158.733338 | -0.782326803 | 0.58560491 | -1.3359294 | 0.18157236 | 0.47998877 |
| TBC1D7 | 19.0208643 | 0.199156524 | 0.46258467 | 0.43052988 | 0.66681023 | 0.86777458 |
| SIRT5 | 50.9677002 | 1.335325426 | 0.35330049 | 3.77957423 | 0.0001571 | 0.00347986 |
| NOL7 | 24.6009755 | -0.565402325 | 0.36458644 | -1.5508046 | 0.12094852 | 0.38441929 |
| RANBP9 | 138.610461 | 0.178105693 | 0.26625475 | 0.66892964 | 0.50354036 | 0.77790293 |
| MCUR1 | 29.719455 | 0.182160212 | 0.41904933 | 0.43469874 | 0.66378112 | 0.86588413 |
| CD83 | 41.1336169 | 0.921867913 | 0.68677654 | 1.3423113 | 0.17949508 | 0.47801043 |
| JARID2 | 179.794771 | 0.8614303 | 0.30658438 | 2.80976576 | 0.00495776 | 0.04981263 |
| DTNBP1 | 118.00698 | -0.710303075 | 0.31476991 | -2.2565787 | 0.02403441 | 0.14714173 |
| MYLIP | 35.2538623 | -0.347722253 | 0.35208653 | -0.9876045 | 0.32334636 | 0.63591679 |
| GMPR | 48.1091601 | -0.496330059 | 0.8648312 | -0.573904 | 0.56603279 | 0.81595078 |
| ATXN1 | 761.974914 | 0.568397077 | 0.32929917 | 1.72608112 | 0.08433282 | 0.31297129 |
| CAP2 | 68.9582763 | 0.609100486 | 0.37817862 | 1.61061586 | 0.10726347 | 0.36040527 |
| NUP153 | 108.988983 | 0.25264193 | 0.32029419 | 0.78878087 | 0.43024009 | 0.72410806 |
| KIF13A | 134.221053 | 0.201550633 | 0.30247365 | 0.66634113 | 0.50519308 | 0.77885156 |
| TPMT | 22.0984984 | 0.272910576 | 0.34664958 | 0.78728084 | 0.43111748 | 0.72479975 |
| KDM1B | 16.9221108 | -0.004288623 | 0.42381449 | -0.0101191 | 0.99192626 | 0.9988777 |
| DEK | 73.6720283 | -0.281516553 | 0.39502249 | -0.7126596 | 0.47605645 | 0.76162498 |
| RNF144B | 157.642253 | -1.975898108 | 0.35564408 | -5.5558302 | 2.76E-08 | 2.48E-06 |
| MBOAT1 | 56.4804895 | 1.707720098 | 0.4927232 | 3.46588127 | 0.0005285 | 0.00895946 |
| E2F3 | 58.1974256 | -0.72383572 | 0.29168382 | -2.4815765 | 0.01308026 | 0.09891352 |
| CDKAL1 | 440.975883 | 0.223991166 | 0.16376931 | 1.36772369 | 0.17139858 | 0.46718638 |
| SOX4 | 62.7788451 | -2.682135642 | 0.73516678 | -3.6483363 | 0.00026394 | 0.00525977 |
| CASC15 | 118.075861 | -1.857995813 | 0.8580174 | -2.1654524 | 0.03035306 | 0.17121109 |
| DCDC2 | 1820.37877 | -1.420707704 | 0.52178422 | -2.7227878 | 0.00647336 | 0.05982046 |
| MRS2 | 61.7656729 | 0.254592285 | 0.2872098 | 0.88643314 | 0.37538417 | 0.67888822 |
| ALDH5A1 | 15.7068919 | 0.524205228 | 0.49650821 | 1.0557836 | 0.29106709 | 0.60564025 |
| TDP2 | 31.0257841 | -0.296634618 | 0.30671174 | -0.9671447 | 0.33347172 | 0.64508099 |
| ACOT13 | 16.3475851 | 0.11567753 | 0.5143083 | 0.22491866 | 0.82204256 | 0.94045703 |
| C6orf62 | 90.6283501 | 0.358205056 | 0.35734347 | 1.00241108 | 0.31614509 | 0.63009595 |
| CARMIL1 | 427.201889 | -0.268405303 | 0.51338993 | -0.5228098 | 0.60110659 | 0.83677484 |
| SLC17A1 | 12.9866414 | 1.18841212 | 1.0146329 | 1.171273 | 0.24148906 | 0.55404626 |
| TRIM38 | 73.5127083 | -0.574585449 | 0.29881311 | -1.9228924 | 0.05449357 | 0.24299285 |
| HIST1H4C | 25.2694234 | -3.295781534 | 0.8906263 | -3.7005212 | 0.00021516 | 0.00447796 |
| HIST1H2AC | 26.2654067 | -1.274032972 | 0.65361057 | -1.9492233 | 0.05126876 | 0.23455111 |
| BTN2A1 | 13.6751346 | -0.273517453 | 0.57915818 | -0.4722673 | 0.63673603 | 0.8538689 |
| ZNF322 | 13.8364507 | 0.154815619 | 0.43746962 | 0.35388885 | 0.72342218 | 0.89263661 |
| LINC00240 | 25.1901116 | 0.586608098 | 0.5201442 | 1.12777976 | 0.25941294 | 0.57248906 |
| ZNF165 | 30.8302312 | -3.368603993 | 1.41098481 | -2.3874134 | 0.0169674 | 0.11753644 |
| ZSCAN16-AS | 19.3420531 | 0.262005271 | 0.43141056 | 0.60732234 | 0.54363702 | 0.8036898 |
| ZKSCAN8 | 26.2839672 | 0.061623738 | 0.3439262 | 0.17917721 | 0.85779855 | 0.95266335 |
| ZSCAN9 | 17.7102431 | -0.165583969 | 0.39229912 | -0.422086 | 0.67296224 | 0.8684158 |
| TRIM27 | 39.4075066 | -0.165498408 | 0.34702756 | -0.4769028 | 0.63343135 | 0.8520131 |
| HLA-A | 229.754179 | -0.794211975 | 0.4458661 | -1.7812791 | 0.07486687 | 0.29289497 |
| TRIM26 | 54.7043852 | 0.5511422 | 0.27216032 | 2.02506452 | 0.04286074 | 0.21080021 |
| HCG17 | 135.353673 | -0.152994454 | 0.34151435 | -0.4479884 | 0.6541616 | 0.86217501 |
| HLA-E | 87.8886151 | -1.857942267 | 0.41365555 | -4.4915203 | 7.07E-06 | 0.00028668 |
| GNL1 | 37.1852868 | -0.155290927 | 0.3642978 | -0.4262747 | 0.66990771 | 0.86777458 |
| PRR3 | 15.3056631 | 0.119811916 | 0.51784007 | 0.23136857 | 0.81702848 | 0.93759674 |
| ABCF1 | 34.6688762 | -0.707797201 | 0.50047915 | -1.4142391 | 0.1572917 | 0.44415138 |
| PPP1R10 | 42.6901107 | 0.105262077 | 0.32189311 | 0.32700941 | 0.74366077 | 0.90463949 |
| MRPS18B | 25.1626388 | 0.007439123 | 0.30725209 | 0.02421179 | 0.98068367 | 0.99646326 |
| ATAT1 | 23.6666753 | -0.791739648 | 0.34782386 | -2.2762661 | 0.02283009 | 0.14238646 |
| C6orf136 | 28.6579254 | 0.843976045 | 0.33160158 | 2.54515085 | 0.01092305 | 0.08734696 |
| DHX16 | 23.6011558 | -0.616590354 | 0.3109998 | -1.9826069 | 0.04741135 | 0.22326436 |
| MDC1 | 21.8930662 | -0.226724847 | 0.46553651 | -0.4870184 | 0.62624529 | 0.8471956 |
| TUBB | 30.5077356 | -1.180859278 | 0.55442038 | -2.1298988 | 0.03317997 | 0.18155742 |
| FLOT1 | 75.8451062 | -1.05312083 | 0.37158969 | -2.8340959 | 0.00459555 | 0.04722006 |
| IER3 | 196.749062 | -3.278672393 | 0.61130261 | -5.3634196 | 8.17E-08 | 6.56E-06 |
| DDR1 | 59.4295805 | -1.3227095 | 0.45745374 | -2.8914607 | 0.00383456 | 0.04157372 |
| CCHCR1 | 17.0624057 | -0.114098981 | 0.62234433 | -0.1833374 | 0.8545333 | 0.95170432 |
| POU5F1 | 32.6302507 | -0.061736605 | 0.56026486 | -0.1101918 | 0.91225725 | 0.97144622 |
| HLA-C | 98.9085281 | -1.072843171 | 0.65405058 | -1.6403061 | 0.10094153 | 0.34704236 |
| HLA-B | 156.779117 | -1.715994018 | 0.62008404 | -2.7673572 | 0.00565128 | 0.05420058 |
| MICA | 30.1303891 | 0.418516174 | 0.46868502 | 0.89295829 | 0.37187951 | 0.67534262 |
| NFKBIL1 | 12.4001365 | 0.206366855 | 0.45670292 | 0.45186235 | 0.65136815 | 0.86120237 |
| PRRC2A | 44.5285728 | -0.088849049 | 0.34197613 | -0.2598107 | 0.79500982 | 0.9268423 |
| BAG6 | 25.8976101 | -0.200218052 | 0.44929486 | -0.4456273 | 0.65586649 | 0.86276795 |
| DDAH2 | 22.4514858 | 0.33889671 | 0.54133067 | 0.6260438 | 0.53128621 | 0.79628881 |
| CLIC1 | 67.7920126 | -1.559586434 | 0.37352623 | -4.1753063 | 2.98E-05 | 0.0009535 |
| LSM2 | 15.1417558 | -0.542101883 | 0.44837776 | -1.2090294 | 0.22665156 | 0.53826211 |
| HSPA1A | 39.0442934 | 0.163778995 | 0.70355914 | 0.2327864 | 0.81592728 | 0.93717111 |
| HSPA1B | 26.525739 | 1.147619681 | 0.73133866 | 1.56920418 | 0.11660037 | 0.37568083 |
| C6orf48 | 19.567791 | 0.009892645 | 0.51969547 | 0.01903546 | 0.98481281 | 0.99742652 |
| NEU1 | 19.5243276 | -0.677852769 | 0.67558606 | -1.0033552 | 0.31568952 | 0.63009595 |
| EHMT2 | 16.4776792 | 0.599228811 | 0.51425393 | 1.16523916 | 0.2439222 | 0.55593512 |
| STK19 | 14.0593862 | -0.221828887 | 0.44282187 | -0.5009438 | 0.61641065 | 0.84223226 |
| ATF6B | 40.6104173 | -0.119977015 | 0.31684234 | -0.3786647 | 0.70493686 | 0.88594572 |
| PBX2 | 22.7735502 | -0.008381695 | 0.48187936 | -0.0173938 | 0.98612249 | 0.99746215 |
| HLA-DRA | 20.3089761 | 0.54451035 | 0.58987123 | 0.92310036 | 0.3559549 | 0.66219544 |
| TAP1 | 17.6977379 | -0.956865986 | 0.72304248 | -1.3233883 | 0.18570627 | 0.4858669 |
| BRD2 | 72.7653613 | -0.392692431 | 0.41547045 | -0.9451754 | 0.34456936 | 0.65359197 |
| RXRB | 12.0315727 | -0.219288963 | 0.51040558 | -0.4296367 | 0.66745995 | 0.86777458 |
| SLC39A7 | 17.890782 | -0.803422252 | 0.63386938 | -1.2674887 | 0.20498062 | 0.51143678 |
| RING1 | 22.2286707 | 0.431489121 | 0.45684793 | 0.9444918 | 0.34491839 | 0.65392823 |
| VPS52 | 27.9502698 | 0.168516936 | 0.32490941 | 0.51865822 | 0.6039991 | 0.83754753 |
| RPS18 | 51.596628 | -0.809370794 | 0.46532478 | -1.7393675 | 0.08197014 | 0.30805708 |
| RGL2 | 19.2145492 | 0.043473709 | 0.46032672 | 0.09444099 | 0.92475886 | 0.97735117 |
| TAPBP | 72.9302838 | -0.996060878 | 0.3245685 | -3.0688773 | 0.00214865 | 0.02660558 |
| PHF1 | 40.0829073 | -0.619073247 | 0.48602824 | -1.2737392 | 0.20275584 | 0.50710716 |
| CUTA | 25.3737586 | -0.406102471 | 0.46610982 | -0.8712592 | 0.38361264 | 0.68771472 |
| SYNGAP1 | 26.6533777 | -0.537240265 | 0.49624581 | -1.0826092 | 0.27898194 | 0.59380484 |
| ITPR3 | 59.3848237 | -2.721202898 | 0.62245267 | -4.3717427 | 1.23E-05 | 0.00046341 |
| UQCC2 | 16.810764 | 0.119119641 | 0.38388325 | 0.31030174 | 0.75633151 | 0.91111045 |
| LEMD2 | 29.6813202 | -0.782545249 | 0.43671169 | -1.7919036 | 0.07314841 | 0.28899744 |
| NUDT3 | 26.4666855 | -0.90627383 | 0.54011325 | -1.677933 | 0.09336018 | 0.33326583 |
| C6orf106 | 101.878277 | -0.047856876 | 0.31498919 | -0.1519318 | 0.87924072 | 0.96035935 |
| SNRPC | 17.0433937 | -0.040558491 | 0.37042585 | -0.1094915 | 0.91281264 | 0.97166624 |

| | | | | | | |
|---|---|---|---|---|---|---|
| UHRF1BP1 | 68.4221213 | 0.316469634 | 0.30447721 | 1.03938692 | 0.29862482 | 0.61248963 |
| TAF11 | 41.3083759 | 0.329837769 | 0.32049948 | 1.02913666 | 0.30341546 | 0.61783048 |
| ANKS1A | 394.069267 | -0.627211457 | 0.36130833 | -1.7359452 | 0.08257352 | 0.30885595 |
| SCUBE3 | 34.6902262 | -0.411929731 | 1.06700575 | -0.3860614 | 0.6994512 | 0.8833378 |
| ZNF76 | 53.5190442 | -0.001767138 | 0.29853005 | -0.0059195 | 0.99527698 | 0.99922221 |
| PPARD | 57.1668074 | -0.868021529 | 0.61120502 | -1.4201806 | 0.1555551 | 0.44111793 |
| RPL10A | 55.2963766 | -0.641427363 | 0.53671574 | -1.195097 | 0.23204914 | 0.54485676 |
| TEAD3 | 31.2613986 | -1.195435991 | 0.43520232 | -2.7468511 | 0.00601704 | 0.05689951 |
| SRPK1 | 74.3624855 | -0.360800871 | 0.3218307 | -1.121089 | 0.26224996 | 0.57629475 |
| MAPK14 | 183.181802 | -0.932990058 | 0.39348411 | -2.3710997 | 0.01773525 | 0.12091034 |
| BRPF3 | 19.0985218 | -0.415312648 | 0.45449813 | -0.913783 | 0.3608309 | 0.66644099 |
| KCTD20 | 64.3809253 | -0.071325061 | 0.25641711 | -0.2781603 | 0.78088931 | 0.92083964 |
| STK38 | 108.206926 | -0.295215116 | 0.22357127 | -1.3204519 | 0.18668418 | 0.48675708 |
| SRSF3 | 79.9847533 | 0.252621619 | 0.27640076 | 0.91396862 | 0.36073333 | 0.66643107 |
| CDKN1A | 22.4012992 | -3.382833863 | 0.7978335 | -4.2400249 | 2.23E-05 | 0.00075834 |
| CPNE5 | 22.6579414 | 2.080442039 | 0.76433942 | 2.72188242 | 0.00649112 | 0.0598944 |
| C6orf89 | 71.1195005 | -0.266231709 | 0.24870361 | -1.0704779 | 0.28440426 | 0.59765276 |
| MTCH1 | 75.510518 | -1.18042743 | 0.60474375 | -1.9519465 | 0.05094456 | 0.23364835 |
| TBC1D22B | 59.7802834 | -0.047543667 | 0.28920261 | -0.1643957 | 0.86941965 | 0.9561822 |
| RNF8 | 31.2265732 | -0.096714914 | 0.38104232 | -0.2538167 | 0.79963715 | 0.92922871 |
| CMTR1 | 22.5165241 | 0.351707928 | 0.5624777 | 0.62528333 | 0.53178512 | 0.79676281 |
| ZFAND3 | 1063.19797 | -0.167101574 | 0.37736488 | -0.4428117 | 0.65790196 | 0.86357172 |
| BTBD9 | 717.267233 | 0.016115457 | 0.26831946 | 0.06006071 | 0.95210728 | 0.98738095 |
| GLO1 | 47.7873336 | -0.219330792 | 0.35624 | -0.6156827 | 0.53810398 | 0.80104055 |
| KIF6 | 15.1254363 | -0.244827503 | 1.03065482 | -0.2375456 | 0.81223356 | 0.93566416 |
| MOCS1 | 18.0456163 | -0.136203427 | 0.43484981 | -0.3132195 | 0.75411392 | 0.91026551 |
| UNC5CL | 12.8678561 | -1.641996618 | 0.57828768 | -2.8394114 | 0.00451969 | 0.04658498 |
| OARD1 | 38.9569975 | -0.225278957 | 0.38982081 | -0.5779039 | 0.56332901 | 0.81448184 |
| NFYA | 20.7631137 | -0.072381337 | 0.44483359 | -0.1627155 | 0.87074241 | 0.95629149 |
| FOXP4 | 14.4715432 | -0.303359334 | 0.44783452 | -0.6773916 | 0.49815754 | 0.77558432 |
| TFEB | 75.9538067 | 0.372657987 | 0.49260541 | 0.75650404 | 0.44934705 | 0.74088628 |
| USP49 | 55.2504381 | -0.007271949 | 0.26827032 | -0.0271068 | 0.97837455 | 0.9951912 |
| CCND3 | 123.39036 | -0.04224083 | 0.48553221 | -0.086999 | 0.93067229 | 0.97888844 |
| TAF8 | 44.2576096 | -0.007520502 | 0.27672752 | -0.0271766 | 0.97831891 | 0.9951912 |
| C6orf132 | 17.2936694 | -1.423003149 | 0.42452996 | -3.3519499 | 0.00080245 | 0.01261432 |
| MRPS10 | 12.0198473 | 0.205513118 | 0.49974062 | 0.41123957 | 0.68089688 | 0.87265005 |
| TRERF1 | 37.565797 | -0.72961638 | 0.6358976 | -1.1473803 | 0.25122448 | 0.56353121 |
| UBR2 | 290.452881 | 0.367711064 | 0.16787681 | 2.19036245 | 0.02849796 | 0.16432591 |
| BICRAL | 123.348338 | 0.244533193 | 0.38546878 | 0.63437872 | 0.52583369 | 0.79321685 |
| RPL7L1 | 47.7520143 | -0.041257788 | 0.32482802 | -0.1270142 | 0.89892912 | 0.96637793 |
| PEX6 | 25.5363573 | -0.417654887 | 0.45490305 | -0.9181185 | 0.35855685 | 0.66491331 |
| PPP2R5D | 13.2924202 | 0.968983234 | 0.42059382 | 2.30384566 | 0.02123131 | 0.13661886 |
| KLHDC3 | 24.587828 | -0.181415943 | 0.35128596 | -0.5164338 | 0.60555146 | 0.83802715 |
| RRP36 | 28.0036266 | -0.440492483 | 0.39127822 | -1.1257782 | 0.26025942 | 0.57381386 |
| CUL7 | 15.6953854 | 0.205708397 | 0.45284259 | 0.45426027 | 0.64964151 | 0.86077172 |
| KLC4 | 31.7438943 | 0.395458184 | 0.30085942 | 1.31442846 | 0.18870206 | 0.48919517 |
| MRPL2 | 17.580866 | 0.969356137 | 0.44080936 | 2.19903711 | 0.02787528 | 0.16193716 |
| PTK7 | 72.7017086 | -0.54236677 | 0.46076283 | -1.1771062 | 0.23915313 | 0.55099114 |
| CUL9 | 27.8274124 | -0.220856986 | 0.44366203 | -0.4978046 | 0.61862179 | 0.84283824 |
| DNPH1 | 16.3453118 | -0.021074968 | 0.7267915 | -0.0289973 | 0.97686677 | 0.99479093 |
| ZNF318 | 20.41919 | 0.218783535 | 0.47752868 | 0.4581579 | 0.64683901 | 0.85896013 |
| ABCC10 | 19.5789694 | 0.0654975 | 0.41341035 | 0.15843217 | 0.87411626 | 0.95761805 |
| TJAP1 | 28.1893526 | 0.064345316 | 0.32329106 | 0.19903215 | 0.84223759 | 0.94776435 |
| POLR1C | 33.7583929 | -0.555760879 | 0.31544991 | -1.7618039 | 0.07810243 | 0.29882113 |
| YIPF3 | 22.1300365 | -0.243884511 | 0.44042061 | -0.5537536 | 0.57974747 | 0.82409732 |
| XPO5 | 72.9892274 | -0.188685352 | 0.27681948 | -0.6816188 | 0.49548005 | 0.77475164 |
| POLH | 37.9525294 | 0.370259541 | 0.42816118 | 0.86476673 | 0.38716684 | 0.69063394 |
| GTPBP2 | 35.5836441 | -0.668781178 | 0.47792832 | -1.3993337 | 0.16171295 | 0.45171558 |
| MRPS18A | 12.1345952 | 0.139692614 | 0.44052399 | 0.31710557 | 0.7511635 | 0.9089517 |
| VEGFA | 244.752726 | -0.093565351 | 0.44584211 | -0.2098621 | 0.83377532 | 0.94633245 |
| MRPL14 | 19.3534576 | -1.506293394 | 0.50459458 | -2.9851557 | 0.00283434 | 0.03303423 |
| TMEM63B | 33.299949 | -0.272543344 | 0.28757218 | -0.9477389 | 0.3432624 | 0.65295057 |
| SLC29A1 | 23.0613114 | -0.187477048 | 0.44762064 | -0.4188302 | 0.67534022 | 0.86960234 |
| HSP90AB1 | 130.599472 | -0.712848382 | 0.42309656 | -1.6848362 | 0.09202019 | 0.33050702 |
| CDC5L | 37.5110592 | 0.565896453 | 0.36147052 | 1.56553969 | 0.11745642 | 0.3777729 |
| SUPT3H | 391.105759 | 0.677808499 | 0.21514094 | 3.15053243 | 0.00162973 | 0.02180146 |
| RUNX2 | 19.591744 | -1.044066825 | 0.55327543 | -1.8870652 | 0.05915155 | 0.25416087 |
| ENPP4 | 42.0102683 | -1.27317886 | 0.67916325 | -1.8746286 | 0.06084383 | 0.25784263 |
| ENPP5 | 17.0391521 | -0.551385215 | 0.58965993 | -0.9350902 | 0.34974182 | 0.65681679 |
| RCAN2 | 21.2653253 | 1.753622667 | 0.41780204 | 4.19725735 | 2.70E-05 | 0.00088387 |
| CYP39A1 | 16.3943484 | 0.29554012 | 0.76461108 | 0.38652346 | 0.69910903 | 0.8833378 |
| SLC25A27 | 106.267018 | 1.229733241 | 0.3938002 | 3.12273389 | 0.0017918 | 0.02323604 |
| ADGRF1 | 29.1205071 | -3.218637995 | 0.72177075 | -4.4593633 | 8.22E-06 | 0.00032616 |
| TNFRSF21 | 16.969048 | -1.555000942 | 0.7155835 | -2.1730531 | 0.02977632 | 0.16882975 |
| CD2AP | 534.937659 | -0.770448384 | 0.31226664 | -2.4672773 | 0.01361449 | 0.10171593 |
| PTCHD4 | 71.5252538 | -3.014409117 | 1.03139411 | -2.922655 | 0.00347061 | 0.03879367 |
| MUT | 39.7983385 | 0.485706004 | 0.36187697 | 1.34218544 | 0.17953588 | 0.47801043 |
| TFAP2B | 75.5266039 | -0.34626666 | 0.42941741 | -0.8063638 | 0.4200331 | 0.71847159 |
| PKHD1 | 2367.7482 | 0.818376628 | 0.50556282 | 1.61874371 | 0.10550242 | 0.35685917 |
| MCM3 | 13.4076776 | -0.490315855 | 0.40883243 | -1.1993076 | 0.23040835 | 0.54333016 |
| EFHC1 | 58.7137398 | -0.41364919 | 0.365313 | -1.1323145 | 0.25750225 | 0.57016172 |
| TRAM2 | 16.0258678 | -0.462333157 | 0.38090062 | -1.2137895 | 0.22482813 | 0.53600039 |
| GSTA4 | 22.9810752 | 0.35938942 | 0.56169882 | 0.63982584 | 0.52228583 | 0.78965349 |
| ICK | 23.7068737 | 0.102437157 | 0.56743226 | 0.18052755 | 0.85673842 | 0.95234014 |
| FBXO9 | 94.5426469 | 0.079281531 | 0.30037397 | 0.26394275 | 0.79182403 | 0.92541581 |
| ELOVL5 | 41.8706221 | -0.400892818 | 0.59194969 | -0.6772414 | 0.49825283 | 0.77558432 |
| GCLC | 68.3647231 | -0.564383866 | 0.39841187 | -1.416584 | 0.15660459 | 0.4431506 |
| LRRC1 | 63.1475069 | 0.256353485 | 0.27733216 | 0.92435543 | 0.35530129 | 0.66163487 |
| TINAG | 61.7285932 | -2.440780175 | 0.65206334 | -3.7431642 | 0.00018172 | 0.00390399 |
| FAM83B | 187.253137 | -1.059036294 | 0.36115763 | -2.9323381 | 0.0033642 | 0.038114 |
| HMGCLL1 | 23.5073463 | 0.779707901 | 0.71876945 | 1.08478164 | 0.27801838 | 0.59346231 |
| DST | 1007.29938 | -1.961479469 | 0.71526277 | -2.7423201 | 0.00610069 | 0.05739939 |
| ZNF451 | 121.736177 | -0.075738902 | 0.19121926 | -0.3960841 | 0.69204301 | 0.87814495 |
| PRIM2 | 78.432388 | 0.381735123 | 0.39279462 | 0.97184407 | 0.33112813 | 0.64324815 |
| AL021368.2 | 15.8947701 | 0.366768068 | 0.52116758 | 0.70374305 | 0.4815928 | 0.76614422 |
| PTP4A1 | 85.5434802 | -0.896328329 | 1.02344698 | -0.8757936 | 0.38114224 | 0.68499812 |
| PHF3 | 206.05694 | 0.143564343 | 0.19388515 | 0.74046074 | 0.45902047 | 0.74758199 |
| EYS | 22.9929324 | -0.882863385 | 0.5926906 | -1.4895856 | 0.13633323 | 0.41138221 |
| LMBRD1 | 191.089943 | 0.224082197 | 0.19260777 | 1.16341206 | 0.24466236 | 0.55684312 |
| COL19A1 | 45.3659191 | -0.167960643 | 0.35613692 | -0.4716182 | 0.63719935 | 0.8538689 |
| FAM135A | 443.98015 | -1.282209715 | 0.36764622 | -3.4876184 | 0.00048734 | 0.00847759 |
| SDHAF4 | 15.1657678 | 0.123461888 | 0.47768682 | 0.25845781 | 0.79605361 | 0.92698936 |
| SMAP1 | 260.757387 | -0.209633853 | 0.36567106 | -0.5732853 | 0.56645152 | 0.81595078 |
| OGFRL1 | 36.7513078 | -3.117002895 | 0.83562917 | -3.730127 | 0.00019138 | 0.00408343 |
| AL136164.2 | 202.26091 | 0.033972813 | 0.55930582 | 0.06074103 | 0.95156545 | 0.98724783 |
| LINC00472 | 130.756545 | -0.025778282 | 0.49772936 | -0.0517918 | 0.95869462 | 0.9891274 |
| AC019205.1 | 18.6119145 | 0.538400331 | 0.64246542 | 0.83802227 | 0.4020182 | 0.70116305 |
| OOEP | 49.6366784 | -1.055983899 | 0.38117858 | -2.7703128 | 0.00560025 | 0.05377622 |
| MTO1 | 40.9073734 | 0.484567682 | 0.31229788 | 1.55162013 | 0.12075314 | 0.3841359 |
| EEF1A1 | 255.693877 | -1.099994253 | 0.50015826 | -2.1992924 | 0.02785714 | 0.16193265 |
| SLC17A5 | 27.1658569 | -0.574775384 | 0.38019467 | -1.5117923 | 0.1305867 | 0.40131523 |
| COX7A2 | 39.0488336 | 7.73E-05 | 0.40858288 | 0.00018916 | 0.99984907 | 0.99992389 |
| TMEM30A | 57.178076 | -0.265456066 | 0.3964647 | -0.6695579 | 0.50313967 | 0.7778729 |

| | | | | | | |
|---|---|---|---|---|---|---|
| SENP6 | 435.138977 | 0.105451293 | 0.15509587 | 0.67991037 | 0.49656121 | 0.77475164 |
| MYO6 | 810.90087 | -0.804460019 | 0.37610755 | -2.1389095 | 0.032443 | 0.17920528 |
| MEI4 | 25.0821514 | 0.670029144 | 0.57128375 | 1.17284825 | 0.24085667 | 0.55327012 |
| IRAK1BP1 | 87.4541206 | 0.20833738 | 0.55535376 | 0.37514355 | 0.70755371 | 0.88614996 |
| PHIP | 248.00213 | 0.186534716 | 0.211555 | 0.88173155 | 0.37792199 | 0.6811801 |
| HMGN3 | 107.307883 | 0.192136288 | 0.38230589 | 0.50257214 | 0.61526512 | 0.8420273 |
| LCA5 | 14.7592545 | -0.154639423 | 0.53681173 | -0.2880701 | 0.77329306 | 0.91791018 |
| SH3BGRL2 | 18.6321814 | 0.722852351 | 0.38910915 | 1.85771098 | 0.06321007 | 0.26477174 |
| BCKDHB | 267.62214 | 0.207622439 | 0.39247311 | 0.52901061 | 0.59679809 | 0.83416944 |
| AL359715.1 | 13.893756 | 0.001705341 | 0.58681765 | 0.00290608 | 0.99768128 | 0.99922221 |
| TENT5A | 20.8834031 | -0.162781992 | 0.68084497 | -0.2390882 | 0.8110372 | 0.93552094 |
| IBTK | 196.650047 | -0.575945726 | 0.46737444 | -1.2323004 | 0.21783688 | 0.52742431 |
| UBE3D | 123.799979 | 0.339750651 | 0.35423775 | 0.95910347 | 0.33750662 | 0.64844536 |
| DOPEY1 | 172.88387 | 0.011339898 | 0.22102675 | 0.05130555 | 0.95908205 | 0.98932194 |
| PGM3 | 42.9908264 | -0.296381632 | 0.38174268 | -0.7763911 | 0.43751808 | 0.73016262 |
| CYB5R4 | 54.092458 | -0.176146587 | 0.34153194 | -0.5157544 | 0.60602598 | 0.83811722 |
| CEP162 | 78.0755344 | 0.054860077 | 0.37582627 | 0.14597191 | 0.88394357 | 0.96170562 |
| SNX14 | 144.019287 | 0.270947876 | 0.25209902 | 1.07476766 | 0.28247876 | 0.59642935 |
| SYNCRIP | 60.4938926 | -0.248926789 | 0.35642083 | -0.698407 | 0.48492272 | 0.76770898 |
| ZNF292 | 786.200905 | -1.495973584 | 0.57034314 | -2.6229361 | 0.00871756 | 0.07443458 |
| SMIM8 | 95.9758735 | 0.92687608 | 0.38870939 | 2.38449623 | 0.01710252 | 0.11799351 |
| RARS2 | 129.168242 | 0.25812959 | 0.21680489 | 1.19060778 | 0.2338076 | 0.54598215 |
| ORC3 | 64.2004246 | 0.519307209 | 0.48430488 | 1.07227334 | 0.28359728 | 0.59716825 |
| AKIRIN2 | 46.3655986 | 0.007680614 | 0.46123329 | 0.01665234 | 0.98671397 | 0.99766074 |
| RNGTT | 183.963321 | 0.265662074 | 0.21148962 | 1.25614711 | 0.20906264 | 0.5174675 |
| PNRC1 | 51.0462495 | -0.139709429 | 0.37741479 | -0.3701748 | 0.71125228 | 0.88694881 |
| PM20D2 | 36.9065289 | -0.208669407 | 0.37855079 | -0.5512323 | 0.58147447 | 0.8255928 |
| UBE2J1 | 49.2988438 | 0.036043397 | 0.28837389 | 0.12498842 | 0.90053272 | 0.967235 |
| RRAGD | 67.8942886 | 0.399885347 | 0.41970561 | 0.95277579 | 0.34070368 | 0.65083559 |
| ANKRD6 | 107.631607 | -0.286237762 | 0.36573949 | -0.7826274 | 0.43384593 | 0.7271579 |
| LYRM2 | 26.4932903 | 0.170837286 | 0.41433385 | 0.41231795 | 0.68010639 | 0.87237893 |
| MDN1 | 109.733823 | 0.237097488 | 0.18724889 | 1.26621574 | 0.20543585 | 0.51190533 |
| CASP8AP2 | 47.7944728 | 0.580851409 | 0.24767202 | 2.34524434 | 0.01901461 | 0.1268185 |
| BACH2 | 345.461113 | -1.086688051 | 0.51509026 | -2.1097041 | 0.03488385 | 0.18591921 |
| MAP3K7 | 117.870994 | 0.075949099 | 0.21269534 | 0.35707928 | 0.72103246 | 0.89175045 |
| MANEA | 17.9050707 | 0.777433839 | 0.47160947 | 1.64846953 | 0.09925636 | 0.34378391 |
| UFL1-AS1 | 27.0220541 | -0.295347862 | 0.69393682 | -0.425612 | 0.67039057 | 0.86777458 |
| UFL1 | 69.0313411 | 0.26970722 | 0.28952014 | 0.93156635 | 0.35156068 | 0.65871783 |
| NDUFAF4 | 15.7747535 | 0.452810191 | 0.41784265 | 1.08368589 | 0.27850409 | 0.59359815 |
| KLHL32 | 60.8862949 | 0.85895584 | 0.55207003 | 1.55588204 | 0.11973615 | 0.38168199 |
| MMS22L | 150.539384 | -2.322890518 | 0.84241647 | -2.7574135 | 0.00582606 | 0.0554621 |
| AL589740.1 | 24.3850129 | -0.871204237 | 0.91468001 | -0.9524689 | 0.34085925 | 0.65083559 |
| FBXL4 | 140.363131 | 0.52701881 | 0.26668635 | 1.9761747 | 0.04813499 | 0.22485402 |
| FAXC | 22.79032 | 1.247835981 | 0.37960239 | 3.28721846 | 0.00101182 | 0.01516759 |
| PNISR | 447.68255 | 0.400032112 | 0.17577459 | 2.27582444 | 0.02285652 | 0.14245601 |
| AL513550.1 | 28.8589224 | 0.195560406 | 0.43082978 | 0.45391571 | 0.6498895 | 0.86079784 |
| USP45 | 83.1673421 | -0.09463593 | 0.35257593 | -0.2684129 | 0.78838151 | 0.92377626 |
| TSTD3 | 16.5314638 | 0.17245226 | 0.46130388 | 0.37383657 | 0.70852591 | 0.88614996 |
| CCNC | 116.608933 | -0.676278165 | 0.34212599 | -1.9766934 | 0.0480763 | 0.22469345 |
| SIM1 | 469.794529 | -0.881199005 | 0.30006376 | -2.9367058 | 0.00331719 | 0.03762706 |
| ASCC3 | 213.956704 | -0.055164822 | 0.22790533 | -0.2420515 | 0.80874027 | 0.93433178 |
| HACE1 | 91.0655129 | 0.874653464 | 0.31207619 | 2.80269206 | 0.0050678 | 0.05064544 |
| PREP | 43.4254194 | 0.440586726 | 0.26762304 | 1.64629597 | 0.09970284 | 0.34450385 |
| ATG5 | 129.677645 | 0.134787897 | 0.2287016 | 0.58936141 | 0.55561886 | 0.81104802 |
| CRYBG1 | 158.49442 | 0.110505418 | 0.58042235 | 0.19038794 | 0.84900514 | 0.95003258 |
| RTN4IP1 | 32.5627607 | 0.547448002 | 0.45058595 | 1.21496908 | 0.22437788 | 0.53561172 |
| QRSL1 | 57.0948305 | 0.358877482 | 0.2691682 | 1.33328335 | 0.18243884 | 0.48134117 |
| LINC02532 | 799.154309 | -0.344678401 | 0.60133919 | -0.5731847 | 0.56651966 | 0.81595078 |
| CD24 | 341.603868 | -1.189902191 | 0.46566136 | -2.5552951 | 0.01060978 | 0.08592203 |
| C6orf203 | 28.8111461 | 0.254951435 | 0.30752152 | 0.82905234 | 0.40707479 | 0.70602034 |
| PDSS2 | 223.925465 | 0.071054167 | 0.17098971 | 0.41554645 | 0.67774192 | 0.87114221 |
| SOBP | 74.8108765 | 0.934335217 | 0.48446334 | 1.9285984 | 0.05378074 | 0.24061977 |
| SEC63 | 97.9097147 | -0.067555749 | 0.27505689 | -0.2456065 | 0.80598689 | 0.93308942 |
| OSTM1 | 41.3004431 | -0.574292061 | 0.6031074 | -0.9522219 | 0.34098447 | 0.65083559 |
| SNX3 | 72.7116284 | 0.132300542 | 0.2378981 | 0.55612274 | 0.57812695 | 0.82311488 |
| AFG1L | 108.33184 | 1.095123777 | 0.27563084 | 3.97315406 | 7.09E-05 | 0.00189491 |
| FOXO3 | 349.249835 | -0.203528803 | 0.26969566 | -0.754661 | 0.45045239 | 0.74156119 |
| ARMC2 | 63.4019862 | 0.379180708 | 0.55692212 | 0.68085051 | 0.49596609 | 0.77475164 |
| SESN1 | 117.759422 | 0.035277041 | 0.29395469 | 0.12000843 | 0.90447647 | 0.96879249 |
| CEP57L1 | 23.6446804 | 0.275727532 | 0.37212566 | 0.74095275 | 0.45872209 | 0.747377 |
| CD164 | 73.9101128 | -0.362625062 | 0.32458183 | -1.1172069 | 0.26390587 | 0.57843402 |
| PPIL6 | 21.0877599 | 0.008554576 | 0.48555338 | 0.0176182 | 0.98594344 | 0.99746215 |
| ZBTB24 | 13.4852881 | -0.539293673 | 0.43936895 | -1.2274278 | 0.21966184 | 0.52962456 |
| AK9 | 73.3789913 | -0.590596531 | 0.27651481 | -2.1358585 | 0.03269095 | 0.17981313 |
| FIG4 | 73.2770666 | 0.00658699 | 0.23750973 | 0.02773356 | 0.97787466 | 0.99505657 |
| WASF1 | 32.4764208 | -0.937534943 | 0.39352935 | -2.3823762 | 0.01720132 | 0.11845278 |
| CDC40 | 38.795105 | 0.002589385 | 0.26579615 | 0.009742 | 0.99222713 | 0.99897698 |
| METTL24 | 13.7530804 | 1.74566325 | 0.64049212 | 2.72550309 | 0.00642036 | 0.05956559 |
| CDK19 | 246.533802 | -0.558973937 | 0.22874809 | -2.4436223 | 0.01454064 | 0.1059672 |
| AMD1 | 114.704242 | -1.443112131 | 0.47203869 | -3.0571903 | 0.00223422 | 0.02737438 |
| GTF3C6 | 24.3723994 | -1.054116026 | 0.49698117 | -2.1210381 | 0.03391859 | 0.18339642 |
| RPF2 | 32.3597399 | -0.306495165 | 0.4141926 | -0.7399822 | 0.45931078 | 0.74779356 |
| SLC16A10 | 91.2535652 | 1.123446652 | 0.28264502 | 3.97476195 | 7.04E-05 | 0.00188835 |
| MFSD4B | 27.3968012 | 0.085709374 | 0.3433825 | 0.24960321 | 0.80289422 | 0.93111762 |
| REV3L | 270.661366 | 0.135765838 | 0.20055428 | 0.67695308 | 0.49843572 | 0.77558432 |
| TRAF3IP2-AS | 119.932111 | -0.09863653 | 0.33470672 | -0.2946954 | 0.76822658 | 0.91600052 |
| TRAF3IP2 | 22.8289144 | -0.686974603 | 0.44662998 | -1.5381292 | 0.12401703 | 0.38919278 |
| TUBE1 | 44.5079458 | 0.586743336 | 0.34676697 | 1.6920393 | 0.09063848 | 0.32769026 |
| MARCKS | 17.634207 | -0.639569203 | 0.40841125 | -1.5659931 | 0.11735023 | 0.37756161 |
| HDAC2 | 49.4001044 | -0.098937576 | 0.26233484 | -0.3771423 | 0.70606783 | 0.88599027 |
| FRK | 235.080606 | 0.424881038 | 0.25645142 | 1.65677006 | 0.09756597 | 0.34041372 |
| NT5DC1 | 249.681342 | 0.906912025 | 0.29418029 | 3.08284424 | 0.00205032 | 0.02569519 |
| DSE | 94.366654 | 0.579015889 | 0.46268666 | 1.25142117 | 0.21078086 | 0.51965116 |
| TSPYL1 | 27.2919263 | -1.48647541 | 0.57711219 | -2.5757131 | 0.01000336 | 0.08239519 |
| RWDD1 | 19.7074958 | -0.094028174 | 0.4026361 | -0.2335314 | 0.81534878 | 0.93716865 |
| ZUP1 | 27.3597514 | 0.044325116 | 0.29464658 | 0.15043485 | 0.88042154 | 0.96097414 |
| KPNA5 | 79.9763316 | 0.745227863 | 0.36743688 | 2.0281793 | 0.04254195 | 0.21037417 |
| DCBLD1 | 18.8342994 | -0.419944941 | 0.59751946 | -0.7028138 | 0.48217178 | 0.76628579 |
| GOPC | 44.4967024 | 0.00561365 | 0.38536628 | 0.01456705 | 0.98837759 | 0.9977555 |
| NUS1 | 52.6540945 | 0.256916563 | 0.27018804 | 0.9508806 | 0.34166499 | 0.65120286 |
| SLC35F1 | 21.7221721 | 2.178249295 | 0.69545473 | 3.13212235 | 0.00173548 | 0.02275889 |
| CEP85L | 78.6012262 | -0.806018349 | 0.25977948 | -3.1027022 | 0.00191763 | 0.02449306 |
| MCM9 | 114.112347 | -0.036441926 | 0.25595825 | -0.1423745 | 0.88678421 | 0.96253908 |
| AL365275.1 | 28.393717 | -0.171667962 | 0.62241529 | -0.2758094 | 0.7826945 | 0.92150025 |
| FAM184A | 119.028004 | 1.172010449 | 0.53694933 | 2.18272078 | 0.02905638 | 0.16620962 |
| MAN1A1 | 398.383982 | 0.861744931 | 0.27667291 | 3.11467048 | 0.00184151 | 0.0236504 |
| TBC1D32 | 110.501317 | 0.810350608 | 0.44679093 | 1.8137132 | 0.06972189 | 0.28154479 |
| HSF2 | 19.6037708 | 0.260243954 | 0.38100164 | 0.68305206 | 0.49457395 | 0.77450579 |
| SERINC1 | 69.8373699 | -0.465645599 | 0.32371238 | -1.4384547 | 0.15030508 | 0.4348261 |
| PKIB | 26.402752 | 0.148180314 | 0.71922737 | 0.20602708 | 0.83676976 | 0.94684966 |
| RNF217 | 101.653771 | -1.883791708 | 0.5923503 | -3.1801988 | 0.00147174 | 0.02032964 |
| TPD52L1 | 195.533866 | -0.07989376 | 0.28048371 | -0.2848428 | 0.7757646 | 0.91926749 |
| HDDC2 | 40.4654339 | -0.511756309 | 0.44205249 | -1.1576822 | 0.24699375 | 0.55965241 |

| | | | | | | |
|---|---|---|---|---|---|---|
| NCOA7 | 117.091805 | -2.01849949 | 0.48772441 | -4.1386066 | 3.49E-05 | 0.00107696 |
| HINT3 | 25.6080394 | -0.263153858 | 0.43854474 | -0.6000616 | 0.54846518 | 0.80647995 |
| TRMT11 | 51.0943732 | -0.399663869 | 0.26037748 | -1.5349402 | 0.12479851 | 0.39027762 |
| AC020559.1 | 14.5697984 | -0.691053446 | 0.64261812 | -1.0753719 | 0.28220828 | 0.59642935 |
| RNF146 | 40.1230453 | 0.289190161 | 0.28345483 | 1.02023368 | 0.30761765 | 0.62161058 |
| ECHDC1 | 99.7758135 | -0.000416147 | 0.25712589 | -0.0016185 | 0.99870866 | 0.99924629 |
| PTPRK | 427.844952 | -0.193014229 | 0.25778721 | -0.7487347 | 0.45401712 | 0.74489806 |
| AL356124.1 | 29.9834004 | -1.259130451 | 0.55284423 | -2.2775501 | 0.02275339 | 0.14209822 |
| ARHGAP18 | 199.574626 | 0.338792212 | 0.40875823 | 0.82883276 | 0.40719905 | 0.70610451 |
| L3MBTL3 | 32.8696566 | -0.14512475 | 0.43673639 | -0.3322937 | 0.7396675 | 0.90304252 |
| EPB41L2 | 42.3328376 | -1.799313813 | 0.38113758 | -4.7209037 | 2.35E-06 | 0.00011462 |
| AKAP7 | 46.6307972 | 1.12429442 | 0.57970296 | 1.93943192 | 0.05244877 | 0.23693425 |
| MED23 | 69.6202731 | 0.430852022 | 0.26221918 | 1.64309881 | 0.10036249 | 0.34581665 |
| CTGF | 20.0823036 | -0.195122792 | 0.79412006 | -0.2457094 | 0.80590717 | 0.93308942 |
| STX7 | 103.311274 | -0.003970147 | 0.26063256 | -0.0152327 | 0.98784651 | 0.9977555 |
| SLC18B1 | 47.9495737 | 0.668080356 | 0.46992049 | 1.42168808 | 0.15511682 | 0.44067923 |
| RPS12 | 51.3485874 | -0.134461565 | 0.4862513 | -0.2765269 | 0.7821434 | 0.92137777 |
| TBPL1 | 27.7354359 | 0.533854317 | 0.39111498 | 1.36495494 | 0.17226721 | 0.46801268 |
| SLC2A12 | 26.5761572 | 0.855009893 | 0.50355956 | 1.69793199 | 0.0895206 | 0.32478213 |
| SGK1 | 108.119903 | -2.042488908 | 0.81600273 | -2.5030418 | 0.0123131 | 0.09519681 |
| HBS1L | 70.0885505 | 0.297679414 | 0.26496324 | 1.12347438 | 0.26123608 | 0.57487967 |
| AHI1 | 444.565096 | -0.433935902 | 0.30190554 | -1.4373234 | 0.15062611 | 0.43488337 |
| LINC00271 | 144.539645 | 0.004708092 | 0.69520804 | 0.00677221 | 0.9945966 | 0.9992177 |
| PDE7B | 469.730502 | 0.022227625 | 0.38671675 | 0.05747779 | 0.9541646 | 0.98807538 |
| AL138828.1 | 193.203183 | -4.867237565 | 1.41425223 | -3.4415626 | 0.00057836 | 0.00961261 |
| BCLAF1 | 152.521371 | 0.030041321 | 0.2153647 | 0.13949046 | 0.88906259 | 0.96307703 |
| MAP7 | 327.518135 | -0.054448579 | 0.29502259 | -0.1845573 | 0.85357626 | 0.95137449 |
| MAP3K5 | 118.091316 | -1.404564813 | 0.40325925 | -3.4830318 | 0.00049577 | 0.00856029 |
| PEX7 | 52.5251616 | 0.471736817 | 0.28336798 | 1.6647499 | 0.09596266 | 0.33853797 |
| IFNGR1 | 35.817907 | 0.156274185 | 0.306349 | 0.51011816 | 0.60996869 | 0.84109213 |
| PERP | 32.5602251 | -1.339761114 | 0.48268418 | -2.7756475 | 0.00550919 | 0.05339677 |
| ARFGEF3 | 63.049921 | -0.761964254 | 0.43763696 | -1.7410875 | 0.08166824 | 0.30754226 |
| HEBP2 | 36.8465367 | -0.38622927 | 0.3530056 | -1.0941165 | 0.27390387 | 0.58894058 |
| NHSL1 | 106.065832 | -2.328361755 | 0.69118806 | -3.3686371 | 0.00075541 | 0.01209035 |
| CCDC28A | 20.4340198 | 0.143146107 | 0.46509949 | 0.30777524 | 0.75825336 | 0.91161939 |
| REPS1 | 132.083805 | -0.032018271 | 0.20183763 | -0.1586338 | 0.87395739 | 0.95761805 |
| HECA | 19.9922702 | -1.002308175 | 0.50880793 | -1.9699146 | 0.04884816 | 0.22682281 |
| CITED2 | 21.6302922 | -1.068456705 | 0.44749061 | -2.3876628 | 0.01695589 | 0.11753644 |
| VTA1 | 60.9393363 | 0.046300672 | 0.24654733 | 0.18779629 | 0.85103635 | 0.95042076 |
| HIVEP2 | 364.585414 | -1.0700924 | 0.52683339 | -2.031178 | 0.04223694 | 0.20951607 |
| AIG1 | 227.544598 | 0.703817881 | 0.24925096 | 2.82373193 | 0.00474681 | 0.04831793 |
| ADAT2 | 21.4459521 | 0.328045047 | 0.54489772 | 0.60203051 | 0.54715383 | 0.80594981 |
| PEX3 | 74.6343311 | -0.265576633 | 0.25450526 | -1.0435015 | 0.29671607 | 0.61093809 |
| FUCA2 | 22.7769134 | -0.194277818 | 0.31172229 | -0.62324 | 0.53312679 | 0.79750909 |
| PHACTR2 | 230.023569 | -0.54398318 | 0.21486055 | -2.5317965 | 0.01134798 | 0.08982054 |
| PLAGL1 | 83.4077197 | -0.606991459 | 0.43268246 | -1.4028566 | 0.1606596 | 0.44998201 |
| SF3B5 | 21.6388011 | -0.498103948 | 0.57574858 | -0.8651414 | 0.38696117 | 0.69039914 |
| UTRN | 401.644648 | 0.463315197 | 0.3886853 | 1.19200596 | 0.23325891 | 0.54577316 |
| EPM2A | 85.9449308 | 0.467052541 | 0.42934711 | 1.08782038 | 0.27667441 | 0.59167711 |
| FBXO30 | 15.5614337 | 0.348880819 | 0.43648495 | 0.79929634 | 0.4241186 | 0.72151771 |
| AL356599.1 | 45.9205448 | 0.362825888 | 0.63149317 | 0.57455236 | 0.56559408 | 0.81595078 |
| SHPRH | 211.746384 | 0.531202832 | 0.29979425 | 1.77189134 | 0.0764126 | 0.29596141 |
| STXBP5-AS1 | 30.2239598 | 0.880138708 | 0.59969983 | 1.46763207 | 0.14220419 | 0.4225818 |
| STXBP5 | 164.620977 | -0.304072678 | 0.20904842 | -1.4545562 | 0.14579216 | 0.42828171 |
| SAMD5 | 901.103969 | -1.356110931 | 0.33656132 | -4.0293131 | 5.59E-05 | 0.00156163 |
| AL033504.1 | 527.360001 | -1.327656999 | 0.35614834 | -3.7278203 | 0.00019314 | 0.00411157 |
| AL513164.1 | 25.5183355 | -2.195336535 | 0.58303445 | -3.7653633 | 0.00016631 | 0.00363149 |
| SASH1 | 369.32054 | -1.890836192 | 0.33139516 | -5.7056845 | 1.16E-08 | 1.21E-06 |
| UST | 132.218431 | 1.150672922 | 0.35398969 | 3.25058317 | 0.00115169 | 0.01664856 |
| TAB2 | 204.305223 | 0.128528409 | 0.33720639 | 0.3811565 | 0.70308712 | 0.88513266 |
| PPIL4 | 80.0313245 | 0.052139191 | 0.25496262 | 0.20449739 | 0.83796483 | 0.94728226 |
| GINM1 | 77.9447128 | 0.123203518 | 0.35168912 | 0.35031939 | 0.72609901 | 0.89457547 |
| KATNA1 | 37.2979449 | 0.01185167 | 0.29860161 | 0.03969058 | 0.96833981 | 0.99156233 |
| LATS1 | 103.054761 | 0.396582946 | 0.17429365 | 2.27537226 | 0.02288361 | 0.14252957 |
| NUP43 | 37.6180417 | 0.104272373 | 0.30494367 | 0.34193979 | 0.73239621 | 0.89806184 |
| PCMT1 | 59.9321159 | -0.441562856 | 0.35369207 | -1.2484387 | 0.21187045 | 0.52068532 |
| LRP11 | 24.0375296 | 0.003445617 | 0.49399453 | 0.00697501 | 0.99443479 | 0.9992177 |
| PLEKHG1 | 57.3518221 | -2.358670767 | 0.52840528 | -4.4637532 | 8.05E-06 | 0.00032091 |
| MTHFD1L | 134.488189 | -2.604928443 | 0.61249647 | -4.2529689 | 2.11E-05 | 0.00073121 |
| RMND1 | 40.4988898 | -0.256619202 | 0.32210249 | -0.7967005 | 0.42562502 | 0.72207563 |
| ARMT1 | 20.4832347 | 0.379911171 | 0.45115404 | 0.84208748 | 0.39973899 | 0.69959441 |
| SYNE1 | 389.52572 | 0.181915664 | 0.45201203 | 0.40245757 | 0.68734731 | 0.87556719 |
| MTRF1L | 20.4997235 | -0.120977552 | 0.38749814 | -0.3122016 | 0.75488729 | 0.91031675 |
| IPCEF1 | 165.828783 | 1.743461082 | 0.48904524 | 3.56503027 | 0.00036381 | 0.00683186 |
| CNKSR3 | 293.068596 | 0.502633463 | 0.32260742 | 1.55803442 | 0.11922509 | 0.38057334 |
| SCAF8 | 277.561609 | 0.452249197 | 0.22519543 | 2.00825211 | 0.04461651 | 0.21576988 |
| TIAM2 | 161.362012 | 0.001240161 | 0.26756741 | 0.00463495 | 0.99630186 | 0.99922221 |
| TFB1M | 83.2335381 | 0.070565085 | 0.33360906 | 0.21152029 | 0.8324813 | 0.94520892 |
| ARID1B | 966.604319 | 0.173336994 | 0.14220571 | 1.21891727 | 0.22287559 | 0.53311751 |
| TMEM242 | 33.5798895 | 0.131103397 | 0.27964621 | 0.46881878 | 0.63919917 | 0.85391761 |
| ZDHHC14 | 609.431035 | 0.498019197 | 0.45387699 | 1.09725588 | 0.27252955 | 0.58820961 |
| SNX9 | 380.739152 | -0.935596299 | 0.52813813 | -1.7714992 | 0.07647772 | 0.29596141 |
| SYNJ2 | 118.740482 | -2.62296176 | 0.74729142 | -3.5099584 | 0.00044818 | 0.00792941 |
| SERAC1 | 23.4079921 | 0.136858192 | 0.37255556 | 0.36734975 | 0.71335815 | 0.88813689 |
| GTF2H5 | 18.4503204 | -0.047457192 | 0.57213358 | -0.0829477 | 0.93389309 | 0.98045386 |
| TULP4 | 282.241889 | -0.267680381 | 0.47820376 | -0.5597622 | 0.57564166 | 0.82147339 |
| TMEM181 | 55.8211374 | -0.746314754 | 0.24973927 | -2.9883756 | 0.00280465 | 0.03293516 |
| DYNLT1 | 40.882471 | -0.727713092 | 0.38042946 | -1.9128726 | 0.05576436 | 0.24602764 |
| SYTL3 | 41.583093 | -1.691132427 | 0.46335504 | -3.6497551 | 0.00026249 | 0.00525206 |
| EZR | 163.163942 | -0.547580721 | 0.35068871 | -1.5614438 | 0.11841907 | 0.37913731 |
| RSPH3 | 47.4106516 | -0.46842612 | 0.40327966 | -1.1615416 | 0.2454217 | 0.55758089 |
| AL035530.2 | 47.1092235 | 1.491676817 | 0.64092526 | 2.32738029 | 0.01994503 | 0.13082568 |
| SOD2.1 | 19.8543092 | -0.316009519 | 0.54082612 | -0.5843089 | 0.5590125 | 0.81279041 |
| WTAP | 171.283376 | -0.249403935 | 0.44903009 | -0.5554281 | 0.57860187 | 0.82318213 |
| TCP1 | 32.0868828 | -0.099914493 | 0.29829147 | -0.3349559 | 0.73765834 | 0.90202313 |
| MRPL18 | 16.5862992 | -0.131694224 | 0.36183599 | -0.3639611 | 0.71588706 | 0.88904248 |
| IGF2R | 285.055902 | -0.336452203 | 0.42503561 | -0.7915859 | 0.42860216 | 0.7236272 |
| SLC22A1 | 15.5318641 | -0.531789118 | 0.82382826 | -0.6455097 | 0.51859695 | 0.78726765 |
| SLC22A2 | 16.0810579 | -0.056169223 | 0.54055229 | -0.1039108 | 0.91724013 | 0.97460959 |
| SLC22A3 | 85.725073 | -0.114355857 | 0.79231911 | -0.1443306 | 0.88523945 | 0.96233796 |
| MAP3K4 | 106.612879 | 0.521929773 | 0.22113689 | 2.36021123 | 0.01826453 | 0.12340471 |
| AGPAT4 | 45.966017 | 0.302301938 | 0.29321731 | 1.03098259 | 0.30254898 | 0.61691783 |
| PRKN | 573.738096 | 0.570615307 | 0.47345517 | 1.20521508 | 0.22812028 | 0.54025743 |
| PACRG | 81.2493133 | 0.095281778 | 0.52342198 | 0.18203626 | 0.85555427 | 0.95215899 |
| QKI | 587.363295 | 0.041064164 | 0.28957582 | 0.14180799 | 0.88723168 | 0.96259578 |
| PDE10A | 706.740629 | 0.372620597 | 0.37364535 | 0.99725743 | 0.31863957 | 0.63093651 |
| SFT2D1 | 43.0420189 | -0.257565302 | 0.31768388 | -0.8107598 | 0.41750365 | 0.71572054 |
| MPC1 | 184.465335 | 0.912294957 | 0.39857883 | 2.28886957 | 0.02208693 | 0.13937617 |
| RPS6KA2 | 86.7018149 | -0.182231145 | 0.26779972 | -0.6804755 | 0.49620344 | 0.77475164 |
| Z94721.1 | 26.384406 | -0.544939227 | 0.42160623 | -1.2925313 | 0.19617323 | 0.49845696 |
| FGFR1OP | 92.3141164 | 0.148034518 | 0.37131489 | 0.39867649 | 0.69013159 | 0.87691291 |
| AFDN | 354.322181 | 0.181108408 | 0.22501636 | 0.80486772 | 0.42089602 | 0.71917261 |
| WDR27 | 123.078895 | 0.704701096 | 0.4108474 | 1.71523807 | 0.08630159 | 0.31751647 |

| | | | | | | |
|---|---|---|---|---|---|---|
| PHF10 | 54.4177026 | 0.324137934 | 0.28058393 | 1.15522628 | 0.24799778 | 0.56056515 |
| ERMARD | 22.653273 | 1.0344792 | 0.36133664 | 2.86292358 | 0.00419752 | 0.04426547 |
| FAM120B | 181.826481 | -0.112109159 | 0.36202665 | -0.309671 | 0.75681115 | 0.91122251 |
| PSMB1 | 48.4594059 | -0.02759866 | 0.28060458 | -0.0983543 | 0.92165098 | 0.9757579 |
| TBP | 24.4629968 | -0.176783546 | 0.35987176 | -0.4912404 | 0.62325643 | 0.84597807 |
| PDCD2 | 16.4910112 | -0.382711105 | 0.46115752 | -0.8298924 | 0.40659964 | 0.70558999 |
| FAM20C | 104.08467 | -2.076631493 | 0.34326631 | -6.0496223 | 1.45E-09 | 2.26E-07 |
| PDGFA | 27.6443492 | -1.305721257 | 0.37307603 | -3.4998797 | 0.00046547 | 0.00818873 |
| PRKAR1B | 84.7529734 | 0.339374483 | 0.41491526 | 0.81793686 | 0.41339324 | 0.71143975 |
| DNAAF5 | 38.2832131 | -0.193012199 | 0.33303898 | -0.5795484 | 0.56221922 | 0.81388399 |
| SUN1 | 95.8242274 | 0.295610737 | 0.27558597 | 1.07266251 | 0.28342257 | 0.59716825 |
| COX19 | 40.3388067 | -0.178395748 | 0.3672959 | -0.4857004 | 0.62717963 | 0.84775629 |
| C7orf50 | 61.6679873 | 0.038330055 | 0.28586658 | 0.13408372 | 0.89333637 | 0.96472879 |
| INTS1 | 32.8845317 | 0.444884578 | 0.44640181 | 0.99660119 | 0.31895813 | 0.63100017 |
| MAFK | 26.5393083 | 0.04860521 | 0.63581948 | 0.07644498 | 0.93906508 | 0.98358154 |
| MAD1L1 | 48.8214245 | -0.303207657 | 0.34952324 | -0.8674893 | 0.38567399 | 0.68921713 |
| SNX8 | 17.5124791 | -0.844820939 | 0.55409622 | -1.5246827 | 0.12733825 | 0.3952403 |
| EIF3B | 52.2633368 | -0.136414929 | 0.34149923 | -0.399459 | 0.68955505 | 0.87665821 |
| CHST12 | 18.8115202 | 0.185514688 | 0.37984208 | 0.48839952 | 0.62526688 | 0.8471532 |
| BRAT1 | 18.2979638 | 0.580142872 | 0.44197424 | 1.31261693 | 0.18931206 | 0.48954068 |
| IQCE | 32.740213 | 0.023297234 | 0.3432307 | 0.06787631 | 0.9458841 | 0.98574808 |
| TTYH3 | 17.4720038 | -1.380914852 | 0.60889126 | -2.267917 | 0.02333427 | 0.14456391 |
| GNA12 | 44.3113525 | -0.168417761 | 0.28867486 | -0.5834168 | 0.55961276 | 0.81314937 |
| SDK1 | 83.2579705 | -1.454056561 | 0.44654908 | -3.2562077 | 0.00112911 | 0.01644974 |
| FOXK1 | 112.039387 | -0.108852121 | 0.26715246 | -0.4074532 | 0.68367517 | 0.87343121 |
| AP5Z1 | 26.4973558 | -1.595625433 | 0.65689793 | -2.429031 | 0.01513924 | 0.10908675 |
| RBAK-RBAKI | 71.9254667 | -0.150981728 | 0.26381444 | -0.5723028 | 0.56711688 | 0.81614413 |
| WIPI2 | 48.3755834 | 0.070865548 | 0.31595486 | 0.2242901 | 0.82253158 | 0.94066053 |
| TNRC18 | 239.235903 | 0.153304675 | 0.38816893 | 0.39494319 | 0.69288482 | 0.87849579 |
| FBXL18 | 34.1226222 | 0.182451749 | 0.32598604 | 0.55969191 | 0.5756896 | 0.82147339 |
| ACTB | 239.26961 | -0.772036562 | 0.56246656 | -1.372591 | 0.16987952 | 0.46436723 |
| RNF216 | 176.889115 | 0.238349526 | 0.19423093 | 1.22714509 | 0.21976806 | 0.52962456 |
| CCZ1 | 24.9686374 | -0.184163191 | 0.42325603 | -0.4351106 | 0.66348214 | 0.8658233 |
| PMS2 | 41.8829038 | 0.379339667 | 0.34232996 | 1.1081112 | 0.26781379 | 0.58378751 |
| AIMP2 | 21.6728713 | -0.278265235 | 0.38579124 | -0.7212845 | 0.4707345 | 0.75739922 |
| EIF2AK1 | 54.5117711 | -0.066894131 | 0.2586318 | -0.2586462 | 0.79590824 | 0.92698936 |
| USP42 | 63.1986919 | 0.078985176 | 0.25233631 | 0.3130155 | 0.75426888 | 0.91026551 |
| CYTH3 | 73.3832785 | 0.591767223 | 0.31026573 | 1.90729162 | 0.05648284 | 0.247484 |
| RAC1 | 111.799749 | -0.402721027 | 0.2414726 | -1.6677711 | 0.09536117 | 0.33742332 |
| DAGLB | 19.3007516 | -0.263528156 | 0.41688973 | -0.6321292 | 0.52730248 | 0.79403853 |
| C7orf26 | 15.5888816 | 0.723177636 | 0.40142899 | 1.80150826 | 0.07162281 | 0.28587802 |
| ZNF316 | 21.143852 | -0.01343063 | 0.56687348 | -0.0236925 | 0.98109792 | 0.99653318 |
| AC073343.2 | 17.472493 | -0.24683255 | 0.63914482 | -0.3861919 | 0.69935455 | 0.8833378 |
| ZNF12 | 29.5335948 | 0.129388694 | 0.35056052 | 0.36909089 | 0.71205998 | 0.88721964 |
| CCZ1B | 30.8840789 | 0.071777142 | 0.42404519 | 0.16926767 | 0.8655861 | 0.9555677 |
| C1GALT1 | 71.3689689 | 0.043021426 | 0.36687779 | 0.11726364 | 0.90665113 | 0.96989619 |
| COL28A1 | 45.7310053 | -0.029584798 | 0.41758452 | -0.0708474 | 0.94351917 | 0.98536881 |
| AC004982.2 | 11.2655782 | 0.544475144 | 0.60046463 | 0.90675639 | 0.36453563 | 0.67012584 |
| MIOS | 55.762872 | 0.038095662 | 0.30226247 | 0.12603504 | 0.89970419 | 0.96656779 |
| RPA3 | 29.0670581 | -0.894497612 | 0.5880749 | -1.5210607 | 0.12824461 | 0.39699626 |
| UMAD1 | 340.828808 | 0.086560282 | 0.22072749 | 0.39215904 | 0.6949407 | 0.87978644 |
| GLCCI1 | 127.470808 | 1.043240302 | 0.39707848 | 2.62728997 | 0.0086068 | 0.07374644 |
| ICA1 | 473.676787 | 1.44778996 | 0.28174573 | 5.13864034 | 2.77E-07 | 1.97E-05 |
| NDUFA4 | 70.7672547 | 0.023491157 | 0.46092264 | 0.05096551 | 0.959353 | 0.98938253 |
| PHF14 | 141.02055 | -0.06163336 | 0.2778888 | -0.2217915 | 0.82447623 | 0.94153354 |
| THSD7A | 598.817119 | -1.536999974 | 0.62337791 | -2.465599 | 0.01367844 | 0.10211187 |
| TMEM106B | 69.2044734 | -0.129994605 | 0.27672826 | -0.4697554 | 0.63852976 | 0.8538689 |
| SCIN | 232.862331 | -0.037151375 | 0.31854824 | -0.1166272 | 0.90715552 | 0.9700616 |
| AC011287.1 | 23.1632471 | -2.3133994 | 0.50576753 | -4.574037 | 4.78E-06 | 0.00020942 |
| ISPD | 61.9839594 | 0.297454718 | 0.30575064 | 0.97286705 | 0.33061939 | 0.6426298 |
| ANKMY2 | 41.0875983 | 0.167028263 | 0.33544452 | 0.49793111 | 0.6185326 | 0.84283824 |
| BZW2 | 55.2214682 | -2.097095286 | 0.37374756 | -5.610994 | 2.01E-08 | 1.88E-06 |
| AC073332.1 | 20.1738725 | -0.813205946 | 0.89721993 | -0.9063619 | 0.36474434 | 0.67012584 |
| SNX13 | 195.292148 | -0.418205225 | 0.21943434 | -1.9058331 | 0.05667187 | 0.24796269 |
| HDAC9 | 34.6786738 | -0.431878442 | 0.58277315 | -0.7410747 | 0.45864814 | 0.747377 |
| MACC1 | 100.203029 | -1.452766672 | 0.64687143 | -2.2458353 | 0.02471456 | 0.14944135 |
| ITGB8 | 74.5476202 | -1.933421429 | 0.57445551 | -3.3656591 | 0.00076361 | 0.01215 |
| SP4 | 61.5438567 | 0.288767425 | 0.28559381 | 1.01111235 | 0.31196266 | 0.62688359 |
| RAPGEF5 | 232.495536 | -1.916423407 | 0.64562276 | -2.9683331 | 0.0029942 | 0.03459466 |
| STEAP1B | 54.3117506 | -1.202713416 | 0.53448362 | -2.2502344 | 0.02443407 | 0.14863223 |
| TOMM7 | 34.4068609 | -0.198101795 | 0.54190699 | -0.3655642 | 0.71469029 | 0.88904248 |
| KLHL7 | 48.653573 | -0.01020957 | 0.26066688 | -0.0391671 | 0.96875715 | 0.99156233 |
| NUPL2 | 56.5097889 | -0.086613617 | 0.30189963 | -0.2868954 | 0.77419241 | 0.91862688 |
| GPNMB | 28.9063134 | -2.169267781 | 0.81123932 | -2.6740171 | 0.00749486 | 0.0666814 |
| MALSU1 | 35.8260523 | 0.360076021 | 0.3085895 | 1.1668447 | 0.24327309 | 0.55555126 |
| TRA2A | 265.969264 | 0.31585449 | 0.26112949 | 1.20957037 | 0.2264438 | 0.53792661 |
| CCDC126 | 26.4107763 | -0.115738683 | 0.45690287 | -0.2533113 | 0.80002763 | 0.92941161 |
| FAM221A | 16.3709792 | 0.306131056 | 0.45256244 | 0.67643938 | 0.49876172 | 0.77558432 |
| MPP6 | 29.6389738 | -0.788872217 | 0.40713679 | -1.9376097 | 0.05267085 | 0.23770718 |
| GSDME | 44.9322803 | 1.526032523 | 0.51111342 | 2.9857023 | 0.00282928 | 0.03302226 |
| OSBPL3 | 591.96416 | 0.782041552 | 0.36936759 | 2.11724467 | 0.03423909 | 0.18418162 |
| CYCS | 37.6459731 | 0.443316119 | 0.50021842 | 0.88624509 | 0.37548547 | 0.67888822 |
| NFE2L3 | 20.2830189 | -1.226856579 | 0.57624114 | -2.129068 | 0.03324863 | 0.1818242 |
| HNRNPA2B1 | 468.694253 | 0.225972391 | 0.23644009 | 0.95572791 | 0.33920972 | 0.64971064 |
| CBX3 | 78.4930781 | -0.635056043 | 0.29134211 | -2.1797606 | 0.02927521 | 0.16684724 |
| SNX10 | 467.00234 | -0.23529582 | 0.22680939 | -1.0374166 | 0.29954176 | 0.61356712 |
| AC004540.1 | 54.3938141 | -0.152263469 | 0.79452674 | -0.1916405 | 0.84802385 | 0.95001494 |
| SKAP2 | 118.179256 | -0.386987302 | 0.39337977 | -0.9837499 | 0.32523851 | 0.63715682 |
| HOXA3 | 17.2366245 | -0.016738152 | 0.81986096 | -0.0204158 | 0.98371165 | 0.99742652 |
| HOXA9 | 12.5479709 | 1.449902846 | 0.66860055 | 2.16856365 | 0.03011583 | 0.170285 |
| HIBADH | 277.322594 | 1.363605752 | 0.34475859 | 3.95524809 | 7.65E-05 | 0.00199691 |
| TAX1BP1 | 275.623057 | 0.084348406 | 0.15375994 | 0.54857207 | 0.58329916 | 0.82696794 |
| JAZF1 | 80.4518004 | -0.221983845 | 0.54230927 | -0.4093307 | 0.68229697 | 0.87274146 |
| CREB5 | 42.2082335 | -0.9282818 | 0.8872253 | -1.0462752 | 0.29543401 | 0.61055359 |
| AC005162.3 | 21.9805374 | 0.956489049 | 0.67877494 | 1.4091402 | 0.15879372 | 0.44717385 |
| CPVL | 55.2400936 | -0.219580997 | 0.50144811 | -0.4378938 | 0.6614633 | 0.86492722 |
| AC007255.1 | 22.2718329 | 0.051949392 | 0.72825707 | 0.07133387 | 0.94313204 | 0.98507485 |
| SCRN1 | 56.8963504 | -0.2685086 | 0.29740523 | -0.9028375 | 0.36661214 | 0.67113782 |
| FKBP14 | 19.137332 | -0.494257676 | 0.41902548 | -1.1795409 | 0.23818288 | 0.54997949 |
| PLEKHA8 | 65.8399053 | -0.592456911 | 0.29624556 | -1.9998845 | 0.04551274 | 0.21869588 |
| MTURN | 122.028475 | 0.733233462 | 0.33009495 | 2.22128047 | 0.02633197 | 0.15607861 |
| ZNRF2 | 108.752614 | -0.822524519 | 0.29633068 | -2.7756982 | 0.00550833 | 0.05339677 |
| AC006978.2 | 90.4668932 | -1.04666499 | 0.33167405 | -3.1557036 | 0.00160111 | 0.0215734 |
| NOD1 | 18.610678 | -0.407730698 | 0.45500052 | -0.8961104 | 0.3701938 | 0.67442107 |
| AC005154.6 | 41.7756971 | 0.876003976 | 0.41296115 | 2.12127452 | 0.03389871 | 0.18339642 |
| GARS | 43.7248048 | -0.593371599 | 0.50039175 | -1.1858141 | 0.23569572 | 0.54835802 |
| AQP1 | 22.4175797 | -0.737230348 | 0.55709204 | -1.3233547 | 0.18571746 | 0.4858669 |
| PDE1C | 49.69614 | -2.50068845 | 0.72881084 | -3.43119 | 0.00060094 | 0.00997004 |
| LSM5 | 14.7173126 | -0.227284405 | 0.44605173 | -0.5095472 | 0.61036873 | 0.84119857 |
| AVL9 | 227.25216 | -0.497030195 | 0.25056759 | -1.9836173 | 0.04729852 | 0.22318392 |
| KBTBD2 | 53.612881 | -0.418129113 | 0.25943183 | -1.6117109 | 0.10702486 | 0.35973315 |
| FKBP9 | 33.9094894 | 0.452566668 | 0.46453829 | 0.97422898 | 0.32994286 | 0.64198398 |
| NT5C3A | 64.5477932 | 0.385835498 | 0.41167744 | 0.93722769 | 0.34864145 | 0.65587927 |

| | | | | | | |
|---|---|---|---|---|---|---|
| RP9 | 16.4800978 | 0.136958621 | 0.39624501 | 0.34564125 | 0.72961234 | 0.89712587 |
| BBS9 | 218.17093 | 0.538138789 | 0.47445773 | 1.13421861 | 0.25670286 | 0.56935259 |
| DPY19L1 | 121.443894 | -0.529502492 | 0.3301785 | -1.6036855 | 0.10878338 | 0.3628967 |
| HERPUD2 | 74.3378106 | -0.044819992 | 0.24824956 | -0.1805441 | 0.85672544 | 0.95234014 |
| SEPT7-AS1 | 23.1812756 | 0.47138766 | 0.58088308 | 0.81150179 | 0.41707757 | 0.7151216 |
| 7-Sep | 164.889954 | -0.002081717 | 0.19171295 | -0.0108585 | 0.99133633 | 0.9988777 |
| EEPD1 | 49.6678015 | -2.392250347 | 0.7537915 | -3.1736234 | 0.00150549 | 0.02065826 |
| KIAA0895 | 37.381835 | -0.052885061 | 0.36193282 | -0.1461184 | 0.88382788 | 0.96170562 |
| AOAH | 20.5587193 | -3.155169559 | 0.99915384 | -3.1578416 | 0.00158942 | 0.02147789 |
| ELMO1 | 26.3882596 | -0.780562381 | 0.97555805 | -0.8001188 | 0.42364194 | 0.72151771 |
| EPDR1 | 21.8964625 | 0.101634772 | 0.68979186 | 0.14734122 | 0.8828627 | 0.96161232 |
| STARD3NL | 30.6175302 | -0.53233211 | 0.36842196 | -1.4448979 | 0.14848657 | 0.4322475 |
| AMPH | 37.8599471 | 2.466976416 | 0.90605021 | 2.72278113 | 0.00647349 | 0.05982046 |
| VPS41 | 151.013231 | -0.281230327 | 0.22793916 | -1.2337956 | 0.21727908 | 0.52689471 |
| YAE1D1 | 17.8777942 | 0.43117034 | 0.34753294 | 1.24066035 | 0.21473125 | 0.52343899 |
| RALA | 80.9046788 | -0.744766387 | 0.38284271 | -1.9453587 | 0.05173182 | 0.23500769 |
| LINC00265 | 14.209792 | 0.318239211 | 0.49002937 | 0.64942885 | 0.51606122 | 0.78456789 |
| CDK13 | 351.322155 | -0.037606303 | 0.22614965 | -0.1662895 | 0.86792916 | 0.95588054 |
| MPLKIP | 27.0300179 | 0.390741208 | 0.40562545 | 0.96330546 | 0.33539425 | 0.64662829 |
| SUGCT | 103.451977 | 1.292987095 | 0.36432285 | 3.54901451 | 0.00038668 | 0.00719634 |
| MRPL32 | 17.8217552 | 0.241718004 | 0.39533841 | 0.6114205 | 0.54092124 | 0.80234642 |
| STK17A | 52.6329122 | -0.91685382 | 0.56183615 | -1.6318883 | 0.10270302 | 0.35025713 |
| COA1 | 204.338546 | 0.375543362 | 0.25684676 | 1.46213002 | 0.14370558 | 0.42455984 |
| BLVRA | 20.8139014 | -0.412058961 | 0.39594512 | -1.0406972 | 0.29801612 | 0.61151019 |
| URGCP | 39.1775808 | 0.253547108 | 0.44188014 | 0.57379159 | 0.56610884 | 0.81595078 |
| UBE2D4 | 11.0331242 | 0.131078576 | 0.44680835 | 0.29336644 | 0.76924207 | 0.916134 |
| AC004951.1 | 10.4734888 | 0.52755902 | 0.45875089 | 1.14999018 | 0.25014792 | 0.56269702 |
| DBNL | 23.9760446 | -0.209966444 | 0.30390388 | -0.6908975 | 0.48962993 | 0.77129743 |
| POLM | 21.6688377 | -0.089767566 | 0.52704663 | -0.1703219 | 0.86475701 | 0.95499164 |
| YKT6 | 14.4738666 | -0.114437672 | 0.61861478 | -0.1849902 | 0.85323672 | 0.95137449 |
| CAMK2B | 10.6176023 | 2.935158386 | 0.65062423 | 4.51129587 | 6.44E-06 | 0.00026358 |
| NUDCD3 | 114.157253 | 0.196069643 | 0.23423985 | 0.83704648 | 0.40256644 | 0.70169281 |
| DDX56 | 19.6130495 | -0.183002361 | 0.36941796 | -0.4953802 | 0.62033173 | 0.84412917 |
| TMED4 | 33.7257278 | -0.13921696 | 0.36463817 | -0.3817948 | 0.70261357 | 0.88469533 |
| OGDH | 317.483623 | 0.480471308 | 0.31622782 | 1.51938343 | 0.12866601 | 0.39737957 |
| ZMIZ2 | 54.3234096 | 0.006230007 | 0.3058796 | 0.02036751 | 0.9837502 | 0.99742652 |
| PPIA | 109.508798 | -1.188834098 | 0.42687751 | -2.7849537 | 0.00535354 | 0.05253187 |
| H2AFV | 58.8910812 | 0.085642711 | 0.33982301 | 0.25202152 | 0.80102443 | 0.93010607 |
| PURB | 44.7068422 | -0.308777576 | 0.2683181 | -1.1507892 | 0.24981896 | 0.5624056 |
| CCM2 | 55.152093 | 0.062819012 | 0.2962361 | 0.21205725 | 0.83206237 | 0.9450226 |
| TBRG4 | 16.698891 | -0.140948354 | 0.51645093 | -0.2729172 | 0.78491684 | 0.92276669 |
| IGFBP3 | 125.243396 | -0.589246525 | 0.9593164 | -0.6142358 | 0.53905949 | 0.80149747 |
| AC004870.4 | 34.9297203 | -0.024796981 | 0.92647923 | -0.0267647 | 0.97864737 | 0.9951912 |
| TNS3 | 520.695403 | -1.325478936 | 0.42059211 | -3.1514593 | 0.00162457 | 0.02180146 |
| HUS1 | 31.9294133 | -0.115152946 | 0.38717595 | -0.2974176 | 0.7661477 | 0.91564296 |
| UPP1 | 146.105197 | -3.069382148 | 0.72755542 | -4.2187606 | 2.46E-05 | 0.00080934 |
| GRB10 | 65.6006304 | -1.309684693 | 0.77023776 | -1.7003642 | 0.08906245 | 0.32396747 |
| COBL | 197.353118 | 0.272196702 | 0.5229031 | 0.52054903 | 0.60268096 | 0.83691476 |
| SEC61G | 19.1841357 | -0.150337854 | 0.3894108 | -0.3860649 | 0.69944858 | 0.8833378 |
| EGFR | 50.0912251 | -1.671082453 | 0.55142915 | -3.0304572 | 0.00244184 | 0.02955001 |
| LANCL2 | 39.1067828 | 0.000189166 | 0.29117977 | 0.00064965 | 0.99948165 | 0.99969609 |
| VOPP1 | 166.938235 | -0.410601001 | 0.21593669 | -1.9014879 | 0.05723813 | 0.24962036 |
| NIPSNAP2 | 112.292348 | 0.255631509 | 0.24047846 | 1.06301211 | 0.28777646 | 0.6024252 |
| PSPH | 16.528292 | -0.122362288 | 0.58940009 | -0.2076048 | 0.83553756 | 0.94654164 |
| CCT6A | 40.5256768 | -0.415495039 | 0.38801532 | -1.0708212 | 0.28424982 | 0.59746288 |
| SUMF2 | 43.8232383 | -0.365373506 | 0.30597658 | -1.1941224 | 0.23243007 | 0.54506955 |
| CHCHD2 | 65.2872608 | -0.421132688 | 0.45323706 | -0.9291665 | 0.35280281 | 0.65972795 |
| ZNF727 | 15.476178 | 0.412826295 | 0.60090701 | 0.68700529 | 0.49207939 | 0.77284018 |
| ZNF736 | 79.5235421 | -0.098417328 | 0.32754088 | -0.3004734 | 0.76381607 | 0.91479214 |
| ZNF680 | 94.1389153 | 0.343341105 | 0.35635564 | 0.96347881 | 0.33530729 | 0.64662829 |
| ZNF107 | 22.6984182 | 0.711740686 | 0.46266061 | 1.53836456 | 0.1239595 | 0.38919278 |
| ZNF138 | 62.8143733 | 0.328387283 | 0.36144687 | 0.90853541 | 0.36359541 | 0.66933142 |
| ZNF117 | 26.4568537 | 0.077391397 | 0.47224222 | 0.16388072 | 0.86982504 | 0.95621002 |
| VKORC1L1 | 86.4248244 | 0.137944691 | 0.27403977 | 0.50337472 | 0.61470084 | 0.84174925 |
| GUSB | 21.6155124 | 0.232532393 | 0.3274926 | 0.71003863 | 0.47768018 | 0.7631764 |
| ASL | 17.9415844 | -0.564392863 | 0.49233786 | -1.1463528 | 0.25164922 | 0.56401021 |
| AC068533.4 | 22.5932259 | -1.030139558 | 0.36114707 | -2.8524101 | 0.00433891 | 0.04525277 |
| CRCP | 12.4235945 | -0.597088904 | 0.49045278 | -1.2174238 | 0.223443 | 0.53397237 |
| TPST1 | 475.691894 | -0.626932376 | 0.26168864 | -2.3957187 | 0.01658781 | 0.115594 |
| AC027644.4 | 89.0882658 | -0.030138895 | 0.31203417 | -0.0965884 | 0.92305323 | 0.97624187 |
| TMEM248 | 55.6775051 | -0.043868776 | 0.33617783 | -0.1304928 | 0.89617657 | 0.96522472 |
| SBDS | 24.8865046 | -0.746284392 | 0.36594251 | -2.0393487 | 0.04141524 | 0.20705398 |
| TYW1 | 147.284094 | 0.295500681 | 0.22645452 | 1.30490079 | 0.19192663 | 0.49284602 |
| AC006480.2 | 19.4981344 | 0.537077171 | 0.44723013 | 1.20089667 | 0.22979129 | 0.54242378 |
| AUTS2 | 219.086541 | 0.965745851 | 0.23057086 | 4.18849918 | 2.81E-05 | 0.00090596 |
| GALNT17 | 215.152097 | -0.6003277 | 0.93801734 | -0.6399964 | 0.52217496 | 0.78965349 |
| CALN1 | 19.0940518 | 1.323289641 | 0.51242102 | 2.58242655 | 0.00981082 | 0.0812399 |
| TYW1B | 113.103505 | 0.243560666 | 0.28085464 | 0.86721254 | 0.38582556 | 0.68921713 |
| POM121 | 111.053153 | -0.103655438 | 0.2582305 | -0.4014066 | 0.68812076 | 0.87567053 |
| NSUN5 | 23.2986025 | 0.39679713 | 0.41448323 | 0.95732976 | 0.33840084 | 0.64922828 |
| TRIM50 | 74.0475331 | 2.021288092 | 1.50948312 | 1.33905975 | 0.18055122 | 0.47907217 |
| BAZ1B | 96.4148206 | 0.070133494 | 0.367253 | 0.19096779 | 0.84855083 | 0.95003258 |
| BCL7B | 17.2585864 | 0.089029088 | 0.4805422 | 0.18526799 | 0.85301884 | 0.95137449 |
| MLXIPL | 31.0278622 | 0.476525342 | 0.52028066 | 0.91590055 | 0.35971906 | 0.66572082 |
| BUD23 | 32.5136334 | -0.395865127 | 0.42762448 | -0.9257307 | 0.35458593 | 0.66121742 |
| ABHD11 | 22.4806918 | -1.155430635 | 0.56106737 | -2.0593438 | 0.03946131 | 0.20080297 |
| CLDN3 | 22.4319019 | -2.345940486 | 0.49194253 | -4.7687288 | 1.85E-06 | 9.50E-05 |
| CLDN4 | 52.9811925 | -1.893723054 | 0.59715418 | -3.1712464 | 0.00151786 | 0.02077605 |
| LIMK1 | 15.4200957 | -1.324010285 | 0.53333387 | -2.4825168 | 0.01304579 | 0.09881314 |
| EIF4H | 45.8108101 | -0.440231431 | 0.32637133 | -1.3488667 | 0.17737978 | 0.47467647 |
| CLIP2 | 85.8602375 | -0.658461699 | 0.5345038 | -1.2319121 | 0.21798192 | 0.52750155 |
| GTF2IRD1 | 740.612562 | -1.745283598 | 0.56515202 | -3.0881666 | 0.00201396 | 0.02537584 |
| GTF2I | 357.607284 | 0.326391831 | 0.23426647 | 1.39325033 | 0.1635441 | 0.45491803 |
| GTF2IRD2 | 41.4198162 | -0.074719821 | 0.45039507 | -0.1658984 | 0.86823691 | 0.95588054 |
| CASTOR2 | 56.3946624 | -0.519311842 | 0.43274991 | -1.2000276 | 0.23012861 | 0.54292511 |
| RCC1L | 38.9043489 | -0.359564082 | 0.37219153 | -0.9660727 | 0.3340078 | 0.64554754 |
| GTF2IRD2B | 72.4479721 | 0.21651667 | 0.42895312 | 0.50475602 | 0.61373021 | 0.84174925 |
| TRIM73 | 40.873538 | -0.229117569 | 0.39282694 | -0.5832532 | 0.55972286 | 0.81314937 |
| POM121C | 96.1390949 | 0.177413666 | 0.26262596 | 0.67553744 | 0.49933438 | 0.77558432 |
| SPDYE5 | 21.2659968 | -0.720239115 | 0.70371728 | -1.023478 | 0.30608192 | 0.61974111 |
| HIP1 | 449.110129 | 0.517372844 | 0.36099942 | 1.43316806 | 0.15180981 | 0.43620176 |
| RHBDD2 | 20.7065535 | -0.334864859 | 0.46547896 | -0.7193985 | 0.47189543 | 0.75795917 |
| POR | 66.0549753 | -1.952267813 | 0.76227532 | -2.5611059 | 0.01043395 | 0.08482906 |
| STYXL1 | 49.6625375 | 0.612365844 | 0.33540119 | 1.82577125 | 0.06788473 | 0.27788783 |
| MDH2 | 24.8293173 | 0.041607388 | 0.48321948 | 0.08610453 | 0.93138332 | 0.97916542 |
| HSPB1 | 72.7916326 | -0.296204339 | 0.51750422 | -0.5723709 | 0.56707075 | 0.81614413 |
| YWHAG | 59.1886154 | -0.35637988 | 0.39069932 | -0.9121589 | 0.36168505 | 0.66686799 |
| SSC4D | 35.1607076 | 1.108286141 | 0.59987323 | 1.84753391 | 0.0646698 | 0.268828 |
| DTX2 | 35.4429528 | 0.38551835 | 0.37364447 | 1.03177856 | 0.30217587 | 0.61638324 |
| POMZP3 | 23.9840918 | -0.226944887 | 0.3878233 | -0.585176 | 0.55842938 | 0.81255224 |
| CCDC146 | 104.263866 | 0.337905612 | 0.37842142 | 0.89293469 | 0.37189215 | 0.67534262 |
| GSAP | 49.9763988 | 0.3039799 | 0.63739551 | 0.47690938 | 0.63342665 | 0.8520131 |
| PTPN12 | 176.109087 | -0.806269925 | 0.30885665 | -2.6104988 | 0.00904103 | 0.07670478 |

| | | | | | | |
|---|---|---|---|---|---|---|
| APTR | 29.9260601 | -0.109381929 | 0.41241513 | -0.2652229 | 0.79083777 | 0.92484276 |
| RSBN1L | 91.9760387 | 0.120050511 | 0.20520047 | 0.58504013 | 0.55852073 | 0.81255224 |
| PHTF2 | 88.9606767 | 0.202459682 | 0.21845407 | 0.92678373 | 0.35403884 | 0.6607207 |
| MAGI2 | 1012.25843 | 0.175993589 | 0.43431182 | 0.40522403 | 0.68531284 | 0.87424503 |
| GNAI1 | 237.059319 | -0.696723076 | 0.68766637 | -1.0131702 | 0.31097887 | 0.62585086 |
| SEMA3C | 77.7095519 | 0.382393133 | 0.96459072 | 0.39643045 | 0.69178751 | 0.87808891 |
| CACNA2D1 | 101.686607 | -1.175792534 | 0.26037442 | -4.5157758 | 6.31E-06 | 0.00026027 |
| PCLO | 213.560098 | 0.622709203 | 0.53750187 | 1.15852472 | 0.24664997 | 0.55931971 |
| KIAA1324L | 28.9899483 | 1.09640561 | 0.69249191 | 1.58327569 | 0.11335864 | 0.37119783 |
| DMTF1 | 155.737028 | 0.054009329 | 0.32147851 | 0.16800292 | 0.86658098 | 0.95578708 |
| TMEM243 | 34.832759 | -1.173001787 | 0.32342007 | -3.6268676 | 0.00028688 | 0.00563131 |
| CROT | 32.6531783 | 0.252679358 | 0.47614737 | 0.53067469 | 0.59564423 | 0.83416944 |
| SLC25A40 | 45.0528652 | 0.634809146 | 0.43612608 | 1.45556335 | 0.14551336 | 0.42813715 |
| DBF4 | 15.5946551 | -0.748482935 | 0.37526966 | -1.9945202 | 0.04609523 | 0.22049887 |
| ADAM22 | 63.4338249 | 0.73433421 | 0.53077269 | 1.38351921 | 0.16650572 | 0.45864087 |
| SRI | 37.7051809 | -0.341216358 | 0.4758485 | -0.7170693 | 0.47333133 | 0.75948052 |
| STEAP2 | 33.9386462 | 1.306328048 | 0.49319458 | 2.64870723 | 0.00808003 | 0.07056452 |
| CFAP69 | 62.6741983 | 0.397375221 | 0.41125156 | 0.96625828 | 0.33391496 | 0.64554754 |
| GTPBP10 | 47.6397969 | -0.15155641 | 0.26231885 | -0.5777565 | 0.56342856 | 0.81448184 |
| CLDN12 | 55.4575233 | -0.352569444 | 0.26140516 | -1.3487471 | 0.17741822 | 0.47467647 |
| CDK14 | 675.38874 | 1.240463415 | 0.21395491 | 5.79777962 | 6.72E-09 | 7.74E-07 |
| AC079760.1 | 39.0365111 | 0.769978748 | 0.61945567 | 1.2429925 | 0.2138706 | 0.52235998 |
| MTERF1 | 165.716852 | 0.098504931 | 0.32954958 | 0.29890778 | 0.76501041 | 0.91549415 |
| AKAP9 | 376.176242 | 0.410502343 | 0.18396215 | 2.23145007 | 0.02565133 | 0.15373119 |
| KRIT1 | 149.51168 | 0.185187755 | 0.20967989 | 0.88319274 | 0.37713214 | 0.68029685 |
| ANKIB1 | 284.258052 | -0.012301972 | 0.1972694 | -0.0623613 | 0.95027513 | 0.98678754 |
| GATAD1 | 39.0328757 | -0.208969719 | 0.34769506 | -0.6010143 | 0.54783044 | 0.80618229 |
| AC007566.1 | 33.3584274 | -0.698762828 | 0.37020227 | -1.8875163 | 0.05909091 | 0.25401736 |
| PEX1 | 55.0644191 | 0.190817897 | 0.35896676 | 0.53157539 | 0.59502011 | 0.83416944 |
| RBM48 | 15.9953244 | -0.457246838 | 0.42317809 | -1.0805069 | 0.27991652 | 0.59397852 |
| FAM133B | 67.3526722 | 0.197931629 | 0.27613001 | 0.71680594 | 0.47349385 | 0.75961057 |
| CDK6 | 210.430018 | 0.857678743 | 0.45482863 | 1.88571844 | 0.05933291 | 0.25458077 |
| SAMD9L | 23.5571864 | 0.08297384 | 1.09690122 | 0.07564386 | 0.93970244 | 0.98358617 |
| VPS50 | 151.442876 | 0.675907182 | 0.18413588 | 3.67069784 | 0.00024189 | 0.00493517 |
| CALCR | 238.054564 | 3.271858088 | 0.97827102 | 3.34453134 | 0.00082422 | 0.01285117 |
| GNGT1 | 152.669321 | 0.672565094 | 0.74097155 | 0.90768005 | 0.36404729 | 0.6696613 |
| BET1 | 37.0311122 | -0.262006692 | 0.424797 | -0.6167809 | 0.53737923 | 0.80078458 |
| CASD1 | 80.8940665 | 0.310907219 | 0.24786738 | 1.25432891 | 0.20972248 | 0.51841262 |
| SGCE | 111.144451 | 0.093266834 | 0.47862134 | 0.1948656 | 0.84549817 | 0.94907846 |
| PPP1R9A | 323.28495 | -0.214265779 | 0.32298701 | -0.6633882 | 0.50708194 | 0.7795601 |
| AC002429.2 | 23.1319822 | -1.11432995 | 0.80998779 | -1.3757367 | 0.16890316 | 0.46258659 |
| PON2 | 68.6644176 | -1.009323184 | 0.45265048 | -2.2298069 | 0.02576026 | 0.15406587 |
| PDK4 | 120.183149 | 0.431393613 | 0.79810434 | 0.54052283 | 0.58883652 | 0.83025227 |
| SLC25A13 | 226.89928 | -0.288214642 | 0.24474742 | -1.1776003 | 0.23895599 | 0.55079752 |
| SEM1 | 86.8730989 | -0.159104006 | 0.26633993 | -0.597372 | 0.55025908 | 0.80796749 |
| SDHAF3 | 18.3454426 | -0.099277244 | 0.4418857 | -0.2246673 | 0.82223814 | 0.94056539 |
| LMTK2 | 91.6550514 | -0.498510768 | 0.27542872 | -1.8099447 | 0.07030436 | 0.28267264 |
| TECPR1 | 15.3810028 | 0.335153867 | 0.48102138 | 0.69675461 | 0.48595638 | 0.76833754 |
| BRI3 | 102.072192 | -0.283556924 | 0.34690439 | -0.8173921 | 0.41370437 | 0.71143975 |
| BAIAP2L1 | 180.097418 | -0.466992551 | 0.42680271 | -1.0941649 | 0.27388266 | 0.58894058 |
| TRRAP | 140.382363 | 0.015090602 | 0.22499523 | 0.06707076 | 0.94652537 | 0.98574808 |
| SMURF1 | 253.607539 | -0.712197905 | 0.3649355 | -1.951572 | 0.05098904 | 0.23373736 |
| PDAP1 | 23.7161939 | -0.652865447 | 0.38644423 | -1.689417 | 0.09113955 | 0.32873701 |
| BUD31 | 13.6648278 | -0.231674296 | 0.49729576 | -0.4658682 | 0.64130982 | 0.85468927 |
| CPSF4 | 14.6808446 | -0.431275491 | 0.44773226 | -0.9632442 | 0.33542501 | 0.64662829 |
| ZNF789 | 38.7084033 | -0.145307038 | 0.37116686 | -0.3914871 | 0.69543722 | 0.8802955 |
| ZNF394 | 28.0897234 | -0.272635895 | 0.44351681 | -0.6147138 | 0.53874375 | 0.80128358 |
| ZKSCAN5 | 21.6049609 | 0.236943319 | 0.36025797 | 0.65770459 | 0.51072797 | 0.78194213 |
| FAM200A | 15.7716115 | 0.096387355 | 0.54830827 | 0.17579045 | 0.86045858 | 0.95363316 |
| ZSCAN25 | 14.7196981 | 0.081308861 | 0.39645134 | 0.20509165 | 0.83750052 | 0.94710186 |
| TRIM4 | 44.9371375 | -0.608569425 | 0.33044945 | -1.8416415 | 0.0655276 | 0.27130523 |
| ZKSCAN1 | 132.310708 | -0.375493167 | 0.23987831 | -1.5653485 | 0.11750121 | 0.37778664 |
| ZNF3 | 27.3411628 | -0.806955112 | 0.55059776 | -1.4655982 | 0.14275777 | 0.4236389 |
| COPS6 | 24.4754241 | -0.094006136 | 0.32315693 | -0.2908993 | 0.77112831 | 0.91720887 |
| AP4M1 | 19.6881835 | -0.59658938 | 0.51840521 | -1.1508167 | 0.24980765 | 0.5624056 |
| LAMTOR4 | 37.856223 | -0.536191409 | 0.34733281 | -1.5437396 | 0.12265143 | 0.38713673 |
| STAG3 | 15.1600263 | -0.101360829 | 0.53039594 | -0.1911041 | 0.84844405 | 0.95003258 |
| CASTOR3 | 92.6634972 | -1.137075653 | 0.30715804 | -3.7019238 | 0.00021397 | 0.00446323 |
| SPDYE3 | 36.1224981 | -0.235764541 | 0.59649122 | -0.3952523 | 0.69265668 | 0.87849579 |
| PILRB | 86.4086214 | -0.20390702 | 0.44649397 | -0.4566848 | 0.6478976 | 0.85943907 |
| PILRA | 37.441334 | 1.245389761 | 0.52863595 | 2.35585522 | 0.01848012 | 0.12417062 |
| ZCWPW1 | 12.5210497 | 0.014705289 | 0.60181536 | 0.02443489 | 0.98050572 | 0.99642892 |
| PPP1R35 | 17.2555071 | -0.950348944 | 0.46958046 | -2.0238256 | 0.0429881 | 0.21114114 |
| TSC22D4 | 46.4447932 | 0.469460661 | 0.32583974 | 1.44077166 | 0.1496492 | 0.433886 |
| AGFG2 | 19.3277951 | -0.610628797 | 0.49332 | -1.2377945 | 0.21579226 | 0.52520152 |
| LRCH4 | 12.9457549 | 0.083507301 | 0.58347409 | 0.14312084 | 0.88619475 | 0.96249798 |
| GNB2 | 44.1089928 | -1.195278576 | 0.41765116 | -2.8619065 | 0.00421101 | 0.04426547 |
| GIGYF1 | 54.3534731 | 0.095685423 | 0.3433993 | 0.27864187 | 0.78051968 | 0.92083964 |
| EPHB4 | 17.0351579 | 1.013301081 | 0.40983534 | 2.472459 | 0.01341871 | 0.10067724 |
| SLC12A9 | 32.4337155 | 0.788903912 | 0.47510846 | 1.66047118 | 0.0968197 | 0.33969491 |
| TRIP6 | 21.8792818 | 0.347147578 | 0.52939536 | 0.65574352 | 0.51198916 | 0.78271634 |
| SRRT | 23.544145 | -0.187293704 | 0.42155421 | -0.4442933 | 0.65683056 | 0.86325042 |
| TRIM56 | 75.4124769 | -1.451375667 | 0.42038015 | -3.4525314 | 0.00055535 | 0.00934677 |
| AP1S1 | 22.3829174 | -0.787556238 | 0.5046892 | -1.5604777 | 0.11864704 | 0.3795077 |
| PLOD3 | 20.5829719 | -1.043967614 | 0.41561954 | -2.5118348 | 0.01201053 | 0.09375173 |
| ZNHIT1 | 22.0173731 | -0.409394112 | 0.45808247 | -0.8937127 | 0.37147563 | 0.67534262 |
| FIS1 | 38.8070159 | -0.140187748 | 0.37397862 | -0.374855 | 0.70776832 | 0.88614996 |
| COL26A1 | 14.0646232 | -1.214687903 | 0.71327329 | -1.7029769 | 0.0885724 | 0.32322859 |
| CUX1 | 633.144182 | -0.248341977 | 0.27794597 | -0.8934901 | 0.37159477 | 0.67534262 |
| PRKRIP1 | 171.517725 | -0.080579431 | 0.22640444 | -0.3559092 | 0.72190854 | 0.89217596 |
| ORAI2 | 24.1714046 | -0.560499297 | 0.4043423 | -1.3862 | 0.16568584 | 0.45785473 |
| AC093668.2 | 10.3228161 | 0.831961661 | 0.6442407 | 1.29138327 | 0.19657081 | 0.49845696 |
| SPDYE2 | 17.1754184 | -0.098208161 | 0.68786048 | -0.1427734 | 0.88646917 | 0.96251467 |
| AC105052.1 | 11.3006823 | 0.605794777 | 0.65180261 | 0.92941447 | 0.35267433 | 0.65972795 |
| AC105052.3 | 44.4271495 | 0.642387413 | 0.57253823 | 1.12199916 | 0.2618628 | 0.57571533 |
| FAM185A | 79.4812386 | 0.673847823 | 0.469107 | 1.43644801 | 0.1508749 | 0.43488337 |
| FBXL13 | 56.185454 | -1.054293133 | 0.39639934 | -2.6596743 | 0.00782162 | 0.06899605 |
| ARMC10 | 43.6082214 | -0.449587344 | 0.32333449 | -1.3904713 | 0.1643858 | 0.45563826 |
| NAPEPLD | 73.2921427 | 0.326741239 | 0.4216778 | 0.77485995 | 0.43842242 | 0.73069774 |
| PMPCB | 42.7247092 | 0.207694328 | 0.2626106 | 0.79088327 | 0.42901211 | 0.7236272 |
| DNAJC2 | 41.5940656 | -0.161613483 | 0.32891995 | -0.4913459 | 0.62318179 | 0.84597807 |
| PSMC2 | 26.5793636 | 0.278002008 | 0.30634256 | 0.90748738 | 0.36414912 | 0.6696613 |
| ORC5 | 52.9391816 | 0.437470887 | 0.2761308 | 1.58428861 | 0.11312806 | 0.37079048 |
| LHFPL3 | 112.983766 | -0.343583725 | 0.52687806 | -0.6521124 | 0.51432866 | 0.7835482 |
| LINC01004 | 159.853207 | 0.24019128 | 0.20983782 | 1.14465203 | 0.25235333 | 0.56479656 |
| KMT2E-AS1 | 12.1504779 | -0.594428011 | 0.64635486 | -0.919662 | 0.35774941 | 0.66447321 |
| KMT2E | 346.557115 | 0.345308168 | 0.17904226 | 1.92864056 | 0.0537755 | 0.24061977 |
| SRPK2 | 540.574176 | 0.36090827 | 0.247489 | 1.45828006 | 0.14476336 | 0.42660354 |
| PUS7 | 27.7766089 | -0.035781948 | 0.36381974 | -0.0983508 | 0.92165377 | 0.9757579 |
| RINT1 | 28.5628994 | 0.183379679 | 0.41660321 | 0.44017827 | 0.659808 | 0.86460873 |
| ATXN7L1 | 172.473367 | -0.256817265 | 0.48276833 | -0.5319679 | 0.59474821 | 0.83416944 |
| CDHR3 | 35.4873701 | -0.860148857 | 0.67399975 | -1.2761857 | 0.20188987 | 0.5060272 |
| SYPL1 | 53.3165073 | 0.022498788 | 0.33039408 | 0.06809682 | 0.94570856 | 0.98574808 |

| | | | | | | |
|---|---|---|---|---|---|---|
| NAMPT | 166.632074 | -1.973656573 | 0.52455111 | -3.7625629 | 0.00016818 | 0.00366383 |
| HBP1 | 89.8217955 | 0.004029638 | 0.26015551 | 0.01548934 | 0.98764179 | 0.9977555 |
| COG5 | 296.117442 | 0.077780806 | 0.26378863 | 0.29486034 | 0.76810057 | 0.91600052 |
| DUS4L | 14.119805 | 0.23587223 | 0.46480135 | 0.5074689 | 0.61182585 | 0.84169124 |
| BCAP29 | 46.579514 | -0.069965251 | 0.25863498 | -0.2705174 | 0.78676227 | 0.92339968 |
| CBLL1 | 21.1242943 | -0.295193879 | 0.39349397 | -0.7501865 | 0.45314236 | 0.74451091 |
| SLC26A3 | 304.21319 | 1.000153361 | 0.77418819 | 1.2918737 | 0.19640089 | 0.49845696 |
| DLD | 65.3387027 | 0.651355016 | 0.25760584 | 2.5284948 | 0.01145528 | 0.09051612 |
| LAMB1 | 564.527963 | -1.019113535 | 0.50615793 | -2.01343 | 0.04406942 | 0.21423527 |
| PNPLA8 | 94.1025133 | -0.826379526 | 0.33589799 | -2.4602098 | 0.01388558 | 0.10308054 |
| IMMP2L | 2419.07997 | -0.035107531 | 0.39137764 | -0.0897024 | 0.92852367 | 0.97829541 |
| DOCK4 | 268.658569 | 0.769918583 | 0.44709397 | 1.72205092 | 0.0850603 | 0.31443489 |
| ZNF277 | 135.393739 | 0.251260001 | 0.27568314 | 0.91140867 | 0.36208009 | 0.66724582 |
| IFRD1 | 124.173136 | 0.048060236 | 0.56224786 | 0.08547874 | 0.9318808 | 0.97938863 |
| LSMEM1 | 17.1609581 | -0.485017896 | 0.68534854 | -0.7076952 | 0.47913453 | 0.76419703 |
| TMEM168 | 86.0442487 | 0.403167555 | 0.27541471 | 1.46385628 | 0.14323322 | 0.4243745 |
| BMT2 | 106.028458 | -0.259087203 | 0.21196917 | -1.2222872 | 0.22159904 | 0.53197463 |
| MDFIC | 40.4600627 | 0.099120368 | 0.57613936 | 0.17204235 | 0.86340424 | 0.95458251 |
| TES | 53.3452401 | -0.654913346 | 0.37298111 | -1.7558888 | 0.0791074 | 0.30106837 |
| MET | 217.781262 | -3.249608513 | 0.84151271 | -3.8616274 | 0.00011263 | 0.00265202 |
| CAPZA2 | 147.34441 | -0.69142778 | 0.44710174 | -1.5464663 | 0.12199199 | 0.38610091 |
| ST7 | 132.57755 | -0.638759318 | 0.26023315 | -2.4545656 | 0.01410549 | 0.10421524 |
| LSM8 | 37.7227913 | -0.088595437 | 0.40467399 | -0.2189304 | 0.82670427 | 0.9422064 |
| ANKRD7 | 54.0578388 | 0.396455491 | 0.62416504 | 0.63517734 | 0.52531276 | 0.79281583 |
| TSPAN12 | 86.4513048 | 0.128299809 | 0.63390876 | 0.20239476 | 0.83960813 | 0.9474978 |
| ING3 | 27.364329 | 0.144639302 | 0.40944646 | 0.35325571 | 0.72389675 | 0.89280599 |
| FAM3C | 110.303118 | -1.980820345 | 0.43908682 | -4.511227 | 6.45E-06 | 0.00026358 |
| AASS | 45.9865936 | 0.008591518 | 0.35812056 | 0.02399057 | 0.98086013 | 0.99646326 |
| CADPS2 | 684.59032 | 1.168038247 | 0.4375692 | 2.66937949 | 0.00759915 | 0.06748049 |
| AC004594.1 | 33.9681843 | -0.01181216 | 0.876876 | -0.0134707 | 0.98925224 | 0.99835349 |
| IQUB | 23.1203792 | -1.340307443 | 0.73166535 | -1.8318586 | 0.06697249 | 0.27545282 |
| NDUFA5 | 69.2423163 | 0.364084611 | 0.30392037 | 1.19796053 | 0.23093238 | 0.54380466 |
| WASL | 108.018247 | 0.092678773 | 0.25453832 | 0.36410539 | 0.71577931 | 0.88904248 |
| POT1 | 61.4186687 | 0.37152725 | 0.23548923 | 1.57768256 | 0.11463855 | 0.37318822 |
| POT1-AS1 | 30.7694825 | 0.326819312 | 0.59027471 | 0.55367324 | 0.57980249 | 0.82409732 |
| ZNF800 | 79.8592107 | -0.193955591 | 0.28553368 | -0.679274 | 0.49696429 | 0.77475164 |
| ARF5 | 14.2628768 | -0.787189087 | 0.5715214 | -1.3773572 | 0.16840185 | 0.46207669 |
| SND1 | 248.269878 | -0.418786142 | 0.25217167 | -1.6607184 | 0.09677001 | 0.33969491 |
| RBM28 | 46.4397568 | 0.467354497 | 0.2482586 | 1.88253095 | 0.05976396 | 0.2554971 |
| HILPDA | 38.0214772 | -3.459548861 | 1.56969188 | -2.2039668 | 0.02752668 | 0.16081374 |
| METTL2B | 37.39398 | -0.212526782 | 0.35053523 | -0.6062922 | 0.54432072 | 0.80393962 |
| FAM71F2 | 14.1617526 | 0.042672547 | 0.45577002 | 0.09362737 | 0.92540517 | 0.97735117 |
| CALU | 51.7910367 | -0.121292313 | 0.31232868 | -0.3883483 | 0.6977583 | 0.8825147 |
| KCP | 23.5415052 | -3.948690206 | 0.81530932 | -4.8431805 | 1.28E-06 | 7.05E-05 |
| ATP6V1F | 29.2715827 | -1.380600934 | 0.46240943 | -2.9856678 | 0.0028296 | 0.03302226 |
| TNPO3 | 136.835894 | -0.181780251 | 0.31487706 | -0.5773055 | 0.56373312 | 0.81450672 |
| TSPAN33 | 29.2743755 | 1.819606409 | 0.55004718 | 3.30809154 | 0.00093934 | 0.01421821 |
| SMO | 27.4894492 | 1.883030359 | 0.4540497 | 4.14718995 | 3.37E-05 | 0.00105666 |
| AHCYL2 | 309.361601 | 0.351573947 | 0.21331948 | 1.64810993 | 0.09933012 | 0.34378396 |
| STRIP2 | 39.6303375 | -2.828461094 | 1.46616427 | -1.929157 | 0.05371137 | 0.24054027 |
| NRF1 | 168.510181 | 0.053133144 | 0.21678689 | 0.2450939 | 0.80638373 | 0.93325451 |
| UBE2H | 864.37545 | -0.759063893 | 0.37773634 | -2.0095072 | 0.04448337 | 0.21534941 |
| ZC3HC1 | 32.3253196 | 0.533077119 | 0.33919365 | 1.57160111 | 0.11604309 | 0.37464881 |
| KLHDC10 | 175.381909 | 0.30029254 | 0.24613247 | 1.22004439 | 0.22244805 | 0.53302786 |
| TMEM209 | 31.5331655 | 0.095203461 | 0.35824775 | 0.26574755 | 0.79043364 | 0.92448611 |
| COPG2 | 55.400575 | -0.297228287 | 0.38016252 | -0.7818453 | 0.43430549 | 0.72766655 |
| AC016831.7 | 971.684588 | -1.393936508 | 0.46148639 | -3.0205366 | 0.00252327 | 0.03031829 |
| LINC00513 | 112.42501 | -2.141159178 | 0.73044652 | -2.9313018 | 0.00337545 | 0.03814869 |
| AC058791.1 | 18.8151438 | -1.339835125 | 0.81948273 | -1.6349766 | 0.10205394 | 0.34893691 |
| MKLN1 | 829.313703 | -0.598216943 | 0.37879543 | -1.5792612 | 0.11427615 | 0.372426 |
| PODXL | 15.7942022 | -0.925105548 | 0.50779859 | -1.8217962 | 0.06848591 | 0.27884832 |
| CHCHD3 | 401.773164 | 0.560969775 | 0.22873816 | 2.45245384 | 0.01418856 | 0.10453042 |
| EXOC4 | 737.787243 | 0.071915478 | 0.28534256 | 0.25203208 | 0.80101627 | 0.93010607 |
| LRGUK | 45.0973511 | 0.42882312 | 0.49873854 | 0.85981548 | 0.38989076 | 0.69292279 |
| SLC35B4 | 111.494103 | 1.650748162 | 0.35259129 | 4.68176103 | 2.84E-06 | 0.0001353 |
| AKR1B1 | 77.9889668 | -0.760506545 | 0.94647359 | -0.8035159 | 0.42167663 | 0.71947943 |
| BPGM | 20.7080471 | 0.233915958 | 0.40362816 | 0.5795333 | 0.56222938 | 0.81388399 |
| CALD1 | 36.9254711 | -0.185266226 | 0.4634117 | -0.3997875 | 0.68931301 | 0.87658953 |
| AGBL3 | 22.2957833 | -0.69925938 | 0.48670532 | -1.4367202 | 0.15079751 | 0.43488337 |
| CYREN | 29.203022 | -0.641995602 | 0.39996067 | -1.6051468 | 0.10846149 | 0.36219464 |
| TMEM140 | 37.8025861 | -1.704187921 | 0.6432064 | -2.6495195 | 0.00806063 | 0.07056452 |
| WDR91 | 135.136706 | -0.385141163 | 0.43125609 | -0.8930683 | 0.37182058 | 0.67534262 |
| AC009542.1 | 15.5136364 | -1.860140341 | 0.81142177 | -2.2924457 | 0.02187994 | 0.13859276 |
| CNOT4 | 234.904824 | -0.111132524 | 0.2678753 | -0.4148666 | 0.67823954 | 0.87130138 |
| NUP205 | 56.2724148 | 0.202295094 | 0.38107067 | 0.53085979 | 0.59551595 | 0.83416944 |
| STMP1 | 60.4971359 | -0.607052376 | 0.41100264 | -1.4770036 | 0.13967467 | 0.41835099 |
| MTPN | 55.9206105 | -0.527945405 | 0.29412023 | -1.7949986 | 0.07265392 | 0.28837569 |
| CREB3L2 | 77.0494179 | 0.598400582 | 0.4189557 | 1.42831469 | 0.1532013 | 0.43862109 |
| TRIM24 | 95.967477 | -0.040401825 | 0.26796789 | -0.1507711 | 0.88015625 | 0.96097414 |
| SVOPL | 63.0901198 | 0.227987579 | 0.62548448 | 0.36449758 | 0.71548648 | 0.88904248 |
| ATP6V0A4 | 447.262718 | 0.692689122 | 0.39004806 | 1.7759071 | 0.07574823 | 0.29465019 |
| TMEM213 | 285.076944 | 0.166499831 | 0.52802298 | 0.31532687 | 0.75251349 | 0.90957166 |
| KIAA1549 | 33.4963694 | 0.052878245 | 0.38656119 | 0.1367914 | 0.89119568 | 0.96328012 |
| ZC3HAV1 | 55.0203356 | -0.726717445 | 0.47296173 | -1.5365248 | 0.12440971 | 0.38965272 |
| TTC26 | 22.4568918 | -0.375469875 | 0.53726844 | -0.6988497 | 0.48464598 | 0.76759625 |
| UBN2 | 112.78377 | 0.257939103 | 0.18921555 | 1.36320249 | 0.1728187 | 0.468183 |
| LUC7L2 | 34.8071376 | 0.775179999 | 0.36170874 | 2.14310553 | 0.03210463 | 0.17765195 |
| HIPK2 | 384.416159 | -1.027473998 | 0.25533037 | -4.0240963 | 5.72E-05 | 0.00158715 |
| TBXAS1 | 34.8747512 | -2.515347774 | 0.52589547 | -4.7829805 | 1.73E-06 | 9.00E-05 |
| PARP12 | 37.1263076 | 0.475288649 | 0.67160819 | 0.7076874 | 0.47913941 | 0.76419703 |
| KDM7A | 195.934108 | -1.66422154 | 0.66593309 | -2.4990822 | 0.01245154 | 0.09610776 |
| SLC37A3 | 79.9152521 | -0.413069853 | 0.2966784 | -1.3923152 | 0.16382696 | 0.45543308 |
| MKRN1 | 69.6314781 | 0.029277673 | 0.31084814 | 0.09418642 | 0.92496107 | 0.97735117 |
| DENND2A | 256.832709 | -0.59833359 | 0.56767131 | -1.0540141 | 0.29187644 | 0.60677688 |
| NDUFB2 | 55.2678836 | 0.020845326 | 0.45606818 | 0.0457066 | 0.9635441 | 0.99076272 |
| BRAF | 525.814551 | 0.07141868 | 0.23490515 | 0.30403199 | 0.76110352 | 0.91320552 |
| MRPS33 | 16.5652858 | 0.284984667 | 0.45006001 | 0.63321481 | 0.52659338 | 0.79346524 |
| TMEM178B | 80.046176 | 1.459136195 | 0.67482349 | 2.16224868 | 0.03059901 | 0.17249407 |
| AGK | 53.8789643 | 0.478548319 | 0.27475367 | 1.74173584 | 0.08155468 | 0.3073629 |
| AC004918.1 | 14.7618042 | -0.026875795 | 0.5386143 | -0.049898 | 0.96020364 | 0.98971244 |
| KIAA1147 | 43.4738522 | 0.101381515 | 0.49378005 | 0.20531715 | 0.83732434 | 0.94706413 |
| SSBP1 | 69.0484313 | 0.031252585 | 0.27881156 | 0.11209214 | 0.91075035 | 0.97128978 |
| MGAM | 16.0946401 | -0.680264891 | 0.91539142 | -0.743141 | 0.45739632 | 0.747377 |
| TRPV6 | 17.414514 | -0.639820278 | 0.69144613 | -0.9253364 | 0.35479094 | 0.66121742 |
| TRPV5 | 64.6570741 | -1.2291221 | 1.36204496 | -0.9024093 | 0.36683946 | 0.67113782 |
| GSTK1 | 49.8598418 | -0.33472032 | 0.44524029 | -0.7517746 | 0.45218664 | 0.74372698 |
| CASP2 | 14.6197286 | -0.478194887 | 0.38478137 | -1.2427704 | 0.21395246 | 0.52235998 |
| TCAF2 | 17.8522292 | -1.54735181 | 0.85109732 | -1.8180668 | 0.06905392 | 0.28006035 |
| TCAF1 | 63.0369126 | 0.064721385 | 0.46912862 | 0.13796086 | 0.89027135 | 0.96323092 |
| AC004889.1 | 30.4642987 | 0.911824938 | 0.43529907 | 2.09470911 | 0.03619686 | 0.19023055 |
| AC074386.1 | 67.9749684 | -0.612474966 | 0.29374477 | -2.0850583 | 0.03706401 | 0.19349655 |
| ARHGEF5 | 15.2960325 | -0.946371597 | 0.59302233 | -1.5958448 | 0.11052345 | 0.36595193 |
| TPK1 | 41.6756555 | 0.086307009 | 0.38522746 | 0.22404169 | 0.82272487 | 0.94066053 |

| | | | | | | |
|---|---|---|---|---|---|---|
| CNTNAP2 | 15.9670664 | 0.206999179 | 0.53483714 | 0.38703217 | 0.69873239 | 0.88302803 |
| CUL1 | 145.338382 | -0.141185635 | 0.2309919 | -0.6112147 | 0.54105748 | 0.80242086 |
| EZH2 | 24.9575181 | -0.617823573 | 0.54814029 | -1.1271267 | 0.2596889 | 0.57296245 |
| PDIA4 | 21.4625378 | -1.031131439 | 0.71217313 | -1.4478662 | 0.14765446 | 0.43117138 |
| ZNF786 | 18.5243973 | -0.177684543 | 0.49439242 | -0.3593998 | 0.71929601 | 0.89102113 |
| ZNF425 | 14.2798163 | 0.433856579 | 0.40672992 | 1.06669453 | 0.2861098 | 0.60028977 |
| ZNF398 | 37.395391 | -0.205705258 | 0.31595302 | -0.6510628 | 0.51500594 | 0.78381327 |
| ZNF282 | 25.047608 | -0.327168996 | 0.39912534 | -0.8197149 | 0.41237865 | 0.71110141 |
| ZNF212 | 17.573893 | 0.226972614 | 0.47762756 | 0.47520837 | 0.63463846 | 0.85244692 |
| ZNF783 | 18.1500094 | 0.571966299 | 0.40406756 | 1.41552145 | 0.15691567 | 0.44349247 |
| ZNF746 | 16.2044141 | 0.205092648 | 0.41803915 | 0.49060632 | 0.62370492 | 0.84597807 |
| ZNF467 | 14.4039782 | 0.773644122 | 0.48067965 | 1.60947968 | 0.10751149 | 0.36071867 |
| ZNF862 | 51.0502101 | 0.016195877 | 0.51020105 | 0.03174411 | 0.97467612 | 0.99361556 |
| ACTR3C | 342.889586 | -0.637356315 | 0.2921269 | -2.181779 | 0.02912585 | 0.16640284 |
| LRRC61 | 17.4837919 | -1.282440263 | 0.44994484 | -2.8502166 | 0.00436895 | 0.04541366 |
| RARRES2 | 16.1115404 | 1.496353505 | 0.60006131 | 2.4936677 | 0.01264308 | 0.0971032 |
| TMEM176B | 32.1488996 | -0.725135438 | 0.54966651 | -1.319228 | 0.18709291 | 0.48725107 |
| TMEM176A | 15.8257982 | -0.618066748 | 0.62821875 | -0.98384 | 0.32519418 | 0.63715682 |
| AOC1 | 12.4162088 | 0.878910282 | 0.80482042 | 1.09205764 | 0.27480777 | 0.58917168 |
| ABCB8 | 15.315575 | 0.248416726 | 0.55702021 | 0.44597435 | 0.65561578 | 0.86268564 |
| SLC4A2 | 28.3502639 | 0.54687471 | 0.53256778 | 1.02686406 | 0.30448448 | 0.61818174 |
| FASTK | 19.9511729 | 0.770444282 | 0.55376384 | 1.39128673 | 0.1641385 | 0.45563826 |
| TMUB1 | 17.3410489 | -0.360083057 | 0.51655412 | -0.6970868 | 0.48574848 | 0.76833754 |
| AGAP3 | 21.3117712 | 0.398665666 | 0.51408217 | 0.77549016 | 0.43805008 | 0.7305377 |
| SMARCD3 | 42.9948386 | -1.45757078 | 0.41249611 | -3.5335382 | 0.00041004 | 0.00746715 |
| NUB1 | 49.9176836 | 0.125474758 | 0.40760343 | 0.30783538 | 0.7582076 | 0.91161939 |
| RHEB | 125.973139 | -0.288828711 | 0.43699478 | -0.6609432 | 0.50864876 | 0.7804435 |
| PRKAG2 | 243.362977 | -0.030397091 | 0.43340592 | -0.0701354 | 0.9440859 | 0.98552865 |
| GALNT11 | 169.316276 | -0.067722353 | 0.46909069 | -0.1443694 | 0.88520876 | 0.96233796 |
| KMT2C | 521.00899 | -0.026790548 | 0.25710042 | -0.1042027 | 0.91700852 | 0.97449139 |
| ACTR3B | 81.548352 | 0.395359618 | 0.23864962 | 1.65665301 | 0.09758965 | 0.34041372 |
| DPP6 | 221.892943 | -0.488168367 | 0.26749174 | -1.8249848 | 0.06800332 | 0.27788783 |
| PAXIP1-AS2 | 19.2351722 | 0.651276161 | 0.49 | 1.32913502 | 0.18380343 | 0.48275582 |
| PAXIP1 | 47.7043864 | -0.249791346 | 0.29077863 | -0.859043 | 0.39031679 | 0.69333469 |
| INSIG1 | 33.2984627 | -1.302323786 | 0.92716386 | -1.4046317 | 0.16013081 | 0.44931077 |
| AC009403.1 | 26.1581326 | -0.587004544 | 0.39839064 | -1.4734396 | 0.14063254 | 0.42027494 |
| RBM33 | 223.917902 | -0.166610671 | 0.27398918 | -0.6080922 | 0.54312635 | 0.80361674 |
| LINC01006 | 33.8593484 | -0.014438191 | 0.48976973 | -0.0294795 | 0.97648213 | 0.99461649 |
| LMBR1 | 216.301548 | 0.26391735 | 0.24200371 | 1.09055086 | 0.27547056 | 0.59004997 |
| NOM1 | 21.9279941 | -0.928663277 | 0.5815929 | -1.5967583 | 0.1103196 | 0.36553657 |
| UBE3C | 289.747842 | 0.088505519 | 0.32947849 | 0.26862306 | 0.78821977 | 0.92377626 |
| DNAJB6 | 123.939661 | -0.155515307 | 0.37243481 | -0.4175638 | 0.67626605 | 0.87020489 |
| ESYT2 | 160.304243 | -0.138406556 | 0.2381706 | -0.5811236 | 0.56115716 | 0.81384822 |
| WDR60 | 160.665149 | -0.361793891 | 0.1833308 | -1.9734484 | 0.04844449 | 0.22559069 |
| GTPBP6 | 33.0195925 | -0.32641099 | 0.47165468 | -0.692055 | 0.48890277 | 0.77060666 |
| LINC00685 | 12.2603578 | -0.9104453 | 0.71221149 | -1.2783356 | 0.20113112 | 0.50511102 |
| PPP2R3B | 12.7580034 | 0.243603548 | 0.40783911 | 0.59730306 | 0.55030506 | 0.80796749 |
| IL3RA | 16.4188225 | -1.181100957 | 0.53000498 | -2.2284715 | 0.02584909 | 0.15415253 |
| SLC25A6 | 80.6484801 | -0.739208103 | 0.55378031 | -1.33484 | 0.18192872 | 0.48053921 |
| LINC00106 | 37.4392231 | -0.094139175 | 0.38044621 | -0.2474441 | 0.80456454 | 0.9324748 |
| ASMTL-AS1 | 24.341851 | 0.120030954 | 0.46048151 | 0.26066401 | 0.79435163 | 0.9267672 |
| ASMTL | 25.8466944 | -0.095688899 | 0.31159218 | -0.3070966 | 0.75876984 | 0.91181467 |
| AKAP17A | 36.6985494 | -0.008084628 | 0.26676745 | -0.0303059 | 0.97582309 | 0.99405381 |
| DHRSX | 204.078664 | 0.214894328 | 0.27276925 | 0.78782461 | 0.4307993 | 0.7245261 |
| CD99 | 36.8679032 | -0.137042225 | 0.39898951 | -0.3434733 | 0.73124246 | 0.89782437 |
| ARSD | 50.8840048 | -0.035918145 | 0.55794164 | -0.0643762 | 0.94867071 | 0.98641632 |
| PRKX | 45.0032648 | -1.500671691 | 0.55494 | -2.7042053 | 0.0068468 | 0.06252649 |
| PUDP | 32.0256188 | -0.337218063 | 0.59850187 | -0.5634369 | 0.57313739 | 0.82088065 |
| TBL1X | 431.389459 | 0.60692448 | 0.34802421 | 1.74391456 | 0.08117399 | 0.30642359 |
| SHROOM2 | 33.0535631 | 0.514741809 | 0.42259354 | 1.21805414 | 0.2232034 | 0.53368772 |
| WWC3 | 29.4498249 | 0.781497887 | 0.43387055 | 1.80122364 | 0.07166764 | 0.28591361 |
| MID1 | 340.079812 | -1.799993931 | 0.30419093 | -5.9173162 | 3.27E-09 | 4.30E-07 |
| AC073529.1 | 31.7422825 | 0.889275639 | 0.41873147 | 2.12373728 | 0.03369212 | 0.18264262 |
| ARHGAP6 | 2051.7259 | 1.433706175 | 0.36610723 | 3.91608263 | 9.00E-05 | 0.00224373 |
| PRPS2 | 33.6051576 | 0.506117233 | 0.40910655 | 1.23712815 | 0.21603951 | 0.52525486 |
| TMSB4X | 117.998277 | -1.587147945 | 0.55309522 | -2.8695745 | 0.00411025 | 0.04359946 |
| RAB9A | 40.7577549 | -0.228762679 | 0.38783941 | -0.5898387 | 0.55529882 | 0.81104802 |
| TRAPPC2 | 23.1809171 | 0.161949768 | 0.38137802 | 0.42464369 | 0.67109644 | 0.86777458 |
| OFD1 | 131.815194 | -0.376690219 | 0.33325501 | -1.1303363 | 0.25833456 | 0.57145203 |
| GEMIN8 | 35.7845633 | 0.324597707 | 0.38385398 | 0.8456281 | 0.39776025 | 0.69815955 |
| MOSPD2 | 21.2460019 | 0.165915211 | 0.3937901 | 0.42132905 | 0.67351482 | 0.86846218 |
| PIR | 23.914181 | 0.533833353 | 0.95068922 | 0.56152246 | 0.57444142 | 0.8213605 |
| CLTRN | 17.2855929 | -1.602671351 | 0.79195734 | -2.0236839 | 0.04300268 | 0.21114114 |
| CA5B | 62.5386435 | -0.207044032 | 0.35513302 | -0.5830042 | 0.55989049 | 0.81314937 |
| ZRSR2 | 28.9308092 | 0.013876901 | 0.34271143 | 0.0404915 | 0.96770128 | 0.99156233 |
| AP1S2 | 27.6685525 | -1.767512463 | 0.54156209 | -3.26373 | 0.00109956 | 0.01609464 |
| CTPS2 | 50.4278636 | -0.212888856 | 0.30857933 | -0.6898999 | 0.49025712 | 0.77196791 |
| SYAP1 | 48.3471565 | -0.434903309 | 0.25889765 | -1.6798272 | 0.09299094 | 0.3326491 |
| TXLNG | 45.1875123 | -0.384867022 | 0.52471031 | -0.7334848 | 0.46326281 | 0.75108023 |
| RBBP7 | 40.5800434 | -0.44407348 | 0.26663471 | -1.6654752 | 0.095818 | 0.33841174 |
| REPS2 | 23.6727649 | -0.089154716 | 0.60923673 | -0.1463384 | 0.88365427 | 0.96170562 |
| NHS | 1027.26458 | 0.196013053 | 0.39156398 | 0.5005901 | 0.61665963 | 0.84223226 |
| SCML1 | 137.046716 | 0.222088234 | 0.32105677 | 0.69174131 | 0.4890998 | 0.77072275 |
| RAI2 | 29.1443095 | -0.80438219 | 0.45653462 | -1.7619303 | 0.07808107 | 0.29882113 |
| CDKL5 | 69.2943501 | 0.568566374 | 0.23467351 | 2.42279747 | 0.01540151 | 0.11036477 |
| PHKA2 | 35.5576786 | 0.313658803 | 0.2937786 | 1.06767071 | 0.28566907 | 0.59963495 |
| PDHA1 | 63.7242324 | 0.161396539 | 0.43653694 | 0.36972023 | 0.71159096 | 0.88701525 |
| MAP3K15 | 95.2136023 | 1.588004114 | 0.55591202 | 2.85657453 | 0.00428239 | 0.04471338 |
| BCLAF3 | 89.5292567 | 0.168400605 | 0.2964348 | 0.56808649 | 0.56997624 | 0.81849045 |
| EIF1AX | 60.5044611 | -0.419960471 | 0.31311861 | -1.3412185 | 0.17984952 | 0.47857217 |
| RPS6KA3 | 157.868265 | -1.849557763 | 0.58330903 | -3.1708026 | 0.00152018 | 0.02077605 |
| MBTPS2 | 40.117781 | 0.066984574 | 0.36342914 | 0.18431261 | 0.85376822 | 0.95142045 |
| SMS | 193.661683 | -0.796879769 | 0.61967608 | -1.2859618 | 0.19845639 | 0.50146542 |
| PTCHD1-AS | 127.334277 | 2.147121642 | 0.85481272 | 2.51180357 | 0.01201159 | 0.09375173 |
| ACOT9 | 30.7399399 | -0.701294323 | 0.52321931 | -1.3403449 | 0.18013324 | 0.47864023 |
| SAT1 | 1198.91712 | -1.04431569 | 0.28588394 | -3.6529358 | 0.00025926 | 0.00520977 |
| APOO | 199.949287 | -0.080179122 | 0.35066179 | -0.2286509 | 0.81914029 | 0.93840325 |
| EIF2S3 | 58.8014317 | -0.122227025 | 0.35638338 | -0.342965 | 0.73162479 | 0.89782437 |
| ZFX | 86.7773538 | 0.118067661 | 0.41775531 | 0.28262396 | 0.77746511 | 0.91968524 |
| PDK3 | 38.062795 | 0.573793749 | 0.55944423 | 1.02564959 | 0.30505678 | 0.61860579 |
| POLA1 | 82.3161599 | 0.04931669 | 0.28624242 | 0.17228994 | 0.86320959 | 0.95458251 |
| GK | 36.4308077 | -0.440594646 | 0.54057408 | -0.8150495 | 0.41504396 | 0.712552 |
| TAB3 | 79.6434207 | -0.499163484 | 0.38968092 | -1.2809544 | 0.20020968 | 0.50411966 |
| DMD | 37.796641 | 0.298789877 | 0.30878534 | 0.96762973 | 0.33322932 | 0.64501354 |
| TMEM47 | 40.812489 | -1.960850424 | 0.80322389 | -2.4412252 | 0.01463752 | 0.10644591 |
| CFAP47 | 37.9471202 | 0.163620236 | 0.60908832 | 0.26863138 | 0.78821336 | 0.92377626 |
| AF241726.2 | 47.1507814 | -1.542121091 | 0.40104559 | -3.8452513 | 0.00012043 | 0.0028072 |
| DYNLT3 | 26.6153285 | -1.557632403 | 0.50794553 | -3.0665343 | 0.00216556 | 0.02673639 |
| RPGR | 54.2283762 | -0.397935861 | 0.29549731 | -1.3466649 | 0.17808818 | 0.47550635 |
| BCOR | 70.9215615 | -0.504445296 | 0.40838823 | -1.2352102 | 0.21675229 | 0.52616463 |
| ATP6AP2 | 40.052272 | -0.393145088 | 0.35600866 | -1.1043133 | 0.26945727 | 0.58564558 |
| CXorf38 | 21.0966221 | -0.452559124 | 0.37933699 | -1.1930266 | 0.23285895 | 0.54525758 |
| MED14 | 106.781481 | -0.8084398 | 0.33979248 | -2.3792163 | 0.01734949 | 0.11929696 |

| | | | | | | |
|---|---|---|---|---|---|---|
| USP9X | 230.848729 | -0.342710642 | 0.31911833 | -1.0739297 | 0.28285421 | 0.59668285 |
| DDX3X | 92.0751947 | -0.276854911 | 0.3244099 | -0.8534108 | 0.39343154 | 0.69595061 |
| CASK | 143.498597 | -0.201657895 | 0.2112829 | -0.9544449 | 0.33985847 | 0.65027011 |
| MAOA | 45.0487655 | -0.950611463 | 0.64701727 | -1.4692212 | 0.14177281 | 0.42192458 |
| MAOB | 32.9440193 | 0.343776911 | 0.64834339 | 0.53023894 | 0.59594628 | 0.83416944 |
| EFHC2 | 18.7363124 | -0.65319268 | 0.60547454 | -1.0788111 | 0.28067193 | 0.5950926 |
| KDM6A | 294.24203 | 0.192856109 | 0.39036518 | 0.49404025 | 0.62127776 | 0.84470972 |
| MIR222HG | 111.385669 | -0.835949023 | 0.73026481 | -1.1447204 | 0.252325 | 0.56479656 |
| KRBOX4 | 40.4753337 | 0.22070328 | 0.38747137 | 0.56959893 | 0.56894975 | 0.81764604 |
| ZNF674 | 13.2395828 | 0.917701773 | 0.49034887 | 1.87152829 | 0.06127189 | 0.25921013 |
| SLC9A7 | 58.5568953 | 0.064242711 | 0.34035645 | 0.18875127 | 0.85028777 | 0.95026767 |
| RP2 | 17.4534315 | -1.123200597 | 0.49074636 | -2.2887599 | 0.02209331 | 0.13937617 |
| JADE3 | 75.1775368 | 0.101357743 | 0.3517585 | 0.28814583 | 0.77323512 | 0.91791018 |
| RGN | 36.0359936 | -0.498197549 | 0.84596424 | -0.5889109 | 0.55592106 | 0.81104802 |
| NDUFB11 | 28.3301736 | -0.10204521 | 0.75660126 | -0.1348732 | 0.89271215 | 0.96427805 |
| RBM10 | 48.4990152 | -0.141225857 | 0.33023122 | -0.4276575 | 0.6689005 | 0.86777458 |
| UBA1 | 82.3811627 | 0.149006967 | 0.33664656 | 0.44262139 | 0.65803961 | 0.86357172 |
| CDK16 | 35.0472717 | 0.101170926 | 0.57502124 | 0.17594294 | 0.86033877 | 0.95363316 |
| USP11 | 26.6911175 | -0.151252027 | 0.52946229 | -0.285671 | 0.77513011 | 0.91892094 |
| ZNF41 | 26.1513371 | 0.137164693 | 0.33044901 | 0.4150858 | 0.67807909 | 0.87130138 |
| ARAF | 18.6980887 | 0.371684961 | 0.43687825 | 0.85077469 | 0.39489452 | 0.69642725 |
| TIMP1 | 22.3931884 | -3.369361628 | 0.58669326 | -5.7429697 | 9.30E-09 | 9.97E-07 |
| UXT | 12.7404729 | -0.077001169 | 0.46329323 | -0.166204 | 0.86799644 | 0.95588054 |
| ZNF81 | 49.1953215 | -0.01760349 | 0.3311638 | -0.0531564 | 0.95760726 | 0.98859719 |
| ZNF630 | 26.9551485 | 0.524907959 | 0.55504746 | 0.94569924 | 0.34430201 | 0.65359197 |
| EBP | 15.1316892 | 0.636559378 | 0.57509253 | 1.10688166 | 0.26834509 | 0.5846368 |
| TBC1D25 | 16.7898282 | 0.217964982 | 0.38740144 | 0.56263338 | 0.57368456 | 0.82110314 |
| RBM3 | 29.7095035 | -0.134632767 | 0.37536027 | -0.3586761 | 0.7198374 | 0.89121815 |
| WDR13 | 23.8354531 | 0.388568777 | 0.41125104 | 0.94484569 | 0.34473766 | 0.65371852 |
| HDAC6 | 20.0026444 | 0.524412466 | 0.34332004 | 1.52747409 | 0.12664316 | 0.39439575 |
| PCSK1N | 58.1598224 | -1.554591838 | 0.72808293 | -2.1351851 | 0.03274589 | 0.17981313 |
| OTUD5 | 48.4882468 | -0.284797109 | 0.33798349 | -0.8426361 | 0.39943197 | 0.69959441 |
| GRIPAP1 | 49.9873635 | -0.799713955 | 0.4038191 | -1.9803768 | 0.04766121 | 0.22360104 |
| TFE3 | 22.245414 | -1.643647467 | 0.48360038 | -3.3987721 | 0.00067689 | 0.0110338 |
| CCDC120 | 14.5403147 | -0.675692625 | 0.46786571 | -1.4442021 | 0.14868214 | 0.43254673 |
| WDR45 | 21.795421 | -0.731783739 | 0.39354841 | -1.8594504 | 0.06296333 | 0.26432691 |
| CLCN5 | 405.893414 | 0.513258635 | 0.40691058 | 1.26135486 | 0.20718102 | 0.51472312 |
| SHROOM4 | 184.011404 | 0.071358644 | 0.27265207 | 0.26172053 | 0.79353691 | 0.92649882 |
| AL158055.1 | 13.5454224 | 0.713626472 | 0.82696422 | 0.86294721 | 0.38816649 | 0.69173015 |
| MAGED1 | 225.680907 | 0.751606662 | 0.18334618 | 4.09938552 | 4.14E-05 | 0.00121461 |
| FAM156A | 14.9460316 | 0.094650983 | 0.42296291 | 0.22378081 | 0.82292786 | 0.9407002 |
| KANTR | 89.3295647 | 0.086161185 | 0.28149667 | 0.30608244 | 0.75954188 | 0.91236458 |
| KDM5C | 51.8300796 | -0.289189823 | 0.45514834 | -0.6353749 | 0.52518395 | 0.79274974 |
| IQSEC2 | 15.7449301 | -0.238036481 | 0.55304596 | -0.4304099 | 0.66689746 | 0.86777458 |
| SMC1A | 46.8437389 | -0.584956497 | 0.50987242 | -1.1472605 | 0.25127396 | 0.56353121 |
| HUWE1 | 209.680442 | 0.092378956 | 0.26064312 | 0.35442698 | 0.72301892 | 0.89243161 |
| PHF8 | 63.3615471 | 0.071188283 | 0.27386217 | 0.25994201 | 0.7949085 | 0.9268423 |
| FAM120C | 37.2512644 | -0.398741166 | 0.28051599 | -1.4214561 | 0.15518421 | 0.44073639 |
| WNK3 | 40.551657 | -0.033935694 | 0.3839261 | -0.0883912 | 0.92956574 | 0.97876742 |
| FGD1 | 13.4841838 | -0.301852389 | 0.45602677 | -0.6619181 | 0.50802369 | 0.78010753 |
| GNL3L | 15.8327785 | -0.408384157 | 0.3802791 | -1.0739064 | 0.28286463 | 0.59668285 |
| MAGED2 | 24.917182 | -0.56408466 | 0.49432298 | -1.1411257 | 0.25381761 | 0.5657651 |
| PFKFB1 | 14.2216801 | 1.11462534 | 0.94977381 | 1.17356925 | 0.2405676 | 0.55302079 |
| RRAGB | 72.5588883 | -0.009438191 | 0.24523404 | -0.0384865 | 0.96929982 | 0.99156233 |
| AL445472.1 | 80.2656197 | -0.36651822 | 0.36791748 | -0.9961968 | 0.31915453 | 0.63100017 |
| UBQLN2 | 17.5269787 | -0.343965583 | 0.46164383 | -0.7450887 | 0.45621813 | 0.74627681 |
| NBDY | 101.218712 | 0.053670761 | 0.26924463 | 0.19933828 | 0.84199814 | 0.94776435 |
| SPIN3 | 29.142181 | -0.15618251 | 0.29438105 | -0.5305454 | 0.59573384 | 0.83416944 |
| FAAH2 | 372.816032 | -0.837948324 | 0.39508936 | -2.1209083 | 0.03392952 | 0.18339642 |
| ARHGEF9 | 79.9931442 | 0.011710303 | 0.30579739 | 0.03829432 | 0.96945302 | 0.99156233 |
| MTMR8 | 36.3440272 | 0.194320862 | 0.54328272 | 0.35767908 | 0.72058349 | 0.89154232 |
| ZC4H2 | 60.9251915 | 0.76223912 | 0.51073238 | 1.49244331 | 0.13558297 | 0.41064171 |
| ZC3H12B | 26.9310063 | -0.096256911 | 0.61995193 | -0.1552651 | 0.87661231 | 0.95879545 |
| LAS1L | 16.0265819 | -0.254909969 | 0.51373931 | -0.4961854 | 0.61976358 | 0.84360228 |
| MSN | 121.304018 | -1.095465827 | 0.33471219 | -3.2728591 | 0.00106466 | 0.01568222 |
| OPHN1 | 366.00045 | 1.409206736 | 0.17819704 | 7.90813768 | 2.61E-15 | 1.74E-12 |
| YIPF6 | 52.814372 | -0.620867155 | 0.33770575 | -1.8384856 | 0.06599088 | 0.27225103 |
| STARD8 | 20.5485678 | 1.129188099 | 0.51513295 | 2.19203238 | 0.02837717 | 0.16420667 |
| EDA | 94.0811958 | 0.757520556 | 0.43514814 | 1.7408337 | 0.08171273 | 0.30758559 |
| IGBP1 | 43.0984719 | -0.377046283 | 0.25254316 | -1.4929974 | 0.13543786 | 0.41064171 |
| DLG3 | 58.5704763 | -1.109694496 | 0.3936601 | -2.8189154 | 0.00481862 | 0.04888883 |
| ZMYM3 | 12.6201242 | 0.646811151 | 0.58570146 | 1.10433589 | 0.26944746 | 0.58564558 |
| NONO | 43.07963 | -0.435310006 | 0.35842634 | -1.2145034 | 0.22455555 | 0.53576151 |
| TAF1 | 112.796958 | -0.184582171 | 0.37517612 | -0.4919881 | 0.62272778 | 0.84578498 |
| OGT | 345.091014 | 0.599719238 | 0.38879979 | 1.54248857 | 0.12295491 | 0.38755404 |
| GCNA | 44.3458475 | 0.414493573 | 0.56498582 | 0.73363536 | 0.46317101 | 0.75106199 |
| PIN4 | 37.9758065 | 0.04667368 | 0.26617228 | 0.17535139 | 0.86080353 | 0.95378872 |
| RPS4X | 85.9016093 | -1.201307642 | 0.51486017 | -2.3332697 | 0.01963399 | 0.12983501 |
| HDAC8 | 314.661127 | -0.012020618 | 0.26794545 | -0.0448622 | 0.96421716 | 0.99082982 |
| PHKA1 | 28.7098909 | 0.753401536 | 0.45073927 | 1.67147969 | 0.09462697 | 0.33551172 |
| CHIC1 | 100.866407 | 0.643376675 | 0.32423459 | 1.98429376 | 0.0472231 | 0.22304735 |
| XIST | 394.694639 | -0.143468132 | 4.20911994 | -0.0340851 | 0.97280932 | 0.99287068 |
| JPX | 245.809756 | 0.210107475 | 0.45775078 | 0.45899971 | 0.64623437 | 0.85869877 |
| FTX | 1729.81101 | -0.015615974 | 0.32908911 | -0.0474521 | 0.96215289 | 0.99029318 |
| SLC16A2 | 32.9424268 | -0.63937454 | 0.38255881 | -1.6713105 | 0.09466037 | 0.33551172 |
| RLIM | 39.1868268 | -0.455862467 | 0.3283222 | -1.3884607 | 0.16499681 | 0.45691425 |
| NEXMIF | 126.040367 | 0.234745225 | 0.349425 | 0.67180432 | 0.50170828 | 0.776256 |
| ABCB7 | 122.056069 | 0.49818794 | 0.2118695 | 2.35139055 | 0.01870339 | 0.12519054 |
| UPRT | 26.8261371 | 0.553062691 | 0.33327484 | 1.65947926 | 0.09701926 | 0.33969491 |
| ZDHHC15 | 26.5444064 | -0.090076581 | 0.38722204 | -0.2326226 | 0.81605451 | 0.93717111 |
| ATRX | 417.967854 | 0.225867009 | 0.22168377 | 1.01887031 | 0.30826454 | 0.62213389 |
| MAGT1 | 70.9524992 | -0.494884301 | 0.3581853 | -1.3816432 | 0.16708127 | 0.45967184 |
| COX7B | 66.5165385 | 0.427034097 | 0.45759601 | 0.93321201 | 0.35071051 | 0.65782032 |
| PGK1 | 155.041197 | -0.91967661 | 0.55288892 | -1.6634021 | 0.09623197 | 0.33897502 |
| BRWD3 | 121.764792 | -0.143734151 | 0.38730006 | -0.3711183 | 0.7105494 | 0.88654658 |
| HMGN5 | 14.685922 | 0.343419126 | 0.4261184 | 0.80592419 | 0.42028656 | 0.71864146 |
| SH3BGRL | 132.049276 | 0.414912156 | 0.48864314 | 0.84911077 | 0.39581966 | 0.69660673 |
| RPS6KA6 | 134.230014 | 0.970781659 | 0.35155221 | 2.76141535 | 0.00575514 | 0.05486805 |
| HDX | 56.5278311 | -0.647010605 | 0.32392926 | -1.9973824 | 0.04578366 | 0.21947908 |
| APOOL | 72.5114195 | 0.175649403 | 0.24479447 | 0.71753828 | 0.47304202 | 0.75927764 |
| ZNF711 | 19.4596736 | 1.433509864 | 0.43687971 | 3.28124614 | 0.0010335 | 0.01533546 |
| CHM | 110.762064 | 0.276379531 | 0.24087724 | 1.1473875 | 0.25122151 | 0.56353121 |
| DIAPH2 | 127.420918 | -1.078748915 | 0.39535566 | -2.7285531 | 0.00636128 | 0.05910136 |
| TSPAN6 | 15.1317015 | -0.122721504 | 0.42561081 | -0.2883421 | 0.7730849 | 0.91791018 |
| SYTL4 | 33.538906 | 0.02003245 | 0.49735445 | 0.04027801 | 0.96787148 | 0.99156233 |
| TRMT2B | 52.7021488 | -0.022952173 | 0.29811646 | -0.0769906 | 0.938631 | 0.98331142 |
| ARMCX4 | 141.816246 | 0.305341516 | 0.48910583 | 0.62428517 | 0.53244032 | 0.79707272 |
| ARMCX3 | 33.6550878 | -0.714225136 | 0.3536402 | -2.0196378 | 0.04342097 | 0.21196707 |
| ZMAT1 | 136.489743 | 0.731219638 | 0.55249249 | 1.32349244 | 0.18567167 | 0.4858669 |
| GPRASP1 | 17.5141768 | 0.866584013 | 0.56554105 | 1.53230966 | 0.12544604 | 0.39171429 |
| BHLHB9 | 22.3388143 | 0.186188323 | 0.4203292 | 0.44295833 | 0.65779587 | 0.86357172 |
| LINC00630 | 60.5523638 | 0.346226968 | 0.33149996 | 1.04442538 | 0.29628863 | 0.61093809 |
| BEX4 | 23.4294545 | -0.377572607 | 0.32738059 | -1.153314 | 0.24878153 | 0.56124825 |

| | | | | | | |
|---|---|---|---|---|---|---|
| BEX2 | 25.6243592 | -0.840193381 | 0.46537646 | -1.8054059 | 0.07101119 | 0.28453304 |
| TCEAL9 | 29.5960098 | -0.747974333 | 0.42331436 | -1.7669477 | 0.07723699 | 0.29758583 |
| BEX3 | 54.1673593 | 0.158045547 | 0.36957466 | 0.42764173 | 0.66891198 | 0.86777458 |
| TCEAL4 | 35.3595503 | -0.392210228 | 0.28209974 | -1.3903247 | 0.1644303 | 0.45563826 |
| TCEAL3 | 20.8520936 | -0.368961199 | 0.47284831 | -0.780295 | 0.43521723 | 0.72827249 |
| MORF4L2 | 92.2732529 | -0.058174889 | 0.31687257 | -0.1835908 | 0.85433448 | 0.95170432 |
| SLC25A53 | 52.9405204 | 0.341316779 | 0.28090204 | 1.21507404 | 0.22433785 | 0.53561172 |
| FAM199X | 66.9010538 | 0.327760491 | 0.33264553 | 0.98531458 | 0.32446957 | 0.63678264 |
| IL1RAPL2 | 216.644821 | 5.032661538 | 0.5829581 | 8.63297298 | 5.98E-18 | 6.97E-15 |
| NRK | 145.452182 | 1.488915189 | 0.35613218 | 4.18079378 | 2.90E-05 | 0.00093399 |
| RNF128 | 28.5329783 | -1.174162048 | 0.65200026 | -1.8008613 | 0.07172474 | 0.2859177 |
| TBC1D8B | 31.9718383 | -0.121374092 | 0.65560478 | -0.185133 | 0.8531247 | 0.95137449 |
| MORC4 | 30.4363791 | -0.228624431 | 0.55467486 | -0.4121774 | 0.68020941 | 0.87237893 |
| RBM41 | 84.5505996 | 0.253594592 | 0.35814245 | 0.70808304 | 0.47889369 | 0.76419703 |
| NUP62CL | 20.5707225 | 0.598894238 | 0.36325831 | 1.64867318 | 0.09921461 | 0.343767 |
| MID2 | 23.4651769 | -0.586588639 | 0.53264663 | -1.1012717 | 0.27077844 | 0.58722969 |
| COL4A6 | 59.5840295 | 2.72042775 | 0.50300051 | 5.40839957 | 6.36E-08 | 5.20E-06 |
| COL4A5 | 294.665493 | 1.529483205 | 0.54397363 | 2.81168629 | 0.00492825 | 0.04959478 |
| ACSL4 | 1543.17495 | -1.23834126 | 0.68430745 | -1.809627 | 0.07035365 | 0.28274888 |
| TMEM164 | 203.179526 | -0.056790962 | 0.20467709 | -0.2774661 | 0.7814222 | 0.92103406 |
| AMMECR1 | 60.7936062 | 0.696677745 | 0.35215258 | 1.97834057 | 0.0478903 | 0.22427381 |
| PAK3 | 13.3839888 | 0.616202615 | 0.47935816 | 1.28547435 | 0.19862657 | 0.50162356 |
| ALG13 | 77.8097122 | 0.003237748 | 0.27000863 | 0.01199128 | 0.99043257 | 0.99857194 |
| AMOT | 29.0171464 | -1.69865654 | 0.66597832 | -2.5506184 | 0.0107532 | 0.08656845 |
| LRCH2 | 14.5084375 | -0.282038088 | 0.53739907 | -0.5248206 | 0.59970793 | 0.83582612 |
| PLS3-AS1 | 34.4326807 | 1.109678043 | 0.42060458 | 2.63829284 | 0.00833246 | 0.07247374 |
| PLS3 | 139.019484 | 0.994208735 | 0.31970155 | 3.10980271 | 0.00187212 | 0.02397758 |
| DANT2 | 687.439687 | -0.136155133 | 0.20360973 | -0.6687064 | 0.50368277 | 0.77790293 |
| KLHL13 | 652.913277 | -0.332090478 | 0.4560287 | -0.7282228 | 0.46647724 | 0.75300195 |
| WDR44 | 69.5783964 | -0.192980697 | 0.32105687 | -0.6010795 | 0.54778705 | 0.80618229 |
| IL13RA1 | 58.2930253 | -1.904189115 | 0.38587911 | -4.9346778 | 8.03E-07 | 4.83E-05 |
| LONRF3 | 41.6621061 | -4.69781865 | 1.11072509 | -4.2295062 | 2.34E-05 | 0.0007799 |
| PGRMC1 | 38.0999101 | -1.127464554 | 0.53008342 | -2.1269568 | 0.03342367 | 0.18214045 |
| SLC25A43 | 37.7860889 | -0.139042202 | 0.53602607 | -0.2593945 | 0.79533089 | 0.9268423 |
| SLC25A5 | 38.732466 | -0.48345527 | 0.611814 | -0.7901997 | 0.42941112 | 0.7236272 |
| UBE2A | 25.9906099 | -0.11045337 | 0.39906775 | -0.2767785 | 0.7819502 | 0.92126657 |
| NKRF | 16.0784182 | 0.178565913 | 0.44009086 | 0.40574783 | 0.6849279 | 0.87408051 |
| 6-Sep | 149.294991 | -0.154629979 | 0.45736176 | -0.3380912 | 0.73529447 | 0.90007689 |
| UPF3B | 53.4905835 | -0.090578851 | 0.49445149 | -0.1831906 | 0.85464849 | 0.95171892 |
| NDUFA1 | 38.692976 | -0.05226972 | 0.41228006 | -0.1267821 | 0.89911289 | 0.96637793 |
| NKAP | 18.743173 | -0.337015887 | 0.37496878 | -0.8987839 | 0.3687678 | 0.67287495 |
| RHOXF1-AS1 | 28.2810366 | 0.893828228 | 0.80149506 | 1.11520117 | 0.26476422 | 0.57977023 |
| LAMP2 | 63.4894889 | -0.406989922 | 0.28201732 | -1.443138 | 0.14898159 | 0.43272656 |
| CUL4B | 33.1810017 | 0.048194614 | 0.31101817 | 0.15495755 | 0.87685478 | 0.95879545 |
| MCTS1 | 29.7600017 | 0.134289639 | 0.37241983 | 0.3605867 | 0.71840843 | 0.8907358 |
| THOC2 | 180.230091 | 0.09932141 | 0.22805571 | 0.4355138 | 0.66318953 | 0.86556259 |
| XIAP | 129.720354 | -0.310872178 | 0.26568509 | -1.1700776 | 0.24196973 | 0.55444462 |
| STAG2 | 208.178041 | 0.096681417 | 0.19357887 | 0.49944199 | 0.61746805 | 0.84244543 |
| TENM1 | 35.6714847 | -0.738365131 | 0.65956258 | -1.119477 | 0.2629367 | 0.57700771 |
| SMARCA1 | 73.4314287 | -0.066830911 | 0.33666872 | -0.1985064 | 0.84264884 | 0.94799096 |
| OCRL | 17.9258777 | 0.278492085 | 0.47121902 | 0.59100349 | 0.55451809 | 0.81072612 |
| ZDHHC9 | 25.1189042 | -1.515809727 | 0.43959203 | -3.4482193 | 0.0005643 | 0.00948017 |
| UTP14A | 12.3611292 | -0.095324058 | 0.5071924 | -0.1879446 | 0.8509201 | 0.95042076 |
| BCORL1 | 14.7999671 | -0.57784345 | 0.67692233 | -0.8536333 | 0.3933082 | 0.69595061 |
| AIFM1 | 31.7494376 | 0.162919486 | 0.37733188 | 0.43176708 | 0.66591071 | 0.86741428 |
| ZNF280C | 13.571477 | 0.327105481 | 0.44746132 | 0.73102515 | 0.46476379 | 0.75233638 |
| SLC25A14 | 14.2254267 | 0.064860005 | 0.52909163 | 0.12258747 | 0.90243378 | 0.96803829 |
| ENOX2 | 63.6514789 | -0.388339841 | 0.28514744 | -1.3618914 | 0.17323217 | 0.46899442 |
| RAP2C | 71.8646272 | 1.897635795 | 0.39984368 | 4.74594415 | 2.08E-06 | 0.00010404 |
| RAP2C-AS1 | 104.05955 | 1.18529282 | 0.52483305 | 2.2584188 | 0.02391956 | 0.14711477 |
| MBNL3 | 56.7012516 | 0.776746174 | 0.39424978 | 1.97018798 | 0.04881683 | 0.2267903 |
| HS6ST2 | 409.394184 | 0.953929937 | 0.34124801 | 2.79541539 | 0.00518331 | 0.05146875 |
| PHF6 | 32.469776 | 0.444568134 | 0.31292936 | 1.42066609 | 0.15541385 | 0.44098563 |
| HPRT1 | 143.851879 | 0.33966824 | 0.20908232 | 1.62456705 | 0.10425484 | 0.35360937 |
| MIR503HG | 38.0254421 | 1.473729757 | 0.64203168 | 2.29541593 | 0.0217093 | 0.13798058 |
| FAM122B | 18.436616 | -0.006254675 | 0.37959944 | -0.016477 | 0.98685382 | 0.99766074 |
| FAM122C | 14.1205007 | 0.312924522 | 0.57357256 | 0.54557095 | 0.5853609 | 0.82884893 |
| RTL8C | 37.7757755 | -0.017997392 | 0.52841979 | -0.0340589 | 0.97283019 | 0.99287068 |
| ZNF75D | 62.7132481 | -0.373483429 | 0.24466611 | -1.5265025 | 0.12688478 | 0.39482486 |
| INTS6L | 27.3631542 | 0.277104986 | 0.43851894 | 0.6319111 | 0.52744497 | 0.79403853 |
| SLC9A6 | 19.9172468 | 0.093124431 | 0.36515861 | 0.2550246 | 0.7987041 | 0.92914747 |
| HTATSF1 | 13.1298769 | 0.242330195 | 0.51333348 | 0.47207167 | 0.63687563 | 0.8538689 |
| AL683813.1 | 27.283698 | -0.238971076 | 0.53123294 | -0.4498424 | 0.65282412 | 0.86183588 |
| RBMX | 135.310745 | 0.176628139 | 0.19737588 | 0.89488207 | 0.37085013 | 0.67515476 |
| FGF13 | 1199.89651 | -0.166653211 | 0.62434903 | -0.2669232 | 0.78952833 | 0.92428059 |
| ATP11C | 34.9668276 | -0.929047415 | 0.43503144 | -2.1355868 | 0.03271311 | 0.17981313 |
| FMR1 | 79.2910603 | -0.315210517 | 0.29592654 | -1.0651648 | 0.28680138 | 0.60119965 |
| IDS | 38.0638176 | -0.327082373 | 0.42718872 | -0.7656625 | 0.44387716 | 0.73655644 |
| TMEM185A | 19.9146424 | 0.249317399 | 0.44293916 | 0.56287052 | 0.57352305 | 0.82110314 |
| LINC00894 | 145.425846 | 0.153700521 | 0.41654442 | 0.36898951 | 0.71213554 | 0.88721964 |
| MAMLD1 | 28.5457899 | 1.034649357 | 0.90659839 | 1.14124332 | 0.25376868 | 0.5657651 |
| MTM1 | 63.7241605 | 0.121280864 | 0.24061759 | 0.50403989 | 0.61423335 | 0.84174925 |
| MTMR1 | 56.904207 | -0.094920466 | 0.25226534 | -0.3762723 | 0.70671445 | 0.88603006 |
| CD99L2 | 38.6495928 | 0.470938453 | 0.4261197 | 1.1051788 | 0.26908212 | 0.58551265 |
| VMA21 | 17.3412734 | -0.525578507 | 0.53376573 | -0.9846614 | 0.32479042 | 0.6371441 |
| DUSP9 | 33.1011426 | 2.834681866 | 0.68506642 | 4.13782046 | 3.51E-05 | 0.00107696 |
| SLC6A8 | 66.1074895 | -0.625700312 | 0.4672244 | -1.3391859 | 0.18051017 | 0.47907217 |
| BCAP31 | 32.4210123 | -0.788360534 | 0.44904454 | -1.7556399 | 0.07914991 | 0.30109904 |
| IDH3G | 16.3281995 | 0.271554221 | 0.57139766 | 0.4752456 | 0.63461193 | 0.85244692 |
| SSR4 | 16.578311 | -0.512466275 | 0.4785221 | -1.0709354 | 0.28419846 | 0.59746288 |
| ARHGAP4 | 12.8880095 | -0.391301116 | 0.49368845 | -0.7926074 | 0.4280066 | 0.7233521 |
| HCFC1 | 13.4580513 | -0.048267484 | 0.5149309 | -0.0937358 | 0.92531899 | 0.97735117 |
| IRAK1 | 14.8766444 | -1.847829817 | 0.55352744 | -3.3382804 | 0.00084299 | 0.01305649 |
| MECP2 | 129.345537 | -0.102202562 | 0.35355001 | -0.2890753 | 0.77252378 | 0.91791018 |
| FLNA | 38.4371359 | -0.594783101 | 0.63254118 | -0.9403073 | 0.34705995 | 0.6553345 |
| EMD | 23.5504804 | -0.211081057 | 0.56740558 | -0.3720109 | 0.70988474 | 0.88614996 |
| RPL10 | 113.041162 | -0.993787774 | 0.46756833 | -2.1254386 | 0.03355003 | 0.18250255 |
| TAZ | 28.6907635 | 0.529159395 | 0.31914611 | 1.65804745 | 0.09730789 | 0.34011144 |
| ATP6AP1 | 25.5352514 | -0.258606208 | 0.51328377 | -0.503827 | 0.61438297 | 0.84174925 |
| GDI1 | 22.487879 | -0.267404001 | 0.4466399 | -0.5987015 | 0.54937192 | 0.80698344 |
| FAM50A | 16.9009054 | -0.027966965 | 0.5264985 | -0.0531188 | 0.95763726 | 0.98859719 |
| FAM3A | 15.0344952 | -0.168132792 | 0.36918543 | -0.4554156 | 0.6488103 | 0.8599651 |
| DKC1 | 16.8405291 | -0.220786646 | 0.40787933 | -0.5413038 | 0.58829817 | 0.83024609 |
| MPP1 | 17.4580028 | 1.646115362 | 0.43595776 | 3.77585977 | 0.00015946 | 0.00349829 |
| F8 | 18.4998629 | -0.037270155 | 0.50640805 | -0.0735971 | 0.94133099 | 0.9844067 |
| FUNDC2 | 24.8775286 | 0.334098755 | 0.43900537 | 0.76103568 | 0.44663575 | 0.73948328 |
| BRCC3 | 41.1774021 | -0.137243425 | 0.29262962 | -0.4690005 | 0.63906931 | 0.85391761 |
| VBP1 | 13.5773733 | 0.010383979 | 0.46183225 | 0.02248431 | 0.98206163 | 0.99659802 |
| TMLHE-AS1 | 28.0019893 | -0.618615881 | 0.74182402 | -0.8339119 | 0.40433063 | 0.70347327 |
| TMLHE | 81.6237199 | 0.394182056 | 0.41804955 | 0.94290751 | 0.34572821 | 0.65414698 |
| VAMP7 | 39.3112849 | -0.335353248 | 0.34532528 | -0.9711228 | 0.33148714 | 0.64324815 |
| ZNF596 | 20.6212265 | 0.20859589 | 0.33159297 | 0.62907211 | 0.52930185 | 0.79493876 |
| FBXO25 | 47.535352 | -0.01654252 | 0.41257447 | -0.0400958 | 0.96801672 | 0.99156233 |

| | | | | | | |
|---|---|---|---|---|---|---|
| TDRP | 68.4990294 | 0.024649398 | 0.31693361 | 0.07777464 | 0.93800732 | 0.98313627 |
| ERICH1 | 238.246403 | -0.595532836 | 0.62901497 | -0.9467705 | 0.34375572 | 0.65306073 |
| AC246817.2 | 27.6728929 | 0.141767416 | 0.61515485 | 0.2304581 | 0.81773582 | 0.93805075 |
| AC246817.1 | 59.2008075 | 0.515250138 | 0.51599745 | 0.99855172 | 0.3180119 | 0.63093651 |
| CSMD1 | 50.2034951 | 1.27663778 | 0.95567323 | 1.33585177 | 0.18159773 | 0.47998877 |
| MCPH1 | 180.204332 | -0.21982079 | 0.27826425 | -0.7899714 | 0.42954448 | 0.72372114 |
| ANGPT2 | 23.5520075 | -0.775808201 | 0.65279825 | -1.1884349 | 0.23466212 | 0.54658747 |
| MCPH1-AS1 | 41.0008253 | -0.378569419 | 0.47168553 | -0.8025886 | 0.42221257 | 0.72008598 |
| AGPAT5 | 56.4592581 | -1.060625808 | 0.56922357 | -1.8632851 | 0.06242216 | 0.2627649 |
| AF233439.1 | 228.790142 | -0.273454532 | 0.65945236 | -0.4146691 | 0.67838414 | 0.87136709 |
| DEFB1 | 479.144038 | 1.616610019 | 0.45746305 | 3.53385919 | 0.00040954 | 0.00746715 |
| FAM85B | 109.686212 | 0.029753056 | 0.4840153 | 0.06147131 | 0.95098386 | 0.98684281 |
| MFHAS1 | 729.042821 | -1.409549227 | 0.46973915 | -3.0007063 | 0.00269354 | 0.03178263 |
| ERI1 | 53.8606491 | -1.576930014 | 0.28012513 | -5.6293771 | 1.81E-08 | 1.70E-06 |
| PPP1R3B | 19.8010448 | -0.743617927 | 0.40455504 | -1.8381131 | 0.06604574 | 0.27225103 |
| TNKS | 326.176582 | -0.254498161 | 0.21235101 | -1.1984787 | 0.23073071 | 0.54354156 |
| MSRA | 141.019825 | -0.513503163 | 0.33087371 | -1.5519612 | 0.12067151 | 0.38413018 |
| XKR6 | 555.912651 | -0.376829024 | 0.25317775 | -1.4883971 | 0.1366462 | 0.41192666 |
| MTMR9 | 54.2140979 | -0.166012067 | 0.27602335 | -0.6014421 | 0.54754556 | 0.80618229 |
| FDFT1 | 82.6783732 | -0.012988527 | 0.24214232 | -0.0536401 | 0.95722194 | 0.98859719 |
| CTSB | 62.3887043 | -1.687049212 | 0.48987564 | -3.4438316 | 0.00057353 | 0.0095823 |
| FAM66D | 54.1386313 | 1.512664622 | 0.80495467 | 1.8791923 | 0.06021824 | 0.25638122 |
| AC145124.1 | 12.6256398 | -0.379418412 | 0.62500596 | -0.6070637 | 0.54380866 | 0.80380646 |
| LONRF1 | 77.0626127 | 0.044864144 | 0.44513144 | 0.10078853 | 0.91971833 | 0.97520237 |
| TRMT9B | 233.067435 | -0.723792009 | 0.50359928 | -1.437238 | 0.15065039 | 0.43488337 |
| TUSC3 | 133.029719 | -0.49558655 | 0.30841821 | -1.6068654 | 0.1080839 | 0.36175274 |
| MSR1 | 169.041248 | -0.052986411 | 0.61204912 | -0.0865722 | 0.9310116 | 0.97899539 |
| AC068992.1 | 70.9882196 | 0.194889579 | 0.63331738 | 0.30772814 | 0.75828921 | 0.91161939 |
| MICU3 | 104.485544 | 0.804412444 | 0.31419951 | 2.56019638 | 0.0104613 | 0.08496619 |
| ZDHHC2 | 130.930153 | -0.299967744 | 0.238667 | -1.2568464 | 0.20880928 | 0.51711493 |
| CNOT7 | 36.6738731 | 0.08466698 | 0.32236899 | 0.26263996 | 0.7928281 | 0.92624097 |
| VPS37A | 60.4986857 | -0.183107003 | 0.28657069 | -0.6389593 | 0.52284942 | 0.79024931 |
| MTMR7 | 25.5170024 | 1.608887428 | 0.90652387 | 1.77478772 | 0.07593294 | 0.29512246 |
| MTUS1 | 247.987778 | 0.068344265 | 0.35222083 | 0.19403811 | 0.84614604 | 0.94909204 |
| AC027117.2 | 42.0773456 | -0.777392368 | 0.73864307 | -1.0524601 | 0.2925885 | 0.60745829 |
| PCM1 | 313.639015 | 0.221416663 | 0.19651742 | 1.12670245 | 0.25986831 | 0.57322264 |
| ASAH1 | 56.3961346 | -0.920337253 | 0.41287398 | -2.2290997 | 0.02580727 | 0.15410988 |
| PSD3 | 27.3442276 | -0.695901195 | 0.40474023 | -1.7193774 | 0.08554568 | 0.31589223 |
| SH2D4A | 29.0918653 | -1.364170138 | 0.58666855 | -2.3252825 | 0.02005686 | 0.13114315 |
| CSGALNACT1 | 727.946667 | -1.662248401 | 0.63065188 | -2.6357622 | 0.00839485 | 0.07273124 |
| INTS10 | 148.192537 | -0.47477884 | 0.27323059 | -1.7376489 | 0.08227269 | 0.30864976 |
| LPL | 19.4867373 | 3.638719058 | 0.80278972 | 4.53259298 | 5.83E-06 | 0.00024582 |
| ATP6V1B2 | 36.4586035 | -0.829899648 | 0.50723312 | -1.6361306 | 0.10181225 | 0.34880722 |
| XPO7 | 101.935794 | 0.357960581 | 0.24829393 | 1.44168075 | 0.14939246 | 0.43339616 |
| DMTN | 44.343571 | 1.049706185 | 0.48037946 | 2.18516044 | 0.02887709 | 0.1657943 |
| FAM160B2 | 27.0737194 | -0.766512753 | 0.38707748 | -1.9802567 | 0.0476747 | 0.22360104 |
| SLC39A14 | 72.6627308 | -0.797822205 | 0.7126875 | -1.1194559 | 0.26294571 | 0.57700771 |
| PPP3CC | 154.897974 | 0.319989255 | 0.27318658 | 1.17132129 | 0.24146966 | 0.55404626 |
| CCAR2 | 16.4346421 | 0.17640149 | 0.42744995 | 0.41268338 | 0.6798386 | 0.87237893 |
| BIN3 | 75.1858244 | -0.881056671 | 0.31603327 | -2.7878605 | 0.00530574 | 0.05229461 |
| PEBP4 | 19.9051005 | 2.450279925 | 0.62971765 | 3.89107707 | 9.98E-05 | 0.00241698 |
| RHOBTB2 | 11.9728939 | -1.252194829 | 0.62341586 | -2.0086028 | 0.04457928 | 0.21570169 |
| TNFRSF10B | 97.8749155 | -1.34643879 | 0.56078265 | -2.4009994 | 0.01635036 | 0.11437129 |
| TNFRSF10D | 59.0053458 | -2.305140282 | 0.3523699 | -6.5418195 | 6.08E-11 | 1.53E-08 |
| TNFRSF10A | 16.8801162 | -2.155912202 | 0.62600416 | -3.4439263 | 0.00057333 | 0.0095823 |
| CHMP7 | 14.2597873 | 0.234642546 | 0.39295981 | 0.59711588 | 0.55043002 | 0.80796749 |
| ENTPD4 | 47.1868572 | -0.537002262 | 0.47014956 | -1.1421945 | 0.25337317 | 0.56559484 |
| SLC25A37 | 111.969584 | -0.225024899 | 0.25514008 | -0.8819661 | 0.37779515 | 0.68108313 |
| DOCK5 | 84.8963922 | -0.881600767 | 0.52677208 | -1.6735905 | 0.0942111 | 0.33463783 |
| KCTD9 | 43.2606677 | 0.518070484 | 0.2950162 | 1.7560747 | 0.07907566 | 0.30106837 |
| PPP2R2A | 184.068172 | 0.237185662 | 0.28818217 | 0.82304071 | 0.41048484 | 0.70916447 |
| BNIP3L | 172.551462 | -1.604968511 | 0.56100472 | -2.8608824 | 0.00422464 | 0.04430424 |
| TRIM35 | 13.6969575 | -0.379319557 | 0.40436476 | -0.9380628 | 0.34821212 | 0.65547624 |
| PTK2B | 61.8957299 | -0.95262211 | 0.31008047 | -3.0721771 | 0.00212504 | 0.02650028 |
| EPHX2 | 90.8929099 | 0.924748626 | 0.64456764 | 1.4346805 | 0.15137816 | 0.43590179 |
| CLU | 95.1545029 | -2.424269722 | 0.53054689 | -4.5693788 | 4.89E-06 | 0.00021313 |
| CCDC25 | 24.7043785 | 0.067419923 | 0.37277505 | 0.18085954 | 0.85647783 | 0.95234014 |
| ELP3 | 113.812737 | -0.729734931 | 0.33546008 | -2.1753257 | 0.02960572 | 0.1681243 |
| ZNF395 | 33.9152478 | -0.925590195 | 0.38346168 | -2.413775 | 0.01578821 | 0.11184941 |
| FBXO16 | 37.2609514 | -0.075808212 | 0.5807977 | -0.1305243 | 0.89615163 | 0.96522472 |
| FZD3 | 27.5715238 | -0.33005524 | 0.42093027 | -0.7841091 | 0.43297611 | 0.72648177 |
| EXTL3 | 32.7233901 | -1.225963007 | 0.54482794 | -2.2501838 | 0.02443728 | 0.14863223 |
| INTS9 | 65.7519782 | -0.016696746 | 0.24593318 | -0.0678914 | 0.94587209 | 0.98574808 |
| HMBOX1 | 188.885117 | -0.485341393 | 0.20634552 | -2.3520811 | 0.0186687 | 0.12504812 |
| KIF13B | 379.833066 | -0.868377593 | 0.30737727 | -2.8251197 | 0.0047263 | 0.04821519 |
| AC108449.1 | 15.9066088 | -1.596041285 | 0.51234098 | -3.1151935 | 0.00183824 | 0.0236504 |
| SARAF | 70.0550151 | -0.509475052 | 0.38776878 | -1.313863 | 0.18889231 | 0.48924896 |
| LEPROTL1 | 25.3771351 | 0.076313864 | 0.35014356 | 0.21795021 | 0.8274679 | 0.9425007 |
| DCTN6 | 13.4655239 | -0.446317631 | 0.51040411 | -0.8744397 | 0.38187883 | 0.68581797 |
| RBPMS | 1318.53496 | -1.399559601 | 0.28916335 | -4.8400311 | 1.30E-06 | 7.12E-05 |
| GTF2E2 | 65.6882525 | 0.1095273 | 0.24886723 | 0.44010334 | 0.65986226 | 0.86460873 |
| GSR | 78.5144892 | 0.519964042 | 0.38878045 | 1.33742331 | 0.1810845 | 0.47966815 |
| PPP2CB | 112.468079 | -0.512931332 | 0.26243207 | -1.9545299 | 0.05063858 | 0.23258823 |
| WRN | 103.611749 | 0.354436474 | 0.26243334 | 1.35057717 | 0.17683092 | 0.47393944 |
| NRG1 | 168.495184 | 0.933738757 | 0.69814298 | 1.33746063 | 0.18107232 | 0.47966815 |
| FUT10 | 58.2315772 | 1.081174564 | 0.40640472 | 2.66033958 | 0.00780619 | 0.06892511 |
| TTI2 | 21.725082 | -0.626671618 | 0.4108834 | -1.5251811 | 0.12721392 | 0.39511746 |
| RNF122 | 18.1446116 | -1.094273164 | 0.55632388 | -1.9669714 | 0.04918651 | 0.22760049 |
| ERLIN2 | 25.9831178 | 0.002905209 | 0.3713898 | 0.00782253 | 0.99375858 | 0.9992177 |
| PLPBP | 26.9201361 | 0.18455199 | 0.31648629 | 0.58312792 | 0.55980719 | 0.81314937 |
| RAB11FIP1 | 100.083104 | -0.49782917 | 0.25117658 | -1.9819888 | 0.0474805 | 0.22347711 |
| ASH2L | 42.6440939 | 0.137691676 | 0.28203406 | 0.4882094 | 0.62540153 | 0.8471532 |
| LSM1 | 18.2806304 | -0.397631792 | 0.39860781 | -0.9975514 | 0.31849693 | 0.63093651 |
| BAG4 | 54.8093795 | 0.25474763 | 0.26082602 | 0.97669561 | 0.32871987 | 0.64027241 |
| DDHD2 | 53.0074768 | 0.580161472 | 0.25341873 | 2.28933933 | 0.02205964 | 0.13935239 |
| PLPP5 | 36.0006955 | -1.087010033 | 0.37681011 | -2.8847688 | 0.00391701 | 0.04212483 |
| NSD3 | 246.097345 | -0.420818504 | 0.19542086 | -2.153396 | 0.03128757 | 0.17510521 |
| LETM2 | 16.3199395 | -1.180738969 | 0.63490782 | -1.8597014 | 0.06292778 | 0.26429668 |
| FGFR1 | 118.202742 | -1.558630712 | 0.65366988 | -2.384431 | 0.01710556 | 0.11799351 |
| TACC1 | 565.357878 | 0.913960939 | 0.5343233 | 1.71050176 | 0.08717313 | 0.31934811 |
| PLEKHA2 | 55.2135759 | -1.155040906 | 0.41358576 | -2.7927483 | 0.00522623 | 0.05178471 |
| ADAM9 | 99.3080576 | -1.209787573 | 0.46166908 | -2.6204648 | 0.008781 | 0.07483916 |
| TCIM | 163.841641 | -1.505191909 | 0.53298291 | -2.8240904 | 0.0047415 | 0.04831667 |
| SFRP1 | 434.978887 | -1.11503708 | 0.51612981 | -2.1603811 | 0.03074318 | 0.17290661 |
| GOLGA7 | 20.9866468 | -0.319426987 | 0.43118259 | -0.7408161 | 0.45880497 | 0.747377 |
| GPAT4 | 28.7849582 | -0.589625947 | 0.44108298 | -1.3367687 | 0.18129813 | 0.47994859 |
| KAT6A | 261.426408 | -0.36281648 | 0.22183962 | -1.63549 | 0.10194637 | 0.34893691 |
| AP3M2 | 19.6012436 | 0.047311099 | 0.37141344 | 0.12738123 | 0.89863867 | 0.96631381 |
| IKBKB | 79.5765503 | -0.721170201 | 0.23478518 | -3.0716172 | 0.00212903 | 0.02650028 |
| POLB | 23.9895385 | 0.417558857 | 0.33036143 | 1.26394556 | 0.20624957 | 0.51322951 |
| VDAC3 | 59.8607144 | 0.539956386 | 0.41657532 | 1.29617948 | 0.19491366 | 0.49736588 |
| SLC20A2 | 225.546134 | -0.360120521 | 0.48575306 | -0.7413654 | 0.45847191 | 0.747377 |
| SMIM19 | 33.1305844 | -0.362292351 | 0.35996517 | -1.006465 | 0.31419194 | 0.62892349 |

| | | | | | | |
|---|---|---|---|---|---|---|
| RNF170 | 70.0115986 | 0.151519704 | 0.38230282 | 0.39633426 | 0.69185846 | 0.87808891 |
| HOOK3 | 119.936262 | -0.221645474 | 0.24585041 | -0.9015461 | 0.36729805 | 0.67137562 |
| FNTA | 32.108589 | -0.570098199 | 0.31373181 | -1.8171514 | 0.06919393 | 0.28038429 |
| HGSNAT | 105.928969 | 0.210866956 | 0.30288951 | 0.69618442 | 0.48631334 | 0.76833754 |
| SPIDR | 539.200684 | -0.572372241 | 0.30508259 | -1.8761222 | 0.0606385 | 0.2573479 |
| CEBPD | 112.602226 | -1.527825542 | 0.47893909 | -3.1900205 | 0.00142263 | 0.01982746 |
| PRKDC | 139.676235 | 0.079168002 | 0.19201325 | 0.4123049 | 0.68011596 | 0.87237893 |
| MCM4 | 15.4607954 | -0.266946165 | 0.53028798 | -0.5033985 | 0.61468414 | 0.84174925 |
| UBE2V2 | 61.7497428 | 0.214553416 | 0.30821072 | 0.69612575 | 0.48635008 | 0.76833754 |
| PXDNL | 69.2608254 | 1.307131697 | 0.40917069 | 3.19458783 | 0.00140031 | 0.0195749 |
| PCMTD1 | 254.918011 | 0.075861405 | 0.22563211 | 0.33621723 | 0.73670706 | 0.90141723 |
| AC064807.1 | 18.9768053 | 0.343305531 | 0.49266225 | 0.6968375 | 0.48590449 | 0.76833754 |
| RB1CC1 | 224.712124 | -0.506943573 | 0.48882602 | -1.0370634 | 0.29970631 | 0.61356712 |
| ATP6V1H | 251.953239 | 0.435783678 | 0.22834365 | 1.90845539 | 0.05633238 | 0.24728962 |
| TCEA1 | 65.8140663 | 0.290960642 | 0.35977836 | 0.8087219 | 0.41867513 | 0.7170696 |
| LYPLA1 | 126.246995 | 0.056663734 | 0.25100151 | 0.22575057 | 0.82139543 | 0.939832 |
| MRPL15 | 27.0002098 | 0.298909627 | 0.35498471 | 0.84203523 | 0.39976824 | 0.69959441 |
| XKR4 | 39.6751996 | -3.88403542 | 0.68957272 | -5.6325247 | 1.78E-08 | 1.70E-06 |
| TMEM68 | 34.874576 | 0.440973019 | 0.33595212 | 1.31260675 | 0.18931549 | 0.48954068 |
| TGS1 | 46.7567896 | -0.380821372 | 0.3525215 | -1.0802784 | 0.28001821 | 0.59405912 |
| LYN | 53.1439943 | -1.429256541 | 0.29647337 | -4.8208598 | 1.43E-06 | 7.62E-05 |
| RPS20 | 48.5785228 | -0.459709303 | 0.34141963 | -1.3464642 | 0.17815286 | 0.47550635 |
| PLAG1 | 45.7531004 | -0.233998592 | 0.43860331 | -0.5335085 | 0.59368163 | 0.83416944 |
| CHCHD7 | 14.7075845 | -0.565175106 | 0.37251424 | -1.5171906 | 0.12921856 | 0.3984239 |
| IMPAD1 | 38.3139062 | -0.158783551 | 0.34121172 | -0.465352 | 0.64167941 | 0.85468927 |
| LINC01606 | 384.988174 | 0.199267975 | 0.5840599 | 0.34117729 | 0.73297012 | 0.89840827 |
| FAM110B | 86.6586034 | -0.835736752 | 0.46376438 | -1.8020719 | 0.07153409 | 0.28564621 |
| UBXN2B | 55.031894 | -0.074595532 | 0.27662536 | -0.2696627 | 0.78741979 | 0.92362588 |
| SDCBP | 61.5117663 | -1.064900773 | 0.40394439 | -2.6362559 | 0.00838265 | 0.07270681 |
| NSMAF | 67.2406674 | -0.075121642 | 0.2356959 | -0.3187227 | 0.74993677 | 0.90857724 |
| TOX | 196.351674 | 1.715287021 | 0.6460933 | 2.65485964 | 0.00793414 | 0.06963542 |
| LINC01301 | 25.3772613 | 0.722346507 | 0.85143488 | 0.84838727 | 0.39622233 | 0.69678933 |
| RAB2A | 204.60494 | -0.301315228 | 0.28494637 | -1.0574454 | 0.29030836 | 0.60499022 |
| CHD7 | 182.991224 | 0.106433152 | 0.23234389 | 0.45808458 | 0.64689168 | 0.85896013 |
| ASPH | 129.087999 | -0.022999817 | 0.20045924 | -0.1147356 | 0.90865467 | 0.97059151 |
| YTHDF3 | 110.752498 | -0.319317162 | 0.34571298 | -0.9236482 | 0.35566953 | 0.66192868 |
| CYP7B1 | 102.181668 | 0.989169206 | 0.34579827 | 2.86053833 | 0.00422922 | 0.04430424 |
| ARMC1 | 34.9392581 | 0.340626973 | 0.29473963 | 1.15568772 | 0.24780891 | 0.56031261 |
| MTFR1 | 64.3445961 | -0.385590352 | 0.31056443 | -1.2415793 | 0.21439184 | 0.52290228 |
| PDE7A | 153.006727 | 1.01287861 | 0.35014506 | 2.89273997 | 0.00381897 | 0.04152772 |
| ADHFE1 | 49.1244784 | 0.878251254 | 0.53009823 | 1.65677078 | 0.09756583 | 0.34041372 |
| VCPIP1 | 29.8517875 | -0.212465516 | 0.36377211 | -0.5840621 | 0.55917851 | 0.81287502 |
| SGK3 | 92.4041404 | 0.359166786 | 0.25371338 | 1.41563991 | 0.15688096 | 0.44349247 |
| TCF24 | 31.7162453 | 2.613202074 | 0.37610853 | 6.94799994 | 3.71E-12 | 1.19E-09 |
| PPP1R42 | 23.6129445 | 0.862331944 | 0.61364938 | 1.40525188 | 0.15994639 | 0.4489284 |
| CSPP1 | 187.319198 | 0.191385706 | 0.1988654 | 0.96238815 | 0.33585467 | 0.6470947 |
| ARFGEF1 | 501.08507 | -0.399201086 | 0.24208116 | -1.6490382 | 0.09913981 | 0.34376333 |
| SULF1 | 14.5846317 | 0.787502698 | 0.60436851 | 1.30301742 | 0.19256881 | 0.49370242 |
| NCOA2 | 710.111919 | -0.095967558 | 0.31963696 | -0.3002392 | 0.76399468 | 0.91479214 |
| TRAM1 | 83.5804717 | -0.516288077 | 0.39805471 | -1.2970279 | 0.19462158 | 0.49702866 |
| LACTB2 | 47.5204615 | -2.255031889 | 0.3837075 | -5.8769554 | 4.18E-09 | 5.34E-07 |
| EYA1 | 34.8885213 | 1.691822382 | 0.58871076 | 2.8737752 | 0.00405598 | 0.04317114 |
| TERF1 | 56.4301302 | 0.137885934 | 0.2828917 | 0.48741598 | 0.62596357 | 0.8471532 |
| RPL7 | 86.5961648 | -0.841230115 | 0.56231012 | -1.4960252 | 0.13464709 | 0.40918252 |
| RDH10 | 79.3718503 | -0.901400009 | 0.57925683 | -1.5561319 | 0.11967674 | 0.38162309 |
| RDH10-AS1 | 23.6834502 | -1.109751127 | 0.66982062 | -1.6567885 | 0.09756223 | 0.34041372 |
| STAU2 | 202.01048 | 0.096774247 | 0.35872193 | 0.26977511 | 0.78733327 | 0.92362588 |
| AC027018.1 | 27.4330261 | -0.146400749 | 0.61166698 | -0.2393472 | 0.81083641 | 0.93552094 |
| ELOC | 36.0143963 | -0.553016977 | 0.33554359 | -1.6481226 | 0.09932751 | 0.34378396 |
| GDAP1 | 63.0724241 | 0.454431072 | 0.48146692 | 0.94384693 | 0.34524788 | 0.65406518 |
| PEX2 | 40.6760062 | 0.039670283 | 0.33174178 | 0.11958181 | 0.90481443 | 0.96904316 |
| ZC2HC1A | 48.0542732 | -0.092562806 | 0.419455 | -0.220674 | 0.82534629 | 0.94180992 |
| IL7 | 23.7342682 | -2.643194994 | 0.6724246 | -3.9308422 | 8.46E-05 | 0.0021424 |
| TPD52 | 230.947974 | 1.197687549 | 0.28043344 | 4.2708442 | 1.95E-05 | 0.00068544 |
| ZBTB10 | 82.6017374 | 0.035049812 | 0.46531752 | 0.07532451 | 0.93995652 | 0.98363127 |
| ZNF704 | 402.886869 | -0.293866059 | 0.32455562 | -0.9054413 | 0.36523166 | 0.67012584 |
| IMPA1 | 40.2742692 | 0.605725926 | 0.34980083 | 1.7316309 | 0.0833393 | 0.31091215 |
| ZFAND1 | 50.0070561 | -0.055190986 | 0.33434589 | -0.1650715 | 0.86888769 | 0.95612718 |
| CHMP4C | 23.0962338 | -1.157424616 | 0.39745198 | -2.9121119 | 0.00358994 | 0.03975369 |
| SNX16 | 30.1114114 | 0.185380972 | 0.58571159 | 0.31650555 | 0.75161881 | 0.90896158 |
| RALYL | 30.9107531 | 2.256891638 | 0.55919989 | 4.03593004 | 5.44E-05 | 0.00152282 |
| LRRCC1 | 17.2488132 | -0.091011514 | 0.49142221 | -0.1852002 | 0.85307198 | 0.95137449 |
| E2F5 | 33.572757 | -0.11804503 | 0.35591108 | -0.33167 | 0.74013845 | 0.90304252 |
| C8orf59 | 26.385449 | -0.000259453 | 0.38679404 | -0.0006708 | 0.9994648 | 0.99969609 |
| CA13 | 41.043693 | -0.640049175 | 0.33731033 | -1.8975084 | 0.05776087 | 0.25049413 |
| CA3 | 55.8863837 | -0.723648712 | 0.30606016 | -2.3644002 | 0.01805929 | 0.1224617 |
| CA2 | 127.916493 | 0.709698657 | 0.56440169 | 1.25743537 | 0.20859603 | 0.51699876 |
| ATP6V0D2 | 154.496717 | 1.147259233 | 0.6601917 | 1.73776682 | 0.08225191 | 0.30864976 |
| AC023194.3 | 29.9822539 | -0.099917757 | 0.89840851 | -0.1112164 | 0.91144475 | 0.97128978 |
| WWP1 | 359.581554 | -0.358465222 | 0.27755068 | -1.2915307 | 0.19651972 | 0.49845696 |
| RMDN1 | 91.2577183 | 0.355639939 | 0.28742638 | 1.23732532 | 0.21596633 | 0.52525486 |
| CPNE3 | 74.5962211 | 0.850845269 | 0.31361632 | 2.71301335 | 0.00666744 | 0.06118822 |
| AF117829.1 | 264.305301 | 0.034582879 | 0.43211759 | 0.08003118 | 0.93621246 | 0.98191732 |
| RIPK2 | 21.9963956 | -0.278060408 | 0.36603486 | -0.7596555 | 0.44746051 | 0.7399821 |
| OSGIN2 | 26.5896084 | 0.170912164 | 0.36738869 | 0.46520802 | 0.6417825 | 0.85468927 |
| NBN | 44.0665247 | 0.05449545 | 0.32767498 | 0.16630946 | 0.86791342 | 0.95588054 |
| DECR1 | 143.846405 | 0.692360458 | 0.35441745 | 1.95351684 | 0.05075838 | 0.23290904 |
| CALB1 | 191.954416 | 3.916306368 | 0.8247435 | 4.74851435 | 2.05E-06 | 0.00010328 |
| LINC00534 | 30.0037184 | 0.82813032 | 0.8778243 | 0.9433896 | 0.34548166 | 0.65406518 |
| PIP4P2 | 45.4385256 | 0.449062724 | 0.37899595 | 1.18487472 | 0.23606699 | 0.54879348 |
| OTUD6B-AS1 | 20.7486642 | 0.214095733 | 0.50376698 | 0.42498961 | 0.67084425 | 0.86777458 |
| OTUD6B | 23.3394708 | 0.342057218 | 0.37155194 | 0.9206175 | 0.35725016 | 0.6639427 |
| LRRC69 | 38.8837215 | -0.959715743 | 0.48011303 | -1.9989371 | 0.04561516 | 0.21889643 |
| TRIQK | 48.929062 | -0.066257839 | 0.44104492 | -0.1502292 | 0.88058376 | 0.96097414 |
| FAM92A | 34.8038101 | 0.332035356 | 0.58161716 | 0.57088302 | 0.56807894 | 0.81689822 |
| RBM12B | 45.7443932 | 0.689468522 | 0.30517434 | 2.25926112 | 0.02386715 | 0.14688929 |
| TMEM67 | 78.5169667 | 0.204530775 | 0.40540017 | 0.50451576 | 0.61389899 | 0.84174925 |
| PDP1 | 19.5253639 | -1.706557744 | 0.72336628 | -2.359189 | 0.01831493 | 0.12347678 |
| GEM | 34.9689009 | 0.463837434 | 1.04200139 | 0.4451409 | 0.65621793 | 0.86294007 |
| VIRMA | 60.2221626 | 0.48687222 | 0.24727907 | 1.96891802 | 0.04896251 | 0.22724062 |
| ESRP1 | 90.9923556 | 0.117034702 | 0.34099012 | 0.34322021 | 0.7314328 | 0.89782437 |
| DPY19L4 | 65.2277752 | 0.27472496 | 0.20535256 | 1.33782095 | 0.18095481 | 0.47966815 |
| INTS8 | 67.3325995 | 0.262570876 | 0.25156292 | 1.04375825 | 0.29659725 | 0.61093809 |
| NDUFAF6 | 58.6519228 | 0.440603092 | 0.32646307 | 1.34962613 | 0.17713594 | 0.47439927 |
| TP53INP1 | 29.4510332 | -0.314734574 | 0.42613247 | -0.7385839 | 0.46015971 | 0.7486528 |
| PLEKHF2 | 18.2733456 | 0.558432857 | 0.44104291 | 1.26616446 | 0.20545421 | 0.51190533 |
| C8orf37-AS1 | 38.2213416 | 0.203967335 | 0.74602976 | 0.27340375 | 0.78454287 | 0.92261923 |
| UQCRB | 58.5490797 | -0.02536147 | 0.54243823 | -0.0467546 | 0.96270883 | 0.99054261 |
| MTERF3 | 24.2468119 | 0.761807716 | 0.34767729 | 2.19113451 | 0.02844206 | 0.16420667 |
| PTDSS1 | 16.9171054 | -1.001915274 | 0.49754948 | -2.0136998 | 0.04404107 | 0.21423527 |
| SDC2 | 139.347992 | -2.213781971 | 0.40454065 | -5.472335 | 4.44E-08 | 3.70E-06 |
| CPQ | 212.281274 | 0.750871388 | 0.62817807 | 1.19531615 | 0.23196353 | 0.54479294 |
| MTDH | 237.562727 | -0.074170371 | 0.20667321 | -0.3588775 | 0.71968672 | 0.89114993 |

| | | | | | | |
|---|---|---|---|---|---|---|
| LAPTM4B | 97.4462823 | -1.52351822 | 0.52431506 | -2.9057304 | 0.00366397 | 0.04033395 |
| RPL30 | 61.7275646 | -0.514844076 | 0.35934206 | -1.4327409 | 0.15193189 | 0.43628364 |
| RIDA | 17.4273143 | 0.453402533 | 0.60961645 | 0.74375049 | 0.45702744 | 0.7472269 |
| POP1 | 14.0623306 | 0.678600715 | 0.4224616 | 1.60630152 | 0.10820768 | 0.36175274 |
| NIPAL2 | 24.0368593 | 0.182078652 | 0.48591394 | 0.37471379 | 0.70787334 | 0.88614996 |
| STK3 | 486.577665 | -1.071807413 | 0.48239961 | -2.2218248 | 0.02629515 | 0.15596435 |
| AC104986.2 | 39.9629698 | 0.524303844 | 0.33086986 | 1.58462255 | 0.11305212 | 0.37079048 |
| VPS13B | 518.196863 | -0.023083367 | 0.19207912 | -0.1201763 | 0.90434345 | 0.9687613 |
| COX6C | 79.7961837 | 0.609813531 | 0.34854273 | 1.74960912 | 0.08018579 | 0.30394212 |
| SPAG1 | 54.9906848 | -0.690109061 | 0.35662113 | -1.9351323 | 0.05297406 | 0.23826828 |
| RNF19A | 373.193784 | -1.718026045 | 0.54961836 | -3.1258527 | 0.0017729 | 0.0231336 |
| AP000424.1 | 40.4612104 | -6.967951924 | 1.04406874 | -6.673844 | 2.49E-11 | 6.83E-09 |
| ANKRD46 | 35.6033255 | 0.2471299 | 0.30284884 | 0.81601732 | 0.41449023 | 0.71211542 |
| PABPC1 | 116.912528 | -0.498681767 | 0.35153329 | -1.4185904 | 0.15601845 | 0.4418943 |
| YWHAZ | 199.410576 | -0.039050813 | 0.33403968 | -0.1169047 | 0.90693556 | 0.9700616 |
| ZNF706 | 55.2706274 | -0.087977274 | 0.33534678 | -0.2623472 | 0.79305381 | 0.92638859 |
| GRHL2 | 181.695639 | 0.073698734 | 0.36507509 | 0.20187281 | 0.84001616 | 0.9474978 |
| RRM2B | 27.5925499 | 0.053052658 | 0.4784989 | 0.1108731 | 0.91171698 | 0.97128978 |
| UBR5 | 304.814166 | 0.212371049 | 0.21658312 | 0.98055219 | 0.32681362 | 0.6384266 |
| AZIN1 | 55.3370433 | 0.024023408 | 0.28667009 | 0.08380159 | 0.93321419 | 0.9798749 |
| AZIN1-AS1 | 134.083507 | 0.403082236 | 0.44488978 | 0.9060272 | 0.36492146 | 0.67012584 |
| AP003354.2 | 31.2491251 | 0.242658799 | 0.62538822 | 0.38801307 | 0.69800636 | 0.88258901 |
| ATP6V1C1 | 89.7243347 | -0.462912543 | 0.35687043 | -1.2971446 | 0.19458144 | 0.49702866 |
| BAALC-AS1 | 64.5055765 | 1.079028474 | 0.46942647 | 2.29861022 | 0.02152708 | 0.1377317 |
| FZD6 | 109.978632 | 0.603663977 | 0.25430025 | 2.37382374 | 0.01760495 | 0.12052024 |
| SLC25A32 | 38.9074226 | -0.116650059 | 0.29080715 | -0.4011251 | 0.68832799 | 0.87569521 |
| DCAF13 | 21.7030248 | -0.258012321 | 0.33461035 | -0.7710829 | 0.44065779 | 0.73302253 |
| LRP12 | 34.2085518 | 0.669873877 | 0.34587092 | 1.93677419 | 0.05277295 | 0.23800665 |
| ZFPM2 | 32.5821633 | -3.256539415 | 1.03791532 | -3.1375772 | 0.0017035 | 0.02246602 |
| AC027031.2 | 29.4826349 | -0.549403987 | 0.72070643 | -0.7623132 | 0.44587313 | 0.7388166 |
| OXR1 | 1085.00781 | 1.229759679 | 0.36455379 | 3.37332852 | 0.00074265 | 0.01191824 |
| AC090579.1 | 81.8874818 | 0.52446913 | 0.75327575 | 0.69625118 | 0.48627154 | 0.76833754 |
| ANGPT1 | 39.7392432 | 2.729731635 | 0.7396682 | 3.69048127 | 0.00022383 | 0.00462748 |
| EIF3E | 95.158494 | 0.318955923 | 0.23201302 | 1.37473285 | 0.16921428 | 0.46322783 |
| EMC2 | 82.1527703 | 0.711921362 | 0.32334668 | 2.20172779 | 0.02768454 | 0.16133167 |
| NUDCD1 | 38.3577554 | 0.273809858 | 0.28348961 | 0.965855 | 0.33411674 | 0.64554754 |
| ENY2 | 46.52586 | -0.406704268 | 0.30391956 | -1.3381971 | 0.18083221 | 0.47954479 |
| EBAG9 | 27.0695017 | 0.649458104 | 0.38104597 | 1.70440879 | 0.08830474 | 0.32275712 |
| SYBU | 23.1050131 | -0.447614041 | 0.52693757 | -0.8494631 | 0.39562363 | 0.69642725 |
| TRPS1 | 584.487102 | -0.035120239 | 0.30354839 | -0.115699 | 0.90789111 | 0.97033204 |
| EIF3H | 124.728606 | -0.041200193 | 0.23075514 | -0.1785451 | 0.85829493 | 0.95271398 |
| UTP23 | 21.2711526 | 0.024723315 | 0.45964533 | 0.05378781 | 0.95710422 | 0.98859719 |
| RAD21 | 62.035806 | 0.125925433 | 0.26686142 | 0.47187576 | 0.63701547 | 0.8538689 |
| MED30 | 14.2173265 | 0.308072446 | 0.68036908 | 0.45280195 | 0.65069136 | 0.86113455 |
| EXT1 | 658.118682 | 0.05970511 | 0.40438532 | 0.14764411 | 0.88262364 | 0.96161232 |
| SAMD12 | 603.695487 | -0.574746997 | 0.26606896 | -2.160143 | 0.0307616 | 0.17290661 |
| AC023590.1 | 46.6042917 | -1.421290877 | 0.73096315 | -1.9444084 | 0.05184622 | 0.235032 |
| SAMD12-AS1 | 40.3089718 | -1.613538951 | 0.52011481 | -3.1022746 | 0.0019204 | 0.02449492 |
| TNFRSF11B | 26.8145474 | -2.092362014 | 0.70785477 | -2.9559199 | 0.00311738 | 0.03579612 |
| MAL2 | 83.8811581 | -0.159701629 | 0.34216597 | -0.4667373 | 0.64068782 | 0.85468927 |
| TAF2 | 107.596962 | 0.018765348 | 0.34640565 | 0.0541716 | 0.95679845 | 0.98859719 |
| DEPTOR | 968.79877 | -2.275784396 | 0.31159182 | -7.3037361 | 2.80E-13 | 1.00E-10 |
| AC091563.1 | 50.0679633 | -2.974938886 | 0.47704363 | -6.2361987 | 4.48E-10 | 8.89E-08 |
| COL14A1 | 55.1320095 | -2.561816949 | 0.57233312 | -4.4760942 | 7.60E-06 | 0.00030553 |
| MRPL13 | 26.9235692 | 0.411069949 | 0.41854002 | 0.98215207 | 0.32602494 | 0.63780184 |
| MTBP | 15.6222399 | 0.236014832 | 0.55616843 | 0.42435856 | 0.67130434 | 0.86777458 |
| SNTB1 | 131.185649 | -2.268236738 | 0.59026926 | -3.8427154 | 0.00012168 | 0.00282714 |
| ZHX2 | 288.748176 | -1.809672807 | 0.47655644 | -3.7973945 | 0.00014622 | 0.00330246 |
| DERL1 | 23.3857265 | -0.027533697 | 0.40697338 | -0.0676548 | 0.94606044 | 0.98574808 |
| TBC1D31 | 16.2360567 | 0.76321067 | 0.45096222 | 1.69240489 | 0.0905688 | 0.32756536 |
| WDYHV1 | 23.5269397 | -0.128451397 | 0.46000854 | -0.279237 | 0.78006296 | 0.9206566 |
| FBXO32 | 22.1011104 | -0.856775577 | 0.38587313 | -2.2203557 | 0.02639463 | 0.15625624 |
| FAM91A1 | 74.3026531 | -0.204701395 | 0.35264404 | -0.580476 | 0.56159368 | 0.81384822 |
| TMEM65 | 106.20407 | -0.504536167 | 0.32634358 | -1.5460276 | 0.12209791 | 0.38617399 |
| RNF139 | 12.283805 | 0.537445743 | 0.49932537 | 1.07634376 | 0.28177355 | 0.59602009 |
| TATDN1 | 71.0083379 | 0.272039241 | 0.29055255 | 0.93628241 | 0.3491278 | 0.65632016 |
| NDUFB9 | 44.2142095 | -0.235512837 | 0.36607808 | -0.6433405 | 0.52000321 | 0.7877576 |
| MTSS1 | 381.521405 | 0.878804848 | 0.32654314 | 2.69123659 | 0.00711877 | 0.06437963 |
| SQLE | 37.1019229 | -0.732908397 | 0.5625485 | -1.3028359 | 0.19263078 | 0.49370242 |
| WASHC5 | 58.935968 | -0.04525007 | 0.28281199 | -0.1600005 | 0.87288065 | 0.95703878 |
| NSMCE2 | 426.639517 | -0.108854268 | 0.38476361 | -0.2829121 | 0.77724424 | 0.91968524 |
| TRIB1 | 22.3326599 | -0.794895292 | 0.40851377 | -1.9458225 | 0.05167606 | 0.23492327 |
| FAM84B | 69.4747788 | -2.026029609 | 0.70465881 | -2.8751923 | 0.00403781 | 0.04311026 |
| PCAT1 | 399.361082 | -0.852062459 | 0.59209553 | -1.4390625 | 0.15013282 | 0.43473243 |
| FAM49B | 392.392763 | -0.193616795 | 0.27084642 | -0.7148582 | 0.47469665 | 0.76075482 |
| ASAP1 | 198.258132 | -0.971984671 | 0.36274493 | -2.6795265 | 0.00737264 | 0.06603502 |
| EFR3A | 160.505206 | -0.635824201 | 0.43263722 | -1.4696475 | 0.14165725 | 0.42176296 |
| KCNQ3 | 37.6167147 | 0.15599617 | 0.92523047 | 0.1686025 | 0.86610931 | 0.95565922 |
| LRRC6 | 51.9703453 | 0.287060378 | 0.45230974 | 0.63465442 | 0.52565382 | 0.79307383 |
| PHF20L1 | 105.823563 | 0.237592625 | 0.25817059 | 0.92029314 | 0.35741959 | 0.66399289 |
| NDRG1 | 361.09699 | -1.416126874 | 0.58014505 | -2.4409876 | 0.01464716 | 0.10644591 |
| ST3GAL1 | 63.1703068 | -2.077907782 | 0.52778476 | -3.9370363 | 8.25E-05 | 0.00210733 |
| ZFAT | 61.0425225 | 0.156201915 | 0.35421364 | 0.44098221 | 0.65922587 | 0.86450914 |
| AC083843.3 | 69.9403094 | 0.53510501 | 0.47565832 | 1.12497773 | 0.26059847 | 0.57426715 |
| KHDRBS3 | 191.078877 | -1.759355002 | 0.50329705 | -3.4956593 | 0.00047289 | 0.00828806 |
| TRAPPC9 | 483.588864 | 0.122281579 | 0.14380781 | 0.85031251 | 0.39515137 | 0.69642725 |
| AGO2 | 122.513334 | -0.584100314 | 0.36559554 | -1.5976681 | 0.11011687 | 0.36538423 |
| PTK2 | 756.236709 | 0.06966196 | 0.24683664 | 0.28221888 | 0.77777568 | 0.91971851 |
| DENND3 | 14.8559717 | -1.332520243 | 0.49430237 | -2.6957594 | 0.00702284 | 0.06382158 |
| SLC45A4 | 78.9506165 | 0.397782251 | 0.34347385 | 1.15811509 | 0.24681707 | 0.55938803 |
| TSNARE1 | 42.9732325 | -0.263147266 | 0.48344274 | -0.5443194 | 0.58622169 | 0.8294442 |
| JRK | 11.6796948 | 0.020929653 | 0.42726925 | 0.04898469 | 0.96093149 | 0.98983463 |
| LY6E | 32.5546921 | -0.567818717 | 0.7966485 | -0.7127594 | 0.47599465 | 0.76162498 |
| GLI4 | 10.5630083 | 1.294338007 | 0.56360448 | 2.29653604 | 0.02164525 | 0.13787238 |
| TOP1MT | 16.3718117 | 0.543146876 | 0.42418226 | 1.28045637 | 0.20038468 | 0.50442407 |
| RHPN1 | 22.6242463 | 0.464886843 | 0.52244243 | 0.88983363 | 0.37355523 | 0.67707283 |
| ZC3H3 | 31.7247832 | 0.066983618 | 0.51213535 | 0.1307928 | 0.89593922 | 0.96519206 |
| GSDMD | 14.6882405 | -0.173356706 | 0.50687017 | -0.342014 | 0.73234034 | 0.89806184 |
| EEF1D | 85.6879655 | -1.057925406 | 0.42958738 | -2.4626548 | 0.01379126 | 0.10262549 |
| ZNF623 | 17.070944 | -0.108105892 | 0.36416088 | -0.296863 | 0.7665711 | 0.91564296 |
| FAM83H | 13.5852454 | -1.17873844 | 0.45835911 | -2.5716483 | 0.01012156 | 0.08292923 |
| IQANK1 | 15.4369922 | -0.746734607 | 0.76402431 | -0.9773702 | 0.3283859 | 0.63988927 |
| SCRIB | 15.1215729 | 0.207855248 | 0.48911504 | 0.42496189 | 0.67086446 | 0.86777458 |
| PUF60 | 14.5707297 | -1.003118066 | 0.50788164 | -1.975102 | 0.04825657 | 0.22503839 |
| NRBP2 | 14.4446788 | -0.622031561 | 0.52316392 | -1.1889802 | 0.23444744 | 0.54657 |
| PLEC | 33.0778189 | -0.357141683 | 0.36513634 | -0.978105 | 0.32802239 | 0.63958193 |
| PARP10 | 15.531019 | 1.879820039 | 0.71397527 | 2.63289238 | 0.00846612 | 0.0729557 |
| GRINA | 22.7277676 | -0.033461373 | 0.60124586 | -0.0556534 | 0.95561793 | 0.98853497 |
| GPAA1 | 20.6832454 | 0.059965088 | 0.52773704 | 0.11362683 | 0.90953362 | 0.97090822 |
| CYC1 | 26.8824506 | -0.008639204 | 0.53619593 | -0.016112 | 0.98714502 | 0.9977555 |
| SHARPIN | 27.9519714 | -0.080525354 | 0.2875325 | -0.2800565 | 0.77943414 | 0.92039563 |
| MAF1 | 18.8786725 | -0.613399947 | 0.66963241 | -0.9160249 | 0.35965385 | 0.66572082 |
| MROH1 | 130.977044 | -0.695784122 | 0.47938641 | -1.4514056 | 0.14666695 | 0.42936346 |

| | | | | | | |
|---|---|---|---|---|---|---|
| HSF1 | 57.7556781 | -0.222369984 | 0.2420812 | -0.918576 | 0.35831738 | 0.6647359 |
| DGAT1 | 13.0009767 | 0.32713849 | 0.48535513 | 0.67401881 | 0.50029936 | 0.77558432 |
| CPSF1 | 11.9810026 | 0.63964534 | 0.58937831 | 1.08528823 | 0.27779402 | 0.59311917 |
| VPS28 | 37.2510154 | -0.412878037 | 0.46924355 | -0.87988 | 0.3789243 | 0.68259083 |
| CYHR1 | 34.043809 | 0.215865021 | 0.49853521 | 0.43299854 | 0.66501583 | 0.86673296 |
| KIFC2 | 18.109221 | 0.747405509 | 0.53885626 | 1.38702205 | 0.16543504 | 0.4575743 |
| PPP1R16A | 30.4982023 | -0.12400493 | 0.40140027 | -0.3089309 | 0.75737412 | 0.91142957 |
| ARHGAP39 | 23.1773826 | -0.112702906 | 0.38356024 | -0.2938336 | 0.76888502 | 0.91600052 |
| ZNF251 | 71.0577023 | -0.003540811 | 0.24401442 | -0.0145107 | 0.98842257 | 0.9977555 |
| RPL8 | 100.376273 | -0.665919153 | 0.48148751 | -1.3830455 | 0.1666509 | 0.4589052 |
| ZNF7 | 24.3022713 | -0.65531194 | 0.32401257 | -2.0224892 | 0.04312583 | 0.21134556 |
| ZNF250 | 17.1672308 | 0.647052972 | 0.39659211 | 1.63153262 | 0.10277798 | 0.35038461 |
| ZNF16 | 18.1372874 | 0.120677143 | 0.41875265 | 0.2881824 | 0.77320712 | 0.91791018 |
| WASHC1 | 101.792986 | 0.46817939 | 0.34500243 | 1.35703216 | 0.17477099 | 0.47124486 |
| CBWD1 | 29.7247747 | 0.341240115 | 0.36564309 | 0.93326012 | 0.35068568 | 0.65782032 |
| DOCK8 | 36.1010819 | 0.827857803 | 0.49535355 | 1.67124633 | 0.09467303 | 0.33551172 |
| KANK1 | 1277.48317 | 2.118265545 | 0.28571005 | 7.41403942 | 1.23E-13 | 5.19E-11 |
| SMARCA2 | 160.651439 | 0.443330197 | 0.33525614 | 1.32236263 | 0.18604743 | 0.48628295 |
| VLDLR-AS1 | 67.5490325 | -1.563894008 | 0.68107012 | -2.2962305 | 0.0216627 | 0.13787238 |
| VLDLR | 38.6038588 | -0.806680431 | 0.38982598 | -2.0693347 | 0.03851469 | 0.19803907 |
| PUM3 | 207.769363 | -0.977639601 | 0.27848304 | -3.5105893 | 0.00044711 | 0.00792566 |
| RFX3 | 262.428783 | 0.278280437 | 0.32269219 | 0.86237115 | 0.38848331 | 0.69192328 |
| RFX3-AS1 | 211.94267 | -0.308115582 | 0.41435754 | -0.7435983 | 0.4571195 | 0.7472269 |
| GLIS3 | 1011.35865 | -0.857618216 | 0.42299458 | -2.0274922 | 0.0426121 | 0.21039469 |
| SLC1A1 | 75.5351805 | 0.419105984 | 0.46706622 | 0.89731598 | 0.36955033 | 0.67364366 |
| SPATA6L | 111.311756 | -0.998121 | 0.37321831 | -2.6743624 | 0.00748715 | 0.0666764 |
| CDC37L1 | 67.0736936 | -0.080570957 | 0.38530532 | -0.2091094 | 0.83436285 | 0.94646712 |
| AK3 | 522.345192 | 1.33725838 | 0.44130247 | 3.03025357 | 0.00244348 | 0.02955001 |
| RCL1 | 119.729135 | -0.009822363 | 0.35343826 | -0.0277909 | 0.97782893 | 0.99505657 |
| JAK2 | 130.148399 | -0.06247873 | 0.21233558 | -0.2942452 | 0.76857052 | 0.91600052 |
| PLGRKT | 55.5584721 | 0.566983266 | 0.30194757 | 1.87775403 | 0.06041482 | 0.25698348 |
| RIC1 | 221.714258 | 0.238725372 | 0.28336661 | 0.8424612 | 0.39952985 | 0.69959441 |
| ERMP1 | 185.372043 | 0.475672511 | 0.31031483 | 1.53287072 | 0.12530771 | 0.39153507 |
| KIAA2026 | 283.343424 | -0.202166952 | 0.22061599 | -0.9163749 | 0.3594703 | 0.66568045 |
| RANBP6 | 15.1526646 | -0.159778192 | 0.58930597 | -0.2711294 | 0.78629148 | 0.92322402 |
| UHRF2 | 144.975375 | 0.60447513 | 0.19526054 | 3.09573628 | 0.00196325 | 0.02487138 |
| GLDC | 176.245745 | 0.900582483 | 0.47935341 | 1.87874431 | 0.06027941 | 0.25652454 |
| AL162411.1 | 27.513749 | -0.28450975 | 0.34791508 | -0.8177563 | 0.41349635 | 0.71143975 |
| KDM4C | 452.913488 | -0.342122877 | 0.2627034 | -1.3023161 | 0.19280834 | 0.4940217 |
| DMAC1 | 16.8425629 | 0.171960134 | 0.38177863 | 0.45041843 | 0.65240876 | 0.86179129 |
| PTPRD | 35.6334634 | -1.26056067 | 0.5595752 | -2.2527101 | 0.02427744 | 0.1479791 |
| LURAP1L | 91.7926003 | -4.425847251 | 0.89091836 | -4.9677361 | 6.77E-07 | 4.23E-05 |
| MPDZ | 141.70127 | 0.517995545 | 0.32823557 | 1.57812132 | 0.11453774 | 0.37301778 |
| NFIB | 257.933603 | -0.212209196 | 0.40335958 | -0.5261043 | 0.59881577 | 0.8356618 |
| ZDHHC21 | 251.671639 | 0.557293139 | 0.27139423 | 2.05344506 | 0.04002943 | 0.20269667 |
| FREM1 | 336.179646 | 1.808566475 | 0.75813898 | 2.3855342 | 0.01705434 | 0.11787593 |
| TTC39B | 93.7785377 | 0.250164758 | 0.33550682 | 0.7456324 | 0.4558895 | 0.74621803 |
| SNAPC3 | 68.511401 | 0.156903107 | 0.2152025 | 0.7290952 | 0.46594343 | 0.75280827 |
| PSIP1 | 69.0641176 | -0.03522542 | 0.38933339 | -0.0904762 | 0.92790878 | 0.97812694 |
| CCDC171 | 153.372057 | 0.556184563 | 0.57033975 | 0.97518113 | 0.32947042 | 0.64146632 |
| BNC2 | 75.0682747 | -2.340952879 | 0.59707665 | -3.9206907 | 8.83E-05 | 0.00221905 |
| CNTLN | 97.1768101 | 0.563845021 | 0.31510404 | 1.78939318 | 0.07355152 | 0.29009913 |
| SH3GL2 | 58.4699214 | -0.63351748 | 0.63472012 | -0.9981052 | 0.31822833 | 0.63093651 |
| HAUS6 | 19.0618484 | 0.001307999 | 0.39571188 | 0.00330543 | 0.99736265 | 0.99922221 |
| PLIN2 | 47.1368139 | -0.643185354 | 0.6318313 | -1.0179701 | 0.30869218 | 0.62245953 |
| DENND4C | 285.352858 | -0.123133956 | 0.16611931 | -0.7412381 | 0.45854911 | 0.747377 |
| RPS6 | 107.229918 | -0.85828168 | 0.53053699 | -1.6177603 | 0.10571427 | 0.35687176 |
| MLLT3 | 480.650272 | -0.372576646 | 0.27790246 | -1.3406742 | 0.18002627 | 0.47863272 |
| FOCAD | 195.906279 | 0.458113831 | 0.4461999 | 1.02670089 | 0.30456133 | 0.61818174 |
| KLHL9 | 17.2250552 | -0.085088035 | 0.44594692 | -0.1908031 | 0.84867989 | 0.95003258 |
| LINC01239 | 24.731759 | -1.103330918 | 0.73271791 | -1.5058058 | 0.13211702 | 0.40375585 |
| CAAP1 | 150.638087 | 0.401198215 | 0.29145719 | 1.37652538 | 0.16865904 | 0.46237864 |
| PLAA | 48.0214667 | -0.109848146 | 0.27118314 | -0.40507 | 0.68542608 | 0.87426987 |
| IFT74 | 116.582651 | 0.23279693 | 0.27456719 | 0.84786872 | 0.39651109 | 0.6969028 |
| MOB3B | 122.11284 | -1.080358718 | 0.33333109 | -3.2410979 | 0.0011907 | 0.01712612 |
| C9orf72 | 37.7425021 | -0.390784549 | 0.36731309 | -1.0639004 | 0.28737382 | 0.60172322 |
| LINGO2 | 104.427008 | 0.501451922 | 0.44615068 | 1.12395194 | 0.26103342 | 0.57470498 |
| ACO1 | 34.8182006 | 0.228613328 | 0.48704073 | 0.46939263 | 0.63878902 | 0.8538689 |
| DDX58 | 77.9071435 | 0.265518492 | 0.93153715 | 0.28503264 | 0.77561914 | 0.91926749 |
| TOPORS | 25.3489115 | -0.002155613 | 0.41033526 | -0.0052533 | 0.99580849 | 0.99922221 |
| NDUFB6 | 39.3960921 | 0.486271983 | 0.3120918 | 1.5581056 | 0.11920822 | 0.38057334 |
| APTX | 66.9588009 | 0.299606859 | 0.35982824 | 0.83263854 | 0.40504862 | 0.70434042 |
| DNAJA1 | 37.553333 | 0.0092017 | 0.34095724 | 0.02698784 | 0.97846943 | 0.9951912 |
| SMU1 | 40.1617694 | 0.513842234 | 0.30188763 | 1.70209765 | 0.08873706 | 0.32344972 |
| B4GALT1 | 106.934303 | -0.432437599 | 0.30078711 | -1.4376866 | 0.15052299 | 0.43488337 |
| BAG1 | 28.4210982 | 0.157716466 | 0.50379171 | 0.31305887 | 0.75423593 | 0.91026551 |
| CHMP5 | 25.8719865 | -0.186619229 | 0.40782132 | -0.4576005 | 0.6472395 | 0.85917726 |
| NFX1 | 149.612615 | -0.425484541 | 0.3650591 | -1.1655224 | 0.24380761 | 0.55580983 |
| UBE2R2 | 236.269396 | -0.253224531 | 0.40166919 | -0.6304306 | 0.52841292 | 0.7943404 |
| UBAP2 | 148.798989 | -0.374395011 | 0.3211889 | -1.1656536 | 0.24375451 | 0.55580983 |
| DCAF12 | 29.4612312 | 0.149037345 | 0.3635411 | 0.4099601 | 0.68183522 | 0.87271442 |
| UBAP1 | 220.465425 | -0.686468845 | 0.34598651 | -1.9840913 | 0.04724566 | 0.22304735 |
| NUDT2 | 14.2115932 | -0.256363879 | 0.64471247 | -0.3976406 | 0.6908951 | 0.87716584 |
| FAM219A | 40.2681439 | -0.363842216 | 0.29130796 | -1.2489951 | 0.21166687 | 0.52063276 |
| VCP | 58.0368 | -0.658016046 | 0.34490646 | -1.9078102 | 0.05641575 | 0.2473063 |
| STOML2 | 15.0201383 | -0.767415938 | 0.63853106 | -1.2018459 | 0.22942324 | 0.54228719 |
| UNC13B | 242.136688 | 0.765270772 | 0.21756185 | 3.51748611 | 0.00043566 | 0.00777862 |
| RUSC2 | 157.35262 | 0.915888302 | 0.39485013 | 2.31958467 | 0.02036336 | 0.13268199 |
| CCDC107 | 16.4334564 | -0.968177436 | 0.39127609 | -2.4744099 | 0.01334565 | 0.10035066 |
| TLN1 | 39.6590778 | -0.272633817 | 0.32872337 | -0.8293716 | 0.40689419 | 0.70596974 |
| GBA2 | 59.5702773 | 0.738847183 | 0.33770453 | 2.18785098 | 0.02868046 | 0.16498593 |
| RGP1 | 18.4099434 | -0.488194328 | 0.51760694 | -0.9431758 | 0.345591 | 0.65413936 |
| AL133410.1 | 20.8313861 | 0.002703704 | 0.41676572 | 0.00648735 | 0.99482388 | 0.9992177 |
| TMEM8B | 18.898896 | 0.884045784 | 0.35757835 | 2.47231347 | 0.01342417 | 0.10067724 |
| RECK | 39.9454352 | 0.461414449 | 0.397463 | 1.16089912 | 0.24568293 | 0.5580384 |
| CLTA | 98.4900935 | -0.616698274 | 0.39042891 | -1.5795405 | 0.11421214 | 0.372426 |
| GNE | 78.4344416 | -2.084237829 | 0.72583 | -2.8715234 | 0.00408499 | 0.04338087 |
| RNF38 | 254.005465 | -0.329872125 | 0.43467058 | -0.7589014 | 0.44791152 | 0.7399821 |
| EBLN3P | 41.1653421 | -0.115172764 | 0.39088277 | -0.2946478 | 0.76826291 | 0.91600052 |
| ZCCHC7 | 476.249845 | 0.174401675 | 0.20837904 | 0.83694442 | 0.40262381 | 0.70169281 |
| GRHPR | 22.6516576 | 0.24534702 | 0.41509335 | 0.59106468 | 0.55447708 | 0.81072612 |
| ZBTB5 | 25.6818751 | 0.029168794 | 0.32464533 | 0.08984818 | 0.92840786 | 0.97829541 |
| AL138752.2 | 63.3772897 | -0.583974227 | 0.30602843 | -1.9082352 | 0.05636082 | 0.24729803 |
| FRMPD1 | 14.6548024 | 2.13030552 | 0.7110196 | 2.99612768 | 0.00273432 | 0.03219039 |
| TRMT10B | 53.5518403 | 0.086383859 | 0.48132523 | 0.17947087 | 0.85756799 | 0.95266335 |
| DCAF10 | 158.613315 | 0.02623325 | 0.20457011 | 0.12823598 | 0.89796222 | 0.96607503 |
| FAM95C | 18.3729303 | 0.279369996 | 0.41851323 | 0.66752967 | 0.50443386 | 0.77792612 |
| ANKRD18A | 20.1306761 | -0.168423511 | 0.44049134 | -0.3823537 | 0.702199 | 0.88441219 |
| GLIDR | 37.2682488 | 0.871531025 | 0.40948006 | 2.12838454 | 0.03330521 | 0.18186117 |
| BMS1P14 | 33.6165879 | -0.684357367 | 0.6212347 | -1.1016084 | 0.27063195 | 0.58722969 |
| CBWD6 | 16.7272429 | 0.243145764 | 0.61134827 | 0.39772054 | 0.6908362 | 0.87716584 |
| CNTNAP3B | 33.4513358 | -0.870591869 | 0.64204178 | -1.3559739 | 0.17510749 | 0.47160665 |
| AL512625.3 | 13.3341207 | 1.39956509 | 0.44624079 | 3.13634505 | 0.00171068 | 0.02249698 |

| | | | | | | |
|---|---|---|---|---|---|---|
| CBWD5 | 46.6607434 | 0.179701767 | 0.32713579 | 0.54931858 | 0.58278684 | 0.82657506 |
| ZNF658 | 17.0176424 | 0.766970049 | 0.59591324 | 1.28704985 | 0.19807691 | 0.50081448 |
| PGM5 | 22.2207128 | -1.421033039 | 0.71261657 | -1.9941061 | 0.04614046 | 0.22049887 |
| AL161457.2 | 11.7835928 | 0.285372491 | 0.60584546 | 0.47103182 | 0.63761801 | 0.8538689 |
| PIP5K1B | 207.7087 | 0.975294441 | 0.3756636 | 2.59619097 | 0.00942637 | 0.07888124 |
| FAM189A2 | 39.4623974 | -0.735252264 | 0.56193324 | -1.3084335 | 0.19072632 | 0.49135864 |
| PTAR1 | 138.449389 | 0.108745578 | 0.17565699 | 0.61907913 | 0.53586423 | 0.79974391 |
| MAMDC2-AS | 69.9593944 | 0.524141843 | 0.65058163 | 0.80565116 | 0.42044402 | 0.71867263 |
| MAMDC2 | 55.9218231 | -0.236123703 | 0.55071266 | -0.4287603 | 0.6680977 | 0.86777458 |
| SMC5-AS1 | 86.5958339 | 0.683990224 | 0.4110946 | 1.66382683 | 0.09614704 | 0.33893195 |
| SMC5 | 189.54772 | 0.010192721 | 0.22647283 | 0.04500637 | 0.96410223 | 0.99082982 |
| KLF9 | 47.7891119 | 0.717620512 | 0.38144187 | 1.88133647 | 0.05992616 | 0.25572153 |
| TRPM3 | 108.204808 | 1.00931762 | 0.68946198 | 1.46392064 | 0.14321563 | 0.4243745 |
| CEMIP2 | 153.137459 | -3.995370486 | 0.47432968 | -8.4231932 | 3.66E-17 | 2.85E-14 |
| ABHD17B | 33.5622594 | -0.291939603 | 0.42922725 | -0.6801516 | 0.49640847 | 0.77475164 |
| C9orf85 | 68.7965277 | -0.052412133 | 0.24147176 | -0.2170529 | 0.82816716 | 0.94268221 |
| ZFAND5 | 136.038429 | 0.030727976 | 0.52747195 | 0.05825518 | 0.95354536 | 0.98771968 |
| ALDH1A1 | 40.8299046 | -0.362636088 | 0.54291487 | -0.6679428 | 0.50417009 | 0.77792612 |
| TRPM6 | 739.203867 | 3.919650174 | 0.644843 | 6.07845658 | 1.21E-09 | 1.95E-07 |
| CARNMT1 | 101.851015 | 0.716442691 | 0.20195936 | 3.54745979 | 0.00038897 | 0.00721016 |
| NMRK1 | 42.0926836 | -0.726639236 | 0.66005412 | -1.1008783 | 0.27094964 | 0.58724651 |
| OSTF1 | 28.627447 | -0.789863461 | 0.35187026 | -2.2447576 | 0.0247837 | 0.14976227 |
| PCSK5 | 46.2619561 | -0.855465641 | 0.8692293 | -0.9841657 | 0.32503406 | 0.63715682 |
| VPS13A | 183.951816 | 0.094289992 | 0.22496807 | 0.4191261 | 0.67512398 | 0.86960234 |
| GNAQ | 553.414471 | -0.498263451 | 0.2355977 | -2.114891 | 0.03443924 | 0.18459472 |
| TLE4 | 26.6758513 | -0.634881578 | 0.53952905 | -1.1767329 | 0.23930214 | 0.55106277 |
| TLE1 | 59.352779 | -0.884420936 | 0.43753634 | -2.0213657 | 0.04324193 | 0.21173421 |
| AL591368.1 | 10.9978533 | 2.183359778 | 0.98787839 | 2.21015036 | 0.02709473 | 0.15969106 |
| RASEF | 26.057963 | -1.409952398 | 0.38920441 | -3.6226527 | 0.0002916 | 0.00567744 |
| FRMD3 | 497.521507 | 2.072266182 | 0.44982467 | 4.60683088 | 4.09E-06 | 0.00018506 |
| IDNK | 16.3603913 | -0.125562976 | 0.35660823 | -0.3521034 | 0.7247607 | 0.89351696 |
| UBQLN1 | 124.458021 | 0.362331305 | 0.27472721 | 1.31887667 | 0.18721035 | 0.48725107 |
| GKAP1 | 49.6164603 | 0.65059671 | 0.32209209 | 2.019909 | 0.04339283 | 0.21196707 |
| KIF27 | 71.807873 | 0.220982948 | 0.33852552 | 0.65278076 | 0.51389763 | 0.78323295 |
| HNRNPK | 135.241297 | -0.138373979 | 0.27657795 | -0.5003073 | 0.61685869 | 0.84223226 |
| RMI1 | 13.83588 | 0.015867043 | 0.50138808 | 0.03164623 | 0.97475418 | 0.99361556 |
| AGTPBP1 | 118.625486 | -0.470063876 | 0.21155282 | -2.2219693 | 0.02628538 | 0.15596435 |
| NAA35 | 132.939095 | 0.213819861 | 0.23510618 | 0.90946083 | 0.36310693 | 0.66856418 |
| GOLM1 | 93.4276225 | 0.174221799 | 0.4177021 | 0.41709582 | 0.67660833 | 0.87028501 |
| C9orf153 | 15.1285497 | 0.880913435 | 0.53203243 | 1.65575139 | 0.09777218 | 0.34066808 |
| ISCA1 | 35.4802685 | 0.188173989 | 0.28039606 | 0.67110068 | 0.50215639 | 0.77659748 |
| TUT7 | 106.069718 | -0.336623857 | 0.23908479 | -1.4079685 | 0.15914039 | 0.44760007 |
| AL513318.2 | 18.6389734 | -0.14449424 | 0.51563154 | -0.2802277 | 0.77930282 | 0.92039563 |
| DAPK1 | 445.013303 | -1.252885196 | 0.60744206 | -2.0625592 | 0.03915453 | 0.20026156 |
| CTSL | 45.9805606 | -0.448682221 | 0.58664131 | -0.7648323 | 0.44437141 | 0.73698311 |
| SPIN1 | 348.969865 | 0.245787193 | 0.24416266 | 1.00665348 | 0.31410132 | 0.62887712 |
| SHC3 | 131.878791 | -0.865378511 | 1.08844327 | -0.7950607 | 0.42657817 | 0.72230448 |
| SECISBP2 | 150.858364 | -0.073981268 | 0.3161386 | -0.2340153 | 0.8149731 | 0.93707884 |
| SEMA4D | 136.98109 | -0.859499535 | 0.39977699 | -2.1499475 | 0.03155937 | 0.17567711 |
| SYK | 53.6230215 | -0.044616972 | 0.36462461 | -0.1223641 | 0.90261065 | 0.96803829 |
| AL158071.1 | 55.0501193 | 1.013575709 | 0.80738385 | 1.25538268 | 0.20933987 | 0.51790437 |
| AL158071.4 | 268.442512 | 1.463139535 | 0.64859775 | 2.25585046 | 0.02408 | 0.14717838 |
| AL158071.5 | 18.3996949 | 2.047083407 | 0.58091465 | 3.52389701 | 0.00042525 | 0.00769399 |
| AUH | 406.818433 | 0.302205762 | 0.25591026 | 1.18090522 | 0.23764038 | 0.54997949 |
| NFIL3 | 54.9390961 | -2.075967526 | 0.79561823 | -2.6092508 | 0.00907407 | 0.07684526 |
| SPTLC1 | 81.1325862 | -0.014308724 | 0.2598636 | -0.0550624 | 0.95608872 | 0.98853497 |
| IARS | 88.6182618 | -1.099814488 | 0.68630091 | -1.6025252 | 0.10903953 | 0.36312941 |
| NOL8 | 29.4813299 | -0.420767209 | 0.32572279 | -1.2917954 | 0.196428 | 0.49845696 |
| CENPP | 49.6250336 | 0.091944363 | 0.72797924 | 0.1263008 | 0.89949382 | 0.96645314 |
| IPPK | 29.6028928 | 0.517838032 | 0.46242509 | 1.11983118 | 0.26278571 | 0.57700771 |
| BICD2 | 27.1385475 | 0.038166693 | 0.39386303 | 0.09690347 | 0.92280305 | 0.97611757 |
| ZNF484 | 18.460873 | -0.050337527 | 0.40406346 | -0.1245783 | 0.90085743 | 0.96747232 |
| NINJ1 | 21.5269754 | -0.149031347 | 0.67907879 | -0.2194611 | 0.82629091 | 0.94219597 |
| FAM120AOS | 12.4988779 | -0.265342798 | 0.44900889 | -0.5909522 | 0.55455244 | 0.81072612 |
| FAM120A | 138.948091 | 0.165698093 | 0.29217851 | 0.56711254 | 0.57063773 | 0.81891359 |
| PHF2 | 42.6211652 | 0.175947695 | 0.31460289 | 0.55926917 | 0.57597803 | 0.82147339 |
| PTPDC1 | 58.8950345 | 0.129342026 | 0.32281308 | 0.40067158 | 0.68866194 | 0.87600053 |
| AL158152.2 | 74.571716 | 0.255585578 | 0.25571979 | 0.99947517 | 0.31756456 | 0.63093651 |
| ZNF169 | 29.9461133 | 0.84482341 | 0.37427669 | 2.2572162 | 0.02399457 | 0.14714173 |
| MFSD14B | 54.3739206 | -0.17796432 | 0.31300495 | -0.5685671 | 0.56964994 | 0.81840001 |
| C9orf3 | 648.396667 | -0.153478476 | 0.23819776 | -0.6443322 | 0.51936007 | 0.78759816 |
| FANCC | 144.616863 | 0.083716281 | 0.28197755 | 0.29688988 | 0.76655058 | 0.91564296 |
| PTCH1 | 48.6151827 | -0.756815219 | 0.36404454 | -2.0789083 | 0.03762578 | 0.19577164 |
| LINC00476 | 89.7625761 | 0.297246813 | 0.45961932 | 0.64672393 | 0.51781063 | 0.78658624 |
| ERCC6L2 | 133.14663 | -0.004254482 | 0.27078458 | -0.0157117 | 0.9874644 | 0.9977555 |
| SLC35D2 | 27.998072 | 0.053293552 | 0.39416146 | 0.13520741 | 0.89244789 | 0.964216 |
| HABP4 | 78.9053804 | 0.526353738 | 0.47840104 | 1.10023536 | 0.27122959 | 0.58758008 |
| CDC14B | 165.318706 | 0.386167803 | 0.50895136 | 0.75875188 | 0.44800099 | 0.7399821 |
| AL589843.1 | 25.2698981 | 0.086810693 | 0.45714483 | 0.18989757 | 0.8493894 | 0.95003258 |
| ZNF510 | 18.4477953 | -0.16105425 | 0.3919807 | -0.4108729 | 0.68116573 | 0.87265585 |
| ZNF782 | 66.7838443 | -0.165316948 | 0.26935759 | -0.6137453 | 0.53938367 | 0.80166126 |
| MFSD14C | 156.579183 | -0.231364125 | 0.24390424 | -0.9485859 | 0.34283126 | 0.65243773 |
| CCDC180 | 14.7396055 | 0.229973917 | 0.60360335 | 0.38100172 | 0.70320197 | 0.88513266 |
| TDRD7 | 32.2448445 | -0.040425711 | 0.43744871 | -0.0924125 | 0.92637034 | 0.97735117 |
| TSTD2 | 37.0620133 | -0.397045285 | 0.27756403 | -1.4304638 | 0.15258396 | 0.43748242 |
| NCBP1 | 40.3474645 | 0.352922975 | 0.36169377 | 0.97575078 | 0.32918798 | 0.64105028 |
| XPA | 54.52131 | 0.811768921 | 0.36086272 | 2.24952283 | 0.02447925 | 0.14869351 |
| ANP32B | 42.8685014 | -0.028249895 | 0.3571109 | -0.0791068 | 0.93694769 | 0.98235694 |
| TRIM14 | 15.6441025 | -1.04801438 | 0.86460388 | -1.2121324 | 0.2254617 | 0.53641361 |
| ANKS6 | 25.9064903 | 0.125258377 | 0.44234026 | 0.283172 | 0.77704499 | 0.91968524 |
| TGFBR1 | 75.6972175 | 0.081491288 | 0.47058343 | 0.17317075 | 0.86251721 | 0.95458251 |
| ALG2 | 16.4002255 | -0.048464854 | 0.48766529 | -0.0993814 | 0.92083546 | 0.9754826 |
| SEC61B | 37.7459763 | -0.713489605 | 0.34384686 | -2.0750214 | 0.03798455 | 0.19654159 |
| AL359710.1 | 69.3232833 | 0.555863607 | 0.47022245 | 1.18212903 | 0.23715451 | 0.54992696 |
| STX17 | 62.825751 | 0.119330818 | 0.36008304 | 0.33139805 | 0.74034384 | 0.90308228 |
| ERP44 | 117.302638 | -0.252376751 | 0.33414906 | -0.7552819 | 0.45007983 | 0.74152951 |
| INVS | 219.875698 | 0.30059188 | 0.30889095 | 0.97313268 | 0.33048736 | 0.64255236 |
| TEX10 | 37.6072063 | -0.02685731 | 0.34729171 | -0.0773336 | 0.93835818 | 0.98331142 |
| PLPPR1 | 183.392239 | 3.081317474 | 0.5145823 | 5.98799741 | 2.12E-09 | 3.19E-07 |
| ZNF189 | 27.9350019 | 0.111082936 | 0.31086715 | 0.3573325 | 0.7208429 | 0.89163428 |
| ALDOB | 50.6100816 | -0.419040606 | 0.89254456 | -0.4694898 | 0.63871954 | 0.8538689 |
| RNF20 | 22.9009305 | 0.322558784 | 0.41271212 | 0.78155878 | 0.43447392 | 0.72781797 |
| SMC2 | 15.3664293 | 0.723222382 | 0.42461273 | 1.70325178 | 0.08852096 | 0.32316736 |
| NIPSNAP3A | 19.4558246 | -0.139513212 | 0.40323433 | -0.3459855 | 0.72935365 | 0.89693557 |
| ABCA1 | 60.8145535 | -1.00184633 | 0.38093168 | -2.6299895 | 0.00853875 | 0.0733037 |
| SLC44A1 | 153.358216 | -0.279230333 | 0.39422503 | -0.7083019 | 0.4787578 | 0.76419703 |
| FSD1L | 95.0372424 | 0.581335907 | 0.27566027 | 2.10888534 | 0.03495448 | 0.18613111 |
| FKTN | 67.7497135 | 0.278019526 | 0.3713973 | 0.74857714 | 0.45411211 | 0.74492283 |
| TMEM38B | 99.6824514 | 0.617063155 | 0.62732404 | 0.98364341 | 0.32529087 | 0.63715682 |
| LINC01505 | 174.990495 | 0.691136729 | 0.5550012 | 1.24528871 | 0.21302565 | 0.52201944 |
| ZNF462 | 107.407815 | -0.704770077 | 0.34436731 | -2.046565 | 0.04070082 | 0.20466781 |
| RAD23B | 132.094574 | -0.44526301 | 0.27505063 | -1.6188402 | 0.10548166 | 0.35685917 |
| ELP1 | 49.4822815 | 0.575591391 | 0.33322398 | 1.72734084 | 0.08410647 | 0.31289963 |

| | | | | | | |
|---|---|---|---|---|---|---|
| FAM206A | 17.0248309 | -0.283678059 | 0.43349394 | -0.6543991 | 0.51285469 | 0.7830488 |
| CTNNAL1 | 110.760482 | 1.034668452 | 0.3644173 | 2.83924078 | 0.0045221 | 0.04658498 |
| TMEM245 | 243.918864 | 0.811085799 | 0.29311962 | 2.76708122 | 0.00565607 | 0.05420058 |
| EPB41L4B | 71.9226371 | 0.47828969 | 0.36052701 | 1.32664038 | 0.18462767 | 0.48423865 |
| PTPN3 | 478.276475 | -1.203944241 | 0.50386426 | -2.3894218 | 0.01687492 | 0.11733164 |
| PALM2 | 62.191866 | -2.137085196 | 0.30811629 | -6.9359695 | 4.03E-12 | 1.25E-09 |
| TXN | 40.2225125 | -0.643844987 | 0.38228114 | -1.6842185 | 0.09213944 | 0.33080792 |
| ECPAS | 187.498658 | 0.238220837 | 0.28378164 | 0.8394512 | 0.40121616 | 0.70114138 |
| ZNF483 | 96.7913982 | -0.264175467 | 0.39927576 | -0.6616366 | 0.50820413 | 0.78013108 |
| PTGR1 | 177.966644 | -0.390814824 | 0.45218597 | -0.864279 | 0.38743466 | 0.69084734 |
| C9orf84 | 33.5682849 | -0.158898225 | 0.70149596 | -0.2265134 | 0.82080215 | 0.93926844 |
| UGCG | 438.151885 | -1.892250601 | 0.57857061 | -3.2705612 | 0.00107334 | 0.01578525 |
| SUSD1 | 140.974808 | -0.435117761 | 0.55043511 | -0.7904978 | 0.42923709 | 0.7236272 |
| PTBP3 | 298.36559 | -0.205877932 | 0.3350926 | -0.6143912 | 0.53895687 | 0.80147271 |
| HSDL2 | 132.275285 | 0.089457671 | 0.26055825 | 0.3433308 | 0.73134961 | 0.89782437 |
| C9orf147 | 30.2383873 | 0.142662236 | 0.43779869 | 0.32586264 | 0.74452828 | 0.90519915 |
| KIAA1958 | 500.984926 | 0.184766133 | 0.31073752 | 0.59460517 | 0.55210742 | 0.80943179 |
| INIP | 39.4638939 | 0.399937335 | 0.26836544 | 1.49027139 | 0.13615289 | 0.41123731 |
| SNX30 | 51.6164251 | 0.221622711 | 0.31690158 | 0.69934241 | 0.48433807 | 0.76759625 |
| FKBP15 | 40.1436072 | 0.080885829 | 0.28290127 | 0.2859154 | 0.77494292 | 0.91881584 |
| SLC31A1 | 28.602101 | -0.016870559 | 0.31238418 | -0.0540058 | 0.95693054 | 0.98859719 |
| CDC26 | 14.0508247 | 0.003796399 | 0.52516536 | 0.00722896 | 0.99423218 | 0.9992177 |
| PRPF4 | 19.5768317 | 0.111093493 | 0.47878955 | 0.23202991 | 0.81651479 | 0.93735334 |
| WDR31 | 15.8792704 | -0.212208194 | 0.56826729 | -0.3734302 | 0.70882825 | 0.88614996 |
| BSPRY | 29.0251982 | -0.659940209 | 0.3019561 | -2.1855502 | 0.02884853 | 0.16573241 |
| HDHD3 | 30.6611489 | -0.168778869 | 0.41368989 | -0.407984 | 0.6832854 | 0.87333147 |
| RGS3 | 128.978952 | -0.247720056 | 0.28243392 | -0.8770903 | 0.38043759 | 0.6843913 |
| ZNF618 | 198.413337 | 0.289488396 | 0.50269302 | 0.57587511 | 0.5646996 | 0.81518178 |
| KIF12 | 80.7656278 | 1.086302407 | 0.51043611 | 2.12818485 | 0.03332176 | 0.18186117 |
| COL27A1 | 93.0685236 | 0.379809962 | 0.56385843 | 0.673591 | 0.50057138 | 0.77569013 |
| WHRN | 48.8764124 | -0.884203224 | 0.45467859 | -1.9446775 | 0.05181381 | 0.235032 |
| ATP6V1G1 | 52.091119 | -0.824563368 | 0.37003373 | -2.2283465 | 0.02585742 | 0.15415253 |
| TMEM268 | 14.4331455 | 0.413203189 | 0.61154674 | 0.67566902 | 0.49925081 | 0.77558432 |
| PAPPA | 25.6638729 | -1.092877931 | 0.48937016 | -2.2332337 | 0.02553354 | 0.15331779 |
| ASTN2 | 105.213359 | -1.168451394 | 0.53146614 | -2.1985434 | 0.0279104 | 0.16193939 |
| CDK5RAP2 | 71.7060713 | 0.412198744 | 0.31250255 | 1.31902523 | 0.18716068 | 0.48725107 |
| MEGF9 | 119.42214 | 0.637462043 | 0.34879189 | 1.82762864 | 0.0676053 | 0.27747209 |
| FBXW2 | 61.0480073 | 0.230124372 | 0.28898187 | 0.79632806 | 0.42584138 | 0.72211376 |
| PSMD5 | 46.5671944 | 0.137530459 | 0.37973831 | 0.36217167 | 0.71722375 | 0.89010971 |
| CNTRL | 35.2857379 | 0.521097626 | 0.31713186 | 1.64315759 | 0.10035033 | 0.34581665 |
| RAB14 | 79.8067071 | 0.1844785 | 0.2822352 | 0.65363392 | 0.51334769 | 0.7830488 |
| GSN | 127.424275 | -0.655463138 | 0.41133233 | -1.5935123 | 0.11104531 | 0.36676814 |
| STOM | 87.9805774 | -2.057870059 | 0.66390364 | -3.0996517 | 0.00193748 | 0.02463849 |
| DAB2IP | 181.440983 | -0.676030942 | 0.40104305 | -1.6856817 | 0.09185712 | 0.33043047 |
| TTLL11 | 33.3056716 | -0.404023477 | 0.60766633 | -0.6648772 | 0.50612905 | 0.77934214 |
| NDUFA8 | 27.3666247 | 0.2189338 | 0.39511076 | 0.55410741 | 0.57950534 | 0.82392616 |
| RBM18 | 17.5084454 | -0.070788477 | 0.4826935 | -0.1466531 | 0.88340587 | 0.96170562 |
| MRRF | 98.0926367 | 0.495540267 | 0.21381225 | 2.31764209 | 0.02046878 | 0.13318277 |
| PDCL | 12.7980412 | -0.197767535 | 0.39732261 | -0.4977505 | 0.6186599 | 0.84283824 |
| RC3H2 | 102.221415 | -0.140172587 | 0.27297579 | -0.5134982 | 0.60760284 | 0.83917773 |
| ZBTB26 | 18.1529088 | 0.020371255 | 0.36389203 | 0.05598159 | 0.95535647 | 0.98853497 |
| RABGAP1 | 267.203307 | -0.178325751 | 0.33476334 | -0.532692 | 0.59424681 | 0.83416944 |
| STRBP | 244.287204 | 0.617037172 | 0.24968641 | 2.47124851 | 0.01346422 | 0.10083567 |
| DENND1A | 298.33091 | 0.181771127 | 0.32624954 | 0.55715367 | 0.57742244 | 0.82246973 |
| NEK6 | 27.3100185 | -2.837183743 | 0.79042524 | -3.5894397 | 0.00033139 | 0.00631876 |
| PSMB7 | 66.6501331 | -0.257248537 | 0.34692068 | -0.7415198 | 0.45837835 | 0.747377 |
| NR6A1 | 515.443862 | 0.011528509 | 0.20974896 | 0.05496337 | 0.95616765 | 0.98853497 |
| MIR181A2HG | 45.334718 | -0.919941946 | 0.42193058 | -2.1803159 | 0.02923406 | 0.16679612 |
| RPL35 | 51.2343725 | -0.699532119 | 0.53944991 | -1.2967508 | 0.19471693 | 0.497136 |
| ARPC5L | 25.8006941 | -1.058704113 | 0.43466695 | -2.4356674 | 0.01486435 | 0.10760494 |
| GOLGA1 | 51.6891231 | 0.662105047 | 0.36299491 | 1.82400644 | 0.0681511 | 0.27829012 |
| SCAI | 66.3302099 | 0.742094549 | 0.41404458 | 1.79230591 | 0.07308398 | 0.28899744 |
| PPP6C | 70.5962795 | 0.226192391 | 0.28819022 | 0.78487184 | 0.43252871 | 0.72599466 |
| RABEPK | 20.534408 | 0.005645301 | 0.43504547 | 0.01297635 | 0.98964666 | 0.99842088 |
| HSPA5 | 82.4411994 | -1.43929804 | 0.74055317 | -1.9435445 | 0.0519504 | 0.23513862 |
| GAPVD1 | 120.175034 | -0.003725929 | 0.28744711 | -0.0129621 | 0.989658 | 0.99842088 |
| MAPKAP1 | 189.272332 | -0.02439366 | 0.2302876 | -0.1059269 | 0.91564032 | 0.97359224 |
| PBX3 | 61.5164469 | -0.955330049 | 0.34447109 | -2.7733243 | 0.00554868 | 0.05355682 |
| MVB12B | 42.0290924 | 0.875500485 | 0.48302713 | 1.81252861 | 0.06990456 | 0.28179425 |
| ZBTB43 | 63.9212383 | -0.499468059 | 0.45578685 | -1.0958369 | 0.27315015 | 0.58894058 |
| ZBTB34 | 11.8693766 | 0.738686555 | 0.43041618 | 1.71621464 | 0.08612277 | 0.31701882 |
| RALGPS1 | 221.921415 | 0.282630056 | 0.24921338 | 1.13408862 | 0.25675738 | 0.56935259 |
| GARNL3 | 24.1120708 | -0.564553089 | 0.39068698 | -1.4450266 | 0.1484504 | 0.4322475 |
| RPL12 | 22.559203 | -0.658566933 | 0.429052 | -1.534935 | 0.1247998 | 0.39027762 |
| LRSAM1 | 58.0300119 | -0.212817814 | 0.31822678 | -0.6687615 | 0.50364763 | 0.77790293 |
| FAM129B | 55.2234506 | -1.470804826 | 0.43148529 | -3.4087021 | 0.00065273 | 0.0107527 |
| STXBP1 | 37.9619209 | -0.718654419 | 0.38694241 | -1.8572646 | 0.06327352 | 0.26477174 |
| CDK9 | 18.8696534 | 0.613585578 | 0.40381987 | 1.51945367 | 0.12864834 | 0.39737957 |
| FPGS | 24.1655205 | -1.177195575 | 0.43572103 | -2.7017185 | 0.00689821 | 0.06287287 |
| DPM2 | 26.208299 | -2.725123281 | 0.88875572 | -3.0662231 | 0.00216782 | 0.02673639 |
| FAM102A | 69.9019101 | -2.611170171 | 0.75053659 | -3.4790711 | 0.00050316 | 0.00865575 |
| SLC25A25 | 45.0245842 | -1.085584875 | 0.62856066 | -1.7270964 | 0.08415034 | 0.31289963 |
| SLC25A25-AS | 19.9521043 | 0.303104394 | 0.57260476 | 0.52934313 | 0.59656744 | 0.83416944 |
| PTGES2 | 25.4543412 | -0.015093626 | 0.37213398 | -0.0405597 | 0.96764695 | 0.99156233 |
| C9orf16 | 20.3630755 | -1.135798942 | 0.40595246 | -2.7978619 | 0.00514421 | 0.05124424 |
| CIZ1 | 25.0219009 | -0.218637668 | 0.40332987 | -0.5420815 | 0.58776235 | 0.83009636 |
| DNM1 | 17.9219257 | -0.281734564 | 0.66098036 | -0.4262374 | 0.66993485 | 0.86777458 |
| GOLGA2 | 64.9230429 | -0.422536731 | 0.38677844 | -1.0924516 | 0.27463464 | 0.58917168 |
| TRUB2 | 12.7692108 | 0.199577628 | 0.44123795 | 0.45231293 | 0.65104356 | 0.86120024 |
| COQ4 | 62.657346 | 0.643226323 | 0.32472786 | 1.98081658 | 0.04761185 | 0.22360104 |
| SLC27A4 | 27.251898 | 0.555611532 | 0.38823056 | 1.43113806 | 0.15239065 | 0.43719706 |
| URM1 | 19.237474 | -0.097574903 | 0.38120695 | -0.2559631 | 0.79797935 | 0.92876787 |
| ODF2 | 39.6888447 | -0.492802878 | 0.34271641 | -1.4379319 | 0.15045337 | 0.43488337 |
| GLE1 | 38.0359716 | -0.039992891 | 0.31037595 | -0.1288531 | 0.89747392 | 0.96607503 |
| SPTAN1 | 99.402775 | 0.200242055 | 0.29181117 | 0.68620421 | 0.49258434 | 0.77284018 |
| WDR34 | 24.8801636 | 0.245769128 | 0.47189242 | 0.52081601 | 0.60249495 | 0.83691476 |
| SET | 54.4057356 | -0.771860093 | 0.39440727 | -1.9570129 | 0.05034596 | 0.23158644 |
| ZER1 | 116.511557 | 0.058255722 | 0.249659 | 0.23334117 | 0.81549649 | 0.93717111 |
| TBC1D13 | 18.3551919 | 0.494863158 | 0.3777549 | 1.31001121 | 0.19019204 | 0.49082497 |
| KYAT1 | 27.511961 | 0.650618594 | 0.34376433 | 1.89262975 | 0.05840712 | 0.25247475 |
| PHYHD1 | 24.881884 | 0.365528416 | 0.41034626 | 0.89078042 | 0.37304698 | 0.67684181 |
| NUP188 | 24.4681277 | 0.094092802 | 0.38887363 | 0.24196241 | 0.80880929 | 0.93433178 |
| SH3GLB2 | 34.8302046 | 0.351135122 | 0.45617458 | 0.76973846 | 0.44145505 | 0.7335817 |
| MIGA2 | 32.1967279 | 0.692239242 | 0.34486124 | 2.00729789 | 0.04471795 | 0.21614837 |
| PTPA | 49.7975382 | -0.818719567 | 0.51718729 | -1.5830234 | 0.11341614 | 0.37119783 |
| IER5L | 27.9645211 | -2.858808002 | 0.56643298 | -5.0470367 | 4.49E-07 | 2.89E-05 |
| LINC01503 | 108.605252 | -2.487746154 | 0.52953426 | -4.697989 | 2.63E-06 | 0.00012628 |
| NTMT1 | 24.7969138 | -0.806222077 | 0.40872592 | -1.972525 | 0.04854971 | 0.22588697 |
| ASB6 | 15.4028368 | 0.247033402 | 0.50445415 | 0.48970437 | 0.62434311 | 0.84662233 |
| TOR1B | 16.108762 | -0.712793525 | 0.45782492 | -1.5569129 | 0.11949117 | 0.38116169 |
| C9orf78 | 17.3131959 | -0.455606369 | 0.40552289 | -1.1235035 | 0.26122373 | 0.57487967 |
| USP20 | 29.713707 | -0.264295977 | 0.31698128 | -0.8337905 | 0.40439905 | 0.70347327 |
| FNBP1 | 96.1086671 | -0.018203344 | 0.24354427 | -0.0747435 | 0.94041882 | 0.98400461 |

| | | | | | | |
|---|---|---|---|---|---|---|
| GPR107 | 89.4410447 | 0.065706504 | 0.23293126 | 0.28208539 | 0.77787804 | 0.9197229 |
| NCS1 | 19.3382689 | 0.424848319 | 0.40431971 | 1.05077321 | 0.29336276 | 0.60865919 |
| ASS1 | 15.4299904 | -0.335161154 | 0.402762 | -0.8321568 | 0.40532042 | 0.70455026 |
| FUBP3 | 84.7671414 | -0.002020552 | 0.34317372 | -0.0058878 | 0.99530221 | 0.99922221 |
| ABL1 | 197.59864 | 0.381095242 | 0.20305716 | 1.87678796 | 0.06054716 | 0.25731164 |
| AIF1L | 129.812349 | 0.123473765 | 0.53119162 | 0.23244675 | 0.81619104 | 0.93721247 |
| NUP214 | 186.564994 | 0.326645158 | 0.2311543 | 1.41310441 | 0.15762503 | 0.44441964 |
| AL157938.2 | 41.1627518 | -0.730788628 | 0.65686984 | -1.1125319 | 0.26590954 | 0.58132252 |
| PRRC2B | 200.442127 | -0.199697468 | 0.3008576 | -0.6637608 | 0.50684345 | 0.77942663 |
| POMT1 | 19.5606434 | 1.217897967 | 0.38830872 | 3.13641674 | 0.00171026 | 0.02249698 |
| RAPGEF1 | 181.100513 | -0.627365859 | 0.56920077 | -1.1021873 | 0.27038025 | 0.58716619 |
| MED27 | 119.873105 | -0.143616727 | 0.32201904 | -0.4459883 | 0.65560569 | 0.86268564 |
| SETX | 179.850419 | -0.32637229 | 0.43198095 | -0.7555247 | 0.4499342 | 0.74152951 |
| TTF1 | 31.7316043 | -0.498727993 | 0.40440011 | -1.2332538 | 0.21748107 | 0.52693657 |
| DDX31 | 36.2585207 | -0.617886943 | 0.36792438 | -1.6793857 | 0.0930769 | 0.3326491 |
| GTF3C4 | 13.7218033 | 0.328089496 | 0.39987109 | 0.82048815 | 0.41193788 | 0.71101607 |
| AK8 | 15.0062988 | 0.674132116 | 0.75762341 | 0.88979843 | 0.37357413 | 0.67707283 |
| TSC1 | 122.962601 | 0.71283799 | 0.2598403 | 2.74336965 | 0.00608122 | 0.05733193 |
| GTF3C5 | 25.8390614 | -0.516725452 | 0.5020379 | -1.0292559 | 0.30335946 | 0.61783048 |
| RALGDS | 17.0285509 | -0.525783484 | 0.55956378 | -0.939631 | 0.34740686 | 0.6553345 |
| ABO | 49.7235243 | 0.343654694 | 0.67794398 | 0.50690722 | 0.61221992 | 0.84174925 |
| RPL7A | 54.5234461 | -0.751500757 | 0.47359605 | -1.5867969 | 0.11255865 | 0.36994129 |
| SURF1 | 12.9255765 | 0.08954163 | 0.48179463 | 0.1858502 | 0.85256223 | 0.95132722 |
| SURF4 | 51.1876703 | -0.434909809 | 0.31273023 | -1.3906868 | 0.16432042 | 0.45563826 |
| REXO4 | 15.002799 | -0.514357699 | 0.39881892 | -1.2897024 | 0.19715402 | 0.49934704 |
| VAV2 | 143.060165 | -0.694422516 | 0.45559679 | -1.5242042 | 0.12745772 | 0.3953479 |
| BRD3 | 30.752463 | 0.238397955 | 0.37333468 | 0.63856365 | 0.52310683 | 0.79051023 |
| WDR5 | 22.5174657 | -0.030917972 | 0.36433679 | -0.084861 | 0.93237192 | 0.9794317 |
| RXRA | 24.5866007 | 0.817470786 | 0.38617266 | 2.11685307 | 0.03427232 | 0.18418162 |
| PPP1R26-AS | 41.0896778 | 0.396868616 | 0.31334153 | 1.26656882 | 0.20530951 | 0.51184648 |
| PPP1R26 | 15.0660768 | 0.299674363 | 0.57695889 | 0.51940332 | 0.60347951 | 0.83748768 |
| CAMSAP1 | 43.5114404 | -0.664375972 | 0.44993507 | -1.4766041 | 0.13978179 | 0.4184541 |
| UBAC1 | 38.9131906 | 0.516055959 | 0.34576282 | 1.49251433 | 0.13556436 | 0.41064171 |
| NACC2 | 40.4582421 | -0.475819436 | 0.37766365 | -1.2599026 | 0.20770449 | 0.51574877 |
| SNAPC4 | 19.2453278 | -0.776025199 | 0.5648081 | -1.3739626 | 0.1694533 | 0.46374597 |
| ENTR1 | 20.8389557 | -0.309847429 | 0.37630727 | -0.8233894 | 0.41028657 | 0.70908471 |
| PMPCA | 24.1087098 | 0.076854384 | 0.39643855 | 0.19386203 | 0.84628391 | 0.94909204 |
| SEC16A | 53.0140477 | -0.432511059 | 0.45156457 | -0.9578056 | 0.33816081 | 0.64897646 |
| NOTCH1 | 33.0539429 | -1.262669191 | 0.3730581 | -3.3846449 | 0.0007127 | 0.0115169 |
| AGPAT2 | 34.6603994 | -0.788360031 | 0.52956097 | -1.4887049 | 0.13656508 | 0.41181527 |
| SNHG7 | 27.1495571 | -0.327958491 | 0.3848207 | -0.8522371 | 0.39408248 | 0.69624829 |
| RABL6 | 64.6881318 | 0.060494651 | 0.36490875 | 0.16578021 | 0.86832992 | 0.95588054 |
| PHPT1 | 35.9072089 | 0.281385445 | 0.57503657 | 0.48933487 | 0.62460464 | 0.84685381 |
| EDF1 | 45.2480673 | -0.284939452 | 0.36108365 | -0.7891231 | 0.43004006 | 0.72390206 |
| FBXW5 | 32.0286323 | -0.674245907 | 0.57122147 | -1.1803581 | 0.23785781 | 0.54997949 |
| ABCA2 | 18.5788815 | 0.804019295 | 0.78726798 | 1.02127778 | 0.30712285 | 0.62090491 |
| UAP1L1 | 12.7941572 | 1.737085166 | 0.5910774 | 2.93884551 | 0.00329437 | 0.03746908 |
| MAN1B1 | 24.6720454 | 0.149967221 | 0.33890677 | 0.44250288 | 0.65812535 | 0.86357172 |
| DPP7 | 22.9391929 | -0.056721064 | 0.54983023 | -0.103161 | 0.91783515 | 0.97481433 |
| SSNA1 | 16.4567644 | -0.669498842 | 0.50570241 | -1.3238989 | 0.18553663 | 0.4858669 |
| TPRN | 15.0658266 | -0.899164732 | 0.48787717 | -1.8430146 | 0.06532688 | 0.27093297 |
| TUBB4B | 33.2637598 | -0.945926018 | 0.48875762 | -1.9353683 | 0.05294511 | 0.238253 |
| NELFB | 31.5383496 | -0.078295963 | 0.37956415 | -0.2062786 | 0.83657329 | 0.94684966 |
| EXD3 | 58.0634741 | -0.212034583 | 0.34665348 | -0.6116615 | 0.54076175 | 0.80223747 |
| NOXA1 | 16.5789057 | -0.268284289 | 0.53443876 | -0.5019926 | 0.61567274 | 0.8422143 |
| PNPLA7 | 25.8269 | 0.24204493 | 0.5805585 | 0.41691738 | 0.67673884 | 0.87033282 |
| MRPL41 | 43.669084 | -0.121294186 | 0.58976354 | -0.2056658 | 0.83705198 | 0.94705408 |
| DPH7 | 27.726656 | 0.462036948 | 0.31562387 | 1.46388469 | 0.14322546 | 0.4243745 |
| ARRDC1 | 20.258175 | -0.545609548 | 0.43546471 | -1.2529363 | 0.21022888 | 0.51886901 |
| EHMT1 | 227.141571 | -0.267024852 | 0.30125157 | -0.8863849 | 0.37541014 | 0.67888822 |
| BET1L | 13.110234 | -0.637944631 | 0.55176587 | -1.1561872 | 0.2476046 | 0.56027856 |
| SIRT3 | 19.0945932 | -0.181493408 | 0.42784677 | -0.4242019 | 0.67141859 | 0.86777458 |
| PSMD13 | 36.2888689 | 0.380891132 | 0.30508535 | 1.248474 | 0.21185752 | 0.52068532 |
| IFITM2 | 104.864161 | -2.764331976 | 0.64773127 | -4.2677142 | 1.97E-05 | 0.00069224 |
| IFITM3 | 163.140897 | -2.60998585 | 0.41336713 | -6.3139656 | 2.72E-10 | 5.76E-08 |
| SIGIRR | 32.4861118 | -0.226254794 | 0.47801765 | -0.4733189 | 0.63598567 | 0.85371875 |
| PTDSS2 | 30.8874759 | -0.200916403 | 0.31722371 | -0.6333587 | 0.52649942 | 0.79346524 |
| RNH1 | 33.9452259 | -0.814127452 | 0.36928407 | -2.2046103 | 0.02748146 | 0.16065021 |
| RASSF7 | 49.9116539 | -0.502138421 | 0.43579825 | -1.1522268 | 0.24922789 | 0.56198328 |
| MIR210HG | 17.6474101 | -2.350780555 | 0.77878719 | -3.0185147 | 0.00254017 | 0.03036691 |
| PHRF1 | 32.985663 | -0.483334948 | 0.41039276 | -1.1777375 | 0.23890126 | 0.55079752 |
| DEAF1 | 87.9113058 | -0.798774607 | 0.31262018 | -2.5550961 | 0.01061585 | 0.08592203 |
| AC131934.1 | 17.2512641 | 0.470817104 | 0.38499868 | 1.22290575 | 0.2213653 | 0.53168728 |
| EPS8L2 | 99.1112683 | -0.474620963 | 0.36425373 | -1.3029955 | 0.19257631 | 0.49370242 |
| AP006621.4 | 17.5531008 | -1.552527431 | 0.64690522 | -2.3999303 | 0.01639819 | 0.11444368 |
| TALDO1 | 40.3945544 | -1.007834853 | 0.37634204 | -2.6779757 | 0.00740686 | 0.06621434 |
| GATD1 | 21.3663112 | -0.698224087 | 0.38511089 | -1.8130469 | 0.06982459 | 0.28168095 |
| AP006621.3 | 21.3651553 | 0.353603338 | 0.43171645 | 0.81906385 | 0.41274998 | 0.71110141 |
| PIDD1 | 10.9528188 | 0.889404743 | 0.50717348 | 1.75364996 | 0.07949051 | 0.30214817 |
| RPLP2 | 59.3040436 | -0.793684681 | 0.38668007 | -2.0525616 | 0.04011511 | 0.20285949 |
| PNPLA2 | 25.4750788 | -0.43384097 | 0.33645538 | -1.2894458 | 0.19724315 | 0.49934704 |
| CD151 | 51.0543554 | -1.19946009 | 0.49882049 | -2.4045927 | 0.0161905 | 0.11387005 |
| POLR2L | 33.7095142 | -1.182737313 | 0.45862504 | -2.5788764 | 0.00991222 | 0.081789 |
| TSPAN4 | 16.7763212 | -0.748498491 | 0.44577048 | -1.6791118 | 0.09313025 | 0.3326491 |
| CHID1 | 16.9363243 | 0.449778846 | 0.38289239 | 1.17468736 | 0.24011982 | 0.55253633 |
| AP2A2 | 213.387094 | -0.032631761 | 0.32356175 | -0.1008517 | 0.91966816 | 0.97520237 |
| TOLLIP | 24.8087093 | -0.312585815 | 0.40863736 | -0.7649467 | 0.44430326 | 0.73698311 |
| MOB2 | 26.8844042 | 0.166791993 | 0.39048544 | 0.4271401 | 0.66927728 | 0.86777458 |
| MRPL23 | 12.3062637 | 0.301931828 | 0.52587012 | 0.57415666 | 0.5658618 | 0.81595078 |
| CD81 | 106.060809 | -1.235486978 | 0.56232668 | -2.1970983 | 0.02801343 | 0.16243608 |
| KCNQ1 | 273.010871 | -0.807011534 | 0.56870739 | -1.4190277 | 0.15589094 | 0.44166732 |
| KCNQ1OT1 | 215.548487 | -1.450210851 | 0.31609391 | -4.5879113 | 4.48E-06 | 0.00019878 |
| NAP1L4 | 248.413208 | 0.292222867 | 0.15376955 | 1.9003949 | 0.05738132 | 0.24970229 |
| CARS | 45.8789544 | -1.238737536 | 0.53111968 | -2.3323134 | 0.01968421 | 0.12997934 |
| OSBPL5 | 20.8979406 | -1.006952568 | 0.52414787 | -1.9211231 | 0.05471619 | 0.24343712 |
| ZNF195 | 31.3777746 | 0.074200271 | 0.31495022 | 0.23559365 | 0.813748 | 0.93633777 |
| AC127526.1 | 16.3952112 | -0.046484917 | 0.50198979 | -0.0926013 | 0.9262203 | 0.97735117 |
| NUP98 | 147.129207 | -0.483478973 | 0.41466989 | -1.165937 | 0.24363991 | 0.55580983 |
| PGAP2 | 18.9140683 | 0.32726911 | 0.67147173 | 0.48739075 | 0.62598145 | 0.8471532 |
| STIM1 | 44.9877195 | -2.41332238 | 0.53416163 | -4.5179628 | 6.24E-06 | 0.0002599 |
| RRM1 | 15.1900694 | 0.163688479 | 0.67576437 | 0.24222715 | 0.80860416 | 0.93433178 |
| AC104389.4 | 39.0289332 | 0.278724907 | 0.77268365 | 0.36072319 | 0.71830639 | 0.8907358 |
| TRIM5 | 15.3830785 | -0.116698814 | 0.91059263 | -0.128157 | 0.89802473 | 0.96607503 |
| TRIM22 | 27.437691 | -0.850058237 | 1.21104115 | -0.7019235 | 0.48272688 | 0.76650978 |
| FAM160A2 | 27.9971609 | 0.722999129 | 0.3216898 | 2.24750404 | 0.02460783 | 0.14891004 |
| SMPD1 | 13.0310287 | 0.89347547 | 0.63515173 | 1.40671186 | 0.15951285 | 0.44798126 |
| APBB1 | 23.1298062 | 0.367689932 | 0.33003793 | 1.11408386 | 0.26524321 | 0.58068272 |
| ARFIP2 | 13.6560651 | 0.410210626 | 0.56217172 | 0.72968919 | 0.46558019 | 0.75274202 |
| AC084337.2 | 34.7696652 | -0.193610982 | 0.59029827 | -0.3279884 | 0.74292044 | 0.90463949 |
| DNHD1 | 47.4385789 | -0.623449032 | 0.44844373 | -1.3902503 | 0.16445289 | 0.45563826 |
| ILK | 35.9615879 | -0.321253775 | 0.35089583 | -0.9155246 | 0.35991628 | 0.66572082 |
| TPP1 | 28.6311946 | -0.822181674 | 0.41372058 | -1.9872873 | 0.04689056 | 0.22193782 |
| AC107884.1 | 29.3519109 | 0.29105861 | 0.43999616 | 0.66150262 | 0.50829004 | 0.78013108 |

| | | | | | | |
|---|---|---|---|---|---|---|
| PPFIBP2 | 26.1192414 | -0.867114702 | 0.47275652 | -1.8341676 | 0.06662911 | 0.27416145 |
| EIF3F | 30.4146591 | -0.65219435 | 0.40919744 | -1.5938378 | 0.11097237 | 0.36676814 |
| TUB | 18.4374249 | 0.716819273 | 0.49755567 | 1.44068155 | 0.14967467 | 0.433886 |
| RIC3 | 99.1903999 | -0.374334521 | 0.45210347 | -0.8279842 | 0.40767946 | 0.70628081 |
| STK33 | 164.80753 | 0.170680349 | 0.64707043 | 0.26377399 | 0.79195408 | 0.92545179 |
| TRIM66 | 70.9742545 | 0.243998544 | 0.52564863 | 0.46418564 | 0.64251475 | 0.85521878 |
| RPL27A | 66.9237513 | -0.841728501 | 0.37706342 | -2.232326 | 0.02559342 | 0.15356053 |
| ST5 | 380.158896 | -2.550281016 | 0.45805604 | -5.5676179 | 2.58E-08 | 2.34E-06 |
| AC026894.1 | 31.4460778 | -3.064661202 | 0.54050829 | -5.6699615 | 1.43E-08 | 1.43E-06 |
| TMEM9B | 27.2895763 | 0.403678678 | 0.41484384 | 0.97308586 | 0.33051063 | 0.64255236 |
| TMEM9B-AS | 24.2763808 | 0.249824624 | 0.66221333 | 0.37725701 | 0.70598261 | 0.88599027 |
| DENND5A | 129.35294 | -0.36049199 | 0.33921209 | -1.0627333 | 0.28790291 | 0.6024252 |
| TMEM41B | 129.042732 | -1.322380784 | 0.61190047 | -2.1611044 | 0.03068728 | 0.17278271 |
| IPO7 | 105.309235 | -0.545971021 | 0.41408757 | -1.3184917 | 0.18733911 | 0.48725107 |
| ZNF143 | 33.7417242 | -0.131490688 | 0.425242 | -0.3092138 | 0.75715891 | 0.9112882 |
| WEE1 | 19.3088186 | -0.367297202 | 0.69877398 | -0.5256309 | 0.59914468 | 0.83579069 |
| SWAP70 | 72.1463408 | -1.332230554 | 0.2556924 | -5.2102861 | 1.89E-07 | 1.41E-05 |
| SBF2-AS1 | 55.8795176 | -1.488415656 | 0.73236516 | -2.0323409 | 0.04211916 | 0.20922698 |
| SBF2 | 718.931018 | 0.474313312 | 0.30738407 | 1.54306404 | 0.12281524 | 0.38745661 |
| AC011092.2 | 27.6509919 | -0.380142642 | 0.55549374 | -0.6843329 | 0.49376498 | 0.7738663 |
| ADM | 41.2731917 | -0.215875771 | 0.74604772 | -0.2893592 | 0.77230651 | 0.91791018 |
| AMPD3 | 73.1511127 | -0.290287103 | 0.54643635 | -0.5312368 | 0.59525469 | 0.83416944 |
| MTRNR2L8 | 50.6403271 | -2.034814588 | 0.75956784 | -2.6789109 | 0.0073862 | 0.06609306 |
| RNF141 | 19.0710426 | 0.736496014 | 0.39291007 | 1.87446461 | 0.06086642 | 0.25784263 |
| CTR9 | 59.9972663 | 0.281312122 | 0.26093795 | 1.07808051 | 0.28099783 | 0.59523128 |
| EIF4G2 | 102.807023 | -0.210168058 | 0.28498278 | -0.7374763 | 0.46083273 | 0.74896364 |
| ZBED5 | 64.6665959 | -0.046611709 | 0.29374387 | -0.1586815 | 0.87391984 | 0.95761805 |
| ZBED5-AS1 | 14.2419243 | 1.219264862 | 0.6240368 | 1.95383488 | 0.05072075 | 0.23285094 |
| GALNT18 | 232.109428 | 0.673322436 | 0.52503159 | 1.28244175 | 0.19968773 | 0.5033686 |
| USP47 | 264.744731 | 0.194119499 | 0.20043776 | 0.9684777 | 0.33280585 | 0.64492217 |
| PARVA | 118.972063 | -0.454204993 | 0.19761513 | -2.2984323 | 0.0215372 | 0.1377317 |
| TEAD1 | 487.901715 | -0.320862384 | 0.60182038 | -0.5331531 | 0.59392763 | 0.83416944 |
| LINC00958 | 68.3505839 | -1.281161784 | 0.43841708 | -2.9222442 | 0.00347519 | 0.03879367 |
| ARNTL | 33.2554854 | 0.14256213 | 0.3700709 | 0.38522924 | 0.70006758 | 0.88363749 |
| BTBD10 | 77.2651232 | 0.102444537 | 0.30037772 | 0.34105238 | 0.73306415 | 0.89840827 |
| FAR1 | 114.650246 | 0.444822159 | 0.25054051 | 1.77545003 | 0.07582361 | 0.2948204 |
| RRAS2 | 92.5328376 | -0.417144491 | 0.47025387 | -0.8870623 | 0.37504535 | 0.67888232 |
| COPB1 | 75.4600479 | 0.449126474 | 0.27730479 | 1.61961311 | 0.10531541 | 0.35668757 |
| PSMA1 | 31.1283288 | -1.098262085 | 0.73101464 | -1.5023804 | 0.13299889 | 0.40565313 |
| PDE3B | 70.144161 | -1.835174348 | 0.72226253 | -2.5408688 | 0.01105774 | 0.08807986 |
| CYP2R1 | 15.1068618 | 0.097786669 | 0.49382264 | 0.19801982 | 0.84302956 | 0.94806508 |
| SOX6 | 772.54488 | 0.742216577 | 0.46693377 | 1.58955431 | 0.1119353 | 0.36854256 |
| C11orf58 | 107.611354 | -0.189235745 | 0.1966784 | -0.9621582 | 0.33597012 | 0.6470947 |
| PLEKHA7 | 512.443447 | -0.471380333 | 0.1919448 | -2.4558119 | 0.01405667 | 0.10405109 |
| RPS13 | 42.4592213 | -1.108940829 | 0.40920967 | -2.7099576 | 0.00672918 | 0.06163349 |
| PIK3C2A | 327.386909 | -0.277737357 | 0.26420973 | -1.0512003 | 0.2931666 | 0.6084367 |
| NUCB2 | 40.2264111 | 0.330876043 | 0.4047991 | 0.81738335 | 0.41370939 | 0.71143975 |
| NCR3LG1 | 13.7865646 | 0.368222287 | 0.72701594 | 0.50648448 | 0.61251659 | 0.84174925 |
| SERGEF | 156.136849 | -0.315627751 | 0.31344473 | -1.0069646 | 0.31395178 | 0.62884777 |
| SAAL1 | 13.1121756 | 0.827240125 | 0.46698472 | 1.77145009 | 0.07648589 | 0.29596141 |
| HPS5 | 32.531982 | -0.14772247 | 0.44259808 | -0.3337621 | 0.73855907 | 0.90241447 |
| GTF2H1 | 60.9653522 | -0.355018341 | 0.36608928 | -0.9697589 | 0.33216667 | 0.64442823 |
| LDHA | 134.896764 | -2.247647399 | 0.33145961 | -6.7810597 | 1.19E-11 | 3.37E-09 |
| TSG101 | 94.4002024 | 0.027532339 | 0.32037011 | 0.08593916 | 0.93151478 | 0.97919321 |
| UEVLD | 78.9865011 | 0.340262204 | 0.22316066 | 1.52474097 | 0.12732372 | 0.3952403 |
| SPTY2D1OS | 34.1685336 | 0.794363625 | 0.3861222 | 2.05728556 | 0.03965877 | 0.20162395 |
| SPTY2D1 | 23.8016502 | 0.047780656 | 0.34152245 | 0.13990488 | 0.88873515 | 0.96307703 |
| ZDHHC13 | 93.6707028 | -0.416761346 | 0.39688042 | -1.050093 | 0.29367536 | 0.60876592 |
| NAV2 | 502.276116 | -0.002051508 | 0.37929892 | -0.0054087 | 0.99568452 | 0.99922221 |
| HTATIP2 | 16.0265121 | -1.55549032 | 0.56972664 | -2.7302397 | 0.00632883 | 0.0589673 |
| PRMT3 | 68.6428297 | 0.232683574 | 0.24762575 | 0.93965824 | 0.34739289 | 0.6553345 |
| NELL1 | 235.669317 | 3.629842225 | 0.64421633 | 5.63450821 | 1.76E-08 | 1.70E-06 |
| ANO5 | 63.5974099 | 0.929507314 | 0.38673901 | 2.40344852 | 0.01624125 | 0.11396363 |
| GAS2 | 22.8955186 | 1.041605648 | 0.76328003 | 1.36464417 | 0.17236491 | 0.4681417 |
| SVIP | 42.9538622 | -0.26392867 | 0.37304426 | -0.7074996 | 0.47925605 | 0.76425234 |
| SLC5A12 | 12.0081829 | 1.641657687 | 0.94543003 | 1.73641374 | 0.08249071 | 0.30885595 |
| BBOX1 | 26.8235872 | -2.247587067 | 0.85330057 | -2.6339922 | 0.00843874 | 0.072922 |
| CCDC34 | 16.1757749 | -1.063615892 | 0.69548492 | -1.5293155 | 0.12618624 | 0.39310409 |
| LGR4 | 1172.95804 | 0.276524573 | 0.3045772 | 0.90789649 | 0.36393291 | 0.6696613 |
| LIN7C | 26.6444149 | -0.448079808 | 0.3964602 | -1.1302012 | 0.25839144 | 0.57145203 |
| BDNF-AS | 105.938953 | 0.271043398 | 0.36312179 | 0.74642558 | 0.45541037 | 0.74574048 |
| METTL15 | 261.668333 | 0.154927246 | 0.41091156 | 0.37703307 | 0.70614903 | 0.88599027 |
| ARL14EP | 30.9142361 | 0.034298994 | 0.33194364 | 0.10332776 | 0.91770284 | 0.97481433 |
| MPPED2 | 121.353186 | -0.843723247 | 0.74358055 | -1.1346763 | 0.25651096 | 0.56935259 |
| DCDC1 | 73.2336927 | -0.861495346 | 0.75065989 | -1.1476507 | 0.25111279 | 0.56353121 |
| DNAJC24 | 45.4075028 | 0.319184387 | 0.26784161 | 1.19169084 | 0.23338249 | 0.54590364 |
| IMMP1L | 127.217453 | 0.608751646 | 0.25262799 | 2.40967621 | 0.01596668 | 0.11269747 |
| ELP4 | 182.056035 | 0.24130751 | 0.26808128 | 0.90012816 | 0.36805205 | 0.67235842 |
| AC131571.1 | 19.2093921 | -0.714059175 | 0.68728649 | -1.0389542 | 0.29882604 | 0.61263281 |
| EIF3M | 44.1577201 | 0.074747346 | 0.24016681 | 0.31123096 | 0.75562505 | 0.91065328 |
| PRRG4 | 37.0300597 | -0.493141087 | 0.43251648 | -1.1401672 | 0.25421667 | 0.5663838 |
| QSER1 | 76.8298587 | 0.446797968 | 0.38315727 | 1.16609551 | 0.24357583 | 0.55580983 |
| TCP11L1 | 33.7063751 | -0.072142815 | 0.43423861 | -0.1661363 | 0.86804966 | 0.95588054 |
| CSTF3 | 80.7365937 | 0.566270761 | 0.32062517 | 1.76614567 | 0.07737141 | 0.29764569 |
| HIPK3 | 211.282332 | -0.57095584 | 0.26933961 | -2.1198362 | 0.03401986 | 0.18367181 |
| CD59 | 17.0488051 | -0.110514802 | 0.37478493 | -0.2948753 | 0.76808917 | 0.91600052 |
| FBXO3 | 50.9906881 | 0.358279075 | 0.27281137 | 1.31328497 | 0.18908694 | 0.48954068 |
| CAPRIN1 | 108.521261 | 0.179624881 | 0.27132003 | 0.66204062 | 0.50794519 | 0.78010753 |
| NAT10 | 41.6109908 | -0.15080247 | 0.33350726 | -0.4521715 | 0.65114546 | 0.86120024 |
| ABTB2 | 365.858053 | 0.072498176 | 0.47456486 | 0.15276769 | 0.87858148 | 0.96013757 |
| CAT | 59.2706136 | 0.966807549 | 0.54758728 | 1.76557706 | 0.07746683 | 0.29785597 |
| EHF | 19.5082558 | -3.023247418 | 0.73458297 | -4.1155969 | 3.86E-05 | 0.00115613 |
| APIP | 38.9950139 | 0.231556384 | 0.39141187 | 0.59159264 | 0.5541234 | 0.81070871 |
| PDHX | 106.613822 | 0.383674799 | 0.20621056 | 1.86059722 | 0.06280108 | 0.26390468 |
| CD44 | 79.5472176 | -5.800640483 | 1.09589464 | -5.293064 | 1.20E-07 | 9.50E-06 |
| TRIM44 | 241.913009 | -0.428215615 | 0.33076566 | -1.2946193 | 0.19545159 | 0.49819317 |
| COMMD9 | 16.9995059 | 0.315518423 | 0.42168932 | 0.74822483 | 0.45432455 | 0.74495384 |
| TRAF6 | 39.3480578 | 0.338533933 | 0.34603212 | 0.97833096 | 0.32791066 | 0.6394978 |
| C11orf74 | 19.5667086 | 0.937709473 | 0.65293266 | 1.43615035 | 0.15095956 | 0.43488337 |
| API5 | 68.6035205 | -0.093963628 | 0.23419753 | -0.4012153 | 0.68826163 | 0.87569521 |
| TTC17 | 352.745638 | 0.243666701 | 0.15218591 | 1.60111211 | 0.10935209 | 0.36375272 |
| HSD17B12 | 307.409285 | 0.280543495 | 0.17198944 | 1.63116693 | 0.1028551 | 0.35039129 |
| ALKBH3 | 30.1738126 | 0.365789368 | 0.38635763 | 0.94676367 | 0.34375922 | 0.65306073 |
| EXT2 | 88.1882375 | -0.261796913 | 0.38023327 | -0.6885166 | 0.49112753 | 0.77215443 |
| PRDM11 | 21.7203348 | 0.578624639 | 0.35455943 | 1.63195386 | 0.10268921 | 0.35025713 |
| SYT13 | 18.0174643 | 1.120021602 | 0.64237926 | 1.74355193 | 0.08123725 | 0.30643842 |
| CRY2 | 40.2887345 | 0.302234243 | 0.2790444 | 1.08310448 | 0.27876205 | 0.59369058 |
| LARGE2 | 25.7103304 | 2.225301062 | 0.50734457 | 4.386173 | 1.15E-05 | 0.00043904 |
| PHF21A | 329.353111 | -0.186894712 | 0.25788486 | -0.7247215 | 0.46862288 | 0.75516139 |
| DGKZ | 24.7940484 | -0.814556499 | 0.32757824 | -2.4866013 | 0.01289698 | 0.09821896 |
| AMBRA1 | 258.537676 | -0.135230494 | 0.26650597 | -0.5074201 | 0.61186006 | 0.84169124 |
| ATG13 | 56.8008328 | -0.422805183 | 0.42227604 | -1.0012531 | 0.31670447 | 0.63070323 |
| ARHGAP1 | 35.70229 | -0.624478742 | 0.36077785 | -1.7309232 | 0.08346547 | 0.31112797 |

| | | | | | | |
|---|---|---|---|---|---|---|
| CKAP5 | 72.2358832 | 0.351533956 | 0.22214176 | 1.5824758 | 0.11354099 | 0.37119783 |
| LRP4 | 45.5269082 | -0.67268915 | 0.52692047 | -1.2766427 | 0.20172844 | 0.5060272 |
| C11orf49 | 235.758015 | -0.161015951 | 0.26405819 | -0.6097745 | 0.5420112 | 0.8031965 |
| ARFGAP2 | 34.0563581 | 0.125064312 | 0.34657928 | 0.3608534 | 0.71820904 | 0.8907358 |
| DDB2 | 19.2936611 | -0.634189306 | 0.59597788 | -1.0641155 | 0.28727637 | 0.60168981 |
| NR1H3 | 21.4632531 | 0.163414311 | 0.50528628 | 0.32340936 | 0.74638524 | 0.90661036 |
| MADD | 31.964264 | 0.444580536 | 0.4471208 | 0.99431863 | 0.32006777 | 0.63212442 |
| SLC39A13 | 26.626737 | -0.194306521 | 0.49531382 | -0.3922897 | 0.69484416 | 0.87978644 |
| CELF1 | 315.379314 | 0.006368799 | 0.29229845 | 0.02178868 | 0.98261652 | 0.99694412 |
| NDUFS3 | 20.7506029 | 0.754851871 | 0.42906441 | 1.75929735 | 0.07852702 | 0.29970982 |
| MTCH2 | 65.0663711 | -0.238059122 | 0.34218736 | -0.6956982 | 0.48661787 | 0.76847808 |
| FNBP4 | 203.417738 | -0.614921386 | 0.37350234 | -1.6463656 | 0.09968852 | 0.34450385 |
| NUP160 | 100.760015 | 0.008103806 | 0.38143796 | 0.02124541 | 0.98304989 | 0.99727529 |
| PTPRJ | 215.272521 | -0.350840915 | 0.41808867 | -0.8391543 | 0.40138271 | 0.70114138 |
| TNKS1BP1 | 27.8587818 | -1.208078839 | 0.41264153 | -2.9276715 | 0.00341511 | 0.03844452 |
| SSRP1 | 16.3455941 | -0.365573493 | 0.53221188 | -0.6868946 | 0.49214912 | 0.77284018 |
| SLC43A1 | 23.5441224 | 2.210395116 | 0.73071574 | 3.02497263 | 0.00248656 | 0.02999308 |
| SERPING1 | 107.010082 | -2.091047855 | 0.31814063 | -6.5727155 | 4.94E-11 | 1.28E-08 |
| ZDHHC5 | 55.3504283 | -0.685581747 | 0.43717826 | -1.5681972 | 0.11683511 | 0.37629379 |
| CTNND1 | 161.214744 | -0.024767421 | 0.2214388 | -0.1118477 | 0.91094416 | 0.97128978 |
| LPXN | 41.4831751 | 0.801128436 | 0.41925959 | 1.91081718 | 0.05602808 | 0.24665053 |
| FAM111A | 36.8309693 | 0.351152816 | 0.5608615 | 0.62609542 | 0.53125235 | 0.79628881 |
| OSBP | 82.4950933 | -0.357250508 | 0.31281108 | -1.1420647 | 0.25342711 | 0.56559484 |
| PATL1 | 58.5841729 | -0.593800884 | 0.32517103 | -1.8261187 | 0.06783239 | 0.27788783 |
| STX3 | 110.651137 | -1.800266387 | 0.52937097 | -3.4007652 | 0.00067198 | 0.0109921 |
| MRPL16 | 16.5678135 | 0.362399098 | 0.61512997 | 0.58914232 | 0.55576581 | 0.81104802 |
| PRPF19 | 20.7543151 | -0.787678669 | 0.50858534 | -1.548764 | 0.12143846 | 0.38487158 |
| VPS37C | 30.6375582 | -0.359113365 | 0.43424458 | -0.8269841 | 0.40824607 | 0.70686841 |
| DDB1 | 65.0515973 | -0.353860395 | 0.39090842 | -0.9052258 | 0.36534577 | 0.67012584 |
| TKFC | 10.6055044 | 0.314038351 | 0.5492973 | 0.57170925 | 0.56751896 | 0.81659673 |
| CPSF7 | 68.0361642 | -0.063134828 | 0.30443794 | -0.2073816 | 0.83571185 | 0.94654164 |
| AP003108.2 | 57.4859471 | 0.126330671 | 0.34617769 | 0.36493013 | 0.71516356 | 0.88904248 |
| SYT7 | 22.9911444 | 0.693145084 | 0.72983402 | 0.94972975 | 0.34224959 | 0.65202651 |
| MYRF | 21.3869004 | -1.268108662 | 0.40660189 | -3.1187968 | 0.00181591 | 0.02351606 |
| TMEM258 | 18.0826864 | -0.606875109 | 0.43444948 | -1.396883 | 0.16244875 | 0.45295218 |
| FTH1 | 437.844526 | -2.032204548 | 0.44167305 | -4.6011514 | 4.20E-06 | 0.00018926 |
| ASRGL1 | 47.5310175 | -0.259198248 | 0.51544953 | -0.5028586 | 0.61506366 | 0.84190945 |
| AHNAK | 137.576044 | -0.366348256 | 0.33572174 | -1.0912259 | 0.2751735 | 0.58956168 |
| B3GAT3 | 24.9679895 | -0.341624103 | 0.39233207 | -0.8707524 | 0.38388934 | 0.68794623 |
| GANAB | 62.3323577 | -0.330056681 | 0.33703014 | -0.9793091 | 0.32742727 | 0.63895602 |
| UBXN1 | 17.4164934 | -0.70373956 | 0.48643397 | -1.4467319 | 0.14797201 | 0.43169307 |
| TTC9C | 32.6197258 | 0.253754606 | 0.32979414 | 0.76943334 | 0.4416361 | 0.73375179 |
| TAF6L | 15.2103234 | 0.300140826 | 0.56131902 | 0.53470632 | 0.59285295 | 0.83349833 |
| TMEM223 | 16.805073 | -0.131954875 | 0.48555954 | -0.2717584 | 0.78580781 | 0.92312864 |
| TMEM179B | 25.6488816 | -0.740215 | 0.41824588 | -1.7698082 | 0.07675909 | 0.29635683 |
| NXF1 | 24.5315221 | 0.483456441 | 0.42415967 | 1.13979824 | 0.25437037 | 0.56645553 |
| STX5 | 60.7540059 | -0.330682709 | 0.40268577 | -0.8211929 | 0.41153638 | 0.71058615 |
| SLC3A2 | 52.3862929 | -1.255752261 | 0.63196402 | -1.9870629 | 0.04691543 | 0.22193782 |
| RARRES3 | 20.8079492 | -2.480387125 | 0.87625479 | -2.8306688 | 0.00464508 | 0.04764655 |
| ATL3 | 57.2124775 | -0.583394027 | 0.25659274 | -2.2736186 | 0.02298893 | 0.1429945 |
| RTN3 | 145.762417 | 0.43184058 | 0.31746868 | 1.36026198 | 0.17374704 | 0.46960235 |
| MARK2 | 90.0120241 | -0.156023422 | 0.27049242 | -0.5768126 | 0.56406609 | 0.81464563 |
| NAA40 | 31.8975778 | 0.614006918 | 0.34830614 | 1.76283691 | 0.077928 | 0.29864393 |
| OTUB1 | 24.9252366 | 0.068776806 | 0.36333742 | 0.18929183 | 0.8498641 | 0.95003258 |
| MACROD1 | 46.7112957 | 0.291752901 | 0.49089906 | 0.5943236 | 0.55229569 | 0.80943179 |
| STIP1 | 34.0161961 | 0.014724394 | 0.34911258 | 0.04217664 | 0.96635789 | 0.99156233 |
| NUDT22 | 16.5053084 | 0.18448355 | 0.40420787 | 0.45640762 | 0.64809689 | 0.85958113 |
| DNAJC4 | 13.2935554 | 1.451559948 | 0.70002877 | 2.07357184 | 0.03811909 | 0.19688489 |
| VEGFB | 41.846796 | -0.087375882 | 0.40320953 | -0.2167009 | 0.82844142 | 0.94280334 |
| FKBP2 | 28.2134539 | 0.229969045 | 0.45579893 | 0.50454055 | 0.61388157 | 0.84174925 |
| PPP1R14B | 17.0652469 | -1.272400715 | 0.71852472 | -1.7708517 | 0.07658537 | 0.29596141 |
| GPR137 | 19.2248265 | 0.040651084 | 0.55990435 | 0.07260362 | 0.94212154 | 0.98479162 |
| ESRRA | 59.635548 | 0.461525928 | 0.42437851 | 1.0875337 | 0.27680101 | 0.59181212 |
| TRMT112 | 20.1991598 | -1.126869928 | 0.55134272 | -2.0438647 | 0.04096691 | 0.20525282 |
| PRDX5 | 28.793549 | 0.089656953 | 0.51200188 | 0.17511059 | 0.86099274 | 0.95388501 |
| SF1 | 170.625939 | -0.078899649 | 0.28079666 | -0.280985 | 0.7787219 | 0.92013725 |
| ATG2A | 20.8852167 | -0.789545606 | 0.65966559 | -1.1968877 | 0.23135034 | 0.54389834 |
| VPS51 | 43.4775457 | 0.132223151 | 0.28562862 | 0.46291983 | 0.64342183 | 0.8563039 |
| FAU | 61.0085393 | -0.248585799 | 0.39788962 | -0.6247607 | 0.53212812 | 0.79691016 |
| SYVN1 | 14.4613276 | -0.074450131 | 0.53887123 | -0.1381594 | 0.89011443 | 0.96323092 |
| CAPN1 | 38.4322325 | -0.812052657 | 0.38213784 | -2.1250255 | 0.03358448 | 0.18250255 |
| DPF2 | 31.4969125 | -0.434045516 | 0.33649011 | -1.2899206 | 0.19707824 | 0.49934704 |
| FRMD8 | 26.846359 | -1.45315777 | 0.50314414 | -2.888154 | 0.0038751 | 0.04191558 |
| NEAT1 | 11322.8536 | -2.206852448 | 0.5336158 | -4.135658 | 3.54E-05 | 0.00107941 |
| MALAT1 | 115875.25 | -0.650375572 | 0.39849013 | -1.6320996 | 0.10265851 | 0.35025713 |
| SCYL1 | 22.5938581 | -0.891402309 | 0.50924128 | -1.7504518 | 0.08004038 | 0.30377247 |
| LTBP3 | 46.8521586 | 0.032660166 | 0.33716066 | 0.09686826 | 0.92283102 | 0.97611757 |
| EHBP1L1 | 18.0200428 | -2.606826173 | 0.63631085 | -4.096781 | 4.19E-05 | 0.0012245 |
| MAP3K11 | 14.1726898 | -0.745559369 | 0.45471755 | -1.6396098 | 0.10108632 | 0.34741202 |
| PCNX3 | 13.9811816 | 0.195109411 | 0.53531809 | 0.36447379 | 0.71550425 | 0.88904248 |
| RELA | 18.0589799 | -0.733383714 | 0.45617797 | -1.6076702 | 0.10790744 | 0.36147286 |
| KAT5 | 24.4037096 | -0.342903064 | 0.43918114 | -0.7807782 | 0.43493296 | 0.72806372 |
| CFL1 | 49.1473857 | -1.353080055 | 0.45478712 | -2.9751943 | 0.00292803 | 0.03404109 |
| SART1 | 20.1335645 | -0.04331559 | 0.45041077 | -0.0961691 | 0.92338629 | 0.97648335 |
| BANF1 | 17.4884904 | -0.786582484 | 0.52450404 | -1.4996691 | 0.13370015 | 0.40702484 |
| SF3B2 | 44.2094673 | -0.303585772 | 0.38609302 | -0.7863021 | 0.4316905 | 0.72537073 |
| PACS1 | 341.29079 | -0.561778602 | 0.19455811 | -2.8874591 | 0.00388367 | 0.04191558 |
| RAB1B | 20.4491074 | -0.22521249 | 0.43453779 | -0.5182806 | 0.60426254 | 0.83754753 |
| SLC29A2 | 18.6369246 | 0.721946234 | 0.65303833 | 1.10551892 | 0.2689348 | 0.58532868 |
| ZDHHC24 | 13.6702739 | -0.549454864 | 0.45909675 | -1.1968171 | 0.23137784 | 0.54389834 |
| CTSF | 14.7176468 | -0.241080866 | 0.70847554 | -0.3402811 | 0.73364482 | 0.89900174 |
| CCS | 23.5212401 | 0.498745074 | 0.43878597 | 1.13664774 | 0.25568558 | 0.56843403 |
| RBM14-RBM | 42.9060876 | 0.447205355 | 0.37213994 | 1.20171288 | 0.22947479 | 0.54228719 |
| RBM4 | 22.2166427 | 0.690439586 | 0.40354667 | 1.71092873 | 0.08709427 | 0.31920872 |
| RBM4B | 30.2179163 | 0.347729678 | 0.34027242 | 1.02191555 | 0.30682087 | 0.62069816 |
| SPTBN2 | 40.9589429 | -0.171353027 | 0.56696817 | -0.3022269 | 0.76247911 | 0.91391634 |
| C11orf80 | 173.354274 | -0.856296263 | 0.46513545 | -1.8409611 | 0.06562726 | 0.27144773 |
| RHOD | 12.7630504 | -0.440988133 | 0.47976916 | -0.9191673 | 0.35800806 | 0.6647359 |
| KDM2A | 344.063699 | -0.211882836 | 0.19280771 | -1.0989334 | 0.27179711 | 0.58786226 |
| GRK2 | 18.3391467 | -0.008591927 | 0.43414207 | -0.0197906 | 0.98421042 | 0.99742652 |
| ANKRD13D | 26.8930915 | -0.38988305 | 0.35482576 | -1.0988014 | 0.27185469 | 0.58786226 |
| RAD9A | 114.131735 | 0.081018702 | 0.45693946 | 0.1773073 | 0.85926702 | 0.95328983 |
| PPP1CA | 23.981482 | -0.76755278 | 0.52276168 | -1.4682652 | 0.1420322 | 0.42241953 |
| RPS6KB2 | 14.5396221 | 0.193950793 | 0.55638906 | 0.34858844 | 0.72739831 | 0.89570283 |
| TMEM134 | 22.3590303 | 0.104203072 | 0.43614481 | 0.23891852 | 0.81116877 | 0.93552094 |
| PITPNM1 | 17.9032449 | -1.028369288 | 0.68251153 | -1.5067427 | 0.13187661 | 0.40351308 |
| CDK2AP2 | 18.0395825 | -1.60160159 | 0.5484673 | -2.9201405 | 0.00349874 | 0.03892866 |
| GSTP1 | 111.106855 | -1.375175793 | 0.58084228 | -2.3675545 | 0.01790608 | 0.12164274 |
| NDUFV1 | 58.1383591 | 0.611396512 | 0.46568697 | 1.3128916 | 0.18921948 | 0.48954068 |
| NDUFS8 | 43.5507922 | 0.032545002 | 0.42206741 | 0.07710854 | 0.9385372 | 0.98331142 |
| CHKA | 150.62549 | -0.2852312 | 0.51744045 | -0.5512348 | 0.58147271 | 0.8255928 |
| KMT5B | 132.371231 | 0.256208604 | 0.18795194 | 1.36316018 | 0.17283203 | 0.468183 |
| LRP5 | 36.2764106 | 0.191680951 | 0.43174605 | 0.44396689 | 0.65706652 | 0.86325042 |

| | | | | | | |
|---|---|---|---|---|---|---|
| PPP6R3 | 443.30396 | 0.076655689 | 0.26599142 | 0.28818858 | 0.77320239 | 0.91791018 |
| CPT1A | 115.499258 | 1.454615661 | 0.50302249 | 2.89175074 | 0.00383102 | 0.04157372 |
| IGHMBP2 | 16.5787444 | -0.337314779 | 0.4899607 | -0.6884527 | 0.49116773 | 0.77215443 |
| TPCN2 | 23.4557908 | -0.083548842 | 0.4119423 | -0.2028169 | 0.83927818 | 0.9474978 |
| CCND1 | 15.7552048 | -0.413646401 | 0.59163138 | -0.6991624 | 0.48445056 | 0.76759625 |
| ORAOV1 | 25.0493168 | 0.164855411 | 0.46308955 | 0.35599035 | 0.72184779 | 0.89217596 |
| PPFIA1 | 164.329528 | -0.502392578 | 0.46423399 | -1.0821969 | 0.27916505 | 0.59380484 |
| AP002336.2 | 57.7901299 | -0.495370964 | 0.43004572 | -1.151903 | 0.24936093 | 0.56214731 |
| AP000487.1 | 14.0014731 | -0.083749063 | 0.42718428 | -0.196049 | 0.84457179 | 0.94905142 |
| CTTN | 155.975128 | -1.376979204 | 0.3840394 | -3.5855155 | 0.00033641 | 0.00638842 |
| SHANK2 | 364.099019 | 0.55578135 | 0.46985182 | 1.18288646 | 0.23685415 | 0.54986214 |
| NADSYN1 | 32.1521214 | -0.651592719 | 0.34653644 | -1.8803007 | 0.0600671 | 0.25597151 |
| AP002495.2 | 58.2082248 | 0.255464541 | 0.38493611 | 0.6636544 | 0.50691153 | 0.77942663 |
| RNF121 | 70.6997073 | -0.071097354 | 0.24959214 | -0.2848541 | 0.77575589 | 0.91926749 |
| NUMA1 | 100.938244 | 0.51904006 | 0.27423358 | 1.89269331 | 0.05839867 | 0.25247475 |
| LAMTOR1 | 29.2250465 | -0.817569581 | 0.29834179 | -2.740379 | 0.00613684 | 0.05756707 |
| INPPL1 | 24.6940455 | 0.145903219 | 0.39207131 | 0.37213439 | 0.70979279 | 0.88614996 |
| CLPB | 18.8945042 | -0.35010999 | 0.45515213 | -0.7692153 | 0.4417655 | 0.73383601 |
| ARAP1 | 35.4468194 | 0.434791884 | 0.4163794 | 1.04422046 | 0.29638341 | 0.61093809 |
| ATG16L2 | 23.9329922 | 0.446707769 | 0.39106153 | 1.14229534 | 0.25333128 | 0.56559484 |
| FCHSD2 | 241.71986 | -0.331761717 | 0.18030081 | -1.8400456 | 0.06576154 | 0.27167064 |
| P2RY6 | 49.8013782 | -3.434587689 | 0.96481611 | -3.5598366 | 0.00037109 | 0.00693387 |
| ARHGEF17 | 39.206409 | 0.124998048 | 0.41197704 | 0.30341023 | 0.76157725 | 0.91342791 |
| FAM168A | 151.174267 | 0.413299166 | 0.33655252 | 1.22803765 | 0.21943283 | 0.52936397 |
| RAB6A | 178.665329 | -0.260332272 | 0.25573706 | -1.0179685 | 0.30869293 | 0.62245953 |
| MRPL48 | 41.8264311 | 0.738185251 | 0.27464698 | 2.68776031 | 0.0071933 | 0.06480226 |
| COA4 | 19.2062487 | -0.774465152 | 0.39301096 | -1.9705943 | 0.0487703 | 0.22668707 |
| PAAF1 | 36.580455 | 0.352819824 | 0.46532974 | 0.75821465 | 0.44832249 | 0.7399821 |
| C2CD3 | 74.2775978 | 0.307951473 | 0.27825959 | 1.1067057 | 0.26842119 | 0.5846368 |
| PPME1 | 86.9042725 | -0.410145107 | 0.44769657 | -0.916123 | 0.3596024 | 0.66572082 |
| PGM2L1 | 23.4344384 | -0.793563106 | 0.5431508 | -1.4610364 | 0.14400545 | 0.42527446 |
| POLD3 | 37.290974 | -0.468364202 | 0.37804615 | -1.2389075 | 0.21537976 | 0.52460839 |
| RNF169 | 129.728189 | -0.801557969 | 0.38873999 | -2.0619385 | 0.03921359 | 0.20036945 |
| XRRA1 | 122.037649 | 0.310112903 | 0.57584089 | 0.53853922 | 0.59020484 | 0.83127944 |
| SPCS2 | 68.4242757 | -0.0723124 | 0.28296412 | -0.2555533 | 0.79829582 | 0.9287884 |
| NEU3 | 23.0460069 | -0.451205816 | 0.42326171 | -1.0660209 | 0.28641421 | 0.60056136 |
| RPS3 | 92.1631414 | -0.840427428 | 0.39683088 | -2.1178478 | 0.03418796 | 0.18404648 |
| UVRAG | 215.872254 | -0.169399784 | 0.41310072 | -0.410069 | 0.68175536 | 0.87271442 |
| THAP12 | 20.1585181 | 0.080966543 | 0.33846136 | 0.23921946 | 0.81093541 | 0.93552094 |
| AP002360.1 | 11.3858335 | 1.193073566 | 0.49252639 | 2.4223546 | 0.0154203 | 0.11036477 |
| EMSY | 130.538907 | 0.183889424 | 0.34029648 | 0.54038003 | 0.58893497 | 0.83025227 |
| ACER3 | 130.603504 | -0.821396536 | 0.42515833 | -1.931978 | 0.05336222 | 0.23909147 |
| PAK1 | 89.7467449 | 0.150400053 | 0.25526199 | 0.58919877 | 0.55572794 | 0.81104802 |
| CLNS1A | 58.1519613 | 0.217102481 | 0.37458988 | 0.5795738 | 0.56220207 | 0.81388399 |
| RSF1 | 206.804046 | 0.172898831 | 0.18164917 | 0.95182838 | 0.34118403 | 0.65083559 |
| AAMDC | 52.9524271 | 0.146035618 | 0.47081095 | 0.31017889 | 0.75642492 | 0.91111045 |
| AP002812.2 | 21.9300296 | -0.279041088 | 0.41336759 | -0.6750435 | 0.49964816 | 0.77558432 |
| INTS4 | 63.1858966 | 0.231148168 | 0.28288595 | 0.81710727 | 0.41386713 | 0.71157977 |
| ALG8 | 28.7435942 | -0.005769316 | 0.49007121 | -0.0117724 | 0.9906072 | 0.99857392 |
| GAB2 | 187.680806 | -0.799253981 | 0.60669036 | -1.3174002 | 0.18770452 | 0.48778065 |
| AP003086.2 | 43.7694211 | -1.299360363 | 0.62880484 | -2.0663969 | 0.03879102 | 0.19898431 |
| NARS2 | 75.0069268 | 0.358904358 | 0.28646052 | 1.25289292 | 0.21024468 | 0.51886901 |
| TENM4 | 26.9376796 | -2.290441339 | 0.8986902 | -2.5486439 | 0.01081427 | 0.08684945 |
| PRCP | 62.908442 | 0.092304111 | 0.47318965 | 0.1950679 | 0.84533979 | 0.94907846 |
| RAB30 | 24.5017262 | 0.510888049 | 0.54904403 | 0.93050469 | 0.35210983 | 0.65925143 |
| RAB30-AS1 | 97.3858882 | 0.317621054 | 0.31622729 | 1.00440748 | 0.31518225 | 0.62968915 |
| PCF11 | 81.4235401 | -0.306953512 | 0.32960324 | -0.9312818 | 0.35170779 | 0.65871783 |
| ANKRD42 | 35.9904634 | -0.648246599 | 0.40541303 | -1.5989782 | 0.10982545 | 0.36480672 |
| CCDC90B | 30.9269614 | -0.771609663 | 0.38574528 | -2.0003088 | 0.04546693 | 0.21867374 |
| AP000446.1 | 34.4646399 | -0.367123508 | 0.49490083 | -0.7418123 | 0.45820109 | 0.747377 |
| DLG2 | 162.28561 | -0.326006575 | 0.3881619 | -0.8398727 | 0.40097978 | 0.70105671 |
| TMEM126B | 20.8378486 | -0.845127723 | 0.37204338 | -2.2715838 | 0.02311166 | 0.14347078 |
| CREBZF | 81.4355358 | -0.097110308 | 0.23681651 | -0.4100656 | 0.68175782 | 0.87271442 |
| SYTL2 | 185.217258 | -0.959874798 | 0.47967575 | -2.001091 | 0.04538259 | 0.21854416 |
| PICALM | 490.635786 | 0.470419952 | 0.30963268 | 1.5192839 | 0.12869105 | 0.39737957 |
| EED | 52.0909815 | 0.659438769 | 0.31369135 | 2.10218982 | 0.03553665 | 0.18804979 |
| HIKESHI | 38.8233138 | 0.453481541 | 0.35214811 | 1.28775798 | 0.19783022 | 0.50042566 |
| AP001831.1 | 51.0681606 | 1.203856742 | 0.56072147 | 2.14697813 | 0.03179502 | 0.17656748 |
| ME3 | 303.000155 | 1.452253084 | 0.42642156 | 3.40567464 | 0.00066001 | 0.01085346 |
| AP003059.1 | 17.0547374 | 1.263810323 | 0.55646059 | 2.27115873 | 0.02313737 | 0.14353483 |
| AP003059.2 | 16.1338139 | 1.876272122 | 0.57154723 | 3.28279456 | 0.00102784 | 0.01528475 |
| PRSS23 | 129.617702 | 1.577049874 | 0.56047619 | 2.81376785 | 0.00489646 | 0.04947544 |
| AP001528.2 | 78.2899701 | 0.132579696 | 0.90530731 | 0.14644717 | 0.88356839 | 0.96170562 |
| FZD4 | 83.9991654 | 1.439854876 | 0.47331318 | 3.04207645 | 0.00234952 | 0.02859914 |
| AP001528.1 | 19.6817569 | 0.265166424 | 0.49586398 | 0.53475637 | 0.59281834 | 0.83349833 |
| TMEM135 | 193.319155 | 0.41438528 | 0.27766005 | 1.49241953 | 0.1355892 | 0.41064171 |
| RAB38 | 32.0555304 | 0.163499957 | 0.47670125 | 0.34298202 | 0.73161198 | 0.89782437 |
| CTSC | 49.1504222 | -0.081585095 | 0.52732015 | -0.1547164 | 0.87704487 | 0.95879545 |
| NOX4 | 18.949581 | -0.146724317 | 0.70790426 | -0.2072658 | 0.83580231 | 0.94654164 |
| CHORDC1 | 44.2633245 | 0.741611103 | 0.4270139 | 1.73673762 | 0.08243349 | 0.30885595 |
| SLC36A4 | 124.128938 | 0.439529622 | 0.26942652 | 1.63135247 | 0.10281597 | 0.350386 |
| SMCO4 | 55.7559787 | -1.224002074 | 0.79716218 | -1.5354492 | 0.12467351 | 0.39027762 |
| CEP295 | 58.6026177 | -0.144300715 | 0.41383148 | -0.3486944 | 0.72731876 | 0.89570283 |
| TAF1D | 134.276522 | -0.450161806 | 0.29870269 | -1.5070564 | 0.1317962 | 0.40351308 |
| MRE11 | 43.666755 | -0.164335862 | 0.31569577 | -0.5205514 | 0.60267934 | 0.83691476 |
| AMOTL1 | 42.1807444 | -1.108548153 | 0.47819105 | -2.3182118 | 0.02043781 | 0.13307413 |
| CWC15 | 22.919875 | -0.393264046 | 0.36701245 | -1.0715278 | 0.28393218 | 0.59743472 |
| SRSF8 | 17.4754101 | -0.766174451 | 0.36070777 | -2.1240864 | 0.03366292 | 0.18264262 |
| ENDOD1 | 33.8996495 | -0.846157505 | 0.43691836 | -1.936649 | 0.05278826 | 0.23800665 |
| AP000787.1 | 28.8366374 | -0.170385838 | 0.43196293 | -0.3944455 | 0.69325215 | 0.87872254 |
| SESN3 | 47.8443188 | -1.045077003 | 0.58244478 | -1.7942937 | 0.0727663 | 0.28837569 |
| FAM76B | 29.3101109 | -0.064658209 | 0.32813779 | -0.1970459 | 0.84379161 | 0.94858974 |
| CEP57 | 97.8880238 | -0.092606046 | 0.2522584 | -0.3671079 | 0.71353855 | 0.88813689 |
| MTMR2 | 68.7800598 | 0.002127601 | 0.25914959 | 0.00820994 | 0.99344949 | 0.9992177 |
| MAML2 | 2323.20742 | 1.374967617 | 0.26625961 | 5.16401121 | 2.42E-07 | 1.73E-05 |
| CCDC82 | 47.5320232 | -0.821979036 | 0.40901678 | -2.0096462 | 0.04446864 | 0.21534941 |
| ARHGAP42 | 21.3056531 | -0.199073689 | 0.41060337 | -0.4848321 | 0.62779546 | 0.84784081 |
| CEP126 | 27.4276933 | 0.231806889 | 0.52347769 | 0.44282095 | 0.65789525 | 0.86357172 |
| YAP1 | 530.522762 | -0.890646698 | 0.23027438 | -3.8677629 | 0.00010984 | 0.00261258 |
| BIRC3 | 81.4538946 | -2.605339576 | 0.60964413 | -4.2735416 | 1.92E-05 | 0.00068208 |
| BIRC2 | 65.4218172 | 0.07539327 | 0.40375235 | 0.18673147 | 0.85187118 | 0.95090866 |
| TMEM123 | 118.016848 | -0.678301235 | 0.32715939 | -2.0733051 | 0.0381439 | 0.19688489 |
| DCUN1D5 | 48.0926636 | -0.344748645 | 0.27834765 | -1.2385542 | 0.21551065 | 0.52479011 |
| DYNC2H1 | 538.996116 | 0.5724636 | 0.4694342 | 1.2194757 | 0.22266369 | 0.53302786 |
| CASP4 | 69.9449728 | -2.284005929 | 0.39548382 | -5.7752196 | 7.69E-09 | 8.55E-07 |
| MSANTD4 | 20.9908768 | -0.403612734 | 0.40205587 | -1.0038723 | 0.31544019 | 0.63006948 |
| KBTBD3 | 21.469079 | 0.524009791 | 0.38746074 | 1.35242036 | 0.17624088 | 0.47349784 |
| AASDHPPT | 30.769408 | -0.148880246 | 0.35030004 | -0.4250078 | 0.670831 | 0.86777458 |
| CWF19L2 | 117.667237 | -0.176296024 | 0.22487365 | -0.7839781 | 0.43305295 | 0.72648177 |
| ALKBH8 | 21.3007769 | 0.69541216 | 0.63633659 | 1.09283698 | 0.27446538 | 0.5891499 |
| SLC35F2 | 70.0569862 | 0.426020713 | 0.28632906 | 1.48787102 | 0.13678491 | 0.41194526 |
| CUL5 | 178.458998 | 0.365993224 | 0.1755885 | 2.08438039 | 0.03712558 | 0.19370952 |
| ACAT1 | 70.36254 | 0.607161094 | 0.28257463 | 2.14867515 | 0.03166016 | 0.17592332 |

| | | | | | | |
|---|---|---|---|---|---|---|
| NPAT | 53.6770544 | 0.019835059 | 0.26328984 | 0.07533545 | 0.93994782 | 0.98363127 |
| ATM | 150.686858 | 0.015442701 | 0.21421496 | 0.07208974 | 0.94253049 | 0.98488741 |
| C11orf65 | 40.8378384 | 0.581908858 | 0.65134707 | 0.89339292 | 0.37164679 | 0.67534262 |
| KDELC2 | 41.9653637 | 1.996050637 | 0.48929107 | 4.07947489 | 4.51E-05 | 0.00130298 |
| EXPH5 | 207.27219 | 1.002156151 | 0.24289089 | 4.12595197 | 3.69E-05 | 0.00111407 |
| DDX10 | 98.2024863 | 0.538110459 | 0.26607511 | 2.02240058 | 0.04313499 | 0.21134556 |
| RDX | 167.43821 | 0.63291703 | 0.29654657 | 2.1342922 | 0.03281887 | 0.18010779 |
| ZC3H12C | 92.3710786 | 0.034443849 | 0.37077967 | 0.09289573 | 0.9259864 | 0.97735117 |
| FDX1 | 94.5811858 | 1.408989119 | 0.37348716 | 3.77252362 | 0.0001616 | 0.00353709 |
| COLCA1 | 20.7037682 | 2.428725985 | 0.69134987 | 3.5130201 | 0.00044304 | 0.00786846 |
| SIK2 | 364.464225 | 0.443760416 | 0.39319916 | 1.12858942 | 0.25907108 | 0.57214086 |
| PPP2R1B | 80.1434992 | 0.095423118 | 0.43233456 | 0.22071592 | 0.82531364 | 0.94180992 |
| CRYAB | 162.703976 | -2.346347546 | 0.83541796 | -2.8085912 | 0.00497588 | 0.0499409 |
| DIXDC1 | 38.9870298 | 0.354974792 | 0.50522209 | 0.70261138 | 0.48229797 | 0.76628579 |
| DLAT | 39.4891402 | 0.798563735 | 0.26699665 | 2.99091293 | 0.00278145 | 0.03270394 |
| NKAPD1 | 17.0662281 | -0.032296142 | 0.39285883 | -0.082208 | 0.9344813 | 0.98085023 |
| AP002884.3 | 26.8973331 | 0.362128927 | 0.46870107 | 0.77262237 | 0.4397459 | 0.73204621 |
| PTS | 55.792089 | -0.57733537 | 0.33596809 | -1.7184232 | 0.08571945 | 0.31615828 |
| AP002884.1 | 33.2762538 | -1.289304988 | 0.44554328 | -2.8937817 | 0.00380633 | 0.04152772 |
| TTC12 | 47.7406626 | 0.676306667 | 0.51212536 | 1.32058812 | 0.18663874 | 0.48675708 |
| ZW10 | 25.1627421 | 0.279124424 | 0.32966615 | 0.84668816 | 0.39716896 | 0.69753312 |
| USP28 | 29.3427332 | 0.001581727 | 0.32506915 | 0.00486582 | 0.99611766 | 0.99922221 |
| NNMT | 82.3861472 | -5.709084845 | 1.42525989 | -4.0056448 | 6.18E-05 | 0.00169113 |
| REXO2 | 42.1286896 | 0.418682094 | 0.52715064 | 0.79423614 | 0.42705797 | 0.72258772 |
| CADM1 | 643.499113 | 1.172703992 | 0.33537299 | 3.49671563 | 0.00047102 | 0.00827086 |
| AP000462.1 | 19.1066638 | 0.388975592 | 0.5821826 | 0.66813332 | 0.50404849 | 0.77792612 |
| BUD13 | 14.704144 | 0.024355982 | 0.49617046 | 0.04908793 | 0.96084922 | 0.98983463 |
| SIK3 | 876.884733 | 0.225308488 | 0.37629452 | 0.5987557 | 0.54933581 | 0.80698344 |
| PAFAH1B2 | 71.9490655 | -0.051716882 | 0.24623382 | -0.2100316 | 0.83364301 | 0.94629747 |
| SIDT2 | 15.0311007 | -0.332157496 | 0.44737997 | -0.7424505 | 0.45781441 | 0.747377 |
| PCSK7 | 68.1457609 | 0.386096908 | 0.27146394 | 1.42227699 | 0.15494586 | 0.44046194 |
| RNF214 | 50.4772521 | 0.537057729 | 0.3510791 | 1.52973426 | 0.12608252 | 0.39291224 |
| BACE1 | 33.6990935 | 0.461028928 | 0.46920874 | 0.98256679 | 0.3258207 | 0.63755555 |
| CEP164 | 41.4129603 | 0.597251158 | 0.44833252 | 1.33216113 | 0.18280725 | 0.48176788 |
| DSCAML1 | 344.390645 | -0.693108968 | 0.67511314 | -1.026656 | 0.30458247 | 0.61818174 |
| FXYD2 | 73.8696694 | 1.159115467 | 0.70411638 | 1.64619871 | 0.09972286 | 0.34450385 |
| FXYD6-FXYD2 | 22.9921119 | 1.230773311 | 1.06889415 | 1.15144546 | 0.24954902 | 0.56229944 |
| AP000757.1 | 26.3037749 | 0.33781316 | 0.49242465 | 0.68602001 | 0.49270049 | 0.77287002 |
| TMPRSS4 | 43.5754327 | -4.547559659 | 0.77798372 | -5.8453147 | 5.06E-09 | 6.04E-07 |
| MPZL3 | 23.1194991 | -0.730582172 | 0.40016575 | -1.8256989 | 0.06789563 | 0.27788783 |
| UBE4A | 65.8817745 | -0.175982274 | 0.28604707 | -0.6152214 | 0.53840853 | 0.80104055 |
| ATP5MG | 78.5680637 | 0.233341598 | 0.34290633 | 0.68048205 | 0.49619929 | 0.77475164 |
| KMT2A | 206.198025 | -0.393339379 | 0.23081233 | -1.7041524 | 0.08835263 | 0.3228056 |
| IFT46 | 20.879554 | 0.982934526 | 0.40422796 | 2.4316342 | 0.01503088 | 0.10847362 |
| ARCN1 | 41.1027898 | -0.125180281 | 0.31146673 | -0.4019058 | 0.68775336 | 0.87565595 |
| PHLDB1 | 63.6709125 | -0.021697029 | 0.5356719 | -0.0405043 | 0.96769106 | 0.99156233 |
| DDX6 | 125.467201 | 0.030215349 | 0.24899689 | 0.1213483 | 0.90341517 | 0.96843447 |
| BCL9L | 24.5419035 | -0.910253695 | 0.65839465 | -1.3825351 | 0.16680747 | 0.45906518 |
| CCDC84 | 109.060404 | 0.383507095 | 0.33497086 | 1.1448969 | 0.25225187 | 0.56479656 |
| RPS25 | 70.0997953 | -0.848776076 | 0.45263091 | -1.8752057 | 0.06076442 | 0.25764779 |
| TRAPPC4 | 13.8092391 | -0.109874344 | 0.55055919 | -0.1995686 | 0.84181796 | 0.94776435 |
| VPS11 | 13.9721046 | -0.449261704 | 0.48059893 | -0.9347955 | 0.34989372 | 0.65681679 |
| HINFP | 15.694854 | -0.024844564 | 0.44529593 | -0.0557934 | 0.95550641 | 0.98853497 |
| CBL | 58.7150639 | -0.329495442 | 0.43395327 | -0.7592878 | 0.44768039 | 0.7399821 |
| USP2 | 28.2862124 | 3.774189846 | 0.61647913 | 6.12216969 | 9.23E-10 | 1.62E-07 |
| USP2-AS1 | 15.8362702 | 3.047035352 | 0.66834732 | 4.55905971 | 5.14E-06 | 0.0002218 |
| OAF | 46.8281925 | 1.63445115 | 0.33448629 | 4.88645181 | 1.03E-06 | 6.02E-05 |
| ARHGEF12 | 468.130192 | -0.204238854 | 0.23623787 | -0.8645475 | 0.38728721 | 0.69071651 |
| SC5D | 17.0407992 | 0.090223686 | 0.42008635 | 0.21477414 | 0.82994341 | 0.94382149 |
| SORL1 | 202.839377 | 0.66698666 | 0.59386975 | 1.12311943 | 0.26138677 | 0.57507557 |
| UBASH3B | 27.394283 | -3.763336704 | 0.76810244 | -4.8995245 | 9.61E-07 | 5.67E-05 |
| JHY | 15.7746187 | -0.519699611 | 0.59149896 | -0.8786146 | 0.37961029 | 0.68343046 |
| HSPA8 | 53.1975376 | -0.101928717 | 0.625269 | -0.1630158 | 0.87050601 | 0.95629149 |
| GRAMD1B | 498.352781 | 1.683793019 | 0.71123443 | 2.36742338 | 0.01791243 | 0.12164274 |
| VWA5A | 21.3450746 | -1.580214223 | 0.39581311 | -3.9923241 | 6.54E-05 | 0.0017786 |
| TBRG1 | 56.8986209 | -0.342340937 | 0.29834739 | -1.1474575 | 0.2511926 | 0.56353121 |
| SIAE | 34.3958844 | 0.397539504 | 0.41580491 | 0.95607216 | 0.33903578 | 0.64971064 |
| MSANTD2 | 72.1771724 | 0.660886982 | 0.33463026 | 1.97497677 | 0.04827078 | 0.22503839 |
| EI24 | 23.8327791 | -0.453254379 | 0.44233362 | -1.024689 | 0.30550997 | 0.61893963 |
| STT3A | 25.218035 | -0.602870465 | 0.41770218 | -1.4433022 | 0.14893537 | 0.43272656 |
| RPUSD4 | 15.1659925 | 0.382027725 | 0.41312349 | 0.9247301 | 0.35510632 | 0.66163487 |
| FAM118B | 24.9239102 | -0.737029126 | 0.46881682 | -1.5721047 | 0.11592627 | 0.37462127 |
| SRPRA | 21.5512387 | -0.703400368 | 0.36210027 | -1.9425569 | 0.05206972 | 0.23556433 |
| FOXRED1 | 23.8771105 | -0.088161738 | 0.50075565 | -0.1760574 | 0.86024885 | 0.95363316 |
| TIRAP | 13.152846 | 0.562873545 | 0.4238816 | 1.32790276 | 0.18421023 | 0.48327975 |
| DCPS | 28.1583185 | 0.021836363 | 0.39030892 | 0.05594636 | 0.95538454 | 0.98853497 |
| ST3GAL4 | 39.772317 | 1.160349202 | 0.722297 | 1.60647103 | 0.10817046 | 0.36175274 |
| ETS1 | 88.4552679 | -0.215717043 | 0.72425284 | -0.2978477 | 0.76581941 | 0.91556612 |
| KCNJ1 | 200.98779 | 0.695564949 | 0.74401157 | 0.93488459 | 0.34984778 | 0.65681679 |
| ARHGAP32 | 192.447511 | 0.1912875 | 0.28575363 | 0.66941407 | 0.50323138 | 0.7778729 |
| BARX2 | 48.1377072 | 0.081594915 | 0.55140466 | 0.14797647 | 0.88236134 | 0.96161232 |
| TMEM45B | 44.0536206 | -0.529671693 | 0.47761305 | -1.1089975 | 0.26743125 | 0.58364201 |
| NFRKB | 37.3642641 | 0.252609966 | 0.3447288 | 0.73277883 | 0.46369334 | 0.75139985 |
| PRDM10 | 54.8533865 | 0.455728693 | 0.25503127 | 1.78695225 | 0.07394521 | 0.29120952 |
| APLP2 | 173.649878 | -0.241183955 | 0.40026982 | -0.6025534 | 0.54680581 | 0.80581909 |
| ST14 | 52.8714017 | -0.224358836 | 0.36634684 | -0.6124219 | 0.54025864 | 0.80166126 |
| ZBTB44 | 113.948609 | 0.190343233 | 0.3221215 | 0.59090509 | 0.55458402 | 0.81072612 |
| AP002986.1 | 17.5450804 | 1.899929 | 0.6806284 | 2.79143363 | 0.00524751 | 0.05193962 |
| ADAMTS15 | 19.3829575 | -0.637773432 | 0.85314669 | -0.7475542 | 0.45472907 | 0.74541031 |
| SNX19 | 54.7802405 | 0.417606112 | 0.37526661 | 1.11282514 | 0.26578354 | 0.58118333 |
| NCAPD3 | 19.9880376 | 0.198050859 | 0.47329501 | 0.41845119 | 0.67561727 | 0.86960234 |
| VPS26B | 20.7202856 | 0.410181674 | 0.44227016 | 0.92744596 | 0.35369504 | 0.6604952 |
| ACAD8 | 15.1219145 | 0.394302856 | 0.43789452 | 0.90045169 | 0.36787992 | 0.67217566 |
| ZMYND11 | 208.735421 | 0.241272057 | 0.28766913 | 0.83871377 | 0.40162995 | 0.70114138 |
| DIP2C | 705.007274 | -2.318595723 | 0.56610394 | -4.0957067 | 4.21E-05 | 0.00122635 |
| LARP4B | 273.407414 | -0.146284401 | 0.20927519 | -0.699005 | 0.48454891 | 0.76759625 |
| GTPBP4 | 28.5278821 | -1.034884045 | 0.51720437 | -2.000919 | 0.04540112 | 0.21854416 |
| IDI1 | 28.11896 | -0.449665864 | 0.52937468 | -0.8494284 | 0.39564298 | 0.69642725 |
| WDR37 | 128.410558 | -0.064738641 | 0.32061524 | -0.20192 | 0.83997924 | 0.9474978 |
| PFKP | 158.687223 | -1.240927327 | 0.389494 | -3.1859986 | 0.00144255 | 0.02007517 |
| PITRM1 | 43.9439199 | -0.487618631 | 0.49660996 | -0.9818946 | 0.32615178 | 0.63780184 |
| KLF6 | 123.812466 | -1.423066581 | 0.60645982 | -2.3465142 | 0.01894994 | 0.12656825 |
| NET1 | 16.0858835 | 0.059449198 | 0.44874665 | 0.13247831 | 0.89460598 | 0.9650125 |
| ASB13 | 13.2394633 | -0.063150632 | 0.47914039 | -0.1317998 | 0.89514261 | 0.96511349 |
| FAM208B | 210.82363 | -0.200967725 | 0.43021825 | -0.4671297 | 0.64040708 | 0.85448707 |
| GDI2 | 120.45017 | -0.594760787 | 0.29913291 | -1.9882827 | 0.04678043 | 0.22193782 |
| ANKRD16 | 18.3615084 | 0.136177737 | 0.35575341 | 0.38278688 | 0.70187779 | 0.88424653 |
| FBH1 | 91.5377019 | -0.083604784 | 0.30035199 | -0.278356 | 0.78073908 | 0.92083964 |
| RBM17 | 60.6568421 | -0.023428068 | 0.25321759 | -0.0925215 | 0.92628372 | 0.97735117 |
| PFKFB3 | 59.0963391 | -2.276179398 | 0.58608434 | -3.8837062 | 0.00010288 | 0.0024786 |
| PRKCQ | 155.501384 | -0.591403043 | 0.39562584 | -1.4948544 | 0.13495244 | 0.4097351 |
| SFMBT2 | 170.246019 | -0.264012738 | 0.68863347 | -0.3833864 | 0.70143325 | 0.88421489 |
| KIN | 31.2274884 | -0.121861317 | 0.33378357 | -0.3650908 | 0.71504361 | 0.88904248 |

| | | | | | | |
|---|---|---|---|---|---|---|
| ATP5F1C | 133.373524 | 0.94850819 | 0.24333316 | 3.89798169 | 9.70E-05 | 0.00235523 |
| TAF3 | 161.499626 | -0.031964042 | 0.2484588 | -0.1286493 | 0.89763518 | 0.96607503 |
| GATA3 | 144.597604 | 0.210238932 | 0.46537996 | 0.45175759 | 0.65144362 | 0.86120237 |
| GATA3-AS1 | 15.183886 | -0.223807159 | 0.69763414 | -0.3208088 | 0.7483553 | 0.90723111 |
| USP6NL | 109.502568 | -0.175619156 | 0.21880001 | -0.8026469 | 0.42217883 | 0.72008598 |
| PROSER2 | 39.2389719 | -0.056521783 | 0.36504192 | -0.1548364 | 0.87695028 | 0.95879545 |
| UPF2 | 116.612778 | -0.050336229 | 0.30100015 | -0.1672299 | 0.86718915 | 0.95588054 |
| DHTKD1 | 45.2010078 | 0.145199208 | 0.28695649 | 0.50599729 | 0.61285855 | 0.84174925 |
| SEC61A2 | 21.7499481 | -0.34070564 | 0.37070943 | -0.9190638 | 0.35806218 | 0.6647359 |
| NUDT5 | 45.8874648 | 0.185471011 | 0.30139509 | 0.61537503 | 0.53830708 | 0.80104055 |
| CDC123 | 44.4425706 | 0.052939793 | 0.37553717 | 0.14097085 | 0.88789297 | 0.96297709 |
| OPTN | 42.3618843 | -0.338479301 | 0.50319937 | -0.6726545 | 0.50116714 | 0.7760974 |
| PHYH | 22.2323364 | 0.907241565 | 0.39432599 | 2.30073995 | 0.02140633 | 0.13717779 |
| SEPHS1 | 26.7898712 | 0.446082935 | 0.33472511 | 1.33268439 | 0.1826354 | 0.4814838 |
| BEND7 | 81.0753473 | -0.172363423 | 0.25466786 | -0.6768166 | 0.49852235 | 0.77558432 |
| PRPF18 | 16.8313559 | -0.476678657 | 0.38579287 | -1.2355818 | 0.21661405 | 0.52610299 |
| AL157392.5 | 28.1514064 | -0.187742434 | 0.31285105 | -0.6001017 | 0.54843849 | 0.80647995 |
| FRMD4A | 361.158794 | 0.530043043 | 0.42057968 | 1.26026784 | 0.20757276 | 0.51555897 |
| FAM107B | 208.200248 | 1.104844067 | 0.3028399 | 3.64827777 | 0.000264 | 0.00525977 |
| HSPA14.1 | 12.695071 | 0.270011151 | 0.40571179 | 0.66552453 | 0.50571506 | 0.77912875 |
| DCLRE1C | 34.0077282 | -0.680261266 | 0.33368669 | -2.0386227 | 0.0414877 | 0.20730509 |
| RPP38 | 34.8606541 | 0.222994705 | 0.32784637 | 0.68018048 | 0.49639019 | 0.77475164 |
| NMT2 | 33.816804 | -0.144845468 | 0.33108202 | -0.4374912 | 0.66175516 | 0.86492722 |
| FAM171A1 | 300.02126 | 0.78544099 | 0.45193252 | 1.73796079 | 0.08221772 | 0.30864976 |
| MINDY3 | 85.5556235 | 0.241530382 | 0.25970555 | 0.93001626 | 0.35236267 | 0.65959235 |
| PTER | 74.8229892 | -0.511181771 | 0.26454775 | -1.9322855 | 0.05332428 | 0.23903633 |
| RSU1 | 75.9642572 | -0.376736039 | 0.2099123 | -1.7947306 | 0.07269663 | 0.28837569 |
| CUBN | 39.9427342 | 0.644331398 | 0.59631925 | 1.08051416 | 0.27991329 | 0.59397852 |
| TRDMT1 | 16.1110129 | -0.602688845 | 0.60731817 | -0.9923774 | 0.32101344 | 0.63269097 |
| VIM | 27.7785233 | -1.48691335 | 0.5000396 | -2.9735912 | 0.00294337 | 0.0341768 |
| STAM | 58.6068576 | 0.09898626 | 0.35034002 | 0.2825434 | 0.77752687 | 0.91968524 |
| CACNB2 | 15.6162337 | 0.955568119 | 0.67838271 | 1.4085974 | 0.15895425 | 0.44748795 |
| AL450384.2 | 28.3046556 | 0.578563399 | 0.46929148 | 1.23284446 | 0.2176338 | 0.52706949 |
| NSUN6 | 88.326766 | 0.231124775 | 0.35692062 | 0.64755232 | 0.51727454 | 0.78615013 |
| ARL5B | 61.2206664 | -1.349312927 | 0.80083802 | -1.6848762 | 0.09201246 | 0.33050702 |
| MALRD1 | 89.7309312 | 1.71173451 | 0.73084799 | 2.34212111 | 0.01917449 | 0.12770211 |
| PLXDC2 | 343.312987 | -0.085172239 | 0.26964363 | -0.3158697 | 0.75210145 | 0.90930937 |
| NEBL | 335.708666 | 0.110193618 | 0.37810469 | 0.29143679 | 0.77071728 | 0.91695393 |
| MLLT10 | 349.975858 | 0.371221261 | 0.18458355 | 2.01112857 | 0.04431188 | 0.21517692 |
| DNAJC1 | 241.587282 | 0.079867455 | 0.17539264 | 0.45536379 | 0.64884754 | 0.8599651 |
| PIP4K2A | 87.6045367 | -0.189698029 | 0.26167092 | -0.7249488 | 0.46848343 | 0.75508029 |
| MSRB2 | 20.8342007 | -0.273513957 | 0.41085743 | -0.665715 | 0.50559327 | 0.77906986 |
| KIAA1217 | 684.964598 | 0.778273129 | 0.29530254 | 2.63551113 | 0.00840107 | 0.07273124 |
| ARHGAP21 | 160.225935 | -0.148532541 | 0.33220107 | -0.4471164 | 0.65479103 | 0.86220471 |
| PRTFDC1 | 32.1162759 | 1.898341791 | 0.60271603 | 3.14964543 | 0.00163469 | 0.02180518 |
| PDSS1 | 28.750263 | 0.488431339 | 0.35384095 | 1.38036973 | 0.16747283 | 0.4603528 |
| ABI1 | 185.093575 | 0.019600293 | 0.26781737 | 0.0731853 | 0.94165867 | 0.98455573 |
| ANKRD26 | 164.348548 | 0.736945497 | 0.2881038 | 2.55791663 | 0.01053013 | 0.0854508 |
| YME1L1 | 103.858618 | -0.179957542 | 0.24737318 | -0.7274739 | 0.46693567 | 0.75315494 |
| MASTL | 25.2256319 | -0.010363833 | 0.53775653 | -0.0192724 | 0.98462384 | 0.99742652 |
| ACBD5 | 48.9106139 | -0.178197268 | 0.29258012 | -0.6090546 | 0.54248826 | 0.80340275 |
| RAB18 | 99.749595 | -0.131927209 | 0.37130983 | -0.3553022 | 0.72236319 | 0.89217596 |
| MPP7 | 95.1507739 | 0.662506141 | 0.23754356 | 2.788988 | 0.0052873 | 0.05216805 |
| WAC-AS1 | 28.2950696 | -0.218318771 | 0.38951173 | -0.5604934 | 0.57514291 | 0.82146824 |
| WAC | 367.513975 | -0.18849891 | 0.23674859 | -0.7961986 | 0.42591658 | 0.72211376 |
| SVIL | 571.218846 | -1.005825336 | 0.57790394 | -1.7404715 | 0.08177626 | 0.30770051 |
| MTPAP | 30.3303111 | 0.733928752 | 0.39946318 | 1.83728761 | 0.06616745 | 0.27250233 |
| MAP3K8 | 33.0974803 | -1.563289195 | 0.46291942 | -3.3770223 | 0.00073275 | 0.01177961 |
| ZNF438 | 80.624302 | 0.436099408 | 0.34014148 | 1.2821118 | 0.19980343 | 0.5033686 |
| ZEB1 | 107.139758 | -3.914197676 | 0.99191222 | -3.946113 | 7.94E-05 | 0.00205154 |
| ARHGAP12 | 194.440872 | 0.217068911 | 0.180643 | 1.20164583 | 0.22950078 | 0.54228719 |
| KIF5B | 112.466149 | -0.183747581 | 0.30113743 | -0.6101785 | 0.54174358 | 0.80301375 |
| EPC1 | 193.296456 | -0.032944174 | 0.22587536 | -0.1458511 | 0.88403892 | 0.96170562 |
| CCDC7 | 142.702771 | 0.418025138 | 0.61249873 | 0.68249144 | 0.49492826 | 0.77475164 |
| ITGB1 | 177.39358 | -1.221771249 | 0.41678114 | -2.9314456 | 0.00337388 | 0.03814869 |
| NRP1 | 240.992134 | -1.844898369 | 0.46296834 | -3.9849342 | 6.75E-05 | 0.00182953 |
| PARD3 | 1291.36463 | -0.149486473 | 0.23728185 | -0.6299954 | 0.5286976 | 0.79434813 |
| CUL2 | 83.494934 | 0.046820137 | 0.23117966 | 0.20252706 | 0.83950471 | 0.9474978 |
| CREM | 196.711245 | -0.851300443 | 0.5125817 | -1.6608093 | 0.09675175 | 0.33969491 |
| CCNY | 376.439699 | 0.276162481 | 0.21946354 | 1.25835244 | 0.20826432 | 0.51645229 |
| AL117336.3 | 55.6245642 | 0.768005355 | 0.33890937 | 2.26610833 | 0.02344475 | 0.14515197 |
| ZNF248 | 155.234021 | -0.154425173 | 0.52968918 | -0.2915392 | 0.77063895 | 0.91695393 |
| ZNF25 | 24.7507539 | 0.700266251 | 0.43285503 | 1.61778472 | 0.105709 | 0.35687176 |
| ZNF33A | 239.712106 | -0.244749581 | 0.25303387 | -0.9672601 | 0.333414 | 0.64508099 |
| AL117339.5 | 28.2438764 | -0.312298762 | 0.35105667 | -0.8895964 | 0.37368262 | 0.67707283 |
| ZNF33B | 124.731457 | -0.135470618 | 0.26662649 | -0.5080914 | 0.61138926 | 0.84169124 |
| BMS1 | 54.2423746 | 0.145381075 | 0.27964751 | 0.51987259 | 0.60315238 | 0.8372477 |
| CSGALNACT2 | 40.9117951 | -0.267010414 | 0.24861635 | -1.0739857 | 0.28282908 | 0.59668285 |
| HNRNPF | 68.0019286 | -1.147258216 | 0.44186496 | -2.5964001 | 0.00942063 | 0.07888124 |
| ZNF487 | 16.1516006 | 0.00897005 | 0.39811801 | 0.02253113 | 0.98202428 | 0.99659802 |
| ZNF32 | 23.7468754 | 0.047996637 | 0.33745576 | 0.1422309 | 0.88689762 | 0.96253908 |
| TMEM72-AS | 32.815253 | 0.188550509 | 0.58712344 | 0.32114288 | 0.74810212 | 0.90723111 |
| TMEM72 | 137.616233 | 1.24029005 | 0.46339777 | 2.67651278 | 0.00743927 | 0.06637682 |
| RASSF4 | 22.9091084 | 0.670061629 | 0.72647534 | 0.92234601 | 0.35634812 | 0.66266252 |
| DEPP1 | 24.5534405 | -1.723882135 | 0.46875799 | -3.6775525 | 0.00023548 | 0.00482558 |
| 8-Mar | 105.911565 | 0.002702586 | 0.23058918 | 0.01172035 | 0.99064873 | 0.99857392 |
| ZFAND4 | 32.6045762 | 0.197688973 | 0.31793377 | 0.62179295 | 0.53407802 | 0.79850222 |
| WASHC2C | 41.7989013 | -0.694947378 | 0.58992354 | -1.1780296 | 0.23878482 | 0.55079752 |
| AGAP4 | 46.2351133 | 0.412479096 | 0.46827531 | 0.88084741 | 0.37840041 | 0.6819106 |
| TIMM23 | 44.5752679 | 0.258273919 | 0.32770749 | 0.78812333 | 0.43062457 | 0.7245261 |
| NCOA4 | 37.2297063 | 0.011728429 | 0.43614581 | 0.02689107 | 0.97854661 | 0.9951912 |
| PTPN20 | 48.5833611 | 0.365749118 | 0.60282401 | 0.60672619 | 0.54403264 | 0.80380646 |
| AGAP9 | 39.7872299 | 0.399560109 | 0.63373328 | 0.63048623 | 0.52837651 | 0.7943404 |
| MAPK8 | 434.23717 | 0.083779081 | 0.36390783 | 0.2302206 | 0.81792035 | 0.93805075 |
| ARHGAP22 | 25.8906051 | -3.910242198 | 0.56830867 | -6.8804902 | 5.96E-12 | 1.74E-09 |
| ERCC6 | 70.0951447 | 0.11359619 | 0.28246874 | 0.4021549 | 0.68757003 | 0.87556719 |
| OGDHL | 99.1726958 | 0.979346728 | 0.66945585 | 1.46289965 | 0.14349484 | 0.42447522 |
| PARG | 133.573613 | 0.214439195 | 0.26842761 | 0.79887159 | 0.42436487 | 0.72151771 |
| TIMM23B | 149.899143 | -0.151746384 | 0.48183012 | -0.3149375 | 0.75280909 | 0.90981102 |
| LINC00843 | 29.8491629 | 0.203528659 | 0.39236005 | 0.51872931 | 0.60394952 | 0.83754753 |
| AGAP6 | 48.9183997 | 0.199877524 | 0.28740442 | 0.69545736 | 0.48676872 | 0.76847808 |
| WASHC2A | 42.3767327 | -0.257496201 | 0.50745714 | -0.5074245 | 0.61185698 | 0.84169124 |
| SGMS1 | 376.62873 | -0.707021081 | 0.21979966 | -3.2166613 | 0.00129692 | 0.01840555 |
| ASAH2B | 15.0559621 | -0.552503155 | 0.53068167 | -1.0411197 | 0.29781999 | 0.61141484 |
| PRKG1 | 261.280009 | -0.440997177 | 0.66784868 | -0.660325 | 0.50904529 | 0.78082426 |
| IPMK | 58.6181757 | -1.171850519 | 0.3751381 | -3.1237843 | 0.00178541 | 0.0231855 |
| CISD1 | 83.7609127 | 0.313544786 | 0.2634754 | 1.19003441 | 0.23403287 | 0.54621338 |
| UBE2D1 | 39.0631245 | -0.229255289 | 0.42051477 | -0.5451777 | 0.58563128 | 0.8289701 |
| TFAM | 20.911792 | 0.595294744 | 0.46643204 | 1.27627327 | 0.20185894 | 0.5060272 |
| BICC1 | 2034.05542 | 0.957193336 | 0.47599424 | 2.0109347 | 0.04433235 | 0.21517692 |
| SLC16A9 | 14.4753494 | 2.648005088 | 0.90635607 | 2.92159471 | 0.00348244 | 0.03879367 |
| CCDC6 | 97.1997246 | -0.091139935 | 0.22074671 | -0.4128711 | 0.67970107 | 0.87237893 |
| ANK3 | 1215.47249 | 0.967003933 | 0.34754608 | 2.78237614 | 0.00539625 | 0.05276491 |

| | | | | | | |
|---|---|---|---|---|---|---|
| ARID5B | 880.636266 | -1.25136167 | 0.63467849 | -1.9716466 | 0.04864997 | 0.22624054 |
| NRBF2 | 22.1051817 | 0.856326515 | 0.50295193 | 1.70260112 | 0.08864274 | 0.32323227 |
| JMJD1C | 850.417105 | 0.158649111 | 0.28779982 | 0.55124812 | 0.58146359 | 0.8255928 |
| REEP3 | 152.807863 | -0.290944138 | 0.3497127 | -0.8319519 | 0.40543611 | 0.70462 |
| AC022387.1 | 27.4184633 | 1.994180634 | 0.57383182 | 3.47520052 | 0.00051047 | 0.00874933 |
| LINC01515 | 33.2064166 | 0.398061572 | 1.02840031 | 0.3870687 | 0.69870535 | 0.88302803 |
| CTNNA3 | 30.2730351 | 2.145656187 | 0.67491169 | 3.17916584 | 0.001477 | 0.02037205 |
| SIRT1 | 55.4316962 | 0.008260664 | 0.4402298 | 0.01876444 | 0.98502903 | 0.99742652 |
| HERC4 | 407.3922 | -0.577430363 | 0.38728413 | -1.4909735 | 0.13596846 | 0.41123731 |
| PBLD | 40.7719897 | -0.048677273 | 0.33936331 | -0.1434371 | 0.88594502 | 0.96249798 |
| HNRNPH3 | 133.661477 | 0.041557638 | 0.37122827 | 0.11194632 | 0.91086597 | 0.97128978 |
| RUFY2 | 100.007641 | 0.505717756 | 0.25964617 | 1.9477189 | 0.05144861 | 0.23455111 |
| DNA2 | 21.5532201 | -0.878253346 | 0.45100861 | -1.9473095 | 0.05149764 | 0.23455111 |
| SLC25A16 | 165.068704 | 0.093834831 | 0.35300559 | 0.26581684 | 0.79038027 | 0.92448611 |
| TET1 | 29.0997975 | 2.127719071 | 0.56765824 | 3.74823954 | 0.00017808 | 0.00383469 |
| CCAR1 | 220.042505 | -0.094024589 | 0.34963118 | -0.2689251 | 0.78798734 | 0.92377626 |
| STOX1 | 18.4230244 | 0.958289569 | 0.62215115 | 1.54028418 | 0.1234911 | 0.38860433 |
| DDX50 | 70.174577 | 0.284106758 | 0.27158151 | 1.04611968 | 0.29550578 | 0.61055359 |
| DDX21 | 46.8779426 | -1.276139355 | 0.43974838 | -2.9019763 | 0.00370817 | 0.0406764 |
| KIF1BP | 33.1726267 | 0.095414336 | 0.30141652 | 0.3165531 | 0.75158272 | 0.90896158 |
| VPS26A | 59.1179486 | 0.065920138 | 0.33121416 | 0.19902572 | 0.84224263 | 0.94776435 |
| SUPV3L1 | 40.619715 | 0.586324422 | 0.40340727 | 1.45343047 | 0.14610427 | 0.42860392 |
| HKDC1 | 179.035863 | -4.759102942 | 0.83447509 | -5.7031096 | 1.18E-08 | 1.22E-06 |
| HK1 | 163.492323 | -0.872979167 | 0.40105821 | -2.1766894 | 0.02950375 | 0.16784194 |
| H2AFY2 | 48.1563379 | 0.467037288 | 0.33166355 | 1.40816584 | 0.15908197 | 0.44758004 |
| AIFM2 | 16.8476279 | -0.467717929 | 0.37614605 | -1.2434477 | 0.21370291 | 0.52235998 |
| TYSND1 | 14.2544952 | 0.370008525 | 0.48693876 | 0.75986665 | 0.4473343 | 0.7399821 |
| SAR1A | 45.774189 | -0.011772702 | 0.34136017 | -0.0344876 | 0.97248831 | 0.99287068 |
| PPA1 | 55.2811282 | -1.340309284 | 0.33058855 | -4.0543125 | 5.03E-05 | 0.00142501 |
| EIF4EBP2 | 41.7555725 | -0.266413937 | 0.3627444 | -0.7344398 | 0.46268072 | 0.75091259 |
| SGPL1 | 76.6970143 | 0.019148258 | 0.29139583 | 0.06571219 | 0.94760697 | 0.98633762 |
| PCBD1 | 23.6311425 | -0.42446365 | 0.40504527 | -1.0479413 | 0.29466567 | 0.60986964 |
| UNC5B | 34.1430404 | -0.288624209 | 0.54287553 | -0.5316582 | 0.59496277 | 0.83416944 |
| PSAP | 142.341462 | -0.844562241 | 0.45708542 | -1.8477121 | 0.06464401 | 0.268828 |
| CHST3 | 33.110727 | -0.484553261 | 0.36727107 | -1.3193341 | 0.18705743 | 0.48725107 |
| ASCC1 | 64.4423779 | -0.15793354 | 0.26571989 | -0.594361 | 0.55227069 | 0.80943179 |
| ANAPC16 | 171.533894 | 0.010555669 | 0.26401116 | 0.03998191 | 0.96810755 | 0.99156233 |
| DDIT4 | 19.1081226 | -1.459879452 | 0.69502843 | -2.10046 | 0.03568839 | 0.18853177 |
| DNAJB12 | 23.865866 | -0.138115951 | 0.34569674 | -0.3995292 | 0.68950329 | 0.87665821 |
| MICU1 | 177.155545 | -0.050957465 | 0.25791837 | -0.1975721 | 0.84337989 | 0.94834468 |
| MCU | 155.440021 | 0.012094271 | 0.32956257 | 0.03669795 | 0.97072584 | 0.99178696 |
| P4HA1 | 187.791998 | -1.263147755 | 0.62101302 | -2.0340117 | 0.04195042 | 0.20900846 |
| ECD | 42.8969776 | 0.258620358 | 0.30081476 | 0.85973293 | 0.38993628 | 0.69292279 |
| FAM149B1 | 68.1539603 | 0.461547551 | 0.43293645 | 1.06608614 | 0.28638471 | 0.60056136 |
| DNAJC9 | 54.2662106 | -0.417214723 | 0.36507816 | -1.1428093 | 0.25311777 | 0.56559484 |
| MRPS16 | 24.8370138 | -0.496302036 | 0.31533676 | -1.5738794 | 0.11551533 | 0.3743642 |
| CFAP70 | 87.1373926 | 0.864906078 | 0.63260325 | 1.36721725 | 0.17155722 | 0.46730924 |
| ANXA7 | 99.4785797 | -0.108996192 | 0.27751941 | -0.3927516 | 0.69450296 | 0.87964032 |
| PPP3CB | 131.352037 | 0.323039133 | 0.19517789 | 1.65510109 | 0.09790399 | 0.34099994 |
| USP54 | 80.750521 | 0.078305854 | 0.28055687 | 0.27910866 | 0.78016143 | 0.9206566 |
| AGAP5 | 16.68439 | 0.410544304 | 0.4385542 | 0.93613128 | 0.3492056 | 0.65632016 |
| SEC24C | 35.5639334 | -0.041220031 | 0.32107203 | -0.1283825 | 0.89784628 | 0.96607503 |
| FUT11 | 24.2851769 | -1.258656642 | 0.60111946 | -2.0938544 | 0.03627295 | 0.19043299 |
| ZSWIM8 | 56.9429073 | -0.307298438 | 0.40762359 | -0.753878 | 0.45092251 | 0.74217149 |
| CAMK2G | 40.651622 | 0.418539442 | 0.38811848 | 1.0783806 | 0.28086394 | 0.59517622 |
| VCL | 177.969204 | -0.705000632 | 0.48190439 | -1.4629471 | 0.14348185 | 0.42447522 |
| AP3M1 | 24.077903 | -0.082698551 | 0.33609293 | -0.2460586 | 0.80563688 | 0.93308942 |
| ADK | 627.245172 | -0.042063049 | 0.11345252 | -0.3707547 | 0.71082027 | 0.88676588 |
| KAT6B | 242.390243 | 0.317784152 | 0.25684995 | 1.23723658 | 0.21599926 | 0.52525486 |
| SAMD8 | 50.7910864 | -0.024741746 | 0.40319569 | -0.0613641 | 0.95106923 | 0.98684281 |
| VDAC2 | 48.1927431 | 0.430591923 | 0.27472477 | 1.56735749 | 0.11703115 | 0.37679506 |
| COMTD1 | 16.8886874 | 0.614363244 | 0.57035368 | 1.07716188 | 0.28140795 | 0.59578741 |
| LRMDA | 1740.16663 | 0.726285331 | 0.37835398 | 1.91959215 | 0.05490944 | 0.24363853 |
| KCNMA1 | 75.0645676 | -5.761264024 | 1.60979802 | -3.5788738 | 0.00034508 | 0.00653965 |
| DLG5 | 128.71518 | -0.530243088 | 0.44907178 | -1.1807525 | 0.23770066 | 0.54997949 |
| POLR3A | 45.7556165 | -0.220120373 | 0.35739148 | -0.6159083 | 0.53795506 | 0.80104055 |
| RPS24 | 117.783366 | -0.448573559 | 0.42254086 | -1.0616099 | 0.28841282 | 0.60295093 |
| ZMIZ1-AS1 | 25.8535399 | -2.923919424 | 0.94289929 | -3.100988 | 0.00192876 | 0.02456799 |
| ZMIZ1 | 273.385023 | -0.591602503 | 0.36224305 | -1.6331645 | 0.1024344 | 0.34985287 |
| PPIF | 20.0116161 | -0.254915561 | 0.47572235 | -0.5358494 | 0.59206261 | 0.83288952 |
| NUTM2B-AS | 210.148196 | -0.498050907 | 0.35081057 | -1.4197147 | 0.15569077 | 0.44123426 |
| TMEM254 | 18.5412442 | 0.174228234 | 0.525781 | 0.33137035 | 0.74036476 | 0.90308228 |
| ANXA11 | 220.448622 | -0.298919713 | 0.22943295 | -1.302863 | 0.19262152 | 0.49370242 |
| FAM213A | 42.7586622 | -0.124282807 | 0.29160584 | -0.4262014 | 0.66996113 | 0.86777458 |
| TSPAN14 | 54.9516728 | -0.853483265 | 0.37076336 | -2.3019623 | 0.0213373 | 0.1369229 |
| NRG3 | 116.972105 | -1.562371099 | 0.61217003 | -2.5521849 | 0.01070497 | 0.08649319 |
| GHITM | 46.2706368 | -0.08157767 | 0.42677676 | -0.1911483 | 0.84840937 | 0.95003258 |
| CCSER2 | 195.108066 | 0.381328506 | 0.25184093 | 1.51416417 | 0.12998421 | 0.40038743 |
| WAPL | 161.604325 | 0.142508463 | 0.25332038 | 0.56256218 | 0.57373305 | 0.82110314 |
| BMPR1A | 277.318673 | 0.328641909 | 0.29616767 | 1.10964815 | 0.26715068 | 0.58335197 |
| GLUD1 | 115.479062 | 0.272073733 | 0.30084164 | 0.90437524 | 0.36579646 | 0.67012584 |
| SHLD2 | 196.323806 | 0.695822614 | 0.20669783 | 3.36637605 | 0.00076163 | 0.01213918 |
| NUTM2A-AS | 228.281409 | 0.106761537 | 0.25513616 | 0.41844926 | 0.67561867 | 0.86960234 |
| MINPP1 | 16.5290042 | -0.033856568 | 0.44025765 | -0.0769017 | 0.93870173 | 0.98331142 |
| ATAD1 | 106.936765 | 0.501831784 | 0.3209414 | 1.56362433 | 0.11790582 | 0.37862775 |
| PTEN | 281.275944 | 0.208465893 | 0.22281398 | 0.93560511 | 0.34947654 | 0.65669474 |
| RNLS | 89.7520411 | 0.210782967 | 0.42104518 | 0.50061841 | 0.61663971 | 0.84223226 |
| LIPA | 47.9891505 | 0.255506983 | 0.42575026 | 0.60013347 | 0.54841729 | 0.80647995 |
| IFIT2 | 23.9538407 | -0.755565014 | 1.13020939 | -0.6685177 | 0.50380317 | 0.77790293 |
| IFIT3 | 49.0434213 | -1.107326734 | 1.60016894 | -0.6920061 | 0.48893347 | 0.77060666 |
| IFIT1 | 37.4763867 | 2.619700369 | 1.06759889 | 2.45382455 | 0.01413459 | 0.10434754 |
| SLC16A12 | 23.6925026 | 0.184888298 | 0.60815846 | 0.30401336 | 0.76111771 | 0.91320552 |
| PANK1 | 52.6187765 | 1.560564342 | 0.51240801 | 3.04555026 | 0.00232255 | 0.02830778 |
| RPP30 | 17.2418613 | 0.606357054 | 0.42474881 | 1.42756623 | 0.15341675 | 0.43892536 |
| PCGF5 | 234.774313 | -0.401382831 | 0.37212716 | -1.0786174 | 0.28075833 | 0.5950926 |
| HECTD2 | 100.334978 | 0.385781845 | 0.31400829 | 1.22857217 | 0.21923226 | 0.52935683 |
| TNKS2 | 174.104567 | -0.289099615 | 0.29043108 | -0.9954156 | 0.3195342 | 0.63148301 |
| BTAF1 | 166.381016 | 0.054899484 | 0.43271514 | 0.12687211 | 0.89904162 | 0.96637793 |
| CPEB3 | 409.809976 | 0.942460371 | 0.37618559 | 2.50530694 | 0.01223452 | 0.09498304 |
| 5-Mar | 83.2655258 | -0.329946757 | 0.31669609 | -1.0418403 | 0.29748572 | 0.61141484 |
| IDE | 67.6473093 | -0.082336119 | 0.34585413 | -0.238066 | 0.81182988 | 0.93552094 |
| EXOC6 | 113.8439 | 0.331692256 | 0.27782048 | 1.19390856 | 0.23251374 | 0.54506955 |
| MYOF | 262.229569 | -1.897292361 | 0.44900708 | -4.2255288 | 2.38E-05 | 0.00078818 |
| FRA10AC1 | 56.9407044 | 0.660058091 | 0.27433014 | 2.40607204 | 0.01612508 | 0.11364345 |
| NOC3L | 21.7993688 | -0.407500052 | 0.3962745 | -1.0283277 | 0.3037957 | 0.61793 |
| TBC1D12 | 60.4342771 | -0.256654972 | 0.27204317 | -0.9434347 | 0.3454586 | 0.65406518 |
| HELLS | 19.0982423 | -0.375913731 | 0.68670544 | -0.5474163 | 0.58409278 | 0.8277977 |
| SORBS1 | 62.1029194 | -2.752945293 | 0.71703904 | -3.8393242 | 0.00012337 | 0.00284736 |
| ALDH18A1 | 53.3808117 | -0.924431944 | 0.42345801 | -2.1830546 | 0.02903179 | 0.16620962 |
| TCTN3 | 57.1724147 | 0.096849041 | 0.27409682 | 0.3533388 | 0.72383446 | 0.89280599 |
| ENTPD1-AS1 | 131.187357 | 0.098595178 | 0.57998421 | 0.16999631 | 0.86501304 | 0.95516124 |
| ZNF518A | 237.577762 | 0.246884462 | 0.30513948 | 0.80908724 | 0.41846497 | 0.71684133 |
| BLNK | 478.039874 | -0.749590428 | 0.50984159 | -1.4702418 | 0.14149627 | 0.42163989 |

| | | | | | | |
|---|---|---|---|---|---|---|
| TM9SF3 | 169.482756 | -0.231746953 | 0.2416389 | -0.9590631 | 0.33752696 | 0.64844536 |
| LCOR | 317.131286 | -0.043980665 | 0.27362473 | -0.1607335 | 0.87230329 | 0.95690422 |
| RRP12 | 26.2264033 | -0.281864381 | 0.46459848 | -0.6066838 | 0.54406076 | 0.80380646 |
| PGAM1 | 53.6304878 | -1.053545591 | 0.44714233 | -2.356175 | 0.01846421 | 0.12417062 |
| EXOSC1 | 28.6723134 | 0.20958274 | 0.35800114 | 0.58542479 | 0.55826213 | 0.81255224 |
| ZDHHC16 | 17.5223935 | -0.260901082 | 0.35718082 | -0.7304454 | 0.46511798 | 0.75251779 |
| MMS19 | 71.1773175 | 0.034441713 | 0.22019783 | 0.15641259 | 0.87570782 | 0.95845753 |
| UBTD1 | 35.5340921 | -0.411424452 | 0.41175541 | -0.9991962 | 0.31769964 | 0.63093651 |
| AL355315.1 | 13.3635749 | 0.004158643 | 0.63612094 | 0.0065375 | 0.99478386 | 0.9992177 |
| MORN4 | 28.5590356 | 0.840229299 | 0.4674433 | 1.79749994 | 0.07225629 | 0.28717717 |
| ZFYVE27 | 35.0207925 | -0.489097662 | 0.31943264 | -1.5311449 | 0.12573358 | 0.39231467 |
| R3HCC1L | 119.039024 | -0.38875396 | 0.18084019 | -2.1497099 | 0.03157817 | 0.17567711 |
| HPS1 | 18.6719506 | -1.162814664 | 0.42060528 | -2.7646221 | 0.00569888 | 0.0544988 |
| GOT1 | 128.383569 | 0.726635067 | 0.2778172 | 2.61551509 | 0.0089093 | 0.07586329 |
| AL391684.1 | 11.4579655 | 0.76242566 | 0.69538145 | 1.09641357 | 0.27289783 | 0.58873191 |
| SLC25A28 | 28.8711928 | -0.171229687 | 0.52225924 | -0.3278634 | 0.74301495 | 0.90463949 |
| CUTC | 33.2684904 | -0.124293949 | 0.33945884 | -0.3661532 | 0.71425074 | 0.88883875 |
| COX15 | 33.1036216 | 0.328388236 | 0.32536371 | 1.00929584 | 0.31283278 | 0.62758402 |
| DNMBP | 85.5824083 | -0.575374329 | 0.52396925 | -1.0981071 | 0.27215774 | 0.58786226 |
| ERLIN1 | 27.6389965 | -0.186044819 | 0.41727877 | -0.4458526 | 0.65570374 | 0.86268564 |
| CHUK | 29.7481959 | -0.257005376 | 0.40860682 | -0.6289797 | 0.52936237 | 0.79493876 |
| CWF19L1 | 41.1906564 | 0.399636465 | 0.44179331 | 0.90457791 | 0.36568904 | 0.67012584 |
| BLOC1S2 | 16.141538 | -0.539667606 | 0.42989272 | -1.2553541 | 0.20935023 | 0.51790437 |
| SEC31B | 18.9355722 | 1.072382003 | 0.42196019 | 2.54142933 | 0.01104003 | 0.08807986 |
| HIF1AN | 33.7728612 | -0.3092524 | 0.32466188 | -0.9525368 | 0.34082479 | 0.65083559 |
| PAX2 | 154.34314 | -0.171335166 | 0.45296715 | -0.3782508 | 0.70524433 | 0.88596489 |
| SLF2 | 80.7819701 | -0.217546315 | 0.28053127 | -0.7754797 | 0.43805623 | 0.7305377 |
| MRPL43 | 17.4841733 | -0.717848461 | 0.43652908 | -1.6444459 | 0.10008413 | 0.34555815 |
| LZTS2 | 25.9642899 | -0.04462651 | 0.36325783 | -0.1228508 | 0.90222526 | 0.96803829 |
| BTRC | 192.704325 | 0.458968935 | 0.25149947 | 1.82493005 | 0.06801159 | 0.27788783 |
| DPCD | 15.0483672 | -0.75229421 | 0.49313538 | -1.5255328 | 0.12712625 | 0.39511746 |
| POLL | 20.9349423 | 0.15646909 | 0.43716998 | 0.35791362 | 0.72040796 | 0.89151237 |
| FBXW4 | 137.577511 | -0.282339251 | 0.22053929 | -1.280222 | 0.20046707 | 0.50442784 |
| OGA | 292.540385 | -0.131679139 | 0.20889194 | -0.6303697 | 0.52845275 | 0.7943404 |
| ARMH3 | 174.766961 | -0.005585261 | 0.19781429 | -0.0282349 | 0.97747483 | 0.99497547 |
| LDB1 | 32.2436563 | 0.645049871 | 0.48136844 | 1.34003357 | 0.18023443 | 0.47864023 |
| PPRC1 | 20.4173213 | -1.030614371 | 0.75030639 | -1.3735914 | 0.16956858 | 0.46392531 |
| GBF1 | 228.104856 | -0.037480903 | 0.26354766 | -0.1422168 | 0.88690877 | 0.96253908 |
| NFKB2 | 25.7753947 | -2.03166069 | 0.74730295 | -2.7186574 | 0.00655475 | 0.06033213 |
| FBXL15 | 16.5054209 | -0.239755195 | 0.68186476 | -0.3516169 | 0.72512557 | 0.89374117 |
| CUEDC2 | 14.5992481 | -0.523641591 | 0.38752903 | -1.3512319 | 0.17662116 | 0.47378082 |
| ACTR1A | 25.396079 | -0.54012699 | 0.32708607 | -1.6513298 | 0.09867124 | 0.34303156 |
| SUFU | 77.2166577 | 0.421745018 | 0.41856008 | 1.00760927 | 0.31364208 | 0.62876773 |
| TRIM8 | 40.8974328 | 0.240882275 | 0.43356215 | 0.5555888 | 0.57849199 | 0.82318213 |
| ARL3 | 38.8840338 | 0.130890882 | 0.38941061 | 0.33612562 | 0.73677614 | 0.90141723 |
| SFXN2 | 18.1962958 | 0.721805532 | 0.42047325 | 1.71665031 | 0.08604308 | 0.3168629 |
| WBP1L | 88.2623753 | -0.046384327 | 0.36481382 | -0.1271452 | 0.89882548 | 0.96637793 |
| CNNM2 | 1090.24453 | 3.385480495 | 0.27061574 | 12.5102867 | 6.56E-36 | 3.06E-32 |
| NT5C2 | 311.050821 | -1.051219411 | 0.46601628 | -2.2557568 | 0.02408586 | 0.14717838 |
| PCGF6 | 28.6590012 | 0.255350112 | 0.4344236 | 0.5877906 | 0.55667285 | 0.81163685 |
| ATP5MD | 37.2539223 | 0.094000328 | 0.41216467 | 0.22806498 | 0.81959572 | 0.9384641 |
| PDCD11 | 23.1256959 | -0.082193958 | 0.44076581 | -0.1864799 | 0.85206845 | 0.95090866 |
| SH3PXD2A | 135.329949 | 1.735559955 | 0.39132391 | 4.43509821 | 9.20E-06 | 0.00036206 |
| STN1 | 43.6321138 | 0.365396417 | 0.38389618 | 0.95181051 | 0.34119309 | 0.65083559 |
| SLK | 116.66724 | 0.074230894 | 0.36481107 | 0.20347763 | 0.83876173 | 0.94741614 |
| CFAP43 | 24.4561493 | -0.644247119 | 0.80076542 | -0.8045391 | 0.42108568 | 0.7192477 |
| GSTO1 | 38.0046567 | -0.880799301 | 0.56032261 | -1.5719503 | 0.11596207 | 0.37462127 |
| GSTO2 | 101.77099 | 0.597006361 | 0.40923998 | 1.45881728 | 0.1446154 | 0.42644619 |
| SORCS3 | 32.2415075 | -5.030159436 | 0.84934593 | -5.9223919 | 3.17E-09 | 4.23E-07 |
| XPNPEP1 | 49.6463698 | -0.143641788 | 0.36071301 | -0.3982163 | 0.69047077 | 0.87710483 |
| ADD3 | 248.73628 | 1.069242387 | 0.34288728 | 3.1183495 | 0.00181867 | 0.02351911 |
| MXI1 | 82.9418962 | -0.695844511 | 0.43855024 | -1.5866928 | 0.11258224 | 0.36994129 |
| SMNDC1 | 26.3750912 | -0.534137203 | 0.45283097 | -1.1795509 | 0.23817886 | 0.54997949 |
| SMC3 | 47.431159 | -0.027400685 | 0.34352412 | -0.0797635 | 0.93642536 | 0.98203015 |
| PDCD4 | 85.9807266 | 0.108788898 | 0.45827072 | 0.23739003 | 0.81235423 | 0.93568756 |
| BBIP1 | 44.4068805 | -0.208492755 | 0.3210359 | -0.6494375 | 0.51605563 | 0.78456789 |
| SHOC2 | 116.79919 | 0.077651421 | 0.28758736 | 0.27000986 | 0.78715267 | 0.92362588 |
| GPAM | 20.9873216 | 0.925769988 | 0.45522273 | 2.03366379 | 0.04198551 | 0.20900846 |
| ZDHHC6 | 28.9255887 | -0.314260234 | 0.28106232 | -1.1181159 | 0.26351752 | 0.57771863 |
| VTI1A | 443.245277 | -0.016125432 | 0.12606496 | -0.1279137 | 0.89821729 | 0.96607503 |
| TCF7L2 | 284.76113 | -0.587041691 | 0.22753922 | -2.5799583 | 0.00988122 | 0.08167778 |
| CASP7 | 27.584089 | -0.54643177 | 0.45377644 | -1.2041872 | 0.22851724 | 0.54092276 |
| NHLRC2 | 93.2448283 | 0.277361758 | 0.22647111 | 1.22471143 | 0.22068397 | 0.53073441 |
| CCDC186 | 69.7428332 | 0.312548527 | 0.29790156 | 1.04916714 | 0.2941012 | 0.60910697 |
| ABLIM1 | 857.584623 | -0.823489752 | 0.34864164 | -2.3619948 | 0.0181769 | 0.12316961 |
| FAM160B1 | 52.2373736 | -0.315635042 | 0.31839348 | -0.9913364 | 0.32152135 | 0.63312885 |
| TRUB1 | 28.2351057 | 0.542725156 | 0.37600858 | 1.44338501 | 0.14891204 | 0.43272656 |
| HSPA12A | 105.081151 | 0.01750983 | 0.50146005 | 0.0349177 | 0.97214537 | 0.99280213 |
| SHTN1 | 111.442431 | -0.757083618 | 0.28039619 | -2.7000495 | 0.00693292 | 0.06312744 |
| PDZD8 | 150.921652 | -0.868976106 | 0.45826339 | -1.8962372 | 0.05792868 | 0.25098839 |
| EMX2OS | 176.590803 | -0.791073371 | 0.4822483 | -1.640386 | 0.10092493 | 0.34704236 |
| EMX2 | 51.4154563 | -0.75737567 | 0.53197324 | -1.4237101 | 0.15453041 | 0.44013998 |
| RAB11FIP2 | 102.216526 | -0.215422968 | 0.42702707 | -0.5044715 | 0.61393011 | 0.84174925 |
| CASC2 | 89.6857721 | 0.317722558 | 0.76541268 | 0.41509968 | 0.67806894 | 0.87130138 |
| FAM204A | 62.0970121 | 0.483925454 | 0.3337068 | 1.45015161 | 0.14701625 | 0.4297124 |
| CACUL1 | 129.218722 | -0.376699464 | 0.29718718 | -1.2675495 | 0.20495888 | 0.51143678 |
| EIF3A | 135.075534 | -0.277125624 | 0.36723472 | -0.7546281 | 0.45047218 | 0.74156119 |
| FAM45A | 24.8327073 | 0.181779002 | 0.42838745 | 0.42433316 | 0.67132286 | 0.86777458 |
| SFXN4 | 38.1544037 | 1.01912973 | 0.4136164 | 2.46394904 | 0.01374156 | 0.10241914 |
| PRDX3 | 34.0857763 | -0.263674112 | 0.46548958 | -0.5664447 | 0.57109151 | 0.81895612 |
| TIAL1 | 85.3222446 | 0.529765819 | 0.23028511 | 2.30047796 | 0.02142116 | 0.13717779 |
| BAG3 | 23.6493737 | 0.125103581 | 0.37240844 | 0.33593111 | 0.73692282 | 0.9014784 |
| INPP5F | 46.1708808 | -0.187324651 | 0.38614939 | -0.4851093 | 0.62759883 | 0.84783129 |
| MCMBP | 43.2900785 | 0.078300496 | 0.32388813 | 0.24175167 | 0.80897259 | 0.93433178 |
| SEC23IP | 80.0378808 | 0.174237538 | 0.3991592 | 0.43651139 | 0.66246573 | 0.86546595 |
| PLPP4 | 14.7027932 | 2.287970672 | 0.47494341 | 4.81735427 | 1.45E-06 | 7.71E-05 |
| WDR11-AS1 | 14.6733784 | 0.423700656 | 0.64524275 | 0.6566531 | 0.511404 | 0.78246322 |
| WDR11 | 113.644424 | 0.495105303 | 0.29849854 | 1.65865237 | 0.09718586 | 0.34011144 |
| FGFR2 | 152.293144 | 0.092423447 | 0.42289877 | 0.21854745 | 0.82700259 | 0.94243121 |
| ATE1 | 170.244291 | 0.141357743 | 0.24663424 | 0.57314728 | 0.56654498 | 0.81595078 |
| NSMCE4A | 19.8147548 | -0.175449271 | 0.56273025 | -0.3117822 | 0.75520606 | 0.91058338 |
| TACC2 | 239.567642 | 1.001326704 | 0.73599667 | 1.36050439 | 0.17367036 | 0.46960235 |
| PLEKHA1 | 560.279109 | -2.951350777 | 0.33499547 | -8.8101214 | 1.25E-18 | 1.94E-15 |
| HTRA1 | 17.8155055 | -2.168724883 | 0.71765221 | -3.021972 | 0.00251134 | 0.03021382 |
| C10orf88 | 23.8756393 | 0.019891798 | 0.41701094 | 0.0477009 | 0.96195462 | 0.99029318 |
| PSTK | 19.5446301 | -0.027559906 | 0.48645149 | -0.056655 | 0.95482003 | 0.98842477 |
| IKZF5 | 25.4304539 | 0.023277635 | 0.3053701 | 0.07622762 | 0.93923801 | 0.98358617 |
| ACADSB | 63.2270431 | -0.359873626 | 0.2792233 | -1.2888381 | 0.19745437 | 0.49974608 |
| BUB3 | 27.9923862 | 0.12860386 | 0.36955739 | 0.34799429 | 0.72784448 | 0.89601556 |
| CPXM2 | 495.176793 | -1.325060117 | 0.62618621 | -2.11608 | 0.034338 | 0.18432213 |
| OAT | 17.0637578 | -0.735323349 | 0.5087659 | -1.4453078 | 0.14837144 | 0.43218222 |
| LHPP | 31.6082045 | 0.588810287 | 0.5027821 | 1.17110432 | 0.24155685 | 0.55406545 |
| ABRAXAS2 | 30.0720857 | -0.17887347 | 0.33036094 | -0.5414486 | 0.58819841 | 0.83024609 |

| | | | | | | |
|---|---|---|---|---|---|---|
| ZRANB1 | 73.2797382 | 0.046113658 | 0.33598857 | 0.1372477 | 0.89083501 | 0.96328012 |
| CTBP2 | 480.403571 | -0.218729802 | 0.39101597 | -0.5593884 | 0.57589667 | 0.82147339 |
| EDRF1 | 52.3838258 | -0.125828843 | 0.30888292 | -0.4073674 | 0.68373814 | 0.87343121 |
| UROS | 21.5862436 | 0.334716278 | 0.41658793 | 0.80347091 | 0.42170261 | 0.71947943 |
| BCCIP | 44.2161105 | -0.108215081 | 0.25613953 | -0.4224849 | 0.67267113 | 0.86833526 |
| DHX32 | 134.476213 | -0.776418032 | 0.267198 | -2.9057779 | 0.00366341 | 0.04033395 |
| FANK1 | 53.6211974 | -1.322946218 | 0.48131494 | -2.7486083 | 0.00598489 | 0.05665288 |
| DOCK1 | 681.432307 | -0.417547074 | 0.23679779 | -1.7633065 | 0.07784882 | 0.29864393 |
| FAM196A | 19.0283625 | -0.695329803 | 0.61200955 | -1.1361421 | 0.25589711 | 0.56876868 |
| MGMT | 96.5134888 | 0.161698875 | 0.46678815 | 0.34640741 | 0.72903656 | 0.89677267 |
| AL157832.3 | 13.7193165 | 0.194084119 | 0.50018778 | 0.38802251 | 0.69799938 | 0.88258901 |
| C10orf143 | 15.8232623 | 0.11506248 | 0.53734321 | 0.21413219 | 0.83044397 | 0.94393022 |
| GLRX3 | 77.2542369 | -0.658534429 | 0.50044645 | -1.3158939 | 0.18820965 | 0.4882768 |
| PPP2R2D | 136.208861 | -0.380578754 | 0.38168592 | -0.9970993 | 0.31871632 | 0.63093651 |
| BNIP3 | 74.6310502 | -1.687420114 | 0.69320202 | -2.43424 | 0.01492309 | 0.10793754 |
| STK32C | 23.7847719 | -0.309147078 | 0.34131569 | -0.9057511 | 0.36506758 | 0.67012584 |
| LRRC27 | 22.1789375 | 0.101576251 | 0.60565902 | 0.16771194 | 0.8668099 | 0.95578708 |
| INPP5A | 95.5152459 | 0.387990322 | 0.3909044 | 0.99254529 | 0.3209316 | 0.63269097 |
| KNDC1 | 22.7329088 | -0.005957729 | 0.59578138 | -0.0099999 | 0.9920214 | 0.9988777 |
| TUBGCP2 | 61.9433866 | -0.15022753 | 0.3132941 | -0.4795096 | 0.63157614 | 0.85110795 |
| ECHS1 | 36.6849465 | -0.292810615 | 0.37750644 | -0.775644 | 0.4379592 | 0.7305377 |
| SCART1 | 22.8200512 | 1.124132453 | 0.49262159 | 2.28193906 | 0.02249294 | 0.14110406 |
| SLC6A12 | 50.5499416 | -0.798053707 | 0.63797059 | -1.2509255 | 0.21096165 | 0.5199594 |
| KDM5A | 160.313196 | 0.091642627 | 0.17541659 | 0.52242852 | 0.60137201 | 0.83679851 |
| B4GALNT3 | 178.252741 | 0.719953685 | 0.40899742 | 1.76028905 | 0.07835881 | 0.29955618 |
| WNK1 | 1504.36638 | 1.827750424 | 0.65635813 | 2.78468466 | 0.00535798 | 0.05253187 |
| RAD52 | 42.3917638 | 0.322417366 | 0.34325261 | 0.93930055 | 0.34757645 | 0.6553345 |
| ERC1 | 343.302817 | -0.218960859 | 0.12263143 | -1.7855199 | 0.07417703 | 0.29129403 |
| ADIPOR2 | 120.162512 | -0.152892035 | 0.38832835 | -0.3937184 | 0.69378892 | 0.87904442 |
| DCP1B | 49.3851011 | 0.325404713 | 0.37388192 | 0.87034086 | 0.38411416 | 0.68808461 |
| CACNA1C | 32.2948827 | 1.454079343 | 0.68336928 | 2.12780907 | 0.03335292 | 0.18186117 |
| FKBP4 | 22.6528531 | 0.218503774 | 0.50441898 | 0.43317913 | 0.66488464 | 0.86668312 |
| ITFG2 | 68.3893374 | 0.045883399 | 0.35896022 | 0.12782307 | 0.89828899 | 0.96607503 |
| TULP3 | 121.098976 | -0.761987288 | 0.29628757 | -2.5717828 | 0.01011763 | 0.08292923 |
| TEAD4 | 56.8291044 | -2.986999029 | 0.72082602 | -4.1438557 | 3.42E-05 | 0.00106143 |
| TSPAN9 | 473.822312 | 0.049109113 | 0.35558912 | 0.13810634 | 0.89015637 | 0.96323092 |
| PARP11 | 24.9508791 | 0.265960929 | 0.36508501 | 0.72849042 | 0.46631343 | 0.75300195 |
| TIGAR | 20.9187173 | -1.02698408 | 0.58840333 | -1.7453744 | 0.08091973 | 0.30571131 |
| C12orf4 | 34.9293436 | 0.52338519 | 0.37353134 | 1.40118146 | 0.16115982 | 0.45084132 |
| CD9 | 70.0388171 | -0.741495911 | 0.42808063 | -1.7321408 | 0.0832485 | 0.31073219 |
| TNFRSF1A | 42.5921509 | -1.893987068 | 0.32261418 | -5.8707496 | 4.34E-09 | 5.47E-07 |
| SCNN1A | 197.615105 | 0.213691563 | 0.35696213 | 0.59863932 | 0.54941343 | 0.80698344 |
| LTBR | 22.7621859 | -1.178304578 | 0.456447 | -2.5814707 | 0.00983803 | 0.08139292 |
| TAPBPL | 11.8907027 | 0.176904046 | 0.48498741 | 0.36476008 | 0.71529051 | 0.88904248 |
| MRPL51 | 18.9414483 | -0.761474932 | 0.51135184 | -1.4891409 | 0.13645027 | 0.41160218 |
| NCAPD2 | 21.9363582 | 0.015081073 | 0.49546923 | 0.03043796 | 0.97571777 | 0.99405381 |
| GAPDH | 167.543287 | -1.386584812 | 0.52781356 | -2.6270352 | 0.00861324 | 0.07374644 |
| CHD4 | 33.1772899 | -0.022135566 | 0.313377 | -0.0706356 | 0.94368779 | 0.98543454 |
| ING4 | 26.9827554 | -0.334482176 | 0.35526501 | -0.9415005 | 0.34644845 | 0.65471653 |
| ZNF384 | 23.3589165 | -0.145535542 | 0.40629427 | -0.3582023 | 0.72019193 | 0.89142036 |
| COPS7A | 18.519654 | 0.095152683 | 0.45747792 | 0.20799405 | 0.83523362 | 0.94654164 |
| MLF2 | 31.3164955 | -0.139875233 | 0.36842939 | -0.3796528 | 0.70420319 | 0.8856427 |
| PTMS | 47.7035631 | -0.741837104 | 0.59499899 | -1.2467872 | 0.21247555 | 0.52134789 |
| USP5 | 14.8196207 | -0.316010251 | 0.50870281 | -0.621208 | 0.53446278 | 0.79873873 |
| TPI1 | 66.9947133 | -0.455636464 | 0.51745016 | -0.8805417 | 0.3785659 | 0.682077 |
| LRRC23 | 23.687885 | -0.885888616 | 0.41790381 | -2.1198386 | 0.03401966 | 0.18367181 |
| ATN1 | 48.7428391 | -0.50713856 | 0.31787262 | -1.5954144 | 0.11061959 | 0.36601032 |
| PTPN6 | 20.9763916 | -0.441546858 | 0.44002078 | -1.0034682 | 0.31563501 | 0.63009595 |
| PHB2 | 30.0547744 | -0.274869641 | 0.37239691 | -0.7381094 | 0.460448 | 0.74872989 |
| LPCAT3 | 111.674944 | -0.289657996 | 0.20915731 | -1.384881 | 0.16608885 | 0.45844062 |
| C1S | 38.2188803 | -0.293220385 | 0.3857188 | -0.7601921 | 0.44713976 | 0.73986356 |
| C1RL | 66.1154964 | -1.366309908 | 0.31713577 | -4.3082806 | 1.65E-05 | 0.00059002 |
| CLSTN3 | 26.3064277 | -0.548145128 | 0.58151273 | -0.9426193 | 0.34587566 | 0.65414698 |
| PEX5 | 17.3925099 | -0.130914585 | 0.42267837 | -0.3097262 | 0.75676914 | 0.91122251 |
| FOXJ2 | 15.0116379 | 0.721240295 | 0.41739194 | 1.7279689 | 0.0839938 | 0.312638 |
| NECAP1 | 20.2581426 | 0.435726177 | 0.39293383 | 1.10890471 | 0.26747129 | 0.58364201 |
| FAM66C | 18.1447139 | 0.433013982 | 0.48404195 | 0.89457945 | 0.37101194 | 0.67522591 |
| RIMKLB | 653.607188 | -2.258991535 | 1.06265015 | -2.1258093 | 0.03351914 | 0.18244746 |
| A2ML1-AS1 | 54.6689218 | -1.185696933 | 0.86560517 | -1.3697896 | 0.1707526 | 0.46658085 |
| PHC1 | 32.726422 | 0.256756675 | 0.35258128 | 0.72821983 | 0.46647903 | 0.75300195 |
| M6PR | 36.8992452 | -0.978678579 | 0.4274097 | -2.2897903 | 0.02203348 | 0.13928144 |
| A2M | 21.4140506 | -0.580099119 | 0.60925457 | -0.9521457 | 0.3410231 | 0.65083559 |
| CLEC2D | 40.3364908 | -0.217477599 | 0.41051603 | -0.5297664 | 0.59627391 | 0.83416944 |
| OLR1 | 63.9081348 | 1.184198771 | 0.79909176 | 1.4819309 | 0.13835868 | 0.41527892 |
| TMEM52B | 488.914749 | 0.957542601 | 0.72758533 | 1.31605539 | 0.18815545 | 0.4882768 |
| GABARAPL1 | 183.2639 | 0.286602298 | 0.27912397 | 1.02679212 | 0.30451836 | 0.61818174 |
| KLRD1 | 152.196988 | -0.540094569 | 0.35709263 | -1.5124775 | 0.13041244 | 0.40117636 |
| YBX3 | 217.030262 | -1.460372721 | 0.32945391 | -4.4327073 | 9.31E-06 | 0.00036304 |
| PRH1 | 14.4455274 | -0.316635414 | 0.72966089 | -0.4339487 | 0.66432567 | 0.8661967 |
| AC244131.2 | 35.7182553 | -0.525549018 | 0.59533838 | -0.8827736 | 0.37735859 | 0.68055929 |
| ETV6 | 351.091836 | -1.543356365 | 0.42432098 | -3.6372379 | 0.00027558 | 0.00543231 |
| BCL2L14 | 49.5836612 | -0.240523782 | 0.60099344 | -0.4002103 | 0.68900161 | 0.87631306 |
| LRP6 | 231.318126 | 0.834282025 | 0.31077318 | 2.68453679 | 0.00726304 | 0.06522614 |
| BORCS5 | 62.0065214 | 0.372732294 | 0.29031465 | 1.28389077 | 0.19918018 | 0.50272636 |
| DUSP16 | 91.671506 | -0.735798191 | 0.41164929 | -1.7874395 | 0.07386649 | 0.29109516 |
| CREBL2 | 31.807494 | -0.36739488 | 0.28822298 | -1.2746898 | 0.20241907 | 0.50680865 |
| CDKN1B | 16.3719226 | 0.202779527 | 0.42355559 | 0.4787554 | 0.63211265 | 0.85146176 |
| APOLD1 | 14.0042572 | 1.182696776 | 0.66484044 | 1.77891823 | 0.07525317 | 0.29345903 |
| GPRC5D-AS1 | 61.5386886 | 0.664446382 | 0.4457772 | 1.4905347 | 0.1360837 | 0.41123731 |
| HEBP1 | 22.9528958 | 0.117679744 | 0.37456561 | 0.31417659 | 0.75338692 | 0.91007188 |
| EMP1 | 38.9825218 | -4.229287827 | 0.73056424 | -5.7890704 | 7.08E-09 | 8.05E-07 |
| ATF7IP | 230.11895 | -0.009998681 | 0.24239958 | -0.0412488 | 0.96709759 | 0.99156233 |
| AC010168.1 | 16.5390459 | -0.076161305 | 0.68603753 | -0.1110162 | 0.91160347 | 0.97128978 |
| H2AFJ | 24.0525495 | -1.277829395 | 0.44908786 | -2.8453884 | 0.00443573 | 0.04598205 |
| WBP11 | 28.1621055 | 0.058775025 | 0.29131571 | 0.20175714 | 0.84010659 | 0.9474978 |
| C12orf60 | 39.2232692 | 0.475853543 | 0.40309214 | 1.18050813 | 0.23779818 | 0.54997949 |
| EPS8 | 96.4656315 | -0.872886318 | 0.61348348 | -1.4228359 | 0.15478373 | 0.44032584 |
| STRAP | 47.9840613 | -0.069873349 | 0.25782663 | -0.271009 | 0.78638408 | 0.92322402 |
| DERA | 130.915758 | -0.388773602 | 0.28318382 | -1.3728666 | 0.16979383 | 0.46436188 |
| MGST1 | 28.367727 | -2.5517987 | 0.69641343 | -3.6642009 | 0.00024811 | 0.00504008 |
| PIK3C2G | 418.728206 | 1.986814758 | 0.91435073 | 2.17292412 | 0.02978603 | 0.16882975 |
| AC087242.1 | 15.8596232 | 1.582501926 | 0.88717388 | 1.78375622 | 0.07446329 | 0.29200098 |
| PLEKHA5 | 1273.01487 | -0.139722587 | 0.31720951 | -0.4404741 | 0.65959374 | 0.86460873 |
| AEBP2 | 399.099943 | 0.684552925 | 0.33478373 | 2.04476165 | 0.04087836 | 0.20496314 |
| AC024901.1 | 54.0405505 | 0.653555649 | 0.43720137 | 1.49486186 | 0.1349505 | 0.4097351 |
| LINC02398 | 24.7411811 | -0.166874719 | 0.72588654 | -0.2298909 | 0.81817654 | 0.9380844 |
| PYROXD1 | 32.8704694 | -0.049937577 | 0.34103124 | -0.1464311 | 0.88358108 | 0.96170562 |
| RECQL | 21.3505657 | -0.603711037 | 0.37045241 | -1.6296588 | 0.10317363 | 0.35088341 |
| GOLT1B | 23.8414443 | -0.212032866 | 0.38138972 | -0.555948 | 0.57824638 | 0.82311488 |
| LDHB | 203.642238 | -0.334878311 | 0.51372856 | -0.6518585 | 0.51449248 | 0.7835482 |
| AC010197.1 | 78.2333462 | -0.93235714 | 0.72562433 | -1.2849034 | 0.19882606 | 0.50199137 |
| ABCC9 | 34.7530909 | -2.217028183 | 1.0067717 | -2.2021161 | 0.02765711 | 0.16127259 |
| AC008250.1 | 77.4198078 | -1.085589875 | 0.83650898 | -1.2977624 | 0.19436901 | 0.49679183 |

| | | | | | | |
|---|---|---|---|---|---|---|
| CMAS | 24.0594128 | -0.153519149 | 0.36105944 | -0.4251908 | 0.67069759 | 0.86777458 |
| C2CD5 | 134.765878 | 0.183924161 | 0.28692971 | 0.64100772 | 0.52151767 | 0.7891319 |
| ETNK1 | 312.350805 | 0.454567789 | 0.37212725 | 1.22153858 | 0.22188217 | 0.53238017 |
| SOX5 | 87.0192066 | -0.293757526 | 0.58276168 | -0.5040783 | 0.61420634 | 0.84174925 |
| CASC1 | 29.3622183 | 0.074876013 | 0.74363193 | 0.10068962 | 0.91979686 | 0.97520237 |
| ETFRF1 | 37.7128607 | 0.166546891 | 0.27172896 | 0.61291549 | 0.53993222 | 0.80166126 |
| KRAS | 79.4298762 | 0.195782647 | 0.3342531 | 0.58573173 | 0.55805581 | 0.81250974 |
| RASSF8-AS1 | 50.6811847 | -0.162950281 | 0.30676542 | -0.5311885 | 0.59528813 | 0.83416944 |
| RASSF8 | 307.081519 | 0.57257396 | 0.58080334 | 0.98583104 | 0.32421603 | 0.63655302 |
| ITPR2 | 778.105213 | 0.023431072 | 0.47277846 | 0.04956036 | 0.96047273 | 0.98977097 |
| INTS13 | 18.7163815 | 0.07167217 | 0.40072598 | 0.17885581 | 0.85805092 | 0.95266335 |
| FGFR1OP2 | 59.4236402 | 0.844275138 | 0.28024801 | 3.01259996 | 0.0025902 | 0.03080489 |
| TM7SF3 | 54.5183391 | 0.489591307 | 0.48971993 | 0.99973735 | 0.31743763 | 0.63093651 |
| MED21 | 17.784639 | -0.468639963 | 0.40043989 | -1.1703129 | 0.24187508 | 0.55444462 |
| STK38L | 78.8224013 | 0.710238675 | 0.29155973 | 2.43599717 | 0.0148508 | 0.10759042 |
| PPFIBP1 | 263.996269 | -0.004524554 | 0.24066206 | -0.0188004 | 0.9850003 | 0.99742652 |
| MRPS35 | 42.1375043 | -0.052504879 | 0.32187914 | -0.1631199 | 0.87042407 | 0.95629149 |
| KLHL42 | 43.994065 | 0.301642562 | 0.34405489 | 0.87672803 | 0.38063438 | 0.68449211 |
| CCDC91 | 307.506669 | 0.195094924 | 0.2456551 | 0.79418226 | 0.42708933 | 0.72258772 |
| AC009318.1 | 17.0995471 | -0.376076316 | 0.67266838 | -0.5590813 | 0.57610623 | 0.82147339 |
| ERGIC2 | 68.915115 | 0.164363754 | 0.286737 | 0.57322129 | 0.56649487 | 0.81595078 |
| TMTC1 | 210.915132 | 1.626110666 | 0.78357325 | 2.07525036 | 0.03796334 | 0.19654159 |
| IPO8 | 78.3431507 | 0.512208777 | 0.2663027 | 1.92340813 | 0.05442881 | 0.24282022 |
| CAPRIN2 | 55.9450559 | -0.070209832 | 0.2315363 | -0.3032347 | 0.76171103 | 0.91342791 |
| DDX11 | 16.2173305 | 0.255736768 | 0.54536478 | 0.46892792 | 0.63912116 | 0.85391761 |
| SINHCAF | 114.058384 | -0.462252065 | 0.20025303 | -2.3083399 | 0.02098024 | 0.13518987 |
| DENND5B | 369.673201 | -1.249440665 | 0.32628521 | -3.8292899 | 0.00012851 | 0.00295867 |
| AMN1 | 84.6509514 | -0.596278814 | 0.35765766 | -1.6671775 | 0.09547911 | 0.3375985 |
| KIAA1551 | 126.502322 | -1.093819818 | 0.42872708 | -2.5513197 | 0.01073158 | 0.08656845 |
| BICD1 | 379.914811 | -2.083792332 | 0.48359189 | -4.3089894 | 1.64E-05 | 0.00059002 |
| FGD4 | 1050.35302 | 0.054813462 | 0.16892088 | 0.32449193 | 0.74556563 | 0.90587099 |
| DNM1L | 161.996471 | 0.336420288 | 0.19487648 | 1.72632577 | 0.08428882 | 0.31297129 |
| YARS2 | 44.9403647 | -0.410554922 | 0.34623722 | -1.185762 | 0.23571632 | 0.54835802 |
| PKP2 | 16.4918528 | -0.633725689 | 0.4545611 | -1.3941485 | 0.16327276 | 0.45454804 |
| ALG10B | 12.3017208 | 0.456712752 | 0.44037649 | 1.03709612 | 0.29969106 | 0.61356712 |
| CPNE8 | 300.548212 | -0.406761703 | 0.51723265 | -0.7864192 | 0.43162192 | 0.72537073 |
| KIF21A | 70.6418045 | -0.265233227 | 0.50142748 | -0.5289563 | 0.59683576 | 0.83416944 |
| SLC2A13 | 82.2213408 | -1.483376365 | 0.44243053 | -3.3527893 | 0.00080002 | 0.01260025 |
| LRRK2 | 123.466016 | -0.641217557 | 0.67188577 | -0.954355 | 0.33990397 | 0.65027011 |
| GXYLT1 | 47.496273 | 0.088477585 | 0.29198142 | 0.30302471 | 0.76187103 | 0.91342791 |
| YAF2 | 126.466539 | -0.099460982 | 0.30806436 | -0.3228578 | 0.74680294 | 0.90661036 |
| PPHLN1 | 159.986276 | 0.090087112 | 0.20930627 | 0.43040809 | 0.66689881 | 0.86777458 |
| ZCRB1 | 21.0835512 | 0.379394965 | 0.48561422 | 0.78126823 | 0.43464475 | 0.72797335 |
| PUS7L | 37.0564419 | 0.416212275 | 0.38499216 | 1.08109285 | 0.27965581 | 0.59383074 |
| IRAK4 | 36.3393857 | -0.616124928 | 0.38577991 | -1.5970892 | 0.11024583 | 0.365422 |
| TWF1 | 17.132792 | -0.437549793 | 0.53411565 | -0.8192042 | 0.41266991 | 0.71110141 |
| TMEM117 | 324.47712 | 0.19254486 | 0.37374935 | 0.51517109 | 0.60643347 | 0.83829217 |
| ANO6 | 192.720117 | -0.829216915 | 0.46604279 | -1.779272 | 0.07519517 | 0.29335556 |
| AC008124.1 | 17.325304 | 0.664554518 | 0.37310153 | 1.78116266 | 0.07488588 | 0.29289497 |
| ARID2 | 285.883028 | 0.398170549 | 0.24141121 | 1.64934573 | 0.09907683 | 0.34368353 |
| SCAF11 | 274.956654 | -0.161156938 | 0.21139271 | -0.7623581 | 0.44584632 | 0.7388166 |
| SLC38A1 | 179.963968 | -1.663555054 | 0.66037245 | -2.5191164 | 0.01176498 | 0.09253427 |
| SLC38A2 | 157.543249 | -1.224831292 | 0.35577725 | -3.4426915 | 0.00057596 | 0.00958967 |
| AC008014.1 | 473.836812 | 0.519229211 | 0.42106162 | 1.23314304 | 0.2175224 | 0.52693657 |
| PCED1B | 57.051132 | -1.751648818 | 0.75807057 | -2.3106672 | 0.02085124 | 0.13454463 |
| RPAP3 | 28.6050131 | 0.560990402 | 0.35292601 | 1.5895411 | 0.11193828 | 0.36854256 |
| RAPGEF3 | 45.4700421 | -2.565163903 | 0.62766637 | -4.086827 | 4.37E-05 | 0.00126974 |
| SLC48A1 | 25.3834608 | -1.914501352 | 0.39341211 | -4.8664017 | 1.14E-06 | 6.54E-05 |
| HDAC7 | 86.0755233 | -2.259199403 | 0.46789873 | -4.828394 | 1.38E-06 | 7.49E-05 |
| VDR | 152.15427 | 1.692911618 | 0.30924199 | 5.47439122 | 4.39E-08 | 3.69E-06 |
| TMEM106C | 15.9436104 | -0.375139777 | 0.54850237 | -0.6839347 | 0.49401643 | 0.7738663 |
| SENP1 | 42.3132203 | -0.491937178 | 0.30060296 | -1.6365015 | 0.10173468 | 0.34880722 |
| PFKM | 20.0009876 | 0.689390448 | 0.46432213 | 1.48472451 | 0.13761682 | 0.413649 |
| ASB8 | 17.1923065 | -0.336655099 | 0.62204467 | -0.5412073 | 0.58836473 | 0.83024609 |
| ZNF641 | 18.0528596 | -0.131462804 | 0.45188457 | -0.2909212 | 0.77111158 | 0.91720887 |
| AC090115.1 | 14.1553371 | 0.427090862 | 0.59802627 | 0.7141674 | 0.47512367 | 0.76125092 |
| KANSL2 | 29.7236735 | 0.430138172 | 0.39030359 | 1.1020605 | 0.27043537 | 0.58716619 |
| CCNT1 | 46.6265207 | -0.215059303 | 0.32657491 | -0.6585298 | 0.51019777 | 0.78158783 |
| ADCY6 | 44.6332319 | 0.325439689 | 0.38109519 | 0.85395907 | 0.39312767 | 0.69595061 |
| DDX23 | 23.5633574 | -0.342434103 | 0.44417347 | -0.7709468 | 0.44073848 | 0.73302253 |
| CCDC65 | 20.9982487 | -1.244973952 | 0.53130988 | -2.3432163 | 0.0191183 | 0.12741887 |
| PRKAG1 | 32.8121089 | -0.076893009 | 0.30597788 | -0.2513025 | 0.80158023 | 0.9304348 |
| KMT2D | 37.0083192 | -0.176957604 | 0.34567449 | -0.5119198 | 0.60870717 | 0.84008076 |
| LMBR1L | 36.8042045 | -0.660936956 | 0.29340638 | -2.2526332 | 0.02428229 | 0.1479791 |
| AC011603.2 | 16.2041129 | 0.830455466 | 0.40962288 | 2.0273659 | 0.04262501 | 0.21039469 |
| TUBA1B | 43.3426503 | -0.835181798 | 0.54758424 | -1.5252115 | 0.12720635 | 0.39511746 |
| TUBA1A | 16.7692278 | -0.772173626 | 0.6178997 | -1.2496747 | 0.2114184 | 0.52053477 |
| TUBA1C | 38.7261573 | -0.500809656 | 0.3517205 | -1.4238853 | 0.15447968 | 0.44013998 |
| SPATS2 | 166.431387 | 0.152198102 | 0.33957873 | 0.4481968 | 0.65401117 | 0.86217501 |
| MCRS1 | 14.8598182 | 0.060168703 | 0.41456745 | 0.1451361 | 0.88460342 | 0.961983 |
| PRPF40B | 88.1992528 | 0.268998165 | 0.3283833 | 0.81915908 | 0.41269566 | 0.71110141 |
| TMBIM6 | 164.767833 | -0.804345569 | 0.41904725 | -1.9194627 | 0.0549258 | 0.24363853 |
| NCKAP5L | 27.3716217 | -0.325123212 | 0.39152227 | -0.830408 | 0.40630817 | 0.70547809 |
| AQP2 | 28.6214269 | 1.28522786 | 0.58272841 | 2.20553494 | 0.02741658 | 0.16065021 |
| SMARCD1 | 22.2755119 | -0.972911383 | 0.43443923 | -2.2394649 | 0.02512568 | 0.15124073 |
| AC074032.1 | 101.16797 | -0.225220215 | 0.30591715 | -0.7362131 | 0.46160102 | 0.74982974 |
| CERS5 | 65.0998748 | -0.758189245 | 0.29992437 | -2.5279348 | 0.01147357 | 0.09058385 |
| LIMA1 | 111.84893 | -1.196918714 | 0.34364002 | -3.4830597 | 0.00049572 | 0.00856029 |
| LARP4 | 173.282124 | 0.39275389 | 0.27448191 | 1.4308917 | 0.15246126 | 0.43726507 |
| DIP2B | 248.907703 | -0.545688331 | 0.44607963 | -1.223298 | 0.22121715 | 0.53160533 |
| ATF1 | 76.0823017 | -0.066651742 | 0.32174745 | -0.2071555 | 0.83588845 | 0.94654164 |
| METTL7A | 26.3916587 | 0.701325672 | 0.60012809 | 1.16862663 | 0.24255409 | 0.55498757 |
| SLC11A2 | 121.064439 | -0.238628164 | 0.22162474 | -1.0767217 | 0.28160462 | 0.59579794 |
| LETMD1 | 85.4306113 | 0.597540252 | 0.32151813 | 1.85849631 | 0.06309857 | 0.26453734 |
| CSRNP2 | 29.6182176 | -0.260379171 | 0.42903713 | -0.6068919 | 0.54392263 | 0.80380646 |
| TFCP2 | 196.116163 | 0.259521901 | 0.22401872 | 1.15848308 | 0.24666695 | 0.55931971 |
| POU6F1 | 27.645433 | 0.718501822 | 0.38923664 | 1.84592546 | 0.06490303 | 0.26967729 |
| DAZAP2 | 78.9196329 | -0.477043499 | 0.28113507 | -1.6968481 | 0.08972539 | 0.32539849 |
| SMAGP | 18.3341148 | 0.568578685 | 0.52589614 | 1.08116156 | 0.27962526 | 0.59383074 |
| ACVR1B | 308.046807 | -0.788408189 | 0.28877846 | -2.7301489 | 0.00633057 | 0.0589673 |
| NR4A1 | 37.7471506 | 0.533623928 | 1.40318088 | 0.3802959 | 0.70372578 | 0.88543615 |
| KRT8 | 56.8886926 | -1.39934315 | 0.3069004 | -4.5596003 | 5.13E-06 | 0.0002218 |
| KRT18 | 49.7127644 | -1.861031346 | 0.50984964 | -3.6501573 | 0.00026208 | 0.00525206 |
| EIF4B | 107.376612 | -0.430807987 | 0.28241721 | -1.5254311 | 0.12715162 | 0.39511746 |
| AC068888.1 | 48.9702301 | 0.312998681 | 0.28867646 | 1.08425426 | 0.27825208 | 0.59355351 |
| TNS2 | 28.4483682 | -0.615960608 | 0.57973105 | -1.0624937 | 0.2880116 | 0.60251743 |
| CSAD | 93.7036543 | 0.429876119 | 0.33336403 | 1.28950961 | 0.19722098 | 0.49934704 |
| ZNF740 | 14.1006812 | 0.334285532 | 0.4966569 | 0.67307136 | 0.50090189 | 0.77607332 |
| RARG | 13.1810038 | -0.218823462 | 0.51284827 | -0.4266827 | 0.66961048 | 0.86777458 |
| PFDN5 | 45.7153529 | 0.124887933 | 0.36886654 | 0.33857214 | 0.73493207 | 0.90007689 |
| AAAS | 11.4988566 | 1.297753267 | 0.47152241 | 2.7522621 | 0.00591851 | 0.05613857 |
| SP1 | 71.4768864 | -0.870004453 | 0.20523957 | -4.2389705 | 2.25E-05 | 0.00075834 |
| PRR13 | 51.6566127 | 0.165844047 | 0.37883162 | 0.43777774 | 0.66154741 | 0.86492722 |

| | | | | | | |
|---|---|---|---|---|---|---|
| PCBP2 | 179.526197 | 0.151947634 | 0.24356812 | 0.62384039 | 0.53273241 | 0.79724684 |
| MAP3K12 | 13.128957 | -0.237396362 | 0.42921337 | -0.5530964 | 0.58019741 | 0.82440729 |
| ATP5MC2 | 78.4635391 | -0.336945329 | 0.34405023 | -0.9793492 | 0.32740745 | 0.63895602 |
| CALCOCO1 | 36.3946381 | 0.20162385 | 0.4110544 | 0.49050405 | 0.62377726 | 0.84597807 |
| SMUG1 | 17.9239112 | 0.631264866 | 0.50172139 | 1.25819802 | 0.20832015 | 0.51645229 |
| LINC02381 | 18.5824309 | -0.177826437 | 0.51566116 | -0.3448513 | 0.73020614 | 0.89731942 |
| CBX5 | 71.8007945 | -0.232339368 | 0.35591682 | -0.6527912 | 0.5138909 | 0.78323295 |
| HNRNPA1 | 48.5859713 | 0.147618665 | 0.32496162 | 0.45426492 | 0.64963816 | 0.86077172 |
| COPZ1 | 35.0762974 | -0.103371581 | 0.31123646 | -0.332132 | 0.7397896 | 0.90304252 |
| PPP1R1A | 27.3447494 | 0.908443245 | 0.54244415 | 1.67472216 | 0.09398876 | 0.3344198 |
| CD63 | 77.6119762 | -1.925382212 | 0.52116647 | -3.6943709 | 0.00022043 | 0.00456735 |
| RAB5B | 45.0282235 | -0.396660888 | 0.3666747 | -1.0817787 | 0.27935087 | 0.59383074 |
| SUOX | 14.7288565 | 0.243215108 | 0.66115797 | 0.36786232 | 0.7129759 | 0.88791068 |
| IKZF4 | 17.7936942 | -0.560780991 | 0.39481983 | -1.4203466 | 0.15550681 | 0.44111516 |
| RPS26 | 15.4566362 | -0.25122871 | 0.40799873 | -0.6157586 | 0.53805388 | 0.80104055 |
| ERBB3 | 27.2742658 | -0.428499061 | 0.42140112 | -1.0168437 | 0.30922781 | 0.62335507 |
| PA2G4 | 15.4589849 | -0.139264046 | 0.4669831 | -0.2982207 | 0.76553469 | 0.91549415 |
| RPL41 | 166.344567 | -0.74461061 | 0.38760394 | -1.9210605 | 0.05472408 | 0.24343712 |
| ESYT1 | 20.9009337 | -1.196582357 | 0.50149419 | -2.3860343 | 0.01703116 | 0.11780307 |
| MYL6B | 13.5273561 | 0.132447161 | 0.55841216 | 0.23718531 | 0.81251303 | 0.93571318 |
| MYL6 | 98.2159828 | 0.222496984 | 0.44674739 | 0.49803756 | 0.61845757 | 0.84283824 |
| SMARCC2 | 82.4089626 | -0.298021183 | 0.28511007 | -1.0452847 | 0.29589142 | 0.61091488 |
| RNF41 | 19.4127313 | -0.11761541 | 0.38000732 | -0.3095083 | 0.75693492 | 0.91125387 |
| ANKRD52 | 27.339878 | -0.076179915 | 0.35090845 | -0.2170934 | 0.82813554 | 0.94268221 |
| COQ10A | 13.9277132 | 1.853861049 | 0.47824583 | 3.8763768 | 0.00010602 | 0.00254784 |
| CS | 43.415813 | 0.455783232 | 0.44380809 | 1.02698271 | 0.30442861 | 0.61818174 |
| AC073896.2 | 16.2633257 | 0.374357363 | 0.5063354 | 0.73934661 | 0.45969655 | 0.74802978 |
| PAN2 | 29.6779126 | 0.472416529 | 0.41444706 | 1.13987184 | 0.2543397 | 0.56645553 |
| STAT2 | 53.4848186 | 0.180451071 | 0.38521666 | 0.46844047 | 0.63946964 | 0.8540918 |
| RBMS2 | 83.677976 | 0.14834562 | 0.34994587 | 0.42391019 | 0.67163131 | 0.86777458 |
| BAZ2A | 70.9332653 | 0.216714354 | 0.21472587 | 1.00926055 | 0.31284969 | 0.62758402 |
| ATP5F1B | 102.978686 | 0.447373397 | 0.45401372 | 0.98537419 | 0.32444031 | 0.63678264 |
| PTGES3 | 145.757426 | 0.206952448 | 0.29935214 | 0.69133445 | 0.48935539 | 0.7709952 |
| NACA | 47.5555094 | -0.565148593 | 0.391745 | -1.4426441 | 0.14912076 | 0.4328578 |
| NEMP1 | 12.7951472 | -0.699229642 | 0.42655168 | -1.6392613 | 0.10115886 | 0.34753323 |
| STAT6 | 61.3236041 | -1.123927288 | 0.31763369 | -3.5384386 | 0.0004025 | 0.00740221 |
| R3HDM2 | 294.938041 | 0.337745444 | 0.18917183 | 1.78538972 | 0.07419813 | 0.29129403 |
| MARS | 35.5946169 | -0.653723245 | 0.55977116 | -1.1678402 | 0.24287124 | 0.55555126 |
| DDIT3 | 44.6463566 | -1.218098112 | 0.74121331 | -1.643384 | 0.10030351 | 0.34581665 |
| MBD6 | 23.5625915 | 0.488606929 | 0.47800761 | 1.02217396 | 0.30669857 | 0.62067717 |
| DCTN2 | 42.0247021 | -0.42244248 | 0.29672579 | -1.4236797 | 0.15453922 | 0.44013998 |
| PIP4K2C | 33.7168744 | -0.073921804 | 0.35299521 | -0.2094131 | 0.8341258 | 0.94640649 |
| DTX3 | 18.8317949 | 0.680920637 | 0.36967404 | 1.84194877 | 0.06548264 | 0.27123953 |
| OS9 | 49.5817797 | -0.109365445 | 0.33881371 | -0.3227893 | 0.74685481 | 0.90661036 |
| 9-Mar | 15.5414089 | -0.903239407 | 0.39325001 | -2.2968579 | 0.02162688 | 0.13787238 |
| TSFM | 17.8973844 | 0.018871042 | 0.44391603 | 0.04251039 | 0.96609183 | 0.99156233 |
| AVIL | 50.5828347 | -1.351352255 | 0.29869923 | -4.5241236 | 6.06E-06 | 0.00025357 |
| CTDSP2 | 57.3102857 | -0.409231517 | 0.24540017 | -1.667609 | 0.09539338 | 0.33742332 |
| LRIG3 | 16.9221685 | 0.650265333 | 0.48068399 | 1.35279175 | 0.17612217 | 0.47338227 |
| SLC16A7 | 318.502641 | 1.113487 | 0.52410896 | 2.12453341 | 0.03362556 | 0.18260033 |
| FAM19A2 | 59.8320569 | 0.887613255 | 0.68225931 | 1.30099105 | 0.19326152 | 0.49477496 |
| USP15 | 216.800187 | 0.270853163 | 0.21772237 | 1.24403003 | 0.21348851 | 0.52235998 |
| MON2 | 324.02956 | 0.314974346 | 0.17390843 | 1.81115052 | 0.07011755 | 0.28228674 |
| AC048341.1 | 97.9582864 | -1.186190541 | 0.50362006 | -2.3553282 | 0.01850635 | 0.12422837 |
| PPM1H | 377.772151 | -1.481287467 | 0.40890622 | -3.6225603 | 0.0002917 | 0.00567744 |
| DPY19L2 | 231.094657 | 0.774034815 | 0.55898393 | 1.38471747 | 0.16613888 | 0.458443 |
| RXYLT1 | 21.5113779 | -0.48867599 | 0.38512925 | -1.2688623 | 0.20449018 | 0.51062303 |
| SRGAP1 | 136.672285 | -1.60791564 | 0.66380044 | -2.4222877 | 0.01542313 | 0.11036477 |
| XPOT | 27.5010399 | -0.25742288 | 0.51442218 | -0.5004117 | 0.61678522 | 0.84223226 |
| TBK1 | 94.442019 | -0.534360174 | 0.30689785 | -1.7411662 | 0.08165445 | 0.30754226 |
| RASSF3 | 22.6146749 | -2.112944917 | 0.58332182 | -3.6222628 | 0.00029204 | 0.00567744 |
| GNS | 79.0516063 | -0.99849254 | 0.38838736 | -2.5708678 | 0.01014441 | 0.08297056 |
| TBC1D30 | 51.3896422 | -0.433217252 | 0.28156122 | -1.5386254 | 0.12389577 | 0.38919278 |
| LINC02389 | 25.6380987 | -0.544342451 | 0.65878147 | -0.8262868 | 0.4086414 | 0.70742154 |
| LEMD3 | 108.222813 | 0.088867807 | 0.24438309 | 0.3636414 | 0.71612581 | 0.88922054 |
| AC090673.1 | 25.0714033 | 0.7064739 | 0.82134775 | 0.86013982 | 0.38971197 | 0.69278827 |
| HELB | 57.7579302 | -0.69003633 | 0.40043913 | -1.7231991 | 0.08485254 | 0.31408459 |
| GRIP1 | 695.990513 | -0.729682107 | 0.32455461 | -2.2482568 | 0.02455982 | 0.14889192 |
| CAND1 | 95.4462785 | 0.266702725 | 0.19809212 | 1.34635708 | 0.17818738 | 0.47550635 |
| MDM1 | 13.5376489 | 0.794249632 | 0.47414567 | 1.67511734 | 0.09391121 | 0.3344198 |
| RAP1B | 204.987389 | -1.337731492 | 0.39315578 | -3.4025482 | 0.00066761 | 0.01095908 |
| NUP107 | 46.8614218 | -0.621907114 | 0.3943947 | -1.5768648 | 0.11482664 | 0.37343691 |
| SLC35E3 | 39.9067132 | -0.086767714 | 0.36402604 | -0.2383558 | 0.81160515 | 0.93552094 |
| MDM2 | 94.6577219 | -0.458563536 | 0.27929597 | -1.6418552 | 0.10062002 | 0.34632707 |
| CPM | 490.17671 | -0.7137359 | 0.65557322 | -1.0887203 | 0.27627723 | 0.59137026 |
| CPSF6 | 169.007286 | 0.65004926 | 0.23749191 | 2.7371427 | 0.00619754 | 0.05790977 |
| FRS2 | 141.19581 | 0.272504674 | 0.22412469 | 1.21586191 | 0.22403753 | 0.53507324 |
| CCT2 | 38.9721403 | -0.296713032 | 0.51486804 | -0.5762895 | 0.56441954 | 0.81490487 |
| RAB3IP | 149.775113 | 0.230689638 | 0.41614228 | 0.5543528 | 0.57933742 | 0.82381304 |
| MYRFL | 59.4468551 | -0.255798044 | 0.49832477 | -0.5133159 | 0.60773034 | 0.83922189 |
| AC025159.1 | 45.1245016 | -0.616331104 | 0.7545556 | -0.8168134 | 0.41403509 | 0.71160612 |
| CNOT2 | 202.754437 | 0.485958087 | 0.27009203 | 1.79923152 | 0.07198207 | 0.2864536 |
| TSPAN8 | 15.7164074 | -0.729851375 | 0.53412153 | -1.3664519 | 0.17179717 | 0.46763314 |
| ZFC3H1 | 175.446196 | -0.43708166 | 0.37891462 | -1.1535096 | 0.24870127 | 0.56120297 |
| RAB21 | 128.180604 | -0.504706355 | 0.3000116 | -1.6822895 | 0.09251272 | 0.33176485 |
| TBC1D15 | 131.120678 | -0.402177388 | 0.35185732 | -1.1430127 | 0.25303332 | 0.56559484 |
| TRHDE | 245.797518 | 0.419387086 | 0.53139516 | 0.78921887 | 0.42998409 | 0.72390206 |
| ATXN7L3B | 24.9050404 | -0.730395715 | 0.33332934 | -2.1912134 | 0.02843635 | 0.16420667 |
| CAPS2 | 112.910722 | 0.832916992 | 0.63803484 | 1.30544124 | 0.19174264 | 0.49284602 |
| KRR1 | 36.8368046 | 0.124906558 | 0.30149146 | 0.41429551 | 0.6786577 | 0.87147102 |
| NAP1L1 | 116.773946 | 0.361744895 | 0.28404849 | 1.27353219 | 0.20282926 | 0.50715474 |
| OSBPL8 | 238.897085 | 0.209567396 | 0.19529754 | 1.07306728 | 0.28324093 | 0.59695715 |
| ZDHHC17 | 117.36989 | 0.769639117 | 0.2092321 | 3.67839884 | 0.0002347 | 0.00482019 |
| AC027288.3 | 56.3144772 | 0.146789438 | 0.57724074 | 0.254295 | 0.79926766 | 0.92922871 |
| PAWR | 366.826679 | 0.563827781 | 0.21757019 | 2.59147531 | 0.00955654 | 0.07955819 |
| AC073569.2 | 26.0185976 | 0.406054561 | 0.51014096 | 0.79596541 | 0.42605214 | 0.72211376 |
| PPP1R12A | 278.746077 | -0.087527264 | 0.21320744 | -0.4105263 | 0.6814199 | 0.87270068 |
| LIN7A | 83.1923117 | 0.297562519 | 0.56654464 | 0.52522343 | 0.59942788 | 0.83581061 |
| ACSS3 | 107.941776 | 0.289343708 | 0.61105451 | 0.47351538 | 0.63584553 | 0.85365352 |
| PPFIA2 | 119.287524 | 0.363606818 | 0.70098093 | 0.51871143 | 0.60396199 | 0.83754753 |
| CCDC59 | 24.7607053 | 0.132943144 | 0.40311815 | 0.32978705 | 0.74156088 | 0.90383185 |
| METTL25 | 107.653566 | 0.593829143 | 0.22360232 | 2.65573786 | 0.00791351 | 0.06954342 |
| TMTC2 | 542.890757 | -1.491830596 | 0.60721727 | -2.4568317 | 0.01401684 | 0.10388949 |
| LRRIQ1 | 35.1930968 | 0.002612207 | 0.90313125 | 0.00289239 | 0.99769221 | 0.99922221 |
| CEP290 | 94.2317543 | 0.334712876 | 0.27927479 | 1.1985073 | 0.23071958 | 0.54354156 |
| TMTC3 | 81.7747877 | 0.19983071 | 0.32076948 | 0.62297294 | 0.5333023 | 0.79764367 |
| KITLG | 386.692042 | -0.200770093 | 0.61319679 | -0.3274154 | 0.7433537 | 0.90463949 |
| DUSP6 | 27.6985125 | -3.153932179 | 0.76775652 | -4.1079849 | 3.99E-05 | 0.00118518 |
| POC1B | 73.5557994 | -0.475881599 | 0.26945441 | -1.7660932 | 0.07738022 | 0.29764569 |
| POC1B-AS1 | 26.6066782 | -0.105402191 | 0.55414506 | -0.1902069 | 0.84914704 | 0.95003258 |
| ATP2B1 | 142.278644 | 0.526250212 | 0.27715673 | 1.89874595 | 0.05759789 | 0.25013758 |
| BTG1 | 74.3239853 | -1.715424682 | 0.37415302 | -4.5848212 | 4.54E-06 | 0.00020077 |
| LINC02391 | 23.6218216 | -0.761967892 | 0.69735387 | -1.092656 | 0.27454487 | 0.5891499 |

| | | | | | | |
|---|---|---|---|---|---|---|
| EEA1 | 164.44061 | 0.035788864 | 0.38773235 | 0.09230301 | 0.9264573 | 0.97735117 |
| AC138123.1 | 148.730096 | 1.373375813 | 0.55156248 | 2.48997325 | 0.01277527 | 0.09763658 |
| AC124947.1 | 56.6964384 | 0.378934904 | 0.37806456 | 1.00230212 | 0.3161977 | 0.63009595 |
| NUDT4 | 215.707591 | 0.735235594 | 0.54367679 | 1.3523395 | 0.17626673 | 0.47349784 |
| UBE2N | 69.0969614 | 0.466731735 | 0.23746318 | 1.96549096 | 0.04935744 | 0.22816499 |
| MRPL42 | 67.3305145 | 0.015610264 | 0.2743675 | 0.05689545 | 0.95462848 | 0.98834154 |
| SOCS2 | 31.8311781 | 0.32519819 | 0.38278629 | 0.84955549 | 0.39557227 | 0.69642725 |
| CRADD | 490.804765 | 0.224902349 | 0.20138769 | 1.11676314 | 0.26409561 | 0.57871387 |
| AC012085.2 | 34.8477279 | -0.916103496 | 0.65893141 | -1.3902866 | 0.16444187 | 0.45563826 |
| CEP83 | 69.7857918 | 0.361641657 | 0.37632217 | 0.96098952 | 0.33655744 | 0.64764721 |
| TMCC3 | 34.1319465 | 0.154448179 | 0.51192153 | 0.30170284 | 0.76287861 | 0.91416015 |
| NDUFA12 | 39.2718629 | 0.75789903 | 0.30437307 | 2.4900331 | 0.01277312 | 0.09763658 |
| NR2C1 | 75.1362391 | 0.399925462 | 0.24198226 | 1.65270571 | 0.09839077 | 0.34231178 |
| FGD6 | 41.142781 | -1.068019984 | 0.42912257 | -2.488846 | 0.01281585 | 0.09786647 |
| VEZT | 143.916095 | -0.033521791 | 0.20391659 | -0.1643897 | 0.86942436 | 0.9561822 |
| METAP2 | 73.3913075 | 0.057867535 | 0.23106323 | 0.25044026 | 0.8022469 | 0.93060267 |
| USP44 | 16.0788888 | 0.155713925 | 0.56250464 | 0.27682247 | 0.78191642 | 0.92126657 |
| NTN4 | 208.636723 | 1.237294063 | 0.48165108 | 2.56885972 | 0.01020338 | 0.08337973 |
| SNRPF | 16.6419405 | -0.606528741 | 0.46076237 | -1.3163591 | 0.18805354 | 0.4882768 |
| LTA4H | 32.6152576 | 0.28180418 | 0.39291292 | 0.7172179 | 0.47323966 | 0.75946412 |
| ELK3 | 38.5791859 | -2.315265809 | 0.72871326 | -3.1771973 | 0.00148706 | 0.02045321 |
| CDK17 | 98.1655066 | -0.116326706 | 0.23551049 | -0.4939343 | 0.6213526 | 0.84470972 |
| CFAP54 | 16.6053167 | -2.312268381 | 0.99351733 | -2.3273559 | 0.01994633 | 0.13082568 |
| NEDD1 | 34.2566925 | 0.500809481 | 0.30079427 | 1.66495685 | 0.09592136 | 0.33852036 |
| TMPO | 38.1141913 | 0.36093547 | 0.31270441 | 1.15423849 | 0.24840241 | 0.56107171 |
| SLC25A3 | 89.5838577 | -0.053815037 | 0.41835614 | -0.1286345 | 0.89764686 | 0.96607503 |
| APAF1 | 33.7664905 | -0.272739889 | 0.37362596 | -0.7299811 | 0.46540173 | 0.75271566 |
| ANKS1B | 71.7516011 | -0.651461377 | 0.69891842 | -0.9320993 | 0.35128521 | 0.65863327 |
| UHRF1BP1L | 56.7340245 | -0.318390359 | 0.33195658 | -0.9591325 | 0.33749198 | 0.64844536 |
| ACTR6 | 29.8942019 | -0.068353542 | 0.33828807 | -0.2020572 | 0.839872 | 0.9474978 |
| SCYL2 | 106.003424 | 0.034724312 | 0.30791836 | 0.11277116 | 0.91021198 | 0.97128978 |
| NR1H4 | 156.01938 | -4.207013753 | 0.73366227 | -5.7342648 | 9.79E-09 | 1.04E-06 |
| UTP20 | 37.8871804 | 0.182076263 | 0.32213885 | 0.5652105 | 0.57193059 | 0.81947131 |
| ARL1 | 31.2793912 | -0.129555697 | 0.4275503 | -0.3030186 | 0.76187568 | 0.91342791 |
| CHPT1 | 75.885336 | 0.566716488 | 0.42675084 | 1.32797979 | 0.18418478 | 0.48327975 |
| GNPTAB | 79.7580848 | -0.133849164 | 0.24198921 | -0.5531204 | 0.58018099 | 0.82440729 |
| DRAM1 | 51.2090811 | -2.637135452 | 0.70605588 | -3.7350237 | 0.0001877 | 0.00401397 |
| WASHC3 | 52.5498536 | 0.741814396 | 0.41684468 | 1.77959424 | 0.07514239 | 0.29327235 |
| NUP37 | 17.789232 | 0.313258131 | 0.53181781 | 0.58903279 | 0.55583928 | 0.81104802 |
| NT5DC3 | 54.6722289 | -0.62376847 | 0.54162346 | -1.1516644 | 0.249459 | 0.56223247 |
| HSP90B1 | 149.004765 | -0.829577615 | 0.68680825 | -1.2078737 | 0.22709585 | 0.53871433 |
| C12orf73 | 12.0203131 | 0.598757244 | 0.48601565 | 1.23197111 | 0.21795988 | 0.52750155 |
| TDG | 51.9423146 | -0.347292246 | 0.34814084 | -0.9975625 | 0.31849155 | 0.63093651 |
| HCFC2 | 26.6475765 | 0.79830721 | 0.31326588 | 2.54833759 | 0.01082377 | 0.08685095 |
| NFYB | 19.0561681 | 0.232078483 | 0.3555444 | 0.65274122 | 0.51392312 | 0.78323295 |
| TXNRD1 | 177.195868 | -1.464958764 | 0.2958088 | -4.9523839 | 7.33E-07 | 4.47E-05 |
| SLC41A2 | 44.628827 | -1.679076061 | 0.30626831 | -5.4823695 | 4.20E-08 | 3.56E-06 |
| C12orf45 | 12.4661981 | 0.142957124 | 0.39890246 | 0.35837614 | 0.72006185 | 0.89137768 |
| WASHC4 | 151.188533 | -0.150140601 | 0.25211093 | -0.5955339 | 0.55148666 | 0.80888181 |
| APPL2 | 102.217962 | -0.338705782 | 0.27676278 | -1.2238126 | 0.22102292 | 0.53127552 |
| C12orf75 | 91.7688096 | -2.42321214 | 0.59515291 | -4.0715791 | 4.67E-05 | 0.00133966 |
| CKAP4 | 16.8657629 | -0.311038273 | 0.61839311 | -0.5029782 | 0.61497958 | 0.84190945 |
| TCP11L2 | 60.8291539 | 0.046074943 | 0.38247408 | 0.12046553 | 0.90411438 | 0.9686272 |
| POLR3B | 37.6633768 | 0.604654491 | 0.29749019 | 2.03251912 | 0.04210113 | 0.20922698 |
| RIC8B | 96.909708 | 0.614038538 | 0.30931474 | 1.98515769 | 0.04712693 | 0.2228253 |
| TMEM263 | 57.2245774 | 0.06946236 | 0.35939619 | 0.19327517 | 0.84674347 | 0.94937904 |
| MTERF2 | 18.0008019 | 0.197243897 | 0.41891652 | 0.47084297 | 0.63775288 | 0.8538689 |
| CRY1 | 77.3461227 | -0.205408175 | 0.69362801 | -0.2961359 | 0.76712628 | 0.9157292 |
| BTBD11 | 310.752584 | 1.024607253 | 0.56826882 | 1.80303269 | 0.07138308 | 0.28516531 |
| PWP1 | 28.4172085 | -0.044094776 | 0.33037481 | -0.1334689 | 0.89382253 | 0.96482804 |
| PRDM4 | 57.1355474 | -0.060868922 | 0.33142479 | -0.1836583 | 0.8542815 | 0.95170432 |
| SART3 | 40.0796508 | -0.073544525 | 0.31403023 | -0.2341957 | 0.81483306 | 0.93703792 |
| ISCU | 54.1197182 | -0.426745362 | 0.41422964 | -1.0302145 | 0.30290934 | 0.61733918 |
| CORO1C | 60.677473 | -2.274419803 | 0.57571726 | -3.9505847 | 7.80E-05 | 0.00203046 |
| SSH1 | 170.648258 | -0.366261725 | 0.34568708 | -1.0595181 | 0.28936389 | 0.60448615 |
| USP30 | 23.4604893 | 0.291987189 | 0.34577621 | 0.84443978 | 0.3984237 | 0.69875231 |
| UNG | 14.492673 | -0.236687963 | 0.43310992 | -0.5464847 | 0.58473277 | 0.82845288 |
| ACACB | 326.742062 | 1.459427122 | 0.41419128 | 3.52355832 | 0.00042579 | 0.00769399 |
| KCTD10 | 22.2728875 | 0.150137473 | 0.34764852 | 0.43186571 | 0.66583902 | 0.86741428 |
| UBE3B | 41.1363561 | 0.171135064 | 0.2732582 | 0.62627604 | 0.5311339 | 0.79628881 |
| MMAB | 25.7660724 | -0.040448221 | 0.44022374 | -0.0918811 | 0.92679254 | 0.97753548 |
| MVK | 20.9451878 | -0.418146398 | 0.42409817 | -0.9859661 | 0.32414977 | 0.63655302 |
| FAM222A | 36.2006031 | -0.002511474 | 0.51565023 | -0.0048705 | 0.99611392 | 0.99922221 |
| TRPV4 | 37.9818068 | 0.480166355 | 0.91128873 | 0.52690913 | 0.59825669 | 0.83505171 |
| GLTP | 34.3384352 | 0.922022436 | 0.30641522 | 3.00906211 | 0.00262056 | 0.03108659 |
| TCHP | 34.2755205 | 0.302377749 | 0.35403933 | 0.85407954 | 0.39306092 | 0.69595061 |
| GIT2 | 75.7464598 | 0.098456274 | 0.25596425 | 0.38464853 | 0.70049784 | 0.88382163 |
| ANKRD13A | 93.3946174 | 0.325983587 | 0.26780922 | 1.21722316 | 0.22351932 | 0.53397237 |
| C12orf76 | 15.939702 | -0.315731577 | 0.54203968 | -0.5824879 | 0.56023806 | 0.81330344 |
| IFT81 | 58.9577465 | 0.595679817 | 0.30738921 | 1.93786831 | 0.05263929 | 0.23767978 |
| ATP2A2 | 185.182168 | 0.672114894 | 0.44796042 | 1.50038903 | 0.13351366 | 0.40681783 |
| ANAPC7 | 41.2390114 | 0.188162115 | 0.34630143 | 0.54334779 | 0.58689037 | 0.8297013 |
| ARPC3 | 78.3313459 | -1.25921187 | 0.29097263 | -4.3275956 | 1.51E-05 | 0.00055337 |
| VPS29 | 48.926194 | -0.312124653 | 0.24806993 | -1.2582124 | 0.20831496 | 0.51645229 |
| PPTC7 | 98.2250195 | 0.455653431 | 0.25519537 | 1.78550822 | 0.07417892 | 0.29129403 |
| TCTN1 | 31.3326383 | 0.579321678 | 0.62485465 | 0.9271303 | 0.35385888 | 0.66066885 |
| PPP1CC | 27.5574587 | 0.045433929 | 0.3324282 | 0.13667291 | 0.89128934 | 0.96328012 |
| ATXN2 | 355.298915 | 0.198750312 | 0.26370102 | 0.75369565 | 0.451032 | 0.74222067 |
| BRAP | 45.6666457 | -0.362081295 | 0.26654141 | -1.3584429 | 0.17432318 | 0.47073576 |
| ACAD10 | 56.7785155 | 0.457039731 | 0.33474482 | 1.36533771 | 0.17214693 | 0.46801268 |
| ALDH2 | 57.8141573 | -0.752524846 | 0.51450851 | -1.4626091 | 0.14357436 | 0.42454311 |
| MAPKAPK5 | 48.7413745 | 0.309691586 | 0.30808816 | 1.00520444 | 0.31479842 | 0.62946385 |
| TMEM116 | 132.794378 | 1.535971291 | 0.28113576 | 5.46345035 | 4.67E-08 | 3.85E-06 |
| ERP29 | 41.7941942 | -0.967087516 | 0.38909594 | -2.4854732 | 0.01293793 | 0.09839584 |
| NAA25 | 99.5303138 | 0.161263876 | 0.33653429 | 0.47919002 | 0.63180345 | 0.85129124 |
| TRAFD1 | 36.7538854 | -0.220959129 | 0.31992557 | -0.6906579 | 0.48978054 | 0.77140435 |
| HECTD4 | 249.80915 | -0.133390874 | 0.16597583 | -0.8036765 | 0.42158383 | 0.71947943 |
| RPL6 | 84.3685862 | -0.633724512 | 0.3922369 | -1.6156677 | 0.10616617 | 0.35826759 |
| PTPN11 | 83.1928642 | -0.310054169 | 0.32417674 | -0.9564356 | 0.33885222 | 0.64971064 |
| RASAL1 | 23.0774362 | -3.877723626 | 0.67436991 | -5.7501433 | 8.92E-09 | 9.67E-07 |
| DDX54 | 17.8903555 | -0.702634083 | 0.45357551 | -1.5491006 | 0.12135755 | 0.38478887 |
| TPCN1 | 90.5007586 | -0.297298923 | 0.3157238 | -0.9416424 | 0.34637574 | 0.65471653 |
| SLC8B1 | 36.0737797 | -1.326747745 | 0.57882314 | -2.292147 | 0.02189716 | 0.13860769 |
| PLBD2 | 14.8289051 | -0.54124142 | 0.46481872 | -1.164414 | 0.24425628 | 0.55622238 |
| RBM19 | 44.6104952 | -0.269601256 | 0.36155395 | -0.7456737 | 0.45586458 | 0.74621803 |
| MED13L | 969.888755 | -0.541968146 | 0.2443544 | -2.2179595 | 0.0265576 | 0.15672344 |
| C12orf49 | 67.3771189 | -2.492784854 | 0.74172747 | -3.3607827 | 0.00077722 | 0.01234548 |
| FBXW8 | 93.495835 | -0.324903237 | 0.22665716 | -1.4334567 | 0.15172735 | 0.43620176 |
| TESC | 108.867569 | -3.155390962 | 0.51603489 | -6.1146854 | 9.67E-10 | 1.64E-07 |
| FBXO21 | 79.6472974 | 0.27238148 | 0.1978978 | 1.3763745 | 0.16870572 | 0.46237864 |
| NOS1 | 69.6412152 | 3.168904565 | 0.76064876 | 4.16605498 | 3.10E-05 | 0.00098623 |
| KSR2 | 101.156552 | 1.769912909 | 0.63334088 | 2.79456605 | 0.00519694 | 0.05154924 |
| WSB2 | 36.2078551 | -0.013563895 | 0.39258241 | -0.0345504 | 0.97243822 | 0.99287068 |

| | | | | | | |
|---|---|---|---|---|---|---|
| VSIG10 | 42.627931 | 0.582286533 | 0.58696177 | 0.99203485 | 0.32118052 | 0.63285867 |
| PEBP1 | 138.730621 | -0.807431931 | 0.43149688 | -1.8712347 | 0.06131256 | 0.25926454 |
| TAOK3 | 359.161655 | 0.342990994 | 0.25973645 | 1.32053471 | 0.18665655 | 0.48675708 |
| SUDS3 | 51.7519991 | -0.046100294 | 0.25503747 | -0.1807589 | 0.85655682 | 0.95234014 |
| SRRM4 | 72.1047004 | 7.649947103 | 0.97316946 | 7.86085816 | 3.82E-15 | 2.37E-12 |
| CCDC60 | 23.0137592 | -1.277500855 | 0.89752074 | -1.4233664 | 0.15462997 | 0.44023507 |
| AC002070.1 | 167.855723 | 0.127837467 | 0.40229576 | 0.31776986 | 0.75065952 | 0.9089517 |
| PRKAB1 | 71.3068113 | 0.108342939 | 0.32922274 | 0.32908704 | 0.7420899 | 0.90424023 |
| CIT | 102.586328 | 0.44141593 | 0.37067094 | 1.19085658 | 0.23370989 | 0.54598215 |
| BICDL1 | 298.300256 | 1.618438082 | 0.39153756 | 4.13354493 | 3.57E-05 | 0.0010849 |
| RAB35 | 29.2644022 | -0.891067576 | 0.57872003 | -1.5397213 | 0.12362829 | 0.38887694 |
| GCN1 | 37.4244941 | -0.185066442 | 0.3263478 | -0.5670835 | 0.57065748 | 0.81891359 |
| RPLP0 | 107.920171 | -1.333589262 | 0.53791691 | -2.4791734 | 0.01316873 | 0.09942124 |
| PXN | 73.3667563 | -0.791017399 | 0.39679217 | -1.9935308 | 0.04620336 | 0.22049887 |
| GATC | 14.24202 | -0.684629135 | 0.48085071 | -1.4237873 | 0.15450805 | 0.44013998 |
| SRSF9 | 42.4633678 | -0.275072909 | 0.26400066 | -1.0419402 | 0.2974394 | 0.61141484 |
| DYNLL1 | 53.9186215 | -0.395010204 | 0.62542337 | -0.6315885 | 0.52765581 | 0.79403853 |
| COQ5 | 34.0338414 | 0.515218086 | 0.65103927 | 0.7913779 | 0.4287235 | 0.7236272 |
| RNF10 | 201.92784 | 0.246939459 | 0.17325947 | 1.42525805 | 0.15408262 | 0.43999572 |
| POP5 | 26.4140924 | 0.369921694 | 0.69983513 | 0.52858406 | 0.59709402 | 0.8342658 |
| MLEC | 77.502442 | -0.609204947 | 0.42402599 | -1.436716 | 0.15079869 | 0.43488337 |
| UNC119B | 16.9322054 | -0.258904728 | 0.44193102 | -0.5858487 | 0.55797717 | 0.81250974 |
| ACADS | 22.939205 | 0.402655602 | 0.49072237 | 0.82053648 | 0.41191034 | 0.71101607 |
| SPPL3 | 302.802515 | -0.141922035 | 0.26136932 | -0.5429942 | 0.58713377 | 0.82983708 |
| P2RX4 | 33.9668189 | -0.712794123 | 0.5272871 | -1.3518141 | 0.1764348 | 0.47354004 |
| CAMKK2 | 18.1569162 | 0.427973993 | 0.3918752 | 1.09211808 | 0.2747812 | 0.58917168 |
| ANAPC5 | 78.417765 | 0.599868394 | 0.31208367 | 1.92213967 | 0.05458819 | 0.24318215 |
| RNF34 | 53.395596 | 0.024940019 | 0.33739995 | 0.07391827 | 0.94107542 | 0.98424983 |
| KDM2B | 65.9884683 | 0.30191994 | 0.2741941 | 1.10111758 | 0.27084549 | 0.58722969 |
| TMEM120B | 114.687529 | 0.674068562 | 0.44323439 | 1.52079481 | 0.12831134 | 0.397071 |
| RHOF | 17.9960718 | -1.346215686 | 0.66224907 | -2.0327936 | 0.04207338 | 0.20922698 |
| SETD1B | 23.4321382 | 0.246173032 | 0.31562808 | 0.77994655 | 0.43542233 | 0.72835986 |
| WDR66 | 35.0969189 | -1.274456455 | 0.54099391 | -2.3557686 | 0.01848443 | 0.12417062 |
| BCL7A | 68.7693027 | 0.583513469 | 0.40238134 | 1.45015043 | 0.14701658 | 0.4297124 |
| MLXIP | 87.0847126 | -1.301704805 | 0.46846425 | -2.7786641 | 0.00545829 | 0.05312435 |
| CLIP1 | 268.013488 | -1.057789669 | 0.53184599 | -1.9889022 | 0.046712 | 0.22187604 |
| ZCCHC8 | 69.3106619 | 0.195142859 | 0.25401097 | 0.7682458 | 0.44234117 | 0.73466139 |
| RSRC2 | 91.677842 | 0.165278551 | 0.31654022 | 0.52214076 | 0.60157233 | 0.83679851 |
| KNTC1 | 14.183261 | 0.152233378 | 0.47844714 | 0.31818223 | 0.75034671 | 0.90883772 |
| DENR | 95.6052695 | -0.097215121 | 0.30107785 | -0.3228903 | 0.74677831 | 0.90661036 |
| HIP1R | 89.4400585 | 0.577596735 | 0.4051409 | 1.42566877 | 0.15396397 | 0.43981621 |
| VPS37B | 142.653421 | -0.51156627 | 0.32318507 | -1.5828895 | 0.11344664 | 0.37119783 |
| ABCB9 | 22.6403975 | 0.301182441 | 0.52121994 | 0.57784136 | 0.56337123 | 0.81448184 |
| ARL6IP4 | 43.5203394 | -0.074870114 | 0.35060011 | -0.2135485 | 0.8308992 | 0.94421744 |
| PITPNM2 | 209.024831 | -0.058253635 | 0.31942809 | -0.1823685 | 0.85529351 | 0.95200547 |
| MPHOSPH9 | 88.1998211 | 0.428297971 | 0.23792929 | 1.80010612 | 0.07184388 | 0.28614796 |
| C12orf65 | 31.0697138 | -0.356122164 | 0.35263658 | -1.0098844 | 0.31255069 | 0.62752426 |
| SBNO1 | 97.7212341 | -0.300028896 | 0.29411179 | -1.0201186 | 0.30767224 | 0.62161058 |
| KMT5A | 17.9583798 | 0.060086085 | 0.47489637 | 0.12652463 | 0.89931666 | 0.96645314 |
| RILPL2 | 23.0456399 | -0.083545736 | 0.38581349 | -0.2165444 | 0.82856345 | 0.94282714 |
| SNRNP35 | 32.5155346 | -0.054161436 | 0.32436765 | -0.1669755 | 0.86738936 | 0.95588054 |
| RILPL1 | 85.9058326 | 0.548236086 | 0.31581855 | 1.7359211 | 0.08257779 | 0.30885595 |
| TMED2 | 31.9460765 | -0.679792957 | 0.35151662 | -1.9338857 | 0.05312718 | 0.23864142 |
| DDX55 | 22.2826851 | 0.107000685 | 0.36874341 | 0.29017654 | 0.77168118 | 0.91763236 |
| EIF2B1 | 28.1141168 | 0.345164031 | 0.28468392 | 1.21244651 | 0.22534151 | 0.53626447 |
| GTF2H3 | 24.9535821 | 0.580643686 | 0.32310716 | 1.79706229 | 0.07232573 | 0.28733069 |
| TCTN2 | 19.8961116 | 0.892048085 | 0.42671407 | 2.09050546 | 0.03657242 | 0.19157373 |
| ATP6V0A2 | 29.9251013 | -0.460870341 | 0.42454694 | -1.085558 | 0.27767457 | 0.59299993 |
| CCDC92 | 82.8314947 | -0.282616251 | 0.20925922 | -1.3505558 | 0.17683776 | 0.47393944 |
| RFLNA | 22.6642146 | 0.879211016 | 0.50068357 | 1.75602131 | 0.07908477 | 0.30106837 |
| NCOR2 | 138.20335 | 0.164488253 | 0.3118587 | 0.5274448 | 0.59788474 | 0.83478247 |
| UBC | 156.243483 | -1.533621347 | 0.36995274 | -4.1454521 | 3.39E-05 | 0.00106113 |
| BRI3BP | 20.4692837 | 0.778667344 | 0.49082865 | 1.58643418 | 0.11264085 | 0.36994129 |
| AACS | 42.5066498 | 1.452158206 | 0.28462931 | 5.10192778 | 3.36E-07 | 2.31E-05 |
| SLC15A4 | 41.7937023 | -0.166004012 | 0.32448582 | -0.511591 | 0.60893731 | 0.84023777 |
| RIMBP2 | 72.0655368 | -2.377287496 | 0.2712636 | -8.763754 | 1.89E-18 | 2.52E-15 |
| STX2 | 17.0546366 | -0.804724144 | 0.53114678 | -1.5150692 | 0.12975487 | 0.3998131 |
| RAN | 23.740978 | -1.17063103 | 0.50967209 | -2.2968317 | 0.02162837 | 0.13787238 |
| SFSWAP | 116.841218 | 0.047897327 | 0.34568151 | 0.13855912 | 0.88979855 | 0.9631927 |
| ULK1 | 58.5475988 | 0.059261845 | 0.33394759 | 0.17745852 | 0.85914824 | 0.95328983 |
| PUS1 | 15.2573236 | 0.025126785 | 0.53243736 | 0.047192 | 0.9623602 | 0.99029318 |
| EP400 | 105.1116 | 0.217429465 | 0.27221239 | 0.79874932 | 0.42443578 | 0.72151771 |
| FBRSL1 | 62.230629 | 0.32117135 | 0.55543533 | 0.57823356 | 0.56310644 | 0.81448184 |
| ANKLE2 | 87.0635595 | -1.486247558 | 0.60832555 | -2.4431779 | 0.01455856 | 0.1059672 |
| GOLGA3 | 58.4785967 | -0.927708665 | 0.4799865 | -1.9327808 | 0.05326321 | 0.23899239 |
| CHFR | 78.7871966 | -0.423166627 | 0.35910692 | -1.1783862 | 0.23864269 | 0.55063213 |
| ZNF26 | 56.7808601 | -0.287046284 | 0.27755684 | -1.0341892 | 0.30104773 | 0.61488917 |
| ZNF84 | 60.6732289 | 0.407358349 | 0.28596117 | 1.42452328 | 0.15429505 | 0.44013998 |
| ZNF140 | 34.4386211 | 0.169152486 | 0.34190023 | 0.49474224 | 0.62078208 | 0.84449549 |
| ZNF891 | 23.1702217 | 0.578713378 | 0.4573779 | 1.26528495 | 0.2057692 | 0.51244445 |
| MPHOSPH8 | 173.851194 | -0.392239267 | 0.20927585 | -1.8742692 | 0.06089334 | 0.25784263 |
| PSPC1 | 241.706352 | -0.04799557 | 0.36238717 | -0.1324428 | 0.89463406 | 0.9650125 |
| ZMYM5 | 83.3355418 | -0.175891054 | 0.28475442 | -0.6176938 | 0.53677717 | 0.80030369 |
| ZMYM2 | 448.36666 | 0.282530511 | 0.25831376 | 1.09374935 | 0.27406492 | 0.58894058 |
| CRYL1 | 123.065685 | 0.325681042 | 0.37965192 | 0.85784115 | 0.39098018 | 0.69372012 |
| IFT88 | 197.27876 | 0.15159554 | 0.31098685 | 0.48746607 | 0.62592808 | 0.8471532 |
| EEF1AKMT1 | 19.783619 | -0.024859473 | 0.35491122 | -0.0700442 | 0.94415848 | 0.98552865 |
| XPO4 | 147.075768 | 0.232227287 | 0.19920795 | 1.16575311 | 0.24371428 | 0.55580983 |
| LATS2 | 113.542395 | 0.747651328 | 0.32681696 | 2.28767601 | 0.0221564 | 0.13967967 |
| SAP18 | 37.2641813 | -0.594059912 | 0.44966783 | -1.3211083 | 0.18646525 | 0.48675708 |
| MRPL57 | 25.5033591 | -0.82152473 | 0.41462124 | -1.981386 | 0.047548 | 0.22356912 |
| ZDHHC20 | 130.07618 | -0.011217959 | 0.25102923 | -0.0446879 | 0.96435611 | 0.99082982 |
| MICU2 | 235.470607 | -0.082696887 | 0.25418475 | -0.3253417 | 0.7449225 | 0.90532552 |
| FGF9 | 78.2287386 | 1.468861145 | 0.5129155 | 2.86374882 | 0.0041866 | 0.0442082 |
| AL136962.1 | 53.3287484 | -0.51068929 | 0.71999871 | -0.7092919 | 0.47814332 | 0.76374064 |
| TNFRSF19 | 127.602739 | 1.935983169 | 0.38322511 | 5.0518171 | 4.38E-07 | 2.83E-05 |
| MIPEP | 119.012865 | 0.355451537 | 0.26393419 | 1.34674306 | 0.178063 | 0.47550635 |
| PARP4 | 64.340481 | -1.753515177 | 0.4965497 | -3.5313991 | 0.00041337 | 0.00751314 |
| MTMR6 | 31.6621517 | -0.086910958 | 0.35544546 | -0.2445128 | 0.8068337 | 0.93325451 |
| NUP58 | 91.2649787 | -0.162171038 | 0.38859391 | -0.4173278 | 0.67643866 | 0.87028412 |
| RNF6 | 16.8121592 | -0.996053428 | 0.57814816 | -1.7228342 | 0.08491851 | 0.31408459 |
| CDK8 | 246.147228 | 0.458088617 | 0.21305856 | 2.15005971 | 0.03155049 | 0.17567711 |
| WASF3 | 43.48525 | 0.049488534 | 0.48483042 | 0.10207391 | 0.91869801 | 0.97506435 |
| USP12 | 122.441689 | 0.411618768 | 0.26768611 | 1.53769191 | 0.12412396 | 0.3892808 |
| LINC02340 | 37.8113324 | 0.902072537 | 0.72413947 | 1.24571656 | 0.21286847 | 0.52201944 |
| RPL21 | 45.0506773 | 0.10664102 | 0.46759126 | 0.22806461 | 0.81959601 | 0.9384641 |
| GTF3A | 21.3589322 | -0.499511773 | 0.43223594 | -1.1556461 | 0.24782595 | 0.56031261 |
| MTIF3 | 45.7702243 | 0.361515218 | 0.29610594 | 1.22089822 | 0.22212456 | 0.53271965 |
| LNX2 | 61.862026 | -0.645431232 | 0.36102956 | -1.7877517 | 0.07381607 | 0.29101946 |
| POLR1D | 41.4808321 | -0.102689742 | 0.34174152 | -0.3004895 | 0.7638038 | 0.91479214 |
| PAN3 | 481.842321 | -0.067396088 | 0.16011245 | -0.4209297 | 0.67380641 | 0.86846218 |
| POMP | 32.4451351 | -0.388705562 | 0.38954502 | -0.997845 | 0.3183545 | 0.63093651 |
| SLC7A1 | 64.728105 | -1.516584977 | 0.38229934 | -3.9670092 | 7.28E-05 | 0.00192239 |

| | | | | | | |
|---|---|---|---|---|---|---|
| UBL3 | 131.068277 | -0.15178528 | 0.31110621 | -0.4878889 | 0.62562851 | 0.8471532 |
| KATNAL1 | 22.7307113 | 0.264765157 | 0.62542661 | 0.42333529 | 0.67205065 | 0.86777458 |
| HMGB1 | 185.959144 | 0.154194664 | 0.29096218 | 0.52994745 | 0.59614837 | 0.83416944 |
| USPL1 | 43.233723 | 0.336403714 | 0.29910227 | 1.12471134 | 0.26071137 | 0.57426715 |
| HSPH1 | 60.6797171 | 0.019071875 | 0.41493621 | 0.04596339 | 0.96333943 | 0.99075412 |
| B3GLCT | 73.690248 | 1.36968041 | 0.3223669 | 4.24882453 | 2.15E-05 | 0.0007421 |
| FRY | 91.3315364 | 0.706889868 | 0.69861236 | 1.01184849 | 0.3116105 | 0.62642275 |
| N4BP2L1 | 52.5359768 | -0.605914669 | 0.44442182 | -1.3633774 | 0.1727636 | 0.468183 |
| N4BP2L2 | 901.012053 | -0.188358351 | 0.22088921 | -0.8527277 | 0.39381031 | 0.69624829 |
| PDS5B | 260.108701 | 0.660386084 | 0.18934283 | 3.48777966 | 0.00048705 | 0.00847759 |
| KL | 258.534174 | 4.162789133 | 0.49326578 | 8.43924171 | 3.19E-17 | 2.71E-14 |
| STARD13 | 186.963527 | 0.23235402 | 0.44338915 | 0.52404083 | 0.60025014 | 0.83602169 |
| AL139383.1 | 54.9214688 | 0.04183684 | 0.65511977 | 0.06386136 | 0.94908062 | 0.98641632 |
| RFC3 | 29.0411232 | 0.537124049 | 0.76560372 | 0.70156927 | 0.48294782 | 0.7665951 |
| LINC02343 | 68.9682689 | -0.701299531 | 0.70579265 | -0.9936339 | 0.32040112 | 0.63212442 |
| NBEA | 409.239483 | -0.55600501 | 0.33835393 | -1.6432645 | 0.10032821 | 0.34581665 |
| SPART | 103.596071 | 0.259780431 | 0.24648818 | 1.05392652 | 0.29191656 | 0.60677688 |
| RFXAP | 14.1816478 | -0.031767393 | 0.55841021 | -0.056889 | 0.95463362 | 0.98834154 |
| ALG5 | 42.3698432 | 0.249308366 | 0.29178494 | 0.85442506 | 0.39286952 | 0.69595061 |
| EXOSC8 | 20.1096385 | 0.470442215 | 0.41133819 | 1.14368717 | 0.2527534 | 0.5654205 |
| SUPT20H | 121.260526 | 0.167936274 | 0.27152025 | 0.61850369 | 0.53624337 | 0.80011732 |
| LINC00571 | 45.120887 | -0.726584718 | 0.47089736 | -1.542979 | 0.12283588 | 0.38745661 |
| UFM1 | 55.3971612 | -0.121341619 | 0.32594997 | -0.3722707 | 0.70969132 | 0.88614996 |
| FREM2 | 233.702561 | -0.132489936 | 0.68105421 | -0.1945365 | 0.84575578 | 0.94909204 |
| PROSER1 | 42.317663 | 0.421832847 | 0.35121326 | 1.20107325 | 0.22972279 | 0.54241242 |
| NHLRC3 | 67.6851628 | 0.246298468 | 0.58178708 | 0.42334812 | 0.67204129 | 0.86777458 |
| COG6 | 106.960476 | 0.004290196 | 0.29624962 | 0.01448169 | 0.98844568 | 0.9977555 |
| LINC00598 | 27.608767 | -1.021640471 | 0.56055716 | -1.8225447 | 0.06837238 | 0.27855451 |
| FOXO1 | 518.174837 | -1.559447826 | 0.6018854 | -2.5909381 | 0.00957147 | 0.0795686 |
| MRPS31 | 82.9838132 | 0.175555139 | 0.26298161 | 0.66755671 | 0.5044166 | 0.77792612 |
| ELF1 | 255.964665 | -0.452553681 | 0.1623821 | -2.7869677 | 0.00532038 | 0.05235679 |
| WBP4 | 40.2755703 | 0.41771371 | 0.37911506 | 1.10181249 | 0.2705432 | 0.58718454 |
| AL354696.2 | 20.3956622 | -0.034038159 | 0.65337788 | -0.0520957 | 0.95845246 | 0.98911021 |
| MTRF1 | 22.9075395 | 0.058163357 | 0.36076665 | 0.16122155 | 0.87191891 | 0.9568935 |
| NAA16 | 75.5117607 | -0.079587966 | 0.27065135 | -0.2940609 | 0.7687114 | 0.91600052 |
| VWA8 | 409.368587 | 1.044029498 | 0.37405735 | 2.7910947 | 0.00525301 | 0.05193962 |
| DGKH | 201.621789 | -1.833479255 | 0.29229381 | -6.272727 | 3.55E-10 | 7.19E-08 |
| AKAP11 | 74.0265703 | 0.112472128 | 0.22568365 | 0.49836187 | 0.618229 | 0.84283824 |
| EPSTI1 | 31.8306005 | 1.08509146 | 1.29693198 | 0.83666027 | 0.40278356 | 0.70169281 |
| DNAJC15 | 53.6364596 | -0.495404413 | 0.3245752 | -1.5263163 | 0.12693111 | 0.39482486 |
| ENOX1 | 18.9800146 | -1.05676711 | 0.77202826 | -1.3688192 | 0.17105579 | 0.46688992 |
| CCDC122 | 43.13153 | -0.513605697 | 0.59499868 | -0.8632048 | 0.38802489 | 0.69163527 |
| TSC22D1 | 768.797562 | 1.280602829 | 0.20661073 | 6.19814278 | 5.71E-10 | 1.09E-07 |
| NUFIP1 | 16.403616 | 0.930926192 | 0.4440631 | 2.09638269 | 0.03604826 | 0.18968056 |
| AL359706.1 | 17.5072399 | -0.524850902 | 0.79699516 | -0.6585371 | 0.51019305 | 0.78158783 |
| GPALPP1 | 52.4457625 | -0.039597669 | 0.226575 | -0.1747663 | 0.86126329 | 0.95407139 |
| GTF2F2 | 101.102974 | -0.44005507 | 0.29599006 | -1.4867224 | 0.13708813 | 0.41253153 |
| TPT1 | 204.083738 | -1.401431796 | 0.38488971 | -3.6411257 | 0.00027145 | 0.00537613 |
| SLC25A30 | 80.7938335 | 0.384917967 | 0.40658841 | 0.94670177 | 0.34379077 | 0.65306073 |
| COG3 | 74.590719 | 0.632162393 | 0.31209568 | 2.02554038 | 0.04281191 | 0.21080021 |
| LINC01055 | 170.711729 | 0.665485924 | 0.64725999 | 1.02815859 | 0.30387523 | 0.61795696 |
| ZC3H13 | 136.777202 | 0.330326411 | 0.16864639 | 1.95869243 | 0.05014882 | 0.23113573 |
| LRCH1 | 646.264764 | 1.27964531 | 0.26887113 | 4.75932585 | 1.94E-06 | 9.84E-05 |
| ESD | 42.13938 | 0.442420696 | 0.37114108 | 1.19205531 | 0.23323956 | 0.54577316 |
| SUCLA2 | 130.653468 | 0.416786173 | 0.25071269 | 1.66240555 | 0.09643149 | 0.33942136 |
| MED4 | 67.4551085 | -0.227284117 | 0.25546443 | -0.8896899 | 0.37363243 | 0.67707283 |
| ITM2B | 435.014874 | -1.134663386 | 0.47836432 | -2.3719649 | 0.01769377 | 0.12083193 |
| RB1 | 242.683678 | -0.698675436 | 0.30564573 | -2.2858995 | 0.02226014 | 0.14014418 |
| RCBTB2 | 27.1388902 | -0.237678046 | 0.43678115 | -0.5441582 | 0.58633261 | 0.82945915 |
| FNDC3A | 601.042546 | -1.014107009 | 0.63388285 | -1.5998335 | 0.10963553 | 0.3644153 |
| CDADC1 | 28.6760135 | 0.067363044 | 0.3031732 | 0.22219326 | 0.82416344 | 0.94149717 |
| CAB39L | 61.9972356 | -0.245123615 | 0.48454711 | -0.5058819 | 0.61293955 | 0.84174925 |
| SETDB2 | 116.824898 | -0.029018824 | 0.41995278 | -0.0691002 | 0.94490986 | 0.98552865 |
| PHF11 | 54.0981129 | -0.176169451 | 0.35728553 | -0.4930775 | 0.62195784 | 0.84535494 |
| RCBTB1 | 21.434423 | -0.645081786 | 0.33931284 | -1.9011417 | 0.05728346 | 0.24969339 |
| EBPL | 16.9856085 | -0.573125454 | 0.4333479 | -1.3225527 | 0.18598416 | 0.48628295 |
| KPNA3 | 89.1560999 | 0.094898123 | 0.21421145 | 0.44301143 | 0.65775746 | 0.86357172 |
| SPRYD7 | 16.1876592 | 0.622660629 | 0.57460484 | 1.08363276 | 0.27852766 | 0.59359815 |
| DLEU2 | 104.393045 | -0.926435045 | 0.48502531 | -1.9100757 | 0.05612347 | 0.24695387 |
| TRIM13 | 36.9494273 | 0.213058919 | 0.30047578 | 0.70907186 | 0.47827988 | 0.76374064 |
| RNASEH2B | 34.5315578 | 0.388854995 | 0.32906996 | 1.18167879 | 0.23733318 | 0.54992696 |
| INTS6 | 178.483353 | 0.184063036 | 0.38032438 | 0.48396328 | 0.62841193 | 0.84817317 |
| INTS6-AS1 | 19.3629278 | 0.537846484 | 0.44756463 | 1.20171803 | 0.22947279 | 0.54228719 |
| WDFY2 | 424.782428 | 1.565711688 | 0.38597914 | 4.05646708 | 4.98E-05 | 0.00141624 |
| ATP7B | 14.3090787 | 0.267893961 | 0.52949426 | 0.50594309 | 0.6128966 | 0.84174925 |
| ALG11 | 15.7362028 | 0.539312257 | 0.47235711 | 1.14174688 | 0.25355925 | 0.56559484 |
| UTP14C | 17.0445754 | -0.20204108 | 0.39499741 | -0.5114998 | 0.60900116 | 0.84023777 |
| NEK3 | 39.3565807 | 0.327900735 | 0.55909381 | 0.58648608 | 0.55754891 | 0.81230403 |
| VPS36 | 89.9359737 | 0.208026297 | 0.24153257 | 0.86127638 | 0.38908584 | 0.69247382 |
| SUGT1 | 59.6282156 | 0.363639718 | 0.2763298 | 1.31596275 | 0.18818655 | 0.4882768 |
| DIAPH3 | 110.542859 | -2.392141599 | 0.81978929 | -2.9179957 | 0.00352289 | 0.03910411 |
| TDRD3 | 230.923442 | 0.49100852 | 0.3178434 | 1.5448127 | 0.12239157 | 0.38683625 |
| PCDH9 | 90.1668986 | -0.236194041 | 0.55967955 | -0.4220166 | 0.67301293 | 0.8684158 |
| DACH1 | 1465.48041 | -0.07678883 | 0.5771745 | -0.1330427 | 0.89415964 | 0.96482804 |
| DIS3 | 41.9333475 | -0.156987623 | 0.41660463 | -0.3768264 | 0.70630262 | 0.88599027 |
| PIBF1 | 255.44541 | -0.092222286 | 0.18520991 | -0.4979339 | 0.61853066 | 0.84283824 |
| KLF5 | 57.9504946 | -2.383826417 | 0.68673145 | -3.4712644 | 0.00051801 | 0.00884093 |
| KLF12 | 380.280689 | 0.907661675 | 0.1902095 | 4.77190509 | 1.82E-06 | 9.40E-05 |
| TBC1D4 | 480.125774 | -0.045938176 | 0.50256641 | -0.0914072 | 0.92716906 | 0.97767557 |
| COMMD6 | 36.9615393 | 0.102887304 | 0.45835576 | 0.22447041 | 0.82239129 | 0.94062517 |
| UCHL3 | 55.1600212 | -0.450695904 | 0.35383183 | -1.2737574 | 0.20274941 | 0.50710716 |
| LMO7-AS1 | 21.2959314 | -0.612432076 | 0.55240577 | -1.1086634 | 0.26757541 | 0.58365544 |
| LMO7 | 366.113183 | 0.869653042 | 0.41637161 | 2.08864635 | 0.03673957 | 0.19223331 |
| AC001226.2 | 24.0208672 | -0.797715333 | 0.61379802 | -1.2996382 | 0.19372501 | 0.49541744 |
| FBXL3 | 65.0972806 | 0.173697157 | 0.30641076 | 0.56687682 | 0.57079788 | 0.81891359 |
| MYCBP2 | 240.202504 | 0.209341344 | 0.24091295 | 0.86895013 | 0.38487441 | 0.68852052 |
| SLAIN1 | 27.2695522 | -0.171035297 | 0.32002191 | -0.5344487 | 0.59303113 | 0.83362313 |
| RNF219 | 19.837755 | -0.482575435 | 0.41111327 | -1.173826 | 0.24046474 | 0.55302079 |
| RBM26 | 249.128073 | 0.533123083 | 0.27379279 | 1.94717722 | 0.05151349 | 0.23455111 |
| NDFIP2 | 140.495432 | -0.847867965 | 0.43317806 | -1.9573197 | 0.05030989 | 0.23153474 |
| GPC5 | 1660.85762 | 1.380596653 | 1.45180072 | 0.95095465 | 0.3416274 | 0.65120286 |
| GPC6 | 58.1452264 | 1.359082933 | 0.99502412 | 1.36587938 | 0.17197683 | 0.46776894 |
| TGDS | 24.507683 | 0.349006474 | 0.35576133 | 0.98101295 | 0.32658635 | 0.63830738 |
| GPR180 | 20.9716401 | -0.002774954 | 0.44716662 | -0.0062056 | 0.99504865 | 0.99922221 |
| ABCC4 | 26.3583692 | -1.737871884 | 0.43896392 | -3.9590312 | 7.53E-05 | 0.00197654 |
| CLDN10 | 31.8792812 | -0.541933709 | 0.66527616 | -0.8145996 | 0.41530152 | 0.71273171 |
| CLDN10-AS1 | 39.479504 | -2.530362867 | 0.86440366 | -2.9272931 | 0.00341926 | 0.03844452 |
| DZIP1 | 37.9395526 | 0.623618545 | 0.30032777 | 2.07645983 | 0.03785144 | 0.19628857 |
| DNAJC3 | 123.605648 | -0.133997472 | 0.43094374 | -0.3109396 | 0.75584654 | 0.91076676 |
| UGGT2 | 252.94102 | 0.065724263 | 0.24223646 | 0.27132275 | 0.78614281 | 0.92322402 |
| MBNL2 | 366.138415 | -0.678143937 | 0.59162932 | -1.1462311 | 0.25169954 | 0.56401021 |
| RAP2A | 13.6687363 | -0.236122254 | 0.50452423 | -0.4680097 | 0.63977762 | 0.85413611 |
| IPO5 | 54.0082463 | -0.071643329 | 0.24390805 | -0.2937309 | 0.76896355 | 0.91600052 |

| | | | | | | |
|---|---|---|---|---|---|---|
| FARP1 | 890.284239 | -0.153741998 | 0.24311434 | -0.6323855 | 0.52713497 | 0.79403853 |
| STK24 | 329.012535 | -1.544813801 | 0.37070225 | -4.167263 | 3.08E-05 | 0.00098438 |
| DOCK9 | 132.365106 | -0.468201788 | 0.27374587 | -1.710352 | 0.0872008 | 0.31934811 |
| UBAC2 | 276.732156 | 0.338182012 | 0.1709899 | 1.97778945 | 0.04795246 | 0.22433958 |
| TM9SF2 | 118.161923 | -0.108124258 | 0.19199585 | -0.5631594 | 0.57332637 | 0.82102521 |
| CLYBL | 151.638273 | 0.309254287 | 0.37744754 | 0.81933052 | 0.41259787 | 0.71110141 |
| AL355338.1 | 20.1198254 | -0.350502541 | 0.40303559 | -0.8696565 | 0.38448813 | 0.68822563 |
| PCCA | 1349.87887 | -0.622532776 | 0.22466602 | -2.7709254 | 0.00558972 | 0.05377622 |
| GGACT | 19.9827271 | -0.656144155 | 0.46089041 | -1.4236446 | 0.15454937 | 0.44013998 |
| TMTC4 | 23.5236283 | -0.307223797 | 0.43642113 | -0.7039618 | 0.48145658 | 0.76608505 |
| FGF14 | 108.831512 | 1.660169 | 0.70202 | 2.36484571 | 0.01803758 | 0.12240351 |
| TPP2 | 132.508563 | 1.113728073 | 0.23501588 | 4.73894812 | 2.15E-06 | 0.00010712 |
| ARGLU1 | 199.163896 | -0.071292805 | 0.21679468 | -0.3288494 | 0.74226951 | 0.9043409 |
| FAM155A | 1354.59069 | -1.562945961 | 0.70408812 | -2.2198158 | 0.02643127 | 0.15637384 |
| LIG4 | 16.9728008 | 0.156673653 | 0.47899041 | 0.32709142 | 0.74359874 | 0.90463949 |
| ABHD13 | 33.1227328 | 0.120626351 | 0.44693854 | 0.26989472 | 0.78724125 | 0.92362588 |
| TNFSF13B | 41.9393857 | -0.035170528 | 0.53631671 | -0.0655779 | 0.94771388 | 0.98633762 |
| IRS2 | 97.4408101 | -1.18772839 | 0.4674099 | -2.5410852 | 0.0110509 | 0.08807986 |
| COL4A1 | 93.9863915 | -0.748561246 | 0.48626893 | -1.5393977 | 0.12370725 | 0.38887694 |
| COL4A2 | 141.248673 | -0.984244045 | 0.4247873 | -2.317028 | 0.02050221 | 0.13330725 |
| RAB20 | 36.6929369 | -0.032915734 | 0.35238768 | -0.0934077 | 0.92557965 | 0.97735117 |
| NAXD | 34.7370411 | -0.250747032 | 0.30693249 | -0.8169452 | 0.41395974 | 0.71160612 |
| CARS2 | 67.4039244 | 0.242784149 | 0.26072401 | 0.93119212 | 0.35175419 | 0.65871783 |
| ANKRD10 | 337.461536 | -0.572025273 | 0.32943427 | -1.7363866 | 0.08249549 | 0.30885595 |
| ARHGEF7 | 245.701917 | -0.132772809 | 0.27442062 | -0.4838296 | 0.62850683 | 0.84817317 |
| TUBGCP3 | 60.1260636 | 0.434040877 | 0.30077339 | 1.44308271 | 0.14899717 | 0.43272656 |
| ATP11A | 94.9240507 | -3.744437557 | 0.79284116 | -4.7228092 | 2.33E-06 | 0.00011444 |
| MCF2L | 69.9139486 | 0.594579636 | 0.42234139 | 1.40781757 | 0.1591851 | 0.44760007 |
| PCID2 | 47.4345498 | 0.027927821 | 0.31467885 | 0.08875023 | 0.92928041 | 0.97876742 |
| CUL4A | 135.876181 | -0.421624749 | 0.28665271 | -1.4708556 | 0.14133017 | 0.42154911 |
| LAMP1 | 70.6616885 | -0.776803154 | 0.39073177 | -1.9880727 | 0.04680365 | 0.22193782 |
| GRTP1 | 28.1605465 | -0.291095805 | 0.36579474 | -0.79579 | 0.42615409 | 0.72211376 |
| DCUN1D2 | 52.1292155 | 0.567622356 | 0.29778599 | 1.90614189 | 0.0566318 | 0.24790371 |
| TMCO3 | 42.0692993 | -0.582060401 | 0.26844032 | -2.1683047 | 0.03013551 | 0.17029303 |
| TFDP1 | 37.2943741 | 0.303566207 | 0.30349854 | 1.00022296 | 0.31720262 | 0.63093651 |
| GAS6 | 90.555162 | 1.685679032 | 0.4241062 | 3.97466253 | 7.05E-05 | 0.00188835 |
| RASA3 | 47.7923399 | -2.14322105 | 0.73783456 | -2.9047447 | 0.00367553 | 0.04037138 |
| CDC16 | 72.017017 | 0.260514386 | 0.26647015 | 0.97764941 | 0.32824775 | 0.63988149 |
| UPF3A | 83.312101 | 0.08916889 | 0.26801033 | 0.33270691 | 0.73935553 | 0.90291434 |
| CHAMP1 | 10.961059 | 0.836400584 | 0.51689918 | 1.61811164 | 0.10563854 | 0.35687176 |
| TTC5 | 24.5537017 | 0.390623624 | 0.43532132 | 0.89732251 | 0.36954685 | 0.67364366 |
| CCNB1IP1 | 72.2124293 | 1.883955119 | 0.37484298 | 5.02598479 | 5.01E-07 | 3.20E-05 |
| TEP1 | 30.3896314 | -1.141358652 | 0.42801344 | -2.6666421 | 0.00766132 | 0.06790319 |
| OSGEP | 22.8195747 | 0.246018285 | 0.42046898 | 0.58510448 | 0.55847747 | 0.81255224 |
| APEX1 | 18.6374831 | -0.340835773 | 0.58859721 | -0.5790645 | 0.56254562 | 0.814089 |
| PNP | 17.4819068 | -0.927225342 | 0.38363312 | -2.4169586 | 0.0156508 | 0.11131049 |
| METTL17 | 23.3857681 | 0.681260097 | 0.3584166 | 1.90074931 | 0.05733486 | 0.24969339 |
| NDRG2 | 31.5764668 | 1.432793355 | 0.51652618 | 2.7739027 | 0.00553882 | 0.05351709 |
| ARHGEF40 | 16.510828 | -0.130725031 | 0.51291105 | -0.2548688 | 0.79882443 | 0.92917153 |
| HNRNPC | 504.339633 | 0.109977442 | 0.33797238 | 0.32540364 | 0.7448756 | 0.90532552 |
| SUPT16H | 53.5817539 | -0.119453204 | 0.28630447 | -0.4172244 | 0.6765143 | 0.87028412 |
| CHD8 | 98.2464257 | 0.124327511 | 0.23679184 | 0.5250498 | 0.59954857 | 0.83582612 |
| RAB2B | 36.2383275 | -0.09282425 | 0.32702044 | -0.2838485 | 0.77652651 | 0.9196152 |
| TOX4 | 44.4063293 | -0.347566387 | 0.29982985 | -1.1592121 | 0.24636975 | 0.55930671 |
| METTL3 | 19.7586969 | 0.462927702 | 0.36166495 | 1.2799905 | 0.20054848 | 0.50442784 |
| DAD1 | 44.1240996 | -0.087693297 | 0.29400944 | -0.2982669 | 0.76549943 | 0.91549415 |
| ABHD4 | 26.0767705 | -0.032383918 | 0.32722674 | -0.0989648 | 0.92116624 | 0.97548809 |
| OXA1L | 22.8103062 | -0.045345763 | 0.44512972 | -0.1018709 | 0.91885915 | 0.97512437 |
| MRPL52 | 16.4839761 | 0.055788023 | 0.4712827 | 0.11837486 | 0.90577065 | 0.96930836 |
| MMP14 | 22.407098 | -0.619128704 | 0.56535943 | -1.0951064 | 0.27347004 | 0.58894058 |
| LRP10 | 54.3020181 | -1.933639156 | 0.42917277 | -4.5055029 | 6.62E-06 | 0.00026961 |
| RBM23 | 70.0958222 | -0.235448224 | 0.33968776 | -0.6931313 | 0.48822715 | 0.77012857 |
| AL132780.1 | 44.4800588 | -1.02776296 | 0.83612644 | -1.2291956 | 0.21899847 | 0.52913753 |
| C14orf93 | 15.9262546 | 0.709054535 | 0.38935623 | 1.8210946 | 0.06859248 | 0.27887855 |
| PSMB5 | 27.2646771 | -0.345557644 | 0.38082986 | -0.9073806 | 0.36420554 | 0.6696613 |
| ACIN1 | 136.706917 | 0.063367802 | 0.31342771 | 0.20217677 | 0.83977853 | 0.9474978 |
| C14orf119 | 13.3544093 | -0.787375118 | 0.39438547 | -1.9964608 | 0.04588379 | 0.21973316 |
| SLC7A8 | 69.1939587 | -0.521667121 | 0.68552467 | -0.760975 | 0.44667201 | 0.73948328 |
| HOMEZ | 14.417681 | 0.953050292 | 0.51766128 | 1.84106931 | 0.0656114 | 0.27144773 |
| PPP1R3E | 10.6782724 | 0.406058101 | 0.76461513 | 0.53106208 | 0.59537576 | 0.83416944 |
| BCL2L2-PABI | 28.0843446 | -0.677002757 | 0.36829418 | -1.8382119 | 0.06603118 | 0.27225103 |
| PABPN1 | 72.3735133 | 0.962663533 | 0.49157715 | 1.95831628 | 0.05019291 | 0.23122466 |
| NGDN | 18.6219756 | 0.047671294 | 0.37269186 | 0.12791075 | 0.89821961 | 0.96607503 |
| ZFHX2 | 32.7209512 | 0.0088109 | 0.55954911 | 0.01574643 | 0.98743669 | 0.9977555 |
| AP1G2 | 15.5403041 | -0.099559199 | 0.49806837 | -0.1998906 | 0.84156612 | 0.94776435 |
| DHRS4-AS1 | 22.7258324 | 0.046911494 | 0.48069176 | 0.09759163 | 0.92225657 | 0.97611757 |
| DHRS4L2 | 13.0100643 | 0.391304889 | 0.64621836 | 0.60553045 | 0.5448266 | 0.80442807 |
| PSME1 | 20.2900223 | -0.790776701 | 0.44758878 | -1.7667483 | 0.0772704 | 0.29759157 |
| AL136295.1 | 18.3113941 | -0.656979698 | 0.46427427 | -1.4150681 | 0.15704854 | 0.44367158 |
| DHRS1 | 15.3304819 | -0.527221979 | 0.41902754 | -1.2582037 | 0.20831811 | 0.51645229 |
| KHNYN | 18.8943649 | -0.427154509 | 0.40838852 | -1.0459513 | 0.29558351 | 0.61055359 |
| SDR39U1 | 21.0363776 | -0.067315565 | 0.41337314 | -0.1628446 | 0.87064083 | 0.95629149 |
| NOVA1 | 91.92041 | 0.290869173 | 0.2955873 | 0.98403814 | 0.32509676 | 0.63715682 |
| LINC00645 | 159.411788 | 0.894539951 | 0.6405616 | 1.39649325 | 0.16256601 | 0.45306004 |
| PRKD1 | 1168.33454 | -0.250894026 | 0.53594738 | -0.4681318 | 0.63969032 | 0.85413611 |
| G2E3 | 40.5406381 | 0.653260601 | 0.27536168 | 2.37237293 | 0.01767424 | 0.12081718 |
| SCFD1 | 170.592968 | 0.182965451 | 0.22111102 | 0.82748228 | 0.40796376 | 0.70664204 |
| STRN3 | 393.279771 | -0.304465609 | 0.41679419 | -0.7304939 | 0.46508835 | 0.75251779 |
| AP4S1 | 24.2299008 | 0.389421322 | 0.46244382 | 0.84209435 | 0.39973515 | 0.69959441 |
| HECTD1 | 394.80346 | -1.221066692 | 0.65990438 | -1.8503691 | 0.06426037 | 0.26796228 |
| HEATR5A | 48.6323103 | 0.118966283 | 0.33004854 | 0.36045087 | 0.71850999 | 0.8907358 |
| AL139353.1 | 12.9413875 | 0.178249896 | 0.51863308 | 0.34369172 | 0.73107814 | 0.89782437 |
| NUBPL | 161.627559 | 0.979390981 | 0.368956 | 2.65449266 | 0.00794278 | 0.06963542 |
| ARHGAP5 | 420.137633 | -0.238601736 | 0.31415711 | -0.7594981 | 0.44755463 | 0.7399821 |
| AKAP6 | 116.657642 | 0.945816368 | 0.4177199 | 2.26423586 | 0.02355961 | 0.14566964 |
| NPAS3 | 206.344343 | -0.146384114 | 0.63817572 | -0.229379 | 0.81857435 | 0.93816073 |
| EGLN3 | 27.1501602 | -1.925408967 | 0.66145797 | -2.9108561 | 0.0036044 | 0.03986646 |
| EAPP | 52.4430854 | -0.466815973 | 0.37268388 | -1.2525789 | 0.21035899 | 0.51888551 |
| SNX6 | 141.03057 | 0.156177811 | 0.27581989 | 0.56623113 | 0.57123667 | 0.81895612 |
| BAZ1A | 74.4358191 | -1.798062373 | 0.54173822 | -3.3190613 | 0.00090321 | 0.01379461 |
| SRP54 | 105.959504 | -0.455308313 | 0.3504633 | -1.2991612 | 0.19388863 | 0.49569991 |
| FAM177A1 | 56.4518571 | -0.487376463 | 0.22717364 | -2.1453918 | 0.03192154 | 0.17712034 |
| PPP2R3C | 45.9099675 | -0.693322828 | 0.45121209 | -1.5365786 | 0.12439654 | 0.38965272 |
| NFKBIA | 38.6742752 | -1.169212164 | 0.51241639 | -2.2817619 | 0.0225034 | 0.14110406 |
| RALGAPA1 | 384.576789 | 0.503793068 | 0.2295925 | 2.19429238 | 0.0282144 | 0.16339816 |
| BRMS1L | 25.1336691 | 0.305954481 | 0.39468582 | 0.77518488 | 0.43823042 | 0.73056686 |
| MBIP | 44.314244 | 0.142568764 | 0.29370837 | 0.48540926 | 0.62738607 | 0.84783129 |
| SLC25A21 | 20.8225605 | 0.779333541 | 0.77536241 | 1.00512164 | 0.31483828 | 0.62946385 |
| MIPOL1 | 170.208944 | 0.796520171 | 0.24613173 | 3.2361539 | 0.00121152 | 0.0173521 |
| SEC23A | 110.08005 | 0.24688136 | 0.24655759 | 1.00131316 | 0.31667543 | 0.63070323 |
| TRAPPC6B | 84.1989402 | 0.427764412 | 0.2984587 | 1.43324491 | 0.15178786 | 0.43620176 |
| PNN | 130.130905 | -0.323316414 | 0.40450711 | -0.7992849 | 0.42412525 | 0.72151771 |
| MIA2 | 181.184544 | 0.036240431 | 0.25931285 | 0.13975563 | 0.88885307 | 0.96307703 |

| | | | | | | |
|---|---|---|---|---|---|---|
| FBXO33 | 84.9887892 | -0.436994583 | 0.25932773 | -1.6851055 | 0.09196822 | 0.33050702 |
| AL049875.1 | 27.8822717 | -0.204418465 | 0.31607757 | -0.6467351 | 0.51780339 | 0.78658624 |
| KLHL28 | 68.7759207 | 0.026350403 | 0.25978747 | 0.10143062 | 0.91920863 | 0.97513464 |
| TOGARAM1 | 123.388257 | 0.217178709 | 0.26543527 | 0.8181984 | 0.41324391 | 0.71143975 |
| PRPF39 | 52.6359328 | 0.270909391 | 0.28543145 | 0.94912245 | 0.34255834 | 0.65223891 |
| FKBP3 | 32.1342799 | 0.341366285 | 0.31072176 | 1.09862367 | 0.27193225 | 0.58786226 |
| FANCM | 32.3327929 | 0.252419901 | 0.3387387 | 0.74517586 | 0.45616541 | 0.74627681 |
| MIS18BP1 | 27.1410613 | -0.628055477 | 0.48934177 | -1.28347 | 0.19932746 | 0.50273976 |
| AL358335.2 | 23.111112 | -0.226451304 | 0.82611474 | -0.274116 | 0.78399544 | 0.92232893 |
| RPS29 | 63.6286526 | 0.002596405 | 0.33732855 | 0.00769696 | 0.99385877 | 0.9992177 |
| RPL36AL | 48.6626565 | -0.926330242 | 0.47841001 | -1.9362685 | 0.05283482 | 0.23809501 |
| DNAAF2 | 16.2933631 | -0.062696962 | 0.4473783 | -0.1401431 | 0.88854696 | 0.96307703 |
| KLHDC1 | 44.4349237 | -0.062219474 | 0.51617441 | -0.1205396 | 0.90405569 | 0.9686272 |
| KLHDC2 | 45.3546953 | -0.094592292 | 0.37267276 | -0.2538213 | 0.7996336 | 0.92922871 |
| NEMF | 213.966247 | 0.065951523 | 0.21965251 | 0.3002539 | 0.76398349 | 0.91479214 |
| VCPKMT | 29.9349349 | 0.463583986 | 0.38841908 | 1.19351495 | 0.23266776 | 0.5452109 |
| SOS2 | 315.713194 | -0.001894992 | 0.35706394 | -0.0053072 | 0.99576553 | 0.99922221 |
| L2HGDH | 81.6108244 | 0.21185953 | 0.28543994 | 0.74222105 | 0.45795342 | 0.747377 |
| ATP5S | 47.6286522 | 0.274162316 | 0.28467981 | 0.96305502 | 0.33551991 | 0.64662829 |
| CDKL1 | 76.5009905 | -0.05797447 | 0.38951705 | -0.1488368 | 0.88168241 | 0.96155389 |
| MAP4K5 | 151.598092 | -0.307641235 | 0.20853693 | -1.4752363 | 0.14014903 | 0.41923311 |
| SAV1 | 47.8085609 | -0.634178735 | 0.29803907 | -2.1278376 | 0.03335055 | 0.18186117 |
| NIN | 55.6763955 | 0.418440391 | 0.39638181 | 1.05564982 | 0.29112823 | 0.60564025 |
| PYGL | 16.6484901 | -2.405477364 | 0.69303551 | -3.4709294 | 0.00051866 | 0.00884093 |
| TMX1 | 26.7389123 | 0.066106686 | 0.33018188 | 0.20021294 | 0.84131405 | 0.94776435 |
| FRMD6 | 18.4696583 | -3.006507936 | 0.46260026 | -6.4991488 | 8.08E-11 | 1.98E-08 |
| RTRAF | 25.5560499 | -0.607032555 | 0.34969179 | -1.7359074 | 0.08258021 | 0.30885595 |
| TXNDC16 | 28.0395831 | 0.695014217 | 0.32934605 | 2.11028556 | 0.03483377 | 0.18591302 |
| ERO1A | 173.014084 | -2.546366213 | 0.67721772 | -3.7600407 | 0.00016989 | 0.00369234 |
| PSMC6 | 68.0381155 | -0.333198491 | 0.31901635 | -1.0444559 | 0.29627453 | 0.61093809 |
| STYX | 36.0504003 | 0.705857555 | 0.36500462 | 1.93383185 | 0.0531338 | 0.23864142 |
| GNPNAT1 | 25.9591582 | -0.362486749 | 0.51217944 | -0.7077339 | 0.47911052 | 0.76419703 |
| FERMT2 | 319.065092 | 0.522645863 | 0.27960434 | 1.86923376 | 0.0615903 | 0.25996738 |
| DDHD1 | 55.4826747 | -0.002947852 | 0.291035 | -0.0101289 | 0.99191848 | 0.9988777 |
| CNIH1 | 23.2257254 | 0.508630823 | 0.34952401 | 1.45520998 | 0.14561114 | 0.4281546 |
| GMFB | 22.6226637 | -0.038219508 | 0.34369408 | -0.1112021 | 0.91145608 | 0.97128978 |
| CGRRF1 | 15.9717471 | 0.576012222 | 0.45219219 | 1.27382169 | 0.20272662 | 0.50710716 |
| SAMD4A | 224.090663 | -1.596200869 | 0.78485016 | -2.0337651 | 0.04197528 | 0.20900846 |
| SOCS4 | 29.6128785 | -0.264807086 | 0.34941079 | -0.7578675 | 0.4485303 | 0.74006308 |
| MAPK1IP1L | 126.955207 | -0.127237839 | 0.25638019 | -0.4962858 | 0.61969281 | 0.84360228 |
| LGALS3 | 69.2178553 | -2.290606021 | 0.38443616 | -5.9583521 | 2.55E-09 | 3.65E-07 |
| FBXO34 | 198.257341 | 0.282906539 | 0.23931246 | 1.18216383 | 0.2371407 | 0.54992696 |
| ATG14 | 57.2843605 | 0.189841651 | 0.36203665 | 0.52437136 | 0.60002027 | 0.83601151 |
| KTN1 | 268.053657 | -0.179737252 | 0.31778096 | -0.5656011 | 0.57166499 | 0.81940114 |
| PELI2 | 215.853002 | 0.059126542 | 0.28686757 | 0.20611093 | 0.83670426 | 0.94684966 |
| TMEM260 | 112.307666 | 0.205312229 | 0.29401485 | 0.69830565 | 0.48498606 | 0.76770898 |
| EXOC5 | 83.8049522 | 0.042580536 | 0.24099025 | 0.17668987 | 0.859752 | 0.95342671 |
| AP5M1 | 73.7048793 | 0.554334579 | 0.31279838 | 1.77217854 | 0.07636492 | 0.29596141 |
| NAA30 | 30.3781819 | 0.433631132 | 0.30941677 | 1.40144676 | 0.16108052 | 0.45075473 |
| CCDC198 | 229.630925 | 0.781094502 | 0.57251361 | 1.36432477 | 0.17246537 | 0.46814182 |
| ARMH4 | 73.8982181 | -0.115600223 | 0.27246193 | -0.4242803 | 0.67136142 | 0.86777458 |
| ACTR10 | 68.8823291 | -0.031448865 | 0.23016737 | -0.1366348 | 0.89131949 | 0.96328012 |
| PSMA3 | 59.6831074 | -0.094418464 | 0.28862532 | -0.3271316 | 0.74356835 | 0.90463949 |
| ARID4A | 113.87769 | -0.057060361 | 0.24499305 | -0.232906 | 0.81583437 | 0.93717111 |
| TIMM9 | 25.5224338 | -0.580373348 | 0.41811908 | -1.3880576 | 0.16511952 | 0.4569826 |
| KIAA0586 | 76.8907034 | 0.080339817 | 0.27888405 | 0.28807606 | 0.77328852 | 0.91791018 |
| DAAM1 | 195.534142 | -0.275613564 | 0.26828444 | -1.0273185 | 0.30427051 | 0.61818174 |
| GPR135 | 20.8711305 | -0.589625801 | 0.45649796 | -1.2916286 | 0.19648581 | 0.49845696 |
| L3HYPDH | 21.9619908 | -0.493868342 | 0.45131149 | -1.094296 | 0.2738252 | 0.58894058 |
| JKAMP | 35.8404246 | 0.305157289 | 0.36982112 | 0.82514836 | 0.40928737 | 0.70777268 |
| AL157911.1 | 26.2357214 | 0.80675202 | 0.68169063 | 1.18345769 | 0.23662781 | 0.54951872 |
| PCNX4 | 163.535923 | -0.008118423 | 0.31647199 | -0.0256529 | 0.9795342 | 0.99576721 |
| DHRS7 | 17.0932425 | -0.528525194 | 0.41320137 | -1.2790984 | 0.20086242 | 0.50467291 |
| PPM1A | 84.3619405 | 0.131521693 | 0.44403579 | 0.29619615 | 0.76708029 | 0.9157292 |
| MNAT1 | 319.937771 | 0.428611901 | 0.19659988 | 2.18012288 | 0.02924835 | 0.16679612 |
| SLC38A6 | 54.0024336 | -0.414968585 | 0.3236559 | -1.2821289 | 0.19979742 | 0.5033686 |
| PRKCH | 144.348461 | -1.954014397 | 0.93918416 | -2.0805445 | 0.03747562 | 0.19520822 |
| TMEM30B | 19.2752878 | -1.360155512 | 0.66503537 | -2.0452378 | 0.04083141 | 0.20490426 |
| HIF1A | 1119.94709 | -1.368974335 | 0.43284665 | -3.1627236 | 0.00156301 | 0.02123122 |
| HIF1A-AS2 | 246.393482 | -2.65232194 | 0.6768202 | -3.9187984 | 8.90E-05 | 0.00223053 |
| SNAPC1 | 146.161724 | -0.556282548 | 0.75584074 | -0.7359785 | 0.46174377 | 0.74992143 |
| SYT16 | 33.6359805 | -3.090430392 | 0.76767176 | -4.0257185 | 5.68E-05 | 0.00158095 |
| PPP2R5E | 209.363236 | 0.11126607 | 0.24173997 | 0.46027171 | 0.6453212 | 0.85797446 |
| WDR89 | 36.6113037 | 0.463736154 | 0.39907824 | 1.16201813 | 0.2452281 | 0.55755248 |
| SGPP1 | 190.03465 | 0.428944748 | 0.49195679 | 0.8719155 | 0.38325448 | 0.68720476 |
| SYNE2 | 1056.65717 | -1.000838714 | 0.44476911 | -2.2502433 | 0.0244335 | 0.14863223 |
| ESR2 | 61.1664498 | -1.429063299 | 0.62210962 | -2.2971246 | 0.02161166 | 0.13787238 |
| MTHFD1 | 27.6567101 | 0.431141577 | 0.31920719 | 1.35066372 | 0.17680318 | 0.47393944 |
| AL122035.1 | 23.9622258 | -1.282836879 | 0.62694773 | -2.0461624 | 0.0407404 | 0.20466781 |
| ZBTB25 | 41.6403747 | -0.499696343 | 0.4234712 | -1.1800008 | 0.23799991 | 0.54997949 |
| ZBTB1 | 112.291847 | -0.39084775 | 0.26886985 | -1.4536689 | 0.14603812 | 0.42860392 |
| AL049869.3 | 13.2711593 | -0.262138806 | 0.71982234 | -0.3641715 | 0.71572992 | 0.88904248 |
| HSPA2 | 23.7861938 | -0.193787266 | 0.62914542 | -0.3080167 | 0.75806966 | 0.91161939 |
| PLEKHG3 | 69.6285249 | -0.280521334 | 0.37490662 | -0.7482432 | 0.45431347 | 0.74495384 |
| CHURC1-FNT | 45.5396333 | -0.679082591 | 0.3550691 | -1.9125364 | 0.05580743 | 0.24602764 |
| RAB15 | 44.1597918 | 0.567870689 | 0.4451975 | 1.2755478 | 0.20211542 | 0.50618429 |
| MAX | 74.056268 | -0.556329271 | 0.27907166 | -1.9934997 | 0.04620676 | 0.22049887 |
| FUT8 | 101.900233 | 0.900535516 | 0.36910706 | 2.43976776 | 0.01469671 | 0.10663976 |
| AL359232.1 | 58.9949246 | 0.829158507 | 0.61089156 | 1.35729246 | 0.1746883 | 0.47119666 |
| GPHN | 1017.0093 | 0.893640979 | 0.2560981 | 3.48944794 | 0.00048402 | 0.0084513 |
| MPP5 | 78.7902434 | -0.472713478 | 0.24353291 | -1.9410661 | 0.05225026 | 0.23626646 |
| ATP6V1D | 56.7548594 | -0.466114473 | 0.56279744 | -0.82821 | 0.40755158 | 0.70619047 |
| EIF2S1 | 35.8902246 | -0.54856188 | 0.35112751 | -1.5622868 | 0.11822044 | 0.37892311 |
| PLEKHH1 | 29.1968432 | -0.484577671 | 0.35726333 | -1.35636 | 0.17498467 | 0.47154827 |
| PIGH | 20.0050665 | 0.37586591 | 0.38451088 | 0.97751696 | 0.32831328 | 0.63988149 |
| ARG2 | 54.2670474 | -0.918482754 | 0.70370302 | -1.3052136 | 0.19182012 | 0.49284602 |
| VTI1B | 23.1681498 | -0.026601613 | 0.48275022 | -0.0551043 | 0.95605537 | 0.98853497 |
| ZFYVE26 | 44.8147777 | -0.787471535 | 0.54470566 | -1.4456827 | 0.14826623 | 0.43214577 |
| RAD51B | 526.795033 | 1.004117825 | 0.37063854 | 2.70915652 | 0.00674545 | 0.06172187 |
| ZFP36L1 | 256.79031 | -0.935656919 | 0.42965445 | -2.1776963 | 0.02942865 | 0.16751695 |
| ACTN1 | 60.3432222 | -1.667233369 | 0.49329176 | -3.3798119 | 0.00072535 | 0.01168084 |
| DCAF5 | 186.404415 | -0.100760493 | 0.21203346 | -0.4752103 | 0.63463706 | 0.85244692 |
| AL359317.2 | 14.4600473 | 0.064773625 | 0.54292409 | 0.11930512 | 0.90503363 | 0.96914626 |
| EXD2 | 41.5366136 | 0.84206363 | 0.35481538 | 2.37324444 | 0.01763259 | 0.12062089 |
| GALNT16 | 33.4942048 | -1.389609742 | 0.95995133 | -1.4475835 | 0.14773356 | 0.43123114 |
| ERH | 32.3481353 | -0.627104394 | 0.30519082 | -2.0547944 | 0.03989887 | 0.20251336 |
| SLC39A9 | 83.948915 | -0.197817394 | 0.23697158 | -0.8347726 | 0.40384574 | 0.70290417 |
| PLEKHD1 | 36.8404481 | 2.972701096 | 0.81652101 | 3.64069148 | 0.00027191 | 0.00537613 |
| SUSD6 | 104.499427 | -0.165138256 | 0.19480459 | -0.8477123 | 0.39659819 | 0.69690885 |
| SRSF5 | 241.315316 | 0.78367255 | 0.22522865 | 3.47945318 | 0.00050244 | 0.00865575 |
| AL357153.2 | 51.2908013 | -0.808455468 | 0.39129829 | -2.0660849 | 0.03882047 | 0.19898957 |
| MED6 | 35.2606101 | -0.106326569 | 0.3104139 | -0.3425316 | 0.73195087 | 0.89787 |
| MAP3K9 | 42.7032766 | 0.219021174 | 0.40728442 | 0.53775976 | 0.59074292 | 0.83191164 |

| | | | | | | |
|---|---|---|---|---|---|---|
| PCNX1 | 269.45688 | -0.084259209 | 0.3054521 | -0.2758508 | 0.78266266 | 0.92150025 |
| SIPA1L1 | 448.68563 | -0.754939867 | 0.41562499 | -1.8163967 | 0.06930954 | 0.28060883 |
| RGS6 | 151.774344 | -4.313999547 | 1.50917462 | -2.8585158 | 0.00425628 | 0.04449052 |
| DPF3 | 14.4485513 | 1.396472314 | 0.45675037 | 3.05740819 | 0.0022326 | 0.02737438 |
| ZFYVE1 | 40.3991439 | -0.188170224 | 0.38235252 | -0.4921381 | 0.62262175 | 0.84576416 |
| RBM25 | 178.518361 | 0.182478747 | 0.25768092 | 0.70815777 | 0.47884729 | 0.76419703 |
| PSEN1 | 123.360501 | -0.410865255 | 0.39455691 | -1.0413333 | 0.29772088 | 0.61141484 |
| PAPLN | 24.2741694 | -0.912848482 | 0.5409561 | -1.6874724 | 0.09151255 | 0.32969979 |
| NUMB | 307.017333 | -0.235741815 | 0.22263809 | -1.0588566 | 0.28966507 | 0.60475529 |
| HEATR4 | 14.2412936 | -1.509931799 | 0.44037771 | -3.4287198 | 0.00060644 | 0.01004334 |
| DNAL1 | 31.6240699 | -0.496694762 | 0.42027409 | -1.1818353 | 0.23727105 | 0.54992696 |
| ELMSAN1 | 56.1650215 | -1.027699917 | 0.41327523 | -2.4867204 | 0.01289267 | 0.09821896 |
| AC005480.1 | 28.575123 | -0.387813878 | 0.48618357 | -0.7976696 | 0.42506222 | 0.72151771 |
| COQ6 | 16.5005388 | 0.815351218 | 0.41551825 | 1.96225127 | 0.04973324 | 0.22967445 |
| ENTPD5 | 45.1623467 | 0.728777474 | 0.39438086 | 1.84790273 | 0.06461642 | 0.268828 |
| BBOF1 | 36.7401157 | -0.100183233 | 0.4916293 | -0.203778 | 0.83852699 | 0.94741614 |
| ALDH6A1 | 41.8106073 | -0.497037432 | 0.46162086 | -1.0767222 | 0.28160439 | 0.59579794 |
| LIN52 | 79.8543826 | 0.641527255 | 0.31680062 | 2.02501891 | 0.04286542 | 0.21080021 |
| ABCD4 | 69.3316836 | -0.71372702 | 0.35904062 | -1.9878726 | 0.04682578 | 0.22193782 |
| NPC2 | 28.2899476 | -1.198234445 | 0.48740347 | -2.4584036 | 0.01395562 | 0.10351809 |
| LTBP2 | 34.2789294 | 2.150977783 | 0.73672051 | 2.91966594 | 0.00350407 | 0.03894151 |
| AREL1 | 57.3883619 | 0.15934181 | 0.30329223 | 0.52537386 | 0.59932332 | 0.83581061 |
| FCF1 | 25.2361227 | 0.043030936 | 0.32824377 | 0.13109445 | 0.89570059 | 0.96519206 |
| YLPM1 | 134.805829 | 0.368693549 | 0.2019055 | 1.82606988 | 0.06783974 | 0.27788783 |
| DLST | 48.2306448 | 0.429746873 | 0.274947 | 1.56301715 | 0.11804856 | 0.37876283 |
| MLH3 | 88.4405873 | -0.071882725 | 0.27503604 | -0.2613575 | 0.79381685 | 0.92658341 |
| ACYP1 | 48.7962532 | 0.415403743 | 0.39553754 | 1.05022583 | 0.2936143 | 0.60876592 |
| ZC2HC1C | 14.6536206 | 0.140260032 | 0.46152635 | 0.30390471 | 0.76120049 | 0.91320552 |
| NEK9 | 72.815945 | 0.284133867 | 0.25054627 | 1.13405746 | 0.25677045 | 0.56935259 |
| TMED10 | 92.2033435 | -0.416384677 | 0.31158701 | -1.3363352 | 0.18143975 | 0.47998877 |
| JDP2 | 20.2700129 | 1.863629178 | 0.6378636 | 2.92167351 | 0.00348156 | 0.03879367 |
| TTLL5 | 148.596004 | -0.077603208 | 0.24095767 | -0.3220616 | 0.74740604 | 0.90692529 |
| IFT43 | 107.947136 | -0.133872068 | 0.36314321 | -0.3686481 | 0.71239001 | 0.88729956 |
| GPATCH2L | 240.618161 | -0.30611728 | 0.29541241 | -1.036237 | 0.30009157 | 0.61374288 |
| AC016526.1 | 38.0871112 | 2.3419398 | 0.56512096 | 4.14413895 | 3.41E-05 | 0.00106143 |
| ESRRB | 397.533861 | 0.852845429 | 0.33874861 | 2.51763521 | 0.01181456 | 0.09280452 |
| ANGEL1 | 19.2648046 | 0.823007927 | 0.40657391 | 2.02425168 | 0.04294426 | 0.2110766 |
| IRF2BPL | 22.8109438 | -0.62902497 | 0.7349703 | -0.8558509 | 0.39208028 | 0.6951429 |
| POMT2 | 18.3486529 | -0.04501371 | 0.36828587 | -0.1222249 | 0.90272089 | 0.96803829 |
| TMED8 | 53.4754777 | 0.148135835 | 0.28367657 | 0.52219975 | 0.60153126 | 0.83679851 |
| VIPAS39 | 23.8825302 | 0.184908094 | 0.49431524 | 0.37406917 | 0.70835285 | 0.88614996 |
| AHSA1 | 19.3402286 | -0.217166505 | 0.40896602 | -0.5310136 | 0.59540938 | 0.83416944 |
| SPTLC2 | 100.168928 | 0.025575559 | 0.25150737 | 0.1016891 | 0.91900345 | 0.97513464 |
| ALKBH1 | 38.4172081 | -0.018472827 | 0.43770902 | -0.0422034 | 0.96633652 | 0.99156233 |
| SLIRP | 33.4379014 | -0.255140372 | 0.31950153 | -0.7985576 | 0.42454699 | 0.72151771 |
| SNW1 | 44.8802472 | -0.292816649 | 0.32888109 | -0.890342 | 0.37328228 | 0.67707283 |
| ADCK1 | 24.4763051 | 0.829202615 | 0.36735132 | 2.25724685 | 0.02399265 | 0.14714173 |
| NRXN3 | 36.1894833 | -1.999406341 | 0.64629542 | -3.0936415 | 0.00197716 | 0.02501365 |
| DIO2 | 38.063948 | 2.673651909 | 0.94345464 | 2.83389555 | 0.00459844 | 0.04722006 |
| CEP128 | 39.5775108 | 0.206692186 | 0.42372661 | 0.4877961 | 0.62569427 | 0.8471532 |
| AC007262.2 | 20.659485 | 0.232452389 | 0.75064174 | 0.30967155 | 0.75681074 | 0.91122251 |
| GTF2A1 | 41.0840768 | -0.062147801 | 0.40046136 | -0.1551905 | 0.87667113 | 0.95879545 |
| STON2 | 96.2659738 | -0.794086568 | 0.49715492 | -1.5972618 | 0.11020736 | 0.365422 |
| SEL1L | 80.1207598 | 0.18128727 | 0.33294418 | 0.54449749 | 0.58609918 | 0.8294442 |
| GALC | 23.5314592 | 0.576211877 | 0.35219406 | 1.63606359 | 0.10182628 | 0.34880722 |
| SPATA7 | 58.0096822 | 0.465707283 | 0.4188448 | 1.11188508 | 0.26618756 | 0.58179391 |
| PTPN21 | 51.4227085 | 0.24360402 | 0.31486091 | 0.77368771 | 0.43911549 | 0.73151917 |
| ZC3H14 | 104.343907 | 0.336665575 | 0.26853674 | 1.25370394 | 0.20994963 | 0.51870552 |
| EML5 | 155.865793 | -4.983272879 | 1.49890354 | -3.3246121 | 0.00088542 | 0.01360643 |
| TTC8 | 46.1160256 | 0.514444965 | 0.44070404 | 1.16732527 | 0.24307904 | 0.55555126 |
| FOXN3 | 237.775629 | 0.974652376 | 0.29671777 | 3.28477924 | 0.00102062 | 0.01522606 |
| EFCAB11 | 46.5951488 | 0.083842189 | 0.59860072 | 0.14006363 | 0.88860972 | 0.96307703 |
| TDP1 | 25.4922666 | -0.574920226 | 0.39353427 | -1.4609153 | 0.14403869 | 0.42527446 |
| PSMC1 | 33.5175625 | -0.150985139 | 0.35833911 | -0.4213471 | 0.67350165 | 0.86846218 |
| NRDE2 | 48.0988447 | 0.246695744 | 0.29012137 | 0.85031912 | 0.39514769 | 0.69642725 |
| CALM1 | 66.2456519 | -0.263261637 | 0.39664886 | -0.6637146 | 0.50687299 | 0.77942663 |
| TTC7B | 170.110166 | 0.522503265 | 0.48278023 | 1.08227974 | 0.27912825 | 0.59380484 |
| RPS6KA5 | 73.4952936 | -1.566010856 | 0.52720208 | -2.9704186 | 0.00297394 | 0.03440328 |
| DGLUCY | 148.313609 | 0.656249624 | 0.43972128 | 1.4924218 | 0.1355886 | 0.41064171 |
| PPP4R3A | 137.253571 | 0.02351649 | 0.3600271 | 0.06531867 | 0.94792028 | 0.98636425 |
| CATSPERB | 25.384052 | -0.192606108 | 0.33971171 | -0.5669693 | 0.57073504 | 0.81891359 |
| TC2N | 32.7843707 | 0.194850151 | 0.44506566 | 0.43780091 | 0.66153061 | 0.86492722 |
| TRIP11 | 152.385977 | 0.19751148 | 0.18678844 | 1.05740739 | 0.2903257 | 0.60499022 |
| ATXN3 | 96.5492316 | -0.001456036 | 0.37947043 | -0.003837 | 0.99693851 | 0.99922221 |
| NDUFB1 | 47.2764279 | 1.006466693 | 0.35734883 | 2.81648243 | 0.00485527 | 0.04915368 |
| CPSF2 | 51.3140196 | 0.227012854 | 0.29361299 | 0.77317033 | 0.43942158 | 0.73176761 |
| LGMN | 29.6696957 | -0.326019415 | 0.43523203 | -0.7490704 | 0.4538148 | 0.74470138 |
| GOLGA5 | 36.0465305 | -0.070158911 | 0.36775962 | -0.1907738 | 0.84870279 | 0.95003258 |
| ITPK1 | 60.0854572 | -0.188135968 | 0.37598184 | -0.5003858 | 0.61680346 | 0.84223226 |
| BTBD7 | 311.061934 | -0.952573605 | 0.21055216 | -4.5241693 | 6.06E-06 | 0.00025357 |
| UNC79 | 30.1053281 | -0.50960574 | 0.69373201 | -0.7345859 | 0.46259174 | 0.75091259 |
| DDX24 | 94.1275496 | -0.628918004 | 0.23745138 | -2.6486181 | 0.00808216 | 0.07056452 |
| IFI27 | 21.5932321 | 0.426761213 | 1.21607279 | 0.35093394 | 0.72563791 | 0.89424371 |
| SERPINA1 | 35.9161243 | -2.388934459 | 0.99041759 | -2.4120477 | 0.01586321 | 0.11205192 |
| DICER1 | 218.62186 | 0.136162082 | 0.1917675 | 0.71003734 | 0.47768098 | 0.7631764 |
| DICER1-AS1 | 23.888174 | 0.498632376 | 0.44998835 | 1.10810064 | 0.26781836 | 0.58378751 |
| CLMN | 514.726716 | 0.407183879 | 0.42587611 | 0.95610875 | 0.33901729 | 0.64971064 |
| GLRX5 | 21.4309741 | -0.241122183 | 0.39355383 | -0.612679 | 0.54008859 | 0.80166126 |
| ATG2B | 110.656359 | 0.354125232 | 0.25898937 | 1.367335 | 0.17152033 | 0.46730924 |
| GSKIP | 19.4994106 | -0.321780418 | 0.35960566 | -0.8948147 | 0.37088617 | 0.67515476 |
| AK7 | 18.1970871 | -1.345826626 | 0.5701843 | -2.3603362 | 0.01825838 | 0.12340471 |
| PAPOLA | 195.14213 | 0.252117825 | 0.27000334 | 0.93375818 | 0.35042865 | 0.65765381 |
| VRK1 | 14.831508 | 0.143534122 | 0.5610874 | 0.2558142 | 0.79809431 | 0.92878574 |
| LINC01550 | 21.3688911 | 0.915004249 | 0.54397162 | 1.68208086 | 0.09255316 | 0.33178227 |
| SETD3 | 102.736401 | 0.100620207 | 0.33754612 | 0.29809321 | 0.76563202 | 0.91549415 |
| CCNK | 38.6975005 | -0.140216445 | 0.27443958 | -0.5109192 | 0.60940764 | 0.84067419 |
| CCDC85C | 20.8444811 | -0.470443511 | 0.50163422 | -0.9378218 | 0.34833599 | 0.65547624 |
| EVL | 107.052802 | -1.338375661 | 0.38863986 | -3.4437426 | 0.00057372 | 0.0095823 |
| YY1 | 118.845457 | -0.463045765 | 0.3627625 | -1.2764433 | 0.20179886 | 0.5060272 |
| SLC25A29 | 144.530092 | -0.834365773 | 0.40169608 | -2.0771071 | 0.03779168 | 0.19628857 |
| WARS | 65.2834835 | -1.228889819 | 0.54635825 | -2.2492381 | 0.02449735 | 0.14870657 |
| WDR25 | 34.297149 | -0.216405928 | 0.36519977 | -0.5925686 | 0.55346987 | 0.81035847 |
| PPP2R5C | 158.692781 | 0.206887593 | 0.245059 | 0.84423586 | 0.39853762 | 0.69875231 |
| DYNC1H1 | 110.840958 | -0.498404046 | 0.36605922 | -1.3615394 | 0.17334329 | 0.46915904 |
| HSP90AA1 | 189.704788 | -0.711095744 | 0.34532737 | -2.0591931 | 0.03947574 | 0.20080297 |
| WDR20 | 140.273348 | -0.117308366 | 0.27694706 | -0.4235769 | 0.67187443 | 0.86777458 |
| MOK | 70.1393238 | 0.325214974 | 0.44540092 | 0.73016233 | 0.46529097 | 0.75266707 |
| ZNF839 | 22.1161096 | 0.585846346 | 0.47467425 | 1.23420713 | 0.21712573 | 0.52665981 |
| TECPR2 | 71.4039154 | -0.009776211 | 0.25949593 | -0.0376739 | 0.96994772 | 0.99156233 |
| ANKRD9 | 40.281903 | 0.930907847 | 0.54653459 | 1.70329174 | 0.08851349 | 0.32316736 |
| RCOR1 | 155.932998 | 0.274296069 | 0.27425343 | 1.00015548 | 0.31723527 | 0.63093651 |
| TRAF3 | 98.3085324 | -1.353683489 | 0.4192775 | -3.2286099 | 0.00124393 | 0.01778903 |
| CDC42BPB | 164.26981 | -0.733166192 | 0.42902862 | -1.7088981 | 0.08746983 | 0.32020757 |

| | | | | | | |
|---|---|---|---|---|---|---|
| EIF5 | 88.6151324 | -1.004364063 | 0.45980147 | -2.1843429 | 0.02893706 | 0.1659343 |
| MARK3 | 339.167015 | -0.134374811 | 0.27618825 | -0.4865334 | 0.62658902 | 0.8472036 |
| CKB | 119.358174 | 0.15709958 | 0.55875844 | 0.28115831 | 0.77858898 | 0.92013725 |
| BAG5 | 29.8293183 | -1.412525085 | 0.3805076 | -3.7122125 | 0.00020546 | 0.0043243 |
| KLC1 | 72.501509 | -0.17776147 | 0.38601556 | -0.4605034 | 0.64515495 | 0.85787574 |
| ZFYVE21 | 16.4879986 | -0.739682557 | 0.54232747 | -1.3639039 | 0.17259781 | 0.468183 |
| PPP1R13B | 100.969492 | -1.117906781 | 0.28179983 | -3.9670243 | 7.28E-05 | 0.00192239 |
| ATP5MPL | 36.1383405 | 0.228961231 | 0.39604899 | 0.57811341 | 0.56318756 | 0.81448184 |
| AL590079.1 | 12.8980729 | 1.598790949 | 0.5758867 | 2.77622481 | 0.00549942 | 0.05339677 |
| SIVA1 | 13.8623148 | -0.05794003 | 0.62160568 | -0.0932103 | 0.92573652 | 0.97735117 |
| AKT1 | 25.1447787 | -0.100385591 | 0.37753449 | -0.2658978 | 0.79031792 | 0.92448611 |
| CEP170B | 23.1320362 | -0.128381093 | 0.48524077 | -0.2645719 | 0.79133924 | 0.92519712 |
| NUDT14 | 10.9185233 | -0.021195292 | 0.72675668 | -0.0291642 | 0.97673362 | 0.99476398 |
| BRF1 | 46.0499584 | 0.344239574 | 0.26275974 | 1.31009256 | 0.19016453 | 0.49082497 |
| PACS2 | 64.5324439 | -0.167362413 | 0.26721133 | -0.6263298 | 0.53109866 | 0.79628881 |
| TEX22 | 12.6895592 | -0.750889983 | 0.53666943 | -1.3991667 | 0.16176301 | 0.45171558 |
| MTA1 | 34.6658654 | 0.783760824 | 0.42397873 | 1.84858526 | 0.06451772 | 0.26879502 |
| CRIP2 | 15.6235833 | 0.431576346 | 0.53461088 | 0.80727191 | 0.41950984 | 0.71797169 |
| AC087386.1 | 13.8103208 | 0.342256557 | 0.5749427 | 0.59528812 | 0.5516509 | 0.80899543 |
| NIPA1 | 62.7745752 | 0.115429645 | 0.28383901 | 0.40667294 | 0.68424822 | 0.87360405 |
| NIPA2 | 52.2362818 | -0.386255081 | 0.38518979 | -1.0027656 | 0.31597396 | 0.63009595 |
| CYFIP1 | 101.683845 | -0.684726133 | 0.32370145 | -2.1153014 | 0.03440427 | 0.18457156 |
| TUBGCP5 | 38.6408871 | 0.110665481 | 0.31141479 | 0.3553636 | 0.72231721 | 0.89217596 |
| SNRPN | 13.83805 | 0.943518733 | 0.80870471 | 1.16670364 | 0.24333007 | 0.55555126 |
| AC124312.1 | 334.109024 | 0.338547851 | 0.25831644 | 1.31059351 | 0.18999513 | 0.49058838 |
| PWAR6 | 79.5507558 | 0.716576285 | 0.34728837 | 2.06334661 | 0.03907971 | 0.19998858 |
| UBE3A | 222.822718 | 0.141064074 | 0.26817152 | 0.52602184 | 0.59887303 | 0.8356618 |
| GABRB3 | 54.5404713 | -0.614974057 | 0.57424347 | -1.0709291 | 0.2842013 | 0.59746288 |
| HERC2 | 235.769782 | 0.4571581 | 0.15138769 | 3.01978385 | 0.00252955 | 0.03035462 |
| FAM189A1 | 121.246956 | 0.813482138 | 0.50893044 | 1.59841518 | 0.10995061 | 0.3650924 |
| TJP1 | 284.134006 | -0.300109057 | 0.40963082 | -0.732633 | 0.4637823 | 0.75139985 |
| FAN1 | 52.3823583 | 0.274785478 | 0.49435623 | 0.55584507 | 0.57831677 | 0.82311488 |
| MTMR10 | 197.035275 | 0.006208195 | 0.37166436 | 0.01670377 | 0.98667294 | 0.99766074 |
| KLF13 | 36.3409583 | 0.090274985 | 0.38597043 | 0.23389094 | 0.81506965 | 0.93707884 |
| OTUD7A | 91.9378977 | 0.29138269 | 0.28358157 | 1.02750924 | 0.30418074 | 0.61818174 |
| LINC02256 | 140.950872 | 0.31153577 | 0.43490371 | 0.71633275 | 0.47378591 | 0.75994836 |
| AC123768.3 | 17.7677619 | 0.828919905 | 0.64936258 | 1.2765132 | 0.20177416 | 0.5060272 |
| GOLGA8N | 19.0251688 | 1.100060554 | 0.45011037 | 2.44397956 | 0.01452625 | 0.1059672 |
| FMN1 | 553.504845 | 1.513959664 | 0.52898288 | 2.86202015 | 0.0042095 | 0.04426547 |
| AVEN | 66.2644208 | 0.31419389 | 0.30602188 | 1.02670402 | 0.30455986 | 0.61818174 |
| EMC7 | 22.0807228 | 0.12655287 | 0.36641582 | 0.34538048 | 0.72980835 | 0.89713026 |
| KATNBL1 | 113.619519 | -0.132118019 | 0.21934253 | -0.6023365 | 0.54695015 | 0.80584949 |
| SLC12A6 | 52.8302205 | -0.20428234 | 0.46353538 | -0.440705 | 0.65942661 | 0.86452386 |
| NOP10 | 23.0535108 | -2.040441777 | 0.50930775 | -4.0063042 | 6.17E-05 | 0.00169113 |
| LPCAT4 | 13.0469881 | 0.104719464 | 0.42209332 | 0.24809553 | 0.8040605 | 0.93223826 |
| GOLGA8A | 80.3801164 | 0.048065975 | 0.48069713 | 0.09999223 | 0.9203505 | 0.9754826 |
| GOLGA8B | 84.9910266 | -0.033360594 | 0.55668084 | -0.0599277 | 0.95221323 | 0.98738095 |
| AQR | 101.770937 | 0.112868107 | 0.1964834 | 0.57444092 | 0.56566947 | 0.81595078 |
| ZNF770 | 51.5610833 | -0.93966101 | 0.43972887 | -2.13691 | 0.03260532 | 0.17956997 |
| AC021231.1 | 94.8060298 | 0.012536799 | 0.7586249 | 0.01652569 | 0.98681501 | 0.99766074 |
| DPH6 | 294.335352 | 0.023368037 | 0.60439061 | 0.0386638 | 0.96915844 | 0.99156233 |
| DPH6-DT | 186.808803 | 0.83307165 | 0.70505661 | 1.18156704 | 0.23737754 | 0.54992696 |
| C15orf41 | 86.9692992 | 0.483246936 | 0.39426551 | 1.22568909 | 0.2203157 | 0.53012221 |
| SPRED1 | 175.101696 | -0.398557272 | 0.28349395 | -1.4058758 | 0.15976102 | 0.44854313 |
| FAM98B | 31.4037241 | 0.080325004 | 0.36091148 | 0.22256151 | 0.8238768 | 0.94138895 |
| RASGRP1 | 19.9388001 | -0.488216828 | 0.48854182 | -0.9993348 | 0.31763254 | 0.63093651 |
| C15orf53 | 110.519961 | 2.946208653 | 0.64087676 | 4.59715323 | 4.28E-06 | 0.00019199 |
| AC013652.1 | 46.1969898 | 0.881196194 | 0.42988681 | 2.04983308 | 0.04038072 | 0.20340889 |
| THBS1 | 17.9302988 | 0.581336509 | 0.54946562 | 1.05800344 | 0.29005388 | 0.60488981 |
| FSIP1 | 15.5934302 | 0.107632711 | 0.55928568 | 0.19244675 | 0.84739228 | 0.94953558 |
| EIF2AK4 | 48.9094932 | 0.317122531 | 0.3040386 | 1.04303377 | 0.29693265 | 0.61093809 |
| SRP14 | 93.2993596 | -1.54931217 | 0.47634657 | -3.2524894 | 0.00114399 | 0.01658872 |
| SRP14-AS1 | 17.6521964 | 0.144617729 | 0.42210169 | 0.34261348 | 0.73188926 | 0.89787 |
| IVD | 39.4150992 | -0.17849927 | 0.3703532 | -0.4819704 | 0.62982699 | 0.84936461 |
| BAHD1 | 20.332087 | 0.264820721 | 0.43864529 | 0.60372407 | 0.54602711 | 0.80553179 |
| CCDC32 | 41.9782092 | -0.90309114 | 0.42903261 | -2.1049475 | 0.03529588 | 0.18752065 |
| RMDN3 | 24.0374597 | 0.036126287 | 0.32404224 | 0.11148635 | 0.91123069 | 0.97128978 |
| DNAJC17 | 39.0954323 | -0.240546325 | 0.27732331 | -0.8673859 | 0.38573061 | 0.68921713 |
| SPINT1-AS1 | 13.8538961 | -0.064947149 | 0.4358116 | -0.1490257 | 0.88153332 | 0.96155389 |
| SPINT1 | 47.0688393 | -0.247609331 | 0.42720656 | -0.579601 | 0.56218374 | 0.81388399 |
| INO80 | 130.323674 | -0.088200704 | 0.30353973 | -0.2905738 | 0.77137727 | 0.91738797 |
| CHP1 | 86.3099263 | -0.500978654 | 0.43885018 | -1.141571 | 0.25363237 | 0.56562263 |
| NDUFAF1 | 13.5955685 | 0.247975919 | 0.48067442 | 0.51589164 | 0.60593009 | 0.83810891 |
| RTF1 | 64.9759981 | -0.539595158 | 0.3013965 | -1.7903166 | 0.07340303 | 0.28975861 |
| RPAP1 | 15.2712999 | -0.122570065 | 0.49907331 | -0.2455953 | 0.80599552 | 0.93308942 |
| MGA | 122.979994 | -0.46910625 | 0.3555764 | -1.319284 | 0.18707419 | 0.48725107 |
| MAPKBP1 | 96.0273279 | -0.791575053 | 0.61182823 | -1.2937864 | 0.1957392 | 0.49824525 |
| EHD4 | 22.7377899 | -0.854574246 | 0.47180536 | -1.8112856 | 0.07009665 | 0.28228674 |
| PLA2G4F | 15.6604933 | 1.031828135 | 0.72990365 | 1.41364978 | 0.15746476 | 0.44423643 |
| VPS39 | 78.2523601 | -0.012800042 | 0.23542778 | -0.0543693 | 0.95664094 | 0.98859719 |
| TMEM87A | 112.973398 | -0.272454048 | 0.26059849 | -1.0454936 | 0.2957949 | 0.61085087 |
| GANC | 63.6048803 | 0.039995943 | 0.36761534 | 0.10879835 | 0.91336243 | 0.97183514 |
| ZNF106 | 80.6215664 | 0.035739389 | 0.24044096 | 0.14864102 | 0.88183691 | 0.96155389 |
| SNAP23 | 57.3132058 | -0.29212228 | 0.24592032 | -1.1878737 | 0.23488318 | 0.54677394 |
| HAUS2 | 16.2666643 | -0.017208625 | 0.42078281 | -0.0408967 | 0.96737826 | 0.99156233 |
| TTBK2 | 161.20442 | -0.528354966 | 0.35116362 | -1.5045834 | 0.13243121 | 0.40428636 |
| UBR1 | 184.155344 | -0.202422938 | 0.23716666 | -0.853505 | 0.39337929 | 0.69595061 |
| AC068724.3 | 18.229566 | -0.493233673 | 0.57543009 | -0.8571566 | 0.39135837 | 0.69425903 |
| TMEM62 | 47.2020218 | 0.209902029 | 0.28110758 | 0.74669643 | 0.45524683 | 0.74560362 |
| CCNDBP1 | 21.089565 | -0.226868213 | 0.37577676 | -0.6037314 | 0.54602226 | 0.80553179 |
| ADAL | 39.5202038 | 1.31951554 | 0.43143959 | 3.05840164 | 0.00222521 | 0.02735153 |
| TUBGCP4 | 28.8413749 | -0.182395726 | 0.3917785 | -0.4655583 | 0.64153171 | 0.85468927 |
| TP53BP1 | 170.875475 | 0.517189407 | 0.21609839 | 2.39330519 | 0.01669735 | 0.11618362 |
| PPIP5K1 | 47.5083557 | 0.284820252 | 0.32197484 | 0.88460408 | 0.37637021 | 0.67982871 |
| CATSPER2 | 61.4838478 | 0.499635784 | 0.50622845 | 0.9869769 | 0.32365396 | 0.63611921 |
| PDIA3 | 80.5807774 | -0.327963109 | 0.31762263 | -1.0325559 | 0.30181179 | 0.61586884 |
| SERF2 | 190.015876 | -0.548028967 | 0.46954365 | -1.1671523 | 0.24314885 | 0.55555126 |
| MFAP1 | 19.3604093 | -0.145699993 | 0.33621256 | -0.4333568 | 0.6647556 | 0.86663607 |
| CASC4 | 201.52397 | -0.333790938 | 0.1766666 | -1.8893834 | 0.05884047 | 0.25354648 |
| CTDSPL2 | 148.928457 | -0.097236178 | 0.28750917 | -0.338202 | 0.73521097 | 0.90007689 |
| EIF3J-DT | 37.0474646 | -0.723174792 | 0.5042096 | -1.4342741 | 0.15149405 | 0.43600335 |
| EIF3J | 64.2406565 | -0.479300929 | 0.31828747 | -1.5058743 | 0.13209944 | 0.40375585 |
| SPG11 | 198.874213 | -0.10534466 | 0.26043619 | -0.4044932 | 0.6858501 | 0.87448517 |
| B2M | 341.798377 | -1.224900881 | 0.35161846 | -3.483608 | 0.0004947 | 0.00856029 |
| TRIM69 | 12.6332776 | -0.105825081 | 0.5648668 | -0.1873452 | 0.85138999 | 0.95058798 |
| GATM | 17.6980868 | 0.687235954 | 0.51972292 | 1.32231218 | 0.18606422 | 0.48628295 |
| SPATA5L1 | 22.4454919 | -0.444556538 | 0.439897 | -1.0105923 | 0.31221158 | 0.6271535 |
| SEMA6D | 168.679076 | 2.448753025 | 0.45298746 | 5.40578552 | 6.45E-08 | 5.23E-06 |
| AC092078.2 | 151.233331 | 4.835705757 | 1.47641617 | 3.27529992 | 0.0010555 | 0.01559662 |
| MYEF2 | 125.635069 | 0.570121033 | 0.21970978 | 2.5948824 | 0.00946233 | 0.07895718 |
| SLC12A1 | 47.8973048 | 2.087276613 | 0.83561792 | 2.49788397 | 0.01249371 | 0.09619433 |
| DUT | 22.6835298 | 0.50649147 | 0.48287265 | 1.04891315 | 0.29421809 | 0.60921374 |
| CEP152 | 56.5989659 | 0.181131141 | 0.34111606 | 0.53099565 | 0.5954218 | 0.83416944 |

| | | | | | | |
|---|---|---|---|---|---|---|
| EID1 | 35.4071247 | -1.172831772 | 0.55631802 | -2.1082038 | 0.03501336 | 0.18623194 |
| SECISBP2L | 220.198433 | 0.116187929 | 0.22615212 | 0.51376006 | 0.60741976 | 0.83917773 |
| COPS2 | 41.2742045 | 0.185985311 | 0.27840919 | 0.66802864 | 0.50411531 | 0.77792612 |
| GALK2 | 131.015061 | 0.331034821 | 0.33478741 | 0.98879114 | 0.32276534 | 0.635042 |
| FAM227B | 84.5469702 | 0.08239175 | 0.67214107 | 0.12258104 | 0.90243887 | 0.96803829 |
| DTWD1 | 63.0088476 | 0.389700324 | 0.35100921 | 1.11022819 | 0.26690071 | 0.58307924 |
| GABPB1 | 59.7774149 | -0.492604989 | 0.41370109 | -1.1907268 | 0.23376084 | 0.54598215 |
| GABPB1-AS1 | 109.897692 | 0.456659814 | 0.38243484 | 1.19408528 | 0.23244461 | 0.54506955 |
| USP8 | 152.132921 | 0.006392111 | 0.18614213 | 0.03433995 | 0.97260607 | 0.99287068 |
| USP50 | 39.5092545 | 0.030413584 | 0.48730811 | 0.06241141 | 0.95023521 | 0.98678754 |
| TRPM7 | 1614.59516 | 2.368617257 | 0.27736267 | 8.53978402 | 1.34E-17 | 1.39E-14 |
| SPPL2A | 340.45207 | -0.020259483 | 0.4035713 | -0.0502005 | 0.95996261 | 0.9895734 |
| AP4E1 | 61.412448 | -0.048092704 | 0.27880031 | -0.1724987 | 0.86304545 | 0.95458251 |
| DMXL2 | 76.8486441 | 1.116131538 | 0.33965862 | 3.28603922 | 0.00101607 | 0.01518241 |
| TMOD3 | 125.069813 | -0.209455695 | 0.19136082 | -1.0945589 | 0.27370992 | 0.58894058 |
| LEO1 | 17.0486103 | 0.084929293 | 0.4175913 | 0.20337898 | 0.83883883 | 0.94741614 |
| MAPK6 | 87.7042225 | -0.137200844 | 0.43651972 | -0.3143062 | 0.7532885 | 0.91007188 |
| GNB5 | 62.7870054 | -0.41331069 | 0.26115583 | -1.582621 | 0.11350788 | 0.37119783 |
| MYO5C | 163.176212 | 0.432377533 | 0.35131032 | 1.23075671 | 0.21841387 | 0.52827262 |
| MYO5A | 215.456168 | 1.053636421 | 0.28772665 | 3.66193544 | 0.00025032 | 0.00506281 |
| ARPP19 | 75.0741209 | -0.480466916 | 0.30411397 | -1.579891 | 0.11413183 | 0.372426 |
| FAM214A | 218.829499 | -0.013691151 | 0.36161196 | -0.0378614 | 0.96979815 | 0.99156233 |
| WDR72 | 552.651238 | 1.247360764 | 0.54647534 | 2.28255636 | 0.02245651 | 0.14099969 |
| UNC13C | 85.3824855 | 4.092457204 | 0.69289027 | 5.90635683 | 3.50E-09 | 4.53E-07 |
| RSL24D1 | 20.0215043 | -0.186282566 | 0.35590795 | -0.523401 | 0.60069525 | 0.8363271 |
| RAB27A | 62.43166 | -0.914174687 | 0.31354833 | -2.9155783 | 0.0035503 | 0.03936148 |
| PIGB | 36.0966443 | -0.364703053 | 0.2719603 | -1.3410158 | 0.17991532 | 0.47860436 |
| CCPG1 | 108.497077 | 0.557140888 | 0.33143342 | 1.68100395 | 0.09276215 | 0.33202083 |
| DNAAF4 | 49.6484644 | 0.173846064 | 0.61933047 | 0.2807 | 0.77894051 | 0.92027893 |
| PYGO1 | 16.9362581 | 0.69679471 | 0.49871375 | 1.39718367 | 0.16235835 | 0.45295218 |
| PRTG | 79.436575 | 0.580050964 | 0.61524935 | 0.94279005 | 0.3457883 | 0.65414698 |
| NEDD4 | 74.4222124 | -1.028488122 | 0.49071474 | -2.0958982 | 0.03609122 | 0.18979954 |
| RFX7 | 182.383711 | 0.416361109 | 0.29151581 | 1.42826252 | 0.15321631 | 0.43862109 |
| TEX9 | 106.273971 | -0.230489482 | 0.4841037 | -0.4761159 | 0.63399179 | 0.85226924 |
| ZNF280D | 72.2817722 | 0.814715712 | 0.47600795 | 1.71155905 | 0.08697796 | 0.31893334 |
| AC090517.4 | 32.8421865 | 0.854221251 | 0.42114586 | 2.02832637 | 0.04252695 | 0.21037417 |
| TCF12 | 733.860242 | 0.118581171 | 0.19503657 | 0.60799455 | 0.54319109 | 0.80361674 |
| CGNL1 | 887.049462 | 0.316127056 | 0.37425448 | 0.84468475 | 0.39828687 | 0.69870683 |
| ALDH1A2 | 322.149254 | -1.346920293 | 0.63788996 | -2.1115245 | 0.03472725 | 0.18545069 |
| LIPC | 21.0806324 | -0.295375975 | 0.53277349 | -0.5544119 | 0.57929697 | 0.82381304 |
| ADAM10 | 352.179843 | -0.87891744 | 0.4286994 | -2.0501952 | 0.04034539 | 0.20334076 |
| MINDY2 | 194.323918 | -0.158194327 | 0.23541768 | -0.671973 | 0.50160089 | 0.776256 |
| RNF111 | 214.680654 | -0.28713662 | 0.27693363 | -1.0368427 | 0.29980915 | 0.61356904 |
| SLTM | 285.639402 | -0.571988236 | 0.33219989 | -1.7218195 | 0.08510223 | 0.31443489 |
| MYO1E | 567.596075 | -1.804408738 | 0.70122094 | -2.5732385 | 0.01007517 | 0.08275141 |
| FAM81A | 37.3551192 | -0.34079255 | 0.50939275 | -0.6690173 | 0.50348446 | 0.77790293 |
| GTF2A2 | 31.6308788 | -0.472992245 | 0.33141399 | -1.4271945 | 0.15352383 | 0.43892689 |
| BNIP2 | 60.7440224 | -0.262360949 | 0.35534142 | -0.7383348 | 0.460311 | 0.74872989 |
| ANXA2 | 98.5656724 | -3.244571744 | 0.51267425 | -6.3287199 | 2.47E-10 | 5.36E-08 |
| ICE2 | 97.1326641 | 0.436446925 | 0.33434802 | 1.30536714 | 0.19176786 | 0.49284602 |
| RORA-AS1 | 60.4050233 | 0.931302222 | 0.6390563 | 1.45730858 | 0.14503122 | 0.42698803 |
| RORA | 1266.701 | 0.585652535 | 0.43600801 | 1.34321507 | 0.17920234 | 0.47753148 |
| AC012404.1 | 31.3558398 | -0.459976618 | 0.6870708 | -0.6694749 | 0.50319262 | 0.7778729 |
| AC009554.1 | 36.2455007 | -1.841712977 | 0.6240793 | -2.9510881 | 0.00316657 | 0.03622707 |
| VPS13C | 362.867333 | -1.206548806 | 0.34305379 | -3.5170834 | 0.00043632 | 0.00777862 |
| C2CD4A | 32.1979316 | -7.187568025 | 1.28251722 | -5.6042663 | 2.09E-08 | 1.93E-06 |
| TLN2 | 158.339224 | 0.654989705 | 0.45924563 | 1.42622958 | 0.15380208 | 0.43948839 |
| TPM1 | 73.389916 | -0.812180128 | 0.44071896 | -1.8428527 | 0.06535052 | 0.27093297 |
| LACTB | 28.2687026 | -1.004333097 | 0.41175698 | -2.4391404 | 0.01472225 | 0.106742 |
| RPS27L | 39.3713784 | -0.4282126 | 0.46258972 | -0.9256855 | 0.35460944 | 0.66121742 |
| RAB8B | 56.6188459 | -0.573450172 | 0.46049931 | -1.2452791 | 0.21302917 | 0.52201944 |
| APH1B | 26.4959861 | -0.700545834 | 0.44532737 | -1.573103 | 0.11569496 | 0.3743642 |
| CA12 | 1030.24944 | 0.807383214 | 0.37800848 | 2.1358865 | 0.03268867 | 0.17981313 |
| USP3 | 245.491067 | 0.304704071 | 0.20860789 | 1.46065455 | 0.14411027 | 0.4253511 |
| USP3-AS1 | 45.0974168 | 0.219889831 | 0.39805559 | 0.55240985 | 0.58066758 | 0.82482397 |
| HERC1 | 310.673322 | -0.522067969 | 0.57670171 | -0.9052652 | 0.36532492 | 0.67012584 |
| FAM96A | 42.844478 | 0.253072077 | 0.27541488 | 0.91887582 | 0.35816053 | 0.6647359 |
| SNX1 | 97.9841051 | -0.097383353 | 0.20750127 | -0.4693145 | 0.63884486 | 0.8538689 |
| PPIB | 54.6351409 | -1.314164729 | 0.40460338 | -3.248032 | 0.00116206 | 0.01677255 |
| CSNK1G1 | 27.7469362 | -0.179224837 | 0.31265341 | -0.5732381 | 0.56648351 | 0.81595078 |
| TRIP4 | 63.3423228 | 0.153967916 | 0.35600133 | 0.43249253 | 0.66538348 | 0.86709093 |
| ZNF609 | 276.648735 | -0.158598401 | 0.22708676 | -0.6984045 | 0.48492429 | 0.76770898 |
| OAZ2 | 27.1988874 | 0.029793474 | 0.45265204 | 0.06581982 | 0.94752128 | 0.98633762 |
| RBPMS2 | 14.9872166 | 0.950356986 | 0.57019434 | 1.66672468 | 0.09556916 | 0.33778881 |
| SPG21 | 43.7042567 | -0.557550107 | 0.44267435 | -1.259504 | 0.20784836 | 0.51588941 |
| MTFMT | 39.1761839 | -0.001113252 | 0.26596922 | -0.0041856 | 0.99666035 | 0.99922221 |
| UBAP1L | 22.3022544 | -0.345930733 | 0.47538825 | -0.7276804 | 0.46680922 | 0.75315494 |
| PDCD7 | 26.4197374 | -0.267266789 | 0.35723771 | -0.7481483 | 0.45437071 | 0.74495384 |
| CLPX | 58.3138329 | 0.001005843 | 0.28084301 | 0.00358151 | 0.99714237 | 0.99922221 |
| PARP16 | 22.4069647 | 0.461314727 | 0.41767669 | 1.10447802 | 0.26938584 | 0.58564558 |
| DPP8 | 94.263105 | -0.260658865 | 0.30067812 | -0.8669033 | 0.38599497 | 0.68921713 |
| HACD3 | 78.8612033 | 0.340390905 | 0.35411083 | 0.96125527 | 0.33642383 | 0.64756726 |
| INTS14 | 24.2687645 | 0.592023463 | 0.30903678 | 1.91570553 | 0.05540259 | 0.24493776 |
| SLC24A1 | 18.4954418 | 0.444002524 | 0.52018706 | 0.85354397 | 0.39335772 | 0.69595061 |
| DENND4A | 255.634926 | -0.152929673 | 0.22659582 | -0.6749007 | 0.49973887 | 0.77558432 |
| RAB11A | 125.169525 | -0.454612314 | 0.4525063 | -1.0046541 | 0.31506343 | 0.62958667 |
| MEGF11 | 200.596825 | -7.478880789 | 0.96392256 | -7.7587984 | 8.57E-15 | 4.44E-12 |
| DIS3L | 29.9309541 | 0.50210692 | 0.32820788 | 1.5298442 | 0.1260553 | 0.39291224 |
| TIPIN | 25.064967 | 0.385960495 | 0.48648769 | 0.79336128 | 0.42756736 | 0.72300292 |
| MAP2K1 | 131.583384 | -0.93155056 | 0.41791767 | -2.2290289 | 0.02581198 | 0.15410988 |
| RPL4 | 64.2216768 | -1.15428084 | 0.5232779 | -2.2058658 | 0.0273934 | 0.16065021 |
| SMAD6 | 17.424941 | 0.017667375 | 0.43908069 | 0.04023719 | 0.96790402 | 0.99156233 |
| SMAD3 | 552.065264 | -1.351361302 | 0.71181371 | -1.8984761 | 0.0576334 | 0.25017402 |
| AAGAB | 98.0498382 | -0.15519345 | 0.2973585 | -0.5219069 | 0.60173516 | 0.83690015 |
| IQCH | 89.5561482 | -0.242764835 | 0.52120668 | -0.4657746 | 0.64137685 | 0.85468927 |
| IQCH-AS1 | 53.6428747 | 0.33455154 | 0.50233308 | 0.66599544 | 0.50541402 | 0.77892236 |
| MAP2K5 | 175.306289 | 0.534533705 | 0.37316293 | 1.43244054 | 0.15201778 | 0.43639587 |
| PIAS1 | 293.974924 | 0.372091282 | 0.25997544 | 1.43125553 | 0.152357 | 0.43719706 |
| FEM1B | 28.6836815 | 0.496308943 | 0.34883345 | 1.42276763 | 0.15480353 | 0.44032584 |
| ANP32A | 50.086906 | -0.165797194 | 0.39955785 | -0.4149517 | 0.67817729 | 0.87130138 |
| SPESP1 | 16.6477226 | 0.472435903 | 0.55592538 | 0.84981892 | 0.39542577 | 0.69642725 |
| GLCE | 112.35046 | 0.737477915 | 0.3546005 | 2.07974299 | 0.03754911 | 0.19548181 |
| AC026992.2 | 15.4226488 | 0.753067361 | 0.55221851 | 1.36371265 | 0.17265802 | 0.468183 |
| PAQR5 | 384.686254 | -0.251890213 | 0.54548701 | -0.4617712 | 0.64424538 | 0.85715524 |
| KIF23 | 41.3496372 | -0.49309416 | 0.75426493 | -0.6537413 | 0.51327847 | 0.7830488 |
| RPLP1 | 126.120625 | -0.756270235 | 0.47276203 | -1.5996848 | 0.10966853 | 0.3644153 |
| TLE3 | 60.2775009 | -0.981596843 | 0.33275522 | -2.9499066 | 0.0031787 | 0.03632132 |
| UACA | 140.687448 | 0.169569717 | 0.277723 | 0.61057138 | 0.54148337 | 0.80292477 |
| LARP6 | 19.7109372 | 1.140646706 | 0.48949678 | 2.33024353 | 0.01979328 | 0.13033374 |
| LRRC49 | 56.6981712 | 0.36246837 | 0.32713014 | 1.10802499 | 0.26785103 | 0.58378751 |
| THSD4 | 25.0503403 | 0.55305036 | 0.47792782 | 1.15718385 | 0.24719725 | 0.55984144 |
| THSD4-AS1 | 28.695457 | -0.839864017 | 0.42940466 | -1.9558801 | 0.05047928 | 0.23198682 |
| MYO9A | 465.299699 | 0.180698158 | 0.27225366 | 0.6637125 | 0.50687434 | 0.77942663 |

| | | | | | | |
|---|---|---|---|---|---|---|
| SENP8 | 19.6091265 | 0.25027458 | 0.3476069 | 0.71999313 | 0.47152922 | 0.75776258 |
| PKM | 299.625541 | -1.364990258 | 0.33006995 | -4.1354576 | 3.54E-05 | 0.00107941 |
| PARP6 | 51.7286339 | -0.221987681 | 0.49008042 | -0.4529617 | 0.65057629 | 0.86113455 |
| AC009690.2 | 58.5807499 | -0.706546432 | 0.5761885 | -1.2262418 | 0.22010769 | 0.52989521 |
| AC009690.1 | 238.816108 | -0.485444988 | 0.53762215 | -0.9029483 | 0.36655336 | 0.67113782 |
| ARIH1 | 473.653863 | -0.051713185 | 0.32234513 | -0.160428 | 0.87254393 | 0.95690422 |
| BBS4 | 71.2345772 | 0.294040646 | 0.26276411 | 1.11902895 | 0.26312779 | 0.57713562 |
| ADPGK | 45.1192776 | 0.109265573 | 0.26677096 | 0.40958571 | 0.68210988 | 0.87274146 |
| NEO1 | 94.5558497 | -0.019378617 | 0.32415388 | -0.0597822 | 0.95232914 | 0.98738095 |
| NPTN | 119.777203 | 0.369156247 | 0.23466802 | 1.57309996 | 0.11569567 | 0.3743642 |
| PML | 19.4263796 | -0.174770808 | 0.54375235 | -0.3214162 | 0.74789502 | 0.90716445 |
| ARID3B | 35.5341086 | 0.034410862 | 0.51123423 | 0.06730939 | 0.94633541 | 0.98574808 |
| CLK3 | 31.9907584 | 0.689917974 | 0.28402311 | 2.42909101 | 0.01513673 | 0.10908675 |
| EDC3 | 41.3403975 | 0.57424681 | 0.27376863 | 2.09756251 | 0.03594381 | 0.18955887 |
| ULK3 | 21.1888346 | 0.781298289 | 0.47429294 | 1.64729059 | 0.09949833 | 0.34423838 |
| SCAMP2 | 30.4567638 | -0.661461303 | 0.4406167 | -1.5012171 | 0.13329943 | 0.40630398 |
| FAM219B | 20.3893959 | 0.02696398 | 0.42174306 | 0.06393461 | 0.94902229 | 0.98641632 |
| COX5A | 47.3596257 | 0.089442318 | 0.31892544 | 0.28044899 | 0.77913305 | 0.92038979 |
| SCAMP5 | 18.9353698 | 0.488905213 | 0.42196517 | 1.15863878 | 0.24660346 | 0.55931971 |
| PPCDC | 55.4343581 | -0.037362845 | 0.3724994 | -0.1003031 | 0.9201037 | 0.97533502 |
| COMMD4 | 20.0051031 | 0.043028387 | 0.45520591 | 0.09452511 | 0.92469204 | 0.97735117 |
| NEIL1 | 21.783478 | 0.047313688 | 0.48769099 | 0.09701571 | 0.92271392 | 0.97611757 |
| MAN2C1 | 49.6603103 | 0.220515673 | 0.3412209 | 0.64625489 | 0.5181143 | 0.78682309 |
| SIN3A | 77.8218175 | -0.262530877 | 0.21689589 | -1.2104004 | 0.22612528 | 0.53730685 |
| PTPN9 | 82.8229849 | -0.30900148 | 0.36406186 | -0.8487609 | 0.39601434 | 0.69668636 |
| SNUPN | 14.5600485 | 0.184720942 | 0.47833188 | 0.38617736 | 0.69936532 | 0.8833378 |
| UBE2Q2 | 55.7408982 | -0.923022682 | 0.36701585 | -2.5149396 | 0.01190528 | 0.09335981 |
| FBXO22 | 76.3762419 | -0.531863669 | 0.28033462 | -1.8972454 | 0.05779556 | 0.25052804 |
| ETFA | 107.535291 | 0.397073925 | 0.19624488 | 2.02335941 | 0.04303611 | 0.21119403 |
| SCAPER | 294.387867 | 0.282460156 | 0.32573679 | 0.86714231 | 0.38586404 | 0.68921713 |
| RCN2 | 59.7478988 | 0.185408394 | 0.30967999 | 0.59870963 | 0.54936653 | 0.80698344 |
| TSPAN3 | 56.9718807 | -0.490570065 | 0.43138075 | -1.137209 | 0.25545093 | 0.5683189 |
| PEAK1 | 158.187507 | -0.310017009 | 0.30558184 | -1.0145138 | 0.31033763 | 0.6252351 |
| HMG20A | 99.5908912 | 0.527585847 | 0.23006748 | 2.29317867 | 0.02183772 | 0.13851354 |
| LINGO1 | 506.323609 | 1.228241425 | 0.6530998 | 1.8806336 | 0.06002178 | 0.25589532 |
| TBC1D2B | 28.9532872 | -0.355349351 | 0.60633308 | -0.586063 | 0.55783321 | 0.81243937 |
| IDH3A | 73.7655061 | 0.402276785 | 0.47352571 | 0.84953526 | 0.39558352 | 0.69642725 |
| DNAJA4 | 22.696746 | -1.646419355 | 0.91119632 | -1.8068767 | 0.07078151 | 0.28397885 |
| WDR61 | 38.0614803 | 0.366191336 | 0.39268119 | 0.93254106 | 0.35105698 | 0.65833774 |
| IREB2 | 94.2819541 | -0.184990252 | 0.27811838 | -0.6651493 | 0.50595499 | 0.77934214 |
| HYKK | 15.729243 | -0.120128873 | 0.60344218 | -0.1990727 | 0.84220586 | 0.94776435 |
| PSMA4 | 29.8902761 | 0.00880408 | 0.38396961 | 0.02292911 | 0.98170682 | 0.99656336 |
| MORF4L1 | 116.336875 | -0.281456235 | 0.2439019 | -1.1539731 | 0.24851119 | 0.56111373 |
| CTSH | 32.5802244 | -1.102955026 | 0.42464345 | -2.5973673 | 0.00939414 | 0.07883975 |
| TMED3 | 27.5573042 | -0.611892943 | 0.34554514 | -1.7708047 | 0.07659319 | 0.29596141 |
| ZFAND6 | 222.663995 | -0.185989206 | 0.27373022 | -0.6794617 | 0.49684535 | 0.77475164 |
| AC016705.2 | 153.985437 | -0.349625646 | 0.64284375 | -0.5438734 | 0.58652856 | 0.82948466 |
| ARNT2 | 332.817563 | -0.413436534 | 0.35127527 | -1.1769588 | 0.23921194 | 0.55099114 |
| ABHD17C | 74.2544627 | 0.100186145 | 0.329047 | 0.30447366 | 0.76076705 | 0.91310141 |
| AC027808.1 | 24.0458715 | 0.286079212 | 0.43379119 | 0.65948598 | 0.50958374 | 0.78121651 |
| MESD | 36.9361944 | -0.14329909 | 0.318673 | -0.4496744 | 0.65294524 | 0.86184334 |
| EFL1 | 52.5209684 | 0.070217832 | 0.28816603 | 0.24367144 | 0.80748529 | 0.93377067 |
| WHAMM | 46.2609371 | -0.379665246 | 0.27489736 | -1.3811164 | 0.16724318 | 0.45985709 |
| HOMER2 | 38.445932 | -0.787921763 | 0.44898831 | -1.7548826 | 0.0792794 | 0.30146864 |
| C15orf40 | 21.9770897 | 0.138594597 | 0.33964602 | 0.408056 | 0.68323256 | 0.87333147 |
| BTBD1 | 80.2408859 | -0.132414076 | 0.29563032 | -0.4479042 | 0.65422231 | 0.86217501 |
| HDGFL3 | 82.0134159 | -0.504998099 | 0.22510699 | -2.2433693 | 0.02487301 | 0.15010741 |
| ADAMTSL3 | 31.6441123 | -1.03567695 | 0.77845007 | -1.3304347 | 0.1833751 | 0.48210665 |
| AC048382.5 | 12.950298 | 0.533048157 | 0.54043338 | 0.98633463 | 0.32396893 | 0.63646555 |
| WDR73 | 65.5561559 | -0.57259764 | 0.30201393 | -1.8959312 | 0.05796914 | 0.25104704 |
| SEC11A | 107.42719 | -0.158872499 | 0.27380511 | -0.5802394 | 0.56175322 | 0.81388399 |
| ZNF592 | 58.5743258 | 0.120485607 | 0.22619178 | 0.53267014 | 0.59426194 | 0.83416944 |
| ALPK3 | 156.377791 | -0.889856831 | 0.62928873 | -1.4140676 | 0.15734204 | 0.44415901 |
| PDE8A | 473.553307 | 0.951387311 | 0.21622999 | 4.39988604 | 1.08E-05 | 0.0004173 |
| AKAP13 | 426.111828 | -1.224447927 | 0.49409516 | -2.4781621 | 0.01320611 | 0.0996228 |
| MRPS11 | 22.9108079 | -0.714381088 | 0.36117735 | -1.9779233 | 0.04793736 | 0.22433958 |
| DET1 | 22.486928 | 0.818453737 | 0.62708114 | 1.30517996 | 0.19183158 | 0.49284602 |
| ABHD2 | 61.8137481 | -1.482640483 | 0.55436112 | -2.674503 | 0.00748401 | 0.0666764 |
| POLG | 19.7752843 | -0.232057406 | 0.3626343 | -0.6399213 | 0.52222377 | 0.78965349 |
| RHCG | 58.309865 | 1.761539323 | 0.80864276 | 2.17839002 | 0.02937701 | 0.16732514 |
| ANPEP | 16.5558138 | -1.225379977 | 0.53414363 | -2.294102 | 0.02178464 | 0.1383651 |
| ZNF710 | 45.3906501 | -0.57957707 | 0.47240495 | -1.2268649 | 0.21987336 | 0.52974139 |
| ZNF710-AS1 | 33.873176 | 0.380026628 | 0.50647442 | 0.75033725 | 0.45305161 | 0.74451091 |
| IDH2 | 253.55186 | 0.258456428 | 0.42455912 | 0.60876429 | 0.54268069 | 0.80340275 |
| IDH2-DT | 43.88773 | 0.366059847 | 0.33246711 | 1.10104079 | 0.27087891 | 0.58722969 |
| SEMA4B | 223.46512 | -2.417425794 | 0.53235901 | -4.540969 | 5.60E-06 | 0.00023841 |
| CIB1 | 70.7903891 | -1.608238004 | 0.60912381 | -2.6402481 | 0.00828453 | 0.07212418 |
| AC091167.6 | 15.0206179 | -0.276758948 | 0.61468958 | -0.4502418 | 0.65253609 | 0.86179129 |
| ZNF774 | 11.7630252 | 0.99556943 | 0.572748 | 1.73823291 | 0.08216978 | 0.30864976 |
| IQGAP1 | 208.01932 | -0.725170858 | 0.20103857 | -3.6071231 | 0.00030961 | 0.00597684 |
| CRTC3 | 109.243033 | 0.155521818 | 0.26661294 | 0.58332433 | 0.55967499 | 0.81314937 |
| CRTC3-AS1 | 19.1372404 | -0.182060128 | 0.41352375 | -0.4402652 | 0.65974502 | 0.86460873 |
| FURIN | 19.2421841 | -1.152048115 | 0.56155668 | -2.051526 | 0.04021575 | 0.20290676 |
| MAN2A2 | 17.1977771 | -0.532616086 | 0.41812443 | -1.273822 | 0.20272652 | 0.50710716 |
| UNC45A | 29.5575121 | -0.015842537 | 0.36632215 | -0.0432476 | 0.9655042 | 0.99156233 |
| AC104035.1 | 37.4968081 | 3.379428474 | 1.52268877 | 2.21938229 | 0.02646073 | 0.15637575 |
| FAM174B | 38.5436953 | -0.229765327 | 0.37991137 | -0.6047867 | 0.54532076 | 0.80477537 |
| AC013394.1 | 145.944595 | 0.432563931 | 0.21831154 | 1.98140666 | 0.04754569 | 0.22356912 |
| CHD2 | 255.158266 | -0.564268633 | 0.404079 | -1.3964315 | 0.16258461 | 0.45306004 |
| MCTP2 | 21.7227093 | -1.523852874 | 0.45413512 | -3.3555055 | 0.0007922 | 0.01251946 |
| AC012409.2 | 62.0312605 | 0.279571742 | 0.74187091 | 0.37684688 | 0.7062874 | 0.88599027 |
| NR2F2-AS1 | 532.726084 | 1.078222076 | 0.50551384 | 2.13292297 | 0.03293105 | 0.18051092 |
| NR2F2 | 86.1641606 | 0.155806322 | 0.33494365 | 0.46517175 | 0.64180847 | 0.85468927 |
| AC087477.2 | 15.05013 | 0.236119555 | 0.57690611 | 0.40928593 | 0.68232984 | 0.87274146 |
| ARRDC4 | 37.611622 | 0.540232927 | 0.4545109 | 1.18860281 | 0.234596 | 0.54657 |
| FAM169B | 20.9791803 | -5.238937603 | 0.85150518 | -6.152561 | 7.62E-10 | 1.42E-07 |
| IGF1R | 817.556143 | -0.122562586 | 0.2422338 | -0.5059681 | 0.61287902 | 0.84174925 |
| TTC23 | 60.6782999 | -0.037265632 | 0.28299143 | -0.1316847 | 0.89523371 | 0.96511349 |
| LRRC28 | 178.076494 | 0.480340118 | 0.41836339 | 1.14814089 | 0.25091041 | 0.56353121 |
| MEF2A | 500.815694 | 0.79830861 | 0.17674934 | 4.5166145 | 6.28E-06 | 0.00026027 |
| LYSMD4 | 12.5403633 | 0.423213777 | 0.47903312 | 0.88347498 | 0.37697969 | 0.68029685 |
| AC090825.1 | 41.7881617 | 1.246297599 | 0.68562626 | 1.81775068 | 0.06910225 | 0.2801345 |
| ADAMTS17 | 561.266879 | 1.139953907 | 0.68560594 | 1.66269549 | 0.09637341 | 0.33934503 |
| LINS1 | 28.1070752 | 0.381085921 | 0.29398923 | 1.29625812 | 0.19488657 | 0.49736588 |
| ASB7 | 37.8051085 | 0.318623401 | 0.29836829 | 1.06788628 | 0.28557181 | 0.59956577 |
| CHSY1 | 179.496594 | -1.823207782 | 0.38923488 | -4.6840812 | 2.81E-06 | 0.00013447 |
| SELENOS | 35.2815022 | -1.029773557 | 0.34107947 | -3.0191602 | 0.00253476 | 0.03036691 |
| SNRPA1 | 33.0088053 | 0.272356668 | 0.28486592 | 0.95608724 | 0.33902816 | 0.64971064 |
| PCSK6 | 64.8925938 | -1.28020659 | 0.61912509 | -2.0677673 | 0.03866191 | 0.19861359 |
| TM2D3 | 40.7288149 | -0.661262329 | 0.55215612 | -1.1976003 | 0.23107267 | 0.54380466 |
| TARSL2 | 76.0713407 | -0.343871913 | 0.45347185 | -0.7583093 | 0.44826585 | 0.7399821 |
| MPG | 14.2919328 | -1.18385842 | 0.37809512 | -3.1311126 | 0.00174145 | 0.02280522 |
| NPRL3 | 65.4664887 | -0.375974988 | 0.30262937 | -1.2423612 | 0.21410333 | 0.52259147 |

| | | | | | | |
|---|---|---|---|---|---|---|
| LUC7L | 134.414311 | 0.367037618 | 0.27979616 | 1.31180363 | 0.1895864 | 0.48980427 |
| FAM234A | 26.0275401 | 0.992976508 | 0.46129144 | 2.15260122 | 0.03135003 | 0.17534955 |
| AXIN1 | 25.6774764 | -0.758128983 | 0.43085649 | -1.7595858 | 0.07847805 | 0.29970982 |
| TMEM8A | 51.9759641 | -0.749503024 | 0.43486129 | -1.7235451 | 0.08479 | 0.31408459 |
| RAB11FIP3 | 152.664807 | 0.749864177 | 0.27064165 | 2.77069027 | 0.00559376 | 0.05377622 |
| CAPN15 | 23.0286785 | -0.687312897 | 0.36764694 | -1.8694917 | 0.06155444 | 0.2599337 |
| PIGQ | 35.3732883 | 0.554173834 | 0.4880414 | 1.13550578 | 0.25616346 | 0.56895381 |
| RAB40C | 62.6300468 | -0.229900449 | 0.27625185 | -0.8322132 | 0.40528859 | 0.70455026 |
| MCRIP2 | 41.0176912 | 0.816343448 | 0.475973 | 1.71510451 | 0.08632607 | 0.31751647 |
| RHOT2 | 30.6932287 | 0.461600596 | 0.37824702 | 1.22036809 | 0.22232537 | 0.53302786 |
| STUB1 | 26.9465772 | -0.52317554 | 0.41850299 | -1.2501118 | 0.2112587 | 0.52029492 |
| METRN | 19.3184199 | -1.066548817 | 0.62315838 | -1.7115213 | 0.08698493 | 0.31893334 |
| FAM173A | 18.8734804 | 0.127925806 | 0.67902705 | 0.18839574 | 0.85056644 | 0.95035128 |
| NARFL | 18.5736323 | 0.075186072 | 0.34180091 | 0.21997037 | 0.82589423 | 0.94197405 |
| LMF1 | 71.6442625 | -0.271754221 | 0.3795824 | -0.7159295 | 0.47403491 | 0.76008624 |
| CACNA1H | 28.1540604 | -1.478071387 | 0.80974267 | -1.8253594 | 0.06794681 | 0.27788783 |
| UBE2I | 88.4027684 | -1.33099288 | 0.58218236 | -2.286213 | 0.0222418 | 0.14012334 |
| BAIAP3 | 21.5054449 | 0.426640874 | 0.34288297 | 1.24427548 | 0.21339819 | 0.52235998 |
| GNPTG | 40.1390919 | -0.223944779 | 0.46163719 | -0.4851099 | 0.62759838 | 0.84783129 |
| UNKL | 35.4233868 | -0.098623102 | 0.36199487 | -0.2724434 | 0.78528113 | 0.92276669 |
| CLCN7 | 51.1544083 | -0.983921265 | 0.61119833 | -1.6098232 | 0.10743646 | 0.36059666 |
| TELO2 | 11.2987079 | -0.19809263 | 0.59284008 | -0.3341418 | 0.73827259 | 0.90241447 |
| IFT140 | 38.4585136 | -0.431268138 | 0.44966077 | -0.9590966 | 0.33751006 | 0.64844536 |
| JPT2 | 33.4998595 | -1.351646143 | 0.39914653 | -3.3863407 | 0.00070831 | 0.01146583 |
| MAPK8IP3 | 88.0940951 | 0.170952448 | 0.21823598 | 0.78333758 | 0.4334289 | 0.7267202 |
| NME3 | 45.3705901 | 0.573626646 | 0.54994392 | 1.04306389 | 0.2969187 | 0.61093809 |
| MRPS34 | 19.8338603 | -0.364456166 | 0.46020811 | -0.7919377 | 0.42839698 | 0.7236272 |
| SPSB3 | 40.7991261 | 0.219803861 | 0.4365348 | 0.50351967 | 0.61459895 | 0.84174925 |
| HAGH | 59.5222098 | -0.311668014 | 0.30570175 | -1.0195166 | 0.30795777 | 0.62178394 |
| FAHD1 | 13.1116421 | -0.898476435 | 0.38527191 | -2.332058 | 0.01969764 | 0.12997934 |
| NDUFB10 | 26.9670259 | 0.219052392 | 0.64043699 | 0.34203582 | 0.73232393 | 0.89806184 |
| RPS2 | 66.6984113 | -0.674787008 | 0.4676301 | -1.4429931 | 0.14902241 | 0.43272656 |
| TSC2 | 54.0187146 | 0.16058438 | 0.44202523 | 0.36329234 | 0.71638651 | 0.88942581 |
| PKD1 | 34.7957207 | 0.370739752 | 0.28058357 | 1.32131666 | 0.1863958 | 0.48675708 |
| SNHG19 | 19.4679609 | 0.709420047 | 0.58408922 | 1.21457481 | 0.2245283 | 0.53576151 |
| PGP | 34.8505731 | -0.419481164 | 0.47262837 | -0.8875497 | 0.37478302 | 0.6785392 |
| E4F1 | 17.1482876 | 0.209013868 | 0.41755369 | 0.50056765 | 0.61667543 | 0.84223226 |
| ECI1 | 67.7656002 | 0.195444642 | 0.29473324 | 0.66312385 | 0.50725124 | 0.77969181 |
| RNPS1 | 49.4305938 | -0.407981724 | 0.32781913 | -1.244533 | 0.21330347 | 0.52227981 |
| ABCA3 | 20.0701046 | -0.080736774 | 0.4721649 | -0.1709927 | 0.86422947 | 0.95486142 |
| TBC1D24 | 17.9665067 | -0.928516849 | 0.66141553 | -1.4038328 | 0.16036863 | 0.44957865 |
| PDPK1 | 114.718938 | -0.574762704 | 0.45490184 | -1.2634873 | 0.2064141 | 0.51350189 |
| AC093525.4 | 11.8258082 | -0.578304991 | 0.5064938 | -1.141781 | 0.25354507 | 0.56559484 |
| KCTD5 | 13.6538582 | -0.205109403 | 0.41159324 | -0.4983303 | 0.61825122 | 0.84283824 |
| SRRM2 | 330.932869 | -0.167667998 | 0.37903852 | -0.4423508 | 0.65823536 | 0.86357172 |
| ELOB | 38.046207 | -0.415431676 | 0.57646338 | -0.7206558 | 0.47112132 | 0.75766284 |
| PRSS22 | 14.6783317 | -3.194107158 | 0.81541093 | -3.9171748 | 8.96E-05 | 0.00223958 |
| FLYWCH1 | 37.9921678 | 0.50149595 | 0.36740593 | 1.36496422 | 0.1722643 | 0.46801268 |
| TNFRSF12A | 48.6929781 | -3.107029161 | 0.80412789 | -3.8638495 | 0.00011161 | 0.00263466 |
| HCFC1R1 | 14.6117556 | 0.054662086 | 0.54159669 | 0.10092766 | 0.91960789 | 0.97520237 |
| ZNF213-AS1 | 20.3145596 | 0.418347342 | 0.43764145 | 0.95591343 | 0.33911597 | 0.64971064 |
| ZNF213 | 32.6225713 | -0.281583846 | 0.41581895 | -0.677179 | 0.49829241 | 0.77558432 |
| ZNF263 | 15.6535251 | -0.064264279 | 0.3762625 | -0.1707964 | 0.86438386 | 0.95487238 |
| TIGD7 | 16.269517 | -0.059038966 | 0.39185373 | -0.1506658 | 0.88023933 | 0.96097414 |
| ZNF75A | 56.6530433 | -0.342888011 | 0.55152494 | -0.621709 | 0.53413324 | 0.79850222 |
| AC025283.3 | 19.3082977 | 0.382893129 | 0.52151413 | 0.73419512 | 0.46282983 | 0.75091259 |
| ZSCAN32 | 12.8495522 | 0.164239387 | 0.42041782 | 0.39065753 | 0.6960504 | 0.88085402 |
| NAA60 | 17.0666495 | -0.031525632 | 0.4681273 | -0.0673441 | 0.94630774 | 0.98574808 |
| CLUAP1 | 41.2140707 | 0.301727318 | 0.42911901 | 0.70313202 | 0.48197348 | 0.76628579 |
| DNASE1 | 145.62838 | 0.691165382 | 0.26799907 | 2.57898422 | 0.00990913 | 0.081789 |
| TRAP1 | 52.5518692 | 0.361774305 | 0.3136241 | 1.15352841 | 0.24869356 | 0.56120297 |
| CREBBP | 238.564056 | 0.123369234 | 0.23396148 | 0.52730574 | 0.59798129 | 0.83479226 |
| ADCY9 | 69.2084525 | -0.262358801 | 0.38598742 | -0.6797082 | 0.49668926 | 0.77475164 |
| GLIS2 | 29.083788 | -0.374421917 | 0.4737221 | -0.790383 | 0.4293041 | 0.7236272 |
| VASN | 17.2571668 | -1.874063544 | 0.5818552 | -3.2208418 | 0.00127815 | 0.01816683 |
| DNAJA3 | 42.7954545 | 0.181577082 | 0.24431416 | 0.74321144 | 0.45735368 | 0.747377 |
| NMRAL1 | 27.7836279 | -0.154186637 | 0.35136655 | -0.4388199 | 0.66079204 | 0.86492722 |
| HMOX2 | 32.370706 | -0.62429449 | 0.40584308 | -1.5382657 | 0.12398366 | 0.38919278 |
| CDIP1 | 28.5886663 | 0.526684999 | 0.40062181 | 1.31466882 | 0.18862123 | 0.48919517 |
| MGRN1 | 64.6781454 | 0.145547996 | 0.31294462 | 0.46509187 | 0.64186567 | 0.85468927 |
| ANKS3 | 39.5844044 | 0.23012124 | 0.25289043 | 0.90996422 | 0.36284138 | 0.66820719 |
| ZNF500 | 16.3387981 | 0.155451481 | 0.36923413 | 0.42101059 | 0.67374735 | 0.86846218 |
| GLYR1 | 113.816802 | -0.391740262 | 0.34527603 | -1.1345713 | 0.25655499 | 0.56935259 |
| UBN1 | 47.63437 | -0.096476014 | 0.28419376 | -0.3394727 | 0.73425368 | 0.89939324 |
| PPL | 22.4785772 | -0.883954276 | 0.38969647 | -2.2683148 | 0.02331002 | 0.14450975 |
| ALG1 | 15.4623861 | 0.847248062 | 0.5311728 | 1.59505168 | 0.11070068 | 0.36614868 |
| RBFOX1 | 19.7606858 | 0.374934879 | 0.6068263 | 0.61786195 | 0.53666634 | 0.80030369 |
| METTL22 | 22.0236353 | -0.221022486 | 0.37354852 | -0.5916835 | 0.55406257 | 0.81070871 |
| ABAT | 20.2276249 | 0.44633855 | 0.46459208 | 0.96071063 | 0.33669769 | 0.64764721 |
| PMM2 | 67.8513334 | -0.571754663 | 0.41795065 | -1.3679956 | 0.17131345 | 0.46718638 |
| CARHSP1 | 32.8734784 | -1.503171351 | 0.35293593 | -4.2590487 | 2.05E-05 | 0.00071693 |
| USP7 | 60.7132182 | 0.075046704 | 0.36283455 | 0.2068345 | 0.83613911 | 0.94671051 |
| C16orf72 | 66.5389939 | -0.223295843 | 0.35599772 | -0.6272395 | 0.53050224 | 0.7958814 |
| ATF7IP2 | 94.4122536 | 0.697589621 | 0.44425763 | 1.57023668 | 0.11636006 | 0.37515255 |
| NUBP1 | 17.4195842 | -0.78043702 | 0.42792332 | -1.8237777 | 0.06818568 | 0.27829012 |
| CIITA | 15.5707295 | -1.380143429 | 0.43157921 | -3.1978914 | 0.00138436 | 0.0193811 |
| DEXI | 15.6105799 | -0.836319771 | 0.40057001 | -2.0878242 | 0.03681369 | 0.19251311 |
| CLEC16A | 268.611028 | -0.646365251 | 0.25543428 | -2.5304562 | 0.01139143 | 0.09008796 |
| AC007220.1 | 24.2759125 | -0.124624458 | 0.45022483 | -0.2768049 | 0.78192989 | 0.92126657 |
| SOCS1 | 17.9155118 | -1.573355079 | 0.77687645 | -2.025232 | 0.04284354 | 0.21080021 |
| AC099489.1 | 45.7816216 | -2.688137061 | 1.14882305 | -2.3399052 | 0.01928863 | 0.12818762 |
| LITAF | 133.043289 | -0.434164764 | 0.23057743 | -1.8829456 | 0.05970774 | 0.25549104 |
| TXNDC11 | 108.097374 | 0.596582771 | 0.24614953 | 2.42366001 | 0.01536498 | 0.11028722 |
| ZC3H7A | 106.055103 | 0.078040054 | 0.22645537 | 0.3446156 | 0.73038338 | 0.89731942 |
| RSL1D1 | 20.067296 | -0.357088448 | 0.36110174 | -0.988886 | 0.32271893 | 0.635042 |
| AC007216.4 | 48.5340381 | -0.225049302 | 0.28235588 | -0.7970413 | 0.42542704 | 0.72187111 |
| GSPT1 | 142.334633 | -0.296048799 | 0.31504128 | -0.9397143 | 0.34736413 | 0.6553345 |
| SNX29 | 1211.78539 | -0.293406181 | 0.52396752 | -0.5599702 | 0.57549978 | 0.82147339 |
| AC010333.2 | 26.1908524 | -2.120367456 | 0.97474614 | -2.1753022 | 0.02960748 | 0.1681243 |
| CPPED1 | 181.214229 | -1.829456235 | 0.70256284 | -2.6039752 | 0.00921494 | 0.07756589 |
| SHISA9 | 57.8593431 | -0.095404862 | 0.82793367 | -0.1152325 | 0.90826084 | 0.97050471 |
| U91319.1 | 64.8329819 | -1.30508364 | 0.89463314 | -1.458792 | 0.14462237 | 0.42644619 |
| ERCC4 | 19.9664825 | -0.043484544 | 0.37880132 | -0.1147951 | 0.90860751 | 0.97059151 |
| MKL2 | 248.950486 | 0.450982143 | 0.2871412 | 1.57059364 | 0.11627707 | 0.37503292 |
| PARN | 113.219526 | 0.196350167 | 0.23367555 | 0.84026832 | 0.40075796 | 0.70092619 |
| BFAR | 34.7665766 | -0.663793441 | 0.27658606 | -2.3999526 | 0.01639719 | 0.11444368 |
| NOMO1 | 18.7280945 | 0.210177498 | 0.41933687 | 0.50121397 | 0.61622054 | 0.84223226 |
| PDXDC1 | 219.537292 | -0.030532039 | 0.25730329 | -0.1186617 | 0.9055434 | 0.96926721 |
| RRN3 | 32.8732019 | 0.276051744 | 0.32345814 | 0.85343884 | 0.39341599 | 0.69595061 |
| C16orf45 | 134.815934 | -0.256100128 | 0.38951927 | -0.6574774 | 0.51087398 | 0.78203726 |
| MARF1 | 59.6427924 | -0.106371113 | 0.34388953 | -0.3093177 | 0.75707988 | 0.9112882 |
| NDE1 | 24.7942521 | -0.14260276 | 0.46429765 | -0.3071365 | 0.75873947 | 0.91181467 |
| FOPNL | 26.5326086 | -0.037583215 | 0.34429585 | -0.1091597 | 0.91307586 | 0.97166624 |

| | | | | | | |
|---|---|---|---|---|---|---|
| ABCC1 | 54.7410089 | -0.082176056 | 0.45895815 | -0.1790491 | 0.85789913 | 0.95266335 |
| ABCC6 | 14.2651014 | 0.55485481 | 0.79724814 | 0.6959625 | 0.48645231 | 0.76836886 |
| RPS15A | 13.5547976 | 0.170201914 | 0.62478558 | 0.27241652 | 0.78530177 | 0.92276669 |
| SMG1 | 284.567667 | -0.127414243 | 0.38231956 | -0.3332663 | 0.73893324 | 0.90263507 |
| COQ7 | 15.0974619 | 0.51298194 | 0.43937121 | 1.16753654 | 0.24299376 | 0.55555126 |
| SYT17 | 31.0126106 | 1.015675059 | 0.89606513 | 1.13348352 | 0.25701126 | 0.56957837 |
| GDE1 | 17.0889484 | -0.356029435 | 0.436246 | -0.8161208 | 0.41443105 | 0.71211542 |
| CCP110 | 26.3270451 | 0.136184985 | 0.40743456 | 0.33424996 | 0.73819095 | 0.90241447 |
| VPS35L | 122.536453 | 0.088778087 | 0.26609546 | 0.33363248 | 0.73865691 | 0.90241575 |
| KNOP1 | 16.5670272 | -0.101630021 | 0.46051299 | -0.2206887 | 0.82533482 | 0.94180992 |
| IQCK | 147.729138 | -0.107478 | 0.45094298 | -0.2383406 | 0.81161697 | 0.93552094 |
| GPRC5B | 100.044326 | -0.213924571 | 0.42088625 | -0.5082717 | 0.61126281 | 0.84169124 |
| UMOD | 117.374312 | 1.24339277 | 0.78911791 | 1.57567424 | 0.1151009 | 0.3738073 |
| ACSM2A | 12.4013065 | 1.053797383 | 1.02222414 | 1.03088681 | 0.3025939 | 0.61691783 |
| ACSM2B | 14.8266194 | 0.829629425 | 1.01465025 | 0.81765064 | 0.4135567 | 0.71143975 |
| ACSM3 | 35.4452725 | -0.507674511 | 0.41473535 | -1.2240927 | 0.22091727 | 0.53115848 |
| THUMPD1 | 44.9140541 | 0.33031395 | 0.29320914 | 1.12654726 | 0.25993395 | 0.57323182 |
| ERI2 | 21.3226884 | -0.263079847 | 0.44129736 | -0.596151 | 0.55107438 | 0.80853147 |
| LYRM1 | 20.8238316 | -0.122464568 | 0.4441175 | -0.2757481 | 0.78274153 | 0.92150025 |
| TMEM159 | 22.431618 | 0.351651784 | 0.36574289 | 0.96147264 | 0.33631458 | 0.64749063 |
| METTL9 | 94.0619604 | -0.385477514 | 0.36713902 | -1.0499497 | 0.29374123 | 0.60876712 |
| NPIPB4 | 37.1500582 | -0.680510506 | 0.7166158 | -0.9496169 | 0.34230692 | 0.65202651 |
| UQCRC2 | 126.293943 | 0.765108829 | 0.22910481 | 3.33955812 | 0.00083912 | 0.0130182 |
| MOSMO | 84.2555487 | 0.237378377 | 0.26212953 | 0.90557661 | 0.36515999 | 0.67012584 |
| EEF2K | 15.796869 | -0.241956736 | 0.37509206 | -0.6450596 | 0.51888856 | 0.78745393 |
| POLR3E | 30.0835329 | 0.299214287 | 0.28734892 | 1.04129255 | 0.29773979 | 0.61141484 |
| CDR2 | 194.548534 | -0.695729281 | 0.47434018 | -1.4667307 | 0.14244934 | 0.42308208 |
| NPIPB5 | 90.6077454 | -0.508171171 | 0.51667763 | -0.9835362 | 0.32534359 | 0.63715682 |
| USP31 | 28.6870933 | -0.84247475 | 0.40099406 | -2.1009657 | 0.03564398 | 0.1884039 |
| COG7 | 94.6438239 | 0.113385464 | 0.26070735 | 0.43491472 | 0.66362433 | 0.86588413 |
| GGA2 | 97.8677204 | -0.240516132 | 0.51703859 | -0.4651802 | 0.64180239 | 0.85468927 |
| EARS2 | 14.7732514 | -0.066177049 | 0.49021147 | -0.1349969 | 0.89261429 | 0.96427805 |
| UBFD1 | 20.4036895 | -0.009682145 | 0.38717386 | -0.0250072 | 0.9800492 | 0.99607355 |
| NDUFAB1 | 27.1918591 | 0.885797722 | 0.39665158 | 2.23318842 | 0.02553652 | 0.15331779 |
| PALB2 | 18.8222721 | -0.340388259 | 0.51579466 | -0.6599298 | 0.50929889 | 0.78103666 |
| DCTN5 | 37.4945972 | -0.34883533 | 0.34472324 | -1.0119287 | 0.31157217 | 0.62642275 |
| RBBP6 | 89.7134004 | 0.483180094 | 0.26085783 | 1.85227368 | 0.06398653 | 0.26717888 |
| TNRC6A | 792.706601 | -0.257861668 | 0.68415232 | -0.3769068 | 0.70624286 | 0.88599027 |
| ARHGAP17 | 72.3276169 | 0.025271426 | 0.29623427 | 0.08530892 | 0.9320158 | 0.97938863 |
| LCMT1 | 57.1423208 | 0.22518107 | 0.30029124 | 0.74987559 | 0.45332964 | 0.74453237 |
| AC008741.2 | 12.1987887 | 0.117204667 | 0.46825658 | 0.2503001 | 0.80235528 | 0.93060837 |
| NSMCE1 | 21.9448915 | 0.290286443 | 0.37208062 | 0.78017083 | 0.43529033 | 0.72827249 |
| IL4R | 32.6007415 | -3.257922864 | 0.66752497 | -4.8806007 | 1.06E-06 | 6.13E-05 |
| GTF3C1 | 51.6848003 | 0.217136943 | 0.28689698 | 0.75684638 | 0.4491419 | 0.74080155 |
| KIAA0556 | 96.2490233 | -0.392148135 | 0.23356031 | -1.6790016 | 0.09315174 | 0.3326491 |
| XPO6 | 106.762571 | 0.250718144 | 0.41807629 | 0.59969472 | 0.54870971 | 0.80661714 |
| SGF29 | 18.7741622 | -0.063949056 | 0.39025167 | -0.1638662 | 0.86983648 | 0.95621002 |
| SULT1A1 | 39.0822495 | 0.94283964 | 0.62335909 | 1.51251446 | 0.13040304 | 0.40117636 |
| ATXN2L | 50.0445396 | -0.20363044 | 0.32157598 | -0.6332265 | 0.52658572 | 0.79346524 |
| TUFM | 44.5238627 | -0.409019014 | 0.54161054 | -0.7551903 | 0.45013482 | 0.74152951 |
| SH2B1 | 67.9764497 | 0.38990159 | 0.36031166 | 1.08212314 | 0.27919781 | 0.59380484 |
| RABEP2 | 57.5506973 | -0.159602752 | 0.48471928 | -0.3292684 | 0.74195281 | 0.90419135 |
| NFATC2IP | 90.4538671 | -0.215601046 | 0.28867146 | -0.7468734 | 0.45513996 | 0.74558223 |
| SPNS1 | 22.6001623 | -0.091230305 | 0.35586765 | -0.2563602 | 0.79767271 | 0.92852689 |
| QPRT | 11.3906083 | -0.681854338 | 0.63249669 | -1.0780362 | 0.2810176 | 0.59523128 |
| KIF22 | 12.1418349 | 0.234905907 | 0.49272495 | 0.47674856 | 0.63354118 | 0.8520131 |
| MAZ | 17.3706144 | -1.288600428 | 0.57548888 | -2.2391404 | 0.02514678 | 0.15127005 |
| AC009133.1 | 19.3968358 | -3.171348247 | 0.72876651 | -4.3516658 | 1.35E-05 | 0.00050592 |
| MVP | 66.5768313 | -1.966392412 | 0.34527561 | -5.6951385 | 1.23E-08 | 1.25E-06 |
| CDIPT | 20.1097644 | 0.636474421 | 0.43056735 | 1.47822268 | 0.13934818 | 0.41770449 |
| KCTD13 | 31.6087314 | -0.181604484 | 0.40580024 | -0.4475219 | 0.6544983 | 0.86217501 |
| TMEM219 | 32.2389279 | -0.01496915 | 0.40193675 | -0.0372426 | 0.97029161 | 0.99156233 |
| TAOK2 | 19.2099337 | -0.085092806 | 0.53235275 | -0.1598429 | 0.87300483 | 0.95703878 |
| INO80E | 23.4255609 | 0.325621057 | 0.48991024 | 0.66465453 | 0.50627148 | 0.77934214 |
| PPP4C | 25.5734521 | -0.404169185 | 0.45256053 | -0.8930721 | 0.37181856 | 0.67534262 |
| YPEL3 | 38.4629484 | -0.189587579 | 0.32379948 | -0.5855092 | 0.55820538 | 0.81255224 |
| NPIPB13 | 23.8463677 | -0.66174131 | 0.72525391 | -0.9124271 | 0.36154394 | 0.66686799 |
| CD2BP2 | 17.7062937 | -0.100350583 | 0.53488609 | -0.1876111 | 0.8511815 | 0.95046902 |
| TBC1D10B | 14.9514005 | -0.454455562 | 0.52424357 | -0.8668787 | 0.3860085 | 0.68921713 |
| SEPHS2 | 31.8529212 | -1.690644496 | 0.70071616 | -2.412738 | 0.0158332 | 0.11192474 |
| ZNF785 | 22.8117212 | -0.044871384 | 0.31357015 | -0.1430984 | 0.88621248 | 0.96249798 |
| FBRS | 23.7691772 | -0.252438126 | 0.30948168 | -0.8156804 | 0.41468298 | 0.71219452 |
| PHKG2 | 15.5308069 | 0.148523584 | 0.47478959 | 0.3128198 | 0.75441757 | 0.91026551 |
| RNF40 | 30.2708298 | 0.053191407 | 0.40343637 | 0.13184584 | 0.89510623 | 0.96511349 |
| BCL7C | 62.5946416 | 0.348121401 | 0.40684632 | 0.85565822 | 0.39218686 | 0.69519967 |
| MIR762HG | 15.8895592 | -0.48122353 | 0.56511223 | -0.8515539 | 0.39446173 | 0.69642725 |
| FBXL19 | 18.167589 | -0.229474101 | 0.45816916 | -0.5008502 | 0.61647658 | 0.84223226 |
| SETD1A | 17.1789561 | -0.371501957 | 0.39346598 | -0.9441781 | 0.34507866 | 0.65406518 |
| STX4 | 15.0769986 | -0.274379866 | 0.45674583 | -0.6007277 | 0.54802138 | 0.80633601 |
| KAT8 | 33.016297 | -0.270555496 | 0.45615982 | -0.5931156 | 0.55310381 | 0.8101068 |
| PRSS8 | 43.1833317 | -1.413249415 | 0.73573936 | -1.9208561 | 0.05474985 | 0.24343712 |
| FUS | 177.688741 | 0.054384092 | 0.29303546 | 0.18558878 | 0.85276726 | 0.95137449 |
| AC026471.4 | 18.2710772 | 0.015233202 | 0.60414502 | 0.02521448 | 0.97988389 | 0.9960141 |
| C16orf58 | 26.6577235 | -0.427056908 | 0.36218587 | -1.1791098 | 0.23835446 | 0.5502394 |
| ZNF720 | 64.1716529 | -0.499534227 | 0.23817639 | -2.0973289 | 0.03596447 | 0.18956063 |
| VPS35 | 49.8206707 | -0.559766603 | 0.28433942 | -1.9686563 | 0.04899257 | 0.22726702 |
| C16orf87 | 14.2825334 | 0.253775112 | 0.4137834 | 0.61330424 | 0.53967519 | 0.80166126 |
| DNAJA2 | 25.3228049 | -0.125433686 | 0.33075093 | -0.3792391 | 0.70451032 | 0.88564841 |
| ITFG1 | 374.595759 | 0.147017034 | 0.17332148 | 0.84823318 | 0.39630812 | 0.69680878 |
| PHKB | 236.910188 | -0.169520496 | 0.24777704 | -0.6841655 | 0.49387068 | 0.7738663 |
| LONP2 | 204.017616 | 0.351850048 | 0.14760145 | 2.3837845 | 0.01713563 | 0.11808769 |
| SIAH1 | 39.7054732 | 0.059794284 | 0.43171568 | 0.13850385 | 0.88984223 | 0.9631927 |
| N4BP1 | 93.3939128 | 0.428982948 | 0.28452807 | 1.50769992 | 0.13163135 | 0.40346176 |
| HEATR3 | 15.3910791 | 0.370991771 | 0.40748245 | 0.91044845 | 0.36258605 | 0.6678689 |
| TENT4B | 58.5164915 | 0.515081342 | 0.34525575 | 1.49188346 | 0.1357297 | 0.41089081 |
| BRD7 | 51.2061566 | -0.264729967 | 0.24411594 | -1.0844436 | 0.27816817 | 0.59352714 |
| CYLD | 57.9499699 | -0.920404654 | 0.71051053 | -1.2954131 | 0.19517776 | 0.49766572 |
| SALL1 | 48.9930441 | -0.335712733 | 0.3400459 | -0.9872571 | 0.32351662 | 0.63611745 |
| AC087564.1 | 165.232589 | -1.072099932 | 0.54164653 | -1.979335 | 0.0477783 | 0.22386175 |
| TOX3 | 501.576126 | 1.765593787 | 0.32134709 | 5.49435125 | 3.92E-08 | 3.35E-06 |
| CHD9 | 628.992967 | 0.137288097 | 0.24282022 | 0.56538989 | 0.5718086 | 0.81947131 |
| RBL2 | 120.800055 | 0.070510929 | 0.24009702 | 0.29367682 | 0.76900487 | 0.91600052 |
| AKTIP | 23.3851157 | 0.78566188 | 0.34129887 | 2.30197623 | 0.02133651 | 0.1369229 |
| RPGRIP1L | 21.666325 | 0.260761161 | 0.75679121 | 0.34456156 | 0.73042401 | 0.89731942 |
| FTO | 336.438779 | -0.032685079 | 0.31988485 | -0.1021776 | 0.91861567 | 0.97506435 |
| LPCAT2 | 62.4350876 | 1.257834594 | 0.60116931 | 2.0923134 | 0.03641049 | 0.19094005 |
| AMFR | 105.761296 | 0.107946594 | 0.50214613 | 0.21497048 | 0.82979033 | 0.94376251 |
| NUDT21 | 60.7949197 | -0.082305201 | 0.31449105 | -0.2617092 | 0.79354565 | 0.92649882 |
| OGFOD1 | 28.7195334 | -0.163655946 | 0.38653472 | -0.4233926 | 0.67200884 | 0.86777458 |
| BBS2 | 57.3959583 | 0.396482277 | 0.25132863 | 1.57754523 | 0.11467012 | 0.37318822 |
| MT2A | 210.649825 | -0.295310281 | 0.4725361 | -0.6249475 | 0.53200548 | 0.79685448 |
| MT1E | 67.6468538 | 0.01702549 | 0.65423438 | 0.02602353 | 0.97923857 | 0.99567616 |
| MT1F | 15.9909302 | 0.267777493 | 0.60297391 | 0.44409466 | 0.65697415 | 0.86325042 |

| | | | | | | |
|---|---|---|---|---|---|---|
| MT1G | 160.327644 | 0.699400463 | 0.79571416 | 0.87895943 | 0.37942328 | 0.68335766 |
| MT1H | 52.7184693 | 1.049978529 | 0.97954246 | 1.07190711 | 0.28376176 | 0.59724483 |
| MT1X | 40.0404212 | 1.109561441 | 0.81028253 | 1.3693513 | 0.17088949 | 0.46658085 |
| NUP93 | 44.5312119 | 0.264415532 | 0.29232346 | 0.90453067 | 0.36571408 | 0.67012584 |
| SLC12A3 | 5322.16201 | 3.339291818 | 0.93077111 | 3.58766164 | 0.00033366 | 0.00634901 |
| HERPUD1 | 76.3590596 | -0.713926827 | 0.62232836 | -1.1471867 | 0.25130446 | 0.56353121 |
| NLRC5 | 58.4836493 | -4.297407604 | 1.49269016 | -2.8789683 | 0.00398978 | 0.04274723 |
| FAM192A | 68.9074914 | -0.029448901 | 0.26728291 | -0.1101788 | 0.9122676 | 0.97144622 |
| RSPRY1 | 78.9193963 | -0.061058022 | 0.29585941 | -0.2063751 | 0.8364979 | 0.94684966 |
| ARL2BP | 27.1316546 | -0.241331525 | 0.38472231 | -0.6272876 | 0.53047075 | 0.7958814 |
| PLLP | 19.078401 | -1.562227678 | 0.71291368 | -2.1913279 | 0.02842807 | 0.16420667 |
| CIAPIN1 | 41.4336883 | -0.110260809 | 0.34125016 | -0.3231084 | 0.74661312 | 0.90661036 |
| COQ9 | 66.0423089 | 0.530807314 | 0.24686672 | 2.15017768 | 0.03154116 | 0.17567711 |
| POLR2C | 15.3040884 | -0.297384843 | 0.46133509 | -0.6446179 | 0.51917485 | 0.78759816 |
| ADGRG1 | 157.42236 | -0.521695575 | 0.41912564 | -1.2447236 | 0.21323337 | 0.52224532 |
| KIFC3 | 184.935546 | 0.960235756 | 0.24347636 | 3.94385629 | 8.02E-05 | 0.00206523 |
| CSNK2A2 | 50.503938 | -0.666678721 | 0.40312657 | -1.6537702 | 0.09817421 | 0.34181341 |
| SETD6 | 12.5922118 | 0.129926662 | 0.55260217 | 0.2351179 | 0.81411723 | 0.93633777 |
| CNOT1 | 167.212577 | 0.076842275 | 0.25992585 | 0.29563153 | 0.76751149 | 0.91600052 |
| SLC38A7 | 20.118547 | 0.040929326 | 0.45625143 | 0.08970783 | 0.92851939 | 0.97829541 |
| GOT2 | 41.3450432 | 0.562624885 | 0.34776943 | 1.61781005 | 0.10570354 | 0.35687176 |
| AC092378.1 | 23.6560309 | 1.083647625 | 0.51575332 | 2.10109675 | 0.03563247 | 0.1884039 |
| AC138305.1 | 843.417789 | 0.868168766 | 0.27357619 | 3.17340768 | 0.00150661 | 0.02065826 |
| AC010542.3 | 27.1303867 | -0.029010107 | 0.61301049 | -0.047324 | 0.962255 | 0.99029318 |
| TK2 | 19.0698823 | -0.076306881 | 0.39088017 | -0.1952181 | 0.84522222 | 0.94907846 |
| CMTM4 | 244.521205 | 1.125257394 | 0.27434168 | 4.10166394 | 4.10E-05 | 0.0012065 |
| DYNC1LI2 | 122.62575 | -0.103269266 | 0.37535703 | -0.2751228 | 0.78322192 | 0.92183302 |
| NAE1 | 41.3504981 | 0.510866531 | 0.31769952 | 1.60801794 | 0.10783125 | 0.36142331 |
| PDP2 | 31.308225 | -0.242246239 | 0.4578436 | -0.5291026 | 0.59673428 | 0.83416944 |
| CDH16 | 231.743953 | 0.391297707 | 0.52897509 | 0.73972804 | 0.45946503 | 0.7479141 |
| RRAD | 111.480928 | -2.468569613 | 0.68735272 | -3.5914161 | 0.00032889 | 0.00628388 |
| FAM96B | 29.1864725 | -0.772386388 | 0.4141265 | -1.8650977 | 0.06216771 | 0.26193029 |
| CES2 | 30.0144537 | -1.058002216 | 0.43184165 | -2.4499773 | 0.01428652 | 0.10505327 |
| CBFB | 161.196718 | -0.098850384 | 0.25013675 | -0.3951854 | 0.69270609 | 0.87849579 |
| C16orf70 | 21.3973087 | 1.080475016 | 0.42566445 | 2.53832572 | 0.01113843 | 0.08853767 |
| NOL3 | 27.8955615 | -0.940597152 | 0.75889153 | -1.2394356 | 0.21518424 | 0.5242691 |
| KIAA0895L | 21.9909461 | 0.062814531 | 0.44022663 | 0.1426868 | 0.88653754 | 0.96251467 |
| E2F4 | 21.8995044 | -0.420375632 | 0.46834398 | -0.8975788 | 0.36941014 | 0.67364366 |
| LRRC29 | 16.5228413 | 0.868961571 | 0.41928339 | 2.07249223 | 0.03821956 | 0.19710132 |
| TMEM208 | 16.8957957 | -1.160755121 | 0.50982494 | -2.276772 | 0.02279985 | 0.14229302 |
| FHOD1 | 10.8913618 | 0.485414185 | 0.47590621 | 1.01997868 | 0.30773857 | 0.62161058 |
| LRRC36 | 15.9465646 | 0.761475883 | 0.78663378 | 0.96801829 | 0.33303524 | 0.64492217 |
| ZDHHC1 | 25.4566023 | -0.128264349 | 0.37242003 | -0.3444078 | 0.73053966 | 0.89731942 |
| HSD11B2 | 217.057938 | 0.927858321 | 1.04355102 | 0.88913556 | 0.37393023 | 0.67738983 |
| ATP6V0D1 | 50.6282328 | -0.316297434 | 0.49145482 | -0.6435941 | 0.51983866 | 0.7877576 |
| AC009061.2 | 41.6534622 | 0.57992751 | 0.54500466 | 1.06407808 | 0.28729333 | 0.60168981 |
| RIPOR1 | 43.2939336 | 0.250891606 | 0.37759389 | 0.66444826 | 0.50640345 | 0.7794166 |
| AC027682.4 | 15.9315046 | 0.123305466 | 0.45576696 | 0.27054499 | 0.78674101 | 0.92339968 |
| CTCF | 91.3465188 | 0.248706385 | 0.26597786 | 0.93506423 | 0.3497552 | 0.65681679 |
| GFOD2 | 22.7639418 | -0.148701251 | 0.31437485 | -0.4730062 | 0.63620876 | 0.85375696 |
| RANBP10 | 94.4374594 | 0.127486532 | 0.34535084 | 0.36915078 | 0.71201534 | 0.88721964 |
| PSKH1 | 26.7772676 | 0.079491898 | 0.33135654 | 0.23989838 | 0.81040903 | 0.93552094 |
| DUS2 | 17.6889171 | 0.158935077 | 0.60846852 | 0.26120509 | 0.79393435 | 0.92660457 |
| NFATC3 | 143.305035 | 0.142265261 | 0.20980918 | 0.67806978 | 0.49772745 | 0.77553656 |
| PLA2G15 | 20.9559788 | 0.600180404 | 0.4424592 | 1.35646498 | 0.17495127 | 0.47154827 |
| SLC7A6 | 63.7938755 | 1.083111838 | 0.37020384 | 2.92571749 | 0.00343663 | 0.03851337 |
| SLC7A6OS | 34.2109345 | -0.758568868 | 0.41263772 | -1.8383411 | 0.06601216 | 0.27225103 |
| PRMT7 | 36.8306829 | 0.896716321 | 0.57013412 | 1.57281645 | 0.11576133 | 0.37438731 |
| ZFP90 | 43.3132372 | 0.174617066 | 0.47836904 | 0.36502585 | 0.71509211 | 0.88904248 |
| CDH3 | 59.6909509 | -2.76622666 | 0.63919798 | -4.3276524 | 1.51E-05 | 0.00055337 |
| CDH1 | 76.4520532 | -0.722600466 | 0.39623079 | -1.8236858 | 0.06819958 | 0.27829012 |
| TANGO6 | 106.187165 | 0.098828329 | 0.36912154 | 0.26773926 | 0.78890002 | 0.92402584 |
| CHTF8 | 32.0328833 | -0.118985872 | 0.28346217 | -0.4197593 | 0.67466132 | 0.86921959 |
| UTP4 | 24.0639077 | -0.349634772 | 0.31798167 | -1.0995438 | 0.27153094 | 0.58786226 |
| SNTB2 | 30.6662523 | -1.381144616 | 0.44998888 | -3.0692861 | 0.00214571 | 0.02660452 |
| VPS4A | 19.2602061 | -0.079952894 | 0.38714619 | -0.2065186 | 0.83638583 | 0.94684966 |
| TERF2 | 42.651635 | 0.017737747 | 0.3056019 | 0.05804201 | 0.95371517 | 0.98771968 |
| CYB5B | 85.7218769 | 0.187888472 | 0.27039401 | 0.69486922 | 0.48713727 | 0.76866947 |
| NFAT5 | 696.293025 | 0.024454303 | 0.18269966 | 0.13384975 | 0.89352139 | 0.96481684 |
| NQO1 | 28.6900251 | -1.869266884 | 0.67309219 | -2.7771335 | 0.00548406 | 0.05331952 |
| WWP2 | 203.287162 | -0.463024583 | 0.27924159 | -1.6581505 | 0.09728709 | 0.34011144 |
| PDPR | 72.2797541 | 0.481730285 | 0.35858862 | 1.3434065 | 0.17914038 | 0.47750284 |
| AC009060.1 | 25.7760293 | 0.023424622 | 0.484507 | 0.04834734 | 0.96143943 | 0.99021995 |
| EXOSC6 | 20.8624345 | -0.103793303 | 0.43301065 | -0.2397015 | 0.81056167 | 0.93552094 |
| AARS | 53.3282934 | -0.941302662 | 0.65631272 | -1.4342289 | 0.15150696 | 0.43600335 |
| ST3GAL2 | 38.1709963 | -0.211669627 | 0.3269392 | -0.6474281 | 0.51735491 | 0.78615013 |
| FUK | 20.0044058 | 0.151902619 | 0.44722283 | 0.33965757 | 0.73411442 | 0.89939324 |
| COG4 | 49.36424 | 0.219383602 | 0.293745 | 0.74685051 | 0.4551538 | 0.74558223 |
| SF3B3 | 64.9418462 | 0.197053301 | 0.25567979 | 0.77070348 | 0.44088271 | 0.73302253 |
| MTSS1L | 55.6209869 | -0.166849147 | 0.57660465 | -0.2893649 | 0.77230215 | 0.91791018 |
| VAC14 | 51.4952157 | -0.544884883 | 0.59995523 | -0.9082092 | 0.36376768 | 0.66951636 |
| HYDIN | 94.2289399 | -1.846093298 | 0.56770146 | -3.2518734 | 0.00114647 | 0.0165989 |
| CMTR2 | 78.3422428 | -2.023132232 | 0.56280048 | -3.5947592 | 0.00032469 | 0.00621648 |
| ZNF19 | 28.9302736 | -0.539312253 | 0.4259827 | -1.2660426 | 0.20549782 | 0.51190533 |
| MARVELD3 | 16.147078 | -0.655380432 | 0.43812112 | -1.4958887 | 0.13468266 | 0.40918252 |
| PHLPP2 | 38.4319937 | 0.272762599 | 0.43615666 | 0.62537759 | 0.53172327 | 0.79676281 |
| AP1G1 | 176.848583 | -0.151177183 | 0.33762772 | -0.4477629 | 0.65432429 | 0.86217501 |
| IST1 | 114.210038 | -0.058092494 | 0.40670507 | -0.1428369 | 0.88641899 | 0.96251467 |
| ZNF821 | 37.8900877 | 0.219371771 | 0.3476735 | 0.63097064 | 0.52805972 | 0.7943404 |
| TXNL4B | 17.8358838 | -0.28430244 | 0.44642599 | -0.6368411 | 0.52422832 | 0.79185875 |
| DHX38 | 18.7814856 | -0.504888061 | 0.46091166 | -1.0954118 | 0.27333628 | 0.58894058 |
| LINC01572 | 48.4487848 | 0.628866956 | 0.64711198 | 0.97180547 | 0.33114734 | 0.64324815 |
| AC004943.2 | 31.3680628 | 0.27880695 | 0.61983213 | 0.44981042 | 0.65284715 | 0.86183588 |
| ZFHX3 | 229.568859 | -0.74186629 | 0.19855382 | -3.7363486 | 0.00018671 | 0.00400207 |
| AC138627.1 | 51.6075104 | 0.847879588 | 0.71002651 | 1.19415201 | 0.23241851 | 0.54506955 |
| PSMD7 | 35.2463144 | -0.50850243 | 0.35967485 | -1.4137837 | 0.15742543 | 0.44423643 |
| NPIPB15 | 18.4917233 | -1.740676891 | 0.93373521 | -1.8642083 | 0.06229246 | 0.26233736 |
| GLG1 | 308.174309 | 0.246847404 | 0.18259342 | 1.35189646 | 0.17640844 | 0.47354004 |
| RFWD3 | 42.1254488 | -0.635137767 | 0.29515041 | -2.1519122 | 0.03140427 | 0.17538139 |
| FA2H | 77.3321211 | 0.696242818 | 0.41803078 | 1.66553005 | 0.09580705 | 0.33841174 |
| WDR59 | 132.114692 | 0.175545896 | 0.2367037 | 0.74162716 | 0.45831326 | 0.747377 |
| ZNRF1 | 81.1795209 | -0.060301445 | 0.45619468 | -0.1321836 | 0.89483909 | 0.96511349 |
| BCAR1 | 79.1247066 | -1.052164217 | 0.45508656 | -2.312009 | 0.02077719 | 0.1342526 |
| CFDP1 | 204.181101 | -0.142151429 | 0.24437987 | -0.5816822 | 0.56078074 | 0.81368185 |
| GABARAPL2 | 33.8624443 | -0.773560922 | 0.45164305 | -1.7127706 | 0.08675476 | 0.31843215 |
| ADAT1 | 16.3976908 | -0.318110105 | 0.4544966 | -0.6999175 | 0.48397885 | 0.76719123 |
| KARS | 24.7793948 | 0.141128176 | 0.43435954 | 0.32491096 | 0.74524846 | 0.90560363 |
| TERF2IP | 118.880791 | -0.23778621 | 0.37349256 | -0.6366558 | 0.52434906 | 0.79185875 |
| WWOX | 1200.64208 | 0.415678657 | 0.3877853 | 1.07192989 | 0.28375152 | 0.59724483 |
| CDYL2 | 70.5204943 | -0.366664586 | 0.8610431 | -0.4258377 | 0.67022614 | 0.86777458 |
| CMC2 | 78.3706415 | 0.851757 | 0.31121899 | 2.73684139 | 0.00620322 | 0.05790977 |
| ATMIN | 15.0969308 | -0.400072106 | 0.44151379 | -0.9061373 | 0.36486319 | 0.67012584 |
| GAN | 48.0853806 | 0.20089053 | 0.45525188 | 0.44127337 | 0.6590151 | 0.86447057 |

| | | | | | | |
|---|---|---|---|---|---|---|
| CMIP | 281.055931 | -0.816955306 | 0.48212532 | -1.6944875 | 0.09017269 | 0.32651269 |
| PLCG2 | 248.23392 | -0.136728795 | 0.22702518 | -0.6022627 | 0.5469993 | 0.80584949 |
| MPHOSPH6 | 25.0485267 | -0.094760248 | 0.33073764 | -0.2865118 | 0.77448613 | 0.91864687 |
| HSBP1 | 30.2472588 | -0.778115373 | 0.45644311 | -1.7047368 | 0.08824352 | 0.32265984 |
| MBTPS1 | 77.2921087 | -0.215893975 | 0.25733751 | -0.8389526 | 0.40149591 | 0.70114138 |
| HSDL1 | 15.3830931 | 0.024867361 | 0.41744397 | 0.05957054 | 0.95249769 | 0.98738742 |
| TAF1C | 18.8945396 | 0.473628049 | 0.47438858 | 0.99839681 | 0.31808698 | 0.63093651 |
| TLDC1 | 30.1894664 | 0.084541069 | 0.39879661 | 0.21199044 | 0.83211449 | 0.9450226 |
| KLHL36 | 45.3465124 | -0.102677054 | 0.27425294 | -0.3743882 | 0.70811554 | 0.88614996 |
| USP10 | 112.452066 | 0.040249773 | 0.32961628 | 0.122111 | 0.90281111 | 0.96803829 |
| ZDHHC7 | 94.8443621 | 0.107948227 | 0.43064012 | 0.25066923 | 0.80206985 | 0.93060267 |
| KIAA0513 | 37.1295663 | -1.304423853 | 0.53256413 | -2.4493273 | 0.01431233 | 0.10507732 |
| GSE1 | 162.510453 | -0.000831903 | 0.36328699 | -0.0022899 | 0.9981729 | 0.99922221 |
| EMC8 | 22.8434097 | -0.668886594 | 0.54946342 | -1.2173451 | 0.22347295 | 0.53397237 |
| COX4I1 | 102.123525 | -0.19616342 | 0.52386959 | -0.3744509 | 0.70806891 | 0.88614996 |
| MTHFSD | 15.3098427 | 0.633806075 | 0.41957599 | 1.51058708 | 0.13089369 | 0.40199365 |
| FBXO31 | 22.4755707 | 0.313289137 | 0.3878991 | 0.80765627 | 0.41928848 | 0.71772459 |
| MAP1LC3B | 38.5266726 | -0.745818813 | 0.46424311 | -1.6065264 | 0.1081583 | 0.36175274 |
| ZCCHC14 | 188.108847 | -1.568009239 | 0.65309405 | -2.4008935 | 0.0163551 | 0.11437129 |
| KLHDC4 | 56.128539 | -0.012185785 | 0.52881492 | -0.0230436 | 0.98161552 | 0.99656336 |
| SLC7A5 | 49.2664828 | -2.9134656 | 0.74462312 | -3.9126714 | 9.13E-05 | 0.00225159 |
| BANP | 76.6135132 | -0.612372663 | 0.56838386 | -1.0773928 | 0.28130483 | 0.59570436 |
| ZFPM1 | 25.9406431 | -0.946694627 | 0.30161518 | -3.1387499 | 0.0016967 | 0.022408 |
| ZC3H18 | 42.1615325 | -0.284690295 | 0.35166749 | -0.809544 | 0.41820233 | 0.71665475 |
| CYBA | 46.8653218 | -2.206400701 | 0.72768402 | -3.0320862 | 0.0024287 | 0.02946992 |
| MVD | 28.4772514 | -0.011968712 | 0.53808119 | -0.0222433 | 0.98225386 | 0.99668462 |
| PIEZO1 | 32.6243558 | 0.424382182 | 0.60943197 | 0.69635694 | 0.48620532 | 0.76833754 |
| APRT | 21.1681163 | -1.194464044 | 0.44874819 | -2.6617691 | 0.00777312 | 0.06869816 |
| GALNS | 30.3261906 | -0.175071931 | 0.31390762 | -0.557718 | 0.57703696 | 0.82229751 |
| ACSF3 | 34.5183864 | -0.003678908 | 0.30583162 | -0.0120292 | 0.99040232 | 0.99857194 |
| ZNF778 | 15.9470386 | -0.127457647 | 0.43666592 | -0.2918882 | 0.77037208 | 0.91677718 |
| ANKRD11 | 370.13334 | -0.46307439 | 0.35431785 | -1.3069463 | 0.19123096 | 0.49200813 |
| SPG7 | 150.031736 | -0.398980843 | 0.37573621 | -1.0618642 | 0.28829733 | 0.60284465 |
| RPL13 | 163.104471 | -0.747574191 | 0.41053856 | -1.8209597 | 0.06861298 | 0.27887855 |
| CHMP1A | 24.007659 | -0.310210438 | 0.46796835 | -0.6628876 | 0.50740252 | 0.77979579 |
| CDK10 | 38.8749835 | -0.236158654 | 0.36544485 | -0.6462224 | 0.51813533 | 0.78682309 |
| VPS9D1 | 38.6900545 | 0.36638925 | 0.3671305 | 0.99798097 | 0.31828859 | 0.63093651 |
| ZNF276 | 24.7264453 | 0.408297778 | 0.48292969 | 0.84546009 | 0.39785401 | 0.69815955 |
| SPIRE2 | 10.0881846 | 0.505771585 | 0.67265578 | 0.75190253 | 0.45210967 | 0.74372698 |
| TCF25 | 275.814196 | -0.908287059 | 0.48408315 | -1.8763038 | 0.06061357 | 0.2573479 |
| DEF8 | 15.0855137 | -0.088809138 | 0.40932051 | -0.2169672 | 0.82823388 | 0.94268221 |
| GAS8 | 49.8152118 | 0.106303374 | 0.34859343 | 0.30494945 | 0.76040465 | 0.91286466 |
| FAM157C | 46.5613295 | 0.302527876 | 0.47884804 | 0.63178263 | 0.52752892 | 0.79403853 |
| RPH3AL | 71.1486216 | -0.871140227 | 0.31326407 | -2.7808495 | 0.00542169 | 0.05287845 |
| VPS53 | 194.160801 | -0.466263065 | 0.33663109 | -1.3850862 | 0.16602612 | 0.45840319 |
| GLOD4 | 21.0801911 | -0.266782936 | 0.36580974 | -0.7292942 | 0.46582169 | 0.75274202 |
| NXN | 27.1775231 | -1.029717696 | 0.64247152 | -1.6027445 | 0.10899107 | 0.36312941 |
| ABR | 22.0842042 | -1.113543745 | 0.45208642 | -2.4631214 | 0.01377333 | 0.10257388 |
| YWHAE | 206.372219 | -0.094201615 | 0.2875915 | -0.3275535 | 0.74324925 | 0.90463949 |
| CRK | 88.6865761 | -0.477431302 | 0.41222284 | -1.1581874 | 0.24678757 | 0.55938803 |
| MYO1C | 62.7235421 | -0.542705367 | 0.28227385 | -1.92262 | 0.05452779 | 0.24302919 |
| INPP5K | 14.2199213 | -0.604027235 | 0.43053285 | -1.4029759 | 0.16062401 | 0.44998201 |
| PITPNA | 105.921947 | -0.443078149 | 0.24990395 | -1.7729938 | 0.07622973 | 0.29596141 |
| SLC43A2 | 94.2304958 | 0.667816193 | 0.48412845 | 1.37941942 | 0.16776547 | 0.46088547 |
| SCARF1 | 15.089714 | 0.230660986 | 0.50321895 | 0.45837102 | 0.64668591 | 0.85893154 |
| PRPF8 | 82.3122824 | -0.152866506 | 0.33948566 | -0.4502886 | 0.65250239 | 0.86179129 |
| MIR22HG | 21.3922248 | -0.423893039 | 0.70300263 | -0.602975 | 0.54652529 | 0.80581909 |
| WDR81 | 10.5284536 | -0.699257842 | 0.78760916 | -0.8878234 | 0.37463575 | 0.6784043 |
| SMYD4 | 21.1115574 | 0.338315978 | 0.35540323 | 0.95192149 | 0.3411368 | 0.65083559 |
| RPA1 | 32.38114 | 0.118953335 | 0.3584211 | 0.33188151 | 0.73997874 | 0.90304252 |
| SMG6 | 275.824333 | -0.128214266 | 0.29715923 | -0.4314665 | 0.66612918 | 0.86745649 |
| TSR1 | 19.1476584 | -0.851452251 | 0.40472627 | -2.1037731 | 0.03539824 | 0.18774357 |
| SGSM2 | 47.3752276 | 0.244049463 | 0.26778542 | 0.91136201 | 0.36210466 | 0.66724582 |
| MNT | 17.6710015 | -0.900015312 | 0.53067538 | -1.6959809 | 0.0898895 | 0.32574025 |
| METTL16 | 82.0010784 | 0.324580245 | 0.21446798 | 1.51342057 | 0.13017286 | 0.40070379 |
| PAFAH1B1 | 311.212206 | -0.075137218 | 0.27066016 | -0.2776072 | 0.78131388 | 0.92103406 |
| CLUH | 65.0696245 | 0.891657301 | 0.40786869 | 2.18613816 | 0.0288055 | 0.16558722 |
| RAP1GAP2 | 180.704527 | 1.123829686 | 0.25379149 | 4.42816151 | 9.50E-06 | 0.00036923 |
| CTNS | 21.3412048 | -0.915946343 | 0.39600812 | -2.3129484 | 0.02072548 | 0.13401134 |
| ITGAE | 16.3535402 | -0.49898018 | 0.44182319 | -1.1293662 | 0.25874338 | 0.57194663 |
| NCBP3 | 66.2514607 | 0.309209483 | 0.2404816 | 1.28579271 | 0.19851541 | 0.50147866 |
| ZZEF1 | 124.822995 | 0.123245868 | 0.39232503 | 0.31414225 | 0.753413 | 0.91007188 |
| CYB5D2 | 36.6085326 | -0.429926837 | 0.38074084 | -1.129185 | 0.25881981 | 0.57194663 |
| ANKFY1 | 107.790699 | 0.561766915 | 0.26698533 | 2.10411153 | 0.03536872 | 0.18769376 |
| UBE2G1 | 149.385707 | 0.023156715 | 0.34919696 | 0.06631419 | 0.94712768 | 0.98626519 |
| SPNS2 | 19.7428116 | -2.817715155 | 0.73334236 | -3.8422916 | 0.00012189 | 0.00282714 |
| GGT6 | 14.915343 | 0.702690805 | 0.57085483 | 1.23094484 | 0.2183435 | 0.52823943 |
| SMTNL2 | 34.5156133 | 1.136751543 | 0.34460849 | 3.29867534 | 0.00097142 | 0.01463253 |
| PELP1 | 23.8741826 | 0.529201914 | 0.32569798 | 1.62482405 | 0.10420006 | 0.35355215 |
| PSMB6 | 17.5644198 | -0.92888635 | 0.59903393 | -1.5506406 | 0.12098783 | 0.38441929 |
| MINK1 | 70.9201837 | -0.793133048 | 0.32423961 | -2.4461325 | 0.01443979 | 0.10559734 |
| SLC25A11 | 15.8392331 | 0.855587647 | 0.52079932 | 1.64283556 | 0.10041696 | 0.34587652 |
| RNF167 | 15.4567195 | 0.030318897 | 0.49322513 | 0.06147071 | 0.95098434 | 0.98684281 |
| PFN1 | 49.3705301 | -1.227681766 | 0.46357702 | -2.64828 | 0.00809025 | 0.07056452 |
| SPAG7 | 24.6496816 | -0.42372223 | 0.51748091 | -0.8188171 | 0.41289076 | 0.71121253 |
| CAMTA2 | 20.4570122 | 0.404532551 | 0.43597311 | 0.92788417 | 0.35346765 | 0.66036212 |
| KIF1C | 27.5378685 | -0.096208222 | 0.43072331 | -0.2233643 | 0.82325197 | 0.94091706 |
| ZNF594 | 23.3285508 | 0.121064479 | 0.45815705 | 0.26424231 | 0.79159321 | 0.92533536 |
| AC087500.1 | 49.9421471 | 0.755430783 | 0.48287834 | 1.56443295 | 0.11771593 | 0.37822242 |
| RABEP1 | 186.445862 | -0.257901341 | 0.379681 | -0.679258 | 0.49697443 | 0.77475164 |
| NUP88 | 124.791017 | -0.105847095 | 0.22629007 | -0.4677496 | 0.63996365 | 0.85413987 |
| RPAIN | 58.1645291 | -0.201867066 | 0.28790035 | -0.7011699 | 0.48319698 | 0.76664595 |
| DERL2 | 19.6988738 | 0.500021278 | 0.38685725 | 1.29252141 | 0.19617664 | 0.49845696 |
| MIS12 | 15.9865879 | -0.433332347 | 0.44611514 | -0.9713464 | 0.3313758 | 0.64324815 |
| KIAA0753 | 56.2467098 | -0.315474959 | 0.40504918 | -0.7788559 | 0.43606458 | 0.72891111 |
| XAF1 | 51.5743608 | -0.188322389 | 2.21525965 | -0.0850114 | 0.9322523 | 0.9794317 |
| ALOX12-AS1 | 36.9587374 | 0.096817799 | 0.67504609 | 0.14342398 | 0.88595534 | 0.96249798 |
| ACADVL | 130.958861 | 0.07059876 | 0.48072449 | 0.14685909 | 0.88324325 | 0.96170562 |
| DVL2 | 16.3696742 | 0.283424847 | 0.39012066 | 0.72650562 | 0.46752887 | 0.75393275 |
| ELP5 | 15.5932602 | 0.105713975 | 0.47456158 | 0.22276134 | 0.82372126 | 0.94133803 |
| EIF5A | 21.5237744 | -1.208357798 | 0.65156805 | -1.8545381 | 0.06366221 | 0.26606295 |
| ZBTB4 | 59.486375 | 0.211993019 | 0.32578329 | 0.6507179 | 0.51522861 | 0.78381327 |
| POLR2A | 97.3733744 | -0.575161683 | 0.39077421 | -1.4718517 | 0.14106093 | 0.42122951 |
| EIF4A1 | 21.4500171 | -0.865578717 | 0.64499372 | -1.3419956 | 0.17959743 | 0.4780378 |
| MPDU1 | 23.8328904 | 0.083689019 | 0.35110319 | 0.23836018 | 0.81160175 | 0.93552094 |
| FXR2 | 20.8788956 | 0.104193926 | 0.33270091 | 0.31317595 | 0.75414699 | 0.91026551 |
| SAT2 | 50.1406613 | 0.656829238 | 0.38910778 | 1.68803932 | 0.09140368 | 0.32943484 |
| TP53 | 28.5600576 | -0.151001631 | 0.32127232 | -0.4700113 | 0.63834692 | 0.8538689 |
| KDM6B | 46.0312929 | -2.106239357 | 0.76228868 | -2.7630469 | 0.00572645 | 0.05465042 |
| NAA38 | 50.4346661 | -0.222402677 | 0.32404927 | -0.6863237 | 0.49250904 | 0.77284018 |
| CHD3 | 24.5568775 | -0.298225983 | 0.43240343 | -0.6896938 | 0.49038674 | 0.77196791 |
| CNTROB | 15.8029364 | 0.531190211 | 0.44487501 | 1.19402124 | 0.23246966 | 0.54506955 |
| PER1 | 36.6442419 | 1.025843234 | 0.5159678 | 1.98819234 | 0.04679042 | 0.22193782 |

| | | | | | | |
|---|---|---|---|---|---|---|
| CTC1 | 24.5000676 | 0.257752887 | 0.34356738 | 0.75022515 | 0.45311911 | 0.74451091 |
| NDEL1 | 78.4377249 | 0.311256394 | 0.26845544 | 1.15943409 | 0.24627929 | 0.55925672 |
| MYH10 | 112.607106 | 0.217869393 | 0.3579444 | 0.60866826 | 0.54274435 | 0.80340275 |
| NTN1 | 26.1836557 | 0.265312645 | 0.58819967 | 0.45105882 | 0.65194716 | 0.86162372 |
| STX8 | 229.105421 | 0.314448516 | 0.21222477 | 1.48167682 | 0.13842631 | 0.41527892 |
| USP43 | 116.236082 | -0.007217882 | 0.68461165 | -0.010543 | 0.99158803 | 0.9988777 |
| SCO1 | 30.0783959 | -0.522107541 | 0.40167686 | -1.2998198 | 0.19366273 | 0.49539404 |
| TMEM220 | 28.7648247 | -0.353073333 | 0.57116509 | -0.6181634 | 0.53646765 | 0.80030143 |
| TMEM220-A! | 21.8636058 | -0.029536712 | 0.39660571 | -0.0744737 | 0.94063343 | 0.98411873 |
| SHISA6 | 24.4850714 | -1.358589408 | 0.67995157 | -1.9980679 | 0.0457093 | 0.21923535 |
| MAP2K4 | 260.628753 | -0.027617029 | 0.30804043 | -0.0896539 | 0.92856224 | 0.97829541 |
| ARHGAP44 | 136.438195 | -0.994296535 | 0.42676278 | -2.3298576 | 0.01981368 | 0.13037595 |
| ELAC2 | 17.3174463 | -0.160208982 | 0.35818565 | -0.4472792 | 0.6546735 | 0.86217501 |
| COX10 | 81.0201816 | 0.394009457 | 0.24436739 | 1.61236511 | 0.10688251 | 0.35938426 |
| TRIM16 | 19.3258043 | 0.519563671 | 0.60836315 | 0.85403541 | 0.39308537 | 0.69595061 |
| ZSWIM7 | 48.1415171 | 0.202994645 | 0.35821251 | 0.56668777 | 0.57092633 | 0.81895612 |
| TTC19 | 129.597922 | 0.182470421 | 0.25017703 | 0.72936521 | 0.46577829 | 0.75274202 |
| NCOR1 | 348.755315 | 0.287115709 | 0.16239488 | 1.76800956 | 0.07705931 | 0.2971468 |
| PIGL | 110.174347 | 0.055458491 | 0.29184306 | 0.19002847 | 0.84928682 | 0.95003258 |
| UBB | 79.4101596 | -1.270099582 | 0.568717 | -2.2332717 | 0.02553103 | 0.15331779 |
| AC098850.3 | 63.7660256 | 0.34336873 | 0.64818741 | 0.52973681 | 0.59629443 | 0.83416944 |
| MPRIP | 182.675461 | -0.222338556 | 0.44418898 | -0.5005495 | 0.61668824 | 0.84223226 |
| COPS3 | 21.8177997 | 0.434937537 | 0.55698609 | 0.78087684 | 0.43487493 | 0.72806372 |
| MED9 | 49.1850242 | 0.003525292 | 0.43632941 | 0.00807943 | 0.99355362 | 0.9992177 |
| RASD1 | 120.26871 | -2.650528222 | 0.76299752 | -3.473836 | 0.00051307 | 0.00877781 |
| PEMT | 19.6319711 | 0.356867214 | 0.45566227 | 0.7831836 | 0.43351931 | 0.7267411 |
| RAI1 | 101.520825 | -0.635502529 | 0.31265957 | -2.0325702 | 0.04209596 | 0.20922698 |
| SREBF1 | 57.0760969 | -0.824112183 | 0.55785219 | -1.4772949 | 0.13959661 | 0.41825155 |
| TOM1L2 | 327.707223 | 1.081376432 | 0.41522577 | 2.60430952 | 0.00920596 | 0.07756589 |
| DRC3 | 16.8335488 | -0.67435274 | 0.70656345 | -0.9544121 | 0.33987506 | 0.65027011 |
| ATPAF2 | 19.2144156 | 0.358626271 | 0.42786394 | 0.8381783 | 0.40193057 | 0.70114138 |
| GID4 | 13.125176 | -0.402178514 | 0.41508753 | -0.9689005 | 0.33259483 | 0.64485636 |
| DRG2 | 12.5899791 | -0.057594097 | 0.45595194 | -0.1263162 | 0.89948167 | 0.96645314 |
| ALKBH5 | 42.7387087 | -0.85935845 | 0.3355479 | -2.5610604 | 0.01043532 | 0.08482906 |
| FLII | 25.9297042 | -0.516711034 | 0.49547848 | -1.0428526 | 0.29701656 | 0.61093809 |
| TOP3A | 34.2054861 | -0.445485332 | 0.38888984 | -1.1455309 | 0.2519893 | 0.56438823 |
| SMCR8 | 14.1840903 | -0.622890787 | 0.49746652 | -1.252126 | 0.21052394 | 0.51915504 |
| SHMT1 | 30.5452187 | -1.056249499 | 0.46444851 | -2.2742015 | 0.02295387 | 0.14287177 |
| TVP23B | 31.552963 | -0.031130336 | 0.33445199 | -0.0930786 | 0.92584109 | 0.97735117 |
| PRPSAP2 | 76.0317554 | 0.323129513 | 0.27029205 | 1.19548284 | 0.23189844 | 0.54477728 |
| EPN2 | 142.086659 | -0.436367078 | 0.23930759 | -1.8234569 | 0.06823422 | 0.27830964 |
| B9D1 | 23.1415538 | -0.278072017 | 0.45345476 | -0.6132299 | 0.53972434 | 0.80166126 |
| SLC47A1 | 35.216964 | 1.936104764 | 0.38122738 | 5.07860894 | 3.80E-07 | 2.53E-05 |
| ALDH3A2 | 46.9732634 | 0.223687574 | 0.28802024 | 0.77663839 | 0.43737214 | 0.73004974 |
| ULK2 | 66.4515997 | 0.756404734 | 0.4544927 | 1.66428358 | 0.09605577 | 0.33873828 |
| AKAP10 | 158.223567 | 0.394907287 | 0.24813527 | 1.59150005 | 0.11149708 | 0.36760999 |
| SPECC1 | 21.5456434 | -0.887272835 | 0.40946462 | -2.1669097 | 0.03024174 | 0.17078983 |
| CCDC144NL-/ | 14.7602763 | -0.990551081 | 0.46614529 | -2.1249835 | 0.03358799 | 0.18250255 |
| USP22 | 134.729683 | -0.097848043 | 0.29885423 | -0.3274106 | 0.74335735 | 0.90463949 |
| DHRS7B | 100.495148 | -0.192790168 | 0.22678602 | -0.8500972 | 0.39527104 | 0.69642725 |
| AC087294.1 | 16.3249449 | 0.107156827 | 0.44168531 | 0.24260899 | 0.80830832 | 0.93433178 |
| MAP2K3 | 35.8942308 | -1.910664297 | 0.63063322 | -3.0297552 | 0.00244752 | 0.02956046 |
| WSB1 | 925.858233 | 0.318977026 | 0.40699511 | 0.78373676 | 0.43319459 | 0.72658865 |
| KSR1 | 111.536747 | -0.081539315 | 0.29526506 | -0.2761563 | 0.782428 | 0.92148019 |
| AC005697.1 | 13.8218987 | 1.896831019 | 1.04202411 | 1.82033314 | 0.06870829 | 0.27897062 |
| NLK | 307.991537 | 0.071269775 | 0.48814287 | 0.14600188 | 0.88391991 | 0.96170562 |
| IFT20 | 13.7400138 | 0.517569498 | 0.42665756 | 1.2130794 | 0.22509947 | 0.53616259 |
| TNFAIP1 | 29.5001222 | -0.806912924 | 0.38342229 | -2.1045019 | 0.03533469 | 0.18761997 |
| POLDIP2 | 19.5029676 | 0.126957303 | 0.40605254 | 0.31266226 | 0.75453727 | 0.91026551 |
| SARM1 | 22.5813991 | -0.545962932 | 0.54311347 | -1.0052465 | 0.31477816 | 0.62946385 |
| AC005726.1 | 22.6802623 | -0.561172011 | 0.36981677 | -1.5174326 | 0.12915751 | 0.39836738 |
| KIAA0100 | 56.2392517 | -0.250278493 | 0.2718497 | -0.9206502 | 0.35723307 | 0.6639427 |
| SDF2 | 21.9849916 | -1.107642663 | 0.4349874 | -2.5463788 | 0.0108847 | 0.08716404 |
| SUPT6H | 86.0933989 | -0.385475871 | 0.36820028 | -1.046919 | 0.29513693 | 0.61025281 |
| RAB34 | 20.6308382 | -0.828811619 | 0.41394691 | -2.0022172 | 0.04526138 | 0.21809669 |
| RPL23A | 61.4283458 | -0.917879946 | 0.38172417 | -2.4045633 | 0.0161918 | 0.11387005 |
| NEK8 | 16.6143264 | -0.548389283 | 0.79749569 | -0.6876392 | 0.49168003 | 0.77269924 |
| TRAF4 | 41.4358983 | -0.408520527 | 0.47548304 | -0.8591695 | 0.39024701 | 0.69333469 |
| FAM222B | 191.499776 | -0.007167404 | 0.24896528 | -0.0287888 | 0.97703306 | 0.99485162 |
| FLOT2 | 73.6897373 | -1.184391234 | 0.28129545 | -4.2104885 | 2.55E-05 | 0.0008366 |
| PHF12 | 60.563028 | -0.093354703 | 0.25082955 | -0.3721838 | 0.70975598 | 0.88614996 |
| MYO18A | 49.4260552 | -1.370814191 | 0.64198494 | -2.1352747 | 0.03273858 | 0.17981313 |
| NUFIP2 | 160.179225 | -0.606714057 | 0.33310238 | -1.8214042 | 0.06854544 | 0.27887855 |
| TAOK1 | 267.556534 | -0.141478723 | 0.31582416 | -0.4479668 | 0.65417719 | 0.86217501 |
| ABHD15-AS1 | 16.9481969 | -1.050527214 | 0.47628303 | -2.2056784 | 0.02740652 | 0.16065021 |
| GIT1 | 14.6603303 | -0.714588459 | 0.49618297 | -1.4401713 | 0.14981895 | 0.43409319 |
| SSH2 | 427.917598 | -0.84151357 | 0.29047462 | -2.8970296 | 0.00376714 | 0.04122633 |
| AC023389.1 | 15.0875297 | -1.360596915 | 0.48357066 | -2.8136465 | 0.00489831 | 0.04947544 |
| EFCAB5 | 15.0619249 | -2.607397289 | 0.60722717 | -4.2939404 | 1.76E-05 | 0.00062706 |
| NSRP1 | 179.900765 | -0.010438805 | 0.17919147 | -0.058255 | 0.95354548 | 0.98771968 |
| BLMH | 29.2554348 | 0.323618114 | 0.34327062 | 0.94274923 | 0.34580919 | 0.65414698 |
| CPD | 168.934469 | -1.255969989 | 0.4558612 | -2.7551588 | 0.00586637 | 0.05571813 |
| GOSR1 | 105.942033 | 0.140566084 | 0.33147921 | 0.42405701 | 0.67152423 | 0.86777458 |
| CRLF3 | 80.1201375 | 0.064253724 | 0.40657482 | 0.15803665 | 0.87442792 | 0.95761805 |
| ATAD5 | 16.7850858 | 0.113535824 | 0.61413303 | 0.18487171 | 0.85332966 | 0.95137449 |
| TEFM | 38.9908564 | 0.077130394 | 0.29769914 | 0.2590884 | 0.79556703 | 0.92698936 |
| RNF135 | 16.5091137 | 0.007018774 | 0.38645566 | 0.01816191 | 0.98550969 | 0.99746215 |
| NF1 | 552.923278 | -0.05674797 | 0.16402325 | -0.3459752 | 0.72936137 | 0.89693557 |
| RAB11FIP4 | 30.3114663 | 0.429510419 | 0.47171018 | 0.9105388 | 0.36253843 | 0.6678689 |
| COPRS | 18.1810472 | -0.056695393 | 0.57263093 | -0.0990086 | 0.92113143 | 0.97548809 |
| UTP6 | 37.000868 | 0.380042037 | 0.40263214 | 0.94389394 | 0.34522385 | 0.65406518 |
| SUZ12 | 130.440373 | 0.181131664 | 0.2268736 | 0.79838141 | 0.42464919 | 0.72151771 |
| LRRC37B | 37.8220063 | -0.336826291 | 0.33774569 | -0.9972778 | 0.31862966 | 0.63093651 |
| RHOT1 | 121.643858 | 0.378526947 | 0.28459782 | 1.33004162 | 0.18350456 | 0.48210665 |
| C17orf75 | 16.2534401 | 1.31857281 | 0.41811677 | 3.15359946 | 0.0016127 | 0.02169818 |
| ZNF207 | 212.213067 | 0.131738866 | 0.24933923 | 0.52835194 | 0.59725509 | 0.8342658 |
| PSMD11 | 56.8702048 | -0.093616436 | 0.46106106 | -0.2030456 | 0.83909936 | 0.94741614 |
| MYO1D | 216.663679 | -0.223750043 | 0.23178254 | -0.9653447 | 0.33437219 | 0.64554754 |
| LIG3 | 16.9290531 | 0.419625265 | 0.38256425 | 1.09687527 | 0.27269592 | 0.58843248 |
| SLFN5 | 19.448667 | -2.396995763 | 0.60391352 | -3.9691043 | 7.21E-05 | 0.0019219 |
| LINC02001 | 13.6145585 | -0.281993518 | 0.47596308 | -0.5924693 | 0.55353637 | 0.81035847 |
| AP2B1 | 163.380175 | 0.236470931 | 0.25260509 | 0.93612893 | 0.34920681 | 0.65632016 |
| TAF15 | 235.734039 | 0.072520745 | 0.18852408 | 0.38467629 | 0.70047726 | 0.88382163 |
| ZNHIT3 | 14.8873105 | -0.746832382 | 0.47235644 | -1.581078 | 0.1138602 | 0.37195243 |
| MYO19 | 84.676608 | 0.426760825 | 0.25127196 | 1.69840208 | 0.0894319 | 0.32467397 |
| GGNBP2 | 106.106869 | -0.448277202 | 0.4357731 | -1.0286941 | 0.30362346 | 0.61784922 |
| LHX1-DT | 180.687292 | 0.387611313 | 0.43864904 | 0.88364792 | 0.3768863 | 0.68029685 |
| LHX1 | 215.205334 | 0.393990681 | 0.54574945 | 0.72192593 | 0.47034002 | 0.75702578 |
| AATF | 113.558381 | -0.447745692 | 0.22237907 | -2.0134345 | 0.04406893 | 0.21423527 |
| ACACA | 125.474898 | -0.043749125 | 0.2407899 | -0.18169 | 0.85582599 | 0.9521686 |
| TADA2A | 42.5152014 | -0.284780875 | 0.3156051 | -0.9023329 | 0.36688004 | 0.67113782 |
| SYNRG | 143.237061 | -0.104383849 | 0.17809918 | -0.5860996 | 0.55780862 | 0.81243937 |
| DDX52 | 108.927498 | -0.853417537 | 0.64154215 | -1.3302595 | 0.18343278 | 0.48210665 |

| | | | | | | |
|---|---|---|---|---|---|---|
| HNF1B | 291.25177 | -0.573593648 | 0.4932869 | -1.1627993 | 0.24491095 | 0.55723517 |
| MRPL45 | 32.2478684 | 0.228504386 | 0.28809476 | 0.79315704 | 0.42768633 | 0.72307296 |
| SOCS7 | 33.1106502 | 0.567752014 | 0.34207395 | 1.65973471 | 0.09696783 | 0.33969491 |
| MLLT6 | 157.114332 | -0.282723587 | 0.28791125 | -0.9819817 | 0.32610884 | 0.63780184 |
| PCGF2 | 28.0717946 | 0.085185146 | 0.42400974 | 0.20090375 | 0.84077384 | 0.94770364 |
| PSMB3 | 24.2033153 | -0.48145949 | 0.40085376 | -1.2010851 | 0.22971819 | 0.54241242 |
| PIP4K2B | 39.2186621 | 0.368521407 | 0.27908009 | 1.32048622 | 0.18667274 | 0.48675708 |
| CWC25 | 29.6617519 | -0.62020636 | 0.30774434 | -2.0153299 | 0.0438701 | 0.21382372 |
| RPL23 | 52.2939219 | -0.635511349 | 0.40622688 | -1.5644246 | 0.11771788 | 0.37822242 |
| LASP1 | 35.075766 | -0.79408901 | 0.36016383 | -2.2047995 | 0.02746817 | 0.16065021 |
| RPL19 | 69.4136122 | -0.91961983 | 0.57320031 | -1.6043603 | 0.10863465 | 0.36253024 |
| FBXL20 | 201.160194 | -0.117440992 | 0.18412768 | -0.6378237 | 0.52358848 | 0.79110987 |
| MED1 | 41.4175829 | 0.084219233 | 0.30065097 | 0.28012294 | 0.77938319 | 0.92039563 |
| CDK12 | 154.757382 | -0.041741106 | 0.23625474 | -0.1766784 | 0.85976102 | 0.95342671 |
| STARD3 | 30.6471728 | -0.450788183 | 0.30660151 | -1.4702739 | 0.1414876 | 0.42163989 |
| PGAP3 | 18.5393583 | 1.178362263 | 0.51310063 | 2.29655196 | 0.02164434 | 0.13787238 |
| ERBB2 | 108.796944 | 0.741516255 | 0.54683489 | 1.3560149 | 0.17509443 | 0.47160665 |
| GRB7 | 97.3345597 | -0.856859983 | 0.33112602 | -2.5877157 | 0.00966147 | 0.08007426 |
| PSMD3 | 23.9122692 | -0.27810829 | 0.48191974 | -0.5770842 | 0.56388255 | 0.81450672 |
| MED24 | 36.6377385 | 0.261057687 | 0.37010774 | 0.70535593 | 0.48058876 | 0.76551071 |
| THRA | 17.2329393 | 0.118861109 | 0.51833885 | 0.2293116 | 0.81862674 | 0.93816073 |
| MSL1 | 48.1303122 | -0.064683333 | 0.28248151 | -0.2289825 | 0.81888249 | 0.93822319 |
| CASC3 | 50.4679033 | 0.279007738 | 0.28421207 | 0.98168855 | 0.32625331 | 0.63786661 |
| WIPF2 | 156.333307 | -0.333993482 | 0.41655932 | -0.8017909 | 0.42267391 | 0.72047744 |
| RARA | 25.6812686 | -0.845428124 | 0.67443046 | -1.2535438 | 0.21000787 | 0.51870552 |
| IGFBP4 | 19.62078 | -0.771796707 | 0.45025183 | -1.7141445 | 0.08650219 | 0.31781798 |
| KRT10 | 20.227872 | -1.156584276 | 0.54589908 | -2.1186778 | 0.0341177 | 0.18384695 |
| KRT19 | 70.8595357 | -3.41161635 | 0.88259865 | -3.8654221 | 0.0001109 | 0.00262438 |
| EIF1 | 132.68272 | -0.978326915 | 0.37429238 | -2.613804 | 0.00895404 | 0.07612871 |
| JUP | 104.233377 | -0.104834791 | 0.372181 | -0.2816769 | 0.77819126 | 0.91990415 |
| NT5C3B | 22.4574242 | 0.014337677 | 0.61647847 | 0.02325738 | 0.98144496 | 0.99656336 |
| ACLY | 33.5853636 | -0.68193931 | 0.33325748 | -2.0462836 | 0.04072848 | 0.20466781 |
| DNAJC7 | 86.4736713 | -0.0317067 | 0.24209264 | -0.1309693 | 0.89579961 | 0.96519206 |
| KAT2A | 25.4513876 | 1.449464328 | 0.46534404 | 3.11482304 | 0.00184055 | 0.0236504 |
| STAT5B | 171.864814 | -0.454934656 | 0.37351318 | -1.2179882 | 0.22322845 | 0.53368772 |
| STAT5A | 22.5180092 | -0.548863306 | 0.52838947 | -1.0387476 | 0.29892212 | 0.61269507 |
| STAT3 | 552.679316 | -1.364759035 | 0.36635008 | -3.7252865 | 0.00019509 | 0.00413421 |
| ATP6V0A1 | 76.136943 | -0.013018956 | 0.3376871 | -0.0385533 | 0.96924653 | 0.99156233 |
| NAGLU | 23.3469371 | -1.163865695 | 0.6415363 | -1.8141853 | 0.06964921 | 0.28149511 |
| COASY | 16.9333559 | -0.29192576 | 0.53047149 | -0.5503138 | 0.58210419 | 0.82598616 |
| MLX | 16.1770928 | -0.27219062 | 0.38149872 | -0.7134772 | 0.47555055 | 0.76153417 |
| RETREG3 | 57.4174818 | -0.015739117 | 0.26448563 | -0.0595084 | 0.95254717 | 0.98738742 |
| TUBG2 | 28.365489 | -0.029429812 | 0.30378371 | -0.0968775 | 0.92282366 | 0.97611757 |
| EZH1 | 74.0423665 | -0.05981844 | 0.30379568 | -0.1969035 | 0.84390304 | 0.94858974 |
| VPS25 | 18.5939731 | -1.068145728 | 0.43605373 | -2.4495736 | 0.01430255 | 0.10507732 |
| WNK4 | 200.727067 | 1.148886767 | 0.66039886 | 1.73968619 | 0.08191414 | 0.30797073 |
| COA3 | 32.6016401 | -0.310832107 | 0.45422994 | -0.6843056 | 0.49378219 | 0.7738663 |
| BECN1 | 55.332447 | 0.198998887 | 0.22779712 | 0.87357946 | 0.38234732 | 0.68644992 |
| PSME3 | 21.6299012 | -0.002714451 | 0.34686454 | -0.0078257 | 0.99375607 | 0.9992177 |
| LINC00671 | 12.9615169 | 3.300063579 | 0.7618834 | 4.33145492 | 1.48E-05 | 0.00054807 |
| RUNDC1 | 27.9541007 | -1.192155152 | 0.52850159 | -2.2557267 | 0.02408775 | 0.14717838 |
| RPL27 | 33.4298098 | -0.74744556 | 0.34621773 | -2.1588887 | 0.0308588 | 0.1733298 |
| BRCA1 | 21.9427904 | -0.004479291 | 0.68135459 | -0.0065741 | 0.99475467 | 0.9992177 |
| NBR1 | 74.7700465 | -0.164784887 | 0.22239528 | -0.740955 | 0.45872074 | 0.747377 |
| LINC00854 | 23.3537506 | -0.262533026 | 0.68234207 | -0.3847528 | 0.70042057 | 0.88382163 |
| DHX8 | 36.703509 | -0.134882715 | 0.33127351 | -0.4071642 | 0.68388739 | 0.87350219 |
| DUSP3 | 36.5008849 | -0.431023395 | 0.40503404 | -1.0641658 | 0.28725358 | 0.60168981 |
| PYY | 21.320565 | -0.637832828 | 0.92934162 | -0.6863276 | 0.49250653 | 0.77284018 |
| TMEM101 | 293.905931 | -0.654865963 | 0.88951766 | -0.7362034 | 0.4616069 | 0.74982974 |
| LSM12 | 41.5132894 | -0.071186803 | 0.3139665 | -0.2267338 | 0.82063078 | 0.9391876 |
| HDAC5 | 51.4249683 | 0.44092078 | 0.27889765 | 1.58094119 | 0.11389148 | 0.37195243 |
| ASB16-AS1 | 10.7353825 | 0.255576078 | 0.44862737 | 0.56968455 | 0.56889167 | 0.81764604 |
| TMUB2 | 24.6169412 | 0.160450182 | 0.317927 | 0.50467617 | 0.6137863 | 0.84174925 |
| UBTF | 29.8731206 | 0.398149196 | 0.32434093 | 1.22756384 | 0.21961074 | 0.52962456 |
| SLC25A39 | 32.0296927 | -0.122470994 | 0.51432204 | -0.2381212 | 0.81178707 | 0.93552094 |
| GRN | 37.4863791 | -1.445953667 | 0.53867205 | -2.6842931 | 0.00726834 | 0.06522614 |
| GPATCH8 | 323.421744 | -0.080015545 | 0.18657428 | -0.428867 | 0.66802004 | 0.86777458 |
| DBF4B | 12.2139044 | 0.454985414 | 0.58251493 | 0.78107082 | 0.43476084 | 0.72803702 |
| EFTUD2 | 44.0796976 | -0.137182389 | 0.3638182 | -0.377063 | 0.70612677 | 0.88599027 |
| DCAKD | 55.9162395 | -0.027171778 | 0.28978484 | -0.0937654 | 0.92529556 | 0.97735117 |
| NMT1 | 69.7019008 | -0.098877122 | 0.2606329 | -0.3793731 | 0.70441079 | 0.8856427 |
| ACBD4 | 12.4537973 | 1.052013702 | 0.52524287 | 2.00290907 | 0.04518705 | 0.21785111 |
| HEXIM1 | 18.3510583 | -1.336825482 | 0.5520504 | -2.4215642 | 0.01545387 | 0.11049992 |
| AC138150.1 | 57.4529228 | 0.448372879 | 0.47753797 | 0.93892613 | 0.34776867 | 0.6553345 |
| MAP3K14 | 45.6270409 | -1.298669524 | 0.45562023 | -2.8503333 | 0.00436734 | 0.04541366 |
| PLEKHM1 | 42.5490993 | -1.369515864 | 0.5850202 | -2.3409719 | 0.01923361 | 0.12791313 |
| LINC02210-C | 36.8654608 | 0.879843395 | 0.54542495 | 1.61313374 | 0.10671545 | 0.35932624 |
| KANSL1 | 710.715127 | 0.189679356 | 0.31194541 | 0.60805304 | 0.5431523 | 0.80361674 |
| ARL17B | 154.891923 | 1.455244269 | 1.00821018 | 1.44339374 | 0.14890959 | 0.43272656 |
| LRRC37A2 | 33.4150953 | 0.757180482 | 0.47548533 | 1.59243711 | 0.11128652 | 0.36720011 |
| ARL17A | 40.8630963 | 0.03280059 | 0.65215594 | 0.05029562 | 0.95988681 | 0.9895734 |
| NSF | 163.829545 | 0.372757778 | 0.35290103 | 1.05626718 | 0.29084617 | 0.60545873 |
| AC005670.2 | 32.8495761 | -0.327976216 | 0.43735356 | -0.7499109 | 0.45330835 | 0.74453237 |
| CDC27 | 164.416344 | -0.726437206 | 0.28893481 | -2.5141906 | 0.01193059 | 0.0934797 |
| AC068234.1 | 40.0827974 | -4.470676794 | 0.76270414 | -5.8616134 | 4.58E-09 | 5.70E-07 |
| EFCAB13 | 63.7273399 | -1.330361554 | 0.36868333 | -3.6084126 | 0.00030808 | 0.00595955 |
| NPEPPS | 344.308641 | -0.431531557 | 0.35761489 | -1.2066935 | 0.22755021 | 0.53959262 |
| AC025682.1 | 38.6989193 | 0.091678289 | 0.35862173 | 0.25564064 | 0.79822833 | 0.9287884 |
| KPNB1 | 134.000206 | -0.538732043 | 0.25974275 | -2.0740984 | 0.03807017 | 0.19687535 |
| TBKBP1 | 14.5195436 | -0.381449022 | 0.6075035 | -0.627896 | 0.53007207 | 0.79562009 |
| SCRN2 | 13.4914754 | 0.364978428 | 0.50971324 | 0.7160466 | 0.47396258 | 0.76008624 |
| SP2 | 18.6755352 | -0.107175269 | 0.40175752 | -0.2667661 | 0.78964929 | 0.92428059 |
| CDK5RAP3 | 33.5954125 | -0.048485331 | 0.30330801 | -0.1598551 | 0.87299522 | 0.95703878 |
| NFE2L1 | 60.4168596 | -0.175794952 | 0.29773217 | -0.5904466 | 0.55489126 | 0.8109414 |
| CBX1 | 45.3434579 | 0.025576985 | 0.28008705 | 0.09131798 | 0.92723993 | 0.97767557 |
| SNX11 | 12.7803531 | -0.176469755 | 0.40116786 | -0.4398901 | 0.66001674 | 0.86468962 |
| SKAP1 | 428.01927 | 0.283532778 | 0.36003274 | 0.78751942 | 0.43097786 | 0.72469569 |
| HOXB2 | 17.9596474 | -0.985724301 | 0.52293023 | -1.8850016 | 0.05942963 | 0.25476866 |
| HOXB-AS1 | 15.3386329 | 0.137769647 | 0.46430074 | 0.29672502 | 0.76667645 | 0.9156515 |
| HOXB3 | 229.034874 | -0.235694917 | 0.43617249 | -0.5403709 | 0.58894128 | 0.83025227 |
| HOXB-AS3 | 44.8406572 | -0.252123863 | 0.53141599 | -0.4744379 | 0.6351877 | 0.85291155 |
| HOXB7 | 27.2558474 | -1.314539969 | 0.46458437 | -2.8294967 | 0.00466213 | 0.04771644 |
| LINC02086 | 16.3696109 | 0.023448072 | 0.74406507 | 0.03151347 | 0.97486005 | 0.99361556 |
| CALCOCO2 | 42.1568402 | 0.383709988 | 0.32703428 | 1.17330206 | 0.2406747 | 0.55313061 |
| ATP5MC1 | 33.8301964 | 0.883735062 | 0.54243993 | 1.62918512 | 0.10327384 | 0.35104821 |
| UBE2Z | 81.6502445 | -0.458673335 | 0.30648152 | -1.4965775 | 0.13450324 | 0.40890387 |
| SNF8 | 26.4841719 | -0.134341786 | 0.36098601 | -0.3721523 | 0.70977944 | 0.88614996 |
| ZNF652 | 122.159574 | 0.031138266 | 0.20697521 | 0.15044443 | 0.88041399 | 0.96097414 |
| PHB | 32.1133009 | -0.550734325 | 0.47878249 | -1.1502808 | 0.25002822 | 0.56269702 |
| SPOP | 82.8845946 | 0.408786487 | 0.22815421 | 1.79171139 | 0.07317921 | 0.28899744 |
| KAT7 | 45.52253 | 0.011520096 | 0.30924099 | 0.03725281 | 0.97028343 | 0.99156233 |
| ITGA3 | 372.474866 | -3.364218769 | 0.70232971 | -4.7900847 | 1.67E-06 | 8.73E-05 |
| PDK2 | 13.557227 | 1.075293597 | 0.61478116 | 1.74906727 | 0.08027939 | 0.30415484 |

| | | | | | | |
|---|---|---|---|---|---|---|
| XYLT2 | 16.5884588 | 0.402755228 | 0.47109153 | 0.8549405 | 0.39258409 | 0.69577153 |
| MRPL27 | 23.6182299 | 0.32146628 | 0.38947441 | 0.82538486 | 0.40915313 | 0.70777268 |
| ACSF2 | 23.5611136 | 0.979616188 | 0.58934178 | 1.66222083 | 0.0964685 | 0.33942352 |
| RSAD1 | 15.2029111 | 0.462713221 | 0.46422627 | 0.99674072 | 0.31889038 | 0.63100017 |
| SPATA20 | 15.4104537 | 0.18669059 | 0.57230732 | 0.32620689 | 0.74426783 | 0.90500173 |
| ABCC3 | 37.6793158 | -3.120789852 | 0.72424071 | -4.3090506 | 1.64E-05 | 0.00059002 |
| ANKRD40 | 16.1449112 | 0.121354135 | 0.38278417 | 0.31703018 | 0.7512207 | 0.9089517 |
| LUC7L3 | 267.710894 | 0.238657744 | 0.31830917 | 0.74976711 | 0.45339498 | 0.74453237 |
| TOB1 | 50.9444821 | 0.220262282 | 0.55583666 | 0.3962716 | 0.69190468 | 0.87808891 |
| SPAG9 | 283.954075 | -0.886685422 | 0.56272005 | -1.5757132 | 0.11509191 | 0.3738073 |
| AC005920.2 | 21.4443706 | -0.724157517 | 0.61259849 | -1.1821079 | 0.23716289 | 0.54992696 |
| MBTD1 | 160.152628 | 0.088532539 | 0.23769796 | 0.37245814 | 0.70955177 | 0.88614996 |
| UTP18 | 36.2449327 | -0.429544713 | 0.35534404 | -1.2088136 | 0.22673446 | 0.53826211 |
| CA10 | 436.179175 | 0.126732366 | 0.59180814 | 0.21414434 | 0.8304345 | 0.94393022 |
| TOM1L1 | 226.606374 | 0.395974637 | 0.33078302 | 1.19708271 | 0.23127432 | 0.54389834 |
| COX11 | 50.0465694 | -0.442415385 | 0.38073157 | -1.1620139 | 0.24522981 | 0.55755248 |
| STXBP4 | 253.507488 | 0.165213736 | 0.38057355 | 0.43411776 | 0.66420293 | 0.86615778 |
| HLF | 206.655837 | -1.432548142 | 0.53516271 | -2.676846 | 0.00743188 | 0.06637437 |
| PCTP | 15.7182648 | 0.106680707 | 0.37740408 | 0.28266973 | 0.77743002 | 0.91968524 |
| DGKE | 33.8728807 | 0.203780042 | 0.38520565 | 0.52901623 | 0.59679419 | 0.83416944 |
| TRIM25 | 45.1414997 | -1.829370956 | 0.89162024 | -2.0517378 | 0.04019515 | 0.20290676 |
| COIL | 19.6079349 | -0.203485874 | 0.37849289 | -0.5376214 | 0.59083846 | 0.83192054 |
| SCPEP1 | 56.3284279 | 1.268296499 | 0.45623445 | 2.77992267 | 0.00543718 | 0.0529742 |
| AKAP1 | 56.9975396 | 0.834840215 | 0.41008011 | 2.03579787 | 0.04177066 | 0.20827251 |
| MSI2 | 1834.07305 | -0.092163681 | 0.33975673 | -0.2712637 | 0.78618819 | 0.92322402 |
| MRPS23 | 17.7973671 | 0.117921553 | 0.39228405 | 0.30060247 | 0.76371765 | 0.91479214 |
| CUEDC1 | 122.95385 | -0.847973556 | 0.41295 | -2.0534534 | 0.04002863 | 0.20269667 |
| VEZF1 | 56.4835038 | -0.118190213 | 0.29697875 | -0.3979753 | 0.69064837 | 0.8771239 |
| AC015813.2 | 43.3966104 | 0.436040246 | 0.32216779 | 1.353457 | 0.17590968 | 0.47294748 |
| SRSF1 | 19.4182778 | 0.073518884 | 0.44949797 | 0.16355777 | 0.8700793 | 0.95629149 |
| DYNLL2 | 25.3990113 | 0.165211256 | 0.43545983 | 0.37939494 | 0.70439461 | 0.8856427 |
| MKS1 | 14.414477 | 0.566295686 | 0.4183653 | 1.35359143 | 0.17586676 | 0.47294748 |
| TSPOAP1 | 28.3913509 | 0.212460028 | 0.81866466 | 0.25952021 | 0.79523389 | 0.9268423 |
| TSPOAP1-AS | 12.4402972 | -0.131242141 | 0.71777271 | -0.1828464 | 0.85491856 | 0.95179231 |
| RNF43 | 79.4654693 | 0.678504064 | 0.46344703 | 1.46403801 | 0.14318356 | 0.4243745 |
| MTMR4 | 27.0498149 | 0.208182018 | 0.37444657 | 0.55597256 | 0.57822961 | 0.82311488 |
| RAD51C | 16.6862005 | 0.553235183 | 0.46190197 | 1.1977329 | 0.23102102 | 0.54380466 |
| TRIM37 | 53.2769807 | -0.006703363 | 0.39305474 | -0.0170545 | 0.98639311 | 0.99762766 |
| SKA2 | 29.5762536 | 1.067470274 | 0.37785026 | 2.82511456 | 0.00472637 | 0.04821519 |
| GDPD1 | 19.7754152 | 0.891968989 | 0.42230473 | 2.11214539 | 0.03467398 | 0.18537855 |
| YPEL2 | 99.168177 | -0.349095497 | 0.36556383 | -0.9549509 | 0.33960255 | 0.65027011 |
| DHX40 | 63.2344209 | -0.027401242 | 0.29993034 | -0.0913587 | 0.92720759 | 0.97767557 |
| CLTC | 222.925992 | 0.239943881 | 0.30328018 | 0.79116242 | 0.42884922 | 0.7236272 |
| PTRH2 | 33.4992639 | -0.485790185 | 0.32352881 | -1.5015361 | 0.13321696 | 0.4061854 |
| VMP1 | 629.645525 | -1.587336305 | 0.62586977 | -2.5362086 | 0.01120599 | 0.08892315 |
| TUBD1 | 47.3716393 | -0.023094873 | 0.37117197 | -0.0622215 | 0.95038645 | 0.98679324 |
| RPS6KB1 | 64.1610585 | 0.302197682 | 0.32611234 | 0.92666742 | 0.35409924 | 0.6607207 |
| RNFT1 | 15.9840544 | 0.132294441 | 0.75324644 | 0.17563235 | 0.86058279 | 0.95365746 |
| HEATR6 | 16.5502211 | 0.594714158 | 0.43427977 | 1.36942635 | 0.17086604 | 0.46658085 |
| USP32 | 237.68277 | -1.089208065 | 0.48120718 | -2.2634909 | 0.02360545 | 0.14585631 |
| APPBP2 | 169.405482 | -0.100266538 | 0.22932811 | -0.4372187 | 0.66195275 | 0.86503818 |
| AC011921.1 | 14.5631764 | 0.341846843 | 0.58886414 | 0.58051903 | 0.56156466 | 0.81384822 |
| PPM1D | 62.0483937 | 0.272778837 | 0.37613855 | 0.72520839 | 0.46832419 | 0.75495415 |
| BCAS3 | 534.690059 | 0.335519724 | 0.29603514 | 1.13337804 | 0.25705553 | 0.56957837 |
| TBX2 | 21.9288761 | 0.817401393 | 0.60871528 | 1.34283041 | 0.17932689 | 0.47772685 |
| INTS2 | 37.4547986 | 0.514300367 | 0.30283701 | 1.69827449 | 0.08945597 | 0.32467397 |
| MED13 | 355.029352 | -0.13961113 | 0.34773127 | -0.4014915 | 0.68805833 | 0.87567053 |
| METTL2A | 14.8829424 | -0.294181233 | 0.4559072 | -0.6452656 | 0.51875508 | 0.78737952 |
| TLK2 | 160.505584 | 0.198438801 | 0.27454748 | 0.72278501 | 0.46981198 | 0.75656768 |
| TANC2 | 249.050446 | -1.583584035 | 0.21614205 | -7.3265894 | 2.36E-13 | 8.80E-11 |
| CYB561 | 15.0840376 | -0.718486425 | 0.46524939 | -1.5443039 | 0.12251474 | 0.38683625 |
| DCAF7 | 106.880058 | -0.312333903 | 0.22105734 | -1.412909 | 0.15768249 | 0.44444727 |
| MAP3K3 | 114.632889 | -0.476893859 | 0.32589026 | -1.4633572 | 0.14336967 | 0.42447522 |
| CCDC47 | 28.2872587 | 0.051705336 | 0.31856021 | 0.16230946 | 0.87106216 | 0.95629149 |
| DDX42 | 84.5929248 | 0.133854685 | 0.33105451 | 0.40432824 | 0.68597137 | 0.87448688 |
| FTSJ3 | 15.1581894 | -0.155842463 | 0.52849282 | -0.2948809 | 0.76808483 | 0.91600052 |
| PSMC5 | 13.8676801 | 0.322196651 | 0.52257558 | 0.61655512 | 0.53752821 | 0.80078458 |
| SMARCD2 | 17.0139047 | 0.06751267 | 0.49020113 | 0.13772443 | 0.89045821 | 0.96323092 |
| ERN1 | 57.9460516 | -2.132334273 | 0.64939782 | -3.2835563 | 0.00102506 | 0.01526785 |
| SNHG25 | 26.02168 | 0.140947175 | 0.4989053 | 0.28251288 | 0.77755027 | 0.91968524 |
| TEX2 | 121.67521 | -0.36772749 | 0.55063994 | -0.6678184 | 0.50424951 | 0.77792612 |
| POLG2 | 28.7035927 | 0.176792214 | 0.28639449 | 0.61730312 | 0.53703481 | 0.80052959 |
| DDX5 | 400.956596 | -0.257305933 | 0.2548949 | -1.0094589 | 0.31275459 | 0.62758402 |
| CEP95 | 150.995042 | 0.057423506 | 0.32401285 | 0.17722601 | 0.85933086 | 0.95328983 |
| SMURF2 | 195.372185 | 0.545882748 | 0.1922851 | 2.83892376 | 0.0045266 | 0.04658498 |
| LRRC37A3 | 146.392049 | 0.531388697 | 0.37397265 | 1.42092929 | 0.15533732 | 0.44090263 |
| GNA13 | 86.8273365 | -0.180129543 | 0.37451965 | -0.4809615 | 0.63054384 | 0.85008542 |
| CEP112 | 144.754543 | 0.103322841 | 0.57758164 | 0.17888872 | 0.85802508 | 0.95266335 |
| PRKCA | 397.203441 | -0.821600572 | 0.25655583 | -3.2024241 | 0.00136276 | 0.01922299 |
| HELZ | 279.081632 | 0.085483447 | 0.29857865 | 0.28630128 | 0.77464738 | 0.91864687 |
| PSMD12 | 58.1284476 | -0.304598075 | 0.39204109 | -0.7769545 | 0.43718563 | 0.72999979 |
| PITPNC1 | 426.084259 | -1.316983315 | 0.73157173 | -1.8002108 | 0.07182736 | 0.28614796 |
| NOL11 | 26.1707227 | 0.358259781 | 0.38115755 | 0.93992572 | 0.34725566 | 0.6553345 |
| BPTF | 358.638068 | -0.239979457 | 0.23243509 | -1.0324579 | 0.30185764 | 0.61586884 |
| AC005332.7 | 16.9267985 | -0.043545294 | 0.40949413 | -0.1063392 | 0.9153132 | 0.97341689 |
| ARSG | 44.9910089 | 0.383298371 | 0.47520918 | 0.80658874 | 0.41990346 | 0.71838163 |
| WIPI1 | 43.8825243 | -1.248109502 | 0.76330965 | -1.6351287 | 0.10202208 | 0.34893691 |
| PRKAR1A | 146.794283 | -0.548323898 | 0.32588825 | -1.6825519 | 0.09246187 | 0.33171007 |
| FAM20A | 81.3801172 | -3.408311862 | 0.58258415 | -5.850334 | 4.91E-09 | 6.02E-07 |
| LINC01482 | 139.091682 | -1.257902533 | 0.84687436 | -1.4853473 | 0.13745185 | 0.41328639 |
| ABCA9 | 75.8222178 | 1.893877061 | 0.55966431 | 3.38395181 | 0.00071451 | 0.01152604 |
| ABCA6 | 68.199887 | 1.830706911 | 0.47103381 | 3.88657219 | 0.00010167 | 0.00245588 |
| ABCA10 | 234.90249 | 1.585342414 | 0.37257478 | 4.25509852 | 2.09E-05 | 0.00072698 |
| ABCA5 | 662.9753 | 1.727944251 | 0.34878858 | 4.95413075 | 7.27E-07 | 4.46E-05 |
| MAP2K6 | 106.873722 | 2.668409402 | 0.68990895 | 3.86777038 | 0.00010984 | 0.00261258 |
| AC005208.1 | 323.775493 | 0.142553452 | 0.74065918 | 0.19246835 | 0.84737535 | 0.94953558 |
| KCNJ16 | 1225.1092 | 1.070317024 | 0.48018881 | 2.22895038 | 0.02581721 | 0.15410988 |
| SOX9 | 51.7438069 | -5.000202321 | 0.81896728 | -6.105497 | 1.02E-09 | 1.68E-07 |
| LINC00511 | 104.890347 | -5.662942634 | 1.0251527 | -5.5239991 | 3.31E-08 | 2.93E-06 |
| SLC39A11 | 134.355356 | -0.173954439 | 0.33707086 | -0.5160768 | 0.6058008 | 0.83805439 |
| COG1 | 18.3253014 | -0.149159727 | 0.37149241 | -0.4015149 | 0.68804109 | 0.87567053 |
| FAM104A | 26.6276116 | 0.068907155 | 0.32353024 | 0.21298521 | 0.8313385 | 0.94460153 |
| C17orf80 | 17.0987025 | -0.136655402 | 0.39836398 | -0.3430416 | 0.7315672 | 0.89782437 |
| SDK2 | 55.342124 | -0.035587322 | 0.57981578 | -0.0613769 | 0.95105901 | 0.98684281 |
| RPL38 | 26.5844283 | -0.664553439 | 0.48642533 | -1.3661983 | 0.17187675 | 0.46763314 |
| GPRC5C | 52.985125 | -1.05334065 | 0.32303339 | -3.2607794 | 0.00111106 | 0.01623756 |
| SLC9A3R1 | 15.4467199 | -1.400502866 | 0.46886852 | -2.9869842 | 0.00281744 | 0.0330023 |
| NAT9 | 14.4345014 | -0.534877894 | 0.47807074 | -1.1188258 | 0.26321445 | 0.57718992 |
| KCTD2 | 26.7612077 | -0.515100644 | 0.35050675 | -1.4695884 | 0.14167327 | 0.42176296 |
| ATP5PD | 26.8995788 | 0.545578949 | 0.36212082 | 1.50662132 | 0.13190775 | 0.40351308 |
| SLC16A5 | 51.8940393 | -1.697556459 | 0.61856197 | -2.7443596 | 0.00606291 | 0.05727514 |
| ARMC7 | 18.144184 | -0.903489373 | 0.40324907 | -2.2405244 | 0.0250569 | 0.15102168 |
| NT5C | 26.4041634 | -0.106345141 | 0.34767647 | -0.3058739 | 0.75970069 | 0.91236458 |

| | | | | | | |
|---|---|---|---|---|---|---|
| JPT1 | 15.1022013 | -0.895784384 | 0.43600118 | -2.0545458 | 0.0399229 | 0.20252509 |
| SUMO2 | 23.9617978 | -0.398910849 | 0.49265977 | -0.8097086 | 0.41810768 | 0.71662427 |
| NUP85 | 28.1327882 | -0.090825503 | 0.34804286 | -0.2609607 | 0.79412283 | 0.92670854 |
| GGA3 | 36.5974228 | -0.294237221 | 0.37769767 | -0.7790284 | 0.43596297 | 0.72891111 |
| MRPS7 | 18.4538176 | 0.022954753 | 0.42896483 | 0.05351197 | 0.95732399 | 0.98859719 |
| GRB2 | 100.560204 | -0.097393242 | 0.2582128 | -0.3771821 | 0.70603829 | 0.88599027 |
| TMEM94 | 32.5904459 | 0.458343941 | 0.29083272 | 1.57597105 | 0.11503248 | 0.3738073 |
| CASKIN2 | 13.3345498 | 0.220590281 | 0.53744528 | 0.41044231 | 0.68148152 | 0.87270068 |
| TSEN54 | 24.4811586 | 0.25887707 | 0.4796051 | 0.5397713 | 0.58935476 | 0.83058401 |
| LLGL2 | 359.697866 | -0.562309002 | 0.38250922 | -1.4700534 | 0.14154728 | 0.42165714 |
| MYO15B | 40.0416124 | 0.327498786 | 0.49449443 | 0.66229015 | 0.50778529 | 0.78008138 |
| RECQL5 | 21.4622836 | 0.280128625 | 0.37055206 | 0.75597643 | 0.44966333 | 0.74127668 |
| SAP30BP | 52.1369458 | -0.149006273 | 0.28107919 | -0.530122 | 0.59602732 | 0.83416944 |
| H3F3B | 121.772088 | -1.151689189 | 0.35329379 | -3.2598625 | 0.00111466 | 0.01626465 |
| UNK | 72.6076335 | 0.228325373 | 0.30481339 | 0.74906608 | 0.45381738 | 0.74470138 |
| WBP2 | 61.0825084 | -1.416592046 | 0.45002026 | -3.1478406 | 0.00164481 | 0.0218465 |
| ACOX1 | 53.0001955 | 0.021093499 | 0.28475426 | 0.07407615 | 0.94094979 | 0.98422884 |
| EVPL | 12.6283881 | -0.295099757 | 0.691465 | -0.4267747 | 0.66954345 | 0.86777458 |
| SRP68 | 28.1164165 | 0.196411881 | 0.33413415 | 0.58782343 | 0.55665082 | 0.81163685 |
| EXOC7 | 73.1437404 | -0.152462845 | 0.33729506 | -0.4520162 | 0.65125728 | 0.86120024 |
| RNF157 | 18.2171013 | 0.420354268 | 0.52976728 | 0.79346966 | 0.42750424 | 0.72300292 |
| UBALD2 | 27.7390095 | -0.442343262 | 0.3457088 | -1.2795256 | 0.20071203 | 0.50462148 |
| PRPSAP1 | 38.8813705 | -0.260087793 | 0.39739832 | -0.6544763 | 0.51280497 | 0.7830488 |
| UBE2O | 63.873829 | 0.360148306 | 0.34009004 | 1.05897929 | 0.28960921 | 0.60475529 |
| RHBDF2 | 18.7238793 | -0.21229884 | 0.43176864 | -0.4916958 | 0.62293439 | 0.84581917 |
| MXRA7 | 25.378383 | 0.570322697 | 0.40805646 | 1.39765632 | 0.1622163 | 0.45271019 |
| JMJD6 | 22.0654827 | -0.424489901 | 0.47761381 | -0.8887723 | 0.37412549 | 0.6776119 |
| SRSF2 | 84.4587595 | 0.706406937 | 0.33036694 | 2.13824947 | 0.0324965 | 0.17931872 |
| MFSD11 | 53.4700439 | -0.066952169 | 0.35372088 | -0.1892797 | 0.84987363 | 0.95003258 |
| SEC14L1 | 111.239083 | -0.223943281 | 0.27842991 | -0.8043076 | 0.42121937 | 0.7192477 |
| 9-Sep | 242.575365 | -1.314075297 | 0.2591057 | -5.07158 | 3.95E-07 | 2.61E-05 |
| TNRC6C | 120.114783 | -0.158610974 | 0.37294899 | -0.4252887 | 0.67062626 | 0.86777458 |
| SYNGR2 | 21.3585477 | -0.164546271 | 0.61816339 | -0.2661857 | 0.79009618 | 0.92448611 |
| AFMID | 71.2364045 | 2.215762204 | 0.45746508 | 4.84356576 | 1.28E-06 | 7.05E-05 |
| PGS1 | 42.7259033 | -0.818612235 | 0.40727508 | -2.0099738 | 0.04443396 | 0.21534941 |
| CYTH1 | 101.033254 | 0.397402897 | 0.2870817 | 1.384285 | 0.16627121 | 0.45864087 |
| USP36 | 97.5185832 | -1.770206062 | 0.76942568 | -2.3006849 | 0.02140945 | 0.13717779 |
| TIMP2 | 32.3106344 | -1.942022986 | 0.61392014 | -3.1633153 | 0.00155983 | 0.02123122 |
| LGALS3BP | 31.5118732 | -0.420707445 | 0.73002958 | -0.5762882 | 0.56442039 | 0.81490487 |
| CANT1 | 10.8080984 | -0.201879991 | 0.51680018 | -0.3906345 | 0.6960674 | 0.88085402 |
| ENGASE | 12.825683 | 0.306985221 | 0.52070053 | 0.58956196 | 0.55548436 | 0.81104802 |
| TBC1D16 | 49.862539 | -0.773731378 | 0.34220102 | -2.2610434 | 0.02375657 | 0.14631883 |
| CCDC40 | 21.1425055 | -0.180981461 | 0.43262656 | -0.4183318 | 0.67570452 | 0.86960234 |
| GAA | 20.3980315 | 0.164232145 | 0.40129972 | 0.40925058 | 0.68235578 | 0.87274146 |
| EIF4A3 | 15.6976328 | -0.188222042 | 0.38678315 | -0.4866345 | 0.62651734 | 0.8472036 |
| AC087741.1 | 15.2276444 | -0.731841407 | 0.60933008 | -1.201059 | 0.2297283 | 0.54241242 |
| SLC26A11 | 37.5425463 | -0.338111054 | 0.44575434 | -0.7585143 | 0.44814314 | 0.7399821 |
| RNF213 | 411.65506 | -1.365391068 | 0.52602853 | -2.5956597 | 0.00944095 | 0.07888124 |
| AC124319.2 | 276.375018 | 0.510076583 | 0.34013519 | 1.49962896 | 0.13371054 | 0.40702484 |
| ENDOV | 28.3717818 | -0.35781182 | 0.29687248 | -1.2052711 | 0.22809866 | 0.54025743 |
| RPTOR | 153.980767 | 0.085309021 | 0.27725426 | 0.30769238 | 0.75831642 | 0.91161939 |
| BAIAP2 | 14.9207084 | -0.215882813 | 0.47619788 | -0.4533469 | 0.650299 | 0.86113455 |
| TEPSIN | 13.7306589 | 0.148344298 | 0.49953726 | 0.29696343 | 0.76649443 | 0.91564296 |
| NDUFAF8 | 17.5609012 | -0.168814751 | 0.5973863 | -0.2825889 | 0.77749197 | 0.91968524 |
| SLC38A10 | 63.2416124 | 0.075413141 | 0.34384051 | 0.21932593 | 0.82639616 | 0.94220076 |
| BAHCC1 | 16.7051917 | 0.585988972 | 0.44052873 | 1.33019469 | 0.18345413 | 0.48210665 |
| ACTG1 | 159.07964 | -0.516707008 | 0.58406106 | -0.8846798 | 0.37632937 | 0.67982871 |
| NPLOC4 | 116.10128 | -0.563700553 | 0.29294411 | -1.9242597 | 0.05432204 | 0.24245986 |
| CCDC137 | 19.1243901 | -0.453342764 | 0.54192316 | -0.8365444 | 0.40284874 | 0.70169281 |
| ARL16 | 19.711302 | 0.082582213 | 0.4117782 | 0.20055023 | 0.84105028 | 0.94776435 |
| HGS | 45.0765097 | -0.954295528 | 0.46316932 | -2.06036 | 0.03936414 | 0.20050645 |
| GCGR | 54.1805303 | 2.156496677 | 0.59008426 | 3.6545572 | 0.00025763 | 0.00518814 |
| MCRIP1 | 21.9098855 | -0.094753414 | 0.41526157 | -0.2281777 | 0.81950813 | 0.9384641 |
| P4HB | 85.2551624 | -0.67635548 | 0.39959477 | -1.6926034 | 0.09053097 | 0.32755561 |
| ARHGDIA | 33.2642858 | -0.984916477 | 0.58435775 | -1.6854683 | 0.09189825 | 0.33045095 |
| ANAPC11 | 22.86228 | -0.467168121 | 0.46498889 | -1.0046866 | 0.31504777 | 0.62958667 |
| SIRT7 | 16.456418 | -0.333028394 | 0.45154026 | -0.7375386 | 0.46079485 | 0.74896364 |
| MAFG | 17.5027308 | -2.394818343 | 0.76867608 | -3.1155104 | 0.00183627 | 0.0236504 |
| ASPSCR1 | 25.2365383 | -0.679234844 | 0.54636819 | -1.2431815 | 0.21380095 | 0.52235998 |
| DCXR | 13.9312729 | 0.769201008 | 0.57428226 | 1.33941279 | 0.18043633 | 0.47903996 |
| RFNG | 15.8186421 | 0.440144893 | 0.48196981 | 0.91322088 | 0.36112639 | 0.66662887 |
| GPS1 | 15.8315438 | -0.006663343 | 0.61357126 | -0.0108599 | 0.9913352 | 0.9988777 |
| DUS1L | 20.0437567 | -0.268744028 | 0.49706132 | -0.5406657 | 0.58873799 | 0.83025227 |
| CCDC57 | 114.356595 | -0.016469708 | 0.26802047 | -0.0614494 | 0.95100128 | 0.98684281 |
| SLC16A3 | 23.6367116 | -1.405847497 | 0.45569001 | -3.0850962 | 0.00203486 | 0.02557017 |
| CSNK1D | 87.8520637 | -0.937525758 | 0.5414543 | -1.7314956 | 0.0833634 | 0.31091215 |
| OGFOD3 | 22.1967363 | -0.187424764 | 0.44538081 | -0.4208191 | 0.67388716 | 0.86846218 |
| HEXDC | 47.6733748 | 0.3799074 | 0.39325446 | 0.96605999 | 0.33401416 | 0.64554754 |
| CYBC1 | 16.7798416 | -1.26927364 | 0.48608174 | -2.611235 | 0.00902159 | 0.07660955 |
| NARF | 44.2542946 | -1.049593024 | 0.42382886 | -2.4764548 | 0.01326944 | 0.09993884 |
| FOXK2 | 134.105733 | -0.310818417 | 0.29527239 | -1.0526498 | 0.29250153 | 0.60745829 |
| WDR45B | 95.1013249 | -0.228608822 | 0.47828439 | -0.4779768 | 0.63266675 | 0.85171597 |
| RAB40B | 59.4441319 | -0.365327454 | 0.32863938 | -1.1116363 | 0.26629457 | 0.58189139 |
| FN3KRP | 16.0418421 | -0.245594749 | 0.50888076 | -0.4826175 | 0.62936737 | 0.84886754 |
| FN3K | 43.5778235 | 0.816430813 | 0.62734786 | 1.3014005 | 0.1931214 | 0.49455204 |
| TBCD | 57.1951813 | 0.022680417 | 0.323432 | 0.07012422 | 0.94409479 | 0.98552865 |
| METRNL | 26.0787633 | -0.763990004 | 0.70648371 | -1.0813979 | 0.27952016 | 0.59383074 |
| USP14 | 101.934721 | 0.30043752 | 0.24903163 | 1.20642315 | 0.22765437 | 0.53970236 |
| THOC1 | 91.8029111 | 0.10983086 | 0.29091445 | 0.37753662 | 0.70577485 | 0.88599027 |
| ENOSF1 | 339.095206 | -0.150867287 | 0.37171954 | -0.4058632 | 0.68484311 | 0.87408051 |
| YES1 | 106.715694 | -0.952177717 | 0.39868638 | -2.3882876 | 0.01692709 | 0.11753077 |
| METTL4 | 33.7003736 | 0.205665808 | 0.36186968 | 0.5683422 | 0.56980263 | 0.81849045 |
| SMCHD1 | 216.966351 | 0.35269465 | 0.26979938 | 1.3072478 | 0.19112857 | 0.49188042 |
| AP001011.1 | 28.7160347 | -0.336683711 | 0.66207905 | -0.5085249 | 0.61108525 | 0.84169124 |
| LPIN2 | 99.0827078 | -1.166549179 | 0.42204886 | -2.7640145 | 0.0057095 | 0.05454445 |
| MYOM1 | 38.7513557 | 2.564554298 | 0.48747402 | 5.26090462 | 1.43E-07 | 1.11E-05 |
| MYL12A | 53.9922919 | -0.794028793 | 0.33538854 | -2.3674893 | 0.01790924 | 0.12164274 |
| MYL12B | 103.944079 | -1.097058044 | 0.32683747 | -3.3565859 | 0.00078911 | 0.01249182 |
| TGIF1 | 43.8036931 | -2.384755255 | 0.71939969 | -3.3149239 | 0.00091668 | 0.01392041 |
| DLGAP1 | 133.583735 | -3.61091112 | 0.87948759 | -4.1056988 | 4.03E-05 | 0.00119316 |
| DLGAP1-AS1 | 17.1977111 | -1.608898139 | 0.41677359 | -3.8603649 | 0.00011322 | 0.00265447 |
| LINC00667 | 18.6641951 | -0.154355821 | 0.41328458 | -0.3734856 | 0.70878709 | 0.88614996 |
| EPB41L3 | 225.650913 | 0.411949034 | 0.36524955 | 1.12785637 | 0.25938058 | 0.57248906 |
| L3MBTL4 | 232.197717 | 0.513312216 | 0.30932076 | 1.65948192 | 0.09701872 | 0.33969491 |
| LAMA1 | 254.980962 | -0.49637044 | 0.35754442 | -1.3882763 | 0.16505293 | 0.45693395 |
| PTPRM | 458.964043 | -1.018803796 | 0.31314397 | -3.2534677 | 0.00114006 | 0.01655746 |
| RAB12 | 68.4162598 | -0.039219169 | 0.23512126 | -0.166804 | 0.86752426 | 0.95588054 |
| MTCL1 | 39.6483347 | 0.396198641 | 0.29397123 | 1.3477463 | 0.17774 | 0.47521885 |
| NDUFV2 | 91.7101791 | -0.051886318 | 0.32761825 | -0.1583743 | 0.87416185 | 0.95761805 |
| ANKRD12 | 455.063416 | -0.197695978 | 0.24860287 | -0.795228 | 0.42648086 | 0.72230448 |
| TWSG1 | 59.6716674 | 0.028363006 | 0.3575897 | 0.07931718 | 0.93678034 | 0.98229194 |
| RALBP1 | 83.4524365 | 0.046468186 | 0.29097162 | 0.15970006 | 0.87311735 | 0.95703878 |
| PPP4R1 | 141.457044 | -0.054902873 | 0.32924459 | -0.1667541 | 0.86756356 | 0.95588054 |

| | | | | | | |
|---|---|---|---|---|---|---|
| RAB31 | 84.0582747 | 0.117096464 | 0.8330589 | 0.14056205 | 0.88821593 | 0.96307703 |
| VAPA | 243.54687 | 0.763706567 | 0.2783506 | 2.74368572 | 0.00607537 | 0.05733193 |
| NAPG | 60.0108911 | 0.099166205 | 0.24647781 | 0.40233319 | 0.68743883 | 0.87556719 |
| GNAL | 33.6092174 | -1.965090968 | 0.57807412 | -3.3993754 | 0.0006754 | 0.01102876 |
| MPPE1 | 47.6404365 | -0.069353252 | 0.40266732 | -0.1722346 | 0.86325308 | 0.95458251 |
| IMPA2 | 112.785183 | -0.073962934 | 0.31453957 | -0.2351467 | 0.81409489 | 0.93633777 |
| AFG3L2 | 116.063878 | 0.090203854 | 0.31675522 | 0.28477464 | 0.7758168 | 0.91926749 |
| SPIRE1 | 491.025228 | -0.95654602 | 0.44740481 | -2.1379878 | 0.03251773 | 0.17931872 |
| PSMG2 | 65.5698461 | -0.222486638 | 0.33518764 | -0.6637674 | 0.50683918 | 0.77942663 |
| CEP76 | 19.724143 | -0.21367396 | 0.48097143 | -0.444255 | 0.65685823 | 0.86325042 |
| PTPN2 | 171.229068 | -0.470049267 | 0.3170487 | -1.4825775 | 0.1381867 | 0.41496063 |
| SEH1L | 30.3034553 | -0.630409314 | 0.33645185 | -1.8736985 | 0.060972 | 0.2580585 |
| CEP192 | 158.217401 | 0.874106544 | 0.3003711 | 2.91008865 | 0.00361326 | 0.03987226 |
| LDLRAD4 | 74.6956451 | 1.31613367 | 0.53642898 | 2.45350962 | 0.01414698 | 0.10435633 |
| FAM210A | 64.2489432 | -0.709172315 | 0.4458888 | -1.590469 | 0.11172912 | 0.36824403 |
| RNMT | 100.962185 | -0.257894658 | 0.42071396 | -0.6129929 | 0.53988105 | 0.80166126 |
| ROCK1 | 215.819897 | -0.139671246 | 0.22240662 | -0.6279995 | 0.53000428 | 0.79562009 |
| ESCO1 | 95.0018386 | -0.242893413 | 0.23953817 | -1.0140071 | 0.31057935 | 0.62531676 |
| SNRPD1 | 16.7309531 | -0.250367377 | 0.36768826 | -0.6809229 | 0.49592025 | 0.77475164 |
| ABHD3 | 101.126332 | -2.967187381 | 0.96351814 | -3.0795345 | 0.00207324 | 0.02594754 |
| MIB1 | 197.726178 | -0.420670456 | 0.50612285 | -0.8311627 | 0.4058817 | 0.70505833 |
| AC090912.1 | 24.6123446 | -0.848235355 | 0.5386777 | -1.5746621 | 0.11533446 | 0.37430509 |
| RBBP8 | 429.426732 | -0.900042634 | 0.3625417 | -2.4825907 | 0.01304309 | 0.09881314 |
| CABLES1 | 195.555523 | -0.051182035 | 0.41925984 | -0.1220771 | 0.90283794 | 0.96803829 |
| TMEM241 | 53.2422837 | -0.665318811 | 0.43831815 | -1.5178902 | 0.12904209 | 0.39814311 |
| RIOK3 | 110.630633 | -0.617267485 | 0.26451972 | -2.3335405 | 0.01961979 | 0.12983501 |
| RMC1 | 32.2520894 | -0.214611785 | 0.38293665 | -0.5604368 | 0.57518155 | 0.82146824 |
| NPC1 | 177.428907 | -2.912055952 | 0.81702132 | -3.564235 | 0.00036492 | 0.00683234 |
| TTC39C | 66.9014614 | -0.015189089 | 0.38746462 | -0.0392012 | 0.96872995 | 0.99156233 |
| OSBPL1A | 79.0687865 | 0.695806722 | 0.33554797 | 2.0736431 | 0.03811247 | 0.19688489 |
| IMPACT | 15.1127236 | 0.194528477 | 0.5250333 | 0.37050693 | 0.71100481 | 0.88677671 |
| SS18 | 158.351567 | 0.058951865 | 0.21249929 | 0.27742146 | 0.7814565 | 0.92103406 |
| TAF4B | 61.8065471 | 1.207678416 | 0.31774882 | 3.80073302 | 0.00014427 | 0.0032729 |
| KCTD1 | 501.972353 | 0.280932776 | 0.51911642 | 0.5411749 | 0.58838704 | 0.83024609 |
| AQP4-AS1 | 45.5980512 | -0.005050179 | 0.67432764 | -0.0074892 | 0.99402453 | 0.9992177 |
| CHST9 | 54.786834 | 0.937208139 | 0.65615551 | 1.42833234 | 0.15319622 | 0.43862109 |
| DSC2 | 16.4928483 | -0.236038205 | 0.42019341 | -0.561737 | 0.57429519 | 0.82127736 |
| DSG2 | 52.439812 | -0.178752168 | 0.43717709 | -0.4088782 | 0.68262907 | 0.87297126 |
| B4GALT6 | 29.4742691 | 1.025283723 | 0.43933591 | 2.33371255 | 0.01961078 | 0.12983501 |
| TRAPPC8 | 139.309247 | 0.048001257 | 0.25315468 | 0.18961236 | 0.8496129 | 0.95003258 |
| RNF138 | 54.2205175 | 0.328673084 | 0.38645869 | 0.85047403 | 0.39506159 | 0.69642725 |
| GAREM1 | 638.28719 | -0.355013426 | 0.49152424 | -0.7222704 | 0.47012823 | 0.75694623 |
| KLHL14 | 37.3212907 | -0.542101616 | 0.46180889 | -1.1738657 | 0.24044882 | 0.55302079 |
| CCDC178 | 294.946367 | -1.262527176 | 0.80323089 | -1.571811 | 0.11599438 | 0.37462127 |
| ASXL3 | 125.785149 | 0.828251543 | 0.47166491 | 1.75601688 | 0.07908553 | 0.30106837 |
| DTNA | 50.5038888 | 0.839743491 | 0.54188388 | 1.54967424 | 0.12121972 | 0.3845705 |
| MAPRE2 | 43.7441009 | -0.665218894 | 0.56125172 | -1.1852416 | 0.23592193 | 0.54869945 |
| ZNF397 | 57.5437864 | -0.023171345 | 0.29368284 | -0.0788992 | 0.93711279 | 0.98241957 |
| ZSCAN30 | 43.1800249 | -0.166438281 | 0.37331228 | -0.445842 | 0.6557114 | 0.86268564 |
| ZNF24 | 45.9784277 | -0.180743975 | 0.34404507 | -0.5253497 | 0.59934011 | 0.83581061 |
| GALNT1 | 68.5380714 | -0.136201444 | 0.28099743 | -0.4847071 | 0.62788414 | 0.84784081 |
| RPRD1A | 168.053368 | -0.249275 | 0.23414196 | -1.0646319 | 0.28704255 | 0.60156995 |
| SLC39A6 | 26.9240633 | -1.315527066 | 0.49996913 | -2.6312166 | 0.00850798 | 0.07324875 |
| ELP2 | 97.8953796 | 0.152184421 | 0.18600423 | 0.81817722 | 0.413256 | 0.71143975 |
| MOCOS | 26.0843193 | -2.044035618 | 0.62958875 | -3.2466203 | 0.00116784 | 0.0168299 |
| FHOD3 | 829.39836 | 1.45658889 | 0.39772334 | 3.66231689 | 0.00024994 | 0.00506281 |
| TPGS2 | 43.0733334 | -1.401942321 | 0.39853354 | -3.5177524 | 0.00043522 | 0.00777862 |
| KIAA1328 | 194.377851 | -0.177410892 | 0.33799529 | -0.5248916 | 0.59965854 | 0.83582612 |
| PIK3C3 | 164.081496 | 0.035213149 | 0.21936978 | 0.1605196 | 0.87247178 | 0.95690422 |
| LINC00907 | 35.3407536 | -0.273115476 | 0.4524583 | -0.6036257 | 0.54609249 | 0.80553179 |
| SETBP1 | 597.932043 | 0.126396768 | 0.30517318 | 0.41418045 | 0.67874196 | 0.87147102 |
| SLC14A2 | 17.0227508 | 0.265337221 | 0.52366332 | 0.50669431 | 0.61236932 | 0.84174925 |
| EPG5 | 103.981424 | -0.436449472 | 0.44386845 | -0.9832856 | 0.32546687 | 0.63726441 |
| PSTPIP2 | 28.9659132 | -0.3032971 | 0.56638982 | -0.5354918 | 0.59230983 | 0.83298595 |
| ATP5F1A | 98.898459 | 0.463994513 | 0.41854084 | 1.10860033 | 0.26760264 | 0.58365544 |
| C18orf25 | 74.7119064 | 0.102635444 | 0.31055133 | 0.3304943 | 0.7410265 | 0.90375649 |
| ST8SIA5 | 18.0139415 | 0.084989844 | 0.36363517 | 0.23372284 | 0.81520016 | 0.93711333 |
| PIAS2 | 247.890538 | 0.476784466 | 0.2914274 | 1.63603169 | 0.10183296 | 0.34880722 |
| KATNAL2 | 112.415045 | 0.364594915 | 0.46099319 | 0.79089003 | 0.42900817 | 0.7236272 |
| SMAD2 | 268.162279 | -0.546286289 | 0.4076553 | -1.3400692 | 0.18022286 | 0.47864023 |
| CTIF | 100.879802 | 0.394919395 | 0.25476884 | 1.55010872 | 0.12111542 | 0.38450126 |
| SMAD7 | 38.781539 | 0.29424052 | 0.52492266 | 0.56054071 | 0.57511068 | 0.82146824 |
| DYM | 365.996004 | 0.078441564 | 0.24444774 | 0.32089298 | 0.74829149 | 0.90723111 |
| LIPG | 89.2066448 | -3.279665784 | 0.88580196 | -3.702482 | 0.0002135 | 0.00446323 |
| ACAA2 | 28.4401492 | -0.322739821 | 0.49996794 | -0.645521 | 0.51858959 | 0.78726765 |
| SNHG22 | 30.1270905 | -0.224484636 | 0.31093659 | -0.7219628 | 0.47031738 | 0.75702578 |
| MYO5B | 135.580155 | -0.832562878 | 0.418955 | -1.987237 | 0.04689614 | 0.22193782 |
| MBD1 | 28.9067498 | -0.126292255 | 0.29441649 | -0.4289578 | 0.66795392 | 0.86777458 |
| CXXC1 | 14.9286669 | 0.021038037 | 0.46301714 | 0.04543684 | 0.96375911 | 0.99082982 |
| MAPK4 | 56.0761276 | 0.072698134 | 0.62576353 | 0.11617509 | 0.90751378 | 0.9700616 |
| ME2 | 39.1979868 | 0.228005835 | 0.31518798 | 0.72339635 | 0.46943642 | 0.75609348 |
| AC091551.1 | 18.7552744 | -0.605861688 | 0.68536141 | -0.8840032 | 0.37669448 | 0.68015091 |
| SMAD4 | 65.7082873 | -0.015751682 | 0.24877767 | -0.0633163 | 0.94951463 | 0.98641632 |
| MEX3C | 17.4890973 | -0.861391204 | 0.38755341 | -2.2226387 | 0.02624017 | 0.15583654 |
| MBD2 | 153.769052 | -0.948498439 | 0.44532662 | -2.1298939 | 0.03318038 | 0.18155742 |
| POLI | 53.0294835 | -0.069151989 | 0.38343325 | -0.1803495 | 0.85687821 | 0.95238198 |
| TCF4 | 26.234686 | 1.786423476 | 0.45802083 | 3.90031051 | 9.61E-05 | 0.0023449 |
| TXNL1 | 64.6985517 | -0.063088985 | 0.32750059 | -0.1926378 | 0.84724266 | 0.94953558 |
| WDR7 | 187.994924 | 0.544385125 | 0.1888002 | 2.88339268 | 0.00393417 | 0.04221193 |
| FECH | 37.6701076 | 1.095156154 | 0.47275751 | 2.31652832 | 0.02052944 | 0.13339129 |
| NARS | 69.3813539 | -0.394534301 | 0.31326192 | -1.2594391 | 0.20787178 | 0.51588941 |
| AC027097.2 | 52.1058287 | -0.009220211 | 0.39008674 | -0.0236363 | 0.98114271 | 0.99653318 |
| ATP8B1 | 102.286387 | -0.359921693 | 0.30849887 | -1.1666872 | 0.2433367 | 0.55555126 |
| NEDD4L | 991.339804 | 0.00460465 | 0.56958768 | 0.00808418 | 0.99354983 | 0.9992177 |
| MALT1 | 54.8373534 | 0.062192921 | 0.36184004 | 0.1718796 | 0.86353219 | 0.95458251 |
| ZNF532 | 198.904882 | -0.508342268 | 0.35733808 | -1.4225807 | 0.15485774 | 0.4403457 |
| SEC11C | 13.0930776 | 0.220867928 | 0.4600312 | 0.48011511 | 0.63114555 | 0.85065063 |
| LMAN1 | 63.8245675 | 0.522019638 | 0.34730585 | 1.50305454 | 0.13282498 | 0.40525528 |
| RNF152 | 125.005681 | 0.458118829 | 0.50217555 | 0.91226828 | 0.36162751 | 0.66686799 |
| PIGN | 136.848207 | 0.232456411 | 0.20409519 | 1.13896074 | 0.25471953 | 0.56709764 |
| KIAA1468 | 154.564794 | -0.243749483 | 0.18326401 | -1.3300456 | 0.18350326 | 0.48210665 |
| ZCCHC2 | 71.5747917 | 0.512372343 | 0.232849 | 2.20044898 | 0.02777506 | 0.16155622 |
| PHLPP1 | 107.309737 | -0.38225131 | 0.24833718 | -1.5392432 | 0.12374495 | 0.38887694 |
| BCL2 | 130.721131 | 0.245320109 | 0.49808745 | 0.49252418 | 0.62234884 | 0.84559756 |
| KDSR | 63.7467229 | -1.48377056 | 0.42968811 | -3.4531339 | 0.00055411 | 0.00934278 |
| VPS4B | 67.0571196 | -0.221942775 | 0.32201277 | -0.6892359 | 0.4906748 | 0.77203035 |
| TMX3 | 76.3401515 | 0.151572841 | 0.29335348 | 0.51669011 | 0.6053725 | 0.83802715 |
| CCDC102B | 35.8905843 | 0.916404045 | 0.63187992 | 1.45028196 | 0.14697991 | 0.4297124 |
| RTTN | 58.3317101 | 0.227983744 | 0.37661257 | 0.60535351 | 0.54494413 | 0.80447421 |
| SOCS6 | 35.7565611 | 0.484366681 | 0.35951095 | 1.34729326 | 0.1778858 | 0.47524562 |
| TIMM21 | 13.9062255 | 0.368109275 | 0.43299586 | 0.85014502 | 0.39524446 | 0.69642725 |
| CYB5A | 90.8127135 | 0.534699025 | 0.50949395 | 1.04947079 | 0.29396149 | 0.60895289 |
| CNDP2 | 86.1481809 | -1.090991902 | 0.35034946 | -3.1140105 | 0.00184563 | 0.02367076 |

| | | | | | | |
|---|---|---|---|---|---|---|
| LINC00909 | 12.9688489 | -0.136436338 | 0.53993561 | -0.25269 | 0.80050776 | 0.92985354 |
| ZNF407 | 300.036067 | -0.065002708 | 0.3018771 | -0.2153284 | 0.8295113 | 0.94356025 |
| ZADH2 | 30.3690879 | -0.382625679 | 0.52449701 | -0.7295097 | 0.46568991 | 0.75274202 |
| TSHZ1 | 80.2176098 | -0.288377426 | 0.3046026 | -0.9467333 | 0.3437747 | 0.65306073 |
| ZNF516 | 190.935182 | -0.190567268 | 0.38448016 | -0.4956492 | 0.62014196 | 0.84399411 |
| ZNF236 | 66.511629 | -0.01018151 | 0.28475554 | -0.0357553 | 0.9714775 | 0.99233745 |
| MBP | 188.223007 | -1.260004123 | 0.51040638 | -2.4686293 | 0.01356317 | 0.10141376 |
| LINC01896 | 73.0762692 | 1.042163849 | 0.44327917 | 2.35103278 | 0.01872138 | 0.12522108 |
| SALL3 | 113.318949 | 0.998914493 | 0.36257504 | 2.75505588 | 0.00586821 | 0.05571813 |
| ATP9B | 353.221873 | 0.386827506 | 0.2046696 | 1.89000956 | 0.05875668 | 0.25348085 |
| NFATC1 | 31.2576387 | 0.783612986 | 0.3933598 | 1.99210238 | 0.04635983 | 0.22110436 |
| CTDP1 | 30.2549134 | -0.795128028 | 0.42600925 | -1.8664572 | 0.06197743 | 0.26124663 |
| PQLC1 | 48.7550487 | 0.382135696 | 0.37230691 | 1.02639969 | 0.30470322 | 0.61829224 |
| HSBP1L1 | 58.5481748 | 0.58834423 | 0.25681567 | 2.29092025 | 0.02196803 | 0.13896193 |
| TXNL4A | 59.5646939 | 0.406155413 | 0.29352998 | 1.38369313 | 0.16645244 | 0.45864087 |
| ADNP2 | 91.3323503 | -1.051905883 | 0.60266409 | -1.7454265 | 0.08091065 | 0.30571131 |
| PARD6G-AS1 | 16.9897849 | -0.908861772 | 0.71731341 | -1.2670358 | 0.20514249 | 0.51160585 |
| C20orf96 | 13.2886312 | -0.33094984 | 0.69402104 | -0.4768585 | 0.63346288 | 0.8520131 |
| NRSN2-AS1 | 87.7565417 | -0.10894197 | 0.57646367 | -0.1889832 | 0.85010596 | 0.95017837 |
| SOX12 | 13.9908767 | 0.823740121 | 0.51653197 | 1.59475146 | 0.11076782 | 0.36624085 |
| RBCK1 | 45.4363105 | -0.240539594 | 0.35332969 | -0.6807795 | 0.49601105 | 0.77475164 |
| TBC1D20 | 29.2986658 | -0.024720045 | 0.32601704 | -0.0758244 | 0.93955881 | 0.98358617 |
| CSNK2A1 | 191.887165 | 0.376308673 | 0.21403855 | 1.75813501 | 0.07872454 | 0.30021579 |
| PSMF1 | 72.4590541 | -0.000513912 | 0.25763917 | -0.0019947 | 0.99840846 | 0.99922221 |
| AL136531.2 | 41.2376986 | -1.283245284 | 0.55895485 | -2.2957942 | 0.02168765 | 0.13793703 |
| SDCBP2-AS1 | 48.0662817 | 0.999610978 | 0.36630152 | 2.72892937 | 0.00635403 | 0.05910136 |
| NSFL1C | 35.1508999 | -0.327427902 | 0.2835724 | -1.1546536 | 0.24823229 | 0.56095925 |
| STK35 | 32.3280777 | -0.642093554 | 0.323849 | -1.9826942 | 0.04740159 | 0.22326436 |
| SNRPB | 25.3787788 | -0.319504772 | 0.45960907 | -0.6951664 | 0.48695105 | 0.76863578 |
| NOP56 | 19.6898449 | -0.506658372 | 0.64111219 | -0.7902804 | 0.42936405 | 0.7236272 |
| IDH3B | 15.3063305 | 0.155761645 | 0.55667353 | 0.27980789 | 0.7796249 | 0.920472 |
| EBF4 | 20.4602817 | -1.670990287 | 0.83562834 | -1.9996812 | 0.0455347 | 0.21869588 |
| PCED1A | 22.525317 | 0.217999996 | 0.37538734 | 0.58073349 | 0.56142009 | 0.81384822 |
| PTPRA | 187.5914 | -0.036907347 | 0.18598743 | -0.19844 | 0.84270082 | 0.94799096 |
| MRPS26 | 16.4559202 | -0.299037287 | 0.45650047 | -0.6550646 | 0.51242618 | 0.7830488 |
| UBOX5-AS1 | 14.0824187 | 0.488713383 | 0.59635461 | 0.81950131 | 0.41250045 | 0.71110141 |
| UBOX5 | 21.6279575 | 0.374253716 | 0.40734948 | 0.9187534 | 0.35822457 | 0.6647359 |
| LZTS3 | 19.2048881 | -0.362681763 | 0.34549834 | -1.0497352 | 0.29383988 | 0.60883624 |
| DDRGK1 | 21.6262797 | -1.000239811 | 0.42512836 | -2.3527949 | 0.01863291 | 0.1248981 |
| ITPA | 35.6720842 | -0.351367977 | 0.43796808 | -0.8022685 | 0.42239768 | 0.72013823 |
| C20orf194 | 544.797231 | 1.080205886 | 0.30302171 | 3.56478053 | 0.00036416 | 0.00683186 |
| ATRN | 257.665954 | 1.044121751 | 0.21686284 | 4.81466412 | 1.47E-06 | 7.77E-05 |
| MAVS | 76.1874478 | 0.017354766 | 0.27105448 | 0.06402685 | 0.94894884 | 0.98641632 |
| PANK2 | 63.9644371 | -0.234065446 | 0.32833078 | -0.7128952 | 0.4759106 | 0.76162498 |
| RNF24 | 225.17304 | -0.757748228 | 0.41461781 | -1.8275824 | 0.06761225 | 0.27747209 |
| PRNP | 19.066482 | -1.980784742 | 0.79664641 | -2.4864039 | 0.01290414 | 0.09821896 |
| SLC23A2 | 45.7359261 | 0.60720186 | 0.30864965 | 1.96728513 | 0.04915035 | 0.22760049 |
| TMEM230 | 48.7913295 | -0.22015299 | 0.37314938 | -0.5899862 | 0.55519989 | 0.81104802 |
| CDS2 | 65.293708 | -0.182187818 | 0.40584798 | -0.4489065 | 0.65349907 | 0.86199934 |
| GPCPD1 | 116.724815 | 0.030886079 | 0.37282391 | 0.08284361 | 0.93397589 | 0.98045386 |
| SHLD1 | 41.4265103 | 0.247850409 | 0.28892243 | 0.85784412 | 0.39097854 | 0.69372012 |
| MCM8 | 15.466511 | 0.58193218 | 0.43812244 | 1.32824097 | 0.18409851 | 0.4832586 |
| CRLS1 | 70.5072514 | 0.359231928 | 0.36467212 | 0.98508197 | 0.32458381 | 0.63687278 |
| TMX4 | 43.0265497 | -0.30462118 | 0.42861078 | -0.7107175 | 0.47725932 | 0.76307714 |
| PLCB1 | 113.4183 | -1.967845591 | 0.77327331 | -2.5448255 | 0.01093324 | 0.08735347 |
| PLCB4 | 264.092159 | -0.339868292 | 0.50522511 | -0.6727067 | 0.50113393 | 0.7760974 |
| MKKS | 16.4918961 | 0.690807229 | 0.39611785 | 1.7439437 | 0.08116891 | 0.30642359 |
| SLX4IP | 152.777137 | 0.528595409 | 0.35924455 | 1.47140831 | 0.14118073 | 0.42136906 |
| JAG1 | 101.877012 | 0.74046325 | 0.53598653 | 1.38149601 | 0.1671265 | 0.45967184 |
| AL050403.2 | 41.5406546 | -0.369232978 | 0.66239348 | -0.5574224 | 0.57723885 | 0.82233385 |
| BTBD3 | 145.940244 | -0.258205421 | 0.25455867 | -1.0143258 | 0.31042732 | 0.62528069 |
| SPTLC3 | 73.0636512 | -0.841429667 | 0.4530836 | -1.8571179 | 0.06329438 | 0.26477174 |
| TASP1 | 179.434022 | 0.094897512 | 0.22450681 | 0.42269324 | 0.67251909 | 0.86825921 |
| ESF1 | 66.144778 | 0.10038794 | 0.27901613 | 0.3597926 | 0.71900224 | 0.89077556 |
| NDUFAF5 | 51.3462942 | 0.447100216 | 0.39497118 | 1.13198188 | 0.25764205 | 0.57033582 |
| MACROD2 | 613.066182 | -0.001273505 | 0.44760914 | -0.0028451 | 0.99772992 | 0.99922221 |
| FLRT3 | 19.1801859 | -2.24457334 | 0.58788438 | -3.8180524 | 0.00013451 | 0.00307025 |
| KIF16B | 187.738506 | 0.35385845 | 0.38087086 | 0.92907724 | 0.35284906 | 0.65972795 |
| SNRPB2 | 22.004021 | 0.719242785 | 0.33569182 | 2.14256872 | 0.03214775 | 0.17776472 |
| DSTN | 228.149506 | -0.458748645 | 0.40359556 | -1.1366543 | 0.25568281 | 0.56843403 |
| RRBP1 | 121.170987 | -1.155223913 | 0.37616814 | -3.0710307 | 0.00213321 | 0.02650028 |
| SNX5 | 36.728729 | 0.01411182 | 0.40103703 | 0.03518832 | 0.97192957 | 0.99269048 |
| OVOL2 | 67.1814134 | 0.439674954 | 0.39184653 | 1.12205907 | 0.26183733 | 0.57571533 |
| MGME1 | 64.7799279 | 0.099860923 | 0.30089069 | 0.33188439 | 0.73997656 | 0.90304252 |
| KAT14 | 14.1394907 | 0.770079786 | 0.49282623 | 1.56257874 | 0.11815171 | 0.37883307 |
| ZNF133 | 49.0423904 | 0.505514914 | 0.43734183 | 1.15588054 | 0.24773002 | 0.56031261 |
| DZANK1 | 27.1894169 | -0.063807937 | 0.37805623 | -0.168779 | 0.86597051 | 0.95565922 |
| POLR3F | 15.5317255 | 0.172617872 | 0.46031373 | 0.37500048 | 0.70766011 | 0.88614996 |
| SEC23B | 72.9427092 | 0.06980492 | 0.237557 | 0.29384493 | 0.7688764 | 0.91600052 |
| SLC24A3 | 189.642376 | 0.961117855 | 0.62604887 | 1.53521219 | 0.12473172 | 0.39027762 |
| AL121761.1 | 38.5543464 | -0.200743404 | 0.44852324 | -0.4475652 | 0.65446701 | 0.86217501 |
| RIN2 | 175.639275 | -1.148571659 | 0.60009049 | -1.9139974 | 0.05562048 | 0.24566811 |
| NAA20 | 47.3970841 | -0.520431196 | 0.26691212 | -1.9498223 | 0.0511973 | 0.23446152 |
| CRNKL1 | 12.3059953 | 0.240224316 | 0.42862042 | 0.56045933 | 0.57516617 | 0.82146824 |
| RALGAPA2 | 493.637865 | -1.501157581 | 0.30731669 | -4.8847252 | 1.04E-06 | 6.04E-05 |
| KIZ | 96.9843019 | 0.091038843 | 0.43681839 | 0.20841348 | 0.83490613 | 0.94654164 |
| KIZ-AS1 | 18.5776194 | -0.986537178 | 0.63830258 | -1.5455635 | 0.12221002 | 0.3863975 |
| XRN2 | 115.245549 | -0.46080284 | 0.31773552 | -1.4502717 | 0.14698276 | 0.4297124 |
| NAPB | 67.7809343 | 0.526740331 | 0.32863113 | 1.60283151 | 0.10897186 | 0.36312941 |
| CST3 | 83.399389 | -1.911774817 | 0.61991572 | -3.083927 | 0.00204288 | 0.02563632 |
| APMAP | 20.0599149 | -0.459509688 | 0.45514267 | -1.0095948 | 0.31268944 | 0.62758402 |
| ACSS1 | 132.071635 | 0.505734399 | 0.40441605 | 1.25053 | 0.211106 | 0.52017769 |
| ENTPD6 | 36.9902168 | 0.304676136 | 0.38038854 | 0.80096034 | 0.42315461 | 0.72111072 |
| PYGB | 47.2902906 | -0.744845523 | 0.41480659 | -1.7956454 | 0.07255094 | 0.28810261 |
| ABHD12 | 66.7668648 | 0.329905116 | 0.21161173 | 1.55901148 | 0.11899366 | 0.38048591 |
| NINL | 264.522266 | 0.120474 | 0.26274112 | 0.45852739 | 0.64657359 | 0.8589047 |
| ZNF337-AS1 | 26.9371511 | 0.93025271 | 0.47485385 | 1.95902953 | 0.05010933 | 0.23113573 |
| ZNF337 | 33.7811098 | 0.228529282 | 0.43614616 | 0.52397408 | 0.60029657 | 0.83602169 |
| FAM182B | 34.4454318 | -0.269498815 | 0.64020721 | -0.4209556 | 0.6737875 | 0.86846218 |
| HM13 | 76.5094791 | -0.173493833 | 0.22272538 | -0.7789585 | 0.43600414 | 0.72891111 |
| ID1 | 17.2625609 | -0.223553245 | 0.83310526 | -0.2683373 | 0.78843967 | 0.92377626 |
| BCL2L1 | 362.747405 | 0.229012061 | 0.52567651 | 0.43565207 | 0.66308919 | 0.86556259 |
| AL117381.1 | 70.8363482 | 0.265637359 | 0.59538734 | 0.4461589 | 0.65548247 | 0.86268564 |
| DUSP15 | 32.4444249 | 2.038582853 | 0.61552022 | 3.31196734 | 0.00092642 | 0.01404549 |
| TTLL9 | 16.2088315 | 0.575854834 | 0.56780541 | 1.01417637 | 0.3104986 | 0.62528919 |
| TM9SF4 | 78.9327384 | 0.070909773 | 0.2456114 | 0.28870717 | 0.77280548 | 0.91791018 |
| PLAGL2 | 13.3095175 | -0.221710635 | 0.46344555 | -0.4783963 | 0.63236817 | 0.85152594 |
| POFUT1 | 21.219467 | -0.23940797 | 0.36654249 | -0.653152 | 0.51365827 | 0.78323295 |
| KIF3B | 71.5323119 | -0.015029689 | 0.28128646 | -0.053432 | 0.95738773 | 0.98859719 |
| ASXL1 | 224.039837 | 0.023885215 | 0.22386552 | 0.10669448 | 0.91503136 | 0.97327771 |
| NOL4L | 17.5005312 | -0.59368646 | 0.44524557 | -1.3333911 | 0.18240351 | 0.48134117 |
| MAPRE1 | 25.6442302 | 0.496960363 | 0.37809719 | 1.31437201 | 0.18872104 | 0.48919517 |
| CDK5RAP1 | 38.2629439 | 0.350779936 | 0.32009102 | 1.09587559 | 0.27313322 | 0.58894058 |

| | | | | | | |
|---|---|---|---|---|---|---|
| SNTA1 | 17.0660396 | 1.211077695 | 0.49509184 | 2.44616775 | 0.01443838 | 0.10559734 |
| CBFA2T2 | 326.922203 | 0.504922218 | 0.18999558 | 2.65754719 | 0.00787116 | 0.06926755 |
| NECAB3 | 11.5951455 | 1.342932707 | 0.55876603 | 2.40339005 | 0.01624385 | 0.11396363 |
| CHMP4B | 55.3514683 | -0.921365562 | 0.51345518 | -1.7944421 | 0.07274264 | 0.28837569 |
| RALY | 112.31247 | -0.820474327 | 0.44571367 | -1.8408103 | 0.06564936 | 0.27144773 |
| EIF2S2 | 60.9581763 | -1.278216987 | 0.36318422 | -3.5194728 | 0.00043241 | 0.00775956 |
| ITCH | 274.43527 | -0.207664854 | 0.30808578 | -0.6740488 | 0.50028028 | 0.77558432 |
| DYNLRB1 | 51.5020747 | -0.24078798 | 0.27498072 | -0.875654 | 0.38121814 | 0.68500249 |
| MAP1LC3A | 24.7290991 | -0.485661142 | 0.74549535 | -0.651461 | 0.51474895 | 0.78372292 |
| PIGU | 56.1269162 | 0.961738345 | 0.32688796 | 2.94210389 | 0.00325991 | 0.03720362 |
| NCOA6 | 162.40728 | 0.458980239 | 0.25911993 | 1.77130427 | 0.07651012 | 0.29596141 |
| TP53INP2 | 21.2381889 | -0.502087376 | 0.63911933 | -0.7855925 | 0.43210623 | 0.7258077 |
| GGT7 | 67.6628883 | 0.886217918 | 0.49129462 | 1.8038421 | 0.07125606 | 0.28479321 |
| ACSS2 | 51.1217393 | 0.187925576 | 0.37026806 | 0.50753926 | 0.61177649 | 0.84169124 |
| GSS | 15.8845294 | -0.44446677 | 0.47222996 | -0.9412083 | 0.34659811 | 0.65471653 |
| TRPC4AP | 155.501129 | -0.107861915 | 0.2902008 | -0.3716803 | 0.71013091 | 0.88614996 |
| EDEM2 | 38.9099572 | 0.629954872 | 0.36494822 | 1.72614864 | 0.08432067 | 0.31297129 |
| PROCR | 16.8080818 | -0.202084877 | 0.63972309 | -0.3158943 | 0.75208274 | 0.90930937 |
| MMP24OS | 47.2274443 | 0.241117127 | 0.42066607 | 0.5731794 | 0.56652323 | 0.81595078 |
| MMP24 | 43.4108326 | 0.03706889 | 0.33851213 | 0.10950535 | 0.91280168 | 0.97166624 |
| UQCC1 | 105.024537 | 0.585198166 | 0.20911016 | 2.79851623 | 0.0051338 | 0.05119523 |
| CEP250 | 40.6379838 | -2.54563881 | 0.98073498 | -2.595644 | 0.00944138 | 0.07888124 |
| ERGIC3 | 35.4264476 | -0.013050574 | 0.33414637 | -0.0390565 | 0.96884537 | 0.99156233 |
| CPNE1 | 38.5956659 | 0.316793348 | 0.31833172 | 0.99516738 | 0.31965487 | 0.63158762 |
| NFS1 | 12.5484777 | 0.811860713 | 0.41618457 | 1.95072275 | 0.05109004 | 0.23408525 |
| ROMO1 | 19.6356684 | 0.327991946 | 0.63673284 | 0.51511706 | 0.60647122 | 0.83829217 |
| RBM39 | 366.884509 | 0.080028255 | 0.25107204 | 0.31874619 | 0.74991898 | 0.90857724 |
| PHF20 | 158.370658 | -0.213700533 | 0.30828082 | -0.6932009 | 0.48818349 | 0.77012857 |
| SCAND1 | 27.4280401 | -0.770892614 | 0.35342961 | -2.1811772 | 0.02917031 | 0.16655479 |
| NORAD | 71.4765482 | -0.788177327 | 0.33194676 | -2.3744089 | 0.01757707 | 0.12050636 |
| EPB41L1 | 299.425229 | -0.300604324 | 0.36971339 | -0.8130739 | 0.41617566 | 0.71390709 |
| AAR2 | 18.4067044 | -0.631186699 | 0.43913206 | -1.4373505 | 0.15061841 | 0.43488337 |
| DLGAP4 | 94.9030171 | -0.363890378 | 0.37122084 | -0.9802531 | 0.3269612 | 0.63858111 |
| DLGAP4-AS1 | 33.427585 | 0.305949765 | 0.44838774 | 0.68233303 | 0.49502839 | 0.77475164 |
| NDRG3 | 28.4400691 | 1.065224213 | 0.48086567 | 2.21522201 | 0.02674483 | 0.15772852 |
| SOGA1 | 37.9056443 | -0.002377031 | 0.41789467 | -0.0056881 | 0.99546157 | 0.99922221 |
| RBL1 | 28.3736684 | 0.977860493 | 0.40944625 | 2.38825119 | 0.01692877 | 0.11753077 |
| RPN2 | 56.9819044 | -0.16643223 | 0.29126191 | -0.5714178 | 0.56771649 | 0.81673499 |
| MANBAL | 14.3578953 | -0.604587258 | 0.45260595 | -1.3357917 | 0.18161737 | 0.47998877 |
| SRC | 25.3316428 | -0.715284999 | 0.36044323 | -1.9844595 | 0.04720464 | 0.22304735 |
| BLCAP | 74.9215678 | -0.125714637 | 0.33564978 | -0.3745411 | 0.70800178 | 0.88614996 |
| CTNNBL1 | 56.1071402 | 0.245409638 | 0.24943978 | 0.98384324 | 0.3251926 | 0.63715682 |
| TTI1 | 33.5996409 | -0.025928347 | 0.39728038 | -0.0652646 | 0.94796332 | 0.98636425 |
| RPRD1B | 73.6761169 | 0.205665433 | 0.30618242 | 0.67170883 | 0.50176908 | 0.776256 |
| TGM2 | 59.8127758 | -4.304701114 | 0.72085247 | -5.9716811 | 2.35E-09 | 3.42E-07 |
| RALGAPB | 127.470204 | -0.023128345 | 0.26418315 | -0.0875466 | 0.93023702 | 0.97884324 |
| DHX35 | 29.9345909 | 0.982433458 | 0.34148371 | 2.8769556 | 0.00401532 | 0.04293446 |
| TOP1 | 240.95597 | -0.827748431 | 0.44472948 | -1.8612403 | 0.06271025 | 0.26385847 |
| PLCG1 | 32.6430156 | -0.93239456 | 0.32289654 | -2.8875954 | 0.00388199 | 0.04191558 |
| ZHX3 | 249.948301 | 0.049093758 | 0.48448575 | 0.10133169 | 0.91928716 | 0.97513464 |
| LPIN3 | 35.8141606 | 0.038653694 | 0.60943084 | 0.06342589 | 0.94942737 | 0.98641632 |
| CHD6 | 229.97403 | 0.236797684 | 0.20639287 | 1.14731521 | 0.25125137 | 0.56353121 |
| SRSF6 | 58.6829563 | 0.735293051 | 0.28300201 | 2.59819024 | 0.00937166 | 0.07872191 |
| L3MBTL1 | 28.2632956 | 0.033184308 | 0.35206431 | 0.09425638 | 0.9249055 | 0.97735117 |
| SGK2 | 93.6616376 | 1.230806778 | 0.47245589 | 2.60512525 | 0.00918407 | 0.07756348 |
| IFT52 | 35.8403648 | -0.215538176 | 0.29545674 | -0.7295084 | 0.46569073 | 0.75274202 |
| OSER1 | 23.7829012 | -0.654348324 | 0.42320257 | -1.5461823 | 0.12206054 | 0.38617399 |
| SERINC3 | 48.5390685 | -0.239831647 | 0.26254163 | -0.9134995 | 0.3609799 | 0.66649041 |
| PKIG | 70.0277014 | -0.0605167 | 0.29013958 | -0.2085779 | 0.83477778 | 0.94654164 |
| YWHAB | 73.5309745 | -1.092649771 | 0.39245517 | -2.7841391 | 0.005367 | 0.05256505 |
| PABPC1L | 27.4090279 | 0.456942648 | 0.35365882 | 1.29204371 | 0.19634201 | 0.49845696 |
| STK4 | 55.1220167 | -0.536849798 | 0.2391889 | -2.2444595 | 0.02480285 | 0.14978095 |
| SLPI | 67.3003207 | -5.13405307 | 0.98412948 | -5.2168472 | 1.82E-07 | 1.38E-05 |
| SDC4 | 180.074615 | -0.327536392 | 0.43620274 | -0.7508811 | 0.45272422 | 0.74447982 |
| PIGT | 14.5833667 | 0.073780954 | 0.55476337 | 0.13299536 | 0.89419705 | 0.96482804 |
| WFDC2 | 496.574447 | -4.908756469 | 0.79969984 | -6.1382487 | 8.34E-10 | 1.53E-07 |
| DNTTIP1 | 15.6364976 | -0.394403 | 0.38339726 | -1.0287058 | 0.30361792 | 0.61784922 |
| SNX21 | 17.0952317 | 0.710100128 | 0.42102841 | 1.68658481 | 0.09168321 | 0.33018706 |
| ACOT8 | 18.0140942 | 0.224502046 | 0.36267702 | 0.61901371 | 0.53590733 | 0.79974391 |
| CTSA | 37.5578693 | -0.371705604 | 0.54194983 | -0.6858672 | 0.49279686 | 0.77289116 |
| ZNF335 | 21.6399922 | -0.632655631 | 0.48894061 | -1.2939315 | 0.1956891 | 0.49824525 |
| NCOA5 | 49.4367404 | -0.43543357 | 0.39864096 | -1.0922951 | 0.27470341 | 0.58917168 |
| SLC35C2 | 25.7649105 | -0.217168466 | 0.40277864 | -0.5391757 | 0.58976561 | 0.83091183 |
| ELMO2 | 23.2929194 | 0.094747059 | 0.43531901 | 0.21764972 | 0.82770204 | 0.94254892 |
| SLC13A3 | 24.5482311 | 0.219993347 | 0.43673892 | 0.5037182 | 0.61445941 | 0.84174925 |
| EYA2 | 123.666474 | 1.446092304 | 0.75105465 | 1.92541557 | 0.05417739 | 0.24204599 |
| ZMYND8 | 367.813446 | -0.411485859 | 0.26714959 | -1.5402826 | 0.12349149 | 0.38860433 |
| NCOA3 | 233.006842 | -0.556149601 | 0.35930831 | -1.5478339 | 0.12166228 | 0.38531899 |
| SULF2 | 71.2722612 | -0.325771486 | 0.43915213 | -0.7418192 | 0.45819688 | 0.747377 |
| ARFGEF2 | 285.958047 | 0.131368209 | 0.18371052 | 0.71508269 | 0.47455795 | 0.76075482 |
| CSE1L | 50.0008043 | -0.357748996 | 0.26460991 | -1.3519864 | 0.17637967 | 0.47354004 |
| STAU1 | 138.000801 | -0.291727219 | 0.20143987 | -1.4482099 | 0.14755836 | 0.43102573 |
| DDX27 | 27.9336521 | -0.102524958 | 0.44067367 | -0.2326551 | 0.81602927 | 0.93717111 |
| ZNFX1 | 35.7796573 | -0.192935388 | 0.51755817 | -0.3727801 | 0.70931211 | 0.88614996 |
| ZFAS1 | 38.0795068 | -0.162901517 | 0.35042457 | -0.464869 | 0.64202528 | 0.85468927 |
| B4GALT5 | 196.18307 | -0.900688096 | 0.38158714 | -2.3603733 | 0.01825655 | 0.12340471 |
| SLC9A8 | 74.5851167 | -0.436949939 | 0.50233608 | -0.8698359 | 0.3843901 | 0.68822563 |
| RNF114 | 71.5391502 | -0.596703594 | 0.23955735 | -2.490859 | 0.01274347 | 0.09755344 |
| CEBPB | 33.6017932 | -3.419860348 | 0.82268104 | -4.15697 | 3.22E-05 | 0.00102277 |
| PTPN1 | 187.206745 | -1.249715709 | 0.4826512 | -2.589273 | 0.00961788 | 0.07978392 |
| PARD6B | 130.614126 | 0.233578232 | 0.30201991 | 0.77338686 | 0.43929347 | 0.73168494 |
| BCAS4 | 35.0662804 | -0.316984211 | 0.40346951 | -0.785646 | 0.4320749 | 0.7258077 |
| ADNP | 240.802163 | 0.060069655 | 0.18211995 | 0.32983566 | 0.74152414 | 0.90383185 |
| DPM1 | 59.8531333 | -0.073283296 | 0.30907839 | -0.2371026 | 0.81257718 | 0.93571318 |
| NFATC2 | 18.0533016 | -0.072727819 | 0.39004808 | -0.1864586 | 0.85208514 | 0.95090866 |
| ATP9A | 77.9453924 | -0.923218937 | 0.28258088 | -3.2670963 | 0.00108657 | 0.01592948 |
| ZFP64 | 42.5221898 | 0.557990713 | 0.38606491 | 1.44532874 | 0.14836557 | 0.43218222 |
| ZNF217 | 52.4462475 | -0.250144439 | 0.35124527 | -0.7121646 | 0.47636287 | 0.76198446 |
| DOK5 | 25.4970212 | -1.847276716 | 0.66007442 | -2.7985886 | 0.00513265 | 0.05119523 |
| RTF2 | 33.0521979 | -0.610287519 | 0.37644088 | -1.6212042 | 0.10497386 | 0.35578926 |
| RAE1 | 20.3681537 | -0.214166032 | 0.40353045 | -0.5307308 | 0.59560535 | 0.83416944 |
| PCK1 | 133.539541 | -1.09886385 | 1.63096446 | -0.6737509 | 0.50046967 | 0.77566143 |
| PMEPA1 | 26.5912528 | -3.499803676 | 1.07066661 | -3.2688081 | 0.00108002 | 0.01585837 |
| RAB22A | 65.309414 | -0.319378544 | 0.37430551 | -0.8532563 | 0.39351717 | 0.69597005 |
| VAPB | 88.5109091 | -0.101500923 | 0.19873026 | -0.5107472 | 0.60952809 | 0.84071597 |
| NPEPL1 | 18.3297707 | 0.018021855 | 0.58759057 | 0.03067077 | 0.9755321 | 0.99397457 |
| GNAS | 242.2012 | 0.023354039 | 0.25320717 | 0.09223293 | 0.92651298 | 0.97735117 |
| NELFCD | 38.2126511 | 0.401065075 | 0.29090289 | 1.37869058 | 0.16799018 | 0.46123098 |
| CTSZ | 17.6529634 | -1.303070604 | 0.38062009 | -3.4235466 | 0.0006181 | 0.01021832 |
| ATP5F1E | 75.6725419 | -0.383436195 | 0.46798142 | -0.8193406 | 0.41259209 | 0.71110141 |
| MIR646HG | 339.876404 | 0.598515641 | 0.9950549 | 0.60149007 | 0.54751362 | 0.80618229 |
| TAF4 | 57.6563295 | 0.222732922 | 0.34012301 | 0.65485991 | 0.51255796 | 0.7830488 |
| PSMA7 | 40.0781329 | -0.913604447 | 0.34954467 | -2.6136987 | 0.0089568 | 0.07612871 |

| | | | | | | |
|---|---|---|---|---|---|---|
| SS18L1 | 55.2735589 | -0.179759161 | 0.37782633 | -0.4757719 | 0.63423687 | 0.85235293 |
| MTG2 | 42.8393987 | 0.372064523 | 0.30672549 | 1.2130212 | 0.22512172 | 0.53616259 |
| OSBPL2 | 71.9807388 | -0.054755142 | 0.32340966 | -0.1693058 | 0.86555609 | 0.9555677 |
| LAMA5 | 60.3410032 | -0.089532461 | 0.37280342 | -0.24016 | 0.81020624 | 0.93552094 |
| SLCO4A1 | 60.460068 | -3.969012584 | 0.75388902 | -5.2647173 | 1.40E-07 | 1.10E-05 |
| OGFR | 29.6641389 | -0.92813082 | 0.42056869 | -2.2068472 | 0.02732474 | 0.16053929 |
| TCFL5 | 13.3368134 | 0.112564818 | 0.4459182 | 0.25243378 | 0.80070579 | 0.92996771 |
| DIDO1 | 112.95365 | 0.034795813 | 0.21238134 | 0.16383649 | 0.86985986 | 0.95621002 |
| GID8 | 26.0144358 | 0.156770771 | 0.35786455 | 0.43807293 | 0.66133342 | 0.86492722 |
| YTHDF1 | 35.2573656 | -0.001249475 | 0.34558937 | -0.0036155 | 0.99711526 | 0.99922221 |
| ARFGAP1 | 31.4261148 | -0.058555711 | 0.425341 | -0.1376677 | 0.89050306 | 0.96323092 |
| PPDPF | 76.2350912 | -0.746281022 | 0.56104293 | -1.3301674 | 0.18346312 | 0.48210665 |
| GMEB2 | 22.6740545 | -0.323687205 | 0.51927025 | -0.6233502 | 0.53305443 | 0.79750909 |
| ARFRP1 | 17.8382584 | -0.257714525 | 0.43475807 | -0.5927769 | 0.55333049 | 0.81031153 |
| SLC2A4RG | 27.465767 | -0.489998819 | 0.53343442 | -0.9185737 | 0.3583186 | 0.6647359 |
| ZBTB46 | 30.5997623 | -0.205326605 | 0.4000946 | -0.5131951 | 0.60781482 | 0.83922189 |
| TPD52L2 | 43.4262944 | 0.136733714 | 0.40965071 | 0.33378122 | 0.73854466 | 0.90241447 |
| DNAJC5 | 110.728148 | -0.484676832 | 0.37443075 | -1.2944365 | 0.19551469 | 0.49821781 |
| UCKL1 | 25.4711937 | 1.130372752 | 0.42361886 | 2.66837211 | 0.00762198 | 0.06761878 |
| ZNF512B | 15.9278177 | 0.453868757 | 0.48010628 | 0.94535059 | 0.34447992 | 0.65359197 |
| PRPF6 | 57.5055093 | 0.012098114 | 0.28475868 | 0.0424855 | 0.96611167 | 0.99156233 |
| PCMTD2 | 78.711501 | 0.195733691 | 0.48713606 | 0.40180497 | 0.68782756 | 0.87565595 |
| PLPP2 | 22.5865471 | -3.260824637 | 0.67065979 | -4.8621144 | 1.16E-06 | 6.60E-05 |
| MIER2 | 17.6937146 | 0.174908754 | 0.39945692 | 0.43786638 | 0.66148315 | 0.86492722 |
| BSG | 71.9532883 | -0.271805257 | 0.43031234 | -0.6316464 | 0.52761793 | 0.79403853 |
| RNF126 | 21.4166223 | -0.275053405 | 0.33711888 | -0.8158944 | 0.41456054 | 0.71211542 |
| PALM | 58.4029715 | 1.683396263 | 0.54553758 | 3.08575673 | 0.00203035 | 0.02554787 |
| PTBP1 | 58.0947122 | -0.053312233 | 0.30572078 | -0.1743821 | 0.86156518 | 0.95417909 |
| MED16 | 14.1739956 | 0.395482794 | 0.50480327 | 0.78343945 | 0.4333691 | 0.7267202 |
| R3HDM4 | 23.8034873 | -1.472854797 | 0.50705885 | -2.9047019 | 0.00367603 | 0.04037138 |
| ARID3A | 113.149793 | -0.398741473 | 0.52927862 | -0.7533678 | 0.45122893 | 0.74241371 |
| WDR18 | 13.8328339 | 0.11836056 | 0.58167478 | 0.20348237 | 0.83875803 | 0.94741614 |
| TMEM259 | 69.9300022 | -0.073737126 | 0.37855157 | -0.1947875 | 0.84555928 | 0.94907846 |
| CNN2 | 15.0660429 | -0.603566449 | 0.47530284 | -1.2698566 | 0.20413571 | 0.50987446 |
| POLR2E | 18.8658811 | -0.542710609 | 0.40696458 | -1.3335573 | 0.18234897 | 0.48134117 |
| GPX4 | 65.6979418 | -0.296332059 | 0.42044057 | -0.7048132 | 0.4809265 | 0.76563273 |
| SBNO2 | 115.229712 | -2.579169539 | 0.35181029 | -7.3311372 | 2.28E-13 | 8.80E-11 |
| STK11 | 44.8244627 | 0.121843502 | 0.37306629 | 0.32660014 | 0.74397034 | 0.904876 |
| ATP5F1D | 49.622304 | 0.204198947 | 0.51194108 | 0.39887197 | 0.68998755 | 0.87691291 |
| MIDN | 78.7179275 | -2.273744599 | 0.57771945 | -3.9357245 | 8.29E-05 | 0.00211309 |
| CIRBP | 139.802482 | 0.710415394 | 0.35528934 | 1.99954042 | 0.04554991 | 0.21869588 |
| MUM1 | 49.3333017 | -0.017562248 | 0.44284188 | -0.0396581 | 0.96836574 | 0.99156233 |
| NDUFS7 | 20.6001787 | 0.88385222 | 0.42429495 | 2.08310801 | 0.03724138 | 0.19409649 |
| DAZAP1 | 79.8043997 | -0.023776388 | 0.40078689 | -0.0593243 | 0.95269383 | 0.98742967 |
| RPS15 | 107.630724 | -1.057040581 | 0.49674503 | -2.1279339 | 0.03334257 | 0.18186117 |
| C19orf25 | 44.4575961 | -0.846257083 | 0.40236031 | -2.103232 | 0.03544549 | 0.18774924 |
| MBD3 | 17.6614994 | -0.847793825 | 0.54286001 | -1.5617172 | 0.11835462 | 0.37909258 |
| TCF3 | 20.7549879 | -0.215153894 | 0.35246396 | -0.6104281 | 0.54157828 | 0.80293781 |
| REXO1 | 21.6227553 | 0.148430942 | 0.38487823 | 0.3856569 | 0.69975079 | 0.88347683 |
| ABHD17A | 20.0821718 | -0.398721686 | 0.45062765 | -0.8848141 | 0.37625692 | 0.67982871 |
| SCAMP4 | 31.2469339 | -0.686398216 | 0.42766418 | -1.6049935 | 0.10849524 | 0.36219464 |
| CSNK1G2 | 38.5769706 | -0.633688547 | 0.30736537 | -2.0616784 | 0.03923837 | 0.20036945 |
| BTBD2 | 48.2146987 | -0.343963625 | 0.33039389 | -1.0410714 | 0.29784241 | 0.61141484 |
| MKNK2 | 21.9202417 | -0.462717613 | 0.49216951 | -0.940159 | 0.34713599 | 0.6553345 |
| MOB3A | 25.1348202 | -1.299976146 | 0.40443496 | -3.2143021 | 0.00130762 | 0.01850114 |
| AP3D1 | 77.9134399 | -0.821807483 | 0.44468321 | -1.848074 | 0.06459163 | 0.268828 |
| DOT1L | 84.911558 | -0.496954239 | 0.65541354 | -0.7582301 | 0.44831327 | 0.7399821 |
| PLEKHJ1 | 32.8317188 | 0.056710309 | 0.50477759 | 0.11234712 | 0.91054818 | 0.97128978 |
| SF3A2 | 12.0157151 | -0.102600892 | 0.53127029 | -0.1931237 | 0.84686208 | 0.94939787 |
| LSM7 | 27.8384086 | -0.494463236 | 0.41629017 | -1.1877851 | 0.23491811 | 0.54677394 |
| SPPL2B | 45.8307188 | 0.38796149 | 0.34299464 | 1.13110075 | 0.25801269 | 0.57102072 |
| TIMM13 | 21.5386482 | -0.401549533 | 0.50495653 | -0.795216 | 0.42648784 | 0.72230448 |
| GADD45B | 42.1011485 | -1.682799994 | 0.50618221 | -3.3244946 | 0.00088579 | 0.01360643 |
| GNG7 | 102.562404 | 0.441342784 | 0.44412996 | 0.99372441 | 0.32035706 | 0.63212442 |
| SGTA | 25.281313 | -0.231134955 | 0.4474636 | -0.5165447 | 0.60547402 | 0.83802715 |
| ZNF554 | 9.94005284 | 0.260503102 | 0.55323428 | 0.47087303 | 0.63773141 | 0.8538689 |
| ZNF555 | 13.3635579 | 0.316017588 | 0.52005946 | 0.60765664 | 0.54341522 | 0.80361674 |
| AES | 82.5808137 | -0.322309557 | 0.33789147 | -0.9538849 | 0.34014193 | 0.65054327 |
| GNA11 | 22.6943957 | -0.122996404 | 0.38176818 | -0.3221756 | 0.74731965 | 0.90692529 |
| NCLN | 20.5949895 | 0.249491361 | 0.43568054 | 0.57264748 | 0.5668834 | 0.81606004 |
| NFIC | 200.565878 | -0.12031041 | 0.28750812 | -0.4184592 | 0.67561143 | 0.86960234 |
| FZR1 | 21.2896644 | -0.437151592 | 0.50136795 | -0.8719177 | 0.38325328 | 0.68720476 |
| MFSD12 | 31.0603651 | -0.84219822 | 0.47164043 | -1.7856786 | 0.07415131 | 0.29129403 |
| HMG20B | 20.5761463 | -0.380422769 | 0.43335551 | -0.8778538 | 0.38002308 | 0.68404154 |
| PIP5K1C | 21.3066719 | 0.433351902 | 0.3982747 | 1.0880729 | 0.27656293 | 0.59165557 |
| TJP3 | 38.4166408 | 0.624717188 | 0.33467417 | 1.86664298 | 0.06195146 | 0.26124663 |
| DAPK3 | 24.9615964 | -1.220642527 | 0.49870063 | -2.4476459 | 0.01437929 | 0.10540292 |
| EEF2 | 98.7880845 | -1.05878834 | 0.48505373 | -2.1828269 | 0.02904856 | 0.16620962 |
| PIAS4 | 25.7972798 | 0.184365976 | 0.43084096 | 0.42792119 | 0.6687085 | 0.86777458 |
| ZBTB7A | 52.9717885 | -0.459484361 | 0.35057948 | -1.3106425 | 0.18997858 | 0.49058838 |
| MAP2K2 | 43.5471616 | -0.082923505 | 0.33846857 | -0.2449962 | 0.8064594 | 0.93325451 |
| YJU2 | 11.4227808 | -0.380223687 | 0.61801518 | -0.6152336 | 0.53840048 | 0.80104055 |
| SH3GL1 | 31.1376001 | -0.607742468 | 0.32000762 | -1.89915 | 0.05754476 | 0.25013758 |
| UBXN6 | 23.5046935 | -0.506011459 | 0.46074632 | -1.0982431 | 0.27209836 | 0.58786226 |
| HDGFL2 | 15.8073864 | 0.157472875 | 0.43103112 | 0.36533992 | 0.71485769 | 0.88904248 |
| PLIN5 | 56.2224145 | 0.156720984 | 0.44143185 | 0.35502872 | 0.72256808 | 0.89222947 |
| MYDGF | 22.2989392 | -0.600552505 | 0.42780242 | -1.4038081 | 0.160376 | 0.44957865 |
| DPP9 | 41.1685497 | -0.264285963 | 0.2927953 | -0.9026305 | 0.36672204 | 0.67113782 |
| PLIN3 | 31.5351365 | -2.034803212 | 0.43868596 | -4.6384051 | 3.51E-06 | 0.00016127 |
| KDM4B | 102.177781 | -0.910541836 | 0.29808006 | -3.0546888 | 0.00225294 | 0.02753136 |
| PTPRS | 66.7245451 | 0.456687177 | 0.3768213 | 1.21194628 | 0.22553294 | 0.53644622 |
| SAFB2 | 98.1890641 | 0.097873473 | 0.33487999 | 0.29226432 | 0.77008454 | 0.91655198 |
| SAFB | 115.886371 | -0.055664333 | 0.33794573 | -0.1647138 | 0.86916923 | 0.95612718 |
| RPL36 | 63.8235656 | -0.612579663 | 0.46955818 | -1.3045873 | 0.1920334 | 0.49298442 |
| C19orf70 | 29.4947551 | -0.206183015 | 0.53241982 | -0.3872565 | 0.69856636 | 0.88302803 |
| LONP1 | 58.3766371 | -1.147762563 | 0.56500663 | -2.0314143 | 0.04221298 | 0.20951607 |
| RANBP3 | 31.9294868 | 0.299588702 | 0.33569302 | 0.89244842 | 0.37215263 | 0.6755422 |
| RFX2 | 71.3609009 | -0.21675302 | 0.51431977 | -0.4214363 | 0.67343652 | 0.86846218 |
| MLLT1 | 30.0692811 | 0.369943524 | 0.34528303 | 1.07142111 | 0.28398012 | 0.59743472 |
| GTF2F1 | 16.3909962 | 0.0349887 | 0.43235233 | 0.08092636 | 0.93550052 | 0.98139141 |
| KHSRP | 38.1362634 | -0.432685908 | 0.48486378 | -0.8923865 | 0.37218579 | 0.6755422 |
| SLC25A23 | 27.0590069 | -0.341402402 | 0.56048092 | -0.609124 | 0.54244222 | 0.80340275 |
| TRIP10 | 30.9947536 | -1.390472183 | 0.49826247 | -2.790642 | 0.00526036 | 0.05195722 |
| SH2D3A | 47.5513099 | -1.371840159 | 0.51472706 | -2.6651798 | 0.00769472 | 0.06806978 |
| INSR | 344.089279 | 0.82336136 | 0.32826174 | 2.5082465 | 0.0121332 | 0.09443235 |
| PNPLA6 | 26.6679203 | 0.42552113 | 0.39263437 | 1.08375924 | 0.27847156 | 0.59359815 |
| CAMSAP3 | 13.3592425 | -0.204080452 | 0.39374505 | -0.5183061 | 0.60424473 | 0.83754753 |
| EVI5L | 14.4521174 | -0.289782112 | 0.39317786 | -0.7370255 | 0.46110684 | 0.74927852 |
| MAP2K7 | 42.025498 | -0.847483653 | 0.52800851 | -1.6050568 | 0.1084813 | 0.36219464 |
| TIMM44 | 26.4679995 | -0.320472111 | 0.70123535 | -0.4570108 | 0.6476633 | 0.85925051 |
| ELAVL1 | 77.6673733 | -0.745072112 | 0.6225844 | -1.1967407 | 0.23140763 | 0.54389834 |
| CERS4 | 62.898234 | -0.468671855 | 0.39676729 | -1.181226 | 0.23751294 | 0.54997949 |
| AC022146.2 | 17.4758891 | -0.661339501 | 0.59406648 | -1.1132416 | 0.26560469 | 0.58092849 |

| | | | | | | |
|---|---|---|---|---|---|---|
| RPS28 | 27.7279494 | -0.200928715 | 0.36789252 | -0.5461614 | 0.58495496 | 0.82864176 |
| ANGPTL4 | 35.0691888 | -5.138490369 | 1.22671787 | -4.1888119 | 2.80E-05 | 0.00090596 |
| RAB11B | 27.2444516 | -0.019763968 | 0.38625306 | -0.0511684 | 0.9591913 | 0.98932518 |
| 2-Mar | 14.9603553 | 0.015709753 | 0.42184663 | 0.03724044 | 0.9702933 | 0.99156233 |
| HNRNPM | 152.898197 | -0.05665814 | 0.28411471 | -0.1994199 | 0.84193426 | 0.94776435 |
| ZNF558 | 15.6081401 | -0.226099405 | 0.39170807 | -0.5772141 | 0.56379486 | 0.81450672 |
| ZNF317 | 20.849951 | 0.153600012 | 0.37976788 | 0.40445762 | 0.68587624 | 0.87448517 |
| ZNF559-ZNF | 31.536053 | 0.424765879 | 0.43912961 | 0.96729045 | 0.33339886 | 0.64508099 |
| ZNF266 | 34.2592155 | 0.036516726 | 0.35452568 | 0.10300164 | 0.91796167 | 0.97482633 |
| ZNF426 | 17.1987217 | -0.228685281 | 0.49501373 | -0.4619777 | 0.64409735 | 0.85708058 |
| ZNF121 | 30.6172781 | -0.046411718 | 0.29793452 | -0.1557783 | 0.87620781 | 0.95866717 |
| ZNF561 | 25.4344143 | -0.350592918 | 0.38321146 | -0.9148811 | 0.36025405 | 0.66607353 |
| ZNF562 | 58.2440476 | -0.465886898 | 0.36038427 | -1.2927504 | 0.19609741 | 0.49845696 |
| ZNF846 | 51.2794037 | -0.02418384 | 0.37931033 | -0.0637574 | 0.9491634 | 0.98641632 |
| FBXL12 | 20.3794044 | -0.153129374 | 0.37775947 | -0.4053621 | 0.68521136 | 0.87423519 |
| UBL5 | 14.8772515 | -0.095632434 | 0.5215954 | -0.183346 | 0.85452654 | 0.95170432 |
| PIN1 | 17.9095908 | -0.31580644 | 0.36311142 | -0.8697232 | 0.38445168 | 0.68822563 |
| C19orf66 | 30.2451041 | -0.070758342 | 0.55044655 | -0.1285472 | 0.89771598 | 0.96607503 |
| EIF3G | 13.5198537 | -0.35017506 | 0.51823321 | -0.6757094 | 0.49922515 | 0.77558432 |
| DNMT1 | 19.9115257 | -0.403294649 | 0.55084449 | -0.7321389 | 0.46408382 | 0.75166353 |
| MRPL4 | 22.9739897 | 0.080059993 | 0.4123675 | 0.19414719 | 0.84606063 | 0.94909204 |
| AC011511.2 | 15.872706 | 2.087386944 | 0.72606211 | 2.87494268 | 0.00404101 | 0.04311026 |
| TYK2 | 40.736388 | 0.429058918 | 0.25980598 | 1.65145896 | 0.09864489 | 0.34303156 |
| CDC37 | 56.4710016 | -0.395921968 | 0.3655401 | -1.083115 | 0.27875738 | 0.59369058 |
| KEAP1 | 13.9401602 | -0.242682456 | 0.47768211 | -0.5080418 | 0.61142406 | 0.84169124 |
| ATG4D | 23.0580888 | 0.065027492 | 0.33345376 | 0.19501202 | 0.84538353 | 0.94907846 |
| KRI1 | 16.3684511 | 0.070081238 | 0.44882888 | 0.15614244 | 0.87592074 | 0.95857805 |
| AP1M2 | 29.7011717 | -0.531008964 | 0.43237513 | -1.228121 | 0.21940155 | 0.52936397 |
| SLC44A2 | 40.3856692 | 0.220375846 | 0.56118347 | 0.39269839 | 0.69454226 | 0.87964032 |
| ILF3-DT | 21.9483706 | -0.042652504 | 0.47908296 | -0.0890295 | 0.92905849 | 0.97870764 |
| ILF3 | 94.263629 | 0.092615449 | 0.37575406 | 0.24647891 | 0.80531153 | 0.93299263 |
| QTRT1 | 33.8013012 | 1.062577551 | 0.42407972 | 2.505608 | 0.01222411 | 0.0949813 |
| DNM2 | 211.922763 | -0.4284449 | 0.27403991 | -1.5634398 | 0.11794919 | 0.37862775 |
| CARM1 | 32.6926726 | -0.201764761 | 0.37284286 | -0.5411523 | 0.58840263 | 0.83024609 |
| YIPF2 | 19.0494387 | 0.50515458 | 0.41711635 | 1.21106395 | 0.2258709 | 0.53705916 |
| TIMM29 | 14.1333586 | -0.198442714 | 0.45309994 | -0.4379668 | 0.66141038 | 0.86492722 |
| SMARCA4 | 145.157523 | 0.08682057 | 0.25907552 | 0.33511685 | 0.73753695 | 0.90199298 |
| LDLR | 58.3124341 | -2.004641495 | 0.7518404 | -2.6663126 | 0.00766884 | 0.06790525 |
| KANK2 | 28.0679119 | 0.607024138 | 0.51034002 | 1.1894504 | 0.23426248 | 0.54647569 |
| DOCK6 | 80.0862317 | -0.275924954 | 0.33133798 | -0.8327598 | 0.40498021 | 0.70434042 |
| EPOR | 28.8082178 | 0.29428225 | 0.57016045 | 0.51613936 | 0.60575708 | 0.83805439 |
| RGL3 | 272.030691 | 0.220004229 | 0.35245524 | 0.62420474 | 0.53249313 | 0.79707272 |
| PRKCSH | 25.5681258 | 0.078657044 | 0.45496185 | 0.17288712 | 0.86274016 | 0.95458251 |
| ECSIT | 29.886856 | 0.704378348 | 0.32079153 | 2.19575108 | 0.02810977 | 0.16289338 |
| ZNF627 | 37.6347799 | 0.374573567 | 0.33206813 | 1.12800215 | 0.25931901 | 0.57248906 |
| AC008543.1 | 25.6003724 | -0.043923071 | 0.3970385 | -0.1106267 | 0.91191235 | 0.97128978 |
| ZNF823 | 29.3041742 | 0.042803293 | 0.40855171 | 0.10476836 | 0.91655962 | 0.97412537 |
| ZNF441 | 27.8726928 | 0.193459368 | 0.41117662 | 0.47050187 | 0.6379965 | 0.8538689 |
| ZNF440 | 39.5544721 | -0.360276939 | 0.32908711 | -1.0947768 | 0.27361443 | 0.58894058 |
| ZNF439 | 53.8356081 | 0.09523412 | 0.28825456 | 0.330382 | 0.74111134 | 0.90375649 |
| ZNF69 | 69.949927 | -0.371766002 | 0.37211558 | -0.9990606 | 0.31776535 | 0.63093651 |
| ZNF433 | 11.7849457 | 0.073113464 | 0.43957047 | 0.16632934 | 0.86789778 | 0.95588054 |
| ZNF844 | 35.5854321 | 0.771797349 | 0.35508027 | 2.17358554 | 0.02973628 | 0.16875291 |
| ZNF788P | 16.7720178 | 0.659883862 | 0.39565064 | 1.66784482 | 0.09534654 | 0.33742332 |
| ZNF136 | 18.1723553 | 0.252648142 | 0.39445887 | 0.640493 | 0.52185214 | 0.78938179 |
| ZNF44 | 241.299671 | 0.350776621 | 0.16251267 | 2.15845707 | 0.03089231 | 0.173407 |
| ZNF563 | 14.9977533 | -0.975572473 | 0.58722956 | -1.6613136 | 0.09665047 | 0.33969491 |
| AC008758.5 | 21.5066116 | -0.892854432 | 0.77377138 | -1.1538995 | 0.24854136 | 0.56111373 |
| AC008758.1 | 16.9812077 | -1.401063297 | 0.5601327 | -2.501306 | 0.01237362 | 0.09558545 |
| ZNF490 | 33.1886442 | -0.048197148 | 0.41495639 | -0.1161499 | 0.90753373 | 0.9700616 |
| ZNF791 | 56.9342986 | -0.532493826 | 0.3791889 | -1.404297 | 0.16023044 | 0.44945506 |
| DHPS | 13.2130799 | 0.426463592 | 0.44713696 | 0.95376503 | 0.3402026 | 0.65054327 |
| TNPO2 | 42.4001615 | -0.243294066 | 0.33783923 | -0.7201475 | 0.4714342 | 0.75776258 |
| TRIR | 37.101596 | -0.627068796 | 0.32247748 | -1.9445352 | 0.05183095 | 0.235032 |
| HOOK2 | 124.608627 | -1.045584277 | 0.37519291 | -2.7867912 | 0.00532328 | 0.05235679 |
| JUNB | 34.4704504 | -1.550702509 | 0.55028419 | -2.818003 | 0.00483234 | 0.04897467 |
| GCDH | 16.6228506 | -0.11795574 | 0.43506397 | -0.2711228 | 0.78629662 | 0.92322402 |
| FARSA | 19.3335753 | -0.298212091 | 0.42800723 | -0.6967454 | 0.48596211 | 0.76833754 |
| CALR | 56.4212051 | -1.170291521 | 0.49358437 | -2.3710061 | 0.01773974 | 0.12091034 |
| RAD23A | 28.8486635 | -0.348781759 | 0.30993765 | -1.1253288 | 0.26044973 | 0.5740977 |
| GADD45GIP1 | 19.0060073 | -1.022173604 | 0.49362423 | -2.0707525 | 0.03838193 | 0.19771996 |
| NFIX | 18.3032397 | 0.141024933 | 0.46249764 | 0.30492033 | 0.76042683 | 0.91286466 |
| NACC1 | 17.983004 | -0.687361521 | 0.41043559 | -1.6747123 | 0.0939907 | 0.3344198 |
| IER2 | 40.5743312 | -0.912625315 | 0.55315658 | -1.6498499 | 0.09897364 | 0.34359002 |
| CCDC130 | 70.1579583 | -0.003878439 | 0.24814121 | -0.01563 | 0.9875296 | 0.9977555 |
| MRI1 | 26.6700659 | 0.197870576 | 0.43426808 | 0.45564154 | 0.64864777 | 0.8599651 |
| C19orf53 | 22.1774983 | -0.942734995 | 0.49998443 | -1.8855287 | 0.05935849 | 0.25458077 |
| ZSWIM4 | 26.0825379 | -1.769068783 | 0.5045034 | -3.5065547 | 0.00045395 | 0.00801631 |
| CC2D1A | 36.1845413 | -0.258979488 | 0.28374749 | -0.9127111 | 0.3613945 | 0.66677568 |
| DCAF15 | 12.8118774 | -0.360977371 | 0.42124709 | -0.8569255 | 0.39148606 | 0.69435343 |
| RFX1 | 28.0571875 | -0.255938108 | 0.38099984 | -0.6717539 | 0.50174041 | 0.776256 |
| PRKACA | 30.3445116 | -0.06997638 | 0.46066504 | -0.151903 | 0.87926348 | 0.96035935 |
| ADGRL1 | 27.3768577 | -0.229419108 | 0.38749708 | -0.5920538 | 0.55381459 | 0.81056645 |
| PKN1 | 19.9228892 | -0.221856915 | 0.37809626 | -0.5867736 | 0.55735575 | 0.81230403 |
| GIPC1 | 58.366406 | -0.173959674 | 0.2667576 | -0.6521264 | 0.51431964 | 0.7835482 |
| DNAJB1 | 15.7238256 | -0.816181319 | 0.37506362 | -2.1761143 | 0.02954671 | 0.16798388 |
| TECR | 22.5777665 | 0.335432773 | 0.42343842 | 0.79216423 | 0.42826492 | 0.7236272 |
| NDUFB7 | 35.1205052 | 0.1719483 | 0.41081475 | 0.41855434 | 0.67554186 | 0.86960234 |
| ZNF333 | 12.8190979 | 0.087466228 | 0.55661486 | 0.15713958 | 0.87513484 | 0.95805533 |
| ILVBL | 17.7773956 | 0.499249659 | 0.43862637 | 1.13821169 | 0.2550321 | 0.56765799 |
| NOTCH3 | 27.415883 | -0.985256429 | 0.56313852 | -1.7495809 | 0.08019065 | 0.30394212 |
| BRD4 | 220.148675 | -0.518204772 | 0.24865537 | -2.0840281 | 0.03715761 | 0.19376823 |
| AKAP8 | 57.9693978 | -0.457337803 | 0.32212396 | -1.4197572 | 0.15567839 | 0.44123426 |
| AKAP8L | 110.923449 | 0.141643338 | 0.33550424 | 0.42218047 | 0.6728933 | 0.8684158 |
| TPM4 | 20.8092157 | -0.641940943 | 0.33465935 | -1.9181922 | 0.05508665 | 0.24411972 |
| RAB8A | 14.4164671 | 0.336026068 | 0.46379272 | 0.72451777 | 0.46874792 | 0.75516139 |
| AP1M1 | 17.1321384 | 0.309307996 | 0.39721877 | 0.77868425 | 0.43616574 | 0.72894951 |
| EPS15L1 | 86.1699053 | 0.343336857 | 0.34901709 | 0.98372505 | 0.32525072 | 0.63715682 |
| C19orf44 | 18.0076853 | 0.00354812 | 0.46346967 | 0.00765556 | 0.99389181 | 0.9992177 |
| TMEM38A | 32.8507715 | 0.88093075 | 0.41579601 | 2.11866091 | 0.03411913 | 0.18384695 |
| SIN3B | 67.9147418 | 0.674412641 | 0.28999789 | 2.32557773 | 0.02004109 | 0.13113201 |
| MYO9B | 84.3834296 | -1.270838562 | 0.50743051 | -2.5044583 | 0.01226391 | 0.09511578 |
| NR2F6 | 16.3574886 | -0.556873689 | 0.60671832 | -0.9178455 | 0.35869975 | 0.66504602 |
| MRPL34 | 21.8235591 | -0.692488298 | 0.58006163 | -1.1938185 | 0.23254898 | 0.54506955 |
| BST2 | 22.3967082 | -1.964321567 | 1.10920951 | -1.7709202 | 0.07657397 | 0.29596141 |
| MVB12A | 17.3591765 | -0.237359484 | 0.38000878 | -0.6246158 | 0.53222326 | 0.79692462 |
| SLC27A1 | 38.8951529 | 0.962683497 | 0.46589233 | 2.06632185 | 0.0387981 | 0.19898431 |
| AC010618.3 | 28.798323 | -0.867358705 | 0.41578819 | -2.086059 | 0.03697328 | 0.19323926 |
| PGLS | 20.6266721 | -0.701139076 | 0.42149807 | -1.6634455 | 0.09622329 | 0.33897502 |
| JUND | 89.5497812 | -1.165082444 | 0.7168177 | -1.6253539 | 0.10408717 | 0.3532977 |
| LSM4 | 22.094252 | -0.045828725 | 0.45919975 | -0.0998013 | 0.92050209 | 0.9754826 |
| PGPEP1 | 54.44801 | -0.934513941 | 0.3193882 | -2.9259501 | 0.00343406 | 0.03851337 |
| GDF15 | 192.902623 | -3.290000598 | 0.72214124 | -4.5558963 | 5.22E-06 | 0.0002231 |

| | | | | | | |
|---|---|---|---|---|---|---|
| SSBP4 | 23.7530074 | -0.949686021 | 0.39352278 | -2.4132936 | 0.01580908 | 0.11184941 |
| ISYNA1 | 24.3552787 | 1.842531214 | 0.62179154 | 2.96326195 | 0.00304397 | 0.03503952 |
| ELL | 58.848809 | 0.0292087 | 0.37354342 | 0.07819359 | 0.93767406 | 0.98289747 |
| FKBP8 | 27.7601721 | -0.132237047 | 0.53549635 | -0.2469429 | 0.80495238 | 0.93269243 |
| KXD1 | 20.1018255 | -0.66743097 | 0.46565202 | -1.4333256 | 0.1517648 | 0.43620176 |
| UBA52 | 37.2317112 | -0.307257387 | 0.43100406 | -0.7128874 | 0.47591541 | 0.76162498 |
| REX1BD | 12.6030995 | 0.634257731 | 0.7318218 | 0.8666833 | 0.38611556 | 0.68921713 |
| CRTC1 | 86.7706587 | 0.01461566 | 0.32713985 | 0.0446771 | 0.96436469 | 0.99082982 |
| UPF1 | 39.7280182 | -0.34606971 | 0.34601125 | -1.000169 | 0.31722875 | 0.63093651 |
| COPE | 22.6533352 | -0.421100542 | 0.4391606 | -0.958876 | 0.33762123 | 0.64844536 |
| DDX49 | 12.5164606 | -0.687292828 | 0.54672808 | -1.2571018 | 0.20871679 | 0.51711493 |
| SUGP2 | 110.836033 | 0.099291573 | 0.21914832 | 0.45307931 | 0.65049163 | 0.86113455 |
| SLC25A42 | 21.600332 | -0.222958775 | 0.5115909 | -0.4358146 | 0.66297126 | 0.86556259 |
| TMEM161A | 18.5818984 | 1.077984028 | 0.56186497 | 1.91858201 | 0.05503725 | 0.24401681 |
| SUGP1 | 28.0611424 | 0.237743203 | 0.3522366 | 0.67495316 | 0.49970553 | 0.77558432 |
| MAU2 | 71.2569459 | -0.184849514 | 0.27903119 | -0.662469 | 0.5076707 | 0.78007936 |
| GATAD2A | 155.305164 | -0.744075134 | 0.34927946 | -2.1303146 | 0.03314565 | 0.18155742 |
| ATP13A1 | 17.0328521 | -0.445681825 | 0.53270294 | -0.8366423 | 0.40279365 | 0.70169281 |
| ZNF101 | 13.6363729 | -0.437885882 | 0.41984362 | -1.0429738 | 0.29696044 | 0.61093809 |
| ZNF14 | 29.2794914 | 0.4591828 | 0.32294154 | 1.42187591 | 0.15506228 | 0.44065854 |
| AC011477.4 | 14.2834183 | -0.709092024 | 0.53098747 | -1.3354214 | 0.18173847 | 0.4801727 |
| ZNF253 | 37.6631882 | 0.28359337 | 0.32565166 | 0.87084883 | 0.3838367 | 0.68794623 |
| ZNF682 | 16.5234479 | -0.861505157 | 0.46558051 | -1.8503892 | 0.06425747 | 0.26796228 |
| ZNF90 | 41.7840696 | -1.002116438 | 0.51666534 | -1.9395852 | 0.05243012 | 0.23693425 |
| AC011447.3 | 18.2713901 | -1.463809445 | 0.49339664 | -2.9668006 | 0.00300916 | 0.0346816 |
| ZNF737 | 35.8590636 | -0.022750224 | 0.39069125 | -0.0582307 | 0.95356487 | 0.98771968 |
| ZNF85 | 15.8063624 | 0.241637235 | 0.57329697 | 0.42148703 | 0.67339948 | 0.86846218 |
| ZNF430 | 34.4459108 | -0.772218264 | 0.45421093 | -1.7001314 | 0.08910621 | 0.32396747 |
| ZNF714 | 22.1001773 | -0.422845882 | 0.47873709 | -0.8832528 | 0.37709969 | 0.68029685 |
| ZNF431 | 77.89282 | -0.290989707 | 0.40788495 | -0.7134112 | 0.47559132 | 0.76153417 |
| ZNF708 | 60.1284174 | -0.400569947 | 0.31336247 | -1.2782958 | 0.20114513 | 0.50511102 |
| ZNF493 | 37.4493734 | 1.018307111 | 0.40764941 | 2.49799727 | 0.01248972 | 0.09619433 |
| AC010615.4 | 58.419703 | -0.287355527 | 0.80856044 | -0.3553915 | 0.7222963 | 0.89217596 |
| ZNF429 | 48.7395399 | -0.187089354 | 0.33997003 | -0.5503113 | 0.58210586 | 0.82598616 |
| AC123912.1 | 19.2164243 | -0.299188732 | 0.56101848 | -0.5332957 | 0.59382892 | 0.83416944 |
| ZNF100 | 37.3653657 | -1.162971435 | 0.49777313 | -2.3363484 | 0.01947309 | 0.12922927 |
| ZNF43 | 57.707846 | -1.04996876 | 0.34314737 | -3.0598187 | 0.00221471 | 0.02727867 |
| ZNF208 | 72.1062048 | 0.114888356 | 0.58732821 | 0.19561185 | 0.84491399 | 0.94907846 |
| ZNF676 | 16.7887363 | 0.611444945 | 0.86853924 | 0.70399231 | 0.48143756 | 0.76608505 |
| ZNF98 | 86.9415929 | 0.63838394 | 0.36740851 | 1.73753171 | 0.08229336 | 0.30864976 |
| ZNF91 | 263.489214 | -0.569897108 | 0.49051842 | -1.1618261 | 0.2453061 | 0.55758089 |
| ZNF675 | 17.9458281 | 0.183285336 | 0.45573781 | 0.40217276 | 0.68755689 | 0.87556719 |
| ZNF254 | 154.918459 | -0.355780368 | 0.29672322 | -1.1990311 | 0.23051585 | 0.54344623 |
| LINC00662 | 58.8539851 | 0.413761055 | 0.24921466 | 1.66025971 | 0.09686222 | 0.33969491 |
| AC006504.5 | 46.5437557 | -0.11735796 | 0.50978238 | -0.2302119 | 0.81792713 | 0.93805075 |
| UQCRFS1 | 36.214405 | 0.028210789 | 0.40385853 | 0.06985315 | 0.94431055 | 0.98552865 |
| C19orf12 | 33.5659339 | -0.336181977 | 0.28941785 | -1.16158 | 0.24540613 | 0.55758089 |
| URI1 | 182.134945 | 0.31512454 | 0.19100514 | 1.64982234 | 0.09897928 | 0.34359002 |
| ZNF507 | 49.8084528 | 0.224372178 | 0.24507713 | 0.9155166 | 0.35992049 | 0.66572082 |
| DPY19L3 | 74.5638954 | -1.251759199 | 0.37896092 | -3.3031353 | 0.0009561 | 0.01442508 |
| PDCD5 | 19.051026 | 0.029032179 | 0.42807751 | 0.06781991 | 0.94592899 | 0.98574808 |
| ANKRD27 | 38.0073141 | 0.234214751 | 0.29025883 | 0.80691689 | 0.41971437 | 0.7181899 |
| NUDT19 | 14.4673769 | 0.933871632 | 0.42342207 | 2.20553368 | 0.02741667 | 0.16065021 |
| CEP89 | 34.7049094 | 0.047721899 | 0.39369219 | 0.12121627 | 0.90351974 | 0.96843523 |
| RHPN2 | 124.114875 | -1.906366398 | 0.40864386 | -4.6651047 | 3.08E-06 | 0.0001438 |
| GPATCH1 | 19.2411223 | -0.075974429 | 0.38320432 | -0.1982609 | 0.84284095 | 0.94799096 |
| PEPD | 33.2310912 | -0.23336558 | 0.46476692 | -0.5021131 | 0.61558793 | 0.8422143 |
| LSM14A | 241.25125 | 0.000954337 | 0.23487013 | 0.00406325 | 0.996758 | 0.99922221 |
| KIAA0355 | 141.334199 | -0.330949825 | 0.32263029 | -1.0257866 | 0.30499219 | 0.61860579 |
| GPI | 203.99804 | -0.498041944 | 0.33466474 | -1.4881817 | 0.13670298 | 0.41194526 |
| UBA2 | 94.5374746 | -0.094991144 | 0.26220332 | -0.3622805 | 0.71714244 | 0.89010971 |
| WTIP | 22.4802471 | 0.061778479 | 0.55237931 | 0.11184068 | 0.91094973 | 0.97128978 |
| SCGB2B2 | 74.7953067 | 0.162160957 | 0.38984829 | 0.41595914 | 0.67743991 | 0.87099416 |
| ZNF302 | 78.2459683 | -0.335988415 | 0.3627834 | -0.9261405 | 0.35437295 | 0.66109911 |
| ZNF181 | 13.2069918 | 0.223467095 | 0.39333598 | 0.56813285 | 0.56994476 | 0.81849045 |
| GRAMD1A | 55.3790465 | -1.610580442 | 0.36331594 | -4.4330025 | 9.29E-06 | 0.00036304 |
| HPN | 228.520086 | 0.922823812 | 0.47466509 | 1.94415774 | 0.05187643 | 0.235032 |
| LSR | 32.3946939 | -1.462502169 | 0.38707415 | -3.7783514 | 0.00015787 | 0.00347986 |
| USF2 | 25.3576215 | -0.042971576 | 0.49559086 | -0.0867078 | 0.9309038 | 0.97899245 |
| DMKN | 47.8155775 | -0.514200054 | 0.49375893 | -1.041399 | 0.29769041 | 0.61141484 |
| TMEM147 | 16.6036453 | 0.091436334 | 0.55677115 | 0.16422606 | 0.86955318 | 0.95621002 |
| COX6B1 | 59.0946463 | 0.459481835 | 0.44141111 | 1.04093853 | 0.29790408 | 0.61141484 |
| KMT2B | 30.4206816 | -0.704983151 | 0.38035312 | -1.8534964 | 0.06381124 | 0.2665663 |
| PROSER3 | 17.6541703 | -0.417360489 | 0.4143591 | -1.0072434 | 0.3138178 | 0.62884777 |
| TBCB | 16.8571725 | -0.526192528 | 0.40452647 | -1.3007617 | 0.19334004 | 0.49484012 |
| CAPNS1 | 47.5054356 | -0.7563406 | 0.43221603 | -1.7499134 | 0.08013326 | 0.30394212 |
| ZNF565 | 60.0902181 | 0.053397686 | 0.40127845 | 0.13306891 | 0.89413888 | 0.96482804 |
| ZNF146 | 77.737052 | 0.090827772 | 0.37967811 | 0.23922309 | 0.8109326 | 0.93552094 |
| LINC00665 | 29.3897702 | -0.118682729 | 0.54195015 | -0.218992 | 0.82665631 | 0.9422064 |
| ZFP14 | 56.9224146 | -0.377254712 | 0.42993717 | -0.8774648 | 0.38023425 | 0.68428956 |
| ZFP82 | 31.0851098 | -0.023842437 | 0.34619891 | -0.0688692 | 0.94509375 | 0.98552865 |
| ZNF566 | 31.789058 | 0.226125553 | 0.41420242 | 0.54593006 | 0.58511401 | 0.82874116 |
| ZNF260 | 25.4027701 | 0.094848561 | 0.34110139 | 0.27806559 | 0.78096201 | 0.92083964 |
| ZNF529 | 46.9169663 | 0.625945175 | 0.28326062 | 2.20978541 | 0.02712006 | 0.15973937 |
| ZNF529-AS1 | 37.1942038 | -0.187731973 | 0.35532142 | -0.5283441 | 0.59726053 | 0.8342658 |
| ZNF382 | 14.9733077 | 0.392051335 | 0.46729956 | 0.8389722 | 0.40148491 | 0.70114138 |
| ZNF461 | 28.1272013 | -0.10954228 | 0.30789848 | -0.355774 | 0.72200981 | 0.89217596 |
| ZNF567 | 25.7513504 | -0.003016251 | 0.38496376 | -0.0078352 | 0.99374852 | 0.9992177 |
| AC020928.1 | 48.3093532 | 0.50144736 | 0.28721084 | 1.74592075 | 0.08082472 | 0.30571131 |
| ZNF790 | 43.1914736 | -0.542470019 | 0.3747731 | -1.4474625 | 0.14776743 | 0.43123114 |
| ZNF345 | 71.9094794 | 0.083581527 | 0.37983882 | 0.22004472 | 0.82583633 | 0.94197405 |
| ZNF568 | 42.5760593 | -0.172467547 | 0.32611753 | -0.5288509 | 0.59690889 | 0.83416944 |
| ZNF420 | 74.3534174 | 0.374321895 | 0.32972836 | 1.13524324 | 0.25627342 | 0.56906248 |
| ZNF383 | 23.49809 | 0.679779919 | 0.43320531 | 1.56918647 | 0.11660449 | 0.37568083 |
| LINC01535 | 15.3426611 | 0.078538691 | 0.42778012 | 0.18359594 | 0.85433045 | 0.95170432 |
| HKR1 | 79.8602023 | 0.488907456 | 0.33609985 | 1.45464941 | 0.14576634 | 0.42828171 |
| ZNF569 | 25.5256305 | -0.376602861 | 0.38910626 | -0.9678664 | 0.33311111 | 0.64492217 |
| ZNF793 | 39.3043509 | 0.473811358 | 0.31844521 | 1.48788972 | 0.13677998 | 0.41194526 |
| ZNF540 | 29.8291087 | 0.327892289 | 0.51172682 | 0.64075651 | 0.52168089 | 0.7892508 |
| ZNF571 | 33.305068 | 0.103922232 | 0.43382429 | 0.23954913 | 0.8106798 | 0.93552094 |
| ZFP30 | 37.576269 | -0.485163686 | 0.41336059 | -1.1737057 | 0.24051292 | 0.55302079 |
| ZNF573 | 50.337284 | 0.612027716 | 0.3941626 | 1.55272903 | 0.12048788 | 0.38381586 |
| SIPA1L3 | 253.284065 | -0.110274694 | 0.22321642 | -0.4940259 | 0.62128789 | 0.84470972 |
| SPINT2 | 23.6602488 | 0.308256364 | 0.31009514 | 0.99407028 | 0.32018865 | 0.63212442 |
| CATSPERG | 26.0777397 | -0.881560319 | 0.61474588 | -1.4340239 | 0.15156543 | 0.43603704 |
| PSMD8 | 33.7675559 | -0.490730148 | 0.44342227 | -1.1066881 | 0.2684288 | 0.5846368 |
| SPRED3 | 19.0852142 | -4.961581817 | 0.67446254 | -7.356349 | 1.89E-13 | 7.66E-11 |
| EIF3K | 66.0806103 | 0.898552062 | 0.2540008 | 3.5375954 | 0.00040379 | 0.00741126 |
| ACTN4 | 291.827401 | -0.622222604 | 0.43601279 | -1.4270742 | 0.15355851 | 0.43892689 |
| ECH1 | 19.077063 | -0.586702624 | 0.61978428 | -0.9466239 | 0.34383045 | 0.65306073 |
| HNRNPL | 70.1387156 | -0.199496211 | 0.38556372 | -0.5174144 | 0.60486693 | 0.83778698 |
| SIRT2 | 32.2153061 | 0.199722551 | 0.5114828 | 0.39047755 | 0.69618345 | 0.88088133 |
| MRPS12 | 18.7295694 | 1.073552753 | 0.43426788 | 2.47209796 | 0.01343227 | 0.10067724 |

| | | | | | | |
|---|---|---|---|---|---|---|
| FBXO17 | 139.572225 | -0.001339453 | 0.45600517 | -0.0029374 | 0.99765633 | 0.99922221 |
| PAK4 | 72.0650983 | 0.502279574 | 0.37065088 | 1.35512852 | 0.17537661 | 0.47219507 |
| SAMD4B | 86.5594988 | -0.191842633 | 0.24055965 | -0.7974847 | 0.4251696 | 0.72156559 |
| MED29 | 13.6765353 | -0.578226472 | 0.42793034 | -1.3512163 | 0.17662614 | 0.47378082 |
| ZFP36 | 27.8159453 | -1.638097844 | 0.81049921 | -2.0210974 | 0.04326968 | 0.21173421 |
| RPS16 | 48.9988862 | -0.817791079 | 0.43323666 | -1.8876313 | 0.05907546 | 0.25401736 |
| SUPT5H | 64.0277673 | -0.241968234 | 0.27889332 | -0.8676014 | 0.38561258 | 0.68921713 |
| TIMM50 | 13.5719041 | -0.062816936 | 0.54636596 | -0.1149723 | 0.9084671 | 0.97059151 |
| FBL | 11.6074793 | 0.149627604 | 0.45759708 | 0.32698549 | 0.74367886 | 0.90463949 |
| PSMC4 | 18.521518 | -0.327490907 | 0.43022419 | -0.7612099 | 0.44653172 | 0.73948328 |
| ZNF546 | 19.1204267 | -0.164974006 | 0.4395951 | -0.3752863 | 0.70744757 | 0.88614996 |
| ZNF780B | 119.701257 | 0.482348923 | 0.33498472 | 1.43991319 | 0.14989196 | 0.43416982 |
| ZNF780A | 13.9546094 | 0.426575911 | 0.52395158 | 0.81415139 | 0.41555823 | 0.71304102 |
| MAP3K10 | 14.1952651 | 0.407427943 | 0.53981687 | 0.75475215 | 0.45039771 | 0.74156119 |
| AKT2 | 66.5614194 | 0.271080913 | 0.30724004 | 0.88230985 | 0.37760927 | 0.68087968 |
| PLD3 | 90.1761405 | 0.013016965 | 0.25829763 | 0.05039521 | 0.95980745 | 0.9895734 |
| BLVRB | 42.5023299 | -0.843435711 | 0.3452062 | -2.4432809 | 0.0145544 | 0.1059672 |
| SPTBN4 | 29.8822703 | -0.229895053 | 0.90460955 | -0.2541373 | 0.79938947 | 0.92922871 |
| COQ8B | 17.2240014 | -0.793770374 | 0.4575909 | -1.7346725 | 0.08279882 | 0.30942534 |
| ITPKC | 31.586813 | -1.658176271 | 0.46039034 | -3.6016748 | 0.00031617 | 0.00607836 |
| SNRPA | 19.258146 | -0.067987086 | 0.43398047 | -0.1566593 | 0.87551335 | 0.95835718 |
| AC008537.1 | 27.0243176 | -1.014555089 | 0.78563202 | -1.2913871 | 0.19656947 | 0.49845696 |
| HNRNPUL1 | 54.0675958 | -0.414685375 | 0.31117431 | -1.3326466 | 0.18264781 | 0.4814838 |
| TGFB1 | 21.2221015 | -1.32637852 | 0.44631956 | -2.9718136 | 0.00296046 | 0.03430713 |
| DMAC2 | 11.7713637 | 0.575505063 | 0.55874485 | 1.02999619 | 0.30301179 | 0.61741302 |
| RPS19 | 133.771834 | -1.125571494 | 0.45748769 | -2.4603317 | 0.01388086 | 0.10308054 |
| ARHGEF1 | 26.9419412 | -0.201852961 | 0.33885724 | -0.5956873 | 0.55138413 | 0.80885866 |
| RABAC1 | 23.1138169 | -0.455587718 | 0.57575168 | -0.791292 | 0.4287736 | 0.7236272 |
| GRIK5 | 54.4882508 | 0.343969181 | 0.44624287 | 0.7708116 | 0.44081861 | 0.73302253 |
| CIC | 20.5152528 | -0.826749842 | 0.3909378 | -2.1147861 | 0.03444818 | 0.18459472 |
| MEGF8 | 29.1565584 | -0.250042587 | 0.36133833 | -0.6919902 | 0.48894348 | 0.77060666 |
| LIPE-AS1 | 20.7636028 | -0.380295651 | 0.50087574 | -0.7592615 | 0.44769616 | 0.7399821 |
| PHLDB3 | 13.5659743 | 0.029540494 | 0.40382017 | 0.0731526 | 0.94168469 | 0.98455573 |
| ETHE1 | 18.8510169 | -0.524387798 | 0.38602711 | -1.3584222 | 0.17432975 | 0.47073576 |
| ZNF428 | 18.6449291 | -0.489102089 | 0.42886372 | -1.1404604 | 0.25409455 | 0.56624703 |
| SMG9 | 13.8618578 | -0.147443269 | 0.38813366 | -0.3798776 | 0.7040363 | 0.8856427 |
| ZNF283 | 26.248665 | 0.605788506 | 0.36840414 | 1.64435855 | 0.10010216 | 0.34555815 |
| ZNF45 | 12.9716334 | -0.001357687 | 0.4659533 | -0.0029138 | 0.99767514 | 0.99922221 |
| ZNF155 | 26.7833568 | -0.245650487 | 0.48662421 | -0.5048053 | 0.61369558 | 0.84174925 |
| ZNF224 | 37.8377214 | -0.162569078 | 0.28978511 | -0.5609987 | 0.57479841 | 0.82146824 |
| ZNF234 | 18.4616804 | 0.248553969 | 0.49875481 | 0.49834902 | 0.61823806 | 0.84283824 |
| ZNF226 | 28.8527458 | 0.703385039 | 0.29600606 | 2.37625217 | 0.01748951 | 0.11999422 |
| ZNF227 | 22.3894571 | 0.397679242 | 0.33911607 | 1.1726936 | 0.2409187 | 0.55327012 |
| ZNF235 | 87.5469089 | 0.108245037 | 0.44759921 | 0.24183474 | 0.80890822 | 0.93433178 |
| ZNF229 | 16.7371822 | -0.491134926 | 0.47259677 | -1.0392262 | 0.29869957 | 0.61250819 |
| ZNF180 | 14.5089699 | 0.334410972 | 0.43599318 | 0.76700963 | 0.44307579 | 0.73535754 |
| AC243964.2 | 31.6552417 | -1.124136601 | 0.4646375 | -2.4193842 | 0.01554681 | 0.11082452 |
| CEACAM19 | 27.6718503 | -2.911694052 | 0.74080381 | -3.9304523 | 8.48E-05 | 0.0021424 |
| BCL3 | 19.7450622 | -3.43432807 | 0.67134503 | -5.1155932 | 3.13E-07 | 2.16E-05 |
| CBLC | 18.9404415 | -0.933042447 | 0.44933526 | -2.076495 | 0.03784819 | 0.19628857 |
| BCAM | 111.855283 | -1.285939019 | 0.52590859 | -2.4451759 | 0.01447815 | 0.10579489 |
| NECTIN2 | 78.4587433 | -0.169628231 | 0.35085527 | -0.4834707 | 0.62876158 | 0.84817317 |
| TOMM40 | 15.6252276 | -0.560177094 | 0.54980018 | -1.018874 | 0.3082628 | 0.62213389 |
| APOE | 66.5300739 | -0.04422031 | 0.69959149 | -0.0632088 | 0.94960027 | 0.98641632 |
| CLPTM1 | 38.6063868 | -0.394716699 | 0.3586889 | -1.100443 | 0.27113914 | 0.58752064 |
| RELB | 63.1330864 | -2.620853019 | 0.67671777 | -3.8728893 | 0.00010755 | 0.00257134 |
| CLASRP | 65.9059129 | -0.173188801 | 0.28737425 | -0.6026594 | 0.54673528 | 0.80581909 |
| GEMIN7-AS1 | 20.3868014 | -0.359016675 | 0.44640475 | -0.8042403 | 0.42125823 | 0.7192477 |
| MARK4 | 58.1084218 | 0.05230221 | 0.30659702 | 0.17058943 | 0.86454661 | 0.95487238 |
| TRAPPC6A | 15.5199939 | 0.386684925 | 0.44906309 | 0.86109265 | 0.38918701 | 0.69247382 |
| PPP1R13L | 38.6129214 | -1.446952979 | 0.35833946 | -4.0379393 | 5.39E-05 | 0.00151439 |
| ERCC1 | 24.6342111 | -1.186878607 | 0.68200424 | -1.7402804 | 0.08180979 | 0.30770251 |
| VASP | 31.3692082 | -1.037757757 | 0.62660161 | -1.6561684 | 0.09768773 | 0.34062841 |
| OPA3 | 64.0303766 | -0.168574179 | 0.39395988 | -0.4278968 | 0.66872624 | 0.86777458 |
| EML2 | 43.4925941 | -0.2258628 | 0.35570642 | -0.6349697 | 0.52544817 | 0.79289185 |
| SNRPD2 | 24.8736224 | -0.688017857 | 0.43366743 | -1.5865103 | 0.11262359 | 0.36994129 |
| FBXO46 | 27.9580696 | -0.295874565 | 0.46023493 | -0.6428773 | 0.52030375 | 0.78804331 |
| DMPK | 16.3184792 | -0.638589192 | 0.36482331 | -1.7504068 | 0.08004815 | 0.30377247 |
| SYMPK | 88.6416342 | -0.138829586 | 0.32369349 | -0.4288921 | 0.66800174 | 0.86777458 |
| MYPOP | 26.2735026 | 0.632520293 | 0.48189341 | 1.31257302 | 0.18932687 | 0.48954068 |
| PPP5C | 41.7997782 | 0.426513263 | 0.27485071 | 1.55179974 | 0.12071015 | 0.38413018 |
| PPP5D1 | 21.2594905 | 0.587665175 | 0.54382269 | 1.08061908 | 0.27986659 | 0.59397852 |
| CALM3 | 19.2916599 | -0.465242234 | 0.61348022 | -0.7583655 | 0.44823221 | 0.7399821 |
| PRKD2 | 40.4677107 | -0.662091066 | 0.31535807 | -2.0994898 | 0.03577374 | 0.18887564 |
| STRN4 | 18.150745 | -0.145365726 | 0.39548893 | -0.3675595 | 0.71320169 | 0.88807326 |
| FKRP | 17.7066168 | -0.126188212 | 0.5166113 | -0.2442614 | 0.80702836 | 0.93335803 |
| SLC1A5 | 17.5574533 | -0.19523632 | 0.54558538 | -0.3578474 | 0.7204575 | 0.89151237 |
| AP2S1 | 20.9336006 | -0.23436985 | 0.49156109 | -0.4767868 | 0.63351393 | 0.8520131 |
| ARHGAP35 | 108.255168 | -0.081816135 | 0.32129889 | -0.2546418 | 0.79899976 | 0.92922871 |
| TMEM160 | 22.0038462 | -0.561964615 | 0.62346282 | -0.9013603 | 0.3673968 | 0.6714245 |
| ZC3H4 | 29.4018437 | 0.111554357 | 0.36488546 | 0.30572431 | 0.75981456 | 0.91236458 |
| SAE1 | 30.7559158 | -0.379850757 | 0.39246348 | -0.9678627 | 0.33311295 | 0.64492217 |
| CCDC9 | 14.2529974 | -0.845098163 | 0.47329282 | -1.7855715 | 0.07416867 | 0.29129403 |
| C5AR1 | 38.1670529 | 1.230995717 | 0.4475184 | 2.75071534 | 0.00594653 | 0.05634699 |
| DHX34 | 33.6413571 | -0.453802276 | 0.38740938 | -1.1713766 | 0.24144744 | 0.55404626 |
| NAPA-AS1 | 15.3809805 | 0.447609534 | 0.54247373 | 0.82512665 | 0.40929969 | 0.70777268 |
| NAPA | 73.0635082 | -0.559586585 | 0.42043364 | -1.3309748 | 0.1831973 | 0.48210665 |
| NOP53 | 12.8806978 | -0.284704696 | 0.5118205 | -0.5562589 | 0.57803389 | 0.82311488 |
| SELENOW | 60.5987746 | -0.157844833 | 0.43867625 | -0.3598208 | 0.71898116 | 0.89077556 |
| PLA2G4C | 38.6237312 | -4.709046503 | 1.42782319 | -3.29806 | 0.00097355 | 0.01464099 |
| ZSWIM9 | 15.2574601 | -0.112591108 | 0.43623721 | -0.2580961 | 0.79633277 | 0.92719864 |
| CARD8 | 38.5771447 | 0.207367374 | 0.41395228 | 0.50094512 | 0.61640975 | 0.84223226 |
| TMEM143 | 21.5793927 | 0.803759963 | 0.42023809 | 1.91262995 | 0.05579544 | 0.24602764 |
| KDELR1 | 27.759229 | -0.394864039 | 0.31602016 | -1.24949 | 0.2114859 | 0.52056351 |
| CYTH2 | 29.1706671 | -0.941604111 | 0.37084347 | -2.5390878 | 0.01111419 | 0.08842043 |
| LMTK3 | 16.2684772 | -0.015545144 | 0.38055251 | -0.0408489 | 0.96741637 | 0.99156233 |
| RPL18 | 61.1754226 | -0.396960843 | 0.47781565 | -0.8307824 | 0.40609655 | 0.70524199 |
| BCAT2 | 25.8705942 | -0.355854844 | 0.52451607 | -0.6784441 | 0.49749014 | 0.77542596 |
| HSD17B14 | 27.3300956 | -0.646091049 | 0.68070057 | -0.949156 | 0.34254126 | 0.65223891 |
| NUCB1 | 48.921442 | -0.320545493 | 0.36851138 | -0.8698388 | 0.38438851 | 0.68822563 |
| BAX | 17.1435964 | -0.51941214 | 0.50165817 | -1.0353906 | 0.30048654 | 0.61414654 |
| FTL | 629.435705 | -1.58796198 | 0.57075212 | -2.782227 | 0.00539873 | 0.05276491 |
| GYS1 | 59.9210039 | -0.266319657 | 0.3660758 | -0.7274987 | 0.46692053 | 0.75315494 |
| SNRNP70 | 187.50541 | 0.456496888 | 0.35317245 | 1.29256088 | 0.19616298 | 0.49845696 |
| TRPM4 | 42.6126681 | 0.805995214 | 0.5410672 | 1.48963977 | 0.13631898 | 0.41138221 |
| SLC6A16 | 72.994305 | 0.467671473 | 0.27243548 | 1.71663204 | 0.08604642 | 0.3168629 |
| TEAD2 | 13.7388389 | -1.272792953 | 0.44002606 | -2.89254 | 0.00382141 | 0.04152772 |
| PIH1D1 | 20.5706028 | -0.274067391 | 0.41788508 | -0.6558439 | 0.51192455 | 0.78271634 |
| RPL13A | 83.2717139 | -0.634780628 | 0.46461228 | -1.3662588 | 0.17185774 | 0.46763314 |
| RPS11 | 68.1814374 | -0.64664829 | 0.45777875 | -1.4125782 | 0.15777978 | 0.44458709 |
| FCGRT | 22.940994 | 0.093267707 | 0.51007794 | 0.18284991 | 0.85491578 | 0.95179231 |
| NOSIP | 32.9190129 | -0.164727207 | 0.38707573 | -0.4255684 | 0.67042236 | 0.86777458 |
| PRR12 | 24.9750748 | 0.426286408 | 0.47273522 | 0.90174454 | 0.36719259 | 0.67131445 |

| | | | | | | |
|---|---|---|---|---|---|---|
| SCAF1 | 20.126751 | 0.254664538 | 0.40413831 | 0.63014204 | 0.52860166 | 0.79434813 |
| IRF3 | 42.1061635 | -0.755271354 | 0.40593735 | -1.8605614 | 0.06280614 | 0.26390468 |
| PRMT1 | 32.6902068 | 0.576148433 | 0.41970155 | 1.37275747 | 0.16982776 | 0.46436188 |
| AP2A1 | 26.0473418 | -0.393700239 | 0.4828657 | -0.8153411 | 0.4148771 | 0.7123967 |
| MED25 | 34.3189644 | -0.091396719 | 0.4029765 | -0.2268041 | 0.82057608 | 0.9391876 |
| PTOV1 | 37.5519947 | -0.156277857 | 0.26736715 | -0.5845066 | 0.55887954 | 0.81279041 |
| PNKP | 37.2812408 | -1.218400177 | 0.52365599 | -2.3267187 | 0.01998024 | 0.13082568 |
| TBC1D17 | 29.7995479 | -0.529596795 | 0.45296335 | -1.1691824 | 0.24233014 | 0.55461125 |
| VRK3 | 39.6957245 | -0.278821567 | 0.34943908 | -0.7979118 | 0.42492167 | 0.72151771 |
| MYH14 | 37.5865059 | -0.484662226 | 0.3288795 | -1.4736772 | 0.14056852 | 0.42021831 |
| NR1H2 | 23.5520033 | -0.294344202 | 0.33583291 | -0.8764603 | 0.38077987 | 0.68461077 |
| EMC10 | 45.876518 | -0.469940132 | 0.30790729 | -1.5262391 | 0.12695034 | 0.39482486 |
| ETFB | 30.2500187 | -0.247657974 | 0.60881865 | -0.4067845 | 0.68416629 | 0.87360405 |
| ZNF175 | 19.6014354 | 0.458770319 | 0.35812758 | 1.28102484 | 0.20018494 | 0.50411966 |
| SPACA6 | 28.5356647 | -0.143847213 | 0.39767273 | -0.3617226 | 0.71755934 | 0.89028919 |
| ZNF577 | 57.1134103 | 0.662102868 | 0.43487367 | 1.52251771 | 0.12787941 | 0.39626043 |
| ZNF350-AS1 | 45.8514242 | 0.134736744 | 0.65796264 | 0.20477872 | 0.83774502 | 0.94722424 |
| ZNF350 | 21.7345465 | 0.445381705 | 0.41781414 | 1.06598046 | 0.28643247 | 0.60056136 |
| ZNF615 | 18.9180145 | 0.862977688 | 0.45807967 | 1.88390307 | 0.05957809 | 0.25511226 |
| ZNF432 | 38.8161338 | -0.52872729 | 0.44165656 | -1.1971458 | 0.23124974 | 0.54389834 |
| ZNF616 | 13.1230983 | -0.098670649 | 0.42688263 | -0.2311423 | 0.81720422 | 0.93768302 |
| PPP2R1A | 77.5801322 | -0.412033508 | 0.35071024 | -1.1748545 | 0.24005292 | 0.55251875 |
| ZNF766 | 37.0726354 | -0.19271693 | 0.35031697 | -0.5501216 | 0.58223598 | 0.82604508 |
| ZNF480 | 29.084487 | -0.480517785 | 0.49028821 | -0.9800721 | 0.32705054 | 0.63862184 |
| ZNF528 | 25.4888994 | 0.31217665 | 0.36374406 | 0.85823161 | 0.39076459 | 0.69372012 |
| ZNF808 | 28.6466956 | 0.350272428 | 0.34781034 | 1.00707883 | 0.31389689 | 0.62884777 |
| ZNF701 | 43.625858 | 0.330513248 | 0.38821744 | 0.85136116 | 0.39456876 | 0.69642725 |
| ZNF83 | 357.934802 | -1.599195739 | 0.82121132 | -1.947362 | 0.05149135 | 0.23455111 |
| ZNF611 | 151.007403 | -1.394058212 | 0.70149883 | -1.9872566 | 0.04689396 | 0.22193782 |
| ZNF28 | 19.1197709 | -0.069923542 | 0.48177472 | -0.1451374 | 0.88460237 | 0.961983 |
| AC010487.3 | 17.6972256 | -0.517781848 | 0.592147 | -0.8744144 | 0.38189263 | 0.68581797 |
| ZNF160 | 87.5930236 | -0.571509438 | 0.34808969 | -1.6418454 | 0.10062206 | 0.34632707 |
| ZNF347 | 29.9963864 | -0.570471622 | 0.39491174 | -1.4445547 | 0.148583 | 0.43239322 |
| ZNF677 | 16.5172943 | -0.428075328 | 0.50813234 | -0.8424485 | 0.39953695 | 0.69959441 |
| ZNF525 | 25.8179603 | -0.054291222 | 0.30292513 | -0.1792232 | 0.85776242 | 0.95266335 |
| ZNF765 | 32.167822 | -0.393347785 | 0.35552997 | -1.1063702 | 0.2685663 | 0.58479966 |
| ZNF761 | 36.9281785 | -0.430481911 | 0.32890848 | -1.3088197 | 0.19059542 | 0.49135864 |
| ZNF331 | 43.8425991 | 0.084991846 | 0.2851547 | 0.29805521 | 0.76566102 | 0.91549415 |
| PRPF31 | 26.6540045 | -0.306663773 | 0.30781206 | -0.9962695 | 0.31911921 | 0.63100017 |
| CNOT3 | 22.9178583 | -0.310115292 | 0.4928281 | -0.6292565 | 0.52918113 | 0.79492265 |
| TMC4 | 53.3344131 | 0.27404843 | 0.33499107 | 0.81807683 | 0.41331332 | 0.71143975 |
| MBOAT7 | 45.0080382 | -1.016685083 | 0.41770321 | -2.4339891 | 0.01493344 | 0.10793754 |
| RPS9 | 120.617356 | -0.765235406 | 0.48055394 | -1.5924027 | 0.11129424 | 0.36720011 |
| LENG8-AS1 | 16.749606 | 0.034609111 | 0.37501567 | 0.09228711 | 0.92646993 | 0.97735117 |
| LENG8 | 125.776523 | 0.405505878 | 0.4948087 | 0.8195205 | 0.41248951 | 0.71110141 |
| AC011476.3 | 18.501048 | -0.868075148 | 0.69293558 | -1.2527501 | 0.21029665 | 0.51886901 |
| RDH13 | 41.1825758 | 0.543587144 | 0.35312826 | 1.53934761 | 0.12371946 | 0.38887694 |
| EPS8L1 | 16.4389858 | 0.259823259 | 0.67827417 | 0.38306524 | 0.70167139 | 0.88424653 |
| PPP1R12C | 45.1751088 | 0.953200205 | 0.39569359 | 2.40893519 | 0.01599914 | 0.11284113 |
| PPP6R1 | 31.2036454 | -0.429597232 | 0.39426513 | -1.0896151 | 0.27588273 | 0.59079711 |
| HSPBP1 | 12.5815383 | 0.504864908 | 0.45974754 | 1.0981351 | 0.2721455 | 0.58786226 |
| RPL28 | 91.1871711 | -0.966717266 | 0.46675802 | -2.0711315 | 0.0383465 | 0.19764666 |
| U2AF2 | 25.8868257 | 0.014650812 | 0.38606407 | 0.03794917 | 0.96972821 | 0.99156233 |
| EPN1 | 55.5016615 | -0.027935169 | 0.34101923 | -0.0819167 | 0.93471296 | 0.98085023 |
| ZNF787 | 21.8228162 | -0.487438624 | 0.47519237 | -1.0257711 | 0.30499947 | 0.61860579 |
| ZNF444 | 22.8372828 | -0.166159477 | 0.37422072 | -0.4440146 | 0.657032 | 0.86325042 |
| ZSCAN5A | 96.9214591 | 0.216124577 | 0.27208715 | 0.79432113 | 0.4270085 | 0.72258772 |
| ZNF667 | 25.5532556 | -0.006396358 | 0.33642814 | -0.0190126 | 0.98483109 | 0.99742652 |
| ZNF667-AS1 | 22.511465 | 0.232209063 | 0.34426566 | 0.67450544 | 0.49999003 | 0.77558432 |
| ZNF471 | 26.7147235 | 0.031930418 | 0.37404978 | 0.08536409 | 0.93197194 | 0.97938863 |
| ZNF264 | 36.6646116 | -0.247458726 | 0.30763914 | -0.8043799 | 0.42117763 | 0.7192477 |
| ZNF460-AS1 | 17.8226758 | 0.450124995 | 0.48032206 | 0.93713162 | 0.34869086 | 0.65587927 |
| ZNF460 | 18.9798877 | -1.114851348 | 0.7643088 | -1.45864 | 0.14466423 | 0.42644619 |
| AC005261.1 | 59.2660103 | -1.065737198 | 0.52760291 | -2.0199608 | 0.04338745 | 0.21196707 |
| AC003002.2 | 13.9541076 | 0.834560062 | 0.45697649 | 1.82626476 | 0.06781039 | 0.27788783 |
| AC004076.1 | 28.096686 | -0.350175892 | 0.42446073 | -0.8249901 | 0.40937722 | 0.70777549 |
| ZNF549 | 19.9215511 | -0.257786282 | 0.3223678 | -0.7996651 | 0.42390486 | 0.72151771 |
| ZNF550 | 10.6525132 | 0.75327013 | 0.42642896 | 1.76646101 | 0.07731854 | 0.29764569 |
| AC003006.1 | 14.2951392 | 0.422843576 | 0.43016624 | 0.98297713 | 0.3256187 | 0.63742782 |
| ZNF154 | 14.9291656 | 0.934715236 | 0.46965737 | 1.9902067 | 0.04656817 | 0.22165096 |
| ZNF586 | 147.797234 | -0.077871486 | 0.28918925 | -0.2692752 | 0.78771793 | 0.92377626 |
| ZNF552 | 65.1611829 | 0.646281259 | 0.44504434 | 1.45217273 | 0.14645358 | 0.42900824 |
| ZNF814 | 80.1959571 | -0.045892846 | 0.37071372 | -0.1237959 | 0.90147686 | 0.96791458 |
| ZNF587 | 38.8728163 | 0.501845012 | 0.28801906 | 1.74240212 | 0.08143811 | 0.30704767 |
| ZNF418 | 15.2252637 | 0.285110927 | 0.44253921 | 0.64426139 | 0.51940595 | 0.78759816 |
| C19orf18 | 14.6896998 | 0.434406156 | 0.47144592 | 0.9214337 | 0.35682404 | 0.66341523 |
| ZNF606 | 19.9874885 | 0.223097178 | 0.39679757 | 0.56224432 | 0.57394957 | 0.82117002 |
| ZNF135 | 39.9096302 | 0.452319061 | 0.39482444 | 1.14562072 | 0.25195212 | 0.56438823 |
| ZSCAN18 | 66.0790309 | -0.574446604 | 0.37047684 | -1.5505601 | 0.12100714 | 0.38441929 |
| ZNF329 | 30.4407044 | -0.050129506 | 0.33895279 | -0.1478952 | 0.88242544 | 0.96161232 |
| ZNF274 | 31.621118 | -0.687376076 | 0.49204795 | -1.3969697 | 0.16242268 | 0.45295218 |
| AC020915.3 | 19.9361606 | 0.333329553 | 0.35328554 | 0.94351315 | 0.34541849 | 0.65406518 |
| RPS5 | 36.9503927 | -0.906451292 | 0.51527094 | -1.7591741 | 0.07854794 | 0.29970982 |
| TRIM28 | 27.5241967 | -0.611254986 | 0.51372066 | -1.1898587 | 0.23410194 | 0.54623787 |
| CHMP2A | 16.7201531 | -0.477442122 | 0.42106311 | -1.1338968 | 0.25683783 | 0.56936661 |
| UBE2M | 15.7235784 | -0.084392469 | 0.46462698 | -0.1816349 | 0.85586927 | 0.9521686 |
| MZF1-AS1 | 15.4231221 | -0.434318988 | 0.47574946 | -0.9129154 | 0.36128706 | 0.66677568 |
| MZF1 | 64.8237459 | -0.399334503 | 0.26237572 | -1.5219949 | 0.12801036 | 0.39653442 |
| RPS4Y1 | 50.5808917 | -0.550072867 | 2.01289895 | -0.273274 | 0.78464263 | 0.92261923 |
| ZFY | 30.60898 | 0.494095616 | 2.38453999 | 0.20720794 | 0.83584747 | 0.94654164 |
| LINC00278 | 97.5651367 | 0.170576038 | 2.65479455 | 0.06425207 | 0.94876952 | 0.98641632 |
| TBL1Y | 59.8463455 | 0.604545911 | 2.49381843 | 0.24241777 | 0.80845646 | 0.93433178 |
| TTTY15 | 35.5959816 | 0.282087408 | 2.40365299 | 0.11735779 | 0.90657653 | 0.96989619 |
| USP9Y | 161.435456 | 0.248993654 | 2.50655304 | 0.09933708 | 0.92087064 | 0.9754826 |
| DDX3Y | 36.2594854 | -0.368167092 | 2.40907869 | -0.1528249 | 0.8785364 | 0.96013757 |
| UTY | 352.431842 | 0.272521869 | 2.93556427 | 0.09283458 | 0.92603498 | 0.97735117 |
| NLGN4Y | 208.933368 | -1.768008457 | 2.52334427 | -0.7006608 | 0.48351471 | 0.76671619 |
| TTTY14 | 244.755253 | 0.170421672 | 1.39510281 | 0.12215707 | 0.90277462 | 0.96803829 |
| KDM5D | 41.6802319 | 0.696262252 | 2.41332245 | 0.28850776 | 0.77295809 | 0.91791018 |
| TTTY10 | 20.8252938 | 0.750776694 | 1.51164494 | 0.49666206 | 0.61942738 | 0.84360228 |
| EIF1AY | 13.690566 | -0.126017206 | 1.83529414 | -0.0686632 | 0.94525769 | 0.98552865 |
| IL17RA | 29.9104065 | -0.988147065 | 0.28228916 | -3.5004782 | 0.00046442 | 0.00818581 |
| HDHD5 | 25.3945044 | -0.112063215 | 0.36629687 | -0.3059355 | 0.75965376 | 0.91236458 |
| ADA2 | 15.2768889 | 0.230406948 | 0.64438759 | 0.35755957 | 0.72067294 | 0.89154232 |
| CECR2 | 236.15723 | 0.653042794 | 0.30319103 | 2.15389881 | 0.03124811 | 0.17498941 |
| ATP6V1E1 | 61.0839864 | -0.795175053 | 0.32582496 | -2.4404976 | 0.01466704 | 0.1065074 |
| BCL2L13 | 110.110527 | 0.157029618 | 0.23355211 | 0.67235368 | 0.50135857 | 0.776256 |
| MICAL3 | 237.495155 | 0.01941783 | 0.48919167 | 0.0396937 | 0.96833732 | 0.99156233 |
| PEX26 | 39.5698711 | -0.647901013 | 0.33880693 | -1.9123015 | 0.05583753 | 0.24604401 |
| DGCR2 | 69.6463253 | 0.197491762 | 0.29290166 | 0.67425962 | 0.50014627 | 0.77558432 |
| SLC25A1 | 22.9707337 | -0.357389107 | 0.51590556 | -0.6927413 | 0.48847189 | 0.77038429 |
| CLTCL1 | 20.4261285 | -0.099393729 | 0.43636606 | -0.227776 | 0.81982036 | 0.93860573 |
| HIRA | 28.9673519 | 0.320024103 | 0.34496349 | 0.92770428 | 0.35356099 | 0.66037714 |

| | | | | | | |
|---|---|---|---|---|---|---|
| UFD1 | 23.310977 | -0.039892909 | 0.37959797 | -0.1050925 | 0.91630239 | 0.97396301 |
| TXNRD2 | 29.309112 | -0.527053341 | 0.38274617 | -1.377031 | 0.16850267 | 0.46209378 |
| COMT | 66.0515253 | -0.659102189 | 0.43325363 | -1.5212849 | 0.12818836 | 0.39695394 |
| TANGO2 | 58.6748514 | 0.227646014 | 0.48689206 | 0.46754924 | 0.64010697 | 0.85420887 |
| DGCR8 | 34.7298154 | -0.129377148 | 0.48501626 | -0.2667481 | 0.78966315 | 0.92428059 |
| ZDHHC8 | 13.797065 | 0.739284391 | 0.54228919 | 1.36326595 | 0.17279871 | 0.468183 |
| KLHL22 | 66.1087754 | 0.430593768 | 0.25721627 | 1.67405337 | 0.09412011 | 0.33444204 |
| MED15 | 96.1797728 | -0.491689028 | 0.34731928 | -1.4156687 | 0.15687253 | 0.44349247 |
| PI4KA | 162.621135 | -0.313749526 | 0.32044405 | -0.9791086 | 0.32752633 | 0.63901559 |
| SNAP29 | 23.5183199 | 0.023258265 | 0.35879258 | 0.06482371 | 0.94831437 | 0.98641632 |
| CRKL | 33.4054316 | 0.336096716 | 0.31265408 | 1.07497947 | 0.28238392 | 0.59642935 |
| LZTR1 | 12.7332258 | -0.061230663 | 0.52499206 | -0.1166316 | 0.907152 | 0.9700616 |
| GGT2 | 132.960986 | -4.225237674 | 0.94441702 | -4.473911 | 7.68E-06 | 0.00030734 |
| HIC2 | 34.0870396 | 0.523416819 | 0.41401058 | 1.26425952 | 0.20613689 | 0.51308606 |
| UBE2L3 | 67.3556327 | -0.416214731 | 0.33480131 | -1.2431694 | 0.21380543 | 0.52235998 |
| PPIL2 | 58.1548184 | 0.475610674 | 0.32971203 | 1.44250324 | 0.14916046 | 0.4328578 |
| YPEL1 | 15.8931212 | -0.0261248 | 0.58509368 | -0.0446506 | 0.96438579 | 0.99082982 |
| MAPK1 | 153.952346 | -0.434183628 | 0.30485111 | -1.4242481 | 0.15437466 | 0.44013998 |
| PPM1F | 19.2343641 | -0.336998041 | 0.47428429 | -0.7105402 | 0.47736922 | 0.76307714 |
| TOP3B | 16.8535056 | 0.360741605 | 0.53358597 | 0.67607026 | 0.49899604 | 0.77558432 |
| AC245060.5 | 47.1764397 | 1.20477967 | 0.51063998 | 2.35935242 | 0.01830686 | 0.12347678 |
| BCR | 314.74246 | -1.633349579 | 0.51039466 | -3.2001698 | 0.00137347 | 0.01930973 |
| CHCHD10 | 76.5295986 | 0.189858173 | 0.44771313 | 0.42406211 | 0.67152052 | 0.86777458 |
| SMARCB1 | 41.2098643 | -0.078724311 | 0.33986341 | -0.2316351 | 0.81682141 | 0.93753535 |
| DDT | 36.4801425 | -0.173224299 | 0.53870745 | -0.3215554 | 0.74778953 | 0.90715451 |
| CABIN1 | 133.972661 | 0.002456722 | 0.32843516 | 0.00748008 | 0.99403181 | 0.9992177 |
| GUCD1 | 20.6537073 | -0.87595554 | 0.43224497 | -2.0265257 | 0.04271094 | 0.21070732 |
| SNRPD3 | 34.6794333 | -0.076313618 | 0.27909626 | -0.2734312 | 0.78452179 | 0.92261923 |
| GGT1 | 16.4686987 | -2.240200256 | 0.67374344 | -3.3250049 | 0.00088417 | 0.01360643 |
| SGSM1 | 143.912713 | 0.830045508 | 0.39460158 | 2.10350275 | 0.03542185 | 0.18774924 |
| KIAA1671 | 200.652314 | -0.748492758 | 0.31791445 | -2.3543842 | 0.01855342 | 0.12445475 |
| AL022323.4 | 14.6974308 | -0.477339145 | 0.46210124 | -1.0329752 | 0.30161548 | 0.61564421 |
| LRP5L | 51.2140506 | 0.241293438 | 0.34455774 | 0.700299 | 0.4837406 | 0.76694394 |
| GRK3 | 45.6217459 | -0.53842633 | 0.49445414 | -1.0889308 | 0.27618441 | 0.59130734 |
| HPS4 | 55.004128 | -0.072273891 | 0.32490727 | -0.2224447 | 0.82396774 | 0.94138895 |
| TFIP11 | 17.8364348 | 0.402559209 | 0.48601321 | 0.82828861 | 0.40750707 | 0.70619047 |
| TPST2 | 137.61269 | 1.170059726 | 0.38900159 | 3.00785336 | 0.002631 | 0.03113128 |
| MIATNB | 40.8419081 | -0.186976488 | 0.42232708 | -0.4427291 | 0.65796169 | 0.86357172 |
| PITPNB | 125.937788 | -0.022114503 | 0.24467497 | -0.0903832 | 0.92798272 | 0.97812694 |
| TTC28 | 289.75735 | -0.653689624 | 0.30182222 | -2.1658101 | 0.0303257 | 0.17116031 |
| HSCB | 39.5890253 | -0.521139553 | 0.33782955 | -1.5426109 | 0.12292521 | 0.38755404 |
| CCDC117 | 15.3911578 | 0.050517903 | 0.36938214 | 0.13676325 | 0.89121793 | 0.96328012 |
| XBP1 | 28.4224275 | -1.040803092 | 0.38403176 | -2.7102006 | 0.00672425 | 0.06163349 |
| Z93930.2 | 66.8305469 | -0.06264793 | 0.5496673 | -0.1139743 | 0.90925819 | 0.97090281 |
| ZNRF3 | 274.034743 | -0.364377217 | 0.35671942 | -1.0214673 | 0.3070331 | 0.62085808 |
| KREMEN1 | 64.4763257 | 1.016738064 | 0.37613618 | 2.70311157 | 0.00686937 | 0.06267122 |
| RHBDD3 | 14.862589 | -0.160490133 | 0.46222509 | -0.3472121 | 0.728432 | 0.89654656 |
| EWSR1 | 123.053461 | 0.274773797 | 0.2874789 | 0.9558051 | 0.33917071 | 0.64971064 |
| AP1B1 | 37.6399594 | -0.075818297 | 0.31006583 | -0.2445232 | 0.80682562 | 0.93325451 |
| THOC5 | 16.9368277 | 0.354935613 | 0.39888896 | 0.88981056 | 0.37356761 | 0.67707283 |
| NIPSNAP1 | 25.7663276 | 0.037250493 | 0.43223858 | 0.0861804 | 0.93132301 | 0.97916542 |
| NF2 | 60.1247621 | 0.378689243 | 0.37678193 | 1.0050621 | 0.31486695 | 0.62946385 |
| UQCR10 | 54.049489 | -0.645296134 | 0.54665188 | -1.1804517 | 0.23782061 | 0.54997949 |
| ASCC2 | 33.972139 | 0.110638912 | 0.29400767 | 0.37631301 | 0.70668421 | 0.88603006 |
| MTMR3 | 225.90467 | -0.072749987 | 0.22035327 | -0.3301516 | 0.74128541 | 0.90383185 |
| TBC1D10A | 19.400448 | -0.678862848 | 0.46591969 | -1.4570383 | 0.1451058 | 0.42707275 |
| SF3A1 | 16.4747286 | 0.132122268 | 0.50923206 | 0.25945395 | 0.79528501 | 0.9268423 |
| RNF215 | 23.4089896 | 0.09273764 | 0.30707145 | 0.30200671 | 0.76264695 | 0.91400002 |
| SEC14L6 | 22.0571235 | 1.094202257 | 0.5629107 | 1.94382922 | 0.05191605 | 0.23509725 |
| GAL3ST1 | 17.8376068 | 1.473579431 | 0.5740034 | 2.56719636 | 0.01025245 | 0.08363419 |
| PES1 | 16.9109118 | -0.060524207 | 0.41483332 | -0.1459001 | 0.88400028 | 0.96170562 |
| TCN2 | 13.5245773 | 0.53866212 | 0.60446835 | 0.8911337 | 0.37285745 | 0.67662959 |
| DUSP18 | 19.3063845 | -0.33588192 | 0.53417415 | -0.6287873 | 0.52948832 | 0.79499985 |
| OSBP2 | 105.086025 | 0.973367052 | 0.28710712 | 3.39025744 | 0.00069827 | 0.01134263 |
| MORC2 | 33.9250378 | -0.409337768 | 0.38862286 | -1.0533034 | 0.29220198 | 0.60706133 |
| SMTN | 23.8016186 | 0.518210635 | 0.45058034 | 1.15009596 | 0.25010435 | 0.56269702 |
| SELENOM | 35.0119387 | -0.297873491 | 0.67582599 | -0.4407547 | 0.65939058 | 0.86452386 |
| INPP5J | 47.173693 | 2.288672395 | 0.73253371 | 3.12432366 | 0.00178214 | 0.02317532 |
| RNF185 | 56.2586108 | -0.105746328 | 0.39507975 | -0.2676582 | 0.78896244 | 0.92402584 |
| LIMK2 | 47.8226896 | -1.461273999 | 0.36471218 | -4.0066498 | 6.16E-05 | 0.00169113 |
| PIK3IP1 | 18.0218409 | -1.190333669 | 0.56517272 | -2.1061414 | 0.03519207 | 0.18707575 |
| PATZ1 | 24.0304908 | -0.059641787 | 0.60283662 | -0.0989352 | 0.92118969 | 0.97548809 |
| DRG1 | 60.3693234 | -0.045408982 | 0.28147142 | -0.1613272 | 0.87183574 | 0.9568935 |
| EIF4ENIF1 | 52.6661168 | -0.216135034 | 0.2585293 | -0.8360176 | 0.40314502 | 0.70207773 |
| SFI1 | 50.4801276 | -0.475617687 | 0.35230074 | -1.3500332 | 0.17700535 | 0.47425226 |
| PISD | 151.799889 | -0.357484615 | 0.4479909 | -0.7979729 | 0.4248862 | 0.72151771 |
| PRR14L | 114.679388 | -0.258800047 | 0.23095291 | -1.120575 | 0.26246882 | 0.5766398 |
| RTCB | 47.99919 | -0.273488282 | 0.40229364 | -0.6798225 | 0.49661684 | 0.77475164 |
| FBXO7 | 55.779903 | 0.143247095 | 0.26717875 | 0.53614704 | 0.59185693 | 0.83276878 |
| TIMP3 | 110.268544 | 0.208840263 | 0.42798063 | 0.48796662 | 0.62557349 | 0.8471532 |
| LARGE1 | 319.550568 | -0.360406508 | 0.57525657 | -0.6265144 | 0.53097762 | 0.79628881 |
| HMGXB4 | 43.1296776 | -0.196394654 | 0.43720951 | -0.4492003 | 0.65328715 | 0.86199934 |
| TOM1 | 60.8816429 | -0.518744886 | 0.47222202 | -1.0985191 | 0.2719779 | 0.58786226 |
| HMOX1 | 19.4092164 | -3.873519592 | 0.89860052 | -4.3106136 | 1.63E-05 | 0.00059002 |
| APOL6 | 14.3666096 | -1.847544475 | 0.87404337 | -2.1137904 | 0.03453317 | 0.18483773 |
| RBFOX2 | 368.972861 | 0.182346203 | 0.27720136 | 0.65781135 | 0.51065936 | 0.78194213 |
| APOL4 | 17.4143547 | -1.866719827 | 1.35257021 | -1.3801279 | 0.16754728 | 0.46042171 |
| APOL2 | 39.6933602 | -0.581853845 | 0.63016339 | -0.9233381 | 0.35583105 | 0.66209714 |
| MYH9 | 192.419002 | -0.344145219 | 0.44242096 | -0.7778682 | 0.43664669 | 0.7294918 |
| TXN2 | 23.0526996 | -0.595441138 | 0.55010333 | -1.0824169 | 0.27906732 | 0.59380484 |
| EIF3D | 34.3990636 | -0.720065162 | 0.35094308 | -2.0518004 | 0.04018906 | 0.20290676 |
| IFT27 | 15.0790177 | 0.81120366 | 0.56921596 | 1.42512458 | 0.15412119 | 0.43999572 |
| PVALB | 14.0232436 | 3.0850191 | 0.80775692 | 3.81924192 | 0.00013386 | 0.00306667 |
| TST | 14.972062 | 0.187683754 | 0.63379835 | 0.29612534 | 0.76713436 | 0.9157292 |
| MPST | 24.1462165 | -0.241946951 | 0.48191957 | -0.5020484 | 0.61563347 | 0.8422143 |
| CARD10 | 14.4011426 | -0.984904025 | 0.48896267 | -2.0142724 | 0.04398094 | 0.21414009 |
| GGA1 | 56.391753 | 0.065643256 | 0.24025492 | 0.27322336 | 0.78468152 | 0.92261923 |
| TRIOBP | 62.1435072 | -0.587023226 | 0.39852205 | -1.4730006 | 0.14075087 | 0.42049378 |
| H1F0 | 25.9962099 | -1.131134083 | 0.64591864 | -1.7512021 | 0.0799111 | 0.30349944 |
| ANKRD54 | 15.8250592 | 0.316701107 | 0.41851691 | 0.75672236 | 0.44921622 | 0.74080155 |
| EIF3L | 66.380341 | 0.061172176 | 0.38739343 | 0.15790711 | 0.87453 | 0.95761805 |
| MICALL1 | 15.83986 | -0.767370067 | 0.66210665 | -1.1589826 | 0.24646329 | 0.55930671 |
| POLR2F | 25.2865319 | 0.419410471 | 0.49342535 | 0.84999781 | 0.3953263 | 0.69642725 |
| PICK1 | 26.6973796 | 0.235736307 | 0.35730631 | 0.65975971 | 0.50940804 | 0.78107558 |
| PLA2G6 | 26.5716038 | 0.514729261 | 0.43995683 | 1.16995401 | 0.24201948 | 0.55444462 |
| TMEM184B | 42.8836956 | -0.056144247 | 0.3753774 | -0.1495675 | 0.88110588 | 0.9612999 |
| CSNK1E | 15.8907707 | 0.260703078 | 0.51589184 | 0.50534445 | 0.61331693 | 0.84174925 |
| TPTEP2-CSNI | 145.523826 | 0.375052671 | 0.3417059 | 1.09758909 | 0.27238396 | 0.5880315 |
| DDX17 | 841.181413 | 0.657873592 | 0.16599296 | 3.96326194 | 7.39E-05 | 0.00194731 |
| FAM227A | 24.278509 | -0.411353029 | 0.58643918 | -0.7014419 | 0.48302726 | 0.7665951 |
| CBY1 | 31.0494734 | 0.127594462 | 0.40843978 | 0.31239479 | 0.75474051 | 0.91026551 |
| JOSD1 | 34.501059 | -0.738159881 | 0.41415493 | -1.7823279 | 0.07469578 | 0.29263171 |
| GTPBP1 | 23.8699891 | -0.220313562 | 0.39480634 | -0.5580294 | 0.57682428 | 0.82212008 |

| | | | | | | |
|---|---|---|---|---|---|---|
| SUN2 | 19.2425773 | -0.323280688 | 0.47390633 | -0.6821616 | 0.49513679 | 0.77475164 |
| AL021707.1 | 14.7798458 | -0.359406285 | 0.51201027 | -0.7019513 | 0.48270952 | 0.76650978 |
| CBX6 | 31.4482829 | -0.454568174 | 0.43890645 | -1.0356835 | 0.30034981 | 0.61413633 |
| CBX7 | 24.7512419 | 0.184621833 | 0.37458568 | 0.49286943 | 0.62210485 | 0.84543151 |
| RPL3 | 67.2696255 | -0.574903257 | 0.57988143 | -0.9914152 | 0.32148289 | 0.63312885 |
| SYNGR1 | 22.0903792 | 0.598922922 | 0.63846651 | 0.93806475 | 0.34821114 | 0.65547624 |
| TAB1 | 25.3816545 | 0.333378214 | 0.38746344 | 0.86041206 | 0.38956194 | 0.69267928 |
| MIEF1 | 12.2498132 | -0.594432389 | 0.47551012 | -1.2500941 | 0.21126518 | 0.52029492 |
| ATF4 | 46.8122672 | -0.564777596 | 0.66414393 | -0.8503843 | 0.39511144 | 0.69642725 |
| RPS19BP1 | 21.5317782 | -0.528921313 | 0.42656162 | -1.2399646 | 0.21498848 | 0.52392906 |
| FAM83F | 29.9012457 | -1.339751818 | 0.73602433 | -1.8202548 | 0.06872021 | 0.27897062 |
| TNRC6B | 719.073661 | 0.187571169 | 0.19431816 | 0.96527866 | 0.33440526 | 0.64554754 |
| MKL1 | 296.754729 | -0.202122921 | 0.41156208 | -0.4911116 | 0.6233475 | 0.84597807 |
| SLC25A17 | 55.7663582 | 0.157525567 | 0.26229637 | 0.60056327 | 0.54813091 | 0.80636993 |
| ST13 | 81.1044513 | -0.025515987 | 0.25679668 | -0.0993626 | 0.92085037 | 0.9754826 |
| XPNPEP3 | 71.9183334 | 0.194958752 | 0.36173715 | 0.53895142 | 0.58992037 | 0.83100432 |
| RBX1 | 50.1988066 | 0.022908523 | 0.42922039 | 0.0533724 | 0.95743519 | 0.98859719 |
| EP300 | 176.876491 | -0.043629934 | 0.27583605 | -0.1581734 | 0.87432014 | 0.95761805 |
| L3MBTL2 | 15.3620946 | 0.1944531 | 0.3982742 | 0.48823926 | 0.62538038 | 0.8471532 |
| RANGAP1 | 29.4446391 | -0.968198352 | 0.49709782 | -1.9477019 | 0.05145065 | 0.23455111 |
| ZC3H7B | 47.8231674 | -0.190236688 | 0.29068911 | -0.6544335 | 0.51283256 | 0.7830488 |
| TEF | 15.3376673 | 1.09640575 | 0.58134025 | 1.88599662 | 0.05929541 | 0.25454438 |
| TOB2 | 35.0819269 | 0.757950228 | 0.32118237 | 2.35987494 | 0.0182811 | 0.12342719 |
| ACO2 | 133.21895 | 0.655285654 | 0.2952645 | 2.21931744 | 0.02646514 | 0.15637575 |
| CSDC2 | 26.2377091 | 0.636765517 | 0.83581082 | 0.76185364 | 0.44614736 | 0.73900835 |
| PMM1 | 20.0455684 | 1.230441294 | 0.55860635 | 2.20269837 | 0.02761602 | 0.16113375 |
| DESI1 | 28.6248241 | -0.315848116 | 0.39534057 | -0.7989266 | 0.42433295 | 0.72151771 |
| XRCC6 | 93.4696449 | -0.250169072 | 0.28789189 | -0.8689688 | 0.38486419 | 0.68852052 |
| SNU13 | 50.5194469 | -0.639607856 | 0.30944845 | -2.0669286 | 0.03874088 | 0.19890966 |
| CCDC134 | 14.7207999 | 0.059581541 | 0.36657367 | 0.16253634 | 0.87088352 | 0.95629149 |
| SREBF2 | 284.976663 | -0.000528564 | 0.23293029 | -0.0022692 | 0.99818945 | 0.99922221 |
| WBP2NL | 36.5507727 | 0.332031282 | 0.64108904 | 0.51791758 | 0.60451578 | 0.8377683 |
| SMDT1 | 20.2351274 | -0.394791046 | 0.40350976 | -0.9783928 | 0.32788009 | 0.6394978 |
| NDUFA6 | 24.1541255 | 0.134953955 | 0.40205152 | 0.33566334 | 0.73712476 | 0.90160714 |
| TCF20 | 131.889791 | -0.423406029 | 0.30597682 | -1.3837847 | 0.1664244 | 0.45864087 |
| RRP7A | 21.6604223 | 0.166973651 | 0.47659996 | 0.3503434 | 0.726081 | 0.89457547 |
| POLDIP3 | 31.253388 | -0.372380726 | 0.40735155 | -0.9141508 | 0.36063763 | 0.66638631 |
| CYB5R3 | 49.1448398 | -1.02203532 | 0.37560296 | -2.7210523 | 0.00650745 | 0.05995596 |
| A4GALT | 35.1979822 | -3.285652499 | 0.58347343 | -5.6311947 | 1.79E-08 | 1.70E-06 |
| ARFGAP3 | 153.563283 | -1.965087018 | 0.38403956 | -5.116887 | 3.11E-07 | 2.16E-05 |
| PACSIN2 | 207.457338 | -0.375194153 | 0.37025973 | -1.0133269 | 0.31090403 | 0.62583531 |
| TSPO | 34.1595048 | -0.906096711 | 0.50894726 | -1.7803352 | 0.07502112 | 0.29304439 |
| EFCAB6 | 111.365061 | -0.233786273 | 0.49201528 | -0.4751606 | 0.63467251 | 0.85244692 |
| SAMM50 | 20.7143347 | 0.515458169 | 0.34158178 | 1.50903298 | 0.13129036 | 0.40268134 |
| PHF21B | 203.750689 | 0.50268874 | 0.25985592 | 1.93449021 | 0.05305288 | 0.23850776 |
| NUP50-DT | 74.4792036 | -0.168506345 | 0.32698863 | -0.5153278 | 0.60632394 | 0.83829217 |
| NUP50 | 25.2131639 | -0.347009795 | 0.32926678 | -1.0538864 | 0.29193491 | 0.60677688 |
| KIAA0930 | 25.1394811 | -3.723108635 | 0.86332955 | -4.3124999 | 1.61E-05 | 0.00058792 |
| FAM118A | 26.0949564 | 1.362436098 | 0.53794605 | 2.53266309 | 0.01131997 | 0.08967493 |
| ATXN10 | 184.722488 | -0.074600493 | 0.25293918 | -0.2949345 | 0.76804391 | 0.91600052 |
| MIRLET7BHC | 82.2766927 | -0.273269723 | 0.25449189 | -1.0737856 | 0.2829188 | 0.59668285 |
| PPARA | 205.27747 | 1.011703251 | 0.19878486 | 5.08943808 | 3.59E-07 | 2.43E-05 |
| TTC38 | 31.82672 | 0.296541973 | 0.43262738 | 0.6854443 | 0.4930636 | 0.77312609 |
| TRMU | 43.6832294 | 0.336147689 | 0.27758484 | 1.21097278 | 0.22590584 | 0.53705916 |
| CELSR1 | 46.7330846 | 0.236482454 | 0.52608423 | 0.44951443 | 0.6530606 | 0.86187361 |
| GRAMD4 | 39.9111048 | 1.033629507 | 0.50039639 | 2.06562141 | 0.03886424 | 0.1991045 |
| CERK | 28.0655255 | -0.042234597 | 0.38560722 | -0.1095275 | 0.91278411 | 0.97166624 |
| TBC1D22A | 302.741434 | -0.6786687 | 0.40180071 | -1.6890679 | 0.09120641 | 0.32885096 |
| BRD1 | 65.5210883 | -0.452247108 | 0.39606761 | -1.1418432 | 0.2535192 | 0.56559484 |
| ZBED4 | 29.2384111 | -0.237445734 | 0.52462823 | -0.4525981 | 0.65083818 | 0.86113455 |
| PIM3 | 65.5057692 | -0.557908717 | 0.43506659 | -1.2823525 | 0.19971903 | 0.5033686 |
| TRABD | 29.6515187 | 0.4181497 | 0.43143345 | 0.9692102 | 0.33244032 | 0.644708 |
| SELENOO | 61.1934466 | -0.203085068 | 0.27438679 | -0.7401416 | 0.4592141 | 0.74776673 |
| TUBGCP6 | 18.3284937 | 0.972510522 | 0.47363339 | 2.05329805 | 0.04004368 | 0.20269667 |
| PLXNB2 | 67.1627946 | -0.878017577 | 0.31738326 | -2.7664268 | 0.00566743 | 0.05425372 |
| DENND6B | 13.8445539 | 0.200972956 | 0.52967747 | 0.37942515 | 0.70437218 | 0.8856427 |
| PPP6R2 | 285.475134 | 0.468235203 | 0.19280465 | 2.42854728 | 0.01515945 | 0.10914803 |
| MIOX | 17.0159684 | -0.156650003 | 0.79898389 | -0.1960615 | 0.84456201 | 0.94905142 |
| RABL2B | 27.3544264 | 0.132423883 | 0.37279998 | 0.35521429 | 0.72242906 | 0.89217596 |
| CU633967.1 | 15.5671853 | -0.073104027 | 0.82851458 | -0.0882351 | 0.92968986 | 0.97876742 |
| FP671120.1 | 482.139949 | -1.037409631 | 0.65291517 | -1.5888888 | 0.11208549 | 0.36863672 |
| FP236383.1 | 268.096977 | -0.995651199 | 0.68850377 | -1.4461086 | 0.14814675 | 0.43194383 |
| CR381653.1 | 40.0082935 | 1.020172625 | 0.57291406 | 1.78067305 | 0.07496588 | 0.29304439 |
| NRIP1 | 134.929073 | -1.187884787 | 0.54712245 | -2.1711498 | 0.02991985 | 0.16937218 |
| USP25 | 379.236414 | 0.847603 | 0.29394924 | 2.88350126 | 0.00393281 | 0.04221193 |
| MIR99AHG | 121.680468 | 0.815055472 | 0.48484055 | 1.68107943 | 0.09274749 | 0.33202083 |
| CXADR | 54.061292 | -0.161785867 | 0.35381532 | -0.4572608 | 0.64748362 | 0.85921615 |
| BTG3 | 28.2783535 | -0.412087133 | 0.37926828 | -1.086532 | 0.27724368 | 0.59235107 |
| C21orf91 | 15.9761328 | -0.149606994 | 0.43302567 | -0.3454922 | 0.72972437 | 0.89713026 |
| CHODL | 56.5202404 | 0.095881033 | 0.9893689 | 0.09691131 | 0.92279683 | 0.97611757 |
| MRPL39 | 21.0431482 | 0.078426416 | 0.35385631 | 0.22163351 | 0.8245992 | 0.94153354 |
| ATP5PF | 55.8258423 | 0.272456247 | 0.41386242 | 0.65832565 | 0.5103289 | 0.78158783 |
| GABPA | 34.4836941 | 0.135330373 | 0.3735365 | 0.36229491 | 0.71713166 | 0.89010971 |
| APP | 916.818723 | -1.437486246 | 0.30711497 | -4.6806127 | 2.86E-06 | 0.00013537 |
| CYYR1 | 39.2811969 | -1.333575156 | 0.5287555 | -2.5221017 | 0.0116656 | 0.09194423 |
| ADAMTS1 | 350.633162 | -6.519530917 | 0.51908621 | -12.55963 | 3.52E-36 | 3.06E-32 |
| LINC00113 | 22.2697292 | 1.409548892 | 1.00787438 | 1.39853628 | 0.16195209 | 0.45210816 |
| LINC01695 | 27.4457327 | -1.462691229 | 0.71404998 | -2.0484438 | 0.04051654 | 0.20398282 |
| AF165147.1 | 33.9275575 | 0.476021008 | 0.54555275 | 0.87254809 | 0.38290944 | 0.68698242 |
| N6AMT1 | 12.6540887 | 0.18293094 | 0.52027353 | 0.35160532 | 0.72513428 | 0.89374117 |
| LTN1 | 91.9108814 | 0.4341971 | 0.23791968 | 1.82497343 | 0.06800504 | 0.27788783 |
| USP16 | 55.2542415 | -0.106787926 | 0.49788383 | -0.2144836 | 0.83016994 | 0.94393022 |
| CCT8 | 31.0306691 | -0.262665951 | 0.33759091 | -0.7780599 | 0.4365337 | 0.72943373 |
| MAP3K7CL | 37.0642287 | 0.110818088 | 0.54484379 | 0.20339424 | 0.8388269 | 0.94741614 |
| BACH1 | 354.530291 | -0.954437561 | 0.30874975 | -3.0912983 | 0.00199283 | 0.02517775 |
| GRIK1 | 23.2787032 | -2.18716678 | 0.80162591 | -2.7284133 | 0.00636398 | 0.05910136 |
| CLDN8 | 24.6518478 | 1.963143897 | 0.89199649 | 2.20084263 | 0.02774717 | 0.16155622 |
| SOD1 | 46.6688304 | -0.596399521 | 0.35509887 | -1.6795309 | 0.09304862 | 0.3326491 |
| SCAF4 | 77.6541952 | 0.275384246 | 0.29122439 | 0.94560846 | 0.34434833 | 0.65359197 |
| HUNK | 29.4689078 | -1.738769556 | 0.84347542 | -2.0614348 | 0.03926158 | 0.20036945 |
| URB1 | 24.4126782 | 0.075657745 | 0.41890755 | 0.18060726 | 0.85667586 | 0.95234014 |
| EVA1C | 28.0460873 | -0.28293799 | 0.46376384 | -0.6100907 | 0.54180175 | 0.80301375 |
| SYNJ1 | 61.6974883 | 0.130936261 | 0.30740418 | 0.42594171 | 0.67015032 | 0.86777458 |
| PAXBP1 | 112.216765 | -0.034542684 | 0.18385103 | -0.1878841 | 0.85096751 | 0.95042076 |
| C21orf62-AS | 30.5227517 | 0.870465855 | 0.71412628 | 1.21892426 | 0.22287294 | 0.53311751 |
| AP000282.1 | 40.9499541 | 1.77821106 | 0.70593089 | 2.51895912 | 0.01177023 | 0.09253427 |
| IFNAR1 | 55.0093853 | -0.86046256 | 0.25553123 | -3.3673479 | 0.00075895 | 0.01211719 |
| IFNGR2 | 82.0790408 | -0.886676427 | 0.3917766 | -2.2632194 | 0.02362216 | 0.14586296 |
| TMEM50B | 69.9224719 | 0.443416751 | 0.41554226 | 1.0670798 | 0.2859358 | 0.60005974 |
| GART | 47.601996 | -0.081271503 | 0.2372469 | -0.3425609 | 0.73192885 | 0.89787 |
| SON | 222.455043 | -0.379266413 | 0.27873048 | -1.3606923 | 0.17361096 | 0.46960235 |
| AP000311.1 | 53.3814314 | -0.150005211 | 0.61660772 | -0.2432749 | 0.80779241 | 0.93400997 |
| ITSN1 | 204.030097 | 0.24791493 | 0.28784639 | 0.8612751 | 0.38908654 | 0.69247382 |

| | | | | | | |
|---|---|---|---|---|---|---|
| MRPS6 | 362.58063 | 1.590796627 | 0.48882388 | 3.25433492 | 0.00113658 | 0.01653275 |
| LINC00310 | 39.9033503 | 0.901600011 | 0.64837708 | 1.39054886 | 0.16436227 | 0.45563826 |
| AP000317.1 | 51.0923341 | 0.694122274 | 0.6560376 | 1.05805258 | 0.29003147 | 0.60488981 |
| RCAN1 | 56.5273386 | 0.583217259 | 0.60403459 | 0.96553619 | 0.33427632 | 0.64554754 |
| RUNX1 | 397.182499 | -2.25336067 | 1.02996021 | -2.1878133 | 0.0286832 | 0.16498593 |
| SETD4 | 31.2159273 | 0.122786065 | 0.3462224 | 0.35464506 | 0.72285551 | 0.89240027 |
| CBR1 | 38.6743113 | -1.27101391 | 0.44224637 | -2.8739951 | 0.00405315 | 0.04317114 |
| CBR3 | 31.0505259 | -0.530281778 | 0.55016823 | -0.9638539 | 0.33511919 | 0.64662829 |
| DOPEY2 | 34.6951686 | -1.294834651 | 0.5191868 | -2.4939668 | 0.01263244 | 0.09710209 |
| MORC3 | 98.1368282 | -0.001877606 | 0.28455221 | -0.0065985 | 0.99473523 | 0.9992177 |
| AP000695.2 | 22.1011347 | -2.838301181 | 0.9430331 | -3.0097578 | 0.00261456 | 0.03105499 |
| CLDN14 | 69.7358164 | -2.605599701 | 0.81808137 | -3.1850129 | 0.00144748 | 0.02011366 |
| HLCS | 175.124555 | 0.081592422 | 0.34287632 | 0.23796459 | 0.81190856 | 0.93552094 |
| PIGP | 11.9970286 | -0.022477979 | 0.70329226 | -0.0319611 | 0.97450309 | 0.99357756 |
| TTC3 | 262.595786 | 0.56944254 | 0.28078059 | 2.02806944 | 0.04255316 | 0.21037417 |
| VPS26C | 55.3246362 | -0.342761408 | 0.28779193 | -1.1910043 | 0.23365191 | 0.54598215 |
| DYRK1A | 466.936452 | -0.155210283 | 0.40783173 | -0.3805743 | 0.70351914 | 0.88535733 |
| KCNJ15 | 20.8646921 | 0.067946045 | 0.48846331 | 0.13910164 | 0.88936984 | 0.96308489 |
| ETS2 | 108.567958 | -3.024229262 | 0.60888066 | -4.966867 | 6.80E-07 | 4.23E-05 |
| BRWD1 | 350.018917 | 0.42592002 | 0.1980718 | 2.15033141 | 0.03152901 | 0.17567711 |
| HMGN1 | 72.210942 | 0.001360104 | 0.34031376 | 0.00399662 | 0.99681117 | 0.99922221 |
| WRB | 25.2227929 | 0.178743331 | 0.36297493 | 0.49243988 | 0.62240841 | 0.84559756 |
| LCA5L | 13.8616098 | 0.179426801 | 0.46558474 | 0.38537947 | 0.69995629 | 0.88361663 |
| SH3BGR | 14.7281174 | 1.899631738 | 0.50990288 | 3.72547756 | 0.00019495 | 0.00413421 |
| PCP4 | 32.9065017 | 3.152368669 | 1.12050738 | 2.8133404 | 0.00490297 | 0.04947544 |
| BACE2 | 949.636725 | -1.265435359 | 0.78598731 | -1.6099946 | 0.10739902 | 0.36059666 |
| FAM3B | 17.3611168 | -0.467908469 | 0.46499683 | -1.0062616 | 0.31428973 | 0.62898422 |
| MX1 | 88.0205837 | -0.337511227 | 1.0092625 | -0.3344137 | 0.73806739 | 0.90240497 |
| TMPRSS2 | 266.867091 | 1.851291051 | 0.38171545 | 4.84992427 | 1.24E-06 | 6.90E-05 |
| RIPK4 | 26.4216403 | -0.450013556 | 0.36076611 | -1.2473831 | 0.21225705 | 0.52134789 |
| PRDM15 | 16.772608 | -0.214494557 | 0.54439998 | -0.3940018 | 0.69357973 | 0.87889881 |
| ZBTB21 | 24.479817 | -0.658379276 | 0.39055636 | -1.6857472 | 0.09184451 | 0.33043047 |
| ABCG1 | 27.0871455 | -0.582515367 | 0.75135552 | -0.7752859 | 0.43817072 | 0.73056686 |
| SLC37A1 | 26.2888408 | -2.138359892 | 0.32219254 | -6.6369007 | 3.20E-11 | 8.53E-09 |
| PDE9A | 148.700352 | -0.072829013 | 0.36595417 | -0.1990113 | 0.84225391 | 0.94776435 |
| WDR4 | 22.9696446 | -0.559836881 | 0.51107725 | -1.0954056 | 0.27333898 | 0.58894058 |
| NDUFV3 | 70.2116312 | 0.558133038 | 0.23120887 | 2.41397764 | 0.01577943 | 0.11184941 |
| PKNOX1 | 70.9883664 | -0.261262031 | 0.23726986 | -1.1011177 | 0.27084545 | 0.58722969 |
| RRP1B | 25.6895377 | 0.365466001 | 0.30117407 | 1.21347101 | 0.22494979 | 0.53615334 |
| PDXK | 114.986376 | 0.168491644 | 0.27939562 | 0.60305758 | 0.54647039 | 0.80581909 |
| CSTB | 47.2500683 | -1.902212664 | 0.39115142 | -4.8631108 | 1.16E-06 | 6.60E-05 |
| AGPAT3 | 122.678912 | 0.228063785 | 0.26488789 | 0.86098229 | 0.3892478 | 0.69247382 |
| TRAPPC10 | 126.625257 | -0.065839986 | 0.31632988 | -0.2081371 | 0.83512191 | 0.94654164 |
| PFKL | 140.030924 | -0.222288315 | 0.34908243 | -0.6367789 | 0.52426889 | 0.79185875 |
| UBE2G2 | 101.48811 | 0.015905639 | 0.26596195 | 0.05980419 | 0.95231159 | 0.98738095 |
| SUMO3 | 18.9709634 | -1.791250025 | 0.59542941 | -3.0083332 | 0.00262685 | 0.03112166 |
| PTTG1IP | 129.972437 | -1.361146142 | 0.40109105 | -3.3936088 | 0.00068978 | 0.0112243 |
| ADARB1 | 55.2690981 | -0.990060618 | 0.63328663 | -1.5633689 | 0.11796585 | 0.37862775 |
| POFUT2 | 25.4744495 | -0.215956012 | 0.33209086 | -0.650292 | 0.51550366 | 0.78410377 |
| COL18A1 | 242.329294 | -1.046054284 | 0.36117361 | -2.8962645 | 0.00377634 | 0.04127854 |
| COL6A1 | 67.4815602 | 1.552626107 | 0.55297486 | 2.80776978 | 0.00498859 | 0.05001461 |
| LSS | 38.2122981 | -0.484676729 | 0.46419187 | -1.0441302 | 0.29642517 | 0.61093809 |
| MCM3AP-AS | 15.3403345 | 0.686309023 | 0.5393136 | 1.2725602 | 0.20317415 | 0.5078809 |
| MCM3AP | 76.8638619 | 0.112183546 | 0.24420571 | 0.45938134 | 0.64596034 | 0.85857936 |
| YBEY | 14.0790158 | 0.247504982 | 0.56784536 | 0.43586687 | 0.66293332 | 0.86556259 |
| PCNT | 32.7093427 | -0.205025138 | 0.33473047 | -0.6125081 | 0.54020161 | 0.80166126 |
| DIP2A | 125.797517 | 0.186215083 | 0.21943412 | 0.84861499 | 0.39609556 | 0.6966978 |
| PRMT2 | 99.7750724 | -0.666773736 | 0.24543545 | -2.7166969 | 0.0065937 | 0.06063078 |
| AC011043.1 | 19.3678039 | 1.002583942 | 0.56371412 | 1.7785326 | 0.07531642 | 0.29358291 |
| AL592183.1 | 68.4249855 | -0.086011697 | 0.57818094 | -0.1487626 | 0.88174097 | 0.96155389 |

**Connecting tubules**

*This means that positive fold changes imply higher mean value in controls than in AKI and vice versa.

| | baseMean | log2 fold change control/AKI* | lfcSE | stat | pvalue | padj |
|---|---|---|---|---|---|---|
| AL669831.5 | 44.0210642 | 0.15105774 | 0.29399963 | 0.51380249 | 0.60739009 | 0.81184603 |
| SAMD11 | 20.726368 | 2.373495209 | 0.55224643 | 4.29789147 | 1.72E-05 | 0.00057938 |
| NOC2L | 29.2722437 | 0.25525373 | 0.30973125 | 0.82411358 | 0.40987502 | 0.6878209 |
| HES4 | 27.2135181 | -1.039115482 | 0.74490987 | -1.3949546 | 0.16302952 | 0.43120156 |
| ISG15 | 141.642756 | 2.076302502 | 1.93049113 | 1.07553071 | 0.28213719 | 0.5750055 |
| AGRN | 112.768069 | -0.722214294 | 0.44891249 | -1.6088086 | 0.10765819 | 0.34161321 |
| C1orf159 | 32.282756 | -0.62768536 | 0.32760955 | -1.9159556 | 0.05537074 | 0.23225616 |
| LINC01342 | 11.2489136 | 0.278346579 | 0.55798308 | 0.49884412 | 0.6178892 | 0.81766147 |
| AL390719.2 | 15.3804117 | -0.83485809 | 0.4898748 | -1.7042275 | 0.0883386 | 0.30586777 |
| SDF4 | 53.8263123 | -0.489769671 | 0.36168171 | -1.3541455 | 0.17568995 | 0.44967898 |
| UBE2J2 | 43.8341854 | -0.034512904 | 0.27865826 | -0.1238539 | 0.90143098 | 0.9603456 |
| ACAP3 | 16.1378878 | 0.648734284 | 0.43045304 | 1.50709652 | 0.13178592 | 0.38082988 |
| INTS11 | 37.3101774 | 0.414193387 | 0.2669134 | 1.55178942 | 0.12071262 | 0.363514 |
| CPTP | 11.9249236 | -0.008497679 | 0.43723586 | -0.019435 | 0.98449409 | 0.99388031 |
| DVL1 | 13.8374438 | 0.555646199 | 0.42662902 | 1.30241069 | 0.19277603 | 0.47341565 |
| AURKAIP1 | 57.154339 | -0.452716654 | 0.42200387 | -1.0727784 | 0.28337053 | 0.57613642 |
| CCNL2 | 101.280223 | 0.248787868 | 0.26136973 | 0.95186184 | 0.34116706 | 0.62956442 |
| MRPL20 | 57.843837 | 0.080576437 | 0.29607752 | 0.27214642 | 0.78550944 | 0.90802494 |
| VWA1 | 22.2996216 | -2.085687093 | 0.77627037 | -2.686805 | 0.00721391 | 0.06177694 |
| ATAD3C | 11.2319531 | 0.07661076 | 0.47454464 | 0.16144058 | 0.87174641 | 0.94844301 |
| ATAD3B | 27.3281183 | 0.435502083 | 0.37231155 | 1.16972487 | 0.24211171 | 0.5328423 |
| ATAD3A | 15.6381034 | 0.379485691 | 0.50523918 | 0.75110108 | 0.45259183 | 0.72046894 |
| SSU72 | 59.4391194 | -0.306616196 | 0.34326024 | -0.893247 | 0.37172489 | 0.65540131 |
| MIB2 | 20.6421402 | 0.553444725 | 0.36376136 | 1.52145 | 0.12814695 | 0.37540371 |
| CDK11B | 28.8836561 | 0.290428146 | 0.4504982 | 0.64468214 | 0.51913319 | 0.75991242 |
| SLC35E2B | 30.5154438 | -0.191730875 | 0.35630881 | -0.5381031 | 0.59050588 | 0.80004855 |
| CDK11A | 18.5704888 | -0.063648166 | 0.38781196 | -0.1641212 | 0.86963573 | 0.94832921 |
| SLC35E2A | 31.3236577 | -0.222447892 | 0.28964238 | -0.7680088 | 0.44248196 | 0.71265656 |
| NADK | 39.4955212 | -0.068834683 | 0.24824401 | -0.2772864 | 0.78156022 | 0.90539001 |
| GNB1 | 275.592998 | -0.307465186 | 0.32757714 | -0.9386039 | 0.34793414 | 0.63501969 |
| PRKCZ | 63.1323577 | 0.12539008 | 0.29093251 | 0.43099371 | 0.66647295 | 0.84452697 |
| FAAP20 | 32.3955021 | -0.298204827 | 0.32729116 | -0.9111301 | 0.36222683 | 0.64639889 |
| SKI | 74.9409621 | -0.012649206 | 0.31182891 | -0.0405646 | 0.96764303 | 0.987279 |
| MORN1 | 31.1670492 | 0.466220002 | 0.2967249 | 1.57121967 | 0.11613163 | 0.35543425 |
| RER1 | 71.3177887 | -0.304510604 | 0.24380595 | -1.2489876 | 0.21166961 | 0.49785853 |
| PEX10 | 19.6798616 | -0.597297627 | 0.36245767 | -1.64791 | 0.09937114 | 0.32555183 |
| TNFRSF14-A | 13.8215883 | -1.180842967 | 0.49575141 | -2.3819256 | 0.01722238 | 0.10980752 |
| TNFRSF14 | 33.2217397 | -0.708774992 | 0.45553638 | -1.555913 | 0.11972878 | 0.36203283 |
| FAM213B | 11.6918607 | -1.539390784 | 0.60479521 | -2.5453091 | 0.0109181 | 0.08115639 |
| LINC00982 | 108.111997 | 2.699022667 | 0.34374855 | 7.85173546 | 4.10E-15 | 3.02E-12 |
| PRDM16 | 321.223399 | 0.88258804 | 0.28240948 | 3.12520682 | 0.0017768 | 0.02248723 |
| ARHGEF16 | 23.627701 | -0.600714076 | 0.36516908 | -1.6450299 | 0.09996364 | 0.32641058 |
| TPRG1L | 13.2126936 | -0.7064714 | 0.49195591 | -1.4360462 | 0.1509892 | 0.41211643 |
| WRAP73 | 11.1572594 | 0.160372783 | 0.41986479 | 0.38196292 | 0.70248887 | 0.86526068 |
| LRRC47 | 17.9846091 | 0.088940785 | 0.32928058 | 0.27010638 | 0.78707841 | 0.908717 |
| CEP104 | 66.2682141 | -0.418904477 | 0.45736526 | -0.915908 | 0.35971517 | 0.64369184 |
| DFFB | 13.476099 | 0.30158304 | 0.46883871 | 0.64325542 | 0.52005838 | 0.76059646 |
| C1orf174 | 12.6580701 | -0.433831411 | 0.43547064 | -0.9962357 | 0.31913563 | 0.611915 |
| NPHP4 | 20.8977575 | -0.120219844 | 0.37999062 | -0.3163758 | 0.75171727 | 0.89132635 |
| RPL22 | 43.754267 | -0.149460786 | 0.36292782 | -0.4118196 | 0.68047166 | 0.8543838 |
| RNF207 | 34.371161 | 0.151854289 | 0.35566904 | 0.42695391 | 0.6694129 | 0.84645464 |
| ICMT | 28.6273691 | -0.804230619 | 0.40565844 | -1.9825315 | 0.04741979 | 0.21024184 |
| ESPN | 18.2956137 | 0.053661701 | 0.6346577 | 0.08455219 | 0.93261741 | 0.97356162 |
| NOL9 | 34.5607552 | 0.102895948 | 0.30480734 | 0.33757701 | 0.73568197 | 0.88426038 |
| ZBTB48 | 13.908182 | -0.554271795 | 0.37870527 | -1.4635967 | 0.14330417 | 0.40052076 |
| KLHL21 | 24.62217 | -0.008978025 | 0.41812049 | -0.0214723 | 0.98286887 | 0.99353746 |
| PHF13 | 13.1075208 | -0.14790361 | 0.47479852 | -0.3115082 | 0.75541434 | 0.89343649 |
| THAP3 | 11.0317388 | 0.417068237 | 0.40114434 | 1.03969619 | 0.29848107 | 0.59122323 |
| DNAJC11 | 96.5753594 | 0.696700863 | 0.28164762 | 2.47366147 | 0.01337364 | 0.09373482 |
| CAMTA1 | 371.935075 | -0.346420104 | 0.26576635 | -1.3034762 | 0.19241225 | 0.47297327 |
| VAMP3 | 24.195392 | -0.751225602 | 0.31539116 | -2.3818854 | 0.01722426 | 0.10980752 |
| PER3 | 90.9762286 | -0.288684386 | 0.59400964 | -0.4859928 | 0.62697229 | 0.82354566 |
| PARK7 | 88.6123018 | -0.603575716 | 0.33841135 | -1.7835564 | 0.07449578 | 0.27636138 |
| ERRFI1 | 78.1494281 | -1.657641026 | 0.5030339 | -3.2952869 | 0.00098321 | 0.01451154 |
| RERE | 1128.23004 | 0.174736329 | 0.19752613 | 0.88462386 | 0.37635954 | 0.65957178 |
| ENO1 | 362.380156 | -1.042984461 | 0.29397963 | -3.547812 | 0.00038845 | 0.00714126 |
| GPR157 | 29.6188511 | 1.491564155 | 0.32301588 | 4.61761861 | 3.88E-06 | 0.00017507 |
| MIR34AHG | 16.8815704 | -0.90574203 | 0.67736198 | -1.337161 | 0.18117008 | 0.45756661 |
| H6PD | 40.6060445 | -0.320091289 | 0.26549504 | -1.2056394 | 0.22795655 | 0.5177793 |
| SPSB1 | 42.8087218 | -2.712955056 | 0.59039633 | -4.5951421 | 4.32E-06 | 0.00018862 |
| SLC25A33 | 59.6447987 | 0.907787902 | 0.36003044 | 2.52141985 | 0.01168823 | 0.08527596 |
| PIK3CD | 14.301563 | -0.670613244 | 0.44646476 | -1.5020519 | 0.1330837 | 0.3830758 |
| CLSTN1 | 110.345495 | 0.211909603 | 0.23634002 | 0.89663021 | 0.36991627 | 0.65432818 |
| CTNNBIP1 | 30.9609797 | -0.281832802 | 0.29953619 | -0.9408973 | 0.34675748 | 0.63406026 |
| LZIC | 28.7808292 | 0.137520699 | 0.39645443 | 0.34687644 | 0.72868415 | 0.88042244 |
| NMNAT1 | 49.3366757 | -0.053350248 | 0.30973117 | -0.1722469 | 0.86324339 | 0.94635064 |
| UBE4B | 165.808087 | -0.036142355 | 0.27222456 | -0.1327667 | 0.8943779 | 0.95838164 |
| KIF1B | 177.69987 | 0.206819213 | 0.17019829 | 1.21516621 | 0.2243027 | 0.5134 |
| PGD | 37.4606466 | -0.099939802 | 0.2594609 | -0.3851825 | 0.70010219 | 0.86376951 |
| DFFA | 38.0876339 | -0.179215547 | 0.32701122 | -0.548041 | 0.58366376 | 0.79598942 |
| PEX14 | 121.010175 | 0.24024569 | 0.24526208 | 0.9795468 | 0.32730988 | 0.61700756 |
| CASZ1 | 238.710087 | 0.267419574 | 0.25111053 | 1.06494769 | 0.2868996 | 0.57931001 |
| SRM | 13.6740638 | -0.278305442 | 0.52819264 | -0.5269014 | 0.59826206 | 0.8059995 |
| EXOSC10 | 32.6390192 | -0.255016942 | 0.44161213 | -0.5774682 | 0.56362324 | 0.78465819 |
| MTOR | 71.519547 | 0.290079639 | 0.26507613 | 1.09432577 | 0.27381213 | 0.56620083 |
| FBXO2 | 27.2785392 | -0.742233572 | 0.38748622 | -1.9155096 | 0.05542755 | 0.23225616 |
| FBXO44 | 23.5850546 | 0.598208316 | 0.37164159 | 1.60963772 | 0.10747697 | 0.34161321 |
| MAD2L2 | 9.08338931 | -0.783895181 | 0.63442182 | -1.2356056 | 0.21660518 | 0.50348976 |
| AGTRAP | 30.3711459 | -1.783453277 | 0.61760061 | -2.8877129 | 0.00388054 | 0.03930906 |
| MTHFR | 27.1082671 | -0.589705511 | 0.31780664 | -1.8555481 | 0.063518 | 0.25100207 |
| CLCN6 | 30.7387379 | -0.654203157 | 0.55050431 | -1.1883706 | 0.23468742 | 0.52479917 |
| KIAA2013 | 19.7849889 | -0.767234192 | 0.57781838 | -1.327812 | 0.18424021 | 0.46080181 |
| PLOD1 | 14.8727839 | 0.063233863 | 0.45596652 | 0.13868093 | 0.88970229 | 0.95702609 |
| MFN2 | 49.9077597 | 0.363983067 | 0.29591597 | 1.2300217 | 0.21868898 | 0.50674964 |
| VPS13D | 348.254052 | 0.259343883 | 0.2389098 | 1.08553056 | 0.27768674 | 0.56997988 |
| DHRS3 | 53.3409941 | -0.237381918 | 0.58797175 | -0.4037301 | 0.68641117 | 0.85766014 |
| PRDM2 | 136.549902 | -0.444851481 | 0.22766773 | -1.9539505 | 0.05070707 | 0.21949971 |
| KAZN | 2972.47758 | -1.227545161 | 0.27251788 | -4.5044573 | 6.65E-06 | 0.00026883 |
| TMEM51-AS | 51.0659576 | -1.728679316 | 0.41675692 | -4.1479318 | 3.35E-05 | 0.00100815 |
| TMEM51 | 342.652194 | -0.398630721 | 0.51992268 | -0.7667115 | 0.44325304 | 0.71338741 |
| EFHD2 | 12.8933645 | -1.191347521 | 0.40909518 | -2.9121524 | 0.00358947 | 0.03723944 |
| CASP9 | 28.6560633 | 1.032193108 | 0.54244465 | 1.90285425 | 0.05705957 | 0.23570501 |
| DNAJC16 | 62.8842557 | 0.214554167 | 0.20775879 | 1.03270802 | 0.30174056 | 0.59542937 |
| DDI2 | 56.6269048 | 0.40307857 | 0.24484171 | 1.64628226 | 0.09970566 | 0.32592604 |
| PLEKHM2 | 57.6428297 | -0.462001603 | 0.32043551 | -1.4417928 | 0.14936084 | 0.4098869 |
| FBLIM1 | 15.9263663 | -2.144355429 | 0.59827488 | -3.5842311 | 0.00033807 | 0.00643522 |
| UQCRHL | 13.3737285 | 0.067311142 | 0.52504121 | 0.12820163 | 0.89798941 | 0.95917897 |
| SPEN | 126.82479 | 0.07583422 | 0.21872931 | 0.34670351 | 0.72881407 | 0.88042244 |
| ZBTB17 | 17.8150059 | 0.374152597 | 0.4158927 | 0.89963733 | 0.36831328 | 0.65369102 |
| CLCNKA | 103.580627 | 3.023631276 | 0.48527721 | 6.23073006 | 4.64E-10 | 8.86E-08 |
| CLCNKB | 132.156684 | 2.101030507 | 0.59730658 | 3.51750773 | 0.00043562 | 0.0078038 |

| | | | | | | |
|---|---|---|---|---|---|---|
| EPHA2 | 20.8920497 | -0.073928229 | 0.65217716 | -0.1133561 | 0.90974828 | 0.9631309 |
| FBXO42 | 143.733772 | 0.275586761 | 0.24978941 | 1.10327642 | 0.26990714 | 0.56184753 |
| SZRD1 | 45.1059142 | -0.438412172 | 0.33735873 | -1.299543 | 0.19375765 | 0.47413095 |
| NECAP2 | 24.9071802 | -0.898158298 | 0.30389217 | -2.9555164 | 0.00312146 | 0.03365503 |
| NBPF1 | 35.2554079 | -0.727003663 | 0.29724394 | -2.4458149 | 0.01445252 | 0.09763268 |
| CROCC | 39.0172438 | 0.351715759 | 0.33420354 | 1.05239985 | 0.29261614 | 0.58421152 |
| ATP13A2 | 18.6606133 | -0.068070564 | 0.45410466 | -0.1499006 | 0.88084303 | 0.95349847 |
| SDHB | 84.5009944 | 0.656483856 | 0.25200009 | 2.60509376 | 0.00918491 | 0.0724525 |
| RCC2 | 39.1533293 | -0.108108488 | 0.29277986 | -0.3692484 | 0.7119426 | 0.87086591 |
| ARHGEF10L | 370.310747 | 0.205447085 | 0.28444718 | 0.72226796 | 0.47012975 | 0.73188482 |
| KLHDC7A | 12.8435265 | 0.944989139 | 0.61591201 | 1.53429244 | 0.12495772 | 0.37009151 |
| UBR4 | 143.013927 | 0.430785464 | 0.29819717 | 1.44463298 | 0.148561 | 0.40889005 |
| AL035413.1 | 31.0877525 | 0.044472966 | 0.32539183 | 0.13667511 | 0.8912876 | 0.95716094 |
| EMC1 | 19.705369 | 0.07425579 | 0.33990274 | 0.21846187 | 0.82706926 | 0.92808376 |
| MRTO4 | 11.6446446 | -0.775365114 | 0.43661014 | -1.7758752 | 0.07575349 | 0.27917785 |
| AKR7A2 | 24.0303452 | -0.206497524 | 0.41851168 | -0.4934092 | 0.62172347 | 0.82030381 |
| CAPZB | 154.966016 | 0.076112 | 0.23418799 | 0.32500385 | 0.74517816 | 0.88808711 |
| NBL1 | 16.342794 | -1.69013107 | 0.49848209 | -3.3905552 | 0.00069751 | 0.01124533 |
| TMCO4 | 130.983853 | -0.167398887 | 0.26361457 | -0.6350138 | 0.52541942 | 0.76221787 |
| OTUD3 | 31.7006795 | -0.354014891 | 0.47560373 | -0.7443484 | 0.45666571 | 0.72302753 |
| VWA5B1 | 25.6904984 | 0.456952306 | 0.54192752 | 0.8431982 | 0.3991176 | 0.67968972 |
| CAMK2N1 | 177.684046 | -0.054583759 | 0.36249744 | -0.150577 | 0.88030944 | 0.95334202 |
| MUL1 | 14.6407906 | 0.604587087 | 0.36539603 | 1.65460771 | 0.0980041 | 0.32348372 |
| PINK1 | 34.1955657 | -0.118641933 | 0.3565766 | -0.332725 | 0.73934191 | 0.88614269 |
| DDOST | 29.4203343 | -0.678221947 | 0.3660835 | -1.8526428 | 0.06393358 | 0.25196776 |
| HP1BP3 | 203.023268 | 0.253751329 | 0.1466857 | 1.72989823 | 0.08364846 | 0.29623459 |
| EIF4G3 | 730.830471 | -0.448555591 | 0.44618667 | -1.0053093 | 0.31474796 | 0.60818748 |
| ECE1 | 38.4244836 | -0.258809448 | 0.38308846 | -0.6755866 | 0.49930312 | 0.75031683 |
| NBPF3 | 17.5801131 | 0.152466491 | 0.34352362 | 0.44383118 | 0.65716464 | 0.83965151 |
| RAP1GAP | 77.5270057 | 0.77816525 | 0.36117957 | 2.15451069 | 0.03120014 | 0.16176341 |
| USP48 | 123.980607 | 0.344741016 | 0.2231142 | 1.54513259 | 0.12231419 | 0.36640908 |
| HSPG2 | 82.6950299 | -0.029923067 | 0.50992819 | -0.0586809 | 0.95320624 | 0.98143529 |
| CDC42 | 136.364376 | -0.487268707 | 0.20480711 | -2.3791591 | 0.01735219 | 0.11021098 |
| ZBTB40 | 44.6093357 | 0.218532438 | 0.30251219 | 0.72239218 | 0.47005339 | 0.73188482 |
| EPHB2 | 40.927607 | -0.825387843 | 0.37183898 | -2.2197453 | 0.02643606 | 0.14524018 |
| KDM1A | 88.8385344 | 0.236123579 | 0.22362646 | 1.05588389 | 0.29102127 | 0.58260806 |
| AL031428.1 | 19.3723305 | -0.234201667 | 0.51074322 | -0.4585507 | 0.64655684 | 0.83458571 |
| LUZP1 | 122.92579 | 0.801462882 | 0.21766185 | 3.68214685 | 0.00023128 | 0.00480368 |
| LINC01355 | 16.6447042 | 0.4029237 | 0.47031351 | 0.856713 | 0.39160352 | 0.6727196 |
| HNRNPR | 128.027657 | 0.11915498 | 0.20174224 | 0.59062981 | 0.55476849 | 0.77938428 |
| TCEA3 | 68.4306432 | 1.115808832 | 0.42382189 | 2.63273052 | 0.00847015 | 0.06865421 |
| ASAP3 | 64.3657933 | 0.374822508 | 0.3464762 | 1.08181313 | 0.27933557 | 0.57211035 |
| RPL11 | 180.837291 | -0.208622696 | 0.43316667 | -0.4816222 | 0.63007434 | 0.82524878 |
| ELOA | 21.9996572 | -0.567436562 | 0.32580883 | -1.7416243 | 0.08157421 | 0.2922036 |
| ELOA-AS1 | 23.3061766 | 0.121788242 | 0.52612924 | 0.23147971 | 0.81694215 | 0.92248811 |
| PITHD1 | 21.9097806 | 0.198275639 | 0.32867163 | 0.60326363 | 0.54633333 | 0.77463416 |
| LYPLA2 | 24.4306873 | -0.335965619 | 0.44665017 | -0.7521896 | 0.45193704 | 0.71982927 |
| GALE | 34.0361412 | -1.562734848 | 0.32572609 | -4.7976963 | 1.61E-06 | 8.23E-05 |
| HMGCL | 43.4375953 | -0.602639629 | 0.31797129 | -1.8952643 | 0.0580574 | 0.23811563 |
| FUCA1 | 45.7865094 | -0.603789778 | 0.37479035 | -1.6110067 | 0.10717826 | 0.34111768 |
| PNRC2 | 48.0953422 | -0.032085379 | 0.25195302 | -0.1273467 | 0.89866602 | 0.95935794 |
| SRSF10 | 103.378962 | 0.131475355 | 0.18942347 | 0.69408166 | 0.487631 | 0.74443126 |
| IFNLR1 | 18.8288368 | -0.148860244 | 0.37070672 | -0.401558 | 0.68800935 | 0.85851167 |
| STPG1 | 17.8001762 | -0.496068328 | 0.41624131 | -1.1917806 | 0.23334727 | 0.52327897 |
| NIPAL3 | 68.1325523 | -0.631184077 | 0.36794382 | -1.715436 | 0.08626532 | 0.30190826 |
| RCAN3 | 33.0483053 | -1.910482219 | 0.42693377 | -4.4748913 | 7.65E-06 | 0.00030061 |
| NCMAP | 52.4179984 | -3.942264348 | 0.78591974 | -5.0161157 | 5.27E-07 | 3.39E-05 |
| SRRM1 | 101.839085 | 0.211617207 | 0.19492594 | 1.08562874 | 0.27764328 | 0.56997988 |
| CLIC4 | 40.1771811 | -1.374353807 | 0.52214188 | -2.6321463 | 0.00848473 | 0.06871832 |
| SYF2 | 35.9666303 | -0.475247565 | 0.34651381 | -1.3715112 | 0.17021566 | 0.44081492 |
| RSRP1 | 241.920978 | -0.2176572 | 0.30236385 | -0.7198519 | 0.47161618 | 0.73286947 |
| TMEM50A | 48.887853 | -1.200920372 | 0.25836867 | -4.6480882 | 3.35E-06 | 0.00015272 |
| MACO1 | 79.2569403 | 0.102415394 | 0.31888387 | 0.32116831 | 0.74808284 | 0.88941136 |
| LDLRAP1 | 14.3701954 | 0.169541056 | 0.42402112 | 0.39984106 | 0.68927358 | 0.85868974 |
| MAN1C1 | 47.128605 | 0.436560273 | 0.45172356 | 0.96643237 | 0.33382787 | 0.62250943 |
| MTFR1L | 34.9870504 | 0.455923387 | 0.43770937 | 1.04161213 | 0.29759154 | 0.59014207 |
| PAQR7 | 24.3496438 | 0.194175293 | 0.42290754 | 0.45914361 | 0.64613104 | 0.83434971 |
| STMN1 | 21.2727114 | -1.448093387 | 0.54864757 | -2.6393872 | 0.00830561 | 0.06774691 |
| PAFAH2 | 23.8057706 | 0.130722055 | 0.40063511 | 0.32628707 | 0.74420717 | 0.88798073 |
| PDIK1L | 11.5218368 | -0.433483104 | 0.49262699 | -0.8799419 | 0.37889081 | 0.66214302 |
| ZNF593 | 15.769816 | -0.723583661 | 0.47979293 | -1.5081166 | 0.1315247 | 0.38039077 |
| CNKSR1 | 22.2431472 | -0.334037994 | 0.31941426 | -1.045783 | 0.29566124 | 0.58721846 |
| CEP85 | 15.659103 | -0.075192574 | 0.44868495 | -0.1675843 | 0.86691029 | 0.94728419 |
| SH3BGRL3 | 45.4686445 | -1.91726662 | 0.47974906 | -3.9963948 | 6.43E-05 | 0.00172972 |
| UBXN11 | 10.5796904 | -0.482865177 | 0.45084187 | -1.07103 | 0.28415593 | 0.5765312 |
| DHDDS | 42.8529707 | -0.238091032 | 0.34289509 | -0.6943553 | 0.48745941 | 0.74443126 |
| HMGN2 | 64.6000193 | -0.192078896 | 0.33088872 | -0.5804939 | 0.56158157 | 0.78361672 |
| RPS6KA1 | 43.7961898 | 0.239103892 | 0.30958148 | 0.77234558 | 0.43990977 | 0.7107827 |
| ARID1A | 178.530833 | 0.051377143 | 0.20442848 | 0.25132087 | 0.80156604 | 0.91498431 |
| ZDHHC18 | 15.1092053 | -0.06965899 | 0.37194092 | -0.1872851 | 0.85143711 | 0.94012264 |
| SFN | 19.2962862 | -3.818159565 | 0.76484926 | -4.9920419 | 5.97E-07 | 3.73E-05 |
| GPN2 | 20.5742474 | -0.27653557 | 0.35467769 | -0.7796813 | 0.43557848 | 0.70733441 |
| NUDC | 51.7318895 | -0.257442093 | 0.30625861 | -0.8406036 | 0.40057004 | 0.68010059 |
| NR0B2 | 34.9177932 | 3.037141142 | 0.72763557 | 4.17398663 | 2.99E-05 | 0.000915 |
| KDF1 | 10.852572 | -0.584672266 | 0.42567676 | -1.3735123 | 0.16959314 | 0.44032355 |
| TRNP1 | 97.0832575 | -2.763922579 | 0.56931349 | -4.8548342 | 1.20E-06 | 6.46E-05 |
| SLC9A1 | 45.3278239 | 0.764717028 | 0.29775102 | 2.56831037 | 0.01021956 | 0.07837003 |
| WDTC1 | 60.6655803 | 0.272500016 | 0.24027669 | 1.13410925 | 0.25674872 | 0.54774827 |
| TMEM222 | 22.8907377 | -0.55579508 | 0.4281066 | -1.2982633 | 0.19419687 | 0.47464204 |
| WASF2 | 131.775803 | -0.195366153 | 0.1512845 | -1.2913825 | 0.19657109 | 0.47806311 |
| AHDC1 | 48.3316168 | 0.178478104 | 0.4702219 | 0.37956144 | 0.70427099 | 0.86621296 |
| IFI6 | 127.248864 | -1.564593546 | 1.83831944 | -0.8510999 | 0.39471385 | 0.67534117 |
| FAM76A | 21.5899041 | -0.724942378 | 0.36029607 | -2.0120741 | 0.04421213 | 0.20044552 |
| STX12 | 105.890121 | -0.001020536 | 0.28269368 | -0.00361 | 0.99711961 | 0.99876774 |
| PPP1R8 | 18.2798846 | -0.242247619 | 0.39278724 | -0.61674 | 0.53740621 | 0.76945357 |
| RPA2 | 30.0219564 | -0.563554569 | 0.3816139 | -1.4767664 | 0.13973828 | 0.39478933 |
| EYA3 | 463.941907 | 0.975773769 | 0.20546584 | 4.74908018 | 2.04E-06 | 0.00010072 |
| PTAFR | 49.0822447 | -0.600826602 | 0.37124879 | -1.6183934 | 0.10557784 | 0.33778352 |
| DNAJC8 | 41.0217909 | -0.442488969 | 0.27703195 | -1.5972489 | 0.11021025 | 0.34604875 |
| ATP5IF1 | 121.740341 | -0.099660146 | 0.40062817 | -0.2487597 | 0.80354666 | 0.91633138 |
| SESN2 | 12.9389225 | -1.033801108 | 0.48009022 | -2.1533476 | 0.03129137 | 0.16199182 |
| PHACTR4 | 208.361791 | -0.089085518 | 0.19198188 | -0.4640309 | 0.64262563 | 0.83318468 |
| RCC1 | 32.5759664 | -0.87010245 | 0.31841206 | -2.7326304 | 0.00628308 | 0.05594494 |
| TRNAU1AP | 44.2589971 | -0.955657732 | 0.30544573 | -3.1287317 | 0.00175563 | 0.02227395 |
| SNHG12 | 13.289271 | -0.794809417 | 0.51788352 | -1.5347262 | 0.12485109 | 0.37001028 |
| TAF12 | 27.2447012 | 0.111693154 | 0.35034543 | 0.31880865 | 0.74987162 | 0.89051036 |
| GMEB1 | 30.6590113 | -0.069353556 | 0.43716126 | -0.1586452 | 0.87394838 | 0.94952712 |
| YTHDF2 | 76.0818556 | -0.186328772 | 0.2420923 | -0.7696601 | 0.44150156 | 0.71223757 |
| EPB41 | 191.196762 | 0.207189936 | 0.19703231 | 1.0515531 | 0.29300464 | 0.5846943 |
| SRSF4 | 227.155288 | -0.01408085 | 0.27339872 | -0.051503 | 0.95892472 | 0.98365454 |
| MECR | 27.7938165 | 0.261313344 | 0.38650893 | 0.67608617 | 0.49898594 | 0.75027574 |
| PTPRU | 17.4642555 | -0.987272339 | 0.46677246 | -2.1151041 | 0.03442108 | 0.1718052 |
| PUM1 | 394.233467 | 0.311808282 | 0.19069752 | 1.63509349 | 0.10202945 | 0.33070354 |

| | | | | | | |
|---|---|---|---|---|---|---|
| SNRNP40 | 34.0839394 | 0.571559944 | 0.27257256 | 2.09690934 | 0.0360016 | 0.17602651 |
| ZCCHC17 | 63.2795774 | -0.166343483 | 0.29076635 | -0.5720864 | 0.56726342 | 0.78656476 |
| SERINC2 | 107.691887 | -0.438083845 | 0.34562797 | -1.2675011 | 0.20497616 | 0.48960454 |
| TINAGL1 | 41.9692584 | 0.039229204 | 0.47904596 | 0.08189027 | 0.93473397 | 0.97513509 |
| PEF1 | 30.9243017 | -0.017238366 | 0.39547289 | -0.0435892 | 0.96523182 | 0.98660829 |
| PTP4A2 | 59.4615971 | -0.35329203 | 0.40891927 | -0.8639652 | 0.387607 | 0.66965073 |
| KHDRBS1 | 148.666651 | -0.433028373 | 0.27408811 | -1.5798875 | 0.11413264 | 0.3526678 |
| TMEM39B | 11.1899399 | 0.482932746 | 0.46923871 | 1.02918351 | 0.30339345 | 0.59727868 |
| KPNA6 | 124.591859 | 0.090183308 | 0.21576128 | 0.41797727 | 0.67596374 | 0.85173672 |
| TXLNA | 40.1461935 | -0.589408219 | 0.26667674 | -2.2101973 | 0.02709147 | 0.14689281 |
| TMEM234 | 13.8513271 | -0.318107551 | 0.40663305 | -0.7822964 | 0.43404043 | 0.70597357 |
| EIF3I | 28.7839193 | -0.827089112 | 0.51385742 | -1.6095693 | 0.10749192 | 0.34161321 |
| HDAC1 | 47.7545373 | 0.237258816 | 0.31331864 | 0.75724449 | 0.4489034 | 0.71802342 |
| BSDC1 | 59.1320721 | -0.289091669 | 0.30257845 | -0.9554272 | 0.33936172 | 0.62811794 |
| ZBTB8OS | 90.5302037 | 0.356326694 | 0.2159085 | 1.65035975 | 0.09886938 | 0.32479347 |
| RBBP4 | 92.7274355 | 0.113185684 | 0.18497388 | 0.61190089 | 0.54060333 | 0.7714342 |
| SYNC | 20.4281386 | -0.862188681 | 0.49772104 | -1.7322729 | 0.08322498 | 0.29554766 |
| KIAA1522 | 112.602138 | -0.186281378 | 0.26795291 | -0.6952019 | 0.48692877 | 0.74426412 |
| YARS | 52.413763 | -0.880851174 | 0.54263639 | -1.6232807 | 0.10452941 | 0.33555699 |
| S100PBP | 85.875865 | -0.150654579 | 0.25002579 | -0.6025562 | 0.54680399 | 0.77463416 |
| TMEM54 | 13.4652685 | -1.025868417 | 0.66387222 | -1.5452799 | 0.12227857 | 0.3664089 |
| RNF19B | 34.3904919 | -0.516717762 | 0.6182918 | -0.8357183 | 0.40331341 | 0.68158378 |
| AL020995.1 | 22.3207829 | -0.453390639 | 0.66786246 | -0.6788683 | 0.49722132 | 0.74900919 |
| AK2 | 108.319278 | 0.312227345 | 0.24815429 | 1.25819845 | 0.20832 | 0.49402397 |
| TRIM62 | 12.3302778 | -0.56887717 | 0.39424177 | -1.4429652 | 0.14903027 | 0.4093068 |
| ZNF362 | 23.6835894 | 0.130098399 | 0.4375353 | 0.29734378 | 0.76620406 | 0.89892202 |
| AL513327.1 | 21.3749936 | -0.317812823 | 0.6361286 | -0.4996047 | 0.61735347 | 0.81758265 |
| PHC2 | 154.799123 | -0.584334248 | 0.37887682 | -1.5422803 | 0.1230055 | 0.36741162 |
| SMIM12 | 32.5956803 | -0.582471677 | 0.46739759 | -1.2462017 | 0.21269035 | 0.49888203 |
| ZMYM1 | 38.96879 | 0.28575644 | 0.30904905 | 0.92463134 | 0.35515771 | 0.64025612 |
| SFPQ | 203.835672 | 0.255992158 | 0.20055241 | 1.27643521 | 0.20180172 | 0.48515376 |
| ZMYM4 | 341.564126 | 0.314935244 | 0.17261309 | 1.82451537 | 0.06807419 | 0.26216836 |
| KIAA0319L | 186.759314 | -0.242501158 | 0.16934925 | -1.4319588 | 0.1521556 | 0.4136994 |
| PSMB2 | 40.3779764 | -0.260928851 | 0.34991991 | -0.7456816 | 0.45585975 | 0.72261381 |
| AGO4 | 70.2576393 | -0.675868006 | 0.22756613 | -2.969985 | 0.00297814 | 0.03236374 |
| AGO1 | 26.9434803 | 0.181332341 | 0.27105258 | 0.66899322 | 0.5034998 | 0.75108109 |
| AGO3 | 221.446063 | 0.135452969 | 0.21436017 | 0.6318943 | 0.52745595 | 0.76221787 |
| ADPRHL2 | 21.2402506 | -1.626800771 | 0.49181677 | -3.3077375 | 0.00094053 | 0.01400192 |
| TRAPPC3 | 30.1909509 | -0.332512749 | 0.294274 | -1.1299427 | 0.25850039 | 0.54999401 |
| MAP7D1 | 20.3127871 | -0.4351605 | 0.41687057 | -1.0438744 | 0.29654352 | 0.58863032 |
| THRAP3 | 159.587415 | 0.01858582 | 0.26936512 | 0.06899861 | 0.94499072 | 0.97858537 |
| SH3D21 | 12.2497538 | 1.02867563 | 0.4951905 | 2.07733312 | 0.03777082 | 0.18216996 |
| EVA1B | 15.4696032 | -0.580469844 | 0.45694617 | -1.2703244 | 0.20396912 | 0.4882179 |
| STK40 | 47.5393277 | -0.374017457 | 0.37029211 | -1.0100606 | 0.31246628 | 0.60551329 |
| OSCP1 | 11.3593147 | 0.118149978 | 0.48394452 | 0.24413951 | 0.80712277 | 0.91776808 |
| MRPS15 | 20.0474747 | -0.253566555 | 0.32761497 | -0.7739773 | 0.43894421 | 0.71027377 |
| LINC01137 | 19.3013827 | 0.452960648 | 0.48641024 | 0.93123173 | 0.35173371 | 0.63801604 |
| ZC3H12A | 25.787307 | 1.250228848 | 0.54469539 | 2.29528076 | 0.02171704 | 0.12733577 |
| MEAF6 | 35.9828081 | -0.138779231 | 0.30731929 | -0.45158 | 0.65157161 | 0.83781115 |
| SNIP1 | 14.82084 | -0.298234129 | 0.4147846 | -0.7190096 | 0.47213498 | 0.73329331 |
| DNALI1 | 12.9960135 | -0.852333298 | 0.48095548 | -1.7721667 | 0.07636688 | 0.28042437 |
| GNL2 | 14.4732727 | -0.13027608 | 0.49060142 | -0.2655436 | 0.79059071 | 0.91080793 |
| C1orf122 | 33.0824031 | -1.005870763 | 0.42348331 | -2.3752312 | 0.01753796 | 0.11098039 |
| MTF1 | 28.7234817 | -0.406269705 | 0.3528553 | -1.1513777 | 0.2495769 | 0.54083745 |
| INPP5B | 80.4334998 | 0.085272817 | 0.31274553 | 0.27265878 | 0.78511552 | 0.90787519 |
| SF3A3 | 34.8814675 | -0.092677975 | 0.27468369 | -0.3373989 | 0.73581621 | 0.88431855 |
| RRAGC | 27.2921114 | -1.50146627 | 0.48169782 | -3.1170294 | 0.00182683 | 0.02295127 |
| AL139260.1 | 42.9812536 | -1.169902249 | 0.51653322 | -2.2649119 | 0.02351809 | 0.13454711 |
| AKIRIN1 | 78.5252921 | 0.328140598 | 0.33284277 | 0.9858727 | 0.32419559 | 0.61460758 |
| NDUFS5 | 113.314627 | -0.74323292 | 0.43564447 | -1.7060538 | 0.08799806 | 0.30539182 |
| MACF1 | 967.094779 | -0.240716341 | 0.24552374 | -0.9804198 | 0.32687893 | 0.61675948 |
| PABPC4 | 39.472452 | 0.084923705 | 0.2860382 | 0.29689638 | 0.76654562 | 0.89911488 |
| PPIE | 23.7640233 | 0.144331351 | 0.32211679 | 0.4480715 | 0.6541016 | 0.83885315 |
| TRIT1 | 56.3153891 | 0.324330233 | 0.26086407 | 1.24329206 | 0.21376023 | 0.50026184 |
| CAP1 | 65.8545967 | -0.674330838 | 0.30335622 | -2.222901 | 0.02622248 | 0.14453931 |
| PPT1 | 25.9407399 | -0.033077746 | 0.35756263 | -0.092509 | 0.92629368 | 0.97111125 |
| RLF | 174.9686 | 0.089272839 | 0.29330604 | 0.30436755 | 0.76084789 | 0.89648567 |
| ZMPSTE24 | 53.9855693 | -0.096799237 | 0.26291394 | -0.3681784 | 0.71274021 | 0.87134026 |
| COL9A2 | 11.5400777 | 0.973738603 | 0.69491935 | 1.40122534 | 0.1611467 | 0.42820049 |
| SMAP2 | 30.5792904 | 0.670052471 | 0.3046946 | 2.1990953 | 0.02787115 | 0.14970206 |
| AL603839.3 | 26.0751417 | 0.264548145 | 0.33413907 | 0.79173066 | 0.42851774 | 0.70284823 |
| ZNF684 | 45.1853898 | 0.751029503 | 0.41904076 | 1.79225883 | 0.07309151 | 0.27311889 |
| NFYC | 161.693192 | 1.046933406 | 0.26539097 | 3.94487196 | 7.98E-05 | 0.00204612 |
| CITED4 | 17.0073572 | -2.40946182 | 0.61445215 | -3.9213173 | 8.81E-05 | 0.00221077 |
| SLFNL1-AS1 | 16.7178439 | -1.087237012 | 0.46072857 | -2.3598211 | 0.01828375 | 0.113881 |
| SCMH1 | 502.642061 | -0.131940379 | 0.1376067 | -0.9588224 | 0.33764824 | 0.62621892 |
| AC093151.3 | 16.4383335 | -2.122659698 | 0.64765458 | -3.2774565 | 0.00104747 | 0.01524156 |
| HIVEP3 | 104.114557 | -0.983360088 | 0.27280947 | -3.6045673 | 0.00031267 | 0.00606622 |
| FOXJ3 | 203.649604 | 0.126070857 | 0.2997064 | 0.42064787 | 0.67401223 | 0.85019422 |
| PPCS | 26.2349538 | -0.698295062 | 0.41717154 | -1.6738799 | 0.09415421 | 0.3160115 |
| CCDC30 | 149.693308 | 0.358639959 | 0.30776417 | 1.1653077 | 0.24389446 | 0.53459591 |
| PPIH | 15.266307 | 0.257064892 | 0.39094068 | 0.65755472 | 0.5108243 | 0.7549696 |
| YBX1 | 99.3558033 | -0.510222198 | 0.3516313 | -1.4510148 | 0.14677575 | 0.40603754 |
| ERMAP | 23.6086158 | 0.216664243 | 0.34244733 | 0.63269363 | 0.52693373 | 0.76221787 |
| SLC2A1 | 120.943809 | -1.661217849 | 0.67197409 | -2.4721457 | 0.01343048 | 0.09373482 |
| SLC2A1-AS1 | 20.1180537 | -0.28625438 | 0.49835292 | -0.5744009 | 0.56569653 | 0.7859225 |
| EBNA1BP2 | 18.9216276 | -0.388556044 | 0.38762971 | -1.0023897 | 0.31615539 | 0.60951749 |
| TMEM125 | 11.8038228 | -1.271496114 | 0.51907088 | -2.4495616 | 0.01430302 | 0.09720405 |
| ELOVL1 | 17.1302198 | -0.596876129 | 0.44907531 | -1.3291226 | 0.18380753 | 0.46016652 |
| SZT2 | 42.8921361 | -0.201500736 | 0.29939966 | -0.6730159 | 0.50093717 | 0.75071921 |
| PTPRF | 145.630186 | -0.410308913 | 0.40684436 | -1.0085157 | 0.31320697 | 0.60594519 |
| IPO13 | 10.444541 | 0.186568497 | 0.51363676 | 0.36323042 | 0.71643276 | 0.8732017 |
| ATP6V0B | 109.322116 | -0.201993858 | 0.33893436 | -0.5959675 | 0.55119695 | 0.77690942 |
| B4GALT2 | 14.4536453 | -0.499300692 | 0.58401404 | -0.8549464 | 0.39258083 | 0.67372442 |
| CCDC24 | 26.6157233 | -0.637706881 | 0.32677313 | -1.9515279 | 0.05099428 | 0.21990081 |
| DMAP1 | 17.3958409 | 0.541362175 | 0.40097625 | 1.35011031 | 0.1769806 | 0.45185765 |
| ERI3 | 90.6675649 | 0.245110312 | 0.35509302 | 0.6902707 | 0.49002397 | 0.74556088 |
| RNF220 | 20.777935 | -0.498144026 | 0.35902859 | -1.3874773 | 0.16529626 | 0.43418718 |
| TMEM53 | 17.8790116 | -0.261467452 | 0.41376393 | -0.6319242 | 0.52743639 | 0.76221787 |
| ARMH1 | 19.7422496 | 0.687209228 | 0.48083475 | 1.42920043 | 0.15294663 | 0.41497399 |
| RPS8 | 145.113195 | -1.002494411 | 0.29252439 | -3.4270456 | 0.00061019 | 0.01007395 |
| EIF2B3 | 59.1666292 | 0.256841938 | 0.29936493 | 0.857956 | 0.39091676 | 0.67223049 |
| HECTD3 | 12.5975089 | -0.694495459 | 0.48315257 | -1.4374248 | 0.15059734 | 0.4115262 |
| ZSWIM5 | 93.9160183 | 0.236225986 | 0.34240729 | 0.68989766 | 0.49025855 | 0.7455837 |
| TESK2 | 30.2333691 | -0.162264002 | 0.33545005 | -0.4837203 | 0.62858437 | 0.82482184 |
| PRDX1 | 172.090696 | -1.178671906 | 0.42773076 | -2.7556398 | 0.00585775 | 0.05330963 |
| AKR1A1 | 21.7192157 | -0.491189087 | 0.34928945 | -1.4062523 | 0.15964922 | 0.42727956 |
| NASP | 61.6790039 | 0.438169825 | 0.34195885 | 1.28135249 | 0.20006988 | 0.48370812 |
| GPBP1L1 | 208.762033 | 0.36566165 | 0.14812593 | 2.4685864 | 0.01356479 | 0.09448578 |
| TMEM69 | 12.3712023 | -0.376045838 | 0.45815104 | -0.8207901 | 0.41176584 | 0.68902299 |
| IPP | 46.0859131 | 0.33221259 | 0.26574707 | 1.2501082 | 0.21126002 | 0.49712778 |
| MAST2 | 58.8371542 | -0.295453508 | 0.38572097 | -0.7659773 | 0.44368981 | 0.71343168 |
| PIK3R3 | 12.2720481 | -0.349284924 | 0.57848462 | -0.6037929 | 0.54598132 | 0.77463416 |

| | | | | | | |
|---|---|---|---|---|---|---|
| TSPAN1 | 290.296938 | -2.99697585 | 0.47957233 | -6.2492676 | 4.12E-10 | 8.02E-08 |
| POMGNT1 | 53.8614751 | -0.258562752 | 0.32707595 | -0.7905282 | 0.42921938 | 0.70310352 |
| LURAP1 | 60.8927814 | 0.826297302 | 0.23786055 | 3.47387289 | 0.000513 | 0.00876446 |
| LRRC41 | 45.1656527 | -0.425387703 | 0.25069366 | -1.6968427 | 0.0897264 | 0.30853184 |
| UQCRH | 131.514804 | 0.530384075 | 0.31417486 | 1.68818116 | 0.09137646 | 0.31120975 |
| NSUN4 | 31.441604 | -0.08966968 | 0.31232646 | -0.2871024 | 0.7740339 | 0.90184317 |
| FAAH | 13.5223075 | 0.609968625 | 0.39610741 | 1.53990711 | 0.123583 | 0.36843999 |
| MKNK1 | 45.3070998 | 0.408323092 | 0.27587021 | 1.48012753 | 0.13883921 | 0.39312191 |
| ATPAF1 | 57.0128605 | 0.596104827 | 0.28738279 | 2.07425375 | 0.03805575 | 0.18276383 |
| EFCAB14 | 150.185979 | 0.302300503 | 0.30718089 | 0.98411233 | 0.32506028 | 0.61535668 |
| PDZK1IP1 | 75.5879573 | -2.147332393 | 0.56966317 | -3.7694773 | 0.00016359 | 0.0036243 |
| CMPK1 | 71.9941481 | -0.704062444 | 0.28388769 | -2.4800739 | 0.01313552 | 0.0924963 |
| TRABD2B | 936.711774 | -0.707750446 | 0.24635009 | -2.8729458 | 0.00406664 | 0.04049316 |
| SPATA6 | 164.314842 | 0.54906457 | 0.24737842 | 2.21953298 | 0.02645049 | 0.14524018 |
| AGBL4 | 778.119499 | 0.970866806 | 0.43207892 | 2.24696637 | 0.02464218 | 0.13752463 |
| FAF1 | 610.965659 | 0.375122353 | 0.1705782 | 2.19912241 | 0.02786922 | 0.14970206 |
| RNF11 | 80.4650149 | 0.060122409 | 0.29828993 | 0.20155695 | 0.8402631 | 0.93552258 |
| TTC39A | 79.7684766 | -2.539306844 | 0.41985004 | -6.0481281 | 1.47E-09 | 2.43E-07 |
| EPS15 | 245.365856 | 0.090792888 | 0.23269586 | 0.39017836 | 0.69640466 | 0.86247999 |
| OSBPL9 | 201.315741 | 0.399014368 | 0.17328455 | 2.30265402 | 0.02129832 | 0.12559546 |
| NRDC | 168.917197 | -0.011483024 | 0.17842327 | -0.0643583 | 0.9486849 | 0.97969618 |
| TXNDC12 | 37.4235878 | -0.454130487 | 0.29329441 | -1.5483776 | 0.12153142 | 0.36544561 |
| BTF3L4 | 63.9151708 | -0.048292865 | 0.24877853 | -0.1941199 | 0.84608199 | 0.93757726 |
| ZFYVE9 | 139.875399 | 0.455341583 | 0.18060667 | 2.52117805 | 0.01169627 | 0.08527596 |
| CC2D1B | 25.9593935 | -0.305609327 | 0.4019752 | -0.7602691 | 0.44709375 | 0.71665782 |
| PRPF38A | 24.9750788 | 0.258634172 | 0.30130616 | 0.85837666 | 0.39068451 | 0.67223049 |
| TUT4 | 218.079423 | 0.243639453 | 0.2310682 | 1.05440496 | 0.29169755 | 0.58328187 |
| ZYG11B | 95.6476055 | 0.353466879 | 0.24075423 | 1.46816476 | 0.14205948 | 0.39812207 |
| ECHDC2 | 193.432647 | 0.384926599 | 0.33739826 | 1.14086717 | 0.2539252 | 0.5452549 |
| SCP2 | 142.116299 | 0.070527653 | 0.25901779 | 0.27228884 | 0.78539994 | 0.90802494 |
| CPT2 | 20.5207549 | 0.043062958 | 0.47067049 | 0.09149279 | 0.92710103 | 0.97111125 |
| C1orf123 | 10.8764507 | 0.271044863 | 0.40211043 | 0.67405578 | 0.50027585 | 0.75052305 |
| MAGOH | 23.3444859 | 0.236674271 | 0.30458373 | 0.77704173 | 0.43713415 | 0.70829758 |
| LRP8 | 13.6634801 | -1.141552854 | 0.40152971 | -2.8430097 | 0.00446897 | 0.04372207 |
| NDC1 | 34.8834814 | 0.355350119 | 0.33435539 | 1.06279166 | 0.28787644 | 0.5800319 |
| YIPF1 | 25.9142551 | -0.469291597 | 0.31950648 | -1.4688015 | 0.14188664 | 0.39807085 |
| HSPB11 | 30.5295408 | -0.239439084 | 0.42789264 | -0.5595775 | 0.57576768 | 0.79171629 |
| LRRC42 | 18.5192928 | -0.408764972 | 0.36454805 | -1.1212924 | 0.2621634 | 0.55389814 |
| TMEM59 | 160.65356 | -0.174195098 | 0.28981321 | -0.6010599 | 0.5478001 | 0.77516644 |
| TCEANC2 | 29.8284601 | -0.188906871 | 0.39707892 | -0.4757414 | 0.63425864 | 0.82785872 |
| MRPL37 | 16.1020805 | 0.165586089 | 0.3723865 | 0.4446619 | 0.65656411 | 0.83951126 |
| SSBP3 | 239.619672 | -0.097964492 | 0.28317172 | -0.3459544 | 0.72937701 | 0.88058619 |
| ACOT11 | 229.270454 | 0.967931232 | 0.34717802 | 2.78799685 | 0.00530351 | 0.0495796 |
| MROH7-TTC4 | 21.5679454 | 1.538766149 | 0.57070838 | 2.69623892 | 0.00701273 | 0.06045623 |
| TTC22 | 11.4042834 | -1.391548949 | 0.44137543 | -3.1527558 | 0.00161737 | 0.02101154 |
| DHCR24 | 41.7955165 | -1.021767848 | 0.51946981 | -1.9669437 | 0.04918971 | 0.21527288 |
| USP24 | 380.826391 | 0.458458803 | 0.30929283 | 1.48228075 | 0.13826561 | 0.39196816 |
| PLPP3 | 167.206959 | -0.79786183 | 0.53909947 | -1.47999 | 0.13887591 | 0.39312191 |
| PRKAA2 | 288.91245 | 0.48382799 | 0.26481168 | 1.8270644 | 0.06769009 | 0.26107947 |
| FYB2 | 315.595842 | -0.697285305 | 0.44291085 | -1.5743243 | 0.1154125 | 0.35464129 |
| DAB1 | 254.020468 | 1.974536611 | 1.09426805 | 1.80443595 | 0.07116299 | 0.26923288 |
| TACSTD2 | 171.089832 | -0.627229451 | 0.48526988 | -1.2925374 | 0.19617112 | 0.47765419 |
| MYSM1 | 121.79729 | 0.655469976 | 0.29644998 | 2.21106436 | 0.02703138 | 0.14670533 |
| AL136985.3 | 14.2060695 | 0.062732549 | 0.48101855 | 0.13041607 | 0.89623725 | 0.95900099 |
| JUN | 79.9237643 | -0.215296427 | 0.55697248 | -0.3865477 | 0.69909109 | 0.86334649 |
| LINC01135 | 38.1159131 | 0.680546043 | 0.56334269 | 1.20804983 | 0.2270281 | 0.51677386 |
| FGGY | 166.070114 | 0.116017451 | 0.26976992 | 0.43006074 | 0.66715146 | 0.84497102 |
| HOOK1 | 129.477062 | 0.268969237 | 0.28435346 | 0.94589753 | 0.34420085 | 0.63200319 |
| AC097066.1 | 15.0012111 | -1.204145031 | 0.89222279 | -1.3496013 | 0.17714391 | 0.4519407 |
| NFIA | 137.427851 | 0.018164197 | 0.58811763 | 0.03088531 | 0.975361 | 0.99023703 |
| TM2D1 | 84.3908124 | 0.106542403 | 0.24204439 | 0.44017713 | 0.65980882 | 0.8409935 |
| PATJ | 1210.34399 | 0.39466989 | 0.16066892 | 2.4564171 | 0.01403302 | 0.09623211 |
| USP1 | 18.8506389 | -0.25353456 | 0.33611292 | -0.7543137 | 0.45066091 | 0.71913083 |
| DOCK7 | 364.050106 | 0.384173348 | 0.1964254 | 1.95582313 | 0.05048599 | 0.21890013 |
| AC103923.1 | 14.7389028 | -0.215651205 | 0.43351119 | -0.4974524 | 0.61887003 | 0.81841635 |
| ATG4C | 42.0603425 | 0.375098937 | 0.35817489 | 1.04725079 | 0.29498393 | 0.58668972 |
| ALG6 | 22.4702553 | 0.721653024 | 0.33094639 | 2.18057379 | 0.02921496 | 0.15474163 |
| ITGB3BP | 63.3736577 | -1.170897791 | 0.48147579 | -2.4318934 | 0.01502013 | 0.10021849 |
| EFCAB7 | 35.4970281 | -0.185815948 | 0.38444195 | -0.4833394 | 0.62885475 | 0.82494825 |
| PGM1 | 66.7556577 | -0.598200744 | 0.24079786 | -2.4842444 | 0.01298267 | 0.09198586 |
| ROR1 | 55.1422887 | 0.844152063 | 0.54036376 | 1.56219221 | 0.11824271 | 0.35954441 |
| CACHD1 | 216.141259 | 0.663042051 | 0.2005191 | 3.30662783 | 0.00094426 | 0.01400282 |
| RAVER2 | 67.0021233 | 0.535459749 | 0.29255997 | 1.8302564 | 0.06721161 | 0.26017544 |
| JAK1 | 230.953277 | -0.369005108 | 0.31331377 | -1.1777494 | 0.23889652 | 0.52912013 |
| AK4 | 26.3804389 | -2.037888352 | 0.66261287 | -3.0755339 | 0.00210126 | 0.02536445 |
| DNAJC6 | 19.3523588 | -0.027315447 | 0.61732967 | -0.0442477 | 0.96470692 | 0.98660829 |
| LEPR | 88.5042095 | -0.932813228 | 0.34286249 | -2.7206628 | 0.00651512 | 0.05721973 |
| SGIP1 | 34.4692748 | 0.25284334 | 0.84570574 | 0.29897318 | 0.76496051 | 0.89797438 |
| WDR78 | 54.5874123 | -0.426020006 | 0.44143761 | -0.9650741 | 0.33450769 | 0.62316424 |
| MIER1 | 132.687782 | -0.343693449 | 0.27690071 | -1.2412155 | 0.21452616 | 0.50091761 |
| SLC35D1 | 43.1703574 | 0.281207591 | 0.24608452 | 1.1427277 | 0.25315167 | 0.54473281 |
| SERBP1 | 120.93739 | -0.320999461 | 0.28294615 | -1.1344896 | 0.25658923 | 0.54762425 |
| GADD45A | 28.5796567 | -1.020005987 | 0.4054623 | -2.5156617 | 0.01188092 | 0.08631678 |
| GNG12 | 193.779423 | -0.410598271 | 0.1633485 | -2.5136336 | 0.01194945 | 0.08669243 |
| GNG12-AS1 | 69.7047492 | -0.640329035 | 0.29814904 | -2.147681 | 0.0317391 | 0.16311062 |
| WLS | 105.87396 | 0.452157104 | 0.24776227 | 1.82496354 | 0.06800653 | 0.26210374 |
| LRRC40 | 44.6975787 | 0.5085374 | 0.31642904 | 1.60711357 | 0.10802946 | 0.34264762 |
| SRSF11 | 206.218189 | 0.013362461 | 0.23471091 | 0.05693157 | 0.95459971 | 0.98226508 |
| ANKRD13C | 133.700484 | 0.494213903 | 0.24882811 | 1.98616586 | 0.04701491 | 0.20913109 |
| LINC01788 | 56.2908164 | 2.165562427 | 0.48531176 | 4.46220881 | 8.11E-06 | 0.00031182 |
| PTGER3 | 634.303121 | 1.085368314 | 0.52108066 | 2.08291807 | 0.03725869 | 0.18041044 |
| ZRANB2-AS1 | 17.1027671 | 0.021143678 | 0.59553318 | 0.03550378 | 0.97167803 | 0.988268 |
| ZRANB2 | 211.621966 | 0.361832995 | 0.20996023 | 1.72334064 | 0.08482694 | 0.29907158 |
| NEGR1 | 15.4807554 | -0.349012107 | 0.50607939 | -0.6896391 | 0.4904212 | 0.7455837 |
| LRRIQ3 | 13.9793412 | 0.589993155 | 0.87285541 | 0.67593458 | 0.49908218 | 0.75027574 |
| CRYZ | 105.413829 | 0.14172201 | 0.33273749 | 0.42592739 | 0.67016076 | 0.84711609 |
| TYW3 | 30.9098255 | 0.034052825 | 0.34941423 | 0.09745689 | 0.92236357 | 0.96885417 |
| ACADM | 81.2051037 | 0.590252637 | 0.27327667 | 2.15990861 | 0.03077975 | 0.16030988 |
| RABGGTB | 23.0968603 | 0.237160084 | 0.33544785 | 0.70699539 | 0.47956934 | 0.7392673 |
| ST6GALNAC3 | 29.4666473 | 0.370509913 | 0.32865463 | 1.12735339 | 0.2595931 | 0.55072654 |
| PIGK | 46.5347042 | -0.045575976 | 0.25449368 | -0.1790849 | 0.85787104 | 0.94359207 |
| AC118549.1 | 241.92711 | 0.435869112 | 0.18314236 | 2.37994706 | 0.01731513 | 0.11006868 |
| USP33 | 177.295672 | 0.327121298 | 0.17705783 | 1.8475393 | 0.06466902 | 0.2538949 |
| MIGA1 | 104.843231 | 0.150652401 | 0.21131848 | 0.71291635 | 0.47589752 | 0.73616591 |
| NEXN | 25.8451268 | 0.403690136 | 0.52682661 | 0.76626756 | 0.44351712 | 0.71343168 |
| FUBP1 | 132.041651 | 0.431221816 | 0.20456814 | 2.1079618 | 0.03503429 | 0.1740015 |
| GIPC2 | 23.3107404 | 1.131149488 | 0.41891944 | 2.70015995 | 0.00693061 | 0.05984844 |
| PTGFR | 61.552397 | 0.296375536 | 0.42397069 | 0.69904723 | 0.48452252 | 0.74250561 |
| IFI44L | 102.662232 | -0.391920901 | 1.81491374 | -0.2159446 | 0.8290309 | 0.92883918 |
| IFI44 | 40.2897682 | -0.029358857 | 1.3999576 | -0.0209712 | 0.98326859 | 0.99368608 |
| LINC01781 | 33.6738283 | -3.072797924 | 0.78142412 | -3.9323049 | 8.41E-05 | 0.00212963 |
| ADGRL2 | 438.166923 | 1.475087434 | 0.30325394 | 4.86419868 | 1.15E-06 | 6.22E-05 |
| LINC01362 | 19.0445223 | -1.013741011 | 0.76532897 | -1.324582 | 0.18530982 | 0.46213065 |

| | | | | | | |
|---|---|---|---|---|---|---|
| LINC01725 | 298.6756 | 0.436450364 | 0.46590892 | 0.93677185 | 0.34887593 | 0.63556496 |
| PRKACB | 110.115088 | 0.556465657 | 0.3311895 | 1.68020319 | 0.09291779 | 0.31376482 |
| RPF1 | 15.2310482 | 0.099928883 | 0.36958505 | 0.27038129 | 0.78686693 | 0.90871373 |
| GNG5 | 58.5974727 | -0.387885601 | 0.31321983 | -1.2383814 | 0.21557469 | 0.5022276 |
| SPATA1 | 22.8331178 | -0.787230492 | 0.64331654 | -1.2237063 | 0.22106305 | 0.5089143 |
| CTBS | 39.1165349 | -0.497482445 | 0.29011268 | -1.7147904 | 0.08638366 | 0.30207891 |
| SSX2IP | 22.3350676 | -0.415940297 | 0.34666711 | -1.1998263 | 0.23020683 | 0.52042807 |
| SYDE2 | 21.9479039 | 1.379882699 | 0.45903896 | 3.00602526 | 0.00264687 | 0.029932 |
| C1orf52 | 35.7036809 | 0.127926386 | 0.35067549 | 0.3647999 | 0.71526078 | 0.87305883 |
| BCL10 | 31.6518623 | -0.483188125 | 0.35628474 | -1.3561853 | 0.17504022 | 0.44812733 |
| AL078459.1 | 50.1852191 | -0.494003703 | 0.46480328 | -1.0628232 | 0.28786214 | 0.5800319 |
| DDAH1 | 245.551024 | 0.379378022 | 0.31572381 | 1.20161361 | 0.22951327 | 0.51974407 |
| ZNHIT6 | 56.2143357 | -0.033922235 | 0.26151615 | -0.1297137 | 0.89679292 | 0.95900099 |
| ODF2L | 130.438896 | -0.274422351 | 0.28854219 | -0.9510649 | 0.34157143 | 0.62971883 |
| SH3GLB1 | 99.3683334 | -0.660662569 | 0.18633916 | -3.5454844 | 0.00039189 | 0.00718376 |
| SELENOF | 101.209272 | 0.197927722 | 0.17695253 | 1.11853568 | 0.26333828 | 0.55501451 |
| HS2ST1 | 109.230799 | -0.024602332 | 0.27072209 | -0.0908767 | 0.92759056 | 0.97114207 |
| LMO4 | 25.4958291 | -1.093194733 | 0.35640507 | -3.0672816 | 0.00216015 | 0.02581658 |
| PKN2-AS1 | 21.1239921 | 0.438401359 | 0.77746352 | 0.56388673 | 0.57283122 | 0.78986357 |
| PKN2 | 282.675235 | 0.127991332 | 0.17200602 | 0.74410962 | 0.45681016 | 0.72302753 |
| GTF2B | 35.4537906 | -0.502360228 | 0.30243821 | -1.6610343 | 0.09670657 | 0.32117055 |
| KYAT3 | 54.7922802 | -0.4859484 | 0.2609972 | -1.8618912 | 0.06261843 | 0.24887925 |
| GBP3 | 28.4330631 | -1.481490414 | 0.58703844 | -2.5236685 | 0.01161374 | 0.08503535 |
| GBP2 | 53.4408446 | -5.104704626 | 0.7189218 | -7.1005006 | 1.24E-12 | 5.12E-10 |
| LRRC8B | 73.8563175 | -0.230460241 | 0.23071048 | -0.9989154 | 0.31783569 | 0.61101185 |
| LRRC8D | 10.1812693 | -0.285745059 | 0.51970134 | -0.5498255 | 0.58243905 | 0.7955793 |
| ZNF326 | 63.8664449 | 0.014164281 | 0.23940217 | 0.05916521 | 0.95282052 | 0.98135593 |
| ZNF644 | 229.473223 | 0.288048234 | 0.24035732 | 1.19841674 | 0.23075482 | 0.52105604 |
| HFM1 | 17.2517985 | 0.931537334 | 0.65531181 | 1.42151769 | 0.15516632 | 0.41868082 |
| TGFBR3 | 29.042097 | -0.583223528 | 0.48731797 | -1.1968028 | 0.23138342 | 0.52111057 |
| BTBD8 | 36.2103222 | -0.364622232 | 0.54263081 | -0.6719527 | 0.50161381 | 0.75108109 |
| GLMN | 42.7159566 | 0.174629666 | 0.2376002 | 0.73497273 | 0.4623561 | 0.72720497 |
| RPAP2 | 136.565783 | 0.458947163 | 0.24667018 | 1.86057007 | 0.06280491 | 0.24926305 |
| EVI5 | 272.768501 | 0.230705183 | 0.19833311 | 1.16322072 | 0.24473996 | 0.53553761 |
| RPL5 | 106.338239 | -0.569313702 | 0.34587628 | -1.6460039 | 0.09976296 | 0.32596194 |
| FAM69A | 28.81521 | 0.435573794 | 0.30826701 | 1.41297571 | 0.15766286 | 0.42342097 |
| MTF2 | 138.773931 | 0.307173132 | 0.21917896 | 1.40147179 | 0.16107304 | 0.42820049 |
| TMED5 | 61.2218494 | -0.340341848 | 0.24575478 | -1.3848839 | 0.16608796 | 0.43493598 |
| CCDC18 | 23.9740662 | -0.988807169 | 0.41007057 | -2.4113098 | 0.01589534 | 0.10383881 |
| DR1 | 29.3506577 | -0.111494818 | 0.33760157 | -0.3302556 | 0.74120682 | 0.88665962 |
| FNBP1L | 407.919571 | 0.338771767 | 0.2087271 | 1.62303683 | 0.10458153 | 0.33555699 |
| BCAR3 | 228.446245 | 0.715312606 | 0.35212723 | 2.03140386 | 0.04221404 | 0.19607262 |
| DNTTIP2 | 12.3005697 | -0.921931313 | 0.39479635 | -2.3352073 | 0.01953259 | 0.1186813 |
| GCLM | 44.6942187 | -1.269337326 | 0.36522626 | -3.4754821 | 0.00050994 | 0.0087556 |
| ARHGAP29 | 651.605593 | 0.05785949 | 0.44491608 | 0.13004585 | 0.89653015 | 0.95900099 |
| ABCD3 | 202.249028 | 0.760042876 | 0.21054523 | 3.6098794 | 0.00030634 | 0.00598844 |
| SLC44A3 | 111.450624 | 0.400728085 | 0.39307596 | 1.01946729 | 0.30798117 | 0.6015969 |
| CNN3 | 172.250143 | -0.194379236 | 0.24148033 | -0.8049485 | 0.42084938 | 0.69661173 |
| AC105942.1 | 12.8912892 | -0.278726198 | 0.57487114 | -0.4848499 | 0.62778284 | 0.82428562 |
| ALG14 | 32.5338247 | -0.086528362 | 0.30087314 | -0.2875909 | 0.77365995 | 0.90184317 |
| PTBP2 | 215.588273 | 0.05853731 | 0.21224456 | 0.27580123 | 0.78270074 | 0.90630359 |
| DPYD | 201.937127 | -0.480358412 | 0.3644607 | -1.3179978 | 0.18750437 | 0.46523845 |
| SNX7 | 49.2750457 | -0.345237141 | 0.2597014 | -1.3293619 | 0.1837286 | 0.46016652 |
| FRRS1 | 17.6310654 | 0.165632764 | 0.45650596 | 0.36282717 | 0.71673399 | 0.8732017 |
| AGL | 107.624309 | 0.592516792 | 0.18496198 | 3.2034518 | 0.00135791 | 0.01835848 |
| TRMT13 | 54.9300363 | 0.82545657 | 0.21990127 | 3.75375985 | 0.0001742 | 0.00377022 |
| DBT | 66.9936091 | 0.600337573 | 0.23153801 | 2.59282515 | 0.00951912 | 0.0744616 |
| RTCA | 36.3431385 | 0.261958423 | 0.30414008 | 0.86130844 | 0.38906819 | 0.67095971 |
| CDC14A | 60.3616461 | 0.412449544 | 0.3862567 | 1.06781202 | 0.28560531 | 0.57805618 |
| VCAM1 | 15.4008927 | -1.305065604 | 0.85783605 | -1.5213462 | 0.12817298 | 0.37540371 |
| EXTL2 | 28.4638648 | 0.604760529 | 0.326695 | 1.85114719 | 0.06414838 | 0.2521767 |
| SLC30A7 | 101.824741 | -0.279644718 | 0.25069074 | -1.1154968 | 0.2646376 | 0.55622662 |
| DPH5 | 26.3122242 | 0.177530795 | 0.28496917 | 0.62298246 | 0.53329605 | 0.76689222 |
| AC093157.1 | 27.9455342 | -0.073918978 | 0.32894205 | -0.2247173 | 0.82219918 | 0.9253109 |
| RNPC3 | 168.648222 | 0.515485489 | 0.21140313 | 2.43840051 | 0.01475242 | 0.09920328 |
| AMY2B | 45.024985 | 0.475234539 | 0.31381003 | 1.51440203 | 0.12992391 | 0.37778043 |
| NTNG1 | 25.5124086 | -4.075095831 | 1.01818593 | -4.0023101 | 6.27E-05 | 0.00169835 |
| VAV3 | 733.237085 | 0.296699385 | 0.2751721 | 1.07823208 | 0.2809302 | 0.57400692 |
| SLC25A24 | 42.5027741 | -0.313668555 | 0.27354721 | -1.1466707 | 0.25151776 | 0.54321509 |
| PRPF38B | 70.0070163 | 0.070242702 | 0.2599418 | 0.27022472 | 0.78698737 | 0.90871373 |
| STXBP3 | 87.6661776 | 0.578531321 | 0.25022648 | 2.31203073 | 0.02077599 | 0.12343382 |
| AL449266.1 | 17.8271088 | -0.606248237 | 0.3725371 | -1.62735 | 0.10366279 | 0.33425166 |
| GPSM2 | 20.1280717 | -1.294928641 | 0.44641549 | -2.9007251 | 0.003723 | 0.03812562 |
| WDR47 | 22.420537 | 0.240266076 | 0.41634832 | 0.57707949 | 0.56388576 | 0.7848083 |
| TAF13 | 15.4944186 | 0.148197892 | 0.38777089 | 0.38217901 | 0.70232859 | 0.86516671 |
| TMEM167B | 21.1683683 | -0.475016531 | 0.37140334 | -1.2789775 | 0.20090496 | 0.48443708 |
| SARS | 49.9177795 | -0.735901052 | 0.29501542 | -2.4944494 | 0.01261527 | 0.09018912 |
| CELSR2 | 22.4386708 | 0.301697005 | 0.46376064 | 0.65054465 | 0.51534047 | 0.75713618 |
| SORT1 | 104.906124 | -0.013192184 | 0.24717369 | -0.0533721 | 0.95743542 | 0.9831804 |
| PSMA5 | 53.2097864 | -0.225355291 | 0.35098154 | -0.6420716 | 0.52082667 | 0.76092013 |
| SYPL2 | 15.863572 | 0.850805694 | 0.46182656 | 1.84226238 | 0.06543678 | 0.25554575 |
| CYB561D1 | 9.73521464 | -0.942060176 | 0.4915828 | -1.9163815 | 0.05531656 | 0.23212675 |
| GNAI3 | 83.9867179 | -0.142231011 | 0.25454523 | -0.5587652 | 0.576322 | 0.79184705 |
| AMPD2 | 15.928367 | 0.124420083 | 0.52898155 | 0.23520685 | 0.81404819 | 0.92107668 |
| GSTM4 | 25.701981 | 0.193494503 | 0.30496513 | 0.63448075 | 0.52576713 | 0.76221787 |
| GSTM3 | 97.1679422 | -0.508141993 | 0.53844269 | -0.9437253 | 0.34531004 | 0.63256941 |
| AHCYL1 | 184.27587 | -0.301913612 | 0.2410307 | -1.252594 | 0.2103535 | 0.49651968 |
| STRIP1 | 26.9393625 | 0.520337923 | 0.38839566 | 1.33971098 | 0.18033933 | 0.45647562 |
| KCNC4 | 106.028717 | 0.092006595 | 0.42947574 | 0.21423001 | 0.83036769 | 0.92993238 |
| RBM15 | 28.6211954 | 0.522217622 | 0.34516028 | 1.51297136 | 0.13028693 | 0.37846669 |
| SLC16A4 | 98.7113561 | -0.52389255 | 0.42711473 | -1.2265851 | 0.21997859 | 0.50818193 |
| AL355488.1 | 12.5500126 | 1.12038129 | 0.43919846 | 2.55096817 | 0.01074241 | 0.08056236 |
| LAMTOR5 | 89.3097184 | -0.717358449 | 0.35458757 | -2.0230784 | 0.04306507 | 0.19806087 |
| LRIF1 | 20.687793 | -0.39934237 | 0.36655489 | -1.0894477 | 0.27595652 | 0.56798847 |
| DRAM2 | 49.2300071 | -0.23235537 | 0.25295333 | -0.9185701 | 0.35832047 | 0.64216076 |
| CEPT1 | 58.9624409 | -0.056365769 | 0.25182261 | -0.2238313 | 0.82288861 | 0.92597982 |
| DENND2D | 33.6271539 | -0.937224362 | 0.28631961 | -3.2733502 | 0.00106281 | 0.01531334 |
| WDR77 | 13.1379107 | 0.098410688 | 0.48373293 | 0.20344012 | 0.83879104 | 0.93540001 |
| ATP5PB | 87.1696378 | -0.028654749 | 0.3865553 | -0.0741285 | 0.94090817 | 0.97727535 |
| RAP1A | 156.313695 | 0.068386345 | 0.27830458 | 0.24572483 | 0.80589525 | 0.91718215 |
| KCND3 | 15.3198264 | -2.877229551 | 0.6540581 | -4.3990427 | 1.09E-05 | 0.00039581 |
| CTTNBP2NL | 70.0159847 | -0.935078533 | 0.24699435 | -3.7858296 | 0.0001532 | 0.00344592 |
| ST7L | 68.9693551 | 0.444524425 | 0.25086329 | 1.77197875 | 0.07639808 | 0.28042437 |
| CAPZA1 | 109.190747 | -0.440821012 | 0.20453941 | -2.1551886 | 0.03114708 | 0.16170525 |
| MOV10 | 31.1394898 | -0.280947618 | 0.45053257 | -0.62359 | 0.53289686 | 0.76685339 |
| RHOC | 54.434503 | -1.003534342 | 0.42269388 | -2.3741398 | 0.01758989 | 0.11119448 |
| LRIG2 | 63.113711 | 0.123677845 | 0.26673616 | 0.46367108 | 0.64288342 | 0.83318468 |
| MAGI3 | 414.769584 | 0.503177158 | 0.280357 | 1.79477295 | 0.07268988 | 0.27219482 |
| PHTF1 | 56.4366663 | -0.121169431 | 0.29755678 | -0.4072145 | 0.68385046 | 0.85606265 |
| RSBN1 | 59.322297 | -0.126550893 | 0.26979896 | -0.4690563 | 0.63902942 | 0.83119926 |
| HIPK1 | 61.086583 | -0.001570636 | 0.2986337 | -0.0052594 | 0.99580362 | 0.99868531 |
| TRIM33 | 198.100796 | 0.212918296 | 0.2841763 | 0.74924721 | 0.45370822 | 0.72126672 |
| BCAS2 | 20.9085439 | 0.334124241 | 0.38279721 | 0.8728492 | 0.38274528 | 0.6661669 |

| | | | | | | |
|---|---|---|---|---|---|---|
| DENND2C | 13.0286164 | 0.660284735 | 0.45474522 | 1.45198828 | 0.14650486 | 0.40573108 |
| NRAS | 14.487058 | -0.925120193 | 0.44603762 | -2.0740856 | 0.03807136 | 0.18276383 |
| CSDE1 | 164.430633 | -0.009415126 | 0.21612132 | -0.0435641 | 0.96525188 | 0.98660829 |
| SIKE1 | 29.1344214 | 0.187708422 | 0.29293147 | 0.64079296 | 0.52165721 | 0.76104628 |
| SLC22A15 | 123.698842 | 0.316709521 | 0.26280905 | 1.20509367 | 0.22816714 | 0.5178625 |
| ATP1A1 | 1384.36523 | -0.101392449 | 0.3198497 | -0.3170003 | 0.75124338 | 0.89132635 |
| LINC01762 | 371.333737 | -0.554987822 | 0.48973713 | -1.1332361 | 0.2571151 | 0.54817875 |
| CD58 | 64.0112697 | -1.517301289 | 0.29555431 | -5.1337478 | 2.84E-07 | 2.02E-05 |
| IGSF3 | 97.6136255 | -0.776034053 | 0.35040664 | -2.2146671 | 0.02678292 | 0.14614285 |
| PTGFRN | 23.9450447 | -0.523476798 | 0.3386854 | -1.5456137 | 0.12219789 | 0.3664089 |
| TTF2 | 33.0934531 | -0.46491074 | 0.32317272 | -1.4385829 | 0.15026874 | 0.41132271 |
| VTCN1 | 72.3188336 | -1.03582649 | 1.12128232 | -0.9237874 | 0.35559701 | 0.64045528 |
| MAN1A2 | 189.049804 | 0.431417041 | 0.20869179 | 2.06724492 | 0.03871108 | 0.18480147 |
| GDAP2 | 68.6710669 | 0.30086154 | 0.27051432 | 1.1121834 | 0.2660593 | 0.55721548 |
| WDR3 | 16.6722691 | -0.079900881 | 0.35327216 | -0.2261737 | 0.8210663 | 0.92474309 |
| SPAG17 | 83.4561697 | 0.059429908 | 0.36370556 | 0.16340115 | 0.8702026 | 0.94835789 |
| WARS2 | 48.8738217 | 0.157719545 | 0.27669376 | 0.57001482 | 0.56866765 | 0.78763298 |
| AL359915.2 | 113.344677 | 0.259989477 | 0.31749651 | 0.8188735 | 0.41285859 | 0.68990579 |
| ZNF697 | 16.8841839 | 0.936771707 | 0.45058597 | 2.07900771 | 0.03761664 | 0.18151131 |
| PHGDH | 24.2848416 | -0.175473514 | 0.51166163 | -0.3429484 | 0.73163732 | 0.88245488 |
| HMGCS2 | 98.3040796 | 2.97699106 | 0.85843063 | 3.467946 | 0.00052445 | 0.00890656 |
| NOTCH2 | 59.5295351 | 0.588202728 | 0.26233826 | 2.24215381 | 0.02495143 | 0.1388707 |
| SEC22B | 70.8481426 | -0.216899811 | 0.25090263 | -0.864478 | 0.38732534 | 0.66959234 |
| NBPF26 | 14.1489209 | 0.782641317 | 0.49505066 | 1.58093179 | 0.11389364 | 0.35248764 |
| LINC00623 | 102.665923 | -1.025985932 | 0.49326965 | -2.0799697 | 0.03752831 | 0.18135358 |
| SRGAP2C | 29.5237106 | -0.042397483 | 0.36952826 | -0.1147341 | 0.9086559 | 0.96256793 |
| AC239800.3 | 13.3621379 | -1.565700973 | 0.47751195 | -3.2788729 | 0.00104223 | 0.01518672 |
| NBPF15 | 24.4678232 | -0.56879292 | 0.35822638 | -1.587803 | 0.11233089 | 0.34961718 |
| FP700111.1 | 23.0119297 | 0.201816563 | 0.35050583 | 0.57578661 | 0.56475943 | 0.78515099 |
| SRGAP2B | 24.1474105 | -0.873782848 | 0.3619823 | -2.4138828 | 0.01578354 | 0.10356817 |
| GPR89A | 17.9068463 | 0.116630069 | 0.37743808 | 0.30900451 | 0.7573181 | 0.89457492 |
| PDZK1 | 48.4017386 | 1.104195621 | 0.57117392 | 1.93320386 | 0.05321109 | 0.22614713 |
| RNF115 | 188.677096 | 0.259462877 | 0.2139506 | 1.2127233 | 0.22523563 | 0.51392634 |
| POLR3C | 20.7993817 | -0.371453343 | 0.37623365 | -0.9872943 | 0.32349837 | 0.61431894 |
| PEX11B | 17.2359745 | -0.289139949 | 0.3908588 | -0.7397555 | 0.45944836 | 0.72484487 |
| RBM8A | 33.3174407 | -0.265390989 | 0.31692687 | -0.8373887 | 0.4023741 | 0.68128594 |
| TXNIP | 801.450077 | 1.148953497 | 0.58360013 | 1.96873412 | 0.04898363 | 0.2147359 |
| NBPF12 | 40.0132301 | 0.040472663 | 0.27996493 | 0.14456334 | 0.88505565 | 0.95619452 |
| PRKAB2 | 29.2116298 | 0.534975212 | 0.367774 | 1.45463032 | 0.14577163 | 0.40412792 |
| AC242426.2 | 52.3864204 | -0.414766938 | 0.32398001 | -1.2802239 | 0.20046641 | 0.48422157 |
| FMO5 | 34.871861 | 0.085878885 | 0.44195446 | 0.19431614 | 0.84592835 | 0.93757726 |
| CHD1L | 52.6772747 | 0.521391819 | 0.24716385 | 2.10949867 | 0.03490156 | 0.17369849 |
| BCL9 | 74.4243533 | 0.600081621 | 0.2146905 | 2.79510091 | 0.00518835 | 0.04891142 |
| ACP6 | 60.6350134 | 0.103300166 | 0.21909068 | 0.47149503 | 0.63728727 | 0.82989931 |
| GPR89B | 24.5110021 | 0.449189155 | 0.30370548 | 1.4790288 | 0.13913262 | 0.39355965 |
| NBPF11 | 39.6375216 | 0.154265887 | 0.28820239 | 0.53526928 | 0.59246367 | 0.80151815 |
| AC239809.3 | 22.8474568 | -0.505459798 | 0.41676751 | -1.21281 | 0.22520248 | 0.51392634 |
| NBPF14 | 16.8911841 | -0.142812667 | 0.36022655 | -0.3964524 | 0.69177134 | 0.85976938 |
| PDE4DIP | 156.066658 | 0.516401357 | 0.27245983 | 1.89533022 | 0.05804867 | 0.23811563 |
| NBPF9 | 17.3212184 | -1.103162458 | 0.41022393 | -2.6891714 | 0.00716296 | 0.06149404 |
| AC245297.3 | 212.385283 | -0.420238863 | 0.33408265 | -1.2578889 | 0.20843193 | 0.49402397 |
| NBPF19 | 27.7649264 | 0.221369024 | 0.66909605 | 0.3308479 | 0.74075938 | 0.88653614 |
| SF3B4 | 13.9973623 | -1.027713191 | 0.38577354 | -2.6640324 | 0.00772101 | 0.06419577 |
| OTUD7B | 103.957312 | -0.341821345 | 0.29563416 | -1.1562309 | 0.24758674 | 0.53836051 |
| VPS45 | 101.35166 | 0.302815029 | 0.20320384 | 1.49020327 | 0.1361708 | 0.38868467 |
| ANP32E | 43.3403272 | 0.087085552 | 0.37211045 | 0.23403146 | 0.81496056 | 0.92163473 |
| APH1A | 48.0532715 | -0.432090348 | 0.41029651 | -1.0531173 | 0.29228724 | 0.58378114 |
| C1orf54 | 32.6263572 | 0.364436914 | 0.46224868 | 0.78840012 | 0.4304627 | 0.7041707 |
| MRPS21 | 69.6923742 | -0.644240008 | 0.23748543 | -2.7127559 | 0.00667262 | 0.0583543 |
| PRPF3 | 55.9930921 | 0.128378358 | 0.27830536 | 0.46128596 | 0.64459345 | 0.83381793 |
| RPRD2 | 213.453905 | 0.491347075 | 0.19830127 | 2.47778081 | 0.01322023 | 0.09290235 |
| TARS2 | 17.6018363 | 0.282551775 | 0.37840926 | 0.74668303 | 0.45525491 | 0.72210564 |
| MCL1 | 77.0721141 | -0.586356756 | 0.37157272 | -1.5780404 | 0.11455633 | 0.35349258 |
| ENSA | 174.171753 | -0.129009189 | 0.31320964 | -0.4118941 | 0.68041707 | 0.8543838 |
| GOLPH3L | 64.4438631 | -0.31796952 | 0.24708576 | -1.2868792 | 0.1981364 | 0.48050875 |
| CTSS | 18.2002767 | -1.91603099 | 0.61442866 | -3.1183946 | 0.00181839 | 0.02287311 |
| ARNT | 80.73055 | 0.383657526 | 0.27804386 | 1.37984537 | 0.16763426 | 0.43776126 |
| SETDB1 | 56.0062837 | -0.02011048 | 0.32377652 | -0.0621122 | 0.95047346 | 0.98025604 |
| CERS2 | 37.7033112 | -0.725603379 | 0.37589313 | -1.9303449 | 0.05356412 | 0.22717891 |
| ANXA9 | 10.736788 | 1.344807168 | 0.59855738 | 2.2467473 | 0.02465618 | 0.13752463 |
| MINDY1 | 20.8769938 | 0.318703437 | 0.4044838 | 0.78792634 | 0.43073979 | 0.70420575 |
| PRUNE1 | 26.8365303 | -0.142840241 | 0.3029472 | -0.4715021 | 0.63728222 | 0.82989931 |
| C1orf56 | 16.0579933 | -0.956542952 | 0.68150535 | -1.4035737 | 0.16044584 | 0.42803048 |
| CDC42SE1 | 44.2173235 | -0.615100252 | 0.32756086 | -1.8778198 | 0.06040582 | 0.24332668 |
| GABPB2 | 69.8791503 | 0.834546863 | 0.25576019 | 3.26300533 | 0.00110237 | 0.01564278 |
| SCNM1 | 13.4779249 | -1.03784689 | 0.41807149 | -2.4824628 | 0.01304777 | 0.09213032 |
| VPS72 | 17.1416917 | 0.005506247 | 0.39151627 | 0.0140639 | 0.988779 | 0.9963225 |
| PIP5K1A | 220.636187 | 0.339435391 | 0.21207246 | 1.60056327 | 0.10947368 | 0.34481257 |
| PSMD4 | 44.0023441 | -0.09681528 | 0.23557108 | -0.4109812 | 0.68108634 | 0.8546627 |
| PI4KB | 53.1721978 | 0.473889288 | 0.26226051 | 1.80694105 | 0.07077147 | 0.26863953 |
| SELENBP1 | 15.8576448 | 1.120221253 | 0.51713256 | 2.16621684 | 0.03029462 | 0.15858495 |
| PSMB4 | 28.9021899 | -0.382623369 | 0.4649609 | -0.8229152 | 0.41055624 | 0.68825899 |
| POGZ | 318.778039 | 0.821435774 | 0.21578365 | 3.80675631 | 0.0001408 | 0.00320915 |
| CGN | 40.1472103 | -0.519515278 | 0.25510494 | -2.0364767 | 0.04170251 | 0.19500656 |
| TUFT1 | 53.3294325 | 0.947661571 | 0.36273895 | 2.61251669 | 0.00898783 | 0.07117035 |
| SNX27 | 87.6905461 | 0.356193628 | 0.2905655 | 1.22586343 | 0.22025007 | 0.50852 |
| MRPL9 | 14.0726695 | -0.97340224 | 0.43627806 | -2.231151 | 0.02567113 | 0.14210852 |
| RORC | 24.935893 | -0.15994259 | 0.57075205 | -0.2802313 | 0.77930006 | 0.90427469 |
| THEM4 | 92.7455751 | 0.469022507 | 0.49587039 | 0.94585706 | 0.3442215 | 0.63200319 |
| S100A10 | 206.331582 | -1.248814141 | 0.39808519 | -3.1370525 | 0.00170656 | 0.02188887 |
| S100A11 | 586.161926 | -2.352654333 | 0.41187202 | -5.7121004 | 1.12E-08 | 1.40E-06 |
| FLG-AS1 | 115.190375 | 1.450793322 | 0.33006774 | 4.39544119 | 1.11E-05 | 0.0003982 |
| S100A6 | 795.213806 | -1.896859748 | 0.60775955 | -3.1210694 | 0.00180196 | 0.02274969 |
| S100A4 | 22.692235 | -0.580949888 | 0.80413702 | -0.7224514 | 0.47001702 | 0.73188482 |
| S100A2 | 245.095159 | 0.648434198 | 1.0487166 | 0.61831213 | 0.53636961 | 0.76895069 |
| S100A16 | 55.8574884 | -1.750965174 | 0.49039252 | -3.5705381 | 0.00035625 | 0.00667286 |
| S100A14 | 31.4601853 | -2.69709737 | 0.63744985 | -4.2310738 | 2.33E-05 | 0.00073724 |
| S100A13 | 59.4506743 | -0.586392439 | 0.400989 | -1.4623654 | 0.1436411 | 0.40102728 |
| CHTOP | 40.3403621 | 0.181782699 | 0.25731171 | 0.70646881 | 0.47989664 | 0.73953305 |
| ILF2 | 31.2369422 | -0.357169782 | 0.37864157 | -0.9432926 | 0.34553127 | 0.63256941 |
| INTS3 | 62.3312813 | 0.047282359 | 0.25105993 | 0.18833097 | 0.85061721 | 0.93973818 |
| GATAD2B | 240.89454 | 0.116226321 | 0.27334828 | 0.425195 | 0.67069453 | 0.84746193 |
| DENND4B | 27.8676609 | -0.425523106 | 0.4140515 | -1.0277057 | 0.30408827 | 0.59822456 |
| CRTC2 | 16.7752708 | -0.202852183 | 0.41378136 | -0.49024 | 0.62396407 | 0.82223405 |
| SLC39A1 | 40.352123 | -1.370433133 | 0.41293741 | -3.318743 | 0.00090424 | 0.01363901 |
| JTB | 38.5621289 | -0.354941404 | 0.36728968 | -0.96638 | 0.33385407 | 0.62250943 |
| RAB13 | 53.7608351 | -0.897811197 | 0.3232939 | -2.7770743 | 0.00548506 | 0.05085757 |
| RPS27 | 45.763811 | -0.275268822 | 0.35888055 | -0.7670207 | 0.44306918 | 0.71326991 |
| TPM3 | 73.9449894 | -0.893118728 | 0.35411722 | -2.5220991 | 0.01166568 | 0.08523395 |
| C1orf43 | 60.3434321 | -0.497888836 | 0.26511765 | -1.877992 | 0.06038226 | 0.24332668 |
| UBAP2L | 67.3132928 | -0.013142563 | 0.21439419 | -0.0613009 | 0.95111955 | 0.98061757 |
| HAX1 | 18.0390256 | -0.262877191 | 0.43588458 | -0.603089 | 0.54644949 | 0.77463416 |
| IL6R | 22.7643072 | -1.427838329 | 0.47919265 | -2.9796749 | 0.00288554 | 0.03172559 |

| | | | | | | |
|---|---|---|---|---|---|---|
| UBE2Q1 | 22.2019661 | -0.113542253 | 0.38063511 | -0.2982968 | 0.76547661 | 0.89836575 |
| ADAR | 115.716105 | -0.492604238 | 0.37507927 | -1.3133337 | 0.18907054 | 0.46792323 |
| KCNN3 | 19.9507267 | 2.081573118 | 0.59725672 | 3.48522345 | 0.00049173 | 0.0084996 |
| PMVK | 23.1981643 | -0.021166355 | 0.51781482 | -0.0408763 | 0.96739451 | 0.98723113 |
| PBXIP1 | 24.9847162 | -0.001123616 | 0.51036506 | -0.0022016 | 0.99824338 | 0.99911623 |
| PYGO2 | 10.5386471 | 0.209650588 | 0.4897217 | 0.42810148 | 0.66857724 | 0.84584093 |
| FLAD1 | 11.1952427 | 0.022764533 | 0.43241988 | 0.05264451 | 0.95801515 | 0.98321101 |
| ZBTB7B | 15.0604574 | 0.178600496 | 0.37626337 | 0.47466883 | 0.63502304 | 0.82850241 |
| EFNA1 | 53.4906059 | -1.706902532 | 0.38273318 | -4.4597715 | 8.20E-06 | 0.00031211 |
| SLC50A1 | 17.0412461 | -1.272320902 | 0.45123213 | -2.8196593 | 0.00480747 | 0.04626289 |
| DPM3 | 31.6766847 | -1.074662998 | 0.54573577 | -1.9692002 | 0.0489301 | 0.21468395 |
| MUC1 | 18.7177182 | -1.176444281 | 0.43140023 | -2.7270367 | 0.00639059 | 0.05653552 |
| THBS3 | 13.5530985 | 0.293441082 | 0.53513674 | 0.54834785 | 0.58345308 | 0.79598942 |
| MTX1 | 11.4977161 | -0.48504841 | 0.48356515 | -1.0030673 | 0.31582837 | 0.60918628 |
| GBA | 17.5132587 | -0.290872032 | 0.44862739 | -0.64836 | 0.51675216 | 0.7580209 |
| FAM189B | 11.251277 | -0.898100288 | 0.47039681 | -1.9092398 | 0.05623117 | 0.23424727 |
| SCAMP3 | 27.1325427 | -0.37235821 | 0.40855994 | -0.9113919 | 0.36208893 | 0.64626475 |
| CLK2 | 21.1814679 | -0.182537694 | 0.33924014 | -0.5380781 | 0.59052312 | 0.80004855 |
| FDPS | 70.9136059 | -0.208429681 | 0.37701105 | -0.5528477 | 0.58036772 | 0.79465023 |
| RUSC1 | 15.99733 | 0.0864857 | 0.39919082 | 0.21665253 | 0.82847915 | 0.92852395 |
| ASH1L | 473.483612 | 0.13556595 | 0.17617745 | 0.76948528 | 0.44160528 | 0.71227192 |
| YY1AP1 | 78.748128 | 0.06341402 | 0.23193539 | 0.27341243 | 0.7845362 | 0.90765805 |
| DAP3 | 96.0441284 | 0.12632729 | 0.21147779 | 0.59735489 | 0.55027046 | 0.77615521 |
| GON4L | 106.851603 | -0.065764933 | 0.27110169 | -0.242584 | 0.80832767 | 0.91848612 |
| RIT1 | 33.4893328 | -1.054273872 | 0.4307074 | -2.4477728 | 0.01437423 | 0.09745114 |
| KHDC4 | 93.546956 | 0.268739724 | 0.26408807 | 1.01761404 | 0.30886141 | 0.60263073 |
| ARHGEF2 | 29.3178481 | -0.830782388 | 0.27437686 | -3.027888 | 0.00246269 | 0.02834712 |
| SSR2 | 50.0183648 | -0.749692286 | 0.28485856 | -2.6318053 | 0.00849325 | 0.06873329 |
| UBQLN4 | 19.7808968 | -0.492869687 | 0.32890075 | -1.4985362 | 0.13399399 | 0.38462138 |
| LAMTOR2 | 18.1988155 | -0.905665623 | 0.58014087 | -1.5611133 | 0.11849702 | 0.35989279 |
| RAB25 | 35.3197204 | -1.150467983 | 0.48614189 | -2.3665271 | 0.01795586 | 0.11278482 |
| LMNA | 92.0430671 | -0.526298982 | 0.28587226 | -1.8410285 | 0.06561738 | 0.25605693 |
| SEMA4A | 23.4377374 | -0.040379943 | 0.33377387 | -0.1209799 | 0.90370692 | 0.96126516 |
| SLC25A44 | 20.5978871 | 0.498522329 | 0.30464196 | 1.63642042 | 0.10175163 | 0.33046825 |
| SMG5 | 53.0071777 | -0.343439862 | 0.25936484 | -1.3241573 | 0.1854508 | 0.4623703 |
| GLMP | 12.6108451 | -0.896324473 | 0.58209056 | -1.5398368 | 0.12360014 | 0.36843999 |
| CCT3 | 73.568022 | 0.247267569 | 0.30127449 | 0.82073849 | 0.41179524 | 0.68902299 |
| RHBG | 103.615037 | 2.539663254 | 0.42656378 | 5.95377138 | 2.62E-09 | 3.86E-07 |
| MEF2D | 38.2152876 | 0.117377808 | 0.31186482 | 0.37637399 | 0.70663888 | 0.86798542 |
| NAXE | 20.4510508 | -0.170723465 | 0.5133694 | -0.3325548 | 0.73947036 | 0.88614269 |
| GPATCH4 | 11.0263215 | -0.936921568 | 0.46677567 | -2.0072202 | 0.04472622 | 0.20164968 |
| ISG20L2 | 14.189264 | -0.704931708 | 0.42806671 | -1.64678 | 0.09960327 | 0.32585356 |
| RRNAD1 | 20.5267951 | 0.253813365 | 0.33348023 | 0.76110468 | 0.44659455 | 0.71619194 |
| MRPL24 | 12.5666332 | -0.346736928 | 0.53473057 | -0.648433 | 0.51670496 | 0.7580209 |
| HDGF | 58.7978148 | -0.764600754 | 0.31113781 | -2.4574344 | 0.01399334 | 0.09620193 |
| PRCC | 45.9274048 | 0.486906302 | 0.25171642 | 1.93434464 | 0.05307077 | 0.22595384 |
| ARHGEF11 | 119.275756 | 0.314417461 | 0.18450381 | 1.70412445 | 0.08835784 | 0.30586777 |
| ETV3 | 10.9467534 | -0.596646792 | 0.47662983 | -1.2518033 | 0.21064155 | 0.49657419 |
| KIRREL1 | 30.8324795 | -0.930243069 | 0.52293393 | -1.7788922 | 0.07525744 | 0.27848498 |
| IFI16 | 17.0796431 | -3.312396738 | 1.40019689 | -2.365665 | 0.01799772 | 0.11284999 |
| DUSP23 | 41.0577533 | -1.824206401 | 0.55707897 | -3.2745921 | 0.00105815 | 0.01528426 |
| TAGLN2 | 90.0170378 | -2.122488923 | 0.48833805 | -4.3463517 | 1.38E-05 | 0.00048339 |
| KCNJ10 | 14.7410225 | 3.529383282 | 0.54288402 | 6.50117364 | 7.97E-11 | 1.91E-08 |
| IGSF8 | 14.8891614 | -0.119193506 | 0.58227832 | -0.2047019 | 0.837805 | 0.93487753 |
| PEA15 | 15.2573021 | -1.345078075 | 0.49541461 | -2.7150553 | 0.00662647 | 0.05803619 |
| DCAF8 | 33.1026341 | 0.424125505 | 0.34275445 | 1.23740334 | 0.21593738 | 0.50279115 |
| COPA | 133.534661 | -0.113813407 | 0.24587559 | -0.4628902 | 0.64344306 | 0.83338577 |
| NCSTN | 30.6069729 | -0.025528318 | 0.39029395 | -0.0654079 | 0.94784921 | 0.979567 |
| VANGL2 | 9.67963375 | 0.422448246 | 0.58489238 | 0.72226663 | 0.47013057 | 0.73188482 |
| F11R | 73.9075829 | -0.359777713 | 0.41503915 | -0.8668525 | 0.38602285 | 0.66811344 |
| AL591806.3 | 25.0018295 | 0.240403099 | 0.4189575 | 0.57381262 | 0.56609461 | 0.78607719 |
| NECTIN4 | 29.9535394 | -1.523903329 | 0.45775093 | -3.3291103 | 0.00087124 | 0.01331677 |
| PFDN2 | 43.961515 | -0.83951031 | 0.28440491 | -2.9518137 | 0.00315913 | 0.03390146 |
| NIT1 | 32.946332 | -0.876996519 | 0.35088708 | -2.4993696 | 0.01244145 | 0.08925861 |
| DEDD | 15.8624534 | -0.422624919 | 0.40142111 | -1.0528219 | 0.29242265 | 0.58393838 |
| UFC1 | 29.9844458 | -0.579551832 | 0.45801526 | -1.2653549 | 0.20574415 | 0.48999801 |
| B4GALT3 | 12.7977181 | 0.149612529 | 0.5063562 | 0.29546894 | 0.76763568 | 0.89979146 |
| NDUFS2 | 50.6076919 | 0.315925584 | 0.3071967 | 1.02841464 | 0.30375482 | 0.59776164 |
| SDHC | 98.881769 | 0.089590332 | 0.2342029 | 0.38253297 | 0.70206608 | 0.86515367 |
| AL592295.4 | 16.5769713 | -0.061105526 | 0.50834851 | -0.120204 | 0.90432155 | 0.96126516 |
| DUSP12 | 12.3290113 | 0.689420182 | 0.40050979 | 1.72135665 | 0.08518613 | 0.2997225 |
| ATF6 | 218.969874 | 0.572188523 | 0.31595077 | 1.8110053 | 0.07014003 | 0.26694834 |
| NOS1AP | 380.842686 | -0.638078446 | 0.44467305 | -1.4349384 | 0.15130465 | 0.41251912 |
| UHMK1 | 89.378358 | -0.205541786 | 0.28555436 | -0.7197991 | 0.47164867 | 0.73286947 |
| UAP1 | 21.2394947 | -0.258115101 | 0.4416384 | -0.584449 | 0.5589183 | 0.78244004 |
| HSD17B7 | 29.5409094 | 0.301593256 | 0.41927822 | 0.71931533 | 0.47194665 | 0.7332219 |
| PBX1 | 192.76913 | 1.056350691 | 0.35351913 | 2.98810051 | 0.00280717 | 0.03126431 |
| MGST3 | 214.917945 | -0.682984292 | 0.18946506 | -3.6048033 | 0.00031239 | 0.00606622 |
| ALDH9A1 | 38.6677809 | 0.053376274 | 0.40087851 | 0.13314826 | 0.89407613 | 0.95815794 |
| TMCO1 | 74.7359473 | -0.706906642 | 0.31100534 | -2.2729727 | 0.02302783 | 0.1323843 |
| UCK2 | 31.8467479 | 0.283405302 | 0.28470778 | 0.99542522 | 0.3195295 | 0.611915 |
| POGK | 38.686797 | 0.098331244 | 0.31277516 | 0.31438316 | 0.75323005 | 0.89205725 |
| TADA1 | 20.0521339 | 0.249145495 | 0.37444759 | 0.66536813 | 0.50581506 | 0.75258619 |
| POU2F1 | 208.859892 | 0.397462006 | 0.18978689 | 2.09425431 | 0.03623733 | 0.17659271 |
| CREG1 | 33.8000805 | -0.08351082 | 0.40084765 | -0.2083356 | 0.83496697 | 0.93320871 |
| MPZL1 | 73.0754475 | -0.956530878 | 0.31483495 | -3.0381979 | 0.00237998 | 0.02751797 |
| MPC2 | 58.9865067 | 0.559714983 | 0.32918891 | 1.70028507 | 0.08907732 | 0.30722281 |
| DCAF6 | 258.830307 | 0.037820092 | 0.22685073 | 0.16671797 | 0.86759197 | 0.94761794 |
| TIPRL | 31.7824228 | -0.220161079 | 0.30046305 | -0.7327393 | 0.46371745 | 0.72812334 |
| SFT2D2 | 44.2083393 | -0.362509126 | 0.24083427 | -1.5052223 | 0.13226692 | 0.38157766 |
| LINC00970 | 154.163599 | 2.619560993 | 0.57081032 | 4.58919701 | 4.45E-06 | 0.00019213 |
| ATP1B1 | 2345.9703 | 1.896370737 | 0.35636552 | 5.32142038 | 1.03E-07 | 8.77E-06 |
| NME7 | 222.492952 | 0.319425176 | 0.18222778 | 1.75288962 | 0.07962096 | 0.28770785 |
| BLZF1 | 46.8876306 | -0.228719972 | 0.28144605 | -0.8126601 | 0.41641297 | 0.69303497 |
| CCDC181 | 15.030632 | -0.059844513 | 0.45189667 | -0.1324296 | 0.89464447 | 0.9584679 |
| SLC19A2 | 86.3363159 | 0.85608664 | 0.41643922 | 2.05573011 | 0.03980854 | 0.18872874 |
| C1orf112 | 11.555576 | -0.267490077 | 0.41805182 | -0.6398491 | 0.52227071 | 0.76155853 |
| SCYL3 | 40.1039295 | 0.017695905 | 0.29943007 | 0.05909862 | 0.95287355 | 0.98135593 |
| KIFAP3 | 87.6010749 | 0.464349169 | 0.23732369 | 1.95660692 | 0.0503937 | 0.218779 |
| GORAB | 14.9453496 | -0.547166997 | 0.36109257 | -1.5153095 | 0.12969405 | 0.37739114 |
| FMO4 | 58.7479107 | 1.175194026 | 0.38761412 | 3.03186588 | 0.00243047 | 0.02803888 |
| PRRC2C | 277.895099 | 0.062471284 | 0.217924 | 0.28666545 | 0.77436849 | 0.90202041 |
| VAMP4 | 26.2652111 | -0.110851627 | 0.34676362 | -0.3196749 | 0.74921479 | 0.89014078 |
| METTL13 | 10.8323614 | 0.027836935 | 0.46024816 | 0.06048245 | 0.95177139 | 0.98065459 |
| DNM3 | 64.8108377 | 0.125311637 | 0.32901334 | 0.38087099 | 0.70329898 | 0.86585351 |
| PIGC | 31.2600048 | -0.180244634 | 0.39150438 | -0.4603898 | 0.64523643 | 0.83387847 |
| SUCO | 85.2507613 | -0.478377518 | 0.26907885 | -1.777834 | 0.07543112 | 0.27878882 |
| AL645568.1 | 42.6973998 | 0.617119506 | 0.39006738 | 1.58208437 | 0.11363031 | 0.35217192 |
| PRDX6 | 107.22173 | -0.421001316 | 0.42311821 | -0.9949969 | 0.31973776 | 0.611915 |
| KLHL20 | 69.2356374 | -0.11065608 | 0.21892081 | -0.5054617 | 0.61323461 | 0.81464125 |
| DARS2 | 15.4299585 | -0.026464092 | 0.41651016 | -0.0635377 | 0.94933835 | 0.9799683 |
| ZBTB37 | 37.6733668 | -0.028743351 | 0.29096308 | -0.0987869 | 0.92130744 | 0.9686919 |
| RC3H1 | 143.254236 | -0.021237658 | 0.22432531 | -0.0946735 | 0.92457418 | 0.97014393 |

| | | | | | | |
|---|---|---|---|---|---|---|
| RABGAP1L | 748.587699 | 0.540445094 | 0.24796582 | 2.17951446 | 0.02929347 | 0.15507777 |
| CACYBP | 17.4878335 | -0.092929402 | 0.3921981 | -0.2369451 | 0.81269941 | 0.92054639 |
| MRPS14 | 20.3235887 | -0.627539854 | 0.36076099 | -1.7394892 | 0.08194875 | 0.29293408 |
| KIAA0040 | 24.6253828 | -0.413636945 | 0.34617414 | -1.1948811 | 0.23213347 | 0.52192035 |
| COP1 | 315.223436 | -0.047568916 | 0.19160229 | -0.248269 | 0.80392625 | 0.91643337 |
| PAPPA2 | 46.8282012 | -0.55431826 | 0.54834429 | -1.0108946 | 0.3120669 | 0.6052171 |
| RASAL2 | 322.871043 | -0.604642355 | 0.54228103 | -1.1149982 | 0.2648512 | 0.55622662 |
| RALGPS2 | 270.866343 | 0.311783403 | 0.24409495 | 1.27730379 | 0.20149502 | 0.4848871 |
| ANGPTL1 | 18.9712913 | -1.379114375 | 0.54764554 | -2.518261 | 0.01179359 | 0.08580337 |
| FAM20B | 47.01696 | -0.158063497 | 0.28487567 | -0.5548508 | 0.57899671 | 0.79372244 |
| TOR3A | 14.0614628 | -0.586535349 | 0.5107909 | -1.1482886 | 0.25084945 | 0.54232519 |
| ABL2 | 145.263786 | 0.227418915 | 0.25775707 | 0.88229941 | 0.37761491 | 0.66066194 |
| SOAT1 | 66.9509384 | -1.21830806 | 0.31088576 | -3.9188288 | 8.90E-05 | 0.00221419 |
| TDRD5 | 14.8787311 | 0.599738465 | 0.56941445 | 1.05325473 | 0.29222426 | 0.58378114 |
| TOR1AIP2 | 99.9220131 | -0.30327298 | 0.23971528 | -1.2651383 | 0.20582176 | 0.48999801 |
| TOR1AIP1 | 46.1762001 | 0.192556931 | 0.29546304 | 0.6517124 | 0.51458672 | 0.75684546 |
| CEP350 | 205.050595 | 0.291777843 | 0.222137 | 1.31350404 | 0.18901316 | 0.46792323 |
| QSOX1 | 46.6792307 | -0.849348687 | 0.38183409 | -2.224392 | 0.02612208 | 0.14414018 |
| ACBD6 | 199.508291 | 0.24758099 | 0.17570255 | 1.40909161 | 0.15880808 | 0.42571971 |
| XPR1 | 184.764472 | -0.392368884 | 0.31339608 | -1.2519904 | 0.21057335 | 0.49657419 |
| KIAA1614 | 30.4787734 | 0.642341489 | 0.40306428 | 1.59364527 | 0.11101551 | 0.34762364 |
| STX6 | 46.3365381 | 0.362424105 | 0.26191924 | 1.38372462 | 0.16644279 | 0.43531192 |
| MR1 | 24.9555264 | -1.050374982 | 0.44463272 | -2.362343 | 0.01815983 | 0.11324612 |
| GLUL | 70.0790574 | -1.508196525 | 0.32116181 | -4.6960643 | 2.65E-06 | 0.00012591 |
| RNASEL | 12.8611018 | -0.731203436 | 0.55951026 | -1.3068633 | 0.19125914 | 0.47160163 |
| DHX9 | 122.942163 | 0.239759598 | 0.2904332 | 0.82552406 | 0.40907413 | 0.6869518 |
| LAMC1 | 177.23871 | -0.682070307 | 0.43514775 | -1.5674453 | 0.11701064 | 0.35699765 |
| LAMC2 | 30.0922993 | -2.823799711 | 0.69510665 | -4.0623978 | 4.86E-05 | 0.00137467 |
| SMG7 | 149.32787 | 0.312344626 | 0.20117621 | 1.55259228 | 0.12052057 | 0.36346292 |
| ARPC5 | 43.7292005 | -1.141445093 | 0.28739156 | -3.9717419 | 7.13E-05 | 0.0018751 |
| RGL1 | 27.3181186 | 0.429867699 | 0.40184409 | 1.06973752 | 0.28473747 | 0.57686567 |
| TSEN15 | 43.573538 | 0.09299605 | 0.281399 | 0.33047755 | 0.74103916 | 0.88665962 |
| EDEM3 | 81.6768379 | 0.079264022 | 0.20310421 | 0.39026282 | 0.69634221 | 0.86247999 |
| RNF2 | 44.0319561 | 0.189931149 | 0.27510507 | 0.69039494 | 0.48994586 | 0.7455572 |
| TRMT1L | 37.1479506 | -0.316297048 | 0.43700441 | -0.7237846 | 0.46919801 | 0.73168094 |
| SWT1 | 57.9924958 | -0.169612586 | 0.31620809 | -0.5363955 | 0.59168526 | 0.80067536 |
| IVNS1ABP | 352.406585 | 0.635937065 | 0.60971841 | 1.04300126 | 0.29694771 | 0.58931907 |
| HMCN1 | 54.1427891 | 1.685075111 | 0.40164079 | 4.19547808 | 2.72E-05 | 0.00084664 |
| TPR | 125.289697 | -0.153257866 | 0.19502297 | -0.7858452 | 0.43195817 | 0.70467666 |
| ODR4 | 72.2118698 | 0.322051178 | 0.2006078 | 1.60537713 | 0.10841083 | 0.34332875 |
| AL596220.1 | 10.581527 | -0.121493415 | 0.74870036 | -0.1622724 | 0.87109134 | 0.94844301 |
| ERVMER61-1 | 116.633497 | 1.472768616 | 0.37420095 | 3.93576932 | 8.29E-05 | 0.00210878 |
| BRINP3 | 46.0554177 | 0.409671394 | 0.59281117 | 0.69106558 | 0.48952433 | 0.7455572 |
| AL390957.1 | 21.1531967 | -0.91551121 | 0.47431973 | -1.9301563 | 0.05358747 | 0.22718443 |
| AL136454.1 | 42.4679341 | -0.710385365 | 0.36992745 | -1.920337 | 0.05481535 | 0.23078465 |
| UCHL5 | 42.0947393 | -0.012978612 | 0.25628266 | -0.0506418 | 0.95961096 | 0.98416248 |
| TROVE2 | 73.7228148 | -0.459736904 | 0.24354554 | -1.8876835 | 0.05906845 | 0.24052299 |
| GLRX2 | 10.5898277 | -0.658789354 | 0.42123747 | -1.5639382 | 0.11783209 | 0.35871933 |
| CDC73 | 167.938465 | 0.257150173 | 0.2687581 | 0.956809 | 0.33866367 | 0.62755046 |
| ZBTB41 | 41.7607541 | -0.241403947 | 0.26930232 | -0.896405 | 0.3700365 | 0.65432818 |
| CRB1 | 20.369947 | -0.107216213 | 0.79174708 | -0.1354173 | 0.89228199 | 0.95772963 |
| DENND1B | 222.21419 | 0.567115442 | 0.22016057 | 2.57591734 | 0.00999745 | 0.07716229 |
| NEK7 | 202.092345 | -0.043390704 | 0.17837098 | -0.243261 | 0.80780322 | 0.91823775 |
| ZNF281 | 14.7180473 | -0.304930127 | 0.37761387 | -0.8075183 | 0.41936791 | 0.69548104 |
| DDX59 | 23.1730899 | 0.311029011 | 0.31716122 | 0.98066534 | 0.3267578 | 0.61667091 |
| CAMSAP2 | 142.307951 | -0.15775441 | 0.20838517 | -0.7570328 | 0.44903022 | 0.71802342 |
| TMEM9 | 30.4606513 | -0.744176967 | 0.51504266 | -1.4448841 | 0.14849043 | 0.40880504 |
| LAD1 | 17.9546587 | -0.498345454 | 0.53297486 | -0.9350262 | 0.34977481 | 0.63669603 |
| CSRP1 | 40.9633483 | 0.52311972 | 0.51398147 | 1.01777934 | 0.30878283 | 0.60259154 |
| AC096677.1 | 40.3197461 | 0.862321433 | 0.32321149 | 2.66797889 | 0.00763091 | 0.06391579 |
| NAV1 | 46.5519657 | -1.108400959 | 0.39580925 | -2.8003412 | 0.00510486 | 0.04833666 |
| IPO9-AS1 | 13.0569213 | -0.873200607 | 0.49651515 | -1.7586585 | 0.07863552 | 0.28554923 |
| IPO9 | 72.8864114 | -0.127423822 | 0.25470181 | -0.5002863 | 0.6168735 | 0.81715711 |
| TIMM17A | 25.3615892 | -0.267357312 | 0.34218825 | -0.7813165 | 0.4346164 | 0.70632878 |
| RNPEP | 34.05096 | -0.164012652 | 0.3272389 | -0.5012016 | 0.61622927 | 0.81691188 |
| ELF3 | 239.923971 | -0.94057591 | 0.38981783 | -2.4128602 | 0.01582789 | 0.10366112 |
| ARL8A | 23.8641901 | -0.496451875 | 0.38266567 | -1.2973515 | 0.19451028 | 0.47506992 |
| PPP1R12B | 218.400433 | 0.329220829 | 0.17235602 | 1.91012088 | 0.05611765 | 0.2339636 |
| KDM5B | 165.863052 | -0.42753729 | 0.23498577 | -1.8194178 | 0.06884771 | 0.26425823 |
| KLHL12 | 28.0389956 | -0.348793232 | 0.34181365 | -1.0204192 | 0.30752967 | 0.60139222 |
| ADIPOR1 | 19.537317 | -0.264386857 | 0.38920675 | -0.6792967 | 0.4969499 | 0.74891426 |
| CYB5R1 | 17.5683955 | 0.153636491 | 0.47328686 | 0.324616 | 0.74547171 | 0.88808711 |
| TMEM183A | 13.4241849 | -0.114706925 | 0.42582769 | -0.269374 | 0.78764186 | 0.90916375 |
| BTG2 | 148.692199 | 1.353058417 | 0.58572256 | 2.31006712 | 0.02088444 | 0.12372138 |
| ATP2B4 | 133.897615 | 0.156220125 | 0.26832759 | 0.58219926 | 0.56043247 | 0.7832825 |
| SNRPE | 17.4355255 | -0.555690963 | 0.38138529 | -1.457033 | 0.14510726 | 0.40304022 |
| SOX13 | 74.5301858 | 0.035929164 | 0.30133116 | 0.11923481 | 0.90508933 | 0.96157183 |
| ETNK2 | 12.1447256 | 0.004824557 | 0.46965979 | 0.01027245 | 0.99180391 | 0.99761413 |
| GOLT1A | 18.0171155 | -0.313183748 | 0.39582037 | -0.791227 | 0.42881157 | 0.70288254 |
| PLEKHA6 | 129.645316 | -0.072559746 | 0.20308109 | -0.3572944 | 0.72087139 | 0.87596332 |
| PPP1R15B | 32.3396796 | -0.490136418 | 0.43408933 | -1.1291142 | 0.25884967 | 0.55002264 |
| PIK3C2B | 27.575702 | -0.409410545 | 0.64367019 | -0.6360564 | 0.52473966 | 0.76221787 |
| MDM4 | 162.205629 | 0.538895011 | 0.19730479 | 2.73128201 | 0.00630885 | 0.05602908 |
| LRRN2 | 101.245526 | 0.984083619 | 0.38456221 | 2.55897119 | 0.01049824 | 0.07934916 |
| RBBP5 | 38.4190826 | 0.081924217 | 0.24672566 | 0.33204579 | 0.73985469 | 0.88614269 |
| DSTYK | 75.3736272 | 0.324085905 | 0.25563822 | 1.26775217 | 0.20488647 | 0.48953841 |
| NUAK2 | 69.9671867 | 0.651022026 | 0.45200635 | 1.440294 | 0.14978424 | 0.41046846 |
| CDK18 | 63.5830492 | -0.197947457 | 0.44516368 | -0.4446622 | 0.6565639 | 0.83951126 |
| MFSD4A | 37.461215 | 0.880463063 | 0.49762338 | 1.76933622 | 0.07683778 | 0.28130164 |
| ELK4 | 66.7787414 | -0.3981227 | 0.3245 | -1.2268804 | 0.21986754 | 0.50818193 |
| NUCKS1 | 233.814256 | 0.035223687 | 0.21002802 | 0.16770946 | 0.86681185 | 0.94728419 |
| RAB29 | 20.5928127 | -0.938426517 | 0.41949536 | -2.2370367 | 0.02528394 | 0.14026664 |
| SLC41A1 | 13.994409 | 0.056551801 | 0.42049894 | 0.13448738 | 0.89301719 | 0.95782021 |
| RHEX | 61.662417 | -3.162905176 | 0.68549116 | -4.6140714 | 3.95E-06 | 0.00017534 |
| SRGAP2 | 46.5958759 | -0.8535528 | 0.3402605 | -2.5085274 | 0.01212356 | 0.08770847 |
| EIF2D | 19.5489113 | -0.002841417 | 0.43496207 | -0.0065326 | 0.99478781 | 0.99837898 |
| MAPKAPK2 | 117.545757 | 0.174840809 | 0.29530495 | 0.59206867 | 0.55380461 | 0.77872693 |
| PIGR | 291.033091 | -2.499179951 | 0.69530776 | -3.5943507 | 0.0003252 | 0.00623878 |
| PFKFB2 | 68.3448572 | -0.357466029 | 0.43810442 | -0.815938 | 0.41453561 | 0.69139449 |
| CD55 | 54.1376898 | -1.567282063 | 0.52216339 | -3.0015166 | 0.00268638 | 0.0303123 |
| CD46 | 199.833893 | -0.014546943 | 0.16401554 | -0.0886925 | 0.92932632 | 0.97220251 |
| MIR29B2CHG | 199.687061 | -0.06501988 | 0.39669903 | -0.1639023 | 0.86980806 | 0.94832921 |
| PLXNA2 | 20.7391177 | -0.62690069 | 0.61855305 | -1.0134954 | 0.31082358 | 0.60394276 |
| LAMB3 | 20.0796175 | -7.764371574 | 1.17416754 | -6.6126607 | 3.77E-11 | 1.05E-08 |
| G0S2 | 65.2702038 | -2.664653158 | 1.18239057 | -2.253615 | 0.0242204 | 0.13597631 |
| IRF6 | 21.9988987 | -0.059357971 | 0.42356029 | -0.1401405 | 0.88854894 | 0.95671313 |
| UTP25 | 10.6317451 | -0.408824048 | 0.45578093 | -0.8969749 | 0.36973231 | 0.65423949 |
| HHAT | 65.1724219 | 0.562078209 | 0.42674391 | 1.31713234 | 0.18779427 | 0.46584555 |
| RCOR3 | 168.787772 | 0.292547698 | 0.18974728 | 1.54177543 | 0.12312817 | 0.36767143 |
| TRAF5 | 34.5878705 | 0.095641082 | 0.37283792 | 0.25652188 | 0.79754789 | 0.91302793 |
| LINC00467 | 38.317842 | -0.182468505 | 0.40980673 | -0.445255 | 0.65613546 | 0.83951126 |
| SLC30A1 | 16.3719443 | -0.90411605 | 0.51341603 | -1.7609814 | 0.07824156 | 0.28462025 |
| LPGAT1 | 151.751766 | -1.003082906 | 0.2380262 | -4.2141701 | 2.51E-05 | 0.00078501 |

| | | | | | | |
|---|---|---|---|---|---|---|
| INTS7 | 26.9392006 | -0.247030702 | 0.3633543 | -0.6798618 | 0.49659198 | 0.74859388 |
| PPP2R5A | 146.48406 | -0.025653722 | 0.23822372 | -0.1076875 | 0.91424357 | 0.96462448 |
| AL360091.3 | 19.1273601 | -1.087087735 | 0.61707865 | -1.761668 | 0.0781254 | 0.28439855 |
| TMEM206 | 11.479141 | 0.064020027 | 0.49614831 | 0.12903405 | 0.89733071 | 0.95917897 |
| NENF | 35.5383011 | -0.357262304 | 0.43779949 | -0.8160409 | 0.41447673 | 0.69139449 |
| ATF3 | 32.0723948 | 2.923737825 | 0.8017115 | 3.64687024 | 0.00026545 | 0.0053727 |
| NSL1 | 59.2301795 | 0.060571883 | 0.25504483 | 0.23749504 | 0.81227277 | 0.92034001 |
| TATDN3 | 41.6477587 | 0.02502129 | 0.324721 | 0.07705473 | 0.93858 | 0.97662903 |
| FLVCR1 | 36.1889935 | -0.169567856 | 0.301311 | -0.5627669 | 0.57359363 | 0.79073486 |
| ANGEL2 | 41.5673528 | 0.063792578 | 0.2484146 | 0.25679883 | 0.79733407 | 0.91298606 |
| RPS6KC1 | 135.544464 | 0.045041899 | 0.27354362 | 0.16466075 | 0.86921101 | 0.94827256 |
| SMYD2 | 31.3553157 | -0.941272786 | 0.32297662 | -2.9143682 | 0.00356409 | 0.03701339 |
| PTPN14 | 334.350968 | -0.196483043 | 0.36606934 | -0.5367372 | 0.59144914 | 0.80046098 |
| KCTD3 | 79.8986505 | 0.427019318 | 0.21776855 | 1.9608861 | 0.04989231 | 0.21717085 |
| ESRRG | 3810.07894 | 0.931946716 | 0.17177451 | 5.4254076 | 5.78E-08 | 5.23E-06 |
| GPATCH2 | 194.204549 | 0.110358919 | 0.17301857 | 0.63784438 | 0.52357499 | 0.76218158 |
| SPATA17 | 53.1011954 | 0.402305654 | 0.41959944 | 0.95878501 | 0.33766706 | 0.62621892 |
| RRP15 | 20.2668407 | -0.31702254 | 0.35467091 | -0.8938498 | 0.37140224 | 0.65494451 |
| LYPLAL1 | 190.174977 | 0.40567319 | 0.31474858 | 1.28888011 | 0.19743977 | 0.47953293 |
| EPRS | 64.1389466 | -0.413839142 | 0.32037478 | -1.2917345 | 0.19644912 | 0.47795702 |
| BPNT1 | 16.5482723 | -0.341917584 | 0.38041497 | -0.8988016 | 0.36875835 | 0.65369102 |
| IARS2 | 60.9123942 | -0.32017105 | 0.22107523 | -1.4482448 | 0.14754859 | 0.40740971 |
| RAB3GAP2 | 182.650686 | 0.182792326 | 0.21077729 | 0.8672297 | 0.38581616 | 0.66811344 |
| MARK1 | 78.9484272 | -0.071540487 | 0.25527651 | -0.280247 | 0.77928798 | 0.90427469 |
| C1orf115 | 12.0390967 | -0.232510789 | 0.51578724 | -0.4507882 | 0.65214222 | 0.83811991 |
| 2-Mar | 81.5957947 | 0.819327118 | 0.31373853 | 2.61149661 | 0.00901469 | 0.0713282 |
| 1-Mar | 12.3201897 | 0.554903631 | 0.4303081 | 1.2895496 | 0.19720709 | 0.47915741 |
| AL513314.2 | 18.643524 | 0.918684736 | 0.35881869 | 2.56030346 | 0.01045808 | 0.07916175 |
| MIA3 | 48.1658818 | 0.280580053 | 0.2471364 | 1.13532465 | 0.25623932 | 0.54710413 |
| AIDA | 20.6329879 | -0.415697134 | 0.37436998 | -1.1103912 | 0.26683048 | 0.55814977 |
| BROX | 47.3847594 | -0.48721311 | 0.25915599 | -1.8799994 | 0.06010815 | 0.24312297 |
| FAM177B | 33.8525749 | -3.337367118 | 0.77138803 | -4.326444 | 1.52E-05 | 0.00052211 |
| AC093152.1 | 44.6917259 | 0.247693949 | 0.28570256 | 0.8669644 | 0.38596152 | 0.66811344 |
| TLR5 | 17.1915635 | -0.352135828 | 0.51310666 | -0.6862819 | 0.49253534 | 0.74674012 |
| SUSD4 | 45.7757602 | 0.79945935 | 0.39501547 | 2.02386846 | 0.04298369 | 0.1979517 |
| CAPN2 | 145.983288 | 0.163779697 | 0.22259444 | 0.73577621 | 0.4618669 | 0.72676972 |
| TP53BP2 | 54.5254592 | -0.325139836 | 0.34211328 | -0.9503865 | 0.34191591 | 0.62990732 |
| FBXO28 | 85.3285008 | -0.388338383 | 0.21072527 | -1.8428658 | 0.06534861 | 0.25530899 |
| DEGS1 | 20.7539711 | -2.003927767 | 0.57305214 | -3.4969379 | 0.00047063 | 0.00823165 |
| NVL | 76.1427569 | 0.555458287 | 0.28564336 | 1.9445867 | 0.05182474 | 0.22230553 |
| CNIH4 | 49.5939658 | -0.436882476 | 0.2490913 | -1.753905 | 0.07944679 | 0.28728003 |
| WDR26 | 151.362427 | 0.288940971 | 0.24295069 | 1.18929881 | 0.2343221 | 0.52443761 |
| CNIH3 | 46.7476083 | 0.633776273 | 0.40948313 | 1.54774698 | 0.12168323 | 0.36568863 |
| DNAH14 | 253.818591 | 0.017168989 | 0.18155573 | 0.09456594 | 0.9246596 | 0.97014393 |
| LBR | 51.7203677 | 0.441534353 | 0.25168121 | 1.75433978 | 0.0793723 | 0.28711147 |
| ENAH | 77.4495285 | 0.173045278 | 0.26357143 | 0.65654036 | 0.51147651 | 0.75522875 |
| SRP9 | 71.649489 | -0.489536382 | 0.3240802 | -1.5105408 | 0.13090548 | 0.37977703 |
| EPHX1 | 23.9039771 | -0.227260552 | 0.35820743 | -0.6344384 | 0.52579475 | 0.76221787 |
| AL591895.1 | 25.7525554 | -0.398934758 | 0.29539035 | -1.3505342 | 0.1768447 | 0.45162472 |
| TMEM63A | 59.9312713 | -0.706339609 | 0.22205681 | -3.180896 | 0.0014682 | 0.01944143 |
| SDE2 | 11.6396422 | -0.338289822 | 0.39614194 | -0.8539611 | 0.39312652 | 0.67385918 |
| H3F3A | 15.7358948 | -0.656769952 | 0.4297034 | -1.5284262 | 0.12640674 | 0.37228594 |
| ACBD3 | 87.1295049 | 0.124270375 | 0.23546183 | 0.52777291 | 0.59765696 | 0.80551365 |
| LIN9 | 12.2191722 | 0.568053616 | 0.48092218 | 1.18117576 | 0.23753291 | 0.52778809 |
| PARP1 | 46.9263805 | -0.155624497 | 0.26256839 | -0.5927008 | 0.5533814 | 0.77828467 |
| STUM | 21.1298195 | -0.369898771 | 0.82518895 | -0.4482595 | 0.65396594 | 0.83885315 |
| ITPKB | 106.544876 | 0.648813924 | 0.41512293 | 1.56294408 | 0.11806575 | 0.35932447 |
| COQ8A | 29.7271058 | 0.223181129 | 0.40455956 | 0.55166445 | 0.58117827 | 0.79484057 |
| CDC42BPA | 316.175993 | -0.222631577 | 0.19555752 | -1.1384455 | 0.25493451 | 0.54590216 |
| ZNF678 | 47.2105139 | -0.040815328 | 0.26929147 | -0.1515656 | 0.87952955 | 0.95279412 |
| SNAP47 | 25.3718425 | 0.370572603 | 0.36438235 | 1.01698836 | 0.30915897 | 0.6027062 |
| ARF1 | 107.25066 | -0.771084101 | 0.28836546 | -2.6739822 | 0.00749564 | 0.06329518 |
| C1orf35 | 14.8906315 | -0.075361202 | 0.37454467 | -0.2012075 | 0.84053632 | 0.93552346 |
| MRPL55 | 15.642766 | 0.168235364 | 0.43265179 | 0.38884704 | 0.6973893 | 0.86275485 |
| GUK1 | 108.514902 | -0.477166438 | 0.40460903 | -1.1793272 | 0.2382679 | 0.52855279 |
| OBSCN | 106.37047 | 0.463183213 | 0.23022254 | 2.01189338 | 0.04423118 | 0.20044552 |
| TRIM11 | 13.0226036 | -0.641922525 | 0.45802958 | -1.4014871 | 0.16106848 | 0.42820049 |
| RNF187 | 33.3303346 | -0.476820739 | 0.50848733 | -0.9377239 | 0.3483863 | 0.6354277 |
| RAB4A | 36.6133176 | 0.000541697 | 0.31855178 | 0.0017005 | 0.9986432 | 0.9994193 |
| NUP133 | 53.0712598 | 0.393066468 | 0.26632925 | 1.47586666 | 0.13997969 | 0.39497967 |
| ABCB10 | 27.0676123 | 0.261925106 | 0.31404359 | 0.83404059 | 0.40425811 | 0.68217314 |
| TAF5L | 16.4418308 | -0.230791026 | 0.40487595 | -0.570029 | 0.56865804 | 0.78763298 |
| GALNT2 | 96.9263335 | -0.967162101 | 0.31038831 | -3.1159747 | 0.00183338 | 0.02300546 |
| COG2 | 37.2600438 | 0.606980473 | 0.2717892 | 2.23327665 | 0.02553071 | 0.14140717 |
| C1orf198 | 14.6202919 | -0.550806747 | 0.48820926 | -1.1282186 | 0.25922763 | 0.55035288 |
| TTC13 | 31.2701101 | 0.398565937 | 0.30773641 | 1.29515364 | 0.19526723 | 0.47624125 |
| ARV1 | 25.920654 | 0.227524262 | 0.35794784 | 0.63563525 | 0.52501419 | 0.76221787 |
| C1orf131 | 17.5879802 | 0.041970525 | 0.35228194 | 0.11913902 | 0.90516522 | 0.96157183 |
| GNPAT | 49.3651771 | 0.332355308 | 0.23353673 | 1.42313931 | 0.15469577 | 0.41795853 |
| EGLN1 | 140.938652 | -0.742709433 | 0.2935079 | -2.5304581 | 0.01139137 | 0.08400422 |
| DISC1 | 25.6872618 | -1.440714144 | 0.4254116 | -3.3866358 | 0.00070755 | 0.0113539 |
| SIPA1L2 | 35.2309447 | -0.066305295 | 0.54642078 | -0.1213448 | 0.90341797 | 0.96126516 |
| NTPCR | 31.8167362 | 0.240846268 | 0.28520043 | 0.84448072 | 0.39840083 | 0.67938457 |
| PCNX2 | 175.186848 | -0.160648698 | 0.22106451 | -0.7267051 | 0.46740664 | 0.73024314 |
| MAP3K21 | 270.307037 | 0.321109479 | 0.39312587 | 0.81681085 | 0.41403653 | 0.69086562 |
| KCNK1 | 81.2445754 | -1.059602077 | 0.31695821 | -3.343034 | 0.00082868 | 0.01274184 |
| AL355472.1 | 12.1191968 | 0.615402108 | 0.61827521 | 0.99535304 | 0.31956459 | 0.611915 |
| COA6 | 13.3323431 | -0.803053016 | 0.42743366 | -1.8787781 | 0.0602748 | 0.24330674 |
| TARBP1 | 62.2371626 | 0.363402292 | 0.29189144 | 1.24499126 | 0.21313496 | 0.49959417 |
| IRF2BP2 | 75.2425858 | -0.338186559 | 0.34632697 | -0.976495 | 0.32881922 | 0.6179306 |
| TOMM20 | 65.900281 | -0.490965629 | 0.34216476 | -1.4348808 | 0.15132108 | 0.41251912 |
| ARID4B | 373.86977 | 0.163136742 | 0.20851468 | 0.78237532 | 0.43399404 | 0.70597357 |
| B3GALNT2 | 60.470619 | 0.493541076 | 0.28509555 | 1.73114268 | 0.08342632 | 0.29605854 |
| LYST | 140.877991 | 0.134812723 | 0.21862203 | 0.61664747 | 0.53746728 | 0.76945357 |
| GPR137B | 38.0599331 | 0.391128853 | 0.35881597 | 1.09005419 | 0.27568927 | 0.56780306 |
| ERO1B | 36.7835202 | 0.754247334 | 0.42576739 | 1.77150095 | 0.07647744 | 0.2804628 |
| LGALS8 | 33.4399701 | -0.973553327 | 0.28114769 | -3.4627826 | 0.00053462 | 0.00899944 |
| HEATR1 | 43.8546006 | 0.159188857 | 0.29941057 | 0.53167414 | 0.59495171 | 0.80330177 |
| MTR | 513.130088 | 0.907283008 | 0.35500323 | 2.55570351 | 0.01059734 | 0.07992223 |
| CHRM3 | 279.006716 | 1.266189544 | 0.43304693 | 2.92390835 | 0.00345667 | 0.03615286 |
| RGS7 | 57.0233165 | 3.322059525 | 0.54286468 | 6.11949838 | 9.39E-10 | 1.70E-07 |
| FH | 22.4617664 | 0.893575287 | 0.38382598 | 2.32807403 | 0.01990817 | 0.12028973 |
| OPN3 | 12.1245736 | -0.594345065 | 0.42908707 | -1.3851386 | 0.16601009 | 0.4348426 |
| CEP170 | 74.9500376 | -0.077033862 | 0.29302132 | -0.2628951 | 0.79263145 | 0.91124908 |
| SDCCAG8 | 287.134088 | 0.225734378 | 0.22574358 | 0.99995925 | 0.31733023 | 0.61071101 |
| AKT3 | 427.399949 | 0.352601503 | 0.26610514 | 1.32504583 | 0.18515595 | 0.46185874 |
| ZBTB18 | 19.3448027 | 0.30653908 | 0.39885624 | 0.76854528 | 0.4421633 | 0.71263553 |
| AL645465.1 | 12.5269693 | -0.82239308 | 0.42304484 | -1.9439856 | 0.05189718 | 0.22230553 |
| ADSS | 71.3576287 | 0.087366334 | 0.2722925 | 0.32085472 | 0.74832049 | 0.8894886 |
| CATSPERE | 49.4607985 | -0.026518583 | 0.32565499 | -0.0814315 | 0.93509878 | 0.97519355 |
| DESI2 | 70.0728706 | -0.2813235 | 0.27312361 | -1.0300226 | 0.30299938 | 0.59684506 |
| COX20 | 77.8840713 | 0.439225045 | 0.31310129 | 1.40282095 | 0.16067024 | 0.42803048 |
| HNRNPU | 311.939217 | 0.1746102 | 0.18694525 | 0.93401784 | 0.35029469 | 0.63689665 |

| | | | | | | |
|---|---|---|---|---|---|---|
| EFCAB2 | 161.786449 | 0.677036818 | 0.42723616 | 1.58468987 | 0.11303681 | 0.35086028 |
| KIF26B | 195.296244 | -0.065238124 | 0.58735848 | -0.1110704 | 0.91156054 | 0.96399248 |
| SMYD3 | 649.420184 | 0.160341014 | 0.14093744 | 1.13767512 | 0.25525617 | 0.54647736 |
| TFB2M | 12.0214475 | -0.425617252 | 0.48140017 | -0.8841236 | 0.3766295 | 0.65964588 |
| CNST | 72.1523359 | 0.159293451 | 0.2361352 | 0.6745858 | 0.49993896 | 0.75045478 |
| SCCPDH | 68.4104162 | -0.447721613 | 0.42348523 | -1.0572308 | 0.29040628 | 0.58205555 |
| AHCTF1 | 100.388565 | 0.64756672 | 0.26873533 | 2.40968215 | 0.01596642 | 0.10410512 |
| ZNF124 | 58.0388698 | -0.158275408 | 0.36661574 | -0.4317202 | 0.6659448 | 0.84396154 |
| AL390728.6 | 19.6037915 | 0.012790303 | 0.34449417 | 0.03712778 | 0.97038312 | 0.98816027 |
| ZNF496 | 26.1085489 | 0.290714415 | 0.31582876 | 0.92048112 | 0.35732139 | 0.64174726 |
| ZNF672 | 19.2798789 | 0.169328872 | 0.33248958 | 0.50927573 | 0.61055897 | 0.81376433 |
| ZNF692 | 18.0573561 | 0.356456137 | 0.382928 | 0.93086988 | 0.35192088 | 0.63801604 |
| FAM110C | 63.8041328 | -3.054793516 | 0.47514318 | -6.4292063 | 1.28E-10 | 2.81E-08 |
| SH3YL1 | 108.141165 | -0.073326111 | 0.24193095 | -0.3030869 | 0.76182361 | 0.89684685 |
| ACP1 | 33.5926607 | -0.410364459 | 0.30541435 | -1.3436319 | 0.17906746 | 0.45449239 |
| EIPR1 | 118.013397 | -0.21417345 | 0.30104508 | -0.7114332 | 0.47681586 | 0.73678733 |
| TRAPPC12 | 123.825123 | 0.107493039 | 0.18675364 | 0.57558738 | 0.56489411 | 0.78515099 |
| ADI1 | 66.3753798 | -0.82143732 | 0.4575197 | -1.7954141 | 0.07258775 | 0.27211179 |
| AC108488.2 | 12.0284673 | 0.358066497 | 0.43291112 | 0.82711318 | 0.40817291 | 0.68657965 |
| RPS7 | 110.48542 | -0.377863257 | 0.40303952 | -0.937534 | 0.34848393 | 0.6354277 |
| RSAD2 | 20.0979944 | -2.79231922 | 2.22978182 | -1.2522836 | 0.21046654 | 0.49657419 |
| RNF144A | 50.9662761 | 0.051636146 | 0.34302244 | 0.15053285 | 0.88034423 | 0.95334202 |
| ID2 | 76.7813071 | 1.321787631 | 0.426685 | 3.09780667 | 0.00194959 | 0.0241112 |
| KIDINS220 | 155.262383 | 0.432286432 | 0.19478 | 2.21935744 | 0.02646242 | 0.14524018 |
| MBOAT2 | 139.310307 | -0.225975024 | 0.33141895 | -0.681841 | 0.49533951 | 0.74770824 |
| ASAP2 | 620.240406 | 1.342900212 | 0.22808162 | 5.88780538 | 3.91E-09 | 5.52E-07 |
| ITGB1BP1 | 53.8621599 | -0.396931567 | 0.22497495 | -1.7643367 | 0.0776753 | 0.28326052 |
| CPSF3 | 25.0092584 | -0.059176634 | 0.33295752 | -0.1777303 | 0.8589348 | 0.94406767 |
| IAH1 | 31.7022197 | -0.408136472 | 0.29612806 | -1.3782432 | 0.16812823 | 0.43857328 |
| ADAM17 | 109.20094 | 0.044050538 | 0.27006952 | 0.16310814 | 0.87043329 | 0.94840865 |
| YWHAQ | 89.6704692 | -0.449771287 | 0.26651106 | -1.6876271 | 0.09148283 | 0.31120975 |
| TAF1B | 105.540208 | 0.018308195 | 0.26433938 | 0.06926018 | 0.94478252 | 0.97858537 |
| GRHL1 | 25.1157706 | 0.619877113 | 0.43470683 | 1.42596589 | 0.15387819 | 0.4169593 |
| KLF11 | 24.6665781 | 0.820358155 | 0.46737041 | 1.75526335 | 0.07921427 | 0.28684198 |
| CYS1 | 243.217455 | 0.531061051 | 0.32720892 | 1.62300298 | 0.10458877 | 0.33555699 |
| HPCAL1 | 110.637979 | 0.705473452 | 0.26219219 | 2.69067299 | 0.00713081 | 0.06131331 |
| ODC1 | 21.4512105 | -0.325966237 | 0.47595129 | -0.6848731 | 0.493424 | 0.74700665 |
| NOL10 | 105.184226 | 0.596959263 | 0.22046734 | 2.70769936 | 0.00677514 | 0.05915039 |
| PDIA6 | 90.7362863 | -0.551617858 | 0.41341807 | -1.3342858 | 0.18211021 | 0.45876773 |
| PQLC3 | 16.347092 | -0.068570317 | 0.46822119 | -0.1464486 | 0.8835673 | 0.95564413 |
| ROCK2 | 251.156877 | 0.553228636 | 0.34293183 | 1.61323211 | 0.10669409 | 0.34029982 |
| E2F6 | 18.8359032 | -0.483632349 | 0.35238004 | -1.3724737 | 0.16991601 | 0.44052905 |
| LPIN1 | 77.8215319 | 0.447430309 | 0.24666176 | 1.81394277 | 0.06968654 | 0.26599137 |
| MIR3681HG | 47.5034235 | -0.892411315 | 0.53048048 | -1.6822698 | 0.09251653 | 0.31323946 |
| TRIB2 | 53.2001436 | -0.771019108 | 0.27735464 | -2.7799034 | 0.00543751 | 0.05051145 |
| LINC00276 | 30.6528134 | -0.688997083 | 0.54546116 | -1.263146 | 0.20653671 | 0.49128174 |
| FAM84A | 76.0636264 | -2.122590121 | 0.37590595 | -5.6465988 | 1.64E-08 | 1.97E-06 |
| NBAS | 240.598649 | 0.06283595 | 0.2231837 | 0.28154363 | 0.77829347 | 0.90417143 |
| DDX1 | 68.5933153 | 0.332500504 | 0.19939865 | 1.66751629 | 0.09541179 | 0.31810105 |
| SMC6 | 61.3343242 | 0.092756399 | 0.28424015 | 0.3263311 | 0.74417386 | 0.88798073 |
| GEN1 | 18.5381611 | -0.686283356 | 0.44214867 | -1.5521552 | 0.1206251 | 0.36346292 |
| KCNS3 | 13.870177 | -0.011815392 | 0.68671106 | -0.0172058 | 0.98627246 | 0.9949057 |
| RDH14 | 13.5432833 | -0.690327906 | 0.46978051 | -1.469469 | 0.14170562 | 0.39788385 |
| LINC01376 | 69.1312886 | -0.018317736 | 0.41909578 | -0.0437078 | 0.96513736 | 0.98660829 |
| TTC32 | 18.2704383 | 0.090479251 | 0.38586786 | 0.23448248 | 0.81461044 | 0.92150178 |
| WDR35 | 61.471406 | -0.053990099 | 0.27150086 | -0.1988579 | 0.84237388 | 0.93584986 |
| LAPTM4A | 136.012857 | -0.809149466 | 0.33117282 | -2.4432846 | 0.01455425 | 0.09806274 |
| SDC1 | 39.8724457 | -0.993064459 | 0.5600316 | -1.7732293 | 0.07619071 | 0.28022732 |
| PUM2 | 306.990139 | 0.224356615 | 0.25878024 | 0.86697737 | 0.38595441 | 0.66811344 |
| RHOB | 35.3264266 | -0.947132892 | 0.44451194 | -2.1307254 | 0.03311177 | 0.16803815 |
| HS1BP3 | 13.6075333 | -0.590641426 | 0.42120257 | -1.402274 | 0.16083343 | 0.42803048 |
| LDAH | 100.762373 | 0.227225867 | 0.1958506 | 1.16019999 | 0.24596739 | 0.53633686 |
| KLHL29 | 36.1412151 | -3.782289597 | 0.75442957 | -5.013443 | 5.35E-07 | 3.40E-05 |
| ATAD2B | 197.150578 | 0.737854048 | 0.24048484 | 3.0681936 | 0.00215357 | 0.02581658 |
| UBXN2A | 71.5525746 | 0.001019276 | 0.34099286 | 0.00298914 | 0.99761501 | 0.99900003 |
| SF3B6 | 35.433607 | -0.847794936 | 0.36252936 | -2.3385552 | 0.01935846 | 0.11819385 |
| FAM228B | 14.0806409 | -0.517342211 | 0.41920991 | -1.2340887 | 0.21716986 | 0.50468845 |
| TP53I3 | 23.7145409 | -0.344994296 | 0.42496336 | -0.8118213 | 0.4168942 | 0.69338933 |
| ITSN2 | 230.322108 | -0.269523845 | 0.22494037 | -1.1982013 | 0.23083865 | 0.52105604 |
| NCOA1 | 459.204201 | 0.391688394 | 0.19880633 | 1.97020086 | 0.04881536 | 0.21436308 |
| DNAJC27 | 35.681907 | 0.539678397 | 0.26497772 | 2.03669351 | 0.04168077 | 0.19500656 |
| DNAJC27-AS | 21.3646432 | 1.01299371 | 0.61957033 | 1.6349939 | 0.10205033 | 0.33070354 |
| DNMT3A | 115.091503 | 0.400062614 | 0.2546073 | 1.57129279 | 0.11611466 | 0.35543425 |
| DTNB | 178.784769 | 0.310331963 | 0.31179565 | 0.99530562 | 0.31958764 | 0.611915 |
| ASXL2 | 143.358683 | 0.013134851 | 0.24150922 | 0.05438654 | 0.9566272 | 0.98286826 |
| RAB10 | 358.551291 | -0.034829983 | 0.31005835 | -0.1123336 | 0.91055887 | 0.96369195 |
| HADHA | 103.301424 | -0.082710877 | 0.22320181 | -0.3705654 | 0.71096122 | 0.87028547 |
| HADHB | 127.47448 | 0.126703237 | 0.23818082 | 0.53196238 | 0.59475205 | 0.80313746 |
| SELENOI | 79.3724625 | 0.287804941 | 0.28990099 | 0.99276977 | 0.32082217 | 0.6120836 |
| MAPRE3 | 26.4429573 | -0.019764246 | 0.31016886 | -0.0637209 | 0.94919244 | 0.97991588 |
| TMEM214 | 15.6885671 | -0.205991604 | 0.37210507 | -0.5535845 | 0.57986323 | 0.79438178 |
| AGBL5 | 25.4083913 | -0.627307425 | 0.36629925 | -1.7125545 | 0.08679453 | 0.30289881 |
| OST4 | 51.1560704 | -0.71617133 | 0.57234443 | -1.2512943 | 0.21082712 | 0.49667071 |
| ATRAID | 39.3879479 | -0.753032663 | 0.39063005 | -1.9277387 | 0.05388764 | 0.2282691 |
| CAD | 11.5303542 | -0.731610415 | 0.55580795 | -1.3163007 | 0.18807312 | 0.46620051 |
| MPV17 | 17.3573598 | -1.272947066 | 0.41896434 | -3.0383184 | 0.00237902 | 0.02751797 |
| GTF3C2 | 53.2395438 | 0.273930746 | 0.22444319 | 1.22049035 | 0.22227905 | 0.5106866 |
| EIF2B4 | 13.2915636 | -0.624028719 | 0.44797378 | -1.3930028 | 0.16361895 | 0.43198422 |
| SNX17 | 21.2336414 | -0.693255154 | 0.35078239 | -1.9763112 | 0.04811955 | 0.21221214 |
| PPM1G | 65.0575724 | -0.635040693 | 0.30602625 | -2.0751184 | 0.03797557 | 0.18260776 |
| NRBP1 | 46.8180186 | -0.281661916 | 0.29492547 | -0.9550274 | 0.33956382 | 0.62815343 |
| KRTCAP3 | 10.7325026 | -1.291882485 | 0.51167044 | -2.5248332 | 0.01157532 | 0.08487163 |
| IFT172 | 55.6531626 | 0.527746969 | 0.33880349 | 1.55767867 | 0.11930944 | 0.36172038 |
| FNDC4 | 14.0572579 | -3.504823205 | 0.72497755 | -4.8343886 | 1.34E-06 | 7.09E-05 |
| ZNF512 | 51.0481035 | -0.261751585 | 0.28208887 | -0.9279047 | 0.35345701 | 0.63947209 |
| GPN1 | 12.3864037 | -0.383210373 | 0.40394455 | -0.9486707 | 0.34278811 | 0.63072032 |
| SUPT7L | 20.1373261 | -0.402901245 | 0.40086667 | -1.0050754 | 0.31486053 | 0.60823049 |
| SLC4A1AP | 26.9860285 | 0.168205217 | 0.29987245 | 0.56092255 | 0.57485034 | 0.79134063 |
| MRPL33 | 162.120808 | -0.017539478 | 0.26077422 | -0.0672592 | 0.94637532 | 0.97905916 |
| RBKS | 103.576652 | 0.719703875 | 0.33857819 | 2.1256652 | 0.03353115 | 0.16924933 |
| BABAM2 | 777.678848 | 0.37213335 | 0.17149049 | 2.16999404 | 0.0300073 | 0.15756125 |
| FOSL2 | 257.296198 | 0.323089763 | 0.61143554 | 0.52841181 | 0.59721354 | 0.80508949 |
| PPP1CB | 144.175646 | 0.344242817 | 0.50227922 | 0.68536145 | 0.49311587 | 0.74685087 |
| TRMT61B | 37.7288206 | 0.818594784 | 0.30979766 | 2.64235303 | 0.00823322 | 0.06735772 |
| WDR43 | 64.0904513 | 0.072005965 | 0.33029796 | 0.21800306 | 0.82742673 | 0.92808376 |
| CLIP4 | 107.691147 | -0.053370037 | 0.31605458 | -0.1688634 | 0.86590412 | 0.94694132 |
| ALK | 21.2974486 | -2.066142989 | 0.81920937 | -2.5221183 | 0.01166505 | 0.08523395 |
| YPEL5 | 61.1547217 | -0.704057218 | 0.28463035 | -2.4735845 | 0.01337652 | 0.09373482 |
| LCLAT1 | 88.9723628 | 0.085376773 | 0.19309776 | 0.44214275 | 0.65838591 | 0.84037209 |
| GALNT14 | 246.038603 | 0.591419816 | 0.39652301 | 1.49151448 | 0.13582647 | 0.38804335 |
| DPY30 | 42.5861601 | 0.080579635 | 0.28514463 | 0.28259215 | 0.7774895 | 0.90366576 |
| SPAST | 73.4440482 | 0.502574574 | 0.28875329 | 1.74049818 | 0.08177158 | 0.29260535 |
| SLC30A6 | 62.2600565 | 0.008293989 | 0.21569901 | 0.03845168 | 0.96932756 | 0.98781534 |

| | | | | | | |
|---|---|---|---|---|---|---|
| YIPF4 | 85.2431026 | 0.093179899 | 0.24524095 | 0.37995245 | 0.70398071 | 0.86617744 |
| BIRC6 | 546.91703 | 0.347744408 | 0.16371156 | 2.12412861 | 0.03365939 | 0.16959564 |
| TTC27 | 67.7521261 | 0.178156787 | 0.27549098 | 0.64668829 | 0.5178337 | 0.75866548 |
| LINC00486 | 104.402646 | 0.63952247 | 0.20899089 | 3.06004952 | 0.002213 | 0.02619294 |
| LTBP1 | 72.5766274 | 0.672896668 | 0.75256468 | 0.89413799 | 0.37124807 | 0.65494451 |
| RASGRP3 | 21.9152547 | 0.001323757 | 0.51550387 | 0.00256789 | 0.99795112 | 0.99900003 |
| FAM98A | 19.5438261 | -0.04528877 | 0.40205005 | -0.1126446 | 0.91031232 | 0.96353 |
| LINC01317 | 20.1756978 | 1.557553814 | 0.60467396 | 2.57585729 | 0.00999919 | 0.07716229 |
| AC068599.1 | 202.032053 | 3.104429778 | 0.69460379 | 4.46935338 | 7.85E-06 | 0.00030726 |
| LINC01320 | 321.907169 | -1.566826578 | 0.56054905 | -2.7951641 | 0.00518734 | 0.04891142 |
| AC012593.1 | 90.3099232 | -1.872186456 | 1.25711453 | -1.4892728 | 0.13641555 | 0.38897122 |
| CRIM1 | 245.245495 | 0.298597229 | 0.23943746 | 1.24707818 | 0.21236883 | 0.49847886 |
| FEZ2 | 94.8989819 | 0.185078144 | 0.22716571 | 0.81472747 | 0.41522833 | 0.69186606 |
| STRN | 156.142454 | 0.335459488 | 0.2677297 | 1.25297822 | 0.21021363 | 0.49648138 |
| HEATR5B | 85.8689418 | -0.306320176 | 0.27664051 | -1.107286 | 0.26817028 | 0.56003648 |
| EIF2AK2 | 93.3113858 | 0.033003606 | 0.60458632 | 0.05458874 | 0.95646611 | 0.98286826 |
| CEBPZOS | 52.2803751 | -0.008161622 | 0.33247163 | -0.0245483 | 0.98041524 | 0.99299415 |
| CEBPZ | 58.3715048 | -0.068949577 | 0.25876242 | -0.266459 | 0.78988573 | 0.91080793 |
| NDUFAF7 | 40.5193625 | 0.213402657 | 0.33513064 | 0.63677453 | 0.52427172 | 0.76221787 |
| PRKD3 | 64.1302221 | -0.547223282 | 0.25222802 | -2.1695578 | 0.03004036 | 0.1576545 |
| CDC42EP3 | 20.6260788 | -0.584063839 | 0.3490731 | -1.6731849 | 0.0942909 | 0.31610311 |
| RMDN2 | 45.9446018 | 0.382677293 | 0.30643704 | 1.24879583 | 0.21173976 | 0.49790984 |
| CYP1B1-AS1 | 12.4805914 | 0.270958624 | 0.55076984 | 0.49196344 | 0.62274519 | 0.82102905 |
| ATL2 | 125.191617 | 0.999146342 | 0.27555164 | 3.6259858 | 0.00028786 | 0.00574718 |
| HNRNPLL | 60.2647189 | 0.452646309 | 0.29517408 | 1.53348934 | 0.12515533 | 0.37032604 |
| GALM | 47.3940769 | -0.146768112 | 0.39358613 | -0.3728996 | 0.70922316 | 0.8691907 |
| SRSF7 | 69.4965063 | -0.295438014 | 0.38714046 | -0.7631287 | 0.44538662 | 0.71514541 |
| DHX57 | 52.638825 | -0.106455711 | 0.26935333 | -0.395227 | 0.69267536 | 0.86052545 |
| MORN2 | 14.4558591 | -0.502899765 | 0.43188273 | -1.1644359 | 0.2442474 | 0.53491427 |
| SOS1 | 310.428281 | 0.183276741 | 0.18508639 | 0.9902227 | 0.32206528 | 0.61306662 |
| MAP4K3 | 743.583159 | 1.08367849 | 0.20350973 | 5.32494671 | 1.01E-07 | 8.67E-06 |
| AC007388.1 | 366.865125 | 0.805005343 | 0.27765848 | 2.89926436 | 0.00374039 | 0.03826568 |
| TMEM178A | 177.832083 | 2.908726833 | 0.52455423 | 5.54514039 | 2.94E-08 | 3.03E-06 |
| THUMPD2 | 67.7116551 | 0.182365467 | 0.24465342 | 0.74540331 | 0.45602795 | 0.72261911 |
| SLC8A1-AS1 | 291.003153 | 2.326674387 | 0.51564046 | 4.51220289 | 6.42E-06 | 0.00026125 |
| SLC8A1 | 9040.06803 | 3.21579902 | 0.43136192 | 7.45499058 | 8.99E-14 | 5.45E-11 |
| EML4 | 267.1023 | 0.342742505 | 0.31773845 | 1.07869382 | 0.28072424 | 0.57381372 |
| COX7A2L | 74.7567238 | -0.215625543 | 0.32717991 | -0.6590427 | 0.50986833 | 0.75480149 |
| MTA3 | 133.164781 | 0.360864542 | 0.17836883 | 2.02313676 | 0.04305905 | 0.19806087 |
| ZFP36L2 | 296.109709 | -1.463955901 | 0.31576577 | -4.6362083 | 3.55E-06 | 0.00016105 |
| THADA | 250.117897 | -0.048794078 | 0.24074424 | -0.2026802 | 0.83938504 | 0.93543959 |
| PLEKHH2 | 122.69501 | -0.536186632 | 0.34458934 | -1.5560163 | 0.11970422 | 0.36203283 |
| DYNC2LI1 | 100.825373 | 0.143982069 | 0.29763007 | 0.48376183 | 0.6285549 | 0.82482184 |
| LRPPRC | 149.694582 | 0.416757267 | 0.16023815 | 2.60086174 | 0.00929899 | 0.07314246 |
| PPM1B | 176.969118 | 0.330968387 | 0.18385083 | 1.80020066 | 0.07182896 | 0.27051259 |
| SLC3A1 | 139.878976 | -0.206483319 | 0.67985216 | -0.303718 | 0.76134277 | 0.89648567 |
| PREPL | 181.591886 | 0.554462245 | 0.27790645 | 1.99513992 | 0.04602762 | 0.2059846 |
| CAMKMT | 551.657766 | 0.079232035 | 0.1978872 | 0.4003899 | 0.68886936 | 0.85864878 |
| SRBD1 | 109.539087 | 0.207285715 | 0.24780067 | 0.83650185 | 0.40287264 | 0.68128594 |
| PRKCE | 531.701446 | 0.361036049 | 0.18928329 | 1.90738468 | 0.05647079 | 0.23458149 |
| EPAS1 | 59.2839505 | -1.127041581 | 0.35029468 | -3.21741 | 0.00129354 | 0.01772076 |
| RHOQ | 77.5742943 | -0.903136338 | 0.35795084 | -2.5230737 | 0.0116334 | 0.08511883 |
| PIGF | 35.1923183 | 0.319806441 | 0.29421765 | 1.08697232 | 0.27704904 | 0.56927037 |
| CRIPT | 12.8267516 | -1.098526145 | 0.4733726 | -2.3206374 | 0.02030642 | 0.12219438 |
| SOCS5 | 52.35474 | 0.215891482 | 0.22343989 | 0.96621727 | 0.33393547 | 0.62254854 |
| MCFD2 | 62.3550158 | -0.251336175 | 0.30892268 | -0.8135893 | 0.41588029 | 0.69248404 |
| TTC7A | 110.161799 | 1.109593753 | 0.27453292 | 4.04175125 | 5.31E-05 | 0.00146924 |
| AC106869.1 | 18.7417621 | -0.519106697 | 0.53668295 | -0.9672502 | 0.33341897 | 0.62214857 |
| EPCAM | 230.557689 | -0.266567181 | 0.32979113 | -0.8082909 | 0.41892312 | 0.69496715 |
| MSH2 | 83.2171301 | 0.358355709 | 0.31771544 | 1.12791406 | 0.25935621 | 0.5504507 |
| MSH6 | 23.2136244 | -0.268242825 | 0.34789424 | -0.771047 | 0.44067905 | 0.71135625 |
| FBXO11 | 260.50664 | -0.124508826 | 0.23360956 | -0.5329783 | 0.59404859 | 0.80267231 |
| FOXN2 | 27.962218 | -0.421328078 | 0.30020643 | -1.4034612 | 0.16047935 | 0.42803048 |
| PPP1R21 | 141.77943 | 0.033653737 | 0.1934492 | 0.17396679 | 0.86189156 | 0.94550174 |
| GPR75-ASB3 | 43.2643492 | -0.394379113 | 0.41395264 | -0.9527155 | 0.34073422 | 0.62907597 |
| ERLEC1 | 31.7609196 | -0.255493847 | 0.33648552 | -0.7593012 | 0.44767243 | 0.71693398 |
| PSME4 | 180.440479 | 0.491118374 | 0.30604089 | 1.60474754 | 0.10854937 | 0.34348281 |
| ACYP2 | 230.235303 | 0.158449843 | 0.31469711 | 0.50349952 | 0.61461312 | 0.81594643 |
| SPTBN1 | 644.101118 | -1.549277913 | 0.3757436 | -4.1232317 | 3.74E-05 | 0.00110597 |
| AC093110.1 | 21.5107609 | -2.741306274 | 0.65546746 | -4.1822157 | 2.89E-05 | 0.00088776 |
| EML6 | 134.961939 | 1.122452368 | 0.31079923 | 3.611503 | 0.00030443 | 0.00596238 |
| RTN4 | 639.481614 | -0.509707732 | 0.35812508 | -1.4232673 | 0.15465867 | 0.41795853 |
| CLHC1 | 42.7560696 | 0.594885316 | 0.27509706 | 2.16245607 | 0.03058304 | 0.15968904 |
| RPS27A | 218.094067 | -0.229544868 | 0.4443631 | -0.5165705 | 0.60545602 | 0.8104748 |
| MTIF2 | 36.1086423 | 0.208437718 | 0.33273868 | 0.62643069 | 0.53103249 | 0.7650254 |
| AC012358.3 | 31.3697761 | 0.027022514 | 0.49619208 | 0.05445978 | 0.95656885 | 0.98286826 |
| CCDC88A | 35.4548783 | -0.213406856 | 0.31961392 | -0.667702 | 0.50432383 | 0.75134405 |
| CFAP36 | 65.6885689 | -0.153757507 | 0.24962861 | -0.6159451 | 0.53793078 | 0.76983624 |
| PPP4R3B | 132.851327 | 0.022564408 | 0.19241517 | 0.11726938 | 0.90664659 | 0.96157183 |
| PNPT1 | 61.9978737 | -0.027930827 | 0.52877731 | -0.0528215 | 0.95787411 | 0.98321101 |
| EFEMP1 | 52.1463064 | -2.726526933 | 0.63781337 | -4.2748037 | 1.91E-05 | 0.00062171 |
| VRK2 | 105.951557 | -0.030806226 | 0.19970473 | -0.1542589 | 0.87740562 | 0.95177769 |
| FANCL | 73.0188472 | 0.456351037 | 0.30209378 | 1.51062707 | 0.13088349 | 0.37977703 |
| PAPOLG | 39.3462038 | 0.549369958 | 0.35097123 | 1.56528485 | 0.11751613 | 0.35807489 |
| REL | 53.4179011 | -0.234174126 | 0.34133711 | -0.6860494 | 0.49268195 | 0.74680729 |
| PUS10 | 21.7931998 | 0.288858347 | 0.45692774 | 0.63217512 | 0.52727246 | 0.76221787 |
| PEX13 | 36.5693053 | -0.359537224 | 0.28870021 | -1.2453653 | 0.21299751 | 0.4993856 |
| KIAA1841 | 21.4274327 | -1.518727824 | 0.41853441 | -3.6286809 | 0.00028487 | 0.00570966 |
| C2orf74 | 16.9474806 | 0.134258545 | 0.39683672 | 0.33832187 | 0.73512064 | 0.88399823 |
| USP34 | 570.264875 | 0.286313149 | 0.23112101 | 1.23880191 | 0.21541887 | 0.50209938 |
| AC016727.1 | 15.0462023 | 0.28943001 | 0.40489896 | 0.71482033 | 0.47472007 | 0.7352452 |
| XPO1 | 239.534253 | 0.338957971 | 0.34121283 | 0.99339165 | 0.32051913 | 0.6120836 |
| FAM161A | 18.2306257 | 0.640596644 | 0.3263397 | 1.96297491 | 0.04964909 | 0.21654738 |
| CCT4 | 55.3496458 | 0.099214899 | 0.25362208 | 0.39119188 | 0.69565541 | 0.86236044 |
| COMMD1 | 221.941964 | 0.463076952 | 0.26722241 | 1.73292707 | 0.08310863 | 0.29523625 |
| AC018462.1 | 14.299221 | -1.072404493 | 0.5313462 | -2.0182783 | 0.04356229 | 0.19928006 |
| B3GNT2 | 46.4697502 | -0.124722461 | 0.27520286 | -0.4532019 | 0.65040335 | 0.83735207 |
| EHBP1 | 267.461173 | 0.311885764 | 0.23630674 | 1.31983438 | 0.18689032 | 0.46438594 |
| WDPCP | 223.982796 | 0.008731788 | 0.26031341 | 0.03354336 | 0.97324129 | 0.98879011 |
| MDH1 | 75.2668558 | 0.432132449 | 0.35706942 | 1.21021972 | 0.22619459 | 0.51520157 |
| UGP2 | 116.733871 | -0.368778047 | 0.18768723 | -1.9648542 | 0.04943111 | 0.21605779 |
| VPS54 | 156.595417 | 0.420616326 | 0.19443731 | 2.16324908 | 0.03052203 | 0.15953218 |
| PELI1 | 33.6905465 | -0.692282913 | 0.42597493 | -1.6251729 | 0.10412572 | 0.33490577 |
| AC008074.1 | 15.1267669 | -0.139652307 | 0.52873769 | -0.264124 | 0.79168438 | 0.91107378 |
| AFTPH | 130.037299 | 0.01856965 | 0.25571539 | 0.07261843 | 0.94210976 | 0.97799423 |
| SERTAD2 | 41.9251118 | 0.238372821 | 0.35550302 | 0.67052262 | 0.50252469 | 0.75108109 |
| CEP68 | 23.7623703 | -0.355139751 | 0.36650417 | -0.9689924 | 0.33254898 | 0.62174935 |
| RAB1A | 232.267763 | 0.375759485 | 0.29792477 | 1.26125628 | 0.20721653 | 0.49241084 |
| ACTR2 | 154.858448 | -0.361640354 | 0.24060478 | -1.5030473 | 0.13282685 | 0.38265722 |
| SPRED2 | 162.968967 | 0.177251715 | 0.27415269 | 0.64654378 | 0.51792725 | 0.75866548 |
| ETAA1 | 21.5116083 | -0.070349362 | 0.33336522 | -0.2110279 | 0.8328655 | 0.93212171 |
| C1D | 32.432234 | 0.051446938 | 0.26886886 | 0.19134584 | 0.84825465 | 0.93871321 |
| PNO1 | 13.4563261 | -0.477751683 | 0.42451372 | -1.1254093 | 0.26041564 | 0.55146957 |

| | | | | | | |
|---|---|---|---|---|---|---|
| APLF | 56.9786732 | -0.149037891 | 0.29439133 | -0.5062578 | 0.61267572 | 0.81450903 |
| ANTXR1 | 31.2482457 | -0.290098245 | 0.35710636 | -0.8123581 | 0.41658618 | 0.69318598 |
| GFPT1 | 124.996884 | -1.223321095 | 0.26992514 | -4.5320754 | 5.84E-06 | 0.00024116 |
| NFU1 | 45.2134332 | 0.50313976 | 0.23346957 | 2.15505495 | 0.03115753 | 0.16170525 |
| AAK1 | 459.631141 | 0.45795583 | 0.31020506 | 1.47630033 | 0.13986329 | 0.39486752 |
| ANXA4 | 271.441341 | -1.01039429 | 0.30472898 | -3.3157145 | 0.00091409 | 0.01372737 |
| GMCL1 | 60.7167377 | -0.452464732 | 0.2738769 | -1.6520733 | 0.0985196 | 0.32413171 |
| MXD1 | 35.5340866 | 0.026690455 | 0.474931 | 0.05619859 | 0.9551836 | 0.98253384 |
| PCBP1 | 62.4276573 | -1.002780203 | 0.4099343 | -2.4461974 | 0.01443719 | 0.09759315 |
| C2orf42 | 22.8315982 | 0.528342675 | 0.38769513 | 1.36277871 | 0.17295226 | 0.44566139 |
| PCYOX1 | 60.8607771 | -1.037450295 | 0.55959567 | -1.8539284 | 0.06374941 | 0.2515306 |
| SNRPG | 29.5760592 | -0.490968631 | 0.34919335 | -1.406008 | 0.15972176 | 0.42727956 |
| FAM136A | 26.0841707 | -0.365069824 | 0.31425357 | -1.1617046 | 0.24535546 | 0.53608738 |
| TGFA | 72.7900376 | -0.049787509 | 0.3808812 | -0.1307166 | 0.89599947 | 0.95900099 |
| ATP6V1B1 | 9.37461384 | 2.265592119 | 0.9028414 | 2.50940213 | 0.01209357 | 0.08755304 |
| AC007040.2 | 52.9381135 | 0.286308928 | 0.80626532 | 0.3551051 | 0.72251086 | 0.87650915 |
| NAGK | 22.9025744 | -0.460432945 | 0.37474174 | -1.2286674 | 0.21919655 | 0.50733831 |
| MCEE | 38.5236767 | 0.088905613 | 0.35563172 | 0.24999349 | 0.80259239 | 0.91564859 |
| MPHOSPH10 | 23.7891766 | -0.602235735 | 0.3974882 | -1.5151034 | 0.12974621 | 0.37739114 |
| PAIP2B | 15.5196629 | -0.417881156 | 0.47048806 | -0.8881865 | 0.37444043 | 0.65806152 |
| ZNF638 | 476.467264 | 0.083861914 | 0.17193643 | 0.48774954 | 0.62572726 | 0.82311785 |
| EXOC6B | 541.447551 | -0.067172763 | 0.3637617 | -0.1846614 | 0.85349459 | 0.94089901 |
| SPR | 19.1243192 | -0.752076094 | 0.53795137 | -1.3980373 | 0.16210186 | 0.42953067 |
| EMX1 | 243.838265 | 1.61895342 | 0.40407149 | 4.00660144 | 6.16E-05 | 0.00168326 |
| SFXN5 | 199.419217 | 0.28325757 | 0.25800645 | 1.09787011 | 0.27226121 | 0.56475837 |
| RAB11FIP5 | 41.2771443 | 0.235717552 | 0.32343679 | 0.72879017 | 0.46613002 | 0.72924396 |
| SMYD5 | 9.71955169 | -0.084280842 | 0.499322 | -0.1687906 | 0.86596138 | 0.94694132 |
| CCT7 | 47.7693968 | -0.20996749 | 0.38255632 | -0.5488538 | 0.58310579 | 0.79586061 |
| ALMS1 | 97.1191265 | -0.049092507 | 0.19997247 | -0.2454963 | 0.80607215 | 0.91723667 |
| NAT8 | 10.2180889 | -0.229952143 | 1.05836989 | -0.2172701 | 0.82799785 | 0.92818649 |
| DUSP11 | 17.2612289 | -0.511348096 | 0.43336844 | -1.1799385 | 0.23802468 | 0.52824866 |
| STAMBP | 54.4471501 | -0.247522138 | 0.29221389 | -0.8470581 | 0.39696275 | 0.67785683 |
| DGUOK | 50.5223951 | -0.320605252 | 0.38148972 | -0.8404034 | 0.40068225 | 0.68010059 |
| TET3 | 47.1080682 | -0.236625849 | 0.34226027 | -0.6913623 | 0.48933789 | 0.74551301 |
| BOLA3 | 16.6717642 | 0.455225344 | 0.60199039 | 0.75620035 | 0.44952908 | 0.71810337 |
| MOB1A | 78.8165967 | -0.125731498 | 0.26046634 | -0.4827169 | 0.62929678 | 0.82502105 |
| AC006030.1 | 24.4881636 | -0.431871047 | 0.45780696 | -0.9433475 | 0.34550319 | 0.63256941 |
| DCTN1 | 10.8972356 | -0.335390225 | 0.51741493 | -0.6482036 | 0.51685327 | 0.75806127 |
| RTKN | 30.9032085 | -0.737854683 | 0.36582227 | -2.0169758 | 0.04369804 | 0.19956366 |
| MOGS | 23.4097925 | -0.669620186 | 0.30881549 | -2.1683504 | 0.03013204 | 0.1578151 |
| CCDC142 | 11.4097881 | 0.262154771 | 0.46267716 | 0.56660409 | 0.5709832 | 0.78905469 |
| TTC31 | 13.2988958 | -0.091880232 | 0.46533351 | -0.1974503 | 0.84347519 | 0.93643755 |
| AUP1 | 18.1924239 | 0.159959219 | 0.37628632 | 0.42509975 | 0.67076396 | 0.84746193 |
| POLE4 | 26.380703 | -0.378374677 | 0.34469454 | -1.0977101 | 0.2723311 | 0.56475837 |
| TACR1 | 24.6457913 | -0.684711121 | 0.58932841 | -1.1618498 | 0.24529647 | 0.53608738 |
| AC007681.1 | 23.0826734 | 0.488677483 | 0.54382597 | 0.89859167 | 0.3688702 | 0.65369102 |
| EVA1A | 52.2358154 | 0.626605513 | 0.4342254 | 1.44304207 | 0.14900862 | 0.4093068 |
| MRPL19 | 77.6107097 | 0.209671083 | 0.29274033 | 0.71623572 | 0.47384581 | 0.73451091 |
| GCFC2 | 98.5293185 | 0.656437929 | 0.32320758 | 2.03101033 | 0.04225395 | 0.19616952 |
| CTNNA2 | 11.9850856 | 1.057883359 | 0.571132 | 1.85225718 | 0.0639889 | 0.25208935 |
| SUCLG1 | 92.0533932 | 0.728393296 | 0.27469917 | 2.6516036 | 0.00801105 | 0.0658687 |
| DNAH6 | 36.1964534 | -0.049029235 | 0.4439438 | -0.1104402 | 0.91206029 | 0.96419139 |
| TMSB10 | 221.341937 | -1.929577677 | 0.4715886 | -4.0916546 | 4.28E-05 | 0.00124293 |
| KCMF1 | 130.911593 | 0.007588505 | 0.24067656 | 0.03152989 | 0.97484696 | 0.98983573 |
| TCF7L1 | 57.1907491 | -0.426718385 | 0.677579 | -0.6297692 | 0.5288456 | 0.76326245 |
| TGOLN2 | 166.384003 | -0.144887842 | 0.3412325 | -0.4246015 | 0.67112718 | 0.84747013 |
| RETSAT | 16.9519465 | 0.226848357 | 0.39088538 | 0.58034496 | 0.56168202 | 0.78364682 |
| ELMOD3 | 49.2234664 | 0.239156826 | 0.24334678 | 0.98278197 | 0.32571476 | 0.6159877 |
| CAPG | 40.0943692 | -0.12468814 | 0.43126386 | -0.2891226 | 0.77248754 | 0.90119866 |
| MAT2A | 70.5577213 | 0.168103226 | 0.29217696 | 0.5753473 | 0.56505644 | 0.78527066 |
| GGCX | 25.4108431 | -0.144987804 | 0.29824946 | -0.4861293 | 0.62687548 | 0.82354566 |
| VAMP8 | 65.9888586 | -0.789117965 | 0.37570985 | -2.1003388 | 0.03569905 | 0.17539507 |
| RNF181 | 11.3402082 | -0.765963646 | 0.48571014 | -1.5769974 | 0.11479612 | 0.35365717 |
| USP39 | 52.0091923 | 0.002801472 | 0.23380663 | 0.011982 | 0.99043997 | 0.99684874 |
| C2orf68 | 25.0025795 | 0.154361121 | 0.3447962 | 0.447688 | 0.65437838 | 0.83900511 |
| POLR1A | 36.712359 | -0.102913955 | 0.27847488 | -0.3695628 | 0.71170828 | 0.87074117 |
| PTCD3 | 83.7791071 | 0.233393414 | 0.20553196 | 1.13555777 | 0.25614169 | 0.54710413 |
| IMMT | 117.251542 | 0.548944683 | 0.22358563 | 2.45518768 | 0.01408111 | 0.09623211 |
| MRPL35 | 18.9547322 | 0.120576807 | 0.41243033 | 0.29235679 | 0.77001384 | 0.900418 |
| KDM3A | 83.0738349 | -0.709377635 | 0.30198309 | -2.3490641 | 0.01882067 | 0.11636543 |
| CHMP3 | 32.831445 | -0.912855933 | 0.47812195 | -1.9092534 | 0.05622942 | 0.23424727 |
| RMND5A | 55.922439 | 0.112912188 | 0.29834779 | 0.37845827 | 0.70509019 | 0.86679461 |
| PLGLB1 | 14.9108799 | 0.543632963 | 0.56136391 | 0.96841452 | 0.33283739 | 0.6218518 |
| CYTOR | 56.0351348 | -0.528372782 | 0.57570828 | -0.9177787 | 0.35873475 | 0.64254425 |
| KRCC1 | 60.489218 | -0.390583387 | 0.25476427 | -1.5331168 | 0.12524707 | 0.37039755 |
| THNSL2 | 16.8352264 | 0.806572772 | 0.92671023 | 0.87036135 | 0.38410296 | 0.66701235 |
| EIF2AK3 | 108.91452 | 0.412830361 | 0.52221864 | 0.79053165 | 0.42921734 | 0.70310352 |
| RPIA | 28.6696487 | 0.331394212 | 0.3026829 | 1.09485609 | 0.27357969 | 0.56583376 |
| MAL | 376.798015 | 1.692223846 | 0.51018821 | 3.31686192 | 0.00091035 | 0.01369107 |
| MRPS5 | 96.4372518 | 0.092415735 | 0.21325707 | 0.43335367 | 0.66475785 | 0.84348349 |
| ZNF514 | 10.4570182 | 0.074305429 | 0.52156105 | 0.14246736 | 0.88671086 | 0.95619452 |
| AC092835.1 | 16.2943087 | -0.141399271 | 0.44744803 | -0.3160127 | 0.75199285 | 0.89148796 |
| PROM2 | 60.0488684 | 0.800125254 | 0.50964549 | 1.56996436 | 0.1164234 | 0.35579765 |
| KCNIP3 | 13.296345 | 0.164634286 | 0.4781585 | 0.34430902 | 0.7306139 | 0.88166621 |
| FAHD2A | 15.9745253 | 0.62126797 | 0.39845389 | 1.55919664 | 0.11894985 | 0.36084255 |
| ANKRD36C | 37.1566667 | -0.262754615 | 0.45853957 | -0.573025 | 0.5666278 | 0.78647713 |
| STARD7 | 67.4406983 | 0.359986315 | 0.24488276 | 1.47003535 | 0.14155218 | 0.39756559 |
| TMEM127 | 43.1801813 | -0.363076976 | 0.29137487 | -1.246082 | 0.2127343 | 0.49888203 |
| CIAO1 | 35.908706 | -0.29654515 | 0.25666819 | -1.1553638 | 0.24794146 | 0.538841 |
| SNRNP200 | 66.219293 | -0.413155864 | 0.27007712 | -1.52977 | 0.12607368 | 0.37183253 |
| NEURL3 | 17.2024334 | 0.211820935 | 0.68579579 | 0.30886882 | 0.75742132 | 0.89457492 |
| KANSL3 | 105.644885 | 0.627701996 | 0.22073762 | 2.84365659 | 0.00445991 | 0.04367489 |
| FER1L5 | 18.8243733 | -0.221716671 | 0.31238437 | -0.709756 | 0.47785547 | 0.73761868 |
| LMAN2L | 33.5283864 | 0.228569113 | 0.27354228 | 0.83558972 | 0.40338575 | 0.68159423 |
| CNNM4 | 24.9202371 | 0.15354257 | 0.32782214 | 0.46837157 | 0.63951889 | 0.83121032 |
| ANKRD23 | 18.5427434 | -0.249852178 | 0.41193133 | -0.6065384 | 0.54415728 | 0.77376236 |
| ANKRD36 | 57.1664912 | -0.217307126 | 0.34076244 | -0.6377086 | 0.52366341 | 0.76218158 |
| AC092683.1 | 55.9458706 | -0.741277159 | 0.4451171 | -1.6653531 | 0.09584232 | 0.31922651 |
| ANKRD36B | 16.4937536 | -0.441969419 | 0.64602978 | -0.6841317 | 0.49389204 | 0.74700665 |
| COX5B | 109.348112 | -0.318130551 | 0.44316754 | -0.7178562 | 0.47284598 | 0.73375471 |
| ACTR1B | 27.5362291 | -0.405483751 | 0.40044241 | -1.0125894 | 0.31125631 | 0.60432766 |
| TMEM131 | 265.556823 | 0.157749976 | 0.20310716 | 0.77668349 | 0.43734553 | 0.70852865 |
| INPP4A | 52.2105891 | -0.165961696 | 0.29587336 | -0.5609214 | 0.57485113 | 0.79134063 |
| COA5 | 27.2611976 | 0.098679863 | 0.31723492 | 0.31106243 | 0.75575317 | 0.89366123 |
| UNC50 | 30.0038069 | -0.160442748 | 0.27187978 | -0.5901239 | 0.55510761 | 0.77954179 |
| MGAT4A | 118.724115 | 0.513013835 | 0.51641978 | 0.9934047 | 0.32051277 | 0.6120836 |
| TSGA10 | 132.885619 | 0.094985404 | 0.3361816 | 0.28254195 | 0.77752799 | 0.90366576 |
| AC079447.1 | 56.3130487 | 0.501242567 | 0.26447301 | 1.89525034 | 0.05805924 | 0.23811563 |
| MITD1 | 39.2703272 | -0.274123367 | 0.3066686 | -0.8938749 | 0.37138882 | 0.65494451 |
| TXNDC9 | 26.9050482 | -0.385782411 | 0.28411919 | -1.3578189 | 0.17452116 | 0.44724303 |
| EIF5B | 115.027715 | -0.534150941 | 0.2504441 | -2.1328151 | 0.0329399 | 0.16741334 |
| REV1 | 189.4834 | 0.634341473 | 0.20997205 | 3.02107582 | 0.00251878 | 0.02883167 |
| LONRF2 | 33.2093079 | -0.22134125 | 0.45162629 | -0.4900982 | 0.62406438 | 0.82224214 |

| | | | | | | |
|---|---|---|---|---|---|---|
| NPAS2 | 282.213915 | -0.766863053 | 0.29642104 | -2.5870736 | 0.00967949 | 0.07548684 |
| RPL31 | 74.0388561 | 0.068814753 | 0.27797947 | 0.24755337 | 0.80447999 | 0.91643337 |
| TBC1D8 | 250.752093 | -0.264119785 | 0.27249397 | -0.9692684 | 0.33241131 | 0.62173226 |
| CNOT11 | 54.7936388 | -0.64400984 | 0.32162548 | -2.0023595 | 0.04524608 | 0.20345923 |
| RNF149 | 254.505173 | -0.460574685 | 0.32369983 | -1.422845 | 0.15478109 | 0.41807041 |
| MAP4K4 | 121.832954 | -1.329737122 | 0.37143322 | -3.5800167 | 0.00034357 | 0.0065064 |
| IL1R1 | 86.5878648 | -0.273231248 | 0.43094215 | -0.6340323 | 0.52605974 | 0.76221787 |
| IL1RL2 | 12.8591235 | 0.029682968 | 0.75215836 | 0.03946372 | 0.96852068 | 0.98759774 |
| SLC9A4 | 90.3677744 | 3.423583256 | 0.59410396 | 5.76259963 | 8.28E-09 | 1.05E-06 |
| SLC9A2 | 85.6632318 | 1.443815809 | 0.41555412 | 3.47443508 | 0.00051193 | 0.00876064 |
| MFSD9 | 12.7197004 | -0.5272143 | 0.41926294 | -1.2574789 | 0.20858027 | 0.49402397 |
| TMEM182 | 31.3107046 | -0.906486232 | 0.35906192 | -2.5245959 | 0.01158314 | 0.08487163 |
| POU3F3 | 126.814961 | -0.43156595 | 0.31434921 | -1.372887 | 0.16978747 | 0.44037022 |
| MRPS9 | 67.5643101 | 0.726748699 | 0.22236821 | 3.26822214 | 0.00108225 | 0.01544593 |
| TGFBRAP1 | 24.9301689 | 0.066508434 | 0.3087948 | 0.21538068 | 0.82947053 | 0.92907256 |
| C2orf49 | 22.6122624 | -0.114040744 | 0.38594195 | -0.2954868 | 0.76762205 | 0.89979146 |
| NCK2 | 95.3834241 | -0.128725605 | 0.22096771 | -0.5825539 | 0.56019364 | 0.78316119 |
| UXS1 | 94.4288995 | 0.755868685 | 0.2304157 | 3.28045659 | 0.00103639 | 0.0151231 |
| LINC01594 | 29.2021002 | 0.408611539 | 0.61002461 | 0.66982795 | 0.50296747 | 0.75108109 |
| SULT1C2 | 310.105231 | 0.65075078 | 0.30977032 | 2.1007525 | 0.0356627 | 0.17536855 |
| SULT1C4 | 10.9058872 | -0.29354556 | 0.684501 | -0.4288461 | 0.66803525 | 0.84558183 |
| GCC2 | 255.701112 | -0.447106025 | 0.19437991 | -2.3001658 | 0.02143883 | 0.12612104 |
| GCC2-AS1 | 29.8888227 | -0.368623653 | 0.50137048 | -0.7352321 | 0.46219818 | 0.72706759 |
| LIMS1 | 290.374751 | -0.556898009 | 0.33938878 | -1.6408851 | 0.10082126 | 0.32827453 |
| RANBP2 | 222.563732 | 1.608099022 | 0.65815868 | 2.44333028 | 0.01455241 | 0.09806274 |
| CCDC138 | 46.7423985 | 0.008396829 | 0.37826131 | 0.02219849 | 0.98228962 | 0.99337859 |
| 10-Sep | 206.15182 | -1.308165523 | 0.55282468 | -2.3663298 | 0.01796543 | 0.11278482 |
| SOWAHC | 29.7076456 | -2.210308461 | 0.74032331 | -2.9855989 | 0.00283024 | 0.03141928 |
| NPHP1 | 47.9683976 | 0.175517817 | 0.40514492 | 0.43322231 | 0.66485327 | 0.84348349 |
| SMIM37 | 33.6073591 | -0.567677932 | 0.43371251 | -1.3088807 | 0.19057477 | 0.47036446 |
| LINC01106 | 11.6720704 | 0.863940107 | 0.54312905 | 1.59067187 | 0.11168344 | 0.34846418 |
| ACOXL | 14.3229332 | -1.988712265 | 0.77669102 | -2.5604934 | 0.01045236 | 0.07916175 |
| MIR4435-2HG | 316.006461 | -0.79526522 | 0.33950758 | -2.3424078 | 0.01915976 | 0.11762443 |
| BCL2L11 | 29.8145085 | -0.605157338 | 0.33392109 | -1.8122765 | 0.06994349 | 0.26667573 |
| AC017002.5 | 23.0851891 | 0.571549068 | 0.87965857 | 0.64973967 | 0.51586039 | 0.75735981 |
| ANAPC1 | 88.7456367 | 0.04857453 | 0.24703654 | 0.19662893 | 0.84411793 | 0.93684082 |
| MERTK | 45.2177231 | 0.230456878 | 0.38401749 | 0.60012079 | 0.54842574 | 0.77527341 |
| TMEM87B | 79.2641761 | -0.709834679 | 0.22720455 | -3.1242098 | 0.00178283 | 0.02253588 |
| ZC3H8 | 55.1991465 | 0.534800177 | 0.24494339 | 2.18336236 | 0.02900914 | 0.15396812 |
| ZC3H6 | 86.1863026 | 0.439386512 | 0.43277981 | 1.01526573 | 0.30997918 | 0.60321222 |
| TTL | 33.6884948 | -0.02385641 | 0.30430587 | -0.0783962 | 0.93751293 | 0.97619962 |
| POLR1B | 18.6987246 | 0.181160645 | 0.33585435 | 0.53940241 | 0.58960922 | 0.79960282 |
| CHCHD5 | 13.3654876 | -0.049871316 | 0.52368062 | -0.0952323 | 0.92413031 | 0.96988492 |
| SLC20A1 | 99.7023409 | 1.444334242 | 0.60638522 | 2.38187572 | 0.01722471 | 0.10980752 |
| PSD4 | 33.9861325 | -0.556342841 | 0.42623029 | -1.3052635 | 0.19180314 | 0.47205703 |
| PAX8 | 1363.01732 | -0.276908485 | 0.31041318 | -0.8920642 | 0.37235853 | 0.65606936 |
| CBWD2 | 32.8524381 | 0.319193899 | 0.32171776 | 0.99215504 | 0.3211219 | 0.6120836 |
| RABL2A | 24.9243782 | 0.112960251 | 0.33827439 | 0.33393084 | 0.73843175 | 0.88591171 |
| AC017074.1 | 23.3407015 | -0.74386887 | 0.47353621 | -1.5708806 | 0.11621038 | 0.35543425 |
| SLC35F5 | 338.467823 | -0.583628986 | 0.19418061 | -3.0055987 | 0.00265059 | 0.02994115 |
| ACTR3 | 202.040851 | -0.230404689 | 0.23530541 | -0.9791729 | 0.32749455 | 0.61719253 |
| DDX18 | 45.2158815 | -0.78190228 | 0.24406153 | -3.2037097 | 0.00135669 | 0.01835848 |
| CCDC93 | 103.386952 | -0.194172954 | 0.25667284 | -0.7564998 | 0.44934958 | 0.7180393 |
| INSIG2 | 79.9753853 | -0.876624888 | 0.55899228 | -1.5682236 | 0.11682896 | 0.3567196 |
| C2orf76 | 20.7419868 | 0.065798785 | 0.32611457 | 0.20176586 | 0.84009978 | 0.93552258 |
| DBI | 116.877356 | -1.448176977 | 0.42654411 | -3.39514 | 0.00068594 | 0.01110064 |
| TMEM37 | 39.3844968 | -1.15978935 | 0.56668109 | -2.046635 | 0.04069394 | 0.19186679 |
| SCTR | 25.8846609 | -2.641497182 | 0.94935224 | -2.7824205 | 0.00539551 | 0.05024962 |
| CFAP221 | 175.84298 | -0.270826902 | 0.42085007 | -0.6435235 | 0.51988448 | 0.76055807 |
| PTPN4 | 198.650888 | 0.975060621 | 0.18777776 | 5.19263089 | 2.07E-07 | 1.61E-05 |
| EPB41L5 | 325.106842 | 0.295599093 | 0.18603071 | 1.58898006 | 0.11206489 | 0.34889469 |
| RALB | 28.9483522 | -0.20299994 | 0.30709135 | -0.6610409 | 0.50858609 | 0.7543124 |
| TFCP2L1 | 618.51039 | 1.933059078 | 0.2950339 | 6.5519897 | 5.68E-11 | 1.50E-08 |
| CLASP1 | 503.606172 | 0.576041417 | 0.26386132 | 2.18312188 | 0.02902684 | 0.15398273 |
| NIFK-AS1 | 24.5377057 | -0.078857786 | 0.46114858 | -0.171003 | 0.86422142 | 0.94663434 |
| NIFK | 12.8664396 | -0.73995908 | 0.50846542 | -1.4552791 | 0.14559202 | 0.40395611 |
| TSN | 29.1685806 | -0.736878415 | 0.28626639 | -2.5741003 | 0.01005011 | 0.0774972 |
| BIN1 | 17.7650546 | -0.534171423 | 0.46018553 | -1.1607741 | 0.24573379 | 0.53627476 |
| ERCC3 | 35.6470793 | 0.155388769 | 0.26766495 | 0.58053461 | 0.56155415 | 0.78361672 |
| MAP3K2 | 206.467904 | 0.329267772 | 0.1990745 | 1.65399275 | 0.09812899 | 0.32365239 |
| IWS1 | 144.297526 | -0.202076139 | 0.25444082 | -0.794197 | 0.42708075 | 0.70162975 |
| AC010976.1 | 15.5941322 | -1.234045266 | 0.76066324 | -1.622328 | 0.10473314 | 0.33560212 |
| WDR33 | 166.76282 | 0.279441705 | 0.20812712 | 1.34264914 | 0.17938561 | 0.45495583 |
| POLR2D | 14.5936637 | -0.019732406 | 0.34960981 | -0.0564412 | 0.95499032 | 0.98245559 |
| AMMECR1L | 20.448653 | 0.041301672 | 0.32784446 | 0.12597947 | 0.89974817 | 0.95974381 |
| SAP130 | 54.8128938 | 0.190351616 | 0.27871708 | 0.68295641 | 0.49463439 | 0.74750235 |
| UGGT1 | 58.7132676 | 0.235480209 | 0.30010673 | 0.78465489 | 0.43265594 | 0.70503345 |
| HS6ST1 | 226.182644 | -0.763024943 | 0.44273542 | -1.7234332 | 0.08481021 | 0.29907158 |
| SMPD4 | 39.0863832 | 0.063133893 | 0.30854167 | 0.20462031 | 0.83786878 | 0.93487753 |
| MZT2B | 50.4149554 | -0.534388227 | 0.37763741 | -1.4150829 | 0.1570442 | 0.42219972 |
| CCDC115 | 17.5231185 | -0.658001527 | 0.49615206 | -1.3262094 | 0.18477035 | 0.46134371 |
| IMP4 | 13.0173656 | -0.46973455 | 0.44897556 | -1.0462363 | 0.29545193 | 0.58702908 |
| PTPN18 | 37.4057405 | -0.218992126 | 0.31133892 | -0.7033882 | 0.48181385 | 0.74068045 |
| FAM168B | 107.124666 | 0.314348082 | 0.27075272 | 1.16101539 | 0.24563564 | 0.53627476 |
| PLEKHB2 | 127.944673 | -0.167879005 | 0.22811356 | -0.7359449 | 0.46176425 | 0.72676972 |
| MZT2A | 34.4447664 | -0.85042675 | 0.35826378 | -2.3737447 | 0.01760872 | 0.1112232 |
| ANKRD30BL | 9.9390733 | -0.270947191 | 0.50093939 | -0.5408782 | 0.58859155 | 0.79900779 |
| GPR39 | 119.141662 | 0.468030229 | 0.30099783 | 1.55492894 | 0.11996299 | 0.36263462 |
| NCKAP5 | 80.1230948 | 1.902370226 | 0.43334371 | 4.38997997 | 1.13E-05 | 0.0004055 |
| MGAT5 | 101.04974 | -0.016148356 | 0.23844018 | -0.067725 | 0.94600457 | 0.97897931 |
| CCNT2-AS1 | 32.5286655 | -0.907064272 | 0.66092557 | -1.3724152 | 0.16993423 | 0.44052905 |
| ACMSD | 15.8789514 | -0.883516225 | 0.8057735 | -1.0964821 | 0.27286785 | 0.56514644 |
| CCNT2 | 117.751174 | -0.047867703 | 0.25704087 | -0.186226 | 0.85226751 | 0.94037032 |
| RAB3GAP1 | 193.655048 | 0.096489457 | 0.17741461 | 0.54386421 | 0.58653492 | 0.79752689 |
| ZRANB3 | 98.2028317 | 0.06199713 | 0.24984047 | 0.24814686 | 0.80402078 | 0.91643337 |
| R3HDM1 | 172.987093 | 0.315764035 | 0.17379876 | 1.81683709 | 0.06924206 | 0.26510127 |
| UBXN4 | 166.01968 | -0.667706562 | 0.210186 | -3.1767414 | 0.0014894 | 0.01967151 |
| DARS | 143.239204 | -0.607624353 | 0.23559838 | -2.5790685 | 0.00990671 | 0.07673607 |
| THSD7B | 31.4981711 | 0.234878363 | 0.80105094 | 0.29321277 | 0.76935953 | 0.90034488 |
| HNMT | 64.0615373 | 0.030084271 | 0.36974706 | 0.08136446 | 0.93515211 | 0.97519355 |
| AC114763.1 | 26.4377217 | 0.00016032 | 0.46479935 | 0.00034492 | 0.99972479 | 0.99975018 |
| SPOPL | 72.8002776 | 0.083944868 | 0.21594579 | 0.38873121 | 0.69747499 | 0.86275485 |
| LRP1B | 1009.09209 | -1.195744123 | 0.49138625 | -2.4334098 | 0.01495736 | 0.09993327 |
| GTDC1 | 97.8487232 | 0.197062277 | 0.23552234 | 0.83670313 | 0.40275946 | 0.68128594 |
| TEX41 | 1104.98071 | 2.624542722 | 0.3110965 | 8.43642651 | 3.27E-17 | 3.36E-14 |
| ACVR2A | 93.2040628 | 0.907304198 | 0.33420071 | 2.71484823 | 0.00663061 | 0.05803619 |
| ORC4 | 144.626304 | 0.840556557 | 0.3002492 | 2.79952968 | 0.00511771 | 0.04841383 |
| MBD5 | 767.661352 | 0.188458664 | 0.13695897 | 1.37602277 | 0.16881459 | 0.43928463 |
| EPC2 | 173.072655 | 0.525331628 | 0.27209835 | 1.93066819 | 0.0535241 | 0.22710266 |
| AC105402.3 | 15.0551055 | 0.214952983 | 1.65753318 | 0.12968246 | 0.89681766 | 0.95900099 |
| LYPD6B | 58.6316751 | -0.430861271 | 0.57641045 | -0.7474904 | 0.4547676 | 0.7219935 |
| MMADHC | 24.4253757 | -0.209901112 | 0.32042141 | -0.6550783 | 0.51241734 | 0.75562371 |
| NMI | 21.184089 | -1.509061734 | 0.75609742 | -1.9958562 | 0.04594958 | 0.20581414 |

| | | | | | | |
|---|---|---|---|---|---|---|
| RIF1 | 127.452981 | 0.078705997 | 0.29840217 | 0.26375813 | 0.7919663 | 0.9111703 |
| NEB | 11.5761134 | 0.572909456 | 0.41825633 | 1.36975681 | 0.17076283 | 0.44178771 |
| ARL5A | 37.8845529 | 0.335743595 | 0.33168947 | 1.01222265 | 0.31143161 | 0.60444017 |
| CACNB4 | 310.214779 | 1.967703218 | 0.21943519 | 8.96712701 | 3.04E-19 | 4.48E-16 |
| STAM2 | 116.270596 | 0.505287862 | 0.24643418 | 2.05039685 | 0.04032572 | 0.19065423 |
| FMNL2 | 388.46487 | -1.47827352 | 0.29858993 | -4.9508486 | 7.39E-07 | 4.43E-05 |
| PRPF40A | 115.098546 | 0.317842791 | 0.1755091 | 1.81097611 | 0.07014455 | 0.26694834 |
| ARL6IP6 | 47.8295118 | -0.312133066 | 0.26129926 | -1.1945425 | 0.23226583 | 0.52210399 |
| KCNJ3 | 317.405947 | 1.667357596 | 0.3388037 | 4.92130877 | 8.60E-07 | 4.95E-05 |
| LINC01876 | 94.3419194 | -0.203066824 | 1.08863892 | -0.1865328 | 0.85202699 | 0.94037032 |
| GPD2 | 106.515872 | -0.073071402 | 0.25157157 | -0.2904597 | 0.77146458 | 0.9008444 |
| ACVR1 | 110.215396 | 0.048556338 | 0.30221843 | 0.16066637 | 0.87235618 | 0.94869771 |
| UPP2 | 36.360216 | 0.242793469 | 0.69990183 | 0.34689646 | 0.7286691 | 0.88042244 |
| CCDC148 | 170.434364 | 0.584760108 | 0.472786 | 1.23683888 | 0.2161469 | 0.50287837 |
| PKP4 | 632.784761 | -0.019564625 | 0.2803854 | -0.0697776 | 0.94437066 | 0.97856665 |
| PKP4-AS1 | 17.9913143 | -1.003184749 | 0.74649894 | -1.3438529 | 0.17899597 | 0.45449239 |
| TANC1 | 304.194833 | 0.219681591 | 0.18064947 | 1.21606553 | 0.22395996 | 0.51317516 |
| WDSUB1 | 88.7126946 | 0.432246516 | 0.27965748 | 1.5456283 | 0.12219436 | 0.3664089 |
| BAZ2B | 584.91728 | 0.289713839 | 0.1437059 | 2.01601909 | 0.04379798 | 0.19976647 |
| AC009961.1 | 38.6045349 | 0.169124867 | 0.45124743 | 0.37479408 | 0.70781362 | 0.86882712 |
| 7-Mar | 199.766347 | 0.296849818 | 0.26124946 | 1.1362696 | 0.25584375 | 0.54693968 |
| PLA2R1 | 34.6477371 | -0.781349252 | 0.28425707 | -2.7487417 | 0.00598245 | 0.05415749 |
| ITGB6 | 462.059327 | -1.424314075 | 0.65359865 | -2.1791876 | 0.02931774 | 0.15512651 |
| RBMS1 | 485.52909 | -0.033012179 | 0.32700586 | -0.1009529 | 0.91958788 | 0.96763053 |
| TANK | 159.497534 | -0.421558935 | 0.17675093 | -2.3850451 | 0.01707703 | 0.10940767 |
| PSMD14 | 68.7959442 | 0.144651979 | 0.32530963 | 0.44465938 | 0.65656593 | 0.83951126 |
| DPP4 | 16.1913273 | -3.18486812 | 0.60557482 | -5.259248 | 1.45E-07 | 1.17E-05 |
| IFIH1 | 35.1779711 | -0.736920744 | 0.86271766 | -0.8541853 | 0.39300232 | 0.67377433 |
| GCA | 40.7660516 | -0.956596381 | 0.39887907 | -2.3982115 | 0.01647535 | 0.10641319 |
| KCNH7 | 63.7422168 | 1.879779784 | 0.75743762 | 2.48176183 | 0.01307346 | 0.0922485 |
| FIGN | 19.7642315 | -0.508272063 | 0.68307958 | -0.7440891 | 0.45682258 | 0.72302753 |
| GRB14 | 214.70462 | 1.247202499 | 0.35637973 | 3.49964488 | 0.00046588 | 0.00817628 |
| COBLL1 | 3251.57532 | 1.267293107 | 0.28238617 | 4.48780155 | 7.20E-06 | 0.00028846 |
| AC019197.1 | 8368.61065 | 0.587006087 | 0.27714345 | 2.11805868 | 0.0341701 | 0.17088366 |
| SLC38A11 | 165.988872 | 1.29276772 | 0.5074363 | 2.54764533 | 0.01084527 | 0.08095937 |
| SCN3A | 296.61585 | -0.949389093 | 0.7354006 | -1.2909822 | 0.19670984 | 0.47828766 |
| SCN2A | 1187.37198 | 0.58239362 | 0.30985647 | 1.87955936 | 0.06016816 | 0.24327015 |
| CSRNP3 | 98.3516113 | 1.033123107 | 0.45310151 | 2.28011402 | 0.02260093 | 0.13102687 |
| GALNT3 | 53.0693665 | -1.07943109 | 0.3351304 | -3.2209286 | 0.00127776 | 0.01757474 |
| TTC21B | 60.1875865 | 0.657203532 | 0.30829525 | 2.13173424 | 0.0330287 | 0.16778189 |
| SCN7A | 36.1948615 | 2.343140858 | 0.52614581 | 4.45340592 | 8.45E-06 | 0.00031894 |
| STK39 | 401.145007 | -0.693135784 | 0.27114102 | -2.5563663 | 0.01057717 | 0.07982857 |
| CERS6 | 37.2734364 | -0.599434467 | 0.53475077 | -1.1209604 | 0.2623047 | 0.55396946 |
| LRP2 | 42.2537188 | 0.690573761 | 0.5905079 | 1.16945728 | 0.24221944 | 0.53285174 |
| FASTKD1 | 96.0708973 | 1.00680429 | 0.25627764 | 3.92856858 | 8.55E-05 | 0.00215769 |
| PPIG | 134.076798 | 0.384485956 | 0.22977857 | 1.67328902 | 0.09427041 | 0.31610311 |
| KLHL23 | 33.4717142 | -0.046807836 | 0.3474718 | -0.1347097 | 0.89284137 | 0.95782021 |
| SSB | 41.3832827 | -0.494402401 | 0.41133393 | -1.201949 | 0.2293833 | 0.51970678 |
| METTL5 | 23.9079186 | -0.03171069 | 0.37078091 | -0.0855241 | 0.93184477 | 0.97351838 |
| UBR3 | 281.252687 | 0.638334191 | 0.1927078 | 3.31244606 | 0.00092484 | 0.0138484 |
| MYO3B | 237.883714 | 0.849127466 | 0.46444153 | 1.82827636 | 0.06750809 | 0.26098543 |
| GORASP2 | 55.1613256 | -0.260424369 | 0.2792223 | -0.9326776 | 0.35098647 | 0.63760583 |
| TLK1 | 362.997987 | 0.533842061 | 0.2229162 | 2.39481047 | 0.01662896 | 0.10727084 |
| METTL8 | 51.1746216 | 0.254748748 | 0.27549349 | 0.92469971 | 0.35512213 | 0.64025612 |
| DCAF17 | 32.5538393 | 0.10026764 | 0.32968742 | 0.30412941 | 0.7610293 | 0.89648567 |
| CYBRD1 | 29.9988528 | 0.353398074 | 0.49304329 | 0.71676885 | 0.47351673 | 0.73418512 |
| DYNC1I2 | 132.473079 | -0.04660766 | 0.19541385 | -0.2385074 | 0.81148754 | 0.9199895 |
| SLC25A12 | 195.898409 | 0.589111451 | 0.2738286 | 2.1513876 | 0.03144562 | 0.16230099 |
| HAT1 | 50.7533934 | 0.272196707 | 0.34329889 | 0.79288549 | 0.42784454 | 0.70230313 |
| METAP1D | 24.7987198 | 0.271607311 | 0.35223797 | 0.77109038 | 0.44065336 | 0.71135625 |
| ITGA6 | 164.204904 | 0.05750747 | 0.24770387 | 0.23216218 | 0.81641205 | 0.92222335 |
| AC078883.1 | 13.8190844 | -2.544424832 | 0.46731119 | -5.444819 | 5.19E-08 | 4.73E-06 |
| PDK1 | 60.5305126 | -2.264796567 | 0.44893428 | -5.0448288 | 4.54E-07 | 2.96E-05 |
| MAP3K20 | 25.5676455 | -1.337870559 | 0.53673143 | -2.4926257 | 0.01268024 | 0.09046528 |
| SP3 | 96.6682673 | -0.176358615 | 0.24226033 | -0.7279715 | 0.46663104 | 0.72991695 |
| OLA1 | 193.154899 | 0.332929635 | 0.21008731 | 1.58472031 | 0.11302989 | 0.35086028 |
| CIR1 | 59.0850715 | -0.58519618 | 0.27634796 | -2.1176063 | 0.03420843 | 0.17099234 |
| SCRN3 | 23.9904774 | 0.274247564 | 0.44065946 | 0.62235714 | 0.53370706 | 0.76708723 |
| GPR155 | 25.0274003 | -0.133728154 | 0.64146749 | -0.2084722 | 0.83486027 | 0.93320871 |
| AC010894.3 | 89.2341685 | -1.293833727 | 0.6076906 | -2.1290994 | 0.03324603 | 0.16847056 |
| WIPF1 | 27.0210194 | -1.262030248 | 0.55837133 | -2.2601989 | 0.02380891 | 0.13541585 |
| ATF2 | 104.330223 | 0.151720215 | 0.2050208 | 0.74002353 | 0.45928571 | 0.72478689 |
| ATP5MC3 | 128.342098 | -0.325637762 | 0.48865708 | -0.6663932 | 0.5051598 | 0.75184725 |
| LNPK | 52.9356567 | -0.414556215 | 0.3192582 | -1.2984982 | 0.19411618 | 0.47456322 |
| HOXD10 | 108.899741 | 0.716064458 | 0.35318969 | 2.02742174 | 0.0426193 | 0.19708211 |
| HOXD-AS2 | 26.7365921 | 1.303174179 | 0.42093885 | 3.0958753 | 0.00196233 | 0.02423597 |
| HOXD9 | 42.0947668 | 1.162901238 | 0.29740603 | 3.91014682 | 9.22E-05 | 0.00227334 |
| HOXD8 | 68.0273215 | 0.940260887 | 0.44002787 | 2.13682123 | 0.03261253 | 0.16624163 |
| HOXD3 | 126.835057 | 0.674135387 | 0.25839685 | 2.60891491 | 0.00908298 | 0.07175835 |
| HAGLR | 30.6745557 | -0.762561635 | 0.32555888 | -2.3423156 | 0.0191645 | 0.11762443 |
| MTX2 | 273.569673 | 0.672380375 | 0.22820683 | 2.946364 | 0.00321534 | 0.03421536 |
| LINC01116 | 37.2091745 | 0.757506937 | 0.44078729 | 1.71853173 | 0.08569967 | 0.30113433 |
| LINC01117 | 22.6923641 | -0.111098107 | 0.4624699 | -0.2402278 | 0.8101537 | 0.91928664 |
| AC079305.3 | 14.8705038 | 0.757857682 | 0.75679369 | 1.00140592 | 0.3166306 | 0.61004834 |
| HNRNPA3 | 186.120244 | 0.191636454 | 0.26263372 | 0.72967192 | 0.46559075 | 0.72890726 |
| NFE2L2 | 1618.84316 | 2.812753208 | 0.28331616 | 9.92796608 | 3.15E-23 | 1.08E-19 |
| AGPS | 79.0346271 | 0.321020747 | 0.248413 | 1.29228642 | 0.19625797 | 0.47775275 |
| RBM45 | 28.6756866 | 0.488824955 | 0.29201445 | 1.67397524 | 0.09413547 | 0.3160115 |
| AC009948.1 | 14.5320298 | -0.367268285 | 0.69159852 | -0.5310426 | 0.59538925 | 0.80357658 |
| PRKRA | 58.5137898 | -0.174374382 | 0.31399034 | -0.5553495 | 0.57865562 | 0.7935467 |
| PJVK | 9.944113 | -0.625169971 | 0.56888438 | -1.0989403 | 0.27179411 | 0.5642932 |
| PLEKHA3 | 39.2306989 | -0.136489112 | 0.29776879 | -0.4583728 | 0.64668464 | 0.83464609 |
| TTN-AS1 | 171.039488 | 0.456950112 | 0.26680137 | 1.7126978 | 0.08676815 | 0.30289881 |
| TTN | 13.7445698 | -1.530721579 | 0.91935961 | -1.6649868 | 0.0959154 | 0.31926347 |
| SESTD1 | 219.094878 | -1.566015222 | 0.39321892 | -3.9825531 | 6.82E-05 | 0.00180128 |
| ZNF385B | 48.8309575 | 0.552085221 | 0.43094442 | 1.28110539 | 0.20015665 | 0.48370812 |
| CWC22 | 50.0286254 | 0.485631215 | 0.25874403 | 1.87687893 | 0.06053469 | 0.24360483 |
| UBE2E3 | 171.900661 | 0.470703128 | 0.1578232 | 2.98247111 | 0.00285932 | 0.031572 |
| CERKL | 18.2157129 | -0.435688419 | 0.47108926 | -0.9248532 | 0.35504226 | 0.64025612 |
| SSFA2 | 40.8924029 | -0.998646285 | 0.2905752 | -3.4367912 | 0.00058865 | 0.00979687 |
| PDE1A | 462.292012 | 0.259377829 | 0.43604697 | 0.59483919 | 0.55195097 | 0.77741728 |
| DNAJC10 | 71.3216758 | -0.554917428 | 0.26321269 | -2.1082472 | 0.03500961 | 0.1740015 |
| NCKAP1 | 272.952518 | 0.037348957 | 0.26148638 | 0.14283328 | 0.88642186 | 0.95619452 |
| NUP35 | 23.8859393 | 0.143310702 | 0.30463845 | 0.4704288 | 0.63804869 | 0.83057589 |
| FSIP2 | 15.6165421 | 0.03739108 | 0.77211902 | 0.04842657 | 0.96137628 | 0.98499239 |
| FSIP2-AS1 | 12.3414185 | 0.799584698 | 0.50784787 | 1.57445713 | 0.1153818 | 0.35464129 |
| LINC01473 | 34.8574719 | 1.043190715 | 0.47946955 | 2.17571841 | 0.02957632 | 0.15609389 |
| ZC3H15 | 67.8725354 | 0.366530627 | 0.25596005 | 1.43198372 | 0.15214848 | 0.4136994 |
| ITGAV | 332.014033 | 0.545451794 | 0.27720204 | 1.96770486 | 0.04910201 | 0.2150718 |
| AC017101.1 | 31.2384591 | 0.052689523 | 0.74817137 | 0.0704244 | 0.94385587 | 0.97842656 |
| AC007319.1 | 56.2072934 | -0.360917282 | 0.60976838 | -0.5918924 | 0.55392263 | 0.77872693 |
| TFPI | 32.1450133 | -2.298426905 | 0.70254219 | -3.2715856 | 0.00106946 | 0.01538771 |
| LINC01090 | 16.0382789 | -0.637960412 | 0.69764282 | -0.9144513 | 0.36047973 | 0.64473302 |

| | | | | | | |
|---|---|---|---|---|---|---|
| GULP1 | 350.050248 | -0.119764684 | 0.49808963 | -0.2404481 | 0.80998293 | 0.91928664 |
| COL5A2 | 16.2357226 | 1.226687212 | 0.61535765 | 1.99345408 | 0.04621175 | 0.20636039 |
| WDR75 | 32.718138 | 0.397327482 | 0.28297792 | 1.40409359 | 0.16029098 | 0.42791337 |
| SLC40A1 | 22.8055218 | 0.631787649 | 0.47480554 | 1.330624 | 0.18331276 | 0.46004581 |
| ASNSD1 | 16.6606905 | -0.330730021 | 0.3910507 | -0.8457472 | 0.3976938 | 0.67867271 |
| ANKAR | 70.0164192 | 0.026575479 | 0.34462506 | 0.07711418 | 0.93853271 | 0.97662903 |
| OSGEPL1 | 10.6240005 | 0.572505662 | 0.40377793 | 1.4178726 | 0.15622796 | 0.42055408 |
| ORMDL1 | 51.4695875 | -0.201490375 | 0.34457285 | -0.5847541 | 0.5587131 | 0.78244004 |
| PMS1 | 61.8451482 | -0.163960956 | 0.24989228 | -0.6561265 | 0.51174271 | 0.75544396 |
| C2orf88 | 26.6303185 | -0.722543944 | 0.37957126 | -1.9035792 | 0.05696501 | 0.23559369 |
| HIBCH | 215.583583 | 0.624413628 | 0.27433395 | 2.27610773 | 0.02283956 | 0.13197761 |
| INPP1 | 43.1381403 | -0.422284207 | 0.35577288 | -1.1869489 | 0.2352478 | 0.52547582 |
| MFSD6 | 51.6865934 | -0.660692412 | 0.27080625 | -2.4397236 | 0.0146985 | 0.09890526 |
| NAB1 | 35.6013497 | 0.533566102 | 0.41573017 | 1.28344331 | 0.19933681 | 0.48246591 |
| GLS | 1368.86657 | -0.209053514 | 0.51966229 | -0.4022872 | 0.68747264 | 0.85804981 |
| STAT1 | 296.426661 | -0.80987122 | 0.81729374 | -0.9909182 | 0.32172554 | 0.6126463 |
| MYO1B | 473.075116 | 0.983845864 | 0.46998314 | 2.09336416 | 0.03631666 | 0.17689561 |
| NABP1 | 28.8515199 | -2.396192786 | 0.8109741 | -2.9547094 | 0.00312964 | 0.03365503 |
| SLC39A10 | 83.0795826 | 0.757621274 | 0.43784424 | 1.73034428 | 0.08356878 | 0.29623459 |
| DNAH7 | 121.525348 | 0.186642756 | 0.29443044 | 0.63391122 | 0.52613876 | 0.76221787 |
| GTF3C3 | 37.4047182 | 0.576883095 | 0.29598368 | 1.94903687 | 0.05129102 | 0.22053427 |
| PGAP1 | 83.6037519 | 2.104857426 | 0.73489459 | 2.86416238 | 0.00418114 | 0.04137758 |
| ANKRD44 | 74.3168025 | -0.09462156 | 0.24560004 | -0.3852669 | 0.7000397 | 0.86376951 |
| SF3B1 | 137.892435 | 0.250186717 | 0.2196337 | 1.13910897 | 0.25465771 | 0.54564969 |
| COQ10B | 34.3752325 | -0.09422025 | 0.2972923 | -0.316928 | 0.75129825 | 0.89132635 |
| HSPD1 | 109.949734 | 0.131849278 | 0.25194337 | 0.52332903 | 0.6007453 | 0.8074205 |
| MOB4 | 13.5084785 | -0.115217409 | 0.44964347 | -0.2562417 | 0.79776421 | 0.9131741 |
| AC011997.1 | 12.8729548 | -0.493215736 | 0.5676288 | -0.8689054 | 0.38489887 | 0.66701235 |
| PLCL1 | 504.02698 | 2.258441222 | 0.2700484 | 8.36309795 | 6.11E-17 | 5.24E-14 |
| FTCDNL1 | 19.6786217 | 0.981948677 | 0.60405264 | 1.62560118 | 0.10403452 | 0.33490577 |
| C2orf69 | 17.7512467 | -0.535680655 | 0.42826556 | -1.2508142 | 0.21100226 | 0.49696965 |
| TYW5 | 36.4802607 | -0.171235984 | 0.31041751 | -0.5516312 | 0.58120106 | 0.79484057 |
| MAIP1 | 18.2810225 | 0.263565695 | 0.40372667 | 0.65283201 | 0.51386459 | 0.75647799 |
| SPATS2L | 243.628132 | -0.910463598 | 0.43434265 | -2.0961874 | 0.03606557 | 0.17602651 |
| KCTD18 | 16.8437457 | -0.20525787 | 0.34523537 | -0.5945447 | 0.55214783 | 0.77741728 |
| BZW1 | 31.173011 | -0.66952043 | 0.40351856 | -1.659206 | 0.09707428 | 0.32146212 |
| CLK1 | 96.1493458 | 0.575523185 | 0.3088788 | 1.86326539 | 0.06242494 | 0.24830181 |
| PPIL3 | 26.6530297 | -0.494498723 | 0.39734558 | -1.2445054 | 0.21331361 | 0.4997935 |
| NIF3L1 | 14.6582719 | -0.032334064 | 0.43159141 | -0.0749182 | 0.94027977 | 0.97688203 |
| ORC2 | 117.928722 | 0.33586901 | 0.32886913 | 1.02128469 | 0.30711958 | 0.60139222 |
| AC005037.1 | 23.8014683 | -0.263195754 | 0.51746714 | -0.5086231 | 0.61101642 | 0.81376433 |
| FAM126B | 94.1739108 | 0.625069651 | 0.24553158 | 2.54578104 | 0.01090336 | 0.08115639 |
| NDUFB3 | 51.3905605 | -0.497817664 | 0.46089755 | -1.0801048 | 0.2800955 | 0.57312294 |
| CFLAR | 216.258596 | -0.453878728 | 0.28492042 | -1.5930017 | 0.1111598 | 0.34769355 |
| CFLAR-AS1 | 17.6475031 | -0.993950888 | 0.59579209 | -1.6682848 | 0.09525921 | 0.31779803 |
| CASP10 | 52.0865393 | -1.172314449 | 0.26791568 | -4.3756843 | 1.21E-05 | 0.00042855 |
| CASP8 | 32.0482396 | -0.912515728 | 0.41404259 | -2.2039175 | 0.02753015 | 0.14844218 |
| ALS2CR12 | 15.4405246 | -0.83504468 | 0.41213803 | -2.0261287 | 0.0427516 | 0.19741236 |
| TRAK2 | 136.214554 | -0.384185961 | 0.24354854 | -1.5774513 | 0.11469172 | 0.35355161 |
| STRADB | 63.320696 | -0.980337998 | 0.38301407 | -2.5595352 | 0.01048122 | 0.07927869 |
| TMEM237 | 14.1755265 | 0.346353321 | 0.47174398 | 0.73419765 | 0.46282828 | 0.72750336 |
| ALS2 | 69.3316502 | 0.327325078 | 0.24566245 | 1.33241801 | 0.18272287 | 0.45952112 |
| SUMO1 | 62.607225 | -0.131390482 | 0.33720527 | -0.3896454 | 0.69679877 | 0.86247999 |
| NOP58 | 72.0033771 | 0.329577775 | 0.28209497 | 1.16832205 | 0.24267688 | 0.53317493 |
| BMPR2 | 330.198823 | 0.017563631 | 0.19932111 | 0.08811727 | 0.92978347 | 0.97253199 |
| FAM117B | 137.693926 | 0.218489009 | 0.2329981 | 0.9377287 | 0.34838385 | 0.6354277 |
| ICA1L | 46.4429369 | -0.983391111 | 0.36043636 | -2.728335 | 0.00636549 | 0.05638634 |
| WDR12 | 41.6233667 | 0.104300107 | 0.33652081 | 0.30993658 | 0.75660919 | 0.89428497 |
| CARF | 62.5899187 | 0.174746123 | 0.36146909 | 0.4834331 | 0.62878825 | 0.82494825 |
| NBEAL1 | 522.090446 | 0.432441724 | 0.24385639 | 1.77334589 | 0.07617141 | 0.28022732 |
| CYP20A1 | 103.120727 | -0.119456992 | 0.40448688 | -0.2953297 | 0.76774203 | 0.89979146 |
| ABI2 | 210.666552 | 0.147380383 | 0.26003068 | 0.56678075 | 0.57086315 | 0.78905469 |
| RAPH1 | 224.501381 | -0.463830837 | 0.44233238 | -1.0486025 | 0.29436111 | 0.58644521 |
| PARD3B | 767.793225 | 0.857184162 | 0.15904224 | 5.38966339 | 7.06E-08 | 6.22E-06 |
| INO80D | 236.587321 | -0.332731446 | 0.64840105 | -0.5131569 | 0.6078416 | 0.81187554 |
| NDUFS1 | 167.924235 | 0.46380696 | 0.24232785 | 1.91396476 | 0.05562466 | 0.23268234 |
| EEF1B2 | 39.3336191 | -0.471307671 | 0.36217309 | -1.3013326 | 0.19314462 | 0.47364338 |
| ZDBF2 | 37.5735225 | 0.142066916 | 0.48758636 | 0.2913677 | 0.77077012 | 0.90076568 |
| FASTKD2 | 43.1406841 | 0.298259434 | 0.27325834 | 1.09149252 | 0.27505622 | 0.56696009 |
| KLF7 | 24.2311938 | -1.309618772 | 0.654822 | -1.9999615 | 0.04550443 | 0.20435335 |
| CREB1 | 103.575088 | -0.07990718 | 0.21240062 | -0.3762097 | 0.70676098 | 0.86803191 |
| METTL21A | 25.9033167 | -0.351922846 | 0.30717271 | -1.145684 | 0.25192594 | 0.54334972 |
| CCNYL1 | 56.108153 | 0.284875833 | 0.36806989 | 0.77397211 | 0.43894728 | 0.71027377 |
| PLEKHM3 | 65.5416315 | 0.637100984 | 0.32108963 | 1.98418423 | 0.04723531 | 0.20965882 |
| IDH1 | 24.4534627 | 0.0985199 | 0.40863902 | 0.24109274 | 0.80948324 | 0.9191701 |
| PIKFYVE | 111.142586 | 0.437715323 | 0.23375102 | 1.87257077 | 0.06112768 | 0.24535775 |
| PTH2R | 1217.66906 | 0.959975206 | 0.28469533 | 3.3719387 | 0.00074641 | 0.01186655 |
| RPE | 54.1441598 | 0.594280185 | 0.27374829 | 2.17090008 | 0.02993873 | 0.15744194 |
| KANSL1L | 513.487067 | -0.060612337 | 0.26189948 | -0.2314336 | 0.81697797 | 0.92248811 |
| AC007038.1 | 53.9948045 | -0.120972644 | 0.32413188 | -0.3732204 | 0.70898438 | 0.86910485 |
| ACADL | 41.0239275 | 0.203673906 | 0.43681238 | 0.4662732 | 0.64101996 | 0.83230084 |
| LANCL1 | 65.3765597 | 0.398917107 | 0.27198255 | 1.46670113 | 0.14245738 | 0.39902009 |
| ERBB4 | 3062.64568 | 0.303465408 | 0.18765858 | 1.61711448 | 0.10585357 | 0.33835045 |
| IKZF2 | 91.4161542 | 0.230932693 | 0.37736831 | 0.61195571 | 0.54056706 | 0.7714342 |
| SPAG16 | 371.850455 | 0.207834303 | 0.35403241 | 0.58704881 | 0.55717093 | 0.7813742 |
| AC068051.1 | 57.2885535 | 0.017677548 | 0.5881708 | 0.03005513 | 0.97602309 | 0.99034668 |
| BARD1 | 23.1927619 | 0.436791613 | 0.36495969 | 1.19682154 | 0.23137612 | 0.52111057 |
| AC016708.1 | 24.9310011 | 0.136700295 | 0.41665324 | 0.32809128 | 0.74284265 | 0.88740176 |
| ATIC | 35.4277547 | 0.076430977 | 0.37815236 | 0.20211688 | 0.83982535 | 0.93543959 |
| MREG | 34.7933745 | 0.358540866 | 0.32433999 | 1.10544762 | 0.26896568 | 0.56124862 |
| PECR | 53.2903482 | 0.708607769 | 0.26405676 | 2.68354341 | 0.00728465 | 0.06217604 |
| XRCC5 | 229.208619 | -0.0741028 | 0.19345314 | -0.383053 | 0.70168049 | 0.86488543 |
| SMARCAL1 | 35.592022 | -0.226751577 | 0.32252832 | -0.7030439 | 0.48202838 | 0.74068045 |
| RPL37A | 126.293091 | -0.214671158 | 0.29112946 | -0.7373735 | 0.46089522 | 0.72601568 |
| IGFBP2 | 18.0910373 | -0.85163837 | 0.64573374 | -1.3188693 | 0.18721283 | 0.4649409 |
| IGFBP5 | 35.3184837 | -0.399113065 | 0.62640428 | -0.6371493 | 0.52402758 | 0.76218158 |
| DIRC3 | 54.0259697 | 0.101403477 | 0.62545912 | 0.16212647 | 0.87120627 | 0.94844301 |
| TNS1 | 195.293633 | 1.064674598 | 0.3302426 | 3.2239166 | 0.0012645 | 0.01746233 |
| ARPC2 | 129.56022 | -0.668783548 | 0.30700982 | -2.1783784 | 0.02937788 | 0.15536495 |
| AAMP | 28.4445861 | -0.229104184 | 0.34780745 | -0.6587098 | 0.51008213 | 0.75482254 |
| PNKD | 78.125039 | -0.358271024 | 0.31927718 | -1.1221316 | 0.26180648 | 0.55337102 |
| TMBIM1 | 73.671036 | -1.033834483 | 0.27776943 | -3.7219161 | 0.00019772 | 0.0041825 |
| CATIP-AS2 | 14.2078462 | 0.277865761 | 0.56883419 | 0.48848288 | 0.62520785 | 0.82285278 |
| CATIP | 16.9417577 | 1.890748735 | 0.48367741 | 3.90911109 | 9.26E-05 | 0.00227766 |
| CTDSP1 | 51.6764028 | -0.347725533 | 0.24938409 | -1.3943373 | 0.16321577 | 0.43145254 |
| USP37 | 91.9807515 | -0.064901623 | 0.32718908 | -0.1983612 | 0.84276246 | 0.93600812 |
| CNOT9 | 47.294268 | -0.101242924 | 0.23401397 | -0.4326362 | 0.66527905 | 0.84348349 |
| PLCD4 | 19.4454529 | -0.945663987 | 0.67460605 | -1.4018018 | 0.16097444 | 0.42818783 |
| ZNF142 | 17.2498492 | -0.705111649 | 0.4400847 | -1.602218 | 0.1091074 | 0.34458138 |
| STK36 | 16.4581668 | 0.273842232 | 0.41928045 | 0.65312426 | 0.51367618 | 0.75647799 |
| TTLL4 | 18.0234306 | -1.309942936 | 0.37608155 | -3.4831354 | 0.00049558 | 0.00855183 |
| CYP27A1 | 16.6758089 | -1.389813396 | 0.46911897 | -2.9626033 | 0.00305049 | 0.03297607 |
| CNPPD1 | 27.504921 | -0.090534836 | 0.33154819 | -0.2730669 | 0.78480179 | 0.90771539 |

| | | | | | | |
|---|---|---|---|---|---|---|
| RETREG2 | 39.9066857 | -0.682039276 | 0.38047809 | -1.7925849 | 0.07303932 | 0.2730229 |
| ZFAND2B | 14.5619158 | -0.016711419 | 0.45059387 | -0.0370875 | 0.97041521 | 0.98816027 |
| ANKZF1 | 28.4984296 | -0.349350997 | 0.29186129 | -1.1969761 | 0.23131586 | 0.52111057 |
| STK16 | 12.0092553 | -0.662655164 | 0.43248035 | -1.5322203 | 0.12546809 | 0.37047069 |
| TUBA4A | 31.9546518 | -0.204637282 | 0.49045289 | -0.4172415 | 0.6765018 | 0.85189115 |
| DNAJB2 | 31.7642668 | 0.12117688 | 0.40168212 | 0.30167358 | 0.76290091 | 0.89739726 |
| DNPEP | 21.425765 | -0.443757629 | 0.40179056 | -1.1044501 | 0.26939793 | 0.56160046 |
| CHPF | 21.5673687 | -0.781555738 | 0.5089233 | -1.5357044 | 0.1246109 | 0.36974123 |
| OBSL1 | 29.3748539 | 0.132940338 | 0.43721562 | 0.30406127 | 0.76108121 | 0.89648567 |
| SGPP2 | 273.862536 | 0.201137747 | 0.3135189 | 0.64154904 | 0.52116603 | 0.76092013 |
| FARSB | 40.8283385 | 0.735499636 | 0.34966985 | 2.10341168 | 0.0354298 | 0.17464009 |
| ACSL3 | 136.303388 | 0.900887794 | 0.28673609 | 3.14187096 | 0.00167872 | 0.02164477 |
| AP1S3 | 52.8109653 | 0.037192935 | 0.51804696 | 0.07179452 | 0.94276543 | 0.9781624 |
| WDFY1 | 82.5279524 | -0.049431205 | 0.22955983 | -0.2153304 | 0.82950975 | 0.92907256 |
| MRPL44 | 12.4508866 | -0.781217259 | 0.55622532 | -1.4044978 | 0.16017066 | 0.42791337 |
| CUL3 | 247.151942 | 0.482712755 | 0.20833474 | 2.31700554 | 0.02050343 | 0.1227566 |
| IRS1 | 24.2943156 | -0.741176671 | 0.45006137 | -1.6468347 | 0.09959203 | 0.32585356 |
| RHBDD1 | 128.849506 | 0.110537266 | 0.26668882 | 0.41448032 | 0.67852238 | 0.85318412 |
| COL4A4 | 304.888211 | -0.810248444 | 0.33028908 | -2.4531494 | 0.01416115 | 0.09655076 |
| COL4A3 | 455.103627 | -0.348545958 | 0.39600228 | -0.8801615 | 0.37877183 | 0.66204741 |
| AC097662.1 | 66.7665494 | -0.669850398 | 0.56718385 | -1.1810111 | 0.23759833 | 0.52778809 |
| MFF | 132.795281 | 0.165607161 | 0.26343631 | 0.62864213 | 0.52958337 | 0.7638395 |
| AGFG1 | 249.478745 | -0.444538114 | 0.31413277 | -1.4151281 | 0.15703093 | 0.42219972 |
| SLC19A3 | 49.74776 | -1.792764758 | 0.87744331 | -2.0431688 | 0.04103573 | 0.19294849 |
| DNER | 25.5392159 | -1.841560709 | 0.68613943 | -2.6839453 | 0.0072759 | 0.06215285 |
| TRIP12 | 287.03261 | 0.127793568 | 0.18792907 | 0.68000957 | 0.4964984 | 0.74859388 |
| FBXO36 | 34.4596826 | -0.028218145 | 0.3161524 | -0.0892549 | 0.92887935 | 0.97199747 |
| SP110 | 48.6605321 | -1.163781317 | 0.72528142 | -1.6045928 | 0.10858346 | 0.34348281 |
| SP140L | 39.2434059 | -1.041475191 | 0.4724718 | -2.2043118 | 0.02750242 | 0.14841799 |
| SP100 | 144.689372 | -0.857704575 | 0.48287246 | -1.7762549 | 0.07569091 | 0.27917785 |
| CAB39 | 236.186537 | 0.618376763 | 0.27860505 | 2.21954612 | 0.02644959 | 0.14524018 |
| ITM2C | 115.64945 | -1.081848797 | 0.52412383 | -2.0641092 | 0.03900737 | 0.18612965 |
| PSMD1 | 124.748247 | 0.238742904 | 0.26497723 | 0.90099403 | 0.36759149 | 0.65303113 |
| ARMC9 | 131.177857 | -0.68493861 | 0.26574394 | -2.5774383 | 0.00995356 | 0.07698319 |
| NCL | 86.1238524 | -0.905037189 | 0.32516706 | -2.7832991 | 0.00538092 | 0.05021215 |
| PTMA | 383.766537 | -0.724687721 | 0.26411471 | -2.743837 | 0.00607257 | 0.05478073 |
| PDE6D | 30.7198761 | -0.17579608 | 0.28114975 | -0.6252756 | 0.53179019 | 0.76558169 |
| COPS7B | 29.9767395 | 0.505077792 | 0.25941119 | 1.94701624 | 0.05153279 | 0.22141969 |
| DIS3L2 | 206.206832 | 0.405157128 | 0.18860185 | 2.14821392 | 0.03169677 | 0.1630255 |
| EIF4E2 | 48.6021377 | -0.371412964 | 0.26585287 | -1.3970621 | 0.16239491 | 0.42996461 |
| EFHD1 | 81.2675981 | 1.395627953 | 0.44475716 | 3.13795496 | 0.00170131 | 0.02188127 |
| GIGYF2 | 258.438 | 0.182225517 | 0.23127444 | 0.78791897 | 0.4307441 | 0.70420575 |
| ATG16L1 | 19.7746392 | -0.004797286 | 0.38404585 | -0.0124914 | 0.99003353 | 0.99680663 |
| DGKD | 38.5861048 | 0.204353392 | 0.34134907 | 0.59866397 | 0.54939699 | 0.77592905 |
| USP40 | 146.784518 | 0.625219764 | 0.29053604 | 2.15195254 | 0.03140109 | 0.16223373 |
| AC114812.2 | 56.9076918 | -1.288642527 | 0.85429016 | -1.5084366 | 0.13144283 | 0.38026509 |
| ARL4C | 72.7220582 | -1.8364266 | 0.71166859 | -2.580452 | 0.00986711 | 0.07654439 |
| SH3BP4 | 120.830738 | 0.648988364 | 0.38146871 | 1.70128859 | 0.08888881 | 0.30697614 |
| AGAP1 | 747.26154 | -0.889640721 | 0.39322626 | -2.2624143 | 0.02367182 | 0.13490566 |
| IQCA1 | 36.3909154 | -0.976901614 | 0.35257536 | -2.7707598 | 0.00559257 | 0.05141788 |
| AC105760.2 | 13.758433 | 0.251341977 | 0.43869116 | 0.57293605 | 0.566688 | 0.78647713 |
| COPS8 | 20.6405405 | -0.359624472 | 0.33893758 | -1.0610345 | 0.28867422 | 0.58064041 |
| RAB17 | 103.32881 | 0.584123928 | 0.3357985 | 1.73950727 | 0.08194558 | 0.29293408 |
| LRRFIP1 | 636.223696 | 0.025015172 | 0.39834396 | 0.06279792 | 0.94992742 | 0.98003064 |
| ILKAP | 74.6437506 | 0.202913179 | 0.21350699 | 0.95038189 | 0.34191824 | 0.62990732 |
| PER2 | 26.4353057 | -0.058357578 | 0.42315365 | -0.1379111 | 0.89031069 | 0.95702609 |
| TRAF3IP1 | 51.1740204 | 0.156416335 | 0.25813474 | 0.6059484 | 0.54454902 | 0.77395839 |
| ASB1 | 14.6896149 | -0.502261068 | 0.4442846 | -1.130494 | 0.25826813 | 0.54961335 |
| HDAC4 | 362.386873 | -0.575838179 | 0.33216904 | -1.7335697 | 0.08299446 | 0.29503415 |
| NDUFA10 | 141.428652 | 0.473971214 | 0.19503882 | 2.43013785 | 0.01509308 | 0.10047582 |
| COPS9 | 28.0302447 | -0.352361702 | 0.44468868 | -0.7923784 | 0.42814008 | 0.70245247 |
| GPC1 | 31.42709 | -1.27427023 | 0.5264538 | -2.4204787 | 0.01550009 | 0.10209839 |
| ANKMY1 | 45.7550456 | -0.069903383 | 0.29630863 | -0.2359141 | 0.81349933 | 0.92085157 |
| RNPEPL1 | 30.4102068 | -1.162096315 | 0.37991209 | -3.0588559 | 0.00222184 | 0.02624931 |
| CAPN10 | 12.4748117 | -0.445602634 | 0.47972265 | -0.9288755 | 0.3529536 | 0.63892716 |
| SNED1 | 33.4897759 | -1.167760774 | 0.46592952 | -2.5063035 | 0.01220009 | 0.08795331 |
| MTERF4 | 43.1145515 | 0.040903211 | 0.354518 | 0.11537696 | 0.90814634 | 0.96232499 |
| PASK | 11.5074769 | 0.201609933 | 0.41304052 | 0.48811175 | 0.62547069 | 0.82303813 |
| PPP1R7 | 34.3245075 | -0.013756031 | 0.27126151 | -0.0507113 | 0.95955555 | 0.98416248 |
| HDLBP | 187.378266 | -0.290872008 | 0.25479491 | -1.1415927 | 0.25362336 | 0.54524749 |
| 2-Sep | 222.483654 | -0.073032038 | 0.15289117 | -0.4776734 | 0.63288271 | 0.82656664 |
| FARP2 | 89.2790385 | 0.047163607 | 0.20778783 | 0.22697964 | 0.82043957 | 0.92413825 |
| STK25 | 34.5165374 | -0.346820867 | 0.30393977 | -1.1410842 | 0.25383488 | 0.54524749 |
| BOK | 13.4253177 | -0.751447247 | 0.66074264 | -1.1372768 | 0.25542261 | 0.54662971 |
| THAP4 | 58.0376968 | 0.120443988 | 0.2354411 | 0.51156739 | 0.60895381 | 0.81272732 |
| ATG4B | 92.2658082 | 0.343477127 | 0.20992526 | 1.63618772 | 0.10180031 | 0.33051935 |
| DTYMK | 12.7665291 | -0.053075935 | 0.43144397 | -0.1230193 | 0.90209182 | 0.9608509 |
| ING5 | 43.1636121 | 0.048726303 | 0.33549258 | 0.14523809 | 0.88452289 | 0.95617574 |
| D2HGDH | 62.6763063 | 1.00597279 | 0.32257211 | 3.1185982 | 0.00181714 | 0.02287311 |
| CHL1 | 45.6363423 | -0.924272389 | 0.7444323 | -1.2415802 | 0.2143915 | 0.50088196 |
| LINC01266 | 122.081171 | 0.365224618 | 0.53360585 | 0.68444643 | 0.4936933 | 0.74700665 |
| CNTN4 | 148.963443 | 1.450508585 | 0.43788182 | 3.31255725 | 0.00092447 | 0.0138484 |
| TRNT1 | 40.4221636 | -0.234284909 | 0.25567932 | -0.9163233 | 0.35949737 | 0.64342285 |
| CRBN | 50.5326606 | -0.156096979 | 0.24213877 | -0.6446592 | 0.51914809 | 0.75991242 |
| AC034195.1 | 97.5763444 | 0.552798262 | 0.38837003 | 1.42338033 | 0.15462593 | 0.41795853 |
| SUMF1 | 457.225842 | 0.121913704 | 0.29998646 | 0.40639735 | 0.68445067 | 0.85614507 |
| SETMAR | 16.1762888 | 0.858940661 | 0.48352821 | 1.77640239 | 0.07566662 | 0.27917785 |
| ITPR1 | 657.351101 | 0.347747557 | 0.2943324 | 1.18147901 | 0.23741249 | 0.52778809 |
| EGOT | 35.3969743 | 0.940214028 | 0.54807952 | 1.71547008 | 0.08625908 | 0.30190826 |
| AC018816.1 | 24.5411129 | -0.008140348 | 0.3514389 | -0.0231629 | 0.98152032 | 0.99318267 |
| BHLHE40 | 325.795943 | 1.089599844 | 0.46605353 | 2.33792852 | 0.01939096 | 0.11820452 |
| ARL8B | 120.942661 | -0.356752741 | 0.3535312 | -1.0091125 | 0.3129207 | 0.60561883 |
| AC026202.2 | 15.9487501 | -1.151375376 | 0.57177788 | -2.013676 | 0.04404356 | 0.20016628 |
| EDEM1 | 30.4844145 | -0.212892412 | 0.29827268 | -0.713751 | 0.47538119 | 0.73587124 |
| AC087857.1 | 21.4423589 | 2.765697497 | 0.72168303 | 3.83228841 | 0.00012696 | 0.00295951 |
| AC026167.1 | 61.1456354 | 3.233154944 | 0.64813441 | 4.98840197 | 6.09E-07 | 3.78E-05 |
| AC069277.1 | 37.5674299 | 1.692123951 | 1.40832412 | 1.20151599 | 0.22955111 | 0.51974407 |
| LMCD1-AS1 | 109.787137 | 1.157044106 | 0.29397653 | 3.93583839 | 8.29E-05 | 0.00210878 |
| LMCD1 | 13.7719948 | 0.040066274 | 0.47167492 | 0.08494468 | 0.93230537 | 0.97356162 |
| RAD18 | 22.394974 | -0.526219357 | 0.42329498 | -1.2431505 | 0.21381239 | 0.50026184 |
| SRGAP3 | 222.278844 | 0.894056983 | 0.2796148 | 3.19745945 | 0.00138644 | 0.01871965 |
| THUMPD3-A! | 89.7530875 | -0.098576613 | 0.25126365 | -0.3923234 | 0.69481927 | 0.86161604 |
| THUMPD3 | 28.1936724 | -0.456406497 | 0.27924644 | -1.6344219 | 0.10217029 | 0.33078515 |
| SETD5 | 278.595611 | -0.075419379 | 0.16752462 | -0.4501988 | 0.65256711 | 0.83835221 |
| MTMR14 | 42.7247791 | 0.215518979 | 0.29925703 | 0.72018018 | 0.47141406 | 0.73283653 |
| OGG1 | 20.6550305 | -0.315226901 | 0.41245212 | -0.7642751 | 0.44470329 | 0.71449366 |
| CAMK1 | 11.8700217 | 0.18650504 | 0.41225953 | 0.45239716 | 0.65098289 | 0.83746575 |
| TADA3 | 26.6127917 | -0.519144653 | 0.32309627 | -1.6067801 | 0.10810262 | 0.34264936 |
| RPUSD3 | 14.3615689 | 0.072057977 | 0.37859422 | 0.19033037 | 0.84905026 | 0.93897371 |
| JAGN1 | 12.0006935 | -1.141010901 | 0.51996843 | -2.194385 | 0.02820775 | 0.15111607 |
| IL17RE | 16.6914508 | 1.00761175 | 0.53633928 | 1.87868349 | 0.06028772 | 0.24330674 |
| IL17RC | 14.4876217 | 0.099504593 | 0.45050031 | 0.22087575 | 0.82518918 | 0.92675231 |
| CRELD1 | 27.3105016 | 0.463227599 | 0.35821809 | 1.29314408 | 0.19596123 | 0.4776213 |

| | | | | | | |
|---|---|---|---|---|---|---|
| EMC3 | 44.9603747 | -0.127152543 | 0.29632768 | -0.4290944 | 0.66785454 | 0.84558183 |
| BRK1 | 46.2286612 | -0.66877564 | 0.32441778 | -2.0614642 | 0.03925877 | 0.18700317 |
| VHL | 27.8354181 | -0.171040106 | 0.32877118 | -0.5202406 | 0.60289592 | 0.80891 |
| IRAK2 | 18.903945 | -1.630122703 | 0.70327177 | -2.3179129 | 0.02045405 | 0.12275018 |
| SEC13 | 46.8322944 | -0.510155236 | 0.30180205 | -1.6903637 | 0.0909584 | 0.3106941 |
| ATG7 | 242.388207 | -0.142810762 | 0.26950558 | -0.5298991 | 0.59618189 | 0.80412393 |
| VGLL4 | 277.171111 | -0.08866522 | 0.17273806 | -0.5132929 | 0.60774645 | 0.81187554 |
| TAMM41 | 75.679626 | 0.111504174 | 0.3008121 | 0.37067716 | 0.710878 | 0.87028547 |
| PPARG | 257.205194 | -0.269195239 | 0.45479763 | -0.5919011 | 0.55391678 | 0.77872693 |
| TSEN2 | 62.3054867 | 0.233230845 | 0.2768427 | 0.84246701 | 0.3995266 | 0.67979406 |
| MKRN2OS | 10.6011261 | -0.936878023 | 0.6931831 | -1.3515592 | 0.17651636 | 0.4510098 |
| MKRN2 | 16.3497969 | 0.187844931 | 0.41608316 | 0.45146007 | 0.651658 | 0.83781115 |
| RAF1 | 130.35775 | 0.413337342 | 0.19700083 | 2.09815023 | 0.03589188 | 0.17574055 |
| RPL32 | 156.979886 | -0.573196359 | 0.40634636 | -1.4106103 | 0.15835955 | 0.42480987 |
| IQSEC1 | 58.8585236 | 0.695487861 | 0.40662087 | 1.71040867 | 0.08719033 | 0.30375002 |
| NUP210 | 39.8706937 | 0.610475431 | 0.34562681 | 1.76628494 | 0.07734806 | 0.28226698 |
| HDAC11 | 20.2097536 | 0.485446344 | 0.49351052 | 0.98365957 | 0.32528293 | 0.61555193 |
| XPC | 47.3681438 | -0.185387607 | 0.28298564 | -0.6551131 | 0.51239492 | 0.75562371 |
| LSM3 | 32.3698999 | -0.253270528 | 0.2853558 | -0.8875605 | 0.37477723 | 0.65806152 |
| SLC6A6 | 103.182542 | -0.398591788 | 0.40078149 | -0.9945364 | 0.31996178 | 0.6120836 |
| CCDC174 | 31.737063 | 0.088602685 | 0.27026254 | 0.32783932 | 0.74303316 | 0.88740176 |
| FGD5-AS1 | 62.4077435 | -0.268567175 | 0.32322364 | -0.830902 | 0.40602899 | 0.68404099 |
| NR2C2 | 107.88746 | -0.001685544 | 0.32047741 | -0.0052595 | 0.99580356 | 0.99868531 |
| MRPS25 | 108.344577 | 0.082375706 | 0.25167619 | 0.3273083 | 0.74343471 | 0.88754632 |
| RBSN | 39.2441694 | 0.433290732 | 0.2987344 | 1.45042129 | 0.14694107 | 0.40616768 |
| CAPN7 | 139.774795 | 0.257243638 | 0.26059889 | 0.98712486 | 0.32358143 | 0.61436342 |
| SH3BP5 | 40.5103708 | -1.087000334 | 0.88736893 | -1.22497 | 0.22058652 | 0.5089143 |
| METTL6 | 31.0187689 | 0.251469762 | 0.34153915 | 0.73628386 | 0.46155797 | 0.72669738 |
| EAF1 | 12.6852024 | -0.374575764 | 0.43556014 | -0.8599863 | 0.38979659 | 0.67185619 |
| COLQ | 16.6917615 | -0.317637793 | 0.37799464 | -0.8403235 | 0.40072701 | 0.68010059 |
| HACL1 | 47.9688464 | 0.128250486 | 0.27198502 | 0.4715351 | 0.63725866 | 0.82989931 |
| BTD | 15.2782877 | -0.225570299 | 0.43034773 | -0.5241582 | 0.6001685 | 0.80738812 |
| ANKRD28 | 360.265208 | 0.311372767 | 0.1891914 | 1.64580829 | 0.09980324 | 0.32599015 |
| OXNAD1 | 124.510872 | 0.4962688 | 0.29593666 | 1.67694264 | 0.0935537 | 0.31496412 |
| RFTN1 | 15.3112642 | -1.628734924 | 0.65408684 | -2.4900897 | 0.01277108 | 0.09086168 |
| PLCL2 | 133.745727 | 0.837775564 | 0.53148182 | 1.57630146 | 0.11495634 | 0.35399477 |
| TBC1D5 | 539.157293 | 0.286193093 | 0.2037932 | 1.40433092 | 0.16022033 | 0.42791337 |
| RAB5A | 105.823208 | -0.307214522 | 0.23080475 | -1.3310581 | 0.18316991 | 0.46004581 |
| KAT2B | 85.6703724 | 1.334402444 | 0.39527933 | 3.37584674 | 0.00073589 | 0.01171735 |
| ZNF385D | 194.515263 | -1.641793792 | 0.79662369 | -2.0609402 | 0.03930874 | 0.18704789 |
| UBE2E2 | 365.257866 | 0.288181261 | 0.21547737 | 1.33740846 | 0.18108934 | 0.45756661 |
| UBE2E1 | 189.849087 | 0.034188854 | 0.26915112 | 0.12702475 | 0.89892081 | 0.95935794 |
| NKIRAS1 | 19.5615676 | 0.167891884 | 0.38050456 | 0.44123487 | 0.65904297 | 0.84059189 |
| RPL15 | 254.759228 | -1.126471869 | 0.46519559 | -2.4215016 | 0.01545653 | 0.101998 |
| NR1D2 | 56.0254087 | 0.204472482 | 0.47175231 | 0.43343186 | 0.66470106 | 0.84348349 |
| THRB | 290.846725 | 0.979484333 | 0.28060151 | 3.49065952 | 0.00048183 | 0.00837068 |
| RARB | 46.1963566 | 0.861148678 | 0.27201614 | 3.1657999 | 0.00154657 | 0.02021926 |
| TOP2B | 166.276806 | 0.606169511 | 0.21747173 | 2.78734855 | 0.00531413 | 0.04963387 |
| NGLY1 | 114.230395 | 0.056814146 | 0.40866304 | 0.13902443 | 0.88943085 | 0.95702609 |
| OXSM | 13.7437455 | -0.682329451 | 0.47264998 | -1.4436253 | 0.14884441 | 0.40923276 |
| NEK10 | 30.6393872 | -0.07923344 | 0.29113813 | -0.2721507 | 0.78550616 | 0.90802494 |
| SLC4A7 | 46.6485061 | -1.249686135 | 0.60185353 | -2.0763958 | 0.03785736 | 0.18238254 |
| CMC1 | 136.261788 | 0.601148555 | 0.25485703 | 2.3587678 | 0.01833572 | 0.11413573 |
| AZI2 | 61.4037238 | -0.011383441 | 0.34848213 | -0.0326658 | 0.97394111 | 0.98930599 |
| TGFBR2 | 78.605553 | 0.356249898 | 0.39586071 | 0.89993751 | 0.36815351 | 0.65369102 |
| STT3B | 130.880105 | 0.433295172 | 0.22727677 | 1.90646483 | 0.05658992 | 0.23469783 |
| OSBPL10 | 45.2258339 | -0.636728348 | 0.32542568 | -1.9566014 | 0.05039435 | 0.218779 |
| GPD1L | 25.3361836 | 0.448430139 | 0.37520871 | 1.19514852 | 0.23202901 | 0.52184597 |
| CMTM8 | 66.9776428 | -0.082726452 | 0.28905944 | -0.2861918 | 0.77473119 | 0.90205458 |
| CMTM7 | 55.2407737 | -2.628866158 | 0.28220128 | -9.3155715 | 1.21E-20 | 2.08E-17 |
| CMTM6 | 63.4826933 | -1.168537636 | 0.31995746 | -3.6521656 | 0.00026004 | 0.00530905 |
| DYNC1LI1 | 36.8301338 | -0.071498875 | 0.30033039 | -0.2380674 | 0.81182882 | 0.92017389 |
| CNOT10 | 88.2838084 | -0.071844581 | 0.27745039 | -0.2589457 | 0.79567715 | 0.91195838 |
| GLB1 | 86.7357348 | 0.254611806 | 0.27508259 | 0.92558314 | 0.35466266 | 0.63999557 |
| CRTAP | 52.1785439 | -0.01557635 | 0.29704129 | -0.0524383 | 0.95817943 | 0.98328165 |
| FBXL2 | 135.119501 | -0.815736714 | 0.27289053 | -2.9892452 | 0.00279668 | 0.03121198 |
| UBP1 | 64.2607492 | 0.232978004 | 0.23611579 | 0.98671081 | 0.32378443 | 0.61448136 |
| CLASP2 | 268.529588 | 0.468619377 | 0.19165495 | 2.4451201 | 0.01448039 | 0.09775687 |
| PDCD6IP | 157.245913 | -0.095393019 | 0.17183433 | -0.5551453 | 0.5787953 | 0.79355192 |
| STAC | 99.9372724 | 0.796211065 | 0.31281677 | 2.54529531 | 0.01091854 | 0.08115639 |
| TRANK1 | 24.4608666 | -2.568564875 | 1.29444535 | -1.9842977 | 0.04722267 | 0.20965882 |
| EPM2AIP1 | 50.8513154 | 0.057220726 | 0.3086885 | 0.18536721 | 0.85294103 | 0.94049004 |
| MLH1 | 31.6310252 | 0.139771801 | 0.27697638 | 0.50463438 | 0.61381566 | 0.81511796 |
| LRRFIP2 | 233.25671 | -0.061638902 | 0.21656267 | -0.2846239 | 0.77593234 | 0.90266675 |
| GOLGA4 | 454.743001 | -0.06351201 | 0.27601754 | -0.2301013 | 0.81801302 | 0.92281757 |
| ITGA9 | 20.8715904 | -1.881278172 | 0.64736131 | -2.9060714 | 0.00365998 | 0.03766744 |
| CTDSPL | 289.44867 | -0.220039765 | 0.27674175 | -0.7951087 | 0.42655029 | 0.70129605 |
| PLCD1 | 15.0172084 | -0.616547133 | 0.42736386 | -1.4426749 | 0.14911206 | 0.40942229 |
| ACAA1 | 40.9572304 | -0.427055288 | 0.35056306 | -1.2181982 | 0.22314867 | 0.51188545 |
| OXSR1 | 155.585404 | 0.219471952 | 0.28529726 | 0.76927466 | 0.44173028 | 0.71227192 |
| ACVR2B | 25.4854711 | -0.403441098 | 0.35848752 | -1.1253979 | 0.26042046 | 0.55146957 |
| EXOG | 26.273192 | -0.677189552 | 0.31428268 | -2.1547148 | 0.03118416 | 0.16176193 |
| WDR48 | 59.4867136 | 0.331366129 | 0.2424269 | 1.36687032 | 0.17166596 | 0.44345604 |
| GORASP1 | 17.1201952 | -0.001675282 | 0.35478141 | -0.004722 | 0.99623239 | 0.99868531 |
| TTC21A | 15.2789901 | -0.167638248 | 0.49922778 | -0.3357951 | 0.73702538 | 0.88493487 |
| CSRNP1 | 14.3113424 | 0.055687393 | 0.48924482 | 0.11382316 | 0.90937798 | 0.96308593 |
| SLC25A38 | 15.189836 | -0.276002633 | 0.42831187 | -0.6443964 | 0.51931841 | 0.76002222 |
| RPSA | 115.39192 | -0.467851321 | 0.44685931 | -1.0469768 | 0.29511028 | 0.58668972 |
| MYRIP | 13.4347598 | -0.475271456 | 0.67974317 | -0.6991927 | 0.48443162 | 0.74250561 |
| EIF1B | 27.7270897 | -0.940223699 | 0.30002284 | -3.1338404 | 0.00172535 | 0.02204834 |
| ENTPD3-AS1 | 34.1535124 | 0.206433967 | 0.49897543 | 0.4137157 | 0.67908233 | 0.85351679 |
| RPL14 | 173.809619 | -0.894550993 | 0.40418155 | -2.2132405 | 0.02688107 | 0.14636826 |
| ZNF621 | 41.0328611 | -0.213869145 | 0.30427842 | -0.7028732 | 0.48213478 | 0.74068045 |
| CTNNB1 | 121.024105 | 0.042413618 | 0.22229147 | 0.19080183 | 0.84868086 | 0.93891735 |
| ULK4 | 292.386453 | -0.107747535 | 0.16114267 | -0.6686468 | 0.5037208 | 0.75108109 |
| TRAK1 | 409.147773 | 0.583340285 | 0.22778321 | 2.56094508 | 0.01043879 | 0.07913199 |
| SEC22C | 24.6626175 | -0.70397232 | 0.35722465 | -1.9706712 | 0.0487615 | 0.21425029 |
| SS18L2 | 12.3987603 | -0.811682863 | 0.459489 | -1.7664903 | 0.07731363 | 0.28224131 |
| NKTR | 283.969713 | 0.181088391 | 0.16798867 | 1.07797981 | 0.28104277 | 0.57412306 |
| AC006059.1 | 32.5761514 | -1.257680226 | 0.56885401 | -2.2109016 | 0.02704265 | 0.14670533 |
| AC099329.3 | 16.4666975 | -0.807840337 | 0.46174393 | -1.7495419 | 0.0801974 | 0.28903174 |
| SNRK | 126.31217 | 0.083026461 | 0.32700597 | 0.25389891 | 0.79957365 | 0.91433098 |
| ANO10 | 261.399331 | 0.670092485 | 0.2496639 | 2.68397824 | 0.00727518 | 0.06215285 |
| ABHD5 | 36.3431425 | -0.385296905 | 0.37736469 | -1.02102 | 0.30724495 | 0.60139222 |
| TCAIM | 160.315035 | 0.468268216 | 0.30821933 | 1.51926945 | 0.12869469 | 0.37611281 |
| ZNF445 | 35.0306704 | 0.050276875 | 0.27033377 | 0.18598074 | 0.85245986 | 0.94037032 |
| ZNF852 | 9.89994681 | -0.07460018 | 0.41844729 | -0.1782786 | 0.85850422 | 0.94389652 |
| AC099669.1 | 80.2322157 | 0.43228916 | 0.26501716 | 1.63117423 | 0.10285356 | 0.33237057 |
| ZNF197 | 34.6307311 | -0.333863276 | 0.32617719 | -1.0235641 | 0.30604119 | 0.60076912 |
| KIAA1143 | 16.8362962 | -0.579144119 | 0.36401111 | -1.5910067 | 0.11160806 | 0.34842008 |
| ZDHHC3 | 104.623875 | 0.244041005 | 0.2699287 | 0.90409432 | 0.36594539 | 0.65142886 |
| EXOSC7 | 34.1189649 | 0.124596922 | 0.30461922 | 0.40902515 | 0.68252121 | 0.85544388 |
| CDCP1 | 70.7588118 | -1.878409378 | 0.36643767 | -5.1261361 | 2.96E-07 | 2.09E-05 |

| | | | | | | |
|---|---|---|---|---|---|---|
| LARS2 | 78.1389512 | 0.24998007 | 0.23563574 | 1.06087504 | 0.28874669 | 0.58064041 |
| LIMD1 | 60.8135556 | 0.404598867 | 0.25523186 | 1.58522087 | 0.11291616 | 0.35086028 |
| SACM1L | 77.7643846 | 0.46979301 | 0.25454451 | 1.84562228 | 0.06494707 | 0.25461247 |
| SLC6A20 | 17.4294605 | -4.380568772 | 0.79588531 | -5.5040201 | 3.71E-08 | 3.54E-06 |
| LZTFL1 | 19.9064654 | -0.058391377 | 0.32524895 | -0.1795283 | 0.85752293 | 0.94359207 |
| FYCO1 | 34.7471507 | -0.041650528 | 0.3522178 | -0.1182522 | 0.90586784 | 0.96157183 |
| LTF | 324.668173 | -5.992732032 | 1.23645221 | -4.8467154 | 1.26E-06 | 6.70E-05 |
| ALS2CL | 18.8726611 | -0.980066664 | 0.38442639 | -2.54927 | 0.01079487 | 0.08071931 |
| PTH1R | 132.61025 | 1.71316602 | 0.47603238 | 3.59884349 | 0.00031964 | 0.00617802 |
| CCDC12 | 106.163679 | 0.392518745 | 0.20615269 | 1.90401951 | 0.05690764 | 0.23559369 |
| NBEAL2 | 53.9937161 | 0.041937132 | 0.45103473 | 0.09297983 | 0.92591959 | 0.97111125 |
| SETD2 | 327.640462 | 0.387521833 | 0.17454468 | 2.22018702 | 0.02640607 | 0.14524018 |
| KIF9-AS1 | 109.86031 | 0.558594461 | 0.30050384 | 1.85885962 | 0.06304704 | 0.24979235 |
| KIF9 | 11.1315988 | -0.79641712 | 0.43322378 | -1.8383504 | 0.06601078 | 0.25681385 |
| KLHL18 | 41.485204 | 0.231437687 | 0.31133102 | 0.74338139 | 0.45725081 | 0.72348301 |
| PTPN23 | 25.7786424 | -0.187921789 | 0.35886571 | -0.5236549 | 0.60051858 | 0.80738812 |
| SCAP | 60.2450193 | 0.462490505 | 0.25619799 | 1.80520739 | 0.07104223 | 0.26907809 |
| ELP6 | 21.2975108 | -0.200983263 | 0.44982953 | -0.4467987 | 0.65502039 | 0.83931753 |
| SMARCC1 | 275.190154 | -0.313242552 | 0.26674982 | -1.1742934 | 0.24027752 | 0.53107467 |
| DHX30 | 53.6975193 | 0.446322263 | 0.22399868 | 1.9925218 | 0.04631384 | 0.20672666 |
| MAP4 | 530.282174 | 0.255600226 | 0.26642475 | 0.95937117 | 0.3373718 | 0.62600941 |
| ZNF589 | 19.2955347 | 0.445109928 | 0.31821516 | 1.39877034 | 0.16188186 | 0.42937872 |
| NME6 | 15.5120564 | -0.652827844 | 0.41276312 | -1.5816041 | 0.11373998 | 0.35229984 |
| PLXNB1 | 101.928165 | -0.029329496 | 0.29271164 | -0.1001993 | 0.92018612 | 0.96801362 |
| TMA7 | 64.8779665 | 0.095320987 | 0.42860109 | 0.22240024 | 0.82400232 | 0.92635145 |
| SHISA5 | 92.2681227 | -1.452456764 | 0.41866792 | -3.4692335 | 0.00052195 | 0.00888774 |
| PFKFB4 | 40.4163759 | -2.651820539 | 0.70874733 | -3.7415598 | 0.00018288 | 0.00390072 |
| UQCRC1 | 52.8289917 | 0.36278009 | 0.35662417 | 1.01726163 | 0.30902898 | 0.6027062 |
| NCKIPSD | 16.5291496 | -0.230998644 | 0.37639045 | -0.6137208 | 0.53939985 | 0.77077724 |
| IP6K2 | 79.4950132 | -0.146830528 | 0.2409765 | -0.6093147 | 0.54231585 | 0.77243948 |
| PRKAR2A | 124.806442 | 0.358848299 | 0.26843802 | 1.3368013 | 0.18128751 | 0.45775095 |
| SLC25A20 | 14.3806389 | 1.468587492 | 0.53674671 | 2.73609037 | 0.00621739 | 0.05545593 |
| ARIH2 | 103.022549 | 0.249694131 | 0.2326256 | 1.07337341 | 0.28310361 | 0.57570734 |
| P4HTM | 34.2370627 | -0.143970636 | 0.34715266 | -0.4147185 | 0.67834797 | 0.85316374 |
| WDR6 | 65.9660942 | 0.316769976 | 0.34484091 | 0.91859744 | 0.35830617 | 0.64216076 |
| DALRD3 | 16.8351161 | 0.253976586 | 0.36126775 | 0.70301482 | 0.48204651 | 0.74068045 |
| NDUFAF3 | 21.7738019 | -0.648916569 | 0.52972471 | -1.2250072 | 0.22057252 | 0.5089143 |
| IMPDH2 | 12.9267618 | 0.238536377 | 0.39300297 | 0.6069582 | 0.54387865 | 0.77347293 |
| QRICH1 | 71.584189 | 0.042527831 | 0.2233488 | 0.19040994 | 0.84898791 | 0.93897371 |
| QARS | 21.4909736 | 0.320284282 | 0.37649164 | 0.85070756 | 0.39493182 | 0.67539635 |
| USP19 | 12.5374909 | 0.526150894 | 0.43013465 | 1.22322369 | 0.22124522 | 0.50910615 |
| LAMB2 | 35.8226414 | 0.54075054 | 0.28504605 | 1.89706379 | 0.05781952 | 0.23769222 |
| USP4 | 88.5097606 | 0.177808479 | 0.26183453 | 0.67908722 | 0.4970826 | 0.74900468 |
| RHOA | 159.828136 | -0.800140195 | 0.32285507 | -2.4783263 | 0.01320004 | 0.09282375 |
| TCTA | 18.7186539 | -0.127639209 | 0.44636781 | -0.2859507 | 0.77491584 | 0.90205458 |
| DAG1 | 68.6352927 | 0.204968501 | 0.26094674 | 0.78548022 | 0.43217206 | 0.70480237 |
| APEH | 23.1003189 | 0.205552161 | 0.54130784 | 0.37973247 | 0.70414402 | 0.86617744 |
| MST1 | 17.7129146 | 0.79983807 | 0.39184687 | 2.04120061 | 0.04123089 | 0.1934247 |
| RNF123 | 16.5563764 | -0.066844036 | 0.37385591 | -0.1787963 | 0.85809769 | 0.94359207 |
| IP6K1 | 118.135543 | 0.157458991 | 0.30465066 | 0.51685098 | 0.60526019 | 0.81044694 |
| RBM6 | 418.211879 | 0.268272999 | 0.19882614 | 1.34928437 | 0.17724564 | 0.45197639 |
| RBM5 | 122.047449 | 0.407217726 | 0.20600005 | 1.97678463 | 0.04806598 | 0.21206669 |
| SEMA3F-AS1 | 14.6094584 | -0.845284872 | 0.8506676 | -0.9936724 | 0.32038241 | 0.6120836 |
| SEMA3F | 65.058006 | -0.504591633 | 0.49652874 | -1.0162385 | 0.30951581 | 0.60299393 |
| GNAI2 | 49.7055047 | -1.198595058 | 0.36459212 | -3.2874958 | 0.00101083 | 0.01485526 |
| SEMA3B | 81.5371508 | -0.653650093 | 0.57873874 | -1.129439 | 0.25871269 | 0.55002264 |
| IFRD2 | 14.4899962 | -0.366400779 | 0.38056055 | -0.9627923 | 0.33565175 | 0.62382904 |
| HYAL1 | 20.5959371 | 0.634154612 | 0.53878572 | 1.17700709 | 0.23919268 | 0.52947206 |
| HYAL2 | 12.5891243 | -0.152759174 | 0.60377094 | -0.2530085 | 0.80026165 | 0.91455697 |
| TUSC2 | 15.4348284 | -0.031512242 | 0.49859015 | -0.0632027 | 0.9496051 | 0.98003064 |
| TMEM115 | 14.8476081 | -0.421377556 | 0.42948632 | -0.9811199 | 0.32653364 | 0.61655967 |
| CACNA2D2 | 33.8860779 | 0.138326823 | 0.62004818 | 0.22309044 | 0.82346512 | 0.92608229 |
| HEMK1 | 16.4751608 | 0.593060044 | 0.35267016 | 1.68162808 | 0.09264098 | 0.31323946 |
| CISH | 39.5924965 | -0.163706018 | 0.38125324 | -0.4293892 | 0.66764002 | 0.84548586 |
| MAPKAPK3 | 37.5795916 | -0.037356721 | 0.42138566 | -0.0886521 | 0.92935841 | 0.97220251 |
| DOCK3 | 54.2013392 | -0.213767219 | 0.70189505 | -0.3045572 | 0.76070339 | 0.89645005 |
| MANF | 30.9351272 | -0.267469786 | 0.57821546 | -0.4625781 | 0.64366684 | 0.83339931 |
| RBM15B | 11.5798567 | -1.045593886 | 0.52682235 | -1.9847182 | 0.04717584 | 0.20957546 |
| DCAF1 | 56.7650123 | 0.170123197 | 0.36543927 | 0.46553069 | 0.64155146 | 0.8325919 |
| RAD54L2 | 70.7005811 | 0.802792149 | 0.28583403 | 2.80859545 | 0.00497581 | 0.04737599 |
| TEX264 | 38.4723644 | -0.561476511 | 0.45877269 | -1.2238665 | 0.2210026 | 0.5089143 |
| RPL29 | 217.310898 | -1.172628329 | 0.42959832 | -2.7295924 | 0.00634127 | 0.0562685 |
| ALAS1 | 35.5136977 | -0.925393052 | 0.56584344 | -1.6354224 | 0.10196053 | 0.33062554 |
| WDR82 | 62.9622494 | -0.247759259 | 0.20772802 | -1.1927099 | 0.23298302 | 0.52282195 |
| BAP1 | 19.7711138 | 0.004611831 | 0.34759487 | 0.01326784 | 0.98941411 | 0.99637773 |
| PHF7 | 18.7116072 | 0.560933164 | 0.49169229 | 1.14082156 | 0.25394419 | 0.5452549 |
| NISCH | 72.1095523 | 0.390308004 | 0.23671273 | 1.64886783 | 0.09917471 | 0.32535676 |
| SMIM4 | 39.5012788 | -0.256894043 | 0.32232851 | -0.7969945 | 0.42545424 | 0.70016446 |
| PBRM1 | 173.765664 | -0.093010289 | 0.27424675 | -0.3391482 | 0.7344981 | 0.88376541 |
| GNL3 | 23.8404572 | -0.493191652 | 0.36880991 | -1.3372516 | 0.18114052 | 0.45756661 |
| GLT8D1 | 18.3820335 | -0.084618795 | 0.38249243 | -0.22123 | 0.82491336 | 0.92670693 |
| SPCS1 | 68.6811629 | -0.818974318 | 0.41755124 | -1.9613744 | 0.04983536 | 0.2170841 |
| NEK4 | 45.5712416 | 0.383618606 | 0.23402885 | 1.63919367 | 0.10117293 | 0.32900364 |
| SFMBT1 | 37.6303001 | 0.602975064 | 0.29059339 | 2.07497859 | 0.03798852 | 0.18260776 |
| AC096887.1 | 10.5475884 | 0.793960326 | 0.68601085 | 1.15735827 | 0.24712602 | 0.53778881 |
| PRKCD | 14.5075977 | 0.410041694 | 0.47179423 | 0.8691113 | 0.38478626 | 0.66701235 |
| TKT | 93.2002652 | -1.333967786 | 0.37391449 | -3.5675744 | 0.0003603 | 0.00673652 |
| DCP1A | 110.054315 | 0.007875419 | 0.20787639 | 0.03788511 | 0.96977929 | 0.98784364 |
| CACNA1D | 172.860661 | -2.100703208 | 0.57375776 | -3.6613068 | 0.00025093 | 0.00515988 |
| CHDH | 34.2100362 | -0.923504839 | 0.38224371 | -2.4160105 | 0.01569161 | 0.10320828 |
| ACTR8 | 13.0320172 | -0.069660413 | 0.4690252 | -0.1485217 | 0.88193107 | 0.95437541 |
| SELENOK | 28.2020321 | -0.698999811 | 0.37207993 | -1.8786281 | 0.06029529 | 0.24330674 |
| CACNA2D3 | 188.358524 | -1.024823779 | 0.60062247 | -1.7062695 | 0.08795792 | 0.30539182 |
| ERC2 | 68.7361306 | -0.654728612 | 0.34260346 | -1.9110391 | 0.05599955 | 0.23384977 |
| CCDC66 | 108.000013 | 0.38644372 | 0.24300714 | 1.59025666 | 0.11177696 | 0.34846418 |
| FAM208A | 123.044129 | 0.221902869 | 0.25322567 | 0.8763048 | 0.38086435 | 0.66423981 |
| ARHGEF3 | 832.392286 | -0.699537042 | 0.24167972 | -2.8944797 | 0.00379787 | 0.03881518 |
| SPATA12 | 18.2528998 | -1.794969588 | 0.63306641 | -2.8353575 | 0.00457744 | 0.04461383 |
| APPL1 | 70.4230455 | -0.022072367 | 0.24227081 | -0.0911062 | 0.92740823 | 0.97111125 |
| PDE12 | 18.0658236 | 0.004886954 | 0.42861529 | 0.01140173 | 0.99090294 | 0.99690254 |
| ARF4 | 82.6007433 | -0.365382724 | 0.37123666 | -0.9842313 | 0.32500182 | 0.61535668 |
| DENND6A | 87.639841 | -0.173203933 | 0.23743531 | -0.7294784 | 0.46570908 | 0.72890726 |
| DENND6A-A! | 19.0830323 | -0.804425152 | 0.52980852 | -1.5183318 | 0.12893076 | 0.37627329 |
| SLMAP | 292.653423 | 0.207189146 | 0.20674867 | 1.00213051 | 0.31628056 | 0.60951749 |
| FLNB | 267.747504 | 0.519581237 | 0.28918334 | 1.79671915 | 0.07238022 | 0.27172106 |
| RPP14 | 10.1947894 | -0.338338747 | 0.49143475 | -0.6884714 | 0.49115601 | 0.74574638 |
| PXK | 85.6162625 | 0.817086007 | 0.33856315 | 2.41339318 | 0.01580476 | 0.1036414 |
| PDHB | 26.0531422 | -0.225039647 | 0.29043795 | -0.7748287 | 0.43844092 | 0.70985672 |
| KCTD6 | 20.8768132 | 0.175605624 | 0.51758659 | 0.33927777 | 0.73440049 | 0.88376541 |
| FAM107A | 75.3821748 | 0.154381846 | 0.65639377 | 0.235197 | 0.81405583 | 0.92107668 |
| FAM3D-AS1 | 18.7701069 | 1.186410116 | 0.79084498 | 1.50018037 | 0.13356769 | 0.38393256 |
| C3orf67 | 18.0453918 | 0.033315743 | 0.62894976 | 0.05297044 | 0.95775546 | 0.98321101 |
| FHIT | 1838.25288 | 0.66225775 | 0.31554964 | 2.09874349 | 0.03583952 | 0.17565115 |

| | | | | | | |
|---|---|---|---|---|---|---|
| PTPRG | 1154.92131 | -0.876864093 | 0.2927514 | -2.9952516 | 0.00274219 | 0.03080697 |
| THOC7 | 34.7609372 | -0.040593909 | 0.33408939 | -0.1215061 | 0.90329016 | 0.96126516 |
| ATXN7 | 21.7442808 | -0.820522387 | 0.33383717 | -2.4578521 | 0.01397707 | 0.09618691 |
| PSMD6-AS2 | 15.9185353 | -0.093370975 | 0.50268296 | -0.1857453 | 0.85264454 | 0.9404651 |
| PSMD6 | 19.6964917 | 0.735748194 | 0.35710032 | 2.06034035 | 0.03936601 | 0.18723393 |
| PRICKLE2 | 345.274815 | 0.592769111 | 0.25043424 | 2.36696508 | 0.01793462 | 0.1127288 |
| ADAMTS9 | 506.097547 | -0.336431666 | 0.34057003 | -0.9878487 | 0.32322674 | 0.61400627 |
| ADAMTS9-A! | 77.5880461 | 0.252530241 | 0.5235873 | 0.48230781 | 0.6295873 | 0.82508693 |
| ADAMTS9-A! | 1486.48657 | 0.508458622 | 0.37682843 | 1.34931069 | 0.17723719 | 0.45197639 |
| MAGI1 | 1595.51061 | 0.159163315 | 0.16798717 | 0.94747303 | 0.34339779 | 0.63107829 |
| MAGI1-AS1 | 24.1365913 | -0.993697939 | 0.74721785 | -1.3298638 | 0.18356316 | 0.46016652 |
| SLC25A26 | 90.4996416 | 0.338481016 | 0.27323653 | 1.23878393 | 0.21542553 | 0.50209938 |
| LRIG1 | 27.5304687 | -0.190317254 | 0.34777659 | -0.54724 | 0.58421387 | 0.79631799 |
| SUCLG2 | 264.170914 | 0.374398491 | 0.15504017 | 2.41484827 | 0.01574177 | 0.10342581 |
| SUCLG2-AS1 | 145.604868 | 1.027058713 | 0.39089615 | 2.6274465 | 0.00860284 | 0.06929353 |
| FAM19A1 | 47.6007554 | 1.290054007 | 0.47044708 | 2.74218729 | 0.00610315 | 0.05491239 |
| EOGT | 13.695152 | -1.005654168 | 0.42233952 | -2.381151 | 0.01725864 | 0.10994649 |
| TMF1 | 88.5226904 | -0.412428904 | 0.22584813 | -1.8261338 | 0.06783011 | 0.26152164 |
| UBA3 | 53.7118418 | 0.030842457 | 0.23570804 | 0.13085026 | 0.89589377 | 0.95900099 |
| ARL6IP5 | 54.2715197 | -1.123531991 | 0.35826045 | -3.136076 | 0.00171225 | 0.02191252 |
| FRMD4B | 68.2874929 | -0.057846749 | 0.30448879 | -0.1899799 | 0.84932489 | 0.93897371 |
| MITF | 532.28882 | 1.260734311 | 0.26825048 | 4.69983989 | 2.60E-06 | 0.00012418 |
| FOXP1 | 722.361961 | 0.223979364 | 0.18439306 | 1.21468438 | 0.22448649 | 0.51340497 |
| EIF4E3 | 36.2331235 | -0.694574685 | 0.32419692 | -2.1424469 | 0.03215754 | 0.16473741 |
| RYBP | 139.436547 | -0.099808409 | 0.2589603 | -0.3854197 | 0.69992646 | 0.86376951 |
| SHQ1 | 61.4772266 | 0.082685382 | 0.20169257 | 0.40995751 | 0.68183712 | 0.85494315 |
| GXYLT2 | 17.0953731 | -0.856955793 | 0.53909518 | -1.5896187 | 0.11192078 | 0.34868346 |
| PPP4R2 | 125.180417 | 0.20658031 | 0.2390527 | 0.86416223 | 0.38749878 | 0.66964123 |
| PDZRN3 | 253.101624 | 1.583326919 | 0.38141573 | 4.15118409 | 3.31E-05 | 0.00099926 |
| CNTN3 | 57.5272465 | -0.972377766 | 0.46576734 | -2.0876899 | 0.03682582 | 0.17886825 |
| ZNF717 | 77.0970018 | 0.181070608 | 0.2950522 | 0.61369007 | 0.53942015 | 0.77077724 |
| ROBO2 | 21.1754067 | 0.96465386 | 0.58348223 | 1.65327033 | 0.09827586 | 0.32387648 |
| ROBO1 | 92.7034167 | 0.116822911 | 0.37906295 | 0.30818868 | 0.75793877 | 0.89462479 |
| GBE1 | 549.006447 | -0.792256333 | 0.54798684 | -1.445758 | 0.1482451 | 0.40845709 |
| VGLL3 | 14.3275898 | 0.061691724 | 0.52597768 | 0.11728962 | 0.90663055 | 0.96157183 |
| CHMP2B | 54.5928943 | -0.229183763 | 0.22224293 | -1.0312308 | 0.30243258 | 0.59607049 |
| CGGBP1 | 87.2169721 | -0.441804441 | 0.29774761 | -1.483822 | 0.13785616 | 0.39134587 |
| ZNF654 | 130.417811 | -0.119175479 | 0.22128397 | -0.5385635 | 0.59018805 | 0.79991018 |
| C3orf38 | 21.4212927 | -0.140926308 | 0.35171578 | -0.4006824 | 0.688654 | 0.8585881 |
| PROS1 | 15.0931321 | -3.138917171 | 0.6515445 | -4.8176559 | 1.45E-06 | 7.60E-05 |
| ARL13B | 45.5065054 | -0.348126533 | 0.25960924 | -1.3409636 | 0.17993228 | 0.45599996 |
| NSUN3 | 39.461464 | -0.098668819 | 0.25808571 | -0.3823103 | 0.70223123 | 0.86516671 |
| MTRNR2L12 | 236.239612 | -1.058849474 | 0.70718981 | -1.4972635 | 0.13432471 | 0.38514144 |
| ARL6 | 31.077059 | -0.220894892 | 0.34834893 | -0.6341196 | 0.52600276 | 0.76221787 |
| CRYBG3 | 210.747909 | -0.809236249 | 0.30251617 | -2.6750182 | 0.00747252 | 0.06323003 |
| RIOX2 | 52.452021 | -0.212833669 | 0.28336419 | -0.7510959 | 0.45259496 | 0.72046894 |
| CLDND1 | 26.8270455 | 0.10051051 | 0.34133451 | 0.29446337 | 0.76840385 | 0.89996549 |
| DCBLD2 | 59.5885512 | 0.285169138 | 0.24580499 | 1.16014381 | 0.24599026 | 0.53633686 |
| CMSS1 | 151.57897 | 0.114176013 | 0.21826604 | 0.5231048 | 0.60090133 | 0.80752484 |
| FILIP1L | 44.9937359 | -0.87487798 | 0.51808437 | -1.6886786 | 0.09128104 | 0.31117713 |
| TBC1D23 | 77.8736049 | -0.088748518 | 0.27339174 | -0.3246203 | 0.74546844 | 0.88808711 |
| NIT2 | 16.7947296 | 0.323173994 | 0.37954031 | 0.85148794 | 0.39449836 | 0.67532771 |
| TOMM70 | 49.8922688 | 0.645173563 | 0.23549975 | 2.73959339 | 0.00615152 | 0.05517739 |
| TFG | 104.154347 | 0.214813447 | 0.24792731 | 0.86643722 | 0.38625044 | 0.66831576 |
| SENP7 | 132.32161 | 0.190124342 | 0.2212624 | 0.8592709 | 0.39019107 | 0.67208635 |
| TRMT10C | 15.8712611 | -1.044176261 | 0.59245437 | -1.7624585 | 0.07799186 | 0.28421369 |
| PCNP | 86.9369852 | -0.18368959 | 0.23268108 | -0.7894479 | 0.42985027 | 0.70346584 |
| ZBTB11 | 36.2002582 | -0.137996572 | 0.34221106 | -0.4032499 | 0.6867644 | 0.8578934 |
| RPL24 | 86.0812047 | -0.368742109 | 0.34617453 | -1.0651913 | 0.28678936 | 0.57922771 |
| CEP97 | 20.9252049 | 0.759311808 | 0.3932645 | 1.93079166 | 0.05350882 | 0.22710266 |
| NFKBIZ | 277.728081 | -1.842864725 | 0.47314408 | -3.8949335 | 9.82E-05 | 0.00238099 |
| ZPLD1 | 249.622478 | -5.865840227 | 1.05894628 | -5.5393181 | 3.04E-08 | 3.10E-06 |
| ALCAM | 67.1043764 | -0.716844609 | 0.4280159 | -1.6748084 | 0.09397184 | 0.31595883 |
| CBLB | 305.298876 | 1.428527487 | 0.26199636 | 5.45247069 | 4.97E-08 | 4.61E-06 |
| LINC00882 | 54.5838066 | 0.651453451 | 0.50884304 | 1.28026406 | 0.20045228 | 0.48422157 |
| BBX | 426.043745 | -0.26990316 | 0.23463232 | -1.1503239 | 0.2500105 | 0.54143539 |
| CD47 | 104.447399 | -1.029580722 | 0.41645498 | -2.4722498 | 0.01342657 | 0.09373482 |
| IFT57 | 51.2436369 | -0.109304797 | 0.28781996 | -0.379768 | 0.70411767 | 0.86617744 |
| DZIP3 | 125.724777 | 1.382833038 | 0.52691132 | 2.62441323 | 0.00867984 | 0.06975017 |
| AC078980.1 | 33.1450764 | 3.345350907 | 0.67102869 | 4.98540664 | 6.18E-07 | 3.81E-05 |
| NECTIN3 | 29.0658441 | -0.491524282 | 0.45930146 | -1.0701561 | 0.28454903 | 0.57686567 |
| PHLDB2 | 417.909872 | 0.862043891 | 0.18757763 | 4.5956646 | 4.31E-06 | 0.00018862 |
| ABHD10 | 20.8301761 | -0.080023088 | 0.42460468 | -0.1884649 | 0.85051221 | 0.93972295 |
| C3orf52 | 60.2451506 | 0.348488867 | 0.38583422 | 0.90320882 | 0.36641508 | 0.65172793 |
| SLC9C1 | 38.6986927 | -1.533805642 | 0.78231166 | -1.9606069 | 0.0499249 | 0.2171986 |
| ATG3 | 31.299976 | 0.150519126 | 0.31987993 | 0.47054883 | 0.63796296 | 0.83056924 |
| SLC35A5 | 17.8766734 | -0.393749034 | 0.49183971 | -0.8005637 | 0.42338425 | 0.69817048 |
| GTPBP8 | 17.7651191 | -0.193270343 | 0.35957127 | -0.5375022 | 0.59092078 | 0.80019 |
| NEPRO | 42.8555589 | -0.184843556 | 0.29207355 | -0.6328665 | 0.52682085 | 0.76221787 |
| CFAP44 | 64.6601936 | -0.180064539 | 0.24901387 | -0.7231105 | 0.46961201 | 0.73178441 |
| SPICE1 | 50.6198009 | -0.292491063 | 0.2885932 | -1.0135064 | 0.31081832 | 0.60394276 |
| SIDT1 | 40.9572309 | -0.083989978 | 0.76848483 | -0.109293 | 0.91297013 | 0.96446045 |
| USF3 | 57.0108201 | 0.226058656 | 0.25161981 | 0.89841357 | 0.3689651 | 0.65369102 |
| NAA50 | 131.302383 | -0.063486879 | 0.26178147 | -0.2425186 | 0.80837834 | 0.91848612 |
| ATP6V1A | 257.01579 | 0.265551693 | 0.28669393 | 0.92625504 | 0.35431346 | 0.63963412 |
| GRAMD1C | 147.8664 | 0.481710983 | 0.32559647 | 1.47947237 | 0.13901411 | 0.39333242 |
| ZDHHC23 | 26.4571794 | -0.170963202 | 0.31099757 | -0.5497252 | 0.58250787 | 0.7955793 |
| CCDC191 | 58.4544249 | -0.386088257 | 0.41440571 | -0.9316673 | 0.35150848 | 0.63801604 |
| QTRT2 | 32.8006776 | -0.554756332 | 0.39673965 | -1.3982881 | 0.16202657 | 0.42953067 |
| ZBTB20 | 2839.63486 | 0.77424102 | 0.18803776 | 4.11747619 | 3.83E-05 | 0.00112836 |
| ZBTB20-AS5 | 51.2770931 | -0.709531861 | 0.64214574 | -1.104939 | 0.26918603 | 0.56148096 |
| LSAMP | 381.757718 | 1.914261818 | 0.6921863 | 2.76552975 | 0.00568304 | 0.05208781 |
| AC092691.1 | 131.638514 | 1.632986432 | 0.78086511 | 2.0912529 | 0.0365054 | 0.17761982 |
| AC068633.1 | 52.884024 | -1.94331318 | 1.25975056 | -1.5426174 | 0.12292363 | 0.36727355 |
| IGSF11 | 23.1183576 | 1.281876023 | 0.39787761 | 3.22178478 | 0.00127395 | 0.01754574 |
| B4GALT4 | 17.2557188 | -0.265672862 | 0.36378851 | -0.7302948 | 0.46521 | 0.72863709 |
| TMEM39A | 30.5412135 | -0.296101459 | 0.38663093 | -0.7658504 | 0.44376531 | 0.71343168 |
| POGLUT1 | 19.9225837 | 0.068854817 | 0.40057061 | 0.17189183 | 0.86352257 | 0.94635064 |
| TIMMDC1 | 46.8124908 | 0.147709771 | 0.37223576 | 0.39681779 | 0.69150183 | 0.85976938 |
| COX17 | 24.1209677 | 0.33821387 | 0.31605255 | 1.07011911 | 0.2845657 | 0.57686567 |
| MAATS1 | 43.8608344 | -0.129967831 | 0.34143718 | -0.3806493 | 0.70346348 | 0.86585351 |
| GSK3B | 353.735807 | 0.162978816 | 0.17389644 | 0.93721766 | 0.34864661 | 0.63548432 |
| LRRC58 | 49.9237915 | -0.233104541 | 0.28967522 | -0.80471 | 0.42098705 | 0.69661173 |
| FSTL1 | 12.3499547 | -0.621085895 | 0.64941773 | -0.9563735 | 0.33888358 | 0.62768404 |
| NDUFB4 | 44.5234439 | -0.078992671 | 0.30502047 | -0.258975 | 0.79565454 | 0.91195838 |
| HGD | 53.3024501 | -0.624858019 | 0.53033343 | -1.1782362 | 0.23870247 | 0.52895523 |
| RABL3 | 34.3281974 | -0.145542238 | 0.27685225 | -0.5257037 | 0.59909413 | 0.80678009 |
| GTF2E1 | 10.9160692 | 0.08721613 | 0.42619844 | 0.20463737 | 0.83785545 | 0.93487753 |
| GOLGB1 | 215.182829 | -0.429275875 | 0.24593045 | -1.7455174 | 0.08089485 | 0.29075313 |
| IQCB1 | 90.0128404 | 0.044217721 | 0.25557449 | 0.17301305 | 0.86264117 | 0.94612258 |
| EAF2 | 14.0065638 | 0.541465498 | 0.40271756 | 1.34452914 | 0.17877734 | 0.45419584 |
| SLC15A2 | 35.1997081 | 1.605906071 | 0.53301985 | 3.01284479 | 0.00258811 | 0.02942907 |
| ILDR1 | 17.9413199 | 0.034518461 | 0.40731043 | 0.08474731 | 0.93246229 | 0.97356162 |

| | | | | | | |
|---|---|---|---|---|---|---|
| CCDC58 | 15.9415694 | 0.120110878 | 0.4276682 | 0.28085062 | 0.77882498 | 0.90427469 |
| FAM162A | 104.843823 | -0.507047448 | 0.47934115 | -1.0578008 | 0.29014627 | 0.58187402 |
| AC083798.2 | 30.2348665 | 0.517462895 | 0.44379318 | 1.16600011 | 0.2436144 | 0.53443687 |
| KPNA1 | 110.236858 | -0.143161166 | 0.29837777 | -0.4797984 | 0.63137077 | 0.82625173 |
| PARP9 | 76.5261279 | -1.511694402 | 0.67051399 | -2.2545307 | 0.0241628 | 0.13587618 |
| DTX3L | 29.0511744 | -1.96734144 | 0.66941928 | -2.9388778 | 0.00329403 | 0.03481518 |
| PARP14 | 199.561462 | -1.050275134 | 0.78246613 | -1.3422627 | 0.17951082 | 0.45516133 |
| HSPBAP1 | 39.066594 | -0.30719138 | 0.37140615 | -0.8271037 | 0.40817831 | 0.68657965 |
| DIRC2 | 107.691602 | 0.271106583 | 0.53529308 | 0.50646383 | 0.61253108 | 0.81444181 |
| PDIA5 | 23.4991663 | -0.963553565 | 0.41692737 | -2.3110825 | 0.0208283 | 0.12353087 |
| SEC22A | 94.7824694 | -0.120406092 | 0.21868864 | -0.5505823 | 0.58192006 | 0.7952959 |
| ADCY5 | 14.0905596 | -1.27580179 | 0.53341011 | -2.391784 | 0.01676671 | 0.10800461 |
| HACD2 | 66.6994158 | 0.174679898 | 0.25624668 | 0.68168649 | 0.49543721 | 0.74770824 |
| MYLK-AS1 | 16.6293247 | 0.197945153 | 0.35919442 | 0.55108081 | 0.58157828 | 0.79525079 |
| MYLK | 23.9436526 | -0.680590397 | 0.66594005 | -1.0219995 | 0.30678114 | 0.60139222 |
| CCDC14 | 83.3701822 | 0.164955146 | 0.2696079 | 0.6118335 | 0.54064792 | 0.7714342 |
| KALRN | 150.87765 | -2.097818891 | 0.43690644 | -4.8015288 | 1.57E-06 | 8.11E-05 |
| UMPS | 19.8553871 | 0.157780015 | 0.34000072 | 0.46405789 | 0.64260627 | 0.83318468 |
| ITGB5 | 89.611579 | 0.68208156 | 0.23512239 | 2.90096393 | 0.00372017 | 0.03812562 |
| SLC12A8 | 27.4735478 | 0.86940116 | 0.45756138 | 1.90007549 | 0.05742321 | 0.23672427 |
| ZNF148 | 309.431 | 0.206840568 | 0.17429381 | 1.18673505 | 0.23533214 | 0.52547582 |
| SNX4 | 118.921973 | 0.274328318 | 0.24413111 | 1.1236926 | 0.26114346 | 0.55242297 |
| OSBPL11 | 42.7820193 | 0.150365652 | 0.27985086 | 0.53730638 | 0.591056 | 0.80019 |
| SLC41A3 | 19.6329473 | -0.035469821 | 0.38527627 | -0.0920633 | 0.92664771 | 0.97111125 |
| ALDH1L1 | 139.250295 | 0.138216818 | 0.43968561 | 0.31435375 | 0.75325238 | 0.89205725 |
| ZXDC | 138.354379 | -0.064766629 | 0.21490373 | -0.3013751 | 0.76312849 | 0.8974069 |
| TXNRD3NB | 11.222533 | 0.158184541 | 0.42973546 | 0.36809749 | 0.71280055 | 0.87134026 |
| TXNRD3 | 21.4461371 | 0.903315199 | 0.33149678 | 2.7249592 | 0.00643094 | 0.05672224 |
| CHCHD6 | 50.2596854 | 0.214469074 | 0.30989907 | 0.69206105 | 0.48889899 | 0.7453955 |
| PLXNA1 | 12.1739105 | -0.453062266 | 0.47664878 | -0.9505159 | 0.34185016 | 0.62990732 |
| TPRA1 | 16.226889 | 0.382560094 | 0.47050609 | 0.81308212 | 0.41617098 | 0.69285607 |
| PODXL2 | 9.60630978 | -0.185373689 | 0.70783117 | -0.2618897 | 0.79340648 | 0.91142199 |
| ABTB1 | 14.9206896 | 0.431514214 | 0.43854508 | 0.98396774 | 0.32513138 | 0.61537818 |
| MGLL | 249.571969 | -1.530355333 | 0.37174916 | -4.1166343 | 3.84E-05 | 0.00112836 |
| SEC61A1 | 77.2464531 | -0.525415994 | 0.34200769 | -1.5362695 | 0.12447229 | 0.36974123 |
| RUVBL1 | 35.9661608 | -0.055945906 | 0.38887719 | -0.1438652 | 0.88560689 | 0.95619452 |
| EEFSEC | 75.7525993 | 0.301779883 | 0.19095764 | 1.58034986 | 0.11402677 | 0.35265799 |
| GATA2 | 72.7641699 | 1.468119891 | 0.41962598 | 3.49863919 | 0.00046764 | 0.00819323 |
| GATA2-AS1 | 23.7733295 | -1.264550896 | 0.43373839 | -2.9154691 | 0.00355154 | 0.03699316 |
| RPN1 | 54.5101218 | -0.212321361 | 0.30006013 | -0.707596 | 0.47919615 | 0.73880257 |
| RAB7A | 197.117358 | -0.134612252 | 0.3085402 | -0.4362876 | 0.6626281 | 0.84255674 |
| ACAD9 | 41.2280661 | 0.264646933 | 0.24007106 | 1.10236916 | 0.27030121 | 0.56232696 |
| CNBP | 92.429742 | -0.728893827 | 0.43587634 | -1.6722491 | 0.09447521 | 0.3161038 |
| COPG1 | 35.1649162 | -0.168350771 | 0.45407554 | -0.370755 | 0.71082003 | 0.87028547 |
| HMCES | 26.7610591 | -0.107550722 | 0.37139514 | -0.2895857 | 0.77213321 | 0.90105761 |
| H1FX | 33.4963833 | -1.250133566 | 0.63529655 | -1.9677953 | 0.0490916 | 0.2150718 |
| MBD4 | 33.4001758 | -0.24562772 | 0.266516 | -0.9216247 | 0.35672439 | 0.64141201 |
| IFT122 | 36.9704035 | -0.165438438 | 0.3446802 | -0.4799766 | 0.63124402 | 0.82625173 |
| TMCC1 | 388.154116 | 0.280749941 | 0.14859723 | 1.889335 | 0.05884695 | 0.24000052 |
| PIK3R4 | 45.2284442 | 0.307487515 | 0.3014414 | 1.02005735 | 0.30770126 | 0.60139222 |
| ATP2C1 | 181.769849 | 0.188868699 | 0.21390102 | 0.8829724 | 0.37725118 | 0.66022373 |
| NEK11 | 125.391442 | -0.760092407 | 0.30633419 | -2.4812523 | 0.01309217 | 0.09231725 |
| NUDT16 | 20.7316346 | -0.503689719 | 0.45135392 | -1.1159529 | 0.26444229 | 0.55608991 |
| MRPL3 | 44.1400916 | 0.212664425 | 0.30840523 | 0.68956166 | 0.49046988 | 0.7455837 |
| DNAJC13 | 102.893826 | -0.091495704 | 0.24263989 | -0.3770843 | 0.70611093 | 0.86764728 |
| UBA5 | 86.1083651 | 0.474851058 | 0.32568371 | 1.45801292 | 0.14483698 | 0.40273129 |
| CDV3 | 94.1630486 | -1.14825618 | 0.43182506 | -2.6590772 | 0.0078355 | 0.06478436 |
| TOPBP1 | 38.76503 | -0.208100779 | 0.28629714 | -0.7268699 | 0.46730564 | 0.73023858 |
| SRPRB | 22.3925731 | 0.158882784 | 0.36938842 | 0.43012389 | 0.66710552 | 0.84497102 |
| SLCO2A1 | 47.7677702 | -1.020035089 | 0.77760136 | -1.3117712 | 0.18959734 | 0.46862566 |
| RYK | 168.924215 | -0.027245514 | 0.1889178 | -0.1442189 | 0.88532763 | 0.95619452 |
| AMOTL2 | 29.2024137 | 0.724003232 | 0.44481861 | 1.62763702 | 0.10360188 | 0.33425166 |
| ANAPC13 | 19.478147 | -0.511777002 | 0.46872157 | -1.0918572 | 0.27489589 | 0.56696009 |
| CEP63 | 52.227256 | 0.28087986 | 0.25358259 | 1.10764647 | 0.26801452 | 0.55994436 |
| EPHB1 | 34.1503668 | -0.372779766 | 0.44591097 | -0.835996 | 0.40315718 | 0.68156125 |
| PPP2R3A | 864.103323 | -0.271490088 | 0.18985898 | -1.4299565 | 0.1527295 | 0.41460325 |
| MSL2 | 54.6302219 | 0.028395028 | 0.28770674 | 0.09869434 | 0.92138096 | 0.9686919 |
| PCCB | 42.8930858 | 0.304690864 | 0.30500459 | 0.99897141 | 0.31780854 | 0.61101185 |
| STAG1 | 510.776116 | 0.197409707 | 0.35213373 | 0.56061004 | 0.5750634 | 0.79149007 |
| NCK1-DT | 12.9961566 | -0.441853249 | 0.41015907 | -1.0772729 | 0.28135837 | 0.57465383 |
| NCK1 | 54.8342673 | -0.338429163 | 0.33805706 | -1.0011007 | 0.31677812 | 0.61010436 |
| DZIP1L | 21.1676342 | 0.518563118 | 0.3883772 | 1.33520483 | 0.18180932 | 0.45839441 |
| DBR1 | 13.4781632 | 0.393524395 | 0.54838701 | 0.71760342 | 0.47300185 | 0.73375471 |
| ARMC8 | 122.30918 | 0.300242594 | 0.18612011 | 1.61316583 | 0.10670849 | 0.34029982 |
| NME9 | 19.0082243 | -1.045173493 | 0.57805141 | -1.8080978 | 0.07059128 | 0.26815316 |
| MRAS | 39.3063164 | -0.553663636 | 0.30034924 | -1.8433995 | 0.0652707 | 0.25528428 |
| CEP70 | 137.599997 | 0.423748448 | 0.27508163 | 1.54044621 | 0.12345163 | 0.36821038 |
| FAIM | 15.7343257 | -0.680882256 | 0.3810149 | -1.7870227 | 0.07393383 | 0.2750634 |
| PIK3CB | 164.593533 | 0.572790735 | 0.19504672 | 2.93668482 | 0.00331741 | 0.03494475 |
| MRPS22 | 11.8642839 | 0.266338347 | 0.4955493 | 0.53746084 | 0.59094933 | 0.80019 |
| COPB2 | 74.5432759 | -0.019351477 | 0.21157837 | -0.0914625 | 0.92712514 | 0.97111125 |
| AC097103.2 | 77.1036996 | 0.392334793 | 0.41261305 | 0.95085406 | 0.34167846 | 0.62980346 |
| NMNAT3 | 45.2976276 | 0.752729545 | 0.34113464 | 2.20654682 | 0.02734574 | 0.14777597 |
| CLSTN2 | 31.4280056 | 0.213412614 | 0.77775918 | 0.27439421 | 0.78378169 | 0.90714343 |
| SLC25A36 | 165.565349 | 0.118704763 | 0.24887583 | 0.47696382 | 0.63338789 | 0.82712156 |
| PXYLP1 | 20.9290597 | -0.601411768 | 0.38207072 | -1.5740849 | 0.11546782 | 0.35464129 |
| AC117383.1 | 19.1625344 | -1.803910274 | 0.51466152 | -3.5050421 | 0.00045654 | 0.00807435 |
| ZBTB38 | 220.73325 | -1.256452477 | 0.38370363 | -3.274539 | 0.00105835 | 0.01528426 |
| RASA2 | 203.416636 | -0.598778561 | 0.30576734 | -1.9582816 | 0.05019698 | 0.21810598 |
| RNF7 | 38.8620755 | -0.847741199 | 0.34025175 | -2.4915117 | 0.01272008 | 0.09058011 |
| ATP1B3 | 409.369715 | 2.120165535 | 0.76741356 | 2.76274184 | 0.00573181 | 0.05239493 |
| TFDP2 | 337.258426 | 0.97483149 | 0.29772367 | 3.27428282 | 0.00105931 | 0.01528426 |
| GK5 | 101.916422 | 0.451343907 | 0.20064239 | 2.24949422 | 0.02448107 | 0.13684427 |
| XRN1 | 236.589222 | 0.248674543 | 0.37264833 | 0.66731695 | 0.5045697 | 0.75160166 |
| ATR | 155.11533 | -0.258495404 | 0.30376137 | -0.8509818 | 0.39477946 | 0.67534117 |
| PLS1 | 70.5600131 | 0.104786262 | 0.42575844 | 0.2461167 | 0.8055919 | 0.91703953 |
| TRPC1 | 77.5040138 | -0.048596083 | 0.36281506 | -0.1339417 | 0.89344864 | 0.95800753 |
| U2SURP | 137.247909 | 0.385161056 | 0.19913423 | 1.93417804 | 0.05309124 | 0.22595384 |
| SLC9A9 | 15.8071396 | -0.529927108 | 0.72533135 | -0.7306 | 0.46502349 | 0.72850852 |
| C3orf58 | 21.1107675 | -0.169185037 | 0.41553262 | -0.4071522 | 0.68389618 | 0.85606265 |
| PLOD2 | 149.857195 | -0.862394327 | 0.42064628 | -2.0501651 | 0.04034832 | 0.19067357 |
| PLSCR4 | 37.5311135 | -0.331625284 | 0.43081066 | -0.7697704 | 0.44143612 | 0.71223757 |
| PLSCR2 | 46.153944 | -0.298449402 | 0.46440256 | -0.6426524 | 0.52044969 | 0.76073677 |
| PLSCR1 | 211.330994 | -0.817863714 | 0.43435628 | -1.8829328 | 0.05970947 | 0.24198543 |
| GYG1 | 14.2249255 | -1.124285692 | 0.4577844 | -2.4559284 | 0.01405212 | 0.09623211 |
| HLTF | 36.9931838 | 0.379141392 | 0.40243238 | 0.94212446 | 0.34612892 | 0.63324812 |
| HPS3 | 43.6575204 | -0.165059193 | 0.32508406 | -0.5077431 | 0.61163351 | 0.81385911 |
| WWTR1 | 526.316266 | 0.619086126 | 0.44574376 | 1.38888344 | 0.1648682 | 0.43375609 |
| COMMD2 | 26.313973 | -0.230528317 | 0.2894513 | -0.7964321 | 0.42578091 | 0.70025461 |
| ANKUB1 | 55.8692991 | -0.498992303 | 0.4963891 | -1.0052443 | 0.31477923 | 0.60818748 |
| RNF13 | 138.918937 | 0.077084547 | 0.17430638 | 0.44223594 | 0.65831848 | 0.84037209 |
| AC117386.2 | 71.014078 | 2.548106669 | 0.79870644 | 3.19029187 | 0.00142129 | 0.01901578 |
| TSC22D2 | 272.69625 | 0.859457804 | 0.27895487 | 3.08099233 | 0.00206312 | 0.02511125 |

| | | | | | | |
|---|---|---|---|---|---|---|
| SERP1 | 109.283935 | -0.396969199 | 0.33504684 | -1.184817 | 0.2360898 | 0.52644622 |
| EIF2A | 42.8177597 | -0.183533847 | 0.31095304 | -0.5902301 | 0.55503639 | 0.77954179 |
| SELENOT | 55.3317523 | -0.11152995 | 0.33920002 | -0.3288029 | 0.74230467 | 0.88720442 |
| SIAH2 | 21.3040907 | -0.006369119 | 0.46989491 | -0.0135543 | 0.98918553 | 0.99637773 |
| MED12L | 30.0674963 | 0.082479129 | 0.47627115 | 0.17317683 | 0.86251243 | 0.9460821 |
| SUCNR1 | 41.1165405 | 1.035845704 | 0.72022756 | 1.43822002 | 0.15037164 | 0.41132271 |
| MBNL1 | 523.850606 | 0.538403957 | 0.22403428 | 2.4032213 | 0.01625135 | 0.10549553 |
| ARHGEF26 | 18.7768893 | -0.083608886 | 0.46774769 | -0.1787478 | 0.85813571 | 0.94359207 |
| DHX36 | 141.112139 | 0.449320997 | 0.21931105 | 2.0487841 | 0.04048323 | 0.19104821 |
| PLCH1 | 82.6158947 | -0.679308263 | 0.58487791 | -1.1614531 | 0.24545769 | 0.53619701 |
| GMPS | 56.6417558 | 0.222070774 | 0.23380689 | 0.94980422 | 0.34221174 | 0.63022262 |
| SSR3 | 46.7587813 | 0.007991832 | 0.28876951 | 0.02767547 | 0.97792099 | 0.99119855 |
| TIPARP | 21.5697587 | -0.427875156 | 0.36538382 | -1.1710293 | 0.24158701 | 0.5324541 |
| LINC00886 | 46.1883962 | 0.351168671 | 0.5076169 | 0.69179863 | 0.4890638 | 0.74542614 |
| LEKR1 | 26.759775 | 1.145345708 | 0.52351908 | 2.18778218 | 0.02868547 | 0.15286634 |
| CCNL1 | 159.578931 | 0.459124318 | 0.3629048 | 1.26513708 | 0.20582219 | 0.48999801 |
| VEPH1 | 152.539413 | 1.951292959 | 0.63107989 | 3.0919904 | 0.00198819 | 0.02447116 |
| RSRC1 | 170.258595 | 0.292807696 | 0.21084528 | 1.38873249 | 0.16491411 | 0.43375609 |
| MLF1 | 22.3909674 | -0.462243965 | 0.46034687 | -1.004121 | 0.31532029 | 0.6086621 |
| GFM1 | 46.6704194 | 0.429384076 | 0.36098304 | 1.18948545 | 0.23424869 | 0.52443761 |
| LXN | 11.8551459 | 0.087129349 | 0.55537614 | 0.15688349 | 0.87533667 | 0.95023376 |
| RARRES1 | 13.6168031 | -1.216584077 | 0.56765614 | -2.1431708 | 0.03209938 | 0.16452131 |
| MFSD1 | 22.2777605 | -0.094406047 | 0.33978875 | -0.2778375 | 0.78113712 | 0.90520525 |
| IQCJ-SCHIP1 | 80.5216105 | 0.733111808 | 0.63666457 | 1.15148831 | 0.2495314 | 0.54083745 |
| SMC4 | 26.6015192 | -0.297478109 | 0.3011668 | -0.987752 | 0.32327412 | 0.61400627 |
| KPNA4 | 130.089057 | -0.035447062 | 0.32478899 | -0.1091387 | 0.91309245 | 0.96449076 |
| PPM1L | 134.064595 | 1.339555194 | 0.3725345 | 3.59578827 | 0.00032341 | 0.006216 |
| B3GALNT1 | 13.7229709 | 0.909474914 | 0.41546071 | 2.1890756 | 0.02859135 | 0.15245758 |
| NMD3 | 106.023168 | 0.655960896 | 0.37916542 | 1.73001243 | 0.08362805 | 0.29623459 |
| LINC01322 | 17.1553522 | 0.587343005 | 0.52469983 | 1.1193886 | 0.26297439 | 0.55458796 |
| PDCD10 | 72.8820516 | -0.07541825 | 0.21414312 | -0.3521862 | 0.72469863 | 0.8779216 |
| GOLIM4 | 57.6323722 | 0.508515339 | 0.36448244 | 1.39517101 | 0.16296428 | 0.4311397 |
| MECOM | 5670.2243 | 0.464379136 | 0.27213976 | 1.70639943 | 0.08793374 | 0.30539182 |
| AC074033.1 | 22.3314022 | -0.567581428 | 0.62867221 | -0.9028257 | 0.36661841 | 0.65175201 |
| MYNN | 17.4389336 | 0.195149618 | 0.40398285 | 0.48306412 | 0.62905021 | 0.8249932 |
| LRRC34 | 10.7964591 | -0.02649264 | 0.47765066 | -0.0554645 | 0.95576844 | 0.98276539 |
| SEC62 | 138.800306 | 0.100390373 | 0.22410314 | 0.44796503 | 0.65417843 | 0.83885315 |
| GPR160 | 103.554389 | -0.071848395 | 0.24669656 | -0.291242 | 0.77086626 | 0.90076568 |
| PHC3 | 235.137458 | 0.367588361 | 0.16293624 | 2.25602579 | 0.02406902 | 0.13578849 |
| PRKCI | 135.090378 | 0.018894081 | 0.30094587 | 0.06278232 | 0.94993984 | 0.98003064 |
| SKIL | 82.9897879 | -0.067661085 | 0.42557761 | -0.1589865 | 0.87367953 | 0.94943205 |
| TNIK | 67.6040464 | -2.602909436 | 0.83577634 | -3.1143612 | 0.00184344 | 0.02310351 |
| PLD1 | 60.6042507 | -0.575677944 | 0.36195876 | -1.5904518 | 0.11173301 | 0.34846418 |
| FNDC3B | 1261.9181 | -0.423635831 | 0.20987546 | -2.0185105 | 0.04353812 | 0.19925797 |
| TNFSF10 | 139.061181 | -3.404049337 | 0.62865338 | -5.4148271 | 6.13E-08 | 5.50E-06 |
| NCEH1 | 126.627718 | -0.998473434 | 0.36896075 | -2.7061779 | 0.00680626 | 0.0591532 |
| ECT2 | 27.636719 | -1.662662405 | 0.40403837 | -4.1151102 | 3.87E-05 | 0.00113262 |
| NLGN1 | 1083.19283 | -3.265086562 | 0.57791415 | -5.6497779 | 1.61E-08 | 1.97E-06 |
| NAALADL2 | 1641.35317 | 0.325447826 | 0.29653923 | 1.09748658 | 0.27242874 | 0.56479948 |
| TBL1XR1 | 656.374365 | -1.006560753 | 0.42544875 | -2.3658801 | 0.01798727 | 0.11284999 |
| ZMAT3 | 47.7935058 | -0.325810375 | 0.36310584 | -0.8972876 | 0.36956546 | 0.65416897 |
| PIK3CA | 110.277076 | -0.143654555 | 0.22566515 | -0.6365828 | 0.52439663 | 0.76221787 |
| KCNMB3 | 18.5386776 | -0.257353462 | 0.43278968 | -0.5946386 | 0.55208506 | 0.77741728 |
| ZNF639 | 15.7132018 | 0.037619346 | 0.36406818 | 0.1033305 | 0.91770067 | 0.9663858 |
| MFN1 | 121.540429 | 0.355664704 | 0.19894821 | 1.78772511 | 0.07382037 | 0.27474619 |
| GNB4 | 17.9280516 | 0.008594413 | 0.53122265 | 0.01617855 | 0.98709195 | 0.99523554 |
| ACTL6A | 23.0747868 | -0.113723099 | 0.33484619 | -0.3396279 | 0.73413679 | 0.88374353 |
| MRPL47 | 23.9670055 | -0.211762688 | 0.38936581 | -0.5438656 | 0.58653393 | 0.79752689 |
| NDUFB5 | 58.2210799 | 0.132619431 | 0.34696492 | 0.38222721 | 0.70229284 | 0.86516671 |
| USP13 | 14.9527938 | -0.061134447 | 0.41942704 | -0.145757 | 0.88411319 | 0.9560338 |
| TTC14 | 148.796194 | 0.730111761 | 0.19279104 | 3.78706269 | 0.00015244 | 0.00343637 |
| FXR1 | 143.21555 | 0.697184527 | 0.26237892 | 2.65716668 | 0.00788005 | 0.06510044 |
| DNAJC19 | 20.4214987 | -0.185037079 | 0.39013214 | -0.4742934 | 0.63529073 | 0.82866107 |
| ATP11B | 160.468139 | 0.1691437 | 0.25190756 | 0.67145146 | 0.50193297 | 0.75108109 |
| DCUN1D1 | 96.2071326 | 0.24467272 | 0.23359021 | 1.04744424 | 0.29489474 | 0.58668972 |
| MCCC1 | 125.283215 | 0.442945379 | 0.23094012 | 1.91800961 | 0.05510979 | 0.23163651 |
| KLHL24 | 159.667709 | -0.383923258 | 0.29535039 | -1.2998908 | 0.19363839 | 0.4740909 |
| YEATS2 | 83.968181 | -0.400094726 | 0.36656023 | -1.0914843 | 0.27505984 | 0.56696009 |
| ABCC5 | 245.592528 | 0.003738253 | 0.28035572 | 0.01333397 | 0.98936135 | 0.99637773 |
| EIF2B5 | 20.7574299 | 0.128535704 | 0.36076464 | 0.35628687 | 0.72162574 | 0.87625492 |
| DVL3 | 32.0809547 | -0.16564737 | 0.35094954 | -0.4719977 | 0.63692844 | 0.82989931 |
| AP2M1 | 68.5232363 | -0.357001178 | 0.37714541 | -0.9465876 | 0.34384895 | 0.63176955 |
| PSMD2 | 29.6649961 | -0.076617573 | 0.30930533 | -0.2477085 | 0.80435992 | 0.91643337 |
| EIF4G1 | 69.947217 | -0.517090357 | 0.3364395 | -1.536949 | 0.12430579 | 0.36936783 |
| FAM131A | 14.3837138 | 0.176994205 | 0.55494 | 0.31894296 | 0.74976977 | 0.89049206 |
| POLR2H | 44.7320062 | 0.254459622 | 0.29286701 | 0.86885723 | 0.38492523 | 0.66701235 |
| MAGEF1 | 16.5671705 | -0.644994278 | 0.44080322 | -1.463225 | 0.14340583 | 0.4006962 |
| VPS8 | 149.250323 | 0.127188873 | 0.26385546 | 0.48203996 | 0.62977756 | 0.82512634 |
| EHHADH | 41.2459326 | 0.591367788 | 0.40215042 | 1.47051393 | 0.14142262 | 0.39752682 |
| MAP3K13 | 189.695948 | 0.298087168 | 0.25809164 | 1.15496637 | 0.2481042 | 0.53889299 |
| TMEM41A | 27.3391302 | -0.720016867 | 0.33980517 | -2.1189109 | 0.034098 | 0.17085487 |
| LIPH | 53.4456726 | 0.915445431 | 0.40238281 | 2.27506097 | 0.02290227 | 0.13210482 |
| SENP2 | 68.683378 | 0.267855639 | 0.28142134 | 0.95179578 | 0.34120056 | 0.62956442 |
| IGF2BP2 | 273.335376 | 0.31485348 | 0.32948079 | 0.95560498 | 0.33927184 | 0.62810099 |
| TRA2B | 78.8243571 | 0.155409424 | 0.25318604 | 0.61381513 | 0.5393375 | 0.77077724 |
| TBCCD1 | 11.828223 | 0.067281748 | 0.43421576 | 0.15495004 | 0.87686071 | 0.95134725 |
| AC068631.2 | 12.957701 | -1.220635565 | 0.66771367 | -1.8280823 | 0.0675372 | 0.26098543 |
| KNG1 | 72.6659686 | 3.332481043 | 0.69387591 | 4.80270465 | 1.57E-06 | 8.11E-05 |
| EIF4A2 | 108.307795 | 0.011763056 | 0.3196877 | 0.03679546 | 0.97064809 | 0.98820206 |
| ST6GAL1 | 436.18296 | -0.999274216 | 0.33573142 | -2.9764096 | 0.00291645 | 0.03186137 |
| BCL6 | 99.1692347 | -2.241399449 | 0.35561893 | -6.3028125 | 2.92E-10 | 5.79E-08 |
| LPP | 1112.30652 | -0.16951507 | 0.30066109 | -0.5638078 | 0.57288494 | 0.78986357 |
| TPRG1 | 16.3637995 | -0.738110575 | 0.46079864 | -1.6018072 | 0.10919825 | 0.34465697 |
| P3H2 | 241.64444 | 1.105147563 | 0.41432921 | 2.66731752 | 0.00764594 | 0.06391579 |
| CCDC50 | 125.211936 | 0.573818943 | 0.20471748 | 2.80297975 | 0.00506328 | 0.0479874 |
| FGF12 | 30.534891 | 1.151245277 | 0.7572823 | 1.52023266 | 0.12845251 | 0.37583578 |
| MB21D2 | 20.7551688 | -0.283984559 | 0.37225003 | -0.7628866 | 0.44553103 | 0.71524207 |
| OPA1 | 141.336371 | 0.433225987 | 0.17857625 | 2.42599997 | 0.01526627 | 0.10133578 |
| HES1 | 151.739217 | 1.699281587 | 0.44615528 | 3.80872237 | 0.00013969 | 0.0031908 |
| ATP13A3 | 181.033176 | 0.724311079 | 0.27688919 | 2.61588785 | 0.00889958 | 0.07080883 |
| LINC00884 | 14.7244605 | 0.792159851 | 0.4266973 | 1.85649134 | 0.06338356 | 0.25066311 |
| LSG1 | 29.9719671 | 0.096343132 | 0.3695199 | 0.26072515 | 0.79430447 | 0.91176607 |
| XXYLT1 | 26.8500336 | -0.220314862 | 0.28265583 | -0.7794457 | 0.43571724 | 0.70739432 |
| ACAP2 | 213.68551 | 0.287690984 | 0.23737709 | 1.21195768 | 0.22552858 | 0.51436694 |
| PPP1R2 | 49.9149055 | -0.197478683 | 0.25146437 | -0.7853148 | 0.43226904 | 0.70484895 |
| MUC20-OT1 | 587.630919 | 0.563656589 | 0.40595793 | 1.3884606 | 0.16499683 | 0.43379411 |
| TNK2 | 12.9115169 | 0.268063696 | 0.40087346 | 0.66869904 | 0.50368748 | 0.75108109 |
| TFRC | 87.4359852 | 0.672964961 | 0.31467348 | 2.13861356 | 0.03246698 | 0.16569809 |
| LINC00885 | 12.8542356 | 1.567411192 | 0.92574634 | 1.69313247 | 0.09043025 | 0.30986868 |
| PCYT1A | 34.7103859 | -0.149016124 | 0.27291031 | -0.546026 | 0.58504806 | 0.79681419 |
| UBXN7 | 66.9811907 | 0.012151105 | 0.26594907 | 0.04568959 | 0.96355766 | 0.98644251 |
| RNF168 | 60.8923653 | -0.2092738 | 0.27600889 | -0.758214 | 0.44832289 | 0.71763328 |
| PIGX | 24.4422317 | -0.259726255 | 0.42185724 | -0.6156733 | 0.53811014 | 0.76994592 |

| | | | | | | |
|---|---|---|---|---|---|---|
| PAK2 | 131.972974 | -0.418779474 | 0.18272205 | -2.2918935 | 0.02191179 | 0.12825868 |
| SENP5 | 147.499654 | -0.154402393 | 0.35435185 | -0.4357319 | 0.66303128 | 0.84275734 |
| NCBP2 | 27.9713538 | -0.604325875 | 0.35494389 | -1.7025955 | 0.08864379 | 0.30665156 |
| NCBP2-AS2 | 12.9646304 | -0.992993214 | 0.55918839 | -1.7757758 | 0.07576988 | 0.27917785 |
| DLG1 | 409.425497 | 0.024258816 | 0.21859543 | 0.11097586 | 0.91163549 | 0.96399248 |
| BDH1 | 59.7513117 | 0.681102478 | 0.34446182 | 1.97729455 | 0.04800834 | 0.21199399 |
| RUBCN | 43.3054413 | 0.184453705 | 0.40869245 | 0.45132643 | 0.6517543 | 0.83783039 |
| FYTTD1 | 48.3956862 | -0.014742556 | 0.33247494 | -0.0443419 | 0.96463191 | 0.98660829 |
| LRCH3 | 152.168338 | 0.024261559 | 0.25157482 | 0.09643874 | 0.92317213 | 0.96927428 |
| IQCG | 27.9575817 | -0.345808366 | 0.29592371 | -1.1685727 | 0.24257583 | 0.53317493 |
| RPL35A | 93.6299036 | -0.391385882 | 0.37099026 | -1.0549762 | 0.29143622 | 0.58292358 |
| LMLN | 39.9039412 | 0.269357661 | 0.35690667 | 0.75470056 | 0.45042867 | 0.71901805 |
| ZNF595 | 33.5054141 | 0.100963074 | 0.4078269 | 0.24756355 | 0.80447212 | 0.91643337 |
| ZNF718 | 102.460993 | 0.119049481 | 0.18584881 | 0.64057166 | 0.52180102 | 0.76109219 |
| ZNF141 | 92.094288 | 0.417189897 | 0.3527908 | 1.18254188 | 0.23699076 | 0.52743022 |
| AC092574.2 | 11.8341439 | -0.595081881 | 0.84724534 | -0.7023726 | 0.48244685 | 0.74082091 |
| ZNF721 | 187.958829 | 0.061871684 | 0.20368614 | 0.30375991 | 0.76131081 | 0.89648567 |
| PIGG | 42.4790576 | 0.064256645 | 0.24170365 | 0.26584888 | 0.7903556 | 0.91080793 |
| ATP5ME | 36.3929085 | -0.062275049 | 0.39424785 | -0.1579591 | 0.874489 | 0.94991414 |
| PCGF3 | 69.3324761 | 0.095683267 | 0.2016181 | 0.47457676 | 0.63508867 | 0.82850241 |
| AC139887.2 | 113.980055 | 0.319178274 | 0.25380487 | 1.25757347 | 0.20854605 | 0.49402397 |
| GAK | 91.1963522 | -0.319817173 | 0.39120776 | -0.8175123 | 0.4136357 | 0.69042044 |
| TMEM175 | 13.5515095 | 0.462872593 | 0.46336922 | 0.99892822 | 0.31782946 | 0.61101185 |
| IDUA | 14.6567622 | 0.870894209 | 0.38675619 | 2.25179127 | 0.02433547 | 0.13647467 |
| FGFRL1 | 14.2370157 | -0.176694744 | 0.39008669 | -0.4529628 | 0.65057555 | 0.83735207 |
| RNF212 | 30.7394992 | 1.793052525 | 0.40790114 | 4.39580171 | 1.10E-05 | 0.0003982 |
| AC092535.3 | 91.6075168 | 1.211169947 | 0.33188402 | 3.6493771 | 0.00026288 | 0.00534154 |
| CTBP1-AS | 18.4426284 | -0.010973857 | 0.51286766 | -0.0213971 | 0.98292892 | 0.99353746 |
| CTBP1 | 52.5964334 | 0.289721214 | 0.25445134 | 1.13861147 | 0.25486524 | 0.5458673 |
| CTBP1-DT | 14.7458078 | 0.336575999 | 0.40028452 | 0.8408419 | 0.40043651 | 0.68010059 |
| MAEA | 77.9423018 | 0.237971245 | 0.24349572 | 0.97731181 | 0.32841481 | 0.61762127 |
| UVSSA | 43.1657265 | 0.024712472 | 0.28719232 | 0.08604851 | 0.93142785 | 0.97334817 |
| SLBP | 57.7162878 | 0.387659706 | 0.34615094 | 1.11991521 | 0.26274989 | 0.55434704 |
| TMEM129 | 23.2504568 | 0.376637982 | 0.4514228 | 0.83433531 | 0.40409206 | 0.68217314 |
| LETM1 | 37.2562779 | 0.617159288 | 0.27775588 | 2.22194858 | 0.02628678 | 0.14481627 |
| NSD2 | 116.703996 | -0.244427983 | 0.19643583 | -1.2443146 | 0.21338378 | 0.49980394 |
| NELFA | 75.1516477 | 0.339791867 | 0.23958078 | 1.41827681 | 0.15610996 | 0.42034628 |
| C4orf48 | 20.1289612 | -1.372843145 | 0.81045876 | -1.6939087 | 0.09028263 | 0.30972084 |
| NAT8L | 24.575189 | 1.594736076 | 0.58203577 | 2.73992797 | 0.00614527 | 0.05517739 |
| POLN | 11.5865549 | 0.848713799 | 0.43882115 | 1.93407678 | 0.05310369 | 0.22595384 |
| MXD4 | 45.0492025 | -0.421278297 | 0.35091222 | -1.2005233 | 0.22993616 | 0.52015862 |
| ZFYVE28 | 74.0706067 | 0.103329524 | 0.27268059 | 0.37893979 | 0.70473257 | 0.86657376 |
| RNF4 | 59.8451602 | -0.048288757 | 0.30327056 | -0.1592267 | 0.87349031 | 0.94942996 |
| FAM193A | 148.887329 | 0.15381001 | 0.29632304 | 0.51906193 | 0.60371756 | 0.80940894 |
| TNIP2 | 20.7121109 | -1.354976842 | 0.36401689 | -3.7222912 | 0.00019742 | 0.0041825 |
| SH3BP2 | 74.7192094 | -2.391577202 | 0.4080109 | -5.8615522 | 4.59E-09 | 6.22E-07 |
| ADD1 | 146.763322 | 0.08686877 | 0.22453837 | 0.38687717 | 0.69884715 | 0.86334649 |
| MFSD10 | 35.3490314 | -0.701210482 | 0.597433 | -1.1737056 | 0.24051295 | 0.53136703 |
| NOP14 | 18.1240813 | -0.245948718 | 0.3751334 | -0.65563 | 0.5120622 | 0.75555211 |
| GRK4 | 26.6857373 | 0.215513147 | 0.54241694 | 0.39732009 | 0.69113143 | 0.85970007 |
| HTT | 143.990491 | 0.417897809 | 0.18893273 | 2.21188681 | 0.02697449 | 0.14648981 |
| RGS12 | 67.3678027 | -0.409496184 | 0.42467454 | -0.9642589 | 0.33491616 | 0.62350658 |
| LRPAP1 | 66.3161116 | -0.384931057 | 0.37818601 | -1.0178353 | 0.30875625 | 0.60259154 |
| TMEM128 | 16.3233737 | 0.21642456 | 0.40717342 | 0.5315292 | 0.59505211 | 0.80333205 |
| ZBTB49 | 12.4454823 | 0.177607808 | 0.45721998 | 0.38845154 | 0.69768191 | 0.86275485 |
| STX18 | 170.664533 | 0.353788581 | 0.1996789 | 1.7717875 | 0.07642984 | 0.28042437 |
| STX18-AS1 | 105.249332 | 0.662138902 | 0.33193191 | 1.99480342 | 0.04606432 | 0.20605934 |
| STK32B | 35.9722509 | 2.48897203 | 0.37346541 | 6.66453158 | 2.66E-11 | 7.60E-09 |
| EVC2 | 49.5216242 | -0.749237254 | 0.329063 | -2.2768809 | 0.02279334 | 0.13197761 |
| EVC | 55.4882147 | -0.054564192 | 0.29877638 | -0.1826255 | 0.85509186 | 0.94225653 |
| CRMP1 | 18.1064141 | -1.017546426 | 0.68970683 | -1.4753318 | 0.14012336 | 0.39525538 |
| WFS1 | 10.2075451 | 0.069329384 | 0.7100839 | 0.09763548 | 0.92222175 | 0.96885417 |
| MAN2B2 | 19.2251351 | -0.309323087 | 0.57512064 | -0.5378404 | 0.59068727 | 0.80016571 |
| MRFAP1 | 57.3584767 | -0.452404578 | 0.41055941 | -1.1019223 | 0.27049544 | 0.56240594 |
| LINC02482 | 11.3366322 | 0.466417208 | 0.49942257 | 0.93391295 | 0.3503488 | 0.63689665 |
| MRFAP1L1 | 18.0244191 | -0.284103036 | 0.46691665 | -0.6084663 | 0.54287826 | 0.77258349 |
| KIAA0232 | 207.919463 | 0.128765489 | 0.21315654 | 0.60408885 | 0.54578457 | 0.77463416 |
| TBC1D14 | 126.721242 | 0.251388689 | 0.21242089 | 1.18344617 | 0.23663237 | 0.52697507 |
| TADA2B | 15.6283424 | -0.299955339 | 0.42767135 | -0.7013688 | 0.48307289 | 0.74111942 |
| GRPEL1 | 19.7050063 | -0.70928508 | 0.41102453 | -1.7256515 | 0.08441014 | 0.29792391 |
| SORCS2 | 95.6352925 | 0.256086149 | 0.36728826 | 0.69723478 | 0.48565587 | 0.74309027 |
| AFAP1 | 22.9080758 | -1.363979224 | 0.55860283 | -2.4417693 | 0.01461548 | 0.09841093 |
| SH3TC1 | 86.2629885 | 0.619494012 | 0.28121394 | 2.20292779 | 0.02759984 | 0.14863227 |
| ACOX3 | 30.2523914 | 0.407871491 | 0.3245765 | 1.2566267 | 0.20888884 | 0.49459271 |
| TRMT44 | 25.2801901 | 0.128039999 | 0.33224534 | 0.38537787 | 0.69995748 | 0.86376951 |
| WDR1 | 86.8349803 | -0.091729621 | 0.3166103 | -0.2897241 | 0.77202735 | 0.90105761 |
| ZNF518B | 13.3359929 | -1.325181985 | 0.51513726 | -2.5724833 | 0.01009718 | 0.07773495 |
| HS3ST1 | 66.5896351 | -2.205473896 | 0.37681921 | -5.8528701 | 4.83E-09 | 6.46E-07 |
| AC005699.1 | 44.5164786 | -0.577304419 | 0.43837092 | -1.3169314 | 0.18786163 | 0.46590046 |
| RAB28 | 198.324147 | 0.625392526 | 0.24903445 | 2.51126913 | 0.01202979 | 0.08721389 |
| BOD1L1 | 124.060675 | -0.41911622 | 0.23233472 | -1.8039328 | 0.07124185 | 0.26933339 |
| LINC01182 | 75.7122918 | -0.577842647 | 0.59040843 | -0.9787168 | 0.32771993 | 0.61729352 |
| CPEB2 | 222.296934 | -0.645003942 | 0.52360347 | -1.2318558 | 0.21800297 | 0.50593977 |
| AC098829.1 | 69.8708111 | -0.767973043 | 0.57759059 | -1.3296149 | 0.18364519 | 0.46016652 |
| CC2D2A | 41.3926004 | -0.376667623 | 0.41043126 | -0.9177362 | 0.35875699 | 0.64254425 |
| FBXL5 | 109.132271 | -0.069925621 | 0.21701151 | -0.3222208 | 0.74728544 | 0.8891386 |
| FAM200B | 37.8803972 | 0.298424597 | 0.33641221 | 0.88708015 | 0.37503575 | 0.65806152 |
| PROM1 | 126.852695 | -1.35018592 | 0.55562246 | -2.430042 | 0.01509707 | 0.10047582 |
| TAPT1 | 88.6465192 | 0.375776858 | 0.25593837 | 1.4682318 | 0.14204127 | 0.39812207 |
| TAPT1-AS1 | 203.427462 | 1.018027801 | 0.42709501 | 2.38360965 | 0.01714378 | 0.10969887 |
| AC097515.1 | 34.3349153 | 1.363796465 | 0.74254544 | 1.83665051 | 0.06626151 | 0.2574031 |
| QDPR | 17.4361735 | 0.228003709 | 0.47711303 | 0.47788196 | 0.63273422 | 0.82649065 |
| LAP3 | 43.6348059 | -0.610357406 | 0.73602833 | -0.829258 | 0.40695841 | 0.6854947 |
| MED28 | 46.6642974 | -0.168902221 | 0.28103776 | -0.6009948 | 0.54784348 | 0.77516644 |
| FAM184B | 11.790174 | -0.775163142 | 0.57812015 | -1.340834 | 0.17997436 | 0.45599996 |
| DCAF16 | 55.4512653 | 0.443721598 | 0.27398552 | 1.61950748 | 0.10533812 | 0.33712125 |
| LCORL | 267.527364 | -0.628364551 | 0.30557034 | -2.0563663 | 0.03974722 | 0.18852482 |
| AC024230.1 | 22.6170759 | -0.258739393 | 0.51577141 | -0.5016552 | 0.6159101 | 0.81666507 |
| SLIT2 | 370.423466 | 1.544813628 | 0.39484578 | 3.91244812 | 9.14E-05 | 0.00225718 |
| PACRGL | 56.9921955 | -0.157987298 | 0.29222882 | -0.5406287 | 0.58876351 | 0.79913593 |
| KCNIP4 | 7250.11125 | -1.109041133 | 0.36025724 | -3.0784701 | 0.00208066 | 0.02524735 |
| ADGRA3 | 86.0131037 | 0.099526338 | 0.35585234 | 0.27968437 | 0.77971967 | 0.90427469 |
| PPARGC1A | 854.21757 | 1.189267539 | 0.31772175 | 3.7431103 | 0.00018176 | 0.00388476 |
| DHX15 | 145.740641 | 0.333472742 | 0.25737062 | 1.2956908 | 0.19508203 | 0.47613296 |
| CCDC149 | 28.1094242 | -0.097613704 | 0.37094689 | -0.2631474 | 0.79243698 | 0.91124908 |
| SEPSECS | 26.702386 | 0.102303301 | 0.35229204 | 0.29039345 | 0.77151525 | 0.9008444 |
| AC104662.2 | 9.73694889 | 0.376511774 | 0.59070399 | 0.63739501 | 0.52386758 | 0.76218158 |
| ZCCHC4 | 17.3800912 | 0.327774454 | 0.39104714 | 0.83819679 | 0.40192019 | 0.68091593 |
| ANAPC4 | 53.6167718 | 0.027610912 | 0.3618488 | 0.07630511 | 0.93917636 | 0.97662903 |
| SEL1L3 | 20.2029101 | -2.556269399 | 0.67328792 | -3.7966958 | 0.00014664 | 0.00332013 |
| SMIM20 | 20.2026103 | -0.734284154 | 0.43396729 | -1.6920265 | 0.09064092 | 0.31018641 |
| RBPJ | 159.457328 | 0.222171245 | 0.18302515 | 1.21388368 | 0.22479214 | 0.51358476 |
| TBC1D19 | 120.625732 | 0.222602714 | 0.2439595 | 0.91245768 | 0.36152784 | 0.64571079 |

| | | | | | | |
|---|---|---|---|---|---|---|
| STIM2 | 363.454563 | -0.800954623 | 0.45860855 | -1.7464886 | 0.08072608 | 0.29058005 |
| PCDH7 | 449.339247 | 0.780611671 | 0.3724923 | 2.09564514 | 0.03611368 | 0.17615679 |
| ARAP2 | 105.462137 | -0.562134233 | 0.56161311 | -1.0009279 | 0.31686167 | 0.6101512 |
| C4orf19 | 628.135912 | 0.023937422 | 0.24149921 | 0.09912008 | 0.92104293 | 0.9686919 |
| RELL1 | 79.9435956 | -0.219692048 | 0.32422638 | -0.6775884 | 0.49803268 | 0.74971166 |
| PGM2 | 25.5559456 | -0.138929364 | 0.28427835 | -0.4887089 | 0.62504779 | 0.82285278 |
| TBC1D1 | 521.128608 | 0.733112128 | 0.19267462 | 3.80492327 | 0.00014185 | 0.00322587 |
| KLF3 | 76.0335484 | -1.353341367 | 0.31972683 | -4.2328051 | 2.31E-05 | 0.00073384 |
| FAM114A1 | 27.9208656 | -1.088749239 | 0.41350378 | -2.632985 | 0.00846381 | 0.06865421 |
| KLHL5 | 138.029199 | 0.33971308 | 0.31612894 | 1.07460291 | 0.28255255 | 0.57535985 |
| WDR19 | 67.641403 | -0.481419159 | 0.30806007 | -1.5627444 | 0.11811272 | 0.35936125 |
| RFC1 | 69.7407089 | 0.386416098 | 0.22154258 | 1.74420688 | 0.08112302 | 0.2912964 |
| RPL9 | 14.8475065 | 0.105106356 | 0.4086124 | 0.25722752 | 0.79700314 | 0.91270858 |
| LIAS | 30.7027323 | 0.729149803 | 0.43080273 | 1.69253757 | 0.09054352 | 0.30999647 |
| UGDH | 63.05471 | -0.54519148 | 0.47262126 | -1.1535484 | 0.24868538 | 0.5398139 |
| UGDH-AS1 | 53.3357037 | 0.14732556 | 0.35031374 | 0.42055319 | 0.67408138 | 0.85019422 |
| SMIM14 | 164.766986 | -0.190700519 | 0.2100814 | -0.9077458 | 0.36401252 | 0.64890415 |
| UBE2K | 355.807292 | 0.348332101 | 0.25293199 | 1.37717691 | 0.16845756 | 0.43890991 |
| PDS5A | 228.549825 | -0.098643258 | 0.18839852 | -0.5235883 | 0.60056493 | 0.80738812 |
| N4BP2 | 48.4628175 | 0.55060609 | 0.24866875 | 2.21421504 | 0.02681399 | 0.1462349 |
| RBM47 | 821.12199 | -0.094148914 | 0.30525534 | -0.3084268 | 0.75775763 | 0.89457492 |
| NSUN7 | 56.2250341 | -0.485498636 | 0.31424296 | -1.5449785 | 0.12235145 | 0.36641415 |
| APBB2 | 576.608103 | 1.208480639 | 0.24842368 | 4.86459528 | 1.15E-06 | 6.22E-05 |
| UCHL1 | 54.9054494 | -1.486129303 | 0.60963242 | -2.4377465 | 0.01477913 | 0.09928814 |
| LIMCH1 | 1088.98277 | 1.073524432 | 0.28131497 | 3.81609419 | 0.00013558 | 0.00311081 |
| TMEM33 | 54.9628211 | -0.265556243 | 0.3473946 | -0.7644225 | 0.44461551 | 0.71446405 |
| SLC30A9 | 202.387567 | 0.278670733 | 0.16304649 | 1.7091489 | 0.08742337 | 0.30416601 |
| AC024022.1 | 21.6060954 | 0.046982241 | 0.86394811 | 0.05438086 | 0.95663173 | 0.98286826 |
| ATP8A1 | 113.102027 | 0.870143302 | 0.24897689 | 3.49487576 | 0.00047428 | 0.00826193 |
| KCTD8 | 328.714537 | 0.587526718 | 0.3522957 | 1.66770905 | 0.0953735 | 0.31807633 |
| GUF1 | 34.6286807 | 0.409974626 | 0.28876894 | 1.41973242 | 0.1556856 | 0.41953257 |
| GNPDA2 | 27.2395079 | -0.093462181 | 0.34426331 | -0.2714846 | 0.78601836 | 0.90831284 |
| GABRA2 | 78.8252166 | 1.088633568 | 0.54653813 | 1.99187121 | 0.04638519 | 0.20695549 |
| ATP10D | 77.9762079 | 0.632462698 | 0.27348591 | 2.31259702 | 0.02074481 | 0.12339087 |
| AC107068.2 | 15.8734058 | -0.981395401 | 0.68601276 | -1.4305789 | 0.15255093 | 0.41441825 |
| NFXL1 | 65.7878068 | 0.026994136 | 0.36092953 | 0.0747906 | 0.94038132 | 0.97688902 |
| NIPAL1 | 58.7383167 | 0.239350783 | 0.3910835 | 0.61201964 | 0.54052476 | 0.7714342 |
| CNGA1 | 40.4268555 | -0.123863885 | 0.39159471 | -0.3163063 | 0.75177001 | 0.89132635 |
| TXK | 33.3901628 | -1.290057884 | 0.2917824 | -4.4213013 | 9.81E-06 | 0.00036097 |
| TEC | 54.9213961 | -0.422133775 | 0.27126142 | -1.556188 | 0.11966338 | 0.36203283 |
| SLAIN2 | 101.90412 | 0.380496718 | 0.20052015 | 1.89754854 | 0.05775558 | 0.237619 |
| FRYL | 316.015757 | 0.126072326 | 0.16130313 | 0.78158635 | 0.43445771 | 0.70618229 |
| OCIAD1 | 164.741473 | 0.345762645 | 0.16493444 | 2.09636412 | 0.0360499 | 0.17602651 |
| OCIAD2 | 32.7281116 | -0.580726429 | 0.27072694 | -2.1450633 | 0.03194778 | 0.1639074 |
| CWH43 | 65.073132 | 1.908987068 | 0.36814151 | 5.18547088 | 2.15E-07 | 1.65E-05 |
| DCUN1D4 | 123.549766 | 0.065054215 | 0.27160714 | 0.23951585 | 0.8107056 | 0.91960902 |
| SGCB | 13.8657574 | -0.64534339 | 0.4475683 | -1.4418881 | 0.14933395 | 0.4098869 |
| USP46 | 38.3229307 | 0.825758681 | 0.28743778 | 2.87282586 | 0.00406818 | 0.04049316 |
| SCFD2 | 129.569618 | 0.166566742 | 0.29335078 | 0.5678074 | 0.57016575 | 0.78843592 |
| AC058822.1 | 56.4630089 | 0.034216332 | 0.57783935 | 0.05921426 | 0.95278145 | 0.98135593 |
| LNX1 | 136.260502 | 0.794667727 | 0.20354237 | 3.90418821 | 9.45E-05 | 0.00231143 |
| CHIC2 | 87.3078076 | -0.115504883 | 0.25329577 | -0.4560079 | 0.64838427 | 0.83558354 |
| TMEM165 | 163.127208 | -0.195770514 | 0.26183041 | -0.7476997 | 0.45464132 | 0.72190427 |
| CLOCK | 112.393454 | 0.13941107 | 0.23589293 | 0.59099299 | 0.55452512 | 0.77922789 |
| EXOC1 | 120.056186 | 0.080144643 | 0.25336085 | 0.31632607 | 0.75175502 | 0.89132635 |
| CEP135 | 31.6839008 | 0.057443964 | 0.38631107 | 0.14869873 | 0.88179137 | 0.95432447 |
| AASDH | 51.312176 | 0.402862221 | 0.29181134 | 1.3805571 | 0.16741518 | 0.4373 |
| PPAT | 12.9751541 | -0.316616684 | 0.42230284 | -0.7497385 | 0.45341224 | 0.72117537 |
| PAICS | 23.1765935 | -0.467458855 | 0.32796114 | -1.4253483 | 0.15405654 | 0.41706471 |
| SRP72 | 54.0011749 | 0.078424523 | 0.26062106 | 0.30091399 | 0.76348008 | 0.8974069 |
| REST | 77.9827757 | -0.214080464 | 0.31903051 | -0.6710344 | 0.50219859 | 0.75108109 |
| NOA1 | 14.2269117 | 0.256344289 | 0.37937939 | 0.67569376 | 0.4992351 | 0.75031683 |
| POLR2B | 73.465096 | 0.262906379 | 0.25940279 | 1.01350636 | 0.31081836 | 0.60394276 |
| IGFBP7 | 1054.422 | 0.125443868 | 0.18037762 | 0.69545141 | 0.48677245 | 0.74422309 |
| AC104806.2 | 19.8473428 | -0.717563656 | 0.5586591 | -1.2844392 | 0.19898832 | 0.48187682 |
| CENPC | 101.888002 | 0.880979726 | 0.32642509 | 2.69887257 | 0.00695748 | 0.06003013 |
| UBA6 | 138.059115 | -0.252476326 | 0.29721607 | -0.8494706 | 0.39561946 | 0.67612337 |
| UBA6-AS1 | 303.162813 | 0.270728049 | 0.35592916 | 0.7606234 | 0.44688204 | 0.71645983 |
| YTHDC1 | 164.612255 | 0.264855547 | 0.20655057 | 1.28227946 | 0.19974463 | 0.48315783 |
| UGT2B7 | 137.941513 | -1.582064295 | 0.68447035 | -2.3113701 | 0.02081242 | 0.12350782 |
| ENAM | 11.626195 | 0.098083777 | 0.40690067 | 0.24105091 | 0.80951566 | 0.9191701 |
| UTP3 | 11.679838 | -1.665060091 | 0.50356064 | -3.3065732 | 0.00094445 | 0.01400282 |
| RUFY3 | 416.345607 | -0.74319715 | 0.32850704 | -2.2623477 | 0.02367593 | 0.13490566 |
| GRSF1 | 54.9386818 | -0.330613882 | 0.27411228 | -1.2061258 | 0.227769 | 0.5177793 |
| MOB1B | 250.53046 | 0.561779278 | 0.30834416 | 1.82192285 | 0.06846669 | 0.26308984 |
| SLC4A4 | 159.84277 | -0.00302279 | 0.34845241 | -0.0086749 | 0.99307852 | 0.9979411 |
| ADAMTS3 | 166.295189 | -0.50227236 | 0.60404343 | -0.831517 | 0.40568165 | 0.68390318 |
| COX18 | 14.4292342 | -0.717850485 | 0.39565536 | -1.8143328 | 0.06962651 | 0.26586074 |
| ANKRD17 | 645.989719 | 0.641130227 | 0.21042172 | 3.04688234 | 0.00231228 | 0.02700808 |
| AC053527.1 | 96.2174865 | 0.680693192 | 0.43347486 | 1.57031759 | 0.11634124 | 0.35565208 |
| RASSF6 | 63.8923179 | -0.565406998 | 0.3437779 | -1.6446869 | 0.1000344 | 0.32653815 |
| CXCL2 | 23.8156542 | -1.199982855 | 0.638505 | -1.8793633 | 0.06019491 | 0.24328283 |
| MTHFD2L | 75.0571704 | -0.18791382 | 0.26331014 | -0.7136596 | 0.47543768 | 0.73587124 |
| BTC | 21.4280931 | -0.755584821 | 0.59949367 | -1.2603716 | 0.20753532 | 0.49286051 |
| PARM1 | 62.1955959 | 1.274791774 | 0.38156026 | 3.3409972 | 0.00083478 | 0.01279749 |
| RCHY1 | 57.3179309 | 0.748363458 | 0.24835208 | 3.0133167 | 0.00258409 | 0.02942907 |
| THAP6 | 35.7453635 | -0.113410985 | 0.31178552 | -0.3637468 | 0.71604709 | 0.87318675 |
| CDKL2 | 39.1205478 | -0.73978264 | 0.34466658 | -2.1463719 | 0.03184332 | 0.16345287 |
| G3BP2 | 73.9425938 | 0.012804211 | 0.2543284 | 0.05034519 | 0.95984732 | 0.98430689 |
| USO1 | 147.439282 | 0.348263762 | 0.26256796 | 1.32637568 | 0.18471529 | 0.46134371 |
| NAAA | 26.4540708 | 0.631162478 | 0.48990406 | 1.28833896 | 0.197628 | 0.47972753 |
| SDAD1 | 39.3943939 | 0.06983665 | 0.27036436 | 0.25830568 | 0.796171 | 0.91195838 |
| NUP54 | 60.88698 | 0.188171249 | 0.23793938 | 0.79083693 | 0.42903916 | 0.70310352 |
| SCARB2 | 142.66903 | -0.314305807 | 0.29981076 | -1.0483473 | 0.29447861 | 0.5865342 |
| SHROOM3 | 564.870521 | 0.076947825 | 0.32674494 | 0.23549814 | 0.81382212 | 0.92101456 |
| 11-Sep | 216.011727 | 0.382005166 | 0.31964406 | 1.19509546 | 0.23204974 | 0.52184597 |
| CCNI | 228.433534 | -0.484304449 | 0.27936214 | -1.733608 | 0.08298766 | 0.29503415 |
| CCNG2 | 88.3904761 | -1.173886545 | 0.35712766 | -3.2870222 | 0.00101253 | 0.01485907 |
| CNOT6L | 162.370771 | 0.891692733 | 0.24849565 | 3.58836357 | 0.00033276 | 0.0063601 |
| MRPL1 | 73.6695972 | 0.65565688 | 0.20398168 | 3.21429302 | 0.00130766 | 0.01786675 |
| FRAS1 | 1060.21366 | 2.402001726 | 0.39783091 | 6.03774535 | 1.56E-09 | 2.52E-07 |
| ANXA3 | 40.4738498 | -0.085070998 | 0.53475932 | -0.1590828 | 0.87360366 | 0.94943205 |
| BMP2K | 85.7544279 | -0.067793596 | 0.33881254 | -0.2000918 | 0.84140882 | 0.93566718 |
| PAQR3 | 15.7334755 | 0.218614392 | 0.44361952 | 0.49279705 | 0.62215599 | 0.82046224 |
| RASGEF1B | 447.513177 | 0.095119305 | 0.73824343 | 0.12884545 | 0.89747994 | 0.95917897 |
| HNRNPD | 155.273363 | 0.007290403 | 0.20487718 | 0.03558426 | 0.97161386 | 0.988268 |
| HNRNPDL | 132.209041 | 0.090516621 | 0.19334287 | 0.46816634 | 0.63966564 | 0.83121032 |
| ENOPH1 | 28.4297679 | -0.067980791 | 0.30135826 | -0.2255813 | 0.82152708 | 0.92485761 |
| TMEM150C | 34.5712541 | 0.129185042 | 0.32560476 | 0.39675415 | 0.69154876 | 0.85976938 |
| SCD5 | 122.346799 | 0.542688933 | 0.57589028 | 0.9423478 | 0.34601461 | 0.63315142 |
| SEC31A | 210.884867 | 0.015592773 | 0.28313891 | 0.0550711 | 0.95608182 | 0.98286826 |
| LIN54 | 36.5805633 | -0.394349567 | 0.30137434 | -1.3085041 | 0.19070237 | 0.47056667 |
| COPS4 | 34.9375917 | 0.091592018 | 0.2821915 | 0.32457398 | 0.74550352 | 0.88808711 |

| | | | | | | |
|---|---|---|---|---|---|---|
| HELQ | 41.1420799 | 0.165082785 | 0.25696699 | 0.64242798 | 0.52059534 | 0.7608417 |
| MRPS18C | 18.670173 | -0.817260875 | 0.4263923 | -1.9166877 | 0.05527762 | 0.23212675 |
| ABRAXAS1 | 19.8195281 | -0.023715202 | 0.43445304 | -0.0545863 | 0.95646802 | 0.98286826 |
| GPAT3 | 165.1488 | 3.58826777 | 0.69350932 | 5.17407288 | 2.29E-07 | 1.71E-05 |
| CDS1 | 80.4196131 | 0.054879879 | 0.32034766 | 0.1713135 | 0.86397727 | 0.94658022 |
| WDFY3 | 505.86684 | 0.284195629 | 0.12343382 | 2.30241297 | 0.02131189 | 0.12560362 |
| ARHGAP24 | 581.773423 | 0.873616949 | 0.25196457 | 3.46722146 | 0.00052587 | 0.00891036 |
| MAPK10 | 938.522102 | 0.688220344 | 0.37005528 | 1.85977711 | 0.06291707 | 0.24948869 |
| AC104827.1 | 14.4490617 | -0.377500214 | 0.68150499 | -0.5539214 | 0.57963262 | 0.79429575 |
| PTPN13 | 784.647162 | 0.812587204 | 0.30848512 | 2.6341212 | 0.00843554 | 0.0685895 |
| AC093827.5 | 21.744871 | 0.291289691 | 0.45646705 | 0.63813957 | 0.52338283 | 0.76218158 |
| AFF1 | 950.148447 | 0.497787286 | 0.25198342 | 1.97547636 | 0.04821411 | 0.21253821 |
| KLHL8 | 88.2324118 | -0.578583268 | 0.30559385 | -1.893308 | 0.05831693 | 0.23878417 |
| HSD17B11 | 18.6940776 | -0.85518944 | 0.41173024 | -2.0770625 | 0.03779579 | 0.18220508 |
| NUDT9 | 42.8797556 | 0.099968945 | 0.28030619 | 0.35664194 | 0.72135987 | 0.87614353 |
| SPP1 | 7134.45954 | -4.585751986 | 0.53637897 | -8.5494628 | 1.24E-17 | 1.59E-14 |
| PKD2 | 230.959974 | -0.023774834 | 0.19593279 | -0.1213418 | 0.90342033 | 0.96126516 |
| PPM1K | 82.6918382 | 0.166504063 | 0.38132931 | 0.43664114 | 0.66237162 | 0.84233458 |
| HERC6 | 39.5302532 | -0.075295462 | 0.95582633 | -0.0787753 | 0.93721139 | 0.97615526 |
| HERC5 | 34.2623831 | -0.760574299 | 0.84559024 | -0.8994596 | 0.36840788 | 0.65369102 |
| PYURF | 21.9336665 | -0.754450687 | 0.38918002 | -1.9385648 | 0.05255436 | 0.22446724 |
| HERC3 | 157.776596 | 0.05656264 | 0.318562 | 0.17755614 | 0.85907157 | 0.94407755 |
| FAM13A-AS1 | 59.7577116 | 0.242622926 | 0.26418481 | 0.91838335 | 0.35841821 | 0.64216076 |
| FAM13A | 220.225536 | 0.140888156 | 0.38796704 | 0.36314465 | 0.71649683 | 0.8732017 |
| GPRIN3 | 165.035192 | -0.185575944 | 0.5431042 | -0.3416949 | 0.7325805 | 0.88300506 |
| CCSER1 | 2895.01609 | 1.389235604 | 0.29038652 | 4.78409126 | 1.72E-06 | 8.76E-05 |
| SMARCAD1 | 111.058547 | 0.409953742 | 0.19926293 | 2.05735079 | 0.03965249 | 0.18824885 |
| PDLIM5 | 242.761155 | -0.347705674 | 0.23087424 | -1.5060393 | 0.13205709 | 0.38129264 |
| BMPR1B | 718.53482 | 1.087010551 | 0.19431482 | 5.5940693 | 2.22E-08 | 2.51E-06 |
| UNC5C | 44.7639551 | 0.913486608 | 0.40184362 | 2.27323904 | 0.02301178 | 0.1323843 |
| STPG2 | 246.613297 | 0.849990173 | 0.47087132 | 1.80514321 | 0.07105228 | 0.26907809 |
| RAP1GDS1 | 158.300886 | 0.05722664 | 0.18488661 | 0.30952291 | 0.75692379 | 0.89445158 |
| TSPAN5 | 105.493685 | -0.655634058 | 0.53942662 | -1.2154277 | 0.22420301 | 0.5134 |
| EIF4E | 85.0894258 | 0.277010422 | 0.27439219 | 1.00954192 | 0.31271481 | 0.6055915 |
| METAP1 | 54.9700766 | 0.129258265 | 0.25558954 | 0.50572595 | 0.61304905 | 0.81464125 |
| ADH5 | 48.9110873 | 0.017370737 | 0.35404951 | 0.04906302 | 0.96086908 | 0.98496251 |
| TRMT10A | 9.25974876 | 0.231218942 | 0.55175639 | 0.41905984 | 0.6751724 | 0.85125763 |
| LAMTOR3 | 29.7555264 | -0.533293372 | 0.38349076 | -1.390629 | 0.16433795 | 0.43321638 |
| DNAJB14 | 121.61174 | -0.039277937 | 0.20127718 | -0.1951435 | 0.84528059 | 0.93715892 |
| H2AFZ | 37.7622662 | -1.314956367 | 0.50132127 | -2.6229814 | 0.0087164 | 0.06982612 |
| PPP3CA | 299.74622 | -0.059761097 | 0.33874263 | -0.1764204 | 0.85996371 | 0.94469462 |
| BANK1 | 43.5240571 | -0.359048553 | 0.43876381 | -0.8183185 | 0.41317533 | 0.69014748 |
| AP002075.1 | 15.0995556 | -0.127572573 | 0.67868214 | -0.187971 | 0.85089937 | 0.9399491 |
| SLC39A8 | 13.6787906 | 0.943507473 | 0.57911947 | 1.6292104 | 0.10326849 | 0.33360675 |
| NFKB1 | 132.963983 | 0.158510069 | 0.23586496 | 0.67203738 | 0.5015599 | 0.75108109 |
| MANBA | 83.0252593 | 0.29967223 | 0.2346909 | 1.27688046 | 0.20164445 | 0.48504517 |
| UBE2D3 | 297.195203 | -0.05393912 | 0.18954072 | -0.284578 | 0.77596746 | 0.90266675 |
| CISD2 | 26.2550839 | -0.926471701 | 0.36992841 | -2.5044621 | 0.01226377 | 0.08835062 |
| SLC9B1 | 10.250159 | 0.594100794 | 0.62968626 | 0.94348699 | 0.34543187 | 0.63256941 |
| SLC9B2 | 10.0214116 | 0.003075621 | 0.60654423 | 0.00507073 | 0.99595416 | 0.99868531 |
| BDH2 | 10.7227911 | -0.00604769 | 0.51962407 | -0.0116386 | 0.99071396 | 0.99684874 |
| TET2 | 193.667371 | 0.477603024 | 0.26306949 | 1.81550138 | 0.06944689 | 0.26541135 |
| PPA2 | 203.990864 | 0.237549153 | 0.23934618 | 0.99249194 | 0.32095761 | 0.6120836 |
| ARHGEF38 | 388.577299 | -1.033276569 | 0.33504311 | -3.0840108 | 0.0020423 | 0.02492802 |
| INTS12 | 50.1568008 | -0.320568616 | 0.30634475 | -1.0464309 | 0.29536213 | 0.58696387 |
| GSTCD | 69.1248982 | -7.01E-05 | 0.22403848 | -0.0003131 | 0.99975018 | 0.99975018 |
| NPNT | 150.325307 | 0.338835976 | 0.35326867 | 0.95914528 | 0.33748556 | 0.62610773 |
| TBCK | 316.656663 | 0.509658382 | 0.22392029 | 2.27607057 | 0.02284179 | 0.13197761 |
| AIMP1 | 41.6344838 | -0.165922523 | 0.3150355 | -0.5266788 | 0.59841665 | 0.8059995 |
| PAPSS1 | 95.5264083 | 0.518810686 | 0.22395663 | 2.31656763 | 0.0205273 | 0.12280616 |
| SGMS2 | 114.864252 | 0.225079244 | 0.51691721 | 0.4354261 | 0.66325317 | 0.84278344 |
| AC096564.1 | 12.6523109 | -0.310182099 | 0.69465872 | -0.4465244 | 0.65521846 | 0.83931753 |
| HADH | 30.0947838 | 0.280330565 | 0.36004213 | 0.77860489 | 0.4362125 | 0.70775777 |
| RPL34-AS1 | 51.3853173 | 0.986593974 | 0.54085203 | 1.82414769 | 0.06812975 | 0.26218628 |
| RPL34 | 104.150257 | -0.352707944 | 0.42726822 | -0.8254954 | 0.4090904 | 0.6869518 |
| OSTC | 37.4727955 | -0.768211875 | 0.36507256 | -2.1042717 | 0.03535475 | 0.17452067 |
| SEC24B-AS1 | 14.3197579 | -0.593095127 | 0.70264536 | -0.8440889 | 0.39861975 | 0.67938457 |
| SEC24B | 196.52012 | 0.276841333 | 0.23854383 | 1.16054704 | 0.24582615 | 0.53627476 |
| MCUB | 16.140318 | -2.891824627 | 0.56211323 | -5.1445589 | 2.68E-07 | 1.95E-05 |
| CASP6 | 22.6254886 | -1.08772317 | 0.370098 | -2.9390139 | 0.00329258 | 0.03481518 |
| PLA2G12A | 32.4440764 | -0.061722597 | 0.34110378 | -0.1809496 | 0.85640713 | 0.94330228 |
| AC126283.2 | 18.3251461 | -0.69226487 | 0.69983523 | -0.9891826 | 0.32257379 | 0.6132414 |
| GAR1 | 15.6846675 | -0.560535978 | 0.4823387 | -1.1621211 | 0.24518628 | 0.53608738 |
| EGF | 226.576007 | 3.372209604 | 0.52663453 | 6.40332035 | 1.52E-10 | 3.20E-08 |
| ELOVL6 | 90.941087 | -0.051890097 | 0.22292532 | -0.232769 | 0.81594081 | 0.92187043 |
| ENPEP | 13.2029037 | -0.132423689 | 0.57803112 | -0.2290944 | 0.81879555 | 0.9234319 |
| FAM241A | 42.2000405 | 0.445579535 | 0.27974726 | 1.59279322 | 0.11120659 | 0.34769355 |
| AP1AR | 41.9054509 | -0.236550402 | 0.39089241 | -0.6051548 | 0.54507617 | 0.77442763 |
| ALPK1 | 164.15321 | -0.935797876 | 0.35083601 | -2.667337 | 0.0076455 | 0.06391579 |
| LARP7 | 60.3203709 | 0.049495748 | 0.20847535 | 0.23741774 | 0.81233273 | 0.92034001 |
| ANK2 | 12.8460158 | 0.745256489 | 0.72823825 | 1.02336906 | 0.30613338 | 0.60083561 |
| CAMK2D | 117.212703 | -0.47810859 | 0.18932609 | -2.525318 | 0.01155936 | 0.08487163 |
| ARSJ | 85.3158462 | 0.681683279 | 0.3137187 | 2.17291249 | 0.0297869 | 0.15672354 |
| UGT8 | 96.6747899 | 0.058036943 | 0.28968393 | 0.20034574 | 0.84121019 | 0.93566718 |
| SNHG8 | 21.5871713 | -0.3038022 | 0.43786479 | -0.6938265 | 0.48779101 | 0.74443126 |
| METTL14 | 43.8766172 | 0.792382933 | 0.26578326 | 2.98131245 | 0.00287016 | 0.0316574 |
| SEC24D | 63.9955599 | -1.104194038 | 0.36983949 | -2.9856034 | 0.00283019 | 0.03141928 |
| USP53 | 346.507859 | -0.267768454 | 0.21593337 | -1.2400513 | 0.21495643 | 0.50124063 |
| C4orf3 | 68.3882032 | -1.61932066 | 0.5397164 | -3.0003177 | 0.00269698 | 0.03039858 |
| PDE5A | 128.40563 | -0.557817922 | 0.44875666 | -1.2430298 | 0.21385684 | 0.50026184 |
| AC108866.1 | 53.7464596 | 0.180510125 | 0.40537965 | 0.44528659 | 0.65611265 | 0.83951126 |
| PRDM5 | 54.1007011 | 0.218848855 | 0.26290432 | 0.83242775 | 0.40516754 | 0.68337198 |
| ANXA5 | 92.6668243 | -0.75909076 | 0.27735191 | -2.7369228 | 0.00620168 | 0.05537004 |
| EXOSC9 | 12.365456 | 0.157505866 | 0.40452395 | 0.38936104 | 0.69700909 | 0.86249983 |
| BBS7 | 18.923913 | -0.21119038 | 0.36077721 | -0.5853762 | 0.55829481 | 0.78233632 |
| KIAA1109 | 332.120714 | 0.448600037 | 0.18823443 | 2.38319863 | 0.01716293 | 0.10975327 |
| FGF2 | 20.2822676 | -0.309188708 | 0.79620845 | -0.3883263 | 0.69777455 | 0.86275485 |
| NUDT6 | 11.293461 | 0.266081987 | 0.56713948 | 0.46916499 | 0.63895171 | 0.83119926 |
| SPATA5 | 135.016186 | 0.176710167 | 0.18547166 | 0.95276101 | 0.34071118 | 0.62907597 |
| SPRY1 | 22.0280612 | -2.919432594 | 0.65070232 | -4.486587 | 7.24E-06 | 0.00028899 |
| LINC01091 | 20.7504585 | -0.807200808 | 0.62082886 | -1.3001986 | 0.19353292 | 0.4740909 |
| ANKRD50 | 24.5544114 | 0.066105716 | 0.3337746 | 0.19805497 | 0.84300206 | 0.93604303 |
| FAT4 | 81.20589 | 1.613791743 | 0.41008514 | 3.93526021 | 8.31E-05 | 0.00210878 |
| INTU | 67.8878386 | 0.022848113 | 0.30090043 | 0.07593247 | 0.93947282 | 0.97673318 |
| HSPA4L | 99.9540623 | 0.831163265 | 0.46141556 | 1.80133343 | 0.07165034 | 0.27018368 |
| MFSD8 | 46.4679763 | -0.029685466 | 0.26932446 | -0.1102219 | 0.91223336 | 0.9642754 |
| ABHD18 | 161.715536 | 0.163520456 | 0.28560185 | 0.5725469 | 0.56695152 | 0.78656476 |
| LARP1B | 135.512793 | -0.073707256 | 0.25849578 | -0.2851391 | 0.77553757 | 0.90237046 |
| PGRMC2 | 44.9924979 | -0.575653748 | 0.3094045 | -1.8605216 | 0.06281177 | 0.24926305 |
| AC108062.1 | 47.0253618 | 0.486517683 | 0.36121841 | 1.34687952 | 0.17801904 | 0.45293953 |
| JADE1 | 146.894338 | -0.000345747 | 0.38742143 | -0.0008924 | 0.99928794 | 0.99957903 |
| SCLT1 | 121.727779 | 0.246916488 | 0.28312916 | 0.87209841 | 0.38315469 | 0.66642912 |
| C4orf33 | 18.7507897 | 0.024183428 | 0.50632122 | 0.04776302 | 0.96190511 | 0.98525601 |

| | | | | | | |
|---|---|---|---|---|---|---|
| AC096711.2 | 74.6536264 | 0.204196596 | 0.36131102 | 0.56515463 | 0.5719686 | 0.78923392 |
| NOCT | 11.6826269 | -1.098747897 | 0.41165535 | -2.6690966 | 0.00760556 | 0.06389755 |
| ELF2 | 364.936871 | -0.259921003 | 0.21461355 | -1.2111118 | 0.22585255 | 0.51476393 |
| NDUFC1 | 62.7239137 | -0.462146298 | 0.46767933 | -0.9881692 | 0.3230698 | 0.61384454 |
| NAA15 | 126.959825 | 0.415056743 | 0.24167296 | 1.71743145 | 0.08590037 | 0.30120681 |
| AC097376.2 | 151.907695 | -0.963182458 | 0.43914158 | -2.1933301 | 0.0282836 | 0.15134187 |
| SETD7 | 21.0724339 | -0.842740703 | 0.33932733 | -2.4835627 | 0.01300754 | 0.09200733 |
| MGST2 | 45.2365879 | 0.113696454 | 0.34212999 | 0.33231946 | 0.73964804 | 0.88614269 |
| MAML3 | 353.916175 | 0.118659226 | 0.25345369 | 0.46816927 | 0.63966355 | 0.83121032 |
| SCOC | 105.87415 | -0.81224923 | 0.30886481 | -2.6297888 | 0.00854379 | 0.06903384 |
| SCOC-AS1 | 14.6161808 | -0.696690241 | 0.60785336 | -1.1461485 | 0.2517337 | 0.54333973 |
| ELMOD2 | 39.0573296 | 0.266463126 | 0.29675833 | 0.89791288 | 0.369232 | 0.65369102 |
| TBC1D9 | 470.722308 | 0.312145825 | 0.31464343 | 0.99206212 | 0.32116722 | 0.6120836 |
| RNF150 | 252.168144 | 1.076682428 | 0.3644835 | 2.95399496 | 0.00313689 | 0.03369787 |
| ZNF330 | 26.0333538 | -0.202210909 | 0.32715446 | -0.61809 | 0.53651599 | 0.76905353 |
| IL15 | 25.0243915 | -0.474095658 | 0.4978732 | -0.9522418 | 0.34097438 | 0.62940657 |
| INPP4B | 106.299165 | 0.49067239 | 0.45613513 | 1.07571717 | 0.28205377 | 0.5750055 |
| AC104596.1 | 29.8837225 | 0.65507471 | 0.61163892 | 1.0710154 | 0.2841625 | 0.5765312 |
| USP38 | 33.6789991 | -0.081068644 | 0.29306054 | -0.2766276 | 0.78206604 | 0.90577227 |
| GAB1 | 363.462945 | 0.21037312 | 0.33198161 | 0.63368909 | 0.52628375 | 0.76221787 |
| SMARCA5 | 89.9445524 | -0.094249558 | 0.19391346 | -0.4860393 | 0.62693932 | 0.82354566 |
| ANAPC10 | 98.6163564 | 0.558976984 | 0.26101557 | 2.1415465 | 0.03222999 | 0.16502654 |
| ABCE1 | 36.6052707 | 0.224402735 | 0.33895013 | 0.66205236 | 0.50793767 | 0.75382716 |
| OTUD4 | 91.0491501 | -0.25003997 | 0.22258849 | -1.1233284 | 0.26129805 | 0.5525231 |
| SMAD1 | 133.003655 | 0.169999462 | 0.26345918 | 0.64525922 | 0.51875922 | 0.75966701 |
| MMAA | 23.5845058 | -0.305482615 | 0.35109216 | -0.8700924 | 0.38424991 | 0.66701235 |
| ZNF827 | 213.941726 | 0.307892124 | 0.24518346 | 1.25576223 | 0.20920219 | 0.4951566 |
| LSM6 | 27.2138549 | 0.311948231 | 0.33773063 | 0.92365988 | 0.35566343 | 0.64045528 |
| SLC10A7 | 107.339823 | 0.695238798 | 0.23240686 | 2.99147279 | 0.00277635 | 0.03108912 |
| TMEM184C | 48.5879202 | 0.260648464 | 0.35210509 | 0.74025758 | 0.45914372 | 0.72469719 |
| PRMT9 | 65.6251953 | 1.425097041 | 0.60899576 | 2.34007712 | 0.01927976 | 0.11811449 |
| ARHGAP10 | 32.0452475 | -0.998948285 | 0.4651448 | -2.1476071 | 0.03174498 | 0.16311062 |
| NR3C2 | 1684.6065 | 0.955520166 | 0.22322573 | 4.2805108 | 1.86E-05 | 0.00060983 |
| AC069272.1 | 22.2359178 | 0.360024485 | 0.67991878 | 0.52951102 | 0.596451 | 0.8043773 |
| DCLK2 | 22.3311458 | 0.638235365 | 0.64486594 | 0.9897179 | 0.32231202 | 0.61319341 |
| LRBA | 1248.87031 | 0.67571358 | 0.18306956 | 3.69102102 | 0.00022336 | 0.00466736 |
| RPS3A | 68.2688847 | 0.012132285 | 0.43823903 | 0.02768417 | 0.97791405 | 0.99119855 |
| SH3D19 | 148.575242 | 0.1840539 | 0.28138602 | 0.65409753 | 0.51304897 | 0.75603355 |
| FAM160A1 | 273.873225 | 0.101483015 | 0.39254825 | 0.25852367 | 0.79600278 | 0.91195838 |
| GATB | 30.0710791 | -0.052053559 | 0.30899276 | -0.1684621 | 0.86621979 | 0.94702879 |
| FBXW7 | 273.24818 | 0.390887716 | 0.26507215 | 1.47464652 | 0.1403076 | 0.39525538 |
| ARFIP1 | 169.738737 | 0.062670042 | 0.20121951 | 0.31145111 | 0.7554577 | 0.89343649 |
| FHDC1 | 14.7025667 | -2.320859021 | 0.4749649 | -4.88638 | 1.03E-06 | 5.75E-05 |
| TRIM2 | 352.546651 | 0.089670018 | 0.17738544 | 0.50550947 | 0.61320105 | 0.81464125 |
| TMEM131L | 25.3994727 | -0.012946495 | 0.42718174 | -0.0303068 | 0.9758224 | 0.99024058 |
| PLRG1 | 25.9269027 | 0.495061494 | 0.32789106 | 1.50983527 | 0.13108546 | 0.38008512 |
| FGB | 10.8385322 | -5.874322281 | 1.92763633 | -3.0474225 | 0.00230813 | 0.02700808 |
| MAP9 | 33.5200739 | -0.03736229 | 0.32424078 | -0.1152301 | 0.90826276 | 0.96234937 |
| GUCY1A1 | 17.3765326 | -1.286783739 | 0.46882787 | -2.7446827 | 0.00605694 | 0.05468766 |
| GUCY1B1 | 19.6502763 | -2.079916441 | 0.54234495 | -3.8350434 | 0.00012554 | 0.00294609 |
| CTSO | 19.6391236 | -0.933305939 | 0.35747858 | -2.6108024 | 0.00903301 | 0.07141829 |
| PDGFC | 130.001943 | -0.341931672 | 0.4175487 | -0.8189025 | 0.41284205 | 0.68990579 |
| TMEM144 | 12.1426282 | -0.410445148 | 0.47824234 | -0.8582367 | 0.39076175 | 0.67223049 |
| ETFDH | 79.7653453 | 1.209541643 | 0.31094988 | 3.88982833 | 0.00010032 | 0.00242593 |
| PPID | 35.7283436 | 0.42245108 | 0.30417007 | 1.38886475 | 0.16487388 | 0.43375609 |
| FNIP2 | 623.487183 | -1.561938594 | 0.62659226 | -2.4927512 | 0.01267576 | 0.09046528 |
| C4orf45 | 17.8347226 | -3.11402932 | 0.73273408 | -4.2498764 | 2.14E-05 | 0.00068859 |
| RAPGEF2 | 350.646554 | 0.500911978 | 0.237645 | 2.10781616 | 0.03504689 | 0.1740015 |
| NAF1 | 38.934115 | 0.401679288 | 0.37458312 | 1.07233687 | 0.28356875 | 0.57631195 |
| TMA16 | 19.6068124 | -0.298700524 | 0.34461381 | -0.8667689 | 0.38606866 | 0.66811344 |
| TMEM192 | 43.2452704 | -0.339794148 | 0.37978277 | -0.8947066 | 0.37094395 | 0.65494451 |
| KLHL2 | 173.897214 | -0.444869872 | 0.35978598 | -1.2364847 | 0.21627843 | 0.50307076 |
| MSMO1 | 112.818938 | 1.749919893 | 0.5206059 | 3.36131402 | 0.00077573 | 0.01221946 |
| DDX60 | 53.2606708 | -0.195403345 | 1.22993205 | -0.1588733 | 0.87376871 | 0.94943205 |
| DDX60L | 59.1162151 | -0.551275934 | 1.38101556 | -0.3991816 | 0.68975944 | 0.85887516 |
| PALLD | 119.831058 | 0.337542055 | 0.39554249 | 0.85336484 | 0.39345701 | 0.67418407 |
| CBR4 | 326.405041 | 0.281856011 | 0.26757287 | 1.05338039 | 0.29216669 | 0.58378114 |
| SH3RF1 | 656.283134 | 0.596174466 | 0.5550199 | 1.0741497 | 0.28275559 | 0.5754607 |
| NEK1 | 214.293627 | 0.735794042 | 0.32724056 | 2.24848056 | 0.02454556 | 0.13713036 |
| CLCN3 | 211.01363 | 0.932034257 | 0.46742076 | 1.99399415 | 0.04615269 | 0.20618606 |
| HPF1 | 19.544748 | -0.019889481 | 0.32912332 | -0.0604317 | 0.95181181 | 0.98065459 |
| MFAP3L | 59.7656171 | 0.154800264 | 0.32702333 | 0.47336154 | 0.63595527 | 0.82931783 |
| GALNT7 | 121.209358 | -0.319831221 | 0.28686136 | -1.1149331 | 0.2648791 | 0.55622662 |
| HMGB2 | 31.3466358 | -0.261234523 | 0.4258906 | -0.6133841 | 0.53962239 | 0.77077724 |
| FBXO8 | 23.5564469 | 0.158291328 | 0.32869992 | 0.48156789 | 0.63011294 | 0.82524878 |
| CEP44 | 42.3966695 | 0.106694344 | 0.32437195 | 0.32892592 | 0.74221168 | 0.88720442 |
| HPGD | 22.7262341 | 1.82116652 | 0.69641232 | 2.61506937 | 0.00892093 | 0.07085849 |
| WDR17 | 25.3449471 | 0.498865243 | 0.46924469 | 1.06312389 | 0.28772577 | 0.5800319 |
| SPCS3 | 61.984593 | -0.106541672 | 0.26264366 | -0.405651 | 0.68499903 | 0.85635073 |
| VEGFC | 65.2818722 | 2.315661147 | 0.38491239 | 6.01607327 | 1.79E-09 | 2.83E-07 |
| AC078881.1 | 55.9224933 | -0.016379759 | 0.39396034 | -0.0415772 | 0.96683577 | 0.98695423 |
| LINC01098 | 180.359362 | 1.251745791 | 1.06682831 | 1.17333387 | 0.24066195 | 0.53146825 |
| LINC01099 | 697.907665 | 2.303423889 | 0.61325942 | 3.75603509 | 0.00017263 | 0.003744 |
| DCTD | 21.7950806 | -1.353804533 | 0.4379201 | -3.0914418 | 0.00199187 | 0.02448716 |
| WWC2 | 212.710312 | 0.69885035 | 0.23381187 | 2.988943 | 0.00279944 | 0.03121198 |
| CDKN2AIP | 17.9725891 | 0.425608885 | 0.66295792 | 0.64198477 | 0.52088307 | 0.76092013 |
| RWDD4 | 18.964774 | -0.021387471 | 0.3512827 | -0.0608839 | 0.95145165 | 0.98061757 |
| TRAPPC11 | 66.9428565 | 0.047683109 | 0.21882467 | 0.21790555 | 0.8275027 | 0.92808376 |
| STOX2 | 332.770364 | -0.553354729 | 0.27555705 | -2.0081313 | 0.04462934 | 0.20147742 |
| IRF2 | 66.8649453 | -0.357981303 | 0.24301421 | -1.473088 | 0.14072732 | 0.39622104 |
| CASP3 | 27.2021748 | -0.121637194 | 0.46465402 | -0.2617801 | 0.79349095 | 0.91142199 |
| PRIMPOL | 27.8265455 | 0.30666086 | 0.47860886 | 0.64073377 | 0.52169567 | 0.76104628 |
| ACSL1 | 172.920158 | -0.134081586 | 0.36706857 | -0.3652767 | 0.71490489 | 0.87272783 |
| SLC25A4 | 43.7112666 | 0.392442368 | 0.42407447 | 0.9254091 | 0.35475315 | 0.64004675 |
| CFAP97 | 59.9533838 | -1.056836512 | 0.28171585 | -3.7514272 | 0.00017583 | 0.00378956 |
| SNX25 | 144.013379 | -0.478364218 | 0.31382238 | -1.5243152 | 0.12742999 | 0.37443918 |
| LRP2BP | 15.5515632 | -0.823943663 | 0.52046021 | -1.583106 | 0.11339732 | 0.3516614 |
| AC112722.1 | 19.0133275 | -0.691022737 | 0.41176827 | -1.6781835 | 0.09331128 | 0.31466211 |
| ANKRD37 | 37.325802 | -3.532208213 | 0.96562367 | -3.6579553 | 0.00025424 | 0.00520702 |
| UFSP2 | 48.9693139 | 0.445016406 | 0.2551219 | 1.7443285 | 0.08110182 | 0.2912964 |
| C4orf47 | 52.683581 | -2.169728864 | 0.59450151 | -3.6496608 | 0.00026259 | 0.00534154 |
| SORBS2 | 319.47729 | 0.251995522 | 0.44119517 | 0.57116564 | 0.56788737 | 0.78718726 |
| TLR3 | 12.0550114 | -0.897404492 | 0.60018382 | -1.4952161 | 0.13485806 | 0.38627205 |
| FAM149A | 54.5904618 | -0.624214099 | 0.26714531 | -2.3366089 | 0.01945953 | 0.11848232 |
| CYP4V2 | 37.971758 | 0.214002251 | 0.28981235 | 0.73841661 | 0.46026132 | 0.72529637 |
| KLKB1 | 22.7698839 | 0.765175771 | 0.47291026 | 1.61801475 | 0.10565942 | 0.33784616 |
| F11 | 10.7288993 | 1.600006713 | 0.68427377 | 2.33825523 | 0.01937401 | 0.11819385 |
| FAT1 | 226.896946 | 0.533779629 | 0.36291426 | 1.47081471 | 0.14134123 | 0.39740649 |
| FRG1-DT | 42.1733647 | 0.885547322 | 0.56193903 | 1.57587794 | 0.11505394 | 0.35402797 |
| FRG1 | 18.0203337 | 0.089386549 | 0.33535885 | 0.26654001 | 0.78982335 | 0.91080793 |
| SDHA | 106.75558 | 0.197918874 | 0.2112415 | 0.93693177 | 0.34879366 | 0.63556496 |
| PDCD6 | 53.6261493 | -0.106451329 | 0.29974992 | -0.3551338 | 0.72248935 | 0.87650915 |
| EXOC3 | 51.1438344 | -0.329939595 | 0.27334503 | -1.2070444 | 0.22741504 | 0.51718095 |

| | | | | | | |
|---|---|---|---|---|---|---|
| SLC9A3 | 12.3363679 | -0.000531049 | 0.4036219 | -0.0013157 | 0.99895022 | 0.99953184 |
| ZDHHC11B | 28.2998597 | 1.407600826 | 0.59305337 | 2.37348085 | 0.01762131 | 0.1112345 |
| BRD9 | 28.7672926 | -0.61266863 | 0.38312388 | -1.5991398 | 0.10978955 | 0.34537939 |
| SLC12A7 | 48.5807786 | -1.635812847 | 0.38061952 | -4.2977639 | 1.73E-05 | 0.00057938 |
| CLPTM1L | 30.8598997 | 0.142045892 | 0.34224814 | 0.41503773 | 0.67811428 | 0.85308748 |
| LPCAT1 | 20.2380755 | -0.903003646 | 0.37815599 | -2.3879131 | 0.01694435 | 0.10882834 |
| NDUFS6 | 52.0638074 | 0.007173267 | 0.44671134 | 0.01605795 | 0.98718816 | 0.99523554 |
| AC022424.1 | 42.9311334 | 0.007489867 | 0.64053232 | 0.01169319 | 0.9906704 | 0.99684874 |
| ADAMTS16 | 369.372624 | 1.023716509 | 0.50954813 | 2.00906735 | 0.04452999 | 0.20147034 |
| AC022424.2 | 17.2001676 | 0.77495994 | 0.68463409 | 1.13193304 | 0.25766259 | 0.54877816 |
| ICE1 | 88.8985022 | 0.167747922 | 0.20085234 | 0.83518033 | 0.40361619 | 0.68187175 |
| MED10 | 25.3659418 | -0.26142722 | 0.33374149 | -0.7833225 | 0.43343775 | 0.70594011 |
| NSUN2 | 45.1451363 | 0.568202122 | 0.30691166 | 1.85135399 | 0.06411864 | 0.2521767 |
| SRD5A1 | 11.4992658 | -0.17838465 | 0.41995717 | -0.4247687 | 0.67100532 | 0.84746193 |
| TENT4A | 67.8003518 | 0.013328252 | 0.28231943 | 0.04720983 | 0.96234599 | 0.98539792 |
| MTRR | 39.4971977 | -0.250411257 | 0.33344011 | -0.7509932 | 0.45265674 | 0.72046894 |
| SEMA5A | 36.6694004 | -0.856053728 | 0.53765259 | -1.5922061 | 0.1113384 | 0.34789452 |
| FAM173B | 35.075938 | 0.28151091 | 0.29427142 | 0.95663692 | 0.33875055 | 0.62755046 |
| CCT5 | 38.4377333 | -0.650953942 | 0.35846926 | -1.8159268 | 0.0693816 | 0.26541135 |
| CMBL | 34.6391659 | -0.851577813 | 0.56170579 | -1.5160567 | 0.12950502 | 0.37720122 |
| 6-Mar | 256.755735 | 0.113305072 | 0.2234784 | 0.50700681 | 0.61215004 | 0.81404024 |
| DAP | 104.342792 | -0.286487719 | 0.30613575 | -0.9358192 | 0.34936627 | 0.63612076 |
| DNAH5 | 14.1631001 | -1.678748177 | 0.7673875 | -2.1876147 | 0.02869768 | 0.15286634 |
| TRIO | 488.886789 | -0.601428195 | 0.28560844 | -2.1057788 | 0.03522357 | 0.1742352 |
| OTULINL | 11.1187364 | 1.671205749 | 0.63204693 | 2.64411657 | 0.00819044 | 0.06712646 |
| OTULIN | 36.9219862 | 0.002104983 | 0.30741835 | 0.00684729 | 0.9945367 | 0.99837898 |
| ANKH | 21.7148645 | -1.199910474 | 0.36716903 | -3.2680056 | 0.00108308 | 0.01544593 |
| FBXL7 | 24.8478013 | -0.236890009 | 0.34305366 | -0.6905334 | 0.48985881 | 0.7455572 |
| RETREG1 | 123.094319 | -1.600766959 | 0.40847381 | -3.9188974 | 8.90E-05 | 0.00221419 |
| MYO10 | 429.828417 | 0.404083369 | 0.32385451 | 1.24773116 | 0.21212952 | 0.49814414 |
| LINC02241 | 37.4886037 | 0.333977346 | 0.27803541 | 1.20120437 | 0.22967193 | 0.51986597 |
| CDH12 | 14.5999199 | -0.514361519 | 0.55952648 | -0.91928 | 0.35794912 | 0.64214377 |
| C5orf17 | 50.7945364 | 0.482712017 | 0.80415649 | 0.60027124 | 0.54832548 | 0.77527341 |
| CDH6 | 17.6691281 | -0.303187564 | 0.83234431 | -0.3642574 | 0.71566582 | 0.87313942 |
| DROSHA | 79.713451 | 0.240441365 | 0.21849331 | 1.10045183 | 0.27113531 | 0.56326597 |
| C5orf22 | 22.0490862 | 0.186543499 | 0.31356982 | 0.59490259 | 0.55190859 | 0.77741728 |
| PDZD2 | 37.7084348 | 0.332353692 | 0.46556635 | 0.71386967 | 0.47530778 | 0.73587124 |
| GOLPH3 | 128.457689 | 0.058799708 | 0.25141372 | 0.23387629 | 0.81508102 | 0.92163473 |
| MTMR12 | 86.051751 | 0.285279874 | 0.22358487 | 1.27593549 | 0.20197832 | 0.48525669 |
| ZFR | 217.551701 | 0.042299494 | 0.14724485 | 0.28727317 | 0.77390317 | 0.90184317 |
| SUB1 | 96.4399307 | 0.314723153 | 0.29777236 | 1.05692533 | 0.29054567 | 0.58210841 |
| NPR3 | 33.7978189 | 2.752158931 | 0.52781695 | 5.21422988 | 1.85E-07 | 1.44E-05 |
| TARS | 36.8945449 | -0.219560274 | 0.31210842 | -0.7034744 | 0.48176018 | 0.74068045 |
| C1QTNF3-AN | 111.574115 | -0.011056148 | 0.17289585 | -0.0639469 | 0.94901253 | 0.97983806 |
| RAI14 | 167.762023 | 0.251878482 | 0.52200128 | 0.48252465 | 0.62943329 | 0.82508693 |
| RAD1 | 16.949126 | -0.239057163 | 0.36210911 | -0.6601799 | 0.50913838 | 0.7544798 |
| BRIX1 | 17.1395132 | -0.709443551 | 0.37269078 | -1.9035715 | 0.05696601 | 0.23559369 |
| DNAJC21 | 54.058739 | -0.0890091 | 0.23771783 | -0.3744317 | 0.70808314 | 0.86882712 |
| PRLR | 80.9835817 | 1.644948633 | 0.40521158 | 4.0594808 | 4.92E-05 | 0.00138796 |
| SPEF2 | 55.8371415 | 0.237678404 | 0.46867395 | 0.50712954 | 0.61206393 | 0.8140308 |
| UGT3A1 | 31.0334809 | 0.165773496 | 0.8217787 | 0.20172523 | 0.84013154 | 0.93552258 |
| LMBRD2 | 45.0992694 | 0.602884217 | 0.27201249 | 2.2163843 | 0.02666519 | 0.14596993 |
| SKP2 | 11.6249496 | 0.486468046 | 0.42922632 | 1.13336024 | 0.257063 | 0.54817875 |
| NADK2 | 42.2956859 | 0.387907802 | 0.33751477 | 1.14930616 | 0.25042976 | 0.54211544 |
| RANBP3L | 24.6152127 | 0.962662867 | 0.85559826 | 1.1251342 | 0.26053217 | 0.55146957 |
| NIPBL | 503.451502 | 0.415152888 | 0.18110252 | 2.292364 | 0.02188465 | 0.12817262 |
| CPLANE1 | 202.075495 | 0.499704084 | 0.21146479 | 2.36306049 | 0.01812471 | 0.11309556 |
| NUP155 | 32.7384644 | 0.210337122 | 0.30207112 | 0.69631656 | 0.4862306 | 0.74374873 |
| WDR70 | 271.969239 | 0.395609084 | 0.20071966 | 1.97095334 | 0.04872921 | 0.21425029 |
| LIFR | 212.140239 | 1.071834916 | 0.20679478 | 5.18308495 | 2.18E-07 | 1.65E-05 |
| LIFR-AS1 | 12.458543 | 0.306496616 | 0.54980259 | 0.55746667 | 0.57720863 | 0.79249085 |
| OSMR-AS1 | 171.758088 | -1.229307627 | 0.30696725 | -4.0046866 | 6.21E-05 | 0.0016919 |
| OSMR | 212.413901 | -2.160255281 | 0.38858333 | -5.5593102 | 2.71E-08 | 2.88E-06 |
| RICTOR | 267.659278 | 0.183994512 | 0.2818317 | 0.65285245 | 0.51385141 | 0.75647799 |
| FYB1 | 36.2305199 | -2.711407846 | 0.60143051 | -4.5082646 | 6.54E-06 | 0.00026509 |
| DAB2 | 11.4530153 | -2.451563437 | 0.66045028 | -3.7119576 | 0.00020566 | 0.00432395 |
| TTC33 | 46.4629377 | 0.09704698 | 0.25877516 | 0.37502433 | 0.70764238 | 0.86880369 |
| PRKAA1 | 67.7520015 | 0.013815147 | 0.22709069 | 0.06083537 | 0.95149032 | 0.98061757 |
| RPL37 | 79.5350095 | -0.236176949 | 0.35030519 | -0.6742034 | 0.50018201 | 0.75049157 |
| OXCT1 | 33.3266807 | 0.460333839 | 0.41198986 | 1.11734265 | 0.26384785 | 0.55534058 |
| C5orf51 | 16.9262696 | -0.431651962 | 0.36350941 | -1.1874575 | 0.23504723 | 0.52531003 |
| FBXO4 | 11.6367351 | -0.794981174 | 0.50680909 | -1.5686009 | 0.11674097 | 0.35655663 |
| GHR | 162.511646 | 1.523189704 | 0.37625904 | 4.04824749 | 5.16E-05 | 0.00144068 |
| SELENOP | 60.8049997 | 0.147012668 | 0.40414474 | 0.36376242 | 0.71603543 | 0.87318675 |
| ZNF131 | 111.057598 | 0.101229962 | 0.24607298 | 0.41138187 | 0.68079255 | 0.85457838 |
| HMGCS1 | 134.154922 | 1.555307118 | 0.45594618 | 3.41116385 | 0.00064686 | 0.0105276 |
| TMEM267 | 49.6379384 | -0.390424221 | 0.26381905 | -1.479894 | 0.13890155 | 0.39312191 |
| PAIP1 | 51.8220551 | -0.159941634 | 0.26264425 | -0.6089668 | 0.54254644 | 0.77243948 |
| NNT-AS1 | 89.8179986 | 0.35779549 | 0.23475029 | 1.52415355 | 0.12747036 | 0.374451 |
| NNT | 221.496617 | 0.640221502 | 0.20083399 | 3.18781444 | 0.00143353 | 0.01910501 |
| MRPS30-DT | 31.9143831 | -0.179707985 | 0.44035027 | -0.4081024 | 0.68319853 | 0.85592986 |
| MRPS30 | 23.2783688 | -0.158836436 | 0.37504909 | -0.4235084 | 0.67192438 | 0.8481392 |
| AC093297.2 | 10.091392 | 0.078745965 | 0.46120856 | 0.1707383 | 0.86442955 | 0.94667303 |
| PARP8 | 122.140371 | -1.856297524 | 0.5193541 | -3.5742425 | 0.00035124 | 0.00661552 |
| ITGA1 | 71.6343279 | 0.841423107 | 0.32017389 | 2.62801916 | 0.00858837 | 0.06925119 |
| ITGA2 | 749.109461 | -0.366738977 | 0.65533121 | -0.5596239 | 0.57573603 | 0.79171629 |
| MOCS2 | 29.3410571 | 0.296885806 | 0.31308718 | 0.94825284 | 0.34300075 | 0.63099889 |
| NDUFS4 | 143.724621 | 0.605020748 | 0.21572406 | 2.80460482 | 0.00503783 | 0.04787798 |
| ARL15 | 2358.65772 | 2.311195379 | 0.2888619 | 8.00103911 | 1.23E-15 | 9.78E-13 |
| SNX18 | 54.5786784 | 0.494059687 | 0.40495125 | 1.22004733 | 0.22244693 | 0.51073063 |
| DHX29 | 63.0508558 | 0.361426237 | 0.25086807 | 1.44070242 | 0.14966877 | 0.41046846 |
| MTREX | 118.676624 | 0.385554045 | 0.22244871 | 1.73322674 | 0.08305538 | 0.29514884 |
| PLPP1 | 194.262864 | 0.582174918 | 0.19514675 | 2.98326729 | 0.00285189 | 0.03152377 |
| SLC38A9 | 66.2084427 | -0.033770047 | 0.26291319 | -0.1284456 | 0.89779633 | 0.95917897 |
| IL6ST | 148.568593 | 0.099625252 | 0.23924397 | 0.41641699 | 0.6771049 | 0.85212981 |
| AC008892.1 | 23.4642893 | 1.278716535 | 0.56168158 | 2.27658621 | 0.02281095 | 0.13197761 |
| LINC01948 | 20.9278694 | -0.576391547 | 0.49022331 | -1.1757734 | 0.23968543 | 0.53016333 |
| MAP3K1 | 254.103896 | 0.285102798 | 0.29923969 | 0.95275731 | 0.34071305 | 0.62907597 |
| MIER3 | 26.3773701 | 0.899054708 | 0.42461697 | 2.11733106 | 0.03423176 | 0.17102599 |
| GPBP1 | 257.960168 | 0.385018393 | 0.27937407 | 1.37814646 | 0.16815807 | 0.43857328 |
| PDE4D | 8929.63419 | 1.636566561 | 0.31120795 | 5.25875572 | 1.45E-07 | 1.17E-05 |
| DEPDC1B | 43.2209905 | 1.037835511 | 0.72817189 | 1.4252617 | 0.15408157 | 0.41706471 |
| ELOVL7 | 191.83662 | 0.812417878 | 0.42008651 | 1.93392994 | 0.05312174 | 0.22595384 |
| ERCC8 | 59.489553 | 0.93318574 | 0.27814771 | 3.35500057 | 0.00079365 | 0.01238813 |
| NDUFAF2 | 100.386675 | 0.03442642 | 0.34977839 | 0.09842352 | 0.921596 | 0.9686919 |
| SMIM15 | 19.140116 | -0.505198279 | 0.41032205 | -1.2312238 | 0.21823917 | 0.50614586 |
| ZSWIM6 | 543.432255 | 0.654452714 | 0.47716795 | 1.37153536 | 0.17020813 | 0.44081492 |
| KIF2A | 77.0240828 | -0.369608956 | 0.2786899 | -1.3262374 | 0.18476109 | 0.46134371 |
| DIMT1 | 12.7779249 | -0.352109997 | 0.39544503 | -0.8904145 | 0.37324334 | 0.65706646 |
| IPO11 | 50.1302645 | 0.075273526 | 0.30526146 | 0.24658706 | 0.80522782 | 0.91682769 |
| RNF180 | 154.535161 | 0.248102946 | 0.26580828 | 0.9333906 | 0.35061833 | 0.63716177 |
| SREK1IP1 | 48.5932462 | -0.153007187 | 0.28461998 | -0.5375841 | 0.59086418 | 0.80019 |
| CWC27 | 131.653457 | 0.762809492 | 0.22732897 | 3.35553142 | 0.00079213 | 0.01238314 |

| | | | | | | | |
|---|---|---|---|---|---|---|---|
| PPWD1 | 48.7952255 | | 0.183380634 | 0.30565065 | 0.59996807 | 0.54852751 | 0.77527341 |
| TRIM23 | 35.5966631 | | 0.088633977 | 0.28275528 | 0.31346533 | 0.75392715 | 0.89238252 |
| TRAPPC13 | 51.525258 | | 0.363683103 | 0.24910753 | 1.45994427 | 0.14430539 | 0.40211906 |
| NLN | 35.3765223 | | -0.951006147 | 0.37666162 | -2.5248289 | 0.01157546 | 0.08487163 |
| ERBIN | 450.347616 | | 0.125059822 | 0.166222 | 0.75236626 | 0.45183083 | 0.71982927 |
| SREK1 | 129.836531 | | 0.548415927 | 0.22120855 | 2.47918053 | 0.01316846 | 0.09266497 |
| MAST4 | 389.653237 | | -0.926544383 | 0.55797852 | -1.6605377 | 0.09680632 | 0.32139824 |
| PIK3R1 | 95.794406 | | 0.254620467 | 0.29133275 | 0.87398505 | 0.38212639 | 0.66549938 |
| SLC30A5 | 59.7380674 | | 0.194138968 | 0.27773878 | 0.69899842 | 0.48455302 | 0.74250561 |
| MRPS36 | 42.7715202 | | 0.220041147 | 0.26540417 | 0.82907948 | 0.40705944 | 0.68555277 |
| CDK7 | 39.2671808 | | -0.337602209 | 0.28113623 | -1.2008492 | 0.22980971 | 0.51998673 |
| CCDC125 | 68.7250647 | | 0.218696194 | 0.37500851 | 0.58317661 | 0.55977441 | 0.78316119 |
| AK6 | 23.2443139 | | -0.003828344 | 0.34217565 | -0.0111882 | 0.99107326 | 0.99697654 |
| RAD17 | 49.3512488 | | 0.331218731 | 0.2523414 | 1.3125818 | 0.1893239 | 0.46828688 |
| MARVELD2 | 32.7175195 | | -0.094861853 | 0.27400906 | -0.3461997 | 0.72919265 | 0.88046679 |
| OCLN | 94.6213676 | | 0.034945456 | 0.22563025 | 0.1548793 | 0.87691647 | 0.95134725 |
| GTF2H2C | 29.9291862 | | -0.031654508 | 0.32152093 | -0.0984524 | 0.92157306 | 0.9686919 |
| NAIP | 107.730655 | | 0.747453036 | 0.39129296 | 1.91021334 | 0.05610575 | 0.2339636 |
| GTF2H2 | 27.9671197 | | 0.400322497 | 0.4368065 | 0.9164756 | 0.3594175 | 0.64339168 |
| BDP1 | 173.419713 | | 0.383820909 | 0.23857464 | 1.60880849 | 0.10765822 | 0.34161321 |
| MCCC2 | 84.4625431 | | 0.073603049 | 0.20381089 | 0.36113403 | 0.71799925 | 0.87412751 |
| MRPS27 | 72.3356673 | | 0.381702416 | 0.20788321 | 1.83613874 | 0.06633714 | 0.2574031 |
| PTCD2 | 16.9991095 | | 0.907576514 | 0.39335144 | 2.30729172 | 0.02103856 | 0.12441979 |
| TNPO1 | 189.459405 | | 0.307238034 | 0.19965726 | 1.53882724 | 0.12384648 | 0.36885413 |
| FCHO2 | 145.377022 | | -0.439798605 | 0.18722949 | -2.3489815 | 0.01882484 | 0.11636543 |
| BTF3 | 60.7281739 | | -0.875873151 | 0.36871716 | -2.3754608 | 0.01752706 | 0.11097955 |
| ANKRA2 | 25.6063801 | | -0.146846548 | 0.38025869 | -0.3861754 | 0.69936677 | 0.86349933 |
| ARHGEF28 | 305.300116 | | -0.227513205 | 0.31663096 | -0.7185438 | 0.47242207 | 0.73351804 |
| HEXB | 23.0106438 | | -0.014232205 | 0.37341454 | -0.0381137 | 0.96959704 | 0.98781534 |
| GFM2 | 40.7155065 | | 0.551027192 | 0.30205993 | 1.82423136 | 0.0681171 | 0.26218628 |
| NSA2 | 30.230981 | | -0.645602349 | 0.38596827 | -1.6726825 | 0.09438982 | 0.3161038 |
| FAM169A | 73.4077958 | | 0.965919474 | 0.31487771 | 3.06760199 | 0.00215784 | 0.02581658 |
| HMGCR | 95.7920905 | | 1.87147419 | 0.46251285 | 4.04631829 | 5.20E-05 | 0.00144867 |
| COL4A3BP | 182.341272 | | 0.11090599 | 0.27313299 | 0.40605124 | 0.68470495 | 0.85625521 |
| POLK | 117.368546 | | 0.202683046 | 0.23799699 | 0.85162023 | 0.39442491 | 0.67531418 |
| ANKDD1B | 25.6896279 | | -0.174869265 | 0.64858343 | -0.2696172 | 0.78745475 | 0.90904962 |
| POC5 | 22.6266567 | | -0.250767617 | 0.3202149 | -0.7831229 | 0.43355495 | 0.70594011 |
| IQGAP2 | 175.006488 | | 1.221609817 | 0.47622852 | 2.56517571 | 0.01031236 | 0.07859336 |
| F2RL1 | 21.8926875 | | -0.007536707 | 0.38729965 | -0.0194596 | 0.98447444 | 0.99388031 |
| ZBED3 | 27.0193563 | | -0.328238922 | 0.36355835 | -0.9028507 | 0.36660512 | 0.65175201 |
| AC022414.1 | 21.8691241 | | 1.602957201 | 0.56547716 | 2.83469839 | 0.0045869 | 0.0446334 |
| PDE8B | 15.6086598 | | 1.116920821 | 0.502332 | 2.22347135 | 0.02618403 | 0.14440466 |
| WDR41 | 85.5804009 | | 0.002852548 | 0.2317742 | 0.01230744 | 0.99018033 | 0.99684874 |
| TBCA | 159.414632 | | 0.052896915 | 0.22636169 | 0.23368316 | 0.81523096 | 0.92163473 |
| AP3B1 | 310.032282 | | 0.546800448 | 0.17616072 | 3.10398624 | 0.00190932 | 0.02378457 |
| SCAMP1 | 80.0388797 | | 0.113329517 | 0.23223093 | 0.48800355 | 0.62554732 | 0.82303813 |
| LHFPL2 | 12.6453533 | | -0.228081051 | 0.40792749 | -0.5591215 | 0.57607877 | 0.79182968 |
| ARSB | 24.1812205 | | -0.446492649 | 0.49863323 | -0.895433 | 0.37055566 | 0.65494451 |
| DMGDH | 110.292512 | | 0.657042756 | 0.33200325 | 1.97902506 | 0.04781318 | 0.21140405 |
| BHMT2 | 96.1290965 | | 0.303014283 | 0.51361033 | 0.58996922 | 0.55521129 | 0.77958112 |
| BHMT | 27.6676763 | | 1.183275568 | 0.40110069 | 2.95007117 | 0.00317701 | 0.03391661 |
| JMY | 89.7759782 | | 0.329745921 | 0.27040675 | 1.21944411 | 0.22267568 | 0.51114189 |
| HOMER1 | 145.616272 | | 1.82290464 | 1.10229839 | 1.65373066 | 0.09818225 | 0.32367153 |
| TENT2 | 318.088922 | | 0.515488348 | 0.18709441 | 2.75523114 | 0.00586507 | 0.05332916 |
| CMYA5 | 159.086513 | | -0.174902587 | 0.34069482 | -0.5133703 | 0.60769235 | 0.81187554 |
| MTX3 | 22.3076294 | | -0.178895654 | 0.34148732 | -0.523872 | 0.60036756 | 0.80738812 |
| THBS4 | 24.7352153 | | 0.500031077 | 0.53725018 | 0.93072296 | 0.35199689 | 0.63801604 |
| SERINC5 | 97.3649447 | | -0.191258387 | 0.21296839 | -0.89806 | 0.36915358 | 0.65369102 |
| ZFYVE16 | 181.82668 | | 0.679616222 | 0.23985135 | 2.83348925 | 0.00460429 | 0.04474844 |
| AC008771.1 | 16.2586159 | | -0.12993345 | 0.49790166 | -0.2609621 | 0.79412175 | 0.91176607 |
| FAM151B | 37.2439579 | | -0.539992698 | 0.37690447 | -1.4327044 | 0.15194232 | 0.41333768 |
| ANKRD34B | 16.4868037 | | -0.358102233 | 1.08739164 | -0.3293222 | 0.74191215 | 0.88719431 |
| DHFR | 379.975814 | | -1.143946339 | 0.5171797 | -2.2118934 | 0.02697403 | 0.14648981 |
| MSH3 | 164.382175 | | 0.186629518 | 0.17038691 | 1.09532782 | 0.27337304 | 0.5655199 |
| RASGRF2 | 140.873303 | | -0.276178394 | 0.25928299 | -1.065162 | 0.28680262 | 0.57922771 |
| CKMT2-AS1 | 83.3553292 | | -0.002371935 | 0.24991214 | -0.0094911 | 0.99242733 | 0.99770844 |
| CKMT2 | 98.3811097 | | -0.074098615 | 0.3838583 | -0.1930364 | 0.84693048 | 0.93758141 |
| ZCCHC9 | 23.6346739 | | -0.121545035 | 0.30196641 | -0.4025118 | 0.68730743 | 0.85804981 |
| SSBP2 | 294.135867 | | 0.209158017 | 0.19446216 | 1.07557179 | 0.28211881 | 0.5750055 |
| ATG10 | 163.358284 | | 0.410123225 | 0.2087881 | 1.96430361 | 0.04949489 | 0.21605779 |
| RPS23 | 204.621105 | | -0.782406635 | 0.44176276 | -1.7711014 | 0.07654384 | 0.28052461 |
| ATP6AP1L | 62.2364396 | | 0.528493477 | 0.41236984 | 1.28160071 | 0.19998275 | 0.48362026 |
| TMEM167A | 42.5547821 | | -0.475429405 | 0.28078475 | -1.6932166 | 0.09041425 | 0.30986868 |
| XRCC4 | 72.4659506 | | -0.058396119 | 0.30408399 | -0.1920394 | 0.84771131 | 0.93823828 |
| COX7C | 124.875032 | | -0.385736485 | 0.38926346 | -0.9909394 | 0.32171519 | 0.6126463 |
| RASA1 | 214.969634 | | 0.182145175 | 0.24356713 | 0.7478233 | 0.45456674 | 0.72190427 |
| CCNH | 260.680976 | | 0.644466825 | 0.37281801 | 1.7286365 | 0.08387417 | 0.29653069 |
| TMEM161B | 87.8389361 | | 0.878498727 | 0.22511131 | 3.90250812 | 9.52E-05 | 0.00231858 |
| TMEM161B-. | 286.300737 | | 0.534939557 | 0.27399549 | 1.95236629 | 0.05089473 | 0.21990081 |
| MEF2C | 22.5938433 | | -0.294372483 | 0.82852272 | -0.355298 | 0.72236634 | 0.87650915 |
| CETN3 | 10.6303116 | | 0.716494543 | 0.58509471 | 1.22457873 | 0.22073398 | 0.5089143 |
| LYSMD3 | 17.6493413 | | -0.891898598 | 0.47067042 | -1.8949536 | 0.05809855 | 0.238174 |
| ARRDC3 | 94.3519739 | | -0.698035572 | 0.39550199 | -1.7649357 | 0.07757458 | 0.28299338 |
| ARRDC3-AS1 | 12.5660915 | | -0.489297137 | 0.57563376 | -0.8500147 | 0.39531693 | 0.67590514 |
| FAM172A | 367.874483 | | 0.414218042 | 0.16999246 | 2.43668476 | 0.0148226 | 0.09936964 |
| KIAA0825 | 114.514624 | | 0.334868781 | 0.35762916 | 0.93635759 | 0.34908911 | 0.6357564 |
| SLF1 | 17.8980465 | | 0.20425892 | 0.35183963 | 0.58054552 | 0.5615468 | 0.78361672 |
| MCTP1 | 69.1933661 | | 0.330998248 | 0.45528955 | 0.72700603 | 0.46722226 | 0.73023858 |
| TTC37 | 141.970097 | | 0.160404625 | 0.2057087 | 0.77976586 | 0.4355287 | 0.70733441 |
| ARSK | 19.1042471 | | 0.305727755 | 0.43538611 | 0.70219914 | 0.48255498 | 0.74087651 |
| RHOBTB3 | 562.885777 | | 1.310961384 | 0.26562816 | 4.93532531 | 8.00E-07 | 4.68E-05 |
| GLRX | 72.5614514 | | 0.060194152 | 0.56502948 | 0.10653276 | 0.91515966 | 0.96466317 |
| ELL2 | 112.743307 | | 0.272182995 | 0.75228903 | 0.36180641 | 0.7174967 | 0.8738209 |
| AC104123.1 | 55.1598472 | | -1.274537765 | 0.30815977 | -4.1359642 | 3.53E-05 | 0.00105243 |
| CAST | 191.836319 | | -0.451485656 | 0.19834684 | -2.2762432 | 0.02283146 | 0.13197761 |
| ERAP1 | 44.256156 | | -0.513392411 | 0.39524213 | -1.2989314 | 0.19396748 | 0.47453169 |
| AC009126.1 | 12.8764666 | | 0.08172815 | 0.59535625 | 0.13727604 | 0.89081261 | 0.95702609 |
| ERAP2 | 26.2524382 | | -0.891433641 | 0.67009689 | -1.3303056 | 0.1834176 | 0.46014503 |
| LNPEP | 169.099214 | | 0.18380682 | 0.26331585 | 0.69804692 | 0.48514785 | 0.74285379 |
| LIX1-AS1 | 38.4878488 | | 0.21399828 | 0.31991515 | 0.66892199 | 0.50354524 | 0.75108109 |
| RIOK2 | 14.0673889 | | -0.217029885 | 0.39279948 | -0.5525208 | 0.58059159 | 0.7947038 |
| RGMB | 12.1027721 | | -0.432032644 | 0.45726996 | -0.9448087 | 0.34475654 | 0.63253461 |
| CHD1 | 162.866483 | | 0.187138598 | 0.31988708 | 0.58501456 | 0.55853793 | 0.78243918 |
| LINC02062 | 43.6094019 | | -0.071070612 | 0.58664122 | -0.1211483 | 0.90357354 | 0.96126516 |
| AC021086.1 | 12.8628223 | | 0.683137693 | 0.57922716 | 1.17939514 | 0.23824087 | 0.52855279 |
| FAM174A | 27.3810286 | | 0.193840896 | 0.31245562 | 0.62037896 | 0.53500832 | 0.76806796 |
| SLCO4C1 | 47.6228914 | | 1.686469732 | 0.3836072 | 4.39634536 | 1.10E-05 | 0.0003982 |
| PAM | 365.598204 | | -0.677370566 | 0.34906538 | -1.9405264 | 0.05231575 | 0.22381931 |
| GIN1 | 17.9289541 | | -0.002783027 | 0.33023243 | -0.0084275 | 0.99327592 | 0.9979411 |
| PPIP5K2 | 110.445425 | | 0.338282891 | 0.20719957 | 1.63264285 | 0.10254413 | 0.33168278 |
| C5orf30 | 15.3234952 | | 0.635474548 | 0.49650277 | 1.27990131 | 0.20057985 | 0.48426847 |
| NUDT12 | 21.8687976 | | 0.266620025 | 0.3580715 | 0.74459997 | 0.45651359 | 0.72298278 |
| EFNA5 | 1053.92066 | | -0.224096874 | 0.28412518 | -0.7887259 | 0.43027225 | 0.70404459 |

| | | | | | | |
|---|---|---|---|---|---|---|
| FBXL17 | 694.927732 | 0.63413101 | 0.20455624 | 3.10003262 | 0.00193499 | 0.02399848 |
| FER | 419.256578 | 0.200830343 | 0.22704036 | 0.88455791 | 0.37639511 | 0.65957178 |
| PJA2 | 134.029189 | -0.353518212 | 0.186617 | -1.8943516 | 0.05817835 | 0.23840629 |
| MAN2A1 | 287.131473 | 0.203180638 | 0.19124364 | 1.06241776 | 0.28804608 | 0.5802602 |
| TMEM232 | 147.091144 | 0.499701607 | 0.4671768 | 1.06961992 | 0.28479043 | 0.57686567 |
| SLC25A46 | 63.3906347 | -0.097228408 | 0.22333799 | -0.435342 | 0.6633142 | 0.84278344 |
| WDR36 | 48.7356833 | 0.36668256 | 0.24831396 | 1.47668928 | 0.13975895 | 0.39478933 |
| NREP | 55.8032534 | -0.356881424 | 0.59208131 | -0.6027575 | 0.54667006 | 0.77463416 |
| EPB41L4A | 234.218897 | -0.043228859 | 0.20372119 | -0.2121962 | 0.83195398 | 0.93120273 |
| APC | 90.5330762 | 0.190836818 | 0.3186206 | 0.59894689 | 0.5492083 | 0.77580474 |
| SRP19 | 32.6325668 | 0.180217873 | 0.29594456 | 0.60895822 | 0.54255213 | 0.77243948 |
| REEP5 | 154.586247 | -0.518342855 | 0.23698954 | -2.1871972 | 0.02872813 | 0.15294948 |
| DCP2 | 106.868011 | -0.390350383 | 0.30791733 | -1.2677116 | 0.20490095 | 0.48953841 |
| MCC | 126.967816 | 0.598128626 | 0.24305946 | 2.46083254 | 0.01386151 | 0.09564717 |
| YTHDC2 | 127.252159 | 0.868661711 | 0.26597953 | 3.26589685 | 0.00109118 | 0.01552671 |
| PGGT1B | 44.1765245 | -0.022361967 | 0.22954144 | -0.0974202 | 0.92239273 | 0.96885417 |
| CCDC112 | 13.4313364 | -0.358475565 | 0.38683073 | -0.9266988 | 0.35408295 | 0.63957058 |
| FEM1C | 30.5973525 | -0.986110181 | 0.51522907 | -1.9139257 | 0.05562965 | 0.23268234 |
| TMED7-TICAI | 16.043585 | -0.011375628 | 0.48721705 | -0.0233482 | 0.98137255 | 0.99318267 |
| ATG12 | 39.5185849 | -0.122017258 | 0.29685935 | -0.4110272 | 0.68105261 | 0.8546627 |
| AP3S1 | 68.1215475 | 0.145985486 | 0.26049693 | 0.56041154 | 0.57519876 | 0.79151817 |
| COMMD10 | 281.370848 | 0.266783797 | 0.21630844 | 1.233349 | 0.21744558 | 0.50510133 |
| SEMA6A | 89.5401589 | 0.809894502 | 0.74626773 | 1.08525998 | 0.27780653 | 0.56999857 |
| SEMA6A-AS1 | 36.1708847 | 0.459904171 | 0.35715588 | 1.28768473 | 0.19785573 | 0.48016719 |
| DTWD2 | 82.3044848 | 0.604822059 | 0.18785651 | 3.21959604 | 0.00128371 | 0.01763222 |
| AC008629.1 | 38.7959656 | 0.857864156 | 0.49702043 | 1.72601388 | 0.08434491 | 0.29788184 |
| DMXL1 | 235.388329 | 0.264596014 | 0.21041411 | 1.25750131 | 0.20857217 | 0.49402397 |
| TNFAIP8 | 72.7256088 | -3.114723274 | 0.47667339 | -6.5342923 | 6.39E-11 | 1.58E-08 |
| HSD17B4 | 120.127281 | 0.337315286 | 0.3079273 | 1.09543808 | 0.27332476 | 0.5655199 |
| SRFBP1 | 45.2474452 | -0.247208725 | 0.30036439 | -0.8230294 | 0.41049128 | 0.68825899 |
| SNX2 | 165.463925 | 0.681797351 | 0.3339838 | 2.04140842 | 0.04121025 | 0.19341593 |
| SNX24 | 226.31803 | 0.599287508 | 0.2722177 | 2.20150089 | 0.02770058 | 0.14909687 |
| PPIC | 37.9934354 | -0.688187548 | 0.30130847 | -2.2839967 | 0.02237172 | 0.13006405 |
| PRDM6 | 14.8745342 | 0.592260733 | 0.51835597 | 1.14257531 | 0.25321496 | 0.54473281 |
| CEP120 | 44.0323704 | 0.103569538 | 0.26949919 | 0.38430371 | 0.70075336 | 0.86415622 |
| CSNK1G3 | 101.191892 | 0.265721361 | 0.29568225 | 0.89867202 | 0.36882739 | 0.65369102 |
| ZNF608 | 225.747944 | -0.453308938 | 0.32321415 | -1.4025034 | 0.16076498 | 0.42803048 |
| GRAMD2B | 311.470428 | 0.155055002 | 0.25591778 | 0.60587818 | 0.54459565 | 0.77395839 |
| AC093535.1 | 16.9056339 | -0.422834884 | 0.70723373 | -0.5978715 | 0.54992568 | 0.77596691 |
| ALDH7A1 | 43.8111086 | 0.434052127 | 0.37039912 | 1.17184979 | 0.24125737 | 0.53187105 |
| PHAX | 23.3920926 | -0.261927472 | 0.30076778 | -0.8708628 | 0.38382907 | 0.66701235 |
| C5orf63 | 21.4156033 | -0.46312732 | 0.2954771 | -1.5673882 | 0.11702398 | 0.35699765 |
| PRRC1 | 50.6423771 | -0.306575144 | 0.34722894 | -0.8829193 | 0.37727986 | 0.66022373 |
| LINC01184 | 115.437599 | 0.456919429 | 0.28466906 | 1.60508986 | 0.10847403 | 0.34342331 |
| SLC12A2 | 96.0999684 | 1.202995175 | 0.33305534 | 3.6119979 | 0.00030385 | 0.00596235 |
| SLC27A6 | 17.3945024 | -1.345853403 | 1.84120933 | -0.7309616 | 0.46480258 | 0.72848544 |
| ISOC1 | 14.67593 | 0.201529831 | 0.44463377 | 0.45324904 | 0.65036942 | 0.83735207 |
| HINT1 | 150.223307 | -1.162723648 | 0.37491305 | -3.1013155 | 0.00192663 | 0.02394226 |
| LYRM7 | 31.0342506 | 0.532515112 | 0.3129052 | 1.70184166 | 0.08878505 | 0.30687325 |
| CDC42SE2 | 294.084159 | -1.090045325 | 0.32385771 | -3.3658156 | 0.00076318 | 0.01211441 |
| P4HA2 | 13.9249012 | -1.135289986 | 0.37957695 | -2.990935 | 0.00278125 | 0.0311101 |
| SLC22A5 | 40.4652109 | 0.176629764 | 0.34061953 | 0.51855442 | 0.6040715 | 0.8095859 |
| AC116366.3 | 113.788786 | 0.165150849 | 0.55108648 | 0.29968227 | 0.76441952 | 0.89764618 |
| IRF1 | 23.179989 | -2.008017109 | 0.46138726 | -4.3521295 | 1.35E-05 | 0.00047243 |
| KIF3A | 62.2542074 | 0.036045142 | 0.19981213 | 0.18039516 | 0.85684235 | 0.94359207 |
| 8-Sep | 37.2842844 | 0.614557448 | 0.38638779 | 1.59051984 | 0.11171768 | 0.34846418 |
| SHROOM1 | 15.3427615 | -1.102826798 | 0.57329409 | -1.9236668 | 0.05439636 | 0.22976275 |
| UQCRQ | 57.8842147 | -0.494497264 | 0.44147291 | -1.1201078 | 0.26266782 | 0.55434704 |
| AFF4 | 298.35784 | -0.116847235 | 0.27923937 | -0.4184483 | 0.67561939 | 0.85167196 |
| ZCCHC10 | 34.9633937 | -0.301092573 | 0.29653109 | -1.0153828 | 0.30992338 | 0.60321222 |
| HSPA4 | 79.0759668 | 0.238435811 | 0.3484113 | 0.68435154 | 0.49375321 | 0.74700665 |
| FSTL4 | 233.287043 | 2.531116995 | 0.69670993 | 3.63295666 | 0.00028019 | 0.0056488 |
| C5orf15 | 75.6213238 | 0.388655514 | 0.39427904 | 0.9857372 | 0.32426209 | 0.61460758 |
| VDAC1 | 129.267604 | 0.266446777 | 0.28992014 | 0.91903507 | 0.35807723 | 0.64216076 |
| SKP1 | 136.216437 | -0.287349158 | 0.35706495 | -0.8047532 | 0.42096212 | 0.69661173 |
| CDKL3 | 42.9735158 | 0.095414458 | 0.32397769 | 0.29450934 | 0.76836873 | 0.89996549 |
| UBE2B | 121.520459 | -0.534141216 | 0.25678587 | -2.0801036 | 0.03751603 | 0.18135358 |
| JADE2 | 35.2732817 | -0.765699872 | 0.36528619 | -2.0961643 | 0.03606762 | 0.17602651 |
| SAR1B | 53.5713816 | -0.058117989 | 0.25719984 | -0.2259643 | 0.82122916 | 0.92482541 |
| SEC24A | 70.0651638 | -0.278738438 | 0.31160194 | -0.8945337 | 0.37103641 | 0.65494451 |
| CAMLG | 26.9364629 | -0.530162337 | 0.31190051 | -1.6997803 | 0.08917226 | 0.30744734 |
| DDX46 | 117.384193 | 0.224816533 | 0.26677286 | 0.84272641 | 0.39938148 | 0.67979406 |
| C5orf24 | 37.336745 | -0.255032947 | 0.36908749 | -0.6909824 | 0.48957663 | 0.7455572 |
| TXNDC15 | 27.8951442 | 0.091498627 | 0.37351136 | 0.24496879 | 0.8064806 | 0.91724035 |
| PCBD2 | 102.743271 | 0.323811066 | 0.27469609 | 1.1787975 | 0.23847882 | 0.52868706 |
| H2AFY | 80.7287561 | -0.126193984 | 0.2728859 | -0.4624423 | 0.64376417 | 0.83339931 |
| CXCL14 | 19.2099919 | -1.177028802 | 0.64979515 | -1.8113844 | 0.07008137 | 0.26694834 |
| SMAD5 | 92.1060026 | 0.036248472 | 0.21277276 | 0.17036237 | 0.86472516 | 0.94689611 |
| KLHL3 | 165.079367 | 2.204439178 | 0.4130861 | 5.33651263 | 9.48E-08 | 8.20E-06 |
| HNRNPA0 | 48.3222238 | -0.864877246 | 0.37757922 | -2.2905848 | 0.02198744 | 0.1285195 |
| AC106791.1 | 29.5969191 | 0.022578068 | 0.46613831 | 0.04843641 | 0.96136844 | 0.98499239 |
| FAM13B | 221.56271 | 0.275973489 | 0.22003843 | 1.25420589 | 0.20976718 | 0.49598841 |
| NME5 | 47.8761498 | 0.468258515 | 0.44143631 | 1.06076122 | 0.28879843 | 0.58064041 |
| BRD8 | 40.6390457 | 0.03436527 | 0.286248 | 0.12005418 | 0.90444023 | 0.96126516 |
| CDC23 | 22.1321692 | 0.151585462 | 0.35043643 | 0.43256194 | 0.66533304 | 0.84348349 |
| FAM53C | 15.7251316 | -0.473647507 | 0.40342269 | -1.1740725 | 0.24036597 | 0.5311562 |
| KDM3B | 105.435812 | 0.136823446 | 0.25160945 | 0.54379295 | 0.58658396 | 0.79752689 |
| EGR1 | 66.3805632 | 1.290026761 | 1.17074765 | 1.10188286 | 0.2705126 | 0.56240594 |
| ETF1 | 72.6147707 | -0.079356428 | 0.34721258 | -0.2285529 | 0.81921647 | 0.92352826 |
| HSPA9 | 140.28453 | 0.095850063 | 0.29225002 | 0.32797282 | 0.74293221 | 0.88740176 |
| CTNNA1 | 415.41285 | 0.130389087 | 0.26130336 | 0.49899507 | 0.61778286 | 0.81764942 |
| SIL1 | 172.94147 | 0.33877777 | 0.17017001 | 1.99081942 | 0.04650074 | 0.20727463 |
| PAIP2 | 75.7502658 | -0.161547119 | 0.26874701 | -0.6011123 | 0.54776522 | 0.77516644 |
| UBE2D2 | 285.986816 | -0.723178338 | 0.19270688 | -3.7527375 | 0.00017491 | 0.0037777 |
| CXXC5 | 121.488543 | 0.032718464 | 0.30232777 | 0.10822183 | 0.91381973 | 0.96462448 |
| PURA | 98.5316022 | -0.335392461 | 0.26233595 | -1.2784846 | 0.20107863 | 0.48456421 |
| IGIP | 15.3765833 | -0.334184675 | 0.58199772 | -0.5742027 | 0.56583063 | 0.7859225 |
| CYSTM1 | 296.778409 | 0.436710996 | 0.39837393 | 1.09623387 | 0.27297644 | 0.56515339 |
| PFDN1 | 49.1849309 | 0.250887823 | 0.34233385 | 0.73287471 | 0.46363485 | 0.72812334 |
| AC008438.1 | 12.9274138 | 0.376327638 | 0.50935145 | 0.73883688 | 0.46000605 | 0.72516945 |
| ANKHD1-EIF4 | 14.2747852 | -0.13727278 | 0.43808279 | -0.3133489 | 0.75401557 | 0.89238252 |
| SRA1 | 23.7264055 | -0.356152482 | 0.38841404 | -0.9169403 | 0.35917393 | 0.6430674 |
| APBB3 | 10.1213058 | 0.226054813 | 0.4663516 | 0.48473044 | 0.62786757 | 0.82428562 |
| SLC35A4 | 24.7800999 | -0.091839425 | 0.46327825 | -0.1982382 | 0.84285874 | 0.93600812 |
| NDUFA2 | 40.8978592 | -0.189037126 | 0.43003938 | -0.439581 | 0.66024063 | 0.84139027 |
| IK | 19.0465437 | -0.836422758 | 0.50735446 | -1.6485964 | 0.09923034 | 0.32535676 |
| WDR55 | 16.3305557 | -0.427849216 | 0.39648885 | -1.0790952 | 0.28054529 | 0.57367559 |
| HARS | 30.1986808 | 0.012754438 | 0.28290332 | 0.04508409 | 0.96404029 | 0.98660829 |
| ZMAT2 | 19.7571515 | -0.241198197 | 0.4421411 | -0.5455231 | 0.58539378 | 0.79697723 |
| TAF7 | 30.114146 | -0.678767879 | 0.36001769 | -1.8853737 | 0.0593794 | 0.24140749 |
| DIAPH1 | 301.609151 | -0.019790252 | 0.22523399 | -0.0878653 | 0.92998374 | 0.97253199 |
| HDAC3 | 33.0046905 | 0.439649907 | 0.32365751 | 1.35838004 | 0.17434312 | 0.44712045 |
| PCDH1 | 44.0343514 | 0.290480337 | 0.40031247 | 0.725634 | 0.46806317 | 0.73071477 |

| | | | | | | |
|---|---|---|---|---|---|---|
| DELE1 | 37.3582103 | 0.035800471 | 0.25492662 | 0.14043441 | 0.88831677 | 0.95661336 |
| RNF14 | 34.1379919 | -0.04663901 | 0.36801022 | -0.1267329 | 0.89915179 | 0.95950505 |
| GNPDA1 | 27.5709634 | -1.470295079 | 0.40698398 | -3.6126608 | 0.00030307 | 0.00595946 |
| NDFIP1 | 187.347031 | -0.100643821 | 0.17669135 | -0.5696024 | 0.56894739 | 0.78780861 |
| SPRY4-AS1 | 49.1528983 | -0.371595141 | 0.48051454 | -0.7733276 | 0.43932854 | 0.71051219 |
| ARHGAP26 | 95.3411209 | -1.747964819 | 0.42455057 | -4.1172123 | 3.83E-05 | 0.00112836 |
| YIPF5 | 22.775642 | -0.690097324 | 0.34153073 | -2.020601 | 0.04332109 | 0.19870608 |
| PRELID2 | 107.161911 | -1.005784595 | 0.44318108 | -2.2694664 | 0.02323998 | 0.13345498 |
| LARS | 91.4835574 | 0.345967852 | 0.19166244 | 1.80508947 | 0.07106068 | 0.26907809 |
| TCERG1 | 114.409666 | 0.361425694 | 0.23366552 | 1.54676518 | 0.12191988 | 0.36611339 |
| PPP2R2B | 130.473295 | -1.915289429 | 0.8017046 | -2.3890214 | 0.01689332 | 0.10870392 |
| STK32A | 39.7155949 | 0.541840232 | 0.8281821 | 0.65425252 | 0.51294913 | 0.75599752 |
| AC091948.1 | 46.2502842 | 1.075290663 | 0.4366209 | 2.46275583 | 0.01378738 | 0.09526329 |
| FBXO38 | 55.883232 | 0.177610109 | 0.27955966 | 0.63532095 | 0.52521911 | 0.76221787 |
| CSNK1A1 | 429.036832 | 0.155196713 | 0.19541888 | 0.79417461 | 0.42709379 | 0.70162975 |
| ARHGEF37 | 14.9614972 | -0.857896267 | 0.58156114 | -1.4751609 | 0.14016929 | 0.39525538 |
| SLC26A2 | 26.5000705 | 0.132231221 | 0.55136396 | 0.23982565 | 0.81046542 | 0.91943781 |
| HMGXB3 | 46.0966549 | -0.444531563 | 0.23767511 | -1.8703328 | 0.06143761 | 0.24564181 |
| TCOF1 | 17.0935647 | -0.329461658 | 0.44613417 | -0.7384811 | 0.46022214 | 0.72529637 |
| CD74 | 316.562729 | -2.557099812 | 0.6545824 | -3.9064598 | 9.37E-05 | 0.0022973 |
| RPS14 | 180.008743 | -0.593929081 | 0.35888803 | -1.6549147 | 0.0979418 | 0.3233963 |
| NDST1 | 35.1110151 | 0.166756579 | 0.31900224 | 0.52274423 | 0.60115226 | 0.80754603 |
| SYNPO | 15.9796269 | -0.560587363 | 0.40124425 | -1.3971225 | 0.16237674 | 0.42996461 |
| RBM22 | 15.2181157 | -0.503755594 | 0.44613924 | -1.1291443 | 0.25883695 | 0.55002264 |
| DCTN4 | 88.1517794 | 0.223287407 | 0.18656673 | 1.19682327 | 0.23137545 | 0.52111057 |
| GPX3 | 256.000626 | -0.639331731 | 0.69295421 | -0.9226176 | 0.35620654 | 0.64081257 |
| TNIP1 | 96.9297022 | -0.851280519 | 0.33181729 | -2.5655098 | 0.01030243 | 0.07859336 |
| ANXA6 | 53.5584226 | 0.453615185 | 0.46070353 | 0.98461408 | 0.32481367 | 0.61522898 |
| ATOX1 | 40.8296703 | 0.006410618 | 0.33931261 | 0.01889296 | 0.9849265 | 0.99403006 |
| G3BP1 | 125.543679 | 0.2427819 | 0.22060858 | 1.10050976 | 0.27111008 | 0.56326597 |
| FAM114A2 | 25.5538354 | 0.13292112 | 0.31096919 | 0.42744145 | 0.66905782 | 0.84624109 |
| MFAP3 | 34.915394 | 0.084094395 | 0.29896212 | 0.28128779 | 0.77848967 | 0.90417143 |
| GALNT10 | 82.9320796 | 0.593549924 | 0.31766185 | 1.86849606 | 0.06169296 | 0.24624197 |
| SAP30L-AS1 | 23.1639205 | 0.065708932 | 0.42854653 | 0.15332975 | 0.87813824 | 0.9522716 |
| SAP30L | 16.6245553 | -0.117704894 | 0.35778707 | -0.3289803 | 0.74217059 | 0.88720442 |
| LARP1 | 175.129372 | 0.024008055 | 0.26289737 | 0.09132102 | 0.92723752 | 0.97111125 |
| FAXDC2 | 28.1683584 | 0.2990841 | 0.47053609 | 0.63562415 | 0.52502143 | 0.76221787 |
| CNOT8 | 26.7173348 | -0.386403408 | 0.33856489 | -1.1412979 | 0.25374596 | 0.54524749 |
| GEMIN5 | 24.4800301 | 0.326416009 | 0.3162802 | 1.03204694 | 0.30205013 | 0.59577262 |
| MRPL22 | 23.2436822 | -0.091633455 | 0.34445365 | -0.2660255 | 0.79021957 | 0.91080793 |
| SGCD | 153.176125 | 1.325315653 | 0.80915218 | 1.63790655 | 0.10144119 | 0.32966787 |
| HAVCR2 | 11.6633087 | -0.362729413 | 0.49987692 | -0.7256374 | 0.46806106 | 0.73071477 |
| CYFIP2 | 200.860863 | 1.049544069 | 0.31620766 | 3.3191608 | 0.00090288 | 0.01363901 |
| CLINT1 | 277.291661 | -0.356996627 | 0.20048565 | -1.7806593 | 0.07496813 | 0.27777682 |
| RNF145 | 85.170259 | -1.167717892 | 0.36557193 | -3.194222 | 0.00140208 | 0.01887257 |
| UBLCP1 | 29.3275208 | 0.173850628 | 0.4456164 | 0.39013516 | 0.6964366 | 0.86247999 |
| AC008691.1 | 15.8458014 | -1.461230415 | 0.77360228 | -1.8888652 | 0.05890989 | 0.24008773 |
| TTC1 | 64.1430353 | -0.154864025 | 0.20874958 | -0.7418651 | 0.45816909 | 0.72410401 |
| PWWP2A | 103.050171 | 0.378465921 | 0.3070333 | 1.2326543 | 0.21770477 | 0.50547544 |
| CCNJL | 26.8096708 | 0.888908204 | 0.35019829 | 2.53829969 | 0.01113925 | 0.08252559 |
| SLU7 | 20.5827002 | -0.53895999 | 0.34911887 | -1.5437722 | 0.12264353 | 0.36686228 |
| AC113414.1 | 106.686714 | -1.105283492 | 0.5193116 | -2.1283628 | 0.03330702 | 0.16856421 |
| CCNG1 | 42.1239668 | -1.267000309 | 0.39172675 | -3.2343982 | 0.00121899 | 0.01692463 |
| NUDCD2 | 15.9236062 | -0.20009235 | 0.39211157 | -0.5102944 | 0.60984521 | 0.81344383 |
| MAT2B | 34.9990188 | -0.349843637 | 0.43795271 | -0.798816 | 0.42439711 | 0.69892168 |
| WWC1 | 624.225216 | 0.249309877 | 0.44566776 | 0.55940748 | 0.57588366 | 0.79171629 |
| RARS | 53.7124399 | -0.148048007 | 0.2387725 | -0.6200379 | 0.53523281 | 0.76808755 |
| PANK3 | 58.436146 | -0.037333795 | 0.28587896 | -0.130593 | 0.89609727 | 0.95900099 |
| KCNIP1 | 210.913843 | 1.368956645 | 0.8117596 | 1.68640647 | 0.09171754 | 0.311704 |
| RANBP17 | 68.658964 | -0.301544942 | 0.41806395 | -0.721289 | 0.4707317 | 0.73210719 |
| NPM1 | 75.3307619 | -0.097579198 | 0.24154826 | -0.4039739 | 0.6862319 | 0.85756108 |
| FBXW11 | 248.771074 | 0.083930657 | 0.28747098 | 0.29196219 | 0.77031553 | 0.90066855 |
| STK10 | 23.2068499 | -0.868180675 | 0.31131032 | -2.7887951 | 0.00529045 | 0.04950733 |
| UBTD2 | 64.9900247 | -0.139854648 | 0.31335804 | -0.4463094 | 0.65537374 | 0.83931753 |
| DUSP1 | 145.422527 | 1.322780638 | 0.69563082 | 1.90155555 | 0.05722928 | 0.23601925 |
| ERGIC1 | 151.165685 | -0.44253064 | 0.23501037 | -1.883026 | 0.05969685 | 0.24198543 |
| ATP6V0E1 | 234.204439 | -0.382774635 | 0.21521866 | -1.7785383 | 0.07531549 | 0.27859969 |
| CREBRF | 150.614903 | -0.478481264 | 0.29021622 | -1.6487061 | 0.09920785 | 0.32535676 |
| BOD1 | 18.7250223 | -0.287067349 | 0.3643652 | -0.7878561 | 0.43078088 | 0.70420575 |
| CPEB4 | 450.875246 | 1.141777088 | 0.3522367 | 3.24150515 | 0.001189 | 0.01657525 |
| SFXN1 | 124.025539 | 0.315386854 | 0.21628476 | 1.4582019 | 0.1447849 | 0.40273129 |
| THOC3 | 55.7145349 | -0.150517585 | 0.37017895 | -0.4066076 | 0.68429619 | 0.85613081 |
| SIMC1 | 62.114446 | 0.066854946 | 0.2706692 | 0.24699872 | 0.80490921 | 0.91666756 |
| KIAA1191 | 45.4213813 | -0.524297937 | 0.36131728 | -1.4510735 | 0.14675939 | 0.40603754 |
| ARL10 | 17.3985898 | -0.320545584 | 0.40043878 | -0.8004859 | 0.42342935 | 0.69817048 |
| CLTB | 35.1170005 | -0.279062305 | 0.3520886 | -0.7925911 | 0.42801608 | 0.70236088 |
| FAF2 | 84.918048 | 0.063710599 | 0.32413509 | 0.1965557 | 0.84417524 | 0.93684082 |
| RNF44 | 24.237708 | 0.230115824 | 0.34097034 | 0.67488516 | 0.49974873 | 0.75045478 |
| TSPAN17 | 15.3579372 | -0.369698188 | 0.51219523 | -0.7217916 | 0.47042264 | 0.73195802 |
| UIMC1 | 106.609716 | -0.055972127 | 0.21571211 | -0.2594761 | 0.79526796 | 0.91195838 |
| ZNF346 | 57.7156359 | 0.370457404 | 0.26022665 | 1.42359517 | 0.15456369 | 0.41793049 |
| NSD1 | 297.557948 | 0.259649989 | 0.18889246 | 1.3745916 | 0.16925809 | 0.44032355 |
| PRELID1 | 28.5085325 | -1.007835083 | 0.43103337 | -2.3381835 | 0.01937773 | 0.11819385 |
| LMAN2 | 60.1037878 | -0.540124032 | 0.30859154 | -1.7502879 | 0.08006865 | 0.28876788 |
| DBN1 | 25.3675591 | -0.627126014 | 0.55184929 | -1.1364081 | 0.2557858 | 0.54693968 |
| PDLIM7 | 13.3140395 | -1.452350336 | 0.63573926 | -2.2845063 | 0.02234179 | 0.12998973 |
| DDX41 | 11.9258338 | -1.074834852 | 0.51404622 | -2.0909304 | 0.0365343 | 0.17761982 |
| FAM193B | 61.6376307 | 0.035893733 | 0.25291945 | 0.14191765 | 0.88714505 | 0.95619452 |
| TMED9 | 60.9685367 | -1.389929325 | 0.45192286 | -3.0755898 | 0.00210087 | 0.02536445 |
| B4GALT7 | 22.1755439 | -0.312354259 | 0.47156817 | -0.6623735 | 0.50773188 | 0.75382716 |
| FAM153A | 13.9269984 | -2.561264392 | 1.0234497 | -2.5025797 | 0.01232919 | 0.08863593 |
| AC106795.3 | 20.5210031 | -0.860129719 | 0.4881428 | -1.7620453 | 0.07806165 | 0.28436743 |
| RMND5B | 42.9938481 | -0.049935074 | 0.29577511 | -0.1688278 | 0.86593206 | 0.94694132 |
| NHP2 | 21.9097377 | -0.958549558 | 0.42471299 | -2.2569348 | 0.02401215 | 0.13578849 |
| HNRNPAB | 29.1640557 | -0.949692602 | 0.36519954 | -2.6004759 | 0.00930946 | 0.07315486 |
| PHYKPL | 104.811131 | -0.307046656 | 0.25646061 | -1.1972468 | 0.23121036 | 0.52111057 |
| CLK4 | 49.194204 | 1.008526432 | 0.30649269 | 3.29053992 | 0.00099995 | 0.01473751 |
| ZNF354A | 20.3757298 | -0.324707344 | 0.32736266 | -0.9918888 | 0.3212518 | 0.6120836 |
| ZNF354B | 17.7306628 | 0.041935413 | 0.36450792 | 0.11504664 | 0.90840815 | 0.96240444 |
| ZFP2 | 19.0260223 | 0.723018889 | 0.4513569 | 1.60187845 | 0.10918249 | 0.34465697 |
| ZNF354C | 13.9276469 | 0.414126904 | 0.37939313 | 1.09155087 | 0.27503056 | 0.56696009 |
| RUFY1 | 86.1850825 | 0.179639768 | 0.2676861 | 0.67108366 | 0.50216723 | 0.75108109 |
| HNRNPH1 | 224.124174 | 0.86600762 | 0.25581373 | 3.38530551 | 0.00071099 | 0.01139133 |
| CANX | 225.138488 | -0.601048826 | 0.28633734 | -2.0990934 | 0.03580867 | 0.17558348 |
| MAML1 | 36.0019211 | 0.235230496 | 0.27983186 | 0.84061371 | 0.40056437 | 0.68010059 |
| MGAT4B | 51.3211215 | -1.684439504 | 0.46519467 | -3.6209347 | 0.00029354 | 0.00584924 |
| SQSTM1 | 239.273779 | -0.449220024 | 0.46524797 | -0.9655497 | 0.33426957 | 0.62283326 |
| MRNIP | 36.7439255 | -0.22067704 | 0.29286931 | -0.7535001 | 0.45114948 | 0.71947224 |
| TBC1D9B | 50.4877657 | -0.576800558 | 0.28553871 | -2.0200433 | 0.0433789 | 0.19888268 |
| RNF130 | 157.385352 | -0.400953629 | 0.41080596 | -0.9760171 | 0.329056 | 0.6182628 |
| MAPK9 | 55.128699 | 0.317900306 | 0.23910205 | 1.32955911 | 0.18366358 | 0.46016652 |
| GFPT2 | 65.8807714 | -2.017691567 | 0.54461479 | -3.7048049 | 0.00021155 | 0.00442972 |
| CNOT6 | 76.6906553 | 0.230808763 | 0.23726106 | 0.97280506 | 0.3306502 | 0.62024005 |

| | | | | | | |
|---|---|---|---|---|---|---|
| MGAT1 | 36.8262123 | -0.413369019 | 0.33328619 | -1.2402825 | 0.21487093 | 0.50124063 |
| ZFP62 | 16.2407099 | -0.520182063 | 0.342074 | -1.5206712 | 0.12834238 | 0.37572697 |
| TRIM41 | 19.5322922 | -0.065304873 | 0.37489281 | -0.1741961 | 0.86171133 | 0.94550174 |
| RACK1 | 124.988838 | -0.629740996 | 0.37540691 | -1.6774891 | 0.09344688 | 0.31481026 |
| TRIM52 | 22.1369558 | 0.269433354 | 0.3220318 | 0.83666692 | 0.40277982 | 0.68128594 |
| AC138035.1 | 10.5137933 | 0.819094694 | 0.45902006 | 1.78444204 | 0.07435187 | 0.27592684 |
| DUSP22 | 32.4856125 | -0.631694256 | 0.2774227 | -2.27701 | 0.02278563 | 0.13197761 |
| EXOC2 | 120.573702 | -0.189074448 | 0.22888233 | -0.8260771 | 0.40876036 | 0.6869518 |
| GMDS | 309.359404 | -0.480706444 | 0.25688143 | -1.8713164 | 0.06130123 | 0.24544318 |
| GMDS-DT | 481.115005 | -0.332552292 | 0.26166645 | -1.2709015 | 0.20376368 | 0.48807721 |
| WRNIP1 | 34.960131 | 0.005957944 | 0.28653787 | 0.02079287 | 0.98341089 | 0.99368608 |
| SERPINB1 | 38.823405 | -1.600928981 | 0.32585237 | -4.91305 | 8.97E-07 | 5.10E-05 |
| SERPINB6 | 70.5257216 | -0.679449677 | 0.26077536 | -2.605498 | 0.00917408 | 0.07242252 |
| NQO2 | 44.0602848 | 0.305398641 | 0.35318061 | 0.86470954 | 0.38719823 | 0.66950996 |
| RIPK1 | 51.0097032 | -0.485744715 | 0.33553426 | -1.4476755 | 0.14770783 | 0.40745374 |
| BPHL | 19.5879418 | -0.773274607 | 0.40961766 | -1.8877961 | 0.05905334 | 0.24052299 |
| PSMG4 | 65.1159189 | 0.095747763 | 0.4400958 | 0.21756119 | 0.82777103 | 0.92813421 |
| SLC22A23 | 122.503203 | -0.538125101 | 0.70865021 | -0.7593663 | 0.44763345 | 0.71693398 |
| PXDC1 | 13.4241914 | -0.816638348 | 0.4453563 | -1.8336742 | 0.06670237 | 0.25852816 |
| PRPF4B | 128.484773 | -0.150963639 | 0.22010729 | -0.6858639 | 0.49279896 | 0.74680729 |
| ECI2 | 36.9131989 | -0.130037313 | 0.26756079 | -0.4860103 | 0.62695983 | 0.82354566 |
| CDYL | 184.876626 | -0.188787019 | 0.26568767 | -0.71056 | 0.47735696 | 0.73707006 |
| LYRM4 | 59.598 | -0.028032102 | 0.35880595 | -0.0781261 | 0.93772776 | 0.97619962 |
| FARS2 | 251.999539 | 0.478072347 | 0.23069883 | 2.07227902 | 0.03823943 | 0.18314393 |
| LY86-AS1 | 14.1991552 | -0.174501556 | 0.86336669 | -0.2021175 | 0.83982483 | 0.93543959 |
| RREB1 | 198.859833 | 0.106879244 | 0.17948765 | 0.59546852 | 0.55153034 | 0.77722877 |
| SSR1 | 76.7380714 | -0.006745822 | 0.29581838 | -0.0228039 | 0.98180667 | 0.99318267 |
| RIOK1 | 18.543424 | -0.136496049 | 0.35287575 | -0.3868105 | 0.69889649 | 0.86334649 |
| DSP | 137.504656 | -0.669848481 | 0.27628065 | -2.4245219 | 0.01532856 | 0.10149612 |
| SNRNP48 | 22.4643979 | -0.062630216 | 0.37011294 | -0.1692192 | 0.86562423 | 0.94694132 |
| BLOC1S5-TXI | 24.9564038 | 0.307165281 | 0.3503239 | 0.87680368 | 0.38059328 | 0.66412918 |
| SLC35B3 | 33.778963 | -0.467961761 | 0.30311444 | -1.5438452 | 0.12262585 | 0.36686228 |
| TFAP2A | 102.709303 | 1.020728104 | 0.24330638 | 4.19523777 | 2.73E-05 | 0.00084664 |
| GCNT2 | 178.583357 | 0.219779401 | 0.32111663 | 0.68442237 | 0.4937085 | 0.74700665 |
| TMEM14C | 35.4484619 | -0.488314801 | 0.39246653 | -1.2442202 | 0.21341851 | 0.49980394 |
| MAK | 41.9080938 | -0.51486209 | 0.48309336 | -1.0657611 | 0.28653167 | 0.57913465 |
| SMIM13 | 13.3068336 | 0.414268447 | 0.37951131 | 1.09158392 | 0.27501603 | 0.56696009 |
| AL139807.1 | 20.9890717 | 0.225328011 | 0.5049682 | 0.44622218 | 0.65543677 | 0.83931753 |
| NEDD9 | 58.2430164 | 0.68301009 | 0.24804932 | 2.75352529 | 0.00589572 | 0.0535606 |
| TMEM170B | 21.0295769 | 0.775712382 | 0.43164299 | 1.79711566 | 0.07231726 | 0.27172106 |
| HIVEP1 | 111.408328 | -0.864697895 | 0.37119704 | -2.3294849 | 0.01983339 | 0.12004913 |
| PHACTR1 | 91.6014063 | -0.511909239 | 0.30611298 | -1.6722886 | 0.09446743 | 0.3161038 |
| TBC1D7 | 35.4618965 | 0.970736505 | 0.44997506 | 2.15731178 | 0.03098138 | 0.16111569 |
| SIRT5 | 51.1698528 | 1.000433207 | 0.35089492 | 2.85109056 | 0.00435696 | 0.0428543 |
| NOL7 | 35.7676851 | -0.387865788 | 0.32611196 | -1.1893639 | 0.23429651 | 0.52443761 |
| RANBP9 | 152.765253 | 0.397443466 | 0.19166758 | 2.07360819 | 0.03811571 | 0.18280637 |
| MCUR1 | 29.5449382 | -0.159318775 | 0.34355896 | -0.4637305 | 0.64284083 | 0.83318468 |
| JARID2 | 218.212381 | 1.728929411 | 0.46014243 | 3.75737881 | 0.0001717 | 0.00373181 |
| DTNBP1 | 123.888747 | -0.818015124 | 0.38887904 | -2.1035207 | 0.03542028 | 0.17464009 |
| MYLIP | 32.5707324 | -1.191815222 | 0.42644848 | -2.7947461 | 0.00519405 | 0.04891142 |
| GMPR | 59.910236 | -2.090202823 | 0.73356199 | -2.8493881 | 0.00438034 | 0.0429774 |
| ATXN1 | 688.374596 | 0.915146496 | 0.30663392 | 2.98449205 | 0.0028405 | 0.03143157 |
| CAP2 | 97.7785546 | 1.169818871 | 0.2578732 | 4.53641116 | 5.72E-06 | 0.00023769 |
| FAM8A1 | 17.966387 | -0.519806196 | 0.39562578 | -1.3138835 | 0.1888854 | 0.4677638 |
| NUP153 | 117.922682 | 0.610508014 | 0.26494662 | 2.30426803 | 0.0212076 | 0.12513214 |
| KIF13A | 153.773708 | 0.08626785 | 0.20129898 | 0.42855582 | 0.6682465 | 0.84573166 |
| TPMT | 28.3391766 | 0.178217682 | 0.2731794 | 0.6523833 | 0.51415393 | 0.75668769 |
| KDM1B | 22.3088395 | -0.182396136 | 0.35618057 | -0.512089 | 0.60858875 | 0.81261032 |
| DEK | 71.8992833 | -0.365584382 | 0.21912257 | -1.6684013 | 0.09523609 | 0.31779803 |
| RNF144B | 204.457936 | -1.987388219 | 0.40863621 | -4.8634658 | 1.15E-06 | 6.22E-05 |
| AL022068.1 | 24.6337799 | -0.857665537 | 0.68725286 | -1.2479621 | 0.21204493 | 0.49805901 |
| ID4 | 15.4547291 | -0.758541647 | 0.43232916 | -1.7545466 | 0.07933689 | 0.28711147 |
| MBOAT1 | 101.863576 | 0.092194249 | 0.36539749 | 0.25231221 | 0.80079975 | 0.9148089 |
| E2F3 | 67.0161843 | -0.665582757 | 0.31903275 | -2.0862521 | 0.03695579 | 0.17933046 |
| CDKAL1 | 455.642577 | 0.342084419 | 0.20103481 | 1.70161786 | 0.08882703 | 0.30687325 |
| SOX4 | 101.044994 | -1.706180469 | 0.61545549 | -2.772224 | 0.00556747 | 0.05134834 |
| CASC15 | 53.971436 | 0.354365701 | 0.48415497 | 0.73192618 | 0.46421362 | 0.72835095 |
| DCDC2 | 2099.70376 | -0.096682485 | 0.28013731 | -0.3451253 | 0.73000014 | 0.88123523 |
| MRS2 | 57.652866 | 0.504722456 | 0.24808274 | 2.03449245 | 0.04190197 | 0.19532764 |
| ALDH5A1 | 18.7786457 | 0.108206518 | 0.44734632 | 0.24188534 | 0.80886901 | 0.91863836 |
| TDP2 | 28.8557237 | -0.487065663 | 0.32409254 | -1.5028598 | 0.13287519 | 0.38268946 |
| ACOT13 | 20.022776 | -0.146593692 | 0.45024878 | -0.3255838 | 0.7447393 | 0.88808711 |
| C6orf62 | 111.136702 | 0.133150568 | 0.29269038 | 0.45491953 | 0.64916713 | 0.83648778 |
| GMNN | 44.0979435 | 1.137553772 | 0.36532413 | 3.11382053 | 0.00184682 | 0.02311776 |
| CARMIL1 | 354.919678 | 0.212186991 | 0.37606288 | 0.56423275 | 0.57259574 | 0.78978194 |
| SLC17A1 | 16.0111241 | 0.405550333 | 0.81588367 | 0.49706882 | 0.61914052 | 0.81858132 |
| TRIM38 | 89.4060788 | -0.155284535 | 0.37818194 | -0.410608 | 0.68135998 | 0.85466584 |
| HIST1H1C | 14.9883727 | -1.598272243 | 0.59911671 | -2.6677143 | 0.00763692 | 0.06391579 |
| HIST1H4C | 28.5377642 | -2.370630594 | 0.96509258 | -2.4563764 | 0.01403461 | 0.09623211 |
| HIST1H2AC | 46.530773 | 1.638336803 | 0.66972776 | 2.4462728 | 0.01443417 | 0.09759315 |
| BTN3A2 | 18.8538168 | -1.437895418 | 0.9373173 | -1.5340541 | 0.12501635 | 0.37009151 |
| BTN3A1 | 16.1247682 | -1.161426617 | 1.01323345 | -1.1462577 | 0.25168855 | 0.54333973 |
| BTN2A1 | 18.244571 | -0.64218088 | 0.44897261 | -1.4303342 | 0.15262112 | 0.41441825 |
| HMGN4 | 13.8189765 | -1.016239103 | 0.52883047 | -1.9216727 | 0.05464695 | 0.23034896 |
| ZNF322 | 14.1476922 | -0.394535514 | 0.47982041 | -0.8222566 | 0.41093085 | 0.68858322 |
| LINC00240 | 23.4484568 | 1.021960937 | 0.54394747 | 1.87878608 | 0.06027371 | 0.24330674 |
| ZSCAN16-AS | 19.8967794 | -0.389395592 | 0.39131508 | -0.9950948 | 0.31969017 | 0.611915 |
| ZKSCAN8 | 20.1143503 | -0.551413194 | 0.39883015 | -1.3825765 | 0.16679475 | 0.43585929 |
| ZSCAN9 | 14.8167513 | -0.853007601 | 0.40387918 | -2.1120366 | 0.03468331 | 0.17300048 |
| ZSCAN23 | 11.9449694 | 0.583712106 | 0.41935874 | 1.3919159 | 0.16394787 | 0.43270826 |
| TRIM27 | 52.1071922 | -0.282161394 | 0.22066018 | -1.2787146 | 0.20099757 | 0.48448221 |
| GABBR1 | 26.524296 | -0.026997283 | 0.41394099 | -0.0652201 | 0.94799874 | 0.979567 |
| HLA-F | 20.1764323 | -1.58152811 | 1.01545005 | -1.5574652 | 0.11936008 | 0.36176744 |
| HLA-A | 451.108514 | -1.837560692 | 0.61475261 | -2.989106 | 0.00279795 | 0.03121198 |
| PPP1R11 | 18.1454275 | -1.17033786 | 0.35663552 | -3.2816077 | 0.00103217 | 0.01508287 |
| TRIM26 | 55.2109373 | 0.428736271 | 0.38035404 | 1.12720315 | 0.2596566 | 0.55074784 |
| HCG17 | 126.778943 | 0.346778198 | 0.23249569 | 1.49154679 | 0.135818 | 0.38804335 |
| HLA-E | 167.117444 | -2.258469478 | 0.5316159 | -4.2483106 | 2.15E-05 | 0.00069126 |
| GNL1 | 44.6155566 | -0.206061531 | 0.27325153 | -0.7541094 | 0.45078356 | 0.71921516 |
| PRR3 | 12.0203626 | -0.497591427 | 0.46455721 | -1.071109 | 0.2841204 | 0.5765312 |
| ABCF1 | 38.5930638 | -0.467783051 | 0.30850542 | -1.516288 | 0.12944656 | 0.37713756 |
| PPP1R10 | 44.8132088 | 0.482647883 | 0.26941255 | 1.79148257 | 0.07321589 | 0.27338532 |
| MRPS18B | 39.0006625 | -0.590086186 | 0.37384977 | -1.5784046 | 0.11447269 | 0.35348341 |
| ATAT1 | 22.641168 | -0.160133799 | 0.39361702 | -0.4068264 | 0.68413549 | 0.85606265 |
| C6orf136 | 34.9461994 | 0.30453344 | 0.26566636 | 1.14630036 | 0.25167089 | 0.54333973 |
| DHX16 | 19.4436215 | -0.605339037 | 0.38841506 | -1.558485 | 0.11911833 | 0.36124727 |
| MDC1 | 23.4888597 | -0.096922713 | 0.34348618 | -0.2821735 | 0.77781044 | 0.90389207 |
| TUBB | 82.1518534 | -1.052032088 | 0.52169049 | -2.0165829 | 0.04373906 | 0.19964547 |
| FLOT1 | 87.2589379 | -0.634240707 | 0.29853435 | -2.124515 | 0.0336271 | 0.16956751 |
| IER3 | 218.535017 | -0.452785987 | 0.77918388 | -0.5811029 | 0.56117113 | 0.78361672 |
| DDR1 | 120.635465 | -0.425354334 | 0.34815627 | -1.221734 | 0.22180825 | 0.51005995 |
| VARS2 | 12.1396653 | 0.500907171 | 0.41734633 | 1.20021941 | 0.23005414 | 0.52031125 |
| CCHCR1 | 17.6368499 | 0.003725162 | 0.4177701 | 0.00891678 | 0.99288554 | 0.9979411 |

| | | | | | | |
|---|---|---|---|---|---|---|
| POU5F1 | 38.6992567 | -0.106227995 | 0.52440867 | -0.2025672 | 0.83947333 | 0.93543959 |
| HLA-C | 228.226222 | -2.007928103 | 0.71581365 | -2.8050989 | 0.00503011 | 0.04784878 |
| HLA-B | 373.367621 | -2.975088009 | 0.65424952 | -4.5473293 | 5.43E-06 | 0.00023129 |
| MICA | 24.2620148 | -0.057979084 | 0.48254166 | -0.1201535 | 0.90436153 | 0.96126516 |
| NFKBIL1 | 14.95118 | -0.634609173 | 0.39035355 | -1.6257292 | 0.10400727 | 0.33490577 |
| PRRC2A | 54.7524527 | 0.025382499 | 0.31249796 | 0.08122453 | 0.93526339 | 0.97521088 |
| BAG6 | 28.5360785 | -0.303620832 | 0.39302567 | -0.7725216 | 0.43980554 | 0.71072575 |
| DDAH2 | 33.6871783 | -0.208659892 | 0.32876494 | -0.6346781 | 0.52563841 | 0.76221787 |
| CLIC1 | 114.245631 | -1.850993689 | 0.41465951 | -4.4638882 | 8.05E-06 | 0.00031166 |
| VARS | 13.9414722 | -0.837509596 | 0.53964537 | -1.5519629 | 0.1206711 | 0.36349523 |
| LSM2 | 23.8696513 | -0.951032385 | 0.43082372 | -2.2074745 | 0.02728093 | 0.14774125 |
| HSPA1A | 95.8871017 | 1.214291027 | 0.63157225 | 1.92264786 | 0.05452429 | 0.22997772 |
| HSPA1B | 70.1100791 | 1.437238062 | 0.64578188 | 2.22557817 | 0.02604245 | 0.14385486 |
| C6orf48 | 25.3285989 | -0.011529828 | 0.32689681 | -0.0352705 | 0.97186401 | 0.988268 |
| NEU1 | 27.5525031 | -0.843768964 | 0.38052997 | -2.2173522 | 0.02659904 | 0.14581205 |
| SLC44A4 | 25.1413766 | -3.236635479 | 0.62972763 | -5.1397387 | 2.75E-07 | 1.97E-05 |
| EHMT2 | 22.061118 | 0.126929625 | 0.3538848 | 0.35867498 | 0.71983824 | 0.87522408 |
| NELFE | 12.3431012 | -0.398464837 | 0.47114966 | -0.8457288 | 0.39770405 | 0.67867271 |
| STK19 | 19.6917571 | -0.647924221 | 0.44541961 | -1.4546378 | 0.14576955 | 0.40412792 |
| ATF6B | 36.4210544 | -0.204322742 | 0.3129523 | -0.6528878 | 0.51382861 | 0.75647799 |
| AGPAT1 | 15.1651974 | -0.296133091 | 0.5249925 | -0.5640711 | 0.57270575 | 0.78982793 |
| RNF5 | 18.7449195 | -0.776478164 | 0.4127945 | -1.8810284 | 0.05996806 | 0.2428146 |
| PBX2 | 26.2394329 | 0.208102427 | 0.35717271 | 0.58263809 | 0.56013696 | 0.78316119 |
| AL662796.1 | 21.608568 | 0.542913241 | 0.5320235 | 1.02046853 | 0.30750631 | 0.60139222 |
| HLA-DRA | 95.7429591 | -2.152569692 | 0.71832198 | -2.9966641 | 0.00272951 | 0.03069808 |
| HLA-DRB5 | 23.8838559 | -0.774235135 | 0.97808362 | -0.7915838 | 0.42860341 | 0.70287684 |
| HLA-DRB1 | 58.9296258 | -0.912927844 | 0.81598376 | -1.1188064 | 0.26322274 | 0.55488453 |
| HLA-DQB1 | 17.6421015 | -0.798314393 | 0.76185618 | -1.0478545 | 0.29470567 | 0.58661907 |
| PSMB8 | 31.428674 | -1.974712696 | 0.60568437 | -3.2602999 | 0.00111294 | 0.01574939 |
| PSMB9 | 16.3904661 | -2.968154733 | 1.16380676 | -2.5503845 | 0.01076042 | 0.08056236 |
| TAP1 | 27.3134362 | -1.848845013 | 0.84143282 | -2.197258 | 0.02800202 | 0.15017014 |
| HLA-DMA | 8.20225407 | 1.275699184 | 0.59985601 | 2.12667567 | 0.03344704 | 0.16897307 |
| BRD2 | 74.7036694 | -0.125092724 | 0.28308235 | -0.4418952 | 0.65856507 | 0.84037209 |
| HLA-DPA1 | 28.4089843 | -2.419844052 | 0.92423265 | -2.6182196 | 0.00883899 | 0.07053391 |
| HLA-DPB1 | 19.5489383 | -1.825260094 | 0.858196 | -2.1268569 | 0.03343197 | 0.16897307 |
| RXRB | 15.2763897 | 0.317336959 | 0.36511434 | 0.86914406 | 0.38476834 | 0.66701235 |
| SLC39A7 | 29.8194381 | -0.875695828 | 0.59632205 | -1.4684948 | 0.14196987 | 0.39809147 |
| RING1 | 23.7255079 | 0.059118381 | 0.41600678 | 0.14210918 | 0.88699377 | 0.95619452 |
| VPS52 | 27.142225 | 0.293717988 | 0.33739764 | 0.87053954 | 0.38400562 | 0.66701235 |
| RPS18 | 104.600506 | -0.574160456 | 0.41868514 | -1.3713419 | 0.17026841 | 0.44084071 |
| WDR46 | 12.1337824 | -0.265127984 | 0.51067294 | -0.5191737 | 0.60363959 | 0.80940894 |
| PFDN6 | 15.0313585 | -0.575279944 | 0.3832951 | -1.50088 | 0.13338661 | 0.38373327 |
| RGL2 | 25.9517197 | -0.445248357 | 0.42380018 | -1.0506092 | 0.29343812 | 0.58494573 |
| TAPBP | 120.65474 | -1.312690327 | 0.41254244 | -3.1819522 | 0.00146286 | 0.0193956 |
| DAXX | 11.6965072 | -0.534354929 | 0.54995996 | -0.9716251 | 0.33123707 | 0.6208887 |
| PHF1 | 28.4927773 | 0.233773397 | 0.27689212 | 0.84427608 | 0.39851515 | 0.67938457 |
| CUTA | 38.8771679 | -0.264529987 | 0.4266245 | -0.6200534 | 0.53522261 | 0.76808755 |
| SYNGAP1 | 17.6026152 | -0.207775776 | 0.38464972 | -0.5401688 | 0.58908063 | 0.79925045 |
| ITPR3 | 69.0185245 | -2.117450967 | 0.42892635 | -4.9366306 | 7.95E-07 | 4.68E-05 |
| UQCC2 | 26.3903688 | -0.261927328 | 0.32927875 | -0.7954577 | 0.42634728 | 0.70107418 |
| LEMD2 | 24.5541408 | -0.234649953 | 0.31754475 | -0.7389508 | 0.45993686 | 0.72516945 |
| HMGA1 | 30.124846 | -1.29555236 | 0.61989241 | -2.0899633 | 0.0366211 | 0.17795781 |
| SMIM29 | 17.4782234 | -0.948728865 | 0.57921591 | -1.6379537 | 0.10143135 | 0.32966787 |
| NUDT3 | 56.4824844 | -0.261303986 | 0.45834537 | -0.5701028 | 0.56860796 | 0.78763298 |
| C6orf106 | 123.165627 | 0.42678405 | 0.31543588 | 1.35299779 | 0.17605634 | 0.45038707 |
| SNRPC | 21.3726586 | -0.883224369 | 0.43675777 | -2.0222293 | 0.04315267 | 0.1982867 |
| UHRF1BP1 | 59.2668081 | -0.039389688 | 0.25869075 | -0.1522655 | 0.8789775 | 0.95273407 |
| TAF11 | 32.0850621 | 0.130755319 | 0.29626209 | 0.44135015 | 0.65895953 | 0.84059189 |
| ANKS1A | 367.959743 | -0.967289904 | 0.35010811 | -2.762832 | 0.00573022 | 0.05239493 |
| SCUBE3 | 22.9876647 | 0.792467102 | 0.67858526 | 1.16782244 | 0.24287839 | 0.53327647 |
| ZNF76 | 53.5983709 | -0.031016767 | 0.28973768 | -0.1070512 | 0.91474836 | 0.96464134 |
| PPARD | 77.0553994 | 0.49723098 | 0.36967453 | 1.34505067 | 0.17860887 | 0.4541038 |
| RPL10A | 114.413418 | -0.365625546 | 0.37128409 | -0.9847595 | 0.32474221 | 0.61520673 |
| TEAD3 | 23.97536 | -0.706991338 | 0.3357491 | -2.1057132 | 0.03522927 | 0.1742352 |
| FKBP5 | 21.4017895 | -2.510848306 | 0.945073 | -2.6567771 | 0.00788916 | 0.06512349 |
| SRPK1 | 70.4210038 | -0.517114699 | 0.22743813 | -2.2736499 | 0.02298704 | 0.1323843 |
| MAPK14 | 231.666944 | -1.255389213 | 0.34993528 | -3.5874897 | 0.00033388 | 0.00636963 |
| MAPK13 | 19.5119126 | -0.97174661 | 0.37838326 | -2.5681543 | 0.01022416 | 0.07837003 |
| BRPF3 | 17.3965266 | -0.087062061 | 0.34588514 | -0.251708 | 0.80126676 | 0.91484195 |
| KCTD20 | 69.1803212 | 0.340840648 | 0.33233667 | 1.02558845 | 0.30508561 | 0.59961934 |
| STK38 | 100.775651 | -0.116828936 | 0.20633705 | -0.5662044 | 0.57125487 | 0.78905469 |
| SRSF3 | 81.6633031 | 0.080194748 | 0.27021114 | 0.2967855 | 0.76663027 | 0.89911488 |
| CDKN1A | 23.6163598 | -1.881243597 | 0.50165923 | -3.7500428 | 0.0001768 | 0.00380259 |
| CPNE5 | 35.8379623 | 1.666712194 | 0.83722931 | 1.99074755 | 0.04650865 | 0.20727463 |
| C6orf89 | 77.0931843 | -0.202170031 | 0.27422103 | -0.7372521 | 0.46096905 | 0.72602097 |
| MTCH1 | 79.9591788 | -0.302700417 | 0.30471103 | -0.9934016 | 0.3205143 | 0.6120836 |
| PIM1 | 18.7693332 | 0.255289289 | 0.7418101 | 0.34414372 | 0.7307382 | 0.88171293 |
| TBC1D22B | 58.0368513 | 0.15981608 | 0.21106032 | 0.75720571 | 0.44892663 | 0.71802342 |
| RNF8 | 35.1560006 | 0.004490433 | 0.29564282 | 0.01518871 | 0.98788163 | 0.99571045 |
| CMTR1 | 29.1904833 | 0.256725281 | 0.38922237 | 0.65958512 | 0.50952011 | 0.75461129 |
| ZFAND3 | 1053.87412 | 0.274238909 | 0.27475607 | 0.99811774 | 0.31822227 | 0.61132712 |
| BTBD9 | 673.394142 | 0.243334153 | 0.29765099 | 0.817515 | 0.41363418 | 0.69042044 |
| GLO1 | 75.7644714 | -0.014169778 | 0.21830998 | -0.0649067 | 0.94824829 | 0.97962835 |
| SAYSD1 | 11.3590798 | -0.60183045 | 0.45547887 | -1.3213137 | 0.1863968 | 0.46383088 |
| KCNK5 | 19.0632552 | -2.111154023 | 0.56037938 | -3.7673656 | 0.00016498 | 0.00363944 |
| KIF6 | 17.522885 | -0.111877802 | 0.78080922 | -0.1432844 | 0.88606555 | 0.95619452 |
| MOCS1 | 21.212439 | -0.6724519 | 0.3732555 | -1.801586 | 0.07161056 | 0.27013255 |
| UNC5CL | 29.199579 | -2.137585079 | 0.60316761 | -3.5439322 | 0.00039421 | 0.00721335 |
| OARD1 | 44.6048355 | -0.471360316 | 0.36058074 | -1.3072254 | 0.19113616 | 0.47141123 |
| NFYA | 26.381426 | -0.19286394 | 0.3687799 | -0.5229785 | 0.60098925 | 0.80753766 |
| FOXP4 | 14.3486658 | -0.629842555 | 0.37446999 | -1.6819574 | 0.09257711 | 0.31323946 |
| TFEB | 83.8111538 | 1.012628972 | 0.40447754 | 2.50354809 | 0.0122955 | 0.08851727 |
| USP49 | 39.7274664 | -0.047080577 | 0.33255418 | -0.1415727 | 0.88741757 | 0.95619452 |
| CCND3 | 106.102602 | -0.533894242 | 0.47211302 | -1.1308611 | 0.25811357 | 0.54951146 |
| TAF8 | 32.5262262 | -0.040820465 | 0.3790882 | -0.1076807 | 0.91424902 | 0.96462448 |
| C6orf132 | 21.3389516 | -1.277683324 | 0.47824497 | -2.6716085 | 0.00754887 | 0.06358824 |
| MRPS10 | 16.9964056 | 0.113183544 | 0.36004081 | 0.3143631 | 0.75324528 | 0.89205725 |
| TRERF1 | 79.7070595 | 0.639205637 | 0.48716563 | 1.31209098 | 0.18948944 | 0.46847137 |
| UBR2 | 272.636654 | 0.590332677 | 0.22261246 | 2.65184022 | 0.00800544 | 0.0658687 |
| BICRAL | 112.330926 | 0.203249806 | 0.27498156 | 0.73913977 | 0.45982213 | 0.72516945 |
| RPL7L1 | 53.9003507 | -0.687483337 | 0.29438777 | -2.3352986 | 0.01952783 | 0.1186813 |
| CNPY3 | 14.1477915 | -0.713407897 | 0.44267576 | -1.6115811 | 0.10705312 | 0.34111768 |
| PEX6 | 21.7182419 | -0.423378736 | 0.39561468 | -1.0701795 | 0.28453851 | 0.57686567 |
| PPP2R5D | 14.1335723 | 0.825458185 | 0.52441546 | 1.57405386 | 0.115475 | 0.35464129 |
| KLHDC3 | 21.629431 | -0.020115085 | 0.37103098 | -0.054214 | 0.95676464 | 0.98290679 |
| RRP36 | 28.9978202 | -0.308765921 | 0.31429746 | -0.9824003 | 0.32590267 | 0.6159877 |
| CUL7 | 16.7689642 | -0.723716688 | 0.49471839 | -1.4628862 | 0.14349853 | 0.40084649 |
| KLC4 | 26.3523845 | 0.545841144 | 0.3409444 | 1.60096818 | 0.10938397 | 0.34481257 |
| MRPL2 | 18.571771 | 0.857769369 | 0.39638796 | 2.16396426 | 0.03046709 | 0.15932588 |
| PTK7 | 24.3579132 | -0.221874045 | 0.47891501 | -0.4632848 | 0.64316024 | 0.83328507 |
| CUL9 | 23.3097824 | 0.483427828 | 0.38777346 | 1.24667589 | 0.21251636 | 0.49867221 |
| DNPH1 | 27.50868 | -0.489186416 | 0.54431728 | -0.8987156 | 0.36880418 | 0.65369102 |
| ZNF318 | 22.0804201 | 0.546301554 | 0.32642562 | 1.67358664 | 0.09421187 | 0.31604384 |

| | | | | | | |
|---|---|---|---|---|---|---|
| ABCC10 | 18.6159057 | -0.12640638 | 0.43096584 | -0.2933095 | 0.76928559 | 0.90034488 |
| TJAP1 | 23.5068455 | 0.235164262 | 0.34323359 | 0.6851435 | 0.49325337 | 0.74694932 |
| POLR1C | 28.2339864 | -0.125097547 | 0.32094615 | -0.3897774 | 0.69670117 | 0.86247999 |
| YIPF3 | 31.7899625 | -0.675369783 | 0.34378567 | -1.9645083 | 0.04947118 | 0.21605779 |
| XPO5 | 99.9204617 | -0.524785469 | 0.28086624 | -1.8684533 | 0.06169891 | 0.24624197 |
| POLH | 31.9478356 | 0.032302361 | 0.40824987 | 0.079124 | 0.936934 | 0.97613368 |
| GTPBP2 | 27.9722424 | -0.811069217 | 0.34563341 | -2.3466169 | 0.01894472 | 0.11674788 |
| MRPS18A | 15.9835314 | -0.806183001 | 0.48910695 | -1.6482755 | 0.09929615 | 0.32546896 |
| VEGFA | 157.234403 | -0.246375826 | 0.49459236 | -0.4981392 | 0.61838596 | 0.81810866 |
| MRPL14 | 26.107473 | -0.835450471 | 0.51391281 | -1.6256658 | 0.10402077 | 0.33490577 |
| TMEM63B | 33.5709424 | -0.01622358 | 0.32338462 | -0.0501681 | 0.95998846 | 0.98435365 |
| SLC29A1 | 33.5125367 | -0.498908987 | 0.39819676 | -1.2529208 | 0.21023455 | 0.49648138 |
| HSP90AB1 | 257.671019 | -0.202475509 | 0.27709947 | -0.7306961 | 0.46496479 | 0.72850852 |
| SLC35B2 | 17.970732 | -0.588229694 | 0.47380217 | -1.2415091 | 0.21441775 | 0.50088196 |
| CDC5L | 38.704303 | -0.147740972 | 0.2849454 | -0.5184887 | 0.60411733 | 0.8095859 |
| SUPT3H | 341.967695 | 0.687882934 | 0.21702607 | 3.16958661 | 0.00152656 | 0.02005946 |
| RUNX2 | 18.6775082 | -1.324878085 | 0.56900333 | -2.3284189 | 0.01988987 | 0.12024969 |
| ENPP4 | 39.1303297 | -0.354557825 | 0.46886523 | -0.7562041 | 0.44952682 | 0.71810337 |
| ENPP5 | 17.3843689 | -1.01393687 | 0.44084359 | -2.2999923 | 0.02144866 | 0.12612104 |
| RCAN2 | 76.5352793 | 1.018986851 | 0.50733254 | 2.00851863 | 0.04458821 | 0.20147742 |
| CYP39A1 | 14.8361212 | -0.124905791 | 0.59274345 | -0.2107249 | 0.83310196 | 0.93218382 |
| SLC25A27 | 78.5983067 | 0.694127833 | 0.38983791 | 1.78055497 | 0.07498518 | 0.27777682 |
| ADGRF1 | 128.314332 | -0.277193795 | 0.40032783 | -0.692417 | 0.4886755 | 0.74516503 |
| TNFRSF21 | 26.2969929 | -1.569764263 | 0.46687949 | -3.3622472 | 0.00077311 | 0.01221559 |
| AL355353.1 | 11.2289793 | -0.387935418 | 0.52830968 | -0.7342955 | 0.46276867 | 0.72750336 |
| CD2AP | 891.306479 | -0.297325098 | 0.27259809 | -1.0907087 | 0.2754011 | 0.56732296 |
| PTCHD4 | 30.4865123 | -1.067985634 | 1.19927718 | -0.8905244 | 0.37318435 | 0.65706646 |
| MUT | 36.2275079 | 0.111602221 | 0.28726877 | 0.3884941 | 0.69765042 | 0.86275485 |
| PKHD1 | 2437.16735 | 0.976074251 | 0.31816226 | 3.06785052 | 0.00215604 | 0.02581658 |
| MCM3 | 19.9774646 | -0.659044179 | 0.49524262 | -1.3307501 | 0.18327125 | 0.46004581 |
| EFHC1 | 63.8206919 | -0.144251724 | 0.24822485 | -0.5811333 | 0.56115061 | 0.78361672 |
| TRAM2 | 15.5311691 | 0.438805144 | 0.375477 | 1.16866052 | 0.24254043 | 0.53317493 |
| TMEM14A | 9.90296495 | 0.335282987 | 0.4946472 | 0.67782247 | 0.49788426 | 0.74971166 |
| GSTA4 | 21.6057161 | 0.214092319 | 0.36102923 | 0.5930055 | 0.55317747 | 0.77810408 |
| ICK | 22.2123717 | -0.189636814 | 0.44424833 | -0.4268712 | 0.66947315 | 0.84645464 |
| FBXO9 | 76.8649158 | 0.147790768 | 0.22287681 | 0.66310519 | 0.50726318 | 0.75376105 |
| ELOVL5 | 55.3513179 | 1.069592498 | 0.68485599 | 1.56177724 | 0.11834047 | 0.35967154 |
| GCLC | 66.75286 | -0.513000014 | 0.32830517 | -1.5625706 | 0.11815362 | 0.35937957 |
| LRRC1 | 89.8010225 | 0.366058034 | 0.32763026 | 1.11729005 | 0.26387033 | 0.55534058 |
| TINAG | 19.5983664 | -0.37486058 | 0.55389717 | -0.6767693 | 0.49855236 | 0.75012216 |
| FAM83B | 127.720833 | -0.57474819 | 0.30773091 | -1.8676973 | 0.06180428 | 0.2464039 |
| DST | 160.295642 | -1.921594323 | 0.56284917 | -3.4140484 | 0.00064005 | 0.01046637 |
| ZNF451 | 107.891231 | 0.059184453 | 0.19595024 | 0.30203818 | 0.76262296 | 0.89732835 |
| ZNF451-AS1 | 14.93785 | -1.178848936 | 0.65847795 | -1.7902633 | 0.07341159 | 0.2738183 |
| PRIM2 | 81.9532101 | -0.118199697 | 0.42289044 | -0.2795043 | 0.77985784 | 0.90433312 |
| AL021368.2 | 13.7232923 | 0.177708035 | 0.40776569 | 0.43580919 | 0.66297517 | 0.84275734 |
| PTP4A1 | 100.364811 | -0.239152129 | 1.07545144 | -0.2223737 | 0.82402297 | 0.92635145 |
| PHF3 | 191.099159 | -0.052816881 | 0.225513 | -0.2342077 | 0.81482374 | 0.92163473 |
| EYS | 20.4532584 | -0.869812302 | 0.70621422 | -1.231655 | 0.21807798 | 0.50599984 |
| LMBRD1 | 158.04552 | 0.480366518 | 0.19391676 | 2.47717892 | 0.01324255 | 0.09299574 |
| COL19A1 | 43.0391978 | 0.8472926 | 0.38610745 | 2.19444771 | 0.02820324 | 0.15111607 |
| FAM135A | 338.311579 | -0.674385795 | 0.24174451 | -2.7896633 | 0.00527629 | 0.0494598 |
| SDHAF4 | 14.9259536 | -0.015300342 | 0.36970578 | -0.0413852 | 0.96698882 | 0.98701267 |
| SMAP1 | 217.717859 | 0.243888386 | 0.29805443 | 0.81826796 | 0.4132042 | 0.69014748 |
| OGFRL1 | 21.4942109 | -1.034708372 | 0.56255867 | -1.8392897 | 0.06587259 | 0.25655016 |
| AL136164.2 | 132.443814 | 0.521446303 | 0.3466837 | 1.50409811 | 0.13255611 | 0.38212969 |
| LINC00472 | 60.716645 | -0.155850302 | 0.426316 | -0.3655746 | 0.71468253 | 0.87255978 |
| AC019205.1 | 22.0950729 | 0.64423558 | 0.38051786 | 1.69304952 | 0.09044604 | 0.30986868 |
| OOEP | 106.846884 | -0.65212112 | 0.41176812 | -1.5837096 | 0.11325982 | 0.35144659 |
| MTO1 | 40.9789318 | 0.336835572 | 0.33204819 | 1.01441773 | 0.31038347 | 0.60377086 |
| EEF1A1 | 584.797577 | -0.504361216 | 0.44816697 | -1.1253869 | 0.26042514 | 0.55146957 |
| SLC17A5 | 33.9896284 | -0.539861948 | 0.40614673 | -1.3292288 | 0.18377249 | 0.46016652 |
| COX7A2 | 68.5977002 | 0.08124271 | 0.36607932 | 0.22192652 | 0.82437108 | 0.92643003 |
| TMEM30A | 78.7064637 | -0.500597975 | 0.30622082 | -1.6347614 | 0.10209907 | 0.33070354 |
| SENP6 | 393.496172 | 0.415860461 | 0.16966757 | 2.4510309 | 0.01424477 | 0.09699249 |
| MYO6 | 777.835584 | -0.060232896 | 0.32326927 | -0.1863242 | 0.85219052 | 0.94037032 |
| MEI4 | 27.6155733 | 0.827900949 | 0.43339295 | 1.91027782 | 0.05609745 | 0.2339636 |
| IRAK1BP1 | 80.4213855 | 0.251413625 | 0.32760454 | 0.76743022 | 0.44282577 | 0.71303393 |
| PHIP | 301.439484 | 0.179231953 | 0.2010296 | 0.89156995 | 0.37262348 | 0.65642393 |
| HMGN3 | 158.161217 | -0.644209834 | 0.30503654 | -2.1119104 | 0.03469414 | 0.17300048 |
| LCA5 | 26.355783 | -0.244634932 | 0.37929801 | -0.6449676 | 0.51894818 | 0.75983572 |
| SH3BGRL2 | 54.6717274 | 1.096428027 | 0.28106392 | 3.9009917 | 9.58E-05 | 0.00232766 |
| BCKDHB | 231.534726 | 0.255166758 | 0.21368183 | 1.19414344 | 0.23242186 | 0.52234076 |
| AL359715.1 | 11.04186 | 0.782027952 | 0.4586193 | 1.7051789 | 0.08816106 | 0.30539182 |
| TENT5A | 27.227579 | -1.236452004 | 0.51242919 | -2.4129227 | 0.01582518 | 0.10366112 |
| IBTK | 183.63343 | 0.099593439 | 0.29351048 | 0.33931818 | 0.73437005 | 0.88376541 |
| TPBG | 14.2350939 | -2.608029355 | 0.61238918 | -4.2587776 | 2.06E-05 | 0.0006659 |
| UBE3D | 103.93963 | 0.382189663 | 0.22629972 | 1.68886494 | 0.09124532 | 0.31117713 |
| DOPEY1 | 203.976264 | 0.182457182 | 0.20409428 | 0.8939848 | 0.37133002 | 0.65494451 |
| PGM3 | 37.3808627 | -0.577994417 | 0.33038474 | -1.7494586 | 0.08021178 | 0.28903174 |
| CYB5R4 | 51.7168458 | 0.299237909 | 0.34348821 | 0.87117373 | 0.3836593 | 0.66701235 |
| MRAP2 | 16.4118388 | 0.452962187 | 0.68930008 | 0.65713352 | 0.51109507 | 0.7549696 |
| CEP162 | 78.2231608 | -0.003922256 | 0.2291269 | -0.0171183 | 0.98634226 | 0.9949057 |
| SNX14 | 133.090073 | 0.118742245 | 0.17610684 | 0.67426254 | 0.50014441 | 0.75049157 |
| SYNCRIP | 50.9398251 | 0.253012046 | 0.23255676 | 1.08795826 | 0.27661353 | 0.56884726 |
| ZNF292 | 569.321132 | -1.435738947 | 0.33104845 | -4.3369451 | 1.44E-05 | 0.00049946 |
| SMIM8 | 92.5998269 | 0.641540672 | 0.27926163 | 2.29727472 | 0.0216031 | 0.12673985 |
| RARS2 | 105.439852 | 0.257787229 | 0.27698705 | 0.93068333 | 0.35201739 | 0.63801604 |
| ORC3 | 64.7290225 | -0.100024125 | 0.31507792 | -0.3174584 | 0.75089581 | 0.89131567 |
| AKIRIN2 | 23.0187769 | -0.205113333 | 0.30068167 | -0.6821611 | 0.49513711 | 0.74770824 |
| RNGTT | 182.556774 | 0.557924696 | 0.26288979 | 2.12227604 | 0.03381456 | 0.1700965 |
| PNRC1 | 74.2073224 | -0.993693468 | 0.32302892 | -3.0761749 | 0.00209675 | 0.02536445 |
| PM20D2 | 36.8030154 | -0.528543073 | 0.3559956 | -1.4846899 | 0.13762599 | 0.39109388 |
| UBE2J1 | 39.6724517 | -0.112443077 | 0.28035482 | -0.4010742 | 0.68836552 | 0.8585881 |
| RRAGD | 73.3581314 | 0.405881515 | 0.33159336 | 1.2240339 | 0.22093944 | 0.5089143 |
| ANKRD6 | 146.695 | 0.104869043 | 0.22644447 | 0.46311153 | 0.64328443 | 0.83328507 |
| LYRM2 | 35.6673792 | -0.937236182 | 0.28197246 | -3.3238572 | 0.00088782 | 0.0134901 |
| MDN1 | 107.305298 | 0.188058474 | 0.19430002 | 0.96787675 | 0.33310592 | 0.62190756 |
| CASP8AP2 | 45.706784 | 0.477457578 | 0.27499518 | 1.73623981 | 0.08252144 | 0.29436838 |
| BACH2 | 242.706275 | 0.092805039 | 0.3591991 | 0.25836657 | 0.79612402 | 0.91195838 |
| MAP3K7 | 96.1316905 | 0.170233897 | 0.22019603 | 0.77310157 | 0.43946227 | 0.71061691 |
| MANEA | 20.2435629 | 0.470596917 | 0.36040693 | 1.30573769 | 0.19164178 | 0.47198029 |
| UFL1-AS1 | 30.8361739 | -0.777924448 | 0.44288452 | -1.756495 | 0.07900392 | 0.28648307 |
| UFL1 | 60.604788 | 0.028326676 | 0.27405444 | 0.1033615 | 0.91767607 | 0.9663858 |
| NDUFAF4 | 21.0071624 | 0.27005538 | 0.39097768 | 0.69071815 | 0.48974268 | 0.7455572 |
| KLHL32 | 49.336428 | 0.969149598 | 0.43071074 | 2.25011711 | 0.02444151 | 0.13669732 |
| MMS22L | 62.4975879 | -0.987307558 | 0.43219005 | -2.2844292 | 0.02234632 | 0.12998973 |
| AL589740.1 | 15.201334 | 0.659812805 | 0.59065806 | 1.11708084 | 0.26395976 | 0.55541533 |
| FBXL4 | 133.90743 | 0.189637021 | 0.2249212 | 0.84312647 | 0.39915772 | 0.67968972 |
| FAXC | 14.7756459 | -0.017216865 | 0.44312075 | -0.0388537 | 0.96900706 | 0.98770265 |
| PNISR | 408.239601 | 0.582541243 | 0.20606465 | 2.82698292 | 0.00469888 | 0.04553894 |
| AL513550.1 | 23.0428814 | 0.521355979 | 0.47362387 | 1.10078062 | 0.27099215 | 0.5631957 |
| USP45 | 72.1604313 | -0.114900963 | 0.33748924 | -0.3404581 | 0.73351153 | 0.88350705 |

| | | | | | | |
|---|---|---|---|---|---|---|
| TSTD3 | 14.5975248 | -0.127641752 | 0.3949328 | -0.3231987 | 0.7465448 | 0.88870184 |
| CCNC | 106.242884 | -0.187868908 | 0.28085414 | -0.6689199 | 0.5035466 | 0.75108109 |
| SIM1 | 274.251625 | 0.626896145 | 0.26502807 | 2.36539529 | 0.01801084 | 0.11286352 |
| ASCC3 | 185.622263 | -0.34518699 | 0.18057025 | -1.9116493 | 0.05592119 | 0.23361723 |
| GRIK2 | 29.6500616 | -1.398220739 | 0.52196498 | -2.6787635 | 0.00738946 | 0.06273624 |
| HACE1 | 76.419824 | 0.553749772 | 0.24092844 | 2.29839934 | 0.02153907 | 0.12650827 |
| PREP | 51.2794537 | -0.102280169 | 0.24477047 | -0.4178616 | 0.67604834 | 0.85173672 |
| ATG5 | 130.570148 | 0.148710611 | 0.25714186 | 0.57832129 | 0.56304723 | 0.78438303 |
| CRYBG1 | 397.158481 | 0.747912608 | 0.3231568 | 2.31439542 | 0.02064604 | 0.1230165 |
| RTN4IP1 | 34.4861278 | 0.413909544 | 0.27890394 | 1.4840577 | 0.13779361 | 0.39127613 |
| QRSL1 | 53.9989929 | 0.246507444 | 0.24219199 | 1.01781833 | 0.3087643 | 0.60259154 |
| LINC02532 | 741.723447 | -0.266117698 | 0.34645565 | -0.7681148 | 0.442419 | 0.71265656 |
| CD24 | 641.392044 | -1.181477476 | 0.32694452 | -3.6136941 | 0.00030187 | 0.00595946 |
| C6orf203 | 35.4181116 | 0.294194954 | 0.3134438 | 0.93858916 | 0.34794172 | 0.63501969 |
| PDSS2 | 238.82393 | 0.295926866 | 0.21503438 | 1.37618395 | 0.16876469 | 0.43928463 |
| SOBP | 76.9216541 | 1.23062733 | 0.47045282 | 2.61583578 | 0.00890094 | 0.07080883 |
| SEC63 | 112.687789 | -0.275054134 | 0.24816252 | -1.1083629 | 0.26770511 | 0.55952485 |
| OSTM1 | 39.7918017 | -0.178633872 | 0.3831693 | -0.4662009 | 0.6410717 | 0.83230084 |
| SNX3 | 95.2227185 | -0.136528396 | 0.19859447 | -0.6874733 | 0.49178451 | 0.74615082 |
| AFG1L | 98.8947052 | 0.868112197 | 0.240648 | 3.6073942 | 0.00030929 | 0.00603462 |
| FOXO3 | 246.20343 | -0.14497284 | 0.2078385 | -0.6975264 | 0.48547342 | 0.74292145 |
| ARMC2 | 56.4887892 | -0.089659634 | 0.26370244 | -0.3400031 | 0.73385423 | 0.88360989 |
| SESN1 | 133.83316 | 0.022705441 | 0.23633363 | 0.09607368 | 0.92346206 | 0.96947989 |
| CEP57L1 | 22.7830756 | -0.095296859 | 0.37690743 | -0.2528389 | 0.80039271 | 0.91455697 |
| CD164 | 131.542683 | -0.362030652 | 0.23863841 | -1.5170678 | 0.12924957 | 0.3768834 |
| PPIL6 | 27.1457091 | -0.270133571 | 0.55459812 | -0.4870799 | 0.62620175 | 0.8231524 |
| MICAL1 | 11.4090086 | -0.950134071 | 0.4392856 | -2.1629074 | 0.0305483 | 0.15958855 |
| ZBTB24 | 12.7493361 | -0.751864173 | 0.47530114 | -1.5818691 | 0.11367946 | 0.35221829 |
| AK9 | 70.9500251 | -0.375159741 | 0.29654491 | -1.2651026 | 0.20583455 | 0.48999801 |
| FIG4 | 57.3343642 | -0.046226273 | 0.29320679 | -0.1576576 | 0.87472663 | 0.94993692 |
| WASF1 | 21.052254 | -1.810779398 | 0.51108753 | -3.5429927 | 0.00039561 | 0.00721347 |
| CDC40 | 36.6287336 | 0.176704739 | 0.25829358 | 0.68412363 | 0.4938971 | 0.74700665 |
| CDK19 | 232.767089 | -0.051442631 | 0.35219091 | -0.1460646 | 0.88387038 | 0.95587158 |
| AMD1 | 107.586894 | -0.247331609 | 0.44540393 | -0.5552973 | 0.57869131 | 0.7935467 |
| GTF3C6 | 47.7664193 | -0.662890581 | 0.40722502 | -1.6278238 | 0.10356225 | 0.33425166 |
| RPF2 | 26.0298335 | 0.407675315 | 0.38108974 | 1.06976197 | 0.28472647 | 0.57686567 |
| SLC16A10 | 176.34085 | 1.383603253 | 0.33264192 | 4.15943745 | 3.19E-05 | 0.00096667 |
| MFSD4B | 25.3049025 | 0.085599557 | 0.32584469 | 0.26270048 | 0.79278145 | 0.91124908 |
| REV3L | 282.276396 | 0.478415779 | 0.15869143 | 3.01475502 | 0.00257187 | 0.02934152 |
| TRAF3IP2-AS | 102.446467 | -0.234409494 | 0.29815927 | -0.7861888 | 0.43175686 | 0.70457139 |
| TRAF3IP2 | 33.1275774 | -0.056440602 | 0.37518094 | -0.1504357 | 0.88042089 | 0.95334202 |
| TUBE1 | 40.3919275 | 0.658063732 | 0.40671478 | 1.61799808 | 0.10566301 | 0.33784616 |
| FAM229B | 13.6197692 | 0.05638281 | 0.53357701 | 0.10566949 | 0.91584459 | 0.96518083 |
| MARCKS | 25.5981806 | -1.092312432 | 0.39962635 | -2.7333344 | 0.00626966 | 0.05587378 |
| HDAC2 | 51.5779837 | -0.199122779 | 0.25303103 | -0.78695 | 0.43131112 | 0.70454366 |
| FRK | 247.702204 | 0.525188294 | 0.25115251 | 2.09111304 | 0.03651793 | 0.17761982 |
| NT5DC1 | 265.909777 | 0.585906177 | 0.20025547 | 2.92579369 | 0.00343579 | 0.03597101 |
| COL10A1 | 12.8696641 | -1.010553787 | 0.82167468 | -1.2298709 | 0.21874545 | 0.50674964 |
| TSPYL4 | 16.4787549 | 0.516582162 | 0.43184875 | 1.19621085 | 0.23161429 | 0.52132191 |
| DSE | 93.2185235 | 0.692132855 | 0.3991354 | 1.73408035 | 0.08290383 | 0.29503415 |
| TSPYL1 | 50.9459359 | -0.273969566 | 0.41024084 | -0.6678262 | 0.50424457 | 0.75133461 |
| RWDD1 | 29.8319523 | 0.187963774 | 0.32588035 | 0.57678769 | 0.56408288 | 0.78488324 |
| ZUP1 | 22.1372668 | -0.235979609 | 0.38718002 | -0.6094829 | 0.54220439 | 0.77243948 |
| KPNA5 | 61.0461765 | 0.229472384 | 0.24371148 | 0.94157395 | 0.34641081 | 0.63353882 |
| DCBLD1 | 34.5646841 | 0.132028643 | 0.33209831 | 0.39755891 | 0.69095535 | 0.85967888 |
| GOPC | 49.9134921 | 0.048331335 | 0.29767417 | 0.16236322 | 0.87101984 | 0.94844301 |
| NUS1 | 59.6811919 | 0.232122079 | 0.26873186 | 0.86376838 | 0.38771514 | 0.66966967 |
| CEP85L | 83.3300436 | -0.257436034 | 0.1895783 | -1.3579404 | 0.1744826 | 0.44724303 |
| MCM9 | 102.751926 | -0.130213577 | 0.21568966 | -0.603708 | 0.54603779 | 0.77463416 |
| AL365275.1 | 29.9141565 | -0.426959987 | 0.4977985 | -0.8576964 | 0.39106012 | 0.67223049 |
| FAM184A | 49.4320794 | 1.267303359 | 0.43608066 | 2.90612143 | 0.00365939 | 0.03766744 |
| MAN1A1 | 443.267826 | 1.237272402 | 0.27229225 | 4.54391346 | 5.52E-06 | 0.00023292 |
| TBC1D32 | 148.330322 | 0.191805021 | 0.25776921 | 0.74409593 | 0.45681844 | 0.72302753 |
| HSF2 | 19.1206931 | 0.092012437 | 0.36564576 | 0.25164366 | 0.80131651 | 0.91484195 |
| SERINC1 | 82.6595172 | -0.752450882 | 0.27627927 | -2.7235156 | 0.00645912 | 0.0568659 |
| PKIB | 17.8945353 | 0.482789708 | 0.60227225 | 0.80161373 | 0.42277643 | 0.69778363 |
| RNF217 | 101.412155 | -1.103583913 | 0.35947831 | -3.0699597 | 0.00214088 | 0.02573549 |
| TPD52L1 | 194.036681 | -0.072220434 | 0.32398586 | -0.2229123 | 0.82360376 | 0.92608229 |
| HDDC2 | 42.7231841 | 0.03483213 | 0.2727878 | 0.12768947 | 0.89839472 | 0.95935794 |
| NCOA7 | 160.242392 | -1.386138539 | 0.34077288 | -4.0676316 | 4.75E-05 | 0.00134787 |
| HINT3 | 30.7195019 | -0.207594995 | 0.34432156 | -0.6029102 | 0.5465684 | 0.77463416 |
| TRMT11 | 43.196193 | -0.305294114 | 0.28797676 | -1.0601346 | 0.28908339 | 0.58094924 |
| AC020559.1 | 16.7162829 | -0.471606884 | 0.43677182 | -1.0797557 | 0.28025096 | 0.57321005 |
| RNF146 | 32.5694678 | -0.18642413 | 0.32293904 | -0.5772734 | 0.56375475 | 0.78473199 |
| ECHDC1 | 84.1561529 | 0.013021095 | 0.19945903 | 0.06528205 | 0.94794943 | 0.979567 |
| PTPRK | 545.50504 | -0.460872812 | 0.27308601 | -1.6876471 | 0.09147899 | 0.31120975 |
| AL356124.1 | 18.0560135 | -0.707924244 | 0.41256454 | -1.7159115 | 0.08617824 | 0.3018729 |
| ARHGAP18 | 168.241694 | 0.565397772 | 0.30874815 | 1.83125885 | 0.06706192 | 0.25972627 |
| L3MBTL3 | 35.6053945 | -0.331053709 | 0.5315248 | -0.6228377 | 0.53339115 | 0.7669018 |
| EPB41L2 | 53.4686093 | -1.079763017 | 0.37810907 | -2.8556919 | 0.00429432 | 0.04233497 |
| AKAP7 | 55.6815265 | 0.95998113 | 0.37811787 | 2.53884094 | 0.01112204 | 0.08249045 |
| MED23 | 62.6441253 | 0.24857236 | 0.23431836 | 1.06083175 | 0.28876637 | 0.58064041 |
| CTGF | 13.9424581 | -0.303895382 | 0.67838635 | -0.447968 | 0.65417628 | 0.83885315 |
| STX7 | 120.594136 | -0.085844177 | 0.19740209 | -0.4348696 | 0.66365705 | 0.84292874 |
| SLC18B1 | 59.2410078 | 0.418345471 | 0.34125702 | 1.22589557 | 0.22023798 | 0.50852 |
| RPS12 | 94.0721191 | -0.421998294 | 0.45140938 | -0.9348461 | 0.34986762 | 0.63669603 |
| EYA4 | 163.904674 | 1.056224728 | 0.6295145 | 1.6778402 | 0.0933783 | 0.31478509 |
| TBPL1 | 20.164614 | -0.007599607 | 0.33195755 | -0.0228933 | 0.98173538 | 0.99318267 |
| SLC2A12 | 12.3696072 | 1.866983835 | 0.60600958 | 3.08078268 | 0.00206457 | 0.02511125 |
| SGK1 | 160.527112 | 1.028749922 | 0.78811303 | 1.30533297 | 0.19177949 | 0.47205703 |
| HBS1L | 71.1359396 | 0.438511437 | 0.22462676 | 1.95217811 | 0.05091706 | 0.21990081 |
| AHI1 | 297.720493 | 0.091464633 | 0.25912644 | 0.35297298 | 0.7241087 | 0.87742017 |
| LINC00271 | 123.338648 | -0.117934711 | 0.42500435 | -0.2774906 | 0.78140343 | 0.90531018 |
| PDE7B | 268.664872 | -0.017991882 | 0.47137889 | -0.0381686 | 0.96955324 | 0.98781534 |
| AL138828.1 | 44.5888006 | -2.213984634 | 0.55592656 | -3.9825128 | 6.82E-05 | 0.00180128 |
| BCLAF1 | 140.027494 | 0.190364912 | 0.17849863 | 1.06647831 | 0.28620748 | 0.57870647 |
| MAP7 | 333.217418 | 0.158516818 | 0.2292229 | 0.69154006 | 0.48922621 | 0.74545311 |
| MAP3K5 | 166.367395 | -1.079186833 | 0.38887688 | -2.7751375 | 0.00551784 | 0.05102761 |
| PEX7 | 54.4223962 | 0.280854768 | 0.24251238 | 1.15810486 | 0.24682125 | 0.53769348 |
| IFNGR1 | 42.6632457 | -0.317964956 | 0.24445643 | -1.3007019 | 0.1933605 | 0.4740563 |
| PERP | 58.0295355 | -1.573620474 | 0.49008054 | -3.2109426 | 0.001323 | 0.01802855 |
| ARFGEF3 | 57.9631293 | -1.799553622 | 0.28414001 | -6.3333341 | 2.40E-10 | 4.85E-08 |
| HEBP2 | 49.1230582 | -1.196991032 | 0.35755262 | -3.3477339 | 0.00081475 | 0.01262192 |
| NHSL1 | 169.174016 | -0.504132847 | 0.40239283 | -1.2528376 | 0.21026483 | 0.49648138 |
| CCDC28A | 24.6322997 | 0.01980426 | 0.36640618 | 0.05405002 | 0.95689531 | 0.98294302 |
| REPS1 | 136.28979 | 0.315954013 | 0.19224994 | 1.64345444 | 0.10028894 | 0.32695465 |
| HECA | 18.4346328 | -0.206864279 | 0.39603385 | -0.5223399 | 0.6014337 | 0.80759541 |
| CITED2 | 26.93539 | 0.041143356 | 0.44666355 | 0.09211263 | 0.92660855 | 0.97111125 |
| VTA1 | 58.1564534 | 0.125974335 | 0.25114375 | 0.50160251 | 0.61594716 | 0.81666507 |
| ADGRG6 | 36.6630719 | -1.243661064 | 0.61692645 | -2.0158984 | 0.0438106 | 0.19976647 |
| HIVEP2 | 307.16891 | 0.648153348 | 0.50721491 | 1.27786729 | 0.20129622 | 0.4848871 |
| AIG1 | 212.035521 | 0.41715932 | 0.15677455 | 2.66088667 | 0.00779352 | 0.06449472 |
| ADAT2 | 22.4971148 | 0.354786345 | 0.38492332 | 0.92170655 | 0.35668166 | 0.64141201 |

| | | | | | | |
|---|---|---|---|---|---|---|
| PEX3 | 64.4328083 | -0.15211081 | 0.28250476 | -0.5384363 | 0.59027589 | 0.799924 |
| FUCA2 | 27.2280034 | -0.831156793 | 0.34805942 | -2.3879738 | 0.01694155 | 0.10882834 |
| PHACTR2 | 348.981171 | -0.105283441 | 0.32950008 | -0.3195248 | 0.7493286 | 0.89017335 |
| PLAGL1 | 81.6917743 | -0.418679098 | 0.32732018 | -1.2791118 | 0.20085769 | 0.48443708 |
| SF3B5 | 46.8504934 | -0.897272499 | 0.52236711 | -1.7177048 | 0.08585047 | 0.30113433 |
| UTRN | 393.1947 | 0.443992268 | 0.19267023 | 2.30441551 | 0.02119933 | 0.12513214 |
| EPM2A | 61.8985682 | 0.454658063 | 0.45456168 | 1.00021204 | 0.3172079 | 0.61058965 |
| FBXO30 | 13.7844368 | -0.130372512 | 0.48860144 | -0.2668279 | 0.78960165 | 0.91080793 |
| AL356599.1 | 51.4229902 | -0.124199807 | 0.47845161 | -0.259587 | 0.79518238 | 0.91195838 |
| SHPRH | 217.489268 | 0.454482699 | 0.22566113 | 2.01400523 | 0.04400898 | 0.20016628 |
| STXBP5-AS1 | 17.7940563 | -0.682143083 | 0.41179094 | -1.6565277 | 0.09761501 | 0.32252399 |
| STXBP5 | 199.453368 | -0.47991995 | 0.18069867 | -2.655913 | 0.0079094 | 0.06518612 |
| SAMD5 | 402.768701 | -0.985022738 | 0.41732296 | -2.3603368 | 0.01825835 | 0.11379161 |
| AL033504.1 | 259.945986 | -0.893745792 | 0.45390827 | -1.9690009 | 0.04895299 | 0.21469294 |
| AL513164.1 | 14.871533 | -1.727872202 | 0.71729945 | -2.4088576 | 0.01600254 | 0.10420871 |
| SASH1 | 160.61695 | -1.20660045 | 0.26212768 | -4.603102 | 4.16E-06 | 0.00018325 |
| UST | 78.8428242 | 1.407700416 | 0.35499261 | 3.9654358 | 7.33E-05 | 0.00190598 |
| TAB2 | 197.153296 | 0.447394969 | 0.26035271 | 1.71841875 | 0.08572026 | 0.30113433 |
| PPIL4 | 71.3571666 | 0.224444306 | 0.24896801 | 0.90149857 | 0.36732329 | 0.65277742 |
| GINM1 | 89.0811647 | -0.116686914 | 0.22361397 | -0.521823 | 0.60179357 | 0.80777556 |
| KATNA1 | 45.2880066 | 0.588970743 | 0.30458442 | 1.93368639 | 0.0531517 | 0.22598795 |
| LATS1 | 96.2290566 | 0.036375448 | 0.26646148 | 0.13651297 | 0.89141577 | 0.9571988 |
| NUP43 | 33.6075885 | 0.368803955 | 0.31743326 | 1.16183149 | 0.24530392 | 0.53608738 |
| PCMT1 | 62.8496441 | 0.056007844 | 0.26729331 | 0.20953702 | 0.83402904 | 0.93291717 |
| LRP11 | 21.7022 | -0.369244139 | 0.41187622 | -0.896493 | 0.36998953 | 0.65432818 |
| PPP1R14C | 31.7118963 | -3.698654084 | 0.86093997 | -4.296065 | 1.74E-05 | 0.00057938 |
| PLEKHG1 | 67.6348757 | -1.209897392 | 0.57420585 | -2.1070795 | 0.03511068 | 0.17415035 |
| MTHFD1L | 194.682599 | -1.441908159 | 0.31737371 | -4.5432501 | 5.54E-06 | 0.00023292 |
| ZBTB2 | 11.361809 | -0.445897345 | 0.54525733 | -0.8177741 | 0.41348616 | 0.69042044 |
| RMND1 | 32.2873237 | -0.315219642 | 0.36098405 | -0.8732232 | 0.38254143 | 0.66592461 |
| ARMT1 | 18.7446994 | 0.454746961 | 0.45802431 | 0.99284461 | 0.32078569 | 0.6120836 |
| SYNE1 | 424.314822 | 0.295987935 | 0.33785253 | 0.87608619 | 0.38098318 | 0.66433459 |
| MTRF1L | 22.7109593 | -0.655450945 | 0.38601829 | -1.697979 | 0.08951173 | 0.30789644 |
| OPRM1 | 14.0856067 | -0.151698971 | 0.65081594 | -0.2330904 | 0.81569118 | 0.92171225 |
| IPCEF1 | 209.751353 | 1.746458799 | 0.44978654 | 3.88286143 | 0.00010323 | 0.00249068 |
| CNKSR3 | 253.823795 | 0.642232289 | 0.30931865 | 2.07628054 | 0.03786801 | 0.18238254 |
| SCAF8 | 255.285754 | 0.589741728 | 0.22878359 | 2.57772744 | 0.00994524 | 0.07697661 |
| TIAM2 | 134.941619 | 0.31051073 | 0.2887943 | 1.07519687 | 0.2822866 | 0.57518127 |
| TFB1M | 67.1669578 | 0.216326461 | 0.25612466 | 0.84461393 | 0.39832643 | 0.67938457 |
| ARID1B | 1071.89804 | 0.480448902 | 0.11879058 | 4.04450325 | 5.24E-05 | 0.001456 |
| TMEM242 | 30.3016814 | -0.153157592 | 0.31350772 | -0.4885289 | 0.62517524 | 0.82285278 |
| ZDHHC14 | 150.660679 | -0.11215234 | 0.3012962 | -0.3722328 | 0.70971949 | 0.86953263 |
| SNX9 | 207.378476 | -0.189458345 | 0.31831113 | -0.5951986 | 0.55171072 | 0.77731453 |
| SYNJ2 | 43.6790138 | -2.642777725 | 0.480155 | -5.5040096 | 3.71E-08 | 3.54E-06 |
| SERAC1 | 21.3882633 | -0.152498633 | 0.37937187 | -0.4019767 | 0.68770121 | 0.85823112 |
| GTF2H5 | 27.541574 | -0.755015921 | 0.47162247 | -1.6008905 | 0.10940118 | 0.34481257 |
| TULP4 | 227.395949 | 0.049487562 | 0.27450203 | 0.18028122 | 0.8569318 | 0.94359207 |
| TMEM181 | 55.1340849 | -0.536795846 | 0.24659389 | -2.1768417 | 0.02949238 | 0.15581053 |
| DYNLT1 | 76.8158071 | -1.008769258 | 0.44390303 | -2.2724991 | 0.02305638 | 0.13247453 |
| SYTL3 | 56.2828134 | -0.810714387 | 0.81272038 | -0.9975318 | 0.31850647 | 0.61144847 |
| EZR | 227.558111 | -0.315957399 | 0.37310113 | -0.8468412 | 0.39708365 | 0.67795091 |
| RSPH3 | 55.0491548 | -0.481510163 | 0.34416359 | -1.3990735 | 0.16179094 | 0.42935864 |
| AL035530.2 | 46.0386615 | 1.082500025 | 0.4637478 | 2.33424292 | 0.01958301 | 0.11888281 |
| SOD2.1 | 22.83831 | -0.884664006 | 0.55653923 | -1.5895807 | 0.11192935 | 0.34868346 |
| WTAP | 137.968987 | -0.985533597 | 0.32153397 | -3.0650995 | 0.00217598 | 0.02585572 |
| ACAT2 | 22.0842115 | -0.045861199 | 0.68146155 | -0.0672983 | 0.94634424 | 0.97905916 |
| TCP1 | 35.0594774 | 0.105660132 | 0.30326403 | 0.3484097 | 0.72753252 | 0.87980279 |
| MRPL18 | 22.9459868 | -0.544244558 | 0.40620096 | -1.3398407 | 0.18029715 | 0.45647562 |
| IGF2R | 246.741654 | 0.14902265 | 0.25729979 | 0.57917905 | 0.56246837 | 0.78400069 |
| SLC22A2 | 26.8803894 | 0.635563792 | 0.72558945 | 0.87592756 | 0.38106942 | 0.66437251 |
| SLC22A3 | 34.9749058 | 0.420084887 | 0.91970908 | 0.45675844 | 0.64784468 | 0.83530612 |
| MAP3K4 | 103.434372 | 0.316896509 | 0.19529719 | 1.62263729 | 0.10466696 | 0.33556952 |
| AGPAT4 | 41.7839352 | -0.452330231 | 0.32801512 | -1.378992 | 0.16789722 | 0.43833683 |
| PRKN | 854.382885 | 0.734964651 | 0.24153209 | 3.04292756 | 0.00234289 | 0.0272113 |
| PACRG | 149.698716 | 0.329829359 | 0.28917829 | 1.14057442 | 0.25404707 | 0.54530511 |
| QKI | 401.146623 | 0.123443641 | 0.21795304 | 0.56637724 | 0.57113737 | 0.78905469 |
| PDE10A | 574.301372 | -1.718322978 | 0.82625217 | -2.0796593 | 0.0375568 | 0.18135358 |
| SFT2D1 | 43.7157281 | -0.382537549 | 0.2625699 | -1.4568979 | 0.14514456 | 0.40304022 |
| MPC1 | 173.788843 | 0.785428913 | 0.22705856 | 3.45914687 | 0.00054189 | 0.00907731 |
| RPS6KA2 | 116.186003 | -0.571866404 | 0.24600463 | -2.3246164 | 0.02009248 | 0.12104838 |
| Z94721.1 | 24.4806558 | -0.872267123 | 0.44024571 | -1.9813189 | 0.04755553 | 0.21044546 |
| FGFR1OP | 80.6819569 | -0.059470246 | 0.22190572 | -0.2679978 | 0.78870101 | 0.90997848 |
| AFDN | 352.716975 | 0.215114801 | 0.18865469 | 1.14025687 | 0.2541793 | 0.54530511 |
| WDR27 | 117.809307 | 0.399705996 | 0.25394755 | 1.57397068 | 0.11549423 | 0.35464129 |
| PHF10 | 47.9070636 | 0.408641571 | 0.37089233 | 1.1017795 | 0.27055754 | 0.56240594 |
| ERMARD | 24.5999919 | 0.47658715 | 0.43155321 | 1.10435315 | 0.26943997 | 0.56160046 |
| FAM120B | 170.937521 | 0.104750081 | 0.23358197 | 0.44845105 | 0.65382771 | 0.83885315 |
| PSMB1 | 66.5812396 | -0.306909716 | 0.32195574 | -0.9532668 | 0.34045491 | 0.62907597 |
| TBP | 23.5505575 | -0.034990635 | 0.31930215 | -0.1095847 | 0.91273873 | 0.96446045 |
| PDCD2 | 26.9783207 | -0.515657017 | 0.32638553 | -1.5799016 | 0.1141294 | 0.3526678 |
| FAM20C | 193.296063 | -1.443600308 | 0.41077212 | -3.5143581 | 0.00044082 | 0.00785693 |
| PDGFA | 32.5709511 | -1.734714601 | 0.51825659 | -3.3472119 | 0.00081629 | 0.01262672 |
| PRKAR1B | 54.1610343 | 0.001682806 | 0.31930862 | 0.00527016 | 0.99579504 | 0.99868531 |
| DNAAF5 | 38.578187 | -0.066941984 | 0.33545415 | -0.1995563 | 0.84182762 | 0.93566718 |
| SUN1 | 89.7724252 | 0.494698427 | 0.18903974 | 2.61690172 | 0.00887319 | 0.07069728 |
| COX19 | 37.5160754 | 0.010750226 | 0.39323167 | 0.02733815 | 0.97819003 | 0.99127618 |
| C7orf50 | 79.4691882 | 0.127151414 | 0.27074771 | 0.46963062 | 0.63861895 | 0.83100321 |
| MICALL2 | 13.1519634 | -1.038172941 | 0.49246479 | -2.1081161 | 0.03502095 | 0.1740015 |
| INTS1 | 36.4372135 | 0.318726909 | 0.35624888 | 0.89467483 | 0.37096094 | 0.65494451 |
| MAFK | 32.0374284 | 1.045181851 | 0.52726337 | 1.98227662 | 0.04744829 | 0.21024184 |
| TMEM184A | 14.5989488 | -0.168562958 | 0.43533454 | -0.3872033 | 0.69860573 | 0.86334649 |
| PSMG3 | 11.8963298 | -0.598243754 | 0.47552049 | -1.258082 | 0.20836211 | 0.49402397 |
| MAD1L1 | 58.4685925 | -0.229986066 | 0.29945149 | -0.7680244 | 0.44247266 | 0.71265656 |
| SNX8 | 21.8329464 | -0.176503274 | 0.52893536 | -0.3336954 | 0.73860945 | 0.88591857 |
| EIF3B | 58.7322952 | -0.255489443 | 0.30941773 | -0.8257104 | 0.40896838 | 0.6869518 |
| CHST12 | 19.9231201 | 0.047147009 | 0.42369256 | 0.11127646 | 0.91139712 | 0.96399248 |
| BRAT1 | 16.3523873 | 0.578703105 | 0.36913697 | 1.56771917 | 0.11694669 | 0.35697326 |
| IQCE | 34.8364177 | -0.259456849 | 0.31642379 | -0.8199663 | 0.41223531 | 0.6893115 |
| TTYH3 | 22.0493412 | -0.643572043 | 0.39171247 | -1.6429705 | 0.10038903 | 0.32707393 |
| GNA12 | 55.97232 | -0.3223865 | 0.28271537 | -1.1403218 | 0.25415226 | 0.54530511 |
| SDK1 | 550.479436 | -0.597860657 | 0.35755922 | -1.6720605 | 0.09451238 | 0.31612552 |
| FOXK1 | 116.039163 | 0.045158343 | 0.2802646 | 0.16112753 | 0.87199296 | 0.94846188 |
| AP5Z1 | 18.7860013 | -0.906804076 | 0.38099613 | -2.3800874 | 0.01730853 | 0.11006868 |
| RBAK-RBAKI | 73.2518338 | -0.262967964 | 0.30691059 | -0.8568227 | 0.39154287 | 0.6727196 |
| WIPI2 | 57.5108558 | -0.09878535 | 0.32558284 | -0.3034108 | 0.76157681 | 0.89665878 |
| TNRC18 | 233.331169 | 0.611062664 | 0.23846474 | 2.5624865 | 0.01039256 | 0.0788977 |
| FBXL18 | 27.6988542 | -0.03028923 | 0.31220212 | -0.097018 | 0.92271209 | 0.96898878 |
| ACTB | 638.639471 | 0.505120943 | 0.46962796 | 1.0755768 | 0.28211657 | 0.5750055 |
| RNF216 | 161.916369 | 0.275325445 | 0.24867561 | 1.10716707 | 0.2682217 | 0.56003648 |
| CCZ1 | 20.2804902 | -0.399318467 | 0.44525808 | -0.8968248 | 0.36981244 | 0.65426888 |
| PMS2 | 39.4575023 | 0.193112633 | 0.29990831 | 0.64390558 | 0.51963666 | 0.7603035 |
| AIMP2 | 20.7333548 | -0.666464464 | 0.43038436 | -1.5485332 | 0.12149398 | 0.36543969 |
| EIF2AK1 | 54.1858766 | -0.15398996 | 0.24249128 | -0.635033 | 0.5254069 | 0.76221787 |

| | | | | | | |
|---|---|---|---|---|---|---|
| USP42 | 59.9277148 | 0.070142516 | 0.30759249 | 0.22803716 | 0.81961736 | 0.92374663 |
| CYTH3 | 78.8431502 | 0.751073664 | 0.25377518 | 2.95960257 | 0.00308036 | 0.03326403 |
| RAC1 | 145.797656 | -0.564849986 | 0.22422468 | -2.519125 | 0.01176469 | 0.08565358 |
| DAGLB | 19.8138668 | -0.073796975 | 0.43607543 | -0.1692298 | 0.86561586 | 0.94694132 |
| ZDHHC4 | 16.4618004 | -0.151240564 | 0.38314237 | -0.3947372 | 0.69303681 | 0.86061538 |
| C7orf26 | 14.7545938 | 0.13668206 | 0.45715101 | 0.29898667 | 0.76495021 | 0.89797438 |
| ZNF316 | 21.4105828 | 0.183383981 | 0.35373497 | 0.51842197 | 0.60416389 | 0.8095859 |
| AC073343.2 | 17.1709404 | -0.085284274 | 0.58137495 | -0.1466941 | 0.88337348 | 0.95561187 |
| ZNF12 | 39.748531 | -0.171542143 | 0.37493971 | -0.4575193 | 0.64729786 | 0.83478232 |
| CCZ1B | 25.3240009 | -0.163444142 | 0.3641215 | -0.4488725 | 0.65352361 | 0.83881384 |
| C1GALT1 | 106.999285 | -0.376612063 | 0.36916638 | -1.0201689 | 0.30764836 | 0.60139222 |
| COL28A1 | 79.3144002 | 0.832767263 | 0.2983611 | 2.79113884 | 0.00525229 | 0.04936964 |
| AC004982.2 | 16.9227009 | 0.512812513 | 0.38846366 | 1.32010421 | 0.18680023 | 0.46437575 |
| MIOS | 61.8591268 | 0.092198608 | 0.28450867 | 0.32406256 | 0.74589067 | 0.88824016 |
| RPA3 | 31.064261 | -0.940253546 | 0.61768545 | -1.5222206 | 0.12795381 | 0.37522918 |
| UMAD1 | 315.892382 | 0.211737113 | 0.17236086 | 1.22845239 | 0.21927719 | 0.50741097 |
| GLCCI1 | 56.5965164 | 0.094033936 | 0.28916994 | 0.32518572 | 0.74504051 | 0.88808711 |
| ICA1 | 239.624503 | -0.047445052 | 0.20713425 | -0.2290546 | 0.81882648 | 0.9234319 |
| NDUFA4 | 153.262414 | -0.238534552 | 0.43298509 | -0.5509071 | 0.58169737 | 0.79526454 |
| PHF14 | 153.991776 | 0.058091658 | 0.27787924 | 0.2090536 | 0.83440639 | 0.93313642 |
| THSD7A | 1006.9009 | 0.591791695 | 0.51591741 | 1.14706673 | 0.25135404 | 0.54308921 |
| TMEM106B | 76.6140306 | -0.358355976 | 0.22803953 | -1.5714643 | 0.11607484 | 0.35543425 |
| SCIN | 444.850318 | 1.217980339 | 0.35122786 | 3.46777828 | 0.00052478 | 0.00890656 |
| AC011287.1 | 26.9280364 | -1.706935794 | 0.5917307 | -2.8846497 | 0.00391849 | 0.03957679 |
| ISPD | 81.7219407 | 1.044227074 | 0.31584303 | 3.30615839 | 0.00094585 | 0.01400282 |
| ANKMY2 | 50.9265151 | 0.170292468 | 0.23238065 | 0.7328169 | 0.46367011 | 0.72812334 |
| BZW2 | 109.21117 | -0.638256633 | 0.3861662 | -1.652803 | 0.09837097 | 0.32402704 |
| AC073332.1 | 17.0837068 | -0.543786378 | 0.531877 | -1.0223912 | 0.30659577 | 0.60139222 |
| SNX13 | 168.116808 | 0.173360859 | 0.20723977 | 0.83652311 | 0.40286068 | 0.68128594 |
| TWISTNB | 13.9944813 | -0.136047408 | 0.39203676 | -0.3470272 | 0.72857091 | 0.88042244 |
| MACC1 | 265.368485 | -0.549215209 | 0.24986237 | -2.1980709 | 0.02794405 | 0.14993732 |
| ITGB8 | 200.644135 | -0.449357918 | 0.46301046 | -0.9705135 | 0.33179059 | 0.62147393 |
| SP4 | 49.1325363 | 0.312708034 | 0.30940824 | 1.01066485 | 0.31217687 | 0.60527563 |
| RAPGEF5 | 195.736171 | -3.27118891 | 0.6598926 | -4.9571535 | 7.15E-07 | 4.31E-05 |
| STEAP1B | 105.097429 | -0.886366426 | 0.40876458 | -2.1684032 | 0.03012802 | 0.1578151 |
| TOMM7 | 54.3505096 | -0.235933583 | 0.37429922 | -0.6303342 | 0.52847598 | 0.76311655 |
| KLHL7 | 50.9336099 | 0.039138234 | 0.24203689 | 0.16170359 | 0.87153928 | 0.94844301 |
| NUPL2 | 50.0185744 | -0.343062505 | 0.38154495 | -0.8991405 | 0.36857785 | 0.65369102 |
| AC005082.1 | 10.2192477 | -1.118541736 | 0.51571068 | -2.1689327 | 0.0300878 | 0.1578151 |
| GPNMB | 29.0045673 | -1.985566201 | 0.42002101 | -4.7273021 | 2.28E-06 | 0.00011056 |
| MALSU1 | 30.4011016 | 0.071777225 | 0.31810389 | 0.22564083 | 0.82148078 | 0.92485761 |
| TRA2A | 209.784808 | 0.02697614 | 0.21973759 | 0.12276525 | 0.90229299 | 0.96086649 |
| CCDC126 | 28.2339866 | -0.208509503 | 0.42533888 | -0.4902197 | 0.62397843 | 0.82223405 |
| FAM221A | 18.1902068 | 0.474737782 | 0.38958116 | 1.21858505 | 0.22300173 | 0.51166231 |
| MPP6 | 33.0214601 | -1.709250177 | 0.61261264 | -2.7900994 | 0.00526919 | 0.04943821 |
| OSBPL3 | 423.398296 | 0.193650127 | 0.31790595 | 0.60914281 | 0.54242978 | 0.77243948 |
| CYCS | 81.4515593 | -0.024690858 | 0.40578695 | -0.0608469 | 0.95148118 | 0.98061757 |
| NFE2L3 | 14.4610159 | -0.993444272 | 0.54903917 | -1.8094233 | 0.07038527 | 0.26766667 |
| HNRNPA2B1 | 404.14164 | 0.295850845 | 0.17494977 | 1.69106169 | 0.09082502 | 0.31034142 |
| CBX3 | 76.2952545 | -0.766902725 | 0.22641419 | -3.3871673 | 0.00070618 | 0.01134961 |
| SNX10 | 324.288297 | -1.134232613 | 0.37274166 | -3.0429457 | 0.00234275 | 0.0272113 |
| AC004540.1 | 71.2494012 | 0.323045642 | 0.61629477 | 0.52417392 | 0.60015758 | 0.80738812 |
| SKAP2 | 259.061609 | -0.077354107 | 0.25707703 | -0.3008985 | 0.76349186 | 0.8974069 |
| HOTAIRM1 | 40.0084743 | -0.270239771 | 0.39748691 | -0.6798709 | 0.49658622 | 0.74859388 |
| HOXA3 | 61.9452948 | 0.3710848 | 0.30440973 | 1.21903068 | 0.22283255 | 0.51138804 |
| HOXA-AS2 | 15.9947542 | -0.230830116 | 0.41489093 | -0.5563634 | 0.57796248 | 0.79336035 |
| HOXA-AS3 | 9.69075047 | -1.156524448 | 0.65870204 | -1.7557627 | 0.07912894 | 0.28668189 |
| HOXA7 | 10.9919257 | -1.035505079 | 0.70321526 | -1.4725293 | 0.140878 | 0.39647869 |
| HOXA9 | 52.119952 | 0.634870525 | 0.32106154 | 1.97741069 | 0.04799523 | 0.21199399 |
| HIBADH | 216.178316 | 0.728230821 | 0.18572797 | 3.92095387 | 8.82E-05 | 0.00221077 |
| TAX1BP1 | 259.638622 | -0.075484706 | 0.17438657 | -0.4328585 | 0.66511759 | 0.84348349 |
| JAZF1 | 57.1434421 | -1.073916391 | 0.56401019 | -1.9040727 | 0.05690072 | 0.23559369 |
| CREB5 | 29.8411106 | 0.018639504 | 0.83783216 | 0.0222473 | 0.98225069 | 0.99337859 |
| AC005162.3 | 27.8559453 | 0.701042154 | 0.55704087 | 1.25851117 | 0.20820695 | 0.49402397 |
| CPVL | 123.799875 | 0.336416163 | 0.29883153 | 1.12577199 | 0.26026204 | 0.55146957 |
| AC007255.1 | 29.9852572 | 0.591647613 | 0.46308901 | 1.27761101 | 0.20138662 | 0.4848871 |
| PRR15 | 19.263232 | -0.239759664 | 0.36436598 | -0.6580188 | 0.51052605 | 0.7549696 |
| SCRN1 | 71.4886281 | -0.024242289 | 0.22503296 | -0.1077277 | 0.91421167 | 0.96462448 |
| FKBP14 | 20.8924218 | -0.254594888 | 0.36099919 | -0.7052506 | 0.48065431 | 0.73988033 |
| PLEKHA8 | 70.3121049 | -0.101724093 | 0.36527609 | -0.2784855 | 0.7806397 | 0.90503603 |
| MTURN | 109.829793 | 0.632854943 | 0.24668265 | 2.56546189 | 0.01030385 | 0.07859336 |
| ZNRF2 | 129.409544 | -0.47147333 | 0.24347864 | -1.9364053 | 0.05281808 | 0.22512695 |
| AC006978.2 | 110.028378 | -0.770862866 | 0.30410053 | -2.5348948 | 0.01124811 | 0.08307399 |
| NOD1 | 27.5183956 | -0.210573622 | 0.59154905 | -0.3559698 | 0.72186315 | 0.87625492 |
| GGCT | 12.2271423 | 0.043236018 | 0.52303421 | 0.08266384 | 0.93411884 | 0.97470802 |
| AC005154.6 | 34.1953314 | 0.464975479 | 0.33189926 | 1.40095365 | 0.16122793 | 0.42830587 |
| GARS | 55.2795517 | -0.431653201 | 0.36211046 | -1.1920484 | 0.23324226 | 0.52315737 |
| AQP1 | 19.2763899 | 0.131233902 | 0.57017701 | 0.23016344 | 0.81796477 | 0.92281757 |
| PDE1C | 258.60781 | 0.229447827 | 0.66935928 | 0.34278725 | 0.73175852 | 0.88245488 |
| LSM5 | 20.7312239 | -0.120309251 | 0.44466317 | -0.2705627 | 0.78672742 | 0.90871373 |
| AVL9 | 192.250203 | -0.115142531 | 0.24330236 | -0.4732487 | 0.63603574 | 0.82931783 |
| KBTBD2 | 43.8047193 | -0.485362066 | 0.25730465 | -1.8863323 | 0.05925019 | 0.24104615 |
| FKBP9 | 38.9578356 | 0.212591463 | 0.30891624 | 0.6881848 | 0.49133642 | 0.74580047 |
| NT5C3A | 74.3309346 | 0.134341325 | 0.47398962 | 0.28342672 | 0.77684975 | 0.90335572 |
| RP9 | 15.8459999 | 0.28552227 | 0.43114296 | 0.66224501 | 0.50781421 | 0.75382716 |
| BBS9 | 208.137694 | 0.567853156 | 0.2285432 | 2.48466439 | 0.01296736 | 0.09194477 |
| DPY19L1 | 80.2400944 | -0.359962641 | 0.33160855 | -1.0855047 | 0.27769817 | 0.56997988 |
| HERPUD2 | 59.9828948 | 0.069418461 | 0.26273696 | 0.26421277 | 0.79161597 | 0.91107378 |
| SEPT7-AS1 | 25.1354125 | 0.466222026 | 0.51635038 | 0.90291796 | 0.36656944 | 0.65175201 |
| 7-Sep | 173.417616 | -0.076140239 | 0.30626218 | -0.2486113 | 0.80366147 | 0.91636087 |
| EEPD1 | 75.3144268 | 0.060116878 | 0.46299238 | 0.12984421 | 0.89668969 | 0.95900099 |
| KIAA0895 | 44.1749666 | -0.094603784 | 0.27140407 | -0.3485717 | 0.72741086 | 0.87975894 |
| AOAH | 45.3789014 | -3.271433157 | 0.68705904 | -4.7615022 | 1.92E-06 | 9.52E-05 |
| EPDR1 | 14.8516987 | 0.357063273 | 0.52210249 | 0.68389498 | 0.49404149 | 0.74700665 |
| STARD3NL | 32.7629289 | -0.034246811 | 0.30346316 | -0.1128533 | 0.91014688 | 0.96345388 |
| VPS41 | 129.70348 | -0.053186188 | 0.21757238 | -0.2444529 | 0.80688011 | 0.91759343 |
| YAE1D1 | 15.5492729 | 0.517480788 | 0.40998626 | 1.26219055 | 0.20688023 | 0.49179557 |
| RALA | 74.684288 | -0.211516561 | 0.26419523 | -0.800607 | 0.42335918 | 0.69817048 |
| LINC00265 | 16.4853751 | 0.324879862 | 0.34822232 | 0.9329668 | 0.35083711 | 0.6374469 |
| CDK13 | 318.012163 | 0.185017147 | 0.20411134 | 0.90645209 | 0.36469661 | 0.64979324 |
| MPLKIP | 27.9573916 | 0.154519426 | 0.30853582 | 0.50081519 | 0.6165012 | 0.81691188 |
| SUGCT | 69.6015088 | 1.207094124 | 0.31261953 | 3.8612243 | 0.00011282 | 0.0026719 |
| MRPL32 | 23.0915051 | -0.252716841 | 0.3649672 | -0.6924371 | 0.48866286 | 0.74516503 |
| STK17A | 33.8243165 | 0.038536385 | 0.45284385 | 0.08509862 | 0.93218299 | 0.97356162 |
| COA1 | 177.229837 | 0.115970029 | 0.17933609 | 0.64666307 | 0.51785003 | 0.75866548 |
| BLVRA | 24.5261549 | -0.479208281 | 0.31135145 | -1.5391233 | 0.1237742 | 0.36874545 |
| URGCP | 47.2852709 | 0.183246535 | 0.41925966 | 0.43707171 | 0.66205935 | 0.84204141 |
| UBE2D4 | 14.6452532 | -0.229232146 | 0.37962495 | -0.6038385 | 0.54595104 | 0.77463416 |
| AC004951.1 | 10.6802138 | 0.915199274 | 0.48739705 | 1.87772839 | 0.06041833 | 0.24332668 |
| DBNL | 26.4599255 | -0.306903837 | 0.42248919 | -0.7264182 | 0.46758244 | 0.73035905 |
| POLM | 21.6341156 | -0.572700169 | 0.41696964 | -1.3734817 | 0.16960264 | 0.44032355 |
| POLD2 | 16.1550233 | -0.125045135 | 0.41667809 | -0.3001001 | 0.7641008 | 0.89757885 |
| YKT6 | 16.5232923 | -0.214455075 | 0.4031247 | -0.531982 | 0.59473848 | 0.80313746 |

| | | | | | | |
|---|---|---|---|---|---|---|
| NUDCD3 | 116.204091 | -0.154111912 | 0.26367337 | -0.5844804 | 0.55889716 | 0.78244004 |
| DDX56 | 21.848334 | 0.330919948 | 0.34775431 | 0.95159122 | 0.34130434 | 0.62956442 |
| TMED4 | 58.5774969 | -0.429410025 | 0.37816287 | -1.1355161 | 0.25615915 | 0.54710413 |
| OGDH | 281.556694 | 0.707764109 | 0.20646667 | 3.42798236 | 0.00060808 | 0.01005536 |
| ZMIZ2 | 52.9024662 | -0.068924836 | 0.26153715 | -0.2635375 | 0.79213636 | 0.91118678 |
| PPIA | 187.323883 | -0.773520641 | 0.37831654 | -2.0446387 | 0.04089049 | 0.19261721 |
| H2AFV | 80.6712587 | -0.117233235 | 0.31301647 | -0.3745274 | 0.708012 | 0.86882712 |
| PURB | 49.2173905 | -0.297295228 | 0.29347221 | -1.0130268 | 0.31104734 | 0.60414964 |
| CCM2 | 57.4306253 | -0.137544889 | 0.29831635 | -0.4610706 | 0.64474797 | 0.83381793 |
| TBRG4 | 16.3569854 | -0.221546014 | 0.40936801 | -0.5411903 | 0.58837639 | 0.79897585 |
| ADCY1 | 21.3155562 | 1.957627949 | 0.37765231 | 5.18367794 | 2.18E-07 | 1.65E-05 |
| IGFBP3 | 127.27307 | -1.699020747 | 0.73705103 | -2.3051603 | 0.0211576 | 0.12498027 |
| AC004870.4 | 68.1735074 | -0.965311274 | 0.7950643 | -1.2141298 | 0.22469816 | 0.5134961 |
| TNS3 | 452.853889 | -0.774132398 | 0.33520511 | -2.309429 | 0.02091979 | 0.12385957 |
| HUS1 | 29.1525239 | 0.21271424 | 0.33084876 | 0.64293499 | 0.52026629 | 0.76073677 |
| UPP1 | 99.9869357 | -1.744110934 | 0.79527803 | -2.1930833 | 0.02830138 | 0.15134187 |
| GRB10 | 75.6961267 | -1.20286998 | 0.513212 | -2.3438072 | 0.01908804 | 0.11739997 |
| COBL | 117.861134 | -0.163385794 | 0.29984763 | -0.5448961 | 0.585825 | 0.79711254 |
| SEC61G | 28.4717056 | -0.432920549 | 0.47453733 | -0.9123003 | 0.36161065 | 0.64574674 |
| EGFR | 200.738906 | -0.171609772 | 0.17184833 | -0.9986118 | 0.31798277 | 0.61109152 |
| LANCL2 | 35.9580085 | 0.260272255 | 0.34120866 | 0.76279498 | 0.44558568 | 0.71524207 |
| VOPP1 | 145.397803 | -0.8348399 | 0.34988574 | -2.3860358 | 0.01703109 | 0.10924074 |
| NIPSNAP2 | 120.591606 | -0.039691513 | 0.23348073 | -0.1699991 | 0.86501085 | 0.94694132 |
| PSPH | 17.9429608 | 0.613026282 | 0.44132788 | 1.38904952 | 0.16481769 | 0.43375609 |
| CCT6A | 50.1723697 | -0.015836373 | 0.27030031 | -0.0585881 | 0.95328022 | 0.98143529 |
| SUMF2 | 55.1423421 | -0.562678256 | 0.24892022 | -2.2604763 | 0.0237917 | 0.13541585 |
| PHKG1 | 17.3720503 | -0.365797134 | 0.4258791 | -0.8589225 | 0.39038328 | 0.67215855 |
| CHCHD2 | 110.650911 | -0.427442594 | 0.372125 | -1.1486532 | 0.25069899 | 0.54232519 |
| NUPR2 | 15.659437 | -0.114011967 | 0.48162768 | -0.2367222 | 0.81287231 | 0.92054639 |
| ZNF727 | 17.4879739 | -0.177674932 | 0.41894588 | -0.4241 | 0.67149291 | 0.8477598 |
| ZNF736 | 80.0498319 | 0.005554561 | 0.2693488 | 0.02062219 | 0.98354704 | 0.99368608 |
| ZNF680 | 100.436219 | 0.340961243 | 0.28301651 | 1.20473974 | 0.22830379 | 0.51805851 |
| ZNF107 | 25.8499569 | 0.085854605 | 0.32300705 | 0.26579793 | 0.79039484 | 0.91080793 |
| ZNF138 | 62.3373606 | 0.111483495 | 0.31527069 | 0.35361198 | 0.72362969 | 0.87731847 |
| ZNF273 | 13.4137805 | -0.122972027 | 0.59666719 | -0.2060982 | 0.83671421 | 0.93419636 |
| ZNF117 | 36.0264905 | -0.296458691 | 0.34692899 | -0.8545227 | 0.39281545 | 0.67377433 |
| VKORC1L1 | 85.1842543 | 0.201718702 | 0.33718395 | 0.5982434 | 0.54967754 | 0.7759356 |
| GUSB | 21.4725376 | -0.279397702 | 0.36918514 | -0.7567956 | 0.44917231 | 0.71802342 |
| ASL | 14.5724568 | -0.150198928 | 0.41959062 | -0.3579654 | 0.7203692 | 0.87576628 |
| AC068533.4 | 24.188978 | 0.413266195 | 0.44870267 | 0.92102459 | 0.35703759 | 0.64169596 |
| CRCP | 17.1103078 | -0.390133535 | 0.45648639 | -0.8546444 | 0.39274804 | 0.67377433 |
| AC068533.3 | 16.2396778 | -0.355512812 | 0.52923746 | -0.6717454 | 0.50174582 | 0.75108109 |
| TPST1 | 461.5758 | 0.587753054 | 0.20714961 | 2.83733607 | 0.00454917 | 0.04438026 |
| AC027644.4 | 83.9195979 | 0.008161011 | 0.40080089 | 0.02036176 | 0.98375479 | 0.99368608 |
| TMEM248 | 72.7798848 | 0.160714157 | 0.24434214 | 0.65774228 | 0.51070375 | 0.7549696 |
| SBDS | 42.3949811 | -0.472497522 | 0.33181286 | -1.423988 | 0.15444995 | 0.41780983 |
| TYW1 | 141.129811 | 0.308113993 | 0.20329321 | 1.51561382 | 0.12961703 | 0.37737742 |
| AC006480.2 | 13.2196355 | 0.347110062 | 0.40467418 | 0.85775193 | 0.39102946 | 0.67223049 |
| AUTS2 | 125.800071 | 0.066312227 | 0.31092375 | 0.21327488 | 0.83111256 | 0.93046312 |
| GALNT17 | 78.7620474 | 3.290517986 | 0.62877528 | 5.23321784 | 1.67E-07 | 1.32E-05 |
| CALN1 | 17.3959471 | 0.382072909 | 0.45442736 | 0.84077884 | 0.40047184 | 0.68010059 |
| TYW1B | 107.419742 | 0.785434064 | 0.22241815 | 3.53133975 | 0.00041346 | 0.00746211 |
| POM121 | 105.504441 | 0.267287072 | 0.23220412 | 1.15108668 | 0.24969658 | 0.54098299 |
| NSUN5 | 18.960353 | 0.009121024 | 0.4136423 | 0.02205051 | 0.98240766 | 0.99340045 |
| BAZ1B | 105.092735 | 0.166809779 | 0.26079309 | 0.63962499 | 0.52241643 | 0.76155853 |
| BCL7B | 25.9702965 | -0.030637706 | 0.34902552 | -0.0877807 | 0.93005097 | 0.97253199 |
| MLXIPL | 23.9904955 | 0.810003133 | 0.43432101 | 1.86498723 | 0.06218319 | 0.24743579 |
| DNAJC30 | 11.3010216 | -0.373911419 | 0.53966008 | -0.6928647 | 0.48839446 | 0.74516503 |
| BUD23 | 38.538894 | -0.198168815 | 0.31985125 | -0.6195655 | 0.53554386 | 0.76808755 |
| ABHD11 | 28.8757695 | -0.969465077 | 0.40422669 | -2.3983203 | 0.01647046 | 0.10641319 |
| CLDN3 | 52.9260049 | -2.865773266 | 0.50499635 | -5.6748396 | 1.39E-08 | 1.72E-06 |
| CLDN4 | 110.822499 | -1.627118321 | 0.32998651 | -4.9308631 | 8.19E-07 | 4.74E-05 |
| LIMK1 | 17.302532 | -0.513065001 | 0.47686269 | -1.0759177 | 0.28196407 | 0.5750055 |
| EIF4H | 55.9050417 | -0.360707371 | 0.23571734 | -1.5302539 | 0.1259539 | 0.37158563 |
| RFC2 | 11.0638861 | -0.308280775 | 0.45780725 | -0.6733855 | 0.50070206 | 0.75071921 |
| CLIP2 | 64.184205 | -1.091691794 | 0.57145048 | -1.9103874 | 0.05608335 | 0.2339636 |
| GTF2IRD1 | 604.958066 | -0.924901443 | 0.58644966 | -1.5771199 | 0.11476793 | 0.35365717 |
| GTF2I | 356.185662 | 0.197614807 | 0.18605755 | 1.06211659 | 0.28818276 | 0.5803086 |
| GTF2IRD2 | 42.2141345 | -0.456381691 | 0.38035843 | -1.1998727 | 0.23018879 | 0.52042807 |
| CASTOR2 | 44.3858642 | -0.254772854 | 0.4242247 | -0.6005611 | 0.54813236 | 0.77516644 |
| RCC1L | 41.7827785 | 0.127678248 | 0.24852203 | 0.51375022 | 0.60742663 | 0.81184603 |
| GTF2IRD2B | 63.9511382 | 0.159539305 | 0.41211689 | 0.38712149 | 0.69866627 | 0.86334649 |
| TRIM73 | 31.0620334 | -0.116749779 | 0.53213896 | -0.2193972 | 0.82634068 | 0.92774212 |
| POM121C | 92.1896713 | 0.22512159 | 0.2624379 | 0.857809 | 0.39099794 | 0.67223049 |
| SPDYE5 | 17.335068 | 0.023431871 | 0.50457227 | 0.04643908 | 0.96296029 | 0.98592893 |
| HIP1 | 192.686904 | 0.067800257 | 0.36242825 | 0.18707222 | 0.85160401 | 0.94012264 |
| RHBDD2 | 29.0830653 | -0.63461453 | 0.45226413 | -1.4031945 | 0.16055885 | 0.42803048 |
| POR | 81.8355116 | -0.650629365 | 0.482795 | -1.3476307 | 0.17777719 | 0.45243593 |
| TMEM120A | 12.3695776 | 0.005374147 | 0.40262701 | 0.01334771 | 0.98935039 | 0.99637773 |
| STYXL1 | 42.0459455 | 0.407182688 | 0.28766963 | 1.41545247 | 0.15693588 | 0.42212882 |
| MDH2 | 45.1641638 | -0.027034665 | 0.36516833 | -0.0740334 | 0.94098378 | 0.97727535 |
| HSPB1 | 194.7878 | 0.738354813 | 0.56668109 | 1.30294592 | 0.19259323 | 0.4733052 |
| YWHAG | 61.9084376 | -0.309736354 | 0.3187033 | -0.9718643 | 0.33111807 | 0.62077859 |
| SSC4D | 45.4962927 | 0.917270582 | 0.39605196 | 2.31603596 | 0.02055631 | 0.12280939 |
| DTX2 | 33.7689766 | 0.053978757 | 0.34335646 | 0.15720909 | 0.87508006 | 0.95005531 |
| POMZP3 | 20.2595432 | 0.07854424 | 0.38295042 | 0.2051029 | 0.83749173 | 0.93487753 |
| CCDC146 | 96.4448821 | -0.058579955 | 0.30194948 | -0.1940058 | 0.84617133 | 0.93757726 |
| FGL2 | 62.4045482 | -1.243787742 | 0.81047356 | -1.5346432 | 0.12487149 | 0.37001028 |
| GSAP | 67.4280595 | 0.193708072 | 0.44504742 | 0.43525266 | 0.66337904 | 0.84278344 |
| PTPN12 | 177.430684 | -0.775359087 | 0.29828107 | -2.5994244 | 0.00933802 | 0.07332342 |
| APTR | 31.7760344 | 0.301397526 | 0.32128005 | 0.93811467 | 0.34818548 | 0.63531825 |
| RSBN1L | 104.55555 | 0.243277073 | 0.18164971 | 1.33926484 | 0.18048447 | 0.45661862 |
| TMEM60 | 10.8203395 | -0.603453159 | 0.62364073 | -0.9676295 | 0.33322944 | 0.62190756 |
| PHTF2 | 126.075779 | 1.058818862 | 0.30676436 | 3.45157064 | 0.00055733 | 0.00930576 |
| MAGI2 | 830.089497 | 1.111554294 | 0.30606637 | 3.63174271 | 0.00028151 | 0.00566124 |
| GNAI1 | 97.3606327 | -1.572901891 | 0.69179047 | -2.2736681 | 0.02298595 | 0.1323843 |
| SEMA3C | 22.7345474 | -0.181063806 | 0.75272257 | -0.2405452 | 0.80990762 | 0.91928664 |
| CACNA2D1 | 182.236255 | 0.124286455 | 0.28969704 | 0.42902218 | 0.66790708 | 0.84558183 |
| PCLO | 228.900055 | 0.794476736 | 0.41025785 | 1.93653025 | 0.05280279 | 0.22512695 |
| KIAA1324L | 28.1032362 | -0.202862501 | 0.44329569 | -0.4576234 | 0.647223 | 0.83478232 |
| DMTF1 | 126.383897 | 0.126624942 | 0.28304615 | 0.44736501 | 0.65461153 | 0.8391996 |
| TMEM243 | 44.2667075 | -0.953090837 | 0.32773049 | -2.9081543 | 0.00363569 | 0.03751236 |
| TP53TG1 | 11.0092605 | 0.538762719 | 0.44855146 | 1.20111686 | 0.22970587 | 0.51986597 |
| CROT | 70.8535282 | 0.661225273 | 0.32876498 | 2.01123996 | 0.04430012 | 0.20060651 |
| ABCB1 | 99.3188021 | 1.288199867 | 0.42568116 | 3.0262083 | 0.00247642 | 0.02847326 |
| SLC25A40 | 50.4308527 | 0.128797613 | 0.36039667 | 0.35737737 | 0.72080932 | 0.87596332 |
| DBF4 | 17.504685 | -0.408236811 | 0.43273377 | -0.9433902 | 0.34548134 | 0.63256941 |
| ADAM22 | 229.561048 | 1.44829448 | 0.35536446 | 4.07551868 | 4.59E-05 | 0.00131751 |
| SRI | 46.9559128 | -0.751823733 | 0.33142297 | -2.268472 | 0.02330045 | 0.13357889 |
| AC003991.1 | 19.8348276 | 0.845375293 | 0.56036541 | 1.50861435 | 0.13139736 | 0.38024035 |
| STEAP2 | 26.0740901 | 0.900217961 | 0.53664311 | 1.67749842 | 0.09344506 | 0.31481026 |
| CFAP69 | 48.4772598 | 0.164416911 | 0.33045529 | 0.49754661 | 0.61880364 | 0.81841635 |
| GTPBP10 | 44.4691443 | -0.169829367 | 0.32247066 | -0.5266506 | 0.59843624 | 0.8059995 |

| | | | | | | |
|---|---|---|---|---|---|---|
| CLDN12 | 43.1979929 | -0.489413473 | 0.26912581 | -1.8185304 | 0.0689831 | 0.26458076 |
| CDK14 | 334.054845 | 0.58839578 | 0.20513172 | 2.8683803 | 0.00412579 | 0.0409479 |
| FZD1 | 10.530972 | -0.887403598 | 0.74849454 | -1.1855846 | 0.23578639 | 0.52588686 |
| AC079760.1 | 32.3058273 | 0.76740168 | 0.48023928 | 1.59795692 | 0.11005257 | 0.345659 |
| MTERF1 | 154.813041 | 0.169600906 | 0.43233896 | 0.39228689 | 0.69484625 | 0.86161604 |
| AKAP9 | 359.742353 | 0.112599381 | 0.16979705 | 0.66314096 | 0.50724028 | 0.75376105 |
| KRIT1 | 130.347607 | -0.096349893 | 0.20877582 | -0.4614993 | 0.64444044 | 0.83381793 |
| ANKIB1 | 266.06453 | -0.230154007 | 0.1738721 | -1.3236971 | 0.18560365 | 0.46252753 |
| GATAD1 | 44.8848174 | -0.746884607 | 0.33890173 | -2.2038383 | 0.02753572 | 0.14844218 |
| AC007566.1 | 33.0268167 | -0.466121451 | 0.54119303 | -0.861285 | 0.38908108 | 0.67095971 |
| PEX1 | 51.7059828 | 0.431584684 | 0.27315214 | 1.58001574 | 0.11410326 | 0.3526678 |
| RBM48 | 11.6005888 | -0.248923275 | 0.46013764 | -0.5409757 | 0.58852434 | 0.79900779 |
| FAM133B | 66.1879008 | 0.213987262 | 0.24589976 | 0.87022152 | 0.38417936 | 0.66701235 |
| CDK6 | 369.025814 | 1.040648795 | 0.31078711 | 3.34842968 | 0.00081271 | 0.01260923 |
| SAMD9 | 15.004754 | -2.192662198 | 0.93736409 | -2.3391788 | 0.01932618 | 0.11815925 |
| SAMD9L | 41.1112188 | -1.426720423 | 0.8859744 | -1.6103405 | 0.10732355 | 0.3413545 |
| VPS50 | 127.143835 | 0.610145012 | 0.22882453 | 2.66643184 | 0.00766611 | 0.06391579 |
| GNGT1 | 81.4190455 | 0.689001572 | 0.4885033 | 1.41043382 | 0.15841162 | 0.42480987 |
| GNG11 | 15.5652184 | -0.631557008 | 0.5158606 | -1.2242784 | 0.22084721 | 0.5089143 |
| BET1 | 44.9245827 | -0.563204822 | 0.36748278 | -1.5326019 | 0.12537396 | 0.37039755 |
| CASD1 | 68.4626809 | 0.354076418 | 0.2528121 | 1.40055168 | 0.16134818 | 0.42831015 |
| SGCE | 72.1820096 | -0.084158256 | 0.50566995 | -0.1664292 | 0.86781918 | 0.94766517 |
| PPP1R9A | 440.664272 | 0.102491083 | 0.17553117 | 0.58389106 | 0.55929361 | 0.78278988 |
| AC002429.2 | 35.0023098 | -0.278577169 | 0.61960187 | -0.4496067 | 0.65299404 | 0.83850297 |
| PON2 | 82.2431756 | -0.681411779 | 0.30169718 | -2.2585951 | 0.02390858 | 0.13541585 |
| PDK4 | 334.614482 | 0.205921275 | 1.21916741 | 0.1689032 | 0.86587278 | 0.94694132 |
| SLC25A13 | 223.430445 | -0.167102356 | 0.22842022 | -0.7315568 | 0.46443914 | 0.7284793 |
| SEM1 | 87.8856015 | 0.293491986 | 0.22616168 | 1.29770873 | 0.19438744 | 0.47488249 |
| SDHAF3 | 24.5918628 | -0.006877024 | 0.31535929 | -0.0218069 | 0.98260195 | 0.99349939 |
| ASNS | 20.4645382 | -0.981300372 | 0.53426887 | -1.8367164 | 0.06625178 | 0.2574031 |
| LMTK2 | 96.6114407 | 0.43024072 | 0.27463267 | 1.56660431 | 0.11720721 | 0.35744016 |
| TECPR1 | 19.6032446 | -0.006146014 | 0.36074744 | -0.0170369 | 0.98640719 | 0.9949057 |
| BRI3 | 114.352213 | -0.662207571 | 0.29562975 | -2.2399896 | 0.0250916 | 0.13948057 |
| BAIAP2L1 | 227.340577 | 0.200519085 | 0.35879224 | 0.55887241 | 0.5762488 | 0.79184705 |
| TRRAP | 142.88383 | 0.245770603 | 0.18877567 | 1.30191888 | 0.19294412 | 0.47358843 |
| SMURF1 | 205.842262 | -0.383265638 | 0.28742896 | -1.3334274 | 0.1823916 | 0.45912471 |
| ARPC1B | 25.6108386 | -1.426689894 | 0.50621718 | -2.8183356 | 0.00482733 | 0.04630755 |
| PDAP1 | 30.0544244 | -0.713862296 | 0.32653564 | -2.1861696 | 0.0288032 | 0.15311173 |
| BUD31 | 21.6929067 | -0.565357018 | 0.47760089 | -1.1837436 | 0.23651456 | 0.52682659 |
| CPSF4 | 15.8743283 | 0.420223581 | 0.33564955 | 1.25197122 | 0.21058035 | 0.49657419 |
| ZNF789 | 32.2755996 | 0.075788205 | 0.31488811 | 0.24068297 | 0.80980084 | 0.91928664 |
| ZNF394 | 22.8322975 | -0.024369033 | 0.33340424 | -0.0730915 | 0.94173327 | 0.97782671 |
| ZKSCAN5 | 20.3541795 | 0.370139473 | 0.37943036 | 0.97551358 | 0.32930557 | 0.6185061 |
| FAM200A | 17.2276721 | -0.447600449 | 0.53254853 | -0.8404876 | 0.40063504 | 0.68010059 |
| ZSCAN25 | 12.4646291 | 0.025129744 | 0.44789525 | 0.0561063 | 0.95525713 | 0.98253384 |
| TRIM4 | 41.9565334 | -0.072992651 | 0.26284742 | -0.2776997 | 0.78124288 | 0.90522598 |
| ZKSCAN1 | 142.943071 | -0.142643789 | 0.21332189 | -0.6686786 | 0.50370049 | 0.75108109 |
| ZNF3 | 22.9405169 | -0.696372958 | 0.35727944 | -1.9490989 | 0.05128362 | 0.22053427 |
| COPS6 | 32.0761632 | -0.034636761 | 0.29963212 | -0.1155976 | 0.90797145 | 0.96223864 |
| MCM7 | 11.9189891 | -0.490306353 | 0.42411251 | -1.1560761 | 0.24765001 | 0.53836051 |
| AP4M1 | 20.6885549 | -0.50950353 | 0.33492286 | -1.5212563 | 0.12819552 | 0.37540371 |
| TAF6 | 15.6566858 | -0.546857022 | 0.50495318 | -1.0829856 | 0.27881481 | 0.57144989 |
| LAMTOR4 | 65.4374447 | -0.208616429 | 0.33608741 | -0.6207207 | 0.53478338 | 0.76795907 |
| CASTOR3 | 76.1027178 | -0.790600113 | 0.47853332 | -1.6521318 | 0.09850769 | 0.32413171 |
| SPDYE3 | 33.2654943 | 0.305338304 | 0.50812235 | 0.60091493 | 0.54789665 | 0.77516644 |
| PILRB | 59.9657573 | -0.099985317 | 0.33355172 | -0.2997596 | 0.76436056 | 0.89764618 |
| PILRA | 17.1325418 | 0.193020566 | 0.72585464 | 0.26592179 | 0.79029944 | 0.91080793 |
| MEPCE | 15.3695794 | -0.56915279 | 0.50658294 | -1.1235135 | 0.26121945 | 0.55247029 |
| PPP1R35 | 20.8261696 | -0.87274277 | 0.37308144 | -2.3392822 | 0.01932083 | 0.11815925 |
| TSC22D4 | 36.4387801 | 0.159806532 | 0.27174117 | 0.58808361 | 0.55647617 | 0.78071064 |
| AGFG2 | 20.4421652 | -0.11868157 | 0.46018079 | -0.2579021 | 0.7964825 | 0.91221376 |
| LRCH4 | 15.0690813 | 0.38695505 | 0.37951499 | 1.01960411 | 0.30791625 | 0.60158415 |
| GNB2 | 69.1218282 | -0.881128988 | 0.39102052 | -2.2534085 | 0.0242334 | 0.13597631 |
| GIGYF1 | 45.3779771 | -0.122988454 | 0.35942182 | -0.3421842 | 0.73221231 | 0.88276784 |
| EPHB4 | 16.1049012 | -0.385982627 | 0.45774604 | -0.8432244 | 0.39910295 | 0.67968972 |
| SLC12A9 | 30.6352188 | 0.865924202 | 0.52302413 | 1.65561041 | 0.09780074 | 0.32303406 |
| TRIP6 | 24.5786259 | 0.130199508 | 0.37077451 | 0.3511555 | 0.72547169 | 0.87840733 |
| SRRT | 21.9459961 | -0.092869808 | 0.3563197 | -0.2606362 | 0.79437308 | 0.91176607 |
| TRIM56 | 104.677037 | -1.071134073 | 0.38107411 | -2.8108288 | 0.00494141 | 0.04713554 |
| AP1S1 | 18.6069731 | -0.555947652 | 0.48612482 | -1.1436315 | 0.2527765 | 0.54433601 |
| PLOD3 | 18.2207641 | -0.708382564 | 0.38268594 | -1.8510807 | 0.06415794 | 0.2521767 |
| ZNHIT1 | 35.3687204 | -0.775744119 | 0.35172552 | -2.2055383 | 0.02741634 | 0.14803099 |
| FIS1 | 55.9191281 | -0.225625327 | 0.39363231 | -0.573188 | 0.56651739 | 0.78647713 |
| IFT22 | 14.1435483 | -0.845817475 | 0.47340599 | -1.7866641 | 0.0739918 | 0.27508608 |
| COL26A1 | 17.0200634 | 0.502753231 | 0.58538629 | 0.85884012 | 0.39042873 | 0.67215855 |
| CUX1 | 528.091248 | 0.092381251 | 0.17764045 | 0.52004626 | 0.60303133 | 0.80891 |
| PRKRIP1 | 150.700083 | -0.07653725 | 0.196326 | -0.3898478 | 0.69664912 | 0.86247999 |
| ORAI2 | 30.2765029 | -0.429631456 | 0.44587736 | -0.9635642 | 0.33526447 | 0.62355923 |
| ALKBH4 | 12.7716878 | -0.011526556 | 0.41163878 | -0.0280016 | 0.97766085 | 0.99112991 |
| POLR2J | 18.3727557 | -0.80510959 | 0.46745903 | -1.7223105 | 0.08501329 | 0.29941397 |
| AC093668.2 | 10.1571002 | 0.980626914 | 0.65087313 | 1.50663297 | 0.13190476 | 0.38098929 |
| SPDYE2 | 11.7889979 | 0.505036902 | 0.61377915 | 0.82283164 | 0.41060374 | 0.68825899 |
| AC105052.1 | 10.7278601 | 0.488052695 | 0.79104771 | 0.61696999 | 0.53725452 | 0.76945357 |
| AC105052.3 | 36.4809153 | 0.710970573 | 0.41870176 | 1.69803579 | 0.08950101 | 0.30789644 |
| FAM185A | 79.5739714 | 0.596291579 | 0.32796949 | 1.81813126 | 0.06904407 | 0.26461757 |
| FBXL13 | 56.8916872 | -0.678356011 | 0.38163489 | -1.7775 | 0.07548601 | 0.27883002 |
| ARMC10 | 43.0055984 | -0.394473198 | 0.2647495 | -1.4899866 | 0.13622776 | 0.38868467 |
| NAPEPLD | 70.1360571 | 0.482467704 | 0.29715997 | 1.6235959 | 0.10446208 | 0.33555699 |
| PMPCB | 40.7174765 | 0.142531811 | 0.28330506 | 0.50310365 | 0.6148914 | 0.81621069 |
| DNAJC2 | 36.0064258 | -0.268583307 | 0.29148008 | -0.9214465 | 0.35681734 | 0.64141201 |
| PSMC2 | 24.7178782 | 0.134050568 | 0.36364926 | 0.36862599 | 0.71240652 | 0.8712265 |
| ORC5 | 55.4168813 | 0.362102625 | 0.25387915 | 1.42627952 | 0.15378768 | 0.41692648 |
| LHFPL3 | 136.402364 | -0.75085881 | 0.47028591 | -1.5966007 | 0.11035474 | 0.34618593 |
| LINC01004 | 146.877172 | 0.407497564 | 0.23555864 | 1.72991986 | 0.08364459 | 0.29623459 |
| KMT2E-AS1 | 9.23988354 | 0.251906374 | 0.60284672 | 0.4178614 | 0.67604846 | 0.85173672 |
| KMT2E | 341.663533 | 0.621843867 | 0.17758244 | 3.50171937 | 0.00046227 | 0.00813175 |
| SRPK2 | 478.291294 | 0.194975622 | 0.21026262 | 0.92729569 | 0.35377303 | 0.63957058 |
| PUS7 | 30.4570064 | -0.58228367 | 0.3888643 | -1.4973955 | 0.13429037 | 0.38514144 |
| RINT1 | 26.9638192 | -0.024210837 | 0.46535284 | -0.0520268 | 0.95850731 | 0.98335155 |
| ATXN7L1 | 159.449347 | 0.031615274 | 0.26720795 | 0.11831712 | 0.9058164 | 0.96157183 |
| CDHR3 | 27.8000666 | 0.404676179 | 0.66093882 | 0.6122748 | 0.54035596 | 0.7714342 |
| SYPL1 | 82.5295012 | -0.124325109 | 0.33784071 | -0.3679992 | 0.71287383 | 0.87134026 |
| NAMPT | 221.711141 | -2.497820858 | 0.4156222 | -6.0098351 | 1.86E-09 | 2.90E-07 |
| HBP1 | 81.2678007 | -0.080693726 | 0.24119174 | -0.3345626 | 0.73795509 | 0.88573617 |
| COG5 | 278.969198 | 0.148782798 | 0.23302422 | 0.6384864 | 0.5231571 | 0.76209905 |
| DUS4L | 12.8377155 | -0.143844643 | 0.36799838 | -0.3908839 | 0.69588306 | 0.86236044 |
| BCAP29 | 50.1459728 | -0.302774946 | 0.27211053 | -1.112691 | 0.26584115 | 0.55721171 |
| CBLL1 | 16.1293287 | -0.077166811 | 0.39903614 | -0.193383 | 0.84665902 | 0.93757726 |
| SLC26A3 | 350.944498 | -0.095731104 | 0.72732629 | -0.1316206 | 0.89528441 | 0.95885435 |
| DLD | 81.4817688 | 0.418702922 | 0.33673899 | 1.24340493 | 0.21371866 | 0.50026184 |
| LAMB1 | 874.803024 | -0.061195372 | 0.50653771 | -0.1208111 | 0.90384067 | 0.96126516 |
| NRCAM | 303.68516 | 1.290236925 | 0.30412856 | 4.24240631 | 2.21E-05 | 0.0007075 |
| PNPLA8 | 82.5427312 | -0.184868828 | 0.28116437 | -0.6575116 | 0.51085204 | 0.7549696 |

| | | | | | | |
|---|---|---|---|---|---|---|
| DNAJB9 | 19.469242 | -1.561255754 | 0.44546103 | -3.5048088 | 0.00045694 | 0.00807435 |
| IMMP2L | 2440.1663 | 0.398554807 | 0.27125053 | 1.46932358 | 0.14174504 | 0.39788385 |
| LRRN3 | 13.5937219 | -1.25243556 | 0.74435247 | -1.682584 | 0.09245565 | 0.31321214 |
| DOCK4 | 99.7210561 | 0.299768428 | 0.40413969 | 0.74174459 | 0.4582421 | 0.72410401 |
| ZNF277 | 106.921696 | 0.242037862 | 0.19713514 | 1.22777638 | 0.21953093 | 0.50777 |
| IFRD1 | 109.409629 | 1.589807534 | 0.54946475 | 2.89337493 | 0.00381126 | 0.03891338 |
| TMEM168 | 101.061737 | 0.713413023 | 0.25980213 | 2.74598601 | 0.00603293 | 0.05451867 |
| BMT2 | 121.437239 | 0.188044684 | 0.16657371 | 1.12889777 | 0.25894096 | 0.55002264 |
| SMIM30 | 16.4526394 | -0.200875176 | 0.39089391 | -0.5138867 | 0.60733122 | 0.81184603 |
| MDFIC | 46.2085542 | 0.142769175 | 0.48223499 | 0.29605727 | 0.76718635 | 0.89966462 |
| TES | 67.2749372 | -0.426675689 | 0.41681534 | -1.0236564 | 0.3059976 | 0.60076912 |
| CAV2 | 29.3518527 | -0.89361418 | 0.49686466 | -1.7985062 | 0.07209682 | 0.2711725 |
| MET | 176.06809 | -2.046876539 | 0.70752963 | -2.8929906 | 0.00381593 | 0.03892246 |
| CAPZA2 | 148.082224 | -0.191037095 | 0.29329602 | -0.6513457 | 0.51482337 | 0.75684546 |
| ST7 | 213.336856 | -0.01362416 | 0.25440289 | -0.0535535 | 0.95729092 | 0.98315333 |
| ST7-AS2 | 12.2025149 | -0.921177359 | 0.70327652 | -1.3098366 | 0.1902511 | 0.46973504 |
| CFTR | 28.5964318 | -4.91645968 | 0.88927408 | -5.5286214 | 3.23E-08 | 3.20E-06 |
| CTTNBP2 | 37.4009622 | -2.490175406 | 0.84014486 | -2.9639834 | 0.00303685 | 0.03286302 |
| LSM8 | 36.3845638 | -0.313115332 | 0.28783758 | -1.0878195 | 0.2766748 | 0.56884726 |
| ANKRD7 | 57.0641482 | 0.507803782 | 0.37247548 | 1.36332137 | 0.17278125 | 0.44544355 |
| TSPAN12 | 71.2642813 | -0.356239324 | 0.45293018 | -0.7865215 | 0.43156203 | 0.70454366 |
| ING3 | 23.2397556 | 0.191651107 | 0.38729932 | 0.49483977 | 0.62071323 | 0.81942374 |
| FAM3C | 74.1510513 | -1.141927583 | 0.33635948 | -3.3949618 | 0.00068638 | 0.01110064 |
| AASS | 47.7064617 | 0.323074151 | 0.26335948 | 1.22674204 | 0.21991956 | 0.50818193 |
| CADPS2 | 2371.71549 | 1.079402212 | 0.24203876 | 4.4596255 | 8.21E-06 | 0.00031211 |
| AC004594.1 | 109.355192 | 0.052403713 | 0.58120526 | 0.09016387 | 0.928157 | 0.97143893 |
| SLC13A1 | 15.9515852 | 0.926617523 | 0.69665406 | 1.33009707 | 0.18348629 | 0.46014503 |
| IQUB | 114.376525 | 0.529734431 | 0.37775948 | 1.40230611 | 0.16082386 | 0.42803048 |
| NDUFA5 | 108.484531 | 0.354639388 | 0.34802159 | 1.01901547 | 0.30819562 | 0.60178758 |
| WASL | 120.84253 | -0.09183175 | 0.20361997 | -0.4509958 | 0.65199258 | 0.83803214 |
| POT1 | 55.5243736 | 0.072687675 | 0.27413627 | 0.26515161 | 0.79089267 | 0.91080793 |
| POT1-AS1 | 31.0563174 | -0.235046205 | 0.35553841 | -0.6610993 | 0.50854862 | 0.7543124 |
| GRM8 | 69.5145738 | -0.532228808 | 0.75823818 | -0.7019283 | 0.4827239 | 0.74093072 |
| ZNF800 | 76.5990357 | -0.099130866 | 0.31497329 | -0.3147279 | 0.7529683 | 0.89205725 |
| ARF5 | 22.8742804 | -0.422906403 | 0.59454881 | -0.7113064 | 0.47689436 | 0.73678733 |
| SND1 | 265.387858 | -0.188546484 | 0.30489895 | -0.6183901 | 0.53631824 | 0.76895069 |
| RBM28 | 45.2678233 | -0.079660976 | 0.34864286 | -0.2284888 | 0.81926629 | 0.92352826 |
| IMPDH1 | 15.1791651 | 0.402214421 | 0.38750737 | 1.03795295 | 0.29929196 | 0.59237383 |
| HILPDA | 54.2162711 | -4.127919786 | 0.93092428 | -4.4342165 | 9.24E-06 | 0.00034244 |
| METTL2B | 40.0298405 | 0.283397636 | 0.29415802 | 0.96341971 | 0.33533694 | 0.62358143 |
| CALU | 67.2095527 | -0.136330197 | 0.37098793 | -0.3674788 | 0.7132619 | 0.87144499 |
| KCP | 57.9368177 | -0.471512059 | 0.34893732 | -1.3512801 | 0.17660572 | 0.45112623 |
| ATP6V1F | 81.1456317 | -0.860516244 | 0.49557877 | -1.7363864 | 0.08249553 | 0.29436838 |
| TNPO3 | 137.157495 | 0.052650212 | 0.26976338 | 0.19517182 | 0.84525844 | 0.93715892 |
| TSPAN33 | 31.0178581 | 0.825483736 | 0.34599356 | 2.38583559 | 0.01704037 | 0.10924074 |
| SMO | 20.6104647 | 0.748400174 | 0.41224005 | 1.81544753 | 0.06945516 | 0.26541135 |
| AHCYL2 | 331.102938 | 0.846909686 | 0.23303593 | 3.63424516 | 0.0002788 | 0.00563167 |
| STRIP2 | 18.4973023 | -2.670418637 | 0.96992662 | -2.7532172 | 0.00590127 | 0.05356381 |
| NRF1 | 165.75809 | 0.137616462 | 0.21193292 | 0.64933972 | 0.51611881 | 0.7575233 |
| UBE2H | 654.514384 | 0.098973762 | 0.29273587 | 0.3380992 | 0.73528843 | 0.88409681 |
| ZC3HC1 | 29.4369447 | 0.37379767 | 0.33155465 | 1.12740892 | 0.25956963 | 0.55072654 |
| KLHDC10 | 174.350278 | 0.081221914 | 0.17391758 | 0.46701381 | 0.64049 | 0.8320413 |
| TMEM209 | 25.4753831 | 0.355759061 | 0.33644263 | 1.05741374 | 0.29032281 | 0.58200489 |
| CEP41 | 12.1925016 | 0.397821173 | 0.40559625 | 0.98083049 | 0.32667634 | 0.61667091 |
| MEST | 38.6770261 | -0.849962551 | 1.28939771 | -0.6591935 | 0.50977154 | 0.75476666 |
| COPG2 | 56.6492719 | -0.107596844 | 0.2722765 | -0.3951749 | 0.6927138 | 0.86052545 |
| AC016831.7 | 740.142406 | -0.761149792 | 0.30853689 | -2.4669653 | 0.01362636 | 0.09485049 |
| LINC00513 | 62.0739223 | -1.535966925 | 0.60213676 | -2.5508606 | 0.01074573 | 0.08056236 |
| AC058791.1 | 14.9606006 | -2.259113032 | 0.78619847 | -2.873464 | 0.00405997 | 0.04048968 |
| MKLN1 | 596.935274 | -0.250068769 | 0.28184447 | -0.887258 | 0.37494 | 0.65806152 |
| PODXL | 11.3891408 | -0.904240681 | 0.47116439 | -1.9191618 | 0.05496387 | 0.23130627 |
| CHCHD3 | 476.361759 | 0.558111886 | 0.1668241 | 3.34551115 | 0.00082131 | 0.01268537 |
| EXOC4 | 702.349091 | 0.364045482 | 0.16276029 | 2.23669714 | 0.02530614 | 0.14031424 |
| LRGUK | 31.0796459 | 0.137522691 | 0.329891 | 0.41687313 | 0.67677121 | 0.85191801 |
| SLC35B4 | 102.61492 | 1.430316162 | 0.35304058 | 4.05142136 | 5.09E-05 | 0.00142901 |
| AKR1B1 | 54.4859526 | 0.133093438 | 0.38018718 | 0.35007345 | 0.72628358 | 0.87880825 |
| BPGM | 31.3113514 | -1.217862727 | 0.38517438 | -3.1618477 | 0.00156772 | 0.02046972 |
| CALD1 | 25.1364085 | -0.552100134 | 0.47565597 | -1.1607131 | 0.24575858 | 0.53627476 |
| AGBL3 | 23.512902 | -0.237858784 | 0.3447 | -0.6900458 | 0.49016539 | 0.74556088 |
| CYREN | 31.7450107 | -0.169916995 | 0.3095012 | -0.5490027 | 0.5830036 | 0.79586061 |
| TMEM140 | 33.1468426 | -1.887251031 | 0.72740236 | -2.5945077 | 0.00947265 | 0.07421082 |
| WDR91 | 86.7050049 | 0.455239717 | 0.35806449 | 1.2713903 | 0.20358983 | 0.48787682 |
| CNOT4 | 219.364835 | 0.142730456 | 0.21398829 | 0.66700126 | 0.50477133 | 0.7517933 |
| NUP205 | 52.0248312 | 0.976959237 | 0.25367188 | 3.85127132 | 0.00011751 | 0.00277014 |
| STMP1 | 76.8218725 | -0.177613042 | 0.29239001 | -0.6074525 | 0.54355066 | 0.77321995 |
| SLC13A4 | 11.4161137 | -2.121939465 | 0.80327572 | -2.6416079 | 0.00825135 | 0.06735772 |
| MTPN | 69.4506717 | -0.456883662 | 0.30111643 | -1.517299 | 0.12919121 | 0.37682602 |
| CREB3L2 | 89.4336352 | 0.861769753 | 0.18829821 | 4.57662208 | 4.73E-06 | 0.00020284 |
| TRIM24 | 124.704283 | 0.184256004 | 0.26259218 | 0.70168124 | 0.48287797 | 0.74093072 |
| SVOPL | 46.1081968 | 1.136101077 | 0.7694739 | 1.47646473 | 0.13981918 | 0.3948512 |
| ATP6V0A4 | 315.103939 | 1.202541796 | 0.30131701 | 3.99095227 | 6.58E-05 | 0.00176093 |
| TMEM213 | 245.799702 | 0.717437364 | 0.36915071 | 1.94348095 | 0.05195807 | 0.22247385 |
| KIAA1549 | 46.9107893 | -0.078819161 | 0.34664489 | -0.2273772 | 0.82013041 | 0.92406023 |
| ZC3HAV1 | 91.9775174 | -0.373048653 | 0.47018862 | -0.7934021 | 0.42754357 | 0.70214472 |
| TTC26 | 28.9078627 | -0.311663789 | 0.34323966 | -0.9080063 | 0.36387488 | 0.64877795 |
| UBN2 | 124.41921 | 0.063870719 | 0.17235947 | 0.37056692 | 0.71096013 | 0.87028547 |
| LUC7L2 | 38.44431 | 0.272944753 | 0.32603103 | 0.83717417 | 0.40249467 | 0.68128594 |
| HIPK2 | 375.165944 | -0.363698858 | 0.19698791 | -1.8463004 | 0.0648486 | 0.25440605 |
| TBXAS1 | 28.0933151 | -1.666562927 | 0.45243429 | -3.6835469 | 0.00023001 | 0.00478702 |
| PARP12 | 44.7866041 | -0.516879268 | 0.758264 | -0.6816614 | 0.49545311 | 0.74770824 |
| KDM7A | 133.244757 | -1.051957327 | 0.25848203 | -4.0697503 | 4.71E-05 | 0.0013423 |
| SLC37A3 | 64.3733172 | -0.173022569 | 0.33669178 | -0.5138901 | 0.60732884 | 0.81184603 |
| MKRN1 | 62.177191 | -0.090207257 | 0.2585578 | -0.3488862 | 0.72717472 | 0.8796799 |
| DENND2A | 18.7615723 | -1.524435345 | 0.64402624 | -2.3670392 | 0.01793103 | 0.1127288 |
| NDUFB2 | 90.3792809 | -0.222439025 | 0.35488584 | -0.6267904 | 0.53079666 | 0.7648996 |
| BRAF | 535.250788 | 0.824814441 | 0.31159415 | 2.64707935 | 0.00811903 | 0.06664721 |
| MRPS33 | 24.4249205 | -0.184495975 | 0.42047133 | -0.4387837 | 0.66081826 | 0.84186478 |
| TMEM178B | 80.747069 | -0.106697154 | 0.65278892 | -0.1634482 | 0.87016558 | 0.94835789 |
| AGK | 55.607315 | 0.606917791 | 0.22619269 | 2.68318924 | 0.00729237 | 0.06219041 |
| AC004918.1 | 15.3299894 | -0.085467617 | 0.39232293 | -0.2178502 | 0.82754585 | 0.92808376 |
| KIAA1147 | 50.9611635 | -0.183372602 | 0.28310121 | -0.6477281 | 0.51716084 | 0.75818855 |
| SSBP1 | 68.9254792 | 0.095562658 | 0.2559473 | 0.3733685 | 0.7088742 | 0.86910485 |
| TRPV6 | 50.7642594 | -0.150389036 | 0.46782837 | -0.321462 | 0.74786032 | 0.88935207 |
| TRPV5 | 227.427665 | -0.239256183 | 0.71272246 | -0.3356933 | 0.73710213 | 0.88493487 |
| GSTK1 | 72.8060952 | -0.070796038 | 0.37482265 | -0.1888788 | 0.85018784 | 0.93946532 |
| TMEM139 | 11.2054917 | -0.312694035 | 0.52261431 | -0.5983266 | 0.54962204 | 0.7759356 |
| CASP2 | 13.9751204 | -0.528316698 | 0.37340296 | -1.4148701 | 0.15710657 | 0.42221043 |
| ZYX | 17.181682 | -1.084731756 | 0.51749044 | -2.0961387 | 0.03606989 | 0.17602651 |
| TCAF2 | 34.2206482 | 0.29554605 | 0.61653449 | 0.47936661 | 0.63167784 | 0.82625173 |
| TCAF1 | 67.8704038 | -0.064860405 | 0.29215492 | -0.2220069 | 0.82430852 | 0.92643003 |
| AC004889.1 | 59.1577519 | 1.147817615 | 0.32716683 | 3.5083557 | 0.00045089 | 0.00800866 |
| AC074386.1 | 78.2193038 | -0.164222749 | 0.36443352 | -0.4506247 | 0.65226006 | 0.83816466 |
| ARHGEF5 | 19.9847026 | -0.316279199 | 0.35886482 | -0.8813324 | 0.37813793 | 0.66119336 |

| | | | | | | |
|---|---|---|---|---|---|---|
| TPK1 | 76.8691773 | -0.396834009 | 0.25257787 | -1.5711353 | 0.11615122 | 0.35543425 |
| CNTNAP2 | 12.0326987 | 0.179764008 | 0.46705395 | 0.38488917 | 0.70031953 | 0.86382804 |
| CUL1 | 138.830483 | 0.108257919 | 0.20310883 | 0.5330045 | 0.59403046 | 0.80267231 |
| EZH2 | 18.7046525 | -0.545194423 | 0.47069041 | -1.1582867 | 0.24674707 | 0.53764558 |
| PDIA4 | 39.2971266 | -1.178537331 | 0.58225695 | -2.0240846 | 0.04296145 | 0.19793776 |
| ZNF786 | 18.4721592 | -0.061422667 | 0.44450415 | -0.1381824 | 0.89009623 | 0.95702609 |
| ZNF425 | 10.9991385 | -0.362913106 | 0.4164786 | -0.8713848 | 0.38354411 | 0.66699382 |
| ZNF398 | 42.121338 | 0.046154844 | 0.28657178 | 0.16105858 | 0.87204726 | 0.94846188 |
| ZNF282 | 22.2098093 | -0.293585009 | 0.29263141 | -1.0032587 | 0.31573606 | 0.60912226 |
| ZNF212 | 11.2457021 | -0.287205545 | 0.42685801 | -0.6728363 | 0.50105147 | 0.75071921 |
| ZNF783 | 16.8123759 | 0.112827117 | 0.3700826 | 0.30487009 | 0.76046509 | 0.89627175 |
| ZNF777 | 11.6847002 | -0.373981509 | 0.38838908 | -0.9629043 | 0.33559556 | 0.62382904 |
| ZNF746 | 18.003958 | 0.297757886 | 0.36688883 | 0.81157523 | 0.41703541 | 0.69338933 |
| ZNF467 | 12.6266969 | 0.631845296 | 0.4226304 | 1.49503041 | 0.13490651 | 0.38627205 |
| ZNF862 | 48.889031 | -0.101426203 | 0.30611642 | -0.3313321 | 0.74039363 | 0.88639183 |
| ATP6V0E2 | 18.4785749 | -0.046629564 | 0.59911886 | -0.0778302 | 0.93796309 | 0.97626283 |
| AC092666.1 | 13.2225379 | -1.51157728 | 0.82196496 | -1.8389802 | 0.0659181 | 0.25655016 |
| ACTR3C | 269.470159 | -0.390383625 | 0.23235211 | -1.6801381 | 0.09293045 | 0.31376482 |
| LRRC61 | 18.3334749 | -1.244487039 | 0.45130738 | -2.7575154 | 0.00582425 | 0.05309858 |
| TMEM176B | 69.2297038 | -1.61698125 | 0.76625384 | -2.1102423 | 0.03483749 | 0.17346343 |
| TMEM176A | 31.9551747 | -1.349809846 | 0.65539128 | -2.0595481 | 0.03944177 | 0.18750765 |
| AOC1 | 37.1836427 | -0.594835849 | 1.37493785 | -0.4326274 | 0.66528545 | 0.84348349 |
| ABCB8 | 19.861447 | -0.361568506 | 0.35786086 | -1.0103606 | 0.31232256 | 0.60537103 |
| SLC4A2 | 23.581301 | 0.20810867 | 0.40373174 | 0.51546274 | 0.6062297 | 0.81114147 |
| FASTK | 25.5826829 | 0.646810598 | 0.44351252 | 1.45838182 | 0.14473533 | 0.40273129 |
| TMUB1 | 27.0599526 | -0.737771351 | 0.47889331 | -1.5405756 | 0.12342011 | 0.36821038 |
| AGAP3 | 21.6349859 | 0.2383554 | 0.52106784 | 0.45743641 | 0.6473574 | 0.83478232 |
| CHPF2 | 13.7218497 | -1.077032706 | 0.57586992 | -1.8702708 | 0.06144622 | 0.24564181 |
| SMARCD3 | 98.2448107 | -1.317260856 | 0.34504398 | -3.8176607 | 0.00013472 | 0.00309857 |
| NUB1 | 59.4888905 | -0.376968762 | 0.37082328 | -1.0165725 | 0.30935683 | 0.60291223 |
| RHEB | 119.471264 | -0.073076457 | 0.31983276 | -0.2284833 | 0.81927051 | 0.92352826 |
| PRKAG2 | 579.651372 | 0.096016525 | 0.40603128 | 0.23647568 | 0.81306357 | 0.92060876 |
| GALNT11 | 233.985935 | 0.744756401 | 0.46748677 | 1.59310692 | 0.11113621 | 0.34769355 |
| KMT2C | 580.823139 | 0.318363025 | 0.21418257 | 1.4864096 | 0.13717081 | 0.39028091 |
| ACTR3B | 74.9105549 | 0.281007597 | 0.2481537 | 1.13239335 | 0.2574691 | 0.54861387 |
| DPP6 | 165.858681 | -0.324503504 | 0.24802874 | -1.3083303 | 0.19076131 | 0.47059938 |
| PAXIP1-AS2 | 23.8794233 | 0.015176589 | 0.42293039 | 0.03588437 | 0.97137456 | 0.988268 |
| PAXIP1 | 40.3230349 | 0.013662256 | 0.33769957 | 0.04045684 | 0.96772892 | 0.987279 |
| PAXIP1-AS1 | 11.6149355 | -0.677016081 | 0.55180282 | -1.2269167 | 0.21985391 | 0.50818193 |
| INSIG1 | 70.6195298 | 1.091260975 | 0.86366708 | 1.2635204 | 0.20640222 | 0.49107522 |
| AC009403.1 | 24.2565595 | -0.718329084 | 0.45496965 | -1.5788506 | 0.11437034 | 0.35329632 |
| RBM33 | 193.506174 | 0.032654862 | 0.18431313 | 0.17717057 | 0.85937441 | 0.94424863 |
| LINC01006 | 58.2431686 | 0.009520566 | 0.31878987 | 0.02986471 | 0.97617495 | 0.99040323 |
| LMBR1 | 226.716345 | 0.432631372 | 0.25820763 | 1.67551741 | 0.09383276 | 0.31579038 |
| NOM1 | 21.6268582 | -0.264387092 | 0.33085657 | -0.7990988 | 0.42423312 | 0.69882468 |
| UBE3C | 252.791388 | 0.368962818 | 0.30676985 | 1.20273494 | 0.22907891 | 0.51924553 |
| DNAJB6 | 131.224819 | -0.003336246 | 0.35078543 | -0.0095108 | 0.9924116 | 0.99770844 |
| PTPRN2 | 11.2280392 | -0.615473964 | 0.57405295 | -1.0721554 | 0.28365024 | 0.57636386 |
| ESYT2 | 325.331896 | 1.174021218 | 0.68000738 | 1.72648306 | 0.08426054 | 0.29778804 |
| WDR60 | 162.121315 | -0.247026492 | 0.24860947 | -0.9936327 | 0.32040174 | 0.6120836 |
| GTPBP6 | 35.833986 | -0.264358692 | 0.38829004 | -0.6808279 | 0.49598039 | 0.74800029 |
| LINC00685 | 20.6582503 | -0.361030129 | 0.69718166 | -0.5178423 | 0.60456833 | 0.81002248 |
| PPP2R3B | 15.6402727 | 0.203115186 | 0.3676982 | 0.55239646 | 0.58067676 | 0.7947038 |
| IL3RA | 15.7597177 | -0.017867648 | 0.74852604 | -0.0238704 | 0.98095595 | 0.99318267 |
| SLC25A6 | 147.054053 | -0.457905154 | 0.42206596 | -1.0849137 | 0.27795987 | 0.57019963 |
| LINC00106 | 50.1605333 | 1.000440176 | 0.44437341 | 2.25135022 | 0.02436336 | 0.1365442 |
| ASMTL-AS1 | 24.347216 | 1.11526035 | 0.46388744 | 2.40416154 | 0.01620961 | 0.10535734 |
| ASMTL | 27.6561702 | 0.244841505 | 0.29920545 | 0.81830563 | 0.41318269 | 0.69014748 |
| AKAP17A | 29.9068379 | -0.533705798 | 0.32934514 | -1.6205061 | 0.10512361 | 0.33670353 |
| DHRSX | 172.129665 | 0.500670735 | 0.36349887 | 1.37736532 | 0.16839933 | 0.43888762 |
| CD99 | 34.8405361 | -0.112328758 | 0.3586557 | -0.3131938 | 0.7541334 | 0.89238252 |
| ARSD | 76.8552611 | -0.062146064 | 0.32682129 | -0.190153 | 0.8491892 | 0.93897371 |
| MXRA5 | 53.4400536 | 0.24102475 | 0.39988037 | 0.60274215 | 0.54668024 | 0.77463416 |
| PRKX | 35.7478987 | -0.790951771 | 0.33740178 | -2.344243 | 0.01906574 | 0.11735971 |
| BX890604.1 | 11.8063591 | 0.005971849 | 0.44538615 | 0.01340825 | 0.98930209 | 0.99637773 |
| PUDP | 38.4925274 | -0.41997373 | 0.32587803 | -1.2887452 | 0.1974867 | 0.47953293 |
| STS | 15.3897161 | -0.252942274 | 0.71662058 | -0.3529654 | 0.72411438 | 0.87742017 |
| PNPLA4 | 15.4812861 | -0.053381682 | 0.42537851 | -0.1254922 | 0.90013391 | 0.95980075 |
| TBL1X | 256.086788 | 0.194099945 | 0.32958759 | 0.58891764 | 0.55591652 | 0.78025232 |
| SHROOM2 | 43.3825941 | 0.222004547 | 0.29326762 | 0.75700328 | 0.4490479 | 0.71802342 |
| WWC3 | 27.5728608 | 0.048726542 | 0.3226509 | 0.15101939 | 0.87996042 | 0.95314397 |
| MID1 | 592.835023 | -0.304354887 | 0.2745312 | -1.108635 | 0.26758768 | 0.55939292 |
| AC073529.1 | 37.9221338 | 0.540978074 | 0.35164522 | 1.53842009 | 0.12394593 | 0.36885651 |
| ARHGAP6 | 829.699061 | 0.982922585 | 0.27067454 | 3.63138179 | 0.00028191 | 0.00566124 |
| PRPS2 | 48.8666613 | 1.577861749 | 0.47403519 | 3.32857512 | 0.00087291 | 0.01332262 |
| TMSB4X | 468.215026 | -1.470765686 | 0.57405091 | -2.5620823 | 0.01040467 | 0.07893141 |
| RAB9A | 45.9224221 | -0.464430866 | 0.29175274 | -1.5918646 | 0.11141512 | 0.34792318 |
| TRAPPC2 | 21.7515078 | -0.191880753 | 0.34350536 | -0.5585961 | 0.57643741 | 0.79190001 |
| OFD1 | 116.758411 | -0.447413885 | 0.30155847 | -1.4836721 | 0.13789593 | 0.39135092 |
| GPM6B | 32.0736969 | -2.147630056 | 0.58807622 | -3.6519587 | 0.00026025 | 0.00530905 |
| GEMIN8 | 32.693359 | -0.187005575 | 0.34262039 | -0.5458098 | 0.58519667 | 0.79681419 |
| MOSPD2 | 24.282358 | 0.357845688 | 0.32661028 | 1.09563509 | 0.2732385 | 0.5655199 |
| PIR | 23.2980437 | 0.229404659 | 0.77285688 | 0.29682683 | 0.76659872 | 0.89911488 |
| CLTRN | 15.9767796 | -0.443790678 | 0.8507998 | -0.5216159 | 0.60193781 | 0.8078639 |
| CA5B | 51.8198056 | -0.364847593 | 0.28684696 | -1.2719242 | 0.20340004 | 0.48753543 |
| ZRSR2 | 25.4795773 | -0.162987203 | 0.33335286 | -0.488933 | 0.62488915 | 0.82285278 |
| AP1S2 | 20.0562751 | -1.614342752 | 0.58414331 | -2.7636074 | 0.00571663 | 0.05234907 |
| CTPS2 | 61.6025908 | -0.119461103 | 0.24076174 | -0.4961798 | 0.61976759 | 0.81899807 |
| SYAP1 | 64.6592172 | -0.001712648 | 0.39368162 | -0.0043503 | 0.99652894 | 0.99868531 |
| TXLNG | 41.5728663 | -0.025542441 | 0.33363168 | -0.0765588 | 0.93897454 | 0.97662903 |
| RBBP7 | 53.9221018 | 0.022055459 | 0.3096739 | 0.07122156 | 0.94322142 | 0.97824588 |
| REPS2 | 105.419487 | -0.59163255 | 0.46767619 | -1.2650474 | 0.20585434 | 0.48999801 |
| NHS | 512.989023 | -0.568408696 | 0.3045275 | -1.8665267 | 0.06196772 | 0.24676931 |
| SCML1 | 113.252066 | 0.634564817 | 0.3029005 | 2.09496128 | 0.03617443 | 0.17636963 |
| RAI2 | 31.3045133 | -0.399753468 | 0.3913171 | -1.0215589 | 0.30698972 | 0.60139222 |
| CDKL5 | 84.0983292 | 0.404255583 | 0.18385806 | 2.19873732 | 0.02789661 | 0.14976073 |
| PHKA2 | 34.7042052 | 0.432655726 | 0.26856208 | 1.61100825 | 0.10717792 | 0.34111768 |
| PDHA1 | 108.119931 | 0.114143645 | 0.37859316 | 0.30149421 | 0.76303767 | 0.8974069 |
| MAP3K15 | 88.1180066 | 1.480799098 | 0.34473706 | 4.29544511 | 1.74E-05 | 0.00057938 |
| SH3KBP1 | 43.418386 | 0.168975046 | 0.4123009 | 0.40983429 | 0.68192751 | 0.85494315 |
| BCLAF3 | 61.3303403 | -0.112609923 | 0.25030874 | -0.4498841 | 0.65279401 | 0.8384346 |
| EIF1AX | 85.7897045 | -0.198931163 | 0.31423805 | -0.6330588 | 0.52669524 | 0.76221787 |
| RPS6KA3 | 132.03546 | -0.195299028 | 0.28111752 | -0.6947238 | 0.48722842 | 0.74434227 |
| MBTPS2 | 33.6911429 | -0.108798552 | 0.30403471 | -0.3578491 | 0.72045623 | 0.87576875 |
| SMS | 127.354075 | -0.279641288 | 0.29767405 | -0.9394211 | 0.34751457 | 0.63465611 |
| PTCHD1-AS | 54.5254407 | 0.29347277 | 0.60869257 | 0.48213628 | 0.62970914 | 0.82512634 |
| PRDX4 | 12.9037952 | -1.202658228 | 0.50037636 | -2.4035073 | 0.01623864 | 0.10547951 |
| ACOT9 | 27.2905908 | 0.521968186 | 0.4218938 | 1.23720279 | 0.21601181 | 0.50279115 |
| SAT1 | 638.018156 | -0.485498511 | 0.3170733 | -1.531187 | 0.12572319 | 0.37111757 |
| APOO | 307.409531 | -0.181203975 | 0.33046238 | -0.5483347 | 0.58346214 | 0.79598942 |
| EIF2S3 | 62.2071284 | -0.181000647 | 0.2677841 | -0.6759201 | 0.4990914 | 0.75027574 |
| ZFX | 69.6777206 | -0.056444223 | 0.32864872 | -0.1717464 | 0.86363693 | 0.94635064 |
| PDK3 | 28.5107907 | -0.178911053 | 0.52480915 | -0.3409069 | 0.73317368 | 0.88344324 |

| | | | | | | |
|---|---|---|---|---|---|---|
| POLA1 | 88.3417594 | 0.294683594 | 0.24627128 | 1.19658126 | 0.23146981 | 0.52111057 |
| IL1RAPL1 | 125.882226 | 1.505985909 | 0.57950607 | 2.59874052 | 0.00935665 | 0.07341369 |
| GK | 32.0304937 | 0.700203667 | 0.35074198 | 1.99634973 | 0.04589587 | 0.205663 |
| TAB3 | 80.0895662 | 0.247876751 | 0.25293154 | 0.98001518 | 0.32707863 | 0.61686572 |
| DMD | 34.4584717 | 0.525371738 | 0.33232662 | 1.58088971 | 0.11390326 | 0.35248764 |
| TMEM47 | 22.4144846 | -2.454673533 | 0.64044893 | -3.8327389 | 0.00012672 | 0.00295951 |
| CFAP47 | 28.1789248 | 0.005197779 | 0.51609257 | 0.01007141 | 0.99196431 | 0.99767806 |
| AF241726.2 | 40.836714 | -1.017353652 | 0.45092819 | -2.2561323 | 0.02406235 | 0.13578849 |
| DYNLT3 | 41.5711581 | -0.968699994 | 0.48888579 | -1.9814443 | 0.04754146 | 0.21044546 |
| RPGR | 33.4259638 | -0.057420259 | 0.3009591 | -0.1907909 | 0.84868942 | 0.93891735 |
| MID1IP1 | 14.0268008 | -0.407018932 | 0.47834853 | -0.8508836 | 0.394834 | 0.67534117 |
| BCOR | 67.1728876 | 0.220026985 | 0.29751897 | 0.73953934 | 0.45957956 | 0.72494085 |
| ATP6AP2 | 70.5426926 | -0.287287391 | 0.36375197 | -0.7897892 | 0.42965086 | 0.70325122 |
| CXorf38 | 29.5985818 | -0.951172134 | 0.33468146 | -2.8420222 | 0.00448284 | 0.04381612 |
| MED14 | 91.81103 | -0.030646695 | 0.30321424 | -0.1010727 | 0.91949271 | 0.96763053 |
| USP9X | 240.728971 | 0.210058601 | 0.25608425 | 0.82027145 | 0.41206138 | 0.68913252 |
| DDX3X | 90.9729815 | -0.231034517 | 0.29860318 | -0.7737175 | 0.43909785 | 0.71036213 |
| CASK | 126.469009 | 0.129479842 | 0.19514734 | 0.66349785 | 0.50701176 | 0.75373237 |
| MAOA | 60.7041686 | -0.859442618 | 0.55650762 | -1.5443501 | 0.12250354 | 0.36665643 |
| MAOB | 15.4927848 | 1.044519087 | 0.70116373 | 1.48969356 | 0.13630483 | 0.38876311 |
| EFHC2 | 16.4093075 | 0.379449042 | 0.46191553 | 0.82146847 | 0.41137948 | 0.68899849 |
| FUNDC1 | 13.4112451 | -0.211705563 | 0.45966486 | -0.460565 | 0.6451107 | 0.83387847 |
| KDM6A | 269.742879 | 0.153626479 | 0.38825624 | 0.39568322 | 0.69233874 | 0.860267 |
| MIR222HG | 79.3228386 | 1.282860582 | 0.5679029 | 2.25894354 | 0.0238869 | 0.13541585 |
| KRBOX4 | 33.6728263 | -0.111254902 | 0.30729961 | -0.3620405 | 0.71732177 | 0.87371114 |
| ZNF674 | 11.4539951 | 0.687621327 | 0.50826929 | 1.35286814 | 0.17609776 | 0.45038707 |
| SLC9A7 | 50.9865379 | -0.801781873 | 0.35235248 | -2.2755108 | 0.0228753 | 0.13202318 |
| RP2 | 14.7066144 | -0.617369316 | 0.35934514 | -1.71804 | 0.08578932 | 0.30113433 |
| JADE3 | 68.9642832 | 0.002399937 | 0.33197129 | 0.00722935 | 0.99423186 | 0.99837898 |
| RGN | 30.7536065 | -0.52659841 | 0.63563699 | -0.8284578 | 0.4074113 | 0.6859211 |
| NDUFB11 | 58.8219752 | -0.219950186 | 0.52804483 | -0.416537 | 0.67701713 | 0.85212346 |
| RBM10 | 46.3886871 | 0.256982511 | 0.23554001 | 1.09103549 | 0.27525727 | 0.56725352 |
| UBA1 | 89.815165 | 0.106001601 | 0.25870396 | 0.40974093 | 0.681996 | 0.85494315 |
| CDK16 | 44.0924202 | -0.001039673 | 0.42115315 | -0.0024686 | 0.99803032 | 0.99900003 |
| USP11 | 38.9385818 | 0.038554091 | 0.33101117 | 0.11647369 | 0.90727714 | 0.96180152 |
| ZNF41 | 21.5907914 | 0.124790032 | 0.34238997 | 0.36446755 | 0.7155089 | 0.87311962 |
| ARAF | 20.5106444 | 0.304061075 | 0.39067048 | 0.77830574 | 0.43638879 | 0.70775777 |
| TIMP1 | 55.5273876 | -3.369716867 | 0.53097811 | -6.3462444 | 2.21E-10 | 4.55E-08 |
| UXT | 14.7431306 | -0.170279473 | 0.40476353 | -0.4206888 | 0.67398236 | 0.85019422 |
| ZNF81 | 59.6515311 | 0.258419604 | 0.33138799 | 0.77980981 | 0.43550283 | 0.70733441 |
| ZNF630 | 24.4900744 | 0.336688136 | 0.56932887 | 0.59137724 | 0.55426768 | 0.7791057 |
| EBP | 22.8118706 | 0.565366458 | 0.42663684 | 1.32517028 | 0.18511468 | 0.46185874 |
| TBC1D25 | 14.226997 | 0.292817261 | 0.47995811 | 0.61008921 | 0.54180272 | 0.77222629 |
| RBM3 | 31.3220804 | -0.692234964 | 0.42346427 | -1.6346951 | 0.10211298 | 0.33070354 |
| WDR13 | 30.3281361 | 0.210509721 | 0.37755265 | 0.55756389 | 0.57714222 | 0.79249085 |
| HDAC6 | 23.0263223 | 0.462717449 | 0.35632561 | 1.29858039 | 0.19408797 | 0.47456322 |
| PCSK1N | 104.492188 | -0.475893501 | 0.52215599 | -0.911401 | 0.36208412 | 0.64626475 |
| TIMM17B | 16.2411326 | -0.293455588 | 0.45765891 | -0.6412103 | 0.52138609 | 0.76102571 |
| OTUD5 | 42.2529019 | -0.397906647 | 0.26191808 | -1.5192027 | 0.12871149 | 0.37611281 |
| GRIPAP1 | 40.3509422 | -0.039402411 | 0.31559081 | -0.1248528 | 0.90064005 | 0.95980075 |
| TFE3 | 25.6444729 | -0.700789208 | 0.39631954 | -1.7682429 | 0.07702031 | 0.28159053 |
| CCDC120 | 12.6646127 | -0.539330329 | 0.40595316 | -1.3285531 | 0.18399545 | 0.4603014 |
| WDR45 | 25.3257459 | -0.466825759 | 0.37018595 | -1.2610575 | 0.20728814 | 0.49241084 |
| PLP2 | 19.666585 | -2.817435868 | 0.57128676 | -4.9317367 | 8.15E-07 | 4.74E-05 |
| PPP1R3F | 9.86386115 | -0.275820776 | 0.48329299 | -0.5707113 | 0.56819535 | 0.78740227 |
| CLCN5 | 211.026039 | 0.797370171 | 0.26851029 | 2.96960745 | 0.00298181 | 0.0323694 |
| SHROOM4 | 146.437485 | -0.051376685 | 0.19776963 | -0.2597805 | 0.79503313 | 0.91195838 |
| AL158055.1 | 14.0842978 | 0.285544735 | 0.55367516 | 0.5157261 | 0.60604572 | 0.81114147 |
| MAGED1 | 185.663522 | 0.59367894 | 0.18587437 | 3.19397954 | 0.00140326 | 0.01887257 |
| FAM156A | 13.166969 | -0.12223707 | 0.63911788 | -0.191259 | 0.84832265 | 0.93871321 |
| TSPYL2 | 14.3717245 | -1.304139672 | 0.6156539 | -2.1183 | 0.03414967 | 0.17086443 |
| KANTR | 83.5578042 | 0.013132646 | 0.32451909 | 0.04046802 | 0.96772 | 0.987279 |
| KDM5C | 42.9604271 | -0.254140767 | 0.33738799 | -0.7532597 | 0.45129389 | 0.71947224 |
| IQSEC2 | 23.3874654 | 0.261040281 | 0.49545319 | 0.52687173 | 0.59828267 | 0.8059995 |
| SMC1A | 45.5612236 | -0.102199417 | 0.33509276 | -0.3049884 | 0.76037496 | 0.89627175 |
| HSD17B10 | 23.0589843 | -0.511031807 | 0.50526165 | -1.0114201 | 0.31181539 | 0.60484318 |
| HUWE1 | 229.705678 | 0.405468231 | 0.19026964 | 2.13101903 | 0.03308757 | 0.16799812 |
| PHF8 | 61.8336864 | 0.39208982 | 0.25386294 | 1.54449413 | 0.12246867 | 0.36665643 |
| FAM120C | 31.6331089 | 0.061244479 | 0.44533389 | 0.13752486 | 0.89061595 | 0.95702609 |
| WNK3 | 39.1390422 | 0.257716172 | 0.30149348 | 0.85479849 | 0.39266271 | 0.67375271 |
| TSR2 | 16.1523449 | -0.458095431 | 0.4109243 | -1.1147928 | 0.26493924 | 0.55622662 |
| FGD1 | 17.519312 | -0.208700813 | 0.38413341 | -0.543303 | 0.58692123 | 0.7977916 |
| GNL3L | 11.9899837 | -0.422717724 | 0.45929201 | -0.9203681 | 0.35738042 | 0.64174726 |
| MAGED2 | 36.911041 | -0.95343597 | 0.49771363 | -1.9156316 | 0.055412 | 0.23225616 |
| TRO | 10.5730089 | 0.555645911 | 0.48099226 | 1.15520759 | 0.24800543 | 0.538841 |
| PFKFB1 | 47.8502759 | 2.614711868 | 0.76584036 | 3.41417351 | 0.00063976 | 0.01046637 |
| APEX2 | 10.5380877 | 0.039230718 | 0.48092819 | 0.08157292 | 0.93498633 | 0.97519355 |
| FAM104B | 11.1046672 | -0.02415163 | 0.55094969 | -0.0438364 | 0.96503484 | 0.98660829 |
| RRAGB | 82.1526802 | 0.048196916 | 0.26911633 | 0.17909324 | 0.85786449 | 0.94359207 |
| AL445472.1 | 97.4930262 | -0.303288407 | 0.23356984 | -1.2984913 | 0.19411856 | 0.47456322 |
| KLF8 | 14.1383225 | -0.737595399 | 0.58888524 | -1.2525282 | 0.21037744 | 0.49651968 |
| UBQLN2 | 20.4118951 | -0.79045399 | 0.46084167 | -1.7152398 | 0.08630126 | 0.30190826 |
| NBDY | 111.1959 | 0.366327564 | 0.18904013 | 1.93782966 | 0.05264401 | 0.22475697 |
| SPIN3 | 26.5613139 | 0.130047569 | 0.36785255 | 0.35353179 | 0.7236898 | 0.87731847 |
| FAAH2 | 267.53691 | -0.60526297 | 0.31661681 | -1.9116577 | 0.05592011 | 0.23361723 |
| ARHGEF9 | 57.9076604 | -0.071033372 | 0.3512741 | -0.2022164 | 0.83974758 | 0.93543959 |
| MTMR8 | 36.2423038 | 0.746721219 | 0.27773817 | 2.68857974 | 0.00717567 | 0.06151922 |
| ZC4H2 | 66.142873 | 0.838966865 | 0.31515448 | 2.66208138 | 0.00776591 | 0.06439823 |
| ZC3H12B | 40.9814865 | 0.3017118 | 0.30587476 | 0.98638997 | 0.32394178 | 0.61448136 |
| LAS1L | 12.8538394 | 0.127986373 | 0.43901106 | 0.29153337 | 0.77064343 | 0.90076568 |
| MSN | 179.139647 | 0.064098189 | 0.27712461 | 0.23129736 | 0.8170838 | 0.92248811 |
| OPHN1 | 350.65656 | 1.528801635 | 0.206095 | 7.41794636 | 1.19E-13 | 6.81E-11 |
| YIPF6 | 54.5246864 | -0.32115131 | 0.29879102 | -1.0748359 | 0.28244822 | 0.57528302 |
| STARD8 | 14.0302025 | 1.092278565 | 0.53112995 | 2.05651849 | 0.03973257 | 0.18852482 |
| EFNB1 | 17.1314346 | -0.489199184 | 0.52523374 | -0.9313933 | 0.35165016 | 0.63801604 |
| EDA | 53.5027431 | -0.528905889 | 0.24794199 | -2.133184 | 0.03290963 | 0.16734207 |
| IGBP1 | 39.4231026 | -0.462079487 | 0.28113248 | -1.6436361 | 0.1002514 | 0.32693571 |
| DLG3 | 57.0432007 | -0.108535887 | 0.28888541 | -0.3757057 | 0.70713572 | 0.86828512 |
| SNX12 | 14.4535009 | -0.569496678 | 0.55285425 | -1.0301027 | 0.30296178 | 0.59684506 |
| MED12 | 11.4764264 | 0.35892348 | 0.48351693 | 0.74231832 | 0.45789449 | 0.72394553 |
| ZMYM3 | 16.0241379 | 0.348717798 | 0.49574008 | 0.70342869 | 0.48178863 | 0.74068045 |
| NONO | 57.819025 | -0.378032974 | 0.27202091 | -1.3897203 | 0.16461383 | 0.43373261 |
| TAF1 | 114.406027 | 0.019211244 | 0.22891117 | 0.08392445 | 0.9331165 | 0.9738594 |
| OGT | 230.668325 | 0.802704113 | 0.31473215 | 2.55043574 | 0.01075884 | 0.08056236 |
| GCNA | 28.3018985 | 0.978398167 | 0.46507662 | 2.10373545 | 0.03540153 | 0.17464009 |
| NHSL2 | 38.02617 | 0.891632275 | 0.28000043 | 3.18439606 | 0.00145056 | 0.01928221 |
| PIN4 | 42.7363429 | -0.184020803 | 0.32898746 | -0.5593551 | 0.57591942 | 0.79171629 |
| RPS4X | 198.548918 | -0.87839215 | 0.4393726 | -1.9991965 | 0.0455871 | 0.20454629 |
| HDAC8 | 293.03415 | 0.079953227 | 0.18381726 | 0.43496039 | 0.66359118 | 0.84292874 |
| PHKA1 | 26.9080732 | 0.45267451 | 0.4011512 | 1.12843864 | 0.25913472 | 0.55032073 |
| NAP1L2 | 11.3852777 | 1.645919001 | 0.70732433 | 2.32696505 | 0.01996713 | 0.12057522 |
| CHIC1 | 93.2107896 | 0.649608518 | 0.23713018 | 2.73945945 | 0.00615403 | 0.05517739 |
| XIST | 409.855067 | -3.23475393 | 3.77162346 | -0.8576556 | 0.39108267 | 0.67223049 |

| | | | | | | |
|---|---|---|---|---|---|---|
| JPX | 254.321206 | 0.05511203 | 0.39615082 | 0.13911881 | 0.88935626 | 0.95702609 |
| FTX | 1456.39912 | -0.06362734 | 0.24362247 | -0.2611719 | 0.79395995 | 0.91176607 |
| SLC16A2 | 15.1055909 | -0.936293292 | 0.44896159 | -2.0854641 | 0.03702719 | 0.17950781 |
| RLIM | 41.9288239 | -0.254053224 | 0.29704465 | -0.8552695 | 0.39240198 | 0.67352969 |
| NEXMIF | 117.15783 | -0.203004313 | 0.31940652 | -0.6355672 | 0.52505854 | 0.76221787 |
| ABCB7 | 107.906857 | 0.646626023 | 0.18880292 | 3.42487304 | 0.00061509 | 0.0101386 |
| UPRT | 20.3876671 | 0.149184191 | 0.32610735 | 0.45746958 | 0.64733356 | 0.83478232 |
| ZDHHC15 | 23.8970858 | 0.131886728 | 0.31439834 | 0.41948926 | 0.6748586 | 0.85096613 |
| PBDC1 | 10.9138255 | -0.630240332 | 0.52553231 | -1.1992418 | 0.23043393 | 0.52060957 |
| ATRX | 397.57622 | 0.41404601 | 0.17475714 | 2.36926518 | 0.01782347 | 0.11217891 |
| MAGT1 | 78.76512 | 0.114782058 | 0.26903494 | 0.42664368 | 0.66963888 | 0.84656028 |
| COX7B | 129.247349 | -0.012954068 | 0.36655333 | -0.0353402 | 0.97180847 | 0.988268 |
| PGK1 | 185.936616 | -1.272925977 | 0.440716 | -2.8883135 | 0.00387314 | 0.03930906 |
| BRWD3 | 104.770264 | 0.057846393 | 0.28413977 | 0.20358429 | 0.83867837 | 0.93537562 |
| HMGN5 | 20.1933373 | 0.091690366 | 0.33591978 | 0.27295317 | 0.78488922 | 0.90771539 |
| SH3BGRL | 196.892612 | 0.062497876 | 0.23826725 | 0.26230158 | 0.79308894 | 0.91136668 |
| RPS6KA6 | 153.731146 | 0.997245668 | 0.27900999 | 3.57422927 | 0.00035126 | 0.00661552 |
| HDX | 48.1026466 | -0.224066671 | 0.38599589 | -0.5804898 | 0.56158437 | 0.78361672 |
| APOOL | 67.0433895 | 0.154583486 | 0.22321874 | 0.69252019 | 0.48861071 | 0.74516503 |
| ZNF711 | 17.7014451 | 0.773213754 | 0.42857626 | 1.8041451 | 0.07120856 | 0.2693064 |
| CHM | 94.6987894 | -0.077680223 | 0.21922641 | -0.3543379 | 0.72308568 | 0.87679245 |
| DIAPH2 | 206.082512 | -0.407087806 | 0.24613605 | -1.6539138 | 0.09814503 | 0.32365239 |
| DIAPH2-AS1 | 15.6059353 | -1.38712321 | 0.56236459 | -2.4665906 | 0.01364062 | 0.09488567 |
| TSPAN6 | 20.1862581 | -0.199227137 | 0.47427049 | -0.4200707 | 0.67443381 | 0.85053459 |
| SYTL4 | 24.9739943 | 0.091292147 | 0.51790754 | 0.17627113 | 0.86008094 | 0.94469532 |
| TRMT2B | 47.4507268 | 0.449714562 | 0.33151466 | 1.35654503 | 0.17492582 | 0.44794576 |
| HNRNPH2 | 28.2042622 | -0.774358773 | 0.49433335 | -1.5664708 | 0.11723843 | 0.35744016 |
| ARMCX4 | 150.58756 | 0.418353242 | 0.34651812 | 1.20730554 | 0.2273145 | 0.51714387 |
| ARMCX6 | 15.2287999 | -0.9729001 | 0.54483164 | -1.7856894 | 0.07414956 | 0.27538206 |
| ARMCX3 | 44.0541567 | -0.4005849 | 0.25999717 | -1.5407279 | 0.12338302 | 0.36821038 |
| ARMCX2 | 21.7593692 | -0.540230422 | 0.44373224 | -1.2174694 | 0.22342568 | 0.51229275 |
| ZMAT1 | 122.974997 | 0.894841498 | 0.39075376 | 2.29003936 | 0.02201904 | 0.12859416 |
| GPRASP1 | 18.5445853 | 0.824123347 | 0.5139937 | 1.60337247 | 0.10885244 | 0.34398708 |
| BHLHB9 | 20.3208728 | 0.104525847 | 0.38054136 | 0.2746767 | 0.78356463 | 0.90699807 |
| LINC00630 | 61.5748211 | 0.664377911 | 0.29993942 | 2.21504031 | 0.0267573 | 0.14614285 |
| BEX4 | 40.635828 | -0.281880292 | 0.35755265 | -0.7883602 | 0.43048603 | 0.7041707 |
| TCEAL8 | 19.389114 | -0.450679119 | 0.45547175 | -0.9894776 | 0.3224295 | 0.61319341 |
| BEX2 | 34.2805068 | -0.161100441 | 0.42693111 | -0.3773453 | 0.70591702 | 0.86753179 |
| TCEAL9 | 50.3859515 | -0.638206499 | 0.3892944 | -1.6393929 | 0.10113145 | 0.32897259 |
| BEX3 | 84.9653596 | 0.380859974 | 0.39356955 | 0.9677069 | 0.33319077 | 0.62190756 |
| TCEAL4 | 59.2667354 | -0.767862874 | 0.31698015 | -2.4224321 | 0.01541701 | 0.10194225 |
| TCEAL3 | 27.6328381 | -0.673806359 | 0.35823509 | -1.8809055 | 0.05998478 | 0.2428146 |
| MORF4L2 | 92.6052981 | -0.270818602 | 0.28341175 | -0.9555659 | 0.3392916 | 0.62810099 |
| SLC25A53 | 51.1194205 | 0.297285549 | 0.2230726 | 1.33268519 | 0.18263514 | 0.45952112 |
| FAM199X | 89.1619805 | 0.181594226 | 0.24916263 | 0.72881808 | 0.46611295 | 0.72924396 |
| IL1RAPL2 | 52.7723002 | 0.689097915 | 0.71964227 | 0.9575562 | 0.3382866 | 0.62725495 |
| NRK | 53.8021634 | 2.187432553 | 0.31364005 | 6.97434062 | 3.07E-12 | 1.09E-09 |
| RNF128 | 74.2206418 | -0.979884606 | 0.53616984 | -1.8275638 | 0.06761504 | 0.26098543 |
| TBC1D8B | 36.1282155 | -0.175434916 | 0.3814225 | -0.4599491 | 0.64555279 | 0.83387847 |
| MORC4 | 28.4458446 | -0.073385994 | 0.46612944 | -0.1574369 | 0.8749005 | 0.94996047 |
| RBM41 | 94.692206 | 0.175657738 | 0.23716448 | 0.74065787 | 0.45890091 | 0.72461104 |
| NUP62CL | 28.9093063 | 0.671326731 | 0.4200321 | 1.59827483 | 0.10998183 | 0.34554217 |
| TSC22D3 | 38.4822957 | 1.32201893 | 0.4214604 | 3.13675717 | 0.00170828 | 0.02188887 |
| MID2 | 26.6503455 | 0.19958051 | 0.35250167 | 0.56618317 | 0.57126927 | 0.78905469 |
| ATG4A | 12.3163697 | 0.550658637 | 0.43512488 | 1.26551861 | 0.20568548 | 0.48999801 |
| COL4A6 | 94.9759048 | 2.183414523 | 0.56989441 | 3.8312615 | 0.00012749 | 0.00295951 |
| COL4A5 | 290.316715 | 1.114741138 | 0.40230736 | 2.77086934 | 0.00559069 | 0.05141788 |
| ACSL4 | 740.459127 | -0.6799702 | 0.55026271 | -1.2357192 | 0.21656296 | 0.50348976 |
| TMEM164 | 164.137215 | 0.75345718 | 0.21750729 | 3.46405491 | 0.0005321 | 0.00898635 |
| AMMECR1 | 54.6063226 | 0.545956928 | 0.30300719 | 1.80179528 | 0.07157762 | 0.27011684 |
| ALG13 | 100.798249 | 0.629320599 | 0.43834316 | 1.43568022 | 0.15109336 | 0.41211643 |
| AMOT | 21.5593549 | -1.052499936 | 0.39473711 | -2.6663314 | 0.00766841 | 0.06391579 |
| PLS3-AS1 | 40.1250526 | 1.752057201 | 0.4397786 | 3.98395288 | 6.78E-05 | 0.00179962 |
| PLS3 | 193.611763 | 1.006950752 | 0.31383842 | 3.20850057 | 0.00133429 | 0.01815834 |
| DANT2 | 646.471327 | 0.214600857 | 0.27341691 | 0.7848851 | 0.43252094 | 0.70501676 |
| KLHL13 | 355.806507 | 0.140190481 | 0.35167493 | 0.39863655 | 0.69016102 | 0.8592192 |
| WDR44 | 104.976573 | 0.893870978 | 0.49521089 | 1.80503095 | 0.07106984 | 0.26907809 |
| IL13RA1 | 66.4353401 | -1.383399444 | 0.28371444 | -4.8760276 | 1.08E-06 | 5.96E-05 |
| PGRMC1 | 66.4483705 | -0.889715507 | 0.38134958 | -2.3330706 | 0.01964444 | 0.1190453 |
| SLC25A43 | 29.4516113 | -0.300706069 | 0.53530431 | -0.5617479 | 0.5742878 | 0.7910707 |
| SLC25A5 | 70.1649231 | -0.514793232 | 0.58990235 | -0.8726753 | 0.38284007 | 0.66617807 |
| CXorf56 | 11.1705034 | -0.151340575 | 0.45939857 | -0.329432 | 0.74182921 | 0.88719431 |
| UBE2A | 31.6301412 | -0.592817562 | 0.40323319 | -1.4701606 | 0.14151825 | 0.39756559 |
| NKRF | 17.0508793 | -0.225809706 | 0.44292348 | -0.5098165 | 0.61018001 | 0.81373132 |
| 6-Sep | 102.6494 | 0.393858232 | 0.32097254 | 1.22707766 | 0.21979341 | 0.50818193 |
| UPF3B | 48.8931505 | -0.235271184 | 0.51865271 | -0.4536199 | 0.65010244 | 0.83735207 |
| NDUFA1 | 66.8808737 | 0.001050513 | 0.38889008 | 0.00270131 | 0.99784467 | 0.99900003 |
| NKAP | 16.9851731 | -0.127418566 | 0.35321904 | -0.3607353 | 0.71829735 | 0.87415486 |
| RHOXF1-AS1 | 33.9887766 | -0.507026661 | 0.6319675 | -0.8022986 | 0.42238024 | 0.69769678 |
| LAMP2 | 85.0505276 | -0.713429349 | 0.30829591 | -2.3141058 | 0.02066192 | 0.12303993 |
| CUL4B | 30.743077 | -0.010004852 | 0.28164795 | -0.0355225 | 0.97166307 | 0.988268 |
| MCTS1 | 28.8248988 | -0.245118241 | 0.30270162 | -0.8097685 | 0.41807323 | 0.6943399 |
| C1GALT1C1 | 12.3612645 | -0.77986713 | 0.46205287 | -1.687831 | 0.09144368 | 0.31120975 |
| THOC2 | 181.730676 | 0.437284273 | 0.20802424 | 2.10208326 | 0.03554598 | 0.17504528 |
| XIAP | 140.590391 | -0.132245095 | 0.18088564 | -0.7310978 | 0.46471941 | 0.72848544 |
| STAG2 | 202.569118 | 0.313400115 | 0.17125922 | 1.82997512 | 0.06725366 | 0.26017544 |
| TENM1 | 22.3973634 | -0.137496977 | 0.6973972 | -0.1971573 | 0.84370441 | 0.93652008 |
| SMARCA1 | 77.2328021 | -0.246460263 | 0.29125493 | -0.8462012 | 0.39744053 | 0.67844777 |
| OCRL | 17.9633639 | -0.438422004 | 0.36742511 | -1.1932282 | 0.23278001 | 0.5228035 |
| ZDHHC9 | 25.7264501 | -1.393842504 | 0.2915801 | -4.7803074 | 1.75E-06 | 8.83E-05 |
| UTP14A | 11.1915236 | 0.138335301 | 0.55831627 | 0.24777229 | 0.80431059 | 0.91643337 |
| BCORL1 | 19.1176681 | -0.602815529 | 0.45710752 | -1.3187609 | 0.18724906 | 0.4649409 |
| AIFM1 | 38.3210652 | 0.045076992 | 0.26542493 | 0.16982953 | 0.8651442 | 0.94694132 |
| ZNF280C | 18.4893058 | 0.311584342 | 0.37630894 | 0.82800144 | 0.40766968 | 0.68624397 |
| SLC25A14 | 12.8048183 | 0.560413563 | 0.43025978 | 1.30250045 | 0.19274536 | 0.47341565 |
| RBMX2 | 9.79156834 | 0.052411758 | 0.44201165 | 0.11857551 | 0.90561167 | 0.96157183 |
| ENOX2 | 75.4271825 | -0.602828654 | 0.33073152 | -1.822713 | 0.06834687 | 0.26278462 |
| RAP2C | 32.177412 | 0.44133855 | 0.38175068 | 1.15609106 | 0.24764391 | 0.53836051 |
| RAP2C-AS1 | 89.1561424 | 0.756624717 | 0.27419156 | 2.75947484 | 0.00578943 | 0.05287478 |
| MBNL3 | 56.7388946 | 0.623436453 | 0.36785266 | 1.69479935 | 0.09011349 | 0.30924357 |
| GPC4 | 22.5779468 | -1.005887367 | 0.473722 | -2.1233706 | 0.03372281 | 0.16980077 |
| PHF6 | 41.2153625 | 0.433861062 | 0.27799355 | 1.56068749 | 0.1185975 | 0.35998571 |
| HPRT1 | 118.835297 | -0.051228649 | 0.20773148 | -0.24661 | 0.8052101 | 0.91682769 |
| FAM122B | 21.1524168 | -0.373573216 | 0.36131144 | -1.0339369 | 0.30116568 | 0.59482532 |
| FAM122C | 16.9199169 | 0.491213551 | 0.64702024 | 0.75919349 | 0.44773682 | 0.71693398 |
| RTL8C | 37.0120447 | -0.698684493 | 0.47724087 | -1.4640081 | 0.14319174 | 0.40031513 |
| RTL8A | 21.7643931 | -1.136453192 | 0.46929676 | -2.4216089 | 0.01545197 | 0.101998 |
| ZNF75D | 53.3039588 | -0.265203513 | 0.2679454 | -0.989767 | 0.32228801 | 0.61319341 |
| INTS6L | 22.8774917 | 0.076287392 | 0.37559124 | 0.20311281 | 0.83904686 | 0.93543959 |
| MMGT1 | 12.8176245 | 0.203844019 | 0.40834543 | 0.49919505 | 0.61764198 | 0.81764942 |
| SLC9A6 | 23.0822569 | 0.451519573 | 0.35337951 | 1.2777186 | 0.20134866 | 0.4848871 |
| HTATSF1 | 26.0640048 | -0.189094686 | 0.34707987 | -0.5448161 | 0.58588004 | 0.79711254 |
| AL683813.1 | 24.22411 | -0.000160773 | 0.47432047 | -0.000339 | 0.99972955 | 0.99975018 |

| | | | | | | |
|---|---|---|---|---|---|---|
| RBMX | 150.484052 | 0.290376723 | 0.19241845 | 1.50908978 | 0.13127584 | 0.38024035 |
| FGF13 | 477.357124 | -0.099834947 | 0.48736907 | -0.2048447 | 0.8376935 | 0.93487753 |
| ATP11C | 82.0247416 | -0.048922418 | 0.38132789 | -0.1282949 | 0.89791561 | 0.95917897 |
| FMR1 | 66.5528823 | 0.095005558 | 0.2921817 | 0.32515917 | 0.74506061 | 0.88808711 |
| IDS | 61.1865804 | -0.60263631 | 0.33693802 | -1.7885672 | 0.07368454 | 0.2745382 |
| TMEM185A | 18.8873631 | -0.180605277 | 0.39897436 | -0.4526739 | 0.65078358 | 0.83735207 |
| LINC00894 | 130.916329 | -0.046252158 | 0.30411125 | -0.1520896 | 0.87911627 | 0.95273407 |
| MTM1 | 68.3363238 | 0.113098143 | 0.25598757 | 0.44181108 | 0.65862592 | 0.84037209 |
| MTMR1 | 66.2498675 | 0.330287577 | 0.27385254 | 1.2060782 | 0.22778734 | 0.5177793 |
| CD99L2 | 43.4937603 | 0.099229922 | 0.24835226 | 0.39955313 | 0.68948568 | 0.85868974 |
| VMA21 | 28.6829288 | -0.393105969 | 0.45831689 | -0.8577165 | 0.39104901 | 0.67223049 |
| CETN2 | 18.9020144 | -0.851946253 | 0.56455417 | -1.5090602 | 0.13128339 | 0.38024035 |
| ZNF185 | 12.6007642 | 0.338256281 | 0.6113937 | 0.55325444 | 0.58008919 | 0.79448004 |
| DUSP9 | 17.0037982 | 2.887576951 | 0.73145371 | 3.94772341 | 7.89E-05 | 0.0020291 |
| SLC6A8 | 42.2506459 | -0.528569287 | 0.51665668 | -1.0230571 | 0.30628085 | 0.60101053 |
| BCAP31 | 59.231942 | -0.657901235 | 0.37879613 | -1.7368214 | 0.08241869 | 0.29420561 |
| IDH3G | 17.9248849 | -0.068390891 | 0.41314765 | -0.1655362 | 0.86852196 | 0.94813111 |
| SSR4 | 29.5212135 | -1.055470242 | 0.4660996 | -2.2644736 | 0.023545 | 0.13454711 |
| L1CAM | 38.0611331 | -0.816565902 | 0.60063971 | -1.3594937 | 0.1739902 | 0.44651726 |
| ARHGAP4 | 13.7390875 | 0.310100349 | 0.39015601 | 0.79481116 | 0.42672336 | 0.70146865 |
| HCFC1 | 22.123157 | 0.13355086 | 0.418565 | 0.31906839 | 0.74967465 | 0.89048176 |
| IRAK1 | 20.9715188 | -1.18859047 | 0.617138 | -1.9259719 | 0.05410788 | 0.2290137 |
| MECP2 | 117.67322 | 0.106073352 | 0.20827234 | 0.5093012 | 0.61054112 | 0.81376433 |
| FLNA | 70.7412909 | 0.140136281 | 0.61652614 | 0.22729982 | 0.82019061 | 0.92406023 |
| EMD | 39.0142832 | 0.297920876 | 0.60995862 | 0.48842801 | 0.62524671 | 0.82285278 |
| RPL10 | 230.148954 | -0.770856704 | 0.50231641 | -1.5346039 | 0.12488116 | 0.37001028 |
| DNASE1L1 | 12.3896992 | -0.594691005 | 0.54718685 | -1.0868152 | 0.27711847 | 0.56927037 |
| TAZ | 25.5159751 | 0.251272696 | 0.30978165 | 0.8111284 | 0.41729194 | 0.69360141 |
| ATP6AP1 | 52.3159754 | -0.511526431 | 0.48658553 | -1.051257 | 0.29314058 | 0.5846943 |
| GDI1 | 28.2698985 | -0.598233844 | 0.44776082 | -1.3360567 | 0.18153074 | 0.45814055 |
| FAM50A | 23.0221485 | -0.414460493 | 0.433886 | -0.955229 | 0.3394619 | 0.62815343 |
| PLXNA3 | 12.825278 | -1.290614664 | 0.45112356 | -2.8608895 | 0.00422454 | 0.04176702 |
| FAM3A | 18.1334119 | 0.067307265 | 0.44364434 | 0.15171447 | 0.87941214 | 0.95279412 |
| IKBKG | 11.6438225 | -0.030305872 | 0.42772986 | -0.0708528 | 0.94351488 | 0.97836843 |
| DKC1 | 20.9136295 | -0.382495474 | 0.38861996 | -0.9842404 | 0.32499731 | 0.61535668 |
| MPP1 | 24.3360406 | 1.223407343 | 0.47313757 | 2.5857328 | 0.00971722 | 0.07566654 |
| F8 | 16.753391 | -0.207225359 | 0.40206265 | -0.5154056 | 0.60626959 | 0.81114147 |
| FUNDC2 | 37.3526932 | 0.021912319 | 0.34972402 | 0.06265603 | 0.95004041 | 0.98003064 |
| BRCC3 | 40.7821701 | -0.11608919 | 0.28530195 | -0.4068994 | 0.68408188 | 0.85606265 |
| VBP1 | 17.6748037 | 0.103997187 | 0.37371359 | 0.27828046 | 0.78079708 | 0.90511663 |
| TMLHE-AS1 | 34.9230396 | -0.723897786 | 0.69439229 | -1.0424911 | 0.29718405 | 0.58944745 |
| TMLHE | 87.7627159 | -0.218118346 | 0.28022388 | -0.7783717 | 0.4363499 | 0.70775777 |
| VAMP7 | 46.2409603 | 0.190939895 | 0.26935557 | 0.70887673 | 0.47840098 | 0.7380184 |
| ZNF596 | 20.0089576 | -0.18110171 | 0.33888544 | -0.5344039 | 0.5930621 | 0.80210249 |
| FBXO25 | 49.8532654 | -0.376714378 | 0.27941986 | -1.3482019 | 0.17759346 | 0.45230354 |
| TDRP | 70.7361529 | 0.464810284 | 0.46318449 | 1.00351004 | 0.31561484 | 0.60911653 |
| ERICH1 | 139.413559 | -0.019042141 | 0.42885322 | -0.0444025 | 0.9645836 | 0.98660829 |
| AC246817.2 | 19.8121085 | 0.388828411 | 0.55888082 | 0.69572688 | 0.48659989 | 0.74420309 |
| AC246817.1 | 46.6929003 | 0.339534235 | 0.51452497 | 0.65989846 | 0.50931899 | 0.75461129 |
| CSMD1 | 52.1034862 | 0.110764147 | 1.17626971 | 0.0941656 | 0.92497761 | 0.97032668 |
| MCPH1 | 165.318347 | 0.021985219 | 0.20992102 | 0.10473091 | 0.91658934 | 0.96570908 |
| ANGPT2 | 24.0094621 | -0.124693773 | 0.74344234 | -0.1677249 | 0.86679973 | 0.94728419 |
| MCPH1-AS1 | 36.8829013 | -0.630748523 | 0.33758902 | -1.8683917 | 0.06170749 | 0.24624197 |
| AGPAT5 | 64.6748381 | -0.87147118 | 0.34756011 | -2.507397 | 0.0121624 | 0.08788108 |
| AF233439.1 | 424.103605 | 1.348417425 | 0.7469552 | 1.80521861 | 0.07104048 | 0.26907809 |
| DEFB1 | 1095.19534 | 0.836358405 | 0.48866622 | 1.71151263 | 0.08698652 | 0.30315803 |
| FAM85B | 108.212346 | 0.477282119 | 0.38270529 | 1.24712706 | 0.21235091 | 0.49847886 |
| MFHAS1 | 661.165105 | -1.202014431 | 0.33272445 | -3.6126423 | 0.00030309 | 0.00595946 |
| ERI1 | 41.4003637 | -1.072779778 | 0.28425213 | -3.774043 | 0.00016062 | 0.00357395 |
| PPP1R3B | 20.8114721 | -1.8082921 | 0.45160558 | -4.0041403 | 6.22E-05 | 0.0016919 |
| TNKS | 300.394129 | 0.071043225 | 0.19323112 | 0.36765933 | 0.71312727 | 0.87138384 |
| MSRA | 106.449534 | -0.00426806 | 0.35932519 | -0.011878 | 0.99052296 | 0.99684874 |
| XKR6 | 368.628136 | 0.053357223 | 0.21114075 | 0.25270926 | 0.80049289 | 0.91457001 |
| MTMR9 | 45.4157795 | -0.020920674 | 0.24786889 | -0.0844022 | 0.93273668 | 0.97356162 |
| FDFT1 | 100.37924 | 0.446090414 | 0.36769584 | 1.21320495 | 0.22505147 | 0.51390457 |
| CTSB | 83.4780294 | -1.508761457 | 0.37077005 | -4.0692646 | 4.72E-05 | 0.0013423 |
| FAM66D | 37.134793 | 1.189060959 | 0.74380627 | 1.59861648 | 0.10990585 | 0.34550245 |
| AC145124.1 | 10.8972218 | 0.593075725 | 0.70951239 | 0.83589199 | 0.40321567 | 0.68156125 |
| LONRF1 | 57.3816853 | 0.093472576 | 0.4178061 | 0.22372238 | 0.82297333 | 0.92597982 |
| TRMT9B | 116.746782 | -0.616290173 | 0.3312162 | -1.8606885 | 0.06278818 | 0.24926305 |
| TUSC3 | 119.333984 | -0.663066873 | 0.26309895 | -2.5202186 | 0.0117282 | 0.08544828 |
| MSR1 | 274.985721 | 1.031397823 | 0.27592979 | 3.73789948 | 0.00018556 | 0.00394161 |
| AC011586.1 | 18.3557917 | -0.380971589 | 0.6046307 | -0.6300897 | 0.52863588 | 0.76311655 |
| AC068992.1 | 117.717006 | 0.767233293 | 0.33506331 | 2.28981586 | 0.02203199 | 0.12859694 |
| MICU3 | 96.0736876 | 0.300595158 | 0.35228125 | 0.85328174 | 0.39350309 | 0.67418407 |
| ZDHHC2 | 133.696925 | -0.368808139 | 0.28287469 | -1.3037863 | 0.19230646 | 0.47282604 |
| CNOT7 | 37.9037477 | -0.121306526 | 0.25904717 | -0.4682797 | 0.6395846 | 0.83121032 |
| VPS37A | 59.26638 | -0.071903162 | 0.24698684 | -0.2911214 | 0.77095845 | 0.90076568 |
| MTUS1 | 271.187576 | 0.382670243 | 0.18970764 | 2.01715783 | 0.04367905 | 0.19956366 |
| AC027117.2 | 29.8550103 | -0.255485121 | 0.41796614 | -0.6112579 | 0.54102884 | 0.77176392 |
| PCM1 | 317.3166 | 0.311666011 | 0.171712 | 1.81505085 | 0.0695161 | 0.26553757 |
| ASAH1 | 110.410631 | -0.449652234 | 0.37310661 | -1.2051575 | 0.22814249 | 0.5178625 |
| PSD3 | 137.900816 | -1.699777377 | 0.64343537 | -2.641722 | 0.00824857 | 0.06735772 |
| SH2D4A | 45.5589204 | -1.224979185 | 0.39991758 | -3.0630791 | 0.00219072 | 0.02597102 |
| CSGALNACT1 | 711.108596 | -0.827170351 | 0.74245952 | -1.1140949 | 0.26523848 | 0.55640131 |
| INTS10 | 120.942868 | -0.213663142 | 0.27697775 | -0.771409 | 0.44046452 | 0.71123283 |
| ATP6V1B2 | 41.226011 | -0.169793817 | 0.33274531 | -0.5102816 | 0.60985418 | 0.81344383 |
| XPO7 | 92.7319205 | 0.228791634 | 0.22179683 | 1.03153697 | 0.30228908 | 0.59594663 |
| DMTN | 50.650354 | 0.700652824 | 0.23931526 | 2.92773986 | 0.00341436 | 0.03581944 |
| FAM160B2 | 24.753015 | -0.319362406 | 0.30670382 | -1.041273 | 0.29774886 | 0.59034041 |
| SLC39A14 | 49.1926915 | -0.459540396 | 0.4499124 | -1.0213997 | 0.3070651 | 0.60139222 |
| PPP3CC | 118.568875 | 0.19809948 | 0.32279964 | 0.61369177 | 0.53941903 | 0.77077724 |
| CCAR2 | 18.8014253 | 0.022263545 | 0.36204886 | 0.06149321 | 0.95096642 | 0.98061757 |
| BIN3 | 52.0198149 | -0.239551667 | 0.28079664 | -0.8531144 | 0.39359585 | 0.67423086 |
| RHOBTB2 | 12.2302718 | -0.655033228 | 0.56432856 | -1.1607302 | 0.24575162 | 0.53627476 |
| TNFRSF10B | 70.4709085 | -0.792758025 | 0.34080217 | -2.3261531 | 0.02001039 | 0.12076566 |
| TNFRSF10D | 36.7190171 | -1.414164197 | 0.32187521 | -4.393517 | 1.12E-05 | 0.00040035 |
| TNFRSF10A | 15.6691618 | -2.371097599 | 0.43110748 | -5.500015 | 3.80E-08 | 3.59E-06 |
| CHMP7 | 15.6949181 | 0.007969028 | 0.41078186 | 0.01939966 | 0.98452228 | 0.99388031 |
| R3HCC1 | 11.2971943 | 0.478743325 | 0.41204889 | 1.16186049 | 0.24529214 | 0.53608738 |
| ENTPD4 | 46.6430512 | -0.13505368 | 0.33570016 | -0.4023045 | 0.68745995 | 0.85804981 |
| SLC25A37 | 133.48312 | -0.350925608 | 0.28249002 | -1.2422584 | 0.21414123 | 0.50069971 |
| STC1 | 81.6996346 | 1.107842238 | 0.58037833 | 1.9088277 | 0.05628432 | 0.23427923 |
| DOCK5 | 100.625891 | -0.694705063 | 0.24107864 | -2.8816534 | 0.00395595 | 0.03979899 |
| KCTD9 | 49.3287781 | -0.020014675 | 0.28963604 | -0.0691028 | 0.94490775 | 0.97858537 |
| PPP2R2A | 199.080673 | 0.326986504 | 0.28653316 | 1.14118206 | 0.25379417 | 0.54524749 |
| BNIP3L | 146.908699 | -1.73976656 | 0.42027658 | -4.1395753 | 3.48E-05 | 0.00103901 |
| TRIM35 | 13.14148 | -0.009962273 | 0.42824174 | -0.0232632 | 0.98144033 | 0.99318267 |
| PTK2B | 56.0490621 | -1.442614648 | 0.32359205 | -4.4581276 | 8.27E-06 | 0.00031315 |
| EPHX2 | 52.0794848 | 0.737521641 | 0.42139094 | 1.75020763 | 0.08008249 | 0.28876788 |
| CLU | 484.947623 | -2.394884685 | 0.63031557 | -3.7995011 | 0.00014499 | 0.00329001 |
| CCDC25 | 28.3846334 | -0.491151701 | 0.30177803 | -1.6275264 | 0.10362535 | 0.33425166 |
| ELP3 | 89.7523706 | 0.086536357 | 0.24920213 | 0.34725369 | 0.72840074 | 0.88042244 |

| | | | | | | |
|---|---|---|---|---|---|---|
| ZNF395 | 28.697101 | -0.779414934 | 0.48957404 | -1.5920267 | 0.11137871 | 0.34791494 |
| FBXO16 | 38.6062242 | -0.094064189 | 0.36337124 | -0.2588653 | 0.79573921 | 0.91195838 |
| FZD3 | 32.6013723 | -0.407761269 | 0.34855395 | -1.1698656 | 0.24205507 | 0.5328423 |
| EXTL3 | 27.8351444 | -0.414297468 | 0.42720747 | -0.9697805 | 0.33215592 | 0.6217267 |
| INTS9 | 62.1720223 | 0.129775365 | 0.20818456 | 0.62336692 | 0.53304343 | 0.76689222 |
| HMBOX1 | 177.942999 | -0.188435111 | 0.19343395 | -0.9741574 | 0.3299784 | 0.61943102 |
| KIF13B | 407.993209 | -0.278256097 | 0.24040076 | -1.1574676 | 0.24708135 | 0.53778881 |
| AC108449.1 | 19.7954891 | -1.061795281 | 0.61424221 | -1.7286264 | 0.08387599 | 0.29653069 |
| SARAF | 98.3464993 | -0.409764976 | 0.28582175 | -1.4336382 | 0.15167554 | 0.41304821 |
| LEPROTL1 | 36.2672085 | -0.538163527 | 0.29623472 | -1.8166794 | 0.06926621 | 0.26510127 |
| DCTN6 | 15.5060355 | -0.049946082 | 0.34556474 | -0.1445347 | 0.88507829 | 0.95619452 |
| RBPMS | 2091.33129 | -0.282009594 | 0.34886774 | -0.8083568 | 0.41888524 | 0.69496715 |
| GTF2E2 | 65.560234 | 0.101170603 | 0.23135529 | 0.4372954 | 0.66189713 | 0.84203637 |
| GSR | 135.068847 | 0.658173308 | 0.26027373 | 2.52877349 | 0.01144619 | 0.08428779 |
| PPP2CB | 142.814494 | -0.145247318 | 0.20822377 | -0.697554 | 0.48545618 | 0.74292145 |
| WRN | 96.4442524 | 0.403595343 | 0.26154581 | 1.54311531 | 0.1228028 | 0.36712549 |
| NRG1 | 68.9267492 | 1.140622502 | 0.51463558 | 2.2163693 | 0.02666622 | 0.14596993 |
| FUT10 | 41.4429593 | -0.068865169 | 0.46973644 | -0.1466038 | 0.88344472 | 0.95561187 |
| TTI2 | 20.2174752 | -0.260839067 | 0.3947307 | -0.6608026 | 0.50873892 | 0.75441538 |
| UNC5D | 14.7281491 | -1.485298501 | 0.44345837 | -3.3493527 | 0.00081001 | 0.01258625 |
| ERLIN2 | 40.6114813 | -0.391308034 | 0.35836218 | -1.0919345 | 0.27486189 | 0.56696009 |
| PLPBP | 29.6741163 | -0.036316295 | 0.3337221 | -0.108822 | 0.9133437 | 0.96461883 |
| RAB11FIP1 | 147.635082 | -0.450022593 | 0.25067126 | -1.79527 | 0.07261069 | 0.27211179 |
| ASH2L | 39.3829259 | 0.076238739 | 0.29798643 | 0.25584634 | 0.79806949 | 0.91332058 |
| LSM1 | 30.5633757 | -0.528369581 | 0.37921444 | -1.3933266 | 0.16352104 | 0.43183638 |
| BAG4 | 50.7946911 | 0.08142935 | 0.22435422 | 0.36294993 | 0.71664229 | 0.8732017 |
| DDHD2 | 56.5839365 | 0.164747682 | 0.27571783 | 0.59752277 | 0.55015841 | 0.77615521 |
| PLPP5 | 50.3017226 | 0.012901111 | 0.31444025 | 0.04102882 | 0.96727292 | 0.98720484 |
| NSD3 | 231.939134 | -0.062166658 | 0.21085431 | -0.2948323 | 0.76812201 | 0.89983999 |
| LETM2 | 11.6490751 | -0.823186475 | 0.51306996 | -1.6044332 | 0.1086186 | 0.34348281 |
| FGFR1 | 71.5238963 | -0.685061262 | 0.50667451 | -1.3520737 | 0.17635175 | 0.45081284 |
| TACC1 | 142.77477 | -0.050528788 | 0.43067986 | -0.1173233 | 0.90660385 | 0.96157183 |
| PLEKHA2 | 170.376123 | 0.037637475 | 0.55937803 | 0.06728451 | 0.94635521 | 0.97905916 |
| TM2D2 | 12.7325531 | -0.714759985 | 0.41558474 | -1.7198899 | 0.08545246 | 0.30045434 |
| ADAM9 | 122.978696 | -0.103127968 | 0.34541739 | -0.2985604 | 0.76527545 | 0.89824174 |
| TCIM | 54.1134617 | -1.102046382 | 0.45560397 | -2.4188691 | 0.01556884 | 0.10248575 |
| AC105999.2 | 54.59445 | 2.474531982 | 0.65682671 | 3.76740461 | 0.00016495 | 0.00363944 |
| GOLGA7 | 25.2991853 | -0.713963529 | 0.35237226 | -2.0261627 | 0.04274812 | 0.19741236 |
| GPAT4 | 43.7602559 | -0.432945107 | 0.44149407 | -0.9806363 | 0.32677213 | 0.61667091 |
| KAT6A | 239.32345 | 0.301528803 | 0.22976212 | 1.3123521 | 0.18940136 | 0.46836602 |
| AP3M2 | 21.0726423 | 0.111518416 | 0.43045187 | 0.25907291 | 0.79557899 | 0.91195838 |
| IKBKB | 84.837942 | -0.069483527 | 0.23009395 | -0.3019789 | 0.76266812 | 0.89732835 |
| POLB | 21.5330499 | 0.296270345 | 0.30840303 | 0.96065964 | 0.33672333 | 0.62525668 |
| VDAC3 | 65.6114856 | -0.056433202 | 0.2803086 | -0.2013253 | 0.84044425 | 0.93552346 |
| SLC20A2 | 199.338695 | 0.35902675 | 0.38901234 | 0.92291866 | 0.35604959 | 0.64081257 |
| SMIM19 | 31.1290549 | -0.407119659 | 0.33499994 | -1.2152828 | 0.22425826 | 0.5134 |
| RNF170 | 70.7449938 | -0.153376854 | 0.25001876 | -0.6134614 | 0.53957131 | 0.77077724 |
| HOOK3 | 121.548947 | 0.156917877 | 0.2910879 | 0.53907385 | 0.5898359 | 0.79960282 |
| FNTA | 29.6096971 | -0.801205059 | 0.4101748 | -1.9533259 | 0.05078099 | 0.21964738 |
| HGSNAT | 133.29876 | 0.463516282 | 0.35967652 | 1.28870322 | 0.19750128 | 0.47953293 |
| SPIDR | 523.06855 | -0.228602578 | 0.22201638 | -1.0296654 | 0.30316711 | 0.59706129 |
| CEBPD | 152.199599 | -1.859170752 | 0.39372016 | -4.7220614 | 2.33E-06 | 0.00011239 |
| PRKDC | 146.814069 | 0.002383771 | 0.20567417 | 0.01159003 | 0.9907527 | 0.99684874 |
| MCM4 | 13.8787348 | -0.343186577 | 0.48190854 | -0.7121405 | 0.47637779 | 0.73644117 |
| UBE2V2 | 55.6382696 | 0.009321203 | 0.32875195 | 0.0283533 | 0.97738037 | 0.99112991 |
| SNTG1 | 543.817179 | 3.942963852 | 0.56187495 | 7.01751132 | 2.26E-12 | 8.31E-10 |
| PXDNL | 24.9600425 | 0.029018165 | 0.37965473 | 0.07643304 | 0.93907458 | 0.97662903 |
| PCMTD1 | 278.682886 | 0.048644055 | 0.18383275 | 0.26461039 | 0.79130961 | 0.91099595 |
| AC064807.1 | 19.769055 | 0.017619678 | 0.52480028 | 0.03357406 | 0.9732168 | 0.98879011 |
| RB1CC1 | 185.604376 | 0.130700457 | 0.26407576 | 0.49493546 | 0.62064568 | 0.81942374 |
| ATP6V1H | 249.084174 | 0.745900219 | 0.18377032 | 4.0588721 | 4.93E-05 | 0.00138796 |
| TCEA1 | 67.9683568 | 0.306868949 | 0.22547093 | 1.36101334 | 0.17350947 | 0.44612537 |
| LYPLA1 | 124.852457 | 0.155745728 | 0.17505437 | 0.88969917 | 0.37362744 | 0.65750816 |
| MRPL15 | 30.1717406 | -0.002462643 | 0.29117743 | -0.0084575 | 0.99325194 | 0.9979411 |
| TMEM68 | 41.2480827 | 0.244945782 | 0.38423234 | 0.63749393 | 0.52380316 | 0.76218158 |
| TGS1 | 68.722827 | 0.622927411 | 0.3917307 | 1.59019298 | 0.11179131 | 0.34846418 |
| LYN | 103.564102 | -0.587524055 | 0.28833284 | -2.0376592 | 0.04158403 | 0.19472668 |
| RPS20 | 84.5059005 | -0.543346543 | 0.35254369 | -1.5412176 | 0.12326383 | 0.36796987 |
| PLAG1 | 43.2021352 | -0.360823579 | 0.42964494 | -0.8398181 | 0.40101041 | 0.6801464 |
| CHCHD7 | 18.4232949 | -0.247241516 | 0.35472625 | -0.6969924 | 0.48580753 | 0.74321193 |
| IMPAD1 | 50.2274092 | -0.137579213 | 0.33527178 | -0.4103513 | 0.68154826 | 0.85479793 |
| LINC01606 | 197.418823 | 0.182419837 | 0.43668942 | 0.41773359 | 0.67614191 | 0.8517503 |
| FAM110B | 326.910932 | 0.313937534 | 0.20923323 | 1.50041911 | 0.13350587 | 0.38393256 |
| AC027698.1 | 23.1709455 | -0.23961245 | 0.53435376 | -0.4484154 | 0.65385344 | 0.83885315 |
| UBXN2B | 54.7722945 | 0.239555772 | 0.24387435 | 0.98229178 | 0.32595612 | 0.6159877 |
| SDCBP | 93.5999061 | -0.998426565 | 0.25240695 | -3.9556224 | 7.63E-05 | 0.00197288 |
| NSMAF | 63.4213056 | 0.391777439 | 0.30967904 | 1.26510801 | 0.20583261 | 0.48999801 |
| TOX | 476.33688 | 0.066337621 | 0.45862004 | 0.14464614 | 0.88499026 | 0.95619452 |
| LINC01301 | 31.234044 | 0.428085014 | 0.62567218 | 0.68420017 | 0.49384877 | 0.74700665 |
| RAB2A | 246.87258 | -0.052581129 | 0.21423827 | -0.2454329 | 0.80612122 | 0.91723667 |
| CHD7 | 165.869542 | -0.061636595 | 0.21964323 | -0.2806214 | 0.77900079 | 0.90427469 |
| ASPH | 159.352303 | -0.599995464 | 0.30923738 | -1.9402424 | 0.05235024 | 0.22387388 |
| YTHDF3 | 98.3059386 | 0.297683469 | 0.26360846 | 1.12926371 | 0.2587866 | 0.55002264 |
| CYP7B1 | 109.131628 | 1.145448297 | 0.25549377 | 4.48327287 | 7.35E-06 | 0.00029126 |
| ARMC1 | 33.7274155 | 0.155991238 | 0.26868946 | 0.5805633 | 0.56153481 | 0.78361672 |
| MTFR1 | 59.1154966 | -0.40102371 | 0.38295535 | -1.0471814 | 0.29501593 | 0.58668972 |
| PDE7A | 60.3892383 | 0.175669399 | 0.41355366 | 0.42478018 | 0.67099693 | 0.84746193 |
| ADHFE1 | 50.7413936 | 0.608393081 | 0.40403377 | 1.5057976 | 0.13211914 | 0.38136491 |
| VCPIP1 | 31.7611867 | -0.304655455 | 0.28167243 | -1.0815949 | 0.27943258 | 0.57219529 |
| SGK3 | 76.1996128 | 0.346333772 | 0.28505957 | 1.21495225 | 0.2243843 | 0.51340497 |
| PPP1R42 | 11.2486527 | -0.613300933 | 0.64988525 | -0.9437065 | 0.34531965 | 0.63256941 |
| CSPP1 | 163.210102 | 0.515098231 | 0.20741426 | 2.48342725 | 0.01301249 | 0.09200733 |
| ARFGEF1 | 477.687128 | 0.014359918 | 0.21802851 | 0.06586257 | 0.94748724 | 0.979567 |
| NCOA2 | 664.783022 | 0.237794117 | 0.13940594 | 1.70576751 | 0.08805137 | 0.30539182 |
| TRAM1 | 131.037896 | -0.670831132 | 0.33224986 | -2.0190562 | 0.04348138 | 0.19908674 |
| LACTB2 | 43.3551196 | -1.343946419 | 0.30944468 | -4.343091 | 1.40E-05 | 0.00048732 |
| XKR9 | 14.4263317 | -0.057618238 | 0.74048505 | -0.0778115 | 0.93797801 | 0.97626283 |
| MSC-AS1 | 107.727207 | -2.760087937 | 0.76283958 | -3.6181761 | 0.00029669 | 0.00590051 |
| TERF1 | 64.7005348 | 0.405489944 | 0.27747546 | 1.46135426 | 0.14391825 | 0.40158336 |
| SBSPON | 17.6040384 | 0.314650872 | 0.573174 | 0.54896222 | 0.58303138 | 0.79586061 |
| RPL7 | 169.587835 | -0.264893404 | 0.46399755 | -0.570894 | 0.56807151 | 0.78733657 |
| RDH10 | 142.496273 | -0.114226451 | 0.56358553 | -0.2026781 | 0.83938663 | 0.93543959 |
| RDH10-AS1 | 44.208454 | -0.640569924 | 0.7487548 | -0.8555136 | 0.39226687 | 0.67340998 |
| STAU2 | 220.942838 | 0.188912619 | 0.17774143 | 1.06285076 | 0.28784963 | 0.5800319 |
| AC027018.1 | 33.4443877 | -0.39155258 | 0.5185336 | -0.7551152 | 0.45017989 | 0.71892004 |
| ELOC | 38.5451723 | -0.072789203 | 0.27520666 | -0.2644893 | 0.79140295 | 0.91099595 |
| GDAP1 | 64.0378222 | 0.119889497 | 0.32328763 | 0.37084467 | 0.71075323 | 0.87028547 |
| HNF4G | 20.7513197 | -1.574241786 | 0.7070559 | -2.2264743 | 0.02598243 | 0.14360031 |
| PEX2 | 39.5483468 | -0.039104968 | 0.28241115 | -0.1384682 | 0.88987039 | 0.95702609 |
| PKIA | 32.2406284 | -0.427777395 | 0.48199751 | -0.8875096 | 0.37480462 | 0.65806152 |
| ZC2HC1A | 67.5815886 | -0.374413498 | 0.29126762 | -1.2854621 | 0.19863084 | 0.48136789 |
| IL7 | 37.7332444 | -1.869329867 | 0.46577783 | -4.0133509 | 5.99E-05 | 0.00164655 |
| TPD52 | 139.071022 | 0.057915365 | 0.26710868 | 0.21682322 | 0.82834611 | 0.92847586 |

| | | | | | | |
|---|---|---|---|---|---|---|
| ZBTB10 | 42.8567384 | -0.295042159 | 0.30170802 | -0.9779063 | 0.32812069 | 0.61729352 |
| ZNF704 | 438.090411 | 0.623407186 | 0.22194555 | 2.80882939 | 0.0049722 | 0.04737599 |
| IMPA1 | 32.3281074 | -0.02794101 | 0.36396776 | -0.0767678 | 0.93880825 | 0.97662903 |
| ZFAND1 | 44.3051013 | 0.273815748 | 0.25668638 | 1.06673267 | 0.28609257 | 0.57870647 |
| CHMP4C | 19.3442537 | -0.826173244 | 0.39123904 | -2.111684 | 0.03471356 | 0.17301358 |
| SNX16 | 33.1340819 | -0.048121297 | 0.44814477 | -0.1073789 | 0.91448838 | 0.96463895 |
| AC015522.1 | 111.682659 | -0.091674028 | 0.6577236 | -0.1393808 | 0.88914926 | 0.95702609 |
| RALYL | 751.794036 | 1.216372499 | 0.36771777 | 3.30789698 | 0.00093999 | 0.01400192 |
| LRRCC1 | 19.7812424 | -0.032461736 | 0.35523658 | -0.0913806 | 0.92719017 | 0.97111125 |
| E2F5 | 26.0777814 | 0.041705666 | 0.4131553 | 0.10094428 | 0.91959469 | 0.96763053 |
| C8orf59 | 30.6145103 | -0.036299191 | 0.39517187 | -0.0918567 | 0.92681188 | 0.97111125 |
| CA13 | 35.6126581 | -0.320685785 | 0.27784942 | -1.1541712 | 0.24843 | 0.53948691 |
| CA3 | 49.8486906 | -0.576555999 | 0.27788268 | -2.0748181 | 0.0380034 | 0.18260776 |
| CA2 | 185.971023 | 0.751724135 | 0.39040591 | 1.92549376 | 0.05416762 | 0.22907832 |
| ATP6V0D2 | 55.392494 | 0.857837854 | 0.77020553 | 1.11377784 | 0.2653745 | 0.55657332 |
| WWP1 | 410.663246 | -0.3548989 | 0.17361904 | -2.0441243 | 0.04094127 | 0.19268019 |
| RMDN1 | 85.2947902 | 0.431647963 | 0.26163233 | 1.64982656 | 0.09897841 | 0.32503708 |
| CPNE3 | 73.1565595 | 0.142356871 | 0.21734445 | 0.65498276 | 0.51247885 | 0.75562371 |
| AF117829.1 | 230.278864 | 0.455565337 | 0.3851153 | 1.18293231 | 0.23683597 | 0.5273145 |
| RIPK2 | 24.7672404 | -0.599211528 | 0.34778212 | -1.7229509 | 0.0848974 | 0.29921758 |
| OSGIN2 | 26.1038794 | -0.499122023 | 0.39027408 | -1.2789013 | 0.20093182 | 0.48443708 |
| NBN | 54.3069074 | 0.022808237 | 0.33263413 | 0.06856854 | 0.94533306 | 0.97858537 |
| DECR1 | 141.95714 | 0.304605051 | 0.27583447 | 1.1043038 | 0.26946138 | 0.56160046 |
| CALB1 | 797.947432 | 3.198121869 | 0.75776647 | 4.2204584 | 2.44E-05 | 0.00076576 |
| LINC00534 | 29.8511008 | 0.651394409 | 0.7669857 | 0.84929146 | 0.39571913 | 0.67616559 |
| TMEM64 | 21.9321164 | -0.296885302 | 0.59692422 | -0.4973584 | 0.6189363 | 0.81841635 |
| PIP4P2 | 42.7552193 | 0.31854713 | 0.30796104 | 1.03437476 | 0.300961 | 0.59482532 |
| OTUD6B-AS1 | 24.8047287 | -0.151739447 | 0.42940686 | -0.3533699 | 0.72381117 | 0.87736236 |
| OTUD6B | 28.1271019 | 0.175819559 | 0.31639652 | 0.55569371 | 0.57842026 | 0.7935467 |
| LRRC69 | 44.9905274 | -0.63905549 | 0.31184915 | -2.0492456 | 0.04043811 | 0.19098451 |
| SLC26A7 | 48.5553734 | 2.205935235 | 0.64240681 | 3.43386028 | 0.00059505 | 0.00988744 |
| TRIQK | 68.8403923 | 0.016836706 | 0.27078492 | 0.06217741 | 0.95042155 | 0.98025604 |
| FAM92A | 36.9672551 | 0.276698988 | 0.43861507 | 0.63084697 | 0.52814059 | 0.76288619 |
| RBM12B | 48.3536639 | 0.368374691 | 0.30384184 | 1.21238962 | 0.22536327 | 0.51410372 |
| TMEM67 | 72.6244497 | 0.382757913 | 0.29218561 | 1.3099821 | 0.19020189 | 0.46973504 |
| PDP1 | 21.8414293 | -0.265917537 | 0.48020542 | -0.5537579 | 0.57974455 | 0.79432483 |
| VIRMA | 58.6469744 | 0.311169944 | 0.20944209 | 1.48570872 | 0.13735618 | 0.39057229 |
| ESRP1 | 82.8469653 | -0.571574164 | 0.26336486 | -2.170275 | 0.02998602 | 0.15755636 |
| DPY19L4 | 76.9405411 | 0.835760022 | 0.23587069 | 3.54329744 | 0.00039516 | 0.00721347 |
| INTS8 | 60.8976255 | 0.227911501 | 0.20393592 | 1.11756431 | 0.26375312 | 0.55532079 |
| NDUFAF6 | 50.4820707 | 0.124040906 | 0.29917089 | 0.41461557 | 0.67842335 | 0.85316374 |
| TP53INP1 | 20.7585572 | -0.221298447 | 0.32483568 | -0.6812627 | 0.49570525 | 0.74780429 |
| PLEKHF2 | 19.2370194 | -0.084233988 | 0.44340127 | -0.1899724 | 0.84933078 | 0.93897371 |
| C8orf37-AS1 | 50.0028384 | -0.645509885 | 0.59856757 | -1.0784244 | 0.28084439 | 0.57394543 |
| UQCRB | 131.059267 | -0.051233726 | 0.47162764 | -0.1086317 | 0.91349459 | 0.96461883 |
| MTERF3 | 19.4935928 | 0.051336001 | 0.36500715 | 0.14064382 | 0.88815133 | 0.95661336 |
| PTDSS1 | 19.7609841 | -0.230626929 | 0.34187149 | -0.6746012 | 0.49992914 | 0.75045478 |
| SDC2 | 53.3144397 | -2.478687151 | 0.34095123 | -7.2699171 | 3.60E-13 | 1.76E-10 |
| CPQ | 148.465724 | 0.660883532 | 0.31871066 | 2.07361603 | 0.03811499 | 0.18280637 |
| MTDH | 241.567089 | 0.188339295 | 0.16270408 | 1.1575573 | 0.24704474 | 0.53778881 |
| LAPTM4B | 115.632003 | -0.514666722 | 0.2647055 | -1.9442994 | 0.05185936 | 0.22230553 |
| MATN2 | 18.623941 | -0.679829505 | 0.66234501 | -1.0263979 | 0.30470408 | 0.59905753 |
| RPL30 | 108.295811 | -0.482486908 | 0.3876966 | -1.2444961 | 0.21331703 | 0.4997935 |
| RIDA | 17.1068621 | 0.106996957 | 0.40357614 | 0.26512211 | 0.7909154 | 0.91080793 |
| POP1 | 13.563664 | -0.214443219 | 0.36290409 | -0.5909088 | 0.55458153 | 0.77922789 |
| NIPAL2 | 17.5632384 | 0.068037333 | 0.53788158 | 0.12649129 | 0.89934305 | 0.95958837 |
| STK3 | 479.64094 | -0.245961764 | 0.26057214 | -0.9439296 | 0.34520561 | 0.63256941 |
| AC104986.2 | 32.8965068 | 0.142441504 | 0.36218671 | 0.39328198 | 0.69411123 | 0.86101552 |
| VPS13B | 439.084108 | 0.2320415 | 0.15378189 | 1.50890004 | 0.13132433 | 0.38024035 |
| COX6C | 126.072169 | 0.253593143 | 0.22746149 | 1.11488386 | 0.26490019 | 0.55622662 |
| POLR2K | 22.4691215 | -0.391873262 | 0.46349237 | -0.8454794 | 0.39784322 | 0.67879776 |
| SPAG1 | 51.0447197 | -0.577600543 | 0.35206266 | -1.6406186 | 0.10087662 | 0.328351 |
| RNF19A | 319.673381 | -0.065846736 | 0.48038444 | -0.1370709 | 0.89097474 | 0.95702609 |
| AP000424.1 | 40.9306355 | -2.952220125 | 0.65255084 | -4.5241228 | 6.06E-06 | 0.00024892 |
| ANKRD46 | 35.7724112 | 0.347601476 | 0.28062671 | 1.23866141 | 0.21547092 | 0.50209938 |
| PABPC1 | 158.154512 | -0.36466008 | 0.29641842 | -1.2302207 | 0.21861447 | 0.50674964 |
| YWHAZ | 231.901061 | -0.286423894 | 0.35407711 | -0.8089308 | 0.41855493 | 0.69466558 |
| ZNF706 | 63.5085782 | 0.069153098 | 0.2239856 | 0.30873904 | 0.75752004 | 0.89457492 |
| AP001207.3 | 10.9427642 | 0.174341728 | 0.55857511 | 0.31211868 | 0.75495033 | 0.89304149 |
| GRHL2 | 244.691193 | 0.361506567 | 0.25157569 | 1.43696937 | 0.1507267 | 0.41166131 |
| RRM2B | 34.4156764 | -0.282725501 | 0.28978145 | -0.9756508 | 0.32923754 | 0.6184911 |
| UBR5 | 292.742078 | 0.484795993 | 0.19806569 | 2.44765256 | 0.01437902 | 0.09745114 |
| KLF10 | 24.393829 | 0.916432572 | 0.60582009 | 1.51271406 | 0.13035231 | 0.37849196 |
| AZIN1 | 68.4921513 | 0.009059031 | 0.29675676 | 0.03052679 | 0.97564693 | 0.99024058 |
| AZIN1-AS1 | 141.854038 | 0.150911547 | 0.25995632 | 0.58052655 | 0.56155959 | 0.78361672 |
| AP003354.2 | 33.1239686 | 0.049428649 | 0.34750637 | 0.14223811 | 0.88689193 | 0.95619452 |
| ATP6V1C1 | 73.0766044 | -0.270120306 | 0.27737777 | -0.9738354 | 0.33013825 | 0.61950534 |
| BAALC-AS1 | 26.6622533 | -0.157804938 | 0.30673681 | -0.5144636 | 0.60692787 | 0.81160059 |
| FZD6 | 73.7774557 | 0.186042539 | 0.24831949 | 0.74920636 | 0.45373284 | 0.72126672 |
| SLC25A32 | 35.136906 | 0.192703003 | 0.2958716 | 0.65130618 | 0.51484886 | 0.75684546 |
| DCAF13 | 19.3429625 | -0.165947215 | 0.36079859 | -0.4599442 | 0.64555628 | 0.83387847 |
| LRP12 | 46.5644358 | 0.631329215 | 0.29779507 | 2.12001232 | 0.03400501 | 0.17063788 |
| ZFPM2 | 25.2563708 | 0.002737138 | 0.54459697 | 0.00502599 | 0.99598986 | 0.99868531 |
| AC027031.2 | 13.4845927 | 0.063185636 | 0.50292024 | 0.12563749 | 0.90001889 | 0.95980075 |
| OXR1 | 489.124732 | 0.302043916 | 0.17898741 | 1.68751488 | 0.09150439 | 0.31120975 |
| AC090579.1 | 34.2335898 | -0.596747514 | 0.54425368 | -1.096451 | 0.27288144 | 0.56514644 |
| ANGPT1 | 177.168864 | 3.336183381 | 0.48502146 | 6.87842424 | 6.05E-12 | 2.08E-09 |
| EIF3E | 92.4461259 | 0.234658649 | 0.21275507 | 1.10295207 | 0.27004798 | 0.56202713 |
| EMC2 | 75.838949 | 0.679431689 | 0.32855887 | 2.06791459 | 0.03864805 | 0.18467173 |
| NUDCD1 | 38.0808343 | 0.281005684 | 0.27745649 | 1.01279191 | 0.31115957 | 0.60425369 |
| ENY2 | 45.0187953 | -0.325079215 | 0.23075904 | -1.4087388 | 0.15891241 | 0.42588856 |
| EBAG9 | 28.7641238 | 0.362987108 | 0.3172812 | 1.1440549 | 0.25260087 | 0.54418531 |
| TRPS1 | 558.537338 | 0.093857066 | 0.2910619 | 0.32246428 | 0.74710099 | 0.88907744 |
| EIF3H | 122.272754 | 0.039644714 | 0.20934428 | 0.18937567 | 0.84979839 | 0.9391357 |
| UTP23 | 18.7155332 | -0.157301161 | 0.47675566 | -0.3299408 | 0.74144467 | 0.88684117 |
| RAD21 | 69.6410076 | -0.303241575 | 0.24946552 | -1.2155651 | 0.22415065 | 0.5134 |
| MED30 | 11.2401855 | 0.013430552 | 0.47284042 | 0.02840398 | 0.97733995 | 0.99112991 |
| EXT1 | 598.348043 | -0.140168711 | 0.31963123 | -0.4385326 | 0.66100025 | 0.84192458 |
| SAMD12 | 745.21306 | -0.5133237 | 0.16172655 | -3.1740225 | 0.00150342 | 0.01983129 |
| AC023590.1 | 59.2243484 | -1.216837996 | 0.55553432 | -2.1903921 | 0.02849581 | 0.15204283 |
| SAMD12-AS1 | 49.658075 | -1.567633856 | 0.29741143 | -5.2709267 | 1.36E-07 | 1.12E-05 |
| TNFRSF11B | 62.9512998 | -2.846091128 | 0.66394818 | -4.2866164 | 1.81E-05 | 0.00059711 |
| COLEC10 | 23.4579946 | -2.308764161 | 0.48696315 | -4.7411476 | 2.13E-06 | 0.00010425 |
| MAL2 | 159.092043 | -0.619512378 | 0.34608425 | -1.7900623 | 0.07344389 | 0.27383966 |
| TAF2 | 95.4681139 | 0.186525583 | 0.25154228 | 0.74152775 | 0.45837351 | 0.72410401 |
| DEPTOR | 502.676174 | -0.997453925 | 0.28703568 | -3.4750172 | 0.00051082 | 0.0087562 |
| AC091563.1 | 32.8640313 | -1.688344149 | 0.32944462 | -5.1248194 | 2.98E-07 | 2.09E-05 |
| COL14A1 | 21.1908831 | -0.947952171 | 0.39009947 | -2.4300268 | 0.01509771 | 0.10047582 |
| MRPL13 | 29.9282673 | -0.132814038 | 0.38617441 | -0.3439224 | 0.73090463 | 0.88180426 |
| MTBP | 14.2451151 | 0.122066963 | 0.41725845 | 0.29254521 | 0.76986979 | 0.90035176 |
| SNTB1 | 129.850807 | -0.877348382 | 0.37760419 | -2.3234604 | 0.02015443 | 0.12135064 |
| ZHX2 | 327.559485 | -0.679057204 | 0.28262183 | -2.4027061 | 0.01627426 | 0.10557772 |
| DERL1 | 37.3038998 | -1.106529015 | 0.3383279 | -3.2705816 | 0.00107327 | 0.0154209 |

| | | | | | | |
|---|---|---|---|---|---|---|
| TBC1D31 | 19.9503163 | 0.092130064 | 0.39105896 | 0.23559124 | 0.81374987 | 0.92101456 |
| ATAD2 | 17.1198984 | -0.400037673 | 0.47375461 | -0.8443985 | 0.39844678 | 0.67938457 |
| WDYHV1 | 19.8179388 | -0.276173787 | 0.46277148 | -0.5967822 | 0.5506528 | 0.77635488 |
| FBXO32 | 23.0976841 | -0.763500657 | 0.32134885 | -2.3759247 | 0.01750504 | 0.11090829 |
| FAM91A1 | 81.2451366 | 0.066614384 | 0.24073224 | 0.27671567 | 0.78199844 | 0.90577227 |
| TMEM65 | 106.646069 | -0.167506795 | 0.20804506 | -0.8051467 | 0.42073502 | 0.69661173 |
| TATDN1 | 71.2423797 | -0.030898697 | 0.24977568 | -0.1237058 | 0.90154823 | 0.9603712 |
| NDUFB9 | 60.4154199 | -0.469245186 | 0.34474754 | -1.361127 | 0.17347356 | 0.44612537 |
| MTSS1 | 280.039218 | 0.677598595 | 0.2873501 | 2.35809418 | 0.01836903 | 0.11417077 |
| SQLE | 60.0845387 | 0.846814219 | 0.51065469 | 1.65829127 | 0.09725869 | 0.32180307 |
| WASHC5 | 47.8951372 | -0.13006207 | 0.29674537 | -0.4382952 | 0.66117231 | 0.84192458 |
| NSMCE2 | 395.52146 | 0.216467298 | 0.30284652 | 0.71477558 | 0.47474772 | 0.7352452 |
| TRIB1 | 36.3076299 | 0.626237533 | 0.43377115 | 1.44370489 | 0.148822 | 0.40923276 |
| FAM84B | 71.0919336 | -0.766341084 | 0.57367005 | -1.3358569 | 0.18159606 | 0.45819314 |
| PCAT1 | 369.53378 | -0.407651948 | 0.3494836 | -1.1664409 | 0.24343625 | 0.53427435 |
| PVT1 | 24.8587583 | -2.001184828 | 0.7481407 | -2.6748777 | 0.00747565 | 0.06323003 |
| FAM49B | 427.715453 | 0.528713396 | 0.29383458 | 1.79935729 | 0.07196218 | 0.27082841 |
| ASAP1 | 222.495343 | -0.214466328 | 0.32627218 | -0.6573234 | 0.51097302 | 0.7549696 |
| EFR3A | 187.907733 | 0.352395649 | 0.32304064 | 1.09087095 | 0.27532967 | 0.56728925 |
| LRRC6 | 61.3075333 | 0.25523446 | 0.33377848 | 0.76468219 | 0.44446082 | 0.71432689 |
| PHF20L1 | 96.0645984 | 0.52779466 | 0.29053443 | 1.81663376 | 0.06927321 | 0.26510127 |
| NDRG1 | 636.639778 | -0.082776952 | 0.48882283 | -0.1693394 | 0.86552971 | 0.94694132 |
| ST3GAL1 | 44.1563742 | -1.574587344 | 0.44038277 | -3.5754972 | 0.00034956 | 0.0066077 |
| ZFAT | 44.2875166 | -0.157837259 | 0.35976051 | -0.4387287 | 0.66085813 | 0.84186478 |
| AC083843.3 | 41.6545719 | 0.973361581 | 0.3385962 | 2.87469729 | 0.00404415 | 0.04041012 |
| KHDRBS3 | 133.100037 | -0.93795217 | 0.35115599 | -2.6710413 | 0.00756163 | 0.06364375 |
| TRAPPC9 | 543.651551 | 0.429107664 | 0.18074795 | 2.37406652 | 0.01759338 | 0.11119448 |
| CHRAC1 | 12.7628661 | -1.141978681 | 0.50604146 | -2.25669 | 0.02402745 | 0.13578849 |
| AGO2 | 121.041696 | 0.133907912 | 0.34265518 | 0.39079494 | 0.69594881 | 0.86236044 |
| PTK2 | 770.148999 | 0.190894126 | 0.18695597 | 1.02106464 | 0.30722382 | 0.60139222 |
| DENND3 | 34.0110411 | -0.772996712 | 0.39574263 | -1.9532814 | 0.05078626 | 0.21964738 |
| SLC45A4 | 137.006131 | 1.159360795 | 0.57226765 | 2.02590658 | 0.04277436 | 0.19741547 |
| PTP4A3 | 12.602008 | -2.17907757 | 0.64935657 | -3.3557489 | 0.0007915 | 0.01238314 |
| TSNARE1 | 39.2734863 | -0.004096423 | 0.25402852 | -0.0161258 | 0.987134 | 0.99523554 |
| JRK | 14.3988461 | 0.045938491 | 0.37027522 | 0.1240658 | 0.90126318 | 0.96026613 |
| LY6E | 51.2532284 | -2.767937016 | 1.43296421 | -1.9316163 | 0.05340688 | 0.22679213 |
| TOP1MT | 14.0566607 | 0.401952009 | 0.40338812 | 0.99643989 | 0.31903646 | 0.611915 |
| RHPN1 | 23.5846451 | -0.314928571 | 0.42405491 | -0.7426599 | 0.45768762 | 0.72377441 |
| ZC3H3 | 28.0643126 | -0.236889282 | 0.39315933 | -0.6025274 | 0.54682311 | 0.77463416 |
| GSDMD | 22.8021317 | -0.145321263 | 0.48301455 | -0.3008631 | 0.76351888 | 0.8974069 |
| NAPRT | 18.060239 | -0.223373936 | 0.45240634 | -0.4937463 | 0.62148539 | 0.82030381 |
| EEF1D | 108.273797 | -0.537220186 | 0.28066883 | -1.9140714 | 0.05561103 | 0.23268234 |
| TSTA3 | 13.5642282 | -1.599461677 | 0.52971188 | -3.0194937 | 0.00253198 | 0.02895052 |
| ZNF623 | 18.3953301 | -0.36198203 | 0.36280985 | -0.9977183 | 0.31841598 | 0.61144847 |
| FAM83H | 23.4387386 | -0.469949602 | 0.52556342 | -0.8941825 | 0.37122426 | 0.65494451 |
| IQANK1 | 22.7107778 | 0.065292027 | 0.52272073 | 0.12490805 | 0.90059634 | 0.95980075 |
| SCRIB | 16.9483779 | -0.182372621 | 0.47626441 | -0.382923 | 0.70177682 | 0.86490068 |
| PUF60 | 22.3446037 | -0.862703762 | 0.49973785 | -1.7263126 | 0.08429118 | 0.29779416 |
| NRBP2 | 16.9063275 | -0.538429645 | 0.42263973 | -1.2739684 | 0.20267463 | 0.48624919 |
| PLEC | 50.3828441 | -0.869530732 | 0.30135099 | -2.8854417 | 0.00390865 | 0.03951608 |
| PARP10 | 19.9648291 | 0.310880129 | 0.80053973 | 0.38833817 | 0.6977658 | 0.86275485 |
| GRINA | 33.0190857 | -0.651120824 | 0.48301118 | -1.348045 | 0.17764389 | 0.45232017 |
| OPLAH | 9.43911344 | 0.91897621 | 0.67388677 | 1.36369528 | 0.17266349 | 0.44525138 |
| GPAA1 | 26.6278184 | 0.040279789 | 0.61212206 | 0.06580352 | 0.94753425 | 0.979567 |
| CYC1 | 41.8148175 | -0.281440587 | 0.48078714 | -0.5853746 | 0.55829586 | 0.78233632 |
| SHARPIN | 25.2739566 | -0.470808669 | 0.34281188 | -1.3733733 | 0.16963634 | 0.44032355 |
| MAF1 | 27.0081596 | -0.765500358 | 0.45975859 | -1.665005 | 0.09591176 | 0.31926347 |
| MROH1 | 120.465974 | -0.216868597 | 0.50063855 | -0.433184 | 0.66488112 | 0.84348349 |
| HSF1 | 63.7212598 | 0.145015311 | 0.27599979 | 0.5254182 | 0.5992925 | 0.80694176 |
| DGAT1 | 18.1056883 | 0.205480717 | 0.44788735 | 0.45877767 | 0.64639383 | 0.83447986 |
| CPSF1 | 15.1335917 | -0.305301623 | 0.53975141 | -0.5656338 | 0.57164277 | 0.78910141 |
| VPS28 | 59.5184588 | -0.872879297 | 0.35420634 | -2.4643243 | 0.01372718 | 0.09507992 |
| CYHR1 | 39.845375 | 0.283241222 | 0.31037763 | 0.91256969 | 0.3614689 | 0.64571079 |
| KIFC2 | 17.6220086 | 0.758203366 | 0.39583958 | 1.9154309 | 0.05543757 | 0.23225616 |
| PPP1R16A | 33.5013211 | -0.350828705 | 0.32846551 | -1.0680839 | 0.28548269 | 0.57792152 |
| ARHGAP39 | 22.5070416 | 0.492820233 | 0.33520584 | 1.47020181 | 0.1415071 | 0.39756559 |
| ZNF251 | 65.2696843 | 0.146334472 | 0.23089736 | 0.63376416 | 0.52623475 | 0.76221787 |
| RPL8 | 211.050892 | -0.690736408 | 0.39392861 | -1.7534558 | 0.0795238 | 0.28745761 |
| ZNF7 | 16.7415461 | -0.660365311 | 0.40676074 | -1.6234736 | 0.10448821 | 0.33555699 |
| ZNF250 | 16.2198671 | 0.35435033 | 0.48393694 | 0.73222418 | 0.46403175 | 0.72835095 |
| ZNF16 | 15.1329841 | -0.556999958 | 0.45506937 | -1.2239891 | 0.22095633 | 0.5089143 |
| C8orf33 | 14.7125323 | -0.882210614 | 0.4758554 | -1.8539468 | 0.06374677 | 0.2515306 |
| WASHC1 | 124.011857 | 1.020525311 | 0.25852651 | 3.9474687 | 7.90E-05 | 0.0020291 |
| CBWD1 | 28.8412946 | 0.300066436 | 0.33541863 | 0.89460278 | 0.37099947 | 0.65494451 |
| DOCK8 | 20.5159274 | 0.045345832 | 0.49164098 | 0.09223363 | 0.92651242 | 0.97111125 |
| KANK1 | 612.944627 | 1.366264451 | 0.24494521 | 5.57783696 | 2.44E-08 | 2.67E-06 |
| SMARCA2 | 304.886148 | 0.737664756 | 0.37619793 | 1.96084217 | 0.04989744 | 0.21717085 |
| VLDLR-AS1 | 27.4008938 | -2.695047632 | 0.71305571 | -3.7795751 | 0.0001571 | 0.00351827 |
| VLDLR | 33.1223518 | -3.427602179 | 0.47452437 | -7.2232375 | 5.08E-13 | 2.30E-10 |
| PUM3 | 223.516888 | -0.792992649 | 0.28106443 | -2.821391 | 0.00478159 | 0.04612352 |
| RFX3 | 211.249077 | 0.679814075 | 0.19534425 | 3.48008243 | 0.00050126 | 0.00863541 |
| RFX3-AS1 | 178.474041 | -0.055631625 | 0.40637266 | -0.1368981 | 0.89111138 | 0.95707146 |
| GLIS3 | 1583.52597 | 0.583928966 | 0.42938276 | 1.35992642 | 0.17385321 | 0.44644175 |
| SLC1A1 | 49.7954987 | 0.027994743 | 0.33134596 | 0.08448796 | 0.93266848 | 0.97356162 |
| SPATA6L | 108.43936 | -1.278510307 | 0.41693148 | -3.0664758 | 0.00216598 | 0.02583292 |
| CDC37L1 | 54.9404488 | 0.066439679 | 0.26747356 | 0.24839718 | 0.80382712 | 0.91643337 |
| AK3 | 292.684422 | 1.180193332 | 0.23357328 | 5.05277544 | 4.35E-07 | 2.86E-05 |
| RCL1 | 91.1052177 | -0.090478258 | 0.3334204 | -0.2713639 | 0.78611117 | 0.90831284 |
| JAK2 | 121.101327 | -0.397145046 | 0.18942973 | -2.0965296 | 0.03603524 | 0.17602651 |
| PLGRKT | 62.8013461 | 0.125198379 | 0.24236496 | 0.51656963 | 0.60545662 | 0.8104748 |
| RIC1 | 194.838704 | 0.315739221 | 0.3055118 | 1.03347635 | 0.30138103 | 0.59513655 |
| ERMP1 | 207.017667 | 0.922723553 | 0.23328263 | 3.95538908 | 7.64E-05 | 0.00197288 |
| KIAA2026 | 215.850201 | -0.130850886 | 0.26903441 | -0.4863723 | 0.62670323 | 0.82354566 |
| RANBP6 | 18.3088221 | -0.232729066 | 0.380594 | -0.611489 | 0.54087587 | 0.77165256 |
| UHRF2 | 107.068498 | 0.184586357 | 0.28292248 | 0.65242732 | 0.51412555 | 0.75668769 |
| GLDC | 256.279775 | 1.679311445 | 0.40566209 | 4.13968047 | 3.48E-05 | 0.00103901 |
| AL162411.1 | 26.8758228 | 0.215080558 | 0.33254503 | 0.64677123 | 0.51778002 | 0.75866548 |
| KDM4C | 399.537983 | -0.317239883 | 0.20955006 | -1.5139098 | 0.13004872 | 0.37803666 |
| DMAC1 | 25.3396303 | -0.011172724 | 0.40818648 | -0.0273716 | 0.97816334 | 0.99127618 |
| PTPRD | 75.5572951 | -1.854906199 | 0.73361844 | -2.5284345 | 0.01145724 | 0.08430895 |
| LURAP1L | 75.3649063 | -2.802477947 | 0.79352495 | -3.5316822 | 0.00041293 | 0.00746211 |
| MPDZ | 97.6911398 | 0.529998042 | 0.25786448 | 2.05533558 | 0.0398466 | 0.18882231 |
| NFIB | 577.058662 | 0.016900826 | 0.37134847 | 0.04551204 | 0.96369918 | 0.98648936 |
| ZDHHC21 | 261.712602 | 0.292021937 | 0.34683954 | 0.84195112 | 0.39981531 | 0.68002268 |
| FREM1 | 361.716736 | 2.182935529 | 0.43142756 | 5.05979625 | 4.20E-07 | 2.79E-05 |
| TTC39B | 87.5209176 | -0.348568876 | 0.33792809 | -1.0314883 | 0.30231189 | 0.59594663 |
| SNAPC3 | 71.9384604 | 0.115969992 | 0.2071346 | 0.55987745 | 0.57556303 | 0.79171629 |
| PSIP1 | 59.5971226 | 0.212842464 | 0.26421502 | 0.80556535 | 0.42049351 | 0.69661173 |
| CCDC171 | 165.540406 | 0.388895147 | 0.28261028 | 1.37608279 | 0.168796 | 0.43928463 |
| BNC2 | 93.8534246 | -1.325363544 | 0.61651272 | -2.1497749 | 0.03157303 | 0.16263267 |
| CNTLN | 64.792099 | -0.212838717 | 0.33630062 | -0.6328823 | 0.52681049 | 0.76221787 |
| SH3GL2 | 73.3160801 | 1.54799637 | 0.50101692 | 3.08970875 | 0.00200353 | 0.02460113 |
| RRAGA | 22.2027764 | -0.881808017 | 0.60048638 | -1.4684896 | 0.14197127 | 0.39809147 |

| | | | | | | |
|---|---|---|---|---|---|---|
| HAUS6 | 15.535494 | -0.093301408 | 0.42041225 | -0.2219284 | 0.82436964 | 0.92643003 |
| PLIN2 | 43.2996522 | -2.691257529 | 0.70246882 | -3.8311416 | 0.00012755 | 0.00295951 |
| DENND4C | 265.710197 | -0.289116677 | 0.22462344 | -1.2871171 | 0.19805347 | 0.48045318 |
| RPS6 | 214.833606 | -0.539949628 | 0.4219906 | -1.27953 | 0.20071049 | 0.48430583 |
| MLLT3 | 713.959011 | -0.346632753 | 0.2591814 | -1.3374137 | 0.18108763 | 0.45756661 |
| FOCAD | 228.015536 | 0.188908769 | 0.20983244 | 0.90028391 | 0.36796918 | 0.65347673 |
| KLHL9 | 23.2904331 | -0.520249437 | 0.53089833 | -0.9799417 | 0.32711487 | 0.61686572 |
| LINC01239 | 21.5162501 | 0.189381179 | 0.50190581 | 0.37732414 | 0.70593273 | 0.86753179 |
| CAAP1 | 154.695542 | 0.679686106 | 0.23841049 | 2.85090684 | 0.00435947 | 0.0428543 |
| PLAA | 42.5442396 | 0.200844288 | 0.30348302 | 0.66179745 | 0.50810104 | 0.75391861 |
| IFT74 | 112.634315 | 0.337227527 | 0.25271097 | 1.33443963 | 0.18205982 | 0.45876773 |
| LRRC19 | 19.1352332 | 0.058889519 | 0.49961332 | 0.11787019 | 0.90617051 | 0.96157183 |
| MOB3B | 185.195156 | -0.716381218 | 0.32141261 | -2.2288523 | 0.02582374 | 0.14286094 |
| C9orf72 | 36.5472604 | -0.299968284 | 0.37493152 | -0.8000615 | 0.42367515 | 0.6983711 |
| LINGO2 | 102.467186 | 0.252303107 | 0.73235039 | 0.34451147 | 0.73046167 | 0.88159862 |
| ACO1 | 47.5210476 | -0.258785787 | 0.31131897 | -0.8312561 | 0.40582898 | 0.68402935 |
| DDX58 | 81.0079734 | -0.971601585 | 1.12491823 | -0.8637086 | 0.38774797 | 0.66966967 |
| TOPORS | 25.5946068 | -0.634500898 | 0.31602383 | -2.0077629 | 0.0446685 | 0.20156586 |
| NDUFB6 | 46.7645112 | 0.47800778 | 0.2914117 | 1.64031772 | 0.10093912 | 0.32845068 |
| APTX | 69.0154783 | 0.232337277 | 0.33636526 | 0.69072911 | 0.48973579 | 0.7455572 |
| DNAJA1 | 61.2564197 | -0.345162323 | 0.26611051 | -1.2970638 | 0.19460922 | 0.47519891 |
| SMU1 | 38.9299839 | 0.045103208 | 0.27145227 | 0.16615521 | 0.86803481 | 0.94780017 |
| B4GALT1 | 188.992274 | -0.001018217 | 0.2573741 | -0.0039562 | 0.99684344 | 0.99868531 |
| BAG1 | 61.567315 | 0.100031651 | 0.35303524 | 0.28334749 | 0.77691047 | 0.90335572 |
| CHMP5 | 40.6364973 | -0.495386206 | 0.42736059 | -1.1591762 | 0.24638438 | 0.53708251 |
| NFX1 | 156.032084 | 0.010076439 | 0.34120038 | 0.02953232 | 0.97644004 | 0.99049082 |
| AQP3 | 186.839873 | -1.062872916 | 0.79860297 | -1.3309153 | 0.18321688 | 0.46004581 |
| UBE2R2 | 217.201037 | 0.05744286 | 0.32021694 | 0.17938733 | 0.85763358 | 0.94359207 |
| UBAP2 | 147.295178 | 0.115141085 | 0.27969694 | 0.41166373 | 0.68058591 | 0.85442311 |
| DCAF12 | 33.0078769 | -0.01109448 | 0.30579562 | -0.0362807 | 0.97105854 | 0.988268 |
| UBAP1 | 166.29416 | -0.125421804 | 0.28352064 | -0.4423727 | 0.6582195 | 0.84037209 |
| NUDT2 | 11.7167789 | -0.351656336 | 0.4275255 | -0.8225389 | 0.41077028 | 0.68842614 |
| FAM219A | 41.9917239 | -0.449784591 | 0.4021857 | -1.1183505 | 0.26341731 | 0.55504737 |
| DNAI1 | 16.5637463 | 0.538956583 | 0.55874765 | 0.9645796 | 0.33475542 | 0.62346575 |
| RPP25L | 12.6412227 | -0.393836432 | 0.50085095 | -0.7863346 | 0.43167148 | 0.70454366 |
| DCTN3 | 14.6893993 | -0.028811698 | 0.6092331 | -0.0472917 | 0.96228071 | 0.98539792 |
| VCP | 73.3950605 | -0.630443089 | 0.25078554 | -2.5138734 | 0.01194133 | 0.08669243 |
| PIGO | 10.263438 | -0.068008355 | 0.52273145 | -0.1301019 | 0.89648581 | 0.95900099 |
| STOML2 | 25.4381713 | -0.264279958 | 0.3933313 | -0.6719017 | 0.50164629 | 0.75108109 |
| FAM214B | 10.8671276 | 0.29783397 | 0.41193434 | 0.72301322 | 0.46967176 | 0.73178441 |
| UNC13B | 183.991599 | 0.575833383 | 0.29344904 | 1.96229431 | 0.04972823 | 0.21680077 |
| RUSC2 | 84.3660393 | 0.204858285 | 0.32691067 | 0.62664913 | 0.53088925 | 0.76492603 |
| TESK1 | 14.4586934 | -0.93090743 | 0.44694164 | -2.0828389 | 0.03726591 | 0.18041044 |
| CCDC107 | 21.3762535 | -0.849204568 | 0.44262196 | -1.9185776 | 0.05503782 | 0.2315229 |
| TLN1 | 43.1967608 | -0.767781452 | 0.30627444 | -2.5068414 | 0.01218154 | 0.08788108 |
| CREB3 | 12.7603062 | -0.673714652 | 0.50891939 | -1.3238141 | 0.1855648 | 0.46252753 |
| GBA2 | 51.3947782 | 0.5018262 | 0.25712355 | 1.9516929 | 0.05097468 | 0.21990081 |
| RGP1 | 25.2994521 | -0.457453609 | 0.31379404 | -1.4578149 | 0.14489158 | 0.40277201 |
| AL133410.1 | 25.308547 | -0.013319092 | 0.38246945 | -0.0348239 | 0.97222013 | 0.98833746 |
| HINT2 | 9.80863302 | 0.134876261 | 0.67352163 | 0.20025528 | 0.84128094 | 0.93566718 |
| TMEM8B | 26.283306 | 0.310420611 | 0.40517071 | 0.76614771 | 0.44358842 | 0.71343168 |
| RECK | 26.8113989 | 0.183838856 | 0.31656499 | 0.58073023 | 0.56142229 | 0.78361672 |
| CLTA | 124.110859 | -1.009645387 | 0.30232539 | -3.3395984 | 0.000839 | 0.012843 |
| GNE | 34.5350853 | -1.096820324 | 0.38818367 | -2.825519 | 0.00472041 | 0.04561881 |
| RNF38 | 217.823656 | 0.003123619 | 0.22067578 | 0.01415479 | 0.98870649 | 0.9963225 |
| EBLN3P | 41.3763545 | -0.422540733 | 0.2688358 | -1.5717428 | 0.11601021 | 0.35543425 |
| ZCCHC7 | 415.095479 | 0.006864601 | 0.19656014 | 0.03492367 | 0.97214061 | 0.98833746 |
| GRHPR | 28.1623913 | -0.283098084 | 0.42182358 | -0.6711291 | 0.50213828 | 0.75108109 |
| ZBTB5 | 22.9618234 | -0.363465466 | 0.39160816 | -0.9281356 | 0.35333724 | 0.6393958 |
| AL138752.2 | 55.2509738 | -0.232258333 | 0.31250779 | -0.7432081 | 0.45735568 | 0.72353781 |
| FRMPD1 | 14.0316131 | 2.131392264 | 0.60990977 | 3.49460261 | 0.00047477 | 0.00826193 |
| TRMT10B | 45.6045227 | -0.234631778 | 0.36814722 | -0.6373314 | 0.52390901 | 0.76218158 |
| DCAF10 | 139.377701 | -0.022958244 | 0.21306322 | -0.1077532 | 0.91419146 | 0.96462448 |
| FAM95C | 21.2083209 | 0.0942438 | 0.37521142 | 0.25117519 | 0.80167867 | 0.91501148 |
| ANKRD18A | 19.7117079 | 0.214250301 | 0.5319969 | 0.40272848 | 0.68714798 | 0.85804981 |
| GLIDR | 34.661622 | 1.005503026 | 0.32790997 | 3.06639966 | 0.00216654 | 0.02583292 |
| BMS1P14 | 25.9751407 | 0.540628042 | 0.52048094 | 1.03870863 | 0.29894027 | 0.59190517 |
| CBWD6 | 14.4365833 | -0.14017554 | 0.49926791 | -0.2807622 | 0.77889282 | 0.90427469 |
| CNTNAP3B | 82.2215497 | -0.434875412 | 0.35384222 | -1.2290094 | 0.21906828 | 0.50726937 |
| AL391987.2 | 10.9763371 | -0.750236907 | 0.43761712 | -1.7143683 | 0.08646111 | 0.30214464 |
| AL512625.3 | 13.2053559 | 1.334113855 | 0.58425224 | 2.28345527 | 0.02240356 | 0.1301757 |
| CBWD5 | 38.2298712 | 0.064085779 | 0.39000816 | 0.16431907 | 0.86947997 | 0.94827256 |
| ZNF658 | 14.4722534 | -0.198026014 | 0.54150828 | -0.3656934 | 0.71459386 | 0.87255493 |
| PGM5 | 82.8983848 | 0.965425911 | 0.38089678 | 2.53461294 | 0.01125716 | 0.08307399 |
| AL161457.2 | 14.0491507 | 0.066011149 | 0.40809178 | 0.16175565 | 0.87149828 | 0.94844301 |
| PIP5K1B | 127.269639 | 1.216290049 | 0.49527313 | 2.45579656 | 0.01405727 | 0.09623211 |
| FAM189A2 | 20.2764718 | 0.82865563 | 0.39624396 | 2.09127638 | 0.0365033 | 0.17761982 |
| PTAR1 | 102.396054 | 0.355612341 | 0.21407877 | 1.6611285 | 0.09668764 | 0.32117055 |
| MAMDC2-AS | 55.2191877 | 0.234689947 | 0.3338765 | 0.70292442 | 0.48210285 | 0.74068045 |
| MAMDC2 | 100.96613 | -0.298008927 | 0.46672191 | -0.638515 | 0.52313852 | 0.76209905 |
| SMC5-AS1 | 82.0295036 | 0.550016679 | 0.22453938 | 2.44953328 | 0.01430415 | 0.09720405 |
| SMC5 | 192.355198 | -0.077310287 | 0.21771171 | -0.355104 | 0.72251172 | 0.87650915 |
| KLF9 | 43.8748803 | 1.013937817 | 0.31111102 | 3.25908681 | 0.00111771 | 0.01579519 |
| TRPM3 | 922.631488 | 2.279257279 | 0.45800344 | 4.9765069 | 6.47E-07 | 3.97E-05 |
| CEMIP2 | 174.500886 | -1.298381432 | 0.30220416 | -4.2963719 | 1.74E-05 | 0.00057938 |
| ABHD17B | 25.8663696 | 0.113689883 | 0.36836831 | 0.308631 | 0.75760223 | 0.89457492 |
| C9orf85 | 58.5136768 | 0.249586135 | 0.25896375 | 0.96378791 | 0.33515227 | 0.62351515 |
| GDA | 42.3154961 | -2.61057702 | 0.79870186 | -3.268525 | 0.0010811 | 0.01544593 |
| ZFAND5 | 191.211504 | -0.535508385 | 0.39560283 | -1.3536516 | 0.17584757 | 0.4499706 |
| ALDH1A1 | 87.2765217 | 0.333383051 | 0.55243212 | 0.60348238 | 0.54618783 | 0.77463416 |
| TRPM6 | 278.74792 | 3.086856576 | 0.70390445 | 4.38533464 | 1.16E-05 | 0.0004114 |
| C9orf41-AS1 | 12.496636 | -0.599528337 | 0.59757127 | -1.003275 | 0.31572818 | 0.60912226 |
| CARNMT1 | 106.863456 | 1.173609808 | 0.23407307 | 5.01386081 | 5.33E-07 | 3.40E-05 |
| NMRK1 | 47.8463724 | -0.510377219 | 0.51531571 | -0.9904166 | 0.32197055 | 0.61299956 |
| OSTF1 | 37.4690003 | -0.074838931 | 0.37188223 | -0.2012436 | 0.84050807 | 0.93552346 |
| PCSK5 | 73.9105272 | -1.467564622 | 0.64977562 | -2.2585714 | 0.02391006 | 0.13541585 |
| RFK | 18.4179886 | -0.527382901 | 0.50367706 | -1.0470655 | 0.29506935 | 0.58668972 |
| VPS13A | 173.778828 | -0.000199696 | 0.21088432 | -0.0009469 | 0.99924445 | 0.99957903 |
| GNAQ | 593.215394 | -0.352932174 | 0.18890232 | -1.8683316 | 0.06171586 | 0.24624197 |
| CEP78 | 15.1330525 | 0.067367422 | 0.36002457 | 0.18711895 | 0.85156737 | 0.94012264 |
| TLE4 | 20.4002551 | -0.430924167 | 0.39466663 | -1.0918688 | 0.27489078 | 0.56696009 |
| TLE1 | 21.7083523 | -0.96346542 | 0.42063994 | -2.2904753 | 0.02199378 | 0.1285195 |
| AL591368.1 | 10.5259782 | 2.180206655 | 0.79277556 | 2.75009316 | 0.00595783 | 0.05398205 |
| RASEF | 50.7860243 | -0.547256908 | 0.30937249 | -1.7689256 | 0.07690629 | 0.28145244 |
| FRMD3 | 148.283267 | 1.37603085 | 0.46209898 | 2.97778381 | 0.00290341 | 0.03178629 |
| IDNK | 14.6946696 | -0.31377248 | 0.4008828 | -0.7827038 | 0.43380109 | 0.70597357 |
| UBQLN1 | 102.859092 | 0.221600355 | 0.19300177 | 1.14817785 | 0.25089515 | 0.54232519 |
| GKAP1 | 51.8749012 | 0.565342119 | 0.26050132 | 2.17020826 | 0.02999107 | 0.15755636 |
| KIF27 | 65.4585016 | 0.205447493 | 0.28412894 | 0.72307839 | 0.46963173 | 0.73178441 |
| HNRNPK | 174.797143 | -0.2450049 | 0.26663745 | -0.918869 | 0.35816408 | 0.64216076 |
| RMI1 | 15.4985754 | -0.484621701 | 0.37481036 | -1.2929784 | 0.19601853 | 0.4776213 |
| AGTPBP1 | 98.4976735 | -0.098232485 | 0.19702761 | -0.4985722 | 0.6180808 | 0.81780997 |
| NAA35 | 120.620402 | 0.280894543 | 0.25710768 | 1.09251713 | 0.27460586 | 0.56696009 |

| | | | | | | |
|---|---|---|---|---|---|---|
| GOLM1 | 55.6426822 | -0.639263665 | 0.25868296 | -2.4712245 | 0.01346513 | 0.09385503 |
| ISCA1 | 31.5970431 | -0.37596222 | 0.30697228 | -1.2247432 | 0.22067199 | 0.5089143 |
| TUT7 | 86.9027566 | -0.416998222 | 0.20650719 | -2.0192915 | 0.04345693 | 0.19906327 |
| DAPK1 | 369.976633 | -0.641563057 | 0.40074922 | -1.6009091 | 0.10939706 | 0.34481257 |
| CTSL | 50.1416358 | -0.1992209 | 0.34858398 | -0.5715148 | 0.56765074 | 0.78696513 |
| SPIN1 | 292.112408 | 0.423097006 | 0.1590078 | 2.66085691 | 0.00779421 | 0.06449472 |
| SHC3 | 16.7163298 | 1.736326742 | 0.76100981 | 2.28160889 | 0.02251244 | 0.13071121 |
| SECISBP2 | 141.840311 | -0.048320588 | 0.26999792 | -0.1789665 | 0.85796399 | 0.94359207 |
| SEMA4D | 147.154386 | -0.587490568 | 0.35099384 | -1.6737917 | 0.09417155 | 0.3160115 |
| LINC01508 | 57.5416954 | 1.059653379 | 0.52194811 | 2.03018912 | 0.04233732 | 0.19633566 |
| SYK | 122.54838 | 0.11325181 | 0.2444904 | 0.46321577 | 0.64320971 | 0.83328507 |
| AL158071.4 | 40.729314 | 0.082986012 | 0.44285996 | 0.18738658 | 0.85135754 | 0.94012264 |
| AUH | 299.493761 | 0.240626579 | 0.24501999 | 0.98206918 | 0.32606577 | 0.6159877 |
| NFIL3 | 25.5811415 | -1.277054531 | 0.71083519 | -1.796555 | 0.07240629 | 0.27172106 |
| ROR2 | 107.952294 | -0.653850164 | 0.31014991 | -2.1081746 | 0.03501589 | 0.1740015 |
| SPTLC1 | 75.5858029 | -0.084702921 | 0.20528423 | -0.4126129 | 0.67989024 | 0.85407015 |
| IARS | 75.0280248 | -0.437575487 | 0.37332734 | -1.172096 | 0.24115851 | 0.53187105 |
| NOL8 | 28.7183884 | -0.296468557 | 0.35157997 | -0.8432464 | 0.39909063 | 0.67968972 |
| CENPP | 55.6584168 | 0.286827741 | 0.42856788 | 0.66927027 | 0.50332309 | 0.75108109 |
| IPPK | 23.5834228 | 0.589931673 | 0.37433654 | 1.57593931 | 0.11503979 | 0.35402797 |
| BICD2 | 28.5236867 | 0.027626253 | 0.36775681 | 0.07512098 | 0.94011845 | 0.97685071 |
| ZNF484 | 13.7069981 | 0.248099073 | 0.40942249 | 0.60597324 | 0.54453252 | 0.77395839 |
| FGD3 | 51.1432026 | 2.231582481 | 0.56331702 | 3.96150372 | 7.45E-05 | 0.00193271 |
| CARD19 | 22.3199949 | -0.345593528 | 0.48619331 | -0.7108151 | 0.47719885 | 0.73693637 |
| NINJ1 | 23.7234822 | -0.427691926 | 0.40096529 | -1.0666557 | 0.28612732 | 0.57870647 |
| FAM120AOS | 19.3846801 | -0.424603411 | 0.39119334 | -1.0854055 | 0.27774209 | 0.56997988 |
| FAM120A | 170.565217 | 0.182220827 | 0.17135101 | 1.06343597 | 0.28758429 | 0.57989692 |
| PHF2 | 50.1327341 | 0.436973966 | 0.24642406 | 1.77326016 | 0.07618561 | 0.28022732 |
| PTPDC1 | 46.2098246 | 0.225011234 | 0.27775762 | 0.81009922 | 0.41788315 | 0.69413613 |
| AL158152.2 | 63.9345695 | 0.199132139 | 0.3015734 | 0.6603107 | 0.50905447 | 0.75446398 |
| ZNF169 | 30.5870137 | -0.005225002 | 0.27717603 | -0.0188508 | 0.9849601 | 0.99403006 |
| MFSD14B | 62.024186 | 0.277669265 | 0.27903018 | 0.99512268 | 0.3196766 | 0.611915 |
| FBP1 | 23.067904 | 0.514701553 | 0.50992619 | 1.00936481 | 0.31279971 | 0.60559801 |
| C9orf3 | 578.46971 | -0.114760656 | 0.24840982 | -0.4619812 | 0.64409483 | 0.83360113 |
| FANCC | 143.069115 | 0.119966466 | 0.25088824 | 0.47816695 | 0.63253138 | 0.82649065 |
| PTCH1 | 32.4385686 | -0.278525381 | 0.35124504 | -0.792966 | 0.42779762 | 0.70230313 |
| LINC00476 | 74.1338184 | 0.294933501 | 0.43031086 | 0.68539637 | 0.49309383 | 0.74685087 |
| ERCC6L2 | 115.249687 | 0.453421548 | 0.22515826 | 2.01379042 | 0.04403154 | 0.20016628 |
| SLC35D2 | 22.6619526 | 0.124981399 | 0.31313871 | 0.39912472 | 0.68980131 | 0.85887516 |
| HABP4 | 58.0509205 | 0.7191663 | 0.39165864 | 1.83620692 | 0.06632706 | 0.2574031 |
| CDC14B | 142.027989 | 0.638878939 | 0.36690862 | 1.74124809 | 0.0816401 | 0.29233796 |
| AL589843.1 | 22.2445823 | 0.193827225 | 0.34285914 | 0.56532612 | 0.57185197 | 0.7891787 |
| ZNF510 | 16.4322269 | 0.618626582 | 0.37650127 | 1.64309295 | 0.1003637 | 0.32707393 |
| ZNF782 | 58.3399983 | 0.303670551 | 0.27235697 | 1.11497258 | 0.26486217 | 0.55622662 |
| MFSD14C | 144.576494 | 0.017359601 | 0.2296681 | 0.0755856 | 0.93974879 | 0.97674567 |
| CCDC180 | 14.3563025 | 0.382643042 | 0.44794229 | 0.85422396 | 0.39298091 | 0.67377433 |
| TDRD7 | 30.2657669 | -0.171307812 | 0.47011876 | -0.3643926 | 0.71556484 | 0.87311962 |
| TSTD2 | 32.2909944 | -0.295564863 | 0.26780487 | -1.1036575 | 0.26974172 | 0.56184375 |
| NCBP1 | 38.9532281 | 0.360996324 | 0.30441007 | 1.18588825 | 0.23566643 | 0.525847 |
| XPA | 40.2828057 | 0.588369375 | 0.27550464 | 2.13560608 | 0.03271154 | 0.16658143 |
| ANP32B | 58.0611459 | -0.052806601 | 0.26321009 | -0.2006253 | 0.84099158 | 0.93566718 |
| NANS | 14.4036844 | -1.004926972 | 0.39202868 | -2.5634016 | 0.01036521 | 0.07880617 |
| TRIM14 | 20.1428878 | -1.860300304 | 0.99644964 | -1.8669286 | 0.06191156 | 0.24673614 |
| ANKS6 | 35.1734256 | 0.314332434 | 0.36687742 | 0.85677782 | 0.39156769 | 0.6727196 |
| TGFBR1 | 89.2546209 | 0.604763352 | 0.55338606 | 1.09284167 | 0.27446332 | 0.56696009 |
| ALG2 | 21.4910327 | -0.348447888 | 0.39360771 | -0.8852669 | 0.37601268 | 0.65935023 |
| SEC61B | 62.3108817 | -0.925411329 | 0.37694153 | -2.4550527 | 0.01408639 | 0.09623211 |
| AL359710.1 | 72.0958106 | 0.84827477 | 0.41925706 | 2.02328083 | 0.0430442 | 0.19806087 |
| STX17 | 62.2315906 | 0.168273847 | 0.29747895 | 0.56566639 | 0.57162058 | 0.78910141 |
| ERP44 | 104.48139 | 0.080040867 | 0.24111125 | 0.33196654 | 0.73991452 | 0.88614269 |
| INVS | 204.513273 | 0.225089499 | 0.26040519 | 0.86438175 | 0.38737821 | 0.66959234 |
| TEX10 | 31.3865749 | 0.243154557 | 0.29638899 | 0.82038999 | 0.41199382 | 0.68913252 |
| PLPPR1 | 44.5858739 | 3.294605971 | 0.44108216 | 7.46937029 | 8.06E-14 | 5.19E-11 |
| MRPL50 | 14.2854871 | -0.032796371 | 0.467641 | -0.0701315 | 0.94408898 | 0.97856665 |
| ZNF189 | 24.159816 | -0.396595539 | 0.34792107 | -1.1399009 | 0.25432759 | 0.54535536 |
| ALDOB | 40.4037311 | -0.263911425 | 0.86821381 | -0.3039705 | 0.76115033 | 0.89648567 |
| RNF20 | 19.8674873 | 0.144987957 | 0.3616785 | 0.40087524 | 0.68851198 | 0.8585881 |
| SMC2 | 16.6745007 | -0.09932242 | 0.48062942 | -0.2066507 | 0.83628264 | 0.93411946 |
| NIPSNAP3A | 18.9139088 | -0.411516716 | 0.37562384 | -1.0955554 | 0.27327341 | 0.5655199 |
| ABCA1 | 36.6015098 | -1.180792904 | 0.36273418 | -3.2552568 | 0.0011329 | 0.01596597 |
| SLC44A1 | 241.639848 | -1.043629403 | 0.51988953 | -2.0074061 | 0.04470644 | 0.20164877 |
| FSD1L | 85.5572992 | -0.725386935 | 0.23122221 | -3.1371854 | 0.00170578 | 0.02188887 |
| FKTN | 57.1953865 | 0.262132911 | 0.25562528 | 1.02545768 | 0.30514728 | 0.59961934 |
| TMEM38B | 36.2415927 | -0.164995228 | 0.59241193 | -0.2785144 | 0.78061755 | 0.90503603 |
| LINC01505 | 55.8900842 | -0.299110673 | 0.75116911 | -0.3981935 | 0.69048754 | 0.85952183 |
| ZNF462 | 73.8117799 | 0.399614725 | 0.38025438 | 1.0509142 | 0.293298 | 0.58477956 |
| RAD23B | 138.1942 | 0.065923633 | 0.21206939 | 0.31085879 | 0.75590797 | 0.89366123 |
| ELP1 | 45.2352781 | 0.528877283 | 0.24000382 | 2.20362024 | 0.02755106 | 0.1484472 |
| FAM206A | 14.9722269 | 0.119689657 | 0.3798538 | 0.31509401 | 0.75269028 | 0.89205725 |
| CTNNAL1 | 81.4681379 | 1.413781191 | 0.4018891 | 3.51783908 | 0.00043508 | 0.0078038 |
| TMEM245 | 234.018633 | 0.364003896 | 0.21624721 | 1.68327672 | 0.09232153 | 0.31296362 |
| EPB41L4B | 54.9660362 | 0.783284622 | 0.28888419 | 2.71141393 | 0.00669969 | 0.05854134 |
| PTPN3 | 335.974119 | -0.366331018 | 0.29304098 | -1.2501017 | 0.21126242 | 0.49712778 |
| PALM2 | 32.3696431 | -0.952056226 | 0.46419044 | -2.0510035 | 0.0402666 | 0.19054963 |
| TXN | 65.5065453 | -0.908214182 | 0.31572481 | -2.8766006 | 0.00401984 | 0.04032366 |
| ECPAS | 180.666669 | 0.242142588 | 0.27356423 | 0.88513981 | 0.37608123 | 0.65935821 |
| ZNF483 | 70.4976075 | 0.280913602 | 0.35791137 | 0.78486919 | 0.43253027 | 0.70501676 |
| PTGR1 | 299.050599 | 1.213974339 | 0.40888622 | 2.96897839 | 0.00298792 | 0.03240159 |
| C9orf84 | 44.4086758 | 0.855495771 | 0.52725907 | 1.622534 | 0.10468906 | 0.33556952 |
| UGCG | 231.755077 | -1.779999043 | 0.34877022 | -5.103644 | 3.33E-07 | 2.30E-05 |
| SUSD1 | 192.99957 | 0.662851245 | 0.32733022 | 2.02502304 | 0.042865 | 0.19767019 |
| PTBP3 | 608.242271 | 0.421703271 | 0.33244054 | 1.26850737 | 0.20461682 | 0.48931349 |
| HSDL2 | 122.057681 | 0.384436222 | 0.2591907 | 1.48321763 | 0.1380166 | 0.39158551 |
| C9orf147 | 20.8404272 | 0.08041361 | 0.45783554 | 0.17563864 | 0.86057785 | 0.945045 |
| KIAA1958 | 328.150078 | 0.264188654 | 0.2001457 | 1.31998165 | 0.18684114 | 0.46437575 |
| INIP | 45.5226935 | 0.816089837 | 0.26686743 | 3.05803462 | 0.00222794 | 0.0262912 |
| SNX30 | 43.0729415 | -0.058070923 | 0.27824302 | -0.2087058 | 0.83467794 | 0.93313642 |
| FKBP15 | 41.5438781 | 0.032757379 | 0.23641052 | 0.13856143 | 0.88979672 | 0.95702609 |
| SLC31A1 | 36.3393562 | 0.096317987 | 0.34634069 | 0.27810185 | 0.78093417 | 0.9051737 |
| CDC26 | 14.5575588 | -0.53238742 | 0.46479304 | -1.145429 | 0.2520315 | 0.54334972 |
| PRPF4 | 18.1840051 | -0.235773301 | 0.49244606 | -0.47878 | 0.63209518 | 0.82642568 |
| WDR31 | 16.0404111 | -0.333185273 | 0.52224207 | -0.6379901 | 0.52348012 | 0.76218158 |
| BSPRY | 28.9077059 | -0.69172964 | 0.34784047 | -1.9886405 | 0.0467409 | 0.20818189 |
| HDHD3 | 33.3942316 | -0.531589057 | 0.38557973 | -1.3786748 | 0.16799504 | 0.43848111 |
| ALAD | 15.6486588 | -0.331126793 | 0.46795419 | -0.7076051 | 0.4791905 | 0.73880257 |
| RGS3 | 89.8644984 | -0.341030001 | 0.35483074 | -0.9611061 | 0.3364988 | 0.62506503 |
| ZNF618 | 271.517152 | 0.207884611 | 0.33706603 | 0.61674744 | 0.53740132 | 0.76945357 |
| KIF12 | 81.4267743 | 0.860587788 | 0.28250565 | 3.04626751 | 0.00231702 | 0.02703273 |
| COL27A1 | 51.1252153 | -0.969089214 | 0.36350774 | -2.6659384 | 0.00767738 | 0.06393883 |
| AKNA | 13.9203312 | -0.528800789 | 0.43646047 | -1.2115663 | 0.22567844 | 0.51459481 |
| WHRN | 33.0425757 | -0.579499306 | 0.38972977 | -1.4869259 | 0.13703437 | 0.39008789 |
| ATP6V1G1 | 78.371762 | -1.139466641 | 0.35436146 | -3.215549 | 0.00130195 | 0.01781237 |
| TMEM268 | 24.232688 | 0.221154028 | 0.42731333 | 0.51754536 | 0.60477551 | 0.81008937 |

| | | | | | | |
|---|---|---|---|---|---|---|
| TNFSF15 | 19.67718 | -2.182292955 | 0.77057758 | -2.8320224 | 0.00462546 | 0.04491188 |
| PAPPA | 327.078961 | -0.284690797 | 1.17625529 | -0.2420315 | 0.80875578 | 0.91861103 |
| ASTN2 | 187.452541 | 0.253037049 | 0.69010314 | 0.36666555 | 0.71386851 | 0.87208269 |
| CDK5RAP2 | 74.0374751 | 0.363427499 | 0.25614551 | 1.41883222 | 0.15594794 | 0.42001978 |
| MEGF9 | 121.157805 | 0.683974134 | 0.25275538 | 2.70607148 | 0.00680844 | 0.0591532 |
| FBXW2 | 59.319732 | 0.276599294 | 0.23423802 | 1.18084713 | 0.23766346 | 0.52778809 |
| PSMD5 | 48.3833926 | -0.100400193 | 0.30198199 | -0.3324708 | 0.73953378 | 0.88614269 |
| CNTRL | 36.2483223 | -0.236187578 | 0.33124983 | -0.7130195 | 0.47583372 | 0.73616591 |
| RAB14 | 96.8304189 | -0.318617666 | 0.23856097 | -1.3355817 | 0.18168604 | 0.45819573 |
| GSN | 113.478476 | -0.865367003 | 0.27979807 | -3.0928269 | 0.0019826 | 0.02443148 |
| STOM | 65.629111 | -2.096252854 | 0.48574171 | -4.315571 | 1.59E-05 | 0.00054304 |
| AL359644.1 | 10.8819575 | -0.112826043 | 0.49007357 | -0.2302227 | 0.81791875 | 0.92281757 |
| DAB2IP | 141.801269 | -0.342640418 | 0.34892914 | -0.9819771 | 0.32611114 | 0.6159877 |
| TTLL11 | 30.6484972 | -0.039185344 | 0.34936588 | -0.1121613 | 0.91069548 | 0.96371076 |
| NDUFA8 | 31.5100917 | -0.202037469 | 0.34966103 | -0.5778095 | 0.56339273 | 0.78465819 |
| RBM18 | 16.3353818 | -0.390224215 | 0.42914695 | -0.9093021 | 0.3631907 | 0.6477823 |
| MRRF | 90.430153 | 0.536671792 | 0.25266388 | 2.12405426 | 0.03366561 | 0.16959564 |
| PDCL | 19.7391596 | -0.640958255 | 0.33794359 | -1.8966427 | 0.05787511 | 0.23782584 |
| RC3H2 | 77.484462 | -0.133133527 | 0.21144358 | -0.6296409 | 0.52892956 | 0.76327669 |
| ZBTB26 | 13.7238227 | 0.207256537 | 0.3821821 | 0.54229786 | 0.58761333 | 0.79838927 |
| RABGAP1 | 219.361071 | 0.268050653 | 0.20275079 | 1.32206958 | 0.18614498 | 0.46365223 |
| STRBP | 219.086227 | 0.438819717 | 0.18009387 | 2.43661664 | 0.01482539 | 0.09936964 |
| DENND1A | 275.657529 | 0.433036539 | 0.24640154 | 1.75744249 | 0.07884242 | 0.28599809 |
| NEK6 | 11.8610529 | -3.85499252 | 0.70643446 | -5.4569712 | 4.84E-08 | 4.54E-06 |
| PSMB7 | 62.6245969 | -0.264209542 | 0.31459817 | -0.8398318 | 0.40100271 | 0.6801464 |
| NR6A1 | 329.758826 | 0.304713135 | 0.20960252 | 1.45376657 | 0.14601103 | 0.40468272 |
| MIR181A2HG | 38.368002 | 0.136515971 | 0.40285095 | 0.33887464 | 0.73470417 | 0.88380093 |
| RPL35 | 93.2292701 | -0.150122355 | 0.38003743 | -0.3950199 | 0.69282819 | 0.86056378 |
| ARPC5L | 35.3215077 | -0.919721779 | 0.31906989 | -2.8825088 | 0.00394522 | 0.03976875 |
| GOLGA1 | 50.1315963 | 0.229534267 | 0.31561108 | 0.72726937 | 0.46706096 | 0.73023858 |
| SCAI | 70.6794829 | 0.358915266 | 0.24600427 | 1.45897981 | 0.14457066 | 0.40242285 |
| PPP6C | 75.8007563 | 0.444879021 | 0.24786627 | 1.79483487 | 0.07268001 | 0.27219482 |
| RABEPK | 24.8670982 | 0.238799679 | 0.29896177 | 0.79876327 | 0.42442769 | 0.69892168 |
| HSPA5 | 140.629071 | -0.99377839 | 0.63121922 | -1.5743792 | 0.11539982 | 0.35464129 |
| GAPVD1 | 107.617883 | 0.347986051 | 0.2349684 | 1.48099083 | 0.13860902 | 0.39280881 |
| MAPKAP1 | 190.884522 | 0.252202367 | 0.25519495 | 0.98827336 | 0.32301879 | 0.61384454 |
| PBX3 | 134.064847 | 0.184201997 | 0.28109623 | 0.6552987 | 0.51227546 | 0.75562371 |
| MVB12B | 37.2319489 | 0.541886767 | 0.3125531 | 1.73374307 | 0.08296368 | 0.29503415 |
| ZBTB43 | 67.7222886 | -0.149963543 | 0.25607256 | -0.5856291 | 0.55812479 | 0.78233632 |
| ZBTB34 | 12.1078697 | 0.354170493 | 0.41613512 | 0.85109494 | 0.39471662 | 0.67534117 |
| RALGPS1 | 211.090991 | 0.41065312 | 0.2351212 | 1.74655934 | 0.08071381 | 0.29058005 |
| GARNL3 | 26.1402638 | 0.128326565 | 0.2962829 | 0.43312174 | 0.66492633 | 0.84348349 |
| RPL12 | 43.3566474 | -0.075592508 | 0.41370543 | -0.1827206 | 0.85501724 | 0.94225653 |
| LRSAM1 | 65.3297772 | -0.172201841 | 0.24499977 | -0.7028653 | 0.48213969 | 0.74068045 |
| FAM129B | 74.5284447 | -1.286277218 | 0.44835768 | -2.8688641 | 0.00411949 | 0.04092475 |
| STXBP1 | 46.1781275 | -0.140615709 | 0.34221246 | -0.4109018 | 0.68114455 | 0.8546627 |
| PTRH1 | 12.9558264 | -0.522682078 | 0.46035476 | -1.1353897 | 0.25621205 | 0.54710413 |
| CDK9 | 21.4800878 | 0.062380633 | 0.38587765 | 0.1616591 | 0.87157432 | 0.94844301 |
| FPGS | 28.4391282 | -0.524404988 | 0.33678447 | -1.5570937 | 0.11944823 | 0.36182172 |
| ENG | 21.6645597 | -0.857584074 | 0.6230679 | -1.3763894 | 0.1687011 | 0.43928463 |
| DPM2 | 36.0389367 | -2.271453037 | 0.89491111 | -2.5381884 | 0.0111428 | 0.08252559 |
| FAM102A | 177.563327 | -0.844231459 | 0.49377949 | -1.7097337 | 0.08731512 | 0.30389202 |
| SLC25A25 | 57.8136537 | 0.522573101 | 0.4094638 | 1.2762376 | 0.20187154 | 0.48515376 |
| SLC25A25-AS | 23.8045108 | 0.867129669 | 0.43278225 | 2.00361652 | 0.04511115 | 0.20302976 |
| PTGES2 | 36.3656858 | -0.115145547 | 0.2881349 | -0.3996237 | 0.68943367 | 0.85868974 |
| LCN2 | 53.4263314 | -7.735818748 | 1.39375415 | -5.5503467 | 2.85E-08 | 2.98E-06 |
| C9orf16 | 55.7568667 | -0.77939605 | 0.465628 | -1.6738599 | 0.09415813 | 0.3160115 |
| CIZ1 | 23.9392407 | -0.304741467 | 0.32974402 | -0.9241759 | 0.35539475 | 0.64037833 |
| DNM1 | 20.6943829 | -0.141373992 | 0.42805536 | -0.3302703 | 0.7411957 | 0.88665962 |
| GOLGA2 | 70.9506866 | -0.233904031 | 0.38441004 | -0.6084753 | 0.54287226 | 0.77258349 |
| SWI5 | 21.9795077 | -0.303986836 | 0.34501934 | -0.8810719 | 0.37827892 | 0.66129805 |
| TRUB2 | 17.1263979 | -0.170039873 | 0.43120826 | -0.3943335 | 0.69333481 | 0.8606282 |
| COQ4 | 67.8846121 | 0.370979757 | 0.23880187 | 1.5535044 | 0.12030268 | 0.36323511 |
| SLC27A4 | 32.226168 | -0.015594059 | 0.42254218 | -0.0369053 | 0.97056049 | 0.98820206 |
| URM1 | 20.1511766 | -0.10537903 | 0.38549327 | -0.2733615 | 0.78457532 | 0.90765805 |
| CERCAM | 18.4851113 | 0.425835853 | 0.4468813 | 0.95290596 | 0.34063772 | 0.62907597 |
| ODF2 | 36.7137324 | 0.272270929 | 0.48846334 | 0.55740299 | 0.57725212 | 0.79249085 |
| GLE1 | 32.7795619 | 0.02998631 | 0.38298733 | 0.07829583 | 0.93759274 | 0.97619962 |
| SPTAN1 | 118.36501 | 0.000681037 | 0.25269172 | 0.00269513 | 0.9978496 | 0.99900003 |
| WDR34 | 29.0068286 | 0.097615566 | 0.44558365 | 0.21907349 | 0.8265928 | 0.92792405 |
| SET | 104.620924 | -0.836059369 | 0.3121555 | -2.6783426 | 0.00739875 | 0.06273624 |
| ZDHHC12 | 16.8715285 | -1.299344951 | 0.63874378 | -2.0342193 | 0.04192949 | 0.19536751 |
| ZER1 | 96.2576126 | 0.128235385 | 0.24548057 | 0.52238508 | 0.60140225 | 0.80759541 |
| TBC1D13 | 22.07849 | 0.06355096 | 0.36518647 | 0.17402331 | 0.86184714 | 0.94550174 |
| ENDOG | 15.4244383 | -0.43989207 | 0.54621855 | -0.8053408 | 0.42062305 | 0.69661173 |
| KYAT1 | 28.5537896 | 0.934610302 | 0.36694437 | 2.54700818 | 0.01086509 | 0.08095937 |
| LRRC8A | 16.0178492 | -0.820244332 | 0.39998461 | -2.0506897 | 0.04029717 | 0.19060673 |
| PHYHD1 | 23.5961082 | 0.190258522 | 0.45871114 | 0.4147676 | 0.67831204 | 0.85316374 |
| NUP188 | 24.1747436 | 0.169213297 | 0.37995829 | 0.44534704 | 0.65606898 | 0.83951126 |
| SH3GLB2 | 54.2665047 | 0.149512482 | 0.40171588 | 0.37218465 | 0.70975538 | 0.86953263 |
| MIGA2 | 32.4736196 | 0.795101683 | 0.3224247 | 2.46600739 | 0.01366285 | 0.09491213 |
| PTPA | 61.4635868 | -0.473221283 | 0.43104993 | -1.097834 | 0.27227697 | 0.56475837 |
| IER5L | 38.9700987 | -1.930790542 | 0.41150296 | -4.6920453 | 2.70E-06 | 0.00012724 |
| LINC01503 | 125.64129 | -1.442097964 | 0.48520945 | -2.9721143 | 0.00295757 | 0.03217406 |
| NTMT1 | 19.9152058 | -0.785660077 | 0.34558146 | -2.273444 | 0.02299944 | 0.1323843 |
| ASB6 | 13.9000659 | -0.533796693 | 0.47591635 | -1.1216187 | 0.26202459 | 0.55371842 |
| TOR1B | 17.1385469 | -0.276156673 | 0.33925485 | -0.8140095 | 0.41563951 | 0.69220102 |
| C9orf78 | 23.1656751 | -0.377599349 | 0.40308093 | -0.936783 | 0.34887021 | 0.63556496 |
| USP20 | 30.9266095 | 0.282472275 | 0.27550264 | 1.02529787 | 0.30522266 | 0.59961934 |
| FNBP1 | 87.4220195 | 0.058150339 | 0.23720975 | 0.24514313 | 0.80634561 | 0.91724035 |
| GPR107 | 85.1835253 | 0.04540608 | 0.24265812 | 0.18711956 | 0.85156689 | 0.94012264 |
| NCS1 | 17.4972413 | -0.22826803 | 0.50474026 | -0.4522485 | 0.65108996 | 0.83749891 |
| ASS1 | 21.1079367 | 0.167399697 | 0.33304638 | 0.50263179 | 0.61522318 | 0.81632065 |
| FUBP3 | 85.977496 | 0.171061929 | 0.19680235 | 0.86920675 | 0.38473406 | 0.66701235 |
| ABL1 | 212.234951 | 0.55102612 | 0.18503962 | 2.97788174 | 0.00290248 | 0.03178629 |
| AIF1L | 275.545618 | 1.00678316 | 0.40370096 | 2.49388347 | 0.0126354 | 0.0902704 |
| NUP214 | 207.494572 | 0.812828528 | 0.18301042 | 4.44143303 | 8.94E-06 | 0.00033355 |
| AL157938.2 | 37.9671128 | -0.270139821 | 0.63793139 | -0.4234622 | 0.67195808 | 0.8481392 |
| AL157938.3 | 17.2541582 | 0.592441198 | 0.46444576 | 1.27558749 | 0.20210138 | 0.48543912 |
| PRRC2B | 246.585514 | -0.018145968 | 0.23004899 | -0.0788787 | 0.93712909 | 0.97615526 |
| POMT1 | 20.8836764 | 0.539648496 | 0.31504505 | 1.71292483 | 0.08672637 | 0.30289881 |
| RAPGEF1 | 101.317881 | 0.068831701 | 0.38251465 | 0.17994527 | 0.85719554 | 0.94359207 |
| MED27 | 108.910016 | 0.204246501 | 0.26596639 | 0.76794101 | 0.44252223 | 0.71265656 |
| SETX | 189.90258 | 0.007598077 | 0.36887722 | 0.02059785 | 0.98356646 | 0.99368608 |
| TTF1 | 28.6823002 | -0.158833875 | 0.2667672 | -0.5954026 | 0.55157441 | 0.77722877 |
| DDX31 | 36.0229225 | -0.10848528 | 0.34120049 | -0.3179517 | 0.75052156 | 0.89107677 |
| GTF3C4 | 18.4818593 | -0.328950066 | 0.34580676 | -0.951254 | 0.34147545 | 0.62965456 |
| AK8 | 16.5550436 | -0.089592169 | 0.57224016 | -0.1565639 | 0.87558853 | 0.95030689 |
| TSC1 | 85.3782752 | 0.329549422 | 0.32468503 | 1.01498189 | 0.31011446 | 0.6033615 |
| GTF3C5 | 21.0347844 | -0.053154022 | 0.44082881 | -0.1205775 | 0.90402572 | 0.96126516 |
| RALGDS | 12.7846893 | -0.853969379 | 0.45614152 | -1.8721588 | 0.06118463 | 0.24535775 |
| ABO | 123.969168 | 0.327339887 | 0.31937398 | 1.02494227 | 0.30539043 | 0.59983455 |
| SURF6 | 15.1012571 | 0.002102816 | 0.43099354 | 0.004879 | 0.99610714 | 0.99868531 |

| | | | | | | |
|---|---|---|---|---|---|---|
| MED22 | 12.5755724 | -0.526627301 | 0.41938792 | -1.2557045 | 0.20922313 | 0.4951566 |
| RPL7A | 128.672253 | -0.606452827 | 0.43815398 | -1.3841089 | 0.16632511 | 0.4351146 |
| SURF1 | 15.2900009 | -0.380977517 | 0.46259686 | -0.8235627 | 0.41018808 | 0.68796444 |
| SURF4 | 60.6637407 | -0.20957341 | 0.32760673 | -0.6397103 | 0.52236094 | 0.76155853 |
| REXO4 | 15.7181733 | -0.937662942 | 0.35401551 | -2.6486493 | 0.00808141 | 0.06639133 |
| CACFD1 | 20.9913342 | -0.192296964 | 0.39976867 | -0.4810206 | 0.63050186 | 0.82555036 |
| VAV2 | 206.61365 | 0.121079198 | 0.6078754 | 0.19918424 | 0.84211863 | 0.93566718 |
| BRD3OS | 10.4445315 | -0.859675982 | 0.43899466 | -1.9582835 | 0.05019676 | 0.21810598 |
| BRD3 | 40.210929 | -0.364462412 | 0.33866658 | -1.0761688 | 0.28185176 | 0.5750055 |
| WDR5 | 21.7703395 | 0.196223298 | 0.34660209 | 0.5661342 | 0.57130256 | 0.78905469 |
| RXRA | 24.8653834 | 1.107509949 | 0.47898394 | 2.31220687 | 0.02076629 | 0.12343382 |
| PPP1R26-AS: | 31.0034045 | 0.149243741 | 0.32972372 | 0.45263271 | 0.65081324 | 0.83735207 |
| PPP1R26 | 16.7406581 | -0.596438231 | 0.36739764 | -1.6234133 | 0.10450107 | 0.33555699 |
| MRPS2 | 12.7315595 | 0.085798408 | 0.60573698 | 0.14164301 | 0.88736199 | 0.95619452 |
| CAMSAP1 | 49.3833155 | -0.277825849 | 0.33839676 | -0.8210062 | 0.41164272 | 0.68899849 |
| UBAC1 | 48.476221 | 0.426614959 | 0.22239048 | 1.9183148 | 0.05507111 | 0.23156838 |
| NACC2 | 37.7993458 | -0.573024082 | 0.43123363 | -1.3288019 | 0.18391332 | 0.46027002 |
| TMEM250 | 14.1622511 | -0.722933009 | 0.5924071 | -1.2203314 | 0.22233926 | 0.51071105 |
| QSOX2 | 11.4273817 | 0.12483945 | 0.39992474 | 0.31215736 | 0.75492094 | 0.89304149 |
| SNAPC4 | 17.5654526 | 0.000490144 | 0.38464284 | 0.00127428 | 0.99898327 | 0.99953184 |
| ENTR1 | 19.1663535 | -0.140787588 | 0.38011902 | -0.3703776 | 0.71110113 | 0.87035331 |
| PMPCA | 23.3346383 | -0.164105888 | 0.36811829 | -0.4457966 | 0.65574416 | 0.83939816 |
| INPP5E | 12.898725 | -0.051690241 | 0.3936689 | -0.1313038 | 0.89553495 | 0.95900099 |
| SEC16A | 54.346847 | -0.011432888 | 0.23507965 | -0.0486341 | 0.96121089 | 0.98499239 |
| NOTCH1 | 20.4989696 | -1.11173209 | 0.43336343 | -2.5653574 | 0.01030696 | 0.07859336 |
| AGPAT2 | 45.6288645 | -0.662596286 | 0.48251288 | -1.3732199 | 0.16968399 | 0.44032355 |
| SNHG7 | 35.0766674 | -0.021387009 | 0.31185859 | -0.0685792 | 0.94532459 | 0.97858537 |
| RABL6 | 68.3209728 | 0.262312351 | 0.37790547 | 0.6941216 | 0.48760596 | 0.74443126 |
| PHPT1 | 60.3191337 | -0.286240822 | 0.42497224 | -0.6735518 | 0.5005963 | 0.75067585 |
| EDF1 | 71.4105554 | -0.193212845 | 0.33408557 | -0.5783334 | 0.56303904 | 0.78438303 |
| TRAF2 | 15.3157144 | -0.91287263 | 0.38530417 | -2.3692259 | 0.01782536 | 0.11217891 |
| FBXW5 | 34.8844078 | -0.640181409 | 0.36882736 | -1.7357211 | 0.08261317 | 0.2945637 |
| LCN12 | 14.7378214 | 0.01907134 | 0.53910262 | 0.03537608 | 0.97177985 | 0.988268 |
| PAXX | 13.1866146 | -0.297520729 | 0.48973621 | -0.6075122 | 0.54351104 | 0.77321995 |
| ABCA2 | 12.456633 | 0.434464217 | 0.73181699 | 0.59367878 | 0.55272698 | 0.77787735 |
| MAN1B1 | 34.9276615 | -0.381229437 | 0.35509367 | -1.0736025 | 0.28300089 | 0.57561209 |
| DPP7 | 30.1917051 | 0.057985089 | 0.39281093 | 0.14761577 | 0.88264601 | 0.95504875 |
| SSNA1 | 22.7898619 | -0.608549245 | 0.43916925 | -1.3856827 | 0.1658438 | 0.4346416 |
| TPRN | 17.1584849 | -0.416280202 | 0.44047096 | -0.9450798 | 0.34461814 | 0.63250598 |
| TMEM203 | 15.7848628 | -0.349289496 | 0.46786419 | -0.7465617 | 0.45532816 | 0.72210564 |
| NDOR1 | 10.742927 | -0.495808287 | 0.43498845 | -1.1398194 | 0.25436153 | 0.54535536 |
| TUBB4B | 81.6902494 | -1.182771679 | 0.44885371 | -2.6350939 | 0.0084114 | 0.06844728 |
| NELFB | 44.3143607 | -0.089290914 | 0.24757953 | -0.3606555 | 0.71835701 | 0.87415486 |
| EXD3 | 64.1317152 | -0.300321044 | 0.26673585 | -1.1259118 | 0.26020285 | 0.55146957 |
| NOXA1 | 17.4366665 | -0.407209845 | 0.35814681 | -1.1369914 | 0.25554188 | 0.54674817 |
| NSMF | 14.1435512 | 0.010206625 | 0.51352497 | 0.01987562 | 0.9841426 | 0.99378916 |
| PNPLA7 | 30.8285586 | -0.459564901 | 0.49291948 | -0.9323326 | 0.35116467 | 0.63773431 |
| MRPL41 | 77.9004334 | -0.673291535 | 0.31606623 | -2.130223 | 0.03315321 | 0.16816561 |
| DPH7 | 22.5328387 | 0.274315562 | 0.34418253 | 0.79700606 | 0.42544751 | 0.70016446 |
| ARRDC1 | 28.0406842 | -0.780221424 | 0.43150735 | -1.8081301 | 0.07058626 | 0.26815316 |
| EHMT1 | 233.243204 | -0.076690258 | 0.22431889 | -0.3418805 | 0.73244081 | 0.88294 |
| AC069287.3 | 9.91772888 | 0.239971973 | 0.73678245 | 0.32570262 | 0.74464936 | 0.88808711 |
| BET1L | 18.8544698 | -0.154652119 | 0.34364425 | -0.4500355 | 0.65268483 | 0.83839889 |
| RIC8A | 15.1137397 | -0.740964697 | 0.48296728 | -1.5341923 | 0.12498235 | 0.37009151 |
| SIRT3 | 19.2501339 | -0.045155049 | 0.41554609 | -0.1086644 | 0.91346871 | 0.96461883 |
| PSMD13 | 38.6118632 | 0.053578893 | 0.37721116 | 0.14203952 | 0.88704879 | 0.95619452 |
| PGGHG | 25.1439066 | -0.591887753 | 0.50914756 | -1.1625073 | 0.24502946 | 0.53594343 |
| IFITM2 | 204.140224 | -2.256864704 | 0.47674976 | -4.7338559 | 2.20E-06 | 0.00010756 |
| IFITM3 | 267.802052 | -2.999885201 | 0.55060772 | -5.4483166 | 5.08E-08 | 4.68E-06 |
| SIGIRR | 38.3378064 | 0.042356674 | 0.25898919 | 0.16354611 | 0.87008847 | 0.94835789 |
| ANO9 | 10.8872938 | -0.584112008 | 0.4476338 | -1.3048881 | 0.19193096 | 0.47212817 |
| PTDSS2 | 28.8810773 | 0.465757232 | 0.29686754 | 1.56890591 | 0.11666987 | 0.35644512 |
| RNH1 | 42.8124989 | -1.011314259 | 0.41771395 | -2.4210689 | 0.01547494 | 0.101998 |
| HRAS | 21.4944654 | -0.678126429 | 0.40260944 | -1.6843282 | 0.09211826 | 0.31254072 |
| RASSF7 | 56.2064582 | -0.784827429 | 0.34753344 | -2.2582789 | 0.02392828 | 0.13544458 |
| MIR210HG | 14.0332138 | -3.574448601 | 0.64804112 | -5.5157744 | 3.47E-08 | 3.37E-06 |
| PHRF1 | 34.2643763 | -0.080402907 | 0.30647115 | -0.2623507 | 0.79305111 | 0.91136668 |
| IRF7 | 12.7590034 | -1.38518117 | 1.48833362 | -0.9306927 | 0.35201257 | 0.63801604 |
| DEAF1 | 64.4463231 | 0.095850243 | 0.28860747 | 0.33211283 | 0.73980406 | 0.88614269 |
| AC131934.1 | 13.9645995 | 0.872787934 | 0.50955656 | 1.71283818 | 0.08674231 | 0.30289881 |
| EPS8L2 | 72.7558942 | 0.16545094 | 0.24719662 | 0.66930907 | 0.50329835 | 0.75108109 |
| TALDO1 | 73.5273014 | -0.672573231 | 0.27606891 | -2.4362512 | 0.01484037 | 0.09940541 |
| GATD1 | 23.5274288 | -0.095072552 | 0.41848246 | -0.2271841 | 0.82028061 | 0.92406023 |
| AP006621.3 | 20.8660809 | -0.201754376 | 0.37859002 | -0.5329099 | 0.59409594 | 0.80267231 |
| RPLP2 | 97.6376071 | -0.715099549 | 0.41816856 | -1.7100749 | 0.08725203 | 0.30377507 |
| PNPLA2 | 31.9416374 | -0.33183358 | 0.3352155 | -0.9899112 | 0.32221752 | 0.61319341 |
| CD151 | 79.9941299 | -1.027507746 | 0.46453373 | -2.211912 | 0.02697274 | 0.14648981 |
| POLR2L | 59.2453537 | -0.609554879 | 0.50551295 | -1.2058146 | 0.227889 | 0.5177793 |
| TSPAN4 | 15.8855823 | -0.534167143 | 0.39597507 | -1.3489918 | 0.17733959 | 0.45201718 |
| CHID1 | 25.7037657 | 0.165901142 | 0.37293261 | 0.44485556 | 0.65642414 | 0.83951126 |
| AP2A2 | 210.388171 | -0.08109722 | 0.28213473 | -0.2874415 | 0.77377431 | 0.90184317 |
| TOLLIP | 32.1953668 | 0.092592841 | 0.36621135 | 0.2528399 | 0.80039194 | 0.91455697 |
| MOB2 | 26.113365 | 0.184282539 | 0.31738891 | 0.58062061 | 0.56149618 | 0.78361672 |
| MRPL23 | 12.7618071 | 0.411588968 | 0.44943183 | 0.91579843 | 0.35977263 | 0.64369184 |
| CD81-AS1 | 15.0006505 | 0.174477965 | 0.44351246 | 0.39340037 | 0.6940238 | 0.86101552 |
| CD81 | 210.775168 | -1.252252641 | 0.40304362 | -3.1069904 | 0.00189003 | 0.02360127 |
| TSSC4 | 14.712294 | -1.050295114 | 0.43313384 | -2.4248743 | 0.01531369 | 0.10149612 |
| KCNQ1 | 428.7332 | 0.4657601 | 0.19279368 | 2.4158474 | 0.01569864 | 0.10320828 |
| KCNQ1OT1 | 291.16304 | -1.21363207 | 0.3982522 | -3.0473958 | 0.00230834 | 0.02700808 |
| CDKN1C | 37.7330368 | -2.552124577 | 0.57454186 | -4.4420167 | 8.91E-06 | 0.00033355 |
| SLC22A18 | 26.9687102 | -0.639895822 | 0.61062917 | -1.0479287 | 0.29467146 | 0.58661907 |
| NAP1L4 | 239.74243 | 0.376742994 | 0.19758742 | 1.90671549 | 0.05655744 | 0.23465757 |
| CARS | 46.0299643 | -1.236424641 | 0.445457 | -2.7756319 | 0.00550946 | 0.05099587 |
| OSBPL5 | 19.1628043 | 0.193729892 | 0.35186188 | 0.55058505 | 0.58191816 | 0.7952959 |
| ZNF195 | 31.4365786 | -0.100330842 | 0.26932991 | -0.3725202 | 0.70950555 | 0.86943335 |
| AC127526.1 | 28.4685518 | 0.858106391 | 0.45430638 | 1.88882751 | 0.05891495 | 0.24008773 |
| NUP98 | 143.936913 | 0.493660767 | 0.32444327 | 1.52156264 | 0.1281187 | 0.37540371 |
| PGAP2 | 26.8679164 | 0.837478027 | 0.41198661 | 2.03277974 | 0.04207478 | 0.19577886 |
| RHOG | 10.4639871 | -0.481748403 | 0.53941542 | -0.8930935 | 0.37180711 | 0.6554341 |
| STIM1 | 92.6185076 | -0.079553456 | 0.25353575 | -0.3137761 | 0.75369111 | 0.89238252 |
| RRM1 | 21.0256452 | 0.177193503 | 0.36204544 | 0.48942338 | 0.62454199 | 0.82266098 |
| TRIM68 | 12.3068181 | 0.066893854 | 0.45951031 | 0.14557639 | 0.8842558 | 0.95603676 |
| AC104389.4 | 51.7686561 | 0.094879896 | 0.60193359 | 0.15762519 | 0.87475215 | 0.94993692 |
| TRIM5 | 31.0597319 | -1.181114151 | 0.58593617 | -2.0157727 | 0.04382375 | 0.19976647 |
| TRIM22 | 51.8383552 | -1.776328853 | 1.14372725 | -1.5531053 | 0.12039798 | 0.36341634 |
| FAM160A2 | 21.8685876 | 0.128458176 | 0.37803347 | 0.33980636 | 0.73400236 | 0.88368497 |
| SMPD1 | 17.7476482 | 0.000761445 | 0.63732161 | 0.00119476 | 0.99904672 | 0.99953184 |
| APBB1 | 20.4611794 | 0.295435491 | 0.36564869 | 0.80797635 | 0.4191042 | 0.6951556 |
| ARFIP2 | 15.9669924 | -0.562963184 | 0.37157445 | -1.5150751 | 0.12975339 | 0.37739114 |
| AC084337.2 | 27.8079043 | 0.256805107 | 0.55759107 | 0.46056173 | 0.64511307 | 0.83387847 |
| DNHD1 | 41.4183597 | -0.163797031 | 0.44407039 | -0.3688538 | 0.71223674 | 0.87112227 |
| RRP8 | 9.93214976 | -0.02600269 | 0.53432068 | -0.048665 | 0.96118631 | 0.98499239 |
| ILK | 37.4935749 | -0.250528352 | 0.30006664 | -0.8349091 | 0.40376892 | 0.68201793 |

| | | | | | | |
|---|---|---|---|---|---|---|
| TAF10 | 19.7761622 | -0.876279647 | 0.40635951 | -2.1564147 | 0.03105129 | 0.1613978 |
| TPP1 | 29.2465589 | -0.753923485 | 0.37300141 | -2.0212349 | 0.04325545 | 0.19853735 |
| MRPL17 | 20.5598387 | -0.560028942 | 0.44199687 | -1.2670428 | 0.20513999 | 0.48964616 |
| AC087280.2 | 19.2833127 | 0.489606254 | 0.80297664 | 0.6097391 | 0.54203465 | 0.77234315 |
| ZNF214 | 13.8422411 | 0.331128317 | 0.38383699 | 0.86267954 | 0.38831368 | 0.67030958 |
| AC107884.1 | 22.7381071 | 0.332878832 | 0.48455462 | 0.68697896 | 0.49209599 | 0.74651345 |
| PPFIBP2 | 59.0520295 | -0.139243722 | 0.34728661 | -0.4009476 | 0.68845872 | 0.8585881 |
| OVCH2 | 23.6775772 | 0.745205506 | 0.65646427 | 1.13518061 | 0.25629965 | 0.54711957 |
| AC044810.3 | 28.1493651 | 0.141991849 | 0.73371944 | 0.19352336 | 0.84654912 | 0.93757726 |
| EIF3F | 40.9520726 | -0.167583419 | 0.34951981 | -0.4794676 | 0.63160602 | 0.82625173 |
| RIC3 | 62.8792529 | -0.131235531 | 0.3684422 | -0.3561903 | 0.72169807 | 0.87625492 |
| STK33 | 363.441346 | 0.481162876 | 0.27047828 | 1.77893353 | 0.07525066 | 0.27848498 |
| TRIM66 | 73.1783472 | 0.394071638 | 0.35483487 | 1.11057754 | 0.26675023 | 0.55809523 |
| RPL27A | 115.156858 | -0.637483884 | 0.35893091 | -1.7760629 | 0.07572255 | 0.27917785 |
| ST5 | 277.12289 | -2.31690955 | 0.47451321 | -4.8827082 | 1.05E-06 | 5.80E-05 |
| AC026894.1 | 20.853993 | -2.230350961 | 0.58209551 | -3.8315893 | 0.00012732 | 0.00295951 |
| AKIP1 | 11.4631067 | -0.905957378 | 0.4152159 | -2.1818947 | 0.0291173 | 0.15438315 |
| TMEM9B | 34.7316498 | -0.421438921 | 0.35829566 | -1.1762323 | 0.23950207 | 0.52992919 |
| TMEM9B-AS | 22.4162456 | 0.566097658 | 0.56881627 | 0.99522059 | 0.31962899 | 0.611915 |
| DENND5A | 101.009221 | 0.222771248 | 0.32988966 | 0.67529017 | 0.49949143 | 0.75032964 |
| TMEM41B | 96.1283962 | -0.084779701 | 0.48695097 | -0.1741032 | 0.86178439 | 0.94550174 |
| IPO7 | 114.029007 | 0.011076317 | 0.3073325 | 0.03604017 | 0.97125033 | 0.988268 |
| ZNF143 | 26.7768443 | 0.073706089 | 0.39829991 | 0.18505173 | 0.85318846 | 0.94066219 |
| WEE1 | 16.38216 | -0.519076912 | 0.42746589 | -1.2143119 | 0.22462865 | 0.51345116 |
| SWAP70 | 77.8985328 | -1.154571774 | 0.26924105 | -4.2882456 | 1.80E-05 | 0.00059464 |
| SBF2-AS1 | 41.4554667 | -0.204188567 | 0.33177654 | -0.61544 | 0.53826415 | 0.77005934 |
| SBF2 | 618.103744 | 0.126339175 | 0.18686813 | 0.67608734 | 0.49898519 | 0.75027574 |
| AC011092.2 | 24.5142559 | -0.646731798 | 0.55066137 | -1.1744637 | 0.24020934 | 0.5310379 |
| ADM | 98.65119 | 0.813709866 | 0.71306935 | 1.14113706 | 0.25381289 | 0.54524749 |
| AMPD3 | 61.6343439 | -0.34888155 | 0.39865762 | -0.8751408 | 0.3814973 | 0.66478098 |
| MTRNR2L8 | 128.503042 | -1.218394725 | 0.72453139 | -1.6816314 | 0.09264034 | 0.31323946 |
| RNF141 | 24.6174074 | 0.179226428 | 0.35689468 | 0.50218297 | 0.61553882 | 0.8164389 |
| CTR9 | 56.1847503 | 0.505816419 | 0.26651239 | 1.89790961 | 0.05770799 | 0.23751806 |
| EIF4G2 | 134.75641 | -0.360941541 | 0.25401335 | -1.420955 | 0.15532984 | 0.41890262 |
| ZBED5 | 49.4632736 | 0.196028899 | 0.26942017 | 0.72759549 | 0.46686124 | 0.73016616 |
| ZBED5-AS1 | 15.558445 | 0.698619622 | 0.47264277 | 1.47811343 | 0.13937742 | 0.39404853 |
| GALNT18 | 246.049382 | 0.703594229 | 0.35086267 | 2.00532655 | 0.04492815 | 0.20229449 |
| USP47 | 238.27234 | 0.328965099 | 0.16167414 | 2.0347416 | 0.04187688 | 0.19529906 |
| PARVA | 99.2312478 | -0.013714133 | 0.35755595 | -0.0383552 | 0.96940447 | 0.98781534 |
| TEAD1 | 392.38992 | 0.405176604 | 0.41747229 | 0.9705473 | 0.33177376 | 0.62147393 |
| LINC00958 | 47.3737295 | -0.45747429 | 0.27555866 | -1.6601702 | 0.09688021 | 0.32146212 |
| ARNTL | 30.7682057 | 0.530648916 | 0.39755529 | 1.33478017 | 0.18194831 | 0.4586326 |
| BTBD10 | 69.5698936 | 0.092680424 | 0.29583923 | 0.31327969 | 0.75406818 | 0.89238252 |
| FAR1 | 120.653608 | 0.552114578 | 0.22558762 | 2.44745069 | 0.01438708 | 0.09745114 |
| RRAS2 | 76.51751 | -0.256309266 | 0.30619766 | -0.8370713 | 0.4025525 | 0.68128594 |
| COPB1 | 85.3534315 | 0.020936197 | 0.25592005 | 0.08180757 | 0.93479974 | 0.97513509 |
| PSMA1 | 88.1013836 | 0.214670368 | 0.23363164 | 0.91884117 | 0.35817866 | 0.64216076 |
| PDE3B | 249.324487 | 0.188463248 | 0.37985685 | 0.49614282 | 0.61979365 | 0.81899807 |
| CYP2R1 | 15.811177 | 0.00169358 | 0.38913473 | 0.00435217 | 0.99652748 | 0.99868531 |
| AC073172.1 | 39.4758617 | 2.317649775 | 0.7003418 | 3.30931234 | 0.00093525 | 0.01396375 |
| SOX6 | 576.959123 | 1.078298645 | 0.34037469 | 3.16797535 | 0.00153505 | 0.02014527 |
| C11orf58 | 105.92404 | -0.16513066 | 0.16846823 | -0.9801887 | 0.32699296 | 0.61686165 |
| PLEKHA7 | 534.643606 | 0.188860874 | 0.19927711 | 0.9477299 | 0.34326698 | 0.6310379 |
| RPS13 | 91.2156311 | -0.724423142 | 0.37124779 | -1.9513197 | 0.05101902 | 0.21991547 |
| PIK3C2A | 241.068481 | 0.027952261 | 0.22114321 | 0.12639891 | 0.89941616 | 0.95958837 |
| NUCB2 | 50.231332 | -0.128266702 | 0.35518249 | -0.361129 | 0.71800301 | 0.87412751 |
| SERGEF | 109.412482 | -0.59651708 | 0.29316398 | -2.0347557 | 0.04187546 | 0.19529906 |
| HPS5 | 28.3284487 | 0.486156764 | 0.35701323 | 1.36173319 | 0.1732821 | 0.44612537 |
| GTF2H1 | 55.4941512 | -0.207478519 | 0.30347278 | -0.6836808 | 0.49417674 | 0.74700665 |
| LDHA | 216.353023 | -2.339479494 | 0.3501443 | -6.6814725 | 2.37E-11 | 7.17E-09 |
| TSG101 | 87.4558499 | -0.161326753 | 0.23073483 | -0.6991868 | 0.48443529 | 0.74250561 |
| UEVLD | 67.7683097 | -0.067849428 | 0.22441095 | -0.3023445 | 0.76238942 | 0.89720537 |
| SPTY2D1OS | 23.6921326 | -0.146192598 | 0.34959609 | -0.4181757 | 0.67581865 | 0.85173672 |
| SPTY2D1 | 23.6089307 | -0.079193214 | 0.35184059 | -0.2250827 | 0.82191497 | 0.92509211 |
| ZDHHC13 | 65.5903374 | -0.86831075 | 0.22501234 | -3.8589472 | 0.00011388 | 0.00269072 |
| NAV2 | 235.705383 | 1.011487902 | 0.19772816 | 5.11554813 | 3.13E-07 | 2.18E-05 |
| HTATIP2 | 24.1458506 | -1.266736901 | 0.46275102 | -2.7374049 | 0.0061926 | 0.05537004 |
| PRMT3 | 79.7936222 | 0.271854882 | 0.21390926 | 1.2708888 | 0.20376821 | 0.48807721 |
| NELL1 | 71.8803263 | 4.014452331 | 0.5713807 | 7.02588023 | 2.13E-12 | 8.12E-10 |
| ANO5 | 45.0926723 | -0.210462704 | 0.41477734 | -0.5074113 | 0.61186627 | 0.81387299 |
| GAS2 | 24.6073978 | 0.340752728 | 0.47182318 | 0.7222043 | 0.47016888 | 0.73188482 |
| SVIP | 52.9355184 | -0.209334456 | 0.29268246 | -0.7152272 | 0.47446866 | 0.73514456 |
| AC006299.1 | 15.6571145 | -0.248088669 | 0.64049503 | -0.3873389 | 0.6985053 | 0.86334649 |
| SLC5A12 | 10.9401542 | 1.885527984 | 0.78998336 | 2.38679455 | 0.01699599 | 0.109092 |
| FIBIN | 37.0259124 | -1.772621158 | 0.823831 | -2.1516806 | 0.03142252 | 0.16226306 |
| BBOX1 | 48.8280561 | -2.628723479 | 0.66036133 | -3.980735 | 6.87E-05 | 0.00181016 |
| LGR4 | 580.430291 | 0.521724293 | 0.2760911 | 1.88968167 | 0.05880055 | 0.24000052 |
| LIN7C | 31.3076263 | -0.051568136 | 0.29568301 | -0.1744034 | 0.8615484 | 0.94550174 |
| BDNF-AS | 116.927603 | 0.348617209 | 0.21853047 | 1.59527967 | 0.11064971 | 0.34689997 |
| METTL15 | 254.183954 | 0.077900769 | 0.26360895 | 0.2955164 | 0.76759942 | 0.89979146 |
| ARL14EP | 34.5790833 | 0.083149442 | 0.28976448 | 0.28695526 | 0.77414658 | 0.90187244 |
| MPPED2 | 661.009519 | 0.808156131 | 0.27578614 | 2.93037259 | 0.00338556 | 0.03555354 |
| AL136088.1 | 33.9454146 | 0.907692807 | 0.51340078 | 1.76800044 | 0.07706083 | 0.28161784 |
| DCDC1 | 95.467935 | -0.731733103 | 0.41535141 | -1.7617205 | 0.07811653 | 0.28439855 |
| DNAJC24 | 41.5127218 | -0.349158601 | 0.31923099 | -1.0937491 | 0.27406505 | 0.56626917 |
| IMMP1L | 117.022436 | 0.541018921 | 0.24457489 | 2.21207879 | 0.02696122 | 0.14648981 |
| ELP4 | 176.893543 | 0.556386682 | 0.1826075 | 3.04689937 | 0.00231215 | 0.02700808 |
| AC131571.1 | 17.3834798 | -0.689476842 | 0.68026339 | -1.013544 | 0.31080041 | 0.60394276 |
| RCN1 | 26.1310161 | -1.381329813 | 0.48025738 | -2.8762282 | 0.00402459 | 0.04032406 |
| EIF3M | 49.3472601 | 0.14059327 | 0.28851237 | 0.48730413 | 0.62604282 | 0.8231524 |
| PRRG4 | 45.6466929 | -0.711206494 | 0.39801872 | -1.7868669 | 0.073959 | 0.2750634 |
| QSER1 | 83.4701817 | 0.468341271 | 0.2292007 | 2.04336757 | 0.04101606 | 0.19294406 |
| TCP11L1 | 40.8130273 | 0.233539021 | 0.30806416 | 0.75808566 | 0.4483997 | 0.71763457 |
| CSTF3 | 76.3176757 | 0.064377355 | 0.2286866 | 0.28150908 | 0.77831996 | 0.90417143 |
| HIPK3 | 206.43535 | -0.375003252 | 0.2067952 | -1.813404 | 0.06976953 | 0.26620951 |
| FBXO3 | 48.4700086 | 0.70358682 | 0.23882124 | 2.94608146 | 0.00321828 | 0.03421536 |
| CAPRIN1 | 128.695641 | 0.045545126 | 0.20532301 | 0.22182183 | 0.82445259 | 0.92643003 |
| NAT10 | 33.275019 | 0.050053394 | 0.26996176 | 0.1854092 | 0.85290809 | 0.94049004 |
| ABTB2 | 672.630911 | 1.707236288 | 0.57295462 | 2.97970593 | 0.00288525 | 0.03172559 |
| CAT | 80.9182455 | 0.311735232 | 0.46154398 | 0.67541827 | 0.49941006 | 0.75031683 |
| ELF5 | 21.3097699 | 1.893269721 | 0.80484056 | 2.35235377 | 0.01865502 | 0.11556467 |
| EHF | 99.2123967 | -0.974850636 | 0.51745091 | -1.8839481 | 0.059572 | 0.24171358 |
| APIP | 45.9306132 | 0.599503441 | 0.43265219 | 1.38564754 | 0.16585456 | 0.4346416 |
| PDHX | 90.6008912 | 0.322981452 | 0.22085098 | 1.46244063 | 0.1436205 | 0.40102728 |
| CD44 | 77.840161 | -4.837610675 | 0.62228194 | -7.7739854 | 7.61E-15 | 5.22E-12 |
| TRIM44 | 203.726469 | -0.098078832 | 0.18542856 | -0.5289305 | 0.59685363 | 0.8047096 |
| COMMD9 | 18.0973661 | -0.276223276 | 0.35125684 | -0.7863855 | 0.43164169 | 0.70454366 |
| PRR5L | 37.9961378 | 0.15823124 | 0.51520016 | 0.30712576 | 0.75874765 | 0.8951693 |
| TRAF6 | 34.1387014 | 0.041484883 | 0.32431185 | 0.12791664 | 0.89821494 | 0.95930026 |
| C11orf74 | 19.1456063 | 0.469095926 | 0.47976341 | 0.97776511 | 0.32819051 | 0.61731215 |
| LRRC4C | 44.5479635 | -3.057675747 | 0.80093826 | -3.8176173 | 0.00013475 | 0.00309857 |
| API5 | 66.6176385 | -0.14707543 | 0.21849098 | -0.6731419 | 0.50085702 | 0.75071921 |
| TTC17 | 337.815866 | 0.2191193 | 0.15588548 | 1.40564276 | 0.15983023 | 0.42745873 |

| | | | | | | |
|---|---|---|---|---|---|---|
| HSD17B12 | 293.219173 | 0.374572169 | 0.15843673 | 2.36417507 | 0.01807027 | 0.11306142 |
| ALKBH3 | 25.5685378 | 0.161335243 | 0.33081351 | 0.48769243 | 0.62576772 | 0.82311785 |
| ACCS | 12.8957476 | -0.198423508 | 0.66145502 | -0.2999803 | 0.76419215 | 0.8975838 |
| EXT2 | 84.0970445 | -0.047927544 | 0.24743832 | -0.1936949 | 0.84641477 | 0.93757726 |
| CD82 | 40.1356773 | -0.707082773 | 0.51819457 | -1.3645121 | 0.17240644 | 0.44481121 |
| TSPAN18 | 17.0951247 | -0.688431372 | 0.79125019 | -0.8700552 | 0.38427022 | 0.66701235 |
| PRDM11 | 27.8165594 | 0.32004921 | 0.31491607 | 1.01630002 | 0.30948654 | 0.60299393 |
| SYT13 | 22.4149151 | 1.002936329 | 0.37111656 | 2.70248333 | 0.00688236 | 0.05958161 |
| CRY2 | 46.8899991 | 0.359751511 | 0.29902903 | 1.20306551 | 0.22895097 | 0.51906974 |
| LARGE2 | 12.9930307 | 1.431717685 | 0.56672299 | 2.52630954 | 0.01152679 | 0.08469969 |
| PHF21A | 293.949445 | -0.163987914 | 0.2377836 | -0.6896519 | 0.49041311 | 0.7455837 |
| CREB3L1 | 15.1229663 | -1.650359019 | 0.71308226 | -2.314402 | 0.02064568 | 0.1230165 |
| DGKZ | 17.7527996 | -0.290404816 | 0.4438748 | -0.6542494 | 0.51295115 | 0.75599752 |
| MDK | 16.8584571 | -4.001318407 | 1.27215206 | -3.1453146 | 0.00165908 | 0.02144527 |
| AMBRA1 | 367.985733 | -0.027112199 | 0.20347486 | -0.1332459 | 0.89399888 | 0.95815794 |
| ATG13 | 45.3854782 | -0.410555787 | 0.29896588 | -1.373253 | 0.16967372 | 0.44032355 |
| ARHGAP1 | 51.0825742 | -0.861082291 | 0.29363796 | -2.9324625 | 0.00336286 | 0.03535116 |
| CKAP5 | 82.3298906 | 0.438970268 | 0.23122146 | 1.89848412 | 0.05763234 | 0.2373015 |
| LRP4 | 43.0617526 | -0.372553743 | 0.54937652 | -0.6781392 | 0.49768345 | 0.74958112 |
| C11orf49 | 261.466141 | 0.149007188 | 0.20096541 | 0.74145689 | 0.45841646 | 0.72410401 |
| ARFGAP2 | 34.4415108 | -0.078990877 | 0.30994334 | -0.2548559 | 0.79883443 | 0.91379784 |
| DDB2 | 24.6170372 | -0.390428811 | 0.5711444 | -0.6835904 | 0.49423387 | 0.74700665 |
| ACP2 | 17.3756042 | -0.125812268 | 0.48486219 | -0.2594805 | 0.79526455 | 0.91195838 |
| NR1H3 | 18.8006529 | 0.668283179 | 0.4236338 | 1.57750202 | 0.11468006 | 0.35355161 |
| MADD | 29.221526 | -0.136145081 | 0.33431666 | -0.4072339 | 0.6838362 | 0.85606265 |
| SLC39A13 | 29.5690392 | 0.045567208 | 0.40099426 | 0.11363556 | 0.9095267 | 0.96308593 |
| PSMC3 | 14.1481603 | -0.461588942 | 0.42371015 | -1.0893979 | 0.27597846 | 0.56798847 |
| CELF1 | 302.074678 | 0.140396567 | 0.24258474 | 0.57875267 | 0.56275607 | 0.78429559 |
| NDUFS3 | 31.0871045 | 0.331730211 | 0.32490598 | 1.02100372 | 0.30725268 | 0.60139222 |
| MTCH2 | 67.2029729 | -0.092968203 | 0.28935383 | -0.3212959 | 0.74798615 | 0.88939905 |
| FNBP4 | 170.700725 | -0.311407482 | 0.24034205 | -1.2956846 | 0.19508418 | 0.47613296 |
| NUP160 | 94.2206736 | -0.030164226 | 0.27995025 | -0.1077485 | 0.91419518 | 0.96462448 |
| PTPRJ | 334.912685 | 0.3220173 | 0.3290989 | 0.97848184 | 0.32783607 | 0.61729352 |
| TNKS1BP1 | 47.6747254 | -1.013467251 | 0.29555094 | -3.429078 | 0.00060564 | 0.01003096 |
| SSRP1 | 24.257403 | -0.81049236 | 0.48771934 | -1.6618008 | 0.09655273 | 0.32086653 |
| TIMM10 | 17.0155142 | -0.738049284 | 0.43730815 | -1.6877099 | 0.09146693 | 0.31120975 |
| UBE2L6 | 20.1976869 | -1.557302348 | 0.85645636 | -1.8183091 | 0.06901691 | 0.26461191 |
| SERPING1 | 160.419358 | -2.80529862 | 0.50756658 | -5.5269569 | 3.26E-08 | 3.20E-06 |
| ZDHHC5 | 61.5347806 | -0.423153457 | 0.28335043 | -1.4933927 | 0.13533443 | 0.38728203 |
| CTNND1 | 170.606275 | 0.039505669 | 0.22348465 | 0.17677128 | 0.85968805 | 0.94449251 |
| LPXN | 30.6071506 | 0.805919776 | 0.31770668 | 2.53667872 | 0.01119096 | 0.08282276 |
| FAM111A | 42.902591 | 0.506536329 | 0.3569401 | 1.41910737 | 0.15586772 | 0.4199135 |
| OSBP | 84.7144864 | -0.165498138 | 0.28731645 | -0.5760134 | 0.5646061 | 0.78515099 |
| PATL1 | 54.41333 | -0.237327097 | 0.35720363 | -0.6644028 | 0.50643253 | 0.75306985 |
| STX3 | 91.5599996 | -0.432717633 | 0.34857552 | -1.2413885 | 0.21446227 | 0.50088196 |
| MRPL16 | 23.4576453 | -0.21874973 | 0.35183884 | -0.6217328 | 0.53411761 | 0.76732389 |
| PRPF19 | 22.3968938 | 0.111677864 | 0.3815398 | 0.29270305 | 0.76974914 | 0.90034488 |
| TMEM109 | 18.2235159 | -0.573094146 | 0.54712518 | -1.0474644 | 0.29488545 | 0.58668972 |
| VPS37C | 35.1720939 | -0.096014121 | 0.31012421 | -0.3095989 | 0.75686598 | 0.89445158 |
| DDB1 | 60.545773 | -0.221681958 | 0.34667302 | -0.6394555 | 0.52252667 | 0.76161146 |
| TKFC | 16.0905683 | -0.038344075 | 0.4687996 | -0.081792 | 0.93481209 | 0.97513509 |
| TMEM138 | 10.7037561 | 0.400503624 | 0.41336458 | 0.96888714 | 0.3326015 | 0.62174935 |
| CPSF7 | 59.6076274 | 0.18398044 | 0.21146335 | 0.87003463 | 0.38428148 | 0.66701235 |
| AP003108.2 | 53.8253333 | 0.211194275 | 0.38128486 | 0.55390154 | 0.57964623 | 0.79429575 |
| SYT7 | 20.6730059 | 1.28900926 | 0.73831493 | 1.74587998 | 0.08083181 | 0.29075313 |
| MYRF | 35.7262818 | -1.588620952 | 0.56401889 | -2.8166095 | 0.00485335 | 0.04642453 |
| TMEM258 | 33.7717048 | -0.978646771 | 0.4226565 | -2.315466 | 0.02058744 | 0.12280939 |
| FADS1 | 16.8366013 | -0.645474304 | 0.78823286 | -0.8188878 | 0.41285041 | 0.68990579 |
| FTH1 | 575.418765 | -1.788436438 | 0.45078105 | -3.9674171 | 7.27E-05 | 0.00189975 |
| ASRGL1 | 34.549164 | -0.180606356 | 0.53330722 | -0.3386535 | 0.73487078 | 0.88380093 |
| AHNAK | 183.425085 | -0.223778946 | 0.28574178 | -0.7831509 | 0.43353848 | 0.70594011 |
| B3GAT3 | 26.9561365 | -0.532310036 | 0.34491283 | -1.5433176 | 0.12275373 | 0.36708533 |
| GANAB | 75.6087321 | -0.091468987 | 0.27585199 | -0.3315872 | 0.74020099 | 0.88638272 |
| UQCC3 | 20.1901407 | -0.406548146 | 0.38783206 | -1.0482582 | 0.29451965 | 0.5865342 |
| UBXN1 | 21.5628927 | -1.107132136 | 0.42556422 | -2.601563 | 0.00928 | 0.07312742 |
| TTC9C | 33.0956306 | -0.309427743 | 0.35578392 | -0.8697069 | 0.38446058 | 0.66701235 |
| POLR2G | 11.4835128 | -0.158177811 | 0.51201683 | -0.3089309 | 0.7573741 | 0.89457492 |
| TAF6L | 15.4984049 | -0.65143526 | 0.50849839 | -1.281096 | 0.20015994 | 0.48370812 |
| TMEM223 | 19.8316142 | -0.842181791 | 0.38164658 | -2.206706 | 0.02733461 | 0.14777597 |
| TMEM179B | 29.8747407 | -1.185645811 | 0.37907668 | -3.1277203 | 0.00176168 | 0.02232325 |
| NXF1 | 16.3382111 | 0.378792837 | 0.39101436 | 0.96874406 | 0.3326729 | 0.62177 |
| STX5 | 61.5126343 | 0.073358943 | 0.33188332 | 0.22103836 | 0.82506257 | 0.92671114 |
| SLC3A2 | 56.6510017 | -1.141987062 | 0.46863475 | -2.4368382 | 0.01481631 | 0.09936964 |
| RARRES3 | 34.538269 | -1.921422495 | 0.95680726 | -2.0081605 | 0.04462624 | 0.20147742 |
| PLA2G16 | 17.2486483 | -1.917749649 | 0.62454828 | -3.0706187 | 0.00213616 | 0.02570875 |
| ATL3 | 65.2261415 | -0.553282143 | 0.21723747 | -2.5469001 | 0.01086845 | 0.08095937 |
| RTN3 | 180.470052 | 0.320713665 | 0.25628361 | 1.25140139 | 0.21078807 | 0.49667071 |
| MARK2 | 98.9369366 | 0.048975557 | 0.19129268 | 0.2560242 | 0.79793215 | 0.91326486 |
| NAA40 | 28.9336966 | 0.457022771 | 0.33011282 | 1.38444416 | 0.1662225 | 0.43507299 |
| OTUB1 | 29.7691457 | -0.172754548 | 0.34913103 | -0.4948129 | 0.62073218 | 0.81942374 |
| MACROD1 | 58.6635034 | 0.915376676 | 0.43205286 | 2.11866823 | 0.03411851 | 0.17086443 |
| AP000721.2 | 17.8141071 | 1.693109175 | 0.54046533 | 3.13268788 | 0.00173214 | 0.02204834 |
| STIP1 | 37.1547479 | 0.45582329 | 0.32529244 | 1.4012723 | 0.16113266 | 0.42820049 |
| TRPT1 | 11.2049651 | 0.110757334 | 0.48078772 | 0.2303664 | 0.81780707 | 0.92281757 |
| NUDT22 | 23.7185853 | -0.061385971 | 0.39444471 | -0.1556263 | 0.87632759 | 0.95100883 |
| DNAJC4 | 15.8606557 | 0.30461333 | 0.52107628 | 0.58458491 | 0.55882685 | 0.78244004 |
| VEGFB | 41.6607728 | -0.497491674 | 0.33020976 | -1.5065929 | 0.13191504 | 0.38098929 |
| FKBP2 | 49.7654311 | -0.264665701 | 0.33230125 | -0.7964631 | 0.42576289 | 0.70025461 |
| PPP1R14B | 34.9830647 | -1.22315421 | 0.45666058 | -2.6784756 | 0.00739581 | 0.06273624 |
| BAD | 25.4799084 | -0.749016472 | 0.37434705 | -2.0008612 | 0.04540735 | 0.20400635 |
| GPR137 | 18.8490335 | -0.261677413 | 0.41380283 | -0.6323722 | 0.52714369 | 0.76221787 |
| ESRRA | 70.5625489 | 0.400569175 | 0.36613577 | 1.09404544 | 0.27393505 | 0.56626457 |
| TRMT112 | 35.6282148 | -0.62174853 | 0.3344595 | -1.8589651 | 0.06303209 | 0.24979235 |
| PRDX5 | 62.8724083 | -0.696823134 | 0.50196314 | -1.3881958 | 0.16507742 | 0.43383357 |
| RPS6KA4 | 10.604081 | -0.391997476 | 0.56146147 | -0.6981734 | 0.48506875 | 0.74285379 |
| SF1 | 170.806344 | -0.090277804 | 0.34238831 | -0.2636708 | 0.79203359 | 0.9111703 |
| MAP4K2 | 10.8833888 | 0.724731517 | 0.40424873 | 1.79278615 | 0.07300713 | 0.2730016 |
| MEN1 | 11.8558343 | -0.263354118 | 0.45349388 | -0.5807225 | 0.56142747 | 0.78361672 |
| EHD1 | 19.4234773 | -0.257237965 | 0.40990243 | -0.627559 | 0.53029286 | 0.76428051 |
| ATG2A | 17.9492532 | 0.00129853 | 0.50776385 | 0.00255735 | 0.99795953 | 0.99900003 |
| PPP2R5B | 11.7292007 | -0.106544605 | 0.49996904 | -0.2131024 | 0.83124709 | 0.93051261 |
| VPS51 | 42.25587 | 0.099405094 | 0.29989079 | 0.33147098 | 0.74028876 | 0.88638479 |
| TM7SF2 | 14.9778717 | 0.396175806 | 0.50865055 | 0.77887619 | 0.43605265 | 0.70765823 |
| FAU | 118.623652 | -0.530840555 | 0.43159449 | -1.2299521 | 0.21871504 | 0.50674964 |
| SYVN1 | 14.9715742 | 0.058414695 | 0.45953283 | 0.12711757 | 0.89884735 | 0.95935794 |
| MRPL49 | 15.4399424 | -0.438934134 | 0.36596865 | -1.1993763 | 0.23038165 | 0.52060957 |
| CAPN1 | 59.3292767 | -0.706496182 | 0.29877116 | -2.3646733 | 0.01804598 | 0.11301502 |
| POLA2 | 11.1173474 | 0.089460884 | 0.53554562 | 0.16704624 | 0.86733366 | 0.94753673 |
| DPF2 | 30.4478261 | 0.052938482 | 0.26949195 | 0.19643809 | 0.84426729 | 0.93684205 |
| FRMD8 | 28.7671595 | -0.371588205 | 0.41893329 | -0.8869866 | 0.37508612 | 0.65806152 |
| NEAT1 | 9036.78831 | -1.32171363 | 0.49620886 | -2.6636236 | 0.0077304 | 0.06419577 |
| MALAT1 | 89533.2753 | -0.543953653 | 0.37887178 | -1.4357196 | 0.15108214 | 0.41211643 |
| SCYL1 | 25.5541042 | -0.611813327 | 0.50884723 | -1.2023517 | 0.2292273 | 0.51946758 |

| | | | | | | |
|---|---|---|---|---|---|---|
| LTBP3 | 54.5820518 | -0.102000698 | 0.31569894 | -0.3230948 | 0.74662342 | 0.88870184 |
| EHBP1L1 | 19.8311536 | -1.448710725 | 0.46976092 | -3.083932 | 0.00204284 | 0.02492802 |
| MAP3K11 | 17.9377073 | -0.211039023 | 0.3899515 | -0.541193 | 0.58837456 | 0.79897585 |
| PCNX3 | 17.628284 | 0.401707587 | 0.45966767 | 0.87390872 | 0.38216796 | 0.66549938 |
| SIPA1 | 15.1746798 | -2.071883417 | 0.48204047 | -4.2981524 | 1.72E-05 | 0.00057938 |
| RELA | 23.8865871 | -0.434836115 | 0.40486867 | -1.0740177 | 0.28281475 | 0.5754607 |
| KAT5 | 25.381085 | 0.227230471 | 0.34912093 | 0.65086466 | 0.51513386 | 0.75713618 |
| RNASEH2C | 19.7739998 | -0.269031777 | 0.4948181 | -0.5436983 | 0.58664908 | 0.79752689 |
| CFL1 | 123.109532 | -1.146081449 | 0.38946548 | -2.9427036 | 0.0032536 | 0.03448412 |
| MUS81 | 12.9443641 | -0.025688402 | 0.37533185 | -0.0684418 | 0.94543392 | 0.97858537 |
| EFEMP2 | 14.7859619 | -0.442550587 | 0.50324528 | -0.8793934 | 0.379188 | 0.66243764 |
| CCDC85B | 11.3232017 | -0.594678408 | 0.56158404 | -1.0589304 | 0.28963148 | 0.58130538 |
| DRAP1 | 20.1526385 | -0.492528817 | 0.41709728 | -1.1808488 | 0.2376628 | 0.52778809 |
| SART1 | 18.9398222 | 0.253741565 | 0.39877075 | 0.63630937 | 0.5245748 | 0.76221787 |
| BANF1 | 35.7977508 | -0.904551076 | 0.40586738 | -2.2286863 | 0.02583478 | 0.14286094 |
| SF3B2 | 47.3965697 | -0.388033142 | 0.38038101 | -1.020117 | 0.30767297 | 0.60139222 |
| PACS1 | 319.711703 | -0.235106399 | 0.20047114 | -1.1727693 | 0.24088833 | 0.53151245 |
| KLC2 | 13.888805 | -0.270800581 | 0.43347345 | -0.6247224 | 0.53215326 | 0.76599732 |
| RAB1B | 33.4984933 | -0.785803764 | 0.48664784 | -1.6147277 | 0.10636967 | 0.33968392 |
| YIF1A | 14.6906058 | -0.463197977 | 0.5687611 | -0.8143981 | 0.41541691 | 0.69204803 |
| BRMS1 | 12.0006399 | -0.227009851 | 0.62965338 | -0.3605315 | 0.71844974 | 0.87415486 |
| SLC29A2 | 28.2424129 | 0.562062291 | 0.30536849 | 1.84060342 | 0.0656797 | 0.25610607 |
| MRPL11 | 13.0027069 | -0.310116728 | 0.39611885 | -0.7828881 | 0.43369283 | 0.70597357 |
| ZDHHC24 | 21.1851176 | -0.444752932 | 0.44418305 | -1.001283 | 0.31669001 | 0.61004871 |
| CTSF | 22.8253115 | -0.351370683 | 0.55167548 | -0.6369155 | 0.52417987 | 0.76221787 |
| CCS | 29.5144136 | 0.225716418 | 0.38969576 | 0.57921189 | 0.56244621 | 0.78400069 |
| RBM14-RBM | 32.5986095 | 0.508096311 | 0.28199702 | 1.80177904 | 0.07158018 | 0.27011684 |
| RBM4 | 22.4349932 | 0.105338454 | 0.42335297 | 0.24881945 | 0.80350045 | 0.91633138 |
| RBM4B | 25.9715946 | 0.115807613 | 0.35754873 | 0.32389323 | 0.74601887 | 0.88829015 |
| SPTBN2 | 127.550612 | 0.689664311 | 0.40146508 | 1.71786874 | 0.08582056 | 0.30113433 |
| C11orf80 | 469.954027 | 0.401645764 | 0.27505551 | 1.46023531 | 0.14422541 | 0.40202041 |
| RCE1 | 13.0704916 | 0.417908557 | 0.44947868 | 0.9297628 | 0.35249391 | 0.63843043 |
| PC | 12.2508287 | 0.023670393 | 0.44175118 | 0.05358309 | 0.95726733 | 0.98315333 |
| RHOD | 18.9153626 | -0.132115732 | 0.43309888 | -0.3050475 | 0.76032997 | 0.89627175 |
| KDM2A | 327.122663 | -0.021963476 | 0.22211783 | -0.0988821 | 0.92123189 | 0.9686919 |
| GRK2 | 19.2280843 | -0.267589623 | 0.37535898 | -0.7128899 | 0.4759139 | 0.73616591 |
| ANKRD13D | 24.9637733 | -0.636949716 | 0.36470048 | -1.7465009 | 0.08072395 | 0.29058005 |
| SSH3 | 17.7443793 | -0.959791936 | 0.41765626 | -2.2980427 | 0.02155936 | 0.12655527 |
| RAD9A | 130.952448 | 0.556677037 | 0.32324327 | 1.72216126 | 0.08504031 | 0.29941397 |
| PPP1CA | 38.8297303 | -0.930397851 | 0.50003498 | -1.8606655 | 0.06279142 | 0.24926305 |
| RPS6KB2 | 14.0001115 | 0.511850965 | 0.42754546 | 1.19718488 | 0.2312345 | 0.52111057 |
| CORO1B | 24.9683045 | -1.279609306 | 0.35274859 | -3.6275391 | 0.00028614 | 0.00572382 |
| TMEM134 | 27.8478273 | -0.570051961 | 0.36859087 | -1.546571 | 0.12196672 | 0.36611339 |
| AIP | 12.733279 | -0.358358397 | 0.42954287 | -0.8342785 | 0.40412404 | 0.68217314 |
| PITPNM1 | 22.5284451 | 0.004922011 | 0.6379514 | 0.00771534 | 0.99384411 | 0.99810704 |
| CDK2AP2 | 41.5289912 | -0.639263693 | 0.54274223 | -1.1778403 | 0.23886026 | 0.52912013 |
| GSTP1 | 270.745117 | -1.343147943 | 0.54331398 | -2.4721395 | 0.01343071 | 0.09373482 |
| NDUFV1 | 75.3228131 | 0.201838148 | 0.33594406 | 0.6008088 | 0.54796734 | 0.77516644 |
| UNC93B1 | 16.2980778 | -0.955066269 | 0.74204184 | -1.2870787 | 0.19806685 | 0.48045318 |
| NDUFS8 | 64.3960377 | -0.080261072 | 0.34702777 | -0.2312814 | 0.81709619 | 0.92248811 |
| TCIRG1 | 10.6169754 | 0.060220197 | 0.7562475 | 0.07963028 | 0.93653132 | 0.97594028 |
| CHKA | 112.490446 | 0.035264643 | 0.30277495 | 0.11647147 | 0.9072789 | 0.96180152 |
| KMT5B | 113.064707 | 0.169123616 | 0.20275168 | 0.83414162 | 0.40420118 | 0.68217314 |
| LRP5 | 32.7304539 | 0.058735504 | 0.37504722 | 0.15660829 | 0.87555357 | 0.95030689 |
| PPP6R3 | 413.406665 | 0.578939719 | 0.16392264 | 3.53178621 | 0.00041276 | 0.00746211 |
| TESMIN | 11.0557294 | -0.194505461 | 0.4221218 | -0.4607804 | 0.64495615 | 0.83387847 |
| CPT1A | 119.042626 | 1.184676349 | 0.28971643 | 4.08908934 | 4.33E-05 | 0.00124972 |
| MRPL21 | 13.2992326 | -0.407881745 | 0.40467744 | -1.0079182 | 0.31349375 | 0.60638615 |
| IGHMBP2 | 16.8688822 | -0.00334222 | 0.40374166 | -0.0082781 | 0.99339509 | 0.99794795 |
| TPCN2 | 29.0785071 | 0.568013861 | 0.3724941 | 1.52489357 | 0.12728565 | 0.37415717 |
| CCND1 | 36.2443285 | -0.258974439 | 0.35354137 | -0.7325152 | 0.46385415 | 0.72822697 |
| ORAOV1 | 23.1337148 | -0.291709193 | 0.32791395 | -0.8895907 | 0.37368572 | 0.65750816 |
| PPFIA1 | 146.932339 | 0.174218475 | 0.35653357 | 0.48864536 | 0.62509279 | 0.82285278 |
| AP002336.2 | 53.6458815 | -0.363223022 | 0.54309478 | -0.6688023 | 0.50362162 | 0.75108109 |
| AP000487.1 | 12.6058407 | 0.143683005 | 0.40023077 | 0.3590004 | 0.71959479 | 0.87503134 |
| CTTN | 167.836458 | -1.076088772 | 0.28603881 | -3.7620377 | 0.00016853 | 0.00369414 |
| SHANK2 | 469.437055 | 0.5648633 | 0.28100451 | 2.0101574 | 0.04441453 | 0.20103626 |
| DHCR7 | 14.4059373 | 0.304650973 | 0.55572375 | 0.54820578 | 0.58355061 | 0.79598942 |
| NADSYN1 | 35.3150774 | -0.709622738 | 0.24999714 | -2.8385235 | 0.00453228 | 0.04425738 |
| AP002495.2 | 64.6602455 | 0.684401114 | 0.27774119 | 2.46416856 | 0.01373315 | 0.09507992 |
| RNF121 | 77.1992962 | 0.166145518 | 0.2555734 | 0.65008924 | 0.51563458 | 0.75713618 |
| NUMA1 | 99.9569513 | 0.272703583 | 0.22274119 | 1.22430691 | 0.22083647 | 0.5089143 |
| LAMTOR1 | 45.5553919 | -1.116998692 | 0.27838421 | -4.0124355 | 6.01E-05 | 0.00164655 |
| ANAPC15 | 10.6005169 | -0.617911113 | 0.55594268 | -1.1114655 | 0.26636803 | 0.55763532 |
| FOLR3 | 14.2884867 | -0.925060824 | 0.66905078 | -1.3826467 | 0.16677324 | 0.43585929 |
| FOLR1 | 24.5450889 | -1.345857677 | 0.62270912 | -2.1612943 | 0.03067261 | 0.15992999 |
| INPPL1 | 25.3785023 | 0.592217357 | 0.36547167 | 1.62041933 | 0.10514223 | 0.33670353 |
| CLPB | 28.2572812 | 0.300606469 | 0.32243501 | 0.93230097 | 0.35118101 | 0.63773431 |
| ARAP1 | 32.0180797 | -0.227436395 | 0.39305951 | -0.5786309 | 0.56283822 | 0.78430399 |
| ATG16L2 | 24.8670641 | 0.241643319 | 0.35420892 | 0.68220563 | 0.49510894 | 0.74770824 |
| FCHSD2 | 216.733094 | -0.274799011 | 0.21275165 | -1.2916422 | 0.19648107 | 0.47795702 |
| P2RY6 | 55.2540642 | -1.935836616 | 0.70570433 | -2.743127 | 0.00608571 | 0.05480335 |
| ARHGEF17 | 34.4233515 | 0.159817253 | 0.25432775 | 0.62839093 | 0.52974788 | 0.76392254 |
| FAM168A | 159.12215 | 0.3798757 | 0.22268968 | 1.70585227 | 0.08803559 | 0.30539182 |
| RAB6A | 215.16023 | 0.009959046 | 0.27496459 | 0.03621938 | 0.97110744 | 0.988268 |
| MRPL48 | 45.6208906 | 0.726561412 | 0.32208168 | 2.25582964 | 0.0240813 | 0.13578849 |
| COA4 | 26.8480598 | -1.066246364 | 0.32629225 | -3.2677649 | 0.001084 | 0.01544593 |
| PAAF1 | 37.9877506 | -0.053689913 | 0.30636428 | -0.1752486 | 0.86088429 | 0.94510123 |
| UCP2 | 16.5706319 | -1.132677631 | 0.8454527 | -1.3397292 | 0.18033342 | 0.45647562 |
| C2CD3 | 68.9712449 | 0.286203884 | 0.26835456 | 1.06651397 | 0.28619137 | 0.57870647 |
| PPME1 | 79.4460718 | -0.581017629 | 0.28543903 | -2.0355227 | 0.0417983 | 0.19510926 |
| P4HA3 | 13.0315103 | -0.697811773 | 0.67786949 | -1.0294191 | 0.3032828 | 0.59717496 |
| PGM2L1 | 30.4159872 | -1.263198386 | 0.39688253 | -3.1828017 | 0.00145857 | 0.01936371 |
| POLD3 | 41.6686939 | -0.250777403 | 0.35585701 | -0.704714 | 0.48098826 | 0.74012562 |
| RNF169 | 115.69291 | -0.317123516 | 0.27699253 | -1.1448811 | 0.25225841 | 0.54367492 |
| XRRA1 | 102.317985 | 0.115300273 | 0.43680605 | 0.26396217 | 0.79180907 | 0.91111549 |
| SPCS2 | 88.7028816 | -0.267550216 | 0.24563371 | -1.0892243 | 0.27605497 | 0.56798847 |
| NEU3 | 15.9739881 | -0.340191148 | 0.36569753 | -0.9302528 | 0.35224022 | 0.63819534 |
| ARRB1 | 51.6239882 | 0.801039646 | 0.31603345 | 2.53466726 | 0.01125542 | 0.08307399 |
| RPS3 | 206.340973 | -0.361479568 | 0.3820691 | -0.9461105 | 0.34409225 | 0.63199115 |
| SERPINH1 | 22.7681799 | -0.196205051 | 0.54447827 | -0.3603542 | 0.71858224 | 0.87421284 |
| UVRAG | 268.246244 | 0.542649831 | 0.25602649 | 2.11950658 | 0.03404768 | 0.17076883 |
| THAP12 | 20.9264638 | 0.111065286 | 0.34527282 | 0.32167399 | 0.74769968 | 0.88935207 |
| AP002360.1 | 9.29383511 | 1.044353972 | 0.63037143 | 1.65672795 | 0.09757449 | 0.32249355 |
| EMSY | 110.92496 | 0.331421614 | 0.28856656 | 1.14851013 | 0.25075804 | 0.54232519 |
| ACER3 | 131.356729 | 0.107185421 | 0.31779438 | 0.33727915 | 0.73590647 | 0.88432217 |
| CAPN5 | 26.3793656 | 0.067849543 | 0.61173416 | 0.11091344 | 0.91168498 | 0.96399248 |
| PAK1 | 105.72285 | 0.899353658 | 0.44482576 | 2.02181112 | 0.04319587 | 0.19839672 |
| CLNS1A | 63.4806697 | -0.158130494 | 0.24603548 | -0.6427142 | 0.52040957 | 0.76073677 |
| RSF1 | 218.102328 | 0.069116578 | 0.18596677 | 0.3716609 | 0.71014534 | 0.86990693 |
| AAMDC | 53.9627019 | -0.10204624 | 0.390789 | -0.2611287 | 0.79399323 | 0.91176607 |
| AP002812.2 | 22.799678 | -0.79687543 | 0.36101318 | -2.2073306 | 0.02729097 | 0.14774125 |
| INTS4 | 57.5916287 | -0.042854898 | 0.30825415 | -0.1390246 | 0.88943074 | 0.95702609 |

| | | | | | | |
|---|---|---|---|---|---|---|
| ALG8 | 28.6282489 | -0.226794162 | 0.29506538 | -0.7686234 | 0.4421169 | 0.71263553 |
| GAB2 | 131.613737 | -1.181286725 | 0.54745661 | -2.1577723 | 0.03094554 | 0.16101056 |
| AP003086.2 | 34.2640988 | -1.456974863 | 0.64736065 | -2.2506386 | 0.02440844 | 0.13661343 |
| NARS2 | 81.0205695 | 0.516802749 | 0.20900609 | 2.47266843 | 0.01341085 | 0.09373482 |
| PRCP | 60.4688081 | 0.381605396 | 0.3076209 | 1.24050541 | 0.21478851 | 0.50124063 |
| RAB30 | 31.0614984 | 0.99421265 | 0.32664544 | 3.04370586 | 0.00233683 | 0.0272113 |
| RAB30-AS1 | 87.8261199 | 0.209601432 | 0.33837013 | 0.61944425 | 0.53562374 | 0.76809518 |
| PCF11 | 82.6388712 | 0.26080876 | 0.28077939 | 0.9288743 | 0.35295423 | 0.63892716 |
| ANKRD42 | 38.7741184 | -0.265886872 | 0.27439155 | -0.9690053 | 0.33254254 | 0.62174935 |
| CCDC90B | 34.6151299 | -0.677319158 | 0.33341275 | -2.0314735 | 0.04220699 | 0.19607262 |
| AP000446.1 | 26.8741619 | 0.792186812 | 0.4811224 | 1.64653905 | 0.09965283 | 0.3259122 |
| DLG2 | 636.582955 | 0.58416774 | 0.50144228 | 1.16497505 | 0.24402909 | 0.5346132 |
| TMEM126B | 21.4108442 | -0.272132409 | 0.32892849 | -0.82733 | 0.40805005 | 0.68657965 |
| CREBZF | 71.2782477 | -0.134751101 | 0.24396404 | -0.55234 | 0.58071541 | 0.7947038 |
| SYTL2 | 442.877718 | -0.661020322 | 0.34941329 | -1.8918007 | 0.05851754 | 0.23922527 |
| PICALM | 481.706531 | 0.745955143 | 0.19760136 | 3.77505065 | 0.00015998 | 0.00356724 |
| EED | 41.902121 | 0.624953444 | 0.31833581 | 1.96318924 | 0.04962419 | 0.21653045 |
| HIKESHI | 43.1240611 | 0.364157796 | 0.28509693 | 1.27731223 | 0.20149204 | 0.4848871 |
| AP001831.1 | 37.3705955 | 0.021695367 | 0.49107886 | 0.04417899 | 0.96476173 | 0.98660829 |
| ME3 | 146.268316 | 1.435618566 | 0.24201073 | 5.93204512 | 2.99E-09 | 4.34E-07 |
| AP003059.1 | 17.7893208 | 0.832419124 | 0.49222687 | 1.69112898 | 0.09081217 | 0.31034142 |
| AP003059.2 | 13.2374498 | 0.573942047 | 0.55564898 | 1.03292199 | 0.30164041 | 0.59542937 |
| PRSS23 | 92.4897773 | 1.171378222 | 0.45079713 | 2.5984598 | 0.0093643 | 0.07341783 |
| AP001528.2 | 57.5645433 | 0.698190582 | 0.6273591 | 1.1129042 | 0.26574957 | 0.55713311 |
| FZD4 | 75.1049426 | 0.371509686 | 0.51836753 | 0.71669167 | 0.47356437 | 0.73418512 |
| AP001528.1 | 17.2045398 | 0.702507046 | 0.48283281 | 1.45496957 | 0.14567768 | 0.40408495 |
| TMEM135 | 165.891928 | 0.380269481 | 0.24207467 | 1.57087677 | 0.11621128 | 0.35543425 |
| RAB38 | 21.1008919 | 0.667448887 | 0.50501105 | 1.32165204 | 0.18628404 | 0.46377434 |
| CTSC | 49.1255011 | 0.239753623 | 0.38676889 | 0.61988859 | 0.53533114 | 0.76808755 |
| NOX4 | 12.6793996 | 1.374058118 | 0.5527577 | 2.48582359 | 0.0129252 | 0.09183133 |
| CHORDC1 | 41.7853341 | 0.707807543 | 0.42905523 | 1.64968864 | 0.09900664 | 0.32503708 |
| SLC36A4 | 106.890333 | 0.288180224 | 0.26617961 | 1.08265327 | 0.27896235 | 0.57145957 |
| SMCO4 | 48.7224768 | -0.521731508 | 0.3948095 | -1.3214766 | 0.1863425 | 0.4638078 |
| CEP295 | 53.0875067 | -0.037276741 | 0.27045848 | -0.137828 | 0.89037638 | 0.95702609 |
| TAF1D | 112.31504 | -0.001236149 | 0.23326849 | -0.0052993 | 0.99577183 | 0.99868531 |
| PANX1 | 13.2988063 | -0.804662982 | 0.38885178 | -2.0693308 | 0.03851505 | 0.1841216 |
| MRE11 | 46.494938 | -0.183293055 | 0.27878143 | -0.6574794 | 0.51087269 | 0.7549696 |
| ANKRD49 | 12.0781697 | -0.219111975 | 0.45498704 | -0.4815785 | 0.6301054 | 0.82524878 |
| AMOTL1 | 76.3183426 | -0.73329541 | 0.28601709 | -2.5638167 | 0.01035282 | 0.07880617 |
| CWC15 | 24.3835942 | -0.261363688 | 0.29199496 | -0.8950966 | 0.37073547 | 0.65494451 |
| SRSF8 | 21.9054664 | -0.381874366 | 0.46496589 | -0.8212954 | 0.411478 | 0.68899849 |
| ENDOD1 | 35.3558281 | -1.100652867 | 0.33931695 | -3.2437309 | 0.00117975 | 0.01646856 |
| AP000787.1 | 27.7047333 | -0.317714637 | 0.34504037 | -0.9208042 | 0.35715264 | 0.64174726 |
| SESN3 | 50.7787427 | -0.672771945 | 0.32484899 | -2.0710298 | 0.03835601 | 0.18353163 |
| FAM76B | 30.7319564 | -0.232888819 | 0.33789167 | -0.6892411 | 0.49067157 | 0.7456686 |
| CEP57 | 102.707147 | 0.095622115 | 0.20589138 | 0.46442991 | 0.64233976 | 0.83318468 |
| MTMR2 | 67.1136318 | 0.19175959 | 0.28902173 | 0.66347811 | 0.50702439 | 0.75373237 |
| MAML2 | 1169.41882 | 0.886885042 | 0.27070178 | 3.27624384 | 0.00105198 | 0.01524257 |
| CCDC82 | 43.2954343 | -0.620602134 | 0.30483516 | -2.0358614 | 0.04176428 | 0.1950388 |
| JRKL | 13.0679135 | -1.324655929 | 0.53560945 | -2.473175 | 0.01339185 | 0.09373482 |
| CNTN5 | 11.9999257 | 0.143294139 | 0.5391445 | 0.26578058 | 0.7904082 | 0.91080793 |
| ARHGAP42 | 41.2584575 | 0.554351002 | 0.34137553 | 1.62387445 | 0.10440261 | 0.33555699 |
| CEP126 | 32.8521521 | -0.218963534 | 0.38719855 | -0.5655071 | 0.57172889 | 0.78911455 |
| CFAP300 | 20.4900647 | -0.742065092 | 0.52814337 | -1.4050448 | 0.16000796 | 0.42782299 |
| YAP1 | 533.798695 | -0.13259245 | 0.19694215 | -0.6732558 | 0.50078455 | 0.75071921 |
| BIRC3 | 104.841787 | -1.738270069 | 0.66146219 | -2.6279205 | 0.00859086 | 0.06925119 |
| BIRC2 | 51.8885747 | -0.040445248 | 0.24848024 | -0.1627705 | 0.87069915 | 0.94844301 |
| TMEM123 | 137.211627 | -0.615276988 | 0.2742234 | -2.2437071 | 0.02485125 | 0.1383879 |
| MMP7 | 220.484378 | -2.992837629 | 0.56675954 | -5.2806127 | 1.29E-07 | 1.07E-05 |
| DCUN1D5 | 40.1034174 | -0.152004727 | 0.31454925 | -0.4832462 | 0.62892094 | 0.82494825 |
| DYNC2H1 | 397.900918 | 0.698872039 | 0.31899712 | 2.19084122 | 0.02846328 | 0.15201076 |
| PDGFD | 25.6875536 | -0.366720338 | 0.5411199 | -0.6777063 | 0.49795796 | 0.74971166 |
| CASP4 | 61.5024381 | -1.507077387 | 0.34553481 | -4.361579 | 1.29E-05 | 0.00045557 |
| MSANTD4 | 15.0863904 | -0.500440591 | 0.40489394 | -1.2359795 | 0.2164662 | 0.50339385 |
| KBTBD3 | 17.335482 | 0.693603365 | 0.41131687 | 1.68629933 | 0.09173816 | 0.311704 |
| AASDHPPT | 31.7976212 | -0.034113218 | 0.32340677 | -0.1054808 | 0.91599427 | 0.96518083 |
| CWF19L2 | 106.42752 | 0.23213412 | 0.23549635 | 0.98572279 | 0.32426916 | 0.61460758 |
| ALKBH8 | 20.7813901 | 0.621870488 | 0.38117526 | 1.63145554 | 0.10279423 | 0.33228309 |
| SLC35F2 | 31.7016769 | 0.253401125 | 0.39982189 | 0.63378502 | 0.52622113 | 0.76221787 |
| CUL5 | 175.560967 | 0.47968952 | 0.21634872 | 2.21720521 | 0.02660907 | 0.14581205 |
| ACAT1 | 69.7997372 | 0.56999836 | 0.22940805 | 2.48464844 | 0.01296794 | 0.09194477 |
| NPAT | 53.4197498 | 0.18033395 | 0.25832434 | 0.6980912 | 0.48512016 | 0.74285379 |
| ATM | 138.055119 | -0.220381045 | 0.22434112 | -0.982348 | 0.32592845 | 0.6159877 |
| C11orf65 | 43.9783583 | 0.683763954 | 0.57723216 | 1.18455624 | 0.23619295 | 0.5264517 |
| KDELC2 | 24.447681 | 1.510671523 | 0.51780112 | 2.9174744 | 0.00352879 | 0.0367951 |
| EXPH5 | 126.521087 | 0.709644875 | 0.35274692 | 2.01176773 | 0.04424443 | 0.20044552 |
| DDX10 | 112.835154 | 0.449861664 | 0.25387619 | 1.77197266 | 0.0763991 | 0.28042437 |
| RDX | 146.762431 | 0.20208599 | 0.35814792 | 0.56425287 | 0.57258205 | 0.78978194 |
| ZC3H12C | 124.838092 | 0.680825731 | 0.57590779 | 1.18217836 | 0.23713494 | 0.52752411 |
| FDX1 | 88.3934602 | 0.571192453 | 0.30412253 | 1.87816556 | 0.06035852 | 0.24332668 |
| COLCA1 | 21.3560385 | 2.06598583 | 0.41847542 | 4.93693474 | 7.94E-07 | 4.68E-05 |
| COLCA2 | 10.900664 | 0.623895993 | 0.49243298 | 1.26696631 | 0.20516734 | 0.48964616 |
| SIK2 | 322.397327 | 0.776172296 | 0.52956151 | 1.46568865 | 0.14273313 | 0.39968379 |
| PPP2R1B | 49.2378967 | -0.385381084 | 0.27797139 | -1.3864056 | 0.1656231 | 0.43438116 |
| C11orf1 | 14.06023 | 0.445452104 | 0.45908134 | 0.97031193 | 0.33189103 | 0.62154906 |
| CRYAB | 220.518828 | -1.550709332 | 0.6300159 | -2.4613813 | 0.01384032 | 0.095565 |
| DIXDC1 | 54.9747754 | 0.010275518 | 0.30063882 | 0.03417895 | 0.97273446 | 0.98856766 |
| DLAT | 43.3335737 | 0.654944921 | 0.29570867 | 2.21483162 | 0.02677162 | 0.14614285 |
| NKAPD1 | 15.423902 | -0.173344777 | 0.46760694 | -0.3707062 | 0.71085639 | 0.87028547 |
| TIMM8B | 24.4986796 | -0.118042954 | 0.41193 | -0.2865607 | 0.7744487 | 0.90202041 |
| AP002884.3 | 28.5692793 | 0.245018306 | 0.73401942 | 0.33380357 | 0.73852779 | 0.88591857 |
| PTS | 52.1728013 | -0.500415192 | 0.40452254 | -1.2370515 | 0.21606798 | 0.5028083 |
| AP002884.1 | 37.1858639 | -1.143429573 | 0.54263182 | -2.1071923 | 0.03510092 | 0.17415035 |
| TTC12 | 47.641328 | 0.19932425 | 0.30515595 | 0.65318816 | 0.51363498 | 0.75647799 |
| ZW10 | 22.3482731 | -0.15713748 | 0.35173342 | -0.4467516 | 0.6550544 | 0.83931753 |
| USP28 | 36.2738286 | -0.37891304 | 0.33940518 | -1.1164032 | 0.26424957 | 0.55579808 |
| ZBTB16 | 31.6395098 | 0.079895863 | 0.79325037 | 0.1007196 | 0.91977305 | 0.96767789 |
| NNMT | 44.0166613 | -6.549775459 | 1.09905229 | -5.9594757 | 2.53E-09 | 3.78E-07 |
| REXO2 | 34.0896364 | -0.078352403 | 0.31097346 | -0.2519585 | 0.80107315 | 0.91484195 |
| CADM1 | 189.930979 | -0.330407902 | 0.44372589 | -0.7446216 | 0.45650048 | 0.72298278 |
| BUD13 | 11.7223697 | 0.185403403 | 0.45642452 | 0.40620824 | 0.6845896 | 0.85621489 |
| SIK3 | 778.494419 | 0.902421952 | 0.35679145 | 2.52927009 | 0.01143 | 0.08422882 |
| PAFAH1B2 | 87.6143763 | -0.178414462 | 0.22713255 | -0.7855081 | 0.4321557 | 0.70480237 |
| SIDT2 | 21.0035214 | 0.06002157 | 0.33434954 | 0.17951743 | 0.85753143 | 0.94359207 |
| PCSK7 | 68.1597669 | 0.529732067 | 0.24770848 | 2.13853017 | 0.03247374 | 0.16569809 |
| RNF214 | 53.1223153 | -0.212550931 | 0.33523041 | -0.6340443 | 0.52605191 | 0.76221787 |
| BACE1 | 35.6509677 | 0.340592471 | 0.32248991 | 1.05613374 | 0.29090712 | 0.58249897 |
| CEP164 | 41.9430818 | 0.303204462 | 0.28071397 | 1.08011888 | 0.28008924 | 0.57312294 |
| DSCAML1 | 243.847948 | -0.635372585 | 0.70751804 | -0.8980302 | 0.36916944 | 0.65369102 |
| FXYD2 | 68.2859308 | 1.308679921 | 0.49464234 | 2.64570948 | 0.00815198 | 0.0668644 |
| AP000757.1 | 21.4081638 | 1.271230412 | 0.53056076 | 2.3960129 | 0.01657451 | 0.10698657 |
| TMPRSS4 | 25.3173933 | -5.092278014 | 0.92096898 | -5.5292612 | 3.22E-08 | 3.20E-06 |
| MPZL3 | 21.3888946 | -0.559541383 | 0.37346731 | -1.4982339 | 0.1340725 | 0.38473951 |

| | | | | | | |
|---|---|---|---|---|---|---|
| MPZL2 | 48.4805511 | -2.889474465 | 0.61142119 | -4.7258331 | 2.29E-06 | 0.00011084 |
| UBE4A | 61.2361523 | 0.233295038 | 0.23437517 | 0.99539142 | 0.31954593 | 0.611915 |
| ATP5MG | 121.486395 | 0.123291401 | 0.36483631 | 0.33793621 | 0.73541126 | 0.8841413 |
| KMT2A | 211.714108 | -0.367836421 | 0.16085221 | -2.2867974 | 0.02220765 | 0.12932913 |
| TMEM25 | 13.3435844 | 0.156751099 | 0.45591641 | 0.34381544 | 0.73098509 | 0.88180426 |
| IFT46 | 19.5937272 | 0.39490322 | 0.3453019 | 1.14364624 | 0.25277038 | 0.54433601 |
| ARCN1 | 35.7557438 | -0.095862343 | 0.31442768 | -0.3048788 | 0.76045844 | 0.89627175 |
| PHLDB1 | 47.4006978 | 0.306458081 | 0.37114426 | 0.82571149 | 0.40896777 | 0.6869518 |
| AP002954.1 | 16.2844288 | -0.330325653 | 0.4200083 | -0.7864741 | 0.43158978 | 0.70454366 |
| DDX6 | 112.860111 | 0.074001375 | 0.16997359 | 0.43536985 | 0.66329399 | 0.84278344 |
| BCL9L | 20.3888475 | -0.866246848 | 0.45684713 | -1.8961416 | 0.05794132 | 0.23800299 |
| CCDC84 | 80.1250498 | 0.246868991 | 0.32996951 | 0.74815698 | 0.45436548 | 0.72190427 |
| RPS25 | 116.361488 | -0.620970212 | 0.41786197 | -1.4860654 | 0.13726182 | 0.39041173 |
| TRAPPC4 | 25.7685851 | -0.578586494 | 0.38586757 | -1.4994432 | 0.1337587 | 0.38416007 |
| HYOU1 | 23.2216063 | -0.23769688 | 0.44759013 | -0.5310593 | 0.59537772 | 0.80357658 |
| VPS11 | 14.5322574 | 0.04392416 | 0.35936744 | 0.12222632 | 0.90271978 | 0.96112228 |
| HINFP | 18.7438059 | -0.254263329 | 0.34081706 | -0.7460405 | 0.45564296 | 0.72243279 |
| NLRX1 | 15.0292249 | -0.584222536 | 0.38703347 | -1.5094884 | 0.13117402 | 0.38023487 |
| CBL | 73.037525 | -0.295778321 | 0.40319387 | -0.7335883 | 0.46319967 | 0.72797604 |
| USP2 | 12.4939188 | 3.535829366 | 0.53034942 | 6.66698074 | 2.61E-11 | 7.60E-09 |
| THY1 | 10.7774387 | 3.22197789 | 0.90983869 | 3.54126279 | 0.00039822 | 0.00724811 |
| NECTIN1 | 11.4478685 | 0.778379239 | 0.53992707 | 1.44163774 | 0.1494046 | 0.40989778 |
| OAF | 50.6615381 | 0.506795501 | 0.36971181 | 1.37078526 | 0.17044191 | 0.44117902 |
| TMEM136 | 13.0470438 | -0.918084376 | 0.43748576 | -2.0985469 | 0.03585686 | 0.17565259 |
| ARHGEF12 | 502.322672 | 0.027945337 | 0.17485116 | 0.15982357 | 0.87302006 | 0.94911911 |
| SC5D | 25.8999575 | 0.482654249 | 0.46965987 | 1.02766764 | 0.3041062 | 0.59822456 |
| SORL1 | 190.896363 | 0.845998159 | 0.35795819 | 2.36339935 | 0.01810814 | 0.11306142 |
| UBASH3B | 23.9686417 | -2.114426586 | 0.65240737 | -3.2409606 | 0.00119128 | 0.0165845 |
| JHY | 18.0770651 | -0.9730127 | 0.67829155 | -1.4345051 | 0.15142817 | 0.41270185 |
| HSPA8 | 100.890794 | 0.664378903 | 0.47360076 | 1.40282481 | 0.16066909 | 0.42803048 |
| GRAMD1B | 41.3177994 | 1.663151043 | 0.50073428 | 3.3214244 | 0.00089559 | 0.01356822 |
| VWA5A | 26.7458067 | -1.15254729 | 0.42612906 | -2.7046907 | 0.0068368 | 0.05928681 |
| TBRG1 | 51.7758373 | 0.047030104 | 0.24314928 | 0.1934207 | 0.8466295 | 0.93757726 |
| SIAE | 43.4085945 | 0.261343075 | 0.36839882 | 0.70940258 | 0.47807469 | 0.7378465 |
| MSANTD2 | 63.8033827 | 0.588165492 | 0.32739645 | 1.7964932 | 0.07241611 | 0.27172106 |
| EI24 | 45.3354087 | -0.512763511 | 0.36948255 | -1.3877882 | 0.16520155 | 0.43404907 |
| STT3A | 30.6618519 | -0.228147346 | 0.41748781 | -0.5464767 | 0.58473833 | 0.79661125 |
| RPUSD4 | 11.409078 | -0.121317147 | 0.45119638 | -0.2688788 | 0.78802294 | 0.90950172 |
| FAM118B | 23.1130596 | -0.632214555 | 0.46452202 | -1.3610002 | 0.17351363 | 0.44612537 |
| SRPRA | 29.5074054 | -0.502669546 | 0.32946142 | -1.5257311 | 0.12707683 | 0.37382797 |
| FOXRED1 | 21.3942 | 0.038955989 | 0.35090489 | 0.11101581 | 0.91160381 | 0.96399248 |
| TIRAP | 10.4565693 | 0.768823894 | 0.51790018 | 1.48450208 | 0.13767577 | 0.39109388 |
| DCPS | 36.3962278 | -0.435762442 | 0.54909307 | -0.793604 | 0.42742602 | 0.70206359 |
| ST3GAL4 | 37.8245396 | -0.782289156 | 0.68293467 | -1.1454817 | 0.25200968 | 0.54334972 |
| ETS1 | 87.9584439 | -0.324163718 | 0.43346399 | -0.7478446 | 0.45455388 | 0.72190427 |
| KCNJ1 | 265.479011 | 0.829379814 | 0.44247502 | 1.87441049 | 0.06087387 | 0.24477853 |
| ARHGAP32 | 225.619435 | 0.20274261 | 0.25910863 | 0.78246183 | 0.43394321 | 0.70597357 |
| BARX2 | 144.832807 | 0.278228742 | 0.40775164 | 0.68234855 | 0.49501858 | 0.74770824 |
| TMEM45B | 86.9895954 | -0.226821264 | 0.29926781 | -0.7579207 | 0.44849846 | 0.71768114 |
| NFRKB | 35.736858 | 0.37562235 | 0.33073659 | 1.13571453 | 0.25607606 | 0.54710413 |
| PRDM10 | 36.1437726 | 0.283822432 | 0.31581137 | 0.89870872 | 0.36880783 | 0.65369102 |
| APLP2 | 229.530994 | -0.772919175 | 0.37331886 | -2.0703995 | 0.03841495 | 0.18372833 |
| ST14 | 98.8921252 | -0.512261126 | 0.29394585 | -1.7427058 | 0.08138502 | 0.2916273 |
| ZBTB44 | 146.559693 | 0.304176828 | 0.23310409 | 1.30489701 | 0.19192792 | 0.47212817 |
| AP002986.1 | 24.6121646 | 1.546605212 | 0.76747457 | 2.01518757 | 0.043885 | 0.19995722 |
| SNX19 | 56.9767297 | -0.064136597 | 0.38453328 | -0.1667908 | 0.86753469 | 0.94761794 |
| JAM3 | 18.8582805 | 0.398876152 | 0.42555123 | 0.93731641 | 0.34859582 | 0.63548432 |
| NCAPD3 | 19.6086798 | -0.118647906 | 0.32907614 | -0.3605485 | 0.718437 | 0.87415486 |
| VPS26B | 22.9555597 | 0.011356802 | 0.37283338 | 0.0304608 | 0.97569956 | 0.99024058 |
| ACAD8 | 17.3631948 | 0.20678781 | 0.40580891 | 0.50956942 | 0.61035315 | 0.81376433 |
| ZMYND11 | 201.994088 | 0.561511592 | 0.14880681 | 3.77342667 | 0.00016102 | 0.00357507 |
| DIP2C | 391.968949 | -0.854197916 | 0.39455312 | -2.1649757 | 0.03038954 | 0.15900106 |
| LARP4B | 262.523651 | 0.047246675 | 0.21363246 | 0.2211587 | 0.82496887 | 0.92670693 |
| GTPBP4 | 26.2384012 | -0.062879534 | 0.32115336 | -0.1957929 | 0.8447723 | 0.93689787 |
| IDI1 | 58.0378368 | 0.831067051 | 0.50120033 | 1.65815343 | 0.0972865 | 0.32180307 |
| WDR37 | 111.935418 | 0.355334257 | 0.24546063 | 1.44762218 | 0.14772275 | 0.40745374 |
| PFKP | 144.483507 | -0.031775521 | 0.49381902 | -0.0643465 | 0.94869434 | 0.97969618 |
| PITRM1 | 40.0377661 | 0.271377342 | 0.2969486 | 0.91388659 | 0.36077644 | 0.64494385 |
| KLF6 | 219.814505 | -0.218323213 | 0.67229175 | -0.3247447 | 0.74537427 | 0.88808711 |
| AKR1C3 | 20.8569453 | -1.777819997 | 0.75865328 | -2.3433893 | 0.01910943 | 0.11746143 |
| NET1 | 12.7059962 | -0.242564215 | 0.44534006 | -0.5446719 | 0.58597919 | 0.79714217 |
| ASB13 | 14.6249884 | -0.334549275 | 0.44925706 | -0.7446723 | 0.45646986 | 0.72298278 |
| FAM208B | 169.378609 | 0.035571865 | 0.33296306 | 0.10683427 | 0.91492046 | 0.96464134 |
| GDI2 | 113.21098 | -0.537825672 | 0.20537746 | -2.6187181 | 0.00882608 | 0.07049178 |
| ANKRD16 | 17.1429343 | 0.085858858 | 0.42359942 | 0.2026888 | 0.83937828 | 0.93543959 |
| FBH1 | 80.5209238 | 0.14952137 | 0.22264897 | 0.67155653 | 0.50186606 | 0.75108109 |
| IL15RA | 15.3256437 | -1.641369897 | 0.59281421 | -2.7687762 | 0.00562673 | 0.05161757 |
| RBM17 | 63.2665019 | 0.304616713 | 0.23513329 | 1.29550655 | 0.19514554 | 0.47616992 |
| PFKFB3 | 289.418912 | -0.950858404 | 0.36035223 | -2.6386916 | 0.00832267 | 0.06783236 |
| AL137145.2 | 56.6059883 | -1.165198819 | 0.37830929 | -3.0800164 | 0.00206989 | 0.02514626 |
| PRKCQ | 58.2150624 | -0.487941843 | 0.34311905 | -1.4220774 | 0.15500379 | 0.41835185 |
| SFMBT2 | 96.1796266 | 0.290331081 | 0.26462869 | 1.09712626 | 0.2725862 | 0.56479948 |
| KIN | 25.0320948 | -0.034144375 | 0.30820733 | -0.1107838 | 0.91178781 | 0.96400225 |
| ATP5F1C | 146.621745 | 0.740050171 | 0.22022794 | 3.3603827 | 0.00077835 | 0.01224201 |
| TAF3 | 178.374265 | 0.318580635 | 0.21942666 | 1.45187755 | 0.14653566 | 0.40573108 |
| GATA3 | 297.105486 | 0.460853111 | 0.29685403 | 1.55245697 | 0.12055292 | 0.36346292 |
| GATA3-AS1 | 34.9728391 | 0.440505146 | 0.39793789 | 1.1069696 | 0.26830707 | 0.56010121 |
| USP6NL | 104.899444 | -0.295053343 | 0.2110503 | -1.3980238 | 0.16210593 | 0.42953067 |
| ECHDC3 | 15.5146081 | -0.209609744 | 0.50099845 | -0.418384 | 0.67566637 | 0.85167196 |
| PROSER2 | 38.6952547 | -0.542355901 | 0.41312121 | -1.3128251 | 0.18924189 | 0.46819643 |
| UPF2 | 103.375295 | 0.129578169 | 0.27934251 | 0.46386842 | 0.64274202 | 0.83318468 |
| DHTKD1 | 41.8144227 | -0.204018968 | 0.26361545 | -0.7739265 | 0.43897428 | 0.71027377 |
| SEC61A2 | 18.492536 | 0.158135792 | 0.33502255 | 0.47201537 | 0.63691582 | 0.82989931 |
| NUDT5 | 40.1185085 | -0.249086569 | 0.26276192 | -0.9479554 | 0.34315218 | 0.6310379 |
| CDC123 | 43.3217419 | 0.02777143 | 0.23964105 | 0.11588762 | 0.90774161 | 0.96209404 |
| OPTN | 39.2005417 | -1.386085331 | 0.42232097 | -3.2820661 | 0.00103049 | 0.01507976 |
| PHYH | 33.121866 | 0.556863969 | 0.31471526 | 1.76942156 | 0.07682355 | 0.28130164 |
| SEPHS1 | 32.7591461 | 0.243733858 | 0.32374638 | 0.7528543 | 0.45153747 | 0.71974919 |
| BEND7 | 107.52375 | -0.044254596 | 0.22940139 | -0.1929134 | 0.84702681 | 0.93758141 |
| PRPF18 | 15.7064895 | -0.289683569 | 0.37342673 | -0.7757441 | 0.43790011 | 0.70931556 |
| AL157392.5 | 30.3041636 | -0.251321974 | 0.32024144 | -0.784789 | 0.43257726 | 0.70501676 |
| FRMD4A | 144.622253 | 0.057864635 | 0.31974778 | 0.18096962 | 0.85639142 | 0.94330228 |
| FAM107B | 347.013887 | 0.482920326 | 0.26600825 | 1.81543368 | 0.06945729 | 0.26541135 |
| DCLRE1C | 32.584151 | -0.246722312 | 0.28123266 | -0.877289 | 0.38032969 | 0.66406325 |
| RPP38 | 42.7611391 | 0.295555717 | 0.34269078 | 0.86245598 | 0.38843665 | 0.6704095 |
| NMT2 | 48.1175609 | 0.225777753 | 0.33685126 | 0.67025948 | 0.50269239 | 0.75108109 |
| FAM171A1 | 389.865943 | 0.779264576 | 0.31614011 | 2.46493421 | 0.01370384 | 0.09506867 |
| MINDY3 | 75.2047174 | 0.131953817 | 0.23471187 | 0.5621949 | 0.57398323 | 0.79097021 |
| PTER | 77.5560595 | -0.404515807 | 0.29066717 | -1.3916804 | 0.1640192 | 0.43270826 |
| RSU1 | 91.0851043 | -0.480239044 | 0.34457152 | -1.3937282 | 0.1633997 | 0.43173729 |
| CUBN | 43.0226791 | 0.996384785 | 0.44033008 | 2.26281335 | 0.0236472 | 0.13490566 |
| TRDMT1 | 14.9213052 | 0.258764383 | 0.40694223 | 0.63587497 | 0.52485792 | 0.76221787 |
| VIM | 32.3933776 | -2.179795341 | 0.43360662 | -5.0271266 | 4.98E-07 | 3.23E-05 |

| | | | | | | |
|---|---|---|---|---|---|---|
| HACD1 | 34.6282421 | -1.734285686 | 0.76486504 | -2.2674401 | 0.02336335 | 0.13386499 |
| STAM | 105.906745 | -0.666819372 | 0.57049018 | -1.1688534 | 0.24246271 | 0.53317493 |
| CACNB2 | 13.6689376 | 0.435117819 | 0.59079422 | 0.73649641 | 0.46142865 | 0.72663375 |
| AL450384.2 | 22.5248964 | 0.13127809 | 0.39045004 | 0.3362225 | 0.73670309 | 0.88466199 |
| NSUN6 | 68.0314673 | 0.222759217 | 0.29838066 | 0.74656051 | 0.45532889 | 0.72210564 |
| ARL5B | 34.1485105 | -0.608661685 | 0.53118923 | -1.1458472 | 0.25185838 | 0.54334972 |
| MALRD1 | 71.8034874 | 1.899190043 | 0.61705773 | 3.07781583 | 0.00208524 | 0.02527308 |
| PLXDC2 | 336.839912 | 0.431644042 | 0.27727866 | 1.55671572 | 0.119538 | 0.36193716 |
| NEBL | 537.932021 | 1.069186939 | 0.36979223 | 2.891318 | 0.0038363 | 0.0390529 |
| MLLT10 | 332.851488 | 0.233793499 | 0.19977043 | 1.17031085 | 0.2418759 | 0.53271231 |
| DNAJC1 | 298.571604 | 0.371090317 | 0.21360448 | 1.73727782 | 0.08233815 | 0.29401996 |
| BMI1 | 18.3497603 | 0.313785029 | 0.51171347 | 0.61320455 | 0.5397411 | 0.77078082 |
| PIP4K2A | 146.026562 | 0.279045907 | 0.24378887 | 1.14462119 | 0.25236611 | 0.54379328 |
| MSRB2 | 36.0227038 | -0.233199008 | 0.28220638 | -0.8263421 | 0.40861007 | 0.68692901 |
| OTUD1 | 19.3430196 | 0.684780546 | 0.41646335 | 1.64427563 | 0.10011928 | 0.326628 |
| KIAA1217 | 599.521729 | 0.752485129 | 0.22505282 | 3.34359337 | 0.00082701 | 0.01273519 |
| ARHGAP21 | 174.725535 | 0.001987967 | 0.23454777 | 0.00847575 | 0.99323741 | 0.9979411 |
| PRTFDC1 | 27.6635188 | 1.218763769 | 0.35672633 | 3.41652313 | 0.00063426 | 0.01040474 |
| PDSS1 | 24.7560243 | 0.609986633 | 0.37360112 | 1.63272162 | 0.10252756 | 0.33168278 |
| ABI1 | 167.286844 | 0.171590143 | 0.22678651 | 0.75661531 | 0.44928037 | 0.7180393 |
| ANKRD26 | 157.819106 | 0.388183404 | 0.25435393 | 1.52615453 | 0.12697138 | 0.3737312 |
| YME1L1 | 109.654772 | 0.009969877 | 0.22300632 | 0.0447067 | 0.96434109 | 0.98660829 |
| MASTL | 20.8029871 | -0.205558094 | 0.38333632 | -0.5362343 | 0.59179662 | 0.80072087 |
| ACBD5 | 60.7842066 | -0.067185942 | 0.22760059 | -0.2951923 | 0.76784698 | 0.89979146 |
| RAB18 | 105.880304 | -0.40673128 | 0.23322581 | -1.7439377 | 0.08116996 | 0.29136339 |
| MPP7 | 121.991975 | 0.916093554 | 0.26605937 | 3.4431922 | 0.00057489 | 0.00958337 |
| WAC-AS1 | 39.7317967 | -0.438626512 | 0.48520491 | -0.9040026 | 0.36599401 | 0.65142886 |
| WAC | 375.152857 | 0.399843624 | 0.217739 | 1.83634362 | 0.06630685 | 0.2574031 |
| SVIL | 244.801669 | -1.688971947 | 0.3036756 | -5.5617637 | 2.67E-08 | 2.88E-06 |
| MTPAP | 24.1433622 | 0.311110681 | 0.33844502 | 0.91923551 | 0.3579724 | 0.64214377 |
| MAP3K8 | 37.2106301 | -1.190783242 | 0.37248064 | -3.1968997 | 0.00138913 | 0.01873147 |
| ZNF438 | 85.8446363 | 0.223864648 | 0.18252094 | 1.22651485 | 0.22000499 | 0.50818193 |
| ZEB1 | 59.0987146 | -2.188871999 | 0.76764113 | -2.8514262 | 0.00435236 | 0.0428543 |
| ARHGAP12 | 200.199352 | 0.005432381 | 0.19318882 | 0.02811954 | 0.97756681 | 0.99112991 |
| KIF5B | 115.984365 | -0.082744159 | 0.18502714 | -0.4472001 | 0.65473059 | 0.83924779 |
| EPC1 | 187.092947 | 0.428159869 | 0.179827 | 2.38095432 | 0.01726785 | 0.10994649 |
| CCDC7 | 148.359455 | 0.368330316 | 0.3013939 | 1.2220895 | 0.22167378 | 0.50986455 |
| ITGB1 | 203.478684 | -0.558208588 | 0.34826609 | -1.6028221 | 0.10897393 | 0.34426539 |
| NRP1 | 375.972782 | -1.013410184 | 0.36941102 | -2.7433134 | 0.00608226 | 0.05480335 |
| PARD3 | 1076.21746 | 0.296354226 | 0.18347655 | 1.61521581 | 0.10626397 | 0.33945159 |
| CUL2 | 73.470813 | 0.341717074 | 0.23676667 | 1.44326512 | 0.1489458 | 0.40929305 |
| CREM | 179.102901 | 0.641807314 | 0.54766326 | 1.17190135 | 0.24123667 | 0.53187105 |
| CCNY | 347.831933 | 0.165065775 | 0.23831237 | 0.69264459 | 0.48853262 | 0.74516503 |
| AL117336.3 | 38.1690685 | 0.253220155 | 0.31046794 | 0.81560805 | 0.41472434 | 0.69145334 |
| FZD8 | 18.9561965 | -0.408089303 | 0.50874286 | -0.8021524 | 0.42246481 | 0.69769678 |
| ZNF248 | 135.709373 | 0.085059625 | 0.33382169 | 0.25480557 | 0.79887327 | 0.91379784 |
| ZNF25 | 22.5660858 | 0.35471825 | 0.3041033 | 1.16644 | 0.2434366 | 0.53427435 |
| ZNF33A | 220.172076 | -0.278617555 | 0.23421009 | -1.1896052 | 0.23420159 | 0.52443761 |
| AL117339.5 | 19.1822127 | -0.140404827 | 0.38172098 | -0.3678206 | 0.71300703 | 0.87134026 |
| ZNF33B | 125.803802 | -0.376708588 | 0.30853933 | -1.2209419 | 0.22210803 | 0.51049171 |
| BMS1 | 60.0462003 | 0.122042966 | 0.25882827 | 0.471521 | 0.63726873 | 0.82989931 |
| CSGALNACT2 | 39.0961259 | -0.514360383 | 0.35711387 | -1.440326 | 0.1497752 | 0.41046846 |
| FXYD4 | 61.077251 | -0.532746945 | 0.70106145 | -0.7599148 | 0.44730554 | 0.71688576 |
| HNRNPF | 87.9065731 | -0.988777299 | 0.26298821 | -3.7597781 | 0.00017006 | 0.00371187 |
| ZNF487 | 19.0925991 | -0.096563627 | 0.36266251 | -0.2662631 | 0.79003662 | 0.91080793 |
| ZNF32 | 23.5950417 | 0.058245914 | 0.36411723 | 0.15996473 | 0.87290886 | 0.94909837 |
| TMEM72-AS | 28.9850591 | 0.371838372 | 0.49885333 | 0.74538617 | 0.4560383 | 0.72261911 |
| TMEM72 | 116.084416 | 1.5485201 | 0.50216853 | 3.08366617 | 0.00204467 | 0.02492802 |
| RASSF4 | 25.8521835 | 0.432655144 | 0.47825257 | 0.90465826 | 0.36564646 | 0.65103522 |
| DEPP1 | 20.0394906 | -1.313083094 | 0.3718462 | -3.5312532 | 0.0004136 | 0.00746211 |
| ZNF22 | 18.2652647 | -1.391888621 | 0.48133281 | -2.8917385 | 0.00383117 | 0.03903925 |
| 8-Mar | 99.516752 | -0.249924297 | 0.23603659 | -1.0588371 | 0.28967398 | 0.58130538 |
| ZFAND4 | 28.7039548 | 0.096989083 | 0.39188071 | 0.24749645 | 0.80452404 | 0.91643337 |
| WASHC2C | 31.5244134 | -0.778213095 | 0.4364105 | -1.7832135 | 0.07455156 | 0.27646874 |
| AGAP4 | 47.874834 | 0.605255445 | 0.31012864 | 1.95162707 | 0.0509825 | 0.21990081 |
| TIMM23 | 41.7668627 | 0.045446873 | 0.35002718 | 0.12983813 | 0.8966945 | 0.95900099 |
| NCOA4 | 55.0436849 | 0.314141538 | 0.33413108 | 0.94017455 | 0.34712803 | 0.6345126 |
| PTPN20 | 56.8604627 | 0.477807833 | 0.46840279 | 1.02007897 | 0.30769101 | 0.60139222 |
| AGAP9 | 45.7413114 | 0.590209173 | 0.50859512 | 1.16046959 | 0.24585767 | 0.53627476 |
| MAPK8 | 415.194759 | 0.514840532 | 0.28923807 | 1.77998881 | 0.07507779 | 0.27801992 |
| ARHGAP22 | 20.2785552 | -4.240474996 | 0.57421305 | -7.3848461 | 1.53E-13 | 8.28E-11 |
| ERCC6 | 56.2647243 | 0.284666653 | 0.28265299 | 1.00712415 | 0.31387512 | 0.60700985 |
| OGDHL | 140.299751 | 1.607946237 | 0.40549124 | 3.96542785 | 7.33E-05 | 0.00190598 |
| PARG | 129.764939 | 0.051467596 | 0.2090602 | 0.24618552 | 0.80553862 | 0.91703953 |
| TIMM23B | 134.423635 | 0.078310668 | 0.48522522 | 0.16139035 | 0.87178597 | 0.94844301 |
| LINC00843 | 28.0867011 | 0.442334376 | 0.29788892 | 1.48489703 | 0.1375711 | 0.39107547 |
| AGAP6 | 37.7562058 | 0.366418736 | 0.31493316 | 1.16348095 | 0.24463442 | 0.53553417 |
| WASHC2A | 33.8545184 | -0.392438932 | 0.34786069 | -1.1281497 | 0.2592567 | 0.55035288 |
| SGMS1 | 558.54914 | -0.415690525 | 0.13970498 | -2.9754882 | 0.00292523 | 0.0318896 |
| ASAH2B | 14.8829796 | -0.606474765 | 0.50793199 | -1.1940078 | 0.23247492 | 0.52234604 |
| PRKG1 | 2036.17323 | 1.73593599 | 0.29028247 | 5.98016126 | 2.23E-09 | 3.38E-07 |
| IPMK | 37.1342927 | -0.685783697 | 0.34888831 | -1.9656253 | 0.0493419 | 0.21584726 |
| CISD1 | 87.1978267 | 0.314231846 | 0.26016866 | 1.20780054 | 0.22712399 | 0.51686136 |
| UBE2D1 | 31.7882245 | -0.342271482 | 0.36643465 | -0.9340587 | 0.35027361 | 0.63689665 |
| TFAM | 30.8275471 | -0.125544692 | 0.32650235 | -0.3845139 | 0.7005976 | 0.86406757 |
| BICC1 | 2038.15406 | 1.149931928 | 0.36396645 | 3.15944485 | 0.0015807 | 0.02058707 |
| PHYHIPL | 18.2214419 | 1.387651909 | 0.43051584 | 3.22323078 | 0.00126753 | 0.01748077 |
| CCDC6 | 99.5737905 | 0.017881194 | 0.18808359 | 0.09507046 | 0.92425886 | 0.96992103 |
| ANK3 | 1656.87893 | 1.183809821 | 0.24079496 | 4.91625664 | 8.82E-07 | 5.05E-05 |
| RHOBTB1 | 14.7734265 | -0.743892239 | 0.52493788 | -1.4171053 | 0.15645215 | 0.42099911 |
| ARID5B | 523.558929 | -0.934290577 | 0.47763471 | -1.9560776 | 0.05045601 | 0.21890013 |
| NRBF2 | 12.1963728 | -0.62541706 | 0.44343862 | -1.4103802 | 0.15842744 | 0.42480987 |
| JMJD1C | 823.21649 | 0.389062089 | 0.23412034 | 1.66180385 | 0.09655211 | 0.32086653 |
| REEP3 | 157.461721 | 0.100590808 | 0.39450969 | 0.25497677 | 0.79874104 | 0.91379784 |
| LINC01515 | 27.5051545 | -0.340805044 | 0.51753152 | -0.6585204 | 0.51020382 | 0.7548518 |
| CTNNA3 | 17.2071025 | 0.78930537 | 0.57051304 | 1.38350101 | 0.1665113 | 0.43538055 |
| SIRT1 | 48.4560873 | -0.19914444 | 0.42117003 | -0.4728362 | 0.63633004 | 0.82959657 |
| HERC4 | 314.18011 | -0.136005073 | 0.24860267 | -0.5470781 | 0.58432509 | 0.79636421 |
| PBLD | 55.0232115 | 0.574801994 | 0.32507588 | 1.76820866 | 0.07702603 | 0.28159053 |
| HNRNPH3 | 112.27884 | 0.551894808 | 0.23786354 | 2.3202161 | 0.02032919 | 0.12225996 |
| RUFY2 | 104.920109 | 0.538250325 | 0.20487995 | 2.62714984 | 0.00861034 | 0.06929979 |
| DNA2 | 13.9530029 | 0.493373003 | 0.46030263 | 1.07184485 | 0.28378972 | 0.57640034 |
| SLC25A16 | 116.70524 | 0.309433079 | 0.2620202 | 1.18095121 | 0.2376221 | 0.52778809 |
| TET1 | 38.4226877 | 1.945826544 | 0.40712204 | 4.77946751 | 1.76E-06 | 8.83E-05 |
| CCAR1 | 198.526364 | 0.260563952 | 0.32186317 | 0.80954883 | 0.41819953 | 0.6944377 |
| STOX1 | 23.8894122 | 1.551783309 | 0.44639904 | 3.47622456 | 0.00050853 | 0.00874597 |
| DDX50 | 59.9380032 | 0.099536835 | 0.27531792 | 0.36153416 | 0.71770017 | 0.87396539 |
| DDX21 | 48.665127 | -0.472036315 | 0.43889356 | -1.0755143 | 0.28214452 | 0.5750055 |
| KIF1BP | 29.3078086 | -0.475628039 | 0.32847689 | -1.4479802 | 0.14762259 | 0.40743891 |
| VPS26A | 66.1091292 | 0.05766506 | 0.32870592 | 0.17543055 | 0.86074134 | 0.945045 |
| SUPV3L1 | 36.3401669 | 0.711552202 | 0.3060421 | 2.32501415 | 0.0200712 | 0.12099095 |
| HKDC1 | 95.6281822 | -0.78253453 | 0.74874615 | -1.0451266 | 0.29596445 | 0.58770736 |
| HK1 | 155.034897 | -0.48066922 | 0.28412164 | -1.6917727 | 0.09068933 | 0.31018641 |

| | | | | | | |
|---|---|---|---|---|---|---|
| TSPAN15 | 55.851501 | -1.286476085 | 0.32869037 | -3.9139452 | 9.08E-05 | 0.00224862 |
| H2AFY2 | 44.7703895 | -0.142086106 | 0.27917616 | -0.5089478 | 0.61078878 | 0.81376433 |
| AIFM2 | 22.8919205 | -0.220054375 | 0.31920818 | -0.6893757 | 0.49058685 | 0.74565148 |
| TYSND1 | 16.087338 | 0.445816182 | 0.43090455 | 1.03460542 | 0.30085322 | 0.59477833 |
| SAR1A | 46.4825931 | -0.169921236 | 0.31163713 | -0.5452535 | 0.58557918 | 0.79701899 |
| PPA1 | 76.1995636 | -1.55143404 | 0.27477798 | -5.6461367 | 1.64E-08 | 1.97E-06 |
| EIF4EBP2 | 56.5215173 | -0.259738649 | 0.27487391 | -0.9449374 | 0.34469082 | 0.63252668 |
| SGPL1 | 101.860365 | 0.134754013 | 0.20093865 | 0.67062266 | 0.50246094 | 0.75108109 |
| PCBD1 | 43.2090659 | -0.411057145 | 0.48891282 | -0.8407575 | 0.40048377 | 0.68010059 |
| PSAP | 262.222933 | -0.960155331 | 0.38793431 | -2.4750462 | 0.01332189 | 0.09348921 |
| CHST3 | 42.1820058 | -0.008926575 | 0.28199595 | -0.031655 | 0.9747472 | 0.98983573 |
| ASCC1 | 71.2936676 | 0.211409195 | 0.29238768 | 0.72304411 | 0.46965278 | 0.73178441 |
| ANAPC16 | 180.39896 | -0.141948179 | 0.21785224 | -0.6515801 | 0.51467211 | 0.75684546 |
| DDIT4 | 108.360228 | -0.193469464 | 0.63681118 | -0.3038098 | 0.76127281 | 0.89648567 |
| DNAJB12 | 21.9562095 | 0.101862148 | 0.34511624 | 0.29515316 | 0.76787688 | 0.89979146 |
| MICU1 | 208.54501 | 0.115323157 | 0.26375872 | 0.43722974 | 0.66194475 | 0.84203637 |
| MCU | 256.946938 | 0.224740832 | 0.46945533 | 0.47872677 | 0.63213302 | 0.82642568 |
| P4HA1 | 131.452453 | -1.447419832 | 0.44330554 | -3.2650615 | 0.0010944 | 0.01555109 |
| ECD | 39.4204606 | 0.254238915 | 0.303149 | 0.83865992 | 0.40166018 | 0.68068813 |
| FAM149B1 | 59.939001 | -0.030592263 | 0.25410884 | -0.1203904 | 0.9041739 | 0.96126516 |
| DNAJC9 | 45.0804199 | -0.500243733 | 0.32221229 | -1.5525284 | 0.12053584 | 0.36346292 |
| MRPS16 | 32.9169789 | -0.319199786 | 0.30865227 | -1.0341728 | 0.30105539 | 0.59482532 |
| CFAP70 | 84.174624 | 0.961209982 | 0.57474115 | 1.6724224 | 0.09444106 | 0.3161038 |
| ANXA7 | 104.549778 | -0.290311459 | 0.21166943 | -1.3715323 | 0.17020908 | 0.44081492 |
| PPP3CB | 107.383223 | 0.334297515 | 0.22137397 | 1.51010306 | 0.13101713 | 0.37999394 |
| USP54 | 81.4344672 | -0.352701634 | 0.27302024 | -1.2918516 | 0.19640854 | 0.47795702 |
| AGAP5 | 18.1222979 | 0.030131998 | 0.33850826 | 0.08901407 | 0.92907073 | 0.97209899 |
| SEC24C | 31.7387006 | -0.103401146 | 0.30330907 | -0.3409102 | 0.73317121 | 0.88344324 |
| FUT11 | 34.7530908 | -2.085412751 | 0.50117365 | -4.1610583 | 3.17E-05 | 0.00096266 |
| CHCHD1 | 26.0806859 | -0.463819341 | 0.4607653 | -1.0066282 | 0.31411348 | 0.60719808 |
| ZSWIM8 | 47.4834498 | -0.359193913 | 0.31414733 | -1.1433932 | 0.25287539 | 0.54443516 |
| CAMK2G | 39.987713 | 0.28735461 | 0.32035423 | 0.89699023 | 0.36972413 | 0.65423949 |
| VCL | 243.539267 | 1.035469482 | 0.37910409 | 2.7313593 | 0.00630737 | 0.05602908 |
| AP3M1 | 25.9211261 | -0.239011936 | 0.30335616 | -0.7878922 | 0.43075979 | 0.70420575 |
| ADK | 711.463153 | 0.003591623 | 0.17653446 | 0.02034517 | 0.98376803 | 0.99368608 |
| KAT6B | 241.754421 | 0.175411341 | 0.21673217 | 0.80934611 | 0.41831609 | 0.69451932 |
| SAMD8 | 59.1645299 | -0.398598949 | 0.25759549 | -1.5473833 | 0.12177085 | 0.36584524 |
| VDAC2 | 52.1596081 | -0.129069868 | 0.31416268 | -0.4108377 | 0.68119155 | 0.8546627 |
| COMTD1 | 20.9401828 | -0.327645409 | 0.46762885 | -0.7006527 | 0.48351979 | 0.74169459 |
| LRMDA | 1721.68553 | 0.490028665 | 0.16518505 | 2.96654371 | 0.00301168 | 0.0326249 |
| DLG5 | 82.5103366 | -0.117941284 | 0.33044668 | -0.3569147 | 0.72115565 | 0.87614353 |
| POLR3A | 34.7466741 | -0.045634304 | 0.31991897 | -0.1426433 | 0.88657188 | 0.95619452 |
| RPS24 | 239.278264 | -0.27537237 | 0.43783808 | -0.6289365 | 0.5293906 | 0.76383501 |
| ZMIZ1-AS1 | 13.6514232 | -2.363412099 | 0.70304142 | -3.3616968 | 0.00077465 | 0.01221946 |
| ZMIZ1 | 261.52095 | -0.046513866 | 0.38035699 | -0.12229 | 0.90266933 | 0.96112228 |
| PPIF | 27.750695 | -0.065587017 | 0.39244156 | -0.1671256 | 0.86727125 | 0.94753673 |
| NUTM2B-AS | 208.387427 | -0.381923035 | 0.34296015 | -1.1136076 | 0.26544756 | 0.55661322 |
| TMEM254 | 18.6570629 | -0.00849202 | 0.34678801 | -0.0244876 | 0.98046364 | 0.99299415 |
| ANXA11 | 260.420138 | 0.371510736 | 0.21586693 | 1.72101735 | 0.08524767 | 0.29983665 |
| FAM213A | 62.4955667 | -0.906454762 | 0.29677416 | -3.0543588 | 0.00225542 | 0.02658507 |
| TSPAN14 | 56.9172613 | -1.068231736 | 0.28382118 | -3.7637492 | 0.00016738 | 0.00368461 |
| NRG3 | 62.3534248 | -3.64059396 | 0.55676847 | -6.5387933 | 6.20E-11 | 1.58E-08 |
| GHITM | 72.343298 | 0.051667264 | 0.36086645 | 0.14317558 | 0.88615151 | 0.95619452 |
| CCSER2 | 180.905914 | 0.220435454 | 0.27753906 | 0.7942502 | 0.42704979 | 0.70162975 |
| WAPL | 151.424748 | 0.299350537 | 0.21652999 | 1.38248995 | 0.16682131 | 0.43585929 |
| BMPR1A | 200.279928 | 0.284515998 | 0.17006337 | 1.67299987 | 0.09432732 | 0.3161038 |
| ADIRF | 17.9631243 | -0.349206541 | 0.69202871 | -0.5046128 | 0.61383082 | 0.81511796 |
| GLUD1 | 100.37472 | 0.130632746 | 0.26123733 | 0.5000539 | 0.61703712 | 0.81726877 |
| SHLD2 | 155.260183 | 0.630713018 | 0.18870569 | 3.3423105 | 0.00083084 | 0.01275607 |
| NUTM2A-AS | 219.224994 | 0.203949589 | 0.28186034 | 0.72358384 | 0.46932126 | 0.73168094 |
| LINC00864 | 22.3683205 | 1.397210526 | 0.49830131 | 2.80394713 | 0.00504812 | 0.04793153 |
| MINPP1 | 19.166287 | 0.289088791 | 0.39552411 | 0.73090056 | 0.4648399 | 0.72848544 |
| PAPSS2 | 76.4727671 | 0.009697545 | 0.40984153 | 0.02366169 | 0.98112246 | 0.99318267 |
| ATAD1 | 116.879531 | 0.398900086 | 0.30197491 | 1.32097097 | 0.18651104 | 0.46400309 |
| PTEN | 267.932607 | -0.187161443 | 0.24646038 | -0.7593977 | 0.44761469 | 0.71693398 |
| RNLS | 92.4909469 | 0.226684814 | 0.36304756 | 0.62439427 | 0.53236869 | 0.76620037 |
| STAMBPL1 | 12.1733892 | -0.225850935 | 0.42359834 | -0.5331724 | 0.59391426 | 0.80267231 |
| LIPA | 42.2187618 | -0.264428609 | 0.35036487 | -0.7547235 | 0.45041489 | 0.71901805 |
| IFIT2 | 29.3865877 | -1.012267436 | 1.6466334 | -0.6147497 | 0.53872001 | 0.7706045 |
| IFIT3 | 59.2979362 | -2.071604881 | 1.37404674 | -1.507667 | 0.13163978 | 0.38051432 |
| IFIT1 | 47.6560821 | 1.334873879 | 1.71009486 | 0.7805847 | 0.4350468 | 0.70680525 |
| PANK1 | 35.5780438 | 1.074350003 | 0.44775042 | 2.39943941 | 0.0164202 | 0.10625683 |
| RPP30 | 16.0769507 | 0.114479713 | 0.41910556 | 0.27315246 | 0.78473602 | 0.90771539 |
| PCGF5 | 203.110576 | 0.01045238 | 0.30340877 | 0.03444983 | 0.97251845 | 0.98854318 |
| HECTD2 | 85.6792952 | 0.182228261 | 0.24954768 | 0.73023424 | 0.46524702 | 0.72863709 |
| PPP1R3C | 15.2744841 | -0.513191147 | 0.71019497 | -0.722606 | 0.46992198 | 0.73188482 |
| TNKS2 | 159.774748 | -0.029065948 | 0.20959633 | -0.1386758 | 0.88970631 | 0.95702609 |
| BTAF1 | 150.319493 | 0.681917049 | 0.36200613 | 1.88371684 | 0.05960329 | 0.2417453 |
| CPEB3 | 366.510816 | 1.455949855 | 0.27526799 | 5.28920868 | 1.23E-07 | 1.03E-05 |
| 5-Mar | 80.2999028 | 0.047739456 | 0.24638263 | 0.19376145 | 0.84636267 | 0.93757726 |
| IDE | 81.3732804 | 0.332356537 | 0.29792159 | 1.11558394 | 0.26460027 | 0.55622662 |
| EXOC6 | 107.108154 | 0.32172574 | 0.27441066 | 1.17242436 | 0.24102673 | 0.53170393 |
| MYOF | 161.405033 | -0.966399038 | 0.32356931 | -2.9866833 | 0.00282022 | 0.03137568 |
| FRA10AC1 | 55.7311981 | 0.447577931 | 0.27585225 | 1.62252776 | 0.10469039 | 0.33556952 |
| SLC35G1 | 14.5849925 | -0.627659156 | 0.49156806 | -1.276851 | 0.20165487 | 0.48504517 |
| PLCE1 | 13.5574352 | -0.881091198 | 0.63178143 | -1.394614 | 0.16313229 | 0.43136263 |
| NOC3L | 20.6990876 | 0.141130742 | 0.34259278 | 0.41194896 | 0.68037683 | 0.8543838 |
| TBC1D12 | 70.5878724 | 0.336081109 | 0.2122791 | 1.58320397 | 0.11337498 | 0.3516614 |
| HELLS | 58.2548234 | -0.260857432 | 0.27741617 | -0.9403108 | 0.34705814 | 0.63449742 |
| SORBS1 | 45.4288353 | -1.587335584 | 0.3972503 | -3.9958072 | 6.45E-05 | 0.00172972 |
| ALDH18A1 | 51.0037136 | -0.348689447 | 0.37890554 | -0.9202543 | 0.35743989 | 0.64174726 |
| TCTN3 | 53.417069 | 0.104929814 | 0.27032863 | 0.3881565 | 0.69790022 | 0.86280668 |
| ENTPD1-AS1 | 148.863393 | 0.002472622 | 0.35319318 | 0.00700076 | 0.99441425 | 0.99837898 |
| ZNF518A | 202.175837 | 0.584854986 | 0.22889592 | 2.55511321 | 0.01061533 | 0.07997578 |
| BLNK | 371.231628 | -0.477407843 | 0.31084396 | -1.5358441 | 0.12457663 | 0.36974123 |
| TM9SF3 | 196.019001 | -0.072749769 | 0.15686302 | -0.463779 | 0.64280611 | 0.83318468 |
| LCOR | 330.382472 | 0.240601229 | 0.23557188 | 1.02134951 | 0.30708888 | 0.60139222 |
| RRP12 | 24.9858536 | -0.191350062 | 0.42836615 | -0.4466974 | 0.65509353 | 0.83931753 |
| PGAM1 | 87.6458896 | -0.882666182 | 0.36641245 | -2.4089416 | 0.01599886 | 0.10420871 |
| EXOSC1 | 24.9751786 | -0.00520152 | 0.32401019 | -0.0160536 | 0.98719166 | 0.99523554 |
| ZDHHC16 | 18.8043509 | -0.280880778 | 0.39685044 | -0.7077749 | 0.47908506 | 0.73880257 |
| MMS19 | 54.4838258 | 0.219256817 | 0.25374199 | 0.86409356 | 0.3875365 | 0.66964123 |
| UBTD1 | 38.719299 | -0.343654753 | 0.31982693 | -1.0745023 | 0.28259764 | 0.57535985 |
| MORN4 | 22.2692574 | 0.319249182 | 0.3069147 | 1.04018863 | 0.29825227 | 0.59099729 |
| ZFYVE27 | 31.0989274 | -0.094740134 | 0.27347443 | -0.3464314 | 0.72901851 | 0.88046679 |
| R3HCC1L | 106.049173 | -0.136832606 | 0.19925399 | -0.6867245 | 0.49225633 | 0.74664675 |
| HPS1 | 19.2521565 | -0.508851281 | 0.34429413 | -1.4779551 | 0.13941978 | 0.39404853 |
| GOT1 | 163.925919 | 0.927048682 | 0.31457025 | 2.94703229 | 0.0032084 | 0.03418088 |
| AL391684.1 | 11.1769075 | 1.375519088 | 0.52876301 | 2.60139052 | 0.00928467 | 0.07312742 |
| SLC25A28 | 34.9034944 | -0.443862723 | 0.55244754 | -0.8034477 | 0.42171599 | 0.69752044 |
| ENTPD7 | 11.5760294 | -0.53772391 | 0.46041626 | -1.1679082 | 0.24284381 | 0.53327647 |
| CUTC | 39.1255572 | -0.448147257 | 0.27732423 | -1.6159686 | 0.1061011 | 0.33903646 |
| COX15 | 35.3169392 | -0.151363489 | 0.24807538 | -0.6101512 | 0.54176166 | 0.77222629 |

| | | | | | | |
|---|---|---|---|---|---|---|
| DNMBP | 91.2251447 | 0.200748413 | 0.23444152 | 0.85628352 | 0.39184097 | 0.67301529 |
| ERLIN1 | 25.5607918 | 0.236448435 | 0.35321172 | 0.66942409 | 0.50322499 | 0.75108109 |
| CHUK | 28.8543938 | 0.457150176 | 0.421933 | 1.08346627 | 0.27860152 | 0.57131913 |
| CWF19L1 | 33.5490523 | 0.165013489 | 0.3034706 | 0.54375445 | 0.58661045 | 0.79752689 |
| BLOC1S2 | 20.9666987 | -0.158188265 | 0.34546425 | -0.4579005 | 0.6470239 | 0.83477022 |
| SEC31B | 15.898343 | 0.658465079 | 0.37976895 | 1.73385706 | 0.08294345 | 0.29503415 |
| HIF1AN | 33.5496592 | -0.595732826 | 0.38461973 | -1.5488878 | 0.12140868 | 0.3652898 |
| PAX2 | 334.947742 | 0.006277476 | 0.31006915 | 0.02024541 | 0.98384761 | 0.99368608 |
| SLF2 | 76.1514629 | -0.209620548 | 0.33338391 | -0.6287662 | 0.5295021 | 0.7638395 |
| MRPL43 | 19.1933028 | -0.983505217 | 0.45738119 | -2.1502966 | 0.03153176 | 0.16250135 |
| LZTS2 | 28.3851557 | -0.730732168 | 0.34688025 | -2.1065833 | 0.03515371 | 0.17419602 |
| BTRC | 190.637611 | 0.447578226 | 0.17526744 | 2.55368731 | 0.01065889 | 0.08015176 |
| DPCD | 19.8838904 | -1.019441818 | 0.42791486 | -2.3823473 | 0.01720266 | 0.10980752 |
| POLL | 21.3274743 | -0.092071436 | 0.39458864 | -0.2333352 | 0.8155011 | 0.92165337 |
| FBXW4 | 133.29854 | 0.05825838 | 0.20551724 | 0.28347199 | 0.77681505 | 0.90335572 |
| OGA | 246.61626 | -0.123668613 | 0.18139856 | -0.6817508 | 0.49539655 | 0.74770824 |
| ARMH3 | 175.084198 | 0.208359172 | 0.20580057 | 1.01243241 | 0.31133135 | 0.60435944 |
| LDB1 | 26.5098714 | 0.37799309 | 0.37210863 | 1.01581382 | 0.30971806 | 0.60315981 |
| PPRC1 | 20.4062469 | -0.790594241 | 0.53606519 | -1.4748099 | 0.14026366 | 0.39525538 |
| NOLC1 | 15.5456684 | -0.199154058 | 0.43957248 | -0.4530631 | 0.65050332 | 0.83735207 |
| GBF1 | 242.495438 | 0.237852745 | 0.20858331 | 1.14032494 | 0.25415095 | 0.54530511 |
| NFKB2 | 20.0308026 | -0.635984182 | 0.43193154 | -1.4724189 | 0.14090778 | 0.39647869 |
| FBXL15 | 23.7554121 | 0.072412442 | 0.42671418 | 0.16969776 | 0.86524783 | 0.94694132 |
| CUEDC2 | 20.1580938 | -0.262400659 | 0.32734892 | -0.8015932 | 0.42278834 | 0.69778363 |
| ACTR1A | 29.2003674 | -0.452381821 | 0.38152632 | -1.1857159 | 0.23573452 | 0.52588501 |
| SUFU | 65.8988794 | 0.414880633 | 0.25013972 | 1.65859557 | 0.09719731 | 0.32176309 |
| TRIM8 | 66.9411877 | -0.114376881 | 0.37083361 | -0.3084318 | 0.75775378 | 0.89457492 |
| ARL3 | 45.3388672 | -0.053505824 | 0.27390887 | -0.1953417 | 0.84512545 | 0.93715892 |
| SFXN2 | 15.2732188 | 0.584222998 | 0.51414733 | 1.13629493 | 0.25583315 | 0.54693968 |
| WBP1L | 64.3414875 | -0.144563876 | 0.30185485 | -0.4789185 | 0.6319966 | 0.82642568 |
| CNNM2 | 255.181949 | 1.88901104 | 0.2016287 | 9.3687607 | 7.34E-21 | 1.51E-17 |
| NT5C2 | 349.611549 | -0.406136298 | 0.29107835 | -1.3952817 | 0.1629309 | 0.4311397 |
| PCGF6 | 30.9088111 | 0.440211042 | 0.394207 | 1.11670023 | 0.26412252 | 0.55564431 |
| ATP5MD | 41.3830422 | 0.381287057 | 0.25590739 | 1.48994157 | 0.1362396 | 0.38868467 |
| PDCD11 | 26.0245442 | -0.059485489 | 0.38789676 | -0.1533539 | 0.87811918 | 0.9522716 |
| SH3PXD2A | 54.8433205 | 0.792719354 | 0.278537 | 2.84601096 | 0.00442707 | 0.04339452 |
| STN1 | 34.9834138 | -0.091493929 | 0.45110299 | -0.2028227 | 0.83927361 | 0.93543959 |
| SLK | 112.365273 | 0.181057121 | 0.29118295 | 0.6217985 | 0.53407437 | 0.76732389 |
| CFAP43 | 31.3073615 | -0.597241206 | 0.43918048 | -1.3598993 | 0.17386181 | 0.44644175 |
| GSTO1 | 43.215608 | -0.844751668 | 0.28304305 | -2.9845342 | 0.0028401 | 0.03143157 |
| GSTO2 | 83.5236135 | 0.157653537 | 0.33763592 | 0.46693355 | 0.64054742 | 0.8320413 |
| LINC01435 | 70.7325829 | 2.782471185 | 0.89542171 | 3.10744218 | 0.00188714 | 0.02359382 |
| XPNPEP1 | 44.9907787 | -0.350008882 | 0.33656574 | -1.0399421 | 0.2983668 | 0.59111054 |
| ADD3 | 215.917973 | 0.544088106 | 0.28542285 | 1.90625282 | 0.05661741 | 0.23471734 |
| MXI1 | 61.3413874 | -1.10671991 | 0.33262721 | -3.327208 | 0.00087721 | 0.01336835 |
| SMNDC1 | 27.5300849 | 0.019075486 | 0.32535702 | 0.0586294 | 0.95324729 | 0.98143529 |
| SMC3 | 46.8489347 | -0.369454492 | 0.27193631 | -1.3586067 | 0.17427125 | 0.4470474 |
| PDCD4 | 60.9130952 | -0.332167577 | 0.43809825 | -0.7582034 | 0.44832924 | 0.71763328 |
| BBIP1 | 35.8011463 | 0.109774721 | 0.30944745 | 0.35474431 | 0.72278115 | 0.87662955 |
| SHOC2 | 119.510923 | -0.067906913 | 0.23782899 | -0.2855283 | 0.77523941 | 0.90226394 |
| GPAM | 24.8592495 | 0.933860456 | 0.36011775 | 2.59320864 | 0.00950851 | 0.07443513 |
| ZDHHC6 | 29.5825572 | -0.165738079 | 0.26614083 | -0.6227458 | 0.53345159 | 0.7669018 |
| VTI1A | 412.93118 | 0.138272667 | 0.16759744 | 0.8250285 | 0.40935541 | 0.68723933 |
| TCF7L2 | 350.899908 | 0.170323362 | 0.20188048 | 0.84368416 | 0.39884592 | 0.67960812 |
| CASP7 | 30.2375465 | -0.649507607 | 0.32959778 | -1.9706068 | 0.04876887 | 0.21425029 |
| NHLRC2 | 84.5351084 | 0.186163739 | 0.21018536 | 0.88571219 | 0.37577264 | 0.65904149 |
| CCDC186 | 82.3130642 | -0.080630467 | 0.21443795 | -0.3760084 | 0.70691066 | 0.86811224 |
| AFAP1L2 | 105.981536 | 1.158841774 | 0.38000667 | 3.04953011 | 0.002292 | 0.02692377 |
| ABLIM1 | 480.616253 | -0.110245061 | 0.29137757 | -0.3783581 | 0.70516459 | 0.86679461 |
| FAM160B1 | 48.4447289 | -0.001405399 | 0.28468073 | -0.0049368 | 0.99606106 | 0.99868531 |
| TRUB1 | 27.3683928 | 0.630201086 | 0.37815513 | 1.66651472 | 0.09561094 | 0.31866189 |
| HSPA12A | 148.543648 | 0.74920426 | 0.28448551 | 2.63354104 | 0.00844996 | 0.06865261 |
| SHTN1 | 98.5102295 | -0.319933963 | 0.32228999 | -0.9926897 | 0.32086118 | 0.6120836 |
| PDZD8 | 120.956431 | -0.295358778 | 0.29746463 | -0.9929207 | 0.32074862 | 0.6120836 |
| AC005871.2 | 14.8805775 | -1.229870436 | 0.59141034 | -2.0795552 | 0.03756635 | 0.18135358 |
| EMX2OS | 315.537225 | -0.156158939 | 0.23499463 | -0.6645213 | 0.50635671 | 0.75306985 |
| EMX2 | 118.928434 | 0.101123848 | 0.25467673 | 0.39706748 | 0.6913177 | 0.85976137 |
| RAB11FIP2 | 217.177383 | 0.497514344 | 0.28546571 | 1.74281646 | 0.08136568 | 0.2916273 |
| AC022395.1 | 17.5715524 | -0.517843072 | 0.64472599 | -0.8031987 | 0.4218599 | 0.69753545 |
| CASC2 | 103.93494 | 0.077401853 | 0.46700924 | 0.16573945 | 0.868362 | 0.94805695 |
| FAM204A | 61.4798904 | 0.11482138 | 0.20174377 | 0.56914461 | 0.56925801 | 0.787921 |
| CACUL1 | 137.342019 | -0.214814905 | 0.27504172 | -0.7810266 | 0.43478683 | 0.70649431 |
| EIF3A | 148.214774 | -0.197856491 | 0.29953926 | -0.6605361 | 0.50890988 | 0.75445608 |
| FAM45A | 25.2054894 | -0.477642253 | 0.30906453 | -1.545445 | 0.12223865 | 0.3664089 |
| SFXN4 | 45.795003 | 0.897056418 | 0.34971469 | 2.56510935 | 0.01031433 | 0.07859336 |
| PRDX3 | 60.3555861 | -0.23037908 | 0.37562531 | -0.6133215 | 0.53966378 | 0.77077724 |
| TIAL1 | 78.464444 | 0.333617516 | 0.21739361 | 1.53462431 | 0.12487614 | 0.37001028 |
| BAG3 | 35.9882661 | 0.954965872 | 0.30330741 | 3.14850826 | 0.00164106 | 0.02126567 |
| INPP5F | 62.6117642 | -0.115649038 | 0.25547979 | -0.4526739 | 0.65078356 | 0.83735207 |
| MCMBP | 47.6512639 | -0.004007882 | 0.25574373 | -0.0156715 | 0.98749648 | 0.99541965 |
| SEC23IP | 76.6061106 | -0.07619751 | 0.33419454 | -0.2280035 | 0.81964356 | 0.92374663 |
| PLPP4 | 103.908616 | 2.836561431 | 0.46810073 | 6.05972444 | 1.36E-09 | 2.34E-07 |
| WDR11-AS1 | 22.5021854 | 0.268141657 | 0.46723466 | 0.57389077 | 0.56604172 | 0.78607719 |
| WDR11 | 108.796893 | 0.339144168 | 0.17778397 | 1.90761954 | 0.05644041 | 0.23454985 |
| FGFR2 | 160.642912 | 0.527657102 | 0.33808107 | 1.56074133 | 0.1185848 | 0.35998571 |
| ATE1 | 155.782857 | 0.101299867 | 0.19948822 | 0.50779873 | 0.6115945 | 0.81385911 |
| NSMCE4A | 18.4272236 | 0.625062172 | 0.33383468 | 1.87237042 | 0.06115537 | 0.24535775 |
| TACC2 | 159.374415 | 0.823797405 | 0.29217616 | 2.819523 | 0.00480951 | 0.04626289 |
| PLEKHA1 | 387.196401 | -1.606651565 | 0.36218939 | -4.4359432 | 9.17E-06 | 0.00034093 |
| C10orf88 | 18.7055887 | -0.017865379 | 0.34360814 | -0.0519935 | 0.9585339 | 0.98335155 |
| PSTK | 12.9130253 | -0.177650307 | 0.42711244 | -0.4159333 | 0.67745878 | 0.85247103 |
| IKZF5 | 22.8414368 | 0.331160072 | 0.33163393 | 0.99857113 | 0.31800249 | 0.61109152 |
| ACADSB | 51.5025994 | 0.091859525 | 0.24564422 | 0.37395354 | 0.70843889 | 0.868892 |
| BUB3 | 31.0610517 | -0.116742186 | 0.35849381 | -0.3256463 | 0.74469197 | 0.88808711 |
| CPXM2 | 71.8027106 | 0.296013473 | 0.69165028 | 0.42798143 | 0.66866464 | 0.84584764 |
| OAT | 31.4094546 | -0.081585172 | 0.39917471 | -0.2043846 | 0.83805295 | 0.93498175 |
| LHPP | 38.2820089 | 0.105737603 | 0.28994608 | 0.36468022 | 0.71535012 | 0.87306444 |
| ABRAXAS2 | 32.1169956 | -0.06372959 | 0.31868892 | -0.1999743 | 0.84150069 | 0.93566718 |
| ZRANB1 | 70.1475269 | 0.241073196 | 0.25795912 | 0.93454032 | 0.35002525 | 0.63675792 |
| CTBP2 | 391.32175 | -0.287606252 | 0.26326803 | -1.0924466 | 0.27463686 | 0.56696009 |
| EDRF1 | 48.7333936 | 0.093189036 | 0.22854842 | 0.40774307 | 0.68346231 | 0.85606265 |
| UROS | 26.5848105 | -0.112151742 | 0.33509158 | -0.3346898 | 0.73785908 | 0.88573617 |
| BCCIP | 44.8393436 | 0.267726712 | 0.25892353 | 1.03399914 | 0.30113656 | 0.59482532 |
| DHX32 | 108.564769 | -0.663198033 | 0.22572768 | -2.9380448 | 0.00330289 | 0.03486312 |
| FANK1 | 55.8131953 | -1.045375965 | 0.38855321 | -2.6904319 | 0.00713596 | 0.06131331 |
| DOCK1 | 590.154248 | -0.119217937 | 0.25721769 | -0.4634904 | 0.64301288 | 0.83324763 |
| FAM196A | 22.1937777 | -1.133990604 | 0.51194965 | -2.2150432 | 0.0267571 | 0.14614285 |
| MGMT | 151.104686 | 0.009173216 | 0.2189689 | 0.04189278 | 0.96658417 | 0.98695423 |
| AL157832.3 | 23.1906095 | -0.376151393 | 0.39317228 | -0.9567088 | 0.33871424 | 0.62755046 |
| C10orf143 | 19.4361554 | 0.230875006 | 0.34181134 | 0.67544572 | 0.49939263 | 0.75031683 |
| GLRX3 | 67.119138 | -0.091196848 | 0.29669458 | -0.3073762 | 0.75855706 | 0.8951693 |
| PPP2R2D | 139.230621 | 0.356669297 | 0.26659956 | 1.33784654 | 0.18094647 | 0.45756661 |
| BNIP3 | 83.048924 | -2.132821288 | 0.59377277 | -3.5919823 | 0.00032817 | 0.00628407 |

| | | | | | | |
|---|---|---|---|---|---|---|
| STK32C | 22.3941565 | -0.359299613 | 0.33072363 | -1.0864044 | 0.2773001 | 0.56952664 |
| LRRC27 | 21.0115552 | -0.352151851 | 0.37814467 | -0.9312622 | 0.35171793 | 0.63801604 |
| PWWP2B | 16.9166842 | -0.215639862 | 0.47760767 | -0.4515 | 0.65162923 | 0.83781115 |
| INPP5A | 85.3577111 | 0.357096036 | 0.25392746 | 1.40629155 | 0.15963757 | 0.42727956 |
| TUBGCP2 | 60.0313616 | 0.069760883 | 0.34602036 | 0.20160919 | 0.84022226 | 0.93552258 |
| ECHS1 | 58.3246041 | -0.619830685 | 0.48193154 | -1.2861385 | 0.19839474 | 0.48097216 |
| SCART1 | 18.081803 | 1.144943736 | 0.51012886 | 2.24442065 | 0.02480535 | 0.13820698 |
| SLC6A12 | 44.7425446 | -0.324218393 | 0.46354511 | -0.6994322 | 0.48428195 | 0.74250561 |
| KDM5A | 150.367581 | -0.1307711 | 0.18052401 | -0.7243973 | 0.46882189 | 0.73156666 |
| CCDC77 | 13.718907 | -0.156313802 | 0.41057258 | -0.3807215 | 0.70340993 | 0.86585351 |
| B4GALNT3 | 286.576064 | 1.859448601 | 0.41662513 | 4.4631216 | 8.08E-06 | 0.00031166 |
| WNK1 | 996.846815 | 2.337735158 | 0.42051246 | 5.55925306 | 2.71E-08 | 2.88E-06 |
| RAD52 | 42.430973 | 0.215262622 | 0.28618135 | 0.75218956 | 0.45193707 | 0.71982927 |
| ERC1 | 663.497734 | 0.442650565 | 0.19929715 | 2.22105815 | 0.02634702 | 0.14499307 |
| FBXL14 | 20.025739 | -1.146220847 | 0.42599046 | -2.6907195 | 0.00712981 | 0.06131331 |
| ADIPOR2 | 115.395692 | 0.428782397 | 0.32850792 | 1.30524217 | 0.1918104 | 0.47205703 |
| DCP1B | 44.6438453 | -0.002064881 | 0.31153513 | -0.0066281 | 0.99471159 | 0.99837898 |
| FKBP4 | 34.8444307 | -0.198524959 | 0.52196417 | -0.3803421 | 0.70369148 | 0.86601715 |
| ITFG2 | 68.610574 | 0.05546539 | 0.38869195 | 0.14269755 | 0.88652905 | 0.95619452 |
| TULP3 | 91.9749641 | -0.181689555 | 0.28590962 | -0.635479 | 0.52511608 | 0.76221787 |
| TEAD4 | 42.6868012 | -2.164985537 | 0.49672136 | -4.3585513 | 1.31E-05 | 0.00046034 |
| TSPAN9 | 318.623415 | 0.139687312 | 0.19231336 | 0.72635263 | 0.46762262 | 0.73035905 |
| PARP11 | 24.3970768 | -0.303098826 | 0.34109244 | -0.888612 | 0.37421166 | 0.65806152 |
| TIGAR | 27.4953104 | 0.135496922 | 0.55801586 | 0.24281912 | 0.80814552 | 0.91842538 |
| C12orf4 | 31.8685801 | 0.216946918 | 0.39737018 | 0.54595671 | 0.58509569 | 0.79681419 |
| CD9 | 134.30131 | -0.224971138 | 0.27763293 | -0.8103186 | 0.41775707 | 0.69403859 |
| TNFRSF1A | 54.7522338 | -1.225563282 | 0.391248 | -3.1324462 | 0.00173356 | 0.02204834 |
| SCNN1A | 255.017249 | 1.066608364 | 0.19727144 | 5.40680566 | 6.42E-08 | 5.70E-06 |
| LTBR | 29.6633139 | -0.727610416 | 0.3922749 | -1.8548483 | 0.0636179 | 0.25120413 |
| TAPBPL | 17.0056896 | -0.301256852 | 0.48170845 | -0.6253925 | 0.53171349 | 0.76557825 |
| MRPL51 | 36.641415 | -0.876048485 | 0.46670048 | -1.8771107 | 0.06050292 | 0.24357213 |
| NCAPD2 | 21.0840264 | 0.094660252 | 0.43557844 | 0.21732079 | 0.82795836 | 0.92818649 |
| GAPDH | 386.464706 | -1.603391014 | 0.48497884 | -3.3061051 | 0.00094603 | 0.01400282 |
| CHD4 | 35.888682 | -0.561950616 | 0.31973252 | -1.7575648 | 0.07882159 | 0.28599809 |
| AC006064.2 | 14.2796721 | -0.159898902 | 0.61030246 | -0.2619994 | 0.79332188 | 0.91142199 |
| ING4 | 24.3257976 | -0.382622669 | 0.31605566 | -1.210618 | 0.22604184 | 0.51508141 |
| ZNF384 | 16.9210528 | 0.146563087 | 0.34746531 | 0.42180639 | 0.67316634 | 0.84956017 |
| COPS7A | 29.3406467 | 0.150085488 | 0.42147528 | 0.35609559 | 0.72176898 | 0.87625492 |
| MLF2 | 44.3366241 | -0.421291556 | 0.37067293 | -1.1365587 | 0.2557228 | 0.54693968 |
| PTMS | 89.6111351 | -0.228558244 | 0.43621781 | -0.5239544 | 0.60031025 | 0.80738812 |
| USP5 | 19.4882654 | -0.092751269 | 0.56549975 | -0.1640165 | 0.86971818 | 0.94832921 |
| TPI1 | 116.471048 | -0.974627913 | 0.52056955 | -1.8722338 | 0.06117426 | 0.24535775 |
| LRRC23 | 24.0553937 | -0.36578928 | 0.32302984 | -1.1323699 | 0.25747894 | 0.54861387 |
| ATN1 | 52.8868261 | -0.303431535 | 0.32293044 | -0.9396189 | 0.34741309 | 0.63465611 |
| C12orf57 | 29.2969802 | -0.314674897 | 0.51283724 | -0.613596 | 0.5394823 | 0.77077724 |
| PTPN6 | 18.5926821 | -0.212856606 | 0.38464774 | -0.5533806 | 0.58000281 | 0.79446735 |
| PHB2 | 41.6436737 | -0.195297564 | 0.37646516 | -0.5187666 | 0.60392349 | 0.80957967 |
| LPCAT3 | 93.4130333 | -0.464467257 | 0.2071987 | -2.2416514 | 0.02498391 | 0.13897638 |
| C1S | 34.2964775 | -0.360183085 | 0.5337609 | -0.6748023 | 0.49980138 | 0.75045478 |
| C1R | 20.3771596 | -2.355630452 | 0.79847495 | -2.950162 | 0.00317607 | 0.03391661 |
| C1RL | 67.2015367 | -1.170701287 | 0.23692742 | -4.9411811 | 7.77E-07 | 4.62E-05 |
| C1RL-AS1 | 11.2636443 | -1.598276444 | 0.65326459 | -2.4465989 | 0.01442112 | 0.09759315 |
| CLSTN3 | 24.5898426 | 0.382852054 | 0.38928429 | 0.98347676 | 0.32537285 | 0.61560901 |
| PEX5 | 15.2483003 | 0.139949494 | 0.36179511 | 0.38681975 | 0.69888965 | 0.86334649 |
| FOXJ2 | 18.0434135 | 0.586266984 | 0.35003383 | 1.67488663 | 0.09395648 | 0.31595883 |
| NECAP1 | 23.9128205 | 0.491887216 | 0.39590211 | 1.24244656 | 0.21407184 | 0.50065111 |
| FAM66C | 15.3627036 | 0.460325033 | 0.44337981 | 1.03821829 | 0.29916844 | 0.59224313 |
| LINC00937 | 26.9890695 | -0.488374586 | 0.32386051 | -1.5079782 | 0.13156011 | 0.38039077 |
| RIMKLB | 125.698432 | -1.524600207 | 0.5490315 | -2.7768902 | 0.00548817 | 0.05085757 |
| A2ML1-AS1 | 15.3996098 | -1.637355468 | 0.63482341 | -2.5792298 | 0.00990209 | 0.07673607 |
| PHC1 | 33.1678009 | -0.355654876 | 0.3009421 | -1.181805 | 0.23728309 | 0.5276258 |
| M6PR | 53.1594234 | -0.405710165 | 0.29786852 | -1.3620444 | 0.17318386 | 0.44603504 |
| KLRG1 | 11.5394083 | -0.311258444 | 0.479643 | -0.6489377 | 0.51637862 | 0.75768873 |
| CLEC2D | 36.3552387 | -0.324094585 | 0.35501051 | -0.9129155 | 0.361287 | 0.64557855 |
| TMEM52B | 237.2275 | 1.91353386 | 0.68615855 | 2.78876342 | 0.00529097 | 0.04950733 |
| GABARAPL1 | 170.06882 | 0.493012857 | 0.34225711 | 1.44047513 | 0.14973302 | 0.41046846 |
| KLRD1 | 59.5735849 | 0.065895799 | 0.46942568 | 0.14037536 | 0.88836342 | 0.95661336 |
| MAGOHB | 14.3558619 | 0.02608827 | 0.44577455 | 0.05852346 | 0.95333167 | 0.98143529 |
| YBX3 | 215.123281 | -1.522372085 | 0.34702667 | -4.3869022 | 1.15E-05 | 0.00040986 |
| PRH1 | 13.4503617 | 0.167258306 | 0.54659688 | 0.30599938 | 0.75960512 | 0.8957706 |
| SMIM10L1 | 17.9726664 | -0.753308586 | 0.3743793 | -2.0121534 | 0.04420377 | 0.20044552 |
| AC244131.2 | 25.3220326 | -0.152334853 | 0.74808792 | -0.2036323 | 0.83864086 | 0.93537562 |
| ETV6 | 390.698342 | -0.988037299 | 0.31195965 | -3.1671958 | 0.00153917 | 0.02017365 |
| BCL2L14 | 42.7104384 | -0.288896129 | 0.43212328 | -0.6685503 | 0.50378242 | 0.75108109 |
| LRP6 | 147.674293 | 0.750570903 | 0.2258978 | 3.32261271 | 0.00089179 | 0.01353046 |
| MANSC1 | 16.4288025 | -0.507666088 | 0.42377727 | -1.197955 | 0.23093455 | 0.52111057 |
| BORCS5 | 53.6243856 | 0.170550284 | 0.23773944 | 0.71738321 | 0.47313768 | 0.73385491 |
| DUSP16 | 87.6546008 | -0.059178886 | 0.20356659 | -0.2907102 | 0.77127296 | 0.90076568 |
| CREBL2 | 35.4619584 | -0.503249569 | 0.31201045 | -1.6129254 | 0.10676073 | 0.34029982 |
| CDKN1B | 23.6557716 | -0.240948415 | 0.50732883 | -0.4749354 | 0.63483303 | 0.82837871 |
| GPRC5A | 50.1416517 | -2.303563459 | 0.65618761 | -3.5105257 | 0.00044722 | 0.0079573 |
| GPRC5D-AS1 | 49.2050774 | 0.084741894 | 0.41112162 | 0.20612366 | 0.83669431 | 0.93419636 |
| HEBP1 | 37.6755969 | 0.208015448 | 0.30812525 | 0.67510029 | 0.49961205 | 0.75040142 |
| FAM234B | 14.7292006 | 0.998997256 | 0.38071651 | 2.62399245 | 0.00869057 | 0.06977484 |
| EMP1 | 30.5458685 | -2.018873653 | 0.38067678 | -5.30338 | 1.14E-07 | 9.60E-06 |
| ATF7IP | 229.136528 | 0.174635495 | 0.22799046 | 0.76597722 | 0.44368986 | 0.71343168 |
| PLBD1 | 22.0474169 | -0.191416666 | 0.48113463 | -0.3978443 | 0.69074496 | 0.85967888 |
| AC010168.1 | 21.0012901 | -0.358378495 | 0.62581297 | -0.5726607 | 0.56687445 | 0.78656476 |
| H2AFJ | 53.0470356 | -1.126127657 | 0.3662164 | -3.0750334 | 0.00210479 | 0.02536445 |
| WBP11 | 35.9959311 | -0.169904102 | 0.31335234 | -0.5422143 | 0.58767092 | 0.79838927 |
| C12orf60 | 28.0477397 | 0.751406935 | 0.3830978 | 1.96139717 | 0.04983271 | 0.2170841 |
| MGP | 69.4695152 | -1.78585568 | 0.64189158 | -2.7821765 | 0.00539957 | 0.05024962 |
| ERP27 | 11.652194 | -0.914198242 | 0.91780536 | -0.9960698 | 0.31921621 | 0.611915 |
| ARHGDIB | 14.0942528 | -2.047176926 | 0.74420003 | -2.7508423 | 0.00594423 | 0.05390617 |
| RERG | 47.0649765 | 0.169381319 | 0.4229918 | 0.40043642 | 0.68883511 | 0.85864878 |
| EPS8 | 164.657641 | -0.180398102 | 0.31414299 | -0.5742547 | 0.56579544 | 0.7859225 |
| STRAP | 52.7005681 | -0.162083434 | 0.29424726 | -0.5508409 | 0.58174273 | 0.79526454 |
| DERA | 131.068257 | -0.143752974 | 0.20449235 | -0.7029748 | 0.48207145 | 0.74068045 |
| MGST1 | 52.1187808 | -0.535517933 | 0.67224391 | -0.7966125 | 0.42567609 | 0.70025461 |
| RERGL | 24.9306045 | 0.098315345 | 0.61240721 | 0.16053917 | 0.87245637 | 0.94870651 |
| PIK3C2G | 921.676957 | 1.8793629 | 0.55630558 | 3.37829238 | 0.00072937 | 0.01164964 |
| PLCZ1 | 13.851797 | -0.238669034 | 0.82786342 | -0.2882952 | 0.7731208 | 0.90149298 |
| AC087242.1 | 35.7649663 | 1.798533375 | 0.5721608 | 3.14340545 | 0.00166994 | 0.02155859 |
| PLEKHA5 | 889.229147 | 0.146045475 | 0.28525257 | 0.51198654 | 0.60866043 | 0.81261032 |
| AEBP2 | 245.070897 | 0.122613219 | 0.26662153 | 0.45987741 | 0.64560422 | 0.83387847 |
| PYROXD1 | 29.3995228 | -0.257143138 | 0.31780862 | -0.8091132 | 0.41845005 | 0.69462979 |
| RECQL | 24.7314149 | -0.785638471 | 0.36139175 | -2.1739247 | 0.02971079 | 0.15648289 |
| GOLT1B | 31.06351 | -0.494538879 | 0.3073823 | -1.6088723 | 0.10764426 | 0.34161321 |
| LDHB | 330.56483 | -0.467670466 | 0.42605607 | -1.0976735 | 0.27234708 | 0.56475837 |
| AC010197.1 | 147.321999 | 0.348274667 | 0.54623231 | 0.6375944 | 0.52373774 | 0.76218158 |
| AC010185.1 | 19.9355068 | -0.387061933 | 0.65772411 | -0.5884868 | 0.55620562 | 0.78054429 |
| ABCC9 | 75.9076485 | -2.572333877 | 1.20864261 | -2.1282833 | 0.0333136 | 0.16856421 |
| AC008250.1 | 134.233911 | -0.546390607 | 0.66973944 | -0.8158256 | 0.41459987 | 0.69139449 |

| | | | | | | |
|---|---|---|---|---|---|---|
| CMAS | 28.1562007 | -0.226627546 | 0.26677292 | -0.8495148 | 0.3955949 | 0.67612337 |
| C2CD5 | 120.256165 | 0.485724745 | 0.22696532 | 2.14008351 | 0.03234802 | 0.16554858 |
| ETNK1 | 272.011962 | 0.642019526 | 0.33432948 | 1.92031981 | 0.05481752 | 0.23078465 |
| SOX5 | 912.873468 | 0.766873065 | 0.41658529 | 1.84085489 | 0.06564283 | 0.25605922 |
| CASC1 | 31.8213813 | 0.24664242 | 0.64057772 | 0.38503122 | 0.70021429 | 0.86380165 |
| ETFRF1 | 30.279462 | -0.139433028 | 0.37722631 | -0.369627 | 0.71166043 | 0.87074117 |
| KRAS | 89.655665 | -0.309545912 | 0.28882509 | -1.0717418 | 0.28383604 | 0.57640034 |
| RASSF8-AS1 | 26.9399613 | -0.634725089 | 0.35814804 | -1.7722423 | 0.07635435 | 0.28042437 |
| RASSF8 | 160.431242 | -0.375378236 | 0.34173055 | -1.0984626 | 0.27200253 | 0.56461214 |
| AC024145.1 | 13.0783103 | -0.471893818 | 0.73532009 | -0.6417529 | 0.52103362 | 0.76092013 |
| ITPR2 | 782.447772 | 0.613379359 | 0.2715132 | 2.25911435 | 0.02387627 | 0.13541585 |
| AC023051.1 | 17.4064433 | -0.631004572 | 0.59118501 | -1.0673555 | 0.28581134 | 0.57835954 |
| INTS13 | 20.4602798 | -0.196894819 | 0.31863887 | -0.6179247 | 0.536625 | 0.76910277 |
| FGFR1OP2 | 58.0447206 | 0.592540207 | 0.29214266 | 2.02825635 | 0.04253409 | 0.19693762 |
| TM7SF3 | 66.9520708 | -0.056619276 | 0.28278538 | -0.20022 | 0.84130856 | 0.93566718 |
| MED21 | 24.0117136 | -0.368397674 | 0.32157195 | -1.1456151 | 0.25195446 | 0.54334972 |
| STK38L | 48.5223753 | 0.802741255 | 0.26545786 | 3.02398749 | 0.00249467 | 0.02865113 |
| ARNTL2 | 17.8270039 | -1.018556456 | 0.72527127 | -1.40438 | 0.16020573 | 0.42791337 |
| PPFIBP1 | 382.187434 | 0.423572577 | 0.29200752 | 1.45055365 | 0.14690419 | 0.40616768 |
| MRPS35 | 48.0529803 | 0.049462126 | 0.27868265 | 0.17748549 | 0.85912706 | 0.94407755 |
| KLHL42 | 38.6257946 | -0.556192558 | 0.40180858 | -1.3842227 | 0.16629027 | 0.4351146 |
| CCDC91 | 300.334791 | 0.122133604 | 0.15455622 | 0.7902212 | 0.4293986 | 0.70316149 |
| FAR2 | 33.1385603 | -1.342384928 | 0.49598437 | -2.7065065 | 0.00679953 | 0.0591532 |
| AC009318.1 | 19.0216054 | -0.05275014 | 0.53650767 | -0.0983213 | 0.92167715 | 0.9686919 |
| ERGIC2 | 69.1231605 | 0.343988882 | 0.23820391 | 1.44409416 | 0.14871249 | 0.4090884 |
| TMTC1 | 23.3999574 | 0.124585054 | 0.72124798 | 0.1727354 | 0.86285942 | 0.94620532 |
| IPO8 | 77.3841656 | 0.586899048 | 0.24239006 | 2.4213 | 0.01546511 | 0.101998 |
| CAPRIN2 | 56.4980132 | 0.038833457 | 0.23312677 | 0.16657657 | 0.86770323 | 0.947639 |
| DDX11 | 11.7286476 | 0.392539067 | 0.59266569 | 0.66232797 | 0.50776105 | 0.75382716 |
| SINHCAF | 106.356611 | -0.454421156 | 0.19115012 | -2.3772998 | 0.01743991 | 0.11063172 |
| DENND5B | 239.663403 | 0.020045476 | 0.33330141 | 0.06014219 | 0.95204239 | 0.98079407 |
| AMN1 | 58.6909171 | 0.148106002 | 0.25842163 | 0.57311766 | 0.56656503 | 0.78647713 |
| KIAA1551 | 90.3853408 | -0.328925954 | 0.24093498 | -1.3652063 | 0.17218823 | 0.44469368 |
| BICD1 | 449.345742 | -0.368412826 | 0.28894687 | -1.2750193 | 0.20230242 | 0.4856498 |
| FGD4 | 955.020134 | 0.484004929 | 0.1964777 | 2.46340895 | 0.01376228 | 0.0951632 |
| DNM1L | 141.963997 | 0.05389013 | 0.16221742 | 0.33220926 | 0.73973125 | 0.88614269 |
| YARS2 | 42.4358222 | -0.010285384 | 0.29442464 | -0.0349338 | 0.9721325 | 0.98833746 |
| PKP2 | 14.8607119 | -0.673700134 | 0.40478475 | -1.6643417 | 0.09604416 | 0.31958881 |
| ALG10B | 14.543675 | 0.530233895 | 0.39402881 | 1.3456729 | 0.17840803 | 0.45374495 |
| CPNE8 | 203.500112 | -0.329850904 | 0.46892539 | -0.7034187 | 0.48179483 | 0.74068045 |
| KIF21A | 56.3750668 | 0.773407912 | 0.33500681 | 2.30863338 | 0.02096393 | 0.12404965 |
| SLC2A13 | 78.3945307 | -1.837639189 | 0.3612849 | -5.0863991 | 3.65E-07 | 2.47E-05 |
| LRRK2 | 140.749034 | -0.982469705 | 0.51727693 | -1.8993109 | 0.05752361 | 0.23694853 |
| GXYLT1 | 55.1286721 | -0.026094888 | 0.24405428 | -0.1069225 | 0.91485048 | 0.96464134 |
| YAF2 | 119.608833 | 0.120927477 | 0.21510528 | 0.56217809 | 0.57399469 | 0.79097021 |
| PPHLN1 | 135.954497 | 0.129263246 | 0.17975116 | 0.7191233 | 0.47206495 | 0.73329331 |
| ZCRB1 | 24.6638681 | -0.48119658 | 0.33854329 | -1.4213738 | 0.15520811 | 0.41868392 |
| PRICKLE1 | 75.8939319 | -3.466497758 | 0.80665068 | -4.2973964 | 1.73E-05 | 0.00057938 |
| PUS7L | 44.0762473 | 0.367643514 | 0.30307959 | 1.2130263 | 0.22511977 | 0.51390457 |
| IRAK4 | 42.1545299 | -0.302497303 | 0.37578958 | -0.8049646 | 0.42084012 | 0.69661173 |
| TWF1 | 32.5213892 | -0.418995487 | 0.30178656 | -1.3883835 | 0.1650203 | 0.43379411 |
| TMEM117 | 382.666561 | 0.477831475 | 0.20363131 | 2.34655214 | 0.01894802 | 0.11674788 |
| ANO6 | 202.09161 | -0.24513395 | 0.29087053 | -0.8427597 | 0.39936287 | 0.67979406 |
| AC008124.1 | 16.7863179 | 0.909585677 | 0.37488601 | 2.42629934 | 0.01525368 | 0.1013175 |
| ARID2 | 283.559434 | 0.516045227 | 0.22784188 | 2.264927 | 0.02351716 | 0.13454711 |
| SCAF11 | 275.652082 | -0.047692469 | 0.14346894 | -0.3324236 | 0.73956939 | 0.88614269 |
| SLC38A1 | 366.317632 | 0.151447496 | 0.50334347 | 0.30088301 | 0.76350371 | 0.8974069 |
| SLC38A2 | 190.275893 | -0.178649504 | 0.20563768 | -0.8687586 | 0.38497917 | 0.66701235 |
| AC008014.1 | 508.506433 | 0.785497611 | 0.28700421 | 2.73688534 | 0.00620239 | 0.05537004 |
| SLC38A4 | 26.2194865 | 1.135074642 | 0.77732311 | 1.4602353 | 0.14422542 | 0.40202041 |
| PCED1B | 122.790825 | -2.019694905 | 0.66006476 | -3.0598436 | 0.00221453 | 0.02619294 |
| RPAP3 | 27.401129 | 0.453414067 | 0.31778715 | 1.42678538 | 0.15364177 | 0.41664056 |
| RAPGEF3 | 109.660199 | 1.042592797 | 0.24002213 | 4.34373604 | 1.40E-05 | 0.00048732 |
| SLC48A1 | 41.8526486 | 0.169443515 | 0.30957859 | 0.54733603 | 0.58414789 | 0.79631799 |
| HDAC7 | 108.362797 | -0.946374339 | 0.44948363 | -2.1054701 | 0.03525041 | 0.1742561 |
| VDR | 35.2900484 | 0.396575441 | 0.52098706 | 0.76120017 | 0.44653752 | 0.71619194 |
| TMEM106C | 24.8743962 | -0.095240454 | 0.47606352 | -0.2000583 | 0.84143499 | 0.93566718 |
| SENP1 | 36.9474596 | 0.359050661 | 0.3266972 | 1.09903194 | 0.27175413 | 0.5642932 |
| PFKM | 24.3150659 | 0.307782118 | 0.4053546 | 0.75929105 | 0.44767847 | 0.71693398 |
| ASB8 | 24.3947351 | -0.403265201 | 0.34595573 | -1.1656555 | 0.24375376 | 0.53451494 |
| ZNF641 | 14.4719222 | -0.213245489 | 0.41926029 | -0.5086232 | 0.6110164 | 0.81376433 |
| AC090115.1 | 16.5732922 | 0.349465488 | 0.58885229 | 0.59346885 | 0.55286743 | 0.77787735 |
| KANSL2 | 22.5029237 | 0.722360627 | 0.32096891 | 2.25056263 | 0.02441325 | 0.13661343 |
| CCNT1 | 48.057109 | 0.00949118 | 0.22790165 | 0.04164595 | 0.96678094 | 0.98695423 |
| ADCY6 | 43.6583434 | 0.317586184 | 0.25607202 | 1.2402221 | 0.21489326 | 0.50124063 |
| DDX23 | 24.3332219 | -0.839979341 | 0.41376361 | -2.0300948 | 0.04234691 | 0.19633566 |
| CCDC65 | 23.3643159 | -0.554391162 | 0.43963374 | -1.2610296 | 0.20729818 | 0.49241084 |
| PRKAG1 | 31.1498185 | -0.362698431 | 0.27553505 | -1.3163422 | 0.1880592 | 0.46620051 |
| KMT2D | 31.8763511 | -0.431699183 | 0.32689719 | -1.3205962 | 0.18663605 | 0.46408993 |
| LMBR1L | 36.7055104 | 0.045606932 | 0.33847576 | 0.13474209 | 0.89281579 | 0.95782021 |
| AC011603.2 | 15.322265 | 0.741091551 | 0.36939211 | 2.00624629 | 0.04482998 | 0.20202906 |
| TUBA1B | 89.5113411 | -1.584429882 | 0.56330221 | -2.8127528 | 0.00491194 | 0.04689786 |
| TUBA1A | 38.803284 | -1.073821719 | 0.57104546 | -1.8804488 | 0.06004694 | 0.24297078 |
| TUBA1C | 64.1733397 | -0.624973961 | 0.29217608 | -2.1390319 | 0.03243308 | 0.16565474 |
| SPATS2 | 268.137882 | 0.810871012 | 0.3361455 | 2.412262 | 0.01585388 | 0.1036995 |
| MCRS1 | 16.1660806 | -0.12563558 | 0.42905609 | -0.2928185 | 0.76966085 | 0.90034488 |
| PRPF40B | 87.2082743 | 0.119664136 | 0.22285827 | 0.53695175 | 0.59130094 | 0.80036557 |
| FMNL3 | 12.8518197 | -0.448009769 | 0.62421824 | -0.7177134 | 0.47293404 | 0.73375471 |
| TMBIM6 | 294.251233 | -0.704264278 | 0.39023826 | -1.8047033 | 0.07112112 | 0.26917332 |
| NCKAP5L | 24.2166076 | -0.435321775 | 0.36873123 | -1.1805937 | 0.23776415 | 0.52789791 |
| AQP2 | 591.177486 | -0.072417387 | 0.79428998 | -0.0911725 | 0.92735554 | 0.97111125 |
| SMARCD1 | 24.2902174 | -0.731355213 | 0.32059438 | -2.2812478 | 0.02253379 | 0.13071121 |
| AC074032.1 | 79.5010428 | -0.073919258 | 0.29210336 | -0.2530586 | 0.80022295 | 0.91455697 |
| CERS5 | 60.2986283 | -0.287106575 | 0.28650528 | -1.0020987 | 0.31629591 | 0.60951749 |
| LIMA1 | 266.649594 | -1.072250004 | 0.37129279 | -2.8878827 | 0.00387845 | 0.03930906 |
| LARP4 | 166.618703 | 0.88549046 | 0.23035693 | 3.84399317 | 0.00012105 | 0.00284712 |
| DIP2B | 235.260002 | 0.008924906 | 0.2608293 | 0.03421742 | 0.97270377 | 0.98856766 |
| ATF1 | 70.1423612 | 0.451905551 | 0.21952858 | 2.05852716 | 0.03953956 | 0.18788584 |
| METTL7A | 51.2068722 | 0.889918935 | 0.51135097 | 1.74032902 | 0.08180126 | 0.29260993 |
| SLC11A2 | 232.240716 | -0.627553648 | 0.27816032 | -2.2560862 | 0.02406523 | 0.13578849 |
| LETMD1 | 92.446289 | 0.335374553 | 0.24695333 | 1.35804832 | 0.17444835 | 0.44724303 |
| CSRNP2 | 26.4895707 | -0.929798296 | 0.32192414 | -2.8882528 | 0.00387388 | 0.03930906 |
| TFCP2 | 171.829886 | 0.760386545 | 0.21592811 | 3.52148016 | 0.00042914 | 0.00771597 |
| POU6F1 | 24.5901395 | 0.595033732 | 0.32647257 | 1.8226148 | 0.06836175 | 0.26278462 |
| DAZAP2 | 127.224273 | -0.206365252 | 0.31842329 | -0.6480847 | 0.51693019 | 0.75806617 |
| SMAGP | 16.0712097 | 0.175388682 | 0.51565427 | 0.34012844 | 0.73375981 | 0.88360989 |
| ACVR1B | 248.208086 | 0.062082467 | 0.17072605 | 0.36363792 | 0.71612841 | 0.87318675 |
| NR4A1 | 109.622728 | 4.26657238 | 1.08867439 | 3.91905278 | 8.89E-05 | 0.00221419 |
| ATG101 | 14.9861461 | -0.808228152 | 0.60152924 | -1.3436224 | 0.17907053 | 0.45449239 |
| AC078864.1 | 31.6426144 | 0.074855215 | 0.52345065 | 0.14300339 | 0.88628751 | 0.95619452 |
| KRT7 | 61.5621236 | -2.19893805 | 0.65078823 | -3.3788842 | 0.00072781 | 0.01164265 |
| KRT8 | 144.830037 | -0.724771863 | 0.42590646 | -1.701716 | 0.08880861 | 0.30687325 |
| KRT18 | 152.879784 | -1.263617934 | 0.46341195 | -2.7267703 | 0.00639576 | 0.05653552 |

| | | | | | | |
|---|---|---|---|---|---|---|
| EIF4B | 121.320031 | -0.031931244 | 0.23823197 | -0.1340343 | 0.89337549 | 0.95800753 |
| AC068888.1 | 40.6517349 | -0.157110133 | 0.33755332 | -0.465438 | 0.64161785 | 0.8325919 |
| TNS2 | 29.4525239 | -0.826872655 | 0.45862221 | -1.8029494 | 0.07139616 | 0.26971882 |
| CSAD | 85.6911381 | 0.83266395 | 0.1833486 | 4.54142517 | 5.59E-06 | 0.00023305 |
| ZNF740 | 15.5418699 | -0.442020524 | 0.39849161 | -1.1092342 | 0.26732916 | 0.55907938 |
| RARG | 16.205395 | -0.249142506 | 0.48507532 | -0.5136161 | 0.60752041 | 0.81186603 |
| PFDN5 | 79.3472078 | -0.172333902 | 0.43003053 | -0.4007481 | 0.68860562 | 0.8585881 |
| C12orf10 | 11.2219619 | -0.52388077 | 0.49352947 | -1.0614984 | 0.28846344 | 0.58064041 |
| AAAS | 14.9260634 | 0.517466996 | 0.37195166 | 1.39122111 | 0.16415839 | 0.4329646 |
| SP1 | 77.9279777 | -0.688640208 | 0.27922487 | -2.4662567 | 0.01365334 | 0.09491009 |
| PRR13 | 70.0921825 | -0.219524943 | 0.37535114 | -0.5848522 | 0.55864709 | 0.78244004 |
| PCBP2 | 179.47173 | 0.185428351 | 0.18796227 | 0.986519 | 0.3238785 | 0.61448136 |
| MAP3K12 | 18.15695 | -0.823238667 | 0.4114248 | -2.0009457 | 0.04539824 | 0.20400635 |
| ATP5MC2 | 122.473806 | -0.14356441 | 0.24890919 | -0.5767742 | 0.56409197 | 0.78488324 |
| CALCOCO1 | 32.6135353 | 0.050757992 | 0.39883514 | 0.12726559 | 0.89873019 | 0.95935794 |
| HOXC4 | 13.1807205 | 1.111024197 | 0.38983709 | 2.84997044 | 0.00437233 | 0.04293969 |
| SMUG1 | 23.6990418 | 0.545682877 | 0.30376083 | 1.79642282 | 0.0724273 | 0.27172106 |
| LINC02381 | 35.1903273 | 0.314025157 | 0.3360021 | 0.93459285 | 0.34999817 | 0.63675792 |
| CBX5 | 108.268745 | -0.217633366 | 0.27319794 | -0.7966142 | 0.42567512 | 0.70025461 |
| HNRNPA1 | 66.1563637 | -0.254743793 | 0.24844995 | -1.0253324 | 0.30520636 | 0.59961934 |
| COPZ1 | 51.9193337 | -0.324140387 | 0.23711665 | -1.3670081 | 0.17162276 | 0.44345564 |
| CD63 | 128.044465 | -1.574585822 | 0.41876909 | -3.7600335 | 0.00016989 | 0.00371187 |
| ORMDL2 | 12.8778016 | -1.177052116 | 0.44865775 | -2.6234967 | 0.00870323 | 0.06977484 |
| RAB5B | 48.1963537 | -0.118701998 | 0.26348118 | -0.4505141 | 0.65233976 | 0.83816466 |
| SUOX | 18.8540128 | -0.33534139 | 0.48696679 | -0.688633 | 0.49105426 | 0.74574113 |
| IKZF4 | 14.3737801 | -0.246414441 | 0.44653762 | -0.5518336 | 0.5810624 | 0.79484057 |
| RPS26 | 21.8389242 | -0.993095361 | 0.44337878 | -2.2398351 | 0.02510163 | 0.13948057 |
| ERBB3 | 22.2282112 | -1.014850714 | 0.45005365 | -2.254955 | 0.02413615 | 0.13583965 |
| PA2G4 | 23.5123438 | -0.734107921 | 0.42876475 | -1.7121462 | 0.08686974 | 0.30295601 |
| RPL41 | 338.234232 | -0.610131473 | 0.38599071 | -1.5806895 | 0.11394904 | 0.35252343 |
| ESYT1 | 22.324144 | -0.621507959 | 0.31922024 | -1.9469566 | 0.05153993 | 0.22141969 |
| MYL6B | 14.2379571 | -0.305262214 | 0.43707259 | -0.6984245 | 0.48491175 | 0.74285379 |
| MYL6 | 166.4437 | 0.03703299 | 0.39350953 | 0.09410951 | 0.92502217 | 0.97032668 |
| SMARCC2 | 72.9513788 | -0.099464355 | 0.26131196 | -0.3806345 | 0.70347446 | 0.86585351 |
| RNF41 | 18.6810275 | -0.753627214 | 0.32104419 | -2.3474252 | 0.01890367 | 0.116684 |
| ANKRD52 | 26.3065472 | -0.127743941 | 0.28951178 | -0.4412392 | 0.65903985 | 0.84059189 |
| COQ10A | 8.53385393 | 0.049368866 | 0.64414684 | 0.07664225 | 0.93890814 | 0.97662903 |
| CS | 48.2982773 | 0.077406883 | 0.30300314 | 0.25546561 | 0.7983635 | 0.91355557 |
| AC073896.2 | 13.240312 | 0.305791806 | 0.35729758 | 0.85584628 | 0.39208281 | 0.67329816 |
| PAN2 | 30.5830961 | 0.036009632 | 0.29533859 | 0.12192661 | 0.90295714 | 0.96126516 |
| STAT2 | 48.9689338 | -0.08078279 | 0.49829584 | -0.1621181 | 0.87121283 | 0.94844301 |
| SPRYD4 | 13.2197035 | -0.513148763 | 0.45601857 | -1.1252804 | 0.26047022 | 0.55146957 |
| RBMS2 | 89.0276396 | 0.853186516 | 0.34244742 | 2.49143801 | 0.01272272 | 0.09058011 |
| BAZ2A | 75.7275416 | -0.147627142 | 0.21593136 | -0.6836763 | 0.49417962 | 0.74700665 |
| ATP5F1B | 156.950944 | 0.134254476 | 0.42660494 | 0.31470446 | 0.75298606 | 0.89205725 |
| PTGES3 | 195.538347 | 0.528939421 | 0.27304507 | 1.93718726 | 0.05272246 | 0.22490549 |
| NACA | 74.4339448 | -0.662591786 | 0.29333374 | -2.2588325 | 0.0238938 | 0.13541585 |
| NEMP1 | 13.8880623 | -0.643102704 | 0.42076286 | -1.5284208 | 0.1264081 | 0.37228594 |
| STAT6 | 77.6661021 | -0.967274607 | 0.26989804 | -3.5838519 | 0.00033856 | 0.00643522 |
| R3HDM2 | 257.706643 | 0.501428957 | 0.23807985 | 2.10613772 | 0.03519239 | 0.1742352 |
| MARS | 31.9386086 | -0.563921858 | 0.55860962 | -1.0095098 | 0.31273023 | 0.6055915 |
| DDIT3 | 50.5460416 | -0.994481999 | 0.83749966 | -1.1874417 | 0.23505345 | 0.52531003 |
| MBD6 | 19.3173169 | -0.242806121 | 0.44963373 | -0.5400087 | 0.58919104 | 0.79929498 |
| DCTN2 | 47.2555997 | -0.45325899 | 0.27508749 | -1.6476903 | 0.09941624 | 0.32555183 |
| PIP4K2C | 34.2576619 | 0.266207505 | 0.31348394 | 0.84919026 | 0.39577543 | 0.67616559 |
| DTX3 | 15.2825511 | 0.251174502 | 0.44824193 | 0.56035476 | 0.57523748 | 0.79151817 |
| OS9 | 65.7832581 | -0.499721021 | 0.37807101 | -1.321765 | 0.18624642 | 0.46377434 |
| TSPAN31 | 19.1816598 | -1.224758808 | 0.47462295 | -2.580488 | 0.00986608 | 0.07654439 |
| CDK4 | 16.8201527 | -0.224963416 | 0.40206485 | -0.5595202 | 0.57580673 | 0.79171629 |
| 9-Mar | 18.1960105 | -0.493541754 | 0.35283446 | -1.3987913 | 0.16187559 | 0.42937872 |
| TSFM | 23.7110601 | -0.719973801 | 0.44987805 | -1.6003755 | 0.10951531 | 0.34481257 |
| AVIL | 35.9955869 | -1.19764311 | 0.36107594 | -3.3168732 | 0.00091031 | 0.01369107 |
| CTDSP2 | 72.2426381 | -0.266823995 | 0.18538999 | -1.4392579 | 0.15007748 | 0.41097772 |
| LRIG3 | 18.4745695 | -0.232156231 | 0.53938146 | -0.430412 | 0.666896 | 0.84492213 |
| SLC16A7 | 370.101239 | 0.975338678 | 0.45355527 | 2.1504296 | 0.03152125 | 0.16250135 |
| FAM19A2 | 68.8708543 | 0.302271706 | 0.41358606 | 0.73085564 | 0.46486733 | 0.72848544 |
| USP15 | 208.052708 | 0.324675457 | 0.23150926 | 1.40242966 | 0.16078698 | 0.42803048 |
| MON2 | 286.419012 | 0.441092525 | 0.18379141 | 2.39996272 | 0.01639674 | 0.10617174 |
| AC048341.1 | 45.5534569 | -0.728357511 | 0.5524079 | -1.3185139 | 0.18733166 | 0.46503392 |
| PPM1H | 228.099883 | -0.405277342 | 0.27982282 | -1.4483355 | 0.14752323 | 0.40740971 |
| AVPR1A | 28.6357104 | 0.250566683 | 0.5188013 | 0.48297234 | 0.62911537 | 0.8249932 |
| AC026116.1 | 21.644983 | 2.520700475 | 0.6496319 | 3.8801981 | 0.00010437 | 0.00250055 |
| DPY19L2 | 521.730721 | 1.661248869 | 0.2842708 | 5.84389551 | 5.10E-09 | 6.74E-07 |
| RXYLT1 | 26.1278898 | -0.404484469 | 0.2944329 | -1.3737747 | 0.16951164 | 0.44032355 |
| SRGAP1 | 118.802352 | -0.90055991 | 0.38865177 | -2.3171383 | 0.0204962 | 0.1227566 |
| XPOT | 32.8573857 | -0.601451038 | 0.39122117 | -1.5373683 | 0.12420314 | 0.36916928 |
| TBK1 | 76.3196246 | -0.382884064 | 0.30791471 | -1.2434744 | 0.21369306 | 0.50026184 |
| RASSF3 | 28.4990996 | -1.405437078 | 0.48904835 | -2.8738203 | 0.0040554 | 0.04048323 |
| GNS | 135.165036 | -0.603904043 | 0.2777716 | -2.1741029 | 0.02969741 | 0.15648289 |
| TBC1D30 | 50.9879872 | -0.088661141 | 0.2532068 | -0.3501531 | 0.72622382 | 0.87880825 |
| LINC02389 | 12.5656671 | -0.579826607 | 0.4262574 | -1.3602734 | 0.17374342 | 0.44636027 |
| LEMD3 | 100.699013 | 0.427977012 | 0.23768536 | 1.80060318 | 0.07176545 | 0.27041976 |
| HMGA2 | 26.3618493 | -0.126450251 | 0.76885073 | -0.1644666 | 0.86936385 | 0.94827256 |
| AC090673.1 | 41.1849592 | 1.452949409 | 0.72350561 | 2.00820753 | 0.04462124 | 0.20147742 |
| LLPH | 10.7248702 | -0.908959804 | 0.48538215 | -1.8726684 | 0.06111419 | 0.24535775 |
| HELB | 40.6332568 | -0.616620482 | 0.34781328 | -1.7728491 | 0.07625371 | 0.28035892 |
| GRIP1 | 1232.33117 | 0.868497342 | 0.18509635 | 4.69213644 | 2.70E-06 | 0.00012724 |
| CAND1 | 99.1342327 | -0.079452951 | 0.22321284 | -0.3559515 | 0.72187687 | 0.87625492 |
| DYRK2 | 23.2693453 | -2.391313427 | 0.57207811 | -4.1800471 | 2.91E-05 | 0.0008936 |
| MDM1 | 13.3555578 | 0.394904614 | 0.44937633 | 0.87878374 | 0.37951854 | 0.66290268 |
| RAP1B | 318.502845 | -1.253653189 | 0.27425561 | -4.5711123 | 4.85E-06 | 0.00020738 |
| NUP107 | 54.1006497 | -0.679722799 | 0.38744918 | -1.7543534 | 0.07936997 | 0.28711147 |
| SLC35E3 | 46.0541486 | -0.619236295 | 0.37660799 | -1.6442463 | 0.10012534 | 0.326628 |
| MDM2 | 82.3552542 | -0.23764404 | 0.31819601 | -0.746848 | 0.45515534 | 0.72210564 |
| CPM | 75.4041933 | 0.109079404 | 0.7269193 | 0.1500571 | 0.88071956 | 0.953465 |
| CPSF6 | 151.272744 | 0.750775691 | 0.25557041 | 2.93764716 | 0.00330713 | 0.03487213 |
| YEATS4 | 20.5050803 | -0.063111278 | 0.46914712 | -0.1345234 | 0.89298869 | 0.95782021 |
| FRS2 | 134.633974 | 0.429305011 | 0.23931136 | 1.79391819 | 0.07282623 | 0.27252299 |
| CCT2 | 39.3219184 | -0.18357266 | 0.35067999 | -0.5234763 | 0.60064284 | 0.80738812 |
| RAB3IP | 136.899333 | 0.617693828 | 0.23542689 | 2.62371825 | 0.00869757 | 0.06977484 |
| MYRFL | 61.2373515 | 1.117727334 | 0.62873764 | 1.77773249 | 0.0754478 | 0.27878882 |
| AC025159.1 | 43.9617017 | -0.546121539 | 0.62844772 | -0.8690008 | 0.38484672 | 0.66701235 |
| CNOT2 | 189.264151 | 0.241150217 | 0.21282355 | 1.1330993 | 0.25717256 | 0.54818781 |
| KCNMB4 | 17.1313895 | 0.919426007 | 0.48655143 | 1.88967896 | 0.05880091 | 0.24000052 |
| ZFC3H1 | 139.798606 | -0.049777626 | 0.27751435 | -0.1793696 | 0.85764753 | 0.94359207 |
| RAB21 | 124.143569 | 0.02953817 | 0.2405752 | 0.12278144 | 0.90228017 | 0.96086649 |
| TBC1D15 | 106.952692 | -0.677587804 | 0.26294068 | -2.5769607 | 0.00996733 | 0.07703182 |
| TRHDE | 368.078712 | 1.228623284 | 0.26267199 | 4.67740502 | 2.91E-06 | 0.00013605 |
| ATXN7L3B | 42.8111297 | -0.796086937 | 0.32396598 | -2.4573165 | 0.01399793 | 0.09620193 |
| CAPS2 | 112.24987 | 1.034727547 | 0.5661774 | 1.82756773 | 0.06761445 | 0.26098543 |
| KRR1 | 39.7830391 | 0.016412727 | 0.24510688 | 0.06696151 | 0.94661234 | 0.9791165 |
| PHLDA1 | 20.1748437 | -0.771425234 | 0.61700167 | -1.2502806 | 0.21119705 | 0.49712778 |
| NAP1L1 | 132.084727 | 0.016222343 | 0.22531865 | 0.07199734 | 0.94260403 | 0.9781624 |

| | | | | | | |
|---|---|---|---|---|---|---|
| OSBPL8 | 204.360819 | 0.346891415 | 0.21983314 | 1.57797599 | 0.11457113 | 0.35349258 |
| ZDHHC17 | 97.1020782 | 0.32380397 | 0.2607765 | 1.24169155 | 0.21435039 | 0.50088196 |
| SYT1 | 25.2423822 | -1.477489668 | 0.77415785 | -1.9085122 | 0.05632505 | 0.23435408 |
| AC027288.3 | 27.3974647 | 0.437632165 | 0.6141845 | 0.71254185 | 0.4761293 | 0.73631255 |
| PAWR | 302.030631 | 0.469072154 | 0.22640423 | 2.07183476 | 0.03828085 | 0.18325713 |
| AC073569.2 | 18.7147124 | -0.236689744 | 0.51197035 | -0.4623114 | 0.643858 | 0.83339931 |
| PPP1R12A | 269.205851 | 0.317212682 | 0.17764283 | 1.78567679 | 0.0741516 | 0.27538206 |
| LIN7A | 127.627842 | 0.785093084 | 0.40375073 | 1.94449947 | 0.05183525 | 0.22230553 |
| ACSS3 | 127.127937 | 0.70052923 | 0.59038272 | 1.18656798 | 0.23539807 | 0.52547582 |
| PPFIA2 | 60.826963 | 1.042218664 | 0.79775296 | 1.30644287 | 0.19140199 | 0.47171515 |
| CCDC59 | 21.3039095 | -0.280616656 | 0.49032239 | -0.5723105 | 0.56711163 | 0.78656476 |
| METTL25 | 91.6350115 | 0.641995788 | 0.20739208 | 3.09556563 | 0.00196438 | 0.02423597 |
| TMTC2 | 1135.40106 | 0.712251224 | 0.3613005 | 1.97135411 | 0.04868339 | 0.21414869 |
| LRRIQ1 | 47.8685624 | -0.161198371 | 0.57554233 | -0.2800808 | 0.7794155 | 0.90427469 |
| RASSF9 | 54.7805007 | -0.177081022 | 0.37692692 | -0.469802 | 0.63849649 | 0.83094881 |
| CEP290 | 90.0047544 | 0.247140133 | 0.2163056 | 1.14255079 | 0.25322515 | 0.54473281 |
| TMTC3 | 69.6248761 | 0.16813529 | 0.21853088 | 0.76938915 | 0.44166233 | 0.71227192 |
| KITLG | 1053.51218 | -0.161848834 | 0.49499951 | -0.3269677 | 0.74369234 | 0.88767449 |
| AC024941.2 | 9.19893786 | 0.685500488 | 0.78284953 | 0.87564783 | 0.38122152 | 0.66452522 |
| DUSP6 | 127.889886 | -0.006960781 | 0.72798305 | -0.0095617 | 0.99237096 | 0.99770844 |
| POC1B | 64.4519354 | -0.491234886 | 0.22245465 | -2.2082474 | 0.02722703 | 0.14755019 |
| POC1B-AS1 | 25.5680316 | -0.237368291 | 0.40741063 | -0.5826266 | 0.56014466 | 0.78316119 |
| ATP2B1 | 106.003789 | 0.617773689 | 0.35434196 | 1.74343926 | 0.08125692 | 0.29141278 |
| BTG1 | 83.5431483 | -1.194035536 | 0.3787002 | -3.1529836 | 0.00161611 | 0.02101154 |
| LINC02391 | 30.8633644 | 0.874388146 | 0.45198983 | 1.93453059 | 0.05304792 | 0.22595384 |
| PLEKHG7 | 14.3879124 | -1.146322623 | 0.9489183 | -1.2080309 | 0.22703538 | 0.51677386 |
| EEA1 | 132.527475 | 0.002465279 | 0.30707922 | 0.00802815 | 0.99359453 | 0.99805098 |
| AC138123.1 | 146.3178 | 2.175851492 | 0.38658856 | 5.62833908 | 1.82E-08 | 2.13E-06 |
| AC124947.1 | 42.3292354 | 1.893896863 | 0.50938702 | 3.71799199 | 0.00020081 | 0.00423062 |
| NUDT4 | 209.722025 | 2.031240265 | 0.35135521 | 5.78115877 | 7.42E-09 | 9.55E-07 |
| UBE2N | 63.1268662 | -0.122231218 | 0.23029524 | -0.5307588 | 0.59558595 | 0.8037044 |
| MRPL42 | 63.2753524 | 0.199558532 | 0.21852035 | 0.9132263 | 0.36112353 | 0.64543627 |
| SOCS2 | 30.9381303 | 0.529964119 | 0.39947446 | 1.32665331 | 0.18462339 | 0.46134371 |
| CRADD | 361.425878 | 0.390665619 | 0.28960476 | 1.34896132 | 0.17734939 | 0.45201718 |
| AC012085.2 | 25.4610714 | -0.640705049 | 0.72219118 | -0.8871682 | 0.37498835 | 0.65806152 |
| CEP83 | 77.4326458 | 0.71581179 | 0.25587 | 2.7975604 | 0.00514901 | 0.04866527 |
| TMCC3 | 42.2935355 | 0.451675477 | 0.37505488 | 1.2042917 | 0.22847685 | 0.51822293 |
| NDUFA12 | 50.8979805 | 0.26368503 | 0.27666377 | 0.95308841 | 0.34054528 | 0.62907597 |
| NR2C1 | 80.295004 | 0.202001454 | 0.20315206 | 0.99433625 | 0.32005919 | 0.6120836 |
| FGD6 | 86.7585528 | -1.204388781 | 0.35091622 | -3.4321263 | 0.00059887 | 0.00993485 |
| VEZT | 187.258584 | -0.366072819 | 0.2144648 | -1.7069133 | 0.08783816 | 0.30529985 |
| METAP2 | 204.697506 | -0.897583671 | 0.47955541 | -1.8716996 | 0.06124817 | 0.24542151 |
| USP44 | 13.0696031 | -0.688396727 | 0.49077889 | -1.4026617 | 0.16071776 | 0.42803048 |
| NTN4 | 189.747219 | 0.544330374 | 0.45886015 | 1.1862664 | 0.23551711 | 0.52562766 |
| SNRPF | 19.2774275 | -0.246898077 | 0.37967469 | -0.6502885 | 0.51550589 | 0.75713618 |
| LTA4H | 38.0886096 | -0.103742592 | 0.3451375 | -0.3005834 | 0.76373222 | 0.89745287 |
| ELK3 | 30.0699017 | -1.98293235 | 0.38993442 | -5.085297 | 3.67E-07 | 2.47E-05 |
| CDK17 | 92.3981903 | -0.097490021 | 0.32210093 | -0.3026692 | 0.76214199 | 0.89711915 |
| CFAP54 | 14.8136026 | -1.123027955 | 0.71778857 | -1.5645665 | 0.11768459 | 0.358386 |
| NEDD1 | 39.3223605 | 0.379367398 | 0.2584406 | 1.46790944 | 0.14212882 | 0.39820809 |
| TMPO | 52.5710196 | -0.105120705 | 0.25179728 | -0.4174815 | 0.67632626 | 0.85177422 |
| SLC25A3 | 131.239313 | -0.006813677 | 0.42528328 | -0.0160215 | 0.98721724 | 0.99523554 |
| IKBIP | 14.7565418 | -2.020814066 | 0.54498345 | -3.7080283 | 0.00020888 | 0.00438264 |
| APAF1 | 38.6670186 | -0.553901419 | 0.2961285 | -1.8704766 | 0.06141767 | 0.24564181 |
| ANKS1B | 91.1701086 | 0.814740486 | 0.70866638 | 1.1496813 | 0.25027516 | 0.54189464 |
| UHRF1BP1L | 62.8503588 | 0.304264042 | 0.25374715 | 1.1990836 | 0.23049544 | 0.52062355 |
| ACTR6 | 26.5165922 | 0.10965414 | 0.34615656 | 0.31677615 | 0.75141347 | 0.89132635 |
| SCYL2 | 116.581234 | 0.413504258 | 0.32605282 | 1.26821247 | 0.20472209 | 0.48945159 |
| NR1H4 | 169.556969 | -2.166159351 | 0.55010079 | -3.93775 | 8.22E-05 | 0.00210256 |
| GAS2L3 | 21.7439171 | -2.933134692 | 0.64543552 | -4.5444271 | 5.51E-06 | 0.00023292 |
| UTP20 | 44.4422048 | 0.171064776 | 0.24943924 | 0.68579737 | 0.4928409 | 0.74680729 |
| ARL1 | 47.0102286 | -0.41151279 | 0.32481341 | -1.2669206 | 0.2051837 | 0.48964616 |
| CHPT1 | 221.827463 | 0.974204591 | 0.30104133 | 3.23611576 | 0.00121168 | 0.01684582 |
| GNPTAB | 93.2446297 | 0.031930404 | 0.2476632 | 0.12892672 | 0.89741563 | 0.95917897 |
| DRAM1 | 44.6905541 | -1.910217397 | 0.43188874 | -4.4229386 | 9.74E-06 | 0.00035953 |
| WASHC3 | 53.2959952 | 0.358071441 | 0.29993315 | 1.19383751 | 0.23254154 | 0.52238179 |
| NUP37 | 21.2545716 | 0.280918204 | 0.35031441 | 0.80190307 | 0.42260903 | 0.69778363 |
| NT5DC3 | 63.9220583 | 0.365986317 | 0.39982684 | 0.91536206 | 0.36000159 | 0.64398965 |
| HSP90B1 | 182.179483 | -0.28292349 | 0.54122005 | -0.5227513 | 0.60114733 | 0.80754603 |
| C12orf73 | 11.7764628 | 0.030018426 | 0.37978991 | 0.07903956 | 0.93700116 | 0.97613368 |
| TDG | 47.3967412 | 0.149096717 | 0.36117217 | 0.41281341 | 0.67974332 | 0.85407015 |
| HCFC2 | 29.3034006 | 0.517706441 | 0.30321216 | 1.70740661 | 0.0877465 | 0.30508419 |
| NFYB | 18.2955398 | 0.401034935 | 0.36578532 | 1.09636695 | 0.27291822 | 0.56514644 |
| TXNRD1 | 226.551831 | -0.99827183 | 0.25768545 | -3.8739937 | 0.00010707 | 0.00255915 |
| CHST11 | 14.5365382 | -1.993932573 | 0.56178275 | -3.5492948 | 0.00038626 | 0.0071186 |
| SLC41A2 | 32.763836 | -1.599926713 | 0.61961533 | -2.582129 | 0.00981929 | 0.07640355 |
| C12orf45 | 11.4647916 | -0.849561822 | 0.48389198 | -1.7556849 | 0.07914223 | 0.28668189 |
| WASHC4 | 147.552279 | 0.14253779 | 0.26698707 | 0.53387526 | 0.59342784 | 0.80219047 |
| APPL2 | 129.203813 | -0.640439358 | 0.31267039 | -2.0482891 | 0.04053168 | 0.19118926 |
| C12orf75 | 193.244447 | -2.698373409 | 0.52371206 | -5.1523988 | 2.57E-07 | 1.91E-05 |
| NUAK1 | 17.0665647 | 0.168491681 | 0.58187342 | 0.28956759 | 0.77214707 | 0.90105761 |
| CKAP4 | 20.4516273 | -0.742547496 | 0.49544549 | -1.4987471 | 0.13393925 | 0.38457139 |
| TCP11L2 | 30.9536245 | -0.168174371 | 0.37461336 | -0.4489279 | 0.6534837 | 0.83881384 |
| POLR3B | 30.1961479 | 0.285699244 | 0.30904139 | 0.92446919 | 0.35524209 | 0.64025612 |
| RIC8B | 71.723022 | 0.223010861 | 0.2467759 | 0.9036979 | 0.36615561 | 0.6515728 |
| TMEM263 | 54.3453312 | -0.449175794 | 0.352226 | -1.2752488 | 0.20222118 | 0.48561366 |
| MTERF2 | 19.1443946 | 0.711688905 | 0.55413419 | 1.28432592 | 0.19902794 | 0.48187682 |
| CRY1 | 74.645757 | 0.606057215 | 0.64235674 | 0.94349009 | 0.34543028 | 0.63256941 |
| BTBD11 | 115.239743 | 1.817435369 | 0.65584189 | 2.77114865 | 0.00558589 | 0.05141788 |
| PWP1 | 30.3544146 | -0.076961459 | 0.30383324 | -0.2533016 | 0.80003512 | 0.91455697 |
| PRDM4 | 46.3520645 | -0.281172554 | 0.29546276 | -0.9516345 | 0.34128238 | 0.62956442 |
| SART3 | 37.8436696 | 0.337670065 | 0.2598216 | 1.29962275 | 0.1937303 | 0.47413095 |
| ISCU | 83.014425 | -1.068373482 | 0.40059981 | -2.6669345 | 0.00765466 | 0.06391579 |
| CORO1C | 29.4714078 | -1.455036771 | 0.62826799 | -2.3159492 | 0.02056104 | 0.12280939 |
| SSH1 | 137.687622 | 0.131573344 | 0.3223474 | 0.4081725 | 0.68314704 | 0.85592986 |
| USP30 | 22.2110446 | 0.189522504 | 0.35971639 | 0.52686647 | 0.59828632 | 0.8059995 |
| UNG | 17.4426124 | -0.347707722 | 0.43549628 | -0.7984172 | 0.42462842 | 0.69914048 |
| ACACB | 246.002303 | 1.020039692 | 0.28719131 | 3.55177773 | 0.00038264 | 0.00707608 |
| KCTD10 | 23.7571136 | -0.435096082 | 0.4371565 | -0.9952868 | 0.31959681 | 0.611915 |
| UBE3B | 41.1749058 | -0.147023991 | 0.30660957 | -0.4795153 | 0.63157206 | 0.82625173 |
| MMAB | 34.2715796 | -0.264801933 | 0.37859356 | -0.6994359 | 0.48427965 | 0.74250561 |
| MVK | 26.9045558 | -0.328017271 | 0.28396013 | -1.1551526 | 0.24802796 | 0.538841 |
| FAM222A | 23.1056061 | 0.867450126 | 0.42848244 | 2.02447068 | 0.04292175 | 0.19784332 |
| TRPV4 | 134.400421 | 1.078137972 | 0.4337705 | 2.48550319 | 0.01293684 | 0.09185069 |
| GLTP | 50.2802977 | 0.600527767 | 0.33006578 | 1.81941844 | 0.06884762 | 0.26425823 |
| TCHP | 39.497782 | -0.009003447 | 0.37858328 | -0.0237819 | 0.98102654 | 0.99318267 |
| GIT2 | 90.023566 | 0.138600855 | 0.27683523 | 0.50066191 | 0.61660909 | 0.81691188 |
| ANKRD13A | 132.064059 | 0.100492163 | 0.30389775 | 0.33067755 | 0.74088806 | 0.88658715 |
| C12orf76 | 19.235241 | 0.151936267 | 0.40622924 | 0.37401608 | 0.70839236 | 0.868892 |
| IFT81 | 55.9494982 | 0.055277761 | 0.24952899 | 0.22152841 | 0.82468102 | 0.92658565 |
| ATP2A2 | 138.674651 | 0.537036989 | 0.3717912 | 1.44445859 | 0.14861002 | 0.40891571 |
| ANAPC7 | 45.1863134 | -0.153529487 | 0.40432971 | -0.3797136 | 0.70415803 | 0.86617744 |
| ARPC3 | 111.312191 | -0.651378165 | 0.22560724 | -2.8872219 | 0.0038866 | 0.03933178 |

| | | | | | | |
|---|---|---|---|---|---|---|
| GPN3 | 12.6942448 | -0.607156982 | 0.48280468 | -1.2575623 | 0.20855008 | 0.49402397 |
| VPS29 | 58.1446648 | -0.60768276 | 0.22828891 | -2.6619022 | 0.00777004 | 0.06439823 |
| PPTC7 | 69.4555395 | 0.203854205 | 0.27583058 | 0.73905586 | 0.45987308 | 0.72516945 |
| TCTN1 | 31.5034058 | 0.050098587 | 0.42307607 | 0.11841508 | 0.90573878 | 0.96157183 |
| PPP1CC | 32.9883988 | 0.015794709 | 0.31578772 | 0.05001685 | 0.96010896 | 0.98437923 |
| SH2B3 | 17.5217951 | -0.861427934 | 0.35974116 | -2.3945771 | 0.01663955 | 0.10727197 |
| ATXN2 | 322.773026 | -0.016303031 | 0.23698611 | -0.0687932 | 0.94515424 | 0.97858537 |
| BRAP | 42.6234419 | -0.140801393 | 0.28010868 | -0.502667 | 0.61519841 | 0.81632065 |
| ACAD10 | 53.7312356 | 0.39945513 | 0.41415368 | 0.96450944 | 0.33479058 | 0.62346575 |
| ALDH2 | 91.7341067 | -1.109916652 | 0.37233176 | -2.9809884 | 0.0028732 | 0.0316574 |
| MAPKAPK5 | 45.1493624 | 0.708788519 | 0.27400719 | 2.58675156 | 0.00968854 | 0.07550027 |
| TMEM116 | 155.246244 | 1.401342882 | 0.21872163 | 6.40696978 | 1.48E-10 | 3.19E-08 |
| ERP29 | 86.7554676 | -0.90500838 | 0.35392319 | -2.5570756 | 0.01055563 | 0.07972439 |
| NAA25 | 97.1424671 | 0.338128081 | 0.29891998 | 1.13116586 | 0.25798529 | 0.54935189 |
| TRAFD1 | 43.42206 | -0.969819094 | 0.3287096 | -2.9503826 | 0.00317381 | 0.03391661 |
| HECTD4 | 257.834783 | 0.164273032 | 0.19499119 | 0.84246388 | 0.39952835 | 0.67979406 |
| RPL6 | 137.676391 | -0.45698883 | 0.43392755 | -1.0531455 | 0.29227433 | 0.58378114 |
| PTPN11 | 88.2133194 | -0.023913398 | 0.25022194 | -0.0955687 | 0.92386309 | 0.96970326 |
| OAS1 | 19.9455526 | -2.411975439 | 3.00500914 | -0.8026516 | 0.42217613 | 0.69766738 |
| OAS3 | 16.1805679 | 0.011445407 | 2.74107553 | 0.00417552 | 0.99666843 | 0.99868531 |
| DTX1 | 11.2119867 | 0.888114344 | 0.4859238 | 1.82768233 | 0.06759724 | 0.26098543 |
| RASAL1 | 105.554023 | -0.923972053 | 0.57359867 | -1.6108337 | 0.10721598 | 0.34111768 |
| DDX54 | 20.3059866 | -0.234382078 | 0.41945192 | -0.5587818 | 0.57631068 | 0.79184705 |
| TPCN1 | 148.752526 | -0.30727226 | 0.28448534 | -1.0800988 | 0.28009821 | 0.57312294 |
| SLC8B1 | 45.1047188 | -0.391966585 | 0.26370504 | -1.4863826 | 0.13717794 | 0.39028091 |
| PLBD2 | 17.4246523 | -0.375443882 | 0.40695501 | -0.9225685 | 0.35623211 | 0.64081257 |
| SDSL | 20.315851 | 0.052374134 | 0.50927456 | 0.10284067 | 0.91808943 | 0.96669637 |
| RBM19 | 40.5997775 | -0.282883363 | 0.29965649 | -0.9440255 | 0.34515662 | 0.63256941 |
| MED13L | 1227.94164 | 0.463636094 | 0.268677 | 1.72562629 | 0.08441467 | 0.29792391 |
| MAP1LC3B2 | 15.4672302 | 0.152675801 | 0.38201305 | 0.39966122 | 0.68940606 | 0.85868974 |
| C12orf49 | 48.1352302 | -1.652670743 | 0.4874124 | -3.3907031 | 0.00069714 | 0.01124533 |
| RNFT2 | 12.0328966 | 0.187164733 | 0.50875529 | 0.36788754 | 0.71295709 | 0.87134026 |
| FBXW8 | 77.6215198 | -0.138732927 | 0.27109055 | -0.5117586 | 0.60881995 | 0.81265394 |
| TESC | 30.8022756 | -2.619784875 | 0.65284541 | -4.0128717 | 6.00E-05 | 0.00164655 |
| FBXO21 | 74.5768069 | 0.249344725 | 0.22439363 | 1.1111934 | 0.26648511 | 0.55774161 |
| NOS1 | 27.2139135 | 3.838729341 | 0.59518267 | 6.44966582 | 1.12E-10 | 2.57E-08 |
| KSR2 | 183.913985 | -0.096208569 | 0.72036453 | -0.1335554 | 0.89375416 | 0.95813441 |
| WSB2 | 33.4088215 | -0.663712845 | 0.34129276 | -1.9447024 | 0.05181081 | 0.22230553 |
| VSIG10 | 56.3479524 | 0.640044562 | 0.38203034 | 1.67537627 | 0.09386043 | 0.31579038 |
| PEBP1 | 237.027918 | -0.551366899 | 0.46333531 | -1.1899954 | 0.23404819 | 0.52443761 |
| TAOK3 | 216.707064 | 0.352762078 | 0.18256649 | 1.93223895 | 0.05333002 | 0.22655912 |
| SUDS3 | 49.7472306 | 0.067477354 | 0.26118994 | 0.25834591 | 0.79613995 | 0.91195838 |
| AC002070.1 | 68.2983286 | -0.605661925 | 0.54363151 | -1.1141038 | 0.26523466 | 0.55640131 |
| PRKAB1 | 59.7157108 | -0.634518092 | 0.30261767 | -2.0967649 | 0.03601439 | 0.17602651 |
| CIT | 45.7398169 | -0.782638894 | 0.46661108 | -1.6772831 | 0.09348713 | 0.31484288 |
| BICDL1 | 86.1555303 | 1.120018103 | 0.31486729 | 3.55711167 | 0.00037495 | 0.00695997 |
| RAB35 | 24.9352207 | -0.707560104 | 0.42766448 | -1.6544748 | 0.09803108 | 0.32348372 |
| GCN1 | 42.9809995 | -0.241127022 | 0.29254467 | -0.82424 | 0.4098032 | 0.6878124 |
| RPLP0 | 201.514982 | -1.077245038 | 0.47485967 | -2.2685545 | 0.02329543 | 0.13357889 |
| PXN | 69.7117365 | 0.197575168 | 0.22345992 | 0.88416376 | 0.37660782 | 0.65964588 |
| GATC | 18.1681352 | -0.070798213 | 0.35529149 | -0.199268 | 0.84205314 | 0.93566718 |
| SRSF9 | 59.5473563 | -0.417991516 | 0.33941785 | -1.2314954 | 0.21813765 | 0.50602433 |
| DYNLL1 | 96.9898006 | -0.711168684 | 0.58135909 | -1.2232864 | 0.22122153 | 0.50910615 |
| COQ5 | 25.4539385 | 0.735673527 | 0.33163566 | 2.2183185 | 0.02653312 | 0.14555072 |
| RNF10 | 187.808986 | 0.875426985 | 0.18967078 | 4.61550791 | 3.92E-06 | 0.00017534 |
| POP5 | 20.180581 | 0.155144573 | 0.39378285 | 0.3939851 | 0.69359204 | 0.86078608 |
| MLEC | 107.711383 | -0.49198365 | 0.39215781 | -1.2545553 | 0.20964024 | 0.49580206 |
| UNC119B | 26.1196959 | 0.318028441 | 0.35323276 | 0.90033676 | 0.36794106 | 0.65347673 |
| ACADS | 32.628942 | 0.175427859 | 0.27945693 | 0.6277456 | 0.53017061 | 0.76428051 |
| SPPL3 | 312.815575 | 0.173681802 | 0.21677694 | 0.80120055 | 0.42301555 | 0.69793501 |
| P2RX4 | 36.3740279 | -0.446500124 | 0.37036031 | -1.2055831 | 0.22797828 | 0.5177793 |
| CAMKK2 | 23.8929104 | -0.370408604 | 0.39073773 | -0.9479724 | 0.34314348 | 0.6310379 |
| ANAPC5 | 82.8864439 | 0.624597669 | 0.19059814 | 3.27703976 | 0.00104902 | 0.01524254 |
| RNF34 | 42.8624926 | -0.340471738 | 0.28446445 | -1.1968868 | 0.23135069 | 0.52111057 |
| KDM2B | 74.4787839 | 0.198582235 | 0.28773671 | 0.69015259 | 0.49009823 | 0.74556088 |
| TMEM120B | 96.8241893 | 1.199592506 | 0.26890677 | 4.46099785 | 8.16E-06 | 0.00031211 |
| RHOF | 19.0681066 | -0.54835015 | 0.48387143 | -1.1332559 | 0.25710681 | 0.54817875 |
| SETD1B | 26.3225171 | 0.295223978 | 0.31859935 | 0.92663083 | 0.35411824 | 0.63957058 |
| WDR66 | 31.0578797 | -0.717036038 | 0.48637573 | -1.474243 | 0.14041617 | 0.39545307 |
| BCL7A | 71.5905777 | 0.659018858 | 0.34656302 | 1.90158445 | 0.0572255 | 0.23601925 |
| MLXIP | 98.4134178 | -1.033769222 | 0.31145961 | -3.3191117 | 0.00090304 | 0.01363901 |
| CLIP1 | 228.738771 | -0.267297566 | 0.39947903 | -0.6691154 | 0.50342187 | 0.75108109 |
| ZCCHC8 | 57.3984055 | 0.103558425 | 0.28837941 | 0.35910478 | 0.71951671 | 0.87503134 |
| RSRC2 | 85.590246 | -0.104531811 | 0.20875486 | -0.5007395 | 0.61655444 | 0.81691188 |
| KNTC1 | 15.5708723 | -0.090061954 | 0.40970418 | -0.2198219 | 0.82600986 | 0.92747178 |
| DENR | 88.3004328 | -0.169213853 | 0.2197514 | -0.770024 | 0.44128564 | 0.71211227 |
| HIP1R | 142.71101 | 0.681042001 | 0.31699145 | 2.14845543 | 0.03167759 | 0.1630255 |
| VPS37B | 198.349735 | 1.20869721 | 0.51725763 | 2.33674118 | 0.01945265 | 0.11848232 |
| ABCB9 | 29.3420898 | 1.273747568 | 0.52346518 | 2.4332995 | 0.01496192 | 0.09993327 |
| ARL6IP4 | 63.5726983 | -0.187152203 | 0.30326119 | -0.6171321 | 0.53714762 | 0.76945357 |
| PITPNM2 | 149.31132 | -0.018822358 | 0.23630447 | -0.079653 | 0.93651325 | 0.97594028 |
| MPHOSPH9 | 81.5534865 | 0.299033485 | 0.28236802 | 1.05902036 | 0.28959051 | 0.58130538 |
| C12orf65 | 37.3271178 | -0.840185224 | 0.30676724 | -2.7388362 | 0.00616571 | 0.05523403 |
| SBNO1 | 95.3489972 | 0.024841315 | 0.23149275 | 0.10730926 | 0.91454364 | 0.96463895 |
| KMT5A | 22.2240066 | -0.290185816 | 0.37069179 | -0.7828223 | 0.43373144 | 0.70597357 |
| RILPL2 | 28.5556141 | -0.273003919 | 0.28422043 | -0.9605359 | 0.33678559 | 0.62525668 |
| SNRNP35 | 27.9488461 | -0.013204343 | 0.27520051 | -0.0479808 | 0.96173154 | 0.98525601 |
| RILPL1 | 73.3685973 | 0.275658159 | 0.2256859 | 1.22142396 | 0.22192554 | 0.51021578 |
| TMED2 | 44.9000419 | -0.462196009 | 0.28763728 | -1.606871 | 0.10808266 | 0.34264936 |
| DDX55 | 16.9751759 | 0.066740138 | 0.35126227 | 0.19000087 | 0.84930845 | 0.93897371 |
| EIF2B1 | 23.4797052 | 0.359263445 | 0.3063052 | 1.17289374 | 0.24083842 | 0.53151245 |
| GTF2H3 | 25.1959475 | 0.322960262 | 0.29681337 | 1.08809202 | 0.27655449 | 0.56884726 |
| TCTN2 | 19.8167068 | -0.143737413 | 0.44222971 | -0.3250288 | 0.74515924 | 0.88808711 |
| ATP6V0A2 | 29.9228884 | -0.053884779 | 0.35371466 | -0.1523397 | 0.87891903 | 0.95273407 |
| CCDC92 | 75.1008007 | -0.499579339 | 0.20532119 | -2.4331601 | 0.01496768 | 0.09993327 |
| RFLNA | 16.7043713 | 0.32366587 | 0.51194929 | 0.63222252 | 0.52724149 | 0.76221787 |
| NCOR2 | 127.575085 | 0.439859553 | 0.27517321 | 1.59848246 | 0.10993565 | 0.34550245 |
| UBC | 311.711781 | -0.590924455 | 0.31913058 | -1.8516698 | 0.06407325 | 0.2521767 |
| DHX37 | 13.4262038 | -0.423477323 | 0.47882735 | -0.8844051 | 0.3764776 | 0.65960412 |
| BRI3BP | 27.0528816 | 0.581768236 | 0.48777868 | 1.19268893 | 0.23299122 | 0.52282195 |
| AACS | 29.1508814 | 0.254882252 | 0.31606701 | 0.80641841 | 0.42000163 | 0.69630783 |
| SLC15A4 | 39.7324198 | -1.149049103 | 0.34567141 | -3.324108 | 0.00088702 | 0.0134901 |
| RIMBP2 | 156.080166 | -0.330241157 | 0.3798676 | -0.8693586 | 0.38465104 | 0.66701235 |
| STX2 | 39.5628419 | 0.213885338 | 0.35584227 | 0.60106782 | 0.54779482 | 0.77516644 |
| RAN | 52.4305859 | -0.795360969 | 0.46388524 | -1.7145641 | 0.08642518 | 0.30212156 |
| SFSWAP | 101.587882 | 0.296668244 | 0.21411931 | 1.38552772 | 0.16589116 | 0.4346416 |
| ULK1 | 52.662924 | 0.191837764 | 0.2304074 | 0.83260244 | 0.40506898 | 0.68331761 |
| PUS1 | 16.0433892 | -0.117241766 | 0.40301993 | -0.2909081 | 0.7711216 | 0.90076568 |
| EP400 | 99.3923229 | 0.179122175 | 0.24831513 | 0.72135022 | 0.47069407 | 0.73210719 |
| DDX51 | 12.6344327 | -0.052425816 | 0.4787997 | -0.1094943 | 0.91281048 | 0.96446045 |
| FBRSL1 | 49.8088419 | 0.484629676 | 0.31940464 | 1.51729068 | 0.12919331 | 0.37682602 |
| POLE | 9.60802261 | 0.070956158 | 0.49348083 | 0.14378706 | 0.88566862 | 0.95619452 |
| ANKLE2 | 61.7900404 | -0.501532701 | 0.41854612 | -1.1982735 | 0.23081058 | 0.52105604 |

| | | | | | | |
|---|---|---|---|---|---|---|
| GOLGA3 | 66.9602867 | -0.436081026 | 0.37829 | -1.1527691 | 0.24900517 | 0.54005289 |
| CHFR | 73.4697684 | -0.261872223 | 0.33025282 | -0.7929447 | 0.42781004 | 0.70230313 |
| ZNF26 | 46.3364821 | 0.024354347 | 0.3719268 | 0.06548156 | 0.94779059 | 0.979567 |
| ZNF84 | 65.1990031 | 0.088261768 | 0.27454034 | 0.32148924 | 0.74783966 | 0.88935207 |
| ZNF140 | 31.2127561 | -0.133339579 | 0.31666659 | -0.4210725 | 0.67370217 | 0.85002814 |
| ZNF891 | 31.7031069 | 1.244369777 | 0.39720232 | 3.13283615 | 0.00173126 | 0.02204834 |
| ZNF268 | 14.0628162 | -0.163665861 | 0.45440569 | -0.3601756 | 0.71871579 | 0.87427206 |
| MPHOSPH8 | 160.446294 | -0.009180083 | 0.26994052 | -0.0340078 | 0.97287093 | 0.98860883 |
| PSPC1 | 217.30765 | 0.490884181 | 0.29584096 | 1.65928402 | 0.09705857 | 0.32146212 |
| ZMYM5 | 65.2202344 | -0.138221237 | 0.27088038 | -0.5102667 | 0.60986462 | 0.81344383 |
| ZMYM2 | 415.996695 | 0.291600964 | 0.26655682 | 1.09395423 | 0.27397505 | 0.56626457 |
| CRYL1 | 115.660815 | 0.180835329 | 0.28557018 | 0.63324304 | 0.52657494 | 0.76221787 |
| IFT88 | 165.570836 | 0.126652289 | 0.2344191 | 0.54028145 | 0.58900295 | 0.79925045 |
| EEF1AKMT1 | 19.3352831 | -0.30104442 | 0.54129958 | -0.5561512 | 0.57810748 | 0.79345367 |
| XPO4 | 133.330822 | 0.490358946 | 0.21427655 | 2.28843962 | 0.02211193 | 0.12891745 |
| LATS2 | 108.079458 | 0.861532032 | 0.35242988 | 2.44454878 | 0.01450335 | 0.09784772 |
| SAP18 | 53.6966837 | -0.827221147 | 0.40369744 | -2.0491117 | 0.04045119 | 0.19098451 |
| MRPL57 | 43.1833209 | -1.061259594 | 0.36494641 | -2.9079875 | 0.00363763 | 0.03751236 |
| ZDHHC20 | 125.785204 | 0.004854582 | 0.25726838 | 0.01886972 | 0.98494504 | 0.99403006 |
| MICU2 | 200.686651 | 0.343374131 | 0.17940229 | 1.91398963 | 0.05562148 | 0.23268234 |
| FGF9 | 60.9034959 | 1.274473712 | 0.3972898 | 3.20791958 | 0.00133699 | 0.01817105 |
| AL136962.1 | 59.2425598 | 0.655372935 | 0.67581053 | 0.9697584 | 0.33216693 | 0.6217267 |
| TNFRSF19 | 84.2549311 | 0.870783102 | 0.31445004 | 2.7692256 | 0.00561897 | 0.05159237 |
| MIPEP | 119.536222 | 0.021169598 | 0.24052385 | 0.08801455 | 0.92986512 | 0.97253199 |
| PARP4 | 103.488707 | -1.309240747 | 0.3426081 | -3.8213947 | 0.0001327 | 0.00307206 |
| ATP12A | 29.8317898 | 4.479966977 | 0.79938484 | 5.60426809 | 2.09E-08 | 2.39E-06 |
| MTMR6 | 33.8069946 | 0.045726891 | 0.26241577 | 0.17425359 | 0.86166617 | 0.94550174 |
| NUP58 | 77.3072997 | -0.141423676 | 0.37533932 | -0.3767889 | 0.70633052 | 0.8678136 |
| RNF6 | 14.132045 | -0.662550292 | 0.45504457 | -1.4560118 | 0.14538934 | 0.40350243 |
| CDK8 | 226.151489 | 0.671175686 | 0.23856263 | 2.81341506 | 0.00490183 | 0.04684479 |
| WASF3 | 39.2264372 | 0.028387932 | 0.23699137 | 0.11978466 | 0.90465373 | 0.9613929 |
| USP12 | 157.147181 | -0.113115918 | 0.27887995 | -0.4056079 | 0.68503071 | 0.85635073 |
| LINC02340 | 26.5236748 | -0.282025033 | 0.50752919 | -0.5556824 | 0.578428 | 0.7935467 |
| RPL21 | 72.0002721 | -0.209345356 | 0.43704113 | -0.4790061 | 0.63193431 | 0.82642568 |
| RASL11A | 16.3847689 | -0.544438307 | 0.59374665 | -0.9169539 | 0.35916679 | 0.6430674 |
| GTF3A | 24.3679479 | -0.459114335 | 0.34926437 | -1.3145181 | 0.18867192 | 0.46745986 |
| MTIF3 | 48.0365733 | 0.089163543 | 0.25457214 | 0.35024862 | 0.72615212 | 0.87880825 |
| LNX2 | 63.0792587 | -0.246899526 | 0.26537157 | -0.9303918 | 0.35216828 | 0.63817724 |
| POLR1D | 44.1357548 | -0.06394678 | 0.24339434 | -0.2627291 | 0.79275937 | 0.91124908 |
| PAN3 | 439.172815 | 0.132311209 | 0.19042561 | 0.69481834 | 0.48716915 | 0.74434227 |
| POMP | 40.1976172 | -0.583310972 | 0.30436307 | -1.9164972 | 0.05530184 | 0.23212675 |
| SLC46A3 | 17.5779191 | -0.448890911 | 0.53378121 | -0.8409642 | 0.40036796 | 0.68010059 |
| SLC7A1 | 124.704427 | 0.198899259 | 0.57362778 | 0.34673924 | 0.72878723 | 0.88042244 |
| UBL3 | 176.407822 | 0.461204835 | 0.25147293 | 1.83401384 | 0.06665194 | 0.25842991 |
| KATNAL1 | 26.3739235 | -0.544771272 | 0.33288524 | -1.6365138 | 0.10173211 | 0.33046825 |
| HMGB1 | 208.425886 | -0.083794063 | 0.24604036 | -0.3405704 | 0.73342702 | 0.88350705 |
| USPL1 | 39.4092148 | -0.004607603 | 0.32350405 | -0.0142428 | 0.98863627 | 0.9963225 |
| HSPH1 | 87.0188595 | 1.026566903 | 0.50999616 | 2.01289144 | 0.04412605 | 0.200435 |
| B3GLCT | 45.2386369 | 0.337308402 | 0.29867584 | 1.12934611 | 0.25875186 | 0.55002264 |
| FRY | 330.891869 | 0.63873122 | 0.37907208 | 1.68498618 | 0.09199124 | 0.31235786 |
| N4BP2L1 | 29.4697091 | -0.555604514 | 0.46579686 | -1.1928043 | 0.23294602 | 0.52282195 |
| N4BP2L2 | 719.599435 | 0.116054153 | 0.2310991 | 0.5021835 | 0.61553845 | 0.8164389 |
| PDS5B | 264.399177 | 0.737229999 | 0.20490847 | 3.59785037 | 0.00032086 | 0.00619004 |
| KL | 148.227061 | 4.039745001 | 0.40467518 | 9.98268542 | 1.81E-23 | 9.35E-20 |
| STARD13 | 475.966685 | 1.289878016 | 0.39910496 | 3.23192677 | 0.00122959 | 0.01704871 |
| AL139383.1 | 81.1672999 | 0.446774222 | 0.35736007 | 1.25020745 | 0.21122378 | 0.49712778 |
| RFC3 | 30.4235486 | 0.170107304 | 0.38147417 | 0.4459209 | 0.65565439 | 0.83938753 |
| LINC02343 | 179.577502 | 0.804455018 | 0.48736874 | 1.65060856 | 0.09881853 | 0.32472998 |
| NBEA | 427.388545 | -1.071016657 | 0.36699077 | -2.9183749 | 0.00351861 | 0.03672616 |
| SPART | 84.4943859 | -0.138808832 | 0.3055061 | -0.454357 | 0.64957191 | 0.83690468 |
| RFXAP | 15.1605389 | 0.778033432 | 0.57112433 | 1.36228383 | 0.17310833 | 0.44595199 |
| ALG5 | 49.8548711 | 0.261509253 | 0.23326353 | 1.12108933 | 0.26224984 | 0.55396716 |
| EXOSC8 | 22.722818 | -0.589586039 | 0.33981751 | -1.7350078 | 0.08273942 | 0.29473773 |
| SUPT20H | 107.607744 | 0.142828077 | 0.22591178 | 0.63222944 | 0.52723696 | 0.76221787 |
| LINC00571 | 32.6702586 | -1.298070871 | 0.3874457 | -3.3503298 | 0.00080715 | 0.01257988 |
| UFM1 | 61.5339743 | -0.273173517 | 0.27139486 | -1.0065538 | 0.31414927 | 0.60719808 |
| FREM2 | 195.775382 | 0.015086702 | 0.34625389 | 0.04357121 | 0.9652462 | 0.98660829 |
| PROSER1 | 35.9793289 | 0.30737847 | 0.37323017 | 0.82356276 | 0.41018804 | 0.68796444 |
| NHLRC3 | 58.6117438 | -0.047708024 | 0.37532593 | -0.1271109 | 0.89885261 | 0.95935794 |
| COG6 | 107.150231 | 0.251178108 | 0.29030375 | 0.86522516 | 0.38691522 | 0.66935358 |
| LINC00598 | 23.0678815 | -0.480845114 | 0.40778782 | -1.1791552 | 0.23833639 | 0.52855279 |
| FOXO1 | 343.10403 | -0.655073757 | 0.52236509 | -1.2540535 | 0.20982257 | 0.49600553 |
| MRPS31 | 75.9063452 | 0.381538802 | 0.28527889 | 1.33742387 | 0.18108432 | 0.45756661 |
| ELF1 | 272.117221 | -0.295736247 | 0.15651888 | -1.8894606 | 0.05883014 | 0.24000052 |
| WBP4 | 39.050885 | 0.123110838 | 0.33415409 | 0.36842535 | 0.7125561 | 0.87130598 |
| AL354696.2 | 18.5285517 | -0.237967848 | 0.45843531 | -0.5190871 | 0.60370001 | 0.80940894 |
| MTRF1 | 20.6203335 | -0.295571153 | 0.35881364 | -0.8237456 | 0.41008413 | 0.68796444 |
| NAA16 | 59.6674023 | 0.328895741 | 0.27574441 | 1.19275578 | 0.23296503 | 0.52282195 |
| VWA8 | 321.880545 | 1.031318437 | 0.20400337 | 5.05539897 | 4.29E-07 | 2.84E-05 |
| DGKH | 155.088294 | -1.535260988 | 0.47898002 | -3.2052714 | 0.00134935 | 0.01829082 |
| AKAP11 | 89.2944776 | -0.255087136 | 0.25446943 | -1.0024274 | 0.31613719 | 0.60951749 |
| LINC02341 | 32.2277789 | -1.363143081 | 0.8641463 | -1.5774448 | 0.11469321 | 0.35355161 |
| EPSTI1 | 73.4146355 | -1.353834201 | 1.28760399 | -1.0514368 | 0.29305803 | 0.5846943 |
| DNAJC15 | 75.5993291 | -0.332727467 | 0.34540651 | -0.9632924 | 0.3354008 | 0.62358764 |
| ENOX1 | 19.4522318 | -0.127141817 | 0.73905544 | -0.1720329 | 0.8634117 | 0.94635064 |
| CCDC122 | 59.3662128 | -0.539341947 | 0.42354702 | -1.2733933 | 0.20287851 | 0.48651172 |
| SMIM2-AS1 | 16.2736423 | -1.529217097 | 0.68312862 | -2.2385493 | 0.02518526 | 0.13986981 |
| SERP2 | 13.4038374 | -0.934465924 | 0.434413 | -2.1511003 | 0.03146829 | 0.16233665 |
| TSC22D1 | 594.752793 | 0.212408072 | 0.18372571 | 1.15611511 | 0.24763408 | 0.53836051 |
| NUFIP1 | 15.34593 | 0.092990354 | 0.37841623 | 0.24573564 | 0.80588688 | 0.91718215 |
| GPALPP1 | 49.1878729 | 0.597359227 | 0.27987288 | 2.13439482 | 0.03281048 | 0.16692026 |
| GTF2F2 | 109.820195 | -0.177938426 | 0.31105196 | -0.5720537 | 0.56728559 | 0.78656476 |
| TPT1 | 395.009249 | -1.339988675 | 0.43150773 | -3.1053642 | 0.00190045 | 0.02370269 |
| SLC25A30 | 67.6404138 | 0.715565642 | 0.34407448 | 2.0796824 | 0.03755467 | 0.18135358 |
| COG3 | 73.4748387 | 0.139134203 | 0.25488 | 0.54588121 | 0.58514759 | 0.79681419 |
| LINC01055 | 44.0678095 | 1.94269188 | 0.6127566 | 3.17041364 | 0.00152222 | 0.02002799 |
| ZC3H13 | 116.384478 | 0.316754959 | 0.21230914 | 1.49195155 | 0.13571185 | 0.3880387 |
| AL138686.1 | 25.4678757 | -0.100204765 | 0.75578017 | -0.1325845 | 0.89452196 | 0.95843632 |
| LRCH1 | 444.555262 | 0.499046263 | 0.23639753 | 2.11104689 | 0.03476828 | 0.17320253 |
| ESD | 46.1844429 | -0.081091193 | 0.33290482 | -0.2435867 | 0.80755091 | 0.91805225 |
| SUCLA2 | 112.641994 | 0.173037605 | 0.27651507 | 0.62578001 | 0.53145925 | 0.76542614 |
| MED4 | 64.5712352 | -0.050738445 | 0.27159702 | -0.1868152 | 0.85180555 | 0.94024437 |
| ITM2B | 669.951422 | -0.77440477 | 0.38109356 | -2.0320594 | 0.04214764 | 0.19602933 |
| RB1 | 244.607503 | -0.443329885 | 0.24721971 | -1.7932627 | 0.07293093 | 0.2728157 |
| LPAR6 | 13.1615506 | -1.704809061 | 0.62687471 | -2.7195372 | 0.00653733 | 0.05736595 |
| RCBTB2 | 30.2114152 | -0.232313455 | 0.3258178 | -0.7130165 | 0.47583557 | 0.73616591 |
| FNDC3A | 518.079979 | 0.26561565 | 0.60304691 | 0.44045604 | 0.65960685 | 0.84097773 |
| CDADC1 | 32.6184197 | 0.473683398 | 0.36310022 | 1.30455276 | 0.19204518 | 0.47229637 |
| CAB39L | 64.2151644 | 1.014357705 | 0.41035137 | 2.47192472 | 0.01343878 | 0.09373482 |
| SETDB2 | 72.1179558 | 0.304146943 | 0.34287946 | 0.88703751 | 0.3750587 | 0.65806152 |
| PHF11 | 46.9395083 | -0.383352852 | 0.3946827 | -0.9712938 | 0.33140201 | 0.62108486 |
| RCBTB1 | 19.8531702 | -0.688750807 | 0.43207728 | -1.5940454 | 0.11092587 | 0.34744854 |
| EBPL | 21.5658708 | -0.335003248 | 0.447189 | -0.7491312 | 0.45377812 | 0.72126672 |

| | | | | | | |
|---|---|---|---|---|---|---|
| KPNA3 | 87.6088598 | 0.019193271 | 0.24449061 | 0.0785031 | 0.93742786 | 0.97619962 |
| SPRYD7 | 17.1930007 | 0.272233522 | 0.39709062 | 0.68557027 | 0.49298413 | 0.74685087 |
| DLEU2 | 104.610384 | -1.914616699 | 0.31986774 | -5.9856511 | 2.16E-09 | 3.31E-07 |
| TRIM13 | 40.5915253 | -0.567649431 | 0.32560423 | -1.7433724 | 0.08126858 | 0.29141278 |
| RNASEH2B | 43.8370713 | -0.158635386 | 0.26102721 | -0.607735 | 0.54336321 | 0.77316683 |
| INTS6 | 191.401678 | 0.930151641 | 0.30871029 | 3.01302445 | 0.00258658 | 0.02942907 |
| INTS6-AS1 | 20.2662922 | 0.549521119 | 0.48536168 | 1.13218893 | 0.25755501 | 0.54866248 |
| WDFY2 | 229.542954 | 1.431712592 | 0.22221605 | 6.44288576 | 1.17E-10 | 2.63E-08 |
| ATP7B | 20.3472869 | 0.753591298 | 0.46925983 | 1.60591477 | 0.10829264 | 0.34305988 |
| ALG11 | 13.465081 | 0.179898098 | 0.44643236 | 0.40296832 | 0.68697153 | 0.85794408 |
| UTP14C | 18.3544082 | -0.209988953 | 0.44947592 | -0.4671862 | 0.64036666 | 0.83201631 |
| NEK5 | 10.567407 | -1.024373035 | 0.54675942 | -1.8735352 | 0.06099451 | 0.24507233 |
| NEK3 | 42.1439715 | 0.135593101 | 0.33054244 | 0.41021389 | 0.68164905 | 0.85482028 |
| VPS36 | 98.3303109 | 0.65694193 | 0.1925064 | 3.41257195 | 0.00064353 | 0.01048994 |
| SUGT1 | 62.3035154 | 0.301782912 | 0.28075148 | 1.07491121 | 0.28241448 | 0.57528302 |
| OLFM4 | 674.312208 | -3.536900844 | 0.77868937 | -4.5421204 | 5.57E-06 | 0.00023305 |
| DIAPH3 | 25.7467522 | -1.85439591 | 0.50544267 | -3.6688551 | 0.00024364 | 0.00503 |
| TDRD3 | 215.958267 | 0.520238039 | 0.21860003 | 2.37986257 | 0.0173191 | 0.11006868 |
| PCDH9 | 437.890693 | 1.107381944 | 0.65077505 | 1.70163554 | 0.08882371 | 0.30687325 |
| DACH1 | 1025.13921 | 0.42931613 | 0.43940871 | 0.97703144 | 0.32855359 | 0.61768835 |
| MZT1 | 12.1907443 | 0.1346406 | 0.41300709 | 0.3260007 | 0.74442383 | 0.88808711 |
| DIS3 | 36.854921 | -0.226274151 | 0.2722933 | -0.8309942 | 0.40597691 | 0.68404099 |
| PIBF1 | 240.676825 | -0.10727778 | 0.19689799 | -0.5448394 | 0.58586399 | 0.79711254 |
| KLF5 | 67.9863136 | -2.352995646 | 0.50962779 | -4.6170866 | 3.89E-06 | 0.00017507 |
| KLF12 | 314.628518 | 0.923914933 | 0.20633917 | 4.47765159 | 7.55E-06 | 0.00029788 |
| TBC1D4 | 375.398096 | 0.848002742 | 0.27881875 | 3.04141217 | 0.00235471 | 0.02731785 |
| COMMD6 | 51.9145913 | 0.383197548 | 0.47634479 | 0.80445416 | 0.42113473 | 0.69672877 |
| UCHL3 | 56.6578131 | -0.115956295 | 0.30104478 | -0.3851796 | 0.70010439 | 0.86376951 |
| LMO7-AS1 | 12.7578108 | -0.280567709 | 0.59658078 | -0.4702929 | 0.63814577 | 0.83059731 |
| LMO7 | 395.22379 | 0.715085642 | 0.35110699 | 2.03666025 | 0.0416841 | 0.19500656 |
| KCTD12 | 59.0919557 | -0.662209551 | 0.63061214 | -1.0501059 | 0.29366941 | 0.58529352 |
| AC001226.2 | 19.9748037 | -0.386196916 | 0.58778189 | -0.6570412 | 0.51115443 | 0.7549696 |
| FBXL3 | 58.9237357 | -0.078839097 | 0.31657245 | -0.2490397 | 0.8033301 | 0.91628728 |
| MYCBP2 | 264.384275 | 0.146239537 | 0.24391967 | 0.59953974 | 0.54881302 | 0.77536011 |
| SLAIN1 | 27.9091098 | 0.281040243 | 0.37511243 | 0.74921603 | 0.45372701 | 0.72126672 |
| EDNRB | 41.6206378 | 0.68746874 | 0.55531627 | 1.23797696 | 0.21572461 | 0.50246323 |
| RNF219 | 22.0043391 | -0.512163102 | 0.42166322 | -1.214626 | 0.22450877 | 0.51340497 |
| RBM26 | 220.069645 | 0.449978359 | 0.15914979 | 2.82738902 | 0.00469293 | 0.04552405 |
| NDFIP2 | 125.223638 | -0.160450605 | 0.3222012 | -0.4979826 | 0.61849628 | 0.81814955 |
| GPC5 | 824.53645 | 5.784486369 | 1.10612667 | 5.22949725 | 1.70E-07 | 1.34E-05 |
| GPC6 | 99.3662503 | 0.129029003 | 0.84887103 | 0.15200071 | 0.87918637 | 0.95273407 |
| TGDS | 28.7980864 | 0.195597203 | 0.41767703 | 0.46829772 | 0.6395717 | 0.83121032 |
| GPR180 | 24.846761 | -0.763784004 | 0.46813124 | -1.6315596 | 0.1027723 | 0.33228309 |
| ABCC4 | 66.7600927 | -1.939723451 | 0.54678337 | -3.5475173 | 0.00038888 | 0.00714126 |
| CLDN10 | 107.225923 | 0.529577952 | 0.40739692 | 1.29990661 | 0.19363298 | 0.4740909 |
| CLDN10-AS1 | 22.835977 | -0.40418343 | 0.5698693 | -0.7092564 | 0.4781654 | 0.73786502 |
| DZIP1 | 24.1694925 | -0.308265119 | 0.38219663 | -0.8065616 | 0.41991911 | 0.69628307 |
| DNAJC3-DT | 17.2427775 | -0.752105533 | 0.51000479 | -1.4747029 | 0.14029244 | 0.39525538 |
| DNAJC3 | 119.269747 | 0.007162959 | 0.29126554 | 0.02459254 | 0.98037997 | 0.99299415 |
| UGGT2 | 228.809299 | 0.013678372 | 0.20723423 | 0.0660044 | 0.94737432 | 0.979567 |
| MBNL2 | 235.426831 | 0.424215214 | 0.28508176 | 1.48804756 | 0.13673835 | 0.38956817 |
| RAP2A | 15.6632105 | -0.308470434 | 0.4825941 | -0.6391923 | 0.52269784 | 0.76175316 |
| IPO5 | 68.9877359 | 0.31866999 | 0.22753507 | 1.40053132 | 0.16135427 | 0.42831015 |
| FARP1 | 637.01564 | -0.054339596 | 0.19317193 | -0.2813017 | 0.77847899 | 0.90417143 |
| STK24 | 292.644222 | -0.818505576 | 0.32088121 | -2.5508056 | 0.01074743 | 0.08056236 |
| DOCK9 | 143.631662 | 0.013794252 | 0.23787525 | 0.05798944 | 0.95375704 | 0.98167699 |
| UBAC2 | 296.417434 | 0.327256972 | 0.23305082 | 1.40423008 | 0.16025035 | 0.42791337 |
| TM9SF2 | 122.392429 | 0.148801736 | 0.18579208 | 0.8009046 | 0.42318688 | 0.69810588 |
| CLYBL | 185.079286 | 0.501092836 | 0.25564316 | 1.96012616 | 0.04998105 | 0.21735109 |
| AL355338.1 | 20.3725046 | -0.022359198 | 0.38530063 | -0.0580305 | 0.95372431 | 0.98167699 |
| PCCA | 1012.06105 | 0.177422853 | 0.22084242 | 0.80339117 | 0.42174867 | 0.69752044 |
| GGACT | 30.7181009 | -0.886614869 | 0.47005184 | -1.8862066 | 0.05926712 | 0.24104615 |
| TMTC4 | 28.8697018 | -0.638636505 | 0.36938395 | -1.7289233 | 0.08382283 | 0.29653069 |
| FGF14 | 52.2236988 | 0.804891432 | 0.64902984 | 1.2401455 | 0.21492158 | 0.50124063 |
| TPP2 | 83.3347495 | 0.212108995 | 0.24841195 | 0.85385987 | 0.39318264 | 0.67385918 |
| EFNB2 | 32.668901 | -0.899590718 | 0.675075 | -1.3325789 | 0.18267002 | 0.45952112 |
| ARGLU1 | 211.519339 | 0.010434874 | 0.2342045 | 0.04455454 | 0.96446238 | 0.98660829 |
| FAM155A | 610.910333 | -0.887472652 | 1.04122132 | -0.8523381 | 0.39402644 | 0.67474408 |
| LIG4 | 17.7616936 | -0.631186992 | 0.35344918 | -1.7857928 | 0.07413282 | 0.27538206 |
| ABHD13 | 27.4742861 | 0.188912093 | 0.34170553 | 0.55285055 | 0.58036575 | 0.79465023 |
| TNFSF13B | 35.3986111 | -0.029827685 | 0.40890692 | -0.0729449 | 0.94184994 | 0.97782671 |
| MYO16 | 28.6474023 | 0.120856045 | 0.85300073 | 0.1416834 | 0.88733009 | 0.95619452 |
| IRS2 | 64.5958608 | -0.940999251 | 0.45635027 | -2.0620109 | 0.03920669 | 0.18699415 |
| COL4A1 | 140.375329 | 0.657977714 | 0.29569405 | 2.2251977 | 0.02606797 | 0.14391866 |
| COL4A2 | 174.778099 | 0.492009997 | 0.30583298 | 1.60875387 | 0.10767017 | 0.34161321 |
| RAB20 | 34.0432397 | -0.778567431 | 0.32279107 | -2.4119857 | 0.01586591 | 0.10371228 |
| NAXD | 33.0223487 | 0.318729651 | 0.27908517 | 1.14205155 | 0.25343259 | 0.54506525 |
| CARS2 | 61.0646785 | 0.314746681 | 0.22628989 | 1.3909003 | 0.16425566 | 0.43311027 |
| ANKRD10 | 302.458732 | -0.169260683 | 0.20161959 | -0.8395052 | 0.40118589 | 0.68033203 |
| ARHGEF7 | 182.621743 | 0.015221907 | 0.1672142 | 0.09103238 | 0.92746686 | 0.97111125 |
| TUBGCP3 | 68.2342697 | 0.78072334 | 0.35700981 | 2.18684003 | 0.0287542 | 0.1530092 |
| ATP11A | 62.1917278 | -2.503019632 | 0.63766853 | -3.9252676 | 8.66E-05 | 0.00218214 |
| MCF2L | 61.9801021 | 0.401825281 | 0.27985441 | 1.43583685 | 0.15104877 | 0.41211643 |
| PCID2 | 44.8464934 | 0.288150132 | 0.31120399 | 0.92592044 | 0.35448733 | 0.63979125 |
| CUL4A | 131.948543 | -0.316106071 | 0.21102448 | -1.4979593 | 0.13414383 | 0.38483703 |
| LAMP1 | 112.254041 | -0.893846083 | 0.31716006 | -2.8182807 | 0.00482816 | 0.04630755 |
| GRTP1 | 38.151721 | 0.275197011 | 0.30612562 | 0.89896758 | 0.36866993 | 0.65369102 |
| DCUN1D2 | 42.8228264 | 0.626014696 | 0.3743369 | 1.67232966 | 0.09445933 | 0.3161038 |
| TMCO3 | 50.6459422 | -0.672532396 | 0.26214856 | -2.5654629 | 0.01030382 | 0.07859336 |
| TFDP1 | 44.9388756 | 0.057467877 | 0.40519393 | 0.14182808 | 0.8872158 | 0.95619452 |
| GAS6 | 70.5452086 | 1.017930915 | 0.32084479 | 3.17265843 | 0.0015105 | 0.01989921 |
| RASA3 | 33.7860771 | -1.668262857 | 0.64955862 | -2.5683022 | 0.0102198 | 0.07837003 |
| CDC16 | 63.9184469 | -0.019504398 | 0.27267458 | -0.0715299 | 0.94297599 | 0.97820347 |
| UPF3A | 67.1159364 | -0.074720144 | 0.26669413 | -0.2801717 | 0.77934579 | 0.90427469 |
| CHAMP1 | 15.7109005 | -0.406255598 | 0.38466551 | -1.0561269 | 0.29091023 | 0.58249897 |
| TTC5 | 24.0601373 | 0.09262302 | 0.34174829 | 0.27102702 | 0.78637025 | 0.90849219 |
| CCNB1IP1 | 52.4430805 | 1.298771699 | 0.29439421 | 4.41167537 | 1.03E-05 | 0.00037472 |
| PARP2 | 12.358428 | 0.605543671 | 0.44497804 | 1.36083946 | 0.17356443 | 0.44612537 |
| TEP1 | 25.2025002 | -0.702882524 | 0.31856276 | -2.2064177 | 0.02735476 | 0.14777597 |
| OSGEP | 21.815539 | 0.241205131 | 0.36696931 | 0.65728965 | 0.51099469 | 0.7549696 |
| APEX1 | 27.6559736 | -0.626918434 | 0.40273891 | -1.5566374 | 0.11955661 | 0.36193716 |
| PIP4P1 | 14.8803316 | -0.371959804 | 0.44269363 | -0.8402195 | 0.40078535 | 0.68010059 |
| PNP | 49.2039752 | 0.853897918 | 0.51334777 | 1.66339073 | 0.09623425 | 0.32011792 |
| METTL17 | 22.0415898 | 0.143123361 | 0.37348339 | 0.38321211 | 0.7015625 | 0.86484347 |
| NDRG2 | 34.8360453 | 1.114473461 | 0.40055754 | 2.78230556 | 0.00539742 | 0.05024962 |
| ARHGEF40 | 16.4857513 | -0.713700337 | 0.40290944 | -1.7713666 | 0.07649975 | 0.2804628 |
| HNRNPC | 427.349919 | 0.068406873 | 0.24275974 | 0.28178837 | 0.77810579 | 0.9041333 |
| SUPT16H | 58.2371339 | -0.306441729 | 0.28860323 | -1.0618098 | 0.28832206 | 0.58047564 |
| CHD8 | 94.395609 | -0.308222211 | 0.23260058 | -1.3251137 | 0.18513345 | 0.46185874 |
| RAB2B | 35.6892055 | 0.59928906 | 0.38152504 | 1.57077256 | 0.11623549 | 0.35543425 |
| TOX4 | 40.594625 | -0.663374164 | 0.28253175 | -2.3479632 | 0.01887639 | 0.11658545 |
| METTL3 | 14.5454278 | 0.228576833 | 0.42394799 | 0.53916243 | 0.58977478 | 0.79960282 |
| DAD1 | 72.0066448 | -0.82585746 | 0.35459836 | -2.328994 | 0.01985938 | 0.12013584 |

| | | | | | | |
|---|---|---|---|---|---|---|
| ABHD4 | 26.0071547 | -0.21556123 | 0.32330551 | -0.6667416 | 0.50493721 | 0.75182297 |
| OXA1L | 22.4777554 | -0.087561382 | 0.33267696 | -0.2632024 | 0.79239456 | 0.91124908 |
| MRPL52 | 19.6571722 | 0.102484343 | 0.34938014 | 0.29333191 | 0.76926846 | 0.90034488 |
| MMP14 | 18.5977916 | -1.157267464 | 0.66858098 | -1.7309309 | 0.08346409 | 0.2960906 |
| LRP10 | 72.3770544 | -1.704610748 | 0.3189486 | -5.3444685 | 9.07E-08 | 7.92E-06 |
| RBM23 | 57.2251159 | 0.171051213 | 0.2831556 | 0.6040891 | 0.54578441 | 0.77463416 |
| PRMT5 | 11.1324893 | -0.151598373 | 0.44757988 | -0.3387069 | 0.73483058 | 0.88380093 |
| AL132780.1 | 26.1018947 | -0.232103666 | 0.67029027 | -0.3462734 | 0.72913728 | 0.88046679 |
| C14orf93 | 14.0504849 | -0.168397183 | 0.54807023 | -0.3072548 | 0.75864947 | 0.8951693 |
| PSMB5 | 41.0077655 | -0.867411597 | 0.320432 | -2.7070068 | 0.00678929 | 0.0591532 |
| ACIN1 | 117.069708 | 0.16450226 | 0.19641632 | 0.83751829 | 0.4023013 | 0.68128594 |
| C14orf119 | 17.250505 | -1.001140904 | 0.42456245 | -2.3580533 | 0.01837105 | 0.11417077 |
| SLC7A8 | 71.92336 | 0.730428107 | 0.61324352 | 1.19108981 | 0.23361833 | 0.52365884 |
| HOMEZ | 15.0858627 | -0.086824485 | 0.38942144 | -0.2229576 | 0.82356847 | 0.92608229 |
| PPP1R3E | 14.8853092 | 0.079409591 | 0.59299078 | 0.1339137 | 0.89347082 | 0.95800753 |
| BCL2L2-PABF | 19.951915 | 0.232354305 | 0.35736618 | 0.65018548 | 0.51557242 | 0.75713618 |
| PABPN1 | 70.5279624 | 0.703756506 | 0.41506469 | 1.6955345 | 0.08997407 | 0.30907252 |
| SLC22A17 | 14.2869205 | 0.216942899 | 0.52210742 | 0.41551392 | 0.67776573 | 0.85275312 |
| NGDN | 20.1572513 | -0.011323305 | 0.36772031 | -0.0307933 | 0.97543442 | 0.99023703 |
| ZFHX2 | 26.4186099 | -0.006249429 | 0.44985977 | -0.013892 | 0.98891618 | 0.99636328 |
| AP1G2 | 21.0936552 | -0.47207637 | 0.4146066 | -1.1386128 | 0.2548647 | 0.5458673 |
| DHRS4-AS1 | 21.1263038 | -0.057932492 | 0.539274 | -0.1074268 | 0.91445038 | 0.96463895 |
| DHRS4L2 | 17.5209483 | 0.162600017 | 0.45828199 | 0.35480342 | 0.72273687 | 0.87662955 |
| NRL | 10.8887271 | -0.034341362 | 0.45159887 | -0.076044 | 0.93938413 | 0.97673318 |
| PSME1 | 40.1957557 | -1.732562762 | 0.45808959 | -3.7821483 | 0.00015548 | 0.00348968 |
| PSME2 | 11.6102584 | -0.134604152 | 0.56326992 | -0.2389692 | 0.81112948 | 0.91988726 |
| AL136295.1 | 23.7742595 | -0.202429334 | 0.41018044 | -0.4935129 | 0.62165023 | 0.82030381 |
| GMPR2 | 14.3919917 | -0.734575697 | 0.53822467 | -1.3648124 | 0.17231202 | 0.44479039 |
| TINF2 | 13.2586827 | -1.348440376 | 0.46669703 | -2.8893271 | 0.00386067 | 0.03926224 |
| DHRS1 | 17.5324266 | -0.371641171 | 0.4230676 | -0.8784439 | 0.37970284 | 0.66311217 |
| KHNYN | 18.6507444 | -0.673234947 | 0.33433748 | -2.0136389 | 0.04404746 | 0.20016628 |
| SDR39U1 | 22.0606985 | 0.109573733 | 0.33710896 | 0.32503952 | 0.74515116 | 0.88808711 |
| NOVA1 | 101.276804 | 0.175763771 | 0.23928665 | 0.7345323 | 0.46262438 | 0.72740482 |
| LINC00645 | 63.235476 | 1.154361939 | 0.56645293 | 2.03787795 | 0.04156214 | 0.19472668 |
| PRKD1 | 526.097654 | -0.724846507 | 0.22933749 | -3.1606106 | 0.00157439 | 0.02053082 |
| G2E3 | 37.4536365 | 0.707817401 | 0.27063199 | 2.61542396 | 0.00891168 | 0.07083957 |
| SCFD1 | 152.322589 | 0.182853036 | 0.24272479 | 0.75333482 | 0.45124874 | 0.71947224 |
| COCH | 11.9183615 | -1.389197846 | 0.46643658 | -2.978321 | 0.00289832 | 0.03178629 |
| STRN3 | 375.295989 | -0.379737627 | 0.35996808 | -1.0549203 | 0.29146179 | 0.58292358 |
| AP4S1 | 26.3277289 | 0.343626905 | 0.37185032 | 0.92410008 | 0.35543421 | 0.64037833 |
| HECTD1 | 293.544902 | -0.156612123 | 0.39942104 | -0.3920978 | 0.69498593 | 0.86168553 |
| HEATR5A | 43.499863 | -0.282786222 | 0.31870046 | -0.8873104 | 0.37491183 | 0.65806152 |
| AL139353.1 | 11.0285075 | 0.582973848 | 0.39370506 | 1.48073752 | 0.13867653 | 0.39280881 |
| NUBPL | 146.80839 | 0.586729067 | 0.24997293 | 2.34717045 | 0.0189166 | 0.11669389 |
| ARHGAP5 | 360.875005 | -0.156949785 | 0.21858546 | -0.7180248 | 0.47274197 | 0.73375471 |
| AKAP6 | 107.154844 | 0.769406574 | 0.35084686 | 2.19299829 | 0.0283075 | 0.15134187 |
| NPAS3 | 106.795781 | -0.354785663 | 0.64788885 | -0.5476027 | 0.58396475 | 0.7962935 |
| EGLN3 | 37.9685779 | -0.419002633 | 0.36490551 | -1.1482497 | 0.25086551 | 0.54232519 |
| EAPP | 49.3578682 | -0.265192845 | 0.33680489 | -0.7873783 | 0.43106047 | 0.70454366 |
| SNX6 | 152.716799 | 0.205314041 | 0.23831629 | 0.86151912 | 0.38895219 | 0.67095971 |
| CFL2 | 12.0038317 | -0.156710606 | 0.53536766 | -0.2927159 | 0.76973934 | 0.90034488 |
| BAZ1A | 63.20223 | -0.890750619 | 0.40220306 | -2.2146789 | 0.02678211 | 0.14614285 |
| SRP54 | 88.5119275 | -0.267095237 | 0.30578603 | -0.873471 | 0.3824064 | 0.66580205 |
| FAM177A1 | 62.0223052 | -0.23011486 | 0.26054459 | -0.8832072 | 0.37712432 | 0.66022373 |
| PPP2R3C | 34.5215739 | -0.715957497 | 0.31102938 | -2.3018967 | 0.021341 | 0.12570324 |
| NFKBIA | 43.3752324 | -0.484873055 | 0.48540627 | -0.9989015 | 0.31784241 | 0.61101185 |
| RALGAPA1 | 354.703194 | 0.723390015 | 0.29704813 | 2.43526198 | 0.01488101 | 0.09961285 |
| BRMS1L | 21.682951 | 0.460499035 | 0.43567047 | 1.05698932 | 0.29051646 | 0.58210841 |
| PTCSC3 | 11.6680229 | 2.083384355 | 0.77381832 | 2.69234304 | 0.00709519 | 0.06111595 |
| MBIP | 53.1801503 | 0.233485411 | 0.3215692 | 0.72608138 | 0.46778888 | 0.73050797 |
| MIPOL1 | 132.186143 | 0.548755409 | 0.2369865 | 2.31555554 | 0.02058255 | 0.12280939 |
| SEC23A | 115.323141 | 0.221835432 | 0.34246073 | 0.64776897 | 0.51713439 | 0.75818855 |
| GEMIN2 | 13.7372148 | 0.283160958 | 0.43458946 | 0.65155965 | 0.51468529 | 0.75684546 |
| TRAPPC6B | 65.2709396 | 0.855709623 | 0.29353691 | 2.91516874 | 0.00355496 | 0.03699316 |
| PNN | 99.5995748 | -0.0338098 | 0.2611185 | -0.1294807 | 0.89697731 | 0.95907216 |
| MIA2 | 190.744083 | 0.081239588 | 0.21449416 | 0.37874965 | 0.70487378 | 0.86664395 |
| FBXO33 | 71.339867 | 0.00820409 | 0.29064923 | 0.02822677 | 0.97748128 | 0.99112991 |
| AL049875.1 | 22.6046126 | 0.923942201 | 0.41300842 | 2.23710257 | 0.02527964 | 0.14026664 |
| KLHL28 | 59.2825988 | -0.324696587 | 0.31477529 | -1.0315187 | 0.30229766 | 0.59594663 |
| TOGARAM1 | 107.053441 | -0.234037532 | 0.25352482 | -0.9231346 | 0.35593707 | 0.64072404 |
| PRPF39 | 46.2213463 | 0.205850858 | 0.29630446 | 0.69472751 | 0.48722608 | 0.74434227 |
| FKBP3 | 35.6597444 | 0.029266122 | 0.24783219 | 0.11808846 | 0.90599756 | 0.96157183 |
| FANCM | 26.0665274 | 0.091585456 | 0.35099378 | 0.26093185 | 0.79414507 | 0.91176607 |
| MIS18BP1 | 32.4462781 | -0.362471287 | 0.35737006 | -1.0142744 | 0.31045186 | 0.60378989 |
| RPS29 | 92.4915968 | -0.043924492 | 0.36127431 | -0.1215821 | 0.90322998 | 0.96126516 |
| RPL36AL | 90.2163681 | -1.490213342 | 0.40502234 | -3.6793361 | 0.00023384 | 0.00484716 |
| DNAAF2 | 16.0300304 | -0.088690635 | 0.36977591 | -0.2398497 | 0.81044679 | 0.91943781 |
| POLE2 | 14.9135468 | -1.116162827 | 0.59640904 | -1.871472 | 0.06127968 | 0.24544318 |
| KLHDC1 | 47.3091277 | 0.640599237 | 0.39550571 | 1.61969656 | 0.10529748 | 0.33709589 |
| KLHDC2 | 47.5264293 | -0.381512917 | 0.28634694 | -1.332345 | 0.18274685 | 0.45952112 |
| NEMF | 213.297411 | 0.242302418 | 0.24853488 | 0.97492319 | 0.32959836 | 0.61894319 |
| ARF6 | 39.7997963 | -1.312953837 | 0.58117913 | -2.2591208 | 0.02387587 | 0.13541585 |
| VCPKMT | 25.8496199 | 0.003089929 | 0.39999835 | 0.00772485 | 0.99383652 | 0.99810704 |
| SOS2 | 300.27586 | 0.660805892 | 0.21185526 | 3.11913841 | 0.00181381 | 0.02287129 |
| L2HGDH | 108.545888 | 0.445516605 | 0.20489161 | 2.17440137 | 0.02967501 | 0.1564544 |
| ATP5S | 46.3905809 | -0.102859951 | 0.25150963 | -0.4089702 | 0.68256151 | 0.85544388 |
| CDKL1 | 135.177133 | 0.901786223 | 0.34788301 | 2.59221116 | 0.00953612 | 0.07448152 |
| MAP4K5 | 193.760995 | 0.13712554 | 0.25877894 | 0.52989451 | 0.59618507 | 0.80412393 |
| ATL1 | 20.4495864 | 0.037865376 | 0.50018789 | 0.0757023 | 0.93965594 | 0.97674567 |
| SAV1 | 44.7001575 | -0.670641176 | 0.26138009 | -2.5657699 | 0.01029471 | 0.07859336 |
| NIN | 60.1478631 | 0.124490848 | 0.24503691 | 0.50804936 | 0.61141873 | 0.81385718 |
| PYGL | 24.8019376 | -0.9733012 | 0.42541834 | -2.2878685 | 0.02214518 | 0.12903828 |
| TMX1 | 38.3635973 | -0.119420228 | 0.28846768 | -0.4139813 | 0.67888779 | 0.85343527 |
| FRMD6 | 18.2053033 | -2.70946201 | 0.5878107 | -4.6094126 | 4.04E-06 | 0.00017854 |
| RTRAF | 40.3717136 | -0.641786532 | 0.43114166 | -1.4885746 | 0.13659942 | 0.38928001 |
| TXNDC16 | 29.7608289 | 0.011965292 | 0.30071023 | 0.03979011 | 0.96826046 | 0.9875279 |
| GPR137C | 16.7720252 | -0.394827592 | 0.42763308 | -0.9232859 | 0.35585823 | 0.64069408 |
| ERO1A | 198.517333 | -2.695630876 | 0.5520542 | -4.8829099 | 1.05E-06 | 5.80E-05 |
| AL133453.1 | 25.8956807 | -4.458245318 | 0.86032216 | -5.182065 | 2.19E-07 | 1.65E-05 |
| PSMC6 | 74.3916208 | -0.11733491 | 0.27862686 | -0.4211184 | 0.6736686 | 0.85002814 |
| STYX | 33.2501991 | 0.171950312 | 0.30320846 | 0.56710262 | 0.57064446 | 0.78888611 |
| GNPNAT1 | 22.4084418 | -0.445200293 | 0.45155771 | -0.9859211 | 0.32417181 | 0.61460758 |
| FERMT2 | 242.342756 | 0.616003079 | 0.23193445 | 2.65593609 | 0.00790886 | 0.06518612 |
| DDHD1 | 66.80456 | 0.175010438 | 0.27749901 | 0.63067049 | 0.528256 | 0.76294593 |
| CNIH1 | 29.5882343 | -0.246217712 | 0.3168029 | -0.7771953 | 0.43704358 | 0.70829758 |
| GMFB | 24.9739996 | -0.471730341 | 0.34438874 | -1.3697612 | 0.17076145 | 0.44178771 |
| CGRRF1 | 17.132666 | 0.166260449 | 0.38327232 | 0.43379195 | 0.66443953 | 0.84348349 |
| SAMD4A | 152.638045 | -0.69278528 | 0.47566989 | -1.4564413 | 0.14527066 | 0.40328169 |
| GCH1 | 15.8171482 | -0.761333704 | 0.66122053 | -1.1514066 | 0.24956499 | 0.54083745 |
| WDHD1 | 14.600021 | 0.182798765 | 0.48812162 | 0.3744943 | 0.70803659 | 0.86882712 |
| SOCS4 | 28.9645499 | -0.342830755 | 0.36614395 | -0.9363278 | 0.34910444 | 0.6357564 |
| MAPK1IP1L | 126.897751 | -0.138340939 | 0.30069813 | -0.4600659 | 0.64546895 | 0.83387847 |
| LGALS3 | 156.771366 | -1.531497855 | 0.37813882 | -4.0500942 | 5.12E-05 | 0.00143324 |

| | | | | | | |
|---|---|---|---|---|---|---|
| FBXO34 | 191.505584 | 0.06870457 | 0.2382528 | 0.28836837 | 0.77306478 | 0.90149298 |
| AL158801.3 | 13.752609 | -0.555774366 | 0.55305953 | -1.0049088 | 0.31494078 | 0.60827146 |
| ATG14 | 38.9634112 | -0.450127411 | 0.28384423 | -1.5858255 | 0.11277891 | 0.35069372 |
| KTN1-AS1 | 11.5885884 | -1.259046409 | 0.64371919 | -1.9558938 | 0.05047766 | 0.21890013 |
| KTN1 | 317.872866 | -0.107496475 | 0.22074161 | -0.4869787 | 0.62627339 | 0.8231524 |
| PELI2 | 226.43173 | 0.704949112 | 0.27444843 | 2.56860319 | 0.01021093 | 0.07837003 |
| TMEM260 | 122.139731 | 0.339317463 | 0.18165431 | 1.86792955 | 0.06177189 | 0.24637012 |
| EXOC5 | 87.4312433 | 0.103721752 | 0.18868212 | 0.54971691 | 0.58251356 | 0.7955793 |
| AP5M1 | 77.1454916 | 0.596621214 | 0.27700137 | 2.15385658 | 0.03125142 | 0.16189143 |
| NAA30 | 31.5685388 | 0.012558819 | 0.29240631 | 0.04294989 | 0.96574148 | 0.98661741 |
| CCDC198 | 268.62169 | 1.142257886 | 0.4739682 | 2.40998843 | 0.01595303 | 0.10408365 |
| ARMH4 | 57.678407 | 0.107711292 | 0.29150662 | 0.36949862 | 0.71175611 | 0.87074117 |
| ACTR10 | 70.1202221 | 0.132663624 | 0.22816322 | 0.58144176 | 0.56094276 | 0.78361672 |
| PSMA3 | 61.6191015 | -0.157932403 | 0.33727537 | -0.4682595 | 0.63959901 | 0.83121032 |
| ARID4A | 98.8726587 | 0.092715109 | 0.22468074 | 0.41265268 | 0.6798611 | 0.85407015 |
| TIMM9 | 27.6053158 | -0.112837102 | 0.31818967 | -0.3546221 | 0.72287268 | 0.87663736 |
| KIAA0586 | 75.6598139 | -0.070258949 | 0.24150318 | -0.2909235 | 0.77110983 | 0.90076568 |
| DAAM1 | 119.03782 | -0.145506233 | 0.25788008 | -0.5642399 | 0.57259088 | 0.78978194 |
| GPR135 | 23.5914908 | -0.417815179 | 0.39472064 | -1.0585086 | 0.28982365 | 0.58136876 |
| L3HYPDH | 22.6109388 | -0.28425428 | 0.44117227 | -0.6443158 | 0.51937065 | 0.76002222 |
| JKAMP | 35.6157284 | 0.020625614 | 0.3694094 | 0.05583403 | 0.95547403 | 0.98258246 |
| AL157911.1 | 12.3049963 | 0.608660423 | 0.52040796 | 1.16958322 | 0.24216873 | 0.53285174 |
| PCNX4 | 142.840821 | -0.355369719 | 0.2982559 | -1.1914927 | 0.23346022 | 0.52341833 |
| DHRS7 | 32.0692996 | -0.516390221 | 0.39519271 | -1.3066795 | 0.19132158 | 0.47164272 |
| PPM1A | 86.7865062 | 0.059529151 | 0.25479043 | 0.23363966 | 0.81526473 | 0.92163473 |
| MNAT1 | 267.354765 | 0.589213784 | 0.18806078 | 3.13310287 | 0.00172969 | 0.02204834 |
| SLC38A6 | 45.5583552 | -0.023481551 | 0.3174661 | -0.0739655 | 0.94103781 | 0.97727535 |
| PRKCH | 73.110405 | -0.125735901 | 0.32519665 | -0.3866457 | 0.69901849 | 0.86334649 |
| TMEM30B | 35.7581626 | -0.671532535 | 0.45655253 | -1.4708768 | 0.14132443 | 0.39740649 |
| HIF1A | 1095.66565 | -1.372309788 | 0.417782 | -3.2847509 | 0.00102073 | 0.01495806 |
| HIF1A-AS2 | 164.891486 | -2.216722028 | 0.54475102 | -4.0692389 | 4.72E-05 | 0.0013423 |
| SNAPC1 | 98.396044 | -0.246714536 | 0.37848104 | -0.6518544 | 0.5144951 | 0.75684546 |
| SYT16 | 44.0062881 | -1.991575172 | 0.83533937 | -2.384151 | 0.01711858 | 0.10960571 |
| PPP2R5E | 269.017507 | 0.135662759 | 0.30396765 | 0.44630657 | 0.65537581 | 0.83931753 |
| WDR89 | 41.9176777 | 0.361470404 | 0.32100835 | 1.12604673 | 0.26014574 | 0.55146957 |
| SGPP1 | 311.30871 | 0.431362344 | 0.44752464 | 0.96388512 | 0.33510352 | 0.62351515 |
| SYNE2 | 947.851962 | -1.044904465 | 0.22142658 | -4.7189658 | 2.37E-06 | 0.00011358 |
| ESR2 | 55.964918 | -1.169101226 | 0.54134379 | -2.1596281 | 0.03080147 | 0.16034199 |
| MTHFD1 | 27.2810642 | -0.045399301 | 0.33459159 | -0.1356857 | 0.89206973 | 0.95760158 |
| AL122035.1 | 23.6313848 | -1.312040095 | 0.38997842 | -3.3643915 | 0.00076713 | 0.01213969 |
| ZBTB25 | 40.8251776 | -0.713602805 | 0.35935069 | -1.9858117 | 0.04705423 | 0.20921567 |
| ZBTB1 | 102.988754 | -0.116052256 | 0.24361923 | -0.4763674 | 0.63381267 | 0.82746649 |
| AL049869.3 | 11.0258018 | -0.580825336 | 0.68337521 | -0.8499362 | 0.39536055 | 0.67590514 |
| HSPA2 | 23.3993718 | -0.234938931 | 0.44518509 | -0.5277331 | 0.59768456 | 0.80551365 |
| PLEKHG3 | 84.779319 | 0.191215999 | 0.33788093 | 0.56592716 | 0.57144329 | 0.78910141 |
| CHURC1-FNT | 40.4610995 | -0.261067626 | 0.41593832 | -0.6276595 | 0.53022704 | 0.76428051 |
| GPX2 | 36.6499462 | -1.358389166 | 0.85073012 | -1.5967334 | 0.11032515 | 0.34618593 |
| RAB15 | 50.2814957 | 0.376210676 | 0.32147694 | 1.17025712 | 0.24189751 | 0.53271231 |
| MAX | 76.8874174 | -0.674956048 | 0.32643886 | -2.0676339 | 0.03867447 | 0.18471226 |
| FUT8 | 74.264571 | -0.332558081 | 0.31049571 | -1.0710553 | 0.28414456 | 0.5765312 |
| AL359232.1 | 67.3588078 | 0.279279501 | 0.42925156 | 0.65061965 | 0.51529205 | 0.75713618 |
| GPHN | 1135.07581 | 1.147061115 | 0.2356799 | 4.86702988 | 1.13E-06 | 6.21E-05 |
| MPP5 | 115.898225 | 0.11602094 | 0.18548826 | 0.6254894 | 0.5316499 | 0.76557825 |
| ATP6V1D | 60.6519281 | -0.096297679 | 0.32759878 | -0.2939501 | 0.76879607 | 0.90022017 |
| EIF2S1 | 33.5435756 | -0.0604474 | 0.25849673 | -0.233842 | 0.81510762 | 0.92163473 |
| PLEKHH1 | 39.2913866 | -0.217783802 | 0.28687238 | -0.7591661 | 0.44775318 | 0.71693398 |
| PIGH | 24.6844066 | 0.304750629 | 0.49015072 | 0.62174881 | 0.53410705 | 0.76732389 |
| ARG2 | 111.293575 | 1.079963157 | 0.57322407 | 1.88401573 | 0.05956285 | 0.24171358 |
| VTI1B | 35.0122023 | 0.146539875 | 0.36321061 | 0.40345703 | 0.68661204 | 0.85780709 |
| ZFYVE26 | 30.4768826 | -0.504175627 | 0.45246666 | -1.1142824 | 0.26515807 | 0.55640131 |
| RAD51B | 1041.3788 | 1.289470251 | 0.28661125 | 4.49902182 | 6.83E-06 | 0.00027472 |
| ZFP36L1 | 423.025798 | -0.420605492 | 0.32106116 | -1.3100479 | 0.19017963 | 0.46973504 |
| ACTN1 | 49.4152586 | -1.091854843 | 0.34676926 | -3.1486494 | 0.00164027 | 0.02126567 |
| DCAF5 | 177.026756 | -0.050029822 | 0.1974483 | -0.2533819 | 0.79997312 | 0.91455697 |
| EXD2 | 42.184757 | 0.329023464 | 0.31217186 | 1.0539818 | 0.29189125 | 0.58344269 |
| GALNT16 | 39.9324636 | 0.251470827 | 0.78059054 | 0.32215459 | 0.74733558 | 0.8891386 |
| ERH | 51.5005404 | -0.161646298 | 0.2252366 | -0.7176733 | 0.47295874 | 0.73375471 |
| SLC39A9 | 69.065041 | 0.040564141 | 0.21812052 | 0.18597123 | 0.85246733 | 0.94037032 |
| PLEKHD1 | 19.3723658 | 2.734615353 | 0.9848057 | 2.776807 | 0.00548958 | 0.05085757 |
| SUSD6 | 113.7882 | -0.092035327 | 0.17293015 | -0.532211 | 0.59457987 | 0.80311549 |
| SRSF5 | 230.188485 | 0.924390227 | 0.23283814 | 3.97009803 | 7.18E-05 | 0.00188328 |
| AL357153.2 | 42.0818621 | -0.458816956 | 0.40265165 | -1.1394886 | 0.25449943 | 0.54553747 |
| MED6 | 32.18043 | -0.132842576 | 0.28856179 | -0.4603609 | 0.64525716 | 0.83387847 |
| MAP3K9 | 42.5949054 | 0.056615163 | 0.39021946 | 0.14508544 | 0.88464341 | 0.95619452 |
| PCNX1 | 250.304676 | 0.157157204 | 0.23998192 | 0.654871 | 0.51255081 | 0.75562371 |
| SIPA1L1 | 487.879476 | -0.157713631 | 0.2353801 | -0.6700381 | 0.50283351 | 0.75108109 |
| RGS6 | 699.719603 | -0.078398323 | 1.04053499 | -0.0753442 | 0.93994082 | 0.97682541 |
| DPF3 | 14.8931507 | 0.591476599 | 0.45568702 | 1.2979887 | 0.19429122 | 0.47476 |
| ZFYVE1 | 42.7171423 | -0.25631773 | 0.34836435 | -0.7357749 | 0.46186773 | 0.72676972 |
| RBM25 | 158.309226 | 0.061879264 | 0.1855251 | 0.33353582 | 0.73872986 | 0.88595984 |
| PSEN1 | 114.212847 | -0.177272665 | 0.3430034 | -0.5168248 | 0.60527847 | 0.81044694 |
| PAPLN | 20.7253016 | -1.373100343 | 0.38513448 | -3.565249 | 0.00036351 | 0.00678422 |
| NUMB | 280.210335 | -0.332639127 | 0.21103558 | -1.576223 | 0.11497442 | 0.35399477 |
| HEATR4 | 13.5491129 | -1.176748785 | 0.62983902 | -1.8683326 | 0.06171572 | 0.24624197 |
| DNAL1 | 42.90299 | 0.130959344 | 0.24932734 | 0.52525064 | 0.59940897 | 0.8069931 |
| PNMA1 | 15.11972 | -0.314193428 | 0.4752692 | -0.6610852 | 0.50855768 | 0.7543124 |
| ELMSAN1 | 51.5703511 | -0.430784025 | 0.31352736 | -1.3739918 | 0.16944422 | 0.44032355 |
| AC005480.1 | 24.7275292 | 0.235567966 | 0.42849912 | 0.54975133 | 0.58248994 | 0.7955793 |
| COQ6 | 12.0801058 | 0.065645006 | 0.4287448 | 0.15310974 | 0.87831173 | 0.95235949 |
| ENTPD5 | 45.7233138 | 0.040176421 | 0.30088198 | 0.13352884 | 0.89377516 | 0.95813441 |
| BBOF1 | 40.1745449 | -0.072499922 | 0.2562596 | -0.2829159 | 0.77724128 | 0.90363838 |
| ALDH6A1 | 42.1214946 | -0.320891584 | 0.47079656 | -0.6815929 | 0.49549642 | 0.74770824 |
| LIN52 | 63.5735114 | 0.037271443 | 0.31538808 | 0.11817645 | 0.90592785 | 0.96157183 |
| ABCD4 | 64.5997498 | -0.152052983 | 0.30681832 | -0.4955799 | 0.62019086 | 0.81932299 |
| NPC2 | 60.5535611 | -1.11356796 | 0.3425698 | -3.2506308 | 0.00115149 | 0.01616169 |
| ISCA2 | 16.8951972 | -0.811306959 | 0.43619915 | -1.8599462 | 0.06289314 | 0.24948869 |
| AREL1 | 57.2306156 | 0.434632149 | 0.24829554 | 1.75046296 | 0.08003846 | 0.28876788 |
| FCF1 | 23.7167845 | 0.17219613 | 0.30356314 | 0.56724981 | 0.57054447 | 0.78885373 |
| YLPM1 | 131.996766 | 0.410251728 | 0.1819096 | 2.25525062 | 0.0241176 | 0.13583965 |
| DLST | 54.381636 | 0.155867035 | 0.25154432 | 0.61964044 | 0.53549454 | 0.76808755 |
| EIF2B2 | 12.9523328 | -0.63136796 | 0.39585734 | -1.5949381 | 0.11072608 | 0.34697143 |
| MLH3 | 89.9262341 | -0.009056691 | 0.23097535 | -0.0392106 | 0.96872245 | 0.98770265 |
| ACYP1 | 50.1547354 | 0.583682505 | 0.2853878 | 2.04522582 | 0.0408326 | 0.1924325 |
| ZC2HC1C | 15.9043346 | 0.422203829 | 0.57393068 | 0.73563558 | 0.4619525 | 0.7267921 |
| NEK9 | 64.591549 | 0.222952453 | 0.24055438 | 0.92682766 | 0.35401602 | 0.63957058 |
| TMED10 | 118.42142 | -0.452607691 | 0.24417562 | -1.8536154 | 0.0637942 | 0.25161098 |
| FOS | 46.5033122 | 1.36187798 | 0.91743374 | 1.48444287 | 0.13769147 | 0.39109388 |
| JDP2 | 11.9030321 | -0.77724975 | 0.42146644 | -1.8441557 | 0.06516045 | 0.25514365 |
| TTLL5 | 151.639411 | -0.041688983 | 0.20331781 | -0.2050434 | 0.83753819 | 0.93487753 |
| ERG28 | 18.1298529 | -0.613305637 | 0.52751394 | -1.162634 | 0.24497804 | 0.53594343 |
| IFT43 | 104.614369 | 0.041235476 | 0.19665882 | 0.20968028 | 0.83391722 | 0.93289339 |
| GPATCH2L | 146.377915 | -0.154864069 | 0.29574496 | -0.5236406 | 0.60052853 | 0.80738812 |
| ESRRB | 97.9480675 | 0.836457633 | 0.58665151 | 1.4258169 | 0.1539212 | 0.4169593 |

| | | | | | | |
|---|---|---|---|---|---|---|
| ANGEL1 | 19.9882503 | 0.346028783 | 0.34882303 | 0.9919895 | 0.32120265 | 0.6120836 |
| IRF2BPL | 41.1221367 | -0.460025988 | 0.37807569 | -1.2167563 | 0.22369694 | 0.5128006 |
| POMT2 | 18.3346425 | 0.128568437 | 0.438477 | 0.29321592 | 0.76935712 | 0.90034488 |
| GSTZ1 | 10.8604949 | -0.759236122 | 0.44795984 | -1.6948754 | 0.09009906 | 0.30924357 |
| TMED8 | 57.0289839 | 0.032309481 | 0.23774104 | 0.13590199 | 0.89189876 | 0.95751782 |
| VIPAS39 | 23.4587827 | -0.001494409 | 0.3434487 | -0.0043512 | 0.99652827 | 0.99868531 |
| AHSA1 | 25.1484591 | 0.296096471 | 0.35857482 | 0.82575923 | 0.40894068 | 0.6869518 |
| SPTLC2 | 127.995096 | -0.283636668 | 0.29284945 | -0.9685409 | 0.3327743 | 0.6218467 |
| ALKBH1 | 34.7678201 | 0.303790776 | 0.33967152 | 0.89436634 | 0.37112592 | 0.65494451 |
| SLIRP | 40.8319899 | 0.036392173 | 0.37204833 | 0.09781571 | 0.92207863 | 0.96885417 |
| SNW1 | 43.5208833 | 0.126150346 | 0.28283853 | 0.44601542 | 0.65558611 | 0.83938753 |
| ADCK1 | 22.870662 | 0.395562385 | 0.46604319 | 0.84876765 | 0.39601059 | 0.67634303 |
| NRXN3 | 14.1897781 | -1.219743697 | 0.63444047 | -1.9225503 | 0.05453655 | 0.22997772 |
| CEP128 | 50.628563 | 0.075723518 | 0.31773297 | 0.23832439 | 0.8116295 | 0.9200492 |
| AC007262.2 | 29.0441913 | 0.600281401 | 0.55577677 | 1.08007645 | 0.28010814 | 0.57312294 |
| GTF2A1 | 43.4487244 | 0.228383493 | 0.41116112 | 0.55545984 | 0.57858017 | 0.7935467 |
| STON2 | 139.57353 | 0.179398785 | 0.31547741 | 0.56865809 | 0.56958819 | 0.78795455 |
| SEL1L | 98.2243197 | -0.001262284 | 0.34773901 | -0.00363 | 0.99710371 | 0.99876774 |
| FLRT2 | 14.4887406 | 2.737933582 | 0.75946686 | 3.60507315 | 0.00031207 | 0.00606622 |
| GALC | 26.7589936 | 0.319254047 | 0.41935296 | 0.76130152 | 0.44647699 | 0.71619194 |
| SPATA7 | 65.5139879 | 0.541223982 | 0.22714559 | 2.38271843 | 0.01718533 | 0.10980752 |
| PTPN21 | 78.0107052 | 0.768997698 | 0.30908776 | 2.48795904 | 0.01284785 | 0.09134478 |
| ZC3H14 | 113.287343 | 0.560335997 | 0.26617375 | 2.10515125 | 0.03527814 | 0.17430956 |
| EML5 | 62.8463416 | -2.470774948 | 0.82132554 | -3.0082773 | 0.00262733 | 0.02974372 |
| TTC8 | 49.5885271 | -0.122850202 | 0.23411413 | -0.5247449 | 0.59976053 | 0.8073609 |
| FOXN3 | 246.902734 | 0.929037468 | 0.18722131 | 4.96224224 | 6.97E-07 | 4.22E-05 |
| EFCAB11 | 51.4181026 | -0.181705257 | 0.37989734 | -0.4783009 | 0.63243603 | 0.82649065 |
| TDP1 | 26.6156919 | -0.851550692 | 0.35032951 | -2.4307136 | 0.01506912 | 0.10047582 |
| KCNK13 | 117.843799 | 1.190408777 | 0.47988811 | 2.48059651 | 0.01311627 | 0.09242398 |
| PSMC1 | 37.5959462 | -0.211879267 | 0.27857145 | -0.7605922 | 0.44690069 | 0.71645983 |
| NRDE2 | 47.6682866 | 0.330222969 | 0.26762533 | 1.23390029 | 0.21724006 | 0.50473773 |
| CALM1 | 113.626245 | -0.721904267 | 0.33402109 | -2.1612536 | 0.03067576 | 0.15992999 |
| TTC7B | 150.697585 | 0.202164925 | 0.3361534 | 0.60140676 | 0.5475691 | 0.77516644 |
| RPS6KA5 | 62.453162 | -1.302198823 | 0.58007493 | -2.2448804 | 0.02477581 | 0.13811712 |
| DGLUCY | 173.759497 | 0.377651432 | 0.39065176 | 0.96672143 | 0.33368331 | 0.62241635 |
| CCDC88C | 44.3736348 | -0.229633565 | 0.41628104 | -0.5516311 | 0.58120113 | 0.79484057 |
| PPP4R3A | 144.197899 | 0.470887519 | 0.2239781 | 2.10238199 | 0.03551983 | 0.17500013 |
| CATSPERB | 52.4738939 | -0.246480075 | 0.41838981 | -0.5891159 | 0.55578355 | 0.78017197 |
| TC2N | 60.0545772 | -0.101796868 | 0.35918197 | -0.2834131 | 0.7768602 | 0.90335572 |
| TRIP11 | 148.205743 | 0.165078978 | 0.15300156 | 1.07893656 | 0.28061601 | 0.57370631 |
| ATXN3 | 87.1418037 | -0.521193512 | 0.32313325 | -1.6129368 | 0.10675824 | 0.34029982 |
| NDUFB1 | 53.3684989 | 0.260026038 | 0.26180345 | 0.99321089 | 0.32060719 | 0.6120836 |
| CPSF2 | 48.3198694 | 0.352630074 | 0.26538952 | 1.32872645 | 0.18393823 | 0.46027002 |
| LGMN | 39.4137707 | -0.297586755 | 0.47879415 | -0.6215338 | 0.53424845 | 0.76740485 |
| GOLGA5 | 35.4720688 | -0.691804613 | 0.34845052 | -1.985374 | 0.04710287 | 0.20934159 |
| ITPK1 | 98.5382645 | 0.183328365 | 0.21112078 | 0.86835776 | 0.38519851 | 0.66728015 |
| MOAP1 | 18.561823 | -1.250304327 | 0.50917805 | -2.4555346 | 0.01406752 | 0.09623211 |
| UBR7 | 14.4448969 | -0.295003339 | 0.43996134 | -0.670521 | 0.50252569 | 0.75108109 |
| BTBD7 | 393.051945 | -0.565099921 | 0.17108937 | -3.3029517 | 0.00095673 | 0.01414092 |
| UNC79 | 35.4743334 | -1.214581282 | 0.65809646 | -1.8455977 | 0.06495065 | 0.25461247 |
| DDX24 | 94.096417 | -0.985330039 | 0.29279985 | -3.3651999 | 0.00076488 | 0.01212279 |
| IFI27 | 26.9498754 | 0.633337509 | 1.57824248 | 0.40129291 | 0.68820449 | 0.85854718 |
| SERPINA1 | 42.7345444 | -2.547442779 | 0.45173737 | -5.6392119 | 1.71E-08 | 2.02E-06 |
| DICER1 | 233.513239 | 0.214062347 | 0.21004323 | 1.0191347 | 0.30813902 | 0.60178758 |
| DICER1-AS1 | 22.8227599 | 0.143953854 | 0.54080144 | 0.26618615 | 0.79009585 | 0.91080793 |
| CLMN | 485.607177 | 0.600972801 | 0.28410262 | 2.11533702 | 0.03440123 | 0.17178939 |
| GLRX5 | 38.7488558 | -0.591485097 | 0.39028322 | -1.5155279 | 0.12963878 | 0.37737742 |
| ATG2B | 103.214814 | 0.560750252 | 0.21643679 | 2.59082682 | 0.00957457 | 0.07472514 |
| GSKIP | 22.8789328 | -0.631788705 | 0.34283299 | -1.8428469 | 0.06535137 | 0.25530899 |
| AK7 | 21.8268625 | -1.089229201 | 0.33480579 | -3.2533165 | 0.00114066 | 0.01605344 |
| PAPOLA | 185.003636 | 0.267350713 | 0.21607827 | 1.23728645 | 0.21598076 | 0.50279115 |
| VRK1 | 15.8634988 | -0.112788076 | 0.39257749 | -0.2873014 | 0.77388152 | 0.90184317 |
| LINC01550 | 11.6729843 | 0.918330639 | 0.65682822 | 1.39812908 | 0.16207431 | 0.42953067 |
| SETD3 | 120.044424 | 0.336153737 | 0.21558016 | 1.5592981 | 0.11892584 | 0.36084255 |
| CCNK | 36.1265798 | 0.028222377 | 0.28662758 | 0.09846358 | 0.92156419 | 0.9686919 |
| CCDC85C | 21.1962138 | 0.514651071 | 0.30558051 | 1.68417504 | 0.09214785 | 0.31254072 |
| EVL | 122.858577 | -0.613775469 | 0.34200602 | -1.7946335 | 0.07271211 | 0.27219482 |
| YY1 | 122.037251 | -0.337215924 | 0.19846828 | -1.6990923 | 0.0893018 | 0.307688 |
| SLC25A29 | 292.325315 | -0.318319711 | 0.4110039 | -0.7744932 | 0.43863921 | 0.71006618 |
| WARS | 81.9893858 | -2.163264515 | 0.66428207 | -3.2565451 | 0.00112777 | 0.01591546 |
| WDR25 | 40.059955 | 0.20754471 | 0.31412648 | 0.66070427 | 0.50880198 | 0.75441538 |
| PPP2R5C | 154.734955 | 0.073399717 | 0.20257002 | 0.36234245 | 0.71709614 | 0.8735396 |
| DYNC1H1 | 121.926697 | -0.470942997 | 0.24128265 | -1.9518311 | 0.05095826 | 0.21990081 |
| HSP90AA1 | 430.457893 | 0.901225844 | 0.54306512 | 1.65951708 | 0.09701164 | 0.32146212 |
| WDR20 | 123.697351 | 0.252079077 | 0.20107479 | 1.2536583 | 0.20996623 | 0.49623127 |
| MOK | 72.246902 | 0.184442803 | 0.37245729 | 0.49520525 | 0.62045525 | 0.81942374 |
| ZNF839 | 23.5441225 | 0.107226993 | 0.37337085 | 0.2871863 | 0.77396968 | 0.90184317 |
| CINP | 14.2854014 | -0.146764716 | 0.43104481 | -0.340486 | 0.73349058 | 0.88350705 |
| TECPR2 | 67.5793298 | 0.015417476 | 0.28015504 | 0.05503194 | 0.95611302 | 0.98286826 |
| ANKRD9 | 41.9294897 | 0.17458385 | 0.32694291 | 0.53398879 | 0.59334929 | 0.80218956 |
| RCOR1 | 180.205673 | 0.436323739 | 0.3684527 | 1.18420558 | 0.23633169 | 0.526647 |
| TRAF3 | 114.460199 | -0.470274567 | 0.27579188 | -1.7051792 | 0.08816102 | 0.30539182 |
| CDC42BPB | 201.678195 | -0.164017956 | 0.34420864 | -0.4765074 | 0.63371295 | 0.82744117 |
| TNFAIP2 | 23.5766398 | -1.124985018 | 1.0045325 | -1.119909 | 0.26275253 | 0.55434704 |
| EIF5 | 91.4791284 | -0.457565076 | 0.27207716 | -1.6817475 | 0.09261781 | 0.31323946 |
| MARK3 | 381.734439 | 0.412543063 | 0.18264248 | 2.25874648 | 0.02389916 | 0.13541585 |
| CKB | 164.836425 | -0.946945724 | 0.46558838 | -2.0338689 | 0.04196482 | 0.19544373 |
| BAG5 | 41.9875347 | -1.103689545 | 0.38337341 | -2.8788891 | 0.00399079 | 0.04011032 |
| KLC1 | 75.0512248 | -0.204047011 | 0.25012288 | -0.8157871 | 0.41462193 | 0.69139449 |
| ZFYVE21 | 29.5396189 | -0.618407214 | 0.34650058 | -1.784722 | 0.07430641 | 0.27585754 |
| PPP1R13B | 64.6872259 | -0.721875155 | 0.25132169 | -2.8723154 | 0.00407476 | 0.04051948 |
| ATP5MPL | 55.3314697 | 0.182235755 | 0.30477602 | 0.59793338 | 0.54988439 | 0.77596691 |
| INF2 | 14.1380465 | -1.137681994 | 0.50246153 | -2.2642171 | 0.02356076 | 0.13454711 |
| SIVA1 | 24.3555772 | -0.321679573 | 0.38907014 | -0.8267907 | 0.40835571 | 0.68672552 |
| AKT1 | 34.9217204 | -0.264460107 | 0.29983834 | -0.882009 | 0.37777196 | 0.66074817 |
| CEP170B | 23.1061404 | -0.237857684 | 0.39315859 | -0.6049917 | 0.54518451 | 0.77447474 |
| CLBA1 | 13.3663726 | -0.500428431 | 0.43577533 | -1.1483634 | 0.25081858 | 0.54232519 |
| GPR132 | 25.7759118 | 0.510875598 | 0.87326808 | 0.58501577 | 0.55853711 | 0.78243918 |
| NUDT14 | 12.6917049 | -0.79338703 | 0.53511442 | -1.4826493 | 0.1381676 | 0.39179815 |
| BRF1 | 43.8144947 | 0.552532694 | 0.2827802 | 1.95392994 | 0.0507095 | 0.21949971 |
| BTBD6 | 15.509852 | -0.835010694 | 0.54760356 | -1.5248453 | 0.1272977 | 0.37415717 |
| PACS2 | 67.1398388 | -0.263912348 | 0.29215718 | -0.9033231 | 0.36635441 | 0.65172793 |
| TEX22 | 10.6888247 | -1.181944917 | 0.50002463 | -2.3637734 | 0.01808987 | 0.11306142 |
| MTA1 | 33.6781834 | 0.400668496 | 0.32838241 | 1.22012776 | 0.22241645 | 0.51073063 |
| CRIP2 | 32.6280999 | -0.830584119 | 0.54683682 | -1.5188884 | 0.12879058 | 0.37613516 |
| AC087386.1 | 12.3022447 | 0.177826305 | 0.56566264 | 0.31436813 | 0.75324146 | 0.89205725 |
| NIPA1 | 45.6915044 | 0.0719868 | 0.29940536 | 0.24043257 | 0.80999493 | 0.91928664 |
| NIPA2 | 48.7482869 | -0.264457522 | 0.24724578 | -1.0696139 | 0.28479313 | 0.57686567 |
| CYFIP1 | 103.624032 | -0.355731137 | 0.2230462 | -1.5948765 | 0.11073986 | 0.34697143 |
| TUBGCP5 | 40.5524867 | 0.16714674 | 0.27403244 | 0.60995238 | 0.54189336 | 0.77224863 |
| AC087463.4 | 11.6967863 | 2.268260044 | 0.71108712 | 3.18984829 | 0.00142348 | 0.01902028 |
| PWRN3 | 39.6113823 | 2.053010872 | 0.81887556 | 2.50710971 | 0.01217229 | 0.08788108 |
| PWRN1 | 312.498638 | 1.917073129 | 0.44745104 | 4.28443102 | 1.83E-05 | 0.00060108 |

| | | | | | | |
|---|---|---|---|---|---|---|
| NPAP1 | 9.15892915 | 2.191845688 | 0.75542351 | 2.9014793 | 0.00371405 | 0.03810974 |
| SNRPN | 33.3217944 | 1.895722106 | 0.63302271 | 2.99471423 | 0.00274702 | 0.03082768 |
| AC124312.1 | 319.708406 | 0.693091909 | 0.23452166 | 2.95534286 | 0.00312322 | 0.03365503 |
| PWAR6 | 80.95726 | 0.535284685 | 0.31386279 | 1.70547357 | 0.08810614 | 0.30539182 |
| UBE3A | 213.0888 | 0.409281861 | 0.25814108 | 1.58549683 | 0.1128535 | 0.35081977 |
| GABRB3 | 95.2694941 | -0.245666671 | 0.38992425 | -0.6300369 | 0.52867044 | 0.76311655 |
| HERC2 | 221.406419 | 0.394274279 | 0.2465584 | 1.59911112 | 0.10979591 | 0.34537939 |
| FAM189A1 | 99.808694 | 0.395835621 | 0.44017107 | 0.89927678 | 0.36850525 | 0.65369102 |
| TJP1 | 247.84526 | 0.119549897 | 0.29407688 | 0.40652599 | 0.68435616 | 0.85613081 |
| FAN1 | 54.8155373 | 0.14957386 | 0.33791061 | 0.44264327 | 0.65802378 | 0.84031559 |
| MTMR10 | 328.876076 | 0.339427603 | 0.2719786 | 1.24799379 | 0.21203333 | 0.49805901 |
| KLF13 | 62.8398402 | 0.074252412 | 0.32232408 | 0.2303657 | 0.81780761 | 0.92281757 |
| OTUD7A | 87.6613013 | 0.470882006 | 0.32899794 | 1.43126128 | 0.15235535 | 0.41413319 |
| LINC02256 | 128.655111 | 0.273158553 | 0.27926631 | 0.97812927 | 0.3280104 | 0.61729352 |
| AC123768.3 | 18.5590676 | 0.327635434 | 0.41472741 | 0.79000189 | 0.42952666 | 0.70316149 |
| GOLGA8N | 15.6422523 | 1.008966609 | 0.44047762 | 2.29061945 | 0.02198543 | 0.1285195 |
| FMN1 | 266.54555 | -0.132902362 | 0.37358635 | -0.3557474 | 0.72202977 | 0.87633726 |
| AVEN | 51.5080667 | 0.942139043 | 0.26788625 | 3.51693697 | 0.00043656 | 0.0078038 |
| EMC7 | 28.522336 | 0.062569452 | 0.35151412 | 0.17799982 | 0.85872311 | 0.94403644 |
| KATNBL1 | 118.153455 | 0.13428625 | 0.20379914 | 0.6589147 | 0.50995055 | 0.75481473 |
| EMC4 | 12.2039168 | -0.762288295 | 0.51239489 | -1.4876969 | 0.13683085 | 0.38961843 |
| SLC12A6 | 52.1839738 | 0.058947061 | 0.43780045 | 0.13464368 | 0.89289361 | 0.95782021 |
| NOP10 | 35.3927018 | -1.226539203 | 0.35028415 | -3.5015549 | 0.00046255 | 0.00813175 |
| LPCAT4 | 11.5689152 | -0.660668162 | 0.45116901 | -1.4643474 | 0.14309905 | 0.40016462 |
| GOLGA8A | 73.0443569 | -0.068532507 | 0.3305572 | -0.2073242 | 0.83575668 | 0.93378849 |
| GOLGA8B | 70.5902352 | 0.0457191 | 0.36657023 | 0.12472126 | 0.90074423 | 0.95981248 |
| AQR | 91.9807326 | 0.361515721 | 0.22465987 | 1.60916909 | 0.10757937 | 0.34161321 |
| ZNF770 | 65.0782713 | -0.898067488 | 0.35992007 | -2.4951859 | 0.01258912 | 0.09012722 |
| AC021231.1 | 87.5412534 | 0.076168525 | 0.34540276 | 0.22052089 | 0.8254655 | 0.92696159 |
| DPH6 | 280.662306 | -0.011146744 | 0.23347038 | -0.0477437 | 0.96192049 | 0.98525601 |
| DPH6-DT | 246.863855 | 1.882775442 | 0.60701431 | 3.10169859 | 0.00192414 | 0.02394018 |
| C15orf41 | 89.6002191 | 0.281045465 | 0.23625628 | 1.18957883 | 0.23421197 | 0.52443761 |
| SPRED1 | 126.963701 | -0.09366836 | 0.37488706 | -0.2498575 | 0.80269752 | 0.91566713 |
| FAM98B | 38.5875185 | 0.399844955 | 0.26320652 | 1.51913015 | 0.12872974 | 0.37611281 |
| RASGRP1 | 107.254617 | 0.053670531 | 0.28832875 | 0.18614353 | 0.85233221 | 0.94037032 |
| C15orf53 | 247.889599 | 3.057782073 | 0.59985274 | 5.09755457 | 3.44E-07 | 2.35E-05 |
| AC013652.1 | 73.8636806 | 1.055404971 | 0.55321288 | 1.90777367 | 0.05642048 | 0.23454985 |
| THBS1 | 67.0075906 | 0.479359159 | 0.61158224 | 0.78380164 | 0.43315651 | 0.70562593 |
| FSIP1 | 23.3724126 | 0.201934405 | 0.4548188 | 0.44398869 | 0.65705076 | 0.83961014 |
| EIF2AK4 | 56.8804692 | 0.145411455 | 0.21757721 | 0.66832117 | 0.50392861 | 0.75108109 |
| SRP14 | 191.475447 | -1.259216039 | 0.38792495 | -3.24603 | 0.00117026 | 0.0163583 |
| SRP14-AS1 | 17.4122974 | -0.19721002 | 0.46379954 | -0.4252053 | 0.67068703 | 0.84746193 |
| IVD | 60.3627179 | 0.030557878 | 0.36687293 | 0.08329281 | 0.93361871 | 0.97428485 |
| BAHD1 | 20.8578932 | 0.054965124 | 0.33421227 | 0.16446172 | 0.86936768 | 0.94827256 |
| CCDC32 | 46.5489961 | -0.538121997 | 0.33860253 | -1.5892439 | 0.11200534 | 0.3488147 |
| RMDN3 | 26.8145708 | -0.304761023 | 0.37293364 | -0.8171991 | 0.41381468 | 0.69060729 |
| GCHFR | 13.1154175 | -2.190296831 | 0.75318443 | -2.9080485 | 0.00363692 | 0.03751236 |
| DNAJC17 | 41.2028763 | 0.16675161 | 0.24197489 | 0.68912775 | 0.49074288 | 0.7456686 |
| ZFYVE19 | 18.0678551 | 0.302367075 | 0.39424575 | 0.76695075 | 0.4431108 | 0.71326991 |
| SPINT1-AS1 | 31.7829291 | 0.096130839 | 0.45582572 | 0.21089385 | 0.83297011 | 0.93213752 |
| SPINT1 | 71.5721925 | -1.014489722 | 0.39572606 | -2.5636161 | 0.0103588 | 0.07880617 |
| VPS18 | 13.8244706 | -0.242568973 | 0.46855383 | -0.5176971 | 0.6046696 | 0.81005282 |
| INO80 | 135.923633 | 0.559952047 | 0.33071066 | 1.69317812 | 0.09042157 | 0.30986868 |
| CHP1 | 83.0794254 | 0.47704071 | 0.4581992 | 1.04112079 | 0.29781949 | 0.59036683 |
| NDUFAF1 | 14.7506736 | -0.171719896 | 0.39207265 | -0.4379798 | 0.66140093 | 0.84192458 |
| RTF1 | 70.6607021 | -0.505746605 | 0.33263273 | -1.5204355 | 0.12840156 | 0.37579343 |
| RPAP1 | 15.4781308 | 0.238822833 | 0.3768828 | 0.63367932 | 0.52629013 | 0.76221787 |
| MGA | 113.041901 | -0.033125471 | 0.25160646 | -0.1316559 | 0.89525648 | 0.95885435 |
| MAPKBP1 | 66.0828895 | 0.460836188 | 0.38499475 | 1.19699342 | 0.23130912 | 0.52111057 |
| EHD4 | 60.0816693 | -0.670776074 | 0.39050018 | -1.7177356 | 0.08584485 | 0.30113433 |
| PLA2G4F | 23.2921461 | 0.736736573 | 0.63655354 | 1.15738351 | 0.24711571 | 0.53778881 |
| VPS39 | 73.9217362 | 0.189762591 | 0.21785628 | 0.87104484 | 0.38372967 | 0.66701235 |
| TMEM87A | 118.379245 | -0.242035682 | 0.18686255 | -1.2952605 | 0.19523038 | 0.47624125 |
| GANC | 50.5084411 | -0.334861035 | 0.34078072 | -0.982629 | 0.32579005 | 0.6159877 |
| ZNF106 | 92.7061744 | 0.104785507 | 0.22974443 | 0.45609596 | 0.64832097 | 0.83558354 |
| SNAP23 | 68.3085266 | -0.193017364 | 0.26141483 | -0.7383566 | 0.46029778 | 0.72529637 |
| LRRC57 | 12.3805022 | -0.183617948 | 0.42318656 | -0.4338936 | 0.66436569 | 0.84348349 |
| HAUS2 | 18.09665 | -0.174494219 | 0.36147007 | -0.4827349 | 0.62928398 | 0.82502105 |
| TTBK2 | 122.733568 | -0.027414236 | 0.26289716 | -0.1042774 | 0.9169492 | 0.96589066 |
| UBR1 | 180.960341 | 0.178423162 | 0.20790113 | 0.8582116 | 0.39077563 | 0.67223049 |
| AC068724.3 | 26.4906121 | -0.216873491 | 0.53260908 | -0.4071908 | 0.68386789 | 0.85606265 |
| TMEM62 | 50.1368124 | -0.324545892 | 0.39870347 | -0.8140032 | 0.41564313 | 0.69220102 |
| CCNDBP1 | 19.8371555 | -0.544621305 | 0.42437041 | -1.283363 | 0.19936492 | 0.48246591 |
| ADAL | 37.6909019 | 1.109640706 | 0.37758433 | 2.93878909 | 0.00329497 | 0.03481518 |
| ZSCAN29 | 11.0296218 | -0.456915819 | 0.44811475 | -1.0196402 | 0.30789912 | 0.60158415 |
| TUBGCP4 | 26.4379048 | -0.397375637 | 0.34161495 | -1.1632267 | 0.24473753 | 0.53553761 |
| TP53BP1 | 153.272156 | 0.401581474 | 0.21470633 | 1.87037561 | 0.06143168 | 0.24564181 |
| PPIP5K1 | 58.8894673 | 0.194440986 | 0.24317682 | 0.79958685 | 0.42395021 | 0.69858206 |
| CATSPER2 | 68.5618401 | 0.611248236 | 0.35459887 | 1.7237738 | 0.08474869 | 0.29900033 |
| PDIA3 | 138.935593 | -0.083833189 | 0.43104345 | -0.194489 | 0.84579305 | 0.93757726 |
| SERF2 | 358.507382 | -0.516294223 | 0.4132836 | -1.2492493 | 0.21157392 | 0.4977471 |
| MFAP1 | 25.9033062 | -0.290975037 | 0.35844374 | -0.8117732 | 0.41692176 | 0.69338933 |
| FRMD5 | 13.1269685 | -1.179305103 | 0.57450848 | -2.0527201 | 0.04009973 | 0.18984716 |
| CASC4 | 249.238914 | -0.214469733 | 0.14369571 | -1.492527 | 0.13556104 | 0.38771511 |
| CTDSPL2 | 134.306462 | 0.097112928 | 0.2242276 | 0.43309979 | 0.66494228 | 0.84348349 |
| EIF3J-DT | 34.2908014 | -0.801255107 | 0.3729701 | -2.1483092 | 0.0316892 | 0.1630255 |
| EIF3J | 67.259915 | -0.20881794 | 0.20871444 | -1.0004959 | 0.31707058 | 0.61043937 |
| SPG11 | 195.722794 | -0.267911926 | 0.2076487 | -1.2902172 | 0.19697526 | 0.47881999 |
| B2M | 644.519896 | -2.019431378 | 0.44007531 | -4.5888314 | 4.46E-06 | 0.00019213 |
| TRIM69 | 19.493062 | -0.649991844 | 0.43420115 | -1.4969832 | 0.13439762 | 0.38524327 |
| SORD | 19.4227597 | 0.638075567 | 0.49066221 | 1.30043756 | 0.19345105 | 0.4740563 |
| DUOX1 | 15.630118 | -1.485258791 | 0.41715412 | -3.5604557 | 0.00037021 | 0.00688433 |
| GATM | 18.3505943 | 1.293914425 | 0.57248109 | 2.26018718 | 0.02380964 | 0.13541585 |
| SPATA5L1 | 23.139131 | -0.263167318 | 0.34931439 | -0.7533824 | 0.45122015 | 0.71947224 |
| SLC30A4 | 12.8927215 | -0.451625867 | 0.51048693 | -0.8846962 | 0.37632048 | 0.65957178 |
| SQOR | 17.4882757 | -1.275096626 | 0.42339673 | -3.0115882 | 0.00259885 | 0.02951856 |
| SEMA6D | 37.5716559 | 0.856589593 | 0.49480494 | 1.73116622 | 0.08342212 | 0.29605854 |
| MYEF2 | 89.44782 | -0.351192937 | 0.24299653 | -1.4452591 | 0.14838513 | 0.40873359 |
| SLC12A1 | 46.0800454 | 3.364505931 | 0.67227522 | 5.00465559 | 5.60E-07 | 3.52E-05 |
| DUT | 41.3529988 | -0.061222718 | 0.35019497 | -0.1748247 | 0.86121741 | 0.94536619 |
| CEP152 | 40.7752729 | 0.686814875 | 0.30269756 | 2.26898054 | 0.02326951 | 0.13355013 |
| EID1 | 76.2455629 | -1.133001861 | 0.4245664 | -2.6686093 | 0.0076166 | 0.06389755 |
| SECISBP2L | 176.715883 | -0.064657942 | 0.1659275 | -0.3896759 | 0.69677624 | 0.86247999 |
| COPS2 | 43.2872876 | 0.263847956 | 0.34134684 | 0.77296146 | 0.43954519 | 0.71063944 |
| GALK2 | 107.689232 | 0.401524832 | 0.25826017 | 1.55472998 | 0.12001039 | 0.36267147 |
| FAM227B | 70.0897932 | 0.29219404 | 0.45554712 | 0.64141343 | 0.52125411 | 0.76094088 |
| DTWD1 | 59.3439991 | 0.219344184 | 0.27909487 | 0.78591263 | 0.43191868 | 0.70467666 |
| SLC27A2 | 12.0943189 | -1.967910115 | 0.7514871 | -2.6186878 | 0.00882687 | 0.07049178 |
| GABPB1 | 51.0441139 | 0.271908407 | 0.34329654 | 0.79205112 | 0.42833087 | 0.7026536 |
| GABPB1-AS1 | 95.1728424 | 0.382366769 | 0.39437522 | 0.96955071 | 0.33227049 | 0.62173066 |
| USP8 | 133.642926 | 0.310619929 | 0.21662081 | 1.43393392 | 0.15159112 | 0.41303669 |
| USP50 | 13.5192907 | -0.266578985 | 0.55007902 | -0.4846194 | 0.62794632 | 0.82428562 |
| TRPM7 | 228.229664 | 0.118695751 | 0.20554633 | 0.57746471 | 0.56362557 | 0.78465819 |

| | | | | | | |
|---|---|---|---|---|---|---|
| SPPL2A | 158.389748 | -0.274507236 | 0.29849915 | -0.9196248 | 0.35776884 | 0.64208096 |
| AP4E1 | 50.2010719 | -0.087425131 | 0.27604536 | -0.3167057 | 0.75146695 | 0.89132635 |
| DMXL2 | 102.909918 | 0.89137557 | 0.24949584 | 3.57270712 | 0.00035331 | 0.00664197 |
| TMOD2 | 15.4243405 | -0.310239236 | 0.40175223 | -0.7722153 | 0.4399869 | 0.71079583 |
| TMOD3 | 122.635026 | -0.482529686 | 0.20601517 | -2.3422046 | 0.0191702 | 0.11762443 |
| LEO1 | 18.4396575 | 0.361659052 | 0.34139835 | 1.05934621 | 0.28944214 | 0.58125398 |
| MAPK6 | 75.7522774 | -0.43477678 | 0.36094854 | -1.2045395 | 0.22838112 | 0.51811987 |
| GNB5 | 74.5957177 | -0.134180699 | 0.31318019 | -0.4284457 | 0.66832668 | 0.84573166 |
| MYO5C | 116.352962 | -0.074372616 | 0.3116833 | -0.238616 | 0.81140338 | 0.9199895 |
| MYO5A | 276.042882 | 0.994607588 | 0.34188241 | 2.90920963 | 0.00362344 | 0.03751236 |
| ARPP19 | 102.025133 | -0.335017984 | 0.26537432 | -1.2624356 | 0.2067921 | 0.49177567 |
| FAM214A | 141.698787 | -0.03162428 | 0.22498569 | -0.1405613 | 0.88821653 | 0.95661336 |
| WDR72 | 562.490766 | 0.995014385 | 0.4106437 | 2.42306013 | 0.01539038 | 0.10183152 |
| UNC13C | 78.9607151 | 2.241142836 | 0.70235388 | 3.19090261 | 0.00141829 | 0.0190003 |
| RSL24D1 | 27.7701021 | -0.690680498 | 0.38591957 | -1.7897006 | 0.07350205 | 0.27395735 |
| RAB27A | 69.3942706 | -0.94181556 | 0.22999918 | -4.0948649 | 4.22E-05 | 0.0012293 |
| PIGB | 31.5307008 | -0.38188829 | 0.30430939 | -1.2549343 | 0.20950261 | 0.49570416 |
| CCPG1 | 123.650651 | -0.194403508 | 0.21011797 | -0.9252112 | 0.35485604 | 0.64012028 |
| DNAAF4 | 56.1376444 | -0.253127593 | 0.31929801 | -0.7927628 | 0.42791602 | 0.70230856 |
| PYGO1 | 23.2046587 | 0.493286409 | 0.41836607 | 1.17907844 | 0.23836694 | 0.52855279 |
| PRTG | 116.730409 | 0.342881142 | 0.61039883 | 0.56173296 | 0.57429798 | 0.7910707 |
| NEDD4 | 47.4404112 | -1.151329239 | 0.33265448 | -3.4610364 | 0.0005381 | 0.00902851 |
| RFX7 | 157.559513 | 0.087680977 | 0.19203176 | 0.45659623 | 0.64796129 | 0.83535192 |
| TEX9 | 102.288424 | -0.018232974 | 0.32189714 | -0.0566422 | 0.95483019 | 0.98238895 |
| MNS1 | 35.2974844 | -0.430682706 | 0.48355168 | -0.8906653 | 0.37310875 | 0.65706383 |
| ZNF280D | 65.6684022 | 0.554991609 | 0.34451817 | 1.61092118 | 0.1071969 | 0.34111768 |
| AC090517.4 | 39.1602609 | 0.208299415 | 0.38109209 | 0.54658551 | 0.58466353 | 0.79661125 |
| TCF12 | 762.207935 | 0.538614586 | 0.19778181 | 2.72327664 | 0.00646379 | 0.0568659 |
| CGNL1 | 321.598667 | 0.993105008 | 0.37285661 | 2.66350381 | 0.00773315 | 0.06419577 |
| POLR2M | 17.8199903 | -0.665565625 | 0.38662971 | -1.7214549 | 0.0851683 | 0.2997225 |
| ALDH1A2 | 146.145692 | -2.147353024 | 0.67264942 | -3.1923807 | 0.00141105 | 0.01895262 |
| LIPC | 55.7698851 | -0.911868252 | 0.50573602 | -1.8030518 | 0.07138008 | 0.26971882 |
| ADAM10 | 474.314251 | -0.340683962 | 0.38088415 | -0.8944556 | 0.37107817 | 0.65494451 |
| MINDY2 | 257.790773 | -0.149613913 | 0.29226148 | -0.511918 | 0.60870839 | 0.81261032 |
| RNF111 | 191.509722 | 0.033453661 | 0.27862945 | 0.12006506 | 0.90443161 | 0.96126516 |
| SLTM | 238.564096 | -0.000604715 | 0.19131443 | -0.0031608 | 0.99747802 | 0.99900003 |
| MYO1E | 631.136795 | 0.502114745 | 0.4676826 | 1.0736229 | 0.28299173 | 0.57561209 |
| FAM81A | 75.6233569 | 0.276291131 | 0.29671835 | 0.93115619 | 0.35177278 | 0.63801604 |
| GCNT3 | 51.3620161 | -4.979875435 | 0.69054551 | -7.2115094 | 5.53E-13 | 2.38E-10 |
| GTF2A2 | 43.745665 | -0.442692016 | 0.3451205 | -1.2827172 | 0.19959116 | 0.48289997 |
| BNIP2 | 60.9103194 | -0.068508684 | 0.28214034 | -0.2428178 | 0.80814658 | 0.91842538 |
| ANXA2 | 159.178375 | -2.069244169 | 0.48636447 | -4.2545134 | 2.10E-05 | 0.00067659 |
| ICE2 | 93.0609883 | 0.515476181 | 0.23262083 | 2.21595025 | 0.02669491 | 0.14604937 |
| RORA-AS1 | 58.688051 | 0.694631345 | 0.41346662 | 1.68001797 | 0.09295382 | 0.31376482 |
| RORA | 1126.35285 | 0.976377365 | 0.39328875 | 2.4825967 | 0.01304287 | 0.09213032 |
| AC012404.1 | 31.8005279 | 0.311162817 | 0.61182322 | 0.50858288 | 0.61104463 | 0.81376433 |
| AC009554.1 | 43.8180595 | -1.436958823 | 0.57408434 | -2.5030448 | 0.012313 | 0.08858134 |
| VPS13C | 369.323139 | -0.854104662 | 0.23626624 | -3.6150093 | 0.00030034 | 0.00595013 |
| C2CD4A | 33.719514 | -5.095955206 | 0.83589607 | -6.0963981 | 1.08E-09 | 1.93E-07 |
| TLN2 | 312.422339 | 1.653050619 | 0.3368617 | 4.90720859 | 9.24E-07 | 5.23E-05 |
| TPM1 | 115.587942 | -1.13121361 | 0.38528528 | -2.9360416 | 0.0033243 | 0.03498152 |
| LACTB | 27.721331 | -1.09930201 | 0.39677119 | -2.7706196 | 0.00559498 | 0.05141788 |
| RPS27L | 43.1768834 | -0.5604854 | 0.36806124 | -1.5228047 | 0.12780757 | 0.37490705 |
| RAB8B | 50.9647054 | -0.073504413 | 0.26448058 | -0.2779199 | 0.78107386 | 0.90520525 |
| APH1B | 25.0850698 | -0.854705289 | 0.40705622 | -2.099723 | 0.03575322 | 0.17539507 |
| CA12 | 1006.50152 | 0.611662186 | 0.29947785 | 2.04242881 | 0.04110901 | 0.19311676 |
| LINC02568 | 51.2542801 | -1.513521844 | 0.49775058 | -3.0407234 | 0.00236011 | 0.02734961 |
| USP3 | 240.345257 | 0.313058676 | 0.20480531 | 1.52856721 | 0.12637177 | 0.37228594 |
| USP3-AS1 | 42.0183462 | -0.109224202 | 0.4760467 | -0.2294401 | 0.81852687 | 0.92329616 |
| HERC1 | 295.300141 | -0.171504473 | 0.40301227 | -0.4255565 | 0.67043107 | 0.84735381 |
| FAM96A | 40.958433 | 0.10957218 | 0.32527218 | 0.33686306 | 0.73622013 | 0.88437124 |
| SNX1 | 91.8804595 | -0.0498542 | 0.21426107 | -0.2326797 | 0.81601014 | 0.92187043 |
| PPIB | 107.686413 | -1.496691202 | 0.32173858 | -4.651886 | 3.29E-06 | 0.0001506 |
| CSNK1G1 | 29.8400323 | -0.391683458 | 0.35235519 | -1.1116154 | 0.26630354 | 0.55761364 |
| TRIP4 | 61.7021499 | 0.027388901 | 0.35897886 | 0.0762967 | 0.93918305 | 0.97662903 |
| ZNF609 | 304.218947 | -0.171087661 | 0.23864387 | -0.7169162 | 0.47342579 | 0.73418512 |
| OAZ2 | 36.3768433 | -0.040124285 | 0.37256461 | -0.1076975 | 0.91423563 | 0.96462448 |
| RBPMS2 | 13.6647566 | 1.468402395 | 0.55707976 | 2.63589256 | 0.00839163 | 0.06834037 |
| SPG21 | 50.4707054 | -0.232946278 | 0.31155033 | -0.7477003 | 0.45464094 | 0.72190427 |
| MTFMT | 36.8541389 | -0.075641636 | 0.31910445 | -0.2370435 | 0.81262304 | 0.92054639 |
| UBAP1L | 19.5261096 | 0.428254715 | 0.37761721 | 1.13409744 | 0.25675368 | 0.54774827 |
| PDCD7 | 35.0827056 | -0.194222934 | 0.27263556 | -0.7123903 | 0.47622312 | 0.73631255 |
| CLPX | 62.6299823 | 0.051935756 | 0.19973691 | 0.26002083 | 0.79484771 | 0.91195838 |
| PARP16 | 24.8989598 | -0.184917686 | 0.32301149 | -0.5724802 | 0.56699669 | 0.78656476 |
| DPP8 | 114.517398 | -0.090360001 | 0.20098555 | -0.4495846 | 0.65301003 | 0.83850297 |
| HACD3 | 135.660558 | 0.106180676 | 0.28406215 | 0.37379381 | 0.70855772 | 0.868892 |
| INTS14 | 30.5382044 | 0.103970358 | 0.31393992 | 0.33117916 | 0.74050916 | 0.88639183 |
| SLC24A1 | 18.6962277 | -0.353475633 | 0.52918179 | -0.6679664 | 0.50415507 | 0.75130993 |
| DENND4A | 227.531517 | -0.292773691 | 0.21999462 | -1.3308221 | 0.18324757 | 0.46004581 |
| RAB11A | 144.167454 | -0.658711479 | 0.35741146 | -1.843006 | 0.06532813 | 0.25530899 |
| MEGF11 | 258.574597 | -6.912923055 | 0.53008682 | -13.041115 | 7.14E-39 | 7.36E-35 |
| DIS3L | 29.8028676 | -0.162013428 | 0.3474289 | -0.4663211 | 0.64098568 | 0.83230084 |
| TIPIN | 18.8461349 | 0.935291851 | 0.49015787 | 1.90814412 | 0.05637259 | 0.23445718 |
| MAP2K1 | 102.099552 | -0.457066842 | 0.30232724 | -1.5118282 | 0.13057758 | 0.37903922 |
| SNAPC5 | 15.7493718 | -0.545814813 | 0.38158955 | -1.4303715 | 0.15261043 | 0.41441825 |
| RPL4 | 126.699569 | -0.60058305 | 0.45246175 | -1.3273676 | 0.18438712 | 0.46094543 |
| SMAD6 | 24.2778644 | 0.028355719 | 0.39827111 | 0.07119703 | 0.94324095 | 0.97824588 |
| SMAD3 | 488.302997 | -0.943232758 | 0.5333795 | -1.7684083 | 0.07699266 | 0.28159053 |
| AAGAB | 102.085001 | 0.018854667 | 0.24953287 | 0.07555985 | 0.93976928 | 0.97674567 |
| IQCH | 110.201631 | 0.087366043 | 0.34913771 | 0.25023376 | 0.80240658 | 0.9156394 |
| IQCH-AS1 | 58.825074 | 0.265603515 | 0.38731725 | 0.68575183 | 0.49286961 | 0.74680729 |
| MAP2K5 | 188.227705 | 0.172756777 | 0.2549752 | 0.67754345 | 0.49806121 | 0.74971166 |
| PIAS1 | 286.574826 | 0.405044338 | 0.18786556 | 2.15603297 | 0.03108109 | 0.1614712 |
| FEM1B | 36.9594022 | -0.09462851 | 0.29548886 | -0.3202439 | 0.74878343 | 0.88983353 |
| CORO2B | 23.3019026 | -0.574676911 | 0.58513736 | -0.9821231 | 0.32603921 | 0.6159877 |
| ANP32A | 64.522962 | -0.0883814 | 0.26212934 | -0.3371671 | 0.73599091 | 0.88432217 |
| SPESP1 | 14.9484137 | -0.112961711 | 0.51756862 | -0.2182546 | 0.82723078 | 0.92808376 |
| GLCE | 107.019449 | 0.70153607 | 0.25028244 | 2.80297761 | 0.00506332 | 0.0479874 |
| AC026992.2 | 15.3720322 | 1.938764653 | 0.5709177 | 3.39587414 | 0.0006841 | 0.01109855 |
| PAQR5 | 391.645324 | 0.43390598 | 0.49503743 | 0.87651145 | 0.38075205 | 0.66415639 |
| KIF23 | 32.1769433 | 0.917655899 | 0.64500263 | 1.42271653 | 0.15481835 | 0.41807041 |
| RPLP1 | 242.030981 | -0.414463136 | 0.43396469 | -0.9550619 | 0.3395464 | 0.62815343 |
| TLE3 | 60.4928121 | -0.328616951 | 0.30041423 | -1.0938794 | 0.27400785 | 0.56626457 |
| UACA | 176.367968 | 0.452824817 | 0.24191642 | 1.87182343 | 0.06123104 | 0.24542151 |
| LARP6 | 23.290905 | 0.736487639 | 0.36664613 | 2.00871519 | 0.04456735 | 0.20147742 |
| LRRC49 | 53.1705772 | 0.218674741 | 0.30218504 | 0.72364516 | 0.46928361 | 0.73168094 |
| THSD4 | 206.743749 | 0.730634911 | 0.38464862 | 1.89948662 | 0.05750052 | 0.23694816 |
| THSD4-AS1 | 72.0766383 | -0.649478767 | 0.27383262 | -2.3718094 | 0.01770122 | 0.11160934 |
| MYO9A | 468.195838 | 0.244251538 | 0.21601171 | 1.13073288 | 0.25816754 | 0.54951281 |
| AC022872.1 | 16.8782513 | -0.683697119 | 0.6889907 | -0.9923169 | 0.32104296 | 0.6120836 |
| SENP8 | 14.2219052 | -0.38793519 | 0.45511499 | -0.8523894 | 0.39399801 | 0.67474408 |
| PKM | 465.381823 | -1.30828182 | 0.27244062 | -4.8020806 | 1.57E-06 | 8.11E-05 |
| PARP6 | 46.1121354 | 0.089419029 | 0.27004905 | 0.33112144 | 0.74055276 | 0.88639183 |

| | | | | | | |
|---|---|---|---|---|---|---|
| AC009690.2 | 21.2978892 | -1.415941968 | 0.67416246 | -2.1002978 | 0.03570265 | 0.17539507 |
| AC009690.1 | 76.8238839 | -0.542999105 | 0.35309711 | -1.537818 | 0.12409311 | 0.36905521 |
| ARIH1 | 405.226015 | 0.401050205 | 0.33542619 | 1.19564368 | 0.23183564 | 0.52159221 |
| BBS4 | 63.174858 | 0.252082091 | 0.22663824 | 1.11226638 | 0.26602364 | 0.55721548 |
| ADPGK | 43.1889644 | -0.242568485 | 0.25844544 | -0.9385675 | 0.34795286 | 0.63501969 |
| NEO1 | 117.520801 | 0.021561872 | 0.23674412 | 0.09107669 | 0.92743165 | 0.97111125 |
| NPTN | 150.227233 | 0.230164846 | 0.26310503 | 0.87480214 | 0.38168157 | 0.66492276 |
| CD276 | 18.4789275 | -1.40014692 | 0.48202213 | -2.9047358 | 0.00367563 | 0.03779078 |
| LOXL1 | 12.2041584 | -0.958376982 | 0.50706539 | -1.8900461 | 0.05875179 | 0.23999245 |
| PML | 32.5948542 | -1.190598564 | 0.50552239 | -2.3551846 | 0.01851351 | 0.114757 |
| ARID3B | 25.0473028 | 0.43469761 | 0.31195406 | 1.39346674 | 0.1634787 | 0.43183526 |
| CLK3 | 28.0888705 | 0.486330599 | 0.33818817 | 1.43804735 | 0.15042062 | 0.41132271 |
| EDC3 | 33.5691824 | 0.449067532 | 0.32175474 | 1.39568272 | 0.16281006 | 0.43095304 |
| CSK | 12.9955053 | -0.266505093 | 0.48865935 | -0.5453801 | 0.58549212 | 0.79700578 |
| ULK3 | 20.0792374 | 0.971939326 | 0.40691965 | 2.38852884 | 0.01691598 | 0.10878179 |
| SCAMP2 | 42.4199198 | -0.228407635 | 0.42514061 | -0.537252 | 0.59109357 | 0.80019 |
| FAM219B | 18.1986613 | -0.246453011 | 0.38393738 | -0.6419094 | 0.52093201 | 0.76092013 |
| COX5A | 72.160218 | -0.009877367 | 0.23102836 | -0.0427539 | 0.9658977 | 0.98661741 |
| SCAMP5 | 40.7896009 | -0.033386355 | 0.3200624 | -0.104312 | 0.91692174 | 0.96589066 |
| PPCDC | 65.2384714 | -0.349807949 | 0.39902405 | -0.8766588 | 0.38067199 | 0.66412918 |
| COMMD4 | 22.8840135 | -0.007048644 | 0.39012041 | -0.0180679 | 0.98558471 | 0.99437976 |
| NEIL1 | 22.7660184 | -0.111618452 | 0.3900306 | -0.2861787 | 0.77474125 | 0.90205458 |
| MAN2C1 | 42.5779377 | 0.443508089 | 0.32985004 | 1.34457491 | 0.17876255 | 0.45419584 |
| SIN3A | 80.1871144 | -0.034360845 | 0.27482116 | -0.1250298 | 0.90049993 | 0.95980075 |
| PTPN9 | 86.5277678 | -0.5399064 | 0.30044126 | -1.7970448 | 0.07232851 | 0.27172106 |
| SNUPN | 18.2324515 | -0.243051075 | 0.40458679 | -0.600739 | 0.54801382 | 0.77516644 |
| IMP3 | 14.5157038 | -1.015681873 | 0.39868564 | -2.5475757 | 0.01084743 | 0.08095937 |
| UBE2Q2 | 38.2065931 | -1.297479972 | 0.28937623 | -4.4837131 | 7.34E-06 | 0.00029126 |
| FBXO22 | 64.9893362 | -0.095275414 | 0.23031936 | -0.4136665 | 0.67911834 | 0.85351679 |
| ETFA | 95.9693486 | 0.275627939 | 0.21618897 | 1.2749399 | 0.20233051 | 0.4856498 |
| SCAPER | 307.983565 | 0.297611146 | 0.25416555 | 1.17093424 | 0.24162521 | 0.5324541 |
| RCN2 | 63.6970927 | 0.20415979 | 0.23897486 | 0.85431494 | 0.39293051 | 0.67377433 |
| TSPAN3 | 66.0319246 | -0.552406833 | 0.27127873 | -2.0363072 | 0.04171952 | 0.19500656 |
| PEAK1 | 141.02551 | 0.156107784 | 0.20788298 | 0.75094068 | 0.45268836 | 0.72046894 |
| HMG20A | 95.4481625 | 0.346254801 | 0.19235157 | 1.80011426 | 0.0718426 | 0.27051259 |
| LINGO1 | 589.740485 | 0.522307873 | 0.6358089 | 0.82148563 | 0.4113697 | 0.68899849 |
| TBC1D2B | 36.3479298 | -0.904899992 | 0.34414446 | -2.629419 | 0.00855309 | 0.06905481 |
| IDH3A | 67.5384988 | 0.383495384 | 0.28366421 | 1.35193434 | 0.17639632 | 0.45081491 |
| DNAJA4 | 16.8657534 | -0.666592218 | 0.64787586 | -1.0288888 | 0.30353193 | 0.59743713 |
| WDR61 | 38.1549183 | 0.314608386 | 0.32027498 | 0.98230711 | 0.32594858 | 0.6159877 |
| IREB2 | 93.8011598 | -0.139746072 | 0.20096937 | -0.6953601 | 0.48682969 | 0.74422309 |
| PSMA4 | 43.5914352 | -0.532582788 | 0.35454752 | -1.5021478 | 0.13305894 | 0.3830758 |
| MORF4L1 | 141.965959 | -0.137640265 | 0.15695819 | -0.8769231 | 0.38052839 | 0.66412918 |
| CTSH | 139.020324 | -0.519716379 | 0.38779017 | -1.3402 | 0.18018034 | 0.45640961 |
| TMED3 | 33.2320481 | -0.763075255 | 0.4493459 | -1.6981912 | 0.08947168 | 0.30789644 |
| ZFAND6 | 211.224867 | 0.130019459 | 0.2489044 | 0.52236706 | 0.60141479 | 0.80759541 |
| FAH | 11.6138928 | 0.161633193 | 0.44926852 | 0.35976968 | 0.71901938 | 0.87453809 |
| AC016705.2 | 44.8468755 | 0.30594509 | 0.50769103 | 0.60262063 | 0.54676109 | 0.77463416 |
| ARNT2 | 150.591423 | 0.089669365 | 0.33908987 | 0.2644413 | 0.79143989 | 0.91099595 |
| ABHD17C | 62.4112132 | 0.579356995 | 0.29307676 | 1.97680973 | 0.04806314 | 0.21206669 |
| CEMIP | 13.9899339 | -1.16738986 | 0.85235681 | -1.3696023 | 0.17081107 | 0.44180158 |
| AC027808.1 | 21.8328799 | 0.029063239 | 0.41180022 | 0.07057607 | 0.94373516 | 0.97839988 |
| MESD | 44.7161072 | -0.496111873 | 0.30031656 | -1.6519631 | 0.09854208 | 0.32413171 |
| EFL1 | 55.8073394 | 0.134383939 | 0.27911905 | 0.48145742 | 0.63019143 | 0.82524878 |
| WHAMM | 35.9081185 | -0.570698249 | 0.29682744 | -1.92266 | 0.05452276 | 0.22997772 |
| HOMER2 | 30.6057845 | -0.488197556 | 0.28747311 | -1.6982373 | 0.08946299 | 0.30789644 |
| C15orf40 | 27.0621416 | -0.621461623 | 0.30562697 | -2.0333992 | 0.04201221 | 0.19557605 |
| BTBD1 | 73.7622576 | 0.109565123 | 0.22552565 | 0.48582112 | 0.627094 | 0.82360051 |
| HDGFL3 | 82.6977909 | 0.005141929 | 0.28481618 | 0.0180535 | 0.98559617 | 0.99437976 |
| ADAMTSL3 | 99.7910212 | -0.258627414 | 0.4875804 | -0.5304303 | 0.59581363 | 0.80383342 |
| WDR73 | 60.7705859 | -0.22098143 | 0.24574138 | -0.8992439 | 0.36852278 | 0.65369102 |
| SEC11A | 124.96647 | 0.37350989 | 0.23759946 | 1.57201487 | 0.1159471 | 0.35543425 |
| ZNF592 | 61.019411 | 0.213400622 | 0.30381166 | 0.70241091 | 0.48242294 | 0.74082091 |
| ALPK3 | 62.8945767 | -0.943599725 | 0.53072862 | -1.7779326 | 0.07541492 | 0.27878882 |
| PDE8A | 360.015991 | 0.742336516 | 0.23149778 | 3.20666799 | 0.00134282 | 0.01822624 |
| AKAP13 | 447.173826 | -0.4300108 | 0.27776293 | -1.5481216 | 0.12159302 | 0.36552415 |
| KLHL25 | 22.7728904 | 0.054231224 | 0.39538238 | 0.13716146 | 0.89090318 | 0.95702609 |
| MRPL46 | 15.1430302 | -0.222680394 | 0.35893711 | -0.6203883 | 0.53500215 | 0.76806796 |
| MRPS11 | 25.3341157 | -0.569693661 | 0.38091565 | -1.4955901 | 0.13476051 | 0.38617602 |
| DET1 | 23.8292046 | 0.231045375 | 0.44136582 | 0.52347818 | 0.60064153 | 0.80738812 |
| ABHD2 | 90.9255309 | -0.261723782 | 0.39936289 | -0.6553533 | 0.51224032 | 0.75562371 |
| POLG | 18.5919112 | -0.367541526 | 0.33443292 | -1.0989992 | 0.2717684 | 0.5642932 |
| RHCG | 216.49326 | 3.166478214 | 0.54514816 | 5.80847269 | 6.30E-09 | 8.22E-07 |
| PEX11A | 11.3887628 | -0.805602837 | 0.39986827 | -2.0146706 | 0.04393918 | 0.20006796 |
| ANPEP | 15.1784524 | -0.834493911 | 0.46980015 | -1.7762742 | 0.07568773 | 0.27917785 |
| ZNF710 | 54.4272117 | -0.741781526 | 0.33392605 | -2.2213946 | 0.02632425 | 0.14494516 |
| ZNF710-AS1 | 38.9753541 | 0.10031539 | 0.48192277 | 0.20815657 | 0.83510672 | 0.9332107 |
| IDH2 | 238.706825 | -0.063726538 | 0.2916915 | -0.2184724 | 0.82706107 | 0.92808376 |
| IDH2-DT | 31.9105953 | 0.081773984 | 0.50611126 | 0.16157314 | 0.87164201 | 0.94844301 |
| SEMA4B | 240.454793 | -1.895917321 | 0.32076171 | -5.9106723 | 3.41E-09 | 4.88E-07 |
| CIB1 | 98.5683015 | -1.311949162 | 0.48017101 | -2.732254 | 0.00629026 | 0.05596053 |
| AC091167.6 | 9.50415294 | -0.192022189 | 0.56089348 | -0.3423505 | 0.7320871 | 0.88272019 |
| ZNF774 | 10.0943542 | 0.436024707 | 0.43616833 | 0.99967072 | 0.31746989 | 0.61075159 |
| IQGAP1 | 250.227178 | -0.253807953 | 0.16681919 | -1.5214554 | 0.1281456 | 0.37540371 |
| CRTC3 | 175.883449 | -0.082977507 | 0.18749918 | -0.4425487 | 0.65809223 | 0.84031559 |
| CRTC3-AS1 | 24.5885649 | 0.20208697 | 0.40483421 | 0.49918452 | 0.6176494 | 0.81764942 |
| FURIN | 31.2934795 | -1.287979222 | 0.52383013 | -2.4587727 | 0.01394129 | 0.09600476 |
| MAN2A2 | 17.8645258 | -0.098985149 | 0.35912088 | -0.2756318 | 0.78283085 | 0.90635238 |
| UNC45A | 31.5981258 | -0.07089147 | 0.31491261 | -0.2251147 | 0.82189002 | 0.92509211 |
| AC104035.1 | 60.9586968 | -1.640346102 | 0.86197548 | -1.9030079 | 0.05703952 | 0.23570501 |
| SLCO3A1 | 93.2710931 | -2.507648939 | 0.58098055 | -4.3162356 | 1.59E-05 | 0.00054304 |
| FAM174B | 34.5138716 | -0.65583182 | 0.3362119 | -1.9506502 | 0.05109867 | 0.22007461 |
| AC013394.1 | 131.210186 | 0.282073084 | 0.24298957 | 1.16084441 | 0.24570518 | 0.53627476 |
| CHD2 | 258.314677 | 0.318856197 | 0.34414604 | 0.92651421 | 0.35417881 | 0.63957058 |
| MCTP2 | 51.4196333 | -1.037722242 | 0.51077438 | -2.0316646 | 0.04218762 | 0.19607262 |
| AC012409.2 | 49.2289805 | -1.253698421 | 0.65089425 | -1.9261169 | 0.05408978 | 0.2290137 |
| NR2F2-AS1 | 327.302746 | -0.488492709 | 0.36122408 | -1.352326 | 0.17627105 | 0.45071839 |
| NR2F2 | 97.1085105 | -0.833782028 | 0.32295614 | -2.5817191 | 0.00983096 | 0.07643662 |
| ARRDC4 | 40.3184107 | 0.263635124 | 0.47997318 | 0.54927053 | 0.58281981 | 0.79578657 |
| FAM169B | 14.8679117 | -4.300280805 | 0.8351557 | -5.1490768 | 2.62E-07 | 1.93E-05 |
| IGF1R | 550.901399 | -0.282600393 | 0.23616131 | -1.1966414 | 0.23144637 | 0.52111057 |
| TTC23 | 63.7399921 | 0.104541721 | 0.24173479 | 0.43246452 | 0.66540384 | 0.84348349 |
| LRRC28 | 132.875535 | 0.700934923 | 0.24802764 | 2.82603554 | 0.0047128 | 0.04559989 |
| MEF2A | 380.124585 | 0.489619284 | 0.16591886 | 2.95095613 | 0.00316792 | 0.03391661 |
| AC090825.1 | 25.8377143 | 0.401438468 | 0.45251753 | 0.88712247 | 0.37501297 | 0.65806152 |
| ADAMTS17 | 53.692837 | 1.070237585 | 0.50030837 | 2.13915587 | 0.03242305 | 0.16565474 |
| LINS1 | 24.8617248 | 0.299103886 | 0.32845632 | 0.9106352 | 0.36248762 | 0.64675224 |
| ASB7 | 31.0029305 | -0.307839025 | 0.32980513 | -0.9333967 | 0.35061517 | 0.63716177 |
| CHSY1 | 257.170126 | -0.491433407 | 0.4882074 | -1.0066079 | 0.31412325 | 0.60719808 |
| SELENOS | 53.0937145 | -1.110099353 | 0.28740472 | -3.8624953 | 0.00011223 | 0.00266415 |
| SNRPA1 | 32.8897596 | -0.251362286 | 0.3243978 | -0.7748582 | 0.43842347 | 0.70985672 |
| PCSK6 | 25.7190964 | 0.208370078 | 0.41018427 | 0.50799139 | 0.61145938 | 0.81385718 |
| TM2D3 | 38.9745265 | -0.18064031 | 0.27646637 | -0.6533898 | 0.513505 | 0.75647799 |

| | | | | | | |
|---|---|---|---|---|---|---|
| TARSL2 | 71.2430969 | 0.253886327 | 0.34486142 | 0.73619812 | 0.46161014 | 0.72669738 |
| SNRNP25 | 11.1877749 | -0.161517265 | 0.48590659 | -0.3324039 | 0.73958427 | 0.88614269 |
| MPG | 30.5132076 | -0.990091572 | 0.29556402 | -3.3498379 | 0.00080859 | 0.0125832 |
| NPRL3 | 64.4621978 | -0.183618387 | 0.28619911 | -0.6415757 | 0.52114873 | 0.76092013 |
| LUC7L | 135.170543 | 0.540182625 | 0.25474877 | 2.12045232 | 0.03396792 | 0.17053484 |
| FAM234A | 22.390201 | 0.325198375 | 0.47247048 | 0.68829353 | 0.49126796 | 0.74580047 |
| AXIN1 | 26.1490453 | -0.363256828 | 0.40942669 | -0.8872329 | 0.37495352 | 0.65806152 |
| MRPL28 | 21.8474294 | -0.286300896 | 0.45377534 | -0.6309309 | 0.52808569 | 0.76288619 |
| TMEM8A | 60.7872729 | -0.362918474 | 0.29626692 | -1.2249713 | 0.22058603 | 0.5089143 |
| NME4 | 14.7536453 | -0.221286185 | 0.43799509 | -0.5052253 | 0.61340063 | 0.81475674 |
| RAB11FIP3 | 183.200535 | 0.792375109 | 0.23462963 | 3.37713143 | 0.00073246 | 0.01168082 |
| CAPN15 | 29.6269171 | -0.300573245 | 0.41535189 | -0.7236593 | 0.46927494 | 0.73168094 |
| PRR35 | 17.5240668 | 2.499905291 | 0.701306 | 3.56464267 | 0.00036435 | 0.00678762 |
| PIGQ | 73.3634626 | 1.475895111 | 0.38912881 | 3.79281896 | 0.00014895 | 0.00336502 |
| RAB40C | 78.1379746 | 0.553841365 | 0.32154276 | 1.72245009 | 0.08498802 | 0.29941397 |
| METTL26 | 12.8202423 | -0.328475371 | 0.58564082 | -0.560882 | 0.57487801 | 0.79134063 |
| MCRIP2 | 50.4794461 | 0.748128776 | 0.32277069 | 2.31783366 | 0.02045836 | 0.12275018 |
| RHOT2 | 40.115664 | 0.282326108 | 0.2845513 | 0.99217999 | 0.32110973 | 0.6120836 |
| STUB1 | 35.2594335 | -0.29668637 | 0.40585824 | -0.7310098 | 0.46477314 | 0.72848544 |
| JMJD8 | 17.5962907 | -0.267159281 | 0.49518 | -0.5395195 | 0.58952843 | 0.79960282 |
| METRN | 23.9835554 | 0.077691083 | 0.50095182 | 0.15508694 | 0.87675278 | 0.95134725 |
| FAM173A | 24.5757204 | -0.030872577 | 0.51055086 | -0.0604692 | 0.95178199 | 0.98065459 |
| NARFL | 15.6844959 | 0.515592864 | 0.35019058 | 1.47232076 | 0.14093428 | 0.39647869 |
| LMF1 | 58.2698299 | 0.080338686 | 0.25496657 | 0.31509498 | 0.75268954 | 0.89205725 |
| CACNA1H | 18.247143 | 1.173326258 | 0.82402911 | 1.42388932 | 0.15447851 | 0.41780983 |
| UBE2I | 87.9663285 | -0.268237948 | 0.332758 | -0.8061052 | 0.4201822 | 0.6964951 |
| BAIAP3 | 10.3410387 | -0.006172468 | 0.52382472 | -0.0117835 | 0.99059838 | 0.99684874 |
| TSR3 | 21.2920732 | -0.411592188 | 0.50520323 | -0.8147062 | 0.41524052 | 0.69186606 |
| GNPTG | 38.2143936 | -0.236714947 | 0.34417861 | -0.6877678 | 0.49159904 | 0.74602485 |
| UNKL | 25.2352587 | -0.041936883 | 0.39220262 | -0.1069266 | 0.91484723 | 0.96464134 |
| CLCN7 | 43.2277811 | -0.4419902 | 0.3722328 | -1.1874026 | 0.23506884 | 0.52531003 |
| TELO2 | 11.6344707 | 0.39536593 | 0.53392778 | 0.74048578 | 0.45900529 | 0.72461104 |
| IFT140 | 45.3937555 | -0.085807304 | 0.26681415 | -0.3215995 | 0.7477561 | 0.88935207 |
| JPT2 | 31.3131632 | -1.204582031 | 0.34858577 | -3.4556259 | 0.00054902 | 0.00918177 |
| MAPK8IP3 | 71.2298014 | -0.201328034 | 0.18646129 | -1.079731 | 0.28026199 | 0.57321005 |
| NME3 | 64.0270161 | 0.206800786 | 0.42869889 | 0.48239169 | 0.62952772 | 0.82508693 |
| MRPS34 | 29.5534316 | -0.760782024 | 0.47535011 | -1.6004667 | 0.10949508 | 0.34481257 |
| SPSB3 | 39.0608691 | 0.31083213 | 0.28594336 | 1.08704092 | 0.27701873 | 0.56927037 |
| NUBP2 | 11.3620592 | -0.040572233 | 0.46741235 | -0.0868018 | 0.93082906 | 0.97314806 |
| HAGH | 53.4646205 | -0.362564741 | 0.35106653 | -1.0327522 | 0.30171986 | 0.59542937 |
| FAHD1 | 19.6729534 | -1.038326792 | 0.41805814 | -2.4836899 | 0.0130029 | 0.09200733 |
| NDUFB10 | 42.5938521 | -0.552884108 | 0.47784494 | -1.1570367 | 0.24725739 | 0.53796106 |
| RPS2 | 145.218784 | -0.910644097 | 0.42280912 | -2.1537948 | 0.03125627 | 0.16189143 |
| TSC2 | 55.908746 | 0.207724898 | 0.3428286 | 0.60591473 | 0.54457138 | 0.77395839 |
| PKD1 | 33.0924283 | -0.41679808 | 0.3190619 | -1.3063236 | 0.19144254 | 0.47171515 |
| SNHG19 | 17.4167252 | 0.402542399 | 0.38815447 | 1.03706755 | 0.29970438 | 0.59284841 |
| TRAF7 | 16.3521437 | -0.506894501 | 0.41788184 | -1.2130092 | 0.22512632 | 0.51390457 |
| PGP | 43.84617 | -0.128108179 | 0.39462468 | -0.324633 | 0.74545888 | 0.88808711 |
| E4F1 | 17.2713459 | 0.316040179 | 0.3474099 | 0.909704 | 0.36297863 | 0.64751617 |
| ECI1 | 96.1894664 | 0.360537253 | 0.20854542 | 1.72881886 | 0.08384152 | 0.29653069 |
| RNPS1 | 51.2945509 | -0.291062305 | 0.23696201 | -1.2283079 | 0.21933142 | 0.50742247 |
| ABCA3 | 22.7344352 | -0.201214751 | 0.3707506 | -0.5427227 | 0.58732077 | 0.79812407 |
| AMDHD2 | 11.2675951 | -1.319609105 | 0.52797886 | -2.4993597 | 0.01244179 | 0.08925861 |
| PDPK1 | 103.871652 | -0.15494054 | 0.25232732 | -0.6140458 | 0.53918503 | 0.77077724 |
| AC093525.4 | 12.5469028 | -0.660087161 | 0.45507591 | -1.450499 | 0.14691941 | 0.40616768 |
| KCTD5 | 12.0278391 | -0.40149721 | 0.50005454 | -0.8029068 | 0.42202859 | 0.69753545 |
| SRRM2 | 288.070936 | 0.090584301 | 0.24283345 | 0.37303057 | 0.70912569 | 0.86917464 |
| ELOB | 60.6524955 | -0.757205335 | 0.40167435 | -1.8851225 | 0.05941331 | 0.24143066 |
| PRSS22 | 22.1857896 | -1.265133712 | 0.46355127 | -2.7292207 | 0.00634842 | 0.0562835 |
| FLYWCH2 | 18.412197 | -0.470652249 | 0.42100274 | -1.1179316 | 0.26359623 | 0.55510391 |
| FLYWCH1 | 39.669732 | 0.540867191 | 0.28578268 | 1.89258212 | 0.05841346 | 0.23897216 |
| TNFRSF12A | 78.5108784 | -1.348088263 | 0.7647035 | -1.7628901 | 0.07791902 | 0.28404875 |
| HCFC1R1 | 31.3191169 | -0.376153933 | 0.50108232 | -0.7506829 | 0.45284351 | 0.72060455 |
| BICDL2 | 17.0478504 | -0.793788255 | 0.46934773 | -1.6912583 | 0.09078747 | 0.31034142 |
| IL32 | 16.1983826 | -0.983642243 | 0.47717161 | -2.0614015 | 0.03926476 | 0.18700317 |
| ZNF213-AS1 | 15.5727491 | -0.079099585 | 0.52195064 | -0.1515461 | 0.87954495 | 0.95279412 |
| ZNF213 | 33.0510456 | -0.148246307 | 0.4434039 | -0.3343369 | 0.73812531 | 0.88573617 |
| ZNF263 | 21.1045588 | -0.15784003 | 0.40906743 | -0.3858533 | 0.6996053 | 0.86367092 |
| TIGD7 | 17.7674975 | -0.271408615 | 0.37490339 | -0.7239428 | 0.46910084 | 0.73168094 |
| ZNF75A | 55.4393923 | -0.404760115 | 0.34410206 | -1.1762793 | 0.2394833 | 0.52992919 |
| AC025283.3 | 22.7108589 | 0.245333057 | 0.35737622 | 0.686484 | 0.49240795 | 0.74674012 |
| ZSCAN32 | 10.4019801 | -0.307109814 | 0.50470708 | -0.6084912 | 0.54286174 | 0.77258349 |
| NAA60 | 27.0399987 | -0.057149189 | 0.42493682 | -0.1344887 | 0.89301617 | 0.95782021 |
| CLUAP1 | 49.513225 | -0.189320466 | 0.31181277 | -0.6071607 | 0.54374426 | 0.77338856 |
| DNASE1 | 84.4661328 | 1.360804965 | 0.36034899 | 3.7763529 | 0.00015914 | 0.00355635 |
| TRAP1 | 52.5096649 | 0.78077822 | 0.25391234 | 3.07499126 | 0.00210509 | 0.02536445 |
| CREBBP | 210.858594 | 0.433383391 | 0.24185228 | 1.79193431 | 0.07314349 | 0.273214 |
| ADCY9 | 98.9617564 | 0.011048511 | 0.2544566 | 0.04342002 | 0.96536672 | 0.98661741 |
| GLIS2 | 43.5973358 | 0.115187727 | 0.3314546 | 0.34752188 | 0.72819928 | 0.88029911 |
| VASN | 58.5625559 | 0.25307164 | 0.35887128 | 0.70518778 | 0.48069338 | 0.73988033 |
| DNAJA3 | 38.2024769 | 0.437915511 | 0.28213099 | 1.55217088 | 0.12062134 | 0.36346292 |
| NMRAL1 | 29.4476224 | -0.598072858 | 0.27015156 | -2.213842 | 0.02683965 | 0.14623587 |
| HMOX2 | 49.6105997 | -0.358199888 | 0.28306673 | -1.2654256 | 0.2057188 | 0.48999801 |
| CDIP1 | 26.3598678 | -0.275704217 | 0.31342777 | -0.879642 | 0.3790533 | 0.66231463 |
| UBALD1 | 11.6262198 | 0.176725319 | 0.43594949 | 0.40538026 | 0.68519802 | 0.85645596 |
| MGRN1 | 64.3143916 | -0.08883036 | 0.20539541 | -0.4324846 | 0.66538921 | 0.84348349 |
| NUDT16L1 | 16.0232334 | -0.066660333 | 0.56187295 | -0.1186395 | 0.90556096 | 0.96157183 |
| ANKS3 | 42.4726371 | 0.236059907 | 0.27462294 | 0.85957826 | 0.39002156 | 0.67201909 |
| ZNF500 | 16.558892 | 0.272260792 | 0.37883819 | 0.71867304 | 0.4723424 | 0.73350488 |
| ROGDI | 11.7918372 | 0.422654779 | 0.6500058 | 0.65023232 | 0.51554217 | 0.75713618 |
| GLYR1 | 113.976903 | 0.007693731 | 0.20333175 | 0.03783832 | 0.96981659 | 0.98784364 |
| UBN1 | 42.4571403 | -0.138533495 | 0.23727518 | -0.5838516 | 0.55932016 | 0.78278988 |
| PPL | 44.0691507 | -0.501864651 | 0.27051226 | -1.8552381 | 0.06356223 | 0.25108056 |
| ALG1 | 16.5191255 | 0.501055426 | 0.35771811 | 1.40069906 | 0.16130408 | 0.42831015 |
| RBFOX1 | 20.7082543 | 0.44401914 | 0.41340215 | 1.07406104 | 0.28279532 | 0.5754607 |
| METTL22 | 16.7264719 | -0.384219638 | 0.41096991 | -0.9349094 | 0.34983499 | 0.63669603 |
| ABAT | 20.0704436 | 0.340531006 | 0.3517666 | 0.96805953 | 0.33301464 | 0.62190756 |
| PMM2 | 72.7500507 | -0.785131397 | 0.41535872 | -1.890249 | 0.05872467 | 0.23997681 |
| CARHSP1 | 41.8228933 | -1.551957904 | 0.34247028 | -4.5316572 | 5.85E-06 | 0.00024116 |
| USP7 | 68.7566308 | 0.443707619 | 0.20108033 | 2.20661875 | 0.02734071 | 0.14777597 |
| C16orf72 | 71.5417808 | 0.210929596 | 0.23488941 | 0.89799535 | 0.36918803 | 0.65369102 |
| ATF7IP2 | 81.0628319 | 0.921564061 | 0.29740145 | 3.09872079 | 0.00194358 | 0.02406583 |
| NUBP1 | 14.4393104 | -0.550984903 | 0.40429437 | -1.362831 | 0.17293578 | 0.44566139 |
| CIITA | 32.2474155 | -2.307186866 | 0.61590296 | -3.7460233 | 0.00017966 | 0.00385595 |
| DEXI | 23.0377555 | -0.470178349 | 0.34963407 | -1.3447727 | 0.17869865 | 0.45419584 |
| CLEC16A | 267.759406 | -0.334082557 | 0.2866652 | -1.1654102 | 0.24385298 | 0.53459591 |
| AC007014.1 | 15.8458045 | -1.618996465 | 0.7149933 | -2.2643519 | 0.02355248 | 0.13454711 |
| AC007220.1 | 17.5565953 | 0.295298629 | 0.42462966 | 0.69542629 | 0.48678819 | 0.74422309 |
| SOCS1 | 59.9883319 | -2.957067493 | 0.52877609 | -5.5922867 | 2.24E-08 | 2.51E-06 |
| AC099489.1 | 13.8798475 | -0.438523586 | 0.54874967 | -0.7991323 | 0.4242137 | 0.69882468 |
| LITAF | 128.502338 | -0.78016247 | 0.19398329 | -4.0218025 | 5.78E-05 | 0.00159514 |
| TXNDC11 | 99.3279935 | 0.225116692 | 0.1916336 | 1.17472455 | 0.24010494 | 0.53092102 |
| ZC3H7A | 97.3931731 | 0.223222161 | 0.21799626 | 1.02397245 | 0.30584829 | 0.60061934 |

| | | | | | | | |
|---|---|---|---|---|---|---|---|
| RSL1D1 | 22.9204351 | | -0.503482432 | 0.30938592 | -1.6273605 | 0.10366056 | 0.33425166 |
| AC007216.4 | 43.9042423 | | 0.226882259 | 0.39148408 | 0.57954402 | 0.56222215 | 0.78397572 |
| GSPT1 | 172.102806 | | 0.147195858 | 0.24432945 | 0.60244829 | 0.54687577 | 0.77463416 |
| SNX29 | 1001.74751 | | 0.662651998 | 0.20152108 | 3.28825151 | 0.00100812 | 0.01483661 |
| CPPED1 | 131.861733 | | -0.854781164 | 0.49254286 | -1.7354452 | 0.08266197 | 0.2945637 |
| SHISA9 | 61.6980483 | | 0.749004221 | 0.70963904 | 1.05547211 | 0.29120946 | 0.58287154 |
| U91319.1 | 69.0830613 | | 1.376288152 | 0.82208763 | 1.67413801 | 0.09410348 | 0.3160115 |
| ERCC4 | 22.2673864 | | -0.258966319 | 0.3533884 | -0.7328093 | 0.46367472 | 0.72812334 |
| MKL2 | 260.779683 | | 0.444341582 | 0.19775935 | 2.24688029 | 0.02464768 | 0.13752463 |
| PARN | 114.854614 | | 0.078415324 | 0.19887389 | 0.39429673 | 0.69336198 | 0.8606282 |
| BFAR | 33.5977105 | | -0.627227485 | 0.27695513 | -2.264726 | 0.0235295 | 0.13454711 |
| NOMO1 | 15.8344661 | | 0.230443763 | 0.47101608 | 0.48924819 | 0.624666 | 0.82271914 |
| PDXDC1 | 236.606376 | | -0.093072639 | 0.25644292 | -0.3629371 | 0.7166519 | 0.8732017 |
| NTAN1 | 12.777131 | | -0.393905064 | 0.39997201 | -0.9848316 | 0.3247068 | 0.61520673 |
| RRN3 | 32.6381472 | | 0.06050817 | 0.29918788 | 0.20224138 | 0.83972803 | 0.93543959 |
| C16orf45 | 76.2651407 | | 0.347003041 | 0.27190548 | 1.27618993 | 0.20188839 | 0.48515376 |
| MARF1 | 50.771609 | | -0.199881627 | 0.45162876 | -0.4425795 | 0.65806992 | 0.84031559 |
| NDE1 | 25.2440273 | | 0.117869869 | 0.36716161 | 0.32102994 | 0.7481877 | 0.88943338 |
| FOPNL | 20.1843917 | | 0.091199956 | 0.34705378 | 0.26278335 | 0.79271757 | 0.91124908 |
| ABCC1 | 75.9518043 | | -0.479910896 | 0.29732912 | -1.614073 | 0.1065116 | 0.33992642 |
| XYLT1 | 19.1066973 | | 0.695446374 | 0.49276814 | 1.41130547 | 0.15815456 | 0.42452014 |
| RPS15A | 24.2922911 | | 0.069127953 | 0.58425539 | 0.11831804 | 0.90581566 | 0.96157183 |
| SMG1 | 216.418747 | | 0.113463303 | 0.24349551 | 0.46597699 | 0.64123197 | 0.83230084 |
| COQ7 | 20.0615707 | | 0.06126315 | 0.34440724 | 0.17787997 | 0.85881724 | 0.94403918 |
| GDE1 | 25.8921578 | | -0.700620519 | 0.33800009 | -2.0728412 | 0.03818706 | 0.18297817 |
| CCP110 | 28.8749979 | | -0.247956457 | 0.3805083 | -0.6516453 | 0.51463 | 0.75684546 |
| VPS35L | 114.627729 | | 0.121379429 | 0.21055519 | 0.57647323 | 0.56429536 | 0.7850602 |
| KNOP1 | 24.4292557 | | -0.282453431 | 0.49357098 | -0.5722651 | 0.56714241 | 0.78656476 |
| IQCK | 173.060028 | | -0.023198751 | 0.20400894 | -0.1137144 | 0.90946421 | 0.96308593 |
| GPRC5B | 118.98469 | | -0.099638292 | 0.31066027 | -0.3207307 | 0.74841446 | 0.88949767 |
| UMOD | 63.0895026 | | 1.138377543 | 0.54175466 | 2.10127873 | 0.03561651 | 0.17524194 |
| ACSM2B | 12.5247033 | | 1.042164573 | 0.87969268 | 1.18469165 | 0.23613939 | 0.52644622 |
| ACSM3 | 51.2499397 | | -0.155599969 | 0.50460762 | -0.3083583 | 0.75780968 | 0.89457492 |
| THUMPD1 | 50.5497184 | | 0.115114445 | 0.35006058 | 0.3288415 | 0.7422755 | 0.88720442 |
| ERI2 | 23.0597765 | | -0.021074038 | 0.48792456 | -0.0431912 | 0.96554914 | 0.98661741 |
| LYRM1 | 24.6353509 | | -0.482143175 | 0.35830443 | -1.3456244 | 0.17842368 | 0.45374495 |
| TMEM159 | 16.9420689 | | -1.121093211 | 0.46504761 | -2.4107063 | 0.01592166 | 0.10394485 |
| METTL9 | 144.916486 | | 0.172270566 | 0.24422389 | 0.70537966 | 0.480574 | 0.73988033 |
| NPIPB4 | 30.8907156 | | 0.410119356 | 0.5015515 | 0.81770138 | 0.41352772 | 0.69042044 |
| UQCRC2 | 134.481672 | | 0.523600205 | 0.21180483 | 2.47208815 | 0.01343264 | 0.09373482 |
| MOSMO | 83.4742881 | | -0.047671743 | 0.24153672 | -0.1973685 | 0.84353917 | 0.93643755 |
| EEF2K | 25.4637656 | | -1.386857623 | 0.38492106 | -3.6029663 | 0.00031461 | 0.00609224 |
| POLR3E | 37.8090868 | | 0.373596192 | 0.35356852 | 1.05664439 | 0.29067391 | 0.58225212 |
| CDR2 | 201.311703 | | 0.016796109 | 0.29507042 | 0.05692237 | 0.95460703 | 0.98226508 |
| NPIPB5 | 82.2736766 | | 0.278318043 | 0.30233341 | 0.92056661 | 0.35727674 | 0.64174726 |
| USP31 | 40.7554629 | | -1.309395776 | 0.3624507 | -3.6126176 | 0.00030312 | 0.00595946 |
| SCNN1G | 365.584483 | | 2.114667555 | 0.37827248 | 5.5903289 | 2.27E-08 | 2.51E-06 |
| AC099482.1 | 35.1803331 | | 1.906491856 | 0.68260731 | 2.79295553 | 0.00522289 | 0.04913807 |
| SCNN1B | 185.855852 | | 2.128070681 | 0.45581833 | 4.66868164 | 3.03E-06 | 0.00014054 |
| COG7 | 94.3635826 | | 0.091310349 | 0.18832712 | 0.48484972 | 0.62778295 | 0.82428562 |
| GGA2 | 110.756809 | | 0.059666858 | 0.23657711 | 0.25220892 | 0.80087959 | 0.9148089 |
| EARS2 | 16.6256243 | | 0.001860144 | 0.3925239 | 0.00473893 | 0.99621889 | 0.99868531 |
| UBFD1 | 20.1807728 | | -0.696417027 | 0.5400121 | -1.2896323 | 0.19717837 | 0.47915741 |
| NDUFAB1 | 38.9980814 | | 0.218833279 | 0.30094166 | 0.72716181 | 0.46712684 | 0.73023858 |
| PALB2 | 20.5779914 | | -0.064142293 | 0.44635825 | -0.1437014 | 0.88573629 | 0.95619452 |
| DCTN5 | 49.3047097 | | -0.14948874 | 0.25807846 | -0.5792376 | 0.56242888 | 0.78400069 |
| RBBP6 | 88.3042953 | | -0.175891172 | 0.23485523 | -0.7489344 | 0.45389672 | 0.72127781 |
| TNRC6A | 468.122968 | | 0.71392979 | 0.25186142 | 2.83461351 | 0.00458812 | 0.0446334 |
| AC008731.1 | 12.7866853 | | 0.11035454 | 0.56290568 | 0.19604446 | 0.84457537 | 0.93689787 |
| ARHGAP17 | 93.7372127 | | 0.243290817 | 0.26226925 | 0.92763758 | 0.3535956 | 0.63949806 |
| LCMT1 | 57.5166739 | | -0.085767669 | 0.29500474 | -0.2907332 | 0.77125539 | 0.90076568 |
| AC008741.2 | 11.2936422 | | 0.301119293 | 0.42096895 | 0.71530048 | 0.47442339 | 0.73514456 |
| NSMCE1 | 28.2691812 | | 0.124221934 | 0.3158563 | 0.39328624 | 0.69410808 | 0.86101552 |
| IL4R | 28.1883401 | | -3.418059726 | 0.81524065 | -4.1927003 | 2.76E-05 | 0.00085279 |
| GTF3C1 | 57.2305489 | | 0.413409329 | 0.28334448 | 1.45903435 | 0.14455565 | 0.40242285 |
| KIAA0556 | 87.7443014 | | -0.24856887 | 0.32911783 | -0.755258 | 0.4500942 | 0.71889465 |
| XPO6 | 99.6463476 | | 0.013120088 | 0.29622176 | 0.04429144 | 0.9646721 | 0.98660829 |
| NUPR1 | 44.5111112 | | -2.600107437 | 1.14390264 | -2.2730146 | 0.0230253 | 0.1323843 |
| SGF29 | 16.1727889 | | 0.408253815 | 0.3935185 | 1.03744505 | 0.29952849 | 0.59261428 |
| SULT1A1 | 29.5480891 | | 1.096205989 | 0.42616387 | 2.57226401 | 0.01010358 | 0.07773495 |
| ATXN2L | 48.5441723 | | -0.364516176 | 0.32713481 | -1.114269 | 0.26516381 | 0.55640131 |
| TUFM | 74.7348237 | | -0.327449947 | 0.43396854 | -0.7545477 | 0.45052044 | 0.71901805 |
| SH2B1 | 52.1487776 | | 0.568593699 | 0.26798174 | 2.12176288 | 0.03385766 | 0.17023014 |
| RABEP2 | 57.7529082 | | 0.062180975 | 0.36384402 | 0.17090009 | 0.86430233 | 0.94663434 |
| NFATC2IP | 91.8059429 | | -0.129369344 | 0.19319957 | -0.6696151 | 0.50310319 | 0.75108109 |
| SPNS1 | 18.2446414 | | -0.635192483 | 0.40419575 | -1.5714972 | 0.11606721 | 0.35543425 |
| QPRT | 29.8172294 | | -0.506010623 | 0.53852871 | -0.9396168 | 0.34741416 | 0.63465611 |
| KIF22 | 13.6580728 | | -0.422073921 | 0.35769812 | -1.1799724 | 0.23801117 | 0.52824866 |
| MAZ | 20.3984503 | | -1.261120121 | 0.44772107 | -2.816754 | 0.00485117 | 0.04642453 |
| AC009133.1 | 16.61536 | | -2.329903669 | 0.54489274 | -4.2758941 | 1.90E-05 | 0.00062064 |
| MVP | 84.7084806 | | -2.006098928 | 0.2946298 | -6.8088799 | 9.84E-12 | 3.27E-09 |
| CDIPT | 30.2790318 | | 0.236933085 | 0.31031138 | 0.76353334 | 0.44514538 | 0.71498094 |
| KCTD13 | 32.0579535 | | -0.142934811 | 0.3260579 | -0.4383725 | 0.66111629 | 0.84192458 |
| TMEM219 | 53.1159391 | | -0.256672989 | 0.44004536 | -0.5832876 | 0.55969973 | 0.78316119 |
| TAOK2 | 17.2188834 | | 0.347021151 | 0.52315125 | 0.66332854 | 0.50712016 | 0.75376105 |
| INO80E | 21.4638798 | | 0.033609246 | 0.3657879 | 0.09188179 | 0.92679196 | 0.97111125 |
| PPP4C | 35.5246801 | | -0.4965422 | 0.38803923 | -1.2796186 | 0.20067932 | 0.48430583 |
| YPEL3 | 36.6879199 | | -0.678626934 | 0.33224973 | -2.0425207 | 0.04109991 | 0.19311676 |
| MAPK3 | 12.1909119 | | 0.068586893 | 0.47348126 | 0.14485662 | 0.88482408 | 0.95619452 |
| NPIPB13 | 20.3946876 | | -0.075107661 | 0.54131077 | -0.1387515 | 0.88964655 | 0.95702609 |
| CD2BP2 | 20.7243684 | | -0.820823233 | 0.43975984 | -1.8665261 | 0.06196779 | 0.24676931 |
| TBC1D10B | 14.0840753 | | -0.72803686 | 0.51479902 | -1.4142157 | 0.15729858 | 0.42255279 |
| DCTPP1 | 13.5299916 | | -1.41007474 | 0.57548322 | -2.4502448 | 0.01427591 | 0.09714033 |
| SEPHS2 | 47.9194947 | | -1.281843973 | 0.48342486 | -2.6515888 | 0.0080114 | 0.0658687 |
| ZNF768 | 19.247647 | | -0.07952945 | 0.40044993 | -0.1986002 | 0.84257547 | 0.93597288 |
| ZNF785 | 23.5734282 | | -0.049773607 | 0.30257894 | -0.1644979 | 0.86933918 | 0.94827256 |
| PRR14 | 13.2748817 | | 0.044175546 | 0.43225921 | 0.10219689 | 0.9186004 | 0.96713554 |
| FBRS | 26.7978944 | | -0.113940432 | 0.29878651 | -0.381344 | 0.70294803 | 0.86561925 |
| PHKG2 | 16.3094209 | | 0.50263772 | 0.39270983 | 1.27992141 | 0.20057278 | 0.48426847 |
| RNF40 | 34.2280986 | | 0.223391218 | 0.29519856 | 0.75674901 | 0.44920025 | 0.71802342 |
| ZNF629 | 13.4698961 | | -0.736251934 | 0.47603233 | -1.5466427 | 0.12194942 | 0.36611339 |
| BCL7C | 67.4843715 | | 0.354193427 | 0.24772665 | 1.42977521 | 0.15278155 | 0.41463527 |
| MIR762HG | 14.1104586 | | -0.254659227 | 0.39585059 | -0.6433216 | 0.52001546 | 0.76059646 |
| FBXL19 | 20.3326587 | | -0.407966952 | 0.4874607 | -0.8369228 | 0.40263599 | 0.68128594 |
| SETD1A | 14.2123392 | | 0.26733518 | 0.40230291 | 0.66451217 | 0.50636256 | 0.75306985 |
| STX4 | 17.9659911 | | -0.279265998 | 0.34357344 | -0.8128277 | 0.41631687 | 0.69298698 |
| BCKDK | 21.746183 | | -0.786614598 | 0.50777455 | -1.5491414 | 0.12134772 | 0.36521304 |
| KAT8 | 31.958726 | | 0.154561049 | 0.30413306 | 0.50820206 | 0.61131165 | 0.81385718 |
| PRSS8 | 42.9380632 | | -0.742414952 | 0.50814668 | -1.4610249 | 0.14400861 | 0.4017267 |
| FUS | 145.247901 | | 0.373917621 | 0.20332792 | 1.8389881 | 0.06591694 | 0.25655016 |
| AC026471.4 | 19.4591928 | | 0.356335212 | 0.44282085 | 0.80469385 | 0.42099637 | 0.69661173 |
| C16orf58 | 26.1208341 | | -0.15166461 | 0.38844496 | -0.3904404 | 0.69621091 | 0.86247999 |
| ZNF720 | 53.8295292 | | -0.030730446 | 0.29119401 | -0.1055325 | 0.91595324 | 0.96518083 |

| | | | | | | |
|---|---|---|---|---|---|---|
| ZNF267 | 12.5559062 | -0.745914862 | 0.40279388 | -1.8518525 | 0.06404701 | 0.2521767 |
| SHCBP1 | 20.8178678 | 2.763193591 | 0.39170751 | 7.05422682 | 1.74E-12 | 6.88E-10 |
| VPS35 | 62.9476501 | -0.64638151 | 0.27438089 | -2.3557818 | 0.01848377 | 0.11464166 |
| C16orf87 | 16.9354845 | 0.108593151 | 0.40466289 | 0.2683546 | 0.78842638 | 0.9097635 |
| GPT2 | 15.8867623 | -0.579201796 | 0.48984694 | -1.1824138 | 0.23704154 | 0.52743022 |
| DNAJA2 | 27.9326251 | -0.239734381 | 0.29965233 | -0.8000418 | 0.4236866 | 0.6983711 |
| ITFG1 | 388.474303 | 0.087922819 | 0.18484688 | 0.47565216 | 0.63432221 | 0.82785872 |
| PHKB | 262.958439 | 0.174843242 | 0.14405007 | 1.21376713 | 0.22483666 | 0.51358476 |
| LONP2 | 175.948342 | 0.180771254 | 0.17252541 | 1.04779495 | 0.29473309 | 0.58661907 |
| SIAH1 | 35.9170325 | -0.498820855 | 0.33168445 | -1.5039018 | 0.13260664 | 0.38212969 |
| N4BP1 | 88.5989545 | -0.175404201 | 0.32372913 | -0.541824 | 0.58793978 | 0.79864921 |
| CNEP1R1 | 13.7465754 | 0.328760073 | 0.39750388 | 0.82706128 | 0.40820233 | 0.68657965 |
| HEATR3 | 19.0255335 | 0.324111843 | 0.32103358 | 1.00958861 | 0.31269243 | 0.6055915 |
| TENT4B | 86.0222116 | 0.902925814 | 0.26407826 | 3.41915995 | 0.00062815 | 0.01032086 |
| BRD7 | 57.1227378 | 0.146747071 | 0.27989924 | 0.52428535 | 0.60008008 | 0.80738812 |
| CYLD | 42.8789785 | -0.107784018 | 0.29450645 | -0.3659819 | 0.71437861 | 0.8724728 |
| SALL1 | 26.7367625 | 0.083459894 | 0.29530094 | 0.28262657 | 0.77746311 | 0.90366576 |
| AC087564.1 | 90.4409645 | -0.067462863 | 0.35969869 | -0.1875538 | 0.85122643 | 0.94012264 |
| TOX3 | 773.937192 | 1.584555011 | 0.42164413 | 3.75803887 | 0.00017125 | 0.00372985 |
| AC007906.2 | 11.4524923 | -0.67855382 | 0.71077601 | -0.9546662 | 0.33974654 | 0.62837861 |
| CHD9 | 553.310804 | 0.202566414 | 0.21956923 | 0.92256286 | 0.35623506 | 0.64081257 |
| RBL2 | 107.792419 | -0.162145279 | 0.23885183 | -0.678853 | 0.49723101 | 0.74900919 |
| AKTIP | 22.601537 | 0.769614507 | 0.33372815 | 2.30611208 | 0.02110437 | 0.12473736 |
| RPGRIP1L | 22.7952922 | -0.326376632 | 0.44077342 | -0.7404635 | 0.4590188 | 0.72461104 |
| FTO | 301.729189 | 0.183961895 | 0.14522708 | 1.266719 | 0.20525578 | 0.48970474 |
| LPCAT2 | 85.6249849 | 0.364318438 | 0.33645234 | 1.08282332 | 0.27888685 | 0.57144989 |
| AMFR | 124.012513 | 0.41207696 | 0.35734927 | 1.15314903 | 0.24884922 | 0.53982832 |
| NUDT21 | 71.3813809 | 0.354623383 | 0.24676337 | 1.43709898 | 0.15068987 | 0.41166131 |
| OGFOD1 | 25.5718764 | -0.021502151 | 0.30868214 | -0.0696579 | 0.94446595 | 0.97856665 |
| BBS2 | 56.0234735 | 0.385776632 | 0.23396733 | 1.64884831 | 0.09917872 | 0.32535676 |
| MT2A | 234.385598 | -1.631009709 | 0.70930171 | -2.2994583 | 0.02147893 | 0.126227 |
| MT1E | 92.6283809 | -1.008833347 | 0.77610973 | -1.2998592 | 0.19364924 | 0.4740909 |
| MT1F | 19.9110896 | -0.744655846 | 0.69976659 | -1.0641489 | 0.28726125 | 0.5796994 |
| MT1G | 150.930429 | -0.099808023 | 0.851762 | -0.1171783 | 0.90671876 | 0.96157183 |
| MT1H | 45.2062286 | -1.031969219 | 1.28635452 | -0.8022432 | 0.42241227 | 0.69769678 |
| MT1X | 48.0171658 | -0.433103465 | 0.72394032 | -0.5982585 | 0.54966745 | 0.7759356 |
| NUP93 | 40.0781042 | 0.141083646 | 0.2858268 | 0.49359838 | 0.62158984 | 0.82030381 |
| SLC12A3 | 256.297294 | 4.801447847 | 0.66488593 | 7.22146108 | 5.14E-13 | 2.30E-10 |
| HERPUD1 | 60.589277 | -0.753485493 | 0.38219756 | -1.9714555 | 0.04867179 | 0.21414869 |
| NLRC5 | 24.3732611 | -2.962284417 | 0.83388491 | -3.55239 | 0.00038175 | 0.00707333 |
| FAM192A | 56.3360175 | -0.265945613 | 0.24719112 | -1.0758704 | 0.28198522 | 0.5750055 |
| RSPRY1 | 78.1424542 | 0.2126847 | 0.26088193 | 0.8152527 | 0.41492768 | 0.69168041 |
| ARL2BP | 40.2746134 | -0.764049925 | 0.31730387 | -2.4079439 | 0.01604264 | 0.10440387 |
| PLLP | 61.7682247 | 0.819123166 | 0.62921611 | 1.30181531 | 0.19297953 | 0.47358843 |
| CX3CL1 | 22.645322 | -0.747927233 | 0.47586848 | -1.57171 | 0.11601783 | 0.35543425 |
| CIAPIN1 | 40.9141814 | -0.025935966 | 0.28584858 | -0.0907332 | 0.92770456 | 0.97116273 |
| COQ9 | 77.0610835 | 0.63193569 | 0.21911749 | 2.88400383 | 0.00392654 | 0.03961921 |
| POLR2C | 20.0774002 | -0.318305281 | 0.44674677 | -0.712496 | 0.47615769 | 0.73631255 |
| DOK4 | 23.615816 | -0.517655119 | 0.52325558 | -0.9892969 | 0.3225179 | 0.6132414 |
| ADGRG1 | 231.219729 | 0.19081971 | 0.30174492 | 0.63238748 | 0.52713372 | 0.76221787 |
| KIFC3 | 189.33426 | 0.952593004 | 0.21572047 | 4.41586755 | 1.01E-05 | 0.00036884 |
| USB1 | 14.7197695 | 0.090425673 | 0.38466052 | 0.23507916 | 0.81414729 | 0.921079 |
| MMP15 | 12.816733 | -1.013669159 | 0.65908779 | -1.5379881 | 0.12405153 | 0.36903808 |
| CSNK2A2 | 53.1040264 | -0.232845201 | 0.25040834 | -0.929862 | 0.35244254 | 0.63843043 |
| SETD6 | 11.4950421 | -0.185092969 | 0.44399909 | -0.4168769 | 0.67676844 | 0.85191801 |
| CNOT1 | 169.234496 | 0.398042512 | 0.20519267 | 1.93984765 | 0.05239821 | 0.22398603 |
| SLC38A7 | 22.4142917 | -0.320070087 | 0.50442305 | -0.6345271 | 0.5257369 | 0.76221787 |
| GOT2 | 76.4139343 | 0.502690812 | 0.30296148 | 1.65925651 | 0.09706411 | 0.32146212 |
| AC092378.1 | 22.0426968 | 0.466977955 | 0.65782569 | 0.70988099 | 0.47777794 | 0.73760952 |
| AC138305.1 | 695.657996 | 1.548792215 | 0.30480461 | 5.08126247 | 3.75E-07 | 2.51E-05 |
| AC010542.3 | 21.4069254 | -0.165068404 | 0.47243795 | -0.349397 | 0.72679128 | 0.87931929 |
| TK2 | 20.7614391 | -0.283225323 | 0.32572905 | -0.869512 | 0.38456713 | 0.66701235 |
| CMTM4 | 236.635502 | 0.950959922 | 0.23762139 | 4.00199631 | 6.28E-05 | 0.00169835 |
| DYNC1LI2 | 116.856977 | 0.089024906 | 0.27608875 | 0.32245032 | 0.74711157 | 0.88907744 |
| NAE1 | 42.0139776 | 0.226855489 | 0.31601661 | 0.71785939 | 0.47284399 | 0.73375471 |
| PDP2 | 25.9728034 | 0.05220598 | 0.47610366 | 0.10965255 | 0.91268493 | 0.96446045 |
| CDH16 | 545.065085 | 0.263041421 | 0.39355386 | 0.66837464 | 0.50389448 | 0.75108109 |
| RRAD | 208.084361 | -1.440477543 | 0.7236141 | -1.9906709 | 0.04651708 | 0.20727463 |
| FAM96B | 42.8293466 | -1.07251851 | 0.3541457 | -3.0284668 | 0.00245798 | 0.02832451 |
| CES2 | 33.5997921 | -0.581493647 | 0.35085135 | -1.657379 | 0.09744288 | 0.32216192 |
| CBFB | 164.703201 | 0.062409043 | 0.26745661 | 0.23334269 | 0.81549531 | 0.92165337 |
| C16orf70 | 23.5978189 | -0.59084459 | 0.42516756 | -1.3896747 | 0.16462769 | 0.43373261 |
| TRADD | 15.243634 | -0.849852174 | 0.45619113 | -1.8629301 | 0.0624721 | 0.24839352 |
| NOL3 | 26.4198358 | -1.167483021 | 0.48623632 | -2.4010609 | 0.01634762 | 0.10592021 |
| KIAA0895L | 20.2367541 | -0.285209767 | 0.36685099 | -0.777454 | 0.43689097 | 0.70829758 |
| E2F4 | 19.3569212 | -0.11402178 | 0.34696369 | -0.3286274 | 0.74243733 | 0.88720442 |
| ELMO3 | 14.1135004 | 0.01841645 | 0.37641303 | 0.04892618 | 0.96097813 | 0.98497629 |
| LRRC29 | 14.5163137 | 0.452453395 | 0.38574412 | 1.17293659 | 0.24082124 | 0.53151245 |
| TMEM208 | 26.7833339 | -0.737230245 | 0.40686716 | -1.8119679 | 0.06999115 | 0.26675872 |
| ZDHHC1 | 31.140216 | 0.425439703 | 0.28785728 | 1.4779536 | 0.1394202 | 0.39404853 |
| HSD11B2 | 724.825019 | 2.690707083 | 0.6271733 | 4.29021309 | 1.79E-05 | 0.00059129 |
| ATP6V0D1 | 68.4918433 | 0.289845481 | 0.29606754 | 0.97898433 | 0.32758772 | 0.61719253 |
| AC009061.2 | 46.1947681 | 0.52735088 | 0.37604684 | 1.40235424 | 0.16080949 | 0.42803048 |
| AC027682.6 | 9.87796682 | 1.499231116 | 0.61063192 | 2.45521248 | 0.01408013 | 0.09623211 |
| RIPOR1 | 45.9177952 | 0.122476518 | 0.32540462 | 0.37638224 | 0.70663275 | 0.86798542 |
| AC027682.4 | 12.0768596 | 0.114759032 | 0.52438434 | 0.21884527 | 0.82677058 | 0.9280225 |
| CTCF | 80.0228728 | 0.071639862 | 0.30990288 | 0.23116875 | 0.8171837 | 0.92248811 |
| ENKD1 | 14.2452424 | -0.172544478 | 0.44312391 | -0.389382 | 0.69699358 | 0.86249983 |
| GFOD2 | 25.4999513 | -0.58457715 | 0.34371301 | -1.7007711 | 0.08898598 | 0.30711342 |
| RANBP10 | 89.8906601 | 0.494208767 | 0.28228914 | 1.75071834 | 0.07999444 | 0.28875358 |
| NUTF2 | 16.5194665 | -0.546805804 | 0.38071445 | -1.4362623 | 0.15092773 | 0.41210109 |
| PSKH1 | 25.4243817 | 0.242276 | 0.32006224 | 0.75696526 | 0.44907067 | 0.71802342 |
| DUS2 | 18.5923753 | 0.483644465 | 0.39958971 | 1.21035264 | 0.22614361 | 0.51519935 |
| NFATC3 | 136.619082 | 0.261429507 | 0.16991756 | 1.53856674 | 0.1239101 | 0.36885651 |
| ESRP2 | 11.0926451 | -0.149599345 | 0.51456105 | -0.290732 | 0.77125632 | 0.90076568 |
| PLA2G15 | 23.1436655 | 0.409903223 | 0.35542908 | 1.15326305 | 0.24880243 | 0.53982832 |
| SLC7A6 | 57.2400583 | 0.894462518 | 0.32626098 | 2.74155532 | 0.00611491 | 0.05497013 |
| SLC7A6OS | 29.1102458 | -0.697019969 | 0.43975806 | -1.5850078 | 0.11296456 | 0.35086028 |
| PRMT7 | 37.2631874 | 0.86990154 | 0.37034579 | 2.34889002 | 0.01882947 | 0.11636543 |
| ZFP90 | 43.3820299 | -0.32444304 | 0.26405475 | -1.2286961 | 0.21918577 | 0.50733831 |
| CDH3 | 19.4326303 | -1.728659243 | 0.62179836 | -2.7800962 | 0.00543428 | 0.05051145 |
| CDH1 | 257.934294 | 0.202910195 | 0.26917168 | 0.75383189 | 0.45095017 | 0.71936957 |
| TANGO6 | 77.0562124 | 0.522889524 | 0.23195351 | 2.2542859 | 0.02417819 | 0.13588854 |
| CHTF8 | 33.4660385 | -0.094585995 | 0.39550886 | -0.2391501 | 0.81098917 | 0.9198294 |
| UTP4 | 23.2120453 | -0.090730411 | 0.33429596 | -0.2714074 | 0.78607768 | 0.90831284 |
| SNTB2 | 40.082923 | 0.104797288 | 0.38772019 | 0.27029103 | 0.78693637 | 0.90871373 |
| VPS4A | 24.9483287 | -0.612576791 | 0.35237341 | -1.7384308 | 0.08213493 | 0.29349776 |
| COG8 | 13.1464167 | 0.40220324 | 0.42505578 | 0.94623637 | 0.34402804 | 0.63198588 |
| TERF2 | 36.1263437 | -0.059500216 | 0.25167075 | -0.2364209 | 0.81310611 | 0.92060876 |
| CYB5B | 120.151086 | 0.393446591 | 0.21361665 | 1.84183487 | 0.06549931 | 0.25569302 |
| NFAT5 | 1080.79479 | 0.416501545 | 0.30512385 | 1.36502456 | 0.17224533 | 0.44472968 |
| NQO1 | 38.0859157 | -1.771907838 | 0.57762594 | -3.0675697 | 0.00215807 | 0.02581658 |
| WWP2 | 181.92459 | 0.080801489 | 0.26470996 | 0.30524537 | 0.76017928 | 0.89627175 |

| | | | | | | |
|---|---|---|---|---|---|---|
| PDPR | 58.8934741 | 0.059249864 | 0.35853436 | 0.16525575 | 0.86874269 | 0.94827158 |
| AC009060.1 | 15.9028382 | 0.314103612 | 0.5286333 | 0.59418053 | 0.55239138 | 0.77752916 |
| EXOSC6 | 24.0459309 | -0.196325858 | 0.35859181 | -0.5474912 | 0.58404132 | 0.7962935 |
| AARS | 45.2905681 | -0.98232366 | 0.47733787 | -2.0579211 | 0.03959771 | 0.18807544 |
| ST3GAL2 | 39.0922899 | 0.022674478 | 0.28438364 | 0.07973201 | 0.9364504 | 0.97594028 |
| FUK | 21.6810312 | 0.63281548 | 0.37325912 | 1.69537849 | 0.09000364 | 0.30907252 |
| COG4 | 41.4927081 | 0.309681328 | 0.25339711 | 1.22211864 | 0.22166277 | 0.50986455 |
| SF3B3 | 72.915129 | 0.312815792 | 0.19247142 | 1.62525838 | 0.10410752 | 0.33490577 |
| MTSS1L | 27.0204171 | -0.876869751 | 0.60891381 | -1.4400556 | 0.14985167 | 0.41046846 |
| VAC14 | 43.7528137 | -0.527583095 | 0.40391737 | -1.3061659 | 0.19149614 | 0.4717344 |
| HYDIN | 127.829883 | -0.803155479 | 0.39413952 | -2.0377441 | 0.04157553 | 0.19472668 |
| CMTR2 | 39.0334144 | -1.169180029 | 0.45098382 | -2.5925099 | 0.00952784 | 0.07447332 |
| ZNF19 | 29.5147697 | -0.674865095 | 0.34611924 | -1.9498052 | 0.05119934 | 0.22032398 |
| MARVELD3 | 16.7779951 | -1.217613607 | 0.36270341 | -3.3570504 | 0.00078779 | 0.01235832 |
| PHLPP2 | 40.1623187 | -0.602100091 | 0.31080756 | -1.9372118 | 0.05271945 | 0.22490549 |
| AP1G1 | 184.804611 | 0.478544051 | 0.2941342 | 1.62695823 | 0.10374597 | 0.3344152 |
| IST1 | 106.498912 | 0.034289729 | 0.19182814 | 0.17875234 | 0.85813217 | 0.94359207 |
| ZNF821 | 35.0341744 | 0.40151072 | 0.30842702 | 1.30180136 | 0.1929843 | 0.47358843 |
| TXNL4B | 19.4254271 | -0.261474847 | 0.37004499 | -0.7066029 | 0.47981331 | 0.73953273 |
| DHX38 | 17.574351 | -0.264588047 | 0.37511676 | -0.7053485 | 0.48059338 | 0.73988033 |
| LINC01572 | 55.7858036 | 0.059731657 | 0.44281994 | 0.13488927 | 0.89269943 | 0.95782021 |
| AC004943.2 | 33.7441431 | 0.412750739 | 0.40653201 | 1.01529702 | 0.30996427 | 0.60321222 |
| ZFHX3 | 276.503039 | -0.50394125 | 0.24870081 | -2.0262951 | 0.04273455 | 0.19741236 |
| AC138627.1 | 24.2608901 | 0.565114922 | 0.72721674 | 0.77709284 | 0.437104 | 0.70829758 |
| PSMD7 | 42.710095 | -0.378166811 | 0.26632462 | -1.4199469 | 0.15562314 | 0.41947399 |
| NPIPB15 | 14.8013042 | -1.811472953 | 0.93088424 | -1.9459702 | 0.05165832 | 0.22183579 |
| GLG1 | 296.862635 | 0.196646482 | 0.16830572 | 1.16838861 | 0.24265005 | 0.53317493 |
| RFWD3 | 43.7216022 | 0.217536724 | 0.4065001 | 0.53514556 | 0.59254921 | 0.80152862 |
| FA2H | 39.3838132 | 1.460735531 | 0.34902 | 4.18524877 | 2.85E-05 | 0.00087861 |
| WDR59 | 130.434138 | 0.202762456 | 0.17829177 | 1.1372508 | 0.25543345 | 0.54662971 |
| ZNRF1 | 118.703566 | -0.556761156 | 0.47784446 | -1.1651514 | 0.2439577 | 0.5346132 |
| ZFP1 | 13.283947 | 0.114816074 | 0.38905512 | 0.29511518 | 0.76790589 | 0.89979146 |
| BCAR1 | 70.4666236 | -0.919189848 | 0.33731435 | -2.7250244 | 0.00642967 | 0.05672224 |
| CFDP1 | 184.330585 | -0.126666778 | 0.32220703 | -0.3931223 | 0.69422914 | 0.8610581 |
| GABARAPL2 | 55.9013057 | -0.108154199 | 0.39776965 | -0.2719016 | 0.78569769 | 0.90814065 |
| ADAT1 | 19.5944323 | -0.194988156 | 0.38800906 | -0.5025351 | 0.61529119 | 0.81632065 |
| KARS | 29.2099788 | -0.476858667 | 0.33999373 | -1.4025514 | 0.16075066 | 0.42803048 |
| TERF2IP | 135.01974 | 0.024021168 | 0.36462196 | 0.06587965 | 0.94747364 | 0.979567 |
| NUDT7 | 12.067569 | 0.709192262 | 0.50127042 | 1.41478976 | 0.15713015 | 0.42221043 |
| WWOX | 1212.5945 | 0.613338074 | 0.23359457 | 2.62565207 | 0.00864832 | 0.0695511 |
| MAF | 11.9374332 | 0.020094275 | 0.47417657 | 0.0423772 | 0.96619801 | 0.98679209 |
| CDYL2 | 32.3840763 | -0.228110731 | 0.48130096 | -0.4739461 | 0.63553832 | 0.82887907 |
| CMC2 | 80.8724562 | 0.517316104 | 0.29197908 | 1.77175744 | 0.07643483 | 0.28042437 |
| ATMIN | 24.2704887 | -0.358197704 | 0.3352798 | -1.0683546 | 0.28536061 | 0.57790151 |
| GAN | 48.5317493 | 0.151636352 | 0.28936386 | 0.52403348 | 0.60025525 | 0.80738812 |
| CMIP | 251.368236 | -0.213271426 | 0.2474206 | -0.8619793 | 0.38869893 | 0.67063748 |
| PLCG2 | 233.096341 | 0.164282508 | 0.23680034 | 0.69375959 | 0.48783299 | 0.74443126 |
| SDR42E1 | 11.8987617 | -0.163947215 | 0.48362308 | -0.3389979 | 0.7346113 | 0.88379839 |
| AC092142.1 | 16.8107323 | 0.317703065 | 0.56086143 | 0.56645554 | 0.57108415 | 0.78905469 |
| MPHOSPH6 | 37.440134 | -0.125816097 | 0.31655321 | -0.3974564 | 0.69103094 | 0.85967888 |
| HSBP1 | 48.9241729 | -0.213522717 | 0.30429951 | -0.701686 | 0.48287498 | 0.74093072 |
| MBTPS1 | 83.2743643 | -0.060940166 | 0.21346019 | -0.2854873 | 0.77527086 | 0.90226394 |
| HSDL1 | 16.5181329 | -0.153511381 | 0.43806484 | -0.3504307 | 0.72601547 | 0.87879349 |
| TAF1C | 18.0732519 | 0.279026184 | 0.36975819 | 0.754618 | 0.45047822 | 0.71901805 |
| TLDC1 | 29.2637963 | 0.151141964 | 0.37141549 | 0.406935 | 0.68405573 | 0.85606265 |
| COTL1 | 23.2700385 | -1.139634131 | 0.61933169 | -1.840103 | 0.06575311 | 0.25622183 |
| KLHL36 | 44.85967 | -0.360694008 | 0.2970152 | -1.2143958 | 0.22459663 | 0.51345116 |
| USP10 | 89.1224623 | -0.154648368 | 0.25078597 | -0.6166548 | 0.53746245 | 0.76945357 |
| ZDHHC7 | 62.2373341 | 0.107965844 | 0.31064053 | 0.34755879 | 0.72817156 | 0.88029911 |
| KIAA0513 | 17.2082803 | 0.002328743 | 0.54378318 | 0.00428248 | 0.99658308 | 0.99868531 |
| GSE1 | 135.309726 | -0.113463586 | 0.24016639 | -0.4724374 | 0.63661462 | 0.82975756 |
| EMC8 | 23.3464656 | -0.024216109 | 0.38668071 | -0.0626256 | 0.95006465 | 0.98003064 |
| COX4I1 | 194.705816 | -0.352369162 | 0.39058405 | -0.9021596 | 0.36697208 | 0.65226818 |
| MTHFSD | 15.6248106 | 0.396102314 | 0.39014137 | 1.01527893 | 0.30997289 | 0.60321222 |
| FBXO31 | 22.179377 | 0.091863818 | 0.32833588 | 0.27978611 | 0.77964161 | 0.90427469 |
| MAP1LC3B | 49.1002881 | -0.7802141 | 0.31764464 | -2.4562483 | 0.01403961 | 0.09623211 |
| ZCCHC14 | 162.444987 | -0.535676816 | 0.55346286 | -0.9678641 | 0.33311226 | 0.62190756 |
| KLHDC4 | 51.0848135 | 0.223817716 | 0.3666537 | 0.61043354 | 0.54157466 | 0.77222175 |
| BANP | 80.4145107 | 0.452944188 | 0.52662939 | 0.86008148 | 0.38974413 | 0.67185619 |
| ZFPM1 | 34.6371211 | -0.083447647 | 0.39977692 | -0.2087355 | 0.8346547 | 0.93313642 |
| ZC3H18 | 42.256957 | -0.280304822 | 0.30195072 | -0.9283132 | 0.35324515 | 0.63934145 |
| CYBA | 56.0715247 | -2.371079934 | 0.6501491 | -3.6469787 | 0.00026534 | 0.0053727 |
| MVD | 29.4763078 | 0.951212962 | 0.50140317 | 1.89710201 | 0.05781448 | 0.23769222 |
| PIEZO1 | 75.2807256 | 1.213973749 | 0.3177923 | 3.82002248 | 0.00013344 | 0.00308227 |
| APRT | 38.7245082 | -0.970635105 | 0.6092253 | -1.5932285 | 0.11110894 | 0.34769355 |
| GALNS | 22.8333927 | -0.029728916 | 0.34965107 | -0.0850245 | 0.93224189 | 0.97356162 |
| TRAPPC2L | 12.2765328 | 0.205305319 | 0.48059618 | 0.42718883 | 0.6692418 | 0.84636987 |
| ACSF3 | 32.1419993 | -0.104940182 | 0.298729 | -0.3512889 | 0.72537162 | 0.87840733 |
| ZNF778 | 12.9879912 | 0.244752573 | 0.41812805 | 0.58535316 | 0.55831028 | 0.78233632 |
| ANKRD11 | 379.387702 | -0.22450803 | 0.31155048 | -0.7206153 | 0.47114626 | 0.73253076 |
| SPG7 | 134.809855 | -0.314842266 | 0.3534952 | -0.890655 | 0.37311429 | 0.65706383 |
| RPL13 | 328.681532 | -0.51825138 | 0.42347883 | -1.2237952 | 0.22102948 | 0.5089143 |
| CHMP1A | 33.8700575 | -0.359762883 | 0.33225581 | -1.0827888 | 0.27890216 | 0.57144989 |
| SPATA33 | 11.3464463 | -0.248878153 | 0.40531184 | -0.6140412 | 0.53918813 | 0.77077724 |
| CDK10 | 35.7039049 | -0.00613132 | 0.26833009 | -0.0228499 | 0.98176999 | 0.99318267 |
| VPS9D1 | 36.8658881 | 0.038889974 | 0.27226834 | 0.14283693 | 0.88641897 | 0.95619452 |
| ZNF276 | 20.3326316 | 0.113604499 | 0.32332941 | 0.35135838 | 0.7253195 | 0.87840733 |
| SPIRE2 | 22.195877 | 1.378149778 | 0.42880929 | 3.2138991 | 0.00130946 | 0.01786758 |
| TCF25 | 215.928298 | -0.272746495 | 0.28184732 | -0.9677101 | 0.33318917 | 0.62190756 |
| DEF8 | 19.4673274 | -0.062077661 | 0.31130471 | -0.1994112 | 0.84194106 | 0.93566718 |
| GAS8 | 43.9273502 | -0.167626381 | 0.2891231 | -0.5797751 | 0.56206628 | 0.78386446 |
| FAM157C | 58.478495 | 1.234928806 | 0.43082426 | 2.86643281 | 0.00415126 | 0.04116103 |
| LINC02193 | 10.8206639 | -0.232759252 | 0.52776962 | -0.4410243 | 0.65919537 | 0.84068219 |
| DOC2B | 51.1079246 | -1.475587333 | 0.35405277 | -4.1677046 | 3.08E-05 | 0.00093779 |
| RPH3AL | 117.912851 | -0.290462362 | 0.32736428 | -0.8872757 | 0.3749305 | 0.65806152 |
| AC129507.2 | 16.818488 | 0.265609803 | 0.56305156 | 0.47173264 | 0.63711763 | 0.82989931 |
| C17orf97 | 9.3968333 | 0.157039096 | 0.54250782 | 0.28946882 | 0.77222264 | 0.90105761 |
| VPS53 | 186.868076 | -0.331477737 | 0.21449242 | -1.5454054 | 0.12224824 | 0.3664089 |
| GLOD4 | 35.0234445 | -0.350131379 | 0.37771707 | -0.9269673 | 0.3539435 | 0.63957058 |
| NXN | 22.7470195 | -0.967267971 | 0.63790374 | -1.5163228 | 0.12943776 | 0.37713756 |
| TIMM22 | 12.3460478 | 0.003586422 | 0.42654505 | 0.00840807 | 0.99329141 | 0.9979411 |
| ABR | 39.2505595 | -1.152096241 | 0.24175374 | -4.7655777 | 1.88E-06 | 9.37E-05 |
| YWHAE | 230.20413 | -0.102125858 | 0.2096972 | -0.4870158 | 0.62624711 | 0.8231524 |
| CRK | 81.4883792 | -0.210788888 | 0.2908292 | -0.7247859 | 0.46858342 | 0.73133883 |
| MYO1C | 79.5382793 | -0.496091968 | 0.27432936 | -1.8083809 | 0.07054724 | 0.26815316 |
| INPP5K | 13.8173366 | 0.032105003 | 0.40216219 | 0.07983098 | 0.93637168 | 0.97594028 |
| PITPNA | 113.803789 | 0.019471327 | 0.2295345 | 0.08482963 | 0.93239684 | 0.97356162 |
| SLC43A2 | 111.74944 | 0.941086268 | 0.43716194 | 2.15271776 | 0.03134087 | 0.16200382 |
| SCARF1 | 16.9765767 | 0.371941212 | 0.54342039 | 0.68444471 | 0.49369439 | 0.74700665 |
| RILP | 11.0670384 | 0.744962459 | 0.47943514 | 1.55383367 | 0.1202241 | 0.36310426 |
| PRPF8 | 90.042239 | 0.271465377 | 0.23660992 | 1.14731189 | 0.25125274 | 0.54298421 |
| MIR22HG | 25.8501486 | 0.174713835 | 0.70593062 | 0.24749434 | 0.80452567 | 0.91643337 |
| WDR81 | 14.1875575 | -0.890817195 | 0.5773431 | -1.5429598 | 0.12284053 | 0.36713175 |

| | | | | | | |
|---|---|---|---|---|---|---|
| SERPINF2 | 13.6633067 | 1.156654321 | 0.47037545 | 2.45900231 | 0.01393237 | 0.09600476 |
| SMYD4 | 21.6824661 | -0.08846636 | 0.63597775 | -0.1391029 | 0.88936882 | 0.95702609 |
| RPA1 | 40.1720727 | -0.085985896 | 0.2587781 | -0.3322766 | 0.73968044 | 0.88614269 |
| SMG6 | 227.218216 | -0.020201359 | 0.23447472 | -0.0861558 | 0.93134256 | 0.97334817 |
| TSR1 | 21.2080263 | -0.238634927 | 0.37762521 | -0.6319359 | 0.52742877 | 0.76221787 |
| SGSM2 | 44.3962096 | 0.180219787 | 0.25583724 | 0.70443141 | 0.48116415 | 0.7402857 |
| MNT | 18.6687949 | -0.659325772 | 0.43222596 | -1.5254192 | 0.12715458 | 0.37394989 |
| METTL16 | 82.2601617 | -0.009219851 | 0.21581529 | -0.042721 | 0.96592392 | 0.98661741 |
| PAFAH1B1 | 302.272814 | 0.219745771 | 0.22852678 | 0.96157557 | 0.33626285 | 0.62473938 |
| CLUH | 65.4764497 | 0.442959904 | 0.35216223 | 1.25782911 | 0.20845357 | 0.49402397 |
| RAP1GAP2 | 143.028474 | 1.02069207 | 0.28984844 | 3.52146823 | 0.00042916 | 0.00771597 |
| SPATA22 | 12.6916851 | 0.826819048 | 0.8463088 | 0.97697087 | 0.32858357 | 0.61768835 |
| ASPA | 41.7300737 | 1.811352533 | 0.60158417 | 3.01097108 | 0.00260414 | 0.02954605 |
| CTNS | 19.9810052 | 0.020516724 | 0.52775508 | 0.03887547 | 0.96898968 | 0.98770265 |
| TAX1BP3 | 20.7132364 | -1.454861796 | 0.55686591 | -2.6125891 | 0.00898593 | 0.07117035 |
| EMC6 | 18.2048086 | -1.049562188 | 0.42029364 | -2.4972117 | 0.01251742 | 0.08973867 |
| ITGAE | 17.9628127 | 0.234709854 | 0.36568149 | 0.64184232 | 0.52097557 | 0.76092013 |
| NCBP3 | 65.8157023 | 0.333229815 | 0.2316595 | 1.43844657 | 0.15030739 | 0.41132271 |
| CAMKK1 | 10.6561496 | -0.291268903 | 0.48314509 | -0.6028601 | 0.54660175 | 0.77463416 |
| ZZEF1 | 108.569303 | 0.420124459 | 0.24542606 | 1.71181685 | 0.08693043 | 0.30306506 |
| CYB5D2 | 38.232471 | -0.425680551 | 0.47108795 | -0.9036116 | 0.36620137 | 0.6515728 |
| ANKFY1 | 100.564318 | -0.070449474 | 0.32621787 | -0.2159584 | 0.82902021 | 0.92883918 |
| UBE2G1 | 141.665789 | 0.307268756 | 0.26974479 | 1.13910912 | 0.25465764 | 0.54564969 |
| SPNS2 | 31.3318714 | -1.127468456 | 0.63264083 | -1.782162 | 0.07472282 | 0.27700412 |
| MYBBP1A | 11.4199802 | -0.528181097 | 0.53168614 | -0.9934077 | 0.32051132 | 0.6120836 |
| GGT6 | 16.6898356 | -0.165511328 | 0.64977933 | -0.2547193 | 0.79893992 | 0.91379784 |
| SMTNL2 | 18.892344 | 0.985935685 | 0.55812036 | 1.76652878 | 0.07730718 | 0.28224131 |
| PELP1 | 25.4359861 | 0.039840156 | 0.32962215 | 0.12086614 | 0.90379706 | 0.96126516 |
| MED11 | 11.5066864 | -0.722540022 | 0.47425254 | -1.5235343 | 0.12762508 | 0.37458507 |
| CXCL16 | 15.9598345 | -2.01391239 | 0.63197433 | -3.1866997 | 0.00143906 | 0.01915401 |
| PSMB6 | 30.3906433 | -0.801185824 | 0.43626261 | -1.836476 | 0.06628729 | 0.2574031 |
| MINK1 | 59.1163547 | -0.463089455 | 0.2329385 | -1.9880331 | 0.04680803 | 0.20830078 |
| SLC25A11 | 25.0090457 | 0.083308169 | 0.52866728 | 0.15758147 | 0.8747866 | 0.94993692 |
| RNF167 | 26.6331611 | -0.61605281 | 0.3686928 | -1.6709109 | 0.09473928 | 0.3166788 |
| PFN1 | 106.948532 | -1.539200485 | 0.39663095 | -3.8806868 | 0.00010416 | 0.00250055 |
| SPAG7 | 39.1246959 | -0.126245766 | 0.38580893 | -0.3272235 | 0.74349881 | 0.88754632 |
| CAMTA2 | 19.7792668 | 0.223357532 | 0.37987313 | 0.58797929 | 0.5565462 | 0.78071064 |
| KIF1C | 31.8457092 | 0.288650665 | 0.37687682 | 0.76590188 | 0.44373469 | 0.71343168 |
| ZNF594 | 23.1403245 | -0.203921469 | 0.33088534 | -0.6162904 | 0.53770286 | 0.76968387 |
| AC087500.1 | 50.987622 | 0.696137505 | 0.3203625 | 2.17296813 | 0.02978272 | 0.15672354 |
| RABEP1 | 170.457712 | 0.139383909 | 0.31651829 | 0.44036606 | 0.65967201 | 0.84097773 |
| NUP88 | 110.398805 | 0.448794168 | 0.28320619 | 1.58469053 | 0.11303666 | 0.35086028 |
| RPAIN | 53.9411799 | -0.057110011 | 0.26485345 | -0.2156287 | 0.82927717 | 0.92901407 |
| C1QBP | 34.9362393 | -0.988148554 | 0.68245879 | -1.4479241 | 0.14763827 | 0.40743891 |
| DERL2 | 25.4336476 | -0.472561587 | 0.3440279 | -1.3736141 | 0.16956151 | 0.44032355 |
| MIS12 | 13.9304421 | -0.347282006 | 0.43314704 | -0.8017647 | 0.42268908 | 0.69778363 |
| KIAA0753 | 58.3110685 | -0.334901163 | 0.24299858 | -1.3782021 | 0.16814089 | 0.43857328 |
| TXNDC17 | 16.7114495 | -1.154598649 | 0.43261966 | -2.6688539 | 0.00761106 | 0.06389755 |
| MED31 | 13.7907511 | -0.396381208 | 0.48316882 | -0.8203783 | 0.41200049 | 0.68913252 |
| XAF1 | 57.2567996 | -1.067456008 | 1.84095924 | -0.5798369 | 0.56202464 | 0.78386446 |
| ALOX12-AS1 | 42.1821259 | 0.083864797 | 0.64265541 | 0.1304973 | 0.89617299 | 0.95900099 |
| DLG4 | 11.7511573 | 0.450522174 | 0.63110862 | 0.71385837 | 0.47531476 | 0.73587124 |
| ACADVL | 179.168982 | 0.1904891 | 0.31854152 | 0.59800399 | 0.54983727 | 0.77596691 |
| DVL2 | 14.0790552 | 0.074965068 | 0.4451069 | 0.16842037 | 0.86625258 | 0.94702879 |
| ELP5 | 18.0692068 | 0.030501232 | 0.40627523 | 0.07507529 | 0.94015481 | 0.97685071 |
| EIF5A | 30.7680394 | -0.636231771 | 0.50515936 | -1.2594674 | 0.20786154 | 0.49352146 |
| GPS2 | 25.0006619 | -0.972809115 | 0.48105558 | -2.0222385 | 0.04315172 | 0.1982867 |
| ZBTB4 | 79.3655059 | 0.454950611 | 0.23046793 | 1.97402998 | 0.04837833 | 0.21317091 |
| POLR2A | 90.91011 | -0.080107869 | 0.25572372 | -0.3132594 | 0.75408356 | 0.89238252 |
| EIF4A1 | 32.7859458 | -1.188747817 | 0.44216373 | -2.6884788 | 0.00717784 | 0.06151922 |
| MPDU1 | 20.7351336 | 0.027070741 | 0.40313519 | 0.06715053 | 0.94646187 | 0.97905916 |
| FXR2 | 21.2262024 | 0.272318674 | 0.38622079 | 0.7050855 | 0.48075703 | 0.73988033 |
| SAT2 | 44.5619193 | -0.037684032 | 0.34448044 | -0.1093938 | 0.91289013 | 0.96446045 |
| TP53 | 23.8108933 | -0.851948376 | 0.34809882 | -2.4474325 | 0.01438781 | 0.09745114 |
| KDM6B | 32.5696305 | -0.572771818 | 0.47667434 | -1.2015999 | 0.22951859 | 0.51974407 |
| NAA38 | 71.4753395 | -0.14249469 | 0.31104128 | -0.4581215 | 0.64686517 | 0.83466996 |
| CYB5D1 | 10.867662 | -0.162214033 | 0.49359823 | -0.3286358 | 0.74243101 | 0.88720442 |
| CHD3 | 48.968764 | 0.291423595 | 0.39281043 | 0.74189372 | 0.45815173 | 0.72410401 |
| TRAPPC1 | 13.2298758 | -1.373737974 | 0.62091628 | -2.2124367 | 0.0269365 | 0.14648981 |
| CNTROB | 16.2457372 | 0.220198944 | 0.39432222 | 0.55842389 | 0.57655497 | 0.7919559 |
| PER1 | 43.7073583 | 1.083243327 | 0.56254786 | 1.92560208 | 0.05415408 | 0.22907832 |
| TMEM107 | 11.2848653 | -1.980046727 | 0.58218458 | -3.4010635 | 0.00067124 | 0.01090716 |
| CTC1 | 20.236635 | -0.119579597 | 0.35507223 | -0.3367754 | 0.73628621 | 0.88437124 |
| SLC25A35 | 10.7747913 | 0.497322381 | 0.62185063 | 0.79974572 | 0.42385814 | 0.69854207 |
| NDEL1 | 70.8787329 | 0.21245923 | 0.35493336 | 0.59858907 | 0.54944695 | 0.77592905 |
| MYH10 | 276.114639 | 0.488528632 | 0.26496089 | 1.84377635 | 0.06521574 | 0.25526313 |
| NTN1 | 25.6508304 | 0.525509132 | 0.3299201 | 1.59283758 | 0.11119663 | 0.34769355 |
| STX8 | 206.626712 | 0.353471365 | 0.27964184 | 1.26401458 | 0.20622479 | 0.49076642 |
| USP43 | 44.4804039 | -0.42905529 | 0.37413976 | -1.1467781 | 0.25147333 | 0.54321509 |
| SCO1 | 30.6668731 | -0.507150199 | 0.29812345 | -1.7011416 | 0.0889164 | 0.30697614 |
| TMEM220 | 34.4581369 | 0.002769671 | 0.42442406 | 0.00652572 | 0.99479327 | 0.99837898 |
| TMEM220-A! | 22.0663305 | -0.517760706 | 0.32073779 | -1.6142803 | 0.10646664 | 0.33988823 |
| SHISA6 | 25.2352904 | -0.499677215 | 0.68992332 | -0.7242504 | 0.46891205 | 0.73159653 |
| DNAH9 | 12.6863334 | -0.849657211 | 0.4553103 | -1.8661059 | 0.06202656 | 0.24690788 |
| MAP2K4 | 238.702928 | 0.196074396 | 0.26477589 | 0.74052966 | 0.45897867 | 0.72461104 |
| ARHGAP44 | 113.611752 | -0.647248098 | 0.26376708 | -2.4538623 | 0.01413311 | 0.09648727 |
| ELAC2 | 20.8813625 | -0.158914936 | 0.31728366 | -0.5008608 | 0.61646912 | 0.81691188 |
| COX10 | 79.8947675 | 0.769639515 | 0.30125326 | 2.55479233 | 0.01062512 | 0.07997578 |
| TRIM16 | 22.404584 | 0.362354434 | 0.3838905 | 0.94390049 | 0.3452205 | 0.63256941 |
| ZSWIM7 | 48.0475708 | -0.010344683 | 0.24779272 | -0.0417473 | 0.96670013 | 0.98695423 |
| TTC19 | 128.642223 | 0.49027718 | 0.324922 | 1.50890729 | 0.13132248 | 0.38024035 |
| NCOR1 | 406.127482 | 0.525160833 | 0.13173967 | 3.9863531 | 6.71E-05 | 0.00178612 |
| PIGL | 102.468423 | 0.220626541 | 0.28213784 | 0.7819814 | 0.4342255 | 0.70613909 |
| UBB | 174.107246 | -0.956849847 | 0.44829196 | -2.1344346 | 0.03280722 | 0.16692026 |
| LRRC75A | 15.8447312 | -0.763551342 | 0.3967809 | -1.9243652 | 0.05430883 | 0.2294871 |
| AC098850.3 | 50.429587 | 0.355844695 | 0.73053603 | 0.48710081 | 0.62618689 | 0.8231524 |
| MPRIP | 177.923388 | 0.247165873 | 0.30488056 | 0.81069739 | 0.41753948 | 0.69378898 |
| COPS3 | 27.1889997 | 0.381619948 | 0.32386122 | 1.17834408 | 0.23865946 | 0.52895523 |
| NT5M | 18.7030686 | 1.125255809 | 0.37291356 | 3.01747087 | 0.00254894 | 0.02911212 |
| MED9 | 46.198882 | 0.020716934 | 0.31522421 | 0.06572127 | 0.94759974 | 0.979567 |
| RASD1 | 201.885142 | -0.290550017 | 0.39121178 | -0.7426924 | 0.45766794 | 0.72377441 |
| PEMT | 22.9318098 | 0.334131453 | 0.34448136 | 0.96995509 | 0.33206888 | 0.6217267 |
| RAI1 | 111.917872 | -0.174296702 | 0.17153823 | -1.0160808 | 0.30959092 | 0.60302622 |
| SREBF1 | 57.1086498 | -0.972791001 | 0.49765626 | -1.9547448 | 0.0506132 | 0.21935934 |
| TOM1L2 | 176.891945 | 0.975121969 | 0.26511837 | 3.67806261 | 0.00023501 | 0.00486164 |
| DRC3 | 16.6438793 | -0.260157808 | 0.51397672 | -0.5061665 | 0.61273976 | 0.81450903 |
| ATPAF2 | 17.80748 | -0.24582621 | 0.39595787 | -0.6208393 | 0.53470536 | 0.76795408 |
| GID4 | 14.1652401 | -0.062456825 | 0.40842855 | -0.1529198 | 0.8784615 | 0.95242163 |
| DRG2 | 11.180171 | -0.421808343 | 0.48642639 | -0.8671576 | 0.38585566 | 0.66811344 |
| ALKBH5 | 52.5217325 | -0.55440668 | 0.26579408 | -2.0858504 | 0.03699218 | 0.17942253 |
| FLII | 37.7717858 | -0.554832958 | 0.38009325 | -1.4597285 | 0.1443647 | 0.40217555 |
| MIEF2 | 13.9593662 | -0.616388655 | 0.36289531 | -1.6985302 | 0.08940773 | 0.30789644 |
| TOP3A | 27.1026802 | -0.154603063 | 0.34061005 | -0.4539005 | 0.64990046 | 0.83722328 |

| | | | | | | |
|---|---|---|---|---|---|---|
| SMCR8 | 13.9949257 | -0.376616492 | 0.3907896 | -0.9637321 | 0.33518025 | 0.62351515 |
| SHMT1 | 51.1352193 | -0.429901314 | 0.4633481 | -0.927815 | 0.35350355 | 0.63947209 |
| TRIM16L | 11.7566529 | 0.31653625 | 0.42562439 | 0.74369857 | 0.45705886 | 0.72329037 |
| TVP23B | 25.6893321 | -0.223821352 | 0.33800617 | -0.6621813 | 0.50785506 | 0.75382716 |
| PRPSAP2 | 65.9942985 | 0.560630174 | 0.29792328 | 1.8817938 | 0.05986402 | 0.24251636 |
| EPN2 | 150.032874 | -0.169420421 | 0.2411967 | -0.702416 | 0.48241977 | 0.74082091 |
| B9D1 | 26.9954424 | -0.515717937 | 0.33631438 | -1.53344 | 0.12516747 | 0.37032604 |
| AC004448.2 | 15.4430185 | -1.341141207 | 0.46207285 | -2.9024454 | 0.00370262 | 0.03803027 |
| SLC47A1 | 21.8023943 | 0.983649401 | 0.3908102 | 2.51694914 | 0.01183759 | 0.08606271 |
| ALDH3A2 | 47.7766664 | 0.176048731 | 0.35687591 | 0.49330517 | 0.62179698 | 0.82030381 |
| ULK2 | 57.7561239 | 0.172699842 | 0.2997415 | 0.5761626 | 0.56450528 | 0.78515099 |
| AKAP10 | 141.750417 | 0.219142624 | 0.23173094 | 0.94567702 | 0.34431335 | 0.63205918 |
| SPECC1 | 18.0928984 | 0.270212142 | 0.39291007 | 0.68772008 | 0.49162908 | 0.74602485 |
| CCDC144NL-/ | 22.2039212 | -0.910978294 | 0.42314956 | -2.1528518 | 0.03133033 | 0.16200382 |
| USP22 | 127.937763 | 0.106933262 | 0.21748387 | 0.49168365 | 0.622943 | 0.82118474 |
| DHRS7B | 86.6939021 | -0.131208577 | 0.26619864 | -0.4928972 | 0.6220852 | 0.82046224 |
| TMEM11 | 12.4905564 | -0.026118516 | 0.42707829 | -0.0611563 | 0.95123476 | 0.98061757 |
| AC087294.1 | 14.4842542 | -0.040850279 | 0.44668632 | -0.0914518 | 0.92713358 | 0.97111125 |
| MAP2K3 | 41.188356 | -0.159271452 | 0.50973489 | -0.3124594 | 0.75469142 | 0.89294028 |
| WSB1 | 1084.92906 | 0.825040967 | 0.41324844 | 1.99647689 | 0.04588204 | 0.205663 |
| KSR1 | 91.8663516 | 0.351065267 | 0.25314331 | 1.3868242 | 0.16549538 | 0.43437793 |
| NLK | 433.108672 | 0.774975376 | 0.39112828 | 1.98138415 | 0.04754821 | 0.21044546 |
| IFT20 | 15.7142915 | -0.13773148 | 0.46847041 | -0.2940025 | 0.76875598 | 0.90022017 |
| TNFAIP1 | 30.1730416 | 0.296655237 | 0.51536532 | 0.57562126 | 0.56487121 | 0.78515099 |
| POLDIP2 | 30.678096 | -0.232707514 | 0.39968172 | -0.5822321 | 0.56041038 | 0.7832825 |
| SARM1 | 26.5938125 | -0.277255808 | 0.31695287 | -0.874754 | 0.38170775 | 0.66492276 |
| AC005726.1 | 19.9255883 | 0.020269716 | 0.38769532 | 0.05228259 | 0.95830352 | 0.98331105 |
| KIAA0100 | 54.8873183 | 0.060425334 | 0.30112471 | 0.20066548 | 0.84096015 | 0.93566718 |
| SDF2 | 25.5215268 | -0.659459565 | 0.35838985 | -1.8400621 | 0.06575913 | 0.25622183 |
| SUPT6H | 71.5395688 | -0.336879532 | 0.26223237 | -1.2846604 | 0.198911 | 0.48182015 |
| RAB34 | 38.2111819 | -0.68080117 | 0.41888043 | -1.6252876 | 0.10410129 | 0.33490577 |
| RPL23A | 115.136952 | -0.323914543 | 0.33980574 | -0.9532345 | 0.34047129 | 0.62907597 |
| NEK8 | 20.0893643 | 0.290039875 | 0.38849773 | 0.74656776 | 0.45532452 | 0.72210564 |
| TRAF4 | 57.5038033 | 0.053629884 | 0.3296947 | 0.16266529 | 0.87078198 | 0.94844301 |
| FAM222B | 198.437171 | -0.176002286 | 0.29523173 | -0.5961496 | 0.55107527 | 0.7768442 |
| ERAL1 | 12.0773486 | 0.003744008 | 0.39113699 | 0.00957211 | 0.99236267 | 0.99770844 |
| FLOT2 | 89.4679552 | -1.185355497 | 0.31505228 | -3.7624089 | 0.00016828 | 0.00369414 |
| DHRS13 | 14.3438051 | -1.305931038 | 0.5595651 | -2.3338322 | 0.01960451 | 0.11894327 |
| PHF12 | 59.900731 | -0.133579456 | 0.22512409 | -0.5933592 | 0.55294077 | 0.77787735 |
| MYO18A | 43.7653421 | -0.391396075 | 0.47061681 | -0.8316662 | 0.40559741 | 0.68389162 |
| NUFIP2 | 150.99397 | -0.240166194 | 0.17952583 | -1.3377807 | 0.18096792 | 0.45756661 |
| TAOK1 | 269.728019 | 0.095178795 | 0.24182472 | 0.39358588 | 0.69388681 | 0.86101552 |
| ABHD15-AS1 | 30.143293 | -0.261072268 | 0.51260882 | -0.5093012 | 0.61054114 | 0.81376433 |
| GIT1 | 20.3866062 | -0.657486436 | 0.36062242 | -1.823199 | 0.06827326 | 0.26264045 |
| SSH2 | 575.440729 | -0.064753675 | 0.21955381 | -0.294933 | 0.76804503 | 0.89983999 |
| AC023389.1 | 18.5700504 | -1.092271113 | 0.42449185 | -2.5731262 | 0.01007844 | 0.07765754 |
| EFCAB5 | 18.5693097 | -1.515548691 | 0.50151194 | -3.0219593 | 0.00251144 | 0.02877963 |
| NSRP1 | 169.23043 | 0.170886599 | 0.17604474 | 0.97069982 | 0.33169778 | 0.62147393 |
| AC104984.4 | 19.0886526 | 0.478639401 | 0.67271486 | 0.71150413 | 0.47677189 | 0.73678733 |
| BLMH | 43.6315752 | -0.071399713 | 0.3462478 | -0.2062099 | 0.83662698 | 0.93419636 |
| CPD | 122.345276 | -1.000879578 | 0.36975104 | -2.7069013 | 0.00679144 | 0.0591532 |
| GOSR1 | 96.6036884 | 0.033394792 | 0.19928409 | 0.16757379 | 0.86691859 | 0.94728419 |
| CRLF3 | 65.6334512 | -0.175751918 | 0.31651986 | -0.5552635 | 0.57871446 | 0.7935467 |
| ATAD5 | 15.2786923 | -0.330489768 | 0.48341008 | -0.6836634 | 0.49418776 | 0.74700665 |
| TEFM | 31.1664214 | -0.06200312 | 0.34439716 | -0.1800338 | 0.85712606 | 0.94359207 |
| RNF135 | 21.3613267 | 0.410085244 | 0.34836936 | 1.17715646 | 0.23913306 | 0.52945386 |
| NF1 | 531.388191 | 0.132243631 | 0.20169061 | 0.65567569 | 0.51203281 | 0.75555211 |
| RAB11FIP4 | 40.226954 | 0.216625574 | 0.3280053 | 0.66043315 | 0.50897591 | 0.75445608 |
| COPRS | 27.6589112 | -0.160065903 | 0.40579077 | -0.3944543 | 0.69324568 | 0.8606282 |
| UTP6 | 33.041179 | 0.155205135 | 0.3986937 | 0.38928414 | 0.69706597 | 0.86249983 |
| SUZ12 | 130.173128 | 0.280572811 | 0.19677181 | 1.42587911 | 0.15390324 | 0.4169593 |
| LRRC37B | 38.8005138 | -0.240197404 | 0.40230289 | -0.5970561 | 0.55046991 | 0.77630952 |
| RHOT1 | 114.026239 | 0.336639724 | 0.19123606 | 1.76033598 | 0.07835086 | 0.28477085 |
| C17orf75 | 16.0507029 | 0.381144156 | 0.37764808 | 1.0092575 | 0.31285116 | 0.60559801 |
| ZNF207 | 187.507869 | 0.287168565 | 0.18848021 | 1.52360058 | 0.12760852 | 0.37458507 |
| PSMD11 | 50.417544 | 0.077341549 | 0.32730621 | 0.23629723 | 0.81320204 | 0.9206162 |
| MYO1D | 239.387438 | -0.328610177 | 0.19629148 | -1.6740929 | 0.09411234 | 0.3160115 |
| ASIC2 | 26.9658986 | 1.534815163 | 0.49714712 | 3.08724538 | 0.00202021 | 0.0247764 |
| LIG3 | 16.1207505 | 0.616519593 | 0.34892754 | 1.76689863 | 0.07724521 | 0.28219153 |
| SLFN5 | 52.9541889 | -3.135166102 | 0.56491713 | -5.5497805 | 2.86E-08 | 2.98E-06 |
| LINC02001 | 16.760869 | -0.022089517 | 0.3515936 | -0.0628268 | 0.94990439 | 0.98003064 |
| AP2B1 | 133.507397 | 0.052854695 | 0.18327611 | 0.28838835 | 0.77304949 | 0.90149298 |
| TAF15 | 214.919065 | 0.09191149 | 0.17043267 | 0.53928327 | 0.58969141 | 0.79960282 |
| AC015849.3 | 12.325295 | -0.141486489 | 0.565935 | -0.2500048 | 0.80258361 | 0.91564859 |
| ZNHIT3 | 14.8657852 | -0.30090593 | 0.40171022 | -0.7490622 | 0.45381973 | 0.72126672 |
| MYO19 | 83.8423279 | 0.304966792 | 0.20277612 | 1.5039581 | 0.13259215 | 0.38212969 |
| GGNBP2 | 105.138037 | -0.06887102 | 0.27232739 | -0.2528979 | 0.80034713 | 0.91455697 |
| DHRS11 | 14.6747755 | 0.831686233 | 0.45022242 | 1.84727857 | 0.06470678 | 0.25394637 |
| LHX1-DT | 158.673896 | 0.400614614 | 0.37758168 | 1.06100121 | 0.28868935 | 0.58064041 |
| LHX1 | 225.100109 | 0.646306474 | 0.34819813 | 1.85614576 | 0.06343279 | 0.25076153 |
| AATF | 121.190918 | 0.179466756 | 0.24546308 | 0.73113543 | 0.46469644 | 0.72848544 |
| ACACA | 194.235804 | 0.237632096 | 0.21941533 | 1.08302414 | 0.27879771 | 0.57144989 |
| TADA2A | 46.3724345 | 0.253071343 | 0.26538207 | 0.9536113 | 0.34028043 | 0.62907597 |
| SYNRG | 113.878649 | -0.250185887 | 0.22233853 | -1.1252475 | 0.26048416 | 0.55146957 |
| DDX52 | 61.0798209 | -0.611953896 | 0.34502848 | -1.773633 | 0.07612388 | 0.28022732 |
| HNF1B | 276.931115 | -0.24274616 | 0.34774323 | -0.6980615 | 0.48513874 | 0.74285379 |
| MRPL45 | 33.3000149 | 0.283440874 | 0.26315054 | 1.07710542 | 0.28143317 | 0.57469267 |
| SOCS7 | 33.608607 | 0.61316151 | 0.28661911 | 2.13929043 | 0.03241215 | 0.16565474 |
| MLLT6 | 117.132158 | -0.230932721 | 0.22794513 | -1.0131066 | 0.31100923 | 0.60414964 |
| CISD3 | 18.9994209 | -0.409562468 | 0.4509439 | -0.9082338 | 0.36375473 | 0.648676 |
| PCGF2 | 29.1803872 | -0.205875034 | 0.31720469 | -0.649029 | 0.51631965 | 0.75768873 |
| PSMB3 | 35.4650701 | -0.596497759 | 0.36459885 | -1.6360385 | 0.10183153 | 0.33051935 |
| PIP4K2B | 39.611928 | 0.076923623 | 0.26251945 | 0.29302067 | 0.76950636 | 0.90034488 |
| CWC25 | 26.9549588 | -0.398007457 | 0.35253347 | -1.128992 | 0.25890122 | 0.55002264 |
| RPL23 | 91.3205546 | -0.182880062 | 0.39309669 | -0.4652292 | 0.64176732 | 0.83268097 |
| LASP1 | 58.4689853 | -0.537758926 | 0.31081441 | -1.7301609 | 0.08360154 | 0.29623459 |
| RPL19 | 161.964914 | -0.778347258 | 0.45022624 | -1.7287914 | 0.08384643 | 0.29653069 |
| FBXL20 | 181.380788 | 0.051082533 | 0.31315334 | 0.16312307 | 0.87042154 | 0.94840865 |
| MED1 | 37.1650573 | 0.010388352 | 0.29176383 | 0.03560535 | 0.97159705 | 0.988268 |
| CDK12 | 118.456053 | 0.044756105 | 0.23616746 | 0.18951004 | 0.84969308 | 0.93912007 |
| PPP1R1B | 14.0836341 | -1.798019553 | 0.79770385 | -2.2539938 | 0.02419656 | 0.13591762 |
| STARD3 | 26.0838056 | -0.460213908 | 0.32729121 | -1.4061297 | 0.15968561 | 0.42727956 |
| PGAP3 | 19.9707073 | 1.06830527 | 0.45332525 | 2.35659778 | 0.01844321 | 0.11445902 |
| ERBB2 | 133.877067 | 0.921294991 | 0.31543129 | 2.92074701 | 0.00349193 | 0.03648468 |
| MIEN1 | 18.1969676 | -1.295436235 | 0.42652967 | -3.0371539 | 0.00238824 | 0.02758251 |
| GRB7 | 73.4830026 | -0.643722217 | 0.26254018 | -2.4518998 | 0.01421042 | 0.09682259 |
| ORMDL3 | 15.270611 | -0.517262312 | 0.46256545 | -1.1182467 | 0.26346162 | 0.55504737 |
| PSMD3 | 27.5179246 | 0.045720426 | 0.3352353 | 0.13638309 | 0.89151844 | 0.95720928 |
| MED24 | 38.094028 | 0.167350756 | 0.27254565 | 0.61402835 | 0.53919659 | 0.77077724 |
| THRA | 32.7518285 | 0.169711905 | 0.3094753 | 0.54838594 | 0.58342693 | 0.79598942 |
| NR1D1 | 15.4963542 | 1.111215499 | 0.71630228 | 1.55132201 | 0.12082454 | 0.3637447 |
| MSL1 | 39.9886162 | -0.525029322 | 0.29014692 | -1.8095292 | 0.07036883 | 0.26766667 |
| CASC3 | 50.1956938 | 0.174273074 | 0.26542422 | 0.65658317 | 0.51144897 | 0.75522875 |

| | | | | | | |
|---|---|---|---|---|---|---|
| WIPF2 | 136.110603 | 0.062703987 | 0.2871966 | 0.21833123 | 0.82717104 | 0.92808376 |
| RARA | 24.8981028 | -0.03020758 | 0.41941786 | -0.0720226 | 0.94258389 | 0.9781624 |
| IGFBP4 | 14.0530776 | -0.459936404 | 0.52533224 | -0.8755153 | 0.3812936 | 0.66453843 |
| TNS4 | 11.0914713 | 2.949203583 | 1.51881473 | 1.94177969 | 0.05216378 | 0.22326183 |
| KRT10 | 30.3584445 | -0.787622871 | 0.40091438 | -1.9645663 | 0.04946446 | 0.21605779 |
| KRT19 | 590.811324 | -1.258840185 | 0.4403887 | -2.8584752 | 0.00425682 | 0.04204583 |
| EIF1 | 201.229621 | -0.941383439 | 0.34390627 | -2.7373256 | 0.0061941 | 0.05537004 |
| JUP | 170.063875 | 0.0550093 | 0.33458487 | 0.1644106 | 0.86940792 | 0.94827256 |
| P3H4 | 14.8608136 | -0.39571979 | 0.4140419 | -0.9557482 | 0.33919948 | 0.62810099 |
| NT5C3B | 24.0008291 | -0.100463498 | 0.30252606 | -0.3320821 | 0.73982724 | 0.88614269 |
| ACLY | 38.9736838 | -0.597719673 | 0.35152098 | -1.7003812 | 0.08905924 | 0.30722281 |
| TTC25 | 11.4193269 | -0.343585939 | 0.44518283 | -0.7717861 | 0.4402411 | 0.71109498 |
| CNP | 15.801734 | -1.343154034 | 0.61449165 | -2.185797 | 0.02883046 | 0.1531776 |
| DNAJC7 | 83.2923907 | 0.089189046 | 0.24134484 | 0.36955024 | 0.71171763 | 0.87074117 |
| NKIRAS2 | 11.8987259 | -1.041550945 | 0.46157284 | -2.2565256 | 0.02403773 | 0.13578849 |
| KAT2A | 20.2592407 | 0.991914715 | 0.41309968 | 2.40115102 | 0.01634359 | 0.10592021 |
| STAT5B | 114.179594 | 0.125311535 | 0.21603916 | 0.58004082 | 0.56188709 | 0.78382678 |
| STAT5A | 16.5228316 | -0.153143096 | 0.44454076 | -0.3444973 | 0.73047232 | 0.88159862 |
| STAT3 | 475.103643 | -0.808648708 | 0.17523079 | -4.6147638 | 3.94E-06 | 0.00017534 |
| ATP6V0A1 | 90.099885 | 0.520215201 | 0.21433107 | 2.42715717 | 0.01521766 | 0.10114345 |
| NAGLU | 24.946384 | -0.67258755 | 0.46521047 | -1.4457705 | 0.14824159 | 0.40845709 |
| COASY | 28.1278127 | -0.29248638 | 0.44790156 | -0.6530149 | 0.51374669 | 0.75647799 |
| MLX | 20.2727591 | -0.505952951 | 0.35946788 | -1.4075053 | 0.1592776 | 0.42671782 |
| RETREG3 | 61.6704383 | -0.279503162 | 0.26274249 | -1.0637912 | 0.28742328 | 0.57979922 |
| TUBG2 | 26.3085549 | -0.296253328 | 0.35819527 | -0.8270721 | 0.40819618 | 0.68657965 |
| PLEKHH3 | 12.5894687 | -0.333738675 | 0.42239652 | -0.7901075 | 0.42946496 | 0.70316149 |
| EZH1 | 66.2649858 | -0.272089482 | 0.3441727 | -0.7905609 | 0.42920026 | 0.70310352 |
| VPS25 | 24.9041562 | -0.606148907 | 0.40260789 | -1.5055564 | 0.13218108 | 0.38143682 |
| WNK4 | 62.9904246 | 1.993661178 | 0.49989514 | 3.98815877 | 6.66E-05 | 0.00177718 |
| COA3 | 60.7333219 | 0.063423897 | 0.44702944 | 0.14187857 | 0.88717592 | 0.95619452 |
| BECN1 | 38.3642594 | 0.115251237 | 0.26664837 | 0.4322218 | 0.66558022 | 0.84360327 |
| PSME3 | 26.7151016 | -0.093971031 | 0.30619823 | -0.3068961 | 0.75892248 | 0.89527304 |
| RUNDC1 | 22.760013 | -0.547133854 | 0.34208227 | -1.5994218 | 0.1097269 | 0.34537323 |
| RPL27 | 61.169945 | -0.90959562 | 0.42099074 | -2.1606072 | 0.0307257 | 0.16010932 |
| IFI35 | 16.7739377 | -2.498977378 | 1.39764256 | -1.7879946 | 0.07377687 | 0.27468354 |
| VAT1 | 18.5320844 | -1.97679875 | 0.44249609 | -4.4673813 | 7.92E-06 | 0.00030783 |
| BRCA1 | 21.407671 | -0.724376055 | 0.47678823 | -1.5192826 | 0.12869138 | 0.37611281 |
| NBR1 | 70.9462054 | -0.23882628 | 0.2575778 | -0.9272006 | 0.35382241 | 0.63957058 |
| LINC00854 | 36.1261617 | -0.604913395 | 0.66647284 | -0.907634 | 0.36407163 | 0.64890415 |
| ARL4D | 54.3288991 | 0.525207325 | 0.47597364 | 1.10343785 | 0.26983707 | 0.56184753 |
| DHX8 | 29.6922716 | -0.427121126 | 0.32817188 | -1.3015165 | 0.19308171 | 0.47361903 |
| DUSP3 | 46.2310322 | -0.66042613 | 0.30759652 | -2.1470533 | 0.03178903 | 0.16325555 |
| TMEM101 | 126.062976 | -2.009330412 | 0.61478776 | -3.2683318 | 0.00108183 | 0.01544593 |
| LSM12 | 43.9737275 | 0.106444351 | 0.35399926 | 0.30069088 | 0.76365022 | 0.89745287 |
| G6PC3 | 20.4145252 | -0.965090424 | 0.63122919 | -1.5289065 | 0.12628762 | 0.37228594 |
| HDAC5 | 44.0614034 | 0.338958638 | 0.33218998 | 1.02037588 | 0.30755023 | 0.60139222 |
| ASB16-AS1 | 11.3943724 | 0.451853778 | 0.38604464 | 1.17047027 | 0.24181177 | 0.53271231 |
| TMUB2 | 24.9106756 | -0.028114358 | 0.40306484 | -0.0697515 | 0.94439149 | 0.97856665 |
| ATXN7L3 | 16.3370712 | -0.687975781 | 0.40110144 | -1.7152164 | 0.08630556 | 0.30190826 |
| UBTF | 49.6843054 | 0.113589844 | 0.30312857 | 0.37472497 | 0.70786502 | 0.86882712 |
| SLC25A39 | 40.1670634 | -0.144334244 | 0.35145659 | -0.4106745 | 0.68131126 | 0.85466584 |
| GRN | 85.924677 | -1.563004344 | 0.43459558 | -3.596457 | 0.00032258 | 0.00621164 |
| GPATCH8 | 333.805133 | 0.373920693 | 0.1517431 | 2.46416927 | 0.01373312 | 0.09507992 |
| DBF4B | 15.0886996 | -0.236007386 | 0.46680779 | -0.5055772 | 0.61315347 | 0.81464125 |
| EFTUD2 | 35.1860027 | 0.029979756 | 0.3354751 | 0.08936507 | 0.92879178 | 0.97199747 |
| DCAKD | 50.0826465 | -0.209692059 | 0.28838716 | -0.72712 | 0.46715247 | 0.73023858 |
| NMT1 | 76.8112011 | 0.111028968 | 0.21538582 | 0.51548875 | 0.60621153 | 0.81114147 |
| ACBD4 | 14.3286197 | 0.490680143 | 0.38632675 | 1.27011693 | 0.20404298 | 0.48826352 |
| HEXIM1 | 19.1714565 | -1.846113306 | 0.44665548 | -4.133193 | 3.58E-05 | 0.00106214 |
| AC138150.1 | 50.7458306 | 0.324850329 | 0.46056744 | 0.7053263 | 0.4806072 | 0.73988033 |
| MAP3K14 | 86.5715459 | 0.154565047 | 0.46213127 | 0.33446135 | 0.73803146 | 0.88573617 |
| ARHGAP27 | 18.6575487 | -0.356560143 | 0.34273065 | -1.0403509 | 0.2981769 | 0.59096161 |
| PLEKHM1 | 34.4971549 | -0.824962642 | 0.53003405 | -1.5564333 | 0.11960509 | 0.36197757 |
| LINC02210-C | 33.0273942 | 0.698199412 | 0.41342839 | 1.68880372 | 0.09125705 | 0.31117713 |
| KANSL1 | 608.119325 | 0.401845842 | 0.24329876 | 1.65165593 | 0.09860471 | 0.32423419 |
| ARL17B | 131.221057 | 1.449702337 | 1.08534021 | 1.33571237 | 0.18164331 | 0.45819573 |
| LRRC37A2 | 35.596559 | 0.378481378 | 0.37761543 | 1.00229319 | 0.31620201 | 0.60951749 |
| ARL17A | 38.2058734 | -0.05618582 | 0.52779387 | -0.1064541 | 0.91522207 | 0.96466317 |
| NSF | 173.123097 | 0.235158483 | 0.34957665 | 0.67269505 | 0.50114132 | 0.7507282 |
| AC005670.2 | 28.608801 | 0.153707232 | 0.35472887 | 0.43330906 | 0.66479026 | 0.84348349 |
| CDC27 | 162.298527 | -0.289723184 | 0.30166355 | -0.9604183 | 0.33684474 | 0.62525668 |
| AC068234.1 | 35.8752844 | -3.847663359 | 0.76762553 | -5.0124224 | 5.37E-07 | 3.40E-05 |
| EFCAB13 | 55.653552 | -1.041500504 | 0.36929515 | -2.8202387 | 0.00479879 | 0.04624618 |
| NPEPPS | 389.545906 | -0.08561562 | 0.2612807 | -0.3276768 | 0.74315606 | 0.88744566 |
| AC025682.1 | 34.6095053 | -0.021612361 | 0.34296532 | -0.0630162 | 0.94975363 | 0.98003064 |
| KPNB1 | 137.704851 | -0.148718591 | 0.23360123 | -0.6366344 | 0.52436298 | 0.76221787 |
| MRPL10 | 13.0154566 | -0.472256899 | 0.43588956 | -1.0834325 | 0.27861651 | 0.57131913 |
| SCRN2 | 17.6127457 | 0.321512216 | 0.36168317 | 0.8889333 | 0.37403893 | 0.65801725 |
| SP2 | 14.5186271 | 0.148173896 | 0.36818955 | 0.40243917 | 0.68736085 | 0.85804981 |
| PNPO | 22.5775224 | -0.326921833 | 0.51305096 | -0.6372112 | 0.52398726 | 0.76218158 |
| PRR15L | 11.0483914 | -1.205217431 | 0.78288136 | -1.5394637 | 0.12369114 | 0.36860459 |
| CDK5RAP3 | 36.198564 | 0.242683374 | 0.28201746 | 0.86052607 | 0.38949912 | 0.67156819 |
| NFE2L1 | 86.3780453 | -0.741950634 | 0.25798154 | -2.8759834 | 0.00402771 | 0.04032406 |
| CBX1 | 40.8640756 | -0.349964744 | 0.426226 | -0.8210779 | 0.4116019 | 0.68899849 |
| SNX11 | 15.1525652 | -0.07502036 | 0.37555105 | -0.1997608 | 0.8416677 | 0.93566718 |
| SKAP1 | 390.872936 | 0.062731172 | 0.21396333 | 0.29318656 | 0.76937956 | 0.90034488 |
| HOXB2 | 21.140311 | -0.970444131 | 0.45756912 | -2.1208689 | 0.03393283 | 0.1705249 |
| HOXB-AS1 | 15.6635896 | 0.518659969 | 0.36875028 | 1.40653445 | 0.15956548 | 0.42727956 |
| HOXB3 | 235.314652 | -0.181867362 | 0.28134638 | -0.646418 | 0.51800869 | 0.7586758 |
| HOXB-AS3 | 57.8625327 | -0.352622253 | 0.4619538 | -0.763328 | 0.44526783 | 0.71506613 |
| HOXB7 | 54.3367865 | -0.969986308 | 0.3784966 | -2.5627345 | 0.01038514 | 0.0788977 |
| AC103702.2 | 24.1234391 | -1.145139207 | 0.35246507 | -3.2489438 | 0.00115834 | 0.01623572 |
| LINC02086 | 59.1354759 | -0.227095264 | 0.39974046 | -0.5681068 | 0.56996247 | 0.78836645 |
| TTLL6 | 11.2367465 | 0.872421348 | 0.642751 | 1.35732398 | 0.17467829 | 0.44753439 |
| CALCOCO2 | 58.6546229 | 0.315525686 | 0.28588435 | 1.10368294 | 0.2697307 | 0.56184375 |
| ATP5MC1 | 70.1902783 | 0.085375497 | 0.4407323 | 0.19371282 | 0.84640075 | 0.93757726 |
| UBE2Z | 90.2416067 | -0.522342685 | 0.22369196 | -2.3350982 | 0.01953829 | 0.1186813 |
| SNF8 | 30.9011726 | -0.078599511 | 0.28673972 | -0.2741145 | 0.78399664 | 0.90719234 |
| ZNF652 | 138.162044 | -0.22198772 | 0.19472521 | -1.140005 | 0.25428421 | 0.54535536 |
| PHB | 52.4243874 | -0.474639134 | 0.50096685 | -0.9474462 | 0.34341146 | 0.63107829 |
| SPOP | 83.1512415 | 0.209574344 | 0.18923178 | 1.10750079 | 0.26807747 | 0.55996231 |
| SLC35B1 | 12.7491825 | -0.045806843 | 0.46708398 | -0.0980698 | 0.92187685 | 0.96880295 |
| FAM117A | 10.5063604 | -0.103706236 | 0.44983777 | -0.2305414 | 0.81767108 | 0.92281757 |
| KAT7 | 39.8648061 | -0.231186387 | 0.2779863 | -0.8316467 | 0.4056084 | 0.68389162 |
| ITGA3 | 257.239984 | -2.615949832 | 0.60829269 | -4.3004788 | 1.70E-05 | 0.00057938 |
| PDK2 | 21.451403 | 0.293919723 | 0.43071043 | 0.68240679 | 0.49498177 | 0.74770824 |
| PPP1R9B | 12.9891397 | -1.170038502 | 0.50196042 | -2.3309378 | 0.01975664 | 0.11965486 |
| XYLT2 | 21.1912772 | 0.367942814 | 0.39708961 | 0.92659894 | 0.35413481 | 0.63957058 |
| MRPL27 | 34.4565827 | -0.190016535 | 0.32586785 | -0.5831092 | 0.5598198 | 0.78316119 |
| LRRC59 | 15.230327 | -0.705621016 | 0.37985773 | -1.8575929 | 0.06322685 | 0.25020219 |
| ACSF2 | 22.5838001 | 0.353548776 | 0.54682981 | 0.64654262 | 0.517928 | 0.75866548 |
| RSAD1 | 19.0802362 | 0.412654284 | 0.38315906 | 1.07697906 | 0.28148962 | 0.57469403 |
| EPN3 | 17.1551373 | 1.041464809 | 0.60651407 | 1.71713216 | 0.08595503 | 0.30129591 |

| | | | | | | |
|---|---|---|---|---|---|---|
| SPATA20 | 16.2057666 | 0.27657546 | 0.43151295 | 0.6409436 | 0.52155933 | 0.76104628 |
| ABCC3 | 85.3774105 | -2.435425072 | 0.78964687 | -3.0841952 | 0.00204104 | 0.02492802 |
| ANKRD40 | 21.4387343 | -0.31109745 | 0.32780487 | -0.9490324 | 0.34260411 | 0.63071973 |
| LUC7L3 | 226.69458 | 0.546832712 | 0.21300225 | 2.56726265 | 0.01025049 | 0.07851345 |
| TOB1 | 66.5885164 | -0.172512021 | 0.3475116 | -0.4964209 | 0.61959748 | 0.81899807 |
| SPAG9 | 273.222602 | 0.089626373 | 0.38421277 | 0.23327276 | 0.81554961 | 0.92165337 |
| AC005920.2 | 23.2637142 | 0.084130565 | 0.66632554 | 0.12626045 | 0.89952576 | 0.95960592 |
| MBTD1 | 130.559057 | -0.068187145 | 0.22704724 | -0.3003214 | 0.76393201 | 0.89757885 |
| UTP18 | 30.4289116 | -0.030068907 | 0.34502001 | -0.0871512 | 0.93055133 | 0.97295644 |
| CA10 | 84.6581135 | -1.02870742 | 0.78430789 | -1.3116117 | 0.18965117 | 0.4686463 |
| TOM1L1 | 219.193163 | 0.358719055 | 0.1708378 | 2.09976397 | 0.03574961 | 0.17539507 |
| COX11 | 41.4369468 | -0.182058105 | 0.29007749 | -0.6276189 | 0.53025364 | 0.76428051 |
| STXBP4 | 241.630792 | 0.765874458 | 0.24985482 | 3.06527791 | 0.00217468 | 0.02585572 |
| HLF | 134.830657 | -0.545525235 | 0.40986281 | -1.3309947 | 0.18319075 | 0.46004581 |
| MMD | 20.7454284 | -0.250322088 | 0.44071455 | -0.5679914 | 0.57004079 | 0.78836893 |
| PCTP | 19.8754072 | -0.216542942 | 0.36993775 | -0.5853497 | 0.55831262 | 0.78233632 |
| DGKE | 35.4143685 | 0.304734726 | 0.37675469 | 0.80884123 | 0.41860647 | 0.69466558 |
| TRIM25 | 36.7495319 | -0.548051755 | 0.49734814 | -1.1019479 | 0.27048431 | 0.56240594 |
| COIL | 23.1978898 | -0.050646543 | 0.29348994 | -0.1725665 | 0.86299216 | 0.94620532 |
| SCPEP1 | 151.99763 | 0.967266241 | 0.35745246 | 2.70599971 | 0.00680991 | 0.0591532 |
| AKAP1 | 61.7852357 | 0.223167793 | 0.33125976 | 0.67369423 | 0.50050573 | 0.7506493 |
| MSI2 | 1187.41282 | 0.518620099 | 0.1498218 | 3.46157966 | 0.00053702 | 0.00902501 |
| MRPS23 | 21.6117875 | -0.433552553 | 0.35512494 | -1.2208451 | 0.22214466 | 0.51049171 |
| CUEDC1 | 110.84524 | -0.69662223 | 0.42316705 | -1.646211 | 0.09972033 | 0.32592604 |
| VEZF1 | 65.9792639 | -0.322152831 | 0.22062012 | -1.4602151 | 0.14423097 | 0.40202041 |
| AC015813.2 | 31.386521 | 0.199930232 | 0.33323781 | 0.59996262 | 0.54853115 | 0.77527341 |
| SRSF1 | 14.9581941 | 0.30910716 | 0.51225903 | 0.60341964 | 0.54622956 | 0.77463416 |
| DYNLL2 | 44.4477536 | -0.142739466 | 0.37646928 | -0.379153 | 0.70457423 | 0.86648248 |
| MKS1 | 13.1154444 | 0.241394512 | 0.41927775 | 0.57573891 | 0.56479167 | 0.78515099 |
| TSPOAP1 | 28.8666974 | 0.742762573 | 0.48746427 | 1.52372721 | 0.12757687 | 0.37458507 |
| TSPOAP1-AS | 11.5313852 | -0.85907744 | 0.69685595 | -1.2327906 | 0.21765391 | 0.50547128 |
| SUPT4H1 | 16.0959986 | -0.857013851 | 0.41290544 | -2.0755693 | 0.0379338 | 0.18260776 |
| RNF43 | 76.724433 | 0.432028484 | 0.44359896 | 0.9739168 | 0.33009785 | 0.61950534 |
| MTMR4 | 29.1931432 | 0.366974884 | 0.34496721 | 1.06379642 | 0.28742093 | 0.57979922 |
| RAD51C | 19.1152265 | 0.329982719 | 0.40493658 | 0.81489974 | 0.4151297 | 0.69186606 |
| TRIM37 | 55.9782669 | 0.322655386 | 0.31203976 | 1.0340201 | 0.30112677 | 0.59482532 |
| SKA2 | 26.274969 | 0.89936239 | 0.33531744 | 2.68212231 | 0.00731567 | 0.0623375 |
| GDPD1 | 19.854771 | 0.615997516 | 0.42432961 | 1.45169581 | 0.1465862 | 0.40573108 |
| YPEL2 | 145.643667 | -0.349944611 | 0.39666422 | -0.8822187 | 0.37765853 | 0.66066194 |
| DHX40 | 53.8253676 | -0.232443234 | 0.29531578 | -0.7871006 | 0.43122297 | 0.70454366 |
| CLTC | 249.255619 | 0.143545994 | 0.21381039 | 0.67137053 | 0.50198451 | 0.75108109 |
| PTRH2 | 33.9365835 | -0.359660985 | 0.32128245 | -1.1194542 | 0.26294643 | 0.55458796 |
| VMP1 | 507.576981 | -0.796651944 | 0.46765534 | -1.7035023 | 0.08847411 | 0.30616737 |
| TUBD1 | 34.3416124 | 0.153899991 | 0.41994612 | 0.36647557 | 0.71401024 | 0.87215242 |
| RPS6KB1 | 50.9931195 | 0.61875208 | 0.29201356 | 2.11891554 | 0.03409761 | 0.17085487 |
| RNFT1 | 11.7391635 | -0.127782167 | 0.42940861 | -0.2975771 | 0.76602596 | 0.89881543 |
| HEATR6 | 15.7772898 | 0.315838314 | 0.39332087 | 0.80300422 | 0.4219723 | 0.69753545 |
| USP32 | 199.21357 | -0.591437239 | 0.302613 | -1.9544344 | 0.05064987 | 0.21942597 |
| APPBP2 | 133.954577 | -0.10631124 | 0.17261045 | -0.6159027 | 0.53795875 | 0.76983624 |
| AC011921.1 | 10.8910187 | 0.148376027 | 0.60565834 | 0.24498305 | 0.80646956 | 0.91724035 |
| PPM1D | 56.3849953 | 0.406150251 | 0.38238284 | 1.06215606 | 0.28816484 | 0.5803086 |
| BCAS3 | 682.156833 | 0.60062066 | 0.17599858 | 3.41264487 | 0.00064336 | 0.01048994 |
| AC005746.2 | 25.3830229 | 1.343747172 | 0.55352321 | 2.4276257 | 0.01519802 | 0.10107811 |
| TBX2 | 164.05819 | 1.29240274 | 0.31671058 | 4.08070598 | 4.49E-05 | 0.00129204 |
| INTS2 | 33.5498444 | 0.262510475 | 0.29123114 | 0.90138188 | 0.36738531 | 0.65277742 |
| MED13 | 312.508608 | 0.33663318 | 0.24280639 | 1.38642639 | 0.16561674 | 0.43438116 |
| METTL2A | 17.3213192 | -0.140934607 | 0.35251267 | -0.3998001 | 0.68930374 | 0.85868974 |
| TLK2 | 156.196301 | 0.372797825 | 0.22082982 | 1.68816799 | 0.09137899 | 0.31120975 |
| TANC2 | 491.192728 | -0.425177078 | 0.23778722 | -1.7880569 | 0.07376683 | 0.27468354 |
| AC015923.1 | 30.3362087 | -1.419724604 | 0.6741902 | -2.1058221 | 0.03521981 | 0.1742352 |
| CYB561 | 23.6183011 | -1.120164475 | 0.43847085 | -2.5547068 | 0.01062773 | 0.07997578 |
| DCAF7 | 116.623773 | -0.395401515 | 0.22782271 | -1.7355667 | 0.08264048 | 0.2945637 |
| MAP3K3 | 98.5577994 | -0.292251044 | 0.31106648 | -0.9395131 | 0.34746736 | 0.63465611 |
| CCDC47 | 23.969349 | -0.596756489 | 0.3134453 | -1.9038616 | 0.05692821 | 0.23559369 |
| DDX42 | 87.3268703 | 0.048867718 | 0.2665106 | 0.18336126 | 0.85451457 | 0.94192265 |
| FTSJ3 | 16.4783054 | -0.337447034 | 0.38038271 | -0.8871251 | 0.37501158 | 0.65806152 |
| PSMC5 | 17.9044767 | 0.164227198 | 0.48781687 | 0.33665748 | 0.73637512 | 0.88437124 |
| SMARCD2 | 27.3063383 | -0.030912333 | 0.42380205 | -0.0729405 | 0.94185347 | 0.97782671 |
| ERN1 | 60.2599882 | -1.746801507 | 0.43674599 | -3.9995822 | 6.35E-05 | 0.00171128 |
| SNHG25 | 19.7360377 | 0.20101679 | 0.50282613 | 0.39977395 | 0.68932302 | 0.85868974 |
| TEX2 | 73.0783394 | -0.453442339 | 0.3756096 | -1.2072171 | 0.22734854 | 0.51714387 |
| POLG2 | 26.3415605 | 0.351766652 | 0.33195923 | 1.05966824 | 0.28929556 | 0.58107288 |
| DDX5 | 360.863728 | -0.121174511 | 0.23539721 | -0.5147661 | 0.60671645 | 0.81143061 |
| CEP95 | 126.182523 | 0.333665491 | 0.27468395 | 1.21472509 | 0.22447096 | 0.51340497 |
| SMURF2 | 130.970477 | 0.274874835 | 0.24049966 | 1.14293231 | 0.2530667 | 0.54473281 |
| LRRC37A3 | 134.876373 | 0.277923972 | 0.3044619 | 0.91283663 | 0.36132847 | 0.64557855 |
| GNA13 | 100.882159 | -0.406647312 | 0.42492686 | -0.9569819 | 0.33857639 | 0.62755046 |
| CEP112 | 158.406986 | 0.121787447 | 0.38330419 | 0.31773054 | 0.75068934 | 0.89117327 |
| PRKCA | 836.401913 | 0.041086614 | 0.32814773 | 0.12520768 | 0.90035914 | 0.95980075 |
| PRKCA-AS1 | 34.0161454 | 0.799016681 | 0.63011382 | 1.26805136 | 0.20477961 | 0.48947553 |
| HELZ | 248.959375 | 0.202402067 | 0.19183364 | 1.0550916 | 0.29138343 | 0.58292358 |
| PSMD12 | 47.6523592 | -0.067669744 | 0.23720473 | -0.2852799 | 0.7754297 | 0.90234686 |
| PITPNC1 | 184.654253 | -1.262372165 | 0.39560451 | -3.1909954 | 0.00141784 | 0.0190003 |
| NOL11 | 28.2132073 | 0.031396151 | 0.36546615 | 0.08590714 | 0.93154024 | 0.97334817 |
| BPTF | 349.869325 | -0.060984827 | 0.19146038 | -0.3185245 | 0.75008709 | 0.89066358 |
| AC005332.7 | 14.3096104 | -0.373408214 | 0.41948009 | -0.8901691 | 0.37337509 | 0.65718609 |
| AMZ2 | 13.2762029 | -0.059434675 | 0.44585718 | -0.1333043 | 0.89395274 | 0.95815794 |
| ARSG | 50.8507025 | 0.181138319 | 0.41222599 | 0.43941509 | 0.66036079 | 0.84143932 |
| WIPI1 | 40.0717236 | -0.575588674 | 0.41922535 | -1.3729815 | 0.16975808 | 0.44037022 |
| PRKAR1A | 197.734694 | -0.137864875 | 0.32148783 | -0.4288339 | 0.66804412 | 0.84558183 |
| FAM20A | 192.530542 | -2.970730311 | 0.43792738 | -6.783614 | 1.17E-11 | 3.66E-09 |
| LINC01482 | 147.87783 | -0.59832285 | 0.66935005 | -0.8938863 | 0.37138272 | 0.65494451 |
| ABCA10 | 48.5264819 | 0.523514681 | 0.35748696 | 1.46443013 | 0.14307646 | 0.40016462 |
| ABCA5 | 160.273387 | 0.672966886 | 0.27607541 | 2.43761982 | 0.01478431 | 0.09928814 |
| MAP2K6 | 18.2601093 | 2.047687769 | 0.74217664 | 2.75903021 | 0.00579732 | 0.05289988 |
| AC005208.1 | 246.613788 | 0.589990155 | 0.66470966 | 0.88759077 | 0.37476092 | 0.65806152 |
| KCNJ16 | 863.984139 | 1.15415262 | 0.32676191 | 3.53209046 | 0.00041229 | 0.00746211 |
| SOX9 | 70.1093218 | -3.736873319 | 0.60628127 | -6.1635968 | 7.11E-10 | 1.31E-07 |
| LINC00511 | 92.1495568 | -4.994976209 | 0.82573397 | -6.0491349 | 1.46E-09 | 2.43E-07 |
| SLC39A11 | 150.944001 | -0.207819416 | 0.28022198 | -0.7416242 | 0.45831503 | 0.72410401 |
| COG1 | 18.761246 | -0.706769974 | 0.35818683 | -1.973188 | 0.04847414 | 0.21350176 |
| FAM104A | 22.0994222 | -0.624782825 | 0.36979207 | -1.6895517 | 0.09111375 | 0.3111216 |
| C17orf80 | 16.8505891 | -0.233637638 | 0.35557794 | -0.6570645 | 0.51113945 | 0.7549696 |
| CDC42EP4 | 26.6240793 | -0.822053699 | 0.39518356 | -2.0801819 | 0.03750885 | 0.18135358 |
| RPL38 | 46.554864 | -0.277171475 | 0.44477062 | -0.6231785 | 0.53316725 | 0.76689222 |
| GPRC5C | 155.615083 | -0.519382045 | 0.23229758 | -2.2358478 | 0.02536174 | 0.14054689 |
| SLC9A3R1 | 23.3049188 | -1.198060896 | 0.47031968 | -2.5473331 | 0.01085498 | 0.08095937 |
| NAT9 | 13.3786151 | -0.633069631 | 0.41844973 | -1.5128929 | 0.13030687 | 0.37846669 |
| TMEM104 | 11.5224074 | 0.31121741 | 0.45675302 | 0.68136914 | 0.49563795 | 0.74780429 |
| CDR2L | 10.3621763 | -0.93116814 | 0.46573706 | -1.999343 | 0.04557125 | 0.20454629 |
| KCTD2 | 30.2985187 | -0.246603954 | 0.29340265 | -0.8404967 | 0.40062996 | 0.68010059 |
| ATP5PD | 40.6664053 | 0.295057812 | 0.35741049 | 0.82554324 | 0.40906325 | 0.6869518 |
| SLC16A5 | 113.38139 | -1.013898085 | 0.40555316 | -2.5000375 | 0.01241802 | 0.08921229 |

| | | | | | | |
|---|---|---|---|---|---|---|
| ARMC7 | 31.6451914 | -0.658832093 | 0.40279897 | -1.635635 | 0.101916 | 0.33058521 |
| NT5C | 32.2666178 | -0.607504473 | 0.38902661 | -1.5616014 | 0.11838193 | 0.35967154 |
| JPT1 | 37.2423472 | -1.344771053 | 0.45209229 | -2.9745499 | 0.00293419 | 0.03195349 |
| SUMO2 | 27.2763187 | -0.336517113 | 0.33353155 | -1.0089514 | 0.31299796 | 0.60565457 |
| NUP85 | 31.3357962 | 0.376596771 | 0.28910995 | 1.30260742 | 0.19270882 | 0.47341565 |
| GGA3 | 35.0166192 | -0.196461203 | 0.31073649 | -0.6322437 | 0.52722761 | 0.76221787 |
| MRPS7 | 25.6807366 | -0.351371652 | 0.41709443 | -0.8424271 | 0.39954893 | 0.67979406 |
| GRB2 | 109.795308 | -0.01935973 | 0.22366977 | -0.086555 | 0.93102526 | 0.97325441 |
| AC011933.4 | 13.802925 | -0.423127807 | 0.59901591 | -0.7063716 | 0.4799571 | 0.73953305 |
| TMEM94 | 29.716313 | 0.386413012 | 0.39753392 | 0.97202525 | 0.33103799 | 0.62074142 |
| CASKIN2 | 17.0855709 | -0.064605342 | 0.40795645 | -0.1583633 | 0.87417051 | 0.94966831 |
| TSEN54 | 23.7249207 | -0.226843844 | 0.30435443 | -0.7453279 | 0.45607352 | 0.72261911 |
| LLGL2 | 371.785717 | -0.158821146 | 0.24884173 | -0.6382416 | 0.52331642 | 0.76218158 |
| MYO15B | 44.8340018 | 0.229277121 | 0.34596466 | 0.66271832 | 0.50751097 | 0.75382716 |
| RECQL5 | 41.4620356 | 0.339511698 | 0.33960093 | 0.99973724 | 0.31743769 | 0.61075159 |
| SMIM5 | 48.0578122 | 2.044292733 | 0.52229718 | 3.91404129 | 9.08E-05 | 0.00224862 |
| SAP30BP | 48.5474908 | -0.047650449 | 0.34526791 | -0.1380101 | 0.89023246 | 0.95702609 |
| ITGB4 | 16.6147732 | -1.939952399 | 0.65650373 | -2.9549754 | 0.00312694 | 0.03365503 |
| H3F3B | 212.101713 | -0.806270045 | 0.42740598 | -1.8864267 | 0.05923748 | 0.24104615 |
| UNK | 70.4445852 | 0.240001678 | 0.29804551 | 0.80525178 | 0.4206744 | 0.69661173 |
| WBP2 | 54.6333324 | -0.647511927 | 0.32467395 | -1.9943452 | 0.04611434 | 0.2061041 |
| TRIM47 | 17.3744225 | -1.716697552 | 0.55378996 | -3.0999073 | 0.00193581 | 0.02399848 |
| TRIM65 | 11.7121547 | -0.413702633 | 0.39029874 | -1.059964 | 0.28916096 | 0.58094924 |
| FBF1 | 8.96380413 | -0.020050503 | 0.50358193 | -0.0398158 | 0.96824 | 0.9875279 |
| ACOX1 | 59.4342259 | -0.0292227 | 0.23273368 | -0.1255628 | 0.90007799 | 0.95980075 |
| EVPL | 19.4333821 | 0.168409883 | 0.39669966 | 0.42452742 | 0.67118121 | 0.84747013 |
| SRP68 | 31.9469962 | 0.176415654 | 0.28347924 | 0.62232301 | 0.5337295 | 0.76708723 |
| EXOC7 | 74.420555 | 0.290334563 | 0.30829901 | 0.94173043 | 0.34633067 | 0.63350472 |
| RNF157 | 10.8777914 | -0.327451485 | 0.47324739 | -0.6919246 | 0.48898471 | 0.74541588 |
| UBALD2 | 30.4551543 | -0.962439724 | 0.42053729 | -2.2885954 | 0.02210287 | 0.12891745 |
| PRPSAP1 | 41.2992884 | 0.611856599 | 0.28772675 | 2.12651967 | 0.03346001 | 0.16897307 |
| UBE2O | 51.5790864 | 0.399088455 | 0.23884314 | 1.67092284 | 0.09473692 | 0.3166788 |
| RHBDF2 | 16.791004 | -0.877593401 | 0.48417063 | -1.8125705 | 0.06989809 | 0.26660131 |
| MXRA7 | 24.2008685 | 0.29146064 | 0.415256 | 0.70188182 | 0.48275286 | 0.74093072 |
| JMJD6 | 15.6151721 | -0.286385049 | 0.46712578 | -0.6130791 | 0.53982405 | 0.77079243 |
| SRSF2 | 75.9145362 | 0.43045458 | 0.28915744 | 1.48865126 | 0.13657923 | 0.38928001 |
| MFSD11 | 46.7821637 | 0.294571099 | 0.25105271 | 1.17334365 | 0.24065803 | 0.53146825 |
| SEC14L1 | 96.3509023 | 0.27618736 | 0.27329425 | 1.01058606 | 0.31221459 | 0.60527563 |
| 9-Sep | 370.079058 | -0.342299628 | 0.34349933 | -0.9965074 | 0.31900366 | 0.611915 |
| TNRC6C | 89.6905658 | 0.308364755 | 0.3192445 | 0.96592032 | 0.33408405 | 0.62271285 |
| TMC6 | 15.6936583 | -1.7019294 | 0.46495622 | -3.6604078 | 0.00025181 | 0.00516771 |
| SYNGR2 | 41.7801641 | -0.511872483 | 0.38454944 | -1.3310967 | 0.1831572 | 0.46004581 |
| AFMID | 57.5292093 | 1.246377786 | 0.2802532 | 4.44732766 | 8.69E-06 | 0.0003269 |
| SOCS3 | 38.3825536 | -3.743256795 | 0.55017648 | -6.8037383 | 1.02E-11 | 3.28E-09 |
| PGS1 | 40.8980034 | -0.605305503 | 0.31507862 | -1.9211253 | 0.05471592 | 0.23054535 |
| CYTH1 | 107.17323 | 0.44468157 | 0.26468532 | 1.68003864 | 0.0929498 | 0.31376482 |
| USP36 | 90.5711008 | 0.654538338 | 0.59252447 | 1.10466044 | 0.26930675 | 0.56160046 |
| TIMP2 | 46.4569696 | -1.209977493 | 0.45277638 | -2.6723512 | 0.00753218 | 0.06349958 |
| LGALS3BP | 95.521811 | -0.87639036 | 0.64927823 | -1.3497917 | 0.17708281 | 0.45189674 |
| CANT1 | 25.4459567 | -0.370133318 | 0.47354267 | -0.7816261 | 0.43443433 | 0.70618229 |
| ENGASE | 14.6305717 | 0.681116984 | 0.43617441 | 1.56157024 | 0.11838926 | 0.35967154 |
| TBC1D16 | 126.335966 | -0.012420256 | 0.33175385 | -0.0374382 | 0.97013564 | 0.98807092 |
| CCDC40 | 40.2293705 | -0.233470077 | 0.3747373 | -0.6230233 | 0.5332692 | 0.76689222 |
| GAA | 47.7768065 | -0.2779165 | 0.44992946 | -0.617689 | 0.53678033 | 0.76921839 |
| EIF4A3 | 20.3402475 | -0.55530069 | 0.3861703 | -1.4379684 | 0.15044302 | 0.41132271 |
| AC087741.1 | 17.2314947 | 0.042206654 | 0.48893009 | 0.08632452 | 0.93120845 | 0.97334714 |
| SGSH | 13.3869434 | -0.250017073 | 0.41562597 | -0.6015434 | 0.54747809 | 0.77516644 |
| SLC26A11 | 37.6912592 | -0.516033271 | 0.30551488 | -1.689061 | 0.09120774 | 0.31117713 |
| RNF213 | 633.065969 | -2.116022944 | 0.51732348 | -4.0903284 | 4.31E-05 | 0.00124655 |
| AC124319.2 | 139.95584 | 0.90633895 | 0.33528166 | 2.70321661 | 0.0068672 | 0.05950031 |
| ENDOV | 19.5685268 | -0.313729497 | 0.35786286 | -0.8766752 | 0.38066307 | 0.66412918 |
| RPTOR | 146.921598 | 0.052185888 | 0.26636269 | 0.19592041 | 0.84467246 | 0.93689787 |
| CHMP6 | 11.0253994 | -0.169114021 | 0.4373189 | -0.3867064 | 0.69897357 | 0.86334649 |
| BAIAP2-DT | 11.0108507 | -0.49046385 | 0.58358674 | -0.8404301 | 0.40066729 | 0.68010059 |
| BAIAP2 | 25.9473328 | 0.499345991 | 0.36388946 | 1.37224637 | 0.16998675 | 0.44055435 |
| TEPSIN | 11.3288058 | 0.082014101 | 0.48298931 | 0.16980521 | 0.86516333 | 0.94694132 |
| NDUFAF8 | 26.3694735 | 0.278281501 | 0.4010598 | 0.69386535 | 0.48776666 | 0.74443126 |
| SLC38A10 | 69.372487 | 0.337373689 | 0.22013404 | 1.53258301 | 0.12537863 | 0.37039755 |
| BAHCC1 | 16.8174948 | 0.792109173 | 0.39938996 | 1.98329768 | 0.04733419 | 0.21000724 |
| ACTG1 | 435.72274 | 0.355145621 | 0.5382563 | 0.65980764 | 0.50937728 | 0.75461129 |
| FAAP100 | 10.8923029 | 0.119156435 | 0.45806349 | 0.26013083 | 0.79476286 | 0.91195838 |
| NPLOC4 | 103.361736 | 0.051447618 | 0.24631033 | 0.20887316 | 0.83454725 | 0.93313642 |
| OXLD1 | 16.294178 | -0.782276738 | 0.3689038 | -2.120544 | 0.03396019 | 0.17053484 |
| CCDC137 | 21.5640273 | -0.907717119 | 0.46504422 | -1.9518942 | 0.05095077 | 0.21990081 |
| ARL16 | 19.9616382 | -0.275565837 | 0.37655555 | -0.7318066 | 0.46428661 | 0.72835095 |
| HGS | 43.8814086 | -0.452786651 | 0.37441837 | -1.2093067 | 0.22654503 | 0.5158857 |
| GCGR | 31.4129979 | 0.976089036 | 0.55151498 | 1.76983231 | 0.07675508 | 0.28119873 |
| MCRIP1 | 26.6861136 | -0.674360412 | 0.34687146 | -1.9441219 | 0.05188075 | 0.22230553 |
| P4HB | 113.69438 | -0.852837986 | 0.24014521 | -3.5513429 | 0.00038327 | 0.00707608 |
| ARHGDIA | 50.918287 | -0.912480827 | 0.49472537 | -1.8444189 | 0.06512211 | 0.2550905 |
| ANAPC11 | 35.7979312 | -0.45671576 | 0.43943286 | -1.03933 | 0.29865128 | 0.59144665 |
| PCYT2 | 13.9100638 | -0.296543939 | 0.67829553 | -0.4371899 | 0.66197365 | 0.84203637 |
| SIRT7 | 16.9055489 | -0.453358284 | 0.40335394 | -1.1239714 | 0.26102517 | 0.55228615 |
| MAFG | 15.7161012 | -1.481473437 | 0.39572495 | -3.7436948 | 0.00018133 | 0.00388379 |
| ASPSCR1 | 25.9506788 | -0.615389779 | 0.37846172 | -1.6260291 | 0.10394346 | 0.33490577 |
| CENPX | 12.9666564 | -0.2569327 | 0.59704676 | -0.4303393 | 0.66694882 | 0.84492213 |
| DCXR | 19.7167195 | 0.413622631 | 0.43809472 | 0.94413973 | 0.34509825 | 0.63256941 |
| RFNG | 18.4702613 | 0.086166806 | 0.43085238 | 0.19999148 | 0.84148724 | 0.93566718 |
| GPS1 | 20.71524 | 0.128068138 | 0.41831293 | 0.30615391 | 0.75948746 | 0.89575355 |
| DUS1L | 23.4093686 | 0.003404108 | 0.48856116 | 0.00696762 | 0.99444069 | 0.99837898 |
| CCDC57 | 122.271278 | 0.05393595 | 0.25896524 | 0.20827486 | 0.83501436 | 0.93320871 |
| SLC16A3 | 44.597908 | -1.444161419 | 0.52398165 | -2.7561297 | 0.00584898 | 0.05327691 |
| CSNK1D | 86.1545697 | -0.406148287 | 0.34226599 | -1.1866452 | 0.23536761 | 0.52547582 |
| OGFOD3 | 26.3808197 | -0.205503754 | 0.34277371 | -0.5995318 | 0.54881828 | 0.77536011 |
| HEXDC | 40.7579582 | 0.123474791 | 0.4114105 | 0.30012552 | 0.76408141 | 0.89757885 |
| CYBC1 | 22.2604025 | -1.295134999 | 0.39527779 | -3.2765185 | 0.00105095 | 0.01524257 |
| NARF | 37.2267213 | -0.693229544 | 0.34370359 | -2.0169401 | 0.04370177 | 0.19956366 |
| FOXK2 | 129.343934 | 0.05814715 | 0.27828566 | 0.2089477 | 0.83448906 | 0.93313642 |
| WDR45B | 81.5304441 | -0.277121444 | 0.32383479 | -0.8557495 | 0.39213638 | 0.67329816 |
| RAB40B | 52.9565122 | -1.36208546 | 0.50439158 | -2.7004524 | 0.00692452 | 0.05984601 |
| FN3KRP | 16.7652121 | -0.639192063 | 0.48435436 | -1.3196785 | 0.18694237 | 0.46440324 |
| FN3K | 41.7642445 | 0.960362831 | 0.49807945 | 1.9281318 | 0.05383873 | 0.22815576 |
| TBCD | 54.2487093 | 0.216511709 | 0.34246667 | 0.63221251 | 0.52724803 | 0.76221787 |
| METRNL | 50.358582 | 0.262869495 | 0.43720762 | 0.60124637 | 0.5476759 | 0.77516644 |
| USP14 | 95.7269609 | 0.177162683 | 0.34691793 | 0.51067606 | 0.60957791 | 0.81344383 |
| THOC1 | 78.0071189 | 0.028400375 | 0.24253461 | 0.11709824 | 0.90678221 | 0.96157183 |
| ENOSF1 | 274.278614 | 0.127973325 | 0.29213978 | 0.43805512 | 0.66134633 | 0.84192458 |
| YES1 | 98.9241004 | -0.487633074 | 0.30977208 | -1.5741673 | 0.11544878 | 0.35464129 |
| METTL4 | 27.9023777 | -0.167938682 | 0.35033589 | -0.4793648 | 0.63167915 | 0.82625173 |
| SMCHD1 | 217.673117 | 0.667897185 | 0.32724176 | 2.04099008 | 0.04125181 | 0.19343477 |
| AP001011.1 | 30.1124712 | -0.158667672 | 0.65497049 | -0.2422516 | 0.80858519 | 0.91854784 |
| LPIN2 | 141.895007 | -0.664963266 | 0.39485283 | -1.6840788 | 0.09216645 | 0.31254072 |
| MYL12A | 96.3425313 | -1.11949523 | 0.35649362 | -3.1402953 | 0.00168778 | 0.02173434 |

| | | | | | | |
|---|---|---|---|---|---|---|
| MYL12B | 202.869526 | -1.051664348 | 0.40242161 | -2.6133396 | 0.00896622 | 0.0711085 |
| TGIF1 | 42.4446809 | -0.763523276 | 0.54250085 | -1.407414 | 0.15930467 | 0.42671782 |
| DLGAP1 | 237.337582 | -4.083286269 | 0.7803291 | -5.2327745 | 1.67E-07 | 1.32E-05 |
| DLGAP1-AS1 | 21.3295744 | -0.584390098 | 0.49705296 | -1.1757099 | 0.23971081 | 0.53016333 |
| LINC00667 | 14.1748134 | 0.133300443 | 0.35653863 | 0.37387377 | 0.70849823 | 0.868892 |
| EPB41L3 | 324.649008 | 1.687586049 | 0.36127726 | 4.67116597 | 2.99E-06 | 0.00013961 |
| L3MBTL4 | 182.391401 | -0.022814078 | 0.35387324 | -0.0644696 | 0.94859629 | 0.97969618 |
| ARHGAP28 | 21.9017706 | 0.82945146 | 0.61926822 | 1.33940583 | 0.18043859 | 0.45661469 |
| LAMA1 | 67.9089494 | -0.002326107 | 0.57215446 | -0.0040655 | 0.99675619 | 0.99868531 |
| PTPRM | 722.007561 | 0.105347216 | 0.15970192 | 0.65964902 | 0.50947909 | 0.75461129 |
| RAB12 | 62.0926071 | -0.056256218 | 0.24612541 | -0.2285673 | 0.81920525 | 0.92352826 |
| MTCL1 | 16.9975324 | -0.334335471 | 0.43272021 | -0.7726366 | 0.43973748 | 0.71072575 |
| NDUFV2 | 93.2545176 | -0.257357396 | 0.30586848 | -0.8413989 | 0.40012451 | 0.68010059 |
| ANKRD12 | 396.088914 | -0.481237602 | 0.20765124 | -2.3175282 | 0.02047498 | 0.1227566 |
| TWSG1 | 159.399223 | 0.749705223 | 0.35280625 | 2.12497719 | 0.03358852 | 0.16945587 |
| AP005432.2 | 12.4713295 | 2.684812387 | 1.0403307 | 2.58072974 | 0.00985917 | 0.07654439 |
| RALBP1 | 391.116798 | 1.032697171 | 0.40211277 | 2.56817801 | 0.01022346 | 0.07837003 |
| PPP4R1 | 137.684269 | -0.167153184 | 0.32882766 | -0.5083307 | 0.61122147 | 0.81385596 |
| RAB31 | 53.4410244 | -0.356745989 | 0.37954278 | -0.9399362 | 0.3472503 | 0.63462349 |
| VAPA | 204.312281 | 0.350009351 | 0.24556674 | 1.42531254 | 0.15406688 | 0.41706471 |
| NAPG | 62.7958643 | 0.094333419 | 0.27941872 | 0.33760594 | 0.73566017 | 0.88426038 |
| GNAL | 28.4834939 | -0.774299359 | 0.51610055 | -1.5002878 | 0.13353987 | 0.38393256 |
| MPPE1 | 42.0302175 | 0.065434544 | 0.31570894 | 0.20726225 | 0.83580506 | 0.93378849 |
| IMPA2 | 95.6700576 | -0.082214206 | 0.3679522 | -0.2234372 | 0.82319526 | 0.92607552 |
| AFG3L2 | 125.824144 | 0.273431983 | 0.26487475 | 1.03230673 | 0.30192845 | 0.59564667 |
| SPIRE1 | 680.557982 | 0.438381965 | 0.30490469 | 1.43776719 | 0.15050012 | 0.41136966 |
| PSMG2 | 98.1497027 | 0.415928737 | 0.23022138 | 1.80664685 | 0.07081736 | 0.26866108 |
| CEP76 | 23.96227 | -0.166498409 | 0.48075101 | -0.3463298 | 0.72909486 | 0.88046679 |
| PTPN2 | 147.394276 | -0.408363817 | 0.35095429 | -1.1635812 | 0.24459377 | 0.53553417 |
| SEH1L | 32.895236 | 0.228733221 | 0.34336115 | 0.66615929 | 0.50530929 | 0.75194227 |
| CEP192 | 138.742506 | 0.440324212 | 0.31742277 | 1.38718535 | 0.16538525 | 0.43431019 |
| LDLRAD4 | 29.9837225 | 0.160108569 | 0.59575867 | 0.26874736 | 0.78812411 | 0.90951658 |
| FAM210A | 54.5509619 | -0.402426843 | 0.38451047 | -1.0465953 | 0.29528628 | 0.58692634 |
| RNMT | 86.4887991 | -0.152941037 | 0.28262886 | -0.5411374 | 0.58841291 | 0.79897585 |
| ROCK1 | 209.177599 | -0.031078946 | 0.18305168 | -0.1697824 | 0.8651813 | 0.94694132 |
| ESCO1 | 79.4062948 | -0.224699066 | 0.29082209 | -0.7726341 | 0.43973895 | 0.71072575 |
| SNRPD1 | 23.0165421 | -0.389573589 | 0.32314221 | -1.2055794 | 0.2279797 | 0.5177793 |
| ABHD3 | 84.014506 | -1.032290156 | 0.59143474 | -1.7453999 | 0.08091528 | 0.29075313 |
| MIB1 | 172.896737 | 0.022304913 | 0.30809248 | 0.07239681 | 0.94228612 | 0.97807875 |
| AC090912.1 | 25.8384198 | -0.727602872 | 0.54658184 | -1.3311874 | 0.18312735 | 0.46004581 |
| RBBP8 | 491.182402 | 0.067022075 | 0.51914578 | 0.12910068 | 0.89727798 | 0.95917897 |
| CABLES1 | 29.7418655 | -2.536360891 | 0.48210875 | -5.2609726 | 1.43E-07 | 1.17E-05 |
| TMEM241 | 31.2074475 | -0.765427694 | 0.28101292 | -2.7238167 | 0.00645323 | 0.0568659 |
| RIOK3 | 103.021837 | -0.466588381 | 0.19672451 | -2.3717857 | 0.01770236 | 0.11160934 |
| RMC1 | 36.5403961 | 0.01377856 | 0.25860256 | 0.05328083 | 0.95750815 | 0.9831804 |
| NPC1 | 131.606146 | -0.792541143 | 0.52030358 | -1.5232283 | 0.1277016 | 0.3747029 |
| TTC39C | 104.708443 | 0.998497202 | 0.32757225 | 3.04817395 | 0.00230237 | 0.02700808 |
| OSBPL1A | 73.0540788 | 0.437261603 | 0.30190635 | 1.44833521 | 0.14752333 | 0.40740971 |
| IMPACT | 18.2948543 | -0.043799833 | 0.38580794 | -0.1135276 | 0.90961232 | 0.96308593 |
| SS18 | 145.341949 | 0.13473371 | 0.23684845 | 0.56886044 | 0.56945086 | 0.78795455 |
| TAF4B | 21.7885199 | 0.164078416 | 0.30228204 | 0.54279908 | 0.58726815 | 0.79812407 |
| KCTD1 | 243.818488 | 0.166536015 | 0.21080509 | 0.78999997 | 0.42952778 | 0.70316149 |
| CHST9 | 24.9585615 | -1.480966095 | 0.60119435 | -2.4633733 | 0.01376365 | 0.0951632 |
| DSC2 | 14.9964547 | -1.243807603 | 0.42676712 | -2.9144879 | 0.00356273 | 0.03701339 |
| DSG2 | 57.721871 | -0.181326129 | 0.36124559 | -0.501947 | 0.61570482 | 0.81655395 |
| B4GALT6 | 22.9951767 | 0.660496259 | 0.43148616 | 1.53074728 | 0.12583186 | 0.37133196 |
| TRAPPC8 | 141.339931 | 0.357358508 | 0.29031473 | 1.23093482 | 0.21834724 | 0.50628253 |
| RNF138 | 46.2313478 | -0.135086666 | 0.2788042 | -0.4845216 | 0.62801571 | 0.82428562 |
| GAREM1 | 769.873416 | 0.222249149 | 0.51005409 | 0.43573643 | 0.66302797 | 0.84275734 |
| KLHL14 | 41.5825846 | 0.503343409 | 0.51525803 | 0.97687639 | 0.32863035 | 0.61768835 |
| CCDC178 | 436.055751 | 1.108747537 | 0.52106018 | 2.12786847 | 0.03334799 | 0.16865537 |
| ASXL3 | 138.396139 | 1.114727365 | 0.30830468 | 3.61566797 | 0.00029957 | 0.00594646 |
| DTNA | 31.5752069 | -1.538327154 | 0.55038928 | -2.7949802 | 0.00519029 | 0.04891142 |
| MAPRE2 | 27.4072378 | 0.16117407 | 0.39717009 | 0.40580617 | 0.68488503 | 0.85635073 |
| ZNF397 | 49.3169813 | -0.156682723 | 0.28376967 | -0.5521475 | 0.58084728 | 0.79477868 |
| ZSCAN30 | 39.3607573 | -0.198004319 | 0.30394712 | -0.6514433 | 0.51476035 | 0.75684546 |
| ZNF24 | 46.4066756 | -0.558238504 | 0.3003374 | -1.8587046 | 0.06306903 | 0.24979235 |
| GALNT1 | 64.1365983 | -0.086486895 | 0.28889585 | -0.2993705 | 0.76465736 | 0.89782313 |
| RPRD1A | 142.048068 | 0.203140934 | 0.2300616 | 0.88298495 | 0.3772444 | 0.66022373 |
| SLC39A6 | 51.1435847 | -0.316031723 | 0.41260267 | -0.7659469 | 0.44370791 | 0.71343168 |
| ELP2 | 95.93948 | 0.423198956 | 0.22119497 | 1.91323947 | 0.0557174 | 0.23295482 |
| MOCOS | 97.9713577 | 0.031496891 | 0.45071334 | 0.06988231 | 0.94428733 | 0.97856665 |
| FHOD3 | 823.322571 | 1.683085959 | 0.19975386 | 8.42579925 | 3.58E-17 | 3.36E-14 |
| TPGS2 | 44.3753791 | -1.084909368 | 0.29008354 | -3.7399894 | 0.00018403 | 0.00391706 |
| KIAA1328 | 167.62801 | 0.037358629 | 0.25975311 | 0.14382361 | 0.88563976 | 0.95619452 |
| AC016205.1 | 25.4466871 | -1.477774184 | 0.52292959 | -2.8259525 | 0.00471403 | 0.04559989 |
| PIK3C3 | 153.506605 | 0.275510063 | 0.20953665 | 1.31485381 | 0.18855904 | 0.46729257 |
| LINC00907 | 32.6482771 | -0.560245951 | 0.33316215 | -1.6816014 | 0.09264615 | 0.31323946 |
| SETBP1 | 985.040337 | 0.88940083 | 0.14726865 | 6.03930859 | 1.55E-09 | 2.52E-07 |
| SLC14A2 | 161.448113 | -0.865030571 | 1.05302706 | -0.8214704 | 0.41137837 | 0.68899849 |
| EPG5 | 92.1506202 | 0.166116379 | 0.21600721 | 0.76903163 | 0.44187453 | 0.71239303 |
| PSTPIP2 | 14.0582333 | -2.086946249 | 0.59519591 | -3.5063182 | 0.00045435 | 0.00805634 |
| ATP5F1A | 134.508675 | 0.382937303 | 0.33185794 | 1.15391936 | 0.24853323 | 0.53959734 |
| HAUS1 | 13.5441487 | 0.135475847 | 0.46759212 | 0.28973082 | 0.77202218 | 0.90105761 |
| C18orf25 | 70.9715036 | 0.389151901 | 0.28978467 | 1.34290022 | 0.17930429 | 0.45495583 |
| ST8SIA5 | 21.2946135 | 0.283352159 | 0.59576916 | 0.4756073 | 0.63435418 | 0.82785872 |
| PIAS2 | 195.818767 | 0.208193426 | 0.2839324 | 0.73324997 | 0.46340598 | 0.72812334 |
| KATNAL2 | 149.169047 | 1.391668961 | 0.31143371 | 4.46858806 | 7.87E-06 | 0.00030726 |
| HDHD2 | 11.0773407 | 0.502251222 | 0.46830782 | 1.07248095 | 0.28350406 | 0.57629417 |
| SMAD2 | 237.125145 | -0.357744302 | 0.37664316 | -0.9498229 | 0.34220224 | 0.63022262 |
| ZBTB7C | 47.3731797 | -1.864048138 | 0.76864716 | -2.4251025 | 0.01530407 | 0.10149612 |
| CTIF | 83.5506464 | 0.45115757 | 0.30799217 | 1.46483456 | 0.14296606 | 0.40000987 |
| SMAD7 | 46.0868924 | 1.703533756 | 0.26265847 | 6.48573702 | 8.83E-11 | 2.07E-08 |
| DYM | 326.196078 | 0.356830906 | 0.2125223 | 1.67902805 | 0.09314658 | 0.31431249 |
| LIPG | 109.537878 | -2.745181956 | 0.78390843 | -3.5019166 | 0.00046192 | 0.00813175 |
| ACAA2 | 43.3790899 | -0.394712956 | 0.27985504 | -1.4104193 | 0.1584159 | 0.42480987 |
| SNHG22 | 28.621335 | -0.6377011 | 0.3384735 | -1.8840503 | 0.05955817 | 0.24171358 |
| MYO5B | 167.388954 | -0.342136639 | 0.25141277 | -1.3608562 | 0.17355913 | 0.44612537 |
| MBD1 | 29.1484569 | 0.115714387 | 0.33759907 | 0.34275683 | 0.73178141 | 0.88245488 |
| CXXC1 | 18.0946209 | 0.30653457 | 0.49644554 | 0.6174586 | 0.53693228 | 0.76932911 |
| MAPK4 | 69.362746 | 2.149508819 | 0.41794525 | 5.14303918 | 2.70E-07 | 1.95E-05 |
| ME2 | 37.676768 | -0.218709005 | 0.27799762 | -0.7867298 | 0.43144006 | 0.70454366 |
| AC091551.1 | 14.7314836 | -0.622436832 | 0.52633836 | -1.1825793 | 0.23697593 | 0.52743022 |
| SMAD4 | 56.0230459 | 0.089828775 | 0.28671302 | 0.31330553 | 0.75404855 | 0.89238252 |
| MEX3C | 17.9811406 | -0.375603098 | 0.38748562 | -0.9693343 | 0.33237842 | 0.62173226 |
| MBD2 | 148.20318 | -1.044588498 | 0.29704336 | -3.5166196 | 0.00043708 | 0.0078038 |
| POLI | 54.8381154 | 0.10974368 | 0.27437366 | 0.39997891 | 0.68917205 | 0.85868974 |
| TCF4 | 75.8472443 | 0.493005727 | 0.46327065 | 1.0641851 | 0.28724485 | 0.5796994 |
| TXNL1 | 54.3211788 | 0.110476195 | 0.26677303 | 0.41412056 | 0.67878582 | 0.85341122 |
| WDR7 | 172.835404 | 0.523614734 | 0.15009743 | 3.48849888 | 0.00048574 | 0.00841026 |
| FECH | 24.4495465 | 0.872160747 | 0.36241202 | 2.40654476 | 0.01610423 | 0.1047385 |
| NARS | 82.5094335 | -0.659918162 | 0.32521417 | -2.0291803 | 0.04243993 | 0.19659002 |
| AC027097.2 | 41.5123337 | -0.561659079 | 0.43404665 | -1.2940063 | 0.19566326 | 0.47709417 |

| | | | | | | |
|---|---|---|---|---|---|---|
| ATP8B1 | 179.831358 | 0.17924147 | 0.27075433 | 0.66200778 | 0.50796624 | 0.75382716 |
| NEDD4L | 729.096819 | 1.195030856 | 0.25068224 | 4.76711407 | 1.87E-06 | 9.35E-05 |
| MALT1 | 129.124514 | -0.642188597 | 0.41834993 | -1.5350513 | 0.12477123 | 0.37001028 |
| AC104365.1 | 14.7771027 | -0.159828117 | 0.66508009 | -0.2403141 | 0.81008677 | 0.91928664 |
| ZNF532 | 127.066113 | -0.134199219 | 0.28401341 | -0.4725102 | 0.6365627 | 0.82975756 |
| SEC11C | 20.4505489 | 0.262418492 | 0.36897904 | 0.71120163 | 0.4769593 | 0.73678733 |
| LMAN1 | 93.1653843 | -0.174059343 | 0.37629739 | -0.4625579 | 0.64368126 | 0.83339931 |
| CDH20 | 24.9096079 | 0.517666018 | 0.91516159 | 0.56565531 | 0.57162812 | 0.78910141 |
| RNF152 | 103.04132 | 0.924426908 | 0.48472694 | 1.90710856 | 0.05650653 | 0.23463534 |
| PIGN | 117.88091 | 0.330721458 | 0.18172858 | 1.8198649 | 0.06877958 | 0.2641936 |
| KIAA1468 | 168.281271 | 0.36740387 | 0.17261051 | 2.12851394 | 0.0332945 | 0.16856421 |
| ZCCHC2 | 73.7044336 | 0.303809115 | 0.25333692 | 1.19922951 | 0.23043871 | 0.52060957 |
| PHLPP1 | 124.031587 | 0.089583462 | 0.24675879 | 0.36304061 | 0.71657455 | 0.8732017 |
| BCL2 | 224.497629 | 0.814933787 | 0.40196231 | 2.02738857 | 0.04262269 | 0.19708211 |
| KDSR | 62.0520719 | -1.014272569 | 0.29286828 | -3.463238 | 0.00053372 | 0.00899892 |
| VPS4B | 69.7536766 | -0.346189176 | 0.20750381 | -1.6683509 | 0.09524609 | 0.31779803 |
| LINC01924 | 164.087587 | -2.200168221 | 0.42762463 | -5.1450924 | 2.67E-07 | 1.95E-05 |
| TMX3 | 77.3402017 | 0.278570834 | 0.22899858 | 1.21647408 | 0.22380438 | 0.51293275 |
| CCDC102B | 35.8078643 | 0.97581544 | 0.53881312 | 1.81104618 | 0.0701337 | 0.26694834 |
| RTTN | 64.3108951 | 0.36639055 | 0.24564196 | 1.49156337 | 0.13581365 | 0.38804335 |
| SOCS6 | 34.0240361 | 0.002969514 | 0.31501364 | 0.00942662 | 0.99247876 | 0.99770844 |
| TIMM21 | 15.4410319 | 0.512433211 | 0.40894355 | 1.25306588 | 0.21018173 | 0.49648138 |
| CYB5A | 175.264421 | -0.048243344 | 0.44768961 | -0.1077607 | 0.91418552 | 0.96462448 |
| CNDP2 | 125.929486 | -0.935892748 | 0.33541909 | -2.7902191 | 0.00526724 | 0.04943821 |
| LINC00909 | 13.4980654 | -0.556350494 | 0.43038296 | -1.2926871 | 0.1961193 | 0.47764091 |
| ZNF407 | 257.073687 | 0.30986072 | 0.22104526 | 1.40179765 | 0.16097568 | 0.42818783 |
| ZADH2 | 33.5230048 | -0.047548813 | 0.34420622 | -0.1381405 | 0.89012939 | 0.95702609 |
| TSHZ1 | 50.6697097 | 0.031633374 | 0.32994899 | 0.09587353 | 0.92362102 | 0.96954797 |
| ZNF516 | 172.951383 | 0.330670477 | 0.2927093 | 1.12968901 | 0.25860729 | 0.55002264 |
| ZNF236-DT | 11.4424809 | 0.374614042 | 0.41884976 | 0.89438763 | 0.37111453 | 0.65494451 |
| ZNF236 | 85.6378904 | 0.358598487 | 0.21335781 | 1.68073759 | 0.0928139 | 0.31370366 |
| MBP | 226.55563 | -0.374556077 | 0.21304755 | -1.7580868 | 0.07873274 | 0.28580152 |
| LINC01896 | 25.2899729 | 1.653153393 | 0.53967428 | 3.06324288 | 0.00218952 | 0.02597102 |
| SALL3 | 33.6441844 | 1.51584155 | 0.46482989 | 3.26106728 | 0.00110994 | 0.01572843 |
| ATP9B | 281.304933 | 0.551900947 | 0.20697788 | 2.66647307 | 0.00766517 | 0.06391579 |
| NFATC1 | 23.3336743 | 0.091886146 | 0.37935255 | 0.24221835 | 0.80861098 | 0.91854784 |
| CTDP1 | 28.6057932 | -0.120842378 | 0.30898364 | -0.3910964 | 0.69572601 | 0.86236044 |
| PQLC1 | 36.3977048 | 0.129926396 | 0.27704733 | 0.46896823 | 0.63909234 | 0.83119926 |
| HSBP1L1 | 43.2269611 | -0.336047482 | 0.29830392 | -1.1265272 | 0.25994243 | 0.55124061 |
| TXNL4A | 61.4464762 | -0.239572951 | 0.248493 | -0.9641034 | 0.33499407 | 0.62350658 |
| ADNP2 | 56.5022981 | 0.094993105 | 0.31030055 | 0.30613257 | 0.75950371 | 0.89575355 |
| C20orf96 | 17.185022 | 0.517725741 | 0.43565484 | 1.18838516 | 0.2346817 | 0.52479917 |
| ZCCHC3 | 15.0681947 | -0.630857958 | 0.54908311 | -1.1489298 | 0.25058492 | 0.54232519 |
| NRSN2-AS1 | 94.3425123 | 0.429024922 | 0.45222422 | 0.94869957 | 0.34277344 | 0.63072032 |
| SOX12 | 19.6343624 | 0.091595965 | 0.50154975 | 0.18262588 | 0.85509157 | 0.94225653 |
| NRSN2 | 17.8303093 | -0.714370841 | 0.47840506 | -1.4932343 | 0.13537587 | 0.38729303 |
| RBCK1 | 65.8349617 | -0.568218989 | 0.34165063 | -1.663158 | 0.09628081 | 0.32016943 |
| TBC1D20 | 29.0826209 | -0.304459685 | 0.30523919 | -0.9974463 | 0.31854795 | 0.61144847 |
| CSNK2A1 | 199.93846 | 0.550366042 | 0.18476957 | 2.97866164 | 0.0028951 | 0.03178629 |
| SLC52A3 | 67.5501065 | -1.072814692 | 0.3727542 | -2.8780754 | 0.00400109 | 0.04017474 |
| PSMF1 | 78.5311793 | -0.236451834 | 0.23821997 | -0.9925777 | 0.32091579 | 0.6120836 |
| AL136531.2 | 37.1606554 | -0.707312448 | 0.38162267 | -1.853434 | 0.06382017 | 0.25161708 |
| SDCBP2-AS1 | 52.9751868 | 0.332587488 | 0.24137882 | 1.37786526 | 0.16824489 | 0.43868866 |
| NSFL1C | 36.5599708 | -0.822332054 | 0.29178493 | -2.8182814 | 0.00482815 | 0.04630755 |
| SIRPA | 10.1068177 | -1.440336817 | 0.65758271 | -2.1903508 | 0.02849881 | 0.15204283 |
| STK35 | 29.7917759 | -0.73496248 | 0.31400188 | -2.3406308 | 0.01925119 | 0.11805107 |
| SNRPB | 38.1764292 | -1.082731647 | 0.40029757 | -2.7048169 | 0.0068342 | 0.05928681 |
| NOP56 | 19.9780975 | -0.632126031 | 0.4389233 | -1.4401743 | 0.14981811 | 0.41046846 |
| IDH3B | 18.7906482 | 0.339755376 | 0.35990692 | 0.94400902 | 0.34516504 | 0.63256941 |
| EBF4 | 18.6247192 | 0.093420627 | 0.45145097 | 0.20693416 | 0.83606128 | 0.93397346 |
| PCED1A | 21.1061579 | 1.008549133 | 0.33026766 | 3.05373261 | 0.00226013 | 0.02661018 |
| PTPRA | 203.040318 | 0.095561604 | 0.21712451 | 0.44012352 | 0.65984765 | 0.8409935 |
| MRPS26 | 30.0198425 | -0.380649726 | 0.41763335 | -0.9114447 | 0.36206109 | 0.64626475 |
| UBOX5-AS1 | 15.1645355 | -0.092910767 | 0.58345963 | -0.1592411 | 0.87347891 | 0.94942996 |
| UBOX5 | 17.8693071 | 0.513832577 | 0.33834499 | 1.51866468 | 0.12884692 | 0.37613516 |
| LZTS3 | 21.0604818 | -0.526717572 | 0.35123397 | -1.4996202 | 0.1337128 | 0.38413532 |
| DDRGK1 | 30.0121066 | -0.60991547 | 0.2970089 | -2.0535259 | 0.0400216 | 0.18956438 |
| ITPA | 43.3977855 | -0.588777509 | 0.27648066 | -2.1295432 | 0.03320935 | 0.16836747 |
| C20orf194 | 438.29426 | 0.377497175 | 0.22863792 | 1.65106986 | 0.09872431 | 0.32452388 |
| ATRN | 178.959907 | 0.551472546 | 0.18523816 | 2.97710017 | 0.00290989 | 0.03182344 |
| CENPB | 26.7587738 | -0.894050935 | 0.47230619 | -1.8929477 | 0.05836482 | 0.23888533 |
| CDC25B | 12.9875392 | -0.867514974 | 0.5076385 | -1.7089227 | 0.08746526 | 0.30420903 |
| MAVS | 75.3160825 | -0.052715609 | 0.30679629 | -0.1718261 | 0.86357425 | 0.94635064 |
| PANK2 | 67.7559085 | -0.332422472 | 0.23237407 | -1.4305489 | 0.15255954 | 0.41441825 |
| RNF24 | 232.280138 | -1.289953651 | 0.36972352 | -3.4889683 | 0.00048489 | 0.00840963 |
| PRNP | 21.3331775 | -0.953175264 | 0.34958212 | -2.7266133 | 0.0063988 | 0.05653552 |
| SLC23A2 | 77.3174981 | 0.457287651 | 0.31377859 | 1.45735774 | 0.14501766 | 0.40290504 |
| TMEM230 | 59.3350252 | -0.117972792 | 0.25291385 | -0.4664545 | 0.64089024 | 0.83230084 |
| PCNA | 16.7173068 | -0.223863388 | 0.51642198 | -0.4334893 | 0.66465936 | 0.84348349 |
| CDS2 | 75.98684 | -0.426584372 | 0.30699064 | -1.389568 | 0.16466009 | 0.43373261 |
| GPCPD1 | 137.950997 | 0.424704389 | 0.38339516 | 1.10774582 | 0.2679716 | 0.55994436 |
| SHLD1 | 44.9589869 | 0.850141612 | 0.36150369 | 2.3516817 | 0.01868876 | 0.11570409 |
| TRMT6 | 11.806459 | 0.219411524 | 0.48449175 | 0.45286948 | 0.65064273 | 0.83735207 |
| MCM8 | 13.9506019 | 0.246334282 | 0.38424032 | 0.64109431 | 0.52146141 | 0.76102783 |
| CRLS1 | 71.6007112 | 0.113249734 | 0.25863505 | 0.43787466 | 0.66147715 | 0.84192458 |
| TMX4 | 48.3929518 | -0.255150124 | 0.35887376 | -0.7109746 | 0.47709998 | 0.73689415 |
| PLCB1 | 61.5126328 | -0.37914181 | 0.63876319 | -0.5935561 | 0.55280902 | 0.77787735 |
| PLCB4 | 151.285196 | -0.02991102 | 0.462049 | -0.0647356 | 0.94838452 | 0.97967084 |
| ANKEF1 | 10.6171885 | -0.271255782 | 0.44402215 | -0.610906 | 0.54126183 | 0.77188253 |
| SNAP25 | 22.6410234 | -2.668858771 | 1.0101251 | -2.6421072 | 0.0082392 | 0.06735772 |
| MKKS | 22.0717778 | 0.621268756 | 0.32828986 | 1.89243965 | 0.05843243 | 0.23897216 |
| SLX4IP | 147.86707 | 0.723519788 | 0.26572558 | 2.7228082 | 0.00647296 | 0.056898 |
| JAG1 | 61.2964324 | -0.309457958 | 0.38952096 | -0.7944578 | 0.42692898 | 0.70162975 |
| AL050403.2 | 43.3166356 | 0.515257065 | 0.40713766 | 1.26555981 | 0.20567072 | 0.48999801 |
| BTBD3 | 90.6322203 | -0.224109032 | 0.22573247 | -0.9928081 | 0.32080346 | 0.6120836 |
| SPTLC3 | 91.9149631 | -0.740937618 | 0.23016002 | -3.2192281 | 0.00128536 | 0.01763222 |
| TASP1 | 184.677426 | -0.066956746 | 0.19027548 | -0.3518937 | 0.72491797 | 0.87808406 |
| ESF1 | 72.0930051 | 0.286596415 | 0.28050481 | 1.02171659 | 0.30691506 | 0.60139222 |
| NDUFAF5 | 49.985211 | 0.743279715 | 0.3403396 | 2.18393546 | 0.02896699 | 0.15382368 |
| MACROD2 | 666.319209 | 0.323748627 | 0.31283086 | 1.0348999 | 0.30071566 | 0.59462049 |
| FLRT3 | 23.3620527 | -1.524278218 | 0.44966341 | -3.3898204 | 0.00069938 | 0.0112579 |
| KIF16B | 208.975575 | 0.622343488 | 0.24935102 | 2.49585298 | 0.01256547 | 0.09002047 |
| SNRPB2 | 25.5143398 | 0.188653736 | 0.37808655 | 0.49896971 | 0.61780073 | 0.81764942 |
| DSTN | 242.968122 | -0.508205267 | 0.30020405 | -1.6928662 | 0.09048095 | 0.30988521 |
| RRBP1 | 107.925125 | -0.920067621 | 0.2218196 | -4.1478194 | 3.36E-05 | 0.00100815 |
| SNX5 | 39.619328 | 0.05370004 | 0.33289056 | 0.16131439 | 0.87184579 | 0.94844301 |
| OVOL2 | 37.5464115 | -0.402598211 | 0.41852414 | -0.9619474 | 0.33607603 | 0.62450491 |
| MGME1 | 41.2203864 | 0.320341206 | 0.27105264 | 1.1818413 | 0.23726868 | 0.5276258 |
| KAT14 | 13.3679499 | 0.385364762 | 0.44546395 | 0.8650863 | 0.38699142 | 0.66937301 |
| ZNF133 | 48.6165752 | 0.19816449 | 0.27974436 | 0.70837706 | 0.47871113 | 0.7383863 |
| DZANK1 | 28.6134831 | -0.543481348 | 0.30634662 | -1.7740733 | 0.07605103 | 0.2801136 |
| POLR3F | 12.2279362 | -0.233647967 | 0.40095413 | -0.5827299 | 0.56007513 | 0.78316119 |
| RBBP9 | 11.0498636 | 0.064088066 | 0.58263506 | 0.10999693 | 0.91241181 | 0.96436509 |

| | | | | | | |
|---|---|---|---|---|---|---|
| SEC23B | 68.9807805 | 0.086928677 | 0.24304522 | 0.35766463 | 0.72059431 | 0.87583324 |
| DTD1 | 14.348339 | -0.648899654 | 0.36947154 | -1.7562913 | 0.07903868 | 0.28650827 |
| SLC24A3 | 65.5198989 | -0.221404058 | 0.73387426 | -0.3016921 | 0.7628868 | 0.89739726 |
| AL121761.1 | 50.2129291 | 0.150785458 | 0.42273335 | 0.35669165 | 0.72132266 | 0.87614353 |
| RIN2 | 213.136441 | -0.256521376 | 0.24447 | -1.0492959 | 0.29404194 | 0.58592264 |
| NAA20 | 72.4535474 | -0.049532857 | 0.28697217 | -0.1726051 | 0.86296185 | 0.94620532 |
| CRNKL1 | 18.3395851 | -0.259753723 | 0.33369227 | -0.778423 | 0.43631971 | 0.70775777 |
| RALGAPA2 | 336.398616 | -1.19754099 | 0.20357162 | -5.882652 | 4.04E-09 | 5.62E-07 |
| KIZ | 189.222994 | 0.448601031 | 0.33272481 | 1.34826445 | 0.17757334 | 0.45230354 |
| KIZ-AS1 | 38.2468917 | -0.334510692 | 0.48997839 | -0.682705 | 0.49479328 | 0.74763279 |
| XRN2 | 112.969268 | -0.155628198 | 0.27694998 | -0.5619361 | 0.57415954 | 0.7910707 |
| NAPB | 51.2192515 | 0.289807578 | 0.29746743 | 0.97424977 | 0.32993254 | 0.61943102 |
| CST3 | 129.446957 | -1.430199609 | 0.54649135 | -2.6170581 | 0.00886912 | 0.07069728 |
| APMAP | 33.2707268 | -0.729355425 | 0.47712702 | -1.52864 | 0.12635372 | 0.37228594 |
| ACSS1 | 110.276711 | 0.419746183 | 0.30873328 | 1.35957545 | 0.17396431 | 0.44651726 |
| ENTPD6 | 36.7086658 | -0.163614075 | 0.28739262 | -0.5693051 | 0.56914913 | 0.78790646 |
| PYGB | 50.7879338 | -0.214498053 | 0.32398022 | -0.6620714 | 0.50792543 | 0.75382716 |
| ABHD12 | 79.2521458 | 0.134563397 | 0.22291677 | 0.60364862 | 0.54607728 | 0.77463416 |
| NINL | 232.549377 | 0.530910804 | 0.33897143 | 1.56624055 | 0.11729231 | 0.35749864 |
| ZNF337-AS1 | 28.0170977 | 0.952859263 | 0.40087836 | 2.37692868 | 0.01745746 | 0.11067494 |
| ZNF337 | 34.654942 | -0.128034055 | 0.31695431 | -0.4039511 | 0.68624865 | 0.85756108 |
| FAM182B | 44.1749038 | -0.025876083 | 0.40472462 | -0.063935 | 0.94902196 | 0.97983806 |
| HM13 | 99.4569869 | -0.576677994 | 0.23391355 | -2.4653467 | 0.01368807 | 0.09502327 |
| ID1 | 74.2034002 | -0.788403279 | 0.70448967 | -1.1191126 | 0.2630921 | 0.55472264 |
| BCL2L1 | 284.184922 | 0.790544946 | 0.3505985 | 2.25484408 | 0.02414312 | 0.13583965 |
| AL117381.1 | 49.8311751 | -0.070154373 | 0.59695772 | -0.1175198 | 0.90644813 | 0.96157183 |
| DUSP15 | 36.9574107 | 1.088688312 | 0.60507876 | 1.79925059 | 0.07197905 | 0.27082841 |
| TTLL9 | 12.6461853 | 0.576527101 | 0.43655357 | 1.32063313 | 0.18662372 | 0.46408993 |
| PDRG1 | 11.840912 | -0.942158215 | 0.46425144 | -2.0294136 | 0.04241618 | 0.19656835 |
| TM9SF4 | 77.0292867 | 0.136391316 | 0.22621431 | 0.60292964 | 0.54655549 | 0.77463416 |
| POFUT1 | 20.4144665 | -0.298767398 | 0.47733032 | -0.6259133 | 0.53137181 | 0.76540721 |
| KIF3B | 94.1198342 | 0.033317483 | 0.23380845 | 0.14249905 | 0.88668583 | 0.95619452 |
| ASXL1 | 215.319757 | 0.062035236 | 0.22896914 | 0.27093274 | 0.78644277 | 0.90849219 |
| NOL4L | 40.9311822 | -0.353806305 | 0.42274304 | -0.8369299 | 0.40263195 | 0.68128594 |
| MAPRE1 | 33.8260206 | -0.186662892 | 0.35063084 | -0.532363 | 0.59447459 | 0.80307858 |
| AL035071.1 | 10.0463439 | 0.433562078 | 0.50947625 | 0.85099566 | 0.39477176 | 0.67534117 |
| CDK5RAP1 | 38.697338 | 0.456665428 | 0.23922696 | 1.90892128 | 0.05627225 | 0.23427923 |
| CBFA2T2 | 307.641166 | 0.218080246 | 0.20466157 | 1.06556519 | 0.28662024 | 0.57920003 |
| NECAB3 | 13.6479457 | 0.990194465 | 0.4201225 | 2.35691842 | 0.01842729 | 0.11442917 |
| PXMP4 | 15.6924569 | 0.608319396 | 0.44707353 | 1.36066968 | 0.1736181 | 0.44614958 |
| CHMP4B | 56.9966873 | -0.418500748 | 0.24845995 | -1.6843791 | 0.09210842 | 0.31254072 |
| RALY | 103.480895 | -0.13263461 | 0.22013072 | -0.6025266 | 0.54682368 | 0.77463416 |
| EIF2S2 | 72.9631201 | -1.175494662 | 0.32082281 | -3.6639996 | 0.00024831 | 0.00511612 |
| AHCY | 16.7707008 | 0.072437561 | 0.40381865 | 0.17938142 | 0.85763822 | 0.94359207 |
| ITCH | 254.684319 | 0.305283465 | 0.19916375 | 1.53282646 | 0.12531862 | 0.37039755 |
| DYNLRB1 | 83.4449386 | -0.743013974 | 0.37823678 | -1.9644149 | 0.049482 | 0.21605779 |
| MAP1LC3A | 26.5014847 | 0.626272683 | 0.587433 | 1.06611765 | 0.28637046 | 0.57892239 |
| PIGU | 57.5095068 | 0.845292775 | 0.31618369 | 2.6734231 | 0.00750815 | 0.06334885 |
| NCOA6 | 171.060463 | 0.337992905 | 0.25036954 | 1.34997615 | 0.17702363 | 0.45185765 |
| TP53INP2 | 31.8183125 | 0.867893662 | 0.59221576 | 1.46550247 | 0.14278388 | 0.39971726 |
| GGT7 | 60.7935367 | 0.449787578 | 0.28135308 | 1.59865879 | 0.10989644 | 0.34550245 |
| ACSS2 | 66.6426511 | 0.619591362 | 0.2833932 | 2.1863311 | 0.02879139 | 0.15311173 |
| GSS | 24.4956996 | -0.7400681 | 0.29476699 | -2.5106885 | 0.0120496 | 0.08729603 |
| TRPC4AP | 142.964414 | 0.131099691 | 0.24267232 | 0.54023339 | 0.58903609 | 0.79925045 |
| EDEM2 | 34.2966063 | 0.030616764 | 0.30309363 | 0.10101421 | 0.91953918 | 0.96763053 |
| PROCR | 15.4121412 | -0.360329593 | 0.64064166 | -0.5624511 | 0.57380873 | 0.79092554 |
| MMP24OS | 50.2896685 | 0.244022233 | 0.33217787 | 0.73461316 | 0.46257512 | 0.72740482 |
| MMP24 | 44.0115588 | 0.532718349 | 0.38360723 | 1.38870779 | 0.16492163 | 0.43375609 |
| UQCC1 | 96.5614844 | 0.558899476 | 0.18780053 | 2.97602717 | 0.00292009 | 0.03186734 |
| CEP250 | 24.2159912 | -1.283531545 | 0.53504799 | -2.3989092 | 0.01644399 | 0.10634401 |
| ERGIC3 | 45.885273 | 0.248265566 | 0.31923869 | 0.77768008 | 0.43675764 | 0.70824448 |
| CPNE1 | 40.4251521 | 0.231799981 | 0.28098375 | 0.82495868 | 0.40939505 | 0.68723933 |
| NFS1 | 10.7498449 | 0.624597551 | 0.423503 | 1.47483618 | 0.14025659 | 0.39525538 |
| ROMO1 | 31.4118357 | -0.787369401 | 0.51919718 | -1.5165132 | 0.12938962 | 0.37713756 |
| RBM39 | 334.67541 | 0.218487167 | 0.19454087 | 1.12309135 | 0.2613987 | 0.55262249 |
| PHF20 | 155.657216 | -0.028652349 | 0.31015227 | -0.0923816 | 0.9263949 | 0.97111125 |
| SCAND1 | 57.1119145 | -1.165592039 | 0.38893244 | -2.9969011 | 0.00272739 | 0.03069808 |
| NORAD | 164.986863 | -0.267699052 | 0.24262666 | -1.1033373 | 0.26988071 | 0.56184753 |
| EPB41L1 | 386.925239 | 0.414568456 | 0.24146221 | 1.71690822 | 0.08599594 | 0.3013368 |
| AAR2 | 20.1291776 | 0.057403077 | 0.39447274 | 0.14551849 | 0.88430152 | 0.95603676 |
| DLGAP4 | 82.9007798 | -0.107057381 | 0.26912778 | -0.3977939 | 0.69078213 | 0.85967888 |
| DLGAP4-AS1 | 40.674843 | 0.314353765 | 0.30900389 | 1.0173133 | 0.30900441 | 0.6027062 |
| MYL9 | 13.0481762 | 0.281416809 | 0.56796296 | 0.49548444 | 0.6202582 | 0.81932299 |
| RAB5IF | 15.3739868 | -0.720827488 | 0.45881783 | -1.5710538 | 0.11617016 | 0.35543425 |
| NDRG3 | 38.0079404 | 0.482679706 | 0.25486782 | 1.89384327 | 0.05824582 | 0.23858783 |
| SOGA1 | 51.36429 | 0.336312923 | 0.31402475 | 1.07097583 | 0.2841803 | 0.5765312 |
| SAMHD1 | 19.5643254 | -0.984162446 | 0.90347574 | -1.089307 | 0.27601853 | 0.56798847 |
| RBL1 | 27.1629358 | 0.529819258 | 0.34086942 | 1.55431737 | 0.12010873 | 0.36286222 |
| RPN2 | 78.275623 | 0.040645167 | 0.31951127 | 0.12721043 | 0.89877385 | 0.95935794 |
| MANBAL | 20.6831994 | -0.44736187 | 0.3521551 | -1.2703547 | 0.20395833 | 0.4882179 |
| SRC | 32.050193 | -0.007383673 | 0.28375866 | -0.026021 | 0.97924062 | 0.99214564 |
| BLCAP | 87.2687843 | -0.044477114 | 0.27564666 | -0.1613555 | 0.87181339 | 0.94844301 |
| CTNNBL1 | 57.3329752 | 0.262186271 | 0.22507123 | 1.16490355 | 0.24405804 | 0.5346132 |
| TTI1 | 30.6215301 | -0.00891826 | 0.36663987 | -0.0243243 | 0.98059393 | 0.99302847 |
| RPRD1B | 73.1988939 | 0.381489021 | 0.22280649 | 1.71219887 | 0.08686003 | 0.30295601 |
| TGM2 | 90.9645897 | -3.950689656 | 0.59848618 | -6.6011376 | 4.08E-11 | 1.11E-08 |
| RALGAPB | 135.736451 | 0.365407592 | 0.25722121 | 1.42059664 | 0.15543405 | 0.41907397 |
| DHX35 | 32.8764512 | 0.630472099 | 0.30883198 | 2.04147285 | 0.04120385 | 0.19341593 |
| TOP1 | 245.160332 | 0.324176313 | 0.28993481 | 1.1181007 | 0.26352399 | 0.55506525 |
| PLCG1 | 28.7346737 | -0.109409928 | 0.34766708 | -0.3146974 | 0.75299141 | 0.89205725 |
| ZHX3 | 211.634555 | 0.61728537 | 0.33460555 | 1.84481511 | 0.06506444 | 0.25496153 |
| LPIN3 | 35.1012707 | 0.049172886 | 0.37488827 | 0.13116677 | 0.89564338 | 0.95900099 |
| CHD6 | 212.001335 | 0.246111184 | 0.14893206 | 1.65250638 | 0.09843137 | 0.32407796 |
| SRSF6 | 51.6804349 | 0.602535688 | 0.30458677 | 1.97820699 | 0.04790536 | 0.21172073 |
| L3MBTL1 | 26.7176602 | 0.057985487 | 0.32911502 | 0.17618609 | 0.86014775 | 0.94469532 |
| SGK2 | 91.8149864 | 1.311671888 | 0.25712028 | 5.10139417 | 3.37E-07 | 2.32E-05 |
| IFT52 | 36.1253483 | 0.160019479 | 0.34922149 | 0.45821773 | 0.64679602 | 0.83466996 |
| OSER1 | 27.1173255 | -0.462311173 | 0.51594273 | -0.8960513 | 0.37022535 | 0.65454977 |
| TTPAL | 17.1317265 | 0.307131284 | 0.43309633 | 0.70915236 | 0.47822993 | 0.73786502 |
| SERINC3 | 53.6680224 | -0.348672673 | 0.2516307 | -1.3856523 | 0.16585309 | 0.4346416 |
| PKIG | 79.0634743 | 0.20496473 | 0.231347 | 0.88596235 | 0.37563782 | 0.65891721 |
| YWHAB | 152.252805 | -1.006685133 | 0.41727361 | -2.4125301 | 0.01584223 | 0.10368911 |
| PABPC1L | 28.597011 | 0.290364098 | 0.2869671 | 1.0118376 | 0.31161571 | 0.60468356 |
| TOMM34 | 15.9007106 | 0.131162045 | 0.4000579 | 0.32785765 | 0.7430193 | 0.88740176 |
| STK4 | 65.4565122 | 0.012015777 | 0.27392067 | 0.0438659 | 0.9650113 | 0.98660829 |
| SLPI | 258.303652 | -6.782010909 | 0.92948084 | -7.2965581 | 2.95E-13 | 1.52E-10 |
| SDC4 | 257.009601 | 0.327380829 | 0.35442185 | 0.92370385 | 0.35564053 | 0.64045528 |
| PIGT | 28.3743142 | 0.048384739 | 0.4968408 | 0.09738479 | 0.92242082 | 0.96885417 |
| WFDC2 | 1513.67598 | -2.90186655 | 0.60179292 | -4.822035 | 1.42E-06 | 7.47E-05 |
| DNTTIP1 | 18.0651158 | -0.532491821 | 0.37578098 | -1.4170271 | 0.15647501 | 0.42099911 |
| SNX21 | 21.0512966 | 0.616566707 | 0.40145911 | 1.53581448 | 0.12458389 | 0.36974123 |
| ACOT8 | 17.4303474 | 0.144686807 | 0.35559806 | 0.406883 | 0.68409392 | 0.85606265 |
| CTSA | 48.6535821 | -0.894988489 | 0.35301534 | -2.535268 | 0.01123613 | 0.08307399 |

| | | | | | | |
|---|---|---|---|---|---|---|
| ZNF335 | 21.095275 | -0.319754461 | 0.40656632 | -0.7864755 | 0.43158895 | 0.70454366 |
| NCOA5 | 38.2839677 | -0.493908305 | 0.31115814 | -1.5873225 | 0.11243963 | 0.34984991 |
| CD40 | 17.8343854 | -1.123190141 | 0.5104969 | -2.2001899 | 0.02779342 | 0.14951846 |
| SLC35C2 | 28.8393783 | -0.230165026 | 0.30232946 | -0.7613053 | 0.44647472 | 0.71619194 |
| ELMO2 | 28.0170044 | -0.488933252 | 0.34380811 | -1.4221109 | 0.15499406 | 0.41835185 |
| ZNF334 | 18.6227148 | 0.655292412 | 0.57212153 | 1.14537276 | 0.25205478 | 0.54334972 |
| SLC13A3 | 26.2864402 | -0.50104527 | 0.63318882 | -0.7913047 | 0.42876623 | 0.70288254 |
| EYA2 | 51.5292115 | 1.888182962 | 0.59991651 | 3.14740955 | 0.00164724 | 0.02131894 |
| ZMYND8 | 320.429698 | -0.184352868 | 0.26560113 | -0.6940967 | 0.48762157 | 0.74443126 |
| NCOA3 | 178.918434 | 0.087739785 | 0.24902106 | 0.35233881 | 0.72458418 | 0.87788619 |
| SULF2 | 18.6223363 | -0.833996826 | 0.56765771 | -1.4691896 | 0.14178136 | 0.39788385 |
| ARFGEF2 | 217.991296 | -0.008932108 | 0.22243549 | -0.0401559 | 0.9679688 | 0.98742594 |
| CSE1L | 54.5142341 | 0.042473838 | 0.22378758 | 0.18979533 | 0.84946951 | 0.93897371 |
| STAU1 | 130.904165 | -0.039155789 | 0.18322697 | -0.213701 | 0.83078023 | 0.93019214 |
| DDX27 | 23.3021536 | -0.4285733 | 0.35006735 | -1.2242596 | 0.22085433 | 0.5089143 |
| ZNFX1 | 45.0762766 | -0.775287137 | 0.53620368 | -1.4458818 | 0.14821036 | 0.40845709 |
| ZFAS1 | 42.1458449 | -0.664114655 | 0.33398805 | -1.9884384 | 0.04676323 | 0.20819134 |
| B4GALT5 | 174.586771 | -0.455929514 | 0.21713163 | -2.0997839 | 0.03574785 | 0.17539507 |
| SLC9A8 | 69.8978163 | -0.052452156 | 0.28761896 | -0.1823668 | 0.85529486 | 0.94237943 |
| RNF114 | 87.194897 | 0.249826876 | 0.39327145 | 0.63525302 | 0.52526341 | 0.76221787 |
| CEBPB | 42.5758851 | -4.619068185 | 0.5450697 | -8.4742706 | 2.37E-17 | 2.71E-14 |
| PTPN1 | 144.223 | -0.691702694 | 0.45513895 | -1.5197616 | 0.12857091 | 0.37607538 |
| RIPOR3 | 16.3711988 | -0.245744111 | 0.64368698 | -0.3817758 | 0.70262767 | 0.86532818 |
| PARD6B | 211.537405 | 0.54816184 | 0.36966289 | 1.48286953 | 0.13810908 | 0.39174001 |
| BCAS4 | 30.1167906 | -0.568818235 | 0.38144982 | -1.4912007 | 0.13590882 | 0.38817095 |
| ADNP | 244.371832 | 0.346000729 | 0.14284652 | 2.42218524 | 0.01542748 | 0.10194608 |
| DPM1 | 50.3771435 | 0.371664192 | 0.22555877 | 1.64774882 | 0.09940423 | 0.32555183 |
| ATP9A | 81.1923611 | -0.369943325 | 0.25467841 | -1.45259 | 0.14633761 | 0.40547877 |
| ZFP64 | 41.6674767 | -0.018058844 | 0.25119324 | -0.0718922 | 0.94268767 | 0.9781624 |
| ZNF217 | 64.7073751 | -0.795357345 | 0.28168801 | -2.82354 | 0.00474965 | 0.04585839 |
| PFDN4 | 11.2826 | -0.235086978 | 0.41203745 | -0.5705476 | 0.56830635 | 0.78745017 |
| DOK5 | 36.4857173 | -0.088119575 | 0.67773801 | -0.1300201 | 0.89655051 | 0.95900099 |
| FAM210B | 19.1759675 | -0.994385072 | 0.42526587 | -2.3382668 | 0.01937341 | 0.11819385 |
| RTF2 | 34.0993349 | -0.255136588 | 0.26888651 | -0.9488635 | 0.34269005 | 0.63072032 |
| RAE1 | 18.5763065 | 0.09651041 | 0.34482156 | 0.27988508 | 0.77956568 | 0.90427469 |
| PCK1 | 42.066121 | -0.423778025 | 1.1879169 | -0.3567405 | 0.72128611 | 0.87614353 |
| PMEPA1 | 24.58607 | -0.805227684 | 0.73378957 | -1.097355 | 0.27248622 | 0.56479948 |
| RAB22A | 65.1413878 | -0.117081861 | 0.25125006 | -0.4659973 | 0.6412174 | 0.83230084 |
| VAPB | 90.9914867 | 0.040899549 | 0.20215916 | 0.20231361 | 0.83967156 | 0.93543959 |
| APCDD1L | 55.2238838 | 0.467377621 | 0.63164942 | 0.739932 | 0.45934126 | 0.72478689 |
| STX16-NPEPI | 10.516302 | 0.643645017 | 0.50841769 | 1.26597684 | 0.20552137 | 0.48999801 |
| NPEPL1 | 28.6006688 | 0.542021498 | 0.39722003 | 1.36453718 | 0.17239856 | 0.44481121 |
| GNAS | 294.474925 | -0.383695188 | 0.28044981 | -1.3681421 | 0.17126761 | 0.442649 |
| NELFCD | 33.6665908 | 0.039318732 | 0.27226974 | 0.14441095 | 0.88517597 | 0.95619452 |
| CTSZ | 24.1404291 | -1.409069684 | 0.43682465 | -3.2257101 | 0.00125661 | 0.01737658 |
| ATP5F1E | 129.435572 | -0.939447461 | 0.53964942 | -1.7408477 | 0.08171028 | 0.29248759 |
| PRELID3B | 10.897364 | -0.391285066 | 0.46896816 | -0.8343532 | 0.40408201 | 0.68217314 |
| FAM217B | 10.1963743 | -0.269008573 | 0.5829052 | -0.4614963 | 0.6444426 | 0.83381793 |
| MIR646HG | 238.105609 | 1.312124593 | 0.7691838 | 1.70586613 | 0.08803301 | 0.30539182 |
| TAF4 | 51.2660304 | 0.296528414 | 0.27947449 | 1.06102141 | 0.28868017 | 0.58064041 |
| LSM14B | 12.596778 | -0.121787928 | 0.39172287 | -0.3109033 | 0.75587414 | 0.89366123 |
| PSMA7 | 71.2707285 | -0.787799081 | 0.37111476 | -2.122791 | 0.03377137 | 0.1699622 |
| SS18L1 | 46.071138 | 0.269871479 | 0.37373188 | 0.72209916 | 0.47023351 | 0.73188482 |
| MTG2 | 39.4242021 | 0.074910763 | 0.28256231 | 0.26511237 | 0.7909229 | 0.91080793 |
| OSBPL2 | 74.887579 | -0.139602219 | 0.23246584 | -0.6005279 | 0.54815449 | 0.77516644 |
| ADRM1 | 10.0417671 | -0.901364258 | 0.54117888 | -1.665557 | 0.09580168 | 0.31919436 |
| LAMA5 | 64.464545 | -0.215082477 | 0.3049691 | -0.7052599 | 0.48064851 | 0.73988033 |
| RPS21 | 13.4764366 | -0.544223437 | 0.54190291 | -1.0042822 | 0.31524262 | 0.6086621 |
| SLCO4A1 | 39.7880253 | -2.398741828 | 0.70052879 | -3.4241874 | 0.00061664 | 0.01014798 |
| OGFR | 29.9793435 | -1.187891138 | 0.39303665 | -3.0223419 | 0.00250827 | 0.02877528 |
| TCFL5 | 17.9947055 | 0.29054051 | 0.42982048 | 0.6759578 | 0.49906744 | 0.75027574 |
| DIDO1 | 106.82649 | 0.198722904 | 0.17919858 | 1.10895359 | 0.2674502 | 0.559219 |
| GID8 | 37.7731818 | -0.424832754 | 0.35630499 | -1.192329 | 0.23313228 | 0.52302456 |
| YTHDF1 | 36.9175639 | -0.016342165 | 0.26662823 | -0.061292 | 0.9511267 | 0.98061757 |
| ARFGAP1 | 30.0921528 | -0.455145107 | 0.43928163 | -1.0361123 | 0.30014975 | 0.59361542 |
| PPDPF | 169.418328 | -0.942693174 | 0.45425236 | -2.0752631 | 0.03796215 | 0.18260776 |
| HELZ2 | 16.620562 | -1.335600698 | 0.95795291 | -1.3942237 | 0.16325005 | 0.43145254 |
| GMEB2 | 23.4525098 | 0.07583663 | 0.38260503 | 0.19821127 | 0.84287977 | 0.93600812 |
| MHENCR | 9.47785003 | 0.179806698 | 0.44796984 | 0.40138126 | 0.68813944 | 0.85854718 |
| ARFRP1 | 21.1337902 | -0.279985042 | 0.38631902 | -0.7247509 | 0.46860489 | 0.73133883 |
| SLC2A4RG | 31.6449554 | -0.565536784 | 0.52569552 | -1.0757877 | 0.2820222 | 0.5750055 |
| ZBTB46 | 29.2082169 | -0.9924708 | 0.52943825 | -1.8745733 | 0.06085145 | 0.24477853 |
| TPD52L2 | 41.1957978 | -0.086407657 | 0.31250943 | -0.2764962 | 0.78216701 | 0.90578738 |
| DNAJC5 | 109.145002 | -0.091750489 | 0.30479393 | -0.3010247 | 0.76339569 | 0.8974069 |
| UCKL1 | 21.9510193 | 0.787259763 | 0.33643782 | 2.33998594 | 0.01928447 | 0.11811449 |
| ZNF512B | 14.745448 | 0.510387687 | 0.37419902 | 1.36394716 | 0.1725842 | 0.44515833 |
| PRPF6 | 53.8512889 | -0.028243652 | 0.2259495 | -0.1249998 | 0.90052367 | 0.95980075 |
| TCEA2 | 11.1604846 | -0.076224568 | 0.46305379 | -0.1646128 | 0.86924878 | 0.94827256 |
| PCMTD2 | 77.6218411 | 0.374761281 | 0.25684439 | 1.45909859 | 0.14453797 | 0.40242285 |
| PLPP2 | 20.7867472 | -3.912303484 | 0.6966093 | -5.6162091 | 1.95E-08 | 2.26E-06 |
| MIER2 | 17.8509204 | 0.595116029 | 0.44288681 | 1.34372038 | 0.17903882 | 0.45449239 |
| TPGS1 | 21.4340282 | -0.52543623 | 0.39501983 | -1.3301515 | 0.18346835 | 0.46014503 |
| CDC34 | 16.1390517 | -0.197385448 | 0.44424213 | -0.4443195 | 0.6568116 | 0.83951711 |
| BSG | 114.883589 | -0.689632232 | 0.35572761 | -1.9386525 | 0.05254367 | 0.22446724 |
| RNF126 | 18.9710855 | -0.58643906 | 0.39100789 | -1.4998139 | 0.13366262 | 0.38409828 |
| PALM | 26.0295375 | 2.140781343 | 0.36486975 | 5.86724814 | 4.43E-09 | 6.09E-07 |
| PTBP1 | 65.4441913 | -0.25314293 | 0.23939903 | -1.05741 | 0.29032451 | 0.58200489 |
| MED16 | 23.1161836 | 0.025418749 | 0.36007311 | 0.0705933 | 0.94372144 | 0.97839988 |
| R3HDM4 | 22.100358 | -0.787796173 | 0.46729307 | -1.6858717 | 0.09182051 | 0.31188094 |
| ARID3A | 80.5732882 | 0.309858382 | 0.29088596 | 1.06522287 | 0.28677508 | 0.57922771 |
| WDR18 | 16.6296286 | 0.775198467 | 0.41268752 | 1.8784151 | 0.0603244 | 0.24332668 |
| TMEM259 | 59.5024717 | 0.259377564 | 0.33911906 | 0.76485693 | 0.44435674 | 0.71427105 |
| CNN2 | 19.191035 | -0.244435439 | 0.41355733 | -0.5910558 | 0.55448306 | 0.77922789 |
| ABCA7 | 13.8627526 | 0.05944247 | 0.58787521 | 0.1011141 | 0.91945989 | 0.96763053 |
| POLR2E | 26.7405175 | -0.24745457 | 0.37134521 | -0.6663734 | 0.50517246 | 0.75184725 |
| GPX4 | 107.895386 | -0.411067801 | 0.35501914 | -1.157875 | 0.24691504 | 0.53778408 |
| SBNO2 | 83.7669762 | -1.864814004 | 0.28543895 | -6.5331449 | 6.44E-11 | 1.58E-08 |
| STK11 | 49.0879923 | 0.486726528 | 0.2884528 | 1.68736974 | 0.09153228 | 0.31120975 |
| ATP5F1D | 102.222029 | 0.128689355 | 0.41695904 | 0.30863788 | 0.757597 | 0.89457492 |
| MIDN | 126.821491 | -1.043327768 | 0.54865595 | -1.9016066 | 0.05722261 | 0.23601925 |
| CIRBP | 135.777593 | 0.447669189 | 0.27969075 | 1.60058632 | 0.10946857 | 0.34481257 |
| C19orf24 | 18.364811 | -0.246786974 | 0.5004593 | -0.493121 | 0.62192712 | 0.82037044 |
| MUM1 | 54.6678428 | -0.033630321 | 0.28949803 | -0.1161677 | 0.90751963 | 0.96195774 |
| NDUFS7 | 34.4699446 | 0.577127467 | 0.39986162 | 1.44331797 | 0.14893092 | 0.40929305 |
| DAZAP1 | 77.3156476 | 0.314319142 | 0.2804397 | 1.1208083 | 0.26236947 | 0.55399268 |
| RPS15 | 199.056845 | -1.041633197 | 0.44523458 | -2.3395155 | 0.01930877 | 0.11815925 |
| C19orf25 | 40.2469409 | -0.602337856 | 0.30071597 | -2.0030125 | 0.04517594 | 0.20323256 |
| MBD3 | 23.6581626 | -0.28976308 | 0.44565116 | -0.6502016 | 0.51556203 | 0.75713618 |
| TCF3 | 24.3642131 | -0.251517278 | 0.44141752 | -0.5697945 | 0.56881708 | 0.78773405 |
| REXO1 | 20.5288237 | -0.353865737 | 0.40663497 | -0.8702295 | 0.38417501 | 0.66701235 |
| ABHD17A | 28.2809105 | -0.326700741 | 0.39307106 | -0.8311493 | 0.40588928 | 0.68402935 |
| SCAMP4 | 46.6061024 | 0.059402087 | 0.30333712 | 0.19582861 | 0.84474432 | 0.93689787 |
| CSNK1G2 | 46.7712447 | -0.718914619 | 0.303168 | -2.3713407 | 0.01772368 | 0.11167548 |

| | | | | | | |
|---|---|---|---|---|---|---|
| BTBD2 | 49.1578291 | -0.22635156 | 0.34332364 | -0.659295 | 0.50970635 | 0.75476666 |
| MKNK2 | 39.7116658 | 0.163104961 | 0.31401779 | 0.51941312 | 0.60347268 | 0.80939664 |
| MOB3A | 29.0111768 | -0.834550013 | 0.36019519 | -2.3169382 | 0.0205071 | 0.1227566 |
| AP3D1 | 75.7366127 | -0.431175254 | 0.33526744 | -1.2860636 | 0.19842086 | 0.48097216 |
| DOT1L | 79.906053 | 0.983488592 | 0.4961263 | 1.98233513 | 0.04744175 | 0.21024184 |
| PLEKHJ1 | 52.9339204 | -0.391476931 | 0.46930618 | -0.834161 | 0.40419024 | 0.68217314 |
| SF3A2 | 10.2289334 | -0.110259227 | 0.43823733 | -0.2515971 | 0.80135253 | 0.91484195 |
| LSM7 | 48.6099783 | -0.178150913 | 0.40095542 | -0.444316 | 0.65681413 | 0.83951711 |
| SPPL2B | 45.2073076 | 0.135565151 | 0.2970241 | 0.45641128 | 0.64809425 | 0.8354188 |
| TIMM13 | 29.5331438 | -0.672558152 | 0.48421702 | -1.3889602 | 0.16484487 | 0.43375609 |
| LMNB2 | 11.2452659 | 0.091855158 | 0.46789151 | 0.19631721 | 0.84436189 | 0.93684611 |
| GADD45B | 99.3053535 | 1.838377983 | 0.43485616 | 4.22755418 | 2.36E-05 | 0.00074656 |
| GNG7 | 76.7722066 | 1.357589367 | 0.29088909 | 4.66703428 | 3.06E-06 | 0.00014054 |
| SGTA | 29.0769149 | 0.217231242 | 0.32803512 | 0.66221947 | 0.50783058 | 0.75382716 |
| THOP1 | 10.9119729 | 0.430480668 | 0.53486815 | 0.80483511 | 0.42091484 | 0.69661173 |
| ZNF555 | 14.3015563 | -0.506494723 | 0.51693398 | -0.9798054 | 0.32718217 | 0.6168797 |
| TLE2 | 46.8093932 | 0.436023872 | 0.31656897 | 1.37734243 | 0.1684064 | 0.43888762 |
| AES | 150.695766 | -0.201091812 | 0.31990828 | -0.6285921 | 0.52961615 | 0.7638395 |
| GNA11 | 38.7245005 | -0.389417544 | 0.38145213 | -1.0208818 | 0.30731044 | 0.60139222 |
| NCLN | 23.7341073 | 0.369333373 | 0.3542291 | 1.04263984 | 0.29711513 | 0.58943269 |
| NFIC | 221.802933 | -0.062670267 | 0.27765057 | -0.2257163 | 0.82142206 | 0.92485761 |
| SMIM24 | 19.9863488 | 0.453548607 | 0.54467957 | 0.83268886 | 0.40502023 | 0.68331761 |
| FZR1 | 22.2235032 | 0.385320535 | 0.39391686 | 0.97817732 | 0.32798663 | 0.61729352 |
| MFSD12 | 36.4778134 | -0.537940742 | 0.42447175 | -1.2673181 | 0.20504156 | 0.48964616 |
| HMG20B | 23.9897065 | -0.143230612 | 0.36125739 | -0.396478 | 0.69175243 | 0.85976938 |
| PIP5K1C | 27.3054431 | -0.280060014 | 0.31926475 | -0.8772031 | 0.38037636 | 0.66406325 |
| TJP3 | 47.5509373 | 0.133946715 | 0.2988711 | 0.44817554 | 0.65402651 | 0.83885315 |
| MRPL54 | 26.3791883 | -1.016658757 | 0.56275627 | -1.8065703 | 0.0708293 | 0.26866108 |
| DAPK3 | 20.007036 | -0.823086084 | 0.32288021 | -2.5491996 | 0.01079705 | 0.08071931 |
| EEF2 | 178.425523 | -0.723435715 | 0.45090768 | -1.6043987 | 0.10862619 | 0.34348281 |
| PIAS4 | 21.3761963 | -0.192573229 | 0.402788 | -0.4781007 | 0.63257853 | 0.82649065 |
| ZBTB7A | 67.4400545 | -0.385432428 | 0.34648532 | -1.1124062 | 0.26596353 | 0.55721548 |
| MAP2K2 | 49.9578478 | 0.00769619 | 0.26286812 | 0.02927776 | 0.97664306 | 0.99058549 |
| SIRT6 | 11.4802642 | -0.549983846 | 0.45404203 | -1.211306 | 0.22577814 | 0.51470821 |
| YJU2 | 12.6256539 | -0.130213694 | 0.50743639 | -0.2566109 | 0.79747918 | 0.91302793 |
| STAP2 | 14.0157216 | -0.702399617 | 0.37814628 | -1.8574812 | 0.06324272 | 0.25020219 |
| SH3GL1 | 43.8229123 | -0.204777716 | 0.29738621 | -0.6885918 | 0.49108016 | 0.74574113 |
| UBXN6 | 34.6160364 | -0.105822136 | 0.40866765 | -0.2589442 | 0.79567826 | 0.91195838 |
| HDGFL2 | 20.0592134 | 0.185961462 | 0.35964724 | 0.51706629 | 0.60510989 | 0.81043188 |
| PLIN5 | 56.7255708 | 0.039711436 | 0.43863704 | 0.0905337 | 0.92786311 | 0.97123001 |
| MYDGF | 30.4596789 | -1.066813154 | 0.32799262 | -3.2525523 | 0.00114374 | 0.01607471 |
| DPP9 | 33.3771867 | -0.473610391 | 0.32087987 | -1.4759742 | 0.13995083 | 0.39497967 |
| TICAM1 | 11.1140467 | -0.571811222 | 0.57458957 | -0.9951646 | 0.31965619 | 0.611915 |
| PLIN3 | 37.7332696 | -1.650001928 | 0.35913061 | -4.5944341 | 4.34E-06 | 0.00018862 |
| KDM4B | 101.020521 | -0.565494043 | 0.27428774 | -2.0616818 | 0.03923804 | 0.18700317 |
| PTPRS | 57.0440817 | 0.741307412 | 0.25784164 | 2.87504928 | 0.00403964 | 0.04040429 |
| TINCR | 11.358723 | -0.503797324 | 0.49679421 | -1.0140966 | 0.31053666 | 0.60384082 |
| SAFB2 | 88.3322551 | 0.448387929 | 0.24655273 | 1.81862891 | 0.06896806 | 0.26458076 |
| SAFB | 88.1100074 | 0.177357685 | 0.22874834 | 0.77533976 | 0.43813892 | 0.70959081 |
| RPL36 | 136.487862 | -0.942703295 | 0.54867502 | -1.7181451 | 0.08577015 | 0.30113433 |
| C19orf70 | 41.22627 | -0.166016135 | 0.4175051 | -0.3976386 | 0.69089661 | 0.85967888 |
| LONP1 | 49.4069085 | -0.410209575 | 0.34249676 | -1.1977035 | 0.23103246 | 0.52111057 |
| CAPS | 39.8756469 | -0.507971728 | 0.56019342 | -0.9067792 | 0.36452359 | 0.6495973 |
| RANBP3 | 35.9790663 | 0.136086178 | 0.25643697 | 0.53068082 | 0.59563998 | 0.8037044 |
| RFX2 | 34.8166242 | -0.796179457 | 0.34439864 | -2.3117962 | 0.02078892 | 0.12343944 |
| MLLT1 | 34.8427529 | 0.416920083 | 0.31191218 | 1.33665857 | 0.18133411 | 0.45775644 |
| CLPP | 18.3500996 | -0.040746939 | 0.41878561 | -0.0972978 | 0.92248986 | 0.96885417 |
| ALKBH7 | 22.6515571 | -0.377660872 | 0.43944806 | -0.8593982 | 0.39012086 | 0.67207778 |
| GTF2F1 | 23.0868159 | -0.732633785 | 0.37241524 | -1.9672498 | 0.04915442 | 0.21520989 |
| KHSRP | 39.8966293 | -0.257748159 | 0.33545022 | -0.7683649 | 0.44227045 | 0.71265656 |
| SLC25A23 | 43.4979794 | -0.139269637 | 0.43930294 | -0.3170241 | 0.75122528 | 0.89132635 |
| AC010503.4 | 16.2631684 | -0.900917113 | 0.55074489 | -1.6358157 | 0.10187818 | 0.3305666 |
| C3 | 51.1813688 | -3.995655511 | 1.35555855 | -2.9476082 | 0.00320243 | 0.03415259 |
| TRIP10 | 33.5787058 | -0.727767689 | 0.31192427 | -2.333155 | 0.01964001 | 0.1190453 |
| SH2D3A | 40.2319458 | -0.637443805 | 0.43720131 | -1.4580098 | 0.14483784 | 0.40273129 |
| ZNF557 | 11.2386983 | -0.188362459 | 0.42180911 | -0.4465585 | 0.65519383 | 0.83931753 |
| INSR | 268.443661 | 0.197609186 | 0.31234512 | 0.63266295 | 0.52695377 | 0.76221787 |
| ZNF358 | 21.4559222 | -0.367576978 | 0.54985316 | -0.6685003 | 0.50381432 | 0.75108109 |
| PNPLA6 | 26.1186016 | -0.211397725 | 0.4317963 | -0.4895774 | 0.62443295 | 0.82262254 |
| CAMSAP3 | 20.4044421 | -0.02957388 | 0.38687384 | -0.0764432 | 0.93906649 | 0.97662903 |
| XAB2 | 12.2982198 | 0.380621182 | 0.44139077 | 0.86232248 | 0.38851009 | 0.67042394 |
| STXBP2 | 17.1483556 | -0.247157207 | 0.42097141 | -0.5871116 | 0.55712874 | 0.7813742 |
| EVI5L | 20.0115476 | -0.07094324 | 0.49502253 | -0.1433132 | 0.88604287 | 0.95619452 |
| MAP2K7 | 43.1286972 | -0.948490124 | 0.50615085 | -1.8739278 | 0.06094038 | 0.24495038 |
| TIMM44 | 25.7640835 | 0.218056443 | 0.41210148 | 0.52913288 | 0.59671328 | 0.80462568 |
| ELAVL1 | 69.4156256 | 0.064781273 | 0.25048078 | 0.25862772 | 0.79592249 | 0.91195838 |
| CERS4 | 56.6270212 | 0.016577314 | 0.38314613 | 0.0432663 | 0.96548926 | 0.98661741 |
| AC022146.2 | 11.2709603 | -0.562029971 | 0.47246385 | -1.1895724 | 0.23421449 | 0.52443761 |
| CD320 | 15.0397307 | -1.895073424 | 0.68952336 | -2.7483817 | 0.00598902 | 0.05416938 |
| RPS28 | 47.867176 | -0.406478195 | 0.42888564 | -0.9477543 | 0.34325457 | 0.6310379 |
| ANGPTL4 | 59.0726203 | -3.243615348 | 1.36417733 | -2.377708 | 0.01742062 | 0.11057743 |
| RAB11B-AS1 | 13.8650454 | -0.677068404 | 0.409665 | -1.6527368 | 0.09838445 | 0.32402704 |
| RAB11B | 43.7047485 | -0.377575218 | 0.32275622 | -1.1698464 | 0.24206277 | 0.5328423 |
| 2-Mar | 16.5416331 | -0.327494353 | 0.34348626 | -0.9534424 | 0.34036599 | 0.62907597 |
| HNRNPM | 140.839787 | 0.311756789 | 0.20771565 | 1.50088255 | 0.13338594 | 0.38373327 |
| ZNF558 | 14.6067091 | -0.169704108 | 0.35373503 | -0.4797492 | 0.63140572 | 0.82625173 |
| ZNF317 | 14.7302729 | -0.259995562 | 0.40686137 | -0.6390274 | 0.52280511 | 0.76180173 |
| ZNF559-ZNF | 23.5504631 | 0.728398312 | 0.33695688 | 2.16169591 | 0.03064162 | 0.15991388 |
| ZNF266 | 29.8413801 | -0.107042424 | 0.28676106 | -0.3732809 | 0.7089394 | 0.86910485 |
| ZNF426 | 16.7447926 | -0.160636774 | 0.36162992 | -0.4442021 | 0.65689647 | 0.83951711 |
| ZNF121 | 30.3553285 | 0.094595205 | 0.33070946 | 0.28603719 | 0.77484964 | 0.90205458 |
| ZNF561 | 22.3481504 | -0.364720051 | 0.32427448 | -1.1247263 | 0.26070502 | 0.55172208 |
| ZNF562 | 55.7747839 | -0.05195199 | 0.27366044 | -0.1898411 | 0.84943367 | 0.93897371 |
| ZNF846 | 47.0737488 | 0.386264341 | 0.25669972 | 1.50473223 | 0.13239293 | 0.38183426 |
| FBXL12 | 17.6986052 | 0.043758896 | 0.368019 | 0.1189039 | 0.90535149 | 0.96157183 |
| UBL5 | 34.4684542 | -0.433569176 | 0.43440611 | -0.9980734 | 0.31824377 | 0.61132712 |
| PIN1 | 20.6453294 | -0.436145052 | 0.38407839 | -1.1355626 | 0.25613967 | 0.54710413 |
| C19orf66 | 32.1407779 | -0.819114823 | 0.65105751 | -1.2581297 | 0.20834484 | 0.49402397 |
| EIF3G | 19.6269325 | -0.724690543 | 0.41347831 | -1.7526688 | 0.07965887 | 0.28774394 |
| DNMT1 | 22.709814 | 0.077767833 | 0.38174903 | 0.20371455 | 0.83857657 | 0.93537562 |
| MRPL4 | 28.3918104 | -0.013482618 | 0.34046493 | -0.0396006 | 0.96841154 | 0.98758421 |
| AC011511.2 | 37.8164028 | 1.977830111 | 0.69010317 | 2.86599192 | 0.00415705 | 0.04117876 |
| ICAM1 | 22.1737196 | -0.983436367 | 0.48815828 | -2.014585 | 0.04394815 | 0.20006796 |
| TYK2 | 39.6258168 | 0.346335134 | 0.32421452 | 1.06822832 | 0.28541753 | 0.57790319 |
| CDC37 | 62.6801736 | -0.396413328 | 0.25072259 | -1.5810834 | 0.11385897 | 0.35248764 |
| PDE4A | 39.1347226 | 0.136169615 | 0.4303663 | 0.316404 | 0.75169588 | 0.89132635 |
| KEAP1 | 20.4203123 | -0.355778407 | 0.48611423 | -0.7318823 | 0.4642404 | 0.72835095 |
| ATG4D | 19.4674665 | -0.112892395 | 0.35277482 | -0.3200126 | 0.74895876 | 0.88993924 |
| KRI1 | 17.875673 | -0.531216124 | 0.35707353 | -1.4876939 | 0.13683163 | 0.38961843 |
| AP1M2 | 40.4589035 | -0.720323485 | 0.41441993 | -1.7381488 | 0.0821846 | 0.29357343 |
| SLC44A2 | 52.8306397 | 0.21774916 | 0.41720651 | 0.52192176 | 0.6017248 | 0.80777556 |
| ILF3-DT | 27.0530161 | -0.264916111 | 0.38988195 | -0.6794778 | 0.4968352 | 0.74885094 |
| ILF3 | 88.5752078 | 0.095580823 | 0.24074143 | 0.3970269 | 0.69134762 | 0.85976137 |

| | | | | | | |
|---|---|---|---|---|---|---|
| QTRT1 | 30.9086212 | 1.288494288 | 0.3491838 | 3.69001733 | 0.00022424 | 0.00467633 |
| DNM2 | 294.683485 | 0.236067362 | 0.22566824 | 1.04608145 | 0.29552343 | 0.58705792 |
| TMED1 | 13.679387 | -1.610607815 | 0.55885031 | -2.8820022 | 0.00395157 | 0.03979382 |
| CARM1 | 24.6716992 | -0.284737209 | 0.32928816 | -0.8647053 | 0.38720057 | 0.66950996 |
| YIPF2 | 21.9404709 | -0.069448477 | 0.34839655 | -0.1993374 | 0.84199879 | 0.93566718 |
| TIMM29 | 11.9407605 | -0.749963549 | 0.57666652 | -1.3005152 | 0.19342447 | 0.4740563 |
| SMARCA4 | 150.148433 | 0.077248717 | 0.29500161 | 0.26185863 | 0.79343044 | 0.91142199 |
| LDLR | 86.7137573 | 0.418503498 | 0.73515456 | 0.5692728 | 0.56917102 | 0.78790646 |
| KANK2 | 17.8691224 | 0.064917943 | 0.38397677 | 0.16906737 | 0.86574365 | 0.94694132 |
| DOCK6 | 74.5391354 | -0.121864426 | 0.245543 | -0.4963058 | 0.61967865 | 0.81899807 |
| RAB3D | 21.5769868 | -0.606475098 | 0.48453306 | -1.2516692 | 0.21069043 | 0.49657581 |
| EPOR | 38.385921 | 1.034223668 | 0.39561974 | 2.6141862 | 0.00894403 | 0.07098719 |
| RGL3 | 270.418966 | 0.506768675 | 0.20601468 | 2.45986677 | 0.01389886 | 0.09584074 |
| PRKCSH | 39.2206205 | -0.229491622 | 0.41428661 | -0.5539441 | 0.5796171 | 0.79429575 |
| ECSIT | 34.6490618 | 0.712740006 | 0.32505194 | 2.19269573 | 0.02832931 | 0.15137372 |
| ELOF1 | 12.1879063 | 0.056471184 | 0.46971606 | 0.12022409 | 0.90430564 | 0.96126516 |
| ZNF627 | 34.5006552 | 0.346783096 | 0.31357257 | 1.10591017 | 0.2687654 | 0.56094432 |
| AC008543.1 | 15.9276461 | 0.479159063 | 0.61479089 | 0.77938543 | 0.43575271 | 0.70739432 |
| ZNF823 | 39.969146 | 0.175960149 | 0.30185025 | 0.58293855 | 0.55993467 | 0.78316119 |
| ZNF441 | 28.45351 | -0.430274882 | 0.33824038 | -1.2720979 | 0.20333833 | 0.48750092 |
| ZNF440 | 35.6418385 | -0.219591242 | 0.42219711 | -0.5201155 | 0.60298311 | 0.80891 |
| ZNF439 | 47.2732839 | 0.019695427 | 0.28753098 | 0.06849845 | 0.94538885 | 0.97858537 |
| ZNF69 | 59.0212565 | -0.384115455 | 0.36442965 | -1.0540181 | 0.29187461 | 0.58344269 |
| ZNF433 | 13.1555268 | 0.287893192 | 0.41182241 | 0.69907121 | 0.48450753 | 0.74250561 |
| ZNF844 | 26.7388839 | 0.325931099 | 0.35189859 | 0.92620745 | 0.35433818 | 0.63963412 |
| ZNF788P | 14.2635179 | 0.068456726 | 0.44544571 | 0.15368141 | 0.87786094 | 0.95217134 |
| ZNF136 | 18.3647637 | 0.294871423 | 0.36302265 | 0.81226728 | 0.41663828 | 0.69318598 |
| ZNF44 | 205.320747 | 0.417013395 | 0.17854147 | 2.33566688 | 0.01950861 | 0.1186813 |
| ZNF563 | 11.2475428 | -0.62708658 | 0.64111685 | -0.9781159 | 0.32801701 | 0.61729352 |
| AC008758.5 | 19.6995956 | -0.704617226 | 0.58462113 | -1.2052545 | 0.22810509 | 0.5178625 |
| ZNF490 | 27.9881406 | -0.334646646 | 0.42805147 | -0.7817907 | 0.4343376 | 0.70618229 |
| ZNF791 | 48.3510036 | -0.317591198 | 0.25013019 | -1.2697036 | 0.20419024 | 0.48840674 |
| MAN2B1 | 11.7699685 | -0.361069322 | 0.38848503 | -0.9294292 | 0.35266671 | 0.63863112 |
| DHPS | 15.1542537 | 0.358852557 | 0.38688298 | 0.92754805 | 0.35364206 | 0.63949806 |
| TNPO2 | 43.4510507 | -0.16633365 | 0.32765348 | -0.5076511 | 0.61169804 | 0.81385911 |
| TRIR | 54.8543953 | -0.530236197 | 0.36865905 | -1.4382834 | 0.15035366 | 0.41132271 |
| ASNA1 | 17.2046902 | -0.708173551 | 0.44999965 | -1.5737202 | 0.11555214 | 0.3547134 |
| HOOK2 | 127.330364 | -0.670501008 | 0.35378241 | -1.8952356 | 0.0580612 | 0.23811563 |
| JUNB | 65.97608 | 0.297508854 | 0.57146456 | 0.52060771 | 0.60264008 | 0.80870107 |
| DNASE2 | 9.54337348 | -0.503552596 | 0.53421182 | -0.9426085 | 0.34588119 | 0.63301973 |
| GCDH | 19.9513136 | -0.321761504 | 0.48091744 | -0.6690577 | 0.50345869 | 0.75108109 |
| FARSA | 16.8739086 | -0.605898122 | 0.48529113 | -1.248525 | 0.21183885 | 0.49802917 |
| CALR | 103.707211 | -0.848791251 | 0.55261808 | -1.5359455 | 0.12455175 | 0.36974123 |
| AC092069.1 | 17.9280309 | -0.164051706 | 0.64430709 | -0.2546173 | 0.79901873 | 0.91379784 |
| RAD23A | 38.1797606 | -0.768132601 | 0.28720188 | -2.6745389 | 0.00748321 | 0.06324203 |
| GADD45GIP1 | 47.6045321 | -1.315302092 | 0.43079122 | -3.053224 | 0.00226397 | 0.02662489 |
| NFIX | 18.863061 | -0.301025176 | 0.47773904 | -0.6301038 | 0.52862668 | 0.76311655 |
| TRMT1 | 10.3945144 | 0.860679625 | 0.48453971 | 1.77628296 | 0.07568629 | 0.27917785 |
| NACC1 | 21.7346717 | -0.112071577 | 0.38695607 | -0.2896235 | 0.77210428 | 0.90105761 |
| STX10 | 12.9851917 | -0.87764526 | 0.47608771 | -1.8434529 | 0.06526291 | 0.25528428 |
| IER2 | 68.77771 | 0.780121466 | 0.58102918 | 1.34265454 | 0.17938386 | 0.45495583 |
| CCDC130 | 61.4803544 | -0.112661807 | 0.21090038 | -0.5341944 | 0.59320703 | 0.80210249 |
| MRI1 | 25.1863891 | 0.062249905 | 0.36979054 | 0.16833829 | 0.86631716 | 0.94702879 |
| C19orf53 | 48.7785928 | -1.044053589 | 0.43248451 | -2.4140832 | 0.01577486 | 0.10356817 |
| ZSWIM4 | 22.4149763 | -1.478565452 | 0.38279937 | -3.8625076 | 0.00011223 | 0.00266415 |
| CC2D1A | 42.4124865 | -0.207420148 | 0.35907157 | -0.5776568 | 0.56349582 | 0.78465819 |
| DCAF15 | 12.3281179 | -0.227173238 | 0.4070091 | -0.5581527 | 0.5767401 | 0.79210458 |
| RFX1 | 31.0791694 | -0.243637037 | 0.34256354 | -0.7112171 | 0.47694974 | 0.73678733 |
| PALM3 | 32.892755 | 0.861125679 | 0.28847994 | 2.98504526 | 0.00283536 | 0.03143157 |
| SAMD1 | 12.8430422 | -1.397194323 | 0.60245363 | -2.3191732 | 0.02038565 | 0.12252796 |
| PRKACA | 33.1100659 | -0.049812302 | 0.30433725 | -0.1636747 | 0.86998725 | 0.94835789 |
| ADGRL1 | 34.208699 | -0.050360886 | 0.33509845 | -0.1502868 | 0.88053832 | 0.95336897 |
| PKN1 | 34.3789903 | -0.401600302 | 0.31063986 | -1.2928164 | 0.19607457 | 0.47764091 |
| PTGER1 | 31.8115922 | 0.600458617 | 0.86926229 | 0.69076805 | 0.48971131 | 0.7455572 |
| GIPC1 | 96.3361849 | -0.831747894 | 0.351929 | -2.3633969 | 0.01810827 | 0.11306142 |
| DNAJB1 | 41.721283 | 0.193475602 | 0.67257661 | 0.28766329 | 0.77360449 | 0.90184317 |
| TECR | 37.9469195 | 0.032315137 | 0.38080057 | 0.08486105 | 0.93237186 | 0.97356162 |
| NDUFB7 | 67.4027271 | -0.204354848 | 0.41486046 | -0.4925869 | 0.62230447 | 0.82055301 |
| ZNF333 | 11.3708048 | -0.275301517 | 0.50050534 | -0.5500471 | 0.58228706 | 0.7955793 |
| ILVBL | 25.7466448 | -0.444363526 | 0.35494974 | -1.2519055 | 0.2106043 | 0.49657419 |
| NOTCH3 | 48.5726925 | 0.350668691 | 0.41546606 | 0.84403691 | 0.39864878 | 0.67938457 |
| BRD4 | 218.314015 | -0.039711035 | 0.26131161 | -0.1519681 | 0.87921208 | 0.95273407 |
| AKAP8 | 45.08944 | -0.029371772 | 0.26207184 | -0.1120753 | 0.91076373 | 0.96371076 |
| AKAP8L | 123.913244 | 0.242875996 | 0.23103445 | 1.05125448 | 0.29314173 | 0.5846943 |
| WIZ | 12.6575208 | -0.641992017 | 0.41355196 | -1.5523854 | 0.12057004 | 0.36346292 |
| TPM4 | 150.04238 | 0.102028821 | 0.42921802 | 0.23770862 | 0.8121071 | 0.92034001 |
| RAB8A | 24.5553439 | -0.083375701 | 0.29663901 | -0.2810679 | 0.77865833 | 0.90426537 |
| FAM32A | 17.1373726 | -0.30710649 | 0.36696349 | -0.8368857 | 0.40265683 | 0.68128594 |
| AP1M1 | 18.3160838 | 0.05013778 | 0.36268005 | 0.13824245 | 0.89004881 | 0.95702609 |
| EPS15L1 | 74.9719556 | 0.314727093 | 0.23603911 | 1.33336841 | 0.18241093 | 0.45912471 |
| C19orf44 | 14.6120068 | 0.208286594 | 0.41988323 | 0.49605838 | 0.61985322 | 0.81899807 |
| TMEM38A | 24.8374158 | 0.190991075 | 0.5407774 | 0.35317873 | 0.72395445 | 0.87742017 |
| SIN3B | 49.0992738 | 0.206585264 | 0.29739081 | 0.6946592 | 0.4872689 | 0.74434227 |
| MYO9B | 75.3384077 | -0.711295904 | 0.39606315 | -1.7959154 | 0.07250798 | 0.27192471 |
| NR2F6 | 32.5215663 | -0.35653017 | 0.40993062 | -0.869733 | 0.38444635 | 0.66701235 |
| ABHD8 | 15.0405686 | -0.015235784 | 0.36368585 | -0.0418927 | 0.96658423 | 0.98695423 |
| MRPL34 | 33.2602636 | -0.728632523 | 0.34585432 | -2.1067614 | 0.03513827 | 0.17419602 |
| DDA1 | 15.9420633 | -0.670938678 | 0.39342241 | -1.7053901 | 0.08812169 | 0.30539182 |
| BST2 | 65.789054 | -1.291542313 | 0.84277588 | -1.5324861 | 0.12540253 | 0.37039755 |
| MVB12A | 22.1046105 | -0.545668145 | 0.34266689 | -1.5924157 | 0.11129133 | 0.34785293 |
| SLC27A1 | 27.8445768 | 0.35705526 | 0.46871106 | 0.76178117 | 0.44619062 | 0.71599 |
| AC010618.3 | 21.2701231 | 0.040691011 | 0.41121599 | 0.09895289 | 0.92117567 | 0.9686919 |
| PGLS | 38.6590292 | -1.145174388 | 0.37354029 | -3.0657319 | 0.00217138 | 0.02585572 |
| COLGALT1 | 16.0349734 | -0.748980787 | 0.49504303 | -1.512961 | 0.13028957 | 0.37846669 |
| RPL18A | 12.4681678 | -1.318550504 | 0.60582952 | -2.1764382 | 0.02952251 | 0.15588975 |
| CCDC124 | 18.2108679 | -0.679562456 | 0.4434436 | -1.5324665 | 0.12540736 | 0.37039755 |
| ARRDC2 | 13.7094536 | -1.939330628 | 0.52152826 | -3.7185533 | 0.00020037 | 0.00422988 |
| IQCN | 11.4953672 | 0.572611318 | 0.74224433 | 0.77145934 | 0.44043472 | 0.71123283 |
| JUND | 155.306836 | -0.493219731 | 0.60075558 | -0.820999 | 0.41164683 | 0.68899849 |
| LSM4 | 50.0973379 | -1.001536239 | 0.39378874 | -2.5433339 | 0.01098003 | 0.08149584 |
| PGPEP1 | 53.0590275 | -0.181276831 | 0.36317368 | -0.4991464 | 0.61767626 | 0.81764942 |
| GDF15 | 210.885181 | -0.629969495 | 0.73326424 | -0.8591303 | 0.39026866 | 0.67210761 |
| SSBP4 | 29.5266878 | -0.440770863 | 0.39537953 | -1.1148045 | 0.26493422 | 0.55622662 |
| ISYNA1 | 14.9988479 | 0.6688872 | 0.56501286 | 1.18384421 | 0.23647474 | 0.52682659 |
| ELL | 58.2308499 | 0.051747921 | 0.30139121 | 0.17169685 | 0.86367586 | 0.94635064 |
| FKBP8 | 47.865281 | -0.132386478 | 0.43783478 | -0.3023663 | 0.76237285 | 0.89720537 |
| KXD1 | 17.8342837 | -0.55047357 | 0.40222832 | -1.36856 | 0.17113687 | 0.44242209 |
| UBA52 | 74.3303311 | -0.695088219 | 0.47436082 | -1.4653154 | 0.14283488 | 0.39975142 |
| REX1BD | 23.616595 | 0.269637109 | 0.40075727 | 0.67281901 | 0.50106244 | 0.75071921 |
| CRTC1 | 94.5560289 | 0.335950569 | 0.25274054 | 1.32923104 | 0.18377176 | 0.46016652 |
| UPF1 | 42.7843662 | -0.034173448 | 0.25880249 | -0.1320445 | 0.89494909 | 0.95869455 |
| COPE | 30.3113176 | -0.079445418 | 0.35634788 | -0.2229434 | 0.82357954 | 0.92608229 |
| DDX49 | 12.7836141 | -0.436557929 | 0.47534916 | -0.9183942 | 0.3584125 | 0.64216076 |

| | | | | | | |
|---|---|---|---|---|---|---|
| SUGP2 | 100.734667 | 0.453184325 | 0.22684252 | 1.99779272 | 0.04573914 | 0.20513913 |
| SLC25A42 | 24.7661421 | 0.396893921 | 0.4053704 | 0.97908956 | 0.32753573 | 0.61719253 |
| TMEM161A | 18.466375 | 0.629550344 | 0.35153556 | 1.79085819 | 0.07331605 | 0.27366014 |
| RFXANK | 12.3901687 | -0.271754516 | 0.41449058 | -0.655635 | 0.51205902 | 0.75555211 |
| SUGP1 | 24.6000691 | 0.18547932 | 0.33753395 | 0.54951308 | 0.58265339 | 0.7956648 |
| MAU2 | 63.4978507 | 0.001179006 | 0.2783963 | 0.00423499 | 0.99662097 | 0.99868531 |
| GATAD2A | 151.151946 | -0.077402914 | 0.23157976 | -0.3342387 | 0.73819946 | 0.88573617 |
| PBX4 | 13.596815 | -0.618898557 | 0.40533225 | -1.526892 | 0.12678788 | 0.37329772 |
| ATP13A1 | 21.2076291 | -0.255322509 | 0.4022862 | -0.6346788 | 0.52563794 | 0.76221787 |
| ZNF101 | 10.6562684 | -0.336529799 | 0.40604357 | -0.8288022 | 0.40721635 | 0.68570495 |
| ZNF14 | 27.525959 | 0.393000109 | 0.32642444 | 1.20395432 | 0.22860723 | 0.51840452 |
| AC011477.4 | 11.3438139 | -0.717895234 | 0.50102647 | -1.4328489 | 0.15190102 | 0.41333768 |
| ZNF253 | 37.5900376 | -0.007021575 | 0.31105579 | -0.0225734 | 0.98199059 | 0.99327119 |
| ZNF93 | 14.7754629 | -0.739247128 | 0.46296168 | -1.5967782 | 0.11031515 | 0.34618593 |
| ZNF682 | 30.0149496 | 0.132868394 | 0.40961459 | 0.32437418 | 0.74565477 | 0.88816458 |
| ZNF90 | 90.6452696 | -0.308107989 | 0.28225909 | -1.0915786 | 0.27501836 | 0.56696009 |
| AC011447.3 | 50.4878999 | -0.336690464 | 0.41530544 | -0.8107056 | 0.41753473 | 0.69378898 |
| ZNF486 | 57.2673961 | -0.869823463 | 0.55861254 | -1.5571141 | 0.1194434 | 0.36182172 |
| AC008554.1 | 20.2566748 | -0.478019654 | 0.89468445 | -0.5342885 | 0.59314192 | 0.80210249 |
| ZNF737 | 40.7864925 | 0.078241882 | 0.32094794 | 0.24378372 | 0.80739833 | 0.91798009 |
| AC010636.2 | 12.9920206 | -0.494189913 | 0.36665214 | -1.3478441 | 0.17770854 | 0.45237296 |
| ZNF66 | 11.7643969 | -0.181695488 | 0.42434336 | -0.4281803 | 0.66851982 | 0.84584093 |
| ZNF85 | 18.3301609 | 0.54979464 | 0.45982718 | 1.19565494 | 0.23183124 | 0.52159221 |
| ZNF430 | 55.1284326 | 0.053218622 | 0.32716213 | 0.16266743 | 0.8707803 | 0.94844301 |
| ZNF714 | 22.6864271 | -0.241768188 | 0.43095182 | -0.5610098 | 0.57479086 | 0.79134063 |
| ZNF431 | 90.3205435 | 0.452701128 | 0.21026767 | 2.15297548 | 0.03132061 | 0.16200382 |
| ZNF708 | 65.2092563 | 0.146823036 | 0.19567487 | 0.75034179 | 0.45304888 | 0.72082001 |
| ZNF738 | 15.5415302 | 0.143060123 | 0.44227702 | 0.32346271 | 0.74634485 | 0.88857559 |
| ZNF493 | 39.1430829 | 0.958596868 | 0.29512782 | 3.24807351 | 0.00116189 | 0.01626334 |
| AC010615.4 | 59.3768368 | -0.213265731 | 0.63343087 | -0.3366835 | 0.73635549 | 0.88437124 |
| ZNF429 | 61.5390388 | 0.039287543 | 0.34557326 | 0.11368803 | 0.9094851 | 0.96308593 |
| AC123912.1 | 22.5284909 | -0.113192887 | 0.58354608 | -0.1939742 | 0.84619608 | 0.93757726 |
| AC123912.4 | 13.3153223 | 0.415056845 | 0.57492475 | 0.72193247 | 0.470336 | 0.73193375 |
| ZNF100 | 70.4655298 | 0.226152147 | 0.33394629 | 0.67721114 | 0.498272 | 0.74991938 |
| ZNF43 | 102.256722 | 0.060395854 | 0.25511884 | 0.23673616 | 0.81286148 | 0.92054639 |
| ZNF208 | 181.455218 | 0.516768741 | 0.37543151 | 1.37646608 | 0.16867738 | 0.43928463 |
| ZNF257 | 12.879406 | -0.199607895 | 0.44188377 | -0.4517204 | 0.65147045 | 0.83781115 |
| ZNF676 | 27.8060414 | 1.198662434 | 0.59306186 | 2.02114235 | 0.04326503 | 0.19853735 |
| ZNF98 | 140.358803 | 0.754026277 | 0.42811179 | 1.76128359 | 0.07819042 | 0.28453467 |
| ZNF99 | 17.9712314 | 0.879940229 | 0.64029934 | 1.37426383 | 0.16935979 | 0.44032355 |
| ZNF730 | 13.3680786 | -0.227680158 | 0.51186359 | -0.4448063 | 0.65645973 | 0.83951126 |
| ZNF91 | 254.814177 | -0.418109501 | 0.3014542 | -1.3869752 | 0.16544932 | 0.43436771 |
| ZNF675 | 36.253809 | 0.594620851 | 0.25217365 | 2.35798166 | 0.0183746 | 0.11417077 |
| AC139769.2 | 14.315313 | -0.264828895 | 0.68953928 | -0.3840664 | 0.70092921 | 0.86426962 |
| ZNF726 | 10.8116712 | 0.015393517 | 0.58641272 | 0.02625031 | 0.97905769 | 0.99205786 |
| ZNF254 | 150.705066 | -0.106273041 | 0.23990593 | -0.442978 | 0.65778167 | 0.84023147 |
| LINC00662 | 51.9066756 | 0.008051189 | 0.43124687 | 0.01866956 | 0.98510471 | 0.99407863 |
| AC006504.5 | 48.7388322 | 0.438998999 | 0.36127018 | 1.21515427 | 0.22430726 | 0.5134 |
| UQCRFS1 | 58.8427618 | -0.114511845 | 0.46316273 | -0.2472389 | 0.80472334 | 0.9165572 |
| C19orf12 | 23.3141743 | -0.587955361 | 0.30667858 | -1.9171712 | 0.05521618 | 0.23198903 |
| URI1 | 182.915935 | 0.189016345 | 0.20529129 | 0.92072268 | 0.35719523 | 0.64174726 |
| ZNF507 | 48.7435722 | 0.369842115 | 0.29024012 | 1.27426255 | 0.20257038 | 0.4861123 |
| DPY19L3 | 58.000034 | -0.665376843 | 0.33066512 | -2.0122378 | 0.04419488 | 0.20044552 |
| PDCD5 | 21.3873714 | -0.025833529 | 0.36323536 | -0.0711206 | 0.94330175 | 0.97824588 |
| ANKRD27 | 38.5636116 | 0.016293739 | 0.36038935 | 0.04521149 | 0.96393874 | 0.98660829 |
| NUDT19 | 13.8372367 | 0.633632404 | 0.38188996 | 1.65920151 | 0.09707519 | 0.32146212 |
| CEP89 | 40.5764745 | -0.220073827 | 0.34026621 | -0.6467696 | 0.5177811 | 0.75866548 |
| RHPN2 | 135.658367 | -1.073405854 | 0.38696932 | -2.7738785 | 0.00553924 | 0.05113371 |
| GPATCH1 | 20.6402278 | 0.408540384 | 0.43528743 | 0.93855315 | 0.34796022 | 0.63501969 |
| LRP3 | 12.8676999 | 0.133847899 | 0.50957872 | 0.26266383 | 0.7928097 | 0.91124908 |
| PEPD | 34.8468732 | -0.440230158 | 0.33824932 | -1.3014961 | 0.19308871 | 0.47361903 |
| LSM14A | 233.452449 | 0.247066973 | 0.17498281 | 1.41194995 | 0.1579647 | 0.42412101 |
| KIAA0355 | 117.591324 | 0.129002584 | 0.23591923 | 0.54680825 | 0.58451047 | 0.7965115 |
| GPI | 179.543438 | -0.195862289 | 0.22224218 | -0.8813011 | 0.37815485 | 0.66119336 |
| UBA2 | 92.5626186 | 0.162874895 | 0.21135773 | 0.77061242 | 0.4409367 | 0.71166064 |
| WTIP | 26.0103492 | 0.483935347 | 0.47346346 | 1.02211761 | 0.30672524 | 0.60139222 |
| SCGB2B2 | 57.6410976 | 0.130739718 | 0.4750607 | 0.27520634 | 0.78315772 | 0.90662892 |
| ZNF302 | 68.1119946 | -0.131928563 | 0.35279803 | -0.3739493 | 0.70844207 | 0.868892 |
| ZNF30 | 11.8291268 | 0.872021539 | 0.68662861 | 1.27000467 | 0.20408297 | 0.48826352 |
| GRAMD1A | 40.5625385 | -1.486768493 | 0.34420548 | -4.3194213 | 1.56E-05 | 0.00053721 |
| HPN | 127.224381 | 0.195541272 | 0.33966731 | 0.5756847 | 0.56482832 | 0.78515099 |
| LSR | 47.8358816 | -1.073741691 | 0.29981838 | -3.5813071 | 0.00034188 | 0.00648627 |
| USF2 | 31.0901144 | -0.150458444 | 0.41719495 | -0.360643 | 0.71836633 | 0.87415486 |
| DMKN | 69.1957052 | -0.701692962 | 0.33394239 | -2.1012396 | 0.03561994 | 0.17524194 |
| TMEM147 | 25.2338655 | -0.220845226 | 0.434114 | -0.5087263 | 0.61094406 | 0.81376433 |
| RBM42 | 13.1465005 | -0.755583059 | 0.56631555 | -1.3342086 | 0.18213551 | 0.45876773 |
| COX6B1 | 106.355337 | -0.113136644 | 0.34674005 | -0.3262866 | 0.7442075 | 0.88798073 |
| KMT2B | 28.7265565 | -0.532955069 | 0.31502689 | -1.6917764 | 0.09068861 | 0.31018641 |
| PROSER3 | 15.9525311 | -0.588163733 | 0.42971936 | -1.3687159 | 0.17108809 | 0.44240699 |
| SYNE4 | 11.297934 | 0.367389559 | 0.43804921 | 0.83869471 | 0.40164065 | 0.68068813 |
| TBCB | 26.2152118 | -0.721003291 | 0.35945523 | -2.0058222 | 0.04487522 | 0.20214451 |
| CAPNS1 | 88.5720777 | -0.871130319 | 0.3766035 | -2.3131233 | 0.02071586 | 0.12328989 |
| COX7A1 | 9.31761887 | 1.621608059 | 0.75867424 | 2.13742339 | 0.03256357 | 0.16607421 |
| ZNF565 | 57.6003472 | -0.100700248 | 0.43444125 | -0.2317926 | 0.81669914 | 0.9224465 |
| ZNF146 | 69.8583245 | 0.133806347 | 0.29108502 | 0.45968133 | 0.64574497 | 0.83395571 |
| LINC00665 | 32.0511194 | -0.339967544 | 0.33854588 | -1.0041993 | 0.31528254 | 0.6086621 |
| ZFP14 | 59.2463369 | -0.017609455 | 0.3094097 | -0.0569131 | 0.95461445 | 0.98226508 |
| ZFP82 | 21.5593919 | -0.552078404 | 0.38079972 | -1.4497868 | 0.14711798 | 0.40654759 |
| ZNF566 | 35.1631361 | 0.103359358 | 0.26938196 | 0.38369072 | 0.70120769 | 0.86450953 |
| ZNF260 | 29.5091368 | -0.195913894 | 0.28327154 | -0.6916116 | 0.48918125 | 0.74545311 |
| ZNF529 | 45.4072069 | -0.157111248 | 0.30879039 | -0.5087958 | 0.61089538 | 0.81376433 |
| ZNF529-AS1 | 41.8928241 | -0.290560507 | 0.33368411 | -0.8707652 | 0.38388238 | 0.66701235 |
| ZNF382 | 11.6672714 | -0.181098767 | 0.40972191 | -0.4420041 | 0.65848623 | 0.84037209 |
| ZNF461 | 24.8218949 | -0.255032521 | 0.41745453 | -0.6109229 | 0.54125064 | 0.77188253 |
| ZNF567 | 26.0374168 | -0.221861493 | 0.36716396 | -0.6042573 | 0.54567261 | 0.77463416 |
| AC020928.1 | 47.7222684 | 0.611043805 | 0.23207265 | 2.63298499 | 0.00846381 | 0.06865421 |
| ZNF790 | 32.6243403 | -0.378924603 | 0.39451303 | -0.9604869 | 0.33681021 | 0.62525668 |
| ZNF345 | 58.7158843 | 0.059625566 | 0.41371084 | 0.14412377 | 0.88540274 | 0.95619452 |
| ZNF568 | 44.4046945 | -0.019183415 | 0.35172719 | -0.0545406 | 0.95650445 | 0.98286826 |
| ZNF420 | 85.8792757 | 0.478717651 | 0.28690894 | 1.66853516 | 0.09520954 | 0.31779803 |
| ZNF383 | 23.508214 | 0.469947169 | 0.37698627 | 1.2465896 | 0.21254802 | 0.49867221 |
| LINC01535 | 15.8731859 | 0.249111617 | 0.4451172 | 0.55965399 | 0.57571547 | 0.79171629 |
| HKR1 | 69.8652357 | 0.374323505 | 0.23925272 | 1.56455276 | 0.11768781 | 0.358386 |
| ZNF569 | 23.2127363 | 0.018446914 | 0.34758394 | 0.05307183 | 0.95767468 | 0.98321101 |
| ZNF793 | 39.1676179 | 0.502017087 | 0.29906249 | 1.67863607 | 0.09322299 | 0.31446734 |
| ZNF540 | 31.3605575 | 0.464784397 | 0.49276897 | 0.94320954 | 0.34557374 | 0.63256941 |
| ZNF571 | 31.257816 | 0.30971749 | 0.39592085 | 0.78227123 | 0.43405519 | 0.70597357 |
| ZFP30 | 34.5205731 | 0.259148368 | 0.307929 | 0.8415848 | 0.4000204 | 0.68010059 |
| ZNF573 | 40.6776273 | 0.268651318 | 0.28655591 | 0.93751797 | 0.34849218 | 0.6354277 |
| SIPA1L3 | 273.178042 | 0.255795409 | 0.25686981 | 0.99581733 | 0.31933891 | 0.611915 |
| SPINT2 | 17.9034249 | -0.231163017 | 0.33556647 | -0.688874 | 0.49090256 | 0.74574113 |
| CATSPERG | 19.8601528 | -0.472320241 | 0.51683554 | -0.9138695 | 0.36078542 | 0.64494385 |
| PSMD8 | 38.9963686 | -0.472422205 | 0.31598779 | -1.4950647 | 0.13489755 | 0.38627205 |

| | | | | | | |
|---|---|---|---|---|---|---|
| EIF3K | 72.7971436 | 0.446677796 | 0.28154664 | 1.58651438 | 0.11262267 | 0.35031364 |
| ACTN4 | 316.883899 | 0.50600772 | 0.3149446 | 1.60665629 | 0.10812979 | 0.34264936 |
| ECH1 | 40.4763642 | -0.362405393 | 0.54534221 | -0.6645467 | 0.50634044 | 0.75306985 |
| HNRNPL | 54.3217608 | 0.322201429 | 0.23890138 | 1.34867962 | 0.17743989 | 0.45213598 |
| SIRT2 | 31.6712155 | 0.623248736 | 0.37544341 | 1.66003374 | 0.09690766 | 0.32146212 |
| NFKBIB | 14.664173 | -0.761343447 | 0.36485635 | -2.0866937 | 0.03691583 | 0.17922097 |
| MRPS12 | 24.5575348 | -0.214624666 | 0.41673068 | -0.5150201 | 0.60653899 | 0.81139653 |
| FBXO17 | 113.583688 | 0.045501542 | 0.33100972 | 0.13746286 | 0.89066495 | 0.95702609 |
| PAK4 | 74.4418975 | 0.863592603 | 0.27078705 | 3.18919465 | 0.0014267 | 0.01903865 |
| SAMD4B | 92.2295056 | 0.247968237 | 0.30880398 | 0.80299559 | 0.42197729 | 0.69753545 |
| PAF1 | 16.176074 | -0.241335303 | 0.42437816 | -0.5686798 | 0.56957343 | 0.78795455 |
| MED29 | 21.6969144 | -0.718539539 | 0.33038365 | -2.1748641 | 0.0296403 | 0.15635146 |
| ZFP36 | 32.8132908 | 0.290903201 | 0.65478315 | 0.44427411 | 0.65684442 | 0.83951711 |
| RPS16 | 77.4821969 | -0.372670188 | 0.41496326 | -0.89808 | 0.36914292 | 0.65369102 |
| SUPT5H | 63.0838417 | -0.350116492 | 0.25474946 | -1.3743561 | 0.16933114 | 0.44032355 |
| TIMM50 | 17.6620631 | 0.423427112 | 0.35620708 | 1.18871055 | 0.23455359 | 0.52479917 |
| DYRK1B | 10.9709601 | -0.457439609 | 0.62413275 | -0.7329204 | 0.463607 | 0.72812334 |
| FBL | 14.4222965 | -0.631467056 | 0.54885033 | -1.1505269 | 0.24992693 | 0.54136822 |
| PSMC4 | 20.6517196 | -0.287167341 | 0.38435173 | -0.7471473 | 0.45497467 | 0.72210564 |
| ZNF546 | 15.9770918 | -0.512788499 | 0.62576276 | -0.8194615 | 0.41252315 | 0.68968086 |
| ZNF780B | 97.4012614 | 0.501509275 | 0.21867395 | 2.29341113 | 0.02182435 | 0.12789216 |
| ZNF780A | 16.1993736 | 0.126618189 | 0.34878639 | 0.36302502 | 0.71658619 | 0.8732017 |
| MAP3K10 | 16.0025369 | -0.109709869 | 0.42100848 | -0.2605883 | 0.79441004 | 0.91176607 |
| AKT2 | 80.7162081 | -0.10379592 | 0.23731943 | -0.437368 | 0.6618445 | 0.84203637 |
| PLD3 | 90.0497388 | -0.404140609 | 0.30308903 | -1.3334056 | 0.18239873 | 0.45912471 |
| SERTAD1 | 14.483126 | -0.963497749 | 0.48283426 | -1.9955041 | 0.04598793 | 0.2058964 |
| BLVRB | 81.7401175 | -0.492886002 | 0.24887149 | -1.980484 | 0.04764917 | 0.21076931 |
| SPTBN4 | 20.1571189 | 1.426927197 | 0.45059116 | 3.16678916 | 0.00154132 | 0.02017622 |
| SHKBP1 | 13.8847738 | -0.157152506 | 0.44811865 | -0.350694 | 0.72581795 | 0.87865765 |
| COQ8B | 19.0227782 | -0.389741901 | 0.46405811 | -0.8398558 | 0.40098923 | 0.6801464 |
| ITPKC | 28.5990149 | -1.339676379 | 0.35162434 | -3.8099648 | 0.00013899 | 0.00318187 |
| SNRPA | 20.2661685 | -0.456742421 | 0.3585742 | -1.2737738 | 0.20274359 | 0.48630139 |
| AC008537.1 | 17.899614 | 0.092691533 | 0.56129507 | 0.16513869 | 0.86883482 | 0.94827167 |
| AXL | 13.00563 | -0.526972913 | 0.67220864 | -0.7839425 | 0.43307386 | 0.70560286 |
| HNRNPUL1 | 49.9002237 | -0.285395819 | 0.24421727 | -1.1686144 | 0.24255903 | 0.53317493 |
| TGFB1 | 37.316486 | -1.40800401 | 0.58872374 | -2.3916209 | 0.01677416 | 0.10800461 |
| DMAC2 | 13.8099287 | -0.080192722 | 0.37070607 | -0.2163243 | 0.82873499 | 0.92870965 |
| RPS19 | 267.005342 | -1.071567685 | 0.407293 | -2.6309504 | 0.00851465 | 0.06885234 |
| ARHGEF1 | 25.6189686 | 0.224784381 | 0.38203832 | 0.58838176 | 0.55627608 | 0.78054429 |
| RABAC1 | 50.2014265 | -0.850637585 | 0.46617467 | -1.8247186 | 0.06804351 | 0.26214816 |
| GRIK5 | 41.3069285 | 0.300173773 | 0.30336239 | 0.98948907 | 0.32242392 | 0.61319341 |
| DEDD2 | 14.0324933 | -0.090576965 | 0.37960262 | -0.23861 | 0.81140804 | 0.9199895 |
| GSK3A | 12.7810143 | 0.170977569 | 0.6111804 | 0.27974976 | 0.7796695 | 0.90427469 |
| CIC | 22.6640281 | -0.702265675 | 0.48373691 | -1.4517513 | 0.14657077 | 0.40573108 |
| PAFAH1B3 | 11.529169 | -1.333261796 | 0.45836253 | -2.9087495 | 0.00362877 | 0.03751236 |
| MEGF8 | 30.7302733 | 0.154626931 | 0.30705987 | 0.50357258 | 0.61456177 | 0.81594643 |
| LIPE-AS1 | 34.9056478 | -0.322448814 | 0.33344657 | -0.9670179 | 0.33353507 | 0.6222525 |
| PHLDB3 | 16.3329502 | -0.725630887 | 0.42427775 | -1.7102733 | 0.08721535 | 0.30375002 |
| ETHE1 | 28.9341642 | -1.249429236 | 0.42690389 | -2.9267226 | 0.00342554 | 0.03590025 |
| ZNF428 | 28.6986172 | -0.987220878 | 0.39060479 | -2.5274162 | 0.01149052 | 0.08449349 |
| SMG9 | 19.0331153 | -0.144067591 | 0.35740996 | -0.4030878 | 0.68688364 | 0.85793832 |
| ZNF283 | 26.5231941 | 0.870238069 | 0.43139478 | 2.01726611 | 0.04366775 | 0.19956366 |
| ZNF45 | 12.8487471 | -0.274058671 | 0.40929773 | -0.6695827 | 0.50312386 | 0.75108109 |
| ZNF155 | 23.2744505 | 0.020894343 | 0.44066714 | 0.04741525 | 0.96218227 | 0.98539792 |
| ZNF224 | 30.827152 | -0.175789087 | 0.35504214 | -0.4951217 | 0.62051418 | 0.81942374 |
| ZNF234 | 16.4639341 | 0.388508222 | 0.39725714 | 0.97797668 | 0.32808586 | 0.61729352 |
| ZNF226 | 29.4098772 | 0.23628481 | 0.32509151 | 0.72682553 | 0.46733284 | 0.73023858 |
| ZNF227 | 17.9915646 | 0.515033315 | 0.35946015 | 1.43279668 | 0.15191595 | 0.41333768 |
| ZNF235 | 74.0288164 | 0.531025298 | 0.43614512 | 1.21754267 | 0.22339781 | 0.51229275 |
| ZNF229 | 17.1710241 | -0.410683429 | 0.36962015 | -1.1110959 | 0.26652708 | 0.55774161 |
| ZNF180 | 12.0961627 | -0.284470815 | 0.45039535 | -0.6316025 | 0.52764667 | 0.76238654 |
| AC243964.2 | 21.54566 | -0.248814268 | 0.49473303 | -0.5029263 | 0.61501607 | 0.816271 |
| CEACAM19 | 16.0913757 | -1.395014836 | 0.61727678 | -2.2599503 | 0.02382434 | 0.13541585 |
| BCL3 | 23.2653816 | -2.20278884 | 0.52146987 | -4.224192 | 2.40E-05 | 0.00075548 |
| CBLC | 22.3017939 | -0.555592677 | 0.47560012 | -1.1681929 | 0.24272896 | 0.53317565 |
| BCAM | 196.78794 | -0.452639639 | 0.43054601 | -1.0513154 | 0.29311377 | 0.5846943 |
| NECTIN2 | 90.3801571 | -0.006858054 | 0.32715292 | -0.0209628 | 0.9832753 | 0.99368608 |
| TOMM40 | 21.228571 | -0.240624463 | 0.38609922 | -0.6232192 | 0.53314046 | 0.76689222 |
| APOE | 57.6194269 | -0.386863119 | 0.75820287 | -0.5102369 | 0.60988547 | 0.81344383 |
| CLPTM1 | 43.5843189 | 0.088091972 | 0.3877503 | 0.22718737 | 0.82027805 | 0.92406023 |
| RELB | 48.329272 | -1.285147905 | 0.4166228 | -3.0846797 | 0.00203771 | 0.02492802 |
| CLASRP | 51.0225188 | -0.114944373 | 0.26273043 | -0.4374993 | 0.66174931 | 0.84203637 |
| GEMIN7-AS1 | 18.108645 | -0.437919775 | 0.41316014 | -1.0599274 | 0.28917761 | 0.58094924 |
| MARK4 | 55.7847406 | -0.038269234 | 0.29426303 | -0.1300511 | 0.89652599 | 0.95900099 |
| TRAPPC6A | 25.1908265 | 0.433261138 | 0.39297673 | 1.10251092 | 0.27023962 | 0.56231236 |
| ERCC2 | 12.0795961 | 0.887084611 | 0.47729872 | 1.85855224 | 0.06309063 | 0.24979235 |
| PPP1R13L | 39.2383703 | -0.735100548 | 0.27542412 | -2.6689767 | 0.00760827 | 0.06389755 |
| ERCC1 | 18.931013 | 0.312960727 | 0.46402021 | 0.67445495 | 0.50002212 | 0.75047026 |
| VASP | 33.6520505 | -0.602047124 | 0.39135029 | -1.5383842 | 0.12395469 | 0.36885651 |
| OPA3 | 58.6692574 | 0.131170571 | 0.43988414 | 0.29819345 | 0.76555552 | 0.89836575 |
| EML2 | 42.9686197 | -0.363023517 | 0.36975445 | -0.9817962 | 0.32620026 | 0.61604309 |
| SNRPD2 | 48.2676119 | -0.962005029 | 0.41641135 | -2.3102277 | 0.02087555 | 0.12372138 |
| FBXO46 | 20.955902 | 0.363079508 | 0.43279789 | 0.83891239 | 0.40151848 | 0.68067193 |
| DMPK | 15.9282891 | -0.26959387 | 0.45132936 | -0.5973329 | 0.55028515 | 0.77615521 |
| SYMPK | 89.7365248 | 0.033304129 | 0.27555784 | 0.12086076 | 0.90380133 | 0.96126516 |
| MYPOP | 18.8569985 | 0.713447088 | 0.39043156 | 1.82732947 | 0.06765025 | 0.26102354 |
| PPP5C | 49.0620931 | 0.055218381 | 0.28618022 | 0.19294968 | 0.84699838 | 0.93758141 |
| CCDC8 | 13.8618037 | -0.052879837 | 0.41179517 | -0.128413 | 0.89782217 | 0.95917897 |
| PNMA8A | 10.7845083 | -0.46657989 | 0.66112325 | -0.7057381 | 0.480351 | 0.73988033 |
| PPP5D1 | 30.7923349 | 0.882175962 | 0.32914298 | 2.68022114 | 0.00735735 | 0.06264087 |
| CALM3 | 41.1263559 | -0.259449747 | 0.57809994 | -0.4487974 | 0.65357782 | 0.83881384 |
| PRKD2 | 50.3150647 | -0.952173733 | 0.33341376 | -2.8558321 | 0.00429242 | 0.04233497 |
| STRN4 | 19.4309357 | 0.141598904 | 0.34273973 | 0.41313828 | 0.6795053 | 0.85389896 |
| FKRP | 18.6936049 | -0.329839458 | 0.41111029 | -0.8023138 | 0.42237146 | 0.69769678 |
| SLC1A5 | 14.4730468 | -0.599050783 | 0.60712708 | -0.9866975 | 0.32379095 | 0.61448136 |
| AP2S1 | 34.1614861 | -0.524579957 | 0.32565123 | -1.6108644 | 0.10720929 | 0.34111768 |
| ARHGAP35 | 130.673938 | 0.345821161 | 0.32263447 | 1.07186677 | 0.28377988 | 0.57640034 |
| TMEM160 | 31.3360554 | -0.579947751 | 0.56301984 | -1.0300663 | 0.30297889 | 0.59684506 |
| ZC3H4 | 31.4372787 | -0.532906838 | 0.39160241 | -1.3608365 | 0.17356537 | 0.44612537 |
| SAE1 | 34.7546794 | -0.055171254 | 0.40111191 | -0.1375458 | 0.8905994 | 0.95702609 |
| CCDC9 | 11.4514388 | -1.029528198 | 0.60576489 | -1.6995508 | 0.08921546 | 0.30749335 |
| INAFM1 | 12.0344152 | 0.010921312 | 0.54467795 | 0.02005095 | 0.98400273 | 0.99374533 |
| C5AR1 | 42.0140159 | 1.011489619 | 0.40110796 | 2.5217391 | 0.01167763 | 0.08526076 |
| DHX34 | 26.2887099 | -0.239297242 | 0.32802668 | -0.7295054 | 0.46569255 | 0.72890726 |
| NAPA-AS1 | 11.2395005 | -0.45482662 | 0.57780216 | -0.7871667 | 0.43118428 | 0.70454366 |
| NAPA | 79.9618566 | 0.240869936 | 0.26194587 | 0.91954088 | 0.35781273 | 0.64208096 |
| NOP53 | 18.4887848 | -0.156350821 | 0.54085248 | -0.2890822 | 0.77251848 | 0.90119866 |
| SELENOW | 79.3925069 | -0.435486041 | 0.39958313 | -1.0898509 | 0.27577882 | 0.56787396 |
| ZSWIM9 | 17.3621802 | -0.017383702 | 0.35100159 | -0.049526 | 0.96050012 | 0.98468228 |
| CARD8 | 36.868843 | 0.340138498 | 0.33134701 | 1.02653257 | 0.30464062 | 0.59905753 |
| TMEM143 | 25.0691178 | 1.291947666 | 0.41267243 | 3.13068571 | 0.00174399 | 0.02215358 |
| KDELR1 | 51.1149213 | -0.68141366 | 0.27344725 | -2.4919382 | 0.01270481 | 0.09057784 |
| CYTH2 | 25.5875881 | -0.715135554 | 0.38941491 | -1.836436 | 0.0662932 | 0.2574031 |

| | | | | | | |
|---|---|---|---|---|---|---|
| LMTK3 | 17.8536607 | -0.344082336 | 0.35686881 | -0.9641704 | 0.33496051 | 0.62350658 |
| RPL18 | 121.78239 | -0.403225418 | 0.52123783 | -0.773592 | 0.4391721 | 0.7103707 |
| SPHK2 | 12.1481463 | -0.319101063 | 0.45720223 | -0.6979429 | 0.48521289 | 0.74285379 |
| BCAT2 | 24.9178946 | 0.027715479 | 0.40825533 | 0.06788761 | 0.9458751 | 0.97894367 |
| HSD17B14 | 20.4720841 | -0.986410337 | 0.61508094 | -1.6037082 | 0.10877839 | 0.34385854 |
| PLEKHA4 | 22.1309212 | -1.349701029 | 0.75377946 | -1.7905782 | 0.07336102 | 0.27372879 |
| NUCB1 | 78.6904324 | -0.505263247 | 0.28933039 | -1.7463193 | 0.08075548 | 0.29058434 |
| BAX | 24.4649038 | -1.339041986 | 0.38317397 | -3.4946058 | 0.00047476 | 0.00826193 |
| FTL | 808.076448 | -1.55461903 | 0.52799569 | -2.9443783 | 0.00323604 | 0.03436877 |
| GYS1 | 58.7411159 | -0.555929216 | 0.35375381 | -1.5715144 | 0.1160632 | 0.35543425 |
| RUVBL2 | 10.5737607 | -0.067440533 | 0.4920188 | -0.137069 | 0.89097624 | 0.95702609 |
| SNRNP70 | 176.343455 | 0.614187604 | 0.22140912 | 2.7739942 | 0.00553727 | 0.05113371 |
| TRPM4 | 45.2544275 | 0.639836259 | 0.45565222 | 1.40422065 | 0.16025315 | 0.42791337 |
| SLC6A16 | 39.1172615 | -0.139115183 | 0.45289531 | -0.3071685 | 0.75871511 | 0.8951693 |
| TEAD2 | 19.4983301 | -1.492345172 | 0.643632 | -2.3186311 | 0.02041505 | 0.12263313 |
| PIH1D1 | 20.986658 | -0.502674473 | 0.34386026 | -1.4618568 | 0.14378044 | 0.40130754 |
| RPL13A | 156.075336 | -0.698951488 | 0.48331516 | -1.4461609 | 0.14813207 | 0.40845709 |
| RPS11 | 125.052364 | -0.384923979 | 0.38834965 | -0.9911789 | 0.32159823 | 0.61263036 |
| FCGRT | 43.7339327 | -0.160832538 | 0.49604785 | -0.3242279 | 0.74576552 | 0.8881938 |
| NOSIP | 30.6703326 | -0.393093687 | 0.28974012 | -1.3567113 | 0.17487297 | 0.44792176 |
| PRRG2 | 11.2901527 | -1.051781656 | 0.44493019 | -2.3639251 | 0.01808247 | 0.11306142 |
| PRR12 | 29.5903622 | -0.14688072 | 0.36066605 | -0.4072485 | 0.68382546 | 0.85606265 |
| SCAF1 | 18.8046078 | -0.412077519 | 0.38744346 | -1.0635811 | 0.28751853 | 0.57987781 |
| IRF3 | 44.5691468 | -0.396126856 | 0.33772304 | -1.172934 | 0.24082226 | 0.53151245 |
| PRMT1 | 37.4295005 | 0.02298172 | 0.33110143 | 0.06940991 | 0.94466334 | 0.97858537 |
| AP2A1 | 29.9444832 | -0.214805483 | 0.38876642 | -0.552531 | 0.58058463 | 0.7947038 |
| MED25 | 33.0978381 | 0.069076357 | 0.32296649 | 0.21388088 | 0.83063995 | 0.93013617 |
| PTOV1 | 32.5195793 | 0.040799668 | 0.40437896 | 0.10089464 | 0.9196341 | 0.96763053 |
| PNKP | 32.0928215 | -0.755891758 | 0.41968211 | -1.8011055 | 0.07168625 | 0.27022019 |
| TBC1D17 | 29.8934772 | -0.077635811 | 0.35633613 | -0.2178724 | 0.82752853 | 0.92808376 |
| IL4I1 | 17.2168822 | -3.106616787 | 0.9506815 | -3.2677787 | 0.00108395 | 0.01544593 |
| ATF5 | 15.0358562 | -0.990712962 | 0.44008679 | -2.2511763 | 0.02437437 | 0.1365442 |
| VRK3 | 39.3161004 | -0.233272856 | 0.2790814 | -0.8358596 | 0.40323392 | 0.68156125 |
| MYH14 | 64.9661729 | -0.292488037 | 0.29598239 | -0.9881941 | 0.32305762 | 0.61384454 |
| KCNC3 | 33.8440723 | 0.988890481 | 0.40304422 | 2.45355331 | 0.01414526 | 0.09650625 |
| NR1H2 | 41.6797229 | 0.129869759 | 0.29280447 | 0.44353748 | 0.65737702 | 0.8398187 |
| EMC10 | 72.0900337 | -0.203775101 | 0.2850836 | -0.7147907 | 0.4747384 | 0.7352452 |
| JOSD2 | 12.1675032 | -0.464711779 | 0.4569256 | -1.0170404 | 0.30913422 | 0.6027062 |
| CLEC11A | 16.0084522 | -0.363035216 | 0.48665103 | -0.7459867 | 0.45567543 | 0.72243279 |
| C19orf48 | 15.988775 | 0.293288551 | 0.51002536 | 0.57504699 | 0.56525952 | 0.78544693 |
| KLK1 | 68.5823637 | 4.743351886 | 1.43277931 | 3.31059491 | 0.00093098 | 0.01392009 |
| ETFB | 51.8126594 | 0.164394563 | 0.48343861 | 0.34005262 | 0.7338169 | 0.88360989 |
| ZNF175 | 14.0238565 | 0.675881355 | 0.41615337 | 1.62411603 | 0.10435105 | 0.33552575 |
| SPACA6 | 34.2549654 | 0.260207223 | 0.42984774 | 0.60534742 | 0.54494818 | 0.77435257 |
| ZNF577 | 57.8931811 | 0.426267002 | 0.31357096 | 1.35939568 | 0.17402124 | 0.44651726 |
| ZNF613 | 13.1813422 | -0.45031502 | 0.42849361 | -1.0509259 | 0.29329264 | 0.58477956 |
| ZNF350-AS1 | 80.1747476 | 0.167268096 | 0.49070905 | 0.34087021 | 0.73320129 | 0.88344324 |
| ZNF350 | 33.9139092 | -0.327058391 | 0.42075955 | -0.7773047 | 0.436979 | 0.70829758 |
| ZNF615 | 20.7098612 | 0.363348764 | 0.39752079 | 0.91403713 | 0.36069733 | 0.64494385 |
| ZNF614 | 12.3973899 | 0.271862013 | 0.42970032 | 0.63267817 | 0.52694383 | 0.76221787 |
| ZNF432 | 41.2721137 | 0.14755384 | 0.33683377 | 0.4380613 | 0.66134185 | 0.84192458 |
| ZNF616 | 13.8930279 | 0.557627781 | 0.50516059 | 1.1038624 | 0.26965283 | 0.56184375 |
| PPP2R1A | 115.156034 | -0.372788391 | 0.36658661 | -1.0169176 | 0.30919261 | 0.6027062 |
| ZNF766 | 32.7995792 | -0.263535624 | 0.27527868 | -0.9573412 | 0.33839506 | 0.62734315 |
| ZNF480 | 31.3911284 | -0.311028948 | 0.43466366 | -0.7155624 | 0.47426163 | 0.73504488 |
| ZNF610 | 11.187004 | 0.38784876 | 0.40852726 | 0.94938281 | 0.34242595 | 0.6305044 |
| ZNF880 | 12.2240614 | -0.192282767 | 0.41362798 | -0.4648689 | 0.64202537 | 0.83291089 |
| ZNF528 | 25.274433 | 0.169439471 | 0.36087989 | 0.46951763 | 0.63869969 | 0.83100331 |
| ZNF808 | 26.8551223 | 0.15871246 | 0.38049835 | 0.41711734 | 0.67659258 | 0.85190134 |
| ZNF701 | 33.7448638 | 0.245847661 | 0.36347062 | 0.67638936 | 0.49879347 | 0.75027574 |
| ZNF83 | 222.910803 | -0.23657353 | 0.26517764 | -0.8921323 | 0.37232203 | 0.65606936 |
| ZNF611 | 100.514287 | -0.17728654 | 0.383389 | -0.4624195 | 0.64378053 | 0.83339931 |
| ZNF600 | 14.3393986 | -0.324507696 | 0.36547705 | -0.8879017 | 0.37459362 | 0.65806152 |
| ZNF28 | 17.8815556 | -0.455206635 | 0.35578126 | -1.2794565 | 0.20073635 | 0.48430583 |
| AC010487.3 | 14.6019851 | -0.205230665 | 0.43781391 | -0.4687623 | 0.63923954 | 0.83121032 |
| ZNF160 | 78.3701717 | -0.463955433 | 0.23780471 | -1.9509934 | 0.05105783 | 0.21999069 |
| ZNF347 | 27.5857892 | -0.465176501 | 0.35164144 | -1.3228717 | 0.18587805 | 0.46309931 |
| ZNF677 | 20.6108753 | -0.54565569 | 0.34572637 | -1.5782877 | 0.11449953 | 0.35348341 |
| ZNF525 | 20.5379385 | -0.441394361 | 0.40577623 | -1.0877778 | 0.27669323 | 0.56884726 |
| ZNF765 | 22.9239714 | -0.108699665 | 0.33190815 | -0.3274992 | 0.74329032 | 0.88750312 |
| ZNF761 | 33.2226834 | -0.131177744 | 0.35851 | -0.365897 | 0.71444191 | 0.8724728 |
| ZNF331 | 120.174605 | 2.811417534 | 1.1053244 | 2.54352254 | 0.0109741 | 0.08149584 |
| NDUFA3 | 21.7164736 | -0.608364762 | 0.69718613 | -0.8726002 | 0.38288103 | 0.66617807 |
| PRPF31 | 33.0202993 | -0.172216151 | 0.32976378 | -0.5222409 | 0.60150258 | 0.80759541 |
| CNOT3 | 20.848703 | -0.126045631 | 0.34639723 | -0.363876 | 0.71595062 | 0.87318675 |
| TMC4 | 75.4413014 | -0.030600567 | 0.24486604 | -0.1249686 | 0.90054841 | 0.95980075 |
| MBOAT7 | 69.8001082 | -1.109238857 | 0.34372989 | -3.2270654 | 0.00125067 | 0.01731773 |
| TSEN34 | 24.3063895 | -0.536267393 | 0.46971276 | -1.1416922 | 0.25358198 | 0.54524749 |
| RPS9 | 251.949287 | -0.671081724 | 0.36276934 | -1.8498854 | 0.06433006 | 0.25266043 |
| LENG8-AS1 | 16.234631 | -0.258396476 | 0.3877074 | -0.6664729 | 0.50510888 | 0.75184725 |
| LENG8 | 113.457248 | 0.664320319 | 0.39161395 | 1.69636529 | 0.08981672 | 0.30870579 |
| AC011476.3 | 18.4892583 | -1.108593993 | 0.6890573 | -1.608856 | 0.10764782 | 0.34161321 |
| RDH13 | 53.2791163 | 0.274787301 | 0.31825191 | 0.86342703 | 0.38790272 | 0.66971239 |
| EPS8L1 | 22.9201796 | -0.041268887 | 0.44411174 | -0.0929246 | 0.9259635 | 0.97111125 |
| PPP1R12C | 33.8682402 | 0.291213254 | 0.33379725 | 0.87242556 | 0.38297626 | 0.66623125 |
| PPP6R1 | 33.1535319 | -0.372374824 | 0.34263051 | -1.0868116 | 0.27712007 | 0.56927037 |
| HSPBP1 | 18.5084233 | 0.23806508 | 0.5254234 | 0.45309189 | 0.65048258 | 0.83735207 |
| KMT5C | 10.3076743 | -0.671173332 | 0.46450186 | -1.4449314 | 0.14847715 | 0.40880504 |
| TMEM238 | 24.7517584 | -0.240217505 | 0.37968476 | -0.6326762 | 0.52694512 | 0.76221787 |
| RPL28 | 185.913372 | -1.012300686 | 0.49717895 | -2.0360892 | 0.0417414 | 0.19502037 |
| ISOC2 | 19.8426115 | -0.722770107 | 0.47371129 | -1.5257608 | 0.12706945 | 0.37382797 |
| NAT14 | 11.9685895 | -0.834664461 | 0.79912595 | -1.0444717 | 0.29626719 | 0.58819515 |
| ZNF524 | 10.5831668 | -0.374495551 | 0.56855607 | -0.6586783 | 0.51010236 | 0.75482254 |
| CCDC106 | 10.1223666 | 0.81951943 | 0.62367408 | 1.31401875 | 0.1888399 | 0.46776355 |
| U2AF2 | 22.9143371 | 0.056996832 | 0.31442279 | 0.1812745 | 0.85615212 | 0.9432231 |
| EPN1 | 57.6860155 | 0.240516344 | 0.29206846 | 0.82349303 | 0.41022768 | 0.68796444 |
| ZNF787 | 19.2944432 | -0.403807333 | 0.36537073 | -1.1051989 | 0.2690734 | 0.5613597 |
| ZNF444 | 29.7247452 | 0.012429608 | 0.29101043 | 0.0427119 | 0.96593119 | 0.98661741 |
| ZSCAN5A | 81.6244226 | 0.058052821 | 0.29068609 | 0.19970967 | 0.84170765 | 0.93566718 |
| ZNF583 | 11.2903976 | -0.437987924 | 0.47530177 | -0.9214944 | 0.35679235 | 0.64141201 |
| ZNF667 | 22.2118532 | -0.975735128 | 0.35939675 | -2.7149247 | 0.00662908 | 0.05803619 |
| ZNF667-AS1 | 31.7418634 | -0.371252872 | 0.33809327 | -1.0980783 | 0.27217031 | 0.56475837 |
| ZNF471 | 27.6781228 | 0.027341002 | 0.33813729 | 0.0808577 | 0.93555512 | 0.97531764 |
| AC005498.3 | 13.8880277 | 0.098442859 | 0.41457141 | 0.23745694 | 0.81230232 | 0.92034001 |
| ZNF264 | 37.9912218 | -0.070465615 | 0.26562157 | -0.2652857 | 0.79078935 | 0.91080793 |
| ZNF460-AS1 | 16.0860805 | 0.028415353 | 0.53902438 | 0.05271627 | 0.95795798 | 0.98321101 |
| ZNF460 | 15.5075769 | -0.097531351 | 0.36749648 | -0.265394 | 0.79070598 | 0.91080793 |
| AC005261.1 | 55.3324919 | -0.294217661 | 0.30075596 | -0.9782605 | 0.32794552 | 0.61729352 |
| AC003002.2 | 10.3796603 | 0.961531729 | 0.53481122 | 1.79788997 | 0.07219445 | 0.27144058 |
| AC004076.1 | 28.8738655 | 0.326727375 | 0.44268585 | 0.73805697 | 0.46047983 | 0.72547228 |
| ZNF549 | 19.1879324 | -0.051220087 | 0.34762862 | -0.1473414 | 0.88286256 | 0.95518274 |
| ZNF134 | 10.4660631 | -0.300294856 | 0.50942575 | -0.5894772 | 0.55554121 | 0.77993807 |
| AC003006.1 | 12.6858178 | -0.019059046 | 0.48945941 | -0.038939 | 0.96893905 | 0.98770265 |

| | | | | | | |
|---|---|---|---|---|---|---|
| ZNF154 | 14.9674675 | 0.712174508 | 0.40455953 | 1.76037012 | 0.07834507 | 0.28477085 |
| ZNF586 | 113.392792 | 0.160017444 | 0.22939847 | 0.69755235 | 0.48545719 | 0.74292145 |
| ZNF552 | 42.7808148 | 0.540300562 | 0.39691299 | 1.36125693 | 0.17343251 | 0.44612537 |
| ZNF814 | 75.6733514 | 0.296833709 | 0.32776567 | 0.90562781 | 0.36513288 | 0.65045806 |
| ZNF587 | 37.1728463 | 0.564933594 | 0.30518547 | 1.85111561 | 0.06415292 | 0.2521767 |
| ZNF418 | 14.1317732 | -0.009077321 | 0.39567469 | -0.0229414 | 0.98169704 | 0.99318267 |
| C19orf18 | 12.0666746 | 0.539793161 | 0.42257929 | 1.27737723 | 0.2014691 | 0.4848871 |
| ZNF606 | 19.1984297 | -0.070018795 | 0.33254765 | -0.2105527 | 0.83323635 | 0.93223294 |
| ZNF135 | 35.9693095 | -0.235937476 | 0.32344959 | -0.7294413 | 0.46573178 | 0.72890726 |
| ZSCAN18 | 71.5246303 | -0.234257599 | 0.24414589 | -0.9594984 | 0.3373077 | 0.62600324 |
| ZNF329 | 28.5971622 | 0.069121246 | 0.31761661 | 0.21762479 | 0.82772147 | 0.92813421 |
| ZNF274 | 37.8813801 | -0.421842983 | 0.33548229 | -1.2574225 | 0.20860069 | 0.49402397 |
| AC020915.3 | 18.8533143 | 0.622696235 | 0.39006338 | 1.59639757 | 0.11040006 | 0.34622265 |
| RPS5 | 70.5330573 | -0.923468347 | 0.42343658 | -2.1808894 | 0.0291916 | 0.15469745 |
| ZNF584 | 9.48496727 | -0.330714599 | 0.48185939 | -0.6863301 | 0.49250498 | 0.74674012 |
| SLC27A5 | 11.696629 | 0.130186637 | 0.48953856 | 0.26593745 | 0.79028738 | 0.91080793 |
| TRIM28 | 35.5556345 | -0.101454041 | 0.41362952 | -0.2452776 | 0.80624153 | 0.91724035 |
| CHMP2A | 33.0483578 | -0.801981679 | 0.48849695 | -1.6417332 | 0.1006453 | 0.32780522 |
| UBE2M | 26.5707672 | -0.467892419 | 0.32769218 | -1.4278413 | 0.15333756 | 0.41592509 |
| MZF1-AS1 | 12.9958663 | -0.057494787 | 0.49199514 | -0.1168605 | 0.90697062 | 0.96167263 |
| MZF1 | 54.520444 | -0.30967997 | 0.33152858 | -0.9340973 | 0.35025368 | 0.63689665 |
| RPS4Y1 | 66.5642906 | 0.526175701 | 1.9872961 | 0.26476965 | 0.79118692 | 0.91099595 |
| ZFY | 24.705633 | 1.165743846 | 1.77164554 | 0.65800061 | 0.51053774 | 0.7549696 |
| LINC00278 | 168.915305 | 0.862189439 | 2.47210118 | 0.34876786 | 0.72726359 | 0.87968411 |
| TBL1Y | 45.5594598 | 0.952942614 | 1.95445322 | 0.48757504 | 0.62585088 | 0.82312214 |
| TTTY15 | 34.7154845 | 1.167403576 | 2.10070666 | 0.55571946 | 0.57840265 | 0.7935467 |
| USP9Y | 127.13777 | 1.102486701 | 2.39652708 | 0.46003515 | 0.64549099 | 0.83387847 |
| DDX3Y | 36.8837691 | 0.991262792 | 2.07434521 | 0.4778678 | 0.6327443 | 0.82649065 |
| UTY | 303.318669 | 1.215755712 | 2.05977167 | 0.5902381 | 0.55503103 | 0.77954179 |
| NLGN4Y | 50.4974739 | -1.136505631 | 2.02519793 | -0.5611825 | 0.57467313 | 0.79134063 |
| TTTY14 | 339.616502 | 0.917191884 | 1.14443565 | 0.80143596 | 0.4228793 | 0.69782197 |
| KDM5D | 34.562307 | 1.21887801 | 2.07735889 | 0.58674407 | 0.5573756 | 0.78155485 |
| TTTY10 | 19.6290661 | 1.145682926 | 1.52209193 | 0.75270284 | 0.4516285 | 0.71978292 |
| EIF1AY | 13.2667999 | 0.355506772 | 1.84205785 | 0.19299436 | 0.84696339 | 0.93758141 |
| IL17RA | 23.4694963 | -0.463989584 | 0.35425312 | -1.3097685 | 0.19027415 | 0.46973504 |
| HDHD5 | 23.7789427 | -0.404020552 | 0.38158547 | -1.0587944 | 0.28969344 | 0.58130538 |
| HDHD5-AS1 | 9.57096203 | -0.61541438 | 0.64527731 | -0.9537208 | 0.340225 | 0.62907597 |
| ADA2 | 16.4428513 | 1.13257816 | 0.55402441 | 2.04427482 | 0.04092641 | 0.19268019 |
| CECR2 | 69.5398583 | 1.044356151 | 0.35478779 | 2.94360795 | 0.00324411 | 0.03441894 |
| ATP6V1E1 | 67.5899742 | -0.507402291 | 0.25125742 | -2.0194519 | 0.04344027 | 0.19906327 |
| BCL2L13 | 126.29814 | 0.153602379 | 0.20150313 | 0.76228286 | 0.4458912 | 0.71562099 |
| MICAL3 | 135.814322 | -0.225076441 | 0.33757178 | -0.6667513 | 0.50493098 | 0.75182297 |
| PEX26 | 43.9885307 | -0.186227449 | 0.27815979 | -0.6694981 | 0.5031778 | 0.75108109 |
| USP18 | 13.8238994 | -1.331540619 | 1.14945537 | -1.15841 | 0.24669675 | 0.53764558 |
| DGCR2 | 80.138799 | 0.017139715 | 0.26184863 | 0.06545658 | 0.94781048 | 0.979567 |
| SLC25A1 | 35.3228686 | -0.352894697 | 0.43484185 | -0.8115472 | 0.41705149 | 0.69338933 |
| CLTCL1 | 24.4688469 | -0.445401385 | 0.37820783 | -1.177663 | 0.23893097 | 0.52912013 |
| HIRA | 25.9191088 | -0.130352049 | 0.45527144 | -0.2863172 | 0.77463518 | 0.90205458 |
| MRPL40 | 21.3831743 | -0.890724634 | 0.51349921 | -1.7346174 | 0.0828086 | 0.29488221 |
| UFD1 | 27.1337177 | -0.073149767 | 0.39421308 | -0.185559 | 0.85279064 | 0.94049004 |
| TXNRD2 | 27.7154498 | -0.011695207 | 0.39624161 | -0.0295153 | 0.97645358 | 0.99049082 |
| COMT | 82.0309196 | 0.037788829 | 0.2948157 | 0.1281778 | 0.89800827 | 0.95917897 |
| TANGO2 | 45.5974123 | -0.012367743 | 0.40792216 | -0.0303189 | 0.97581274 | 0.99024058 |
| DGCR8 | 27.6521529 | -0.493755009 | 0.48811097 | -1.011563 | 0.31174703 | 0.60482447 |
| RANBP1 | 15.4021314 | -0.519511463 | 0.50867671 | -1.0212999 | 0.30711238 | 0.60139222 |
| ZDHHC8 | 17.649663 | 1.176980016 | 0.44877622 | 2.62264347 | 0.00872505 | 0.06984109 |
| DGCR6L | 11.3823319 | -0.326553881 | 0.59579921 | -0.5480938 | 0.58362747 | 0.79598942 |
| KLHL22 | 58.7617455 | 0.336201338 | 0.26160221 | 1.28516244 | 0.19873553 | 0.48150832 |
| MED15 | 92.1292958 | -0.22737184 | 0.22512911 | -1.009962 | 0.31251351 | 0.60551329 |
| PI4KA | 125.366674 | 0.006629267 | 0.2001426 | 0.03312272 | 0.97357673 | 0.98903337 |
| SNAP29 | 27.3211119 | -0.350959238 | 0.31555372 | -1.1122012 | 0.26605163 | 0.55721548 |
| CRKL | 30.7779705 | 0.160168261 | 0.34905969 | 0.45885636 | 0.64633732 | 0.83447986 |
| LZTR1 | 12.9728452 | 0.443927613 | 0.41487171 | 1.07003588 | 0.28460316 | 0.57686567 |
| GGT2 | 97.0432577 | -2.178307219 | 0.81158263 | -2.684024 | 0.00727419 | 0.06215285 |
| HIC2 | 28.0403008 | 0.832967711 | 0.36930489 | 2.25550141 | 0.02410187 | 0.13583014 |
| UBE2L3 | 85.240742 | -0.243187454 | 0.23836317 | -1.0202392 | 0.30761502 | 0.60139222 |
| SDF2L1 | 21.3732533 | -1.112200869 | 0.58453888 | -1.9026978 | 0.05707998 | 0.23570501 |
| PPIL2 | 62.021236 | 0.350993108 | 0.35192899 | 0.9973407 | 0.31859916 | 0.61144847 |
| YPEL1 | 11.5316879 | -1.106603427 | 0.61235316 | -1.8071327 | 0.07074159 | 0.26862509 |
| MAPK1 | 162.291098 | -0.330396785 | 0.2779794 | -1.1885657 | 0.23461061 | 0.52479917 |
| PPM1F | 20.7367506 | -0.421709291 | 0.50249809 | -0.8392257 | 0.40134269 | 0.68048591 |
| TOP3B | 13.9956632 | 0.0311599 | 0.45351132 | 0.0687081 | 0.94522197 | 0.97858537 |
| AC245060.5 | 40.1173395 | 0.536937952 | 0.40513246 | 1.32533927 | 0.18505865 | 0.46185874 |
| RSPH14 | 10.6864082 | -0.061422732 | 0.47710767 | -0.1287398 | 0.89756356 | 0.95917897 |
| RAB36 | 13.2417343 | 0.034398777 | 0.52937825 | 0.06497958 | 0.94819026 | 0.97962835 |
| BCR | 196.476616 | -0.481416938 | 0.38567734 | -1.2482376 | 0.21194407 | 0.49805901 |
| CHCHD10 | 154.174224 | -0.18732803 | 0.36909107 | -0.5075388 | 0.61177683 | 0.81385911 |
| SMARCB1 | 47.7441994 | -0.056481563 | 0.32176536 | -0.1755365 | 0.8606581 | 0.945045 |
| DDT | 70.7053142 | -0.152376552 | 0.54340119 | -0.2804126 | 0.77916095 | 0.90427469 |
| CABIN1 | 158.67924 | 0.188680058 | 0.21141193 | 0.89247593 | 0.37213789 | 0.65590496 |
| GUCD1 | 24.8455263 | -1.523959922 | 0.39037575 | -3.9038284 | 9.47E-05 | 0.00231143 |
| SNRPD3 | 44.6849858 | -0.560282389 | 0.26186658 | -2.1395719 | 0.03238938 | 0.16565474 |
| SGSM1 | 101.651013 | 0.434555769 | 0.41678784 | 1.04263064 | 0.29711939 | 0.58943269 |
| KIAA1671 | 152.500434 | -0.272739005 | 0.28052304 | -0.9722517 | 0.33092534 | 0.62064315 |
| LRP5L | 55.2823457 | 0.373426521 | 0.37886456 | 0.98564648 | 0.32430662 | 0.61460758 |
| GRK3 | 41.8645284 | 0.242031232 | 0.37649047 | 0.64286151 | 0.52031397 | 0.76073677 |
| HPS4 | 47.7060567 | -0.358423355 | 0.25851719 | -1.3864585 | 0.16560695 | 0.43438116 |
| TFIP11 | 14.6695025 | -0.315227206 | 0.45765811 | -0.6887832 | 0.49095972 | 0.74574113 |
| TPST2 | 62.2065684 | 0.670464478 | 0.3690056 | 1.81694932 | 0.06922487 | 0.26510127 |
| MIATNB | 39.9147823 | 0.047814087 | 0.40567176 | 0.11786398 | 0.90617544 | 0.96157183 |
| PITPNB | 118.350655 | 0.113292038 | 0.25230041 | 0.44903628 | 0.65340549 | 0.83881384 |
| TTC28 | 199.930381 | -0.292572382 | 0.23792148 | -1.2297014 | 0.21880893 | 0.50678272 |
| HSCB | 39.3227894 | -0.735915328 | 0.27624906 | -2.663956 | 0.00772277 | 0.06419577 |
| CCDC117 | 17.7167083 | 0.257501842 | 0.38212534 | 0.67386749 | 0.50039556 | 0.75059335 |
| XBP1 | 58.4653409 | -1.388135858 | 0.35761134 | -3.8816886 | 0.00010373 | 0.00249688 |
| Z93930.2 | 78.244553 | 0.127732705 | 0.38938494 | 0.3280371 | 0.74288361 | 0.88740176 |
| ZNRF3 | 221.625666 | 0.128474624 | 0.23835532 | 0.53900464 | 0.58988366 | 0.79960282 |
| KREMEN1 | 56.838151 | 0.666384435 | 0.28957737 | 2.30123103 | 0.02137858 | 0.12585262 |
| RHBDD3 | 14.8282357 | -0.372985742 | 0.38341158 | -0.9728077 | 0.33064887 | 0.62024005 |
| EWSR1 | 115.50507 | 0.544504346 | 0.28618003 | 1.90266364 | 0.05708445 | 0.23570501 |
| GAS2L1 | 9.65104979 | 0.125646337 | 0.60958905 | 0.20611646 | 0.83669994 | 0.93419636 |
| AP1B1 | 46.8765142 | 0.08756594 | 0.27617308 | 0.31706906 | 0.7511912 | 0.89132635 |
| THOC5 | 15.4273486 | -0.744114581 | 0.3774583 | -1.9713822 | 0.04868018 | 0.21414869 |
| NIPSNAP1 | 26.6974314 | -0.167321207 | 0.36283054 | -0.4611553 | 0.64468722 | 0.83381793 |
| NF2 | 71.6885631 | 0.683955122 | 0.33181908 | 2.06122905 | 0.03928119 | 0.18700317 |
| ZMAT5 | 11.2085499 | 0.353764865 | 0.45404732 | 0.77913655 | 0.43589928 | 0.70752078 |
| UQCR10 | 116.256855 | -0.896716179 | 0.54027758 | -1.6597324 | 0.09696831 | 0.32146212 |
| ASCC2 | 38.0856425 | -0.233278516 | 0.23647522 | -0.9864819 | 0.32389672 | 0.61448136 |
| MTMR3 | 211.96325 | 0.291164911 | 0.22915186 | 1.2706199 | 0.2038639 | 0.48819292 |
| TBC1D10A | 17.843772 | -0.193734355 | 0.33531646 | -0.5777657 | 0.5634223 | 0.78465819 |
| SF3A1 | 18.6782013 | -0.405521679 | 0.4086336 | -0.9923846 | 0.32100997 | 0.6120836 |
| RNF215 | 23.1084043 | -0.557439958 | 0.52664328 | -1.0584773 | 0.28983789 | 0.58136876 |
| SEC14L6 | 21.8210579 | -0.274380661 | 0.57405836 | -0.4779665 | 0.63267405 | 0.82649065 |

| | | | | | | |
|---|---|---|---|---|---|---|
| GAL3ST1 | 11.4948496 | -0.852411559 | 0.44725266 | -1.9058837 | 0.05666529 | 0.23482134 |
| PES1 | 17.4514873 | -0.635111991 | 0.41073929 | -1.5462655 | 0.12204046 | 0.36622803 |
| TCN2 | 18.1971057 | -0.074130129 | 0.711882 | -0.1041326 | 0.91706411 | 0.96591294 |
| DUSP18 | 23.8623384 | -0.811422985 | 0.37849793 | -2.1437977 | 0.0320491 | 0.16434537 |
| OSBP2 | 62.8593329 | 0.552729999 | 0.30154371 | 1.83300126 | 0.06680238 | 0.25881841 |
| MORC2 | 29.6759836 | -0.016958049 | 0.30386679 | -0.0558075 | 0.95549515 | 0.98258246 |
| SMTN | 12.4013106 | 1.48166708 | 0.52578365 | 2.81801663 | 0.00483213 | 0.04630755 |
| SELENOM | 66.6676488 | -0.381963668 | 0.58812284 | -0.6494624 | 0.51603954 | 0.75751487 |
| INPP5J | 62.1609854 | 1.770376547 | 0.49567585 | 3.57164172 | 0.00035475 | 0.0066569 |
| RNF185 | 57.570324 | 0.287899045 | 0.29695251 | 0.96951209 | 0.33228975 | 0.62173066 |
| LIMK2 | 63.9653969 | -0.576655512 | 0.25341692 | -2.275521 | 0.0228747 | 0.13202318 |
| PIK3IP1 | 25.5287144 | -1.233254395 | 0.41795656 | -2.950676 | 0.00317079 | 0.03391661 |
| PIK3IP1-AS1 | 14.5838081 | -0.927199013 | 0.51434526 | -1.8026783 | 0.07143876 | 0.26978083 |
| PATZ1 | 35.0951895 | 0.156110463 | 0.30697686 | 0.50854147 | 0.61107366 | 0.81376433 |
| DRG1 | 56.527752 | -0.02602208 | 0.30311856 | -0.0858479 | 0.93158736 | 0.97334817 |
| EIF4ENIF1 | 56.3139922 | 0.074885887 | 0.24111807 | 0.31057766 | 0.75612171 | 0.89381134 |
| SFI1 | 54.3054389 | -0.178100387 | 0.37237798 | -0.4782785 | 0.63245199 | 0.82649065 |
| PISD | 123.967103 | 0.299721524 | 0.25244704 | 1.18726494 | 0.23512313 | 0.5253174 |
| PRR14L | 104.262018 | -0.141878298 | 0.23773874 | -0.5967824 | 0.55065266 | 0.77635488 |
| YWHAH | 27.4461606 | -1.63437063 | 0.54290118 | -3.0104385 | 0.00260871 | 0.02956535 |
| RTCB | 49.5372151 | -0.477239725 | 0.22527923 | -2.1184364 | 0.03413812 | 0.17086443 |
| FBXO7 | 55.6476353 | -0.079000182 | 0.26993177 | -0.2926672 | 0.76977652 | 0.90034488 |
| SYN3 | 19.1149782 | 1.47867037 | 0.63591017 | 2.32528184 | 0.02005689 | 0.12097549 |
| TIMP3 | 222.702024 | 1.482936069 | 0.44175407 | 3.35692675 | 0.00078814 | 0.01235832 |
| LARGE1 | 305.3289 | -0.386071293 | 0.39979835 | -0.965665 | 0.33421182 | 0.62283326 |
| HMGXB4 | 41.8472311 | -0.184447836 | 0.31031831 | -0.5943827 | 0.55225618 | 0.77744509 |
| TOM1 | 53.3284598 | -0.031295094 | 0.27980455 | -0.1118463 | 0.9109453 | 0.96380389 |
| APOL6 | 34.0779538 | -1.964852216 | 0.75548114 | -2.6007958 | 0.00930078 | 0.07314246 |
| RBFOX2 | 365.423961 | 0.377063042 | 0.18906278 | 1.99438004 | 0.04611054 | 0.2061041 |
| APOL2 | 42.0030934 | -1.265728573 | 0.64275122 | -1.9692356 | 0.04892604 | 0.21468395 |
| APOL1 | 25.2330608 | -2.413438133 | 0.72159466 | -3.3445898 | 0.00082404 | 0.01270853 |
| MYH9 | 171.510663 | 0.135162696 | 0.2814675 | 0.48020711 | 0.63108013 | 0.82620251 |
| TXN2 | 35.3651655 | -0.19128284 | 0.3818975 | -0.5008748 | 0.61645923 | 0.81691188 |
| EIF3D | 40.838714 | -0.821364321 | 0.37454216 | -2.1929823 | 0.02830866 | 0.15134187 |
| IFT27 | 18.7105315 | 0.334009743 | 0.38854787 | 0.85963602 | 0.38998971 | 0.67201909 |
| PVALB | 33.3136002 | 2.226301047 | 0.47697351 | 4.66755701 | 3.05E-06 | 0.00014054 |
| TST | 21.8394809 | -1.093302132 | 0.53926001 | -2.0274119 | 0.04262031 | 0.19708211 |
| MPST | 25.2860617 | -0.882416791 | 0.38105235 | -2.3157364 | 0.02057267 | 0.12280939 |
| CARD10 | 21.4087378 | 0.006029834 | 0.46192597 | 0.01305368 | 0.98958497 | 0.99645238 |
| CDC42EP1 | 20.1634132 | -1.073101857 | 0.55745188 | -1.9250126 | 0.05422779 | 0.22923868 |
| GGA1 | 52.6300315 | 0.17093295 | 0.2370413 | 0.7211104 | 0.47084159 | 0.73216756 |
| LGALS1 | 22.2082621 | -2.339445712 | 0.56880446 | -4.1129173 | 3.91E-05 | 0.0011402 |
| TRIOBP | 73.0226952 | -0.664391928 | 0.26502999 | -2.5068556 | 0.01218105 | 0.08788108 |
| H1F0 | 53.0319764 | -1.783016529 | 0.72810053 | -2.4488603 | 0.0143309 | 0.09732164 |
| GCAT | 11.244394 | 0.070785604 | 0.50108561 | 0.14126449 | 0.887661 | 0.95635679 |
| ANKRD54 | 15.7333369 | -0.161067174 | 0.41203123 | -0.3909101 | 0.69586368 | 0.86236044 |
| EIF3L | 73.3225478 | 0.344971843 | 0.33608461 | 1.02644344 | 0.30468261 | 0.59905753 |
| MICALL1 | 14.7488897 | -0.569913855 | 0.44074326 | -1.2930745 | 0.1959853 | 0.4776213 |
| POLR2F | 26.8735688 | 0.377413166 | 0.32374947 | 1.16575687 | 0.24371276 | 0.53451494 |
| PICK1 | 21.315395 | -0.1227999 | 0.35291276 | -0.3479611 | 0.72786943 | 0.88010691 |
| PLA2G6 | 22.7520358 | 0.40596279 | 0.36244005 | 1.12008258 | 0.26267857 | 0.55434704 |
| MAFF | 18.6815935 | -1.707891194 | 0.70138502 | -2.4350266 | 0.01489069 | 0.09961294 |
| TMEM184B | 43.8353281 | -0.318629196 | 0.34463364 | -0.9245447 | 0.35520281 | 0.64025612 |
| CSNK1E | 15.3132521 | 0.015713772 | 0.43229373 | 0.03634976 | 0.97100348 | 0.988268 |
| TPTEP2-CSNI | 113.40594 | 0.142480183 | 0.2335336 | 0.61010571 | 0.54179179 | 0.77222629 |
| DDX17 | 781.66943 | 0.717996697 | 0.11579342 | 6.20066937 | 5.62E-10 | 1.05E-07 |
| FAM227A | 24.9353074 | -0.767783805 | 0.43621992 | -1.7600842 | 0.07839353 | 0.28477085 |
| CBY1 | 32.0528459 | -0.067822738 | 0.2695064 | -0.2516554 | 0.80130744 | 0.91484195 |
| TOMM22 | 13.6919185 | -0.663698223 | 0.64270403 | -1.0326654 | 0.30176051 | 0.59542937 |
| JOSD1 | 34.4506477 | -0.490909422 | 0.57820331 | -0.8490256 | 0.39586704 | 0.67620996 |
| GTPBP1 | 24.3542702 | -0.542493768 | 0.41308069 | -1.3132876 | 0.18908604 | 0.46792323 |
| SUN2 | 28.7071078 | -0.50648379 | 0.40580972 | -1.248082 | 0.21200104 | 0.49805901 |
| AL021707.1 | 17.5709997 | -0.854026437 | 0.38826978 | -2.1995697 | 0.02783744 | 0.1496771 |
| CBX6 | 43.8587543 | -0.338152969 | 0.4527952 | -0.7468122 | 0.45517694 | 0.72210564 |
| CBX7 | 28.4766094 | 0.130551776 | 0.30735278 | 0.42476199 | 0.67101019 | 0.84746193 |
| RPL3 | 156.142458 | -0.21857744 | 0.4962488 | -0.4404594 | 0.65960442 | 0.84097773 |
| SYNGR1 | 26.3691161 | 0.163848903 | 0.41557478 | 0.39427056 | 0.69338129 | 0.8606282 |
| TAB1 | 24.9932606 | 0.188512509 | 0.28631069 | 0.65841939 | 0.51026868 | 0.7548518 |
| MIEF1 | 17.8869263 | -0.24010681 | 0.34777813 | -0.6904023 | 0.48994125 | 0.7455572 |
| ATF4 | 55.7509272 | -0.788317713 | 0.46473912 | -1.6962586 | 0.08983692 | 0.30870579 |
| RPS19BP1 | 29.4758049 | -0.572070918 | 0.38288898 | -1.4940908 | 0.13515188 | 0.38686708 |
| FAM83F | 29.2265254 | -0.737324093 | 0.42242109 | -1.7454718 | 0.08090278 | 0.29075313 |
| TNRC6B | 689.763582 | 0.335036142 | 0.21173816 | 1.58231343 | 0.11357804 | 0.35211584 |
| MKL1 | 298.695827 | 0.483968311 | 0.34957775 | 1.38443682 | 0.16622474 | 0.43507299 |
| SLC25A17 | 54.2381538 | -0.017850548 | 0.22042022 | -0.0809842 | 0.93545454 | 0.97531148 |
| ST13 | 100.858919 | -0.092074947 | 0.23217364 | -0.396578 | 0.69167871 | 0.85976938 |
| XPNPEP3 | 70.5060883 | 0.03269471 | 0.23165191 | 0.14113723 | 0.88776153 | 0.95636508 |
| RBX1 | 53.8729493 | 0.156816462 | 0.30559265 | 0.51315521 | 0.60784275 | 0.81187554 |
| EP300 | 164.435797 | -0.086206876 | 0.2211615 | -0.3897915 | 0.69669072 | 0.86247999 |
| L3MBTL2 | 18.2616631 | -0.095389002 | 0.36638979 | -0.2603484 | 0.79459503 | 0.91187679 |
| RANGAP1 | 29.7242927 | -0.420725571 | 0.30109904 | -1.3972996 | 0.16232349 | 0.42996461 |
| ZC3H7B | 60.5995692 | 0.227099589 | 0.30226749 | 0.75131993 | 0.45246014 | 0.72046894 |
| TEF | 23.7599774 | 1.243188747 | 0.4868735 | 2.55341222 | 0.01066732 | 0.0801566 |
| TOB2 | 42.4883541 | 0.342255043 | 0.45480003 | 0.75253962 | 0.45172661 | 0.71982793 |
| ACO2 | 150.752358 | 0.641410723 | 0.24488405 | 2.61924258 | 0.00881253 | 0.07048652 |
| POLR3H | 13.2558733 | -0.012825442 | 0.40534687 | -0.0316407 | 0.97475862 | 0.98983573 |
| PMM1 | 13.7011918 | 0.38111439 | 0.40887653 | 0.9321014 | 0.35128412 | 0.63780914 |
| DESI1 | 41.2457598 | 0.554368288 | 0.44185772 | 1.25463075 | 0.20961284 | 0.49580206 |
| XRCC6 | 104.074699 | -0.06399557 | 0.25513578 | -0.2508295 | 0.80194597 | 0.91521517 |
| SNU13 | 80.327318 | -0.518978245 | 0.28051215 | -1.8501097 | 0.06429774 | 0.25262981 |
| CCDC134 | 13.6638902 | 0.170833732 | 0.41801562 | 0.40867787 | 0.68277608 | 0.8556087 |
| SREBF2 | 354.165301 | 0.680505757 | 0.35689191 | 1.90675592 | 0.0565522 | 0.23465757 |
| WBP2NL | 35.9307167 | 0.901361829 | 0.54362071 | 1.65807117 | 0.0973031 | 0.32180307 |
| SMDT1 | 28.2684195 | -0.595586158 | 0.44402113 | -1.3413464 | 0.179808 | 0.45580267 |
| NDUFA6 | 42.4222205 | -0.251090571 | 0.49633953 | -0.5058847 | 0.6129376 | 0.81464125 |
| TCF20 | 132.602732 | -0.387621733 | 0.27851494 | -1.3917448 | 0.16399968 | 0.43270826 |
| RRP7A | 28.039002 | -0.201193568 | 0.39085063 | -0.5147582 | 0.606722 | 0.81143061 |
| SERHL2 | 17.5343319 | 0.029182734 | 0.36902542 | 0.07908055 | 0.93696855 | 0.97613368 |
| POLDIP3 | 31.7818319 | -0.166358503 | 0.27307652 | -0.6092011 | 0.54239118 | 0.77243948 |
| CYB5R3 | 83.0948733 | -1.050364167 | 0.30300006 | -3.4665477 | 0.00052719 | 0.00891805 |
| A4GALT | 88.2389009 | -1.39501847 | 0.36029535 | -3.8718748 | 0.000108 | 0.00257553 |
| ARFGAP3 | 146.7914 | -0.953498452 | 0.19752907 | -4.8271297 | 1.39E-06 | 7.32E-05 |
| PACSIN2 | 213.750998 | 0.220380012 | 0.29676292 | 0.74261302 | 0.457716 | 0.72377441 |
| TTLL1 | 10.9499924 | -0.271590603 | 0.52949984 | -0.5129191 | 0.60800789 | 0.81199083 |
| MCAT | 12.7039804 | 0.335288209 | 0.45969351 | 0.72937338 | 0.46577329 | 0.72890726 |
| TSPO | 66.4037203 | -1.035476861 | 0.3867767 | -2.6771956 | 0.00742413 | 0.06289751 |
| EFCAB6 | 106.561345 | -0.246026003 | 0.29216379 | -0.8420824 | 0.3997418 | 0.68000992 |
| SAMM50 | 16.7353024 | 0.75415769 | 0.371483 | 2.03012708 | 0.04234362 | 0.19633566 |
| PARVB | 13.122584 | 1.578583661 | 0.90078329 | 1.75245664 | 0.07969532 | 0.28777469 |
| NUP50-DT | 63.6624126 | -0.372541516 | 0.29190821 | -1.2762283 | 0.20187484 | 0.48515376 |
| NUP50 | 32.137708 | -0.239138278 | 0.27695734 | -0.8634481 | 0.38789116 | 0.66971239 |
| KIAA0930 | 23.3603094 | -0.949738719 | 0.48693094 | -1.9504587 | 0.05112147 | 0.22008079 |
| FAM118A | 24.559923 | 0.823560481 | 0.4723111 | 1.74368224 | 0.08121451 | 0.29141278 |

| | | | | | | |
|---|---|---|---|---|---|---|
| ATXN10 | 202.862392 | 0.129924861 | 0.17320446 | 0.75012424 | 0.45317988 | 0.7209171 |
| MIRLET7BHG | 83.2629086 | -0.077701868 | 0.26932815 | -0.2885026 | 0.77296205 | 0.90149298 |
| PPARA | 181.148459 | 0.571419685 | 0.27151492 | 2.10456088 | 0.03532955 | 0.17447988 |
| TTC38 | 29.5968482 | 0.078222194 | 0.35017224 | 0.22338206 | 0.82323817 | 0.92607552 |
| TRMU | 38.7858722 | -0.050216768 | 0.29348225 | -0.1711067 | 0.8641399 | 0.94663434 |
| CELSR1 | 50.0145531 | 0.141374158 | 0.24858897 | 0.56870649 | 0.56955534 | 0.78795455 |
| GRAMD4 | 26.0558311 | 0.269667176 | 0.37462105 | 0.71983989 | 0.47162358 | 0.73286947 |
| CERK | 32.7027506 | 0.056214436 | 0.31291576 | 0.17964719 | 0.85742956 | 0.94359207 |
| TBC1D22A | 270.556293 | -0.316444917 | 0.31886668 | -0.9924051 | 0.32099995 | 0.6120836 |
| BRD1 | 56.3717619 | -0.035769506 | 0.32054093 | -0.1115911 | 0.91114765 | 0.96391899 |
| ZBED4 | 29.0369712 | -0.505831161 | 0.38136427 | -1.3263727 | 0.18471627 | 0.46134371 |
| ALG12 | 13.6442543 | 0.04290439 | 0.40238392 | 0.10662551 | 0.91508608 | 0.96466317 |
| PIM3 | 87.3212395 | 0.73102217 | 0.43439372 | 1.68285623 | 0.09240293 | 0.3131365 |
| TRABD | 35.9289843 | 0.137169554 | 0.37686496 | 0.36397534 | 0.71587642 | 0.87318675 |
| SELENOO | 54.0691403 | -0.2429843 | 0.2281923 | -1.0648225 | 0.28695625 | 0.57931085 |
| TUBGCP6 | 16.1926647 | 0.350977915 | 0.38787067 | 0.90488386 | 0.36552692 | 0.65093488 |
| PLXNB2 | 108.379275 | -0.255693039 | 0.3050556 | -0.838185 | 0.40192679 | 0.68091593 |
| DENND6B | 17.6466854 | -0.377622808 | 0.44033409 | -0.8575825 | 0.39112304 | 0.67223049 |
| PPP6R2 | 218.007551 | 0.585601739 | 0.21684386 | 2.70056872 | 0.0069221 | 0.05984601 |
| SBF1 | 11.3935242 | -0.451493995 | 0.41330164 | -1.092408 | 0.27465383 | 0.56696009 |
| MIOX | 14.3195222 | 1.452056828 | 0.85986549 | 1.6887023 | 0.09127649 | 0.31117713 |
| LMF2 | 16.7497903 | -0.900312596 | 0.49194574 | -1.8301055 | 0.06723417 | 0.26017544 |
| KLHDC7B | 23.2817378 | -5.767322343 | 0.94829614 | -6.0817735 | 1.19E-09 | 2.08E-07 |
| ARSA | 14.1700595 | -0.394885874 | 0.47768376 | -0.826668 | 0.40842527 | 0.6867304 |
| RABL2B | 24.736866 | -0.098398316 | 0.358746 | -0.2742841 | 0.78386632 | 0.90714343 |
| CU633967.1 | 32.3438182 | -0.377134184 | 0.39639788 | -0.9514031 | 0.34139977 | 0.62962773 |
| FP671120.1 | 506.551147 | -0.753000321 | 0.50848323 | -1.4808754 | 0.13863977 | 0.39280881 |
| FP236383.1 | 290.726029 | -0.989175667 | 0.47718732 | -2.0729295 | 0.03817884 | 0.18297817 |
| CR381653.1 | 42.3289359 | 0.853810099 | 0.48505398 | 1.76023729 | 0.07836758 | 0.28477085 |
| RBM11 | 11.5962983 | -0.687926029 | 0.65211167 | -1.0549206 | 0.29146164 | 0.58292358 |
| HSPA13 | 14.5837407 | 0.010610509 | 0.45819342 | 0.02315727 | 0.98152482 | 0.99318267 |
| NRIP1 | 94.498655 | -0.291793931 | 0.31347205 | -0.9308451 | 0.35193368 | 0.63801604 |
| USP25 | 316.40353 | 0.651946381 | 0.29941612 | 2.17739236 | 0.02945131 | 0.15567334 |
| MIR99AHG | 125.921461 | 0.440418302 | 0.34894378 | 1.26214687 | 0.20689594 | 0.49179557 |
| CXADR | 72.0278688 | -0.421319079 | 0.2940292 | -1.4329158 | 0.1518819 | 0.41333768 |
| BTG3 | 52.735998 | 0.559428119 | 0.4540596 | 1.23205878 | 0.21792713 | 0.50587771 |
| C21orf91 | 16.86096 | -0.637826982 | 0.38225099 | -1.6686078 | 0.09519513 | 0.31779803 |
| CHODL | 55.4894768 | 0.079584106 | 0.66929314 | 0.1189077 | 0.90534848 | 0.96157183 |
| MRPL39 | 22.9095172 | 0.256991265 | 0.32237201 | 0.79718852 | 0.42534155 | 0.70016446 |
| ATP5PF | 89.6371589 | -0.008609329 | 0.39796822 | -0.0216332 | 0.98274054 | 0.99353746 |
| GABPA | 32.5024849 | -0.384380538 | 0.38540906 | -0.9973313 | 0.31860371 | 0.61144847 |
| APP | 804.421499 | -0.113454303 | 0.18307752 | -0.6197064 | 0.53545113 | 0.76808755 |
| CYYR1 | 51.7792722 | -0.081186668 | 0.54531806 | -0.1488795 | 0.88164873 | 0.95427036 |
| ADAMTS1 | 376.204314 | -4.867228045 | 0.51420148 | -9.4656049 | 2.92E-21 | 7.52E-18 |
| ADAMTS5 | 14.997443 | 0.05456779 | 0.69830217 | 0.07814352 | 0.93771389 | 0.97619962 |
| AF165147.1 | 20.1850293 | -1.197175357 | 0.65501439 | -1.8277085 | 0.06759331 | 0.26098543 |
| N6AMT1 | 12.3173717 | 0.316811385 | 0.46524297 | 0.68095899 | 0.49589744 | 0.74798468 |
| LTN1 | 78.0951909 | 0.308112535 | 0.23374868 | 1.31813596 | 0.18745814 | 0.46523578 |
| USP16 | 46.2804914 | 0.286400161 | 0.31130638 | 0.9199945 | 0.35757563 | 0.6418791 |
| CCT8 | 35.1209489 | -0.150160375 | 0.30587427 | -0.4909219 | 0.62348169 | 0.82178971 |
| MAP3K7CL | 36.8237754 | 0.150273385 | 0.35992435 | 0.4175138 | 0.67630263 | 0.85177422 |
| BACH1 | 378.961619 | -1.211221538 | 0.26780493 | -4.5227753 | 6.10E-06 | 0.00024951 |
| GRIK1 | 21.3309227 | -2.03831347 | 0.68118274 | -2.9923152 | 0.0027687 | 0.03103718 |
| CLDN8 | 71.290138 | 1.552421358 | 0.57936096 | 2.67954084 | 0.00737232 | 0.06271647 |
| SOD1 | 92.3267835 | -0.284835857 | 0.47909813 | -0.5945251 | 0.55216097 | 0.77741728 |
| SCAF4 | 71.8335777 | 0.412597925 | 0.28305298 | 1.4576703 | 0.14493144 | 0.40277414 |
| HUNK | 178.163962 | 0.015942652 | 0.4158706 | 0.03833561 | 0.9694201 | 0.98781534 |
| URB1 | 27.4812846 | 0.6057488 | 0.39400935 | 1.537397 | 0.12419612 | 0.36916928 |
| EVA1C | 53.8618163 | -0.990479685 | 0.4474116 | -2.2137997 | 0.02684256 | 0.14623587 |
| SYNJ1 | 75.6235121 | 0.436566645 | 0.23797818 | 1.83448181 | 0.06658251 | 0.2582579 |
| PAXBP1 | 108.941097 | 0.20914245 | 0.20152558 | 1.03779602 | 0.29936503 | 0.59240463 |
| C21orf62-AS | 31.055202 | 0.724394661 | 0.58323903 | 1.2420202 | 0.21422911 | 0.50079153 |
| AP000282.1 | 17.1256973 | 1.632364279 | 0.56059798 | 2.91182689 | 0.00359322 | 0.03724077 |
| IFNAR1 | 69.8377482 | -0.570268725 | 0.22860122 | -2.4946006 | 0.0126099 | 0.09018912 |
| IFNGR2 | 96.065609 | -0.543095162 | 0.38139406 | -1.4239738 | 0.15445405 | 0.41780983 |
| TMEM50B | 94.6899898 | 0.220166447 | 0.33557569 | 0.6560858 | 0.51176892 | 0.75544396 |
| GART | 43.926542 | 0.249762489 | 0.27589316 | 0.90528698 | 0.36531337 | 0.65066708 |
| SON | 186.03528 | -0.304323848 | 0.26098833 | -1.1660439 | 0.2435967 | 0.53443687 |
| AP000311.1 | 57.9087747 | -0.217820094 | 0.46254084 | -0.4709208 | 0.63769731 | 0.83032832 |
| ITSN1 | 233.376388 | 0.69195391 | 0.25835264 | 2.6783311 | 0.007399 | 0.06273624 |
| MRPS6 | 145.508372 | 0.6495716 | 0.40587609 | 1.6004185 | 0.10950577 | 0.34481257 |
| LINC00310 | 16.5718293 | 0.469302312 | 0.58262952 | 0.8054901 | 0.42053691 | 0.69661173 |
| AP000317.1 | 25.1179483 | 0.791236567 | 0.37952918 | 2.08478456 | 0.03708886 | 0.17972222 |
| RCAN1 | 76.6354303 | 1.265136463 | 0.55453868 | 2.28142147 | 0.02252352 | 0.13071121 |
| CLIC6 | 26.4076721 | -0.874652114 | 0.43476934 | -2.0117613 | 0.04424511 | 0.20044552 |
| RUNX1 | 580.998556 | -0.32325026 | 0.72639151 | -0.4450083 | 0.65631375 | 0.83951126 |
| SETD4 | 32.6774513 | 0.203913672 | 0.33995797 | 0.59982024 | 0.54862605 | 0.77530117 |
| CBR1 | 81.5883103 | -1.001982315 | 0.37903908 | -2.6434802 | 0.00820586 | 0.06719931 |
| AP000688.2 | 14.4973849 | -0.794478081 | 0.4877265 | -1.6289418 | 0.10332534 | 0.33368578 |
| CBR3 | 42.6202501 | -0.12872508 | 0.44258672 | -0.2908471 | 0.77116824 | 0.90076568 |
| DOPEY2 | 66.4790335 | -0.223381891 | 0.30524137 | -0.7318205 | 0.46427813 | 0.72835095 |
| MORC3 | 103.939084 | 0.147624421 | 0.23070628 | 0.63988037 | 0.52225038 | 0.76155853 |
| CLDN14 | 29.7059083 | -2.933945931 | 0.61371731 | -4.7806146 | 1.75E-06 | 8.83E-05 |
| HLCS | 211.044456 | 0.012676712 | 0.17680458 | 0.071699 | 0.94284145 | 0.9781624 |
| PIGP | 14.0303909 | -0.247981464 | 0.62801558 | -0.3948651 | 0.69294241 | 0.8606019 |
| TTC3 | 284.845006 | 0.23971401 | 0.22603267 | 1.06052814 | 0.2889044 | 0.58074012 |
| VPS26C | 54.4320213 | -0.103561341 | 0.26818651 | -0.3861542 | 0.69938249 | 0.86349933 |
| DYRK1A | 412.060967 | 0.53595744 | 0.22863539 | 2.34415785 | 0.0190701 | 0.11735971 |
| KCNJ15 | 357.791543 | 0.718391835 | 0.43620565 | 1.64691088 | 0.09957636 | 0.32585356 |
| ETS2 | 59.8704757 | -2.714153185 | 0.64063511 | -4.2366601 | 2.27E-05 | 0.00072359 |
| BRWD1 | 309.175862 | 0.380545015 | 0.15695868 | 2.42449168 | 0.01532983 | 0.10149612 |
| HMGN1 | 92.9850147 | -0.571684423 | 0.26760733 | -2.1362809 | 0.03265653 | 0.16638356 |
| WRB | 27.3450447 | 0.037917553 | 0.32368972 | 0.11714166 | 0.90674779 | 0.96157183 |
| LCA5L | 14.9056484 | -0.1434057 | 0.42280146 | -0.3391798 | 0.73447431 | 0.88376541 |
| SH3BGR | 18.5754767 | 0.749071692 | 0.60367741 | 1.24084763 | 0.21466204 | 0.50112131 |
| BACE2 | 442.369906 | -2.007306128 | 0.65952202 | -3.0435771 | 0.00233783 | 0.0272113 |
| FAM3B | 41.0320957 | -0.46662036 | 0.30725342 | -1.5186824 | 0.12884246 | 0.37613516 |
| MX1 | 101.504519 | -1.044649865 | 1.28735826 | -0.8114679 | 0.41709704 | 0.69338933 |
| TMPRSS2 | 268.082725 | 1.916147899 | 0.39178651 | 4.89079606 | 1.00E-06 | 5.65E-05 |
| RIPK4 | 25.4373997 | -0.259533775 | 0.33980454 | -0.7637737 | 0.4450021 | 0.71486226 |
| PRDM15 | 15.8472625 | -0.650341302 | 0.56481833 | -1.1514168 | 0.24956082 | 0.54083745 |
| C2CD2 | 32.97353 | -0.647008571 | 0.38753654 | -1.6695421 | 0.09500999 | 0.31748068 |
| ZBTB21 | 28.1303162 | -0.181593717 | 0.3915341 | -0.4638005 | 0.64279068 | 0.83318468 |
| RSPH1 | 15.7735643 | -0.942747976 | 0.50013574 | -1.8849842 | 0.05943197 | 0.24143066 |
| SLC37A1 | 49.5208663 | -1.325010222 | 0.23962391 | -5.5295408 | 3.21E-08 | 3.20E-06 |
| LINC01671 | 18.335095 | -0.686286955 | 0.47866322 | -1.4337574 | 0.15164149 | 0.41304821 |
| PDE9A | 166.227287 | 0.116707691 | 0.29462909 | 0.39611735 | 0.69201846 | 0.85997276 |
| WDR4 | 20.0620337 | -0.014814514 | 0.46399231 | -0.0319284 | 0.97452918 | 0.98980574 |
| NDUFV3 | 68.8057685 | 0.425472209 | 0.19621954 | 2.16834778 | 0.03013224 | 0.1578151 |
| PKNOX1 | 71.5510019 | -0.069012554 | 0.266707 | -0.2587579 | 0.79582201 | 0.91195838 |
| RRP1B | 27.0038798 | 0.206731008 | 0.33346063 | 0.61995627 | 0.53528658 | 0.76808755 |
| PDXK | 135.562761 | 0.49508785 | 0.3281555 | 1.50869894 | 0.13137574 | 0.38024035 |
| CSTB | 112.087862 | -1.507142419 | 0.35926914 | -4.1950234 | 2.73E-05 | 0.00084664 |

| | | | | | | |
|---|---|---|---|---|---|---|
| AGPAT3 | 119.275164 | 0.27966894 | 0.18769356 | 1.49002951 | 0.13621648 | 0.38868467 |
| TRAPPC10 | 105.004641 | 0.113726651 | 0.24661244 | 0.46115536 | 0.64468715 | 0.83381793 |
| PFKL | 147.180414 | -0.584393447 | 0.30391138 | -1.9229074 | 0.05449168 | 0.22997772 |
| C21orf2 | 16.659501 | 0.464035711 | 0.54967278 | 0.84420355 | 0.39855567 | 0.67938457 |
| UBE2G2 | 91.2202524 | 0.250317546 | 0.21706008 | 1.1532178 | 0.248821 | 0.53982832 |
| SUMO3 | 33.1650297 | -1.029541039 | 0.45507073 | -2.2623759 | 0.02367419 | 0.13490566 |
| PTTG1IP | 135.650853 | -1.121735981 | 0.22557108 | -4.9728714 | 6.60E-07 | 4.02E-05 |
| FAM207A | 13.2438718 | 0.169770186 | 0.48356108 | 0.35108323 | 0.72552591 | 0.87840733 |
| ADARB1 | 35.0134461 | 0.463438752 | 0.34909181 | 1.3275555 | 0.18432499 | 0.46090196 |
| POFUT2 | 20.0814728 | 0.177285334 | 0.43026112 | 0.41204126 | 0.68030918 | 0.8543838 |
| COL18A1 | 274.767156 | 0.02861726 | 0.29603985 | 0.09666692 | 0.92299091 | 0.96918279 |
| COL6A1 | 56.3747359 | 1.3094709 | 0.41196343 | 3.1786096 | 0.00147983 | 0.01957026 |
| LSS | 29.7565058 | -0.370391639 | 0.51347746 | -0.7213396 | 0.47070058 | 0.73210719 |
| MCM3AP-AS | 14.4500417 | 0.621272601 | 0.38043645 | 1.63305226 | 0.10245801 | 0.33161243 |
| MCM3AP | 65.0221379 | 0.517436964 | 0.25543005 | 2.02574817 | 0.0427906 | 0.19741547 |
| YBEY | 11.8429736 | -0.013024746 | 0.51300988 | -0.0253889 | 0.97974478 | 0.99255883 |
| PCNT | 33.4970516 | -0.389278841 | 0.44073284 | -0.8832535 | 0.3770993 | 0.66022373 |
| DIP2A | 111.486224 | 0.162839831 | 0.20578362 | 0.79131579 | 0.42875974 | 0.70288254 |
| PRMT2 | 104.158109 | -0.794278456 | 0.22890843 | -3.4698523 | 0.00052074 | 0.00888197 |
| AC011043.1 | 33.765574 | 0.507768231 | 0.46276724 | 1.09724324 | 0.27253507 | 0.56479948 |
| AL592183.1 | 68.9485541 | -0.332943967 | 0.49193294 | -0.6768076 | 0.49852802 | 0.75012216 |

**Collecting duct principal cells**

| | baseMean | log2 fold change control/AKI* | lfcSE | stat | pvalue | padj |
|---|---|---|---|---|---|---|
| AL669831.5 | 117.625093 | 0.5368652 | 0.41493114 | 1.29386578 | 0.19571179 | 0.64948524 |
| AL645608.1 | 37.838824 | 0.788673419 | 0.61239835 | 1.28784379 | 0.19780034 | 0.652979 |
| SAMD11 | 118.20368 | 1.029872876 | 0.60412465 | 1.70473573 | 0.08824372 | 0.47036301 |
| NOC2L | 80.3916088 | 0.073414075 | 0.24824249 | 0.29573534 | 0.76743221 | 0.95408212 |
| HES4 | 131.461973 | -0.436871742 | 1.56219353 | -0.2796528 | 0.77974392 | 0.95636474 |
| ISG15 | 634.877613 | 1.881746313 | 1.91064278 | 0.98487605 | 0.32468496 | 0.77020457 |
| AGRN | 470.478214 | -0.231507244 | 0.31063711 | -0.7452659 | 0.45611099 | 0.84372683 |
| C1orf159 | 129.910862 | -0.166584485 | 0.29247319 | -0.5695718 | 0.56896815 | 0.89617938 |
| LINC01342 | 43.7069962 | -0.241886853 | 0.39713435 | -0.6090807 | 0.54247097 | 0.88450129 |
| AL390719.2 | 38.6331177 | -0.1139079 | 0.44281385 | -0.2572365 | 0.79699618 | 0.95961857 |
| SDF4 | 194.743158 | -0.48157749 | 0.3027567 | -1.5906419 | 0.11169019 | 0.52161905 |
| UBE2J2 | 120.117691 | 0.093736352 | 0.23972423 | 0.39101743 | 0.69578435 | 0.93130502 |
| INTS11 | 123.392404 | 0.275197699 | 0.22324299 | 1.23272718 | 0.21767757 | 0.675129 |
| DVL1 | 44.9676192 | 0.26859489 | 0.37207176 | 0.72189002 | 0.4703621 | 0.85236058 |
| AURKAIP1 | 240.795271 | 0.009933161 | 0.52204844 | 0.01902728 | 0.98481934 | 0.9954565 |
| CCNL2 | 274.326611 | -0.021698993 | 0.33286892 | -0.0651878 | 0.94802447 | 0.98977092 |
| MRPL20 | 244.574701 | 0.101847388 | 0.34099182 | 0.29867985 | 0.76518433 | 0.95378284 |
| VWA1 | 109.481148 | -1.38091697 | 0.55354445 | -2.4946813 | 0.01260703 | 0.1792696 |
| ATAD3C | 29.2610394 | -0.747676386 | 0.49450886 | -1.5119575 | 0.13054467 | 0.55395564 |
| ATAD3B | 71.7650765 | 0.248615784 | 0.33624188 | 0.73939565 | 0.45966678 | 0.84632956 |
| ATAD3A | 37.646709 | -0.081896156 | 0.45152952 | -0.181375 | 0.85607326 | 0.97182813 |
| SSU72 | 251.688729 | 0.025899785 | 0.35551268 | 0.07285193 | 0.94192395 | 0.98885525 |
| MIB2 | 64.2940999 | 0.782405185 | 0.3516839 | 2.22473982 | 0.02609871 | 0.27515522 |
| CDK11B | 102.200513 | 1.188114389 | 0.54714591 | 2.17147634 | 0.02989518 | 0.29144328 |
| SLC35E2B | 85.2218091 | -0.16756232 | 0.42077537 | -0.3982227 | 0.69046601 | 0.93073907 |
| CDK11A | 45.9637821 | -0.180266774 | 0.34874 | -0.5169088 | 0.60521982 | 0.90428025 |
| SLC35E2A | 95.2218604 | -0.210201928 | 0.40892917 | -0.5140302 | 0.6072309 | 0.90510401 |
| NADK | 134.31432 | 0.462498583 | 0.31157368 | 1.48439553 | 0.13770402 | 0.56434148 |
| GNB1 | 948.793386 | -0.164432181 | 0.28433277 | -0.5783089 | 0.56305556 | 0.8927792 |
| PRKCZ | 179.441755 | -0.045658347 | 0.23204497 | -0.1967651 | 0.84401138 | 0.96876046 |
| FAAP20 | 177.778053 | -0.06507859 | 0.31802245 | -0.2046352 | 0.83785715 | 0.96757829 |
| SKI | 241.762857 | 0.271088543 | 0.41148303 | 0.65880857 | 0.51001871 | 0.87094607 |
| MORN1 | 122.211515 | -0.031255552 | 0.23844062 | -0.1310832 | 0.89570952 | 0.98272987 |
| RER1 | 305.643726 | 0.028061809 | 0.35368295 | 0.07934171 | 0.93676083 | 0.98822345 |
| PEX10 | 76.0813871 | -0.234382254 | 0.40592688 | -0.5774002 | 0.56366916 | 0.89295783 |
| AL139246.5 | 78.1323222 | -0.711025727 | 0.81175609 | -0.8759106 | 0.38107866 | 0.80315561 |
| TNFRSF14-A | 103.219763 | -1.243169337 | 0.68638348 | -1.8111877 | 0.0701118 | 0.42868822 |
| TNFRSF14 | 258.973109 | -0.651299791 | 0.5354477 | -1.2163649 | 0.22384595 | 0.68115724 |
| FAM213B | 117.683282 | -0.561603778 | 0.52811163 | -1.0634187 | 0.28759212 | 0.73650262 |
| MMEL1 | 37.0811475 | -0.829066866 | 0.62648999 | -1.3233521 | 0.1857183 | 0.63626275 |
| LINC00982 | 377.940021 | 1.573562358 | 0.44526229 | 3.53401215 | 0.0004093 | 0.02177405 |
| PRDM16 | 1017.52816 | 0.20978942 | 0.3924751 | 0.53452925 | 0.59297542 | 0.9025973 |
| ARHGEF16 | 89.1986175 | -0.034581346 | 0.47424392 | -0.0729189 | 0.94187065 | 0.98885525 |
| TPRG1L | 81.2447232 | -0.404113552 | 0.28694588 | -1.4083267 | 0.15903434 | 0.59747329 |
| WRAP73 | 42.0303651 | 0.3346637 | 0.28757463 | 1.16374554 | 0.24452715 | 0.70121778 |
| LRRC47 | 67.7882402 | 0.02181368 | 0.31966242 | 0.06823974 | 0.94559479 | 0.98949015 |
| CEP104 | 234.865246 | -0.555335336 | 0.37561335 | -1.478476 | 0.13928041 | 0.56656585 |
| C1orf174 | 50.188868 | -0.111956273 | 0.46335805 | -0.2416194 | 0.80907512 | 0.96245698 |
| NPHP4 | 69.6948276 | 0.23506245 | 0.27199783 | 0.86420707 | 0.38747415 | 0.8065026 |
| RPL22 | 190.042941 | 0.231272767 | 0.33662315 | 0.68703762 | 0.49205902 | 0.86172738 |
| RNF207 | 81.4088126 | 0.081857864 | 0.23650292 | 0.34611777 | 0.72925421 | 0.94004725 |
| ICMT | 99.8263651 | 0.030738504 | 0.27984596 | 0.1098408 | 0.91253563 | 0.98480378 |
| ESPN | 86.5684673 | 1.145554541 | 0.62787231 | 1.8245024 | 0.06807615 | 0.42365683 |
| NOL9 | 93.1733938 | -0.064559883 | 0.29283038 | -0.2204685 | 0.82550628 | 0.96534651 |
| ZBTB48 | 47.724122 | 0.175986329 | 0.30652569 | 0.5741324 | 0.56587821 | 0.89394085 |
| KLHL21 | 58.7267779 | -0.77541298 | 0.41838512 | -1.8533474 | 0.06383258 | 0.41357646 |
| PHF13 | 47.3358921 | 0.110081763 | 0.29060168 | 0.37880635 | 0.70483167 | 0.93241442 |
| THAP3 | 40.278405 | -0.056669521 | 0.25887291 | -0.2189087 | 0.8267212 | 0.96545294 |
| DNAJC11 | 175.71743 | -0.019729513 | 0.20839438 | -0.0946739 | 0.92457383 | 0.98710257 |
| CAMTA1 | 1006.88619 | -0.359718163 | 0.29419772 | -1.2227089 | 0.22143968 | 0.67915945 |
| VAMP3 | 108.061993 | -0.319268202 | 0.34209613 | -0.9332704 | 0.35068036 | 0.78242416 |
| PER3 | 198.979431 | -1.028884702 | 0.58964124 | -1.7449334 | 0.08099646 | 0.45368408 |
| PARK7 | 422.227972 | -0.326306584 | 0.49049013 | -0.6652664 | 0.50588014 | 0.87068877 |
| ERRFI1 | 328.135005 | 0.520334142 | 1.29986876 | 0.40029744 | 0.68893745 | 0.93036007 |
| RERE | 2945.66642 | 0.089064182 | 0.22619813 | 0.39374411 | 0.69376997 | 0.93105058 |
| ENO1 | 1957.39888 | -0.659518977 | 0.44763575 | -1.4733385 | 0.14065979 | 0.56886692 |
| GPR157 | 36.3476524 | -0.06194486 | 0.29553342 | -0.2096036 | 0.83397709 | 0.96749268 |
| MIR34AHG | 44.3913371 | -0.972807183 | 0.81666833 | -1.19119 | 0.23357899 | 0.69247746 |
| H6PD | 117.65147 | -0.406088774 | 0.22001108 | -1.8457651 | 0.06492632 | 0.41540163 |
| SPSB1 | 127.347321 | -1.386246915 | 0.47534001 | -2.916327 | 0.00354179 | 0.09053163 |
| SLC25A33 | 147.008776 | 0.787960795 | 0.44040946 | 1.789155 | 0.07358986 | 0.4369528 |
| PIK3CD | 37.551523 | -0.701482961 | 0.47660474 | -1.4718338 | 0.14106577 | 0.56895751 |
| CLSTN1 | 260.614526 | 0.186207264 | 0.25149314 | 0.74040693 | 0.45905311 | 0.84594279 |
| CTNNBIP1 | 91.3337699 | -0.594777761 | 0.31653076 | -1.879052 | 0.06023739 | 0.40493126 |
| LZIC | 106.411177 | 0.051248271 | 0.24093778 | 0.21270334 | 0.83155836 | 0.96656674 |
| NMNAT1 | 119.906887 | -0.133068694 | 0.23993823 | -0.5545956 | 0.57917128 | 0.89975022 |
| UBE4B | 407.426611 | -0.175615602 | 0.28089886 | -0.6251916 | 0.53184533 | 0.88045351 |
| KIF1B | 457.333209 | 0.180930576 | 0.19780829 | 0.9146764 | 0.36036154 | 0.78892604 |
| PGD | 89.896907 | -0.537086113 | 0.35693226 | -1.5047284 | 0.13239392 | 0.55543964 |
| DFFA | 134.99283 | -0.589990551 | 0.20195556 | -2.9213881 | 0.00348475 | 0.0899513 |
| PEX14 | 333.793624 | 0.16241398 | 0.21637602 | 0.75060988 | 0.45288747 | 0.84248055 |
| CASZ1 | 639.012269 | -0.24748938 | 0.39003212 | -0.6345359 | 0.52573115 | 0.87791848 |
| SRM | 50.1581133 | -0.118044481 | 0.39320086 | -0.3002142 | 0.76401378 | 0.95378284 |
| EXOSC10 | 97.3655633 | 0.26900635 | 0.25190998 | 1.06786697 | 0.28558052 | 0.73607046 |
| MTOR | 195.624262 | 0.100211298 | 0.29434006 | 0.34046095 | 0.73350942 | 0.94163693 |
| FBXO2 | 120.212283 | -0.166612705 | 0.62646037 | -0.2659589 | 0.79027087 | 0.95812572 |
| FBXO44 | 54.3543407 | 0.177833395 | 0.2887359 | 0.61590331 | 0.53795834 | 0.88265739 |
| AGTRAP | 124.79997 | -0.339983637 | 0.40245327 | -0.8447779 | 0.39823484 | 0.81400298 |
| MTHFR | 71.6660496 | -0.11945191 | 0.30932352 | -0.3861714 | 0.6993697 | 0.93180634 |
| CLCN6 | 93.0908092 | -0.991914626 | 0.49140361 | -2.0185334 | 0.04353573 | 0.35177679 |
| KIAA2013 | 86.9208264 | -0.174850838 | 0.31426528 | -0.5563797 | 0.57795128 | 0.89918786 |
| PLOD1 | 52.4796337 | -0.338047147 | 0.36407772 | -0.9285027 | 0.35314687 | 0.78377365 |
| MFN2 | 126.603999 | 0.061353005 | 0.250393 | 0.24502684 | 0.80643565 | 0.96218371 |
| VPS13D | 780.902363 | 0.035678776 | 0.16622688 | 0.21463902 | 0.83004877 | 0.96643829 |
| DHRS3 | 171.081593 | -2.228987584 | 0.44544176 | -5.0039934 | 5.62E-07 | 0.00021796 |
| PRDM2 | 537.651129 | -0.037748308 | 0.26085607 | -0.1447093 | 0.88494037 | 0.97891513 |
| KAZN | 10345.8817 | -1.346316686 | 0.43823823 | -3.0721114 | 0.0021255 | 0.06513239 |
| TMEM51-AS | 137.149302 | -1.581594816 | 0.41739939 | -3.7891641 | 0.00015116 | 0.01139644 |
| TMEM51 | 942.153407 | -0.077485825 | 0.4235385 | -0.1829487 | 0.85483826 | 0.97122276 |
| C1orf195 | 38.9416262 | 0.030892486 | 0.73013933 | 0.0423104 | 0.96625126 | 0.99284614 |
| FHAD1 | 54.0977 | -0.499586166 | 0.3917999 | -1.2751054 | 0.20227194 | 0.65696752 |
| EFHD2 | 32.3744813 | -0.911285323 | 0.38687249 | -2.3555185 | 0.01849688 | 0.22752025 |
| CASP9 | 208.132488 | 2.451123317 | 0.89923761 | 2.72577935 | 0.00641499 | 0.12805535 |
| DNAJC16 | 193.13378 | 0.398098195 | 0.20124354 | 1.97819118 | 0.04790714 | 0.36948395 |
| DDI2 | 155.113889 | 0.04917204 | 0.18820343 | 0.26127069 | 0.79388377 | 0.95937941 |
| PLEKHM2 | 153.184686 | -0.161819895 | 0.28526517 | -0.5672613 | 0.57053666 | 0.89724252 |
| FBLIM1 | 142.039183 | -1.270869728 | 0.66463958 | -1.9121186 | 0.05586098 | 0.39368206 |
| UQCRHL | 65.2065987 | 0.354140046 | 0.53921773 | 0.65676633 | 0.51133118 | 0.87151901 |
| SPEN | 371.626344 | 0.297189556 | 0.20355238 | 1.46001516 | 0.14428591 | 0.57406643 |
| ZBTB17 | 45.118242 | -0.124466647 | 0.33082533 | -0.3762307 | 0.70674539 | 0.93271523 |
| CLCNKA | 59.9151531 | 1.833337055 | 0.66798901 | 2.7445617 | 0.00605918 | 0.1254944 |

*This means that positive fold changes imply higher mean value in controls than in AKI and vice versa.

| | | | | | | |
|---|---|---|---|---|---|---|
| CLCNKB | 85.0498752 | 0.254579121 | 1.01686468 | 0.25035693 | 0.80231133 | 0.96071123 |
| EPHA2 | 56.3072095 | 0.365282699 | 0.67511542 | 0.54106704 | 0.58846138 | 0.90200952 |
| FBXO42 | 419.64711 | 0.30866245 | 0.27294885 | 1.13084356 | 0.25812094 | 0.71534188 |
| SZRD1 | 140.50314 | -0.210156491 | 0.22695465 | -0.9259845 | 0.35445404 | 0.78551033 |
| NECAP2 | 87.1199448 | -0.467973882 | 0.25911137 | -1.8060723 | 0.07090704 | 0.43078597 |
| NBPF1 | 107.798233 | -0.337274743 | 0.2524684 | -1.3359088 | 0.1815791 | 0.63191037 |
| CROCC | 99.2842336 | 0.390080712 | 0.32731902 | 1.19174472 | 0.23336136 | 0.69247746 |
| ATP13A2 | 55.7764779 | 0.092529685 | 0.30548543 | 0.30289394 | 0.76197069 | 0.95374766 |
| SDHB | 202.678012 | 0.2153468 | 0.19873575 | 1.0835836 | 0.27854947 | 0.72935218 |
| RCC2 | 168.609561 | 0.324668221 | 0.27249818 | 1.19145097 | 0.23347659 | 0.69247746 |
| ARHGEF10L | 1114.38733 | 0.033598063 | 0.43290627 | 0.07761048 | 0.93813791 | 0.98846521 |
| KLHDC7A | 59.0736232 | 0.062201273 | 0.47689331 | 0.13043016 | 0.8962261 | 0.98298792 |
| UBR4 | 452.743077 | 0.611382832 | 0.26637816 | 2.29516872 | 0.02172346 | 0.24935578 |
| AL035413.1 | 67.0274747 | -0.254373439 | 0.25056656 | -1.0151931 | 0.3100138 | 0.75409553 |
| EMC1 | 70.0611332 | 0.176578986 | 0.2831372 | 0.62365166 | 0.53285637 | 0.88114359 |
| MRTO4 | 48.971908 | -0.575206889 | 0.30717553 | -1.8725674 | 0.06112814 | 0.40676596 |
| AKR7A2 | 121.446563 | -0.235851102 | 0.46586583 | -0.506264 | 0.61267132 | 0.90714827 |
| CAPZB | 520.874249 | -0.027182495 | 0.23119003 | -0.1175764 | 0.90640329 | 0.98320957 |
| NBL1 | 190.560919 | -0.53291518 | 0.50509951 | -1.0550697 | 0.29139346 | 0.73816859 |
| TMCO4 | 387.482238 | -0.032223736 | 0.28350275 | -0.1136629 | 0.90950505 | 0.98441374 |
| OTUD3 | 69.6283561 | 0.256586682 | 0.35558309 | 0.72159417 | 0.47054403 | 0.85236058 |
| VWA5B1 | 44.1749799 | -1.185188946 | 0.84712965 | -1.3990644 | 0.16179367 | 0.60170251 |
| CAMK2N1 | 822.682763 | 0.009835782 | 0.45574077 | 0.02158197 | 0.98278142 | 0.99507903 |
| MUL1 | 47.6634072 | -0.133010995 | 0.33843017 | -0.3930235 | 0.69430216 | 0.93105058 |
| PINK1 | 146.357573 | -0.217698786 | 0.39625989 | -0.5493839 | 0.58274205 | 0.90085422 |
| DDOST | 117.3025 | -0.472788244 | 0.31000267 | -1.5251103 | 0.1272316 | 0.54746778 |
| HP1BP3 | 621.034485 | 0.221656047 | 0.15024936 | 1.47525448 | 0.14014414 | 0.56749674 |
| EIF4G3 | 1709.10288 | 0.370985075 | 0.26415779 | 1.40440708 | 0.16019767 | 0.60010648 |
| ECE1 | 45.433246 | -1.174445079 | 0.41194672 | -2.8509635 | 0.0043587 | 0.10311318 |
| NBPF3 | 44.1343552 | -0.73785028 | 0.32381457 | -2.2786198 | 0.02268968 | 0.25189387 |
| RAP1GAP | 178.175856 | -0.502233108 | 0.39357809 | -1.2760698 | 0.20193085 | 0.65666058 |
| USP48 | 371.882542 | 0.356583823 | 0.1938132 | 1.83983245 | 0.06579284 | 0.41788422 |
| HSPG2 | 378.91538 | -0.358016657 | 0.46772246 | -0.7654468 | 0.44400554 | 0.83873671 |
| CDC42 | 632.178369 | 0.081056524 | 0.25314666 | 0.32019591 | 0.74881983 | 0.94686219 |
| ZBTB40 | 105.559977 | 0.125159605 | 0.32003894 | 0.39107617 | 0.69574093 | 0.93130502 |
| EPHB2 | 155.456401 | -1.038132117 | 0.47846085 | -2.1697326 | 0.03002711 | 0.29169628 |
| KDM1A | 334.690617 | 0.261282719 | 0.17140296 | 1.52437697 | 0.12741457 | 0.54746778 |
| AL031428.1 | 76.2179286 | -0.110476868 | 0.52096922 | -0.2120603 | 0.83206002 | 0.96656674 |
| LUZP1 | 257.893076 | 0.232286013 | 0.28159392 | 0.82489711 | 0.40943001 | 0.82136801 |
| LINC01355 | 48.8025773 | 0.312114789 | 0.45178832 | 0.69084298 | 0.48966422 | 0.86055851 |
| HNRNPR | 453.180165 | 0.132481441 | 0.16314969 | 0.81202385 | 0.41677794 | 0.82489761 |
| TCEA3 | 183.846387 | 0.221581476 | 0.41550808 | 0.53327838 | 0.59384089 | 0.90304695 |
| ASAP3 | 136.635425 | -0.142978793 | 0.37342521 | -0.3828847 | 0.70180527 | 0.93215897 |
| ID3 | 260.672505 | 1.597812715 | 0.81548954 | 1.95932951 | 0.05007421 | 0.37753792 |
| RPL11 | 1139.17476 | 0.236235951 | 0.55475708 | 0.42583675 | 0.67022681 | 0.92474338 |
| ELOA | 79.1258634 | -0.242797348 | 0.22930991 | -1.0588175 | 0.28968289 | 0.73754646 |
| ELOA-AS1 | 80.1749033 | -0.265700928 | 0.27730778 | -0.9581445 | 0.3379899 | 0.77836439 |
| PITHD1 | 99.538017 | 0.199293818 | 0.28450817 | 0.70048538 | 0.48362423 | 0.85829223 |
| LYPLA2 | 89.4895438 | -0.374521351 | 0.39056746 | -0.9589159 | 0.33760111 | 0.77827084 |
| GALE | 143.289428 | -1.146852567 | 0.3439837 | -3.3340317 | 0.00085597 | 0.03624114 |
| HMGCL | 160.306143 | -0.876110477 | 0.27126688 | -3.2296994 | 0.0012392 | 0.04561933 |
| FUCA1 | 404.593578 | 0.37650946 | 0.51209052 | 0.73524005 | 0.46219331 | 0.8478533 |
| PNRC2 | 224.389965 | 0.169213718 | 0.35661025 | 0.47450604 | 0.63513909 | 0.91454294 |
| SRSF10 | 356.534344 | 0.215009406 | 0.23883216 | 0.90025317 | 0.36798554 | 0.79498834 |
| IFNLR1 | 61.8167618 | -0.171867896 | 0.34456043 | -0.4988034 | 0.61791793 | 0.90794578 |
| STPG1 | 81.8260301 | -0.152267438 | 0.22966296 | -0.6630039 | 0.50732806 | 0.87094607 |
| NIPAL3 | 255.000562 | -0.287403739 | 0.3110756 | -0.9239032 | 0.35553672 | 0.78607579 |
| RCAN3 | 131.189247 | -0.904569471 | 0.35332025 | -2.560197 | 0.01046128 | 0.16338934 |
| NCMAP | 109.424766 | -3.86073276 | 0.82764446 | -4.6647237 | 3.09E-06 | 0.00080718 |
| SRRM1 | 367.70621 | 0.284347513 | 0.19439738 | 1.46271267 | 0.14354602 | 0.57324898 |
| CLIC4 | 117.072269 | 0.035920514 | 0.65432743 | 0.05489685 | 0.95622064 | 0.9908068 |
| SYF2 | 164.675969 | 0.04283842 | 0.43437223 | 0.09862145 | 0.92143884 | 0.98607457 |
| RSRP1 | 628.684243 | -0.207317461 | 0.33812924 | -0.6131308 | 0.53978983 | 0.8833006 |
| TMEM50A | 221.659194 | -0.381899609 | 0.24081588 | -1.5858573 | 0.11277169 | 0.52363638 |
| MACO1 | 203.295896 | 0.092455035 | 0.23974464 | 0.38563963 | 0.69976358 | 0.93180634 |
| LDLRAP1 | 52.403616 | 0.264400158 | 0.28049758 | 0.94261118 | 0.34587982 | 0.78180304 |
| MAN1C1 | 534.034854 | 1.117785848 | 0.49467613 | 2.25963168 | 0.02384412 | 0.26048096 |
| AL031280.1 | 54.2432632 | 1.209715482 | 0.65876695 | 1.836333 | 0.06630842 | 0.41912682 |
| MTFR1L | 116.670478 | 0.253572042 | 0.40136091 | 0.63178062 | 0.52753024 | 0.87810931 |
| AL033528.2 | 66.1860173 | 0.472867305 | 0.51369697 | 0.92051799 | 0.35730214 | 0.78612469 |
| PAQR7 | 142.561257 | 0.187177293 | 0.32618943 | 0.57383003 | 0.56608282 | 0.89399457 |
| STMN1 | 175.725479 | -1.581224936 | 0.7208158 | -2.1936602 | 0.02825985 | 0.28587181 |
| PAFAH2 | 92.4337536 | -0.19112609 | 0.22599043 | -0.8457265 | 0.39770535 | 0.81357642 |
| PDIK1L | 33.7890669 | -0.428656741 | 0.31366513 | -1.3666063 | 0.17174875 | 0.6187442 |
| ZNF593 | 90.1074409 | 0.688498726 | 0.77670867 | 0.8864311 | 0.37538527 | 0.79878936 |
| CNKSR1 | 78.7576155 | -0.0344961 | 0.28927725 | -0.1192493 | 0.90507788 | 0.98320957 |
| CEP85 | 34.9747699 | -0.079026193 | 0.34130537 | -0.231541 | 0.81689452 | 0.96380902 |
| SH3BGRL3 | 224.632146 | -1.020413387 | 0.48598754 | -2.0996699 | 0.03575789 | 0.31730966 |
| UBXN11 | 47.5477272 | 0.272339149 | 0.35513531 | 0.76686023 | 0.44316462 | 0.83847661 |
| DHDDS | 116.96175 | -0.151262656 | 0.29768792 | -0.5081249 | 0.61136573 | 0.90658678 |
| HMGN2 | 285.687663 | -0.075885965 | 0.42090125 | -0.180294 | 0.85692177 | 0.97191993 |
| RPS6KA1 | 131.294394 | -0.002118246 | 0.27444172 | -0.0077184 | 0.99384168 | 0.99870178 |
| ARID1A | 519.847675 | -0.010337199 | 0.32699914 | -0.0316123 | 0.97478122 | 0.9939318 |
| ZDHHC18 | 38.4641641 | 0.055694047 | 0.39496395 | 0.14101046 | 0.88786168 | 0.98027712 |
| GPN2 | 77.0192507 | 0.014296922 | 0.20782733 | 0.06879231 | 0.94515494 | 0.98949015 |
| NUDC | 220.276074 | -0.100876352 | 0.32257819 | -0.3127191 | 0.7544941 | 0.95002982 |
| NR0B2 | 97.365884 | -0.019867991 | 0.79556811 | -0.0249733 | 0.98007623 | 0.99469867 |
| KDF1 | 37.9525618 | 0.06330123 | 0.33173488 | 0.19081873 | 0.84866761 | 0.96991131 |
| TRNP1 | 94.2973391 | -1.543479494 | 0.50717095 | -3.0433121 | 0.0023399 | 0.06986353 |
| SLC9A1 | 115.052503 | 0.631148718 | 0.34407597 | 1.83432954 | 0.06660509 | 0.41998878 |
| WDTC1 | 169.958294 | 0.17896844 | 0.23696903 | 0.75523979 | 0.45010512 | 0.84143804 |
| TMEM222 | 71.0179272 | -0.598979026 | 0.28200632 | -2.1239915 | 0.03367085 | 0.31035229 |
| WASF2 | 435.698649 | -0.194129532 | 0.21068446 | -0.9214231 | 0.35682956 | 0.78612469 |
| AHDC1 | 115.047607 | 0.027385134 | 0.54733958 | 0.05003317 | 0.96009596 | 0.99121324 |
| IFI6 | 506.653485 | -0.550792698 | 1.6163396 | -0.3407655 | 0.73328016 | 0.94163693 |
| FAM76A | 88.085855 | 0.229249775 | 0.33226791 | 0.6899546 | 0.49022274 | 0.86055851 |
| STX12 | 291.854552 | 0.494559422 | 0.25957853 | 1.90524011 | 0.05674887 | 0.39727338 |
| PPP1R8 | 66.648626 | -0.110122646 | 0.26022923 | -0.4231755 | 0.67216719 | 0.92498948 |
| RPA2 | 116.571271 | 0.186222416 | 0.3535472 | 0.52672575 | 0.59838405 | 0.90369786 |
| EYA3 | 1663.55735 | 0.588880416 | 0.46303378 | 1.27178716 | 0.20344875 | 0.6582233 |
| PTAFR | 49.2511279 | -1.340537534 | 0.7715896 | -1.7373712 | 0.08232167 | 0.45595206 |
| DNAJC8 | 165.55849 | -0.039155937 | 0.23326697 | -0.1678589 | 0.86669428 | 0.97550812 |
| ATP5IF1 | 714.682592 | 0.448950081 | 0.46272907 | 0.97022235 | 0.33193567 | 0.77618491 |
| SESN2 | 39.6966167 | -0.242104111 | 0.31768656 | -0.7620848 | 0.44600937 | 0.83968362 |
| PHACTR4 | 540.120061 | -0.088518142 | 0.19757903 | -0.4480139 | 0.65414319 | 0.92002724 |
| RCC1 | 86.7078187 | -0.558140519 | 0.33365956 | -1.6727844 | 0.09436974 | 0.48561474 |
| TRNAU1AP | 147.244612 | -0.546615936 | 0.28104985 | -1.9449074 | 0.05178612 | 0.38247133 |
| TAF12 | 83.487837 | 0.432862173 | 0.36942592 | 1.17171577 | 0.24131119 | 0.69706209 |
| GMEB1 | 87.2502021 | -0.111683333 | 0.29359501 | -0.3803993 | 0.70364905 | 0.93241442 |
| YTHDF2 | 218.538744 | 0.019528545 | 0.21097075 | 0.09256518 | 0.92624901 | 0.98729553 |
| EPB41 | 391.934336 | -0.173630408 | 0.18657732 | -0.9306083 | 0.3520562 | 0.78314007 |
| SRSF4 | 622.392479 | -0.060713056 | 0.20723625 | -0.2929654 | 0.76954858 | 0.95482817 |
| MECR | 76.1403365 | -0.388597348 | 0.27531205 | -1.4114796 | 0.15810324 | 0.5960151 |

| | | | | | | |
|---|---|---|---|---|---|---|
| PTPRU | 42.3054309 | -0.704097869 | 0.36841149 | -1.9111724 | 0.05598243 | 0.39402004 |
| SDC3 | 74.155773 | -1.761414807 | 0.43735432 | -4.027432 | 5.64E-05 | 0.00609238 |
| PUM1 | 1206.23581 | 0.452861632 | 0.24332062 | 1.86117242 | 0.06271983 | 0.41131726 |
| SNRNP40 | 122.29513 | 0.445940067 | 0.21704548 | 2.05459273 | 0.03991836 | 0.33656026 |
| ZCCHC17 | 185.706914 | -0.114852559 | 0.1914933 | -0.5997732 | 0.54865737 | 0.88854099 |
| SERINC2 | 470.38466 | -0.264725866 | 0.31131556 | -0.8503458 | 0.39513289 | 0.81175661 |
| TINAGL1 | 141.719441 | 0.745329404 | 0.37156863 | 2.00589972 | 0.04486695 | 0.35515061 |
| PEF1 | 127.740521 | -0.005637969 | 0.36359053 | -0.0155064 | 0.9876282 | 0.9964791 |
| PTP4A2 | 390.329501 | 0.646432513 | 0.69315131 | 0.93259943 | 0.35102683 | 0.78254865 |
| KHDRBS1 | 483.386738 | -0.475498122 | 0.28152395 | -1.6890148 | 0.09121659 | 0.47749744 |
| TMEM39B | 41.5369332 | 0.182781262 | 0.31547615 | 0.57938219 | 0.56233132 | 0.89242209 |
| KPNA6 | 387.836786 | 0.158318028 | 0.15929339 | 0.99387694 | 0.32028278 | 0.76529037 |
| TXLNA | 113.643447 | -0.112199371 | 0.20048861 | -0.5596297 | 0.57573207 | 0.89839467 |
| TMEM234 | 42.5973776 | -0.197215806 | 0.31106266 | -0.6340067 | 0.52607646 | 0.87791848 |
| EIF3I | 141.603637 | -0.364196732 | 0.51102575 | -0.7126778 | 0.47604513 | 0.85414749 |
| HDAC1 | 228.359656 | 0.361223287 | 0.27277748 | 1.32424162 | 0.18542281 | 0.63605149 |
| MARCKSL1 | 103.850744 | -2.374435781 | 0.94118548 | -2.5228139 | 0.011642 | 0.17111944 |
| BSDC1 | 163.179509 | -0.226620242 | 0.21878685 | -1.0358038 | 0.30029371 | 0.74729373 |
| ZBTB8OS | 230.57967 | 0.16799444 | 0.22611794 | 0.74295053 | 0.45751163 | 0.84409989 |
| RBBP4 | 297.062143 | -0.053802075 | 0.19054575 | -0.2823578 | 0.77766919 | 0.95601455 |
| SYNC | 79.3060396 | -0.546637627 | 0.32395031 | -1.687412 | 0.09152417 | 0.47836139 |
| KIAA1522 | 461.930942 | -0.421230004 | 0.20541227 | -2.0506565 | 0.04030041 | 0.33814918 |
| YARS | 105.501448 | -0.727758598 | 0.39759555 | -1.8303992 | 0.06719026 | 0.42120786 |
| S100PBP | 219.212996 | -0.312534483 | 0.17969036 | -1.7392947 | 0.08198294 | 0.45535449 |
| TMEM54 | 96.8300194 | -0.022243372 | 0.46189988 | -0.0481563 | 0.96159171 | 0.99171131 |
| RNF19B | 104.671708 | 0.557281184 | 0.80174774 | 0.69508295 | 0.48700332 | 0.85951411 |
| AL020995.1 | 63.8883558 | -0.354872773 | 0.64668474 | -0.548757 | 0.58317223 | 0.90085422 |
| AK2 | 345.087233 | 0.392858117 | 0.26030177 | 1.50924103 | 0.1312372 | 0.55395564 |
| TRIM62 | 36.0729333 | -1.23815715 | 0.311178 | -3.9789353 | 6.92E-05 | 0.00718291 |
| ZNF362 | 100.78087 | 0.263918614 | 0.40352365 | 0.65403505 | 0.51308922 | 0.87178584 |
| AL513327.1 | 58.6898798 | -0.036795013 | 0.67178757 | -0.0547718 | 0.95632027 | 0.9908068 |
| PHC2 | 345.877088 | -0.235943697 | 0.30695325 | -0.7686633 | 0.44209322 | 0.83770294 |
| SMIM12 | 109.238792 | -0.333000915 | 0.34676317 | -0.9603122 | 0.33689809 | 0.77827084 |
| AC114490.3 | 30.6738716 | 0.17433835 | 0.50094979 | 0.34801562 | 0.72782845 | 0.93931348 |
| ZMYM1 | 143.404268 | 0.460395578 | 0.23668991 | 1.9451424 | 0.05175784 | 0.38247133 |
| SFPQ | 565.883419 | 0.060620097 | 0.22231015 | 0.27268255 | 0.78509725 | 0.95702626 |
| ZMYM4 | 1080.5385 | 0.157867281 | 0.15205237 | 1.03824283 | 0.29915702 | 0.74594723 |
| KIAA0319L | 604.871011 | -0.063384851 | 0.2868853 | -0.2209414 | 0.82513803 | 0.96511681 |
| PSMB2 | 152.969838 | 0.095566663 | 0.2478434 | 0.38559293 | 0.69979817 | 0.93180634 |
| AGO4 | 249.834588 | -0.687116984 | 0.20192904 | -3.4027645 | 0.00066708 | 0.03052826 |
| AGO1 | 96.0591099 | 0.169279886 | 0.26489022 | 0.63905677 | 0.522786 | 0.87688817 |
| AGO3 | 656.799001 | 0.242223291 | 0.17090848 | 1.41726896 | 0.15640431 | 0.59366792 |
| AL138787.2 | 34.2457536 | 0.062030363 | 0.4527989 | 0.13699318 | 0.89103618 | 0.98095103 |
| ADPRHL2 | 88.4757974 | -0.892622607 | 0.41082896 | -2.1727353 | 0.02980024 | 0.29105042 |
| TRAPPC3 | 103.32709 | -0.065500779 | 0.25817128 | -0.2537106 | 0.79971918 | 0.96043078 |
| MAP7D1 | 62.4961369 | -0.128491337 | 0.26467086 | -0.4854759 | 0.62733877 | 0.91061758 |
| THRAP3 | 530.052005 | -0.105542954 | 0.17584357 | -0.6002094 | 0.54836671 | 0.88823542 |
| EVA1B | 51.7255514 | -0.653355495 | 0.37301337 | -1.7515605 | 0.07984941 | 0.45063101 |
| STK40 | 125.467527 | -0.002427104 | 0.40808866 | -0.0059475 | 0.99525462 | 0.99908534 |
| LSM10 | 50.1135247 | 0.222415701 | 0.62355391 | 0.35669041 | 0.72132358 | 0.93744589 |
| OSCP1 | 46.5342461 | 0.003294358 | 0.2837627 | 0.01160955 | 0.99073713 | 0.99720756 |
| MRPS15 | 73.6851179 | -0.488303431 | 0.47338689 | -1.0315103 | 0.3023016 | 0.74896472 |
| LINC01137 | 42.3246439 | -1.116221325 | 0.42951601 | -2.5987886 | 0.00935534 | 0.15464341 |
| ZC3H12A | 104.234178 | 0.815772365 | 0.89930181 | 0.90711745 | 0.36434469 | 0.79268955 |
| MEAF6 | 145.188655 | 0.243601214 | 0.29819751 | 0.8169123 | 0.41397855 | 0.82456685 |
| SNIP1 | 54.8561217 | -0.198660309 | 0.29841568 | -0.6657167 | 0.50559218 | 0.87058024 |
| DNALI1 | 58.044687 | 0.019627277 | 0.39222296 | 0.05004112 | 0.96008962 | 0.99121324 |
| GNL2 | 44.0021222 | 0.026766293 | 0.39760975 | 0.067318 | 0.94632855 | 0.98949015 |
| C1orf122 | 162.034531 | -0.350124649 | 0.39812069 | -0.8794435 | 0.37916086 | 0.80274043 |
| MTF1 | 82.1399954 | -0.108196413 | 0.20472264 | -0.5285024 | 0.59715065 | 0.90369786 |
| AL929472.2 | 29.7127187 | -1.189856655 | 0.64693785 | -1.8392132 | 0.06588383 | 0.41813998 |
| INPP5B | 220.186984 | -0.238887285 | 0.36321195 | -0.6577077 | 0.510726 | 0.87138909 |
| SF3A3 | 100.127115 | -0.148878749 | 0.24205176 | -0.6150699 | 0.53850856 | 0.88265739 |
| UTP11 | 32.1309271 | -0.264147909 | 0.36694587 | -0.7198553 | 0.47161412 | 0.85244471 |
| RRAGC | 101.636007 | -0.885501448 | 0.44405917 | -1.9941069 | 0.04614038 | 0.36110228 |
| AL139260.1 | 116.180628 | -1.258082816 | 0.60999475 | -2.0624486 | 0.03916505 | 0.33315722 |
| MYCBP | 33.0828475 | -0.012763051 | 0.5273288 | -0.0242032 | 0.98069052 | 0.99493094 |
| AKIRIN1 | 194.451125 | 0.281991257 | 0.32934063 | 0.85622979 | 0.39187069 | 0.80874455 |
| NDUFS5 | 486.227944 | -0.53623978 | 0.362697 | -1.4784787 | 0.1392797 | 0.56656585 |
| MACF1 | 2184.10625 | -0.232773336 | 0.23854837 | -0.9757909 | 0.32916808 | 0.77573228 |
| PABPC4 | 157.179917 | 0.171490012 | 0.47170426 | 0.36355409 | 0.71619102 | 0.93573028 |
| PPIE | 79.5246979 | -0.081328352 | 0.31058594 | -0.2618546 | 0.79343355 | 0.95906299 |
| TRIT1 | 185.40328 | -0.161007544 | 0.32136724 | -0.501008 | 0.61636553 | 0.90794578 |
| MFSD2A | 221.101566 | -2.975173026 | 1.81023315 | -1.6435303 | 0.10027326 | 0.49569046 |
| CAP1 | 317.031895 | -0.465068496 | 0.3317816 | -1.4017308 | 0.16099564 | 0.60032459 |
| PPT1 | 111.092582 | 0.115758087 | 0.28803121 | 0.40189425 | 0.68776186 | 0.93036007 |
| RLF | 675.565037 | 0.431803627 | 0.3782774 | 1.14149993 | 0.25366194 | 0.71061335 |
| ZMPSTE24 | 217.363795 | 0.459943636 | 0.20635071 | 2.22894143 | 0.0258178 | 0.2733036 |
| COL9A2 | 158.449471 | 0.903709695 | 0.53123482 | 1.70114922 | 0.08891497 | 0.47038308 |
| SMAP2 | 86.9380736 | -0.178176137 | 0.28878663 | -0.616982 | 0.53724663 | 0.88236087 |
| AL603839.3 | 68.2766557 | -0.013694893 | 0.4461455 | -0.030696 | 0.97551196 | 0.9939318 |
| ZNF684 | 124.571708 | 0.261926562 | 0.44452656 | 0.5892259 | 0.55570975 | 0.89081623 |
| NFYC | 306.69665 | 0.410444211 | 0.37975786 | 1.08080505 | 0.27978384 | 0.72938673 |
| SLFNL1-AS1 | 40.9777881 | -1.148757003 | 0.5245497 | -2.189987 | 0.02852518 | 0.28715843 |
| SCMH1 | 1323.74787 | -0.37977217 | 0.30412864 | -1.2487221 | 0.21176672 | 0.66975892 |
| AC093151.3 | 33.4345019 | -3.183217815 | 0.85838639 | -3.708374 | 0.00020859 | 0.01374886 |
| HIVEP3 | 127.687032 | -0.527099843 | 0.46457868 | -1.134576 | 0.25655299 | 0.71374446 |
| FOXJ3 | 524.9843 | 0.323676812 | 0.21612145 | 1.49766166 | 0.13422118 | 0.55858006 |
| RIMKLA | 36.7382704 | -2.424963788 | 0.66669959 | -3.6372661 | 0.00027555 | 0.01640759 |
| PPCS | 118.343045 | 0.045747615 | 0.37209466 | 0.12294617 | 0.90214972 | 0.98320957 |
| CCDC30 | 486.914119 | 0.259868397 | 0.2822103 | 0.92083243 | 0.35713792 | 0.78612469 |
| PPIH | 61.3547535 | 0.455188848 | 0.24385422 | 1.86664334 | 0.06195141 | 0.40910573 |
| YBX1 | 446.06213 | -0.240688006 | 0.50496204 | -0.4766457 | 0.6336144 | 0.91415622 |
| SVBP | 41.6130875 | -0.795101577 | 0.3242965 | -2.4517736 | 0.01421541 | 0.19297602 |
| ERMAP | 67.5376608 | -0.064422851 | 0.33967719 | -0.189659 | 0.84957632 | 0.96991131 |
| SLC2A1 | 675.977665 | -1.517736814 | 0.92338125 | -1.643673 | 0.10024377 | 0.49569046 |
| SLC2A1-AS1 | 75.3709917 | -2.207067942 | 0.7209616 | -3.0612836 | 0.0022039 | 0.06714214 |
| EBNA1BP2 | 90.7587039 | -0.06594047 | 0.27546704 | -0.239377 | 0.81081327 | 0.96288868 |
| TMEM125 | 50.1898641 | 0.3928531 | 0.41978139 | 0.93585163 | 0.34934958 | 0.78180304 |
| C1orf210 | 36.6032379 | -0.43962115 | 0.45730179 | -0.961337 | 0.33638273 | 0.77820994 |
| ELOVL1 | 76.7984395 | -0.235302638 | 0.2933946 | -0.8020006 | 0.42255263 | 0.82757457 |
| MED8 | 40.6126208 | -0.301705503 | 0.35067797 | -0.8603492 | 0.3895966 | 0.80680464 |
| SZT2 | 117.89245 | -0.124706014 | 0.26422683 | -0.4719658 | 0.63695123 | 0.91454294 |
| HYI | 43.6187533 | 0.602761631 | 0.36982533 | 1.62985492 | 0.10313216 | 0.50171849 |
| PTPRF | 592.603403 | -0.157847127 | 0.31386595 | -0.5029125 | 0.61502576 | 0.90792412 |
| IPO13 | 33.427407 | 0.0146635 | 0.37353616 | 0.0392559 | 0.96868636 | 0.99303855 |
| ATP6V0B | 366.475306 | -0.286207938 | 0.38380672 | -0.7457085 | 0.45584352 | 0.84372683 |
| B4GALT2 | 44.8637001 | -0.618518748 | 0.44949159 | -1.3760408 | 0.16880902 | 0.61405939 |
| CCDC24 | 106.546895 | -0.224975187 | 0.28977394 | -0.7763817 | 0.43752364 | 0.83420562 |
| DMAP1 | 63.1790279 | 0.477572255 | 0.24338657 | 1.9621964 | 0.04973962 | 0.37609758 |
| ERI3 | 258.768972 | 0.018385712 | 0.30752364 | 0.05978634 | 0.95232581 | 0.9903694 |
| RNF220 | 76.2152161 | 0.159621059 | 0.27349197 | 0.58364076 | 0.55946203 | 0.89179371 |
| TMEM53 | 45.4225601 | -0.850941488 | 0.35120177 | -2.4229419 | 0.01539539 | 0.20206713 |

| | | | | | | |
|---|---|---|---|---|---|---|
| ARMH1 | 70.6893567 | 0.128552745 | 0.45140372 | 0.28478442 | 0.77580931 | 0.95601455 |
| RPS8 | 771.956117 | -0.134735324 | 0.45448274 | -0.2964586 | 0.76687987 | 0.95408212 |
| EIF2B3 | 140.597869 | 0.190486372 | 0.31216241 | 0.6102156 | 0.541719 | 0.8841637 |
| HECTD3 | 38.3156741 | 0.106761579 | 0.28007539 | 0.38118872 | 0.70306322 | 0.93241442 |
| UROD | 44.1961927 | 0.271487982 | 0.42826669 | 0.63392271 | 0.52613126 | 0.87791848 |
| ZSWIM5 | 221.230743 | 0.0008555 | 0.51340943 | 0.00166631 | 0.99867048 | 0.99962432 |
| TESK2 | 73.7245193 | -0.57146496 | 0.41858349 | -1.3652353 | 0.1721791 | 0.61884138 |
| PRDX1 | 1214.13835 | -0.501642747 | 0.62215817 | -0.8062946 | 0.42007303 | 0.82694802 |
| AKR1A1 | 78.4198128 | -0.534154727 | 0.39055566 | -1.3676789 | 0.17141262 | 0.61817077 |
| NASP | 283.002052 | 0.805926984 | 0.36853471 | 2.18684145 | 0.0287541 | 0.28768102 |
| GPBP1L1 | 693.222781 | 0.210355808 | 0.19883744 | 1.05792858 | 0.29008801 | 0.73754646 |
| TMEM69 | 50.9990297 | 0.218139618 | 0.32941169 | 0.66220971 | 0.50783683 | 0.87094607 |
| IPP | 141.832707 | -0.003146011 | 0.23922156 | -0.013151 | 0.9895073 | 0.9965778 |
| MAST2 | 164.04168 | -0.186589626 | 0.21625914 | -0.8628058 | 0.38824428 | 0.8065026 |
| PIK3R3 | 33.9505263 | -0.422051084 | 0.51179903 | -0.8246422 | 0.40957475 | 0.82150113 |
| TSPAN1 | 1304.56006 | -1.290016655 | 0.36669819 | -3.5179248 | 0.00043494 | 0.02245382 |
| POMGNT1 | 133.678431 | -0.405024646 | 0.29894796 | -1.3548333 | 0.17547068 | 0.62477136 |
| LURAP1 | 140.360903 | 0.058349979 | 0.2265953 | 0.25750745 | 0.79678706 | 0.95961857 |
| LRRC41 | 146.991187 | 0.236837122 | 0.2821877 | 0.83928933 | 0.40130697 | 0.81648166 |
| UQCRH | 611.542952 | 0.796420738 | 0.4814543 | 1.65419801 | 0.09808729 | 0.49265259 |
| NSUN4 | 98.0989004 | 0.176767431 | 0.2322713 | 0.76103862 | 0.446634 | 0.83989311 |
| FAAH | 49.5651106 | 0.352982927 | 0.28159289 | 1.25352215 | 0.21001574 | 0.66797116 |
| MKNK1 | 116.041428 | 0.267951444 | 0.26882788 | 0.99673977 | 0.31889084 | 0.76370567 |
| ATPAF1 | 155.470703 | 0.057544922 | 0.24216761 | 0.23762435 | 0.81217246 | 0.96288868 |
| EFCAB14-AS | 43.6837064 | 0.179691552 | 0.73873306 | 0.24324287 | 0.80781725 | 0.96245698 |
| EFCAB14 | 566.61901 | 0.237559553 | 0.31492387 | 0.75433962 | 0.45064531 | 0.84153105 |
| PDZK1IP1 | 162.05664 | -2.066689405 | 0.79855489 | -2.5880367 | 0.00965247 | 0.15717671 |
| CMPK1 | 228.748195 | -0.449187794 | 0.25177932 | -1.7840536 | 0.07441496 | 0.43775442 |
| TRABD2B | 746.205175 | -0.812352315 | 0.59978024 | -1.3544166 | 0.1756035 | 0.62477136 |
| SPATA6 | 474.78239 | 0.187532632 | 0.25159612 | 0.74537172 | 0.45604704 | 0.84372683 |
| AGBL4 | 2350.09715 | 0.837706363 | 0.42670258 | 1.96320906 | 0.04962189 | 0.37602124 |
| FAF1 | 1284.76968 | 0.023341746 | 0.17495926 | 0.13341247 | 0.89386719 | 0.98194215 |
| RNF11 | 268.201759 | 0.241669083 | 0.18109908 | 1.33445783 | 0.18205386 | 0.63239126 |
| TTC39A | 157.249802 | -1.825776578 | 0.45416586 | -4.0200656 | 5.82E-05 | 0.0062219 |
| EPS15 | 648.44047 | 0.127505329 | 0.21847236 | 0.58362224 | 0.55947449 | 0.89179371 |
| OSBPL9 | 592.962565 | 0.250985639 | 0.1664056 | 1.50827639 | 0.1314838 | 0.55397858 |
| NRDC | 502.024415 | 0.189209155 | 0.24161112 | 0.78311444 | 0.43355991 | 0.83187655 |
| TXNDC12 | 110.351929 | -0.226575774 | 0.24384024 | -0.9291976 | 0.35278667 | 0.78314007 |
| BTF3L4 | 211.674133 | 0.070582184 | 0.19883828 | 0.35497282 | 0.72260996 | 0.93852427 |
| ZFYVE9 | 364.354859 | 0.252623286 | 0.2238533 | 1.1285216 | 0.2590997 | 0.71645511 |
| CC2D1B | 74.0899405 | -0.689210414 | 0.45625643 | -1.5105769 | 0.13089629 | 0.55395564 |
| PRPF38A | 82.230589 | 0.18293371 | 0.25196474 | 0.726029 | 0.46782099 | 0.85161077 |
| TUT4 | 633.014317 | 0.183671785 | 0.22224117 | 0.82645257 | 0.40854741 | 0.82073894 |
| ZYG11B | 270.619629 | 0.472663807 | 0.21528525 | 2.19552346 | 0.02812607 | 0.28491502 |
| ECHDC2 | 597.828245 | 0.196716718 | 0.32745871 | 0.60073746 | 0.54801486 | 0.88807728 |
| SCP2 | 499.574198 | 0.069234595 | 0.20193812 | 0.34285055 | 0.73171089 | 0.94066857 |
| CPT2 | 67.1596139 | -0.271884337 | 0.35219496 | -0.7719711 | 0.44013154 | 0.83682484 |
| C1orf123 | 53.1150534 | -0.111843303 | 0.4007706 | -0.2790706 | 0.78019062 | 0.95639467 |
| MAGOH | 89.701129 | 0.428908658 | 0.36407717 | 1.17807074 | 0.23876841 | 0.69499004 |
| LRP8 | 58.1671802 | -1.281784794 | 0.49234711 | -2.6034169 | 0.00922996 | 0.15397695 |
| NDC1 | 100.952854 | 0.09043541 | 0.312849 | 0.28907048 | 0.77252744 | 0.95543807 |
| YIPF1 | 56.8358815 | 0.072425453 | 0.27439803 | 0.26394305 | 0.7918238 | 0.95839852 |
| HSPB11 | 112.95165 | 0.011447065 | 0.25981448 | 0.04405861 | 0.96485768 | 0.99243116 |
| LRRC42 | 59.8658634 | -0.36009078 | 0.31843766 | -1.1308046 | 0.25813733 | 0.71534188 |
| TMEM59 | 535.176843 | 0.033795296 | 0.40864055 | 0.08270177 | 0.93408868 | 0.98822345 |
| TCEANC2 | 76.0253457 | -0.130160142 | 0.32204539 | -0.4041671 | 0.68608987 | 0.92974759 |
| MRPL37 | 53.7681798 | 0.006071047 | 0.31090292 | 0.01952715 | 0.98442058 | 0.9954565 |
| SSBP3 | 464.006012 | -0.262447684 | 0.53485294 | -0.4906913 | 0.6236448 | 0.91058133 |
| ACOT11 | 534.401997 | 0.309722081 | 0.4117592 | 0.75219226 | 0.45193545 | 0.84234337 |
| MROH7-TTC | 47.881913 | 0.948437097 | 0.52396449 | 1.81011714 | 0.07027762 | 0.42920134 |
| TTC22 | 31.9160757 | -0.508257165 | 0.3979941 | -1.277047 | 0.20158566 | 0.65588877 |
| DHCR24 | 149.888359 | -1.084728259 | 0.64666549 | -1.6774179 | 0.09346079 | 0.4832112 |
| USP24 | 713.964448 | 0.113380862 | 0.24644528 | 0.46006505 | 0.64546953 | 0.91685019 |
| PLPP3 | 235.205851 | -0.238129801 | 0.32619067 | -0.7300325 | 0.4653703 | 0.84991999 |
| PRKAA2 | 703.901679 | 0.083019447 | 0.27502179 | 0.30186498 | 0.76275499 | 0.95378284 |
| FYB2 | 1496.87844 | -0.052528038 | 0.49859854 | -0.1053514 | 0.916097 | 0.98520052 |
| DAB1 | 463.746613 | 0.361834741 | 0.8291557 | 0.43638938 | 0.66255425 | 0.9229791 |
| TACSTD2 | 659.512064 | 0.607750724 | 0.69840191 | 0.87020198 | 0.38419003 | 0.80561637 |
| MYSM1 | 290.686547 | 0.241899062 | 0.21403996 | 1.13015842 | 0.25840948 | 0.71549044 |
| JUN | 302.306899 | 0.563301969 | 0.69889154 | 0.8059934 | 0.42024665 | 0.82694802 |
| LINC01135 | 126.977163 | 0.337229673 | 0.4525158 | 0.74523292 | 0.45613093 | 0.84372683 |
| FGGY | 468.51233 | -0.133630138 | 0.32431734 | -0.4120351 | 0.68031368 | 0.92700898 |
| HOOK1 | 354.135637 | 0.051954658 | 0.30915789 | 0.16805218 | 0.86654223 | 0.97544173 |
| NFIA | 816.862283 | 0.203732335 | 0.42167978 | 0.48314467 | 0.62899302 | 0.9114564 |
| TM2D1 | 278.020816 | 0.246438257 | 0.20170091 | 1.22180044 | 0.2217831 | 0.6794174 |
| PATJ | 3650.1527 | 0.054148224 | 0.2016354 | 0.26854522 | 0.78827967 | 0.95801395 |
| USP1 | 77.816446 | -0.17120553 | 0.22515106 | -0.760403 | 0.44701376 | 0.84014281 |
| DOCK7 | 1123.02204 | 0.347691542 | 0.23652544 | 1.46999639 | 0.14156273 | 0.56994907 |
| AC103923.1 | 48.3170134 | 0.23354798 | 0.46288242 | 0.50455142 | 0.61387394 | 0.90751853 |
| ATG4C | 138.527406 | 0.255988395 | 0.30486146 | 0.83968761 | 0.40108356 | 0.81648166 |
| ALG6 | 59.3838369 | 0.26092999 | 0.24686618 | 1.05696935 | 0.29052558 | 0.73754646 |
| ITGB3BP | 200.177768 | -0.849990779 | 0.47324535 | -1.7960891 | 0.07248035 | 0.43315584 |
| EFCAB7 | 117.151134 | -0.015088129 | 0.26360808 | -0.057237 | 0.95435643 | 0.99060627 |
| PGM1 | 162.36331 | -1.222520958 | 0.38847326 | -3.1469887 | 0.00164961 | 0.05505717 |
| ROR1 | 100.65168 | -1.343565982 | 0.79978875 | -1.6799011 | 0.09297657 | 0.48165814 |
| CACHD1 | 199.829066 | -0.116131182 | 0.68502654 | -0.169528 | 0.86538135 | 0.97497275 |
| RAVER2 | 163.106843 | -0.134659198 | 0.28618963 | -0.4705244 | 0.63798042 | 0.915143 |
| JAK1 | 846.864383 | 0.18259748 | 0.3098026 | 0.58939944 | 0.55559335 | 0.89081623 |
| AK4 | 134.859561 | -2.030214956 | 0.76731701 | -2.6458621 | 0.0081483 | 0.14312705 |
| DNAJC6 | 52.3023997 | -0.741820865 | 0.41468892 | -1.7888611 | 0.07363719 | 0.43698629 |
| LEPR | 234.24322 | -0.820274142 | 0.41739812 | -1.9652081 | 0.04939016 | 0.3755208 |
| PDE4B | 123.38708 | -4.417038824 | 1.38816464 | -3.1819272 | 0.00146299 | 0.05060096 |
| WDR78 | 211.696445 | 0.13306482 | 0.38164609 | 0.34866025 | 0.72734439 | 0.93931348 |
| MIER1 | 435.708116 | 0.268297822 | 0.18971028 | 1.41425033 | 0.15728842 | 0.59486921 |
| SLC35D1 | 135.018543 | 0.572585017 | 0.26422078 | 2.16707031 | 0.03022949 | 0.29256342 |
| SERBP1 | 489.1971 | -0.267908509 | 0.3125648 | -0.8571295 | 0.39137332 | 0.80851416 |
| GADD45A | 102.858025 | -1.277116481 | 0.34836663 | -3.6660127 | 0.00024636 | 0.01527734 |
| GNG12 | 682.259091 | -0.235791643 | 0.15761995 | -1.4959505 | 0.13466655 | 0.55868799 |
| GNG12-AS1 | 224.50603 | -1.093928281 | 0.33891629 | -3.2277241 | 0.00124779 | 0.04572331 |
| WLS | 304.941344 | -0.175172573 | 0.34486264 | -0.5079488 | 0.61148923 | 0.90658678 |
| LRRC40 | 133.813838 | 0.517983371 | 0.23877002 | 2.16938197 | 0.0300537 | 0.29169628 |
| SRSF11 | 689.456573 | 0.224050816 | 0.19262972 | 1.16311654 | 0.24478222 | 0.70130056 |
| ANKRD13C | 398.887495 | 1.025477995 | 0.34518305 | 2.97082374 | 0.00297002 | 0.08063686 |
| HHLA3 | 42.345526 | -0.037356135 | 0.41462914 | -0.0900953 | 0.92821149 | 0.98751845 |
| LINC01788 | 163.892487 | 0.820312762 | 0.67647857 | 1.21262194 | 0.2252744 | 0.68272865 |
| PTGER3 | 884.465057 | -0.189987086 | 0.43498594 | -0.436766 | 0.66228103 | 0.92284375 |
| ZRANB2-AS1 | 46.6181173 | 0.399601458 | 0.66678815 | 0.59929298 | 0.54897753 | 0.88854099 |
| ZRANB2 | 563.890686 | 0.252174419 | 0.19219158 | 1.31209919 | 0.18948667 | 0.64328484 |
| NEGR1 | 46.8095705 | -0.023811384 | 0.51534894 | -0.0462044 | 0.96314734 | 0.99211837 |
| LRRIQ3 | 42.0070767 | -0.405623446 | 0.56262393 | -0.7209495 | 0.47094058 | 0.85236058 |
| CRYZ | 404.303801 | 0.197810288 | 0.27668466 | 0.71493044 | 0.47465203 | 0.85368787 |
| TYW3 | 92.9211729 | 0.266873261 | 0.21063978 | 1.26696512 | 0.20516776 | 0.66097699 |
| SLC44A5 | 97.5813723 | -0.153030744 | 1.31896659 | -0.1160232 | 0.90763414 | 0.9835597 |

| | | | | | | |
|---|---|---|---|---|---|---|
| ACADM | 290.414273 | 0.38642382 | 0.22863855 | 1.69010792 | 0.09100731 | 0.47749744 |
| RABGGTB | 65.7560337 | 0.622150375 | 0.29507929 | 2.10841762 | 0.03499488 | 0.31514522 |
| ST6GALNAC3 | 55.233953 | -0.439244899 | 0.43757187 | -1.0038234 | 0.31546372 | 0.75912793 |
| PIGK | 173.062297 | 0.409179492 | 0.19002521 | 2.1532906 | 0.03129585 | 0.29704233 |
| AC118549.1 | 729.749476 | 0.138503469 | 0.26988758 | 0.51318948 | 0.60781879 | 0.90514346 |
| USP33 | 467.633953 | 0.170860193 | 0.21717702 | 0.78673238 | 0.43143854 | 0.8312408 |
| MIGA1 | 341.68976 | 0.09559521 | 0.18173265 | 0.52602112 | 0.59887354 | 0.90369786 |
| NEXN | 36.8179886 | -0.262138972 | 0.462444 | -0.5668556 | 0.57081229 | 0.8975413 |
| FUBP1 | 370.803236 | 0.379987419 | 0.1826601 | 2.08029788 | 0.03749822 | 0.32424203 |
| PTGFR | 315.240855 | -0.256587312 | 0.50844061 | -0.5046554 | 0.61380088 | 0.90751853 |
| IFI44L | 360.955907 | -0.510280174 | 1.85580137 | -0.2749649 | 0.78334323 | 0.9568898 |
| IFI44 | 123.186765 | 0.483179631 | 1.03090682 | 0.4686938 | 0.63928852 | 0.91521016 |
| AL136234.1 | 104.297624 | 1.967174885 | 0.6499794 | 3.02651883 | 0.00247387 | 0.07262241 |
| LINC01781 | 73.0825915 | -1.635394472 | 0.69221799 | -2.3625426 | 0.01815006 | 0.2244336 |
| ADGRL2 | 961.92353 | 1.050811866 | 0.28300358 | 3.71306922 | 0.00020476 | 0.01374886 |
| LINC01362 | 53.0282692 | -1.03347433 | 0.77500177 | -1.3335122 | 0.18236377 | 0.6328385 |
| LINC01725 | 732.977474 | -0.388069918 | 0.41536696 | -0.9342821 | 0.35015839 | 0.78223232 |
| PRKACB | 350.858025 | 0.165301726 | 0.28754457 | 0.57487341 | 0.56537691 | 0.89382261 |
| SAMD13 | 40.4538776 | -0.061037384 | 0.58485207 | -0.1043638 | 0.91688065 | 0.98520052 |
| RPF1 | 55.0169987 | 0.051737302 | 0.32427576 | 0.15954724 | 0.87323774 | 0.97622753 |
| GNG5 | 292.930632 | -0.072985804 | 0.37957676 | -0.1922821 | 0.84752127 | 0.96991131 |
| SPATA1 | 61.407825 | -0.576062554 | 0.55382881 | -1.0401455 | 0.2982723 | 0.74594723 |
| CTBS | 100.944422 | -0.577568288 | 0.3364364 | -1.7167235 | 0.0860297 | 0.46572214 |
| SSX2IP | 64.8020427 | -0.145051873 | 0.26484312 | -0.5476898 | 0.58390492 | 0.90085422 |
| WDR63 | 58.5709384 | 0.52033513 | 0.46876456 | 1.1100138 | 0.26699308 | 0.72152517 |
| MCOLN3 | 56.0659469 | 0.488877296 | 0.58928996 | 0.82960398 | 0.40676272 | 0.82018309 |
| SYDE2 | 62.3358545 | 0.328778647 | 0.26260602 | 1.25198445 | 0.21057553 | 0.66871998 |
| C1orf52 | 110.751507 | 0.356706665 | 0.28778188 | 1.23950358 | 0.21515906 | 0.67132603 |
| BCL10 | 124.082712 | -0.553964571 | 0.37838689 | -1.4640163 | 0.14318949 | 0.5729912 |
| AL078459.1 | 234.143691 | -0.414918984 | 0.49971362 | -0.8303135 | 0.40636153 | 0.82002585 |
| DDAH1 | 921.460836 | 0.224800722 | 0.20218028 | 1.11188253 | 0.26618866 | 0.72053648 |
| CYR61 | 51.9484265 | -0.662867501 | 0.91745621 | -0.7225059 | 0.46998352 | 0.85232523 |
| ZNHIT6 | 204.275066 | 0.026535278 | 0.27910145 | 0.09507395 | 0.92425609 | 0.98697817 |
| AL136382.1 | 41.2595926 | 0.072405411 | 0.6542793 | 0.11066438 | 0.9118825 | 0.98480302 |
| ODF2L | 542.036972 | 0.127291581 | 0.33593174 | 0.37892097 | 0.70474655 | 0.93241442 |
| SH3GLB1 | 345.744949 | -0.126656631 | 0.18261828 | -0.6935594 | 0.48795854 | 0.86006752 |
| SELENOF | 345.822168 | 0.240732511 | 0.16358358 | 1.47161779 | 0.14112412 | 0.56895751 |
| HS2ST1 | 225.409101 | -0.133856191 | 0.26814035 | -0.499202 | 0.61763711 | 0.90794578 |
| LMO4 | 174.429873 | -0.378487559 | 0.4173781 | -0.9068218 | 0.36450104 | 0.79268955 |
| PKN2-AS1 | 60.0801645 | -0.324650415 | 0.49581822 | -0.6547771 | 0.51261128 | 0.87177451 |
| PKN2 | 812.031449 | 0.201678012 | 0.18057491 | 1.11686616 | 0.26405155 | 0.71989632 |
| GTF2B | 100.904313 | 0.025897313 | 0.25368867 | 0.10208305 | 0.91869076 | 0.98520052 |
| KYAT3 | 179.332074 | -0.673035113 | 0.23674421 | -2.8428788 | 0.00447081 | 0.10481891 |
| GBP3 | 125.157947 | -0.604608184 | 0.40816848 | -1.4812711 | 0.13853434 | 0.56513813 |
| GBP2 | 88.8645168 | -3.011201821 | 0.62587108 | -4.8112174 | 1.50E-06 | 0.00049129 |
| LRRC8B | 202.127395 | -0.098676091 | 0.23829722 | -0.4140883 | 0.67880945 | 0.9268955 |
| LRRC8D | 36.3676859 | -0.80249386 | 0.35744565 | -2.2450794 | 0.02476304 | 0.26640367 |
| ZNF326 | 217.368938 | 0.184542051 | 0.1890563 | 0.97612221 | 0.32900391 | 0.77569425 |
| ZNF644 | 581.254403 | 0.307297013 | 0.16545984 | 1.85723018 | 0.06327841 | 0.41267121 |
| HFM1 | 44.8428067 | 0.355882642 | 0.38793456 | 0.91737802 | 0.35894459 | 0.7881289 |
| TGFBR3 | 240.03009 | 1.162066476 | 0.69026514 | 1.6835074 | 0.09227691 | 0.48040835 |
| BTBD8 | 129.865689 | -0.14198092 | 0.22470373 | -0.6318583 | 0.52747946 | 0.87810931 |
| GLMN | 121.637575 | 0.084737081 | 0.26913563 | 0.31484899 | 0.75287631 | 0.94913313 |
| RPAP2 | 366.897953 | -0.162535297 | 0.18381645 | -0.8842261 | 0.37657419 | 0.80001976 |
| EVI5 | 800.152032 | 0.635270069 | 0.23247243 | 2.73266837 | 0.00628236 | 0.12759512 |
| RPL5 | 544.570179 | -0.344359084 | 0.3603375 | -0.9556571 | 0.33924551 | 0.77836439 |
| FAM69A | 83.2111835 | -0.253983192 | 0.32369301 | -0.7846422 | 0.43266338 | 0.83162989 |
| MTF2 | 456.165574 | 0.192274033 | 0.21957875 | 0.87564956 | 0.38122057 | 0.80315561 |
| TMED5 | 449.557184 | 0.842450914 | 0.36950361 | 2.27995313 | 0.02261047 | 0.25181478 |
| CCDC18 | 116.253877 | 0.325777701 | 0.43273607 | 0.75283233 | 0.45155068 | 0.84234337 |
| DR1 | 113.680114 | 0.467704252 | 0.26662044 | 1.75419505 | 0.07939709 | 0.44953856 |
| FNBP1L | 1483.92658 | -0.155437026 | 0.33238926 | -0.4676355 | 0.64004526 | 0.91559846 |
| BCAR3 | 732.260826 | 0.372560354 | 0.43609676 | 0.85430663 | 0.39293512 | 0.81010924 |
| DNTTIP2 | 50.1179814 | -0.146168521 | 0.44773309 | -0.3264635 | 0.74407369 | 0.9456489 |
| GCLM | 136.349628 | -1.393354769 | 0.35237194 | -3.9542161 | 7.68E-05 | 0.00745109 |
| ABCA4 | 88.3571112 | -0.685450794 | 0.75031906 | -0.9135458 | 0.36095558 | 0.78927419 |
| ARHGAP29 | 2060.67737 | 0.054717895 | 0.45868641 | 0.1192926 | 0.90504355 | 0.98320957 |
| ABCD3 | 565.260758 | 0.474669783 | 0.24584056 | 1.9308034 | 0.05350737 | 0.38854227 |
| SLC44A3 | 257.175071 | 0.110802139 | 0.39580614 | 0.27994042 | 0.77952321 | 0.95636474 |
| CNN3 | 533.100164 | -0.245637192 | 0.35121343 | -0.6993958 | 0.48430472 | 0.85909165 |
| AC105942.1 | 51.4473948 | -0.378399468 | 0.33876984 | -1.116981 | 0.26400246 | 0.71989632 |
| ALG14 | 109.447745 | -0.120993833 | 0.31041168 | -0.389785 | 0.6966955 | 0.93130502 |
| TMEM56-RWDD3 | 33.2226697 | 0.801406043 | 0.42270124 | 1.895916 | 0.05797115 | 0.3985947 |
| AC092802.1 | 59.4015085 | 0.684393771 | 0.52829544 | 1.29547546 | 0.19515626 | 0.64928179 |
| PTBP2 | 683.057643 | -0.007254318 | 0.22603457 | -0.0320938 | 0.97439721 | 0.9939318 |
| DPYD | 556.313727 | -0.456039497 | 0.2375283 | -1.9199375 | 0.0548658 | 0.39224566 |
| SNX7 | 147.560779 | -0.232845276 | 0.23720035 | -0.9816397 | 0.32627739 | 0.77134831 |
| FRRS1 | 114.439575 | 0.069157131 | 0.49296969 | 0.14028678 | 0.88843341 | 0.98051184 |
| AGL | 257.337226 | 0.131027321 | 0.30065999 | 0.43579899 | 0.66298257 | 0.9229791 |
| SASS6 | 37.4204082 | 0.383768247 | 0.34636474 | 1.10798877 | 0.26786667 | 0.72276232 |
| TRMT13 | 147.239516 | 0.182372494 | 0.25352325 | 0.71935213 | 0.47192398 | 0.85244471 |
| DBT | 200.80657 | -0.137322681 | 0.20817472 | -0.6596511 | 0.50947775 | 0.87094607 |
| RTCA | 108.695249 | 0.005904704 | 0.27231961 | 0.02168299 | 0.98270083 | 0.99507903 |
| CDC14A | 111.525575 | -0.340948922 | 0.39560375 | -0.8618445 | 0.38877308 | 0.8065026 |
| EXTL2 | 85.206148 | 0.670192476 | 0.28696042 | 2.33548752 | 0.01951797 | 0.23473196 |
| SLC30A7 | 274.990959 | -0.023395516 | 0.24440995 | -0.0957224 | 0.92374102 | 0.98694936 |
| DPH5 | 64.5952301 | -0.308447422 | 0.23016401 | -1.3401202 | 0.18020628 | 0.63015077 |
| AC093157.1 | 90.0533258 | 0.084469811 | 0.27527055 | 0.30686105 | 0.75894914 | 0.95232124 |
| RNPC3 | 356.64996 | 0.220771616 | 0.30236656 | 0.73014561 | 0.46530119 | 0.84991999 |
| AMY2B | 95.5061295 | 0.089705804 | 0.35051247 | 0.25592757 | 0.79800677 | 0.96000168 |
| PRMT6 | 31.6889994 | -0.123352887 | 0.38109024 | -0.3236842 | 0.74617714 | 0.94594643 |
| VAV3 | 1724.28945 | 0.058166117 | 0.42551804 | 0.13669483 | 0.89127202 | 0.98095103 |
| SLC25A24 | 152.393984 | 0.065391994 | 0.24869228 | 0.26294341 | 0.7925942 | 0.95859644 |
| FAM102B | 82.8298527 | 0.58585191 | 0.56389706 | 1.03893415 | 0.29883536 | 0.74594723 |
| PRPF38B | 246.458005 | 0.342866235 | 0.28107798 | 1.21982602 | 0.22253083 | 0.6798044 |
| STXBP3 | 293.255599 | 0.810725209 | 0.17461954 | 4.6428092 | 3.44E-06 | 0.00085762 |
| AL449266.1 | 42.8219428 | -0.66162661 | 0.4216729 | -1.5690518 | 0.11663588 | 0.53183794 |
| GPSM2 | 65.0829203 | -0.745595695 | 0.46794624 | -1.5933362 | 0.11108478 | 0.51948619 |
| WDR47 | 71.3303453 | 0.494686426 | 0.26095382 | 1.89568572 | 0.05800161 | 0.3985947 |
| TAF13 | 56.9810052 | 0.251029011 | 0.36702123 | 0.6839632 | 0.49399841 | 0.86339027 |
| TMEM167B | 87.1325128 | -0.093295184 | 0.2829887 | -0.3296781 | 0.74164318 | 0.94487568 |
| SARS | 183.061209 | -0.249220001 | 0.18090326 | -1.3776424 | 0.16831373 | 0.6131323 |
| CELSR2 | 143.874565 | 1.148751481 | 0.37158966 | 3.09145166 | 0.0019918 | 0.06287382 |
| SORT1 | 341.391809 | 0.186402584 | 0.21512689 | 0.86647737 | 0.38622843 | 0.80634213 |
| PSMA5 | 153.973659 | -0.364768594 | 0.29576927 | -1.2332877 | 0.21746846 | 0.67507983 |
| GNAI3 | 271.555535 | 0.039910253 | 0.24356989 | 0.16385545 | 0.86984494 | 0.97590997 |
| AMPD2 | 53.910004 | 0.579570713 | 0.39482025 | 1.46793563 | 0.14212171 | 0.57093391 |
| GSTM4 | 70.8293538 | 0.083989775 | 0.35522437 | 0.23644148 | 0.81309012 | 0.96288868 |
| GSTM3 | 637.994537 | -0.064101164 | 0.5443436 | -0.1177586 | 0.9062589 | 0.98320957 |
| AHCYL1 | 490.272107 | -0.706986991 | 0.25254631 | -2.799435 | 0.00511921 | 0.11471549 |
| STRIP1 | 72.8918776 | 0.129596735 | 0.2375947 | 0.54545298 | 0.58544201 | 0.90173902 |
| KCNC4 | 182.659882 | -0.206293217 | 0.47604109 | -0.4333517 | 0.66475928 | 0.92324595 |
| RBM15 | 86.2073164 | 0.47964764 | 0.31494362 | 1.52296352 | 0.12776783 | 0.54791812 |

| | | | | | | |
|---|---|---|---|---|---|---|
| SLC16A4 | 512.67367 | 0.120890707 | 0.41919519 | 0.28838763 | 0.77305004 | 0.95543807 |
| AL355488.1 | 38.5583839 | 0.158981475 | 0.32524351 | 0.48880752 | 0.62497798 | 0.91058133 |
| LAMTOR5 | 487.422757 | 0.178767446 | 0.49409191 | 0.3618101 | 0.71749394 | 0.93573028 |
| LRIF1 | 48.8936316 | -0.220772576 | 0.48952619 | -0.4509924 | 0.65199505 | 0.91889565 |
| DRAM2 | 135.060528 | -0.309170862 | 0.17513237 | -1.7653553 | 0.07750407 | 0.44546179 |
| CEPT1 | 156.270568 | 0.088700219 | 0.23271597 | 0.38115227 | 0.70309027 | 0.93241442 |
| DENND2D | 101.221173 | 0.259846861 | 0.35457326 | 0.73284394 | 0.46365362 | 0.84922793 |
| WDR77 | 45.3452344 | 0.670522085 | 0.41530651 | 1.61452341 | 0.10641394 | 0.50807204 |
| ATP5PB | 310.354266 | -0.049330358 | 0.4873054 | -0.1012309 | 0.91936718 | 0.98520052 |
| RAP1A | 437.816402 | 0.354648262 | 0.16200168 | 2.18916405 | 0.02858492 | 0.28721951 |
| CTTNBP2NL | 186.150856 | -0.296787564 | 0.29122721 | -1.0190928 | 0.3081589 | 0.75313873 |
| ST7L | 191.043689 | 0.133139134 | 0.1990477 | 0.66888055 | 0.50357168 | 0.86946722 |
| CAPZA1 | 377.554835 | -0.008853712 | 0.23342557 | -0.0379295 | 0.9697439 | 0.99337569 |
| MOV10 | 103.246369 | -0.229242376 | 0.33112505 | -0.6923136 | 0.4887404 | 0.86014074 |
| RHOC | 252.177597 | -0.268651052 | 0.5069176 | -0.5299699 | 0.59613283 | 0.90346667 |
| LRIG2 | 179.508628 | 0.155794343 | 0.24416044 | 0.63808184 | 0.52342041 | 0.87725117 |
| MAGI3 | 1234.11247 | 0.492484277 | 0.33840243 | 1.45532135 | 0.14558032 | 0.57618473 |
| PHTF1 | 158.333284 | -0.060185081 | 0.32538427 | -0.1849662 | 0.85325557 | 0.97062426 |
| RSBN1 | 199.964561 | -0.20116017 | 0.18749075 | -1.0729072 | 0.28331277 | 0.73329656 |
| HIPK1 | 248.552295 | 0.264851074 | 0.20643272 | 1.28298981 | 0.19949565 | 0.65475264 |
| TRIM33 | 572.224699 | 0.088423407 | 0.26231708 | 0.33708596 | 0.7360521 | 0.94289524 |
| BCAS2 | 89.9198533 | -0.030021397 | 0.52033455 | -0.0576963 | 0.95399051 | 0.99060627 |
| NRAS | 69.7578055 | -0.427179274 | 0.39545294 | -1.0802278 | 0.28004073 | 0.72945471 |
| CSDE1 | 620.706592 | 0.157957961 | 0.21091424 | 0.74892031 | 0.45390524 | 0.84279368 |
| SIKE1 | 89.5671357 | 0.05063802 | 0.25811915 | 0.1961808 | 0.84446866 | 0.96876046 |
| VANGL1 | 35.545361 | -0.377234025 | 0.3764822 | -1.001997 | 0.31634505 | 0.76056347 |
| SLC22A15 | 244.908474 | -0.421936994 | 0.36973391 | -1.141191 | 0.25379047 | 0.71077608 |
| ATP1A1 | 2055.60026 | -0.119640226 | 0.32797584 | -0.3647837 | 0.71527291 | 0.93573028 |
| LINC01762 | 319.798884 | -0.874933877 | 0.99610787 | -0.8783525 | 0.37975243 | 0.80295536 |
| CD58 | 110.199924 | -0.814765296 | 0.23712354 | -3.4360371 | 0.00059029 | 0.02774786 |
| IGSF3 | 151.79031 | -0.655189625 | 0.41249334 | -1.5883641 | 0.11220402 | 0.5224487 |
| PTGFRN | 65.4090099 | -0.55498037 | 0.38208629 | -1.4525001 | 0.1463626 | 0.57722788 |
| TTF2 | 76.8123376 | -0.662440222 | 0.3003724 | -2.2053965 | 0.02742629 | 0.2806909 |
| VTCN1 | 290.794849 | -1.248839932 | 1.08951523 | -1.1462345 | 0.25169814 | 0.70813507 |
| MAN1A2 | 573.051448 | 0.170065541 | 0.31544606 | 0.53912716 | 0.58979911 | 0.90233654 |
| GDAP2 | 195.125213 | 0.245551919 | 0.23322444 | 1.05285674 | 0.29240666 | 0.73844969 |
| WDR3 | 53.7378582 | -0.007257305 | 0.27043005 | -0.0268362 | 0.97859041 | 0.99469867 |
| SPAG17 | 445.663514 | 0.356786783 | 0.24260793 | 1.47063117 | 0.14139089 | 0.56947599 |
| WARS2 | 159.761388 | 0.170396293 | 0.24183246 | 0.70460473 | 0.48105626 | 0.85600365 |
| AL359915.2 | 345.796083 | -0.006641883 | 0.49017033 | -0.0135502 | 0.98918887 | 0.99651056 |
| PHGDH | 53.7026597 | -0.907972615 | 0.71087576 | -1.2772592 | 0.20151075 | 0.65584864 |
| HMGCS2 | 612.409901 | 1.611812282 | 0.6580586 | 2.44934462 | 0.01431164 | 0.19353036 |
| NOTCH2 | 198.435468 | 0.095183883 | 0.21971796 | 0.43320938 | 0.66486266 | 0.92324595 |
| SEC22B | 204.376146 | 0.121004503 | 0.22790239 | 0.53094882 | 0.59545425 | 0.90323766 |
| NBPF26 | 42.7122275 | 0.419098483 | 0.44926573 | 0.93285211 | 0.35089633 | 0.78242416 |
| LINC00623 | 499.366113 | -0.127637415 | 0.38051754 | -0.3354311 | 0.73729992 | 0.9429136 |
| SRGAP2C | 113.161826 | 0.679742707 | 0.36260375 | 1.87461575 | 0.06084561 | 0.40676596 |
| AC239800.3 | 85.2034806 | -0.241673372 | 0.41474256 | -0.5827069 | 0.56009059 | 0.89233565 |
| NBPF15 | 73.2113098 | -0.556773386 | 0.33754879 | -1.6494605 | 0.09905334 | 0.4933116 |
| FP700111.1 | 66.2046797 | 0.134208502 | 0.37608374 | 0.35685803 | 0.72119809 | 0.93744589 |
| SRGAP2B | 123.000387 | 0.071312126 | 0.29346971 | 0.24299654 | 0.80800808 | 0.96245698 |
| GPR89A | 50.4331264 | 0.164712451 | 0.27282718 | 0.60372448 | 0.54602683 | 0.88669883 |
| PDZK1 | 157.620061 | 0.912879856 | 0.65019054 | 1.40401897 | 0.1603132 | 0.60010648 |
| RNF115 | 486.500655 | 0.207141491 | 0.33835286 | 0.61220552 | 0.54040179 | 0.88342811 |
| POLR3C | 70.4655552 | -0.328019404 | 0.28024432 | -1.1704766 | 0.24180924 | 0.69770754 |
| PIAS3 | 34.202358 | -0.55958692 | 0.32865891 | -1.7026373 | 0.08863597 | 0.47038308 |
| PEX11B | 59.8327465 | -0.443203997 | 0.28972226 | -1.5297547 | 0.12607745 | 0.54580422 |
| RBM8A | 139.896246 | -0.252621776 | 0.42440884 | -0.5952321 | 0.55168832 | 0.88923362 |
| POLR3GL | 38.5022252 | 0.086819208 | 0.42884459 | 0.20244912 | 0.83956563 | 0.96833625 |
| TXNIP | 2661.63171 | 0.09079185 | 0.6850529 | 0.13253261 | 0.89456303 | 0.9821637 |
| NBPF12 | 107.685347 | -0.168111854 | 0.26348394 | -0.6380345 | 0.5234512 | 0.87725117 |
| PRKAB2 | 97.3375914 | 0.240281833 | 0.43760032 | 0.54908971 | 0.58294388 | 0.90085422 |
| AC242426.2 | 209.913406 | -0.165160636 | 0.30375189 | -0.5437353 | 0.58662361 | 0.90196444 |
| FMO5 | 149.164883 | 0.13674415 | 0.43557107 | 0.31394223 | 0.75356492 | 0.94967914 |
| CHD1L | 146.603318 | 0.461192906 | 0.28134306 | 1.63925462 | 0.10116024 | 0.49726048 |
| BCL9 | 191.448064 | -0.184289697 | 0.39237324 | -0.4696796 | 0.63858396 | 0.91521016 |
| ACP6 | 189.821715 | 0.163956246 | 0.27786903 | 0.59004865 | 0.55515803 | 0.89081623 |
| GPR89B | 63.0748305 | 0.296885161 | 0.34754835 | 0.85422694 | 0.39297926 | 0.81010924 |
| NBPF11 | 101.927454 | -0.06575177 | 0.31538675 | -0.2084798 | 0.83485435 | 0.96751891 |
| AC239809.3 | 72.9340394 | -0.467978054 | 0.50572337 | -0.9253637 | 0.35477675 | 0.78589311 |
| NBPF14 | 49.4595219 | -0.026703632 | 0.3357166 | -0.0795422 | 0.93660138 | 0.98822345 |
| PDE4DIP | 432.267491 | 0.794054497 | 0.42904555 | 1.85074638 | 0.06420605 | 0.41357646 |
| NBPF9 | 45.7550977 | -0.072960451 | 0.26654623 | -0.2737253 | 0.78429572 | 0.95697044 |
| AC245297.3 | 567.09025 | -0.37649739 | 0.30236363 | -1.2451808 | 0.21306529 | 0.67054344 |
| NBPF19 | 73.685457 | 0.107960418 | 0.63481715 | 0.17006538 | 0.86495872 | 0.97485201 |
| HIST2H2BF | 65.2363784 | 3.125045865 | 0.84476417 | 3.69931155 | 0.00021619 | 0.01407217 |
| HIST2H2BE | 70.7998257 | 2.146477123 | 0.76278928 | 2.81398438 | 0.00489316 | 0.11130863 |
| SF3B4 | 57.2255286 | -0.31662676 | 0.42571719 | -0.7437491 | 0.45702831 | 0.84408051 |
| OTUD7B | 261.362741 | 0.050156849 | 0.26357806 | 0.1902922 | 0.84908017 | 0.96991131 |
| VPS45 | 288.328347 | 0.263297561 | 0.20736794 | 1.26971198 | 0.20418724 | 0.65923669 |
| ANP32E | 223.025386 | -0.217138017 | 0.2745418 | -0.7909106 | 0.42899618 | 0.82995753 |
| APH1A | 197.286495 | -0.241358974 | 0.30649745 | -0.7874747 | 0.43100405 | 0.83111714 |
| C1orf54 | 38.1873017 | -0.949291997 | 0.62866267 | -1.5100181 | 0.13103881 | 0.55395564 |
| MRPS21 | 282.838576 | -0.111649399 | 0.34717225 | -0.3215966 | 0.74775835 | 0.94666677 |
| PRPF3 | 155.886357 | 0.238229716 | 0.26661 | 0.89355133 | 0.371562 | 0.79539603 |
| RPRD2 | 885.573085 | 1.238574741 | 0.34157131 | 3.6261088 | 0.00028772 | 0.01703587 |
| TARS2 | 57.0182196 | -0.105972636 | 0.289349 | -0.366245 | 0.71418224 | 0.93573028 |
| MCL1 | 237.057571 | -0.477074938 | 0.33526328 | -1.4229859 | 0.15474023 | 0.59091117 |
| ENSA | 442.411354 | -0.535688455 | 0.21029832 | -2.5472788 | 0.01085667 | 0.16452817 |
| GOLPH3L | 178.844829 | -0.421326873 | 0.23224491 | -1.814149 | 0.06965479 | 0.42828237 |
| CTSS | 72.4613685 | -1.192108907 | 0.38703326 | -3.0801201 | 0.00206917 | 0.06434693 |
| ARNT | 249.552899 | 0.546709628 | 0.24716191 | 2.21194934 | 0.02697017 | 0.27874491 |
| SETDB1 | 147.827143 | -0.157279956 | 0.19156501 | -0.8210265 | 0.41163115 | 0.82355334 |
| CERS2 | 125.822965 | -0.230032008 | 0.33989266 | -0.6767784 | 0.49854656 | 0.86603149 |
| ANXA9 | 79.8732817 | 1.085022568 | 0.70229575 | 1.5449653 | 0.12235466 | 0.54052568 |
| MINDY1 | 91.236805 | 0.303713039 | 0.26509811 | 1.14566277 | 0.25193471 | 0.70813507 |
| PRUNE1 | 98.7293897 | 0.464362196 | 0.21638072 | 2.14604242 | 0.03186959 | 0.2995969 |
| BNIPL | 78.0746614 | 0.867088396 | 0.66287701 | 1.30806829 | 0.19085015 | 0.64458858 |
| C1orf56 | 61.2650168 | 0.230648315 | 0.4080868 | 0.56519425 | 0.57194165 | 0.89797476 |
| CDC42SE1 | 142.605824 | 0.276549829 | 0.25430875 | 1.087457 | 0.27683489 | 0.72743146 |
| GABPB2 | 211.297327 | 0.319465318 | 0.24836185 | 1.28628979 | 0.19834194 | 0.65386083 |
| SCNM1 | 45.8408588 | -0.556881781 | 0.42583152 | -1.3077514 | 0.19095763 | 0.64458858 |
| VPS72 | 53.6024799 | 0.014102866 | 0.27819053 | 0.05069499 | 0.95956857 | 0.99119981 |
| PIP5K1A | 593.929692 | 0.309549796 | 0.26459717 | 1.16989082 | 0.24204491 | 0.6978349 |
| PSMD4 | 156.534136 | 0.142167944 | 0.30670542 | 0.46353254 | 0.6429827 | 0.91643619 |
| ZNF687 | 34.8082028 | -0.180703725 | 0.32248385 | -0.5603497 | 0.57524094 | 0.89807676 |
| PI4KB | 162.264538 | 0.478184989 | 0.24654108 | 1.93957533 | 0.05243132 | 0.38418477 |
| SELENBP1 | 51.0686497 | 0.803449298 | 0.64464191 | 1.24634976 | 0.21263602 | 0.6699289 |
| PSMB4 | 120.663915 | 0.068151183 | 0.38500643 | 0.1770131 | 0.8594981 | 0.97244655 |
| POGZ | 918.219102 | 0.394049157 | 0.18688649 | 2.10849464 | 0.03498823 | 0.31514522 |
| CGN | 136.988392 | 0.372718167 | 0.29317825 | 1.27130225 | 0.20362114 | 0.6582233 |
| TUFT1 | 151.993985 | 0.245339374 | 0.45622902 | 0.53775486 | 0.5907463 | 0.90233654 |
| SNX27 | 256.100119 | 0.29358072 | 0.17954804 | 1.63510955 | 0.10202608 | 0.4989423 |

| | | | | | | |
|---|---|---|---|---|---|---|
| MRPL9 | 41.3367863 | -0.780000275 | 0.50267483 | -1.5516995 | 0.12073415 | 0.53842291 |
| TDRKH | 31.8455878 | -0.534723745 | 0.38991508 | -1.3713851 | 0.17025493 | 0.61613093 |
| RORC | 92.9607202 | -1.190218448 | 0.5108286 | -2.3299761 | 0.01980741 | 0.23615663 |
| THEM4 | 252.088539 | -0.266204483 | 0.57816032 | -0.4604337 | 0.64520496 | 0.91685018 |
| S100A10 | 1328.25283 | -0.31580125 | 0.57463088 | -0.5495724 | 0.58261272 | 0.90085422 |
| S100A11 | 3245.8857 | -1.091905372 | 0.58684731 | -1.8606294 | 0.06279653 | 0.41131726 |
| FLG-AS1 | 355.202572 | 1.270015323 | 0.39672533 | 3.20124584 | 0.00136835 | 0.04877646 |
| S100A6 | 4301.60372 | -0.859487148 | 0.72078001 | -1.1924403 | 0.23308864 | 0.69247746 |
| S100A4 | 193.239316 | 0.911224789 | 1.31711482 | 0.69183398 | 0.48904159 | 0.86014074 |
| S100A2 | 818.295799 | -0.469718565 | 0.90148637 | -0.521049 | 0.60233264 | 0.90369786 |
| S100A16 | 196.116123 | -1.416363005 | 0.62677476 | -2.259764 | 0.0238359 | 0.26048096 |
| S100A14 | 124.679602 | -0.083917816 | 0.70174539 | -0.1195844 | 0.90481236 | 0.98320957 |
| S100A13 | 359.530208 | 0.232744028 | 0.59252672 | 0.39279921 | 0.69446779 | 0.93105058 |
| S100A1 | 185.669373 | 0.481184322 | 0.98746391 | 0.48729307 | 0.62605066 | 0.91061758 |
| CHTOP | 147.378416 | 0.130157286 | 0.19656537 | 0.66215776 | 0.50787012 | 0.87094607 |
| SNAPIN | 36.8957756 | -0.256316165 | 0.33633253 | -0.7620915 | 0.44600539 | 0.83968362 |
| ILF2 | 133.750343 | -0.174622519 | 0.42297495 | -0.4128436 | 0.67972118 | 0.92700898 |
| INTS3 | 223.03248 | 0.109467669 | 0.20630237 | 0.5306176 | 0.5956838 | 0.90323766 |
| GATAD2B | 604.773002 | 0.166842974 | 0.26091225 | 0.6394601 | 0.52252366 | 0.87688817 |
| DENND4B | 81.6995273 | -0.263602694 | 0.23625532 | -1.1157535 | 0.26452767 | 0.71989632 |
| CRTC2 | 51.9977489 | 0.339987361 | 0.27421219 | 1.2398696 | 0.21502363 | 0.67132603 |
| SLC39A1 | 129.733837 | -0.934903768 | 0.35212664 | -2.6550214 | 0.00793034 | 0.14172148 |
| JTB | 158.407666 | 0.373208588 | 0.43138339 | 0.86514363 | 0.38695996 | 0.8065026 |
| RAB13 | 254.256411 | -0.439566688 | 0.48219771 | -0.9115902 | 0.36198451 | 0.79033284 |
| RPS27 | 225.980462 | 0.350127706 | 0.32504009 | 1.07718314 | 0.28139845 | 0.73086885 |
| NUP210L | 45.0837913 | -2.063921643 | 1.00952197 | -2.0444544 | 0.04090868 | 0.34079725 |
| TPM3 | 275.47415 | -0.355418388 | 0.35527865 | -1.0003933 | 0.3171202 | 0.76120469 |
| C1orf43 | 188.741126 | -0.382842311 | 0.19805934 | -1.9329677 | 0.05324018 | 0.38756321 |
| UBAP2L | 238.388835 | -0.039314625 | 0.31019715 | -0.1267408 | 0.89914558 | 0.98312332 |
| HAX1 | 65.6744717 | -0.021796261 | 0.47475799 | -0.0459103 | 0.96338178 | 0.99225956 |
| IL6R | 38.9614489 | -1.289276097 | 0.55730375 | -2.3134172 | 0.02069971 | 0.24280431 |
| UBE2Q1 | 85.6343903 | 0.100275794 | 0.36073898 | 0.27797327 | 0.78103288 | 0.9566945 |
| ADAR | 390.753053 | -0.189977878 | 0.32553499 | -0.5835867 | 0.55949844 | 0.89179371 |
| PMVK | 124.376449 | 0.327397891 | 0.45845271 | 0.71413666 | 0.47514268 | 0.85385826 |
| PBXIP1 | 127.621888 | -0.040746386 | 0.32890023 | -0.1238868 | 0.90140493 | 0.98320957 |
| PYGO2 | 47.7386737 | 0.080426051 | 0.30368637 | 0.2648326 | 0.79113842 | 0.95832571 |
| FLAD1 | 33.20632 | 0.020804354 | 0.30881906 | 0.06736745 | 0.94628918 | 0.98949015 |
| ZBTB7B | 54.0272713 | -0.355434674 | 0.34112017 | -1.0419632 | 0.29742873 | 0.74513589 |
| EFNA1 | 139.410547 | -1.021257956 | 0.48442009 | -2.1082073 | 0.03501306 | 0.31514522 |
| SLC50A1 | 56.6655161 | -0.552435235 | 0.41502657 | -1.331084 | 0.18316136 | 0.633903 |
| DPM3 | 137.863669 | -0.372479281 | 0.28043044 | -1.3282412 | 0.18409842 | 0.63486392 |
| MUC1 | 69.6195062 | -1.045462982 | 0.36681006 | -2.8501481 | 0.00436989 | 0.10314511 |
| THBS3 | 44.3707727 | -0.379314977 | 0.31808205 | -1.1925067 | 0.23306262 | 0.69247746 |
| MTX1 | 48.4057963 | -0.28657286 | 0.41778693 | -0.6859306 | 0.49275684 | 0.86213395 |
| GBA | 57.803082 | -0.624175014 | 0.39321224 | -1.5873743 | 0.11242789 | 0.52250301 |
| SCAMP3 | 95.5794928 | -0.261960544 | 0.24803015 | -1.0561641 | 0.29089324 | 0.73772562 |
| CLK2 | 66.9318458 | -0.341134736 | 0.31737892 | -1.07485 | 0.28244188 | 0.73267102 |
| FDPS | 237.105507 | -0.654629195 | 0.55563684 | -1.1781602 | 0.23873276 | 0.69499004 |
| RUSC1 | 39.8810695 | -0.444178617 | 0.28813891 | -1.5415433 | 0.12318461 | 0.54197241 |
| ASH1L | 1188.97471 | 0.052509291 | 0.1540066 | 0.34095482 | 0.7331376 | 0.94163693 |
| YY1AP1 | 220.720083 | 0.00649764 | 0.1958773 | 0.03317199 | 0.97353743 | 0.9939318 |
| DAP3 | 235.063691 | -0.172587165 | 0.19257369 | -0.8962136 | 0.37013869 | 0.79532503 |
| GON4L | 300.279487 | -0.0333507 | 0.22693324 | -0.1469626 | 0.88316154 | 0.97878487 |
| SYT11 | 38.8919344 | 0.117783635 | 0.57370697 | 0.20530278 | 0.83733557 | 0.96757829 |
| RIT1 | 121.324413 | -0.74775075 | 0.44191981 | -1.6920508 | 0.0906363 | 0.4761245 |
| KHDC4 | 276.380659 | 0.116445319 | 0.20671916 | 0.563302 | 0.57322926 | 0.89797476 |
| ARHGEF2 | 97.7337995 | -0.611484732 | 0.29818216 | -2.0507086 | 0.04029533 | 0.33814918 |
| SSR2 | 237.532269 | -0.493869372 | 0.39137144 | -1.2618943 | 0.20698684 | 0.66434881 |
| UBQLN4 | 77.7628717 | 0.636321787 | 0.28805291 | 2.20904479 | 0.02717153 | 0.27917411 |
| LAMTOR2 | 91.1586569 | -0.185457521 | 0.43672545 | -0.4246547 | 0.6710884 | 0.92474338 |
| RAB25 | 157.704009 | -0.341450857 | 0.40967554 | -0.8334665 | 0.40458166 | 0.81914727 |
| LMNA | 482.848967 | 0.239306 | 0.50445381 | 0.47438635 | 0.63522443 | 0.91454294 |
| SEMA4A | 94.9116122 | -0.208206665 | 0.42203147 | -0.4933439 | 0.62176959 | 0.91007616 |
| SLC25A44 | 52.4359847 | 0.530012205 | 0.25457923 | 2.08191455 | 0.03735028 | 0.32376418 |
| SMG5 | 154.745071 | -0.268374866 | 0.29118478 | -0.9216652 | 0.35670326 | 0.78612469 |
| GLMP | 62.6355806 | -0.431093168 | 0.38699314 | -1.1139556 | 0.26529824 | 0.72029159 |
| CCT3 | 242.371796 | -0.082751886 | 0.273749 | -0.3022911 | 0.76243017 | 0.95374766 |
| RHBG | 200.065166 | 0.798709229 | 0.62641653 | 1.27504494 | 0.20229334 | 0.65696752 |
| MEF2D | 126.787235 | 0.31365701 | 0.34949686 | 0.89745301 | 0.36947723 | 0.79532503 |
| NAXE | 88.359842 | 0.200098099 | 0.38650959 | 0.51770539 | 0.60466383 | 0.90416461 |
| GPATCH4 | 45.1380628 | 0.01229813 | 0.3480797 | 0.03533136 | 0.97181551 | 0.9939318 |
| CRABP2 | 47.1994763 | -1.862362015 | 0.72112142 | -2.5825914 | 0.00980614 | 0.15842224 |
| ISG20L2 | 63.6261365 | -0.329603915 | 0.37017359 | -0.8904036 | 0.37324919 | 0.79681088 |
| RRNAD1 | 64.8630836 | -0.147043423 | 0.29731884 | -0.4945648 | 0.62090738 | 0.90957637 |
| MRPL24 | 47.9327475 | 0.017012728 | 0.3206303 | 0.05306026 | 0.95768389 | 0.99106618 |
| HDGF | 262.780255 | -0.292734761 | 0.30996782 | -0.9444037 | 0.34496339 | 0.78180304 |
| PRCC | 153.119889 | 0.322396406 | 0.20526617 | 1.57062609 | 0.11626953 | 0.53070759 |
| ARHGEF11 | 301.370543 | 0.0273014 | 0.28656644 | 0.09527075 | 0.92409977 | 0.98697817 |
| ETV3 | 48.5224023 | 0.323406803 | 0.40635848 | 0.79586577 | 0.42611006 | 0.82774157 |
| KIRREL1 | 64.8954283 | -1.349391865 | 0.67395273 | -2.0022055 | 0.04526263 | 0.35665593 |
| IFI16 | 59.6801299 | -1.667504347 | 0.69528631 | -2.3982988 | 0.01647142 | 0.21098114 |
| DUSP23 | 131.274898 | -1.13616055 | 0.48664507 | -2.3346801 | 0.01956014 | 0.23481129 |
| TAGLN2 | 472.707934 | -0.990744207 | 0.64072912 | -1.5462762 | 0.12203787 | 0.54028588 |
| IGSF8 | 68.2009579 | 0.076724784 | 0.36740659 | 0.208828 | 0.83458251 | 0.96749268 |
| PEA15 | 52.2768746 | -0.986406387 | 0.40506267 | -2.4351945 | 0.01488379 | 0.19869417 |
| DCAF8 | 101.369344 | 0.428857823 | 0.2679966 | 1.60023603 | 0.10954623 | 0.51481816 |
| COPA | 470.903069 | 0.127098482 | 0.26840917 | 0.47352511 | 0.63583859 | 0.91454294 |
| NCSTN | 121.402268 | -0.151031096 | 0.29105303 | -0.5189126 | 0.60382166 | 0.90371579 |
| VANGL2 | 31.7008338 | -0.258698524 | 0.58836244 | -0.4396925 | 0.66015987 | 0.92284375 |
| F11R | 404.88785 | 0.3675475 | 0.31009297 | 1.18528164 | 0.23590611 | 0.69335782 |
| AL591806.3 | 74.9974784 | 0.452011692 | 0.42358473 | 1.06711046 | 0.28592196 | 0.73607046 |
| USF1 | 32.4455948 | 0.408521336 | 0.4180424 | 0.97722464 | 0.32845795 | 0.7745813 |
| NECTIN4 | 122.403778 | -0.763908306 | 0.59696526 | -1.2796529 | 0.20066725 | 0.65487239 |
| PFDN2 | 185.063312 | -0.155026387 | 0.53075841 | -0.2920847 | 0.7702219 | 0.95482817 |
| NIT1 | 121.414157 | -0.070227269 | 0.37563205 | -0.1869576 | 0.85169387 | 0.97053519 |
| DEDD | 44.3539013 | 0.10055587 | 0.31770226 | 0.31650977 | 0.75161561 | 0.94834236 |
| UFC1 | 166.953438 | -0.098616037 | 0.4560247 | -0.2162515 | 0.82879169 | 0.96606563 |
| PPOX | 32.0903013 | 0.289406306 | 0.38578979 | 0.7501658 | 0.45315485 | 0.84248055 |
| B4GALT3 | 52.5659315 | 0.278030219 | 0.40181999 | 0.69192729 | 0.48898299 | 0.86014074 |
| NDUFS2 | 162.576373 | 0.406825625 | 0.31073814 | 1.30922335 | 0.19045871 | 0.64458858 |
| SDHC | 314.595515 | -0.004892482 | 0.22264048 | -0.0219748 | 0.98246805 | 0.99507903 |
| AL592295.4 | 40.9625429 | -0.302302243 | 0.28242045 | -1.0703979 | 0.28444026 | 0.73476311 |
| DUSP12 | 57.5919096 | 0.628561536 | 0.45626435 | 1.37762578 | 0.16831886 | 0.6131323 |
| ATF6 | 855.749944 | 0.687984654 | 0.38696247 | 1.77791054 | 0.07541855 | 0.44057213 |
| NOS1AP | 1112.49666 | 0.493000248 | 0.57320643 | 0.86007453 | 0.38974796 | 0.80680464 |
| UHMK1 | 320.980805 | 0.160529298 | 0.22227314 | 0.72221637 | 0.47016146 | 0.85232523 |
| UAP1 | 90.7677711 | 1.632767467 | 0.68184418 | 2.39463429 | 0.01663695 | 0.21108198 |
| HSD17B7 | 70.3110138 | 0.092550113 | 0.36532086 | 0.25333925 | 0.80000607 | 0.96048386 |
| PBX1 | 610.494478 | 0.459450081 | 0.2686985 | 1.70990938 | 0.08728263 | 0.46735992 |
| MGST3 | 1091.14978 | -0.018691658 | 0.37625178 | -0.0496786 | 0.96037852 | 0.9913918 |
| ALDH9A1 | 188.303017 | -0.161649945 | 0.35376655 | -0.4569396 | 0.64771446 | 0.91763881 |
| TMCO1 | 266.263064 | -0.436139513 | 0.32876069 | -1.326617 | 0.18463542 | 0.63541313 |
| UCK2 | 119.792179 | 0.133733821 | 0.31511669 | 0.4243946 | 0.67127806 | 0.92474338 |

| | | | | | | |
|---|---|---|---|---|---|---|
| POGK | 129.073962 | 0.676309554 | 0.31280594 | 2.16207392 | 0.03061248 | 0.29432916 |
| TADA1 | 52.36667 | 0.004875897 | 0.33629078 | 0.01449905 | 0.98843184 | 0.9964791 |
| POU2F1 | 601.468531 | -0.079664608 | 0.25620151 | -0.3109451 | 0.75584235 | 0.95067145 |
| CREG1 | 122.219194 | -0.14513911 | 0.36409238 | -0.3986326 | 0.6901639 | 0.93073907 |
| MPZL1 | 199.990735 | -0.355762019 | 0.31087366 | -1.1443942 | 0.25246018 | 0.70813507 |
| MPC2 | 202.382291 | 0.419006947 | 0.36036837 | 1.16271844 | 0.24494376 | 0.70148313 |
| DCAF6 | 779.941669 | 0.188619457 | 0.27565748 | 0.684253 | 0.49381542 | 0.86339027 |
| TIPRL | 102.490113 | 0.029675843 | 0.24681819 | 0.12023361 | 0.90429809 | 0.98320957 |
| SFT2D2 | 190.259831 | -0.033605938 | 0.28263801 | -0.118901 | 0.9053538 | 0.98320957 |
| TBX19 | 34.9510003 | 0.130186838 | 0.31969393 | 0.40722337 | 0.68384394 | 0.92844727 |
| LINC00970 | 165.535836 | 1.97739806 | 0.62483396 | 3.16467765 | 0.00155255 | 0.05282697 |
| ATP1B1 | 4443.92114 | 1.049900804 | 0.44673331 | 2.35017356 | 0.01876466 | 0.22823596 |
| NME7 | 678.960494 | 0.328776009 | 0.19716494 | 1.6675176 | 0.09541153 | 0.48704959 |
| BLZF1 | 135.572738 | -0.258013137 | 0.21777764 | -1.184755 | 0.23611434 | 0.69335782 |
| CCDC181 | 107.84852 | 0.343445368 | 0.32215893 | 1.06607435 | 0.28639003 | 0.73607046 |
| SLC19A2 | 338.342772 | 0.410384573 | 0.69671846 | 0.58902497 | 0.55584452 | 0.89081623 |
| C1orf112 | 37.9159551 | -0.21393726 | 0.43178256 | -0.4954745 | 0.6202652 | 0.90914396 |
| SCYL3 | 127.519931 | -0.217622972 | 0.30762032 | -0.7074402 | 0.47929298 | 0.85532469 |
| KIFAP3 | 291.373103 | 0.161946226 | 0.15680183 | 1.03280829 | 0.30169363 | 0.748343 |
| GORAB | 48.6908346 | -0.084125818 | 0.32607843 | -0.2579926 | 0.79641263 | 0.95948555 |
| FMO4 | 87.4043552 | 0.405552017 | 0.35851915 | 1.13118649 | 0.25797661 | 0.71534188 |
| PRRC2C | 884.655718 | 0.030999589 | 0.19873235 | 0.15598663 | 0.87604356 | 0.97622753 |
| VAMP4 | 76.2215287 | 0.288122318 | 0.2707239 | 1.06426628 | 0.28720809 | 0.73638964 |
| METTL13 | 38.5948368 | -0.312111784 | 0.306922 | -1.0169091 | 0.30919666 | 0.75357697 |
| DNM3 | 132.472285 | -0.387514565 | 0.41305006 | -0.9381782 | 0.34815284 | 0.78180304 |
| PIGC | 77.9906777 | -0.322036345 | 0.31452285 | -1.0238885 | 0.30588792 | 0.75110249 |
| SUCO | 260.313964 | -0.012412772 | 0.24444021 | -0.0507804 | 0.95950051 | 0.99119981 |
| AL645568.1 | 84.8161877 | -0.053060695 | 0.45869296 | -0.115678 | 0.90790772 | 0.98375444 |
| PRDX6 | 434.182427 | -0.392176558 | 0.47473424 | -0.826097 | 0.40874904 | 0.82073894 |
| ANKRD45 | 43.3352507 | 0.421361576 | 0.38254103 | 1.10148073 | 0.27068749 | 0.72343657 |
| KLHL20 | 165.651394 | -0.275355332 | 0.29097498 | -0.9463196 | 0.34398559 | 0.78180304 |
| DARS2 | 47.7091827 | -0.199150112 | 0.34448109 | -0.5781162 | 0.56318565 | 0.8927792 |
| ZBTB37 | 143.531683 | -0.091393113 | 0.26136596 | -0.3496749 | 0.7265827 | 0.93906703 |
| RC3H1 | 379.720196 | 0.101142057 | 0.22198242 | 0.45563093 | 0.6486554 | 0.91768694 |
| RABGAP1L | 2249.92191 | 0.391151527 | 0.23226114 | 1.68410232 | 0.0921619 | 0.48040835 |
| CACYBP | 74.151909 | 0.085761248 | 0.36081386 | 0.23768834 | 0.81212283 | 0.96288868 |
| MRPS14 | 89.2683708 | 0.183891218 | 0.40758423 | 0.45117353 | 0.65186448 | 0.91887313 |
| KIAA0040 | 66.7386802 | -0.514774167 | 0.32877612 | -1.5657286 | 0.11741217 | 0.53282158 |
| COP1 | 893.267675 | 0.089816749 | 0.22806514 | 0.3938206 | 0.69371349 | 0.93105058 |
| RASAL2 | 602.105719 | 0.421329877 | 0.31274349 | 1.34720589 | 0.17791393 | 0.62729004 |
| RALGPS2 | 743.86808 | 0.238132388 | 0.25903759 | 0.91929664 | 0.35794044 | 0.78674827 |
| ANGPTL1 | 63.905346 | -0.815724034 | 0.44758549 | -1.8224988 | 0.06837933 | 0.42478683 |
| FAM20B | 139.106307 | 0.101888343 | 0.17444019 | 0.58408755 | 0.55916141 | 0.89179371 |
| TOR3A | 48.8038175 | -0.22766445 | 0.43591584 | -0.522267 | 0.60148446 | 0.90369786 |
| ABL2 | 374.928488 | 0.098424551 | 0.26212686 | 0.37548442 | 0.70730023 | 0.93274945 |
| SOAT1 | 200.420481 | -0.641118163 | 0.29188657 | -2.1964634 | 0.02805879 | 0.28466231 |
| TDRD5 | 57.7897681 | 0.153918613 | 0.66907297 | 0.23004757 | 0.8180548 | 0.96404074 |
| TOR1AIP2 | 395.855729 | 0.133166786 | 0.2293153 | 0.5807148 | 0.56143268 | 0.89242209 |
| TOR1AIP1 | 128.963623 | 0.045950088 | 0.2078784 | 0.22104311 | 0.82505887 | 0.96511681 |
| CEP350 | 561.555453 | 0.369819962 | 0.15957679 | 2.31750477 | 0.02047625 | 0.24101888 |
| QSOX1 | 202.656991 | -0.86241425 | 0.28435561 | -3.0328723 | 0.00242238 | 0.07177223 |
| ACBD6 | 523.690956 | 0.124130738 | 0.19566444 | 0.63440623 | 0.52581575 | 0.87791848 |
| XPR1 | 412.08286 | 0.322909531 | 0.23955001 | 1.34798381 | 0.17766359 | 0.6269762 |
| KIAA1614 | 85.5575486 | 0.177463175 | 0.33694678 | 0.52668013 | 0.59841574 | 0.90369786 |
| STX6 | 117.307589 | -0.120559463 | 0.30358002 | -0.3971258 | 0.69127468 | 0.93073907 |
| MR1 | 83.5171623 | -0.506132456 | 0.26635118 | -1.9002448 | 0.057401 | 0.39785877 |
| GLUL | 190.45199 | -0.31916676 | 0.31992945 | -0.9976161 | 0.31846556 | 0.76321041 |
| RNASEL | 35.2850391 | -0.350391902 | 0.33391887 | -1.0493324 | 0.29402514 | 0.73965037 |
| DHX9 | 403.785325 | 0.648901912 | 0.31135912 | 2.08409478 | 0.03715155 | 0.32359312 |
| LAMC1 | 848.105189 | 0.723200193 | 0.37359749 | 1.93577369 | 0.05289542 | 0.38630247 |
| LAMC2 | 157.731713 | -0.889776724 | 0.69441131 | -1.2813396 | 0.2000744 | 0.65475264 |
| SMG7 | 383.478089 | 0.537909279 | 0.21597566 | 2.49060136 | 0.01275271 | 0.18036223 |
| ARPC5 | 202.809329 | -0.396524485 | 0.33569993 | -1.1811873 | 0.23752833 | 0.6939708 |
| RGL1 | 124.414127 | -0.026365275 | 0.34336833 | -0.0767842 | 0.93879519 | 0.98848592 |
| TSEN15 | 112.502836 | 0.11977057 | 0.27793799 | 0.4309255 | 0.66652254 | 0.92396246 |
| EDEM3 | 255.757775 | 0.059194406 | 0.22065538 | 0.26826632 | 0.78849433 | 0.95801395 |
| RNF2 | 111.949135 | 0.005613801 | 0.28125272 | 0.01995999 | 0.98407529 | 0.99540861 |
| TRMT1L | 130.514047 | 0.170962503 | 0.32175737 | 0.53133981 | 0.59518332 | 0.90323766 |
| SWT1 | 159.393478 | -0.232202676 | 0.3057001 | -0.7595767 | 0.44750765 | 0.84026779 |
| IVNS1ABP | 1792.3199 | 0.42637848 | 0.71430988 | 0.59690968 | 0.55056768 | 0.88916026 |
| HMCN1 | 64.7775178 | 0.380896049 | 0.72273637 | 0.52701934 | 0.59818016 | 0.90369786 |
| TPR | 401.775455 | -0.21449156 | 0.18514768 | -1.1584891 | 0.24666451 | 0.70292817 |
| ODR4 | 210.466969 | -0.091945883 | 0.18812593 | -0.4887465 | 0.62502121 | 0.91058133 |
| BRINP3 | 334.657962 | -0.273963227 | 0.69978675 | -0.3914953 | 0.69543116 | 0.93126994 |
| AL390957.1 | 57.0902677 | -0.500184568 | 0.50742678 | -0.9857276 | 0.32426682 | 0.76970832 |
| AL136454.1 | 282.591721 | -0.038018457 | 0.51755527 | -0.0734578 | 0.94144185 | 0.98876195 |
| RGS2 | 194.725036 | 2.011392454 | 1.54255039 | 1.30393954 | 0.1922542 | 0.64529552 |
| UCHL5 | 137.395793 | 0.161981284 | 0.20868081 | 0.77621552 | 0.43762174 | 0.83420562 |
| TROVE2 | 273.648502 | -0.034389669 | 0.17131812 | -0.2007357 | 0.84090522 | 0.96854218 |
| GLRX2 | 33.7258035 | -0.049518244 | 0.43551844 | -0.1136995 | 0.90947598 | 0.98441374 |
| CDC73 | 487.118272 | 0.496128137 | 0.26700964 | 1.85809076 | 0.06315613 | 0.41264103 |
| ZBTB41 | 154.843088 | -0.000928897 | 0.24600373 | -0.0037759 | 0.99698724 | 0.99937123 |
| CRB1 | 74.6625975 | -1.118334186 | 0.79669521 | -1.4037165 | 0.1604033 | 0.60010648 |
| DENND1B | 731.840528 | 0.532423059 | 0.21593018 | 2.46571857 | 0.01367387 | 0.18905301 |
| NEK7 | 673.109842 | 0.161790011 | 0.19042792 | 0.84961285 | 0.39554037 | 0.81184157 |
| ZNF281 | 62.3197897 | -0.275319473 | 0.28578729 | -0.963372 | 0.33536087 | 0.77820994 |
| DDX59 | 61.2042264 | -0.071228261 | 0.23859756 | -0.2985289 | 0.76529955 | 0.95378284 |
| CAMSAP2 | 420.755846 | 0.099167303 | 0.19608916 | 0.50572558 | 0.61304931 | 0.90751853 |
| INAVA | 32.5165257 | -0.814470204 | 0.42244927 | -1.9279716 | 0.05385865 | 0.38953671 |
| TMEM9 | 123.303529 | -0.270980756 | 0.40442218 | -0.6700442 | 0.50282958 | 0.86900627 |
| LAD1 | 83.5476669 | -0.448312506 | 0.25439953 | -1.7622379 | 0.07802911 | 0.44731172 |
| PHLDA3 | 44.8931307 | -0.699169069 | 0.50067804 | -1.3964444 | 0.1625807 | 0.60211533 |
| CSRP1 | 148.550338 | 0.424054888 | 0.45929184 | 0.92327982 | 0.3558614 | 0.78612469 |
| AC096677.1 | 94.0129456 | 0.168266775 | 0.36296454 | 0.46359012 | 0.64294144 | 0.91643619 |
| NAV1 | 122.15627 | -1.309734456 | 0.29398089 | -4.4551687 | 8.38E-06 | 0.00162687 |
| IPO9-AS1 | 44.1458569 | -1.741587393 | 0.52757759 | -3.3011019 | 0.00096306 | 0.03852235 |
| IPO9 | 211.98666 | -0.308100342 | 0.2481513 | -1.2415826 | 0.21439059 | 0.67132603 |
| TIMM17A | 82.0449648 | 0.168448938 | 0.37500564 | 0.44919041 | 0.65329431 | 0.91973728 |
| RNPEP | 133.807287 | -0.063444129 | 0.22858361 | -0.2775533 | 0.7813553 | 0.95673616 |
| ELF3 | 1160.00513 | -0.238949849 | 0.52640767 | -0.4539255 | 0.64988247 | 0.91803345 |
| ARL8A | 107.216545 | 0.071232771 | 0.2578438 | 0.27626327 | 0.78234586 | 0.95688565 |
| PPP1R12B | 610.675119 | 0.133805808 | 0.22144805 | 0.60423115 | 0.54568997 | 0.88669883 |
| KDM5B | 633.077272 | -0.484695946 | 0.26225485 | -1.8481868 | 0.06457533 | 0.41412378 |
| KLHL12 | 86.5635563 | 0.08585511 | 0.23756528 | 0.36139587 | 0.71780353 | 0.93573028 |
| ADIPOR1 | 78.278407 | -0.020384189 | 0.36773996 | -0.055431 | 0.95579511 | 0.9908068 |
| CYB5R1 | 66.5657765 | -0.230445772 | 0.40177114 | -0.5735747 | 0.56625562 | 0.89413272 |
| TMEM183A | 45.5672774 | -0.034300429 | 0.28276846 | -0.1213022 | 0.9034517 | 0.98320957 |
| BTG2 | 1321.48329 | 1.454050742 | 0.54323371 | 2.67665778 | 0.00743605 | 0.13720044 |
| ATP2B4 | 170.258957 | -0.399427477 | 0.35105895 | -1.1377789 | 0.25521281 | 0.71266224 |
| SNRPE | 73.9447588 | -0.058404291 | 0.36144268 | -0.1615866 | 0.87163142 | 0.97622753 |
| SOX13 | 240.437183 | 0.471565953 | 0.57996268 | 0.81309706 | 0.41616241 | 0.82489761 |
| ETNK2 | 36.2734322 | -0.833760924 | 0.42485448 | -1.9624624 | 0.04970868 | 0.37609758 |
| GOLT1A | 47.4166024 | -0.423918287 | 0.3116878 | -1.3600734 | 0.1738067 | 0.62124632 |

| | | | | | | |
|---|---|---|---|---|---|---|
| PLEKHA6 | 573.066613 | -0.336497809 | 0.37373988 | -0.900353 | 0.36793244 | 0.79498834 |
| PPP1R15B | 111.239245 | 0.277590837 | 0.37790919 | 0.73454376 | 0.4626174 | 0.84818586 |
| PIK3C2B | 62.2556443 | 0.245259786 | 0.48420767 | 0.50651777 | 0.61249322 | 0.90714827 |
| MDM4 | 467.223078 | 0.329036123 | 0.20588707 | 1.59813884 | 0.11001208 | 0.51562013 |
| LRRN2 | 131.723419 | -0.582843762 | 0.67837022 | -0.8591824 | 0.39023989 | 0.80728662 |
| RBBP5 | 99.8323828 | 0.31214422 | 0.2121229 | 1.47152534 | 0.1411491 | 0.56895751 |
| DSTYK | 201.467276 | 0.058219265 | 0.32160707 | 0.18102607 | 0.85634711 | 0.97182813 |
| NUAK2 | 401.741455 | 1.233246053 | 0.45555934 | 2.70710297 | 0.00678732 | 0.1314808 |
| CDK18 | 428.945093 | -0.145307984 | 0.64238436 | -0.226201 | 0.82104509 | 0.96502145 |
| MFSD4A | 54.6940133 | 0.321553828 | 0.66098068 | 0.48647992 | 0.62662694 | 0.91061758 |
| ELK4 | 222.876884 | 0.608790812 | 0.52048392 | 1.16966306 | 0.24213659 | 0.6978349 |
| NUCKS1 | 1050.15491 | 0.25084122 | 0.18366637 | 1.3657439 | 0.17201936 | 0.61884138 |
| RAB29 | 100.72905 | -0.597496101 | 0.31882252 | -1.8740712 | 0.06092062 | 0.40676596 |
| SLC41A1 | 52.4965196 | -0.495142352 | 0.33955044 | -1.4582291 | 0.14477739 | 0.57450476 |
| RHEX | 561.005933 | -0.39421097 | 0.40497852 | -0.973412 | 0.33034856 | 0.77618491 |
| SRGAP2 | 196.129585 | 0.044294499 | 0.30170079 | 0.14681598 | 0.88327727 | 0.97878487 |
| EIF2D | 66.7557175 | 0.045065874 | 0.22001127 | 0.20483438 | 0.83770152 | 0.96757829 |
| DYRK3 | 47.3837411 | -0.125953032 | 0.31423219 | -0.4008279 | 0.68854683 | 0.93036007 |
| MAPKAPK2 | 341.781193 | 0.527310367 | 0.33557335 | 1.5713714 | 0.1160964 | 0.53066791 |
| PIGR | 791.063809 | -1.590375177 | 0.71948891 | -2.2104235 | 0.02707579 | 0.27917411 |
| C1orf116 | 38.8347453 | -0.154472379 | 0.44046596 | -0.3507022 | 0.72581178 | 0.93895336 |
| PFKFB2 | 164.479742 | -0.708100717 | 0.59962717 | -1.1809017 | 0.2376418 | 0.6939708 |
| YOD1 | 61.4569273 | 0.109508983 | 0.78198869 | 0.14003909 | 0.88862911 | 0.98051184 |
| CD55 | 299.655594 | 0.082775177 | 0.7931065 | 0.1043683 | 0.91687708 | 0.98520052 |
| CD46 | 689.28181 | 0.03848698 | 0.20047593 | 0.19197806 | 0.84775939 | 0.96991131 |
| MIR29B2CHC | 558.911946 | -0.233031082 | 0.49717499 | -0.4687104 | 0.63927666 | 0.91521016 |
| IRF6 | 98.7085751 | 0.275378189 | 0.48963082 | 0.56242005 | 0.57382986 | 0.89797476 |
| UTP25 | 40.6450665 | -0.527152967 | 0.27762161 | -1.8988182 | 0.05758838 | 0.39817926 |
| HHAT | 170.19601 | -0.163327969 | 0.33331707 | -0.4900078 | 0.62412837 | 0.91058133 |
| RCOR3 | 504.929892 | 0.280061031 | 0.24196201 | 1.15745869 | 0.24708501 | 0.70327292 |
| TRAF5 | 159.159388 | -0.421654254 | 0.36077873 | -1.1687337 | 0.24251095 | 0.6978349 |
| LINC00467 | 92.8903478 | -0.705186002 | 0.38456153 | -1.8337404 | 0.06669254 | 0.42003475 |
| SLC30A1 | 59.6670747 | -0.876183284 | 0.39019191 | -2.2455188 | 0.02473484 | 0.26640367 |
| LPGAT1 | 372.865827 | 0.062845803 | 0.21434564 | 0.29319842 | 0.76937049 | 0.95482817 |
| INTS7 | 85.1339249 | 0.50912096 | 0.46702857 | 1.09012807 | 0.27565673 | 0.7263609 |
| PPP2R5A | 393.945373 | 0.17856078 | 0.25064173 | 0.7124144 | 0.4762082 | 0.85414749 |
| AL360091.3 | 58.3034416 | -0.593790169 | 0.66691043 | -0.8903597 | 0.37327278 | 0.79681088 |
| TMEM206 | 48.8832556 | 0.864383447 | 0.47853602 | 1.80630801 | 0.07087024 | 0.43078597 |
| NENF | 210.914574 | 0.148835065 | 0.5090672 | 0.29236821 | 0.77000511 | 0.95482817 |
| ATF3 | 394.581128 | 2.996383272 | 1.40422671 | 2.13383155 | 0.03285657 | 0.30588131 |
| NSL1 | 154.064064 | -0.237529978 | 0.21188279 | -1.1210442 | 0.26226905 | 0.71989632 |
| TATDN3 | 118.893902 | -0.566616448 | 0.36893645 | -1.5358104 | 0.12458488 | 0.54288963 |
| FLVCR1 | 117.404913 | -0.367659524 | 0.31959401 | -1.1503956 | 0.24998099 | 0.70674386 |
| ANGEL2 | 127.74713 | 0.180724348 | 0.19815818 | 0.91202063 | 0.36175786 | 0.79000258 |
| RPS6KC1 | 345.273261 | -0.029688369 | 0.25323017 | -0.1172387 | 0.90667092 | 0.98321965 |
| SMYD2 | 104.021163 | -0.777303738 | 0.26575334 | -2.9249067 | 0.00344559 | 0.08938078 |
| PTPN14 | 802.214698 | 0.382682419 | 0.29518272 | 1.29642553 | 0.19482892 | 0.64860454 |
| KCTD3 | 232.168673 | 0.450590649 | 0.18153404 | 2.48212764 | 0.01306005 | 0.18379304 |
| ESRRG | 4685.15676 | 0.394141703 | 0.32727427 | 1.2043162 | 0.22846739 | 0.68678297 |
| GPATCH2 | 558.422677 | 0.105683224 | 0.25499699 | 0.41444891 | 0.67854538 | 0.9268955 |
| SPATA17 | 150.326628 | -0.018383091 | 0.30250387 | -0.0607698 | 0.95154256 | 0.99030211 |
| RRP15 | 76.7199502 | -0.138822785 | 0.24765469 | -0.5605498 | 0.57510449 | 0.89807676 |
| LYPLAL1 | 492.767449 | -0.132554098 | 0.29077884 | -0.4558588 | 0.6484915 | 0.91768694 |
| EPRS | 222.561454 | -0.250997648 | 0.17612514 | -1.4251097 | 0.15412549 | 0.5897171 |
| BPNT1 | 66.4151381 | -0.394427917 | 0.29847953 | -1.3214572 | 0.18634897 | 0.63723614 |
| IARS2 | 216.283733 | -0.44903931 | 0.1890413 | -2.3753502 | 0.01753231 | 0.21933738 |
| RAB3GAP2 | 561.035997 | 0.331146002 | 0.22702191 | 1.45865215 | 0.14466087 | 0.57447743 |
| MARK1 | 302.956524 | -0.013380174 | 0.33597478 | -0.0398249 | 0.9682327 | 0.99303855 |
| C1orf115 | 51.2018643 | 0.147641667 | 0.41936688 | 0.35205848 | 0.72479441 | 0.93895336 |
| 2-Mar | 197.155226 | 0.05998767 | 0.29066864 | 0.20637819 | 0.8364955 | 0.96757829 |
| 1-Mar | 61.1186637 | 0.361080832 | 0.52138082 | 0.6925472 | 0.48859375 | 0.86014074 |
| AL513314.2 | 45.5209922 | 0.247607566 | 0.35133117 | 0.70476971 | 0.48095356 | 0.85600365 |
| MIA3 | 145.510025 | -0.136307009 | 0.2411229 | -0.565301 | 0.57186907 | 0.89797476 |
| AIDA | 79.568725 | 0.218348943 | 0.31555283 | 0.69195686 | 0.48896442 | 0.86014074 |
| BROX | 187.601335 | -0.100058884 | 0.23580149 | -0.4243352 | 0.67132134 | 0.92474338 |
| FAM177B | 86.1211772 | -1.492528163 | 0.76270804 | -1.9568801 | 0.05036157 | 0.37839277 |
| AC093152.1 | 125.048443 | 0.420140198 | 0.29268014 | 1.43549266 | 0.15114676 | 0.58535779 |
| TLR5 | 55.5003259 | 0.119338218 | 0.49266008 | 0.24223237 | 0.80860011 | 0.96245698 |
| SUSD4 | 197.316278 | 0.713305189 | 0.44544771 | 1.60132195 | 0.10930563 | 0.51461693 |
| CAPN2 | 292.743663 | -0.408729945 | 0.18884175 | -2.1644046 | 0.03043331 | 0.29341408 |
| TP53BP2 | 150.541406 | -0.281862981 | 0.31617084 | -0.8914895 | 0.37266663 | 0.7966304 |
| FBXO28 | 199.200685 | -0.259763795 | 0.19028728 | -1.3651138 | 0.17221728 | 0.61884138 |
| NVL | 200.934739 | 0.360871437 | 0.25033061 | 1.44157933 | 0.14942108 | 0.58256434 |
| CNIH4 | 167.264923 | -0.226082602 | 0.2527677 | -0.8944284 | 0.37109274 | 0.79539603 |
| WDR26 | 454.330915 | 0.23748079 | 0.19862029 | 1.1956522 | 0.23183231 | 0.69186326 |
| CNIH3 | 154.381813 | 0.103630942 | 0.37339008 | 0.2775407 | 0.78136496 | 0.95673616 |
| DNAH14 | 711.985737 | -0.380290771 | 0.29582784 | -1.2855138 | 0.1986128 | 0.65403502 |
| LBR | 169.919962 | 0.40485585 | 0.19889992 | 2.0354752 | 0.04180308 | 0.34565724 |
| ENAH | 277.18518 | 0.601109219 | 0.19238855 | 3.12445425 | 0.00178135 | 0.05833929 |
| SRP9 | 284.377017 | -0.06065436 | 0.3385812 | -0.1791427 | 0.85782563 | 0.97220857 |
| EPHX1 | 77.6988403 | -0.840767634 | 0.3083874 | -2.7263359 | 0.00640418 | 0.12805535 |
| AL591895.1 | 86.2907283 | -0.245128588 | 0.24199373 | -1.0129543 | 0.311082 | 0.75435382 |
| TMEM63A | 183.735207 | -0.395633509 | 0.24373793 | -1.6231922 | 0.10454833 | 0.50362646 |
| SDE2 | 39.2140526 | -0.021458077 | 0.3379596 | -0.063493 | 0.94937391 | 0.98977092 |
| H3F3A | 63.5071434 | 0.080418616 | 0.31990128 | 0.25138573 | 0.8015159 | 0.96069141 |
| ACBD3 | 239.454657 | -0.375492779 | 0.16372977 | -2.293369 | 0.02182677 | 0.24999406 |
| PARP1 | 119.963401 | -0.175104016 | 0.34248074 | -0.5112813 | 0.60915406 | 0.90552475 |
| ITPKB | 185.812836 | 0.144327518 | 0.39783275 | 0.3627844 | 0.71676594 | 0.93573028 |
| COQ8A | 82.4677882 | -0.589508819 | 0.37028489 | -1.5920412 | 0.11137544 | 0.52038102 |
| CDC42BPA | 1411.4819 | 0.168011748 | 0.2047329 | 0.82063874 | 0.41185207 | 0.82355334 |
| ZNF678 | 167.453912 | -0.138161604 | 0.26539589 | -0.5205868 | 0.60265463 | 0.90369786 |
| SNAP47 | 70.4914553 | -0.093666391 | 0.26336605 | -0.355651 | 0.72210197 | 0.93798074 |
| ARF1 | 385.657952 | -0.286886245 | 0.23811516 | -1.2048214 | 0.22827226 | 0.68654401 |
| C1orf35 | 50.5621818 | 0.128416824 | 0.30983926 | 0.41446273 | 0.67853526 | 0.9268955 |
| MRPL55 | 72.5436738 | 0.226771914 | 0.38114706 | 0.59497222 | 0.55186204 | 0.88923362 |
| GUK1 | 447.230002 | 0.332420891 | 0.55983147 | 0.59378744 | 0.55265429 | 0.88968003 |
| OBSCN | 210.848264 | -0.247543631 | 0.28759948 | -0.8607235 | 0.38939034 | 0.80680464 |
| RNF187 | 117.760415 | -0.294724184 | 0.33523023 | -0.8791695 | 0.3793094 | 0.80274043 |
| RAB4A | 125.947613 | 0.000211153 | 0.23895367 | 0.00088365 | 0.99929495 | 0.99992293 |
| NUP133 | 165.651686 | -0.01088364 | 0.21708545 | -0.0501353 | 0.96001458 | 0.99121324 |
| ABCB10 | 78.3367057 | 0.412059691 | 0.27606755 | 1.49260459 | 0.13554072 | 0.55916513 |
| TAF5L | 41.2854042 | -0.045915188 | 0.2525385 | -0.1818146 | 0.85572822 | 0.9717401 |
| GALNT2 | 259.612447 | -0.451058605 | 0.39602868 | -1.1389544 | 0.25472217 | 0.71221409 |
| COG2 | 93.2829256 | 0.127012416 | 0.22951269 | 0.5534004 | 0.57998927 | 0.89995373 |
| C1orf198 | 56.8823011 | -0.176078124 | 0.26432799 | -0.666135 | 0.50532482 | 0.87044775 |
| TTC13 | 123.925533 | 0.773413881 | 0.2670452 | 2.89619091 | 0.00377723 | 0.09515702 |
| ARV1 | 70.3488804 | 0.157020816 | 0.25799067 | 0.60862983 | 0.54276983 | 0.88450129 |
| C1orf131 | 51.1111752 | -0.267781199 | 0.36332435 | -0.7370307 | 0.46110368 | 0.84722486 |
| GNPAT | 148.330109 | 0.158882022 | 0.21233075 | 0.74827608 | 0.45429365 | 0.84280358 |
| EGLN1 | 416.727644 | -0.849532232 | 0.42949419 | -1.9779831 | 0.04793062 | 0.36948395 |
| DISC1 | 51.3692416 | -1.392127502 | 0.61029296 | -2.2810807 | 0.02254367 | 0.25166501 |
| SIPA1L2 | 120.213234 | -0.161481728 | 0.54591963 | -0.2957976 | 0.76738464 | 0.95408212 |
| NTPCR | 104.424939 | -0.162317358 | 0.25416813 | -0.638622 | 0.52306887 | 0.87717892 |

| | | | | | | |
|---|---|---|---|---|---|---|
| PCNX2 | 549.392048 | -0.23692963 | 0.20400512 | -1.1613906 | 0.24548309 | 0.70154448 |
| MAP3K21 | 1286.20522 | 0.460327765 | 0.65681672 | 0.7008466 | 0.48339875 | 0.85829223 |
| KCNK1 | 166.646431 | -0.534154485 | 0.5204062 | -1.0264184 | 0.30469443 | 0.74998115 |
| COA6 | 73.2049892 | -0.077919982 | 0.43080742 | -0.1808696 | 0.8564699 | 0.97182813 |
| TARBP1 | 193.228017 | -0.188539183 | 0.29147823 | -0.646838 | 0.51773682 | 0.87526312 |
| IRF2BP2 | 258.697261 | 0.125487672 | 0.4960953 | 0.25295073 | 0.80030628 | 0.96063297 |
| TOMM20 | 310.137666 | -0.064245292 | 0.35985063 | -0.1785332 | 0.85830423 | 0.97237339 |
| ARID4B | 1066.43615 | 0.367357356 | 0.24778819 | 1.48254587 | 0.13819511 | 0.56438077 |
| B3GALNT2 | 134.475882 | 0.06488702 | 0.26318514 | 0.24654515 | 0.80526026 | 0.96218371 |
| LYST | 310.827602 | -0.129778865 | 0.27252732 | -0.476205 | 0.63392836 | 0.91415622 |
| GPR137B | 66.960005 | -0.237227662 | 0.47300174 | -0.5015366 | 0.61599356 | 0.90794578 |
| ERO1B | 89.4106164 | 0.138597896 | 0.47006259 | 0.29484988 | 0.76810856 | 0.95411028 |
| EDARADD | 32.0990232 | -1.1538056 | 0.45210084 | -2.5520979 | 0.01070764 | 0.16452817 |
| LGALS8 | 93.3263875 | -0.614397093 | 0.31036834 | -1.979574 | 0.04775142 | 0.36905226 |
| HEATR1 | 142.579237 | 0.40488979 | 0.33278626 | 1.21666619 | 0.22373125 | 0.68103746 |
| MTR | 1862.55103 | 0.736815651 | 0.35364684 | 2.08347871 | 0.03720761 | 0.32359312 |
| CHRM3 | 225.076653 | -0.900107651 | 0.73908127 | -1.2178737 | 0.22327197 | 0.6809925 |
| FH | 60.2117379 | 0.405328134 | 0.35416813 | 1.14445117 | 0.25243658 | 0.70813507 |
| OPN3 | 30.2697442 | -1.11794115 | 0.39730967 | -2.8137779 | 0.00489631 | 0.11130863 |
| CEP170 | 466.5905 | 0.160570298 | 0.63662269 | 0.25222208 | 0.80086941 | 0.96063297 |
| SDCCAG8 | 1179.93166 | 0.052029956 | 0.35885534 | 0.14498866 | 0.88471982 | 0.97891513 |
| AKT3 | 2882.06145 | 0.593993235 | 0.21187083 | 2.80356314 | 0.00505413 | 0.11390821 |
| ZBTB18 | 66.0607946 | 0.151270619 | 0.31993492 | 0.47281685 | 0.63634386 | 0.91454294 |
| AL645465.1 | 40.2001952 | -1.12445804 | 0.59286628 | -1.896647 | 0.05787455 | 0.3985947 |
| ADSS | 241.290104 | -0.005301495 | 0.2740111 | -0.0193477 | 0.9845637 | 0.9954565 |
| CATSPERE | 180.753478 | -0.765747814 | 0.36311258 | -2.1088441 | 0.03495804 | 0.31514522 |
| DESI2 | 173.107999 | -0.111554226 | 0.23069228 | -0.4835629 | 0.62869612 | 0.9113467 |
| COX20 | 316.804384 | 0.412641442 | 0.37472967 | 1.101171 | 0.27082224 | 0.72343657 |
| HNRNPU | 1158.38699 | 0.378662729 | 0.24198025 | 1.56484975 | 0.11761814 | 0.53282158 |
| EFCAB2 | 312.536954 | 0.340002872 | 0.38418665 | 0.88499397 | 0.37615989 | 0.79946372 |
| KIF26B | 513.04736 | -0.309927086 | 0.6253459 | -0.495609 | 0.62017027 | 0.90914396 |
| SMYD3 | 2044.03155 | 0.144224654 | 0.20268417 | 0.71157336 | 0.47672901 | 0.85414749 |
| TFB2M | 43.0214804 | -0.005639025 | 0.36271171 | -0.0155469 | 0.98759591 | 0.9964791 |
| CNST | 278.505986 | 0.36726515 | 0.19492881 | 1.8840989 | 0.0595516 | 0.40313286 |
| SCCPDH | 670.846096 | 0.705760709 | 0.54294218 | 1.29988189 | 0.19364145 | 0.64723766 |
| AHCTF1 | 288.606022 | 0.526284255 | 0.16014081 | 3.28638427 | 0.00101482 | 0.03983282 |
| ZNF124 | 169.932374 | -0.279724856 | 0.33647013 | -0.8313512 | 0.40577524 | 0.82002585 |
| AL390728.6 | 60.6287968 | 0.041638463 | 0.42035223 | 0.09905612 | 0.92109371 | 0.98581108 |
| ZNF496 | 65.5755634 | -0.02140574 | 0.34003679 | -0.0629512 | 0.94980533 | 0.98977092 |
| ZNF672 | 55.2640072 | -0.072580852 | 0.32168686 | -0.2256258 | 0.82149248 | 0.96502145 |
| ZNF692 | 48.6661866 | 0.643250356 | 0.2978292 | 2.15979613 | 0.03078846 | 0.29520861 |
| FAM110C | 133.121822 | -1.408360201 | 0.65425249 | -2.1526249 | 0.03134817 | 0.29704233 |
| SH3YL1 | 346.593695 | -0.053577813 | 0.22117779 | -0.2422387 | 0.80859524 | 0.96245698 |
| ACP1 | 129.653585 | -0.166160536 | 0.19420816 | -0.8555796 | 0.39223038 | 0.8093275 |
| TMEM18 | 38.8683924 | -0.023639564 | 0.42287435 | -0.0559021 | 0.9554198 | 0.9908068 |
| EIPR1 | 343.487914 | -0.238712489 | 0.33284003 | -0.7171988 | 0.47325141 | 0.85298656 |
| TRAPPC12 | 363.036849 | -0.320462422 | 0.25943873 | -1.2352143 | 0.21675077 | 0.67424993 |
| ADI1 | 219.393011 | -0.255015901 | 0.48868226 | -0.521844 | 0.60177896 | 0.90369786 |
| AC108488.2 | 33.7045294 | -0.072599972 | 0.35523934 | -0.2043692 | 0.83806501 | 0.96760178 |
| RPS7 | 560.381465 | 0.132994175 | 0.5041608 | 0.26379317 | 0.7919393 | 0.95839852 |
| RNF144A | 150.901845 | 0.197836618 | 0.38513302 | 0.51368387 | 0.60747303 | 0.90510401 |
| ID2 | 408.763862 | 0.661029888 | 0.67798438 | 0.9749928 | 0.32956383 | 0.77596696 |
| KIDINS220 | 415.923807 | 0.096282305 | 0.22039863 | 0.43685527 | 0.66221631 | 0.92284375 |
| MBOAT2 | 702.519878 | 0.246553833 | 0.37906026 | 0.6504344 | 0.51541166 | 0.87374866 |
| ASAP2 | 1844.45995 | 0.751770541 | 0.27669296 | 2.71698473 | 0.00658796 | 0.12933064 |
| ITGB1BP1 | 195.502782 | 0.234495382 | 0.39321008 | 0.59636158 | 0.5509337 | 0.88916026 |
| CPSF3 | 71.9430117 | -0.043175093 | 0.34915757 | -0.123655 | 0.90158841 | 0.98320957 |
| IAH1 | 149.749581 | 0.008108653 | 0.30296102 | 0.02676467 | 0.97864743 | 0.99469867 |
| ADAM17 | 352.974644 | 0.085453053 | 0.26274009 | 0.32523797 | 0.74500097 | 0.94591836 |
| YWHAQ | 331.789698 | 0.068417273 | 0.25274606 | 0.27069571 | 0.78662508 | 0.9576974 |
| TAF1B | 331.749766 | -0.086781349 | 0.29317541 | -0.2960049 | 0.76722636 | 0.95408212 |
| GRHL1 | 87.3929267 | 0.219633231 | 0.3818161 | 0.57523302 | 0.56513371 | 0.89382261 |
| KLF11 | 66.4234491 | 0.563045119 | 0.3160044 | 1.78176355 | 0.0747878 | 0.43878029 |
| CYS1 | 812.255191 | 0.413151883 | 0.40981062 | 1.0081532 | 0.31338092 | 0.75717828 |
| HPCAL1 | 296.922102 | 0.489134906 | 0.48879606 | 1.00069323 | 0.31697514 | 0.76103142 |
| ODC1 | 178.501633 | 1.101086645 | 0.51109122 | 2.15438382 | 0.03121008 | 0.29698524 |
| NOL10 | 254.920506 | 0.209906867 | 0.21216169 | 0.98937214 | 0.3224811 | 0.76844064 |
| PDIA6 | 347.153349 | -0.222904374 | 0.3339979 | -0.6673826 | 0.5045278 | 0.86946722 |
| PQLC3 | 109.500371 | 0.720621094 | 0.46893754 | 1.53671018 | 0.1243643 | 0.54275732 |
| ROCK2 | 907.415106 | 0.935331367 | 0.16649295 | 5.61784355 | 1.93E-08 | 1.69E-05 |
| E2F6 | 41.8185768 | -0.3359095 | 0.30035858 | -1.1183616 | 0.26341259 | 0.71989632 |
| LPIN1 | 181.419379 | -1.020801085 | 0.48408721 | -2.1087132 | 0.03496935 | 0.31514522 |
| MIR3681HG | 119.977215 | -1.078298623 | 0.54178172 | -1.9902824 | 0.04655984 | 0.36244837 |
| TRIB2 | 163.631163 | 0.210314655 | 0.45737885 | 0.45982593 | 0.64564117 | 0.91696971 |
| FAM84A | 220.260482 | -1.960702797 | 0.52527817 | -3.7326942 | 0.00018944 | 0.01314806 |
| NBAS | 703.367171 | 0.037423577 | 0.25713491 | 0.14554063 | 0.88428404 | 0.9788678 |
| DDX1 | 205.144906 | 0.181724129 | 0.16746238 | 1.08516393 | 0.27784906 | 0.72814657 |
| SMC6 | 202.154444 | 0.443400795 | 0.30386002 | 1.45922714 | 0.1445026 | 0.57406643 |
| GEN1 | 65.0476542 | 0.447886478 | 0.47469502 | 0.94352471 | 0.34541258 | 0.78180304 |
| KCNS3 | 61.9815187 | 0.562225562 | 0.50076461 | 1.12273421 | 0.26155039 | 0.7198106 |
| RDH14 | 59.5334146 | -0.268964488 | 0.3700145 | -0.7269026 | 0.46728565 | 0.85120743 |
| LINC01376 | 193.322838 | -0.458990725 | 0.3642395 | -1.2601344 | 0.20762088 | 0.66560625 |
| TTC32 | 52.0934663 | 0.013632066 | 0.31417844 | 0.04338957 | 0.965391 | 0.99267049 |
| WDR35 | 221.214424 | 0.488893674 | 0.19496124 | 2.50764551 | 0.01215385 | 0.17590979 |
| LAPTM4A | 565.043441 | -0.302680747 | 0.37814179 | -0.8004425 | 0.42345448 | 0.82774157 |
| SDC1 | 86.4506542 | -0.602339234 | 0.54599741 | -1.1031906 | 0.26994439 | 0.72333985 |
| PUM2 | 922.247787 | 0.257512837 | 0.18392671 | 1.40008394 | 0.16148818 | 0.60099295 |
| RHOB | 198.024956 | -0.399859582 | 0.69803836 | -0.5728333 | 0.5667576 | 0.89454608 |
| HS1BP3 | 55.6431537 | -0.270578571 | 0.24530273 | -1.1030394 | 0.27001007 | 0.72333985 |
| GDF7 | 50.657202 | 0.254299948 | 0.56940877 | 0.4466035 | 0.65516137 | 0.92051095 |
| LDAH | 287.239887 | -0.110793045 | 0.31836918 | -0.3480018 | 0.72783884 | 0.93931348 |
| AC018742.1 | 203.226907 | 0.329137523 | 0.7683035 | 0.42839519 | 0.66836343 | 0.92422713 |
| KLHL29 | 64.1653855 | -2.033443038 | 0.54365881 | -3.7402926 | 0.00018381 | 0.01293366 |
| ATAD2B | 520.743538 | 0.508204482 | 0.26440144 | 1.92209423 | 0.0545939 | 0.39095834 |
| UBXN2A | 202.261665 | -0.240673774 | 0.26084009 | -0.922687 | 0.35617032 | 0.78612469 |
| SF3B6 | 196.997961 | 0.034158773 | 0.52975087 | 0.06448082 | 0.94858737 | 0.98977092 |
| FAM228B | 57.0890769 | -0.531894561 | 0.4702798 | -1.1310172 | 0.25804784 | 0.71534188 |
| TP53I3 | 111.330729 | -0.183514006 | 0.40822121 | -0.4495455 | 0.6530382 | 0.9196238 |
| ITSN2 | 587.044892 | -0.028954032 | 0.14808983 | -0.1955167 | 0.84498849 | 0.96881318 |
| NCOA1 | 1185.22248 | 0.100210715 | 0.23838586 | 0.42037189 | 0.6742138 | 0.92595152 |
| PTRHD1 | 38.903839 | -0.423175072 | 0.54758621 | -0.7728008 | 0.43964027 | 0.83634598 |
| DNAJC27 | 94.9726916 | -0.070979476 | 0.24691643 | -0.2874636 | 0.7737574 | 0.95568386 |
| DNAJC27-AS | 59.3268205 | 0.176063245 | 0.4737278 | 0.37165487 | 0.71014983 | 0.93471656 |
| DNMT3A | 300.569476 | 0.088409614 | 0.29700707 | 0.29766838 | 0.76595628 | 0.95389523 |
| DTNB | 759.464198 | 0.427945427 | 0.32860142 | 1.30232374 | 0.19280574 | 0.64567765 |
| ASXL2 | 418.872992 | 0.046417219 | 0.23822053 | 0.19484979 | 0.84551054 | 0.96884569 |
| KIF3C | 34.6863794 | -0.824637727 | 0.51355026 | -1.6057586 | 0.10832697 | 0.51253572 |
| RAB10 | 917.195203 | 0.088630364 | 0.22810231 | 0.38855531 | 0.69760513 | 0.93156241 |
| HADHA | 401.55045 | -0.304891162 | 0.24029404 | -1.2688253 | 0.20450338 | 0.65976734 |
| HADHB | 451.423601 | -0.023351981 | 0.19103855 | -0.122237 | 0.90271132 | 0.98320957 |
| SELENOI | 189.569965 | 0.264278461 | 0.24055744 | 1.09860856 | 0.27193884 | 0.72372882 |
| MAPRE3 | 76.0245339 | -0.336219635 | 0.31798332 | -1.0573499 | 0.29035192 | 0.73754646 |
| TMEM214 | 59.0746425 | -0.44390008 | 0.33042316 | -1.3434291 | 0.17913308 | 0.62850818 |

| | | | | | | |
|---|---|---|---|---|---|---|
| AGBL5 | 94.9200919 | -0.110415564 | 0.31802462 | -0.3471919 | 0.72844718 | 0.93931348 |
| OST4 | 313.078856 | 0.00297902 | 0.41567205 | 0.00716675 | 0.99428181 | 0.99885672 |
| PREB | 30.4483897 | 0.232250146 | 0.42443596 | 0.54719715 | 0.58424329 | 0.90090543 |
| ATRAID | 198.979152 | -0.345764823 | 0.45165297 | -0.7655542 | 0.44394161 | 0.83873671 |
| CAD | 35.5804407 | -0.513170999 | 0.34077639 | -1.5058878 | 0.13209598 | 0.55448417 |
| MPV17 | 63.0971635 | -0.342916337 | 0.25372346 | -1.3515358 | 0.17652387 | 0.62528328 |
| GTF3C2 | 155.718962 | 0.088955182 | 0.26068063 | 0.34124201 | 0.7329214 | 0.94163693 |
| EIF2B4 | 43.6039707 | 0.15043102 | 0.24819469 | 0.60610087 | 0.54444777 | 0.88654641 |
| SNX17 | 90.8550613 | 0.124860205 | 0.4367931 | 0.28585663 | 0.77498793 | 0.95601455 |
| PPM1G | 255.204172 | -0.573884877 | 0.33467428 | -1.7147565 | 0.08638989 | 0.4662029 |
| NRBP1 | 132.963189 | -0.277119225 | 0.23403716 | -1.1840822 | 0.23638054 | 0.69335782 |
| KRTCAP3 | 53.2773305 | -0.903982226 | 0.42097202 | -2.1473689 | 0.03176392 | 0.2995969 |
| IFT172 | 165.045077 | 0.031330901 | 0.21954197 | 0.14271031 | 0.88651898 | 0.97973087 |
| ZNF512 | 187.813493 | 0.137376173 | 0.25591597 | 0.53680187 | 0.59140448 | 0.90233654 |
| GPN1 | 45.5448134 | -0.005554684 | 0.31138811 | -0.0178385 | 0.98576772 | 0.99587186 |
| SUPT7L | 60.2810088 | 0.204281011 | 0.25707029 | 0.79465041 | 0.42681689 | 0.82788099 |
| SLC4A1AP | 80.7254782 | 0.148135344 | 0.21960493 | 0.67455384 | 0.49995927 | 0.86719185 |
| MRPL33 | 605.699512 | 0.66661853 | 0.355727 | 1.87396102 | 0.0609358 | 0.40676596 |
| RBKS | 451.381183 | 0.677333818 | 0.69672407 | 0.9721694 | 0.33096628 | 0.77618491 |
| BABAM2 | 1635.23234 | -0.145598179 | 0.32138516 | -0.4530333 | 0.65052477 | 0.91817658 |
| FOSL2 | 1205.35802 | 1.013395455 | 0.80478808 | 1.25920784 | 0.20795527 | 0.66589349 |
| PPP1CB | 548.608627 | 0.751588971 | 0.71408788 | 1.05251607 | 0.29256284 | 0.73844969 |
| TRMT61B | 99.0293199 | 0.171762247 | 0.27693543 | 0.62022489 | 0.53510973 | 0.88137638 |
| WDR43 | 209.980761 | 0.078355094 | 0.32634872 | 0.24009622 | 0.81025566 | 0.96275276 |
| CLIP4 | 358.622512 | -0.207070368 | 0.44789663 | -0.4623173 | 0.64385378 | 0.91666725 |
| YPEL5 | 334.217048 | 0.698390788 | 0.58032519 | 1.2034473 | 0.22880327 | 0.68686861 |
| LCLAT1 | 278.239452 | 0.202588528 | 0.25343356 | 0.79937531 | 0.42407282 | 0.82774157 |
| GALNT14 | 372.468318 | -0.04607576 | 0.64594704 | -0.0713306 | 0.94313468 | 0.98885525 |
| MEMO1 | 33.9052865 | -0.090996258 | 0.34237051 | -0.265783 | 0.79040634 | 0.95817912 |
| DPY30 | 193.302449 | 0.305452596 | 0.39670338 | 0.76997729 | 0.44131336 | 0.83770294 |
| SPAST | 203.279813 | 0.574726241 | 0.29477088 | 1.94973888 | 0.05120725 | 0.38114487 |
| SLC30A6 | 167.055032 | -0.029294415 | 0.23233694 | -0.1260859 | 0.89966392 | 0.98312332 |
| YIPF4 | 264.006948 | 0.123873447 | 0.18518408 | 0.66892062 | 0.50354612 | 0.86946722 |
| BIRC6 | 1493.89129 | 0.258023123 | 0.23121266 | 1.11595585 | 0.26444103 | 0.71989632 |
| TTC27 | 215.55311 | 0.127256659 | 0.2363702 | 0.5383786 | 0.59031569 | 0.90233654 |
| LINC00486 | 300.432025 | 0.577897531 | 0.2408324 | 2.39958385 | 0.01641372 | 0.21054564 |
| LTBP1 | 68.1630228 | 0.586390188 | 0.77940768 | 0.75235362 | 0.45183843 | 0.84234337 |
| FAM98A | 60.6222576 | 0.690785011 | 0.26612254 | 2.59574033 | 0.00943874 | 0.15525074 |
| LINC01317 | 56.428322 | 2.61934034 | 0.64458295 | 4.06362025 | 4.83E-05 | 0.00576513 |
| AC068599.1 | 473.166867 | 2.706508054 | 0.83552027 | 3.23930865 | 0.0011982 | 0.04468725 |
| LINC01320 | 700.908043 | -0.926053238 | 0.44711622 | -2.071169 | 0.03834301 | 0.32856477 |
| AC012593.1 | 116.816291 | -1.077353539 | 1.06872653 | -1.0080722 | 0.31341979 | 0.75717828 |
| CRIM1 | 962.397628 | -0.154235481 | 0.31070936 | -0.4963979 | 0.61961369 | 0.90907933 |
| FEZ2 | 328.864526 | 0.14861753 | 0.29893579 | 0.49715537 | 0.61907949 | 0.90854966 |
| STRN | 484.899194 | 0.693342287 | 0.16489256 | 4.20481253 | 2.61E-05 | 0.00365121 |
| HEATR5B | 217.473122 | -0.215266052 | 0.29246511 | -0.7360401 | 0.46170629 | 0.84738158 |
| EIF2AK2 | 333.125301 | -0.216085376 | 0.4854175 | -0.4451537 | 0.65620871 | 0.92099468 |
| CEBPZOS | 170.707623 | -0.092682017 | 0.33137473 | -0.2796894 | 0.77971577 | 0.95636474 |
| CEBPZ | 180.683849 | -0.279537424 | 0.29185024 | -0.9578112 | 0.33815798 | 0.77836439 |
| NDUFAF7 | 103.338157 | 0.44427276 | 0.25455008 | 1.74532554 | 0.08092822 | 0.45354424 |
| PRKD3 | 287.833726 | 0.102528321 | 0.19546159 | 0.52454459 | 0.59989982 | 0.90369786 |
| CDC42EP3 | 51.5844807 | -0.37064824 | 0.3286409 | -1.1278214 | 0.25939536 | 0.7168415 |
| RMDN2 | 138.853622 | 0.10938043 | 0.30222396 | 0.36191846 | 0.71741296 | 0.93573028 |
| CYP1B1-AS1 | 43.5411835 | 0.197879501 | 0.59995535 | 0.32982371 | 0.74153317 | 0.94487568 |
| ATL2 | 233.18124 | 0.798759528 | 0.21481521 | 3.71835642 | 0.00020052 | 0.01361583 |
| HNRNPLL | 184.64837 | 0.668075916 | 0.22680331 | 2.94561805 | 0.0032231 | 0.08616863 |
| GALM | 188.597175 | 0.340945516 | 0.35193518 | 0.96877362 | 0.33265815 | 0.77653541 |
| SRSF7 | 206.787556 | -0.183968325 | 0.3969582 | -0.4634451 | 0.64304537 | 0.91643619 |
| DHX57 | 161.697901 | -0.201763524 | 0.2946052 | -0.6848607 | 0.49343183 | 0.86307787 |
| MORN2 | 90.5599003 | 0.517681147 | 0.48191513 | 1.07421642 | 0.2827257 | 0.73268183 |
| SOS1 | 810.416268 | -0.017680776 | 0.26138756 | -0.067642 | 0.94607062 | 0.98949015 |
| MAP4K3 | 1681.99623 | 0.624011854 | 0.29786772 | 2.09492941 | 0.03617727 | 0.31913954 |
| AC007388.1 | 812.826622 | 0.200944794 | 0.3853627 | 0.52144329 | 0.602058 | 0.90369786 |
| TMEM178A | 133.974867 | 1.229320544 | 0.721643 | 1.70350236 | 0.0884741 | 0.47038308 |
| THUMPD2 | 168.405922 | -0.290754531 | 0.35453822 | -0.8200936 | 0.41216274 | 0.82355334 |
| SLC8A1 | 845.096328 | 0.086103754 | 0.88538943 | 0.09724958 | 0.92252819 | 0.98664103 |
| EML4 | 894.184628 | 0.683721432 | 0.26646729 | 2.56587381 | 0.01029163 | 0.16264592 |
| COX7A2L | 276.704032 | 0.104392272 | 0.26557364 | 0.3930822 | 0.69425877 | 0.93105058 |
| MTA3 | 468.401578 | 0.255831238 | 0.22547658 | 1.13462445 | 0.2565327 | 0.71374446 |
| ZFP36L2 | 757.336618 | -0.301956796 | 0.47700392 | -0.6330279 | 0.52671541 | 0.87791848 |
| THADA | 659.662634 | 0.134292332 | 0.25772238 | 0.52107362 | 0.60231549 | 0.90369786 |
| PLEKHH2 | 536.997015 | -0.46073182 | 0.39598496 | -1.1635084 | 0.2446233 | 0.70121778 |
| DYNC2LI1 | 469.439462 | 0.116194859 | 0.19735025 | 0.58877483 | 0.55601233 | 0.89081623 |
| LRPPRC | 447.344024 | 0.202479406 | 0.19516531 | 1.0374764 | 0.29951389 | 0.74611494 |
| PPM1B | 415.13433 | 0.164037741 | 0.27149318 | 0.60420574 | 0.54570686 | 0.88669883 |
| SLC3A1 | 225.634202 | -0.753379115 | 0.83302314 | -0.9043916 | 0.3657878 | 0.79329304 |
| PREPL | 533.869504 | 0.306515316 | 0.31938716 | 0.95969829 | 0.33720708 | 0.77827084 |
| CAMKMT | 1366.58143 | 0.287911924 | 0.22503793 | 1.2793929 | 0.20075873 | 0.65487239 |
| SRBD1 | 316.190003 | 0.210084089 | 0.28813812 | 0.72910897 | 0.46593501 | 0.85028134 |
| PRKCE | 1317.67186 | 0.173078993 | 0.30018104 | 0.57658202 | 0.56422185 | 0.8930743 |
| EPAS1 | 70.8983863 | -1.250474273 | 0.57734045 | -2.1659218 | 0.03031716 | 0.29256342 |
| RHOQ | 285.507879 | -0.402135878 | 0.39474582 | -1.018721 | 0.30833542 | 0.75313873 |
| AC018682.1 | 49.7560376 | -0.518875518 | 0.72302034 | -0.71765 | 0.47297315 | 0.85298656 |
| PIGF | 119.735088 | 0.289292725 | 0.21247815 | 1.36151751 | 0.1733502 | 0.62003758 |
| CRIPT | 61.2643707 | 0.151299471 | 0.35590992 | 0.42510607 | 0.67075935 | 0.92474338 |
| SOCS5 | 131.731066 | 0.140281952 | 0.27414735 | 0.51170274 | 0.60885907 | 0.90549306 |
| MCFD2 | 194.883748 | 0.019207012 | 0.23586535 | 0.0814321 | 0.93509832 | 0.98822345 |
| TTC7A | 201.190574 | 0.094843754 | 0.32445776 | 0.29231464 | 0.77004607 | 0.95482817 |
| AC106869.1 | 48.2540411 | -0.614954681 | 0.43066444 | -1.4279207 | 0.15331468 | 0.58860244 |
| EPCAM | 1419.62174 | 0.137388275 | 0.50692247 | 0.27102424 | 0.78637239 | 0.9576974 |
| MSH2 | 218.673766 | -0.080211636 | 0.31021002 | -0.258572 | 0.79596547 | 0.95948555 |
| MSH6 | 78.3851895 | -0.232004116 | 0.32493102 | -0.7140104 | 0.47522077 | 0.85385826 |
| FBXO11 | 765.165809 | 0.134804722 | 0.19592803 | 0.68803183 | 0.49143273 | 0.86152811 |
| FOXN2 | 107.433606 | -0.036512207 | 0.25807021 | -0.1414817 | 0.88748943 | 0.98009982 |
| PPP1R21 | 614.758555 | 0.411744606 | 0.20290347 | 2.02926353 | 0.04243146 | 0.34798255 |
| GPR75-ASB3 | 107.513759 | -0.545162594 | 0.36062442 | -1.5117185 | 0.1306055 | 0.55395564 |
| ERLEC1 | 112.236619 | -0.259245618 | 0.18813627 | -1.3779672 | 0.1682134 | 0.6131323 |
| PSME4 | 381.969003 | 0.365190536 | 0.25455399 | 1.43462897 | 0.15139285 | 0.5856763 |
| ACYP2 | 670.723793 | 0.134640928 | 0.32346466 | 0.41624618 | 0.67722987 | 0.9268955 |
| SPTBN1 | 1738.60311 | -0.923876926 | 0.37223375 | -2.4819806 | 0.01306544 | 0.18379304 |
| AC093110.1 | 86.7018687 | -0.737204829 | 0.65419091 | -1.1268956 | 0.25978664 | 0.7168415 |
| EML6 | 812.950245 | 1.330692203 | 0.47283909 | 2.81426013 | 0.00488896 | 0.11130863 |
| RTN4 | 2369.52103 | -0.027838548 | 0.25990791 | -0.1071093 | 0.91470229 | 0.98480378 |
| CLHC1 | 158.447722 | 0.549182004 | 0.21922969 | 2.50505303 | 0.0122433 | 0.17590979 |
| RPS27A | 1293.27595 | 0.633069294 | 0.50613497 | 1.25079146 | 0.21101057 | 0.66930714 |
| MTIF2 | 100.970518 | 0.118363303 | 0.22170266 | 0.5338831 | 0.59342242 | 0.90277205 |
| AC012358.3 | 101.027733 | -0.037008669 | 0.45207936 | -0.0818632 | 0.93475549 | 0.98822345 |
| CCDC88A | 154.735193 | -0.030544535 | 0.36447753 | -0.0838036 | 0.93321257 | 0.98822345 |
| CFAP36 | 236.085207 | 0.265703425 | 0.28830538 | 0.92160411 | 0.35673511 | 0.78612469 |
| PPP4R3B | 410.574084 | -0.000970081 | 0.23853643 | -0.0040668 | 0.99675517 | 0.99932313 |
| PNPT1 | 159.821105 | 0.125653209 | 0.44462955 | 0.28260202 | 0.77748193 | 0.95601455 |
| EFEMP1 | 151.432865 | -2.090257854 | 0.80141067 | -2.6082231 | 0.00910136 | 0.15239051 |
| VRK2 | 268.449411 | -0.316206909 | 0.26832173 | -1.1784618 | 0.23861255 | 0.69499004 |

| | | | | | | |
|---|---|---|---|---|---|---|
| FANCL | 179.05181 | 0.134668793 | 0.26724153 | 0.50392165 | 0.61431644 | 0.90764425 |
| PAPOLG | 107.277796 | 0.394025517 | 0.2695231 | 1.46193599 | 0.14375875 | 0.57324898 |
| REL | 345.438024 | 0.927094629 | 0.54808711 | 1.69150964 | 0.09073951 | 0.47642788 |
| PUS10 | 74.7475413 | 0.291788836 | 0.33255531 | 0.87741445 | 0.38026157 | 0.8029673 |
| PEX13 | 106.951046 | -0.376998178 | 0.31499657 | -1.1968326 | 0.23137179 | 0.69121333 |
| KIAA1841 | 78.5721397 | -0.371650996 | 0.35265311 | -1.0538713 | 0.29194184 | 0.73844969 |
| C2orf74 | 61.6724883 | 0.703304972 | 0.47442045 | 1.48245078 | 0.13822039 | 0.56438077 |
| USP34 | 1601.24102 | 0.220353545 | 0.23100027 | 0.95391035 | 0.34012902 | 0.77836439 |
| AC016727.1 | 38.6398269 | 0.228154197 | 0.36172689 | 0.63073606 | 0.52821312 | 0.87826011 |
| XPO1 | 730.685419 | 0.091144335 | 0.34340064 | 0.26541691 | 0.79068831 | 0.95830066 |
| FAM161A | 56.3912533 | -0.074275639 | 0.30388159 | -0.244423 | 0.80690326 | 0.96218371 |
| CCT4 | 191.045494 | 0.270242878 | 0.22071823 | 1.22437953 | 0.22080909 | 0.67871976 |
| COMMD1 | 559.141738 | 0.403576639 | 0.23009482 | 1.75395792 | 0.07943772 | 0.44953856 |
| AC018462.1 | 38.6489386 | -0.479056601 | 0.48825236 | -0.981166 | 0.3265109 | 0.77145036 |
| B3GNT2 | 98.3700242 | -0.402552213 | 0.2714018 | -1.4832334 | 0.1380124 | 0.56438077 |
| EHBP1 | 697.353487 | 0.414306829 | 0.16120579 | 2.57004925 | 0.01016841 | 0.16136868 |
| WDPCP | 800.714608 | -0.050189381 | 0.22370654 | -0.2243537 | 0.82248213 | 0.96511681 |
| MDH1 | 259.216819 | -0.057460306 | 0.37666152 | -0.1525516 | 0.87875192 | 0.97692415 |
| UGP2 | 304.183298 | -0.693072935 | 0.26045074 | -2.6610519 | 0.0077897 | 0.14099484 |
| VPS54 | 507.685534 | 0.206952012 | 0.23891572 | 0.86621345 | 0.38637312 | 0.80634213 |
| PELI1 | 138.98807 | 0.382014291 | 0.43532168 | 0.87754484 | 0.38019077 | 0.8029673 |
| AC008074.1 | 52.371359 | -0.16475753 | 0.38760834 | -0.4250619 | 0.67079156 | 0.92474338 |
| AFTPH | 366.684155 | 0.400872739 | 0.23917206 | 1.67608514 | 0.09372152 | 0.48360487 |
| SERTAD2 | 116.779304 | -0.450524645 | 0.54560061 | -0.8257407 | 0.40895118 | 0.82087883 |
| CEP68 | 88.8015043 | -0.67101163 | 0.3019815 | -2.222029 | 0.02628135 | 0.27568539 |
| RAB1A | 688.44015 | 0.331880609 | 0.2987012 | 1.11107894 | 0.26653437 | 0.72068431 |
| ACTR2 | 528.007567 | -0.062943755 | 0.24645097 | -0.2554007 | 0.79841361 | 0.96000168 |
| SPRED2 | 390.155343 | 0.481821889 | 0.29541423 | 1.63100434 | 0.1028894 | 0.50161615 |
| ETAA1 | 67.4551322 | -0.236402808 | 0.24381614 | -0.9695946 | 0.33224862 | 0.77618491 |
| C1D | 122.077549 | -0.013585492 | 0.28554772 | -0.047577 | 0.9620534 | 0.99171131 |
| PNO1 | 39.6956252 | -0.517190326 | 0.31521275 | -1.6407659 | 0.10084603 | 0.49726048 |
| APLF | 172.738028 | 0.024811476 | 0.19869938 | 0.12486942 | 0.90062693 | 0.9831844 |
| ANTXR1 | 64.987032 | -0.18153121 | 0.47385147 | -0.3830973 | 0.70164764 | 0.93215897 |
| GFPT1 | 394.918782 | -0.224696827 | 0.26785919 | -0.8388618 | 0.40154689 | 0.81660753 |
| NFU1 | 150.853841 | 0.733300268 | 0.32394656 | 2.26364576 | 0.02359591 | 0.25948074 |
| AAK1 | 1835.02893 | 1.185225254 | 0.40402252 | 2.93356233 | 0.00335096 | 0.08779525 |
| ANXA4 | 785.419559 | -0.941349265 | 0.38050559 | -2.4739433 | 0.01336309 | 0.18614502 |
| GMCL1 | 206.06264 | -0.04361079 | 0.26150564 | -0.1667681 | 0.86755255 | 0.97576204 |
| MXD1 | 92.1258825 | 0.35324716 | 0.59169447 | 0.5970094 | 0.5505011 | 0.88916026 |
| PCBP1 | 319.726851 | -0.730412922 | 0.39456491 | -1.8511857 | 0.06414284 | 0.41357646 |
| C2orf42 | 56.0611426 | 0.249409311 | 0.3747727 | 0.66549488 | 0.50573402 | 0.87058024 |
| PCYOX1 | 293.505707 | -0.468020379 | 0.43900347 | -1.0660972 | 0.28637969 | 0.73607046 |
| SNRPG | 122.548333 | -0.057126628 | 0.37001305 | -0.1543908 | 0.87730157 | 0.97640529 |
| FAM136A | 88.4667065 | -0.357442763 | 0.27323858 | -1.3081709 | 0.19081534 | 0.64458858 |
| TGFA | 240.444538 | 0.167122557 | 0.33420592 | 0.50005864 | 0.61703379 | 0.90794578 |
| NAGK | 111.269254 | 0.090407533 | 0.44427607 | 0.20349404 | 0.8387489 | 0.96796482 |
| MCEE | 87.3639232 | -0.403352036 | 0.33893093 | -1.1900715 | 0.2340183 | 0.69285918 |
| MPHOSPH10 | 73.565617 | 0.030792187 | 0.26080724 | 0.11806492 | 0.90601622 | 0.98320957 |
| PAIP2B | 48.6113931 | -0.506830719 | 0.36527349 | -1.3875376 | 0.16527788 | 0.6066943 |
| ZNF638 | 1304.00868 | 0.0679502 | 0.25117326 | 0.27053118 | 0.78675163 | 0.9576974 |
| EXOC6B | 1152.98202 | 0.215105722 | 0.25497084 | 0.84364832 | 0.39886595 | 0.81451972 |
| SPR | 69.9415091 | -0.595832531 | 0.49048577 | -1.2147805 | 0.22444983 | 0.68200471 |
| EMX1 | 759.045274 | 1.122369925 | 0.58033158 | 1.9340149 | 0.0531113 | 0.3871585 |
| SFXN5 | 450.887724 | 0.838980606 | 0.43630505 | 1.92292205 | 0.05448984 | 0.39095834 |
| RAB11FIP5 | 98.0244257 | 0.183813364 | 0.50604354 | 0.36323626 | 0.7164284 | 0.93573028 |
| CCT7 | 182.996257 | -0.180766701 | 0.43637693 | -0.4142444 | 0.67869513 | 0.9268955 |
| ALMS1 | 271.084976 | 0.086880406 | 0.20873355 | 0.41622637 | 0.67724437 | 0.9268955 |
| TPRKB | 34.7429325 | -0.215208087 | 0.36524381 | -0.5892176 | 0.5557153 | 0.89081623 |
| DUSP11 | 61.5870284 | -0.143352342 | 0.28770074 | -0.4982689 | 0.61829449 | 0.90816066 |
| STAMBP | 131.71573 | -0.079934881 | 0.23497918 | -0.3401786 | 0.73372206 | 0.94163693 |
| DGUOK | 159.689026 | 0.167660241 | 0.36175787 | 0.46345983 | 0.6430348 | 0.91643619 |
| TET3 | 158.652411 | -0.188471261 | 0.38609138 | -0.488152 | 0.62544221 | 0.91061758 |
| BOLA3 | 53.9829385 | 0.328074823 | 0.47683075 | 0.68803202 | 0.49143262 | 0.86152811 |
| MOB1A | 304.326036 | 0.022382572 | 0.27538091 | 0.08127859 | 0.9352204 | 0.98822345 |
| AC006030.1 | 76.4236705 | -0.696501269 | 0.34515966 | -2.0179104 | 0.04360059 | 0.35202942 |
| WDR54 | 48.6914874 | -1.137391054 | 0.44270222 | -2.5692012 | 0.01019333 | 0.16136868 |
| RTKN | 78.8058759 | -0.49072021 | 0.30187367 | -1.6255814 | 0.10403874 | 0.50291791 |
| MOGS | 77.7898831 | -0.392350322 | 0.31114397 | -1.2609928 | 0.20731143 | 0.66481754 |
| TTC31 | 35.5002423 | 0.125217943 | 0.33711177 | 0.3714434 | 0.71030731 | 0.93471656 |
| AUP1 | 56.1191184 | 0.119801043 | 0.37690578 | 0.31785408 | 0.75059562 | 0.94794781 |
| M1AP | 105.778573 | -0.454210841 | 0.44637235 | -1.0175604 | 0.3088869 | 0.75355748 |
| HK2 | 90.9363955 | -5.212149924 | 1.81385643 | -2.8735185 | 0.00405927 | 0.09893299 |
| POLE4 | 83.6385061 | -0.01232844 | 0.44299108 | -0.02783 | 0.97779774 | 0.99469867 |
| TACR1 | 51.876616 | -1.276526758 | 0.62519749 | -2.0417976 | 0.04117161 | 0.34162982 |
| AC007681.1 | 57.6716004 | -0.719208874 | 0.46298692 | -1.5534108 | 0.12032503 | 0.53797196 |
| EVA1A | 77.6558336 | 0.038555296 | 0.46954506 | 0.08211203 | 0.93455763 | 0.98822345 |
| MRPL19 | 295.164355 | 0.307012997 | 0.37104211 | 0.82743438 | 0.4079909 | 0.82036543 |
| GCFC2 | 330.858178 | 0.372007708 | 0.26234897 | 1.41798808 | 0.15619424 | 0.59366792 |
| CTNNA2 | 39.9922541 | 1.417386358 | 0.55666019 | 2.54623267 | 0.01088926 | 0.16452817 |
| SUCLG1 | 292.393534 | 0.282068743 | 0.22446645 | 1.2566187 | 0.20889174 | 0.66655368 |
| DNAH6 | 203.897415 | -0.01628043 | 0.45048881 | -0.0361395 | 0.97117114 | 0.9939318 |
| TMSB10 | 1182.22193 | -1.364699499 | 0.51081322 | -2.6716214 | 0.00754858 | 0.13878787 |
| KCMF1 | 331.684171 | 0.122931446 | 0.24236614 | 0.50721378 | 0.61200483 | 0.90693023 |
| TCF7L1 | 974.132364 | 1.134449779 | 0.36689112 | 3.09206111 | 0.00198772 | 0.06287382 |
| TGOLN2 | 691.902252 | -0.063261396 | 0.29574044 | -0.2139085 | 0.83061841 | 0.96648703 |
| RETSAT | 69.6239264 | 0.218885322 | 0.29037417 | 0.75380438 | 0.4509667 | 0.84184734 |
| ELMOD3 | 161.358428 | 0.366872397 | 0.20941943 | 1.75185462 | 0.07979881 | 0.45058811 |
| CAPG | 206.925873 | 0.341573335 | 0.48200843 | 0.70864598 | 0.4785442 | 0.85532469 |
| MAT2A | 268.350063 | -0.175573691 | 0.53244744 | -0.3297484 | 0.74159008 | 0.94487568 |
| GGCX | 93.2569261 | 0.206396009 | 0.26878665 | 0.76788043 | 0.44255822 | 0.83826789 |
| VAMP8 | 372.172125 | -0.059394633 | 0.44924644 | -0.1322095 | 0.89481861 | 0.9821637 |
| RNF181 | 53.6787812 | -0.592122178 | 0.37891189 | -1.5626909 | 0.11812531 | 0.53360052 |
| USP39 | 171.325309 | -0.045499168 | 0.20638976 | -0.2204526 | 0.82551865 | 0.96534651 |
| C2orf68 | 65.6156472 | -0.388335402 | 0.30777369 | -1.2617564 | 0.20703647 | 0.66434881 |
| POLR1A | 88.8364203 | -0.457885479 | 0.26290935 | -1.7416097 | 0.08157676 | 0.45528236 |
| PTCD3 | 218.94066 | -0.112019711 | 0.18535441 | -0.6043542 | 0.54560819 | 0.88669883 |
| IMMT | 306.16983 | 0.057806009 | 0.2056325 | 0.2811132 | 0.77862358 | 0.95623387 |
| MRPL35 | 77.6798687 | 0.04005845 | 0.40422978 | 0.09909822 | 0.92106029 | 0.98581108 |
| KDM3A | 263.73855 | -0.886376115 | 0.46642545 | -1.9003597 | 0.05738594 | 0.39785877 |
| CHMP3 | 162.476921 | -0.094774428 | 0.31716708 | -0.2988155 | 0.76508085 | 0.95378284 |
| RMND5A | 218.387356 | 0.341800578 | 0.25532373 | 1.3386949 | 0.18067002 | 0.63124852 |
| PLGLB1 | 56.6790579 | 1.186505396 | 0.69362431 | 1.71058798 | 0.0871572 | 0.46735992 |
| CYTOR | 214.472161 | -1.064165538 | 0.69514821 | -1.530847 | 0.12580722 | 0.54504325 |
| KRCC1 | 198.840115 | -0.311208118 | 0.261087 | -1.1919709 | 0.23327264 | 0.69247746 |
| THNSL2 | 78.7870685 | 0.307773089 | 0.84566006 | 0.36394422 | 0.71589966 | 0.93573028 |
| EIF2AK3 | 526.814096 | 1.24743183 | 0.92973683 | 1.34170423 | 0.17969192 | 0.6296126 |
| RPIA | 101.310794 | 0.183396925 | 0.25831271 | 0.70998025 | 0.47771639 | 0.85479418 |
| MAL | 1109.5094 | 0.740761245 | 0.50517193 | 1.46635472 | 0.14255167 | 0.57173423 |
| MRPS5 | 287.368158 | 0.049063101 | 0.20030805 | 0.24493824 | 0.80650425 | 0.96218371 |
| AC092835.1 | 46.4881044 | -0.458417229 | 0.37179018 | -1.2329998 | 0.21757583 | 0.6751218 |
| PROM2 | 161.266745 | 0.523661947 | 0.44032166 | 1.18927139 | 0.23433289 | 0.69333955 |
| KCNIP3 | 47.8360945 | 0.129587239 | 0.6429022 | 0.20156602 | 0.84025601 | 0.96833625 |
| FAHD2A | 53.8570624 | 0.145539232 | 0.26990333 | 0.53922726 | 0.58973005 | 0.90233654 |

| | | | | | | |
|---|---|---|---|---|---|---|
| ANKRD36C | 173.04571 | 0.187896204 | 0.30130137 | 0.6236155 | 0.53288013 | 0.88114359 |
| STARD7 | 220.138031 | 0.683702915 | 0.36882846 | 1.85371519 | 0.06377992 | 0.41357646 |
| TMEM127 | 126.856997 | -0.068762637 | 0.28439909 | -0.2417822 | 0.80894894 | 0.96245698 |
| CIAO1 | 115.011838 | -0.428712852 | 0.25184486 | -1.7022895 | 0.08870112 | 0.47038308 |
| SNRNP200 | 233.846176 | -0.222049156 | 0.22073303 | -1.0059625 | 0.3144336 | 0.75823062 |
| NEURL3 | 71.3208945 | 0.344511197 | 0.72456983 | 0.47546997 | 0.63445203 | 0.91430092 |
| KANSL3 | 261.743984 | 0.141663216 | 0.20274428 | 0.69872856 | 0.48472168 | 0.85909499 |
| FER1L5 | 59.285176 | -0.663622583 | 0.28206263 | -2.3527491 | 0.0186352 | 0.2280158 |
| LMAN2L | 100.002324 | 0.054359073 | 0.2324858 | 0.23381675 | 0.81512724 | 0.96333253 |
| CNNM4 | 74.5471376 | -0.067875252 | 0.4272519 | -0.1588647 | 0.87377546 | 0.97622753 |
| ANKRD23 | 56.3434021 | -0.200928452 | 0.35129127 | -0.5719711 | 0.56734156 | 0.89463429 |
| ANKRD36 | 245.049268 | 0.297764967 | 0.29514462 | 1.00887816 | 0.31303307 | 0.75694198 |
| AC092683.1 | 195.707461 | -0.70478188 | 0.44944788 | -1.5681059 | 0.11685641 | 0.53183794 |
| ANKRD36B | 69.3218223 | -0.477013485 | 0.42605966 | -1.1195932 | 0.26288717 | 0.71989632 |
| COX5B | 464.990953 | 0.106648057 | 0.47895132 | 0.22266993 | 0.82379241 | 0.96511681 |
| ACTR1B | 86.7263249 | -0.073198875 | 0.27129491 | -0.2698129 | 0.78730417 | 0.95791052 |
| TMEM131 | 1065.46296 | 0.395882258 | 0.22032455 | 1.79681407 | 0.07236514 | 0.43315305 |
| INPP4A | 156.753417 | -0.124890433 | 0.23277098 | -0.5365378 | 0.59158691 | 0.90233654 |
| COA5 | 102.924755 | 0.144710464 | 0.37057315 | 0.39050446 | 0.69616356 | 0.93130502 |
| UNC50 | 106.084912 | -0.044026901 | 0.2925135 | -0.1505124 | 0.88036039 | 0.97745279 |
| MGAT4A | 461.390015 | 0.616912012 | 0.46408901 | 1.32929676 | 0.18375008 | 0.63428883 |
| KIAA1211L | 124.796201 | -0.844900084 | 0.31548671 | -2.6780845 | 0.00740445 | 0.13685832 |
| TSGA10 | 481.88565 | 0.334148331 | 0.2839451 | 1.17680611 | 0.2392729 | 0.69526124 |
| AC079447.1 | 138.325538 | 0.18741897 | 0.24368193 | 0.76911311 | 0.44182616 | 0.83770294 |
| MITD1 | 112.422831 | -0.215394787 | 0.31186088 | -0.6906759 | 0.48976926 | 0.86055851 |
| TXNDC9 | 87.2342192 | -0.285499066 | 0.29935766 | -0.9537056 | 0.3402327 | 0.77836439 |
| EIF5B | 576.088184 | -0.113332441 | 0.20835405 | -0.5439416 | 0.58648164 | 0.90196444 |
| REV1 | 635.113339 | 0.472679761 | 0.15529634 | 3.04372755 | 0.00233667 | 0.06986353 |
| LONRF2 | 54.9820512 | -0.446616945 | 0.75775901 | -0.5893918 | 0.55559848 | 0.89081623 |
| NPAS2 | 1265.31355 | 0.035227589 | 0.43930058 | 0.08019017 | 0.93608601 | 0.98822345 |
| RPL31 | 285.404807 | -0.002815349 | 0.34779034 | -0.008095 | 0.99354123 | 0.99870178 |
| TBC1D8 | 668.465358 | 0.215331071 | 0.28687644 | 0.75060563 | 0.45289003 | 0.84248055 |
| CNOT11 | 143.101672 | -0.599966784 | 0.2081185 | -2.8828133 | 0.00394141 | 0.09740819 |
| RNF149 | 841.784237 | -0.052740682 | 0.35990996 | -0.1465385 | 0.88349626 | 0.97878487 |
| MAP4K4 | 403.755735 | -0.443934593 | 0.33743118 | -1.31563 | 0.18829826 | 0.64111947 |
| IL1R1 | 237.421858 | 0.161138765 | 0.39380074 | 0.40918858 | 0.68240127 | 0.92806156 |
| SLC9A4 | 325.240523 | 2.819654419 | 0.70968977 | 3.97308029 | 7.09E-05 | 0.00728967 |
| SLC9A2 | 307.389552 | 1.346096195 | 0.52608906 | 2.558685 | 0.01050689 | 0.16367902 |
| MFSD9 | 51.3371137 | -0.389809382 | 0.35136449 | -1.109416 | 0.26725078 | 0.72166661 |
| TMEM182 | 86.9651002 | -0.577683271 | 0.32573348 | -1.7734845 | 0.07614846 | 0.4426298 |
| POU3F3 | 525.30728 | -0.294720136 | 0.30288286 | -0.9730499 | 0.33052851 | 0.77618491 |
| MRPS9 | 173.583994 | 0.353771771 | 0.15849694 | 2.23204171 | 0.02561221 | 0.27225313 |
| TGFBRAP1 | 76.9468271 | -0.121137983 | 0.23730073 | -0.510483 | 0.60971313 | 0.90583975 |
| C2orf49 | 68.512796 | -0.488602632 | 0.31400762 | -1.5560216 | 0.11970296 | 0.53725356 |
| FHL2 | 67.9243543 | 1.59498824 | 1.35303094 | 1.17882614 | 0.23846741 | 0.69499004 |
| NCK2 | 327.535943 | 0.036222537 | 0.24742924 | 0.14639554 | 0.88360915 | 0.97878487 |
| UXS1 | 285.846523 | 0.466978721 | 0.3828466 | 1.21975411 | 0.2225581 | 0.6798044 |
| SULT1C2 | 372.83687 | -0.778453875 | 0.49111552 | -1.5850729 | 0.11294978 | 0.52423105 |
| GCC2 | 820.058825 | -0.238325417 | 0.24982711 | -0.9539614 | 0.34010319 | 0.77836439 |
| GCC2-AS1 | 74.3238336 | -0.460005986 | 0.45234836 | -1.0169286 | 0.3091874 | 0.75357697 |
| LIMS1 | 775.600699 | -0.341850878 | 0.38731569 | -0.8826156 | 0.37744399 | 0.80119855 |
| RANBP2 | 1106.65406 | 2.50981039 | 0.78324292 | 3.2043831 | 0.00135352 | 0.04844574 |
| CCDC138 | 133.233175 | 0.011575204 | 0.39517583 | 0.02929128 | 0.97663228 | 0.99456869 |
| 10-Sep | 670.76311 | -0.62527712 | 0.52340447 | -1.1946347 | 0.2322298 | 0.69186326 |
| SOWAHC | 119.534201 | -1.210943025 | 0.84101166 | -1.4398647 | 0.14990567 | 0.58336854 |
| NPHP1 | 160.728225 | -0.003627683 | 0.30162028 | -0.0120273 | 0.99040382 | 0.9970636 |
| SMIM37 | 109.858744 | -0.00551198 | 0.39291183 | -0.0140285 | 0.98880721 | 0.9964791 |
| LINC01106 | 38.0014621 | 0.405712717 | 0.5176862 | 0.78370394 | 0.43321385 | 0.83179416 |
| MIR4435-2H( | 829.364086 | -1.725543054 | 0.42039079 | -4.1046167 | 4.05E-05 | 0.00493518 |
| BCL2L11 | 91.1521748 | 0.128816856 | 0.47949939 | 0.26864863 | 0.78820008 | 0.95801395 |
| AC017002.5 | 69.0101794 | -1.350358451 | 0.94024412 | -1.4361786 | 0.15095154 | 0.58535779 |
| ANAPC1 | 265.636794 | -0.017452366 | 0.1675785 | -0.1041444 | 0.91705474 | 0.98520052 |
| MERTK | 129.311738 | 0.397640815 | 0.42760874 | 0.92991741 | 0.35241385 | 0.78314007 |
| TMEM87B | 229.766214 | -0.404302901 | 0.22520504 | -1.7952658 | 0.07261136 | 0.43315584 |
| ZC3H8 | 126.62768 | 0.192570621 | 0.26583082 | 0.72441043 | 0.4688138 | 0.85206059 |
| ZC3H6 | 263.610888 | -0.272724342 | 0.32448214 | -0.8404911 | 0.4006331 | 0.8162653 |
| TTL | 79.6900358 | -0.440131754 | 0.39801712 | -1.1058111 | 0.26880829 | 0.72322263 |
| POLR1B | 44.9248214 | 0.133339972 | 0.32084256 | 0.41559315 | 0.67770774 | 0.9268955 |
| CHCHD5 | 80.1097348 | -0.400585906 | 0.42674171 | -0.9387081 | 0.34788062 | 0.78180304 |
| SLC20A1 | 68.9370511 | 0.364031162 | 0.5563456 | 0.65432559 | 0.51290206 | 0.87178584 |
| PSD4 | 76.0486855 | -0.356285149 | 0.30914781 | -1.1524751 | 0.2491259 | 0.70622072 |
| PAX8 | 4625.23234 | 0.440306725 | 0.25621871 | 1.71847995 | 0.08570911 | 0.46492311 |
| CBWD2 | 102.296564 | 0.017155171 | 0.31210841 | 0.05496542 | 0.95616601 | 0.9908068 |
| RABL2A | 80.3499791 | 0.01901033 | 0.31645156 | 0.06007343 | 0.95209715 | 0.9903694 |
| SLC35F5 | 726.182942 | -0.391141116 | 0.23231599 | -1.6836599 | 0.09224742 | 0.48040835 |
| ACTR3 | 508.980569 | -0.074164584 | 0.2629557 | -0.2820421 | 0.77791121 | 0.95601455 |
| DDX18 | 168.886852 | -0.55485205 | 0.34807181 | -1.5940735 | 0.11091958 | 0.51894517 |
| CCDC93 | 248.018848 | -0.352269149 | 0.23358032 | -1.5081286 | 0.13152163 | 0.55397858 |
| INSIG2 | 274.463432 | 0.049615517 | 0.54009167 | 0.09186499 | 0.92680531 | 0.98729622 |
| C2orf76 | 74.7955925 | -0.269647143 | 0.29864951 | -0.9028883 | 0.36658519 | 0.7935157 |
| DBI | 546.424155 | -1.099846213 | 0.49445434 | -2.2243636 | 0.02612399 | 0.27515522 |
| TMEM37 | 64.0857816 | -1.353172626 | 0.6436307 | -2.1024053 | 0.03551778 | 0.31644624 |
| CFAP221 | 545.485075 | -0.700885033 | 0.52552834 | -1.3336769 | 0.18230975 | 0.6328385 |
| PTPN4 | 441.06628 | 0.514660213 | 0.2460605 | 2.09160026 | 0.03647429 | 0.32067999 |
| EPB41L5 | 1164.40565 | 0.597620696 | 0.29868647 | 2.00082945 | 0.04541077 | 0.35755439 |
| RALB | 101.399587 | -0.247831095 | 0.2684876 | -0.9230635 | 0.35597414 | 0.78612469 |
| TFCP2L1 | 1502.28279 | 1.264419875 | 0.4456788 | 2.83706537 | 0.00455303 | 0.10650836 |
| CLASP1 | 1021.3717 | 0.243874054 | 0.25343953 | 0.96225737 | 0.33592033 | 0.77820994 |
| NIFK-AS1 | 71.6047874 | -0.767748529 | 0.3847383 | -1.9955085 | 0.04598745 | 0.36047296 |
| NIFK | 50.9068367 | -0.283057781 | 0.37393929 | -0.7569619 | 0.44907271 | 0.84115854 |
| TSN | 124.865419 | -0.410039718 | 0.30072274 | -1.3635142 | 0.17272052 | 0.61884138 |
| ERCC3 | 110.556859 | -0.031973049 | 0.31471909 | -0.1015923 | 0.91908026 | 0.98520052 |
| MAP3K2 | 576.573891 | 0.328787169 | 0.17438712 | 1.88538678 | 0.05937764 | 0.40275455 |
| IWS1 | 395.845208 | -0.51821743 | 0.41824268 | -1.2390353 | 0.21533244 | 0.67151527 |
| AC010976.1 | 39.2158188 | -0.480485526 | 0.60213773 | -0.7979662 | 0.42489013 | 0.82774157 |
| WDR33 | 425.620069 | 0.050382584 | 0.24487227 | 0.20575047 | 0.83698583 | 0.96757829 |
| POLR2D | 43.5818669 | -0.073928892 | 0.25678735 | -0.2878993 | 0.77342384 | 0.95561066 |
| AMMECR1L | 58.5814055 | -0.037781706 | 0.23870973 | -0.1582747 | 0.87424036 | 0.97622753 |
| SAP130 | 127.319872 | 0.18607506 | 0.24814954 | 0.74985053 | 0.45334473 | 0.84253454 |
| UGGT1 | 148.101033 | -0.076090285 | 0.25247158 | -0.3013816 | 0.76312354 | 0.95378284 |
| HS6ST1 | 1190.32929 | -0.076862125 | 0.51189473 | -0.1501522 | 0.88064453 | 0.97756113 |
| SMPD4 | 118.581442 | -0.0729626 | 0.2525974 | -0.2888494 | 0.77269665 | 0.95543807 |
| MZT2B | 250.103674 | -0.018331205 | 0.47984956 | -0.038202 | 0.96952664 | 0.99334395 |
| CCDC115 | 101.34767 | -0.504772854 | 0.44770173 | -1.1274758 | 0.25954138 | 0.7168415 |
| IMP4 | 51.9526313 | -0.238117967 | 0.35478092 | -0.671169 | 0.50211285 | 0.86837155 |
| PTPN18 | 125.569197 | -0.253740642 | 0.27453346 | -0.9242613 | 0.3553503 | 0.78607579 |
| FAM168B | 333.451223 | 0.209857725 | 0.24857587 | 0.84424012 | 0.39853524 | 0.81400298 |
| PLEKHB2 | 412.323027 | 0.022428803 | 0.27062637 | 0.08287738 | 0.93394904 | 0.98822345 |
| MZT2A | 174.554198 | -0.254778737 | 0.34304798 | -0.7426913 | 0.45766862 | 0.84413361 |
| GPR39 | 398.642992 | 0.717631647 | 0.27603351 | 2.59979897 | 0.00932784 | 0.15443246 |
| NCKAP5 | 293.035599 | 2.997385602 | 0.43715411 | 6.85658801 | 7.05E-12 | 3.70E-08 |
| MGAT5 | 209.479202 | -0.022342131 | 0.29111546 | -0.0767466 | 0.93882511 | 0.98848592 |
| CCNT2-AS1 | 89.8036212 | -0.354424075 | 0.55093202 | -0.6433173 | 0.52001826 | 0.87631314 |

| | | | | | | |
|---|---|---|---|---|---|---|
| CCNT2 | 318.229745 | 0.233843486 | 0.19240241 | 1.21538751 | 0.22421833 | 0.68187922 |
| RAB3GAP1 | 619.358277 | 0.056550109 | 0.18681031 | 0.30271406 | 0.76210778 | 0.95374766 |
| ZRANB3 | 258.196627 | -0.502901071 | 0.24557052 | -2.0478887 | 0.04057091 | 0.33987462 |
| R3HDM1 | 473.194605 | 0.0390549 | 0.23376493 | 0.16706911 | 0.86731567 | 0.97576204 |
| UBXN4 | 512.482837 | -0.307939759 | 0.23217633 | -1.3263185 | 0.18473422 | 0.63541313 |
| DARS | 479.309964 | -0.75427018 | 0.32397464 | -2.3281766 | 0.01990273 | 0.23692294 |
| HNMT | 222.367928 | -0.452180825 | 0.33784927 | -1.33841 | 0.18076283 | 0.63125441 |
| AC114763.1 | 123.858247 | -0.072912112 | 0.42412212 | -0.171913 | 0.86350591 | 0.97461028 |
| SPOPL | 217.607872 | 0.11909961 | 0.24265469 | 0.49081931 | 0.62355425 | 0.91058133 |
| LRP1B | 842.682545 | -2.763025923 | 0.55014704 | -5.0223408 | 5.10E-07 | 0.00020575 |
| GTDC1 | 265.504496 | -0.280152511 | 0.35400761 | -0.7913743 | 0.42872562 | 0.8296433 |
| TEX41 | 1492.53726 | 0.911935672 | 0.80178036 | 1.13738839 | 0.25537596 | 0.71266224 |
| ACVR2A | 306.291782 | 0.701654623 | 0.45571255 | 1.53968687 | 0.1236367 | 0.54273186 |
| ORC4 | 429.516296 | 0.531748856 | 0.29791965 | 1.78487338 | 0.07428186 | 0.43775442 |
| MBD5 | 2411.65952 | 0.435855789 | 0.22662812 | 1.92322026 | 0.05445239 | 0.39095834 |
| EPC2 | 419.631757 | 0.299703239 | 0.22854134 | 1.31137428 | 0.18973134 | 0.64390689 |
| AC105402.3 | 45.5580766 | 1.726780652 | 1.11705824 | 1.54582867 | 0.12214595 | 0.54028588 |
| KIF5C | 255.436649 | -0.617826902 | 0.71156064 | -0.8682702 | 0.38524643 | 0.80570396 |
| LYPD6B | 693.790863 | 0.6419676 | 0.31117013 | 2.06307592 | 0.03910542 | 0.33313698 |
| MMADHC | 91.5309511 | -0.139653376 | 0.27139812 | -0.5145702 | 0.60685341 | 0.90510401 |
| RBM43 | 42.8115377 | -0.82534437 | 0.427144 | -1.9322392 | 0.05332999 | 0.38758552 |
| NMI | 71.9536005 | -0.818843781 | 0.53140472 | -1.5409042 | 0.1233401 | 0.54197241 |
| RIF1 | 335.434157 | -0.038188844 | 0.18947987 | -0.2015457 | 0.84027193 | 0.96833625 |
| ARL5A | 118.92029 | 0.458395362 | 0.21163383 | 2.16598341 | 0.03031246 | 0.29256342 |
| CACNB4 | 82.3134031 | 1.030749439 | 0.58878324 | 1.75064331 | 0.08000737 | 0.45074086 |
| STAM2 | 280.094544 | 0.33132695 | 0.24528523 | 1.35078232 | 0.17676517 | 0.62542168 |
| FMNL2 | 1464.0061 | -0.359040356 | 0.29409842 | -1.220817 | 0.22215531 | 0.67976281 |
| PRPF40A | 378.951115 | 0.281885394 | 0.13299561 | 2.11950903 | 0.03404747 | 0.3120805 |
| ARL6IP6 | 130.227716 | -0.240226017 | 0.27528159 | -0.8726556 | 0.38285084 | 0.80423549 |
| KCNJ3 | 1223.97177 | 1.155243366 | 0.3626947 | 3.18516748 | 0.0014467 | 0.05037024 |
| LINC01876 | 362.034491 | 1.284481585 | 0.63383247 | 2.02653168 | 0.04271033 | 0.3484612 |
| NR4A2 | 431.712541 | 4.833003916 | 1.45329386 | 3.32555172 | 0.00088244 | 0.03655313 |
| GPD2 | 425.87309 | 0.631913428 | 0.26870584 | 2.35169218 | 0.01868823 | 0.22804166 |
| ACVR1 | 219.560094 | -0.680972309 | 0.47606721 | -1.4304121 | 0.15259877 | 0.58795409 |
| UPP2 | 87.1797168 | -0.519064091 | 0.6495297 | -0.7991383 | 0.4242102 | 0.82774157 |
| CCDC148 | 512.172438 | 0.353583162 | 0.39039373 | 0.90570913 | 0.36508983 | 0.79281836 |
| PKP4 | 1522.2614 | 0.145282088 | 0.22051074 | 0.65884359 | 0.50999621 | 0.87094607 |
| PKP4-AS1 | 54.2796845 | -0.293552349 | 0.57442989 | -0.5110325 | 0.60932829 | 0.90552475 |
| TANC1 | 859.994405 | 0.093650535 | 0.22913796 | 0.40870807 | 0.68275391 | 0.92806156 |
| WDSUB1 | 309.902458 | 0.301241503 | 0.32263516 | 0.93369087 | 0.35046337 | 0.78223232 |
| BAZ2B | 1993.7951 | 0.407270112 | 0.25291466 | 1.61030643 | 0.10733098 | 0.51064149 |
| AC009961.1 | 144.676258 | 0.60127775 | 0.47440932 | 1.26742397 | 0.20500373 | 0.66065163 |
| 7-Mar | 714.508005 | 0.439178848 | 0.25353546 | 1.73221865 | 0.08323464 | 0.45768271 |
| PLA2R1 | 85.8755346 | -0.107802166 | 0.48145946 | -0.223907 | 0.82282963 | 0.96511681 |
| ITGB6 | 1056.35284 | -2.73317448 | 0.82412188 | -3.3164688 | 0.00091163 | 0.03755638 |
| RBMS1 | 1033.24959 | 0.026292968 | 0.26690916 | 0.09850905 | 0.92152809 | 0.98607457 |
| TANK | 377.567401 | -0.269706707 | 0.22163793 | -1.2168798 | 0.22364997 | 0.68103746 |
| PSMD14 | 192.547492 | 0.165614105 | 0.2191152 | 0.75583119 | 0.44975041 | 0.84122512 |
| IFIH1 | 73.9125154 | -0.25700098 | 0.80302912 | -0.3200394 | 0.74893844 | 0.94686219 |
| GCA | 151.443927 | -0.063861307 | 0.26401517 | -0.241885 | 0.80886928 | 0.96245698 |
| FIGN | 85.3974017 | 0.01653875 | 0.48457098 | 0.03413071 | 0.97277292 | 0.9939318 |
| GRB14 | 140.768852 | 0.678665966 | 0.59593963 | 1.13881664 | 0.25477964 | 0.71221409 |
| COBLL1 | 9286.99595 | 0.881924525 | 0.38495385 | 2.29098769 | 0.02196413 | 0.25042784 |
| AC019197.1 | 13234.3471 | 0.257320218 | 0.44397136 | 0.57958742 | 0.56219288 | 0.89242209 |
| SLC38A11 | 576.883719 | 0.673611062 | 0.54243752 | 1.24182239 | 0.2143021 | 0.67132603 |
| SCN3A | 632.036451 | -0.681645066 | 0.79892304 | -0.8532049 | 0.39354568 | 0.81028658 |
| SCN2A | 1790.8389 | 0.367511393 | 0.48227918 | 0.7620304 | 0.44604186 | 0.83968362 |
| CSRNP3 | 410.738715 | 0.700200278 | 0.39154559 | 1.78829819 | 0.07372791 | 0.4370297 |
| GALNT3 | 220.378303 | -0.203033939 | 0.27331591 | -0.7428545 | 0.4575698 | 0.84409989 |
| TTC21B | 202.163243 | 0.239778381 | 0.22683177 | 1.05707581 | 0.29047699 | 0.73754646 |
| STK39 | 387.307735 | -1.383757092 | 0.46355221 | -2.9851159 | 0.00283471 | 0.07817828 |
| CERS6 | 125.568047 | -0.512783642 | 0.37942846 | -1.3514633 | 0.17654706 | 0.62528328 |
| LRP2 | 90.0603257 | 0.744940656 | 0.77461373 | 0.96169307 | 0.3362038 | 0.77820994 |
| FASTKD1 | 173.564939 | 0.640255506 | 0.2920624 | 2.19218738 | 0.02836598 | 0.2862587 |
| PPIG | 440.913135 | 0.305126248 | 0.19914196 | 1.53220468 | 0.12547193 | 0.54417366 |
| KLHL23 | 102.880776 | 0.053261045 | 0.3321578 | 0.16034862 | 0.87260646 | 0.97622753 |
| SSB | 192.084979 | -0.073647963 | 0.27850649 | -0.2644389 | 0.7914417 | 0.95832571 |
| METTL5 | 69.6124053 | -0.303627795 | 0.29453651 | -1.0308664 | 0.30260348 | 0.74899204 |
| UBR3 | 662.982482 | 0.59140539 | 0.18457271 | 3.20418656 | 0.00135445 | 0.04844574 |
| MYO3B | 950.056337 | 0.382335072 | 0.57854812 | 0.66085267 | 0.5087068 | 0.87094607 |
| GORASP2 | 168.28883 | -0.143553741 | 0.20093277 | -0.7144367 | 0.47495721 | 0.85385826 |
| TLK1 | 1105.26139 | 0.108738573 | 0.28805618 | 0.37749085 | 0.70580886 | 0.93241442 |
| METTL8 | 140.186715 | -0.233216591 | 0.28182179 | -0.8275322 | 0.4079355 | 0.82036543 |
| DCAF17 | 102.389187 | 0.082024254 | 0.26352574 | 0.31125709 | 0.75560519 | 0.95067145 |
| CYBRD1 | 164.925015 | 0.554716577 | 0.6797477 | 0.81606246 | 0.41446442 | 0.82461178 |
| DYNC1I2 | 521.743891 | 0.020141558 | 0.12908138 | 0.15603767 | 0.87600333 | 0.97622753 |
| SLC25A12 | 364.765437 | -0.131497683 | 0.26818335 | -0.4903276 | 0.62390213 | 0.91058133 |
| HAT1 | 162.172926 | 0.456022399 | 0.25555785 | 1.78441947 | 0.07435553 | 0.43775442 |
| METAP1D | 78.1513573 | -0.161782177 | 0.26346524 | -0.6140551 | 0.53917892 | 0.88290547 |
| ITGA6 | 382.041604 | 0.327188949 | 0.25712009 | 1.27251413 | 0.20319051 | 0.6582233 |
| PDK1 | 258.524086 | -1.245534632 | 0.92613677 | -1.3448712 | 0.17866684 | 0.6279103 |
| MAP3K20 | 91.8096036 | -1.069455828 | 0.37096166 | -2.8829282 | 0.00393997 | 0.09740819 |
| SP3 | 284.997915 | -0.012440687 | 0.18356968 | -0.0677709 | 0.94596799 | 0.98949015 |
| OLA1 | 482.326317 | 0.292405002 | 0.19842982 | 1.47359405 | 0.14059092 | 0.56886692 |
| CIR1 | 196.950743 | -0.190164227 | 0.23185312 | -0.8201927 | 0.41210628 | 0.82355334 |
| SCRN3 | 70.2518056 | -0.473188275 | 0.27561433 | -1.7168493 | 0.08600671 | 0.46572214 |
| GPR155 | 110.77174 | -0.219465846 | 0.38684898 | -0.5673166 | 0.5704991 | 0.89724252 |
| AC010894.3 | 518.222464 | -0.970456994 | 0.52984117 | -1.8315998 | 0.06701106 | 0.42077646 |
| WIPF1 | 114.568098 | -1.076051422 | 0.36863392 | -2.9190244 | 0.00351129 | 0.09019188 |
| ATF2 | 327.883167 | 0.156812085 | 0.20516471 | 0.76432289 | 0.44467485 | 0.83922067 |
| ATP5MC3 | 541.602713 | -0.176926507 | 0.5709642 | -0.3098732 | 0.75665738 | 0.95072713 |
| LNPK | 164.525995 | -0.269285652 | 0.28209672 | -0.9545862 | 0.33978697 | 0.77836439 |
| HOXD10 | 367.997358 | -0.354199219 | 0.56237786 | -0.6298243 | 0.52880957 | 0.8788295 |
| HOXD-AS2 | 97.8750665 | 0.687744102 | 0.63323981 | 1.08607211 | 0.27744709 | 0.72788245 |
| HOXD9 | 156.077738 | -0.103431099 | 0.51426572 | -0.2011238 | 0.84060174 | 0.96854218 |
| HOXD8 | 278.751419 | 0.039358493 | 0.53627455 | 0.07339243 | 0.94149384 | 0.98876195 |
| HOXD3 | 403.126584 | 0.251714636 | 0.39554422 | 0.63637546 | 0.52453173 | 0.87772379 |
| HAGLR | 65.9503915 | 0.180245114 | 0.31658189 | 0.56934753 | 0.56912032 | 0.89617938 |
| MTX2 | 616.080712 | 0.388681553 | 0.1990394 | 1.95278703 | 0.05084484 | 0.38016512 |
| LINC01116 | 132.305668 | 0.460737809 | 0.38483817 | 1.19722483 | 0.23121893 | 0.69095364 |
| LINC01117 | 62.1182697 | -0.077938428 | 0.42714029 | -0.1824656 | 0.85521731 | 0.97135336 |
| HNRNPA3 | 694.304409 | 0.367360604 | 0.26473742 | 1.38764139 | 0.16524627 | 0.6066943 |
| NFE2L2 | 1195.5949 | -0.281479323 | 0.2638746 | -1.0667162 | 0.28609999 | 0.73607046 |
| AGPS | 260.267427 | 0.573172801 | 0.20503429 | 2.79549733 | 0.00518199 | 0.11481456 |
| RBM45 | 67.1386073 | 0.353307498 | 0.334379 | 1.05660792 | 0.29069057 | 0.73763611 |
| PRKRA | 159.580674 | -0.105927924 | 0.18326591 | -0.5780012 | 0.56326328 | 0.8927792 |
| PJVK | 35.890042 | -0.583883358 | 0.33925886 | -1.7210556 | 0.08524073 | 0.46310155 |
| PLEKHA3 | 128.225839 | 0.454283377 | 0.24702531 | 1.8390155 | 0.06591291 | 0.41813998 |
| TTN-AS1 | 388.963974 | -0.199110405 | 0.34239857 | -0.5815165 | 0.56089243 | 0.89242209 |
| TTN | 46.1856391 | -1.649244712 | 0.75357073 | -2.1885732 | 0.02862788 | 0.28737561 |
| SESTD1 | 784.622494 | -1.460957338 | 0.54960317 | -2.658204 | 0.00785583 | 0.14121635 |
| ZNF385B | 81.7035559 | 0.469190661 | 0.62336233 | 0.75267727 | 0.45164387 | 0.84234337 |
| CWC22 | 158.513244 | 0.542773708 | 0.24494574 | 2.2158936 | 0.02669879 | 0.27785831 |

| | | | | | | |
|---|---|---|---|---|---|---|
| UBE2E3 | 443.31037 | 0.034797284 | 0.2392137 | 0.14546526 | 0.88434354 | 0.9788678 |
| CERKL | 42.1369746 | 0.114008654 | 0.69720366 | 0.16352274 | 0.87010687 | 0.9759113 |
| SSFA2 | 134.443657 | -0.305047803 | 0.37131127 | -0.821542 | 0.41133761 | 0.82314649 |
| PDE1A | 3339.95871 | 0.492310847 | 0.28425241 | 1.73194962 | 0.08328253 | 0.45768271 |
| DNAJC10 | 257.198347 | -0.34902418 | 0.32982701 | -1.0582038 | 0.28996256 | 0.73754646 |
| NCKAP1 | 798.730188 | 0.103357685 | 0.2754621 | 0.37521563 | 0.70750011 | 0.93277156 |
| NUP35 | 74.109932 | 0.264956794 | 0.25088897 | 1.0560719 | 0.29093537 | 0.73772562 |
| FSIP2 | 50.7067098 | -0.290936556 | 0.6739671 | -0.4316777 | 0.66597569 | 0.92385693 |
| FSIP2-AS1 | 37.6338229 | 0.65140757 | 0.35243069 | 1.84832816 | 0.06455488 | 0.41412378 |
| LINC01473 | 103.085373 | 0.307495714 | 0.25865355 | 1.18883237 | 0.23450564 | 0.69335782 |
| ZC3H15 | 222.88347 | 0.113163303 | 0.22297114 | 0.50752444 | 0.61178689 | 0.90680628 |
| ITGAV | 952.07375 | 0.358059724 | 0.47788979 | 0.74925167 | 0.45370553 | 0.84268043 |
| AC017101.1 | 109.882729 | -0.155010675 | 0.74621039 | -0.2077305 | 0.83543938 | 0.96757829 |
| AC007319.1 | 264.818725 | 0.335236706 | 0.90862555 | 0.36894924 | 0.71216556 | 0.9348115 |
| TFPI | 204.845951 | -0.761790546 | 0.9169243 | -0.8308107 | 0.40608056 | 0.82002585 |
| GULP1 | 420.062448 | -1.477713735 | 0.45506237 | -3.2472774 | 0.00116515 | 0.04392356 |
| WDR75 | 100.104202 | 0.112196443 | 0.21291098 | 0.52696411 | 0.59821851 | 0.90369786 |
| SLC40A1 | 66.7935776 | -0.195568435 | 0.24192893 | -0.8083714 | 0.41887679 | 0.82694802 |
| ASNSD1 | 56.1404911 | -0.122571747 | 0.31161509 | -0.3933434 | 0.69406585 | 0.93105058 |
| ANKAR | 216.410849 | -0.012598322 | 0.30242933 | -0.0416571 | 0.96677207 | 0.99291246 |
| ORMDL1 | 166.474338 | -0.223209676 | 0.28653917 | -0.7789849 | 0.43598864 | 0.83364877 |
| PMS1 | 180.815447 | 0.051957332 | 0.24506527 | 0.21201426 | 0.83209591 | 0.96656674 |
| C2orf88 | 85.3237196 | -0.775513471 | 0.46436798 | -1.6700408 | 0.09491129 | 0.48627909 |
| HIBCH | 582.862397 | 0.138563225 | 0.26540861 | 0.52207508 | 0.60161805 | 0.90369786 |
| INPP1 | 101.9883 | -0.228531422 | 0.32270926 | -0.7081651 | 0.47884276 | 0.85532469 |
| MFSD6 | 197.757008 | 0.093916401 | 0.22431352 | 0.41868364 | 0.67544736 | 0.92671696 |
| NAB1 | 102.177231 | 0.292848317 | 0.38827956 | 0.75422027 | 0.45071697 | 0.84153105 |
| GLS | 3668.9224 | 0.622208693 | 0.47263687 | 1.31646245 | 0.18801887 | 0.64079278 |
| STAT1 | 1025.9541 | -0.782047134 | 0.84005904 | -0.9309431 | 0.35188302 | 0.78314007 |
| MYO1B | 1326.4769 | 0.538001633 | 0.42572376 | 1.26373411 | 0.20632548 | 0.66368662 |
| NABP1 | 46.9612065 | -3.443816484 | 0.81203316 | -4.2409801 | 2.23E-05 | 0.0031949 |
| SLC39A10 | 216.236138 | -0.059209305 | 0.36835257 | -0.1607409 | 0.87229751 | 0.97622753 |
| DNAH7 | 487.162815 | 0.22898489 | 0.25074881 | 0.91320428 | 0.36113511 | 0.78946516 |
| STK17B | 42.625696 | 0.319827139 | 0.65715082 | 0.48668757 | 0.62647975 | 0.91061758 |
| GTF3C3 | 107.129846 | 0.139492904 | 0.22095327 | 0.63132309 | 0.52782929 | 0.87817923 |
| PGAP1 | 450.436456 | 1.484161554 | 0.52707688 | 2.81583505 | 0.00486506 | 0.11130863 |
| ANKRD44 | 376.788487 | 0.301625697 | 0.29324448 | 1.02858098 | 0.30367662 | 0.74953627 |
| SF3B1 | 437.729735 | 0.143732874 | 0.15579849 | 0.92255624 | 0.35623851 | 0.78612469 |
| COQ10B | 93.9738639 | 0.038825368 | 0.29887734 | 0.12990402 | 0.89664237 | 0.98302816 |
| HSPD1 | 333.213574 | 0.162682009 | 0.29502373 | 0.55142009 | 0.58134573 | 0.90045866 |
| MOB4 | 61.3099176 | 0.332702555 | 0.36840378 | 0.90309212 | 0.366477 | 0.7935157 |
| PLCL1 | 315.071646 | 2.297121368 | 0.66389461 | 3.46006932 | 0.00054004 | 0.02637847 |
| FTCDNL1 | 87.1208125 | -0.288885105 | 0.52553193 | -0.5497004 | 0.58252489 | 0.90085422 |
| C2orf69 | 59.048889 | 0.192490981 | 0.41417229 | 0.46476065 | 0.64210287 | 0.91643619 |
| TYW5 | 90.126488 | 0.098591779 | 0.2166946 | 0.45498031 | 0.6491234 | 0.91768694 |
| MAIP1 | 51.6931805 | -0.368571361 | 0.29623107 | -1.2442023 | 0.21342513 | 0.67075711 |
| SPATS2L | 849.123895 | 0.123443539 | 0.4129001 | 0.29896708 | 0.76496516 | 0.95378284 |
| KCTD18 | 52.8278746 | -0.446757494 | 0.29730151 | -1.5027084 | 0.13291424 | 0.55601567 |
| BZW1 | 106.297038 | -0.430780311 | 0.43453676 | -0.9913553 | 0.32151212 | 0.7663589 |
| CLK1 | 275.021284 | 0.507110898 | 0.40350907 | 1.25675217 | 0.20884339 | 0.66655368 |
| PPIL3 | 84.1044893 | -0.383182501 | 0.32963123 | -1.1624581 | 0.24504942 | 0.70148313 |
| NIF3L1 | 51.3424596 | -0.401489274 | 0.29199178 | -1.375002 | 0.16913083 | 0.61478828 |
| ORC2 | 300.397274 | -0.19317175 | 0.28477777 | -0.6783245 | 0.49756595 | 0.86550172 |
| AC005037.1 | 65.5154438 | -0.112714249 | 0.32418562 | -0.3476843 | 0.72807729 | 0.93931348 |
| FAM126B | 265.166215 | 0.450458542 | 0.24975351 | 1.80361243 | 0.07129209 | 0.43137474 |
| NDUFB3 | 247.651752 | 0.198688739 | 0.46437567 | 0.42786208 | 0.66875154 | 0.92448438 |
| CFLAR | 524.231459 | -0.347015795 | 0.19195223 | -1.8078238 | 0.07063394 | 0.42987436 |
| CFLAR-AS1 | 44.4551193 | -0.471016211 | 0.61038791 | -0.771667 | 0.44031168 | 0.83701549 |
| CASP10 | 121.732125 | -0.534124086 | 0.36098005 | -1.4796499 | 0.13896671 | 0.56646097 |
| CASP8 | 150.068996 | -0.718824934 | 0.28690376 | -2.5054566 | 0.01222934 | 0.17590979 |
| TRAK2 | 358.449157 | -0.284478745 | 0.22597208 | -1.258911 | 0.20806248 | 0.66589349 |
| STRADB | 167.889135 | -0.532587006 | 0.3781594 | -1.4083664 | 0.15902259 | 0.59747329 |
| TMEM237 | 40.4102636 | 0.405050245 | 0.3515211 | 1.1522786 | 0.24920661 | 0.70622072 |
| ALS2 | 185.324039 | -0.144781473 | 0.31555455 | -0.458816 | 0.64636631 | 0.91743182 |
| SUMO1 | 279.083033 | 0.296829514 | 0.40547435 | 0.73205498 | 0.46413501 | 0.84965121 |
| NOP58 | 217.112247 | 0.201834286 | 0.22874638 | 0.88234966 | 0.37758775 | 0.80119855 |
| BMPR2 | 990.959521 | -0.225001498 | 0.24425713 | -0.9211665 | 0.35696349 | 0.78612469 |
| FAM117B | 513.487808 | 0.142349965 | 0.2728281 | 0.52175698 | 0.60183955 | 0.90369786 |
| ICA1L | 270.885092 | 0.106263166 | 0.28971704 | 0.36678258 | 0.7137812 | 0.93573028 |
| WDR12 | 116.46342 | -0.510682438 | 0.35027708 | -1.4579385 | 0.1448575 | 0.57455768 |
| CARF | 196.327301 | -0.087582848 | 0.3498035 | -0.2503773 | 0.8022956 | 0.96071123 |
| NBEAL1 | 1837.89291 | 0.436638812 | 0.24644223 | 1.77176947 | 0.07643283 | 0.4426298 |
| CYP20A1 | 305.173097 | -0.101264029 | 0.2889849 | -0.3504129 | 0.72602887 | 0.93895336 |
| ABI2 | 617.89375 | 0.113829659 | 0.28576937 | 0.39832701 | 0.69038916 | 0.93073907 |
| RAPH1 | 920.53565 | -0.257493267 | 0.44780427 | -0.575013 | 0.56528252 | 0.89382261 |
| PARD3B | 1913.41926 | 0.757918946 | 0.12599987 | 6.01523578 | 1.80E-09 | 3.14E-06 |
| INO80D | 686.911254 | 0.653843361 | 0.70352822 | 0.92937759 | 0.35269344 | 0.78314007 |
| NDUFS1 | 431.909574 | 0.206540488 | 0.17991696 | 1.14797677 | 0.25097815 | 0.70813507 |
| EEF1B2 | 191.704489 | -0.366955272 | 0.52217215 | -0.7027477 | 0.482213 | 0.85726755 |
| ZDBF2 | 178.090511 | 1.598866111 | 0.72585857 | 2.20272404 | 0.0276142 | 0.28206322 |
| ADAM23 | 122.27126 | -0.024327691 | 0.48549677 | -0.0501089 | 0.96003564 | 0.99121324 |
| FASTKD2 | 94.7530893 | -0.099611205 | 0.25507529 | -0.3905169 | 0.69615438 | 0.93130502 |
| CREB1 | 316.57669 | 0.184352899 | 0.15818776 | 1.16540559 | 0.24385486 | 0.70054794 |
| METTL21A | 81.8403209 | 0.298144152 | 0.34802118 | 0.85668393 | 0.39161959 | 0.80866481 |
| CCNYL1 | 197.491258 | 1.0363127 | 0.45704964 | 2.26739636 | 0.02336602 | 0.25830794 |
| PLEKHM3 | 182.451012 | 0.352858438 | 0.19963314 | 1.76753441 | 0.07713877 | 0.44451877 |
| IDH1 | 73.7795411 | -0.161794471 | 0.41253264 | -0.392198 | 0.69491193 | 0.93125819 |
| PIKFYVE | 329.912343 | 0.388127108 | 0.20350568 | 1.90720526 | 0.05649401 | 0.39629 |
| PTH2R | 3246.7206 | 0.65651128 | 0.33696585 | 1.94830212 | 0.05137882 | 0.38215051 |
| RPE | 213.038556 | 0.400676722 | 0.33784954 | 1.18596203 | 0.2356373 | 0.69335782 |
| KANSL1L | 1668.6619 | 0.086386775 | 0.36856598 | 0.23438619 | 0.81468518 | 0.96321082 |
| AC007038.1 | 169.073168 | -0.360710308 | 0.32746939 | -1.1015085 | 0.27067541 | 0.72343657 |
| ACADL | 253.921639 | 0.431930784 | 0.38519505 | 1.12133006 | 0.26214739 | 0.71989632 |
| LANCL1 | 207.971432 | 0.007153544 | 0.20334498 | 0.03517935 | 0.97193673 | 0.9939318 |
| ERBB4 | 4148.00411 | 0.453607671 | 0.27212701 | 1.6668969 | 0.09553491 | 0.48710328 |
| IKZF2 | 421.028177 | 0.149654782 | 0.44882044 | 0.33344021 | 0.73880202 | 0.94364962 |
| SPAG16 | 1828.96363 | 0.521431767 | 0.39180341 | 1.33085052 | 0.1832382 | 0.633903 |
| AC068051.1 | 412.426308 | -0.245793948 | 0.47995924 | -0.5121142 | 0.60857108 | 0.90549306 |
| VWC2L | 38.2387144 | -0.921180565 | 0.70556346 | -1.3055956 | 0.19169011 | 0.64458858 |
| BARD1 | 73.4349262 | -0.496413753 | 0.42416501 | -1.1703317 | 0.24186751 | 0.69770754 |
| AC016708.1 | 119.35412 | -0.115926666 | 0.24840169 | -0.4666903 | 0.64072145 | 0.91619059 |
| ATIC | 131.150803 | 0.54682261 | 0.26275039 | 2.08114863 | 0.0374203 | 0.3238355 |
| MREG | 78.3175492 | 0.486334982 | 0.24116873 | 2.01657561 | 0.04373982 | 0.35261025 |
| PECR | 173.756107 | 0.54427681 | 0.29544588 | 1.84222169 | 0.06544273 | 0.41750899 |
| XRCC5 | 761.362706 | -0.119712379 | 0.21386601 | -0.5597541 | 0.57564717 | 0.89839467 |
| SMARCAL1 | 103.763313 | -0.206165286 | 0.24149204 | -0.8537146 | 0.39326312 | 0.81010924 |
| RPL37A | 530.26165 | 0.004059372 | 0.29076856 | 0.01396084 | 0.98886123 | 0.9964791 |
| IGFBP2 | 387.581537 | 0.05687612 | 0.69826079 | 0.08145398 | 0.93508092 | 0.98822345 |
| IGFBP5 | 97.5383664 | -0.1441734 | 0.57376095 | -0.2512778 | 0.80159932 | 0.96069141 |
| DIRC3 | 143.686529 | -0.780082876 | 0.80064061 | -0.9743234 | 0.329896 | 0.77618491 |
| TNS1 | 405.698307 | -0.067226368 | 0.46055192 | -0.1459691 | 0.88394575 | 0.97884102 |
| ARPC2 | 558.000775 | -0.089543908 | 0.32402858 | -0.2763457 | 0.78228255 | 0.95688565 |
| AAMP | 116.091984 | 0.054086696 | 0.42745762 | 0.12653113 | 0.89931151 | 0.98312332 |

| | | | | | | |
|---|---|---|---|---|---|---|
| PNKD | 447.266161 | 0.343969838 | 0.36516309 | 0.94196223 | 0.34621198 | 0.78180304 |
| TMBIM1 | 253.478084 | -0.409067667 | 0.32776472 | -1.2480528 | 0.21201173 | 0.66975892 |
| CATIP-AS2 | 62.4531756 | 0.092816869 | 0.52231005 | 0.17770454 | 0.85895502 | 0.97244655 |
| CATIP | 46.7747168 | 0.324642574 | 0.67650386 | 0.47988281 | 0.63131072 | 0.9133034 |
| CTDSP1 | 212.033844 | 0.292151993 | 0.22844271 | 1.27888519 | 0.20093749 | 0.65520376 |
| USP37 | 313.171132 | 0.414609406 | 0.57815059 | 0.71713048 | 0.47329359 | 0.85298656 |
| CNOT9 | 140.384524 | 0.16816665 | 0.30945699 | 0.54342495 | 0.58683725 | 0.90196444 |
| ZNF142 | 48.3162615 | 0.083410335 | 0.2329536 | 0.35805557 | 0.72030173 | 0.93668719 |
| STK36 | 62.4707143 | 0.595811663 | 0.26702021 | 2.23133545 | 0.02565892 | 0.27225313 |
| TTLL4 | 43.8679468 | -0.428466465 | 0.32168672 | -1.3319371 | 0.18288087 | 0.63358396 |
| CYP27A1 | 45.409806 | -0.728485536 | 0.68572776 | -1.0623539 | 0.28807507 | 0.73653438 |
| CNPPD1 | 107.833879 | -0.069097176 | 0.36041043 | -0.191718 | 0.84796309 | 0.96991131 |
| RETREG2 | 135.619146 | -0.506325448 | 0.26253561 | -1.9285972 | 0.05378089 | 0.38935477 |
| ZFAND2B | 50.1749877 | 0.340936629 | 0.34862933 | 0.97793444 | 0.32810675 | 0.77410148 |
| ANKZF1 | 106.467963 | -0.551952087 | 0.3904959 | -1.4134645 | 0.15751919 | 0.59509775 |
| STK16 | 56.1930789 | -0.126282486 | 0.28894894 | -0.4370408 | 0.66208174 | 0.92284375 |
| TUBA4A | 189.01939 | 1.26580167 | 0.78676597 | 1.60886682 | 0.10764547 | 0.51115745 |
| DNAJB2 | 129.801564 | 0.158044137 | 0.31213499 | 0.50633265 | 0.61262314 | 0.90714827 |
| DNPEP | 69.616461 | -0.445647305 | 0.28596874 | -1.5583777 | 0.11914375 | 0.53606632 |
| CHPF | 63.6986503 | -0.274955036 | 0.50642847 | -0.5429297 | 0.58717824 | 0.90196444 |
| OBSL1 | 152.222607 | -0.171737732 | 0.21456041 | -0.8004167 | 0.42346942 | 0.82774157 |
| STK11IP | 31.7968961 | 0.193257161 | 0.30093669 | 0.64218543 | 0.52075279 | 0.87638467 |
| SGPP2 | 649.945548 | 0.175139176 | 0.44274195 | 0.39557845 | 0.69241603 | 0.93105058 |
| FARSB | 104.272345 | 0.294646822 | 0.31306895 | 0.94115632 | 0.34662476 | 0.78180304 |
| ACSL3 | 450.863357 | 0.594831349 | 0.42418111 | 1.40230512 | 0.16082415 | 0.6002217 |
| AP1S3 | 67.2338071 | 0.478808282 | 0.40807151 | 1.17334406 | 0.24065786 | 0.6963541 |
| WDFY1 | 195.851449 | -0.315288588 | 0.26649916 | -1.1830754 | 0.23677928 | 0.69335782 |
| MRPL44 | 50.4032612 | 0.21288129 | 0.35195734 | 0.6048497 | 0.54527887 | 0.88669883 |
| CUL3 | 715.585178 | 0.397832638 | 0.18825994 | 2.11320925 | 0.03458286 | 0.31447717 |
| DOCK10 | 198.355787 | 0.151202224 | 0.61187341 | 0.24711357 | 0.80482033 | 0.96210606 |
| IRS1 | 51.508641 | -1.517593551 | 0.64499361 | -2.3528815 | 0.01862857 | 0.2280158 |
| RHBDD1 | 336.162122 | 0.173596262 | 0.21786736 | 0.79679794 | 0.42556839 | 0.82774157 |
| COL4A4 | 198.473437 | -0.550214783 | 0.97386205 | -0.5649823 | 0.57208583 | 0.89797476 |
| COL4A3 | 195.396082 | -0.789901672 | 0.83703917 | -0.9436854 | 0.34533042 | 0.78180304 |
| AC097662.1 | 55.8429113 | 0.19633423 | 0.45701454 | 0.42960171 | 0.66748539 | 0.92422713 |
| MFF | 390.955655 | 0.462402724 | 0.27540603 | 1.6789855 | 0.09315487 | 0.48210881 |
| AGFG1 | 527.286677 | 0.143321309 | 0.27071173 | 0.52942409 | 0.59651129 | 0.90364821 |
| TRIP12 | 852.239884 | 0.15549799 | 0.19278699 | 0.80657928 | 0.41990891 | 0.82694802 |
| FBXO36 | 100.851837 | 0.291321418 | 0.24620772 | 1.1832343 | 0.2367163 | 0.69335782 |
| SLC16A14 | 47.4715605 | -0.171194232 | 0.72366729 | -0.2365648 | 0.8129944 | 0.96288868 |
| SP110 | 153.711536 | -0.594089862 | 0.4843986 | -1.2264483 | 0.22003 | 0.67801071 |
| SP140L | 167.211188 | -0.447521016 | 0.33803156 | -1.323903 | 0.18553525 | 0.63605149 |
| SP100 | 433.786753 | -0.538944826 | 0.48719868 | -1.1062116 | 0.26863495 | 0.72322263 |
| CAB39 | 788.004298 | 0.724114098 | 0.16190259 | 4.4725294 | 7.73E-06 | 0.0015335 |
| ITM2C | 577.416849 | -0.448594772 | 0.60698968 | -0.7390484 | 0.45987759 | 0.84642028 |
| PSMD1 | 328.425288 | 0.251237671 | 0.30017495 | 0.8369708 | 0.40260898 | 0.81738515 |
| ARMC9 | 477.779714 | -0.078847661 | 0.21968425 | -0.3589136 | 0.71965974 | 0.93636596 |
| NCL | 370.571876 | -0.683720333 | 0.33972484 | -2.0125709 | 0.04415979 | 0.35332084 |
| PTMA | 2137.96196 | -0.247710638 | 0.3410436 | -0.7263313 | 0.4676357 | 0.85157639 |
| PDE6D | 85.2402472 | 0.298018014 | 0.29532799 | 1.0091086 | 0.31292256 | 0.75694198 |
| COPS7B | 89.5048371 | 0.136937804 | 0.2873355 | 0.47657809 | 0.63366259 | 0.91415622 |
| DIS3L2 | 578.287404 | 0.123502688 | 0.24522394 | 0.50363227 | 0.61451981 | 0.90770509 |
| EIF4E2 | 157.5624 | -0.192742901 | 0.19679747 | -0.9793972 | 0.32738375 | 0.77309185 |
| EFHD1 | 221.202613 | 0.504367523 | 0.52496052 | 0.96077229 | 0.33666668 | 0.77827084 |
| GIGYF2 | 717.875931 | 0.23915304 | 0.24655977 | 0.9699597 | 0.33206658 | 0.77618491 |
| ATG16L1 | 55.9259851 | 0.56089034 | 0.27431304 | 2.04470899 | 0.04088356 | 0.34079725 |
| DGKD | 82.1579064 | 0.12755954 | 0.31916619 | 0.39966495 | 0.68940331 | 0.93045031 |
| USP40 | 461.551687 | 0.308109933 | 0.33315739 | 0.92481796 | 0.35506061 | 0.78607579 |
| AC114812.2 | 192.785358 | -2.878798745 | 1.00538977 | -2.8633659 | 0.00419166 | 0.10017148 |
| SH3BP4 | 165.300016 | -0.089054692 | 0.40531241 | -0.2197186 | 0.82609029 | 0.96545294 |
| AGAP1 | 1220.12964 | -0.774881775 | 0.3640473 | -2.1285195 | 0.03329403 | 0.30804159 |
| IQCA1 | 317.634146 | -0.24710775 | 0.22135467 | -1.1163431 | 0.26427531 | 0.71989632 |
| ACKR3 | 65.9500219 | 2.288969289 | 0.69077765 | 3.31361224 | 0.00092099 | 0.03755638 |
| AC105760.2 | 41.0345866 | -0.074389503 | 0.38440898 | -0.1935166 | 0.84655444 | 0.96949956 |
| COPS8 | 83.1935106 | 0.290089708 | 0.30926913 | 0.93798469 | 0.34825228 | 0.78180304 |
| RAB17 | 374.100533 | 0.469755998 | 0.48921163 | 0.96023064 | 0.33693915 | 0.77827084 |
| LRRFIP1 | 2507.12995 | 0.270431953 | 0.43509828 | 0.62154222 | 0.53424293 | 0.88137638 |
| ILKAP | 208.892852 | 0.165435578 | 0.17650816 | 0.93726873 | 0.34862035 | 0.78180304 |
| PER2 | 92.0147979 | -0.190037295 | 0.38604389 | -0.4922686 | 0.62252945 | 0.91030371 |
| TRAF3IP1 | 147.9627 | 0.179660633 | 0.22533908 | 0.79729017 | 0.42528253 | 0.82774157 |
| ASB1 | 44.4608943 | -0.277009489 | 0.27154166 | -1.0201363 | 0.30766384 | 0.75262615 |
| HDAC4 | 1313.43307 | -0.402529527 | 0.48051499 | -0.8377044 | 0.40219673 | 0.81702302 |
| NDUFA10 | 469.939162 | 0.184037196 | 0.24225574 | 0.75968147 | 0.44744501 | 0.84026779 |
| COPS9 | 144.137492 | 0.013354604 | 0.43004913 | 0.03105367 | 0.97522674 | 0.9939318 |
| GPC1 | 89.3886211 | -1.00878872 | 0.39620102 | -2.5461538 | 0.01089172 | 0.16452817 |
| ANKMY1 | 129.976148 | -0.503032724 | 0.3576421 | -1.4065255 | 0.15956814 | 0.59895206 |
| RNPEPL1 | 102.437945 | -0.542713672 | 0.32576021 | -1.6659913 | 0.09571515 | 0.48710328 |
| CAPN10 | 41.9838526 | 0.068558785 | 0.36328059 | 0.1887213 | 0.85031126 | 0.97017003 |
| SNED1 | 251.01957 | -0.413881167 | 0.56588984 | -0.7313812 | 0.46454637 | 0.84978984 |
| AC005237.1 | 107.399125 | -0.458240594 | 0.68331407 | -0.6706149 | 0.50246589 | 0.86852095 |
| MTERF4 | 126.210932 | -0.130079998 | 0.29743636 | -0.4373372 | 0.66186679 | 0.92284375 |
| PASK | 34.5477525 | 0.273164437 | 0.31217246 | 0.87504334 | 0.38155032 | 0.80315561 |
| PPP1R7 | 122.923872 | -0.292101906 | 0.23902005 | -1.2220812 | 0.22167693 | 0.6794174 |
| HDLBP | 611.803897 | 0.007621017 | 0.18463637 | 0.04127582 | 0.96707601 | 0.99291246 |
| 2-Sep | 818.831876 | 0.061967204 | 0.17173519 | 0.36082997 | 0.71822656 | 0.93573028 |
| FARP2 | 292.873424 | 0.07298179 | 0.20647908 | 0.35345853 | 0.72374472 | 0.93893205 |
| STK25 | 127.03663 | 0.071896912 | 0.26577892 | 0.27051397 | 0.78676487 | 0.9576974 |
| THAP4 | 168.5505 | -0.067751568 | 0.23460506 | -0.2887899 | 0.77274218 | 0.95543807 |
| ATG4B | 219.85591 | 0.258779532 | 0.26175795 | 0.98862149 | 0.32284837 | 0.76879139 |
| DTYMK | 59.3866596 | 0.463315851 | 0.4201369 | 1.10277352 | 0.27012553 | 0.72343657 |
| ING5 | 120.378054 | 0.045068123 | 0.32323947 | 0.13942642 | 0.88911319 | 0.98054894 |
| D2HGDH | 173.726574 | 0.523192918 | 0.2707324 | 1.93250946 | 0.05329666 | 0.38758552 |
| CHL1 | 95.7319262 | 0.220572491 | 0.94179494 | 0.23420437 | 0.81482632 | 0.96324591 |
| LINC01266 | 565.761094 | -0.195616526 | 0.50809118 | -0.3850028 | 0.70023534 | 0.93180634 |
| CNTN4 | 198.183022 | -0.196099185 | 0.47599031 | -0.4119815 | 0.68035301 | 0.92700898 |
| TRNT1 | 127.659712 | 0.049879432 | 0.17652031 | 0.2825705 | 0.7775061 | 0.95601455 |
| CRBN | 156.619486 | -0.023922383 | 0.25429052 | -0.094075 | 0.92504957 | 0.98710257 |
| AC034195.1 | 237.913968 | -0.007355366 | 0.5381279 | -0.0136684 | 0.98909451 | 0.99651056 |
| SUMF1 | 1198.19665 | -0.140643936 | 0.38873163 | -0.3618021 | 0.71749989 | 0.93573028 |
| SETMAR | 35.3342327 | -0.423854646 | 0.41845977 | -1.0128922 | 0.31111165 | 0.75435382 |
| ITPR1 | 491.509398 | 0.375986072 | 0.34133388 | 1.10151993 | 0.27067044 | 0.72343657 |
| EGOT | 114.942212 | 0.309924514 | 0.84847801 | 0.36527112 | 0.71490903 | 0.93573028 |
| AC018816.1 | 63.8522581 | 0.000581244 | 0.338412 | 0.00171756 | 0.99862958 | 0.99962432 |
| BHLHE40-AS | 68.811835 | 0.001594478 | 0.59457656 | 0.0026817 | 0.99786031 | 0.99954455 |
| BHLHE40 | 1164.36445 | -0.403241956 | 0.75226296 | -0.5360386 | 0.5919319 | 0.90233654 |
| ARL8B | 372.018752 | -0.084654647 | 0.35458309 | -0.2387442 | 0.81130396 | 0.96288868 |
| AC026202.2 | 68.7829322 | -0.274901278 | 0.5412357 | -0.5079142 | 0.61151354 | 0.90658678 |
| EDEM1 | 95.5129926 | -0.153529569 | 0.3537804 | -0.4339686 | 0.66431127 | 0.92300139 |
| LMCD1-AS1 | 310.907736 | 0.242087479 | 0.47783874 | 0.50663008 | 0.6124144 | 0.90714827 |
| LMCD1 | 40.666427 | 0.123231384 | 0.63095632 | 0.1953089 | 0.84515113 | 0.96881318 |
| RAD18 | 76.9126873 | -0.031643079 | 0.32952334 | -0.0960268 | 0.92349927 | 0.98694936 |
| SRGAP3 | 937.076841 | 0.662416896 | 0.36226309 | 1.82855199 | 0.06746675 | 0.42262495 |
| THUMPD3-A! | 264.202691 | -0.128344367 | 0.20900957 | -0.6140598 | 0.53917583 | 0.88290547 |

| | | | | | | |
|---|---|---|---|---|---|---|
| THUMPD3 | 92.8573891 | -0.374871729 | 0.21579545 | -1.7371624 | 0.08235852 | 0.45595206 |
| SETD5 | 923.894402 | -0.166644021 | 0.16115046 | -1.0340896 | 0.30109427 | 0.74809578 |
| MTMR14 | 156.46338 | 0.268504503 | 0.38311621 | 0.7008435 | 0.48340069 | 0.85829223 |
| OGG1 | 60.148999 | -0.555143263 | 0.32708928 | -1.6972224 | 0.08965462 | 0.47357883 |
| CAMK1 | 46.8856648 | 0.34113855 | 0.28039423 | 1.21663899 | 0.2237416 | 0.68103746 |
| TADA3 | 90.4981537 | -0.728204902 | 0.24042873 | -3.0287765 | 0.00245546 | 0.0722844 |
| RPUSD3 | 50.0907971 | -0.169720674 | 0.25201413 | -0.673457 | 0.50065662 | 0.86743068 |
| JAGN1 | 64.2516252 | -0.592149294 | 0.45310114 | -1.306881 | 0.19125312 | 0.64458858 |
| IL17RE | 55.3790469 | 0.561052419 | 0.36996669 | 1.51649443 | 0.12939438 | 0.55140574 |
| IL17RC | 49.6468701 | 0.1498062 | 0.37049266 | 0.40434323 | 0.68596034 | 0.92974759 |
| CRELD1 | 80.3260401 | -0.046896699 | 0.34292683 | -0.1367542 | 0.89122505 | 0.98095103 |
| EMC3 | 146.312828 | -0.344490329 | 0.22008603 | -1.5652531 | 0.11752358 | 0.53282158 |
| BRK1 | 219.093909 | 0.129859026 | 0.45643873 | 0.28450483 | 0.77602353 | 0.95601455 |
| VHL | 90.9062772 | -0.049203049 | 0.23039299 | -0.2135614 | 0.83088911 | 0.96648703 |
| SEC13 | 154.187161 | -0.130582569 | 0.3073885 | -0.4248128 | 0.67097315 | 0.92474338 |
| ATG7 | 567.924371 | -0.380163684 | 0.25307432 | -1.502182 | 0.13305012 | 0.55601567 |
| VGLL4 | 596.513848 | -0.192325046 | 0.23641831 | -0.8134947 | 0.41593448 | 0.82489761 |
| TAMM41 | 173.901078 | -0.331022282 | 0.26534024 | -1.2475389 | 0.21219996 | 0.66984203 |
| PPARG | 1505.56936 | 0.444551816 | 0.45173272 | 0.98410364 | 0.32506456 | 0.77058828 |
| TSEN2 | 218.353432 | 0.035908455 | 0.26958348 | 0.13319976 | 0.8940354 | 0.982024 |
| MKRN2OS | 40.6826461 | -0.367622307 | 0.50037423 | -0.7346947 | 0.46252543 | 0.84816562 |
| MKRN2 | 48.0674004 | 0.238646992 | 0.28182315 | 0.84679699 | 0.39710828 | 0.81298981 |
| RAF1 | 431.997897 | 0.794774752 | 0.17242851 | 4.60930014 | 4.04E-06 | 0.00092048 |
| RPL32 | 844.374661 | -0.098072726 | 0.47612681 | -0.2059803 | 0.83680633 | 0.96757829 |
| IQSEC1 | 100.236477 | 0.472995629 | 0.34418054 | 1.37426604 | 0.16935911 | 0.61478828 |
| NUP210 | 96.9177639 | 0.206341594 | 0.37638426 | 0.54822058 | 0.58354046 | 0.90085422 |
| HDAC11 | 68.9688822 | 0.283887931 | 0.25373883 | 1.11881944 | 0.26321718 | 0.71989632 |
| XPC | 186.538518 | 0.210197208 | 0.28437735 | 0.73914891 | 0.45981658 | 0.84642028 |
| LSM3 | 185.363791 | 0.398762673 | 0.55512924 | 0.71832403 | 0.47255752 | 0.85283327 |
| SLC6A6 | 248.491399 | -0.304676836 | 0.68936237 | -0.4419691 | 0.6585116 | 0.92225064 |
| CCDC174 | 96.6084104 | -0.212382109 | 0.20605733 | -1.0306943 | 0.30268421 | 0.74899204 |
| FGD5-AS1 | 295.66451 | -0.117156102 | 0.2774841 | -0.4222083 | 0.67287295 | 0.92506804 |
| NR2C2 | 294.826773 | 0.071141278 | 0.28521772 | 0.24942797 | 0.80302975 | 0.96125092 |
| MRPS25 | 285.811136 | -0.15665589 | 0.16809033 | -0.9319744 | 0.35134974 | 0.78276898 |
| RBSN | 102.12909 | 0.146225419 | 0.27043852 | 0.54069745 | 0.58871614 | 0.90200952 |
| CAPN7 | 418.75116 | 0.130581824 | 0.20825487 | 0.62702891 | 0.53064029 | 0.88004345 |
| SH3BP5-AS1 | 36.6609034 | 0.25435226 | 0.43790028 | 0.58084516 | 0.56134481 | 0.89242209 |
| SH3BP5 | 459.514087 | 0.239067871 | 1.07636153 | 0.22210741 | 0.82423027 | 0.96511681 |
| METTL6 | 91.4247232 | -0.227719662 | 0.27567139 | -0.8260547 | 0.40877304 | 0.82073894 |
| EAF1 | 48.2840016 | 0.166380672 | 0.40468584 | 0.41113539 | 0.68097326 | 0.92731286 |
| COLQ | 54.621641 | -0.612762875 | 0.29411893 | -2.0833847 | 0.03721617 | 0.32359312 |
| HACL1 | 174.030356 | 0.142902256 | 0.25614619 | 0.55789336 | 0.57691721 | 0.89869922 |
| BTD | 72.1385746 | -0.220558801 | 0.34213173 | -0.6446605 | 0.51914724 | 0.87597215 |
| ANKRD28 | 838.62589 | 0.093282129 | 0.21335369 | 0.43721826 | 0.66195307 | 0.92284375 |
| DPH3 | 38.3970177 | -0.13665379 | 0.50497607 | -0.2706144 | 0.78668763 | 0.9576974 |
| OXNAD1 | 211.62511 | -0.221042517 | 0.38116334 | -0.5799155 | 0.56197163 | 0.89242209 |
| RFTN1 | 39.2082546 | -1.154476307 | 0.48155869 | -2.397374 | 0.01651306 | 0.21100962 |
| PLCL2 | 2591.27144 | 1.521705775 | 0.40930952 | 3.71773853 | 0.00020101 | 0.01361583 |
| TBC1D5 | 1752.93411 | 0.384127612 | 0.16765249 | 2.29121322 | 0.02195108 | 0.25042784 |
| RAB5A | 315.69223 | 0.059971906 | 0.19721621 | 0.30409217 | 0.76105767 | 0.95354509 |
| KAT2B | 236.02665 | 0.209210124 | 0.31328598 | 0.66779281 | 0.50426586 | 0.86946722 |
| ZNF385D | 53.8558268 | -1.247964799 | 0.5129889 | -2.4327325 | 0.01498537 | 0.19879321 |
| UBE2E2 | 793.837475 | -0.101706305 | 0.34069463 | -0.2985263 | 0.76530151 | 0.95378284 |
| UBE2E1 | 572.965901 | -0.007920529 | 0.23840467 | -0.033223 | 0.97349672 | 0.9939318 |
| NKIRAS1 | 61.5288036 | 0.209120667 | 0.30252647 | 0.6912475 | 0.48941002 | 0.86042896 |
| RPL15 | 1478.5133 | -0.569113879 | 0.48186945 | -1.1810541 | 0.23758124 | 0.6939708 |
| NR1D2 | 163.725299 | -0.354426977 | 0.33491843 | -1.0582487 | 0.2899421 | 0.73754646 |
| THRB | 1791.69811 | 1.38116482 | 0.26989184 | 5.1174753 | 3.10E-07 | 0.00014433 |
| RARB | 251.884691 | 0.447106529 | 0.33639745 | 1.329102 | 0.18381432 | 0.63430163 |
| TOP2B | 526.377907 | 0.58815594 | 0.20600746 | 2.85502249 | 0.00430338 | 0.10203485 |
| NGLY1 | 345.43345 | 0.44814822 | 0.68388558 | 0.65529707 | 0.51227651 | 0.87177451 |
| OXSM | 44.4344443 | -0.428794531 | 0.37742419 | -1.1361077 | 0.25591149 | 0.71330223 |
| NEK10 | 381.700248 | 0.009433779 | 0.35051025 | 0.02691442 | 0.978528 | 0.99469867 |
| SLC4A7 | 305.268932 | 0.317565535 | 0.6802287 | 0.46685113 | 0.64060639 | 0.91615106 |
| CMC1 | 313.768321 | 0.100102511 | 0.18207644 | 0.54978288 | 0.5824683 | 0.90085422 |
| AZI2 | 244.754583 | 0.043839917 | 0.26682331 | 0.16430317 | 0.86949249 | 0.97576204 |
| TGFBR2 | 326.911867 | 1.423902152 | 0.46104618 | 3.08841544 | 0.00201227 | 0.062951 |
| STT3B | 318.912124 | 0.450896855 | 0.22987676 | 1.96147216 | 0.04982397 | 0.37645791 |
| OSBPL10 | 110.566128 | 1.007900153 | 0.55027692 | 1.83162353 | 0.06700753 | 0.42077646 |
| GPD1L | 36.1646103 | -0.111856088 | 0.35980503 | -0.3108797 | 0.75589206 | 0.95067145 |
| CMTM8 | 56.410003 | -0.73477897 | 0.52497785 | -1.399638 | 0.16162174 | 0.60127647 |
| CMTM7 | 153.836752 | -2.114657871 | 0.61372499 | -3.4456115 | 0.00056977 | 0.02714172 |
| CMTM6 | 202.379482 | -0.438231747 | 0.26132509 | -1.6769601 | 0.09355029 | 0.48341808 |
| DYNC1LI1 | 89.1859915 | -0.305271573 | 0.23384609 | -1.305438 | 0.19174374 | 0.64458858 |
| CNOT10 | 265.492627 | 0.103049559 | 0.27244645 | 0.37823785 | 0.70525391 | 0.93241442 |
| GLB1 | 235.287875 | -0.061717496 | 0.29101609 | -0.2120759 | 0.83204783 | 0.96656674 |
| CRTAP | 135.370921 | -0.07357606 | 0.29578022 | -0.2487525 | 0.80355226 | 0.9615369 |
| FBXL2 | 524.223079 | -0.393819869 | 0.2637644 | -1.4930744 | 0.13541772 | 0.55916513 |
| UBP1 | 163.287607 | 0.040996506 | 0.29602386 | 0.13849055 | 0.88985274 | 0.98064269 |
| CLASP2 | 786.524792 | 0.533808509 | 0.28083591 | 1.90078434 | 0.05733027 | 0.39785877 |
| PDCD6IP | 451.098163 | 0.04538886 | 0.14997547 | 0.30264189 | 0.76216279 | 0.95374766 |
| STAC | 181.748257 | 0.153141396 | 0.55261749 | 0.27712006 | 0.78168792 | 0.95688565 |
| EPM2AIP1 | 151.167897 | 0.062908917 | 0.22878014 | 0.27497543 | 0.78333512 | 0.9568898 |
| MLH1 | 95.1141704 | 0.331212387 | 0.23974309 | 1.38153046 | 0.16711592 | 0.61066068 |
| LRRFIP2 | 740.129556 | -0.154684636 | 0.24484252 | -0.631772 | 0.5275359 | 0.87810931 |
| GOLGA4 | 1251.1376 | -0.214506151 | 0.23985015 | -0.894334 | 0.3711432 | 0.79539603 |
| ITGA9 | 69.5473024 | -0.72316521 | 0.49808287 | -1.4518974 | 0.14653015 | 0.57722788 |
| CTDSPL | 551.781073 | -0.001133406 | 0.28653168 | -0.0039556 | 0.99684389 | 0.99932313 |
| PLCD1 | 49.5337436 | -0.413549611 | 0.45069758 | -0.9175767 | 0.35884051 | 0.78806549 |
| ACAA1 | 145.588168 | -0.233454043 | 0.48281952 | -0.4835224 | 0.62872487 | 0.9113467 |
| OXSR1 | 646.329861 | 1.013027465 | 0.5295865 | 1.91286498 | 0.05576534 | 0.39368206 |
| ACVR2B | 69.4550421 | -0.162529254 | 0.29706691 | -0.5471133 | 0.5843009 | 0.90090543 |
| EXOG | 54.0804094 | -0.349667319 | 0.28489346 | -1.2273617 | 0.21968668 | 0.67800045 |
| WDR48 | 196.409716 | 0.232279774 | 0.22468858 | 1.0337854 | 0.3012365 | 0.74814891 |
| GORASP1 | 62.2634128 | 0.046721349 | 0.25045891 | 0.18654297 | 0.85201899 | 0.97053519 |
| TTC21A | 62.3819314 | 0.003409173 | 0.40292553 | 0.00846105 | 0.99324914 | 0.99858509 |
| CSRNP1 | 78.0875936 | 0.461184916 | 0.851276 | 0.54175722 | 0.58798577 | 0.90198646 |
| SLC25A38 | 53.9844736 | -0.104069517 | 0.36091516 | -0.288349 | 0.77307962 | 0.95543807 |
| RPSA | 566.747636 | -0.1279419 | 0.41175347 | -0.3107245 | 0.75601005 | 0.95068219 |
| MYRIP | 60.095204 | -0.159719914 | 0.52287945 | -0.3054622 | 0.76001415 | 0.95297299 |
| EIF1B | 145.913303 | -0.287171532 | 0.51470535 | -0.5579338 | 0.57688956 | 0.89869922 |
| ENTPD3-AS1 | 84.4607322 | -0.46440273 | 0.57683249 | -0.8050911 | 0.42076709 | 0.82694802 |
| RPL14 | 962.514007 | 0.111663608 | 0.57311554 | 0.19483612 | 0.84552124 | 0.96884569 |
| ZNF621 | 129.175161 | -0.341694359 | 0.26620632 | -1.2835697 | 0.19929256 | 0.65475264 |
| CTNNB1 | 380.944811 | 0.307208759 | 0.17250701 | 1.78084794 | 0.0749373 | 0.43922979 |
| ULK4 | 1027.54577 | 0.01406852 | 0.34132846 | 0.04121696 | 0.96712294 | 0.99291246 |
| TRAK1 | 1140.64711 | 0.252345611 | 0.38777313 | 0.65075579 | 0.51520414 | 0.87353816 |
| SEC22C | 96.9850628 | -0.115447853 | 0.34098409 | -0.3385726 | 0.73493176 | 0.94215106 |
| SS18L2 | 63.1296276 | -0.141087805 | 0.42902864 | -0.328854 | 0.74226601 | 0.94508888 |
| NKTR | 732.769937 | 0.037530046 | 0.16497377 | 0.22749099 | 0.82004198 | 0.96502145 |
| AC006059.1 | 110.984423 | -0.353872316 | 0.51254102 | -0.6904273 | 0.48992551 | 0.86055851 |
| CCDC13 | 36.265746 | 0.364600234 | 0.37519091 | 0.97177257 | 0.3311637 | 0.77618491 |
| AC099329.3 | 39.4004537 | -0.320784983 | 0.40785372 | -0.7865197 | 0.43156309 | 0.8312408 |

| | | | | | | |
|---|---|---|---|---|---|---|
| POMGNT2 | 37.8644752 | 0.09036701 | 0.33720115 | 0.26799141 | 0.78870593 | 0.95801395 |
| SNRK | 284.982527 | 1.237180035 | 0.48083315 | 2.5729924 | 0.01008234 | 0.16107157 |
| ANO10 | 604.723295 | -0.041555491 | 0.34184055 | -0.121564 | 0.90324436 | 0.98320957 |
| ABHD5 | 74.1997716 | -0.351955556 | 0.33903598 | -1.0381068 | 0.29922033 | 0.74594723 |
| TCAIM | 548.653208 | 0.102713077 | 0.2552842 | 0.40234797 | 0.68742796 | 0.93030023 |
| ZNF445 | 153.455067 | 0.505303811 | 0.29095662 | 1.73669812 | 0.08244047 | 0.45616479 |
| ZNF852 | 38.0886034 | -0.250976241 | 0.26571639 | -0.9445267 | 0.34490054 | 0.78180304 |
| AC099669.1 | 178.190271 | 0.03204084 | 0.38147511 | 0.08399195 | 0.93306283 | 0.98822345 |
| ZNF197 | 95.717252 | -0.613583978 | 0.35990824 | -1.7048345 | 0.0882253 | 0.47036301 |
| KIAA1143 | 76.2403643 | -0.577811902 | 0.36602259 | -1.5786236 | 0.11442241 | 0.52703872 |
| ZDHHC3 | 303.494781 | 0.153652146 | 0.2136359 | 0.71922438 | 0.47200268 | 0.85244471 |
| EXOSC7 | 95.2602009 | -0.128545959 | 0.28093643 | -0.4575624 | 0.64726684 | 0.91743182 |
| CDCP1 | 204.851737 | -1.632883768 | 0.44007618 | -3.7104571 | 0.00020689 | 0.01374886 |
| LARS2 | 206.461204 | 0.043819797 | 0.25125996 | 0.17440024 | 0.86155093 | 0.97369284 |
| LIMD1 | 148.735213 | -0.064492568 | 0.30405807 | -0.2121061 | 0.83202427 | 0.96656674 |
| SACM1L | 227.122905 | 0.462419249 | 0.21219835 | 2.17918397 | 0.029318 | 0.28931514 |
| SLC6A20 | 43.5895795 | -5.022227806 | 1.08340093 | -4.6356133 | 3.56E-06 | 0.00086736 |
| LZTFL1 | 70.6868679 | 0.203888686 | 0.24199512 | 0.84253224 | 0.3994901 | 0.81547649 |
| FYCO1 | 113.225462 | -0.089065525 | 0.18083462 | -0.4925248 | 0.62234843 | 0.91028772 |
| LTF | 896.092408 | -4.640607064 | 1.4057436 | -3.301176 | 0.0009628 | 0.03852235 |
| ALS2CL | 35.5537642 | -1.21259246 | 0.27799391 | -4.3619389 | 1.29E-05 | 0.00230675 |
| CCDC12 | 316.284078 | 0.140639384 | 0.24493317 | 0.57419492 | 0.5658359 | 0.89394085 |
| NBEAL2 | 200.239995 | 0.433284257 | 0.44162922 | 0.98110413 | 0.32654139 | 0.77145036 |
| SETD2 | 918.377898 | 0.414186526 | 0.17711457 | 2.33852319 | 0.01936012 | 0.23374897 |
| KIF9-AS1 | 241.688533 | 0.018190895 | 0.28918521 | 0.06290396 | 0.94984298 | 0.98977092 |
| KIF9 | 53.210615 | 0.12714391 | 0.40792926 | 0.31168127 | 0.75528276 | 0.95067145 |
| KLHL18 | 101.085008 | 0.207429778 | 0.21875419 | 0.94823225 | 0.34301123 | 0.78096612 |
| PTPN23 | 75.4942687 | -0.424452897 | 0.25314154 | -1.6767414 | 0.09359306 | 0.48341808 |
| SCAP | 158.065056 | 0.185838385 | 0.2476363 | 0.75044887 | 0.4529844 | 0.84248055 |
| ELP6 | 79.5399996 | -0.287563537 | 0.28603045 | -1.0053599 | 0.31472359 | 0.75823062 |
| SMARCC1 | 800.37027 | -0.23016579 | 0.28578483 | -0.8053814 | 0.42059962 | 0.82694802 |
| DHX30 | 164.675145 | 0.31613641 | 0.22748703 | 1.3896898 | 0.16462309 | 0.60598877 |
| MAP4 | 1647.04582 | 0.189860193 | 0.21004992 | 0.9038813 | 0.36605834 | 0.79329304 |
| ZNF589 | 65.0167272 | 0.278766737 | 0.27215402 | 1.0242977 | 0.30569469 | 0.75109981 |
| NME6 | 64.1493465 | -0.740658031 | 0.28208297 | -2.6256744 | 0.00864775 | 0.14712406 |
| PLXNB1 | 370.316853 | -0.063110488 | 0.27804518 | -0.2269793 | 0.82043987 | 0.96502145 |
| TMA7 | 376.272148 | 0.556955442 | 0.50783871 | 1.09671719 | 0.27276504 | 0.72451099 |
| SHISA5 | 287.087954 | -1.030348863 | 0.41135541 | -2.5047656 | 0.01225326 | 0.17590979 |
| PFKFB4 | 199.882039 | -3.188929519 | 0.87074296 | -3.6623087 | 0.00024995 | 0.01531872 |
| UQCRC1 | 165.025059 | 0.197735981 | 0.29057917 | 0.68048918 | 0.49619477 | 0.86452555 |
| NCKIPSD | 49.4030793 | -0.844316628 | 0.4075174 | -2.0718542 | 0.03827904 | 0.32828506 |
| IP6K2 | 219.914004 | 0.288862355 | 0.15664024 | 1.84411331 | 0.06516663 | 0.41617688 |
| PRKAR2A | 383.373607 | 0.187537469 | 0.21632205 | 0.86693645 | 0.38597683 | 0.80634213 |
| SLC25A20 | 34.7265744 | 0.830610444 | 0.29679359 | 2.79861313 | 0.00513226 | 0.11471549 |
| ARIH2 | 314.723224 | 0.286074402 | 0.20317107 | 1.40804694 | 0.15911717 | 0.59747329 |
| P4HTM | 112.489035 | -0.122998692 | 0.29132995 | -0.4221972 | 0.67288107 | 0.92506804 |
| WDR6 | 204.046466 | 0.32803648 | 0.25458365 | 1.2885214 | 0.19756452 | 0.65273525 |
| DALRD3 | 59.2538256 | 0.243432048 | 0.37887178 | 0.64251828 | 0.52053672 | 0.87638467 |
| NDUFAF3 | 137.544509 | -0.01854823 | 0.57276795 | -0.0323835 | 0.97416622 | 0.9939318 |
| IMPDH2 | 56.1574682 | -0.490877923 | 0.3172337 | -1.54737 | 0.12177405 | 0.54028588 |
| QRICH1 | 202.502279 | 0.360666021 | 0.16497801 | 2.18614598 | 0.02880493 | 0.28768102 |
| QARS | 77.238538 | 0.335782697 | 0.24381607 | 1.37719677 | 0.16845142 | 0.61330788 |
| USP19 | 48.755571 | -0.159200461 | 0.29070805 | -0.54763 | 0.58394596 | 0.90085422 |
| LAMB2 | 122.830412 | 0.755849905 | 0.40901842 | 1.84796054 | 0.06460805 | 0.41412378 |
| KLHDC8B | 52.4952796 | 0.126160462 | 0.44042197 | 0.28645361 | 0.77453072 | 0.95586283 |
| C3orf62 | 35.0254852 | -0.15959003 | 0.33807852 | -0.4720502 | 0.63689096 | 0.91454294 |
| USP4 | 199.046849 | -0.147580446 | 0.30807211 | -0.4790451 | 0.63190652 | 0.9133034 |
| RHOA | 569.489337 | -0.19463685 | 0.2354861 | -0.8265322 | 0.40850224 | 0.82073894 |
| TCTA | 57.7255601 | -0.669191178 | 0.34484846 | -1.940537 | 0.05231447 | 0.3839325 |
| DAG1 | 229.384616 | 0.267936367 | 0.28742685 | 0.93218977 | 0.35123846 | 0.78268744 |
| APEH | 98.1398248 | 0.374932186 | 0.38802919 | 0.96624738 | 0.33392041 | 0.77766353 |
| MST1 | 76.1597838 | 1.21287463 | 0.32840772 | 3.69319766 | 0.00022145 | 0.01432601 |
| RNF123 | 57.9327589 | 0.1791164 | 0.22487431 | 0.79651784 | 0.42573111 | 0.82774157 |
| IP6K1 | 249.445479 | 0.17263455 | 0.26433736 | 0.65308418 | 0.51370201 | 0.87226135 |
| RBM6 | 1104.37516 | 0.274562248 | 0.25516326 | 1.0760258 | 0.28191572 | 0.73166833 |
| RBM5 | 401.814096 | 0.523986833 | 0.17933279 | 2.92186858 | 0.00347938 | 0.0899513 |
| SEMA3F-AS1 | 54.4035796 | -0.844168791 | 0.65892675 | -1.2811269 | 0.20014909 | 0.65475264 |
| SEMA3F | 217.878339 | -0.132498966 | 0.36453773 | -0.3634712 | 0.71625293 | 0.93573028 |
| GNAI2 | 248.824433 | -0.372669428 | 0.27310618 | -1.3645588 | 0.17239177 | 0.61884138 |
| SEMA3B | 356.751866 | 0.391567739 | 0.72213138 | 0.54223892 | 0.58765393 | 0.90196444 |
| IFRD2 | 52.2585143 | -0.081702669 | 0.36934628 | -0.2212089 | 0.82492981 | 0.96511681 |
| HYAL1 | 48.1073109 | 0.551548542 | 0.4823637 | 1.14342879 | 0.25286061 | 0.70874008 |
| HYAL2 | 54.8112093 | -0.7692065 | 0.32696924 | -2.3525347 | 0.01864595 | 0.2280158 |
| TUSC2 | 76.711823 | 0.280751773 | 0.48464442 | 0.57929435 | 0.56239057 | 0.89242209 |
| RASSF1 | 41.3647808 | -0.370345877 | 0.49463201 | -0.7487301 | 0.45401989 | 0.84279368 |
| TMEM115 | 54.8386121 | -0.142375016 | 0.4379305 | -0.3251087 | 0.7450988 | 0.94592798 |
| CACNA2D2 | 89.8670547 | -0.529764838 | 0.83453284 | -0.6348041 | 0.52555621 | 0.87791848 |
| HEMK1 | 51.7330867 | 0.12728839 | 0.32270956 | 0.39443637 | 0.6932589 | 0.93105058 |
| CISH | 77.9282485 | -1.420656673 | 0.37293136 | -3.8094321 | 0.00013929 | 0.0106549 |
| MAPKAPK3 | 97.2461093 | -0.224678535 | 0.3162757 | -0.7103882 | 0.47746342 | 0.85462282 |
| DOCK3 | 157.196907 | 0.067735504 | 0.66760696 | 0.10146015 | 0.91918519 | 0.98520052 |
| MANF | 84.9299355 | -0.296089207 | 0.44303214 | -0.6683244 | 0.50392652 | 0.86946722 |
| RBM15B | 50.2671663 | -0.611873842 | 0.6128748 | -0.9983668 | 0.31810153 | 0.76268681 |
| DCAF1 | 162.74177 | 0.183885157 | 0.34358504 | 0.53519547 | 0.5925147 | 0.90245475 |
| RAD54L2 | 153.808306 | 0.242980087 | 0.30216047 | 0.80414253 | 0.42131466 | 0.82694802 |
| TEX264 | 162.195114 | -0.58237098 | 0.38613463 | -1.5082071 | 0.13150154 | 0.55397858 |
| RPL29 | 1258.11328 | -0.660326909 | 0.52912739 | -1.2479545 | 0.21204772 | 0.66975892 |
| ALAS1 | 280.626107 | -1.308925489 | 0.45524972 | -2.8751813 | 0.00403796 | 0.09864282 |
| WDR82 | 195.943891 | -0.079197744 | 0.1999661 | -0.3960558 | 0.69206383 | 0.93093471 |
| DNAH1 | 36.7499772 | -0.035324989 | 0.39189421 | -0.0901391 | 0.92817668 | 0.98751845 |
| BAP1 | 56.7365308 | -0.137503102 | 0.35671147 | -0.3854743 | 0.69988605 | 0.93180634 |
| PHF7 | 34.8432096 | -0.628563221 | 0.42993136 | -1.4620083 | 0.14373893 | 0.57324898 |
| NISCH | 215.089732 | 0.303195481 | 0.22432369 | 1.35159811 | 0.17650392 | 0.62528328 |
| SMIM4 | 116.801818 | -0.501207715 | 0.27453023 | -1.8256922 | 0.06789663 | 0.42329369 |
| PBRM1 | 538.81353 | -0.186436511 | 0.25499927 | -0.7311257 | 0.4647024 | 0.84991999 |
| GNL3 | 92.4525512 | -0.163857568 | 0.47948133 | -0.3417392 | 0.73254716 | 0.94151267 |
| GLT8D1 | 80.1022967 | 0.269698819 | 0.32506968 | 0.82966464 | 0.40672842 | 0.82018309 |
| SPCS1 | 361.887495 | -0.499986469 | 0.49519099 | -1.0096841 | 0.31264666 | 0.75677346 |
| NEK4 | 152.288952 | 0.307160251 | 0.16439716 | 1.86840361 | 0.06170583 | 0.40825577 |
| SFMBT1 | 87.8031725 | 0.079699652 | 0.23076408 | 0.34537286 | 0.72981408 | 0.94030631 |
| PRKCD | 64.8924528 | 0.169578569 | 0.31335055 | 0.54117845 | 0.58838459 | 0.90200952 |
| TKT | 224.827064 | -1.665431901 | 0.38581116 | -4.3167023 | 1.58E-05 | 0.00259343 |
| DCP1A | 285.222627 | 0.019269433 | 0.20477687 | 0.09409966 | 0.92502999 | 0.98710257 |
| CACNA1D | 77.2256565 | -2.220134179 | 0.39874387 | -5.5678202 | 2.58E-08 | 2.08E-05 |
| CHDH | 49.4150599 | -0.567240648 | 0.41893925 | -1.3539926 | 0.17573874 | 0.62477136 |
| ACTR8 | 35.137569 | -0.00826093 | 0.33088437 | -0.0249662 | 0.98008191 | 0.99469867 |
| SELENOK | 125.353647 | 0.278060109 | 0.68991508 | 0.40303527 | 0.68692228 | 0.92992151 |
| CACNA2D3 | 60.4256208 | -0.357530567 | 0.42834123 | -0.8346863 | 0.40389435 | 0.81888426 |
| ERC2 | 215.097114 | -0.355159656 | 0.30887251 | -1.1498584 | 0.25020219 | 0.70696117 |
| CCDC66 | 309.24526 | 0.18450668 | 0.19515387 | 0.94544209 | 0.34443322 | 0.78180304 |
| FAM208A | 372.13904 | 0.175154845 | 0.21838628 | 0.80204144 | 0.42252899 | 0.82757457 |
| ARHGEF3 | 1057.11433 | -0.837102705 | 0.37814586 | -2.2137032 | 0.0268492 | 0.27859368 |
| APPL1 | 241.306544 | -0.042089816 | 0.15912306 | -0.2645111 | 0.79138611 | 0.95832571 |

| | | | | | | |
|---|---|---|---|---|---|---|
| PDE12 | 64.8197196 | 0.155839081 | 0.352363 | 0.44226857 | 0.65829487 | 0.92207034 |
| ARF4 | 263.573591 | 0.189515935 | 0.28591539 | 0.66283922 | 0.50743353 | 0.87094607 |
| DENND6A | 284.857553 | 0.179909201 | 0.21872889 | 0.82252142 | 0.4107802 | 0.82265937 |
| DENND6A-AS | 65.7906743 | 0.025062027 | 0.39399825 | 0.06360949 | 0.94928118 | 0.98977092 |
| SLMAP | 765.183752 | 0.185967585 | 0.27315064 | 0.68082427 | 0.49598269 | 0.86452555 |
| FLNB | 768.735975 | -0.034136366 | 0.1921534 | -0.1776516 | 0.85899657 | 0.97244655 |
| RPP14 | 41.4971039 | -0.052603468 | 0.31845456 | -0.1651836 | 0.86879948 | 0.97576204 |
| PXK | 216.558168 | 0.379032593 | 0.38036298 | 0.99650234 | 0.31900613 | 0.76378437 |
| PDHB | 104.359722 | 0.178938657 | 0.39536034 | 0.45259638 | 0.65083941 | 0.91825484 |
| KCTD6 | 51.2385773 | 0.176879359 | 0.54173408 | 0.32650587 | 0.74404165 | 0.9456489 |
| FAM107A | 93.2145681 | -0.025155682 | 0.76954958 | -0.0326888 | 0.97392272 | 0.9939318 |
| C3orf67 | 64.9461286 | -0.249812361 | 0.3472981 | -0.7193024 | 0.47195461 | 0.85244471 |
| FHIT | 4610.70647 | 0.09223759 | 0.25680285 | 0.35917666 | 0.71946294 | 0.9362935 |
| PTPRG | 1509.98122 | -0.864081252 | 0.48722381 | -1.7734791 | 0.07614935 | 0.4426298 |
| THOC7 | 124.036318 | -0.113825954 | 0.28311396 | -0.40205 | 0.68764726 | 0.93036007 |
| ATXN7 | 57.5541445 | -0.331789896 | 0.25108584 | -1.3214202 | 0.18636131 | 0.63723614 |
| PSMD6-AS2 | 45.0057026 | 0.560747986 | 0.4409067 | 1.27180646 | 0.20344189 | 0.6582233 |
| PSMD6 | 64.1020803 | 0.62126934 | 0.26829902 | 2.31558554 | 0.02058091 | 0.24180261 |
| PRICKLE2 | 629.602764 | -0.270492241 | 0.48925583 | -0.5528646 | 0.58035611 | 0.90025637 |
| ADAMTS9 | 1146.30309 | -0.297211889 | 0.53039879 | -0.5603555 | 0.57523697 | 0.89807676 |
| ADAMTS9-AS | 99.7558157 | 0.478029701 | 0.54556193 | 0.87621528 | 0.38091301 | 0.80308469 |
| ADAMTS9-AS | 4499.34738 | 0.293919525 | 0.45058742 | 0.652303 | 0.51420572 | 0.87226916 |
| MAGI1 | 4163.72096 | 0.488016537 | 0.12485797 | 3.90857335 | 9.28E-05 | 0.00855087 |
| MAGI1-AS1 | 71.3016291 | 0.263557748 | 0.66512856 | 0.39625084 | 0.69191999 | 0.93093471 |
| SLC25A26 | 264.366938 | 0.083381384 | 0.29203812 | 0.28551541 | 0.7752493 | 0.95601455 |
| LRIG1 | 38.1414179 | -0.193844052 | 0.51328722 | -0.3776522 | 0.70568898 | 0.93241442 |
| SUCLG2 | 685.568242 | -0.055324562 | 0.2261376 | -0.24465 | 0.80672745 | 0.96218371 |
| SUCLG2-AS1 | 348.071751 | 0.085690651 | 0.40154429 | 0.21340274 | 0.83101284 | 0.96648703 |
| FAM19A1 | 62.0396522 | -0.543982535 | 0.55832604 | -0.9743098 | 0.32990274 | 0.77618491 |
| EOGT | 73.9618167 | -0.330490392 | 0.23503649 | -1.4061238 | 0.15968738 | 0.59918502 |
| AC109587.1 | 38.7316319 | 0.051100453 | 0.74402596 | 0.06868101 | 0.94524354 | 0.98949015 |
| TMF1 | 322.890986 | 0.323773893 | 0.26121892 | 1.23947338 | 0.21517024 | 0.67132603 |
| UBA3 | 201.120355 | 0.621128488 | 0.17864766 | 3.47683538 | 0.00050737 | 0.02532014 |
| ARL6IP5 | 290.42072 | -0.105185805 | 0.44440473 | -0.2366892 | 0.81289791 | 0.96288868 |
| FRMD4B | 89.9345197 | -0.630566462 | 0.34747279 | -1.8147219 | 0.06956666 | 0.42828237 |
| MITF | 1035.327 | 0.891723477 | 0.27476983 | 3.24534715 | 0.00117308 | 0.0440639 |
| FOXP1 | 1792.95274 | 0.093705206 | 0.21632138 | 0.43317588 | 0.664887 | 0.92324595 |
| EIF4E3 | 84.9775451 | -0.242669311 | 0.30347794 | -0.7996275 | 0.42392664 | 0.82774157 |
| RYBP | 505.490748 | -0.067610522 | 0.50211846 | -0.1346505 | 0.89288818 | 0.98168537 |
| SHQ1 | 172.087473 | -0.121289785 | 0.32172127 | -0.3770027 | 0.70617161 | 0.93242767 |
| GXYLT2 | 83.3474569 | -0.346817873 | 0.45672712 | -0.7593547 | 0.44764042 | 0.84026779 |
| PPP4R2 | 349.239488 | 0.255772422 | 0.20768629 | 1.23153254 | 0.21812375 | 0.67576162 |
| PDZRN3 | 412.975415 | 0.989313431 | 0.63609769 | 1.55528536 | 0.11987812 | 0.53734932 |
| CNTN3 | 252.183686 | -0.51854515 | 0.26679568 | -1.943604 | 0.05194322 | 0.38290999 |
| ZNF717 | 198.395744 | -0.238220941 | 0.25952691 | -0.9179046 | 0.35866881 | 0.78785353 |
| ROBO2 | 1195.50779 | 1.394827095 | 1.26395279 | 1.10354367 | 0.26979114 | 0.72333985 |
| ROBO1 | 114.96236 | 0.192031056 | 0.67514075 | 0.28443114 | 0.77608 | 0.95601455 |
| GBE1 | 1191.98923 | -0.852279775 | 0.69228971 | -1.2311028 | 0.21828443 | 0.67601089 |
| VGLL3 | 156.933399 | 1.052575747 | 0.37722284 | 2.7903288 | 0.00526545 | 0.11544342 |
| CHMP2B | 217.474858 | -0.192869411 | 0.30061902 | -0.6415742 | 0.52114968 | 0.87650871 |
| CGGBP1 | 321.749308 | -0.248181225 | 0.23608219 | -1.0512492 | 0.29314413 | 0.73864467 |
| ZNF654 | 390.441403 | -0.292135608 | 0.27759756 | -1.0523709 | 0.29262939 | 0.73844969 |
| C3orf38 | 83.9922527 | -0.017731692 | 0.18528265 | -0.0957008 | 0.92375823 | 0.98694936 |
| ARL13B | 165.573988 | 0.322984059 | 0.20194677 | 1.59935242 | 0.10974232 | 0.51504679 |
| NSUN3 | 113.906755 | -0.025905506 | 0.31294437 | -0.0827799 | 0.93402655 | 0.98822345 |
| MTRNR2L12 | 767.994788 | -0.648156838 | 0.58640767 | -1.1053008 | 0.26902929 | 0.72322263 |
| ARL6 | 124.578136 | 0.329222266 | 0.2047294 | 1.60808492 | 0.10781658 | 0.51150646 |
| CRYBG3 | 302.270978 | -1.017245911 | 0.27929203 | -3.6422303 | 0.00027029 | 0.01627201 |
| RIOX2 | 95.3499393 | -0.36320833 | 0.24735608 | -1.4683623 | 0.14200585 | 0.57093391 |
| CLDND1 | 86.163137 | 0.41804079 | 0.4294885 | 0.97334571 | 0.33038151 | 0.77618491 |
| DCBLD2 | 144.799325 | -0.231253013 | 0.33210557 | -0.6963238 | 0.48622607 | 0.85912348 |
| CMSS1 | 640.960263 | 0.919303733 | 0.42113621 | 2.18291308 | 0.02904221 | 0.28822195 |
| FILIP1L | 189.15376 | 0.436031771 | 0.73836632 | 0.59053583 | 0.55483147 | 0.89081623 |
| TBC1D23 | 200.024197 | 0.107436624 | 0.17544691 | 0.61235972 | 0.54029978 | 0.88342811 |
| NIT2 | 63.5744395 | 0.212262582 | 0.28494737 | 0.74491855 | 0.45632096 | 0.84372683 |
| TOMM70 | 158.904023 | 0.453968834 | 0.23919599 | 1.89789483 | 0.05770994 | 0.39817926 |
| TFG | 316.37336 | 0.181987862 | 0.16954515 | 1.07338876 | 0.28309673 | 0.73293254 |
| ABI3BP | 54.250466 | 0.30280193 | 0.71826751 | 0.42157264 | 0.67333698 | 0.92545201 |
| SENP7 | 432.842912 | 0.063073004 | 0.24721499 | 0.25513422 | 0.79861943 | 0.96005379 |
| TRMT10C | 65.796392 | -1.222040248 | 0.4523727 | -2.7014014 | 0.00690479 | 0.13279571 |
| PCNP | 278.786728 | 0.085104037 | 0.23066917 | 0.36894413 | 0.71216937 | 0.9348115 |
| ZBTB11 | 99.080838 | 0.388708586 | 0.2370531 | 1.63975325 | 0.10105648 | 0.49726048 |
| RPL24 | 438.638102 | 0.033866864 | 0.43232115 | 0.07833728 | 0.93755976 | 0.98846521 |
| CEP97 | 70.9068166 | 0.291565671 | 0.21023394 | 1.38686301 | 0.16548354 | 0.60681158 |
| NFKBIZ | 685.82635 | -0.974904479 | 0.82513576 | -1.181508 | 0.23740099 | 0.6939708 |
| ZPLD1 | 571.530324 | -3.530785435 | 1.68385495 | -2.0968465 | 0.03600716 | 0.31832076 |
| ALCAM | 237.13238 | -1.275146276 | 0.44463577 | -2.8678446 | 0.00413278 | 0.09955609 |
| CBLB | 593.374278 | 0.430099019 | 0.30631631 | 1.40410094 | 0.16028879 | 0.60010648 |
| LINC00882 | 278.960289 | 1.144999901 | 0.38193171 | 2.99791789 | 0.00271831 | 0.0761708 |
| DUBR | 51.5500039 | -0.321238289 | 0.39322236 | -0.816938 | 0.41396385 | 0.82456685 |
| BBX | 1334.83899 | -0.254573228 | 0.30851976 | -0.825144 | 0.40928984 | 0.82124403 |
| CD47 | 250.668021 | -0.741077696 | 0.28234419 | -2.6247315 | 0.00867173 | 0.14729294 |
| IFT57 | 164.458159 | 0.299403092 | 0.24014721 | 1.24674818 | 0.21248985 | 0.66984203 |
| CIP2A | 38.9093563 | 0.539019972 | 0.48959917 | 1.10094135 | 0.27092219 | 0.72343657 |
| DZIP3 | 517.165478 | 0.686178995 | 0.45308516 | 1.51445923 | 0.12990941 | 0.55282113 |
| NECTIN3 | 72.1467905 | 0.085397195 | 0.27394903 | 0.31172658 | 0.75524833 | 0.95067145 |
| PHLDB2 | 909.442911 | 0.485896624 | 0.16505247 | 2.94389194 | 0.00324113 | 0.08621081 |
| ABHD10 | 74.6485749 | 0.537317852 | 0.30609768 | 1.75538033 | 0.07919427 | 0.44935354 |
| C3orf52 | 258.441721 | 0.50948215 | 0.44298413 | 1.15011377 | 0.25009702 | 0.70685456 |
| AC128688.2 | 38.9097092 | 0.394653263 | 0.56987563 | 0.69252525 | 0.48860753 | 0.86014074 |
| SLC9C1 | 214.28726 | -1.870888654 | 1.00663193 | -1.8585628 | 0.06308913 | 0.41264103 |
| ATG3 | 78.8836447 | 0.093620876 | 0.24847042 | 0.37678882 | 0.70633055 | 0.93243538 |
| SLC35A5 | 54.7112079 | -0.222527777 | 0.28858129 | -0.7711095 | 0.44064202 | 0.83733111 |
| GTPBP8 | 41.5745418 | 0.35758108 | 0.38176074 | 0.93666278 | 0.34893205 | 0.78180304 |
| NEPRO | 97.846333 | -0.271362983 | 0.23694275 | -1.1452681 | 0.2520981 | 0.70813507 |
| CFAP44 | 226.185299 | -0.082547895 | 0.22295285 | -0.3702482 | 0.71119756 | 0.9348115 |
| SPICE1 | 167.022215 | 0.021679327 | 0.24983036 | 0.08677619 | 0.93084941 | 0.98822345 |
| SIDT1 | 97.6297134 | -1.540947436 | 0.87932189 | -1.752427 | 0.07970041 | 0.45027508 |
| USF3 | 177.823686 | 0.061025293 | 0.27320723 | 0.22336631 | 0.82325042 | 0.96511681 |
| NAA50 | 504.530113 | 0.388820671 | 0.26016195 | 1.49453322 | 0.1350363 | 0.55891804 |
| ATP6V1A | 822.711293 | 0.023240615 | 0.3244077 | 0.07164015 | 0.94288829 | 0.98885525 |
| GRAMD1C | 523.601705 | 0.188659902 | 0.32276429 | 0.58451293 | 0.55887527 | 0.89179371 |
| ZDHHC23 | 67.3092636 | -0.219878021 | 0.33200591 | -0.6622714 | 0.5077973 | 0.87094607 |
| CCDC191 | 246.689596 | -0.502109075 | 0.32652307 | -1.5377446 | 0.12411108 | 0.54273186 |
| QTRT2 | 90.2197298 | -0.544035107 | 0.36047879 | -1.5092014 | 0.13124732 | 0.55395564 |
| ZBTB20 | 8194.12004 | 0.536039376 | 0.20325731 | 2.63724525 | 0.00835824 | 0.1442148 |
| ZBTB20-AS1 | 41.2293648 | 0.158101681 | 0.53039862 | 0.29808087 | 0.76564144 | 0.95389523 |
| ZBTB20-AS5 | 173.608363 | 0.203908403 | 0.61429881 | 0.33193683 | 0.73993696 | 0.94440864 |
| ZBTB20-AS2 | 39.5617992 | 0.470041483 | 0.57584234 | 0.81626767 | 0.41434706 | 0.82461178 |
| LSAMP | 242.053193 | 0.941104329 | 0.82251934 | 1.14417289 | 0.25255195 | 0.70813507 |
| AC068633.1 | 314.501437 | 0.651906234 | 0.86612235 | 0.75267223 | 0.4516469 | 0.84234337 |
| IGSF11 | 53.0546198 | 0.049964917 | 0.5323339 | 0.09386011 | 0.92522028 | 0.98710257 |
| B4GALT4 | 62.6926045 | -0.211048193 | 0.28328194 | -0.7450111 | 0.456265 | 0.84372683 |

| | | | | | | |
|---|---|---|---|---|---|---|
| TMEM39A | 79.5394177 | -0.369188207 | 0.34263957 | -1.0774827 | 0.28126467 | 0.73086885 |
| POGLUT1 | 57.0919883 | 0.225734099 | 0.33634391 | 0.67114074 | 0.50213088 | 0.86837155 |
| TIMMDC1 | 158.298325 | -0.087669438 | 0.20622133 | -0.425123 | 0.67074699 | 0.92474338 |
| COX17 | 123.900887 | 0.82109618 | 0.33194108 | 2.47362023 | 0.01337518 | 0.18614502 |
| MAATS1 | 174.13772 | 0.299850676 | 0.28297668 | 1.05963034 | 0.28931281 | 0.73754646 |
| GSK3B | 853.848243 | 0.283956056 | 0.25813363 | 1.10003513 | 0.27131682 | 0.72343657 |
| LRRC58 | 152.910253 | -0.446850679 | 0.24893146 | -1.7950751 | 0.07264173 | 0.43315584 |
| FSTL1 | 40.7392558 | -1.250167906 | 0.66586406 | -1.8775122 | 0.06044793 | 0.40556611 |
| NDUFB4 | 194.478777 | -0.337401686 | 0.28730028 | -1.1743869 | 0.24024009 | 0.6963541 |
| HGD | 216.631502 | -0.974337808 | 0.66217476 | -1.4714209 | 0.14117732 | 0.56895751 |
| RABL3 | 98.7207018 | -0.249902807 | 0.22478333 | -1.1117497 | 0.26624578 | 0.72053648 |
| GOLGB1 | 723.28774 | -0.139011075 | 0.19271045 | -0.7213468 | 0.47069614 | 0.85236058 |
| IQCB1 | 235.203742 | -0.174889917 | 0.16105651 | -1.0858916 | 0.27752694 | 0.72788245 |
| SLC15A2 | 47.8278653 | 0.632866392 | 0.57307433 | 1.10433562 | 0.26944758 | 0.72322263 |
| ILDR1 | 59.3482849 | -0.355869088 | 0.31471338 | -1.130772 | 0.25815105 | 0.71534188 |
| CCDC58 | 58.8617298 | -0.175846566 | 0.31766603 | -0.553558 | 0.57988139 | 0.89995373 |
| FAM162A | 439.742829 | -0.857719362 | 0.56772877 | -1.5107907 | 0.13084178 | 0.55395564 |
| AC083798.2 | 96.6984702 | -0.141079454 | 0.43567507 | -0.3238181 | 0.74607579 | 0.94594643 |
| KPNA1 | 337.664831 | 0.370548587 | 0.25184979 | 1.47130789 | 0.14120787 | 0.56895751 |
| PARP9 | 185.977084 | -0.932773469 | 0.64395367 | -1.4485102 | 0.14747441 | 0.57923594 |
| DTX3L | 82.882122 | -1.135642209 | 0.54010215 | -2.1026434 | 0.03549695 | 0.31644624 |
| PARP14 | 555.580294 | -0.306500367 | 0.73134077 | -0.4190938 | 0.6751476 | 0.92656883 |
| HSPBAP1 | 96.5497458 | 0.050547645 | 0.38174019 | 0.13241374 | 0.89465705 | 0.9821637 |
| DIRC2 | 299.439487 | 1.092894269 | 0.61182011 | 1.78630002 | 0.07405069 | 0.43749352 |
| PDIA5 | 43.4959712 | -0.259572228 | 0.39166557 | -0.6627395 | 0.50749743 | 0.87094607 |
| SEC22A | 240.539977 | -0.032813777 | 0.19794801 | -0.1657697 | 0.86833821 | 0.97576204 |
| ADCY5 | 51.0682238 | -0.191934747 | 0.79708656 | -0.2407954 | 0.80971373 | 0.96247146 |
| HACD2 | 161.219346 | 0.284293486 | 0.18168931 | 1.56472324 | 0.11764782 | 0.53282158 |
| MYLK-AS1 | 41.6737738 | -0.00169316 | 0.36353573 | -0.0046575 | 0.99628388 | 0.99916339 |
| MYLK | 39.9581598 | -1.062482967 | 0.62899069 | -1.6891871 | 0.09118358 | 0.47749744 |
| CCDC14 | 266.47929 | 0.115964772 | 0.1561045 | 0.74286629 | 0.45756264 | 0.84409989 |
| KALRN | 473.381185 | -0.916824287 | 0.50707502 | -1.8080644 | 0.07059648 | 0.42987436 |
| UMPS | 58.7645334 | -0.340343348 | 0.28051071 | -1.2132989 | 0.22501555 | 0.68233881 |
| ITGB5 | 330.174136 | -0.256758975 | 0.26868507 | -0.9556131 | 0.33926773 | 0.77836439 |
| SLC12A8 | 70.0875304 | -0.375646534 | 0.58464214 | -0.6425239 | 0.5205331 | 0.87638467 |
| ZNF148 | 933.853488 | 0.1564898 | 0.2256276 | 0.69357562 | 0.48794839 | 0.86006752 |
| SNX4 | 345.687406 | 0.169008352 | 0.19281812 | 0.87651695 | 0.38074907 | 0.80308469 |
| OSBPL11 | 131.827294 | 0.34860118 | 0.24492582 | 1.42329289 | 0.15465127 | 0.59086594 |
| SLC41A3 | 78.8058983 | -0.179543505 | 0.26744982 | -0.6713166 | 0.50201886 | 0.86837155 |
| ALDH1L1 | 140.01634 | -0.974141333 | 0.62627132 | -1.5554622 | 0.11983604 | 0.53734932 |
| ZXDC | 328.46019 | -0.196128185 | 0.22259664 | -0.8810923 | 0.37826786 | 0.80215442 |
| TXNRD3NB | 43.2365244 | -0.247042268 | 0.61736173 | -0.4001581 | 0.68904011 | 0.93036007 |
| TXNRD3 | 53.2336898 | 0.065104934 | 0.39908198 | 0.16313674 | 0.87041077 | 0.97602235 |
| CHCHD6 | 135.64595 | 0.102071115 | 0.24550031 | 0.41576777 | 0.67757994 | 0.9268955 |
| PLXNA1 | 40.4529607 | 0.298060338 | 0.31837712 | 0.93618641 | 0.34917722 | 0.78180304 |
| TPRA1 | 52.2699344 | 0.024357145 | 0.34652011 | 0.07029071 | 0.94396227 | 0.98949015 |
| ABTB1 | 46.1029953 | 0.250933804 | 0.34029227 | 0.73740671 | 0.46087505 | 0.84706604 |
| MGLL | 503.722367 | -1.486096828 | 0.31547961 | -4.7105955 | 2.47E-06 | 0.00071903 |
| SEC61A1 | 190.167253 | -0.213262346 | 0.30974578 | -0.6885077 | 0.49113314 | 0.86143519 |
| RUVBL1 | 129.075215 | 0.023376451 | 0.24791288 | 0.094293 | 0.92487641 | 0.98710257 |
| EEFSEC | 203.781915 | 0.087343839 | 0.2033358 | 0.42955464 | 0.66751964 | 0.92422713 |
| GATA2 | 535.069708 | 0.968649835 | 0.60264584 | 1.60732851 | 0.10798232 | 0.51162458 |
| GATA2-AS1 | 118.489296 | -0.418113888 | 0.53947399 | -0.7750399 | 0.43831606 | 0.8347986 |
| RPN1 | 161.967536 | -0.450096815 | 0.3135886 | -1.4353099 | 0.15119881 | 0.58535779 |
| RAB7A | 654.358429 | -0.184732637 | 0.3326273 | -0.5553742 | 0.57863871 | 0.89972309 |
| ACAD9 | 144.456747 | 0.227583354 | 0.24527612 | 0.92786593 | 0.35347712 | 0.78434051 |
| CNBP | 384.397577 | -0.453371382 | 0.38201975 | -1.1867747 | 0.23531649 | 0.69335782 |
| COPG1 | 100.920077 | -0.169735551 | 0.28458551 | -0.5964308 | 0.5508875 | 0.88916026 |
| HMCES | 103.435353 | -0.14456388 | 0.31698757 | -0.4560554 | 0.64835017 | 0.91768694 |
| H1FX | 236.762812 | -1.120420676 | 0.49089636 | -2.2823976 | 0.02246588 | 0.25166501 |
| MBD4 | 116.027721 | -0.052816037 | 0.20677401 | -0.2554288 | 0.79839193 | 0.96000168 |
| IFT122 | 174.664863 | -0.099598364 | 0.24545711 | -0.4057669 | 0.6849139 | 0.92881914 |
| TMCC1 | 1049.46768 | -0.053723703 | 0.2750272 | -0.1953396 | 0.8451271 | 0.96881318 |
| PIK3R4 | 127.316179 | 0.221383482 | 0.21895499 | 1.01109126 | 0.31197275 | 0.75559844 |
| ATP2C1 | 460.267042 | 0.231378765 | 0.25095146 | 0.92200605 | 0.35652542 | 0.78612469 |
| NEK11 | 461.158184 | -0.417820356 | 0.30585424 | -1.3660767 | 0.17191488 | 0.61884138 |
| NUDT16 | 96.004867 | -0.221889425 | 0.32627873 | -0.680061 | 0.49646586 | 0.86473156 |
| MRPL3 | 147.662451 | 0.219032931 | 0.2497141 | 0.87713481 | 0.38041342 | 0.8029673 |
| DNAJC13 | 333.595932 | -0.157877516 | 0.33111089 | -0.4768116 | 0.63349628 | 0.91415622 |
| UBA5 | 293.844825 | 0.267009514 | 0.33163498 | 0.80513073 | 0.42074424 | 0.82694802 |
| CDV3 | 326.621263 | -0.235400352 | 0.52924646 | -0.444784 | 0.65647586 | 0.92124626 |
| TOPBP1 | 95.79904 | 0.017316294 | 0.22248893 | 0.07782991 | 0.93796335 | 0.98846521 |
| SRPRB | 56.7053482 | 0.303096274 | 0.33577545 | 0.9026755 | 0.36669814 | 0.7935157 |
| RAB6B | 65.4690525 | 0.176628654 | 0.39635599 | 0.44563135 | 0.65586356 | 0.92098622 |
| SLCO2A1 | 118.924191 | -1.854502493 | 0.82455933 | -2.2490832 | 0.02450721 | 0.26518037 |
| RYK | 457.548425 | 0.214465529 | 0.15313253 | 1.40052229 | 0.16135697 | 0.60092341 |
| AMOTL2 | 108.466716 | 0.564898884 | 0.7669527 | 0.73654983 | 0.46139615 | 0.84722486 |
| ANAPC13 | 95.6300981 | -0.132095816 | 0.33242094 | -0.3973751 | 0.69109084 | 0.93073907 |
| CEP63 | 163.682845 | 0.135446721 | 0.29481376 | 0.45943147 | 0.64592435 | 0.91724759 |
| EPHB1 | 147.708789 | 1.479371784 | 0.57968867 | 2.55201085 | 0.01071032 | 0.16452817 |
| PPP2R3A | 2003.57004 | -0.115596227 | 0.15585816 | -0.7416758 | 0.45828377 | 0.84511947 |
| MSL2 | 127.500159 | 0.425078945 | 0.29479703 | 1.44193767 | 0.14931996 | 0.58238674 |
| PCCB | 142.904329 | 0.381615958 | 0.28490058 | 1.33947062 | 0.18041751 | 0.63067897 |
| STAG1 | 1548.34725 | 0.432348885 | 0.25983136 | 1.66395961 | 0.0961205 | 0.48772364 |
| NCK1 | 111.187214 | -0.618604649 | 0.31478165 | -1.9651865 | 0.04939265 | 0.3755208 |
| DZIP1L | 87.2462189 | 0.962891721 | 0.4705079 | 2.04649426 | 0.04070777 | 0.34074872 |
| DBR1 | 49.9006776 | 0.005858197 | 0.28906349 | 0.02026612 | 0.98383108 | 0.99540861 |
| ARMC8 | 349.791477 | 0.040890152 | 0.20224834 | 0.20217793 | 0.83977762 | 0.96833625 |
| NME9 | 71.4514222 | -0.546444685 | 0.36231066 | -1.5082214 | 0.13149788 | 0.55397858 |
| MRAS | 148.315721 | -0.8267336 | 0.32580217 | -2.5375325 | 0.0111637 | 0.16686755 |
| CEP70 | 417.971519 | 0.206597746 | 0.26566304 | 0.77766837 | 0.43676454 | 0.83393108 |
| FAIM | 67.5581237 | -0.036286984 | 0.24075118 | -0.150724 | 0.88019343 | 0.97740892 |
| PIK3CB | 564.927244 | 0.397474178 | 0.28487557 | 1.39525538 | 0.16293884 | 0.60211533 |
| MRPS22 | 42.8153175 | -0.119425565 | 0.29186573 | -0.4091798 | 0.68240771 | 0.92806156 |
| COPB2 | 240.38433 | -0.101461185 | 0.21963696 | -0.4619495 | 0.64411752 | 0.91682456 |
| AC097103.2 | 285.065758 | -0.190273017 | 0.34910358 | -0.5450331 | 0.58573073 | 0.90196444 |
| NMNAT3 | 123.265987 | 0.07989706 | 0.32093206 | 0.24895319 | 0.80339699 | 0.96152281 |
| SLC25A36 | 622.086315 | 0.058926759 | 0.36882275 | 0.15976986 | 0.87306237 | 0.97622753 |
| PXYLP1 | 48.6888607 | -0.253605543 | 0.4055464 | -0.6253429 | 0.53174606 | 0.88045351 |
| ZBTB38 | 440.649123 | -0.568118081 | 0.3881922 | -1.4634969 | 0.14333147 | 0.57310712 |
| RASA2 | 602.460204 | -0.167338842 | 0.19249025 | -0.8693367 | 0.38466297 | 0.80561637 |
| RNF7 | 168.946187 | 0.125933463 | 0.48023708 | 0.26223186 | 0.79314269 | 0.9588344 |
| ATP1B3 | 1792.54498 | 3.312235094 | 1.02498744 | 3.23148847 | 0.00123147 | 0.04560366 |
| TFDP2 | 988.146646 | 0.424078177 | 0.27631044 | 1.53478885 | 0.1248357 | 0.54322641 |
| GK5 | 406.180708 | 0.348552342 | 0.26001622 | 1.3405023 | 0.1800821 | 0.63015077 |
| XRN1 | 713.130047 | 0.047793961 | 0.37953207 | 0.12592865 | 0.8997884 | 0.98312332 |
| ATR | 393.620291 | -0.100771031 | 0.2587095 | -0.3895142 | 0.69689579 | 0.93130502 |
| PLS1 | 215.207227 | 0.280406817 | 0.3460694 | 0.8102618 | 0.41778972 | 0.82580843 |
| TRPC1 | 261.973906 | 0.24953367 | 0.31492187 | 0.79236691 | 0.42814677 | 0.8293089 |
| PCOLCE2 | 36.8071554 | -0.533737024 | 0.50313374 | -1.0608254 | 0.28876928 | 0.73668501 |
| U2SURP | 409.541857 | 0.442607391 | 0.22065416 | 2.00588738 | 0.04486826 | 0.35515061 |
| C3orf58 | 70.9021565 | -0.176084194 | 0.32494813 | -0.541884 | 0.58789841 | 0.90198646 |
| AC107021.1 | 60.9088826 | -0.713850002 | 0.73788948 | -0.9674213 | 0.33333346 | 0.777038 |

| | | | | | | |
|---|---|---|---|---|---|---|
| PLOD2 | 485.896991 | -0.607371308 | 0.44857081 | -1.3540143 | 0.1757318 | 0.62477136 |
| PLSCR4 | 113.131988 | 0.139691696 | 0.28789891 | 0.48521092 | 0.62752673 | 0.91061758 |
| PLSCR2 | 178.87523 | 0.133398033 | 0.32129612 | 0.4151872 | 0.67800487 | 0.9268955 |
| PLSCR1 | 999.639496 | -0.007045315 | 0.32127603 | -0.0219292 | 0.98250446 | 0.99507903 |
| GYG1 | 45.8184346 | -0.50295827 | 0.37341836 | -1.3469029 | 0.17801151 | 0.62729004 |
| HLTF | 119.146979 | 0.161222084 | 0.33995272 | 0.47424854 | 0.63532268 | 0.91454294 |
| HPS3 | 109.442863 | -0.320527595 | 0.25952973 | -1.2350323 | 0.21681848 | 0.67426045 |
| WWTR1 | 1892.07107 | 0.842850508 | 0.53238085 | 1.58317209 | 0.11338225 | 0.52545147 |
| COMMD2 | 96.859152 | -0.324452186 | 0.31525259 | -1.0291817 | 0.30339431 | 0.74953627 |
| ANKUB1 | 152.628288 | -0.263262173 | 0.46468091 | -0.566544 | 0.57102404 | 0.89773478 |
| RNF13 | 323.660443 | -0.10886536 | 0.18861092 | -0.5771954 | 0.56380744 | 0.89295783 |
| TSC22D2 | 633.188532 | 0.056920486 | 0.32129173 | 0.17716138 | 0.85938163 | 0.97244655 |
| SERP1 | 446.050087 | 0.074458658 | 0.40377832 | 0.18440479 | 0.85369591 | 0.97062426 |
| EIF2A | 147.61664 | -0.068762787 | 0.30907565 | -0.2224788 | 0.82394116 | 0.96511681 |
| SELENOT | 223.160178 | 0.19144214 | 0.25327519 | 0.75586613 | 0.44972946 | 0.84122512 |
| SIAH2 | 79.2729491 | 0.609276502 | 0.52872804 | 1.15234385 | 0.2491798 | 0.70622072 |
| MBNL1 | 1345.0721 | 0.432047938 | 0.22478122 | 1.92208202 | 0.05459544 | 0.39095834 |
| ARHGEF26 | 51.9638214 | -0.541472049 | 0.59915556 | -0.9037253 | 0.36614107 | 0.79329304 |
| DHX36 | 406.911818 | 0.157364099 | 0.19666853 | 0.80014886 | 0.42362455 | 0.82774157 |
| PLCH1 | 257.164874 | -1.180375366 | 0.5921099 | -1.9935072 | 0.04620593 | 0.36110228 |
| GMPS | 126.102907 | 0.047756539 | 0.18346617 | 0.26030161 | 0.79463113 | 0.95937941 |
| SSR3 | 142.902723 | -0.154036373 | 0.32918333 | -0.4679349 | 0.63983114 | 0.91541711 |
| TIPARP | 112.739891 | 2.051306991 | 1.02192844 | 2.00729026 | 0.04471877 | 0.35515061 |
| LINC00886 | 132.460621 | 0.358189569 | 0.5038756 | 0.71086904 | 0.47716539 | 0.85452722 |
| LEKR1 | 124.247713 | 0.491134654 | 0.37039556 | 1.32597337 | 0.18484852 | 0.63541313 |
| CCNL1 | 527.480532 | 1.489109563 | 0.52311215 | 2.8466354 | 0.00441839 | 0.10405565 |
| VEPH1 | 682.341096 | 0.846052182 | 0.62485516 | 1.35399726 | 0.17573725 | 0.62477136 |
| PQLC2L | 134.239692 | -0.502476766 | 0.31638933 | -1.5881596 | 0.11225025 | 0.5224487 |
| RSRC1 | 511.074587 | -0.127557791 | 0.21995137 | -0.5799363 | 0.56195756 | 0.89242209 |
| MLF1 | 115.665537 | 0.252791834 | 0.35084057 | 0.72053194 | 0.47119754 | 0.85236058 |
| GFM1 | 145.987817 | 0.350464413 | 0.30355963 | 1.15451586 | 0.24828874 | 0.70482674 |
| RARRES1 | 35.5918188 | -0.616638477 | 0.6122882 | -1.007105 | 0.31388434 | 0.75763653 |
| MFSD1 | 71.9928281 | 0.349086701 | 0.28783313 | 1.21280931 | 0.22520273 | 0.6827089 |
| IQCJ-SCHIP1 | 287.735687 | 1.647757486 | 0.97144804 | 1.69618695 | 0.08985048 | 0.47358503 |
| SMC4 | 143.395036 | 0.736694725 | 0.37416185 | 1.96891994 | 0.04896229 | 0.3746651 |
| KPNA4 | 308.732787 | 0.158101501 | 0.27302823 | 0.5790665 | 0.5625443 | 0.89244086 |
| PPM1L | 77.6731845 | -0.097596967 | 0.34562074 | -0.2823817 | 0.77765086 | 0.95601455 |
| NMD3 | 194.466915 | -0.396747312 | 0.32809423 | -1.209248 | 0.22656756 | 0.68467102 |
| LINC01322 | 50.3940228 | 0.465854707 | 0.38563796 | 1.20801052 | 0.22704322 | 0.68523226 |
| PDCD10 | 229.970312 | 0.16569471 | 0.22939521 | 0.72231112 | 0.47010322 | 0.85232523 |
| GOLIM4 | 239.233905 | 0.975176973 | 0.32433956 | 3.00665441 | 0.0026414 | 0.07481583 |
| MECOM | 23218.6649 | 0.423578888 | 0.38553757 | 1.09867086 | 0.27191166 | 0.72372882 |
| AC074033.1 | 106.595785 | -0.070251814 | 0.64670889 | -0.1086297 | 0.91349618 | 0.98480378 |
| MYNN | 60.7456558 | 0.338076445 | 0.29673579 | 1.13931806 | 0.25457052 | 0.71200935 |
| LRRC34 | 45.4982107 | 0.17863776 | 0.38139376 | 0.46838145 | 0.63951183 | 0.91521016 |
| SEC62 | 695.3026 | 0.664526117 | 0.29603673 | 2.24474211 | 0.02478469 | 0.26640367 |
| GPR160 | 498.268591 | 0.66736612 | 0.27692243 | 2.40993884 | 0.0159552 | 0.20617811 |
| PHC3 | 748.05664 | 0.276773303 | 0.23188218 | 1.19359454 | 0.23263661 | 0.69218101 |
| PRKCI | 588.550795 | 0.284456271 | 0.41650263 | 0.68296392 | 0.49462964 | 0.86389349 |
| SKIL | 317.960408 | 0.645425498 | 0.46207915 | 1.39678559 | 0.16247806 | 0.60211533 |
| RPL22L1 | 40.9793326 | -0.313802975 | 0.45170046 | -0.6947148 | 0.48723408 | 0.85968086 |
| TNIK | 203.876735 | -0.750639623 | 0.7255317 | -1.0346062 | 0.30085284 | 0.7478505 |
| PLD1 | 115.734011 | -0.735717747 | 0.40772361 | -1.8044522 | 0.07116045 | 0.43107604 |
| FNDC3B | 2708.86165 | -0.380083134 | 0.28908778 | -1.3147672 | 0.18858816 | 0.64159008 |
| TNFSF10 | 257.449112 | -2.349154219 | 0.69328881 | -3.3884208 | 0.00070296 | 0.03203067 |
| NCEH1 | 292.686737 | 1.079420723 | 0.51718643 | 2.08710177 | 0.03687894 | 0.32258577 |
| ECT2 | 72.0702911 | -0.020075565 | 0.38378224 | -0.0523098 | 0.95828186 | 0.99119981 |
| NLGN1 | 2841.82305 | -1.891686229 | 0.78888241 | -2.3979318 | 0.01648793 | 0.21098114 |
| NAALADL2 | 4050.16914 | -0.225998764 | 0.32373619 | -0.6980955 | 0.4851175 | 0.85912348 |
| TBL1XR1 | 2153.28742 | -0.486684402 | 0.44090569 | -1.1038288 | 0.26966741 | 0.72333985 |
| LINC00578 | 61.2976523 | -1.338433662 | 0.54299308 | -2.4649184 | 0.01370444 | 0.18912039 |
| ZMAT3 | 139.80389 | 0.062091522 | 0.40157902 | 0.15461844 | 0.87712213 | 0.97640529 |
| PIK3CA | 363.140472 | 0.243282253 | 0.19494446 | 1.24795675 | 0.2120469 | 0.66975892 |
| KCNMB3 | 55.6554431 | 0.033930482 | 0.34017535 | 0.0997441 | 0.92054749 | 0.98581108 |
| ZNF639 | 64.0285892 | 0.007555971 | 0.281252 | 0.02686549 | 0.97856702 | 0.99469867 |
| MFN1 | 407.112891 | 0.543709324 | 0.21997092 | 2.47173275 | 0.013446 | 0.18664115 |
| GNB4 | 51.6246015 | -0.483675321 | 0.37788538 | -1.2799524 | 0.20056186 | 0.65479387 |
| ACTL6A | 73.5939835 | -0.206396109 | 0.35900825 | -0.5749063 | 0.56535467 | 0.89382261 |
| MRPL47 | 77.680458 | -0.195349458 | 0.36780914 | -0.5311164 | 0.59533814 | 0.90323766 |
| NDUFB5 | 211.766083 | 0.168867949 | 0.40478544 | 0.41717891 | 0.67654755 | 0.9268955 |
| TTC14 | 335.113733 | 0.393759562 | 0.16689602 | 2.3593107 | 0.01830892 | 0.22547294 |
| FXR1 | 400.553686 | 0.537957306 | 0.19763797 | 2.72193301 | 0.00649013 | 0.12862701 |
| DNAJC19 | 78.8024999 | 0.334485773 | 0.30437738 | 1.09891798 | 0.27180384 | 0.72372882 |
| ATP11B | 638.322181 | 0.592078224 | 0.33354428 | 1.77511132 | 0.07587951 | 0.4425249 |
| DCUN1D1 | 653.776073 | 1.507397514 | 0.53864094 | 2.79852013 | 0.00513374 | 0.11471549 |
| MCCC1 | 602.499551 | 0.691568942 | 0.37304533 | 1.85384693 | 0.06376106 | 0.41357646 |
| MCF2L2 | 55.547504 | 0.669067303 | 0.63277838 | 1.05734856 | 0.29035254 | 0.73754646 |
| B3GNT5 | 51.0057272 | 0.249878324 | 0.5479524 | 0.45602195 | 0.64837419 | 0.91768694 |
| KLHL24 | 521.478054 | -0.433477396 | 0.29556028 | -1.4666294 | 0.1424769 | 0.57165311 |
| YEATS2 | 325.234883 | -0.881965821 | 0.424135 | -2.079446 | 0.03757638 | 0.32438258 |
| ABCC5 | 583.851903 | -0.679608726 | 0.27711098 | -2.4524785 | 0.01418759 | 0.19284812 |
| EIF2B5 | 66.4421819 | 0.20245478 | 0.2667258 | 0.75903709 | 0.44783037 | 0.84033344 |
| DVL3 | 113.522213 | -0.03216105 | 0.23272605 | -0.1381927 | 0.8900881 | 0.98066897 |
| AP2M1 | 283.693755 | -0.419180294 | 0.3719141 | -1.127089 | 0.25970485 | 0.7168415 |
| ABCF3 | 33.9063009 | 0.304835565 | 0.38845202 | 0.78474445 | 0.43260342 | 0.83162989 |
| PSMD2 | 121.814356 | -0.306350929 | 0.32466627 | -0.9435872 | 0.34538065 | 0.78180304 |
| EIF4G1 | 266.33382 | -0.386493354 | 0.26234938 | -1.473201 | 0.14069686 | 0.56886692 |
| FAM131A | 44.3967777 | 0.901105264 | 0.48331901 | 1.86441097 | 0.06226401 | 0.40987868 |
| POLR2H | 151.55007 | -0.021162015 | 0.2779502 | -0.076136 | 0.9393109 | 0.98854974 |
| MAGEF1 | 82.9656107 | -0.094429112 | 0.36255944 | -0.2604514 | 0.7945156 | 0.95937941 |
| VPS8 | 399.172021 | 0.097574074 | 0.32403054 | 0.30112616 | 0.76331829 | 0.95378284 |
| EHHADH | 164.326176 | 0.250479882 | 0.32790283 | 0.76388447 | 0.44493609 | 0.8392603 |
| MAP3K13 | 465.753609 | 0.195131225 | 0.21902786 | 0.89089683 | 0.37298452 | 0.79681088 |
| TMEM41A | 105.887283 | -0.543356839 | 0.26989422 | -2.0132214 | 0.04409134 | 0.35332084 |
| LIPH | 236.479356 | 0.909589922 | 0.37858775 | 2.40258678 | 0.01627957 | 0.20933731 |
| SENP2 | 205.040112 | 0.608019146 | 0.24674344 | 2.46417555 | 0.01373288 | 0.18912039 |
| IGF2BP2 | 467.407036 | 0.121327363 | 0.27351216 | 0.44359039 | 0.65733876 | 0.92160948 |
| TRA2B | 253.888703 | 0.301650092 | 0.26287015 | 1.14752508 | 0.25116467 | 0.70813507 |
| AC068631.2 | 52.5041984 | -0.798491722 | 0.41360332 | -1.9305738 | 0.05353579 | 0.38854227 |
| EIF4A2 | 379.408106 | 0.020143817 | 0.34464615 | 0.05844782 | 0.95339192 | 0.99060627 |
| ST6GAL1 | 3796.77654 | 0.430734871 | 0.55589486 | 0.77484953 | 0.43842858 | 0.8347986 |
| BCL6 | 223.719663 | -1.038517522 | 0.49517372 | -2.0972791 | 0.03596887 | 0.31832076 |
| LPP | 3264.74055 | 0.161928351 | 0.26522653 | 0.61052849 | 0.54151177 | 0.88410086 |
| P3H2 | 699.366388 | 0.953813288 | 0.36014731 | 2.64839766 | 0.00808743 | 0.14312705 |
| CCDC50 | 282.696068 | 0.444766808 | 0.14890943 | 2.98682764 | 0.00281889 | 0.07794703 |
| FGF12 | 532.005682 | 1.132877202 | 0.91502293 | 1.23808614 | 0.21568412 | 0.67215996 |
| MB21D2 | 91.8596843 | 0.489522186 | 0.48302866 | 1.01344335 | 0.31084844 | 0.75435382 |
| OPA1 | 380.671823 | 0.205366358 | 0.14174304 | 1.44886379 | 0.14737562 | 0.57911382 |
| HES1 | 787.482102 | 0.992139247 | 0.5726378 | 1.7325773 | 0.08317083 | 0.45768271 |
| ATP13A3 | 696.942929 | 1.070528819 | 0.44842527 | 2.3873071 | 0.01697231 | 0.21364736 |
| LINC00884 | 49.8988747 | -0.041574822 | 0.31161065 | -0.1334191 | 0.89386192 | 0.98194215 |
| LSG1 | 87.0287109 | -0.355289693 | 0.30031364 | -1.1830621 | 0.23678453 | 0.69335782 |
| XXYLT1 | 89.1066205 | -0.064485846 | 0.28811753 | -0.2238179 | 0.82289904 | 0.96511681 |

| | | | | | | |
|---|---|---|---|---|---|---|
| ACAP2 | 578.418628 | 0.264516694 | 0.22742246 | 1.16310719 | 0.24478602 | 0.70130056 |
| PPP1R2 | 179.076899 | -0.14425978 | 0.20475099 | -0.7045621 | 0.48108282 | 0.85600365 |
| MUC20-OT1 | 1528.37035 | 0.310326117 | 0.46928436 | 0.66127521 | 0.50843584 | 0.87094607 |
| LINC01983 | 77.2255987 | 1.508500828 | 0.60397856 | 2.49760656 | 0.01250349 | 0.17852391 |
| TNK2 | 44.7853307 | 0.494011621 | 0.33321284 | 1.48257077 | 0.13818849 | 0.56438077 |
| TFRC | 257.798086 | 0.495280577 | 0.60959219 | 0.81247854 | 0.41651709 | 0.82489761 |
| PCYT1A | 108.300455 | -0.257857815 | 0.22204953 | -1.1612626 | 0.24553512 | 0.70154448 |
| UBXN7 | 221.507514 | 0.134782458 | 0.18137567 | 0.74311211 | 0.45741381 | 0.84409989 |
| RNF168 | 185.742924 | 0.285278138 | 0.23632095 | 1.20716397 | 0.227369 | 0.68590303 |
| PIGX | 81.1018111 | -0.008417853 | 0.22054197 | -0.0381689 | 0.96955299 | 0.99334395 |
| PAK2 | 527.016892 | -0.074971965 | 0.18027212 | -0.4158822 | 0.6774962 | 0.9268955 |
| SENP5 | 385.289145 | -0.232408865 | 0.21871107 | -1.0626296 | 0.28794994 | 0.73653438 |
| NCBP2 | 104.392155 | -0.213342079 | 0.32991243 | -0.6466627 | 0.51785024 | 0.87526312 |
| NCBP2-AS2 | 66.3787952 | -0.462732084 | 0.52602329 | -0.8796798 | 0.37903278 | 0.80274043 |
| DLG1 | 1313.92372 | 0.32921288 | 0.24359396 | 1.35148213 | 0.17654104 | 0.62528328 |
| BDH1 | 113.579409 | 0.111470917 | 0.39082375 | 0.28522043 | 0.77547527 | 0.95601455 |
| RUBCN | 108.948713 | 0.171229821 | 0.31759877 | 0.5391388 | 0.58979108 | 0.90233654 |
| FYTTD1 | 154.73654 | 0.57555334 | 0.38461625 | 1.49643532 | 0.13454025 | 0.55868799 |
| LRCH3 | 424.792329 | 0.125137889 | 0.21938122 | 0.57041294 | 0.56839765 | 0.8957783 |
| IQCG | 97.7196401 | -0.350097739 | 0.2975149 | -1.1767402 | 0.23929923 | 0.69526124 |
| RPL35A | 581.553756 | 0.099271306 | 0.43471678 | 0.22835858 | 0.81936749 | 0.96461147 |
| LMLN | 138.382746 | 0.069603794 | 0.27850208 | 0.24992199 | 0.80264768 | 0.96090332 |
| ZNF595 | 84.4382769 | 0.171439155 | 0.25187445 | 0.68065321 | 0.49609095 | 0.86452555 |
| ZNF718 | 261.274616 | 0.027758855 | 0.25777874 | 0.10768481 | 0.91424572 | 0.98480378 |
| ZNF141 | 417.97102 | 2.058415733 | 0.7419037 | 2.77450529 | 0.00552857 | 0.11921698 |
| AC092574.2 | 39.189185 | -0.171786758 | 0.60928414 | -0.2819485 | 0.77798299 | 0.95601455 |
| ZNF721 | 562.80835 | -0.139199459 | 0.28242069 | -0.4928798 | 0.6220975 | 0.91028772 |
| PIGG | 123.554757 | -0.226488638 | 0.20933378 | -1.0819498 | 0.27927484 | 0.72938673 |
| ATP5ME | 141.071402 | 0.152407064 | 0.28438785 | 0.53591272 | 0.59201887 | 0.90233654 |
| PCGF3 | 191.117279 | 0.120160257 | 0.28799868 | 0.41722503 | 0.67651382 | 0.9268955 |
| AC139887.2 | 277.077055 | 0.022720344 | 0.35521011 | 0.06396311 | 0.9489996 | 0.98977092 |
| GAK | 250.996138 | -0.395437695 | 0.30180982 | -1.3102214 | 0.19012094 | 0.64458858 |
| TMEM175 | 37.6877389 | 0.158270141 | 0.33425432 | 0.47350215 | 0.63585497 | 0.91454294 |
| IDUA | 39.2068422 | 0.416388147 | 0.4194813 | 0.99262624 | 0.32089213 | 0.7656989 |
| FGFRL1 | 43.6319602 | -0.92867987 | 0.29828249 | -3.113424 | 0.0018493 | 0.05926813 |
| RNF212 | 46.3197518 | 1.062419098 | 0.52414498 | 2.02695653 | 0.04266686 | 0.3484612 |
| AC092535.3 | 161.598187 | 0.218463101 | 0.44544475 | 0.49043815 | 0.62382389 | 0.91058133 |
| CTBP1-AS | 59.0279201 | 0.452507152 | 0.48595595 | 0.93116908 | 0.35176611 | 0.78314007 |
| CTBP1 | 154.213986 | 0.240414584 | 0.24210473 | 0.99301893 | 0.32070073 | 0.76568446 |
| CTBP1-DT | 35.2714685 | -0.319071114 | 0.32183261 | -0.9914195 | 0.3214808 | 0.7663589 |
| MAEA | 200.280789 | 0.049929835 | 0.23358816 | 0.21375157 | 0.83074079 | 0.96648703 |
| UVSSA | 142.897941 | 0.018346732 | 0.35950571 | 0.05103322 | 0.95929905 | 0.99119981 |
| SLBP | 223.685555 | 0.001797605 | 0.31334057 | 0.0057369 | 0.99542264 | 0.99908534 |
| TMEM129 | 84.0599311 | 0.02334256 | 0.34352862 | 0.06794939 | 0.94582592 | 0.98949015 |
| FGFR3 | 65.6691053 | 0.436146576 | 0.82221668 | 0.53045211 | 0.5957985 | 0.90323766 |
| LETM1 | 93.7125048 | 0.173968844 | 0.31929575 | 0.54485175 | 0.58585549 | 0.90196444 |
| NSD2 | 267.614149 | -0.564772179 | 0.23527493 | -2.4004775 | 0.0163737 | 0.21028962 |
| NELFA | 237.904608 | 0.070493887 | 0.28634877 | 0.24618191 | 0.80554142 | 0.96218371 |
| C4orf48 | 63.8104006 | -0.273669891 | 0.59022299 | -0.463672 | 0.64288274 | 0.91643619 |
| NAT8L | 34.5694601 | 1.264010544 | 0.60672654 | 2.08332826 | 0.03722131 | 0.32359312 |
| POLN | 51.0961623 | 0.74217107 | 0.49700347 | 1.49329153 | 0.13536089 | 0.55916513 |
| MXD4 | 284.459425 | 0.180085646 | 0.30783878 | 0.58499986 | 0.55854781 | 0.89179371 |
| ZFYVE28 | 176.217693 | 0.115306557 | 0.33022655 | 0.34917409 | 0.72695861 | 0.93928323 |
| RNF4 | 190.99752 | 0.005868467 | 0.21974479 | 0.02670583 | 0.97869436 | 0.99469867 |
| FAM193A | 372.960747 | 0.07633097 | 0.29326793 | 0.26027725 | 0.79464992 | 0.95937941 |
| TNIP2 | 85.9068314 | -0.653049583 | 0.37866472 | -1.7246116 | 0.08459749 | 0.46272529 |
| SH3BP2 | 279.118754 | -1.320922445 | 0.32588475 | -4.0533423 | 5.05E-05 | 0.00576513 |
| ADD1 | 572.700364 | 0.146273609 | 0.16305325 | 0.89709101 | 0.36967035 | 0.79532503 |
| MFSD10 | 147.537602 | -0.011913204 | 0.36503413 | -0.0326359 | 0.97396497 | 0.9939318 |
| NOP14-AS1 | 35.3142208 | -0.371338818 | 0.34315197 | -1.082141 | 0.27918987 | 0.72938673 |
| NOP14 | 52.764545 | -0.179103072 | 0.27708071 | -0.6463932 | 0.51802477 | 0.87526312 |
| GRK4 | 65.4881234 | -0.762553333 | 0.35534136 | -2.145974 | 0.03187505 | 0.2995969 |
| HTT | 398.1139 | 0.264821039 | 0.20400505 | 1.29811022 | 0.19424946 | 0.64791037 |
| RGS12 | 125.880037 | -0.361225111 | 0.36599502 | -0.9869673 | 0.32365867 | 0.76937573 |
| LRPAP1 | 386.149957 | 0.038836921 | 0.56649894 | 0.06855604 | 0.94534302 | 0.98949015 |
| ADRA2C | 35.5710033 | 0.167373727 | 0.75533407 | 0.22158901 | 0.82463384 | 0.96511681 |
| TMEM128 | 62.0561209 | 0.348891212 | 0.26377362 | 1.32269183 | 0.18593788 | 0.63659883 |
| STX18 | 431.084186 | -0.020511117 | 0.30111894 | -0.0681163 | 0.94569303 | 0.98949015 |
| STX18-AS1 | 251.62187 | -0.342272503 | 0.35727156 | -0.9580178 | 0.3380538 | 0.77836439 |
| EVC2 | 138.682474 | -0.327665742 | 0.28289574 | -1.1582562 | 0.24675951 | 0.70292817 |
| EVC | 171.615528 | 0.1967307 | 0.43094522 | 0.45650977 | 0.64802345 | 0.91768694 |
| CRMP1 | 54.5656984 | -0.186226286 | 0.64830477 | -0.2872511 | 0.77392003 | 0.95577208 |
| WFS1 | 54.6943127 | 0.20412157 | 0.46886519 | 0.43535236 | 0.66330668 | 0.9229791 |
| MAN2B2 | 94.2693531 | -0.396374122 | 0.36751614 | -1.0785217 | 0.28080101 | 0.73022198 |
| MRFAP1 | 291.336835 | 0.071710881 | 0.47030249 | 0.15247821 | 0.87880978 | 0.97692415 |
| LINC02482 | 32.1810845 | 0.028337456 | 0.33909548 | 0.08356778 | 0.93340009 | 0.98822345 |
| AC093323.1 | 31.1045093 | -0.271388674 | 0.40676761 | -0.6671836 | 0.50465487 | 0.86946722 |
| MRFAP1L1 | 69.4583647 | -0.175180112 | 0.30528157 | -0.5738313 | 0.56608198 | 0.89399457 |
| BLOC1S4 | 54.9223479 | -0.900220661 | 0.39496692 | -2.2792306 | 0.02265336 | 0.25189387 |
| KIAA0232 | 474.592747 | 0.146875421 | 0.17322612 | 0.84788265 | 0.39650333 | 0.81238609 |
| TBC1D14 | 205.961508 | -0.365741265 | 0.23226546 | -1.5746692 | 0.11533282 | 0.52873491 |
| TADA2B | 54.2730883 | -0.054572677 | 0.30294115 | -0.1801428 | 0.85704044 | 0.97193497 |
| GRPEL1 | 64.4809411 | -0.915872811 | 0.41191101 | -2.2234725 | 0.02618395 | 0.27550988 |
| SORCS2 | 695.497825 | 0.123034657 | 0.42170046 | 0.29175841 | 0.77047134 | 0.95482817 |
| AFAP1 | 258.245794 | 0.053779903 | 0.62227896 | 0.08642411 | 0.93112928 | 0.98822345 |
| SH3TC1 | 302.922198 | 0.031506484 | 0.48484138 | 0.06498308 | 0.94818748 | 0.98977092 |
| ACOX3 | 95.8474457 | 0.106057143 | 0.23034331 | 0.46043074 | 0.64520707 | 0.91685018 |
| TRMT44 | 76.6782427 | 0.503903294 | 0.26966777 | 1.86860777 | 0.0616774 | 0.40825577 |
| WDR1 | 335.163854 | -0.020785728 | 0.28140684 | -0.0738636 | 0.9411189 | 0.98876195 |
| ZNF518B | 51.8921747 | 0.020964322 | 0.27662182 | 0.07578694 | 0.9395886 | 0.98855185 |
| HS3ST1 | 101.772771 | -0.611301941 | 0.76158853 | -0.8026669 | 0.42216726 | 0.82743836 |
| AC005699.1 | 92.8786962 | -0.903841136 | 0.70737199 | -1.2777452 | 0.2013393 | 0.65580391 |
| RAB28 | 464.212268 | 0.441065261 | 0.20924521 | 2.10788698 | 0.03504077 | 0.31514522 |
| BOD1L1 | 344.220821 | -0.307070961 | 0.22371086 | -1.3726243 | 0.16986917 | 0.61569949 |
| LINC01182 | 55.8734586 | -1.771808328 | 0.68068283 | -2.6029866 | 0.00924156 | 0.15397695 |
| CPEB2 | 377.306177 | 0.362246146 | 0.35684339 | 1.01514042 | 0.3100389 | 0.75409553 |
| AC098829.1 | 148.939748 | -0.52702284 | 0.67217281 | -0.7840585 | 0.43300576 | 0.8317266 |
| CC2D2A | 180.233021 | 0.220860659 | 0.40628218 | 0.54361394 | 0.58670717 | 0.90196444 |
| FBXL5 | 381.511418 | 0.33193758 | 0.18624922 | 1.78222269 | 0.07471292 | 0.43878029 |
| FAM200B | 125.688543 | -0.008050743 | 0.26860767 | -0.0299721 | 0.97608928 | 0.99420893 |
| PROM1 | 492.301186 | -0.171301575 | 0.65296488 | -0.2623442 | 0.79305606 | 0.9588344 |
| TAPT1 | 258.201567 | 0.373827164 | 0.24139187 | 1.54863198 | 0.12147021 | 0.53989034 |
| TAPT1-AS1 | 449.734367 | 0.345619591 | 0.48214871 | 0.71683193 | 0.47347781 | 0.85310957 |
| AC097515.1 | 52.6267762 | -1.26741871 | 0.87507344 | -1.448357 | 0.14751725 | 0.57923594 |
| QDPR | 74.9284889 | 0.293361904 | 0.33349545 | 0.8796579 | 0.37904466 | 0.80274043 |
| LAP3 | 105.625351 | 0.045390305 | 0.74916834 | 0.06058759 | 0.95168766 | 0.99033727 |
| MED28 | 167.492336 | 0.191984995 | 0.16581582 | 1.15782071 | 0.24693721 | 0.70304318 |
| DCAF16 | 149.736628 | 0.224824263 | 0.21136707 | 1.06366739 | 0.2874794 | 0.73650262 |
| LCORL | 732.338182 | -0.308299877 | 0.29171874 | -1.0568395 | 0.29058486 | 0.73754646 |
| AC024230.1 | 109.54322 | 0.785897317 | 0.48305591 | 1.62692829 | 0.10375233 | 0.50269275 |
| SLIT2 | 242.554833 | 0.329699077 | 0.63765048 | 0.51705297 | 0.60511918 | 0.90428025 |
| PACRGL | 175.303546 | 0.056963472 | 0.27786768 | 0.20500215 | 0.83757044 | 0.96757829 |
| KCNIP4 | 12670.1722 | -2.573659424 | 0.68568172 | -3.7534316 | 0.00017443 | 0.01269464 |

| | | | | | | |
|---|---|---|---|---|---|---|
| ADGRA3 | 257.734824 | -0.076944556 | 0.44256325 | -0.1738612 | 0.86197458 | 0.9739172 |
| PPARGC1A | 1609.08413 | -0.235784647 | 0.4101337 | -0.574897 | 0.56536093 | 0.89382261 |
| DHX15 | 414.328038 | 0.136534689 | 0.26958337 | 0.50646555 | 0.61252987 | 0.90714827 |
| CCDC149 | 70.5042669 | -0.243203564 | 0.41295273 | -0.588938 | 0.55590286 | 0.89081623 |
| SEPSECS | 90.4122689 | 0.127608286 | 0.29004493 | 0.43996041 | 0.65996578 | 0.92284375 |
| ZCCHC4 | 54.9854381 | 0.371926042 | 0.32641347 | 1.13943228 | 0.2545229 | 0.71200935 |
| ANAPC4 | 145.637437 | 0.000730433 | 0.20420868 | 0.00357689 | 0.99714606 | 0.99938948 |
| SMIM20 | 70.0156908 | -0.175587544 | 0.29865783 | -0.5879221 | 0.55658457 | 0.89094338 |
| RBPJ | 499.248028 | -0.339038094 | 0.26127255 | -1.2976415 | 0.19441056 | 0.64808107 |
| TBC1D19 | 389.069143 | 0.255707537 | 0.24608072 | 1.03912058 | 0.29874866 | 0.74594723 |
| STIM2 | 1244.48145 | -1.267128377 | 0.56474754 | -2.2437077 | 0.02485122 | 0.26654723 |
| PCDH7 | 969.123822 | 0.202205947 | 0.65302127 | 0.3096468 | 0.75682956 | 0.95080003 |
| ARAP2 | 355.230538 | 0.044371831 | 0.36195602 | 0.12258901 | 0.90243255 | 0.98320957 |
| C4orf19 | 1632.27974 | -0.123212198 | 0.34230211 | -0.3599516 | 0.71888331 | 0.9359691 |
| RELL1 | 147.517649 | -1.042062671 | 0.40693962 | -2.5607304 | 0.01044524 | 0.16338221 |
| PGM2 | 84.2001998 | 0.19041638 | 0.4157122 | 0.45804858 | 0.64691754 | 0.91743182 |
| TBC1D1 | 1360.20325 | 0.459589897 | 0.23334835 | 1.96954426 | 0.04889063 | 0.3746651 |
| KLF3 | 323.084495 | -0.099607611 | 0.48448545 | -0.2055946 | 0.83710757 | 0.96757829 |
| FAM114A1 | 73.1830723 | -1.12415236 | 0.44181156 | -2.5444159 | 0.01094607 | 0.16452817 |
| KLHL5 | 320.641601 | 0.104184645 | 0.42166245 | 0.24708068 | 0.80484579 | 0.96210606 |
| WDR19 | 223.031034 | -0.075514371 | 0.25858289 | -0.2920316 | 0.77026247 | 0.95482817 |
| RFC1 | 253.392649 | 0.594967695 | 0.24286052 | 2.4498329 | 0.01429225 | 0.19351784 |
| RPL9 | 56.6943847 | 0.368418863 | 0.27991757 | 1.31616911 | 0.18811729 | 0.64091976 |
| LIAS | 78.1125039 | 0.185557745 | 0.34353866 | 0.54013642 | 0.58910296 | 0.90233654 |
| UGDH | 127.872348 | 0.12802819 | 0.46038607 | 0.27808876 | 0.78094423 | 0.9566945 |
| UGDH-AS1 | 158.559653 | -0.083968098 | 0.30613274 | -0.2742866 | 0.7838644 | 0.9568898 |
| SMIM14 | 600.345693 | 0.390694357 | 0.20521919 | 1.90379059 | 0.05693746 | 0.39727338 |
| UBE2K | 1018.39714 | 0.303651154 | 0.2646941 | 1.14717765 | 0.25130821 | 0.70813507 |
| PDS5A | 740.904687 | 0.234105571 | 0.22360323 | 1.04696864 | 0.29511404 | 0.74132195 |
| N4BP2 | 147.859391 | 0.323904262 | 0.29674537 | 1.09152255 | 0.27504301 | 0.7263609 |
| RBM47 | 2968.33198 | 3.39E-05 | 0.48864019 | 6.95E-05 | 0.99994458 | 0.99994458 |
| NSUN7 | 228.401138 | -0.511825929 | 0.27303453 | -1.8745832 | 0.06085009 | 0.40676596 |
| APBB2 | 1059.97438 | 0.856415357 | 0.19641055 | 4.3603328 | 1.30E-05 | 0.00230675 |
| UCHL1 | 280.37801 | -0.877022719 | 0.45763417 | -1.9164275 | 0.0553107 | 0.39343117 |
| LIMCH1 | 2683.32324 | 0.905292805 | 0.54468361 | 1.6620526 | 0.09650223 | 0.48810007 |
| TMEM33 | 169.629901 | 0.041860472 | 0.29093207 | 0.14388401 | 0.88559207 | 0.97931886 |
| SLC30A9 | 608.531156 | 0.533398931 | 0.14789271 | 3.60666134 | 0.00031016 | 0.01805832 |
| ATP8A1 | 251.975413 | 0.429175894 | 0.27224795 | 1.57641555 | 0.11493007 | 0.52758086 |
| KCTD8 | 1255.41717 | 0.765274043 | 0.46456271 | 1.64729977 | 0.09949644 | 0.49465025 |
| GUF1 | 79.0353377 | 0.205169155 | 0.24283501 | 0.84489116 | 0.39817161 | 0.81400298 |
| GNPDA2 | 77.5392657 | 0.044529393 | 0.26190818 | 0.1700191 | 0.86499511 | 0.97485201 |
| GABRA2 | 313.043536 | -0.143416387 | 0.5867577 | -0.2444218 | 0.80690414 | 0.96218371 |
| COMMD8 | 55.042733 | -0.143842219 | 0.49659776 | -0.2896554 | 0.77207989 | 0.95537275 |
| ATP10D | 261.199411 | 0.532685063 | 0.25015863 | 2.12938913 | 0.03322208 | 0.30784031 |
| AC107068.2 | 78.5005087 | -0.434684524 | 0.54402296 | -0.7990187 | 0.42427956 | 0.82774157 |
| NFXL1 | 248.183906 | 0.68083805 | 0.38775854 | 1.75582994 | 0.07911745 | 0.44916083 |
| NIPAL1 | 363.997679 | 0.804503977 | 0.41364781 | 1.94490087 | 0.05178691 | 0.38247133 |
| CNGA1 | 419.782641 | 0.414145297 | 0.41144182 | 1.00657075 | 0.3141411 | 0.75804713 |
| TXK | 67.9199489 | -0.850698257 | 0.54809916 | -1.5520882 | 0.12064112 | 0.53823708 |
| TEC | 163.162941 | -0.647262493 | 0.31442603 | -2.0585525 | 0.03953713 | 0.3346923 |
| SLAIN2 | 288.515351 | 0.304364527 | 0.14723861 | 2.06715153 | 0.03871988 | 0.33071257 |
| FRYL | 819.892136 | -0.060447211 | 0.24407296 | -0.2476604 | 0.80439716 | 0.96210606 |
| OCIAD1 | 493.955522 | 0.174588785 | 0.15625306 | 1.11734632 | 0.26384628 | 0.71989632 |
| OCIAD2 | 109.378661 | -0.579299474 | 0.39024394 | -1.4844548 | 0.13768832 | 0.56434148 |
| CWH43 | 142.508055 | 0.96477849 | 0.54774455 | 1.76136575 | 0.07817652 | 0.44739666 |
| DCUN1D4 | 244.444358 | -0.462355148 | 0.28439596 | -1.6257444 | 0.10400403 | 0.50291791 |
| SGCB | 60.8151155 | -0.150892318 | 0.43061607 | -0.3504103 | 0.72603078 | 0.93895336 |
| USP46 | 119.762349 | 0.696545557 | 0.22461475 | 3.10106771 | 0.00192824 | 0.06142242 |
| ERVMER34-1 | 47.7409533 | 1.517492966 | 0.5743625 | 2.64204744 | 0.00824065 | 0.14386723 |
| RASL11B | 80.017932 | 2.181354392 | 0.55003523 | 3.9658449 | 7.31E-05 | 0.00734782 |
| SCFD2 | 253.665713 | -0.24033827 | 0.20006479 | -1.2013022 | 0.22963401 | 0.68857352 |
| AC058822.1 | 168.619421 | -0.289012752 | 0.49492651 | -0.5839508 | 0.55925339 | 0.89179371 |
| LNX1 | 249.770152 | -0.179391253 | 0.41015622 | -0.437373 | 0.66184086 | 0.92284375 |
| CHIC2 | 245.791167 | 0.164983544 | 0.22548861 | 0.73167131 | 0.46436921 | 0.84976241 |
| TMEM165 | 475.793199 | -0.021968156 | 0.19254782 | -0.1140919 | 0.90916491 | 0.98441374 |
| CLOCK | 312.189695 | 0.207430535 | 0.19689678 | 1.05349888 | 0.29211241 | 0.73844969 |
| EXOC1 | 337.443645 | -0.007865111 | 0.22926984 | -0.034305 | 0.97263391 | 0.9939318 |
| CEP135 | 125.70019 | 0.178667928 | 0.24839872 | 0.71927878 | 0.47196917 | 0.85244471 |
| AASDH | 157.049245 | 0.25304467 | 0.27679125 | 0.91420763 | 0.36060775 | 0.78927419 |
| PPAT | 41.5734804 | 0.163783444 | 0.40305384 | 0.40635624 | 0.68448087 | 0.92881914 |
| PAICS | 91.1888714 | -0.260938241 | 0.46330462 | -0.563211 | 0.57329124 | 0.89797476 |
| SRP72 | 154.699345 | -0.057898317 | 0.19894257 | -0.2910303 | 0.77102814 | 0.95498333 |
| REST | 270.86016 | -0.470654537 | 0.2209494 | -2.1301462 | 0.03315954 | 0.30780515 |
| NOA1 | 40.893993 | -0.154238331 | 0.36284293 | -0.4250829 | 0.67077623 | 0.92474338 |
| POLR2B | 213.549656 | 0.262351048 | 0.27601512 | 0.95049521 | 0.34186069 | 0.77973697 |
| IGFBP7 | 2812.04252 | 0.112095686 | 0.38379942 | 0.29206841 | 0.77023432 | 0.95482817 |
| AC104806.2 | 38.6296114 | -0.981604606 | 0.49173711 | -1.9961979 | 0.04591238 | 0.3604208 |
| CENPC | 283.753693 | 0.648995607 | 0.31203445 | 2.07988447 | 0.03753613 | 0.32430227 |
| UBA6 | 451.992439 | 0.091328401 | 0.38674365 | 0.23614713 | 0.81331851 | 0.96288868 |
| UBA6-AS1 | 624.846624 | -0.644561157 | 0.30946749 | -2.0828073 | 0.03726879 | 0.32359312 |
| YTHDC1 | 603.796195 | 0.389500514 | 0.19231768 | 2.0252975 | 0.04283683 | 0.3484612 |
| UGT2B7 | 239.099374 | -1.015201539 | 0.58374686 | -1.7391126 | 0.08201495 | 0.45535449 |
| UTP3 | 43.8981976 | -0.964824628 | 0.3636744 | -2.6529902 | 0.00797822 | 0.14172148 |
| RUFY3 | 825.528616 | -0.346354563 | 0.34795129 | -0.9954111 | 0.31953638 | 0.76402949 |
| GRSF1 | 206.296901 | 0.136925294 | 0.22046745 | 0.62106806 | 0.53455484 | 0.88137638 |
| MOB1B | 1331.46939 | 0.901678177 | 0.35538933 | 2.537156 | 0.01117571 | 0.16686755 |
| DCK | 39.5464274 | 0.0372524 | 0.35890718 | 0.10379397 | 0.91733285 | 0.98520052 |
| SLC4A4 | 1661.00365 | 0.950280488 | 0.44647305 | 2.12841624 | 0.03330259 | 0.30804159 |
| ADAMTS3 | 339.179326 | 0.141877568 | 0.57834421 | 0.24531683 | 0.80621112 | 0.96218371 |
| COX18 | 45.6474577 | 0.12995662 | 0.40236976 | 0.3229781 | 0.74671183 | 0.94597987 |
| ANKRD17 | 1583.84071 | 0.374566051 | 0.22668815 | 1.65234069 | 0.09846512 | 0.49303127 |
| AC053527.1 | 226.72021 | -0.066661057 | 0.65873436 | -0.1011957 | 0.91939514 | 0.98520052 |
| RASSF6 | 273.72114 | -0.714168523 | 0.43207918 | -1.6528649 | 0.09835836 | 0.49296777 |
| CXCL2 | 106.544056 | 0.993052469 | 1.36403508 | 0.72802561 | 0.46659791 | 0.85057334 |
| MTHFD2L | 238.519848 | -1.147081776 | 0.46135611 | -2.4863262 | 0.01290696 | 0.18229777 |
| BTC | 108.949713 | 0.468992878 | 0.35984669 | 1.30331302 | 0.19246792 | 0.64546041 |
| PARM1 | 79.7177485 | 0.736596324 | 0.38988624 | 1.88925963 | 0.05885705 | 0.40079394 |
| RCHY1 | 138.620358 | 0.440827732 | 0.2681005 | 1.64426302 | 0.10012189 | 0.49569046 |
| THAP6 | 86.785795 | -0.159518747 | 0.27097828 | -0.5886772 | 0.5560778 | 0.89081623 |
| CDKL2 | 206.909709 | 0.545513312 | 0.2030801 | 2.68619781 | 0.00722703 | 0.13573345 |
| G3BP2 | 306.685011 | 0.33177364 | 0.30334183 | 1.09372861 | 0.27407402 | 0.72590691 |
| USO1 | 383.31823 | -5.18E-05 | 0.20334779 | -0.0002547 | 0.99979675 | 0.99993938 |
| NAAA | 48.3102568 | -0.162323815 | 0.63531586 | -0.255501 | 0.7983362 | 0.96000168 |
| SDAD1 | 130.264116 | 0.152499331 | 0.28562863 | 0.53390772 | 0.59340538 | 0.90277205 |
| NUP54 | 181.503231 | 0.403497394 | 0.19367499 | 2.08337376 | 0.03721717 | 0.32359312 |
| SCARB2 | 482.07626 | -0.032331029 | 0.25436816 | -0.1271033 | 0.89885865 | 0.98312332 |
| SHROOM3 | 1324.83118 | 0.073101025 | 0.49568707 | 0.14747414 | 0.88275779 | 0.97866303 |
| 11-Sep | 905.795268 | 0.539089837 | 0.27175876 | 1.98370727 | 0.04728848 | 0.36682701 |
| CCNI | 862.16023 | -0.171288205 | 0.22792507 | -0.751511 | 0.45234516 | 0.84234337 |
| CCNG2 | 428.738863 | -0.858331608 | 0.36796565 | -2.3326406 | 0.01966702 | 0.23555469 |
| CNOT6L | 541.216256 | 0.802360877 | 0.18664944 | 4.29875866 | 1.72E-05 | 0.0027273 |
| MRPL1 | 234.244864 | 0.667725852 | 0.16755022 | 3.98522806 | 6.74E-05 | 0.00713649 |
| FRAS1 | 3774.90282 | 1.370047909 | 0.42359553 | 3.23433045 | 0.00121928 | 0.04531236 |

| | | | | | | |
|---|---|---|---|---|---|---|
| ANXA3 | 201.39205 | -0.044423853 | 0.46838192 | -0.0948454 | 0.92443766 | 0.98707149 |
| BMP2K | 198.841313 | -0.221222801 | 0.42827049 | -0.5165492 | 0.60547085 | 0.90428025 |
| PAQR3 | 61.7727664 | 0.005095416 | 0.33941428 | 0.01501238 | 0.9880223 | 0.9964791 |
| CFAP299 | 64.2676119 | -0.441672036 | 0.73652555 | -0.5996697 | 0.54872641 | 0.88854099 |
| RASGEF1B | 1445.19302 | 1.289365017 | 0.64230401 | 2.00740615 | 0.04470643 | 0.35515061 |
| HNRNPD | 455.082227 | -0.05687797 | 0.1722647 | -0.3301777 | 0.74126566 | 0.94487568 |
| HNRNPDL | 473.794923 | 0.069810609 | 0.26057493 | 0.26790992 | 0.78876866 | 0.95801395 |
| ENOPH1 | 88.6179764 | -0.604140369 | 0.2410016 | -2.5067898 | 0.01218331 | 0.17590979 |
| TMEM150C | 66.4110228 | -0.880058502 | 0.57382139 | -1.5336802 | 0.12510835 | 0.54337567 |
| SCD5 | 93.7317518 | -1.413754917 | 0.80391224 | -1.7585936 | 0.07864656 | 0.44843088 |
| SEC31A | 564.375087 | 0.004374962 | 0.22920665 | 0.01908741 | 0.98477137 | 0.9954565 |
| LIN54 | 89.4566084 | -0.200194987 | 0.1984181 | -1.0089553 | 0.31299609 | 0.75694198 |
| COPS4 | 120.772014 | 0.327645675 | 0.19103669 | 1.71509291 | 0.08632819 | 0.46616751 |
| HELQ | 109.535835 | 0.044954804 | 0.32157893 | 0.139794 | 0.88882276 | 0.98051184 |
| MRPS18C | 58.4823332 | -0.266141876 | 0.37196825 | -0.7154962 | 0.47430247 | 0.85363042 |
| ABRAXAS1 | 72.9695367 | 0.347311552 | 0.28651015 | 1.21221378 | 0.22543056 | 0.68280317 |
| GPAT3 | 200.875855 | 2.978605542 | 0.77285309 | 3.8540385 | 0.00011619 | 0.00966367 |
| CDS1 | 268.256728 | 0.452017792 | 0.14336621 | 3.15288935 | 0.00161663 | 0.05465255 |
| WDFY3 | 2075.62131 | 0.682674508 | 0.24418187 | 2.7957625 | 0.00517774 | 0.11481456 |
| ARHGAP24 | 807.61499 | 0.367924286 | 0.56403929 | 0.65230258 | 0.514206 | 0.87226916 |
| MAPK10 | 3160.16107 | 0.642663819 | 0.30434542 | 2.11162638 | 0.03471851 | 0.31514522 |
| AC104827.1 | 67.4222003 | -0.094862486 | 0.587402 | -0.161495 | 0.87170355 | 0.97622753 |
| PTPN13 | 2829.25022 | 0.86171477 | 0.33237896 | 2.59256714 | 0.00952626 | 0.15599248 |
| AC093827.5 | 89.0073817 | -0.278966911 | 0.30839176 | -0.9045861 | 0.36568469 | 0.79329304 |
| AFF1 | 4713.87615 | 0.284832473 | 0.250648 | 1.13638439 | 0.25579573 | 0.71330223 |
| KLHL8 | 455.006131 | 0.013382991 | 0.3114141 | 0.0429749 | 0.96572154 | 0.99271817 |
| HSD17B11 | 95.3707051 | -0.116702458 | 0.44264358 | -0.2636488 | 0.79205054 | 0.95839852 |
| NUDT9 | 165.219588 | 0.030731103 | 0.21438794 | 0.14334343 | 0.88601896 | 0.9795842 |
| SPP1 | 2952.15867 | -3.68482569 | 0.7668921 | -4.8048816 | 1.55E-06 | 0.00049174 |
| PKD2 | 1034.34026 | 0.658026053 | 0.33319414 | 1.97490284 | 0.04827918 | 0.37148734 |
| PPM1K | 367.400585 | 1.006717031 | 0.45117315 | 2.23133188 | 0.02565915 | 0.27225313 |
| HERC6 | 132.933495 | -0.22751486 | 0.84342318 | -0.2697517 | 0.78735126 | 0.95791052 |
| HERC5 | 97.9852641 | -0.869608686 | 0.74709712 | -1.1639835 | 0.24443072 | 0.70121778 |
| PYURF | 101.670435 | -0.115403415 | 0.50154288 | -0.2300968 | 0.81801655 | 0.96404074 |
| HERC3 | 324.436429 | -0.462657329 | 0.36268982 | -1.2756281 | 0.20208701 | 0.65670446 |
| FAM13A-AS1 | 95.6123953 | -0.427107127 | 0.25847547 | -1.6524088 | 0.09845125 | 0.49303127 |
| FAM13A | 750.652624 | -0.330686364 | 0.61001922 | -0.5420917 | 0.58775533 | 0.90198646 |
| GPRIN3 | 420.398449 | -0.814460556 | 0.61025543 | -1.3346224 | 0.18199997 | 0.63239126 |
| CCSER1 | 7766.5275 | 0.565728652 | 0.35795972 | 1.58042546 | 0.11400946 | 0.52658404 |
| SMARCAD1 | 307.118403 | 0.244144889 | 0.18034302 | 1.35378065 | 0.17580637 | 0.62477136 |
| PDLIM5 | 664.675028 | 0.055375951 | 0.23384437 | 0.23680686 | 0.81280663 | 0.96288868 |
| BMPR1B | 4439.78201 | 0.997846086 | 0.39921189 | 2.49954001 | 0.01243547 | 0.17817202 |
| STPG2 | 539.910632 | -0.08746195 | 0.41909404 | -0.2086929 | 0.83468798 | 0.96749268 |
| RAP1GDS1 | 511.082348 | -0.124374132 | 0.22310553 | -0.5574677 | 0.57720791 | 0.89873672 |
| TSPAN5 | 1213.06577 | -0.387836137 | 0.62183307 | -0.6236982 | 0.53282584 | 0.88114359 |
| EIF4E | 252.451048 | 0.321081627 | 0.25695586 | 1.24955948 | 0.21146051 | 0.66975892 |
| METAP1 | 211.340772 | -0.020308278 | 0.25509781 | -0.0796098 | 0.93654762 | 0.98822345 |
| ADH5 | 249.702516 | -0.069067459 | 0.33732 | -0.2047535 | 0.8377647 | 0.96757829 |
| AP002026.1 | 52.4829088 | -0.18350822 | 0.7055634 | -0.2600875 | 0.79479628 | 0.95937941 |
| TRMT10A | 44.3632353 | -0.040848511 | 0.36707263 | -0.1112818 | 0.91139287 | 0.98468493 |
| AC083902.1 | 46.9785215 | 0.950480725 | 0.76267965 | 1.24623848 | 0.21267685 | 0.6699289 |
| C4orf54 | 93.6887533 | 1.47633086 | 0.72797948 | 2.02798416 | 0.04256186 | 0.3484612 |
| LAMTOR3 | 121.975953 | -0.084792596 | 0.26664731 | -0.3179953 | 0.75048849 | 0.94794737 |
| DNAJB14 | 382.669642 | 0.227787169 | 0.1574504 | 1.44672339 | 0.14797441 | 0.57952643 |
| H2AFZ | 245.327374 | -0.289719205 | 0.64996252 | -0.4457476 | 0.65577961 | 0.92098622 |
| PPP3CA | 513.103682 | -0.13360861 | 0.29289706 | -0.4561623 | 0.64827324 | 0.91768694 |
| BANK1 | 157.121493 | -0.549601512 | 0.44359786 | -1.2389634 | 0.21535905 | 0.67151527 |
| AP002075.1 | 49.8550219 | 0.194447879 | 0.71164552 | 0.27323699 | 0.78467105 | 0.95702626 |
| SLC39A8 | 57.9277882 | 2.278707531 | 0.52021611 | 4.38030944 | 1.19E-05 | 0.00221785 |
| NFKB1 | 337.434426 | 0.21616611 | 0.39607131 | 0.54577573 | 0.58522011 | 0.90173902 |
| MANBA | 238.260418 | -0.190023915 | 0.30539792 | -0.6222175 | 0.53379889 | 0.88137638 |
| UBE2D3 | 949.916326 | 0.238600669 | 0.18907665 | 1.26192563 | 0.20697555 | 0.66434881 |
| CISD2 | 113.440855 | -0.424092427 | 0.27237694 | -1.5570056 | 0.11946915 | 0.53689397 |
| SLC9B1 | 44.9203267 | -0.76724722 | 0.62052081 | -1.2364569 | 0.21628878 | 0.67323263 |
| SLC9B2 | 35.3077294 | 0.343112609 | 0.27178972 | 1.26241937 | 0.20679792 | 0.66434881 |
| TET2 | 682.978307 | 0.464111346 | 0.38486766 | 1.20589854 | 0.22785662 | 0.68629723 |
| PPA2 | 616.4955 | 0.17747158 | 0.2003546 | 0.8857874 | 0.3757321 | 0.79926035 |
| ARHGEF38 | 911.09103 | -0.950938638 | 0.49161508 | -1.9343154 | 0.05307436 | 0.3871585 |
| INTS12 | 168.617419 | -0.099582993 | 0.28995591 | -0.3434419 | 0.73126608 | 0.94055824 |
| GSTCD | 271.086832 | 0.010491643 | 0.26442134 | 0.03967775 | 0.96835004 | 0.99303855 |
| NPNT | 617.249906 | -0.165696815 | 0.45422797 | -0.3647878 | 0.71526983 | 0.93573028 |
| TBCK | 889.393832 | 0.275599448 | 0.31307477 | 0.88029913 | 0.37869728 | 0.80274043 |
| AIMP1 | 156.189122 | -0.395335131 | 0.22506336 | -1.7565504 | 0.07899447 | 0.44916083 |
| PAPSS1 | 445.212601 | 0.781584759 | 0.1973337 | 3.96072622 | 7.47E-05 | 0.00738763 |
| SGMS2 | 158.497805 | 0.031148783 | 0.30354403 | 0.10261702 | 0.91826693 | 0.98520052 |
| HADH | 100.534293 | -0.050333654 | 0.33023871 | -0.152416 | 0.87885886 | 0.97692415 |
| RPL34-AS1 | 73.9663874 | -0.306964049 | 0.55241315 | -0.5556784 | 0.57843074 | 0.89953318 |
| RPL34 | 558.811811 | 0.39422087 | 0.42064924 | 0.93717243 | 0.34866987 | 0.78180304 |
| OSTC | 127.573197 | -0.33348658 | 0.4087675 | -0.8158344 | 0.41459487 | 0.82461178 |
| COL25A1 | 116.639123 | -0.414054721 | 1.33402491 | -0.3103801 | 0.75627196 | 0.95072713 |
| SEC24B-AS1 | 40.0396106 | -1.022967453 | 0.53301726 | -1.9192013 | 0.05495886 | 0.39224566 |
| SEC24B | 554.266659 | 0.45606887 | 0.23062737 | 1.97751405 | 0.04798355 | 0.36948395 |
| CASP6 | 66.2814227 | -1.074871663 | 0.2667551 | -4.0294324 | 5.59E-05 | 0.00609238 |
| PLA2G12A | 100.914245 | -0.193032697 | 0.21699477 | -0.8895731 | 0.37369518 | 0.79681088 |
| AC126283.2 | 57.0221799 | -1.072646692 | 0.55151047 | -1.9449253 | 0.05178396 | 0.38247133 |
| GAR1 | 50.0709148 | 0.213461442 | 0.38484525 | 0.55466826 | 0.57912159 | 0.89975022 |
| ELOVL6 | 292.344619 | -0.067481836 | 0.30665866 | -0.2200552 | 0.82582816 | 0.96545294 |
| FAM241A | 293.477333 | 1.254204099 | 0.35487773 | 3.53418659 | 0.00040903 | 0.02177405 |
| AP1AR | 200.144467 | 0.601612607 | 0.3561341 | 1.68928674 | 0.09116449 | 0.47749744 |
| TIFA | 31.601756 | -0.538772498 | 0.35827212 | -1.503808 | 0.1326308 | 0.55598831 |
| ALPK1 | 404.262768 | -0.507274779 | 0.35753467 | -1.4188128 | 0.15595359 | 0.5934642 |
| LARP7 | 214.375545 | 0.173044157 | 0.22324215 | 0.77514107 | 0.43825631 | 0.8347986 |
| ANK2 | 79.2600153 | 2.537445561 | 0.71080995 | 3.56979463 | 0.00035726 | 0.02002191 |
| CAMK2D | 424.535409 | -0.082328424 | 0.2466154 | -0.3338333 | 0.73850538 | 0.94349597 |
| ARSJ | 341.630393 | 0.240498106 | 0.38025805 | 0.63246026 | 0.52708617 | 0.87791848 |
| UGT8 | 278.8101 | 0.064254832 | 0.26100229 | 0.24618493 | 0.80553908 | 0.96218371 |
| SNHG8 | 93.9156242 | 0.743502787 | 0.51440453 | 1.44536593 | 0.14835513 | 0.58049694 |
| METTL14 | 126.707766 | 0.471929535 | 0.19077567 | 2.4737407 | 0.01337067 | 0.18614502 |
| SEC24D | 148.991448 | -0.349388444 | 0.4912032 | -0.7112911 | 0.47690389 | 0.85420488 |
| USP53 | 1200.83333 | 0.044151889 | 0.25498627 | 0.17315398 | 0.86253039 | 0.97436475 |
| C4orf3 | 385.583612 | -0.394917086 | 0.49993296 | -0.7899401 | 0.42956276 | 0.82998114 |
| PDE5A | 392.137198 | 0.029142704 | 0.35153038 | 0.08290238 | 0.93392916 | 0.98822345 |
| AC108866.1 | 179.329102 | -0.372919389 | 0.37837232 | -0.9855885 | 0.32433511 | 0.76970832 |
| PRDM5 | 317.451188 | 0.435881059 | 0.33337176 | 1.30749244 | 0.19104552 | 0.64458858 |
| ANXA5 | 329.288146 | -0.418670211 | 0.40063967 | -1.0450044 | 0.29602094 | 0.74305872 |
| EXOSC9 | 35.7652657 | 0.166625388 | 0.3091758 | 0.53893412 | 0.58993231 | 0.90233654 |
| BBS7 | 70.3709504 | 0.841112148 | 0.3188751 | 2.637748 | 0.00834586 | 0.1442148 |
| KIAA1109 | 1431.63416 | 0.735276988 | 0.22897653 | 3.21114567 | 0.00132207 | 0.04777684 |
| FGF2 | 91.7231339 | 0.679752717 | 0.64222555 | 1.058433 | 0.28985808 | 0.73754646 |
| NUDT6 | 49.3064145 | -0.046369741 | 0.37633213 | -0.1232149 | 0.9019369 | 0.98320957 |
| SPATA5 | 458.727617 | 0.453308226 | 0.2525824 | 1.79469443 | 0.0727024 | 0.43315584 |
| SPRY1 | 119.854554 | -2.290962868 | 0.58893385 | -3.8900173 | 0.00010024 | 0.00890241 |
| LINC01091 | 140.944967 | -0.360298366 | 0.5801819 | -0.6210093 | 0.53459349 | 0.88137638 |

| | | | | | | |
|---|---|---|---|---|---|---|
| ANKRD50 | 76.3730449 | -0.581051475 | 0.56361948 | -1.0309287 | 0.30257427 | 0.74899204 |
| FAT4 | 148.142416 | -0.221887616 | 0.49215579 | -0.4508483 | 0.65209887 | 0.9189184 |
| INTU | 262.624163 | 0.120863317 | 0.37722573 | 0.32040052 | 0.74866473 | 0.94678489 |
| HSPA4L | 214.674562 | 0.170236853 | 0.6119243 | 0.2781992 | 0.78085945 | 0.9566945 |
| MFSD8 | 176.130924 | 0.615189686 | 0.37195112 | 1.65395302 | 0.09813706 | 0.49265259 |
| ABHD18 | 477.654729 | 0.318019697 | 0.2664686 | 1.1934603 | 0.23268915 | 0.69218101 |
| LARP1B | 287.749095 | 0.200027715 | 0.24295895 | 0.82329841 | 0.41033832 | 0.82240305 |
| PGRMC2 | 187.876063 | -0.411832149 | 0.26802079 | -1.5365679 | 0.12439915 | 0.54275732 |
| AC108062.1 | 145.246092 | -0.374048611 | 0.34113448 | -1.0964843 | 0.27286688 | 0.72451099 |
| JADE1 | 413.341927 | 0.717370845 | 0.3944039 | 1.81887362 | 0.06893071 | 0.42719921 |
| SCLT1 | 354.225238 | 0.337882417 | 0.3190972 | 1.05886988 | 0.28965904 | 0.73754646 |
| C4orf33 | 70.93439 | 0.46837461 | 0.59115859 | 0.79229942 | 0.42818611 | 0.8293089 |
| AC096711.2 | 236.166225 | 0.266620209 | 0.37640866 | 0.70832644 | 0.47874256 | 0.85532469 |
| ELF2 | 868.669321 | -0.335112513 | 0.28082873 | -1.1932986 | 0.23275247 | 0.69218101 |
| NDUFC1 | 231.329064 | -0.143569542 | 0.37413105 | -0.3837413 | 0.70117019 | 0.93183677 |
| NAA15 | 324.569631 | 0.251825888 | 0.25489671 | 0.98795268 | 0.32317582 | 0.76904692 |
| AC097376.2 | 350.110197 | -0.816986456 | 0.35038249 | -2.3316989 | 0.01971654 | 0.23587823 |
| SETD7 | 67.4044287 | -0.733493724 | 0.39593622 | -1.8525552 | 0.06394613 | 0.41357646 |
| MGST2 | 110.200735 | -0.198955781 | 0.25759764 | -0.772351 | 0.4399066 | 0.83670075 |
| MAML3 | 1357.77111 | -0.031438187 | 0.34320684 | -0.0916013 | 0.92701483 | 0.98730848 |
| SCOC | 414.85506 | -0.372840439 | 0.22546703 | -1.6536362 | 0.09820146 | 0.49265259 |
| SCOC-AS1 | 39.0302714 | -0.881542204 | 0.57822192 | -1.5245742 | 0.12736534 | 0.54746778 |
| ELMOD2 | 126.492853 | 0.182883272 | 0.22955845 | 0.79667412 | 0.42564032 | 0.82774157 |
| TBC1D9 | 824.771215 | 0.574396736 | 0.24914107 | 2.305508 | 0.02113814 | 0.24614189 |
| RNF150 | 555.12123 | 0.117868472 | 0.53262813 | 0.22129599 | 0.82486197 | 0.96511681 |
| ZNF330 | 80.1263448 | -0.388335616 | 0.39135055 | -0.9922961 | 0.32105312 | 0.76590864 |
| IL15 | 78.7333079 | -0.483338218 | 0.49693937 | -0.9726302 | 0.33073715 | 0.77618491 |
| INPP4B | 251.117772 | -0.850989382 | 0.55790864 | -1.5253203 | 0.12717922 | 0.54746778 |
| AC104596.1 | 151.157128 | 0.379786385 | 0.64439597 | 0.58936804 | 0.55561441 | 0.89081623 |
| USP38 | 102.004784 | 0.27460379 | 0.25193518 | 1.08997793 | 0.27572286 | 0.7263609 |
| GAB1 | 1113.54709 | 0.246508361 | 0.30100421 | 0.81895321 | 0.41281311 | 0.82421059 |
| SMARCA5 | 324.973778 | 0.03219944 | 0.17349907 | 0.18558855 | 0.85276744 | 0.97062426 |
| ANAPC10 | 231.917929 | 0.140068917 | 0.19322149 | 0.72491378 | 0.46850493 | 0.85167938 |
| ABCE1 | 99.9412445 | 0.228467756 | 0.24339063 | 0.93868756 | 0.34789118 | 0.78180304 |
| OTUD4 | 287.843122 | 0.163244357 | 0.18770249 | 0.86969734 | 0.38446583 | 0.80561637 |
| SMAD1 | 383.891983 | -0.013887727 | 0.40181943 | -0.0345621 | 0.97242892 | 0.9939318 |
| MMAA | 76.4364274 | -0.066682526 | 0.23888178 | -0.2791445 | 0.78013397 | 0.95639467 |
| ZNF827 | 697.996351 | -0.085246206 | 0.41229818 | -0.2067586 | 0.83619837 | 0.96757829 |
| LSM6 | 94.5157418 | 0.032694018 | 0.28475071 | 0.11481628 | 0.90859074 | 0.98427726 |
| SLC10A7 | 256.696801 | 0.577602522 | 0.17710546 | 3.26134785 | 0.00110884 | 0.04225907 |
| TMEM184C | 173.571884 | 0.973688053 | 0.54200982 | 1.79643987 | 0.07242459 | 0.43315584 |
| PRMT9 | 285.103589 | 2.309020466 | 0.76385795 | 3.02284012 | 0.00250415 | 0.0731015 |
| ARHGAP10 | 64.204251 | -0.392993803 | 0.62565741 | -0.6281294 | 0.52991918 | 0.87951796 |
| NR3C2 | 4193.69502 | 0.660850256 | 0.31414566 | 2.10364279 | 0.03540962 | 0.31644624 |
| AC069272.1 | 58.0144212 | -0.832461194 | 0.67026528 | -1.2419876 | 0.21424112 | 0.67132603 |
| DCLK2 | 52.8860444 | 0.320949552 | 0.42425941 | 0.75649366 | 0.44935327 | 0.84117804 |
| LRBA | 3362.20315 | 1.025112834 | 0.30985295 | 3.30838495 | 0.00093836 | 0.03807006 |
| RPS3A | 393.989469 | 0.321294752 | 0.49923028 | 0.64358026 | 0.51984766 | 0.87631314 |
| SH3D19 | 343.251253 | -0.068265967 | 0.31877423 | -0.2141515 | 0.83042895 | 0.96648703 |
| FAM160A1 | 541.096645 | -0.342735359 | 0.39352145 | -0.8709446 | 0.38378443 | 0.80553992 |
| GATB | 84.5110917 | -0.297735777 | 0.33130405 | -0.8986784 | 0.36882401 | 0.79532503 |
| FBXW7 | 685.458702 | 0.549677914 | 0.21373325 | 2.57179411 | 0.0101173 | 0.16113882 |
| TMEM154 | 37.6984548 | -1.712627861 | 0.6586092 | -2.6003704 | 0.00931232 | 0.15441946 |
| ARFIP1 | 507.912867 | -0.124986428 | 0.20091756 | -0.6220782 | 0.53389047 | 0.88137638 |
| FHDC1 | 70.0489953 | -1.494561628 | 0.35220226 | -4.2434754 | 2.20E-05 | 0.0031949 |
| TRIM2 | 1038.97587 | -0.174030676 | 0.36481865 | -0.4770334 | 0.63333835 | 0.91415622 |
| TMEM131L | 97.9732393 | 0.479719056 | 0.41672508 | 1.15116435 | 0.24966463 | 0.70658528 |
| PLRG1 | 81.164015 | 0.289791612 | 0.269851 | 1.0738949 | 0.28286979 | 0.73287402 |
| MAP9 | 152.527271 | 0.224381395 | 0.27732566 | 0.80908992 | 0.41846343 | 0.82682819 |
| GUCY1A1 | 128.051153 | -0.666555882 | 0.4442956 | -1.5002532 | 0.13354884 | 0.5567191 |
| GUCY1B1 | 132.060161 | -0.546798975 | 0.35630177 | -1.5346513 | 0.1248695 | 0.54322641 |
| CTSO | 73.6124213 | -0.398182132 | 0.39033682 | -1.0200988 | 0.3076816 | 0.75262615 |
| PDGFC | 347.461982 | -0.401343346 | 0.37378874 | -1.0737171 | 0.28294951 | 0.73289938 |
| ETFDH | 202.267622 | 0.459184562 | 0.22912546 | 2.00407478 | 0.04506205 | 0.35614652 |
| PPID | 108.434414 | 0.153380975 | 0.24462802 | 0.62699675 | 0.53066137 | 0.88004345 |
| FNIP2 | 1329.64569 | -1.348081364 | 0.61236901 | -2.20142 | 0.02770631 | 0.2824534 |
| C4orf45 | 38.3509391 | -1.935512385 | 0.67692541 | -2.8592698 | 0.00424617 | 0.10090683 |
| RAPGEF2 | 898.822193 | 0.583080455 | 0.15879835 | 3.67182954 | 0.00024082 | 0.0150226 |
| NAF1 | 146.338943 | 1.072282055 | 0.50163332 | 2.13758142 | 0.03255073 | 0.30458184 |
| TMA16 | 75.2162032 | -0.088436157 | 0.25176143 | -0.3512697 | 0.72538604 | 0.93895336 |
| 1-Mar | 39.977174 | -0.713509454 | 0.39624985 | -1.8006555 | 0.0717572 | 0.43222361 |
| TMEM192 | 156.673511 | -0.498077031 | 0.38143579 | -1.3057952 | 0.19162221 | 0.64458858 |
| KLHL2 | 567.999783 | -0.117269981 | 0.36293673 | -0.3231141 | 0.74660882 | 0.94597987 |
| MSMO1 | 364.916026 | 0.284479729 | 0.64709552 | 0.43962556 | 0.66020833 | 0.92284375 |
| DDX60 | 187.449365 | -0.353794816 | 0.95429423 | -0.3707398 | 0.71083137 | 0.93475719 |
| DDX60L | 135.390964 | -0.195651772 | 0.95453538 | -0.2049707 | 0.83759503 | 0.96757829 |
| PALLD | 551.78751 | 0.431386596 | 0.2397056 | 1.7996517 | 0.07191565 | 0.43230175 |
| CBR4 | 943.505842 | 0.084386276 | 0.25587661 | 0.32979284 | 0.7415565 | 0.94487568 |
| SH3RF1 | 1663.55366 | 0.83163806 | 0.38380647 | 2.16681613 | 0.03024888 | 0.29256342 |
| AC096741.1 | 42.4519703 | 1.546860602 | 0.7290653 | 2.12170379 | 0.03386262 | 0.31129847 |
| NEK1 | 658.002958 | 0.788169544 | 0.38022192 | 2.07291979 | 0.03817974 | 0.32797026 |
| CLCN3 | 774.879528 | 0.880652108 | 0.45361871 | 1.94139284 | 0.05221065 | 0.38343909 |
| HPF1 | 69.2980719 | 0.461901859 | 0.32910292 | 1.40351798 | 0.16046243 | 0.60010648 |
| MFAP3L | 182.605616 | 0.180726975 | 0.40831348 | 0.4426182 | 0.65804192 | 0.92185868 |
| AC105285.1 | 42.1898194 | 0.242222817 | 0.46617197 | 0.5195997 | 0.60334261 | 0.90369786 |
| GALNT7 | 523.691073 | 0.292056091 | 0.26164745 | 1.11621989 | 0.26432802 | 0.71989632 |
| HMGB2 | 144.914196 | -0.843463497 | 0.40131994 | -2.1017233 | 0.03557752 | 0.31651305 |
| FBXO8 | 91.0939967 | 0.330114562 | 0.29743885 | 1.10985689 | 0.2670607 | 0.72152517 |
| CEP44 | 141.947427 | 0.312671735 | 0.3005909 | 1.04019027 | 0.29825151 | 0.74594723 |
| HPGD | 141.832577 | 0.386200441 | 0.59490037 | 0.64918507 | 0.51621876 | 0.87455102 |
| WDR17 | 111.200366 | 0.453754786 | 0.32198949 | 1.40922233 | 0.15876944 | 0.59723751 |
| SPCS3 | 169.340288 | -0.171768922 | 0.20345789 | -0.844248 | 0.39853082 | 0.81400298 |
| AGA | 36.7537212 | -0.205949992 | 0.42980581 | -0.4791699 | 0.63181779 | 0.9133034 |
| AC078881.1 | 149.586132 | -0.18192984 | 0.67018164 | -0.2714635 | 0.78603458 | 0.95753138 |
| DCTD | 88.4358853 | -0.386489876 | 0.28190476 | -1.3709945 | 0.17037666 | 0.61613093 |
| WWC2 | 820.16334 | 0.578607628 | 0.28466342 | 2.03260269 | 0.04209268 | 0.34673083 |
| CDKN2AIP | 113.132819 | 1.834440798 | 0.75759694 | 2.42139415 | 0.0154611 | 0.20228759 |
| ING2 | 39.9947963 | -1.277377159 | 0.47151366 | -2.7090989 | 0.00674662 | 0.13117736 |
| RWDD4 | 72.9860634 | 0.079350784 | 0.24890398 | 0.31880079 | 0.74987758 | 0.9474041 |
| TRAPPC11 | 210.372147 | -0.109338359 | 0.23109136 | -0.4731391 | 0.63611394 | 0.91454294 |
| STOX2 | 889.996558 | -0.085750481 | 0.3100896 | -0.2765345 | 0.78213755 | 0.95688565 |
| IRF2 | 192.72968 | -0.188693937 | 0.27898883 | -0.6763494 | 0.49881882 | 0.86607376 |
| CASP3 | 70.4183869 | 0.013959852 | 0.33481894 | 0.04169373 | 0.96674285 | 0.99291246 |
| PRIMPOL | 97.6064568 | -0.048484529 | 0.38021602 | -0.1275184 | 0.89853013 | 0.98312332 |
| ACSL1 | 304.004918 | -0.399880548 | 0.44119492 | -0.906358 | 0.3647464 | 0.79269964 |
| SLC25A4 | 138.464237 | -0.066893329 | 0.54436655 | -0.1228829 | 0.90219984 | 0.98320957 |
| CFAP97 | 216.501536 | -0.614495284 | 0.29703866 | -2.0687384 | 0.03857064 | 0.32997578 |
| SNX25 | 347.181847 | -0.183621739 | 0.38997252 | -0.4708581 | 0.63774204 | 0.91505156 |
| LRP2BP | 39.788453 | -1.09590961 | 0.5321756 | -2.0593008 | 0.03946544 | 0.3346923 |
| ANKRD37 | 547.444763 | -0.171707962 | 1.67347351 | -0.1026057 | 0.9182759 | 0.98520052 |
| UFSP2 | 162.832403 | 0.018656094 | 0.32154637 | 0.05801992 | 0.95373276 | 0.99060627 |
| C4orf47 | 190.345375 | -1.895755851 | 0.71346171 | -2.6571235 | 0.00788106 | 0.14142719 |
| SORBS2 | 861.232673 | -0.087337454 | 0.37809273 | -0.2309948 | 0.81731884 | 0.96383504 |

| | | | | | | |
|---|---|---|---|---|---|---|
| TLR3 | 58.5052682 | -0.652135864 | 0.46083235 | -1.415126 | 0.15703154 | 0.59475627 |
| FAM149A | 366.456627 | -0.11437763 | 0.28057757 | -0.4076507 | 0.68353017 | 0.92827258 |
| CYP4V2 | 94.2908382 | -0.221579773 | 0.23520524 | -0.9420699 | 0.34615687 | 0.78180304 |
| KLKB1 | 78.8302159 | 1.177657919 | 0.44095361 | 2.67070706 | 0.00756917 | 0.13892272 |
| FAT1 | 609.547128 | 0.332910837 | 0.35378531 | 0.94099678 | 0.34670651 | 0.78180304 |
| FRG1-DT | 126.141132 | 0.098370704 | 0.52733165 | 0.18654428 | 0.85201796 | 0.97053519 |
| FRG1 | 59.3623112 | -0.103720707 | 0.24378756 | -0.4254553 | 0.67050481 | 0.92474338 |
| PLEKHG4B | 66.0140424 | -0.22721695 | 0.82806349 | -0.2743956 | 0.78378065 | 0.9568898 |
| CCDC127 | 37.3757189 | -0.114651239 | 0.29083734 | -0.3942109 | 0.69342536 | 0.93105058 |
| SDHA | 371.453173 | 0.049933552 | 0.20829221 | 0.23972837 | 0.81054084 | 0.96287327 |
| PDCD6 | 143.197705 | -0.307591798 | 0.3029305 | -1.0153873 | 0.30992123 | 0.75409553 |
| EXOC3 | 145.839874 | -0.387577655 | 0.2392006 | -1.6203039 | 0.10516702 | 0.50534174 |
| SLC9A3 | 37.8313569 | -0.30447662 | 0.336863 | -0.9038589 | 0.36607023 | 0.79329304 |
| BRD9 | 79.7123632 | -0.48860137 | 0.2983119 | -1.6378876 | 0.10144513 | 0.49779302 |
| SLC12A7 | 189.23044 | -0.821947772 | 0.69832114 | -1.1770341 | 0.23918191 | 0.69526124 |
| CLPTM1L | 100.607526 | -0.342856094 | 0.26170446 | -1.3100888 | 0.19016578 | 0.64458858 |
| LPCAT1 | 42.3442231 | -0.427187989 | 0.46648519 | -0.915759 | 0.35979334 | 0.78892604 |
| MRPL36 | 50.2931851 | -0.928307388 | 0.53892741 | -1.7225091 | 0.08497733 | 0.46310155 |
| NDUFS6 | 196.971929 | 0.164734688 | 0.34898405 | 0.47204074 | 0.63689771 | 0.91454294 |
| AC022424.1 | 60.6968761 | -0.025938842 | 0.98541852 | -0.0263227 | 0.97899998 | 0.99469867 |
| ADAMTS16 | 514.741841 | 0.775631818 | 0.99414671 | 0.78019855 | 0.43527402 | 0.83348651 |
| ICE1 | 257.005998 | 0.139078338 | 0.19414578 | 0.71636036 | 0.47376886 | 0.85310957 |
| MED10 | 94.6874614 | -0.298466738 | 0.30771249 | -0.9699533 | 0.33206979 | 0.77618491 |
| NSUN2 | 109.075406 | 0.407008803 | 0.23888223 | 1.70380526 | 0.08841748 | 0.47036301 |
| SRD5A1 | 36.4551565 | -0.383640288 | 0.29032814 | -1.3214023 | 0.18636725 | 0.63723614 |
| TENT4A | 161.328364 | -0.086151628 | 0.23066637 | -0.3734902 | 0.70878364 | 0.93375896 |
| MTRR | 89.9883943 | -0.029746396 | 0.23403233 | -0.1271038 | 0.89885825 | 0.98312332 |
| SEMA5A | 55.3968281 | -1.099433394 | 0.53532415 | -2.0537713 | 0.03999783 | 0.33668856 |
| FAM173B | 97.721459 | 0.060344529 | 0.30758557 | 0.19618778 | 0.8444632 | 0.96876046 |
| CCT5 | 144.13889 | -0.089619526 | 0.41122755 | -0.2179317 | 0.82748231 | 0.96545294 |
| CMBL | 102.872281 | -1.059557889 | 0.52384062 | -2.0226723 | 0.04310695 | 0.34966009 |
| 6-Mar | 671.573132 | 0.263690102 | 0.20593351 | 1.28046235 | 0.20038258 | 0.65475264 |
| DAP | 488.50891 | 0.715285503 | 0.5315505 | 1.3456586 | 0.17841264 | 0.62759322 |
| DNAH5 | 141.71548 | -0.818559156 | 0.46478052 | -1.7611735 | 0.07820903 | 0.44739666 |
| TRIO | 1233.78242 | -0.317823441 | 0.29394041 | -1.0812513 | 0.27958535 | 0.72938673 |
| OTULINL | 82.0933738 | 1.036989033 | 0.39535722 | 2.62291664 | 0.00871806 | 0.14784024 |
| OTULIN | 96.5889649 | -0.074218459 | 0.2239967 | -0.3313373 | 0.74038973 | 0.94454656 |
| ANKH | 90.5358442 | -0.585354483 | 0.56532943 | -1.0354219 | 0.30047192 | 0.74743548 |
| FBXL7 | 94.8189114 | 0.760419876 | 0.64178324 | 1.18485469 | 0.23607491 | 0.69335782 |
| ZNF622 | 35.0687396 | -0.147133128 | 0.37943093 | -0.3877732 | 0.69818391 | 0.93177271 |
| RETREG1 | 323.36588 | -0.804797551 | 0.31381957 | -2.5645231 | 0.01033177 | 0.16264592 |
| MYO10 | 2177.13278 | 0.313613167 | 0.51789291 | 0.60555602 | 0.54480962 | 0.88669883 |
| LINC02241 | 94.6928599 | 0.296191081 | 0.25247204 | 1.17316388 | 0.24073009 | 0.6963541 |
| CDH12 | 42.645191 | 0.144676458 | 0.40589324 | 0.35643968 | 0.72151132 | 0.93744589 |
| C5orf17 | 177.491562 | -0.782879768 | 1.47616745 | -0.5303462 | 0.59587193 | 0.90323766 |
| DROSHA | 224.124632 | 0.192219921 | 0.23152849 | 0.83022147 | 0.40641358 | 0.82002585 |
| C5orf22 | 57.1750667 | -0.011469509 | 0.24045566 | -0.0476991 | 0.96195608 | 0.99171131 |
| PDZD2 | 149.567645 | -0.017417066 | 0.4087951 | -0.0426059 | 0.96601573 | 0.99284614 |
| GOLPH3 | 329.914852 | 0.386297921 | 0.23294282 | 1.65833799 | 0.09724927 | 0.49030463 |
| MTMR12 | 265.925063 | 0.727593071 | 0.21699695 | 3.35301055 | 0.00079938 | 0.03447517 |
| ZFR | 709.539656 | 0.276532223 | 0.1462976 | 1.89020336 | 0.05873077 | 0.40066749 |
| SUB1 | 434.567545 | 0.26237132 | 0.33263294 | 0.78877131 | 0.43024568 | 0.83047752 |
| TARS | 95.8893786 | -0.305016637 | 0.1974459 | -1.5448112 | 0.12239193 | 0.54052568 |
| C1QTNF3-AM | 291.368332 | 0.137308074 | 0.24139158 | 0.56881881 | 0.56947911 | 0.89626255 |
| RAI14 | 663.711772 | 0.449241565 | 0.39247029 | 1.14465114 | 0.2523537 | 0.70813507 |
| TTC23L | 45.1359533 | -0.097399342 | 0.48127041 | -0.2023797 | 0.83961993 | 0.96833625 |
| RAD1 | 42.8369041 | -0.320834563 | 0.32288113 | -0.9936615 | 0.32038767 | 0.76536648 |
| BRIX1 | 56.4985202 | 0.089128099 | 0.34978203 | 0.2548104 | 0.79886954 | 0.96006966 |
| DNAJC21 | 154.089955 | -0.229151095 | 0.1980556 | -1.1570039 | 0.24727078 | 0.70341959 |
| SPEF2 | 199.461306 | -0.092344737 | 0.4275308 | -0.2159955 | 0.82899124 | 0.96606563 |
| UGT3A1 | 423.170439 | 0.430027154 | 0.47740342 | 0.90076262 | 0.36771455 | 0.79489448 |
| LMBRD2 | 121.569745 | 0.33715709 | 0.24414049 | 1.38099618 | 0.16728013 | 0.61104769 |
| SKP2 | 33.8664936 | 0.240997788 | 0.33213779 | 0.72559581 | 0.46808659 | 0.85161077 |
| NADK2 | 157.278778 | 0.608662411 | 0.34191864 | 1.78013814 | 0.07505336 | 0.43941853 |
| NIPBL | 1385.0864 | 0.260422092 | 0.20643497 | 1.26152122 | 0.20712112 | 0.66441057 |
| CPLANE1 | 646.20285 | 0.862273913 | 0.24614704 | 3.50308466 | 0.0004599 | 0.02351115 |
| NUP155 | 96.0921481 | 0.404260841 | 0.19792379 | 2.0425076 | 0.04110121 | 0.34131622 |
| WDR70 | 703.245631 | 0.254274953 | 0.21747626 | 1.16920784 | 0.2423199 | 0.6978349 |
| LIFR | 500.853882 | 0.56397277 | 0.2888888 | 1.95221401 | 0.0509128 | 0.3802749 |
| LIFR-AS1 | 37.4394704 | -0.045405101 | 0.56275559 | -0.0806835 | 0.93569365 | 0.98822345 |
| OSMR-AS1 | 416.681579 | -1.363250087 | 0.50458186 | -2.7017422 | 0.00689772 | 0.13279571 |
| OSMR | 525.720535 | -1.593899326 | 0.67471381 | -2.3623339 | 0.01816028 | 0.2244336 |
| RICTOR | 638.46856 | 0.04259463 | 0.23128154 | 0.18416787 | 0.85388176 | 0.97067804 |
| DAB2 | 37.4201052 | -2.141080382 | 0.53055713 | -4.0355322 | 5.45E-05 | 0.00600986 |
| TTC33 | 153.325591 | 0.046968246 | 0.26048376 | 0.18031161 | 0.85690794 | 0.97191993 |
| PRKAA1 | 248.344539 | 0.422228364 | 0.20838458 | 2.02619774 | 0.04274453 | 0.3484612 |
| RPL37 | 364.283066 | -0.263168139 | 0.33032969 | -0.7966833 | 0.42563499 | 0.82774157 |
| CARD6 | 53.004368 | -0.980820031 | 0.43532906 | -2.2530543 | 0.02425572 | 0.26396676 |
| OXCT1 | 50.2202723 | 0.899765448 | 0.41343067 | 2.17633937 | 0.02952989 | 0.29105042 |
| C5orf51 | 55.1966394 | -0.071918447 | 0.32520312 | -0.2211493 | 0.82497618 | 0.96511681 |
| FBXO4 | 38.2243157 | -0.806271777 | 0.40378358 | -1.9967919 | 0.0458478 | 0.3601836 |
| GHR | 443.375426 | 0.411401734 | 0.35556716 | 1.15702962 | 0.24726026 | 0.70341959 |
| SELENOP | 347.284099 | 0.144065659 | 0.37815943 | 0.3809654 | 0.70322892 | 0.93241442 |
| ZNF131 | 337.506167 | 0.018735521 | 0.18382212 | 0.10192201 | 0.91881858 | 0.98520052 |
| HMGCS1 | 448.546705 | -0.072110744 | 0.74036057 | -0.0973995 | 0.92240915 | 0.9866345 |
| CCL28 | 70.0332648 | -1.821679546 | 0.62154417 | -2.9308932 | 0.00337989 | 0.08828112 |
| TMEM267 | 137.495214 | -0.431292327 | 0.27088447 | -1.5921634 | 0.111348 | 0.52038102 |
| PAIP1 | 154.158338 | -0.015585075 | 0.25622927 | -0.0608247 | 0.9514988 | 0.99030211 |
| NNT-AS1 | 179.136959 | -0.503113255 | 0.27958568 | -1.7994958 | 0.07194029 | 0.43230175 |
| NNT | 367.121164 | 0.011407474 | 0.2561227 | 0.0445391 | 0.96447469 | 0.99243116 |
| MRPS30-DT | 92.4142388 | -0.13933986 | 0.37091181 | -0.3756684 | 0.7071634 | 0.93274945 |
| MRPS30 | 72.9485641 | -0.294808511 | 0.29956531 | -0.984121 | 0.32505602 | 0.77058828 |
| PARP8 | 655.070571 | -0.781670879 | 0.42640243 | -1.8331764 | 0.06677634 | 0.42005766 |
| ITGA1 | 101.439695 | 0.155340719 | 0.37463886 | 0.41464124 | 0.67840456 | 0.9268955 |
| ITGA2 | 1347.63525 | -0.519703055 | 0.63700081 | -0.8158593 | 0.4145806 | 0.82461178 |
| MOCS2 | 105.493114 | 0.524379948 | 0.2570306 | 2.04014599 | 0.04133579 | 0.34244985 |
| NDUFS4 | 367.714842 | 0.288402967 | 0.20349091 | 1.41727689 | 0.15640199 | 0.59366792 |
| ARL15 | 4975.36033 | 1.122336756 | 0.23296787 | 4.81756032 | 1.45E-06 | 0.00049129 |
| SNX18 | 363.807916 | 0.675178389 | 0.53316763 | 1.26635293 | 0.20538676 | 0.66147916 |
| CCNO | 47.2450106 | 0.145509056 | 0.5143504 | 0.28289869 | 0.7772545 | 0.95601455 |
| AC026704.1 | 99.2081446 | -0.602572061 | 0.44875096 | -1.3427761 | 0.17934449 | 0.62889415 |
| DHX29 | 246.216007 | 0.446077866 | 0.19466997 | 2.29145697 | 0.021937 | 0.25042784 |
| MTREX | 408.492454 | 0.286008069 | 0.24150367 | 1.18428043 | 0.23630207 | 0.69335782 |
| PLPP1 | 488.019824 | 0.497322718 | 0.3421501 | 1.45352204 | 0.14607887 | 0.57722788 |
| SLC38A9 | 235.227341 | 0.205232166 | 0.2345125 | 0.87514382 | 0.38149566 | 0.80315561 |
| IL6ST | 454.341986 | 0.395362287 | 0.29935867 | 1.32069764 | 0.1866022 | 0.6378314 |
| LINC01948 | 38.3451511 | -0.062360839 | 0.37900915 | -0.1645365 | 0.86930882 | 0.97576204 |
| MAP3K1 | 575.712206 | 0.287101051 | 0.40144069 | 0.71517676 | 0.47449982 | 0.85368787 |
| MIER3 | 76.6613108 | 0.625077651 | 0.24020115 | 2.6023092 | 0.00925983 | 0.15403658 |
| GPBP1 | 741.789626 | 0.604423787 | 0.21489289 | 2.81267462 | 0.00491313 | 0.11144945 |
| PDE4D | 26196.0279 | 1.307841629 | 0.44017161 | 2.97120847 | 0.0029663 | 0.08063686 |
| AC092343.1 | 38.5766198 | 0.637723645 | 0.7689926 | 0.8292975 | 0.40693608 | 0.82018309 |
| ELOVL7 | 283.976057 | 0.179445842 | 0.36518255 | 0.49138669 | 0.62315298 | 0.91044796 |

| | | | | | | |
|---|---|---|---|---|---|---|
| ERCC8 | 154.154727 | 0.213511523 | 0.27323226 | 0.78142867 | 0.43455042 | 0.83266948 |
| NDUFAF2 | 243.895699 | -0.043262178 | 0.22374192 | -0.1933575 | 0.846679 | 0.96953627 |
| SMIM15 | 103.586844 | 0.021577095 | 0.32734623 | 0.06591521 | 0.94744533 | 0.98977092 |
| ZSWIM6 | 1524.22094 | 0.988864364 | 0.54557515 | 1.81251724 | 0.06990631 | 0.42843167 |
| KIF2A | 196.323017 | -0.090927671 | 0.27209777 | -0.3341728 | 0.73824918 | 0.94349597 |
| DIMT1 | 37.5814758 | 0.192351956 | 0.36303446 | 0.5298449 | 0.59621947 | 0.90346733 |
| IPO11 | 142.379832 | 0.237806372 | 0.21243356 | 1.1194388 | 0.26295299 | 0.71989632 |
| RNF180 | 391.579894 | 0.36909212 | 0.21336241 | 1.72988353 | 0.08365109 | 0.45946718 |
| SREK1IP1 | 165.727399 | -0.084882668 | 0.19780855 | -0.4291153 | 0.66783934 | 0.92422713 |
| CWC27 | 331.174505 | 0.274240884 | 0.21415087 | 1.28059664 | 0.20033538 | 0.65475264 |
| PPWD1 | 127.16546 | 0.084216645 | 0.27174819 | 0.30990693 | 0.75663174 | 0.95072713 |
| TRIM23 | 97.7649997 | 0.134087408 | 0.33812368 | 0.3965632 | 0.69168959 | 0.93077901 |
| TRAPPC13 | 159.690069 | 0.085566174 | 0.23557511 | 0.36322247 | 0.7164387 | 0.93573028 |
| NLN | 119.000731 | -1.268837418 | 0.41492823 | -3.0579684 | 0.00222843 | 0.06749698 |
| ERBIN | 1350.85021 | -0.004827507 | 0.29471618 | -0.0163802 | 0.98693108 | 0.99643909 |
| SREK1 | 393.857059 | 0.321339663 | 0.22328543 | 1.43914299 | 0.15011001 | 0.5835137 |
| MAST4 | 261.695301 | 0.416197624 | 0.451951 | 0.92089104 | 0.35710732 | 0.78612469 |
| PIK3R1 | 160.901369 | -0.197452552 | 0.36370917 | -0.5428858 | 0.58720841 | 0.90196444 |
| SLC30A5 | 149.875868 | 0.232073458 | 0.25894055 | 0.89624223 | 0.37012341 | 0.79532503 |
| MRPS36 | 143.491483 | 0.047502466 | 0.29509046 | 0.16097594 | 0.87211235 | 0.97622753 |
| CDK7 | 117.321971 | 0.019039923 | 0.27591275 | 0.06900704 | 0.94498401 | 0.98949015 |
| CCDC125 | 187.280403 | -0.277066095 | 0.34015517 | -0.8145286 | 0.41534222 | 0.82486005 |
| AK6 | 87.576498 | -0.087766097 | 0.37733254 | -0.2325962 | 0.81607502 | 0.96333253 |
| RAD17 | 157.919723 | 0.119025788 | 0.18712716 | 0.63606902 | 0.52473144 | 0.87790318 |
| MARVELD2 | 115.617663 | -0.016826271 | 0.26565624 | -0.0633385 | 0.94949695 | 0.98977092 |
| OCLN | 311.16926 | 0.21384936 | 0.26139049 | 0.81812217 | 0.41328743 | 0.82437234 |
| GTF2H2C | 86.5242053 | 0.006303203 | 0.27975142 | 0.02253144 | 0.98202403 | 0.99507903 |
| NAIP | 281.85368 | 0.527225055 | 0.44517788 | 1.184302 | 0.23629354 | 0.69335782 |
| GTF2H2 | 71.8766682 | -0.048488853 | 0.52729979 | -0.0919569 | 0.92673228 | 0.98729622 |
| BDP1 | 515.038851 | 0.324053801 | 0.20954303 | 1.54647856 | 0.12198903 | 0.54028588 |
| MCCC2 | 237.716695 | 0.037433862 | 0.21668018 | 0.17276089 | 0.86283938 | 0.97441343 |
| MRPS27 | 216.674387 | 0.114910062 | 0.18162885 | 0.63266415 | 0.52695299 | 0.87791848 |
| PTCD2 | 39.7281481 | -0.173174515 | 0.37563661 | -0.4610161 | 0.64478707 | 0.91682456 |
| TNPO1 | 542.806262 | 0.292661156 | 0.23267015 | 1.25783712 | 0.20845067 | 0.66602531 |
| FCHO2 | 477.621143 | -0.007999078 | 0.23318622 | -0.0343034 | 0.97263522 | 0.9939318 |
| BTF3 | 335.77621 | -0.189989449 | 0.51772926 | -0.3669668 | 0.71364378 | 0.93573028 |
| ANKRA2 | 71.3935981 | -0.048766269 | 0.25624348 | -0.1903122 | 0.84906447 | 0.96991131 |
| ARHGEF28 | 799.198378 | -0.214678497 | 0.29086687 | -0.7380644 | 0.46047529 | 0.84677681 |
| HEXB | 73.4840337 | -0.247900564 | 0.36320974 | -0.6825273 | 0.49490559 | 0.86389349 |
| GFM2 | 102.566564 | 0.487782075 | 0.23268526 | 2.09631702 | 0.03605408 | 0.31832076 |
| NSA2 | 133.558524 | -0.118014028 | 0.40905506 | -0.288504 | 0.77296095 | 0.95543807 |
| FAM169A | 241.747857 | 0.627612904 | 0.49735848 | 1.26189243 | 0.2069875 | 0.66434881 |
| AC116337.3 | 59.8872605 | 0.823687118 | 0.67686185 | 1.21692058 | 0.22363443 | 0.68103746 |
| GCNT4 | 69.8533301 | 1.307100972 | 0.68716232 | 1.90217207 | 0.05714866 | 0.39761441 |
| LINC01336 | 35.6918988 | 1.205989767 | 0.76471008 | 1.57705488 | 0.1147829 | 0.52713618 |
| HMGCR | 206.345229 | 0.374157641 | 0.570542 | 0.65579333 | 0.51195711 | 0.87155791 |
| COL4A3BP | 557.541195 | 0.484912128 | 0.2481539 | 1.95407823 | 0.05069197 | 0.3799001 |
| POLK | 438.708192 | 0.216965016 | 0.27153876 | 0.79902042 | 0.42427857 | 0.82774157 |
| ANKDD1B | 101.254572 | 0.022698248 | 0.48100432 | 0.04718928 | 0.96236237 | 0.99171131 |
| POC5 | 72.5895839 | -0.105531403 | 0.28294411 | -0.3729761 | 0.7091662 | 0.93391076 |
| IQGAP2 | 730.866795 | 0.318036403 | 0.49158954 | 0.64695519 | 0.51766095 | 0.87526312 |
| F2RL1 | 101.534744 | 0.097748309 | 0.31160961 | 0.31368837 | 0.75375773 | 0.94970964 |
| AC008581.2 | 40.866331 | -0.335497418 | 0.46904718 | -0.7152744 | 0.47443953 | 0.85368787 |
| ZBED3 | 110.659919 | -0.395289346 | 0.32569983 | -1.2136615 | 0.22487701 | 0.68211607 |
| AC022414.1 | 47.0225109 | 0.6780303 | 0.43588838 | 1.55551359 | 0.1198238 | 0.53734932 |
| PDE8B | 53.8394532 | 0.794158138 | 0.49645095 | 1.59967091 | 0.10967161 | 0.51494555 |
| WDR41 | 252.496696 | 0.077528941 | 0.24548753 | 0.3158162 | 0.75214202 | 0.94854974 |
| TBCA | 586.175268 | 0.085325448 | 0.20000998 | 0.42660594 | 0.66966637 | 0.92474338 |
| AP3B1 | 812.018946 | -0.097670646 | 0.30803586 | -0.3170756 | 0.75118626 | 0.94814309 |
| SCAMP1 | 316.272298 | 0.06886795 | 0.21698682 | 0.31738309 | 0.75095293 | 0.94796275 |
| LHFPL2 | 42.6903048 | -1.433636557 | 0.42650844 | -3.3613322 | 0.00077567 | 0.03373058 |
| ARSB | 134.642334 | -1.231416694 | 0.63339523 | -1.9441521 | 0.05187711 | 0.38286767 |
| DMGDH | 330.589434 | 0.014756043 | 0.25853694 | 0.05707518 | 0.95448531 | 0.99060627 |
| BHMT2 | 310.553533 | -0.728969834 | 0.48481805 | -1.5035947 | 0.13268577 | 0.55599634 |
| BHMT | 55.8736602 | 0.400191656 | 0.62730985 | 0.63794894 | 0.52350692 | 0.87725117 |
| JMY | 383.799582 | 0.94177608 | 0.49902487 | 1.88723276 | 0.05912903 | 0.4016022 |
| HOMER1 | 761.821788 | 3.424618971 | 0.78781621 | 4.34697706 | 1.38E-05 | 0.00237801 |
| TENT2 | 824.980562 | 0.235653583 | 0.2129895 | 1.10640937 | 0.26854937 | 0.72322263 |
| CMYA5 | 398.455079 | -0.488339246 | 0.50759434 | -0.962066 | 0.33601646 | 0.77820994 |
| MTX3 | 83.8247065 | -0.115148741 | 0.2278907 | -0.5052806 | 0.61336179 | 0.90751853 |
| THBS4 | 55.4559436 | -0.322303627 | 0.56214156 | -0.5733496 | 0.56640802 | 0.89423863 |
| SERINC5 | 300.988969 | -0.159932818 | 0.24079036 | -0.6641994 | 0.50656267 | 0.87094607 |
| ZFYVE16 | 583.743633 | 0.909216585 | 0.34468605 | 2.6378108 | 0.00834431 | 0.1442148 |
| AC008771.1 | 59.143821 | -0.208371422 | 0.39092063 | -0.5330274 | 0.59401458 | 0.90304695 |
| FAM151B | 89.4828319 | 0.39386734 | 0.35678528 | 1.10393382 | 0.26962185 | 0.72333985 |
| DHFR | 1121.93966 | -0.674522901 | 0.4752918 | -1.4191764 | 0.15584759 | 0.5934642 |
| MSH3 | 466.325958 | 0.065681053 | 0.23282297 | 0.28210727 | 0.77786126 | 0.95601455 |
| RASGRF2 | 367.512119 | -0.041517693 | 0.40611674 | -0.1022309 | 0.91857338 | 0.98520052 |
| CKMT2-AS1 | 264.418514 | 0.283585723 | 0.28739009 | 0.98676235 | 0.32375916 | 0.76937573 |
| CKMT2 | 178.374119 | -0.856888756 | 0.43661367 | -1.9625789 | 0.04969513 | 0.37609758 |
| ZCCHC9 | 112.00006 | -0.14838697 | 0.23367429 | -0.6350162 | 0.52541782 | 0.87791848 |
| ACOT12 | 64.1637569 | 2.01377871 | 0.70121152 | 2.8718563 | 0.00408068 | 0.09922408 |
| SSBP2 | 1367.08197 | 0.790773281 | 0.29065989 | 2.72061374 | 0.00651609 | 0.12862701 |
| ATG10 | 654.325871 | 0.366209589 | 0.23166597 | 1.5807656 | 0.11393164 | 0.52645663 |
| RPS23 | 1251.57151 | -0.127818371 | 0.51663944 | -0.2474034 | 0.80459602 | 0.96210606 |
| ATP6AP1L | 150.304854 | -0.190816331 | 0.54063282 | -0.35295 | 0.72412595 | 0.93893205 |
| TMEM167A | 152.386879 | 0.088530013 | 0.29996416 | 0.29513531 | 0.76789052 | 0.95411028 |
| XRCC4 | 192.251052 | -0.403556913 | 0.26089876 | -1.546795 | 0.12191267 | 0.54028588 |
| COX7C | 569.048625 | -0.418232562 | 0.38723018 | -1.0800619 | 0.28011462 | 0.72945471 |
| RASA1 | 593.042915 | 0.253705601 | 0.17675647 | 1.43533979 | 0.15119029 | 0.58535779 |
| CCNH | 698.231323 | 0.825022259 | 0.44229909 | 1.86530398 | 0.06213881 | 0.40942507 |
| TMEM161B | 224.217542 | 0.302980643 | 0.21902325 | 1.38332642 | 0.1665648 | 0.60907157 |
| TMEM161B-/ | 850.223817 | 0.071894131 | 0.28715729 | 0.25036498 | 0.80230511 | 0.96071123 |
| LINC00461 | 68.558947 | 0.114898807 | 0.69274327 | 0.16586059 | 0.86826666 | 0.97576204 |
| CETN3 | 50.9742601 | 0.457118032 | 0.33486814 | 1.36506875 | 0.17223144 | 0.61884138 |
| LYSMD3 | 59.3764514 | 0.148509216 | 0.45291141 | 0.32789904 | 0.742988 | 0.94519474 |
| ARRDC3 | 539.076436 | -0.12170636 | 0.68353156 | -0.1780552 | 0.85867961 | 0.97237339 |
| ARRDC3-AS1 | 38.7845257 | 0.252459871 | 0.35153024 | 0.71817397 | 0.47265002 | 0.85285335 |
| FAM172A | 1041.3053 | 0.169262369 | 0.16527001 | 1.02415661 | 0.30576131 | 0.75109981 |
| KIAA0825 | 424.667758 | 0.357395848 | 0.20497505 | 1.74360657 | 0.08122772 | 0.45449358 |
| SLF1 | 59.6036244 | 0.038837358 | 0.35797581 | 0.10849157 | 0.91360577 | 0.98480378 |
| MCTP1 | 530.13326 | 0.040445812 | 0.38835231 | 0.10414722 | 0.91705252 | 0.98520052 |
| FAM81B | 47.1009579 | -0.612114612 | 0.60123866 | -1.0180892 | 0.30863555 | 0.7532605 |
| TTC37 | 490.490595 | 0.101221134 | 0.27368341 | 0.36984753 | 0.7114961 | 0.9348115 |
| ARSK | 85.9984463 | 0.24548885 | 0.32062285 | 0.76566236 | 0.44387723 | 0.83873671 |
| RHOBTB3 | 1165.51151 | 0.44117493 | 0.38214171 | 1.15447992 | 0.24830347 | 0.70482674 |
| GLRX | 269.640375 | 0.2724247 | 0.54431699 | 0.50048906 | 0.61673076 | 0.90794578 |
| ELL2 | 627.329785 | 1.150214407 | 0.85131168 | 1.35110845 | 0.17666069 | 0.62541921 |
| AC104123.1 | 240.885661 | -0.382668119 | 0.27766099 | -1.3781847 | 0.16814628 | 0.6131323 |
| CAST | 695.151262 | -0.18256127 | 0.19824217 | -0.9209003 | 0.35710249 | 0.78612469 |
| ERAP1 | 144.719949 | -0.4580691 | 0.33906663 | -1.3509708 | 0.1767048 | 0.62541921 |
| AC009126.1 | 58.2558605 | -0.290154532 | 0.33142146 | -0.875485 | 0.38131005 | 0.80315561 |
| ERAP2 | 93.659909 | -0.839297755 | 0.514267 | -1.6320272 | 0.10267375 | 0.50117415 |

| | | | | | | |
|---|---|---|---|---|---|---|
| LNPEP | 452.03445 | 0.001503541 | 0.16443553 | 0.00914365 | 0.99270452 | 0.99832487 |
| LIX1-AS1 | 88.8310066 | -0.171083514 | 0.39213557 | -0.4362867 | 0.66262876 | 0.9229791 |
| RIOK2 | 42.2588186 | 0.296141947 | 0.4161085 | 0.71169406 | 0.47665425 | 0.85414749 |
| RGMB | 57.3118025 | -0.011810965 | 0.35661887 | -0.0331193 | 0.97357946 | 0.9939318 |
| CHD1 | 491.517575 | 0.953096044 | 0.41753287 | 2.28268508 | 0.02244893 | 0.25166501 |
| LINC02062 | 127.141074 | 0.021117405 | 0.53525848 | 0.03945272 | 0.96852944 | 0.99303855 |
| FAM174A | 79.1478865 | 0.54561343 | 0.35399636 | 1.54129671 | 0.12324459 | 0.54197241 |
| SLCO4C1 | 128.59842 | 1.469537547 | 0.3463347 | 4.24311381 | 2.20E-05 | 0.0031949 |
| PAM | 932.220981 | -0.551081271 | 0.34341587 | -1.6047053 | 0.10855867 | 0.51254482 |
| GIN1 | 55.5843912 | 0.064859001 | 0.27630252 | 0.23473909 | 0.81441125 | 0.96310425 |
| PPIP5K2 | 331.575207 | 0.239543938 | 0.17830938 | 1.3434175 | 0.17913682 | 0.62850818 |
| NUDT12 | 75.838246 | -0.059943512 | 0.31704151 | -0.1890715 | 0.85003679 | 0.97017003 |
| EFNA5 | 2275.27865 | 0.12700894 | 0.42874417 | 0.29623479 | 0.76705078 | 0.95408212 |
| FBXL17 | 1532.21245 | 0.325069941 | 0.21749306 | 1.49462214 | 0.13501308 | 0.55891804 |
| FER | 1638.80967 | 0.560887244 | 0.30256964 | 1.85374593 | 0.06377552 | 0.41357646 |
| PJA2 | 474.953875 | 0.183583707 | 0.28667999 | 0.64037851 | 0.52192655 | 0.87670505 |
| MAN2A1 | 736.702288 | 0.496829493 | 0.27605704 | 1.79973492 | 0.0719025 | 0.43230175 |
| TMEM232 | 481.129651 | 0.175504711 | 0.43229171 | 0.40598676 | 0.68475233 | 0.92881914 |
| SLC25A46 | 198.050623 | 0.33491255 | 0.2084132 | 1.60696417 | 0.10806223 | 0.51174521 |
| WDR36 | 124.339891 | 0.135927184 | 0.20650135 | 0.65823873 | 0.51038474 | 0.87133207 |
| NREP | 114.259401 | 0.337908606 | 0.5006295 | 0.67496743 | 0.49969646 | 0.86702299 |
| EPB41L4A | 975.536384 | 0.376034288 | 0.29317672 | 1.28261989 | 0.19962528 | 0.65475264 |
| EPB41L4A-AS | 36.1676445 | 0.035790736 | 0.33077629 | 0.10820224 | 0.91383527 | 0.98480378 |
| APC | 205.787239 | 0.171277595 | 0.26634191 | 0.64307413 | 0.520176 | 0.87638467 |
| SRP19 | 111.828468 | 0.218127906 | 0.32487131 | 0.67142865 | 0.5019475 | 0.86837155 |
| REEP5 | 539.944904 | -0.025943579 | 0.26943146 | -0.0962901 | 0.92329019 | 0.98694936 |
| DCP2 | 237.285354 | 0.005609639 | 0.16144641 | 0.03474614 | 0.97228217 | 0.9939318 |
| MCC | 509.460163 | 0.684716295 | 0.48527628 | 1.4109824 | 0.1582498 | 0.59635309 |
| YTHDC2 | 366.508836 | 0.76786108 | 0.16826047 | 4.56352637 | 5.03E-06 | 0.00107584 |
| PGGT1B | 149.180905 | 0.117232252 | 0.18359621 | 0.63853306 | 0.52312674 | 0.87717892 |
| CCDC112 | 65.3270599 | -0.465093543 | 0.33398078 | -1.3925758 | 0.16374811 | 0.60340373 |
| FEM1C | 94.1687324 | -0.275250177 | 0.54707188 | -0.5031335 | 0.61487043 | 0.90792412 |
| TMED7-TICA | 57.7394586 | 0.040095328 | 0.28097248 | 0.14270198 | 0.88652555 | 0.97973087 |
| ATG12 | 128.618248 | 0.100531901 | 0.23290872 | 0.43163648 | 0.66600564 | 0.92385693 |
| AP3S1 | 256.065126 | 0.114474609 | 0.31592415 | 0.36234839 | 0.7170917 | 0.93573028 |
| COMMD10 | 739.98867 | 0.126122019 | 0.25801087 | 0.48882445 | 0.62496599 | 0.91058133 |
| SEMA6A | 384.846099 | 2.182402826 | 0.85735797 | 2.5454978 | 0.01091221 | 0.16452817 |
| SEMA6A-AS1 | 123.795287 | 0.789404847 | 0.50942225 | 1.54960811 | 0.12123561 | 0.53989034 |
| DTWD2 | 221.480574 | 0.353209578 | 0.22754141 | 1.55228703 | 0.12059356 | 0.53823708 |
| AC008629.1 | 97.5471402 | 0.397399262 | 0.62309369 | 0.63778412 | 0.52361422 | 0.87725117 |
| DMXL1 | 759.617852 | 0.317321993 | 0.20228912 | 1.56865576 | 0.11672817 | 0.53183794 |
| TNFAIP8 | 216.212087 | -1.253539342 | 0.43079329 | -2.9098395 | 0.00361614 | 0.09198346 |
| HSD17B4 | 375.126151 | 0.219231519 | 0.19579184 | 1.11971733 | 0.26283424 | 0.71989632 |
| SRFBP1 | 127.531404 | -0.199660652 | 0.20909893 | -0.9548621 | 0.33964743 | 0.77836439 |
| SNX2 | 599.140185 | 0.970772697 | 0.4312334 | 2.25115379 | 0.0243758 | 0.26499831 |
| SNX24 | 532.280157 | 0.721451181 | 0.31607231 | 2.28255103 | 0.02245683 | 0.25166501 |
| PPIC | 132.837067 | 0.027466078 | 0.42128627 | 0.06519576 | 0.94801814 | 0.98977092 |
| CEP120 | 154.146755 | 0.638879773 | 0.28008871 | 2.28099077 | 0.02254899 | 0.25166501 |
| CSNK1G3 | 345.340819 | 0.69452536 | 0.23436258 | 2.96346518 | 0.00304196 | 0.08216441 |
| ZNF608 | 566.511827 | 0.067087901 | 0.59964414 | 0.11187953 | 0.91091893 | 0.98468493 |
| GRAMD2B | 1006.19448 | 0.333162536 | 0.1981232 | 1.68159271 | 0.09264784 | 0.48138295 |
| AC093535.1 | 56.9275236 | 0.080656779 | 0.57816276 | 0.13950532 | 0.88905085 | 0.98054894 |
| ALDH7A1 | 153.639939 | 0.208543002 | 0.31014448 | 0.67240598 | 0.50132528 | 0.86804029 |
| PHAX | 86.0119446 | -0.133932303 | 0.29071498 | -0.4606997 | 0.64501407 | 0.91682456 |
| C5orf63 | 68.5932445 | -0.421432729 | 0.29694165 | -1.4192443 | 0.15582782 | 0.5934642 |
| PRRC1 | 147.40806 | -0.172431733 | 0.26893231 | -0.6411715 | 0.52141126 | 0.87650871 |
| LINC01184 | 301.815539 | -0.289984447 | 0.34168132 | -0.8486986 | 0.39604905 | 0.8120904 |
| SLC12A2 | 96.5206685 | 0.403533321 | 0.30467626 | 1.32446593 | 0.18534835 | 0.63605149 |
| SLC27A6 | 53.2478693 | 3.125436555 | 1.6423524 | 1.90302431 | 0.05703737 | 0.39754595 |
| ISOC1 | 44.6297846 | -0.100605181 | 0.4362346 | -0.2306217 | 0.81760868 | 0.96404074 |
| HINT1 | 812.973088 | -0.382120685 | 0.50114911 | -0.762489 | 0.4457682 | 0.83968362 |
| LYRM7 | 85.5598126 | -0.06963184 | 0.23399894 | -0.2975733 | 0.76602884 | 0.95389523 |
| CDC42SE2 | 977.958626 | -0.128694041 | 0.39215157 | -0.3281742 | 0.74277993 | 0.94508888 |
| P4HA2 | 50.4581791 | -0.607288668 | 0.42371183 | -1.4332587 | 0.15178392 | 0.58653964 |
| SLC22A4 | 55.9763908 | 0.104192391 | 0.50709685 | 0.20546843 | 0.83720616 | 0.96757829 |
| SLC22A5 | 224.476069 | 0.607552659 | 0.33681522 | 1.80381594 | 0.07126017 | 0.43137474 |
| AC116366.3 | 283.129129 | -0.19782669 | 0.50361229 | -0.3928155 | 0.69445579 | 0.93105058 |
| IRF1 | 71.7915026 | -1.102473358 | 0.41645308 | -2.6472931 | 0.0081139 | 0.14312705 |
| AC004039.1 | 44.8221869 | 0.031753333 | 0.68728645 | 0.04620102 | 0.96315003 | 0.99211837 |
| KIF3A | 251.864726 | 0.447757731 | 0.22452927 | 1.99420654 | 0.04612949 | 0.36110228 |
| 8-Sep | 148.509762 | 0.567490827 | 0.31572359 | 1.79742927 | 0.0722675 | 0.43302651 |
| SOWAHA | 34.6776175 | -0.166319753 | 0.60920883 | -0.2730094 | 0.78484597 | 0.95702626 |
| SHROOM1 | 94.1288329 | -0.508492709 | 0.57552467 | -0.883529 | 0.37695054 | 0.80049477 |
| UQCRQ | 277.368064 | -0.171913541 | 0.40144315 | -0.4282388 | 0.66847726 | 0.92422713 |
| AFF4 | 959.983481 | 0.145690049 | 0.33056753 | 0.44072705 | 0.65941061 | 0.92284375 |
| ZCCHC10 | 93.1220596 | -0.294705002 | 0.23880436 | -1.2340855 | 0.21717103 | 0.67475613 |
| HSPA4 | 231.318808 | 0.413256529 | 0.26597418 | 1.55374677 | 0.12024483 | 0.53784287 |
| FSTL4 | 221.492103 | 3.199097373 | 0.96749974 | 3.30656149 | 0.00094449 | 0.03807006 |
| C5orf15 | 278.918483 | 0.827592999 | 0.34526361 | 2.39698879 | 0.01653043 | 0.21100962 |
| VDAC1 | 381.886915 | 0.074133376 | 0.32626764 | 0.22721645 | 0.82025543 | 0.96502145 |
| SKP1 | 739.19565 | 0.215599637 | 0.49613839 | 0.43455544 | 0.66388514 | 0.92300139 |
| CDKL3 | 124.274474 | 0.184196039 | 0.32382487 | 0.56881375 | 0.56948254 | 0.89626255 |
| UBE2B | 362.61157 | -0.260659584 | 0.32776212 | -0.7952706 | 0.4264561 | 0.8277744 |
| JADE2 | 127.788469 | -0.176269682 | 0.28153503 | -0.6261021 | 0.53124796 | 0.88023377 |
| SAR1B | 164.296992 | 0.175157261 | 0.22317784 | 0.78483269 | 0.43255167 | 0.83162989 |
| SEC24A | 170.112393 | -0.091603509 | 0.46189134 | -0.1983226 | 0.84279264 | 0.96876046 |
| CAMLG | 112.087655 | 0.014867759 | 0.35459844 | 0.04192844 | 0.96655574 | 0.99291246 |
| DDX46 | 342.612566 | 0.041887444 | 0.26693835 | 0.15691804 | 0.87530944 | 0.97622753 |
| C5orf24 | 275.271637 | 0.984301107 | 0.46862229 | 2.10041461 | 0.03569238 | 0.31699677 |
| TXNDC15 | 103.135501 | 0.194502009 | 0.21129722 | 0.92051382 | 0.35730432 | 0.78612469 |
| PCBD2 | 264.061858 | 0.202935888 | 0.3512734 | 0.57771493 | 0.5634566 | 0.89294196 |
| H2AFY | 255.467696 | -0.245972405 | 0.23965498 | -1.0263605 | 0.30472169 | 0.74998115 |
| CXCL14 | 48.6430069 | -1.203357806 | 0.75710672 | -1.5894164 | 0.11196643 | 0.52244352 |
| SMAD5 | 293.40913 | 0.100786304 | 0.15804623 | 0.63770139 | 0.52366808 | 0.87725117 |
| KLHL3 | 140.836433 | 0.308938575 | 0.43302234 | 0.7134472 | 0.47556908 | 0.85401927 |
| HNRNPA0 | 212.535823 | -0.265516258 | 0.34875024 | -0.7613364 | 0.44645615 | 0.83980766 |
| AC106791.1 | 109.388332 | -0.018079104 | 0.44502096 | -0.0406253 | 0.96759463 | 0.99292814 |
| FAM13B | 660.363108 | 0.309141116 | 0.19637229 | 1.57426039 | 0.11542726 | 0.52893648 |
| NME5 | 150.876114 | 0.12337262 | 0.34877178 | 0.35373452 | 0.72353785 | 0.93893205 |
| BRD8 | 132.970903 | 0.173461994 | 0.22714714 | 0.76365477 | 0.445073 | 0.83929814 |
| CDC23 | 71.6610368 | 0.298685464 | 0.21259724 | 1.40493577 | 0.16004038 | 0.59986522 |
| FAM53C | 60.1221428 | 0.074720268 | 0.31973097 | 0.23369731 | 0.81521998 | 0.96333253 |
| KDM3B | 314.31743 | -0.02365635 | 0.1634969 | -0.1446899 | 0.88495571 | 0.97891513 |
| EGR1 | 373.69177 | 2.380034364 | 1.71441966 | 1.38824491 | 0.16506248 | 0.60654096 |
| ETF1 | 264.927265 | 0.60690673 | 0.34563722 | 1.75590675 | 0.07910433 | 0.44916083 |
| HSPA9 | 429.349523 | 0.154123834 | 0.33956656 | 0.45388402 | 0.64991231 | 0.91803345 |
| CTNNA1 | 1378.55555 | 0.281819872 | 0.25855313 | 1.08998823 | 0.27571833 | 0.7263609 |
| SIL1 | 485.44343 | 0.083561119 | 0.24047084 | 0.34748961 | 0.72822352 | 0.93931348 |
| PAIP2 | 249.971415 | 0.088415338 | 0.19290497 | 0.45833625 | 0.64671088 | 0.91743182 |
| TMEM173 | 32.3125504 | -1.913511428 | 0.53837554 | -3.5542318 | 0.00037909 | 0.02099512 |
| UBE2D2 | 763.103175 | -0.141943842 | 0.17889324 | -0.7934556 | 0.42751242 | 0.82876992 |
| CXXC5 | 517.494716 | -0.351629848 | 0.29757324 | -1.1816582 | 0.23734136 | 0.6939708 |
| PURA | 383.40347 | 0.004413204 | 0.212296 | 0.02078797 | 0.98341479 | 0.99540861 |

| | | | | | | |
|---|---|---|---|---|---|---|
| IGIP | 66.7335846 | 0.189545039 | 0.36616521 | 0.51764896 | 0.60470322 | 0.90416461 |
| CYSTM1 | 1545.39421 | 0.743993823 | 0.56489803 | 1.31704093 | 0.1878249 | 0.64035097 |
| PFDN1 | 175.362367 | 0.304863885 | 0.29829844 | 1.02200964 | 0.30677634 | 0.75187465 |
| ANKHD1-EIF4 | 41.0769349 | -0.159712454 | 0.41942285 | -0.380791 | 0.70335833 | 0.93241442 |
| SRA1 | 90.3665465 | -0.452467258 | 0.38545571 | -1.1738502 | 0.24045503 | 0.6963541 |
| SLC35A4 | 91.4828925 | -0.292404329 | 0.33221843 | -0.8801569 | 0.37877434 | 0.80274043 |
| NDUFA2 | 198.290395 | -0.146665319 | 0.37181329 | -0.3944596 | 0.69324175 | 0.93105058 |
| IK | 87.3881852 | -0.582591876 | 0.30952282 | -1.8822259 | 0.05980534 | 0.40360868 |
| WDR55 | 54.079677 | 0.244167333 | 0.43570367 | 0.56039769 | 0.57520821 | 0.89807676 |
| HARS | 96.3219174 | 0.081583183 | 0.18572228 | 0.43927516 | 0.66046217 | 0.92284375 |
| ZMAT2 | 94.3564211 | -0.166534288 | 0.45127244 | -0.3690327 | 0.71210335 | 0.9348115 |
| TAF7 | 128.011653 | -0.352139294 | 0.35854061 | -0.9821462 | 0.32602784 | 0.77110624 |
| DIAPH1 | 613.369959 | -0.25682534 | 0.26740015 | -0.9604533 | 0.33682715 | 0.77827084 |
| HDAC3 | 106.849428 | 0.187838671 | 0.38607303 | 0.48653663 | 0.62658674 | 0.91061758 |
| PCDH1 | 143.00079 | 0.200049885 | 0.57907535 | 0.34546434 | 0.72974531 | 0.94030631 |
| DELE1 | 102.719067 | -0.0179588 | 0.33571746 | -0.0534938 | 0.95733848 | 0.99106618 |
| RNF14 | 109.046946 | 0.261835514 | 0.29408949 | 0.89032599 | 0.37329087 | 0.79681088 |
| GNPDA1 | 96.7290686 | -1.15278989 | 0.38519819 | -2.9927189 | 0.00276504 | 0.0769785 |
| NDFIP1 | 633.019973 | -0.007732509 | 0.17298227 | -0.0447012 | 0.96434551 | 0.99243116 |
| SPRY4-AS1 | 539.769373 | 0.305368108 | 0.49338777 | 0.61892111 | 0.53596833 | 0.88137638 |
| FGF1 | 96.6421623 | -0.105962291 | 0.67489135 | -0.1570064 | 0.87523977 | 0.97622753 |
| ARHGAP26 | 123.606041 | -0.675068436 | 0.36788022 | -1.8350224 | 0.06650238 | 0.41989619 |
| YIPF5 | 71.8699603 | -0.510465618 | 0.23561323 | -2.1665405 | 0.03026991 | 0.29256342 |
| PRELID2 | 346.662282 | -0.866373861 | 0.51100878 | -1.6954187 | 0.08999601 | 0.47371081 |
| LARS | 286.042147 | 0.084423301 | 0.21269965 | 0.39691321 | 0.69143146 | 0.93073907 |
| TCERG1 | 335.161974 | 0.176572823 | 0.22064758 | 0.80024816 | 0.42356703 | 0.82774157 |
| PPP2R2B | 130.208708 | -0.190213669 | 0.92325012 | -0.2060262 | 0.83677048 | 0.96757829 |
| STK32A | 41.4056512 | -1.594891613 | 0.89991647 | -1.7722663 | 0.07635037 | 0.4426298 |
| AC091948.1 | 84.0131596 | 0.145288423 | 0.46678065 | 0.31125631 | 0.75560578 | 0.95067145 |
| FBXO38 | 157.08987 | 0.306321515 | 0.18776916 | 1.63137292 | 0.10281166 | 0.50161366 |
| CSNK1A1 | 1293.17334 | 0.165998081 | 0.20364144 | 0.81514884 | 0.41498712 | 0.82472987 |
| ARHGEF37 | 77.7290334 | -0.87959962 | 0.26386546 | -3.3335156 | 0.00085756 | 0.03624114 |
| SLC26A2 | 56.8813341 | -0.146442546 | 0.40689805 | -0.3598999 | 0.71892203 | 0.9359691 |
| HMGXB3 | 130.09529 | -0.331104247 | 0.28254085 | -1.171881 | 0.24124484 | 0.69706209 |
| TCOF1 | 59.0632145 | -0.251267395 | 0.3084409 | -0.8146371 | 0.41528007 | 0.82486005 |
| CD74 | 813.710538 | -1.35977133 | 0.65566003 | -2.0738969 | 0.03808888 | 0.32772702 |
| RPS14 | 969.174859 | 0.203163347 | 0.47216094 | 0.4302841 | 0.66698899 | 0.92422364 |
| NDST1 | 127.392559 | 0.390172189 | 0.23936704 | 1.63001635 | 0.10309804 | 0.50171849 |
| SYNPO | 45.2807751 | 0.114562616 | 0.43909564 | 0.26090584 | 0.79416512 | 0.95937941 |
| RBM22 | 57.8728608 | -0.199307565 | 0.28073415 | -0.7099513 | 0.47773437 | 0.85479418 |
| DCTN4 | 308.626978 | 0.599752041 | 0.25315895 | 2.36907306 | 0.01783273 | 0.22195607 |
| GPX3 | 677.603539 | -0.76793996 | 0.78729323 | -0.975418 | 0.32935298 | 0.77581911 |
| TNIP1 | 248.88547 | -1.086141998 | 0.27545546 | -3.9430767 | 8.04E-05 | 0.00773433 |
| ANXA6 | 76.6015023 | -0.080879611 | 0.51192159 | -0.1579922 | 0.87446295 | 0.97622753 |
| ATOX1 | 186.403633 | 0.151007986 | 0.22801973 | 0.66225841 | 0.50780562 | 0.87094607 |
| G3BP1 | 371.573631 | 0.40522485 | 0.25493562 | 1.58951837 | 0.11194341 | 0.52244352 |
| FAM114A2 | 87.6579823 | -0.230470314 | 0.31442957 | -0.7329791 | 0.46357115 | 0.84922793 |
| MFAP3 | 106.900829 | -0.390176526 | 0.26160543 | -1.4914695 | 0.13583827 | 0.55980539 |
| GALNT10 | 125.924919 | -0.752579875 | 0.33261555 | -2.2626118 | 0.02365963 | 0.25990871 |
| SAP30L-AS1 | 57.1879863 | 0.056286285 | 0.31349749 | 0.17954301 | 0.85751135 | 0.97201489 |
| SAP30L | 51.1973561 | 0.020529429 | 0.51636795 | 0.03975737 | 0.96828657 | 0.99303855 |
| LARP1 | 503.506834 | -0.203028146 | 0.26954865 | -0.7532152 | 0.45132059 | 0.84220793 |
| FAXDC2 | 96.8825087 | -0.265741713 | 0.43233226 | -0.6146701 | 0.53877261 | 0.88287241 |
| CNOT8 | 108.826917 | -0.095031598 | 0.20625638 | -0.460745 | 0.64498155 | 0.91682456 |
| GEMIN5 | 66.7927835 | -0.059182209 | 0.34228023 | -0.1729057 | 0.86272554 | 0.97438987 |
| MRPL22 | 96.9475527 | -0.139663937 | 0.24085926 | -0.579857 | 0.56201103 | 0.89242209 |
| SGCD | 401.108047 | 1.54998366 | 0.9660787 | 1.60440723 | 0.10862432 | 0.51254482 |
| HAVCR2 | 37.1204823 | -0.54407765 | 0.32594629 | -1.6692249 | 0.09507282 | 0.48627909 |
| CYFIP2 | 1371.94276 | 1.159476599 | 0.41080638 | 2.82244061 | 0.00476596 | 0.10977431 |
| ADAM19 | 64.5273458 | 1.000791803 | 0.39941806 | 2.5056248 | 0.01222353 | 0.17590979 |
| CLINT1 | 751.015732 | 0.229421156 | 0.16729939 | 1.37132094 | 0.17027493 | 0.61613093 |
| RNF145 | 224.385156 | -0.791512178 | 0.24894407 | -3.179478 | 0.00147541 | 0.05086268 |
| UBLCP1 | 80.3794031 | 0.257444199 | 0.31317802 | 0.82203789 | 0.41105533 | 0.82289588 |
| TTC1 | 214.622283 | -0.214675697 | 0.18356967 | -1.1694508 | 0.24222206 | 0.6978349 |
| PWWP2A | 298.711072 | 0.056559214 | 0.37090367 | 0.1524903 | 0.87880024 | 0.97692415 |
| SLU7 | 70.3174487 | -0.185666809 | 0.29317936 | -0.6332874 | 0.52654595 | 0.87791848 |
| AC113414.1 | 252.221605 | -1.067496109 | 0.48028691 | -2.2226217 | 0.02624132 | 0.27568539 |
| CCNG1 | 143.539962 | -1.073502596 | 0.3634109 | -2.9539637 | 0.00313721 | 0.08408684 |
| NUDCD2 | 73.4275509 | -0.413838304 | 0.48036279 | -0.861512 | 0.38895612 | 0.806701 |
| MAT2B | 168.29816 | -0.018822169 | 0.29654824 | -0.0634709 | 0.94939157 | 0.98977092 |
| WWC1 | 1874.09253 | 0.504386317 | 0.39067352 | 1.2910686 | 0.19667989 | 0.65097045 |
| RARS | 146.619851 | -0.450439359 | 0.24986154 | -1.8027559 | 0.07142656 | 0.43193905 |
| PANK3 | 226.521644 | -0.320548363 | 0.29773043 | -1.0766396 | 0.28164132 | 0.7313184 |
| KCNIP1 | 93.5476576 | 0.308553504 | 1.00460757 | 0.30713834 | 0.75873807 | 0.9521704 |
| RANBP17 | 332.182904 | -0.265633092 | 0.38158251 | -0.6961354 | 0.48634404 | 0.85912348 |
| NPM1 | 316.660153 | -0.064137276 | 0.33742879 | -0.1900765 | 0.8492492 | 0.96991131 |
| FBXW11 | 745.233665 | 0.480705405 | 0.25500024 | 1.88511746 | 0.05941398 | 0.40275455 |
| STK10 | 57.5671873 | -0.571044166 | 0.30998981 | -1.8421385 | 0.06545489 | 0.41750899 |
| UBTD2 | 165.304881 | -0.218155887 | 0.21662324 | -1.0070752 | 0.31389865 | 0.75763653 |
| SH3PXD2B | 44.3193452 | -1.928844324 | 0.55972273 | -3.4460711 | 0.0005688 | 0.02714172 |
| AC022217.3 | 56.5086369 | 2.065982625 | 0.77056574 | 2.68112443 | 0.00733752 | 0.13610131 |
| DUSP1 | 1337.87601 | 1.583402654 | 0.90083447 | 1.75770656 | 0.07879745 | 0.44904692 |
| ERGIC1 | 573.973789 | -0.150634914 | 0.35241213 | -0.4274396 | 0.66905914 | 0.92463889 |
| RPL26L1 | 51.2118544 | -0.376412485 | 0.38813667 | -0.9697937 | 0.33214934 | 0.77618491 |
| ATP6V0E1 | 1072.08163 | -0.462438737 | 0.22941436 | -2.0157358 | 0.04382761 | 0.35277527 |
| CREBRF | 555.365394 | -0.286286583 | 0.24444461 | -1.1711716 | 0.24152981 | 0.69750135 |
| BNIP1 | 38.3632377 | 0.403121806 | 0.36129685 | 1.1157634 | 0.26452342 | 0.71989632 |
| BOD1 | 76.5417916 | -0.273137298 | 0.39250317 | -0.6958856 | 0.48650048 | 0.85912348 |
| CPEB4 | 562.187342 | 0.856403598 | 0.34322173 | 2.49519047 | 0.01258895 | 0.17925575 |
| SFXN1 | 374.716465 | -0.030762612 | 0.21742751 | -0.1414845 | 0.88748724 | 0.98009982 |
| THOC3 | 171.378221 | -0.547709785 | 0.46840467 | -1.169309 | 0.24227917 | 0.6978349 |
| SIMC1 | 270.54088 | 0.388031598 | 0.30703265 | 1.26381218 | 0.20629745 | 0.66368662 |
| KIAA1191 | 183.651761 | -0.337269089 | 0.26431317 | -1.2760207 | 0.20194819 | 0.65666058 |
| ARL10 | 63.1685869 | -0.345650417 | 0.43315554 | -0.7979822 | 0.42488082 | 0.82774157 |
| CLTB | 129.976575 | -0.146550467 | 0.42360839 | -0.3459574 | 0.72937471 | 0.94008695 |
| FAF2 | 242.751292 | 0.145329413 | 0.2258367 | 0.64351549 | 0.51988967 | 0.87631314 |
| RNF44 | 80.3423746 | 0.11406829 | 0.24268257 | 0.47003083 | 0.63833299 | 0.91521016 |
| TSPAN17 | 53.1442474 | -0.471677171 | 0.35029593 | -1.3465106 | 0.17813791 | 0.62752447 |
| UIMC1 | 313.931168 | 0.267929847 | 0.21785786 | 1.22983785 | 0.21875783 | 0.67667712 |
| ZNF346 | 150.466877 | 0.015692381 | 0.30533788 | 0.0513935 | 0.95901197 | 0.99119981 |
| NSD1 | 807.232616 | 0.29147302 | 0.31111391 | 0.93686915 | 0.34882587 | 0.78180304 |
| PRELID1 | 149.0939 | 0.366093102 | 0.54209776 | 0.67532672 | 0.49946821 | 0.86691402 |
| LMAN2 | 221.152608 | -0.308874596 | 0.27790124 | -1.1114545 | 0.26637274 | 0.72059534 |
| DBN1 | 93.0908528 | -0.547835189 | 0.37845952 | -1.4475397 | 0.14774582 | 0.57952643 |
| DDX41 | 42.8285263 | -0.653576205 | 0.33049025 | -1.977596 | 0.0479743 | 0.36948395 |
| FAM193B | 163.961177 | -0.081575173 | 0.28912101 | -0.2821489 | 0.77782934 | 0.95601455 |
| TMED9 | 226.823506 | -0.699601502 | 0.41622532 | -1.680824 | 0.09279711 | 0.48153712 |
| B4GALT7 | 73.4281462 | -0.639226707 | 0.32666534 | -1.9568244 | 0.05036812 | 0.37839277 |
| FAM153A | 56.8931363 | -3.055283442 | 1.0673025 | -2.8626218 | 0.00420152 | 0.10017148 |
| AC106795.3 | 63.4020082 | -0.846172617 | 0.37924943 | -2.231177 | 0.0256694 | 0.27225313 |
| RMND5B | 118.986654 | -0.160794556 | 0.28092625 | -0.5723728 | 0.56706941 | 0.89454608 |
| NHP2 | 117.257457 | -0.339326488 | 0.48229193 | -0.7035707 | 0.48170014 | 0.85664644 |
| HNRNPAB | 149.164159 | -0.88078144 | 0.34778474 | -2.5325477 | 0.0113237 | 0.16768662 |

| | | | | | | |
|---|---|---|---|---|---|---|
| PHYKPL | 304.830584 | -0.45759086 | 0.3124735 | -1.4644149 | 0.14308061 | 0.5729912 |
| CLK4 | 118.992443 | 0.435638518 | 0.24742357 | 1.76069938 | 0.0782893 | 0.44761148 |
| ZNF354A | 74.8982148 | -0.351528873 | 0.34214752 | -1.027419 | 0.30422319 | 0.74968437 |
| ZNF354B | 47.2548949 | -0.234789518 | 0.35583412 | -0.6598286 | 0.50936385 | 0.87094607 |
| ZFP2 | 53.6604667 | -0.085720347 | 0.45858415 | -0.1869239 | 0.85172028 | 0.97053519 |
| ZNF354C | 48.5218594 | -0.008069388 | 0.31848994 | -0.0253364 | 0.97978664 | 0.99469867 |
| RUFY1 | 242.755145 | -0.15653859 | 0.23239983 | -0.6735745 | 0.50058189 | 0.86743068 |
| HNRNPH1 | 718.5327 | 0.885018681 | 0.23215258 | 3.81222851 | 0.00013772 | 0.01065488 |
| CANX | 730.302777 | -0.375265196 | 0.21731626 | -1.726816 | 0.08420071 | 0.46162908 |
| MAML1 | 87.0218969 | 0.159090865 | 0.20971739 | 0.75859645 | 0.448094 | 0.8406776 |
| MGAT4B | 173.484464 | -1.360704528 | 0.58253521 | -2.3358323 | 0.01949999 | 0.23473196 |
| SQSTM1 | 1162.24988 | 0.178130017 | 0.67113675 | 0.26541538 | 0.79068949 | 0.95830066 |
| MRNIP | 133.450156 | -0.072955252 | 0.27727683 | -0.2631134 | 0.79246317 | 0.95856579 |
| TBC1D9B | 169.313637 | -0.544442987 | 0.20655785 | -2.6357894 | 0.00839418 | 0.1442148 |
| RNF130 | 462.338562 | -0.225862095 | 0.34505647 | -0.6545656 | 0.51274748 | 0.87177451 |
| MAPK9 | 193.712571 | 0.045291222 | 0.18456631 | 0.24539269 | 0.80615239 | 0.96218371 |
| GFPT2 | 174.694211 | -1.834082591 | 0.99038715 | -1.8518845 | 0.06404241 | 0.41357646 |
| CNOT6 | 223.593089 | 0.229042739 | 0.27941102 | 0.81973409 | 0.41236771 | 0.82364303 |
| MGAT1 | 92.8619843 | -0.233036855 | 0.28676633 | -0.8126367 | 0.41642636 | 0.82489761 |
| ZFP62 | 46.2776172 | -0.376294828 | 0.27349877 | -1.3758556 | 0.16886633 | 0.61405939 |
| TRIM41 | 63.3579246 | 0.120295406 | 0.29323359 | 0.41023746 | 0.68163176 | 0.92760692 |
| RACK1 | 552.650257 | -0.428589659 | 0.37912102 | -1.1304825 | 0.25827299 | 0.71549044 |
| TRIM52 | 64.5613556 | -0.024152012 | 0.25862801 | -0.0933851 | 0.9255976 | 0.98717098 |
| AC138035.1 | 42.2975099 | 0.338022375 | 0.35369982 | 0.95567584 | 0.33923603 | 0.77836439 |
| DUSP22 | 87.7415336 | -0.490337576 | 0.27263509 | -1.7985124 | 0.07209585 | 0.43274028 |
| EXOC2 | 341.801281 | -0.030564435 | 0.30002703 | -0.1018723 | 0.91885806 | 0.98520052 |
| FOXQ1 | 74.1744335 | 1.406988112 | 1.30257827 | 1.08015629 | 0.28007259 | 0.72945471 |
| GMDS | 974.868461 | 0.003922526 | 0.27156315 | 0.01444425 | 0.98847556 | 0.9964791 |
| GMDS-DT | 1267.42186 | -0.536888297 | 0.26978665 | -1.9900477 | 0.04658568 | 0.36244837 |
| WRNIP1 | 120.591467 | 0.326382749 | 0.26151762 | 1.24803349 | 0.21201879 | 0.66975892 |
| SERPINB1 | 115.20738 | -0.609329062 | 0.36573813 | -1.6660255 | 0.09570833 | 0.48710328 |
| SERPINB6 | 248.630438 | -0.423952205 | 0.24286223 | -1.745649 | 0.08087196 | 0.45347147 |
| NQO2 | 91.7834327 | -0.631712034 | 0.34395349 | -1.8366205 | 0.06626594 | 0.41911108 |
| RIPK1 | 122.749068 | -0.415007906 | 0.32376762 | -1.281808 | 0.19991002 | 0.65475264 |
| BPHL | 91.4191875 | -0.309486367 | 0.30552886 | -1.012953 | 0.31108263 | 0.75435382 |
| PSMG4 | 130.790803 | -0.511479855 | 0.3432364 | -1.4901679 | 0.1361801 | 0.5601128 |
| SLC22A23 | 177.494314 | 0.512659646 | 0.35699633 | 1.43603619 | 0.15099204 | 0.58535779 |
| PXDC1 | 34.6132437 | -1.055784243 | 0.58375138 | -1.8086197 | 0.0705101 | 0.42987436 |
| PRPF4B | 370.93988 | -0.160358226 | 0.18778118 | -0.8539632 | 0.39312536 | 0.81010924 |
| ECI2 | 140.674576 | -0.35694813 | 0.32237716 | -1.1072377 | 0.26819119 | 0.72308815 |
| CDYL | 507.469744 | -0.015634531 | 0.28408955 | -0.0550338 | 0.95611153 | 0.9908068 |
| LYRM4 | 173.731905 | -0.116485024 | 0.32519584 | -0.3581996 | 0.72019393 | 0.93668719 |
| FARS2 | 716.430152 | 0.27622055 | 0.20384185 | 1.3550728 | 0.17539436 | 0.62477136 |
| RREB1 | 758.18852 | 0.223496774 | 0.30588138 | 0.73066486 | 0.46498388 | 0.84991999 |
| SSR1 | 250.245308 | -0.02962772 | 0.21327376 | -0.1389187 | 0.88951437 | 0.98064269 |
| AL139095.4 | 46.8734024 | -0.225764753 | 0.59581715 | -0.3789162 | 0.70475011 | 0.93241442 |
| RIOK1 | 44.9240529 | 0.254468427 | 0.32202275 | 0.79021878 | 0.42940001 | 0.82998114 |
| DSP | 518.518748 | -0.464935007 | 0.32537689 | -1.4289122 | 0.15302947 | 0.58860244 |
| SNRNP48 | 67.9945602 | 0.132888698 | 0.32241342 | 0.41216864 | 0.68021582 | 0.92700898 |
| BLOC1S5-TXI | 85.9965943 | -0.052809755 | 0.44552285 | -0.1185343 | 0.90564429 | 0.98320957 |
| SLC35B3 | 127.747398 | 0.018490182 | 0.42081584 | 0.04393889 | 0.96495311 | 0.99243116 |
| TFAP2A | 281.705203 | 0.160009525 | 0.32843294 | 0.48719086 | 0.62612308 | 0.91061758 |
| TFAP2A-AS1 | 32.9000437 | 0.111196222 | 0.33206179 | 0.33486606 | 0.73772613 | 0.94319505 |
| GCNT2 | 142.884102 | -0.213197902 | 0.51058123 | -0.4175592 | 0.67626941 | 0.9268955 |
| TMEM14C | 176.539107 | -0.021469923 | 0.44473271 | -0.048276 | 0.96149627 | 0.99171131 |
| MAK | 133.59252 | -0.199009988 | 0.55823059 | -0.3565014 | 0.7214651 | 0.93744589 |
| SMIM13 | 32.7725749 | 0.209649063 | 0.31806258 | 0.65914407 | 0.50980326 | 0.87094607 |
| AL139807.1 | 61.793219 | 0.412466525 | 0.55860919 | 0.7383812 | 0.46028284 | 0.84657145 |
| NEDD9 | 157.699672 | 0.474401479 | 0.39685862 | 1.19539164 | 0.23193405 | 0.69186326 |
| TMEM170B | 41.8218565 | -0.160664461 | 0.33357132 | -0.4816495 | 0.63005495 | 0.91226525 |
| HIVEP1 | 250.571018 | -0.820670679 | 0.32162945 | -2.5516031 | 0.01072286 | 0.16452817 |
| EDN1 | 62.211549 | -1.105989709 | 0.86951232 | -1.2719655 | 0.20338536 | 0.6582233 |
| PHACTR1 | 540.782573 | -0.150694542 | 0.27866745 | -0.5407684 | 0.58866725 | 0.90200952 |
| TBC1D7 | 166.297015 | 1.13545395 | 0.44285876 | 2.56391893 | 0.01034977 | 0.16264592 |
| SIRT5 | 114.705958 | 0.138454988 | 0.28300637 | 0.48922923 | 0.62467942 | 0.91058133 |
| NOL7 | 165.108758 | -0.05270111 | 0.4327222 | -0.1217897 | 0.90306557 | 0.98320957 |
| RANBP9 | 461.331322 | 0.599786422 | 0.15988227 | 3.75142549 | 0.00017583 | 0.01270841 |
| MCUR1 | 111.087193 | 0.340430289 | 0.4121424 | 0.82600162 | 0.40880317 | 0.82073894 |
| JARID2 | 829.597261 | 2.202130427 | 0.534348 | 4.12115402 | 3.77E-05 | 0.00475993 |
| DTNBP1 | 280.570259 | -0.542051424 | 0.27543764 | -1.9679642 | 0.04907215 | 0.3746651 |
| MYLIP | 83.5844155 | -0.476214851 | 0.51039542 | -0.9330312 | 0.35080386 | 0.78242416 |
| GMPR | 111.52382 | -1.654382942 | 0.80997019 | -2.0425232 | 0.04109966 | 0.34131622 |
| ATXN1 | 1434.37092 | 0.456528984 | 0.38482884 | 1.18631697 | 0.23549715 | 0.69335782 |
| CAP2 | 147.153279 | 0.167027961 | 0.47504575 | 0.35160395 | 0.7251353 | 0.93895336 |
| FAM8A1 | 60.4012245 | 0.211294926 | 0.36352207 | 0.58124373 | 0.5610762 | 0.89242209 |
| NUP153 | 347.77379 | 0.679533685 | 0.23076903 | 2.94464852 | 0.00323322 | 0.08621081 |
| KIF13A | 535.749272 | 0.051289033 | 0.33168745 | 0.15463061 | 0.87711254 | 0.97640529 |
| TPMT | 74.9615861 | 0.458654708 | 0.2983524 | 1.53729185 | 0.12422186 | 0.54273186 |
| KDM1B | 67.2387733 | -0.232775466 | 0.30422167 | -0.7651508 | 0.44418172 | 0.83873671 |
| DEK | 228.265225 | -0.392725755 | 0.25705838 | -1.5277687 | 0.12656997 | 0.54691843 |
| RNF144B | 614.492087 | -1.376535028 | 0.39747521 | -3.4631972 | 0.0005338 | 0.02626382 |
| AL022068.1 | 387.850832 | 0.600535154 | 0.47172094 | 1.27307292 | 0.20299217 | 0.65821717 |
| ID4 | 282.548997 | 0.599737392 | 0.45090437 | 1.33007668 | 0.18349301 | 0.63405547 |
| MBOAT1 | 242.557609 | 0.557840956 | 0.72088826 | 0.77382444 | 0.43903462 | 0.83559755 |
| E2F3 | 133.30516 | -1.044992775 | 0.49380479 | -2.1162062 | 0.03432727 | 0.31309819 |
| CDKAL1 | 1282.24026 | 0.008229869 | 0.37163648 | 0.02214494 | 0.98233234 | 0.99507903 |
| SOX4 | 496.356773 | -1.466392632 | 0.71693441 | -2.0453651 | 0.04081888 | 0.34079725 |
| CASC15 | 86.6369761 | 0.869376031 | 0.60693541 | 1.43240288 | 0.15202856 | 0.58683582 |
| DCDC2 | 4916.3203 | -0.618287245 | 0.45467735 | -1.3598374 | 0.1738814 | 0.62125058 |
| MRS2 | 122.585343 | -0.028033951 | 0.23742381 | -0.1180756 | 0.90600778 | 0.98320957 |
| GPLD1 | 40.6471069 | 0.280454882 | 0.49192976 | 0.57011164 | 0.56860198 | 0.8958131 |
| ALDH5A1 | 79.8366635 | 0.115381204 | 0.34630859 | 0.33317454 | 0.73900254 | 0.94379071 |
| TDP2 | 92.7073686 | 0.00310439 | 0.23766221 | 0.01306219 | 0.98957817 | 0.9965778 |
| ACOT13 | 76.4850989 | -0.29771329 | 0.31276766 | -0.9518673 | 0.34116431 | 0.77954601 |
| C6orf62 | 409.998298 | 0.086253119 | 0.30446712 | 0.28329206 | 0.77695296 | 0.95601455 |
| GMNN | 220.398225 | 1.610550356 | 0.60582833 | 2.65842692 | 0.00785064 | 0.14121635 |
| CARMIL1 | 1453.43022 | 0.664161413 | 0.44079489 | 1.50673573 | 0.13187841 | 0.55448417 |
| TRIM38 | 267.136682 | -0.103956153 | 0.37418146 | -0.2778228 | 0.78114836 | 0.9566945 |
| HIST1H1C | 158.3701 | 0.106524252 | 0.64246763 | 0.16580485 | 0.86831053 | 0.97576204 |
| HFE | 31.7629771 | -0.26549546 | 0.42583398 | -0.6234718 | 0.53297456 | 0.88114425 |
| HIST1H4C | 191.593773 | -3.262754495 | 0.71214092 | -4.5816136 | 4.61E-06 | 0.00102883 |
| HIST1H2BC | 108.485296 | 3.646953424 | 0.68718883 | 5.30706154 | 1.11E-07 | 6.87E-05 |
| HIST1H2AC | 1211.29043 | 2.893902303 | 0.61727792 | 4.68816753 | 2.76E-06 | 0.00078079 |
| HIST1H1E | 97.9109089 | -1.448644629 | 0.66306004 | -2.1847865 | 0.0289045 | 0.28768102 |
| HIST1H2BD | 184.916599 | 1.994235811 | 0.5536579 | 3.60192785 | 0.00031587 | 0.01808965 |
| AL031777.3 | 59.2740957 | 3.445412208 | 0.79271191 | 4.3463611 | 1.38E-05 | 0.00237801 |
| BTN3A2 | 55.7506793 | -0.869704476 | 0.8506355 | -1.0224173 | 0.30658342 | 0.7515776 |
| BTN3A1 | 52.2668311 | -0.221679294 | 0.67374595 | -0.329025 | 0.74213675 | 0.94508888 |
| BTN2A1 | 51.8622384 | -0.184697088 | 0.25360501 | -0.7282864 | 0.46643826 | 0.85057334 |
| HCG11 | 37.631277 | -0.270504678 | 0.43585764 | -0.6206262 | 0.53484559 | 0.88137638 |
| HMGN4 | 54.9814686 | -0.081256408 | 0.42771004 | -0.1899801 | 0.8493247 | 0.96991131 |
| ABT1 | 39.2659935 | -0.238328644 | 0.50300742 | -0.4738074 | 0.63563725 | 0.91454294 |
| ZNF322 | 45.2878185 | -0.287728753 | 0.38490909 | -0.7475239 | 0.45474737 | 0.84290184 |

| | | | | | | |
|---|---|---|---|---|---|---|
| LINC00240 | 51.7236004 | -0.444475253 | 0.56766522 | -0.7829884 | 0.43363395 | 0.83187655 |
| ZSCAN16-AS | 77.8199984 | -0.456576715 | 0.37605656 | -1.2141171 | 0.224703 | 0.68208993 |
| ZKSCAN8 | 67.9835336 | -0.242906979 | 0.28898261 | -0.8405592 | 0.40059495 | 0.8162653 |
| ZSCAN9 | 43.3515297 | -0.467304894 | 0.3131428 | -1.492306 | 0.13561894 | 0.55916513 |
| ZSCAN23 | 38.9962909 | -0.262577498 | 0.340831 | -0.7704038 | 0.4410604 | 0.83752728 |
| TRIM27 | 145.767025 | -0.439970261 | 0.24114034 | -1.8245403 | 0.06807044 | 0.42365683 |
| GABBR1 | 130.867512 | 0.919089545 | 0.29366603 | 3.12971009 | 0.00174979 | 0.057666 |
| HLA-F | 57.6250082 | -2.109416316 | 0.92013298 | -2.2925125 | 0.02187609 | 0.25028539 |
| HLA-A | 1378.48429 | -1.184391658 | 0.59000763 | -2.0074175 | 0.04470523 | 0.35515061 |
| ZNRD1 | 51.7252194 | 0.7522778 | 0.7165473 | 1.04986482 | 0.29378027 | 0.73938934 |
| PPP1R11 | 75.178808 | -0.31210101 | 0.3330257 | -0.937168 | 0.34867217 | 0.78180304 |
| TRIM26 | 166.168844 | 0.357464206 | 0.32214943 | 1.10962235 | 0.2671618 | 0.72161227 |
| HCG17 | 313.697445 | 0.095872137 | 0.25855612 | 0.37079817 | 0.71078786 | 0.93475719 |
| HLA-E | 557.920367 | -1.517578881 | 0.51080452 | -2.9709582 | 0.00296872 | 0.08063686 |
| GNL1 | 163.0327 | 0.239714826 | 0.25604432 | 0.93622397 | 0.34915788 | 0.78180304 |
| PRR3 | 39.3218836 | -0.519868514 | 0.44819389 | -1.1599188 | 0.24608187 | 0.70206841 |
| ABCF1 | 146.47149 | 0.035512812 | 0.27567389 | 0.12882182 | 0.89749864 | 0.98312332 |
| PPP1R10 | 182.157576 | 0.419036947 | 0.36169269 | 1.15854415 | 0.24664205 | 0.70292817 |
| MRPS18B | 143.482656 | -0.761170962 | 0.28597602 | -2.6616601 | 0.00777564 | 0.14098385 |
| ATAT1 | 55.9498553 | -0.622396453 | 0.37209393 | -1.6726864 | 0.09438905 | 0.48561474 |
| C6orf136 | 95.6699062 | 0.341417622 | 0.27259198 | 1.25248595 | 0.21039285 | 0.66835921 |
| DHX16 | 62.7303081 | 0.155948401 | 0.33573025 | 0.46450506 | 0.64228593 | 0.91643619 |
| MDC1 | 87.3550991 | -0.049007816 | 0.22888856 | -0.2141121 | 0.83045962 | 0.96648703 |
| TUBB | 439.133775 | -0.686098158 | 0.66851045 | -1.0263088 | 0.30474605 | 0.74998115 |
| FLOT1 | 354.317003 | -0.279916997 | 0.36720585 | -0.762289 | 0.44588751 | 0.83968362 |
| IER3 | 1262.10078 | -0.212598911 | 0.94610136 | -0.2247105 | 0.82220449 | 0.96502145 |
| DDR1 | 416.249562 | 0.138160245 | 0.21936901 | 0.62980748 | 0.52882055 | 0.8788295 |
| VARS2 | 33.9695765 | 0.110392629 | 0.32907717 | 0.33546122 | 0.73727719 | 0.9429136 |
| CCHCR1 | 72.6870586 | 0.254549717 | 0.2210224 | 1.15169193 | 0.24944769 | 0.70629672 |
| TCF19 | 37.6998712 | 0.093666005 | 0.62735067 | 0.14930406 | 0.88131371 | 0.97768262 |
| POU5F1 | 320.388956 | 0.876001741 | 0.43692545 | 2.00492267 | 0.04497131 | 0.35569764 |
| HLA-C | 899.143165 | -1.185017061 | 0.69067485 | -1.715738 | 0.08621001 | 0.46595201 |
| HLA-B | 1351.27791 | -2.314193708 | 0.67869661 | -3.4097617 | 0.0006502 | 0.03001789 |
| MICA | 64.9182135 | -0.468205326 | 0.32905861 | -1.422863 | 0.15477587 | 0.59091117 |
| NFKBIL1 | 50.969583 | -0.135405646 | 0.31997536 | -0.4231752 | 0.67216742 | 0.92498948 |
| PRRC2A | 192.672748 | 0.085399188 | 0.19552632 | 0.43676569 | 0.66228128 | 0.92284375 |
| BAG6 | 95.917251 | -0.337524306 | 0.26851629 | -1.2569975 | 0.20875454 | 0.66655368 |
| C6orf47 | 39.287563 | -0.386911448 | 0.41634726 | -0.9292998 | 0.35273372 | 0.78314007 |
| GPANK1 | 37.1421694 | -0.322837196 | 0.31247474 | -1.0331625 | 0.30152784 | 0.74814891 |
| DDAH2 | 158.872248 | -0.356758422 | 0.27572228 | -1.293905 | 0.19569825 | 0.64948524 |
| CLIC1 | 501.810502 | -0.946624982 | 0.37440474 | -2.5283467 | 0.01146011 | 0.16915774 |
| VARS | 42.3769479 | -0.338365985 | 0.39007239 | -0.8674441 | 0.38569874 | 0.8060156 |
| LSM2 | 103.174911 | -0.240870062 | 0.32507787 | -0.7409611 | 0.45871702 | 0.84564214 |
| HSPA1A | 469.176785 | 1.427738549 | 0.45299848 | 3.15175132 | 0.00162294 | 0.05468957 |
| HSPA1B | 274.467515 | 1.520354457 | 0.54822408 | 2.77323547 | 0.00555019 | 0.11943745 |
| C6orf48 | 87.0119629 | 0.170478673 | 0.46851798 | 0.36386794 | 0.71595663 | 0.93573028 |
| NEU1 | 95.6017845 | -0.584495815 | 0.41042612 | -1.4241194 | 0.15441191 | 0.59038189 |
| SLC44A4 | 144.813663 | -0.662819911 | 0.56482486 | -1.1734963 | 0.24059682 | 0.6963541 |
| EHMT2 | 64.2351746 | 0.058877304 | 0.28106505 | 0.20947928 | 0.83407411 | 0.96749268 |
| NELFE | 51.6654822 | -0.192947122 | 0.40735139 | -0.4736626 | 0.63574052 | 0.91454294 |
| STK19 | 73.0370909 | 0.302902831 | 0.50613528 | 0.5984622 | 0.54953157 | 0.88888577 |
| ATF6B | 126.825492 | 0.144302412 | 0.20720978 | 0.69640735 | 0.48617376 | 0.85912348 |
| AGPAT1 | 63.3659143 | 0.321910341 | 0.37281374 | 0.86346157 | 0.38788374 | 0.8065026 |
| RNF5 | 114.921342 | 0.156762251 | 0.52413186 | 0.29908934 | 0.76487188 | 0.95378284 |
| PBX2 | 82.142906 | 0.44471976 | 0.24156227 | 1.841015 | 0.06561936 | 0.41788422 |
| AL662796.1 | 86.5302745 | 0.725034349 | 0.71680312 | 1.01148326 | 0.31178519 | 0.75531873 |
| HLA-DRA | 402.005642 | -0.099531448 | 0.68092904 | -0.1461701 | 0.88378713 | 0.97878487 |
| HLA-DRB5 | 65.4841813 | 1.055502989 | 0.89093858 | 1.18470904 | 0.23613251 | 0.69335782 |
| HLA-DRB1 | 250.775874 | 1.022366981 | 1.01767153 | 1.00461391 | 0.3150828 | 0.75892156 |
| HLA-DQB1 | 70.6845642 | 0.607415644 | 1.42217665 | 0.42710281 | 0.66930445 | 0.92463889 |
| PSMB8 | 122.488861 | -0.891185612 | 0.57312759 | -1.5549515 | 0.11995762 | 0.53747577 |
| TAP1 | 67.6844114 | -1.228870112 | 0.91076924 | -1.3492662 | 0.17725149 | 0.62617642 |
| BRD2 | 260.124856 | 0.044016638 | 0.1651305 | 0.26655668 | 0.78981051 | 0.95801395 |
| HLA-DPA1 | 73.541158 | -1.375360868 | 0.91003621 | -1.5113254 | 0.13070556 | 0.55395564 |
| HLA-DPB1 | 82.8702073 | -0.336195046 | 0.80514727 | -0.4175572 | 0.67627089 | 0.9268955 |
| RXRB | 54.2316518 | 0.110524263 | 0.25529014 | 0.43293588 | 0.66506135 | 0.92324595 |
| SLC39A7 | 89.0599268 | -0.447538306 | 0.43540544 | -1.0278657 | 0.30401301 | 0.74966033 |
| RING1 | 75.6017638 | 0.183608267 | 0.21776997 | 0.84312943 | 0.39915606 | 0.81495334 |
| VPS52 | 82.6105669 | 0.599889942 | 0.21702473 | 2.76415474 | 0.00570705 | 0.1213371 |
| RPS18 | 518.752546 | -0.14483767 | 0.44937161 | -0.3223116 | 0.74721667 | 0.94632395 |
| B3GALT4 | 33.1307452 | -0.902194739 | 0.32327889 | -2.7907629 | 0.0052584 | 0.11544342 |
| WDR46 | 43.1995083 | -0.234789488 | 0.33688077 | -0.6969513 | 0.48583328 | 0.85912348 |
| PFDN6 | 72.7270522 | -0.279070931 | 0.32244822 | -0.8654752 | 0.38677804 | 0.8065026 |
| RGL2 | 127.310323 | 0.238876391 | 0.21419708 | 1.11521778 | 0.26475711 | 0.71989632 |
| TAPBP | 341.435843 | -0.324707274 | 0.3899669 | -0.8326534 | 0.40504022 | 0.81946362 |
| DAXX | 45.2958065 | 0.15135926 | 0.3984092 | 0.37990905 | 0.70401293 | 0.93241442 |
| PHF1 | 101.10297 | 0.248422421 | 0.30591166 | 0.81207243 | 0.41675007 | 0.82489761 |
| CUTA | 187.678483 | -0.002697976 | 0.4925087 | -0.005478 | 0.99562919 | 0.99908534 |
| SYNGAP1 | 58.0568225 | 0.044712697 | 0.24229973 | 0.18453466 | 0.85359404 | 0.97062426 |
| BAK1 | 46.0390744 | 0.542884936 | 0.44901874 | 1.2090474 | 0.22664464 | 0.68470332 |
| ITPR3 | 300.284641 | -0.782868074 | 0.56167329 | -1.393814 | 0.16337379 | 0.6028723 |
| UQCC2 | 79.8671521 | 0.131308016 | 0.29922965 | 0.4388202 | 0.66079182 | 0.92284375 |
| LEMD2 | 82.7119426 | -0.436709273 | 0.3245768 | -1.3454728 | 0.17847259 | 0.62759322 |
| HMGA1 | 108.619218 | -1.008848466 | 0.49314859 | -2.0457292 | 0.04078302 | 0.34079725 |
| SMIM29 | 48.8744807 | -0.696382207 | 0.36876519 | -1.8884163 | 0.05897008 | 0.40122703 |
| NUDT3 | 227.058555 | -0.335811486 | 0.38897135 | -0.8633322 | 0.38795485 | 0.8065026 |
| C6orf106 | 409.340949 | 0.718544982 | 0.35032217 | 2.05109765 | 0.04025744 | 0.33814918 |
| SNRPC | 90.9457735 | -0.551045487 | 0.42823551 | -1.2867814 | 0.19817049 | 0.65370687 |
| UHRF1BP1 | 144.855245 | -0.047649822 | 0.22129609 | -0.2153216 | 0.82951661 | 0.96642181 |
| TAF11 | 113.479095 | 0.223230785 | 0.26804932 | 0.83279744 | 0.40495898 | 0.81945744 |
| ANKS1A | 724.38962 | -0.199948987 | 0.5026675 | -0.3977758 | 0.69079543 | 0.93073907 |
| SCUBE3 | 44.1714063 | 1.428878927 | 0.58728987 | 2.43300454 | 0.01497411 | 0.19879321 |
| ZNF76 | 162.351856 | -0.004986656 | 0.28205065 | -0.01768 | 0.98589413 | 0.99587186 |
| PPARD | 198.793181 | 0.262222749 | 0.28521213 | 0.91939551 | 0.35788874 | 0.78674827 |
| RPL10A | 622.279298 | 0.054616417 | 0.51740141 | 0.10555908 | 0.91593219 | 0.98520052 |
| TEAD3 | 74.7643368 | -0.277855056 | 0.30491639 | -0.91125 | 0.36216368 | 0.79051672 |
| SRPK1 | 193.289811 | -0.315948318 | 0.28128497 | -1.1232321 | 0.26133892 | 0.71941999 |
| MAPK14 | 650.364842 | -0.830203693 | 0.40962962 | -2.0267179 | 0.04269127 | 0.3484612 |
| MAPK13 | 63.4962841 | -0.432540857 | 0.25236091 | -1.7139772 | 0.08653291 | 0.46649429 |
| BRPF3 | 42.6681903 | -0.236102033 | 0.32253883 | -0.7320112 | 0.46416173 | 0.84965121 |
| KCTD20 | 238.488303 | 0.567730522 | 0.42118912 | 1.34792303 | 0.17768314 | 0.6269762 |
| STK38 | 379.063902 | 0.33712964 | 0.19365839 | 1.74084705 | 0.08171039 | 0.45528236 |
| SRSF3 | 352.382152 | 0.098096411 | 0.38694508 | 0.25351508 | 0.79987021 | 0.96043078 |
| CDKN1A | 124.270842 | -0.43274921 | 0.89163305 | -0.4853445 | 0.62743198 | 0.91061758 |
| C6orf89 | 222.641678 | -0.289001698 | 0.25084048 | -1.1521334 | 0.24926626 | 0.70622072 |
| MTCH1 | 265.83923 | -0.034397454 | 0.28653583 | -0.1200459 | 0.90444678 | 0.98320957 |
| PIM1 | 57.9290561 | -0.222533466 | 0.5882506 | -0.3782971 | 0.70520993 | 0.93241442 |
| TBC1D22B | 143.619647 | -0.056022307 | 0.22386713 | -0.250248 | 0.80239555 | 0.96071123 |
| RNF8 | 110.088444 | -0.327666793 | 0.32171195 | -1.0185099 | 0.30843572 | 0.75313873 |
| CMTR1 | 88.2070323 | 0.180311304 | 0.25080254 | 0.71893733 | 0.47217954 | 0.85244471 |
| ZFAND3 | 3112.5367 | 0.538505688 | 0.26027439 | 2.06899225 | 0.03854681 | 0.32997578 |
| BTBD9 | 1783.20776 | 0.288026802 | 0.24114459 | 1.19441535 | 0.23231553 | 0.69186326 |
| GLO1 | 340.736474 | -0.060308094 | 0.20917141 | -0.288319 | 0.77310256 | 0.95543807 |

| | | | | | | |
|---|---|---|---|---|---|---|
| SAYSD1 | 48.4526542 | -0.493152054 | 0.43334819 | -1.1380042 | 0.25511873 | 0.71259177 |
| KCNK5 | 40.8711611 | -0.369251546 | 0.47563486 | -0.7763341 | 0.43755176 | 0.83420562 |
| KIF6 | 82.7005009 | 0.121635356 | 0.56980935 | 0.21346676 | 0.83096291 | 0.96648703 |
| MOCS1 | 50.9690111 | -0.929237334 | 0.35176439 | -2.6416469 | 0.0082504 | 0.14386723 |
| UNC5CL | 104.4415 | -1.473501855 | 0.66591285 | -2.2127548 | 0.02691456 | 0.2787199 |
| OARD1 | 163.423614 | -0.347451079 | 0.27772015 | -1.2510834 | 0.21090404 | 0.66917175 |
| NFYA | 93.9386223 | 0.059717995 | 0.34006983 | 0.1756051 | 0.8606042 | 0.97304262 |
| FOXP4 | 36.4824831 | -0.700757306 | 0.36461159 | -1.9219282 | 0.05461479 | 0.39095834 |
| TFEB | 132.300204 | -0.258526132 | 0.59530056 | -0.4342783 | 0.66408634 | 0.92300139 |
| USP49 | 89.8868701 | -0.038235939 | 0.26424827 | -0.144697 | 0.88495009 | 0.97891513 |
| CCND3 | 253.564351 | -0.303920167 | 0.48203158 | -0.6304985 | 0.52836851 | 0.87837912 |
| TAF8 | 94.1253299 | -0.258286556 | 0.34743791 | -0.7434035 | 0.45723743 | 0.84409989 |
| C6orf132 | 85.442489 | 0.239230499 | 0.36246038 | 0.66001834 | 0.50924206 | 0.87094607 |
| MRPS10 | 61.162782 | -0.071626692 | 0.27964481 | -0.2561345 | 0.79784696 | 0.95999536 |
| TRERF1 | 407.658839 | 0.522639943 | 0.48380719 | 1.08026493 | 0.28002422 | 0.72945471 |
| UBR2 | 701.839416 | 0.174808636 | 0.24307152 | 0.71916545 | 0.47203899 | 0.85244471 |
| BICRAL | 333.65748 | -0.143819964 | 0.36520781 | -0.3938031 | 0.69372642 | 0.93105058 |
| RPL7L1 | 198.293041 | -0.281472309 | 0.30470159 | -0.9237638 | 0.35560929 | 0.78607579 |
| C6orf226 | 34.5428638 | -0.575921261 | 0.40345373 | -1.4274778 | 0.15344221 | 0.58860244 |
| CNPY3 | 50.0572851 | -0.459290344 | 0.30201113 | -1.5207729 | 0.12831684 | 0.54843482 |
| PEX6 | 74.7761668 | -0.632042913 | 0.36701085 | -1.7221368 | 0.08504473 | 0.46310155 |
| PPP2R5D | 40.3597724 | 0.519743237 | 0.27075863 | 1.91958142 | 0.05491079 | 0.39224566 |
| MEA1 | 56.0749669 | -0.347448663 | 0.56556351 | -0.6143407 | 0.53899024 | 0.88287241 |
| KLHDC3 | 98.9300556 | -0.05423853 | 0.2512808 | -0.2158483 | 0.82910601 | 0.96609195 |
| RRP36 | 108.687688 | 0.025024152 | 0.32689055 | 0.07655208 | 0.93897988 | 0.98848592 |
| CUL7 | 50.0421931 | -0.393445146 | 0.32493152 | -1.2108556 | 0.22595076 | 0.68375483 |
| KLC4 | 71.4686265 | -0.084038866 | 0.26827897 | -0.3132518 | 0.75408938 | 0.94983584 |
| MRPL2 | 72.3577034 | 0.14031487 | 0.38989421 | 0.35987934 | 0.71893737 | 0.9359691 |
| CUL9 | 73.1932252 | 0.123966837 | 0.29793413 | 0.41608807 | 0.67734556 | 0.9268955 |
| DNPH1 | 146.276341 | -0.062557851 | 0.52150612 | -0.1199561 | 0.90451791 | 0.98320957 |
| ZNF318 | 64.2234155 | 0.957724972 | 0.33452804 | 2.86291386 | 0.00419765 | 0.10017148 |
| ABCC10 | 61.5084902 | 0.486249517 | 0.40920067 | 1.18829111 | 0.23471874 | 0.69335782 |
| TJAP1 | 52.3970953 | 0.062222697 | 0.27412556 | 0.22698612 | 0.82043454 | 0.96502145 |
| POLR1C | 92.3671117 | -0.259585514 | 0.24847765 | -1.0447037 | 0.29615995 | 0.74305872 |
| YIPF3 | 129.309496 | -0.231188712 | 0.32443697 | -0.7125844 | 0.47610299 | 0.85414749 |
| XPO5 | 341.984295 | 0.145941225 | 0.26052359 | 0.5601843 | 0.57535373 | 0.8980797 |
| POLH | 102.336971 | -0.145678779 | 0.28884963 | -0.5043412 | 0.61402161 | 0.90760881 |
| GTPBP2 | 76.5645391 | -0.677325685 | 0.33655593 | -2.0125204 | 0.0441651 | 0.35332084 |
| MRPS18A | 67.7441201 | -0.326135491 | 0.44630517 | -0.7307455 | 0.46493462 | 0.84991999 |
| VEGFA | 884.575293 | -0.681655717 | 0.8534035 | -0.7987496 | 0.42443561 | 0.82774157 |
| MRPL14 | 110.500499 | -0.577239899 | 0.44824685 | -1.2877723 | 0.19782522 | 0.652979 |
| TMEM63B | 81.1050278 | 0.07636197 | 0.28251039 | 0.27029791 | 0.78693107 | 0.95773286 |
| SLC29A1 | 126.068227 | -0.704494735 | 0.39873613 | -1.7668194 | 0.07725847 | 0.44472917 |
| HSP90AB1 | 1056.49456 | -0.123069713 | 0.23128731 | -0.5321075 | 0.59465155 | 0.90323392 |
| SLC35B2 | 56.7334072 | -0.564111826 | 0.51193277 | -1.1019256 | 0.27049402 | 0.72343657 |
| CDC5L | 133.599342 | 0.177206064 | 0.24933516 | 0.71071431 | 0.47726129 | 0.85455294 |
| SUPT3H | 869.03028 | 0.38106962 | 0.2802611 | 1.35969503 | 0.17392645 | 0.62125058 |
| ENPP4 | 109.088592 | -0.149838764 | 0.39298012 | -0.3812884 | 0.70298925 | 0.93241442 |
| ENPP5 | 77.7837416 | -0.500044766 | 0.39237566 | -1.2744031 | 0.20252059 | 0.6574307 |
| RCAN2 | 167.729883 | 0.999729877 | 0.70967133 | 1.40872237 | 0.15891728 | 0.59747329 |
| CYP39A1 | 51.5788481 | -0.579278367 | 0.55045352 | -1.0523656 | 0.29263183 | 0.73844969 |
| SLC25A27 | 200.286296 | 0.486738821 | 0.47040968 | 1.0347126 | 0.30080315 | 0.7478505 |
| ADGRF1 | 495.276723 | 0.138586816 | 0.42943208 | 0.32272115 | 0.74690643 | 0.94604537 |
| TNFRSF21 | 52.6242727 | -1.702172561 | 0.76006228 | -2.2395172 | 0.02512228 | 0.26892903 |
| CD2AP | 2070.65024 | -0.159782423 | 0.32030067 | -0.4988514 | 0.6178841 | 0.90794578 |
| MUT | 124.812537 | 0.103880459 | 0.30136793 | 0.34469647 | 0.73032257 | 0.94048525 |
| TFAP2B | 210.254727 | -0.348385968 | 0.54684346 | -0.6370854 | 0.52406923 | 0.87744743 |
| PKHD1 | 7003.51155 | 0.703447043 | 0.30807621 | 2.28335398 | 0.02240953 | 0.25166501 |
| MCM3 | 86.5062388 | -0.125724242 | 0.33333429 | -0.3771716 | 0.70604605 | 0.93241442 |
| EFHC1 | 184.288594 | 0.017256181 | 0.21101781 | 0.08177594 | 0.93482489 | 0.98822345 |
| TRAM2 | 53.6106891 | 0.257634857 | 0.48108921 | 0.53552409 | 0.59228751 | 0.90233654 |
| TMEM14A | 58.9452828 | 0.186619839 | 0.35081259 | 0.53196449 | 0.59475059 | 0.90323392 |
| GSTA4 | 81.4650924 | 0.098096402 | 0.36589334 | 0.26810109 | 0.78862151 | 0.95801395 |
| ICK | 71.3323437 | -0.114731724 | 0.31648177 | -0.3625224 | 0.7169617 | 0.93573028 |
| FBXO9 | 215.19846 | 0.118780667 | 0.24066881 | 0.49354409 | 0.62162819 | 0.9099963 |
| ELOVL5 | 111.911068 | 0.556615961 | 0.50925261 | 1.09300562 | 0.27439133 | 0.7263609 |
| GCLC | 115.027073 | -1.054019371 | 0.41621352 | -2.5324006 | 0.01132845 | 0.16768662 |
| LRRC1 | 390.500201 | 1.662174591 | 0.44338635 | 3.74881771 | 0.00017767 | 0.01275331 |
| FAM83B | 339.547571 | 0.336898829 | 0.44867924 | 0.75086788 | 0.45273217 | 0.84248055 |
| DST | 559.208721 | -0.466539106 | 0.39288703 | -1.1874637 | 0.23504475 | 0.69335782 |
| ZNF451 | 323.162196 | 0.370414771 | 0.14497935 | 2.55494841 | 0.01062035 | 0.16452817 |
| ZNF451-AS1 | 42.2740003 | -0.326192195 | 0.63933907 | -0.5102022 | 0.60990981 | 0.90587512 |
| PRIM2 | 300.592513 | 0.108174091 | 0.34428137 | 0.31420257 | 0.75336719 | 0.94963774 |
| AL021368.2 | 39.5411637 | 0.22248054 | 0.25614144 | 0.86858472 | 0.38507431 | 0.80565309 |
| PTP4A1 | 680.466116 | 1.265405207 | 0.97801637 | 1.2938487 | 0.19571769 | 0.64948524 |
| PHF3 | 631.113904 | 0.03396543 | 0.21733136 | 0.15628408 | 0.87580911 | 0.97622753 |
| EYS | 55.0979432 | -1.067719312 | 0.74168976 | -1.4395767 | 0.14998719 | 0.5834691 |
| LMBRD1 | 473.484187 | 0.492643011 | 0.15739882 | 3.12990278 | 0.00174864 | 0.057666 |
| COL19A1 | 109.398385 | 0.055770726 | 0.36179831 | 0.15414866 | 0.87749252 | 0.97640529 |
| FAM135A | 453.221931 | -0.399886417 | 0.29639088 | -1.349186 | 0.17727724 | 0.62617642 |
| SDHAF4 | 104.335052 | 0.015402682 | 0.45530949 | 0.03382904 | 0.97301348 | 0.9939318 |
| SMAP1 | 844.568502 | 0.535639875 | 0.3890375 | 1.37683354 | 0.16856372 | 0.61338465 |
| OGFRL1 | 58.0650118 | -1.013038483 | 0.46698975 | -2.1692949 | 0.0300603 | 0.29169628 |
| AL136164.2 | 330.786573 | 1.086746145 | 0.50300775 | 2.16049582 | 0.03073431 | 0.29495928 |
| LINC00472 | 117.955877 | -0.416708824 | 0.48252772 | -0.8635956 | 0.38781007 | 0.8065026 |
| AC019205.1 | 77.8355313 | 0.181175953 | 0.32086849 | 0.56464239 | 0.57231703 | 0.89797476 |
| OOEP | 583.097435 | -0.287507873 | 0.46524516 | -0.6179707 | 0.53659466 | 0.88156639 |
| MTO1 | 98.809049 | 0.287375846 | 0.25940275 | 1.10783656 | 0.26793241 | 0.72276232 |
| EEF1A1 | 3240.38349 | 0.081224324 | 0.47521258 | 0.17092208 | 0.86428503 | 0.97466129 |
| SLC17A5 | 117.460118 | -0.328249657 | 0.29841341 | -1.0999829 | 0.27133955 | 0.72343657 |
| COX7A2 | 266.905879 | 0.057540615 | 0.37327745 | 0.15414972 | 0.87749169 | 0.97640529 |
| TMEM30A | 237.715691 | -0.071007475 | 0.2967937 | -0.2392486 | 0.81091283 | 0.96288868 |
| SENP6 | 1049.66499 | 0.334501269 | 0.20846651 | 1.60458036 | 0.10858619 | 0.51254482 |
| MYO6 | 2330.19756 | 0.321561244 | 0.34426449 | 0.93405289 | 0.35027661 | 0.78223232 |
| MEI4 | 145.409418 | 0.284293564 | 0.26025651 | 1.09235909 | 0.2746753 | 0.7263609 |
| IRAK1BP1 | 245.397344 | -0.245859289 | 0.41783444 | -0.5884132 | 0.556255 | 0.89082402 |
| PHIP | 998.745758 | 0.264956798 | 0.19098623 | 1.38730835 | 0.16534776 | 0.60673827 |
| HMGN3 | 582.633102 | -0.179304355 | 0.26038325 | -0.6886171 | 0.49106426 | 0.86143519 |
| LCA5 | 120.683831 | 0.306243753 | 0.28852022 | 1.06142908 | 0.28849495 | 0.73656893 |
| SH3BGRL2 | 187.974205 | 0.786771698 | 0.20643438 | 3.81124354 | 0.00013827 | 0.01065488 |
| BCKDHB | 613.576134 | -0.03631279 | 0.2745455 | -0.1322651 | 0.8947746 | 0.9821637 |
| TENT5A | 132.14608 | -1.424671203 | 0.58991401 | -2.4150489 | 0.0157331 | 0.20433451 |
| IBTK | 533.364681 | 0.14670395 | 0.36610789 | 0.40071235 | 0.68863192 | 0.93036007 |
| TPBG | 42.0088486 | -0.923273558 | 0.57157748 | -1.6153078 | 0.10624405 | 0.50765436 |
| UBE3D | 290.196937 | -0.239795551 | 0.24348104 | -0.9848633 | 0.3246912 | 0.77020457 |
| DOPEY1 | 589.246112 | 0.095654439 | 0.39879855 | 0.23985654 | 0.81044148 | 0.96286438 |
| PGM3 | 105.906882 | -0.547465478 | 0.32563974 | -1.6811998 | 0.0927241 | 0.48153712 |
| ME1 | 46.689117 | -2.294021861 | 0.60976262 | -3.7621556 | 0.00016846 | 0.01234553 |
| CYB5R4 | 147.594126 | 0.351140756 | 0.33719198 | 1.04136748 | 0.29770503 | 0.74550746 |
| CEP162 | 252.502516 | 0.194153365 | 0.22379781 | 0.86753917 | 0.38564666 | 0.8060156 |
| SNX14 | 405.997074 | 0.140635688 | 0.15926673 | 0.88301989 | 0.37722553 | 0.80091643 |
| SYNCRIP | 139.779391 | -0.26524433 | 0.17895665 | -1.4821709 | 0.13829483 | 0.56438077 |
| ZNF292 | 1531.6656 | -1.206421765 | 0.44255909 | -2.7260128 | 0.00641045 | 0.12805535 |

| | | | | | | |
|---|---|---|---|---|---|---|
| SMIM8 | 218.369411 | 0.198145777 | 0.39417654 | 0.50268282 | 0.61518729 | 0.90792412 |
| SLC35A1 | 33.442596 | 0.185211667 | 0.57774828 | 0.32057502 | 0.74853247 | 0.94678489 |
| RARS2 | 279.497076 | 0.347886299 | 0.25396937 | 1.36979626 | 0.17075051 | 0.61726986 |
| ORC3 | 192.489778 | 0.083466612 | 0.21384286 | 0.39031751 | 0.69630178 | 0.93130502 |
| AKIRIN2 | 62.6266183 | -0.167975105 | 0.31295368 | -0.5367411 | 0.59144645 | 0.90233654 |
| RNGTT | 492.302251 | 0.531670148 | 0.2765234 | 1.92269493 | 0.05451837 | 0.39095834 |
| PNRC1 | 311.833245 | -0.182124418 | 0.38002224 | -0.4792467 | 0.63176311 | 0.9133034 |
| PM20D2 | 111.42854 | -0.319957807 | 0.22343333 | -1.4320057 | 0.15214218 | 0.5870582 |
| UBE2J1 | 140.921241 | 0.134824163 | 0.2593675 | 0.51981902 | 0.60318972 | 0.90369786 |
| RRAGD | 92.9830235 | -1.797185886 | 0.51684157 | -3.4772472 | 0.00050659 | 0.02532014 |
| ANKRD6 | 356.818138 | -0.048325347 | 0.2047855 | -0.2359803 | 0.81344795 | 0.96288868 |
| LYRM2 | 139.055121 | -0.522684632 | 0.3400535 | -1.5370659 | 0.12427718 | 0.54273186 |
| MDN1 | 312.247734 | 0.349623709 | 0.24728205 | 1.41386608 | 0.15740123 | 0.59507808 |
| CASP8AP2 | 142.875892 | 0.254242761 | 0.24151491 | 1.05270004 | 0.29247849 | 0.73844969 |
| BACH2 | 715.295586 | 0.510126035 | 0.64356616 | 0.79265516 | 0.42797876 | 0.8292138 |
| MAP3K7 | 317.403628 | 0.146229083 | 0.25318457 | 0.57755921 | 0.56356175 | 0.89295783 |
| MANEA | 70.5676054 | 0.088071763 | 0.23612076 | 0.37299458 | 0.70915248 | 0.93391076 |
| UFL1-AS1 | 99.3308094 | -0.899297619 | 0.48483301 | -1.8548605 | 0.06361615 | 0.41357646 |
| UFL1 | 196.906623 | -0.026269268 | 0.17433734 | -0.1506807 | 0.88022762 | 0.97740892 |
| NDUFAF4 | 51.4114665 | -0.127934855 | 0.52818319 | -0.2422168 | 0.80861216 | 0.96245698 |
| KLHL32 | 126.264293 | 0.472479849 | 0.49455765 | 0.95535849 | 0.33939644 | 0.77836439 |
| MMS22L | 254.1414 | -1.409059685 | 0.46668133 | -3.0193187 | 0.00253344 | 0.0735219 |
| AL589740.1 | 60.8537199 | -1.031182982 | 0.56204486 | -1.8346987 | 0.06655034 | 0.41989619 |
| FBXL4 | 369.006691 | -0.144537962 | 0.3219575 | -0.4489349 | 0.65347861 | 0.91974964 |
| PNISR | 1032.76889 | 0.4173614 | 0.19780698 | 2.10994268 | 0.03486329 | 0.31514522 |
| AL513550.1 | 58.7164843 | -0.726335714 | 0.51272626 | -1.416615 | 0.15659552 | 0.59374856 |
| USP45 | 205.570842 | -0.15003027 | 0.3451085 | -0.4347336 | 0.66375579 | 0.92300139 |
| TSTD3 | 47.4523557 | -0.068724955 | 0.36152493 | -0.1900974 | 0.8492328 | 0.96991131 |
| CCNC | 303.309127 | -0.162835815 | 0.28441334 | -0.5725323 | 0.56696144 | 0.89454608 |
| SIM1 | 725.462292 | 0.992215461 | 0.36467845 | 2.72079547 | 0.0065125 | 0.12862701 |
| ASCC3 | 339.960792 | -0.922275377 | 0.20624115 | -4.4718301 | 7.76E-06 | 0.0015335 |
| GRIK2 | 153.368133 | -0.991515658 | 0.6538709 | -1.5163783 | 0.12942372 | 0.55140574 |
| HACE1 | 197.284276 | 0.302288152 | 0.29344076 | 1.03015053 | 0.30293935 | 0.74930478 |
| PREP | 134.255413 | -0.057532502 | 0.23541492 | -0.2443877 | 0.8069306 | 0.96218371 |
| ATG5 | 393.014223 | 0.171475294 | 0.21354324 | 0.80300033 | 0.42197454 | 0.8273701 |
| CRYBG1 | 1600.4954 | 0.927789967 | 0.42305782 | 2.19305711 | 0.02830326 | 0.28603492 |
| RTN4IP1 | 82.2373424 | -0.237011632 | 0.38660904 | -0.6130525 | 0.53984164 | 0.8833006 |
| QRSL1 | 135.740654 | 0.080983615 | 0.20228427 | 0.40034558 | 0.688902 | 0.93036007 |
| LINC02532 | 3118.4703 | -0.408202762 | 0.43219756 | -0.9444819 | 0.34492346 | 0.78180304 |
| CD24 | 3068.37802 | -0.479902567 | 0.29692277 | -1.6162538 | 0.10603945 | 0.50765436 |
| C6orf203 | 75.3657877 | -0.294667413 | 0.32991345 | -0.8931658 | 0.37176839 | 0.7954538 |
| PDSS2 | 709.094622 | 0.534527889 | 0.23649441 | 2.26021362 | 0.023808 | 0.26048096 |
| SOBP | 434.35978 | 2.128709487 | 0.65983871 | 3.22610577 | 0.00125487 | 0.04582242 |
| SCML4 | 58.3238506 | -1.026567878 | 0.47681769 | -2.1529568 | 0.03132208 | 0.29704233 |
| SEC63 | 382.472171 | 0.155999549 | 0.18526895 | 0.8420167 | 0.39977861 | 0.81574763 |
| OSTM1 | 122.397043 | -0.222300246 | 0.29236329 | -0.7603562 | 0.44704169 | 0.84014281 |
| SNX3 | 363.979179 | 0.098396237 | 0.33248954 | 0.29593784 | 0.76727755 | 0.95408212 |
| AFG1L | 247.01512 | 0.349865232 | 0.31951134 | 1.09500097 | 0.27351621 | 0.72531626 |
| FOXO3 | 658.850834 | -0.530464656 | 0.27732461 | -1.9127933 | 0.05577452 | 0.39368206 |
| ARMC2 | 186.555636 | -0.112949762 | 0.26709632 | -0.4228803 | 0.67238262 | 0.92498948 |
| SESN1 | 330.815969 | 0.127007472 | 0.27711469 | 0.45832097 | 0.64672186 | 0.91743182 |
| CEP57L1 | 93.0668038 | 0.41022006 | 0.37925079 | 1.08165909 | 0.27940404 | 0.72938673 |
| CD164 | 551.519243 | -0.030628014 | 0.26093483 | -0.117378 | 0.9065605 | 0.98320957 |
| PPIL6 | 125.706731 | -0.186902377 | 0.39592503 | -0.4720651 | 0.63688034 | 0.91454294 |
| MICAL1 | 34.0059309 | 0.223151746 | 0.31928344 | 0.69891424 | 0.48460562 | 0.85909499 |
| ZBTB24 | 38.5536957 | -0.410373425 | 0.41507704 | -0.9886681 | 0.32282556 | 0.76879139 |
| AK9 | 198.006622 | -0.173808366 | 0.23963318 | -0.7253101 | 0.46826182 | 0.85161077 |
| FIG4 | 179.03236 | -0.298181005 | 0.27301471 | -1.0921793 | 0.27475432 | 0.7263609 |
| WASF1 | 40.984607 | -0.889722924 | 0.33050108 | -2.6920424 | 0.00710159 | 0.13409852 |
| CDC40 | 101.127286 | -0.164798672 | 0.25079279 | -0.6571109 | 0.51110961 | 0.87138909 |
| CDK19 | 582.178194 | 0.072206367 | 0.34598181 | 0.2086999 | 0.83468252 | 0.96749268 |
| AMD1 | 262.966278 | 0.178274661 | 0.34715177 | 0.51353523 | 0.60757697 | 0.90510401 |
| GTF3C6 | 201.406498 | -0.094012876 | 0.47584248 | -0.1975714 | 0.84338039 | 0.96876046 |
| RPF2 | 76.5522086 | -0.564089464 | 0.26611296 | -2.1197369 | 0.03402824 | 0.3120805 |
| SLC16A10 | 450.132783 | 0.70758296 | 0.36950587 | 1.91494376 | 0.05549968 | 0.39368206 |
| MFSD4B | 100.003856 | 0.19808517 | 0.25414426 | 0.77942021 | 0.43573222 | 0.83364877 |
| REV3L | 929.339904 | 0.436530415 | 0.28963342 | 1.50718245 | 0.1317639 | 0.55435605 |
| TRAF3IP2-AS | 290.266769 | -0.272032434 | 0.33498778 | -0.8120667 | 0.41675333 | 0.82489761 |
| TRAF3IP2 | 105.785082 | -0.254689042 | 0.3761239 | -0.6771413 | 0.49831629 | 0.86591853 |
| TUBE1 | 107.48822 | 0.103228241 | 0.25266913 | 0.40855106 | 0.68286915 | 0.92808568 |
| FAM229B | 59.7904443 | -0.35445097 | 0.39086838 | -0.9068295 | 0.36449697 | 0.79268955 |
| MARCKS | 155.561469 | -1.253844404 | 0.32953492 | -3.8048909 | 0.00014187 | 0.01077365 |
| HDAC2 | 175.395678 | 0.093111943 | 0.16881388 | 0.55156567 | 0.58124596 | 0.90045866 |
| FRK | 635.326715 | 0.540256682 | 0.2536827 | 2.12965522 | 0.03320009 | 0.30784031 |
| NT5DC1 | 637.842634 | 0.305351674 | 0.22042304 | 1.38529835 | 0.16596126 | 0.60771278 |
| TSPYL4 | 58.6317339 | 0.344164433 | 0.36220986 | 0.95017964 | 0.34202098 | 0.77973697 |
| DSE | 313.160108 | 0.128043039 | 0.30260531 | 0.42313547 | 0.67219643 | 0.92498948 |
| TSPYL1 | 308.710282 | 0.301774281 | 0.40369974 | 0.74752161 | 0.45474876 | 0.84290184 |
| RWDD1 | 104.120208 | 0.189080672 | 0.40944151 | 0.46180143 | 0.64422373 | 0.91682456 |
| ZUP1 | 49.0685669 | 0.004145278 | 0.29432523 | 0.01408401 | 0.98876296 | 0.9964791 |
| KPNA5 | 155.041907 | 0.178951024 | 0.25989613 | 0.68854823 | 0.49110761 | 0.86143519 |
| DCBLD1 | 139.614433 | -0.317858662 | 0.29786802 | -1.0671124 | 0.28592108 | 0.73607046 |
| GOPC | 166.907504 | 0.205945864 | 0.27971728 | 0.73626435 | 0.46156984 | 0.84730283 |
| NUS1 | 183.669764 | 0.388211925 | 0.18074193 | 2.14787969 | 0.03172331 | 0.29951381 |
| SLC35F1 | 46.6194522 | -0.443746754 | 0.31761168 | -1.3971361 | 0.16237264 | 0.60211533 |
| CEP85L | 266.678195 | -0.405930846 | 0.36073609 | -1.1252848 | 0.26046836 | 0.7176381 |
| MCM9 | 240.562749 | -0.080419431 | 0.27539217 | -0.2920179 | 0.77027297 | 0.95482817 |
| ASF1A | 39.4121774 | -0.248190463 | 0.32421356 | -0.7655154 | 0.44396472 | 0.83873671 |
| AL365275.1 | 67.1245852 | -0.122808594 | 0.66921354 | -0.1835118 | 0.85439645 | 0.97115778 |
| FAM184A | 72.1874647 | 0.100674158 | 0.56351921 | 0.17865257 | 0.85821051 | 0.97237339 |
| MAN1A1 | 2158.07567 | 1.111859001 | 0.37890418 | 2.93440683 | 0.00334186 | 0.08777614 |
| TBC1D32 | 507.283107 | -0.162324497 | 0.26629811 | -0.6095593 | 0.54215376 | 0.88450129 |
| HSF2 | 60.944035 | 0.182303207 | 0.23403665 | 0.77895152 | 0.43600827 | 0.83364877 |
| SERINC1 | 348.956307 | -0.177353981 | 0.33307565 | -0.5324736 | 0.59439804 | 0.90305763 |
| PKIB | 52.942757 | 0.089867752 | 0.52336601 | 0.1717111 | 0.86366466 | 0.97461028 |
| RNF217 | 266.001141 | -0.075746251 | 0.39665013 | -0.1909649 | 0.8485531 | 0.96991131 |
| TPD52L1 | 396.937549 | 0.185447477 | 0.38709368 | 0.47907648 | 0.63188422 | 0.9133034 |
| HDDC2 | 168.158007 | 0.010750802 | 0.33227433 | 0.0323552 | 0.97418879 | 0.9939318 |
| NCOA7 | 689.483031 | -0.363416287 | 0.28198869 | -1.2887619 | 0.19748086 | 0.65266459 |
| HINT3 | 141.975697 | 0.086661124 | 0.29250108 | 0.29627625 | 0.76701912 | 0.95408212 |
| TRMT11 | 139.612397 | 0.089501317 | 0.26397758 | 0.33904894 | 0.73457287 | 0.94215106 |
| RNF146 | 89.826986 | -0.17100128 | 0.2705739 | -0.6319947 | 0.52739032 | 0.87810931 |
| ECHDC1 | 222.168234 | -0.554896006 | 0.27907278 | -1.9883559 | 0.04677234 | 0.36336111 |
| PTPRK | 1473.44996 | -0.335588623 | 0.33981628 | -0.987559 | 0.32336868 | 0.76915656 |
| ARHGAP18 | 297.212279 | 0.079140748 | 0.40058478 | 0.19756305 | 0.84338695 | 0.96876046 |
| L3MBTL3 | 96.0163804 | -0.378932014 | 0.46064466 | -0.8226124 | 0.41072844 | 0.82265937 |
| EPB41L2 | 71.6817193 | -0.795204873 | 0.44741239 | -1.7773421 | 0.07551197 | 0.44087212 |
| AKAP7 | 139.493989 | 0.355502527 | 0.44121556 | 0.80573434 | 0.42039604 | 0.82694802 |
| MED23 | 178.791097 | 0.354494524 | 0.18258401 | 1.94154203 | 0.05219257 | 0.38343909 |
| STX7 | 476.562699 | 0.10878694 | 0.27692453 | 0.39283966 | 0.69443791 | 0.93105058 |
| SLC18B1 | 181.691736 | -0.152313005 | 0.43237714 | -0.3522689 | 0.72463663 | 0.93895336 |
| RPS12 | 499.539189 | 0.332211856 | 0.40378401 | 0.82274643 | 0.41065221 | 0.82265937 |
| EYA4 | 2129.66239 | 1.44274197 | 0.33475773 | 4.30980931 | 1.63E-05 | 0.00263444 |

| | | | | | | |
|---|---|---|---|---|---|---|
| TARID | 55.6600659 | 0.535982944 | 0.65729756 | 0.81543425 | 0.41482379 | 0.82461178 |
| TBPL1 | 56.1180904 | 0.123545016 | 0.30844461 | 0.40054198 | 0.68875737 | 0.93036007 |
| SGK1 | 198.832939 | 1.669800304 | 0.7555001 | 2.21019203 | 0.02709184 | 0.27917411 |
| HBS1L | 204.124683 | 0.396545878 | 0.23034525 | 1.72152833 | 0.085155 | 0.46310155 |
| AHI1 | 1105.49054 | 0.354204368 | 0.26554011 | 1.33390159 | 0.18223611 | 0.63281459 |
| LINC00271 | 363.429699 | -0.421002277 | 0.4959034 | -0.8489603 | 0.39590341 | 0.8120904 |
| PDE7B | 1959.59449 | 0.139496301 | 0.50003325 | 0.27897405 | 0.78026474 | 0.95639467 |
| AL138828.1 | 252.833254 | -1.078129262 | 0.51480005 | -2.094268 | 0.03623611 | 0.31938981 |
| BCLAF1 | 456.898232 | 0.263463476 | 0.14599106 | 1.80465489 | 0.0711287 | 0.43107604 |
| MAP7 | 920.815313 | 0.357066829 | 0.17309747 | 2.06280793 | 0.03913088 | 0.33313698 |
| MAP3K5 | 459.41119 | -1.023647479 | 0.49735119 | -2.0581985 | 0.03957108 | 0.3347094 |
| PEX7 | 166.627413 | 0.224655912 | 0.20327036 | 1.10520741 | 0.26906973 | 0.72322263 |
| IFNGR1 | 158.027757 | -0.034660074 | 0.41141807 | -0.0842454 | 0.93286133 | 0.98822345 |
| PERP | 256.596157 | -0.611870995 | 0.35509199 | -1.7231338 | 0.08486434 | 0.46310155 |
| ARFGEF3 | 204.654468 | -1.477316017 | 0.42955766 | -3.4391565 | 0.00058353 | 0.02767146 |
| HEBP2 | 235.340133 | -0.379708887 | 0.40172891 | -0.9451869 | 0.34456348 | 0.78180304 |
| NHSL1 | 693.210837 | 0.118612234 | 0.46438187 | 0.2554196 | 0.79839903 | 0.96000168 |
| CCDC28A | 94.2970194 | -0.153022175 | 0.41950266 | -0.3647705 | 0.71528276 | 0.93573028 |
| REPS1 | 324.356715 | 0.166870462 | 0.2038153 | 0.81873374 | 0.41293834 | 0.82430359 |
| HECA | 74.1678998 | 1.325371197 | 0.55495822 | 2.38823601 | 0.01692947 | 0.21364736 |
| CITED2 | 79.1864937 | 0.069481326 | 0.37641292 | 0.18458805 | 0.85355216 | 0.97062426 |
| VTA1 | 156.248948 | -0.090925539 | 0.24872011 | -0.3655737 | 0.71468318 | 0.93573028 |
| ADGRG6 | 100.170578 | -1.645495442 | 0.64596 | -2.5473643 | 0.01085401 | 0.16452817 |
| HIVEP2 | 907.045303 | 0.937934195 | 0.55878615 | 1.67852084 | 0.09324547 | 0.48233587 |
| AIG1 | 618.719664 | 0.312147826 | 0.12878219 | 2.42384309 | 0.01535724 | 0.20199029 |
| ADAT2 | 63.1434075 | 0.171242668 | 0.26909629 | 0.63636205 | 0.52454047 | 0.87772379 |
| PEX3 | 185.139956 | -0.239604964 | 0.26774078 | -0.894914 | 0.37083308 | 0.79539603 |
| FUCA2 | 77.4179547 | -0.590911486 | 0.23061822 | -2.5622932 | 0.01039835 | 0.16289193 |
| PHACTR2 | 377.380165 | -0.357867554 | 0.54665431 | -0.6546506 | 0.51269276 | 0.87177451 |
| PLAGL1 | 166.691381 | -1.042010984 | 0.59343409 | -1.7559001 | 0.07910547 | 0.44916083 |
| SF3B5 | 213.931391 | -0.597781534 | 0.43852033 | -1.3631786 | 0.17282622 | 0.61900848 |
| UTRN | 1246.70008 | 0.565431933 | 0.22199638 | 2.54703218 | 0.01086434 | 0.16452817 |
| EPM2A | 104.727346 | -0.301347687 | 0.36286549 | -0.8304667 | 0.406275 | 0.82002585 |
| FBXO30 | 37.2840953 | 0.147454947 | 0.36633228 | 0.40251694 | 0.68730363 | 0.9302521 |
| AL356599.1 | 139.861638 | -0.693096143 | 0.41919912 | -1.6533817 | 0.09825321 | 0.49267637 |
| SHPRH | 623.863666 | 0.195806273 | 0.23811016 | 0.82233483 | 0.41088636 | 0.82271476 |
| RAB32 | 41.6884725 | -0.75260949 | 0.37985273 | -1.9813192 | 0.04755549 | 0.36819774 |
| STXBP5 | 869.700403 | 0.045098364 | 0.18015555 | 0.25033015 | 0.80233205 | 0.96071123 |
| SAMD5 | 770.128883 | -0.540843022 | 0.35423687 | -1.5267835 | 0.12681485 | 0.54692166 |
| AL033504.1 | 480.448606 | -0.747144171 | 0.40126562 | -1.8619691 | 0.06260745 | 0.41084915 |
| SASH1 | 547.665738 | -0.308285765 | 0.49632437 | -0.6211377 | 0.53450904 | 0.88137638 |
| TAB2 | 518.589448 | 0.205023164 | 0.1880625 | 1.09018632 | 0.27563108 | 0.7263609 |
| PPIL4 | 237.066849 | 0.591016137 | 0.23558948 | 2.50866951 | 0.01211868 | 0.17590979 |
| GINM1 | 349.733992 | -0.020884673 | 0.2519174 | -0.0829029 | 0.93392878 | 0.98822345 |
| KATNA1 | 144.423053 | 0.502800935 | 0.20535292 | 2.44847226 | 0.01434635 | 0.19356764 |
| LATS1 | 250.001389 | -0.160518248 | 0.26829359 | -0.5982933 | 0.54964426 | 0.88889487 |
| NUP43 | 102.303412 | 0.394431509 | 0.24199086 | 1.62994382 | 0.10311337 | 0.50171849 |
| PCMT1 | 233.592797 | 0.279070338 | 0.23343053 | 1.19551773 | 0.23188481 | 0.69186326 |
| LRP11 | 58.2500656 | -0.292592652 | 0.52654898 | -0.5556798 | 0.57842974 | 0.89953318 |
| PPP1R14C | 216.684624 | -1.196393349 | 0.44601508 | -2.6824056 | 0.00730948 | 0.13610131 |
| PLEKHG1 | 269.749613 | -0.288230719 | 0.33485112 | -0.8607728 | 0.38936321 | 0.80680464 |
| MTHFD1L | 521.037637 | -0.962231874 | 0.51112454 | -1.8825781 | 0.05975756 | 0.40360868 |
| AKAP12 | 109.753079 | 1.176130782 | 0.75957991 | 1.54839637 | 0.12152689 | 0.53989034 |
| RMND1 | 98.8131921 | -0.118034672 | 0.29286115 | -0.4030397 | 0.68691902 | 0.92992151 |
| ARMT1 | 59.7820491 | -0.295840591 | 0.39928538 | -0.7409252 | 0.45873881 | 0.84564214 |
| SYNE1 | 1557.42088 | 0.086820428 | 0.55623484 | 0.15608592 | 0.8759653 | 0.97622753 |
| MTRF1L | 58.7492777 | -0.597876265 | 0.35902812 | -1.665263 | 0.09586029 | 0.48710328 |
| OPRM1 | 47.5646245 | 0.778294332 | 0.84333373 | 0.92287823 | 0.35607067 | 0.78612469 |
| IPCEF1 | 471.333886 | 0.580655083 | 0.66502242 | 0.87313611 | 0.3825889 | 0.80405156 |
| CNKSR3 | 568.981084 | -0.12259148 | 0.36506268 | -0.3358094 | 0.7370146 | 0.9429136 |
| SCAF8 | 673.662711 | 0.529463843 | 0.2315256 | 2.28684792 | 0.0222047 | 0.25157326 |
| TIAM2 | 362.275049 | -0.057903061 | 0.34019508 | -0.1702055 | 0.86484856 | 0.97485201 |
| TFB1M | 192.521051 | -0.044742361 | 0.28499183 | -0.1569952 | 0.8752486 | 0.97622753 |
| ARID1B | 3251.88329 | 0.186517272 | 0.23122371 | 0.80665287 | 0.4198665 | 0.82694802 |
| TMEM242 | 87.3266976 | -0.108893558 | 0.23399767 | -0.4653617 | 0.64167244 | 0.91643619 |
| ZDHHC14 | 290.275109 | 0.442311135 | 0.40190038 | 1.10054918 | 0.27109292 | 0.72343657 |
| SNX9 | 667.529645 | -0.266039431 | 0.4678198 | -0.5686793 | 0.56957381 | 0.89626629 |
| SYNJ2 | 186.869828 | -1.457180778 | 0.78823336 | -1.8486667 | 0.06450595 | 0.41412378 |
| SERAC1 | 58.5143634 | -0.216489366 | 0.31089604 | -0.6963401 | 0.48621589 | 0.85912348 |
| GTF2H5 | 155.513972 | 0.011219502 | 0.44077017 | 0.02545431 | 0.97969259 | 0.99469867 |
| TULP4 | 520.772389 | -0.24325855 | 0.2419417 | -1.0054428 | 0.31468367 | 0.75823062 |
| TMEM181 | 163.22806 | -0.668891945 | 0.26269956 | -2.5462241 | 0.01088952 | 0.16452817 |
| DYNLT1 | 354.052983 | -0.196834678 | 0.59409887 | -0.3313164 | 0.74040553 | 0.94454656 |
| SYTL3 | 296.633044 | 0.362957472 | 1.27760207 | 0.28409274 | 0.77633931 | 0.95601455 |
| EZR | 819.57587 | 0.492664792 | 0.47938545 | 1.02770075 | 0.30409061 | 0.74967528 |
| RSPH3 | 180.409313 | -0.207141323 | 0.26871893 | -0.7708475 | 0.44079731 | 0.83733111 |
| AL035530.2 | 125.649968 | 0.377540688 | 0.46447731 | 0.81282913 | 0.41631603 | 0.82489761 |
| SOD2 | 33.8775365 | -1.886265325 | 0.68908486 | -2.7373484 | 0.00619367 | 0.12652949 |
| SOD2.1 | 90.1358494 | -0.277871765 | 0.55676704 | -0.4990808 | 0.61772243 | 0.90794578 |
| WTAP | 378.015048 | -0.143441515 | 0.37220743 | -0.3853806 | 0.69995547 | 0.93180634 |
| ACAT2 | 83.1544011 | -0.99816925 | 0.70695124 | -1.4119351 | 0.15796908 | 0.59593808 |
| TCP1 | 139.201902 | 0.106144912 | 0.33091234 | 0.32076444 | 0.74838891 | 0.94678489 |
| MRPL18 | 106.854751 | -0.230731487 | 0.50017808 | -0.4612987 | 0.64458433 | 0.91682456 |
| IGF2R | 583.142703 | -0.291893966 | 0.29159132 | -1.0010379 | 0.31680848 | 0.76097935 |
| MAP3K4 | 312.742524 | 0.236470928 | 0.20739291 | 1.1402074 | 0.25419991 | 0.71135248 |
| AGPAT4 | 149.844798 | -0.228741946 | 0.28471018 | -0.8034203 | 0.42173183 | 0.82719689 |
| PRKN | 2904.81475 | 0.585779421 | 0.19997013 | 2.9293346 | 0.00339689 | 0.08833588 |
| PACRG | 584.103346 | 0.195688416 | 0.29176895 | 0.6706965 | 0.50241389 | 0.86852095 |
| QKI | 1021.18887 | 0.309010801 | 0.18730583 | 1.64976606 | 0.09899079 | 0.4933116 |
| PDE10A | 4570.11467 | -1.909677607 | 1.00405633 | -1.9019626 | 0.05717604 | 0.39761441 |
| LINC00473 | 260.90232 | 0.018830877 | 1.70704562 | 0.01103127 | 0.9911985 | 0.9974665 |
| SFT2D1 | 156.543667 | 0.204321666 | 0.22058383 | 0.92627673 | 0.35430218 | 0.78548769 |
| MPC1 | 418.514648 | -0.011996182 | 0.24353948 | -0.0492576 | 0.96071397 | 0.9914606 |
| RPS6KA2 | 364.035089 | -0.956373541 | 0.25268374 | -3.7848639 | 0.00015379 | 0.01151249 |
| Z94721.1 | 89.7361994 | -1.079892787 | 0.45939006 | -2.35071 | 0.01873763 | 0.22823596 |
| FGFR1OP | 249.359619 | 0.138226195 | 0.23283929 | 0.59365495 | 0.55274292 | 0.88968604 |
| AFDN | 1278.32155 | 0.157958705 | 0.24657848 | 0.64060216 | 0.5217812 | 0.87670505 |
| WDR27 | 277.837022 | 0.211309203 | 0.20144724 | 1.04895554 | 0.29419858 | 0.73973155 |
| C6orf120 | 40.3061726 | -0.213018911 | 0.34369311 | -0.619794 | 0.53539345 | 0.88137638 |
| PHF10 | 142.93927 | 0.113141653 | 0.29256869 | 0.38671826 | 0.6989648 | 0.93180634 |
| ERMARD | 80.2963022 | -0.048704655 | 0.26285354 | -0.185292 | 0.85300002 | 0.97062426 |
| FAM120B | 446.165156 | -0.106787199 | 0.26014788 | -0.4104865 | 0.68144908 | 0.92759922 |
| PSMB1 | 278.34928 | -0.319254974 | 0.39842125 | -0.8013001 | 0.42295795 | 0.82774157 |
| TBP | 61.5037146 | -0.14183794 | 0.27446831 | -0.5167735 | 0.6053143 | 0.90428025 |
| PDCD2 | 93.1545939 | -0.131296122 | 0.24878614 | -0.5277469 | 0.59767499 | 0.90369786 |
| FAM20C | 797.758847 | -0.393531441 | 0.44043821 | -0.8934998 | 0.3715896 | 0.79539603 |
| PDGFA | 53.9602868 | -1.016200557 | 0.54743088 | -1.8563084 | 0.06340961 | 0.41300975 |
| PRKAR1B | 134.429704 | -0.436455085 | 0.2571047 | -1.6975773 | 0.08958758 | 0.47350347 |
| DNAAF5 | 108.366631 | -0.046865596 | 0.28052753 | -0.1670624 | 0.86732097 | 0.97576204 |
| SUN1 | 250.807525 | -0.035579848 | 0.17454902 | -0.2038387 | 0.83847954 | 0.96786711 |
| ADAP1 | 44.342013 | 0.488435664 | 0.43722144 | 1.11713566 | 0.26393632 | 0.71989632 |
| COX19 | 107.910716 | -0.233931392 | 0.27815155 | -0.8410214 | 0.40033597 | 0.8162653 |
| C7orf50 | 326.583443 | 0.223560856 | 0.33923549 | 0.65901375 | 0.50988694 | 0.87094607 |

| | | | | | | |
|---|---|---|---|---|---|---|
| MICALL2 | 49.1878605 | -0.669146191 | 0.35635616 | -1.8777456 | 0.06041598 | 0.40556611 |
| INTS1 | 129.461697 | 0.328034815 | 0.29006491 | 1.13090143 | 0.25809658 | 0.71534188 |
| MAFK | 143.211914 | 1.450755109 | 0.59726373 | 2.42900253 | 0.01514043 | 0.20009038 |
| TMEM184A | 53.8701286 | 0.39534845 | 0.42314561 | 0.93430829 | 0.35014489 | 0.78223232 |
| PSMG3 | 53.6812529 | -0.244570308 | 0.34512401 | -0.7086447 | 0.47854497 | 0.85532469 |
| MAD1L1 | 159.065332 | 0.094910917 | 0.31350053 | 0.30274564 | 0.76208371 | 0.95374766 |
| MRM2 | 39.2205033 | -0.233606845 | 0.37168763 | -0.6285031 | 0.52967438 | 0.87929472 |
| SNX8 | 81.1811385 | -0.415924446 | 0.39896202 | -1.0425164 | 0.29717233 | 0.74470732 |
| EIF3B | 198.775556 | -0.195991882 | 0.23047452 | -0.8503842 | 0.39511153 | 0.81175661 |
| CHST12 | 50.6870445 | 0.023638261 | 0.39907891 | 0.05923205 | 0.95276728 | 0.99050799 |
| BRAT1 | 49.4904702 | 0.095383293 | 0.27287247 | 0.34955264 | 0.72667446 | 0.93906703 |
| IQCE | 127.680644 | 0.271216501 | 0.26218812 | 1.03443476 | 0.30093296 | 0.74787229 |
| TTYH3 | 99.860459 | -0.985025122 | 0.54360854 | -1.8120119 | 0.06998436 | 0.42862997 |
| GNA12 | 160.464887 | 0.033683326 | 0.26430509 | 0.12744108 | 0.8985913 | 0.98312332 |
| SDK1 | 2224.78755 | -0.051898943 | 0.33034181 | -0.1571068 | 0.87516068 | 0.97622753 |
| FOXK1 | 310.274388 | -0.13424841 | 0.26870238 | -0.4996175 | 0.61734444 | 0.90794578 |
| AP5Z1 | 51.0760532 | -0.609484897 | 0.39477247 | -1.5438891 | 0.12261521 | 0.54116729 |
| RBAK-RBAKD | 171.529503 | -0.630725435 | 0.20256203 | -3.1137397 | 0.00184732 | 0.05926813 |
| WIPI2 | 212.841273 | 0.139732287 | 0.19245233 | 0.7260618 | 0.46780089 | 0.85161077 |
| TNRC18 | 781.264254 | 0.337712715 | 0.26159943 | 1.29095354 | 0.19671979 | 0.65097045 |
| FBXL18 | 62.6400967 | -0.351015064 | 0.40238591 | -0.8723344 | 0.38302598 | 0.80427015 |
| ACTB | 2795.32504 | 0.218522007 | 0.38782877 | 0.56344971 | 0.5731287 | 0.89797476 |
| RNF216 | 365.282063 | -0.222539973 | 0.21055766 | -1.0569075 | 0.2905538 | 0.73754646 |
| CCZ1 | 65.833248 | 0.517502465 | 0.35244554 | 1.46831896 | 0.1420176 | 0.57093391 |
| PMS2 | 108.289039 | -0.020797377 | 0.32731174 | -0.06354 | 0.94933654 | 0.98977092 |
| AIMP2 | 65.7090925 | -0.577431387 | 0.36345924 | -1.5887102 | 0.11212585 | 0.5224487 |
| EIF2AK1 | 209.690604 | -0.030839009 | 0.18697028 | -0.1649407 | 0.86899066 | 0.97576204 |
| USP42 | 168.910193 | 0.120836079 | 0.29350288 | 0.41170322 | 0.68055697 | 0.92710737 |
| CYTH3 | 172.66848 | -0.123705615 | 0.30346793 | -0.4076398 | 0.68353812 | 0.92827258 |
| RAC1 | 633.68506 | -0.236406193 | 0.26703086 | -0.8853141 | 0.37598723 | 0.79942102 |
| DAGLB | 46.403502 | -0.71046891 | 0.3707718 | -1.9161892 | 0.05534102 | 0.39343117 |
| ZDHHC4 | 60.0722951 | -0.161850271 | 0.27058146 | -0.5981573 | 0.54973499 | 0.88889487 |
| C7orf26 | 42.5895903 | 0.361748404 | 0.28451583 | 1.27145264 | 0.20356767 | 0.6582233 |
| ZNF316 | 60.20809 | 0.220155026 | 0.31562241 | 0.6975266 | 0.4854733 | 0.85912348 |
| AC073343.2 | 50.6335637 | -0.599618129 | 0.47812359 | -1.254107 | 0.20980312 | 0.6679604 |
| ZNF12 | 110.827071 | -0.118962669 | 0.28350184 | -0.4196187 | 0.67476404 | 0.92632004 |
| CCZ1B | 77.3422047 | 0.193987469 | 0.36069685 | 0.53781304 | 0.59070613 | 0.90233654 |
| C1GALT1 | 327.668798 | 0.020681549 | 0.39287378 | 0.05264171 | 0.95801738 | 0.99119981 |
| COL28A1 | 264.023639 | 0.567959926 | 0.4643003 | 1.22325988 | 0.22123156 | 0.67871976 |
| AC004982.2 | 57.9092211 | 0.561513048 | 0.39359017 | 1.42664399 | 0.15368254 | 0.58888227 |
| MIOS | 163.527634 | 0.402522036 | 0.2499979 | 1.61010165 | 0.10737567 | 0.51064149 |
| RPA3 | 134.298061 | 0.146787714 | 0.4893761 | 0.29994868 | 0.76421631 | 0.95378284 |
| UMAD1 | 1020.16366 | 0.249374513 | 0.25767849 | 0.96777388 | 0.33315731 | 0.777038 |
| GLCCI1 | 174.334644 | 0.218687902 | 0.32496429 | 0.67295978 | 0.50097288 | 0.86769956 |
| ICA1 | 622.243555 | -0.599949476 | 0.36799425 | -1.630323 | 0.10303326 | 0.50171849 |
| NDUFA4 | 755.276147 | -0.038673793 | 0.44690488 | -0.086537 | 0.93103957 | 0.98822345 |
| PHF14 | 529.253456 | 0.014062166 | 0.26404865 | 0.05325597 | 0.95752796 | 0.99106618 |
| THSD7A | 6303.15089 | 0.35790953 | 0.66878632 | 0.53516275 | 0.59253732 | 0.90245475 |
| TMEM106B | 255.195424 | 0.147171988 | 0.18565275 | 0.79272722 | 0.42793677 | 0.8292138 |
| VWDE | 47.6157968 | -0.132921387 | 0.63125423 | -0.2105671 | 0.83322507 | 0.96749268 |
| AC013470.2 | 52.6471789 | -0.596203485 | 0.60319241 | -0.9884134 | 0.32295021 | 0.76879947 |
| SCIN | 2169.03331 | 1.08346984 | 0.31016766 | 3.49317474 | 0.00047731 | 0.02428277 |
| ARL4A | 34.4491057 | -0.055443555 | 0.73847387 | -0.0750786 | 0.94015221 | 0.98855185 |
| AC011287.1 | 65.2063937 | -0.747823927 | 0.48247926 | -1.5499608 | 0.12115093 | 0.53989034 |
| AGMO | 251.914964 | 0.00120528 | 0.60105732 | 0.00200527 | 0.99840003 | 0.99954455 |
| ISPD | 144.318997 | -0.153565232 | 0.33162024 | -0.4630756 | 0.6433102 | 0.91643619 |
| ANKMY2 | 155.901087 | 0.188941306 | 0.26541227 | 0.71187855 | 0.47653999 | 0.85414749 |
| BZW2 | 292.00027 | -0.849314588 | 0.55224173 | -1.5379399 | 0.12406331 | 0.54273186 |
| TSPAN13 | 48.823807 | -1.343838512 | 0.35676836 | -3.7666975 | 0.00016542 | 0.01228554 |
| AGR2 | 143.086866 | -2.698886876 | 0.61547198 | -4.3850687 | 1.16E-05 | 0.00220936 |
| AC073332.1 | 52.2954245 | -0.542264339 | 0.42554572 | -1.2742798 | 0.20256426 | 0.6574307 |
| SNX13 | 450.865325 | 0.309060436 | 0.23305682 | 1.32611627 | 0.18480119 | 0.63541313 |
| HDAC9 | 69.3094969 | -1.072812299 | 0.61145961 | -1.7545105 | 0.07934307 | 0.44953856 |
| TWISTNB | 53.8983601 | -0.07731472 | 0.47585672 | -0.1624748 | 0.87093198 | 0.97622753 |
| MACC1 | 1713.24472 | 0.238676725 | 0.25278646 | 0.94418317 | 0.34507605 | 0.78180304 |
| MACC1-AS1 | 39.102979 | -0.658717361 | 0.70356435 | -0.9362574 | 0.34914065 | 0.78180304 |
| ITGB8 | 967.250693 | 0.678960094 | 0.33816947 | 2.00775102 | 0.04466976 | 0.35515061 |
| SP4 | 167.601182 | 0.138104493 | 0.30375071 | 0.45466393 | 0.64935103 | 0.91788058 |
| DNAH11 | 140.137155 | 1.304807657 | 1.4891282 | 0.87622251 | 0.38090908 | 0.80308469 |
| RAPGEF5 | 68.5468706 | -2.913193753 | 0.79030176 | -3.686179 | 0.00022765 | 0.01460392 |
| STEAP1B | 552.447921 | -0.043781098 | 0.43123741 | -0.1015244 | 0.91913423 | 0.98520052 |
| TOMM7 | 300.801368 | 0.495622631 | 0.41429311 | 1.19630914 | 0.23157594 | 0.69142903 |
| KLHL7 | 148.953332 | -0.113426914 | 0.29560166 | -0.3837154 | 0.70118939 | 0.93183677 |
| NUPL2 | 133.684426 | -0.109010287 | 0.30439316 | -0.3581233 | 0.72025103 | 0.93668719 |
| AC005082.1 | 55.7301231 | -0.243492381 | 0.33754012 | -0.7213732 | 0.47067996 | 0.85236058 |
| GPNMB | 55.6451156 | -3.694483766 | 0.73219965 | -5.0457328 | 4.52E-07 | 0.00019728 |
| MALSU1 | 112.406469 | 0.415518684 | 0.24499533 | 1.69602699 | 0.08988077 | 0.47358503 |
| TRA2A | 644.520471 | 0.265700242 | 0.22947479 | 1.15786244 | 0.24692018 | 0.70304318 |
| CCDC126 | 68.9977784 | -0.097520136 | 0.33665873 | -0.2896706 | 0.77206825 | 0.95537275 |
| FAM221A | 59.19666 | 0.213729883 | 0.29588061 | 0.72235177 | 0.47007823 | 0.85232523 |
| OSBPL3 | 873.188569 | 0.313529487 | 0.27722679 | 1.13094946 | 0.25807636 | 0.71534188 |
| CYCS | 331.533549 | 0.268886482 | 0.52605981 | 0.51113291 | 0.60925799 | 0.90552475 |
| HNRNPA2B1 | 1222.20395 | 0.440180768 | 0.25709771 | 1.71211473 | 0.08687553 | 0.46701529 |
| CBX3 | 221.070726 | -0.529038938 | 0.26285635 | -2.0126542 | 0.04415102 | 0.35332084 |
| SNX10 | 160.788137 | -1.610778625 | 0.41526457 | -3.8789214 | 0.00010492 | 0.00908734 |
| AC004540.2 | 57.6809575 | 0.18950209 | 0.34880339 | 0.54329199 | 0.58692878 | 0.90196444 |
| AC004540.1 | 348.270242 | 0.544267363 | 0.53328745 | 1.02058911 | 0.30744915 | 0.75251398 |
| SKAP2 | 159.510795 | -0.65664146 | 0.60550957 | -1.0844444 | 0.27816781 | 0.72879966 |
| HIBADH | 700.017248 | 0.187255061 | 0.25990919 | 0.7204634 | 0.47123973 | 0.85236058 |
| TAX1BP1 | 898.537091 | 0.251551519 | 0.22824834 | 1.10209572 | 0.27042006 | 0.72343657 |
| JAZF1 | 199.342337 | -0.478040581 | 0.57007489 | -0.8385575 | 0.40171766 | 0.81668304 |
| AC005162.3 | 77.6331086 | 0.450949409 | 0.56731715 | 0.79488062 | 0.42668295 | 0.8277744 |
| CPVL | 558.636406 | -0.075842953 | 0.41059465 | -0.1847149 | 0.85345265 | 0.97062426 |
| AC007255.1 | 76.2903464 | -0.102356262 | 0.48637073 | -0.2104491 | 0.83331721 | 0.96749268 |
| PRR15 | 88.2596085 | -0.048917152 | 0.57846198 | -0.0845642 | 0.93260789 | 0.98822345 |
| SCRN1 | 279.402219 | 0.253532727 | 0.23917721 | 1.06002042 | 0.28913531 | 0.73743928 |
| FKBP14 | 63.4703872 | -0.392657841 | 0.29394909 | -1.3358022 | 0.18161394 | 0.63191037 |
| PLEKHA8 | 152.944103 | -0.191920865 | 0.25634392 | -0.7486851 | 0.45404706 | 0.84279368 |
| MTURN | 358.796738 | 0.388771945 | 0.33638586 | 1.15573211 | 0.24779075 | 0.70405328 |
| ZNRF2 | 368.263186 | -0.277947109 | 0.45146145 | -0.6156608 | 0.53811843 | 0.88265739 |
| AC006978.2 | 336.842808 | -0.568707971 | 0.30834324 | -1.8443991 | 0.06512501 | 0.41616467 |
| NOD1 | 128.144166 | -0.089345939 | 0.37917333 | -0.2356335 | 0.81371707 | 0.96288868 |
| GGCT | 83.4851044 | 0.318788772 | 0.55026371 | 0.57933817 | 0.56236101 | 0.89242209 |
| AC005154.6 | 90.635987 | -0.036677245 | 0.27317978 | -0.1342605 | 0.89319661 | 0.98168537 |
| GARS | 182.1536 | 0.146379044 | 0.38642837 | 0.37879994 | 0.70483643 | 0.93241442 |
| AQP1 | 51.9255984 | -0.502192557 | 0.51807895 | -0.969336 | 0.3323776 | 0.77629941 |
| LSM5 | 94.2393704 | 0.241952383 | 0.38005029 | 0.63663255 | 0.52436422 | 0.87770915 |
| AVL9 | 505.340331 | -0.29448432 | 0.26995875 | -1.0908493 | 0.27533919 | 0.7263609 |
| KBTBD2 | 119.358056 | -0.46630633 | 0.21068794 | -2.2132559 | 0.02688001 | 0.27863748 |
| FKBP9 | 128.571628 | 0.178832156 | 0.26370026 | 0.67816451 | 0.49766739 | 0.8655085 |
| NT5C3A | 191.472294 | 0.183111696 | 0.42323759 | 0.43264516 | 0.66527257 | 0.92324595 |
| RP9 | 43.9697682 | 0.046508251 | 0.30169684 | 0.15415558 | 0.87748707 | 0.97640529 |
| BBS9 | 584.125217 | 0.339568384 | 0.17657184 | 1.92311748 | 0.0544653 | 0.39095834 |

| | | | | | | |
|---|---|---|---|---|---|---|
| DPY19L1 | 133.397891 | -0.451821687 | 0.34524607 | -1.3086947 | 0.1906378 | 0.64458858 |
| HERPUD2 | 169.693851 | 0.070237261 | 0.24078328 | 0.29170324 | 0.77051354 | 0.95482817 |
| SEPT7-AS1 | 59.0806824 | -0.268754106 | 0.43897141 | -0.612236 | 0.54038161 | 0.88342811 |
| 7-Sep | 580.382497 | 0.435449195 | 0.27137002 | 1.60463266 | 0.10857467 | 0.51254482 |
| EEPD1 | 130.867434 | -1.018967728 | 0.61020288 | -1.6698835 | 0.09494241 | 0.48627909 |
| KIAA0895 | 149.330715 | 0.039613935 | 0.25228713 | 0.15701925 | 0.87522968 | 0.97622753 |
| AOAH | 82.067537 | -3.525842016 | 0.85430313 | -4.1271557 | 3.67E-05 | 0.00469399 |
| STARD3NL | 95.1928116 | -0.074085 | 0.25061511 | -0.2956127 | 0.7675259 | 0.95408212 |
| VPS41 | 456.11122 | -0.221603924 | 0.26227265 | -0.8449372 | 0.39814589 | 0.81400298 |
| YAE1D1 | 48.9782868 | 0.001838746 | 0.32417879 | 0.00567201 | 0.99547441 | 0.99908534 |
| RALA | 197.935747 | -0.22511066 | 0.32801079 | -0.6862904 | 0.49253 | 0.86200976 |
| LINC00265 | 35.40831 | 0.072141232 | 0.45078527 | 0.16003458 | 0.87285383 | 0.97622753 |
| CDK13 | 785.533825 | 0.239400698 | 0.22897823 | 1.04551727 | 0.29578396 | 0.74264876 |
| MPLKIP | 109.57671 | -0.087453263 | 0.38007199 | -0.2300966 | 0.81801673 | 0.96404074 |
| SUGCT | 94.6244775 | 0.644413773 | 0.37491024 | 1.71884817 | 0.08564202 | 0.46479978 |
| MRPL32 | 90.6074159 | -0.419668555 | 0.37734797 | -1.1121527 | 0.26607251 | 0.72053648 |
| STK17A | 67.8806584 | 0.054342958 | 0.48988934 | 0.11092905 | 0.91167261 | 0.98468493 |
| COA1 | 527.393144 | 0.151036851 | 0.18201139 | 0.82982086 | 0.40664007 | 0.82018309 |
| BLVRA | 91.5173131 | -0.103622167 | 0.33333244 | -0.3108673 | 0.75590148 | 0.95067145 |
| URGCP | 113.636408 | 0.114351732 | 0.3898085 | 0.29335361 | 0.76925188 | 0.95482817 |
| UBE2D4 | 50.7113395 | 0.014806036 | 0.28148288 | 0.05260013 | 0.95805051 | 0.99119981 |
| DBNL | 102.074965 | -0.153314061 | 0.26061322 | -0.588282 | 0.55634304 | 0.89082889 |
| POLM | 61.3466473 | -0.366244829 | 0.31577984 | -1.1598107 | 0.24612589 | 0.70206841 |
| POLD2 | 72.1114509 | -0.026103097 | 0.35556499 | -0.073413 | 0.94147746 | 0.98876195 |
| YKT6 | 50.5520182 | 0.056939081 | 0.35891911 | 0.15864043 | 0.87395218 | 0.97622753 |
| NUDCD3 | 325.964336 | -0.087536711 | 0.232046 | -0.3772386 | 0.70599629 | 0.93241442 |
| DDX56 | 78.0712568 | 0.634347516 | 0.24051587 | 2.63744559 | 0.0083533 | 0.1442148 |
| TMED4 | 278.81233 | -0.05961398 | 0.37998175 | -0.1568864 | 0.87533435 | 0.97622753 |
| OGDH | 635.834937 | -0.046177395 | 0.25728649 | -0.1794785 | 0.85756199 | 0.97201489 |
| ZMIZ2 | 158.615802 | 0.039851303 | 0.27658766 | 0.144082 | 0.88543572 | 0.9792493 |
| PPIA | 917.126129 | -0.311604613 | 0.48883897 | -0.6374382 | 0.52383948 | 0.87725117 |
| H2AFV | 283.547365 | 0.22383361 | 0.32994046 | 0.67840607 | 0.49751426 | 0.86550172 |
| PURB | 166.767254 | -0.139995097 | 0.17716454 | -0.7901982 | 0.42941204 | 0.82998114 |
| CCM2 | 176.504458 | -0.20689956 | 0.28093526 | -0.736467 | 0.46144652 | 0.84722486 |
| TBRG4 | 50.8194852 | 0.216941799 | 0.2741502 | 0.79132461 | 0.42875459 | 0.8296433 |
| ADCY1 | 40.8003994 | -0.046280129 | 0.59900472 | -0.0772617 | 0.93841535 | 0.98848592 |
| IGFBP3 | 260.575085 | -1.464771816 | 1.00479417 | -1.457783 | 0.14490038 | 0.57455768 |
| AC004870.4 | 320.750001 | -1.786747134 | 0.89627113 | -1.9935342 | 0.04620298 | 0.36110228 |
| TNS3 | 1017.74514 | -0.513683734 | 0.30445168 | -1.6872422 | 0.09155678 | 0.47836139 |
| HUS1 | 82.0764648 | 0.191289273 | 0.24251058 | 0.78878733 | 0.43023631 | 0.83047752 |
| UPP1 | 192.46331 | -1.947163634 | 1.05231446 | -1.8503629 | 0.06426126 | 0.41357646 |
| GRB10 | 136.511754 | -1.403960767 | 0.52603136 | -2.6689678 | 0.00760847 | 0.13907288 |
| COBL | 500.941011 | 0.076261795 | 0.22379409 | 0.3407677 | 0.73327847 | 0.94163693 |
| SEC61G | 144.284175 | -0.285603826 | 0.38088105 | -0.7498505 | 0.45334478 | 0.84253454 |
| EGFR | 844.377475 | 0.19497551 | 0.2535552 | 0.76896673 | 0.44191306 | 0.83770294 |
| LANCL2 | 124.80556 | 0.372348835 | 0.34496362 | 1.07938581 | 0.28041578 | 0.72945471 |
| VOPP1 | 410.068798 | -0.749003665 | 0.24941462 | -3.0030463 | 0.00267292 | 0.07550451 |
| NIPSNAP2 | 427.172141 | -0.141630702 | 0.26595403 | -0.5325383 | 0.59435324 | 0.90305763 |
| PSPH | 49.2480263 | 0.193961529 | 0.42683078 | 0.45442255 | 0.64952473 | 0.91788284 |
| CCT6A | 151.974824 | 0.007512435 | 0.32701853 | 0.02297251 | 0.9816722 | 0.99507903 |
| SUMF2 | 215.180808 | -0.237685425 | 0.21376979 | -1.1118756 | 0.26619162 | 0.72053648 |
| PHKG1 | 60.6745363 | 0.045282516 | 0.35785547 | 0.12653856 | 0.89930563 | 0.98312332 |
| CHCHD2 | 604.866974 | 0.206187638 | 0.52618496 | 0.39185391 | 0.69516616 | 0.93126994 |
| NUPR2 | 92.486461 | 0.682578799 | 0.6454104 | 1.05758879 | 0.29024296 | 0.73754646 |
| ZNF727 | 78.9062033 | -0.210536904 | 0.46002796 | -0.4576611 | 0.64719593 | 0.91743182 |
| ZNF736 | 294.550087 | 0.09882607 | 0.27028187 | 0.36564076 | 0.71463316 | 0.93573028 |
| ZNF680 | 273.437233 | 0.114872004 | 0.31273396 | 0.36731541 | 0.71338376 | 0.93573028 |
| ZNF107 | 114.888927 | 0.393845753 | 0.38206427 | 1.03083638 | 0.30261756 | 0.74899204 |
| ZNF138 | 211.947052 | -0.191313072 | 0.30794017 | -0.621267 | 0.53442395 | 0.88137638 |
| ZNF273 | 49.1903406 | -0.273647609 | 0.40933086 | -0.6685243 | 0.503799 | 0.86946722 |
| ZNF117 | 136.109797 | 0.083948067 | 0.27615685 | 0.30398691 | 0.76113786 | 0.95354509 |
| VKORC1L1 | 208.73695 | -0.089622631 | 0.25325245 | -0.3538865 | 0.72342392 | 0.93893205 |
| GUSB | 79.6749061 | -0.145762761 | 0.30430846 | -0.4789967 | 0.63194095 | 0.9133034 |
| ASL | 41.4021891 | -0.33595519 | 0.37431746 | -0.8975141 | 0.36944466 | 0.79532503 |
| AC068533.4 | 55.507312 | -0.67761156 | 0.38872574 | -1.743161 | 0.08130549 | 0.454686 |
| CRCP | 63.072304 | -0.054427822 | 0.3404095 | -0.1598893 | 0.87296831 | 0.97622753 |
| AC068533.3 | 45.5933357 | -0.193082498 | 0.46671832 | -0.4137024 | 0.67909205 | 0.92700898 |
| TPST1 | 1420.25482 | 0.653849052 | 0.26599029 | 2.45816883 | 0.01396475 | 0.19155836 |
| AC027644.4 | 348.67097 | 0.206467348 | 0.31184061 | 0.66209255 | 0.50791191 | 0.87094607 |
| AC027644.3 | 51.370311 | -0.23482786 | 0.46721442 | -0.5026126 | 0.61523666 | 0.90792412 |
| TMEM248 | 248.855978 | 0.207382648 | 0.24393837 | 0.8501436 | 0.39524525 | 0.81175661 |
| SBDS | 198.386469 | 0.098821968 | 0.54045851 | 0.18284839 | 0.85491698 | 0.97122276 |
| TYW1 | 415.86816 | 0.469002454 | 0.24814867 | 1.89000594 | 0.05875717 | 0.40066749 |
| AUTS2 | 285.729752 | 0.267153158 | 0.64758285 | 0.41253896 | 0.67994443 | 0.92700898 |
| TYW1B | 294.35724 | 0.564986655 | 0.24631791 | 2.29372945 | 0.02180604 | 0.24999406 |
| POM121 | 273.34483 | 0.154416076 | 0.26235705 | 0.58857224 | 0.55614826 | 0.89081623 |
| NSUN5 | 54.3127248 | -0.029515292 | 0.37960331 | -0.077753 | 0.93802454 | 0.98846521 |
| BAZ1B | 304.335133 | 0.463934644 | 0.23563191 | 1.96889569 | 0.04896507 | 0.3746651 |
| BCL7B | 83.7218368 | 0.149603981 | 0.27564964 | 0.54273235 | 0.58731409 | 0.90196444 |
| TBL2 | 34.2938167 | -0.364938119 | 0.32160269 | -1.1347483 | 0.25648078 | 0.71374446 |
| MLXIPL | 47.1298668 | -0.331370457 | 0.69190474 | -0.478925 | 0.63199201 | 0.9133034 |
| VPS37D | 48.3985511 | -0.060701817 | 0.41533437 | -0.1461517 | 0.88380165 | 0.97878487 |
| DNAJC30 | 48.8878252 | -0.631138429 | 0.37950936 | -1.6630378 | 0.09630486 | 0.4878081 |
| BUD23 | 145.067922 | 0.109505951 | 0.26166481 | 0.41849706 | 0.67558374 | 0.92671696 |
| ABHD11 | 67.9470496 | -1.021668508 | 0.39218944 | -2.6050383 | 0.0091864 | 0.15354619 |
| CLDN3 | 185.424393 | -1.836622755 | 0.42319726 | -4.3398739 | 1.43E-05 | 0.0024098 |
| CLDN4 | 405.305314 | -0.79140041 | 0.3850028 | -2.0555705 | 0.03982393 | 0.33603445 |
| LIMK1 | 51.1922262 | -0.677219744 | 0.36909167 | -1.834828 | 0.06653118 | 0.41989619 |
| EIF4H | 283.926173 | 0.26368771 | 0.45288622 | 0.58223832 | 0.56040617 | 0.89242209 |
| CLIP2 | 189.348934 | -0.497477611 | 0.47995916 | -1.0364999 | 0.299969 | 0.74689358 |
| GTF2IRD1 | 1185.14719 | -0.808069796 | 0.37501879 | -2.1547448 | 0.03118181 | 0.29698524 |
| GTF2I | 1143.51532 | 0.072391977 | 0.19997215 | 0.3620103 | 0.71734434 | 0.93573028 |
| GTF2IRD2 | 123.77853 | -0.515804019 | 0.37032258 | -1.3928506 | 0.16366498 | 0.60330954 |
| CASTOR2 | 86.0861946 | -0.186112086 | 0.54931883 | -0.3388052 | 0.73475647 | 0.94215106 |
| RCC1L | 141.236627 | -0.42656877 | 0.21916805 | -1.9463091 | 0.05161762 | 0.38247133 |
| GTF2IRD2B | 179.860729 | -0.202036097 | 0.30946872 | -0.6528482 | 0.51385415 | 0.87226916 |
| TRIM73 | 78.3839986 | -0.231077457 | 0.3569236 | -0.6474143 | 0.51736381 | 0.87526312 |
| POM121C | 231.318306 | -0.310146838 | 0.28021022 | -1.1068363 | 0.26836471 | 0.72322263 |
| SPDYE5 | 55.714431 | -0.477188982 | 0.41481251 | -1.1503727 | 0.24999041 | 0.70674386 |
| HIP1 | 737.281204 | 0.40613008 | 0.5139812 | 0.79016525 | 0.42943127 | 0.82998114 |
| RHBDD2 | 125.178204 | -0.37554135 | 0.58083422 | -0.6465551 | 0.5179199 | 0.87526312 |
| POR | 455.680159 | -0.552182862 | 0.65167875 | -0.8473237 | 0.39681471 | 0.81254751 |
| TMEM120A | 55.4405258 | 0.320379939 | 0.4195784 | 0.76357587 | 0.44512003 | 0.83929814 |
| STYXL1 | 126.143468 | 0.22618652 | 0.36411938 | 0.62118781 | 0.53447606 | 0.88137638 |
| MDH2 | 206.522283 | 0.315913053 | 0.438338 | 0.72070651 | 0.47109011 | 0.85236058 |
| HSPB1 | 704.744436 | 0.876973656 | 0.51165122 | 1.71400677 | 0.08652748 | 0.46649429 |
| YWHAG | 203.8976 | -0.232486174 | 0.31065029 | -0.7483855 | 0.45422766 | 0.84280358 |
| SSC4D | 80.3096166 | 0.543784395 | 0.59369543 | 0.91593158 | 0.35970278 | 0.78892604 |
| DTX2 | 82.4385335 | 0.150606746 | 0.35130841 | 0.42870237 | 0.66813984 | 0.92422713 |
| POMZP3 | 58.9688355 | -0.23432704 | 0.33423309 | -0.7010887 | 0.48324767 | 0.85829223 |
| CCDC146 | 293.174348 | 0.033307304 | 0.30778516 | 0.10821608 | 0.91382429 | 0.98480378 |
| FGL2 | 583.705792 | -0.126842925 | 0.64437525 | -0.1968464 | 0.84394777 | 0.96876046 |
| GSAP | 171.846491 | -0.115536747 | 0.30605924 | -0.377498 | 0.70580356 | 0.93241442 |

| | | | | | | |
|---|---|---|---|---|---|---|
| PTPN12 | 521.211489 | -0.558583298 | 0.38017 | -1.4692987 | 0.14175178 | 0.57049105 |
| APTR | 95.0830342 | 0.179026558 | 0.31015339 | 0.57721941 | 0.56379125 | 0.89295783 |
| RSBN1L | 348.324506 | 0.60709715 | 0.19489345 | 3.11502083 | 0.00183932 | 0.05926813 |
| TMEM60 | 52.3450029 | 0.179422153 | 0.58905108 | 0.30459524 | 0.76067445 | 0.95329513 |
| PHTF2 | 607.937073 | 1.440311388 | 0.35547684 | 4.05177277 | 5.08E-05 | 0.00576513 |
| MAGI2 | 1296.44291 | -0.04330707 | 0.32207966 | -0.1344607 | 0.89303825 | 0.98168537 |
| GNAI1 | 531.261588 | 0.620393208 | 0.77086861 | 0.8047976 | 0.42093649 | 0.82694802 |
| SEMA3C | 59.5581822 | 0.666101699 | 0.75911868 | 0.87746714 | 0.38023296 | 0.8029673 |
| CACNA2D1 | 844.147766 | 0.521493073 | 0.31890739 | 1.63524925 | 0.10199681 | 0.4989423 |
| PCLO | 766.034908 | 0.007819233 | 0.50259718 | 0.01555765 | 0.98758729 | 0.9964791 |
| KIAA1324L | 80.3994996 | -0.135863237 | 0.64721863 | -0.2099186 | 0.83373119 | 0.96749268 |
| DMTF1 | 383.117446 | 0.300879696 | 0.25543163 | 1.17792653 | 0.2388259 | 0.69499004 |
| TMEM243 | 148.503609 | -0.354499303 | 0.3930014 | -0.9020306 | 0.3670406 | 0.79392889 |
| TP53TG1 | 59.3577986 | 0.164808851 | 0.33094151 | 0.49799994 | 0.61848409 | 0.90831183 |
| CROT | 314.730861 | 0.35614274 | 0.42990243 | 0.8284269 | 0.40742877 | 0.82018309 |
| ABCB1 | 764.996177 | 0.990887688 | 0.46440307 | 2.13368033 | 0.03286896 | 0.30588131 |
| SLC25A40 | 129.950102 | -0.066961286 | 0.23029435 | -0.2907639 | 0.77123188 | 0.95498333 |
| DBF4 | 42.6801158 | -0.701206441 | 0.28930673 | -2.4237474 | 0.01536128 | 0.20199029 |
| ADAM22 | 571.174526 | 0.645769707 | 0.51729534 | 1.24835786 | 0.21190004 | 0.66975892 |
| SRI | 277.716321 | 0.265698732 | 0.42202584 | 0.62957929 | 0.52896987 | 0.8788295 |
| STEAP2 | 101.999841 | 0.386846251 | 0.3870005 | 0.99960144 | 0.31750343 | 0.76155577 |
| CFAP69 | 188.372912 | 0.316141959 | 0.22450087 | 1.40819928 | 0.15907207 | 0.59747329 |
| GTPBP10 | 124.476829 | -0.148943596 | 0.27899101 | -0.5338652 | 0.59343479 | 0.90277205 |
| CLDN12 | 121.476445 | -0.136139623 | 0.25011173 | -0.5443152 | 0.58622457 | 0.90196444 |
| CDK14 | 687.054271 | 0.420514042 | 0.29133019 | 1.4434276 | 0.14890005 | 0.5817833 |
| FZD1 | 42.8848311 | 0.240602223 | 0.42170374 | 0.57054799 | 0.56830608 | 0.8957783 |
| AC079760.1 | 82.4747946 | -1.006745368 | 0.67532174 | -1.490764 | 0.13602346 | 0.5601128 |
| MTERF1 | 359.537428 | -0.520356072 | 0.43493219 | -1.1964074 | 0.23153763 | 0.69142903 |
| AKAP9 | 1089.1838 | 0.122669985 | 0.14649803 | 0.83734906 | 0.40239639 | 0.81711183 |
| KRIT1 | 367.496795 | -0.141815747 | 0.21454642 | -0.6610026 | 0.50861063 | 0.87094607 |
| ANKIB1 | 796.993393 | 0.115792805 | 0.19016101 | 0.6089198 | 0.5425776 | 0.88450129 |
| GATAD1 | 133.842956 | -0.655450479 | 0.24362474 | -2.6904102 | 0.00713642 | 0.13451389 |
| AC007566.1 | 86.4498321 | -0.882076419 | 0.43754798 | -2.0159536 | 0.04380483 | 0.35277527 |
| PEX1 | 155.393933 | 0.060436393 | 0.21578904 | 0.28007165 | 0.77942254 | 0.95636474 |
| RBM48 | 33.9233757 | -0.283025629 | 0.42637318 | -0.6637979 | 0.50681966 | 0.87094607 |
| FAM133B | 189.508577 | 0.343125027 | 0.22718239 | 1.51035048 | 0.13095402 | 0.55395564 |
| CDK6 | 1062.57092 | 1.072583856 | 0.26448976 | 4.0552944 | 5.01E-05 | 0.00576513 |
| SAMD9 | 72.4499237 | -1.086441002 | 0.67793908 | -1.6025643 | 0.10903087 | 0.51387157 |
| SAMD9L | 165.804253 | -0.871872152 | 0.76812785 | -1.1350612 | 0.25634966 | 0.71374446 |
| VPS50 | 372.118407 | 0.381503548 | 0.273293 | 1.3959507 | 0.16272934 | 0.60211533 |
| GNGT1 | 59.6224909 | -0.207349607 | 0.69266473 | -0.2993506 | 0.76467255 | 0.95378284 |
| TFPI2 | 62.2223104 | 0.042780482 | 1.05313079 | 0.04062219 | 0.96759709 | 0.99292814 |
| GNG11 | 134.968711 | -0.079818185 | 0.58196613 | -0.1371526 | 0.89091015 | 0.98095103 |
| BET1 | 152.865985 | -0.140277753 | 0.31503012 | -0.4452836 | 0.6561148 | 0.92098622 |
| CASD1 | 213.005552 | -0.103321479 | 0.28167827 | -0.3668067 | 0.71376321 | 0.93573028 |
| SGCE | 291.355738 | 0.210159163 | 0.3329963 | 0.63111562 | 0.52796493 | 0.87826011 |
| PEG10 | 36.1966726 | 0.545164323 | 0.67582002 | 0.80667087 | 0.41985613 | 0.82694802 |
| PPP1R9A | 1880.6393 | 0.480660227 | 0.17665457 | 2.72090454 | 0.00651036 | 0.12862701 |
| AC002429.2 | 165.634289 | -0.04910544 | 0.5292164 | -0.092789 | 0.92607122 | 0.98729553 |
| PON3 | 38.0438964 | -0.78159881 | 0.46446915 | -1.6827787 | 0.09241793 | 0.48066498 |
| PON2 | 241.72705 | -0.614086772 | 0.27373668 | -2.2433485 | 0.02487435 | 0.26654723 |
| PDK4 | 3412.80273 | 2.706326533 | 1.43807349 | 1.88191115 | 0.05984808 | 0.40360868 |
| SLC25A13 | 513.393326 | -0.545667753 | 0.35099795 | -1.554618 | 0.12003706 | 0.53751263 |
| SEM1 | 291.466121 | 0.300988009 | 0.13833652 | 2.17576676 | 0.02957271 | 0.29105042 |
| SDHAF3 | 78.6836336 | -0.401054145 | 0.23279885 | -1.7227497 | 0.0849338 | 0.46310155 |
| ASNS | 46.5664721 | -0.41967996 | 0.32156419 | -1.3051203 | 0.19185188 | 0.64458858 |
| LMTK2 | 277.774404 | 0.672059241 | 0.37760775 | 1.77978137 | 0.07511175 | 0.43951486 |
| TECPR1 | 50.3599891 | -0.301930433 | 0.35080823 | -0.8606709 | 0.38941932 | 0.80680464 |
| BRI3 | 448.610748 | -0.389171879 | 0.41978839 | -0.9270668 | 0.35389185 | 0.78491204 |
| BAIAP2L1 | 1070.92166 | 0.781553921 | 0.63030491 | 1.23996165 | 0.21498958 | 0.67132603 |
| TRRAP | 341.180579 | 0.072764333 | 0.17664572 | 0.41192242 | 0.68039629 | 0.92700898 |
| SMURF1 | 422.868293 | -0.068740698 | 0.31704205 | -0.2168189 | 0.82834951 | 0.96596226 |
| ARPC1B | 97.989595 | 0.33532412 | 0.53556513 | 0.62611268 | 0.53124103 | 0.88023377 |
| PDAP1 | 126.990308 | -0.113253421 | 0.37906267 | -0.2987723 | 0.7651138 | 0.95378284 |
| BUD31 | 95.2954769 | -0.217136582 | 0.33084834 | -0.6563025 | 0.51162953 | 0.87151901 |
| CPSF4 | 47.6991049 | -0.010934889 | 0.24843206 | -0.0440156 | 0.96489196 | 0.99243116 |
| ZNF789 | 98.6411929 | -0.153962958 | 0.33382972 | -0.4612021 | 0.64465364 | 0.91682456 |
| ZNF394 | 77.0853014 | -0.447016692 | 0.25699767 | -1.7393803 | 0.08196788 | 0.45535449 |
| ZKSCAN5 | 56.5638842 | -0.166015112 | 0.33517299 | -0.4953117 | 0.6203801 | 0.90918521 |
| FAM200A | 48.521818 | -0.463411598 | 0.43992487 | -1.053388 | 0.29216318 | 0.73844969 |
| ZSCAN25 | 40.4130549 | 0.065732923 | 0.33133316 | 0.19838921 | 0.84274056 | 0.96876046 |
| TRIM4 | 123.776546 | -0.394616329 | 0.22260342 | -1.7727326 | 0.07627303 | 0.4426298 |
| ZKSCAN1 | 424.758342 | -0.089127355 | 0.19160937 | -0.4651513 | 0.64182308 | 0.91643619 |
| ZSCAN21 | 34.0012893 | -0.609473388 | 0.36300176 | -1.6789819 | 0.09315557 | 0.48210881 |
| ZNF3 | 71.8385747 | -0.41407825 | 0.36592904 | -1.1315807 | 0.25781075 | 0.71534188 |
| COPS6 | 117.339004 | 0.180813099 | 0.3076415 | 0.58773963 | 0.55670707 | 0.89100338 |
| MCM7 | 39.3917872 | -0.154010365 | 0.41862872 | -0.3678925 | 0.7129534 | 0.93548912 |
| AP4M1 | 61.8579227 | -0.640670443 | 0.30140361 | -2.125623 | 0.03353466 | 0.30964163 |
| TAF6 | 66.9531523 | 0.165868389 | 0.22538667 | 0.73592811 | 0.46177445 | 0.84738158 |
| LAMTOR4 | 313.701419 | -0.018016802 | 0.46531174 | -0.0387199 | 0.96911374 | 0.99318522 |
| CASTOR3 | 158.600199 | -0.983950538 | 0.48689214 | -2.0208799 | 0.0432922 | 0.35061999 |
| SPDYE3 | 118.888761 | -0.216122471 | 0.45951685 | -0.4703254 | 0.63812252 | 0.91521016 |
| PILRB | 191.635901 | -0.362719043 | 0.44922713 | -0.8074291 | 0.41941932 | 0.82694802 |
| PILRA | 36.355978 | 0.002020363 | 0.46272007 | 0.00436628 | 0.99651623 | 0.99926854 |
| MEPCE | 67.5437774 | -0.292548956 | 0.48840713 | -0.5989858 | 0.54918233 | 0.88873237 |
| PPP1R35 | 97.8541896 | 0.034077457 | 0.44375388 | 0.0767936 | 0.93878774 | 0.98848592 |
| TSC22D4 | 146.760313 | 0.041679546 | 0.3952691 | 0.105446 | 0.91602192 | 0.98520052 |
| AGFG2 | 122.300893 | 0.133449468 | 0.4983072 | 0.26780562 | 0.78884894 | 0.95801395 |
| LRCH4 | 47.4210766 | 0.479120164 | 0.41125124 | 1.16503033 | 0.24400672 | 0.70060011 |
| GNB2 | 315.421954 | -0.513814615 | 0.37120344 | -1.384186 | 0.1663015 | 0.60834438 |
| GIGYF1 | 128.334234 | -0.160836707 | 0.31547311 | -0.509827 | 0.61017267 | 0.90600873 |
| POP7 | 38.2878731 | -0.579541252 | 0.52016855 | -1.1141413 | 0.26521858 | 0.72029159 |
| EPHB4 | 47.3140139 | -0.574274278 | 0.37725251 | -1.5222543 | 0.12794536 | 0.54791812 |
| SLC12A9 | 96.7299454 | -0.246556886 | 0.40503472 | -0.6087302 | 0.54270326 | 0.88450129 |
| TRIP6 | 148.898503 | 0.588011201 | 0.41224635 | 1.42635877 | 0.15376481 | 0.58898217 |
| SRRT | 77.1959564 | -0.057100752 | 0.3034961 | -0.1881433 | 0.85076433 | 0.97017003 |
| TRIM56 | 369.393067 | -1.032246161 | 0.29855277 | -3.4574999 | 0.00054521 | 0.0264529 |
| AP1S1 | 95.3467602 | -0.386048761 | 0.49894168 | -0.7737352 | 0.43908738 | 0.83559755 |
| PLOD3 | 57.3138187 | -0.586959766 | 0.32056093 | -1.8310396 | 0.06709463 | 0.42104891 |
| ZNHIT1 | 162.701533 | 0.005487545 | 0.387811 | 0.01415005 | 0.98871027 | 0.9964791 |
| CLDN15 | 38.3961943 | -0.45913584 | 0.27598166 | -1.6636462 | 0.09618315 | 0.48772364 |
| FIS1 | 314.59783 | 0.237259969 | 0.47585281 | 0.49859949 | 0.61806157 | 0.90794578 |
| IFT22 | 85.4813849 | -0.340925682 | 0.48808123 | -0.6985019 | 0.48486335 | 0.85909499 |
| COL26A1 | 89.9263576 | 1.341274149 | 0.57699396 | 2.32458959 | 0.02009392 | 0.23835055 |
| CUX1 | 1463.02968 | 0.161626236 | 0.16792256 | 0.96250457 | 0.33579621 | 0.77820994 |
| PRKRIP1 | 401.355165 | -0.287393612 | 0.28461443 | -1.0097647 | 0.31260802 | 0.75677346 |
| ORAI2 | 161.014486 | 0.046832241 | 0.27592039 | 0.16973099 | 0.8652217 | 0.97489769 |
| ALKBH4 | 39.7254097 | -0.089854622 | 0.31914905 | -0.2815444 | 0.77829289 | 0.95620223 |
| POLR2J | 98.203159 | 0.305619441 | 0.4774345 | 0.64012853 | 0.52208904 | 0.87670505 |
| SPDYE2 | 43.4721056 | 0.30309843 | 0.53233442 | 0.56937598 | 0.56910101 | 0.89617938 |
| AC105052.3 | 101.122939 | 0.55844606 | 0.47636479 | 1.17230759 | 0.24107359 | 0.69675984 |
| FAM185A | 237.736102 | 0.129672452 | 0.32110097 | 0.403837 | 0.68633259 | 0.92974759 |
| FBXL13 | 182.938043 | -0.365439845 | 0.30461639 | -1.1996723 | 0.23026664 | 0.6900756 |

| | | | | | | |
|---|---|---|---|---|---|---|
| ARMC10 | 140.494855 | -0.126041845 | 0.28799178 | -0.4376578 | 0.66163438 | 0.92284375 |
| NAPEPLD | 257.013688 | 0.754590034 | 0.2731359 | 2.76269076 | 0.00573271 | 0.12137121 |
| PMPCB | 124.995796 | 0.102283225 | 0.23790073 | 0.42994078 | 0.66723872 | 0.92422364 |
| DNAJC2 | 103.474328 | -0.156244238 | 0.25101674 | -0.6224455 | 0.53364897 | 0.88137638 |
| PSMC2 | 97.6722282 | 0.062729896 | 0.21330356 | 0.29408743 | 0.76869109 | 0.95453142 |
| ORC5 | 156.563179 | 0.016914762 | 0.27336006 | 0.06187723 | 0.9506606 | 0.98992048 |
| LHFPL3 | 310.967075 | -0.313673796 | 0.51863841 | -0.6048025 | 0.54531025 | 0.88669883 |
| LINC01004 | 404.997178 | 0.260277156 | 0.34741304 | 0.74918648 | 0.45374482 | 0.84268043 |
| KMT2E | 1188.68748 | 0.381754011 | 0.28798028 | 1.32562552 | 0.18496377 | 0.63541313 |
| SRPK2 | 1216.72821 | 0.198464342 | 0.22172059 | 0.89511013 | 0.37072822 | 0.79539603 |
| PUS7 | 93.2142115 | -0.443575277 | 0.37218972 | -1.1917989 | 0.23334012 | 0.69247746 |
| RINT1 | 65.6579048 | -0.022605701 | 0.35092167 | -0.0644181 | 0.94863733 | 0.98977092 |
| EFCAB10 | 43.958892 | -1.085515071 | 0.5719649 | -1.8978701 | 0.0577132 | 0.39817926 |
| ATXN7L1 | 431.63914 | 0.077897892 | 0.2156965 | 0.36114583 | 0.71799043 | 0.93573028 |
| CDHR3 | 100.774048 | -0.410839745 | 0.55173937 | -0.7446265 | 0.45649756 | 0.84375562 |
| SYPL1 | 386.1813 | 0.270964691 | 0.35585035 | 0.76145687 | 0.44638423 | 0.83980766 |
| NAMPT | 834.552059 | -0.791138678 | 0.91587952 | -0.8638021 | 0.3876966 | 0.8065026 |
| PRKAR2B | 75.9312175 | 0.038735412 | 0.50956042 | 0.07601731 | 0.93940533 | 0.98854984 |
| HBP1 | 293.399157 | 0.027382539 | 0.3355968 | 0.08159356 | 0.93496992 | 0.98822345 |
| COG5 | 741.035728 | 0.358613843 | 0.17705994 | 2.02538098 | 0.04282826 | 0.3484612 |
| DUS4L | 41.4542585 | -0.056858878 | 0.33257772 | -0.1709642 | 0.86425193 | 0.97466129 |
| BCAP29 | 178.737415 | 0.16952873 | 0.28622902 | 0.59228352 | 0.55366075 | 0.89006974 |
| CBLL1 | 47.1076114 | -0.09778136 | 0.24644117 | -0.3967736 | 0.69153439 | 0.93073907 |
| SLC26A3 | 1144.35256 | 1.076873913 | 0.65206697 | 1.65147746 | 0.09864112 | 0.4932056 |
| DLD | 274.232066 | 0.301893806 | 0.29090522 | 1.03777378 | 0.29937539 | 0.74594723 |
| LAMB1 | 2398.95964 | -0.887904334 | 0.50602645 | -1.7546599 | 0.07931749 | 0.44953856 |
| NRCAM | 810.948114 | 0.691681922 | 0.5616624 | 1.23149051 | 0.21813946 | 0.67576162 |
| PNPLA8 | 264.88972 | 0.234122948 | 0.43366942 | 0.53986502 | 0.58929012 | 0.90233654 |
| DNAJB9 | 68.2698845 | -0.408054578 | 0.47231857 | -0.8639393 | 0.38762124 | 0.8065026 |
| IMMP2L | 7403.74566 | 0.420235246 | 0.22257597 | 1.88805307 | 0.05901883 | 0.40122703 |
| LRRN3 | 49.4436396 | -0.418181521 | 0.76106613 | -0.549468 | 0.58268429 | 0.90085422 |
| DOCK4 | 75.4282718 | 0.027828312 | 0.46517505 | 0.05982331 | 0.95229636 | 0.9903694 |
| ZNF277 | 228.017597 | 0.188436662 | 0.20473043 | 0.92041357 | 0.35735668 | 0.78612469 |
| IFRD1 | 297.469206 | 2.000536587 | 0.64710478 | 3.09151879 | 0.00199135 | 0.06287382 |
| TMEM168 | 271.200241 | 0.281336094 | 0.24862703 | 1.13155874 | 0.25781999 | 0.71534188 |
| BMT2 | 355.49198 | 0.016975608 | 0.2293964 | 0.0740012 | 0.94100943 | 0.98876195 |
| SMIM30 | 80.966934 | -0.226474356 | 0.32411892 | -0.6987385 | 0.48471549 | 0.85909499 |
| FOXP2 | 178.42868 | 0.552781669 | 1.61500619 | 0.34227836 | 0.73214142 | 0.9411066 |
| MDFIC | 355.135372 | 0.536280576 | 0.3477311 | 1.5422278 | 0.12301825 | 0.54192149 |
| TES | 243.351268 | 0.18280561 | 0.47314589 | 0.38636204 | 0.69922856 | 0.93180634 |
| CAV2 | 108.69592 | -0.052599405 | 0.67425247 | -0.0780114 | 0.93781895 | 0.98846521 |
| MET | 336.362484 | -2.23835113 | 0.83455403 | -2.6820925 | 0.00731632 | 0.13610131 |
| CAPZA2 | 522.024494 | -0.045473981 | 0.28557982 | -0.1592339 | 0.87348462 | 0.97622753 |
| ST7 | 725.671957 | 0.45824742 | 0.29910173 | 1.5320788 | 0.12550299 | 0.54417366 |
| ST7-AS2 | 56.0138249 | 0.469561649 | 0.56326047 | 0.83364922 | 0.40447868 | 0.81914727 |
| LSM8 | 131.993564 | 0.059240195 | 0.20663824 | 0.28668554 | 0.77435311 | 0.95586283 |
| ANKRD7 | 153.184915 | -0.351389799 | 0.32891448 | -1.0683318 | 0.28537087 | 0.73607046 |
| TSPAN12 | 227.493721 | 0.156494452 | 0.37649719 | 0.41565902 | 0.67765953 | 0.9268955 |
| ING3 | 59.684752 | 0.434876644 | 0.24140293 | 1.80145551 | 0.07163111 | 0.43222361 |
| FAM3C | 185.509642 | -0.352043118 | 0.29924929 | -1.1764209 | 0.23942673 | 0.69526124 |
| AASS | 132.924522 | 0.650829349 | 0.3299478 | 1.97252218 | 0.04855003 | 0.37302367 |
| CADPS2 | 6641.07028 | 0.567263763 | 0.47935759 | 1.1833833 | 0.23665727 | 0.69335782 |
| AC004594.1 | 404.846818 | -0.57014547 | 0.66063464 | -0.8630269 | 0.38812267 | 0.8065026 |
| IQUB | 488.419978 | 0.855564849 | 0.56207636 | 1.52215058 | 0.12797136 | 0.54791812 |
| NDUFA5 | 500.457139 | 0.541944245 | 0.41907947 | 1.29317774 | 0.19594959 | 0.64965255 |
| WASL | 414.034279 | 0.420408526 | 0.17406352 | 2.41525921 | 0.01572402 | 0.20433451 |
| POT1 | 177.554905 | 0.270779249 | 0.17727763 | 1.52743048 | 0.126654 | 0.54691843 |
| POT1-AS1 | 106.106509 | -0.466081101 | 0.21342247 | -2.1838427 | 0.02897381 | 0.28781563 |
| ZNF800 | 205.877634 | 0.208986076 | 0.17011367 | 1.2285084 | 0.21925618 | 0.67731484 |
| ARF5 | 131.538756 | -0.361601889 | 0.42445049 | -0.8519295 | 0.39425323 | 0.81094677 |
| SND1 | 610.619583 | -0.109731175 | 0.28408531 | -0.3862613 | 0.69930313 | 0.93180634 |
| RBM28 | 123.944943 | 0.247600926 | 0.21230436 | 1.16625458 | 0.24351153 | 0.69975346 |
| IMPDH1 | 35.4041283 | -0.536859918 | 0.49565072 | -1.0831416 | 0.27874558 | 0.72938673 |
| HILPDA | 366.542934 | -2.657718264 | 1.41986429 | -1.8718115 | 0.06123269 | 0.40676991 |
| METTL2B | 104.865203 | -0.376501304 | 0.24088211 | -1.5630107 | 0.11805009 | 0.53360052 |
| CALU | 178.879166 | -0.098815581 | 0.27687981 | -0.3568898 | 0.72117431 | 0.93744589 |
| KCP | 379.033239 | 0.231996244 | 0.37030946 | 0.62649289 | 0.5309917 | 0.88022667 |
| ATP6V1F | 453.548002 | -0.549554393 | 0.4916203 | -1.1178432 | 0.26363399 | 0.71989632 |
| IRF5 | 42.6820304 | 0.56801797 | 0.32283894 | 1.75944688 | 0.07850164 | 0.44809213 |
| TNPO3 | 400.899253 | 0.240964921 | 0.28704416 | 0.83946986 | 0.4012057 | 0.81648166 |
| TSPAN33 | 53.4353632 | 0.023747599 | 0.32882224 | 0.07222017 | 0.94242669 | 0.98885525 |
| SMO | 34.9757339 | -0.138344684 | 0.46511844 | -0.2974397 | 0.76613085 | 0.95390891 |
| AHCYL2 | 647.24603 | 0.425328782 | 0.30345037 | 1.401642 | 0.16102218 | 0.60032459 |
| AC078846.1 | 33.5862687 | -0.042958459 | 0.32997357 | -0.1301876 | 0.89641803 | 0.98302816 |
| NRF1 | 454.581628 | 0.285282269 | 0.2430415 | 1.17380066 | 0.24047488 | 0.6963541 |
| UBE2H | 1626.3455 | 0.290046113 | 0.28323638 | 1.02404259 | 0.30581516 | 0.75109981 |
| ZC3HC1 | 67.2500941 | -0.032651301 | 0.25976935 | -0.1256934 | 0.8999746 | 0.98312332 |
| KLHDC10 | 465.343557 | -0.090084557 | 0.23429463 | -0.3844926 | 0.70061336 | 0.93180634 |
| TMEM209 | 87.6981774 | 0.271955846 | 0.27507726 | 0.98865259 | 0.32283315 | 0.76879139 |
| CEP41 | 49.4598764 | -0.118290482 | 0.33697452 | -0.3510369 | 0.72556068 | 0.93895336 |
| MEST | 115.115849 | -1.073839291 | 0.76950011 | -1.3955025 | 0.16286438 | 0.60211533 |
| COPG2 | 174.074602 | -0.280667641 | 0.29127479 | -0.9635837 | 0.33525468 | 0.77820994 |
| AC016831.7 | 2631.27207 | 0.210626845 | 0.52922754 | 0.3979892 | 0.69063815 | 0.93073907 |
| LINC00513 | 182.402953 | -1.248847096 | 0.67274253 | -1.8563522 | 0.06340337 | 0.41300975 |
| AC058791.1 | 49.8658702 | 0.087249496 | 0.74498707 | 0.11711545 | 0.90676857 | 0.98321965 |
| MKLN1 | 1744.37313 | 0.006143458 | 0.29820675 | 0.02060134 | 0.98356367 | 0.99540861 |
| CHCHD3 | 1293.06002 | 0.177803978 | 0.22849182 | 0.77816344 | 0.43647267 | 0.83393108 |
| EXOC4 | 1987.60185 | 0.177911284 | 0.1887083 | 0.94278461 | 0.34579109 | 0.78180304 |
| LRGUK | 95.7609402 | 0.157812856 | 0.30199257 | 0.52257198 | 0.60127214 | 0.90369786 |
| SLC35B4 | 190.472617 | 0.950916996 | 0.40597518 | 2.34230331 | 0.01916513 | 0.23192909 |
| AKR1B1 | 579.221354 | 1.681314796 | 0.79221619 | 2.1222929 | 0.03381315 | 0.31120503 |
| BPGM | 102.695129 | -1.528287331 | 0.54746485 | -2.7915716 | 0.00524527 | 0.11544342 |
| CALD1 | 71.0706126 | -0.311340716 | 0.45587676 | -0.6829493 | 0.49463888 | 0.86389349 |
| AGBL3 | 87.0792135 | -0.301564697 | 0.44414804 | -0.6789734 | 0.49715472 | 0.86519122 |
| CYREN | 117.919923 | -0.47489892 | 0.23351179 | -2.0337257 | 0.04197926 | 0.34641157 |
| TMEM140 | 105.366932 | -1.446152356 | 0.73974414 | -1.9549359 | 0.05059064 | 0.37952033 |
| WDR91 | 142.160524 | -0.150414953 | 0.36781958 | -0.4089368 | 0.68258606 | 0.92806156 |
| CNOT4 | 545.939797 | 0.278651265 | 0.19261038 | 1.44670946 | 0.14797831 | 0.57952643 |
| NUP205 | 134.384433 | 0.440187435 | 0.22609067 | 1.94695093 | 0.05154062 | 0.38247133 |
| STMP1 | 246.496667 | -0.087867214 | 0.26857374 | -0.3271623 | 0.7435451 | 0.94555911 |
| MTPN | 247.328575 | -0.080671922 | 0.30820313 | -0.2617492 | 0.79351481 | 0.95906299 |
| CREB3L2 | 218.497313 | 0.353415517 | 0.23817813 | 1.48382857 | 0.13785441 | 0.56434148 |
| TRIM24 | 374.834586 | 0.302035041 | 0.35449648 | 0.85201141 | 0.39420776 | 0.81094677 |
| SVOPL | 143.600402 | 1.54929378 | 0.58398372 | 2.65297428 | 0.0079786 | 0.14172148 |
| ATP6V0A4 | 562.686261 | 1.099286907 | 0.41349397 | 2.65853191 | 0.00784819 | 0.14121635 |
| TMEM213 | 440.912996 | 0.313048972 | 0.42037707 | 0.74468612 | 0.45646149 | 0.84375562 |
| KIAA1549 | 299.088298 | 0.317464881 | 0.3148875 | 1.0081851 | 0.31336561 | 0.75717828 |
| ZC3HAV1 | 741.517621 | 0.654245933 | 0.62007006 | 1.05511615 | 0.29137221 | 0.73816859 |
| TTC26 | 155.129767 | -0.023445312 | 0.18874927 | -0.1242141 | 0.90114579 | 0.98320957 |
| UBN2 | 380.59672 | 0.109544376 | 0.19502101 | 0.56170552 | 0.57431667 | 0.89807676 |
| LUC7L2 | 137.494569 | 0.144445824 | 0.25844188 | 0.55891029 | 0.57622295 | 0.89869922 |
| HIPK2 | 1058.65016 | -0.094905094 | 0.26989599 | -0.3516358 | 0.72511141 | 0.93895336 |
| TBXAS1 | 46.8661677 | -1.488011761 | 0.43322177 | -3.4347576 | 0.00059308 | 0.02774786 |

| | | | | | | |
|---|---|---|---|---|---|---|
| PARP12 | 131.410486 | -0.195508996 | 0.68237896 | -0.2865109 | 0.77448686 | 0.95586283 |
| KDM7A | 425.45851 | -0.543371176 | 0.34353322 | -1.5817136 | 0.11371496 | 0.52602773 |
| SLC37A3 | 160.645309 | -0.148491461 | 0.24700339 | -0.6011718 | 0.5477256 | 0.88774579 |
| MKRN1 | 217.492977 | 0.112478207 | 0.17577302 | 0.63990597 | 0.52223373 | 0.87670505 |
| NDUFB2 | 436.674996 | 0.038062443 | 0.41492031 | 0.09173434 | 0.92690911 | 0.98729622 |
| BRAF | 1535.65409 | 0.80699992 | 0.23954159 | 3.36893454 | 0.00075459 | 0.03310667 |
| MRPS33 | 113.060395 | 0.210570135 | 0.5152245 | 0.40869589 | 0.68276285 | 0.92806156 |
| AGK | 171.084141 | 0.405135154 | 0.288766 | 1.40298774 | 0.1606205 | 0.60010648 |
| AC004918.1 | 47.8387971 | 0.147060369 | 0.45182138 | 0.32548342 | 0.74481523 | 0.94579712 |
| KIAA1147 | 187.973598 | -0.064134564 | 0.28080893 | -0.2283922 | 0.81934137 | 0.96461147 |
| WEE2-AS1 | 37.4704222 | -0.408286706 | 0.42830707 | -0.953257 | 0.34045988 | 0.77836439 |
| SSBP1 | 226.566078 | -0.142817784 | 0.27465626 | -0.5199874 | 0.60307239 | 0.90369786 |
| EPHB6 | 51.3780354 | -0.153346093 | 0.40200849 | -0.3814499 | 0.70286945 | 0.93241442 |
| TRPV6 | 158.124681 | -0.715317989 | 0.48467716 | -1.4758649 | 0.13998018 | 0.56739811 |
| TRPV5 | 115.30927 | -4.17317076 | 0.81378918 | -5.1280736 | 2.93E-07 | 0.00014433 |
| GSTK1 | 262.054913 | -0.361336177 | 0.31558611 | -1.1449686 | 0.25222217 | 0.70813507 |
| AC073342.1 | 34.1559277 | -0.263809446 | 0.4864484 | -0.5423174 | 0.58759985 | 0.90196444 |
| TMEM139 | 72.5654783 | 0.636926003 | 0.71226857 | 0.89422169 | 0.37120329 | 0.79539603 |
| CASP2 | 55.3212956 | 0.035946794 | 0.29997609 | 0.1198322 | 0.90461608 | 0.98320957 |
| ZYX | 71.8992527 | -0.786185588 | 0.48668882 | -1.6153763 | 0.10622922 | 0.50765436 |
| TCAF2 | 122.418399 | 0.271272993 | 0.57652149 | 0.47053405 | 0.63797351 | 0.915143 |
| TCAF1 | 258.01517 | -0.233270688 | 0.25916647 | -0.9000805 | 0.3680774 | 0.79501778 |
| AC004889.1 | 217.204657 | 0.997624864 | 0.3455307 | 2.88722497 | 0.00388656 | 0.09674863 |
| AC074386.1 | 276.872473 | 0.05373569 | 0.33697795 | 0.15946352 | 0.8733037 | 0.97622753 |
| ARHGEF5 | 76.8804983 | 0.09446477 | 0.3959941 | 0.23855096 | 0.8114538 | 0.96288868 |
| TPK1 | 203.681604 | -0.425413927 | 0.32785507 | -1.297567 | 0.19443616 | 0.64808107 |
| CNTNAP2 | 44.4297586 | 0.469270603 | 0.39764616 | 1.18012104 | 0.23795208 | 0.69406342 |
| CUL1 | 328.458485 | -0.023738927 | 0.18252028 | -0.1300619 | 0.89651749 | 0.98302816 |
| PDIA4 | 121.738925 | -0.707489128 | 0.37141647 | -1.9048405 | 0.05680081 | 0.39727338 |
| ZNF786 | 42.23227 | -0.297647634 | 0.4808317 | -0.6190266 | 0.5358988 | 0.88137638 |
| ZNF398 | 120.600697 | -0.233480563 | 0.34199611 | -0.6826995 | 0.49479676 | 0.86389349 |
| ZNF282 | 68.3740768 | -0.373804308 | 0.26906533 | -1.3892697 | 0.16475075 | 0.60604843 |
| ZNF212 | 32.2941756 | -0.036521281 | 0.33371773 | -0.1094376 | 0.91285538 | 0.98480378 |
| ZNF783 | 62.2086912 | -0.029461722 | 0.29038297 | -0.1014582 | 0.91918677 | 0.98520052 |
| ZNF777 | 38.5623423 | -0.374480138 | 0.35213476 | -1.0634569 | 0.2875748 | 0.73650262 |
| ZNF746 | 43.856302 | -0.258930151 | 0.27599417 | -0.9381725 | 0.34815575 | 0.78180304 |
| ZNF467 | 33.41682 | -0.114280422 | 0.77388631 | -0.1476708 | 0.88260257 | 0.97865059 |
| ZNF862 | 143.600166 | 0.015682047 | 0.34941174 | 0.04488128 | 0.96420194 | 0.99243116 |
| ATP6V0E2 | 51.1027014 | -0.119995499 | 0.4274693 | -0.2807114 | 0.77893177 | 0.95623387 |
| ACTR3C | 457.096878 | -0.433345571 | 0.27692867 | -1.5648274 | 0.11762339 | 0.53282158 |
| LRRC61 | 51.5568876 | -1.270548231 | 0.40757949 | -3.1173017 | 0.00182515 | 0.05903562 |
| TMEM176B | 349.565449 | -0.853575264 | 0.66676705 | -1.2801701 | 0.20048531 | 0.65475264 |
| TMEM176A | 168.517329 | -0.84776764 | 0.67359671 | -1.2585686 | 0.20818621 | 0.66589349 |
| AOC1 | 284.794248 | 0.041268373 | 0.87096883 | 0.04738215 | 0.96220866 | 0.99171131 |
| ABCB8 | 72.7950548 | -0.119707413 | 0.25096419 | -0.47699 | 0.63336923 | 0.91415622 |
| SLC4A2 | 59.4047949 | -0.205143338 | 0.33387252 | -0.6144361 | 0.53892717 | 0.88287241 |
| FASTK | 78.5908871 | 0.276043624 | 0.32323245 | 0.85400961 | 0.39309966 | 0.81010924 |
| TMUB1 | 95.947015 | 0.284117187 | 0.44393582 | 0.63999608 | 0.52217515 | 0.87670505 |
| AGAP3 | 61.1097951 | 1.105973631 | 0.50056206 | 2.20946356 | 0.02714241 | 0.27917411 |
| CHPF2 | 45.7094161 | -0.189241071 | 0.53611163 | -0.3529882 | 0.7240973 | 0.93893205 |
| SMARCD3 | 177.113695 | -1.486642564 | 0.35966263 | -4.1334363 | 3.57E-05 | 0.00462387 |
| NUB1 | 182.724547 | 0.131147712 | 0.33862274 | 0.38729742 | 0.69853604 | 0.93180634 |
| RHEB | 320.956854 | 0.088985365 | 0.34258354 | 0.25974793 | 0.79505822 | 0.95937941 |
| PRKAG2 | 4993.40831 | -0.699267228 | 0.64363919 | -1.0864274 | 0.27728996 | 0.72788245 |
| PRKAG2-AS1 | 63.7517734 | 0.431454633 | 0.88551138 | 0.48723782 | 0.62608981 | 0.91061758 |
| GALNT11 | 886.143011 | 1.084239762 | 0.47068577 | 2.30353204 | 0.02124893 | 0.24688332 |
| KMT2C | 1760.14377 | 0.241153781 | 0.25212972 | 0.95646708 | 0.33883631 | 0.77836439 |
| ACTR3B | 163.44233 | 0.374522221 | 0.23065609 | 1.62372568 | 0.10443437 | 0.50345841 |
| DPP6 | 289.627629 | 0.050905769 | 0.31157742 | 0.16338081 | 0.87021862 | 0.9759113 |
| PAXIP1-AS2 | 65.6985786 | -0.172930912 | 0.32996823 | -0.5240835 | 0.60022045 | 0.90369786 |
| PAXIP1 | 102.805716 | 0.018635531 | 0.25855665 | 0.07207523 | 0.94254204 | 0.98885525 |
| PAXIP1-AS1 | 47.8562 | -0.52588938 | 0.47623044 | -1.104275 | 0.26947385 | 0.72322263 |
| INSIG1 | 281.683368 | 0.082186519 | 1.06153189 | 0.07742256 | 0.93828739 | 0.98846521 |
| AC009403.1 | 64.0330162 | -1.130709213 | 0.37468006 | -3.0177993 | 0.00254617 | 0.0735219 |
| RBM33 | 517.914279 | -0.059223034 | 0.26308814 | -0.2251072 | 0.82189588 | 0.96502145 |
| LINC01006 | 201.86677 | -0.021517602 | 0.40113048 | -0.0536424 | 0.95722007 | 0.99106618 |
| LMBR1 | 486.910264 | 0.211036761 | 0.2083762 | 1.01276808 | 0.31117095 | 0.75435382 |
| NOM1 | 50.8377971 | -0.259573504 | 0.39085037 | -0.664125 | 0.50661031 | 0.87094607 |
| UBE3C | 651.613283 | 0.249584698 | 0.28036607 | 0.89021005 | 0.37335311 | 0.79681088 |
| DNAJB6 | 516.440915 | 0.443533223 | 0.31747815 | 1.39705117 | 0.16239818 | 0.60211533 |
| PTPRN2 | 50.1401678 | -0.342093717 | 0.46764827 | -0.7315193 | 0.46446204 | 0.8497839 |
| NCAPG2 | 44.4089303 | 0.149435323 | 0.35782728 | 0.41761859 | 0.676226 | 0.9268955 |
| ESYT2 | 2331.76522 | 2.129574373 | 0.84357179 | 2.5244732 | 0.01158718 | 0.17055293 |
| WDR60 | 608.198757 | 0.079170147 | 0.25284226 | 0.31312071 | 0.75418895 | 0.94983584 |
| GTPBP6 | 127.766686 | 0.164384202 | 0.33297774 | 0.49367925 | 0.62153272 | 0.9099963 |
| LINC00685 | 58.0919269 | 0.127139616 | 0.64508844 | 0.19708866 | 0.84375816 | 0.96876046 |
| PPP2R3B | 50.149927 | 0.081658024 | 0.36667936 | 0.22269599 | 0.82377212 | 0.96511681 |
| SLC25A6 | 556.365284 | -0.734335651 | 0.46502996 | -1.5791147 | 0.11430975 | 0.52703872 |
| LINC00106 | 108.017606 | 0.357586167 | 0.40227882 | 0.8889013 | 0.37405613 | 0.79709398 |
| ASMTL-AS1 | 54.7814216 | 0.761978724 | 0.34659155 | 2.1984919 | 0.02791407 | 0.28401388 |
| ASMTL | 82.4235873 | -0.123620341 | 0.33436797 | -0.3697135 | 0.711596 | 0.9348115 |
| AKAP17A | 82.0852298 | -0.001948321 | 0.29398133 | -0.0066274 | 0.99471217 | 0.99888869 |
| DHRSX | 526.35023 | 0.472171726 | 0.30498454 | 1.54818251 | 0.12157836 | 0.53989034 |
| CD99 | 134.267094 | -0.265945436 | 0.36800929 | -0.7226596 | 0.46988906 | 0.85232523 |
| ARSD | 290.749109 | -0.338444029 | 0.2453722 | -1.3793088 | 0.16779957 | 0.6125181 |
| MXRA5 | 237.372739 | 0.411583732 | 0.47604195 | 0.8645955 | 0.38726084 | 0.8065026 |
| PRKX | 74.9772145 | -0.53077044 | 0.29613656 | -1.7923165 | 0.07308228 | 0.43517176 |
| BX890604.1 | 35.2408115 | 0.123166853 | 0.34072961 | 0.36147975 | 0.71774084 | 0.93573028 |
| PUDP | 97.9057616 | -0.305758281 | 0.35332068 | -0.8653846 | 0.38682773 | 0.8065026 |
| STS | 46.0089792 | 0.236634597 | 0.57834485 | 0.4091583 | 0.68242349 | 0.92806156 |
| PNPLA4 | 53.6963564 | -0.374341536 | 0.25641231 | -1.4599203 | 0.14431198 | 0.57406643 |
| TBL1X | 912.434414 | 0.526921359 | 0.38352657 | 1.3738849 | 0.16947742 | 0.61478828 |
| SHROOM2 | 125.314886 | 0.952846528 | 0.36010765 | 2.64600465 | 0.00814487 | 0.14312705 |
| WWC3 | 95.2898755 | 0.4047273 | 0.26532079 | 1.52542624 | 0.12715282 | 0.54746778 |
| MID1 | 1365.60404 | -0.723147953 | 0.40580969 | -1.7819879 | 0.07475121 | 0.43878029 |
| AC073529.1 | 82.4376486 | -0.755541597 | 0.32988685 | -2.2903053 | 0.02200362 | 0.25042784 |
| HCCS | 31.1723237 | -0.744284588 | 0.32350695 | -2.3006757 | 0.02140997 | 0.24847891 |
| ARHGAP6 | 1754.16313 | 0.322895583 | 0.33980163 | 0.9502473 | 0.34198661 | 0.77973697 |
| MSL3 | 40.0183892 | 0.087779829 | 0.38348323 | 0.22890135 | 0.81894559 | 0.96443578 |
| PRPS2 | 212.567006 | 1.145147073 | 0.42205138 | 2.7132883 | 0.00666191 | 0.13031298 |
| TMSB4X | 4320.04348 | -0.637561166 | 0.59789652 | -1.0663403 | 0.28626983 | 0.73607046 |
| RAB9A | 146.622302 | 0.101157693 | 0.45372495 | 0.22294937 | 0.82357491 | 0.96511681 |
| TRAPPC2 | 72.2977212 | 0.021232989 | 0.341133 | 0.06224255 | 0.95036968 | 0.98992048 |
| OFD1 | 360.25933 | -0.048079321 | 0.39824637 | -0.1207276 | 0.90390681 | 0.98320957 |
| GPM6B | 306.659407 | 0.114759227 | 0.51934778 | 0.22096797 | 0.82511737 | 0.96511681 |
| GEMIN8 | 129.833652 | -0.118361111 | 0.31221457 | -0.3791018 | 0.70461226 | 0.93241442 |
| MOSPD2 | 91.1713028 | -0.114262495 | 0.29698571 | -0.3847407 | 0.70042952 | 0.93180634 |
| PIR | 41.0188203 | -1.241098875 | 0.72578063 | -1.7100193 | 0.08726231 | 0.46735992 |
| CLTRN | 42.1896018 | -1.614900924 | 0.7630818 | -2.1162881 | 0.03432031 | 0.31309819 |
| CA5B | 154.253094 | -0.105322969 | 0.34543176 | -0.3049024 | 0.7604405 | 0.95327948 |
| ZRSR2 | 75.9714494 | 0.036728651 | 0.29123224 | 0.12611464 | 0.89964118 | 0.98312332 |
| CTPS2 | 192.946082 | 0.154053732 | 0.29271642 | 0.52629002 | 0.59868672 | 0.90369786 |
| SYAP1 | 261.204001 | 0.683891054 | 0.4368418 | 1.56553482 | 0.11745756 | 0.53282158 |

| | | | | | | |
|---|---|---|---|---|---|---|
| TXLNG | 127.615572 | 0.19477847 | 0.26701345 | 0.72947063 | 0.46571383 | 0.85020431 |
| RBBP7 | 202.349341 | -0.021969645 | 0.26232554 | -0.0837495 | 0.93325556 | 0.98822345 |
| REPS2 | 582.435872 | 0.072379082 | 0.36553463 | 0.19800882 | 0.84303816 | 0.96876046 |
| NHS | 299.857893 | -1.197143733 | 0.61534572 | -1.9454815 | 0.05171705 | 0.38247133 |
| SCML1 | 397.247174 | 0.253654452 | 0.40788582 | 0.62187612 | 0.53402333 | 0.88137638 |
| RAI2 | 124.566766 | -0.296331801 | 0.24728084 | -1.1983613 | 0.23077638 | 0.69081304 |
| CDKL5 | 318.940431 | 0.347258286 | 0.19123689 | 1.81585408 | 0.06939276 | 0.42828237 |
| PHKA2 | 118.565337 | -0.129577094 | 0.24759373 | -0.5233456 | 0.60073376 | 0.90369786 |
| PDHA1 | 503.615329 | 0.414145719 | 0.43983656 | 0.94159003 | 0.34640258 | 0.78180304 |
| MAP3K15 | 471.538265 | 0.775071722 | 0.60016589 | 1.29142915 | 0.19655491 | 0.65097045 |
| SH3KBP1 | 207.023157 | 0.191065498 | 0.35535785 | 0.53767069 | 0.59080442 | 0.90233654 |
| BCLAF3 | 191.475873 | -0.049905319 | 0.238063 | -0.2096307 | 0.8339559 | 0.96749268 |
| EIF1AX | 371.990138 | -0.124203183 | 0.35737261 | -0.3475453 | 0.72818167 | 0.93931348 |
| RPS6KA3 | 350.803619 | -0.283067473 | 0.23197968 | -1.2202253 | 0.22237948 | 0.6798044 |
| MBTPS2 | 99.9224081 | -0.160517482 | 0.29962784 | -0.5357228 | 0.59215011 | 0.90233654 |
| SMS | 472.943409 | -0.117946306 | 0.42743392 | -0.2759404 | 0.78259381 | 0.95688565 |
| PTCHD1-AS | 208.540637 | 0.048434433 | 0.83835166 | 0.05777341 | 0.95392912 | 0.99060627 |
| PRDX4 | 60.3878766 | -1.011288851 | 0.37455368 | -2.6999838 | 0.00693429 | 0.13285432 |
| ACOT9 | 92.6115817 | 0.962728568 | 0.44250144 | 2.17565069 | 0.02958139 | 0.29105042 |
| SAT1 | 1694.81145 | 0.18564502 | 0.38258602 | 0.48523733 | 0.627508 | 0.91061758 |
| APOO | 1503.42118 | 0.163324257 | 0.45280682 | 0.36069302 | 0.71832895 | 0.93574734 |
| EIF2S3 | 193.50081 | 0.106344259 | 0.3318281 | 0.32047997 | 0.74860452 | 0.94678489 |
| ZFX | 210.718317 | 0.359299324 | 0.28274664 | 1.27074658 | 0.20381882 | 0.6584529 |
| PDK3 | 132.736781 | -0.947394272 | 0.78411264 | -1.2082375 | 0.22695594 | 0.68523226 |
| POLA1 | 262.13951 | -0.118538211 | 0.24263978 | -0.4885358 | 0.6251704 | 0.91060262 |
| IL1RAPL1 | 72.825927 | 1.569811453 | 0.74596908 | 2.10439211 | 0.03534426 | 0.31644624 |
| GK | 80.0676586 | 0.311627886 | 0.33016457 | 0.94385624 | 0.34524311 | 0.78180304 |
| TAB3 | 259.087835 | 0.308071324 | 0.28145039 | 1.09458483 | 0.27369856 | 0.72561623 |
| DMD | 149.287621 | 0.244696042 | 0.29824274 | 0.82045933 | 0.41195431 | 0.82355334 |
| CFAP47 | 91.535345 | 0.128668965 | 0.51995904 | 0.24745981 | 0.80455239 | 0.96210606 |
| AF241726.2 | 172.302165 | -0.209379628 | 0.46193096 | -0.4532704 | 0.65035405 | 0.91806445 |
| DYNLT3 | 234.47998 | 0.134362003 | 0.42660406 | 0.31495716 | 0.75279418 | 0.94913313 |
| RPGR | 128.847111 | 0.061720585 | 0.27695753 | 0.22285216 | 0.82365057 | 0.96511681 |
| MID1IP1 | 41.1713456 | -0.625538754 | 0.38228554 | -1.6363129 | 0.10177412 | 0.49869583 |
| BCOR | 280.801684 | -0.338285778 | 0.28238199 | -1.1979722 | 0.23092784 | 0.69089594 |
| ATP6AP2 | 300.432573 | 0.031105659 | 0.36982925 | 0.08410816 | 0.93297043 | 0.98822345 |
| CXorf38 | 116.2235 | -0.229006478 | 0.21392892 | -1.0704793 | 0.28440363 | 0.73476311 |
| MED14 | 244.876384 | 0.368482207 | 0.29490554 | 1.24949232 | 0.21148506 | 0.66975892 |
| USP9X | 884.021033 | 0.612726905 | 0.36176444 | 1.69371788 | 0.0903189 | 0.47517175 |
| DDX3X | 337.148019 | 0.422818413 | 0.34645695 | 1.22040679 | 0.2223107 | 0.6798044 |
| CASK | 298.984255 | -0.076930464 | 0.2196315 | -0.3502706 | 0.72613561 | 0.93895336 |
| MAOA | 195.133532 | -0.698020922 | 0.54715493 | -1.2757281 | 0.20205165 | 0.65670446 |
| EFHC2 | 109.900911 | 0.22427127 | 0.32860442 | 0.68249621 | 0.49492524 | 0.86389349 |
| FUNDC1 | 53.8237392 | 0.127981775 | 0.3805139 | 0.33633929 | 0.73661503 | 0.9429136 |
| KDM6A | 756.184899 | 0.494838586 | 0.46499617 | 1.06417778 | 0.28724817 | 0.73638964 |
| MIR222HG | 79.830422 | -0.120816724 | 0.85280852 | -0.1416692 | 0.88734128 | 0.98009982 |
| KRBOX4 | 94.7823441 | -0.291965802 | 0.30413141 | -0.9599989 | 0.33705579 | 0.77827084 |
| SLC9A7 | 93.7744339 | -0.658228844 | 0.27486261 | -2.3947559 | 0.01663143 | 0.21108198 |
| RP2 | 52.4202959 | 0.341481942 | 0.37246246 | 0.91682244 | 0.35923569 | 0.78860286 |
| JADE3 | 157.045371 | -0.080332836 | 0.38249579 | -0.2100228 | 0.83364988 | 0.96749268 |
| RGN | 117.878691 | -1.372307079 | 0.76157794 | -1.801926 | 0.07155705 | 0.43222361 |
| NDUFB11 | 265.220171 | 0.002961398 | 0.50504834 | 0.00586359 | 0.99532156 | 0.99908534 |
| RBM10 | 118.695532 | 0.20903385 | 0.22406338 | 0.93292288 | 0.35085979 | 0.78242416 |
| UBA1 | 306.253202 | 0.165879032 | 0.22215839 | 0.7466701 | 0.45526272 | 0.84340698 |
| CDK16 | 133.959896 | 0.14743943 | 0.32370096 | 0.45548037 | 0.64876369 | 0.91768694 |
| USP11 | 170.608238 | 0.183561159 | 0.28250963 | 0.64975186 | 0.51585252 | 0.87425708 |
| ZNF41 | 75.4179647 | 0.063984128 | 0.30141838 | 0.2122768 | 0.8318911 | 0.96656674 |
| ARAF | 70.7863799 | 0.270028874 | 0.29229462 | 0.9238243 | 0.3555778 | 0.78607579 |
| TIMP1 | 66.9413126 | -3.26252947 | 0.70007204 | -4.6602768 | 3.16E-06 | 0.00080718 |
| ELK1 | 34.8086777 | -0.353140446 | 0.35124842 | -1.0053866 | 0.31471074 | 0.75823062 |
| UXT | 50.8729536 | 0.01787547 | 0.28876974 | 0.06190216 | 0.95064075 | 0.98992048 |
| ZNF81 | 164.320138 | -0.022206157 | 0.3188956 | -0.0696346 | 0.94448452 | 0.98949015 |
| ZNF182 | 35.6645066 | -0.458908521 | 0.30087667 | -1.525238 | 0.12719975 | 0.54746778 |
| ZNF630 | 72.3052341 | -0.131283374 | 0.46289679 | -0.2836126 | 0.77670726 | 0.95601455 |
| EBP | 78.503344 | -0.595460901 | 0.64053093 | -0.9296365 | 0.35255935 | 0.78314007 |
| TBC1D25 | 40.7325417 | 0.335220298 | 0.32512011 | 1.03106602 | 0.30250986 | 0.74899204 |
| RBM3 | 158.34827 | -0.390540421 | 0.44428294 | -0.8790354 | 0.37938209 | 0.80274043 |
| WDR13 | 125.603521 | 0.129258342 | 0.27661774 | 0.46728146 | 0.64029852 | 0.91583574 |
| HDAC6 | 78.4180167 | 0.261254222 | 0.21736421 | 1.2019192 | 0.22939484 | 0.68825018 |
| PCSK1N | 598.989942 | 0.019715227 | 0.4189147 | 0.04706263 | 0.96246331 | 0.99171131 |
| TIMM17B | 64.2821041 | -0.221262777 | 0.49148635 | -0.4501911 | 0.65257266 | 0.91909172 |
| SLC35A2 | 32.8586202 | -0.033132567 | 0.37374421 | -0.0886504 | 0.92935977 | 0.9878984 |
| OTUD5 | 156.979507 | 0.123970167 | 0.1746838 | 0.70968327 | 0.47790057 | 0.85479418 |
| GRIPAP1 | 126.012983 | -0.133173843 | 0.18366076 | -0.7251078 | 0.46838592 | 0.85161077 |
| TFE3 | 83.5337524 | -0.189907387 | 0.20617211 | -0.9211109 | 0.35699251 | 0.78612469 |
| CCDC120 | 47.3601182 | -0.717773627 | 0.328447 | -2.185356 | 0.02886275 | 0.28768102 |
| WDR45 | 84.4465309 | 0.135894007 | 0.34493924 | 0.39396506 | 0.69360684 | 0.93105058 |
| MAGIX | 39.2496674 | -0.092735967 | 0.3682847 | -0.2518051 | 0.80119171 | 0.96063297 |
| PLP2 | 98.3670084 | -0.784419441 | 0.52187645 | -1.503075 | 0.1328197 | 0.55601567 |
| PPP1R3F | 37.3420977 | -0.072929946 | 0.32617016 | -0.2235948 | 0.82307263 | 0.96511681 |
| CLCN5 | 567.477677 | 0.481281828 | 0.30186839 | 1.59434322 | 0.11085918 | 0.51889425 |
| SHROOM4 | 371.71897 | -0.298622477 | 0.2332296 | -1.2803799 | 0.20041158 | 0.65475264 |
| MAGED1 | 366.904386 | -0.284613399 | 0.32251643 | -0.8824772 | 0.3775188 | 0.80119855 |
| FAM156A | 47.5043337 | 0.011268271 | 0.35905041 | 0.03138354 | 0.97496367 | 0.9939318 |
| GPR173 | 68.5827421 | -0.075167404 | 0.48477257 | -0.155057 | 0.87677634 | 0.97640529 |
| TSPYL2 | 45.9447928 | 0.355383332 | 0.47639393 | 0.74598627 | 0.45567571 | 0.84372464 |
| KANTR | 243.790407 | -0.116411891 | 0.24587011 | -0.4734691 | 0.63587857 | 0.91454294 |
| KDM5C | 136.926469 | 0.156929894 | 0.4559353 | 0.34419334 | 0.73070089 | 0.94048525 |
| IQSEC2 | 77.5022288 | 0.021180411 | 0.39744912 | 0.05329088 | 0.95750015 | 0.99106618 |
| SMC1A | 136.17566 | -0.11371291 | 0.23400866 | -0.4859346 | 0.62701351 | 0.91061758 |
| HSD17B10 | 114.755102 | -0.016704549 | 0.52234149 | -0.0319801 | 0.9744879 | 0.9939318 |
| HUWE1 | 802.028103 | 0.57393095 | 0.14786318 | 3.88150016 | 0.00010381 | 0.00906643 |
| PHF8 | 192.039165 | 0.598862036 | 0.26810224 | 2.23370772 | 0.02550231 | 0.27216315 |
| FAM120C | 101.045336 | -0.028271038 | 0.35069634 | -0.080614 | 0.93574892 | 0.98822345 |
| WNK3 | 107.976195 | -0.523691126 | 0.3998331 | -1.3097743 | 0.19027219 | 0.64458858 |
| TSR2 | 83.4298438 | -0.114032771 | 0.36129829 | -0.3156195 | 0.75229137 | 0.94862393 |
| FGD1 | 44.0579742 | 0.007529416 | 0.36953984 | 0.02037511 | 0.98374414 | 0.99540861 |
| MAGED2 | 179.551214 | -0.643621684 | 0.34218612 | -1.8809111 | 0.05998401 | 0.40367252 |
| TRO | 34.1586363 | 0.861383723 | 0.51149459 | 1.68405245 | 0.09217153 | 0.48040835 |
| APEX2 | 33.4107645 | 0.002361388 | 0.50670287 | 0.0046603 | 0.99628163 | 0.99916339 |
| FAM104B | 60.5843731 | 0.152727475 | 0.32602049 | 0.46845974 | 0.63945585 | 0.91521016 |
| RRAGB | 233.672947 | -0.114701785 | 0.28727135 | -0.3992803 | 0.6896867 | 0.93071293 |
| AL445472.1 | 343.850292 | -0.311894259 | 0.24304963 | -1.2832534 | 0.19940331 | 0.65475264 |
| KLF8 | 41.399577 | -0.283739915 | 0.50260438 | -0.5645393 | 0.57238718 | 0.89797476 |
| UBQLN2 | 86.2336078 | -0.449589631 | 0.29535175 | -1.5222176 | 0.12795456 | 0.54791812 |
| NBDY | 344.558872 | 0.10555347 | 0.28333243 | 0.37254284 | 0.70948872 | 0.93421809 |
| SPIN3 | 64.6430432 | 0.023613228 | 0.27509685 | 0.08583606 | 0.93159674 | 0.98822345 |
| FAAH2 | 655.602316 | -0.280451238 | 0.39509572 | -0.7098311 | 0.47780886 | 0.85479418 |
| ARHGEF9 | 131.958597 | -0.388800697 | 0.34431606 | -1.1291971 | 0.25881469 | 0.7159728 |
| MTMR8 | 108.563852 | 0.502638248 | 0.27674993 | 1.81621815 | 0.06933692 | 0.42828237 |
| ZC4H2 | 223.221779 | 0.509613175 | 0.19646108 | 2.59396502 | 0.00948761 | 0.15560281 |
| ZC3H12B | 165.136403 | 0.375112036 | 0.3815857 | 0.98303482 | 0.3255903 | 0.77073504 |
| LAS1L | 54.1883317 | 0.006805545 | 0.34903028 | 0.01949844 | 0.98444348 | 0.9954565 |

| | | | | | | |
|---|---|---|---|---|---|---|
| MSN | 613.37117 | 0.324041199 | 0.49402809 | 0.65591654 | 0.51187782 | 0.87155791 |
| OPHN1 | 1038.05103 | 1.355289022 | 0.34715393 | 3.90400023 | 9.46E-05 | 0.00861114 |
| YIPF6 | 190.905158 | 0.161734555 | 0.21356455 | 0.75730994 | 0.4488642 | 0.84115854 |
| EFNB1 | 89.0963999 | 0.341016083 | 0.43878783 | 0.77717763 | 0.43705398 | 0.83399958 |
| EDA | 122.734079 | -0.296891486 | 0.22496627 | -1.3197155 | 0.18693001 | 0.63853538 |
| IGBP1 | 136.683707 | -0.372804218 | 0.32463505 | -1.1483794 | 0.25081195 | 0.70811133 |
| PDZD11 | 43.8811716 | -0.252679386 | 0.50495671 | -0.5003981 | 0.61679478 | 0.90794578 |
| DLG3 | 155.249788 | -0.100325057 | 0.32355887 | -0.3100674 | 0.7565097 | 0.95072713 |
| SNX12 | 70.1092386 | -0.19229504 | 0.23073703 | -0.8333948 | 0.40462211 | 0.81914727 |
| MED12 | 37.6045916 | 0.260808526 | 0.28628804 | 0.91100044 | 0.36229514 | 0.79051672 |
| ZMYM3 | 64.9318378 | 0.150093282 | 0.28468246 | 0.52723051 | 0.59803352 | 0.90369786 |
| NONO | 218.386793 | -0.199940614 | 0.23519982 | -0.8500883 | 0.39527599 | 0.81175661 |
| TAF1 | 298.495984 | 0.007724537 | 0.26669444 | 0.028964 | 0.97689331 | 0.99460933 |
| OGT | 610.320174 | 1.19814676 | 0.3657602 | 3.27577128 | 0.00105374 | 0.04090067 |
| GCNA | 38.2631764 | 0.499337472 | 0.55877302 | 0.89363204 | 0.3715188 | 0.79539603 |
| NHSL2 | 87.3503129 | 0.146040783 | 0.24363829 | 0.59941638 | 0.54889526 | 0.88854099 |
| PIN4 | 131.939897 | -0.657255265 | 0.41069181 | -1.6003613 | 0.10951846 | 0.51481816 |
| RPS4X | 1024.55867 | -0.117334026 | 0.51710209 | -0.2269069 | 0.82049615 | 0.96502145 |
| HDAC8 | 788.36516 | -0.170907106 | 0.22733717 | -0.7517781 | 0.4521845 | 0.84234337 |
| PHKA1 | 63.8304748 | 0.180007285 | 0.37152737 | 0.48450612 | 0.62802672 | 0.91090895 |
| NAP1L2 | 37.7152719 | -0.04034721 | 0.69942688 | -0.0576861 | 0.95399866 | 0.99060627 |
| CHIC1 | 228.600477 | 0.671287554 | 0.24581011 | 2.73091924 | 0.00631579 | 0.12786365 |
| XIST | 1177.92794 | 0.041458118 | 3.98004077 | 0.01041651 | 0.99168898 | 0.99767558 |
| JPX | 688.232672 | -0.208651894 | 0.45284179 | -0.4607611 | 0.64496999 | 0.91682456 |
| FTX | 4660.79993 | 0.056972262 | 0.28420544 | 0.20046155 | 0.84111963 | 0.96854218 |
| SLC16A2 | 50.0121442 | -1.118342942 | 0.45941231 | -2.4342903 | 0.01492102 | 0.19869417 |
| RLIM | 186.940565 | 0.084946538 | 0.3177417 | 0.26734464 | 0.78920382 | 0.95801395 |
| NEXMIF | 423.807766 | -0.629402572 | 0.38675944 | -1.6273748 | 0.10365752 | 0.50253415 |
| ABCB7 | 283.071469 | 0.319278711 | 0.24459894 | 1.30531516 | 0.19178555 | 0.64458858 |
| UPRT | 60.8782775 | 0.076435773 | 0.30140615 | 0.25359726 | 0.79980671 | 0.96043078 |
| ZDHHC15 | 76.3558672 | -0.512531952 | 0.3673273 | -1.3953005 | 0.16292526 | 0.60211533 |
| PBDC1 | 68.3430793 | 0.042723088 | 0.4672067 | 0.09144365 | 0.92714008 | 0.98732047 |
| ATRX | 1376.34322 | 0.561519042 | 0.17291784 | 3.24731697 | 0.00116499 | 0.04392356 |
| MAGT1 | 200.830742 | -0.118716145 | 0.23999412 | -0.4946627 | 0.62083822 | 0.90957637 |
| COX7B | 534.137458 | -0.059742614 | 0.41209539 | -0.1449728 | 0.88473236 | 0.97891513 |
| PGK1 | 893.829113 | -1.009690776 | 0.57696319 | -1.750009 | 0.08011676 | 0.45074086 |
| TAF9B | 45.5608398 | -0.634940726 | 0.43261255 | -1.4676891 | 0.14218869 | 0.57093391 |
| BRWD3 | 349.169288 | 0.018891551 | 0.36341744 | 0.05198306 | 0.95854219 | 0.99119981 |
| HMGN5 | 135.301804 | 0.215207385 | 0.44265615 | 0.48617281 | 0.62684465 | 0.91061758 |
| SH3BGRL | 720.072991 | -0.07267112 | 0.24028371 | -0.3024388 | 0.76231758 | 0.95374766 |
| POU3F4 | 57.421268 | -0.154242209 | 0.51624284 | -0.2987784 | 0.76510913 | 0.95378284 |
| RPS6KA6 | 374.867101 | 0.508137043 | 0.33418671 | 1.5205184 | 0.12838074 | 0.54843482 |
| HDX | 94.0486357 | -0.319726082 | 0.34402751 | -0.929362 | 0.35270153 | 0.78314007 |
| APOOL | 174.592239 | -0.046950608 | 0.23964339 | -0.1959186 | 0.84467384 | 0.96876046 |
| ZNF711 | 61.0612009 | 0.582393025 | 0.35513706 | 1.63991058 | 0.10102376 | 0.49726048 |
| POF1B | 104.987383 | -0.842235156 | 0.368802 | -2.2837055 | 0.02238884 | 0.25166501 |
| CHM | 292.935971 | -0.004895747 | 0.25889926 | -0.0189099 | 0.98491302 | 0.9954565 |
| PCDH11X | 42.1030528 | -0.494274299 | 0.47554965 | -1.0393747 | 0.29863048 | 0.74594723 |
| DIAPH2 | 683.479111 | -0.038827753 | 0.19024505 | -0.2040934 | 0.83828054 | 0.967744 |
| DIAPH2-AS1 | 54.2324571 | -0.330715552 | 0.6320586 | -0.5232356 | 0.60081032 | 0.90369786 |
| TSPAN6 | 105.81857 | -0.037878979 | 0.36850312 | -0.1027915 | 0.91812847 | 0.98520052 |
| SYTL4 | 63.6098208 | 0.307810887 | 0.5501422 | 0.5595115 | 0.57581269 | 0.89839467 |
| TRMT2B | 113.760522 | 0.055431415 | 0.35002169 | 0.15836566 | 0.87416867 | 0.97622753 |
| GLA | 38.9403345 | -0.070594754 | 0.55361724 | -0.1275155 | 0.89853244 | 0.98312332 |
| HNRNPH2 | 122.923811 | 0.041047865 | 0.39333446 | 0.10435868 | 0.91688471 | 0.98520052 |
| ARMCX4 | 410.032146 | 0.058413629 | 0.35901838 | 0.16270373 | 0.87075171 | 0.97622753 |
| ARMCX1 | 38.3960292 | 0.020556353 | 0.43274625 | 0.04750209 | 0.96211306 | 0.99171131 |
| ARMCX6 | 78.8869162 | -0.310867897 | 0.35106217 | -0.8855067 | 0.37588341 | 0.79936245 |
| ARMCX3 | 217.676552 | 0.148042132 | 0.38192435 | 0.38762161 | 0.69829607 | 0.93177271 |
| ARMCX2 | 170.992103 | 0.113458906 | 0.56796223 | 0.19976488 | 0.84166447 | 0.96866282 |
| ZMAT1 | 343.154138 | 0.039868839 | 0.312138 | 0.12772824 | 0.89836404 | 0.98312332 |
| TCEAL2 | 66.6181897 | -0.19345074 | 0.7082226 | -0.2731496 | 0.7847382 | 0.95702626 |
| BEX5 | 37.6452653 | -0.076426572 | 0.68901959 | -0.1109208 | 0.91167919 | 0.98468493 |
| ARMCX5 | 34.6472906 | -0.18614242 | 0.3306685 | -0.5629276 | 0.5734842 | 0.89797476 |
| GPRASP1 | 61.1926167 | 0.387087464 | 0.62587144 | 0.6184776 | 0.53626056 | 0.88137638 |
| BHLHB9 | 63.1260008 | -0.009665692 | 0.37155835 | -0.0260139 | 0.97924623 | 0.99469867 |
| LINC00630 | 178.763639 | 0.565081063 | 0.2799364 | 2.01860515 | 0.04352827 | 0.35177679 |
| BEX4 | 265.834532 | 0.335057478 | 0.46424472 | 0.72172598 | 0.47046296 | 0.85236058 |
| TCEAL8 | 96.6647534 | -0.628687594 | 0.40100883 | -1.567765 | 0.116936 | 0.53189637 |
| BEX2 | 158.057016 | -0.596790592 | 0.4360509 | -1.368626 | 0.17111622 | 0.6176702 |
| TCEAL9 | 384.957505 | 0.049458905 | 0.44167121 | 0.11198127 | 0.91083825 | 0.98468493 |
| BEX3 | 654.640692 | 0.638672536 | 0.52982216 | 1.205447 | 0.22803079 | 0.68631896 |
| TCEAL4 | 318.318995 | -0.451618221 | 0.38501496 | -1.1729888 | 0.24080031 | 0.6963541 |
| TCEAL3 | 142.662099 | -0.3411913 | 0.4105303 | -0.8310989 | 0.40591774 | 0.82002585 |
| TCEAL1 | 32.0731125 | -0.192207012 | 0.51370127 | -0.3741611 | 0.70828449 | 0.9332187 |
| MORF4L2 | 343.112351 | 0.322457137 | 0.33607097 | 0.9594912 | 0.33731135 | 0.77827084 |
| SLC25A53 | 145.071668 | 0.232591822 | 0.24890767 | 0.93445021 | 0.35007171 | 0.78223232 |
| FAM199X | 324.126228 | 0.225174897 | 0.23197532 | 0.97068471 | 0.33170531 | 0.77618491 |
| IL1RAPL2 | 902.942044 | 0.90523872 | 0.60061881 | 1.50717676 | 0.13176536 | 0.55435605 |
| RNF128 | 208.542185 | -0.035251301 | 0.3105355 | -0.1135178 | 0.90962007 | 0.98441374 |
| TBC1D8B | 122.311678 | -0.31469372 | 0.36175376 | -0.8699114 | 0.38434882 | 0.80561637 |
| MORC4 | 100.867633 | -0.617012159 | 0.48021595 | -1.2848639 | 0.19883983 | 0.65426733 |
| RBM41 | 222.54259 | -0.162453354 | 0.31298024 | -0.5190531 | 0.60372374 | 0.90371579 |
| NUP62CL | 86.6415049 | 0.390112596 | 0.4851097 | 0.80417397 | 0.42129651 | 0.82694802 |
| PRPS1 | 40.3182296 | 0.302813411 | 0.39451564 | 0.76755744 | 0.44275016 | 0.83826789 |
| TSC22D3 | 126.687688 | 1.086997236 | 0.30833505 | 3.52537684 | 0.00042288 | 0.02204871 |
| MID2 | 65.2997236 | 0.103112327 | 0.29488712 | 0.34966711 | 0.72658854 | 0.93906703 |
| PSMD10 | 34.4975715 | -0.08345591 | 0.35315211 | -0.2363172 | 0.81318656 | 0.96288868 |
| ATG4A | 35.6536734 | 0.145683621 | 0.31463494 | 0.4630243 | 0.64334695 | 0.91643619 |
| COL4A6 | 569.887617 | 1.951088715 | 0.4691754 | 4.1585486 | 3.20E-05 | 0.0043032 |
| COL4A5 | 1402.04403 | 1.178305708 | 0.29723084 | 3.96427805 | 7.36E-05 | 0.00734782 |
| ACSL4 | 1214.28609 | -1.13589079 | 0.6218272 | -1.8266984 | 0.06774512 | 0.42296693 |
| TMEM164 | 306.969926 | 0.068020793 | 0.3971416 | 0.17127592 | 0.86400682 | 0.97466129 |
| AMMECR1 | 156.190254 | -0.324881038 | 0.25689604 | -1.2646401 | 0.20600036 | 0.66304785 |
| ALG13 | 412.158543 | 1.110316641 | 0.52254327 | 2.12483195 | 0.03360064 | 0.30997771 |
| AMOT | 41.5177878 | -0.102543419 | 0.38227722 | -0.2682436 | 0.78851181 | 0.95801395 |
| PLS3-AS1 | 97.0571927 | 1.009643821 | 0.59007357 | 1.71104736 | 0.08707237 | 0.46723935 |
| PLS3 | 873.822724 | 0.784566831 | 0.33923893 | 2.31272639 | 0.02073769 | 0.24280431 |
| DANT2 | 1442.11006 | -0.212441462 | 0.29651536 | -0.7164602 | 0.47370721 | 0.85310957 |
| KLHL13 | 498.287297 | -0.387937087 | 0.43798213 | -0.8857372 | 0.37575913 | 0.79926035 |
| WDR44 | 319.759012 | 1.428599891 | 0.47491476 | 3.00811856 | 0.00262871 | 0.07465809 |
| IL13RA1 | 153.621481 | -0.952848848 | 0.31650979 | -3.0104878 | 0.00260828 | 0.0744818 |
| PGRMC1 | 253.119424 | -0.417759261 | 0.27481501 | -1.5201471 | 0.128474 | 0.54843482 |
| SLC25A43 | 102.93804 | -1.060767362 | 0.45506816 | -2.3310076 | 0.01975296 | 0.23590615 |
| SLC25A5 | 309.90276 | -0.192242298 | 0.62585986 | -0.3071651 | 0.75871772 | 0.9521704 |
| CXorf56 | 36.1587554 | 0.061541385 | 0.30015071 | 0.20503495 | 0.83754482 | 0.96757829 |
| UBE2A | 112.507204 | -0.102838692 | 0.31473494 | -0.326747 | 0.74385926 | 0.9456489 |
| NKRF | 52.6789112 | 0.216038899 | 0.39450595 | 0.54761886 | 0.58395363 | 0.90085422 |
| 6-Sep | 264.227439 | 0.036630275 | 0.27545444 | 0.13298125 | 0.89420821 | 0.98211088 |
| RPL39 | 38.6508123 | 0.666515859 | 0.44114442 | 1.51087906 | 0.13081927 | 0.55395564 |
| UPF3B | 117.566265 | -0.539543777 | 0.34558452 | -1.5612498 | 0.11846482 | 0.53417263 |
| NDUFA1 | 288.077608 | -0.407716271 | 0.36673024 | -1.1117607 | 0.26624105 | 0.72053648 |
| NKAP | 62.3447511 | -0.191440289 | 0.24405489 | -0.7844149 | 0.4327967 | 0.83162989 |

| | | | | | | |
|---|---|---|---|---|---|---|
| RHOXF1-AS1 | 76.7219528 | -0.229231517 | 0.53874143 | -0.4254945 | 0.67047623 | 0.92474338 |
| LAMP2 | 289.199955 | -0.265282394 | 0.22955501 | -1.1556376 | 0.24782944 | 0.70405328 |
| CUL4B | 105.654958 | 0.125950986 | 0.215799 | 0.58364954 | 0.55945612 | 0.89179371 |
| MCTS1 | 91.1931907 | -0.065617812 | 0.23845759 | -0.275176 | 0.783181 | 0.9568898 |
| C1GALT1C1 | 54.3670905 | -0.523626638 | 0.44260515 | -1.1830559 | 0.23678699 | 0.69335782 |
| THOC2 | 573.113784 | 0.22197693 | 0.24019054 | 0.92417018 | 0.35539772 | 0.78607579 |
| XIAP | 509.30518 | 0.007980905 | 0.19486965 | 0.04095509 | 0.9673317 | 0.99291246 |
| STAG2 | 720.625269 | 0.291808902 | 0.18887957 | 1.54494689 | 0.12235911 | 0.54052568 |
| SMARCA1 | 294.546106 | 0.257463469 | 0.25845444 | 0.99616578 | 0.31916961 | 0.76378437 |
| OCRL | 71.5803353 | 0.054140506 | 0.32163741 | 0.16832776 | 0.86632543 | 0.9753562 |
| ZDHHC9 | 106.07939 | -0.710641316 | 0.29981429 | -2.3702717 | 0.01777502 | 0.22150083 |
| UTP14A | 40.3652438 | -0.196048509 | 0.25737154 | -0.7617335 | 0.44621911 | 0.83971561 |
| BCORL1 | 66.6624964 | -0.288375799 | 0.40817525 | -0.7065 | 0.47987728 | 0.85561033 |
| ELF4 | 36.0618923 | -0.328408987 | 0.4024733 | -0.8159771 | 0.41451324 | 0.82461178 |
| AIFM1 | 110.811423 | -0.061942237 | 0.23904744 | -0.2591211 | 0.7955418 | 0.95948555 |
| ZNF280C | 44.1305204 | 0.307817494 | 0.26730102 | 1.15157618 | 0.24949527 | 0.70629672 |
| SLC25A14 | 33.2915341 | 0.53360825 | 0.30124006 | 1.77137215 | 0.07649884 | 0.4426298 |
| RBMX2 | 38.4572635 | 0.049621764 | 0.27094386 | 0.18314408 | 0.85468497 | 0.97122276 |
| ENOX2 | 201.745475 | -0.50365955 | 0.25925489 | -1.9427196 | 0.05205005 | 0.38306494 |
| RAP2C | 72.1606697 | 0.348617109 | 0.35167324 | 0.99130974 | 0.32153436 | 0.7663589 |
| RAP2C-AS1 | 258.09991 | 0.18156062 | 0.29820206 | 0.608851 | 0.5426232 | 0.88450129 |
| MBNL3 | 219.736559 | 0.544964348 | 0.32429742 | 1.68044614 | 0.09287055 | 0.48153712 |
| GPC4 | 112.371734 | -0.164617485 | 0.30558886 | -0.5386894 | 0.59010118 | 0.90233654 |
| GPC3 | 43.6823585 | 0.92454703 | 0.73581028 | 1.25650193 | 0.20893405 | 0.66655368 |
| PHF6 | 137.170512 | 0.126646644 | 0.21022443 | 0.60243543 | 0.54688434 | 0.88734291 |
| HPRT1 | 243.231424 | -0.350379124 | 0.3420278 | -1.0244171 | 0.30563831 | 0.75109981 |
| FAM122B | 57.0414313 | 0.354930613 | 0.31122125 | 1.14044465 | 0.2541011 | 0.7112659 |
| FAM122C | 53.634948 | 0.582804671 | 0.45321018 | 1.28594788 | 0.19846125 | 0.65403502 |
| MOSPD1 | 33.471731 | -0.150434936 | 0.30963078 | -0.4858526 | 0.62707164 | 0.91061758 |
| RTL8C | 150.054006 | -0.324839933 | 0.38963125 | -0.8337112 | 0.40444375 | 0.81914727 |
| RTL8A | 97.6236185 | -0.522253817 | 0.32260265 | -1.6188764 | 0.10547386 | 0.50611998 |
| ZNF75D | 132.476704 | -0.497838819 | 0.34839924 | -1.428932 | 0.15302378 | 0.58860244 |
| ZNF449 | 37.0266132 | -0.348491289 | 0.3692522 | -0.9437758 | 0.34528423 | 0.78180304 |
| INTS6L | 68.8261875 | 0.271161914 | 0.43636924 | 0.62140474 | 0.53433335 | 0.88137638 |
| MMGT1 | 48.9649528 | -0.104360339 | 0.31598258 | -0.3302724 | 0.74119411 | 0.94487568 |
| SLC9A6 | 87.3653005 | 0.505471929 | 0.33563951 | 1.5059965 | 0.13206807 | 0.55448417 |
| FHL1 | 149.499166 | 3.092878207 | 0.93920871 | 3.29306806 | 0.000991 | 0.03919144 |
| HTATSF1 | 144.625567 | 0.22514513 | 0.44911902 | 0.50130393 | 0.61615724 | 0.90794578 |
| AL683813.1 | 78.829577 | -0.231297643 | 0.36022498 | -0.6420922 | 0.52081333 | 0.87638467 |
| RBMX | 453.225003 | 0.170696585 | 0.22938401 | 0.74415206 | 0.45678448 | 0.84383948 |
| ATP11C | 415.839642 | 1.159764852 | 0.65097767 | 1.78157393 | 0.07481874 | 0.43878029 |
| LDOC1 | 38.7527757 | -0.40736149 | 0.44976313 | -0.9057245 | 0.36508168 | 0.79281836 |
| FMR1 | 221.757363 | -0.062903272 | 0.22168183 | -0.2837547 | 0.77659834 | 0.95601455 |
| IDS | 257.129648 | 0.524911202 | 0.37130955 | 1.41367548 | 0.15745721 | 0.59507808 |
| TMEM185A | 61.4730347 | -0.156548444 | 0.2837559 | -0.5517011 | 0.58115316 | 0.90045866 |
| LINC00894 | 355.454778 | 0.01841161 | 0.37554341 | 0.04902658 | 0.96089811 | 0.99155299 |
| MTM1 | 168.817877 | -0.174878568 | 0.24482976 | -0.7142864 | 0.47505011 | 0.85385826 |
| MTMR1 | 161.010644 | 0.008442039 | 0.17561735 | 0.04807065 | 0.96165994 | 0.99171131 |
| CD99L2 | 127.038414 | -0.183206543 | 0.22227445 | -0.8242357 | 0.40980563 | 0.8216497 |
| HMGB3 | 47.0079842 | -0.900630098 | 0.63673373 | -1.4144533 | 0.15722886 | 0.59486921 |
| VMA21 | 117.57327 | 0.208994197 | 0.46525808 | 0.44920057 | 0.65328697 | 0.91973728 |
| CETN2 | 120.411695 | -0.04693799 | 0.4421569 | -0.1061569 | 0.91545789 | 0.98520052 |
| ZNF185 | 63.2110924 | 0.212294401 | 0.65339724 | 0.32490863 | 0.74525023 | 0.94594643 |
| BCAP31 | 197.998036 | -0.383259721 | 0.35947208 | -1.0661738 | 0.28634507 | 0.73607046 |
| IDH3G | 66.9447731 | -0.11281246 | 0.38762113 | -0.291038 | 0.77102229 | 0.95498333 |
| SSR4 | 143.248679 | 0.019654883 | 0.43903555 | 0.04476832 | 0.96429198 | 0.99243116 |
| L1CAM | 314.622781 | -0.110515513 | 0.45273391 | -0.244107 | 0.80714796 | 0.96226758 |
| AVPR2 | 118.60423 | 0.425525668 | 0.66071781 | 0.64403541 | 0.51955247 | 0.87623268 |
| HCFC1 | 78.4859478 | 0.17594278 | 0.22351273 | 0.78717118 | 0.43118167 | 0.83111714 |
| IRAK1 | 65.222452 | -0.884742848 | 0.38718989 | -2.285036 | 0.02231072 | 0.25166501 |
| MECP2 | 318.233316 | -0.11056721 | 0.2134482 | -0.5180049 | 0.60445486 | 0.90405122 |
| FLNA | 296.162638 | -0.092260781 | 0.39896965 | -0.2312476 | 0.81712244 | 0.96380902 |
| EMD | 187.886274 | 0.677980518 | 0.54532792 | 1.24325291 | 0.21377465 | 0.67116787 |
| RPL10 | 1364.34063 | 0.114231499 | 0.58223212 | 0.1961958 | 0.84445691 | 0.96876046 |
| DNASE1L1 | 39.4739106 | -0.573068917 | 0.29910643 | -1.9159365 | 0.05537318 | 0.39343117 |
| TAZ | 91.8297492 | -0.288265782 | 0.26952124 | -1.0695475 | 0.28482302 | 0.7352082 |
| ATP6AP1 | 235.306037 | 0.059488532 | 0.40131039 | 0.14823571 | 0.88215675 | 0.97830717 |
| GDI1 | 101.325827 | 0.14805283 | 0.33408383 | 0.44316072 | 0.65764949 | 0.92180056 |
| FAM50A | 81.4161533 | 0.294500745 | 0.45039751 | 0.65386851 | 0.51319652 | 0.87182681 |
| PLXNA3 | 52.0423147 | -0.553372926 | 0.47558804 | -1.1635552 | 0.24460432 | 0.70121778 |
| LAGE3 | 35.7085192 | -0.094088393 | 0.46980394 | -0.2002716 | 0.84126816 | 0.96854218 |
| UBL4A | 40.8475878 | -0.041913381 | 0.44630813 | -0.0939113 | 0.92517961 | 0.98710257 |
| FAM3A | 64.4529201 | 0.024786999 | 0.245732 | 0.10087005 | 0.91965362 | 0.98527601 |
| IKBKG | 35.188024 | -0.260492757 | 0.39461575 | -0.6601175 | 0.50917844 | 0.87094607 |
| DKC1 | 69.1520524 | -0.057926158 | 0.23826105 | -0.2431206 | 0.80791201 | 0.96245698 |
| MPP1 | 71.4306085 | 0.043739735 | 0.49722361 | 0.08796794 | 0.92990216 | 0.98807358 |
| F8 | 52.6951122 | -0.296140439 | 0.3929708 | -0.753594 | 0.45109304 | 0.84193323 |
| FUNDC2 | 161.712404 | 0.199219848 | 0.3788088 | 0.52591135 | 0.5989498 | 0.90369786 |
| BRCC3 | 131.777629 | -0.368811947 | 0.20225856 | -1.8234677 | 0.06823258 | 0.42412661 |
| VBP1 | 59.0689875 | -0.106907032 | 0.32974272 | -0.3242135 | 0.74577642 | 0.94594643 |
| TMLHE-AS1 | 124.186808 | -1.289589862 | 0.70936092 | -1.8179601 | 0.06907023 | 0.42755818 |
| TMLHE | 259.286757 | -0.403199517 | 0.28735353 | -1.403148 | 0.16057272 | 0.60010648 |
| VAMP7 | 139.792948 | 0.115433635 | 0.30924382 | 0.37327709 | 0.70894222 | 0.93385049 |
| ZNF596 | 53.9548076 | -0.38436709 | 0.28790114 | -1.3350663 | 0.18185466 | 0.63211837 |
| FBXO25 | 212.800859 | 0.27298352 | 0.41821827 | 0.65272978 | 0.5139305 | 0.87226916 |
| TDRP | 263.366322 | 0.590016225 | 0.2674146 | 2.20637251 | 0.02735793 | 0.28061519 |
| ERICH1 | 311.145266 | -0.521691155 | 0.39203159 | -1.3307375 | 0.18327539 | 0.633903 |
| AC100810.1 | 49.553699 | -0.0326609 | 0.40762187 | -0.0801255 | 0.93613745 | 0.98822345 |
| AC246817.2 | 50.3613341 | -0.5577944 | 0.68376049 | -0.8157745 | 0.4146291 | 0.82461178 |
| AC246817.1 | 91.5428001 | 0.162851502 | 0.59337034 | 0.27445171 | 0.78373751 | 0.9568898 |
| CSMD1 | 175.27034 | 1.827724462 | 0.98849241 | 1.84900201 | 0.06445752 | 0.41412378 |
| MCPH1 | 427.462222 | 0.102562463 | 0.22147343 | 0.46309149 | 0.64329879 | 0.91643619 |
| ANGPT2 | 67.9304238 | -0.525956083 | 0.66119739 | -0.79546 | 0.42634597 | 0.8277744 |
| MCPH1-AS1 | 154.347206 | -0.121704691 | 0.31165857 | -0.3905065 | 0.69616206 | 0.93130502 |
| AGPAT5 | 405.815959 | -0.261457678 | 0.25998389 | -1.0056688 | 0.31457494 | 0.75823062 |
| AF233439.1 | 703.175892 | -0.9492433 | 0.86459781 | -1.0979016 | 0.27224747 | 0.72405776 |
| DEFB1 | 2065.50345 | -1.508128925 | 0.54574282 | -2.7634425 | 0.00571952 | 0.1213371 |
| FAM85B | 417.242069 | 0.304256327 | 0.46101155 | 0.65997549 | 0.50926956 | 0.87094607 |
| PRAG1 | 48.1843386 | -0.520178866 | 0.57957817 | -0.8975129 | 0.36944531 | 0.79532503 |
| MFHAS1 | 1196.75838 | -0.649051779 | 0.47487529 | -1.3667837 | 0.17169313 | 0.6187442 |
| ERI1 | 117.952488 | 0.117432649 | 0.32090538 | 0.36594166 | 0.71440861 | 0.93573028 |
| PPP1R3B | 112.098081 | -1.408977959 | 0.45039001 | -3.1283508 | 0.0017579 | 0.05775177 |
| TNKS | 856.625126 | 0.045034784 | 0.25628166 | 0.17572379 | 0.86051094 | 0.97304216 |
| MSRA | 226.885975 | -0.57364526 | 0.2920446 | -1.9642386 | 0.04950243 | 0.3755208 |
| XKR6 | 689.382506 | -0.308546304 | 0.34363249 | -0.8978962 | 0.36924089 | 0.79532503 |
| MTMR9 | 118.816892 | -0.022413892 | 0.26454624 | -0.0847258 | 0.93247938 | 0.98822345 |
| FAM167A | 45.6926844 | -0.180065818 | 0.44673945 | -0.4030667 | 0.68689912 | 0.92992151 |
| NEIL2 | 30.0701917 | -0.159684217 | 0.47232171 | -0.3380836 | 0.7353002 | 0.94232801 |
| FDFT1 | 352.8773 | -0.20602434 | 0.56148828 | -0.3669255 | 0.71367463 | 0.93573028 |
| CTSB | 245.653839 | -0.835490211 | 0.32933501 | -2.5369007 | 0.01118387 | 0.16686755 |
| FAM66D | 111.709968 | 0.087891868 | 0.63347561 | 0.13874546 | 0.88965129 | 0.98064269 |
| AC145124.1 | 48.2787777 | -0.115241279 | 0.6405044 | -0.1799227 | 0.85721326 | 0.97193497 |
| LONRF1 | 129.369612 | 0.028083836 | 0.34702825 | 0.08092666 | 0.93550028 | 0.98822345 |

| | | | | | | |
|---|---|---|---|---|---|---|
| TRMT9B | 112.422359 | -0.674885876 | 0.47179415 | -1.4304668 | 0.15258308 | 0.58795409 |
| TUSC3 | 364.98135 | -0.0686619 | 0.26741151 | -0.2567649 | 0.79736024 | 0.95991117 |
| MSR1 | 537.818406 | -0.403699444 | 0.51170109 | -0.7889361 | 0.43014938 | 0.83047752 |
| AC068992.1 | 225.746464 | -0.184472592 | 0.41962269 | -0.4396154 | 0.6602157 | 0.92284375 |
| MICU3 | 268.476839 | 0.625200887 | 0.26585904 | 2.3516255 | 0.01869158 | 0.22804166 |
| ZDHHC2 | 386.721963 | 0.047702724 | 0.1973559 | 0.24170913 | 0.80900555 | 0.96245698 |
| CNOT7 | 124.297046 | 0.053767451 | 0.29865021 | 0.18003487 | 0.85712519 | 0.97193497 |
| VPS37A | 152.843088 | 0.397205822 | 0.28916057 | 1.37365139 | 0.16954994 | 0.61483852 |
| SLC7A2 | 180.728749 | 1.07968804 | 1.67135614 | 0.6459952 | 0.51828247 | 0.87526312 |
| MTUS1 | 1046.57871 | 0.904364409 | 0.40027417 | 2.25936241 | 0.02386085 | 0.26048096 |
| AC027117.2 | 130.862799 | 0.09014563 | 0.41169564 | 0.21896183 | 0.82667978 | 0.96545294 |
| PCM1 | 1327.67818 | 0.534247695 | 0.18611682 | 2.87049666 | 0.00409828 | 0.09942112 |
| ASAH1 | 899.730123 | 0.099278632 | 0.3095667 | 0.32070191 | 0.7484363 | 0.94678489 |
| PSD3 | 675.972565 | -0.27137827 | 0.76818021 | -0.3532742 | 0.72388287 | 0.93893205 |
| SH2D4A | 175.833282 | -0.190892785 | 0.3634802 | -0.5251807 | 0.59945759 | 0.90369786 |
| CSGALNACT1 | 1266.42798 | -0.808133599 | 1.35536147 | -0.5962495 | 0.55100856 | 0.88916026 |
| INTS10 | 257.203723 | -0.15307818 | 0.24832223 | -0.6164497 | 0.53759773 | 0.88239554 |
| ATP6V1B2 | 159.338581 | 0.005956893 | 0.2659338 | 0.02239991 | 0.98212895 | 0.99507903 |
| XPO7 | 262.812465 | 0.23616342 | 0.19185837 | 1.23092582 | 0.21835061 | 0.67601607 |
| DMTN | 128.010536 | 0.25876526 | 0.2723319 | 0.95018342 | 0.34201906 | 0.77973697 |
| FAM160B2 | 69.0305377 | -0.038525892 | 0.2344635 | -0.1643151 | 0.8694831 | 0.97576204 |
| SLC39A14 | 203.314447 | -0.962243947 | 0.71884351 | -1.3386 | 0.18070091 | 0.63124852 |
| PPP3CC | 292.808386 | -0.110767861 | 0.35333177 | -0.3134953 | 0.75390436 | 0.94974369 |
| CCAR2 | 68.1777036 | 0.161399505 | 0.30512717 | 0.52895815 | 0.59683448 | 0.90369786 |
| BIN3 | 131.702493 | -0.124492234 | 0.25510253 | -0.4880086 | 0.62554374 | 0.91061758 |
| TNFRSF10B | 110.326762 | -0.353084645 | 0.56499692 | -0.624932 | 0.5320157 | 0.88045351 |
| TNFRSF10D | 84.9572786 | -1.745988391 | 0.58757447 | -2.9715185 | 0.00296331 | 0.08063686 |
| TNFRSF10A | 61.9944701 | -1.135905794 | 0.41691609 | -2.724543 | 0.00643905 | 0.12829144 |
| CHMP7 | 62.6235825 | 0.308159915 | 0.28283488 | 1.08953999 | 0.27591583 | 0.72653214 |
| R3HCC1 | 32.9679211 | 0.282395612 | 0.45627813 | 0.61891113 | 0.53597491 | 0.88137638 |
| ENTPD4 | 94.438475 | -0.192102536 | 0.26804261 | -0.7166866 | 0.47356752 | 0.85310957 |
| SLC25A37 | 273.677426 | 0.030386987 | 0.24606802 | 0.12349019 | 0.90171894 | 0.98320957 |
| STC1 | 296.987457 | 1.429465426 | 0.54954942 | 2.60115903 | 0.00929094 | 0.15430907 |
| DOCK5 | 371.370578 | -0.491824397 | 0.3225914 | -1.5246048 | 0.1273577 | 0.54746778 |
| KCTD9 | 174.856957 | 0.403730803 | 0.24130531 | 1.67311197 | 0.09430526 | 0.48561474 |
| PPP2R2A | 547.869689 | 0.485466857 | 0.21587764 | 2.2488057 | 0.02452486 | 0.26518037 |
| BNIP3L | 544.485587 | -1.481578783 | 0.53344756 | -2.7773654 | 0.00548015 | 0.11866118 |
| PTK2B | 180.623138 | -0.931358585 | 0.36419808 | -2.5572858 | 0.01054925 | 0.16402988 |
| EPHX2 | 121.011716 | 0.066782101 | 0.41579313 | 0.16061377 | 0.87239761 | 0.97622753 |
| CLU | 1870.04194 | -0.331607175 | 0.57538651 | -0.5763207 | 0.56439841 | 0.89321887 |
| SCARA3 | 48.0394906 | 0.426889678 | 0.99205245 | 0.43030958 | 0.66697046 | 0.92422364 |
| CCDC25 | 128.964622 | -0.015713096 | 0.23069736 | -0.0681113 | 0.94569704 | 0.98949015 |
| ELP3 | 229.272627 | 0.117209185 | 0.26933792 | 0.4351752 | 0.66343526 | 0.9229791 |
| ZNF395 | 103.8567 | -1.085694191 | 0.51512535 | -2.1076311 | 0.03506291 | 0.31514522 |
| FBXO16 | 123.356911 | -0.306123275 | 0.44518269 | -0.6876352 | 0.49168255 | 0.86172738 |
| FZD3 | 104.060297 | -0.061652325 | 0.28175784 | -0.2188132 | 0.82679559 | 0.96545294 |
| EXTL3 | 54.9989418 | -0.545075741 | 0.38178849 | -1.4276903 | 0.15338101 | 0.58860244 |
| INTS9 | 184.879479 | -0.028884503 | 0.23838236 | -0.1211688 | 0.90355734 | 0.98320957 |
| HMBOX1 | 556.162662 | 0.040241394 | 0.23319686 | 0.17256405 | 0.86299411 | 0.9744399 |
| KIF13B | 1612.98219 | 0.655372411 | 0.45656533 | 1.43544059 | 0.15116159 | 0.58535779 |
| AC108449.1 | 67.7533713 | 0.345481629 | 0.57524379 | 0.60058298 | 0.54811778 | 0.88810673 |
| SARAF | 425.189089 | -0.054052579 | 0.39993468 | -0.1351535 | 0.8924905 | 0.98168537 |
| LEPROTL1 | 143.406119 | 0.127716638 | 0.25661531 | 0.49769687 | 0.61869772 | 0.90835482 |
| DCTN6 | 78.7919165 | 0.207820146 | 0.42765757 | 0.48594988 | 0.6270027 | 0.91061758 |
| RBPMS | 7726.74508 | 0.300367918 | 0.28259015 | 1.06291009 | 0.28782272 | 0.73653438 |
| GTF2E2 | 229.204597 | 0.611394037 | 0.24015565 | 2.54582403 | 0.01090201 | 0.16452817 |
| GSR | 425.395473 | 0.408379973 | 0.37771021 | 1.08119919 | 0.27960852 | 0.72938673 |
| UBXN8 | 40.3899545 | -0.100655924 | 0.34685422 | -0.2901966 | 0.77166582 | 0.95523952 |
| PPP2CB | 557.501205 | 0.024382483 | 0.15915964 | 0.15319514 | 0.87824439 | 0.97692415 |
| WRN | 275.470065 | 0.116920301 | 0.23903192 | 0.48914096 | 0.62474191 | 0.91058133 |
| NRG1 | 68.438469 | -2.023937214 | 0.75895227 | -2.6667516 | 0.00765882 | 0.13910656 |
| FUT10 | 114.026276 | -0.125772643 | 0.42549937 | -0.2955883 | 0.7675445 | 0.95408212 |
| TTI2 | 46.0297743 | -0.256849608 | 0.32950931 | -0.7794912 | 0.43569042 | 0.83364877 |
| UNC5D | 53.7611685 | -0.389291436 | 0.41979094 | -0.927346 | 0.35374693 | 0.78477304 |
| ERLIN2 | 152.510048 | -0.11557936 | 0.24280539 | -0.4760164 | 0.63406266 | 0.91415622 |
| PLPBP | 104.067921 | -0.004582717 | 0.32100673 | -0.0142761 | 0.98860973 | 0.9964791 |
| RAB11FIP1 | 478.17657 | -0.108650929 | 0.24869756 | -0.4368798 | 0.66219855 | 0.92284375 |
| EIF4EBP1 | 41.1424191 | -1.469891081 | 0.55678535 | -2.63996 | 0.00829158 | 0.1442148 |
| ASH2L | 128.567157 | 0.230336644 | 0.19604293 | 1.17492962 | 0.24002287 | 0.6963541 |
| LSM1 | 134.752989 | 0.246179616 | 0.45110804 | 0.54572208 | 0.58525699 | 0.90173902 |
| BAG4 | 166.022559 | 0.493830932 | 0.26046813 | 1.89593611 | 0.05796849 | 0.3985947 |
| DDHD2 | 227.536722 | 0.129972515 | 0.24685704 | 0.52650925 | 0.59853443 | 0.90369786 |
| PLPP5 | 187.906177 | 0.125248011 | 0.43105257 | 0.29056319 | 0.77138541 | 0.95500521 |
| NSD3 | 752.192712 | -0.048387544 | 0.14021031 | -0.3451069 | 0.73001402 | 0.9404483 |
| LETM2 | 48.9927798 | 0.444537833 | 0.47248095 | 0.94085874 | 0.34677726 | 0.78180304 |
| FGFR1 | 275.435487 | 0.558418442 | 0.59669474 | 0.9358528 | 0.34934898 | 0.78180304 |
| TACC1 | 374.812639 | 0.695600762 | 0.69304715 | 1.00368461 | 0.31553066 | 0.75912793 |
| PLEKHA2 | 1399.2522 | 0.210143733 | 0.69006438 | 0.30452772 | 0.76072588 | 0.95329513 |
| AC108863.1 | 58.1705022 | 0.907634336 | 0.95184612 | 0.95355154 | 0.3403107 | 0.77836439 |
| TM2D2 | 73.5170363 | -0.295377329 | 0.61602008 | -0.479493 | 0.63158793 | 0.9133034 |
| ADAM9 | 510.126564 | 0.465936699 | 0.5570708 | 0.83640481 | 0.40292721 | 0.81787277 |
| TCIM | 128.238277 | 0.101914322 | 0.87448431 | 0.1165422 | 0.90722285 | 0.98341906 |
| GOLGA7 | 84.0765187 | -0.185030424 | 0.35368831 | -0.5231454 | 0.60087305 | 0.90369786 |
| GPAT4 | 97.700508 | -0.306060487 | 0.44347591 | -0.6901401 | 0.49010612 | 0.86055851 |
| KAT6A | 515.696939 | 0.35083899 | 0.22732174 | 1.54335872 | 0.12274376 | 0.54116729 |
| AP3M2 | 67.0413108 | 0.046248633 | 0.27968769 | 0.16535813 | 0.86866211 | 0.97576204 |
| IKBKB | 258.699822 | -0.056870886 | 0.2320103 | -0.2451223 | 0.80636177 | 0.96218371 |
| POLB | 105.868771 | 1.691717739 | 0.67794054 | 2.49537774 | 0.01258231 | 0.17925575 |
| VDAC3 | 239.862125 | -0.141589371 | 0.2754676 | -0.5139965 | 0.60725446 | 0.90510401 |
| SLC20A2 | 987.339934 | 1.298125255 | 0.47256582 | 2.74697236 | 0.00601482 | 0.12509033 |
| SMIM19 | 146.096777 | -0.131034466 | 0.39705206 | -0.3300184 | 0.74138609 | 0.94487568 |
| RNF170 | 237.149324 | -0.024465935 | 0.28561314 | -0.0856611 | 0.93173583 | 0.98822345 |
| HOOK3 | 482.323201 | 1.174923059 | 0.55973737 | 2.09906133 | 0.03581149 | 0.31751647 |
| FNTA | 104.716101 | -0.612560075 | 0.2765457 | -2.215041 | 0.02675725 | 0.27804646 |
| HGSNAT | 579.909076 | 1.158390604 | 0.62562133 | 1.85158425 | 0.06408555 | 0.41357646 |
| SPIDR | 1435.48935 | 0.029142995 | 0.23934676 | 0.12176056 | 0.90308865 | 0.98320957 |
| CEBPD | 769.05717 | -0.315834358 | 0.78369152 | -0.4030085 | 0.68694196 | 0.92992151 |
| PRKDC | 434.744755 | -0.085380002 | 0.19272218 | -0.4430211 | 0.65775044 | 0.92180056 |
| MCM4 | 37.5658969 | -0.958999083 | 0.45983646 | -2.0855221 | 0.03702194 | 0.32332491 |
| UBE2V2 | 195.030723 | 0.178566856 | 0.3860461 | 0.46255319 | 0.64368467 | 0.91655099 |
| SNTG1 | 310.563397 | 2.874588712 | 0.75095986 | 3.82788598 | 0.00012925 | 0.01044017 |
| PXDNL | 62.2151952 | -7.58E-05 | 0.38757463 | -0.0001956 | 0.99984396 | 0.99993938 |
| PCMTD1 | 889.078914 | 0.065544533 | 0.20253592 | 0.32361931 | 0.74622627 | 0.94594643 |
| AC064807.1 | 50.2384557 | -0.831977435 | 0.48574931 | -1.7127712 | 0.08675464 | 0.46672926 |
| RB1CC1 | 481.162109 | 0.101326547 | 0.25683738 | 0.39451636 | 0.69319985 | 0.93105058 |
| ATP6V1H | 551.019528 | 0.013478561 | 0.30689481 | 0.04391916 | 0.96496885 | 0.99243116 |
| TCEA1 | 219.76576 | 0.609612956 | 0.18011497 | 3.38457684 | 0.00071288 | 0.03220254 |
| LYPLA1 | 328.214033 | 0.11740557 | 0.22662688 | 0.51805667 | 0.60441873 | 0.90405122 |
| MRPL15 | 104.47952 | -0.399446545 | 0.44221299 | -0.9032899 | 0.36637203 | 0.7935157 |
| TMEM68 | 169.472101 | 0.615816034 | 0.26811609 | 2.2968261 | 0.02162869 | 0.24935578 |
| TGS1 | 343.977606 | 0.65499373 | 0.43755571 | 1.49693791 | 0.13440941 | 0.55868799 |
| LYN | 578.080477 | 0.246868834 | 0.34408656 | 0.71746142 | 0.47308943 | 0.85298656 |
| RPS20 | 421.67466 | 0.060757544 | 0.38338958 | 0.15847469 | 0.87408276 | 0.97622753 |

| | | | | | | |
|---|---|---|---|---|---|---|
| PLAG1 | 118.763511 | -0.483798457 | 0.4034612 | -1.1991202 | 0.23048123 | 0.69012664 |
| CHCHD7 | 72.6079072 | -0.144425012 | 0.26944535 | -0.5360086 | 0.59195264 | 0.90233654 |
| IMPAD1 | 169.464288 | 0.004476726 | 0.2956696 | 0.01514097 | 0.98791971 | 0.9964791 |
| LINC01606 | 255.505391 | 0.434268888 | 0.747705 | 0.58080244 | 0.56137361 | 0.89242209 |
| FAM110B | 920.896009 | 0.185029804 | 0.39922954 | 0.46346722 | 0.64302951 | 0.91643619 |
| AC027698.1 | 67.7388069 | -0.781156362 | 0.63009386 | -1.239746 | 0.21506935 | 0.67132603 |
| UBXN2B | 177.798513 | 0.292236421 | 0.17025278 | 1.71648543 | 0.08607323 | 0.46572214 |
| SDCBP | 505.978531 | -0.169918387 | 0.37084073 | -0.4581977 | 0.64681038 | 0.91743182 |
| NSMAF | 174.352508 | 0.220770725 | 0.25366074 | 0.87033856 | 0.38411541 | 0.80561637 |
| TOX | 3955.20487 | 0.9306744 | 0.51134633 | 1.8200471 | 0.06875183 | 0.42634272 |
| LINC01301 | 103.624603 | -0.310260218 | 0.59005689 | -0.5258141 | 0.59901739 | 0.90369786 |
| RAB2A | 731.974987 | 0.172606379 | 0.17911899 | 0.96364088 | 0.33522601 | 0.77820994 |
| CHD7 | 398.158986 | -0.208325834 | 0.28790509 | -0.723592 | 0.46931623 | 0.85226721 |
| ASPH | 497.178411 | -0.616896485 | 0.21428725 | -2.8788297 | 0.00399154 | 0.09782854 |
| YTHDF3 | 279.079855 | 0.379347099 | 0.34289536 | 1.10630573 | 0.26859421 | 0.72322263 |
| CYP7B1 | 320.391673 | 0.664855145 | 0.32162742 | 2.06715941 | 0.03871914 | 0.33071257 |
| ARMC1 | 103.803769 | 0.329321649 | 0.25157757 | 1.30902625 | 0.19052546 | 0.64458858 |
| MTFR1 | 126.976773 | -0.309916393 | 0.38768989 | -0.7993925 | 0.42406286 | 0.82774157 |
| PDE7A | 121.256429 | 0.334070763 | 0.4309396 | 0.77521482 | 0.43821273 | 0.8347986 |
| ADHFE1 | 102.009024 | -0.038708726 | 0.51909417 | -0.0745698 | 0.94055704 | 0.98855185 |
| VCPIP1 | 98.3903433 | -0.480586782 | 0.29303421 | -1.6400364 | 0.10099759 | 0.49726048 |
| SGK3 | 166.718163 | -0.557597872 | 0.24855121 | -2.2433923 | 0.02487153 | 0.26654723 |
| PPP1R42 | 59.6481637 | -0.248477184 | 0.4121137 | -0.6029336 | 0.54655288 | 0.88719155 |
| CSPP1 | 443.292032 | 0.362461843 | 0.29662429 | 1.22195606 | 0.22172424 | 0.6794174 |
| ARFGEF1 | 1485.22398 | 0.030268853 | 0.17083973 | 0.17717689 | 0.85936944 | 0.97244655 |
| NCOA2 | 1785.67129 | 0.363067437 | 0.13241787 | 2.74183111 | 0.00610977 | 0.12629278 |
| TRAM1 | 379.323258 | -0.263871364 | 0.2144378 | -1.2305263 | 0.21850007 | 0.67607934 |
| LACTB2 | 69.6586089 | -1.288306501 | 0.33660127 | -3.8273964 | 0.00012951 | 0.01044017 |
| MSC-AS1 | 434.866312 | -1.46385216 | 0.92207224 | -1.5875678 | 0.11238411 | 0.52250301 |
| TERF1 | 200.718993 | 0.302119945 | 0.1701478 | 1.77563238 | 0.07579353 | 0.44226959 |
| SBSPON | 115.967499 | 0.960573349 | 0.71400253 | 1.34533606 | 0.17851673 | 0.62759322 |
| RPL7 | 972.146465 | 0.380697877 | 0.53485837 | 0.71177324 | 0.47660521 | 0.85414749 |
| RDH10 | 906.156206 | 0.67510213 | 0.75840407 | 0.89016153 | 0.37337916 | 0.79681088 |
| RDH10-AS1 | 302.569083 | 0.935252401 | 0.85864819 | 1.0892149 | 0.27605913 | 0.72654437 |
| STAU2 | 653.000366 | -0.236778709 | 0.27009789 | -0.8766404 | 0.38068201 | 0.80308469 |
| AC027018.1 | 97.8465963 | -0.506488697 | 0.49323262 | -1.0268759 | 0.3044789 | 0.74992687 |
| ELOC | 142.490292 | 0.077967107 | 0.35102017 | 0.22211575 | 0.82422378 | 0.96511681 |
| TMEM70 | 42.6842144 | -0.558914614 | 0.49675331 | -1.1251351 | 0.26053177 | 0.7176381 |
| GDAP1 | 155.324306 | 0.151442951 | 0.38375335 | 0.39463617 | 0.69311142 | 0.93105058 |
| HNF4G | 220.894328 | -0.203619964 | 0.39783772 | -0.5118166 | 0.60877934 | 0.90549306 |
| ZFHX4 | 48.3933537 | 0.100263158 | 1.12336074 | 0.08925286 | 0.92888096 | 0.98761849 |
| PEX2 | 154.866996 | 0.220741318 | 0.25503195 | 0.86554378 | 0.3867404 | 0.8065026 |
| PKIA-AS1 | 50.0825663 | 0.422252068 | 0.60915216 | 0.69317995 | 0.48819663 | 0.86014074 |
| PKIA | 611.026227 | 0.610993438 | 0.43758948 | 1.39627087 | 0.16263294 | 0.60211533 |
| ZC2HC1A | 372.387764 | 0.103583265 | 0.20822155 | 0.49746658 | 0.61886006 | 0.90835482 |
| IL7 | 144.664781 | -0.914153968 | 0.44756463 | -2.0425072 | 0.04110125 | 0.34131622 |
| TPD52 | 457.361445 | -0.161761394 | 0.42801521 | -0.3779338 | 0.70547981 | 0.93241442 |
| ZBTB10 | 131.709364 | 0.507538398 | 0.3814492 | 1.33055306 | 0.18333612 | 0.63390383 |
| ZNF704 | 1120.75772 | 0.267279252 | 0.3324138 | 0.80405583 | 0.42136473 | 0.82694802 |
| PAG1 | 92.4141329 | 1.497089029 | 0.48708041 | 3.07359732 | 0.00211495 | 0.06499896 |
| IMPA1 | 74.9066441 | 0.281166739 | 0.25236872 | 1.1141109 | 0.26523161 | 0.72029159 |
| ZFAND1 | 140.56002 | 0.301407846 | 0.20747015 | 1.45277695 | 0.14628569 | 0.57722788 |
| CHMP4C | 52.7010152 | -0.589169235 | 0.34015589 | -1.7320566 | 0.08326349 | 0.45768271 |
| SNX16 | 110.234234 | -0.231244164 | 0.35328078 | -0.654562 | 0.51274982 | 0.87177451 |
| AC015522.1 | 229.29151 | -1.449128527 | 1.53606947 | -0.9434004 | 0.34547615 | 0.78180304 |
| RALYL | 2412.03299 | 0.489066937 | 0.39389203 | 1.24162689 | 0.21437426 | 0.67132603 |
| LRRCC1 | 64.1130758 | 0.193227037 | 0.27849664 | 0.69382179 | 0.48779398 | 0.86006752 |
| E2F5 | 81.7045756 | 0.304361289 | 0.32128308 | 0.94733059 | 0.34347035 | 0.78148489 |
| C8orf59 | 111.595455 | -0.278040603 | 0.2734856 | -1.0166554 | 0.30931741 | 0.75369599 |
| CA13 | 106.675294 | -0.075147093 | 0.33417285 | -0.2248749 | 0.82207658 | 0.96502145 |
| CA1 | 38.3134012 | -0.779814979 | 0.65448654 | -1.1914912 | 0.2334608 | 0.69247746 |
| CA3 | 119.902805 | -0.367489141 | 0.29942462 | -1.2273177 | 0.2197032 | 0.67800045 |
| CA2 | 823.521408 | 1.047894311 | 0.56268872 | 1.86229841 | 0.06256104 | 0.41084915 |
| WWP1 | 1152.58764 | -0.410469113 | 0.24869543 | -1.6504892 | 0.09884293 | 0.4933116 |
| RMDN1 | 235.911663 | 0.101001499 | 0.27783439 | 0.36353131 | 0.71620803 | 0.93573028 |
| CPNE3 | 232.649245 | 0.320375734 | 0.16894878 | 1.8962891 | 0.05792183 | 0.3985947 |
| CNGB3 | 44.8898322 | 0.044605926 | 0.60884462 | 0.07326323 | 0.94159665 | 0.98877083 |
| AF117829.1 | 671.525865 | -0.065806157 | 0.36638988 | -0.1796069 | 0.85746117 | 0.97201489 |
| RIPK2 | 97.4509702 | -0.311229954 | 0.31626317 | -0.9840854 | 0.32507355 | 0.77058828 |
| OSGIN2 | 101.873164 | -0.324180006 | 0.3388035 | -0.9568378 | 0.33864912 | 0.77836439 |
| NBN | 182.504179 | -0.025627447 | 0.22917056 | -0.111827 | 0.91096061 | 0.98468493 |
| DECR1 | 438.884643 | 0.232106196 | 0.26543073 | 0.87445112 | 0.38187263 | 0.8032969 |
| CALB1 | 432.990686 | 2.355435061 | 0.90962292 | 2.58946318 | 0.00961257 | 0.15716027 |
| TMEM64 | 119.518224 | 1.147447268 | 0.39018744 | 2.94075914 | 0.00327409 | 0.08642938 |
| PIP4P2 | 133.222573 | -0.051067538 | 0.33495129 | -0.1524626 | 0.8788221 | 0.97692415 |
| OTUD6B-AS1 | 99.9881358 | 0.117936269 | 0.41904824 | 0.28143841 | 0.77837416 | 0.95620223 |
| OTUD6B | 99.4262093 | 0.625255508 | 0.30119279 | 2.07593122 | 0.03790031 | 0.32690969 |
| LRRC69 | 141.505593 | -0.336940814 | 0.29257118 | -1.1516541 | 0.24946322 | 0.70629672 |
| SLC26A7 | 235.24274 | 1.08217891 | 0.66167941 | 1.63550337 | 0.10194357 | 0.4989423 |
| AC117834.1 | 62.3127884 | -0.988276567 | 0.59019301 | -1.6744973 | 0.09403291 | 0.48492647 |
| TRIQK | 294.342632 | -0.110693681 | 0.16591879 | -0.6671558 | 0.50467262 | 0.86946722 |
| FAM92A | 96.108735 | 0.327833152 | 0.32653223 | 1.00398405 | 0.3153863 | 0.75912793 |
| RBM12B | 124.138812 | 0.009934822 | 0.24954606 | 0.03981158 | 0.96824335 | 0.99303855 |
| TMEM67 | 250.737304 | 0.302253053 | 0.18752995 | 1.61175882 | 0.10701443 | 0.50977784 |
| PDP1 | 76.0415572 | 0.817946661 | 0.42410156 | 1.92865754 | 0.05377339 | 0.38935477 |
| VIRMA | 189.790889 | 0.013990006 | 0.20646999 | 0.06775806 | 0.94597823 | 0.98949015 |
| ESRP1 | 268.783659 | -0.160187848 | 0.27022 | -0.5928053 | 0.55331146 | 0.88991776 |
| DPY19L4 | 225.613776 | 0.229786181 | 0.1733741 | 1.32537777 | 0.18504589 | 0.63541313 |
| INTS8 | 152.801859 | 0.066204194 | 0.29195449 | 0.22676204 | 0.82060878 | 0.96502145 |
| NDUFAF6 | 150.17164 | 0.110114735 | 0.31671504 | 0.34767763 | 0.7280823 | 0.93931348 |
| TP53INP1 | 69.9655644 | 0.23670102 | 0.25182368 | 0.93994742 | 0.34724453 | 0.78180304 |
| PLEKHF2 | 68.1958017 | 0.250091492 | 0.44405944 | 0.56319372 | 0.57330298 | 0.89797476 |
| C8orf37-AS1 | 173.865744 | -0.140880083 | 0.38126609 | -0.3695059 | 0.71175065 | 0.9348115 |
| UQCRB | 560.583501 | 0.228274629 | 0.43918633 | 0.51976715 | 0.60322587 | 0.90369786 |
| MTERF3 | 39.6592936 | -0.285036229 | 0.29799055 | -0.9565277 | 0.33880567 | 0.77836439 |
| PTDSS1 | 60.3118488 | 0.176527096 | 0.38521215 | 0.45825941 | 0.64676608 | 0.91743182 |
| SDC2 | 99.9161433 | -1.635223462 | 0.5351724 | -3.0555078 | 0.0022468 | 0.06785717 |
| CPQ | 559.351425 | -0.230005339 | 0.26770236 | -0.8591831 | 0.39023951 | 0.80728662 |
| TSPYL5 | 52.7290184 | -0.082877377 | 0.39408547 | -0.2103031 | 0.83343115 | 0.96749268 |
| MTDH | 695.996461 | 0.101729096 | 0.12921886 | 0.78726198 | 0.43112852 | 0.83111714 |
| LAPTM4B | 432.948051 | -0.272734849 | 0.25214639 | -1.0816528 | 0.27940684 | 0.72938673 |
| MATN2 | 100.546017 | 1.262785821 | 0.48393746 | 2.60939878 | 0.00907015 | 0.15233197 |
| RPL30 | 701.987599 | 0.42150586 | 0.4983838 | 0.84574551 | 0.39769473 | 0.81357642 |
| RIDA | 58.8724242 | 0.011138489 | 0.36544451 | 0.03047929 | 0.97568481 | 0.9939318 |
| POP1 | 35.5489972 | -0.153539552 | 0.33798101 | -0.4542846 | 0.64962403 | 0.9178994 |
| NIPAL2 | 44.8136327 | -0.983848954 | 0.46794715 | -2.1024788 | 0.03551135 | 0.31644624 |
| STK3 | 1325.59488 | -0.069442774 | 0.27141346 | -0.255856 | 0.798062 | 0.96000168 |
| AC104986.2 | 89.8355278 | -0.170045748 | 0.34000559 | -0.5001263 | 0.61698613 | 0.90794578 |
| VPS13B | 1276.1304 | 0.211619287 | 0.22552831 | 0.93832693 | 0.34807643 | 0.78180304 |
| COX6C | 466.224506 | 0.249881647 | 0.31914222 | 0.78297897 | 0.43363946 | 0.83187655 |
| POLR2K | 115.968331 | 0.186982479 | 0.53198525 | 0.35148057 | 0.72522784 | 0.93895336 |
| SPAG1 | 176.41171 | -0.242417665 | 0.34031971 | -0.7123233 | 0.47626459 | 0.85414749 |
| RNF19A | 1406.77501 | 0.212531895 | 0.51004721 | 0.41669063 | 0.67690471 | 0.9268955 |

| | | | | | | |
|---|---|---|---|---|---|---|
| AP000424.1 | 235.690655 | -2.054564891 | 0.67139915 | -3.0601243 | 0.00221245 | 0.06720721 |
| ANKRD46 | 146.398824 | 0.525862363 | 0.28425536 | 1.84996462 | 0.06431865 | 0.41357646 |
| PABPC1 | 640.436902 | 0.519451841 | 0.36832827 | 1.41029587 | 0.15845234 | 0.59690169 |
| YWHAZ | 906.533269 | 0.492983805 | 0.35999092 | 1.36943398 | 0.17086366 | 0.61746591 |
| ZNF706 | 243.660884 | 0.136820423 | 0.24486196 | 0.55876553 | 0.57632175 | 0.89869922 |
| AP001207.3 | 42.7051627 | -0.404893118 | 0.58987373 | -0.6864064 | 0.49245686 | 0.86200976 |
| GRHL2 | 830.925684 | 0.066647522 | 0.29130772 | 0.22878735 | 0.8190342 | 0.96443578 |
| NCALD | 67.9365815 | -2.326043263 | 0.90330363 | -2.5750403 | 0.01002284 | 0.16036545 |
| RRM2B | 152.44683 | 0.459187605 | 0.30729425 | 1.49429286 | 0.13509909 | 0.55895716 |
| UBR5 | 875.981148 | 0.530396632 | 0.20230626 | 2.62175098 | 0.00874793 | 0.14810716 |
| KLF10 | 173.040168 | 1.410383613 | 0.86860098 | 1.62374168 | 0.10443095 | 0.50345841 |
| AZIN1 | 256.189782 | 0.277437743 | 0.29834265 | 0.92992988 | 0.35240739 | 0.78314007 |
| AZIN1-AS1 | 382.416283 | -0.58472292 | 0.36062702 | -1.6214063 | 0.10493053 | 0.50459257 |
| AP003354.2 | 75.8851322 | -0.662973226 | 0.41882295 | -1.5829439 | 0.11343426 | 0.52545147 |
| ATP6V1C1 | 258.137178 | -0.204497194 | 0.29521804 | -0.6926988 | 0.48849857 | 0.86014074 |
| BAALC-AS1 | 68.9810223 | -0.716011571 | 0.38231623 | -1.8728255 | 0.06109249 | 0.40676596 |
| FZD6 | 244.924995 | 0.235101738 | 0.17272242 | 1.3611536 | 0.17346515 | 0.62023706 |
| SLC25A32 | 76.6981862 | -0.108884827 | 0.2288506 | -0.47579 | 0.63422399 | 0.91425962 |
| DCAF13 | 55.5947282 | -0.010320269 | 0.31561657 | -0.0326988 | 0.97391482 | 0.9939318 |
| LRP12 | 130.545631 | 0.715200247 | 0.25845863 | 2.76717498 | 0.00565444 | 0.12123309 |
| OXR1 | 985.616974 | -0.050825211 | 0.40773496 | -0.1246526 | 0.90079861 | 0.98320957 |
| AC090579.1 | 81.3423451 | -0.877002988 | 0.52888398 | -1.6582143 | 0.09727422 | 0.49030463 |
| ANGPT1 | 85.1815908 | 1.254511191 | 0.65925962 | 1.90290919 | 0.0570524 | 0.39754595 |
| EIF3E | 307.800447 | 0.372967508 | 0.19942886 | 1.87017823 | 0.06145907 | 0.4077045 |
| EMC2 | 264.925375 | 0.466139428 | 0.37511914 | 1.24264367 | 0.21399917 | 0.67132603 |
| NUDCD1 | 109.37163 | -0.206648166 | 0.23971667 | -0.8620517 | 0.38865906 | 0.8065026 |
| ENY2 | 193.67305 | -0.134610445 | 0.27585765 | -0.4879707 | 0.62557061 | 0.91061758 |
| EBAG9 | 100.67872 | 0.052704343 | 0.21855402 | 0.24115018 | 0.80943872 | 0.96245698 |
| SYBU | 84.1366676 | -0.280501019 | 0.44466755 | -0.6308106 | 0.52816435 | 0.87826011 |
| TRPS1 | 2811.38665 | 1.416264304 | 0.56041008 | 2.52719279 | 0.01149784 | 0.16947585 |
| EIF3H | 419.102567 | 0.273876109 | 0.22066277 | 1.24115233 | 0.21454948 | 0.67132603 |
| UTP23 | 54.1156007 | -0.444548459 | 0.24683802 | -1.8009724 | 0.07170723 | 0.43222361 |
| RAD21 | 271.241555 | 0.278510341 | 0.24626493 | 1.13093788 | 0.25808124 | 0.71534188 |
| MED30 | 43.4053947 | 0.586366099 | 0.30028843 | 1.95267627 | 0.05085797 | 0.38016512 |
| EXT1 | 1452.00162 | 0.15625745 | 0.32372831 | 0.48268083 | 0.62932238 | 0.9114564 |
| SAMD12 | 2693.27401 | -0.318450691 | 0.1812323 | -1.757141 | 0.07889378 | 0.44910748 |
| AC023590.1 | 278.966799 | -0.620878427 | 0.65036045 | -0.9546682 | 0.33974552 | 0.77836439 |
| SAMD12-AS1 | 184.450021 | -1.22363225 | 0.39797941 | -3.074612 | 0.00210777 | 0.06499896 |
| TNFRSF11B | 327.357974 | -1.613510504 | 0.62914894 | -2.5645922 | 0.01032971 | 0.16264592 |
| COLEC10 | 96.334981 | -1.7123919 | 0.44057161 | -3.8867504 | 0.0001016 | 0.0089472 |
| MAL2 | 1136.24785 | 0.205739382 | 0.41345203 | 0.49761367 | 0.61875637 | 0.90835482 |
| MAL2-AS1 | 74.7549785 | 0.530427178 | 0.74324012 | 0.71366865 | 0.47543209 | 0.85401927 |
| NOV | 106.62949 | -1.824698105 | 0.88631011 | -2.0587581 | 0.03951742 | 0.3346923 |
| TAF2 | 280.326646 | 0.248990852 | 0.22761219 | 1.09392585 | 0.2739875 | 0.72590691 |
| DEPTOR | 605.598762 | -1.193335406 | 0.48299407 | -2.4707041 | 0.01348474 | 0.18693126 |
| AC091563.1 | 60.6777062 | -1.171652714 | 0.44546174 | -2.6301983 | 0.00853351 | 0.14565335 |
| MRPL13 | 116.403928 | -0.167522269 | 0.40283942 | -0.4158537 | 0.67751705 | 0.9268955 |
| MTBP | 57.427399 | 0.30806469 | 0.31845336 | 0.96737774 | 0.33335523 | 0.777038 |
| SNTB1 | 172.282796 | -0.42342169 | 0.45237831 | -0.9359903 | 0.3492782 | 0.78180304 |
| ZHX2 | 1015.40152 | 0.036689187 | 0.38161787 | 0.09614117 | 0.92340846 | 0.98694936 |
| AC104316.1 | 42.3209481 | -0.892801928 | 0.49281087 | -1.8116523 | 0.07003994 | 0.42862997 |
| DERL1 | 125.890017 | -0.101539968 | 0.27502299 | -0.3692054 | 0.71197465 | 0.9348115 |
| TBC1D31 | 54.2137132 | 0.025329226 | 0.32413508 | 0.07814404 | 0.93771347 | 0.98846521 |
| ATAD2 | 78.2479493 | -0.105109858 | 0.38027243 | -0.2764067 | 0.78223569 | 0.95688565 |
| WDYHV1 | 54.0799063 | -0.363354239 | 0.29556309 | -1.2293627 | 0.21893585 | 0.67702796 |
| FBXO32 | 89.1872372 | 0.206281245 | 0.68056439 | 0.3031032 | 0.76181122 | 0.95374766 |
| FAM91A1 | 260.942962 | 0.370214632 | 0.22886668 | 1.61759949 | 0.10574894 | 0.50651229 |
| TMEM65 | 273.64325 | -0.19480522 | 0.33853473 | -0.5754364 | 0.56499616 | 0.89382261 |
| RNF139 | 40.7741897 | 0.518201361 | 0.45103975 | 1.14890397 | 0.25059558 | 0.70788186 |
| TATDN1 | 186.228604 | 0.071770721 | 0.22226326 | 0.32290861 | 0.74676445 | 0.94597987 |
| NDUFB9 | 204.990054 | -0.02137366 | 0.38290416 | -0.0558199 | 0.95548531 | 0.9908068 |
| MTSS1 | 600.144937 | 0.683121002 | 0.44075512 | 1.54988784 | 0.12116844 | 0.53989034 |
| SQLE | 200.843979 | -0.314130952 | 0.68137667 | -0.4610239 | 0.64478144 | 0.91682456 |
| WASHC5 | 136.331849 | -0.220589366 | 0.28004845 | -0.7876829 | 0.43088223 | 0.83111714 |
| NSMCE2 | 895.967565 | -0.108604205 | 0.25281821 | -0.4295743 | 0.66750534 | 0.92422713 |
| TRIB1 | 87.4964509 | 0.157175864 | 0.49482834 | 0.31763715 | 0.75076019 | 0.94794781 |
| FAM84B | 209.610466 | -0.612495555 | 0.4570291 | -1.3401675 | 0.18019089 | 0.63015077 |
| PCAT1 | 1161.15295 | 0.134768087 | 0.32641037 | 0.4128793 | 0.67969505 | 0.92700898 |
| PVT1 | 163.335202 | 0.104777867 | 0.77273718 | 0.13559315 | 0.89214292 | 0.98168537 |
| FAM49B | 1290.36985 | 0.670930466 | 0.26176666 | 2.56308601 | 0.01037463 | 0.16276372 |
| ASAP1 | 407.644081 | -0.374326366 | 0.40215829 | -0.9307936 | 0.35196034 | 0.78314007 |
| EFR3A | 754.241251 | 0.823274345 | 0.29068745 | 2.83216336 | 0.00462342 | 0.10762557 |
| LRRC6 | 203.584288 | 0.528662006 | 0.23736674 | 2.22719496 | 0.02593425 | 0.27398277 |
| PHF20L1 | 236.946806 | 0.146936629 | 0.25033749 | 0.58695415 | 0.5572345 | 0.89148045 |
| NDRG1 | 4661.83604 | 0.099807132 | 0.57345853 | 0.1740442 | 0.86183072 | 0.97390403 |
| ZFAT | 98.5452952 | -0.230214873 | 0.39490761 | -0.5829588 | 0.55992103 | 0.89233156 |
| AC083843.3 | 78.442458 | 0.032069099 | 0.44804773 | 0.07157519 | 0.94293999 | 0.98885525 |
| KHDRBS3 | 97.3409384 | -1.6890574 | 0.44295342 | -3.8131716 | 0.00013719 | 0.01065488 |
| COL22A1 | 95.4975039 | 0.206795046 | 0.96261982 | 0.21482525 | 0.82990356 | 0.96643829 |
| TRAPPC9 | 1268.20196 | 0.231686257 | 0.20739786 | 1.11711016 | 0.26394723 | 0.71989632 |
| CHRAC1 | 72.6458887 | -0.060824086 | 0.444996 | -0.1366846 | 0.89128013 | 0.98095103 |
| AGO2 | 345.081534 | 0.258583598 | 0.29988244 | 0.86228323 | 0.38853168 | 0.8065026 |
| PTK2 | 2630.17004 | 0.080147235 | 0.23296354 | 0.34403339 | 0.73082118 | 0.94048525 |
| DENND3 | 142.21838 | -0.14762741 | 0.56598048 | -0.2608348 | 0.7942199 | 0.95937941 |
| SLC45A4 | 1174.04603 | 0.363311238 | 1.04692737 | 0.34702621 | 0.72857163 | 0.93931348 |
| TSNARE1 | 103.307696 | -0.640500279 | 0.38461683 | -1.6652945 | 0.09585403 | 0.48710328 |
| JRK | 44.8747785 | 0.254053318 | 0.29752632 | 0.8538852 | 0.3931686 | 0.81010924 |
| THEM6 | 40.8298605 | 0.186759113 | 0.47569595 | 0.39260185 | 0.69461357 | 0.93112691 |
| LY6E | 146.173729 | -2.056389191 | 1.39402631 | -1.4751437 | 0.1401739 | 0.56749674 |
| GLI4 | 37.6882713 | -0.405836418 | 0.39852528 | -1.0183455 | 0.30851379 | 0.75313873 |
| TOP1MT | 54.5885941 | 0.805799323 | 0.34530546 | 2.33358409 | 0.01961751 | 0.23523056 |
| RHPN1 | 94.9730563 | -0.001193656 | 0.42346623 | -0.0028188 | 0.99775095 | 0.99954455 |
| ZC3H3 | 79.5605791 | -0.176710394 | 0.32735528 | -0.5398123 | 0.58932651 | 0.90233654 |
| GSDMD | 100.708695 | -0.190072216 | 0.44622748 | -0.4259536 | 0.67014164 | 0.92474338 |
| NAPRT | 83.991955 | 0.298704467 | 0.54831317 | 0.54476982 | 0.58591184 | 0.90196444 |
| EEF1D | 425.55393 | -0.228299413 | 0.41013185 | -0.5566488 | 0.57776739 | 0.89916873 |
| TSTA3 | 54.4740809 | -1.012904886 | 0.67695059 | -1.4962759 | 0.13458176 | 0.55868799 |
| ZNF623 | 51.8901919 | -0.359428206 | 0.29376841 | -1.2235087 | 0.22113761 | 0.67871976 |
| FAM83H | 106.005099 | 0.350338695 | 0.34016368 | 1.02991212 | 0.30305126 | 0.74940473 |
| IQANK1 | 99.7432499 | -0.120910783 | 0.37368967 | -0.3235593 | 0.74627168 | 0.94594643 |
| SCRIB | 67.37657 | -0.112770028 | 0.37398286 | -0.301538 | 0.76300431 | 0.95378284 |
| PUF60 | 105.766061 | -0.398080282 | 0.41094291 | -0.9686997 | 0.33269504 | 0.77653541 |
| NRBP2 | 71.5557411 | -0.130407114 | 0.33129584 | -0.3936274 | 0.69385616 | 0.93105058 |
| EPPK1 | 35.5422656 | 0.206588979 | 0.52000508 | 0.39728262 | 0.69115906 | 0.93073907 |
| PLEC | 174.074302 | -0.228851296 | 0.30273168 | -0.7559542 | 0.44967663 | 0.84122512 |
| PARP10 | 68.7125641 | 0.226706229 | 0.64990685 | 0.34882881 | 0.72721784 | 0.93931348 |
| GRINA | 142.948949 | -0.260174559 | 0.37881744 | -0.6868072 | 0.49220421 | 0.86172738 |
| EXOSC4 | 35.3687248 | 0.037706952 | 0.56277748 | 0.06700153 | 0.94658049 | 0.9895481 |
| GPAA1 | 94.2100871 | 0.151045414 | 0.46009037 | 0.3282951 | 0.74268855 | 0.94508888 |
| CYC1 | 168.941387 | -0.181558706 | 0.57282256 | -0.3169545 | 0.7512781 | 0.94814481 |
| SHARPIN | 80.0530419 | -0.322867796 | 0.23246706 | -1.3888755 | 0.16487062 | 0.60604843 |
| MAF1 | 124.981391 | -0.367560007 | 0.33277075 | -1.1045442 | 0.26935714 | 0.72322263 |
| MROH1 | 273.026199 | -0.539030033 | 0.48346279 | -1.1149359 | 0.26487788 | 0.71989632 |

| | | | | | | |
|---|---|---|---|---|---|---|
| HSF1 | 196.768827 | 0.096403349 | 0.24867777 | 0.38766372 | 0.69826491 | 0.93177271 |
| DGAT1 | 61.8957743 | 0.242367414 | 0.37363875 | 0.64866777 | 0.51655314 | 0.87483466 |
| SLC52A2 | 43.4649457 | -0.669595706 | 0.42271541 | -1.5840343 | 0.11318591 | 0.52509446 |
| CPSF1 | 44.2456584 | 0.088993249 | 0.41162198 | 0.2162014 | 0.82883076 | 0.96606563 |
| SLC39A4 | 43.9774383 | -0.638018144 | 0.4247762 | -1.5020101 | 0.13309449 | 0.55601567 |
| VPS28 | 321.631394 | -0.453927155 | 0.38552789 | -1.1774172 | 0.23902904 | 0.69526071 |
| CYHR1 | 168.767825 | -0.02633071 | 0.393884 | -0.0668489 | 0.946702 | 0.98957081 |
| KIFC2 | 62.058143 | 0.387977279 | 0.3651265 | 1.0625832 | 0.28797101 | 0.73653438 |
| PPP1R16A | 125.01867 | 0.041288013 | 0.24877564 | 0.16596485 | 0.86818461 | 0.97576204 |
| MFSD3 | 31.1731029 | 0.477517267 | 0.56450498 | 0.84590444 | 0.39760606 | 0.81357642 |
| LRRC14 | 31.5203112 | -0.15705274 | 0.40311911 | -0.3895939 | 0.69683688 | 0.93130502 |
| ARHGAP39 | 53.8017416 | 0.112626753 | 0.32837047 | 0.34298685 | 0.73160835 | 0.94065213 |
| ZNF251 | 165.982064 | -0.067476005 | 0.3004336 | -0.2245954 | 0.82229404 | 0.96502145 |
| RPL8 | 1106.11385 | 0.132585307 | 0.5172862 | 0.25630938 | 0.79771195 | 0.95999536 |
| ZNF7 | 64.7501584 | -0.38534943 | 0.26880029 | -1.4335901 | 0.15168927 | 0.58639009 |
| COMMD5 | 34.5835414 | -0.466555538 | 0.39450922 | -1.1826226 | 0.23695874 | 0.69347321 |
| ZNF250 | 56.5507119 | -0.02968512 | 0.32948582 | -0.0900953 | 0.92821149 | 0.98751845 |
| ZNF16 | 43.1251335 | -0.279500642 | 0.34146883 | -0.8185246 | 0.4130577 | 0.82437234 |
| C8orf33 | 45.5894598 | -0.569993893 | 0.39939944 | -1.4271274 | 0.15354317 | 0.58860244 |
| WASHC1 | 355.473354 | 0.724712342 | 0.30013622 | 2.41461145 | 0.015752 | 0.20433451 |
| CBWD1 | 85.0879344 | 0.063624754 | 0.28290521 | 0.22489779 | 0.82205879 | 0.96502145 |
| DOCK8 | 128.674376 | -1.693119693 | 1.02376006 | -1.6538247 | 0.09816313 | 0.49265259 |
| KANK1 | 1668.41327 | 0.820890798 | 0.43435424 | 1.88991088 | 0.05876988 | 0.40066749 |
| SMARCA2 | 1270.07307 | 1.790063018 | 0.53776981 | 3.3286789 | 0.00087259 | 0.0365037 |
| VLDLR-AS1 | 46.9324815 | -2.0596681 | 0.65379584 | -3.1503231 | 0.0016309 | 0.05470286 |
| VLDLR | 75.9151142 | -2.959212267 | 0.48013205 | -6.16333 | 7.12E-10 | 1.49E-06 |
| PUM3 | 394.079128 | -0.872691746 | 0.32414124 | -2.6923194 | 0.0070957 | 0.13409852 |
| RFX3 | 625.514624 | 0.731264715 | 0.20290648 | 3.60394959 | 0.00031342 | 0.01808965 |
| RFX3-AS1 | 534.361585 | -0.477663625 | 0.46567423 | -1.0257463 | 0.30501118 | 0.75035614 |
| GLIS3 | 5184.86761 | 0.494377775 | 0.46755918 | 1.05735871 | 0.29034791 | 0.73754646 |
| SLC1A1 | 133.770968 | -1.041987161 | 0.38224647 | -2.7259563 | 0.00641155 | 0.12805535 |
| SPATA6L | 395.92495 | -0.082726587 | 0.3769816 | -0.2194446 | 0.82630371 | 0.96545294 |
| CDC37L1 | 158.214193 | 0.456327601 | 0.27538169 | 1.65707315 | 0.09750469 | 0.49033067 |
| AK3 | 603.364537 | 0.749313212 | 0.23789171 | 3.14980795 | 0.00163378 | 0.05470286 |
| RCL1 | 178.968005 | -0.067084179 | 0.30969525 | -0.2166135 | 0.82850955 | 0.96604139 |
| JAK2 | 350.114116 | 0.344443826 | 0.20584258 | 1.67333611 | 0.09426115 | 0.48561474 |
| PLGRKT | 179.744847 | -0.206617275 | 0.22087455 | -0.9354508 | 0.34955603 | 0.78199953 |
| RIC1 | 585.711035 | 0.812961471 | 0.20730189 | 3.92163075 | 8.80E-05 | 0.0083794 |
| ERMP1 | 443.261682 | 0.465210912 | 0.2837266 | 1.63964505 | 0.10107899 | 0.49726048 |
| KIAA2026 | 527.547992 | 0.099405508 | 0.22650119 | 0.43887411 | 0.66075276 | 0.92284375 |
| RANBP6 | 56.1896995 | 0.180072219 | 0.29781128 | 0.60465211 | 0.54541018 | 0.88669883 |
| UHRF2 | 292.599053 | 0.213271906 | 0.24042146 | 0.88707518 | 0.37503843 | 0.79825864 |
| GLDC | 685.820922 | 1.202770735 | 0.54712434 | 2.19834988 | 0.02792418 | 0.28401388 |
| AL162411.1 | 83.0507824 | -0.255128624 | 0.55055892 | -0.4633993 | 0.64307817 | 0.91643619 |
| KDM4C | 1053.53094 | -0.356538547 | 0.29611259 | -1.2040641 | 0.22856479 | 0.68678297 |
| DMAC1 | 110.720742 | -0.582707603 | 0.39588884 | -1.471897 | 0.14104868 | 0.56895751 |
| PTPRD | 435.337356 | 0.210513949 | 0.94333242 | 0.22315988 | 0.82341108 | 0.96511681 |
| MPDZ | 247.605687 | 0.501553 | 0.22146238 | 2.2647323 | 0.02352911 | 0.25901796 |
| NFIB | 2640.38716 | 0.976306009 | 0.3565438 | 2.73824984 | 0.00617671 | 0.12643442 |
| ZDHHC21 | 957.758899 | 0.947517284 | 0.25906076 | 3.65750984 | 0.00025468 | 0.01551756 |
| FREM1 | 815.607774 | 1.413985085 | 0.32415387 | 4.36207998 | 1.29E-05 | 0.00230675 |
| TTC39B | 246.908506 | -0.29994905 | 0.39402232 | -0.7612489 | 0.44650842 | 0.83980766 |
| SNAPC3 | 237.569108 | 0.046216666 | 0.25872977 | 0.1786291 | 0.85822894 | 0.97237339 |
| PSIP1 | 225.407208 | 0.377440811 | 0.21310151 | 1.77117846 | 0.07653103 | 0.4426298 |
| CCDC171 | 458.379344 | 0.115059376 | 0.28203101 | 0.4079671 | 0.68329783 | 0.92818681 |
| BNC2 | 501.968903 | -0.458592652 | 0.47970917 | -0.9559806 | 0.33908205 | 0.77836439 |
| CNTLN | 170.212866 | -0.160155502 | 0.28460122 | -0.5627365 | 0.57361431 | 0.89797476 |
| SH3GL2 | 229.673301 | 1.871939222 | 0.65399631 | 2.86230854 | 0.00420567 | 0.10017148 |
| RRAGA | 85.1647975 | -0.199155961 | 0.50573894 | -0.393792 | 0.69373459 | 0.93105058 |
| HAUS6 | 63.183519 | -0.24265854 | 0.30591095 | -0.7932326 | 0.42764232 | 0.82886841 |
| PLIN2 | 224.15008 | -1.73480925 | 0.59104901 | -2.935136 | 0.00333401 | 0.08777614 |
| DENND4C | 820.621107 | -0.32173838 | 0.32711779 | -0.9835551 | 0.3253343 | 0.77073504 |
| RPS6 | 1111.89188 | 0.037334095 | 0.52039729 | 0.07174152 | 0.94280761 | 0.98885525 |
| ACER2 | 40.3314149 | -1.394962965 | 0.4846089 | -2.8785335 | 0.00399529 | 0.09782854 |
| MLLT3 | 1485.48113 | -0.470082498 | 0.26406579 | -1.7801719 | 0.07504784 | 0.43941853 |
| FOCAD | 610.326301 | 0.518119032 | 0.30328271 | 1.70836982 | 0.08756774 | 0.46829366 |
| KLHL9 | 97.1127898 | -0.225698402 | 0.27307777 | -0.8264986 | 0.4085213 | 0.82073894 |
| LINC01239 | 76.7648083 | -1.55607748 | 0.57747778 | -2.6946101 | 0.0070471 | 0.13403564 |
| AL391117.1 | 70.8703503 | 0.721235088 | 0.57614739 | 1.25182392 | 0.21063403 | 0.66871998 |
| TUSC1 | 37.6888037 | 0.002153506 | 0.29786985 | 0.00722969 | 0.99423159 | 0.99885672 |
| CAAP1 | 418.264272 | 0.373172587 | 0.22413843 | 1.66492016 | 0.09592869 | 0.48710328 |
| PLAA | 139.227175 | 0.115351519 | 0.23971 | 0.4812128 | 0.63036527 | 0.91258848 |
| IFT74 | 379.602028 | 0.427431972 | 0.25930598 | 1.64836911 | 0.09927695 | 0.49402775 |
| LRRC19 | 87.5442245 | 0.493763365 | 0.58776308 | 0.8400721 | 0.40086796 | 0.8162653 |
| MOB3B | 340.029187 | -0.00011027 | 0.39211114 | -0.0002812 | 0.99977562 | 0.99993938 |
| C9orf72 | 183.063783 | 1.509324379 | 0.65293026 | 2.31161654 | 0.02079882 | 0.24300076 |
| LINGO2 | 112.305023 | -2.134632449 | 0.99029167 | -2.1555593 | 0.03111809 | 0.29675133 |
| ACO1 | 157.261589 | -0.120236772 | 0.21346958 | -0.5632501 | 0.57326457 | 0.89797476 |
| DDX58 | 184.369669 | -0.545782549 | 0.90921454 | -0.6002792 | 0.5483202 | 0.88823542 |
| TOPORS | 78.0953178 | -0.307205997 | 0.30939475 | -0.9929257 | 0.32074617 | 0.76568446 |
| NDUFB6 | 135.886721 | 0.171901477 | 0.2875937 | 0.59772337 | 0.55002453 | 0.88899708 |
| APTX | 229.991527 | 0.269350451 | 0.25548252 | 1.05428131 | 0.29175414 | 0.73844969 |
| DNAJA1 | 337.838872 | 0.6343928 | 0.46488329 | 1.3646281 | 0.17236997 | 0.61884138 |
| SMU1 | 136.771498 | 0.441455166 | 0.19022379 | 2.32071477 | 0.02030224 | 0.24014392 |
| B4GALT1 | 943.179809 | 0.788123578 | 0.3627899 | 2.17239667 | 0.02982576 | 0.29105042 |
| BAG1 | 483.247498 | 1.102138602 | 0.643864 | 1.71175683 | 0.08694149 | 0.46701529 |
| CHMP5 | 184.217517 | 0.073444431 | 0.44213318 | 0.16611382 | 0.86806738 | 0.97576204 |
| NFX1 | 490.671689 | 0.106954253 | 0.30748165 | 0.34783947 | 0.72796075 | 0.93931348 |
| AQP7 | 123.283988 | 1.769222353 | 0.61055708 | 2.8977182 | 0.00375888 | 0.09515236 |
| AQP3 | 3438.93743 | 0.683139512 | 0.72611668 | 0.94081232 | 0.34680105 | 0.78180304 |
| PRSS3 | 52.7886919 | -0.538333459 | 0.75989054 | -0.7084355 | 0.47867483 | 0.85532469 |
| UBE2R2 | 543.532496 | 0.076603791 | 0.29192668 | 0.26240764 | 0.79300718 | 0.9588344 |
| UBAP2 | 418.065773 | -0.065920543 | 0.30465319 | -0.216379 | 0.82869236 | 0.96606563 |
| DCAF12 | 92.3620103 | 0.304356378 | 0.28731371 | 1.05931727 | 0.28945531 | 0.73754646 |
| UBAP1 | 462.290494 | 0.02057684 | 0.3276189 | 0.06280724 | 0.94992 | 0.98977092 |
| KIF24 | 37.1083414 | 0.674948825 | 0.38929148 | 1.73378779 | 0.08295574 | 0.45768271 |
| NUDT2 | 38.7312362 | -0.457707108 | 0.36784062 | -1.2443082 | 0.21338614 | 0.67075711 |
| FAM219A | 94.1518795 | -0.520370756 | 0.43972331 | -1.183405 | 0.23664869 | 0.69335782 |
| DNAI1 | 74.6499215 | 0.522278238 | 0.50186781 | 1.04066892 | 0.29802923 | 0.74594723 |
| RPP25L | 43.197209 | -0.593120339 | 0.39248945 | -1.5111752 | 0.13074381 | 0.55395564 |
| DCTN3 | 96.1120108 | 0.280155976 | 0.60109988 | 0.46607226 | 0.64116378 | 0.91643619 |
| VCP | 287.459247 | -0.503481122 | 0.25822418 | -1.949783 | 0.05120199 | 0.38114487 |
| PIGO | 43.6290758 | 0.025721581 | 0.34448069 | 0.0746677 | 0.94047911 | 0.98855185 |
| STOML2 | 121.688598 | 0.016046457 | 0.44294802 | 0.0362265 | 0.97110176 | 0.9939318 |
| FAM214B | 37.4123954 | -0.651784723 | 0.30713003 | -2.1221784 | 0.03382276 | 0.31120503 |
| UNC13B | 530.783875 | 0.423591234 | 0.31287247 | 1.35387827 | 0.17577522 | 0.62477136 |
| RUSC2 | 220.365866 | -0.015245345 | 0.44617396 | -0.0341691 | 0.97274234 | 0.9939318 |
| TESK1 | 41.1045554 | -0.146669995 | 0.32232753 | -0.455034 | 0.64908475 | 0.91768694 |
| CD72 | 38.9062632 | -0.603104669 | 0.38442746 | -1.5688387 | 0.11668553 | 0.53183794 |
| CCDC107 | 128.961253 | 0.004373753 | 0.66983759 | 0.00652957 | 0.99479019 | 0.99888869 |
| TLN1 | 183.130332 | -0.576745877 | 0.21012061 | -2.7448325 | 0.00605418 | 0.1254944 |
| CREB3 | 64.7540139 | -0.025283024 | 0.38917845 | -0.0649651 | 0.94820177 | 0.98977092 |
| GBA2 | 144.357707 | 0.214235671 | 0.29706598 | 0.72117202 | 0.47080369 | 0.85236058 |

| | | | | | | |
|---|---|---|---|---|---|---|
| RGP1 | 83.1537211 | -0.001200852 | 0.23595742 | -0.0050893 | 0.99593936 | 0.99908534 |
| AL133410.1 | 91.8726444 | 0.193849788 | 0.30644452 | 0.63257711 | 0.52700984 | 0.87791848 |
| HINT2 | 54.7480906 | 0.354925913 | 0.4378224 | 0.81066184 | 0.4175599 | 0.82566561 |
| TMEM8B | 87.2207323 | 0.130627335 | 0.41288369 | 0.31637805 | 0.75171558 | 0.94835431 |
| RECK | 70.1815836 | -0.316930853 | 0.34320281 | -0.9234506 | 0.35577241 | 0.78612469 |
| CLTA | 429.313156 | -0.56582444 | 0.33424399 | -1.6928485 | 0.0904843 | 0.47556444 |
| GNE | 70.0669373 | -0.298520103 | 0.38121743 | -0.7830705 | 0.43358574 | 0.83187655 |
| RNF38 | 625.664552 | 0.039546864 | 0.22207353 | 0.17808004 | 0.85866011 | 0.97237339 |
| EBLN3P | 166.545739 | -0.152287887 | 0.30388681 | -0.5011336 | 0.61627711 | 0.90794578 |
| ZCCHC7 | 1225.35562 | 0.207186745 | 0.25398043 | 0.81575869 | 0.41463816 | 0.82461178 |
| GRHPR | 108.658534 | 0.093201826 | 0.34281501 | 0.27187207 | 0.78572039 | 0.9573712 |
| ZBTB5 | 58.4612916 | -0.25552897 | 0.3220373 | -0.7934763 | 0.42750037 | 0.82876992 |
| AL138752.2 | 186.123814 | -0.069110336 | 0.26857473 | -0.2573226 | 0.79692978 | 0.95961857 |
| TRMT10B | 128.754352 | -0.375468572 | 0.31639414 | -1.1867115 | 0.23534143 | 0.69335782 |
| DCAF10 | 395.530155 | -0.012033087 | 0.21088896 | -0.0570589 | 0.9544983 | 0.99060627 |
| FAM95C | 55.5033409 | 0.429769416 | 0.45506218 | 0.94441911 | 0.34495552 | 0.78180304 |
| ANKRD18A | 61.9602621 | 0.215841571 | 0.43908779 | 0.49156814 | 0.62302467 | 0.91038742 |
| CNTNAP3 | 41.7413073 | 0.189510423 | 0.32428273 | 0.58439875 | 0.55895207 | 0.89179371 |
| GLIDR | 138.481892 | 0.38736725 | 0.43186005 | 0.89697402 | 0.36973278 | 0.79532503 |
| BMS1P14 | 83.1867503 | 0.345033682 | 0.4967287 | 0.69461193 | 0.48729854 | 0.85968086 |
| CBWD6 | 53.9893237 | 0.137119689 | 0.42355564 | 0.32373477 | 0.74613884 | 0.94594643 |
| CNTNAP3B | 838.282311 | -0.130276661 | 0.42203477 | -0.308687 | 0.75755961 | 0.95149422 |
| FAM27C | 33.9630263 | -0.189073165 | 0.53571614 | -0.3529354 | 0.72413691 | 0.93893205 |
| AL512625.3 | 46.1562595 | 0.825325678 | 0.48440009 | 1.70380992 | 0.0884166 | 0.47036301 |
| CBWD5 | 164.162982 | 0.102253639 | 0.33469006 | 0.3055174 | 0.75997212 | 0.95297299 |
| ZNF658 | 48.7968853 | -0.295371298 | 0.44907712 | -0.6577296 | 0.51071192 | 0.87138909 |
| CBWD3 | 47.6018508 | 0.622490209 | 0.3382069 | 1.84056032 | 0.06568602 | 0.41788422 |
| PGM5 | 393.837569 | 0.655488002 | 0.31324419 | 2.0925783 | 0.03638682 | 0.32044859 |
| AL161457.2 | 49.4733796 | 0.187821902 | 0.47704336 | 0.39372081 | 0.69378717 | 0.93105058 |
| PIP5K1B | 89.7339503 | 0.946548151 | 0.62518938 | 1.51401828 | 0.1300212 | 0.55282113 |
| FAM189A2 | 39.6776642 | -0.455373883 | 0.54513225 | -0.8353457 | 0.4035231 | 0.81828987 |
| APBA1 | 119.000617 | 0.794684424 | 0.36956768 | 2.15030824 | 0.03153084 | 0.29823395 |
| PTAR1 | 293.007172 | 0.285686169 | 0.172136 | 1.65965379 | 0.09698412 | 0.48959228 |
| MAMDC2-AS | 119.913945 | -0.399810322 | 0.33743001 | -1.1848689 | 0.23606929 | 0.69335782 |
| MAMDC2 | 529.437754 | 0.265635165 | 0.50600703 | 0.52496339 | 0.59960864 | 0.90369786 |
| SMC5-AS1 | 190.818402 | -0.201249684 | 0.24291917 | -0.8284636 | 0.40740801 | 0.82018309 |
| SMC5 | 547.851997 | 0.362975206 | 0.23869241 | 1.52068182 | 0.1283397 | 0.54843482 |
| KLF9 | 136.91644 | 0.901541111 | 0.36330593 | 2.48149299 | 0.01308333 | 0.18379796 |
| TRPM3 | 3768.17106 | 1.684199881 | 0.5831727 | 2.88799509 | 0.00387706 | 0.09674186 |
| CEMIP2 | 495.180835 | -0.259155608 | 0.32599101 | -0.7949778 | 0.42662643 | 0.8277744 |
| ABHD17B | 64.5933552 | 0.290391044 | 0.34076638 | 0.85217047 | 0.39411948 | 0.81094677 |
| C9orf85 | 149.522788 | -0.020650917 | 0.27054509 | -0.0763308 | 0.93915594 | 0.98848592 |
| GDA | 60.5925684 | -2.34115617 | 0.6044781 | -3.8730206 | 0.00010749 | 0.00923397 |
| ZFAND5 | 725.795732 | 0.279899375 | 0.55430845 | 0.50495238 | 0.61359228 | 0.90751853 |
| ALDH1A1 | 349.618784 | 0.208638257 | 0.52074382 | 0.40065431 | 0.68867466 | 0.93036007 |
| CARNMT1 | 230.753325 | 0.223190523 | 0.2075359 | 1.07543091 | 0.28218185 | 0.73217772 |
| NMRK1 | 159.891472 | 0.066448002 | 0.34835461 | 0.19074816 | 0.8487229 | 0.96991131 |
| OSTF1 | 117.189609 | 0.532395841 | 0.44390916 | 1.1993351 | 0.23039767 | 0.69012664 |
| PCSK5 | 186.755606 | -1.26009995 | 0.73064059 | -1.7246509 | 0.0845904 | 0.46272529 |
| RFK | 76.6409356 | -0.35900157 | 0.40472055 | -0.8870357 | 0.37505969 | 0.79825864 |
| VPS13A | 523.022719 | 0.267228746 | 0.20603519 | 1.29700539 | 0.19462933 | 0.648352 |
| GNAQ | 1856.84582 | -0.020368944 | 0.19684984 | -0.1034745 | 0.91758637 | 0.98520052 |
| CEP78 | 60.4639739 | 0.595052036 | 0.36200154 | 1.64378318 | 0.100221 | 0.49569046 |
| TLE4 | 67.5851438 | 0.212426962 | 0.40414551 | 0.52562 | 0.59915226 | 0.90369786 |
| TLE1 | 85.8890885 | 0.484597091 | 0.64522741 | 0.75104852 | 0.45262346 | 0.84248055 |
| AL591368.1 | 32.9057827 | 2.038228257 | 0.82362522 | 2.47470356 | 0.01333468 | 0.18614502 |
| RASEF | 206.122069 | -0.527607295 | 0.24472576 | -2.1559124 | 0.03109051 | 0.29675133 |
| FRMD3 | 206.144612 | 1.084032712 | 0.27728286 | 3.90948325 | 9.25E-05 | 0.00855087 |
| IDNK | 51.7112614 | -0.57849507 | 0.38407115 | -1.5062185 | 0.13201109 | 0.55448417 |
| UBQLN1 | 294.260365 | 0.287144643 | 0.15625145 | 1.83770861 | 0.06610535 | 0.41858129 |
| GKAP1 | 139.509995 | 0.595781453 | 0.22102107 | 2.69558664 | 0.00702648 | 0.13388639 |
| KIF27 | 198.504379 | 0.136530406 | 0.26207358 | 0.52096212 | 0.60239316 | 0.90369786 |
| HNRNPK | 764.472642 | 0.300209577 | 0.34636088 | 0.8667537 | 0.38607698 | 0.80634213 |
| RMI1 | 66.4516598 | -0.730617663 | 0.3181813 | -2.2962307 | 0.02166269 | 0.24935578 |
| AGTPBP1 | 249.338356 | 0.417793588 | 0.33207588 | 1.25812688 | 0.20834587 | 0.66589349 |
| NAA35 | 314.811589 | 0.184034895 | 0.24876825 | 0.73978449 | 0.45943077 | 0.84619235 |
| GOLM1 | 77.6429028 | -0.474710625 | 0.37031735 | -1.2819022 | 0.19987696 | 0.65475264 |
| ISCA1 | 113.196711 | 0.189389353 | 0.36419553 | 0.52002108 | 0.60304888 | 0.90369786 |
| TUT7 | 275.491221 | -0.29036182 | 0.20016008 | -1.450648 | 0.1468779 | 0.57793161 |
| DAPK1 | 1988.92862 | 1.032201596 | 0.61649464 | 1.67430749 | 0.09407018 | 0.48492647 |
| CTSL | 207.00918 | -0.200209085 | 0.40375574 | -0.4958668 | 0.61998836 | 0.90914396 |
| SPIN1 | 813.193115 | 0.188802092 | 0.22634826 | 0.83412214 | 0.40421216 | 0.81914727 |
| SECISBP2 | 379.935284 | -0.1398084 | 0.23886412 | -0.5853052 | 0.55834255 | 0.89179371 |
| SEMA4D | 506.062329 | -0.253482301 | 0.31130052 | -0.8142688 | 0.41549098 | 0.82487773 |
| LINC01508 | 213.256008 | 0.954514861 | 0.62948542 | 1.51634148 | 0.12943303 | 0.55140574 |
| DIRAS2 | 37.5058596 | -0.399901695 | 0.37160684 | -1.0761419 | 0.2818638 | 0.73166833 |
| SYK | 667.98879 | 0.603984843 | 0.28038261 | 2.15414515 | 0.03122879 | 0.29698524 |
| AL158071.4 | 64.706502 | 0.652145447 | 0.39236432 | 1.66209162 | 0.09649441 | 0.48810007 |
| AUH | 716.387899 | 0.043794002 | 0.25613719 | 0.17097869 | 0.86424052 | 0.97466129 |
| NFIL3 | 126.765468 | 1.328517959 | 1.07059802 | 1.24091203 | 0.21463825 | 0.67132603 |
| ROR2 | 756.841973 | 0.1753685 | 0.24263411 | 0.72276937 | 0.46982159 | 0.85232523 |
| SPTLC1 | 245.321414 | 0.143615326 | 0.16909241 | 0.84933042 | 0.39569746 | 0.81200497 |
| IARS | 198.857188 | -0.266326005 | 0.22602484 | -1.1783041 | 0.23867537 | 0.69499004 |
| NOL8 | 74.9318206 | -0.395787045 | 0.24618934 | -1.6076531 | 0.10791118 | 0.51162458 |
| CENPP | 155.726452 | -0.113677025 | 0.44572829 | -0.2550366 | 0.79869484 | 0.96005379 |
| IPPK | 47.5556783 | 0.331920823 | 0.34722618 | 0.95592108 | 0.33911211 | 0.77836439 |
| BICD2 | 71.6877234 | 0.189490142 | 0.26038037 | 0.72774357 | 0.46677057 | 0.85074011 |
| ZNF484 | 38.9844942 | -0.066737178 | 0.26516945 | -0.2516775 | 0.80129037 | 0.96063297 |
| FGD3 | 228.589675 | 0.979341162 | 0.61013724 | 1.60511618 | 0.10846824 | 0.51254482 |
| CARD19 | 65.3908699 | -0.179427458 | 0.401603 | -0.4467782 | 0.65503522 | 0.92051095 |
| NINJ1 | 77.4116302 | -0.788509053 | 0.29563083 | -2.6672085 | 0.00764842 | 0.13910656 |
| FAM120AOS | 68.4404452 | -0.398669149 | 0.30503103 | -1.306979 | 0.19121984 | 0.64458858 |
| FAM120A | 525.164763 | 0.09027273 | 0.19109129 | 0.4724063 | 0.63663681 | 0.91454294 |
| PHF2 | 117.982825 | 0.40513326 | 0.28573877 | 1.41784489 | 0.15623606 | 0.59366792 |
| PTPDC1 | 104.342451 | -0.062143704 | 0.26678119 | -0.2329388 | 0.81580889 | 0.96333253 |
| AL158152.2 | 136.771076 | -0.070349171 | 0.3121418 | -0.2253757 | 0.82168703 | 0.96502145 |
| ZNF169 | 66.2115564 | -0.023789918 | 0.35827908 | -0.0664005 | 0.94705895 | 0.98977092 |
| MFSD14B | 183.09002 | 0.718036709 | 0.22342689 | 3.21374351 | 0.00131017 | 0.04751054 |
| FBP1 | 164.82255 | 0.542947168 | 0.48781306 | 1.11302303 | 0.26569854 | 0.72053648 |
| C9orf3 | 1149.76024 | -0.290375362 | 0.29159156 | -0.9958291 | 0.31933319 | 0.76378437 |
| FANCC | 447.407728 | -0.241873462 | 0.27172754 | -0.8901323 | 0.37339485 | 0.79681088 |
| PTCH1 | 69.064994 | -1.386391481 | 0.35040987 | -3.9564853 | 7.61E-05 | 0.00744967 |
| LINC00476 | 169.525054 | -0.213047102 | 0.41003709 | -0.5195801 | 0.60335629 | 0.90369786 |
| ERCC6L2 | 291.948299 | 0.142488335 | 0.24112041 | 0.59094266 | 0.55455884 | 0.8907979 |
| SLC35D2 | 47.4258618 | 0.07510084 | 0.35184912 | 0.21344615 | 0.83097899 | 0.96648703 |
| HABP4 | 137.8256 | 0.390893237 | 0.32126205 | 1.21674264 | 0.22370215 | 0.68103746 |
| CDC14B | 387.22337 | 0.403231002 | 0.24914101 | 1.61848506 | 0.1055581 | 0.50629242 |
| AL589843.1 | 60.3386466 | 0.675911098 | 0.36533524 | 1.85011196 | 0.06429741 | 0.41357646 |
| ZNF510 | 37.8637447 | -0.320143742 | 0.3301185 | -0.9697843 | 0.33215402 | 0.77618491 |
| ZNF782 | 137.989509 | -0.031208386 | 0.25782652 | -0.1210441 | 0.90365608 | 0.98320957 |
| MFSD14C | 386.775648 | 0.102495 | 0.18775386 | 0.54590088 | 0.58513407 | 0.90173902 |
| CCDC180 | 38.1380002 | -0.409402901 | 0.47115324 | -0.8689379 | 0.38488111 | 0.80561637 |
| TDRD7 | 78.6169372 | -0.213384634 | 0.4771169 | -0.4472376 | 0.65470349 | 0.92047627 |

| | | | | | | |
|---|---|---|---|---|---|---|
| TSTD2 | 110.625745 | -0.228568605 | 0.2505188 | -0.912381 | 0.36156817 | 0.78981248 |
| NCBP1 | 99.8293825 | 0.132048266 | 0.24953924 | 0.52916834 | 0.59668868 | 0.90369786 |
| XPA | 96.61703 | 0.322994609 | 0.27814409 | 1.16124921 | 0.24554057 | 0.70154448 |
| ANP32B | 196.430629 | -0.156471652 | 0.23801605 | -0.6573996 | 0.51092401 | 0.87138909 |
| NANS | 38.0490828 | -0.344356793 | 0.39230522 | -0.8777777 | 0.38006434 | 0.8029673 |
| TRIM14 | 56.5989667 | -2.166539463 | 0.8627375 | -2.5112383 | 0.01203085 | 0.17535919 |
| ANKS6 | 114.846497 | 0.128140794 | 0.39961559 | 0.32066015 | 0.74846795 | 0.94678489 |
| GALNT12 | 44.8234315 | -0.682011624 | 0.53381082 | -1.277628 | 0.20138064 | 0.65580391 |
| TGFBR1 | 696.54011 | 0.825624631 | 0.70340372 | 1.17375642 | 0.2404926 | 0.6963541 |
| ALG2 | 74.4600291 | -0.109555679 | 0.31226 | -0.3508476 | 0.72570267 | 0.93895336 |
| SEC61B | 260.331387 | -0.043720048 | 0.39964132 | -0.1093982 | 0.91288665 | 0.98480378 |
| AL359710.1 | 220.666289 | 0.774585838 | 0.40564216 | 1.90952989 | 0.05619377 | 0.39444789 |
| NR4A3 | 174.524968 | 5.684142307 | 1.84046823 | 3.08842186 | 0.00201223 | 0.062951 |
| STX17 | 192.383759 | 0.345243027 | 0.28178824 | 1.22518609 | 0.22050512 | 0.67869352 |
| ERP44 | 267.180581 | 0.142971411 | 0.2254796 | 0.63407693 | 0.52603062 | 0.87791848 |
| INVS | 566.834633 | 0.16333878 | 0.17494 | 0.93368455 | 0.35046664 | 0.78223232 |
| TEX10 | 97.3811374 | 0.174364011 | 0.28200485 | 0.61830146 | 0.53637664 | 0.88137638 |
| MRPL50 | 68.1879055 | 0.18225197 | 0.45549634 | 0.40011731 | 0.68907012 | 0.93036007 |
| ZNF189 | 58.7956595 | -0.213848225 | 0.33054213 | -0.6469621 | 0.51765648 | 0.87526312 |
| ALDOB | 110.236773 | -0.760808398 | 0.99139775 | -0.7674098 | 0.44283787 | 0.83826789 |
| TMEM246 | 43.8540377 | 0.031508203 | 0.32315829 | 0.09750084 | 0.92232867 | 0.9866345 |
| RNF20 | 81.6217629 | 0.395899981 | 0.35449201 | 1.11680933 | 0.26407586 | 0.71989632 |
| SMC2 | 59.0945513 | -0.324142942 | 0.33378714 | -0.9711067 | 0.33149512 | 0.77618491 |
| NIPSNAP3A | 83.5854802 | 0.025692486 | 0.29958644 | 0.08575984 | 0.93165733 | 0.98822345 |
| ABCA1 | 157.636979 | -0.238834928 | 0.45869966 | -0.5206782 | 0.60259094 | 0.90369786 |
| SLC44A1 | 1490.3124 | -0.218952069 | 0.70293683 | -0.3114819 | 0.75543433 | 0.95067145 |
| FSD1L | 256.548746 | -0.195068469 | 0.22213866 | -0.8781383 | 0.37986866 | 0.80295536 |
| FKTN | 178.300946 | 0.081181142 | 0.21909201 | 0.37053448 | 0.71098429 | 0.93475719 |
| TMEM38B | 71.9226399 | -0.201468686 | 0.41469343 | -0.4858256 | 0.62709082 | 0.91061758 |
| LINC01505 | 70.9875608 | -0.136048643 | 0.57055338 | -0.2384503 | 0.81153183 | 0.96288868 |
| ZNF462 | 94.5206284 | 0.012742697 | 0.50949164 | 0.02501061 | 0.9800465 | 0.99469867 |
| RAD23B | 429.106295 | 0.197570915 | 0.18789672 | 1.05148676 | 0.29303509 | 0.73864467 |
| ELP1 | 142.165226 | 0.655352861 | 0.19777321 | 3.31365845 | 0.00092084 | 0.03755638 |
| FAM206A | 54.4523583 | 0.126877348 | 0.31019407 | 0.40902571 | 0.68252079 | 0.92806156 |
| CTNNAL1 | 219.465484 | 0.639326808 | 0.33863925 | 1.88792884 | 0.0590355 | 0.40122703 |
| TMEM245 | 1029.55571 | 0.369417277 | 0.32362622 | 1.14149364 | 0.25366456 | 0.71061335 |
| EPB41L4B | 209.42693 | 0.268534681 | 0.32051489 | 0.83782281 | 0.40213023 | 0.81702302 |
| PTPN3 | 966.674468 | -0.350924063 | 0.29106979 | -1.2056355 | 0.22795808 | 0.68629723 |
| PALM2 | 148.867284 | -0.915190295 | 0.45582025 | -2.0077877 | 0.04466585 | 0.35515061 |
| PALM2-AKAF | 40.4183532 | -0.923959327 | 0.49721161 | -1.8582819 | 0.063129 | 0.41264103 |
| TXN | 243.741466 | -0.582683036 | 0.35295697 | -1.6508614 | 0.09876688 | 0.4933116 |
| ECPAS | 449.964841 | 0.392642384 | 0.2407495 | 1.63091672 | 0.10290789 | 0.50161615 |
| ZNF483 | 255.4643 | 0.182632039 | 0.28317125 | 0.64495262 | 0.51895789 | 0.87579366 |
| PTGR1 | 969.51936 | 0.870212706 | 0.52486391 | 1.65797778 | 0.09732195 | 0.49030463 |
| C9orf84 | 98.4766797 | -0.42390248 | 0.59319917 | -0.714604 | 0.47485379 | 0.85385826 |
| UGCG | 288.441768 | -0.623416249 | 0.35811772 | -1.7408137 | 0.08171623 | 0.45528236 |
| SUSD1 | 632.119147 | 0.628833693 | 0.38266677 | 1.64329318 | 0.10032229 | 0.495699 |
| PTBP3 | 2262.31299 | 0.979452824 | 0.37159614 | 2.63579927 | 0.00839394 | 0.1442148 |
| HSDL2 | 366.705772 | 0.479741669 | 0.2875267 | 1.66851172 | 0.09521419 | 0.48627909 |
| C9orf147 | 50.5941267 | 0.004033951 | 0.29173039 | 0.01382767 | 0.98896747 | 0.9964791 |
| KIAA1958 | 598.206929 | -0.025797795 | 0.2057694 | -0.1253724 | 0.90022877 | 0.98312332 |
| INIP | 105.546822 | 0.136993353 | 0.2227148 | 0.61510665 | 0.53848429 | 0.88265739 |
| SNX30 | 76.6074805 | -0.404671293 | 0.25982543 | -1.5574738 | 0.11935804 | 0.53662476 |
| FKBP15 | 105.991323 | -0.053596916 | 0.18436847 | -0.2907054 | 0.77127661 | 0.95498333 |
| SLC31A1 | 86.9876456 | -0.245195799 | 0.24448643 | -1.0029015 | 0.31590841 | 0.75968795 |
| CDC26 | 79.9776011 | 0.279869837 | 0.45377315 | 0.61676157 | 0.53739201 | 0.88239554 |
| PRPF4 | 69.308401 | 0.060263399 | 0.45864641 | 0.13139403 | 0.89546361 | 0.98256294 |
| WDR31 | 57.2054063 | -0.324374422 | 0.42941692 | -0.7553834 | 0.45001896 | 0.84142706 |
| BSPRY | 101.274874 | -0.462811875 | 0.28753458 | -1.6095868 | 0.10748809 | 0.51064149 |
| HDHD3 | 141.511903 | -0.33251364 | 0.32429611 | -1.0253396 | 0.30520298 | 0.75054742 |
| ALAD | 69.7757853 | -0.240947888 | 0.23190737 | -1.0389833 | 0.29881249 | 0.74594723 |
| RGS3 | 138.563203 | -1.232238531 | 0.51934998 | -2.3726554 | 0.01766073 | 0.22033867 |
| ZNF618 | 802.801796 | 0.27123976 | 0.38393407 | 0.70647484 | 0.47989289 | 0.85561033 |
| KIF12 | 320.137869 | 0.48593328 | 0.30700555 | 1.58281593 | 0.11346342 | 0.52545147 |
| COL27A1 | 130.064239 | -1.185270399 | 0.62414173 | -1.8990405 | 0.05755915 | 0.39817926 |
| AKNA | 43.2427387 | -0.427715791 | 0.34306947 | -1.2467323 | 0.21249567 | 0.66984203 |
| WHRN | 67.3991064 | -0.453372441 | 0.279913 | -1.6196906 | 0.10529876 | 0.50574291 |
| ATP6V1G1 | 407.002157 | -0.516155991 | 0.49403816 | -1.0447695 | 0.29612952 | 0.74305872 |
| TMEM268 | 51.0320544 | -0.663948235 | 0.41812435 | -1.5879205 | 0.11230431 | 0.5224487 |
| PAPPA | 3461.5327 | -0.424612643 | 0.91221172 | -0.465476 | 0.64159064 | 0.91643619 |
| AL137024.1 | 126.354687 | 0.094691539 | 0.99099114 | 0.09555236 | 0.92387611 | 0.98697468 |
| ASTN2 | 377.892156 | -0.758572651 | 0.71598218 | -1.0594854 | 0.28937879 | 0.73754646 |
| CDK5RAP2 | 244.105938 | 0.197840888 | 0.23234846 | 0.85148353 | 0.39450081 | 0.81129679 |
| MEGF9 | 406.583681 | 0.804022888 | 0.38247578 | 2.10215373 | 0.03553981 | 0.31644624 |
| FBXW2 | 171.462249 | 0.328312881 | 0.19899013 | 1.6498953 | 0.09896435 | 0.4933116 |
| PSMD5 | 148.069046 | 0.185678995 | 0.23092335 | 0.80407199 | 0.4213554 | 0.82694802 |
| TRAF1 | 35.1376588 | -0.496330745 | 0.47310241 | -1.0490979 | 0.29413306 | 0.73973155 |
| CNTRL | 133.601592 | -0.224707829 | 0.30413981 | -0.7388307 | 0.4600098 | 0.84651496 |
| RAB14 | 393.72945 | -0.101250533 | 0.2131968 | -0.4749158 | 0.63484698 | 0.91454294 |
| GSN | 294.654232 | -0.83207133 | 0.20261383 | -4.1066857 | 4.01E-05 | 0.00493518 |
| STOM | 96.4993973 | -0.675999044 | 0.37264314 | -1.8140655 | 0.06966765 | 0.42828237 |
| DAB2IP | 298.016442 | -0.677783299 | 0.28328408 | -2.3925923 | 0.01672982 | 0.21174942 |
| TTLL11 | 67.7031163 | 0.187806142 | 0.31117387 | 0.60354086 | 0.54614894 | 0.88669883 |
| NDUFA8 | 135.69595 | 0.032459919 | 0.48451406 | 0.06699479 | 0.94658585 | 0.9895481 |
| RBM18 | 57.4566297 | 0.261428348 | 0.29378898 | 0.88985078 | 0.37354602 | 0.79681088 |
| MRRF | 225.31215 | 0.024266609 | 0.23427355 | 0.10358237 | 0.91750078 | 0.98520052 |
| PTGS1 | 119.319234 | 1.988562243 | 0.84754804 | 2.34625313 | 0.01896322 | 0.23028341 |
| PDCL | 76.4030943 | -0.301353432 | 0.25571553 | -1.1784714 | 0.23860873 | 0.69499004 |
| RC3H2 | 259.691335 | -0.106640924 | 0.19806204 | -0.5384218 | 0.59028586 | 0.90233654 |
| ZBTB26 | 32.825053 | -0.229302814 | 0.36850589 | -0.6222501 | 0.53377745 | 0.88137638 |
| RABGAP1 | 657.153707 | 0.215114969 | 0.18211928 | 1.18117625 | 0.23753272 | 0.6939708 |
| STRBP | 588.407444 | 0.108612948 | 0.29848795 | 0.36387716 | 0.71594974 | 0.93573028 |
| DENND1A | 928.0985 | 0.084166792 | 0.30561761 | 0.27539903 | 0.78300969 | 0.9568898 |
| PSMB7 | 209.88261 | -0.051108731 | 0.27869178 | -0.183388 | 0.85449358 | 0.97116286 |
| NR6A1 | 804.269667 | 0.280316494 | 0.20557576 | 1.36356786 | 0.17270361 | 0.61884138 |
| MIR181A2HC | 118.512159 | 0.003506383 | 0.32082688 | 0.01092921 | 0.99127993 | 0.9974665 |
| RPL35 | 421.82524 | 0.25093368 | 0.38388635 | 0.65366658 | 0.51332664 | 0.87190651 |
| ARPC5L | 146.660511 | -0.160567789 | 0.45385421 | -0.3537872 | 0.7234984 | 0.93893205 |
| GOLGA1 | 136.540138 | -0.1077893 | 0.25972653 | -0.4150107 | 0.67813405 | 0.9268955 |
| SCAI | 256.980204 | 0.470398643 | 0.31830681 | 1.47781518 | 0.13945725 | 0.56656585 |
| PPP6C | 195.691573 | 0.35882509 | 0.15300136 | 2.34524116 | 0.01901478 | 0.23064219 |
| RABEPK | 62.3872166 | -0.250238762 | 0.3009712 | -0.8314376 | 0.40572648 | 0.82002585 |
| HSPA5 | 436.139176 | -0.43453849 | 0.46996864 | -0.9246117 | 0.35516794 | 0.78607579 |
| GAPVD1 | 320.398041 | 0.256477484 | 0.22385616 | 1.14572448 | 0.25190917 | 0.70813507 |
| MAPKAP1 | 474.502485 | 0.162252484 | 0.21590764 | 0.75149023 | 0.45235768 | 0.84234337 |
| PBX3 | 370.303098 | 0.139936474 | 0.43478466 | 0.32185237 | 0.74756454 | 0.94664513 |
| MVB12B | 106.283246 | 0.079922847 | 0.3425736 | 0.23330125 | 0.81552749 | 0.96333253 |
| ZBTB43 | 130.589586 | -0.500188481 | 0.24292188 | -2.0590507 | 0.03948938 | 0.3346923 |
| ZBTB34 | 33.2715789 | -0.452590285 | 0.31208572 | -1.4502114 | 0.14699956 | 0.57806944 |
| RALGPS1 | 538.467529 | 0.131058724 | 0.25643621 | 0.51107729 | 0.60929694 | 0.90552475 |
| GARNL3 | 89.2155692 | -0.164257302 | 0.31148885 | -0.5273296 | 0.59796469 | 0.90369786 |
| SLC2A8 | 31.7758476 | -0.270857933 | 0.27933346 | -0.969658 | 0.33221698 | 0.77618491 |
| RPL12 | 157.252194 | -0.025328302 | 0.35540948 | -0.0712651 | 0.94318675 | 0.98885525 |

| | | | | | | |
|---|---|---|---|---|---|---|
| LRSAM1 | 187.573384 | -0.378720115 | 0.24041981 | -1.575245 | 0.11519989 | 0.52858795 |
| FAM129B | 304.451781 | -1.057196661 | 0.46337368 | -2.2815207 | 0.02251765 | 0.25166501 |
| STXBP1 | 117.231754 | -0.31217928 | 0.19840471 | -1.5734469 | 0.11561537 | 0.52956691 |
| PTRH1 | 68.1929266 | -0.326219265 | 0.41880958 | -0.7789202 | 0.4360267 | 0.83364877 |
| CDK9 | 81.7139297 | 0.30435231 | 0.31456641 | 0.9675296 | 0.33327935 | 0.777038 |
| FPGS | 93.811504 | -0.550123955 | 0.35442754 | -1.5521479 | 0.12062685 | 0.53823708 |
| ENG | 63.4086527 | -1.023094688 | 0.48933488 | -2.0907864 | 0.03654722 | 0.32100172 |
| DPM2 | 114.867263 | -1.770933272 | 0.58869649 | -3.008228 | 0.00262776 | 0.07465809 |
| FAM102A | 695.822352 | -0.627195441 | 0.56859411 | -1.1030636 | 0.26999957 | 0.72333985 |
| SLC25A25 | 176.239672 | 1.083343529 | 0.50763894 | 2.1340828 | 0.032836 | 0.30588131 |
| SLC25A25-AS | 70.4266547 | 0.802288453 | 0.46004711 | 1.74392672 | 0.08117187 | 0.45442371 |
| PTGES2 | 118.78739 | -0.209913782 | 0.3031532 | -0.6924347 | 0.48866441 | 0.86014074 |
| LCN2 | 97.4290056 | -7.389548053 | 1.09758329 | -6.7325624 | 1.67E-11 | 5.82E-08 |
| C9orf16 | 275.401313 | -0.095379344 | 0.49144003 | -0.1940814 | 0.84611218 | 0.96931085 |
| CIZ1 | 99.9103976 | -0.056422976 | 0.24869388 | -0.2268772 | 0.82051921 | 0.96502145 |
| DNM1 | 61.273769 | -0.203187046 | 0.50674161 | -0.4009678 | 0.68844387 | 0.93036007 |
| GOLGA2 | 188.75176 | -0.057986263 | 0.28904968 | -0.20061 | 0.84100352 | 0.96854218 |
| SWI5 | 101.593306 | -0.20918635 | 0.33761546 | -0.6195994 | 0.53552154 | 0.88137638 |
| TRUB2 | 68.0085889 | -0.166795845 | 0.34573057 | -0.4824446 | 0.62949015 | 0.91157341 |
| COQ4 | 223.502591 | 0.22171903 | 0.19181431 | 1.1559045 | 0.24772022 | 0.70405328 |
| SLC27A4 | 124.285034 | -0.333877684 | 0.32769192 | -1.0188768 | 0.30826147 | 0.75313873 |
| URM1 | 80.4457973 | -0.462816788 | 0.30814442 | -1.5019477 | 0.13311063 | 0.55601567 |
| CERCAM | 84.1095326 | 0.039649092 | 0.32637552 | 0.12148305 | 0.90330844 | 0.98320957 |
| ODF2 | 80.9131822 | 0.000794386 | 0.3182803 | 0.00249587 | 0.99800859 | 0.99954455 |
| GLE1 | 95.3305369 | 0.008065245 | 0.24447421 | 0.03299017 | 0.97368243 | 0.9939318 |
| SPTAN1 | 473.387185 | 0.110702463 | 0.2343617 | 0.47235731 | 0.63667177 | 0.91454294 |
| WDR34 | 122.308659 | 0.559235774 | 0.35724105 | 1.56542978 | 0.11748217 | 0.53282158 |
| SET | 496.532731 | -0.150292183 | 0.30823236 | -0.4875938 | 0.6258376 | 0.91061758 |
| ZDHHC12 | 70.0331796 | -0.476673764 | 0.37458662 | -1.2725328 | 0.20318387 | 0.6582233 |
| ZER1 | 256.05289 | -0.137960613 | 0.2760613 | -0.4997463 | 0.61725372 | 0.90794578 |
| TBC1D13 | 75.0815403 | 0.197432389 | 0.27975304 | 0.70573814 | 0.480351 | 0.85584469 |
| ENDOG | 73.1052891 | -0.183532311 | 0.40874819 | -0.4490107 | 0.65342393 | 0.91974964 |
| SPOUT1 | 34.2638525 | 0.176493455 | 0.31742039 | 0.55602432 | 0.57819423 | 0.89943232 |
| KYAT1 | 80.1327815 | 0.5713529 | 0.32294286 | 1.76920743 | 0.07685926 | 0.44354906 |
| LRRC8A | 41.9732514 | -0.372002244 | 0.2588324 | -1.4372321 | 0.15065205 | 0.58480232 |
| PHYHD1 | 60.8799109 | -0.305520004 | 0.45288252 | -0.6746121 | 0.49992227 | 0.86719185 |
| NUP188 | 58.5052835 | 0.054798522 | 0.24105128 | 0.22733139 | 0.82016607 | 0.96502145 |
| SH3GLB2 | 245.487094 | 0.23434472 | 0.30462756 | 0.76928273 | 0.44172548 | 0.83770294 |
| MIGA2 | 65.2982174 | 0.618382639 | 0.30714306 | 2.01333749 | 0.04407914 | 0.35332084 |
| PTPA | 155.255067 | -0.163382912 | 0.2962706 | -0.5514652 | 0.58131484 | 0.90045866 |
| IER5L | 271.651809 | -0.496642194 | 0.51170985 | -0.9705543 | 0.33177027 | 0.77618491 |
| LINC01503 | 767.402659 | -0.90438078 | 0.51234002 | -1.7651965 | 0.07753075 | 0.44546179 |
| NTMT1 | 70.7113629 | -0.250223983 | 0.40870955 | -0.6122294 | 0.54038602 | 0.88342811 |
| ASB6 | 51.07482 | 0.162339883 | 0.2340957 | 0.69347657 | 0.48801052 | 0.86006752 |
| TOR1B | 39.2035093 | -0.009590917 | 0.31337301 | -0.0306054 | 0.97558421 | 0.9939318 |
| TOR1A | 37.0348226 | -0.153027178 | 0.29066393 | -0.5264746 | 0.59855849 | 0.90369786 |
| C9orf78 | 107.089014 | 0.049573614 | 0.41569329 | 0.11925527 | 0.90507312 | 0.98320957 |
| USP20 | 78.9140537 | -0.21246401 | 0.27169332 | -0.7819994 | 0.43421492 | 0.83243867 |
| FNBP1 | 242.972754 | -0.194141518 | 0.32639758 | -0.5948007 | 0.55197669 | 0.88923362 |
| GPR107 | 257.206334 | -0.119828822 | 0.15931633 | -0.752144 | 0.45196447 | 0.84234337 |
| NCS1 | 101.422835 | 0.390268487 | 0.33587263 | 1.16195384 | 0.24525422 | 0.70149132 |
| ASS1 | 77.8813294 | 0.398089951 | 0.50408758 | 0.78972378 | 0.4296891 | 0.83007221 |
| FUBP3 | 236.908841 | 0.137936932 | 0.2145483 | 0.64291784 | 0.52027741 | 0.87638467 |
| ABL1 | 481.777267 | 0.257392665 | 0.24929234 | 1.03249328 | 0.30184109 | 0.74853163 |
| AIF1L | 1619.04972 | 0.808553214 | 0.5009598 | 1.61400818 | 0.10652565 | 0.50837378 |
| NUP214 | 759.673468 | 0.768309442 | 0.23518289 | 3.26685946 | 0.00108748 | 0.04205444 |
| AL157938.2 | 204.130346 | 0.366445303 | 0.70316227 | 0.52113903 | 0.60226992 | 0.90369786 |
| AL157938.3 | 55.5559174 | 0.513499735 | 0.55173085 | 0.93070694 | 0.35200518 | 0.78314007 |
| PRRC2B | 800.491517 | -0.083167118 | 0.18790625 | -0.442599 | 0.6580558 | 0.92185868 |
| POMT1 | 65.5732988 | 0.156864282 | 0.24158871 | 0.64930304 | 0.51614252 | 0.87455102 |
| UCK1 | 39.5405263 | -0.154703207 | 0.4502568 | -0.3435888 | 0.73115553 | 0.94053147 |
| RAPGEF1 | 234.945607 | 0.097728322 | 0.3023302 | 0.32325029 | 0.7465057 | 0.94597987 |
| MED27 | 287.491845 | -0.008380019 | 0.23404727 | -0.0358048 | 0.97143799 | 0.9939318 |
| SETX | 470.854772 | 0.134255031 | 0.20267504 | 0.6624152 | 0.50770516 | 0.87094607 |
| TTF1 | 97.6422799 | -0.162080537 | 0.24063409 | -0.673556 | 0.50059363 | 0.86743068 |
| DDX31 | 90.4689006 | -0.065174795 | 0.25699464 | -0.2536037 | 0.79980173 | 0.96043078 |
| GTF3C4 | 63.5153812 | -0.067444284 | 0.27310458 | -0.2469541 | 0.80494378 | 0.96211346 |
| AK8 | 74.7244864 | -0.046521783 | 0.3595611 | -0.1293849 | 0.89705308 | 0.98302816 |
| SPACA9 | 36.7371444 | 0.054916104 | 0.37029524 | 0.14830357 | 0.8821032 | 0.97830717 |
| TSC1 | 249.844004 | -0.061480536 | 0.35847054 | -0.1715079 | 0.86382441 | 0.97466129 |
| GTF3C5 | 59.3924053 | -0.25685519 | 0.40188647 | -0.6391238 | 0.52274243 | 0.87688817 |
| RALGDS | 44.3799728 | -0.16022777 | 0.34413322 | -0.4655981 | 0.64150321 | 0.91643619 |
| ABO | 383.074303 | 0.366115671 | 0.28175372 | 1.29941735 | 0.19380074 | 0.64723766 |
| SURF6 | 67.0690701 | -0.204942655 | 0.25202714 | -0.8131769 | 0.41611663 | 0.82489761 |
| MED22 | 45.0336832 | 0.301514381 | 0.29914721 | 1.00791306 | 0.3134962 | 0.75718834 |
| RPL7A | 664.239902 | 0.141501325 | 0.49938359 | 0.28335197 | 0.77690704 | 0.95601455 |
| SURF1 | 61.7654338 | -0.056389154 | 0.28659567 | -0.1967551 | 0.84401921 | 0.96876046 |
| SURF2 | 43.4324177 | 0.618757522 | 0.55280923 | 1.11929665 | 0.2630136 | 0.71989632 |
| SURF4 | 190.09179 | 0.419533819 | 0.47481369 | 0.88357566 | 0.37692532 | 0.80049477 |
| REXO4 | 60.0930901 | -0.106470042 | 0.28033894 | -0.3797904 | 0.704101 | 0.93241442 |
| ADAMTS13 | 35.4752124 | 0.17169639 | 0.36413297 | 0.47152114 | 0.63726863 | 0.91466334 |
| CACFD1 | 55.1637755 | 0.075659833 | 0.40081778 | 0.18876366 | 0.85027805 | 0.97017003 |
| VAV2 | 930.343974 | -0.490139631 | 0.843863 | -0.5808284 | 0.56135608 | 0.89242209 |
| BRD3OS | 54.9837827 | -0.163971342 | 0.48386914 | -0.3388754 | 0.73470361 | 0.94215106 |
| BRD3 | 141.145179 | -0.023473625 | 0.29237538 | -0.0802859 | 0.93600986 | 0.98822345 |
| WDR5 | 63.3260155 | 0.034214897 | 0.31762798 | 0.10772004 | 0.91421777 | 0.98480378 |
| RXRA | 95.1647106 | 0.237202791 | 0.40634367 | 0.5837492 | 0.55938906 | 0.89179371 |
| PPP1R26-AS: | 77.2824342 | -0.405128323 | 0.43311101 | -0.9353914 | 0.34958662 | 0.78199953 |
| PPP1R26 | 58.0877126 | -0.064871271 | 0.26432193 | -0.2454252 | 0.80612719 | 0.96218371 |
| MRPS2 | 66.7754819 | 0.240135557 | 0.37144349 | 0.64649285 | 0.51796023 | 0.87526312 |
| CAMSAP1 | 97.7170283 | -0.297803691 | 0.34126513 | -0.8726461 | 0.382856 | 0.80423549 |
| UBAC1 | 116.72837 | 0.001820309 | 0.34045675 | 0.00534667 | 0.995734 | 0.99908534 |
| NACC2 | 87.2127338 | 0.117971627 | 0.24297028 | 0.48553933 | 0.62729382 | 0.91061758 |
| TMEM250 | 41.9156423 | -0.372637675 | 0.34847461 | -1.0693395 | 0.2849167 | 0.7352689 |
| QSOX2 | 41.3045684 | 0.273779297 | 0.29731618 | 0.92083552 | 0.3571363 | 0.78612469 |
| SNAPC4 | 51.8081886 | -0.057308828 | 0.36455326 | -0.1572029 | 0.87508495 | 0.97622753 |
| ENTR1 | 54.7309534 | -0.237077927 | 0.39425972 | -0.6013242 | 0.54762405 | 0.88773467 |
| PMPCA | 80.2772955 | 0.090765511 | 0.25968887 | 0.34951637 | 0.72670168 | 0.93906703 |
| INPP5E | 39.1744846 | 0.047986161 | 0.35047106 | 0.13691904 | 0.89109479 | 0.98095103 |
| SEC16A | 154.539858 | -0.073161052 | 0.26441297 | -0.2766924 | 0.78201633 | 0.95688565 |
| NOTCH1 | 43.2712468 | -0.274144117 | 0.4132575 | -0.6633736 | 0.5070913 | 0.87094607 |
| AGPAT2 | 141.81678 | -0.549939675 | 0.36926252 | -1.4892919 | 0.13641052 | 0.56040074 |
| SNHG7 | 118.193628 | 0.336775192 | 0.37757803 | 0.89193535 | 0.37242759 | 0.79637648 |
| RABL6 | 212.575259 | 0.208568839 | 0.26577859 | 0.78474658 | 0.43260217 | 0.83162989 |
| PHPT1 | 228.38733 | -0.258100991 | 0.54183346 | -0.4763475 | 0.63382686 | 0.91415622 |
| EDF1 | 357.806725 | 0.085425094 | 0.45998065 | 0.18571454 | 0.85266863 | 0.97062426 |
| TRAF2 | 38.9484847 | 0.397374519 | 0.39252454 | 1.01235585 | 0.31136794 | 0.7546568 |
| FBXW5 | 110.200152 | -0.325110354 | 0.32007008 | -1.0157474 | 0.30974969 | 0.75409553 |
| LCN12 | 91.8616333 | 0.229951684 | 0.48569377 | 0.47344993 | 0.63589221 | 0.91454294 |
| PAXX | 69.4509047 | 0.058413327 | 0.3652245 | 0.15993814 | 0.8729298 | 0.97622753 |
| ABCA2 | 33.0452626 | -0.082925454 | 0.70587502 | -0.1174789 | 0.90648053 | 0.98320957 |
| MAN1B1 | 110.507855 | -0.215463043 | 0.21421604 | -1.0058212 | 0.31450158 | 0.75823062 |
| DPP7 | 119.042546 | 0.01410042 | 0.38963264 | 0.03618901 | 0.97113165 | 0.9939318 |

| | | | | | | |
|---|---|---|---|---|---|---|
| SSNA1 | 120.584513 | -0.091908737 | 0.45476814 | -0.2021002 | 0.83983838 | 0.96833625 |
| TPRN | 44.3165985 | -0.512856952 | 0.34710987 | -1.4775061 | 0.13954002 | 0.56656585 |
| TMEM203 | 62.872876 | 0.096573573 | 0.50510393 | 0.19119545 | 0.84837247 | 0.96991131 |
| RNF208 | 33.549932 | 0.72874693 | 0.47889854 | 1.52171466 | 0.12808059 | 0.54809499 |
| TUBB4B | 378.910348 | 0.359120274 | 0.61219748 | 0.58660855 | 0.55746664 | 0.89148045 |
| NELFB | 135.194179 | 0.154562645 | 0.26619042 | 0.58064692 | 0.56147845 | 0.89242209 |
| TOR4A | 43.6502878 | -0.466008875 | 0.30570597 | -1.5243696 | 0.12741642 | 0.54746778 |
| EXD3 | 180.439044 | -0.187494546 | 0.27926888 | -0.6713764 | 0.50198076 | 0.86837155 |
| NOXA1 | 76.7437867 | -0.249268701 | 0.27286806 | -0.9135137 | 0.36097244 | 0.78927419 |
| NSMF | 38.8227217 | 0.532685089 | 0.310384 | 1.71621311 | 0.08612304 | 0.46572214 |
| PNPLA7 | 105.434096 | -0.2875827 | 0.50518274 | -0.5692647 | 0.56917652 | 0.89617938 |
| MRPL41 | 316.340791 | -0.334645121 | 0.43743193 | -0.7650222 | 0.44425835 | 0.83873671 |
| DPH7 | 64.4647334 | 0.227604847 | 0.23542887 | 0.96676692 | 0.33366056 | 0.77744565 |
| ARRDC1 | 105.394303 | 0.42636219 | 0.22883882 | 1.86315497 | 0.06244047 | 0.41052452 |
| ARRDC1-AS1 | 31.3819182 | 0.042314104 | 0.37089829 | 0.11408547 | 0.90917004 | 0.98441374 |
| EHMT1 | 627.39191 | 0.088071776 | 0.21149228 | 0.41643022 | 0.67709522 | 0.9268955 |
| AC069287.3 | 44.2212623 | 0.463932227 | 0.70098419 | 0.6618298 | 0.5080803 | 0.87094607 |
| BET1L | 66.6179047 | 0.036922229 | 0.31066281 | 0.11884985 | 0.90539431 | 0.98320957 |
| RIC8A | 65.0825855 | -0.431630908 | 0.39756257 | -1.085693 | 0.27761482 | 0.72788245 |
| SIRT3 | 69.2556685 | -0.133245524 | 0.28289249 | -0.4710112 | 0.63763275 | 0.91502002 |
| PSMD13 | 126.970639 | -0.035001167 | 0.27091059 | -0.1291982 | 0.8972008 | 0.98302816 |
| AC136475.3 | 38.1994481 | 1.805567847 | 1.04142498 | 1.73374739 | 0.08296291 | 0.45768271 |
| PGGHG | 246.063759 | 1.006466064 | 0.68232058 | 1.47506332 | 0.14019552 | 0.56749674 |
| IFITM2 | 651.158259 | -2.01342201 | 0.41077277 | -4.9015469 | 9.51E-07 | 0.00034362 |
| IFITM3 | 1112.39215 | -2.065387538 | 0.62692934 | -3.2944503 | 0.00098614 | 0.03914693 |
| PKP3 | 53.5440523 | -0.233920762 | 0.31270674 | -0.7480515 | 0.45442907 | 0.84290184 |
| SIGIRR | 99.2448245 | -0.336166739 | 0.31972231 | -1.0514335 | 0.29305955 | 0.73864467 |
| ANO9 | 38.7573661 | -0.035748318 | 0.38313361 | -0.0933051 | 0.92566118 | 0.98717098 |
| PTDSS2 | 79.133099 | -0.252480637 | 0.28323093 | -0.8914303 | 0.37269836 | 0.7966304 |
| RNH1 | 177.109333 | -0.449710269 | 0.36855139 | -1.2202105 | 0.22238509 | 0.6798044 |
| HRAS | 97.1299798 | 0.202799296 | 0.44550895 | 0.45520813 | 0.64895951 | 0.91768694 |
| AP006284.1 | 45.5352068 | 0.880646054 | 0.65845932 | 1.33743425 | 0.18108093 | 0.63152351 |
| RASSF7 | 263.284834 | 0.006420263 | 0.35981862 | 0.01784305 | 0.98576406 | 0.99587186 |
| MIR210HG | 83.8403415 | -1.657944508 | 0.6866433 | -2.4145645 | 0.01575403 | 0.20433451 |
| PHRF1 | 108.138277 | 0.255224834 | 0.2761433 | 0.92424779 | 0.35535732 | 0.78607579 |
| DEAF1 | 198.214173 | -0.092106786 | 0.20219531 | -0.4555337 | 0.6487253 | 0.91768694 |
| AC131934.1 | 35.5665652 | 0.033479973 | 0.40225861 | 0.08322997 | 0.93366868 | 0.98822345 |
| EPS8L2 | 467.741274 | 0.509121922 | 0.40435385 | 1.25909997 | 0.20799422 | 0.66589349 |
| AP006621.4 | 65.1828093 | -0.634376237 | 0.62988499 | -1.0071303 | 0.31387218 | 0.75763653 |
| TALDO1 | 273.146939 | -0.923500695 | 0.4272208 | -2.1616473 | 0.03064537 | 0.29437535 |
| GATD1 | 68.2908658 | -0.059511303 | 0.29489515 | -0.201805 | 0.84006921 | 0.96833625 |
| AP006621.3 | 85.6722259 | -0.212316056 | 0.3312529 | -0.6409485 | 0.52155614 | 0.87650871 |
| RPLP2 | 458.584995 | 0.04595302 | 0.35813272 | 0.12831282 | 0.89790142 | 0.98312332 |
| PNPLA2 | 155.948992 | 0.277274768 | 0.34201982 | 0.81069795 | 0.41753915 | 0.82566561 |
| CD151 | 355.850665 | -0.383469919 | 0.3490213 | -1.0987006 | 0.27189867 | 0.72372882 |
| POLR2L | 316.233674 | -0.082663217 | 0.51487586 | -0.1605498 | 0.87244799 | 0.97622753 |
| TSPAN4 | 43.7496413 | -0.663727488 | 0.45607536 | -1.4553022 | 0.14558561 | 0.57618473 |
| CHID1 | 96.5820291 | 0.185722279 | 0.34580837 | 0.53706705 | 0.5912213 | 0.90233654 |
| AP2A2 | 642.654688 | 0.140245396 | 0.32418723 | 0.43260618 | 0.6653009 | 0.92324595 |
| MUC6 | 82.6057908 | 0.377515896 | 0.93038128 | 0.40576472 | 0.68491549 | 0.92881914 |
| TOLLIP | 96.2288976 | -0.083260937 | 0.26522921 | -0.3139207 | 0.75358127 | 0.94967914 |
| MOB2 | 74.1856073 | 0.151961512 | 0.25505742 | 0.59579334 | 0.5513133 | 0.88916026 |
| MRPL23 | 41.8786936 | -0.204582421 | 0.6149747 | -0.332668 | 0.73938489 | 0.94404893 |
| CD81-AS1 | 41.1741237 | 0.028067954 | 0.3659942 | 0.07668961 | 0.93887047 | 0.98848592 |
| CD81 | 1179.15295 | -0.597208763 | 0.32013102 | -1.8655136 | 0.06210945 | 0.40942507 |
| TSSC4 | 57.7318225 | -0.018564088 | 0.38567459 | -0.0481341 | 0.96160939 | 0.99171131 |
| KCNQ1 | 1183.1197 | 0.131020664 | 0.23779325 | 0.55098562 | 0.58164353 | 0.90053894 |
| KCNQ1OT1 | 1165.96078 | -0.977326008 | 0.42414194 | -2.3042428 | 0.02120902 | 0.24669313 |
| CDKN1C | 507.853436 | -1.969060428 | 0.84937792 | -2.3182383 | 0.02043637 | 0.24101888 |
| SLC22A18 | 111.857186 | -0.931638576 | 0.45774284 | -2.0352881 | 0.04182189 | 0.34565724 |
| PHLDA2 | 134.179809 | 0.365215254 | 1.31909939 | 0.27686712 | 0.78188214 | 0.95688565 |
| NAP1L4 | 694.968777 | -0.046583602 | 0.22912749 | -0.2033087 | 0.83889379 | 0.96802542 |
| CARS | 106.117455 | -0.697443869 | 0.34289959 | -2.0339595 | 0.04195568 | 0.34641157 |
| OSBPL5 | 53.3326278 | -0.24718101 | 0.39302362 | -0.6289215 | 0.52940044 | 0.87911847 |
| ZNF195 | 93.72594 | -0.133828167 | 0.3417996 | -0.3915399 | 0.69539823 | 0.93126994 |
| AC127526.1 | 59.3690936 | 0.118840624 | 0.43411489 | 0.27375385 | 0.78427379 | 0.95697044 |
| NUP98 | 452.840345 | 0.577542055 | 0.29541164 | 1.95504165 | 0.05057815 | 0.37952033 |
| PGAP2 | 93.4953221 | 0.025803472 | 0.3511418 | 0.07348448 | 0.94142059 | 0.98876195 |
| RHOG | 44.7082379 | -0.60762477 | 0.39360816 | -1.5437301 | 0.12265373 | 0.54116729 |
| STIM1 | 215.280882 | -0.412754789 | 0.30884739 | -1.3364361 | 0.18140678 | 0.63181008 |
| RRM1 | 78.0175643 | 0.229391818 | 0.28037861 | 0.81815022 | 0.41327142 | 0.82437234 |
| TRIM68 | 28.4810399 | -0.11712384 | 0.40704855 | -0.2877392 | 0.77354635 | 0.95561066 |
| AC104389.4 | 181.843855 | 0.298523476 | 0.43928314 | 0.67956962 | 0.49677701 | 0.86487238 |
| TRIM5 | 74.5564131 | -1.578993973 | 0.57213767 | -2.7598147 | 0.00578341 | 0.1217072 |
| TRIM22 | 148.173748 | -1.329599827 | 0.8059647 | -1.6496998 | 0.09900435 | 0.4933116 |
| FAM160A2 | 70.3910495 | 0.480319535 | 0.33653865 | 1.42723439 | 0.15351234 | 0.58860244 |
| SMPD1 | 49.3267085 | 0.133143393 | 0.54845265 | 0.24276187 | 0.80818987 | 0.96245698 |
| APBB1 | 67.5746921 | 0.000586139 | 0.27086034 | 0.00216399 | 0.99827339 | 0.99954455 |
| ARFIP2 | 59.9844327 | -0.028608524 | 0.3377172 | -0.0847115 | 0.93249077 | 0.98822345 |
| AC084337.2 | 58.7276715 | -0.505082222 | 0.52977846 | -0.9533838 | 0.34039563 | 0.77836439 |
| DNHD1 | 95.2219765 | -0.852049223 | 0.46310068 | -1.839879 | 0.065786 | 0.41788422 |
| RRP8 | 31.7719418 | -0.457933044 | 0.31995201 | -1.4312554 | 0.15235703 | 0.58767084 |
| ILK | 153.144525 | -0.106254857 | 0.30618561 | -0.3470276 | 0.72857058 | 0.93931348 |
| TAF10 | 91.7688046 | 0.226255524 | 0.40165783 | 0.56330415 | 0.57322779 | 0.89797476 |
| TPP1 | 79.2322838 | -0.377399085 | 0.2821766 | -1.3374571 | 0.18107349 | 0.63152351 |
| MRPL17 | 103.889828 | -0.881132027 | 0.45948178 | -1.9176648 | 0.05515353 | 0.3929361 |
| AC087280.2 | 90.2477361 | 0.140755831 | 0.59466901 | 0.23669609 | 0.81289257 | 0.96288868 |
| ZNF215 | 46.9519981 | -0.13308389 | 0.53575594 | -0.2484039 | 0.80382188 | 0.96164992 |
| ZNF214 | 41.1152172 | -0.41293803 | 0.4353476 | -0.9485249 | 0.34286232 | 0.78096003 |
| SYT9 | 166.518954 | 0.623790222 | 0.64797743 | 0.96267277 | 0.33571177 | 0.77820994 |
| AC107884.1 | 73.0929232 | 0.061651609 | 0.50148921 | 0.12293706 | 0.90215694 | 0.98320957 |
| PPFIBP2 | 230.33918 | -0.294996092 | 0.28879289 | -1.0214798 | 0.3070272 | 0.75214355 |
| OVCH2 | 97.1602346 | -0.281588289 | 0.5384377 | -0.5229728 | 0.60099316 | 0.90369786 |
| AC044810.3 | 206.022039 | -1.437467277 | 0.86339715 | -1.6648969 | 0.09593332 | 0.48710328 |
| EIF3F | 191.977584 | -0.007510941 | 0.31779585 | -0.0236345 | 0.98114417 | 0.99507903 |
| TUB | 34.2315877 | 1.883571173 | 0.62481305 | 3.0146156 | 0.00257305 | 0.07369381 |
| RIC3 | 134.905538 | -0.281549426 | 0.42064852 | -0.6693223 | 0.50328992 | 0.86946722 |
| STK33 | 1145.68823 | 0.752846256 | 0.27493828 | 2.73823731 | 0.00617695 | 0.12643442 |
| TRIM66 | 199.907867 | -0.200275572 | 0.29719628 | -0.6738832 | 0.50038559 | 0.86743068 |
| RPL27A | 518.230893 | 0.066152939 | 0.34739197 | 0.19042737 | 0.84897425 | 0.96991131 |
| ST5 | 409.758737 | -1.595354391 | 0.35262817 | -4.5241831 | 6.06E-06 | 0.00127079 |
| AC026894.1 | 39.1158964 | -2.121577506 | 0.62283145 | -3.406343 | 0.00065839 | 0.03026303 |
| AKIP1 | 42.7818488 | -0.423379525 | 0.36382762 | -1.1636817 | 0.24455304 | 0.70121778 |
| TMEM9B | 161.121771 | 0.227962828 | 0.34733092 | 0.65632749 | 0.51161343 | 0.87151901 |
| TMEM9B-AS | 53.7107897 | -0.826283285 | 0.44126234 | -1.8725443 | 0.06113133 | 0.40676596 |
| DENND5A | 247.593023 | -0.627018897 | 0.3893233 | -1.6105353 | 0.10728105 | 0.51064149 |
| TMEM41B | 269.927209 | -0.114805646 | 0.40080847 | -0.2864352 | 0.77454483 | 0.95586283 |
| IPO7 | 332.73799 | 0.004675129 | 0.31540934 | 0.01482242 | 0.98817385 | 0.9964791 |
| ZNF143 | 92.5560567 | 0.241057902 | 0.42567445 | 0.56629638 | 0.57119232 | 0.89773478 |
| WEE1 | 81.8484863 | 0.326273372 | 0.33460541 | 0.9750989 | 0.3295112 | 0.77596696 |
| SWAP70 | 349.568178 | -0.46826023 | 0.26751118 | -1.7504324 | 0.08004373 | 0.45074086 |
| SBF2-AS1 | 107.806654 | -1.171887506 | 0.367451 | -3.1892348 | 0.0014265 | 0.05016683 |
| SBF2 | 1774.48941 | 0.015992073 | 0.24649696 | 0.06487737 | 0.94827164 | 0.98977092 |

| | | | | | | |
|---|---|---|---|---|---|---|
| AC011092.2 | 84.9058383 | -0.281983548 | 0.4819954 | -0.5850337 | 0.55852506 | 0.89179371 |
| ADM | 233.181281 | -2.242893767 | 1.4478523 | -1.5491178 | 0.1213534 | 0.53989034 |
| AMPD3 | 95.8626138 | -2.374488246 | 0.70146333 | -3.3850497 | 0.00071165 | 0.03220254 |
| MTRNR2L8 | 274.671879 | -1.486425191 | 0.60577568 | -2.4537551 | 0.01413732 | 0.19259627 |
| RNF141 | 94.9310413 | 0.169582367 | 0.24668643 | 0.687441 | 0.49180487 | 0.86172738 |
| CTR9 | 150.676887 | 0.034452975 | 0.19086377 | 0.18051081 | 0.85675157 | 0.97191993 |
| EIF4G2 | 567.063012 | -0.027339935 | 0.32214622 | -0.0848681 | 0.93236626 | 0.98822345 |
| ZBED5 | 132.799068 | -0.181204316 | 0.20952801 | -0.8648214 | 0.38713681 | 0.8065026 |
| ZBED5-AS1 | 49.8957404 | 0.34171018 | 0.47177529 | 0.72430707 | 0.46887723 | 0.85206059 |
| GALNT18 | 1035.79625 | 0.569718573 | 0.46650037 | 1.2212607 | 0.22198733 | 0.67955938 |
| USP47 | 691.923734 | 0.330197 | 0.19698595 | 1.67624647 | 0.09368992 | 0.48360487 |
| PARVA | 255.996129 | -0.356048601 | 0.24949238 | -1.4270921 | 0.15355335 | 0.58860244 |
| TEAD1 | 1439.22824 | 0.446198434 | 0.30471147 | 1.46433094 | 0.14310355 | 0.5729912 |
| LINC00958 | 109.606203 | -0.787905426 | 0.22775066 | -3.4595089 | 0.00054116 | 0.02637847 |
| ARNTL | 84.2170048 | 0.232614805 | 0.36740761 | 0.63312463 | 0.52665226 | 0.87791848 |
| BTBD10 | 179.652119 | 0.203628164 | 0.26185293 | 0.77764324 | 0.43677936 | 0.83393108 |
| FAR1 | 420.506334 | 0.304943244 | 0.16116122 | 1.89216262 | 0.05846932 | 0.39982063 |
| RRAS2 | 204.26874 | -0.101573534 | 0.31814085 | -0.3192722 | 0.7495201 | 0.94706664 |
| COPB1 | 268.642233 | 0.069581174 | 0.27525519 | 0.25278787 | 0.80043214 | 0.96063297 |
| PSMA1 | 119.385473 | 1.040657559 | 0.76056839 | 1.36826295 | 0.17122978 | 0.61772396 |
| PDE3B | 473.163542 | 0.798836167 | 0.89771201 | 0.88985795 | 0.37354217 | 0.79681088 |
| AC073172.2 | 46.9564675 | 0.110608971 | 1.09068289 | 0.10141259 | 0.91922295 | 0.98520052 |
| AC073172.1 | 324.51854 | 0.570238904 | 0.89725383 | 0.635538 | 0.52507759 | 0.87791848 |
| AC087379.2 | 546.335085 | 0.214668515 | 0.96882104 | 0.22157706 | 0.82464314 | 0.96511681 |
| SOX6 | 2427.26555 | 0.655231914 | 0.39221527 | 1.67059257 | 0.09480218 | 0.48627909 |
| C11orf58 | 404.317035 | 0.004553759 | 0.19447004 | 0.02341625 | 0.98131824 | 0.99507903 |
| PLEKHA7 | 1710.05719 | 0.078883691 | 0.2164103 | 0.36450987 | 0.7154773 | 0.93573028 |
| RPS13 | 464.423299 | -0.022929986 | 0.55663604 | -0.0411939 | 0.96714135 | 0.99291246 |
| PIK3C2A | 644.999137 | -0.195681371 | 0.20479123 | -0.9555164 | 0.33931664 | 0.77836439 |
| NUCB2 | 165.665552 | -0.438928289 | 0.24833684 | -1.7674715 | 0.07714929 | 0.44451877 |
| SERGEF | 219.678937 | -0.118232406 | 0.26350315 | -0.4486945 | 0.65365207 | 0.91983951 |
| HPS5 | 102.232056 | 0.165133287 | 0.49235251 | 0.33539645 | 0.73732604 | 0.9429136 |
| GTF2H1 | 159.523578 | -0.317684492 | 0.29388174 | -1.0809943 | 0.27969964 | 0.72938673 |
| LDHA | 990.259918 | -0.863979293 | 0.57806139 | -1.4946151 | 0.13501492 | 0.55891804 |
| TSG101 | 242.297093 | 0.02287615 | 0.21603114 | 0.10589284 | 0.91566738 | 0.98520052 |
| UEVLD | 194.462827 | -0.355555681 | 0.25796691 | -1.3782996 | 0.16811082 | 0.6131323 |
| SPTY2D1OS | 61.679994 | 0.214927974 | 0.28468544 | 0.75496652 | 0.45026907 | 0.84147218 |
| SPTY2D1 | 94.876844 | -0.170271985 | 0.33026397 | -0.5155633 | 0.60615945 | 0.90493874 |
| ZDHHC13 | 182.409903 | -0.065781397 | 0.18447615 | -0.3565848 | 0.72140263 | 0.93744589 |
| NAV2 | 769.044816 | 1.074545923 | 0.5031122 | 2.13579777 | 0.0326959 | 0.30539487 |
| HTATIP2 | 114.847721 | -0.371614893 | 0.57001456 | -0.6519393 | 0.51444033 | 0.87252543 |
| PRMT3 | 242.461581 | 0.754749972 | 0.42588502 | 1.77219186 | 0.07636271 | 0.4426298 |
| ANO5 | 82.7415628 | -0.039107815 | 0.41427561 | -0.0944005 | 0.92479104 | 0.98710257 |
| FANCF | 32.5796836 | -0.196581693 | 0.29723087 | -0.6613771 | 0.50837051 | 0.87094607 |
| GAS2 | 75.186458 | 0.004811912 | 0.44743137 | 0.01075452 | 0.9914193 | 0.99751097 |
| SVIP | 229.542146 | -0.096401685 | 0.29299418 | -0.3290225 | 0.74213866 | 0.94508888 |
| MUC15 | 72.6937926 | 1.023052826 | 0.69234712 | 1.47765882 | 0.13949912 | 0.56656585 |
| FIBIN | 51.5727056 | -1.177467576 | 0.87040364 | -1.3527834 | 0.17612485 | 0.62516039 |
| BBOX1 | 89.9161365 | -0.841887339 | 0.58359688 | -1.4425837 | 0.14913778 | 0.58194377 |
| CCDC34 | 53.3128234 | 0.356073838 | 0.46602764 | 0.76406163 | 0.44483052 | 0.83925275 |
| LGR4 | 761.076012 | 0.197214911 | 0.35827957 | 0.55044978 | 0.58201091 | 0.90085422 |
| LIN7C | 104.875295 | 0.540213853 | 0.35144764 | 1.5371105 | 0.12426625 | 0.54273186 |
| BDNF-AS | 415.379849 | -0.087254241 | 0.1891736 | -0.461239 | 0.64462716 | 0.91682456 |
| METTL15 | 861.397104 | -0.012978052 | 0.32431761 | -0.0400165 | 0.96807998 | 0.99303855 |
| ARL14EP | 106.246657 | -0.311443329 | 0.27221489 | -1.144112 | 0.25257718 | 0.70813507 |
| MPPED2 | 1723.36919 | 0.538362334 | 0.3830288 | 1.40554008 | 0.15986074 | 0.59962081 |
| AL136088.1 | 50.2149934 | 0.461891601 | 0.47497434 | 0.9724559 | 0.33082379 | 0.77618491 |
| DCDC1 | 410.172021 | 0.069614117 | 0.25791972 | 0.26990615 | 0.78723246 | 0.95791052 |
| DNAJC24 | 107.779403 | 0.047615201 | 0.30399693 | 0.15663053 | 0.87553604 | 0.97622753 |
| IMMP1L | 263.434763 | 0.068311318 | 0.25117742 | 0.2719644 | 0.78564939 | 0.9573712 |
| ELP4 | 444.78749 | 0.303691235 | 0.21751882 | 1.39616076 | 0.16266609 | 0.60211533 |
| AC131571.1 | 57.4231018 | -0.136331648 | 0.69226484 | -0.1969357 | 0.84387787 | 0.96876046 |
| RCN1 | 99.2217096 | -0.629568308 | 0.32806756 | -1.9190203 | 0.05498176 | 0.39224566 |
| EIF3M | 146.911079 | 0.272566256 | 0.19397559 | 1.40515751 | 0.15997445 | 0.59983263 |
| PRRG4 | 142.646502 | 0.006435849 | 0.38905728 | 0.01654216 | 0.98680186 | 0.99643909 |
| QSER1 | 230.200228 | 0.083104297 | 0.25423082 | 0.32688522 | 0.7437547 | 0.9456489 |
| TCP11L1 | 105.804851 | 0.222906056 | 0.35815621 | 0.62237105 | 0.53369791 | 0.88137638 |
| CSTF3 | 213.466602 | 0.249037862 | 0.24904506 | 0.99997109 | 0.3173245 | 0.76134633 |
| HIPK3 | 595.290402 | 0.217122796 | 0.15316025 | 1.41761842 | 0.1563022 | 0.59366792 |
| FBXO3 | 142.663666 | 0.572661098 | 0.27072696 | 2.11527177 | 0.03440679 | 0.31355058 |
| CAPRIN1 | 430.04368 | 0.164535659 | 0.1819387 | 0.90434667 | 0.3658116 | 0.79329304 |
| NAT10 | 94.9966037 | -0.618343203 | 0.22370877 | -2.7640544 | 0.0057088 | 0.1213371 |
| ABTB2 | 3134.42333 | 1.109861506 | 0.59911449 | 1.85250319 | 0.0639536 | 0.41357646 |
| CAT | 328.284641 | 0.201889265 | 0.38791226 | 0.52045084 | 0.60274938 | 0.90369786 |
| ELF5 | 323.172577 | 1.848207451 | 0.67055658 | 2.75622894 | 0.00584721 | 0.12255743 |
| EHF | 664.125793 | 0.344995359 | 0.40829199 | 0.84497214 | 0.3981264 | 0.81400298 |
| APIP | 163.11281 | -0.062308722 | 0.44699011 | -0.1393962 | 0.88913708 | 0.98054894 |
| PDHX | 251.216044 | 0.145938279 | 0.26207721 | 0.55685224 | 0.57762839 | 0.89916873 |
| CD44 | 121.482935 | -5.717760767 | 1.34311835 | -4.2570789 | 2.07E-05 | 0.00314575 |
| TRIM44 | 506.888303 | -0.087865804 | 0.16500845 | -0.5324928 | 0.59438475 | 0.90305763 |
| COMMD9 | 58.4429387 | -0.300891796 | 0.23334468 | -1.2894736 | 0.1972335 | 0.6524002 |
| PRR5L | 162.238007 | 0.004297952 | 0.44298408 | 0.00970227 | 0.99225883 | 0.99797241 |
| TRAF6 | 76.8563068 | 0.093102765 | 0.27394258 | 0.33986233 | 0.73396021 | 0.94176201 |
| C11orf74 | 74.9488782 | 0.397967178 | 0.27536531 | 1.44523354 | 0.1483923 | 0.58049694 |
| LRRC4C | 85.4616137 | -2.156760033 | 0.76591061 | -2.8159422 | 0.00486344 | 0.11130863 |
| API5 | 214.838718 | 0.122428035 | 0.20868496 | 0.58666438 | 0.55742913 | 0.89148045 |
| TTC17 | 939.633607 | -0.023460127 | 0.19595269 | -0.1197234 | 0.90470225 | 0.98320957 |
| HSD17B12 | 830.811828 | -0.042662369 | 0.22581978 | -0.1889222 | 0.8501538 | 0.97017003 |
| ALKBH3 | 68.3208524 | -0.044317783 | 0.27815341 | -0.1593286 | 0.87341002 | 0.97622753 |
| ACCS | 43.288646 | -0.253385718 | 0.37895704 | -0.6686397 | 0.50372535 | 0.86946722 |
| EXT2 | 216.220113 | 0.041268523 | 0.23974839 | 0.17213264 | 0.86333325 | 0.97455111 |
| CD82 | 281.941312 | -0.082056046 | 0.51432627 | -0.1595408 | 0.87324278 | 0.97622753 |
| TSPAN18 | 55.6227555 | -0.839730803 | 0.62403559 | -1.3456457 | 0.17841681 | 0.62759322 |
| TP53I11 | 57.8248884 | -0.143236418 | 0.53643318 | -0.2670163 | 0.78945659 | 0.95801395 |
| PRDM11 | 83.9227207 | 0.157068871 | 0.24435775 | 0.64278244 | 0.52036528 | 0.87638467 |
| CRY2 | 125.733261 | 0.23805097 | 0.34211711 | 0.6958172 | 0.48654332 | 0.85912348 |
| LARGE2 | 33.0587196 | 0.906278299 | 0.45128905 | 2.0081992 | 0.04462213 | 0.35515061 |
| PHF21A | 872.318133 | 0.076072641 | 0.26867762 | 0.28313724 | 0.77707164 | 0.95601455 |
| CREB3L1 | 117.716732 | -0.843476953 | 1.11510739 | -0.7564087 | 0.44940418 | 0.84117804 |
| DGKZ | 49.2873976 | -0.145060832 | 0.28380865 | -0.511122 | 0.60926566 | 0.90552475 |
| AMBRA1 | 1464.06065 | 0.212979341 | 0.27161556 | 0.78412055 | 0.43296938 | 0.8317266 |
| ATG13 | 145.160423 | -0.095416077 | 0.22140384 | -0.4309595 | 0.66649785 | 0.92396246 |
| ARHGAP1 | 161.932861 | -0.480285831 | 0.26497674 | -1.8125584 | 0.06989995 | 0.42843167 |
| CKAP5 | 210.137472 | 0.01586357 | 0.18464785 | 0.08591256 | 0.93153593 | 0.98822345 |
| LRP4 | 164.409668 | -0.441797594 | 0.45975608 | -0.9609391 | 0.33658279 | 0.77827084 |
| C11orf49 | 856.603633 | 0.292113318 | 0.21136191 | 1.38205281 | 0.16695549 | 0.61028723 |
| ARFGAP2 | 112.272017 | 0.143437183 | 0.26770854 | 0.53579607 | 0.5920995 | 0.90233654 |
| DDB2 | 57.5803033 | -0.77550185 | 0.41577834 | -1.865181 | 0.06215604 | 0.40942507 |
| ACP2 | 58.3503359 | -0.146898466 | 0.32380525 | -0.453663 | 0.65007139 | 0.91803345 |
| NR1H3 | 42.1718625 | -0.270845497 | 0.31624603 | -0.8564392 | 0.39175489 | 0.80866481 |
| MADD | 77.3692957 | 0.089484884 | 0.23724489 | 0.37718361 | 0.70603716 | 0.93241442 |
| SLC39A13 | 91.3805655 | -0.017650061 | 0.25733218 | -0.0685886 | 0.94531707 | 0.98949015 |
| PSMC3 | 51.9044585 | -0.234921666 | 0.54726494 | -0.429265 | 0.66773041 | 0.92422713 |

| | | | | | | |
|---|---|---|---|---|---|---|
| CELF1 | 832.391924 | 0.232346944 | 0.25800638 | 0.9005473 | 0.36782907 | 0.79497806 |
| NDUFS3 | 124.789298 | 0.211813073 | 0.46355185 | 0.45693502 | 0.64771775 | 0.91763881 |
| MTCH2 | 193.877188 | -0.140468001 | 0.2540245 | -0.5529703 | 0.58028375 | 0.90025637 |
| FNBP4 | 459.500975 | -0.166877009 | 0.17562926 | -0.9501663 | 0.34202775 | 0.77973697 |
| NUP160 | 274.691983 | 0.260205049 | 0.21461753 | 1.21241284 | 0.22535439 | 0.68277363 |
| PTPRJ | 1227.68697 | 0.22986359 | 0.34961233 | 0.65748137 | 0.51087145 | 0.87138909 |
| TNKS1BP1 | 176.924607 | -0.400298819 | 0.21738157 | -1.841457 | 0.06555462 | 0.41788422 |
| SSRP1 | 98.6348984 | -0.178875139 | 0.33951618 | -0.5268531 | 0.59829563 | 0.90369786 |
| TIMM10 | 80.2285765 | -0.685291594 | 0.44253462 | -1.5485604 | 0.12148743 | 0.53989034 |
| UBE2L6 | 94.9547934 | -0.740143315 | 0.67021112 | -1.1043435 | 0.26944415 | 0.72322263 |
| SERPING1 | 522.025256 | -1.631057803 | 0.39127221 | -4.1686012 | 3.06E-05 | 0.00417124 |
| ZDHHC5 | 161.816573 | -0.427480716 | 0.22423235 | -1.9064185 | 0.05659593 | 0.39673933 |
| MED19 | 39.6211768 | -0.630825799 | 0.4527925 | -1.3931896 | 0.16356246 | 0.60314377 |
| CTNND1 | 624.600448 | 0.386197511 | 0.1625946 | 2.37521736 | 0.01753862 | 0.21933738 |
| LPXN | 97.6049595 | 0.312500154 | 0.36666965 | 0.85226623 | 0.39406635 | 0.81094677 |
| FAM111A | 153.174088 | 0.011045673 | 0.25305933 | 0.04364855 | 0.96518455 | 0.9925556 |
| OSBP | 295.966239 | 0.133044246 | 0.17573012 | 0.75709416 | 0.44899345 | 0.84115854 |
| PATL1 | 168.563182 | -0.310671172 | 0.36047704 | -0.8618334 | 0.38877918 | 0.8065026 |
| STX3 | 194.308674 | -0.294350155 | 0.43688903 | -0.6737412 | 0.50047584 | 0.86743068 |
| MRPL16 | 98.2217093 | 0.216596466 | 0.38645537 | 0.56046955 | 0.5751592 | 0.89807676 |
| PRPF19 | 87.377977 | -0.231681194 | 0.36081801 | -0.6420999 | 0.52080835 | 0.87638467 |
| TMEM109 | 60.0985466 | -0.251209161 | 0.40958788 | -0.6133218 | 0.53966361 | 0.8833006 |
| VPS37C | 92.9551359 | -0.193426914 | 0.26518006 | -0.7294173 | 0.46574646 | 0.85020431 |
| DDB1 | 231.755365 | 0.099111992 | 0.22545632 | 0.43960618 | 0.66022236 | 0.92284375 |
| TKFC | 50.6567595 | -0.283689324 | 0.27920506 | -1.0160608 | 0.30960043 | 0.75403497 |
| CYB561A3 | 31.3151376 | -0.232861393 | 0.3637082 | -0.6402424 | 0.52201505 | 0.87670505 |
| TMEM138 | 35.8998962 | -0.012957046 | 0.36465144 | -0.0355327 | 0.97165498 | 0.9939318 |
| CPSF7 | 169.249267 | 0.118769351 | 0.16773438 | 0.70807993 | 0.47889562 | 0.85532469 |
| AP003108.2 | 133.119064 | -0.293531343 | 0.28041458 | -1.0467763 | 0.29520275 | 0.74136707 |
| SYT7 | 40.4954521 | 1.508271864 | 0.89262249 | 1.68970856 | 0.09108373 | 0.47749744 |
| MYRF | 170.171025 | -0.665985622 | 0.40379869 | -1.6493011 | 0.09908596 | 0.4933116 |
| TMEM258 | 154.239831 | -0.502085983 | 0.24423152 | -2.0557788 | 0.03980384 | 0.33603445 |
| FADS2 | 54.7298511 | -1.622606734 | 0.98879621 | -1.6409921 | 0.10079906 | 0.49726048 |
| FADS1 | 73.5171536 | -1.501062872 | 0.89335296 | -1.6802573 | 0.09290726 | 0.48153712 |
| FTH1 | 2515.18135 | -1.149673006 | 0.54477506 | -2.1103628 | 0.03482712 | 0.31514522 |
| ASRGL1 | 87.7585163 | -0.796661921 | 0.30788949 | -2.587493 | 0.00966772 | 0.15717671 |
| AHNAK | 822.694267 | -0.000799041 | 0.33870178 | -0.0023591 | 0.99811769 | 0.99954455 |
| B3GAT3 | 87.7555348 | -0.178684683 | 0.35554983 | -0.5025588 | 0.61527453 | 0.90792412 |
| GANAB | 262.533585 | 0.023144419 | 0.31059856 | 0.07451554 | 0.94060018 | 0.98855185 |
| UQCC3 | 86.4532355 | -0.199606521 | 0.35092024 | -0.5688088 | 0.56948591 | 0.89626255 |
| UBXN1 | 144.705994 | -0.189943727 | 0.44592395 | -0.4259554 | 0.67014032 | 0.92474338 |
| TTC9C | 88.587008 | -0.185079878 | 0.25389917 | -0.7289503 | 0.46603207 | 0.85028134 |
| POLR2G | 56.7975599 | 0.075079509 | 0.37676211 | 0.19927564 | 0.84204714 | 0.96876046 |
| TAF6L | 40.827168 | -0.14488889 | 0.37922024 | -0.3820706 | 0.70240902 | 0.93241442 |
| TMEM223 | 88.7392795 | -0.666420705 | 0.32583453 | -2.0452734 | 0.04082791 | 0.34079725 |
| TMEM179B | 124.92911 | -0.702993268 | 0.41426883 | -1.6969495 | 0.08970622 | 0.47358503 |
| NXF1 | 63.7178831 | 0.32359829 | 0.27746714 | 1.16625809 | 0.24351011 | 0.69975346 |
| STX5 | 169.767517 | 0.263188427 | 0.24116409 | 1.09132513 | 0.27512984 | 0.7263609 |
| SLC3A2 | 163.609417 | -1.308028767 | 0.43351207 | -3.0172834 | 0.00255051 | 0.0735219 |
| RARRES3 | 134.382666 | -2.287059283 | 0.84065944 | -2.7205538 | 0.00651727 | 0.12862701 |
| PLA2G16 | 60.25525 | -1.408751741 | 0.64279175 | -2.1916145 | 0.02840735 | 0.2862587 |
| ATL3 | 248.623087 | 0.037929746 | 0.16012333 | 0.23687832 | 0.81275119 | 0.96288868 |
| RTN3 | 782.01783 | 0.65547654 | 0.24805733 | 2.64243975 | 0.00823111 | 0.14386723 |
| MARK2 | 280.366073 | 0.320611032 | 0.19557049 | 1.63936303 | 0.10113768 | 0.49726048 |
| NAA40 | 93.935446 | 0.18400126 | 0.26024889 | 0.70702034 | 0.47955384 | 0.85532469 |
| OTUB1 | 119.351988 | 0.097045958 | 0.27018845 | 0.35917878 | 0.71946135 | 0.9362935 |
| MACROD1 | 169.764855 | 0.191627577 | 0.36536673 | 0.5244801 | 0.59994466 | 0.90369786 |
| AP000721.2 | 47.6889049 | 0.277908895 | 0.61601994 | 0.4511362 | 0.65189139 | 0.91887313 |
| STIP1 | 105.414879 | 0.162482311 | 0.25903063 | 0.62727064 | 0.53048185 | 0.88004345 |
| TRPT1 | 37.9672638 | -0.092689902 | 0.45795723 | -0.2023986 | 0.83960512 | 0.96833625 |
| NUDT22 | 78.9254131 | -0.013288593 | 0.29148271 | -0.0455896 | 0.96363732 | 0.99242523 |
| DNAJC4 | 68.1679777 | 0.329584568 | 0.34483195 | 0.95578316 | 0.3391818 | 0.77836439 |
| VEGFB | 186.929384 | -0.537736884 | 0.3678309 | -1.4619133 | 0.14376497 | 0.57324898 |
| FKBP2 | 209.29935 | 0.384682854 | 0.40945733 | 0.93949436 | 0.34747698 | 0.78180304 |
| PPP1R14B | 146.805245 | -0.207412463 | 0.54895025 | -0.3778347 | 0.70555338 | 0.93241442 |
| PLCB3 | 29.8169693 | 0.160935531 | 0.42937244 | 0.3748157 | 0.70779754 | 0.9330268 |
| BAD | 127.490127 | 0.204577314 | 0.45168859 | 0.45291672 | 0.65060871 | 0.91817658 |
| GPR137 | 54.7525843 | -0.114309974 | 0.32500477 | -0.3517178 | 0.72504989 | 0.93895336 |
| ESRRA | 161.634853 | -0.320918184 | 0.29635064 | -1.0829003 | 0.27885269 | 0.72938673 |
| TRMT112 | 228.435233 | 0.369903162 | 0.61513674 | 0.60133485 | 0.54761698 | 0.88773467 |
| PRDX5 | 340.44528 | 0.133228676 | 0.56673099 | 0.23508274 | 0.81414451 | 0.96303374 |
| SF1 | 506.217239 | -0.087133205 | 0.23853086 | -0.3652911 | 0.71489411 | 0.93573028 |
| MAP4K2 | 38.9373877 | 0.328020125 | 0.35285479 | 0.9296179 | 0.35256896 | 0.78314007 |
| MEN1 | 38.2092933 | -0.144191772 | 0.37410948 | -0.3854267 | 0.69992132 | 0.93180634 |
| EHD1 | 59.5541969 | 0.213130024 | 0.33205828 | 0.64184524 | 0.52097368 | 0.87650871 |
| ATG2A | 62.0808585 | -0.362486056 | 0.30584413 | -1.1851987 | 0.23593891 | 0.69335782 |
| PPP2R5B | 33.8907685 | -0.481764726 | 0.3617948 | -1.3315966 | 0.18299278 | 0.63376217 |
| ZFPL1 | 40.7434383 | -0.525808453 | 0.3250415 | -1.6176656 | 0.10573468 | 0.50651229 |
| VPS51 | 155.164645 | 0.11549123 | 0.22919518 | 0.50389903 | 0.61433233 | 0.90764425 |
| TM7SF2 | 73.578837 | 0.250014023 | 0.58284936 | 0.42895136 | 0.66795862 | 0.92422713 |
| FAU | 624.935209 | 0.182690944 | 0.43179596 | 0.42309553 | 0.67222556 | 0.92498948 |
| SYVN1 | 56.2037345 | 0.268940081 | 0.34688583 | 0.77529856 | 0.43816326 | 0.8347986 |
| MRPL49 | 72.1091832 | -0.022361406 | 0.39178956 | -0.057075 | 0.95448542 | 0.99060627 |
| CAPN1 | 251.786977 | -0.169884364 | 0.29044408 | -0.5849125 | 0.55860658 | 0.89179371 |
| CDC42EP2 | 47.7923626 | 0.330115859 | 0.41473525 | 0.79596768 | 0.42605082 | 0.82774157 |
| DPF2 | 99.606968 | 0.010898395 | 0.19579216 | 0.05566308 | 0.95561021 | 0.9908068 |
| FRMD8 | 61.546989 | -0.692799774 | 0.37806033 | -1.8325112 | 0.0668753 | 0.42042779 |
| NEAT1 | 18827.8969 | -0.972606951 | 0.60217714 | -1.6151509 | 0.10627802 | 0.50765436 |
| MALAT1 | 224543.617 | -0.302888517 | 0.41570602 | -0.7286123 | 0.46623886 | 0.85051057 |
| SCYL1 | 72.8052854 | 0.106946272 | 0.30289225 | 0.35308355 | 0.72402581 | 0.93893205 |
| LTBP3 | 236.141351 | 0.666550691 | 0.3041021 | 2.1918648 | 0.02838927 | 0.2862587 |
| SSSCA1 | 55.0013222 | -0.060595847 | 0.47383645 | -0.1278835 | 0.8982412 | 0.98312332 |
| FAM89B | 44.1169399 | -0.052229462 | 0.52294104 | -0.0998764 | 0.92044246 | 0.98581108 |
| EHBP1L1 | 40.8531187 | -1.074486204 | 0.56200689 | -1.9118737 | 0.05589239 | 0.39368206 |
| MAP3K11 | 73.7873818 | -0.01401732 | 0.29308822 | -0.0478263 | 0.96185469 | 0.99171131 |
| PCNX3 | 58.3388758 | 0.378519299 | 0.28961057 | 1.30699408 | 0.19121472 | 0.64458858 |
| SIPA1 | 59.3129637 | -0.201235316 | 0.52329594 | -0.3845536 | 0.70056821 | 0.93180634 |
| RELA | 95.1858308 | -0.310245925 | 0.32192936 | -0.9637081 | 0.3351923 | 0.77820994 |
| KAT5 | 97.4307568 | 0.058786553 | 0.24945278 | 0.23566205 | 0.81369492 | 0.96288868 |
| RNASEH2C | 82.3983814 | 0.217316368 | 0.51383699 | 0.42292862 | 0.67234734 | 0.92498948 |
| OVOL1 | 34.9653243 | 0.86585592 | 0.50392608 | 1.71822012 | 0.08575647 | 0.46493939 |
| CFL1 | 623.560423 | -0.660896413 | 0.49694468 | -1.3299195 | 0.1835448 | 0.63405547 |
| MUS81 | 51.5745846 | -0.445630924 | 0.2737958 | -1.6276032 | 0.10360905 | 0.50253415 |
| EFEMP2 | 83.8897973 | -0.109736021 | 0.3663644 | -0.299527 | 0.76453799 | 0.95378284 |
| FIBP | 37.9924586 | 0.689345601 | 0.43698277 | 1.57751207 | 0.11467775 | 0.52703872 |
| CCDC85B | 46.2585636 | 0.85475462 | 0.49894525 | 1.71312309 | 0.0866899 | 0.46672926 |
| C11orf68 | 38.7737309 | 0.110099221 | 0.47665621 | 0.23098245 | 0.81732843 | 0.96383504 |
| DRAP1 | 116.013683 | 0.216872822 | 0.51096791 | 0.42443531 | 0.67124838 | 0.92474338 |
| SART1 | 52.8870014 | 0.248939372 | 0.26664538 | 0.93359716 | 0.35051173 | 0.78223232 |
| BANF1 | 161.182147 | -0.377358947 | 0.48102673 | -0.7844864 | 0.43275474 | 0.83162989 |
| SF3B2 | 185.248602 | -0.210272109 | 0.1856471 | -1.1326442 | 0.2573637 | 0.71530073 |
| PACS1 | 640.559957 | -0.452001027 | 0.16732419 | -2.701349 | 0.00690588 | 0.13279571 |
| KLC2 | 46.8852697 | 0.257052612 | 0.27099196 | 0.94856176 | 0.34284355 | 0.78096003 |

| | | | | | | |
|---|---|---|---|---|---|---|
| RAB1B | 124.802807 | -0.317005304 | 0.31062224 | -1.0205493 | 0.30746802 | 0.75251398 |
| YIF1A | 51.6629999 | -0.388897547 | 0.54614147 | -0.7120821 | 0.47641396 | 0.85414749 |
| BRMS1 | 50.3605467 | 0.111437781 | 0.4341792 | 0.25666311 | 0.79743885 | 0.95991117 |
| B4GAT1 | 38.9705039 | 0.145750264 | 0.46996103 | 0.31013266 | 0.75646008 | 0.95072713 |
| SLC29A2 | 101.634922 | 0.497890774 | 0.32388811 | 1.53723078 | 0.1242368 | 0.54273186 |
| MRPL11 | 66.4760952 | -0.19182155 | 0.48935544 | -0.3919882 | 0.69506694 | 0.93125819 |
| ZDHHC24 | 64.2083033 | -0.428203739 | 0.28330827 | -1.511441 | 0.13067612 | 0.55395564 |
| CTSF | 96.2682079 | -0.226225798 | 0.47961114 | -0.4716859 | 0.63715102 | 0.91466334 |
| CCS | 117.2377 | 0.237982493 | 0.35399581 | 0.6722749 | 0.50140871 | 0.86804029 |
| RBM14-RBM | 67.6079391 | 0.427012675 | 0.35790507 | 1.19308921 | 0.23283444 | 0.69222834 |
| RBM4 | 57.3957572 | 0.143737683 | 0.38932321 | 0.36919885 | 0.71197951 | 0.9348115 |
| RBM4B | 92.1269623 | 0.153310348 | 0.27502985 | 0.55743168 | 0.57723253 | 0.89873672 |
| SPTBN2 | 988.286561 | 0.044619068 | 0.51955282 | 0.08587975 | 0.93156201 | 0.98822345 |
| C11orf80 | 2380.26282 | 0.426487531 | 0.41752375 | 1.02146891 | 0.30703233 | 0.75214355 |
| RCE1 | 85.6413523 | 0.873828979 | 0.39553157 | 2.2092522 | 0.0271571 | 0.27917411 |
| LRFN4 | 29.967122 | -0.288355938 | 0.6221481 | -0.4634844 | 0.64301719 | 0.91643619 |
| RHOD | 84.6532079 | 0.203195804 | 0.39276586 | 0.51734589 | 0.60491472 | 0.90428025 |
| KDM2A | 919.638631 | -0.239015022 | 0.27659618 | -0.8641299 | 0.38751656 | 0.8065026 |
| GRK2 | 73.8476833 | -0.23290515 | 0.25747347 | -0.9045792 | 0.36568835 | 0.79329304 |
| ANKRD13D | 94.8092464 | -0.098398975 | 0.2386983 | -0.4122316 | 0.6801697 | 0.92700898 |
| SSH3 | 89.4411122 | -0.583395915 | 0.31536309 | -1.8499182 | 0.06432535 | 0.41357646 |
| RAD9A | 399.135377 | 0.423210822 | 0.22502737 | 1.88070822 | 0.06001162 | 0.40367252 |
| PPP1CA | 171.17088 | -0.452936955 | 0.46062255 | -0.9833148 | 0.32545255 | 0.77073504 |
| RPS6KB2 | 49.4296768 | 0.188924285 | 0.38651104 | 0.48879402 | 0.62498753 | 0.91058133 |
| CORO1B | 112.808922 | -0.518805721 | 0.46702993 | -1.1108618 | 0.26662783 | 0.72072728 |
| TMEM134 | 144.619786 | -0.382412653 | 0.34148102 | -1.119865 | 0.26277128 | 0.71989632 |
| AIP | 81.1832803 | 0.431466697 | 0.49451881 | 0.87249805 | 0.38293673 | 0.80424388 |
| PITPNM1 | 94.9937908 | 0.665517348 | 0.54765124 | 1.21522111 | 0.22428177 | 0.68187922 |
| CDK2AP2 | 216.573843 | 0.898770744 | 0.65412248 | 1.37400986 | 0.16943862 | 0.61478828 |
| GSTP1 | 1591.21378 | -0.702968645 | 0.54098122 | -1.2994326 | 0.1937955 | 0.64723766 |
| NDUFV1 | 228.64654 | 0.307735009 | 0.26141182 | 1.17720387 | 0.23911414 | 0.69526124 |
| NUDT8 | 41.8155364 | -0.144761571 | 0.45610981 | -0.3173832 | 0.75095288 | 0.94796275 |
| UNC93B1 | 64.1387187 | -1.095557853 | 0.51735888 | -2.1175974 | 0.03420917 | 0.31256508 |
| NDUFS8 | 253.77333 | -0.117313973 | 0.44273847 | -0.2649735 | 0.79102986 | 0.95832571 |
| TCIRG1 | 53.5873239 | 0.374587969 | 0.36535269 | 1.02527772 | 0.30523216 | 0.75054742 |
| CHKA | 352.364449 | -0.200321995 | 0.27786966 | -0.7209207 | 0.47095829 | 0.85236058 |
| KMT5B | 350.976802 | 0.172622613 | 0.25016164 | 0.69004428 | 0.49016634 | 0.86055851 |
| LRP5 | 94.1972517 | -0.220900936 | 0.37051553 | -0.5961989 | 0.55104239 | 0.88916026 |
| PPP6R3 | 1240.39828 | 0.610248224 | 0.24771658 | 2.46349369 | 0.01375903 | 0.18923183 |
| TESMIN | 39.4784559 | -0.488848696 | 0.46516477 | -1.0509151 | 0.29329757 | 0.73870669 |
| CPT1A | 329.892351 | 0.315605904 | 0.29443643 | 1.07189827 | 0.28376573 | 0.73392518 |
| MRPL21 | 57.4974695 | -0.017319978 | 0.32596222 | -0.0531349 | 0.9576244 | 0.99106618 |
| IGHMBP2 | 47.1086879 | 0.124835705 | 0.3382089 | 0.36910828 | 0.71204702 | 0.9348115 |
| TPCN2 | 75.3105553 | 0.25873108 | 0.27641293 | 0.93603103 | 0.34925721 | 0.78180304 |
| CCND1 | 148.202442 | 0.015323482 | 0.5360764 | 0.02858451 | 0.97719597 | 0.99469867 |
| ORAOV1 | 68.3023656 | 0.039519034 | 0.29345901 | 0.13466628 | 0.89287573 | 0.98168537 |
| FADD | 37.8164569 | -0.381351217 | 0.37625222 | -1.0135521 | 0.31079654 | 0.75435382 |
| PPFIA1 | 513.924925 | 0.048763989 | 0.28382955 | 0.1718073 | 0.86358903 | 0.97461028 |
| AP002336.2 | 167.244788 | -0.017257647 | 0.43845659 | -0.03936 | 0.96860338 | 0.99303855 |
| AP000487.1 | 44.51998 | 0.468571553 | 0.37879453 | 1.23700717 | 0.21608442 | 0.67297615 |
| CTTN | 658.795311 | -0.612467493 | 0.33156723 | -1.8471895 | 0.06471968 | 0.41458575 |
| SHANK2 | 1517.45476 | -0.09053287 | 0.29837049 | -0.3034243 | 0.7615665 | 0.95366435 |
| DHCR7 | 54.7748678 | -0.613650351 | 0.70846556 | -0.8661682 | 0.38639792 | 0.80634213 |
| NADSYN1 | 94.3022612 | -0.532156273 | 0.22703011 | -2.3439898 | 0.01907869 | 0.23114993 |
| AP002495.2 | 259.071023 | 0.512662515 | 0.23465674 | 2.18473386 | 0.02890837 | 0.28768102 |
| RNF121 | 225.623866 | 0.278775991 | 0.22578115 | 1.23471773 | 0.21693557 | 0.67442445 |
| NUMA1 | 394.888538 | 0.192794053 | 0.24781796 | 0.77796644 | 0.4365888 | 0.83393108 |
| LAMTOR1 | 242.609422 | -0.520759489 | 0.41964056 | -1.2409656 | 0.21461847 | 0.67132603 |
| ANAPC15 | 49.8191832 | 0.012205868 | 0.49998279 | 0.02441258 | 0.98052352 | 0.99485782 |
| FOLR1 | 308.611435 | -0.124512348 | 0.70964476 | -0.1754573 | 0.86072033 | 0.97306893 |
| INPPL1 | 96.9726136 | 0.365131027 | 0.32764614 | 1.11440662 | 0.26510479 | 0.72029159 |
| CLPB | 85.5452889 | 0.131778494 | 0.26993714 | 0.48818215 | 0.62542082 | 0.91061758 |
| ARAP1 | 82.9669603 | -0.359026662 | 0.26868038 | -1.3362593 | 0.18146454 | 0.63181008 |
| ATG16L2 | 63.6293881 | 0.214764336 | 0.34918223 | 0.61504943 | 0.53852207 | 0.88265739 |
| FCHSD2 | 402.214446 | -0.472169681 | 0.21883428 | -2.1576586 | 0.03095438 | 0.29646676 |
| ARHGEF17 | 98.574831 | -0.147691493 | 0.35016059 | -0.4217822 | 0.67318403 | 0.92536314 |
| FAM168A | 455.990974 | 0.361506655 | 0.2788246 | 1.2965379 | 0.19479023 | 0.64860454 |
| PLEKHB1 | 45.1846953 | 0.421625638 | 0.63961428 | 0.65918734 | 0.50977548 | 0.87094607 |
| RAB6A | 836.254837 | 0.642062546 | 0.22277528 | 2.88210852 | 0.00395024 | 0.09740819 |
| MRPL48 | 131.568792 | 0.472394951 | 0.37601504 | 1.25631931 | 0.20900022 | 0.66656188 |
| COA4 | 122.956393 | -0.543915904 | 0.52516906 | -1.0356968 | 0.30034363 | 0.74729373 |
| PAAF1 | 125.923986 | 0.013209057 | 0.34859459 | 0.03789232 | 0.96977354 | 0.99337569 |
| UCP2 | 242.621086 | 0.41110946 | 0.43817452 | 0.93823223 | 0.34812508 | 0.78180304 |
| C2CD3 | 182.425209 | -0.099607171 | 0.17956494 | -0.5547139 | 0.57909037 | 0.89975022 |
| PPME1 | 279.530021 | -0.742607371 | 0.47266279 | -1.5711145 | 0.11615605 | 0.53066791 |
| P4HA3 | 55.0866041 | -0.66088417 | 0.6609026 | -0.9999721 | 0.317324 | 0.76134633 |
| PGM2L1 | 86.8026897 | -0.887649597 | 0.51496865 | -1.7236964 | 0.08476267 | 0.46310155 |
| POLD3 | 148.156689 | 0.025334957 | 0.29830065 | 0.08493095 | 0.93231629 | 0.98822345 |
| RNF169 | 271.704808 | -0.064786802 | 0.24079602 | -0.2690526 | 0.78788919 | 0.95801395 |
| XRRA1 | 272.94439 | 0.081486881 | 0.50680742 | 0.1607847 | 0.87226297 | 0.97622753 |
| SPCS2 | 307.576468 | 0.136543433 | 0.33280468 | 0.41028099 | 0.68159983 | 0.92760692 |
| NEU3 | 39.7413074 | -0.318378786 | 0.28530999 | -1.1159048 | 0.2644629 | 0.71989632 |
| ARRB1 | 103.357512 | 0.476145049 | 0.36459499 | 1.30595608 | 0.19156749 | 0.64458858 |
| RPS3 | 1048.10816 | 0.01071107 | 0.47256621 | 0.02266576 | 0.98191689 | 0.99507903 |
| SERPINH1 | 65.0310611 | -0.20325633 | 0.48567621 | -0.4185017 | 0.67558033 | 0.92671696 |
| MAP6 | 42.9410958 | -0.112864863 | 0.59970158 | -0.1882017 | 0.85071853 | 0.97017003 |
| UVRAG | 594.39936 | 0.463063399 | 0.36802058 | 1.25825409 | 0.20829988 | 0.66589349 |
| THAP12 | 66.9416968 | 0.61404955 | 0.24134739 | 2.54425606 | 0.01095108 | 0.16452817 |
| EMSY | 244.45519 | 0.18680441 | 0.2381435 | 0.78441953 | 0.43279398 | 0.83162989 |
| ACER3 | 275.916858 | -0.546875461 | 0.30389296 | -1.7995661 | 0.07192919 | 0.43230175 |
| CAPN5 | 101.222374 | -0.081909691 | 0.37424636 | -0.2188657 | 0.82675467 | 0.96545294 |
| PAK1 | 328.221599 | 0.841678756 | 0.4642868 | 1.81284232 | 0.06985614 | 0.42843167 |
| CLNS1A | 220.030507 | -0.092440129 | 0.20754494 | -0.4453981 | 0.65603206 | 0.92098622 |
| RSF1 | 687.403339 | 0.220243705 | 0.17665106 | 1.24677263 | 0.21248088 | 0.66984203 |
| AAMDC | 183.06409 | -0.087112427 | 0.27143801 | -0.3209294 | 0.74826392 | 0.94678489 |
| AP002812.2 | 65.8200269 | -0.063352281 | 0.4848324 | -0.1306684 | 0.89603762 | 0.98291229 |
| INTS4 | 152.850736 | 0.025972454 | 0.23612643 | 0.10999384 | 0.91241426 | 0.98480378 |
| ALG8 | 82.893568 | -0.347350956 | 0.28320378 | -1.2265053 | 0.22000856 | 0.67801071 |
| GAB2 | 327.164242 | -1.37117514 | 0.6305197 | -2.1746746 | 0.02965451 | 0.29105042 |
| AP003086.2 | 103.420599 | -0.523418737 | 0.83721253 | -0.6251922 | 0.53184493 | 0.88045351 |
| NARS2 | 210.751347 | 0.402341519 | 0.20895465 | 1.92549684 | 0.05416723 | 0.39095834 |
| PRCP | 141.415427 | -0.144868586 | 0.26829708 | -0.5399559 | 0.58922746 | 0.90233654 |
| RAB30 | 125.802258 | 0.706064361 | 0.40117206 | 1.76000381 | 0.07840716 | 0.44789124 |
| RAB30-AS1 | 251.542512 | -0.155858975 | 0.32274941 | -0.4829102 | 0.62915951 | 0.9114564 |
| PCF11 | 218.13641 | 0.0332525 | 0.26339322 | 0.12624661 | 0.89953672 | 0.98312332 |
| ANKRD42 | 121.849794 | 0.040417504 | 0.21875427 | 0.18476213 | 0.85341562 | 0.97062426 |
| CCDC90B | 108.398668 | -0.335190818 | 0.2563091 | -1.3077601 | 0.19095468 | 0.64458858 |
| AP000446.1 | 64.2039768 | -0.272971503 | 0.29645718 | -0.9207788 | 0.3571659 | 0.78612469 |
| DLG2 | 2470.36525 | 0.840101743 | 0.5484159 | 1.53186978 | 0.12555457 | 0.54417366 |
| TMEM126B | 75.8287177 | -0.555277141 | 0.32631264 | -1.7016722 | 0.08881684 | 0.47038308 |
| TMEM126A | 42.5607351 | 0.541630583 | 0.4556825 | 1.18861397 | 0.23459161 | 0.69335782 |
| CREBZF | 190.41495 | 0.121334954 | 0.23556229 | 0.5150865 | 0.60649257 | 0.90493874 |
| SYTL2 | 1274.72603 | -0.369066992 | 0.55979454 | -0.6592901 | 0.5097095 | 0.87094607 |

| | | | | | | |
|---|---|---|---|---|---|---|
| PICALM | 1571.23473 | 0.850392873 | 0.23952017 | 3.55040197 | 0.00038464 | 0.02099512 |
| EED | 94.1927131 | 0.402615264 | 0.32439682 | 1.24111964 | 0.21456156 | 0.67132603 |
| HIKESHI | 110.707779 | 0.224620558 | 0.22908256 | 0.98052229 | 0.32682837 | 0.77195432 |
| AP001831.1 | 102.297165 | -1.417914474 | 0.60336497 | -2.3500113 | 0.01877284 | 0.22823596 |
| ME3 | 234.673569 | 0.148851293 | 0.39593367 | 0.37595007 | 0.70695401 | 0.93274945 |
| AP003059.1 | 35.7223638 | -0.265929606 | 0.70730444 | -0.3759762 | 0.70693462 | 0.93274945 |
| PRSS23 | 264.027381 | -0.135311905 | 0.48003236 | -0.2818808 | 0.77803491 | 0.95601455 |
| AP001528.2 | 250.32665 | -0.463402522 | 0.577781 | -0.8020383 | 0.42253078 | 0.82757457 |
| FZD4 | 268.708683 | 0.732889191 | 0.541801 | 1.35269074 | 0.17615445 | 0.62516039 |
| AP001528.1 | 45.6401468 | -0.045949744 | 0.43951891 | -0.1045455 | 0.91673643 | 0.98520052 |
| TMEM135 | 457.344751 | 0.394788244 | 0.23696967 | 1.66598636 | 0.09571613 | 0.48710328 |
| RAB38 | 55.6154427 | 0.19547151 | 0.46395634 | 0.42131445 | 0.67352548 | 0.92557344 |
| CTSC | 168.113204 | -0.402286135 | 0.34362253 | -1.1707211 | 0.24171091 | 0.69763987 |
| NOX4 | 98.9245513 | 1.270880108 | 0.58566135 | 2.16999142 | 0.0300075 | 0.29169628 |
| CHORDC1 | 87.5376401 | 0.890121539 | 0.31736794 | 2.80469901 | 0.00503636 | 0.11375224 |
| SLC36A4 | 232.554241 | -0.416859354 | 0.3209196 | -1.2989526 | 0.19396019 | 0.64756381 |
| SMCO4 | 181.294364 | -0.326490477 | 0.39466281 | -0.8272644 | 0.40808724 | 0.82040175 |
| CEP295 | 151.055061 | 0.13506389 | 0.29521951 | 0.45750326 | 0.64730936 | 0.91743182 |
| TAF1D | 299.128454 | -0.167576322 | 0.31885705 | -0.5255531 | 0.59919872 | 0.90369786 |
| PANX1 | 39.4229649 | -0.523515315 | 0.34877029 | -1.5010318 | 0.13334732 | 0.55619258 |
| MRE11 | 179.160796 | 0.268976761 | 0.22389607 | 1.20134651 | 0.22961682 | 0.68857352 |
| ANKRD49 | 35.8311677 | 0.335203135 | 0.43010201 | 0.77935729 | 0.43576928 | 0.83364877 |
| AMOTL1 | 137.012411 | -0.732101747 | 0.54473425 | -1.3439613 | 0.17896091 | 0.62831166 |
| CWC15 | 91.1778147 | -0.103928541 | 0.29674329 | -0.3502305 | 0.72616574 | 0.93895336 |
| SRSF8 | 76.0391114 | -0.344087608 | 0.42729435 | -0.8052707 | 0.42066351 | 0.82694802 |
| ENDOD1 | 51.9154382 | -0.643387088 | 0.30359655 | -2.1192174 | 0.0340721 | 0.3120805 |
| AP000787.1 | 134.45786 | 0.743832382 | 0.57967535 | 1.28318789 | 0.19942626 | 0.65475264 |
| SESN3 | 348.450253 | 1.000113638 | 0.71407741 | 1.40056753 | 0.16134344 | 0.60092341 |
| FAM76B | 85.9257215 | 0.235561159 | 0.29517065 | 0.79805076 | 0.42484103 | 0.82774157 |
| CEP57 | 313.55144 | 0.046860683 | 0.15337076 | 0.30553858 | 0.75995599 | 0.95297299 |
| MTMR2 | 206.670052 | 0.185569319 | 0.2459449 | 0.75451582 | 0.45053955 | 0.84153105 |
| MAML2 | 2870.0755 | 0.604774939 | 0.35511677 | 1.70303121 | 0.08856223 | 0.47038308 |
| CCDC82 | 166.746381 | 0.102061073 | 0.32539775 | 0.31365021 | 0.75378672 | 0.94970964 |
| JRKL | 52.1733259 | -0.559221674 | 0.34897372 | -1.602475 | 0.10905061 | 0.51387157 |
| CNTN5 | 69.1506959 | 1.320899554 | 0.69090521 | 1.91183905 | 0.05589684 | 0.39368206 |
| ARHGAP42 | 146.396084 | 2.399651794 | 0.96238781 | 2.49343536 | 0.01265136 | 0.17941309 |
| CEP126 | 104.734017 | 0.05877274 | 0.38915719 | 0.15102571 | 0.87995544 | 0.97740892 |
| CFAP300 | 93.4926174 | -0.072478005 | 0.48496185 | -0.1494509 | 0.88119782 | 0.97768262 |
| YAP1 | 1428.90631 | 0.02356465 | 0.31467611 | 0.07488541 | 0.94030589 | 0.98855185 |
| BIRC3 | 183.216604 | -0.127824831 | 0.94584968 | -0.1351429 | 0.89249892 | 0.98168537 |
| BIRC2 | 164.165388 | 0.444192229 | 0.3431907 | 1.29430146 | 0.19556131 | 0.64948524 |
| TMEM123 | 519.25002 | -0.49942636 | 0.16876722 | -2.9592616 | 0.00308377 | 0.08286649 |
| MMP7 | 447.803752 | -2.897611325 | 0.77503658 | -3.7386769 | 0.00018499 | 0.01293366 |
| DCUN1D5 | 97.9689193 | -0.048668154 | 0.20015723 | -0.2431496 | 0.8078895 | 0.96245698 |
| DYNC2H1 | 731.83034 | 0.367709498 | 0.27505219 | 1.33687174 | 0.18126451 | 0.63174328 |
| PDGFD | 199.139512 | 1.005019295 | 0.6034555 | 1.66544061 | 0.09582488 | 0.48710328 |
| CASP4 | 158.480091 | -0.992876218 | 0.35570875 | -2.7912618 | 0.0052503 | 0.11544342 |
| MSANTD4 | 49.7417872 | -0.302428292 | 0.27817092 | -1.0872031 | 0.27694705 | 0.72743146 |
| KBTBD3 | 64.0318587 | 0.432944172 | 0.26395403 | 1.64022568 | 0.10095825 | 0.49726048 |
| AASDHPPT | 116.23223 | 0.461383866 | 0.31198544 | 1.47886343 | 0.13917682 | 0.56656585 |
| CWF19L2 | 339.631128 | 0.223608388 | 0.23322859 | 0.95875206 | 0.33768367 | 0.77827084 |
| ALKBH8 | 55.4000799 | 0.159536603 | 0.2520262 | 0.63301595 | 0.52672323 | 0.87791848 |
| SLC35F2 | 35.0067634 | -0.30330821 | 0.35721386 | -0.8490942 | 0.39582889 | 0.8120904 |
| CUL5 | 559.461897 | 0.374866127 | 0.18911819 | 1.98217909 | 0.0474592 | 0.367879 |
| ACAT1 | 229.747277 | 0.247222124 | 0.29587451 | 0.83556412 | 0.40340016 | 0.81828987 |
| NPAT | 171.158569 | 0.000157456 | 0.1824647 | 0.00086294 | 0.99931147 | 0.99992293 |
| ATM | 374.559224 | 0.024523811 | 0.25060438 | 0.09785867 | 0.92204452 | 0.98652645 |
| C11orf65 | 116.120523 | -0.173319296 | 0.44927415 | -0.3857762 | 0.69966239 | 0.93180634 |
| KDELC2 | 70.6315679 | 0.519913372 | 0.40329276 | 1.2891711 | 0.19733861 | 0.6524002 |
| EXPH5 | 273.147655 | -0.159059993 | 0.37286191 | -0.4265922 | 0.66967636 | 0.92474338 |
| DDX10 | 272.961586 | 0.55483825 | 0.31683074 | 1.75121343 | 0.07990915 | 0.45072547 |
| RDX | 500.799086 | 0.769890479 | 0.42420576 | 1.8148987 | 0.06953948 | 0.42828237 |
| ZC3H12C | 313.639648 | 1.394055053 | 0.63892264 | 2.18188396 | 0.0291181 | 0.28842879 |
| FDX1 | 322.250276 | 0.880965387 | 0.37957966 | 2.32089726 | 0.02029239 | 0.24014392 |
| COLCA1 | 73.9167686 | 1.004298725 | 0.50440525 | 1.99105527 | 0.04647481 | 0.36244837 |
| COLCA2 | 59.4676875 | 0.154542113 | 0.32596854 | 0.47410131 | 0.63542766 | 0.91454294 |
| SIK2 | 1051.21931 | 1.37318444 | 0.58125445 | 2.36244976 | 0.0181546 | 0.2244336 |
| PPP2R1B | 150.687711 | 0.234793713 | 0.28237794 | 0.83148744 | 0.40569832 | 0.82002585 |
| C11orf1 | 47.5946676 | 0.035569659 | 0.54868494 | 0.06482711 | 0.94831166 | 0.98977092 |
| CRYAB | 863.436759 | -0.304156637 | 0.93169315 | -0.3264558 | 0.74407952 | 0.9456489 |
| DIXDC1 | 189.749746 | -0.275988272 | 0.27748222 | -0.9946161 | 0.31992303 | 0.76477952 |
| DLAT | 109.12129 | 0.133330852 | 0.26728794 | 0.49882854 | 0.61790018 | 0.90794578 |
| NKAPD1 | 58.7058833 | 0.10078883 | 0.2769504 | 0.36392376 | 0.71591494 | 0.93573028 |
| TIMM8B | 95.6979541 | -0.522700066 | 0.392422 | -1.3319846 | 0.18286525 | 0.63358396 |
| AP002884.3 | 55.805209 | -0.268024045 | 0.68907971 | -0.3889594 | 0.69730617 | 0.93140054 |
| PTS | 198.794858 | -0.025986585 | 0.39248321 | -0.0662107 | 0.94721009 | 0.98977092 |
| AP002884.1 | 115.960833 | -0.253924593 | 0.55863285 | -0.4545465 | 0.64943555 | 0.91788058 |
| TTC12 | 163.496672 | 0.135620743 | 0.29218115 | 0.46416664 | 0.64252836 | 0.91643619 |
| ZW10 | 57.9641941 | -0.161872256 | 0.23927799 | -0.6765029 | 0.49872139 | 0.86605292 |
| USP28 | 106.387366 | -0.031448089 | 0.21355832 | -0.1472576 | 0.88292869 | 0.97874896 |
| ZBTB16 | 129.819137 | 0.926876727 | 0.86694625 | 1.06912825 | 0.28501188 | 0.73533347 |
| NNMT | 64.405721 | -3.943198445 | 0.97082511 | -4.061698 | 4.87E-05 | 0.00576513 |
| REXO2 | 117.543914 | -0.257879095 | 0.43409242 | -0.594065 | 0.55246866 | 0.88951784 |
| CADM1 | 549.829778 | -0.609940679 | 0.55559697 | -1.0978114 | 0.27228686 | 0.72405776 |
| BUD13 | 37.5108492 | 0.263940448 | 0.29510666 | 0.89439001 | 0.37111325 | 0.79539603 |
| ZPR1 | 36.939489 | -0.07195191 | 0.33027806 | -0.2178525 | 0.82754402 | 0.96545294 |
| SIK3 | 2603.0962 | 1.056927714 | 0.3313687 | 3.18958223 | 0.00142479 | 0.05016683 |
| PAFAH1B2 | 315.961112 | 0.179327532 | 0.21731797 | 0.82518499 | 0.40926657 | 0.82124403 |
| SIDT2 | 74.7886947 | 0.128704418 | 0.24251379 | 0.53070968 | 0.59561998 | 0.90323766 |
| PCSK7 | 153.169325 | 0.055139684 | 0.32324938 | 0.1705794 | 0.8645545 | 0.97477474 |
| RNF214 | 138.997973 | -0.114048959 | 0.28647344 | -0.3981136 | 0.69054648 | 0.93073907 |
| BACE1 | 84.556561 | -0.345950014 | 0.33968896 | -1.0184317 | 0.30847283 | 0.75313873 |
| CEP164 | 128.122268 | 0.065115112 | 0.27886755 | 0.23349835 | 0.81537445 | 0.96333253 |
| FXYD2 | 47.4998657 | -0.038655788 | 0.70795892 | -0.0546017 | 0.95645575 | 0.9908068 |
| MPZL3 | 71.3333676 | -0.039173702 | 0.31681641 | -0.123648 | 0.90159402 | 0.98320957 |
| MPZL2 | 194.723634 | -1.78898415 | 0.37534818 | -4.766199 | 1.88E-06 | 0.00057866 |
| UBE4A | 177.259462 | 0.244443722 | 0.19650514 | 1.24395585 | 0.21351581 | 0.67075711 |
| ATP5MG | 500.247101 | 0.050730837 | 0.40095579 | 0.12652476 | 0.89931655 | 0.98312332 |
| KMT2A | 629.226354 | -0.191871878 | 0.18944743 | -1.0127975 | 0.31115692 | 0.75435382 |
| TMEM25 | 51.8907413 | -0.355987342 | 0.34293221 | -1.0380691 | 0.29923788 | 0.74594723 |
| IFT46 | 73.5434508 | 0.312227368 | 0.34734009 | 0.89890969 | 0.36870077 | 0.79532503 |
| ARCN1 | 123.847445 | -0.223350599 | 0.32996126 | -0.6768995 | 0.49846974 | 0.86603149 |
| PHLDB1 | 157.673901 | 0.956327354 | 0.3897943 | 2.45341545 | 0.01415068 | 0.19259627 |
| AP002954.1 | 71.7136977 | -0.200248707 | 0.4822529 | -0.4152359 | 0.67796923 | 0.9268955 |
| DDX6 | 377.607007 | -0.028758926 | 0.15046074 | -0.1911391 | 0.84841664 | 0.96991131 |
| BCL9L | 67.4811834 | -0.244944823 | 0.37310105 | -0.6565107 | 0.5114956 | 0.87151901 |
| CCDC84 | 184.848588 | 0.301009966 | 0.29653163 | 1.01510238 | 0.31005703 | 0.75409553 |
| RPS25 | 624.950265 | -0.052319587 | 0.50373116 | -0.1038641 | 0.91727719 | 0.98520052 |
| TRAPPC4 | 146.611483 | 0.463292454 | 0.51311052 | 0.90290968 | 0.36657383 | 0.7935157 |
| HYOU1 | 64.0574497 | -0.562306941 | 0.32661813 | -1.7216036 | 0.08514136 | 0.46310155 |
| VPS11 | 58.689245 | 0.122140561 | 0.27507274 | 0.44403004 | 0.65702086 | 0.92160948 |
| H2AFX | 34.2059393 | -0.773567913 | 0.47766881 | -1.619465 | 0.10534726 | 0.50574404 |
| DPAGT1 | 36.3628258 | -0.083797053 | 0.42806509 | -0.1957577 | 0.84479979 | 0.96876046 |

| | | | | | | |
|---|---|---|---|---|---|---|
| HINFP | 51.0666698 | -0.052014998 | 0.26558337 | -0.1958519 | 0.84472611 | 0.96876046 |
| NLRX1 | 50.8538302 | -0.450243034 | 0.36766542 | -1.2245999 | 0.22072601 | 0.67871976 |
| CBL | 191.852584 | -0.008750267 | 0.34905791 | -0.0250682 | 0.98000054 | 0.99469867 |
| RNF26 | 39.4762661 | -0.915248282 | 0.29334805 | -3.1200081 | 0.00180846 | 0.05874854 |
| NECTIN1 | 34.3881947 | 0.620420326 | 0.42771822 | 1.4505352 | 0.14690933 | 0.57793161 |
| OAF | 164.378073 | 0.707721661 | 0.63617055 | 1.11247159 | 0.26593544 | 0.72053648 |
| TMEM136 | 64.4765532 | 0.179179689 | 0.39618025 | 0.4522681 | 0.65107585 | 0.91846479 |
| ARHGEF12 | 1313.66969 | 0.421933938 | 0.23291788 | 1.81151371 | 0.07006137 | 0.42862997 |
| SC5D | 77.5136632 | -0.129886156 | 0.36100326 | -0.3597922 | 0.71900254 | 0.9359691 |
| SORL1 | 602.998718 | 0.816559784 | 0.32090697 | 2.54453736 | 0.01094226 | 0.16452817 |
| JHY | 70.5165624 | -0.951547545 | 0.35829735 | -2.6557482 | 0.00791327 | 0.14172148 |
| HSPA8 | 460.815413 | 0.9939113 | 0.49237775 | 2.01859509 | 0.04352932 | 0.35177679 |
| VWA5A | 86.7006699 | -0.872595988 | 0.25484152 | -3.424073 | 0.0006169 | 0.02873385 |
| TBRG1 | 138.188761 | 0.191526885 | 0.27198464 | 0.70418272 | 0.48131899 | 0.85611389 |
| SIAE | 142.085814 | -0.250361241 | 0.3794271 | -0.6598402 | 0.5093564 | 0.87094607 |
| SPA17 | 44.0823678 | -0.410257424 | 0.33573262 | -1.2219766 | 0.22171646 | 0.6794174 |
| MSANTD2 | 190.619906 | 0.512874842 | 0.26192789 | 1.95807645 | 0.05022104 | 0.37837277 |
| TMEM218 | 40.723017 | 0.39081989 | 0.34699264 | 1.12630598 | 0.26003603 | 0.71715199 |
| EI24 | 188.433268 | -0.470344844 | 0.3083495 | -1.5253627 | 0.12716865 | 0.54746778 |
| STT3A | 86.8030252 | -0.109371016 | 0.29144258 | -0.3752747 | 0.7074562 | 0.93277156 |
| FAM118B | 66.0993527 | -0.716842623 | 0.32498417 | -2.205777 | 0.02739962 | 0.2806909 |
| SRPRA | 118.389849 | -0.340198413 | 0.30573213 | -1.1127336 | 0.26582286 | 0.72053648 |
| FOXRED1 | 65.1982595 | -0.328345088 | 0.34762041 | -0.9445507 | 0.34488833 | 0.78180304 |
| TIRAP | 39.7175287 | 0.326802989 | 0.34469226 | 0.94810074 | 0.34307817 | 0.78096612 |
| DCPS | 329.275457 | 0.20813071 | 0.77993747 | 0.26685564 | 0.78958032 | 0.95801395 |
| ST3GAL4 | 485.308978 | -1.16709105 | 0.98520332 | -1.1846195 | 0.23616793 | 0.69335782 |
| ETS1 | 260.169387 | -0.245203716 | 0.55809689 | -0.4393569 | 0.66040295 | 0.92284375 |
| KCNJ1 | 597.333512 | -0.657455617 | 0.61330352 | -1.0719906 | 0.28372425 | 0.73392518 |
| ARHGAP32 | 582.591592 | 0.187530597 | 0.2339979 | 0.80141999 | 0.42288855 | 0.82774157 |
| BARX2 | 100.894809 | -0.425518583 | 0.90088041 | -0.4723364 | 0.63668672 | 0.91454294 |
| TMEM45B | 381.786012 | -0.444042644 | 0.50537422 | -0.8786413 | 0.37959582 | 0.80285856 |
| NFRKB | 101.800922 | 0.234298993 | 0.20797385 | 1.1265791 | 0.25992048 | 0.71702201 |
| PRDM10 | 111.819916 | 0.162300649 | 0.33405272 | 0.4858534 | 0.62707111 | 0.91061758 |
| APLP2 | 843.818097 | -0.551587025 | 0.29370931 | -1.8780032 | 0.06038073 | 0.40556611 |
| ST14 | 270.33243 | -0.104323387 | 0.25286246 | -0.4125697 | 0.67992191 | 0.92700898 |
| ZBTB44 | 510.992683 | 0.403982981 | 0.2981832 | 1.35481469 | 0.17547661 | 0.62477136 |
| AP002986.1 | 44.321867 | 0.054647105 | 0.71528546 | 0.07639902 | 0.93910165 | 0.98848592 |
| SNX19 | 178.302856 | 0.537272034 | 0.36337337 | 1.47856746 | 0.13925596 | 0.56656585 |
| JAM3 | 177.049755 | 1.090758556 | 0.31753099 | 3.43512473 | 0.00059228 | 0.02774786 |
| NCAPD3 | 69.974597 | 0.2444637 | 0.28350298 | 0.86229676 | 0.38852424 | 0.8065026 |
| VPS26B | 112.357964 | 0.583339167 | 0.33792987 | 1.72621369 | 0.08430898 | 0.46162908 |
| THYN1 | 36.9191514 | 0.052723084 | 0.36189635 | 0.14568559 | 0.8841696 | 0.9788678 |
| ACAD8 | 67.9377749 | 0.629641293 | 0.36934085 | 1.70477024 | 0.08823728 | 0.47036301 |
| ZMYND11 | 600.617512 | 0.616238637 | 0.25816478 | 2.38699735 | 0.01698661 | 0.21364736 |
| DIP2C | 1290.97161 | -0.043563683 | 0.48827875 | -0.0892189 | 0.92890796 | 0.98761849 |
| LARP4B | 953.146223 | 0.435220009 | 0.20404216 | 2.13299064 | 0.03292549 | 0.30590353 |
| GTPBP4 | 81.3476043 | -0.036341119 | 0.24238345 | -0.1499323 | 0.88081799 | 0.97765013 |
| IDI1 | 251.98914 | 0.428772312 | 0.71019999 | 0.6037346 | 0.5460201 | 0.88669883 |
| WDR37 | 263.220433 | -0.143473422 | 0.2507489 | -0.5721797 | 0.56720026 | 0.89454608 |
| PFKP | 640.266538 | -0.438379264 | 0.82979368 | -0.5282991 | 0.59729175 | 0.90369786 |
| PITRM1 | 94.9404253 | -0.207737363 | 0.2812763 | -0.7385527 | 0.46017867 | 0.84657145 |
| KLF6 | 1108.94764 | 0.024467817 | 0.72564597 | 0.03371867 | 0.97310149 | 0.9939318 |
| AKR1C1 | 169.834971 | 2.893542275 | 1.53799631 | 1.88137141 | 0.05992141 | 0.40367252 |
| AKR1C3 | 120.514467 | -0.478154105 | 0.67608852 | -0.7072359 | 0.47941986 | 0.85532469 |
| NET1 | 48.3799193 | 0.454253544 | 0.34490718 | 1.31703129 | 0.18782814 | 0.64035097 |
| ASB13 | 43.3780838 | 0.008175749 | 0.40813198 | 0.02003212 | 0.98401775 | 0.99540861 |
| FAM208B | 416.107007 | 0.295068215 | 0.22958137 | 1.2852446 | 0.19870682 | 0.65403502 |
| GDI2 | 322.452808 | -0.307926869 | 0.21381967 | -1.4401242 | 0.14983227 | 0.58336854 |
| ANKRD16 | 42.3599019 | -0.336619547 | 0.34250422 | -0.9828187 | 0.32569668 | 0.77073504 |
| FBH1 | 210.127434 | 0.05961441 | 0.19312588 | 0.30868163 | 0.75756372 | 0.95149422 |
| IL15RA | 52.4774455 | -1.16673234 | 0.43093845 | -2.7074222 | 0.0067808 | 0.1314808 |
| RBM17 | 194.525334 | 0.158751563 | 0.23738041 | 0.66876437 | 0.5036458 | 0.86946722 |
| PFKFB3 | 2322.21366 | -0.139844087 | 0.30348493 | -0.4607942 | 0.64494629 | 0.91682456 |
| AL137145.2 | 380.657653 | -0.61882558 | 0.50140113 | -1.2341926 | 0.21713114 | 0.67475613 |
| AL137145.1 | 43.0030958 | -0.1247419 | 0.59876847 | -0.2083308 | 0.8349707 | 0.96754677 |
| PRKCQ | 160.720933 | -0.244300606 | 0.47744415 | -0.5116842 | 0.60887208 | 0.90549306 |
| SFMBT2 | 272.087861 | 0.673791254 | 0.353907 | 1.9038653 | 0.05692773 | 0.39727338 |
| KIN | 81.0227603 | 0.064804543 | 0.21942047 | 0.29534411 | 0.76773102 | 0.95408765 |
| ATP5F1C | 389.053221 | -0.030719285 | 0.20212713 | -0.15198 | 0.87920269 | 0.97699546 |
| TAF3 | 470.925771 | 0.562917743 | 0.17606473 | 3.19722041 | 0.00138759 | 0.04915682 |
| GATA3 | 2027.51207 | 0.732926707 | 0.27301853 | 2.68453095 | 0.00726317 | 0.13601674 |
| GATA3-AS1 | 238.936081 | 0.348841678 | 0.493396 | 0.7070217 | 0.47955299 | 0.85532469 |
| USP6NL | 386.331819 | 0.252546844 | 0.15193937 | 1.66215537 | 0.09648163 | 0.48810007 |
| ECHDC3 | 87.9688896 | -0.30910301 | 0.55694854 | -0.5549938 | 0.57889887 | 0.89975022 |
| PROSER2 | 157.986669 | -0.262029352 | 0.40371236 | -0.6490496 | 0.5163063 | 0.87455795 |
| UPF2 | 334.986443 | 0.173342309 | 0.19174709 | 0.90401534 | 0.36598727 | 0.79329304 |
| DHTKD1 | 130.084938 | -0.427469359 | 0.21994835 | -1.9434988 | 0.05195592 | 0.38290999 |
| SEC61A2 | 57.7890785 | 0.519805582 | 0.26093664 | 1.99207584 | 0.04636274 | 0.36205774 |
| NUDT5 | 133.240616 | -0.340395395 | 0.24253686 | -1.403479 | 0.16047405 | 0.60010648 |
| CDC123 | 136.734322 | 0.065409495 | 0.21077141 | 0.31033381 | 0.75630713 | 0.95072713 |
| CAMK1D | 100.244939 | -0.368365828 | 0.82229813 | -0.4479711 | 0.65417403 | 0.92002724 |
| OPTN | 136.597028 | -0.498395506 | 0.38836647 | -1.2833124 | 0.19938265 | 0.65475264 |
| PHYH | 109.2179 | 0.391831635 | 0.40262867 | 0.97318365 | 0.33046203 | 0.77618491 |
| SEPHS1 | 116.123137 | 0.123187839 | 0.24404671 | 0.50477157 | 0.61371929 | 0.90751853 |
| BEND7 | 307.2374 | -0.183363465 | 0.2256722 | -0.8125213 | 0.41649259 | 0.82489761 |
| PRPF18 | 39.770121 | -0.735641163 | 0.36365143 | -2.0229294 | 0.04308043 | 0.34966009 |
| AL157392.5 | 93.4287454 | 0.056118411 | 0.31795219 | 0.17649953 | 0.85990152 | 0.97256291 |
| FRMD4A | 347.695138 | 0.377201195 | 0.47294794 | 0.79755331 | 0.42512976 | 0.82774157 |
| FAM107B | 1914.25123 | 0.923067438 | 0.42944835 | 2.14942599 | 0.03160065 | 0.29862469 |
| DCLRE1C | 103.552469 | -0.212290526 | 0.29574948 | -0.7178052 | 0.47287738 | 0.85298656 |
| RPP38 | 152.737965 | 0.165396737 | 0.22849635 | 0.7238485 | 0.46915875 | 0.8522644 |
| NMT2 | 200.526034 | 0.148661477 | 0.39712646 | 0.37434292 | 0.70814921 | 0.9332187 |
| FAM171A1 | 805.499663 | -0.026889337 | 0.41591381 | -0.0646512 | 0.9484517 | 0.98977092 |
| MINDY3 | 220.313317 | 0.500826522 | 0.21151674 | 2.36778669 | 0.01789485 | 0.22246504 |
| PTER | 274.563172 | -0.499087133 | 0.21642388 | -2.3060632 | 0.0211071 | 0.24605388 |
| RSU1 | 253.44206 | -0.310626627 | 0.35998554 | -0.8628864 | 0.38819993 | 0.8065026 |
| CUBN | 114.15157 | 1.135235738 | 0.55637686 | 2.04040789 | 0.04130972 | 0.34244985 |
| TRDMT1 | 57.4648111 | 0.211366896 | 0.302331 | 0.69912412 | 0.48447447 | 0.85909499 |
| VIM-AS1 | 36.1911455 | -0.472165583 | 0.58453121 | -0.807768 | 0.41922417 | 0.82694802 |
| VIM | 169.390309 | -2.0233077 | 0.66589276 | -3.0384888 | 0.00237768 | 0.07079 |
| HACD1 | 312.586652 | 0.022135758 | 0.59180915 | 0.03740354 | 0.97016325 | 0.99348357 |
| STAM | 807.762315 | 0.464373897 | 0.65129429 | 0.71300164 | 0.47584474 | 0.85414749 |
| AL450384.2 | 63.6309485 | -0.396802738 | 0.44182018 | -0.8981091 | 0.36912737 | 0.79532503 |
| NSUN6 | 212.901292 | -0.034124148 | 0.29777967 | -0.1145953 | 0.90876591 | 0.98427726 |
| ARL5B | 104.803556 | 0.039213048 | 0.76103536 | 0.05152592 | 0.95890645 | 0.99119981 |
| MALRD1 | 164.9398 | 0.021471234 | 0.55744142 | 0.03851747 | 0.9692751 | 0.99325345 |
| PLXDC2 | 857.703511 | 0.062050448 | 0.27903793 | 0.22237281 | 0.82402368 | 0.96511681 |
| NEBL | 1355.26891 | 0.137549009 | 0.47151388 | 0.29171784 | 0.77050237 | 0.95482817 |
| MLLT10 | 855.646491 | 0.224675744 | 0.28298523 | 0.79394866 | 0.42722532 | 0.82851986 |
| DNAJC1 | 650.620743 | 0.331515227 | 0.18791702 | 1.76415756 | 0.07770546 | 0.44622092 |
| BMI1 | 48.4528218 | 0.719633541 | 0.35867699 | 2.00635549 | 0.04481833 | 0.35515061 |
| PIP4K2A | 488.982773 | 0.240611982 | 0.29421328 | 0.81781482 | 0.41346294 | 0.82440569 |
| MSRB2 | 147.374194 | 0.114341935 | 0.30295677 | 0.37741997 | 0.70586153 | 0.93241442 |

| | | | | | | |
|---|---|---|---|---|---|---|
| OTUD1 | 63.1224998 | 0.282779778 | 0.32851175 | 0.86079047 | 0.38935346 | 0.80680464 |
| KIAA1217 | 2355.90023 | 0.616470265 | 0.23902281 | 2.57912737 | 0.00990503 | 0.15945418 |
| ARHGAP21 | 452.597883 | -0.242574968 | 0.35225182 | -0.6886408 | 0.49104935 | 0.86143519 |
| PRTFDC1 | 87.0208177 | 0.30492208 | 0.4122582 | 0.7396386 | 0.45951931 | 0.84620671 |
| PDSS1 | 46.9566742 | -0.061250854 | 0.48836335 | -0.1254207 | 0.90019053 | 0.98312332 |
| ABI1 | 425.355121 | 0.235239098 | 0.21080158 | 1.11592664 | 0.26445354 | 0.71989632 |
| ANKRD26 | 383.527279 | 0.037210862 | 0.26736612 | 0.13917568 | 0.88931132 | 0.98063791 |
| YME1L1 | 361.270146 | 0.056260852 | 0.17389179 | 0.32353943 | 0.74628675 | 0.94594643 |
| MASTL | 52.5029706 | -0.082004449 | 0.27542713 | -0.2977356 | 0.76590501 | 0.95389523 |
| ACBD5 | 257.227264 | 0.063263343 | 0.17293958 | 0.36581181 | 0.7145055 | 0.93573028 |
| RAB18 | 341.4094 | -0.001608669 | 0.20229945 | -0.0079519 | 0.99365535 | 0.99870178 |
| MPP7 | 451.813573 | 0.396561705 | 0.26411035 | 1.50150005 | 0.13322627 | 0.55609507 |
| AL358234.1 | 76.6627022 | -0.425030679 | 0.68973679 | -0.6162216 | 0.5377483 | 0.8824933 |
| WAC-AS1 | 209.394094 | -0.65519832 | 0.36990192 | -1.7712758 | 0.07651485 | 0.4426298 |
| WAC | 1255.18474 | 0.429722558 | 0.20439813 | 2.10238009 | 0.03551999 | 0.31644624 |
| SVIL | 468.781859 | -0.920781937 | 0.33752694 | -2.728025 | 0.00637148 | 0.12805535 |
| MTPAP | 71.0401687 | -0.047193351 | 0.22962851 | -0.2055204 | 0.83716553 | 0.96757829 |
| MAP3K8 | 134.510732 | -0.98472935 | 0.37433906 | -2.6305813 | 0.0085239 | 0.14565335 |
| ZNF438 | 324.411759 | 0.181556192 | 0.26055327 | 0.69681026 | 0.48592154 | 0.85912348 |
| ZEB1 | 60.555923 | -2.075316455 | 0.760171 | -2.7300653 | 0.00633218 | 0.12786365 |
| ARHGAP12 | 682.96247 | -0.027314361 | 0.1711366 | -0.1596056 | 0.87319176 | 0.97622753 |
| KIF5B | 442.700171 | 0.234685322 | 0.16841853 | 1.39346497 | 0.16347923 | 0.60304904 |
| EPC1 | 585.654168 | 0.309415579 | 0.24338528 | 1.27129947 | 0.20362213 | 0.6582233 |
| CCDC7 | 477.911157 | 0.07372516 | 0.33531436 | 0.21986878 | 0.82597335 | 0.96545294 |
| ITGB1 | 663.554374 | -0.471678084 | 0.30702375 | -1.5362919 | 0.12446682 | 0.54282656 |
| NRP1 | 399.589457 | -2.721418314 | 0.6617497 | -4.1124587 | 3.91E-05 | 0.00488402 |
| PARD3 | 2953.60663 | 0.350670222 | 0.22220425 | 1.57814364 | 0.11453262 | 0.52703872 |
| CUL2 | 182.424622 | 0.090858333 | 0.17310056 | 0.52488757 | 0.59966135 | 0.90369786 |
| CREM | 978.880953 | 1.455415962 | 0.76673542 | 1.89819843 | 0.05766995 | 0.39817926 |
| CCNY | 1183.92754 | 0.479802123 | 0.21747241 | 2.20626663 | 0.02736534 | 0.28061519 |
| AL117336.3 | 113.22244 | 0.049891207 | 0.32762827 | 0.15227992 | 0.87896616 | 0.9769398 |
| ZNF248 | 338.035266 | -0.104015843 | 0.26215295 | -0.3967754 | 0.69153309 | 0.93073907 |
| ZNF25 | 62.6457103 | -0.387938714 | 0.37479777 | -1.0350614 | 0.30064023 | 0.7476767 |
| ZNF33A | 603.829349 | -0.243862438 | 0.28273961 | -0.8624983 | 0.38841335 | 0.8065026 |
| AL117339.5 | 38.0528339 | -0.1250811 | 0.29284645 | -0.4271218 | 0.66929063 | 0.92463889 |
| ZNF33B | 420.832909 | -0.261333039 | 0.26095835 | -1.0014358 | 0.31661615 | 0.76069171 |
| BMS1 | 154.720673 | 0.092969704 | 0.20596363 | 0.45138895 | 0.65170925 | 0.91887313 |
| CSGALNACT2 | 125.194137 | -0.301832533 | 0.27622391 | -1.0927096 | 0.2745213 | 0.7263609 |
| RASGEF1A | 49.3389884 | -0.126942204 | 0.46241267 | -0.2745215 | 0.78368391 | 0.9568898 |
| FXYD4 | 1628.65458 | 1.105800945 | 0.67033019 | 1.64963621 | 0.09901737 | 0.4933116 |
| HNRNPF | 368.81537 | -0.465370881 | 0.30151108 | -1.543462 | 0.12271873 | 0.54116729 |
| ZNF487 | 83.2117134 | 0.643399647 | 0.43383275 | 1.4830592 | 0.13805868 | 0.56438077 |
| ZNF32 | 94.5394837 | 0.115379355 | 0.28481781 | 0.4050988 | 0.68540489 | 0.92924233 |
| TMEM72-AS | 113.88352 | -0.432343646 | 0.63576046 | -0.6800417 | 0.49647803 | 0.86473156 |
| TMEM72 | 345.66373 | 0.319848341 | 0.83064192 | 0.38506164 | 0.70019174 | 0.93180634 |
| RASSF4 | 40.7960944 | 0.473688529 | 0.60060416 | 0.78868673 | 0.43029513 | 0.83047752 |
| DEPP1 | 52.1170845 | 0.173865407 | 0.71910353 | 0.24178077 | 0.80895004 | 0.96245698 |
| ZNF22 | 85.8568746 | -0.903600394 | 0.3543234 | -2.5502137 | 0.01076569 | 0.16452817 |
| 8-Mar | 288.460638 | -0.199829299 | 0.24189174 | -0.8261105 | 0.40874143 | 0.82073894 |
| ZFAND4 | 79.9859849 | 0.190746783 | 0.26129406 | 0.73000812 | 0.46538522 | 0.84991999 |
| WASHC2C | 106.815641 | -0.504447817 | 0.37483865 | -1.3457732 | 0.17837566 | 0.62759322 |
| AGAP4 | 137.965163 | 0.425773787 | 0.34198317 | 1.24501388 | 0.21312665 | 0.67054344 |
| TIMM23 | 124.27734 | 0.388063139 | 0.26528631 | 1.46280876 | 0.14351971 | 0.57324898 |
| NCOA4 | 196.634786 | 0.048250989 | 0.30841362 | 0.15644896 | 0.87567915 | 0.97622753 |
| PTPN20 | 230.566259 | 0.523854718 | 0.44183563 | 1.18563258 | 0.23576743 | 0.69335782 |
| AGAP9 | 152.123752 | 0.450241789 | 0.3665193 | 1.22842586 | 0.21928714 | 0.67731484 |
| MAPK8 | 945.038763 | 0.16299703 | 0.31700325 | 0.51418095 | 0.60712549 | 0.90510401 |
| ARHGAP22 | 44.4851941 | -3.561578316 | 0.69026769 | -5.1597059 | 2.47E-07 | 0.00012961 |
| ERCC6 | 142.463876 | -0.212165781 | 0.24929362 | -0.8510678 | 0.39473166 | 0.81161229 |
| OGDHL | 231.012834 | 0.272990763 | 0.53535093 | 0.50992863 | 0.61010147 | 0.90600873 |
| PARG | 357.673184 | 0.110504206 | 0.20887965 | 0.52903289 | 0.59678263 | 0.90369786 |
| TIMM23B | 266.117324 | -0.045390483 | 0.40095041 | -0.1132072 | 0.90986627 | 0.98454672 |
| LINC00843 | 73.5656572 | -0.260093912 | 0.35522283 | -0.7321993 | 0.46404692 | 0.84965121 |
| AGAP6 | 111.857384 | 0.228781436 | 0.23803852 | 0.961111 | 0.33649636 | 0.77827084 |
| WASHC2A | 98.2664831 | -0.446328524 | 0.35475625 | -1.2581273 | 0.20834572 | 0.66589349 |
| SGMS1 | 1455.35676 | 0.07596465 | 0.2733981 | 0.27785361 | 0.78112474 | 0.9566945 |
| SGMS1-AS1 | 37.3245096 | -0.505648619 | 0.35822728 | -1.4115302 | 0.15808834 | 0.5960151 |
| ASAH2B | 51.9670686 | -0.577069038 | 0.36911844 | -1.5633709 | 0.11796537 | 0.53360052 |
| PRKG1 | 1636.88218 | 0.620882314 | 0.41450547 | 1.49788691 | 0.13416264 | 0.55858006 |
| CSTF2T | 33.6005989 | -0.259127515 | 0.36132562 | -0.7171579 | 0.47327669 | 0.85298656 |
| IPMK | 111.96959 | -1.066516679 | 0.38592005 | -2.763569 | 0.0057173 | 0.1213371 |
| CISD1 | 244.660974 | 0.042782616 | 0.2847547 | 0.15024376 | 0.8805723 | 0.97756113 |
| UBE2D1 | 83.6314169 | 0.262636234 | 0.3143009 | 0.83562037 | 0.40336851 | 0.81828987 |
| TFAM | 93.0994337 | -0.22184421 | 0.25316186 | -0.876294 | 0.38087025 | 0.80308469 |
| BICC1 | 4333.08272 | 0.438095396 | 0.48472594 | 0.90380019 | 0.36610136 | 0.79329304 |
| PHYHIPL | 41.2720206 | -0.120389154 | 0.68866495 | -0.1748153 | 0.86122478 | 0.97353422 |
| CCDC6 | 240.703793 | -0.092962237 | 0.23733567 | -0.391691 | 0.69528656 | 0.93126994 |
| ANK3 | 8417.15534 | 1.267829862 | 0.234975 | 5.39559469 | 6.83E-08 | 4.47E-05 |
| RHOBTB1 | 64.0100402 | -0.294337589 | 0.50606339 | -0.581622 | 0.56082134 | 0.89242209 |
| CABCOCO1 | 68.6059043 | 0.521663195 | 0.45441749 | 1.14798221 | 0.25097591 | 0.70813507 |
| ARID5B | 1146.41567 | 0.192484682 | 0.59023615 | 0.3261147 | 0.74433758 | 0.9456489 |
| ADO | 33.522221 | 0.044187935 | 0.45566452 | 0.09697471 | 0.92274648 | 0.9866731 |
| NRBF2 | 36.5323641 | -0.481118935 | 0.32551662 | -1.4780165 | 0.13940337 | 0.56656585 |
| JMJD1C | 2830.41985 | 0.731523903 | 0.2760995 | 2.64949373 | 0.00806125 | 0.14294732 |
| REEP3 | 629.241619 | 1.080636859 | 0.53253647 | 2.029226 | 0.04243528 | 0.34798255 |
| LINC01515 | 195.275907 | -0.254244192 | 0.40262585 | -0.6314652 | 0.52773643 | 0.87816414 |
| CTNNA3 | 49.7712739 | -0.290649182 | 0.49689428 | -0.5849316 | 0.55859369 | 0.89179371 |
| SIRT1 | 118.563195 | 0.389401737 | 0.26028127 | 1.4960805 | 0.13463268 | 0.55868799 |
| HERC4 | 848.65544 | -0.047607411 | 0.25701135 | -0.1852347 | 0.85304498 | 0.97062426 |
| PBLD | 212.005811 | -0.083865428 | 0.29064531 | -0.2885491 | 0.77292649 | 0.95543807 |
| HNRNPH3 | 365.085919 | 0.612961281 | 0.18917317 | 3.24021257 | 0.00119441 | 0.04468725 |
| RUFY2 | 322.878104 | 0.421846889 | 0.20777326 | 2.03032325 | 0.04232369 | 0.34798255 |
| DNA2 | 37.5167011 | 0.038062331 | 0.42537698 | 0.08947906 | 0.9287012 | 0.98761086 |
| SLC25A16 | 252.013617 | 0.120044039 | 0.31947764 | 0.37575098 | 0.70710203 | 0.93274945 |
| TET1 | 158.735759 | 0.978879404 | 0.45175563 | 2.16683388 | 0.03024752 | 0.29256342 |
| CCAR1 | 542.217791 | 0.285851607 | 0.24979363 | 1.14435108 | 0.25247807 | 0.70813507 |
| STOX1 | 76.8773213 | 1.006804501 | 0.50116707 | 2.00891991 | 0.04454563 | 0.35515061 |
| DDX50 | 207.12499 | -0.299319005 | 0.18318688 | -1.6339544 | 0.10226842 | 0.49966111 |
| DDX21 | 200.58915 | -0.234307617 | 0.47226385 | -0.4961371 | 0.61979769 | 0.90909444 |
| KIF1BP | 117.246979 | 0.004048868 | 0.25996743 | 0.01557452 | 0.98757383 | 0.9964791 |
| VPS26A | 224.013342 | 0.213919759 | 0.33875916 | 0.63148038 | 0.52772648 | 0.87816414 |
| SUPV3L1 | 109.910235 | 0.517406037 | 0.29802835 | 1.73609672 | 0.08254673 | 0.45651176 |
| HKDC1 | 409.8574 | 0.470863369 | 0.96282438 | 0.48904388 | 0.62481063 | 0.91058133 |
| HK1 | 499.55359 | 0.081654388 | 0.3926314 | 0.20796703 | 0.83525471 | 0.96757829 |
| TSPAN15 | 219.115146 | -0.936211682 | 0.29459762 | -3.1779337 | 0.00148329 | 0.05096671 |
| FAM241B | 31.6918811 | -0.269203615 | 0.34525642 | -0.7797208 | 0.43555522 | 0.83364877 |
| H2AFY2 | 105.506025 | -0.393967032 | 0.41411072 | -0.9513568 | 0.3414233 | 0.77954601 |
| AIFM2 | 72.2724425 | -0.411810268 | 0.33060976 | -1.2456083 | 0.21290823 | 0.67045621 |
| TYSND1 | 50.0754081 | -0.493869549 | 0.30630866 | -1.6123264 | 0.10689093 | 0.50942107 |
| SAR1A | 160.507953 | -0.025298839 | 0.2324607 | -0.1088306 | 0.91333685 | 0.98480378 |
| PPA1 | 247.620761 | -0.741877619 | 0.42743562 | -1.7356476 | 0.08262616 | 0.45670998 |
| EIF4EBP2 | 228.033812 | 0.243274971 | 0.22449567 | 1.08365104 | 0.27851955 | 0.72935218 |
| SGPL1 | 356.470321 | 0.410275191 | 0.33384978 | 1.22892156 | 0.21910122 | 0.67731484 |

| | | | | | | |
|---|---|---|---|---|---|---|
| PCBD1 | 234.305626 | -0.084060656 | 0.58966238 | -0.1425573 | 0.88663985 | 0.97975386 |
| UNC5B | 34.8627213 | -0.994021607 | 0.76569784 | -1.2981904 | 0.19422191 | 0.64791037 |
| PSAP | 1130.34439 | -0.692714111 | 0.30962133 | -2.2372946 | 0.02526709 | 0.27020321 |
| CHST3 | 112.696437 | 0.005847928 | 0.45547053 | 0.01283931 | 0.98975599 | 0.99660288 |
| ASCC1 | 212.044497 | -0.127202018 | 0.28045986 | -0.453548 | 0.65015416 | 0.91803345 |
| ANAPC16 | 604.967561 | 0.013366999 | 0.22833913 | 0.05854012 | 0.95331841 | 0.99060627 |
| DDIT4 | 1153.15052 | -0.186739169 | 0.78485621 | -0.2379279 | 0.81193704 | 0.96288868 |
| DNAJB12 | 76.2102083 | 0.002153109 | 0.30981897 | 0.00694957 | 0.99445509 | 0.99888869 |
| MICU1 | 766.691103 | 0.241326365 | 0.20353265 | 1.18568871 | 0.23574526 | 0.69335782 |
| MCU | 1123.12458 | 1.622757571 | 0.34688757 | 4.67805048 | 2.90E-06 | 0.00079873 |
| P4HA1 | 554.942878 | -1.276323799 | 0.71449137 | -1.786339 | 0.07404439 | 0.43749352 |
| ECD | 113.095572 | 0.258923719 | 0.30118701 | 0.85967758 | 0.3899668 | 0.80704029 |
| FAM149B1 | 173.920221 | 0.021605285 | 0.23871227 | 0.09050764 | 0.92788382 | 0.98751845 |
| DNAJC9 | 140.748033 | -0.517903113 | 0.25856905 | -2.0029586 | 0.04518173 | 0.35634897 |
| MRPS16 | 127.867153 | -0.053168089 | 0.28182867 | -0.1886539 | 0.85036405 | 0.97017003 |
| CFAP70 | 245.361711 | 0.412618065 | 0.44873859 | 0.91950654 | 0.35783069 | 0.78674827 |
| ANXA7 | 326.385681 | -0.089230987 | 0.17267504 | -0.5167567 | 0.605326 | 0.90428025 |
| PPP3CB | 342.224155 | 0.153820089 | 0.23227608 | 0.66222958 | 0.5078241 | 0.87094607 |
| USP54 | 212.9767 | 0.385024671 | 0.2342574 | 1.64359662 | 0.10025955 | 0.49569046 |
| AGAP5 | 40.3777779 | 0.251355443 | 0.27660379 | 0.90872017 | 0.36349786 | 0.79182239 |
| SEC24C | 99.6560093 | -0.301598165 | 0.26894961 | -1.1213928 | 0.26212069 | 0.71989632 |
| FUT11 | 197.398817 | -0.982834064 | 0.74150857 | -1.3254521 | 0.18502126 | 0.63541313 |
| CHCHD1 | 130.816131 | 0.055275468 | 0.47046154 | 0.117492 | 0.90647018 | 0.98320957 |
| ZSWIM8 | 146.35581 | -0.040057536 | 0.20269178 | -0.1976278 | 0.84333626 | 0.96876046 |
| CAMK2G | 104.058165 | -0.126754415 | 0.34768016 | -0.3645719 | 0.715431 | 0.93573028 |
| VCL | 600.41398 | 0.582284636 | 0.26492521 | 2.19792081 | 0.02795475 | 0.28401388 |
| AP3M1 | 80.1457372 | -0.157550842 | 0.25766127 | -0.611465 | 0.5408918 | 0.88391488 |
| ADK | 2198.71613 | 0.236619088 | 0.16132002 | 1.46676824 | 0.14243911 | 0.57165311 |
| KAT6B | 604.69284 | 0.053362047 | 0.28697217 | 0.18594851 | 0.85248514 | 0.97062426 |
| SAMD8 | 182.017954 | 0.269276943 | 0.22391572 | 1.20258171 | 0.22913823 | 0.68767716 |
| VDAC2 | 215.574237 | 0.003759978 | 0.42662521 | 0.0088133 | 0.99296809 | 0.99839831 |
| COMTD1 | 76.9165615 | 0.041979212 | 0.40874331 | 0.10270312 | 0.9181986 | 0.98520052 |
| ZNF503-AS1 | 50.487867 | 0.041394114 | 0.40112215 | 0.10319578 | 0.91780759 | 0.98520052 |
| AC010997.3 | 73.2058347 | -0.201544707 | 0.66414099 | -0.3034667 | 0.76153419 | 0.95366435 |
| ZNF503-AS2 | 45.3771467 | -0.95634219 | 0.44208422 | -2.1632579 | 0.03052135 | 0.2939924 |
| LRMDA | 3673.47355 | 0.023226455 | 0.29394975 | 0.07901505 | 0.93702065 | 0.98822345 |
| DLG5 | 180.703316 | -0.035621739 | 0.32030272 | -0.1112127 | 0.91144766 | 0.98468493 |
| POLR3A | 93.908421 | -0.117684049 | 0.2719379 | -0.4327607 | 0.6651886 | 0.92324595 |
| RPS24 | 1136.32392 | 0.381787678 | 0.48632537 | 0.78504577 | 0.43242673 | 0.83162989 |
| ZMIZ1 | 542.085489 | -0.292022245 | 0.43257714 | -0.6750755 | 0.49962778 | 0.86702299 |
| PPIF | 71.1115501 | -0.466344172 | 0.50582908 | -0.9219402 | 0.35655976 | 0.78612469 |
| NUTM2B-AS | 544.379128 | -0.034903223 | 0.37245953 | -0.0937101 | 0.92533944 | 0.98712921 |
| TMEM254 | 66.0044474 | -0.298352431 | 0.30690341 | -0.9721379 | 0.33098196 | 0.77618491 |
| ANXA11 | 660.16523 | 0.033994564 | 0.2684722 | 0.12662228 | 0.89923936 | 0.98312332 |
| FAM213A | 277.454743 | -0.85427648 | 0.43655329 | -1.9568664 | 0.05036317 | 0.37839277 |
| TSPAN14 | 173.05157 | -0.982404342 | 0.231431 | -4.2449125 | 2.19E-05 | 0.0031949 |
| NRG3 | 125.628403 | -0.48114165 | 0.96322298 | -0.4995122 | 0.61741858 | 0.90794578 |
| GHITM | 256.668218 | -0.130339347 | 0.39028322 | -0.3339609 | 0.73840904 | 0.94349597 |
| CCSER2 | 447.898692 | 0.128392877 | 0.28036348 | 0.4579515 | 0.64698728 | 0.91743182 |
| WAPL | 505.638135 | 0.603778099 | 0.18696426 | 3.2293771 | 0.0012406 | 0.04561933 |
| BMPR1A | 567.376027 | 0.4527237 | 0.14997376 | 3.01868602 | 0.00253873 | 0.0735219 |
| ADIRF | 51.7087352 | 0.216124779 | 0.59793987 | 0.36144902 | 0.71776381 | 0.93573028 |
| GLUD1 | 272.520355 | -0.115243413 | 0.27103514 | -0.4251973 | 0.67069284 | 0.92474338 |
| SHLD2 | 407.502794 | 0.147136856 | 0.29380321 | 0.5008007 | 0.61651139 | 0.90794578 |
| NUTM2A-AS | 562.757403 | -0.007119728 | 0.25739091 | -0.0276611 | 0.97793241 | 0.99469867 |
| MINPP1 | 59.6069686 | 0.014803733 | 0.31996511 | 0.04626671 | 0.96309767 | 0.99211837 |
| PAPSS2 | 429.388384 | 0.135757938 | 0.51988621 | 0.2611301 | 0.79399218 | 0.95937941 |
| ATAD1 | 282.331378 | 0.325181206 | 0.21630513 | 1.50334486 | 0.13275014 | 0.55601567 |
| PTEN | 882.193648 | 0.292162042 | 0.22438774 | 1.30204104 | 0.19290236 | 0.64567765 |
| RNLS | 200.15984 | 0.230575139 | 0.22955246 | 1.0044551 | 0.3151593 | 0.75893141 |
| LIPA | 116.65149 | 0.173415762 | 0.40995653 | 0.42301012 | 0.67228788 | 0.92498948 |
| IFIT2 | 95.6682629 | -0.749447615 | 1.32586876 | -0.5652502 | 0.57190358 | 0.89797476 |
| IFIT3 | 178.340804 | -1.632221965 | 1.27498863 | -1.2801855 | 0.2004799 | 0.65475264 |
| PANK1 | 73.6053751 | 0.779116973 | 0.40487768 | 1.92432682 | 0.05431363 | 0.39095834 |
| KIF20B | 55.2690233 | -0.149478961 | 0.23113419 | -0.6467194 | 0.51781357 | 0.87526312 |
| RPP30 | 53.6250862 | -0.120542652 | 0.29155441 | -0.4134482 | 0.67927825 | 0.92700898 |
| PCGF5 | 869.173068 | 0.59865503 | 0.54140998 | 1.10573327 | 0.26884198 | 0.72322263 |
| HECTD2 | 237.794378 | 0.108405137 | 0.19251914 | 0.56308757 | 0.57337525 | 0.89797476 |
| PPP1R3C | 136.998192 | -2.152624559 | 0.93714012 | -2.2970146 | 0.02161794 | 0.24935578 |
| TNKS2 | 492.593373 | 0.186203097 | 0.17078916 | 1.09025127 | 0.27560248 | 0.7263609 |
| BTAF1 | 446.999132 | 0.415904802 | 0.44431949 | 0.93604898 | 0.34924797 | 0.78180304 |
| CPEB3 | 711.507879 | 0.541751058 | 0.32922381 | 1.64554035 | 0.09985843 | 0.4953766 |
| 5-Mar | 228.490296 | 0.148252741 | 0.24562888 | 0.60356397 | 0.54613357 | 0.88669883 |
| IDE | 180.913869 | 0.103626854 | 0.24521122 | 0.42260242 | 0.67258536 | 0.92502554 |
| EXOC6 | 242.631105 | 0.033634694 | 0.30282404 | 0.11107009 | 0.91156077 | 0.98468493 |
| MYOF | 490.593387 | -1.028317148 | 0.44765256 | -2.2971323 | 0.02161123 | 0.24935578 |
| FRA10AC1 | 173.860427 | 0.335918344 | 0.25392809 | 1.32288766 | 0.18587274 | 0.63658376 |
| SLC35G1 | 36.4288178 | -0.62193234 | 0.43921064 | -1.4160229 | 0.15676879 | 0.59397576 |
| NOC3L | 76.613269 | 0.325370955 | 0.23997114 | 1.35587537 | 0.17513883 | 0.62472937 |
| TBC1D12 | 312.485286 | 0.701108304 | 0.24648987 | 2.84436966 | 0.00444994 | 0.10456358 |
| HELLS | 351.833168 | -0.214690262 | 0.3651511 | -0.5879491 | 0.55656646 | 0.89094338 |
| PDLIM1 | 59.3740615 | -0.793232324 | 0.42416077 | -1.8701218 | 0.0614669 | 0.4077045 |
| SORBS1 | 231.32613 | -0.059083083 | 0.39486756 | -0.1496276 | 0.88105844 | 0.97768262 |
| ALDH18A1 | 149.44701 | -0.809864419 | 0.38295186 | -2.1147943 | 0.03444748 | 0.31364863 |
| TCTN3 | 145.653103 | 0.063571307 | 0.22759953 | 0.27931212 | 0.78000531 | 0.95639467 |
| ENTPD1-AS1 | 484.327032 | -0.107131863 | 0.35636904 | -0.3006206 | 0.76370385 | 0.95378284 |
| AL365273.2 | 38.395824 | -0.210650933 | 0.53492366 | -0.3937963 | 0.69373147 | 0.93105058 |
| CC2D2B | 44.4797779 | -0.32001086 | 0.61512116 | -0.5202404 | 0.60289605 | 0.90369786 |
| ZNF518A | 465.995159 | -0.18839038 | 0.27968281 | -0.6735858 | 0.50057466 | 0.86743068 |
| BLNK | 1049.73766 | -0.281220405 | 0.3530189 | -0.7966157 | 0.42567424 | 0.82774157 |
| TM9SF3 | 624.576814 | -0.053748247 | 0.16584515 | -0.3240869 | 0.74587221 | 0.94594643 |
| LCOR | 930.893146 | 0.322034097 | 0.22660587 | 1.42111982 | 0.15528193 | 0.59198058 |
| RRP12 | 87.9328665 | -0.140623998 | 0.33951537 | -0.4141904 | 0.67873469 | 0.9268955 |
| PGAM1 | 390.08918 | -0.520162972 | 0.3750568 | -1.3868912 | 0.16547494 | 0.60681158 |
| EXOSC1 | 84.2595589 | -0.045218138 | 0.22109974 | -0.2045147 | 0.83795134 | 0.96757829 |
| ZDHHC16 | 56.4893816 | -0.061462523 | 0.25310649 | -0.2428327 | 0.80813503 | 0.96245698 |
| MMS19 | 187.511149 | 0.14396699 | 0.26278653 | 0.54784768 | 0.5837965 | 0.90085422 |
| UBTD1 | 131.962959 | 0.523668949 | 0.36004037 | 1.45447287 | 0.14581525 | 0.57663146 |
| AL355315.1 | 32.9021923 | -0.386573618 | 0.4910772 | -0.7871952 | 0.4311676 | 0.83111714 |
| MORN4 | 71.2573363 | -0.22905657 | 0.32039596 | -0.7149172 | 0.47466023 | 0.85368787 |
| AVPI1 | 62.3303626 | 1.541071799 | 0.97052975 | 1.58786663 | 0.1123165 | 0.5224487 |
| ZFYVE27 | 82.768116 | -0.029003026 | 0.23183777 | -0.1251005 | 0.90044397 | 0.98312332 |
| CRTAC1 | 62.1727138 | -0.409685782 | 0.956347 | -0.4283861 | 0.66837003 | 0.92422713 |
| R3HCC1L | 284.408302 | -0.294847517 | 0.2161077 | -1.3643545 | 0.17245601 | 0.61884138 |
| HPS1 | 71.0352919 | -0.603925234 | 0.32089073 | -1.8820277 | 0.05983226 | 0.40360868 |
| GOT1 | 359.394171 | 0.896185356 | 0.5596556 | 1.60131579 | 0.10930699 | 0.51461693 |
| SLC25A28 | 102.536886 | -0.315056744 | 0.42135118 | -0.7477296 | 0.45462328 | 0.84290184 |
| ENTPD7 | 39.9264682 | 0.119132057 | 0.47342879 | 0.2516367 | 0.80132189 | 0.96063297 |
| CUTC | 135.192861 | -0.361607181 | 0.23751558 | -1.5224567 | 0.12789469 | 0.54791812 |
| COX15 | 124.121655 | -0.415286225 | 0.26596394 | -1.5614381 | 0.11842042 | 0.53417263 |
| DNMBP | 260.267965 | -0.080910671 | 0.336189 | -0.2406702 | 0.80981075 | 0.96247146 |
| ERLIN1 | 82.2865111 | -0.095236151 | 0.22858789 | -0.4166282 | 0.67695041 | 0.9268955 |
| CHUK | 81.7215704 | 0.389237142 | 0.30904452 | 1.25948564 | 0.20785497 | 0.66589349 |

| | | | | | | |
|---|---|---|---|---|---|---|
| CWF19L1 | 96.6328013 | 0.09315148 | 0.26833693 | 0.34714371 | 0.72848335 | 0.93931348 |
| BLOC1S2 | 62.1646302 | 0.022369367 | 0.33960332 | 0.06586911 | 0.94748204 | 0.98977092 |
| SEC31B | 50.9857858 | 0.398045873 | 0.27330966 | 1.45639154 | 0.14528441 | 0.57564486 |
| HIF1AN | 104.923921 | -0.031366401 | 0.2053455 | -0.1527494 | 0.87859591 | 0.97692415 |
| PAX2 | 1473.71141 | 0.149942401 | 0.42958427 | 0.34904071 | 0.72705874 | 0.9392968 |
| SLF2 | 196.140317 | -0.315693608 | 0.28635692 | -1.102448 | 0.27026696 | 0.72343657 |
| MRPL43 | 102.10886 | -0.196205564 | 0.38856231 | -0.5049526 | 0.6135921 | 0.90751853 |
| LZTS2 | 109.181255 | -0.627863081 | 0.34714042 | -1.8086718 | 0.07050201 | 0.42987436 |
| BTRC | 511.031932 | 0.438294156 | 0.22675893 | 1.93286395 | 0.05325296 | 0.38756321 |
| DPCD | 92.8858543 | -0.421729274 | 0.38253565 | -1.1024574 | 0.27026285 | 0.72343657 |
| POLL | 70.1361919 | -0.305902218 | 0.24968115 | -1.2251714 | 0.22051063 | 0.67869352 |
| FBXW4 | 422.073514 | 0.067954837 | 0.20846278 | 0.32598067 | 0.74443898 | 0.94566309 |
| OGA | 823.766282 | -0.258021714 | 0.26819953 | -0.9620513 | 0.33602381 | 0.77820994 |
| ARMH3 | 469.271373 | 0.346730101 | 0.21960838 | 1.57885642 | 0.11436899 | 0.52703872 |
| LDB1 | 83.6508577 | -0.335339365 | 0.41018003 | -0.8175419 | 0.41361881 | 0.82440569 |
| PPRC1 | 43.2740341 | -0.333672899 | 0.41623817 | -0.8016394 | 0.42276161 | 0.82774157 |
| NOLC1 | 58.829126 | -0.415310859 | 0.34889074 | -1.1903751 | 0.23389899 | 0.69285918 |
| GBF1 | 651.56545 | 0.291554967 | 0.16330677 | 1.78532075 | 0.07420931 | 0.43765533 |
| NFKB2 | 46.6447223 | -0.425994621 | 0.41424063 | -1.0283748 | 0.30377356 | 0.749599 |
| FBXL15 | 90.1638365 | 0.12685951 | 0.44325118 | 0.28620231 | 0.77472318 | 0.95586283 |
| CUEDC2 | 84.5528087 | -0.018361529 | 0.35363601 | -0.0519221 | 0.95859075 | 0.99119981 |
| ACTR1A | 92.8538475 | -0.007649591 | 0.2909577 | -0.0262911 | 0.97902517 | 0.99469867 |
| SUFU | 161.980189 | 0.099071935 | 0.40165889 | 0.2466569 | 0.80517377 | 0.96218371 |
| TRIM8 | 320.812776 | -0.149578489 | 0.39077064 | -0.3827782 | 0.70188421 | 0.93215897 |
| ARL3 | 230.513625 | 0.347679108 | 0.36359623 | 0.95622308 | 0.33895954 | 0.77836439 |
| SFXN2 | 42.9234361 | 0.06159291 | 0.3544041 | 0.17379288 | 0.86202824 | 0.9739172 |
| WBP1L | 197.86303 | 0.153194894 | 0.23884738 | 0.64139239 | 0.52126777 | 0.87650871 |
| CNNM2 | 401.334242 | 0.323281763 | 0.39987784 | 0.80845132 | 0.41883082 | 0.82694802 |
| NT5C2 | 1048.13202 | 0.161476398 | 0.35729762 | 0.45193808 | 0.65131359 | 0.91866508 |
| PCGF6 | 81.8964939 | 0.244433583 | 0.38066709 | 0.64211903 | 0.5207959 | 0.87638467 |
| ATP5MD | 168.822507 | 0.445969284 | 0.33477849 | 1.33213245 | 0.18281667 | 0.63358396 |
| PDCD11 | 80.4762258 | 0.001443481 | 0.27585574 | 0.00523274 | 0.9958249 | 0.99908534 |
| SH3PXD2A | 76.7699205 | -0.054990306 | 0.35124767 | -0.1565571 | 0.87559394 | 0.97622753 |
| STN1 | 79.2464869 | -0.218994394 | 0.35734188 | -0.6128428 | 0.5399803 | 0.88338956 |
| SLK | 278.581875 | 0.192330289 | 0.15680254 | 1.22657635 | 0.21998186 | 0.67801071 |
| CFAP43 | 138.197361 | -0.1399013 | 0.4661953 | -0.3000916 | 0.76410727 | 0.95378284 |
| GSTO1 | 145.529131 | -0.330032899 | 0.23210988 | -1.4218822 | 0.15506046 | 0.59139011 |
| GSTO2 | 296.792479 | 0.264950015 | 0.22270692 | 1.18968019 | 0.23417211 | 0.69305952 |
| XPNPEP1 | 131.174247 | -0.288207434 | 0.28557656 | -1.0092125 | 0.31287273 | 0.75694198 |
| ADD3 | 710.528739 | 0.21919094 | 0.29879202 | 0.73359034 | 0.46319845 | 0.84880569 |
| MXI1 | 243.156386 | -0.717929892 | 0.53557798 | -1.3404769 | 0.18009036 | 0.63015077 |
| SMNDC1 | 89.3470383 | 0.273685485 | 0.27477541 | 0.99603341 | 0.31923391 | 0.76378437 |
| SMC3 | 153.182409 | 0.126784779 | 0.2638546 | 0.48051002 | 0.63086478 | 0.91315397 |
| PDCD4 | 217.088522 | -0.031140494 | 0.24900725 | -0.1250586 | 0.90047717 | 0.98312332 |
| BBIP1 | 121.071919 | -0.078435566 | 0.25155639 | -0.3118011 | 0.75519167 | 0.95067145 |
| SHOC2 | 332.821751 | 0.147976464 | 0.19063948 | 0.77621102 | 0.4376244 | 0.83420562 |
| GPAM | 62.4002252 | 0.906310503 | 0.38884091 | 2.33080028 | 0.01976389 | 0.23590615 |
| ACSL5 | 33.265527 | -0.547841554 | 0.37600787 | -1.4569949 | 0.14511779 | 0.57520213 |
| ZDHHC6 | 104.543946 | -0.37842846 | 0.23341699 | -1.621255 | 0.10496296 | 0.50459257 |
| VTI1A | 1042.24401 | -0.034549921 | 0.20206984 | -0.1709801 | 0.86423941 | 0.97466129 |
| TCF7L2 | 1336.39533 | 0.220529795 | 0.3637525 | 0.60626331 | 0.54433992 | 0.88654641 |
| CASP7 | 100.838909 | 0.12735994 | 0.27214072 | 0.46799296 | 0.63978963 | 0.91541711 |
| NHLRC2 | 260.159371 | 0.044961285 | 0.23511648 | 0.19122983 | 0.84834553 | 0.96991131 |
| CCDC186 | 323.414232 | 0.211076481 | 0.23053395 | 0.91559825 | 0.35987765 | 0.78892604 |
| AFAP1L2 | 453.745299 | 0.998389472 | 0.45956444 | 2.17246892 | 0.02982031 | 0.29105042 |
| ABLIM1 | 1353.19725 | -0.32332827 | 0.39059987 | -0.8277736 | 0.40779871 | 0.82029376 |
| FAM160B1 | 135.727374 | 0.148782993 | 0.18214246 | 0.8168496 | 0.41401438 | 0.82456685 |
| TRUB1 | 83.4251139 | 0.112703761 | 0.20037857 | 0.56245416 | 0.57380662 | 0.89797476 |
| HSPA12A | 363.742281 | 0.356354128 | 0.29817028 | 1.19513632 | 0.23203377 | 0.69186326 |
| SHTN1 | 227.808899 | -0.836355693 | 0.40183216 | -2.0813558 | 0.03740135 | 0.3238355 |
| PDZD8 | 259.868612 | 0.1466776 | 0.23102942 | 0.63488711 | 0.52550204 | 0.87791848 |
| EMX2OS | 645.604666 | -0.088987034 | 0.29708413 | -0.2995348 | 0.76453203 | 0.95378284 |
| EMX2 | 285.494285 | -0.036933631 | 0.35759235 | -0.1032842 | 0.91773743 | 0.98520052 |
| RAB11FIP2 | 518.278259 | 0.224966727 | 0.21900179 | 1.02723694 | 0.30430891 | 0.74968437 |
| AC022395.1 | 45.0986665 | -0.227414604 | 0.67043477 | -0.3392047 | 0.73445555 | 0.94215106 |
| CASC2 | 348.204849 | -0.333209224 | 0.43315276 | -0.7692649 | 0.44173606 | 0.83770294 |
| FAM204A | 187.863741 | 0.413839206 | 0.19198824 | 2.15554453 | 0.03111925 | 0.29675133 |
| CACUL1 | 389.863686 | -0.053738829 | 0.28194825 | -0.1905982 | 0.8488404 | 0.96991131 |
| EIF3A | 485.966979 | -0.116338268 | 0.32418542 | -0.3588634 | 0.71969731 | 0.93636596 |
| FAM45A | 104.930764 | -0.448623313 | 0.27945458 | -1.6053532 | 0.10841609 | 0.51254482 |
| SFXN4 | 87.9497228 | -0.17889096 | 0.38597781 | -0.4634747 | 0.64302413 | 0.91643619 |
| PRDX3 | 245.739621 | -0.062330577 | 0.45654357 | -0.1365271 | 0.89140459 | 0.980985 |
| TIAL1 | 216.828206 | 0.207026111 | 0.18050196 | 1.14694661 | 0.25140369 | 0.70813507 |
| BAG3 | 65.0828619 | -0.342830224 | 0.33007954 | -1.0386291 | 0.29897725 | 0.74594723 |
| INPP5F | 148.297063 | -0.170345273 | 0.25435609 | -0.6697118 | 0.50304153 | 0.86922923 |
| MCMBP | 141.817355 | -0.15159192 | 0.17015315 | -0.8909146 | 0.37297501 | 0.79681088 |
| SEC23IP | 219.285641 | 0.156159651 | 0.26173408 | 0.59663477 | 0.55075125 | 0.88916026 |
| PLPP4 | 156.35539 | 1.33707894 | 0.90325864 | 1.48028358 | 0.13879758 | 0.5659917 |
| WDR11-AS1 | 53.3027372 | -0.53544913 | 0.47959838 | -1.1164532 | 0.26422821 | 0.71989632 |
| WDR11 | 304.893845 | 0.190131108 | 0.16961435 | 1.12096123 | 0.26230437 | 0.71989632 |
| FGFR2 | 529.703732 | 0.798616858 | 0.46911222 | 1.70240047 | 0.08868032 | 0.47038308 |
| ATE1 | 465.577815 | 0.210588024 | 0.17298151 | 1.21740193 | 0.22345133 | 0.68103746 |
| NSMCE4A | 61.639604 | 0.697788296 | 0.4012002 | 1.73925212 | 0.08199043 | 0.45535449 |
| TACC2 | 375.759513 | 0.905583879 | 0.51660298 | 1.75295907 | 0.07960903 | 0.45000144 |
| PLEKHA1 | 917.046695 | -0.578119202 | 0.37347612 | -1.5479415 | 0.12163637 | 0.53991917 |
| C10orf88 | 51.443043 | 0.13669088 | 0.28196547 | 0.48477879 | 0.62783327 | 0.91081017 |
| PSTK | 40.9782422 | -0.272062523 | 0.4102378 | -0.6631825 | 0.50721368 | 0.87094607 |
| IKZF5 | 75.7563425 | 0.348634726 | 0.22333036 | 1.56107181 | 0.11850681 | 0.53417263 |
| ACADSB | 141.598166 | 0.180450196 | 0.28520531 | 0.63270279 | 0.52692775 | 0.87791848 |
| BUB3 | 104.061987 | -0.293259073 | 0.26195209 | -1.1195141 | 0.26292086 | 0.71989632 |
| OAT | 147.836263 | 0.017914632 | 0.34408943 | 0.05206388 | 0.95847779 | 0.99119981 |
| LHPP | 126.460218 | -0.121747774 | 0.25399664 | -0.4793283 | 0.6317051 | 0.9133034 |
| ABRAXAS2 | 81.7460769 | -0.046997471 | 0.26924107 | -0.1745554 | 0.86142903 | 0.97366008 |
| ZRANB1 | 209.699219 | 0.556957482 | 0.21438057 | 2.59798491 | 0.00937726 | 0.15476173 |
| CTBP2 | 1041.25562 | -0.299072356 | 0.20750401 | -1.4412847 | 0.14950427 | 0.5826719 |
| EDRF1 | 147.450645 | 0.150383974 | 0.23649138 | 0.63589622 | 0.52484407 | 0.87791848 |
| UROS | 114.073779 | 0.080406639 | 0.21672958 | 0.37099985 | 0.71063765 | 0.93475719 |
| BCCIP | 147.988473 | 0.221986813 | 0.23593652 | 0.94087515 | 0.34676884 | 0.78180304 |
| DHX32 | 342.358692 | -0.563448502 | 0.21376657 | -2.635812 | 0.00839362 | 0.1442148 |
| FANK1 | 270.690415 | -0.332656489 | 0.26801783 | -1.241173 | 0.21454185 | 0.67132603 |
| DOCK1 | 1572.32422 | -0.098053919 | 0.2019314 | -0.4855804 | 0.62726473 | 0.91061758 |
| FAM196A | 68.79994 | -0.131619508 | 0.49852065 | -0.2640202 | 0.79176437 | 0.95839852 |
| MGMT | 766.546107 | 0.02088 | 0.20050228 | 0.10413847 | 0.91705946 | 0.98520052 |
| AL157832.3 | 120.022711 | -0.561847175 | 0.48264188 | -1.1641078 | 0.24438033 | 0.70121778 |
| C10orf143 | 48.8400287 | 0.162313546 | 0.40983605 | 0.39604506 | 0.69207178 | 0.93093471 |
| GLRX3 | 171.320406 | 0.177674303 | 0.28597203 | 0.62129958 | 0.53440253 | 0.88137638 |
| PPP2R2D | 308.038051 | 0.073492789 | 0.22158089 | 0.33167476 | 0.74013486 | 0.94445369 |
| BNIP3 | 412.466808 | -1.623734645 | 0.66555787 | -2.4396596 | 0.01470111 | 0.1970174 |
| STK32C | 85.0999311 | 0.061456511 | 0.28210183 | 0.21785222 | 0.82754425 | 0.96545294 |
| LRRC27 | 81.4231945 | -0.422806265 | 0.29697217 | -1.4237235 | 0.15452652 | 0.59060466 |
| PWWP2B | 68.4796858 | -0.174719773 | 0.44158078 | -0.3956689 | 0.69234931 | 0.93105058 |
| INPP5A | 186.039915 | 0.027677989 | 0.27838948 | 0.09942182 | 0.92080336 | 0.98581108 |
| CFAP46 | 81.2616226 | -0.873708667 | 0.66912334 | -1.3057513 | 0.19163715 | 0.64458858 |

| | | | | | | |
|---|---|---|---|---|---|---|
| TUBGCP2 | 191.255325 | -0.138155147 | 0.22794064 | -0.6061014 | 0.5444474 | 0.88654641 |
| ECHS1 | 311.353484 | -0.495699976 | 0.51709273 | -0.9586288 | 0.33774578 | 0.77827084 |
| SCART1 | 43.6048628 | 0.583721737 | 0.387451 | 1.50656918 | 0.13192112 | 0.55448417 |
| SLC6A12 | 165.065896 | 1.019269238 | 0.67416063 | 1.51190858 | 0.13055712 | 0.55395564 |
| SLC6A13 | 42.675542 | 2.672036331 | 0.47430735 | 5.63355453 | 1.77E-08 | 1.68E-05 |
| KDM5A | 469.809089 | 0.05550444 | 0.18107075 | 0.30653454 | 0.75919769 | 0.95246973 |
| CCDC77 | 36.9232053 | -0.45172424 | 0.34875135 | -1.2952616 | 0.19523 | 0.64932097 |
| B4GALNT3 | 199.106339 | 0.00769646 | 0.32684725 | 0.02354758 | 0.98121349 | 0.99507903 |
| WNK1 | 907.769012 | 0.881991011 | 0.53527876 | 1.64772279 | 0.09940957 | 0.49445291 |
| RAD52 | 137.494117 | 0.114999692 | 0.29182078 | 0.39407644 | 0.69352461 | 0.93105058 |
| ERC1 | 3535.34324 | 0.611948412 | 0.27146213 | 2.25426804 | 0.02417931 | 0.2634087 |
| FBXL14 | 50.3399038 | -0.666253286 | 0.34879966 | -1.9101317 | 0.05611626 | 0.39444789 |
| ADIPOR2 | 326.474309 | 0.170891506 | 0.26907173 | 0.63511506 | 0.52535337 | 0.87791848 |
| DCP1B | 107.715471 | -0.211924646 | 0.33942137 | -0.6243704 | 0.53238438 | 0.88086332 |
| FKBP4 | 165.992504 | -0.219749863 | 0.46521813 | -0.4723588 | 0.63667073 | 0.91454294 |
| ITFG2 | 157.46848 | -0.353329476 | 0.37321363 | -0.9467218 | 0.34378056 | 0.7816924 |
| TULP3 | 248.539946 | -0.05278041 | 0.21227245 | -0.2486447 | 0.80363566 | 0.9615369 |
| TEAD4 | 70.4738687 | -1.979662203 | 0.5496134 | -3.6019177 | 0.00031588 | 0.01808965 |
| TSPAN9 | 780.006489 | -0.056482555 | 0.25079827 | -0.2252111 | 0.82181506 | 0.96502145 |
| PARP11 | 88.0476789 | 0.256485402 | 0.28155553 | 0.91095851 | 0.36231724 | 0.79051672 |
| CCND2 | 135.297648 | 0.28780144 | 0.41898191 | 0.6869066 | 0.49214159 | 0.86172738 |
| TIGAR | 76.5145322 | 0.351798553 | 0.4865831 | 0.72299788 | 0.46968119 | 0.85232523 |
| C12orf4 | 92.7471932 | -0.358672613 | 0.30026944 | -1.1945026 | 0.23228144 | 0.69186326 |
| ANO2 | 66.0914876 | 1.09953252 | 0.45891557 | 2.39593642 | 0.01657797 | 0.21108198 |
| CD9 | 1283.8246 | 1.146805567 | 0.63802054 | 1.79744303 | 0.07226532 | 0.43302651 |
| TNFRSF1A | 173.799067 | -1.335671989 | 0.37908129 | -3.5234448 | 0.00042598 | 0.02210013 |
| SCNN1A | 810.930009 | 0.174931335 | 0.34283068 | 0.51025577 | 0.60987228 | 0.90587512 |
| LTBR | 102.261028 | -0.820770621 | 0.33230033 | -2.4699663 | 0.01351258 | 0.18706977 |
| TAPBPL | 59.1318741 | -0.093200912 | 0.39751344 | -0.2344598 | 0.81462806 | 0.96321082 |
| MRPL51 | 190.766624 | -0.048310921 | 0.52287043 | -0.0923956 | 0.92638374 | 0.98729622 |
| NCAPD2 | 64.8131398 | -0.151771019 | 0.39485269 | -0.3843738 | 0.70070144 | 0.93180634 |
| GAPDH | 2169.01156 | -0.98268119 | 0.62127686 | -1.5817122 | 0.11371529 | 0.52602773 |
| CHD4 | 97.1831164 | -0.782169648 | 0.32941829 | -2.3743965 | 0.01757766 | 0.21956364 |
| AC006064.2 | 42.8714332 | -0.187767986 | 0.36069502 | -0.5205727 | 0.60266446 | 0.90369786 |
| ING4 | 87.0826366 | -0.279467159 | 0.25578833 | -1.092572 | 0.27458178 | 0.7263609 |
| ZNF384 | 64.4720752 | 0.316969865 | 0.25709166 | 1.23290604 | 0.21761082 | 0.6751218 |
| COPS7A | 99.3304652 | -0.185814427 | 0.33303464 | -0.5579432 | 0.57688315 | 0.89869922 |
| MLF2 | 187.558099 | -0.203246352 | 0.35090255 | -0.5792102 | 0.56244732 | 0.89242209 |
| PTMS | 446.576367 | 0.121540893 | 0.45538155 | 0.26689903 | 0.78954691 | 0.95801395 |
| USP5 | 69.2932838 | 0.028862418 | 0.30880396 | 0.09346518 | 0.92553401 | 0.98717098 |
| TPI1 | 571.585601 | -0.482085071 | 0.49780138 | -0.9684285 | 0.33283039 | 0.77667835 |
| LRRC23 | 84.9323462 | -0.367598579 | 0.28714915 | -1.280166 | 0.20048676 | 0.65475264 |
| ENO2 | 75.4899392 | -1.864794227 | 1.00080204 | -1.8632998 | 0.0624201 | 0.41052452 |
| ATN1 | 206.473705 | -0.203813934 | 0.26067592 | -0.7818671 | 0.43429267 | 0.83243867 |
| C12orf57 | 129.684104 | -0.322698303 | 0.54246763 | -0.5948711 | 0.55192965 | 0.88923362 |
| PTPN6 | 66.2216212 | -0.091428336 | 0.33585923 | -0.2722222 | 0.78545118 | 0.95726577 |
| PHB2 | 176.830098 | -0.36714039 | 0.40454058 | -0.907549 | 0.36411656 | 0.79268955 |
| LPCAT3 | 254.106399 | -0.46980466 | 0.2750138 | -1.7082949 | 0.08758164 | 0.46829366 |
| C1S | 90.2526436 | -0.559978653 | 0.39642318 | -1.412578 | 0.15777984 | 0.59566075 |
| C1R | 43.6277424 | -2.215983001 | 0.74688072 | -2.9669838 | 0.00300737 | 0.08143983 |
| C1RL | 210.95572 | -1.040553939 | 0.26910202 | -3.8667637 | 0.00011029 | 0.009397 |
| CLSTN3 | 64.3713901 | -0.04596868 | 0.32628114 | -0.1408867 | 0.88795943 | 0.98028177 |
| PEX5 | 51.854655 | 0.287835227 | 0.30177023 | 0.95382247 | 0.34017351 | 0.77836439 |
| CD163L1 | 38.5327373 | -0.161393171 | 0.49549811 | -0.325719 | 0.74463694 | 0.94575914 |
| FOXJ2 | 48.2022153 | 0.346267922 | 0.28635044 | 1.20924531 | 0.22656862 | 0.68467102 |
| NECAP1 | 56.8747516 | 0.635901228 | 0.28505179 | 2.23082701 | 0.02569259 | 0.27225313 |
| FAM66C | 47.6417875 | -0.145011718 | 0.49175224 | -0.2948878 | 0.76807962 | 0.95411028 |
| LINC00937 | 89.660974 | -0.90890097 | 0.39223597 | -2.31723 | 0.0204912 | 0.24101888 |
| RIMKLB | 268.103472 | -1.882010119 | 0.65634454 | -2.8674118 | 0.00413844 | 0.09955609 |
| PHC1 | 123.802772 | 0.172611607 | 0.23737939 | 0.72715499 | 0.46713101 | 0.85112152 |
| M6PR | 172.878567 | -0.634331985 | 0.24620117 | -2.5764784 | 0.00998124 | 0.16018903 |
| CLEC2D | 147.333305 | -0.532970059 | 0.29335813 | -1.8167898 | 0.06924931 | 0.42828237 |
| GABARAPL1 | 566.738693 | -0.055577492 | 0.49201099 | -0.1129599 | 0.91006238 | 0.98454672 |
| KLRD1 | 164.033303 | -0.47217587 | 0.531838 | -0.887819 | 0.37463813 | 0.79800969 |
| MAGOHB | 52.4233798 | -0.07868872 | 0.29533961 | -0.2664347 | 0.78990445 | 0.95801395 |
| STYK1 | 49.1630661 | -0.687022699 | 0.63498257 | -1.0819552 | 0.27927243 | 0.72938673 |
| YBX3 | 914.921923 | -0.399800255 | 0.5948595 | -0.6720919 | 0.50152519 | 0.86804029 |
| SMIM10L1 | 59.7032597 | -0.77526141 | 0.35571483 | -2.1794464 | 0.02929852 | 0.28931514 |
| AC244131.2 | 66.4952235 | -0.187872224 | 0.76977972 | -0.2440597 | 0.80718457 | 0.96226758 |
| ETV6 | 940.267926 | -0.766276343 | 0.2273977 | -3.3697629 | 0.00075233 | 0.03310667 |
| BCL2L14 | 146.087536 | -0.450821728 | 0.35704379 | -1.2626511 | 0.2067146 | 0.66434881 |
| LRP6 | 359.94435 | 0.440330638 | 0.31230885 | 1.40992048 | 0.15856316 | 0.59710455 |
| MANSC1 | 105.681832 | 0.041128864 | 0.31215122 | 0.13175942 | 0.89517458 | 0.98234865 |
| BORCS5 | 165.83022 | 0.254064917 | 0.2393701 | 1.06138951 | 0.28851292 | 0.73656893 |
| DUSP16 | 275.00522 | 0.600591245 | 0.16156739 | 3.71728003 | 0.00020138 | 0.01361583 |
| CREBL2 | 155.578727 | 0.198003264 | 0.26170492 | 0.75658976 | 0.44929568 | 0.84117804 |
| CDKN1B | 129.508073 | -0.161249759 | 0.44187647 | -0.3649204 | 0.71517079 | 0.93573028 |
| GPRC5A | 524.510886 | -1.358985014 | 1.00062922 | -1.3581305 | 0.17442229 | 0.62259729 |
| GPRC5D-AS1 | 165.065272 | -0.231248725 | 0.44986741 | -0.5140375 | 0.60722576 | 0.90510401 |
| HEBP1 | 175.99334 | 0.373524591 | 0.49846675 | 0.74934706 | 0.45364805 | 0.84268043 |
| FAM234B | 73.736048 | 1.1489593 | 0.32542167 | 3.53067855 | 0.0004145 | 0.02193894 |
| GRIN2B | 128.06352 | 0.771430659 | 0.71610574 | 1.07725803 | 0.281365 | 0.73086885 |
| ATF7IP | 788.211328 | 0.371443481 | 0.33395913 | 1.11224233 | 0.26603397 | 0.72053648 |
| PLBD1 | 100.452384 | -0.850889833 | 0.51244387 | -1.6604547 | 0.09682301 | 0.48901455 |
| AC010168.1 | 61.701819 | -0.259074343 | 0.56936344 | -0.4550246 | 0.64909158 | 0.91768694 |
| H2AFJ | 267.606952 | -0.467967456 | 0.45272751 | -1.0336625 | 0.30129396 | 0.74814891 |
| WBP11 | 154.042277 | 0.24366795 | 0.28206372 | 0.86387553 | 0.38765627 | 0.8065026 |
| C12orf60 | 79.7209376 | -0.148344676 | 0.38632812 | -0.3839862 | 0.70098866 | 0.93180634 |
| MGP | 279.281754 | -2.146404887 | 0.7996128 | -2.6843053 | 0.00726807 | 0.13601674 |
| ERP27 | 229.998626 | -1.775232649 | 0.84137199 | -2.109926 | 0.03486473 | 0.31514522 |
| ARHGDIB | 99.9471095 | -0.921241932 | 0.70062775 | -1.3148807 | 0.18854999 | 0.64159008 |
| RERG | 94.0092113 | -0.794735 | 0.64944783 | -1.2237088 | 0.22106211 | 0.67871976 |
| EPS8 | 604.110956 | 0.102996067 | 0.21551475 | 0.47790727 | 0.63271621 | 0.91415622 |
| STRAP | 178.862953 | 0.42517218 | 0.34866969 | 1.2194125 | 0.22268767 | 0.68000197 |
| DERA | 344.974051 | -0.119936324 | 0.27438674 | -0.4371069 | 0.66203386 | 0.92284375 |
| MGST1 | 365.371746 | -1.157371348 | 0.40928929 | -2.8277587 | 0.00468751 | 0.10823805 |
| LMO3 | 260.188563 | 1.363117232 | 0.41344503 | 3.29697335 | 0.00097733 | 0.03894447 |
| RERGL | 93.9297361 | -0.303391616 | 0.80371392 | -0.3774871 | 0.70581167 | 0.93241442 |
| PIK3C2G | 2909.47011 | 0.759919993 | 0.46724653 | 1.62637912 | 0.10386903 | 0.50279327 |
| PLCZ1 | 60.2488241 | -1.483184921 | 0.98938352 | -1.4991001 | 0.13384767 | 0.55774297 |
| AC087242.1 | 86.1737009 | 0.366870921 | 0.6956168 | 0.52740377 | 0.59791323 | 0.90369786 |
| PLEKHA5 | 2322.07794 | 0.252503534 | 0.32531345 | 0.77618536 | 0.43763955 | 0.83420562 |
| AEBP2 | 623.465089 | 0.500667583 | 0.27197693 | 1.84084577 | 0.06564416 | 0.41788422 |
| AC024901.1 | 76.2546094 | 0.188437031 | 0.73573937 | 0.25611927 | 0.79785874 | 0.95999536 |
| LINC02398 | 60.2139907 | -0.133796367 | 0.85769888 | -0.1559946 | 0.8760373 | 0.97622753 |
| PDE3A | 59.6894747 | 2.297483842 | 0.7436827 | 3.08933344 | 0.00200606 | 0.062951 |
| PYROXD1 | 102.515395 | 0.064066334 | 0.28918882 | 0.22153808 | 0.82467349 | 0.96511681 |
| RECQL | 68.4183901 | 0.068066946 | 0.33990404 | 0.20025342 | 0.84128239 | 0.96854218 |
| GOLT1B | 76.9536309 | -0.146139337 | 0.28921518 | -0.5052962 | 0.6133508 | 0.90751853 |
| LDHB | 1077.14508 | -0.64891144 | 0.42311061 | -1.5336686 | 0.12511121 | 0.54337567 |
| CMAS | 114.398281 | -0.245512591 | 0.20052009 | -1.224379 | 0.22080929 | 0.67871976 |
| C2CD5 | 349.547875 | 0.495999939 | 0.28022208 | 1.77002449 | 0.07672306 | 0.44322095 |
| ETNK1 | 713.957318 | 0.553733142 | 0.25715639 | 2.15329334 | 0.03129563 | 0.29704233 |

| | | | | | | |
|---|---|---|---|---|---|---|
| AC084816.1 | 30.9728802 | -0.066014443 | 0.85224947 | -0.0774591 | 0.93825836 | 0.98846521 |
| SOX5 | 2688.08229 | 1.476276214 | 0.48370127 | 3.05204125 | 0.00227291 | 0.06844851 |
| BCAT1 | 66.1239749 | 2.367700156 | 1.45352807 | 1.62893322 | 0.10332716 | 0.50179268 |
| CASC1 | 153.771758 | 0.085076535 | 0.50853822 | 0.16729625 | 0.86713696 | 0.97576204 |
| ETFRF1 | 96.4643067 | -0.063902109 | 0.30593077 | -0.2088777 | 0.83454373 | 0.96749268 |
| KRAS | 276.11438 | 0.100224244 | 0.26495845 | 0.37826401 | 0.70523448 | 0.93241442 |
| RASSF8-AS1 | 98.1895643 | -0.504914753 | 0.4540554 | -1.1120113 | 0.26613327 | 0.72053648 |
| RASSF8 | 651.642557 | 0.150170553 | 0.46036219 | 0.32620089 | 0.74427237 | 0.9456489 |
| BHLHE41 | 64.6359536 | -1.134632061 | 0.67902994 | -1.6709603 | 0.09472952 | 0.48627909 |
| SSPN | 85.6608814 | -0.699065954 | 0.44261886 | -1.5793858 | 0.11424759 | 0.52703872 |
| AC024145.1 | 37.3878944 | -0.273088318 | 0.65428643 | -0.4173834 | 0.67639797 | 0.9268955 |
| ITPR2 | 1377.00116 | 0.56070956 | 0.24503339 | 2.2882986 | 0.02212014 | 0.25088641 |
| AC023051.1 | 40.1227672 | -0.051967851 | 0.76981578 | -0.0675069 | 0.9461782 | 0.98949015 |
| INTS13 | 63.5983994 | -0.236712491 | 0.3478296 | -0.6805415 | 0.49616163 | 0.86452555 |
| FGFR1OP2 | 181.270756 | 0.734429413 | 0.29905043 | 2.45587145 | 0.01405434 | 0.19221396 |
| TM7SF3 | 326.904646 | 0.390489294 | 0.33002758 | 1.18320201 | 0.2367291 | 0.69335782 |
| MED21 | 128.650046 | 0.835176214 | 0.3841097 | 2.1743169 | 0.02968134 | 0.29105042 |
| STK38L | 154.327614 | 1.294289117 | 0.34375708 | 3.76512715 | 0.00016646 | 0.01228554 |
| PPFIBP1 | 945.206451 | 0.295311459 | 0.49877381 | 0.5920749 | 0.55380043 | 0.89015775 |
| MRPS35 | 162.23055 | -0.021149328 | 0.19373038 | -0.1091689 | 0.91306855 | 0.98480378 |
| KLHL42 | 105.694155 | -0.213627426 | 0.2220383 | -0.9621197 | 0.33598947 | 0.77820994 |
| CCDC91 | 964.597135 | -0.014360986 | 0.19432552 | -0.0739017 | 0.9410886 | 0.98876195 |
| FAR2 | 58.1164572 | -0.585820143 | 0.58341532 | -1.004122 | 0.31531983 | 0.75912793 |
| AC009318.1 | 44.0273028 | -0.55746868 | 0.56502741 | -0.9866224 | 0.3238278 | 0.76937573 |
| ERGIC2 | 221.164723 | 0.099005191 | 0.19835699 | 0.4991263 | 0.61769041 | 0.90794578 |
| OVCH1-AS1 | 45.0420922 | -2.391357326 | 1.07105887 | -2.2327039 | 0.02556848 | 0.27225313 |
| TMTC1 | 172.743733 | 2.054227603 | 0.93898845 | 2.18770275 | 0.02869126 | 0.28768102 |
| IPO8 | 226.688455 | 0.420171831 | 0.20784742 | 2.02153979 | 0.04322392 | 0.35033771 |
| CAPRIN2 | 147.895556 | 0.188453607 | 0.20911375 | 0.90120141 | 0.36748124 | 0.79455404 |
| DDX11 | 35.0666636 | 0.025680994 | 0.35881556 | 0.07157157 | 0.94294286 | 0.98885525 |
| SINHCAF | 326.466017 | -0.200012758 | 0.21826339 | -0.9163825 | 0.3594663 | 0.78892604 |
| DENND5B | 390.012938 | -0.941267589 | 0.37410118 | -2.5160776 | 0.01186691 | 0.17330267 |
| AMN1 | 204.11519 | -0.072671789 | 0.34833349 | -0.208627 | 0.83473939 | 0.96749268 |
| KIAA1551 | 215.811785 | -0.438113686 | 0.32806578 | -1.3354447 | 0.18173086 | 0.63200456 |
| BICD1 | 1543.0822 | 0.084501017 | 0.52052475 | 0.16233813 | 0.87103959 | 0.97622753 |
| FGD4 | 2216.96119 | 0.272999124 | 0.32750519 | 0.83357191 | 0.40452226 | 0.81914727 |
| DNM1L | 434.086498 | 0.384524835 | 0.19242435 | 1.99831696 | 0.04568231 | 0.35915273 |
| YARS2 | 110.093248 | -0.06546214 | 0.25164255 | -0.2601394 | 0.79475625 | 0.95937941 |
| PKP2 | 42.8640794 | -0.720678964 | 0.49933607 | -1.4432744 | 0.14894319 | 0.5817833 |
| ALG10B | 47.3875159 | 0.378551985 | 0.32070204 | 1.18038533 | 0.23784699 | 0.69406342 |
| CPNE8 | 731.130206 | 0.367471039 | 0.61672221 | 0.59584531 | 0.55127857 | 0.88916026 |
| KIF21A | 147.784052 | 0.864878751 | 0.40540386 | 2.13337573 | 0.03289392 | 0.30588131 |
| SLC2A13 | 242.906397 | -1.459799464 | 0.41917971 | -3.4825146 | 0.00049673 | 0.02514836 |
| LRRK2 | 612.609038 | 0.426552912 | 0.31221763 | 1.36620379 | 0.17187501 | 0.61884138 |
| CNTN1 | 125.222335 | 0.841625827 | 0.6218962 | 1.35332204 | 0.17595277 | 0.62486786 |
| GXYLT1 | 192.209619 | 0.610380467 | 0.19149406 | 3.18746425 | 0.00143526 | 0.05030617 |
| YAF2 | 303.53204 | 0.108156393 | 0.22539427 | 0.47985423 | 0.63133105 | 0.9133034 |
| PPHLN1 | 438.541736 | 0.054200181 | 0.19649422 | 0.27583601 | 0.78267402 | 0.95688565 |
| ZCRB1 | 127.535613 | 0.12941682 | 0.33532565 | 0.38594369 | 0.69953837 | 0.93180634 |
| PRICKLE1 | 176.668374 | -0.654337284 | 0.80294867 | -0.8149179 | 0.41511928 | 0.82472987 |
| PUS7L | 117.545269 | 0.054861788 | 0.25237853 | 0.21737898 | 0.82791301 | 0.96569646 |
| IRAK4 | 105.352288 | -0.619420846 | 0.32388711 | -1.9124591 | 0.05581734 | 0.39368206 |
| TWF1 | 123.213648 | -0.094831874 | 0.28810444 | -0.329158 | 0.74203629 | 0.94508888 |
| TMEM117 | 661.184365 | 0.236879149 | 0.27608158 | 0.85800419 | 0.39089015 | 0.80778625 |
| ANO6 | 448.464044 | -0.351198777 | 0.38482091 | -0.9126291 | 0.36143762 | 0.78979697 |
| AC008124.1 | 50.5810192 | -0.016756656 | 0.39471136 | -0.0424529 | 0.96613763 | 0.99284614 |
| ARID2 | 804.120725 | 0.423410639 | 0.17431369 | 2.42901538 | 0.01513989 | 0.20009038 |
| SCAF11 | 809.588481 | 0.083199356 | 0.12576018 | 0.66157153 | 0.50824586 | 0.87094607 |
| SLC38A1 | 1945.66568 | 0.730808009 | 0.40187985 | 1.8184739 | 0.06899173 | 0.42732468 |
| SLC38A2 | 529.338882 | 0.668255429 | 0.30104367 | 2.21979567 | 0.02643264 | 0.27646116 |
| AC008014.1 | 1259.29803 | 0.154882968 | 0.33015638 | 0.46912002 | 0.63898385 | 0.91521016 |
| SLC38A4 | 136.158824 | 1.716572023 | 0.47022444 | 3.65053768 | 0.00026169 | 0.01585278 |
| PCED1B | 166.742873 | -3.381749511 | 0.86863751 | -3.8931654 | 9.89E-05 | 0.00886273 |
| RPAP3 | 85.2358332 | 0.355208929 | 0.33202991 | 1.06981003 | 0.28470483 | 0.73512071 |
| RAPGEF3 | 502.329676 | 0.773501799 | 0.52816777 | 1.46450019 | 0.14305733 | 0.5729912 |
| SLC48A1 | 187.957078 | -0.071101084 | 0.45025148 | -0.1579142 | 0.87452444 | 0.97622753 |
| HDAC7 | 306.436577 | 0.035313247 | 0.36264901 | 0.09737583 | 0.92242794 | 0.9866345 |
| VDR | 39.6798855 | -1.246305746 | 0.42826783 | -2.9101083 | 0.00361304 | 0.09198346 |
| TMEM106C | 98.8483337 | 0.031812519 | 0.42694196 | 0.07451252 | 0.94060258 | 0.98855185 |
| SENP1 | 100.427891 | 0.063172888 | 0.21065341 | 0.29989018 | 0.76426093 | 0.95378284 |
| PFKM | 70.6309347 | -0.007081936 | 0.28755088 | -0.0246285 | 0.98035132 | 0.99482084 |
| ASB8 | 88.783444 | -0.096872696 | 0.33569954 | -0.2885696 | 0.77291073 | 0.95543807 |
| ZNF641 | 50.9247928 | 0.026117343 | 0.29178219 | 0.08950972 | 0.92867683 | 0.98761086 |
| AC090115.1 | 38.9487884 | -0.540541255 | 0.50638725 | -1.0674464 | 0.2857703 | 0.73607046 |
| KANSL2 | 51.7978809 | 0.278724695 | 0.26128239 | 1.06675652 | 0.2860818 | 0.73607046 |
| CCNT1 | 139.422899 | 0.476036229 | 0.22631168 | 2.10345411 | 0.03542609 | 0.31644624 |
| ADCY6 | 170.833657 | 0.576703242 | 0.23968409 | 2.40609728 | 0.01612397 | 0.20785139 |
| DDX23 | 89.8742391 | -0.118773728 | 0.28617142 | -0.415044 | 0.6781097 | 0.9268955 |
| CCDC65 | 76.8284043 | -0.302214677 | 0.35066803 | -0.8618256 | 0.3887835 | 0.8065026 |
| PRKAG1 | 100.417926 | -0.43119988 | 0.26209385 | -1.6452117 | 0.09992616 | 0.4953766 |
| KMT2D | 97.1374617 | -0.107017397 | 0.27858067 | -0.3841523 | 0.70086559 | 0.93180634 |
| LMBR1L | 89.5702932 | -0.315155822 | 0.2098337 | -1.5019314 | 0.13311482 | 0.55601567 |
| AC011603.2 | 34.6474032 | 0.45102342 | 0.44520938 | 1.0130591 | 0.31103193 | 0.75435382 |
| TUBA1B | 457.176001 | -0.866168393 | 0.66903295 | -1.2946573 | 0.19543847 | 0.64948524 |
| TUBA1A | 227.182226 | -1.781787647 | 0.86748357 | -2.0539728 | 0.03997832 | 0.33668856 |
| TUBA1C | 245.360141 | -0.275894693 | 0.40222756 | -0.6859169 | 0.49276549 | 0.86213395 |
| DNAJC22 | 32.1404299 | -0.44137746 | 0.61233395 | -0.7208117 | 0.47102539 | 0.85236058 |
| SPATS2 | 776.658629 | 0.336156052 | 0.31072532 | 1.08184314 | 0.27932223 | 0.72938673 |
| MCRS1 | 55.8586398 | -0.013812024 | 0.28780597 | -0.0479907 | 0.96172362 | 0.99171131 |
| PRPF40B | 228.396552 | -0.243240194 | 0.25611105 | -0.949745 | 0.34224183 | 0.78005531 |
| FMNL3 | 69.3358981 | -0.42847652 | 0.5544081 | -0.772854 | 0.43960879 | 0.83634598 |
| TMBIM6 | 1131.3004 | -0.265269232 | 0.31831807 | -0.8333464 | 0.40464937 | 0.81914727 |
| NCKAP5L | 63.1826084 | 0.429711193 | 0.31714221 | 1.35494797 | 0.17543414 | 0.62477136 |
| AQP2 | 9762.38265 | 1.123941294 | 0.62725337 | 1.79184576 | 0.07315768 | 0.43537334 |
| AC025154.2 | 178.193546 | 1.923398534 | 0.73396015 | 2.62057626 | 0.00877813 | 0.1483787 |
| SMARCD1 | 79.53272 | -0.401989005 | 0.28656248 | -1.4027971 | 0.16067736 | 0.60010648 |
| AC074032.1 | 186.696158 | -0.215566586 | 0.26427458 | -0.8156917 | 0.41467648 | 0.82461178 |
| CERS5 | 154.110796 | 0.059188514 | 0.21845678 | 0.27093924 | 0.78643776 | 0.9576974 |
| LIMA1 | 1008.83766 | -0.544861143 | 0.37771026 | -1.4425373 | 0.14915086 | 0.58194377 |
| LARP4 | 388.282794 | 0.185323507 | 0.16295147 | 1.13729264 | 0.25541597 | 0.71266224 |
| DIP2B | 557.380542 | 0.22074843 | 0.29363031 | 0.75179033 | 0.45217716 | 0.84234337 |
| ATF1 | 167.355925 | 0.07674125 | 0.22050275 | 0.34802854 | 0.72781875 | 0.93931348 |
| METTL7A | 217.480357 | 0.360339463 | 0.43251138 | 0.83313291 | 0.40476977 | 0.81923276 |
| SLC11A2 | 1009.6862 | -0.629492535 | 0.32729152 | -1.923339 | 0.05443749 | 0.39095834 |
| LETMD1 | 225.688989 | -0.156357861 | 0.19206206 | -0.8141007 | 0.41558725 | 0.82487773 |
| CSRNP2 | 68.2796078 | -0.68926462 | 0.36413262 | -1.8928945 | 0.05837191 | 0.39982063 |
| TFCP2 | 451.227678 | 0.332541144 | 0.20463157 | 1.62507255 | 0.1041471 | 0.5032096 |
| POU6F1 | 102.389391 | 0.539183977 | 0.33610891 | 1.60419426 | 0.10867124 | 0.51254482 |
| DAZAP2 | 565.055585 | 0.013426658 | 0.3154562 | 0.04256267 | 0.96605015 | 0.99284614 |
| SMAGP | 37.9979067 | -0.168385962 | 0.40942058 | -0.4112787 | 0.68086819 | 0.92729027 |
| FIGNL2 | 51.4678417 | -0.130152247 | 0.4282626 | -0.3039076 | 0.76119831 | 0.95354509 |
| ACVR1B | 602.916339 | -0.016642502 | 0.42204595 | -0.0394329 | 0.96854524 | 0.99303855 |
| NR4A1 | 651.622634 | 4.723861954 | 1.05068765 | 4.49597173 | 6.93E-06 | 0.00142308 |

| | | | | | | |
|---|---|---|---|---|---|---|
| ATG101 | 62.7952863 | -0.108321792 | 0.42983147 | -0.2520099 | 0.8010334 | 0.96063297 |
| AC078864.2 | 68.0691921 | 0.398421822 | 0.7840209 | 0.50817755 | 0.61132883 | 0.90658678 |
| AC078864.1 | 166.783883 | 0.280129202 | 0.53017176 | 0.52837443 | 0.59723948 | 0.90369786 |
| KRT7 | 227.278188 | -1.324353963 | 0.60325251 | -2.1953559 | 0.02813808 | 0.28491502 |
| KRT8 | 722.124091 | -0.751353493 | 0.56772134 | -1.3234547 | 0.1856842 | 0.63626275 |
| KRT18 | 834.989409 | -0.805354958 | 0.61691342 | -1.3054586 | 0.19173672 | 0.64458858 |
| EIF4B | 427.342437 | -0.006632403 | 0.27842971 | -0.0238207 | 0.98099559 | 0.99507903 |
| AC068888.1 | 146.599711 | -0.230627579 | 0.2895732 | -0.7964397 | 0.42577654 | 0.82774157 |
| TNS2 | 59.6449924 | -1.172594456 | 0.48475616 | -2.4189367 | 0.01556595 | 0.2032348 |
| SPRYD3 | 40.9667994 | -0.241252559 | 0.35428235 | -0.6809613 | 0.495896 | 0.86452555 |
| CSAD | 251.06643 | 0.327747318 | 0.29358719 | 1.1163543 | 0.26427051 | 0.71989632 |
| ZNF740 | 52.1403052 | 0.087417804 | 0.31183213 | 0.2803361 | 0.77921965 | 0.95623735 |
| RARG | 79.7518273 | 0.236443503 | 0.38109639 | 0.62042966 | 0.53497495 | 0.88137638 |
| MFSD5 | 36.7989491 | -0.173358397 | 0.41549607 | -0.4172323 | 0.67650847 | 0.9268955 |
| PFDN5 | 440.933131 | 0.291875653 | 0.47444516 | 0.61519365 | 0.53842684 | 0.88265739 |
| C12orf10 | 47.3596472 | 0.055628132 | 0.40312375 | 0.13799269 | 0.89024619 | 0.98074005 |
| AAAS | 42.8081747 | -0.133548225 | 0.35890434 | -0.3720998 | 0.70981856 | 0.93453499 |
| SP1 | 201.803557 | -0.218574487 | 0.24804487 | -0.8811893 | 0.37821536 | 0.80215442 |
| PRR13 | 235.894336 | -0.416060765 | 0.38494582 | -1.0808294 | 0.279773 | 0.72938673 |
| PCBP2 | 566.013632 | 0.169029244 | 0.21193075 | 0.79756827 | 0.42512107 | 0.82774157 |
| MAP3K12 | 58.0594322 | -0.670338676 | 0.38558133 | -1.7385143 | 0.08212023 | 0.45535449 |
| ATP5MC2 | 532.60026 | 0.203389586 | 0.38754396 | 0.52481681 | 0.59971055 | 0.90369786 |
| CALCOCO1 | 108.875426 | 0.160304002 | 0.29041052 | 0.55199104 | 0.5809545 | 0.90045866 |
| SMUG1 | 58.0997965 | 0.455838483 | 0.32492587 | 1.40289995 | 0.16064668 | 0.60010648 |
| LINC02381 | 89.1206424 | -0.041237969 | 0.35955742 | -0.1146909 | 0.90869012 | 0.98427726 |
| CBX5 | 469.201012 | -0.117521268 | 0.21305189 | -0.5516087 | 0.5812165 | 0.90045866 |
| HNRNPA1 | 319.58324 | 0.411291668 | 0.36495431 | 1.12696755 | 0.2597562 | 0.7168415 |
| COPZ1 | 189.792507 | -0.199173552 | 0.23652922 | -0.8420674 | 0.39975021 | 0.81574763 |
| CD63 | 596.493001 | -1.005574574 | 0.4990901 | -2.0148157 | 0.04392396 | 0.35300853 |
| ORMDL2 | 63.6871063 | -0.572733451 | 0.43139154 | -1.3276418 | 0.18429646 | 0.63497106 |
| PYM1 | 35.5394179 | -0.469470986 | 0.2688709 | -1.7460833 | 0.08079648 | 0.45329073 |
| RAB5B | 180.483371 | -0.068177031 | 0.24815051 | -0.2747406 | 0.7835155 | 0.9568898 |
| SUOX | 87.1475216 | -0.180045724 | 0.40134748 | -0.4486031 | 0.653718 | 0.91983951 |
| IKZF4 | 57.9945298 | -0.278840765 | 0.30361926 | -0.9183896 | 0.35841494 | 0.78746091 |
| RPS26 | 104.936399 | -0.737452554 | 0.42967325 | -1.7163101 | 0.0861053 | 0.46572214 |
| ERBB3 | 69.3689298 | -1.027746909 | 0.39044822 | -2.6322233 | 0.00848281 | 0.14526117 |
| PA2G4 | 83.2556182 | -0.170115878 | 0.3586902 | -0.4742697 | 0.63530763 | 0.91454294 |
| RPL41 | 2120.69781 | 0.226844424 | 0.42355798 | 0.53556877 | 0.59225662 | 0.90233654 |
| ESYT1 | 102.656721 | -0.67517755 | 0.29575662 | -2.2828823 | 0.0224373 | 0.25166501 |
| MYL6B | 64.4171982 | 0.024915999 | 0.43586188 | 0.0571649 | 0.95441384 | 0.99060627 |
| MYL6 | 927.382018 | 0.473882916 | 0.46829933 | 1.0119231 | 0.31157482 | 0.75498361 |
| SMARCC2 | 237.41906 | -0.237280695 | 0.23065154 | -1.028741 | 0.30360141 | 0.74953627 |
| RNF41 | 66.0992722 | -0.444264126 | 0.27848812 | -1.5952714 | 0.11065157 | 0.5181539 |
| NABP2 | 37.0404256 | -0.327479998 | 0.30583769 | -1.070764 | 0.28427554 | 0.73476311 |
| ANKRD52 | 79.1459125 | 0.019159891 | 0.29726401 | 0.06445412 | 0.94860864 | 0.98977092 |
| CS | 124.088126 | 0.128092486 | 0.30523336 | 0.41965428 | 0.67473803 | 0.92632004 |
| PAN2 | 78.786446 | 0.251510589 | 0.23021681 | 1.09249445 | 0.27461583 | 0.7263609 |
| STAT2 | 138.059741 | 0.067821484 | 0.40106987 | 0.16910142 | 0.86571687 | 0.9750363 |
| SPRYD4 | 46.9506708 | -0.279262966 | 0.33688535 | -0.8289555 | 0.40712958 | 0.82018309 |
| RBMS2 | 326.940196 | 0.879139883 | 0.1702074 | 5.16510956 | 2.40E-07 | 0.00012961 |
| BAZ2A | 214.713401 | -0.036223783 | 0.17269904 | -0.2097509 | 0.83386208 | 0.96749268 |
| ATP5F1B | 638.878877 | 0.012162156 | 0.42932712 | 0.02832841 | 0.97740022 | 0.99469867 |
| PTGES3 | 795.022672 | 0.612654254 | 0.23241129 | 2.63607786 | 0.00838705 | 0.1442148 |
| NACA | 463.688074 | 0.065168506 | 0.50757146 | 0.12839277 | 0.89783815 | 0.98312332 |
| NEMP1 | 45.9171928 | -0.981801143 | 0.2664806 | -3.684325 | 0.00022931 | 0.01460392 |
| STAT6 | 299.202064 | -0.436198502 | 0.25908194 | -1.6836315 | 0.09225291 | 0.48040835 |
| LRP1 | 45.2105827 | -0.218026865 | 0.33982117 | -0.6415929 | 0.52113751 | 0.87650871 |
| R3HDM2 | 619.562483 | 0.278373412 | 0.22299535 | 1.24833732 | 0.21190755 | 0.66975892 |
| MARS | 80.3737665 | 0.035302401 | 0.40879413 | 0.08635741 | 0.93118231 | 0.98822345 |
| DDIT3 | 223.257429 | 0.228118231 | 0.93281574 | 0.24454801 | 0.80680643 | 0.96218371 |
| MBD6 | 86.213692 | 0.396244944 | 0.25223985 | 1.57090538 | 0.11620463 | 0.53066791 |
| DCTN2 | 181.127194 | -0.2945302 | 0.33902106 | -0.8687667 | 0.38497476 | 0.8056181 |
| PIP4K2C | 114.682443 | -0.497638677 | 0.2341 | -2.1257526 | 0.03352387 | 0.30964163 |
| DTX3 | 61.7346269 | 0.337922807 | 0.30035418 | 1.12508109 | 0.26055467 | 0.7176381 |
| OS9 | 258.983612 | -0.299030601 | 0.23699656 | -1.2617508 | 0.20703847 | 0.66434881 |
| TSPAN31 | 70.1464777 | -0.194067054 | 0.32225265 | -0.6022202 | 0.54702757 | 0.88743792 |
| CDK4 | 70.2116364 | -0.431293269 | 0.35318214 | -1.2211639 | 0.22202398 | 0.67955938 |
| 9-Mar | 61.5709187 | -0.320725632 | 0.33497926 | -0.9574492 | 0.33834057 | 0.77836439 |
| TSFM | 88.2503758 | -0.622432336 | 0.48723622 | -1.2774755 | 0.20143443 | 0.65580391 |
| AVIL | 47.3403499 | -0.916128908 | 0.59575636 | -1.5377577 | 0.12410788 | 0.54273186 |
| CTDSP2 | 255.071225 | -0.23126362 | 0.25280966 | -0.9147737 | 0.36031046 | 0.78892604 |
| LRIG3 | 75.7107555 | 0.210719008 | 0.41824754 | 0.50381411 | 0.61439201 | 0.90764425 |
| SLC16A7 | 497.954629 | -0.111746584 | 0.31841129 | -0.3509504 | 0.72562552 | 0.93895336 |
| FAM19A2 | 214.135782 | -0.093339841 | 0.4190019 | -0.2227671 | 0.82371677 | 0.96511681 |
| USP15 | 575.810717 | 0.58157725 | 0.26237582 | 2.21658093 | 0.02665174 | 0.27764439 |
| MON2 | 1047.31256 | 0.697614676 | 0.19707543 | 3.53983587 | 0.00040038 | 0.02151764 |
| AC048341.1 | 58.9012419 | 0.232090812 | 0.49115471 | 0.47254115 | 0.63654058 | 0.91454294 |
| PPM1H | 485.56644 | -0.032256018 | 0.29794006 | -0.1082634 | 0.91378672 | 0.98480378 |
| DPY19L2 | 665.799316 | 0.73211582 | 0.30347237 | 2.41246284 | 0.01584515 | 0.2052623 |
| RXYLT1 | 69.8638304 | -0.344763717 | 0.27381722 | -1.2591016 | 0.20799362 | 0.66589349 |
| SRGAP1 | 150.152576 | -1.00687765 | 0.51755509 | -1.9454502 | 0.05172082 | 0.38247133 |
| XPOT | 91.056251 | -0.027393537 | 0.25916534 | -0.1056991 | 0.91582111 | 0.98520052 |
| TBK1 | 228.741775 | -0.088250702 | 0.27639183 | -0.3192956 | 0.74950236 | 0.94706664 |
| RASSF3 | 77.350566 | -0.909480348 | 0.44910889 | -2.0250776 | 0.0428594 | 0.3484612 |
| GNS | 455.075481 | -0.293598378 | 0.27546509 | -1.0658279 | 0.28650144 | 0.73607046 |
| TBC1D30 | 167.605737 | -0.127296535 | 0.26742233 | -0.4760131 | 0.63406504 | 0.91415622 |
| LEMD3 | 269.312142 | 0.656532913 | 0.237269 | 2.76704047 | 0.00565677 | 0.12123309 |
| AC090673.1 | 65.9976936 | -1.243727699 | 1.01354709 | -1.227104 | 0.2197835 | 0.67801071 |
| LLPH | 50.2775489 | -0.136784285 | 0.36503464 | -0.3747159 | 0.70787176 | 0.9330268 |
| HELB | 92.5791583 | -0.285906998 | 0.31522236 | -0.907001 | 0.36440626 | 0.79268955 |
| GRIP1 | 4720.01277 | 1.329616408 | 0.2642191 | 5.03224933 | 4.85E-07 | 0.00020321 |
| CAND1 | 320.319933 | 0.30026671 | 0.16996028 | 1.76668751 | 0.07728058 | 0.44472917 |
| DYRK2 | 46.2915855 | -1.13947393 | 0.47357935 | -2.4060887 | 0.01612435 | 0.20785139 |
| MDM1 | 48.5998799 | 0.061129274 | 0.27120574 | 0.22539816 | 0.82166955 | 0.96502145 |
| RAP1B | 1611.14686 | -0.239070408 | 0.56283109 | -0.424764 | 0.67100869 | 0.92474338 |
| NUP107 | 174.310087 | -0.322446156 | 0.30614656 | -1.0532412 | 0.29223048 | 0.73844969 |
| SLC35E3 | 136.03039 | -0.972503296 | 0.40619239 | -2.3941938 | 0.01665695 | 0.21108198 |
| MDM2 | 274.658345 | -0.308531865 | 0.34191358 | -0.902368 | 0.36686143 | 0.79370516 |
| CPM | 50.0561963 | -0.007675471 | 0.55178062 | -0.0139104 | 0.98890149 | 0.9964791 |
| CPSF6 | 415.114311 | 0.557384079 | 0.20184941 | 2.76138572 | 0.00575567 | 0.12145831 |
| YEATS4 | 82.9664752 | 0.012910377 | 0.42162951 | 0.03062019 | 0.97557244 | 0.9939318 |
| FRS2 | 439.162562 | 0.620864016 | 0.18389407 | 3.37620466 | 0.00073493 | 0.03279545 |
| CCT2 | 152.033865 | 0.000626 | 0.30804202 | 0.00203219 | 0.99837855 | 0.99954455 |
| RAB3IP | 271.102213 | 0.191524607 | 0.19378906 | 0.98831484 | 0.32299848 | 0.76879947 |
| MYRFL | 74.2000275 | 0.969528191 | 0.74377685 | 1.30352026 | 0.1923972 | 0.6454298 |
| AC025159.1 | 186.317266 | -1.488046487 | 0.59676524 | -2.4935207 | 0.01264832 | 0.17941309 |
| CNOT2 | 586.916366 | 0.475542112 | 0.24017719 | 1.97996372 | 0.04770761 | 0.36898576 |
| KCNMB4 | 65.1669968 | 0.515008314 | 0.94839795 | 0.54302976 | 0.58710931 | 0.90196444 |
| ZFC3H1 | 366.679759 | 0.102592132 | 0.2312627 | 0.44361728 | 0.65731931 | 0.92160948 |
| TMEM19 | 35.9833711 | 0.071814588 | 0.33685253 | 0.21319296 | 0.83117646 | 0.96656674 |
| RAB21 | 383.59546 | 0.223278021 | 0.28964554 | 0.77086641 | 0.44078612 | 0.83733111 |
| TBC1D15 | 307.127457 | 0.019042554 | 0.37041488 | 0.05140872 | 0.95899984 | 0.99119981 |
| TRHDE | 721.982401 | 0.801536175 | 0.32526848 | 2.46422946 | 0.01373082 | 0.18912039 |

| | | | | | | |
|---|---|---|---|---|---|---|
| ATXN7L3B | 187.358762 | -0.390457493 | 0.37622292 | -1.0378355 | 0.29934666 | 0.74594723 |
| CAPS2 | 293.602314 | 0.33675262 | 0.31629067 | 1.0646935 | 0.28701465 | 0.73638964 |
| KRR1 | 141.35752 | 0.233675353 | 0.26907015 | 0.86845513 | 0.38514523 | 0.80565309 |
| PHLDA1 | 188.369895 | -0.618664202 | 0.61073154 | -1.0129888 | 0.31106551 | 0.75435382 |
| NAP1L1 | 456.745083 | 0.259407645 | 0.28329156 | 0.9156914 | 0.35982878 | 0.78892604 |
| OSBPL8 | 541.636279 | -0.062735733 | 0.26015374 | -0.2411487 | 0.80943988 | 0.96245698 |
| ZDHHC17 | 329.797688 | 0.359291702 | 0.23594874 | 1.52275321 | 0.12782046 | 0.54791812 |
| AC027288.3 | 78.0846591 | 0.235559186 | 0.64182843 | 0.3670127 | 0.71360954 | 0.93573028 |
| PAWR | 928.838127 | 0.195244701 | 0.27725559 | 0.70420475 | 0.48130528 | 0.85611389 |
| AC073569.2 | 54.7441024 | 0.069973764 | 0.4995142 | 0.14008363 | 0.88859391 | 0.98051184 |
| PPP1R12A | 725.768026 | 0.393184126 | 0.17243346 | 2.28020777 | 0.02259537 | 0.25181478 |
| LIN7A | 327.971724 | 0.010037414 | 0.48749423 | 0.02058981 | 0.98357287 | 0.99540861 |
| ACSS3 | 254.829568 | -0.395490169 | 0.54137797 | -0.7305251 | 0.4650693 | 0.84991999 |
| PPFIA2 | 183.717702 | -0.722336941 | 0.8139394 | -0.8874579 | 0.37483243 | 0.79825864 |
| CCDC59 | 75.0064686 | -0.250370702 | 0.46529526 | -0.5380899 | 0.59051495 | 0.90233654 |
| METTL25 | 195.10881 | 0.309866821 | 0.30344245 | 1.02117163 | 0.30717313 | 0.75214355 |
| TMTC2 | 3622.43225 | 0.711621097 | 0.36754878 | 1.93612693 | 0.05285216 | 0.38625565 |
| LRRIQ1 | 245.973315 | 0.193055863 | 0.2651213 | 0.72817937 | 0.4665038 | 0.85057334 |
| RASSF9 | 104.756076 | 0.255532867 | 0.36584417 | 0.69847462 | 0.48488043 | 0.85909499 |
| C12orf29 | 39.6772812 | -0.324275721 | 0.37261743 | -0.8702645 | 0.38415587 | 0.80561637 |
| CEP290 | 288.828607 | 0.156291286 | 0.20968615 | 0.74535817 | 0.45605522 | 0.84372683 |
| TMTC3 | 187.360352 | 0.16688832 | 0.26952229 | 0.61920045 | 0.53578432 | 0.88137638 |
| KITLG | 2086.92693 | 0.024844745 | 0.46578398 | 0.05333963 | 0.95746131 | 0.99106618 |
| DUSP6 | 607.397 | 0.105921968 | 0.88754288 | 0.11934293 | 0.90500367 | 0.98320957 |
| POC1B | 202.476506 | 0.305732982 | 0.26011594 | 1.17537198 | 0.23984593 | 0.6960912 |
| POC1B-AS1 | 72.5463574 | 0.153612861 | 0.40466519 | 0.37960483 | 0.70423877 | 0.93241442 |
| ATP2B1 | 329.022434 | 0.676088125 | 0.35058214 | 1.92847279 | 0.05379635 | 0.38935477 |
| BTG1 | 398.840983 | -0.238448659 | 0.55310003 | -0.4311131 | 0.66638615 | 0.92396246 |
| PLEKHG7 | 53.3061108 | -2.285098731 | 0.82065776 | -2.7844722 | 0.00536149 | 0.11705923 |
| EEA1 | 296.513733 | -0.034715208 | 0.20869154 | -0.166347 | 0.86788389 | 0.97576204 |
| AC138123.1 | 218.161786 | 0.50562407 | 0.35096294 | 1.44067654 | 0.14967609 | 0.58312467 |
| AC124947.1 | 64.6069698 | 0.255187019 | 0.49743727 | 0.51300342 | 0.60794893 | 0.90514346 |
| NUDT4 | 294.719801 | 1.094014601 | 0.38635936 | 2.83159851 | 0.0046316 | 0.10762557 |
| UBE2N | 195.604131 | 0.011830066 | 0.18404197 | 0.06427917 | 0.94874794 | 0.98977092 |
| MRPL42 | 194.728089 | -0.194893204 | 0.21754302 | -0.8958835 | 0.37031498 | 0.79532503 |
| SOCS2 | 116.382347 | 0.78178876 | 0.52281011 | 1.49535893 | 0.13482079 | 0.55868799 |
| CRADD | 947.805531 | -0.061647855 | 0.25966085 | -0.2374168 | 0.81233344 | 0.96288868 |
| AC012085.2 | 83.8629143 | -0.409003176 | 0.61498423 | -0.6650629 | 0.50601029 | 0.87076975 |
| CEP83 | 228.665231 | 0.248851758 | 0.17770836 | 1.4003379 | 0.16141216 | 0.60092341 |
| TMCC3 | 149.02779 | 0.753266256 | 0.33477955 | 2.25003665 | 0.02444662 | 0.26518037 |
| NDUFA12 | 189.637171 | 0.038199499 | 0.31792135 | 0.12015393 | 0.90436121 | 0.98320957 |
| NR2C1 | 195.81098 | 0.121653986 | 0.23892873 | 0.50916433 | 0.61063704 | 0.9062166 |
| FGD6 | 186.324525 | -1.146635016 | 0.40433457 | -2.835857 | 0.00457029 | 0.10667403 |
| VEZT | 640.979465 | 0.042514386 | 0.28459942 | 0.14938325 | 0.88125123 | 0.97768262 |
| METAP2 | 1011.19976 | -0.940254113 | 0.53741096 | -1.7495998 | 0.08018739 | 0.45074086 |
| USP44 | 78.3164204 | 0.095328902 | 0.40464277 | 0.2355878 | 0.81375254 | 0.96288868 |
| NTN4 | 1492.13477 | 1.104405775 | 0.57334507 | 1.9262497 | 0.0540732 | 0.39065232 |
| SNRPF | 78.4557018 | -0.349496858 | 0.40029959 | -0.8730882 | 0.38261499 | 0.80405156 |
| LTA4H | 223.984521 | 0.012405814 | 0.45037087 | 0.02754577 | 0.97802443 | 0.99469867 |
| ELK3 | 87.1530459 | -1.025201027 | 0.41736042 | -2.4563925 | 0.01403398 | 0.19221396 |
| CDK17 | 183.890731 | -0.02807977 | 0.298923 | -0.0939365 | 0.92515963 | 0.98710257 |
| CFAP54 | 104.429981 | -0.144751791 | 0.5113483 | -0.2830787 | 0.77711654 | 0.95601455 |
| NEDD1 | 160.536622 | 0.761619166 | 0.2253966 | 3.37901794 | 0.00072745 | 0.03271976 |
| AC007656.1 | 55.1944004 | 0.046792333 | 0.65594142 | 0.07133615 | 0.94313023 | 0.98885525 |
| TMPO | 225.671426 | -0.015384169 | 0.27686958 | -0.0555647 | 0.95568861 | 0.9908068 |
| SLC25A3 | 538.77688 | -0.009140261 | 0.51476546 | -0.0177562 | 0.98583337 | 0.99587186 |
| IKBIP | 46.3548067 | -1.09763144 | 0.33744211 | -3.2527993 | 0.00114274 | 0.04339107 |
| APAF1 | 148.555268 | -0.464902314 | 0.24984162 | -1.8607881 | 0.0627741 | 0.41131726 |
| ANKS1B | 252.610823 | 1.505824686 | 0.59378356 | 2.53598244 | 0.01121324 | 0.16686755 |
| UHRF1BP1L | 151.88945 | 0.240415188 | 0.22572602 | 1.06507519 | 0.2868419 | 0.73638964 |
| ACTR6 | 106.088285 | -0.182708424 | 0.33541973 | -0.5447158 | 0.585949 | 0.90196444 |
| DEPDC4 | 41.7530483 | -0.465098758 | 0.40423689 | -1.1505599 | 0.24991334 | 0.70674386 |
| SCYL2 | 357.32192 | 0.329324079 | 0.2297076 | 1.43366643 | 0.15166748 | 0.58639009 |
| NR1H4 | 542.053839 | -0.669550953 | 0.69082899 | -0.9691993 | 0.33244578 | 0.77629941 |
| UTP20 | 115.305475 | -0.00173153 | 0.30938299 | -0.0055967 | 0.99553449 | 0.99908534 |
| ARL1 | 197.86086 | -0.047530478 | 0.28957068 | -0.1641412 | 0.86961999 | 0.97576204 |
| CHPT1 | 614.317615 | 0.974910048 | 0.35348095 | 2.75802714 | 0.00581514 | 0.12212953 |
| GNPTAB | 244.054282 | -0.094799295 | 0.21869743 | -0.4334724 | 0.66467163 | 0.92324595 |
| DRAM1 | 160.035144 | -0.165405049 | 0.37581067 | -0.4401287 | 0.65984392 | 0.92284375 |
| WASHC3 | 195.339981 | 0.019218133 | 0.2392221 | 0.08033594 | 0.93597007 | 0.98822345 |
| NUP37 | 80.0487833 | 0.072629357 | 0.21120845 | 0.34387524 | 0.73094011 | 0.94048525 |
| NT5DC3 | 51.8269232 | -0.639336426 | 0.40934317 | -1.5618593 | 0.11832114 | 0.5340248 |
| HSP90B1 | 567.973933 | -0.369236836 | 0.40902925 | -0.902715 | 0.36667718 | 0.7935157 |
| C12orf73 | 41.1423556 | -0.039022979 | 0.29078537 | -0.1341986 | 0.89324556 | 0.98168537 |
| TDG | 110.678291 | -0.025098197 | 0.19412853 | -0.1292865 | 0.89713096 | 0.98302816 |
| HCFC2 | 82.1936221 | 0.466864054 | 0.20605919 | 2.26567941 | 0.02347102 | 0.25893742 |
| NFYB | 81.6611482 | 0.070998363 | 0.23943372 | 0.29652616 | 0.76682829 | 0.95408212 |
| TXNRD1 | 958.776245 | -0.7707077 | 0.47712353 | -1.6153211 | 0.10624118 | 0.50765436 |
| SLC41A2 | 64.367712 | -1.292740359 | 0.51724656 | -2.499273 | 0.01244484 | 0.17817202 |
| C12orf45 | 42.2004361 | -0.032684579 | 0.3224899 | -0.1013507 | 0.91927207 | 0.98520052 |
| WASHC4 | 437.423449 | 0.275533759 | 0.24929608 | 1.10524704 | 0.26905256 | 0.72322263 |
| APPL2 | 294.127435 | -0.831895368 | 0.31133844 | -2.671997 | 0.00754013 | 0.13878787 |
| C12orf75 | 500.003373 | -1.405513589 | 0.52050236 | -2.700302 | 0.00692766 | 0.13285432 |
| NUAK1 | 41.2479664 | 0.234554175 | 0.49340095 | 0.47538249 | 0.63451437 | 0.91430092 |
| CKAP4 | 84.9340048 | -0.368535525 | 0.34144445 | -1.0793426 | 0.28043505 | 0.72945471 |
| TCP11L2 | 83.0825045 | -0.97508759 | 0.33519005 | -2.9090589 | 0.00362519 | 0.09199019 |
| POLR3B | 88.4399354 | 0.247153797 | 0.30379259 | 0.81356099 | 0.41589649 | 0.82489761 |
| RIC8B | 188.409687 | 0.340672192 | 0.22488412 | 1.51487883 | 0.12980309 | 0.55275758 |
| TMEM263 | 195.328387 | 0.072427389 | 0.27882731 | 0.25975716 | 0.7950511 | 0.95937941 |
| MTERF2 | 64.6507594 | 0.061895195 | 0.3283964 | 0.18847708 | 0.85050268 | 0.97017003 |
| CRY1 | 173.028095 | 0.463458934 | 0.54892407 | 0.84430426 | 0.3984994 | 0.81400298 |
| PWP1 | 92.6832726 | 0.110541804 | 0.24220453 | 0.45639858 | 0.64810339 | 0.91768694 |
| PRDM4 | 141.586511 | -0.327730477 | 0.28564562 | -1.1473324 | 0.25124426 | 0.70813507 |
| SART3 | 107.47794 | 0.27953347 | 0.17102974 | 1.6344144 | 0.10217187 | 0.49942218 |
| ISCU | 377.050539 | -0.211357565 | 0.51699403 | -0.4088201 | 0.68267167 | 0.92806156 |
| CORO1C | 83.140349 | -1.055487185 | 0.5006068 | -2.1084156 | 0.03499506 | 0.31514522 |
| SSH1 | 314.373383 | 0.368053495 | 0.2420925 | 1.52030112 | 0.12843531 | 0.54843482 |
| USP30 | 69.0138912 | 0.462212445 | 0.23233758 | 1.98940026 | 0.04665704 | 0.36273427 |
| UNG | 109.301288 | 0.825481396 | 0.54008374 | 1.52843222 | 0.12640526 | 0.54650459 |
| ACACB | 899.799482 | 0.757297001 | 0.52381789 | 1.44572574 | 0.14825414 | 0.58038977 |
| MYO1H | 36.9735335 | 2.091252155 | 0.62867076 | 3.32646639 | 0.00087955 | 0.03655313 |
| KCTD10 | 85.0930729 | -0.579806186 | 0.27363189 | -2.1189277 | 0.03409658 | 0.3120805 |
| UBE3B | 142.644962 | -0.224177365 | 0.21914544 | -1.0229616 | 0.30632602 | 0.75140194 |
| MMAB | 132.978215 | -0.486644185 | 0.37671855 | -1.2917978 | 0.19642719 | 0.65097045 |
| MVK | 108.267569 | -0.349460324 | 0.42612872 | -0.8200816 | 0.41216958 | 0.82355334 |
| TRPV4 | 445.170552 | 0.443606548 | 0.39288616 | 1.1290969 | 0.25885696 | 0.7159728 |
| GLTP | 197.609169 | 0.699975268 | 0.36086353 | 1.93972292 | 0.05241337 | 0.38418477 |
| TCHP | 141.012141 | 0.385797027 | 0.35119954 | 1.09851233 | 0.27198084 | 0.72372882 |
| GIT2 | 340.666388 | 1.006964903 | 0.46195721 | 2.17977961 | 0.0292738 | 0.28931514 |
| ANKRD13A | 661.947182 | 0.658288624 | 0.51285859 | 1.28356752 | 0.19929332 | 0.65475264 |
| C12orf76 | 89.961479 | 0.682711086 | 0.53865875 | 1.26742782 | 0.20500236 | 0.66065163 |
| IFT81 | 224.134628 | 0.136736343 | 0.18189845 | 0.75171801 | 0.45222066 | 0.84234337 |
| ATP2A2 | 291.381182 | -0.090079869 | 0.33573613 | -0.2683056 | 0.78846413 | 0.95801395 |

| | | | | | | |
|---|---|---|---|---|---|---|
| ANAPC7 | 123.853983 | 0.125695483 | 0.34594563 | 0.36333883 | 0.71635179 | 0.93573028 |
| ARPC3 | 563.448177 | -0.231989785 | 0.46401849 | -0.4999581 | 0.61710461 | 0.90794578 |
| GPN3 | 55.7737972 | -0.235349475 | 0.38548238 | -0.6105324 | 0.5415092 | 0.88410086 |
| VPS29 | 232.781353 | -0.138695958 | 0.32661888 | -0.4246416 | 0.67109797 | 0.92474338 |
| PPTC7 | 116.865141 | -0.434548344 | 0.26703505 | -1.6273083 | 0.10367164 | 0.50253415 |
| TCTN1 | 102.205138 | -0.156023372 | 0.37859064 | -0.4121163 | 0.68025419 | 0.92700898 |
| PPP1CC | 117.864951 | 0.27332665 | 0.27730921 | 0.98563855 | 0.32431051 | 0.76970832 |
| SH2B3 | 57.0922596 | -0.731883531 | 0.41251364 | -1.7742045 | 0.07602934 | 0.4426298 |
| ATXN2 | 902.224375 | 0.166988109 | 0.28323831 | 0.58956752 | 0.55548063 | 0.89081623 |
| BRAP | 119.238577 | 0.012639806 | 0.25991661 | 0.04863024 | 0.96121397 | 0.99171131 |
| ACAD10 | 135.150577 | 0.005091666 | 0.35768355 | 0.01423511 | 0.98864241 | 0.9964791 |
| ALDH2 | 371.312175 | -0.716807086 | 0.35385534 | -2.0257066 | 0.04279486 | 0.3484612 |
| MAPKAPK5 | 131.733365 | 0.390175376 | 0.17455913 | 2.2352047 | 0.02540391 | 0.27138941 |
| TMEM116 | 370.293579 | 0.178710832 | 0.22386838 | 0.79828529 | 0.42470495 | 0.82774157 |
| ERP29 | 456.206875 | -0.161894215 | 0.45147826 | -0.358587 | 0.71990411 | 0.93651875 |
| NAA25 | 220.949283 | 0.300237375 | 0.27806972 | 1.07971976 | 0.28026699 | 0.72945471 |
| TRAFD1 | 110.489079 | 0.0380343 | 0.28345332 | 0.13418188 | 0.89325875 | 0.98168537 |
| HECTD4 | 780.476154 | 0.224869136 | 0.21912339 | 1.0262215 | 0.30478719 | 0.74998115 |
| RPL6 | 633.958003 | -0.4034172 | 0.49337989 | -0.8176604 | 0.41355113 | 0.82440569 |
| PTPN11 | 269.411659 | 0.309776093 | 0.26063733 | 1.18853311 | 0.23462344 | 0.69335782 |
| RASAL1 | 535.947066 | -0.292365906 | 0.53865153 | -0.5427737 | 0.5872856 | 0.90196444 |
| DDX54 | 90.0294536 | -0.289283522 | 0.3240687 | -0.8926611 | 0.37203869 | 0.79586967 |
| IQCD | 42.4037316 | -0.395849237 | 0.26460807 | -1.4959832 | 0.13465802 | 0.55868799 |
| TPCN1 | 686.699745 | -0.26316547 | 0.25111485 | -1.0479885 | 0.29464391 | 0.74067358 |
| SLC8B1 | 137.886408 | -0.193452647 | 0.31183891 | -0.6203608 | 0.53502025 | 0.88137638 |
| PLBD2 | 47.4737983 | -0.431767131 | 0.28225312 | -1.5297161 | 0.12608703 | 0.54580422 |
| SDSL | 77.3714235 | -0.685079134 | 0.540266 | -1.2680404 | 0.20478351 | 0.66034806 |
| RBM19 | 105.663141 | -0.262458687 | 0.32715486 | -0.802246 | 0.42241066 | 0.82757457 |
| TBX3 | 162.08484 | 0.52746133 | 0.6369174 | 0.82814715 | 0.40758717 | 0.82018309 |
| MED13L | 3118.31309 | 0.384481426 | 0.29973521 | 1.28273695 | 0.19958425 | 0.65475264 |
| C12orf49 | 85.0369402 | -0.646712699 | 0.41168779 | -1.5708814 | 0.1162102 | 0.53066791 |
| FBXW8 | 206.833452 | -0.177090022 | 0.25866938 | -0.6846192 | 0.49358425 | 0.86313415 |
| FBXO21 | 254.993312 | 0.251148118 | 0.22168209 | 1.13292019 | 0.25724777 | 0.7152976 |
| KSR2 | 1867.7967 | 0.316676475 | 0.81370971 | 0.38917623 | 0.69714579 | 0.93130502 |
| RFC5 | 42.4200769 | 0.457939447 | 0.41597226 | 1.1008894 | 0.2709448 | 0.72343657 |
| WSB2 | 139.168784 | 0.231577114 | 0.43237712 | 0.53559059 | 0.59224154 | 0.90233654 |
| VSIG10 | 198.015635 | 0.039529511 | 0.32631225 | 0.12114014 | 0.90358004 | 0.98320957 |
| PEBP1 | 1626.9796 | 0.143852847 | 0.57099495 | 0.25193366 | 0.80109234 | 0.96063297 |
| TAOK3 | 679.772473 | 0.514082865 | 0.23176125 | 2.21815709 | 0.02654412 | 0.27684155 |
| SUDS3 | 174.966462 | 0.279337208 | 0.1714498 | 1.62926527 | 0.10325688 | 0.50175883 |
| AC002070.1 | 156.147628 | -0.201892245 | 0.58702198 | -0.3439262 | 0.73090178 | 0.94048525 |
| PRKAB1 | 220.65313 | -0.412029354 | 0.38983177 | -1.0569414 | 0.29053832 | 0.73754646 |
| CIT | 154.227566 | 0.363671898 | 0.65663213 | 0.55384421 | 0.57968547 | 0.89989814 |
| BICDL1 | 89.1091587 | 0.194245556 | 0.36151597 | 0.53730837 | 0.59105463 | 0.90233654 |
| RAB35 | 65.3102304 | -0.113707932 | 0.35684162 | -0.318651 | 0.74999119 | 0.94743342 |
| GCN1 | 122.989356 | 0.01001663 | 0.22510619 | 0.04449735 | 0.96450796 | 0.99243116 |
| RPLP0 | 846.278295 | -0.57740541 | 0.47847125 | -1.2067714 | 0.2275202 | 0.68604702 |
| PXN | 184.018413 | -0.241886488 | 0.15973593 | -1.5142898 | 0.12995236 | 0.55282113 |
| GATC | 56.9182016 | -0.452103508 | 0.26523514 | -1.7045385 | 0.08828053 | 0.47036301 |
| SRSF9 | 319.018242 | 0.314514449 | 0.43541624 | 0.72233054 | 0.47009128 | 0.85232523 |
| DYNLL1 | 506.656271 | -0.350307568 | 0.62136247 | -0.5637733 | 0.57290843 | 0.89797476 |
| COQ5 | 67.3774623 | 0.061300907 | 0.23656922 | 0.25912461 | 0.7955391 | 0.95948555 |
| RNF10 | 485.550096 | 0.223783967 | 0.22535488 | 0.99302916 | 0.32069574 | 0.76568446 |
| POP5 | 80.1259349 | 0.219726412 | 0.35463244 | 0.61958915 | 0.53552831 | 0.88137638 |
| MLEC | 320.983334 | -0.180558101 | 0.22277312 | -0.8105022 | 0.41765161 | 0.82569116 |
| UNC119B | 80.8252343 | 0.420825366 | 0.32508505 | 1.29450853 | 0.19548982 | 0.64948524 |
| ACADS | 89.662606 | -0.013655989 | 0.36391003 | -0.0375257 | 0.97006583 | 0.99348088 |
| SPPL3 | 871.241996 | 0.278930301 | 0.23485098 | 1.18769061 | 0.23495532 | 0.69335782 |
| C12orf43 | 37.7798939 | -0.033643361 | 0.41777255 | -0.0805303 | 0.93581548 | 0.98822345 |
| P2RX4 | 93.9654064 | 0.36067748 | 0.42005261 | 0.85864835 | 0.39053456 | 0.80773675 |
| CAMKK2 | 62.6729394 | -0.064726623 | 0.31632085 | -0.2046233 | 0.83786643 | 0.96757829 |
| ANAPC5 | 231.498637 | 0.288084124 | 0.20044992 | 1.43718751 | 0.15066472 | 0.58480232 |
| RNF34 | 105.193422 | -0.295226853 | 0.25442964 | -1.1603477 | 0.24590727 | 0.70206841 |
| KDM2B | 209.462658 | -0.751489964 | 0.31479189 | -2.3872596 | 0.0169745 | 0.21364736 |
| ORAI1 | 36.3008872 | -1.355288373 | 0.3634434 | -3.7290219 | 0.00019222 | 0.01325337 |
| MORN3 | 36.8209958 | -0.221511535 | 0.46501617 | -0.4763523 | 0.63382339 | 0.91415622 |
| TMEM120B | 158.522462 | 0.366287942 | 0.31041855 | 1.17998083 | 0.23800784 | 0.69406342 |
| RHOF | 73.3763823 | -0.048659093 | 0.71302236 | -0.0682434 | 0.94559185 | 0.98949015 |
| SETD1B | 83.5051758 | -0.30372915 | 0.35962546 | -0.8445708 | 0.39835054 | 0.81400298 |
| WDR66 | 123.018202 | -0.260944408 | 0.41197561 | -0.6333977 | 0.52647396 | 0.87791848 |
| BCL7A | 241.541677 | 0.64077275 | 0.30014387 | 2.13488536 | 0.03277038 | 0.30581793 |
| MLXIP | 199.490668 | -0.406477701 | 0.26955829 | -1.5079399 | 0.13156991 | 0.55397858 |
| CLIP1 | 414.007607 | -0.180433268 | 0.35135385 | -0.5135372 | 0.60757561 | 0.90510401 |
| ZCCHC8 | 137.084771 | 0.131629223 | 0.21243825 | 0.61961169 | 0.53551347 | 0.88137638 |
| RSRC2 | 279.892011 | 0.127654483 | 0.27957002 | 0.45661005 | 0.64795135 | 0.91768694 |
| KNTC1 | 46.9474156 | 0.032912148 | 0.30694414 | 0.1072252 | 0.91461032 | 0.98480378 |
| DENR | 239.315761 | -0.198560032 | 0.20650266 | -0.9615374 | 0.33628203 | 0.77820994 |
| HIP1R | 492.622837 | 0.029218589 | 0.29329211 | 0.09962283 | 0.92064377 | 0.98581108 |
| VPS37B | 612.33814 | 1.51348083 | 0.59031396 | 2.56385742 | 0.01035161 | 0.16264592 |
| ABCB9 | 42.2722554 | 0.329615993 | 0.34278534 | 0.96158135 | 0.33625994 | 0.77820994 |
| ARL6IP4 | 260.00454 | 0.243122579 | 0.40264646 | 0.60381154 | 0.54596895 | 0.88669883 |
| PITPNM2 | 262.832733 | -0.173469125 | 0.31985311 | -0.54234 | 0.58758433 | 0.90196444 |
| MPHOSPH9 | 168.394284 | -0.165937475 | 0.21615077 | -0.7676932 | 0.44266949 | 0.83826789 |
| C12orf65 | 108.25951 | -0.576238237 | 0.37349552 | -1.5428251 | 0.12287323 | 0.54151028 |
| SBNO1 | 282.330514 | 0.095312861 | 0.21438113 | 0.44459538 | 0.65661218 | 0.92131419 |
| KMT5A | 82.0936567 | -0.231203715 | 0.24673433 | -0.9370553 | 0.3487301 | 0.78180304 |
| RILPL2 | 145.509692 | -0.253041471 | 0.40888734 | -0.6188538 | 0.53601269 | 0.88137638 |
| SNRNP35 | 105.48264 | 0.036570829 | 0.24413359 | 0.14979843 | 0.88092365 | 0.97766386 |
| RILPL1 | 193.540678 | -0.371331161 | 0.33110359 | -1.1214954 | 0.26207704 | 0.71989632 |
| TMED2 | 177.413931 | -0.25369682 | 0.25510993 | -0.9944608 | 0.31999859 | 0.76478568 |
| DDX55 | 52.8517523 | -0.082269831 | 0.43927902 | -0.1872838 | 0.85143815 | 0.97053206 |
| EIF2B1 | 79.1014697 | 0.028940423 | 0.25795736 | 0.11219072 | 0.91067218 | 0.98468493 |
| GTF2H3 | 82.3696966 | 0.005821608 | 0.2167354 | 0.02686044 | 0.97857105 | 0.99469867 |
| TCTN2 | 81.1720851 | 0.516929891 | 0.2463565 | 2.09830016 | 0.03587864 | 0.31784288 |
| ATP6V0A2 | 68.7536955 | -0.132164432 | 0.48424585 | -0.2729284 | 0.78490828 | 0.95702626 |
| CCDC92 | 251.490078 | -0.242947365 | 0.17148213 | -1.4167503 | 0.15655595 | 0.59374856 |
| RFLNA | 46.4169526 | -0.365495768 | 0.45648534 | -0.8006736 | 0.42332062 | 0.82774157 |
| NCOR2 | 303.080367 | -0.297538986 | 0.23733916 | -1.2536447 | 0.20997117 | 0.66797116 |
| UBC | 1643.7255 | -0.363156697 | 0.4151755 | -0.8747065 | 0.38173364 | 0.8032969 |
| DHX37 | 35.4751188 | -0.458070291 | 0.36943866 | -1.2399089 | 0.21500909 | 0.67132603 |
| BRI3BP | 77.0941436 | -0.154103413 | 0.27502345 | -0.5603283 | 0.57525554 | 0.89807676 |
| AACS | 83.285733 | -0.392255355 | 0.39712136 | -0.9877468 | 0.32327666 | 0.76911223 |
| SLC15A4 | 128.00908 | -0.703872249 | 0.3130409 | -2.2484993 | 0.02454437 | 0.26518037 |
| RIMBP2 | 492.677205 | -0.224846321 | 0.38469075 | -0.5844859 | 0.55889344 | 0.89179371 |
| STX2 | 170.562747 | 0.6832003 | 0.36553455 | 1.86904441 | 0.06161664 | 0.408181 |
| RAN | 278.098796 | -0.140976643 | 0.54526225 | -0.2585483 | 0.79598376 | 0.95948555 |
| SFSWAP | 255.543422 | -0.038039599 | 0.27440176 | -0.1386274 | 0.8897446 | 0.98064269 |
| ULK1 | 149.806587 | -0.292080173 | 0.31085608 | -0.9395993 | 0.34742311 | 0.78180304 |
| PUS1 | 42.4096671 | -0.039188011 | 0.42898765 | -0.09135 | 0.92721451 | 0.98732047 |
| EP400 | 255.748914 | -0.044764816 | 0.22725512 | -0.1969805 | 0.84384284 | 0.96876046 |
| DDX51 | 37.3175291 | -0.112763832 | 0.41526491 | -0.2715467 | 0.78597056 | 0.95753138 |
| FBRSL1 | 135.972497 | -0.008509791 | 0.32668759 | -0.0260487 | 0.97921848 | 0.99469867 |
| ANKLE2 | 167.054547 | -0.429257003 | 0.34495569 | -1.244383 | 0.21335863 | 0.67075711 |

| | | | | | | |
|---|---|---|---|---|---|---|
| GOLGA3 | 163.617517 | -0.165577344 | 0.34203982 | -0.4840879 | 0.62832347 | 0.91101688 |
| CHFR | 181.521579 | -0.167076964 | 0.27721284 | -0.6027028 | 0.54670639 | 0.88719155 |
| ZNF26 | 120.020117 | -0.104258527 | 0.27432195 | -0.380059 | 0.70390161 | 0.93241442 |
| ZNF84 | 195.682842 | -0.215442811 | 0.26767486 | -0.8048675 | 0.42089612 | 0.82694802 |
| ZNF140 | 99.8154048 | 0.147051052 | 0.26527348 | 0.55433755 | 0.57934786 | 0.89989114 |
| ZNF891 | 110.530342 | 1.435917685 | 0.55054559 | 2.60817217 | 0.00910272 | 0.15239051 |
| ZNF268 | 52.4280818 | -0.145787868 | 0.38727587 | -0.3764445 | 0.70658647 | 0.93262295 |
| TPTE2 | 46.0147155 | -0.223344226 | 0.43581806 | -0.5124713 | 0.60832123 | 0.90531192 |
| MPHOSPH8 | 555.339125 | -0.239527594 | 0.26254513 | -0.9123292 | 0.36159545 | 0.78981248 |
| PSPC1 | 564.471038 | 0.558364961 | 0.28581517 | 1.95358756 | 0.05075001 | 0.3799001 |
| ZMYM5 | 166.049491 | 0.056481905 | 0.23821873 | 0.23710103 | 0.81257842 | 0.96288868 |
| ZMYM2 | 1430.67888 | 0.799256024 | 0.29574378 | 2.70252863 | 0.00688143 | 0.13279571 |
| CRYL1 | 436.655152 | 0.287465633 | 0.26622345 | 1.07979079 | 0.28023536 | 0.72945471 |
| IFT88 | 603.270023 | 0.285580044 | 0.23792739 | 1.20028235 | 0.2300297 | 0.68956272 |
| EEF1AKMT1 | 62.9319318 | -0.543421265 | 0.33175853 | -1.6380024 | 0.10142119 | 0.49779302 |
| XPO4 | 325.721006 | 0.39765394 | 0.25185618 | 1.57889295 | 0.11436061 | 0.52703872 |
| LATS2 | 239.859694 | 0.727136882 | 0.29866716 | 2.43460604 | 0.01490801 | 0.19869417 |
| SAP18 | 339.29321 | 0.063686787 | 0.51850842 | 0.12282691 | 0.90224417 | 0.98320957 |
| MRPL57 | 224.602393 | -0.331848105 | 0.4328409 | -0.7666746 | 0.44327503 | 0.83847661 |
| ZDHHC20 | 383.182021 | 0.047384221 | 0.26577701 | 0.17828563 | 0.85849867 | 0.97237339 |
| MICU2 | 623.363501 | 0.09098372 | 0.19604649 | 0.46409258 | 0.64258142 | 0.91643619 |
| FGF9 | 76.0261139 | 0.200865906 | 0.61171055 | 0.32836757 | 0.74263376 | 0.94508888 |
| AL136962.1 | 73.6110028 | 0.191645543 | 1.36988322 | 0.13989918 | 0.88873965 | 0.98051184 |
| TNFRSF19 | 160.58187 | -0.299822613 | 0.57284489 | -0.5233923 | 0.60070127 | 0.90369786 |
| MIPEP | 340.115971 | -0.138893125 | 0.33816993 | -0.4107199 | 0.68127792 | 0.92759922 |
| PARP4 | 420.262053 | -0.866299868 | 0.3063662 | -2.8276613 | 0.00468894 | 0.10823805 |
| ATP12A | 78.4487913 | 3.880593021 | 0.9290469 | 4.17696137 | 2.95E-05 | 0.00407381 |
| MTMR6 | 103.636832 | 0.166926881 | 0.22606738 | 0.73839438 | 0.46027483 | 0.84657145 |
| NUP58 | 277.213598 | 0.154944551 | 0.51527647 | 0.30070178 | 0.76364191 | 0.95378284 |
| RNF6 | 46.1819655 | -0.232136256 | 0.33898369 | -0.6848007 | 0.49346971 | 0.86307787 |
| CDK8 | 420.726867 | 0.320318916 | 0.25691715 | 1.24677904 | 0.21247853 | 0.66984203 |
| WASF3 | 120.26111 | 0.005089552 | 0.23623312 | 0.02154462 | 0.98281121 | 0.99507903 |
| USP12 | 520.937698 | 0.266638905 | 0.23657884 | 1.12706151 | 0.25971648 | 0.7168415 |
| LINC02340 | 76.0363849 | -0.613430667 | 0.59011676 | -1.0395073 | 0.29856887 | 0.74594723 |
| RPL21 | 394.019541 | 0.580744773 | 0.48146056 | 1.20621464 | 0.22773474 | 0.68629723 |
| RASL11A | 143.583417 | -0.100273126 | 0.63326169 | -0.1583439 | 0.87418582 | 0.97622753 |
| GTF3A | 77.3734787 | -0.323157776 | 0.31279548 | -1.033128 | 0.30154399 | 0.74814891 |
| MTIF3 | 175.040713 | -0.06640862 | 0.28264554 | -0.2349537 | 0.81424465 | 0.96303374 |
| LNX2 | 182.105169 | 0.100783808 | 0.21230932 | 0.4747027 | 0.63499889 | 0.91454294 |
| POLR1D | 162.51203 | -0.285947285 | 0.20923336 | -1.3666429 | 0.17173726 | 0.6187442 |
| PAN3 | 1322.70683 | 0.108205602 | 0.21323135 | 0.50745634 | 0.61183466 | 0.90680628 |
| POMP | 167.897915 | -0.051480739 | 0.33135292 | -0.1553653 | 0.87653335 | 0.97640529 |
| SLC7A1 | 629.687498 | 0.457912761 | 0.73785377 | 0.62060096 | 0.53486221 | 0.88137638 |
| UBL3 | 737.776494 | 0.58861384 | 0.43212672 | 1.36213246 | 0.17315609 | 0.61955472 |
| KATNAL1 | 129.101114 | 0.198123003 | 0.32712665 | 0.60564617 | 0.54474974 | 0.88669883 |
| HMGB1 | 901.696171 | -0.237321045 | 0.24746451 | -0.9590104 | 0.33755349 | 0.77827084 |
| USPL1 | 95.5449998 | -0.167366199 | 0.28065991 | -0.596331 | 0.55095414 | 0.88916026 |
| HSPH1 | 227.702836 | 1.182258057 | 0.45667144 | 2.58885922 | 0.00962945 | 0.15717671 |
| B3GLCT | 136.400377 | 0.48918076 | 0.30025967 | 1.62919234 | 0.10327231 | 0.50175883 |
| FRY | 1912.48795 | 1.009526126 | 0.46944644 | 2.15046071 | 0.03151879 | 0.29823395 |
| N4BP2L1 | 41.9015266 | 0.013939769 | 0.61224589 | 0.02276825 | 0.98183513 | 0.99507903 |
| N4BP2L2 | 1872.18608 | 0.128743299 | 0.21530952 | 0.59794524 | 0.54987647 | 0.88889487 |
| PDS5B | 705.454846 | 0.442302003 | 0.26542693 | 1.66637955 | 0.09563784 | 0.48710328 |
| KL | 112.136416 | 1.885953387 | 0.32643447 | 5.77743327 | 7.58E-09 | 8.56E-06 |
| STARD13 | 1577.76858 | 1.183038904 | 0.24943074 | 4.7429556 | 2.11E-06 | 0.00063066 |
| AL139383.1 | 132.73045 | 0.302415344 | 0.43372453 | 0.6972521 | 0.48564504 | 0.85912348 |
| RFC3 | 98.7186951 | -0.564185936 | 0.39179898 | -1.4399883 | 0.14987072 | 0.58336854 |
| LINC02343 | 357.525064 | -1.136373311 | 0.68656339 | -1.6551615 | 0.09789173 | 0.49204095 |
| NBEA | 1345.16719 | -0.175901245 | 0.31074574 | -0.5660616 | 0.57135188 | 0.89785091 |
| SPART | 301.699875 | -0.467509494 | 0.35413343 | -1.3201507 | 0.18678471 | 0.63824708 |
| RFXAP | 42.6748799 | -0.332178333 | 0.40298502 | -0.8242945 | 0.40977224 | 0.8216497 |
| SMAD9 | 45.6607397 | -1.971212182 | 0.61450292 | -3.2078158 | 0.00133747 | 0.04816735 |
| ALG5 | 113.119631 | -0.072619691 | 0.21284762 | -0.3411816 | 0.73296687 | 0.94163693 |
| EXOSC8 | 86.2396063 | -0.1371233 | 0.32013502 | -0.4283296 | 0.66841118 | 0.92422713 |
| SUPT20H | 362.157628 | 0.273082941 | 0.20795374 | 1.31319082 | 0.18911866 | 0.64237956 |
| LINC00571 | 116.653971 | -1.347492427 | 0.35114156 | -3.8374621 | 0.00012431 | 0.01017808 |
| UFM1 | 226.478617 | 0.098141531 | 0.26610919 | 0.36880174 | 0.71227551 | 0.93483373 |
| FREM2 | 529.25617 | 0.796937823 | 0.26150568 | 3.04749722 | 0.00230756 | 0.06929282 |
| PROSER1 | 84.9675629 | 0.079795824 | 0.38749636 | 0.20592664 | 0.83684822 | 0.96757829 |
| NHLRC3 | 171.059324 | -0.462962172 | 0.34783399 | -1.330986 | 0.18319362 | 0.633903 |
| COG6 | 227.386494 | 0.104057975 | 0.23592813 | 0.44105794 | 0.65917105 | 0.92284375 |
| LINC00598 | 60.574015 | -0.597645648 | 0.43129348 | -1.3857053 | 0.16583692 | 0.60746974 |
| FOXO1 | 1303.99617 | -0.15969754 | 0.68214409 | -0.2341112 | 0.81489869 | 0.96324591 |
| MRPS31 | 211.235656 | 0.061527568 | 0.25915856 | 0.23741283 | 0.81233654 | 0.96288868 |
| ELF1 | 730.964487 | 0.172068199 | 0.16928719 | 1.01642773 | 0.30942574 | 0.75378471 |
| WBP4 | 122.590746 | 0.165297882 | 0.23068531 | 0.71655139 | 0.47365095 | 0.85310957 |
| AL354696.2 | 56.5742345 | -0.380979035 | 0.4468208 | -0.8526439 | 0.39385681 | 0.81076789 |
| MTRF1 | 55.6489324 | -0.022111315 | 0.30860191 | -0.07165 | 0.94288048 | 0.98885525 |
| NAA16 | 150.828978 | 0.111244422 | 0.22534555 | 0.49366149 | 0.62154527 | 0.9099963 |
| VWA8 | 740.358479 | 0.379919417 | 0.24653562 | 1.54103258 | 0.12330886 | 0.54197241 |
| DGKH | 348.730325 | -1.285366495 | 0.52631358 | -2.4422066 | 0.01459779 | 0.19588328 |
| AKAP11 | 383.071374 | 0.077032813 | 0.32507682 | 0.23696803 | 0.81268159 | 0.96288868 |
| EPSTI1 | 248.753441 | 0.003925232 | 0.84718138 | 0.00463328 | 0.99630319 | 0.99916339 |
| DNAJC15 | 305.465707 | 0.144239514 | 0.2520712 | 0.57221736 | 0.56717473 | 0.89454608 |
| CCDC122 | 131.723219 | -0.598671604 | 0.40477505 | -1.479023 | 0.13913417 | 0.56656585 |
| SERP2 | 56.7504231 | 0.088263327 | 0.26729117 | 0.33021416 | 0.74123815 | 0.94487568 |
| TSC22D1 | 1676.4044 | 0.446610348 | 0.32635051 | 1.36849898 | 0.17115594 | 0.6176702 |
| NUFIP1 | 44.0004604 | 0.184038196 | 0.34426298 | 0.53458607 | 0.59293612 | 0.9025973 |
| AL359706.1 | 40.2093632 | -0.423315299 | 0.55028482 | -0.7692658 | 0.44173553 | 0.83770294 |
| GPALPP1 | 151.319775 | 0.068169282 | 0.19280659 | 0.35356303 | 0.72366638 | 0.93893205 |
| GTF2F2 | 244.509095 | 0.242842953 | 0.26765218 | 0.90730795 | 0.36424397 | 0.79268955 |
| TPT1 | 1827.85903 | -0.303700838 | 0.53978488 | -0.5626331 | 0.57368475 | 0.89797476 |
| SLC25A30 | 285.683407 | 0.544473218 | 0.41752823 | 1.30403928 | 0.19222019 | 0.64529552 |
| COG3 | 218.516612 | 0.273019899 | 0.1915178 | 1.42555888 | 0.15399571 | 0.58943574 |
| ZC3H13 | 363.136597 | 0.123643404 | 0.15899219 | 0.77766969 | 0.43676376 | 0.83393108 |
| AL138686.1 | 111.548856 | 1.14024356 | 0.68231433 | 1.67114116 | 0.0946938 | 0.48627909 |
| LRCH1 | 992.759824 | 0.471783485 | 0.25999566 | 1.81458216 | 0.06958815 | 0.42828237 |
| ESD | 178.770144 | -0.055815405 | 0.28225076 | -0.1977511 | 0.84323979 | 0.96876046 |
| SUCLA2 | 271.805612 | -0.057517649 | 0.26357054 | -0.2182249 | 0.8272539 | 0.96545294 |
| MED4 | 195.057289 | -0.036227478 | 0.23513459 | -0.1540712 | 0.87755357 | 0.97640529 |
| ITM2B | 2677.27816 | -0.216347569 | 0.37266736 | -0.580538 | 0.56155186 | 0.89242209 |
| RB1 | 778.896408 | -0.124595116 | 0.21556502 | -0.5779932 | 0.56326871 | 0.8927792 |
| LPAR6 | 41.3331321 | -0.78103708 | 0.50429898 | -1.548758 | 0.12143991 | 0.53989034 |
| RCBTB2 | 150.738364 | -0.138581004 | 0.3550019 | -0.3903669 | 0.69626523 | 0.93130502 |
| CYSLTR2 | 76.3281243 | 0.81199655 | 0.50066658 | 1.62183095 | 0.10483955 | 0.50446211 |
| FNDC3A | 2236.98222 | 1.183777356 | 0.76437704 | 1.54868252 | 0.12145806 | 0.53989034 |
| CDADC1 | 150.59431 | 2.376842524 | 0.78460496 | 3.02934936 | 0.00245081 | 0.0722844 |
| CAB39L | 293.609798 | 1.72644435 | 0.65948041 | 2.61788572 | 0.00884764 | 0.14907283 |
| SETDB2 | 193.869372 | -0.276593254 | 0.40704024 | -0.6795231 | 0.49680646 | 0.86487238 |
| PHF11 | 136.199861 | -0.447334683 | 0.41045721 | -1.0898449 | 0.27578149 | 0.7263609 |
| RCBTB1 | 74.5454059 | -0.186536497 | 0.37145741 | -0.5021747 | 0.61554467 | 0.90794578 |
| EBPL | 76.9514974 | -0.411654355 | 0.36560838 | -1.1259434 | 0.26018946 | 0.71719766 |
| KPNA3 | 264.808017 | 0.22234608 | 0.29126304 | 0.76338583 | 0.44523333 | 0.83929814 |

| | | | | | | |
|---|---|---|---|---|---|---|
| SPRYD7 | 54.8040574 | -0.19319652 | 0.34587659 | -0.5585707 | 0.57645476 | 0.89869922 |
| DLEU2 | 480.557704 | -0.151749003 | 0.36694573 | -0.4135462 | 0.6792065 | 0.92700898 |
| TRIM13 | 137.309181 | 0.114376066 | 0.24573347 | 0.46544765 | 0.64161092 | 0.91643619 |
| RNASEH2B | 113.425869 | 0.396799005 | 0.21365133 | 1.85722692 | 0.06327888 | 0.41267121 |
| FAM124A | 115.969252 | -0.883659881 | 0.537993 | -1.6425118 | 0.10048397 | 0.49626392 |
| INTS6 | 555.592124 | 0.582306504 | 0.34105781 | 1.70735426 | 0.08775623 | 0.46898789 |
| INTS6-AS1 | 56.8337084 | -0.13745041 | 0.40392957 | -0.3402831 | 0.73364333 | 0.94163693 |
| WDFY2 | 921.14649 | 1.34237826 | 0.35135858 | 3.82053648 | 0.00013316 | 0.01065294 |
| ATP7B | 70.2373108 | 0.317206532 | 0.36878361 | 0.8601427 | 0.38971039 | 0.80680464 |
| UTP14C | 80.4532195 | 0.036963553 | 0.25259398 | 0.14633585 | 0.88365627 | 0.97878487 |
| NEK5 | 89.206893 | 0.488037716 | 0.56504229 | 0.86371892 | 0.38774232 | 0.8065026 |
| NEK3 | 142.848706 | -0.311082458 | 0.44715163 | -0.6956979 | 0.48661803 | 0.85912348 |
| VPS36 | 274.028616 | 0.310388741 | 0.18716781 | 1.65834468 | 0.09724792 | 0.49030463 |
| CKAP2 | 33.3618398 | 0.065968132 | 0.52487154 | 0.12568434 | 0.8999818 | 0.98312332 |
| SUGT1 | 273.328956 | 0.755195734 | 0.41572243 | 1.81658645 | 0.06928046 | 0.42828237 |
| OLFM4 | 141.863187 | -5.4887029 | 0.99867849 | -5.4959659 | 3.89E-08 | 2.91E-05 |
| PCDH17 | 71.4574262 | 0.580116177 | 1.00114932 | 0.5794502 | 0.56228544 | 0.89242209 |
| TDRD3 | 565.674159 | 0.373722643 | 0.28757065 | 1.29958547 | 0.19374308 | 0.64723766 |
| PCDH9 | 222.97549 | 0.293560883 | 0.65169283 | 0.45045897 | 0.65237953 | 0.91909172 |
| DACH1 | 789.339609 | -0.148033577 | 0.79557328 | -0.1860716 | 0.85238863 | 0.97062426 |
| MZT1 | 58.4350691 | 0.270323897 | 0.38280011 | 0.70617509 | 0.48007926 | 0.85579702 |
| DIS3 | 108.946388 | 0.045195744 | 0.19660437 | 0.22988169 | 0.8181837 | 0.96407382 |
| PIBF1 | 617.87971 | -0.338092062 | 0.20250523 | -1.6695473 | 0.09500896 | 0.48627909 |
| KLF5 | 229.999878 | 0.371970511 | 0.53229287 | 0.698808 | 0.48467203 | 0.85909499 |
| KLF12 | 601.041563 | 0.346640811 | 0.2952716 | 1.17397276 | 0.24040594 | 0.6963541 |
| TBC1D4 | 723.148086 | 0.194998104 | 0.23674556 | 0.82366108 | 0.41013216 | 0.8221471 |
| COMMD6 | 266.952291 | 0.286658 | 0.39247827 | 0.73037929 | 0.46515838 | 0.84991999 |
| UCHL3 | 140.718068 | -0.017205923 | 0.23726243 | -0.0725185 | 0.94218926 | 0.98885525 |
| LMO7-AS1 | 62.2903644 | -0.69573082 | 0.51766463 | -1.3439798 | 0.17895491 | 0.62831166 |
| LMO7 | 1018.19608 | 0.903750287 | 0.32182537 | 2.8082009 | 0.00498191 | 0.11276556 |
| KCTD12 | 426.839992 | -0.002625996 | 0.8372735 | -0.0031364 | 0.99749755 | 0.99954455 |
| AC001226.2 | 85.4699145 | -0.87132787 | 0.66224892 | -1.3157105 | 0.18827122 | 0.64111947 |
| FBXL3 | 209.695548 | -0.085672476 | 0.39227117 | -0.2184012 | 0.82711657 | 0.96545294 |
| MYCBP2 | 696.23987 | 0.168545307 | 0.17157095 | 0.98236506 | 0.32592003 | 0.77102527 |
| SLAIN1 | 72.1591663 | 0.797615764 | 0.42405443 | 1.88092779 | 0.05998174 | 0.40367252 |
| EDNRB | 56.2834412 | 1.974087119 | 0.71759067 | 2.75099329 | 0.00594149 | 0.1240374 |
| RNF219 | 61.0085908 | -0.001838573 | 0.25504775 | -0.0072087 | 0.99424831 | 0.99885672 |
| RBM26 | 599.514608 | 0.399840476 | 0.21321738 | 1.87527151 | 0.06075538 | 0.40676596 |
| NDFIP2 | 285.967962 | 0.009045786 | 0.15969297 | 0.05664486 | 0.9548281 | 0.99073937 |
| GPC5 | 116.610583 | 1.932990709 | 0.6672717 | 2.89685703 | 0.00376921 | 0.09515702 |
| GPC6 | 325.19696 | 1.041800379 | 0.98096324 | 1.06201775 | 0.28822763 | 0.73653438 |
| TGDS | 70.8700378 | -0.159278481 | 0.27004511 | -0.5898218 | 0.55531015 | 0.89081623 |
| GPR180 | 117.754598 | -0.009297793 | 0.36594019 | -0.025408 | 0.97972956 | 0.99469867 |
| ABCC4 | 73.6138148 | -1.408986825 | 0.47904987 | -2.9412111 | 0.00326932 | 0.08642938 |
| CLDN10 | 156.519743 | 0.348724287 | 0.50635488 | 0.68869542 | 0.49101496 | 0.86143519 |
| DNAJC3-DT | 44.0332121 | -1.457848664 | 0.50840538 | -2.8674926 | 0.00413738 | 0.09955609 |
| DNAJC3 | 306.960789 | 0.096949011 | 0.17005138 | 0.570116 | 0.56859902 | 0.8958131 |
| UGGT2 | 702.525812 | 0.17655027 | 0.21765527 | 0.81114632 | 0.41728165 | 0.8254526 |
| MBNL2 | 694.929756 | 0.02318605 | 0.28289072 | 0.08196115 | 0.93467761 | 0.98822345 |
| RAP2A | 55.5524672 | -0.100203262 | 0.26055548 | -0.3845755 | 0.70055192 | 0.93180634 |
| IPO5 | 209.625222 | -0.039277577 | 0.19998007 | -0.1964075 | 0.84429126 | 0.96876046 |
| FARP1 | 1569.7818 | -0.048331106 | 0.18727595 | -0.2580743 | 0.79634958 | 0.95948555 |
| STK24 | 705.960921 | 0.056197919 | 0.42616603 | 0.1318686 | 0.89508822 | 0.98234865 |
| SLC15A1 | 297.13111 | -3.075745354 | 2.04697474 | -1.502581 | 0.13294712 | 0.55601567 |
| DOCK9 | 544.531723 | 0.424053914 | 0.22518628 | 1.88312496 | 0.05968343 | 0.40353703 |
| UBAC2 | 803.84153 | 0.334198154 | 0.15889706 | 2.10323685 | 0.03544507 | 0.31644624 |
| TM9SF2 | 407.258824 | 0.179164031 | 0.18794946 | 0.95325645 | 0.34046015 | 0.77836439 |
| CLYBL | 462.063435 | 0.158711182 | 0.30388335 | 0.52227667 | 0.60147771 | 0.90369786 |
| AL355338.1 | 46.1168216 | -0.692182306 | 0.3831726 | -1.8064504 | 0.07084801 | 0.43078597 |
| PCCA | 2387.30086 | -0.037210141 | 0.30317887 | -0.1227333 | 0.9023183 | 0.98320957 |
| GGACT | 145.668035 | -0.609921124 | 0.35484224 | -1.7188515 | 0.08564141 | 0.46479978 |
| TMTC4 | 89.085065 | -0.347766355 | 0.38874167 | -0.894595 | 0.37100362 | 0.79539603 |
| FGF14 | 201.198396 | 0.041966001 | 0.65179676 | 0.0643851 | 0.94866359 | 0.98977092 |
| TPP2 | 227.074853 | 0.290126509 | 0.24062348 | 1.20572817 | 0.22792232 | 0.68629723 |
| EFNB2 | 84.9803445 | 0.161306605 | 0.4603497 | 0.35040015 | 0.72603841 | 0.93895336 |
| ARGLU1 | 594.255505 | 0.205046789 | 0.18109621 | 1.13225335 | 0.25752794 | 0.71534188 |
| FAM155A | 1061.09161 | -0.871767485 | 0.97243661 | -0.8964774 | 0.36999782 | 0.79532503 |
| LIG4 | 55.6620841 | 0.275858124 | 0.34177709 | 0.80712878 | 0.41959229 | 0.82694802 |
| ABHD13 | 81.2055018 | 0.673635912 | 0.27535073 | 2.44646497 | 0.01442648 | 0.19433101 |
| TNFSF13B | 63.1217859 | 0.124858556 | 0.47066774 | 0.26527961 | 0.79079407 | 0.95831659 |
| MYO16 | 49.3748785 | -0.789007982 | 0.86196235 | -0.9153625 | 0.36000138 | 0.78892604 |
| IRS2 | 266.123167 | -0.611085622 | 0.38642282 | -1.5813911 | 0.11378863 | 0.52602773 |
| COL4A1 | 456.133857 | 0.277843129 | 0.44872489 | 0.61918369 | 0.53579536 | 0.88137638 |
| COL4A2 | 540.78759 | -0.15114117 | 0.41656659 | -0.362826 | 0.7167349 | 0.93573028 |
| RAB20 | 117.419004 | -0.988938131 | 0.49204095 | -2.0098696 | 0.044445 | 0.35501796 |
| NAXD | 116.498391 | 0.164882701 | 0.30836436 | 0.5347009 | 0.5928567 | 0.9025973 |
| CARS2 | 183.370712 | -0.271805086 | 0.22309708 | -1.2183265 | 0.22309992 | 0.68086406 |
| ING1 | 36.6579108 | -0.257526002 | 0.58905351 | -0.4371861 | 0.66197639 | 0.92284375 |
| ANKRD10 | 883.167844 | 0.327210378 | 0.30006222 | 1.0904751 | 0.27550391 | 0.7263609 |
| ARHGEF7 | 491.838432 | -0.138776581 | 0.24582188 | -0.5645412 | 0.57238586 | 0.89797476 |
| TUBGCP3 | 263.506801 | 1.161394188 | 0.46087598 | 2.51997118 | 0.01173644 | 0.17202509 |
| ATP11A | 349.383039 | -0.741300445 | 0.62476299 | -1.1865307 | 0.2354128 | 0.69335782 |
| MCF2L | 232.22734 | 0.195346962 | 0.31260946 | 0.6248914 | 0.53204233 | 0.88045351 |
| PCID2 | 149.811803 | 0.333848798 | 0.22371944 | 1.4922655 | 0.13562956 | 0.55916513 |
| CUL4A | 387.076538 | -0.521901526 | 0.23039425 | -2.2652541 | 0.02349709 | 0.25893742 |
| LAMP1 | 504.055126 | -0.576373715 | 0.29331614 | -1.9650256 | 0.04941128 | 0.3755208 |
| GRTP1 | 83.3306621 | -0.436316813 | 0.27312351 | -1.5975073 | 0.11015266 | 0.51604824 |
| DCUN1D2 | 106.406344 | 0.187001507 | 0.24393683 | 0.7665981 | 0.44332051 | 0.83847661 |
| TMCO3 | 179.872799 | -0.266483714 | 0.28232767 | -0.943881 | 0.34523048 | 0.78180304 |
| TFDP1 | 248.158709 | 0.958416219 | 0.70684179 | 1.35591334 | 0.17512674 | 0.62472937 |
| GAS6 | 39.7951853 | 1.135112194 | 0.58968501 | 1.92494667 | 0.05423603 | 0.39095834 |
| RASA3 | 39.9168406 | 0.244684642 | 0.74560896 | 0.32816752 | 0.74278501 | 0.94508888 |
| CDC16 | 208.21558 | -0.094761427 | 0.23321825 | -0.4063208 | 0.6845069 | 0.92881914 |
| UPF3A | 222.891937 | 0.003296344 | 0.18947227 | 0.0173975 | 0.9861195 | 0.99590752 |
| CHAMP1 | 51.0068111 | -0.485248077 | 0.3710916 | -1.3076235 | 0.19100106 | 0.64458858 |
| TTC5 | 86.9248305 | 0.224362011 | 0.24670268 | 0.90944295 | 0.36311636 | 0.79148491 |
| CCNB1IP1 | 116.088224 | 0.598363344 | 0.31332629 | 1.90971317 | 0.05617015 | 0.39444789 |
| PARP2 | 47.6168912 | 0.470527034 | 0.28176488 | 1.66992789 | 0.09493363 | 0.48627909 |
| TEP1 | 64.7863938 | -0.079752711 | 0.35811395 | -0.2227021 | 0.82376741 | 0.96511681 |
| OSGEP | 72.609086 | 0.355022811 | 0.2677194 | 1.32610042 | 0.18480644 | 0.63541313 |
| APEX1 | 123.747318 | -0.087329546 | 0.49375897 | -0.1768668 | 0.85961305 | 0.97244655 |
| PIP4P1 | 56.8720752 | 0.246635736 | 0.4393617 | 0.5613501 | 0.57455889 | 0.89807676 |
| PNP | 331.622985 | 1.010399559 | 0.66481774 | 1.51981437 | 0.12855764 | 0.54856841 |
| METTL17 | 59.8809527 | 0.230799018 | 0.27474947 | 0.84003444 | 0.40088907 | 0.8162653 |
| NDRG2 | 102.001372 | -0.04925045 | 0.36616165 | -0.1345047 | 0.89300352 | 0.98168537 |
| ARHGEF40 | 34.9438115 | -0.784112451 | 0.53733809 | -1.4592534 | 0.14449537 | 0.57406643 |
| HNRNPC | 1168.01829 | 0.195411628 | 0.21524841 | 0.90784236 | 0.36396151 | 0.79266764 |
| SUPT16H | 210.130377 | -0.189177367 | 0.19571772 | -0.9665827 | 0.33375267 | 0.77744565 |
| CHD8 | 288.794165 | -0.052857818 | 0.24194546 | -0.21847 | 0.82706295 | 0.96545294 |
| RAB2B | 101.377701 | 0.071782872 | 0.23317418 | 0.30785087 | 0.75819581 | 0.95205992 |
| TOX4 | 156.599859 | -0.265674483 | 0.22978891 | -1.1561676 | 0.24761262 | 0.70405328 |
| METTL3 | 52.3358413 | -0.159924249 | 0.32617644 | -0.4902998 | 0.62392176 | 0.91058133 |
| DAD1 | 305.183069 | -0.222763866 | 0.43461366 | -0.5125561 | 0.60826189 | 0.90531192 |

| | | | | | | |
|---|---|---|---|---|---|---|
| ABHD4 | 80.205 | 0.089153584 | 0.33658523 | 0.2648767 | 0.79110445 | 0.95832571 |
| OXA1L | 77.0897192 | -0.498881477 | 0.37519696 | -1.3296522 | 0.18363288 | 0.63409311 |
| MRPL52 | 75.7738044 | 0.21637305 | 0.27536559 | 0.78576648 | 0.43200431 | 0.83162989 |
| MMP14 | 57.9058307 | -0.57158675 | 0.38412421 | -1.4880259 | 0.13674408 | 0.56155091 |
| LRP10 | 249.714515 | -0.731627587 | 0.2777223 | -2.6343855 | 0.00842897 | 0.14457551 |
| RBM23 | 161.130434 | 0.238539379 | 0.17848814 | 1.33644388 | 0.18140423 | 0.63181008 |
| PRMT5 | 45.9063962 | 0.047980043 | 0.25922499 | 0.18509035 | 0.85315817 | 0.97062426 |
| AL132780.1 | 104.23397 | -0.598080158 | 0.6431969 | -0.9298555 | 0.35244592 | 0.78314007 |
| C14orf93 | 44.1867853 | -0.206730458 | 0.36601788 | -0.5648097 | 0.57220319 | 0.89797476 |
| PSMB5 | 189.409381 | -0.334069456 | 0.43650565 | -0.7653268 | 0.44407699 | 0.83873671 |
| ACIN1 | 327.549624 | 0.036160801 | 0.16176585 | 0.22353792 | 0.82311688 | 0.96511681 |
| C14orf119 | 81.2651854 | -0.601954449 | 0.44672771 | -1.3474751 | 0.17782728 | 0.62727361 |
| SLC7A8 | 147.702189 | -1.141853938 | 0.7338056 | -1.5560714 | 0.11969111 | 0.53725356 |
| HOMEZ | 50.2611621 | -0.737946966 | 0.29838247 | -2.4731579 | 0.01339249 | 0.18614502 |
| PPP1R3E | 45.5707688 | -1.126738523 | 0.43404994 | -2.595873 | 0.00943509 | 0.15525074 |
| BCL2L2-PABI | 56.4222829 | -0.3924892 | 0.34654052 | -1.1325925 | 0.25738541 | 0.71530073 |
| PABPN1 | 232.533162 | 0.96748386 | 0.37503459 | 2.579719 | 0.00988807 | 0.15942618 |
| SLC22A17 | 58.3790921 | 0.142180453 | 0.496102 | 0.2865952 | 0.77442229 | 0.95586283 |
| NGDN | 60.0742697 | 0.034704936 | 0.27613175 | 0.12568253 | 0.89998323 | 0.98312332 |
| ZFHX2 | 97.7501131 | 0.134063362 | 0.45037235 | 0.29767227 | 0.76595331 | 0.95389523 |
| AP1G2 | 63.736377 | -0.266895136 | 0.35221141 | -0.7577697 | 0.44858886 | 0.84115427 |
| DHRS4-AS1 | 84.4471821 | -0.537046688 | 0.38020044 | -1.4125357 | 0.15779229 | 0.59566075 |
| DHRS4L2 | 77.9252234 | -0.233656616 | 0.40818427 | -0.5724292 | 0.56703121 | 0.89454608 |
| PCK2 | 37.1369237 | -0.567229953 | 0.29153886 | -1.9456409 | 0.05169788 | 0.38247133 |
| PSME1 | 235.37395 | -0.329416843 | 0.52209165 | -0.630956 | 0.52806932 | 0.87826011 |
| EMC9 | 46.07675 | -0.565634588 | 0.42189988 | -1.3406844 | 0.18002294 | 0.63015077 |
| REC8 | 64.6221736 | -0.655442698 | 0.47683649 | -1.3745649 | 0.16926638 | 0.61478828 |
| AL136295.1 | 61.4252418 | -0.183365276 | 0.39472311 | -0.4645415 | 0.64225982 | 0.91643619 |
| GMPR2 | 77.3890446 | -0.357791327 | 0.49293821 | -0.725834 | 0.46794052 | 0.85161077 |
| TINF2 | 41.8570001 | -0.017094742 | 0.41692865 | -0.0410016 | 0.96729462 | 0.99291246 |
| DHRS1 | 63.910255 | -0.206448308 | 0.38976205 | -0.5296778 | 0.59633534 | 0.90351227 |
| NFATC4 | 48.3539755 | 0.569250561 | 0.47174153 | 1.20670011 | 0.22754766 | 0.68604702 |
| KHNYN | 66.6992157 | 0.136352649 | 0.29554913 | 0.46135358 | 0.64454495 | 0.91682456 |
| SDR39U1 | 68.8865127 | -0.124543177 | 0.30831255 | -0.4039511 | 0.68624871 | 0.92974759 |
| STXBP6 | 315.541426 | 2.619424868 | 0.63281825 | 4.13930048 | 3.48E-05 | 0.0045636 |
| NOVA1 | 203.471465 | -0.404425798 | 0.54159524 | -0.7467307 | 0.45522613 | 0.84340698 |
| LINC00645 | 74.6764619 | -1.348397314 | 0.94461629 | -1.4274551 | 0.15344877 | 0.58860244 |
| PRKD1 | 1578.75929 | -0.1373918 | 0.20978174 | -0.6549274 | 0.51251453 | 0.87177451 |
| G2E3 | 132.102801 | 0.559751286 | 0.24359576 | 2.29786962 | 0.02156921 | 0.24935578 |
| SCFD1 | 409.782646 | 0.23439685 | 0.2514148 | 0.93231128 | 0.35117568 | 0.78268744 |
| COCH | 45.0082176 | -0.55963973 | 0.46493656 | -1.2036905 | 0.22870922 | 0.68678297 |
| STRN3 | 863.219518 | 0.094229404 | 0.34444485 | 0.27356892 | 0.78441592 | 0.95700569 |
| AP4S1 | 74.0242395 | 0.501685237 | 0.40927609 | 1.22578681 | 0.22027891 | 0.67837878 |
| HECTD1 | 911.77546 | 0.063816907 | 0.39995484 | 0.15956028 | 0.87322747 | 0.97622753 |
| HEATR5A | 106.659452 | -0.275527285 | 0.31529786 | -0.8738635 | 0.38219261 | 0.80380866 |
| AL139353.1 | 32.511718 | 0.111377608 | 0.49017026 | 0.22722229 | 0.8202509 | 0.96502145 |
| NUBPL | 487.482155 | 0.277446255 | 0.27804841 | 0.99783435 | 0.31835969 | 0.76313119 |
| ARHGAP5 | 926.804643 | 0.068213644 | 0.16867204 | 0.40441583 | 0.68590696 | 0.92974759 |
| AKAP6 | 223.163741 | 0.05673625 | 0.25576608 | 0.22182867 | 0.82444726 | 0.96511681 |
| NPAS3 | 1082.74482 | 1.166343307 | 0.53655233 | 2.17377363 | 0.02972214 | 0.29105042 |
| EGLN3 | 146.27035 | -0.905390529 | 0.64920582 | -1.3946125 | 0.16313273 | 0.6021948 |
| EAPP | 182.539306 | -0.357763687 | 0.23702581 | -1.509387 | 0.13119991 | 0.55395564 |
| SNX6 | 439.3222 | 0.08441795 | 0.20561521 | 0.41056278 | 0.68139316 | 0.92759922 |
| CFL2 | 63.2349758 | 0.403628252 | 0.31376353 | 1.28640908 | 0.19830033 | 0.65386083 |
| BAZ1A | 147.691668 | -0.755325772 | 0.29393479 | -2.5697053 | 0.01017851 | 0.16136868 |
| SRP54 | 296.046131 | -0.193370765 | 0.22539651 | -0.8579137 | 0.39094009 | 0.80778625 |
| FAM177A1 | 316.805647 | 0.504773928 | 0.38441135 | 1.3131088 | 0.18914629 | 0.64237956 |
| PPP2R3C | 126.880778 | -0.498338658 | 0.35956977 | -1.3859304 | 0.16576817 | 0.60743022 |
| NFKBIA | 262.942099 | 1.082719829 | 0.71630224 | 1.51154048 | 0.1306508 | 0.55395564 |
| RALGAPA1 | 1133.07387 | 1.021386602 | 0.3533066 | 2.8909355 | 0.00384097 | 0.09653084 |
| BRMS1L | 61.7203975 | 0.356569502 | 0.23891008 | 1.49248411 | 0.13557228 | 0.55916513 |
| LINC00609 | 60.7968524 | 0.677358431 | 0.77010237 | 0.87956934 | 0.37909265 | 0.80274043 |
| PTCSC3 | 229.703533 | 1.970201581 | 0.85004873 | 2.31775134 | 0.02046284 | 0.24101888 |
| MBIP | 195.721455 | 0.34788998 | 0.30686629 | 1.13368589 | 0.25692633 | 0.7145934 |
| MIPOL1 | 446.228417 | 0.100302572 | 0.2341476 | 0.42837327 | 0.66837938 | 0.92422713 |
| SEC23A | 338.426463 | 0.490725612 | 0.29505283 | 1.66317882 | 0.09627665 | 0.4878081 |
| GEMIN2 | 36.8348167 | -0.209716147 | 0.29167073 | -0.7190168 | 0.47213059 | 0.85244471 |
| TRAPPC6B | 151.371846 | 0.475009126 | 0.20186182 | 2.35314004 | 0.01861562 | 0.2280158 |
| PNN | 313.724751 | -0.007892528 | 0.22265514 | -0.0354473 | 0.97172305 | 0.9939318 |
| MIA2 | 426.061662 | 0.013785735 | 0.25266421 | 0.05456149 | 0.95648782 | 0.9908068 |
| FBXO33 | 223.786553 | 0.358540687 | 0.27857387 | 1.28705788 | 0.19807412 | 0.65359469 |
| AL049875.1 | 64.0320957 | 0.064553289 | 0.32598008 | 0.19802833 | 0.8430229 | 0.96876046 |
| KLHL28 | 162.05845 | -0.220649376 | 0.29956794 | -0.7365587 | 0.46139075 | 0.84722486 |
| TOGARAM1 | 299.144172 | 0.045620127 | 0.27744041 | 0.16443217 | 0.86939094 | 0.97576204 |
| PRPF39 | 132.893646 | 0.382636955 | 0.33393828 | 1.14583138 | 0.25186493 | 0.70813507 |
| FKBP3 | 148.366553 | 0.219172899 | 0.29348364 | 0.74679767 | 0.4551857 | 0.84340698 |
| FANCM | 69.5264671 | -0.088768996 | 0.36946356 | -0.2402646 | 0.81012517 | 0.96270686 |
| MIS18BP1 | 92.9573364 | -0.591702052 | 0.47087834 | -1.2565922 | 0.20890134 | 0.66655368 |
| RPS29 | 328.453786 | 0.044412745 | 0.269796 | 0.16461602 | 0.86924622 | 0.97576204 |
| RPL36AL | 484.506431 | -0.541689941 | 0.46698385 | -1.1599758 | 0.24605868 | 0.70206841 |
| DNAAF2 | 44.6935119 | -0.267759054 | 0.32327448 | -0.8282715 | 0.40751673 | 0.82018309 |
| POLE2 | 44.0890959 | -1.007851148 | 0.36944666 | -2.7280018 | 0.00637193 | 0.12805535 |
| KLHDC1 | 140.956873 | 0.412908033 | 0.34697555 | 1.19002056 | 0.23403831 | 0.69285918 |
| KLHDC2 | 192.71911 | 0.132217111 | 0.26888184 | 0.49172942 | 0.62291064 | 0.91034771 |
| NEMF | 645.985896 | 0.331174029 | 0.21571477 | 1.53524039 | 0.12472479 | 0.54319079 |
| ARF6 | 221.926527 | 0.359664177 | 0.41462899 | 0.86743616 | 0.38570308 | 0.8060156 |
| LINC01588 | 51.6358684 | -0.249865411 | 0.31127384 | -0.802719 | 0.42213719 | 0.82743836 |
| VCPKMT | 72.9927219 | 0.307458354 | 0.21928138 | 1.40211793 | 0.16088003 | 0.6002217 |
| SOS2 | 859.333518 | 0.52385069 | 0.29741696 | 1.76133431 | 0.07818184 | 0.44739666 |
| L2HGDH | 384.218946 | 0.414450814 | 0.26427037 | 1.56828333 | 0.11681503 | 0.53183794 |
| ATP5S | 164.117811 | 0.141367035 | 0.19679338 | 0.7183526 | 0.47253991 | 0.85283327 |
| AL359397.2 | 37.2671268 | 1.225336761 | 0.33655049 | 3.64087058 | 0.00027172 | 0.01627201 |
| CDKL1 | 644.591154 | 0.993859962 | 0.3746062 | 2.65307935 | 0.00797611 | 0.14172148 |
| MAP4K5 | 936.660405 | 0.801142507 | 0.32628342 | 2.45535767 | 0.01407445 | 0.19221396 |
| ATL1 | 101.116963 | 0.104051208 | 0.36756843 | 0.28307983 | 0.77711564 | 0.95601455 |
| SAV1 | 148.121361 | -0.648423936 | 0.24302847 | -2.6680987 | 0.00762818 | 0.13907288 |
| NIN | 127.413043 | -0.170754199 | 0.3039128 | -0.5618526 | 0.57421644 | 0.89807676 |
| PYGL | 56.6486229 | -1.738102687 | 0.63530938 | -2.7358367 | 0.00622219 | 0.12678619 |
| TMX1 | 124.947369 | 0.269267149 | 0.25170363 | 1.06977858 | 0.28471898 | 0.73512071 |
| RTRAF | 187.921839 | -0.20540965 | 0.43068201 | -0.4769404 | 0.63340456 | 0.91415622 |
| NID2 | 38.4503393 | 0.506476693 | 1.03583708 | 0.48895401 | 0.62487426 | 0.91058133 |
| TXNDC16 | 71.8613851 | -0.090278967 | 0.25514308 | -0.3538366 | 0.72346132 | 0.93893205 |
| GPR137C | 42.5084293 | -1.080687708 | 0.64723927 | -1.6696881 | 0.09498109 | 0.48627909 |
| ERO1A | 1133.75371 | -2.340889128 | 1.02211991 | -2.2902295 | 0.02200802 | 0.25042784 |
| AL133453.1 | 104.279616 | -2.275821496 | 0.85584498 | -2.6591515 | 0.00783377 | 0.14121635 |
| PSMC6 | 201.309845 | 0.093058127 | 0.2677839 | 0.34751204 | 0.72820668 | 0.93931348 |
| STYX | 78.9705564 | -0.227926804 | 0.39295407 | -0.5800342 | 0.56189155 | 0.89242209 |
| GNPNAT1 | 89.6990379 | -0.106667318 | 0.31750553 | -0.3359542 | 0.7369054 | 0.9429136 |
| FERMT2 | 438.090622 | 0.014298011 | 0.27675197 | 0.05166363 | 0.95879672 | 0.99119981 |
| DDHD1 | 134.183829 | -0.069227448 | 0.28474191 | -0.2431235 | 0.80790974 | 0.96245698 |
| CNIH1 | 112.708954 | -0.192948422 | 0.28651766 | -0.6734259 | 0.50067635 | 0.86743068 |
| GMFB | 120.458247 | -0.030206393 | 0.25132264 | -0.1201897 | 0.90433288 | 0.98320957 |
| CGRRF1 | 60.661146 | 0.621642605 | 0.34773641 | 1.78768337 | 0.07382711 | 0.43712321 |

| | | | | | | |
|---|---|---|---|---|---|---|
| SAMD4A | 504.33887 | -0.024625504 | 0.55938555 | -0.0440224 | 0.96488654 | 0.99243116 |
| GCH1 | 44.8397964 | 0.667579883 | 0.5702292 | 1.17072203 | 0.24171053 | 0.69763987 |
| WDHD1 | 45.4267174 | 0.286856494 | 0.47060311 | 0.60955078 | 0.54215942 | 0.88450129 |
| SOCS4 | 99.3523533 | 0.122315991 | 0.36652402 | 0.33371889 | 0.73859169 | 0.94349597 |
| MAPK1IP1L | 428.588226 | 0.254398516 | 0.3475847 | 0.73190366 | 0.46422737 | 0.84965121 |
| LGALS3 | 581.999307 | -1.889036388 | 0.63166212 | -2.9905805 | 0.00278448 | 0.07719927 |
| FBXO34 | 630.736551 | 0.31927084 | 0.27371563 | 1.16643264 | 0.24343957 | 0.69975346 |
| AL158801.3 | 44.2601526 | 0.30313783 | 0.50083485 | 0.60526505 | 0.5450029 | 0.88669883 |
| ATG14 | 132.091295 | -0.288750926 | 0.30706796 | -0.9403486 | 0.34703878 | 0.78180304 |
| KTN1-AS1 | 42.7616455 | -1.092576206 | 0.45105944 | -2.4222444 | 0.01542497 | 0.20206713 |
| KTN1 | 1158.9562 | -0.018025917 | 0.25120658 | -0.0717573 | 0.94279502 | 0.98885525 |
| PELI2 | 958.194675 | 0.908517123 | 0.31687091 | 2.86715222 | 0.00414184 | 0.09955609 |
| TMEM260 | 386.46363 | 0.380089255 | 0.26465063 | 1.43619253 | 0.15094756 | 0.58535779 |
| EXOC5 | 260.578556 | 0.049460515 | 0.162331 | 0.30468929 | 0.76060281 | 0.95329513 |
| AP5M1 | 179.608707 | 0.136146758 | 0.27657293 | 0.49226349 | 0.62253308 | 0.91030371 |
| NAA30 | 86.9679686 | 0.117135409 | 0.21142257 | 0.55403455 | 0.5795552 | 0.89989814 |
| CCDC198 | 803.787015 | 0.926189123 | 0.50031802 | 1.85120083 | 0.06414067 | 0.41357646 |
| ARMH4 | 137.128281 | -0.10161081 | 0.28814216 | -0.3526412 | 0.72435741 | 0.93893205 |
| ACTR10 | 226.306213 | 0.02258319 | 0.18672884 | 0.1209411 | 0.90373769 | 0.98320957 |
| PSMA3 | 174.649969 | 0.167533914 | 0.27796476 | 0.60271638 | 0.54669738 | 0.88719155 |
| ARID4A | 349.199592 | 0.086598298 | 0.22712606 | 0.38127857 | 0.70299655 | 0.93241442 |
| AL139021.1 | 38.3658745 | -0.333063487 | 0.49004219 | -0.6796629 | 0.49671794 | 0.86487238 |
| TIMM9 | 79.8705354 | -0.226498877 | 0.23343911 | -0.9702696 | 0.33191211 | 0.77618491 |
| KIAA0586 | 179.957869 | -0.063006338 | 0.18639199 | -0.3380314 | 0.73533955 | 0.94232801 |
| DAAM1 | 272.929859 | 0.11405467 | 0.26049255 | 0.43784236 | 0.66150056 | 0.92284375 |
| GPR135 | 49.3435781 | -0.725370568 | 0.53596481 | -1.3533922 | 0.17593038 | 0.62486786 |
| L3HYPDH | 50.1261098 | -0.48959526 | 0.36222224 | -1.3516433 | 0.17648946 | 0.62528328 |
| JKAMP | 115.529631 | -0.082970256 | 0.31924907 | -0.2598919 | 0.79494714 | 0.95937941 |
| CCDC175 | 82.5739556 | 0.424665356 | 0.52355191 | 0.81112369 | 0.41729464 | 0.8254526 |
| PCNX4 | 472.31091 | 0.398736396 | 0.50379575 | 0.79146439 | 0.42867305 | 0.8296433 |
| DHRS7 | 193.742635 | 0.589148762 | 0.61208065 | 0.96253454 | 0.33578116 | 0.77820994 |
| PPM1A | 324.572145 | 0.421681936 | 0.34617089 | 1.21813229 | 0.22317371 | 0.68089096 |
| MNAT1 | 751.739804 | 0.288423477 | 0.23931139 | 1.20522253 | 0.22811741 | 0.68638255 |
| SLC38A6 | 160.205963 | 0.02232913 | 0.23004528 | 0.09706406 | 0.92267552 | 0.9866731 |
| PRKCH | 325.244137 | -0.111101014 | 0.49067377 | -0.2264254 | 0.82087056 | 0.96502145 |
| TMEM30B | 176.602353 | -0.081727609 | 0.35305792 | -0.231485 | 0.81693805 | 0.96380902 |
| AL359220.1 | 35.9729989 | -0.860733024 | 0.73201264 | -1.1758445 | 0.23965703 | 0.69573564 |
| HIF1A | 3115.05291 | -0.877759494 | 0.49107519 | -1.7874238 | 0.07386901 | 0.43712436 |
| HIF1A-AS2 | 872.421697 | -1.266943102 | 0.76812288 | -1.6494016 | 0.09906539 | 0.4933116 |
| SNAPC1 | 433.523945 | -0.143887796 | 0.53705357 | -0.2679208 | 0.78876032 | 0.95801395 |
| SYT16 | 45.0306683 | -2.364835085 | 0.6148563 | -3.846159 | 0.00011998 | 0.00990104 |
| PPP2R5E | 677.30136 | 0.244304479 | 0.2733842 | 0.89363056 | 0.37151959 | 0.79539603 |
| WDR89 | 148.17765 | 0.020554032 | 0.36208912 | 0.05676512 | 0.9547323 | 0.99073937 |
| SGPP1 | 1265.58723 | 0.456889533 | 0.47119813 | 0.96963358 | 0.33222917 | 0.77618491 |
| SYNE2 | 2764.83313 | 0.27047867 | 0.21291361 | 1.27036814 | 0.20395353 | 0.65868504 |
| ESR2 | 221.63662 | 0.810654939 | 0.64910223 | 1.24888639 | 0.21170663 | 0.66975892 |
| MTHFD1 | 66.3919687 | -0.83215213 | 0.24095973 | -3.4534904 | 0.00055338 | 0.02672555 |
| AL122035.1 | 70.0202636 | -1.210821706 | 0.43475281 | -2.7850808 | 0.00535144 | 0.11705923 |
| ZBTB25 | 160.493046 | -0.946405993 | 0.37744679 | -2.5073891 | 0.01216267 | 0.17590979 |
| ZBTB1 | 314.488891 | -0.38934028 | 0.29615011 | -1.3146721 | 0.18862014 | 0.64159008 |
| AL049869.3 | 38.5974472 | -0.335851029 | 0.73422457 | -0.4574228 | 0.64736721 | 0.91743182 |
| HSPA2 | 155.613944 | 0.952978389 | 0.58185669 | 1.63782322 | 0.10145858 | 0.49779302 |
| PPP1R36 | 39.154808 | -0.23517122 | 0.45076408 | -0.5217169 | 0.60186748 | 0.90369786 |
| PLEKHG3 | 385.869959 | -0.157661074 | 0.40914777 | -0.3853402 | 0.6999854 | 0.93180634 |
| SPTB | 35.1116152 | 0.019898089 | 0.6060018 | 0.03283503 | 0.97380614 | 0.9939318 |
| CHURC1-FNT | 93.6845949 | -0.816508433 | 0.40756858 | -2.0033645 | 0.04513818 | 0.35634897 |
| RAB15 | 135.412247 | 0.555468997 | 0.42408548 | 1.30980434 | 0.19026203 | 0.64458858 |
| MAX | 241.549489 | -0.688981219 | 0.27167514 | -2.5360481 | 0.01121113 | 0.16686755 |
| FUT8 | 229.396509 | 0.462126096 | 0.45306558 | 1.01999823 | 0.3077293 | 0.75262615 |
| AL359232.1 | 109.946292 | -0.836688873 | 0.41168337 | -2.0323602 | 0.04211721 | 0.34673083 |
| GPHN | 1957.08997 | 0.384771479 | 0.34393256 | 1.11874107 | 0.26325062 | 0.71989632 |
| MPP5 | 389.387593 | 0.450391591 | 0.14851543 | 3.03262494 | 0.00242437 | 0.07177223 |
| ATP6V1D | 220.393398 | -0.027118409 | 0.22996822 | -0.1179224 | 0.90612913 | 0.98320957 |
| EIF2S1 | 119.558549 | 0.034319108 | 0.25488538 | 0.13464526 | 0.89289236 | 0.98168537 |
| PLEKHH1 | 131.594637 | 0.127100197 | 0.25185949 | 0.50464725 | 0.61380662 | 0.90751853 |
| PIGH | 88.1181447 | 1.076444919 | 0.63484272 | 1.69560881 | 0.08995999 | 0.47371081 |
| ARG2 | 855.487349 | 2.263441421 | 0.84070416 | 2.69231618 | 0.00709576 | 0.13409852 |
| VTI1B | 141.036407 | 0.490584325 | 0.41489514 | 1.18242968 | 0.23703525 | 0.69350348 |
| ZFYVE26 | 95.5564769 | -0.679016362 | 0.39079673 | -1.737518 | 0.08229577 | 0.45595206 |
| RAD51B | 2428.82614 | 0.654792294 | 0.35018996 | 1.86982031 | 0.06150877 | 0.4077242 |
| ZFP36L1 | 2127.96204 | 0.287297411 | 0.41365161 | 0.69453958 | 0.48734389 | 0.85968086 |
| ACTN1 | 139.247971 | -0.609116214 | 0.43571375 | -1.3979734 | 0.16212106 | 0.60211533 |
| DCAF5 | 475.194937 | -0.110473435 | 0.23554865 | -0.4690047 | 0.63906625 | 0.91521016 |
| EXD2 | 115.617866 | 0.245412523 | 0.20913411 | 1.17346962 | 0.24060753 | 0.6963541 |
| ERH | 212.333511 | 0.105749487 | 0.35964056 | 0.29404216 | 0.76872569 | 0.95453142 |
| SLC39A9 | 193.089862 | 0.238062179 | 0.1569914 | 1.51640265 | 0.12941757 | 0.55140574 |
| SUSD6 | 185.875125 | -0.283894584 | 0.29480604 | -0.9629877 | 0.33555371 | 0.77820994 |
| SRSF5 | 640.127204 | 0.731842153 | 0.20751372 | 3.5267169 | 0.00042075 | 0.02204711 |
| AL357153.2 | 109.779295 | -0.561573275 | 0.40084654 | -1.4009683 | 0.16122357 | 0.60086165 |
| MED6 | 99.0849082 | 0.229583919 | 0.25340266 | 0.90600436 | 0.36493354 | 0.79280753 |
| MAP3K9 | 106.752477 | -0.146245507 | 0.42443747 | -0.3445631 | 0.73042282 | 0.94048525 |
| PCNX1 | 657.290321 | 0.315367019 | 0.25567679 | 1.23345969 | 0.21740431 | 0.67507983 |
| SIPA1L1 | 1092.60059 | -0.022902459 | 0.2407764 | -0.0951192 | 0.92422015 | 0.98697817 |
| ZFYVE1 | 98.1212552 | -0.552030866 | 0.33873487 | -1.6296842 | 0.10316826 | 0.50171849 |
| RBM25 | 493.015498 | 0.222294657 | 0.21400873 | 1.03871771 | 0.29893604 | 0.74594723 |
| PSEN1 | 321.662456 | -0.003227643 | 0.24732617 | -0.0130501 | 0.98958779 | 0.9965778 |
| PAPLN | 88.773131 | -1.03490834 | 0.75151773 | -1.377091 | 0.1684841 | 0.61330788 |
| NUMB | 924.841601 | 0.05759283 | 0.2386387 | 0.24133902 | 0.80929237 | 0.96245698 |
| HEATR4 | 43.3521893 | -1.661817545 | 0.41873805 | -3.9686327 | 7.23E-05 | 0.00734782 |
| DNAL1 | 150.122087 | 0.259566016 | 0.27215837 | 0.95373153 | 0.34021955 | 0.77836439 |
| PNMA1 | 68.3290215 | -0.079528194 | 0.3703866 | -0.2147167 | 0.82998819 | 0.96643829 |
| ELMSAN1 | 219.980858 | -0.332692193 | 0.38387772 | -0.8666619 | 0.38612729 | 0.80634213 |
| AC005480.1 | 89.0342427 | 0.32718691 | 0.47870961 | 0.6834768 | 0.49430561 | 0.86367502 |
| COQ6 | 39.19258 | 0.248289254 | 0.29538948 | 0.84054874 | 0.40060079 | 0.8162653 |
| ENTPD5 | 124.034177 | 0.251318414 | 0.32735575 | 0.76772262 | 0.442652 | 0.83826789 |
| BBOF1 | 156.21987 | -0.270101021 | 0.27791617 | -0.9718795 | 0.33111051 | 0.77618491 |
| ALDH6A1 | 167.812999 | -0.284000706 | 0.50686107 | -0.5603127 | 0.57526615 | 0.89807676 |
| LIN52 | 159.422797 | 0.251752741 | 0.32850894 | 0.76634975 | 0.44346822 | 0.83860465 |
| ABCD4 | 193.222375 | -0.313023673 | 0.22084375 | -1.4173988 | 0.15636635 | 0.59366792 |
| NPC2 | 350.545772 | -0.59481 | 0.47574942 | -1.250259 | 0.21120494 | 0.66972097 |
| ISCA2 | 88.8830892 | -0.340117332 | 0.54880497 | -0.6197417 | 0.53542785 | 0.88137638 |
| AREL1 | 139.083838 | 0.437034283 | 0.23098535 | 1.8920433 | 0.05848521 | 0.39982063 |
| FCF1 | 73.7295574 | -0.110556825 | 0.23863289 | -0.4632925 | 0.64315473 | 0.91643619 |
| YLPM1 | 352.713113 | 0.141052604 | 0.24296795 | 0.58053995 | 0.56155055 | 0.89242209 |
| DLST | 156.498382 | -0.120101948 | 0.18407528 | -0.652461 | 0.5141038 | 0.87226916 |
| EIF2B2 | 46.4927692 | -0.215894487 | 0.39098579 | -0.5521799 | 0.58082512 | 0.90045866 |
| MLH3 | 278.247206 | -0.187660798 | 0.16881057 | -1.111665 | 0.26628223 | 0.72053648 |
| ACYP1 | 194.643918 | 0.479931177 | 0.412977 | 1.16212568 | 0.24518442 | 0.70148313 |
| ZC2HC1C | 40.4677777 | -0.013156926 | 0.4814485 | -0.0273278 | 0.97819829 | 0.99469867 |
| NEK9 | 211.603854 | -0.02475143 | 0.18388627 | -0.1346018 | 0.89292669 | 0.98168537 |
| AL049780.2 | 44.8529445 | -0.082123734 | 0.68542451 | -0.1198144 | 0.90463017 | 0.98320957 |
| TMED10 | 517.961065 | -0.040654205 | 0.2588447 | -0.1570602 | 0.87519739 | 0.97622753 |
| FOS | 526.506955 | 2.923116263 | 1.46876728 | 1.99018341 | 0.04657074 | 0.36244837 |

| | | | | | | |
|---|---|---|---|---|---|---|
| JDP2 | 56.5808086 | 1.170331492 | 0.59957323 | 1.95194087 | 0.05094523 | 0.3802749 |
| TTLL5 | 448.162606 | 0.10252214 | 0.26667651 | 0.38444384 | 0.70064951 | 0.93180634 |
| ERG28 | 83.5183836 | -0.481441009 | 0.36947497 | -1.3030409 | 0.19256079 | 0.64556529 |
| IFT43 | 310.480364 | 0.169029813 | 0.21260616 | 0.79503722 | 0.42659185 | 0.8277744 |
| GPATCH2L | 318.057293 | -0.169668667 | 0.25296149 | -0.6707292 | 0.50239304 | 0.86852095 |
| ANGEL1 | 46.112947 | 0.087507089 | 0.42693054 | 0.20496798 | 0.83759714 | 0.96757829 |
| IRF2BPL | 203.817786 | -0.154746327 | 0.38885287 | -0.397956 | 0.69066263 | 0.93073907 |
| POMT2 | 57.6173598 | -0.041091058 | 0.36207284 | -0.1134884 | 0.90964338 | 0.98441374 |
| GSTZ1 | 41.5830766 | -0.348966092 | 0.3595639 | -0.9705259 | 0.33178441 | 0.77618491 |
| TMED8 | 206.119004 | 0.311375254 | 0.16339345 | 1.90567771 | 0.05669203 | 0.39714739 |
| VIPAS39 | 73.490448 | -0.219654124 | 0.23197194 | -0.9468995 | 0.34368998 | 0.78165602 |
| AHSA1 | 94.5718995 | 0.584677653 | 0.41039926 | 1.42465573 | 0.15425674 | 0.59000389 |
| SPTLC2 | 612.474361 | 0.024651806 | 0.30952446 | 0.07964413 | 0.9365203 | 0.98822345 |
| ALKBH1 | 88.9301097 | 0.082480766 | 0.22539886 | 0.36593248 | 0.71441546 | 0.93573028 |
| SLIRP | 129.432979 | -0.368550077 | 0.26413967 | -1.3952848 | 0.16292996 | 0.60211533 |
| SNW1 | 147.524358 | -0.127597539 | 0.2332718 | -0.5469908 | 0.58438503 | 0.90090543 |
| ADCK1 | 56.3234144 | -0.191772419 | 0.46464177 | -0.4127318 | 0.67980314 | 0.92700898 |
| CEP128 | 206.180143 | -0.073557508 | 0.34883281 | -0.2108675 | 0.83299065 | 0.96739162 |
| AC007262.2 | 65.3392466 | -0.096676243 | 0.57320585 | -0.1686589 | 0.86606498 | 0.97521876 |
| GTF2A1 | 141.893513 | 0.00830217 | 0.27857338 | 0.02980245 | 0.9762246 | 0.99425013 |
| STON2 | 448.209745 | 0.688171504 | 0.31823115 | 2.16248945 | 0.03058047 | 0.29429138 |
| SEL1L | 314.409855 | 0.123711942 | 0.19291145 | 0.64128874 | 0.5213351 | 0.87650871 |
| GALC | 80.6667013 | -0.088012037 | 0.21179544 | -0.4155521 | 0.67773779 | 0.9268955 |
| SPATA7 | 229.237525 | -0.009458402 | 0.34516402 | -0.0274026 | 0.9781386 | 0.99469867 |
| PTPN21 | 312.99644 | 1.096408218 | 0.36521121 | 3.0021209 | 0.00268106 | 0.07553085 |
| ZC3H14 | 340.632249 | 0.573232043 | 0.2287324 | 2.50612528 | 0.01220624 | 0.17590979 |
| EML5 | 90.9553767 | -0.591337625 | 0.51616848 | -1.1456291 | 0.25194865 | 0.70813507 |
| TTC8 | 180.280493 | 0.14047623 | 0.29008376 | 0.48426093 | 0.6282007 | 0.9109649 |
| FOXN3 | 675.702735 | -0.127457084 | 0.25969447 | -0.4907963 | 0.62357053 | 0.91058133 |
| EFCAB11 | 180.600182 | -0.378948942 | 0.39318428 | -0.9637947 | 0.33514884 | 0.77820994 |
| TDP1 | 94.1590421 | -0.398283295 | 0.274343 | -1.4517713 | 0.14656521 | 0.57722788 |
| KCNK13 | 455.971689 | 1.648577549 | 0.62104381 | 2.65452699 | 0.00794197 | 0.14172148 |
| PSMC1 | 132.889725 | -0.233241453 | 0.21417 | -1.0890482 | 0.27613264 | 0.72655538 |
| NRDE2 | 127.177774 | -0.301075201 | 0.31035951 | -0.9700853 | 0.33200397 | 0.77618491 |
| CALM1 | 694.441038 | 0.216369971 | 0.51037186 | 0.42394573 | 0.67160539 | 0.92489152 |
| TTC7B | 415.883162 | 0.662049096 | 0.31699148 | 2.08853911 | 0.03674923 | 0.32201668 |
| RPS6KA5 | 198.684627 | -0.716858576 | 0.3745356 | -1.9139932 | 0.05562103 | 0.39368206 |
| DGLUCY | 421.423218 | -0.351208796 | 0.37509178 | -0.9363276 | 0.34910453 | 0.78180304 |
| CCDC88C | 300.945543 | 0.237726724 | 0.46707263 | 0.50897164 | 0.6107721 | 0.9062166 |
| PPP4R3A | 465.747588 | 0.876496343 | 0.3934971 | 2.22745313 | 0.025917 | 0.27398277 |
| CATSPERB | 255.282211 | -0.202882384 | 0.57664015 | -0.3518353 | 0.72496175 | 0.93895336 |
| TC2N | 526.850572 | 1.345933158 | 0.76385862 | 1.76201869 | 0.07806614 | 0.44731172 |
| TRIP11 | 483.500847 | 0.027170555 | 0.19418396 | 0.13992172 | 0.88872184 | 0.98051184 |
| ATXN3 | 265.376523 | 0.027674531 | 0.23651202 | 0.1170111 | 0.90685126 | 0.98321965 |
| NDUFB1 | 156.662623 | 0.174994486 | 0.27023586 | 0.64756204 | 0.51726826 | 0.87526312 |
| CPSF2 | 147.754774 | 0.323227339 | 0.19368309 | 1.66884642 | 0.09514782 | 0.48627909 |
| LGMN | 118.992457 | -0.357024839 | 0.48008493 | -0.7436702 | 0.45707604 | 0.84408051 |
| GOLGA5 | 107.263681 | -0.116546701 | 0.23940542 | -0.4868173 | 0.62638781 | 0.91061758 |
| ITPK1 | 415.811069 | 0.477094222 | 0.31968951 | 1.49236747 | 0.13560284 | 0.55916513 |
| MOAP1 | 117.309851 | -0.898535277 | 0.41419642 | -2.1693459 | 0.03005644 | 0.29169628 |
| GON7 | 48.6700632 | -0.382321293 | 0.48030823 | -0.7959915 | 0.42603695 | 0.82774157 |
| UBR7 | 47.5582962 | 0.070691072 | 0.30311506 | 0.23321531 | 0.81559422 | 0.96333253 |
| BTBD7 | 932.307852 | -0.274165361 | 0.25520916 | -1.0742771 | 0.2826985 | 0.73268183 |
| UNC79 | 47.3320138 | -1.353016864 | 0.6321332 | -2.1403984 | 0.03232259 | 0.30298811 |
| DDX24 | 425.920846 | -0.559128837 | 0.32461013 | -1.7224627 | 0.08498573 | 0.46310155 |
| IFI27L1 | 39.5552547 | -0.027076219 | 0.30089651 | -0.0899852 | 0.92829901 | 0.98751845 |
| IFI27L2 | 59.3402784 | -0.386491982 | 0.346545 | -1.1152721 | 0.26473382 | 0.71989632 |
| SERPINA1 | 84.4563915 | -2.099643027 | 0.81630406 | -2.5721335 | 0.01010739 | 0.16113882 |
| DICER1 | 652.857139 | 0.096146691 | 0.1535965 | 0.62596927 | 0.5313351 | 0.88023899 |
| DICER1-AS1 | 60.677484 | -0.550414657 | 0.47712521 | -1.1536063 | 0.24866161 | 0.70546119 |
| CLMN | 1300.99712 | 0.117393857 | 0.33459193 | 0.35085681 | 0.72569578 | 0.93895336 |
| GLRX5 | 213.78954 | 0.153394442 | 0.56233837 | 0.27277961 | 0.78502263 | 0.95702626 |
| ATG2B | 268.80644 | 0.348710376 | 0.24962178 | 1.39695491 | 0.16242713 | 0.60211533 |
| GSKIP | 60.7249234 | -0.369326967 | 0.31120829 | -1.1867517 | 0.23532558 | 0.69335782 |
| AK7 | 80.9062657 | -1.026884697 | 0.31300043 | -3.2807773 | 0.00103521 | 0.04033103 |
| PAPOLA | 592.466936 | 0.438980832 | 0.16453556 | 2.66799982 | 0.00763043 | 0.13907288 |
| VRK1 | 69.2037427 | 0.123026255 | 0.2489979 | 0.49408552 | 0.62124579 | 0.90994491 |
| LINC01550 | 60.1585732 | -0.052342171 | 0.54654702 | -0.0957688 | 0.92370417 | 0.98694936 |
| SETD3 | 385.615106 | 0.51091665 | 0.16152163 | 3.16314693 | 0.00156074 | 0.05293368 |
| CCNK | 109.902377 | -0.079471803 | 0.18899203 | -0.4205035 | 0.6741177 | 0.92595152 |
| CCDC85C | 59.5244894 | 0.195038575 | 0.24673568 | 0.79047576 | 0.42924998 | 0.82998114 |
| EML1 | 46.1083431 | -0.636068883 | 0.3650455 | -1.7424373 | 0.08143196 | 0.45515037 |
| EVL | 228.71569 | -0.103216145 | 0.39201019 | -0.2632996 | 0.79231963 | 0.9585028 |
| DEGS2 | 69.0749299 | 0.195159355 | 0.76536451 | 0.25498877 | 0.79873177 | 0.96005379 |
| YY1 | 469.768244 | -0.063680841 | 0.27335488 | -0.2329603 | 0.81579222 | 0.96333253 |
| SLC25A29 | 1635.69472 | 0.183326172 | 0.64437933 | 0.28450039 | 0.77602693 | 0.95601455 |
| WARS | 220.032154 | -0.758479924 | 0.74567783 | -1.0171684 | 0.30907333 | 0.75357697 |
| WDR25 | 128.206554 | -0.019509342 | 0.28987921 | -0.0673016 | 0.94634158 | 0.98949015 |
| PPP2R5C | 470.307195 | -0.026102734 | 0.16740236 | -0.1559281 | 0.87608969 | 0.97622753 |
| DYNC1H1 | 409.796666 | -0.41408895 | 0.2104272 | -1.967849 | 0.04908542 | 0.3746651 |
| HSP90AA1 | 1573.49229 | 0.798893477 | 0.43411492 | 1.84028109 | 0.06572698 | 0.41788422 |
| WDR20 | 316.443997 | 0.195949047 | 0.21404491 | 0.91545761 | 0.35995145 | 0.78892604 |
| MOK | 268.693899 | 0.480185088 | 0.26482232 | 1.81323497 | 0.06979559 | 0.42843167 |
| ZNF839 | 77.582291 | 0.060226231 | 0.29272646 | 0.20574236 | 0.83699217 | 0.96757829 |
| CINP | 62.0537057 | -0.479607261 | 0.2473416 | -1.9390481 | 0.05249548 | 0.38418477 |
| TECPR2 | 170.672383 | -0.073383873 | 0.24405617 | -0.3006844 | 0.7636552 | 0.95378284 |
| ANKRD9 | 99.0713812 | -0.440437485 | 0.38994464 | -1.1294872 | 0.25869236 | 0.7159728 |
| RCOR1 | 436.109288 | 0.351988573 | 0.3045603 | 1.15572704 | 0.24779282 | 0.70405328 |
| TRAF3 | 302.677254 | -0.162504963 | 0.42489114 | -0.3824626 | 0.70211828 | 0.93224371 |
| CDC42BPB | 875.310045 | 0.306257868 | 0.36976281 | 0.82825493 | 0.40752614 | 0.82018309 |
| TNFAIP2 | 54.5442756 | -0.648482381 | 0.67404531 | -0.9620753 | 0.33601176 | 0.77820994 |
| EIF5 | 320.418666 | -0.487279405 | 0.35461298 | -1.3741161 | 0.16940564 | 0.61478828 |
| MARK3 | 1153.95913 | 0.327350082 | 0.19335891 | 1.69296614 | 0.09046191 | 0.47556444 |
| CKB | 494.462254 | -0.160635614 | 0.57302223 | -0.2803305 | 0.77922395 | 0.95623735 |
| BAG5 | 136.685313 | -0.641959162 | 0.22218435 | -2.8893087 | 0.0038609 | 0.09660987 |
| KLC1 | 228.568296 | 0.137347369 | 0.27414194 | 0.50100823 | 0.61636533 | 0.90794578 |
| ZFYVE21 | 149.638128 | 0.079664514 | 0.39772446 | 0.20030077 | 0.84124536 | 0.96854218 |
| PPP1R13B | 277.580326 | -0.475449835 | 0.20764773 | -2.2896944 | 0.02203904 | 0.25050883 |
| ATP5MPL | 207.469871 | -0.072873116 | 0.27964315 | -0.2605932 | 0.7944062 | 0.95937941 |
| INF2 | 60.1087475 | -0.425442201 | 0.31690529 | -1.3424901 | 0.17943715 | 0.6289302 |
| SIVA1 | 126.206245 | 0.077532944 | 0.49005992 | 0.15821115 | 0.87429042 | 0.97622753 |
| AKT1 | 105.095376 | -0.078288595 | 0.3377143 | -0.231819 | 0.81667859 | 0.96380902 |
| CEP170B | 74.2132204 | 0.248823475 | 0.35938589 | 0.69235738 | 0.48871292 | 0.86014074 |
| CLBA1 | 53.2993471 | -0.279206496 | 0.24576022 | -1.1360931 | 0.2559176 | 0.71330223 |
| GPR132 | 72.1477299 | -1.083880665 | 1.11534338 | -0.971791 | 0.33115453 | 0.77618491 |
| NUDT14 | 45.8618483 | -0.669303371 | 0.37436208 | -1.7878503 | 0.07380016 | 0.43712321 |
| BRF1 | 124.070168 | 0.098240292 | 0.26075351 | 0.37675539 | 0.70635539 | 0.93243538 |
| BTBD6 | 72.1933696 | -0.113721966 | 0.38791318 | -0.2931635 | 0.76939722 | 0.95482817 |
| PACS2 | 217.158929 | -0.381170009 | 0.35113128 | -1.0855484 | 0.27767882 | 0.72788245 |
| TEX22 | 34.0127707 | -0.606361497 | 0.41895537 | -1.4473176 | 0.14780798 | 0.57952643 |
| MTA1 | 143.844116 | 0.251632222 | 0.24443983 | 1.02942398 | 0.30328049 | 0.74953627 |
| CRIP2 | 291.563798 | 0.179084618 | 0.42805581 | 0.41836745 | 0.67567848 | 0.92672563 |
| NIPA1 | 105.721018 | -0.502579356 | 0.19957838 | -2.5182054 | 0.01179545 | 0.17264852 |

| | | | | | | |
|---|---|---|---|---|---|---|
| NIPA2 | 144.627023 | 0.277591342 | 0.3161153 | 0.8781332 | 0.37987144 | 0.80295536 |
| CYFIP1 | 289.854453 | -0.102560893 | 0.21245143 | -0.4827498 | 0.62927337 | 0.9114564 |
| TUBGCP5 | 117.684371 | 0.226763813 | 0.28229391 | 0.80328977 | 0.42180727 | 0.82719689 |
| NDN | 34.2999914 | -0.199943067 | 0.48048429 | -0.4161282 | 0.67731619 | 0.9268955 |
| AC087463.4 | 122.997134 | 2.495065718 | 0.63732268 | 3.91491749 | 9.04E-05 | 0.00853838 |
| PWRN3 | 387.976753 | 2.496552035 | 0.97908439 | 2.54988443 | 0.01077586 | 0.16452817 |
| PWRN1 | 3243.20903 | 2.223338199 | 0.5202361 | 4.27370998 | 1.92E-05 | 0.00296287 |
| NPAP1 | 103.61937 | 2.225766146 | 0.57038447 | 3.90222079 | 9.53E-05 | 0.00861114 |
| SNRPN | 184.165342 | 2.190089921 | 0.66217782 | 3.30740454 | 0.00094165 | 0.03807006 |
| AC124312.1 | 945.491539 | 0.698767283 | 0.24237342 | 2.88301944 | 0.00393883 | 0.09740819 |
| PWAR6 | 232.035092 | 0.622753255 | 0.24070818 | 2.58717113 | 0.00967675 | 0.15717671 |
| UBE3A | 539.098116 | 0.238395177 | 0.1580107 | 1.50872804 | 0.1313683 | 0.55397858 |
| GABRB3 | 94.2331414 | -0.899457253 | 0.58904202 | -1.5269832 | 0.12676519 | 0.54692166 |
| HERC2 | 577.222752 | 0.039293722 | 0.24668542 | 0.15928676 | 0.87344295 | 0.97622753 |
| FAM189A1 | 56.8139376 | 0.252942763 | 0.37873077 | 0.66786958 | 0.50421685 | 0.86946722 |
| TJP1 | 536.403586 | 0.133096293 | 0.27375288 | 0.48619138 | 0.62683148 | 0.91061758 |
| FAN1 | 155.649452 | -0.066711893 | 0.27063007 | -0.2465058 | 0.80529069 | 0.96218371 |
| MTMR10 | 1123.36812 | 0.15262067 | 0.50777473 | 0.30056768 | 0.76374418 | 0.95378284 |
| KLF13 | 222.630328 | -0.047693115 | 0.37365387 | -0.1276398 | 0.89843401 | 0.98312332 |
| OTUD7A | 198.476007 | 0.008171224 | 0.26282635 | 0.03108982 | 0.97519791 | 0.9939318 |
| LINC02256 | 376.607683 | -0.377354755 | 0.29469409 | -1.2804965 | 0.20037056 | 0.65475264 |
| AC123768.3 | 61.6223982 | -0.148000995 | 0.41011419 | -0.3608775 | 0.718191 | 0.93573028 |
| GOLGA8N | 44.1623628 | 0.285782374 | 0.31383531 | 0.91061256 | 0.36249955 | 0.79063377 |
| FMN1 | 103.591261 | -0.878042148 | 0.54847954 | -1.6008658 | 0.10940664 | 0.51481816 |
| AVEN | 122.379618 | 0.537420455 | 0.29422283 | 1.8265763 | 0.0677635 | 0.42296693 |
| EMC7 | 111.213242 | 0.126124776 | 0.37046761 | 0.34044751 | 0.73351955 | 0.94163693 |
| KATNBL1 | 397.464346 | 0.440507976 | 0.19050295 | 2.31234204 | 0.02075884 | 0.24280431 |
| EMC4 | 67.0198684 | -0.433874597 | 0.39179679 | -1.107397 | 0.26812232 | 0.72308815 |
| SLC12A6 | 116.924946 | -0.254301345 | 0.35791 | -0.7105176 | 0.47738323 | 0.85462282 |
| NOP10 | 191.836099 | -0.162489867 | 0.557884 | -0.291261 | 0.77085169 | 0.95498333 |
| LPCAT4 | 35.626357 | -0.766998358 | 0.32418355 | -2.3659386 | 0.01798443 | 0.22331375 |
| GOLGA8A | 180.228499 | 0.052415957 | 0.28711071 | 0.18256357 | 0.85514047 | 0.97135336 |
| GOLGA8B | 194.642239 | 0.003429104 | 0.38390897 | 0.00893208 | 0.99287333 | 0.99839831 |
| AQR | 288.018795 | 0.254007576 | 0.18169001 | 1.39802717 | 0.16210491 | 0.60211533 |
| ZNF770 | 217.844081 | -0.571126444 | 0.27659406 | -2.0648544 | 0.03893678 | 0.33202399 |
| AC021231.1 | 296.23874 | 0.082657149 | 0.4681964 | 0.17654375 | 0.85986678 | 0.97256291 |
| DPH6 | 840.938414 | 0.045735002 | 0.32351287 | 0.14136996 | 0.88757768 | 0.98009982 |
| DPH6-DT | 569.628415 | 0.731745113 | 0.67099578 | 1.09053609 | 0.27547706 | 0.7263609 |
| C15orf41 | 252.299265 | -0.019588637 | 0.24908474 | -0.0786425 | 0.93731701 | 0.98843654 |
| SPRED1 | 287.565162 | -0.634781368 | 0.41756177 | -1.5202095 | 0.12845833 | 0.54843482 |
| FAM98B | 124.960328 | 0.342917129 | 0.18995808 | 1.80522532 | 0.07103943 | 0.43107604 |
| RASGRP1 | 408.157657 | -0.049083931 | 0.33430324 | -0.1468246 | 0.88327048 | 0.97878487 |
| C15orf53 | 74.1956982 | -0.838917823 | 0.61180638 | -1.3712146 | 0.17030808 | 0.61613093 |
| AC013652.1 | 69.0492967 | 0.3975085 | 0.38231105 | 1.03975154 | 0.29845535 | 0.74594723 |
| THBS1 | 167.722063 | -0.285084046 | 0.7354432 | -0.3876357 | 0.69828564 | 0.93177271 |
| FSIP1 | 128.710677 | 0.509510132 | 0.36531223 | 1.39472508 | 0.16309876 | 0.6021948 |
| EIF2AK4 | 193.529711 | 0.061684963 | 0.17847039 | 0.34563136 | 0.72961978 | 0.94028717 |
| SRP14 | 1192.79645 | -0.466434784 | 0.49510812 | -0.9420867 | 0.34614824 | 0.78180304 |
| SRP14-AS1 | 45.8655785 | -0.632032879 | 0.44175454 | -1.4307332 | 0.1525067 | 0.58795409 |
| BMF | 54.1089125 | 1.557297425 | 0.74791813 | 2.08217632 | 0.03732637 | 0.32376418 |
| PAK6 | 33.8989286 | 0.201255412 | 0.51872283 | 0.38798256 | 0.69802894 | 0.93177271 |
| IVD | 217.397224 | 0.093059255 | 0.31148988 | 0.2987553 | 0.76512676 | 0.95378284 |
| BAHD1 | 70.405815 | -0.336177816 | 0.29024672 | -1.1582484 | 0.24676267 | 0.70292817 |
| CCDC32 | 142.091446 | -0.129263228 | 0.31609491 | -0.408938 | 0.68258515 | 0.92806156 |
| RMDN3 | 88.0670917 | -0.004741577 | 0.22032827 | -0.0215205 | 0.98283044 | 0.99507903 |
| GCHFR | 50.9446309 | -1.385266991 | 0.7314393 | -1.8938919 | 0.05823935 | 0.39970428 |
| DNAJC17 | 125.874132 | 0.011317737 | 0.22400601 | 0.05052425 | 0.95970462 | 0.99119981 |
| ZFYVE19 | 53.6066808 | -0.000267508 | 0.32859904 | -0.0008141 | 0.99935045 | 0.99992293 |
| SPINT1-AS1 | 115.948989 | 0.355858025 | 0.44298197 | 0.80332396 | 0.42178751 | 0.82719689 |
| SPINT1 | 259.044325 | -0.736831745 | 0.32753091 | -2.2496556 | 0.02447082 | 0.26518037 |
| RHOV | 53.3205692 | -1.037533042 | 1.0820022 | -0.958901 | 0.3376086 | 0.77827084 |
| VPS18 | 56.9046037 | -0.392326047 | 0.42882553 | -0.914885 | 0.360252 | 0.78892604 |
| INO80 | 408.962766 | 0.456001995 | 0.33786194 | 1.34966962 | 0.17712198 | 0.62617642 |
| CHP1 | 323.462868 | 0.527909698 | 0.50670866 | 1.04184068 | 0.29748555 | 0.74513589 |
| NDUFAF1 | 41.150476 | -0.26878602 | 0.25221677 | -1.0656945 | 0.28656178 | 0.73607046 |
| RTF1 | 274.278524 | -0.25931844 | 0.2984167 | -0.868981 | 0.38485753 | 0.80561637 |
| RPAP1 | 44.2893619 | -0.15731527 | 0.29312559 | -0.5366821 | 0.5914872 | 0.90233654 |
| MGA | 329.22879 | 0.21430303 | 0.29181234 | 0.73438645 | 0.46271324 | 0.84821318 |
| MAPKBP1 | 173.915333 | 0.155633401 | 0.39138409 | 0.39764876 | 0.69088911 | 0.93073907 |
| EHD4 | 289.508648 | 0.387522969 | 0.35590889 | 1.08882634 | 0.27623048 | 0.72663037 |
| PLA2G4F | 67.7834412 | 0.032294074 | 0.51323596 | 0.06292247 | 0.94982824 | 0.98977092 |
| VPS39 | 205.705503 | 0.093099842 | 0.27036932 | 0.34434322 | 0.73058818 | 0.94048525 |
| TMEM87A | 353.053936 | -0.088969382 | 0.22252149 | -0.3998238 | 0.68928632 | 0.93041766 |
| GANC | 141.852607 | -0.397491001 | 0.32148036 | -1.2364395 | 0.21629526 | 0.67323263 |
| ZNF106 | 292.612752 | 0.012446751 | 0.21785157 | 0.05713409 | 0.95443838 | 0.99060627 |
| SNAP23 | 209.992004 | -0.102352838 | 0.20526571 | -0.4986358 | 0.61803596 | 0.90794578 |
| LRRC57 | 47.7889584 | -0.188088607 | 0.28117941 | -0.6689274 | 0.50354181 | 0.86946722 |
| HAUS2 | 48.1351725 | -0.110713377 | 0.33382453 | -0.3316514 | 0.74015249 | 0.94445369 |
| TTBK2 | 303.332084 | 0.205277643 | 0.21136155 | 0.97121562 | 0.33144092 | 0.77618491 |
| UBR1 | 563.071936 | 0.189258514 | 0.26273731 | 0.72033361 | 0.47131962 | 0.85236058 |
| AC068724.3 | 123.965962 | 0.414086254 | 0.39900287 | 1.03780269 | 0.29936192 | 0.74594723 |
| TMEM62 | 157.714016 | -0.086742677 | 0.32549704 | -0.266493 | 0.78985956 | 0.95801395 |
| CCNDBP1 | 83.795662 | 0.047945968 | 0.45399862 | 0.10560818 | 0.91589323 | 0.98520052 |
| ADAL | 91.9444377 | 0.00886549 | 0.32936207 | 0.02691715 | 0.97852581 | 0.99469867 |
| ZSCAN29 | 33.746607 | -0.338497288 | 0.31832524 | -1.0633693 | 0.28761451 | 0.73650262 |
| TUBGCP4 | 85.23533 | -0.225634639 | 0.26608726 | -0.8479724 | 0.39645336 | 0.81238609 |
| TP53BP1 | 455.10523 | 0.356287723 | 0.16681451 | 2.13583174 | 0.03269313 | 0.30539487 |
| PPIP5K1 | 123.328566 | -0.220552992 | 0.27918397 | -0.7899916 | 0.42953268 | 0.82998114 |
| CATSPER2 | 148.663998 | 0.037430037 | 0.40421299 | 0.09259979 | 0.92622151 | 0.98729553 |
| PDIA3 | 439.608098 | -0.116925791 | 0.1994939 | -0.5861121 | 0.55780017 | 0.89166349 |
| SERF2 | 1586.81829 | 0.021341493 | 0.41284528 | 0.05169368 | 0.95877277 | 0.99119981 |
| MFAP1 | 98.0381595 | -0.496307437 | 0.28827064 | -1.7216718 | 0.08512899 | 0.46310155 |
| FRMD5 | 49.0659834 | -0.830267823 | 0.39859297 | -2.0829967 | 0.03725153 | 0.32359312 |
| CASC4 | 779.125995 | -0.005346999 | 0.24077646 | -0.0222073 | 0.98228258 | 0.99507903 |
| AC025043.1 | 40.1583118 | -0.342351658 | 0.42852545 | -0.7989062 | 0.42434478 | 0.82774157 |
| CTDSPL2 | 403.45643 | -0.054782781 | 0.25123073 | -0.2180576 | 0.8273842 | 0.96545294 |
| EIF3J-DT | 123.635777 | -0.771304071 | 0.39262114 | -1.9644996 | 0.04947218 | 0.3755208 |
| EIF3J | 223.9642 | -0.306825928 | 0.28221843 | -1.0871931 | 0.27695148 | 0.72743146 |
| SPG11 | 585.278687 | -0.254012759 | 0.24417616 | -1.0402849 | 0.29820758 | 0.74594723 |
| B2M | 2851.24279 | -1.133176472 | 0.53903904 | -2.1022159 | 0.03553436 | 0.31644624 |
| TRIM69 | 68.4733634 | -0.593892198 | 0.5162237 | -1.1504551 | 0.24995647 | 0.70674386 |
| SORD | 135.197373 | -0.023304907 | 0.35721266 | -0.065241 | 0.94798213 | 0.98977092 |
| DUOX1 | 47.9971772 | -1.11097811 | 0.41639751 | -2.668071 | 0.00762881 | 0.13907288 |
| GATM | 108.58395 | 0.795510446 | 0.32402825 | 2.45506508 | 0.01408591 | 0.19221396 |
| SPATA5L1 | 74.8078931 | -0.248760774 | 0.27708722 | -0.8977707 | 0.36930783 | 0.79532503 |
| SLC30A4 | 39.295068 | 0.15397131 | 0.29209966 | 0.52711909 | 0.59811089 | 0.90369786 |
| SQOR | 48.9360039 | -0.597432834 | 0.33670896 | -1.7743301 | 0.07600857 | 0.4426298 |
| SEMA6D | 153.5313 | 0.399847999 | 0.29133852 | 1.37245152 | 0.16992292 | 0.61569949 |
| MYEF2 | 296.806612 | 0.000561927 | 0.24533021 | 0.00229049 | 0.99817245 | 0.99954455 |
| SLC12A1 | 110.504699 | 2.874625087 | 0.69302086 | 4.14796331 | 3.35E-05 | 0.00444997 |
| DUT | 164.681575 | 0.132948801 | 0.39389211 | 0.33752593 | 0.73572047 | 0.94270088 |
| FBN1 | 46.9035943 | -1.236752208 | 0.76168532 | -1.6237049 | 0.1044388 | 0.50345841 |
| CEP152 | 100.286436 | 0.672761277 | 0.31947981 | 2.10580215 | 0.03522154 | 0.31629971 |

| | | | | | | |
|---|---|---|---|---|---|---|
| EID1 | 480.680463 | -0.809679767 | 0.43351231 | -1.8677203 | 0.06180106 | 0.40838664 |
| SECISBP2L | 598.318065 | -0.246002748 | 0.14582854 | -1.6869315 | 0.09161653 | 0.47839622 |
| COPS2 | 141.215523 | 0.414715915 | 0.26117709 | 1.58787247 | 0.11231518 | 0.5224487 |
| GALK2 | 282.050046 | 0.087851049 | 0.24289282 | 0.36168648 | 0.71758633 | 0.93573028 |
| FAM227B | 167.345472 | -0.365447844 | 0.42810576 | -0.8536392 | 0.39330495 | 0.81010924 |
| DTWD1 | 202.066932 | 0.097425855 | 0.23495969 | 0.41464924 | 0.6783987 | 0.9268955 |
| GABPB1 | 150.605447 | -0.110884001 | 0.33994467 | -0.3261825 | 0.74428628 | 0.9456489 |
| GABPB1-AS1 | 252.007388 | -0.02222614 | 0.41833146 | -0.0531305 | 0.95762797 | 0.99106618 |
| USP8 | 389.398717 | 0.20739245 | 0.19480716 | 1.06460381 | 0.28705525 | 0.73638964 |
| TRPM7 | 573.921312 | -0.043375667 | 0.21961215 | -0.1975103 | 0.8434282 | 0.96876046 |
| SPPL2A | 382.110784 | -0.263735311 | 0.18987541 | -1.3889914 | 0.16483537 | 0.60604843 |
| AP4E1 | 142.057698 | 0.1363335 | 0.26529177 | 0.51390022 | 0.60732176 | 0.90510401 |
| DMXL2 | 343.999605 | 0.56731064 | 0.26435647 | 2.14600624 | 0.03187248 | 0.2995969 |
| LYSMD2 | 43.3743173 | 0.230596251 | 0.49841643 | 0.4626578 | 0.64360967 | 0.91655099 |
| TMOD2 | 53.7676833 | 0.174065888 | 0.30920698 | 0.56294294 | 0.57347374 | 0.89797476 |
| TMOD3 | 402.747806 | -0.477644402 | 0.1888602 | -2.5290898 | 0.01143588 | 0.16903805 |
| LEO1 | 57.8600522 | 0.24672191 | 0.26105884 | 0.94508162 | 0.34461721 | 0.78180304 |
| MAPK6 | 246.084567 | -0.102685416 | 0.45446111 | -0.2259498 | 0.82124043 | 0.96502145 |
| GNB5 | 286.613767 | 0.4816985 | 0.46146843 | 1.04383847 | 0.29656013 | 0.74370667 |
| CERNA1 | 36.2590488 | -0.827137468 | 0.54929354 | -1.5058205 | 0.13211326 | 0.55448417 |
| MYO5C | 309.508107 | -0.611916221 | 0.18762764 | -3.261333 | 0.0011089 | 0.04225907 |
| MYO5A | 935.167916 | 0.761186213 | 0.36727016 | 2.07255119 | 0.03821407 | 0.32799625 |
| ARPP19 | 413.108212 | 0.313978944 | 0.27588551 | 1.13807698 | 0.25508834 | 0.71259177 |
| FAM214A | 371.518501 | 0.351033682 | 0.17550264 | 2.00016184 | 0.04548279 | 0.35785259 |
| WDR72 | 1820.17789 | 0.461785036 | 0.39566634 | 1.16710721 | 0.24316708 | 0.6993389 |
| UNC13C | 343.434638 | 1.854923499 | 0.82085845 | 2.25973613 | 0.02383763 | 0.26048096 |
| RSL24D1 | 121.919092 | -0.179442361 | 0.49173398 | -0.3649176 | 0.71517295 | 0.93573028 |
| RAB27A | 312.277697 | -0.205409422 | 0.36514471 | -0.5625425 | 0.57374644 | 0.89797476 |
| PIGB | 153.293881 | 0.985612118 | 0.48322718 | 2.03964544 | 0.04138565 | 0.34259213 |
| CCPG1 | 573.818837 | 0.422275352 | 0.31622807 | 1.33535063 | 0.18176162 | 0.63200456 |
| C15orf65 | 41.839754 | 0.478098213 | 0.38568024 | 1.2396233 | 0.21511476 | 0.67132603 |
| DNAAF4 | 227.036603 | 0.202816527 | 0.25187028 | 0.80524201 | 0.42068004 | 0.82694802 |
| PYGO1 | 106.271021 | 1.344017653 | 0.31750139 | 4.233108 | 2.30E-05 | 0.00326415 |
| PRTG | 409.711011 | -0.551979285 | 0.65863787 | -0.8380619 | 0.40199596 | 0.81693186 |
| NEDD4 | 55.3008408 | -1.835571871 | 0.46968094 | -3.9081251 | 9.30E-05 | 0.00855087 |
| RFX7 | 510.863769 | -0.000993185 | 0.28262891 | -0.0035141 | 0.99719616 | 0.99938948 |
| TEX9 | 310.341798 | -0.117488505 | 0.25632341 | -0.4583604 | 0.64669353 | 0.91743182 |
| MNS1 | 341.818092 | 0.470498496 | 0.4572973 | 1.02886785 | 0.30354178 | 0.74953627 |
| ZNF280D | 170.179359 | 0.155483686 | 0.22952562 | 0.67741321 | 0.49814382 | 0.86583246 |
| AC090517.4 | 107.804103 | -0.167869183 | 0.35049581 | -0.4789478 | 0.6319758 | 0.9133034 |
| TCF12 | 1969.32938 | 0.499995764 | 0.23917423 | 2.09050854 | 0.03657214 | 0.32100172 |
| CGNL1 | 648.033903 | 0.733237191 | 0.53480924 | 1.37102566 | 0.17036696 | 0.61613093 |
| POLR2M | 52.4647553 | 0.218340088 | 0.34929713 | 0.62508412 | 0.53191585 | 0.88045351 |
| ALDH1A2 | 280.916103 | -2.162921879 | 0.58821731 | -3.6770796 | 0.00023592 | 0.01482952 |
| LIPC | 205.71047 | -0.290078809 | 0.47385342 | -0.6121699 | 0.54042535 | 0.88342811 |
| ADAM10 | 1321.57511 | 0.295463554 | 0.38962698 | 0.75832416 | 0.44825695 | 0.84083279 |
| MINDY2 | 830.261493 | 0.267575975 | 0.3524109 | 0.7592727 | 0.44768945 | 0.84026779 |
| RNF111 | 562.572565 | 0.053133083 | 0.26825116 | 0.19807214 | 0.84298862 | 0.96876046 |
| SLTM | 751.899499 | 0.142286991 | 0.18978756 | 0.74971715 | 0.45342508 | 0.84253454 |
| MYO1E | 2528.70519 | 1.493310562 | 0.53731922 | 2.77918696 | 0.00544951 | 0.11829207 |
| FAM81A | 475.645451 | 0.711872599 | 0.29773717 | 2.39094296 | 0.01680516 | 0.21244646 |
| GCNT3 | 179.08929 | -3.157554643 | 0.85225948 | -3.7049217 | 0.00021146 | 0.01385037 |
| GTF2A2 | 179.651676 | -0.011020318 | 0.41783764 | -0.0263746 | 0.97895852 | 0.99469867 |
| BNIP2 | 203.603871 | -0.10241672 | 0.18913972 | -0.5414871 | 0.58817189 | 0.90200952 |
| ANXA2 | 711.871371 | -0.295846938 | 0.43615882 | -0.6783009 | 0.4975809 | 0.86550172 |
| ICE2 | 281.470803 | 0.114640874 | 0.23102832 | 0.49622 | 0.6197392 | 0.90909444 |
| RORA-AS1 | 185.276582 | 0.048935848 | 0.4935915 | 0.0991424 | 0.9210252 | 0.98581108 |
| RORA | 2134.01178 | 0.289186848 | 0.45398171 | 0.63700109 | 0.52412413 | 0.87744743 |
| AC012404.1 | 70.6292638 | -0.389644609 | 0.72878141 | -0.5346522 | 0.59289036 | 0.9025973 |
| AC009554.1 | 180.721257 | -0.356043788 | 0.594109 | -0.5992903 | 0.54897929 | 0.88854099 |
| VPS13C | 1233.49382 | -0.13188556 | 0.29979438 | -0.4399201 | 0.65999501 | 0.92284375 |
| TLN2 | 985.947792 | 0.87066362 | 0.24774278 | 3.51438539 | 0.00044077 | 0.02264362 |
| TPM1 | 426.890142 | -0.438865204 | 0.44904514 | -0.9773298 | 0.3284059 | 0.7745813 |
| LACTB | 98.5949192 | -1.2523191 | 0.39848093 | -3.1427328 | 0.00167379 | 0.05568658 |
| RPS27L | 195.878089 | -0.603693214 | 0.25118811 | -2.403351 | 0.01624558 | 0.20915691 |
| RAB8B | 143.036551 | -0.085502336 | 0.2150709 | -0.3975542 | 0.69095883 | 0.93073907 |
| APH1B | 86.1572108 | -0.617234313 | 0.31154812 | -1.9811845 | 0.04757059 | 0.36819774 |
| CA12 | 1203.18607 | -1.415599606 | 0.46795965 | -3.0250463 | 0.00248595 | 0.07277309 |
| LINC02568 | 98.1825754 | -2.503539387 | 0.54094087 | -4.6281202 | 3.69E-06 | 0.00087889 |
| USP3 | 704.931078 | 0.233672877 | 0.2716669 | 0.86014482 | 0.38970922 | 0.80680464 |
| USP3-AS1 | 158.185152 | -0.262591316 | 0.44578972 | -0.5890475 | 0.55582941 | 0.89081623 |
| HERC1 | 708.790961 | -0.077882268 | 0.26951597 | -0.2889709 | 0.77260366 | 0.95543807 |
| AC073167.1 | 36.5664443 | -0.342617623 | 0.64606786 | -0.5303121 | 0.59589554 | 0.90323766 |
| DAPK2 | 76.6121281 | -0.043379027 | 0.84625484 | -0.05126 | 0.95911834 | 0.99119981 |
| FAM96A | 104.926229 | -0.012482785 | 0.22745068 | -0.0548813 | 0.95623304 | 0.9908068 |
| SNX1 | 271.487295 | -0.032593135 | 0.14833946 | -0.2197199 | 0.82608929 | 0.96545294 |
| PPIB | 438.577096 | -0.68506058 | 0.37885559 | -1.8082367 | 0.07056967 | 0.42987436 |
| CSNK1G1 | 107.640389 | -0.201560475 | 0.31773453 | -0.6343675 | 0.52584099 | 0.87791848 |
| TRIP4 | 162.532542 | -0.010105805 | 0.29105031 | -0.0347219 | 0.97230154 | 0.9939318 |
| ZNF609 | 814.616347 | 0.104498183 | 0.24305356 | 0.42993891 | 0.66724008 | 0.92422364 |
| OAZ2 | 133.824764 | 0.08377712 | 0.29846739 | 0.28069104 | 0.77894738 | 0.95623387 |
| SPG21 | 175.624383 | -0.204640194 | 0.25717465 | -0.7957246 | 0.42619213 | 0.82774157 |
| MTFMT | 95.8656211 | 0.195279352 | 0.24477346 | 0.79779628 | 0.42498872 | 0.82774157 |
| UBAP1L | 44.8672636 | -0.108934659 | 0.34486803 | -0.3158735 | 0.75209855 | 0.94854974 |
| PDCD7 | 94.4509088 | -0.421438013 | 0.27595356 | -1.5272063 | 0.12670973 | 0.54691843 |
| CLPX | 200.989863 | 0.20513236 | 0.16366781 | 1.25334577 | 0.2100799 | 0.66797249 |
| PARP16 | 52.5942269 | -0.336501752 | 0.3570949 | -0.9423314 | 0.34602298 | 0.78180304 |
| DPP8 | 330.291923 | 0.116791246 | 0.2124609 | 0.54970701 | 0.58252035 | 0.90085422 |
| HACD3 | 514.559071 | 0.282918022 | 0.29427965 | 0.96139173 | 0.33635524 | 0.77820994 |
| INTS14 | 93.6491244 | -0.193145669 | 0.19481615 | -0.9914254 | 0.32147792 | 0.7663589 |
| SLC24A1 | 61.1710278 | -0.325629183 | 0.33735202 | -0.9652504 | 0.33441939 | 0.77820994 |
| DENND4A | 706.142081 | -0.041636385 | 0.29956292 | -0.1389905 | 0.88945769 | 0.98064269 |
| RAB11A | 542.306947 | -0.287763938 | 0.37150414 | -0.7745915 | 0.4385811 | 0.83493732 |
| MEGF11 | 126.70834 | -6.008486605 | 0.75842566 | -7.9223145 | 2.33E-15 | 2.44E-11 |
| DIS3L | 84.421569 | -0.043303695 | 0.2774166 | -0.1560963 | 0.87595714 | 0.97622753 |
| TIPIN | 76.2925701 | 0.500633533 | 0.43157542 | 1.16001402 | 0.2460431 | 0.70206841 |
| MAP2K1 | 291.981374 | -0.254073568 | 0.24190547 | -1.050301 | 0.29357973 | 0.73906211 |
| SNAPC5 | 59.240706 | -0.023373373 | 0.36639348 | -0.0637931 | 0.94913498 | 0.98977092 |
| RPL4 | 533.259186 | -0.349247464 | 0.4203325 | -0.8308838 | 0.40603928 | 0.82002585 |
| ZWILCH | 35.4266692 | -0.195311703 | 0.3519468 | -0.5549467 | 0.57893114 | 0.89975022 |
| SMAD6 | 105.915789 | 0.477172648 | 0.43576475 | 1.09502352 | 0.27350633 | 0.72531626 |
| SMAD3 | 1613.12777 | -0.809480117 | 0.66620625 | -1.2150593 | 0.22434347 | 0.68187922 |
| AAGAB | 359.840626 | 0.465153519 | 0.23680847 | 1.96426045 | 0.0494999 | 0.3755208 |
| IQCH | 360.143139 | -0.032501009 | 0.29683884 | -0.1094904 | 0.91281352 | 0.98480378 |
| IQCH-AS1 | 159.517655 | -0.161848888 | 0.32712464 | -0.4947621 | 0.62076803 | 0.90957637 |
| MAP2K5 | 576.668468 | 0.239151998 | 0.23878617 | 1.00153205 | 0.31656966 | 0.76069171 |
| PIAS1 | 739.439097 | 0.293434967 | 0.24700768 | 1.18795887 | 0.23484961 | 0.69335782 |
| FEM1B | 117.474692 | 0.099376082 | 0.21035102 | 0.47242976 | 0.63662007 | 0.91454294 |
| CORO2B | 215.178363 | 0.072507403 | 0.42445275 | 0.17082562 | 0.86436088 | 0.97466129 |
| ANP32A | 287.79363 | -0.036728671 | 0.20065469 | -0.1830442 | 0.85476336 | 0.97122276 |
| SPESP1 | 56.47211 | -0.26618339 | 0.58172509 | -0.4575759 | 0.64725715 | 0.91743182 |
| NOX5 | 33.7044407 | -0.086807769 | 0.83714038 | -0.1036956 | 0.91741092 | 0.98520052 |
| GLCE | 231.28595 | 0.364875816 | 0.32706977 | 1.11559017 | 0.2645976 | 0.71989632 |

| | | | | | | |
|---|---|---|---|---|---|---|
| PAQR5 | 420.720664 | -0.387190889 | 0.38053576 | -1.0174888 | 0.30892094 | 0.75355748 |
| RPLP1 | 1089.08306 | 0.326714698 | 0.45899146 | 0.71180997 | 0.47658246 | 0.85414749 |
| TLE3 | 240.905548 | 0.366070457 | 0.40379825 | 0.90656773 | 0.36463544 | 0.79269964 |
| UACA | 816.765497 | 1.017201406 | 0.27202998 | 3.73929886 | 0.00018453 | 0.01293366 |
| LARP6 | 59.88171 | 0.120642244 | 0.29880635 | 0.40374726 | 0.68639858 | 0.92974759 |
| LRRC49 | 147.659301 | 0.315690885 | 0.31547235 | 1.00069271 | 0.31697539 | 0.76103142 |
| THSD4 | 1733.70207 | -0.120271398 | 0.55064707 | -0.2184183 | 0.82710321 | 0.96545294 |
| THSD4-AS1 | 390.238685 | -0.440204551 | 0.3720553 | -1.1831697 | 0.23674191 | 0.69335782 |
| MYO9A | 1592.4132 | 0.274709555 | 0.212756 | 1.29119532 | 0.19663596 | 0.65097045 |
| AC022872.1 | 58.6312907 | 0.112823056 | 0.50553403 | 0.22317599 | 0.82339854 | 0.96511681 |
| SENP8 | 42.8823977 | -0.493093511 | 0.44634208 | -1.1047435 | 0.26927075 | 0.72322263 |
| PKM | 1655.17477 | -1.089866395 | 0.26977323 | -4.0399353 | 5.35E-05 | 0.00596089 |
| PARP6 | 143.914163 | -0.022948662 | 0.22838339 | -0.1004831 | 0.91996083 | 0.98540367 |
| AC009690.2 | 63.311166 | -1.055137077 | 0.60310397 | -1.7495111 | 0.08020272 | 0.45074086 |
| AC009690.1 | 201.075 | -0.82380288 | 0.36645505 | -2.2480326 | 0.02457411 | 0.26522833 |
| ARIH1 | 1192.41948 | 0.788992051 | 0.32568232 | 2.42258176 | 0.01541066 | 0.20206713 |
| BBS4 | 215.244422 | 0.205864305 | 0.22966972 | 0.89634937 | 0.3700662 | 0.79532503 |
| ADPGK | 124.133059 | -0.456613693 | 0.26790566 | -1.7043824 | 0.08830967 | 0.47036301 |
| NEO1 | 418.5561 | -0.535950055 | 0.21028077 | -2.5487354 | 0.01081143 | 0.16452817 |
| NPTN | 626.108948 | 0.48867959 | 0.18126828 | 2.69589134 | 0.00702006 | 0.13388639 |
| CD276 | 52.9904039 | -0.869322296 | 0.50750724 | -1.7129259 | 0.08672618 | 0.46672926 |
| LOXL1-AS1 | 31.0634135 | -0.31215942 | 0.47223593 | -0.6610243 | 0.50859674 | 0.87094607 |
| LOXL1 | 37.3500601 | -0.40211149 | 0.46709828 | -0.8608713 | 0.38930894 | 0.80680464 |
| PML | 107.358342 | -0.575502862 | 0.3705995 | -1.552897 | 0.12044773 | 0.53823708 |
| ARID3B | 78.3883146 | 0.274103038 | 0.41903015 | 0.65413679 | 0.51302368 | 0.87178584 |
| CLK3 | 91.7424588 | 0.093396462 | 0.25195983 | 0.37067996 | 0.71087592 | 0.93475719 |
| EDC3 | 97.1690355 | 0.325775825 | 0.22467085 | 1.45001374 | 0.14705469 | 0.57806944 |
| CSK | 40.1479114 | -0.04036958 | 0.37649236 | -0.1072255 | 0.91461009 | 0.98480378 |
| ULK3 | 56.0771648 | 0.299445085 | 0.31395137 | 0.95379449 | 0.34018768 | 0.77836439 |
| SCAMP2 | 170.523554 | 0.289733305 | 0.2719919 | 1.0652277 | 0.2867729 | 0.73638964 |
| MPI | 45.0948261 | -0.64239961 | 0.39034403 | -1.6457267 | 0.09982003 | 0.4953766 |
| FAM219B | 83.2326466 | 0.251477152 | 0.24113185 | 1.04290308 | 0.29699318 | 0.74443639 |
| COX5A | 253.279228 | 0.002937741 | 0.44085727 | 0.0066637 | 0.99468318 | 0.99888869 |
| SCAMP5 | 243.84427 | 0.08086923 | 0.43704064 | 0.18503824 | 0.85319904 | 0.97062426 |
| PPCDC | 205.655037 | -0.406391965 | 0.41347108 | -0.9828788 | 0.32566708 | 0.77073504 |
| C15orf39 | 48.796486 | -0.238317357 | 0.39294519 | -0.6064901 | 0.54418937 | 0.88653888 |
| COMMD4 | 102.475787 | -0.133430657 | 0.44157141 | -0.3021723 | 0.7625207 | 0.95374766 |
| NEIL1 | 80.817034 | 0.108554362 | 0.24957929 | 0.4349494 | 0.66359915 | 0.92300139 |
| MAN2C1 | 121.689371 | 0.098721946 | 0.33228447 | 0.29710069 | 0.76638964 | 0.95398103 |
| SIN3A | 245.003642 | 0.053012504 | 0.16375285 | 0.32373486 | 0.74613878 | 0.94594643 |
| PTPN9 | 237.286459 | -0.260932062 | 0.24659165 | -1.0581545 | 0.28998501 | 0.73754646 |
| SNUPN | 76.6465523 | -0.410630067 | 0.22514171 | -1.8238738 | 0.06817116 | 0.42399627 |
| IMP3 | 92.7849438 | -0.417045145 | 0.49155169 | -0.8484258 | 0.39620087 | 0.81224279 |
| UBE2Q2 | 138.065333 | -0.976959353 | 0.31183966 | -3.13289 | 0.00173094 | 0.05740596 |
| FBXO22 | 192.403289 | -0.19313035 | 0.20370673 | -0.9480803 | 0.34308855 | 0.78096612 |
| NRG4 | 124.325718 | -1.068759215 | 0.61811666 | -1.7290575 | 0.08379881 | 0.46003748 |
| ETFA | 281.515256 | 0.28452072 | 0.14643752 | 1.94294958 | 0.05202225 | 0.38306494 |
| SCAPER | 973.369311 | -0.06618664 | 0.22593234 | -0.2929489 | 0.76956118 | 0.95482817 |
| RCN2 | 215.060609 | 0.179240061 | 0.22275097 | 0.80466568 | 0.42101263 | 0.82694802 |
| TSPAN3 | 268.54976 | 0.140476278 | 0.22666697 | 0.61974747 | 0.53542406 | 0.88137638 |
| PEAK1 | 478.156176 | 0.382802021 | 0.18260622 | 2.09632521 | 0.03605335 | 0.31832076 |
| HMG20A | 266.137749 | 0.087262979 | 0.23065716 | 0.3783233 | 0.70519044 | 0.93241442 |
| LINGO1 | 2068.12934 | 1.241876836 | 0.62004412 | 2.00288462 | 0.04518967 | 0.35634897 |
| TBC1D2B | 120.279886 | -1.300820608 | 0.53236354 | -2.4434818 | 0.0145463 | 0.19569355 |
| IDH3A | 122.443672 | -0.162348917 | 0.23630093 | -0.6870431 | 0.49205558 | 0.86172738 |
| DNAJA4 | 41.9833767 | 0.259593523 | 0.39560729 | 0.65618994 | 0.51170192 | 0.87151901 |
| WDR61 | 128.661673 | 0.157663887 | 0.22109472 | 0.71310562 | 0.4757804 | 0.85414749 |
| IREB2 | 278.03199 | -0.015388281 | 0.16697273 | -0.0921604 | 0.92657056 | 0.98729622 |
| PSMA4 | 164.914913 | -0.446771996 | 0.29834413 | -1.4975056 | 0.13426175 | 0.55858006 |
| MORF4L1 | 601.617615 | 0.284045947 | 0.26687326 | 1.06434771 | 0.28717121 | 0.73638964 |
| CTSH | 1162.13192 | -0.4817968 | 0.31653671 | -1.5220882 | 0.12798698 | 0.54791812 |
| TMED3 | 99.9080902 | -1.093602265 | 0.33942575 | -3.221919 | 0.00127335 | 0.04633584 |
| ZFAND6 | 610.017468 | 0.358066355 | 0.24157307 | 1.48222792 | 0.13827966 | 0.56438077 |
| AC016705.2 | 66.5520752 | -0.244424068 | 0.44610699 | -0.5479046 | 0.58375741 | 0.90085422 |
| ARNT2 | 221.452077 | -0.264300224 | 0.50838783 | -0.5198791 | 0.60314782 | 0.90369786 |
| ABHD17C | 91.1734859 | -0.051413044 | 0.31313306 | -0.1641891 | 0.86958226 | 0.97576204 |
| AC027808.1 | 53.7433683 | -0.627581656 | 0.43731824 | -1.4350686 | 0.15126756 | 0.58540769 |
| MESD | 184.361789 | -0.477496393 | 0.32050371 | -1.4898311 | 0.13626865 | 0.56025716 |
| EFL1 | 122.57824 | -0.361598974 | 0.35670784 | -1.0137119 | 0.31072025 | 0.75435382 |
| RPS17 | 47.0764224 | 0.495412047 | 0.41865343 | 1.18334643 | 0.23667188 | 0.69335782 |
| WHAMM | 102.657192 | -0.229640074 | 0.22282772 | -1.0305723 | 0.30274144 | 0.74899204 |
| HOMER2 | 52.2792119 | -0.088267787 | 0.42561199 | -0.2073903 | 0.83570508 | 0.96757829 |
| C15orf40 | 103.189696 | -0.408209278 | 0.28104098 | -1.4524902 | 0.14636535 | 0.57722788 |
| BTBD1 | 198.856655 | 0.410060571 | 0.21974835 | 1.86604617 | 0.06203491 | 0.40939913 |
| HDGFL3 | 247.655552 | 0.048978051 | 0.2177461 | 0.22493193 | 0.82203223 | 0.96502145 |
| AC103876.1 | 37.0439173 | -0.079971144 | 0.48315839 | -0.1655174 | 0.86853672 | 0.97576204 |
| ADAMTSL3 | 673.697953 | 0.431115387 | 0.53535583 | 0.80528756 | 0.42065376 | 0.82694802 |
| WDR73 | 193.069021 | -0.047264111 | 0.21978732 | -0.2150448 | 0.82973242 | 0.96642181 |
| NMB | 38.6251348 | -0.140787262 | 0.62319174 | -0.2259132 | 0.8212689 | 0.96502145 |
| SEC11A | 409.191773 | 0.319818502 | 0.29060854 | 1.1005131 | 0.27110863 | 0.72343657 |
| ZNF592 | 138.270037 | 0.036840243 | 0.312251 | 0.11798278 | 0.9060813 | 0.98320957 |
| ALPK3 | 100.486559 | -1.664346795 | 0.62908718 | -2.6456536 | 0.00815333 | 0.14312705 |
| PDE8A | 1002.18721 | 0.309234486 | 0.31018157 | 0.99694668 | 0.31879039 | 0.76363962 |
| AKAP13 | 1035.33135 | -0.2450408 | 0.32380473 | -0.7567548 | 0.44919675 | 0.84117804 |
| AC087286.2 | 48.4492079 | -0.03667709 | 0.59496445 | -0.0616459 | 0.95084486 | 0.98995173 |
| KLHL25 | 43.1283723 | 0.031645944 | 0.38252084 | 0.08272999 | 0.93406624 | 0.98822345 |
| MRPL46 | 64.8647998 | 0.017766296 | 0.32924326 | 0.053961 | 0.95696624 | 0.99106618 |
| MRPS11 | 90.8157611 | -0.362234122 | 0.29654057 | -1.2215331 | 0.22188424 | 0.67952858 |
| DET1 | 70.9207678 | -0.610783825 | 0.45661604 | -1.3376311 | 0.18101672 | 0.63152351 |
| ABHD2 | 241.235718 | -0.283753197 | 0.36334626 | -0.7809443 | 0.43483524 | 0.83295071 |
| POLG | 61.1045875 | -0.048718274 | 0.22780312 | -0.2138613 | 0.83065521 | 0.96648703 |
| PEX11A | 51.6362014 | -0.488883688 | 0.31970091 | -1.5291908 | 0.12621716 | 0.54614195 |
| AP3S2 | 38.8051793 | 0.313529221 | 0.30500491 | 1.02794812 | 0.30397423 | 0.74966033 |
| ZNF710 | 192.540649 | -0.393291489 | 0.30774082 | -1.2779958 | 0.2012509 | 0.65580391 |
| ZNF710-AS1 | 75.3994895 | -0.879205946 | 0.37343986 | -2.3543441 | 0.01855542 | 0.22797284 |
| IDH2 | 484.317039 | -1.153894452 | 0.23285857 | -4.9553445 | 7.22E-07 | 0.00027024 |
| SEMA4B | 825.484961 | -1.512692983 | 0.6217453 | -2.4329786 | 0.01497519 | 0.19879321 |
| CIB1 | 412.033843 | -0.54996711 | 0.49690445 | -1.1067864 | 0.26838627 | 0.72322263 |
| IQGAP1 | 1051.09897 | 0.041410931 | 0.17367914 | 0.23843354 | 0.81154486 | 0.96288868 |
| CRTC3 | 747.098999 | 0.058463377 | 0.19409032 | 0.30121737 | 0.76324874 | 0.95378284 |
| CRTC3-AS1 | 67.4710106 | -0.280402579 | 0.46049167 | -0.60892 | 0.54257748 | 0.88450129 |
| FURIN | 96.1485041 | -0.474840807 | 0.36832238 | -1.2891989 | 0.19732894 | 0.6524002 |
| MAN2A2 | 54.8735269 | -0.129987013 | 0.32077454 | -0.4052286 | 0.6853095 | 0.92923321 |
| HDDC3 | 46.4996782 | -0.844160992 | 0.33675478 | -2.5067528 | 0.01218459 | 0.17590979 |
| UNC45A | 110.689379 | -0.007912174 | 0.25596798 | -0.0309108 | 0.97534068 | 0.9939318 |
| RCCD1 | 33.3727201 | -0.187488494 | 0.3448572 | -0.5436699 | 0.58666862 | 0.90196444 |
| AC104035.1 | 351.273843 | -0.723648283 | 0.79615547 | -0.9089284 | 0.36338794 | 0.79174754 |
| SLCO3A1 | 833.728373 | 0.082436945 | 0.43161557 | 0.19099623 | 0.84852855 | 0.96991131 |
| FAM174B | 126.783199 | -0.268616945 | 0.38830397 | -0.6917698 | 0.48908193 | 0.86014074 |
| AC013394.1 | 437.490464 | 0.820686409 | 0.31231778 | 2.62772873 | 0.0085957 | 0.14647637 |
| CHD2 | 801.814887 | 0.485459295 | 0.41533871 | 1.16882746 | 0.24247315 | 0.6978349 |
| MCTP2 | 211.311745 | -0.258579905 | 0.30823993 | -0.8388917 | 0.40153011 | 0.81660753 |
| AC012409.2 | 245.346537 | -0.333030279 | 0.74338473 | -0.4479918 | 0.6541591 | 0.92002724 |

| | | | | | | |
|---|---|---|---|---|---|---|
| NR2F2-AS1 | 1390.19254 | -0.061406282 | 0.58226693 | -0.1054607 | 0.91601025 | 0.98520052 |
| NR2F2 | 650.361112 | -0.162547903 | 0.64785571 | -0.2509014 | 0.80189034 | 0.96071123 |
| AC087477.2 | 42.9000311 | -0.551590379 | 0.52245804 | -1.0557601 | 0.29107781 | 0.73772562 |
| LINC00923 | 180.071782 | 0.035528641 | 0.59493775 | 0.05971825 | 0.95238004 | 0.9903694 |
| ARRDC4 | 127.094005 | 0.510958772 | 0.48453373 | 1.05453706 | 0.2916371 | 0.73844969 |
| IGF1R | 1168.59772 | -0.548535386 | 0.25560822 | -2.1460005 | 0.03187294 | 0.2995969 |
| TTC23 | 194.750078 | 0.204435546 | 0.24109691 | 0.8479393 | 0.39647177 | 0.81238609 |
| LRRC28 | 452.480693 | 0.647524715 | 0.27669242 | 2.3402329 | 0.01927172 | 0.23294995 |
| MEF2A | 1342.93967 | 0.165598285 | 0.24635982 | 0.67218057 | 0.50146875 | 0.86804029 |
| AC090825.1 | 101.038471 | 0.236066688 | 0.42715183 | 0.55265288 | 0.58050112 | 0.90034805 |
| LINS1 | 56.8289088 | 0.306325303 | 0.2980089 | 1.02790657 | 0.30399378 | 0.74966033 |
| ASB7 | 92.4443493 | -0.138694371 | 0.25865996 | -0.5362035 | 0.59181792 | 0.90233654 |
| AC015712.1 | 34.6430037 | -0.536695114 | 0.53509898 | -1.0029829 | 0.31586913 | 0.75968795 |
| ALDH1A3 | 48.7929581 | 0.882378955 | 1.36128153 | 0.64819726 | 0.51685737 | 0.87520848 |
| CHSY1 | 530.852771 | -1.155362415 | 0.4887354 | -2.3639835 | 0.01807962 | 0.22423005 |
| SELENOS | 181.606836 | -0.515837217 | 0.43299537 | -1.1913227 | 0.23352692 | 0.69247746 |
| SNRPA1 | 111.114016 | -0.260588988 | 0.24676988 | -1.056 | 0.29096822 | 0.73772562 |
| TM2D3 | 129.275311 | -0.224830019 | 0.29385061 | -0.7651167 | 0.44420203 | 0.83873671 |
| TARSL2 | 165.405435 | -0.356143692 | 0.28561521 | -1.2469353 | 0.21242122 | 0.66984203 |
| SNRNP25 | 61.8048654 | 0.231101254 | 0.49252506 | 0.46921725 | 0.63891436 | 0.91521016 |
| MPG | 111.381288 | -0.831739872 | 0.24782615 | -3.3561424 | 0.00079038 | 0.03422796 |
| NPRL3 | 185.239486 | -0.570541597 | 0.27698855 | -2.0598021 | 0.03941746 | 0.3346923 |
| LUC7L | 329.48958 | 0.417297835 | 0.22799027 | 1.8303318 | 0.06720034 | 0.42120786 |
| FAM234A | 43.5459749 | 0.273177424 | 0.52056543 | 0.52477059 | 0.59974268 | 0.90369786 |
| AXIN1 | 76.1049796 | -0.295851543 | 0.3238193 | -0.9136316 | 0.36091046 | 0.78927419 |
| MRPL28 | 113.242664 | -0.096407716 | 0.4462379 | -0.2160456 | 0.82895224 | 0.96606563 |
| TMEM8A | 166.452225 | -0.893233871 | 0.28156781 | -3.172358 | 0.00151207 | 0.05169888 |
| NME4 | 89.114036 | -0.586928826 | 0.51928319 | -1.1302673 | 0.2583636 | 0.71549044 |
| RAB11FIP3 | 385.113049 | 0.057011966 | 0.37752496 | 0.15101509 | 0.87996381 | 0.97740892 |
| CAPN15 | 80.7897483 | -0.063200581 | 0.3207724 | -0.1970262 | 0.843807 | 0.96876046 |
| PRR35 | 135.58046 | 0.658378035 | 0.90762962 | 0.72538183 | 0.46821781 | 0.85161077 |
| PIGQ | 433.389218 | 1.014824874 | 0.68972165 | 1.47135423 | 0.14119534 | 0.56895751 |
| RAB40C | 332.941933 | 0.628705908 | 0.33661994 | 1.86770249 | 0.06180355 | 0.40838664 |
| METTL26 | 62.1425838 | 0.380959248 | 0.53545044 | 0.71147434 | 0.47679035 | 0.85414749 |
| MCRIP2 | 151.742605 | 0.30700337 | 0.3415289 | 0.8989089 | 0.36870118 | 0.79532503 |
| WDR90 | 47.1894921 | 0.382853375 | 0.33901677 | 1.12930512 | 0.25876914 | 0.7159728 |
| RHOT2 | 113.751832 | 0.013692505 | 0.21840176 | 0.06269411 | 0.95001009 | 0.98977092 |
| STUB1 | 206.757262 | 0.129937024 | 0.45329655 | 0.28664905 | 0.77438105 | 0.95586283 |
| JMJD8 | 78.8285602 | 0.0575725 | 0.4714708 | 0.12211255 | 0.90280988 | 0.98320957 |
| METRN | 85.9478704 | -1.014703971 | 0.54749591 | -1.8533544 | 0.06383157 | 0.41357646 |
| FAM173A | 89.3546707 | 0.302303328 | 0.41020944 | 0.73694873 | 0.46115353 | 0.84722486 |
| NARFL | 53.485907 | 0.100235037 | 0.41400755 | 0.2421092 | 0.80869555 | 0.96245698 |
| LMF1 | 167.147391 | -0.392007247 | 0.32044123 | -1.2233359 | 0.22120286 | 0.67871976 |
| UBE2I | 302.197459 | -0.578370627 | 0.3069007 | -1.884553 | 0.05949022 | 0.40301067 |
| TSR3 | 98.5347523 | 0.129872877 | 0.47143123 | 0.27548637 | 0.7829426 | 0.9568898 |
| GNPTG | 109.954656 | -0.392809487 | 0.30308985 | -1.2960166 | 0.19496975 | 0.64886726 |
| UNKL | 70.7062988 | -0.238538803 | 0.2871632 | -0.8306733 | 0.40615821 | 0.82002585 |
| CLCN7 | 122.424167 | -0.450364677 | 0.41209078 | -1.0928773 | 0.27444766 | 0.7263609 |
| TELO2 | 40.7035936 | 0.269746589 | 0.36753737 | 0.73392969 | 0.46299159 | 0.848575 |
| IFT140 | 145.737497 | -0.147874316 | 0.28469359 | -0.5194157 | 0.6034709 | 0.90371579 |
| JPT2 | 143.079488 | -0.544790173 | 0.4362008 | -1.2489436 | 0.21168572 | 0.66975892 |
| MAPK8IP3 | 260.473181 | 0.305061663 | 0.4197358 | 0.72679447 | 0.46735187 | 0.85120743 |
| NME3 | 257.851761 | 0.124204971 | 0.4248112 | 0.29237687 | 0.76999849 | 0.95482817 |
| MRPS34 | 139.025228 | 0.024569376 | 0.57074864 | 0.04304763 | 0.96566357 | 0.99271817 |
| SPSB3 | 151.211275 | 0.08895374 | 0.27510515 | 0.32334451 | 0.74643435 | 0.94597987 |
| NUBP2 | 59.6340716 | 0.169102814 | 0.42131409 | 0.40136994 | 0.68814777 | 0.93036007 |
| HAGH | 163.805382 | -0.232470878 | 0.25936997 | -0.8962906 | 0.37009756 | 0.79532503 |
| FAHD1 | 67.9940143 | -1.06360554 | 0.39643864 | -2.6829008 | 0.00729866 | 0.13610131 |
| MSRB1 | 41.4069324 | -0.733485508 | 0.38745249 | -1.8930979 | 0.05834485 | 0.39982063 |
| NDUFB10 | 220.214287 | -0.01670899 | 0.5490562 | -0.0304322 | 0.97572236 | 0.9939318 |
| RPS2 | 614.382214 | 0.187303604 | 0.40696044 | 0.46025015 | 0.64533668 | 0.91685019 |
| SNHG9 | 34.7232007 | -0.177232832 | 0.33405032 | -0.5305573 | 0.59572561 | 0.90323766 |
| TSC2 | 163.376356 | -0.0775527 | 0.29028965 | -0.2671563 | 0.78934884 | 0.95801395 |
| PKD1 | 94.3550475 | -0.049708857 | 0.23289178 | -0.2134419 | 0.83098233 | 0.96648703 |
| SNHG19 | 50.7991489 | 0.170430538 | 0.3481536 | 0.48952687 | 0.62446874 | 0.91058133 |
| TRAF7 | 51.8340785 | -0.239844974 | 0.31184123 | -0.7691253 | 0.44181893 | 0.83770294 |
| MLST8 | 34.4951873 | -0.224173944 | 0.38205605 | -0.5867567 | 0.55736713 | 0.89148045 |
| PGP | 144.755545 | -0.360406219 | 0.32900458 | -1.0954444 | 0.273322 | 0.72516823 |
| E4F1 | 49.0233043 | -0.036223312 | 0.32469061 | -0.1115625 | 0.91117027 | 0.98468493 |
| ECI1 | 311.74507 | 0.042502273 | 0.30337732 | 0.14009707 | 0.8885833 | 0.98051184 |
| RNPS1 | 172.190798 | -0.145830484 | 0.20440672 | -0.7134329 | 0.47557791 | 0.85401927 |
| ABCA3 | 81.3370274 | 0.19185149 | 0.3500376 | 0.54808824 | 0.58363131 | 0.90085422 |
| PDPK1 | 266.678036 | 0.085709818 | 0.19669733 | 0.4357447 | 0.66302197 | 0.9229791 |
| KCTD5 | 42.431574 | -0.211535764 | 0.35942028 | -0.5885471 | 0.55616513 | 0.89081623 |
| SRRM2 | 870.451883 | 0.044003803 | 0.24288329 | 0.18117262 | 0.85623208 | 0.97182813 |
| ELOB | 284.4664 | -0.201577692 | 0.49372045 | -0.4082831 | 0.68306588 | 0.92818681 |
| PRSS22 | 73.5310029 | -0.233440171 | 0.43980062 | -0.5307864 | 0.59556683 | 0.90323766 |
| FLYWCH2 | 88.0044919 | 0.903461579 | 0.58059827 | 1.55608726 | 0.11968734 | 0.53725356 |
| FLYWCH1 | 147.364691 | 0.457911474 | 0.4448771 | 1.02929882 | 0.30333928 | 0.74953627 |
| TNFRSF12A | 198.698107 | -1.07778287 | 0.78219342 | -1.3778982 | 0.16823474 | 0.6131323 |
| HCFC1R1 | 191.324635 | 0.098479327 | 0.63337217 | 0.15548414 | 0.87643966 | 0.97640529 |
| BICDL2 | 88.9804967 | 0.257174643 | 0.27522896 | 0.93440255 | 0.35009628 | 0.78223232 |
| ZNF213-AS1 | 61.1077243 | 0.191076932 | 0.27072657 | 0.70579306 | 0.48031684 | 0.85584469 |
| ZNF205 | 32.4258622 | -0.422093268 | 0.32533239 | -1.2974216 | 0.19448616 | 0.64808107 |
| ZNF213 | 101.68056 | -0.269500854 | 0.40889083 | -0.6591022 | 0.50983012 | 0.87094607 |
| ZNF263 | 54.3162535 | -0.177511229 | 0.26566443 | -0.6681784 | 0.50401974 | 0.86946722 |
| TIGD7 | 44.7828099 | -0.473984727 | 0.30390858 | -1.5596293 | 0.11884751 | 0.53524793 |
| ZNF75A | 156.730675 | -0.551432382 | 0.39452318 | -1.3977186 | 0.1621976 | 0.60211533 |
| AC025283.3 | 59.9970254 | -0.154477152 | 0.54180703 | -0.2851147 | 0.77555627 | 0.95601455 |
| ZSCAN32 | 34.9696023 | -0.328042322 | 0.35031471 | -0.9364218 | 0.34905608 | 0.78180304 |
| NAA60 | 93.5934947 | 0.025166801 | 0.27928074 | 0.09011291 | 0.92819749 | 0.98751845 |
| CLUAP1 | 184.356943 | 0.069021368 | 0.22434555 | 0.3076565 | 0.75834372 | 0.95213157 |
| DNASE1 | 169.714616 | 0.177400604 | 0.33431724 | 0.53063553 | 0.59567137 | 0.90323766 |
| TRAP1 | 138.597374 | 0.302314976 | 0.21374214 | 1.41439106 | 0.15724711 | 0.59486921 |
| CREBBP | 548.053827 | 0.400047947 | 0.21221093 | 1.88514303 | 0.05941053 | 0.40275455 |
| ADCY9 | 610.372705 | 0.707053584 | 0.45713096 | 1.54671996 | 0.12193079 | 0.54028588 |
| GLIS2 | 133.741658 | -0.509280206 | 0.52474682 | -0.9705256 | 0.33178458 | 0.77618491 |
| VASN | 251.412941 | 0.136239413 | 0.54098985 | 0.25183359 | 0.8011697 | 0.96063297 |
| DNAJA3 | 111.987047 | -0.109231085 | 0.20070715 | -0.5442312 | 0.58628241 | 0.90196444 |
| NMRAL1 | 93.0304736 | -0.558760743 | 0.2472026 | -2.2603352 | 0.02380046 | 0.26048096 |
| HMOX2 | 104.982964 | -0.148963366 | 0.34328102 | -0.43394 | 0.66433201 | 0.92300139 |
| CDIP1 | 85.4748042 | 0.02883463 | 0.3647283 | 0.07905784 | 0.93698662 | 0.98822345 |
| UBALD1 | 50.3322849 | 0.734141783 | 0.52089361 | 1.40938911 | 0.15872014 | 0.59723751 |
| MGRN1 | 193.066713 | -0.163617378 | 0.15329261 | -1.0673533 | 0.28581233 | 0.73607046 |
| NUDT16L1 | 89.8387829 | 0.398266241 | 0.50941941 | 0.78180422 | 0.43432964 | 0.83243867 |
| ANKS3 | 131.897071 | -0.120806276 | 0.2200918 | -0.5488904 | 0.58308066 | 0.90085422 |
| ZNF500 | 52.2952732 | 0.191443413 | 0.37256967 | 0.51384595 | 0.6073597 | 0.90510401 |
| SMIM22 | 114.022488 | -0.225616451 | 0.42105666 | -0.5358339 | 0.59207332 | 0.90233654 |
| ROGDI | 69.6057935 | 0.29211267 | 0.52315864 | 0.55836347 | 0.57659622 | 0.89869922 |
| GLYR1 | 367.887474 | 0.049735922 | 0.1969428 | 0.25253994 | 0.80062375 | 0.96063297 |
| UBN1 | 130.306144 | 0.098323889 | 0.29222915 | 0.3364616 | 0.7365228 | 0.9429136 |
| PPL | 224.852135 | 0.04158537 | 0.21002847 | 0.19799873 | 0.84304606 | 0.96876046 |
| ALG1 | 34.9613904 | -0.015771992 | 0.3266553 | -0.0482833 | 0.96149047 | 0.99171131 |

| | | | | | | |
|---|---|---|---|---|---|---|
| RBFOX1 | 51.5066585 | 0.144096478 | 0.4424994 | 0.3256422 | 0.74469508 | 0.94575914 |
| METTL22 | 48.5010275 | -0.464951253 | 0.35660745 | -1.3038181 | 0.1922956 | 0.64529552 |
| ABAT | 53.0542429 | -0.093049325 | 0.2843707 | -0.3272114 | 0.74350802 | 0.94555911 |
| PMM2 | 156.200077 | -0.853999653 | 0.39060598 | -2.1863456 | 0.02879033 | 0.28768102 |
| CARHSP1 | 226.861192 | -0.285234538 | 0.47658049 | -0.5985023 | 0.5495048 | 0.88888577 |
| USP7 | 213.065392 | 0.220086542 | 0.15478917 | 1.42184721 | 0.15507061 | 0.59139011 |
| C16orf72 | 227.591556 | 0.13949256 | 0.17244952 | 0.80888925 | 0.41857886 | 0.82690036 |
| AC087190.1 | 42.5929762 | -0.104830058 | 0.565922 | -0.1852376 | 0.85304264 | 0.97062426 |
| ATF7IP2 | 185.612758 | -0.009899199 | 0.44756264 | -0.022118 | 0.98235381 | 0.99507903 |
| EMP2 | 168.560416 | 0.846017057 | 0.516951 | 1.63655172 | 0.10172417 | 0.49869583 |
| NUBP1 | 55.7249595 | -0.605168544 | 0.34584834 | -1.749809 | 0.08015127 | 0.45074086 |
| CIITA | 72.6264195 | -1.35272479 | 0.5169869 | -2.6165553 | 0.0088822 | 0.14941483 |
| DEXI | 82.6064026 | -0.531888923 | 0.27726204 | -1.9183619 | 0.05506514 | 0.39257328 |
| CLEC16A | 699.178712 | -0.475037581 | 0.23859672 | -1.9909644 | 0.0464848 | 0.36244837 |
| AC007014.1 | 36.1181734 | -1.304642569 | 0.51180937 | -2.5490791 | 0.01080078 | 0.16452817 |
| SOCS1 | 397.479005 | -2.66939535 | 0.4524678 | -5.8996361 | 3.64E-09 | 4.77E-06 |
| LITAF | 452.471308 | -0.460089587 | 0.31690329 | -1.4518296 | 0.14654898 | 0.57722788 |
| SNN | 44.6394594 | -0.933400873 | 0.65400499 | -1.4272076 | 0.15352007 | 0.58860244 |
| TXNDC11 | 290.420705 | 0.253766866 | 0.2104322 | 1.20593173 | 0.22784382 | 0.68629723 |
| ZC3H7A | 316.264644 | 0.087972506 | 0.20069206 | 0.43834573 | 0.66113568 | 0.92284375 |
| RSL1D1 | 93.6813866 | -0.582109395 | 0.30287713 | -1.9219325 | 0.05461425 | 0.39095834 |
| AC007216.4 | 133.566615 | 0.192694822 | 0.32333391 | 0.59596231 | 0.5512004 | 0.88916026 |
| GSPT1 | 437.844023 | 0.092203313 | 0.22159818 | 0.41608335 | 0.67734902 | 0.9268955 |
| SNX29 | 1952.71874 | 0.470262794 | 0.31573286 | 1.48943253 | 0.13637351 | 0.56040074 |
| CPPED1 | 197.300714 | -0.714501111 | 0.48916282 | -1.4606611 | 0.14410846 | 0.57402382 |
| U91319.1 | 53.9532224 | 0.837411779 | 0.54483008 | 1.53701458 | 0.12428974 | 0.54273186 |
| ERCC4 | 62.448225 | -0.317346099 | 0.25288532 | -1.2549012 | 0.20951461 | 0.66739001 |
| MKL2 | 782.4171 | 0.135926914 | 0.22082616 | 0.61553809 | 0.53819942 | 0.88265739 |
| PARN | 331.942455 | 0.086909178 | 0.2164612 | 0.40150002 | 0.68805202 | 0.93036007 |
| BFAR | 111.014581 | -0.604638484 | 0.22507443 | -2.6863935 | 0.0072228 | 0.13573345 |
| NOMO1 | 52.1460928 | 0.654425643 | 0.31151397 | 2.10079065 | 0.03565935 | 0.31697198 |
| PDXDC1 | 637.606089 | -0.060730131 | 0.16623546 | -0.365326 | 0.71486809 | 0.93573028 |
| NTAN1 | 51.503669 | -0.160230856 | 0.29179729 | -0.549117 | 0.58292515 | 0.90085422 |
| RRN3 | 107.336871 | -0.057800091 | 0.23383454 | -0.2471837 | 0.80476605 | 0.96210606 |
| C16orf45 | 155.126753 | -0.054485543 | 0.39414128 | -0.1382386 | 0.89005184 | 0.98066897 |
| MARF1 | 182.987429 | -0.0508969 | 0.24547866 | -0.2073374 | 0.83574639 | 0.96757829 |
| NDE1 | 58.2504474 | -0.385896131 | 0.29690717 | -1.2997198 | 0.19369701 | 0.64723766 |
| FOPNL | 80.1701129 | 0.151504762 | 0.25893602 | 0.58510502 | 0.55847711 | 0.89179371 |
| ABCC1 | 350.09668 | -0.346806545 | 0.38683967 | -0.8965124 | 0.36997918 | 0.79532503 |
| XYLT1 | 182.799012 | 0.816046144 | 0.4356056 | 1.87336008 | 0.06101868 | 0.40676596 |
| RPS15A | 134.489006 | 0.510220382 | 0.40418831 | 1.26233336 | 0.20682886 | 0.66434881 |
| SMG1 | 684.292297 | 0.136397914 | 0.17717792 | 0.76983584 | 0.44139728 | 0.83770294 |
| COQ7 | 73.207073 | 0.133102932 | 0.35017061 | 0.38010881 | 0.70386465 | 0.93241442 |
| SYT17 | 158.550094 | 0.394502612 | 0.40142168 | 0.98276359 | 0.3257238 | 0.77073504 |
| GDE1 | 74.8704895 | -0.781037621 | 0.29128939 | -2.6813116 | 0.00733342 | 0.13610131 |
| CCP110 | 101.588449 | 0.400656184 | 0.35581745 | 1.12601613 | 0.26015869 | 0.71719766 |
| VPS35L | 326.876307 | 0.027303402 | 0.25086678 | 0.10883626 | 0.91333236 | 0.98480378 |
| KNOP1 | 81.886891 | -0.773136645 | 0.37294243 | -2.0730724 | 0.03816555 | 0.32797026 |
| IQCK | 654.717174 | -0.275965074 | 0.40522937 | -0.6810096 | 0.49586544 | 0.86452555 |
| GPRC5B | 421.951623 | -0.360149045 | 0.39367624 | -0.9148356 | 0.36027793 | 0.78892604 |
| UMOD | 171.570971 | 1.908315795 | 0.52694242 | 3.62148826 | 0.00029291 | 0.01724567 |
| ACSM3 | 239.246449 | -0.191954495 | 0.35184686 | -0.5455626 | 0.58536663 | 0.90173902 |
| THUMPD1 | 218.574096 | -0.006064579 | 0.23544886 | -0.0257575 | 0.97945074 | 0.99469867 |
| ERI2 | 75.1883004 | -0.491164242 | 0.39818513 | -1.2335073 | 0.21738657 | 0.67507983 |
| LYRM1 | 76.9301191 | -0.304895055 | 0.33363941 | -0.913846 | 0.36079775 | 0.78927419 |
| TMEM159 | 48.1946242 | -0.855214862 | 0.37358126 | -2.2892338 | 0.02206577 | 0.25054091 |
| CRYM | 33.1752178 | -1.684310247 | 0.57678597 | -2.9201651 | 0.00349846 | 0.0900832 |
| METTL9 | 470.504439 | 0.472285718 | 0.20697006 | 2.28190358 | 0.02249503 | 0.25166501 |
| NPIPB4 | 88.0940159 | 0.703457099 | 0.41831395 | 1.68164867 | 0.09263699 | 0.48138295 |
| UQCRC2 | 380.282389 | 0.258239968 | 0.17620005 | 1.46560666 | 0.14275547 | 0.57233258 |
| MOSMO | 217.696593 | 0.101419486 | 0.24078146 | 0.42120969 | 0.67360197 | 0.92557344 |
| EEF2K | 201.805476 | 0.673100227 | 0.57539432 | 1.16980686 | 0.2420787 | 0.6978349 |
| POLR3E | 115.137878 | 0.990957765 | 0.47479301 | 2.08713638 | 0.03687581 | 0.32258577 |
| CDR2 | 647.844031 | 0.227622779 | 0.24188588 | 0.94103376 | 0.34668756 | 0.78180304 |
| NPIPB5 | 242.094699 | 0.07732252 | 0.22542729 | 0.34300426 | 0.73159525 | 0.94065213 |
| USP31 | 98.259963 | -0.304158825 | 0.32995106 | -0.9218301 | 0.3566172 | 0.78612469 |
| SCNN1G | 2133.53593 | 0.941831246 | 0.46938881 | 2.00650555 | 0.04480234 | 0.35515061 |
| AC099482.1 | 249.001529 | 1.129865326 | 0.66618559 | 1.69602186 | 0.08988174 | 0.47358503 |
| SCNN1B | 898.792127 | 0.992062393 | 0.43980945 | 2.25566414 | 0.02409167 | 0.26272709 |
| COG7 | 330.306634 | 0.019887589 | 0.27510697 | 0.07229039 | 0.94237082 | 0.98885525 |
| GGA2 | 293.57614 | -0.474663965 | 0.27961921 | -1.6975371 | 0.08959517 | 0.47350347 |
| EARS2 | 49.1513166 | 0.022213702 | 0.25284739 | 0.08785419 | 0.92999257 | 0.98807358 |
| UBFD1 | 83.835962 | 0.068744474 | 0.23160503 | 0.2968177 | 0.76660569 | 0.95404674 |
| NDUFAB1 | 144.943542 | 0.531701302 | 0.42434765 | 1.25298515 | 0.21021111 | 0.66798437 |
| PALB2 | 49.2678433 | -0.403048634 | 0.39786522 | -1.0130281 | 0.31104675 | 0.75435382 |
| DCTN5 | 147.880502 | -0.210402125 | 0.23680781 | -0.8884932 | 0.37427551 | 0.79739934 |
| RBBP6 | 235.496454 | -0.207576121 | 0.24990464 | -0.8306213 | 0.4061876 | 0.82002585 |
| TNRC6A | 1048.42298 | 0.206171539 | 0.22717747 | 0.90753515 | 0.36412387 | 0.79268955 |
| AC008731.1 | 37.5286602 | -0.852416533 | 0.44841175 | -1.9009683 | 0.05730616 | 0.39785877 |
| ARHGAP17 | 241.160171 | 0.162073804 | 0.21131152 | 0.76698992 | 0.44308751 | 0.83847661 |
| LCMT1 | 170.804732 | -0.040634505 | 0.19953126 | -0.2036498 | 0.83862716 | 0.96793091 |
| NSMCE1 | 105.878493 | 0.181473393 | 0.31216771 | 0.58133301 | 0.56101604 | 0.89242209 |
| GTF3C1 | 156.028903 | 0.032603304 | 0.21098561 | 0.15452857 | 0.87719299 | 0.97640529 |
| KIAA0556 | 268.823565 | -0.131587052 | 0.20799128 | -0.6326566 | 0.52695793 | 0.87791848 |
| XPO6 | 274.876984 | -0.076373389 | 0.31678677 | -0.2410877 | 0.80948715 | 0.96245698 |
| NUPR1 | 510.324641 | 0.190444014 | 1.02140224 | 0.18645349 | 0.85208915 | 0.97053519 |
| SGF29 | 49.1753776 | -0.563455492 | 0.4496461 | -1.2531088 | 0.21016611 | 0.66798437 |
| SULT1A1 | 48.6152454 | -0.365654386 | 0.60792449 | -0.6014799 | 0.54752037 | 0.88773467 |
| ATXN2L | 142.995765 | 0.014097471 | 0.24984834 | 0.05642411 | 0.95500395 | 0.99073937 |
| TUFM | 306.593058 | -0.14297275 | 0.44437586 | -0.3217383 | 0.74765093 | 0.94664513 |
| SH2B1 | 153.53659 | 0.194447454 | 0.25980171 | 0.74844563 | 0.45419141 | 0.84280358 |
| RABEP2 | 116.122104 | -0.463172115 | 0.29584671 | -1.5655814 | 0.11744665 | 0.53282158 |
| NFATC2IP | 253.22771 | -0.046928462 | 0.26352849 | -0.1780774 | 0.85866221 | 0.97237339 |
| SPNS1 | 60.5089427 | -0.580663489 | 0.37803339 | -1.5360111 | 0.12453565 | 0.54288963 |
| QPRT | 215.30371 | -0.562356182 | 0.48116064 | -1.1687494 | 0.24250463 | 0.6978349 |
| KIF22 | 40.2635946 | -0.268432278 | 0.42097065 | -0.6376508 | 0.52370101 | 0.87725117 |
| MAZ | 60.9498958 | -0.52863027 | 0.40506899 | -1.3050376 | 0.19188004 | 0.64458858 |
| AC009133.1 | 43.1423209 | -1.596572236 | 0.44549704 | -3.5837999 | 0.00033863 | 0.01918302 |
| MVP | 272.453028 | -1.564969254 | 0.33495939 | -4.6721164 | 2.98E-06 | 0.00080108 |
| CDIPT | 114.063136 | 0.075946854 | 0.32635552 | 0.23271203 | 0.81598503 | 0.96333253 |
| KCTD13 | 77.8879029 | -0.080467041 | 0.24230189 | -0.3320942 | 0.73981816 | 0.94440785 |
| TMEM219 | 246.626963 | 0.029566007 | 0.36847514 | 0.08023881 | 0.93604732 | 0.98822345 |
| TAOK2 | 53.4878532 | 0.283473962 | 0.28962611 | 0.97875831 | 0.32769941 | 0.77348871 |
| INO80E | 75.9494136 | 0.395742129 | 0.26678407 | 1.48337991 | 0.1379735 | 0.56438077 |
| PPP4C | 145.854529 | -0.297920173 | 0.31745484 | -0.9384647 | 0.34800563 | 0.78180304 |
| YPEL3 | 152.172177 | 0.117272155 | 0.44616776 | 0.26284319 | 0.79267145 | 0.95859644 |
| MAPK3 | 63.0539974 | -0.209319113 | 0.37459132 | -0.5587933 | 0.57630281 | 0.89869922 |
| NPIPB13 | 51.3713982 | 0.082059386 | 0.47418838 | 0.17305229 | 0.86261032 | 0.97436475 |
| CD2BP2 | 85.6123154 | 0.042805235 | 0.39112828 | 0.1094404 | 0.91285319 | 0.98480378 |
| TBC1D10B | 51.5502094 | 0.225290915 | 0.25160998 | 0.89539736 | 0.37057471 | 0.79539603 |
| DCTPP1 | 66.8900973 | -0.485178447 | 0.43916115 | -1.1047845 | 0.26925297 | 0.72322263 |
| SEPHS2 | 230.591907 | -1.396199912 | 0.44740418 | -3.120668 | 0.00180441 | 0.05874854 |
| ZNF768 | 63.5150094 | -0.021397054 | 0.36219378 | -0.0590763 | 0.95289137 | 0.99050799 |

| | | | | | | |
|---|---|---|---|---|---|---|
| ZNF688 | 32.6627434 | -0.205188345 | 0.32736402 | -0.6267895 | 0.5307972 | 0.88004345 |
| ZNF785 | 64.8695625 | -0.231015475 | 0.31217912 | -0.7400094 | 0.45929431 | 0.84608972 |
| PRR14 | 40.7538739 | 0.177465396 | 0.30277814 | 0.58612354 | 0.5577925 | 0.89166349 |
| FBRS | 91.7520585 | 0.027412781 | 0.21172008 | 0.12947653 | 0.8969806 | 0.98302816 |
| PHKG2 | 60.0095455 | -0.010365095 | 0.30678752 | -0.0337859 | 0.97304787 | 0.9939318 |
| RNF40 | 109.890035 | 0.039129306 | 0.29258353 | 0.13373721 | 0.89361038 | 0.98186588 |
| ZNF629 | 44.7048363 | -0.843364697 | 0.40318245 | -2.0917694 | 0.03645915 | 0.32067999 |
| BCL7C | 252.197988 | 0.264018534 | 0.31777123 | 0.83084467 | 0.40606139 | 0.82002585 |
| MIR762HG | 44.6321821 | -0.358656976 | 0.35207551 | -1.0186933 | 0.30834857 | 0.75313873 |
| FBXL19 | 67.9477814 | 0.043780145 | 0.37181146 | 0.11774824 | 0.90626714 | 0.98320957 |
| ORAI3 | 33.1685755 | -0.552662812 | 0.37988869 | -1.454802 | 0.14572409 | 0.57651509 |
| SETD1A | 48.8666469 | 0.054855803 | 0.28622073 | 0.19165559 | 0.84801199 | 0.96991131 |
| STX4 | 67.4366077 | 0.168943502 | 0.3807461 | 0.44371696 | 0.65724723 | 0.92160948 |
| BCKDK | 100.061718 | -0.374862497 | 0.40280721 | -0.9306251 | 0.35204753 | 0.78314007 |
| KAT8 | 106.016046 | 0.169588731 | 0.29433429 | 0.57617728 | 0.56449536 | 0.89323741 |
| PRSS8 | 220.454732 | -0.432645503 | 0.38803359 | -1.1149692 | 0.26486362 | 0.71989632 |
| FUS | 414.85916 | 0.301959059 | 0.18591011 | 1.62422074 | 0.10432871 | 0.50345841 |
| PYCARD | 37.9322197 | -0.752519766 | 0.63207074 | -1.1905626 | 0.23382535 | 0.69285918 |
| AC026471.4 | 55.8002338 | -0.480870714 | 0.36499866 | -1.3174589 | 0.18768486 | 0.64035097 |
| C16orf58 | 77.5622184 | -0.506439085 | 0.29621185 | -1.7097192 | 0.08731782 | 0.46735992 |
| ZNF720 | 154.600336 | -0.155749493 | 0.26341931 | -0.5912607 | 0.55434574 | 0.89076101 |
| ZNF267 | 40.5727722 | -0.001548204 | 0.29965006 | -0.0051667 | 0.99587758 | 0.99908534 |
| SHCBP1 | 172.044097 | 2.294119852 | 0.36909511 | 6.21552485 | 5.12E-10 | 1.34E-06 |
| VPS35 | 299.555802 | -0.458928974 | 0.25696421 | -1.7859645 | 0.074105 | 0.43749352 |
| C16orf87 | 113.802643 | 0.652648079 | 0.25739349 | 2.53560443 | 0.01122535 | 0.16686755 |
| GPT2 | 65.9626501 | 0.838474019 | 0.46953667 | 1.78574767 | 0.07414012 | 0.43749352 |
| DNAJA2 | 118.469151 | -0.251221353 | 0.28704897 | -0.8751864 | 0.38147249 | 0.80315561 |
| ITFG1 | 1179.78254 | 0.027999856 | 0.18286419 | 0.15311831 | 0.87830498 | 0.97692415 |
| PHKB | 890.08298 | 0.30235037 | 0.16667599 | 1.8140007 | 0.06967762 | 0.42828237 |
| LONP2 | 558.144194 | 0.226083557 | 0.14727548 | 1.53510656 | 0.12475765 | 0.54319079 |
| SIAH1 | 122.751499 | -0.090587007 | 0.30203629 | -0.2999209 | 0.76423746 | 0.95378284 |
| N4BP1 | 255.556412 | 0.162882493 | 0.27471711 | 0.5929099 | 0.55324145 | 0.88991776 |
| CNEP1R1 | 34.7624321 | 0.183348165 | 0.26335638 | 0.69619793 | 0.48630488 | 0.85912348 |
| HEATR3 | 55.3046749 | 0.370973252 | 0.28427553 | 1.30497777 | 0.19190042 | 0.64458858 |
| TENT4B | 255.728302 | 1.035074496 | 0.2768687 | 3.73850306 | 0.00018512 | 0.01293366 |
| BRD7 | 191.280602 | 0.069992465 | 0.29376952 | 0.23825639 | 0.81168224 | 0.96288868 |
| CYLD | 125.521432 | 0.236298066 | 0.1997737 | 1.18282869 | 0.23687705 | 0.69342778 |
| HNRNPA1P4 | 58.2627096 | -0.462533903 | 0.48808985 | -0.9476409 | 0.3433123 | 0.78130572 |
| TOX3 | 1508.04781 | 0.994822752 | 0.3236304 | 3.07394721 | 0.00211247 | 0.06499896 |
| AC007906.2 | 72.7057415 | 0.323305516 | 0.87120462 | 0.3711017 | 0.71056178 | 0.93475719 |
| CHD9 | 1450.0366 | -0.120541874 | 0.19047149 | -0.6328605 | 0.52682477 | 0.87791848 |
| RBL2 | 415.639566 | -0.070870542 | 0.17056561 | -0.4155031 | 0.67777363 | 0.9268955 |
| AKTIP | 73.1849879 | 0.787290324 | 0.34298112 | 2.29543341 | 0.0217083 | 0.24935578 |
| RPGRIP1L | 85.9773996 | 0.278840679 | 0.3107451 | 0.89732928 | 0.36954324 | 0.79532503 |
| FTO | 952.135781 | 0.261604956 | 0.19566697 | 1.33699089 | 0.18122561 | 0.63174328 |
| MMP2 | 42.2483845 | -0.397116408 | 0.7021928 | -0.5655376 | 0.57170818 | 0.89797476 |
| LPCAT2 | 591.382502 | 0.107995782 | 0.38145116 | 0.28311824 | 0.7770862 | 0.95601455 |
| AMFR | 514.998664 | 0.767240266 | 0.40288289 | 1.9043754 | 0.05686131 | 0.39727338 |
| NUDT21 | 273.089594 | 0.572896621 | 0.24532715 | 2.33523529 | 0.01953113 | 0.23473196 |
| OGFOD1 | 97.4564113 | 0.194803775 | 0.24183923 | 0.80550939 | 0.42052579 | 0.82694802 |
| BBS2 | 171.235188 | 0.230590872 | 0.21456508 | 1.07468966 | 0.28251369 | 0.73267595 |
| MT2A | 663.235469 | -1.655558189 | 1.00621879 | -1.6453263 | 0.09990255 | 0.4953766 |
| MT1E | 310.004878 | -1.878313156 | 1.04090389 | -1.804502 | 0.07115265 | 0.43107604 |
| MT1F | 68.4012928 | -1.062809921 | 0.81347651 | -1.3065035 | 0.19138139 | 0.64458858 |
| MT1G | 451.539241 | -1.039810435 | 1.08464717 | -0.9586624 | 0.33772886 | 0.77827084 |
| MT1H | 132.639091 | -0.64883883 | 1.38488881 | -0.4685133 | 0.63941756 | 0.91521016 |
| MT1X | 160.166167 | -1.926099597 | 0.98065905 | -1.9640869 | 0.04952001 | 0.3755208 |
| NUP93 | 139.937932 | 0.20036289 | 0.35017126 | 0.57218542 | 0.56719637 | 0.89454608 |
| SLC12A3 | 69.0748464 | 2.464127289 | 0.42743748 | 5.76488355 | 8.17E-09 | 8.56E-06 |
| HERPUD1 | 193.170276 | -0.173622581 | 0.59711256 | -0.2907703 | 0.77122702 | 0.95498333 |
| FAM192A | 173.887668 | -0.027082457 | 0.22887415 | -0.118329 | 0.90580695 | 0.98320957 |
| RSPRY1 | 217.422925 | 0.053141991 | 0.21709224 | 0.24478992 | 0.8066191 | 0.96218371 |
| ARL2BP | 150.58516 | 0.108058145 | 0.20930601 | 0.51626872 | 0.60566674 | 0.90444392 |
| PLLP | 269.077543 | 0.549666612 | 0.61205216 | 0.89807152 | 0.36914742 | 0.79532503 |
| CIAPIN1 | 109.835535 | -0.211250244 | 0.26042651 | -0.8111703 | 0.4172679 | 0.8254526 |
| COQ9 | 227.259534 | -0.021616702 | 0.28956471 | -0.0746524 | 0.94049128 | 0.98855185 |
| POLR2C | 62.7989838 | 0.000989766 | 0.39398243 | 0.00251221 | 0.99799555 | 0.99954455 |
| DOK4 | 54.1833619 | -0.24899276 | 0.41972818 | -0.5932238 | 0.55303137 | 0.88991776 |
| ADGRG1 | 640.193277 | -0.690572776 | 0.28357431 | -2.4352445 | 0.01488173 | 0.19869417 |
| KIFC3 | 886.793631 | 1.088615989 | 0.38480217 | 2.82902765 | 0.00466897 | 0.10823805 |
| USB1 | 38.4567851 | -0.49301733 | 0.34059346 | -1.4475244 | 0.14775012 | 0.57952643 |
| MMP15 | 39.4829973 | -0.659351737 | 0.3752268 | -1.7572085 | 0.07888227 | 0.44910748 |
| CSNK2A2 | 149.056851 | 0.110842772 | 0.23944797 | 0.46290962 | 0.64342915 | 0.91643619 |
| CCDC113 | 39.5166936 | -0.776841137 | 0.30504987 | -2.5466038 | 0.01087769 | 0.16452817 |
| SETD6 | 41.1499522 | -0.14526672 | 0.27274649 | -0.5326071 | 0.59430558 | 0.90305763 |
| CNOT1 | 465.194656 | 0.364397652 | 0.16428396 | 2.21809641 | 0.02654826 | 0.27684155 |
| SLC38A7 | 82.8124262 | -0.465273758 | 0.45290328 | -1.0273137 | 0.30427275 | 0.74968437 |
| GOT2 | 286.94093 | 0.403646641 | 0.31814005 | 1.26877029 | 0.204523 | 0.65976734 |
| AC092378.1 | 66.7647461 | 0.003558977 | 0.44243956 | 0.00804399 | 0.9935819 | 0.99870178 |
| AC138305.1 | 615.796582 | 0.491196299 | 0.4204241 | 1.16833526 | 0.24267156 | 0.69810539 |
| AC010542.3 | 39.2919462 | -1.450653588 | 0.59250329 | -2.4483469 | 0.01435134 | 0.19356764 |
| TK2 | 55.9675285 | -0.615893638 | 0.32592416 | -1.8896839 | 0.05880025 | 0.40066749 |
| CMTM4 | 475.142426 | 0.371531724 | 0.21186922 | 1.75358991 | 0.0795008 | 0.44963216 |
| DYNC1LI2 | 329.445572 | 0.030447655 | 0.26170697 | 0.11634254 | 0.90738107 | 0.98348884 |
| NAE1 | 124.493545 | 0.171507562 | 0.25704487 | 0.66722812 | 0.50462643 | 0.86946722 |
| PDP2 | 82.3686408 | 0.438598696 | 0.46830992 | 0.93655649 | 0.34898674 | 0.78180304 |
| CDH16 | 1495.63039 | 0.376325305 | 0.43568675 | 0.86375201 | 0.38772413 | 0.8065026 |
| RRAD | 287.689451 | 0.043461572 | 0.77027864 | 0.05642318 | 0.95500469 | 0.99073937 |
| FAM96B | 210.992063 | -0.074439351 | 0.47449917 | -0.1568798 | 0.87533955 | 0.97622753 |
| CES2 | 152.43788 | -0.706298497 | 0.3615087 | -1.9537524 | 0.0507305 | 0.3799001 |
| CBFB | 492.201771 | 0.016969057 | 0.21189383 | 0.08008283 | 0.93617138 | 0.98822345 |
| C16orf70 | 56.2500168 | -0.415188396 | 0.33112237 | -1.253882 | 0.20988488 | 0.6679604 |
| TRADD | 55.7466901 | -0.547153569 | 0.35498053 | -1.5413622 | 0.12322866 | 0.54197241 |
| NOL3 | 49.7533674 | -2.152060261 | 0.63842812 | -3.3708732 | 0.0007493 | 0.03310667 |
| KIAA0895L | 43.8599779 | -0.08641626 | 0.45955545 | -0.1880432 | 0.85084282 | 0.97017003 |
| E2F4 | 60.7625496 | 0.320751391 | 0.39129042 | 0.81972717 | 0.41237166 | 0.82364303 |
| ELMO3 | 55.1187451 | 0.516940573 | 0.45013451 | 1.14841356 | 0.25079788 | 0.70811133 |
| LRRC29 | 46.3028284 | -0.207079498 | 0.36064998 | -0.5741841 | 0.5658432 | 0.89394085 |
| TMEM208 | 93.6240153 | -0.093211669 | 0.38001708 | -0.2452828 | 0.80623744 | 0.96218371 |
| ZDHHC1 | 103.133827 | -0.179466376 | 0.22860792 | -0.7850401 | 0.43243009 | 0.83162989 |
| HSD11B2 | 2291.87214 | 0.748383612 | 0.77841749 | 0.96141675 | 0.33634266 | 0.77820994 |
| ATP6V0D1 | 208.770896 | 0.182564008 | 0.30130996 | 0.605901 | 0.54458049 | 0.88662476 |
| AC009061.2 | 98.4288273 | -0.057380496 | 0.4585362 | -0.1251384 | 0.90041397 | 0.98312332 |
| RIPOR1 | 120.225219 | -0.415179487 | 0.2846714 | -1.4584517 | 0.14471607 | 0.57447896 |
| AC027682.4 | 40.1453921 | -0.33147065 | 0.47275423 | -0.7011479 | 0.4832107 | 0.85829223 |
| CTCF | 256.356405 | 0.02885552 | 0.25560537 | 0.1128909 | 0.91011705 | 0.98454672 |
| ENKD1 | 61.4482953 | 0.168787028 | 0.38479152 | 0.4386454 | 0.6609185 | 0.92284375 |
| GFOD2 | 84.083065 | -0.264564071 | 0.27562149 | -0.9598819 | 0.33711467 | 0.77827084 |
| RANBP10 | 222.086796 | 0.095582444 | 0.24874354 | 0.38426101 | 0.700785 | 0.93180634 |
| CENPT | 33.6363999 | 0.247732865 | 0.27232025 | 0.90971152 | 0.36297467 | 0.79134066 |
| THAP11 | 39.9922649 | 0.288593973 | 0.49377884 | 0.58445998 | 0.55891088 | 0.89179371 |
| NUTF2 | 51.2134515 | 0.070397173 | 0.32019078 | 0.21986009 | 0.82598012 | 0.96545294 |
| PSKH1 | 63.171998 | 0.194027378 | 0.27804666 | 0.69782309 | 0.48528784 | 0.85912348 |

| | | | | | | |
|---|---|---|---|---|---|---|
| DUS2 | 51.3391191 | 0.251290248 | 0.29538274 | 0.85072759 | 0.39492069 | 0.81175661 |
| NFATC3 | 393.713574 | 0.312118882 | 0.16986445 | 1.83745852 | 0.06614223 | 0.41858129 |
| AC020978.5 | 33.7286827 | 0.590830528 | 0.35581325 | 1.6605074 | 0.09681242 | 0.48901455 |
| ESRP2 | 56.7238459 | 0.270456821 | 0.34798065 | 0.77721798 | 0.43703018 | 0.83399958 |
| PLA2G15 | 87.962995 | -0.055058979 | 0.21962573 | -0.2506946 | 0.80205025 | 0.96071123 |
| SLC7A6 | 196.86345 | 0.827117083 | 0.27807078 | 2.97448403 | 0.00293482 | 0.08051541 |
| SLC7A6OS | 104.520396 | -0.247125497 | 0.40437566 | -0.6111285 | 0.54111451 | 0.88400312 |
| PRMT7 | 137.106857 | 0.531373303 | 0.29824906 | 1.78164286 | 0.07480749 | 0.43878029 |
| ZFP90 | 161.869323 | -0.186999879 | 0.20895344 | -0.8949356 | 0.3708215 | 0.79539603 |
| CDH3 | 55.1906103 | -0.131908656 | 0.45821962 | -0.2878721 | 0.77344463 | 0.95561066 |
| CDH1 | 1035.58982 | 0.478962127 | 0.29683847 | 1.61354463 | 0.10662623 | 0.50849828 |
| TANGO6 | 187.577336 | -0.055147392 | 0.3033486 | -0.1817954 | 0.85574327 | 0.9717401 |
| CHTF8 | 121.255039 | -0.324837609 | 0.37032512 | -0.8771687 | 0.38039501 | 0.8029673 |
| UTP4 | 69.4626912 | 0.165322073 | 0.29327478 | 0.5637105 | 0.57295117 | 0.89797476 |
| SNTB2 | 127.492847 | -0.119396092 | 0.35682884 | -0.3346033 | 0.73792439 | 0.94333345 |
| VPS4A | 99.8891216 | 0.226753649 | 0.24177342 | 0.93787666 | 0.3483078 | 0.78180304 |
| COG8 | 49.1708851 | 0.357383712 | 0.27092149 | 1.31914126 | 0.18712189 | 0.6387744 |
| NIP7 | 35.4734695 | -0.208302243 | 0.56054213 | -0.3716085 | 0.71018433 | 0.93471656 |
| TERF2 | 133.107045 | 0.205315761 | 0.28669038 | 0.71615854 | 0.47389346 | 0.85318734 |
| CYB5B | 369.66994 | 0.264772233 | 0.21864527 | 1.21096714 | 0.225908 | 0.68375483 |
| NFAT5 | 4177.16644 | 0.841093131 | 0.30012611 | 2.80246567 | 0.00507136 | 0.11405122 |
| NQO1 | 158.83977 | -1.955493634 | 0.62935011 | -3.1071634 | 0.00188892 | 0.06035328 |
| NOB1 | 37.3597057 | -0.112345793 | 0.36764811 | -0.3055797 | 0.75992469 | 0.95297299 |
| WWP2 | 551.517528 | -0.163894812 | 0.29398352 | -0.5574966 | 0.57718819 | 0.89873672 |
| PDPR | 150.90518 | 0.14692768 | 0.31291063 | 0.46955158 | 0.63867542 | 0.91521016 |
| AC009060.1 | 46.6408355 | -0.226670068 | 0.41293777 | -0.5489206 | 0.58305991 | 0.90085422 |
| EXOSC6 | 86.3222061 | -0.105648738 | 0.3519656 | -0.3001678 | 0.76404917 | 0.95378284 |
| AARS | 124.654929 | -0.289564589 | 0.28151008 | -1.0286118 | 0.30366214 | 0.74953627 |
| DDX19A | 37.4145478 | 0.09614922 | 0.30714565 | 0.31304113 | 0.75424941 | 0.94983584 |
| ST3GAL2 | 118.005349 | 0.054592864 | 0.32635134 | 0.16728249 | 0.86714778 | 0.97576204 |
| FUK | 65.298963 | 0.013495958 | 0.22837757 | 0.05909494 | 0.95287649 | 0.99050799 |
| COG4 | 123.064263 | -0.145380524 | 0.2712346 | -0.5359955 | 0.59196166 | 0.90233654 |
| SF3B3 | 237.003709 | 0.053506728 | 0.16303951 | 0.32818258 | 0.74277361 | 0.94508888 |
| MTSS1L | 45.4980103 | -0.48562711 | 0.44865426 | -1.0824083 | 0.27907112 | 0.72938673 |
| VAC14 | 120.12421 | -0.645021011 | 0.44059491 | -1.4639775 | 0.14320011 | 0.5729912 |
| HYDIN | 429.515705 | -0.184925043 | 0.31204206 | -0.5926286 | 0.55342975 | 0.88997143 |
| CMTR2 | 87.8747216 | -0.804423643 | 0.37955181 | -2.1194041 | 0.03405633 | 0.3120805 |
| ZNF19 | 89.4626526 | -0.341230262 | 0.36539731 | -0.9338609 | 0.35037565 | 0.78223232 |
| MARVELD3 | 52.7317953 | -0.76140458 | 0.3005886 | -2.5330454 | 0.01130763 | 0.16768662 |
| PHLPP2 | 102.292698 | -0.072402754 | 0.34032439 | -0.2127463 | 0.83152485 | 0.96656674 |
| AP1G1 | 630.2974 | 0.759772064 | 0.21227307 | 3.57922027 | 0.00034462 | 0.01941735 |
| IST1 | 351.370779 | 0.109768225 | 0.1462391 | 0.75060791 | 0.45288866 | 0.84248055 |
| ZNF821 | 120.92653 | -0.029495941 | 0.42034219 | -0.0701713 | 0.94405735 | 0.98949015 |
| TXNL4B | 53.769151 | -0.311798994 | 0.29366743 | -1.0617418 | 0.28835291 | 0.73653438 |
| DHX38 | 60.298105 | -0.099405431 | 0.25976899 | -0.3826686 | 0.70196552 | 0.93215897 |
| LINC01572 | 204.520776 | 0.006605007 | 0.33154725 | 0.01992177 | 0.98410578 | 0.99540861 |
| AC004943.2 | 156.503906 | 0.184473109 | 0.26316561 | 0.70097727 | 0.4833172 | 0.85829223 |
| ZFHX3 | 1509.32034 | -0.086528942 | 0.32418732 | -0.2669103 | 0.78953821 | 0.95801395 |
| AC138627.1 | 46.0457108 | -0.529765462 | 0.67341342 | -0.7866868 | 0.43146522 | 0.8312408 |
| PSMD7 | 145.069571 | -0.410706354 | 0.34297218 | -1.1974917 | 0.23111495 | 0.69089594 |
| NPIPB15 | 39.7491995 | -1.606979697 | 0.9056272 | -1.7744384 | 0.07599066 | 0.4426298 |
| GLG1 | 762.372825 | 0.147704431 | 0.19426861 | 0.76031035 | 0.4470691 | 0.84014281 |
| RFWD3 | 90.5392777 | 0.11045565 | 0.22549635 | 0.48983342 | 0.62425178 | 0.91058133 |
| WDR59 | 404.723163 | 0.096729933 | 0.23005408 | 0.42046606 | 0.67414502 | 0.92595152 |
| ZNRF1 | 410.207774 | 0.2172338 | 0.44092758 | 0.49267456 | 0.62224255 | 0.91028772 |
| ZFP1 | 34.0397565 | 0.100300749 | 0.32548653 | 0.30815637 | 0.75796335 | 0.95188207 |
| BCAR1 | 201.35192 | -0.573579005 | 0.23532016 | -2.437441 | 0.01479163 | 0.19797735 |
| CFDP1 | 574.482214 | 0.139219231 | 0.26951604 | 0.51655268 | 0.60546845 | 0.90428025 |
| GABARAPL2 | 306.587516 | 0.132859647 | 0.57096219 | 0.2326943 | 0.8159988 | 0.96333253 |
| ADAT1 | 75.806757 | -0.376801118 | 0.25967352 | -1.4510572 | 0.14676394 | 0.57779343 |
| KARS | 99.961551 | -0.354130179 | 0.31873225 | -1.1110585 | 0.26654317 | 0.72068431 |
| TERF2IP | 479.05469 | -0.439093669 | 0.43542628 | -1.0084225 | 0.31325166 | 0.75717828 |
| AC010528.1 | 33.8724668 | 0.926078084 | 0.57031432 | 1.62380295 | 0.10441787 | 0.50345841 |
| NUDT7 | 40.9431633 | 0.374678923 | 0.42335381 | 0.8850255 | 0.37614288 | 0.79946372 |
| WWOX | 3224.61534 | -0.073539148 | 0.27508185 | -0.2673355 | 0.78921084 | 0.95801395 |
| CDYL2 | 99.2196669 | 0.123328935 | 0.36748325 | 0.33560424 | 0.73716933 | 0.9429136 |
| CMC2 | 197.297967 | 0.238394632 | 0.27423354 | 0.86931245 | 0.38467625 | 0.80561637 |
| ATMIN | 97.3824613 | -0.035247865 | 0.31609397 | -0.1115107 | 0.91121137 | 0.98468493 |
| GAN | 117.884891 | -0.174875214 | 0.21830483 | -0.8010598 | 0.42309705 | 0.82774157 |
| CMIP | 578.57046 | -0.485519469 | 0.24672354 | -1.9678685 | 0.04908317 | 0.3746651 |
| PLCG2 | 770.296458 | -0.163466325 | 0.28198562 | -0.5796974 | 0.56211871 | 0.89242209 |
| MPHOSPH6 | 86.7872566 | -0.240675544 | 0.29074582 | -0.8277868 | 0.40779127 | 0.82029376 |
| HSBP1 | 219.15001 | -0.217072848 | 0.42838926 | -0.5067187 | 0.61235221 | 0.90714827 |
| MBTPS1 | 258.861022 | 0.016851242 | 0.1585757 | 0.10626623 | 0.91537112 | 0.98520052 |
| HSDL1 | 45.4551469 | -0.192200791 | 0.52675817 | -0.3648748 | 0.71520486 | 0.93573028 |
| TAF1C | 56.0831674 | 0.361713926 | 0.26002071 | 1.3910966 | 0.16419614 | 0.60471773 |
| TLDC1 | 105.522852 | -0.819619167 | 0.33891866 | -2.4183359 | 0.01559168 | 0.2032348 |
| COTL1 | 103.801481 | -0.534888213 | 0.56888607 | -0.9402378 | 0.34709557 | 0.78180304 |
| KLHL36 | 131.627731 | -0.559588157 | 0.19184276 | -2.9169105 | 0.00353517 | 0.09053163 |
| USP10 | 255.906699 | -0.136184074 | 0.20660065 | -0.6591657 | 0.50978934 | 0.87094607 |
| ZDHHC7 | 183.050137 | 0.111310652 | 0.33427546 | 0.3329908 | 0.73914123 | 0.94385282 |
| KIAA0513 | 47.082174 | -0.711065238 | 0.58846969 | -1.2083294 | 0.22692058 | 0.68523226 |
| GSE1 | 388.958235 | -0.307708611 | 0.27114685 | -1.1348412 | 0.25644187 | 0.71374446 |
| EMC8 | 70.4206776 | -0.144349093 | 0.29413385 | -0.4907599 | 0.62359629 | 0.91058133 |
| COX4I1 | 880.499527 | 0.032055153 | 0.50842592 | 0.06304783 | 0.94972841 | 0.98977092 |
| MTHFSD | 51.2761226 | 0.046092247 | 0.32612719 | 0.14133212 | 0.88760758 | 0.98009982 |
| FBXO31 | 46.3532624 | -0.204541066 | 0.31572861 | -0.6478382 | 0.51708958 | 0.87526312 |
| MAP1LC3B | 228.551781 | -0.185870726 | 0.44007115 | -0.4223652 | 0.6727585 | 0.92506804 |
| ZCCHC14 | 239.872099 | -0.450234077 | 0.31828056 | -1.4145824 | 0.15719097 | 0.59486921 |
| KLHDC4 | 123.139763 | 0.302882985 | 0.35034779 | 0.86452089 | 0.38730181 | 0.8065026 |
| BANP | 440.026984 | 0.379234734 | 0.85893568 | 0.44151703 | 0.65883873 | 0.9225855 |
| ZFPM1 | 119.049023 | -0.061876247 | 0.28284869 | -0.2187609 | 0.82683627 | 0.96545294 |
| ZC3H18 | 115.027988 | -0.302823173 | 0.25664137 | -1.1799468 | 0.23802137 | 0.69406342 |
| CYBA | 209.557502 | -1.772460755 | 0.48208483 | -3.6766574 | 0.00023631 | 0.01482952 |
| MVD | 89.5781015 | -0.445630895 | 0.56675635 | -0.786283 | 0.4317017 | 0.83133261 |
| PIEZO1 | 349.642089 | 0.785570213 | 0.47396306 | 1.6574503 | 0.09742847 | 0.49030463 |
| APRT | 189.377901 | -0.535449457 | 0.50782308 | -1.0544016 | 0.2916991 | 0.73844969 |
| GALNS | 62.0688061 | -0.082303044 | 0.30145196 | -0.2730221 | 0.78483624 | 0.95702626 |
| TRAPPC2L | 60.8254146 | 0.145389898 | 0.41107503 | 0.35368214 | 0.72357711 | 0.93893205 |
| ACSF3 | 84.8968902 | -0.230681141 | 0.31362527 | -0.7355311 | 0.4620161 | 0.8476766 |
| ZNF778 | 41.8634868 | -0.421701212 | 0.38452228 | -1.0966886 | 0.27277753 | 0.72451099 |
| ANKRD11 | 1253.13571 | 0.194904328 | 0.31881967 | 0.61133094 | 0.54098051 | 0.883922 |
| SPG7 | 386.512496 | -0.203155803 | 0.37693756 | -0.5389641 | 0.58991164 | 0.90233654 |
| RPL13 | 1740.41321 | 0.069740642 | 0.45018707 | 0.1549148 | 0.87688848 | 0.97640529 |
| CHMP1A | 127.442302 | -0.034378867 | 0.25563005 | -0.1344868 | 0.89301765 | 0.98168537 |
| SPATA33 | 43.6857876 | -0.574242404 | 0.33153303 | -1.732082 | 0.08325896 | 0.45768271 |
| CDK10 | 115.180093 | 0.13376194 | 0.24273091 | 0.5510709 | 0.58158508 | 0.90053894 |
| VPS9D1 | 92.7242038 | 0.201575247 | 0.37894881 | 0.53193265 | 0.59477264 | 0.90323392 |
| ZNF276 | 58.7772684 | 0.294392024 | 0.2593918 | 1.13493191 | 0.25640385 | 0.71374446 |
| SPIRE2 | 98.5848933 | 1.878252865 | 0.56435504 | 3.32814049 | 0.00087428 | 0.0365037 |
| TCF25 | 667.446292 | -0.127721061 | 0.2030502 | -0.6290122 | 0.52934104 | 0.87911847 |
| DEF8 | 63.1341629 | -0.419530498 | 0.24636608 | -1.7028744 | 0.08859157 | 0.47038308 |
| GAS8 | 126.510268 | -0.091585984 | 0.27900261 | -0.3282621 | 0.74271349 | 0.94508888 |

| | | | | | | |
|---|---|---|---|---|---|---|
| FAM157C | 192.603144 | 0.830484906 | 0.39797117 | 2.08679665 | 0.03690652 | 0.32258577 |
| AC133919.2 | 36.6124485 | 0.058521391 | 0.71324843 | 0.0820491 | 0.93460767 | 0.98822345 |
| DOC2B | 295.498107 | 0.038571979 | 0.51671989 | 0.07464775 | 0.94049498 | 0.98855185 |
| RPH3AL | 483.353042 | -0.428635416 | 0.47389035 | -0.9045034 | 0.36572855 | 0.79329304 |
| AC129507.2 | 68.2978587 | -0.870964207 | 0.49211307 | -1.7698457 | 0.07675285 | 0.44322095 |
| C17orf97 | 42.3103836 | 0.330365849 | 0.42022327 | 0.78616743 | 0.43176941 | 0.83133261 |
| VPS53 | 558.088295 | -0.039069318 | 0.25226428 | -0.1548746 | 0.87692021 | 0.97640529 |
| FAM57A | 41.7817972 | -1.867767181 | 0.53930202 | -3.4633046 | 0.00053358 | 0.02626382 |
| GLOD4 | 148.290385 | -0.183113666 | 0.32333972 | -0.5663197 | 0.57117644 | 0.89773478 |
| NXN | 87.0364446 | -0.5529288 | 0.65145704 | -0.8487571 | 0.39601646 | 0.8120904 |
| TIMM22 | 44.7376932 | -0.458849697 | 0.36100333 | -1.2710401 | 0.20371439 | 0.65831849 |
| ABR | 96.5591962 | -0.498212874 | 0.27714602 | -1.7976548 | 0.07223173 | 0.43302651 |
| YWHAE | 892.494076 | 0.157120789 | 0.23929633 | 0.65659507 | 0.51144132 | 0.87151901 |
| CRK | 203.72587 | -0.459168225 | 0.28378738 | -1.6180009 | 0.10566241 | 0.50651229 |
| MYO1C | 294.773947 | -0.388993909 | 0.2143719 | -1.8145751 | 0.06958924 | 0.42828237 |
| INPP5K | 51.0945053 | -0.240582553 | 0.290122 | -0.8292462 | 0.40696512 | 0.82018309 |
| PITPNA | 284.123572 | 0.004722296 | 0.24915899 | 0.01895294 | 0.98487865 | 0.9954565 |
| SLC43A2 | 228.186393 | -0.029548626 | 0.44554457 | -0.0663202 | 0.94712286 | 0.98977092 |
| PRPF8 | 280.986155 | 0.082645747 | 0.18228309 | 0.45339229 | 0.65026628 | 0.91806421 |
| MIR22HG | 58.3080918 | 0.476202524 | 0.66697545 | 0.71397309 | 0.47524382 | 0.85385826 |
| WDR81 | 42.0274378 | -1.021341985 | 0.47014915 | -2.1723787 | 0.02982711 | 0.29105042 |
| SMYD4 | 66.9617305 | -0.334963636 | 0.44080467 | -0.7598913 | 0.44731956 | 0.84026779 |
| RPA1 | 146.893465 | -0.261043017 | 0.21468501 | -1.215935 | 0.22400968 | 0.68145759 |
| SMG6 | 620.935109 | -0.051337303 | 0.22056425 | -0.2327544 | 0.81595211 | 0.96333253 |
| TSR1 | 69.7264964 | -0.61454217 | 0.26647014 | -2.3062328 | 0.02109763 | 0.24605388 |
| SGSM2 | 139.221148 | -0.177399326 | 0.28274661 | -0.6274145 | 0.53038756 | 0.88004345 |
| MNT | 76.0023868 | -0.403874859 | 0.49154985 | -0.8216356 | 0.41128431 | 0.82314649 |
| METTL16 | 234.732744 | 0.127778514 | 0.18424098 | 0.69354015 | 0.48797064 | 0.86006752 |
| PAFAH1B1 | 868.284527 | 0.212110237 | 0.18594003 | 1.14074543 | 0.25397588 | 0.71110531 |
| CLUH | 158.544676 | -0.035317493 | 0.32950049 | -0.1071849 | 0.91464226 | 0.98480378 |
| RAP1GAP2 | 343.417471 | 0.494360492 | 0.4103448 | 1.20474415 | 0.22830209 | 0.68654401 |
| SPATA22 | 89.7952515 | 0.255564622 | 0.79873354 | 0.3199623 | 0.74899691 | 0.94686219 |
| ASPA | 226.995608 | 1.29232685 | 0.43313377 | 2.98366678 | 0.00284817 | 0.0783433 |
| TRPV1 | 36.4301977 | 0.41737202 | 0.31084369 | 1.34270705 | 0.17936685 | 0.62889415 |
| CTNS | 70.5114158 | -0.205738366 | 0.43095657 | -0.4773993 | 0.63307781 | 0.91415622 |
| TAX1BP3 | 110.19279 | 0.038028585 | 0.47825338 | 0.07951556 | 0.93662256 | 0.98822345 |
| EMC6 | 83.1144787 | 0.049060738 | 0.5898863 | 0.08316982 | 0.93371651 | 0.98822345 |
| ITGAE | 67.153934 | 0.407693308 | 0.28220798 | 1.44465551 | 0.14855467 | 0.58091529 |
| NCBP3 | 187.560656 | 0.192975069 | 0.22368969 | 0.86269094 | 0.38830742 | 0.8065026 |
| ZZEF1 | 270.684187 | 0.271530409 | 0.24198524 | 1.12209493 | 0.26182208 | 0.71989632 |
| CYB5D2 | 121.181475 | -0.586267422 | 0.39929608 | -1.4682524 | 0.14203568 | 0.57093391 |
| ANKFY1 | 267.645575 | -0.057305192 | 0.28682583 | -0.1997909 | 0.84164411 | 0.96866282 |
| UBE2G1 | 377.709119 | 0.589474755 | 0.28420846 | 2.07409293 | 0.03807068 | 0.32772702 |
| SPNS2 | 258.634069 | -0.627446139 | 0.80059757 | -0.7837223 | 0.4332031 | 0.83179416 |
| MYBBP1A | 39.4715635 | -0.253695195 | 0.34797558 | -0.7290603 | 0.46596475 | 0.85028134 |
| GGT6 | 65.1863565 | 0.136064665 | 0.64107491 | 0.21224456 | 0.83191624 | 0.96656674 |
| SMTNL2 | 75.7319186 | 0.284374223 | 0.54329335 | 0.52342666 | 0.60067738 | 0.90369786 |
| PELP1 | 82.3218875 | -0.045969647 | 0.23118437 | -0.1988441 | 0.8423847 | 0.96876046 |
| MED11 | 49.5545005 | -0.693940428 | 0.38020952 | -1.8251527 | 0.06797799 | 0.42354894 |
| CXCL16 | 47.6428279 | -0.596082946 | 0.55639395 | -1.0713325 | 0.28401993 | 0.7344014 |
| PSMB6 | 136.580815 | -0.109396063 | 0.57207531 | -0.1912267 | 0.848348 | 0.96991131 |
| MINK1 | 159.59627 | -0.340198196 | 0.20359738 | -1.6709361 | 0.09473431 | 0.48627909 |
| SLC25A11 | 126.433042 | 0.236684974 | 0.50929317 | 0.46473228 | 0.64212319 | 0.91643619 |
| RNF167 | 82.7368625 | -0.118026268 | 0.35129086 | -0.3359788 | 0.73688688 | 0.9429136 |
| PFN1 | 572.042999 | -0.454379601 | 0.50942994 | -0.8919374 | 0.3724265 | 0.79637648 |
| SPAG7 | 161.634915 | 0.033794916 | 0.3852094 | 0.08773129 | 0.93009025 | 0.98807358 |
| CAMTA2 | 67.9750728 | 0.425018123 | 0.24620193 | 1.72629889 | 0.08429365 | 0.46162908 |
| KIF1C | 128.311209 | 0.184135283 | 0.33335466 | 0.55237051 | 0.58069453 | 0.90045866 |
| ZNF594 | 67.0004516 | -0.226026477 | 0.29944901 | -0.7548079 | 0.45036426 | 0.84147218 |
| AC087500.1 | 145.376923 | 0.295340698 | 0.41919373 | 0.70454465 | 0.48109366 | 0.85600365 |
| RABEP1 | 403.751782 | -0.006994481 | 0.22793778 | -0.0306859 | 0.97552002 | 0.9939318 |
| NUP88 | 273.037489 | 0.483549998 | 0.26854594 | 1.80062303 | 0.07176232 | 0.43222361 |
| RPAIN | 154.87332 | 0.093643809 | 0.26551542 | 0.3526869 | 0.72432318 | 0.93893205 |
| C1QBP | 161.804962 | -0.694815675 | 0.4681976 | -1.4840223 | 0.13780301 | 0.56434148 |
| DERL2 | 103.191392 | -0.062917488 | 0.32259417 | -0.195036 | 0.84536474 | 0.96884569 |
| MIS12 | 46.0060697 | -0.095818377 | 0.45605244 | -0.2101039 | 0.83358662 | 0.96749268 |
| PITPNM3 | 36.2875478 | -0.257392663 | 0.47448896 | -0.5424629 | 0.58749965 | 0.90196444 |
| KIAA0753 | 179.422495 | -0.25046828 | 0.19154628 | -1.3076123 | 0.19100483 | 0.64458858 |
| TXNDC17 | 73.7341931 | -0.250680262 | 0.35952233 | -0.6972592 | 0.48564059 | 0.85912348 |
| MED31 | 54.7087923 | -0.128426822 | 0.34031733 | -0.3773737 | 0.70589588 | 0.93241442 |
| XAF1 | 171.329676 | 0.247251765 | 1.71010313 | 0.14458296 | 0.88504016 | 0.97891513 |
| ALOX12-AS1 | 127.555435 | -0.479864216 | 0.57067794 | -0.8408669 | 0.40042248 | 0.8162653 |
| DLG4 | 44.9243858 | 0.874274388 | 0.42328227 | 2.06546421 | 0.0388791 | 0.33180208 |
| ACADVL | 657.849211 | 0.122218905 | 0.29666101 | 0.4119817 | 0.68035284 | 0.92700898 |
| DVL2 | 59.2252735 | -0.254067307 | 0.2872646 | -0.8844365 | 0.37646061 | 0.79994063 |
| PHF23 | 64.4075705 | -0.074555262 | 0.27230549 | -0.2737927 | 0.78424392 | 0.95697044 |
| ELP5 | 69.7992062 | -0.625294964 | 0.32898119 | -1.9007013 | 0.05734115 | 0.39785877 |
| CLDN7 | 42.3940771 | 0.103267497 | 0.50231602 | 0.20558273 | 0.83711687 | 0.96757829 |
| EIF5A | 150.698317 | -0.331508089 | 0.55608503 | -0.5961464 | 0.55107743 | 0.88916026 |
| GPS2 | 104.350495 | -0.199304837 | 0.395495 | -0.5039377 | 0.61430516 | 0.90764425 |
| ZBTB4 | 324.495451 | 0.375169507 | 0.21575646 | 1.73885646 | 0.08206001 | 0.45535449 |
| POLR2A | 342.588586 | 0.21431497 | 0.38418468 | 0.55784361 | 0.57695118 | 0.89869922 |
| EIF4A1 | 134.015605 | -0.747630427 | 0.52176687 | -1.4328821 | 0.15189153 | 0.58670849 |
| MPDU1 | 71.6112497 | -0.129620013 | 0.33026777 | -0.3924695 | 0.69471137 | 0.93113891 |
| FXR2 | 65.8451627 | 0.08006984 | 0.28502969 | 0.28091754 | 0.77877365 | 0.95623387 |
| SAT2 | 150.3662 | -0.601519912 | 0.42455748 | -1.4168162 | 0.15653667 | 0.59374856 |
| TP53 | 67.3749535 | -0.829292161 | 0.5109336 | -1.6230919 | 0.10456977 | 0.50362646 |
| KDM6B | 116.073208 | -0.334988828 | 0.43559713 | -0.7690336 | 0.44187336 | 0.83770294 |
| NAA38 | 296.652631 | -0.056500496 | 0.23714002 | -0.238258 | 0.81168103 | 0.96288868 |
| CYB5D1 | 41.0688645 | 0.07450916 | 0.32375325 | 0.23014182 | 0.81798157 | 0.96404074 |
| CHD3 | 388.034331 | 0.475392818 | 0.33728725 | 1.4094598 | 0.15869925 | 0.59723751 |
| TRAPPC1 | 92.8491537 | -0.815948049 | 0.5191213 | -1.5717869 | 0.11599999 | 0.53066791 |
| CNTROB | 39.5003042 | 0.067987651 | 0.35113007 | 0.19362526 | 0.84646932 | 0.96949956 |
| PER1 | 226.413538 | 0.913216066 | 0.79659021 | 1.14640634 | 0.25162706 | 0.70813507 |
| TMEM107 | 62.3084864 | -0.58771216 | 0.33394515 | -1.7599062 | 0.07842371 | 0.44789124 |
| CTC1 | 77.526932 | 0.080424883 | 0.23589144 | 0.34094023 | 0.73314858 | 0.94163693 |
| PFAS | 33.5233805 | -0.191856492 | 0.32293965 | -0.5940939 | 0.55244928 | 0.88951784 |
| SLC25A35 | 43.9616595 | 0.767053605 | 0.50992402 | 1.50425078 | 0.13251681 | 0.55573274 |
| RANGRF | 39.5249556 | -0.154256918 | 0.32912019 | -0.4686948 | 0.63928781 | 0.91521016 |
| NDEL1 | 194.73047 | 0.367209356 | 0.39438843 | 0.93108551 | 0.35180933 | 0.78314007 |
| MYH10 | 1734.58036 | 0.113789517 | 0.25271359 | 0.45027067 | 0.65251528 | 0.91909172 |
| STX8 | 554.045719 | 0.167783037 | 0.25973998 | 0.64596539 | 0.51830177 | 0.87526312 |
| USP43 | 80.0308531 | -0.547868947 | 0.31362683 | -1.7468816 | 0.08065788 | 0.45299814 |
| SCO1 | 105.008656 | -0.090594122 | 0.26052506 | -0.3477367 | 0.72803794 | 0.93931348 |
| TMEM220 | 147.872665 | 0.23519288 | 0.43488118 | 0.54082101 | 0.58863096 | 0.90200952 |
| TMEM220-A | 98.6111663 | -0.812590191 | 0.46537123 | -1.7461118 | 0.08079153 | 0.45329073 |
| TMEM238L | 108.453998 | 3.359926012 | 1.46359874 | 2.29566064 | 0.02169529 | 0.24935578 |
| SHISA6 | 80.7002628 | -0.578421804 | 0.61464677 | -0.9410638 | 0.34667219 | 0.78180304 |
| DNAH9 | 73.3722016 | -0.701405018 | 0.34499771 | -2.0330715 | 0.0420453 | 0.34668351 |
| MAP2K4 | 669.669608 | 0.307054561 | 0.21039999 | 1.45938487 | 0.14445921 | 0.57406643 |
| ARHGAP44 | 475.014569 | -0.397899991 | 0.33337103 | -1.193565 | 0.23264817 | 0.69218101 |
| ELAC2 | 79.5786651 | -0.247256635 | 0.25050421 | -0.9870358 | 0.32362507 | 0.76937573 |
| COX10 | 159.870663 | 0.502102983 | 0.26067895 | 1.92613552 | 0.05408745 | 0.39065232 |

| | | | | | | |
|---|---|---|---|---|---|---|
| HS3ST3B1 | 44.6772238 | 1.545697418 | 0.8065959 | 1.91632193 | 0.05532413 | 0.39343117 |
| TRIM16 | 58.1946101 | -0.497758955 | 0.36503566 | -1.3635899 | 0.17269666 | 0.61884138 |
| AC015922.4 | 36.3187845 | 0.422246191 | 0.44180597 | 0.95572767 | 0.33920984 | 0.77836439 |
| ZSWIM7 | 173.50217 | 0.115040629 | 0.2712543 | 0.42410619 | 0.67148837 | 0.9248519 |
| TTC19 | 472.040892 | 0.771621744 | 0.49079124 | 1.57219951 | 0.11590429 | 0.53065835 |
| NCOR1 | 1457.86596 | 0.451000545 | 0.25042814 | 1.80091798 | 0.07171581 | 0.43222361 |
| PIGL | 307.53343 | -0.098891977 | 0.31916865 | -0.3098424 | 0.75668082 | 0.95072713 |
| CENPV | 45.9869017 | 0.010068882 | 0.36208462 | 0.02780809 | 0.97781521 | 0.99469867 |
| UBB | 1006.28674 | -0.432981477 | 0.49788394 | -0.8696434 | 0.38449532 | 0.80561637 |
| LRRC75A | 49.2142423 | 0.431145829 | 0.45787493 | 0.94162357 | 0.3463854 | 0.78180304 |
| AC098850.3 | 90.4183456 | 1.037029771 | 0.69586306 | 1.49027853 | 0.13615102 | 0.5601128 |
| MPRIP | 513.969871 | 0.155127024 | 0.2094022 | 0.74080897 | 0.45880928 | 0.84564214 |
| COPS3 | 61.8447074 | 0.215534233 | 0.24726041 | 0.87168922 | 0.38337795 | 0.80484793 |
| NT5M | 41.6581467 | 0.162936422 | 0.31267156 | 0.52111046 | 0.60228983 | 0.90369786 |
| MED9 | 101.896544 | -0.173604822 | 0.29842436 | -0.5817381 | 0.5607431 | 0.89242209 |
| RASD1 | 1098.29402 | 0.573681754 | 0.75787347 | 0.75696244 | 0.44907236 | 0.84115854 |
| PEMT | 48.8204594 | -0.194885082 | 0.32976316 | -0.590985 | 0.55453047 | 0.8907979 |
| RAI1 | 237.896884 | -0.634258596 | 0.19033714 | -3.3322902 | 0.00086134 | 0.03625254 |
| SREBF1 | 213.806562 | -0.726794717 | 0.58238593 | -1.2479606 | 0.21204548 | 0.66975892 |
| TOM1L2 | 321.718597 | 0.182818529 | 0.22713379 | 0.80489359 | 0.42088109 | 0.82694802 |
| DRC3 | 61.1619194 | 0.053321353 | 0.39009026 | 0.13668979 | 0.891276 | 0.98095103 |
| ATPAF2 | 59.7460449 | -0.082903601 | 0.23710171 | -0.3496542 | 0.72659825 | 0.93906703 |
| GID4 | 36.8051159 | -0.552291256 | 0.30772522 | -1.7947546 | 0.0726928 | 0.43315584 |
| DRG2 | 46.7512671 | -0.118187546 | 0.31476594 | -0.3754776 | 0.70730533 | 0.93274945 |
| ALKBH5 | 193.836028 | -0.44267076 | 0.23874321 | -1.8541711 | 0.06371469 | 0.41357646 |
| FLII | 147.389221 | -0.266050981 | 0.27640234 | -0.9625497 | 0.33577357 | 0.77820994 |
| MIEF2 | 41.9473778 | -0.476889501 | 0.35088603 | -1.3591008 | 0.17411464 | 0.62171087 |
| TOP3A | 65.7170034 | -0.419305303 | 0.3018889 | -1.3889391 | 0.16485127 | 0.60604843 |
| SMCR8 | 51.7994171 | -0.34958074 | 0.27876142 | -1.2540499 | 0.20982386 | 0.6679604 |
| SHMT1 | 257.545301 | 0.225383446 | 0.37272626 | 0.60468893 | 0.54538571 | 0.88669883 |
| TVP23B | 70.4561078 | -0.441756528 | 0.33365735 | -1.3239826 | 0.18550882 | 0.63605149 |
| PRPSAP2 | 176.452311 | 0.145844189 | 0.20978041 | 0.69522313 | 0.48691548 | 0.85950383 |
| EPN2 | 480.048968 | -0.364079691 | 0.21805919 | -1.669637 | 0.09499121 | 0.48627909 |
| B9D1 | 116.199056 | 0.272740302 | 0.23406548 | 1.1652308 | 0.24392558 | 0.70055909 |
| AC004448.2 | 59.102014 | -0.28239306 | 0.50599471 | -0.5580949 | 0.57677959 | 0.89869922 |
| SLC47A1 | 92.3946071 | 1.144665692 | 0.28261513 | 4.05026332 | 5.12E-05 | 0.00576513 |
| ALDH3A2 | 199.046753 | 0.054658483 | 0.34745662 | 0.15731023 | 0.87500036 | 0.97622753 |
| ULK2 | 183.491215 | -0.061952657 | 0.24744254 | -0.2503719 | 0.80229976 | 0.96071123 |
| AKAP10 | 414.425072 | 0.093757659 | 0.3173986 | 0.29539405 | 0.76769288 | 0.95408765 |
| CCDC144NL-/ | 108.879766 | -0.660943063 | 0.5281422 | -1.2514491 | 0.21077069 | 0.66895118 |
| USP22 | 412.266276 | -0.048390001 | 0.20921782 | -0.23129 | 0.81708948 | 0.96380902 |
| DHRS7B | 242.244262 | -0.166122971 | 0.23076774 | -0.7198708 | 0.47160452 | 0.85244471 |
| TMEM11 | 45.4292512 | -0.483843273 | 0.3853484 | -1.2555995 | 0.2092612 | 0.66698825 |
| AC087294.1 | 38.6692405 | -0.23689672 | 0.28523131 | -0.8305425 | 0.40623214 | 0.82002585 |
| MAP2K3 | 128.485811 | -0.024386331 | 0.6203619 | -0.0393098 | 0.96864336 | 0.99303855 |
| WSB1 | 3662.47465 | 0.441513519 | 0.5983442 | 0.7378922 | 0.46057996 | 0.8467861 |
| KSR1 | 234.221206 | 0.220845616 | 0.31742398 | 0.69574332 | 0.48658959 | 0.85912348 |
| NLK | 1590.88877 | 1.004585773 | 0.30592175 | 3.28379981 | 0.00102418 | 0.04004989 |
| ERVE-1 | 34.5873982 | 0.83727218 | 0.76087928 | 1.10040081 | 0.27115753 | 0.72343657 |
| IFT20 | 75.8484462 | 0.423426646 | 0.33244898 | 1.27365905 | 0.20278428 | 0.65774658 |
| TNFAIP1 | 126.583342 | 0.266310524 | 0.40590713 | 0.65608733 | 0.51176794 | 0.87151901 |
| POLDIP2 | 119.458365 | -0.340255665 | 0.2304153 | -1.476706 | 0.13975446 | 0.56680602 |
| TMEM199 | 33.4657841 | 0.184810738 | 0.35245889 | 0.52434693 | 0.60003726 | 0.90369786 |
| SARM1 | 64.4420186 | -0.572914173 | 0.38911472 | -1.4723529 | 0.14092561 | 0.56895751 |
| AC005726.1 | 49.6077325 | -0.09160429 | 0.35484668 | -0.2581517 | 0.79628979 | 0.95948555 |
| KIAA0100 | 161.986287 | 0.11322404 | 0.2585698 | 0.43788579 | 0.66146908 | 0.92284375 |
| SDF2 | 105.024281 | -0.312124811 | 0.32400874 | -0.9633222 | 0.33538586 | 0.77820994 |
| SUPT6H | 219.602362 | -0.274254967 | 0.21061058 | -1.3021899 | 0.19285148 | 0.64567765 |
| RAB34 | 139.423276 | -0.662615196 | 0.37391956 | -1.7720795 | 0.07638136 | 0.4426298 |
| RPL23A | 559.72769 | 0.031892311 | 0.46863173 | 0.0680541 | 0.94574257 | 0.98949015 |
| NEK8 | 65.1595737 | -0.488548261 | 0.45803657 | -1.0666141 | 0.28614613 | 0.73607046 |
| TRAF4 | 163.555892 | 0.133981596 | 0.34493227 | 0.3884287 | 0.69769881 | 0.9315688 |
| FAM222B | 506.937081 | -0.16156728 | 0.27240375 | -0.5931169 | 0.5531029 | 0.88991776 |
| ERAL1 | 42.762552 | 0.052499021 | 0.31454511 | 0.16690458 | 0.86744513 | 0.97576204 |
| FLOT2 | 334.97431 | -0.674271579 | 0.26990822 | -2.4981513 | 0.01248429 | 0.17849296 |
| DHRS13 | 56.4924857 | -1.66411619 | 0.49809771 | -3.3409433 | 0.00083494 | 0.03573026 |
| PHF12 | 157.998076 | -0.053584956 | 0.28677708 | -0.1868523 | 0.85177644 | 0.97053519 |
| MYO18A | 109.916731 | -0.5224623 | 0.33776148 | -1.546838 | 0.12190231 | 0.54028588 |
| NUFIP2 | 414.450572 | -0.030212055 | 0.21164378 | -0.1427496 | 0.88648798 | 0.97973087 |
| TAOK1 | 864.253383 | 0.185387493 | 0.23415328 | 0.79173561 | 0.42851486 | 0.8296433 |
| ABHD15-AS1 | 119.665148 | 0.387255977 | 0.47969537 | 0.80729564 | 0.41949618 | 0.82694802 |
| GIT1 | 61.399037 | -0.082229089 | 0.38222532 | -0.2151325 | 0.82966402 | 0.96642181 |
| SSH2 | 2020.49578 | 0.399041188 | 0.21087801 | 1.89228449 | 0.05845309 | 0.39982063 |
| AC023389.1 | 74.7785846 | -0.048313315 | 0.36111044 | -0.133791 | 0.89356787 | 0.98186588 |
| EFCAB5 | 91.4728291 | -0.166860142 | 0.45641613 | -0.3655877 | 0.71467273 | 0.93573028 |
| NSRP1 | 465.646863 | -0.11985716 | 0.18630167 | -0.6433499 | 0.51999708 | 0.87631314 |
| AC104984.4 | 92.0421799 | -0.762998088 | 0.8165798 | -0.9343828 | 0.35010649 | 0.78223232 |
| BLMH | 127.579768 | 0.050167067 | 0.25598829 | 0.19597407 | 0.84463046 | 0.96876046 |
| CPD | 259.128015 | -0.033924606 | 0.23605872 | -0.1437126 | 0.88572744 | 0.97936523 |
| GOSR1 | 267.334007 | 0.219237712 | 0.16504073 | 1.32838551 | 0.18405079 | 0.63486392 |
| CRLF3 | 201.964402 | 0.060651422 | 0.28547554 | 0.21245751 | 0.83175012 | 0.96656674 |
| ATAD5 | 43.6412519 | -0.706454996 | 0.49449504 | -1.4286392 | 0.15310797 | 0.58860244 |
| TEFM | 89.417981 | -0.315426881 | 0.260664 | -1.2100899 | 0.22624439 | 0.68421917 |
| RNF135 | 70.5292971 | 0.096818854 | 0.25480485 | 0.37997257 | 0.70396578 | 0.93241442 |
| NF1 | 1476.51781 | 0.174207292 | 0.2165178 | 0.80458646 | 0.42105836 | 0.82694802 |
| OMG | 35.5972169 | -0.175044612 | 0.67480194 | -0.2594015 | 0.7953255 | 0.95948555 |
| RAB11FIP4 | 119.831752 | 0.028090751 | 0.34988676 | 0.08028527 | 0.93601038 | 0.98822345 |
| COPRS | 129.393766 | 0.712337451 | 0.59163872 | 1.20400748 | 0.22858669 | 0.68678297 |
| UTP6 | 99.3740662 | 0.629794342 | 0.21868528 | 2.87991187 | 0.00397786 | 0.09782854 |
| SUZ12 | 401.39175 | 0.172415648 | 0.19379563 | 0.88967771 | 0.37363897 | 0.79681088 |
| LRRC37B | 109.476972 | -0.426698806 | 0.35447242 | -1.2037574 | 0.22868336 | 0.68678297 |
| RHOT1 | 304.81643 | 0.092178369 | 0.19630411 | 0.46956923 | 0.63866281 | 0.91521016 |
| C17orf75 | 47.6206991 | 0.18096789 | 0.30223084 | 0.59877374 | 0.54932378 | 0.88882402 |
| ZNF207 | 544.618592 | 0.046552717 | 0.16656812 | 0.27948155 | 0.7798753 | 0.95636474 |
| PSMD11 | 144.066075 | 0.139849792 | 0.24000222 | 0.58270207 | 0.56009388 | 0.89233565 |
| MYO1D | 967.956324 | -0.281893108 | 0.26698366 | -1.055844 | 0.29103948 | 0.73772562 |
| TMEM98 | 36.3781729 | -0.709958188 | 0.74658414 | -0.950942 | 0.34163384 | 0.77973697 |
| ASIC2 | 103.881204 | 1.481789124 | 0.69093436 | 2.14461636 | 0.03198353 | 0.30034716 |
| ZNF830 | 38.4414615 | -0.729423101 | 0.46057048 | -1.5837383 | 0.11325329 | 0.52517453 |
| LIG3 | 53.4517875 | -0.11354179 | 0.24035901 | -0.4723842 | 0.63665261 | 0.91454294 |
| SLFN5 | 198.202666 | -1.998189505 | 0.43704555 | -4.5720394 | 4.83E-06 | 0.00105455 |
| LINC02001 | 60.3607588 | -0.48530952 | 0.27122667 | -1.7893134 | 0.07356436 | 0.4369528 |
| AP2B1 | 388.636345 | 0.111158387 | 0.27334253 | 0.40666334 | 0.68425527 | 0.92881914 |
| TAF15 | 695.758744 | -0.017333006 | 0.14861686 | -0.1166288 | 0.90715421 | 0.98341906 |
| AC015849.3 | 44.5642619 | -0.010126598 | 0.49790467 | -0.0203384 | 0.9837734 | 0.99540861 |
| ZNHIT3 | 66.1344448 | -0.087822409 | 0.42418522 | -0.2070379 | 0.83598027 | 0.96757829 |
| MYO19 | 184.212775 | -0.64785587 | 0.28596046 | -2.2655435 | 0.02347934 | 0.25893742 |
| GGNBP2 | 315.986886 | -0.074479753 | 0.22592755 | -0.3296621 | 0.74165528 | 0.94487568 |
| DHRS11 | 34.1711499 | 0.393894689 | 0.70781875 | 0.55649089 | 0.57787532 | 0.89918786 |
| LHX1-DT | 516.167417 | -0.077995182 | 0.54818796 | -0.1422782 | 0.88686029 | 0.97989413 |
| LHX1 | 600.099274 | -0.069910856 | 0.63621874 | -0.1098849 | 0.91250063 | 0.98480378 |
| AATF | 286.510864 | 0.005841389 | 0.2327497 | 0.0250973 | 0.97997735 | 0.99469867 |
| ACACA | 617.132511 | -0.049613966 | 0.30360183 | -0.1634179 | 0.87018943 | 0.9759113 |

| | | | | | | |
|---|---|---|---|---|---|---|
| TADA2A | 124.799365 | -0.059478859 | 0.24889018 | -0.2389763 | 0.81112394 | 0.96288868 |
| SYNRG | 350.05855 | -0.132790569 | 0.18779934 | -0.7070875 | 0.47951208 | 0.85532469 |
| DDX52 | 170.922652 | -0.070518511 | 0.19513239 | -0.361388 | 0.71780939 | 0.93573028 |
| HNF1B | 642.215938 | 0.192095886 | 0.29504727 | 0.65106817 | 0.51500248 | 0.87333754 |
| MRPL45 | 100.686727 | 0.117211106 | 0.19708964 | 0.59470962 | 0.55203759 | 0.88923362 |
| SOCS7 | 91.0158429 | 0.101324618 | 0.31572965 | 0.32092209 | 0.74826943 | 0.94678489 |
| MLLT6 | 356.046923 | -0.24187569 | 0.21515607 | -1.1241872 | 0.26093364 | 0.71849305 |
| CISD3 | 81.1671904 | 0.023363356 | 0.46014294 | 0.05077413 | 0.95950551 | 0.99119981 |
| PCGF2 | 121.897853 | 0.084366831 | 0.24596992 | 0.34299653 | 0.73160107 | 0.94065213 |
| PSMB3 | 162.458648 | -0.334797893 | 0.33722982 | -0.9927885 | 0.32081302 | 0.76568446 |
| PIP4K2B | 120.460573 | 0.011773882 | 0.20585635 | 0.05719465 | 0.95439014 | 0.99060627 |
| CWC25 | 86.9278765 | 0.136115394 | 0.2234607 | 0.60912454 | 0.54244189 | 0.88450129 |
| RPL23 | 432.195856 | 0.024944135 | 0.40361269 | 0.06180216 | 0.95072039 | 0.98992048 |
| LASP1 | 257.798481 | -0.253924496 | 0.26626339 | -0.9536591 | 0.34025624 | 0.77836439 |
| RPL19 | 959.405894 | -0.044362992 | 0.56953118 | -0.0778939 | 0.93791247 | 0.98846521 |
| STAC2 | 100.74122 | -4.797808007 | 0.92720656 | -5.1744759 | 2.29E-07 | 0.00012961 |
| FBXL20 | 543.094164 | -0.042671332 | 0.25851586 | -0.1650627 | 0.86889462 | 0.97576204 |
| MED1 | 124.133022 | 0.331651765 | 0.18714454 | 1.7721691 | 0.07636649 | 0.4426298 |
| CDK12 | 327.037105 | -0.175299069 | 0.2053414 | -0.8536957 | 0.39327361 | 0.81010924 |
| STARD3 | 68.4285485 | -0.120342164 | 0.25228395 | -0.4770108 | 0.63335445 | 0.91415622 |
| PGAP3 | 56.6455528 | 0.350565787 | 0.33280076 | 1.05338036 | 0.2921667 | 0.73844969 |
| ERBB2 | 646.240507 | 0.993975121 | 0.31090364 | 3.19705206 | 0.0013884 | 0.04915682 |
| MIEN1 | 88.7136812 | -0.340919372 | 0.56159788 | -0.6070525 | 0.5438161 | 0.88606852 |
| GRB7 | 233.764442 | -0.11367495 | 0.27631028 | -0.4114033 | 0.68077686 | 0.92728639 |
| ORMDL3 | 63.0405451 | 0.113055583 | 0.44515385 | 0.25396968 | 0.79951898 | 0.96043078 |
| PSMD3 | 95.3453168 | -0.21149228 | 0.28389464 | -0.7449675 | 0.45629137 | 0.84372683 |
| MED24 | 111.662191 | 0.445347869 | 0.24542581 | 1.81459262 | 0.06958654 | 0.42828237 |
| THRA | 175.109107 | 0.516781414 | 0.32584552 | 1.58597058 | 0.11274598 | 0.52363638 |
| NR1D1 | 48.0797188 | 0.184210541 | 0.9435729 | 0.19522661 | 0.84521555 | 0.96881318 |
| MSL1 | 177.341143 | -0.076167752 | 0.22643362 | -0.33638 | 0.7365843 | 0.9429136 |
| CASC3 | 169.866035 | 0.260871556 | 0.26468179 | 0.98560448 | 0.32432724 | 0.76970832 |
| WIPF2 | 415.799756 | -0.082843528 | 0.30160017 | -0.27468 | 0.78356211 | 0.9568898 |
| RARA | 90.788251 | -0.862518113 | 0.66055007 | -1.3057573 | 0.19163509 | 0.64458858 |
| KRT10 | 157.128585 | -0.26830107 | 0.60915165 | -0.4404504 | 0.65961095 | 0.92284375 |
| TMEM99 | 36.3719505 | 0.541206066 | 0.48817966 | 1.10862067 | 0.26759386 | 0.72222087 |
| KRT19 | 3220.16233 | -1.227621183 | 0.40723278 | -3.0145441 | 0.00257366 | 0.07369381 |
| EIF1 | 958.329245 | -0.316285317 | 0.44850252 | -0.705203 | 0.48068393 | 0.85600365 |
| JUP | 856.814574 | -0.098752859 | 0.32143539 | -0.3072246 | 0.75867242 | 0.9521704 |
| P3H4 | 37.2078906 | 0.316104273 | 0.34632606 | 0.91273604 | 0.36138138 | 0.78979697 |
| NT5C3B | 64.7991617 | -0.18428393 | 0.28642706 | -0.6433887 | 0.51997192 | 0.87631314 |
| ACLY | 144.02162 | -0.957407726 | 0.28720818 | -3.3334974 | 0.00085761 | 0.03624114 |
| TTC25 | 43.1778316 | -0.136949205 | 0.25682397 | -0.5332415 | 0.59386641 | 0.90304695 |
| CNP | 83.1343201 | -0.639153088 | 0.30596924 | -2.0889456 | 0.03671262 | 0.32196511 |
| DNAJC7 | 271.37319 | 0.212847594 | 0.17379574 | 1.22469971 | 0.22068839 | 0.67871976 |
| NKIRAS2 | 44.8726041 | -0.483826014 | 0.32604311 | -1.4839326 | 0.13782679 | 0.56434148 |
| DHX58 | 33.76011 | -1.222821796 | 0.52584335 | -2.3254488 | 0.02004797 | 0.23821174 |
| KAT2A | 64.6461638 | 0.110852957 | 0.43066676 | 0.25739845 | 0.7968712 | 0.95961857 |
| GHDC | 33.2926697 | -0.549118714 | 0.31527462 | -1.7417156 | 0.08155822 | 0.45528236 |
| STAT5B | 262.730614 | -0.032103803 | 0.28445845 | -0.1128594 | 0.91014205 | 0.98454672 |
| STAT5A | 50.6143794 | -0.413551695 | 0.35585615 | -1.1621316 | 0.245182 | 0.70148313 |
| STAT3 | 1514.89265 | -0.675049413 | 0.38071992 | -1.7730866 | 0.07621435 | 0.4426298 |
| CAVIN1 | 33.4306493 | -0.657328606 | 0.6191116 | -1.0617288 | 0.28835883 | 0.73653438 |
| ATP6V0A1 | 253.370345 | -0.429218715 | 0.29084617 | -1.4757585 | 0.14000873 | 0.56739811 |
| NAGLU | 78.437669 | -1.394945205 | 0.54832044 | -2.5440328 | 0.01095808 | 0.16452817 |
| COASY | 95.3415224 | 0.093617226 | 0.31520305 | 0.29700609 | 0.76646186 | 0.95398103 |
| MLX | 67.1921033 | -0.363002801 | 0.40638707 | -0.893244 | 0.37172654 | 0.7954538 |
| RETREG3 | 181.720121 | 0.088688793 | 0.22247714 | 0.39864228 | 0.69015681 | 0.93073907 |
| TUBG2 | 68.4002707 | -0.10062697 | 0.27315948 | -0.3683818 | 0.7125886 | 0.93512754 |
| PLEKHH3 | 48.6313948 | 0.545434396 | 0.31842117 | 1.71293383 | 0.08672471 | 0.46672926 |
| EZH1 | 207.543414 | -0.131476295 | 0.27087961 | -0.485368 | 0.62741532 | 0.91061758 |
| VPS25 | 97.3054269 | -0.785192717 | 0.30848682 | -2.545304 | 0.01091826 | 0.16452817 |
| WNK4 | 67.8476704 | -0.768628854 | 0.46672685 | -1.6468494 | 0.099589 | 0.49487561 |
| COA3 | 329.577444 | 0.265580516 | 0.50253525 | 0.52848137 | 0.59716527 | 0.90369786 |
| BECN1 | 103.884973 | 0.234864003 | 0.19398777 | 1.21071551 | 0.22600446 | 0.68375483 |
| PSME3 | 72.9734233 | -0.161504663 | 0.25715601 | -0.6280416 | 0.52997671 | 0.87951796 |
| RUNDC1 | 91.0221949 | -0.007083405 | 0.4037923 | -0.0175422 | 0.98600407 | 0.99588692 |
| RPL27 | 279.803917 | -0.35209617 | 0.38175535 | -0.9223084 | 0.35636772 | 0.78612469 |
| IFI35 | 62.8881921 | -0.487161897 | 0.79984148 | -0.6090731 | 0.54247601 | 0.88450129 |
| VAT1 | 95.1578662 | -1.104426288 | 0.40324822 | -2.7388249 | 0.00616592 | 0.12643442 |
| BRCA1 | 90.8268815 | -0.331562819 | 0.4201542 | -0.7891456 | 0.43002694 | 0.83047752 |
| NBR1 | 262.996148 | 0.123903235 | 0.23897328 | 0.51848153 | 0.60412234 | 0.90381187 |
| LINC00854 | 98.4278971 | 0.080176192 | 0.64077729 | 0.12512334 | 0.90042591 | 0.98312332 |
| ARL4D | 353.29955 | 0.748451261 | 0.7055145 | 1.06085879 | 0.28875408 | 0.73668501 |
| DHX8 | 92.2701601 | -0.112480197 | 0.25247319 | -0.4455134 | 0.65594876 | 0.92098622 |
| DUSP3 | 149.640189 | -0.022346312 | 0.32699703 | -0.068338 | 0.9455166 | 0.98949015 |
| LINC01976 | 54.5570547 | 1.958501451 | 1.07262937 | 1.82588834 | 0.06786708 | 0.42329369 |
| TMEM101 | 427.05998 | -2.449076331 | 0.72699146 | -3.3687828 | 0.00075501 | 0.03310667 |
| LSM12 | 145.28627 | -0.018009721 | 0.2897796 | -0.0621497 | 0.9504436 | 0.98992048 |
| G6PC3 | 75.5647644 | -0.57422282 | 0.43365266 | -1.3241538 | 0.18545197 | 0.63605149 |
| HDAC5 | 104.952016 | 1.025249164 | 0.38075993 | 2.69263931 | 0.00708889 | 0.13409852 |
| TMUB2 | 85.6174112 | -0.057621344 | 0.33404739 | -0.1724945 | 0.86304877 | 0.9744399 |
| ATXN7L3 | 55.9915113 | -0.116179886 | 0.29837604 | -0.389374 | 0.69699947 | 0.93130502 |
| UBTF | 193.851499 | 0.220958012 | 0.27019558 | 0.81777063 | 0.41348817 | 0.82440569 |
| SLC25A39 | 123.281643 | -0.257901166 | 0.30999689 | -0.8319476 | 0.40543852 | 0.82002585 |
| GRN | 416.373522 | -0.703985387 | 0.41891627 | -1.6804919 | 0.09286165 | 0.48153712 |
| GPATCH8 | 855.596906 | 0.258818181 | 0.19748814 | 1.31055052 | 0.19000967 | 0.64458858 |
| CCDC43 | 35.2735581 | -0.389113741 | 0.36645632 | -1.0618284 | 0.28831358 | 0.73653438 |
| DBF4B | 46.5418455 | -0.025912659 | 0.29756948 | -0.087081 | 0.9306071 | 0.98822345 |
| EFTUD2 | 109.703285 | -0.149704519 | 0.24205195 | -0.6184809 | 0.53625835 | 0.88137638 |
| DCAKD | 138.457159 | -0.099718939 | 0.37779897 | -0.2639471 | 0.79182069 | 0.95839852 |
| NMT1 | 203.255735 | -0.068892742 | 0.20565645 | -0.3349895 | 0.73763304 | 0.9431911 |
| ACBD4 | 56.3581066 | 0.538742106 | 0.23727064 | 2.27058058 | 0.02317238 | 0.25643774 |
| HEXIM1 | 95.846955 | 0.436735336 | 0.59293939 | 0.73655983 | 0.46139007 | 0.84722486 |
| AC138150.1 | 136.212261 | -0.021953719 | 0.51992826 | -0.0422245 | 0.96631972 | 0.99284614 |
| MAP3K14 | 217.378447 | -0.086731628 | 0.43911459 | -0.1975148 | 0.84342471 | 0.96876046 |
| ARHGAP27 | 58.0865479 | -0.187579656 | 0.28632604 | -0.6551261 | 0.51238658 | 0.87177451 |
| PLEKHM1 | 106.547787 | -0.597336805 | 0.34144227 | -1.7494518 | 0.08021295 | 0.45074086 |
| LINC02210-C | 94.3759704 | 0.35374821 | 0.43126453 | 0.82025807 | 0.412069 | 0.82355334 |
| KANSL1 | 1770.28817 | 0.447023224 | 0.37558834 | 1.19019463 | 0.2339699 | 0.69285918 |
| ARL17B | 386.087638 | 1.9488877 | 1.02946942 | 1.89309916 | 0.05834468 | 0.39982063 |
| LRRC37A2 | 105.240406 | 0.668162897 | 0.30587925 | 2.18440087 | 0.0289328 | 0.28768102 |
| ARL17A | 119.338759 | -0.043712999 | 0.48860411 | -0.0894651 | 0.92871231 | 0.98761086 |
| NSF | 470.986215 | 0.407635748 | 0.32396784 | 1.2582599 | 0.20829778 | 0.66589349 |
| WNT9B | 123.162769 | 2.319982035 | 1.04880073 | 2.21203319 | 0.02696437 | 0.27874491 |
| AC005670.2 | 73.9160633 | -0.319621729 | 0.36251959 | -0.8816675 | 0.37795666 | 0.80181899 |
| CDC27 | 486.486242 | -0.060488894 | 0.27620252 | -0.219002 | 0.82664852 | 0.96545294 |
| AC068234.1 | 83.4950252 | -2.89890351 | 0.96879947 | -2.9922637 | 0.00276917 | 0.0769785 |
| EFCAB13 | 183.907035 | -0.57573739 | 0.38072898 | -1.5121974 | 0.13048365 | 0.55395564 |
| AC040934.1 | 34.1886232 | -0.132314431 | 0.50845315 | -0.2602293 | 0.79468687 | 0.95937941 |
| NPEPPS | 989.692958 | 0.058434103 | 0.16973092 | 0.34427495 | 0.73063952 | 0.94048525 |
| AC025682.1 | 109.719044 | -0.223794921 | 0.29647689 | -0.7548478 | 0.45034032 | 0.84147218 |
| KPNB1 | 439.2991 | -0.115166998 | 0.19149774 | -0.6014013 | 0.5475727 | 0.88773467 |
| MRPL10 | 53.0693598 | -0.387382061 | 0.41892222 | -0.9247112 | 0.35511616 | 0.78607579 |

| | | | | | | |
|---|---|---|---|---|---|---|
| SCRN2 | 67.209666 | -0.083303662 | 0.34268062 | -0.2430942 | 0.80793245 | 0.96245698 |
| SP2 | 48.3077584 | 0.030835317 | 0.28246153 | 0.10916643 | 0.91307049 | 0.98480378 |
| PNPO | 67.6317358 | -0.227476074 | 0.43844819 | -0.5188209 | 0.60388566 | 0.90371579 |
| CDK5RAP3 | 114.486789 | 0.294072621 | 0.25899563 | 1.1354347 | 0.25619323 | 0.71374446 |
| NFE2L1 | 322.934723 | -0.351988317 | 0.23443271 | -1.5014471 | 0.13323995 | 0.55609507 |
| CBX1 | 174.764173 | -0.203264782 | 0.32069991 | -0.6338162 | 0.52620081 | 0.87791848 |
| SNX11 | 43.4265739 | 0.371835504 | 0.29307894 | 1.26872135 | 0.20454047 | 0.65976734 |
| SKAP1 | 1282.40932 | 0.1455488 | 0.19661664 | 0.74026696 | 0.45913802 | 0.84595051 |
| HOXB2 | 138.867956 | 0.360360811 | 0.46667416 | 0.77218934 | 0.4400023 | 0.83673091 |
| HOXB-AS1 | 35.7445148 | 0.31763269 | 0.40910037 | 0.77641751 | 0.43750251 | 0.83420562 |
| HOXB3 | 907.151453 | -0.106041134 | 0.24496936 | -0.4328751 | 0.66510551 | 0.92324595 |
| HOXB-AS3 | 213.418486 | 0.279538023 | 0.4125377 | 0.67760601 | 0.49802153 | 0.86583246 |
| HOXB7 | 314.250955 | -0.068126178 | 0.36211579 | -0.1881337 | 0.85077186 | 0.97017003 |
| AC103702.2 | 81.0779424 | -0.605885965 | 0.45063454 | -1.3445174 | 0.17878114 | 0.62810137 |
| LINC02086 | 153.170667 | -0.090132527 | 0.43434409 | -0.2075141 | 0.83560837 | 0.96757829 |
| CALCOCO2 | 204.682244 | 0.022936851 | 0.21902864 | 0.10472079 | 0.91659737 | 0.98520052 |
| ATP5MC1 | 253.702771 | -0.362463114 | 0.44446498 | -0.8155043 | 0.41478369 | 0.82461178 |
| UBE2Z | 285.808768 | -0.049048015 | 0.20671533 | -0.2372732 | 0.81244483 | 0.96288868 |
| SNF8 | 120.966298 | 0.316370061 | 0.33658374 | 0.93994456 | 0.347246 | 0.78180304 |
| B4GALNT2 | 160.064734 | 2.2874763 | 0.82845357 | 2.76114 | 0.00576 | 0.12145831 |
| ZNF652 | 421.078833 | -0.066393232 | 0.19876582 | -0.3340274 | 0.73835888 | 0.94349597 |
| AC091180.5 | 46.7709908 | 0.24625132 | 0.52529753 | 0.46878446 | 0.63922371 | 0.91521016 |
| PHB | 230.828747 | -0.404870582 | 0.46135675 | -0.8775651 | 0.38017978 | 0.8029673 |
| SPOP | 196.507136 | 0.2309613 | 0.16953818 | 1.36229671 | 0.17310426 | 0.61955472 |
| SLC35B1 | 42.1320537 | -0.03039951 | 0.35352016 | -0.0859909 | 0.93147366 | 0.98822345 |
| FAM117A | 41.8799451 | 0.066521063 | 0.33968583 | 0.19583114 | 0.84474234 | 0.96876046 |
| KAT7 | 120.889351 | 0.224102356 | 0.27356611 | 0.81918904 | 0.41267857 | 0.82409897 |
| ITGA3 | 553.19487 | -2.677502859 | 0.77009403 | -3.4768519 | 0.00050734 | 0.02532014 |
| PDK2 | 72.9473033 | 0.28505579 | 0.36291286 | 0.78546621 | 0.43218028 | 0.83162989 |
| PPP1R9B | 42.3193158 | -0.594911383 | 0.30993142 | -1.9194936 | 0.05492189 | 0.39224566 |
| XYLT2 | 71.1722102 | 0.029121256 | 0.28390768 | 0.10257298 | 0.91830189 | 0.98520052 |
| MRPL27 | 132.633859 | -0.562311784 | 0.32008598 | -1.7567523 | 0.07896004 | 0.44916083 |
| LRRC59 | 52.4117321 | -0.582475336 | 0.345546 | -1.6856665 | 0.09186004 | 0.47942892 |
| ACSF2 | 65.618969 | -0.058978236 | 0.38770115 | -0.1521229 | 0.87908998 | 0.9769738 |
| RSAD1 | 64.0397287 | 0.130743743 | 0.23261539 | 0.56205973 | 0.57407533 | 0.89807676 |
| EPN3 | 56.9897658 | 0.942258463 | 0.51025274 | 1.84665047 | 0.06479781 | 0.41483268 |
| SPATA20 | 66.7053141 | -0.17267645 | 0.40118703 | -0.4304138 | 0.66689463 | 0.92422364 |
| ABCC3 | 236.983123 | -2.318943027 | 0.84922132 | -2.7306698 | 0.00632058 | 0.12786365 |
| ANKRD40 | 75.2380698 | 0.070507341 | 0.37575468 | 0.18764195 | 0.85115734 | 0.97042184 |
| LUC7L3 | 683.569853 | 0.50767299 | 0.14296153 | 3.55111608 | 0.0003836 | 0.02099512 |
| TOB1 | 357.589654 | 1.086501235 | 0.49620424 | 2.18962507 | 0.02855144 | 0.28715843 |
| SPAG9 | 885.59692 | 0.677247582 | 0.30519721 | 2.21904907 | 0.02648339 | 0.27671574 |
| AC005920.2 | 89.2696896 | 1.060204522 | 0.64699498 | 1.63865957 | 0.10128418 | 0.49763628 |
| MBTD1 | 370.814849 | -0.096453184 | 0.23019431 | -0.4190077 | 0.67521051 | 0.92656883 |
| UTP18 | 96.6782134 | 0.073625431 | 0.27806271 | 0.26477995 | 0.79117898 | 0.95832571 |
| TOM1L1 | 633.594893 | 0.23633447 | 0.20016751 | 1.18068344 | 0.2377285 | 0.69398181 |
| COX11 | 128.854895 | 0.062063879 | 0.22018185 | 0.28187554 | 0.77803894 | 0.95601455 |
| STXBP4 | 469.934093 | 0.130088136 | 0.24352977 | 0.53417755 | 0.5932187 | 0.90277205 |
| HLF | 111.513983 | -0.950395069 | 0.69682665 | -1.3638902 | 0.17260211 | 0.61884138 |
| MMD | 50.6531984 | 0.872374763 | 0.58679354 | 1.48668093 | 0.1370991 | 0.56256796 |
| PCTP | 53.3082178 | -0.281361373 | 0.30024147 | -0.9371169 | 0.3486984 | 0.78180304 |
| DGKE | 71.9978711 | 0.117510125 | 0.34135105 | 0.34425008 | 0.73065822 | 0.94048525 |
| TRIM25 | 80.2874996 | -0.741462147 | 0.32955747 | -2.249872 | 0.02445707 | 0.26518037 |
| COIL | 80.2098731 | -0.117286485 | 0.22314793 | -0.5255997 | 0.59916636 | 0.90369786 |
| SCPEP1 | 353.946473 | 0.358481143 | 0.30902204 | 1.16005042 | 0.24602828 | 0.70206841 |
| AKAP1 | 166.526344 | 0.33480444 | 0.25759795 | 1.29971701 | 0.19369798 | 0.64723766 |
| MSI2 | 2132.88633 | 0.226049135 | 0.20712988 | 1.09134006 | 0.27512328 | 0.7263609 |
| MRPS23 | 100.979298 | -0.186459656 | 0.34560305 | -0.5395197 | 0.5895283 | 0.90233654 |
| CUEDC1 | 280.64232 | -0.308727938 | 0.40046739 | -0.770919 | 0.44075492 | 0.83733111 |
| VEZF1 | 189.358847 | -0.251731422 | 0.25931952 | -0.9707384 | 0.33167856 | 0.77618491 |
| AC015813.2 | 76.7588064 | 0.19784225 | 0.24879676 | 0.79519626 | 0.42649935 | 0.8277744 |
| SRSF1 | 42.4416839 | 0.128234944 | 0.3786007 | 0.33870763 | 0.73483 | 0.94215106 |
| DYNLL2 | 167.360996 | 0.220663865 | 0.28332418 | 0.77883881 | 0.43607467 | 0.83364877 |
| MKS1 | 39.1628551 | -0.097288461 | 0.34477574 | -0.282179 | 0.77780629 | 0.95601455 |
| TSPOAP1 | 109.114917 | 0.11014794 | 0.42628581 | 0.25838988 | 0.79610602 | 0.95948555 |
| TSPOAP1-AS | 35.4856295 | -0.604481255 | 0.60097612 | -1.0058324 | 0.31449619 | 0.75823062 |
| SUPT4H1 | 76.8486166 | -0.188102811 | 0.43869872 | -0.4287745 | 0.66808736 | 0.92422713 |
| RNF43 | 206.670356 | 0.137135265 | 0.49599783 | 0.2764836 | 0.78217666 | 0.95688565 |
| MTMR4 | 68.7773607 | 0.065080157 | 0.25832346 | 0.25193282 | 0.80109299 | 0.96063297 |
| RAD51C | 78.0254796 | 0.566243526 | 0.33281716 | 1.7013652 | 0.08887443 | 0.47038308 |
| TRIM37 | 179.413792 | 0.301118177 | 0.28253349 | 1.06577872 | 0.28652368 | 0.73607046 |
| SKA2 | 81.887359 | 0.411630468 | 0.349486 | 1.17781675 | 0.23886967 | 0.69499004 |
| GDPD1 | 45.2356943 | 0.22996357 | 0.38096724 | 0.60363083 | 0.54608911 | 0.88669883 |
| YPEL2 | 518.026254 | -0.037466359 | 0.49728279 | -0.0753422 | 0.93994248 | 0.98855185 |
| DHX40 | 207.084585 | -0.105638801 | 0.27113345 | -0.3896192 | 0.69681814 | 0.93130502 |
| CLTC | 718.284503 | 0.020425376 | 0.21409175 | 0.09540478 | 0.92399332 | 0.98697817 |
| PTRH2 | 95.7867532 | -0.075534258 | 0.34752707 | -0.2173478 | 0.82793728 | 0.96569646 |
| VMP1 | 972.281599 | -0.219245926 | 0.50734104 | -0.432147 | 0.66563455 | 0.92358667 |
| TUBD1 | 58.9773394 | -0.178369491 | 0.29884881 | -0.5968553 | 0.55060401 | 0.88916026 |
| RPS6KB1 | 136.567917 | 0.69577211 | 0.27604133 | 2.52053599 | 0.01171763 | 0.1719898 |
| HEATR6 | 50.1890464 | 0.242911582 | 0.30065521 | 0.80794071 | 0.41912472 | 0.82694802 |
| USP32 | 483.105332 | -0.476508612 | 0.34367762 | -1.3864988 | 0.16559464 | 0.6070066 |
| APPBP2 | 388.727671 | -0.054227638 | 0.19164385 | -0.2829605 | 0.77720713 | 0.95601455 |
| PPM1D | 131.950944 | 0.519244682 | 0.19111761 | 2.71688556 | 0.00658994 | 0.12933064 |
| BCAS3 | 1727.76718 | 0.394831151 | 0.17327264 | 2.27866997 | 0.02268669 | 0.25189387 |
| AC005746.2 | 78.9563209 | 1.463605736 | 0.51906484 | 2.81969733 | 0.0048069 | 0.11047429 |
| TBX2 | 549.397843 | 0.882522972 | 0.45511897 | 1.93910389 | 0.05248869 | 0.38418477 |
| NACA2 | 69.3134989 | 0.492910178 | 0.63236155 | 0.77947525 | 0.43569981 | 0.83364877 |
| INTS2 | 109.471757 | 0.231656573 | 0.26739658 | 0.86634082 | 0.38630329 | 0.80634213 |
| MED13 | 867.291771 | 0.345638002 | 0.25614138 | 1.34940323 | 0.17720748 | 0.62617642 |
| METTL2A | 56.0286317 | 0.145384401 | 0.31922244 | 0.4554329 | 0.64879783 | 0.91768694 |
| TLK2 | 452.931454 | 0.292623356 | 0.27234256 | 1.07446795 | 0.282613 | 0.73268183 |
| 10-Mar | 34.9447144 | 0.695336036 | 0.51297566 | 1.35549518 | 0.17525984 | 0.62477136 |
| TANC2 | 1737.86277 | 0.115977514 | 0.2218492 | 0.52277635 | 0.6011299 | 0.90369786 |
| AC015923.1 | 142.063389 | -0.273357768 | 0.63984371 | -0.4272258 | 0.66921484 | 0.92463889 |
| CYB561 | 107.802903 | -0.096560052 | 0.32041826 | -0.3013563 | 0.76314283 | 0.95378284 |
| DCAF7 | 388.126425 | -0.37833682 | 0.21417773 | -1.766462 | 0.07731837 | 0.44472917 |
| TACO1 | 35.8442247 | -0.505857945 | 0.45488853 | -1.1120482 | 0.26611745 | 0.72053648 |
| MAP3K3 | 253.847368 | -0.359842457 | 0.30381967 | -1.1843949 | 0.23625679 | 0.69335782 |
| CCDC47 | 104.417813 | 0.01583661 | 0.40579961 | 0.03902569 | 0.96886991 | 0.99312312 |
| DDX42 | 267.38093 | -0.062809054 | 0.19361451 | -0.3244026 | 0.74563323 | 0.94594643 |
| FTSJ3 | 54.0238594 | -0.127775352 | 0.28607054 | -0.4466568 | 0.65512287 | 0.92051095 |
| PSMC5 | 80.1192558 | 0.276257601 | 0.36537979 | 0.75608343 | 0.44959918 | 0.84122512 |
| SMARCD2 | 79.1955621 | 0.032622752 | 0.24652571 | 0.13233002 | 0.89472326 | 0.9821637 |
| ERN1 | 188.407885 | -0.568681615 | 0.51863898 | -1.0964884 | 0.27286511 | 0.72451099 |
| SNHG25 | 51.7075862 | 0.191139576 | 0.48151465 | 0.39695485 | 0.69140075 | 0.93073907 |
| TEX2 | 126.474122 | -0.371286266 | 0.28884389 | -1.2854219 | 0.19864489 | 0.65403502 |
| POLG2 | 66.5304483 | 0.003570492 | 0.28274316 | 0.01262804 | 0.98992455 | 0.99667684 |
| DDX5 | 1063.92328 | -0.065801924 | 0.28712705 | -0.2291735 | 0.81873404 | 0.96440741 |
| CEP95 | 340.82121 | -0.07237287 | 0.18747241 | -0.3860455 | 0.69946301 | 0.93180634 |
| SMURF2 | 341.746175 | 0.160962731 | 0.24990269 | 0.64410164 | 0.51950953 | 0.87623268 |
| LRRC37A3 | 467.981258 | 0.395536402 | 0.32585159 | 1.21385446 | 0.2248033 | 0.68208993 |
| GNA13 | 421.900784 | 0.258584638 | 0.64675535 | 0.39981832 | 0.68929033 | 0.93041766 |

| | | | | | | |
|---|---|---|---|---|---|---|
| CEP112 | 567.291227 | 0.040160793 | 0.27018665 | 0.14864092 | 0.88183698 | 0.97815957 |
| PRKCA | 3531.77386 | 0.213880598 | 0.27054173 | 0.79056418 | 0.42919836 | 0.82998114 |
| PRKCA-AS1 | 164.377907 | 0.333066862 | 0.48192652 | 0.69111544 | 0.489493 | 0.86043049 |
| HELZ | 661.842702 | 0.04965116 | 0.24757811 | 0.20054746 | 0.84105245 | 0.96854218 |
| PSMD12 | 142.814792 | -0.052381958 | 0.22109677 | -0.2369187 | 0.81271987 | 0.96288868 |
| PITPNC1 | 273.25357 | -1.117359367 | 0.59655265 | -1.8730273 | 0.06106462 | 0.40676596 |
| NOL11 | 80.7419835 | 0.28756881 | 0.22229102 | 1.29365914 | 0.19578319 | 0.64948524 |
| BPTF | 1099.06 | -0.100197468 | 0.24657451 | -0.4063578 | 0.68447974 | 0.92881914 |
| AC005332.7 | 44.1441267 | 0.353188968 | 0.31629933 | 1.11662887 | 0.26415304 | 0.71989632 |
| AMZ2 | 57.0083814 | -0.60334878 | 0.30688198 | -1.9660613 | 0.04929153 | 0.3755208 |
| ARSG | 127.48531 | -0.999157792 | 0.46654671 | -2.1416029 | 0.03222545 | 0.302348 |
| WIPI1 | 86.816679 | -1.523123707 | 0.4105633 | -3.709839 | 0.00020739 | 0.01374886 |
| PRKAR1A | 967.785635 | 0.229248423 | 0.42535184 | 0.53896187 | 0.58991317 | 0.90233654 |
| FAM20A | 1212.28652 | -0.808624087 | 0.39971725 | -2.0229902 | 0.04307416 | 0.34966009 |
| AC079210.1 | 49.9741979 | -1.374057868 | 0.56257409 | -2.4424478 | 0.01458804 | 0.19588328 |
| LINC01482 | 1058.0531 | 0.728704798 | 0.58062925 | 1.25502599 | 0.20946933 | 0.66739001 |
| ABCA10 | 69.9216559 | 0.373420443 | 0.48920984 | 0.76331343 | 0.44527649 | 0.83929814 |
| ABCA5 | 324.631129 | 0.468902774 | 0.27791713 | 1.68720358 | 0.09156421 | 0.47836139 |
| AC005208.1 | 878.213906 | 1.139097844 | 0.67675535 | 1.68317524 | 0.09234117 | 0.48050422 |
| KCNJ16 | 2926.83949 | 1.165251034 | 0.41929839 | 2.77904962 | 0.00545182 | 0.11829207 |
| SOX9 | 109.24278 | -4.088940197 | 0.69099913 | -5.9174317 | 3.27E-09 | 4.77E-06 |
| SLC39A11 | 455.063011 | -0.101067719 | 0.23263895 | -0.4344402 | 0.66396879 | 0.92300139 |
| COG1 | 53.522171 | -0.432637986 | 0.31358298 | -1.3796603 | 0.16769126 | 0.61233603 |
| FAM104A | 72.9928106 | -0.194057364 | 0.26921094 | -0.7208376 | 0.47100945 | 0.85236058 |
| C17orf80 | 51.7238018 | -0.290729158 | 0.30721805 | -0.9463284 | 0.34398113 | 0.78180304 |
| CDC42EP4 | 138.375777 | -0.002529471 | 0.32472904 | -0.0077895 | 0.99378495 | 0.99870178 |
| RPL38 | 221.875476 | -0.004485928 | 0.32227534 | -0.0139196 | 0.98889416 | 0.9964791 |
| GPRC5C | 762.150097 | -0.010590936 | 0.18394644 | -0.0575762 | 0.95408621 | 0.99060627 |
| SLC9A3R1 | 45.2679193 | -0.569992345 | 0.572348 | -0.9958842 | 0.3193064 | 0.76378437 |
| NAT9 | 44.1229701 | -0.580886156 | 0.35041045 | -1.6577307 | 0.09737184 | 0.49030463 |
| TMEM104 | 34.7253726 | -0.051108302 | 0.39870856 | -0.1281846 | 0.89800288 | 0.98312332 |
| MRPL58 | 40.224783 | -0.41455113 | 0.43883765 | -0.9446572 | 0.34483393 | 0.78180304 |
| KCTD2 | 79.4488568 | -0.078560306 | 0.22078807 | -0.3558177 | 0.72197708 | 0.93793477 |
| ATP5PD | 177.277764 | 0.299333572 | 0.45479596 | 0.65817114 | 0.51042817 | 0.87133207 |
| SLC16A5 | 607.534667 | -0.745288344 | 0.4127523 | -1.8056552 | 0.07097221 | 0.43093209 |
| ARMC7 | 147.337198 | -0.11472351 | 0.37956975 | -0.3022462 | 0.7624644 | 0.95374766 |
| NT5C | 162.754621 | -0.375838355 | 0.3476486 | -1.0810869 | 0.27965845 | 0.72938673 |
| JPT1 | 337.630363 | 0.406972077 | 0.71292986 | 0.57084448 | 0.56810507 | 0.89570349 |
| SUMO2 | 115.339237 | 0.296387997 | 0.35244096 | 0.84095786 | 0.40037154 | 0.8162653 |
| NUP85 | 83.5807709 | 0.196597 | 0.26827686 | 0.73281386 | 0.46367197 | 0.84922793 |
| GGA3 | 101.347166 | 0.021755923 | 0.1913866 | 0.11367527 | 0.90949522 | 0.98441374 |
| MRPS7 | 121.817025 | -0.167305533 | 0.38286554 | -0.4369825 | 0.66212406 | 0.92284375 |
| MIF4GD | 35.8676186 | -0.706418567 | 0.39598996 | -1.7839305 | 0.07443496 | 0.43775442 |
| GRB2 | 362.305808 | 0.240185953 | 0.19266325 | 1.24666203 | 0.21252145 | 0.66984203 |
| AC011933.4 | 50.9531827 | 0.286437562 | 0.43268888 | 0.66199427 | 0.50797489 | 0.87094607 |
| TMEM94 | 100.102505 | 0.128143302 | 0.18296264 | 0.70037963 | 0.48369026 | 0.85829223 |
| CASKIN2 | 53.4231718 | 0.083065537 | 0.29718603 | 0.27950687 | 0.77985587 | 0.95636474 |
| TSEN54 | 60.607911 | -0.097568477 | 0.30534842 | -0.3195316 | 0.74932342 | 0.94704648 |
| LLGL2 | 1146.0293 | -0.369442608 | 0.26450185 | -1.3967487 | 0.16248916 | 0.60211533 |
| MYO15B | 169.345517 | -0.063785667 | 0.338563 | -0.1884012 | 0.85056218 | 0.97017003 |
| RECQL5 | 169.051469 | 0.500950996 | 0.27415174 | 1.82727635 | 0.06765823 | 0.42296693 |
| SMIM5 | 377.52308 | 1.109881763 | 0.70998145 | 1.5632546 | 0.11799272 | 0.53360052 |
| SMIM6 | 68.1568043 | 0.834402575 | 0.80161952 | 1.04089603 | 0.29792381 | 0.74587709 |
| SAP30BP | 145.426292 | -0.177928646 | 0.26301409 | -0.6764985 | 0.49872418 | 0.86605292 |
| ITGB4 | 83.6794475 | -0.493833077 | 0.40275181 | -1.2261474 | 0.22014322 | 0.67816019 |
| H3F3B | 1690.00429 | 0.24481078 | 0.63206241 | 0.38732058 | 0.69851889 | 0.93180634 |
| UNK | 218.359976 | 0.170831206 | 0.26293383 | 0.64971178 | 0.51587842 | 0.87425708 |
| WBP2 | 188.026044 | -0.646230056 | 0.27378676 | -2.3603407 | 0.01825816 | 0.22511232 |
| TRIM47 | 40.7916774 | -2.337294963 | 0.45711 | -5.1132002 | 3.17E-07 | 0.00014433 |
| TRIM65 | 35.3683634 | -0.31354927 | 0.32956883 | -0.9513924 | 0.34140523 | 0.77954601 |
| FBF1 | 38.7082125 | -0.859153184 | 0.44922318 | -1.9125308 | 0.05580814 | 0.39368206 |
| ACOX1 | 190.298046 | 0.08489555 | 0.17229667 | 0.49272889 | 0.62220416 | 0.91028772 |
| TEN1 | 40.7292374 | -0.216392518 | 0.48841628 | -0.4430494 | 0.65773004 | 0.92180056 |
| EVPL | 51.7217131 | -0.364424301 | 0.27618679 | -1.3194849 | 0.18700705 | 0.63859039 |
| SRP68 | 95.3156274 | 0.058340555 | 0.19218826 | 0.30355941 | 0.76146358 | 0.95366435 |
| EXOC7 | 228.669991 | -0.033685791 | 0.2793442 | -0.1205888 | 0.90401671 | 0.98320957 |
| UBALD2 | 114.346063 | -0.616687857 | 0.33195963 | -1.8577195 | 0.06320886 | 0.41267121 |
| PRPSAP1 | 94.1945617 | 0.105420461 | 0.27952909 | 0.37713591 | 0.7060726 | 0.93241442 |
| UBE2O | 140.400585 | -0.183448435 | 0.20674747 | -0.8873068 | 0.37491375 | 0.79825864 |
| MXRA7 | 111.803402 | 0.468244075 | 0.3619969 | 1.29350299 | 0.19583715 | 0.64948524 |
| JMJD6 | 54.4848665 | 0.01271729 | 0.44775428 | 0.02840239 | 0.97734122 | 0.99469867 |
| SRSF2 | 242.794983 | 0.392190013 | 0.35249181 | 1.11262164 | 0.26587096 | 0.72053648 |
| MFSD11 | 126.841355 | -0.098992479 | 0.25764561 | -0.3842195 | 0.70081574 | 0.93180634 |
| SEC14L1 | 295.151717 | -0.596286695 | 0.5665438 | -1.0524989 | 0.29257074 | 0.73844969 |
| 9-Sep | 1352.22855 | -0.190050732 | 0.61138051 | -0.3108551 | 0.7559108 | 0.95067145 |
| TNRC6C | 186.432473 | -0.083834657 | 0.4048361 | -0.207083 | 0.83594507 | 0.96757829 |
| TMC6 | 70.9618949 | -0.319360927 | 0.38320215 | -0.8334007 | 0.40461879 | 0.81914727 |
| SYNGR2 | 171.229835 | -0.408894737 | 0.40819533 | -1.0017134 | 0.31648203 | 0.76069171 |
| AFMID | 266.203752 | 0.689639896 | 0.49175595 | 1.40240274 | 0.16079502 | 0.6002217 |
| SOCS3 | 226.655715 | -0.784485973 | 0.97456928 | -0.8049566 | 0.42084473 | 0.82694802 |
| PGS1 | 94.4076501 | -0.73269942 | 0.301496 | -2.4302127 | 0.01508996 | 0.19992772 |
| CYTH1 | 273.482706 | 0.304671132 | 0.24839191 | 1.22657431 | 0.21998263 | 0.67801071 |
| USP36 | 387.246517 | 1.333357595 | 1.00596127 | 1.32545619 | 0.18501989 | 0.63541313 |
| TIMP2 | 207.66613 | -0.642179156 | 0.39801843 | -1.6134408 | 0.10664878 | 0.50849828 |
| LGALS3BP | 417.830292 | -0.606455187 | 0.59385333 | -1.0212205 | 0.30714999 | 0.75214355 |
| CANT1 | 68.1421954 | -0.53231832 | 0.28432227 | -1.8722358 | 0.06117399 | 0.40676991 |
| ENGASE | 34.4832288 | 0.175844258 | 0.45136291 | 0.38958508 | 0.69684338 | 0.93130502 |
| CBX4 | 32.9630876 | -0.001615204 | 0.60182667 | -0.0026838 | 0.99785861 | 0.99954455 |
| TBC1D16 | 483.790452 | -0.118604304 | 0.44187815 | -0.2684095 | 0.78838411 | 0.95801395 |
| CCDC40 | 137.216598 | -0.538188216 | 0.67624704 | -0.7958456 | 0.4261218 | 0.82774157 |
| GAA | 181.778987 | -0.191896528 | 0.43334076 | -0.4428306 | 0.6578883 | 0.92185868 |
| EIF4A3 | 109.650986 | -0.224011829 | 0.4366358 | -0.5130405 | 0.60792302 | 0.90514346 |
| AC087741.1 | 47.3240928 | -0.898972756 | 0.40802655 | -2.2032213 | 0.02757916 | 0.28198008 |
| SGSH | 53.1525252 | -0.324079989 | 0.38548637 | -0.8407041 | 0.4005137 | 0.8162653 |
| SLC26A11 | 128.28708 | -0.396453822 | 0.24630154 | -1.6096279 | 0.10747912 | 0.51064149 |
| RNF213 | 1822.34254 | -1.94219075 | 0.54175797 | -3.5849787 | 0.00033711 | 0.01918302 |
| AC124319.2 | 382.264203 | 0.654743634 | 0.28650107 | 2.28530959 | 0.02229468 | 0.25166501 |
| ENDOV | 80.0696601 | 0.383733975 | 0.26255301 | 1.46154854 | 0.14386496 | 0.5732718 |
| RPTOR | 358.379854 | -0.025919646 | 0.28491625 | -0.0909729 | 0.92751416 | 0.98751845 |
| CHMP6 | 43.6005051 | -0.111729128 | 0.30631089 | -0.3647573 | 0.7152926 | 0.93573028 |
| BAIAP2-DT | 43.0505187 | 0.287960392 | 0.38008821 | 0.75761463 | 0.44868172 | 0.84115854 |
| BAIAP2 | 136.42731 | 1.039149958 | 0.57708853 | 1.80067685 | 0.07175383 | 0.43222361 |
| TEPSIN | 30.4549176 | 0.159396638 | 0.31294532 | 0.50934342 | 0.61051153 | 0.9062166 |
| NDUFAF8 | 102.552333 | 0.406017502 | 0.38550771 | 1.05320202 | 0.29224841 | 0.73844969 |
| SLC38A10 | 201.671764 | 0.027828668 | 0.23479437 | 0.11852357 | 0.90565282 | 0.98320957 |
| AC110285.6 | 38.0791264 | 0.942165959 | 0.57038082 | 1.65181915 | 0.09857142 | 0.49309237 |
| BAHCC1 | 58.8634029 | 0.101979583 | 0.45377687 | 0.22473508 | 0.82218537 | 0.96502145 |
| ACTG1 | 2411.33955 | 0.235578444 | 0.4467512 | 0.52731463 | 0.59797512 | 0.90369786 |
| FAAP100 | 35.3533942 | 0.430343096 | 0.32429496 | 1.32701136 | 0.18450493 | 0.63541313 |
| NPLOC4 | 309.174681 | 0.344562579 | 0.27849475 | 1.23723186 | 0.21600102 | 0.67291637 |
| OXLD1 | 79.8233701 | -0.307052323 | 0.29716779 | -1.0332625 | 0.30148108 | 0.74814891 |
| CCDC137 | 70.8320184 | 0.022709517 | 0.27488699 | 0.08261401 | 0.93415846 | 0.98822345 |
| ARL16 | 85.9425479 | 0.123077341 | 0.46054489 | 0.26724287 | 0.78928217 | 0.95801395 |

| | | | | | | |
|---|---|---|---|---|---|---|
| HGS | 155.466533 | -0.425709859 | 0.34591084 | -1.2306925 | 0.21843791 | 0.67607934 |
| MCRIP1 | 122.123658 | -0.436412924 | 0.38337544 | -1.1383435 | 0.25497709 | 0.71257599 |
| P4HB | 392.058954 | -0.680202156 | 0.3587211 | -1.8961866 | 0.05793537 | 0.3985947 |
| ARHGDIA | 222.224582 | -0.669150681 | 0.37606121 | -1.7793664 | 0.07517971 | 0.43966708 |
| ANAPC11 | 186.242694 | -0.041560902 | 0.54290082 | -0.0765534 | 0.93897883 | 0.98848592 |
| PCYT2 | 43.8779314 | -0.606486819 | 0.57698983 | -1.0511222 | 0.29320247 | 0.73864467 |
| SIRT7 | 47.821198 | -0.197676835 | 0.30156274 | -0.6555082 | 0.51214064 | 0.87172874 |
| MAFG | 43.3401454 | -1.360334667 | 0.44230019 | -3.0755914 | 0.00210086 | 0.06499896 |
| ASPSCR1 | 63.9462927 | -0.411942183 | 0.34045514 | -1.2099749 | 0.22628851 | 0.68421917 |
| CENPX | 57.6540857 | -0.325021652 | 0.59242182 | -0.5486321 | 0.58325792 | 0.90085422 |
| DCXR | 58.7689274 | -0.20053763 | 0.35598733 | -0.563328 | 0.57321156 | 0.89797476 |
| RFNG | 66.1280813 | -0.020519065 | 0.24700694 | -0.0830708 | 0.93379524 | 0.98822345 |
| GPS1 | 83.7857403 | -0.264167793 | 0.38280262 | -0.6900888 | 0.49013836 | 0.86055851 |
| DUS1L | 83.1789512 | -0.052750817 | 0.31841718 | -0.1656657 | 0.86842001 | 0.97576204 |
| FASN | 41.0966027 | -1.99110901 | 0.85661944 | -2.3243799 | 0.02010514 | 0.23835055 |
| CCDC57 | 356.02673 | -0.173710986 | 0.24568165 | -0.7070572 | 0.47953091 | 0.85532469 |
| SLC16A3 | 201.190775 | -1.037552242 | 0.55433736 | -1.8716982 | 0.06124837 | 0.40676991 |
| CSNK1D | 264.056827 | -0.30660118 | 0.32718061 | -0.9371007 | 0.34870676 | 0.78180304 |
| OGFOD3 | 86.9766161 | -0.117135323 | 0.3050355 | -0.3840055 | 0.70097434 | 0.93180634 |
| HEXDC | 96.0156284 | -0.243563529 | 0.31431533 | -0.7749019 | 0.43839766 | 0.8347986 |
| CYBC1 | 67.9175004 | -0.773579063 | 0.32306027 | -2.3945348 | 0.01664146 | 0.21108198 |
| NARF | 100.614251 | -0.61597709 | 0.31575933 | -1.9507804 | 0.05108318 | 0.38076225 |
| FOXK2 | 390.275043 | 0.60234387 | 0.40911827 | 1.47229767 | 0.14094051 | 0.56895751 |
| WDR45B | 305.390255 | 0.501826938 | 0.57742063 | 0.86908384 | 0.38480128 | 0.80561637 |
| RAB40B | 149.166346 | -0.398619922 | 0.36712656 | -1.0857834 | 0.27757484 | 0.72788245 |
| FN3KRP | 61.8696206 | 0.191354974 | 0.39434663 | 0.48524561 | 0.62750213 | 0.91061758 |
| FN3K | 228.887253 | 1.891537452 | 0.8514048 | 2.22166641 | 0.02630586 | 0.27568539 |
| TBCD | 178.580711 | 0.204963759 | 0.25405898 | 0.8067566 | 0.41980673 | 0.82694802 |
| METRNL | 202.182091 | 0.371488743 | 0.37944112 | 0.97904186 | 0.3275593 | 0.77333216 |
| AC144831.1 | 43.3066365 | 1.288901268 | 0.4991038 | 2.58243127 | 0.00981069 | 0.15842224 |
| USP14 | 257.76768 | 0.289780789 | 0.22617778 | 1.28120804 | 0.2001206 | 0.65475264 |
| THOC1 | 193.693789 | 0.2621547 | 0.21486761 | 1.22007546 | 0.22243627 | 0.6798044 |
| ENOSF1 | 742.029463 | -0.189107037 | 0.32280859 | -0.5858178 | 0.55799793 | 0.89179371 |
| YES1 | 291.460946 | 0.2352548 | 0.21417505 | 1.09842299 | 0.27201983 | 0.72372882 |
| METTL4 | 80.3050374 | 0.416089951 | 0.27762157 | 1.49876665 | 0.13393418 | 0.55788164 |
| SMCHD1 | 589.088031 | 0.934713925 | 0.29962149 | 3.11964911 | 0.00181067 | 0.05874854 |
| AP001011.1 | 100.890877 | 0.67890578 | 0.64696325 | 1.04937302 | 0.29400647 | 0.73965037 |
| LPIN2 | 840.07737 | 0.732038333 | 0.42909862 | 1.70599087 | 0.08800978 | 0.4701032 |
| AP000919.2 | 81.5642283 | 0.762549937 | 0.3714199 | 2.053067 | 0.04006608 | 0.33699239 |
| MYOM1 | 49.16541 | 0.388568194 | 0.60235525 | 0.64508144 | 0.51887441 | 0.87579366 |
| MYL12A | 587.986297 | -0.132029566 | 0.50966952 | -0.2590494 | 0.79559715 | 0.95948555 |
| MYL12B | 1068.07224 | -0.097320193 | 0.53667114 | -0.1813405 | 0.85610035 | 0.97182813 |
| TGIF1 | 188.548009 | 0.805495377 | 0.5775661 | 1.39463755 | 0.16312517 | 0.6021948 |
| DLGAP1 | 383.003119 | -1.60868961 | 0.92851081 | -1.7325481 | 0.08317603 | 0.45768271 |
| DLGAP1-AS1 | 105.615094 | 1.601716348 | 0.73291202 | 2.18541423 | 0.02885849 | 0.28768102 |
| DLGAP1-AS2 | 43.5511284 | 1.1132644 | 0.72339214 | 1.5389501 | 0.12381648 | 0.54273186 |
| LINC00667 | 60.2844868 | 0.055940221 | 0.33877252 | 0.16512621 | 0.86884465 | 0.97576204 |
| EPB41L3 | 296.915659 | 0.468685654 | 0.62960475 | 0.74441251 | 0.45662695 | 0.84382603 |
| L3MBTL4 | 443.199401 | -0.327181097 | 0.36590248 | -0.8941757 | 0.37122791 | 0.79539603 |
| AP001021.2 | 37.399076 | -0.276402503 | 0.69473165 | -0.3978551 | 0.69073703 | 0.93073907 |
| ARHGAP28 | 109.411566 | 0.76664602 | 0.44836914 | 1.70985455 | 0.08729277 | 0.46735992 |
| PTPRM | 1646.10398 | -0.195629521 | 0.41634067 | -0.4698785 | 0.63844184 | 0.91521016 |
| RAB12 | 162.947747 | 0.215135889 | 0.22877995 | 0.94036163 | 0.3470321 | 0.78180304 |
| MTCL1 | 56.9477479 | 0.771105874 | 0.74732463 | 1.03182184 | 0.30215559 | 0.74896472 |
| NDUFV2 | 299.148565 | 0.215363005 | 0.22586783 | 0.95349126 | 0.34034122 | 0.77836439 |
| ANKRD12 | 1369.10183 | -0.092260904 | 0.22848879 | -0.4037874 | 0.68636903 | 0.92974759 |
| TWSG1 | 604.958536 | 0.872551951 | 0.36559502 | 2.38666254 | 0.01700209 | 0.21364736 |
| AP005432.2 | 57.5286614 | 1.654366074 | 1.33439003 | 1.23979199 | 0.21505234 | 0.67132603 |
| RALBP1 | 1593.3708 | 0.428753295 | 0.40766259 | 1.05173568 | 0.29292083 | 0.73864467 |
| PPP4R1 | 342.700512 | 0.442499067 | 0.34981023 | 1.26496893 | 0.20588247 | 0.66287197 |
| RAB31 | 61.8180248 | -0.298390291 | 0.45096062 | -0.661677 | 0.50817822 | 0.87094607 |
| VAPA | 530.060498 | 0.367765903 | 0.22529166 | 1.63239913 | 0.10259543 | 0.50102523 |
| NAPG | 165.726405 | 0.159106813 | 0.2984278 | 0.53315011 | 0.59392967 | 0.90304695 |
| MPPE1 | 119.763256 | -0.220268253 | 0.28157945 | -0.7822597 | 0.43406197 | 0.83243867 |
| IMPA2 | 160.217242 | -0.250203953 | 0.37677306 | -0.6640707 | 0.50664506 | 0.87094607 |
| AFG3L2 | 377.148681 | 0.172222711 | 0.24061582 | 0.71575806 | 0.47414076 | 0.85348594 |
| SPIRE1 | 1580.98745 | 0.476230034 | 0.31544809 | 1.50969381 | 0.13112157 | 0.55395564 |
| PSMG2 | 260.281325 | 0.380534285 | 0.20828861 | 1.8269568 | 0.06770627 | 0.42296693 |
| CEP76 | 55.8436717 | 0.069403907 | 0.41024456 | 0.16917691 | 0.86565749 | 0.9750363 |
| PTPN2 | 339.299769 | 0.069085575 | 0.29082604 | 0.23754948 | 0.81223054 | 0.96288868 |
| SEH1L | 85.6805644 | 0.079810021 | 0.28274425 | 0.2822693 | 0.77773703 | 0.95601455 |
| CEP192 | 342.429811 | 0.278198478 | 0.30846783 | 0.90187194 | 0.3671249 | 0.79394737 |
| LDLRAD4 | 440.18779 | 0.734039082 | 0.51027103 | 1.43852783 | 0.15028435 | 0.58397478 |
| FAM210A | 158.911575 | -0.594008496 | 0.50492914 | -1.1764195 | 0.23942727 | 0.69526124 |
| RNMT | 345.604604 | -0.397360757 | 0.55838104 | -0.7116301 | 0.47669386 | 0.85414749 |
| ZNF519 | 43.7842932 | 0.630807667 | 0.38800599 | 1.62576787 | 0.10399904 | 0.50291791 |
| ROCK1 | 562.511948 | 0.140102339 | 0.19870242 | 0.70508623 | 0.48075658 | 0.85600365 |
| GREB1L | 391.175758 | 2.655818367 | 1.04457631 | 2.54248381 | 0.01100677 | 0.16502282 |
| ESCO1 | 225.266607 | 0.310003536 | 0.27044304 | 1.14628031 | 0.25167919 | 0.70813507 |
| SNRPD1 | 79.4186729 | -0.342149111 | 0.39666329 | -0.8625681 | 0.38837496 | 0.8065026 |
| ABHD3 | 192.85036 | -0.995871507 | 0.63510698 | -1.5680374 | 0.1168724 | 0.53183794 |
| MIB1 | 502.037668 | 0.161395995 | 0.31542367 | 0.51168003 | 0.60887496 | 0.90549306 |
| AC090912.1 | 76.0026976 | 0.076790436 | 0.55991077 | 0.13714763 | 0.8909141 | 0.98095103 |
| RBBP8 | 1303.84118 | 0.906183941 | 0.53256292 | 1.70155283 | 0.08883923 | 0.47038308 |
| CABLES1 | 147.401249 | -0.51895123 | 0.6123448 | -0.8474821 | 0.39672649 | 0.81254751 |
| TMEM241 | 106.327438 | -0.265974857 | 0.28939713 | -0.9190653 | 0.35806142 | 0.78684917 |
| RIOK3 | 333.689046 | -0.127813753 | 0.38294047 | -0.3337693 | 0.73855369 | 0.94349597 |
| RMC1 | 76.8043362 | 0.05693397 | 0.28959889 | 0.19659596 | 0.84414374 | 0.96876046 |
| NPC1 | 243.291184 | -0.405572917 | 0.47858534 | -0.8474412 | 0.39674927 | 0.81254751 |
| TTC39C | 176.224871 | -0.031433786 | 0.30629461 | -0.102626 | 0.91825982 | 0.98520052 |
| OSBPL1A | 128.036866 | -0.066566106 | 0.40702688 | -0.1635423 | 0.87009148 | 0.9759113 |
| IMPACT | 74.2791773 | 0.124454436 | 0.27050574 | 0.46008057 | 0.64545839 | 0.91685019 |
| SS18 | 397.389065 | 0.440337225 | 0.1826362 | 2.41100737 | 0.01590853 | 0.20582884 |
| TAF4B | 45.20201 | 0.201893732 | 0.2845272 | 0.70957622 | 0.47796697 | 0.85479418 |
| KCTD1 | 721.109027 | -0.942939322 | 0.36854488 | -2.5585468 | 0.01051107 | 0.16367902 |
| AQP4-AS1 | 65.0851716 | -1.621622039 | 0.37840256 | -4.2854415 | 1.82E-05 | 0.0028527 |
| CHST9 | 109.738366 | -0.534202962 | 0.76770031 | -0.6958483 | 0.48652383 | 0.85912348 |
| DSG2 | 99.8638295 | -0.254892537 | 0.51823895 | -0.4918436 | 0.62282988 | 0.91034771 |
| TRAPPC8 | 373.568583 | 0.898894547 | 0.27114928 | 3.31512792 | 0.00091601 | 0.03755638 |
| RNF125 | 65.4102782 | 0.945074831 | 0.79147271 | 1.19407128 | 0.23245009 | 0.69206731 |
| RNF138 | 137.809085 | 0.815479623 | 0.36896304 | 2.21019324 | 0.02709175 | 0.27917411 |
| GAREM1 | 2695.956 | 0.846388339 | 0.42896528 | 1.97309286 | 0.04848498 | 0.37279723 |
| AC025887.2 | 58.6015972 | 0.922734411 | 0.46931183 | 1.96614353 | 0.04928203 | 0.3755208 |
| KLHL14 | 213.892342 | 1.371456735 | 0.63570545 | 2.15737766 | 0.03097625 | 0.29646676 |
| CCDC178 | 362.144145 | 0.268136023 | 0.52052536 | 0.51512577 | 0.60646514 | 0.90493874 |
| ASXL3 | 313.886898 | 0.565855354 | 0.39495346 | 1.43271401 | 0.15193958 | 0.58670849 |
| MAPRE2 | 50.827116 | 0.025149577 | 0.40574188 | 0.06198418 | 0.95057543 | 0.98992048 |
| ZNF397 | 141.178275 | -0.272346758 | 0.25139522 | -1.083341 | 0.27865707 | 0.72935218 |
| ZSCAN30 | 101.355853 | -0.005753192 | 0.2595741 | -0.022164 | 0.98231716 | 0.99507903 |
| ZNF24 | 140.118645 | -0.140145982 | 0.21699273 | -0.6458556 | 0.51837285 | 0.87526312 |
| GALNT1 | 166.347862 | 0.331551506 | 0.24125441 | 1.37428166 | 0.16935426 | 0.61478828 |
| RPRD1A | 348.930048 | 0.036623477 | 0.27638145 | 0.13251062 | 0.89458043 | 0.9821637 |
| SLC39A6 | 366.920071 | 0.251713192 | 0.48007052 | 0.52432545 | 0.6000522 | 0.90369786 |

| | | | | | | |
|---|---|---|---|---|---|---|
| ELP2 | 350.496244 | 0.512604677 | 0.35956361 | 1.42563004 | 0.15397516 | 0.58943574 |
| MOCOS | 220.438913 | 0.198968883 | 0.40651413 | 0.48945134 | 0.6245222 | 0.91058133 |
| FHOD3 | 1782.91728 | 1.177950028 | 0.44492011 | 2.64755402 | 0.00810764 | 0.14312705 |
| TPGS2 | 155.873645 | -1.035124233 | 0.33725878 | -3.0692284 | 0.00214612 | 0.06557255 |
| KIAA1328 | 465.020167 | 0.024335905 | 0.30543959 | 0.07967502 | 0.93649573 | 0.98822345 |
| AC016205.1 | 154.428684 | -0.090687871 | 0.5342137 | -0.1697595 | 0.86519925 | 0.97489769 |
| PIK3C3 | 377.985728 | 0.002894086 | 0.25797718 | 0.01121838 | 0.99104922 | 0.99742589 |
| LINC00907 | 70.0538867 | -0.781333814 | 0.5097443 | -1.5327956 | 0.12532623 | 0.54408404 |
| SETBP1 | 2144.48856 | 0.780697866 | 0.35929502 | 2.17286023 | 0.02979084 | 0.29105042 |
| SLC14A2 | 742.310479 | 4.061469907 | 1.2804541 | 3.17189807 | 0.00151446 | 0.05169888 |
| EPG5 | 234.146233 | 0.161655501 | 0.27640302 | 0.58485432 | 0.55864568 | 0.89179371 |
| ATP5F1A | 424.610975 | 0.232816931 | 0.35589856 | 0.65416655 | 0.51300451 | 0.87178584 |
| HAUS1 | 33.4979641 | 0.133713535 | 0.49440292 | 0.27045458 | 0.78681056 | 0.9576974 |
| C18orf25 | 147.467985 | 0.565550713 | 0.21112512 | 2.67874663 | 0.00738983 | 0.13682934 |
| PIAS2 | 440.335013 | -0.375219883 | 0.34416575 | -1.0902302 | 0.27561177 | 0.7263609 |
| KATNAL2 | 236.809542 | 0.664022717 | 0.44402872 | 1.49544993 | 0.13479706 | 0.55868799 |
| HDHD2 | 39.6447421 | 0.458531927 | 0.35706874 | 1.28415589 | 0.19908742 | 0.65475264 |
| SMAD2 | 1078.22121 | 0.257663152 | 0.63153211 | 0.40799691 | 0.68327594 | 0.92818681 |
| ZBTB7C | 94.6287785 | -0.681264301 | 0.82243197 | -0.8283534 | 0.4074704 | 0.82018309 |
| CTIF | 273.095846 | 0.638097703 | 0.3564988 | 1.78990142 | 0.07346976 | 0.4369528 |
| SMAD7 | 116.152285 | 0.260727256 | 0.34666434 | 0.7521029 | 0.45198918 | 0.84234337 |
| DYM | 787.222834 | 0.277280873 | 0.19413071 | 1.42832054 | 0.15319962 | 0.58860244 |
| LIPG | 169.218326 | -2.607663275 | 1.04994327 | -2.483623 | 0.01300534 | 0.18344006 |
| ACAA2 | 171.68114 | -0.114937963 | 0.32583342 | -0.3527507 | 0.72427536 | 0.93893205 |
| SNHG22 | 76.700141 | -0.980140745 | 0.37083564 | -2.6430597 | 0.00821605 | 0.14386723 |
| MYO5B | 455.969277 | 0.343320648 | 0.31553973 | 1.08804255 | 0.27657632 | 0.72735756 |
| CFAP53 | 35.3914547 | -0.110085183 | 0.41376494 | -0.2660573 | 0.79019508 | 0.95812572 |
| MBD1 | 96.6437382 | 0.03301748 | 0.2555276 | 0.12921297 | 0.89718913 | 0.98302816 |
| CXXC1 | 70.7589721 | 0.412017245 | 0.31989183 | 1.28798925 | 0.1977497 | 0.652979 |
| MAPK4 | 184.168175 | 0.703310499 | 0.52211074 | 1.34705236 | 0.17796337 | 0.62729004 |
| ME2 | 106.86988 | -0.010710705 | 0.22763055 | -0.047053 | 0.96247097 | 0.99171131 |
| AC091551.1 | 49.6901828 | -0.610731237 | 0.35689774 | -1.7112219 | 0.08704015 | 0.46723935 |
| SMAD4 | 184.535979 | 0.45635496 | 0.23597456 | 1.93391593 | 0.05312347 | 0.3871585 |
| MEX3C | 51.7871238 | -0.29270661 | 0.27495105 | -1.0645772 | 0.28706732 | 0.73638964 |
| MBD2 | 446.707977 | -0.267126138 | 0.26097677 | -1.0235629 | 0.30604178 | 0.75130426 |
| POLI | 164.127506 | 0.191517664 | 0.28054874 | 0.68265381 | 0.49482562 | 0.86389349 |
| TCF4 | 260.795714 | 0.208456268 | 0.57771068 | 0.3608316 | 0.71822534 | 0.93573028 |
| TXNL1 | 173.700846 | 0.186028235 | 0.19572152 | 0.95047409 | 0.34187141 | 0.77973697 |
| WDR7 | 399.018196 | 0.383073657 | 0.22376659 | 1.71193411 | 0.08690881 | 0.46701529 |
| FECH | 64.7286007 | 0.193188855 | 0.32459014 | 0.59517783 | 0.55172461 | 0.88923362 |
| NARS | 247.55551 | -0.329270762 | 0.29997378 | -1.0976652 | 0.27235073 | 0.72405776 |
| AC027097.2 | 152.527745 | 0.213925211 | 0.44915132 | 0.47628761 | 0.63386949 | 0.91415622 |
| ATP8B1 | 1204.37303 | 1.025887251 | 0.53561407 | 1.91534783 | 0.05544816 | 0.39368206 |
| NEDD4L | 2661.7568 | 1.203349763 | 0.30215551 | 3.98255113 | 6.82E-05 | 0.00714521 |
| MALT1 | 319.263847 | -0.524775248 | 0.45175628 | -1.1616335 | 0.24538435 | 0.70154448 |
| AC104365.1 | 50.2882625 | -0.652252845 | 0.62919522 | -1.0366462 | 0.29990077 | 0.74689358 |
| ZNF532 | 282.62227 | -0.403377351 | 0.4428545 | -0.9108575 | 0.36237045 | 0.79051672 |
| SEC11C | 61.3492345 | 0.594089933 | 0.37389537 | 1.58892026 | 0.1120784 | 0.5224487 |
| LMAN1 | 314.504632 | 0.317704031 | 0.33075758 | 0.9605344 | 0.33678633 | 0.77827084 |
| RNF152 | 253.678287 | 0.369847686 | 0.64134132 | 0.5766784 | 0.56415673 | 0.8930743 |
| PIGN | 344.263644 | 0.158690489 | 0.18258168 | 0.86914795 | 0.38476621 | 0.80561637 |
| KIAA1468 | 530.799048 | 0.467261484 | 0.24562505 | 1.90233643 | 0.05712719 | 0.39761441 |
| ZCCHC2 | 228.531933 | 0.375469315 | 0.3595409 | 1.04430211 | 0.29634564 | 0.74334665 |
| PHLPP1 | 383.249464 | 0.500472812 | 0.23012506 | 2.17478626 | 0.02964614 | 0.29105042 |
| BCL2 | 811.460611 | 1.163002847 | 0.30504707 | 3.81253576 | 0.00013755 | 0.01065488 |
| KDSR | 148.030857 | -0.335482126 | 0.30380901 | -1.1042534 | 0.26948324 | 0.72322263 |
| VPS4B | 211.511881 | -0.19737678 | 0.17680037 | -1.1163822 | 0.26425858 | 0.71989632 |
| LINC01924 | 519.698105 | -1.362987494 | 0.85279951 | -1.5982508 | 0.10998717 | 0.51562013 |
| TMX3 | 190.013374 | 0.427858691 | 0.27828436 | 1.53748738 | 0.124174 | 0.54273186 |
| CCDC102B | 87.0095581 | -0.329087638 | 0.43562491 | -0.7554381 | 0.44998619 | 0.84142706 |
| RTTN | 181.881882 | 0.014769532 | 0.3935132 | 0.03753249 | 0.97006043 | 0.99348088 |
| SOCS6 | 104.130037 | -0.054977049 | 0.30475515 | -0.1803974 | 0.85684057 | 0.97191993 |
| FBXO15 | 36.3154841 | -0.180391873 | 0.51222474 | -0.3521733 | 0.7247083 | 0.93895336 |
| TIMM21 | 44.0645838 | 0.166580676 | 0.26131471 | 0.63747149 | 0.52381777 | 0.87725117 |
| CYB5A | 1159.06398 | 0.412721941 | 0.52467839 | 0.7866189 | 0.43150499 | 0.8312408 |
| CNDP2 | 496.373131 | -0.52103975 | 0.2587832 | -2.0134219 | 0.04407027 | 0.35332084 |
| LINC00909 | 42.423181 | -0.749889759 | 0.38506525 | -1.9474356 | 0.05148254 | 0.38247133 |
| ZNF407 | 649.589187 | 0.117247611 | 0.27899447 | 0.42025067 | 0.67430234 | 0.92595152 |
| ZADH2 | 79.942865 | -0.202308619 | 0.29436013 | -0.6872827 | 0.4919046 | 0.86172738 |
| TSHZ1 | 68.820972 | 0.206754376 | 0.458194 | 0.45123763 | 0.65181829 | 0.91887313 |
| ZNF516 | 449.569489 | 0.249297432 | 0.39399792 | 0.63273795 | 0.52690479 | 0.87791848 |
| ZNF236-DT | 45.9185892 | 0.715188512 | 0.53867679 | 1.3276765 | 0.184285 | 0.63497106 |
| ZNF236 | 316.319908 | 1.090467964 | 0.48844409 | 2.23253383 | 0.0255797 | 0.27225313 |
| MBP | 962.377646 | -0.192855058 | 0.3524409 | -0.5471983 | 0.58424251 | 0.90090543 |
| ATP9B | 602.439696 | 0.212379238 | 0.20763032 | 1.02287201 | 0.30636837 | 0.75140194 |
| NFATC1 | 78.479445 | -0.082869204 | 0.42675176 | -0.194186 | 0.84603027 | 0.96931085 |
| CTDP1 | 84.1105281 | -0.141039039 | 0.36232299 | -0.3892633 | 0.69708139 | 0.93130502 |
| PQLC1 | 110.224906 | 0.340075051 | 0.19389243 | 1.75393669 | 0.07944136 | 0.44953856 |
| HSBP1L1 | 140.786489 | -0.222795988 | 0.39488858 | -0.5641996 | 0.57261828 | 0.89797476 |
| TXNL4A | 186.486905 | 0.042501865 | 0.35399474 | 0.12006355 | 0.90443281 | 0.98320957 |
| ADNP2 | 146.550033 | 0.163932985 | 0.30757722 | 0.53298156 | 0.59404634 | 0.90304695 |
| C20orf96 | 62.0332208 | -0.111358508 | 0.31236846 | -0.3564973 | 0.72146818 | 0.93744589 |
| ZCCHC3 | 54.2673113 | -0.335827219 | 0.31761674 | -1.0573348 | 0.29035884 | 0.73754646 |
| NRSN2-AS1 | 193.070032 | 0.000123348 | 0.4029904 | 0.00030608 | 0.99975578 | 0.99993938 |
| SOX12 | 67.2993008 | -0.299771378 | 0.44027348 | -0.6808754 | 0.49595034 | 0.86452555 |
| NRSN2 | 62.9033982 | -1.016621922 | 0.33697412 | -3.0169139 | 0.00255362 | 0.0735219 |
| TRIB3 | 35.7749199 | -1.343657447 | 0.39939082 | -3.3642672 | 0.00076747 | 0.03351294 |
| RBCK1 | 215.953821 | -0.293347742 | 0.26458285 | -1.1087179 | 0.26755189 | 0.72222087 |
| TBC1D20 | 94.198619 | 0.012046509 | 0.23820331 | 0.05057238 | 0.95966627 | 0.99119981 |
| CSNK2A1 | 522.16283 | 0.021057414 | 0.21245568 | 0.09911439 | 0.92104745 | 0.98581108 |
| SLC52A3 | 210.811225 | -0.899294386 | 0.40919089 | -2.197738 | 0.02796778 | 0.28401388 |
| PSMF1 | 221.783616 | -0.386627343 | 0.20242704 | -1.909959 | 0.05613849 | 0.39444789 |
| AL136531.2 | 94.6131337 | -0.669149826 | 0.40295769 | -1.6605957 | 0.09679467 | 0.48901455 |
| SDCBP2-AS1 | 152.41371 | 0.181128002 | 0.293736 | 0.61663536 | 0.53747526 | 0.88239554 |
| NSFL1C | 130.838003 | -0.290233668 | 0.21772645 | -1.3330198 | 0.1825253 | 0.63297987 |
| STK35 | 70.2785644 | -0.852452798 | 0.24999017 | -3.4099453 | 0.00064976 | 0.03001789 |
| SNRPB | 164.728898 | -0.517182919 | 0.47735987 | -1.0834235 | 0.27862048 | 0.72935218 |
| NOP56 | 55.4280587 | -0.516341672 | 0.33529727 | -1.5399519 | 0.12357207 | 0.54273186 |
| IDH3B | 66.0862937 | -0.239556069 | 0.34809903 | -0.6881837 | 0.49133712 | 0.86152811 |
| EBF4 | 43.3242377 | 0.210754916 | 0.497685 | 0.4234705 | 0.67195201 | 0.92498948 |
| PCED1A | 54.0911085 | 0.356342784 | 0.28654924 | 1.24356564 | 0.21365947 | 0.67100726 |
| PTPRA | 522.65348 | -0.011185601 | 0.19028218 | -0.0587843 | 0.95312393 | 0.99060627 |
| MRPS26 | 161.264859 | 0.092988344 | 0.56282464 | 0.16521726 | 0.86877298 | 0.97576204 |
| UBOX5-AS1 | 49.4781526 | -0.088892946 | 0.5373093 | -0.1654409 | 0.86859694 | 0.97576204 |
| UBOX5 | 67.9893243 | 0.274931215 | 0.40346253 | 0.68142936 | 0.49559985 | 0.86452555 |
| LZTS3 | 88.4086576 | -0.286806787 | 0.29763168 | -0.9636299 | 0.3352315 | 0.77820994 |
| DDRGK1 | 118.161136 | -0.036768966 | 0.40066357 | -0.0917702 | 0.92688064 | 0.98729622 |
| ITPA | 124.58247 | -0.706209801 | 0.35874253 | -1.9685701 | 0.04900248 | 0.3746651 |
| C20orf194 | 1234.11826 | 0.091259254 | 0.19871807 | 0.45923983 | 0.64606195 | 0.91731869 |
| ATRN | 492.131826 | 0.369601212 | 0.25007997 | 1.47793209 | 0.13942596 | 0.56656585 |
| CENPB | 109.706087 | -0.796613306 | 0.36175769 | -2.2020632 | 0.02766084 | 0.28226449 |
| CDC25B | 38.6893008 | -0.277375612 | 0.36533823 | -0.7592296 | 0.44771521 | 0.84026779 |
| MAVS | 203.936537 | -0.303713015 | 0.23691358 | -1.2819569 | 0.19985775 | 0.65475264 |

| | | | | | | |
|---|---|---|---|---|---|---|
| PANK2 | 284.555633 | 0.328680097 | 0.25720676 | 1.27788279 | 0.20129076 | 0.65580391 |
| RNF24 | 928.073311 | -0.605242689 | 0.3045115 | -1.9875856 | 0.04685753 | 0.36375331 |
| SMOX | 40.4531253 | -0.977616304 | 0.70628587 | -1.3841652 | 0.16630789 | 0.60834438 |
| PRNP | 84.2169301 | -0.433889922 | 0.39895813 | -1.0875575 | 0.27679048 | 0.72743146 |
| SLC23A2 | 450.575469 | 0.531748207 | 0.27250746 | 1.95131612 | 0.05101945 | 0.38055791 |
| TMEM230 | 252.892907 | -0.070628027 | 0.28376397 | -0.2488971 | 0.80344039 | 0.96152281 |
| PCNA | 86.1692971 | -0.07781535 | 0.42211519 | -0.1843462 | 0.85374184 | 0.97062426 |
| CDS2 | 332.979887 | 0.400278984 | 0.33776459 | 1.18508275 | 0.23598473 | 0.69335782 |
| GPCPD1 | 756.273777 | 0.68822738 | 0.56976634 | 1.20791161 | 0.22708126 | 0.68523226 |
| SHLD1 | 142.300135 | 0.178222469 | 0.30901699 | 0.57674004 | 0.56411508 | 0.8930743 |
| CHGB | 315.287755 | 1.838972606 | 1.31781364 | 1.39547243 | 0.16287342 | 0.60211533 |
| TRMT6 | 82.2769137 | 0.778496451 | 0.40701372 | 1.91270323 | 0.05578606 | 0.39368206 |
| MCM8 | 78.4111755 | 0.554320926 | 0.24342815 | 2.27714394 | 0.02277763 | 0.25260273 |
| CRLS1 | 242.571089 | 0.593060287 | 0.33626514 | 1.76366865 | 0.07778778 | 0.44644905 |
| TMX4 | 155.123914 | -0.094265905 | 0.3338089 | -0.2823948 | 0.77764078 | 0.95601455 |
| PLCB1 | 130.303616 | 0.664137984 | 0.59448072 | 1.11717329 | 0.26392024 | 0.71989632 |
| PLCB4 | 169.385072 | -0.376311378 | 0.60527673 | -0.6217179 | 0.53412738 | 0.88137638 |
| ANKEF1 | 40.7076403 | -0.434562228 | 0.26122011 | -1.6635864 | 0.09619511 | 0.48772364 |
| MKKS | 58.0556183 | -0.270068381 | 0.48763851 | -0.5538291 | 0.57969584 | 0.89989814 |
| SLX4IP | 399.849817 | 0.296905572 | 0.31065246 | 0.95574835 | 0.33919939 | 0.77836439 |
| JAG1 | 177.017493 | 0.013556759 | 0.43645085 | 0.03106136 | 0.9752206 | 0.9939318 |
| AL050403.2 | 99.5230541 | 0.2965902 | 0.42898682 | 0.69137368 | 0.48933074 | 0.86042896 |
| BTBD3 | 186.371483 | -0.040749993 | 0.21134377 | -0.1928138 | 0.84710481 | 0.96991131 |
| SPTLC3 | 244.120525 | 0.28916904 | 0.34165599 | 0.84637487 | 0.39734365 | 0.81331279 |
| TASP1 | 633.827127 | 0.278118005 | 0.17797037 | 1.56272081 | 0.11811828 | 0.53360052 |
| ESF1 | 196.271173 | 0.073294975 | 0.26058796 | 0.2812677 | 0.77850508 | 0.95623387 |
| NDUFAF5 | 107.863676 | 0.403183827 | 0.36864107 | 1.09370297 | 0.27408527 | 0.72590691 |
| MACROD2 | 1716.94239 | 0.34769421 | 0.19027138 | 1.8273595 | 0.06764574 | 0.42296693 |
| FLRT3 | 66.3045266 | -0.558154021 | 0.54802788 | -1.0184774 | 0.30845113 | 0.75313873 |
| KIF16B | 572.142837 | 0.684447617 | 0.41434513 | 1.65187804 | 0.09855941 | 0.49309237 |
| SNRPB2 | 100.052253 | -0.149627533 | 0.27911669 | -0.5360752 | 0.59190659 | 0.90233654 |
| DSTN | 1070.72382 | -0.418088173 | 0.40529042 | -1.0315767 | 0.30227044 | 0.74896472 |
| RRBP1 | 265.122876 | -0.91741433 | 0.25889465 | -3.5435817 | 0.00039473 | 0.02132361 |
| SNX5 | 172.944731 | 0.480720687 | 0.33093292 | 1.45262273 | 0.14632853 | 0.57722788 |
| OVOL2 | 127.582564 | -0.45266372 | 0.42664674 | -1.0609801 | 0.28869892 | 0.73668501 |
| MGME1 | 130.199845 | -0.140210313 | 0.31301725 | -0.4479316 | 0.65420257 | 0.92002724 |
| KAT14 | 45.6235634 | 0.25614498 | 0.39571486 | 0.64729685 | 0.51743984 | 0.87526312 |
| ZNF133 | 154.939174 | -0.096772318 | 0.28163764 | -0.3436058 | 0.73114278 | 0.94053147 |
| DZANK1 | 110.886695 | -0.058514152 | 0.26020589 | -0.2248764 | 0.82207546 | 0.96502145 |
| POLR3F | 38.4233936 | 0.109998101 | 0.32628705 | 0.33712064 | 0.73602596 | 0.94289524 |
| RBBP9 | 51.3606493 | -0.472139574 | 0.36275434 | -1.3015408 | 0.19307341 | 0.64604385 |
| SEC23B | 173.435858 | 0.091114969 | 0.22610883 | 0.40296953 | 0.68697064 | 0.92992151 |
| DTD1 | 66.9381166 | -0.633987252 | 0.36837091 | -1.7210568 | 0.08524052 | 0.46310155 |
| SLC24A3 | 98.951366 | -2.730075392 | 1.05193785 | -2.595282 | 0.00945133 | 0.15525074 |
| AL121761.1 | 142.015122 | -0.131458432 | 0.46310533 | -0.2838629 | 0.77651543 | 0.95601455 |
| RIN2 | 425.190147 | -0.092074705 | 0.24034305 | -0.383097 | 0.70164784 | 0.93215897 |
| NAA20 | 271.209317 | -0.292146644 | 0.38936583 | -0.750314 | 0.4530656 | 0.84248055 |
| CRNKL1 | 67.9303505 | -0.425686602 | 0.27654133 | -1.5393236 | 0.12372531 | 0.54273186 |
| CFAP61 | 39.0542355 | -0.177762096 | 0.3448223 | -0.515518 | 0.60619111 | 0.90493874 |
| RALGAPA2 | 878.224843 | -1.073189771 | 0.28113797 | -3.8173064 | 0.00013492 | 0.01065488 |
| KIZ | 800.228981 | 0.881271717 | 0.56424508 | 1.56185982 | 0.11832102 | 0.5340248 |
| KIZ-AS1 | 191.094177 | 0.182925526 | 0.63088307 | 0.28995155 | 0.7718533 | 0.95535876 |
| XRN2 | 412.577631 | -0.13033789 | 0.24333682 | -0.5356275 | 0.59221603 | 0.90233654 |
| NXT1 | 35.4005482 | 0.332273904 | 0.53379764 | 0.62247166 | 0.53363177 | 0.88137638 |
| GZF1 | 59.0660798 | 1.318010953 | 0.82391065 | 1.59970132 | 0.10966486 | 0.51494555 |
| NAPB | 173.00583 | 0.718702208 | 0.25742055 | 2.791938 | 0.00523934 | 0.11544342 |
| CST3 | 618.001326 | -0.87865413 | 0.40762388 | -2.1555511 | 0.03111873 | 0.29675133 |
| AL035661.1 | 52.4474397 | 0.417030122 | 0.57132321 | 0.7299373 | 0.46542851 | 0.84991999 |
| APMAP | 136.300331 | -0.121937431 | 0.36004043 | -0.3386771 | 0.73485303 | 0.94215106 |
| ACSS1 | 168.732674 | -0.522395966 | 0.29517197 | -1.7698021 | 0.07676012 | 0.44322095 |
| ENTPD6 | 97.9138011 | 0.12028371 | 0.23996268 | 0.50126007 | 0.6161881 | 0.90794578 |
| PYGB | 133.364896 | -0.414349198 | 0.33631401 | -1.2320307 | 0.21793761 | 0.67553567 |
| ABHD12 | 227.192776 | 0.017398812 | 0.28640651 | 0.06074866 | 0.95155938 | 0.99030211 |
| NINL | 390.534571 | -0.13144494 | 0.38527114 | -0.3411751 | 0.73297173 | 0.94163693 |
| ZNF337-AS1 | 79.7540579 | 0.171753017 | 0.39457276 | 0.43528859 | 0.66335297 | 0.9229791 |
| ZNF337 | 101.101702 | -0.062689485 | 0.29563295 | -0.2120518 | 0.83206666 | 0.96656674 |
| FAM182B | 126.2486 | -0.574537156 | 0.37453022 | -1.5340208 | 0.12502453 | 0.54337567 |
| HM13 | 248.872611 | -0.360528371 | 0.19722281 | -1.8280257 | 0.06754569 | 0.42286671 |
| ID1 | 1215.32897 | -0.055154106 | 0.79640622 | -0.0692537 | 0.94478765 | 0.98949015 |
| BCL2L1 | 345.215872 | -0.049426348 | 0.35697275 | -0.1384597 | 0.8898771 | 0.98064269 |
| AL117381.1 | 58.7054991 | -0.987604358 | 0.58050702 | -1.7012789 | 0.08889063 | 0.47038308 |
| DUSP15 | 74.6735588 | -0.690331686 | 0.64044034 | -1.0779016 | 0.28107766 | 0.73069157 |
| TTLL9 | 35.3784016 | -0.610848268 | 0.44613687 | -1.3691948 | 0.1709384 | 0.61752307 |
| PDRG1 | 42.3351619 | -0.334704137 | 0.32958199 | -1.0155413 | 0.30984785 | 0.75409553 |
| TM9SF4 | 202.774198 | -0.040240149 | 0.22746853 | -0.1769042 | 0.8595836 | 0.97244655 |
| POFUT1 | 96.7533813 | 0.109407449 | 0.29455307 | 0.37143544 | 0.71031324 | 0.93471656 |
| KIF3B | 335.494697 | 0.205672648 | 0.24611101 | 0.83569055 | 0.40332901 | 0.81828987 |
| ASXL1 | 561.269783 | 0.227701942 | 0.25369262 | 0.89755052 | 0.36942523 | 0.79532503 |
| NOL4L | 183.393799 | -0.211225194 | 0.38877929 | -0.5433036 | 0.58692077 | 0.90196444 |
| MAPRE1 | 156.055903 | 0.332402878 | 0.35761377 | 0.92950245 | 0.35262876 | 0.78314007 |
| AL035071.2 | 35.1451816 | -0.160767663 | 0.49588878 | -0.324201 | 0.74578583 | 0.94594643 |
| AL035071.1 | 32.9942211 | 0.056455467 | 0.44610986 | 0.12655059 | 0.8992961 | 0.98312332 |
| CDK5RAP1 | 112.256274 | 0.360938438 | 0.23351502 | 1.54567545 | 0.12218297 | 0.54028588 |
| CBFA2T2 | 711.467334 | 0.104634995 | 0.28357047 | 0.36899116 | 0.71213431 | 0.9348115 |
| PXMP4 | 49.8367675 | -0.014060888 | 0.444464 | -0.0316356 | 0.97476265 | 0.9939318 |
| CHMP4B | 197.138182 | -0.038487609 | 0.2976868 | -0.1292889 | 0.89712903 | 0.98302816 |
| RALY | 247.540695 | -0.324399273 | 0.29761176 | -1.0900082 | 0.27570951 | 0.7263609 |
| EIF2S2 | 255.470081 | -0.840853964 | 0.37016602 | -2.2715591 | 0.02311316 | 0.25605273 |
| AHCY | 65.2656887 | -0.247154991 | 0.36567086 | -0.6758947 | 0.49910751 | 0.86643145 |
| ITCH | 701.920078 | 0.207395761 | 0.2154159 | 0.96276903 | 0.33566344 | 0.77820994 |
| DYNLRB1 | 432.391182 | -0.049002609 | 0.42449711 | -0.1154369 | 0.90809887 | 0.98385983 |
| MAP1LC3A | 123.073972 | 0.823243952 | 0.55007886 | 1.49659261 | 0.13449929 | 0.55868799 |
| PIGU | 188.770743 | 0.319885715 | 0.24553748 | 1.30279792 | 0.19264376 | 0.64563691 |
| NCOA6 | 435.897277 | 0.13803992 | 0.23811927 | 0.57970916 | 0.56211077 | 0.89242209 |
| TP53INP2 | 96.384839 | 0.528777366 | 0.54787849 | 0.96513621 | 0.33447659 | 0.77820994 |
| GGT7 | 146.504022 | -0.245442903 | 0.2423141 | -1.0129122 | 0.31110211 | 0.75435382 |
| ACSS2 | 150.103192 | 0.042776591 | 0.39615283 | 0.10798002 | 0.91401154 | 0.98480378 |
| GSS | 77.7780833 | -0.000212847 | 0.39864562 | -0.0005339 | 0.99957399 | 0.99993938 |
| TRPC4AP | 461.85497 | -0.024254496 | 0.23336078 | -0.1039356 | 0.91722044 | 0.98520052 |
| EDEM2 | 85.2385596 | -0.221003888 | 0.263915 | -0.8374055 | 0.40236465 | 0.81711183 |
| PROCR | 55.6688629 | -0.625517278 | 0.48436643 | -1.2914134 | 0.19656038 | 0.65097045 |
| MMP24OS | 140.739708 | 0.07192238 | 0.26068949 | 0.27589291 | 0.78263032 | 0.95688565 |
| MMP24 | 106.840936 | -0.544549881 | 0.4070446 | -1.3378138 | 0.18095714 | 0.63152351 |
| UQCC1 | 253.410866 | 0.311801209 | 0.24305325 | 1.28285141 | 0.19954414 | 0.65475264 |
| CEP250 | 65.5851633 | -0.879052687 | 0.31429383 | -2.7969136 | 0.00515933 | 0.11481456 |
| ERGIC3 | 143.90442 | 0.090293508 | 0.27517739 | 0.32812836 | 0.74281461 | 0.94508888 |
| CPNE1 | 123.409779 | 0.282307175 | 0.31742701 | 0.8893609 | 0.37380915 | 0.79689176 |
| ROMO1 | 152.208573 | -0.209649952 | 0.52704925 | -0.3977806 | 0.69079194 | 0.93073907 |
| RBM39 | 915.473265 | 0.261430218 | 0.15342945 | 1.70391161 | 0.0883976 | 0.47036301 |
| PHF20 | 457.522113 | 0.227280876 | 0.25975047 | 0.87499697 | 0.38157555 | 0.80315561 |
| SCAND1 | 229.147071 | -0.638983356 | 0.44294174 | -1.4425901 | 0.14913598 | 0.58194377 |
| NORAD | 608.672782 | 0.028701926 | 0.158624 | 0.18094315 | 0.8564122 | 0.97182813 |
| EPB41L1 | 1045.11239 | 0.109805928 | 0.36849088 | 0.29798819 | 0.76571218 | 0.95389523 |

| | | | | | | |
|---|---|---|---|---|---|---|
| AAR2 | 51.4794892 | 0.203131852 | 0.25798188 | 0.78738805 | 0.43105474 | 0.83111714 |
| DLGAP4 | 224.811239 | -0.25649112 | 0.25068126 | -1.0231763 | 0.3062245 | 0.75140194 |
| DLGAP4-AS1 | 142.88053 | 0.295477077 | 0.29126394 | 1.01446501 | 0.31036092 | 0.75435382 |
| MYL9 | 98.6822381 | 0.447515986 | 0.41692356 | 1.07337658 | 0.28310219 | 0.73293254 |
| RAB5IF | 79.7960732 | -0.197329583 | 0.53269549 | -0.370436 | 0.71105766 | 0.93475719 |
| NDRG3 | 90.1587583 | 0.338227508 | 0.34711953 | 0.97438341 | 0.3298662 | 0.77618491 |
| SOGA1 | 230.729709 | 0.562037322 | 0.38049317 | 1.47712855 | 0.13964118 | 0.56656585 |
| SAMHD1 | 58.5972613 | -0.520216642 | 0.69587963 | -0.747567 | 0.45472138 | 0.84290184 |
| RBL1 | 64.6564638 | -0.021067398 | 0.35150393 | -0.059935 | 0.95220738 | 0.9903694 |
| RPN2 | 244.480756 | 0.123857635 | 0.2531974 | 0.48917419 | 0.62471838 | 0.91058133 |
| MANBAL | 84.7255886 | -0.035241078 | 0.27372245 | -0.1287475 | 0.89755746 | 0.98312332 |
| SRC | 121.097719 | 0.467910722 | 0.24313364 | 1.9245001 | 0.05429193 | 0.39095834 |
| BLCAP | 235.802334 | 0.110663403 | 0.23812074 | 0.46473652 | 0.64212015 | 0.91643619 |
| CTNNBL1 | 191.087308 | 0.445023327 | 0.24443239 | 1.82063977 | 0.06866163 | 0.42628787 |
| TTI1 | 90.4289504 | 0.120443023 | 0.30875963 | 0.39008669 | 0.69647245 | 0.93130502 |
| RPRD1B | 179.760358 | 0.193224835 | 0.21981678 | 0.87902676 | 0.37938677 | 0.80274043 |
| TGM2 | 68.4941402 | -2.497973432 | 0.64640499 | -3.8644092 | 0.00011136 | 0.00941158 |
| RALGAPB | 397.769371 | 0.324300622 | 0.17689868 | 1.83325629 | 0.06676447 | 0.42005766 |
| DHX35 | 88.3010857 | 0.410141908 | 0.34238085 | 1.19791137 | 0.23095152 | 0.69089594 |
| TOP1 | 834.695407 | 0.353104804 | 0.36580976 | 0.96526897 | 0.33441011 | 0.77820994 |
| PLCG1 | 89.4558494 | -0.485455745 | 0.30776358 | -1.5773658 | 0.11471139 | 0.52703872 |
| ZHX3 | 567.181717 | 0.866696702 | 0.39751267 | 2.18029956 | 0.02923526 | 0.28931514 |
| LPIN3 | 101.29379 | 0.19677935 | 0.25902752 | 0.7596851 | 0.44744283 | 0.84026779 |
| CHD6 | 621.747212 | -0.126886038 | 0.23029641 | -0.5509684 | 0.58165535 | 0.90053894 |
| SRSF6 | 148.445761 | 0.658644312 | 0.35089873 | 1.87702109 | 0.0605152 | 0.40575773 |
| L3MBTL1 | 76.8371631 | -0.362339185 | 0.33309605 | -1.0877919 | 0.27668699 | 0.72743146 |
| SGK2 | 215.27606 | 0.647763048 | 0.36005068 | 1.79908853 | 0.07200468 | 0.43244071 |
| IFT52 | 138.981876 | 0.288304537 | 0.26289291 | 1.09666151 | 0.27278939 | 0.72451099 |
| OSER1 | 74.7358527 | -0.228208309 | 0.45506623 | -0.5014837 | 0.61603073 | 0.90794578 |
| FITM2 | 32.5806751 | 0.341121117 | 0.32560227 | 1.04766199 | 0.29479437 | 0.74087409 |
| TTPAL | 48.6633398 | -0.231350676 | 0.35724718 | -0.6475927 | 0.5172484 | 0.87526312 |
| SERINC3 | 159.277531 | 0.052380433 | 0.32441761 | 0.1614599 | 0.8717312 | 0.97622753 |
| PKIG | 285.969329 | 0.331172546 | 0.18059219 | 1.83381433 | 0.06668156 | 0.42003475 |
| YWHAB | 697.523445 | -0.608731348 | 0.37037753 | -1.6435429 | 0.10027066 | 0.49569046 |
| PABPC1L | 84.4699116 | 0.208334902 | 0.25859018 | 0.80565665 | 0.42044085 | 0.82694802 |
| TOMM34 | 46.2947012 | 0.153712324 | 0.2864653 | 0.5365827 | 0.59155589 | 0.90233654 |
| STK4 | 199.886467 | 0.025819606 | 0.23102855 | 0.11175938 | 0.9110142 | 0.98468493 |
| SLPI | 2334.70845 | -3.593804117 | 1.37202095 | -2.6193508 | 0.00880973 | 0.14867308 |
| SDC4 | 644.189925 | -0.160062572 | 0.48489236 | -0.3300992 | 0.74132502 | 0.94487568 |
| PIGT | 112.763575 | 0.028072101 | 0.38723285 | 0.07249411 | 0.94220869 | 0.98885525 |
| WFDC2 | 6791.59647 | -1.85120054 | 0.52474056 | -3.5278396 | 0.00041897 | 0.02204711 |
| DNTTIP1 | 58.1457439 | -0.209361007 | 0.30338575 | -0.6900819 | 0.4901427 | 0.86055851 |
| SNX21 | 60.0430657 | 0.196104738 | 0.45856688 | 0.42764697 | 0.66890816 | 0.92457894 |
| ACOT8 | 45.4908403 | -0.218460754 | 0.26803382 | -0.8150492 | 0.41504413 | 0.82472987 |
| CTSA | 161.473596 | -1.020960961 | 0.34690473 | -2.9430586 | 0.00324987 | 0.08622436 |
| ZNF335 | 62.0306526 | 0.098416354 | 0.26562413 | 0.37050984 | 0.71100265 | 0.93475719 |
| NCOA5 | 101.489667 | -0.23198321 | 0.2866061 | -0.8094148 | 0.41827661 | 0.82661491 |
| CD40 | 71.6893032 | -0.964850925 | 0.55834325 | -1.7280605 | 0.08397737 | 0.46077638 |
| SLC35C2 | 79.1831012 | -0.335419226 | 0.27030621 | -1.2408861 | 0.21464782 | 0.67132603 |
| ELMO2 | 83.3027083 | -0.132482148 | 0.30394744 | -0.4358719 | 0.66292967 | 0.9229791 |
| ZNF334 | 43.1901822 | -0.500206328 | 0.58858033 | -0.8498523 | 0.39540723 | 0.81175661 |
| SLC13A3 | 65.1119499 | -0.128375259 | 0.68782954 | -0.1866382 | 0.85194432 | 0.97053519 |
| ZMYND8 | 976.926235 | -0.479029282 | 0.37665988 | -1.2717821 | 0.20345056 | 0.6582233 |
| NCOA3 | 477.140443 | 0.110949144 | 0.26402129 | 0.42022802 | 0.67431889 | 0.92595152 |
| ARFGEF2 | 556.534808 | -0.130215023 | 0.19482016 | -0.6683858 | 0.50388738 | 0.86946722 |
| CSE1L | 193.342346 | 0.293185567 | 0.21718819 | 1.34991485 | 0.1770433 | 0.62617642 |
| STAU1 | 397.460777 | 0.289439604 | 0.19886211 | 1.45547886 | 0.14553674 | 0.57618473 |
| DDX27 | 93.6194354 | 0.225779392 | 0.42808686 | 0.52741491 | 0.59790549 | 0.90369786 |
| ZNFX1 | 140.46099 | -0.507180141 | 0.5023446 | -1.0096259 | 0.31267453 | 0.75677346 |
| ZFAS1 | 145.996879 | -0.14764018 | 0.25381095 | -0.5816935 | 0.56077315 | 0.89242209 |
| B4GALT5 | 460.099313 | 0.215167909 | 0.28386255 | 0.75800034 | 0.44845078 | 0.84104584 |
| SLC9A8 | 168.974179 | -0.133082072 | 0.2333151 | -0.5703963 | 0.56840894 | 0.8957783 |
| RNF114 | 317.089285 | 0.2985038 | 0.31340145 | 0.95246463 | 0.3408614 | 0.77911176 |
| CEBPB | 123.662788 | -2.645504073 | 0.65189866 | -4.0581524 | 4.95E-05 | 0.00576513 |
| PTPN1 | 280.274899 | 0.02647333 | 0.2427734 | 0.10904543 | 0.91316646 | 0.98480378 |
| RIPOR3 | 61.6393003 | -1.097215787 | 0.70414991 | -1.5582133 | 0.11918268 | 0.53606632 |
| PARD6B | 764.625983 | 0.50461307 | 0.47959806 | 1.05215828 | 0.29272694 | 0.73851187 |
| BCAS4 | 111.141348 | -0.589752045 | 0.38693083 | -1.5241795 | 0.12746387 | 0.54746778 |
| ADNP | 757.864146 | 0.204673616 | 0.28949788 | 0.70699521 | 0.47956945 | 0.85532469 |
| DPM1 | 170.807908 | 0.322893554 | 0.22127702 | 1.45922769 | 0.14450245 | 0.57406643 |
| ATP9A | 298.081573 | -0.22022067 | 0.2504361 | -0.8793487 | 0.37921221 | 0.80274043 |
| ZFP64 | 108.318364 | -0.381212744 | 0.39062301 | -0.9759096 | 0.32910927 | 0.77573228 |
| ZNF217 | 261.170622 | -0.269917142 | 0.36730925 | -0.7348498 | 0.46243096 | 0.84814078 |
| PFDN4 | 53.0014081 | -0.161978747 | 0.36808432 | -0.4400588 | 0.65989451 | 0.92284375 |
| DOK5 | 115.424493 | -0.593199474 | 0.64501992 | -0.9196607 | 0.35775009 | 0.78674827 |
| FAM210B | 79.2892651 | -0.430872858 | 0.32137405 | -1.3407207 | 0.18001114 | 0.63015077 |
| CSTF1 | 42.011984 | -0.292249186 | 0.35971232 | -0.8124525 | 0.41653202 | 0.82489761 |
| RTF2 | 94.6092178 | -0.012691155 | 0.21226214 | -0.05979 | 0.95232289 | 0.9903694 |
| RAE1 | 54.5971836 | 0.040897909 | 0.26825326 | 0.15246006 | 0.87882409 | 0.97692415 |
| PCK1 | 178.34614 | 0.36771047 | 1.31082829 | 0.28051765 | 0.77908039 | 0.95623735 |
| PMEPA1 | 101.700994 | -1.970962747 | 0.76194536 | -2.5867508 | 0.00968856 | 0.15717671 |
| RAB22A | 188.839785 | 0.402234511 | 0.25469872 | 1.57925614 | 0.11427732 | 0.52703872 |
| VAPB | 268.613873 | 0.023770124 | 0.17382226 | 0.1367496 | 0.89122873 | 0.98095103 |
| APCDD1L | 112.847528 | 0.506061863 | 0.60580728 | 0.83535124 | 0.40351997 | 0.81828987 |
| NPEPL1 | 94.4183509 | 0.15048854 | 0.24914178 | 0.60402771 | 0.54582522 | 0.88669883 |
| GNAS | 1019.84681 | -0.401078524 | 0.26911655 | -1.4903525 | 0.13613158 | 0.5601128 |
| NELFCD | 123.728506 | 0.051260193 | 0.19440427 | 0.26367832 | 0.7920278 | 0.95839852 |
| CTSZ | 132.534707 | -0.544974501 | 0.33237241 | -1.6396502 | 0.10107791 | 0.49726048 |
| ATP5F1E | 543.092586 | -0.574661439 | 0.46517988 | -1.2353532 | 0.21669909 | 0.67424993 |
| PRELID3B | 64.4485073 | 0.379848063 | 0.39917148 | 0.95159119 | 0.34130435 | 0.77954601 |
| FAM217B | 31.9054984 | -0.632827214 | 0.40386778 | -1.5669168 | 0.11713414 | 0.53233555 |
| MIR646HG | 198.044832 | 0.398443727 | 1.05392404 | 0.37805735 | 0.70538799 | 0.93241442 |
| TAF4 | 125.894127 | 0.15203233 | 0.26279634 | 0.57851768 | 0.56291466 | 0.8927792 |
| LSM14B | 40.6400833 | -0.5257797 | 0.31561213 | -1.6659046 | 0.09573242 | 0.48710328 |
| PSMA7 | 333.950501 | -0.183881207 | 0.47304334 | -0.3887196 | 0.6974836 | 0.93151881 |
| SS18L1 | 101.57399 | 0.037463787 | 0.24175537 | 0.15496569 | 0.87684837 | 0.97640529 |
| MTG2 | 107.285465 | 0.163104775 | 0.31646526 | 0.51539551 | 0.60627667 | 0.90493874 |
| OSBPL2 | 203.759642 | -0.200642206 | 0.17981317 | -1.1158371 | 0.26449186 | 0.71989632 |
| ADRM1 | 46.1934825 | -0.173621821 | 0.44147694 | -0.3932749 | 0.69411642 | 0.93105058 |
| LAMA5 | 227.778986 | 0.052602938 | 0.32786482 | 0.16044093 | 0.87253375 | 0.97622753 |
| RPS21 | 49.7247394 | 0.222551586 | 0.35356878 | 0.62944355 | 0.52905871 | 0.8788295 |
| SLCO4A1 | 177.88349 | 0.280086755 | 1.29084685 | 0.21697907 | 0.82822465 | 0.96592414 |
| NTSR1 | 42.4217955 | -1.008861139 | 0.61310365 | -1.6454985 | 0.09986705 | 0.4953766 |
| MRGBP | 46.3358005 | -0.151721436 | 0.33452401 | -0.4535442 | 0.6501569 | 0.91803345 |
| OGFR | 119.789309 | -0.569145694 | 0.35357042 | -1.6097096 | 0.10746127 | 0.51064149 |
| TCFL5 | 76.8386274 | -0.478205 | 0.32710963 | -1.4619105 | 0.14376574 | 0.57324898 |
| DIDO1 | 298.352878 | -0.044533953 | 0.15202337 | -0.2929415 | 0.76956687 | 0.95482817 |
| GID8 | 132.18796 | 0.228631558 | 0.2287481 | 0.99949052 | 0.31755713 | 0.76155577 |
| YTHDF1 | 96.7378227 | 0.007233093 | 0.23367451 | 0.03095371 | 0.97530645 | 0.9939318 |
| ARFGAP1 | 89.3558488 | 0.077562666 | 0.37437307 | 0.20718014 | 0.83586918 | 0.96757829 |
| PPDPF | 941.663333 | -0.366746313 | 0.45849066 | -0.7998992 | 0.4237692 | 0.82774157 |
| GMEB2 | 66.8778559 | -0.085907832 | 0.28432844 | -0.3021429 | 0.7625431 | 0.95374766 |
| ARFRP1 | 76.0310023 | 0.17117006 | 0.3584669 | 0.4775059 | 0.63300192 | 0.91415622 |

| | | | | | | |
|---|---|---|---|---|---|---|
| SLC2A4RG | 144.329158 | -0.329194549 | 0.36651732 | -0.8981692 | 0.36909538 | 0.79532503 |
| ZBTB46 | 88.9046496 | -0.19020024 | 0.3656369 | -0.5201889 | 0.60293196 | 0.90369786 |
| TPD52L2 | 131.20813 | 0.007877677 | 0.27134753 | 0.02903169 | 0.97683932 | 0.99460933 |
| DNAJC5 | 308.45155 | -0.271949748 | 0.24703266 | -1.1008656 | 0.27095517 | 0.72343657 |
| UCKL1 | 72.6346848 | 0.882380509 | 0.27730475 | 3.18198846 | 0.00146268 | 0.05060096 |
| ZNF512B | 53.4986063 | 0.323090634 | 0.42287935 | 0.76402557 | 0.444852 | 0.83925275 |
| PRPF6 | 171.093024 | -0.125669175 | 0.20487766 | -0.6133864 | 0.53962086 | 0.8833006 |
| TCEA2 | 61.135082 | 0.383951871 | 0.34890684 | 1.10044237 | 0.27113943 | 0.72343657 |
| PCMTD2 | 234.906202 | 0.010290244 | 0.30822319 | 0.03338569 | 0.97336702 | 0.9939318 |
| PLPP2 | 42.2336446 | -3.317893779 | 0.71764937 | -4.6232798 | 3.78E-06 | 0.00087967 |
| MIER2 | 44.4184983 | 0.077284807 | 0.29880128 | 0.25864952 | 0.79590568 | 0.95948555 |
| TPGS1 | 74.1577767 | -0.303589559 | 0.47445669 | -0.6398678 | 0.52225855 | 0.87670505 |
| CDC34 | 72.3584044 | 0.029947101 | 0.32219211 | 0.09294797 | 0.9259449 | 0.98729553 |
| BSG | 408.367711 | -0.233385032 | 0.37269106 | -0.6262158 | 0.5311734 | 0.88023377 |
| RNF126 | 66.2485765 | -0.198793017 | 0.34998312 | -0.5680074 | 0.57002992 | 0.89684936 |
| PTBP1 | 224.852949 | 0.021146286 | 0.28394315 | 0.07447366 | 0.9406335 | 0.98855185 |
| MED16 | 93.0463996 | -0.035343532 | 0.33503448 | -0.1054922 | 0.91598524 | 0.98520052 |
| R3HDM4 | 80.5975558 | -0.475772663 | 0.35837128 | -1.3275971 | 0.18431126 | 0.63497106 |
| ARID3A | 305.940393 | -0.861174946 | 0.49501752 | -1.7396858 | 0.08191421 | 0.45535449 |
| WDR18 | 53.98206 | 0.101200835 | 0.40152284 | 0.25204253 | 0.80100819 | 0.96063297 |
| TMEM259 | 193.6131 | -0.024305806 | 0.2915537 | -0.0833665 | 0.93356014 | 0.98822345 |
| CNN2 | 61.6716184 | -0.1902661 | 0.50324648 | -0.3780774 | 0.70537313 | 0.93241442 |
| ABCA7 | 34.5967597 | 0.270952018 | 0.45969338 | 0.58941901 | 0.55558023 | 0.89081623 |
| POLR2E | 131.409707 | -0.051097848 | 0.39739811 | -0.128581 | 0.8976892 | 0.98312332 |
| GPX4 | 587.685435 | -0.343362919 | 0.47443708 | -0.723727 | 0.46923336 | 0.8522644 |
| SBNO2 | 191.262112 | -1.394919053 | 0.42750166 | -3.2629559 | 0.00110257 | 0.04225907 |
| STK11 | 160.244359 | 0.055619765 | 0.20897673 | 0.26615291 | 0.79012145 | 0.95812572 |
| ATP5F1D | 481.686315 | 0.234770658 | 0.49794244 | 0.47148152 | 0.63729692 | 0.91466334 |
| MIDN | 574.993342 | 0.166915989 | 0.7763101 | 0.215012 | 0.82975796 | 0.96642181 |
| CIRBP | 564.974638 | 0.145167379 | 0.35564826 | 0.40817683 | 0.68314386 | 0.92818681 |
| C19orf24 | 85.6833985 | -0.14542324 | 0.38415201 | -0.3785565 | 0.70501723 | 0.93241442 |
| MUM1 | 117.330299 | -0.163802272 | 0.22821033 | -0.7177689 | 0.4728998 | 0.85298656 |
| NDUFS7 | 129.984867 | -0.028239684 | 0.44532667 | -0.0634134 | 0.94943731 | 0.98977092 |
| GAMT | 51.2149778 | 0.027028204 | 0.62858087 | 0.04299877 | 0.96570251 | 0.99271817 |
| DAZAP1 | 213.389155 | 0.090335096 | 0.23598014 | 0.38280805 | 0.70186209 | 0.93215897 |
| RPS15 | 1061.01694 | -0.402188088 | 0.46957434 | -0.856495 | 0.39172402 | 0.80866481 |
| C19orf25 | 108.526249 | -0.483903829 | 0.22617333 | -2.1395265 | 0.03239305 | 0.3033773 |
| MBD3 | 83.9225815 | 0.238959793 | 0.42581322 | 0.56118453 | 0.57467175 | 0.89807676 |
| TCF3 | 69.1137384 | -0.180477705 | 0.4269697 | -0.4226944 | 0.67251824 | 0.92502554 |
| REXO1 | 54.6582796 | 0.177801558 | 0.32460518 | 0.54774713 | 0.58386555 | 0.90085422 |
| ABHD17A | 107.952386 | 0.057609177 | 0.38143886 | 0.15103122 | 0.87995109 | 0.97740892 |
| SCAMP4 | 169.57804 | 0.007030892 | 0.28607288 | 0.02457728 | 0.98039214 | 0.99482084 |
| CSNK1G2 | 149.159554 | -0.025138582 | 0.30745121 | -0.0817645 | 0.93483402 | 0.98822345 |
| BTBD2 | 131.545577 | -0.044506024 | 0.34282335 | -0.129822 | 0.89670723 | 0.98302816 |
| MKNK2 | 146.403758 | 0.53267801 | 0.43543637 | 1.22331998 | 0.22120886 | 0.67871976 |
| MOB3A | 82.6925469 | -0.432802513 | 0.36143875 | -1.1974436 | 0.2311337 | 0.69089594 |
| AP3D1 | 205.010965 | -0.217321656 | 0.22220706 | -0.9780142 | 0.32806731 | 0.77410148 |
| DOT1L | 139.685005 | 0.096309221 | 0.5314338 | 0.18122524 | 0.85619078 | 0.97182813 |
| PLEKHJ1 | 185.506039 | -0.185166315 | 0.3683821 | -0.5026474 | 0.61521218 | 0.90792412 |
| SF3A2 | 33.0275086 | -0.426509694 | 0.29473335 | -1.4471036 | 0.14786791 | 0.57952643 |
| LSM7 | 196.91554 | -0.048940538 | 0.38707459 | -0.126437 | 0.89938604 | 0.98312332 |
| SPPL2B | 145.804628 | 0.168627334 | 0.23291877 | 0.72397487 | 0.46908116 | 0.8522644 |
| TIMM13 | 155.524023 | -0.141716404 | 0.46242007 | -0.3064668 | 0.75924925 | 0.95246973 |
| LMNB2 | 36.7854901 | 0.195182806 | 0.44752985 | 0.4361336 | 0.6627398 | 0.9229791 |
| GADD45B | 309.396575 | 0.853829741 | 0.61527047 | 1.38773073 | 0.16521905 | 0.6066943 |
| GNG7 | 245.044747 | 1.086068289 | 0.40269172 | 2.69702166 | 0.00699627 | 0.13379732 |
| SGTA | 94.0185142 | 0.088164873 | 0.20456955 | 0.43097749 | 0.66648474 | 0.92396246 |
| THOP1 | 44.0671287 | 0.738741011 | 0.41282628 | 1.78947184 | 0.07353886 | 0.4369528 |
| ZNF555 | 44.8963569 | 0.151294633 | 0.44470066 | 0.34021679 | 0.73369327 | 0.94163693 |
| TLE2 | 288.340321 | 0.58477473 | 0.41237783 | 1.4180557 | 0.1561745 | 0.59366792 |
| AES | 656.039932 | -0.185437055 | 0.30039466 | -0.6173114 | 0.53702933 | 0.88214222 |
| GNA11 | 164.024053 | 0.060117642 | 0.30058911 | 0.1999994 | 0.84148105 | 0.96866282 |
| NCLN | 76.0238381 | 0.213615526 | 0.26144987 | 0.81704202 | 0.41390442 | 0.82456685 |
| NFIC | 617.941706 | -0.19201915 | 0.23183756 | -0.8282487 | 0.40752968 | 0.82018309 |
| SMIM24 | 57.9904972 | -0.353637064 | 0.48634198 | -0.7271366 | 0.46714227 | 0.85112152 |
| FZR1 | 72.245423 | 0.546356597 | 0.22044611 | 2.4784134 | 0.01319681 | 0.18489654 |
| MFSD12 | 135.349756 | -0.651655878 | 0.56400411 | -1.1554098 | 0.24792265 | 0.7041272 |
| HMG20B | 90.7488399 | -0.256536376 | 0.33504052 | -0.7656876 | 0.4438622 | 0.83873671 |
| PIP5K1C | 88.3374865 | -0.258102188 | 0.28388848 | -0.9091675 | 0.3632617 | 0.79163705 |
| TJP3 | 144.351751 | 0.200750475 | 0.45588767 | 0.44035075 | 0.65968309 | 0.92284375 |
| MRPL54 | 102.627238 | -0.378009068 | 0.51235305 | -0.7377902 | 0.46064194 | 0.8467861 |
| DAPK3 | 54.2523804 | 0.000331277 | 0.3363174 | 0.00098501 | 0.99921407 | 0.99992293 |
| EEF2 | 765.589234 | -0.231578703 | 0.37914201 | -0.6107967 | 0.54133415 | 0.88408632 |
| PIAS4 | 57.6902661 | 0.058386533 | 0.29407352 | 0.198544 | 0.84261947 | 0.96876046 |
| ZBTB7A | 197.660188 | -0.454269027 | 0.36637694 | -1.2398953 | 0.21501414 | 0.67132603 |
| MAP2K2 | 192.205356 | -0.238939265 | 0.29590187 | -0.8074949 | 0.41938139 | 0.82694802 |
| SIRT6 | 33.8948093 | -0.203255156 | 0.30931329 | -0.6571174 | 0.51110541 | 0.87138909 |
| YJU2 | 37.9746872 | 0.062111708 | 0.29697454 | 0.20914826 | 0.83433251 | 0.96749268 |
| STAP2 | 48.0207799 | -0.606729851 | 0.37091785 | -1.6357527 | 0.10189136 | 0.4989423 |
| SH3GL1 | 174.263605 | -0.306929018 | 0.42152735 | -0.7281355 | 0.46653066 | 0.85057334 |
| UBXN6 | 145.440305 | -0.318162055 | 0.40874394 | -0.7783897 | 0.43633933 | 0.83393108 |
| HDGFL2 | 86.4614461 | 0.221944937 | 0.2783666 | 0.79731167 | 0.42527004 | 0.82774157 |
| PLIN5 | 59.5062394 | -0.402794833 | 0.46334814 | -0.8693136 | 0.38467563 | 0.80561637 |
| MYDGF | 150.876381 | -0.675399481 | 0.4046873 | -1.6689416 | 0.09512895 | 0.48627909 |
| DPP9 | 88.5447748 | -0.500230478 | 0.27627769 | -1.8106076 | 0.07020162 | 0.42898715 |
| PLIN3 | 93.7733943 | -1.070394765 | 0.30140462 | -3.5513549 | 0.00038325 | 0.02099512 |
| KDM4B | 258.133659 | -0.925694574 | 0.23971636 | -3.8616245 | 0.00011264 | 0.00944337 |
| PTPRS | 220.26987 | 0.371948944 | 0.16305467 | 2.28113022 | 0.02254074 | 0.25166501 |
| TINCR | 58.5170942 | -0.677259346 | 0.35404844 | -1.9129002 | 0.05576083 | 0.39368206 |
| SAFB2 | 222.261669 | 0.269734674 | 0.24426342 | 1.1042778 | 0.26947265 | 0.72322263 |
| SAFB | 268.122266 | 0.244543387 | 0.1559461 | 1.56812761 | 0.11685135 | 0.53183794 |
| RPL36 | 663.983445 | -0.252035387 | 0.42543049 | -0.5924244 | 0.55356646 | 0.89005469 |
| C19orf70 | 190.667961 | -0.270621172 | 0.39952951 | -0.6773496 | 0.49818414 | 0.86583246 |
| LONP1 | 155.198964 | -0.582403388 | 0.32738093 | -1.7789778 | 0.07524341 | 0.43979415 |
| CAPS | 331.818379 | 0.300874047 | 0.60681072 | 0.4958285 | 0.62001542 | 0.90914396 |
| RANBP3 | 97.991845 | 0.051349948 | 0.22773059 | 0.22548551 | 0.8216016 | 0.96502145 |
| RFX2 | 179.433268 | -0.652731332 | 0.67075394 | -0.9731308 | 0.33048829 | 0.77618491 |
| MLLT1 | 162.334159 | 0.886656181 | 0.38342855 | 2.3124417 | 0.02075335 | 0.24280431 |
| CLPP | 95.4748495 | 0.051454536 | 0.4521875 | 0.11379027 | 0.90940406 | 0.98441374 |
| ALKBH7 | 133.030598 | 0.008445995 | 0.58216605 | 0.01450788 | 0.98842479 | 0.9964791 |
| GTF2F1 | 99.852908 | -0.291034365 | 0.25124306 | -1.1583777 | 0.24670992 | 0.70292817 |
| KHSRP | 121.375057 | -0.127406757 | 0.32566594 | -0.3912192 | 0.69563524 | 0.93130502 |
| SLC25A23 | 127.942519 | -0.242466545 | 0.30317972 | -0.7997453 | 0.4238584 | 0.82774157 |
| CRB3 | 52.5242488 | 0.060986549 | 0.41828649 | 0.14580091 | 0.88407856 | 0.9788678 |
| AC010503.4 | 61.1446588 | 0.059652632 | 0.34630678 | 0.17225372 | 0.86323806 | 0.97454863 |
| TRIP10 | 118.261459 | -0.010256043 | 0.32517835 | -0.0315397 | 0.97483909 | 0.9939318 |
| SH2D3A | 154.404399 | 0.338075365 | 0.48560606 | 0.69619265 | 0.48630819 | 0.85912348 |
| ZNF557 | 34.1527769 | -0.142621406 | 0.33256821 | -0.4288486 | 0.66803342 | 0.92422713 |
| INSR | 859.006138 | -0.040199447 | 0.32950415 | -0.1219998 | 0.90289916 | 0.98320957 |
| PEX11G | 30.8526406 | -0.420527873 | 0.33966699 | -1.2380593 | 0.21569408 | 0.67215996 |
| ZNF358 | 69.4748036 | -0.04620347 | 0.43120486 | -0.1071497 | 0.91467022 | 0.98480378 |
| PNPLA6 | 105.619689 | 0.450516879 | 0.28149035 | 1.60047009 | 0.10949434 | 0.51481816 |
| CAMSAP3 | 81.7793093 | 0.248749878 | 0.24324116 | 1.02264714 | 0.30647471 | 0.7514869 |

| | | | | | | |
|---|---|---|---|---|---|---|
| XAB2 | 41.5615416 | 0.108153963 | 0.35208438 | 0.30718194 | 0.75870489 | 0.9521704 |
| STXBP2 | 61.764148 | 0.065893538 | 0.42143215 | 0.15635622 | 0.87575224 | 0.97622753 |
| EVI5L | 51.7905372 | 0.022178537 | 0.27959726 | 0.07932315 | 0.93677559 | 0.98822345 |
| MAP2K7 | 111.118356 | -0.552756715 | 0.32021034 | -1.7262301 | 0.08430603 | 0.46162908 |
| CTXN1 | 31.7768154 | -1.202204349 | 0.6821835 | -1.7622888 | 0.07802052 | 0.44731172 |
| TIMM44 | 67.9205556 | -0.152494934 | 0.27521574 | -0.5540923 | 0.57951565 | 0.89989814 |
| ELAVL1 | 230.637981 | 0.17157705 | 0.21472092 | 0.79907001 | 0.42424982 | 0.82774157 |
| CERS4 | 127.202819 | -0.389970355 | 0.36364447 | -1.0723946 | 0.28354285 | 0.73371088 |
| CD320 | 53.0095202 | -0.6056975 | 0.46643111 | -1.2985787 | 0.19408856 | 0.64778601 |
| RPS28 | 206.11585 | 0.103288413 | 0.34758611 | 0.29715921 | 0.76634496 | 0.95398103 |
| ANGPTL4 | 498.450023 | -0.930107227 | 1.55683761 | -0.5974337 | 0.55021787 | 0.88916026 |
| RAB11B | 171.802889 | 0.061601538 | 0.34829223 | 0.17686739 | 0.85961255 | 0.97244655 |
| 2-Mar | 62.4949263 | -0.360614146 | 0.31705734 | -1.1373783 | 0.25538017 | 0.71266224 |
| HNRNPM | 508.849956 | 0.482959821 | 0.25636843 | 1.88385057 | 0.05958519 | 0.40313286 |
| ZNF558 | 36.331586 | -0.38758844 | 0.3191598 | -1.2144024 | 0.22459408 | 0.68208993 |
| ZNF317 | 46.5474129 | -0.148762651 | 0.25896029 | -0.5744613 | 0.56565571 | 0.89394085 |
| ZNF559-ZNF | 57.8969215 | 0.085528549 | 0.40871237 | 0.20926342 | 0.83424261 | 0.96749268 |
| ZNF266 | 94.0573034 | -0.28576453 | 0.21235325 | -1.3457036 | 0.17839813 | 0.62759322 |
| ZNF426 | 63.9775832 | 0.021122713 | 0.24804156 | 0.08515796 | 0.93213581 | 0.98822345 |
| ZNF121 | 102.618979 | 0.191848029 | 0.26525179 | 0.72326762 | 0.46951548 | 0.85232523 |
| ZNF561 | 64.9622467 | 0.220990026 | 0.28963103 | 0.76300535 | 0.4454602 | 0.83944393 |
| ZNF562 | 170.886391 | 0.096035294 | 0.23571755 | 0.40741682 | 0.68370188 | 0.92837467 |
| ZNF846 | 171.297409 | 0.432832554 | 0.29490605 | 1.46769643 | 0.1421867 | 0.57093391 |
| FBXL12 | 57.5117495 | 0.153787465 | 0.23273752 | 0.66077641 | 0.50875571 | 0.87094607 |
| UBL5 | 180.658814 | -0.275361974 | 0.50290774 | -0.5475397 | 0.58400797 | 0.90085422 |
| PIN1 | 108.025128 | -0.125028295 | 0.45303096 | -0.2759818 | 0.78256205 | 0.95688565 |
| C19orf66 | 137.575512 | -0.351573405 | 0.48485217 | -0.7251146 | 0.46838171 | 0.85161077 |
| EIF3G | 98.9213141 | -0.260490414 | 0.39873198 | -0.653297 | 0.51356481 | 0.8721697 |
| DNMT1 | 64.8301968 | 0.21912928 | 0.38975312 | 0.56222586 | 0.57396215 | 0.89804767 |
| MRPL4 | 96.0690422 | -0.039061892 | 0.36419423 | -0.1072557 | 0.91458616 | 0.98480378 |
| TYK2 | 69.3573204 | 0.119502884 | 0.2742873 | 0.43568508 | 0.66306523 | 0.9229791 |
| CDC37 | 227.165814 | -0.201656214 | 0.32713062 | -0.6164394 | 0.53760454 | 0.88239554 |
| PDE4A | 68.8699036 | 0.13926061 | 0.57549252 | 0.24198509 | 0.80879172 | 0.96245698 |
| KEAP1 | 57.9511695 | -0.494628939 | 0.41515408 | -1.1914346 | 0.23348302 | 0.69247746 |
| ATG4D | 59.5728649 | 0.383701409 | 0.2541105 | 1.50997858 | 0.13104889 | 0.55395564 |
| KRI1 | 53.5714029 | -0.288664626 | 0.25771387 | -1.1200974 | 0.26267227 | 0.71989632 |
| AP1M2 | 179.677942 | -0.203533931 | 0.30966835 | -0.6572642 | 0.51101103 | 0.87138909 |
| SLC44A2 | 270.283013 | 0.316051099 | 0.42543859 | 0.74288301 | 0.45755251 | 0.84409989 |
| ILF3-DT | 93.9168249 | -0.454967218 | 0.42504396 | -1.0704004 | 0.28443912 | 0.73476311 |
| ILF3 | 287.275242 | 0.251983147 | 0.28173518 | 0.89439717 | 0.37110942 | 0.79539603 |
| QTRT1 | 81.4303853 | 0.72156519 | 0.25802406 | 2.79650355 | 0.00516588 | 0.11481456 |
| DNM2 | 1069.98461 | -0.20663584 | 0.40552543 | -0.5095509 | 0.61036614 | 0.90616761 |
| TMED1 | 74.7801215 | -0.534875844 | 0.37161124 | -1.4393425 | 0.15005351 | 0.5835105 |
| CARM1 | 66.6625978 | -0.12044633 | 0.32644686 | -0.3689615 | 0.7121564 | 0.9348115 |
| YIPF2 | 61.3473741 | -0.212817065 | 0.2958522 | -0.7193358 | 0.47193407 | 0.85244471 |
| TIMM29 | 53.0476405 | 0.121850297 | 0.32562885 | 0.37419994 | 0.70825557 | 0.9332187 |
| SMARCA4 | 433.645834 | -0.335205151 | 0.34467071 | -0.9725374 | 0.33078327 | 0.77618491 |
| LDLR | 193.87897 | -1.45557191 | 0.87826304 | -1.6573303 | 0.09745272 | 0.49030463 |
| KANK2 | 107.929433 | 0.74764635 | 0.38509686 | 1.94145012 | 0.0522037 | 0.38343909 |
| DOCK6 | 250.752562 | -0.710130153 | 0.44221718 | -1.6058403 | 0.10830901 | 0.51253572 |
| RAB3D | 97.3903784 | -0.201807325 | 0.32295598 | -0.6248756 | 0.53205268 | 0.88045351 |
| EPOR | 182.321885 | 0.435362412 | 0.47007847 | 0.92614838 | 0.35436887 | 0.78548769 |
| RGL3 | 780.235815 | 0.036658949 | 0.35315902 | 0.10380295 | 0.91732572 | 0.98520052 |
| CCDC151 | 37.6388413 | -0.483048744 | 0.51698263 | -0.9343617 | 0.35011737 | 0.78223232 |
| PRKCSH | 176.891035 | -0.254490387 | 0.2993128 | -0.8502489 | 0.39518671 | 0.81175661 |
| ECSIT | 94.5044761 | 0.206549831 | 0.31146888 | 0.66314757 | 0.50723605 | 0.87094607 |
| ELOF1 | 53.8087159 | -0.297563869 | 0.42988958 | -0.6921867 | 0.48882007 | 0.86014074 |
| ZNF627 | 86.0644333 | -0.203427881 | 0.39072055 | -0.520648 | 0.60261199 | 0.90369786 |
| AC008543.1 | 42.9292633 | -0.568317168 | 0.59721453 | -0.9516131 | 0.34129324 | 0.77954601 |
| ZNF823 | 108.298804 | -0.068778193 | 0.29948314 | -0.2296563 | 0.81835884 | 0.96407382 |
| ZNF441 | 83.8086777 | -0.520735719 | 0.4225711 | -1.2323032 | 0.21783585 | 0.67542003 |
| ZNF440 | 118.73986 | -1.175846938 | 0.42702526 | -2.753577 | 0.00589479 | 0.12330819 |
| ZNF439 | 118.931611 | -0.386463901 | 0.42242722 | -0.9148651 | 0.36026248 | 0.78892604 |
| ZNF69 | 207.487639 | 0.024717418 | 0.47879316 | 0.05162442 | 0.95882796 | 0.99119981 |
| ZNF433 | 40.0905839 | -0.228815072 | 0.32815165 | -0.6972845 | 0.48562474 | 0.85912348 |
| ZNF844 | 77.3562755 | -0.264296853 | 0.41341519 | -0.6393013 | 0.52262697 | 0.87688817 |
| ZNF788P | 44.5671313 | -0.288886042 | 0.41239991 | -0.7004998 | 0.48361525 | 0.85829223 |
| ZNF136 | 60.7442576 | -0.008623019 | 0.24575819 | -0.0350874 | 0.97201004 | 0.9939318 |
| ZNF44 | 580.826324 | 0.107582013 | 0.22959804 | 0.46856678 | 0.63937933 | 0.91521016 |
| ZNF563 | 37.9188054 | -1.369657314 | 0.50019195 | -2.7382634 | 0.00617646 | 0.12643442 |
| AC008758.5 | 43.3065511 | -0.618309008 | 0.53913942 | -1.1468444 | 0.25144595 | 0.70813507 |
| ZNF490 | 81.0432803 | -0.044301924 | 0.37289776 | -0.1188045 | 0.90543026 | 0.98320957 |
| ZNF791 | 143.335801 | -0.49993038 | 0.20325705 | -2.4595967 | 0.01390932 | 0.19104807 |
| DHPS | 47.4051704 | 0.348349001 | 0.40311239 | 0.86414859 | 0.38750627 | 0.8065026 |
| TNPO2 | 126.206527 | -0.144344202 | 0.2607922 | -0.5534836 | 0.57993232 | 0.89995373 |
| TRIR | 292.75303 | 0.204404396 | 0.48728394 | 0.41947698 | 0.67486757 | 0.92634082 |
| ASNA1 | 82.1842504 | -0.473017105 | 0.47943085 | -0.9866222 | 0.32382789 | 0.76937573 |
| HOOK2 | 373.776022 | -0.540905847 | 0.26064135 | -2.0752879 | 0.03795986 | 0.32715403 |
| JUNB | 553.91758 | 1.450608861 | 0.9522818 | 1.52329789 | 0.12768419 | 0.54791812 |
| DNASE2 | 51.4830939 | -0.430755377 | 0.44512849 | -0.9677102 | 0.33318912 | 0.777038 |
| GCDH | 61.8681584 | -0.243528438 | 0.34522615 | -0.7054171 | 0.48055069 | 0.85600365 |
| FARSA | 64.2548348 | -0.144677967 | 0.32611994 | -0.4436342 | 0.65730706 | 0.92160948 |
| CALR | 424.309794 | -0.093694024 | 0.38804288 | -0.2414528 | 0.80920423 | 0.96245698 |
| AC092069.1 | 77.6175592 | 0.346516139 | 0.45082221 | 0.76863147 | 0.44211212 | 0.83770294 |
| RAD23A | 127.315943 | -0.224797155 | 0.35411438 | -0.6348151 | 0.52554902 | 0.87791848 |
| GADD45GIP1 | 256.197798 | -0.430683764 | 0.51262726 | -0.8401499 | 0.40082432 | 0.8162653 |
| NFIX | 83.178793 | -0.012883908 | 0.44605326 | -0.0288842 | 0.97695692 | 0.99460933 |
| NACC1 | 67.14007 | -0.471222759 | 0.30715141 | -1.5341709 | 0.12498761 | 0.54337567 |
| STX10 | 53.9018217 | -0.340230241 | 0.32755659 | -1.0386915 | 0.29894825 | 0.74594723 |
| IER2 | 429.434472 | 1.811578538 | 0.76725752 | 2.36110888 | 0.01822038 | 0.22491117 |
| CCDC130 | 189.921044 | -0.30128404 | 0.26726871 | -1.1272702 | 0.25962827 | 0.7168415 |
| MRI1 | 108.175728 | -0.069809921 | 0.36813457 | -0.1896315 | 0.84959788 | 0.96991131 |
| C19orf53 | 265.409163 | -0.300959979 | 0.5301619 | -0.5676756 | 0.57025526 | 0.89706922 |
| ZSWIM4 | 68.9535811 | -1.130782284 | 0.31253989 | -3.6180414 | 0.00029684 | 0.01737929 |
| AC020916.1 | 86.0581369 | 1.420295807 | 0.81686933 | 1.73870624 | 0.08208645 | 0.45535449 |
| CC2D1A | 120.667269 | -0.039251198 | 0.23326726 | -0.1682671 | 0.86637318 | 0.9753562 |
| DCAF15 | 44.7735112 | -0.010799949 | 0.34554836 | -0.0312545 | 0.97506656 | 0.9939318 |
| RFX1 | 87.3813873 | 0.146954061 | 0.28340127 | 0.51853706 | 0.60408361 | 0.90381187 |
| PALM3 | 115.083604 | 0.404300401 | 0.19934134 | 2.02818137 | 0.04254174 | 0.3484612 |
| MISP3 | 31.8516344 | -0.640797899 | 0.48648132 | -1.3172097 | 0.18776836 | 0.64035097 |
| SAMD1 | 56.1819039 | -0.097986578 | 0.4160488 | -0.235517 | 0.81380746 | 0.96288868 |
| PRKACA | 97.2939448 | -0.248344386 | 0.25706929 | -0.9660601 | 0.33401411 | 0.77770893 |
| ADGRL1 | 104.598006 | 0.099266819 | 0.29564181 | 0.33576719 | 0.73704643 | 0.9429136 |
| DDX39A | 37.0361627 | -0.206474861 | 0.4561447 | -0.4526521 | 0.65079927 | 0.91825484 |
| PKN1 | 187.200852 | -0.413501054 | 0.34061308 | -1.2139905 | 0.22475134 | 0.68208993 |
| PTGER1 | 656.749057 | 0.643396471 | 0.82108515 | 0.78359287 | 0.43327904 | 0.83179416 |
| GIPC1 | 491.973442 | -0.172084676 | 0.32539784 | -0.5288439 | 0.59691371 | 0.90369786 |
| DNAJB1 | 137.146491 | 0.79625639 | 0.3923338 | 2.0295381 | 0.04240351 | 0.34798255 |
| TECR | 133.263068 | 0.222780705 | 0.24754995 | 0.89994243 | 0.36815089 | 0.79501778 |
| NDUFB7 | 296.28826 | 0.048123892 | 0.39963375 | 0.12041999 | 0.90415046 | 0.98320957 |
| ADGRE3 | 31.5133238 | -0.226160103 | 0.36579282 | -0.6182738 | 0.5363949 | 0.88137638 |
| ZNF333 | 43.2397857 | -0.169164392 | 0.32969351 | -0.5130959 | 0.60788423 | 0.90514346 |
| ILVBL | 106.884786 | -0.224054886 | 0.2936826 | -0.7629151 | 0.44551404 | 0.83944393 |

| | | | | | | |
|---|---|---|---|---|---|---|
| NOTCH3 | 170.73236 | -0.406645345 | 0.47394475 | -0.8580016 | 0.39089159 | 0.80778625 |
| BRD4 | 550.414365 | 0.127121089 | 0.20192399 | 0.62954921 | 0.52898956 | 0.8788295 |
| AKAP8 | 139.84579 | -0.006706186 | 0.1845305 | -0.0363419 | 0.97100975 | 0.9939318 |
| AKAP8L | 373.827475 | 0.301200222 | 0.27945721 | 1.07780445 | 0.28112103 | 0.73069157 |
| WIZ | 47.447396 | 0.07764517 | 0.28250744 | 0.27484292 | 0.78343692 | 0.9568898 |
| TPM4 | 1061.9397 | 0.579679602 | 0.56659219 | 1.02309846 | 0.30626129 | 0.75140194 |
| RAB8A | 93.8120328 | 0.228242738 | 0.29689192 | 0.76877383 | 0.44202759 | 0.83770294 |
| FAM32A | 70.6286326 | -0.362413991 | 0.46384296 | -0.7813291 | 0.43460897 | 0.83266948 |
| AP1M1 | 55.50099 | 0.059743702 | 0.23506351 | 0.25415983 | 0.79937208 | 0.96043078 |
| EPS15L1 | 173.284308 | -0.159802656 | 0.30797253 | -0.5188861 | 0.60384019 | 0.90371579 |
| C19orf44 | 62.442834 | 0.400500173 | 0.30648009 | 1.30677388 | 0.19128952 | 0.64458858 |
| TMEM38A | 39.7433699 | -0.360956382 | 0.4296735 | -0.8400713 | 0.40086841 | 0.8162653 |
| SIN3B | 132.254395 | 0.098240217 | 0.2244896 | 0.4376159 | 0.66166474 | 0.92284375 |
| CPAMD8 | 391.58682 | -0.743858364 | 0.69676901 | -1.0675824 | 0.28570891 | 0.73607046 |
| MYO9B | 209.03692 | -0.241870386 | 0.30553307 | -0.7916341 | 0.42857408 | 0.8296433 |
| USE1 | 37.2554688 | -0.248476324 | 0.44629291 | -0.5567562 | 0.57769404 | 0.89916873 |
| OCEL1 | 41.822116 | -0.380584293 | 0.4435404 | -0.85806 | 0.39085931 | 0.80778625 |
| NR2F6 | 135.777163 | 0.114947682 | 0.39088704 | 0.2940688 | 0.76870533 | 0.95453142 |
| ABHD8 | 50.50281 | -0.265859435 | 0.38341153 | -0.6934049 | 0.48805549 | 0.86006752 |
| MRPL34 | 141.428931 | -0.12985143 | 0.4822386 | -0.269268 | 0.78772345 | 0.95801395 |
| DDA1 | 50.8010178 | -0.097442125 | 0.36425069 | -0.2675139 | 0.78907351 | 0.95801395 |
| BST2 | 112.242384 | -1.852495013 | 1.29727863 | -1.4279855 | 0.15329605 | 0.58860244 |
| MVB12A | 98.5947995 | -0.058697969 | 0.33322218 | -0.1761526 | 0.86017402 | 0.97276613 |
| SLC27A1 | 63.7853262 | -0.331653562 | 0.36956466 | -0.8974169 | 0.36949652 | 0.79532503 |
| AC010618.3 | 61.2844691 | -0.114784574 | 0.36132692 | -0.3176751 | 0.75073138 | 0.94794781 |
| PGLS | 163.829346 | -0.397659988 | 0.421338 | -0.9438028 | 0.34527042 | 0.78180304 |
| COLGALT1 | 49.7204291 | -0.398210735 | 0.45312019 | -0.8788192 | 0.3794993 | 0.80281644 |
| RPL18A | 60.5826442 | -1.433399973 | 0.56264954 | -2.5475893 | 0.01084701 | 0.16452817 |
| CCDC124 | 78.4675552 | -0.082180576 | 0.48558821 | -0.1692392 | 0.86560847 | 0.9750363 |
| ARRDC2 | 46.7100074 | -0.865023463 | 0.43898529 | -1.9705067 | 0.04878033 | 0.37451854 |
| MAST3 | 47.2132053 | -0.583764885 | 0.34418957 | -1.6960563 | 0.08987522 | 0.47358503 |
| IQCN | 50.2602984 | -0.330068135 | 0.60929842 | -0.5417184 | 0.58801255 | 0.90198646 |
| JUND | 1059.24472 | 0.838810337 | 0.73791198 | 1.13673495 | 0.2556491 | 0.71312287 |
| LSM4 | 271.190071 | -0.293612954 | 0.44182969 | -0.6645388 | 0.50634555 | 0.87094607 |
| PGPEP1 | 204.129667 | -0.325101718 | 0.29571871 | -1.0993613 | 0.27161049 | 0.72372882 |
| GDF15 | 1591.01861 | 0.045310477 | 0.82561534 | 0.05488086 | 0.95623338 | 0.9908068 |
| SSBP4 | 186.484475 | -0.077520755 | 0.27588258 | -0.2809918 | 0.77871665 | 0.95623387 |
| ISYNA1 | 75.2781587 | 0.80953372 | 0.66418001 | 1.21884686 | 0.22290232 | 0.68045917 |
| ELL | 148.019923 | -0.078377697 | 0.26294173 | -0.2980801 | 0.76564203 | 0.95389523 |
| FKBP8 | 204.306119 | -0.146112531 | 0.32699865 | -0.4468292 | 0.65499842 | 0.92051095 |
| KXD1 | 76.6464239 | -0.427688416 | 0.30281444 | -1.4123779 | 0.15783873 | 0.59566075 |
| UBA52 | 357.955943 | 0.31656923 | 0.45010621 | 0.70332118 | 0.48185562 | 0.85677755 |
| REX1BD | 94.5575026 | 0.429596796 | 0.453547 | 0.94719355 | 0.34354016 | 0.78148489 |
| CRTC1 | 350.728885 | 0.181054205 | 0.34080914 | 0.53124809 | 0.59524688 | 0.90323766 |
| UPF1 | 164.308099 | -0.14646386 | 0.26016335 | -0.5629688 | 0.57345611 | 0.89797476 |
| COPE | 127.589227 | 0.111621351 | 0.43319319 | 0.25767107 | 0.79666078 | 0.95961857 |
| DDX49 | 48.2324309 | -0.4299109 | 0.30789499 | -1.3962907 | 0.16262699 | 0.60211533 |
| SUGP2 | 316.51293 | -0.012266034 | 0.21321303 | -0.0575295 | 0.95412343 | 0.99060627 |
| SLC25A42 | 59.4907342 | -0.305125542 | 0.29650251 | -1.0290825 | 0.30344091 | 0.74953627 |
| TMEM161A | 50.6562461 | -0.113228626 | 0.33738599 | -0.3356056 | 0.7371683 | 0.9429136 |
| RFXANK | 50.1283359 | 0.286783959 | 0.33056699 | 0.86755172 | 0.38563979 | 0.8060156 |
| SUGP1 | 66.7277579 | 0.23294327 | 0.28610283 | 0.81419421 | 0.4155337 | 0.82487773 |
| MAU2 | 181.542581 | -0.02058562 | 0.21474144 | -0.0958624 | 0.92362989 | 0.98694936 |
| GATAD2A | 398.574386 | -0.234235989 | 0.16106458 | -1.4542986 | 0.14586355 | 0.57663146 |
| PBX4 | 64.6214595 | -0.073333374 | 0.28788164 | -0.2547345 | 0.7989282 | 0.96006966 |
| ATP13A1 | 60.6412705 | 0.133801276 | 0.30484551 | 0.43891502 | 0.66072311 | 0.92284375 |
| ZNF14 | 87.1045286 | 0.398678801 | 0.33360864 | 1.1950494 | 0.23206773 | 0.69186326 |
| AC011477.4 | 35.8987937 | -0.600615095 | 0.38219979 | -1.5714689 | 0.11607377 | 0.53066791 |
| ZNF253 | 93.8157612 | -0.069726077 | 0.34573682 | -0.2016739 | 0.8401717 | 0.96833625 |
| ZNF93 | 55.6738144 | -0.375931476 | 0.37079471 | -1.0138534 | 0.31065271 | 0.75435382 |
| ZNF682 | 85.8410635 | -0.412482061 | 0.48148866 | -0.8566807 | 0.39162135 | 0.80866481 |
| ZNF90 | 368.985419 | -0.197942058 | 0.32758111 | -0.6042536 | 0.54567506 | 0.88669883 |
| AC011447.3 | 281.762512 | -0.260093089 | 0.52853004 | -0.4921065 | 0.62264403 | 0.91033892 |
| ZNF486 | 230.744654 | -0.602763254 | 0.51044448 | -1.1808596 | 0.23765851 | 0.6939708 |
| AC008554.1 | 253.143424 | 0.603503179 | 0.64410105 | 0.93696972 | 0.34877414 | 0.78180304 |
| ZNF737 | 127.967858 | 0.016325942 | 0.32964455 | 0.0495259 | 0.9605002 | 0.9913918 |
| AC010636.2 | 58.5818706 | -1.004141178 | 0.45142465 | -2.2243827 | 0.02612271 | 0.27515522 |
| AC010329.1 | 43.0868877 | 0.274263956 | 0.42027193 | 0.6525869 | 0.51402263 | 0.87226916 |
| ZNF66 | 46.3957365 | 0.486252859 | 0.33482904 | 1.45224218 | 0.14643428 | 0.57722788 |
| ZNF85 | 50.0869781 | -0.07964594 | 0.55637565 | -0.1431514 | 0.88617063 | 0.97964855 |
| ZNF430 | 148.729107 | 0.148607035 | 0.22519906 | 0.6598919 | 0.5093232 | 0.87094607 |
| ZNF714 | 77.9180324 | -0.018300811 | 0.34337691 | -0.0532966 | 0.95749561 | 0.99106618 |
| ZNF431 | 270.90396 | 0.152658935 | 0.25675425 | 0.59457218 | 0.55212948 | 0.88924496 |
| ZNF708 | 177.311737 | -0.086847619 | 0.29762331 | -0.2918038 | 0.77043662 | 0.95482817 |
| ZNF738 | 50.3513358 | 0.29448794 | 0.40339783 | 0.73001864 | 0.46537879 | 0.84991999 |
| ZNF493 | 82.6309207 | 0.333275596 | 0.33807178 | 0.98581312 | 0.32422483 | 0.76970832 |
| AC010615.4 | 153.06871 | -0.789675057 | 0.74266086 | -1.0633051 | 0.28764363 | 0.73650262 |
| ZNF429 | 221.210309 | -0.128160238 | 0.33754329 | -0.3796853 | 0.704179 | 0.93241442 |
| AC123912.1 | 67.0575637 | -0.272024215 | 0.5849243 | -0.4650588 | 0.64188932 | 0.91643619 |
| ZNF100 | 201.416543 | -0.142186775 | 0.53978478 | -0.2634138 | 0.79223164 | 0.9585028 |
| ZNF43 | 306.188032 | -0.110893246 | 0.30574238 | -0.3627016 | 0.71682782 | 0.93573028 |
| ZNF208 | 612.562079 | -0.396055539 | 0.36623554 | -1.081423 | 0.27950899 | 0.72938673 |
| ZNF257 | 38.0725541 | -0.077294009 | 0.30126389 | -0.2565658 | 0.79751398 | 0.95991117 |
| ZNF676 | 110.618827 | -0.023646888 | 0.432093 | -0.0547264 | 0.95635645 | 0.9908068 |
| ZNF98 | 415.289861 | 0.004689697 | 0.36103114 | 0.01298973 | 0.98963599 | 0.9965778 |
| ZNF99 | 46.1054249 | -0.788003231 | 0.55392258 | -1.4225873 | 0.15485583 | 0.59100111 |
| ZNF728 | 42.4913045 | -0.147879605 | 0.55332503 | -0.2672563 | 0.78927182 | 0.95801395 |
| ZNF730 | 69.1943551 | -0.566878026 | 0.61248486 | -0.925538 | 0.35468611 | 0.78585845 |
| ZNF91 | 782.145089 | -0.60163653 | 0.25956865 | -2.317832 | 0.02045845 | 0.24101888 |
| ZNF675 | 182.356566 | 1.51445533 | 0.47534454 | 3.18601606 | 0.00144247 | 0.05037024 |
| AC139769.2 | 68.6632351 | 0.002562448 | 0.60111015 | 0.00426286 | 0.99659874 | 0.99926854 |
| ZNF726 | 41.236394 | -0.021484617 | 0.52302053 | -0.041078 | 0.96723374 | 0.99291246 |
| ZNF254 | 457.235576 | -0.3812714 | 0.26354298 | -1.4467143 | 0.14797695 | 0.57952643 |
| LINC00662 | 139.412509 | -0.408711921 | 0.3366283 | -1.2141342 | 0.22469649 | 0.68208993 |
| AC006504.5 | 141.706986 | 0.100804064 | 0.42813823 | 0.23544747 | 0.81386144 | 0.96288868 |
| UQCRFS1 | 237.927345 | -0.043265 | 0.50861357 | -0.0850646 | 0.93221005 | 0.98822345 |
| POP4 | 39.039043 | -0.165300266 | 0.32754942 | -0.5046575 | 0.61379943 | 0.90751853 |
| C19orf12 | 97.2889856 | -0.188157602 | 0.23613805 | -0.7968119 | 0.42556031 | 0.82774157 |
| URI1 | 627.17358 | 0.373026596 | 0.1851217 | 2.01503446 | 0.04390104 | 0.35300853 |
| ZNF507 | 120.079184 | 0.053871219 | 0.26597993 | 0.20253866 | 0.83949564 | 0.96833625 |
| DPY19L3 | 158.722639 | -0.048101277 | 0.19658018 | -0.2446904 | 0.80669618 | 0.96218371 |
| PDCD5 | 66.1803186 | -0.015471118 | 0.27700172 | -0.0558521 | 0.95545966 | 0.9908068 |
| ANKRD27 | 128.991573 | -0.237690602 | 0.27210834 | -0.8735146 | 0.38238265 | 0.8038857 |
| NUDT19 | 38.3655647 | 0.161507528 | 0.31235223 | 0.5170686 | 0.60510827 | 0.90428025 |
| CEP89 | 150.509552 | -0.338910368 | 0.21720282 | -1.5603405 | 0.11867943 | 0.53472074 |
| RHPN2 | 373.96021 | -0.924767662 | 0.34092658 | -2.7125127 | 0.00667752 | 0.13031298 |
| GPATCH1 | 51.6929855 | 0.193616208 | 0.31177637 | 0.62100989 | 0.53459312 | 0.88137638 |
| LRP3 | 91.3126842 | 0.558406821 | 0.31298377 | 1.78413987 | 0.07440094 | 0.43775442 |
| PEPD | 122.340202 | -0.696739817 | 0.34072011 | -2.0449037 | 0.04086435 | 0.34079725 |
| LSM14A | 677.347922 | 0.197897877 | 0.20989016 | 0.94286402 | 0.34575046 | 0.78180304 |
| KIAA0355 | 337.384632 | -0.12496461 | 0.20998975 | -0.5950986 | 0.55177755 | 0.88923362 |
| GPI | 437.482786 | -1.190529359 | 0.42900483 | -2.7750955 | 0.00551855 | 0.11921698 |
| UBA2 | 271.609291 | 0.09349172 | 0.23944752 | 0.39044764 | 0.69620556 | 0.93130502 |

| | | | | | | |
|---|---|---|---|---|---|---|
| WTIP | 66.2629696 | 0.314350323 | 0.61306738 | 0.51275004 | 0.60812618 | 0.90527875 |
| SCGB2B2 | 126.362265 | -0.293073521 | 0.41822912 | -0.7007487 | 0.48345984 | 0.85829223 |
| ZNF302 | 195.97989 | -0.159936168 | 0.27669464 | -0.5780241 | 0.56324785 | 0.8927792 |
| GRAMD1A | 105.395696 | -1.091612104 | 0.29803097 | -3.6627472 | 0.00024952 | 0.01531872 |
| HPN | 243.184338 | -0.766393704 | 0.41335824 | -1.8540666 | 0.06372963 | 0.41357646 |
| FXYD3 | 59.9409219 | 0.607004183 | 0.58174914 | 1.04341226 | 0.2967574 | 0.74402333 |
| FXYD5 | 58.5556003 | -0.702510142 | 0.64114231 | -1.0957164 | 0.2732029 | 0.72503581 |
| LSR | 157.956993 | -0.519935328 | 0.3218635 | -1.6153908 | 0.1062261 | 0.50765436 |
| USF2 | 127.874458 | 0.130338766 | 0.33240746 | 0.39210542 | 0.69498032 | 0.93125819 |
| DMKN | 348.596673 | 0.008959321 | 0.26821893 | 0.03340302 | 0.97335321 | 0.9939318 |
| TMEM147 | 93.0705089 | -0.055431859 | 0.45095337 | -0.1229215 | 0.90216927 | 0.98320957 |
| HAUS5 | 32.0485245 | 0.066501903 | 0.34520448 | 0.19264496 | 0.84723703 | 0.96991131 |
| RBM42 | 67.1055473 | -0.331874537 | 0.35924171 | -0.9238196 | 0.35558024 | 0.78607579 |
| COX6B1 | 457.607154 | -0.072991268 | 0.38054459 | -0.1918074 | 0.84789308 | 0.96991131 |
| KMT2B | 87.6118241 | -0.23928648 | 0.31894974 | -0.7502326 | 0.45311466 | 0.84248055 |
| PROSER3 | 51.5807342 | 0.143607602 | 0.29198121 | 0.49183851 | 0.62283351 | 0.91034771 |
| SDHAF1 | 39.4083618 | -0.700252748 | 0.31511202 | -2.2222343 | 0.02626747 | 0.27568539 |
| TBCB | 106.46824 | -0.031477568 | 0.40154325 | -0.0783915 | 0.93751665 | 0.98846521 |
| CAPNS1 | 421.651705 | -0.562078724 | 0.35967407 | -1.5627446 | 0.11811267 | 0.53360052 |
| ZNF565 | 183.233826 | -0.127414147 | 0.45078765 | -0.2826478 | 0.77744682 | 0.95601455 |
| ZNF146 | 218.559734 | 0.163759228 | 0.20564956 | 0.79630236 | 0.42585632 | 0.82774157 |
| LINC00665 | 105.821064 | -0.184052932 | 0.27554885 | -0.6679503 | 0.50416533 | 0.86946722 |
| ZFP14 | 162.642727 | -0.206428762 | 0.31015794 | -0.6655601 | 0.50569229 | 0.87058024 |
| ZFP82 | 67.8217699 | -0.351002681 | 0.25964092 | -1.3518773 | 0.17641456 | 0.62528328 |
| ZNF566 | 77.4043352 | 0.109815998 | 0.27405346 | 0.40071014 | 0.68863355 | 0.93036007 |
| ZNF260 | 88.9703151 | -0.117438489 | 0.18832332 | -0.6236003 | 0.53289008 | 0.88114359 |
| ZNF529 | 139.186939 | 0.126115549 | 0.2422658 | 0.52056686 | 0.60266854 | 0.90369786 |
| ZNF529-AS1 | 124.348582 | -0.552103636 | 0.29811746 | -1.8519668 | 0.06403059 | 0.41357646 |
| ZNF461 | 64.7735641 | -0.389011909 | 0.25694526 | -1.5139875 | 0.13002902 | 0.55282113 |
| ZNF567 | 57.9922017 | -0.00550358 | 0.29067315 | -0.0189339 | 0.98489383 | 0.9954565 |
| AC020928.1 | 124.668046 | 0.062441003 | 0.32895611 | 0.1898156 | 0.84945363 | 0.96991131 |
| ZNF790 | 94.2923688 | -0.564745422 | 0.35863645 | -1.5747017 | 0.11532532 | 0.52873491 |
| ZNF345 | 148.513967 | -0.323666033 | 0.36021697 | -0.8985308 | 0.36890265 | 0.79532503 |
| ZNF568 | 103.571572 | -0.302832622 | 0.32670258 | -0.9269367 | 0.35395938 | 0.78491204 |
| ZNF420 | 211.33662 | -0.034747509 | 0.54873437 | -0.063323 | 0.9495093 | 0.98977092 |
| ZNF383 | 56.456271 | 0.010283129 | 0.29796342 | 0.03451138 | 0.97246937 | 0.9939318 |
| LINC01535 | 38.2317816 | -0.082505092 | 0.33378911 | -0.2471773 | 0.80477101 | 0.96210606 |
| HKR1 | 225.977894 | 0.364805665 | 0.15892329 | 2.29548275 | 0.02170547 | 0.24935578 |
| ZNF569 | 67.5802043 | 0.079196124 | 0.39795866 | 0.1990059 | 0.84225813 | 0.96876046 |
| ZNF570 | 35.1873978 | -0.303428056 | 0.29409442 | -1.0317369 | 0.30219541 | 0.74896472 |
| ZNF793 | 102.755692 | 0.154729621 | 0.30393627 | 0.50908574 | 0.61069213 | 0.9062166 |
| ZNF540 | 84.0586151 | 0.456154179 | 0.55439216 | 0.82280056 | 0.41062142 | 0.82265937 |
| ZNF571 | 93.8346608 | 1.014361356 | 0.62362887 | 1.6265465 | 0.10383345 | 0.50279327 |
| ZFP30 | 98.8303029 | -0.177791318 | 0.32659141 | -0.5443846 | 0.58617687 | 0.90196444 |
| ZNF573 | 119.44649 | -0.423868502 | 0.34179644 | -1.2401197 | 0.21493112 | 0.67132603 |
| SIPA1L3 | 1054.53761 | -0.171606162 | 0.44619192 | -0.3846017 | 0.70053255 | 0.93180634 |
| SPINT2 | 58.6866271 | -0.118379553 | 0.32202806 | -0.3676063 | 0.71316679 | 0.93565198 |
| CATSPERG | 48.8153122 | -0.65317994 | 0.42459409 | -1.5383632 | 0.12395982 | 0.54273186 |
| PSMD8 | 165.451513 | -0.621240018 | 0.36769866 | -1.6895357 | 0.09111681 | 0.47749744 |
| FAM98C | 34.7701411 | 0.226277822 | 0.37071272 | 0.61038592 | 0.54160619 | 0.88411727 |
| EIF3K | 231.163256 | 0.696079574 | 0.329423 | 2.11302661 | 0.03459849 | 0.31447717 |
| ACTN4 | 1029.48805 | 0.222358027 | 0.24536298 | 0.90624112 | 0.36480824 | 0.79269964 |
| CAPN12 | 40.7962165 | -0.63539869 | 0.36654969 | -1.7334585 | 0.0830142 | 0.45768271 |
| ECH1 | 227.961169 | 0.381326525 | 0.54025708 | 0.7058242 | 0.48029747 | 0.85584469 |
| HNRNPL | 156.379203 | 0.113018237 | 0.28358886 | 0.39852848 | 0.69024067 | 0.93073907 |
| SIRT2 | 101.2619 | 0.217884891 | 0.27855546 | 0.78219571 | 0.43409956 | 0.83243867 |
| NFKBIB | 42.415857 | -0.041560222 | 0.28156059 | -0.1476067 | 0.88265319 | 0.97865059 |
| MRPS12 | 110.894234 | -0.221863325 | 0.50969503 | -0.4352864 | 0.66335454 | 0.9229791 |
| FBXO17 | 209.70617 | -0.888719949 | 0.57490768 | -1.5458481 | 0.12214125 | 0.54028588 |
| PAK4 | 194.549507 | 0.013187152 | 0.33887315 | 0.03891472 | 0.96895838 | 0.99312312 |
| SAMD4B | 275.764736 | 0.140192703 | 0.22923823 | 0.611559 | 0.54082957 | 0.88391488 |
| PAF1 | 63.079207 | 0.34784246 | 0.40032106 | 0.86890872 | 0.38489706 | 0.80561637 |
| MED29 | 89.7577594 | 0.192007403 | 0.37755982 | 0.50854829 | 0.61106888 | 0.90644046 |
| ZFP36 | 240.719757 | 1.275935471 | 0.87165795 | 1.46380294 | 0.1432478 | 0.5729912 |
| RPS16 | 366.781519 | 0.128797895 | 0.3975519 | 0.32397756 | 0.74595502 | 0.94594643 |
| SUPT5H | 206.319997 | -0.002184762 | 0.18473021 | -0.0118268 | 0.99056382 | 0.99712889 |
| TIMM50 | 46.3827037 | -0.052902749 | 0.28487791 | -0.1857032 | 0.85267749 | 0.97062426 |
| DYRK1B | 41.9980207 | -0.131185787 | 0.32683122 | -0.401387 | 0.68813526 | 0.93036007 |
| FBL | 73.2063644 | 0.00640378 | 0.39056182 | 0.01639633 | 0.98691821 | 0.99643909 |
| FCGBP | 42.0797023 | -0.150364226 | 0.43609527 | -0.3447967 | 0.73024718 | 0.94048525 |
| PSMC4 | 74.4143438 | -0.362570151 | 0.29513227 | -1.2285005 | 0.21925913 | 0.67731484 |
| ZNF546 | 46.3919375 | -0.611036805 | 0.44525801 | -1.3723207 | 0.16996361 | 0.61569949 |
| ZNF780B | 256.997639 | 0.207107779 | 0.29275866 | 0.7074352 | 0.47929607 | 0.85532469 |
| ZNF780A | 55.5115602 | 0.054669405 | 0.29153094 | 0.18752522 | 0.85124885 | 0.97042184 |
| MAP3K10 | 44.9796755 | 0.086559969 | 0.32722076 | 0.2645308 | 0.79137094 | 0.95832571 |
| AKT2 | 298.440935 | -0.024395035 | 0.41263592 | -0.05912 | 0.95285653 | 0.99050799 |
| PLD3 | 293.828316 | -0.500180038 | 0.24396179 | -2.0502393 | 0.04034109 | 0.33821967 |
| SERTAD1 | 58.5206011 | -0.840437776 | 0.64906009 | -1.2948536 | 0.19537075 | 0.64948524 |
| SERTAD3 | 44.9016694 | 0.153202395 | 0.35947058 | 0.42618897 | 0.66997015 | 0.92474338 |
| BLVRB | 375.600937 | -0.780781016 | 0.41313635 | -1.889887 | 0.05877308 | 0.40066749 |
| SPTBN4 | 43.861736 | -0.5810029 | 0.59700895 | -0.9731896 | 0.33045908 | 0.77618491 |
| SHKBP1 | 44.9873645 | -0.265631404 | 0.3797285 | -0.6995298 | 0.48422099 | 0.85908853 |
| LTBP4 | 83.1646452 | -0.4186885 | 0.28830359 | -1.4522486 | 0.14643248 | 0.57722788 |
| COQ8B | 54.0026685 | -0.278797113 | 0.39569774 | -0.7045709 | 0.48107732 | 0.85600365 |
| ITPKC | 70.3533723 | -0.410675255 | 0.35005965 | -1.1731579 | 0.24073248 | 0.6963541 |
| SNRPA | 72.0763436 | -0.243614589 | 0.29021699 | -0.8394222 | 0.40123242 | 0.81648166 |
| AC008537.1 | 72.4154963 | -0.11910299 | 0.5099505 | -0.2335579 | 0.81532818 | 0.96333253 |
| HNRNPUL1 | 163.257767 | -0.317323089 | 0.24155314 | -1.3136782 | 0.18895453 | 0.64237956 |
| TGFB1 | 148.866583 | -0.348644001 | 0.4053888 | -0.8600238 | 0.38977594 | 0.80680464 |
| CCDC97 | 37.8869118 | -0.836454983 | 0.53369373 | -1.567294 | 0.11704599 | 0.5321657 |
| B3GNT8 | 32.7710652 | -1.406310042 | 0.78638964 | -1.788312 | 0.07372568 | 0.4370297 |
| DMAC2 | 45.7134174 | -0.400094798 | 0.30723059 | -1.3022623 | 0.19282675 | 0.64567765 |
| RPS19 | 1220.62199 | -0.454404017 | 0.39658219 | -1.1458004 | 0.25187776 | 0.70813507 |
| ARHGEF1 | 77.9518102 | -0.135125688 | 0.31322387 | -0.4314029 | 0.66617544 | 0.92396246 |
| RABAC1 | 262.841767 | 0.375145037 | 0.4924779 | 0.76174999 | 0.44620923 | 0.83971561 |
| GRIK5 | 132.794683 | -0.239132133 | 0.32128581 | -0.7442972 | 0.45669668 | 0.84382603 |
| DEDD2 | 42.7646228 | 0.024620867 | 0.26559306 | 0.09270147 | 0.92614073 | 0.98729553 |
| GSK3A | 59.5628339 | 0.562325825 | 0.32440516 | 1.73340592 | 0.08302355 | 0.45768271 |
| CIC | 79.1402561 | -0.294493298 | 0.2365388 | -1.2450105 | 0.21312788 | 0.67054344 |
| PAFAH1B3 | 53.5067795 | -1.038091575 | 0.52021495 | -1.9955051 | 0.04598782 | 0.36047296 |
| MEGF8 | 101.214861 | 0.211232209 | 0.27617015 | 0.76486257 | 0.44435339 | 0.83876504 |
| LIPE-AS1 | 75.1331446 | -0.248888851 | 0.35138954 | -0.708299 | 0.47875961 | 0.85532469 |
| AC004784.1 | 53.1817189 | -4.754209724 | 1.10004835 | -4.3218189 | 1.55E-05 | 0.00257422 |
| PHLDB3 | 52.0169148 | 1.039981683 | 0.59740374 | 1.74083557 | 0.0817124 | 0.45528236 |
| ETHE1 | 77.2822023 | -0.33672336 | 0.38259337 | -0.8801077 | 0.37880098 | 0.80274043 |
| XRCC1 | 33.5556646 | 0.13434656 | 0.30641296 | 0.43844934 | 0.66106059 | 0.92284375 |
| PINLYP | 34.7938626 | -0.37266422 | 0.3045772 | -1.223546 | 0.22112354 | 0.67871976 |
| ZNF428 | 137.755164 | -0.273433945 | 0.48754982 | -0.5608328 | 0.57491151 | 0.89807676 |
| PLAUR | 72.2624294 | -0.84386335 | 1.34627624 | -0.6268129 | 0.53078187 | 0.88004345 |
| SMG9 | 48.4162625 | 0.04535478 | 0.3842912 | 0.11802191 | 0.9060503 | 0.98320957 |
| ZNF283 | 54.8006129 | 0.487423291 | 0.34354359 | 1.41881061 | 0.15595424 | 0.5934642 |
| ZNF45 | 35.5140907 | -0.568832654 | 0.38950764 | -1.460389 | 0.1441832 | 0.57406643 |
| ZNF155 | 75.7019546 | -0.226928128 | 0.49671908 | -0.4568541 | 0.64777595 | 0.91763881 |

| | | | | | | |
|---|---|---|---|---|---|---|
| ZNF224 | 89.0971454 | -0.139860354 | 0.29112217 | -0.4804181 | 0.63093014 | 0.91315397 |
| ZNF234 | 39.4007091 | 0.253428085 | 0.39892746 | 0.6352736 | 0.52524999 | 0.87791848 |
| ZNF226 | 84.0319278 | -0.022892233 | 0.26422371 | -0.0866396 | 0.930958 | 0.98822345 |
| ZNF227 | 45.33576 | 0.079243544 | 0.32930523 | 0.24063858 | 0.80983525 | 0.96247146 |
| ZNF235 | 179.650475 | -0.122794408 | 0.41647311 | -0.2948435 | 0.7681134 | 0.95411028 |
| ZNF229 | 42.8584657 | -0.222755388 | 0.43249047 | -0.5150527 | 0.60651619 | 0.90493874 |
| AC243964.2 | 42.3610636 | -0.476791428 | 0.37964567 | -1.2558853 | 0.20915755 | 0.66686071 |
| BCL3 | 66.9891495 | -1.275512948 | 0.63250996 | -2.0165895 | 0.04373837 | 0.35261025 |
| CBLC | 75.1884272 | -0.504046076 | 0.43923442 | -1.147556 | 0.25115192 | 0.70813507 |
| BCAM | 677.203121 | -0.538412765 | 0.52534885 | -1.0248671 | 0.30542589 | 0.7508476 |
| NECTIN2 | 249.100137 | 0.036482412 | 0.23290171 | 0.15664296 | 0.87552624 | 0.97622753 |
| TOMM40 | 62.8339641 | -0.323721506 | 0.29341551 | -1.103287 | 0.26990257 | 0.72333985 |
| APOE | 156.231858 | -0.524648 | 0.91304308 | -0.5746147 | 0.56555189 | 0.89394085 |
| CLPTM1 | 138.8901 | 0.0512421 | 0.24551238 | 0.20871493 | 0.83467078 | 0.96749268 |
| RELB | 73.1950776 | -0.473561692 | 0.65489095 | -0.7231153 | 0.46960903 | 0.85232523 |
| CLASRP | 129.288845 | -0.327022069 | 0.2566669 | -1.2741108 | 0.20262416 | 0.6574307 |
| GEMIN7-AS1 | 55.4125734 | -0.230386049 | 0.33087837 | -0.6962862 | 0.4862496 | 0.85912348 |
| MARK4 | 152.697134 | -0.198507613 | 0.26609247 | -0.7460099 | 0.45566145 | 0.84372464 |
| TRAPPC6A | 109.04673 | 0.092083801 | 0.38891371 | 0.2367718 | 0.81283383 | 0.96288868 |
| ERCC2 | 38.4446085 | 0.284041164 | 0.43162862 | 0.65806842 | 0.51049417 | 0.87133207 |
| PPP1R13L | 122.976751 | -0.702589829 | 0.33171052 | -2.1180812 | 0.03416819 | 0.31246302 |
| ERCC1 | 57.9423684 | 0.098118514 | 0.34270022 | 0.28631004 | 0.77464067 | 0.95586283 |
| VASP | 122.808455 | -0.058760518 | 0.40158241 | -0.1463224 | 0.88366685 | 0.97878487 |
| OPA3 | 156.975177 | -0.12279738 | 0.28271117 | -0.4343563 | 0.66402972 | 0.92300139 |
| EML2 | 122.625274 | -0.490027865 | 0.34504802 | -1.4201729 | 0.15555734 | 0.59281487 |
| SNRPD2 | 237.718801 | -0.305128617 | 0.44939411 | -0.6789778 | 0.49715194 | 0.86519122 |
| FBXO46 | 54.3579945 | 0.16345997 | 0.32553889 | 0.50212117 | 0.61558229 | 0.90794578 |
| DMPK | 51.3605427 | -0.382134668 | 0.43739042 | -0.8736695 | 0.38229827 | 0.80386955 |
| SYMPK | 265.817365 | -0.011317071 | 0.25747801 | -0.0439535 | 0.96494143 | 0.99243116 |
| MYPOP | 39.0828137 | 0.193944285 | 0.38725132 | 0.50082278 | 0.61649585 | 0.90794578 |
| HIF3A | 36.4901228 | 0.597111725 | 0.57637522 | 1.03597744 | 0.30021267 | 0.74729373 |
| PPP5C | 146.686228 | 0.341667031 | 0.18223841 | 1.87483541 | 0.06081537 | 0.40676596 |
| CCDC8 | 62.2015426 | -0.037999768 | 0.41305289 | -0.0919973 | 0.92670015 | 0.98729622 |
| PNMA8A | 63.8005393 | 0.009316923 | 0.52252719 | 0.0178305 | 0.98577407 | 0.99587186 |
| PPP5D1 | 80.6918457 | 0.385098912 | 0.35057389 | 1.09848145 | 0.27199431 | 0.72372882 |
| CALM3 | 216.319175 | 0.005800977 | 0.32422033 | 0.01789208 | 0.98572494 | 0.99587186 |
| PRKD2 | 130.849718 | -0.677612634 | 0.29480145 | -2.298539 | 0.02153113 | 0.24935578 |
| STRN4 | 54.1172341 | 0.197620125 | 0.21805688 | 0.90627788 | 0.36478879 | 0.79269964 |
| FKRP | 39.2052693 | -0.118908311 | 0.37202438 | -0.3196251 | 0.74925258 | 0.94704648 |
| AP2S1 | 159.517602 | -0.414653184 | 0.39585605 | -1.0474848 | 0.29487606 | 0.74090172 |
| ARHGAP35 | 466.557206 | 0.147566922 | 0.40870061 | 0.36106362 | 0.71805189 | 0.93573028 |
| TMEM160 | 183.213681 | 0.023008009 | 0.45531685 | 0.05053186 | 0.95969856 | 0.99119981 |
| ZC3H4 | 103.086907 | -0.116667317 | 0.26871026 | -0.4341751 | 0.66416126 | 0.92300139 |
| SAE1 | 107.893607 | 0.088431266 | 0.32440343 | 0.27259658 | 0.78516334 | 0.95702626 |
| INAFM1 | 35.5345541 | -0.302922988 | 0.45016086 | -0.6729216 | 0.50099714 | 0.86769956 |
| C5AR1 | 144.244665 | 0.718392968 | 0.59904477 | 1.19923085 | 0.23043819 | 0.69012664 |
| DHX34 | 62.5729392 | -0.249319489 | 0.36567544 | -0.6818054 | 0.49536201 | 0.86436794 |
| NAPA-AS1 | 50.1822568 | -0.254341351 | 0.57339043 | -0.4435745 | 0.65735028 | 0.92160948 |
| NAPA | 338.402661 | -0.061030883 | 0.25746792 | -0.2370427 | 0.81262369 | 0.96288868 |
| NOP53 | 85.0190601 | -0.354640326 | 0.32533489 | -1.0900777 | 0.2756789 | 0.7263609 |
| SELENOW | 352.719879 | 0.448530876 | 0.46940893 | 0.95552267 | 0.33931345 | 0.77836439 |
| LIG1 | 36.3971218 | -0.40062835 | 0.5014915 | -0.7988737 | 0.42436368 | 0.82774157 |
| ZSWIM9 | 60.8603112 | -0.102454834 | 0.36867961 | -0.2778967 | 0.78109169 | 0.9566945 |
| CARD8 | 96.3646595 | 0.065366612 | 0.29549395 | 0.22121134 | 0.82492788 | 0.96511681 |
| TMEM143 | 44.3611737 | 0.089772245 | 0.34799119 | 0.25797275 | 0.79642794 | 0.95948555 |
| KDELR1 | 187.755449 | -0.7363023 | 0.28476998 | -2.5856037 | 0.00972086 | 0.15745696 |
| CYTH2 | 82.4125852 | -0.641758992 | 0.46350347 | -1.3845829 | 0.16618003 | 0.60830133 |
| LMTK3 | 38.8629486 | 0.196832265 | 0.54713816 | 0.35974874 | 0.71903504 | 0.9359691 |
| RPL18 | 628.362633 | -0.037162137 | 0.49161044 | -0.0755927 | 0.93974318 | 0.98855185 |
| SPHK2 | 32.3464297 | -0.325819332 | 0.36653889 | -0.888908 | 0.37405255 | 0.79709398 |
| BCAT2 | 76.2167471 | -0.295806036 | 0.31180168 | -0.9486993 | 0.34277356 | 0.78096003 |
| HSD17B14 | 40.2580605 | -1.128871875 | 0.58971086 | -1.9142803 | 0.05558435 | 0.39368206 |
| PLEKHA4 | 49.345312 | -0.969791817 | 0.77956741 | -1.2440128 | 0.21349486 | 0.67075711 |
| PPP1R15A | 45.4773276 | -0.411605439 | 0.4823249 | -0.853378 | 0.39344974 | 0.81024824 |
| NUCB1 | 242.457583 | -0.780339942 | 0.31016455 | -2.5158902 | 0.01187322 | 0.17330267 |
| BAX | 113.357333 | -0.460322139 | 0.4238449 | -1.0860627 | 0.27745125 | 0.72788245 |
| FTL | 3207.66535 | -1.576343718 | 0.64923107 | -2.4280165 | 0.01518165 | 0.20038254 |
| GYS1 | 140.049261 | -0.779397632 | 0.43377576 | -1.7967754 | 0.07237128 | 0.43315305 |
| RUVBL2 | 36.899945 | -0.118014261 | 0.29476686 | -0.4003648 | 0.68888788 | 0.93036007 |
| SNRNP70 | 490.470956 | 0.413390664 | 0.26869477 | 1.53851397 | 0.12392299 | 0.54273186 |
| TRPM4 | 125.682337 | -0.188899418 | 0.36036469 | -0.5241896 | 0.60014668 | 0.90369786 |
| SLC6A16 | 94.642618 | -0.329885561 | 0.43576431 | -0.7570275 | 0.44903339 | 0.84115854 |
| TEAD2 | 50.8469408 | -1.259945564 | 0.64011147 | -1.9683221 | 0.04903099 | 0.3746651 |
| PIH1D1 | 82.1821273 | -0.225115834 | 0.38380581 | -0.5865358 | 0.55751554 | 0.89148045 |
| RPL13A | 784.356689 | -0.042280075 | 0.43878988 | -0.0963561 | 0.92323777 | 0.98694936 |
| RPS11 | 573.543719 | 0.138346368 | 0.41663442 | 0.33205699 | 0.73984623 | 0.94440785 |
| FCGRT | 172.066061 | -0.190387275 | 0.38353784 | -0.4963976 | 0.6196139 | 0.90907933 |
| NOSIP | 119.068862 | -0.209205218 | 0.27264338 | -0.7673218 | 0.4428902 | 0.83826789 |
| PRRG2 | 48.1151451 | -0.556717555 | 0.34188936 | -1.6283559 | 0.10344945 | 0.50215387 |
| PRR12 | 82.3006452 | -0.020542958 | 0.45289143 | -0.0453596 | 0.96382071 | 0.99243116 |
| RRAS | 74.3599433 | 0.106411701 | 0.66441426 | 0.16015866 | 0.87275609 | 0.97622753 |
| SCAF1 | 62.2286798 | 0.023724375 | 0.26331806 | 0.09009779 | 0.92820951 | 0.98751845 |
| IRF3 | 142.481839 | -0.296351482 | 0.21718827 | -1.3644912 | 0.17241302 | 0.61884138 |
| PRMT1 | 149.632584 | 0.020596896 | 0.30154348 | 0.0683049 | 0.94554293 | 0.98949015 |
| AP2A1 | 87.1700123 | -0.379644222 | 0.23601322 | -1.6085718 | 0.10771 | 0.51123225 |
| MED25 | 121.748117 | -0.107020681 | 0.25830681 | -0.4143161 | 0.67864259 | 0.9268955 |
| PTOV1 | 115.705778 | 0.236845159 | 0.23785081 | 0.99577191 | 0.31936098 | 0.76378437 |
| PNKP | 91.0964833 | -0.837658942 | 0.3088124 | -2.7125172 | 0.00667743 | 0.13031298 |
| TBC1D17 | 87.8023377 | -0.185561354 | 0.27983736 | -0.6631043 | 0.50726376 | 0.87094607 |
| ATF5 | 57.9816968 | -0.532440082 | 0.39490108 | -1.3482872 | 0.17756601 | 0.6269762 |
| VRK3 | 103.799198 | -0.095950575 | 0.21830572 | -0.4395239 | 0.66028198 | 0.92284375 |
| MYH14 | 245.194142 | 0.710115303 | 0.32540572 | 2.18224588 | 0.02909139 | 0.28842879 |
| KCNC3 | 197.635976 | 1.110478728 | 0.55334536 | 2.00684566 | 0.0447661 | 0.35515061 |
| NR1H2 | 157.164062 | 0.143599133 | 0.27956417 | 0.51365357 | 0.60749422 | 0.90510401 |
| EMC10 | 209.42477 | 0.13676512 | 0.34097626 | 0.40109865 | 0.68834749 | 0.93036007 |
| JOSD2 | 54.8018985 | 0.171735782 | 0.44103108 | 0.3893961 | 0.69698316 | 0.93130502 |
| CLEC11A | 57.7085493 | 0.01462758 | 0.3324168 | 0.04400373 | 0.96490143 | 0.99243116 |
| C19orf48 | 47.8094181 | -0.476423827 | 0.34924772 | -1.364143 | 0.17252257 | 0.61884138 |
| CTU1 | 35.713269 | 0.03497838 | 0.38748799 | 0.09026958 | 0.92807299 | 0.98751845 |
| ETFB | 215.225426 | 0.210009651 | 0.39845165 | 0.52706433 | 0.59814892 | 0.90369786 |
| ZNF175 | 48.4094956 | -0.293642731 | 0.42932783 | -0.6839592 | 0.49400091 | 0.86339027 |
| SPACA6 | 68.4837595 | 0.188666478 | 0.33591135 | 0.56165556 | 0.57435072 | 0.89807676 |
| ZNF577 | 145.142716 | -0.2150451 | 0.44549918 | -0.4827059 | 0.62930455 | 0.9114564 |
| ZNF613 | 39.1076524 | -0.309978141 | 0.47997265 | -0.6458246 | 0.51839296 | 0.87526312 |
| ZNF350-AS1 | 227.336879 | 0.146318835 | 0.86648574 | 0.16886468 | 0.86590308 | 0.97514123 |
| ZNF350 | 97.2989393 | -0.513291321 | 0.61227582 | -0.8383335 | 0.40184344 | 0.8167803 |
| ZNF615 | 66.8477873 | 0.074312818 | 0.4368512 | 0.17011014 | 0.86492352 | 0.97485201 |
| ZNF614 | 38.1547649 | 0.05930065 | 0.48325178 | 0.1227117 | 0.9023354 | 0.98320957 |
| ZNF432 | 118.88017 | -0.074254876 | 0.30809902 | -0.2410098 | 0.80954754 | 0.96245698 |
| ZNF616 | 41.2564108 | 0.072555496 | 0.55540252 | 0.13063588 | 0.89606336 | 0.98291229 |
| AC011468.3 | 53.740638 | -0.348890653 | 0.44285561 | -0.7878203 | 0.43080181 | 0.83111714 |
| ZNF836 | 67.3101992 | 0.83155453 | 0.84608966 | 0.98282081 | 0.32569564 | 0.77073504 |
| PPP2R1A | 548.949266 | -0.209097725 | 0.4244395 | -0.4926444 | 0.6222639 | 0.91028772 |

| | | | | | | |
|---|---|---|---|---|---|---|
| ZNF766 | 100.099966 | -0.646415116 | 0.26636187 | -2.4268305 | 0.01523137 | 0.20078585 |
| ZNF480 | 76.58877 | -0.479414199 | 0.3503074 | -1.3685529 | 0.17113909 | 0.6176702 |
| ZNF610 | 34.1773688 | -0.217775696 | 0.52800185 | -0.4124525 | 0.68000777 | 0.92700898 |
| ZNF880 | 37.599271 | 0.095914303 | 0.41744146 | 0.22976707 | 0.81827277 | 0.96407382 |
| ZNF528 | 74.5388846 | 0.341235029 | 0.34979715 | 0.97552262 | 0.32930109 | 0.77581911 |
| ZNF808 | 55.1811536 | -0.119752062 | 0.31084255 | -0.3852499 | 0.70005228 | 0.93180634 |
| ZNF701 | 86.9178981 | 0.065210608 | 0.25246825 | 0.25829231 | 0.79618132 | 0.95948555 |
| ZNF83 | 621.414182 | -0.065538793 | 0.40342081 | -0.1624576 | 0.87094548 | 0.97622753 |
| ZNF611 | 273.290335 | -0.289871523 | 0.29759501 | -0.974047 | 0.33003322 | 0.77618491 |
| ZNF600 | 35.6476965 | -0.419638517 | 0.38604209 | -1.0870279 | 0.27702449 | 0.7274409 |
| ZNF28 | 58.3258937 | -0.11539172 | 0.23789119 | -0.4850609 | 0.62763314 | 0.91064589 |
| AC010487.3 | 66.2529163 | 0.589292568 | 0.38458605 | 1.53227753 | 0.12545396 | 0.54417366 |
| ZNF160 | 237.972633 | -0.453445207 | 0.28360868 | -1.5988411 | 0.10985592 | 0.51534917 |
| ZNF347 | 73.7276256 | -0.254729531 | 0.31328083 | -0.8131028 | 0.4161591 | 0.82489761 |
| ZNF677 | 74.612736 | -0.17594651 | 0.30312274 | -0.5804464 | 0.5616136 | 0.89242209 |
| ZNF525 | 48.350818 | -0.941327668 | 0.40303281 | -2.3356105 | 0.01951155 | 0.23473196 |
| ZNF765 | 60.0645007 | -0.236115286 | 0.3363381 | -0.7020177 | 0.48266815 | 0.85778569 |
| ZNF761 | 88.2910421 | -0.196568268 | 0.37744575 | -0.5207855 | 0.60251622 | 0.90369786 |
| ZNF331 | 1301.54198 | 3.387772812 | 1.66907398 | 2.02973197 | 0.04238379 | 0.34798255 |
| MYADM | 58.9958069 | -0.554612022 | 0.35159604 | -1.5774126 | 0.11470062 | 0.52703872 |
| NDUFA3 | 89.2705209 | -0.131331587 | 0.46030066 | -0.285317 | 0.77540132 | 0.95601455 |
| TFPT | 38.1497137 | -0.155416927 | 0.39371356 | -0.3947462 | 0.69303021 | 0.93105058 |
| PRPF31 | 120.712119 | -0.028439052 | 0.33532555 | -0.0848103 | 0.93241222 | 0.98822345 |
| CNOT3 | 59.8575745 | 0.132104652 | 0.26551131 | 0.49754812 | 0.61880258 | 0.90835482 |
| TMC4 | 274.210744 | 0.167479737 | 0.20194081 | 0.82935061 | 0.40690604 | 0.82018309 |
| MBOAT7 | 220.087998 | -0.789236791 | 0.30645002 | -2.5754176 | 0.01001191 | 0.16036545 |
| TSEN34 | 89.1842259 | -0.232603474 | 0.45708284 | -0.5088869 | 0.61083149 | 0.9062166 |
| RPS9 | 1317.89095 | 0.235464526 | 0.4751668 | 0.49554078 | 0.62021845 | 0.90914396 |
| LENG8-AS1 | 50.9665406 | -0.156773649 | 0.32361977 | -0.4844378 | 0.6280752 | 0.91090895 |
| LENG8 | 297.625911 | 1.062015082 | 0.41318015 | 2.57034392 | 0.01015976 | 0.16136868 |
| AC011476.3 | 68.6853006 | -0.985248685 | 0.65964339 | -1.4936081 | 0.13527809 | 0.55916513 |
| RDH13 | 182.365397 | -0.396821443 | 0.43586293 | -0.9104271 | 0.36259729 | 0.7906824 |
| EPS8L1 | 141.354323 | 0.804616435 | 0.45524395 | 1.76744013 | 0.07715455 | 0.44451877 |
| PPP1R12C | 104.638071 | 0.447958202 | 0.23588441 | 1.8990581 | 0.05755684 | 0.39817926 |
| PPP6R1 | 103.400244 | 0.096822596 | 0.2681538 | 0.36107113 | 0.71804628 | 0.93573028 |
| HSPBP1 | 63.2178337 | 0.207503042 | 0.33844961 | 0.61309878 | 0.53981103 | 0.8833006 |
| TMEM238 | 96.6689778 | -1.013503164 | 0.503993 | -2.0109469 | 0.04433107 | 0.35437802 |
| RPL28 | 946.436941 | -0.580909017 | 0.42627676 | -1.3627508 | 0.17296105 | 0.61927974 |
| ISOC2 | 107.659561 | -0.273878482 | 0.54191352 | -0.5053915 | 0.61328389 | 0.90751853 |
| NAT14 | 65.1965578 | -0.056053593 | 0.52058547 | -0.1076741 | 0.91425418 | 0.98480378 |
| ZNF579 | 36.2335258 | -0.325134411 | 0.40689327 | -0.7990656 | 0.42425238 | 0.82774157 |
| ZNF524 | 37.551933 | -0.008215666 | 0.40903419 | -0.0200855 | 0.98397515 | 0.99540861 |
| CCDC106 | 51.7633823 | 0.260087008 | 0.39439517 | 0.65945789 | 0.50960178 | 0.87094607 |
| U2AF2 | 92.1529705 | -0.03814957 | 0.2516524 | -0.1515963 | 0.87950536 | 0.97722818 |
| EPN1 | 284.214673 | 0.411907988 | 0.27746226 | 1.48455499 | 0.13766175 | 0.56434148 |
| ZNF787 | 59.569634 | -0.546818516 | 0.30612867 | -1.7862375 | 0.07406081 | 0.43749352 |
| ZNF444 | 82.6442409 | 0.02357908 | 0.21662363 | 0.10884814 | 0.91332294 | 0.98480378 |
| ZSCAN5A | 182.723344 | -0.212678463 | 0.34607527 | -0.6145439 | 0.53885594 | 0.88287241 |
| ZNF667 | 73.7701004 | -0.892018974 | 0.46984014 | -1.8985585 | 0.05762255 | 0.39817926 |
| ZNF667-AS1 | 102.473062 | -0.092056963 | 0.30563815 | -0.3011959 | 0.76326511 | 0.95378284 |
| ZNF471 | 73.379451 | 0.01232655 | 0.30376377 | 0.0405794 | 0.96763121 | 0.99292814 |
| AC005498.3 | 36.0259436 | -0.566134888 | 0.48271584 | -1.1728119 | 0.24087124 | 0.69636706 |
| ZNF470 | 36.0262921 | -1.417666781 | 0.49068236 | -2.8891741 | 0.00386255 | 0.09660987 |
| ZNF264 | 114.268217 | -0.081847296 | 0.20759534 | -0.3942636 | 0.6933864 | 0.93105058 |
| ZNF460-AS1 | 34.6121704 | -0.398921813 | 0.32604555 | -1.2235156 | 0.22113502 | 0.67871976 |
| ZNF460 | 45.5859966 | -0.39833948 | 0.52393811 | -0.7602796 | 0.44708745 | 0.84014281 |
| AC005261.1 | 203.826032 | -0.176848566 | 0.39699025 | -0.4454733 | 0.65597774 | 0.92098622 |
| AC004076.1 | 75.8165902 | 0.213487326 | 0.43248253 | 0.49363226 | 0.62156591 | 0.9099963 |
| ZNF549 | 56.1615594 | -0.353562569 | 0.31678694 | -1.1160895 | 0.26438384 | 0.71989632 |
| ZNF134 | 36.4285853 | -0.325415618 | 0.36325348 | -0.8958362 | 0.37034026 | 0.79532503 |
| ZNF154 | 43.3960731 | 0.283348804 | 0.34517453 | 0.82088561 | 0.41171143 | 0.82355334 |
| ZNF776 | 34.3093356 | 0.177729401 | 0.32581659 | 0.5454891 | 0.58541717 | 0.90173902 |
| ZNF586 | 311.375981 | 0.051966837 | 0.25889873 | 0.20072264 | 0.84091545 | 0.96854218 |
| ZNF552 | 132.553507 | 0.129755679 | 0.29802229 | 0.43538918 | 0.66327996 | 0.9229791 |
| ZNF814 | 195.601407 | -0.025876836 | 0.29494574 | -0.0877342 | 0.93008792 | 0.98807358 |
| ZNF587 | 103.126636 | 0.42349229 | 0.22343462 | 1.89537453 | 0.0580428 | 0.39861635 |
| ZNF418 | 34.2395057 | 0.010557716 | 0.40455459 | 0.02609714 | 0.97917986 | 0.99469867 |
| ZNF606 | 63.3855267 | -0.250466219 | 0.43419052 | -0.5768579 | 0.56403547 | 0.8930743 |
| ZNF135 | 92.7809073 | -0.375810466 | 0.34286894 | -1.0960761 | 0.27304546 | 0.72480152 |
| ZSCAN18 | 314.426259 | -0.565504361 | 0.28836478 | -1.9610729 | 0.04987051 | 0.37645791 |
| ZNF329 | 81.5533957 | -0.241869448 | 0.36557196 | -0.6616193 | 0.50821526 | 0.87094607 |
| ZNF274 | 120.401665 | -0.386231645 | 0.25732543 | -1.5009463 | 0.13336946 | 0.55619258 |
| AC020915.3 | 56.3327607 | 0.081872903 | 0.34686889 | 0.23603415 | 0.81340617 | 0.96288868 |
| RPS5 | 338.880118 | -0.419082289 | 0.53903053 | -0.7774741 | 0.4368791 | 0.83396957 |
| ZNF584 | 38.132509 | -0.7251977 | 0.40007728 | -1.8126441 | 0.06988674 | 0.42843167 |
| ZNF446 | 34.5229498 | -0.278472702 | 0.30428108 | -0.9151824 | 0.36009586 | 0.78892604 |
| SLC27A5 | 55.5292242 | 0.293007532 | 0.34153768 | 0.85790689 | 0.39094388 | 0.80778625 |
| TRIM28 | 130.947994 | -0.057441358 | 0.38102477 | -0.1507549 | 0.88016905 | 0.97740892 |
| CHMP2A | 166.876349 | 0.005286971 | 0.52298965 | 0.01010913 | 0.99193422 | 0.99774168 |
| UBE2M | 115.659874 | 0.13051408 | 0.44781573 | 0.29144595 | 0.77071028 | 0.95495906 |
| MZF1-AS1 | 42.4692228 | -0.020664364 | 0.45665778 | -0.0452513 | 0.96390699 | 0.99243116 |
| MZF1 | 144.045014 | -0.140060529 | 0.23292595 | -0.6013093 | 0.54763403 | 0.88773467 |
| RPS4Y1 | 274.00365 | 0.509694533 | 2.21225605 | 0.23039581 | 0.81778421 | 0.96404074 |
| ZFY | 77.3103475 | 1.096745184 | 2.0976097 | 0.52285474 | 0.60107534 | 0.90369786 |
| LINC00278 | 649.214111 | 0.231116105 | 2.61591676 | 0.08834995 | 0.92959854 | 0.98804151 |
| PCDH11Y | 123.55098 | -0.084287514 | 0.94983407 | -0.0887392 | 0.92928918 | 0.9878984 |
| TBL1Y | 167.479896 | 0.751790992 | 1.85260062 | 0.40580306 | 0.68488731 | 0.92881914 |
| TTTY15 | 90.8886887 | 0.869541528 | 1.73677545 | 0.50066434 | 0.61660738 | 0.90794578 |
| USP9Y | 333.606469 | 0.63237143 | 2.44581972 | 0.25855194 | 0.79598097 | 0.95948555 |
| DDX3Y | 130.905573 | 0.956643842 | 2.53325584 | 0.37763412 | 0.70570241 | 0.93241442 |
| UTY | 764.361165 | 0.620811573 | 2.15814298 | 0.28766008 | 0.77360695 | 0.95561066 |
| NLGN4Y | 123.718279 | -0.354188552 | 1.5769905 | -0.2245978 | 0.8222922 | 0.96502145 |
| TTTY14 | 1078.81855 | 0.829650999 | 0.92827416 | 0.89375643 | 0.37145223 | 0.79539603 |
| KDM5D | 98.8400507 | 0.856623448 | 2.46869366 | 0.34699463 | 0.72859535 | 0.93931348 |
| TTTY10 | 69.7109273 | 1.25019798 | 1.60196317 | 0.78041618 | 0.43514595 | 0.83339355 |
| EIF1AY | 61.488158 | 1.083844027 | 2.36631149 | 0.458031 | 0.64693017 | 0.91743182 |
| IL17RA | 64.7642947 | -0.456411901 | 0.28930976 | -1.577589 | 0.11466006 | 0.52703872 |
| HDHD5 | 78.5590332 | -0.283539676 | 0.22110169 | -1.2823949 | 0.19970417 | 0.65475264 |
| HDHD5-AS1 | 33.0246642 | -1.223702802 | 0.51477733 | -2.3771498 | 0.017447 | 0.21871363 |
| ATP6V1E1 | 276.42159 | -0.326560632 | 0.21861002 | -1.4938045 | 0.13522673 | 0.55916513 |
| BCL2L13 | 387.108436 | 0.119491508 | 0.21762678 | 0.54906619 | 0.58296003 | 0.90085422 |
| MICAL3 | 250.31445 | -0.552741143 | 0.42095268 | -1.3130719 | 0.18915871 | 0.64237956 |
| PEX26 | 124.066044 | -0.35096932 | 0.24409627 | -1.4378316 | 0.15048185 | 0.58452549 |
| DGCR2 | 210.744035 | 0.311496786 | 0.21992209 | 1.41639606 | 0.15665958 | 0.59377662 |
| SLC25A1 | 144.557601 | -0.384947616 | 0.39157194 | -0.9830827 | 0.32556672 | 0.77073504 |
| CLTCL1 | 89.8462246 | 0.139885795 | 0.29416385 | 0.475537 | 0.63440427 | 0.91430092 |
| HIRA | 80.0978902 | -0.121991771 | 0.34570211 | -0.3528812 | 0.72417752 | 0.93893205 |
| MRPL40 | 115.879162 | -0.863910777 | 0.59080044 | -1.4622717 | 0.14366676 | 0.57324898 |
| UFD1 | 68.3812432 | 0.16321054 | 0.32518733 | 0.50189699 | 0.61573998 | 0.90794578 |
| TXNRD2 | 84.0190625 | 0.329223237 | 0.25733607 | 1.27935129 | 0.20077338 | 0.65487239 |
| COMT | 349.816245 | -0.189641792 | 0.42413669 | -0.4471242 | 0.65478536 | 0.92047627 |
| TANGO2 | 105.817102 | -0.933370615 | 0.40858793 | -2.2843813 | 0.02234913 | 0.25166501 |
| DGCR8 | 77.7511122 | -0.360931818 | 0.31050318 | -1.1624094 | 0.24506921 | 0.70148313 |

| | | | | | | |
|---|---|---|---|---|---|---|
| RANBP1 | 79.8204569 | 0.214014524 | 0.48896017 | 0.43769317 | 0.66160872 | 0.92284375 |
| ZDHHC8 | 52.5935503 | 0.440754418 | 0.39529759 | 1.11499393 | 0.26485302 | 0.71989632 |
| DGCR6L | 65.3009584 | 0.009623017 | 0.4251183 | 0.02263609 | 0.98194055 | 0.99507903 |
| KLHL22 | 135.954163 | -0.035562056 | 0.31751382 | -0.1120016 | 0.91082214 | 0.98468493 |
| MED15 | 281.85942 | -0.368772911 | 0.20537241 | -1.7956302 | 0.07255334 | 0.43315584 |
| PI4KA | 315.283965 | 0.119725702 | 0.17934171 | 0.66758425 | 0.50439901 | 0.86946722 |
| SNAP29 | 83.6515925 | 0.151055236 | 0.2957709 | 0.51071703 | 0.60954921 | 0.90572462 |
| CRKL | 89.3460292 | -0.375020447 | 0.21618177 | -1.734746 | 0.08278581 | 0.45735124 |
| LZTR1 | 48.1067408 | 0.193987781 | 0.25991147 | 0.746361 | 0.45544937 | 0.84360365 |
| THAP7 | 40.6473919 | -0.569276722 | 0.39850386 | -1.428535 | 0.15313792 | 0.58860244 |
| HIC2 | 64.0570323 | 0.218198549 | 0.39312531 | 0.55503561 | 0.5788703 | 0.89975022 |
| UBE2L3 | 294.537559 | -0.046150614 | 0.22839388 | -0.2020659 | 0.83986521 | 0.96833625 |
| YDJC | 29.9426695 | -0.78415328 | 0.43359293 | -1.8085011 | 0.07052855 | 0.42987436 |
| SDF2L1 | 70.7258067 | -0.696714307 | 0.45193335 | -1.5416306 | 0.12316338 | 0.54197241 |
| PPIL2 | 160.504604 | -0.005004898 | 0.32486787 | -0.015406 | 0.98770832 | 0.9964791 |
| YPEL1 | 38.1884953 | -0.650246469 | 0.5442688 | -1.1947157 | 0.23219814 | 0.69186326 |
| MAPK1 | 524.039032 | -0.020145462 | 0.25456491 | -0.0791368 | 0.93692378 | 0.98822345 |
| PPM1F | 53.3119765 | -0.316611698 | 0.41089237 | -0.7705466 | 0.44097575 | 0.83751828 |
| TOP3B | 41.2181029 | 0.076386316 | 0.39724038 | 0.19229243 | 0.84751315 | 0.96991131 |
| AC245060.5 | 94.1421505 | 0.717382118 | 0.46152527 | 1.55437233 | 0.12009563 | 0.53751263 |
| RSPH14 | 35.3731343 | -0.175884527 | 0.35076148 | -0.5014363 | 0.61606412 | 0.90794578 |
| GNAZ | 44.7698391 | 1.046315924 | 0.40582515 | 2.57824321 | 0.00993041 | 0.1596176 |
| RAB36 | 57.0922959 | 0.01153837 | 0.31838832 | 0.03623993 | 0.97109105 | 0.9939318 |
| BCR | 324.807303 | -0.25935053 | 0.270319 | -0.959424 | 0.3373452 | 0.77827084 |
| CHCHD10 | 655.020538 | -0.090478613 | 0.43700659 | -0.2070418 | 0.83597725 | 0.96757829 |
| SMARCB1 | 137.008579 | -0.067179284 | 0.29022714 | -0.2314714 | 0.8169486 | 0.96380902 |
| DDT | 315.800858 | 0.265327614 | 0.47259804 | 0.56142343 | 0.57450891 | 0.89807676 |
| CABIN1 | 557.474682 | -0.09347734 | 0.27790952 | -0.3363589 | 0.73660024 | 0.9429136 |
| GUCD1 | 100.937048 | -0.604469604 | 0.2369404 | -2.5511463 | 0.01073693 | 0.16452817 |
| SNRPD3 | 219.471433 | -0.089182393 | 0.32073895 | -0.2780529 | 0.78097175 | 0.9566945 |
| SGSM1 | 173.292806 | -0.12071186 | 0.44001266 | -0.2743372 | 0.78382547 | 0.9568898 |
| KIAA1671 | 356.835574 | -0.308520307 | 0.29350173 | -1.0511703 | 0.29318037 | 0.73864467 |
| LRP5L | 107.695431 | -0.379079993 | 0.33626502 | -1.1273251 | 0.25960506 | 0.7168415 |
| GRK3 | 115.041657 | -0.06023961 | 0.3808731 | -0.1581619 | 0.87432923 | 0.97622753 |
| ASPHD2 | 35.9021839 | 0.690661612 | 0.65744121 | 1.05052984 | 0.29347458 | 0.73897491 |
| HPS4 | 134.217286 | -0.248171128 | 0.22992867 | -1.0793396 | 0.28043636 | 0.72945471 |
| TFIP11 | 48.0520953 | 0.023195524 | 0.26285744 | 0.08824374 | 0.92968295 | 0.98804151 |
| TPST2 | 84.159154 | -0.203992677 | 0.29674822 | -0.6874268 | 0.49181382 | 0.86172738 |
| MIATNB | 87.4095428 | -0.729300957 | 0.43706339 | -1.6686389 | 0.09518897 | 0.48627909 |
| PITPNB | 286.291168 | 0.227735264 | 0.32622336 | 0.69809612 | 0.48511708 | 0.85912348 |
| TTC28 | 502.583028 | -0.251965889 | 0.21478886 | -1.1730864 | 0.24076116 | 0.6963541 |
| HSCB | 110.444772 | -0.508990296 | 0.25716382 | -1.9792454 | 0.04778839 | 0.36906581 |
| CCDC117 | 60.6255596 | 0.303280835 | 0.31523197 | 0.9620878 | 0.3360055 | 0.77820994 |
| XBP1 | 174.631586 | -0.755996468 | 0.27343256 | -2.764837 | 0.00569512 | 0.1213371 |
| Z93930.2 | 205.543457 | -0.595078867 | 0.47637353 | -1.2491854 | 0.21159726 | 0.66975892 |
| ZNRF3 | 362.434209 | -0.24469418 | 0.40055284 | -0.6108911 | 0.54127166 | 0.88408632 |
| KREMEN1 | 90.5996368 | -0.391305773 | 0.29798928 | -1.3131538 | 0.18913111 | 0.64237956 |
| RHBDD3 | 46.0205079 | -0.451025467 | 0.36744846 | -1.2274523 | 0.21965263 | 0.67800045 |
| EWSR1 | 338.318039 | 0.318769683 | 0.26198544 | 1.21674578 | 0.22370095 | 0.68103746 |
| AP1B1 | 146.092607 | 0.01848979 | 0.26911216 | 0.06870663 | 0.94522314 | 0.98949015 |
| THOC5 | 48.5936347 | -0.063687361 | 0.31132288 | -0.2045701 | 0.837908 | 0.96757829 |
| NIPSNAP1 | 90.2741676 | -0.063694484 | 0.27472627 | -0.2318471 | 0.81665678 | 0.96380902 |
| NF2 | 175.589332 | 0.230433327 | 0.3853361 | 0.59800608 | 0.54983588 | 0.88889487 |
| ZMAT5 | 48.0799965 | -0.010575767 | 0.25500035 | -0.0414735 | 0.96691839 | 0.99291246 |
| UQCR10 | 532.436366 | -0.462007 | 0.46339244 | -0.9970102 | 0.31875955 | 0.76363962 |
| ASCC2 | 105.075444 | 0.060964504 | 0.24140989 | 0.25253524 | 0.80062738 | 0.96063297 |
| MTMR3 | 505.986066 | 0.344716898 | 0.1851199 | 1.86212771 | 0.06258509 | 0.41084915 |
| TBC1D10A | 47.3700245 | -0.225254556 | 0.45172012 | -0.4986596 | 0.61801924 | 0.90794578 |
| SF3A1 | 87.2169645 | 0.020907587 | 0.30338997 | 0.06891324 | 0.94505868 | 0.98949015 |
| RNF215 | 70.617564 | -0.670863425 | 0.32829795 | -2.0434591 | 0.04100701 | 0.34131622 |
| SEC14L6 | 88.4385878 | -0.010409246 | 0.36886898 | -0.0282194 | 0.9774872 | 0.99469867 |
| GAL3ST1 | 65.83879 | 0.29545699 | 0.32842112 | 0.89962847 | 0.368318 | 0.7952148 |
| PES1 | 64.1200205 | -0.563048631 | 0.29195906 | -1.9285191 | 0.05379059 | 0.38935477 |
| TCN2 | 49.617492 | -0.262993445 | 0.48614806 | -0.540974 | 0.58852552 | 0.90200952 |
| DUSP18 | 81.0430312 | -0.177411238 | 0.26852883 | -0.6606785 | 0.50881849 | 0.87094607 |
| OSBP2 | 171.65056 | -0.08819846 | 0.26958449 | -0.3271644 | 0.74354351 | 0.94555911 |
| MORC2 | 77.3355493 | 0.043108553 | 0.24276845 | 0.17757066 | 0.85906017 | 0.97244655 |
| SELENOM | 410.303019 | -0.158883675 | 0.41820383 | -0.3799192 | 0.70400538 | 0.93241442 |
| INPP5J | 200.196783 | 0.858638739 | 0.55243813 | 1.55427132 | 0.12011971 | 0.53751263 |
| RNF185 | 184.594378 | -0.132427058 | 0.24125848 | -0.5489012 | 0.58307329 | 0.90085422 |
| LIMK2 | 237.511951 | -0.488897089 | 0.22381261 | -2.1844036 | 0.0289326 | 0.28768102 |
| PIK3IP1 | 169.162874 | -0.685354041 | 0.35013884 | -1.9573779 | 0.05030305 | 0.37839277 |
| PIK3IP1-AS1 | 54.883795 | -0.806431308 | 0.43697927 | -1.8454681 | 0.06496947 | 0.41542409 |
| PATZ1 | 114.907621 | -0.555253593 | 0.37985045 | -1.4617689 | 0.14380454 | 0.57324898 |
| DRG1 | 148.337495 | -0.144397755 | 0.27830855 | -0.5188405 | 0.60387195 | 0.90371579 |
| EIF4ENIF1 | 181.184574 | 0.316394138 | 0.21221677 | 1.49090073 | 0.13598756 | 0.5601128 |
| SFI1 | 164.361374 | 0.035392811 | 0.32413986 | 0.10918994 | 0.91305184 | 0.98480378 |
| PISD | 389.167033 | 0.465185332 | 0.43571952 | 1.06762564 | 0.28568941 | 0.73607046 |
| PRR14L | 308.892018 | -0.10670051 | 0.18061776 | -0.5907532 | 0.55468583 | 0.8907979 |
| YWHAH | 99.5605996 | -0.101549039 | 0.53114317 | -0.1911896 | 0.84837707 | 0.96991131 |
| RTCB | 162.596414 | 0.051927237 | 0.25736568 | 0.20176442 | 0.8401009 | 0.96833625 |
| FBXO7 | 189.571736 | 0.05572982 | 0.24352439 | 0.22884698 | 0.81898785 | 0.96443578 |
| TIMP3 | 332.235116 | -0.412477702 | 0.78544601 | -0.5251509 | 0.59947828 | 0.90369786 |
| LARGE1 | 553.484148 | -0.425534667 | 0.31998242 | -1.3298689 | 0.18356148 | 0.63405547 |
| HMGXB4 | 117.186016 | -0.387042729 | 0.32662881 | -1.184962 | 0.23603247 | 0.69335782 |
| TOM1 | 118.152518 | -0.491523935 | 0.20606664 | -2.3852669 | 0.01706673 | 0.21420282 |
| APOL6 | 97.6987026 | -0.93117546 | 0.70432377 | -1.3220844 | 0.18614005 | 0.63708286 |
| RBFOX2 | 953.688973 | 0.176318807 | 0.21021303 | 0.8387625 | 0.4016026 | 0.81660753 |
| APOL2 | 119.18544 | -1.472223071 | 0.53742709 | -2.7393912 | 0.00615531 | 0.12643442 |
| APOL1 | 110.597002 | -1.566481838 | 0.46889065 | -3.3408255 | 0.0008353 | 0.03573026 |
| MYH9 | 557.632777 | -0.110582255 | 0.22628808 | -0.4886791 | 0.6250689 | 0.91058133 |
| TXN2 | 146.112936 | -0.202431302 | 0.38080729 | -0.5315846 | 0.59501371 | 0.90323766 |
| EIF3D | 162.46874 | -0.584776455 | 0.26333798 | -2.2206309 | 0.02637597 | 0.27614403 |
| IFT27 | 80.2105165 | 0.345950547 | 0.23264285 | 1.48704567 | 0.13700275 | 0.5623928 |
| TST | 93.9726885 | -0.626607095 | 0.40993044 | -1.5285693 | 0.12637125 | 0.54650459 |
| MPST | 96.1363376 | -0.173188354 | 0.37130337 | -0.4664336 | 0.64090519 | 0.9163283 |
| CARD10 | 46.3127511 | -1.041094992 | 0.49336556 | -2.1101898 | 0.03484201 | 0.31514522 |
| CDC42EP1 | 66.4587737 | -1.268114466 | 0.4283533 | -2.9604405 | 0.00307199 | 0.08276221 |
| GGA1 | 144.069047 | 0.088703456 | 0.23330818 | 0.38019866 | 0.70379795 | 0.93241442 |
| LGALS1 | 145.994595 | -1.942871016 | 0.52743259 | -3.6836386 | 0.00022993 | 0.01460392 |
| TRIOBP | 247.74448 | -0.576647033 | 0.209925 | -2.7469192 | 0.00601579 | 0.12509033 |
| H1F0 | 304.713019 | -0.562457405 | 0.69849303 | -0.8052441 | 0.42067882 | 0.82694802 |
| GCAT | 35.1390126 | -0.486675551 | 0.42198192 | -1.153309 | 0.24878357 | 0.70561619 |
| ANKRD54 | 53.0596918 | -0.149041657 | 0.29623089 | -0.5031267 | 0.61487522 | 0.90792412 |
| EIF3L | 216.511776 | 0.277044935 | 0.29295668 | 0.94568567 | 0.34430893 | 0.78180304 |
| MICALL1 | 39.7920736 | 0.233269916 | 0.3214727 | 0.72562902 | 0.46806623 | 0.85161077 |
| POLR2F | 87.0104288 | 0.236366852 | 0.34885858 | 0.67754346 | 0.4980612 | 0.86583246 |
| PICK1 | 71.0616145 | -0.023190376 | 0.2654439 | -0.0873645 | 0.93038178 | 0.98818294 |
| PLA2G6 | 61.4556893 | 0.386245528 | 0.28970439 | 1.33324015 | 0.18245301 | 0.6329386 |
| MAFF | 119.013122 | -0.994831562 | 1.02910068 | -0.9666999 | 0.33369406 | 0.77744565 |
| TMEM184B | 144.333796 | 0.198448178 | 0.21715259 | 0.9138651 | 0.36078773 | 0.78927419 |
| CSNK1E | 58.78246 | 0.387939274 | 0.34558517 | 1.12255765 | 0.26162541 | 0.7198106 |
| TPTEP2-CSNI | 489.995925 | 0.737911178 | 0.30114827 | 2.45032515 | 0.01427273 | 0.19350346 |

| | | | | | | |
|---|---|---|---|---|---|---|
| DDX17 | 2381.56801 | 0.558837389 | 0.19071 | 2.93029937 | 0.00338636 | 0.08828112 |
| FAM227A | 98.1498455 | 0.01696518 | 0.28257921 | 0.0600369 | 0.95212625 | 0.9903694 |
| CBY1 | 117.46588 | -0.242905409 | 0.31232555 | -0.7777315 | 0.43672733 | 0.83393108 |
| TOMM22 | 57.5693304 | 0.514758344 | 0.47257879 | 1.08925401 | 0.27604189 | 0.72654437 |
| JOSD1 | 134.772865 | 0.716906938 | 0.74977919 | 0.95615742 | 0.33899271 | 0.77836439 |
| GTPBP1 | 91.4665406 | 0.418383541 | 0.38246806 | 1.09390451 | 0.27399686 | 0.72590691 |
| SUN2 | 84.5605582 | -0.474692738 | 0.27559584 | -1.7224234 | 0.08499284 | 0.46310155 |
| AL021707.1 | 40.4662084 | -0.07550264 | 0.33753054 | -0.2236913 | 0.82299753 | 0.96511681 |
| CBX6 | 182.799423 | 0.013834565 | 0.37857602 | 0.03654369 | 0.97084884 | 0.9939318 |
| CBX7 | 103.988952 | -0.030980254 | 0.36506774 | -0.0848617 | 0.93237137 | 0.98822345 |
| RPL3 | 804.368483 | 0.362466218 | 0.44301875 | 0.81817354 | 0.4132581 | 0.82437234 |
| SYNGR1 | 86.7333857 | -0.14646088 | 0.36917588 | -0.3967239 | 0.6915711 | 0.93073907 |
| TAB1 | 69.7815472 | -0.210054058 | 0.30861899 | -0.6806258 | 0.49610827 | 0.86452555 |
| MIEF1 | 58.5967715 | -0.272535418 | 0.31129444 | -0.8754908 | 0.38130692 | 0.80315561 |
| ATF4 | 159.718496 | -0.369337018 | 0.46277846 | -0.798086 | 0.42482058 | 0.82774157 |
| RPS19BP1 | 170.475381 | 0.321194117 | 0.48241303 | 0.66580729 | 0.50553428 | 0.87058024 |
| FAM83F | 73.2511305 | -1.286227203 | 0.51248207 | -2.5097994 | 0.01207998 | 0.17583076 |
| TNRC6B | 1827.66149 | 0.131450398 | 0.23562062 | 0.55789005 | 0.57691947 | 0.89869922 |
| MKL1 | 978.448517 | 0.675763567 | 0.27259495 | 2.47900254 | 0.01317504 | 0.18483853 |
| SLC25A17 | 154.083731 | 0.176193779 | 0.2756205 | 0.63926225 | 0.52265234 | 0.87688817 |
| ST13 | 396.061882 | -0.291142806 | 0.26007794 | -1.1194445 | 0.26295058 | 0.71989632 |
| XPNPEP3 | 222.627798 | 0.025577334 | 0.2298931 | 0.11125751 | 0.91141215 | 0.98468493 |
| RBX1 | 231.824164 | 0.164609959 | 0.27669123 | 0.59492295 | 0.55189498 | 0.88923362 |
| EP300 | 503.201864 | 0.293221673 | 0.19604125 | 1.49571418 | 0.13472815 | 0.55868799 |
| L3MBTL2 | 64.8261152 | 0.152036733 | 0.27992455 | 0.54313469 | 0.58703707 | 0.90196444 |
| RANGAP1 | 73.0982043 | -0.2540645 | 0.4000959 | -0.635009 | 0.52542254 | 0.87791848 |
| ZC3H7B | 172.547841 | -0.011759025 | 0.23774812 | -0.04946 | 0.9605527 | 0.9913918 |
| TEF | 63.1546153 | 0.197248499 | 0.4084428 | 0.48292808 | 0.6291468 | 0.9114564 |
| TOB2 | 277.293555 | 1.227943573 | 0.66826598 | 1.83750722 | 0.06613505 | 0.41858129 |
| PHF5A | 49.7835376 | -0.059512698 | 0.48837421 | -0.1218588 | 0.90301084 | 0.98320957 |
| ACO2 | 379.19431 | 0.174937119 | 0.20002591 | 0.87457231 | 0.38180666 | 0.8032969 |
| POLR3H | 54.0120274 | -0.004946271 | 0.34799927 | -0.0142135 | 0.98865969 | 0.9964791 |
| PMM1 | 62.8346765 | -0.07696509 | 0.31448234 | -0.2447358 | 0.806661 | 0.96218371 |
| DESI1 | 108.15741 | 0.814935996 | 0.51532652 | 1.58139734 | 0.11378721 | 0.52602773 |
| XRCC6 | 394.74532 | 0.209657994 | 0.35698603 | 0.58730028 | 0.55700205 | 0.89133937 |
| SNU13 | 346.144405 | -0.184354646 | 0.43414099 | -0.4246423 | 0.67109742 | 0.92474338 |
| CCDC134 | 37.2323892 | -0.168103267 | 0.37583871 | -0.447275 | 0.6546765 | 0.92047627 |
| SREBF2 | 955.544973 | 0.131692003 | 0.36578098 | 0.36002966 | 0.71882495 | 0.9359691 |
| WBP2NL | 86.3109713 | -0.105744799 | 0.51705566 | -0.2045134 | 0.83795234 | 0.96757829 |
| NAGA | 37.9905549 | -0.362296469 | 0.41349255 | -0.8761862 | 0.38092881 | 0.80308469 |
| SMDT1 | 178.154758 | 0.435889099 | 0.53626782 | 0.8128198 | 0.41632138 | 0.82489761 |
| NDUFA6 | 185.912218 | 0.138777227 | 0.43932648 | 0.31588632 | 0.75208879 | 0.94854974 |
| TCF20 | 370.220564 | 0.01994216 | 0.19838628 | 0.10052187 | 0.91993002 | 0.98540367 |
| RRP7A | 96.8937825 | -0.253980774 | 0.3614828 | -0.7026082 | 0.48229996 | 0.85727672 |
| SERHL2 | 73.6849889 | -0.163130063 | 0.36104488 | -0.4518277 | 0.65139315 | 0.91866508 |
| POLDIP3 | 124.412854 | 0.147080472 | 0.23771251 | 0.61873257 | 0.53609255 | 0.88137638 |
| CYB5R3 | 315.013898 | -0.632199308 | 0.35220902 | -1.7949549 | 0.07266089 | 0.43315584 |
| A4GALT | 130.400298 | -2.624751393 | 0.48336199 | -5.4301982 | 5.63E-08 | 3.93E-05 |
| ARFGAP3 | 383.951982 | -0.957231776 | 0.34421842 | -2.7808848 | 0.0054211 | 0.11811454 |
| PACSIN2 | 616.257945 | 0.325890701 | 0.29628011 | 1.0999412 | 0.27135774 | 0.72343657 |
| TTLL1 | 41.8095365 | -0.056563948 | 0.39980331 | -0.1414794 | 0.8874912 | 0.98009982 |
| MCAT | 39.287372 | -0.226734392 | 0.36831078 | -0.6156062 | 0.53815447 | 0.88265739 |
| TSPO | 405.319195 | -0.842285745 | 0.4887672 | -1.7232862 | 0.08483679 | 0.46310155 |
| EFCAB6 | 335.186777 | -0.30457168 | 0.33998728 | -0.8958326 | 0.37034219 | 0.79532503 |
| SAMM50 | 52.2944859 | 0.114022043 | 0.27603787 | 0.41306667 | 0.67955777 | 0.92700898 |
| RTL6 | 38.1292537 | 0.097634524 | 0.38864477 | 0.2512179 | 0.80164565 | 0.96069141 |
| NUP50-DT | 259.608915 | -0.239845339 | 0.35157603 | -0.6822005 | 0.49511218 | 0.86407588 |
| NUP50 | 145.902474 | 0.40018214 | 0.37988959 | 1.05341696 | 0.29214993 | 0.73844969 |
| KIAA0930 | 84.1001549 | -0.155547109 | 0.51530968 | -0.3018517 | 0.76276511 | 0.95378284 |
| FAM118A | 71.1465193 | 0.733195583 | 0.61221373 | 1.19761375 | 0.23106742 | 0.69089594 |
| ATXN10 | 709.676283 | 0.176931268 | 0.17390883 | 1.01737947 | 0.30897295 | 0.75355748 |
| MIRLET7BHG | 360.063234 | -0.076744712 | 0.36226265 | -0.2118483 | 0.83222542 | 0.96661004 |
| PPARA | 340.50714 | -0.280789377 | 0.31049188 | -0.9043373 | 0.3658166 | 0.79329304 |
| TTC38 | 82.3158725 | -0.081378753 | 0.37345002 | -0.2179107 | 0.82749869 | 0.96545294 |
| TRMU | 104.445242 | -0.406799038 | 0.20745922 | -1.9608627 | 0.04989504 | 0.37645791 |
| CELSR1 | 166.958266 | -0.094843804 | 0.37332484 | -0.2540517 | 0.79945563 | 0.96043078 |
| GRAMD4 | 50.3196308 | -0.034322465 | 0.33493155 | -0.1024761 | 0.91837881 | 0.98520052 |
| CERK | 135.222893 | 0.116165743 | 0.40115346 | 0.28957931 | 0.77213809 | 0.95537275 |
| TBC1D22A | 767.599847 | -0.427909092 | 0.26151668 | -1.6362592 | 0.10178534 | 0.49869583 |
| BRD1 | 153.495991 | 0.034151962 | 0.29415617 | 0.11610146 | 0.90757213 | 0.9835597 |
| ZBED4 | 79.1077116 | -0.163511049 | 0.45061503 | -0.3628619 | 0.71670801 | 0.93573028 |
| ALG12 | 43.5966288 | -0.306595536 | 0.35854724 | -0.855105 | 0.39249302 | 0.80971002 |
| CRELD2 | 37.8447552 | 0.32014853 | 0.28598128 | 1.11947375 | 0.26293808 | 0.71989632 |
| PIM3 | 564.481997 | 0.367611582 | 0.85466468 | 0.43012376 | 0.66710562 | 0.92422364 |
| TRABD | 107.353383 | 0.226910542 | 0.36098248 | 0.62859156 | 0.52961648 | 0.87929472 |
| SELENOO | 146.490371 | -0.318488267 | 0.24775308 | -1.2855068 | 0.19861524 | 0.65403502 |
| TUBGCP6 | 45.1781366 | 0.572750873 | 0.27836387 | 2.05756185 | 0.03963221 | 0.33495611 |
| PLXNB2 | 593.569235 | -0.065326534 | 0.34355431 | -0.1901491 | 0.84919232 | 0.96991131 |
| DENND6B | 74.4799549 | 0.141183189 | 0.31355174 | 0.45027078 | 0.6525152 | 0.91909172 |
| PPP6R2 | 576.158748 | 0.487800046 | 0.28442574 | 1.71503479 | 0.08633885 | 0.46616751 |
| SBF1 | 42.8889231 | 0.275898498 | 0.26697579 | 1.03342142 | 0.30140673 | 0.74814891 |
| LMF2 | 53.108038 | -0.473907846 | 0.32080692 | -1.477237 | 0.13961212 | 0.56656585 |
| ARSA | 42.2967794 | -0.63375997 | 0.307672 | -2.0598559 | 0.03941232 | 0.3346923 |
| RABL2B | 80.132067 | -0.076257048 | 0.25002546 | -0.3049971 | 0.76036834 | 0.95327948 |
| CU633967.1 | 66.4813652 | -0.073268265 | 0.65486739 | -0.1118826 | 0.91091649 | 0.98468493 |
| FP671120.1 | 1488.62304 | -0.776547397 | 0.4078009 | -1.9042317 | 0.05688002 | 0.39727338 |
| FP236383.1 | 915.378242 | -0.97019542 | 0.43197048 | -2.2459762 | 0.02470553 | 0.26637239 |
| CR381653.1 | 97.2669494 | -0.047777196 | 0.63290645 | -0.0754886 | 0.939826 | 0.98855185 |
| AP001347.1 | 86.962113 | -0.170846934 | 0.39317194 | -0.4345349 | 0.66390005 | 0.92300139 |
| HSPA13 | 56.4496489 | 0.392322717 | 0.3690811 | 1.06297157 | 0.28779485 | 0.73653438 |
| SAMSN1 | 222.037666 | 0.797803593 | 0.71079324 | 1.12241303 | 0.26168687 | 0.7198106 |
| NRIP1 | 301.249482 | 0.027571825 | 0.39659064 | 0.06952213 | 0.94457402 | 0.98949015 |
| USP25 | 814.424747 | 0.476157476 | 0.29624626 | 1.60730291 | 0.10798794 | 0.51162458 |
| MIR99AHG | 2261.71451 | 1.0916846 | 0.36470456 | 2.99333963 | 0.00275942 | 0.0769785 |
| CXADR | 379.989847 | 0.00179772 | 0.26644271 | 0.00674712 | 0.99461662 | 0.99888869 |
| BTG3 | 277.406056 | 0.789478344 | 0.28861519 | 2.73540123 | 0.00623043 | 0.12678619 |
| C21orf91 | 38.859694 | -0.124430048 | 0.37611646 | -0.3308285 | 0.74077402 | 0.94487568 |
| CHODL | 154.796164 | 0.138882318 | 0.52102242 | 0.26655728 | 0.78981005 | 0.95801395 |
| NCAM2 | 80.3758188 | 1.776123043 | 0.51195962 | 3.46926395 | 0.00052189 | 0.02592118 |
| MRPL39 | 65.5985906 | 0.003052844 | 0.2960817 | 0.01031082 | 0.99177331 | 0.99767558 |
| ATP5PF | 365.288455 | 0.049915062 | 0.4943803 | 0.10096491 | 0.91957832 | 0.98527601 |
| GABPA | 115.330821 | 0.295428459 | 0.22569647 | 1.30896359 | 0.19054668 | 0.64458858 |
| APP | 2458.10458 | 0.138438608 | 0.25769821 | 0.53721215 | 0.59112108 | 0.90233654 |
| CYYR1 | 95.4167177 | -0.927767555 | 0.60747878 | -1.5272427 | 0.12670066 | 0.54691843 |
| ADAMTS1 | 250.577812 | -3.273168454 | 0.67090686 | -4.8787226 | 1.07E-06 | 0.000373 |
| ADAMTS5 | 92.7343816 | 0.359700743 | 0.56119136 | 0.64095916 | 0.52154922 | 0.87650871 |
| N6AMT1 | 38.8535238 | -0.02550813 | 0.28364381 | -0.0899301 | 0.92834272 | 0.98751845 |
| LTN1 | 209.674612 | -0.085928405 | 0.21743534 | -0.3951906 | 0.69270222 | 0.93105058 |
| RWDD2B | 31.4727471 | -0.017771943 | 0.53809124 | -0.0330277 | 0.97365246 | 0.9939318 |
| USP16 | 148.043288 | 0.351292763 | 0.27611349 | 1.27227669 | 0.20327483 | 0.6582233 |
| CCT8 | 127.338592 | 0.017366554 | 0.21515462 | 0.08071662 | 0.93566732 | 0.98822345 |
| MAP3K7CL | 102.98955 | -0.227528072 | 0.53497604 | -0.4253052 | 0.67061423 | 0.92474338 |
| BACH1 | 1056.23484 | -0.995748167 | 0.40232984 | -2.4749548 | 0.0133253 | 0.18614502 |

| | | | | | | |
|---|---|---|---|---|---|---|
| GRIK1 | 70.9021455 | -2.120187982 | 0.62801185 | -3.3760318 | 0.00073539 | 0.03279545 |
| CLDN8 | 352.473494 | 0.541767323 | 0.79221354 | 0.68386526 | 0.49406025 | 0.86339027 |
| TIAM1 | 80.8250119 | 0.403878921 | 0.3885023 | 1.03957924 | 0.29853543 | 0.74594723 |
| SOD1 | 472.982456 | 0.029858473 | 0.4866983 | 0.06134904 | 0.95108124 | 0.99009947 |
| SCAF4 | 180.026118 | 0.452074956 | 0.18866419 | 2.39618848 | 0.01656657 | 0.21108198 |
| HUNK | 650.161629 | 0.829806664 | 0.44783329 | 1.85293652 | 0.06389146 | 0.41357646 |
| URB1 | 77.6016286 | 0.048283131 | 0.27123484 | 0.17801228 | 0.85871334 | 0.97237339 |
| EVA1C | 294.986768 | 0.520407761 | 0.53001341 | 0.9818766 | 0.32616065 | 0.77124629 |
| SYNJ1 | 229.115864 | 0.522803753 | 0.28721201 | 1.8202712 | 0.06871772 | 0.42634272 |
| PAXBP1 | 339.297896 | 0.237944608 | 0.24870845 | 0.95672103 | 0.33870808 | 0.77836439 |
| C21orf62-AS | 78.8122397 | -0.424967244 | 0.5286697 | -0.8038426 | 0.42148786 | 0.82703478 |
| IFNAR1 | 280.261854 | -0.156068337 | 0.26322854 | -0.5929005 | 0.55324773 | 0.88991776 |
| IFNGR2 | 382.757323 | 0.092043647 | 0.40723906 | 0.22601871 | 0.82118687 | 0.96502145 |
| TMEM50B | 476.437846 | 1.111518452 | 0.3225793 | 3.44572152 | 0.00056954 | 0.02714172 |
| GART | 120.088139 | 0.120360098 | 0.20353788 | 0.59134003 | 0.55429261 | 0.89076101 |
| SON | 661.187192 | -0.3000495 | 0.21996525 | -1.3640769 | 0.17254338 | 0.61884138 |
| AP000311.1 | 122.394 | 0.187035348 | 0.46740357 | 0.40015815 | 0.68904004 | 0.93036007 |
| ITSN1 | 456.687307 | 0.637869581 | 0.34306127 | 1.85934594 | 0.06297812 | 0.41224903 |
| MRPS6 | 810.965615 | 2.222488696 | 0.68126512 | 3.26229633 | 0.00110514 | 0.04225907 |
| LINC00310 | 69.4634575 | 1.862203124 | 0.68667656 | 2.71190722 | 0.00668973 | 0.13031298 |
| AP000317.1 | 85.5821482 | 1.646578972 | 0.50102538 | 3.28641826 | 0.0010147 | 0.03983282 |
| KCNE1 | 66.079562 | 2.908438256 | 0.81692204 | 3.56023969 | 0.00037052 | 0.02065432 |
| RCAN1 | 654.959155 | 2.042196524 | 0.92204472 | 2.21485627 | 0.02676993 | 0.27804646 |
| CLIC6 | 95.8364513 | -0.700425123 | 0.60692761 | -1.1540505 | 0.24847946 | 0.70513532 |
| RUNX1 | 261.458892 | -1.725626748 | 0.63446367 | -2.7198196 | 0.00653175 | 0.12867063 |
| SETD4 | 81.478231 | 0.005598288 | 0.24012294 | 0.02331426 | 0.9813996 | 0.99507903 |
| CBR1 | 178.319339 | -0.811639251 | 0.46145245 | -1.7588795 | 0.07859798 | 0.44839785 |
| CBR3 | 88.6534658 | -0.326527076 | 0.50584369 | -0.6455098 | 0.51859685 | 0.87546633 |
| DOPEY2 | 174.87757 | -0.19388306 | 0.23872019 | -0.812177 | 0.41669005 | 0.82489761 |
| MORC3 | 312.011377 | 0.06567929 | 0.19529999 | 0.3362995 | 0.73664503 | 0.9429136 |
| CLDN14 | 100.767056 | -2.377395781 | 0.79216863 | -3.0011233 | 0.00268986 | 0.07557557 |
| HLCS | 567.042731 | 0.115394832 | 0.25853886 | 0.44633458 | 0.65535558 | 0.92066039 |
| PIGP | 71.3028666 | 0.255422783 | 0.47210435 | 0.54103035 | 0.58848666 | 0.90200952 |
| TTC3 | 1111.31462 | 0.633693698 | 0.39059155 | 1.6223948 | 0.10471884 | 0.50411275 |
| VPS26C | 148.16936 | -0.399117418 | 0.23646485 | -1.6878509 | 0.09143985 | 0.47836139 |
| DYRK1A | 1108.47364 | 0.458589913 | 0.26340689 | 1.74099437 | 0.08168457 | 0.45528236 |
| KCNJ15 | 1435.57648 | 0.23196468 | 0.73264224 | 0.31661385 | 0.75153663 | 0.94834236 |
| ETS2 | 206.921898 | -0.897606353 | 0.84493655 | -1.0623358 | 0.28808326 | 0.73653438 |
| PSMG1 | 37.9501106 | -0.029315116 | 0.33537589 | -0.0874097 | 0.93034583 | 0.98818294 |
| BRWD1 | 978.741863 | 0.361621464 | 0.14515209 | 2.491328 | 0.01272666 | 0.18023698 |
| HMGN1 | 465.4897 | -0.200805601 | 0.28363413 | -0.707974 | 0.47896138 | 0.85532469 |
| WRB | 101.466608 | 0.499063303 | 0.25766339 | 1.93688089 | 0.05275991 | 0.38585053 |
| LCA5L | 43.0106943 | -0.083637613 | 0.35600427 | -0.2349343 | 0.81425973 | 0.96303374 |
| SH3BGR | 40.3669241 | 0.487182367 | 0.40195011 | 1.21204686 | 0.22549444 | 0.68280317 |
| PCP4 | 35.2721234 | 0.144224747 | 0.58776205 | 0.24537948 | 0.80616261 | 0.96218371 |
| BACE2 | 1545.98152 | -2.599463625 | 0.84194315 | -3.0874574 | 0.00201877 | 0.0629663 |
| FAM3B | 210.082354 | -0.740261014 | 0.30609971 | -2.4183656 | 0.0155904 | 0.2032348 |
| MX1 | 352.600715 | -0.691787492 | 1.07346384 | -0.6444441 | 0.51928746 | 0.87606771 |
| TMPRSS2 | 311.075576 | 0.18391714 | 0.52277806 | 0.3518073 | 0.72498278 | 0.93895336 |
| RIPK4 | 91.4505149 | 0.19777071 | 0.36037156 | 0.54879667 | 0.583145 | 0.90085422 |
| PRDM15 | 46.3841185 | -0.361026437 | 0.32312259 | -1.1173048 | 0.26386401 | 0.71989632 |
| C2CD2 | 116.574391 | -0.357847474 | 0.2478797 | -1.4436337 | 0.14884205 | 0.5817833 |
| ZBTB21 | 91.1867848 | 0.279624757 | 0.43295558 | 0.64585092 | 0.51837591 | 0.87526312 |
| RSPH1 | 81.3855767 | -0.069643119 | 0.48057359 | -0.1449167 | 0.88477667 | 0.97891513 |
| SLC37A1 | 134.492279 | -0.67058197 | 0.33108675 | -2.0253966 | 0.04282666 | 0.3484612 |
| LINC01671 | 77.6745951 | -1.381720719 | 0.75161807 | -1.8383282 | 0.06601405 | 0.41852829 |
| PDE9A | 605.013503 | -0.062842169 | 0.50468431 | -0.1245178 | 0.90090533 | 0.98320957 |
| WDR4 | 45.304616 | -0.3222473 | 0.42713292 | -0.7544427 | 0.45058346 | 0.84153105 |
| NDUFV3 | 203.134592 | 0.201620745 | 0.20544044 | 0.98140729 | 0.32639193 | 0.77144508 |
| PKNOX1 | 247.617794 | 0.080696817 | 0.23747594 | 0.33981049 | 0.73399925 | 0.94176201 |
| RRP1B | 77.8437804 | 0.542414937 | 0.28966417 | 1.87256481 | 0.0611285 | 0.40676596 |
| PDXK | 421.625362 | 0.298051484 | 0.31691443 | 0.94047936 | 0.34697173 | 0.78180304 |
| CSTB | 616.643106 | -0.443279064 | 0.55007522 | -0.8058517 | 0.42032835 | 0.82694802 |
| RRP1 | 33.3458837 | 0.110312368 | 0.3984467 | 0.27685602 | 0.78189066 | 0.95688565 |
| AGPAT3 | 300.319028 | -0.111975374 | 0.20288993 | -0.5519021 | 0.58101545 | 0.90045866 |
| TRAPPC10 | 246.925268 | 0.102417019 | 0.25238875 | 0.40579075 | 0.68489635 | 0.92881914 |
| PFKL | 592.122244 | -0.3820514 | 0.39718385 | -0.9619006 | 0.33609951 | 0.77820994 |
| C21orf2 | 66.762933 | 0.543431123 | 0.47477275 | 1.14461312 | 0.25236945 | 0.70813507 |
| UBE2G2 | 279.306907 | 0.209449057 | 0.24646094 | 0.84982657 | 0.39542152 | 0.81175661 |
| SUMO3 | 165.531249 | -0.082066255 | 0.42312288 | -0.1939537 | 0.84621212 | 0.96931938 |
| PTTG1IP | 447.799622 | -0.778890267 | 0.21947279 | -3.548915 | 0.00038682 | 0.02100463 |
| FAM207A | 49.2314787 | -0.06809735 | 0.41405444 | -0.1644647 | 0.86936531 | 0.97576204 |
| ADARB1 | 123.461102 | 0.189537071 | 0.32085824 | 0.59071903 | 0.55470869 | 0.8907979 |
| POFUT2 | 76.3334372 | 0.474726235 | 0.30576883 | 1.55256584 | 0.12052689 | 0.53823708 |
| COL18A1 | 827.424487 | 0.142021308 | 0.28445233 | 0.49927982 | 0.61758227 | 0.90794578 |
| PCBP3 | 198.872135 | 1.390054589 | 0.79887741 | 1.74000989 | 0.08185728 | 0.45535449 |
| COL6A1 | 46.5944288 | 0.285700908 | 0.59885962 | 0.47707492 | 0.63330877 | 0.91415622 |
| LSS | 105.902491 | -0.98369882 | 0.62335762 | -1.578065 | 0.11455069 | 0.52703872 |
| MCM3AP-AS | 51.44215 | -0.581359382 | 0.4179382 | -1.3910176 | 0.1642201 | 0.60471773 |
| MCM3AP | 183.56963 | 0.086074716 | 0.254578 | 0.33810744 | 0.73528222 | 0.94232801 |
| YBEY | 43.5688778 | -0.127038474 | 0.39643233 | -0.3204544 | 0.74862391 | 0.94678489 |
| PCNT | 102.70895 | 0.026995085 | 0.24648342 | 0.1095209 | 0.91278935 | 0.98480378 |
| DIP2A | 295.757193 | 0.058625171 | 0.21405379 | 0.27388055 | 0.78417642 | 0.95697044 |
| PRMT2 | 329.61481 | -0.317721253 | 0.27735603 | -1.1455358 | 0.25198729 | 0.70813507 |
| AC011043.1 | 103.39831 | 1.10649145 | 0.47531545 | 2.32790968 | 0.0199169 | 0.23692294 |
| AL592183.1 | 188.551408 | -0.719751113 | 0.46168652 | -1.5589606 | 0.1190057 | 0.53573012 |

**Collecting duct intercalated cells type A**

*This means that positive fold changes imply higher mean value in controls than in AKI and vice versa.

| | baseMean | log2 fold change control/AKI* | lfcSE | stat | pvalue | padj |
|---|---|---|---|---|---|---|
| AL669831.5 | 23.9983634 | 0.510478665 | 0.29007062 | 1.75984268 | 0.07843448 | 0.44182027 |
| NOC2L | 21.9691869 | 0.336508245 | 0.32238034 | 1.04382372 | 0.29656695 | 0.7415547 |
| ISG15 | 175.761502 | -2.502491493 | 1.57291861 | -1.590986 | 0.11161273 | 0.51407571 |
| AGRN | 45.9738332 | 0.420063179 | 0.356876 | 1.17705641 | 0.23917299 | 0.69356592 |
| C1orf159 | 23.2007535 | 0.210752912 | 0.31904866 | 0.66056668 | 0.50889024 | 0.86281145 |
| AL390719.2 | 15.8393265 | -0.218388646 | 0.35100126 | -0.6221876 | 0.53381849 | 0.86909171 |
| SDF4 | 62.3658481 | -0.136434042 | 0.25746783 | -0.5299071 | 0.59617632 | 0.89360216 |
| UBE2J2 | 49.111668 | 0.016689546 | 0.23755809 | 0.07025459 | 0.94399103 | 0.99086173 |
| ACAP3 | 14.4411895 | 0.404421951 | 0.40968781 | 0.98714666 | 0.32357074 | 0.76304922 |
| INTS11 | 38.8687937 | -0.061412272 | 0.26442463 | -0.2322487 | 0.81634487 | 0.96332667 |
| CPTP | 15.173372 | 0.136054765 | 0.54854792 | 0.24802713 | 0.80411342 | 0.95966218 |
| DVL1 | 25.6079257 | 0.163941556 | 0.32322638 | 0.50720352 | 0.61201203 | 0.90011356 |
| AURKAIP1 | 87.6826146 | 0.08160068 | 0.3642076 | 0.22404991 | 0.82271847 | 0.96521156 |
| CCNL2 | 78.1415717 | -0.076862968 | 0.23974602 | -0.3206017 | 0.74851229 | 0.94674262 |
| MRPL20 | 60.6860931 | 0.067158527 | 0.26286042 | 0.25549122 | 0.79834373 | 0.95848876 |
| VWA1 | 14.0596711 | -1.147221417 | 0.56407595 | -2.0338066 | 0.04197109 | 0.32888549 |
| ATAD3C | 10.3198 | 0.476765811 | 0.50632172 | 0.94162622 | 0.34638404 | NA |
| ATAD3B | 26.8393954 | 0.565049441 | 0.31161354 | 1.81330195 | 0.06978526 | 0.41879921 |
| ATAD3A | 17.6943416 | 0.093027251 | 0.35420064 | 0.26263999 | 0.79282808 | 0.95746995 |
| SSU72 | 79.5260158 | -0.068476518 | 0.31610397 | -0.2166266 | 0.82849938 | 0.96638336 |
| MIB2 | 16.8975317 | 0.963184354 | 0.33352813 | 2.88786541 | 0.00387866 | 0.08178909 |
| CDK11B | 29.3520728 | -0.132645585 | 0.3811406 | -0.3480227 | 0.72782313 | 0.94030592 |
| SLC35E2B | 25.2846838 | -0.021370791 | 0.44717241 | -0.0477909 | 0.96188285 | 0.99289813 |
| CDK11A | 12.9060735 | 0.104556394 | 0.40454652 | 0.25845332 | 0.79605707 | 0.95848876 |
| SLC35E2A | 21.7280457 | 0.237722135 | 0.41265607 | 0.57607813 | 0.56456237 | 0.88191823 |
| NADK | 50.1602058 | 0.244687046 | 0.27355585 | 0.89446835 | 0.37107135 | 0.7900012 |
| GNB1 | 312.9607 | -0.238795666 | 0.22637789 | -1.0548542 | 0.29149202 | 0.73994767 |
| PRKCZ | 92.9317922 | -0.257675476 | 0.403416 | -0.6387339 | 0.52299606 | 0.86740339 |
| FAAP20 | 48.9496007 | -0.324539705 | 0.33442619 | -0.9704375 | 0.33182848 | 0.76685572 |
| SKI | 34.711009 | 0.017260544 | 0.23628918 | 0.07304839 | 0.94176761 | 0.98991882 |
| MORN1 | 15.1996098 | 0.547818094 | 0.42310325 | 1.29476219 | 0.19540228 | 0.6459894 |
| RER1 | 85.0233206 | 0.052210666 | 0.18711118 | 0.27903553 | 0.78021756 | 0.95537097 |
| PEX10 | 17.5634942 | -0.188250364 | 0.38287492 | -0.4916759 | 0.6229485 | 0.90430717 |
| PANK4 | 13.3661344 | 0.460312324 | 0.39082621 | 1.17779287 | 0.23887919 | 0.69356592 |
| TNFRSF14-A | 17.1347379 | -0.649095783 | 0.45175831 | -1.4368209 | 0.15076888 | 0.58759787 |
| TNFRSF14 | 45.9019014 | -0.938964851 | 0.41264653 | -2.2754701 | 0.02287774 | 0.23759542 |
| LINC00982 | 49.9927911 | 0.821919985 | 0.52937697 | 1.55261756 | 0.12051453 | 0.53341392 |
| PRDM16 | 245.376301 | 0.101099757 | 0.28066489 | 0.3602152 | 0.71868621 | 0.93762693 |
| ARHGEF16 | 40.0161729 | -0.252991292 | 0.31866472 | -0.7939106 | 0.42724749 | 0.82375285 |
| TPRG1L | 25.4803409 | -0.179201371 | 0.36710623 | -0.4881458 | 0.62544656 | 0.90447745 |
| WRAP73 | 11.7268162 | 0.345115797 | 0.36098489 | 0.95603945 | 0.33905231 | NA |
| LRRC47 | 21.8441422 | -0.235428786 | 0.43765614 | -0.537931 | 0.59062472 | 0.89149106 |
| CEP104 | 48.8020517 | 0.280587411 | 0.28828029 | 0.97331458 | 0.33039698 | 0.76685572 |
| C1orf174 | 11.2212296 | -0.694211863 | 0.41518306 | -1.6720621 | 0.09451207 | NA |
| NPHP4 | 23.0466631 | -0.126258362 | 0.36629707 | -0.3446884 | 0.73032861 | 0.94155624 |
| RPL22 | 44.7341461 | 0.616877927 | 0.35549736 | 1.7352532 | 0.08269597 | 0.45473406 |
| RNF207 | 21.3646298 | -0.339052609 | 0.34932064 | -0.9706057 | 0.33174466 | 0.76685572 |
| ICMT | 25.514415 | 0.492067999 | 0.31803431 | 1.54721673 | 0.12181099 | 0.53742103 |
| NOL9 | 26.6235704 | 0.419700311 | 0.3967351 | 1.0578855 | 0.29010765 | 0.73979353 |
| ZBTB48 | 15.5919373 | 0.290283773 | 0.36999332 | 0.7845649 | 0.43270872 | 0.8248269 |
| KLHL21 | 25.9406754 | -0.331858316 | 0.60026889 | -0.5528494 | 0.58036652 | 0.89012555 |
| DNAJC11 | 72.1303552 | 0.129949185 | 0.24756728 | 0.52490452 | 0.59964956 | 0.89476262 |
| CAMTA1 | 380.132492 | -0.119653662 | 0.23308967 | -0.5133375 | 0.60771527 | 0.89764552 |
| VAMP3 | 30.2765187 | 0.102338124 | 0.33738762 | 0.30332507 | 0.76164214 | 0.9499444 |
| PER3 | 118.051848 | -0.254057947 | 0.39248954 | -0.6472986 | 0.51743868 | 0.86500999 |
| PARK7 | 118.135707 | -0.448997905 | 0.36445376 | -1.2319749 | 0.21795844 | 0.67202699 |
| ERRFI1 | 21.4685093 | -1.658128286 | 0.63492592 | -2.6115303 | 0.0090138 | 0.13790829 |
| RERE | 1223.53237 | 0.025785832 | 0.2296773 | 0.11226983 | 0.91060946 | 0.98340144 |
| ENO1 | 260.217417 | -0.838545965 | 0.35486094 | -2.363027 | 0.01812635 | 0.20906729 |
| GPR157 | 24.1032696 | 0.096537888 | 0.47322193 | 0.2040013 | 0.83835249 | 0.96748214 |
| H6PD | 47.0041557 | -0.503542553 | 0.26967635 | -1.8672107 | 0.06187217 | 0.39536238 |
| SPSB1 | 16.6124766 | -0.941529911 | 0.73912968 | -1.2738359 | 0.20272159 | 0.65568502 |
| SLC25A33 | 39.4475162 | -0.049000415 | 0.36779858 | -0.1332262 | 0.89401447 | 0.97833263 |
| PIK3CD | 11.3799688 | -0.210700147 | 0.41154863 | -0.511969 | 0.60867269 | NA |
| CLSTN1 | 65.3058163 | -0.082752659 | 0.29247879 | -0.2829356 | 0.77722621 | 0.95512504 |
| CTNNBIP1 | 23.2544558 | 0.240452956 | 0.28638322 | 0.83961958 | 0.40112172 | 0.80857869 |
| LZIC | 28.7232656 | 0.563526557 | 0.25871412 | 2.17818242 | 0.02939246 | 0.27178917 |
| NMNAT1 | 40.9502545 | 0.498339834 | 0.34405822 | 1.44841718 | 0.14750041 | 0.58421969 |
| RBP7 | 11.8498178 | 0.457846138 | 0.57440532 | 0.79707852 | 0.42540543 | NA |
| UBE4B | 186.937593 | -0.120138909 | 0.19218068 | -0.6251352 | 0.53188232 | 0.86846407 |
| KIF1B | 398.451509 | 0.017160978 | 0.2601783 | 0.06595853 | 0.94741084 | 0.99149867 |
| PGD | 47.7708866 | 0.179462427 | 0.32946529 | 0.54470815 | 0.58595426 | 0.89059199 |
| DFFA | 54.9936222 | 0.840163921 | 0.35393546 | 2.37377717 | 0.01760717 | 0.20601879 |
| PEX14 | 119.637217 | 0.136952632 | 0.19191554 | 0.71360886 | 0.47546907 | 0.84609246 |
| CASZ1 | 273.702332 | 0.115369773 | 0.42405064 | 0.27206603 | 0.78557125 | 0.95577371 |
| AL139423.1 | 12.6022701 | -0.726634622 | 0.69535876 | -1.044978 | 0.29603313 | 0.74123516 |
| SRM | 14.0198955 | 0.115528101 | 0.42476653 | 0.27198023 | 0.78563722 | 0.95577371 |
| EXOSC10 | 32.3016013 | 0.272000003 | 0.27667861 | 0.98309011 | 0.32556309 | 0.76334591 |
| MTOR | 127.608258 | 0.347795955 | 0.2153805 | 1.61479782 | 0.10635448 | 0.50162992 |
| UBIAD1 | 12.5945827 | 0.180651612 | 0.53686267 | 0.33649501 | 0.73649761 | 0.94365644 |
| FBXO2 | 18.383391 | -0.339476056 | 0.43241026 | -0.7850786 | 0.43240746 | 0.8248269 |
| FBXO44 | 17.8601803 | 0.818981917 | 0.40050718 | 2.04486203 | 0.04086846 | 0.32473787 |
| AGTRAP | 20.4507002 | -1.05969267 | 0.44539552 | -2.3792172 | 0.01734945 | 0.20451748 |
| MTHFR | 30.5191057 | -0.239792161 | 0.44492604 | -0.5389484 | 0.58992248 | 0.89149106 |
| CLCN6 | 32.9856089 | -1.190838102 | 0.30823384 | -3.8634243 | 0.00011181 | 0.00699705 |
| KIAA2013 | 26.0253803 | 0.821141383 | 0.35425211 | 2.31795762 | 0.02045162 | 0.22365361 |
| MFN2 | 59.7695047 | 0.400715774 | 0.28131305 | 1.42444789 | 0.15431686 | 0.59443382 |
| VPS13D | 343.01607 | -0.140193167 | 0.15388862 | -0.9110041 | 0.36229321 | 0.78714716 |
| DHRS3 | 115.573632 | -0.912558049 | 0.42656936 | -2.1392958 | 0.03241172 | 0.28526325 |
| PRDM2 | 232.797862 | -0.490111281 | 0.2453724 | -1.9974181 | 0.04577978 | 0.34053978 |
| KAZN | 992.31998 | -1.207191202 | 0.45103587 | -2.676486 | 0.00743987 | 0.12466529 |
| TMEM51-AS | 52.8140427 | -0.855758008 | 0.41215586 | -2.0762971 | 0.03786648 | 0.3123341 |
| TMEM51 | 433.819024 | -0.585625887 | 0.40048402 | -1.4622953 | 0.14366031 | 0.57797804 |
| C1orf195 | 17.938098 | -1.062798288 | 0.41433819 | -2.5650503 | 0.01031609 | 0.14935229 |
| CASP9 | 16.6300124 | -0.002212364 | 0.46037603 | -0.0048056 | 0.99616573 | 0.99956632 |
| DNAJC16 | 60.2943817 | 0.130554565 | 0.2272654 | 0.5744586 | 0.56565751 | 0.88294821 |
| DDI2 | 62.169987 | -0.209145114 | 0.2209515 | -0.9465657 | 0.34386013 | 0.77370523 |
| PLEKHM2 | 62.3262527 | -0.563588394 | 0.31630361 | -1.7817957 | 0.07478256 | 0.43178025 |
| UQCRHL | 16.4537425 | 0.817248667 | 0.58074107 | 1.40725137 | 0.15935286 | 0.60024133 |
| SPEN | 116.657918 | 0.082947332 | 0.21326827 | 0.38893424 | 0.69732479 | 0.92989421 |
| ZBTB17 | 21.1938301 | -0.117205593 | 0.31191397 | -0.3757626 | 0.70709342 | 0.93432317 |
| SRARP | 32.6270528 | -0.204147623 | 0.53627112 | -0.3806799 | 0.7034408 | 0.93304742 |
| CLCNKA | 201.227552 | 1.219296648 | 0.53468476 | 2.28040283 | 0.02258381 | 0.23595493 |
| CLCNKB | 336.39926 | 0.99213591 | 0.59885429 | 1.65672339 | 0.09757541 | 0.48339198 |
| FAM131C | 18.2227334 | -0.082636589 | 0.79352081 | -0.1041392 | 0.91705892 | 0.98495314 |
| EPHA2 | 12.7350112 | 0.648678971 | 0.74906802 | 0.8659814 | 0.38650036 | 0.80009555 |
| FBXO42 | 114.544861 | 0.142082186 | 0.20064158 | 0.7081393 | 0.47885876 | 0.84833462 |
| SZRD1 | 61.9413711 | -0.004938936 | 0.28079 | -0.0175894 | 0.98596639 | 0.99804172 |
| NECAP2 | 20.7324652 | -0.349439724 | 0.32652203 | -1.0701873 | 0.28453502 | 0.73638801 |
| NBPF1 | 29.3109943 | 0.365789239 | 0.32108457 | 1.13923021 | 0.25460715 | 0.70793186 |
| CROCC | 26.3791058 | 0.596129614 | 0.30590608 | 1.94873412 | 0.05132719 | 0.36077232 |

| | | | | | | |
|---|---|---|---|---|---|---|
| ATP13A2 | 18.8221856 | -0.037223231 | 0.37562622 | -0.0990965 | 0.92106168 | 0.98614962 |
| SDHB | 84.2649204 | 0.194825433 | 0.23512509 | 0.82860331 | 0.40732891 | 0.81294693 |
| RCC2 | 18.6532187 | 0.365944489 | 0.33982285 | 1.07686838 | 0.28153907 | 0.73311066 |
| ARHGEF10L | 127.309384 | -0.2466972 | 0.37692881 | -0.6544928 | 0.51279435 | 0.86346419 |
| KLHDC7A | 13.4925687 | 0.591652825 | 1.05733589 | 0.55956942 | 0.57577317 | 0.88746465 |
| UBR4 | 131.634152 | 0.226028665 | 0.20661613 | 1.09395458 | 0.2739749 | 0.72510729 |
| AL035413.1 | 18.3873377 | -0.224888182 | 0.50100945 | -0.4488701 | 0.65352534 | 0.9158679 |
| EMC1 | 20.02347 | 0.14626098 | 0.43946842 | 0.3328134 | 0.73927514 | 0.94376279 |
| MRTO4 | 12.776665 | 0.198094876 | 0.40436539 | 0.48989078 | 0.62421118 | 0.90447745 |
| AKR7A2 | 20.1338855 | 0.107248164 | 0.47560858 | 0.2254967 | 0.82159289 | 0.96447429 |
| CAPZB | 204.922011 | -0.201385622 | 0.2437325 | -0.8262567 | 0.40865847 | 0.81362627 |
| NBL1 | 16.0745965 | -0.576996056 | 0.45600836 | -1.265319 | 0.20575699 | 0.65836247 |
| TMCO4 | 223.10674 | 0.197806351 | 0.36709439 | 0.53884329 | 0.58999499 | 0.89149106 |
| OTUD3 | 25.2840262 | 0.379122497 | 0.35448095 | 1.06951446 | 0.28483792 | 0.73638801 |
| VWA5B1 | 220.719665 | 0.272009656 | 0.58620767 | 0.46401586 | 0.64263638 | 0.91146968 |
| CAMK2N1 | 250.152909 | 0.132119657 | 0.21391493 | 0.61762709 | 0.53682118 | 0.86963427 |
| MUL1 | 19.0175074 | 0.028535389 | 0.37733277 | 0.07562394 | 0.93971829 | 0.98885522 |
| CDA | 40.6734775 | -0.213176895 | 0.47460663 | -0.4491654 | 0.65331232 | 0.9158679 |
| PINK1 | 61.817769 | 0.313147955 | 0.38555829 | 0.81219354 | 0.41668059 | 0.81850986 |
| DDOST | 25.0116331 | 0.012138886 | 0.36146455 | 0.03358251 | 0.97321007 | 0.99526459 |
| HP1BP3 | 220.693694 | 0.660020918 | 0.18677482 | 3.53377895 | 0.00040966 | 0.01742867 |
| EIF4G3 | 509.462222 | -0.063384138 | 0.28555771 | -0.2219661 | 0.82434025 | 0.96539788 |
| ECE1 | 130.31039 | -0.574278535 | 0.38054704 | -1.5090869 | 0.13127659 | 0.55812058 |
| NBPF3 | 25.1203345 | 0.226769184 | 0.40850312 | 0.55512229 | 0.57881102 | 0.88935005 |
| RAP1GAP | 197.575599 | 0.375422099 | 0.39558208 | 0.94903718 | 0.3426017 | 0.7728457 |
| USP48 | 141.074019 | 0.072521753 | 0.21290636 | 0.34062747 | 0.73338405 | 0.94273028 |
| HSPG2 | 36.3732965 | 0.907779568 | 0.50724766 | 1.78961806 | 0.07351533 | 0.42788861 |
| CDC42 | 188.461102 | -0.33016306 | 0.19782837 | -1.6689369 | 0.09512989 | 0.48035393 |
| ZBTB40 | 56.1108881 | 0.437662064 | 0.30767098 | 1.4225003 | 0.15488106 | 0.59498864 |
| EPHB2 | 19.8508388 | -2.419551427 | 0.70507711 | -3.4316125 | 0.0006 | 0.0230954 |
| KDM1A | 128.144554 | 0.201288435 | 0.25956916 | 0.77547129 | 0.43806122 | 0.82709089 |
| AL031428.1 | 28.070482 | -0.270332041 | 0.42635261 | -0.6340574 | 0.52604334 | 0.86750588 |
| LUZP1 | 140.698394 | 0.290133354 | 0.15718047 | 1.84586139 | 0.06491233 | 0.40484427 |
| LINC01355 | 12.0548772 | 1.123999754 | 0.48484385 | 2.31827167 | 0.02043456 | NA |
| HNRNPR | 120.998681 | 0.113653582 | 0.17571252 | 0.6468155 | 0.51775136 | 0.86500999 |
| TCEA3 | 33.0076746 | 1.257847947 | 0.51649525 | 2.43535242 | 0.01487729 | 0.18484189 |
| ASAP3 | 51.9194 | 0.164192328 | 0.36112797 | 0.45466523 | 0.6493501 | 0.91417939 |
| ID3 | 29.8817794 | 1.055174102 | 0.47834757 | 2.20587323 | 0.02739288 | 0.26050089 |
| RPL11 | 348.952635 | 0.433994629 | 0.42836738 | 1.0131365 | 0.31099497 | 0.75416279 |
| ELOA | 27.7869937 | -0.0421127 | 0.34604882 | -0.1216958 | 0.90313991 | 0.98016176 |
| ELOA-AS1 | 21.4079812 | 0.329066554 | 0.50545915 | 0.65102502 | 0.51503034 | 0.86463667 |
| PITHD1 | 28.0890999 | 0.441452978 | 0.34304137 | 1.28687971 | 0.19813622 | 0.64973675 |
| LYPLA2 | 30.757692 | -0.111964779 | 0.39114846 | -0.2862462 | 0.77468953 | 0.95470568 |
| GALE | 39.9441778 | -1.499845588 | 0.31546865 | -4.7543411 | 1.99E-06 | 0.00033297 |
| HMGCL | 47.5642846 | -0.624435967 | 0.26214651 | -2.3820114 | 0.01721837 | 0.20451748 |
| FUCA1 | 59.4963331 | -0.234471425 | 0.27501453 | -0.8525783 | 0.39389318 | 0.80438874 |
| PNRC2 | 63.3867833 | 0.331136607 | 0.33048587 | 1.00196902 | 0.31635855 | 0.75864788 |
| SRSF10 | 103.645114 | 0.144336292 | 0.24300998 | 0.59395213 | 0.55254413 | 0.87576439 |
| IFNLR1 | 19.5459155 | 0.215454853 | 0.35461538 | 0.60757334 | 0.54347048 | 0.87233405 |
| STPG1 | 16.3573775 | 0.274459935 | 0.37222008 | 0.73735929 | 0.46090388 | 0.83769303 |
| NIPAL3 | 78.7056864 | 0.238570926 | 0.29696734 | 0.80335746 | 0.42176816 | 0.8205277 |
| RCAN3 | 55.7277606 | 0.203404663 | 0.37903969 | 0.53663157 | 0.59152214 | 0.89151328 |
| SRRM1 | 117.763204 | 0.272795383 | 0.21308819 | 1.28019945 | 0.200475 | 0.65342994 |
| CLIC4 | 21.3220876 | -0.637441383 | 0.58109798 | -1.0969603 | 0.27265877 | 0.72345643 |
| SYF2 | 47.4779667 | -0.334417413 | 0.31959528 | -1.0463778 | 0.29538663 | 0.74111836 |
| RSRP1 | 258.866914 | -0.018932479 | 0.29579154 | -0.0640062 | 0.94896533 | 0.99185581 |
| TMEM50A | 77.1227292 | -0.474563501 | 0.28568117 | -1.6611648 | 0.09668036 | 0.48257296 |
| MACO1 | 73.7200718 | -0.05180571 | 0.31101729 | -0.1665686 | 0.86770952 | 0.97434023 |
| LDLRAP1 | 23.0087878 | 0.386158386 | 0.38202505 | 1.01081955 | 0.31210281 | 0.75484355 |
| MAN1C1 | 326.860318 | 0.051015018 | 0.35938207 | 0.14195204 | 0.88711789 | 0.97729058 |
| AL031280.1 | 29.5523108 | -0.565101238 | 0.41912082 | -1.3483015 | 0.17756143 | 0.6251709 |
| MTFR1L | 38.0550042 | 0.743303965 | 0.37664871 | 1.97346744 | 0.04844233 | 0.35179202 |
| PAQR7 | 10.8645823 | 0.441918466 | 0.43128433 | 1.02465689 | 0.30552511 | NA |
| STMN1 | 13.4964445 | 0.085982957 | 0.64784477 | 0.13272154 | 0.89441361 | 0.97850297 |
| PAFAH2 | 28.5905567 | -0.220768108 | 0.32934708 | -0.6703205 | 0.50265348 | 0.86010152 |
| PDIK1L | 14.2478128 | 0.426114649 | 0.38031759 | 1.12041793 | 0.26253571 | 0.71333232 |
| ZNF593 | 24.9190626 | -0.315492876 | 0.38501314 | -0.819434 | 0.41253882 | 0.81556555 |
| CNKSR1 | 27.4204385 | -0.496726982 | 0.33713922 | -1.4733587 | 0.14065435 | 0.57494615 |
| CEP85 | 99.1872962 | -1.210724693 | 0.29868737 | -4.0534847 | 5.05E-05 | 0.00407887 |
| SH3BGRL3 | 57.1116077 | -0.913328589 | 0.32401709 | -2.8187667 | 0.00482085 | 0.09530573 |
| DHDDS | 38.0679391 | -0.149416172 | 0.30910294 | -0.4833864 | 0.62882137 | 0.90659442 |
| HMGN2 | 43.9609475 | 0.193225152 | 0.29060842 | 0.66489868 | 0.5061153 | 0.86214551 |
| RPS6KA1 | 40.7943899 | -0.674652785 | 0.40087288 | -1.6829594 | 0.09238295 | 0.4764741 |
| ARID1A | 144.316335 | 0.453049488 | 0.22937269 | 1.9751675 | 0.04824914 | 0.35162786 |
| ZDHHC18 | 15.5912539 | 0.834825652 | 0.34363222 | 2.42941612 | 0.01512316 | 0.18663446 |
| GPN2 | 20.6579375 | -0.145366116 | 0.39398515 | -0.3689634 | 0.71215498 | 0.9357596 |
| NUDC | 52.1447005 | 0.031505726 | 0.30767205 | 0.10240035 | 0.91843891 | 0.98521039 |
| NR0B2 | 14.9197282 | -0.492998799 | 0.77346981 | -0.637386 | 0.52387347 | 0.86747074 |
| KDF1 | 9.96923888 | -0.801214688 | 0.52414248 | -1.52862 | 0.12635867 | NA |
| TRNP1 | 27.6948409 | -0.801679046 | 0.39477007 | -2.0307493 | 0.04228043 | 0.32888549 |
| SLC9A1 | 34.7700708 | 0.593184661 | 0.42453088 | 1.39727093 | 0.16233211 | 0.60094226 |
| WDTC1 | 53.892314 | -0.161257984 | 0.22899335 | -0.7042038 | 0.48130587 | 0.85008502 |
| TMEM222 | 21.9451615 | -0.39777736 | 0.37180167 | -1.0698644 | 0.28468037 | 0.73638801 |
| WASF2 | 119.116841 | -0.064819253 | 0.24329125 | -0.2664266 | 0.7899107 | 0.95741689 |
| AHDC1 | 17.7318173 | 0.828542519 | 0.34851257 | 2.3773677 | 0.0174367 | 0.20451748 |
| IFI6 | 181.637526 | -2.340375672 | 1.5191855 | -1.5405463 | 0.12342724 | 0.54049755 |
| FAM76A | 24.4826784 | 0.129400686 | 0.29753099 | 0.43491499 | 0.66362413 | 0.91855484 |
| STX12 | 131.443262 | -0.248274382 | 0.29279307 | -0.8479517 | 0.39646487 | 0.80674902 |
| PPP1R8 | 21.3988322 | 0.173162707 | 0.34588101 | 0.50064243 | 0.6166228 | 0.90160403 |
| RPA2 | 23.8225872 | 0.282579429 | 0.3163197 | 0.89333489 | 0.37167786 | 0.7900012 |
| EYA3 | 282.001285 | -0.208349166 | 0.25274988 | -0.8243294 | 0.40975239 | 0.81413754 |
| DNAJC8 | 45.9072028 | -0.267809653 | 0.31718054 | -0.8443445 | 0.39847692 | 0.80714564 |
| ATP5IF1 | 135.433229 | 0.169801258 | 0.37246624 | 0.45588362 | 0.64847367 | 0.91417939 |
| SESN2 | 12.1313557 | -0.501878066 | 0.64894122 | -0.7733799 | 0.43929761 | NA |
| PHACTR4 | 215.740261 | 0.00062068 | 0.20355032 | 0.00304927 | 0.99756704 | 0.99982181 |
| RCC1 | 30.0834967 | -0.414848429 | 0.30629056 | -1.3544277 | 0.17559996 | 0.62231623 |
| TRNAU1AP | 59.332435 | -0.45934464 | 0.2385343 | -1.9256964 | 0.0541423 | 0.3701059 |
| SNHG12 | 12.8242921 | -0.290231667 | 0.46038218 | -0.6304146 | 0.52842333 | 0.86750588 |
| TAF12 | 25.1069917 | 0.063941803 | 0.36115442 | 0.17704838 | 0.85947039 | 0.97287712 |
| GMEB1 | 26.7185532 | 0.532209457 | 0.52608359 | 1.01164429 | 0.31170816 | 0.75460103 |
| YTHDF2 | 91.3780734 | 0.345116937 | 0.24658294 | 1.39959776 | 0.16163381 | 0.60060549 |
| EPB41 | 257.660586 | -0.203485723 | 0.30349883 | -0.6704663 | 0.50256061 | 0.86010152 |
| SRSF4 | 245.375654 | 0.159308324 | 0.20195453 | 0.78883265 | 0.43020982 | 0.82469033 |
| MECR | 26.5901407 | 0.067990235 | 0.36941645 | 0.18404767 | 0.85397606 | 0.97135895 |
| PTPRU | 11.2849979 | -0.581346609 | 0.44162088 | -1.3163929 | 0.18804219 | NA |
| LINC01648 | 18.8656871 | -7.023343637 | 1.79636258 | -3.9097584 | 9.24E-05 | 0.00605519 |
| PUM1 | 327.861926 | 0.120150155 | 0.15458579 | 0.77723932 | 0.43701759 | 0.82687287 |
| SNRNP40 | 44.0428847 | 0.181487239 | 0.24025785 | 0.75538528 | 0.45001785 | 0.83118301 |
| ZCCHC17 | 54.2659223 | 0.077162569 | 0.23883986 | 0.32307241 | 0.7466404 | 0.94618979 |
| SERINC2 | 118.030869 | -0.049212687 | 0.29240381 | -0.1683038 | 0.86634425 | 0.97434023 |
| TINAGL1 | 32.3380522 | 0.174293046 | 0.45426016 | 0.38368552 | 0.70121155 | 0.93127962 |
| PEF1 | 31.3981212 | -0.299236633 | 0.26997303 | -1.1083946 | 0.26769146 | 0.71922155 |
| PTP4A2 | 64.3151693 | -0.487380155 | 0.32456804 | -1.5016271 | 0.13319345 | 0.56148477 |

| | | | | | | |
|---|---|---|---|---|---|---|
| KHDRBS1 | 136.5728 | -0.23088708 | 0.22659723 | -1.0189316 | 0.30823544 | 0.7512664 |
| TMEM39B | 13.2117413 | 0.498316633 | 0.50362064 | 0.98946825 | 0.32243409 | 0.76283188 |
| KPNA6 | 143.676094 | 0.042876809 | 0.17147674 | 0.25004445 | 0.80255297 | 0.95908975 |
| TXLNA | 38.2790628 | -0.006125381 | 0.31297189 | -0.0195717 | 0.98438507 | 0.99773864 |
| EIF3I | 38.3882493 | -0.448633642 | 0.270272 | -1.6599339 | 0.09692776 | 0.48257296 |
| HDAC1 | 72.3963594 | 0.167704979 | 0.28675501 | 0.58483715 | 0.55865722 | 0.88004847 |
| MARCKSL1 | 49.4972918 | 0.357980146 | 0.56915758 | 0.62896491 | 0.52937203 | 0.86758926 |
| FAM229A | 13.0909113 | 1.020038516 | 0.3939731 | 2.58910699 | 0.00962252 | 0.14462571 |
| BSDC1 | 55.478729 | -0.046572557 | 0.26360093 | -0.1766783 | 0.85976111 | 0.97287712 |
| ZBTB8OS | 77.5998273 | -0.105513488 | 0.17477675 | -0.6037044 | 0.54604021 | 0.87258486 |
| RBBP4 | 73.8071335 | -0.006421036 | 0.21437018 | -0.029953 | 0.97610451 | 0.99556048 |
| SYNC | 17.8591545 | -0.302242946 | 0.41960987 | -0.7202951 | 0.4713433 | 0.84510721 |
| KIAA1522 | 156.299673 | -0.256403279 | 0.25989801 | -0.9865534 | 0.3238616 | 0.76304922 |
| YARS | 64.3130863 | -0.884017645 | 0.32888879 | -2.6878923 | 0.00719046 | 0.1219361 |
| S100PBP | 68.7500215 | -0.334286585 | 0.27082251 | -1.2343383 | 0.21707689 | 0.67156502 |
| TMEM54 | 33.7981424 | -0.424612989 | 0.40998003 | -1.0356919 | 0.30034591 | 0.7445324 |
| RNF19B | 36.0557501 | -0.329105271 | 0.45640392 | -0.7210834 | 0.47085823 | 0.84439356 |
| AL020995.1 | 29.9225966 | -0.537149588 | 0.60526242 | -0.8874656 | 0.37482827 | 0.7926388 |
| AK2 | 132.389044 | -0.23785334 | 0.22448341 | -1.0595587 | 0.28934543 | 0.73917583 |
| ZNF362 | 14.2826409 | 0.556608543 | 0.52341828 | 1.06341059 | 0.28759579 | 0.73803612 |
| AL513327.1 | 22.2297266 | -1.116726073 | 0.64966581 | -1.7189239 | 0.08562824 | 0.46118484 |
| PHC2 | 122.300226 | -0.255414724 | 0.27467121 | -0.9298926 | 0.3524267 | 0.77964174 |
| SMIM12 | 32.7490789 | -0.154429497 | 0.36739002 | -0.4203421 | 0.67423555 | 0.92189158 |
| ZMYM1 | 45.8208819 | 0.415407557 | 0.26575515 | 1.56312138 | 0.11802405 | 0.5279131 |
| SFPQ | 206.290411 | 0.26516953 | 0.20446535 | 1.29689225 | 0.19466826 | 0.6459894 |
| ZMYM4 | 315.856387 | 0.138968159 | 0.15522365 | 0.89527699 | 0.37063904 | 0.7900012 |
| KIAA0319L | 206.429286 | -0.039936022 | 0.22987991 | -0.1737256 | 0.86208113 | 0.97302617 |
| PSMB2 | 48.0799642 | -0.017111436 | 0.28043112 | -0.0610183 | 0.95134462 | 0.99217308 |
| AGO4 | 97.5481493 | -0.851431329 | 0.24868096 | -3.4237898 | 0.00061754 | 0.02349088 |
| AGO1 | 24.4235746 | 0.917979868 | 0.28817989 | 3.18544037 | 0.00144534 | 0.04136384 |
| AGO3 | 225.917826 | 0.342797338 | 0.20796158 | 1.64836858 | 0.09927706 | 0.48586654 |
| ADPRHL2 | 17.8451982 | -0.029608443 | 0.45898177 | -0.064509 | 0.94856496 | 0.99179534 |
| TRAPPC3 | 32.0519794 | -0.338408811 | 0.28746233 | -1.1772284 | 0.23910437 | 0.69356592 |
| MAP7D1 | 16.4020039 | -0.017489165 | 0.43147571 | -0.0405334 | 0.9676679 | 0.99393148 |
| THRAP3 | 150.956946 | -0.102485564 | 0.28702647 | -0.3570596 | 0.72104716 | 0.93818343 |
| EVA1B | 22.1367435 | -0.810586918 | 0.39368364 | -2.0589804 | 0.03949611 | 0.31899441 |
| STK40 | 69.2763342 | -0.339177837 | 0.37665227 | -0.9005065 | 0.36785074 | 0.78877653 |
| LSM10 | 16.9016712 | -0.510771113 | 0.48382579 | -1.0556922 | 0.29110886 | 0.73994767 |
| MRPS15 | 28.6139964 | -0.204467146 | 0.40406944 | -0.5060198 | 0.61284273 | 0.90011356 |
| MEAF6 | 33.124331 | -0.061892396 | 0.24316212 | -0.2545314 | 0.79908505 | 0.95848876 |
| SNIP1 | 15.5709844 | -0.254588793 | 0.33902598 | -0.7509419 | 0.45268765 | 0.83328686 |
| GNL2 | 14.5935614 | 0.088824667 | 0.45836814 | 0.19378456 | 0.84634458 | 0.97027687 |
| C1orf122 | 49.3303793 | -0.012451019 | 0.37004549 | -0.0336473 | 0.97315843 | 0.99526459 |
| MTF1 | 34.9584524 | -0.002670969 | 0.39186599 | -0.006816 | 0.99456164 | 0.99923337 |
| AL929472.2 | 13.8336167 | -0.957408648 | 0.50375211 | -1.9005551 | 0.05736031 | 0.38078905 |
| INPP5B | 99.384039 | -0.259396897 | 0.2784957 | -0.9314215 | 0.35163555 | 0.77927001 |
| SF3A3 | 29.2613836 | 0.517856685 | 0.25738045 | 2.01202806 | 0.04421698 | 0.33403793 |
| UTP11 | 10.5232274 | 0.311838014 | 0.43275547 | 0.72058711 | 0.47116359 | NA |
| RRAGC | 40.2981918 | -0.942970392 | 0.51233641 | -1.8405297 | 0.06569051 | 0.40715521 |
| AL139260.1 | 78.8955372 | -1.030505176 | 0.5952654 | -1.7311693 | 0.08342157 | 0.4560197 |
| RHBDL2 | 23.461865 | -1.397879521 | 0.59876311 | -2.334612 | 0.0195637 | 0.21965565 |
| AKIRIN1 | 89.841048 | 0.409358 | 0.30360285 | 1.34833386 | 0.17755103 | 0.6251709 |
| NDUFS5 | 223.601401 | -0.350658286 | 0.28446927 | -1.2326754 | 0.21769688 | 0.67202699 |
| MACF1 | 1210.32324 | -0.498528772 | 0.23610146 | -2.1115023 | 0.03472916 | 0.29785398 |
| PABPC4 | 37.2536393 | 0.425098323 | 0.26496187 | 1.60437544 | 0.10863132 | 0.50659799 |
| PPIE | 21.9451397 | 0.31659761 | 0.32625532 | 0.9703983 | 0.331848 | 0.76685572 |
| TRIT1 | 33.2897561 | 0.47572698 | 0.30496316 | 1.55994903 | 0.11887193 | 0.52993914 |
| CAP1 | 85.8625161 | -0.281367 | 0.23010112 | -1.2227972 | 0.2214063 | 0.67492938 |
| PPT1 | 22.801651 | -0.022824676 | 0.39768278 | -0.0573942 | 0.9542312 | 0.99228059 |
| RLF | 162.317225 | -0.395034587 | 0.20656126 | -1.9124331 | 0.05582067 | 0.37625808 |
| ZMPSTE24 | 168.753818 | 0.434254204 | 0.27551696 | 1.57614327 | 0.11499279 | 0.51928773 |
| COL9A2 | 50.7580362 | -0.421734986 | 0.73683905 | -0.572357 | 0.56708014 | 0.88417195 |
| SMAP2 | 36.095012 | -0.330200168 | 0.31950399 | -1.0334774 | 0.30138052 | 0.74591425 |
| AL603839.3 | 14.7141143 | 0.561600617 | 0.39785859 | 1.41155836 | 0.15808004 | 0.59870582 |
| ZNF684 | 40.686732 | 0.346960658 | 0.42077863 | 0.82456816 | 0.40961681 | 0.81413754 |
| NFYC | 167.905015 | 0.581185902 | 0.25951351 | 2.23952076 | 0.02512205 | 0.25044595 |
| CITED4 | 95.0889676 | -0.893282576 | 0.52500976 | -1.701459 | 0.08885683 | 0.4687332 |
| CTPS1 | 26.3164105 | -0.213558539 | 0.41124342 | -0.5192996 | 0.60355184 | 0.8956362 |
| SLFNL1-AS1 | 21.8780985 | -1.10889774 | 0.4891647 | -2.266921 | 0.02339505 | 0.23883233 |
| SCMH1 | 491.661835 | -0.443264985 | 0.22656828 | -1.95643 | 0.05041452 | 0.35799473 |
| HIVEP3 | 34.1796528 | -0.089868117 | 0.35308199 | -0.2545248 | 0.79909016 | 0.95848876 |
| FOXJ3 | 173.198557 | 0.202035834 | 0.18789021 | 1.07528668 | 0.2822464 | 0.73374523 |
| PPCS | 37.9807388 | 0.0344444 | 0.32258719 | 0.10677547 | 0.91496711 | 0.98407488 |
| CCDC30 | 49.0099995 | 0.357564043 | 0.40907355 | 0.87408252 | 0.38207331 | 0.79731722 |
| PPIH | 15.6029761 | 0.682262762 | 0.3594377 | 1.89813914 | 0.05767776 | 0.38148784 |
| YBX1 | 116.861617 | -0.096308964 | 0.39807782 | -0.241935 | 0.80883052 | 0.96124185 |
| SVBP | 19.9431761 | -0.220881643 | 0.42046997 | -0.5253209 | 0.59936016 | 0.89476262 |
| ERMAP | 29.1541755 | 0.350685736 | 0.25434941 | 1.37875583 | 0.16797005 | 0.60954339 |
| SLC2A1 | 27.4708722 | -1.451567562 | 0.56822451 | -2.554567 | 0.010632 | 0.15188567 |
| EBNA1BP2 | 22.8198049 | 0.324285196 | 0.3623185 | 0.89502798 | 0.37077213 | 0.7900012 |
| TMEM125 | 11.5005534 | 0.509906344 | 0.5717354 | 0.89185723 | 0.37246946 | NA |
| ELOVL1 | 26.0463493 | -0.117937075 | 0.33924138 | -0.3476494 | 0.72810348 | 0.9404266 |
| MED8 | 14.9248813 | -0.717220327 | 0.45646472 | -1.5712503 | 0.11612452 | 0.52196843 |
| SZT2 | 38.6740412 | 0.128119258 | 0.24123904 | 0.53108842 | 0.59535751 | 0.893081 |
| PTPRF | 127.604024 | 0.278533675 | 0.27211224 | 1.02359847 | 0.30602497 | 0.74897897 |
| IPO13 | 15.3074422 | 0.865472666 | 0.39726116 | 2.17859874 | 0.02936149 | 0.27178917 |
| ATP6V0B | 531.68551 | -0.07013951 | 0.34355049 | -0.2041607 | 0.83822792 | 0.96748214 |
| B4GALT2 | 22.5393472 | 0.22510411 | 0.36616136 | 0.61476751 | 0.53870827 | 0.86972442 |
| CCDC24 | 24.7889306 | -0.243333245 | 0.36267515 | -0.6709399 | 0.50225879 | 0.86010152 |
| DMAP1 | 16.3952519 | 0.534160193 | 0.42595113 | 1.25404102 | 0.2098271 | 0.66164938 |
| ERI3 | 64.4545965 | 0.293359584 | 0.26459363 | 1.10871747 | 0.26755208 | 0.71922155 |
| RNF220 | 29.0040649 | 0.217760474 | 0.25358948 | 0.85871256 | 0.39049912 | 0.8023388 |
| TMEM53 | 46.5340829 | 0.063071906 | 0.36816634 | 0.17131361 | 0.86397718 | 0.97395945 |
| ARMH1 | 12.8470632 | 0.337347496 | 0.39251062 | 0.85946081 | 0.39008633 | 0.80191835 |
| RPS8 | 246.794105 | 0.184619629 | 0.40137702 | 0.45996562 | 0.6455409 | 0.91225913 |
| EIF2B3 | 47.2876845 | 0.803883289 | 0.27957149 | 2.87541223 | 0.004035 | 0.08381056 |
| HECTD3 | 13.2433242 | -0.228801904 | 0.51824225 | -0.441496 | 0.65885393 | 0.91706167 |
| UROD | 13.543191 | 0.610863061 | 0.44250201 | 1.38047522 | 0.16744037 | 0.60898823 |
| ZSWIM5 | 116.338998 | 0.320423685 | 0.2497275 | 1.28309329 | 0.19945939 | 0.65266113 |
| TESK2 | 14.1774966 | 0.363744264 | 0.44077013 | 0.82524709 | 0.40923133 | 0.81413754 |
| PRDX1 | 297.807616 | -0.421314679 | 0.52106509 | -0.8085644 | 0.41876575 | 0.8189355 |
| AKR1A1 | 18.8744774 | 0.190308497 | 0.33629931 | 0.56589024 | 0.5714684 | 0.88578992 |
| NASP | 74.2717054 | 0.54893837 | 0.29704704 | 1.84798464 | 0.06460456 | 0.40351853 |
| GPBP1L1 | 213.943532 | 0.103720969 | 0.18235004 | 0.56880146 | 0.56949088 | 0.88562534 |
| TMEM69 | 13.1441783 | 0.424666896 | 0.37221695 | 1.14091231 | 0.25390642 | 0.70701839 |
| IPP | 44.3657729 | 0.459438096 | 0.30674042 | 1.4978075 | 0.13418327 | 0.56229288 |
| MAST2 | 55.9034701 | -0.313647526 | 0.27289641 | -1.1493282 | 0.25042069 | 0.70453262 |
| TSPAN1 | 309.937517 | -1.660481579 | 0.4876051 | -3.4053819 | 0.00066072 | 0.02464978 |
| POMGNT1 | 63.3393096 | 0.019476927 | 0.25415497 | 0.07663406 | 0.93891466 | 0.98871562 |
| LURAP1 | 48.6882202 | 0.597444343 | 0.36193574 | 1.65069174 | 0.09880153 | 0.48476221 |
| LRRC41 | 43.661885 | -0.254453121 | 0.23500207 | -1.0827697 | 0.27891066 | 0.72981748 |
| UQCRH | 157.174575 | 0.515205291 | 0.3511591 | 1.46715631 | 0.14233354 | 0.57673923 |
| NSUN4 | 42.5739297 | 0.117713951 | 0.24093156 | 0.48857839 | 0.62514022 | 0.90447745 |

| | | | | | | |
|---|---|---|---|---|---|---|
| FAAH | 18.8190814 | 0.666708224 | 0.37266951 | 1.78900662 | 0.07361375 | 0.42788861 |
| MKNK1 | 80.6818358 | -0.020894744 | 0.23015954 | -0.0907837 | 0.92766443 | 0.98690525 |
| ATPAF1 | 61.0008257 | 0.064010073 | 0.24758254 | 0.25854034 | 0.79598992 | 0.95848876 |
| EFCAB14 | 125.384313 | 0.153915945 | 0.18229093 | 0.8443423 | 0.39847816 | 0.80714564 |
| PDZK1IP1 | 28.80599 | -1.785057918 | 0.71781727 | -2.486786 | 0.01289029 | 0.16919599 |
| CMPK1 | 73.6281342 | 0.014314458 | 0.31306265 | 0.04572394 | 0.96353028 | 0.99329511 |
| TRABD2B | 139.339693 | -0.480255391 | 0.68308435 | -0.7030689 | 0.48201279 | 0.85071398 |
| SPATA6 | 113.47569 | -0.180615119 | 0.31559573 | -0.572299 | 0.56711943 | 0.88417195 |
| AGBL4 | 165.87497 | 0.917806234 | 0.53546005 | 1.71405174 | 0.08651922 | 0.46341053 |
| FAF1 | 569.431393 | -0.089780193 | 0.18611306 | -0.482396 | 0.62952467 | 0.90720388 |
| RNF11 | 113.716088 | -0.113750284 | 0.22124461 | -0.5141381 | 0.60715544 | 0.89696793 |
| TTC39A | 38.323504 | -1.860747573 | 0.44439423 | -4.1871552 | 2.82E-05 | 0.0026346 |
| EPS15 | 350.184419 | -0.091193735 | 0.23146124 | -0.3939914 | 0.69358739 | 0.92826395 |
| OSBPL9 | 246.144869 | 0.212054766 | 0.1669747 | 1.26998145 | 0.20409124 | 0.65771647 |
| NRDC | 566.567795 | -0.161137129 | 0.34460914 | -0.4675939 | 0.64007503 | 0.91096398 |
| RAB3B | 49.3007467 | 0.707903335 | 0.50493958 | 1.40195653 | 0.16092823 | 0.60060549 |
| TXNDC12 | 39.9870979 | -0.229979278 | 0.23989918 | -0.9586497 | 0.33773525 | 0.76966704 |
| BTF3L4 | 69.4679026 | -0.07268815 | 0.19314261 | -0.3763444 | 0.70666084 | 0.93432317 |
| ZFYVE9 | 92.6397773 | -0.397838157 | 0.23557614 | -1.688788 | 0.09126007 | 0.47439586 |
| CC2D1B | 36.9761157 | -0.454045088 | 0.40869415 | -1.1109655 | 0.26658322 | 0.71754988 |
| PRPF38A | 22.0621891 | 0.505468857 | 0.38586473 | 1.30996386 | 0.19020806 | 0.6421543 |
| TUT4 | 231.293641 | -0.279011669 | 0.1791153 | -1.557721 | 0.1192994 | 0.53058422 |
| ZYG11B | 129.2276 | 0.377715163 | 0.28100936 | 1.34413731 | 0.17890399 | 0.62639418 |
| ECHDC2 | 185.917337 | 0.322151093 | 0.20559868 | 1.56689284 | 0.11713974 | 0.52482932 |
| SCP2 | 162.542733 | 0.40329325 | 0.27531412 | 1.46484768 | 0.14296248 | 0.57698886 |
| CPT2 | 28.5505532 | -0.218950163 | 0.32217333 | -0.6796036 | 0.49675547 | 0.85814183 |
| C1orf123 | 12.2189638 | 0.234771517 | 0.47144836 | 0.49797928 | 0.61849865 | NA |
| MAGOH | 19.7983921 | 0.073606551 | 0.34108319 | 0.21580234 | 0.82914182 | 0.96669991 |
| NDC1 | 25.4649657 | 0.608846759 | 0.27270722 | 2.23260227 | 0.02557518 | 0.25205612 |
| YIPF1 | 24.2341432 | -0.463313712 | 0.3078817 | -1.5048433 | 0.13236437 | 0.55993651 |
| HSPB11 | 32.6654844 | -0.015820518 | 0.33823492 | -0.0467738 | 0.96269354 | 0.99306991 |
| LRRC42 | 16.2154744 | 0.093577881 | 0.34532875 | 0.27098202 | 0.78640486 | 0.95577371 |
| TMEM59 | 208.399496 | -0.284191237 | 0.29502589 | -0.9632756 | 0.33540925 | 0.76777176 |
| TCEANC2 | 22.366409 | 0.196222179 | 0.32049562 | 0.61224605 | 0.54037498 | 0.87026896 |
| MRPL37 | 23.60467 | 0.092162668 | 0.312992 | 0.29445695 | 0.76840876 | 0.95205353 |
| SSBP3 | 115.652335 | -0.33854147 | 0.38477664 | -0.8798389 | 0.37894657 | 0.79535699 |
| ACOT11 | 91.9218544 | -0.330900393 | 0.44706273 | -0.7401655 | 0.4591996 | 0.83645518 |
| DHCR24 | 73.5879981 | -0.408366662 | 0.50689987 | -0.805616 | 0.42046428 | 0.81938194 |
| TMEM61 | 129.573638 | -0.388866266 | 0.39214743 | -0.9916328 | 0.32137667 | 0.76163052 |
| BSND | 87.5344687 | 0.415965804 | 0.39005272 | 1.06643484 | 0.28622712 | 0.73672385 |
| USP24 | 244.709084 | -0.164475369 | 0.17904363 | -0.9186329 | 0.35828763 | 0.78380469 |
| PLPP3 | 108.928072 | -0.037501212 | 0.36238073 | -0.1034857 | 0.91757752 | 0.98511367 |
| PRKAA2 | 429.250301 | 0.298839184 | 0.14030877 | 2.12986818 | 0.0331825 | 0.28919158 |
| FYB2 | 1025.82632 | -1.357529785 | 0.44282478 | -3.0656139 | 0.00217224 | 0.05402742 |
| DAB1 | 118.96525 | 0.875984312 | 0.57030305 | 1.53599795 | 0.12453888 | 0.54320519 |
| TACSTD2 | 66.8800593 | 0.376226908 | 0.6941682 | 0.54198234 | 0.58783067 | 0.89073775 |
| MYSM1 | 127.41908 | -0.009981427 | 0.22040236 | -0.0452873 | 0.96387831 | 0.99334616 |
| JUN | 68.8402241 | 1.198568259 | 0.42911469 | 2.79311866 | 0.00522025 | 0.09995328 |
| LINC01135 | 66.7847068 | 0.020820172 | 0.44845882 | 0.04642605 | 0.96297067 | 0.99306991 |
| FGGY | 116.185459 | 0.077921197 | 0.32056658 | 0.24307336 | 0.80794856 | 0.96054677 |
| HOOK1 | 81.2955108 | 0.093770243 | 0.30710848 | 0.30533264 | 0.76011282 | 0.94952011 |
| NFIA | 230.604398 | 0.09835418 | 0.43348006 | 0.22689436 | 0.82050589 | 0.96389705 |
| TM2D1 | 96.0275696 | 0.295632632 | 0.18162421 | 1.62771603 | 0.10358511 | 0.4956959 |
| PATJ | 947.684313 | -0.027931124 | 0.15415378 | -0.18119 | 0.85621844 | 0.97166553 |
| USP1 | 22.607273 | 0.407851591 | 0.36383793 | 1.12097051 | 0.26230041 | 0.71321158 |
| DOCK7 | 247.391786 | 0.04770743 | 0.31298295 | 0.15242821 | 0.87884921 | 0.97556614 |
| ATG4C | 41.2545826 | 0.112257954 | 0.47443185 | 0.23661555 | 0.81295505 | 0.96262567 |
| LINC01739 | 28.1845605 | 1.209631504 | 0.73982083 | 1.63503304 | 0.10204212 | 0.49220184 |
| ALG6 | 18.0053026 | 0.591501257 | 0.32930835 | 1.79619269 | 0.07246388 | 0.42448303 |
| ITGB3BP | 54.2933861 | -0.532674721 | 0.4159847 | -1.2805152 | 0.20036401 | 0.65330283 |
| EFCAB7 | 22.8116028 | -0.207156806 | 0.3414898 | -0.6066266 | 0.54409872 | 0.87247631 |
| PGM1 | 126.984264 | -0.711474557 | 0.24933324 | -2.8535086 | 0.00432394 | 0.08811382 |
| CACHD1 | 40.2094262 | 0.170730232 | 0.61365169 | 0.2782201 | 0.78084341 | 0.95537097 |
| RAVER2 | 32.6971025 | -0.186462538 | 0.41002267 | -0.4547615 | 0.64928081 | 0.91417939 |
| JAK1 | 286.678911 | -1.093055693 | 0.23502976 | -4.6507119 | 3.31E-06 | 0.00049364 |
| AK4 | 18.4468464 | -2.055776691 | 0.66418775 | -3.0951741 | 0.00196697 | 0.05060915 |
| DNAJC6 | 12.4259551 | -0.122614573 | 0.50139964 | -0.2445446 | 0.80680907 | NA |
| LEPR | 72.9014727 | -0.683631258 | 0.41499571 | -1.6473212 | 0.09949203 | 0.48642778 |
| WDR78 | 23.3129927 | 0.312825575 | 0.41430272 | 0.75506522 | 0.45020985 | 0.83118301 |
| MIER1 | 120.850914 | -0.220316447 | 0.19625855 | -1.1225827 | 0.26161478 | 0.71281761 |
| SLC35D1 | 32.3168149 | 0.030556687 | 0.30964137 | 0.09868412 | 0.92138908 | 0.98614962 |
| SERBP1 | 116.912488 | 0.060337352 | 0.22804167 | 0.26458915 | 0.79132598 | 0.95746995 |
| GADD45A | 74.6254915 | -1.280370129 | 0.43011666 | -2.9767973 | 0.00291276 | 0.0667939 |
| GNG12 | 408.765594 | 0.501482338 | 0.39446547 | 1.27129591 | 0.2036234 | 0.65771647 |
| GNG12-AS1 | 151.765287 | 0.308367341 | 0.40205834 | 0.76697163 | 0.44309838 | 0.82894008 |
| WLS | 224.753255 | -0.200018099 | 0.32165859 | -0.6218335 | 0.53405133 | 0.86909171 |
| LRRC40 | 52.6531858 | 0.210950505 | 0.26222517 | 0.80446324 | 0.42112949 | 0.81961398 |
| SRSF11 | 214.234112 | 0.102923293 | 0.18039407 | 0.57054698 | 0.56830677 | 0.88512775 |
| ANKRD13C | 102.254289 | -0.044414268 | 0.21162621 | -0.2098713 | 0.83376812 | 0.96701848 |
| CTH | 15.4504603 | -0.78946577 | 0.70182986 | -1.1248677 | 0.26064508 | 0.71118349 |
| PTGER3 | 3332.00678 | -0.180113627 | 0.35149699 | -0.5124187 | 0.60835801 | 0.8980479 |
| ZRANB2-AS1 | 22.7509993 | 0.431849866 | 0.49535405 | 0.87180042 | 0.38331727 | 0.7980695 |
| ZRANB2 | 202.482207 | 0.141247193 | 0.20960137 | 0.67388488 | 0.50038451 | 0.85942996 |
| CRYZ | 122.301025 | 0.265443463 | 0.33419294 | 0.79428207 | 0.42703124 | 0.82375285 |
| TYW3 | 35.6067655 | -0.120440187 | 0.43046002 | -0.2797941 | 0.77963546 | 0.95537097 |
| ACADM | 99.7046179 | 0.306654694 | 0.25610605 | 1.19737389 | 0.23116086 | 0.68570652 |
| RABGGTB | 26.8482441 | 0.195177538 | 0.41420525 | 0.47120972 | 0.63749098 | 0.91003128 |
| PIGK | 45.4252858 | 0.345122641 | 0.24670413 | 1.39893337 | 0.16183296 | 0.60088379 |
| AC118549.1 | 222.016114 | -0.021481583 | 0.1463631 | -0.1467691 | 0.88331426 | 0.97662748 |
| USP33 | 224.364838 | -0.124513552 | 0.20265724 | -0.6144047 | 0.53894795 | 0.86972442 |
| MIGA1 | 113.971006 | 0.012137304 | 0.19733501 | 0.06150609 | 0.95095617 | 0.99217308 |
| NEXN | 37.2867615 | 0.097326813 | 0.34141087 | 0.28507239 | 0.77558868 | 0.95493992 |
| FUBP1 | 135.216674 | 0.115408301 | 0.20044816 | 0.57575136 | 0.56478326 | 0.88191823 |
| GIPC2 | 27.8512075 | -0.337393627 | 0.55176958 | -0.6114756 | 0.54088478 | 0.87065754 |
| PTGFR | 15.4134734 | 0.347313043 | 0.96252346 | 0.36083592 | 0.71822211 | 0.93733389 |
| IFI44L | 101.769402 | -3.59235183 | 2.54799813 | -1.4098722 | 0.15857741 | 0.59881912 |
| IFI44 | 44.1598777 | -1.830048811 | 1.3478164 | -1.3577879 | 0.17453099 | 0.62052871 |
| ADGRL2 | 236.764378 | 0.350729575 | 0.30517033 | 1.1492912 | 0.25043592 | 0.70453262 |
| LINC01725 | 106.11703 | 0.006675705 | 0.29877273 | 0.02234375 | 0.98217375 | 0.99718289 |
| PRKACB | 102.355385 | 0.796937363 | 0.40774922 | 1.95447921 | 0.05064457 | 0.35884028 |
| RPF1 | 20.1519276 | 0.333901921 | 0.30336129 | 1.10067411 | 0.27103852 | 0.72194341 |
| GNG5 | 76.9225415 | -0.121582551 | 0.27923685 | -0.4354101 | 0.66326477 | 0.91843944 |
| SPATA1 | 21.3041834 | -1.036997402 | 0.58429063 | -1.7747973 | 0.07593137 | 0.43402139 |
| CTBS | 46.8725198 | 0.067369514 | 0.26219551 | 0.25694382 | 0.79722215 | 0.95848876 |
| SSX2IP | 25.8437464 | 0.704790493 | 0.42640351 | 1.65287217 | 0.09835688 | 0.48476221 |
| WDR63 | 23.7910404 | -0.54005682 | 0.61035673 | -0.8848216 | 0.37625286 | 0.79422885 |
| MCOLN3 | 29.4985358 | -0.589344906 | 0.60680688 | -0.9712232 | 0.33143715 | 0.76685572 |
| SYDE2 | 19.6444307 | 0.809114072 | 0.40325236 | 2.00647076 | 0.04480605 | 0.33628081 |
| C1orf52 | 31.7468447 | 0.183820009 | 0.26779364 | 0.68642411 | 0.4924457 | 0.85714521 |
| BCL10 | 36.0707076 | -0.043370515 | 0.31867472 | -0.1360965 | 0.89174499 | 0.97793768 |
| AL078459.1 | 35.1802386 | -1.214251214 | 0.39861655 | -3.0461636 | 0.00231782 | 0.05620706 |
| DDAH1 | 98.5332609 | 0.014085072 | 0.41093606 | 0.03427558 | 0.9726574 | 0.99526459 |
| ZNHIT6 | 48.6973964 | 0.014867853 | 0.27480249 | 0.05410378 | 0.95685248 | 0.99236316 |

| | | | | | | |
|---|---|---|---|---|---|---|
| COL24A1 | 180.167199 | 0.003245277 | 0.36129514 | 0.00898234 | 0.99283322 | 0.99923337 |
| ODF2L | 96.3081347 | -0.268401045 | 0.27678213 | -0.9697196 | 0.33218629 | 0.76721253 |
| SH3GLB1 | 104.044169 | -0.375360153 | 0.18126954 | -2.0707294 | 0.03838409 | 0.3141788 |
| SELENOF | 113.89338 | 0.162189584 | 0.19451918 | 0.83379739 | 0.40439517 | 0.81056858 |
| HS2ST1 | 51.9423835 | 0.494021175 | 0.26226024 | 1.88370594 | 0.05960476 | 0.38765901 |
| LMO4 | 32.2939283 | -0.411476122 | 0.41945847 | -0.9809699 | 0.3266076 | 0.76371899 |
| PKN2-AS1 | 14.6976123 | 0.569436881 | 0.55106077 | 1.0333468 | 0.30144164 | 0.74591425 |
| PKN2 | 258.206381 | 0.010560208 | 0.16899219 | 0.06248932 | 0.95017316 | 0.99192022 |
| GTF2B | 33.6451364 | -0.126772814 | 0.36309113 | -0.3491488 | 0.72697764 | 0.94009907 |
| KYAT3 | 64.8753985 | -0.678875384 | 0.25235731 | -2.6901356 | 0.0071423 | 0.12146099 |
| GBP3 | 23.1982353 | -0.968108417 | 0.4882305 | -1.9828921 | 0.04737948 | 0.34764068 |
| GBP2 | 38.6761885 | -1.940019562 | 0.53304816 | -3.6394827 | 0.00027319 | 0.0134513 |
| LRRC8B | 86.6488411 | -0.360042077 | 0.26110034 | -1.3789414 | 0.16791281 | 0.60954339 |
| ZNF326 | 69.2505991 | 0.052083134 | 0.22010028 | 0.23663366 | 0.812941 | 0.96262567 |
| ZNF644 | 253.804081 | -0.081208986 | 0.16748684 | -0.4848679 | 0.62777008 | 0.90603677 |
| HFM1 | 19.9248046 | -0.237829577 | 0.62530071 | -0.3803443 | 0.70368984 | 0.93306088 |
| BTBD8 | 41.687509 | -0.414150814 | 0.47857912 | -0.8653758 | 0.38683253 | 0.80009555 |
| GLMN | 44.8821826 | 0.061533937 | 0.31863999 | 0.19311429 | 0.84686946 | 0.97027687 |
| RPAP2 | 107.513089 | 0.116361607 | 0.15244693 | 0.76329255 | 0.44528894 | 0.83035712 |
| EVI5 | 275.783268 | 0.010536132 | 0.20632934 | 0.05106463 | 0.95927402 | 0.99238848 |
| RPL5 | 165.915756 | -0.128806946 | 0.33419824 | -0.3854208 | 0.69992569 | 0.93127962 |
| FAM69A | 131.015681 | -0.077553875 | 0.30042685 | -0.2581456 | 0.79629453 | 0.95848876 |
| MTF2 | 144.815653 | 0.129234897 | 0.24837967 | 0.5203119 | 0.60284621 | 0.8956362 |
| TMED5 | 120.953103 | 0.077950591 | 0.29103105 | 0.26784287 | 0.78882027 | 0.95715263 |
| CCDC18 | 24.481434 | 0.593631044 | 0.41523132 | 1.42963937 | 0.15282055 | 0.59152407 |
| DR1 | 40.7669639 | 0.456608669 | 0.24208306 | 1.88616531 | 0.05927268 | 0.3871608 |
| FNBP1L | 525.442488 | 0.039343037 | 0.2088294 | 0.18839798 | 0.85056468 | 0.97027687 |
| BCAR3 | 94.048058 | -0.122575367 | 0.36824258 | -0.3328658 | 0.73923557 | 0.94376279 |
| DNTTIP2 | 20.7874551 | -0.245037807 | 0.36164114 | -0.6775717 | 0.49804329 | 0.85839018 |
| GCLM | 57.8828359 | -1.493858059 | 0.33463379 | -4.4641578 | 8.04E-06 | 0.00098271 |
| ARHGAP29 | 99.5513966 | -0.373128509 | 0.28531122 | -1.3077947 | 0.19094294 | 0.6421543 |
| ABCD3 | 101.019766 | 0.396425979 | 0.19393677 | 2.04409914 | 0.04094375 | 0.32473787 |
| SLC44A3 | 82.1746876 | 0.211419432 | 0.23402955 | 0.90338778 | 0.36632012 | 0.78793182 |
| CNN3 | 232.817388 | -0.61096569 | 0.28321144 | -2.1572776 | 0.03098404 | 0.27936455 |
| AC105942.1 | 32.8914254 | -0.895208714 | 0.34724228 | -2.5780522 | 0.0099359 | 0.14718772 |
| ALG14 | 56.1098538 | -0.290179764 | 0.28603677 | -1.0144841 | 0.3103518 | 0.75363506 |
| PTBP2 | 199.477627 | 0.011522109 | 0.21872113 | 0.05267945 | 0.95798731 | 0.99238848 |
| DPYD | 270.937678 | -0.449891583 | 0.22476636 | -2.0015966 | 0.04532814 | 0.33821763 |
| DPYD-AS1 | 12.7485962 | -1.50760636 | 0.56329241 | -2.6764187 | 0.00744136 | 0.12466529 |
| SNX7 | 49.9283449 | -0.201604139 | 0.30331783 | -0.664663 | 0.50626606 | 0.86214551 |
| FRRS1 | 23.2791093 | -0.263335554 | 0.53572821 | -0.4915469 | 0.62303967 | 0.90430717 |
| AGL | 98.8892087 | 0.486866422 | 0.18310496 | 2.65894714 | 0.00783853 | 0.12855183 |
| SASS6 | 11.5258485 | 0.267158901 | 0.41525797 | 0.64335646 | 0.51999283 | NA |
| TRMT13 | 51.0684927 | 0.232055069 | 0.28014542 | 0.82833789 | 0.40747917 | 0.81294693 |
| DBT | 68.1387014 | 0.36360682 | 0.31003609 | 1.17278868 | 0.24088056 | 0.69556484 |
| RTCA | 35.3537264 | 0.207744146 | 0.25650153 | 0.80991386 | 0.41798968 | 0.8188001 |
| CDC14A | 71.5132926 | -0.282808339 | 0.30830616 | -0.9172971 | 0.35898699 | 0.78445006 |
| EXTL2 | 57.954085 | 1.447212812 | 0.35619326 | 4.06299891 | 4.84E-05 | 0.00394898 |
| SLC30A7 | 88.6753435 | 0.125539146 | 0.20875949 | 0.60135778 | 0.54760171 | 0.87309151 |
| DPH5 | 20.8342211 | 0.173488423 | 0.37203837 | 0.46631863 | 0.64098745 | 0.91146968 |
| AC093157.1 | 20.0133503 | 0.63521741 | 0.36303297 | 1.74975129 | 0.08016124 | 0.44789428 |
| RNPC3 | 107.821716 | 0.005018367 | 0.21262714 | 0.02360173 | 0.9811703 | 0.99714727 |
| VAV3 | 276.985566 | 0.129622061 | 0.32304012 | 0.40125685 | 0.68823103 | 0.9267526 |
| SLC25A24 | 61.2439993 | -0.099089351 | 0.28955251 | -0.3422155 | 0.73218874 | 0.94194042 |
| PRPF38B | 72.8697362 | -0.090642944 | 0.32481715 | -0.2790584 | 0.78020003 | 0.95537097 |
| STXBP3 | 108.874606 | 0.10125632 | 0.21038496 | 0.48129067 | 0.63030992 | 0.90764716 |
| AL449266.1 | 21.059978 | -0.182599013 | 0.35861792 | -0.5091743 | 0.61063009 | 0.89989545 |
| GPSM2 | 56.3581426 | -0.387039741 | 0.43208573 | -0.8957476 | 0.3703876 | 0.7900012 |
| WDR47 | 17.1018108 | 0.36071445 | 0.40058044 | 0.90047945 | 0.36786516 | 0.78877653 |
| TAF13 | 20.5103556 | 0.341043693 | 0.3689627 | 0.92433109 | 0.35531396 | 0.78124141 |
| TMEM167B | 31.5026591 | 0.286582167 | 0.30799768 | 0.93046858 | 0.35212852 | 0.77964174 |
| SARS | 61.0803528 | -0.385693564 | 0.21401412 | -1.8021874 | 0.07151592 | 0.42281025 |
| SORT1 | 88.7367542 | 0.145587097 | 0.19780078 | 0.73602894 | 0.46171309 | 0.83846194 |
| PSMA5 | 63.0819878 | -0.257417802 | 0.27481919 | -0.9366806 | 0.34892288 | 0.77868979 |
| SYPL2 | 21.0518958 | 1.13542094 | 0.34487922 | 3.29222778 | 0.00099397 | 0.03257351 |
| CYB561D1 | 11.798385 | 0.420619511 | 0.44174881 | 0.95216897 | 0.3410113 | NA |
| GNAI3 | 74.2856711 | 0.165825501 | 0.30761233 | 0.539073 | 0.58983649 | 0.89149106 |
| GSTM3 | 20.1391418 | 0.864672149 | 0.64434629 | 1.34193704 | 0.17961641 | 0.62717033 |
| LINC01768 | 70.6327151 | 0.654879533 | 0.62578371 | 1.04649501 | 0.29533255 | 0.74111836 |
| AHCYL1 | 174.801454 | -0.360055633 | 0.24484786 | -1.470528 | 0.14141881 | 0.57625424 |
| STRIP1 | 35.0389791 | 0.308923567 | 0.24341904 | 1.26910191 | 0.20440472 | 0.65771647 |
| KCNC4 | 96.1269627 | 0.535676473 | 0.33004728 | 1.6230295 | 0.1045831 | 0.4985042 |
| RBM15 | 34.6854443 | 0.369290687 | 0.31993032 | 1.15428475 | 0.24838345 | 0.7032456 |
| SLC16A4 | 58.8630638 | -0.679520656 | 0.59860431 | -1.135175 | 0.256302 | 0.70971436 |
| AL355488.1 | 12.3328107 | 0.393051245 | 0.60327954 | 0.65152424 | 0.51470814 | NA |
| LAMTOR5 | 100.276293 | -0.381979388 | 0.33833661 | -1.1289922 | 0.25890114 | 0.71022613 |
| LRIF1 | 24.7351969 | -0.373723041 | 0.31522413 | -1.1855788 | 0.23578869 | 0.69035627 |
| DRAM2 | 68.7242713 | -0.158331237 | 0.33240004 | -0.4763274 | 0.63384117 | 0.90805737 |
| CEPT1 | 101.041118 | 0.156615831 | 0.25054917 | 0.6250902 | 0.53191186 | 0.86846407 |
| DENND2D | 48.1945503 | 0.109390784 | 0.23373472 | 0.46801256 | 0.63977561 | 0.91096398 |
| WDR77 | 17.9469493 | 0.459648606 | 0.31820564 | 1.44450175 | 0.14859789 | 0.58569376 |
| ATP5PB | 138.467191 | 0.101371807 | 0.34284458 | 0.2956786 | 0.76747554 | 0.95205353 |
| RAP1A | 192.007247 | -0.161425306 | 0.20510292 | -0.7870454 | 0.43125529 | 0.8248269 |
| KCND3 | 28.7584006 | -0.989328095 | 0.72420978 | -1.3660794 | 0.17191403 | 0.61601998 |
| CTTNBP2NL | 57.4225416 | -0.506455642 | 0.22641178 | -2.2368785 | 0.02529428 | 0.25137888 |
| WNT2B | 23.4634566 | -0.374628133 | 0.62562798 | -0.5988034 | 0.54930402 | 0.8740566 |
| ST7L | 59.0533918 | 0.231671882 | 0.23718562 | 0.97675351 | 0.3286912 | 0.76649357 |
| CAPZA1 | 125.95098 | -0.329300971 | 0.17165501 | -1.9183883 | 0.05506179 | 0.37349819 |
| MOV10 | 30.9914886 | -0.302771008 | 0.40149454 | -0.7541099 | 0.45078323 | 0.83185162 |
| RHOC | 43.7105819 | -0.853326493 | 0.36955965 | -2.3090359 | 0.02094159 | 0.22580589 |
| LRIG2 | 63.5334768 | 0.397729157 | 0.2153137 | 1.84720784 | 0.06471702 | 0.40396083 |
| MAGI3 | 1021.22249 | 0.579260319 | 0.26210289 | 2.21004935 | 0.02710174 | 0.25953238 |
| PHTF1 | 47.8685645 | -0.163348282 | 0.22482477 | -0.7265582 | 0.46749663 | 0.84315561 |
| RSBN1 | 68.8793045 | 0.149087758 | 0.21569213 | 0.6912063 | 0.4894359 | 0.8552564 |
| HIPK1 | 104.097403 | 0.537183348 | 0.23588802 | 2.27728122 | 0.02276944 | 0.23723259 |
| TRIM33 | 181.213853 | 0.029601981 | 0.17408062 | 0.17004754 | 0.86497275 | 0.97424938 |
| BCAS2 | 22.8615595 | 0.325925753 | 0.33052267 | 0.98609198 | 0.32408798 | 0.76304922 |
| DENND2C | 14.866962 | 0.418611901 | 0.47000201 | 0.89065981 | 0.3731117 | 0.7913202 |
| NRAS | 19.0036621 | -0.191179605 | 0.51603612 | -0.3704772 | 0.71102698 | 0.93563937 |
| CSDE1 | 184.620897 | -0.060534787 | 0.18450187 | -0.3280985 | 0.74283718 | 0.94546379 |
| SIKE1 | 32.3986199 | 0.014640273 | 0.34838188 | 0.04202364 | 0.96647986 | 0.99351999 |
| SLC22A15 | 130.225673 | -0.079239547 | 0.26719613 | -0.2965595 | 0.76680284 | 0.95205353 |
| MAB21L3 | 48.7819396 | -1.797883869 | 0.65637929 | -2.7390929 | 0.0061609 | 0.11066794 |
| ATP1A1 | 1291.35166 | -0.716419038 | 0.27303389 | -2.6239198 | 0.00869243 | 0.13512263 |
| LINC01762 | 99.6121782 | 1.091677348 | 1.06069027 | 1.02921407 | 0.30337909 | 0.74787746 |
| CD58 | 48.9228206 | -0.579523326 | 0.2636062 | -2.1984435 | 0.02791752 | 0.26291255 |
| IGSF3 | 99.2750893 | -0.021104815 | 0.31374514 | -0.0672674 | 0.94636884 | 0.99134285 |
| PTGFRN | 20.3033413 | -0.69682438 | 0.52222177 | -1.3343457 | 0.18209058 | 0.63058858 |
| TTF2 | 45.4931253 | -0.403725 | 0.2840891 | -1.4211211 | 0.15528157 | 0.59533243 |
| MAN1A2 | 230.918626 | 0.003029819 | 0.20628469 | 0.01468756 | 0.98828144 | 0.99915789 |
| GDAP2 | 61.3837597 | 0.27709298 | 0.23642613 | 1.1720066 | 0.24119441 | 0.69556484 |
| WDR3 | 13.1112088 | 0.741085124 | 0.39224168 | 1.88935841 | 0.05884382 | 0.38488541 |
| SPAG17 | 56.4988032 | 0.126138346 | 0.5673936 | 0.22231189 | 0.82407109 | 0.96539788 |

| | | | | | | |
|---|---|---|---|---|---|---|
| WARS2 | 50.0063284 | 0.044076659 | 0.27458064 | 0.16052355 | 0.87246867 | 0.97434023 |
| AL359915.2 | 178.500118 | 0.1434149 | 0.37347476 | 0.38400159 | 0.70097727 | 0.93127962 |
| HMGCS2 | 62.0341246 | 1.687004797 | 1.51939978 | 1.11031002 | 0.26686546 | 0.71785772 |
| SEC22B | 67.2545195 | 0.108010201 | 0.2096326 | 0.51523569 | 0.60638833 | 0.89643487 |
| LINC00623 | 167.825164 | 0.319327314 | 0.26582213 | 1.20128192 | 0.22964186 | 0.68523674 |
| AC239800.3 | 19.0597247 | 0.164619723 | 0.31748327 | 0.51851464 | 0.60409925 | 0.8958512 |
| NBPF15 | 15.3089346 | -0.514587041 | 0.5180675 | -0.9932818 | 0.32057263 | 0.76131909 |
| FP700111.1 | 12.0353628 | 0.763335255 | 0.40772868 | 1.87216471 | 0.06118382 | NA |
| SRGAP2B | 13.919997 | -0.268272893 | 0.40980919 | -0.6546288 | 0.51270679 | 0.86346419 |
| GPR89A | 23.858661 | 0.058263574 | 0.27784346 | 0.20969928 | 0.83390239 | 0.96701848 |
| PDZK1 | 18.7657885 | 0.488255921 | 0.4684975 | 1.04217402 | 0.29733101 | 0.74160729 |
| RNF115 | 173.897227 | 0.011590259 | 0.20491281 | 0.0565619 | 0.95489418 | 0.99228059 |
| POLR3C | 20.4629791 | -0.028203546 | 0.3893913 | -0.0724298 | 0.94225985 | 0.98991882 |
| PEX11B | 15.2454145 | 0.44794573 | 0.3496871 | 1.28099015 | 0.20019713 | 0.65330283 |
| RBM8A | 41.2784834 | -0.087739843 | 0.33757918 | -0.2599089 | 0.79493403 | 0.95804683 |
| TXNIP | 1254.00711 | 0.141660041 | 0.37685483 | 0.37590083 | 0.70699063 | 0.93432317 |
| NBPF12 | 32.5888928 | -0.114803787 | 0.26830247 | -0.4278894 | 0.66873164 | 0.92049996 |
| PRKAB2 | 44.6453632 | 0.389456807 | 0.40419186 | 0.96354441 | 0.33527439 | 0.76777176 |
| AC242426.2 | 63.1143543 | 0.071172942 | 0.26280493 | 0.27082042 | 0.78652915 | 0.95577371 |
| FMO5 | 45.2939498 | 0.096041266 | 0.43740434 | 0.2195709 | 0.82620535 | 0.96539788 |
| CHD1L | 90.2867744 | -0.53590186 | 0.3793079 | -1.4128413 | 0.15770239 | 0.59847934 |
| LINC00624 | 23.9805634 | -0.908261625 | 0.52289933 | -1.7369723 | 0.08239207 | 0.45473406 |
| BCL9 | 101.501392 | -0.267123865 | 0.39004229 | -0.6848587 | 0.49343307 | 0.85721581 |
| ACP6 | 37.7077325 | 0.08586951 | 0.31271779 | 0.27459106 | 0.78363044 | 0.95544332 |
| GPR89B | 29.0767424 | -0.239440803 | 0.28196418 | -0.8491887 | 0.39577628 | 0.80624334 |
| NBPF11 | 31.8235001 | -0.040801632 | 0.31394366 | -0.1299648 | 0.89659426 | 0.97885533 |
| AC239809.3 | 26.4235632 | -0.098370664 | 0.40283821 | -0.244194 | 0.80708059 | 0.96016552 |
| PDE4DIP | 126.907256 | -0.730104393 | 0.46336623 | -1.575653 | 0.11510579 | 0.51955616 |
| AC245297.3 | 196.134703 | -0.156436983 | 0.35579883 | -0.4396782 | 0.6601702 | 0.91752855 |
| SF3B4 | 14.6413064 | 0.347830147 | 0.40350202 | 0.86202827 | 0.38867196 | 0.80130032 |
| OTUD7B | 76.5123973 | -0.312338509 | 0.2561365 | -1.2194221 | 0.22268402 | 0.67606266 |
| VPS45 | 106.193395 | 0.291477872 | 0.19417442 | 1.50111363 | 0.13332617 | 0.5615981 |
| ANP32E | 28.6030404 | -0.078512751 | 0.31694884 | -0.2477143 | 0.80435549 | 0.95973006 |
| APH1A | 40.7614336 | 0.038105209 | 0.37170244 | 0.10251536 | 0.91834762 | 0.98521039 |
| MRPS21 | 67.6880252 | -0.314240217 | 0.2493288 | -1.2603446 | 0.20754506 | 0.65919683 |
| PRPF3 | 40.2364188 | 0.244433604 | 0.31160712 | 0.78442882 | 0.43278853 | 0.8248269 |
| RPRD2 | 158.567734 | 0.130393146 | 0.16448695 | 0.79272637 | 0.42793726 | 0.82407405 |
| TARS2 | 16.0078622 | 0.218040949 | 0.41659201 | 0.52339205 | 0.60070146 | 0.89490801 |
| MCL1 | 69.6012673 | -0.806592605 | 0.39334511 | -2.0505978 | 0.04030613 | 0.32205065 |
| ENSA | 140.921739 | -0.087348333 | 0.15809654 | -0.5525 | 0.58060584 | 0.89012555 |
| GOLPH3L | 49.6740117 | 0.211506483 | 0.22573035 | 0.93698737 | 0.34876505 | 0.77868979 |
| CTSS | 24.0151775 | -1.70455311 | 0.52928256 | -3.2204974 | 0.00127968 | 0.03819363 |
| ARNT | 56.171635 | 0.192249151 | 0.24153925 | 0.79593337 | 0.42607077 | 0.82328415 |
| SETDB1 | 42.7556148 | 0.039479372 | 0.31799344 | 0.12415153 | 0.9011953 | 0.979694 |
| CERS2 | 43.1568467 | -0.146034946 | 0.29073902 | -0.5022888 | 0.6154644 | 0.90113279 |
| MINDY1 | 84.1360813 | 0.528942345 | 0.28410356 | 1.86179413 | 0.06263211 | 0.39785954 |
| PRUNE1 | 43.1853572 | 0.376242194 | 0.35710024 | 1.05360388 | 0.29206431 | 0.73999615 |
| C1orf56 | 23.1917273 | 0.023267287 | 0.4703475 | 0.04946829 | 0.96054611 | 0.99238848 |
| CDC42SE1 | 62.4437155 | -0.45490837 | 0.22138046 | -2.0548714 | 0.03989143 | 0.3200553 |
| GABPB2 | 66.4734409 | 0.764912291 | 0.31917734 | 2.39651187 | 0.01655196 | 0.19944595 |
| SCNM1 | 15.2918559 | 0.05658028 | 0.39740556 | 0.14237415 | 0.88678448 | 0.97714523 |
| VPS72 | 21.9670561 | 0.212638963 | 0.29181251 | 0.7286835 | 0.46619529 | 0.84226007 |
| PIP5K1A | 195.496061 | -0.028524134 | 0.18470633 | -0.1544296 | 0.87727098 | 0.97541592 |
| PSMD4 | 46.9371035 | -0.020034399 | 0.22254025 | -0.090026 | 0.92826659 | 0.98719284 |
| ZNF687 | 10.5386166 | -0.043341296 | 0.40429137 | -0.1072031 | 0.91462784 | NA |
| PI4KB | 60.3131855 | 0.457313172 | 0.23254392 | 1.96656684 | 0.04923317 | 0.35506449 |
| SELENBP1 | 37.4158324 | 0.745301003 | 0.65913303 | 1.13072927 | 0.25816906 | 0.71022118 |
| PSMB4 | 41.9734105 | 0.253081472 | 0.34462429 | 0.73436922 | 0.46272374 | 0.83921832 |
| POGZ | 257.321609 | 0.456112146 | 0.16520168 | 2.76094139 | 0.0057635 | 0.10528431 |
| CGN | 19.943396 | -0.086840834 | 0.40818208 | -0.2127502 | 0.83152178 | 0.96701848 |
| TUFT1 | 39.4228909 | -0.076035908 | 0.39881684 | -0.1906537 | 0.84879691 | 0.97027687 |
| SNX27 | 81.5472617 | 0.265902222 | 0.23029514 | 1.15461499 | 0.24824812 | 0.70306768 |
| RORC | 42.8931162 | 0.408787286 | 0.42405341 | 0.96399953 | 0.33504616 | 0.76777176 |
| AL450992.2 | 11.3569996 | -0.702511627 | 0.4887299 | -1.4374231 | 0.15059782 | NA |
| THEM4 | 121.541482 | 0.81883756 | 0.24518851 | 3.33962456 | 0.00083892 | 0.02920851 |
| S100A10 | 411.034825 | -0.509072834 | 0.5511994 | -0.9235729 | 0.35570872 | 0.78153728 |
| S100A11 | 629.921078 | -1.108446046 | 0.5140187 | -2.1564314 | 0.03105 | 0.27952435 |
| FLG-AS1 | 52.0745225 | 1.257682207 | 0.40812605 | 3.08160237 | 0.0020589 | 0.05200859 |
| S100A6 | 585.229974 | -1.005304647 | 0.60020523 | -1.6749348 | 0.09394702 | 0.4781144 |
| S100A2 | 29.2176525 | -1.430879461 | 0.66864995 | -2.139953 | 0.03235857 | 0.28508455 |
| S100A16 | 19.0364371 | -1.420127747 | 0.3914071 | -3.6282626 | 0.00028533 | 0.01390829 |
| S100A13 | 65.5535237 | -0.76364826 | 0.39895335 | -1.9141292 | 0.05560365 | 0.37507329 |
| S100A1 | 27.9635069 | -0.541955641 | 0.91558755 | -0.5919212 | 0.55390339 | 0.87691576 |
| CHTOP | 45.5501151 | 0.190325847 | 0.2438693 | 0.78044202 | 0.43513074 | 0.82565889 |
| SNAPIN | 12.3583175 | -0.405301289 | 0.51137881 | -0.7925657 | 0.42803091 | NA |
| ILF2 | 33.6743339 | -0.05304205 | 0.33133859 | -0.1600841 | 0.8728148 | 0.97434023 |
| INTS3 | 65.1949787 | -0.267736038 | 0.35637449 | -0.7512772 | 0.45248583 | 0.83316488 |
| GATAD2B | 205.512727 | 0.073722266 | 0.21610751 | 0.34113699 | 0.73300045 | 0.94269288 |
| DENND4B | 25.101714 | -0.191464129 | 0.41181534 | -0.4649271 | 0.64198363 | 0.91146968 |
| CRTC2 | 18.4091905 | 0.040202386 | 0.37939932 | 0.10596325 | 0.91561151 | 0.98437981 |
| SLC39A1 | 44.0977272 | -1.328900619 | 0.44512112 | -2.985481 | 0.00283133 | 0.06539018 |
| JTB | 53.6862184 | 0.030533865 | 0.40003829 | 0.07632736 | 0.93915866 | 0.98871562 |
| RAB13 | 56.5064491 | 0.228758083 | 0.36218282 | 0.63160943 | 0.52764212 | 0.86750588 |
| RPS27 | 89.5702669 | -0.042895876 | 0.30695955 | -0.1397444 | 0.88886195 | 0.97765744 |
| TPM3 | 71.1379628 | -0.477977085 | 0.32286892 | -1.480406 | 0.13876493 | 0.57155829 |
| C1orf43 | 70.524264 | -0.295440833 | 0.28425381 | -1.0393558 | 0.29863931 | 0.74296125 |
| UBAP2L | 84.8463036 | 0.023934665 | 0.22277027 | 0.10744102 | 0.91443911 | 0.98407488 |
| HAX1 | 24.455763 | 0.094753139 | 0.33075548 | 0.28647489 | 0.77451443 | 0.95470568 |
| IL6R | 28.4738942 | -0.888910596 | 0.54750197 | -1.6235752 | 0.10446651 | 0.49819329 |
| UBE2Q1 | 26.9497252 | -0.152000507 | 0.4122252 | -0.3687317 | 0.7123277 | 0.9357596 |
| ADAR | 126.143327 | -0.115775085 | 0.36697407 | -0.3154857 | 0.75239288 | 0.94728927 |
| KCNN3 | 26.0281648 | 1.406682291 | 0.79971675 | 1.75897565 | 0.07858164 | 0.4421357 |
| PMVK | 44.9582238 | 0.257133962 | 0.3254917 | 0.78998625 | 0.4295358 | 0.82469033 |
| PBXIP1 | 42.7450808 | 0.173686255 | 0.32467763 | 0.53494987 | 0.59268452 | 0.89198431 |
| PYGO2 | 13.0013571 | 0.286500294 | 0.41064284 | 0.6976873 | 0.48537277 | 0.85356796 |
| FLAD1 | 13.2988596 | -0.013353819 | 0.42246723 | -0.0316091 | 0.97478377 | 0.99543096 |
| ZBTB7B | 15.2336594 | 0.08135237 | 0.43135508 | 0.18859722 | 0.85040851 | 0.97027687 |
| EFNA1 | 72.5508792 | -1.460444632 | 0.38749604 | -3.7689279 | 0.00016395 | 0.0098352 |
| SLC50A1 | 24.5928476 | -0.575105194 | 0.33190707 | -1.7327296 | 0.08314375 | 0.45590411 |
| DPM3 | 40.221026 | -0.153405061 | 0.38065002 | -0.4030081 | 0.68694223 | 0.92610697 |
| MUC1 | 17.4445967 | -0.066693307 | 0.5087636 | -0.131089 | 0.89570491 | 0.97885533 |
| MTX1 | 14.2510495 | -0.026298021 | 0.39518812 | -0.0665456 | 0.94694347 | 0.99149867 |
| GBA | 12.2304211 | -0.268498523 | 0.5043829 | -0.5323307 | 0.59449694 | NA |
| SCAMP3 | 27.0441025 | 0.593919201 | 0.35746869 | 1.66145795 | 0.09662151 | 0.48257296 |
| CLK2 | 20.5959351 | 0.000204094 | 0.37570533 | 0.00054323 | 0.99956657 | 0.99982181 |
| FDPS | 106.672682 | -0.899147412 | 0.42214226 | -2.129963 | 0.03317467 | 0.28919158 |
| RUSC1 | 12.5423353 | 0.324092659 | 0.35963986 | 0.9011589 | 0.36750384 | 0.78877653 |
| ASH1L | 466.941529 | 0.088944887 | 0.13907952 | 0.63952542 | 0.52248119 | 0.86728831 |
| YY1AP1 | 90.1023565 | -0.018165375 | 0.22069249 | -0.0823108 | 0.93439957 | 0.9881638 |
| DAP3 | 105.548897 | -0.040571234 | 0.17708934 | -0.2291004 | 0.8187909 | 0.9635539 |
| GON4L | 76.8175821 | 0.208757375 | 0.29821065 | 0.70003327 | 0.48390653 | 0.85235034 |
| RIT1 | 27.8811866 | -0.315606945 | 0.41554749 | -0.7594967 | 0.44755549 | 0.83066268 |
| KHDC4 | 83.4511184 | -0.036969841 | 0.1907277 | -0.1938357 | 0.84630452 | 0.97027687 |

| | | | | | | |
|---|---|---|---|---|---|---|
| ARHGEF2 | 36.0781979 | -0.174038273 | 0.26421606 | -0.6586968 | 0.51009048 | 0.86281145 |
| SSR2 | 53.0421067 | -0.011171838 | 0.26036486 | -0.0429084 | 0.96577456 | 0.99334616 |
| UBQLN4 | 17.4033593 | 0.492086994 | 0.39235903 | 1.25417527 | 0.20977831 | 0.66164938 |
| LAMTOR2 | 25.5442936 | 0.303885183 | 0.41249611 | 0.7366983 | 0.46130584 | 0.83826652 |
| RAB25 | 36.7536297 | -0.133165538 | 0.36412992 | -0.3657089 | 0.71458234 | 0.93684603 |
| LMNA | 103.236995 | -0.531525976 | 0.26511474 | -2.0048903 | 0.04497477 | 0.33661674 |
| SEMA4A | 15.6803505 | -0.008587906 | 0.41748214 | -0.0205707 | 0.9835881 | 0.99757472 |
| SLC25A44 | 16.3270441 | 0.338557176 | 0.3885471 | 0.8713414 | 0.38356777 | 0.7980695 |
| SMG5 | 60.8332846 | -0.266119904 | 0.30718555 | -0.8663165 | 0.38631663 | 0.80001522 |
| GLMP | 15.8586185 | -1.039815402 | 0.57697474 | -1.8021853 | 0.07151625 | 0.42281025 |
| CCT3 | 66.5779644 | 0.430163976 | 0.28223805 | 1.52411758 | 0.12747934 | 0.55055637 |
| RHBG | 286.814666 | 1.721800531 | 0.52255858 | 3.29494262 | 0.00098442 | 0.03257351 |
| MEF2D | 43.1994622 | -0.161304145 | 0.38911787 | -0.414538 | 0.67848014 | 0.92291026 |
| NAXE | 27.8353025 | 0.155128775 | 0.5076041 | 0.30560978 | 0.75990178 | 0.94952011 |
| GPATCH4 | 13.7796366 | -0.187599301 | 0.45176679 | -0.415257 | 0.67795381 | 0.9226233 |
| ISG20L2 | 15.8790188 | 0.06601358 | 0.40471463 | 0.16311143 | 0.8704307 | 0.97434023 |
| RRNAD1 | 22.5593155 | -0.286365656 | 0.30672024 | -0.9336379 | 0.35049068 | 0.77868979 |
| MRPL24 | 11.3585564 | 0.350524523 | 0.47133613 | 0.74368269 | 0.45706846 | NA |
| HDGF | 49.2985111 | -0.227950962 | 0.29717393 | -0.7670624 | 0.4430444 | 0.82894008 |
| PRCC | 43.1024005 | 0.612880141 | 0.2253969 | 2.71911516 | 0.00654568 | 0.11607515 |
| ARHGEF11 | 109.776999 | 0.113119753 | 0.26424829 | 0.42808131 | 0.66859192 | 0.92049996 |
| ETV3 | 11.3142876 | 0.321170473 | 0.4233338 | 0.75866957 | 0.44805024 | NA |
| DUSP23 | 38.1086315 | -0.735565009 | 0.42288695 | -1.7393892 | 0.08196632 | 0.45345012 |
| TAGLN2 | 70.2776761 | -1.268290634 | 0.39256099 | -3.2308117 | 0.00123439 | 0.03747651 |
| IGSF8 | 18.8013129 | 0.658481905 | 0.38045583 | 1.73077095 | 0.08349263 | 0.4560197 |
| PEA15 | 17.5388836 | -0.18292412 | 0.52209294 | -0.350367 | 0.72606332 | 0.94003188 |
| DCAF8 | 34.3560513 | 0.382599172 | 0.23711471 | 1.61356155 | 0.10662256 | 0.50199866 |
| COPA | 143.374073 | -0.107386163 | 0.18989817 | -0.5654934 | 0.5717382 | 0.88578992 |
| NCSTN | 29.5440897 | 0.211795123 | 0.35619234 | 0.59460886 | 0.55210496 | 0.87576439 |
| F11R | 55.8826765 | -0.160282349 | 0.30955671 | -0.5177802 | 0.6046116 | 0.89593125 |
| AL591806.3 | 12.0971562 | 0.300464552 | 0.53637263 | 0.56017876 | 0.57535751 | NA |
| PFDN2 | 61.2480635 | -0.358910885 | 0.40119983 | -0.8945938 | 0.37100426 | 0.7900012 |
| NIT1 | 35.7219803 | -0.300700187 | 0.3717184 | -0.8089462 | 0.41854609 | 0.81880353 |
| DEDD | 14.7597304 | 0.13094753 | 0.36931788 | 0.35456591 | 0.72291482 | 0.93873366 |
| UFC1 | 39.6103104 | -0.268679144 | 0.4027241 | -0.6671544 | 0.50467353 | 0.86155108 |
| B4GALT3 | 18.1770109 | 0.077831419 | 0.35333673 | 0.22027548 | 0.82565661 | 0.96539788 |
| NDUFS2 | 90.8389044 | 0.199444751 | 0.2928106 | 0.68113912 | 0.49578347 | 0.85814183 |
| SDHC | 111.508912 | 0.076016291 | 0.18970425 | 0.40070948 | 0.68863403 | 0.92684197 |
| AL592295.4 | 10.6160377 | 0.357860535 | 0.41935779 | 0.85335374 | 0.39346317 | NA |
| FCGR2A | 15.6057437 | -1.839282809 | 0.67363104 | -2.7304009 | 0.00632573 | 0.11279343 |
| DUSP12 | 15.5146722 | 0.274499518 | 0.36107146 | 0.76023599 | 0.44711354 | 0.83066268 |
| ATF6 | 195.462812 | -0.293154067 | 0.1773059 | -1.6533802 | 0.09825351 | 0.48476221 |
| NOS1AP | 316.317956 | -0.580954925 | 0.34761092 | -1.6712793 | 0.09466653 | 0.47996672 |
| C1orf226 | 12.2284818 | -0.627735816 | 0.48263179 | -1.3006516 | 0.19337774 | NA |
| UHMK1 | 286.341464 | 0.655270214 | 0.23601576 | 2.77638335 | 0.00549674 | 0.10201977 |
| UAP1 | 248.470107 | 0.446464252 | 0.31049788 | 1.4378979 | 0.15046302 | 0.58759787 |
| DDR2 | 46.3800492 | -0.48665938 | 0.45569635 | -1.0679466 | 0.28554459 | 0.73672385 |
| HSD17B7 | 37.2921696 | -0.43430998 | 0.41483422 | -1.0469483 | 0.29512343 | 0.74111836 |
| PBX1 | 160.226152 | 1.128472153 | 0.37825728 | 2.98334554 | 0.00285116 | 0.06569179 |
| MGST3 | 176.258596 | 0.071945057 | 0.28356281 | 0.25371824 | 0.79971324 | 0.95861074 |
| ALDH9A1 | 37.3260049 | 0.145017952 | 0.35243973 | 0.4114688 | 0.68072882 | 0.92375164 |
| TMCO1 | 77.1306034 | 0.067533185 | 0.18784844 | 0.35950889 | 0.71921443 | 0.93781153 |
| UCK2 | 12.7471963 | 0.367991636 | 0.42764267 | 0.86051197 | 0.38950689 | 0.80191835 |
| POGK | 36.4837269 | 0.825559337 | 0.41668863 | 1.98123799 | 0.04756459 | 0.3482087 |
| TADA1 | 20.2100222 | 0.602566778 | 0.32761483 | 1.83925369 | 0.06587788 | 0.40779544 |
| POU2F1 | 199.187204 | -0.203071361 | 0.20975203 | -0.9681497 | 0.33296962 | 0.76753928 |
| CREG1 | 26.5379682 | 0.203967232 | 0.33473553 | 0.60933846 | 0.54230012 | 0.87177844 |
| MPZL1 | 102.980531 | -0.47918122 | 0.30817716 | -1.5548888 | 0.11997255 | 0.53162804 |
| MPC2 | 56.6817598 | 0.117407582 | 0.43846508 | 0.26776951 | 0.78887674 | 0.95715263 |
| DCAF6 | 257.087523 | -0.094368238 | 0.21198568 | -0.4451633 | 0.65620176 | 0.9158679 |
| TIPRL | 30.2822411 | 0.112041163 | 0.25285061 | 0.4431121 | 0.65768465 | 0.91601353 |
| SFT2D2 | 57.4372951 | -0.050655264 | 0.29429315 | -0.1721252 | 0.86333911 | 0.97368057 |
| TBX19 | 11.4099139 | 0.580540248 | 0.45804346 | 1.26743486 | 0.20499984 | NA |
| LINC00970 | 54.0480212 | 1.57535357 | 0.69394279 | 2.27014904 | 0.02319854 | 0.23879078 |
| ATP1B1 | 1151.22078 | 0.995201469 | 0.27511267 | 3.61743236 | 0.00029754 | 0.0143589 |
| NME7 | 117.131199 | 0.36340923 | 0.23919337 | 1.5193115 | 0.12868411 | 0.55308726 |
| BLZF1 | 42.8543534 | 0.060206512 | 0.27022984 | 0.22279743 | 0.82369317 | 0.96539788 |
| CCDC181 | 11.8901313 | -0.656688159 | 0.49001151 | -1.3401484 | 0.18019709 | NA |
| SLC19A2 | 163.4661 | -0.838110379 | 0.38847244 | -2.1574513 | 0.03097052 | 0.27936455 |
| SCYL3 | 36.0683142 | -0.011827524 | 0.27696572 | -0.0427039 | 0.96593755 | 0.99334616 |
| KIFAP3 | 74.6651374 | -0.066908996 | 0.31081269 | -0.2152711 | 0.82955595 | 0.96669991 |
| GORAB | 14.5983729 | 0.012360427 | 0.40304024 | 0.03066797 | 0.97553433 | 0.99554793 |
| FMO4 | 19.063729 | 0.186487852 | 0.46949378 | 0.39721049 | 0.69121225 | 0.92786587 |
| PRRC2C | 276.929532 | -0.035924165 | 0.18833203 | -0.1907491 | 0.84872217 | 0.97027687 |
| VAMP4 | 21.1566347 | -0.119927789 | 0.30931324 | -0.3877228 | 0.69822119 | 0.93015138 |
| METTL13 | 12.7784954 | 0.15563288 | 0.3781008 | 0.41161744 | 0.68061985 | 0.92375164 |
| DNM3 | 23.0341409 | 0.584005717 | 0.42272305 | 1.38153271 | 0.16711523 | 0.60871863 |
| PIGC | 21.5408469 | -0.228133768 | 0.36298857 | -0.6284875 | 0.52968465 | 0.86758926 |
| SUCO | 99.248634 | -0.210753809 | 0.25867756 | -0.8147356 | 0.41522369 | 0.81726257 |
| AL645568.1 | 25.471815 | 0.207007126 | 0.45703901 | 0.45293098 | 0.65059844 | 0.91487459 |
| PRDX6 | 147.515041 | -0.112392129 | 0.31093508 | -0.3614649 | 0.71775192 | 0.93723455 |
| KLHL20 | 57.3036378 | 0.078798558 | 0.25583741 | 0.30800248 | 0.75808044 | 0.94924107 |
| DARS2 | 19.0010986 | -0.041660547 | 0.32123625 | -0.1296882 | 0.89681313 | 0.97885533 |
| ZBTB37 | 28.92072 | 0.177534524 | 0.41106566 | 0.43188849 | 0.66582246 | 0.91958319 |
| RC3H1 | 166.216206 | 0.149730534 | 0.20164106 | 0.74255973 | 0.45774828 | 0.83603055 |
| RABGAP1L | 659.045356 | 0.095276555 | 0.2048689 | 0.4650611 | 0.6418877 | 0.91146968 |
| CACYBP | 23.8912166 | 0.383456282 | 0.32130992 | 1.19341564 | 0.23270663 | 0.68763908 |
| MRPS14 | 25.0592891 | 0.515651339 | 0.34352134 | 1.50107515 | 0.13333613 | 0.5615981 |
| KIAA0040 | 74.7168657 | -0.956442058 | 0.35404446 | -2.7014744 | 0.00690328 | 0.11936151 |
| COP1 | 281.010437 | -0.136792556 | 0.14684267 | -0.9315586 | 0.35156468 | 0.77927001 |
| PAPPA2 | 47.7592759 | 0.968789894 | 0.69826368 | 1.38742701 | 0.16531159 | 0.60578861 |
| SEC16B | 19.2615235 | 1.126696489 | 0.6468391 | 1.74184968 | 0.08153475 | 0.45193547 |
| RASAL2 | 137.908374 | 0.906436501 | 0.27474316 | 3.29921404 | 0.00096956 | 0.03220797 |
| RALGPS2 | 418.897909 | -0.080235975 | 0.27207966 | -0.2948988 | 0.76807116 | 0.95205353 |
| ANGPTL1 | 53.4543119 | 0.231861805 | 0.50880791 | 0.45569615 | 0.64860849 | 0.91417939 |
| FAM20B | 50.6973091 | 0.280122629 | 0.22049098 | 1.27044937 | 0.20392461 | 0.65771647 |
| TOR3A | 15.3877687 | 0.562419122 | 0.45972753 | 1.22337491 | 0.22118813 | 0.67492938 |
| ABL2 | 175.875498 | -0.06815512 | 0.22304509 | -0.3055666 | 0.75993469 | 0.94952011 |
| SOAT1 | 113.685241 | 0.141845888 | 0.26516325 | 0.53493796 | 0.59269276 | 0.89198431 |
| TOR1AIP2 | 109.802825 | 0.126368477 | 0.2258745 | 0.55946323 | 0.57584562 | 0.88746465 |
| TOR1AIP1 | 57.6342449 | 0.230671489 | 0.27915243 | 0.82632808 | 0.40861801 | 0.81362627 |
| CEP350 | 196.210315 | 0.207328456 | 0.14429931 | 1.43679447 | 0.1507764 | 0.58759787 |
| QSOX1 | 84.0035059 | -0.512847885 | 0.27731447 | -1.8493369 | 0.06440918 | 0.40281695 |
| ACBD6 | 173.439361 | -0.082113808 | 0.217856 | -0.3769178 | 0.70623469 | 0.93407096 |
| XPR1 | 277.583514 | -0.028025939 | 0.25160567 | -0.1113883 | 0.9113084 | 0.9835359 |
| STX6 | 56.7998212 | 0.255636027 | 0.38755871 | 0.65960593 | 0.50950675 | 0.86281145 |
| MR1 | 116.475299 | 0.010009016 | 0.32082522 | 0.03119772 | 0.97511186 | 0.99543096 |
| CACNA1E | 23.7131143 | 0.399326513 | 1.45749703 | 0.27398101 | 0.78409921 | 0.95544332 |
| GLUL | 115.601994 | -0.546714437 | 0.28306754 | -1.9313922 | 0.05343457 | 0.36655965 |
| RNASEL | 29.9852395 | -0.527906453 | 0.41714704 | -1.2655165 | 0.20568625 | 0.65836247 |
| RGS16 | 29.594964 | -0.894237953 | 0.70729434 | -1.2643081 | 0.20611947 | 0.65836247 |
| DHX9 | 112.194165 | -0.077934693 | 0.20822153 | -0.3742874 | 0.70819052 | 0.93436453 |
| LAMC1 | 103.74101 | 0.108812416 | 0.25874959 | 0.42053174 | 0.67409705 | 0.92189158 |

| | | | | | | |
|---|---|---|---|---|---|---|
| SMG7 | 187.449714 | -0.008187128 | 0.24620959 | -0.0332527 | 0.97347309 | 0.99526459 |
| ARPC5 | 65.1006292 | -0.59522526 | 0.28848294 | -2.0632945 | 0.03908466 | 0.31752191 |
| RGL1 | 16.9275161 | -0.106856936 | 0.50666536 | -0.2109024 | 0.83296344 | 0.96701848 |
| TSEN15 | 33.1132284 | -0.044168288 | 0.34366668 | -0.1285207 | 0.8977369 | 0.97892709 |
| C1orf21 | 14.3259238 | -2.088263679 | 0.55555313 | -3.758891 | 0.00017067 | 0.01003322 |
| EDEM3 | 92.1340577 | 0.212730089 | 0.35857045 | 0.59327279 | 0.55299861 | 0.87591223 |
| RNF2 | 33.1560157 | 0.098915925 | 0.28073432 | 0.3523471 | 0.72457797 | 0.93935288 |
| TRMT1L | 35.0801078 | 0.078945786 | 0.29082802 | 0.27145179 | 0.78604358 | 0.95577371 |
| SWT1 | 56.2657792 | -0.16791553 | 0.26005053 | -0.6457035 | 0.51847142 | 0.86552266 |
| IVNS1ABP | 112.845698 | 0.338348144 | 0.38625374 | 0.87597376 | 0.3810443 | 0.79688435 |
| TPR | 132.760049 | 0.027909996 | 0.17655794 | 0.1580784 | 0.87439502 | 0.97434023 |
| ODR4 | 74.0256775 | 0.058706766 | 0.18422315 | 0.31867203 | 0.74997523 | 0.94684415 |
| PLA2G4A | 57.3849597 | 0.029025409 | 0.41956797 | 0.06917928 | 0.94484692 | 0.99121467 |
| BRINP3 | 37.158075 | -0.120602682 | 0.72796433 | -0.1656711 | 0.86841576 | 0.97434023 |
| AL390957.1 | 73.9218806 | 0.834664069 | 0.68847495 | 1.21233759 | 0.22538318 | 0.67852788 |
| AL136454.1 | 71.1719748 | 0.254254234 | 0.32588817 | 0.7801886 | 0.43527987 | 0.82578032 |
| UCHL5 | 46.21622 | -0.209246387 | 0.26533088 | -0.7886243 | 0.43033161 | 0.82469033 |
| TROVE2 | 87.8474288 | -0.029433433 | 0.19713506 | -0.1493059 | 0.88131224 | 0.97630358 |
| GLRX2 | 11.4596479 | -0.342599784 | 0.4168021 | -0.8219723 | 0.41109265 | NA |
| CDC73 | 137.721736 | -0.009673225 | 0.19539042 | -0.0495072 | 0.96051513 | 0.99238848 |
| ZBTB41 | 41.2470155 | 0.305392388 | 0.24522712 | 1.24534506 | 0.21300494 | 0.66628439 |
| DENND1B | 252.722117 | 0.192617242 | 0.25789383 | 0.74688582 | 0.45513249 | 0.83423755 |
| NEK7 | 209.715766 | -0.15135332 | 0.21399989 | -0.7072589 | 0.47940562 | 0.84833462 |
| ATP6V1G3 | 200.728887 | 0.660505309 | 0.60503551 | 1.09168025 | 0.27497367 | 0.72554623 |
| PTPRC | 20.1804007 | 1.351895107 | 0.63297064 | 2.13579434 | 0.03269618 | 0.28675674 |
| ZNF281 | 17.6875325 | 0.73511315 | 0.37217675 | 1.97517214 | 0.04824862 | 0.35162786 |
| DDX59 | 35.2018543 | 0.17445534 | 0.26039219 | 0.66997146 | 0.50287598 | 0.86010152 |
| CAMSAP2 | 382.657111 | 0.34215154 | 0.23804593 | 1.43733415 | 0.15062307 | 0.58759787 |
| TMEM9 | 29.7946218 | -0.03915123 | 0.4076394 | -0.0960438 | 0.9234858 | 0.98672316 |
| LAD1 | 15.1475792 | -0.280894941 | 0.39213329 | -0.7163252 | 0.4737906 | 0.84593347 |
| CSRP1 | 19.7810749 | -0.379070573 | 0.49159443 | -0.7711043 | 0.44064512 | 0.82793978 |
| AC096677.1 | 20.5090107 | 0.207083725 | 0.39230117 | 0.52786925 | 0.59759009 | 0.89363331 |
| NAV1 | 18.9733751 | -0.634317261 | 0.70298564 | -0.9023189 | 0.36688746 | 0.78821891 |
| IPO9 | 71.7532566 | -0.09272628 | 0.2665292 | -0.3479029 | 0.72791311 | 0.94030592 |
| TIMM17A | 40.2709475 | -0.137079804 | 0.33970442 | -0.4035267 | 0.68656079 | 0.92597882 |
| RNPEP | 46.9698796 | -0.169857005 | 0.22638034 | -0.7503169 | 0.45306384 | 0.83359621 |
| ELF3 | 211.348752 | -1.07248407 | 0.3062884 | -3.5015497 | 0.00046256 | 0.01917452 |
| ARL8A | 30.2615882 | 0.207780499 | 0.34138853 | 0.60863351 | 0.54276739 | 0.87210113 |
| PPP1R12B | 521.106898 | 0.379779156 | 0.29101562 | 1.30501296 | 0.19188843 | 0.6421543 |
| KDM5B | 159.358407 | -0.203279572 | 0.19754609 | -1.0290235 | 0.30346863 | 0.74787746 |
| RABIF | 9.94999756 | -0.520752708 | 0.61358512 | -0.8487049 | 0.39604551 | NA |
| KLHL12 | 38.6711105 | -0.185898592 | 0.24735648 | -0.7515412 | 0.45232701 | 0.83316488 |
| ADIPOR1 | 24.8024526 | 0.042509627 | 0.30449932 | 0.139605 | 0.88897209 | 0.97766772 |
| CYB5R1 | 19.047876 | 0.613072365 | 0.44115304 | 1.38970449 | 0.16461863 | 0.60511982 |
| TMEM183A | 14.0748976 | 0.006200774 | 0.44128449 | 0.01405165 | 0.98878878 | 0.99915789 |
| BTG2 | 101.368425 | -0.400573787 | 0.64742204 | -0.6187213 | 0.5361 | 0.86946459 |
| ATP2B4 | 66.2691825 | -0.721851587 | 0.44931988 | -1.6065427 | 0.10815471 | 0.50607849 |
| SNRPE | 28.4614595 | -0.156148394 | 0.30212013 | -0.5168421 | 0.6052664 | 0.89600432 |
| SOX13 | 26.8166779 | -0.047035107 | 0.45187847 | -0.104088 | 0.91709955 | 0.98495314 |
| GOLT1A | 28.5918614 | -0.970981944 | 0.38872508 | -2.4978629 | 0.01249445 | 0.16647826 |
| PLEKHA6 | 124.174524 | 0.338465431 | 0.30146436 | 1.1227378 | 0.26154887 | 0.71281761 |
| PPP1R15B | 37.7333395 | 0.44658335 | 0.33052667 | 1.35112651 | 0.17665491 | 0.62424504 |
| MDM4 | 136.547928 | 0.123754467 | 0.19658936 | 0.62950746 | 0.52901689 | 0.86758926 |
| LRRN2 | 95.5991802 | 0.35734015 | 0.43832248 | 0.81524486 | 0.41493216 | 0.81726257 |
| RBBP5 | 43.6168006 | 0.198991482 | 0.25370109 | 0.78435407 | 0.43283238 | 0.8248269 |
| DSTYK | 56.9497127 | 0.200815309 | 0.30562049 | 0.6570741 | 0.51113327 | 0.86312853 |
| NUAK2 | 49.8777091 | -0.785624305 | 0.50131402 | -1.5671301 | 0.11708428 | 0.52482323 |
| CDK18 | 17.8185356 | -0.002711586 | 0.34761617 | -0.0078005 | 0.99377615 | 0.99923337 |
| MFSD4A | 17.8885704 | -0.031871679 | 0.48635296 | -0.065532 | 0.94775043 | 0.99149867 |
| ELK4 | 148.390215 | -0.12671798 | 0.3692795 | -0.3431492 | 0.73148619 | 0.94155624 |
| NUCKS1 | 205.132484 | 0.302358096 | 0.13552384 | 2.23103259 | 0.02567897 | 0.25211135 |
| RAB29 | 12.6012714 | -0.396080804 | 0.49975758 | -0.7925459 | 0.42804246 | 0.82407405 |
| SLC41A1 | 16.6436472 | 0.257605044 | 0.33007292 | 0.78044889 | 0.4351267 | 0.82565889 |
| RHEX | 18.8473102 | -0.521760466 | 0.71980042 | -0.7248682 | 0.46853286 | 0.84326495 |
| SRGAP2 | 12.5119452 | -1.077855968 | 0.48950911 | -2.201912 | 0.02767153 | NA |
| EIF2D | 20.0412192 | 0.720702015 | 0.32153577 | 2.24143649 | 0.02499781 | 0.24972071 |
| MAPKAPK2 | 129.25088 | 0.190284473 | 0.21100403 | 0.90180492 | 0.3671605 | 0.78856614 |
| PIGR | 46.8061186 | -1.509545896 | 0.55967161 | -2.6971993 | 0.00699254 | 0.12004891 |
| PFKFB2 | 107.362073 | -0.63712879 | 0.2884823 | -2.2085542 | 0.02720567 | 0.25997792 |
| YOD1 | 15.4481175 | -0.933332601 | 0.50312936 | -1.8550549 | 0.06358838 | 0.40032348 |
| CD55 | 86.6038922 | -0.417320185 | 0.44456407 | -0.9387178 | 0.34787567 | 0.77751014 |
| CR1L | 97.9691751 | 0.792997227 | 0.60145691 | 1.31846059 | 0.18734951 | 0.63899094 |
| CD46 | 324.730856 | -0.09064645 | 0.22027837 | -0.4115086 | 0.68069962 | 0.92375164 |
| MIR29B2CH( | 187.754128 | -0.878982041 | 0.28352849 | -3.1001542 | 0.0019342 | 0.05043477 |
| PLXNA2 | 56.3010084 | 0.84927833 | 0.47502479 | 1.78786107 | 0.07379842 | 0.42813679 |
| G0S2 | 134.399858 | -1.806715832 | 0.80382825 | -2.2476391 | 0.02459921 | 0.24624595 |
| IRF6 | 38.5124171 | 0.1643717 | 0.3452597 | 0.47608133 | 0.63401643 | 0.90805737 |
| HHAT | 48.1639963 | -0.349193757 | 0.48951704 | -0.7133434 | 0.47563328 | 0.84622943 |
| RCOR3 | 163.455435 | -0.011409165 | 0.19287748 | -0.0591524 | 0.95283072 | 0.99217529 |
| LINC00467 | 33.5065624 | -0.104836773 | 0.34465634 | -0.3041777 | 0.76099251 | 0.94990484 |
| SLC30A1 | 14.5572249 | 0.368314866 | 0.51510258 | 0.71503207 | 0.47458923 | 0.84593347 |
| LPGAT1 | 261.582417 | -0.242097205 | 0.27428727 | -0.8826411 | 0.37743024 | 0.79432848 |
| INTS7 | 62.3142407 | 0.474124523 | 0.2775063 | 1.70851805 | 0.08754026 | 0.46528247 |
| PPP2R5A | 551.035396 | -0.488877796 | 0.29125243 | -1.6785364 | 0.09324243 | 0.47790456 |
| AL360091.3 | 76.1966827 | -1.415442449 | 0.55662426 | -2.5429047 | 0.01099352 | 0.15544774 |
| NENF | 33.5351096 | -0.161313707 | 0.29942124 | -0.5387517 | 0.59005818 | 0.89149106 |
| NSL1 | 45.3980347 | -0.15775361 | 0.25417458 | -0.6206506 | 0.53482953 | 0.86942289 |
| TATDN3 | 46.6662431 | -0.444200778 | 0.31410978 | -1.4141577 | 0.1573156 | 0.59804205 |
| FLVCR1 | 32.8029817 | -0.295342702 | 0.46372782 | -0.636888 | 0.52419777 | 0.86747074 |
| ANGEL2 | 36.3930902 | 0.115373945 | 0.25735515 | 0.44830634 | 0.65393213 | 0.9158679 |
| RPS6KC1 | 140.56877 | -0.137490436 | 0.19764521 | -0.6956426 | 0.48665265 | 0.85365949 |
| SMYD2 | 36.2747197 | -0.369337359 | 0.32652663 | -1.1311094 | 0.25800903 | 0.71022118 |
| PTPN14 | 117.434823 | 0.263126927 | 0.21754455 | 1.20953122 | 0.22645882 | 0.68070982 |
| KCTD3 | 72.8508916 | -0.348126685 | 0.20938421 | -1.6626215 | 0.09638822 | 0.48243848 |
| ESRRG | 3023.32315 | 0.431649788 | 0.33149486 | 1.30213116 | 0.19287155 | 0.64373732 |
| GPATCH2 | 153.908973 | -0.068762974 | 0.19616236 | -0.3505411 | 0.72593263 | 0.94000087 |
| RRP15 | 27.4319967 | 0.357414282 | 0.27336804 | 1.30744723 | 0.19106087 | 0.6421543 |
| LYPLAL1 | 209.836564 | -0.07291734 | 0.23395676 | -0.3116702 | 0.75529121 | 0.94795939 |
| ZC3H11B | 10.4481885 | 0.651557987 | 0.51572703 | 1.26337761 | 0.20645351 | NA |
| EPRS | 56.6937783 | 0.012648597 | 0.20372166 | 0.06208764 | 0.95049304 | 0.99209822 |
| BPNT1 | 24.0412326 | -0.278672702 | 0.35513039 | -0.7847053 | 0.43262637 | 0.8248269 |
| IARS2 | 71.4183988 | 0.133014405 | 0.21042376 | 0.63212636 | 0.52730431 | 0.86750588 |
| RAB3GAP2 | 194.417958 | -0.043497366 | 0.21656081 | -0.2008552 | 0.8408118 | 0.96748214 |
| MARK1 | 80.7175584 | -0.136687276 | 0.32530415 | -0.420183 | 0.67435177 | 0.92189158 |
| 2-Mar | 55.1330895 | 0.65859812 | 0.30484202 | 2.16045712 | 0.0307373 | 0.27916836 |
| AL513314.2 | 16.9660664 | 0.382449078 | 0.47195057 | 0.81035833 | 0.41773426 | 0.8188001 |
| MIA3 | 42.67005 | 0.236095045 | 0.22727619 | 1.03880239 | 0.29889665 | 0.74296125 |
| AIDA | 18.4505439 | -0.456902878 | 0.33497252 | -1.3640011 | 0.17256722 | 0.61726473 |
| BROX | 56.2137846 | -0.043279999 | 0.23092744 | -0.1874182 | 0.85133278 | 0.97027687 |
| FAM177B | 27.6966519 | -2.639185348 | 0.73824823 | -3.5749295 | 0.00035032 | 0.0158791 |
| AC093152.1 | 23.3649729 | 0.281765697 | 0.29917333 | 0.94181422 | 0.34628776 | 0.77642952 |
| SUSD4 | 11.3998305 | 0.460990368 | 0.64049063 | 0.71974568 | 0.47168159 | NA |
| CAPN2 | 248.173629 | -0.090648853 | 0.20460564 | -0.4430418 | 0.65773549 | 0.91601353 |

| | | | | | | |
|---|---|---|---|---|---|---|
| TP53BP2 | 52.2668678 | -0.385697395 | 0.31694126 | -1.2169365 | 0.22362837 | 0.67606266 |
| FBXO28 | 86.5151069 | 0.064689949 | 0.23519424 | 0.27504903 | 0.78327857 | 0.95544332 |
| DEGS1 | 13.4334769 | -0.601440196 | 0.70993707 | -0.847174 | 0.39689817 | 0.80674902 |
| NVL | 68.0811393 | 0.288560941 | 0.20541567 | 1.40476596 | 0.16009089 | 0.60024417 |
| CNIH4 | 61.2461446 | 0.078310494 | 0.29300988 | 0.2672623 | 0.78926721 | 0.95719943 |
| WDR26 | 168.879664 | 0.162443592 | 0.26070336 | 0.62309743 | 0.5332205 | 0.86909171 |
| CNIH3 | 43.0295348 | 0.616003767 | 0.31772028 | 1.93882421 | 0.05252275 | 0.36358594 |
| DNAH14 | 254.228899 | 0.122020081 | 0.24640582 | 0.49519966 | 0.62045919 | 0.90380166 |
| LBR | 52.7016555 | -0.262466449 | 0.3298927 | -0.7956116 | 0.42625783 | 0.82329929 |
| ENAH | 33.1840993 | 0.546167465 | 0.31331686 | 1.74317928 | 0.0813023 | 0.45116267 |
| SRP9 | 100.345026 | 0.130449704 | 0.3148563 | 0.41431506 | 0.67864339 | 0.92291026 |
| EPHX1 | 21.139246 | 0.158336282 | 0.48431026 | 0.3269315 | 0.74371969 | 0.94546379 |
| AL591895.1 | 23.1163699 | -0.433324666 | 0.38337268 | -1.1302961 | 0.25835146 | 0.71022613 |
| TMEM63A | 64.7521073 | -0.309743614 | 0.32193923 | -0.9621183 | 0.3359902 | 0.76847558 |
| SDE2 | 17.3667492 | 0.176869837 | 0.41957717 | 0.42154304 | 0.67335859 | 0.92189158 |
| H3F3A | 24.931393 | -0.378031927 | 0.31409223 | -1.2035698 | 0.2287559 | 0.68401116 |
| ACBD3 | 103.040916 | -0.061434133 | 0.20805715 | -0.2952753 | 0.7677836 | 0.95205353 |
| LIN9 | 14.605825 | 1.106961767 | 0.381867 | 2.89881494 | 0.00374576 | 0.07967952 |
| PARP1 | 52.3358032 | -0.026352297 | 0.33483192 | -0.0787031 | 0.93726881 | 0.98871562 |
| STUM | 106.557595 | -0.592187022 | 0.41041757 | -1.4428891 | 0.14905172 | 0.58629428 |
| ITPKB | 50.4833383 | -0.045163252 | 0.31547904 | -0.1431577 | 0.88616564 | 0.97693497 |
| PSEN2 | 46.461462 | -0.549898439 | 0.40333391 | -1.3633826 | 0.17276196 | 0.61746168 |
| COQ8A | 42.4268475 | -0.742333344 | 0.41808217 | -1.775568 | 0.07580414 | 0.43380542 |
| CDC42BPA | 450.782517 | -0.259136599 | 0.22127186 | -1.1711231 | 0.24154929 | 0.69556484 |
| ZNF678 | 73.8892878 | -0.034456931 | 0.25548239 | -0.1348701 | 0.8927146 | 0.97823448 |
| SNAP47 | 19.1443132 | 0.524063825 | 0.29614917 | 1.7695941 | 0.07679478 | 0.43580177 |
| ARF1 | 136.103725 | -0.394331207 | 0.2219789 | -1.7764355 | 0.07566116 | 0.43325908 |
| MRPL55 | 18.5361676 | 0.31683139 | 0.38958148 | 0.81326091 | 0.41606849 | 0.81763864 |
| GUK1 | 128.243799 | -0.277511023 | 0.26239067 | -1.0576253 | 0.29022629 | 0.73979353 |
| IBA57 | 13.5023886 | 0.424203845 | 0.62264077 | 0.68129789 | 0.49568302 | 0.85814183 |
| OBSCN | 80.5307159 | -0.04466329 | 0.25703264 | -0.1737651 | 0.86205011 | 0.97302617 |
| TRIM11 | 11.9330403 | -0.404515841 | 0.48655028 | -0.8313958 | 0.40575009 | NA |
| RNF187 | 27.9934814 | -0.300540371 | 0.36567834 | -0.8218709 | 0.4111504 | 0.81457492 |
| RHOU | 24.0880491 | -1.710105155 | 0.42442485 | -4.0292296 | 5.60E-05 | 0.00444927 |
| RAB4A | 43.6529163 | 0.004928867 | 0.25834399 | 0.0190787 | 0.98477832 | 0.99773864 |
| NUP133 | 50.760794 | -0.230658774 | 0.22099996 | -1.043705 | 0.29662188 | 0.7415547 |
| ABCB10 | 26.2009398 | 0.148894982 | 0.32027106 | 0.46490302 | 0.64200091 | 0.91146968 |
| TAF5L | 17.7813993 | 0.590625883 | 0.43351237 | 1.36241991 | 0.1730654 | 0.61786323 |
| GALNT2 | 48.5781396 | -0.305230579 | 0.2837067 | -1.0758667 | 0.28198689 | 0.73351377 |
| COG2 | 33.3132375 | 0.440730627 | 0.28584104 | 1.54187315 | 0.12310442 | 0.5395901 |
| C1orf198 | 17.1156901 | -0.407480309 | 0.39088206 | -1.0424636 | 0.29719681 | 0.74160729 |
| TTC13 | 25.2204729 | 0.511785889 | 0.44038056 | 1.16214459 | 0.24517674 | 0.69883704 |
| ARV1 | 23.8635367 | 0.108796052 | 0.34437224 | 0.31592573 | 0.75205888 | 0.94728927 |
| C1orf131 | 18.6098672 | 0.14785723 | 0.31863981 | 0.46402622 | 0.64262896 | 0.91146968 |
| GNPAT | 59.7262557 | 0.01086249 | 0.24590415 | 0.04417367 | 0.96476597 | 0.99334616 |
| EXOC8 | 11.791216 | 0.82317592 | 0.40634685 | 2.02579622 | 0.04278567 | NA |
| SPRTN | 15.3224501 | 0.40708737 | 0.48185401 | 0.8448355 | 0.39820269 | 0.80714564 |
| EGLN1 | 238.939835 | -0.18997915 | 0.24529759 | -0.7744844 | 0.4386444 | 0.82745796 |
| DISC1 | 52.185804 | -1.807640528 | 0.57943154 | -3.1196792 | 0.00181048 | 0.04811415 |
| SIPA1L2 | 106.961805 | 0.392548922 | 0.40731047 | 0.96375849 | 0.33516702 | 0.76777176 |
| NTPCR | 26.7405479 | 0.651904426 | 0.28505511 | 2.28694175 | 0.02219922 | 0.2338029 |
| PCNX2 | 146.967564 | -0.226424762 | 0.28054303 | -0.8070946 | 0.41961199 | 0.81899879 |
| MAP3K21 | 130.032987 | 0.258449194 | 0.20610071 | 1.25399469 | 0.20984393 | 0.66164938 |
| KCNK1 | 193.430889 | -0.786442371 | 0.30603642 | -2.5697672 | 0.01017669 | 0.14844189 |
| SLC35F3 | 106.575839 | -1.321362217 | 0.40205029 | -3.2865595 | 0.00101419 | 0.03290193 |
| COA6 | 21.767948 | -0.122960771 | 0.48338055 | -0.2543767 | 0.79920452 | 0.95848876 |
| TARBP1 | 59.3376465 | -0.133155071 | 0.42084274 | -0.316401 | 0.75169815 | 0.94719044 |
| IRF2BP2 | 59.9609771 | 0.037223412 | 0.30621171 | 0.12156103 | 0.90324668 | 0.98016176 |
| TOMM20 | 92.2399121 | 0.375833013 | 0.29252008 | 1.284811 | 0.19885834 | 0.65144406 |
| ARID4B | 368.158288 | -0.359119518 | 0.24386923 | -1.4725905 | 0.14086149 | 0.57506583 |
| B3GALNT2 | 54.1888424 | -0.09735744 | 0.27692974 | -0.3515601 | 0.72516822 | 0.93940528 |
| LYST | 137.358917 | 0.486678662 | 0.24137566 | 2.0162707 | 0.04377168 | 0.33268382 |
| GPR137B | 81.0392155 | -0.69330152 | 0.26158627 | -2.6503743 | 0.00804026 | 0.130419 |
| ERO1B | 146.725005 | -0.07178683 | 0.50497453 | -0.1421593 | 0.88695417 | 0.9772212 |
| EDARADD | 140.186563 | 0.650540977 | 0.43189214 | 1.50625797 | 0.13200096 | 0.55916709 |
| LGALS8 | 45.3874262 | -0.604561437 | 0.25283825 | -2.3910996 | 0.016798 | 0.20115876 |
| HEATR1 | 35.7598486 | -0.039884614 | 0.29726967 | -0.1341698 | 0.8932683 | 0.97831181 |
| MTR | 695.705029 | -0.403751901 | 0.39431157 | -1.0239413 | 0.305863 | 0.74897897 |
| CHRM3 | 18.4078542 | 0.761267137 | 0.8044648 | 0.9463026 | 0.34399427 | 0.77382755 |
| FH | 34.1399224 | 0.220914607 | 0.29077974 | 0.75973177 | 0.44741493 | 0.83066268 |
| OPN3 | 21.0671509 | -0.249173996 | 0.35795167 | -0.6961107 | 0.48635949 | 0.85363814 |
| CEP170 | 54.1942024 | 0.012440363 | 0.33354352 | 0.03729757 | 0.97024774 | 0.99496808 |
| SDCCAG8 | 261.873517 | 0.00374239 | 0.19992793 | 0.0187187 | 0.98506551 | 0.99777861 |
| AKT3 | 378.952892 | -0.771890443 | 0.20684212 | -3.7317855 | 0.00019013 | 0.01042206 |
| ZBTB18 | 44.1516442 | 0.393784104 | 0.30581225 | 1.28766621 | 0.19786218 | 0.64914661 |
| AL645465.1 | 15.3597603 | -0.006061099 | 0.45245742 | -0.013396 | 0.98931189 | 0.99923337 |
| ADSS | 55.7937226 | -0.325577294 | 0.32355823 | -1.0062402 | 0.31430004 | 0.75700972 |
| CATSPERE | 35.6016241 | -0.02224816 | 0.4270609 | -0.052096 | 0.95845221 | 0.99238848 |
| DESI2 | 68.6727324 | -0.493005072 | 0.26837788 | -1.8369811 | 0.06621269 | 0.40888394 |
| COX20 | 81.9067869 | -0.117499891 | 0.298257 | -0.3939552 | 0.69361413 | 0.92826395 |
| HNRNPU | 263.328632 | -0.061889613 | 0.20368023 | -0.3038568 | 0.76123702 | 0.9499444 |
| EFCAB2 | 101.528491 | 0.462059745 | 0.43241169 | 1.06856441 | 0.28526599 | 0.73659018 |
| KIF26B | 18.3022415 | 0.312684944 | 0.41787191 | 0.74827941 | 0.45429164 | 0.83364149 |
| SMYD3 | 351.996689 | 0.467941986 | 0.33212572 | 1.40893029 | 0.15885578 | 0.5993392 |
| TFB2M | 11.3509915 | -0.0455012 | 0.41525735 | -0.1095735 | 0.91274763 | NA |
| CNST | 85.9905028 | -0.320698667 | 0.19915938 | -1.6102614 | 0.10734079 | 0.50372797 |
| SCCPDH | 128.800395 | 0.003417231 | 0.48874498 | 0.00699185 | 0.99442136 | 0.99923337 |
| AHCTF1 | 99.7347564 | -0.075678351 | 0.21944133 | -0.3448683 | 0.73019341 | 0.94155624 |
| ZNF124 | 32.0854441 | -0.275689496 | 0.27234405 | -1.0122839 | 0.31140234 | 0.75450562 |
| AL390728.6 | 21.9113058 | -0.163977952 | 0.40893689 | -0.400986 | 0.68843046 | 0.9267526 |
| ZNF496 | 20.7613237 | 0.166112878 | 0.45678252 | 0.36365857 | 0.71611299 | 0.93689626 |
| SH3BP5L | 10.741168 | 0.231766598 | 0.43889021 | 0.5280742 | 0.59744783 | NA |
| ZNF672 | 32.7342144 | -0.296231162 | 0.26972675 | -1.0982639 | 0.27208926 | 0.7232847 |
| ZNF692 | 15.9771473 | 0.491844396 | 0.36949708 | 1.33111848 | 0.18315003 | 0.63200117 |
| FAM110C | 12.8325646 | -2.68000618 | 0.62498813 | -4.2880913 | 1.80E-05 | 0.00183599 |
| SH3YL1 | 116.414303 | -0.358641006 | 0.2729218 | -1.3140797 | 0.18881938 | 0.64021224 |
| ACP1 | 45.9317881 | -0.277781739 | 0.27582674 | -1.0070878 | 0.31389259 | 0.75646164 |
| TMEM18 | 12.8045445 | 0.44655309 | 0.44818497 | 0.99635892 | 0.31907579 | 0.76063778 |
| EIPR1 | 101.739432 | -0.099708871 | 0.37323154 | -0.2671502 | 0.78935354 | 0.95719943 |
| TRAPPC12 | 125.853887 | -0.273904094 | 0.30053011 | -0.9114032 | 0.36208298 | 0.78714716 |
| ADI1 | 91.7243452 | -0.115825559 | 0.43802583 | -0.2644263 | 0.79145143 | 0.95746995 |
| AC108488.2 | 11.1912791 | -0.086693008 | 0.43248509 | -0.2004532 | 0.84112618 | NA |
| RPS7 | 179.534943 | 0.370455823 | 0.45472748 | 0.81467657 | 0.41525747 | 0.81726257 |
| RNF144A | 28.9637705 | -0.299806222 | 0.43438017 | -0.6901932 | 0.49007273 | 0.85590664 |
| ID2 | 20.9131645 | -0.096984194 | 0.46341335 | -0.2092823 | 0.8342279 | 0.96701848 |
| KIDINS220 | 278.688723 | 0.231690583 | 0.17395396 | 1.3319075 | 0.18289058 | 0.63154788 |
| MBOAT2 | 241.470627 | -0.808057537 | 0.58304257 | -1.3859323 | 0.16576758 | 0.60621113 |
| ASAP2 | 224.497767 | 0.327368255 | 0.33920132 | 0.96511491 | 0.33448726 | 0.76777176 |
| ITGB1BP1 | 68.2014416 | 0.446665189 | 0.29491388 | 1.51456145 | 0.1298835 | 0.55598851 |
| CPSF3 | 25.9638641 | 0.216936465 | 0.31655544 | 0.68530322 | 0.4931526 | 0.85714521 |
| IAH1 | 58.3429357 | -0.232008491 | 0.30567877 | -0.7589944 | 0.44785588 | 0.83066268 |
| ADAM17 | 123.05057 | -0.359233516 | 0.23162694 | -1.5509142 | 0.12092224 | 0.53461518 |
| YWHAQ | 147.534 | 0.010453841 | 0.16052718 | 0.06512194 | 0.94807691 | 0.9916267 |

| | | | | | | |
|---|---|---|---|---|---|---|
| TAF1B | 82.8814696 | 0.419128352 | 0.257545 | 1.62739853 | 0.10365248 | 0.49577372 |
| GRHL1 | 39.5306247 | -0.025902863 | 0.27362426 | -0.0946658 | 0.92458027 | 0.98672316 |
| KLF11 | 22.9526235 | 0.205938878 | 0.43290748 | 0.47571107 | 0.63428023 | 0.90805737 |
| CYS1 | 33.9354087 | 0.28462401 | 0.4073044 | 0.69879926 | 0.48467749 | 0.85339838 |
| HPCAL1 | 49.1993385 | 0.308035601 | 0.43142222 | 0.71400032 | 0.47522698 | 0.84597206 |
| ODC1 | 21.1837657 | 0.095836101 | 0.35357585 | 0.27104821 | 0.78635395 | 0.95577371 |
| NOL10 | 98.7599373 | 0.045084967 | 0.23990105 | 0.18793151 | 0.85093034 | 0.97027687 |
| ATP6V1C2 | 550.290121 | 0.655772754 | 0.39697066 | 1.65194265 | 0.09854624 | 0.48476221 |
| PDIA6 | 113.455001 | -0.150950239 | 0.2858275 | -0.5281166 | 0.59741843 | 0.89363331 |
| PQLC3 | 43.1391364 | 0.286308977 | 0.40348079 | 0.70959754 | 0.47795375 | 0.84771464 |
| ROCK2 | 189.30584 | 0.201595856 | 0.19719055 | 1.02234034 | 0.30661984 | 0.74936067 |
| E2F6 | 18.757128 | 0.082673599 | 0.34167927 | 0.24196258 | 0.80880916 | 0.96124185 |
| LPIN1 | 93.9896303 | -0.856085446 | 0.39702635 | -2.1562434 | 0.03106467 | 0.27952435 |
| MIR3681HG | 163.296787 | -0.433810109 | 0.41330972 | -1.0496005 | 0.29390181 | 0.74044476 |
| TRIB2 | 43.7188106 | -0.122790612 | 0.36931423 | -0.3324828 | 0.73952476 | 0.94376279 |
| NBAS | 240.644136 | -0.006645591 | 0.25588609 | -0.0259709 | 0.97928055 | 0.99623022 |
| DDX1 | 85.8505536 | -0.055582656 | 0.23980958 | -0.2317783 | 0.81671021 | 0.96340618 |
| SMC6 | 41.1091186 | 0.364941444 | 0.32537413 | 1.1216056 | 0.26203017 | 0.71302363 |
| GEN1 | 17.0939029 | 0.000257374 | 0.53209192 | 0.0004837 | 0.99961406 | 0.99982181 |
| RDH14 | 25.1905699 | -0.377458444 | 0.43172114 | -0.8743108 | 0.38194904 | 0.79731722 |
| LINC01376 | 58.7860609 | -0.161428131 | 0.27769551 | -0.5813134 | 0.56102922 | 0.88072236 |
| TTC32 | 20.9639887 | 0.294983406 | 0.35663758 | 0.82712374 | 0.40816693 | 0.81361336 |
| WDR35 | 45.1212363 | 0.213058743 | 0.25820197 | 0.82516312 | 0.40927899 | 0.81413754 |
| LAPTM4A | 173.497271 | -0.172464843 | 0.28378658 | -0.6077273 | 0.54336838 | 0.87233405 |
| SDC1 | 29.801949 | 0.021497747 | 0.44036129 | 0.04881843 | 0.961064 | 0.99238848 |
| PUM2 | 292.590532 | 0.143437005 | 0.13160398 | 1.08991387 | 0.27575109 | 0.72554623 |
| RHOB | 23.3858221 | -0.854371103 | 0.49948751 | -1.7104954 | 0.08717429 | 0.46486567 |
| HS1BP3 | 11.9116018 | -0.684458059 | 0.46519599 | -1.4713327 | 0.14120117 | NA |
| LDAH | 74.2996913 | -0.03445999 | 0.21611822 | -0.1594497 | 0.87331456 | 0.97434023 |
| KLHL29 | 12.7455147 | -1.572504388 | 0.6851902 | -2.2949896 | 0.02173372 | 0.23061274 |
| ATAD2B | 221.300874 | 0.019176212 | 0.26080804 | 0.07352615 | 0.94138743 | 0.98985996 |
| UBXN2A | 77.7329625 | -0.064504355 | 0.23728653 | -0.2718416 | 0.7857438 | 0.95577371 |
| SF3B6 | 37.8488751 | 0.023925636 | 0.42436701 | 0.05637959 | 0.95503942 | 0.99228059 |
| FAM228B | 17.3206558 | -0.200488819 | 0.35808497 | -0.5598917 | 0.57555328 | 0.88746465 |
| TP53I3 | 26.6013495 | -0.082457506 | 0.43669632 | -0.1888212 | 0.85023299 | 0.97027687 |
| ITSN2 | 349.29837 | -0.466242013 | 0.17176915 | -2.7143525 | 0.00664055 | 0.11674591 |
| NCOA1 | 469.333775 | -0.226186358 | 0.17849247 | -1.2672039 | 0.20508239 | 0.65805651 |
| PTRHD1 | 12.6150849 | -0.48890675 | 0.62268099 | -0.7851641 | 0.43235737 | 0.8248269 |
| DNAJC27 | 34.0647406 | 0.267124507 | 0.27868029 | 0.9585339 | 0.3377936 | 0.76966704 |
| DNMT3A | 92.4770651 | 0.359062198 | 0.33517793 | 1.07125847 | 0.28405323 | 0.73573199 |
| DTNB | 153.424277 | 0.399802155 | 0.27537502 | 1.45184612 | 0.1465444 | 0.58349206 |
| ASXL2 | 154.431532 | -0.102233424 | 0.3131884 | -0.3264279 | 0.74410065 | 0.94546379 |
| RAB10 | 288.696717 | -0.289887073 | 0.22860138 | -1.2680898 | 0.20476588 | 0.65790959 |
| HADHA | 208.90849 | -0.119839005 | 0.18175451 | -0.6593454 | 0.50967397 | 0.86281145 |
| HADHB | 319.807886 | 0.137218751 | 0.31467424 | 0.43606604 | 0.66278881 | 0.918408 |
| SELENOI | 178.915814 | 0.507493593 | 0.1800554 | 2.81854144 | 0.00482424 | 0.09530573 |
| AC013472.2 | 73.7897044 | 2.608341072 | 0.55128674 | 4.73136917 | 2.23E-06 | 0.00035462 |
| MAPRE3 | 103.611356 | 0.850343595 | 0.29595426 | 2.87322645 | 0.00406303 | 0.08421233 |
| TMEM214 | 22.810919 | 0.233283579 | 0.36418201 | 0.64056865 | 0.52180297 | 0.86703098 |
| AGBL5 | 24.3165802 | 0.421506781 | 0.32074781 | 1.31413769 | 0.18879987 | 0.64021224 |
| OST4 | 73.6289535 | 0.018539246 | 0.41783633 | 0.04436964 | 0.96460976 | 0.99334616 |
| ATRAID | 56.7474196 | -0.105161117 | 0.3410196 | -0.3083726 | 0.75779879 | 0.94918855 |
| MPV17 | 19.8618907 | -0.889566665 | 0.37402562 | -2.3783575 | 0.01738996 | 0.20451748 |
| GTF3C2 | 46.7694232 | 0.308409327 | 0.25542657 | 1.2074285 | 0.22726716 | 0.68208276 |
| EIF2B4 | 10.5843879 | -0.032339404 | 0.42360956 | -0.0763425 | 0.93914663 | NA |
| SNX17 | 24.6885083 | -0.11943821 | 0.31390789 | -0.3804881 | 0.70358315 | 0.93306088 |
| ZNF513 | 16.4519162 | -0.252123555 | 0.38263748 | -0.6589097 | 0.50995374 | 0.86281145 |
| PPM1G | 92.6128714 | -0.535504927 | 0.31563615 | -1.6965893 | 0.08977433 | 0.47198427 |
| NRBP1 | 33.6167576 | 0.418336475 | 0.28524888 | 1.4665666 | 0.14249399 | 0.57673923 |
| KRTCAP3 | 22.3278249 | -0.447073458 | 0.4558975 | -0.9806447 | 0.326768 | 0.76371899 |
| IFT172 | 28.9089837 | 0.324187701 | 0.45318231 | 0.71535824 | 0.47438771 | 0.84593347 |
| ZNF512 | 55.0626993 | 0.036521972 | 0.36920714 | 0.09892 | 0.92120179 | 0.98614962 |
| GPN1 | 12.2459956 | -0.100734474 | 0.45533912 | -0.2212296 | 0.82491369 | NA |
| SUPT7L | 23.5969219 | -0.214260592 | 0.30741384 | -0.6969777 | 0.48581675 | 0.85356796 |
| SLC4A1AP | 23.1756091 | 0.144470169 | 0.28396624 | 0.50875826 | 0.61092168 | 0.90002868 |
| MRPL33 | 84.5220593 | 0.216020196 | 0.33253421 | 0.64961796 | 0.51593903 | 0.86476217 |
| RBKS | 47.9655851 | -0.177472521 | 0.33341682 | -0.5322842 | 0.59452914 | 0.89289841 |
| BABAM2 | 447.465842 | -0.505760494 | 0.31436769 | -1.6088183 | 0.10765607 | 0.50496321 |
| FOSL2 | 121.902912 | -1.015371042 | 0.3544255 | -2.8648363 | 0.00417225 | 0.08574324 |
| PPP1CB | 84.8484955 | -0.233521095 | 0.23201079 | -1.0065096 | 0.31417047 | 0.75694327 |
| TRMT61B | 30.3903999 | 0.708237645 | 0.28449746 | 2.48943401 | 0.01279467 | 0.16857023 |
| WDR43 | 56.8914501 | 0.31058934 | 0.20543192 | 1.5118845 | 0.13056325 | 0.55687707 |
| CLIP4 | 150.077929 | -0.772050649 | 0.23761258 | -3.2491994 | 0.0011573 | 0.03609595 |
| YPEL5 | 72.8911126 | -0.223699995 | 0.25549518 | -0.8755546 | 0.38127219 | 0.79688435 |
| LBH | 17.7638597 | -0.292803166 | 0.74283219 | -0.3941713 | 0.69345454 | 0.92826395 |
| LCLAT1 | 106.482364 | -0.117483188 | 0.36792066 | -0.3193166 | 0.74948642 | 0.94684415 |
| CAPN13 | 45.4153964 | 0.526719723 | 0.53660934 | 0.98157017 | 0.32631165 | 0.76362437 |
| MEMO1 | 19.5567699 | 1.1246852 | 0.3545689 | 3.17197927 | 0.00151404 | 0.04256861 |
| DPY30 | 44.0171964 | 0.002807595 | 0.3444434 | 0.00815111 | 0.99349643 | 0.99923337 |
| SPAST | 48.1664194 | 0.115636263 | 0.2474095 | 0.46738812 | 0.64022222 | 0.91096398 |
| SLC30A6 | 61.1052892 | 0.178834181 | 0.18013758 | 0.99276441 | 0.32082478 | 0.76131909 |
| YIPF4 | 115.848894 | 0.159940979 | 0.21368728 | 0.74848151 | 0.45416978 | 0.83364149 |
| BIRC6 | 546.157275 | -0.00529597 | 0.17689444 | -0.0299386 | 0.97611603 | 0.99556048 |
| TTC27 | 65.0987826 | 0.253107526 | 0.32568775 | 0.77714783 | 0.43707156 | 0.82687287 |
| LINC00486 | 97.4282287 | 0.62490178 | 0.21114897 | 2.95953035 | 0.00308108 | 0.06902196 |
| LTBP1 | 15.3701799 | -0.806586674 | 0.77907911 | -1.0353078 | 0.30052519 | 0.74458947 |
| RASGRP3 | 25.4078689 | -0.485160257 | 0.49884618 | -0.9725649 | 0.33076962 | 0.76685572 |
| FAM98A | 19.7265299 | 0.798146271 | 0.34231332 | 2.33162492 | 0.01972043 | 0.22063228 |
| LINC01317 | 13.6519823 | 1.385198779 | 0.76893361 | 1.80145434 | 0.0716313 | 0.42284325 |
| AC068599.1 | 72.1513171 | 2.31200042 | 0.81625356 | 2.8324537 | 0.00461923 | 0.09276706 |
| LINC01320 | 129.09247 | -1.671521163 | 0.5142474 | -3.2504222 | 0.00115234 | 0.03605702 |
| AC012593.1 | 19.6761973 | -0.903759654 | 0.73024565 | -1.2376105 | 0.21586053 | 0.67024558 |
| CRIM1 | 317.649176 | -0.638247041 | 0.40441213 | -1.5782094 | 0.11451751 | 0.51928773 |
| FEZ2 | 141.58688 | -0.008335556 | 0.24539157 | -0.0339684 | 0.97290236 | 0.99526459 |
| STRN | 222.950911 | 0.371831868 | 0.24872148 | 1.49497291 | 0.13492151 | 0.56411149 |
| HEATR5B | 70.6102611 | -0.080081033 | 0.2009983 | -0.3984165 | 0.69032322 | 0.92757102 |
| EIF2AK2 | 108.629646 | -0.220386879 | 0.52665472 | -0.4184656 | 0.67560674 | 0.92193843 |
| CEBPZOS | 51.1304937 | -0.138034452 | 0.35153412 | -0.392663 | 0.69456838 | 0.92841398 |
| CEBPZ | 60.7312432 | -0.261526305 | 0.23647462 | -1.1059381 | 0.26875329 | 0.72009952 |
| NDUFAF7 | 39.8197228 | 0.039288065 | 0.25236722 | 0.15567817 | 0.87628671 | 0.97532685 |
| PRKD3 | 73.9102123 | -0.161835162 | 0.39179269 | -0.4130632 | 0.67956028 | 0.92360021 |
| CDC42EP3 | 22.2515092 | -0.523167127 | 0.38867009 | -1.3460442 | 0.17828825 | 0.62598987 |
| RMDN2 | 28.4540093 | 0.665820459 | 0.35407936 | 1.88042718 | 0.06004988 | 0.38930355 |
| ATL2 | 212.198437 | 0.288806452 | 0.22768081 | 1.26847078 | 0.20462988 | 0.65790847 |
| HNRNPLL | 54.7874518 | -0.223842203 | 0.26446208 | -0.8464057 | 0.39732647 | 0.80674902 |
| GALM | 35.8050135 | 0.405927552 | 0.37976014 | 1.0689051 | 0.28511243 | 0.73659018 |
| SRSF7 | 71.7739596 | 0.176085545 | 0.37273953 | 0.4724091 | 0.63663481 | 0.90920631 |
| DHX57 | 36.3926163 | 0.059220933 | 0.35588164 | 0.16640626 | 0.86783725 | 0.97434023 |
| SOS1 | 320.821645 | 0.019005114 | 0.19844161 | 0.09577182 | 0.9237018 | 0.98672316 |
| MAP4K3 | 731.793332 | 0.292482022 | 0.16725202 | 1.74875029 | 0.08033419 | 0.44809757 |
| AC007388.1 | 386.667995 | 0.324057848 | 0.24773493 | 1.30808298 | 0.19084517 | 0.6421543 |
| TMEM178A | 25.0992963 | 1.305577729 | 0.34855214 | 3.7457172 | 0.00017988 | 0.01020367 |
| THUMPD2 | 51.1742582 | -0.04677517 | 0.23165276 | -0.2019193 | 0.83997979 | 0.96748214 |

| | | | | | | |
|---|---|---|---|---|---|---|
| SLC8A1 | 83.1480985 | 1.931278041 | 0.59879656 | 3.22526578 | 0.00125856 | 0.0377957 |
| PKDCC | 12.5279806 | 0.65617638 | 0.40503151 | 1.62006253 | 0.10521884 | NA |
| EML4 | 244.569056 | -0.145479418 | 0.28085934 | -0.5179796 | 0.60447248 | 0.89593125 |
| COX7A2L | 136.184341 | -0.213208162 | 0.35562221 | -0.5995356 | 0.5488158 | 0.87371493 |
| KCNG3 | 31.6516161 | -1.601086156 | 0.74330848 | -2.1539996 | 0.0312402 | 0.28032376 |
| MTA3 | 118.388748 | -0.142070496 | 0.23723878 | -0.5988502 | 0.54927277 | 0.8740566 |
| ZFP36L2 | 96.5968665 | -0.198394477 | 0.42781059 | -0.4637437 | 0.64283136 | 0.91148432 |
| THADA | 158.411485 | 0.172995159 | 0.25695945 | 0.67323914 | 0.50079516 | 0.85942996 |
| PLEKHH2 | 88.7315253 | -0.56548072 | 0.41106728 | -1.3756403 | 0.16893302 | 0.6112086 |
| DYNC2LI1 | 80.8237775 | 0.065487005 | 0.32050561 | 0.20432405 | 0.83810027 | 0.96748214 |
| LRPPRC | 151.832215 | 0.297546153 | 0.18018236 | 1.65136118 | 0.09866485 | 0.48476221 |
| PPM1B | 195.221845 | 0.183635566 | 0.20171047 | 0.91039183 | 0.3626159 | 0.78718296 |
| SLC3A1 | 130.568029 | -0.296935996 | 0.33912983 | -0.8755821 | 0.38125728 | 0.79688435 |
| PREPL | 144.148648 | 0.240934476 | 0.28219409 | 0.85378995 | 0.39322138 | 0.80400536 |
| CAMKMT | 332.488747 | -0.006178285 | 0.18104304 | -0.0341261 | 0.97277663 | 0.99526459 |
| SRBD1 | 93.2802937 | 0.348594588 | 0.19115785 | 1.82359548 | 0.06821325 | 0.41547228 |
| PRKCE | 581.871993 | 0.028241231 | 0.15856954 | 0.17809998 | 0.85864446 | 0.97287712 |
| EPAS1 | 130.55453 | -0.405546516 | 0.23839187 | -1.7011759 | 0.08890996 | 0.4687332 |
| RHOQ | 95.6387782 | -0.690916174 | 0.3240505 | -2.132125 | 0.03299657 | 0.28912987 |
| AC018682.1 | 16.9849341 | -1.08029085 | 0.69860134 | -1.5463624 | 0.12201707 | 0.53747904 |
| PIGF | 43.771876 | 0.133848549 | 0.2849094 | 0.46979338 | 0.63850264 | 0.91053744 |
| CRIPT | 14.0984543 | -0.340884517 | 0.41672136 | -0.8180155 | 0.41334835 | 0.81626201 |
| SOCS5 | 50.5013358 | 0.419540484 | 0.23812808 | 1.76182702 | 0.07809853 | 0.44084536 |
| MCFD2 | 67.9493256 | 0.3105834 | 0.31860965 | 0.97480852 | 0.32965525 | 0.76685572 |
| TTC7A | 56.9558899 | 0.444918692 | 0.33444954 | 1.3303014 | 0.18341899 | 0.63239131 |
| AC106869.1 | 31.8586988 | -0.335172135 | 0.51807037 | -0.6469626 | 0.51765617 | 0.86500999 |
| EPCAM | 357.082781 | 0.238689618 | 0.33990676 | 0.70222085 | 0.48254144 | 0.85133721 |
| MSH2 | 42.8260025 | 0.499864638 | 0.28204175 | 1.77230723 | 0.07634357 | 0.43515065 |
| MSH6 | 23.5911814 | -0.21347133 | 0.43123539 | -0.4950228 | 0.62058406 | 0.90380166 |
| FBXO11 | 250.138029 | 0.016831736 | 0.13909102 | 0.12101238 | 0.90368123 | 0.98028494 |
| FOXN2 | 51.5625225 | -0.337192135 | 0.32798861 | -1.0280605 | 0.30392137 | 0.74820946 |
| PPP1R21 | 101.363221 | 0.497202955 | 0.31351634 | 1.5858917 | 0.11276388 | 0.51571293 |
| GPR75-ASB3 | 44.7084243 | -0.709769359 | 0.44749502 | -1.5860944 | 0.11271789 | 0.51571293 |
| ERLEC1 | 44.5896067 | 0.072199826 | 0.22888355 | 0.31544349 | 0.75242495 | 0.94728927 |
| PSME4 | 212.790032 | 0.050801795 | 0.21261266 | 0.23894059 | 0.81115165 | 0.96247289 |
| ACYP2 | 206.677661 | -0.00018636 | 0.17853201 | -0.0010438 | 0.99916713 | 0.99982181 |
| SPTBN1 | 583.022791 | -0.465586437 | 0.30158359 | -1.5438056 | 0.12263543 | 0.53899578 |
| AC093110.1 | 24.2591934 | -0.48589788 | 0.57104916 | -0.8508863 | 0.39483254 | 0.8052114 |
| EML6 | 39.6791609 | 0.811606139 | 0.66641027 | 1.2178776 | 0.22327049 | 0.67606266 |
| RTN4 | 1013.86352 | 0.721865368 | 0.28043034 | 2.5741343 | 0.01004913 | 0.1476917 |
| CLHC1 | 36.7161566 | 0.40818005 | 0.33442356 | 1.22054812 | 0.22225716 | 0.67557645 |
| RPS27A | 331.313438 | 0.465552854 | 0.41425992 | 1.12381825 | 0.26109014 | 0.7121975 |
| MTIF2 | 38.7332112 | 0.05668614 | 0.29393623 | 0.19285183 | 0.84707501 | 0.97027687 |
| AC012358.3 | 30.0442409 | 0.446515247 | 0.37237107 | 1.19911369 | 0.23048374 | 0.68523674 |
| CCDC88A | 38.6839442 | 0.096342088 | 0.34312233 | 0.28078058 | 0.7788787 | 0.95537097 |
| CFAP36 | 35.0814148 | -0.37673346 | 0.29061994 | -1.2963097 | 0.19486879 | 0.6459894 |
| PPP4R3B | 146.162908 | -0.114337646 | 0.2110939 | -0.5416435 | 0.58806409 | 0.89073775 |
| PNPT1 | 66.2868284 | -0.295978428 | 0.6191355 | -0.4780511 | 0.63261381 | 0.90805737 |
| EFEMP1 | 103.928625 | 0.388887661 | 0.61982817 | 0.62741205 | 0.53038917 | 0.86784453 |
| VRK2 | 119.40192 | 0.067283138 | 0.25487455 | 0.26398532 | 0.79179123 | 0.95746995 |
| FANCL | 49.9125522 | 0.265171881 | 0.31406984 | 0.84430865 | 0.39849695 | 0.80714564 |
| PAPOLG | 31.8936341 | -0.196357201 | 0.27599459 | -0.7114531 | 0.47680351 | 0.84706851 |
| REL | 63.308748 | -0.06236078 | 0.3871492 | -0.1610769 | 0.87203286 | 0.97434023 |
| PEX13 | 37.29939 | 0.020529377 | 0.31938732 | 0.06427737 | 0.94874937 | 0.99179534 |
| KIAA1841 | 19.7088333 | 0.084705911 | 0.40348652 | 0.20993492 | 0.83371846 | 0.96701848 |
| C2orf74 | 14.0508103 | 0.376102846 | 0.56629316 | 0.66414867 | 0.50659516 | 0.86235197 |
| USP34 | 687.341247 | -0.261493898 | 0.20539615 | -1.2731197 | 0.20297556 | 0.65623093 |
| AC016727.1 | 15.9384518 | 0.116407898 | 0.32813929 | 0.35475148 | 0.72277578 | 0.93873366 |
| XPO1 | 211.364048 | -0.140861136 | 0.26062984 | -0.5404643 | 0.58887685 | 0.89126313 |
| FAM161A | 13.414897 | 0.249177096 | 0.53964952 | 0.46173875 | 0.64426868 | 0.9119294 |
| CCT4 | 73.9164908 | 0.040486001 | 0.23987616 | 0.16877876 | 0.86597066 | 0.97434023 |
| COMMD1 | 218.265504 | 0.280224229 | 0.21144256 | 1.32529718 | 0.18507261 | 0.63569556 |
| AC018462.1 | 14.6181133 | -0.788808694 | 0.57508208 | -1.3716454 | 0.17017385 | 0.61252734 |
| B3GNT2 | 128.823164 | -0.327627007 | 0.29189402 | -1.1224177 | 0.2616849 | 0.71281761 |
| EHBP1 | 124.374907 | 0.482962272 | 0.22386528 | 2.15737905 | 0.03097614 | 0.27936455 |
| WDPCP | 177.229487 | -0.040764287 | 0.21480955 | -0.1897694 | 0.84948981 | 0.97027687 |
| MDH1 | 103.832701 | -0.178593016 | 0.32608121 | -0.5476949 | 0.58390141 | 0.89036744 |
| UGP2 | 131.221274 | -0.371321266 | 0.2086703 | -1.7794639 | 0.07516373 | 0.43243667 |
| VPS54 | 171.181873 | -0.302888723 | 0.25404595 | -1.1922596 | 0.23315946 | 0.68763965 |
| PELI1 | 41.1208232 | 0.001425911 | 0.58589618 | 0.00243373 | 0.99805817 | 0.99982181 |
| AFTPH | 82.472937 | 0.094288497 | 0.21165709 | 0.44547762 | 0.65597463 | 0.9158679 |
| SERTAD2 | 22.7733347 | 0.065805728 | 0.40995519 | 0.16051932 | 0.872472 | 0.97434023 |
| CEP68 | 43.2099334 | -0.855226911 | 0.29258353 | -2.923018 | 0.00346656 | 0.0754849 |
| RAB1A | 252.10034 | 0.037896745 | 0.17148159 | 0.220996 | 0.82509555 | 0.96539788 |
| ACTR2 | 193.977229 | -0.215944234 | 0.19528298 | -1.1058016 | 0.26881241 | 0.72009952 |
| SPRED2 | 252.130353 | 0.400723043 | 0.25537523 | 1.56915389 | 0.11661209 | 0.52367464 |
| ETAA1 | 18.4207869 | -0.186681365 | 0.35354213 | -0.5280315 | 0.5974775 | 0.89363331 |
| C1D | 40.3684157 | 0.141743493 | 0.29723604 | 0.47687182 | 0.6334534 | 0.90805737 |
| PNO1 | 14.2141236 | -0.078104019 | 0.43062446 | -0.1813739 | 0.85607414 | 0.97166553 |
| APLF | 42.8773614 | -0.098082452 | 0.30801323 | -0.3184358 | 0.75015436 | 0.94684415 |
| GFPT1 | 116.458961 | -0.174066837 | 0.2213968 | -0.7862211 | 0.43173796 | 0.8248269 |
| NFU1 | 49.6539599 | 0.628416363 | 0.24055871 | 2.61232009 | 0.008993 | 0.13780744 |
| AAK1 | 354.026112 | -0.050254199 | 0.18539593 | -0.2710642 | 0.78634166 | 0.95577371 |
| ANXA4 | 520.6698 | 0.444029831 | 0.25860767 | 1.71700176 | 0.08597885 | 0.46168448 |
| GMCL1 | 64.3362781 | 0.290130451 | 0.23235085 | 1.24867396 | 0.21178435 | 0.66386489 |
| MXD1 | 31.8431758 | -0.686622673 | 0.30582884 | -2.2451208 | 0.02476038 | 0.24760381 |
| PCBP1 | 83.5234228 | -0.511944682 | 0.32806015 | -1.5605208 | 0.11863686 | 0.52958009 |
| C2orf42 | 20.1633236 | 0.402095628 | 0.37326017 | 1.07725299 | 0.28136726 | 0.73311066 |
| PCYOX1 | 66.9811046 | 0.754607187 | 0.41671496 | 1.81084735 | 0.07016448 | 0.41948432 |
| SNRPG | 38.3724133 | -0.155949648 | 0.38190285 | -0.408349 | 0.68301747 | 0.92474072 |
| FAM136A | 19.840325 | 0.206015875 | 0.32858265 | 0.62698342 | 0.5306701 | 0.86786739 |
| TGFA | 27.8910428 | 0.495439038 | 0.50016623 | 0.99054875 | 0.32190597 | 0.76248108 |
| ATP6V1B1 | 39.6545437 | 1.998077485 | 0.74391998 | 2.68587689 | 0.00723397 | 0.12224659 |
| AC007040.2 | 119.997313 | 0.305825007 | 1.01155107 | 0.30233274 | 0.76239842 | 0.95030387 |
| NAGK | 24.3086559 | 0.163529122 | 0.48421053 | 0.33772318 | 0.7355718 | 0.94318131 |
| MCEE | 38.8211769 | 0.313157191 | 0.30039852 | 1.04247247 | 0.29719268 | 0.74160729 |
| MPHOSPH10 | 27.7886923 | -0.019257676 | 0.32507157 | -0.0592413 | 0.95275988 | 0.99217529 |
| ZNF638 | 446.046585 | -0.131189989 | 0.17866608 | -0.7342747 | 0.46278132 | 0.83921832 |
| EXOC6B | 399.625312 | -0.05597306 | 0.27517866 | -0.2034063 | 0.83881751 | 0.96748214 |
| SPR | 16.1413345 | -0.270073038 | 0.58043455 | -0.4652946 | 0.64172053 | 0.91146968 |
| EMX1 | 18.1276969 | 1.478709388 | 0.62770815 | 2.35572757 | 0.01848647 | 0.21121173 |
| SFXN5 | 46.0761288 | 0.537154934 | 0.47365648 | 1.13406014 | 0.25676932 | 0.70972499 |
| RAB11FIP5 | 17.1793199 | -0.208342818 | 0.41302121 | -0.5044361 | 0.61395494 | 0.90041774 |
| SMYD5 | 12.4182901 | 0.09210311 | 0.42519354 | 0.21661456 | 0.82850874 | NA |
| PRADC1 | 13.2877947 | 0.221646676 | 0.59244308 | 0.37412316 | 0.70831269 | 0.93439862 |
| CCT7 | 69.578205 | 0.047482062 | 0.29896232 | 0.1588229 | 0.87380841 | 0.97434023 |
| ALMS1 | 105.287499 | 0.050645494 | 0.24995678 | 0.202617 | 0.8394344 | 0.96748214 |
| DUSP11 | 17.5913743 | -0.596079134 | 0.36991372 | -1.6114004 | 0.10709247 | 0.50318505 |
| STAMBP | 42.6869102 | 0.044400709 | 0.25352306 | 0.17513479 | 0.86097372 | 0.97302617 |
| DGUOK | 45.8395992 | -0.461534308 | 0.29840242 | -1.5466842 | 0.12193942 | 0.53747904 |
| TET3 | 51.5610588 | 0.41934838 | 0.36743795 | 1.14127673 | 0.25375478 | 0.70701839 |
| BOLA3 | 20.3333795 | 0.375901288 | 0.57082321 | 0.65852488 | 0.51020092 | 0.86281145 |
| MOB1A | 115.46267 | 0.201457313 | 0.25493334 | 0.79023526 | 0.42939039 | 0.82469033 |

| | | | | | | |
|---|---|---|---|---|---|---|
| AC006030.1 | 31.297854 | -0.117002668 | 0.30556542 | -0.3829055 | 0.70178986 | 0.93149448 |
| DCTN1 | 18.8944269 | 0.146715847 | 0.42152362 | 0.3480608 | 0.72779453 | 0.94030592 |
| RTKN | 32.5664094 | -0.182229276 | 0.28405573 | -0.6415265 | 0.52118067 | 0.8669958 |
| MOGS | 26.8854269 | 0.346299504 | 0.39466146 | 0.87745964 | 0.38023703 | 0.79609308 |
| AUP1 | 26.6398056 | 0.095286529 | 0.44533585 | 0.21396555 | 0.83057392 | 0.96694276 |
| M1AP | 26.0046744 | -0.248526696 | 0.65966623 | -0.3767461 | 0.70636228 | 0.93411235 |
| POLE4 | 29.5756893 | -0.302692156 | 0.32375051 | -0.934955 | 0.34981149 | 0.77868979 |
| TACR1 | 16.8273232 | 0.555108509 | 0.53120952 | 1.04498975 | 0.29602771 | 0.74123516 |
| AC007681.1 | 13.9107694 | 0.696874665 | 0.55754869 | 1.24989024 | 0.21133964 | 0.66321402 |
| MRPL19 | 68.7974648 | 0.296557649 | 0.31989628 | 0.92704314 | 0.35390414 | 0.78037511 |
| GCFC2 | 87.6233385 | 0.02128986 | 0.31653428 | 0.06725926 | 0.94637531 | 0.99134285 |
| SUCLG1 | 119.692204 | 0.289878687 | 0.21421966 | 1.35318435 | 0.17599674 | 0.6225246 |
| DNAH6 | 45.5528212 | 1.804130439 | 0.68699532 | 2.62611749 | 0.0086365 | 0.13511944 |
| TMSB10 | 223.133561 | -1.214840943 | 0.42940728 | -2.8291112 | 0.00466775 | 0.09335495 |
| KCMF1 | 119.91063 | 0.063056813 | 0.23734944 | 0.26567078 | 0.79049277 | 0.95746995 |
| TCF7L1 | 159.530529 | 0.288663579 | 0.31629036 | 0.91265373 | 0.36142469 | 0.78668094 |
| TGOLN2 | 181.109194 | 0.056837435 | 0.29939799 | 0.18983907 | 0.84943524 | 0.97027687 |
| RETSAT | 30.1895995 | 0.246519862 | 0.34883271 | 0.70669938 | 0.47975331 | 0.84857898 |
| ELMOD3 | 47.9896043 | 0.562017327 | 0.27092424 | 2.07444457 | 0.03803804 | 0.31295081 |
| CAPG | 55.7774191 | 0.232171779 | 0.41725976 | 0.55642024 | 0.5779236 | 0.88877358 |
| MAT2A | 51.0953156 | 0.159917116 | 0.29232598 | 0.54705064 | 0.58434394 | 0.89036744 |
| GGCX | 21.8702965 | 0.430017089 | 0.33255035 | 1.29308867 | 0.19598039 | 0.64659037 |
| VAMP8 | 126.007269 | -0.452394488 | 0.38253467 | -1.1826235 | 0.2369584 | 0.69190142 |
| VAMP5 | 10.9938864 | -0.171600461 | 0.44802304 | -0.383017 | 0.70170712 | NA |
| RNF181 | 17.1885435 | 0.119448714 | 0.39774678 | 0.30031347 | 0.76393806 | 0.9513483 |
| USP39 | 50.8532771 | 0.002896265 | 0.2420896 | 0.01196361 | 0.99045465 | 0.99923337 |
| C2orf68 | 20.1328335 | -0.085542593 | 0.31194906 | -0.2742197 | 0.78391575 | 0.95544332 |
| ST3GAL5 | 11.3571549 | -0.07024361 | 0.56552186 | -0.1242102 | 0.90114881 | NA |
| POLR1A | 41.4301001 | 0.387199508 | 0.27461422 | 1.40997618 | 0.15854672 | 0.59881912 |
| PTCD3 | 85.9167946 | 0.236316546 | 0.22076933 | 1.07042289 | 0.284429 | 0.73638801 |
| IMMT | 167.99172 | 0.086490102 | 0.19912873 | 0.43434266 | 0.66403963 | 0.91876908 |
| MRPL35 | 35.6826465 | 0.572549965 | 0.34721524 | 1.64897705 | 0.09915234 | 0.48550109 |
| REEP1 | 10.9544044 | 1.078030672 | 0.63759065 | 1.69078808 | 0.09087729 | NA |
| KDM3A | 114.75045 | 0.016182396 | 0.31974718 | 0.05060997 | 0.95963632 | 0.99238848 |
| CHMP3 | 45.7702292 | -0.435776366 | 0.29591706 | -1.4726301 | 0.14085081 | 0.57506583 |
| RNF103-CHM | 19.2477838 | -3.098062818 | 1.33583171 | -2.3192014 | 0.02038412 | 0.22348867 |
| RMND5A | 65.269176 | 0.003260466 | 0.2963661 | 0.01100148 | 0.99122227 | 0.99923337 |
| CYTOR | 13.6909659 | -0.392697186 | 0.59684778 | -0.657952 | 0.51056898 | 0.86310894 |
| KRCC1 | 63.229798 | 0.172280037 | 0.22669183 | 0.75997462 | 0.44726975 | 0.83066268 |
| THNSL2 | 19.4720374 | -0.073897086 | 0.65229361 | -0.1132881 | 0.90980217 | 0.98329594 |
| EIF2AK3 | 101.461049 | -1.170134657 | 0.33379844 | -3.5055127 | 0.00045573 | 0.0190052 |
| RPIA | 26.2623233 | 0.05588035 | 0.38288863 | 0.14594414 | 0.88396549 | 0.97662748 |
| MAL | 370.049254 | 0.638607559 | 0.35053952 | 1.82178475 | 0.06848765 | 0.41572605 |
| MRPS5 | 153.56079 | -0.020665633 | 0.20692355 | -0.0998709 | 0.92044685 | 0.98612044 |
| ZNF514 | 10.1907726 | -0.763241192 | 0.41720941 | -1.8293959 | 0.06734032 | NA |
| AC092835.1 | 19.4364772 | 0.230363598 | 0.45930009 | 0.50155357 | 0.61598159 | 0.90148181 |
| PROM2 | 20.7748113 | 1.34819031 | 0.5069718 | 2.65930039 | 0.00783031 | 0.12855183 |
| KCNIP3 | 24.2131724 | 0.254906754 | 0.52588326 | 0.48472118 | 0.62787414 | 0.90603677 |
| FAHD2A | 19.3877345 | 0.381316031 | 0.36738817 | 1.03791049 | 0.29931173 | 0.74336229 |
| ANKRD36C | 30.9258677 | -0.29290074 | 0.3318624 | -0.882597 | 0.37745408 | 0.79432848 |
| STARD7 | 107.544926 | 0.594148961 | 0.17793013 | 3.33922628 | 0.00084012 | 0.02920851 |
| TMEM127 | 51.9279137 | 0.032758112 | 0.3554648 | 0.09215572 | 0.92657432 | 0.98672316 |
| CIAO1 | 40.9616241 | -0.350186706 | 0.25118022 | -1.3941651 | 0.16326774 | 0.60228925 |
| SNRNP200 | 65.214764 | -0.185678057 | 0.27002043 | -0.6876445 | 0.49167669 | 0.85654883 |
| KANSL3 | 102.014666 | 0.362041262 | 0.17883521 | 2.02444062 | 0.04292484 | 0.33179556 |
| LMAN2L | 24.8495056 | 0.61142512 | 0.30341488 | 2.01514545 | 0.04388941 | 0.33287289 |
| CNNM4 | 31.3657037 | 0.867815896 | 0.52076586 | 1.66642241 | 0.09562931 | 0.48081904 |
| ANKRD23 | 14.1131075 | -0.011459266 | 0.39584363 | -0.028949 | 0.97690529 | 0.9957152 |
| ANKRD36 | 35.4635883 | -0.055026944 | 0.3498516 | -0.1572865 | 0.87501904 | 0.97469966 |
| AC092683.1 | 14.2980811 | -0.209820975 | 0.39059504 | -0.5371829 | 0.59114129 | 0.89149106 |
| ANKRD36B | 15.6875725 | -0.98605938 | 0.51084806 | -1.93024 | 0.0535771 | 0.36684184 |
| COX5B | 149.307523 | 0.095338378 | 0.45284572 | 0.2105317 | 0.83325272 | 0.96701848 |
| ACTR1B | 22.3900845 | -0.074826707 | 0.31195937 | -0.2398604 | 0.81043846 | 0.9623221 |
| TMEM131 | 186.494379 | 0.118675196 | 0.17062166 | 0.69554588 | 0.48671327 | 0.85365949 |
| INPP4A | 386.803831 | 0.1649711 | 0.28429861 | 0.58027404 | 0.56172983 | 0.88153686 |
| COA5 | 23.9477843 | 0.414534289 | 0.4206524 | 0.98545565 | 0.32440031 | 0.76304922 |
| UNC50 | 44.5019645 | -0.000810375 | 0.30234519 | -0.0026803 | 0.99786143 | 0.99982181 |
| MGAT4A | 63.3717854 | 0.009707132 | 0.28757488 | 0.03375514 | 0.9730724 | 0.99526459 |
| KIAA1211L | 19.2728912 | -0.70985157 | 0.62651838 | -1.13301 | 0.25721006 | 0.71022118 |
| TSGA10 | 102.797837 | 0.176734738 | 0.27327105 | 0.64673787 | 0.51780161 | 0.86500999 |
| AC079447.1 | 33.5127712 | 0.711791304 | 0.38839396 | 1.83265285 | 0.06685421 | 0.41121488 |
| MITD1 | 35.0371669 | -0.263958576 | 0.23936656 | -1.1027379 | 0.270141 | 0.72068562 |
| TXNDC9 | 29.0816411 | -0.343667666 | 0.34384884 | -0.9994731 | 0.31756557 | 0.7596513 |
| EIF5B | 145.694034 | -0.229244561 | 0.2567039 | -0.8930311 | 0.37184053 | 0.7900012 |
| REV1 | 245.731699 | 0.065367553 | 0.16246863 | 0.40233954 | 0.68743416 | 0.92643753 |
| LONRF2 | 160.164085 | 0.155929058 | 0.41612545 | 0.37471647 | 0.70787134 | 0.93432317 |
| PDCL3 | 18.4789129 | -0.636368162 | 0.44619572 | -1.4262086 | 0.15380815 | 0.59345228 |
| RPL31 | 89.9858127 | 0.133784095 | 0.27392736 | 0.4883926 | 0.62527178 | 0.90447745 |
| TBC1D8 | 66.8048465 | -0.399452904 | 0.33774385 | -1.1827096 | 0.23692425 | 0.69190142 |
| CNOT11 | 38.7511928 | -0.056464704 | 0.25503782 | -0.2213974 | 0.82478304 | 0.96539788 |
| RNF149 | 179.367072 | -0.293105203 | 0.27142101 | -1.0798914 | 0.28019056 | 0.73179753 |
| MAP4K4 | 155.744728 | -0.705782453 | 0.3348562 | -2.107718 | 0.03505539 | 0.29932858 |
| IL1R1 | 113.613473 | -0.842898619 | 0.36451231 | -2.312401 | 0.0207556 | 0.22444738 |
| IL1RL2 | 37.7311097 | -0.146560684 | 0.40943844 | -0.3579554 | 0.72037672 | 0.93806337 |
| SLC9A4 | 19.0539636 | 1.412437099 | 0.57571192 | 2.45337476 | 0.01415228 | 0.17828198 |
| SLC9A2 | 62.017599 | 0.318227397 | 0.47373417 | 0.67174254 | 0.50174761 | 0.85942996 |
| MFSD9 | 13.8375037 | -0.184154644 | 0.367448 | -0.501172 | 0.61625008 | 0.90160403 |
| TMEM182 | 21.2910021 | -0.209579411 | 0.33382711 | -0.6278082 | 0.53012956 | 0.86764582 |
| POU3F3 | 54.7897237 | 0.354254793 | 0.31630438 | 1.11998069 | 0.26272199 | 0.71348562 |
| MRPS9 | 53.0238032 | 0.191888647 | 0.27858675 | 0.68879317 | 0.49095344 | 0.85654883 |
| TGFBRAP1 | 26.8820081 | -0.046325007 | 0.35241575 | -0.1314499 | 0.89541944 | 0.97885533 |
| C2orf49 | 23.9303294 | 0.080393413 | 0.27462384 | 0.2927401 | 0.76972081 | 0.95218859 |
| NCK2 | 58.057098 | -0.731876996 | 0.29658509 | -2.4676797 | 0.0135992 | 0.17531992 |
| UXS1 | 47.3715915 | 0.31705541 | 0.27879079 | 1.13725208 | 0.25543292 | 0.70849619 |
| LINC01594 | 14.1986754 | -0.830975727 | 0.63431826 | -1.3100297 | 0.19018581 | 0.6421543 |
| SULT1C2 | 262.146355 | 0.24454301 | 0.32735832 | 0.74701939 | 0.45505185 | 0.83423755 |
| GCC2 | 240.724995 | -0.521682463 | 0.21909114 | -2.3811208 | 0.01726005 | 0.20451748 |
| GCC2-AS1 | 44.9993136 | -0.638819984 | 0.3357844 | -1.9024707 | 0.05710965 | 0.37973269 |
| LIMS1 | 1241.07222 | -1.536993863 | 0.27468211 | -5.5955369 | 2.20E-08 | 8.46E-06 |
| RANBP2 | 208.842814 | 0.05439513 | 0.1601155 | 0.33972432 | 0.73406414 | 0.94277981 |
| CCDC138 | 35.8481036 | 0.439934285 | 0.27739905 | 1.58592573 | 0.11275616 | 0.51571293 |
| 10-Sep | 193.781005 | -1.195350158 | 0.30008516 | -3.9833697 | 6.79E-05 | 0.00510904 |
| SOWAHC | 37.4375054 | -1.457233899 | 0.35611484 | -4.0920336 | 4.28E-05 | 0.00368188 |
| AC013271.1 | 99.7327844 | -0.196798181 | 0.60622901 | -0.3246268 | 0.74546354 | 0.94614106 |
| NPHP1 | 22.2469795 | 0.101821608 | 0.42307979 | 0.24066762 | 0.80981273 | 0.96215121 |
| SMIM37 | 87.1822752 | -1.202083127 | 0.52035988 | -2.3100995 | 0.02088264 | 0.22556976 |
| LINC01106 | 11.8683803 | 0.282928783 | 0.47714119 | 0.59296659 | 0.55320351 | NA |
| AC112229.3 | 203.998796 | -1.416371389 | 0.52177444 | -2.7145281 | 0.00663703 | 0.11674591 |
| MIR4435-2H | 83.4850519 | -0.278137863 | 0.28758254 | -0.9671584 | 0.33346487 | 0.76777176 |
| BCL2L11 | 35.6645343 | -0.550390712 | 0.32652471 | -1.685602 | 0.09187248 | 0.4764741 |
| ANAPC1 | 69.2295536 | 0.22826461 | 0.19629752 | 1.16285022 | 0.24489028 | 0.69845213 |
| MERTK | 62.9244854 | 0.163724433 | 0.33341046 | 0.49105969 | 0.62338423 | 0.90447745 |
| TMEM87B | 85.8172898 | -0.471927007 | 0.25198938 | -1.8728051 | 0.0610953 | 0.39246665 |

| | | | | | | |
|---|---|---|---|---|---|---|
| ZC3H8 | 38.8302159 | 0.537714124 | 0.24682397 | 2.17853283 | 0.02936639 | 0.27178917 |
| ZC3H6 | 79.8526608 | -0.410217712 | 0.31756768 | -1.2917489 | 0.19644411 | 0.64685325 |
| RGPD8 | 11.0009153 | -0.305200891 | 0.48275794 | -0.6322027 | 0.52725441 | NA |
| TTL | 45.7874172 | 0.05723057 | 0.30473717 | 0.18780305 | 0.85103105 | 0.97027687 |
| POLR1B | 11.1787049 | 0.442272361 | 0.4491705 | 0.98464248 | 0.32479971 | NA |
| CHCHD5 | 21.7908913 | -0.468740737 | 0.41050583 | -1.1418613 | 0.25351166 | 0.70701839 |
| SLC20A1 | 90.4865593 | -0.759759947 | 0.42310463 | -1.7956786 | 0.07254564 | 0.42448303 |
| PSD4 | 41.4823585 | -0.28429746 | 0.32218433 | -0.8824062 | 0.37755717 | 0.79432848 |
| PAX8 | 1677.22527 | -0.628492356 | 0.32417956 | -1.9387168 | 0.05253583 | 0.36358594 |
| CBWD2 | 31.7909251 | 0.185315363 | 0.35074547 | 0.52834714 | 0.59725842 | 0.89363331 |
| SLC35F5 | 252.216112 | -0.276633125 | 0.21440425 | -1.2902409 | 0.19696704 | 0.64743488 |
| ACTR3 | 233.079915 | -0.267877065 | 0.23107333 | -1.159273 | 0.24634494 | 0.69972061 |
| DPP10 | 51.5067365 | 0.623313439 | 0.5467529 | 1.14002769 | 0.25427477 | 0.70733158 |
| DDX18 | 53.7102272 | -0.254931898 | 0.24180163 | -1.0543018 | 0.29174476 | 0.73994767 |
| CCDC93 | 114.703937 | -0.31685457 | 0.25341517 | -1.2503378 | 0.21117616 | 0.66313007 |
| INSIG2 | 90.5059254 | -0.108017159 | 0.29322054 | -0.368382 | 0.71258844 | 0.9357596 |
| C2orf76 | 19.3295433 | -0.193643948 | 0.3960644 | -0.4889204 | 0.62489809 | 0.90447745 |
| DBI | 191.83145 | -1.403140559 | 0.39130129 | -3.5858316 | 0.00033601 | 0.0156695 |
| TMEM37 | 19.3108456 | 0.678132885 | 0.51793208 | 1.30930852 | 0.19042987 | 0.6421543 |
| SCTR | 66.4079207 | 1.480755925 | 0.8555506 | 1.73076371 | 0.08349392 | 0.4560197 |
| CFAP221 | 91.6702039 | 0.543190357 | 0.61760682 | 0.87950835 | 0.37912571 | 0.79535699 |
| PTPN4 | 168.142293 | 0.340783142 | 0.24672654 | 1.38121801 | 0.16721194 | 0.60884227 |
| EPB41L5 | 336.11128 | 0.142195122 | 0.17689897 | 0.80382108 | 0.42150031 | 0.82017112 |
| RALB | 31.7190994 | 0.038532522 | 0.27656467 | 0.13932554 | 0.88919291 | 0.97778408 |
| TFCP2L1 | 1106.23603 | 0.182507949 | 0.37671647 | 0.48447032 | 0.62805212 | 0.906154 |
| CLASP1 | 298.879291 | 0.008604385 | 0.25723108 | 0.03345002 | 0.97331572 | 0.99526459 |
| NIFK-AS1 | 22.2245636 | -0.835710864 | 0.42050198 | -1.9874125 | 0.04687671 | 0.34525746 |
| NIFK | 16.3627258 | 0.252695783 | 0.32788649 | 0.77068067 | 0.44089623 | 0.82801422 |
| TSN | 24.5101754 | 0.298238075 | 0.34483973 | 0.86485995 | 0.38711566 | 0.80009555 |
| CNTNAP5 | 283.033813 | -1.099479622 | 0.95315996 | -1.1535101 | 0.24870108 | 0.70352887 |
| BIN1 | 18.0032348 | -0.540287835 | 0.50701609 | -1.0656227 | 0.28659425 | 0.7371955 |
| ERCC3 | 35.0656862 | -0.108964352 | 0.356737 | -0.3054473 | 0.76002551 | 0.94952011 |
| MAP3K2 | 188.459629 | 0.046051083 | 0.19389377 | 0.23750676 | 0.81226367 | 0.96262567 |
| IWS1 | 118.907588 | -0.172794547 | 0.17984219 | -0.9608121 | 0.33664668 | 0.76907036 |
| AC010976.1 | 12.083965 | -0.472725111 | 0.63967169 | -0.7390121 | 0.45989966 | NA |
| WDR33 | 170.668419 | 0.254640134 | 0.18629733 | 1.36684799 | 0.17167296 | 0.61538348 |
| POLR2D | 12.2766819 | 0.329867929 | 0.42603816 | 0.77426851 | 0.43877202 | NA |
| AMMECR1L | 22.0646257 | 0.532280999 | 0.30042701 | 1.77174815 | 0.07643637 | 0.43515065 |
| SAP130 | 42.2956771 | 0.296446228 | 0.24952315 | 1.18805099 | 0.23481332 | 0.68936745 |
| UGGT1 | 65.6589591 | 0.144371106 | 0.21919724 | 0.65863559 | 0.5101298 | 0.86281145 |
| HS6ST1 | 104.917502 | -0.2970037 | 0.34427997 | -0.8626807 | 0.38831302 | 0.80109289 |
| SMPD4 | 35.4713797 | 0.247100398 | 0.32235419 | 0.76654937 | 0.44334949 | 0.82924991 |
| MZT2B | 49.9011223 | -0.042735997 | 0.27964729 | -0.1528211 | 0.87853939 | 0.97548444 |
| CCDC115 | 17.1216425 | -0.010169167 | 0.39162592 | -0.0259665 | 0.97928403 | 0.99623022 |
| IMP4 | 18.1358263 | -0.121893892 | 0.41518384 | -0.2935902 | 0.76907109 | 0.95205353 |
| PTPN18 | 44.1196307 | 0.159697776 | 0.29966849 | 0.53291481 | 0.59409255 | 0.89288778 |
| FAM168B | 131.911395 | 0.093907565 | 0.23954441 | 0.39202571 | 0.69503922 | 0.92863367 |
| PLEKHB2 | 272.915857 | -0.196810118 | 0.32809155 | -0.5998634 | 0.54859727 | 0.87371493 |
| MZT2A | 24.3644612 | 0.495110615 | 0.35801329 | 1.3829392 | 0.16668351 | 0.60783084 |
| GPR39 | 23.7256362 | 0.811531111 | 0.52053876 | 1.55902148 | 0.1189913 | 0.53056597 |
| NCKAP5 | 164.335649 | -1.083700998 | 0.72938013 | -1.4857835 | 0.13733638 | 0.56857145 |
| MGAT5 | 152.568939 | -1.16938767 | 0.39776567 | -2.9398909 | 0.00328328 | 0.07254624 |
| TMEM163 | 86.7810103 | -1.828286587 | 0.55722692 | -3.281045 | 0.00103423 | 0.03332909 |
| CCNT2-AS1 | 18.5696794 | -0.619436659 | 0.54530742 | -1.1359403 | 0.25598156 | 0.70923197 |
| CCNT2 | 201.278907 | -0.217162879 | 0.33056352 | -0.6569475 | 0.51121467 | 0.86312853 |
| RAB3GAP1 | 226.171484 | -0.170407519 | 0.16853042 | -1.011138 | 0.31195037 | 0.75484355 |
| ZRANB3 | 89.9084192 | 0.103620824 | 0.22433462 | 0.46190296 | 0.64415091 | 0.9119294 |
| R3HDM1 | 138.893497 | 0.147469733 | 0.21641534 | 0.68141996 | 0.4956058 | 0.85814183 |
| UBXN4 | 186.32567 | -0.513423845 | 0.20654402 | -2.4857842 | 0.01292663 | 0.16944371 |
| MCM6 | 11.1237697 | -0.345685128 | 0.46975124 | -0.7358898 | 0.46179779 | NA |
| DARS | 173.668406 | -0.61364437 | 0.31560663 | -1.9443329 | 0.05185531 | 0.36134809 |
| THSD7B | 164.748557 | 1.248564978 | 0.67859345 | 1.83993079 | 0.0657784 | 0.40743964 |
| HNMT | 53.7417534 | -0.070213669 | 0.30962805 | -0.2267678 | 0.82060431 | 0.96389705 |
| AC114763.1 | 45.3004749 | -0.235818187 | 0.46827596 | -0.5035881 | 0.61455088 | 0.90054121 |
| SPOPL | 170.630092 | 0.265336726 | 0.34416619 | 0.77095523 | 0.44073347 | 0.82793978 |
| NXPH2 | 417.983523 | 0.202790798 | 0.55359999 | 0.36631287 | 0.71413163 | 0.93655477 |
| LRP1B | 26.4502921 | -1.617246874 | 0.6414044 | -2.5214153 | 0.01168838 | 0.16081885 |
| KYNU | 46.9593979 | 1.424025571 | 0.45908505 | 3.10187749 | 0.00192298 | 0.05027725 |
| GTDC1 | 69.953453 | 0.109820024 | 0.25597348 | 0.42902891 | 0.66790219 | 0.92049996 |
| ACVR2A | 62.4075362 | 0.169289341 | 0.25564358 | 0.66220845 | 0.50783764 | 0.86281145 |
| ORC4 | 136.88463 | 0.125431168 | 0.2470578 | 0.50769968 | 0.61166397 | 0.90011356 |
| MBD5 | 668.083498 | 0.212809219 | 0.21115725 | 1.00782339 | 0.31353925 | 0.75632348 |
| EPC2 | 186.576828 | 0.393840359 | 0.21581455 | 1.82490181 | 0.06801585 | 0.4151827 |
| AC105402.3 | 18.5305982 | 1.161673815 | 1.11100585 | 1.04560549 | 0.29574321 | 0.74123516 |
| LYPD6B | 30.0530844 | 2.37429644 | 0.59890613 | 3.9643883 | 7.36E-05 | 0.00526714 |
| MMADHC | 34.0551997 | -0.220498945 | 0.29089505 | -0.7580017 | 0.44844996 | 0.83118301 |
| NMI | 20.16201 | -1.005913514 | 0.59600199 | -1.6877687 | 0.09145563 | 0.47515782 |
| RIF1 | 113.580851 | -0.023272116 | 0.2288045 | -0.1017118 | 0.91898544 | 0.98542547 |
| ARL5A | 35.0443114 | 0.130747243 | 0.26100289 | 0.50094174 | 0.61641213 | 0.90160403 |
| STAM2 | 91.2004481 | 0.2024005 | 0.17213927 | 1.17579504 | 0.23967679 | 0.69419674 |
| FMNL2 | 499.763142 | -0.297271274 | 0.23830019 | -1.2474655 | 0.21222686 | 0.66453706 |
| PRPF40A | 113.982011 | 0.156616053 | 0.1705516 | 0.9182913 | 0.35846638 | 0.78398112 |
| ARL6IP6 | 43.9404076 | 0.079504038 | 0.3229067 | 0.24621366 | 0.80551685 | 0.96005568 |
| KCNJ3 | 309.104635 | 1.221736902 | 0.50649115 | 2.41215846 | 0.01585839 | 0.19341706 |
| LINC01876 | 110.194188 | 0.257950606 | 0.64919689 | 0.39733802 | 0.69111822 | 0.92786587 |
| NR4A2 | 25.7750381 | 0.775775886 | 0.4663843 | 1.66338336 | 0.09623572 | 0.48192386 |
| GPD2 | 234.324604 | 0.043670114 | 0.31222096 | 0.13986926 | 0.88876329 | 0.97765744 |
| CYTIP | 44.7217408 | 0.282197912 | 0.62948749 | 0.44829789 | 0.65393823 | 0.9158679 |
| ACVR1C | 17.1799065 | -0.925508163 | 0.71285925 | -1.2983042 | 0.19418282 | 0.6455015 |
| ACVR1 | 96.2119087 | -0.129335754 | 0.31616848 | -0.4090723 | 0.68248663 | 0.92474072 |
| UPP2 | 14.2458129 | 0.219340531 | 0.55785449 | 0.39318592 | 0.69418217 | 0.92841398 |
| CCDC148 | 42.5885931 | 0.292398584 | 0.45639078 | 0.64067592 | 0.52173326 | 0.86703098 |
| PKP4 | 274.485131 | -0.086604429 | 0.25987382 | -0.3332557 | 0.73894129 | 0.94376279 |
| DAPL1 | 31.6857664 | -3.406999687 | 0.85290157 | -3.9945989 | 6.48E-05 | 0.00493631 |
| TANC1 | 200.46546 | -0.304809691 | 0.35802895 | -0.8513549 | 0.39457226 | 0.80508012 |
| WDSUB1 | 90.8373955 | -0.236566181 | 0.21906703 | -1.0798803 | 0.28019547 | 0.73179753 |
| BAZ2B | 443.073797 | 0.013864178 | 0.30684871 | 0.04518246 | 0.96396188 | 0.99334616 |
| AC009961.1 | 16.2840558 | 0.129998973 | 0.40476794 | 0.32116914 | 0.74808222 | 0.94656788 |
| 7-Mar | 189.539779 | -0.012248101 | 0.20120944 | -0.0608724 | 0.95146083 | 0.99217308 |
| PLA2R1 | 12.3720803 | -0.653032153 | 0.60749566 | -1.0749577 | 0.28239366 | NA |
| ITGB6 | 82.4212593 | -0.328013704 | 0.43793067 | -0.7490083 | 0.4538522 | 0.83364149 |
| RBMS1 | 164.97372 | 0.117792375 | 0.32902731 | 0.35800182 | 0.72034195 | 0.93806337 |
| TANK | 129.769729 | -0.386879024 | 0.23455481 | -1.6494184 | 0.09906194 | 0.48530345 |
| PSMD14 | 77.737245 | -0.153000274 | 0.22999811 | -0.6652241 | 0.5059072 | 0.86213981 |
| IFIH1 | 30.6343987 | -0.956847812 | 0.56108147 | -1.7053634 | 0.08812667 | 0.46763055 |
| GCA | 37.9245247 | -0.316473521 | 0.29944827 | -1.0568554 | 0.29057758 | 0.73979353 |
| FIGN | 34.5641714 | -0.155275103 | 0.61700641 | -0.2516588 | 0.80130479 | 0.95877735 |
| GRB14 | 670.937814 | 0.889451512 | 0.29638327 | 3.00101794 | 0.00269079 | 0.06304501 |
| COBLL1 | 1965.30172 | -0.032453251 | 0.23828817 | -0.1361933 | 0.89166847 | 0.97793768 |
| AC019197.1 | 2807.52072 | 0.455829169 | 0.36693058 | 1.24227631 | 0.21413463 | 0.66787972 |
| SLC38A11 | 96.163377 | -0.28219757 | 0.61980968 | -0.4552971 | 0.64889549 | 0.91417939 |
| SCN3A | 127.362681 | -0.24812583 | 0.67190415 | -0.3692875 | 0.71191341 | 0.9357596 |
| SCN2A | 411.850455 | 0.968393769 | 0.50050159 | 1.93484655 | 0.05300913 | 0.36519073 |

| | | | | | | |
|---|---|---|---|---|---|---|
| CSRNP3 | 54.1352494 | 0.371866032 | 0.4456294 | 0.83447374 | 0.40401408 | 0.81027913 |
| GALNT3 | 71.0545276 | -0.154247857 | 0.34715654 | -0.4443179 | 0.65681278 | 0.9158679 |
| TTC21B | 77.1499768 | 0.194322608 | 0.27230012 | 0.71363395 | 0.47545356 | 0.84609246 |
| SCN9A | 26.0751825 | -0.253184968 | 0.53806069 | -0.4705509 | 0.63796145 | 0.91025608 |
| STK39 | 496.727536 | -0.964946098 | 0.43382565 | -2.2242716 | 0.02613017 | 0.25383383 |
| CERS6 | 87.2107676 | 0.172063751 | 0.35180972 | 0.48908186 | 0.62478374 | 0.90447745 |
| LRP2 | 25.6994891 | 0.666405426 | 0.74804152 | 0.8908669 | 0.37300058 | 0.7913202 |
| FASTKD1 | 99.7147129 | 0.506673689 | 0.27884255 | 1.81706015 | 0.0692079 | 0.41776205 |
| PPIG | 142.590592 | 0.120471635 | 0.17747462 | 0.67881049 | 0.49725794 | 0.85814183 |
| KLHL23 | 39.4901197 | 0.413816458 | 0.25451787 | 1.6258837 | 0.1039744 | 0.49650823 |
| SSB | 50.9712561 | -0.093468501 | 0.22900571 | -0.4081492 | 0.68316413 | 0.92474072 |
| METTL5 | 30.0834992 | -0.201677471 | 0.36274583 | -0.5559746 | 0.57822822 | 0.88901786 |
| UBR3 | 558.592638 | -0.074028296 | 0.20662494 | -0.3582738 | 0.72013844 | 0.93806337 |
| MYO3B | 191.924398 | 0.298904879 | 0.48566456 | 0.61545541 | 0.53825401 | 0.86972442 |
| AC007405.1 | 13.6668701 | 0.093638405 | 0.55969114 | 0.16730371 | 0.86713108 | 0.97434023 |
| GAD1 | 58.6036333 | -0.698698584 | 0.70642113 | -0.9890681 | 0.32262984 | 0.76292283 |
| GORASP2 | 67.0587797 | 0.32428737 | 0.29536724 | 1.09791245 | 0.27224272 | 0.72345643 |
| TLK1 | 261.340787 | -0.02626785 | 0.15662045 | -0.1677166 | 0.86680623 | 0.97434023 |
| METTL8 | 21.1701619 | 0.386801991 | 0.38690507 | 0.99973357 | 0.31743946 | 0.75958338 |
| DCAF17 | 25.9721787 | 0.079329051 | 0.29283446 | 0.27090067 | 0.78646743 | 0.95577371 |
| DYNC1I2 | 133.413109 | -0.302533479 | 0.18870268 | -1.6032284 | 0.10888425 | 0.50680288 |
| SLC25A12 | 153.244509 | 0.056453532 | 0.31953281 | 0.17667523 | 0.8597635 | 0.97287712 |
| HAT1 | 38.2152713 | 0.214271713 | 0.25609414 | 0.8366912 | 0.40276617 | 0.80903538 |
| METAP1D | 18.751087 | 0.744548111 | 0.30990101 | 2.40253528 | 0.01628186 | 0.19692529 |
| ITGA6 | 1079.88322 | 0.177423514 | 0.3081413 | 0.57578622 | 0.56475969 | 0.88191823 |
| AC078883.1 | 43.6160276 | -1.28181667 | 0.53916252 | -2.3774217 | 0.01743414 | 0.20451748 |
| PDK1 | 74.4575379 | -1.157886028 | 0.416561 | -2.7796314 | 0.00544206 | 0.10201977 |
| MAP3K20 | 61.4945602 | -0.557120658 | 0.49849905 | -1.1175962 | 0.26373948 | 0.71380385 |
| SP3 | 95.6902541 | 0.087324923 | 0.19187092 | 0.45512328 | 0.64902055 | 0.91417939 |
| OLA1 | 151.973563 | -0.009882356 | 0.19230989 | -0.0513877 | 0.95901662 | 0.99238848 |
| CIR1 | 54.290893 | -0.295768706 | 0.20376892 | -1.4514908 | 0.14664325 | 0.58349206 |
| SCRN3 | 22.8770764 | -0.004046981 | 0.36783175 | -0.0110023 | 0.99122164 | 0.99923337 |
| GPR155 | 41.6789737 | 0.504932049 | 0.55557533 | 0.90884534 | 0.36343177 | 0.78724613 |
| AC010894.3 | 158.954819 | -0.864082319 | 0.35137301 | -2.4591596 | 0.01392627 | 0.17681258 |
| WIPF1 | 142.794301 | -0.127122442 | 0.40504685 | -0.3138463 | 0.7536378 | 0.94791062 |
| CHN1 | 12.9896525 | 0.078124489 | 0.56579355 | 0.1380795 | 0.89017759 | 0.97782578 |
| ATF2 | 122.673944 | 0.201274394 | 0.22614118 | 0.89003867 | 0.37344512 | 0.79178529 |
| ATP5MC3 | 203.418308 | -0.055536751 | 0.44218259 | -0.1255969 | 0.90005104 | 0.97940229 |
| LNPK | 75.4412208 | -0.103126194 | 0.26460484 | -0.3897366 | 0.69673133 | 0.92961401 |
| HOXD10 | 14.3541556 | 0.564560179 | 0.70101571 | 0.80534597 | 0.42062006 | 0.81938194 |
| HOXD-AS2 | 21.2314149 | -0.517626416 | 0.42513287 | -1.2175639 | 0.22338976 | 0.67606266 |
| HOXD8 | 38.9239969 | 1.006397912 | 0.36063513 | 2.79062642 | 0.00526062 | 0.10005484 |
| HOXD3 | 49.9535729 | 0.346605642 | 0.39805869 | 0.87074004 | 0.38389611 | 0.7980695 |
| HAGLR | 43.402256 | -0.700218528 | 0.26776521 | -2.6150467 | 0.00892153 | 0.13736318 |
| HOXD1 | 15.543455 | -1.363831901 | 0.53719134 | -2.5388196 | 0.01112272 | 0.15593979 |
| MTX2 | 249.891147 | 0.134022199 | 0.24016272 | 0.55804746 | 0.57681197 | 0.88821715 |
| LINC01116 | 15.2777675 | 0.43105112 | 0.45077439 | 0.95624581 | 0.33894806 | 0.77015605 |
| AC074286.1 | 27.0405632 | 0.631401279 | 0.90589276 | 0.69699341 | 0.48580692 | 0.85356796 |
| HNRNPA3 | 194.128769 | 0.026380769 | 0.20164441 | 0.13082817 | 0.89591124 | 0.97885533 |
| NFE2L2 | 428.889409 | -0.098705188 | 0.19099219 | -0.5168022 | 0.60529422 | 0.89600432 |
| AGPS | 75.7041055 | 0.235111827 | 0.17615645 | 1.33467626 | 0.18198233 | 0.63058858 |
| AC012499.1 | 16.1792967 | 2.383462157 | 0.60047632 | 3.96928585 | 7.21E-05 | 0.00526714 |
| RBM45 | 23.3536201 | 0.335580853 | 0.34730387 | 0.96624565 | 0.33392127 | 0.76777176 |
| AC009948.1 | 37.8825015 | -0.226529857 | 0.554086 | -0.4088352 | 0.68266061 | 0.92474072 |
| PRKRA | 85.9817509 | -0.035665145 | 0.21072577 | -0.1692491 | 0.86560072 | 0.97434023 |
| PLEKHA3 | 52.9870615 | -0.046760754 | 0.22234383 | -0.2103083 | 0.83342706 | 0.96701848 |
| TTN-AS1 | 125.100511 | -0.165007993 | 0.26178144 | -0.6303273 | 0.52848046 | 0.86750588 |
| SESTD1 | 234.077598 | -0.637623326 | 0.35762852 | -1.7829208 | 0.0745992 | 0.43097811 |
| ZNF385B | 67.3737023 | 0.674966748 | 0.44058972 | 1.53196209 | 0.12553179 | 0.54505746 |
| CWC22 | 46.6357351 | 0.278834471 | 0.25383993 | 1.09846575 | 0.27200117 | 0.7232847 |
| UBE2E3 | 123.951892 | 0.187127238 | 0.20596775 | 0.90852687 | 0.36359993 | 0.7873616 |
| SSFA2 | 65.302693 | -0.261952435 | 0.36606223 | -0.7155954 | 0.47424124 | 0.84593347 |
| PPP1R1C | 21.3982337 | -0.061901867 | 0.70275742 | -0.0880843 | 0.92980971 | 0.98733028 |
| PDE1A | 1099.90465 | 2.266312354 | 0.23635917 | 9.5884257 | 8.94E-22 | 8.68E-18 |
| DNAJC10 | 70.3509188 | -0.238996623 | 0.2114824 | -1.1301017 | 0.25843336 | 0.71022613 |
| NCKAP1 | 269.642378 | -0.366291607 | 0.25028182 | -1.4635166 | 0.14332607 | 0.57698886 |
| NUP35 | 22.7563128 | -0.064783151 | 0.27575431 | -0.2349307 | 0.81426253 | 0.96286073 |
| LINC01473 | 29.3895197 | 0.181688069 | 0.3399257 | 0.53449348 | 0.59300016 | 0.89215305 |
| ZC3H15 | 72.4393566 | -0.258003675 | 0.19014691 | -1.3568649 | 0.17482414 | 0.62071528 |
| ITGAV | 313.041804 | 0.478729608 | 0.43699425 | 1.09550552 | 0.27329523 | 0.7244699 |
| AC017101.1 | 26.5661352 | 0.246038973 | 0.60453186 | 0.40699091 | 0.68401466 | 0.92538534 |
| AC007319.1 | 33.2415757 | -0.299571928 | 0.80267865 | -0.3732153 | 0.70898823 | 0.93482669 |
| TFPI | 24.9986264 | -1.826043169 | 0.50162548 | -3.640252 | 0.00027237 | 0.0134513 |
| LINC01090 | 29.5479397 | -1.126495276 | 0.4395938 | -2.5625823 | 0.0103897 | 0.14952531 |
| GULP1 | 703.110847 | -0.989448067 | 0.53873187 | -1.8366243 | 0.06626537 | 0.40888394 |
| WDR75 | 38.3387026 | 0.302053492 | 0.29584992 | 1.02096866 | 0.30726929 | 0.75038069 |
| SLC40A1 | 49.3999251 | 0.859212323 | 0.22316645 | 3.85009635 | 0.00011807 | 0.00729486 |
| ASNSD1 | 16.5281848 | 0.559039902 | 0.59366335 | 0.94167831 | 0.34635736 | 0.77642952 |
| ANKAR | 59.9450292 | 0.281683386 | 0.24104895 | 1.16857338 | 0.24257556 | 0.69621873 |
| ORMDL1 | 47.0978526 | 0.056635766 | 0.31587086 | 0.17930038 | 0.85770185 | 0.97243282 |
| PMS1 | 41.3489923 | 0.526614725 | 0.22683853 | 2.32154002 | 0.02025772 | 0.22348867 |
| C2orf88 | 38.2139701 | 0.227064827 | 0.29063727 | 0.78126534 | 0.43464645 | 0.82565889 |
| HIBCH | 175.889163 | 0.281438895 | 0.31519373 | 0.89290766 | 0.37190662 | 0.7900012 |
| INPP1 | 62.0558366 | -0.502245059 | 0.34712649 | -1.4468647 | 0.14793483 | 0.58498485 |
| MFSD6 | 64.3753428 | -0.294720703 | 0.29902932 | -0.9855913 | 0.3243337 | 0.76304922 |
| NAB1 | 28.3319249 | 0.90511976 | 0.32681052 | 2.76955515 | 0.00561329 | 0.10331862 |
| GLS | 484.794586 | 0.681896436 | 0.39571609 | 1.72319611 | 0.08485307 | 0.45956155 |
| STAT1 | 163.65537 | -1.207671776 | 0.59104377 | -2.0432865 | 0.04102408 | 0.32510914 |
| SLC39A10 | 18.0123109 | -0.647866918 | 0.38845497 | -1.6678044 | 0.09535456 | 0.48081904 |
| DNAH7 | 24.0844656 | -0.420642335 | 0.40950484 | -1.0271975 | 0.30432748 | 0.74820946 |
| GTF3C3 | 49.1354906 | -0.153084504 | 0.24613895 | -0.6219434 | 0.53397907 | 0.86909171 |
| PGAP1 | 50.2135052 | -0.872479266 | 0.33011411 | -2.6429627 | 0.00821841 | 0.13111608 |
| ANKRD44 | 314.152441 | 0.351840455 | 0.38776135 | 0.90736339 | 0.36421466 | 0.78780828 |
| SF3B1 | 170.002391 | -0.090946042 | 0.19440544 | -0.4678163 | 0.63991594 | 0.91096398 |
| COQ10B | 51.4501528 | -0.482834388 | 0.22359509 | -2.1594141 | 0.03081805 | 0.27936455 |
| HSPD1 | 163.74233 | 0.139779443 | 0.26845678 | 0.52067765 | 0.60259135 | 0.8956362 |
| MOB4 | 21.3719938 | 0.215690578 | 0.37407962 | 0.57659002 | 0.56421644 | 0.88191823 |
| PLCL1 | 1571.2662 | 0.770172069 | 0.39300136 | 1.95971858 | 0.05002869 | 0.35682229 |
| FTCDNL1 | 20.4727804 | 1.229709094 | 0.56972506 | 2.15842548 | 0.03089477 | 0.27936455 |
| C2orf69 | 24.5579897 | -0.095331319 | 0.3892424 | -0.244915 | 0.80652221 | 0.96005568 |
| TYW5 | 43.1428183 | -0.123702187 | 0.26655027 | -0.4640858 | 0.6425863 | 0.91146968 |
| MAIP1 | 16.569839 | -0.069900855 | 0.36297464 | -0.1925778 | 0.84728964 | 0.97027687 |
| SPATS2L | 116.363329 | -0.988636794 | 0.55099413 | -1.7942783 | 0.07276876 | 0.42521506 |
| KCTD18 | 14.4587033 | -0.536989852 | 0.41338173 | -1.2990169 | 0.19393813 | 0.64535157 |
| BZW1 | 28.9713272 | -0.147491361 | 0.38808517 | -0.3800489 | 0.70390908 | 0.93306088 |
| CLK1 | 102.630674 | 0.233562704 | 0.22921471 | 1.0189691 | 0.30821764 | 0.7512664 |
| PPIL3 | 21.6074782 | 0.047043578 | 0.4448168 | 0.10575944 | 0.91577322 | 0.98437981 |
| NIF3L1 | 17.7612453 | 0.696135281 | 0.32129044 | 2.1666853 | 0.03025886 | 0.27663612 |
| ORC2 | 66.8990462 | 0.005616035 | 0.30125032 | 0.01864242 | 0.98512636 | 0.99777861 |
| AC005037.1 | 16.8198694 | -0.534631354 | 0.48362992 | -1.1054555 | 0.26896227 | 0.72010324 |
| FAM126B | 106.297989 | 0.422358331 | 0.22099579 | 1.91116005 | 0.05598402 | 0.37674523 |
| NDUFB3 | 78.8930029 | 0.237194565 | 0.53963311 | 0.43954783 | 0.66026463 | 0.91752855 |
| CFLAR | 244.031949 | -0.305234457 | 0.27684831 | -1.1025332 | 0.27022992 | 0.72071219 |

| | | | | | | |
|---|---|---|---|---|---|---|
| CFLAR-AS1 | 25.9444357 | -1.696209161 | 0.62052173 | -2.7335209 | 0.00626611 | 0.11214263 |
| CASP10 | 40.1695566 | -1.294996865 | 0.41137478 | -3.1479734 | 0.00164407 | 0.04479619 |
| CASP8 | 32.9052236 | -0.609654831 | 0.27061333 | -2.2528632 | 0.02426777 | 0.24374605 |
| TRAK2 | 118.817114 | -0.39761868 | 0.30679633 | -1.2960347 | 0.19496354 | 0.6459894 |
| STRADB | 45.1285117 | -0.761432767 | 0.3963256 | -1.9212303 | 0.05470267 | 0.37236207 |
| TMEM237 | 11.0533801 | 0.737438253 | 0.44895642 | 1.64256088 | 0.10047382 | NA |
| ALS2 | 80.7597702 | -0.502659186 | 0.22846794 | -2.2001301 | 0.02779767 | 0.26203825 |
| FZD7 | 96.2843625 | 0.673538639 | 0.40242587 | 1.67369618 | 0.09419032 | 0.47910128 |
| SUMO1 | 57.001044 | -0.371336063 | 0.26552974 | -1.3984726 | 0.16197119 | 0.60094226 |
| NOP58 | 74.2878678 | -0.016119081 | 0.28233638 | -0.0570918 | 0.9544721 | 0.99228059 |
| BMPR2 | 315.660001 | -0.284139171 | 0.25889663 | -1.0975005 | 0.27242268 | 0.72345643 |
| FAM117B | 108.682806 | 0.178485419 | 0.33586391 | 0.53142184 | 0.59512649 | 0.893081 |
| ICA1L | 32.296111 | -0.179901616 | 0.43140004 | -0.4170181 | 0.67666519 | 0.92237057 |
| WDR12 | 28.420698 | 0.336988619 | 0.393925 | 0.8554639 | 0.39229438 | 0.80367402 |
| CARF | 39.6489118 | 0.17449171 | 0.3927753 | 0.44425326 | 0.65685949 | 0.9158679 |
| NBEAL1 | 474.564633 | 0.253273862 | 0.18960534 | 1.33579499 | 0.18161629 | 0.62984555 |
| CYP20A1 | 93.4192207 | 0.462782323 | 0.36525474 | 1.26701252 | 0.20515081 | 0.65805651 |
| ABI2 | 148.978366 | -0.156354606 | 0.27719839 | -0.5640531 | 0.572718 | 0.88578992 |
| RAPH1 | 216.445906 | -0.550890372 | 0.36924733 | -1.4919278 | 0.13571806 | 0.56627586 |
| PARD3B | 537.646668 | 0.787883523 | 0.18989103 | 4.14913493 | 3.34E-05 | 0.00299743 |
| INO80D | 286.259602 | 0.39523881 | 0.42694296 | 0.92574148 | 0.35458035 | 0.78097851 |
| NDUFS1 | 219.823891 | 0.261642024 | 0.16270695 | 1.60805684 | 0.10782273 | 0.50550047 |
| EEF1B2 | 71.4435492 | 0.492754823 | 0.44977147 | 1.0955671 | 0.27326827 | 0.7244699 |
| ZDBF2 | 36.4587713 | 0.05467635 | 0.29493844 | 0.18538224 | 0.85292923 | 0.97107539 |
| FASTKD2 | 43.8507999 | 0.164730876 | 0.319771 | 0.51515265 | 0.60644635 | 0.89643487 |
| KLF7 | 80.0396337 | -0.737751851 | 0.38900632 | -1.8965035 | 0.0578935 | 0.38201831 |
| CREB1 | 118.665193 | -0.187072526 | 0.21054831 | -0.8885017 | 0.37427095 | 0.7923487 |
| METTL21A | 34.4738718 | -0.061060806 | 0.34241289 | -0.1783251 | 0.85846768 | 0.97287712 |
| CCNYL1 | 39.9494376 | -0.609899914 | 0.28342613 | -2.1518832 | 0.03140656 | 0.28099688 |
| PLEKHM3 | 46.8806145 | 0.064899339 | 0.31737932 | 0.20448509 | 0.83797444 | 0.96748214 |
| IDH1 | 13.9261964 | -0.590670645 | 0.43827205 | -1.347726 | 0.17774652 | 0.62542578 |
| PIKFYVE | 128.084668 | -0.264500832 | 0.20962498 | -1.261781 | 0.20702758 | 0.65895397 |
| PTH2R | 76.3101922 | 0.966171287 | 0.5345238 | 1.80753652 | 0.07067867 | 0.42119392 |
| MAP2 | 111.324979 | 0.327006735 | 0.38901107 | 0.84061035 | 0.40056626 | 0.80830282 |
| UNC80 | 21.1649831 | 2.106665881 | 0.59851827 | 3.5198021 | 0.00043187 | 0.01813475 |
| RPE | 48.0992101 | -0.032619232 | 0.27126132 | -0.1202502 | 0.90428494 | 0.98041914 |
| KANSL1L | 494.382009 | 0.20821075 | 0.25766004 | 0.80808319 | 0.41904269 | 0.8189355 |
| AC007038.1 | 49.8475392 | 0.171769582 | 0.41359822 | 0.41530542 | 0.67791834 | 0.9226233 |
| ACADL | 106.276917 | 0.523833827 | 0.2308881 | 2.26877792 | 0.02328183 | 0.23879078 |
| LANCL1 | 51.0586827 | 0.249409304 | 0.23480131 | 1.06221426 | 0.28813843 | 0.73803612 |
| ERBB4 | 1516.42934 | -0.055282746 | 0.2872776 | -0.1924367 | 0.84740017 | 0.97027687 |
| IKZF2 | 48.7174235 | -0.657829222 | 0.54612407 | -1.2045417 | 0.22838027 | 0.68364117 |
| SPAG16 | 177.943562 | 0.160854907 | 0.36144115 | 0.44503762 | 0.65629257 | 0.9158679 |
| BARD1 | 16.0979133 | -0.795243342 | 0.48415287 | -1.642546 | 0.10047691 | 0.48886326 |
| AC016708.1 | 16.8238527 | 0.564315669 | 0.50309636 | 1.12168505 | 0.26199637 | 0.71302363 |
| ATIC | 23.2513184 | -0.302412369 | 0.32962467 | -0.9174446 | 0.35890972 | 0.78445006 |
| MREG | 28.2323578 | 0.499089272 | 0.46477524 | 1.0738293 | 0.2828992 | 0.7344003 |
| PECR | 14.8295755 | -0.43335987 | 0.52856896 | -0.8198739 | 0.41228803 | 0.81556555 |
| XRCC5 | 210.80627 | -0.119447121 | 0.16082866 | -0.742698 | 0.45766457 | 0.83603055 |
| SMARCAL1 | 23.9617756 | 0.408435094 | 0.36213406 | 1.12785607 | 0.25938071 | 0.71066775 |
| RPL37A | 218.430535 | 0.237842252 | 0.34982124 | 0.67989655 | 0.49656997 | 0.85814183 |
| IGFBP5 | 1368.9944 | -0.277032551 | 0.42652108 | -0.6495167 | 0.51600448 | 0.86476217 |
| AC007563.2 | 84.993138 | -0.87965147 | 0.53909038 | -1.6317328 | 0.10273578 | 0.49357954 |
| DIRC3 | 29.7217442 | -1.323989561 | 0.52264164 | -2.5332646 | 0.01130056 | 0.15732016 |
| TNS1 | 139.468069 | 0.00996524 | 0.40652031 | 0.02451351 | 0.98044301 | 0.99687019 |
| ARPC2 | 127.739626 | -0.402023245 | 0.23178234 | -1.7344861 | 0.08283188 | 0.45496557 |
| AAMP | 25.5625911 | 0.235880916 | 0.34332623 | 0.68704601 | 0.49205374 | 0.85689789 |
| PNKD | 94.9173947 | -0.27979418 | 0.36977498 | -0.7566607 | 0.44925318 | 0.83118301 |
| TMBIM1 | 68.2922416 | -1.29209607 | 0.4094145 | -3.1559607 | 0.0015997 | 0.04420835 |
| CATIP-AS2 | 12.4977046 | -0.891126798 | 0.59262705 | -1.503689 | 0.13266145 | NA |
| CTDSP1 | 40.4601862 | 0.05349834 | 0.33340729 | 0.16045942 | 0.87251919 | 0.97434023 |
| USP37 | 74.6048212 | -0.338204544 | 0.29454482 | -1.1482278 | 0.25087454 | 0.70535741 |
| CNOT9 | 49.5619359 | -0.438784813 | 0.27832016 | -1.576547 | 0.1148998 | 0.51928773 |
| PLCD4 | 11.6475732 | 1.06551251 | 0.68158981 | 1.5632753 | 0.11798786 | NA |
| ZNF142 | 15.0757774 | 0.049553697 | 0.38871914 | 0.12747944 | 0.89856095 | 0.97908342 |
| TTLL4 | 15.0685033 | -0.523178971 | 0.39944201 | -1.3097745 | 0.19027212 | 0.6421543 |
| CYP27A1 | 24.109583 | -0.186402154 | 0.28288513 | -0.6589323 | 0.50993923 | 0.86281145 |
| CNPPD1 | 58.4301335 | -0.180149062 | 0.23891396 | -0.7540332 | 0.45082927 | 0.83185162 |
| RETREG2 | 51.2148927 | 0.28256131 | 0.38124104 | 0.74116185 | 0.45859531 | 0.83645518 |
| ZFAND2B | 15.1307188 | 0.590847557 | 0.3611622 | 1.63596181 | 0.10184758 | 0.49220184 |
| ANKZF1 | 25.3413618 | -0.452217357 | 0.35941806 | -1.2581932 | 0.2083219 | 0.65973983 |
| STK16 | 14.7196327 | -0.420352817 | 0.41529065 | -1.0121895 | 0.31144747 | 0.75450562 |
| TUBA4A | 72.2542579 | -0.318785931 | 0.37619698 | -0.847391 | 0.39677722 | 0.80674902 |
| DNAJB2 | 35.2050558 | 0.402007169 | 0.31932124 | 1.25894278 | 0.208051 | 0.65973983 |
| DNPEP | 22.9545752 | 0.209938004 | 0.32561311 | 0.64474677 | 0.5190913 | 0.86618562 |
| CHPF | 20.2193098 | -0.625525786 | 0.41928743 | -1.4918782 | 0.13573107 | 0.56627586 |
| OBSL1 | 14.7280784 | -0.060638867 | 0.45441356 | -0.1334442 | 0.89384207 | 0.97833263 |
| SGPP2 | 243.513866 | 0.372901011 | 0.20692575 | 1.8021006 | 0.07152958 | 0.42281025 |
| FARSB | 40.9702292 | 0.437806224 | 0.31166429 | 1.40473655 | 0.16009963 | 0.60024417 |
| ACSL3 | 199.523743 | 0.350670948 | 0.29744361 | 1.17894933 | 0.23841835 | 0.69324282 |
| AP1S3 | 25.8515795 | -0.413094137 | 0.4415287 | -0.9355997 | 0.34947931 | 0.77868979 |
| WDFY1 | 64.6601316 | -0.3411375 | 0.34394442 | -0.991839 | 0.32127606 | 0.76163052 |
| MRPL44 | 18.6296353 | 0.156028304 | 0.33378102 | 0.46745709 | 0.64017289 | 0.91096398 |
| CUL3 | 203.857319 | -0.086503763 | 0.19334167 | -0.447414 | 0.6545762 | 0.9158679 |
| DOCK10 | 281.151282 | -0.004062043 | 0.67533618 | -0.0060148 | 0.99520088 | 0.99923337 |
| IRS1 | 107.096248 | -0.700986612 | 0.49219758 | -1.4241976 | 0.15438928 | 0.59443382 |
| RHBDD1 | 191.047531 | -0.08485091 | 0.30600389 | -0.277287 | 0.78155972 | 0.95540382 |
| COL4A4 | 294.068355 | 0.396217737 | 0.28308609 | 1.39963692 | 0.16162207 | 0.60060549 |
| COL4A3 | 282.496502 | 0.321771203 | 0.27606734 | 1.16555331 | 0.24379509 | 0.69758478 |
| AC097662.1 | 43.8835026 | -0.292652524 | 0.51484119 | -0.5684326 | 0.56974125 | 0.88562534 |
| MFF | 99.2194005 | 0.290477603 | 0.33892013 | 0.85706801 | 0.3914073 | 0.80332153 |
| AGFG1 | 185.795853 | 0.149493506 | 0.23403958 | 0.63875309 | 0.52298357 | 0.86740339 |
| TRIP12 | 283.96057 | -0.073369928 | 0.17091556 | -0.4292759 | 0.66772246 | 0.92049996 |
| FBXO36 | 20.4236792 | 0.677537683 | 0.30310689 | 2.23530943 | 0.02539704 | 0.25137888 |
| SP110 | 47.9264008 | -1.329949549 | 0.51313754 | -2.5917994 | 0.00954754 | 0.14402982 |
| SP140L | 40.6594906 | -1.053136509 | 0.50218034 | -2.0971281 | 0.03598223 | 0.30345958 |
| SP100 | 135.202254 | -0.902137757 | 0.40576375 | -2.2233079 | 0.02619504 | 0.25409193 |
| CAB39 | 271.436903 | 0.814691806 | 0.25698981 | 3.17013266 | 0.00152369 | 0.04267815 |
| ITM2C | 49.1185991 | -0.097602051 | 0.297235 | -0.3283666 | 0.74263448 | 0.94546379 |
| C2orf72 | 12.0705712 | 0.714602681 | 0.56996499 | 1.25376592 | 0.2099271 | NA |
| PSMD1 | 136.785766 | 0.1126665 | 0.19560705 | 0.57598385 | 0.5646261 | 0.88191823 |
| ARMC9 | 101.088793 | -0.279023888 | 0.23325137 | -1.1962369 | 0.23160413 | 0.68597254 |
| NCL | 93.6147492 | -0.346940079 | 0.38436992 | -0.9026203 | 0.36672747 | 0.78809407 |
| PTMA | 256.895035 | -0.191126901 | 0.31020574 | -0.6161295 | 0.53780907 | 0.86972442 |
| PDE6D | 34.4813789 | 0.705524023 | 0.33668829 | 2.09548135 | 0.03612822 | 0.30394083 |
| COPS7B | 33.7363734 | 0.226747072 | 0.26540992 | 0.85432779 | 0.39292339 | 0.80400536 |
| DIS3L2 | 162.176465 | -0.058728839 | 0.25437533 | -0.2308747 | 0.81741211 | 0.9635539 |
| EIF4E2 | 45.8077926 | -0.453206653 | 0.29265203 | -1.5486196 | 0.1214732 | 0.53631772 |
| EFHD1 | 293.337177 | 0.523059605 | 0.30296872 | 1.72644756 | 0.08426693 | 0.4585327 |
| GIGYF2 | 408.921577 | 0.064345736 | 0.17336515 | 0.37115726 | 0.7105204 | 0.93540281 |
| KCNJ13 | 13.1552583 | 1.929603374 | 0.65594793 | 2.94170204 | 0.00326414 | 0.072288 |
| SNORC | 11.9406874 | -0.228477025 | 0.49577521 | -0.460848 | 0.64490764 | NA |
| ATG16L1 | 18.1009589 | 0.223846933 | 0.33709008 | 0.66405672 | 0.50665401 | 0.86235197 |

| | | | | | | |
|---|---|---|---|---|---|---|
| DGKD | 22.5175863 | 0.642251854 | 0.4163668 | 1.54251456 | 0.1229486 | 0.53921501 |
| USP40 | 110.947764 | -0.118823438 | 0.29555473 | -0.4020353 | 0.68765804 | 0.92655688 |
| SH3BP4 | 213.465623 | -0.113373791 | 0.36845122 | -0.3077037 | 0.75830783 | 0.94924107 |
| AGAP1 | 368.934924 | -0.753049424 | 0.45366958 | -1.6599072 | 0.09693311 | 0.48257296 |
| IQCA1 | 32.2575006 | -0.013113408 | 0.59654806 | -0.0219821 | 0.9824622 | 0.99719767 |
| COPS8 | 26.7023361 | -0.212791425 | 0.30063095 | -0.7078161 | 0.47905947 | 0.84833462 |
| RAB17 | 97.3429201 | 0.108684629 | 0.46491756 | 0.23377183 | 0.81516212 | 0.96300833 |
| LRRFIP1 | 457.018197 | -0.588726454 | 0.24812963 | -2.3726568 | 0.01766067 | 0.20618531 |
| ILKAP | 61.737328 | 0.045918655 | 0.22687442 | 0.20239679 | 0.83960654 | 0.96748214 |
| PER2 | 26.8594522 | -0.048714973 | 0.38439545 | -0.1267314 | 0.89915299 | 0.97931552 |
| TRAF3IP1 | 41.5601143 | 0.40280351 | 0.32867097 | 1.22555246 | 0.22036714 | 0.67406151 |
| ASB1 | 28.701229 | -0.088259223 | 0.46479343 | -0.1898891 | 0.849396 | 0.97027687 |
| HDAC4 | 134.536876 | -0.372280931 | 0.2632765 | -1.4140302 | 0.15735303 | 0.59804205 |
| NDUFA10 | 153.626056 | 0.113822629 | 0.22473983 | 0.50646398 | 0.61253097 | 0.90011356 |
| COPS9 | 39.6995764 | 0.028042446 | 0.41658162 | 0.06731561 | 0.94633045 | 0.99134285 |
| GPC1 | 9.98539265 | 0.17292301 | 0.57189447 | 0.30236874 | 0.76237098 | NA |
| ANKMY1 | 33.2719838 | 0.478662646 | 0.26657328 | 1.79561376 | 0.07255597 | 0.42448303 |
| RNPEPL1 | 42.9286615 | -0.531284429 | 0.34159913 | -1.5552863 | 0.11987791 | 0.53145142 |
| CAPN10 | 22.7549278 | -0.368123342 | 0.33054252 | -1.1136944 | 0.26541033 | 0.71660319 |
| SNED1 | 44.3590441 | -0.690174165 | 0.49572122 | -1.3922627 | 0.16384286 | 0.60359885 |
| AC005237.1 | 12.2145558 | -1.335449459 | 0.57275095 | -2.3316408 | 0.0197196 | NA |
| MTERF4 | 37.0744872 | 0.132635553 | 0.22799069 | 0.58175863 | 0.56072927 | 0.88071368 |
| PPP1R7 | 44.9288056 | -0.40023175 | 0.23436467 | -1.7077307 | 0.08768632 | 0.46580356 |
| HDLBP | 180.542159 | -0.103581558 | 0.18362051 | -0.5641067 | 0.57268153 | 0.88578992 |
| 2-Sep | 200.896761 | -0.060258217 | 0.19438965 | -0.3099868 | 0.75657103 | 0.94828001 |
| FARP2 | 114.972981 | 0.000746838 | 0.19173467 | 0.00389516 | 0.99689212 | 0.99975875 |
| STK25 | 38.1840571 | 0.057530752 | 0.30516738 | 0.18852196 | 0.85046751 | 0.97027687 |
| BOK | 20.0053338 | 0.120622474 | 0.40608249 | 0.29703934 | 0.76643648 | 0.95205353 |
| THAP4 | 64.7865975 | -0.619917559 | 0.20342033 | -3.047471 | 0.00230776 | 0.05610339 |
| ATG4B | 76.9912498 | -0.135321675 | 0.24949699 | -0.542378 | 0.58755814 | 0.89073775 |
| DTYMK | 16.8456489 | 0.606649094 | 0.44864876 | 1.35216934 | 0.17632115 | 0.62329269 |
| ING5 | 48.9930033 | 0.624393933 | 0.32164705 | 1.94123943 | 0.05222924 | 0.36271841 |
| D2HGDH | 32.9816641 | 0.415508281 | 0.32090174 | 1.29481469 | 0.19538416 | 0.6459894 |
| CHL1 | 366.110422 | 1.142453221 | 0.66749563 | 1.7115516 | 0.08697933 | 0.46422617 |
| LINC01266 | 36.1962732 | 0.211808007 | 0.4195354 | 0.50486326 | 0.61365488 | 0.9002782 |
| CNTN4 | 17.6339776 | -0.217787554 | 0.60639698 | -0.3591501 | 0.71948278 | 0.93786761 |
| TRNT1 | 41.3284673 | 0.094945744 | 0.23681411 | 0.40092943 | 0.68847208 | 0.9267526 |
| CRBN | 36.8363946 | -0.084081269 | 0.30155242 | -0.278828 | 0.7803768 | 0.95537097 |
| AC034195.1 | 75.2510761 | 0.337126325 | 0.42432234 | 0.79450525 | 0.42690135 | 0.82375285 |
| SUMF1 | 390.95825 | -0.174349453 | 0.30255892 | -0.5762496 | 0.56444649 | 0.88191823 |
| SETMAR | 24.5172781 | 0.837254668 | 0.33578018 | 2.49346066 | 0.01265046 | 0.16734742 |
| ITPR1 | 509.346086 | 0.899208467 | 0.32495924 | 2.76714229 | 0.00565501 | 0.10388933 |
| EGOT | 17.5717947 | -2.277555003 | 0.66645669 | -3.4174089 | 0.0006322 | 0.02386134 |
| AC018816.1 | 20.6613629 | -0.474939095 | 0.41064127 | -1.1565791 | 0.24744438 | 0.70202121 |
| BHLHE40-AS | 15.1758351 | -1.223345935 | 0.69088325 | -1.7706985 | 0.07661085 | 0.43573154 |
| BHLHE40 | 171.271498 | -1.76538094 | 0.49477939 | -3.5680163 | 0.00035969 | 0.01600474 |
| ARL8B | 116.924152 | -0.279349942 | 0.24771619 | -1.1277016 | 0.25944596 | 0.71066775 |
| AC026202.2 | 20.9622226 | -0.98657505 | 0.45101705 | -2.1874451 | 0.02871004 | 0.26751914 |
| EDEM1 | 33.0901279 | -0.00788456 | 0.28681536 | -0.02749 | 0.9780689 | 0.99609121 |
| LMCD1-AS1 | 71.4192478 | 0.517226242 | 0.36834008 | 1.40420843 | 0.16025679 | 0.60024417 |
| RAD18 | 14.5092567 | 0.817030099 | 0.34679702 | 2.3559317 | 0.01847631 | 0.21121173 |
| SRGAP3 | 139.232208 | 0.103291479 | 0.64261927 | 0.16073511 | 0.87230203 | 0.97434023 |
| THUMPD3-A! | 97.6030354 | -0.002850938 | 0.30448903 | -0.009363 | 0.9925295 | 0.99923337 |
| THUMPD3 | 32.3744019 | -0.115284844 | 0.25861246 | -0.4457823 | 0.65575454 | 0.9158679 |
| SETD5 | 301.14251 | -0.165234683 | 0.19517427 | -0.8466007 | 0.3972177 | 0.80674902 |
| MTMR14 | 28.3599124 | -0.059532404 | 0.29566067 | -0.2013538 | 0.84042193 | 0.96748214 |
| OGG1 | 15.6886022 | -0.070127015 | 0.35403653 | -0.1980785 | 0.84298367 | 0.96906158 |
| CAMK1 | 11.7573308 | 0.573415425 | 0.50733022 | 1.13026072 | 0.25836638 | NA |
| TADA3 | 29.7840129 | -0.201882673 | 0.25171569 | -0.8020266 | 0.42253758 | 0.82138304 |
| JAGN1 | 15.6748376 | 0.118479901 | 0.48556877 | 0.24400231 | 0.80722903 | 0.96016552 |
| IL17RC | 12.5239514 | 0.498936215 | 0.38865196 | 1.28376097 | 0.1992256 | NA |
| CRELD1 | 26.4303943 | 0.21093072 | 0.37221248 | 0.56669437 | 0.57092184 | 0.88578992 |
| EMC3 | 49.2438152 | 0.093412694 | 0.28586479 | 0.32677229 | 0.74384012 | 0.94546379 |
| BRK1 | 43.8215617 | 0.123403524 | 0.38222906 | 0.32285228 | 0.74680712 | 0.94618979 |
| VHL | 26.2500397 | 0.572721415 | 0.37311703 | 1.53496455 | 0.12479254 | 0.54379497 |
| SEC13 | 38.0356703 | -0.039580845 | 0.33247053 | -0.1190507 | 0.9052352 | 0.9810929 |
| ATG7 | 215.894626 | -0.243593904 | 0.21512712 | -1.1323254 | 0.25749766 | 0.71022118 |
| VGLL4 | 263.516358 | 0.034738441 | 0.21663434 | 0.16035519 | 0.87260128 | 0.97434023 |
| TAMM41 | 58.9934191 | 0.010171918 | 0.21758466 | 0.04674924 | 0.96271308 | 0.99306991 |
| PPARG | 283.521728 | 0.057819238 | 0.28475314 | 0.2030504 | 0.83909564 | 0.96748214 |
| TSEN2 | 44.4005185 | 0.090612512 | 0.32903037 | 0.27539255 | 0.78301467 | 0.95544332 |
| MKRN2 | 15.4864254 | 0.020463387 | 0.36559011 | 0.05597358 | 0.95536286 | 0.99228059 |
| RAF1 | 121.460211 | 0.067396642 | 0.22816313 | 0.29538797 | 0.76769752 | 0.95205353 |
| RPL32 | 231.066049 | 0.370435316 | 0.44297416 | 0.83624588 | 0.40301659 | 0.80924894 |
| IQSEC1 | 100.550171 | 0.072527236 | 0.44660845 | 0.16239558 | 0.87099435 | 0.97434023 |
| NUP210 | 31.8440788 | 0.18411614 | 0.44113889 | 0.41736547 | 0.67641111 | 0.92229234 |
| HDAC11 | 17.7520416 | 0.365249668 | 0.4771329 | 0.76550929 | 0.44396833 | 0.83006558 |
| XPC | 48.7804718 | -0.241469011 | 0.23326975 | -1.0351493 | 0.30059921 | 0.74458947 |
| LSM3 | 36.8231951 | -0.439803267 | 0.3878527 | -1.1339441 | 0.25681801 | 0.70972499 |
| SLC6A6 | 78.2420864 | -2.247288597 | 0.49027188 | -4.58376 | 4.57E-06 | 0.00063284 |
| GRIP2 | 54.8071733 | 1.050324846 | 0.73084218 | 1.43714317 | 0.15067732 | 0.58759787 |
| CCDC174 | 40.2712718 | -0.060969706 | 0.21606339 | -0.2821843 | 0.77780216 | 0.95526475 |
| FGD5-AS1 | 80.1562547 | 0.291013403 | 0.22720317 | 1.280851 | 0.20024601 | 0.65330283 |
| NR2C2 | 92.130876 | 0.091021096 | 0.26698519 | 0.3409219 | 0.73316238 | 0.94269288 |
| MRPS25 | 99.4061224 | -0.095465888 | 0.22396179 | -0.4262597 | 0.66991862 | 0.92120933 |
| RBSN | 31.7782742 | -0.137345171 | 0.2984941 | -0.4601269 | 0.64542512 | 0.91225913 |
| CAPN7 | 170.870771 | -0.147498915 | 0.20765052 | -0.7103229 | 0.47750394 | 0.84771464 |
| SH3BP5-AS1 | 11.6283122 | 0.473458048 | 0.49435918 | 0.95772075 | 0.33820359 | NA |
| METTL6 | 18.4092619 | 0.090251378 | 0.34142014 | 0.26434111 | 0.79151709 | 0.95746995 |
| EAF1 | 15.9643856 | 0.273105793 | 0.4228311 | 0.64589807 | 0.51834537 | 0.86552266 |
| COLQ | 14.4343223 | -0.402935757 | 0.62154423 | -0.6482817 | 0.51680276 | 0.86500999 |
| HACL1 | 45.9216113 | -0.42654141 | 0.22997863 | -1.8547002 | 0.06363905 | 0.40032348 |
| BTD | 19.3476376 | 0.167001318 | 0.32175703 | 0.51902928 | 0.60374032 | 0.8956362 |
| ANKRD28 | 300.848349 | -0.266764199 | 0.23068492 | -1.1564007 | 0.2475173 | 0.70202275 |
| DPH3 | 11.8073761 | 0.194400102 | 0.59913698 | 0.32446687 | 0.7455846 | NA |
| OXNAD1 | 99.217619 | 0.410406093 | 0.27708369 | 1.48116292 | 0.13856316 | 0.57119278 |
| PLCL2 | 310.642758 | -0.137328526 | 0.4435634 | -0.3096029 | 0.75686293 | 0.94852331 |
| TBC1D5 | 549.155362 | 0.143231804 | 0.29253716 | 0.48961918 | 0.6244034 | 0.90447745 |
| RAB5A | 96.6638695 | -0.38049161 | 0.19540504 | -1.9471945 | 0.05151142 | 0.36077232 |
| KAT2B | 71.6146062 | 0.546707441 | 0.37673154 | 1.45118574 | 0.14672815 | 0.58349206 |
| UBE2E2 | 272.439775 | 0.03543139 | 0.42455461 | 0.08345544 | 0.93348941 | 0.9881638 |
| UBE2E1 | 214.331442 | -0.207148508 | 0.21671487 | -0.9558574 | 0.33914429 | 0.77042145 |
| NKIRAS1 | 21.0081998 | -0.030921515 | 0.32983703 | -0.0937479 | 0.92530945 | 0.98672316 |
| RPL15 | 422.323987 | -0.3740239 | 0.3622721 | -1.0324391 | 0.30186644 | 0.74658452 |
| NR1D2 | 104.509563 | -0.470423976 | 0.28589427 | -1.6454473 | 0.09987759 | 0.48708529 |
| THRB | 577.656259 | 0.504811869 | 0.15696355 | 3.21610897 | 0.00129941 | 0.03866357 |
| AC092422.1 | 499.248652 | -0.412163483 | 0.64307514 | -0.6409259 | 0.52157086 | 0.86703098 |
| RARB | 344.532632 | -0.022210515 | 0.51531666 | -0.0431007 | 0.96562125 | 0.99334616 |
| TOP2B | 131.415899 | 0.181252115 | 0.22957811 | 0.78950085 | 0.42981933 | 0.82469033 |
| NGLY1 | 80.5840398 | 0.049507723 | 0.24754411 | 0.19999556 | 0.84148406 | 0.9677342 |
| OXSM | 15.7948561 | 0.466992832 | 0.45062672 | 1.03631856 | 0.30005355 | 0.74437099 |
| NEK10 | 63.7340518 | -0.40293713 | 0.49282488 | -0.8176071 | 0.41358156 | 0.81655631 |
| SLC4A7 | 72.3430767 | -1.12248656 | 0.61424198 | -1.8274338 | 0.06763457 | 0.41443799 |

| | | | | | | |
|---|---|---|---|---|---|---|
| CMC1 | 149.288779 | -0.339089717 | 0.26851845 | -1.2628172 | 0.20665487 | 0.65836247 |
| AZI2 | 53.0592918 | -0.190552485 | 0.3275069 | -0.5818274 | 0.56068295 | 0.88071368 |
| TGFBR2 | 50.5253464 | 1.034929486 | 0.28913735 | 3.5793697 | 0.00034442 | 0.0157922 |
| STT3B | 165.841472 | 0.482215619 | 0.22279737 | 2.16436855 | 0.03043608 | 0.27757967 |
| OSBPL10 | 191.184075 | -0.140861579 | 0.5147421 | -0.2736547 | 0.78435002 | 0.95544332 |
| GPD1L | 62.6619001 | 0.599588412 | 0.29185533 | 2.05440282 | 0.03993672 | 0.32015388 |
| CMTM8 | 23.8612243 | -0.812281013 | 0.33059463 | -2.4570303 | 0.01400909 | 0.17763157 |
| CMTM6 | 73.1100669 | -0.222579203 | 0.30430165 | -0.7314427 | 0.46450882 | 0.84133578 |
| DYNC1LI1 | 41.5047242 | -0.183569914 | 0.3376768 | -0.5436261 | 0.58669881 | 0.89059199 |
| CNOT10 | 65.5856418 | -0.101907197 | 0.22729271 | -0.4483523 | 0.65389899 | 0.9158679 |
| GLB1 | 97.3402288 | 0.253138377 | 0.17940602 | 1.41098038 | 0.1582504 | 0.59881912 |
| CRTAP | 57.5400706 | -0.118818208 | 0.30563817 | -0.3887545 | 0.69745778 | 0.9299437 |
| FBXL2 | 117.749125 | -0.198853202 | 0.23938695 | -0.8306769 | 0.40615621 | 0.8115705 |
| UBP1 | 55.1040505 | 0.197173865 | 0.23847791 | 0.8268014 | 0.40834964 | 0.81362627 |
| CLASP2 | 226.892239 | 0.107735336 | 0.19453982 | 0.55379581 | 0.5797186 | 0.88972503 |
| PDCD6IP | 202.341181 | -0.213552521 | 0.16670147 | -1.2810476 | 0.20017695 | 0.65330283 |
| TRANK1 | 15.2775871 | -1.382104032 | 0.82505821 | -1.6751594 | 0.09390296 | 0.4781144 |
| EPM2AIP1 | 44.0039997 | 0.012556716 | 0.30392037 | 0.04131581 | 0.96704413 | 0.99388939 |
| MLH1 | 32.7223404 | 0.040424816 | 0.24383476 | 0.16578775 | 0.86832399 | 0.97434023 |
| LRRFIP2 | 448.505001 | -0.625968245 | 0.30643699 | -2.0427307 | 0.0410791 | 0.32527942 |
| GOLGA4 | 442.4556 | -0.173826774 | 0.2122037 | -0.8191505 | 0.41270052 | 0.81556555 |
| ITGA9 | 34.8433501 | -0.802589847 | 0.47368381 | -1.6943578 | 0.09019732 | 0.47226726 |
| CTDSPL | 357.751328 | 0.099023991 | 0.31887385 | 0.31054284 | 0.75614819 | 0.94796399 |
| ACAA1 | 38.718684 | -0.000182752 | 0.31593984 | -0.0005784 | 0.99953847 | 0.99982181 |
| OXSR1 | 117.549462 | 0.134748237 | 0.27682261 | 0.48676745 | 0.62642314 | 0.90541055 |
| ACVR2B | 19.400995 | -0.062569586 | 0.37643133 | -0.1662178 | 0.86798555 | 0.97434023 |
| EXOG | 31.4002052 | -0.203715561 | 0.28031384 | -0.726741 | 0.46738463 | 0.84315249 |
| WDR48 | 46.5671664 | -0.238242457 | 0.26677254 | -0.8930546 | 0.37182791 | 0.7900012 |
| GORASP1 | 23.1995924 | -0.000536059 | 0.30384055 | -0.0017643 | 0.99859231 | 0.99982181 |
| CSRNP1 | 15.2552195 | -1.294469503 | 0.56820054 | -2.2781913 | 0.02271518 | 0.23692181 |
| SLC25A38 | 15.8614764 | -0.259499154 | 0.36693722 | -0.7072031 | 0.47944025 | 0.84833462 |
| RPSA | 226.907394 | 0.312783368 | 0.30889161 | 1.01259911 | 0.31125169 | 0.75450562 |
| MYRIP | 143.718783 | -0.592923671 | 0.35001512 | -1.6939944 | 0.09026634 | 0.47226726 |
| EIF1B | 52.8586597 | -0.359984278 | 0.4124087 | -0.8728824 | 0.3827272 | 0.79786242 |
| ENTPD3-AS1 | 18.9370938 | -0.110041826 | 0.40843315 | -0.2694243 | 0.78760317 | 0.95640344 |
| RPL14 | 291.299755 | 0.156599852 | 0.51609486 | 0.30343231 | 0.76156043 | 0.9499444 |
| ZNF621 | 35.0603825 | 0.197286945 | 0.23482044 | 0.84016086 | 0.4008182 | 0.80830282 |
| CTNNB1 | 118.433628 | 0.193617206 | 0.22620126 | 0.85595106 | 0.39202485 | 0.80360124 |
| ULK4 | 85.2035755 | 0.083854309 | 0.2822948 | 0.29704518 | 0.76643202 | 0.95205353 |
| TRAK1 | 312.005255 | -0.377803514 | 0.20944375 | -1.8038424 | 0.07125601 | 0.42274211 |
| SEC22C | 42.073614 | 0.265780511 | 0.22588487 | 1.17661934 | 0.23934747 | 0.69386447 |
| SS18L2 | 16.9722882 | 0.167116143 | 0.49496404 | 0.3376329 | 0.73563985 | 0.94318131 |
| NKTR | 287.95176 | 0.025300187 | 0.14492665 | 0.17457236 | 0.86141567 | 0.97302617 |
| AC006059.1 | 41.6714501 | -0.558321196 | 0.48775721 | -1.1446703 | 0.25234576 | 0.70621866 |
| AC099329.3 | 16.4854825 | -0.336861983 | 0.51032866 | -0.6600883 | 0.50919716 | 0.86281145 |
| SNRK | 91.700839 | -0.247397002 | 0.24804794 | -0.9973758 | 0.31858215 | 0.76052272 |
| ANO10 | 132.419777 | -0.370689274 | 0.30111643 | -1.2310496 | 0.2183043 | 0.67239244 |
| ABHD5 | 32.8726571 | -0.474596794 | 0.39588176 | -1.1988347 | 0.23059222 | 0.68523674 |
| TCAIM | 102.786225 | 0.40399248 | 0.24731903 | 1.63348727 | 0.10236656 | 0.49278193 |
| ZNF445 | 39.889596 | 0.712335184 | 0.30469313 | 2.33787738 | 0.01939361 | 0.21848783 |
| AC099669.1 | 53.0165536 | 0.248746133 | 0.41915782 | 0.59344266 | 0.55288494 | 0.87587522 |
| ZNF197 | 36.2571127 | -0.391923793 | 0.28262329 | -1.3867357 | 0.16552238 | 0.60594576 |
| KIAA1143 | 21.9291421 | -0.037970266 | 0.41165089 | -0.092239 | 0.92650815 | 0.98672316 |
| ZDHHC3 | 278.24964 | 0.223703003 | 0.25100253 | 0.89123805 | 0.37280147 | 0.7911123 |
| EXOSC7 | 203.74562 | -0.169976811 | 0.35535939 | -0.4783237 | 0.63241984 | 0.90805737 |
| CLEC3B | 12.1577343 | 1.753478517 | 0.81742457 | 2.14512579 | 0.03194279 | NA |
| CDCP1 | 68.6321449 | -1.394367638 | 0.48029518 | -2.9031473 | 0.00369433 | 0.07910593 |
| LARS2 | 69.8385381 | 0.223774807 | 0.28788315 | 0.77731125 | 0.43697516 | 0.82687287 |
| LIMD1 | 50.1324862 | -0.331446463 | 0.35954861 | -0.9218405 | 0.35661178 | 0.78241164 |
| SACM1L | 119.024477 | 0.385697196 | 0.25390664 | 1.51905127 | 0.12874959 | 0.55308726 |
| LZTFL1 | 20.7292311 | -0.033695361 | 0.43662482 | -0.0771723 | 0.93848644 | 0.98871562 |
| FYCO1 | 71.3760041 | -0.346003865 | 0.32497492 | -1.0647094 | 0.28700744 | 0.73803612 |
| LTF | 22.4861723 | -4.118857019 | 1.04298409 | -3.9491082 | 7.84E-05 | 0.00551374 |
| LRRC2 | 13.8813896 | 1.002505743 | 0.48612742 | 2.06222832 | 0.039186 | 0.3178129 |
| ALS2CL | 70.6007835 | -0.076800966 | 0.41129451 | -0.1867299 | 0.85187244 | 0.97053536 |
| CCDC12 | 97.0363021 | 0.230832004 | 0.19725521 | 1.17022005 | 0.24191242 | 0.69556484 |
| NBEAL2 | 45.2263332 | 0.320148251 | 0.4137464 | 0.77377895 | 0.43906153 | 0.82745796 |
| SETD2 | 372.430907 | 0.282356694 | 0.16845592 | 1.67614587 | 0.09370962 | 0.4781144 |
| KIF9-AS1 | 155.136237 | 0.206941086 | 0.30327908 | 0.6823454 | 0.49502057 | 0.85814183 |
| KIF9 | 11.3656515 | -0.195782307 | 0.39086728 | -0.500892 | 0.61644711 | NA |
| KLHL18 | 52.4391008 | 0.464961509 | 0.36015343 | 1.29100951 | 0.19670038 | 0.64685325 |
| PTPN23 | 30.0802258 | -0.688684108 | 0.40429186 | -1.703433 | 0.08848706 | 0.46851772 |
| SCAP | 70.3598122 | 0.626912679 | 0.21882088 | 2.86495816 | 0.00417064 | 0.08574324 |
| ELP6 | 24.6489778 | 0.141126585 | 0.3198255 | 0.44126121 | 0.65902391 | 0.91714948 |
| SMARCC1 | 244.736547 | -0.310852901 | 0.21506268 | -1.4454061 | 0.14834386 | 0.58517098 |
| DHX30 | 50.1809766 | 0.437361688 | 0.22454909 | 1.94773305 | 0.05144691 | 0.36077232 |
| MAP4 | 362.142175 | -0.34929466 | 0.19617669 | -1.7805106 | 0.07499244 | 0.43243667 |
| ZNF589 | 12.5425134 | 0.38341889 | 0.42414441 | 0.90398194 | 0.36600498 | 0.78793182 |
| NME6 | 16.6997031 | 0.282364381 | 0.36565517 | 0.77221492 | 0.43998714 | 0.82793978 |
| PLXNB1 | 55.9105402 | -0.433874112 | 0.27467698 | -1.5795794 | 0.11420321 | 0.51869892 |
| TMA7 | 82.7886614 | 0.172986768 | 0.42699273 | 0.40512813 | 0.68538333 | 0.92591798 |
| SHISA5 | 57.1336208 | -1.181405569 | 0.37855109 | -3.1208617 | 0.00180323 | 0.04805303 |
| PFKFB4 | 17.2544483 | -2.111978875 | 0.76036571 | -2.777583 | 0.00547648 | 0.10201977 |
| UQCRC1 | 83.9271467 | 0.308987689 | 0.32334383 | 0.95560102 | 0.33927385 | 0.77045469 |
| IP6K2 | 67.7429342 | 0.008892674 | 0.20832601 | 0.04268634 | 0.96595157 | 0.99334616 |
| PRKAR2A | 115.942027 | 0.31078243 | 0.2851626 | 1.08984287 | 0.27578237 | 0.72554623 |
| SLC25A20 | 15.958315 | 0.689414928 | 0.40494398 | 1.70249456 | 0.0886627 | 0.4687332 |
| ARIH2 | 133.64606 | 0.290130389 | 0.17888563 | 1.62187646 | 0.1048298 | 0.49918954 |
| P4HTM | 31.4906239 | 0.07947199 | 0.36859257 | 0.21560931 | 0.8292923 | 0.96669991 |
| WDR6 | 58.4639012 | 0.614256644 | 0.36235834 | 1.69516351 | 0.09004441 | 0.47226726 |
| DALRD3 | 13.9139235 | 0.640570423 | 0.4850965 | 1.32050102 | 0.1866678 | 0.63791877 |
| NDUFAF3 | 29.1436735 | -0.417490504 | 0.46419694 | -0.8993823 | 0.36844908 | 0.78877653 |
| IMPDH2 | 19.3470459 | 0.076636959 | 0.39775057 | 0.19267593 | 0.84721278 | 0.97027687 |
| QRICH1 | 64.2489658 | 0.26096078 | 0.21825075 | 1.19569247 | 0.23181659 | 0.68622616 |
| QARS | 27.6963237 | 0.909279099 | 0.29497068 | 3.0826084 | 0.00205195 | 0.05196888 |
| USP19 | 13.0669238 | 0.145363096 | 0.44427613 | 0.32719087 | 0.74352352 | 0.94546379 |
| LAMB2 | 27.4176655 | 0.369027582 | 0.28300208 | 1.3039748 | 0.19224217 | 0.6421543 |
| C3orf62 | 11.7030281 | -0.439924849 | 0.52101706 | -0.8443579 | 0.39846947 | NA |
| USP4 | 82.8246935 | 0.062137674 | 0.29461531 | 0.21091122 | 0.83295655 | 0.96701848 |
| RHOA | 178.13228 | -0.261233695 | 0.23661867 | -1.1040282 | 0.26958089 | 0.72046011 |
| TCTA | 16.2301608 | 0.399233714 | 0.50186778 | 0.7954958 | 0.42632515 | 0.82329929 |
| DAG1 | 71.3647185 | 0.362427163 | 0.24736956 | 1.46512436 | 0.14288699 | 0.57698886 |
| APEH | 37.4975758 | 0.326414249 | 0.41688443 | 0.78298498 | 0.43363593 | 0.82504445 |
| MST1 | 14.5414148 | 1.05429895 | 0.39996558 | 2.63597419 | 0.00838961 | 0.13244159 |
| RNF123 | 19.4521507 | 0.019616817 | 0.29850989 | 0.0657158 | 0.94760409 | 0.99149867 |
| IP6K1 | 103.850058 | 0.05264471 | 0.24640018 | 0.21365532 | 0.83081585 | 0.96701848 |
| RBM6 | 420.369082 | 0.273355834 | 0.18121584 | 1.50845442 | 0.13143826 | 0.55820979 |
| RBM5 | 148.951582 | 0.113058219 | 0.15652736 | 0.72229044 | 0.47011593 | 0.84415485 |
| SEMA3F | 63.3379298 | -0.439274242 | 0.49978672 | -0.8789234 | 0.37944281 | 0.79535699 |
| GNAI2 | 47.5291611 | -0.619561482 | 0.32853608 | -1.8858248 | 0.05931857 | 0.3871608 |
| SEMA3B | 20.825429 | 0.096190127 | 0.40926602 | 0.23503082 | 0.81418481 | 0.96286073 |
| IFRD2 | 15.6071519 | -0.357472928 | 0.41929211 | -0.852563 | 0.3939017 | 0.80438874 |
| HYAL1 | 22.2272764 | 0.157156782 | 0.47237782 | 0.33269298 | 0.73936605 | 0.94376279 |

| | | | | | | |
|---|---|---|---|---|---|---|
| HYAL2 | 16.6343433 | 0.117081224 | 0.44782425 | 0.26144458 | 0.79374968 | 0.95770265 |
| TUSC2 | 21.3151044 | -0.29060619 | 0.37426901 | -0.7764634 | 0.43747542 | 0.82687287 |
| TMEM115 | 12.8161095 | 0.109055986 | 0.38839831 | 0.28078388 | 0.77887617 | 0.95537097 |
| CACNA2D2 | 20.5172042 | -2.160265725 | 0.81237868 | -2.6591856 | 0.00783298 | 0.12855183 |
| HEMK1 | 16.5249551 | 0.69028862 | 0.3822707 | 1.80575863 | 0.07095605 | 0.42199491 |
| CISH | 37.4023336 | -0.343101995 | 0.3067353 | -1.1185605 | 0.26332768 | 0.71348562 |
| MAPKAPK3 | 35.6963518 | -0.295524502 | 0.25682948 | -1.1506643 | 0.24987038 | 0.70437161 |
| DOCK3 | 19.6298827 | -0.833760663 | 0.70593432 | -1.181074 | 0.23757333 | 0.69207518 |
| MANF | 41.4653747 | -0.536084081 | 0.32452113 | -1.6519235 | 0.09855015 | 0.48476221 |
| RBM15B | 16.1728697 | 0.076766679 | 0.52463571 | 0.14632378 | 0.8836658 | 0.97662748 |
| DCAF1 | 74.8546973 | -0.085285965 | 0.26503184 | -0.3217952 | 0.74760787 | 0.9462156 |
| RAD54L2 | 137.879135 | 0.093307429 | 0.28678584 | 0.32535578 | 0.74491182 | 0.94595436 |
| TEX264 | 66.6900554 | -0.007617268 | 0.3396367 | -0.0224277 | 0.98210679 | 0.99718289 |
| RPL29 | 287.770548 | -0.323055417 | 0.42286619 | -0.7639661 | 0.44488747 | 0.83035712 |
| ALAS1 | 25.2582139 | 0.042372657 | 0.35191091 | 0.12040734 | 0.90416047 | 0.98041914 |
| WDR82 | 66.4560655 | 0.216326522 | 0.21739068 | 0.99510488 | 0.31968526 | 0.76106474 |
| DNAH1 | 12.6613384 | 0.270273394 | 0.36502315 | 0.74042809 | 0.45904028 | 0.83645518 |
| BAP1 | 22.7078916 | 0.080267486 | 0.42327177 | 0.18963581 | 0.84959452 | 0.97027687 |
| PHF7 | 13.4807043 | 0.299596539 | 0.40076941 | 0.7475534 | 0.45472958 | 0.83385406 |
| NISCH | 53.7333227 | 0.242699276 | 0.23690627 | 1.02445271 | 0.30562149 | 0.74897897 |
| SMIM4 | 33.9252082 | 0.07549919 | 0.33693867 | 0.22407398 | 0.82269974 | 0.96521156 |
| PBRM1 | 181.583095 | -0.09845349 | 0.2189115 | -0.4497411 | 0.65289717 | 0.9158679 |
| GNL3 | 27.3800329 | 0.104296157 | 0.38973975 | 0.26760462 | 0.78900367 | 0.95715263 |
| GLT8D1 | 19.0621216 | 0.083186664 | 0.34950862 | 0.23801033 | 0.81187308 | 0.96256453 |
| SPCS1 | 94.6022757 | -0.154777201 | 0.4120545 | -0.3756231 | 0.70719709 | 0.93432317 |
| NEK4 | 40.6761663 | -0.09032742 | 0.30829553 | -0.2929897 | 0.76953001 | 0.95209708 |
| SFMBT1 | 44.3469821 | -0.018390878 | 0.22292532 | -0.0824979 | 0.93425076 | 0.9881638 |
| PRKCD | 68.3980243 | 0.906140607 | 0.39674425 | 2.2839414 | 0.02237498 | 0.23514329 |
| TKT | 87.25246 | -0.740975971 | 0.2566456 | -2.8871564 | 0.00388741 | 0.08179582 |
| DCP1A | 93.6552059 | 0.046423751 | 0.19346289 | 0.23996204 | 0.81035968 | 0.9623221 |
| CACNA1D | 390.422863 | -0.766367502 | 0.53681194 | -1.4276275 | 0.1533991 | 0.59258114 |
| CHDH | 38.0292816 | -0.549659089 | 0.40934939 | -1.3427627 | 0.17934883 | 0.62668718 |
| ACTR8 | 13.5831502 | 0.064649044 | 0.40435546 | 0.15988171 | 0.87297426 | 0.97434023 |
| SELENOK | 35.1265095 | -0.456615074 | 0.31330731 | -1.4574032 | 0.14500512 | 0.58002047 |
| CACNA2D3 | 18.3002045 | 0.032875408 | 0.54713845 | 0.06008609 | 0.95208707 | 0.99217529 |
| ERC2 | 194.8531 | 0.423554091 | 0.38366032 | 1.10398202 | 0.26960094 | 0.72046011 |
| CCDC66 | 102.217412 | 0.235159985 | 0.2394532 | 0.98207074 | 0.326065 | 0.76353952 |
| FAM208A | 111.216806 | 0.055209455 | 0.20165885 | 0.2737765 | 0.78425639 | 0.95544332 |
| ARHGEF3 | 396.634292 | -0.844983898 | 0.32833434 | -2.5735471 | 0.01006619 | 0.14771874 |
| APPL1 | 64.1469713 | 0.030221472 | 0.21856113 | 0.13827469 | 0.89002333 | 0.97782578 |
| PDE12 | 28.506945 | 0.069749984 | 0.29968463 | 0.23274461 | 0.81595972 | 0.96310651 |
| ARF4 | 68.9523292 | -0.074038362 | 0.34116127 | -0.2170187 | 0.8281938 | 0.96636016 |
| DENND6A | 72.6091835 | 0.09672653 | 0.236763 | 0.40853735 | 0.68287922 | 0.92474072 |
| DENND6A-A! | 17.6951257 | -1.323255182 | 0.42728355 | -3.0969018 | 0.00195555 | 0.05060915 |
| SLMAP | 226.634895 | -0.150598298 | 0.16939969 | -0.8890116 | 0.37399683 | 0.7923487 |
| FLNB | 361.305353 | -0.051961587 | 0.23341511 | -0.2226145 | 0.82383555 | 0.96539788 |
| ABHD6 | 38.418873 | 0.486421555 | 0.36914874 | 1.31768445 | 0.1876093 | 0.63942735 |
| RPP14 | 13.4682302 | 0.455943475 | 0.35470692 | 1.28540902 | 0.19864939 | 0.65097942 |
| PXK | 315.309949 | 0.950798436 | 0.28085008 | 3.38543049 | 0.00071067 | 0.0259153 |
| PDHB | 51.6117461 | 0.192178781 | 0.31717457 | 0.60590854 | 0.54457549 | 0.87247631 |
| KCTD6 | 17.7981614 | -0.719562986 | 0.4503824 | -1.5976712 | 0.11011617 | 0.5105769 |
| FHIT | 1353.80637 | 0.110660969 | 0.25315149 | 0.43713338 | 0.66201462 | 0.91810778 |
| PTPRG | 1055.67056 | 0.153615494 | 0.62660283 | 0.24515608 | 0.80633558 | 0.96005568 |
| THOC7 | 33.2137246 | 0.223307597 | 0.26147668 | 0.8540249 | 0.39309119 | 0.80400536 |
| ATXN7 | 23.9847341 | -0.057258566 | 0.31280289 | -0.18305 | 0.8547588 | 0.97152899 |
| PSMD6-AS2 | 14.2064105 | -0.352114339 | 0.54133636 | -0.6504539 | 0.51539908 | 0.86463667 |
| PSMD6 | 22.7503113 | -0.044322947 | 0.38280582 | -0.1157844 | 0.90782341 | 0.98250721 |
| ADAMTS9-A! | 51.2408371 | -0.092662631 | 0.46937748 | -0.197416 | 0.843502 | 0.96954253 |
| MAGI1 | 9871.07419 | -0.511187978 | 0.25762616 | -1.9842239 | 0.04723088 | 0.3468127 |
| MAGI1-AS1 | 175.233841 | -1.387828367 | 0.56668789 | -2.4490172 | 0.01432466 | 0.17998604 |
| SLC25A26 | 70.0400877 | -0.118991653 | 0.27011624 | -0.4405202 | 0.6595604 | 0.91732025 |
| LRIG1 | 22.6168957 | -0.516351736 | 0.66789997 | -0.7730974 | 0.43946473 | 0.82760292 |
| SUCLG2 | 238.425999 | 0.102594177 | 0.22526388 | 0.45543999 | 0.64879273 | 0.91417939 |
| SUCLG2-AS1 | 74.292053 | 0.721807567 | 0.30770605 | 2.34576983 | 0.01898783 | 0.21617599 |
| FAM19A1 | 18.2872449 | 1.03083486 | 0.48884412 | 2.10871894 | 0.03496885 | 0.29911626 |
| EOGT | 22.784742 | -0.261779035 | 0.37806106 | -0.6924253 | 0.4886703 | 0.85461689 |
| TMF1 | 85.2621799 | -0.012667424 | 0.24315488 | -0.0520961 | 0.95845211 | 0.99238848 |
| UBA3 | 84.4395228 | 0.144034249 | 0.24268511 | 0.59350262 | 0.55284483 | 0.87587522 |
| ARL6IP5 | 102.180524 | 0.052791852 | 0.31407492 | 0.16808681 | 0.86651498 | 0.97434023 |
| FRMD4B | 16.7201331 | -0.445106549 | 0.43712917 | -1.0182495 | 0.30855941 | 0.75132915 |
| MITF | 338.770354 | 0.547742069 | 0.21288033 | 2.57300464 | 0.01008199 | 0.14772698 |
| FOXP1 | 1878.42348 | -0.085519995 | 0.23475246 | -0.3642986 | 0.71563503 | 0.93689626 |
| EIF4E3 | 19.7052615 | -0.155241504 | 0.34925586 | -0.4444922 | 0.65668677 | 0.9158679 |
| RYBP | 109.87254 | 0.085663811 | 0.2030143 | 0.4219595 | 0.67305458 | 0.92189158 |
| SHQ1 | 45.3136595 | 0.139984048 | 0.29092262 | 0.48117278 | 0.6303937 | 0.90764716 |
| GXYLT2 | 23.8269586 | -1.096517941 | 0.53187484 | -2.061609 | 0.03924498 | 0.31802531 |
| PPP4R2 | 125.281965 | 0.305011551 | 0.21240098 | 1.43601763 | 0.15099733 | 0.58822251 |
| PDZRN3 | 35.3316284 | 1.52310856 | 0.63696271 | 2.39120523 | 0.01679316 | 0.20115876 |
| CNTN3 | 79.6928348 | -0.370271067 | 0.27063036 | -1.3681801 | 0.17125571 | 0.61479659 |
| ZNF717 | 50.0409278 | 0.073168583 | 0.31393879 | 0.2330664 | 0.81570985 | 0.96300833 |
| GBE1 | 107.991641 | -0.532950913 | 0.45610326 | -1.1684874 | 0.24261022 | 0.69621873 |
| CHMP2B | 47.7674533 | -0.423738769 | 0.33120236 | -1.2793954 | 0.20075786 | 0.65358156 |
| CGGBP1 | 126.757361 | -0.096226344 | 0.24670648 | -0.3900438 | 0.69650413 | 0.9295666 |
| ZNF654 | 126.222272 | -0.381091307 | 0.23579558 | -1.6161936 | 0.10605246 | 0.50162992 |
| C3orf38 | 23.91381 | 0.489623768 | 0.27358623 | 1.78965061 | 0.0735101 | 0.42788861 |
| ARL13B | 36.3696388 | -0.015874558 | 0.2640502 | -0.0601195 | 0.95206048 | 0.99217529 |
| NSUN3 | 48.1891315 | 0.045962396 | 0.23636226 | 0.19445742 | 0.84581773 | 0.97027687 |
| MTRNR2L12 | 317.465453 | 0.224259133 | 0.45700359 | 0.49071635 | 0.62362708 | 0.90447745 |
| ARL6 | 87.6494661 | 0.284467247 | 0.27609352 | 1.03032933 | 0.30285544 | 0.74712557 |
| CRYBG3 | 399.200316 | -0.738673106 | 0.31133634 | -2.3725888 | 0.01766392 | 0.20618531 |
| AC110491.2 | 13.1335178 | -1.308568876 | 0.73249089 | -1.7864644 | 0.0740241 | 0.42893297 |
| RIOX2 | 86.7869359 | -0.599454261 | 0.39118047 | -1.5324238 | 0.12541789 | 0.54505746 |
| CLDND1 | 29.2528774 | 0.349133774 | 0.36866971 | 0.94700964 | 0.34363387 | 0.77370523 |
| ST3GAL6 | 333.248007 | 0.304948722 | 0.37890598 | 0.8048137 | 0.4209272 | 0.81954914 |
| DCBLD2 | 56.2400085 | 0.479016061 | 0.22628962 | 2.11682734 | 0.0342745 | 0.29499795 |
| CMSS1 | 139.144215 | 0.342580499 | 0.26476374 | 1.29391019 | 0.19569644 | 0.64610467 |
| FILIP1L | 35.2426838 | -0.546522135 | 0.43579048 | -1.2540938 | 0.20980792 | 0.66164938 |
| TBC1D23 | 69.0272979 | -0.076062239 | 0.25712569 | -0.2958173 | 0.76736957 | 0.95205353 |
| NIT2 | 18.094055 | -0.300825243 | 0.33367352 | -0.9015556 | 0.36729297 | 0.78856614 |
| TOMM70 | 47.9747591 | 0.275711624 | 0.21265157 | 1.29654171 | 0.19478892 | 0.6459894 |
| TFG | 136.285821 | -0.122343061 | 0.2277489 | -0.537184 | 0.59114054 | 0.89149106 |
| SENP7 | 121.39304 | -0.190406626 | 0.29253919 | -0.6508756 | 0.51512679 | 0.86463667 |
| TRMT10C | 25.8469485 | -0.836094181 | 0.46558748 | -1.7957832 | 0.072529 | 0.42448303 |
| PCNP | 104.238898 | -0.175579229 | 0.22100499 | -0.7944582 | 0.42692874 | 0.82375285 |
| ZBTB11 | 46.1497197 | 0.208550386 | 0.29149114 | 0.71546046 | 0.47432457 | 0.84593347 |
| RPL24 | 138.361794 | 0.236836033 | 0.3948418 | 0.59982512 | 0.54862279 | 0.87371493 |
| CEP97 | 20.2862002 | 0.567104206 | 0.44452328 | 1.27575817 | 0.20204102 | 0.6545025 |
| NFKBIZ | 31.3455995 | -1.241340741 | 0.54325973 | -2.2849857 | 0.02231366 | 0.2347533 |
| ZPLD1 | 80.3561597 | -3.785596568 | 0.68006026 | -5.5665604 | 2.60E-08 | 9.00E-06 |
| ALCAM | 38.1509306 | -1.463104658 | 0.66455296 | -2.2016374 | 0.02769093 | 0.26179535 |
| CBLB | 143.685951 | -0.637740107 | 0.28084113 | -2.2708216 | 0.02315778 | 0.23879078 |
| BBX | 414.980437 | -0.573702261 | 0.25821782 | -2.2217764 | 0.02629842 | 0.25458548 |
| CD47 | 95.9949234 | -0.965825901 | 0.32113284 | -3.0075588 | 0.00263355 | 0.06230597 |

| | | | | | | |
|---|---|---|---|---|---|---|
| IFT57 | 48.5647115 | -0.368490532 | 0.40696662 | -0.9054564 | 0.36522365 | 0.78793182 |
| DZIP3 | 73.0911513 | -0.314500446 | 0.50470147 | -0.6231415 | 0.53319152 | 0.86909171 |
| NECTIN3 | 36.1436108 | 0.251363066 | 0.30130234 | 0.83425526 | 0.40413716 | 0.81027913 |
| PHLDB2 | 393.040178 | 0.094725434 | 0.22721544 | 0.416897 | 0.67675375 | 0.92237057 |
| ABHD10 | 37.284201 | 0.308615698 | 0.33357402 | 0.92517906 | 0.35487277 | 0.7810615 |
| C3orf52 | 31.7013465 | -0.40754883 | 0.58448739 | -0.6972757 | 0.48563029 | 0.85356796 |
| SLC9C1 | 44.7377831 | -0.110215221 | 0.72800094 | -0.1513943 | 0.87966466 | 0.97581687 |
| CD200 | 27.0000434 | -0.54539384 | 0.76981325 | -0.7084755 | 0.47865001 | 0.84833462 |
| AC092894.1 | 69.5716593 | 0.254008381 | 0.45176588 | 0.56225668 | 0.57394115 | 0.88637538 |
| ATG3 | 35.7541707 | 0.039371443 | 0.30855123 | 0.12760099 | 0.89846475 | 0.97908342 |
| SLC35A5 | 24.6613409 | -0.393848732 | 0.35596395 | -1.1064287 | 0.268541 | 0.72009952 |
| GTPBP8 | 21.9174477 | 0.173436712 | 0.37650484 | 0.46064935 | 0.6450502 | 0.9122579 |
| NEPRO | 38.9517375 | -0.591239597 | 0.26005729 | -2.2734975 | 0.02299622 | 0.23831548 |
| CFAP44 | 35.7397665 | 0.644731633 | 0.27235859 | 2.36721611 | 0.01792246 | 0.20770358 |
| SPICE1 | 42.4138201 | 0.349308797 | 0.26829842 | 1.3019413 | 0.19293645 | 0.64373732 |
| SIDT1 | 14.7293892 | -1.489997184 | 0.76173868 | -1.9560477 | 0.05045954 | 0.3580523 |
| USF3 | 80.1744054 | -0.091416332 | 0.28012771 | -0.3263381 | 0.7441686 | 0.94546379 |
| NAA50 | 142.214257 | -0.325581731 | 0.22097862 | -1.473363 | 0.14065318 | 0.57494615 |
| ATP6V1A | 889.306067 | 0.385230085 | 0.25881423 | 1.48844243 | 0.13663425 | 0.56808927 |
| GRAMD1C | 220.721783 | -0.543313105 | 0.22788106 | -2.384196 | 0.01711648 | 0.20380223 |
| ZDHHC23 | 35.6600293 | 0.363391097 | 0.28473711 | 1.27623371 | 0.20187292 | 0.6545025 |
| CCDC191 | 42.7557422 | -0.513455108 | 0.37593687 | -1.3658014 | 0.1720013 | 0.61610509 |
| QTRT2 | 29.4202039 | 0.282931536 | 0.2869157 | 0.98611382 | 0.32407726 | 0.76304922 |
| ZBTB20 | 4900.96819 | -0.016114775 | 0.27825876 | -0.0579129 | 0.953818 | 0.99217529 |
| ZBTB20-AS1 | 24.0149501 | -0.448384041 | 0.64688736 | -0.6931408 | 0.48822118 | 0.85440032 |
| ZBTB20-AS5 | 102.96786 | -0.551297671 | 0.63067318 | -0.8741416 | 0.38204114 | 0.79731722 |
| ZBTB20-AS2 | 30.6284389 | -0.557361494 | 0.64070339 | -0.8699213 | 0.38434344 | 0.79852888 |
| LSAMP | 20.38397 | -0.090357485 | 0.40311264 | -0.2241495 | 0.822641 | 0.96521156 |
| B4GALT4 | 20.4459059 | 0.12270728 | 0.33820707 | 0.36281702 | 0.71674158 | 0.93689626 |
| ARHGAP31 | 21.9494162 | 0.523115259 | 0.50805316 | 1.02964671 | 0.30317588 | 0.74772592 |
| TMEM39A | 27.2961357 | 0.508178526 | 0.29740217 | 1.708725 | 0.0875019 | 0.46528247 |
| POGLUT1 | 15.1264471 | 0.491148339 | 0.4232483 | 1.16042601 | 0.2458754 | 0.69938066 |
| TIMMDC1 | 48.6670858 | -0.043288487 | 0.28485484 | -0.1519668 | 0.8792131 | 0.97567183 |
| COX17 | 40.9280254 | 0.160424736 | 0.32456524 | 0.49427578 | 0.62111143 | 0.90380166 |
| GSK3B | 295.848522 | 0.043072926 | 0.19302596 | 0.22314577 | 0.82342206 | 0.96539788 |
| LRRC58 | 36.1362213 | 0.118650964 | 0.41235529 | 0.28773964 | 0.77354604 | 0.95457609 |
| NDUFB4 | 62.769505 | -0.089964288 | 0.37936994 | -0.2371413 | 0.81254716 | 0.96262567 |
| HGD | 198.977268 | 0.226943067 | 0.54674038 | 0.41508379 | 0.67808057 | 0.9226233 |
| RABL3 | 34.8393124 | 0.201345237 | 0.29070005 | 0.69262196 | 0.48854682 | 0.85461689 |
| GTF2E1 | 11.8931286 | 0.055255 | 0.39733248 | 0.1390649 | 0.88939887 | NA |
| GOLGB1 | 218.757235 | -0.104772584 | 0.23912271 | -0.4381541 | 0.66127461 | 0.91804261 |
| IQCB1 | 80.2967413 | -0.246491141 | 0.20938967 | -1.1771886 | 0.23912023 | 0.69356592 |
| ILDR1 | 22.8518193 | -0.015846813 | 0.41809374 | -0.0379025 | 0.96976539 | 0.99486579 |
| CCDC58 | 24.1329714 | -0.191243204 | 0.35730244 | -0.5352418 | 0.59248266 | 0.89198431 |
| FAM162A | 151.605962 | -0.167659506 | 0.37935267 | -0.4419621 | 0.65851663 | 0.91683813 |
| AC083798.2 | 20.102142 | -0.056930072 | 0.40727015 | -0.1397845 | 0.88883022 | 0.97765744 |
| KPNA1 | 121.971309 | 0.159444904 | 0.20981039 | 0.7599476 | 0.44728591 | 0.83066268 |
| PARP9 | 87.1063025 | -1.088633148 | 0.60096107 | -1.811487 | 0.07006551 | 0.41948432 |
| DTX3L | 42.5495899 | -1.242437581 | 0.41425123 | -2.9992369 | 0.00270657 | 0.06326193 |
| PARP14 | 275.347037 | -0.974444881 | 0.65418213 | -1.4895621 | 0.13633942 | 0.5673498 |
| HSPBAP1 | 27.6590849 | -0.997246793 | 0.40015352 | -2.4921605 | 0.01269687 | 0.16756408 |
| DIRC2 | 59.9068338 | -0.88511701 | 0.39157202 | -2.2604195 | 0.02379523 | 0.2419431 |
| PDIA5 | 12.8123611 | -0.071300479 | 0.35684202 | -0.1998096 | 0.84162945 | 0.9677342 |
| SEC22A | 93.9446731 | -0.064809222 | 0.22729491 | -0.2851327 | 0.77554244 | 0.95493992 |
| ADCY5 | 114.460831 | -1.037815346 | 0.44768829 | -2.3181651 | 0.02044035 | 0.22365361 |
| HACD2 | 80.3215542 | 0.546230645 | 0.28566297 | 1.91215068 | 0.05585687 | 0.37625808 |
| MYLK-AS1 | 12.2087027 | 0.629669663 | 0.4000393 | 1.5740195 | 0.11548294 | NA |
| MYLK | 34.0409346 | -1.448427617 | 0.52211093 | -2.774176 | 0.00553417 | 0.10218578 |
| CCDC14 | 56.8700089 | 0.307112672 | 0.33254534 | 0.9235212 | 0.35573566 | 0.78153728 |
| KALRN | 24.7651797 | 0.07198139 | 0.43593616 | 0.16511911 | 0.86885023 | 0.97434023 |
| UMPS | 22.5844396 | 0.703071222 | 0.35875795 | 1.95973701 | 0.05002654 | 0.35682229 |
| ITGB5 | 39.5082215 | -0.373213335 | 0.34124625 | -1.0936775 | 0.27409646 | 0.72510729 |
| ZNF148 | 260.335197 | 0.262419556 | 0.15733252 | 1.66792951 | 0.09532972 | 0.48081904 |
| SNX4 | 111.314069 | 0.124051105 | 0.19078614 | 0.65021027 | 0.51555641 | 0.86463667 |
| OSBPL11 | 36.860205 | 0.046438541 | 0.30992863 | 0.14983624 | 0.88089382 | 0.97624626 |
| SLC41A3 | 20.1933319 | -0.035760646 | 0.32184423 | -0.1111117 | 0.9115278 | 0.9835359 |
| ALDH1L1 | 44.0770691 | 0.147827923 | 0.56771747 | 0.26038995 | 0.794563 | 0.95786532 |
| ZXDC | 104.503327 | 0.144843016 | 0.22396701 | 0.64671586 | 0.51781586 | 0.86500999 |
| TXNRD3NB | 12.5136203 | 1.088427264 | 0.50345228 | 2.16192738 | 0.03062377 | NA |
| TXNRD3 | 15.6405142 | 0.143407081 | 0.32179945 | 0.44564116 | 0.65585647 | 0.9158679 |
| CHCHD6 | 29.0521893 | 0.205963117 | 0.3269548 | 0.6299437 | 0.52873142 | 0.86750588 |
| TPRA1 | 18.9539851 | 0.55164782 | 0.3293361 | 1.6750299 | 0.09392837 | 0.4781144 |
| ABTB1 | 13.1272279 | -0.158516148 | 0.49025006 | -0.3233373 | 0.74643978 | 0.94614106 |
| MGLL | 266.491849 | -0.461745887 | 0.36132554 | -1.2779221 | 0.2012769 | 0.65374585 |
| KBTBD12 | 149.177452 | -0.022745895 | 0.28428255 | -0.0800116 | 0.93622805 | 0.98871562 |
| SEC61A1 | 72.0205859 | 0.001177487 | 0.34504084 | 0.0034126 | 0.99727714 | 0.99982181 |
| RUVBL1 | 37.6400491 | 0.011079522 | 0.2939808 | 0.03768791 | 0.96993652 | 0.99486579 |
| EEFSEC | 71.4278563 | 0.32022394 | 0.26782304 | 1.19565492 | 0.23183125 | 0.68622616 |
| GATA2 | 55.7307526 | 0.821008244 | 0.49497346 | 1.65869144 | 0.09717798 | 0.48257296 |
| GATA2-AS1 | 48.1172292 | -0.344072534 | 0.31887855 | -1.0790081 | 0.28058412 | 0.73241819 |
| RPN1 | 69.8891532 | 0.233366052 | 0.282897 | 0.82491525 | 0.40941971 | 0.81413754 |
| RAB7A | 204.278724 | -0.271476698 | 0.30038748 | -0.903755 | 0.36612531 | 0.78793182 |
| ACAD9 | 34.2811653 | 0.021421039 | 0.25917121 | 0.08265208 | 0.93412819 | 0.9881638 |
| CNBP | 113.035016 | -0.789492091 | 0.382397 | -2.0645875 | 0.03896204 | 0.31679112 |
| COPG1 | 40.1358486 | -0.027738003 | 0.30083977 | -0.0922019 | 0.92653762 | 0.98672316 |
| HMCES | 30.7123003 | 0.079138993 | 0.26716608 | 0.29621647 | 0.76706477 | 0.95205353 |
| H1FX | 59.267748 | -1.627509554 | 0.62076921 | -2.6217627 | 0.00874763 | 0.13576324 |
| MBD4 | 64.1385982 | -0.260503293 | 0.25819768 | -1.0089296 | 0.31300838 | 0.75583304 |
| IFT122 | 98.564954 | -0.296596601 | 0.4049768 | -0.7323792 | 0.46393713 | 0.84068563 |
| TMCC1 | 445.388386 | -0.113397367 | 0.2355068 | -0.4815036 | 0.63015864 | 0.90764716 |
| PIK3R4 | 38.1506014 | 0.046211247 | 0.24393912 | 0.18943762 | 0.84974984 | 0.97027687 |
| ATP2C1 | 102.712523 | 0.120042585 | 0.29593097 | 0.40564388 | 0.68500428 | 0.92581857 |
| NEK11 | 51.4196997 | -0.503773396 | 0.46960187 | -1.072767 | 0.28337566 | 0.73495826 |
| NUDT16 | 38.8374293 | 0.353691838 | 0.31253058 | 1.13170314 | 0.25775926 | 0.71022118 |
| MRPL3 | 63.0405792 | 0.291653338 | 0.26665806 | 1.0937353 | 0.27407109 | 0.72510729 |
| DNAJC13 | 93.4963164 | -0.057199 | 0.20818844 | -0.2747463 | 0.78351117 | 0.95544332 |
| UBA5 | 58.2717763 | -0.400646349 | 0.30659555 | -1.3067585 | 0.19129474 | 0.6421543 |
| CDV3 | 65.1814234 | -0.957654201 | 0.48935392 | -1.9569767 | 0.05035021 | 0.35799473 |
| TOPBP1 | 28.992432 | -0.010803042 | 0.26242369 | -0.0411664 | 0.96716323 | 0.99390649 |
| SRPRB | 19.7628935 | 0.412482629 | 0.33759558 | 1.22182473 | 0.22177391 | 0.67499434 |
| RYK | 163.026225 | -0.035696245 | 0.2134224 | -0.1672563 | 0.86716837 | 0.97434023 |
| AMOTL2 | 26.6760718 | 1.452447276 | 0.42775104 | 3.3955435 | 0.00068492 | 0.02520723 |
| ANAPC13 | 28.8866582 | 0.391198028 | 0.32001586 | 1.22243325 | 0.22154383 | 0.67492938 |
| CEP63 | 46.4477645 | -0.081833351 | 0.22859123 | -0.3579899 | 0.72035088 | 0.93806337 |
| PPP2R3A | 500.844864 | -0.360891746 | 0.29258678 | -1.233452 | 0.21740717 | 0.67202699 |
| MSL2 | 61.7879185 | 0.427337625 | 0.26791828 | 1.59502977 | 0.11070558 | 0.51233021 |
| PCCB | 58.8884037 | 0.281227221 | 0.26372527 | 1.06636431 | 0.28625899 | 0.73672385 |
| STAG1 | 419.306805 | 0.041832103 | 0.28620663 | 0.1461605 | 0.88379469 | 0.97662748 |
| NCK1 | 43.6458123 | -0.083749689 | 0.3507008 | -0.2388067 | 0.8112555 | 0.96247289 |
| DBR1 | 12.2136416 | 0.013221229 | 0.41347425 | 0.03197594 | 0.97449124 | NA |
| ARMC8 | 114.952407 | 0.262900474 | 0.17067742 | 1.54033541 | 0.12347862 | 0.54049755 |
| NME9 | 14.8776798 | -0.578658062 | 0.60615054 | -0.9546441 | 0.33975769 | 0.77073189 |
| MRAS | 12.3898513 | -1.388898326 | 0.55204364 | -2.5159212 | 0.01187217 | NA |

| | | | | | | |
|---|---|---|---|---|---|---|
| CEP70 | 160.74565 | -0.149328171 | 0.33824525 | -0.441479 | 0.65886627 | 0.91706167 |
| FAIM | 17.9024741 | -0.255859639 | 0.47261719 | -0.5413676 | 0.58825423 | 0.89073775 |
| PIK3CB | 105.145887 | 0.24882927 | 0.19655994 | 1.26592055 | 0.20554153 | 0.65836247 |
| MRPS22 | 15.8840206 | 0.118580029 | 0.38183775 | 0.31055082 | 0.75614212 | 0.94796399 |
| COPB2 | 78.0602367 | -0.131033626 | 0.27453712 | -0.4772893 | 0.63315613 | 0.90805737 |
| AC097103.2 | 63.2858618 | 0.198740474 | 0.40718236 | 0.48808715 | 0.62548811 | 0.90447745 |
| NMNAT3 | 34.1053356 | -0.108922635 | 0.27562316 | -0.3951868 | 0.69270503 | 0.928133 |
| SLC25A36 | 194.714475 | -0.329323123 | 0.24510135 | -1.3436202 | 0.17907124 | 0.62639418 |
| SPSB4 | 34.5653916 | 2.260868109 | 0.82348007 | 2.74550435 | 0.0060418 | 0.10913487 |
| PXYLP1 | 13.5998209 | 0.83204941 | 0.37266786 | 2.23268358 | 0.02556982 | 0.25205612 |
| ZBTB38 | 107.257749 | 0.099080919 | 0.20005653 | 0.49526462 | 0.62041334 | 0.90380166 |
| RASA2 | 154.277608 | -0.312193025 | 0.2128203 | -1.4669325 | 0.14239441 | 0.57673923 |
| RNF7 | 47.8662867 | -0.835476363 | 0.3213333 | -2.6000304 | 0.00932155 | 0.14127974 |
| ATP1B3 | 77.7523687 | -0.012554322 | 0.52196193 | -0.0240522 | 0.98081099 | 0.99705162 |
| TFDP2 | 135.830626 | 0.250329581 | 0.23554334 | 1.06277504 | 0.28788398 | 0.73803612 |
| GK5 | 111.248978 | 0.222599662 | 0.20299074 | 1.09660006 | 0.27281626 | 0.72363077 |
| XRN1 | 244.195694 | -0.254662379 | 0.26179834 | -0.9727425 | 0.33068129 | 0.76685572 |
| ATR | 169.514895 | -0.092535218 | 0.21333071 | -0.4337642 | 0.66445971 | 0.91900848 |
| PLS1 | 45.2805438 | 0.156688706 | 0.36682048 | 0.42715365 | 0.66926742 | 0.92049996 |
| TRPC1 | 60.5968838 | -0.134025046 | 0.56268634 | -0.2381878 | 0.81173541 | 0.96256453 |
| PCOLCE2 | 17.5947042 | 0.295850554 | 0.56933328 | 0.51964388 | 0.60331181 | 0.8956362 |
| U2SURP | 174.031066 | 0.181936267 | 0.18534535 | 0.98160686 | 0.32629357 | 0.76362437 |
| CHST2 | 29.0264702 | 0.512120974 | 0.53491748 | 0.95738314 | 0.33837391 | 0.76998177 |
| SLC9A9 | 19.2842653 | -0.504274963 | 0.5022914 | -1.003949 | 0.31540318 | 0.75786443 |
| C3orf58 | 19.5674383 | 0.320261681 | 0.35603011 | 0.89953537 | 0.36836756 | 0.78877653 |
| AC107021.1 | 52.1360481 | -0.581519329 | 0.7400656 | -0.7857673 | 0.43200384 | 0.8248269 |
| LNCSRLR | 14.2153237 | -0.713202167 | 0.60334968 | -1.182071 | 0.23717753 | 0.69207518 |
| PLOD2 | 384.895949 | -0.346877332 | 0.36241005 | -0.9571405 | 0.33849635 | 0.76998177 |
| PLSCR4 | 15.7474541 | -1.620542461 | 0.50939845 | -3.1812866 | 0.00146623 | 0.04146468 |
| PLSCR2 | 15.770199 | 0.005660092 | 0.48615722 | 0.01164251 | 0.99071083 | 0.99923337 |
| PLSCR1 | 216.105065 | -1.217917372 | 0.43908492 | -2.7737627 | 0.00554121 | 0.10218578 |
| AGTR1 | 26.2850121 | 0.761647109 | 2.09185819 | 0.36410074 | 0.71578279 | 0.93689626 |
| GYG1 | 51.1516688 | -0.407652828 | 0.37534967 | -1.0860615 | 0.27745179 | 0.72678432 |
| HLTF | 42.8989039 | 0.356179441 | 0.3724286 | 0.95636974 | 0.33888547 | 0.77015605 |
| HPS3 | 38.7811576 | -0.460572772 | 0.24130944 | -1.9086397 | 0.05630859 | 0.37709488 |
| WWTR1 | 224.32924 | -0.539809895 | 0.30212133 | -1.7867322 | 0.07398078 | 0.42893297 |
| COMMD2 | 28.5662401 | -0.100629683 | 0.3432917 | -0.2931317 | 0.76942148 | 0.95208424 |
| ANKUB1 | 85.7596727 | 0.056404851 | 0.64023994 | 0.08809955 | 0.92979755 | 0.98733028 |
| RNF13 | 142.422329 | -0.181139643 | 0.19829893 | -0.9134676 | 0.36099667 | 0.78616766 |
| PFN2 | 15.7421977 | 0.579865331 | 0.46496689 | 1.24711103 | 0.21235679 | 0.66468566 |
| AC117386.2 | 53.9601446 | 1.715987442 | 0.75955737 | 2.25919399 | 0.02387132 | 0.24214086 |
| TSC22D2 | 260.759841 | -0.094911217 | 0.28134319 | -0.3373503 | 0.73585283 | 0.94327639 |
| SERP1 | 112.019805 | -0.126595992 | 0.29791063 | -0.4249462 | 0.67087589 | 0.92170876 |
| EIF2A | 48.5903264 | -0.076954282 | 0.29129519 | -0.2641797 | 0.79164144 | 0.95746995 |
| SELENOT | 97.9619339 | 0.440981863 | 0.17091283 | 2.5801566 | 0.00987555 | 0.1471432 |
| SIAH2 | 25.7223222 | -0.514385336 | 0.43104462 | -1.1933459 | 0.23273393 | 0.68763908 |
| MBNL1 | 627.769484 | 0.04389275 | 0.23299425 | 0.18838555 | 0.85057442 | 0.97027687 |
| RAP2B | 28.5289967 | 0.593079087 | 0.6412068 | 0.92494198 | 0.35499609 | 0.7810615 |
| DHX36 | 152.334179 | -0.42616138 | 0.22181643 | -1.9212345 | 0.05470215 | 0.37236207 |
| PLCH1 | 164.40783 | 0.064516426 | 0.48890596 | 0.13196081 | 0.89501529 | 0.97862357 |
| GMPS | 55.9858754 | 0.210070475 | 0.23587541 | 0.89059928 | 0.37314418 | 0.7913202 |
| SSR3 | 46.1444815 | 0.063866561 | 0.31468417 | 0.20295447 | 0.83917062 | 0.96748214 |
| TIPARP | 15.9631524 | -0.553438439 | 0.33608451 | -1.646724 | 0.09961478 | 0.48648077 |
| LINC00886 | 79.2778134 | 0.177869776 | 0.36954138 | 0.48132573 | 0.63028501 | 0.90764716 |
| LEKR1 | 20.9282174 | 1.036063587 | 0.62743965 | 1.65125617 | 0.09868628 | 0.48476221 |
| CCNL1 | 132.959794 | -0.42889513 | 0.28360246 | -1.5123111 | 0.13045475 | 0.55685884 |
| VEPH1 | 13.7661851 | 1.450557693 | 0.40541867 | 3.57792525 | 0.00034633 | 0.0157922 |
| RSRC1 | 176.423433 | -0.23991957 | 0.16213871 | -1.479718 | 0.13894851 | 0.57182884 |
| MLF1 | 13.1911706 | 0.038962394 | 0.50285027 | 0.07748309 | 0.93823924 | 0.98871562 |
| GFM1 | 62.0944451 | 0.271722967 | 0.23662706 | 1.14831739 | 0.25083756 | 0.70535741 |
| MFSD1 | 31.4705088 | 0.156055896 | 0.34736227 | 0.44925978 | 0.65324427 | 0.9158679 |
| IQCJ-SCHIP1 | 77.8911854 | -1.158948043 | 0.75961872 | -1.5256971 | 0.12708533 | 0.54952841 |
| SMC4 | 23.9967202 | 0.390324698 | 0.42121457 | 0.92666477 | 0.35410062 | 0.78045353 |
| KPNA4 | 115.314128 | 0.103752385 | 0.29708495 | 0.34923474 | 0.72691308 | 0.94009907 |
| PPM1L | 314.740507 | -0.337271985 | 0.35474983 | -0.9507319 | 0.34174049 | 0.77203275 |
| B3GALNT1 | 12.744397 | -0.449338748 | 0.48347146 | -0.9294008 | 0.35268143 | 0.779984 |
| NMD3 | 70.3850488 | -0.268379455 | 0.24145108 | -1.1115273 | 0.26634146 | 0.71747544 |
| LINC01324 | 17.0393516 | 0.445098086 | 0.93589558 | 0.4755852 | 0.63436992 | 0.90805737 |
| LINC01322 | 12.8025674 | -0.201836475 | 0.49121615 | -0.4108914 | 0.68115219 | 0.92382219 |
| PDCD10 | 87.8307649 | -0.288718125 | 0.22767473 | -1.2681167 | 0.20475628 | 0.65790959 |
| GOLIM4 | 112.096905 | 1.060164299 | 0.17411258 | 6.08895876 | 1.14E-09 | 6.48E-07 |
| MECOM | 2528.44048 | 0.573522454 | 0.24015661 | 2.38811855 | 0.01693488 | 0.20205205 |
| MYNN | 25.6470238 | 0.313500668 | 0.35344911 | 0.88697541 | 0.37509214 | 0.79285112 |
| SEC62 | 162.662665 | 0.166996064 | 0.20395608 | 0.81878446 | 0.4129094 | 0.81572733 |
| GPR160 | 97.8697507 | -0.631261135 | 0.38455883 | -1.6415203 | 0.10068945 | 0.4893225 |
| PHC3 | 298.941612 | -0.152441333 | 0.23431439 | -0.6505846 | 0.51531468 | 0.86463667 |
| PRKCI | 215.151125 | -0.661738373 | 0.43988046 | -1.5043596 | 0.13248881 | 0.55997449 |
| SKIL | 46.8283901 | 0.185933219 | 0.29116826 | 0.63857654 | 0.52309844 | 0.86740339 |
| RPL22L1 | 13.0915136 | -0.668464601 | 0.55416296 | -1.20626 | 0.22771726 | 0.68265461 |
| TNIK | 93.1400871 | -0.851691169 | 0.8140011 | -1.0463022 | 0.29542152 | 0.74111836 |
| PLD1 | 97.9361396 | -1.591329626 | 0.48188079 | -3.3023305 | 0.00095885 | 0.03196167 |
| FNDC3B | 660.27973 | -0.769334345 | 0.25001578 | -3.0771431 | 0.00208995 | 0.05234522 |
| TNFSF10 | 57.625063 | -0.971339583 | 0.65022636 | -1.4938484 | 0.13521526 | 0.56489756 |
| NCEH1 | 132.779693 | -0.30316263 | 0.31534229 | -0.9613764 | 0.33636295 | 0.76884797 |
| ECT2 | 33.336254 | -0.22258843 | 0.45349552 | -0.4908283 | 0.6235479 | 0.90447745 |
| NLGN1 | 170.770053 | 1.580071606 | 0.47240793 | 3.34471861 | 0.00082366 | 0.028843 |
| NAALADL2 | 422.972452 | 0.332007406 | 0.32562632 | 1.01959635 | 0.30791994 | 0.7512664 |
| TBL1XR1 | 479.72771 | -0.234796049 | 0.25803995 | -0.9099213 | 0.36286402 | 0.78718296 |
| KCNMB2-AS1 | 37.859971 | 0.211057283 | 1.11666529 | 0.18900675 | 0.85008752 | 0.97027687 |
| ZMAT3 | 26.9863418 | -0.016823958 | 0.3310353 | -0.0508222 | 0.95946716 | 0.99238848 |
| PIK3CA | 106.057625 | -0.389694758 | 0.22698241 | -1.71685 | 0.08600658 | 0.46168448 |
| KCNMB3 | 16.319836 | -0.278799869 | 0.45165986 | -0.6172784 | 0.53705112 | 0.86963427 |
| ZNF639 | 20.1089758 | 0.029010027 | 0.3587633 | 0.08086119 | 0.93555235 | 0.98871562 |
| MFN1 | 88.8984668 | -0.351018461 | 0.22039568 | -1.5926739 | 0.11123336 | 0.51305924 |
| ACTL6A | 20.0302535 | 0.100954386 | 0.3686178 | 0.27387279 | 0.78418238 | 0.95544332 |
| MRPL47 | 26.2025917 | -0.099918108 | 0.32081006 | -0.3114557 | 0.75545425 | 0.94795939 |
| NDUFB5 | 79.2409918 | 0.149547773 | 0.38579347 | 0.38763687 | 0.69828478 | 0.93015138 |
| TTC14 | 93.5616668 | 0.644938006 | 0.22287729 | 2.89369106 | 0.00380743 | 0.08063762 |
| FXR1 | 118.986993 | 0.427101862 | 0.16823149 | 2.53877478 | 0.01112414 | 0.15593979 |
| DNAJC19 | 20.3395868 | -0.286269127 | 0.46582725 | -0.6145392 | 0.53885905 | 0.86972442 |
| ATP11B | 160.357285 | -0.163914147 | 0.166892 | -0.982157 | 0.32602252 | 0.76353952 |
| DCUN1D1 | 78.6139983 | 0.178714303 | 0.21320039 | 0.83824566 | 0.40189275 | 0.80890307 |
| MCCC1 | 227.341516 | 0.184950933 | 0.35467594 | 0.52146456 | 0.60204319 | 0.89526582 |
| MCF2L2 | 38.1522991 | -1.06790953 | 0.31182479 | -3.4247102 | 0.00061546 | 0.02349088 |
| B3GNT5 | 29.9246876 | -0.607196515 | 0.32365452 | -1.8760638 | 0.06064653 | 0.39165865 |
| KLHL24 | 181.617463 | -0.229567427 | 0.28446297 | -0.8070204 | 0.41965471 | 0.81899879 |
| YEATS2 | 86.0919141 | -0.362633768 | 0.33439444 | -1.0844492 | 0.27816569 | 0.72806455 |
| ABCC5 | 365.433647 | 0.556732076 | 0.25248451 | 2.20501481 | 0.02745306 | 0.26056227 |
| EIF2B5 | 25.4260061 | 0.130713575 | 0.32591766 | 0.40106318 | 0.68837361 | 0.9267526 |
| DVL3 | 35.6170933 | 0.46687664 | 0.32121597 | 1.45346648 | 0.14609428 | 0.58293481 |
| AP2M1 | 61.7586382 | -0.254517352 | 0.27255477 | -0.9338209 | 0.35039627 | 0.77868979 |
| ABCF3 | 12.0244725 | 0.602244421 | 0.46140879 | 1.30522961 | 0.19181467 | NA |
| PSMD2 | 40.2616277 | -0.229282694 | 0.25521838 | -0.8983785 | 0.36898382 | 0.78889478 |

| | | | | | | |
|---|---|---|---|---|---|---|
| EIF4G1 | 97.7329612 | -0.363069895 | 0.36160472 | -1.0040519 | 0.31535361 | 0.75786443 |
| FAM131A | 14.582046 | 0.487654395 | 0.38229579 | 1.27559448 | 0.20209891 | 0.6545025 |
| POLR2H | 48.9874275 | 0.360719112 | 0.34029395 | 1.06002211 | 0.28913454 | 0.73917583 |
| MAGEF1 | 20.3613695 | 0.276457787 | 0.42197667 | 0.65514947 | 0.51237153 | 0.86346419 |
| VPS8 | 137.7107 | 0.102385787 | 0.17780139 | 0.57584356 | 0.56472093 | 0.88191823 |
| EHHADH | 31.6760802 | 0.358718017 | 0.39366651 | 0.9112231 | 0.36217784 | 0.78714716 |
| MAP3K13 | 110.583225 | -0.113562272 | 0.24718964 | -0.4594136 | 0.64593721 | 0.91242041 |
| TMEM41A | 28.3433589 | 0.327079319 | 0.2850834 | 1.14731098 | 0.25125312 | 0.70545786 |
| LIPH | 116.381897 | -0.017171151 | 0.34313379 | -0.0500421 | 0.9600888 | 0.99238848 |
| SENP2 | 58.8767108 | 0.425526333 | 0.30394092 | 1.40002978 | 0.1615044 | 0.60060549 |
| IGF2BP2 | 196.360665 | -0.232549318 | 0.16392129 | -1.4186645 | 0.15599685 | 0.59597065 |
| TRA2B | 94.1366552 | 0.059008628 | 0.20720447 | 0.28478453 | 0.77580922 | 0.95493992 |
| TBCCD1 | 11.7055414 | 0.152425898 | 0.4432003 | 0.34392102 | 0.73090569 | NA |
| AC068631.2 | 15.6846243 | -0.775916772 | 0.55983313 | -1.3859787 | 0.16575342 | 0.60621113 |
| EIF4A2 | 143.278384 | 0.275643468 | 0.2826853 | 0.9750895 | 0.32951586 | 0.76685572 |
| ST6GAL1 | 573.469819 | -1.030748608 | 0.20397877 | -5.0532152 | 4.34E-07 | 0.00010484 |
| AC072022.1 | 17.6308195 | -1.430034094 | 0.74611709 | -1.9166349 | 0.05528432 | 0.37383273 |
| BCL6 | 215.735107 | -1.557614385 | 0.44933754 | -3.4664684 | 0.00052734 | 0.02087851 |
| LPP | 1521.90532 | -0.260133862 | 0.23846811 | -1.0908539 | 0.27533718 | 0.72554623 |
| TPRG1 | 24.4093397 | 0.183501957 | 0.30801174 | 0.59576287 | 0.55133365 | 0.87527601 |
| P3H2 | 199.578728 | 0.842133196 | 0.62237431 | 1.35309761 | 0.17602445 | 0.6225246 |
| CCDC50 | 198.823584 | -0.537522901 | 0.2369328 | -2.2686724 | 0.02328826 | 0.23879078 |
| FGF12 | 62.7651027 | 0.627792878 | 0.33889364 | 1.85247763 | 0.06395727 | 0.40111702 |
| MB21D2 | 20.5730456 | 0.551696923 | 0.56835637 | 0.97068838 | 0.33170348 | 0.76685572 |
| OPA1 | 144.566325 | 0.175957821 | 0.16881195 | 1.04233034 | 0.29725855 | 0.74160729 |
| HES1 | 21.7157693 | -1.258249987 | 0.55138625 | -2.2819756 | 0.02249078 | 0.23562337 |
| ATP13A3 | 58.6753217 | -0.155945126 | 0.34975188 | -0.4458736 | 0.65568856 | 0.9158679 |
| LSG1 | 30.3676302 | 0.154954699 | 0.3832653 | 0.40430141 | 0.68599109 | 0.92597882 |
| XXYLT1 | 27.3935476 | 0.607027249 | 0.39729078 | 1.52791677 | 0.1265332 | 0.54817868 |
| ACAP2 | 200.098864 | -0.095617082 | 0.15816407 | -0.6045436 | 0.54548227 | 0.87247631 |
| PPP1R2 | 72.5923728 | -0.431925529 | 0.24579752 | -1.7572412 | 0.0788767 | 0.44254372 |
| MUC20-OT1 | 1266.31049 | 0.142666889 | 0.37169497 | 0.38382787 | 0.70110603 | 0.93127962 |
| MUC20 | 15.5613486 | 0.677455513 | 0.33340239 | 2.03194555 | 0.04215917 | 0.32888549 |
| MUC4 | 12.7723852 | 0.006379172 | 0.64716024 | 0.00985718 | 0.99213524 | 0.99923337 |
| TNK2 | 21.044615 | -0.253274106 | 0.41662283 | -0.6079218 | 0.54323933 | 0.87233405 |
| TFRC | 190.036687 | -0.409321553 | 0.32401996 | -1.2632603 | 0.20649564 | 0.65836247 |
| LINC00885 | 27.6576349 | 0.375223088 | 0.82480352 | 0.45492421 | 0.64916377 | 0.91417939 |
| PCYT1A | 35.0630616 | -0.015070628 | 0.28160174 | -0.0535175 | 0.95731957 | 0.99236316 |
| UBXN7 | 78.0205399 | 0.056486383 | 0.22206864 | 0.25436453 | 0.79921395 | 0.95848876 |
| RNF168 | 65.5302872 | -0.367081322 | 0.2267232 | -1.6190726 | 0.10543164 | 0.50058098 |
| WDR53 | 11.2948068 | -0.022071955 | 0.39450695 | -0.0559482 | 0.95538307 | NA |
| PIGX | 19.4261299 | 0.248201068 | 0.2982193 | 0.83227701 | 0.40525261 | 0.81150914 |
| PAK2 | 133.608067 | -0.110596296 | 0.20587725 | -0.5371953 | 0.5911327 | 0.89149106 |
| SENP5 | 137.263485 | -0.26304235 | 0.26641726 | -0.9873322 | 0.32347979 | 0.76304922 |
| NCBP2 | 33.4406577 | -0.09095002 | 0.3058135 | -0.2974035 | 0.76615843 | 0.95205353 |
| NCBP2-AS2 | 17.315928 | -0.05763843 | 0.44886338 | -0.1284097 | 0.89782472 | 0.97892709 |
| DLG1 | 559.211007 | -0.140260798 | 0.23397103 | -0.5994793 | 0.54885329 | 0.87371493 |
| BDH1 | 32.8421481 | 0.714343621 | 0.3929142 | 1.81806517 | 0.06905417 | 0.41751716 |
| AC024560.1 | 59.4134046 | 3.593643909 | 1.46443452 | 2.45394647 | 0.0141298 | 0.17828198 |
| RUBCN | 48.0515359 | -0.136311339 | 0.31584571 | -0.4315757 | 0.66604981 | 0.9196702 |
| FYTTD1 | 67.6688035 | -0.002151193 | 0.25259443 | -0.0085164 | 0.99320499 | 0.99923337 |
| LRCH3 | 199.190361 | -0.086133443 | 0.17263812 | -0.4989248 | 0.61783235 | 0.90214869 |
| IQCG | 25.9952776 | 0.400131762 | 0.30240809 | 1.32315164 | 0.18578496 | 0.63633971 |
| RPL35A | 153.354411 | 0.35241278 | 0.38832791 | 0.9075134 | 0.36413537 | 0.78780828 |
| LMLN | 26.4299815 | 0.697092507 | 0.28541795 | 2.44235689 | 0.01459171 | 0.18286773 |
| ZNF595 | 35.8363355 | 0.070708337 | 0.27443857 | 0.25764723 | 0.79667917 | 0.95848876 |
| ZNF718 | 80.9497571 | 0.349671276 | 0.30255694 | 1.15572058 | 0.24779547 | 0.70240969 |
| ZNF141 | 58.0148085 | 0.353792296 | 0.20671479 | 1.71149965 | 0.08698892 | 0.46422617 |
| AC092574.2 | 15.9310765 | -0.840009951 | 0.52038175 | -1.6142187 | 0.10648 | 0.50162992 |
| ZNF721 | 203.85264 | 0.352714416 | 0.16779481 | 2.10205791 | 0.0355482 | 0.30204646 |
| PIGG | 44.6705076 | 0.229816752 | 0.2175248 | 1.05650826 | 0.29073607 | 0.73979353 |
| ATP5ME | 45.4789136 | 0.488071377 | 0.3180454 | 1.53459654 | 0.12488296 | 0.54394465 |
| SLC49A3 | 11.5017474 | -0.050911059 | 0.56418894 | -0.0902376 | 0.9280984 | NA |
| PCGF3 | 59.0097412 | 0.148773888 | 0.1926264 | 0.77234422 | 0.43991058 | 0.82793978 |
| AC139887.2 | 113.516866 | 0.434306951 | 0.34047981 | 1.27557329 | 0.2021064 | 0.6545025 |
| GAK | 115.0365 | -0.38971248 | 0.27390362 | -1.4228088 | 0.15479159 | 0.59498864 |
| TMEM175 | 17.783861 | 0.973913987 | 0.37985887 | 2.56388374 | 0.01035082 | 0.14940917 |
| FGFRL1 | 26.7619236 | 0.220856643 | 0.36240539 | 0.60941876 | 0.54224691 | 0.87177844 |
| RNF212 | 28.9507327 | 0.007023264 | 0.28491831 | 0.0246501 | 0.98033406 | 0.99687019 |
| AC092535.3 | 73.6317435 | -0.038766344 | 0.23764038 | -0.1631303 | 0.87041585 | 0.97434023 |
| CTBP1-AS | 19.6418997 | 0.672934895 | 0.60026007 | 1.12107223 | 0.26225712 | 0.71321158 |
| CTBP1 | 47.4675391 | 0.202939549 | 0.29597227 | 0.68567081 | 0.49292072 | 0.85714521 |
| MAEA | 77.5950226 | 0.024520814 | 0.18001078 | 0.13621859 | 0.89164848 | 0.97793768 |
| UVSSA | 53.897423 | 0.116097846 | 0.30984461 | 0.374697 | 0.70788583 | 0.93432317 |
| SLBP | 59.677502 | -0.074638767 | 0.26336901 | -0.2834 | 0.77687026 | 0.95512504 |
| TMEM129 | 32.0286536 | 0.764201457 | 0.28226497 | 2.70739036 | 0.00678145 | 0.11790324 |
| LETM1 | 38.4154312 | 0.314199494 | 0.34952762 | 0.89892607 | 0.36869204 | 0.78877653 |
| NSD2 | 120.621644 | 0.395864152 | 0.20065412 | 1.97286827 | 0.04851057 | 0.35179202 |
| NELFA | 54.3240088 | 0.218027528 | 0.29318051 | 0.74366312 | 0.45708031 | 0.83569193 |
| MXD4 | 56.9044247 | 0.715765259 | 0.32067821 | 2.23203587 | 0.02561259 | 0.25205612 |
| ZFYVE28 | 57.782282 | 0.206030556 | 0.51568247 | 0.39952988 | 0.68950281 | 0.92734996 |
| RNF4 | 58.4553112 | -0.127140075 | 0.2918923 | -0.4355719 | 0.66314738 | 0.918408 |
| FAM193A | 120.693909 | -0.038119266 | 0.24751509 | -0.1540078 | 0.87760355 | 0.97542218 |
| TNIP2 | 25.9001773 | -0.817635797 | 0.37630575 | -2.1727965 | 0.02979564 | 0.27473167 |
| SH3BP2 | 38.0913978 | -1.218120255 | 0.37536437 | -3.2451675 | 0.00117382 | 0.03637703 |
| ADD1 | 150.139171 | -0.012948781 | 0.19590138 | -0.0660985 | 0.94729943 | 0.99149867 |
| MFSD10 | 50.4288448 | -0.224497828 | 0.3489293 | -0.6433906 | 0.51997069 | 0.86676674 |
| NOP14-AS1 | 10.4963181 | 0.350870441 | 0.44838643 | 0.78251798 | 0.43391022 | NA |
| NOP14 | 16.6488947 | 0.17088992 | 0.49719123 | 0.34371065 | 0.7310639 | 0.94155624 |
| GRK4 | 10.7497121 | 0.511205863 | 0.51132909 | 0.99975902 | 0.31742714 | NA |
| HTT | 121.299875 | 0.210919258 | 0.20496617 | 1.02904426 | 0.30345888 | 0.74787746 |
| RGS12 | 39.1701359 | -0.314508293 | 0.39544111 | -0.7953353 | 0.42641846 | 0.82329929 |
| LRPAP1 | 65.7884253 | -0.188370305 | 0.29505228 | -0.6384303 | 0.52319364 | 0.86740339 |
| TMEM128 | 17.792554 | 0.39031381 | 0.31740482 | 1.22970349 | 0.21880816 | 0.67250923 |
| ZBTB49 | 11.3758172 | 0.262010075 | 0.45034142 | 0.58180319 | 0.56069926 | NA |
| STX18 | 196.89582 | 0.313019651 | 0.23353136 | 1.34037524 | 0.18012338 | 0.62768048 |
| STX18-AS1 | 66.2392636 | 0.246219866 | 0.30795934 | 0.79952069 | 0.42398855 | 0.82220891 |
| EVC2 | 55.484754 | -0.641325649 | 0.35589752 | -1.8019953 | 0.07154615 | 0.42281025 |
| EVC | 45.7758305 | -0.034655852 | 0.31256179 | -0.1108768 | 0.91171404 | 0.9835359 |
| CRMP1 | 13.2160747 | -0.080317373 | 0.60654172 | -0.1324185 | 0.89465325 | 0.97850297 |
| MAN2B2 | 22.9962094 | 0.007790504 | 0.45742573 | 0.01703119 | 0.98641173 | 0.99810751 |
| MRFAP1 | 64.5603397 | 0.035332134 | 0.38336119 | 0.09216409 | 0.92656767 | 0.98672316 |
| MRFAP1L1 | 19.4558905 | 0.132833964 | 0.36920988 | 0.35977901 | 0.7190124 | 0.93767415 |
| BLOC1S4 | 16.827258 | -0.33374914 | 0.42598404 | -0.783478 | 0.43334644 | 0.82501678 |
| KIAA0232 | 233.53541 | 0.12998317 | 0.21643201 | 0.60057277 | 0.54812458 | 0.87341177 |
| TBC1D14 | 742.538251 | 0.073993464 | 0.25748285 | 0.2873724 | 0.77382719 | 0.95457609 |
| TADA2B | 23.8222523 | 0.204483101 | 0.44158768 | 0.46306342 | 0.64331891 | 0.91192138 |
| GRPEL1 | 29.9012137 | -0.839016378 | 0.3800219 | -2.2078106 | 0.02725748 | 0.25997792 |
| SORCS2 | 69.8089074 | 1.098327467 | 1.09417121 | 1.00379854 | 0.31547573 | 0.75786443 |
| SH3TC1 | 124.268203 | -0.266051816 | 0.23321039 | -1.1408232 | 0.25394351 | 0.70701839 |
| ACOX3 | 27.2047997 | 0.156827742 | 0.28523479 | 0.54981982 | 0.58244296 | 0.89036744 |
| TRMT44 | 15.6015731 | 0.450681553 | 0.41433727 | 1.08771667 | 0.27672021 | 0.72584803 |
| SLC2A9 | 22.761709 | 0.388672221 | 0.41684447 | 0.93241543 | 0.35112187 | 0.77910174 |

| | | | | | | |
|---|---|---|---|---|---|---|
| WDR1 | 198.792557 | -0.23507956 | 0.21606904 | -1.0879836 | 0.27660237 | 0.72584803 |
| ZNF518B | 15.5182598 | -0.318398779 | 0.38839223 | -0.8197867 | 0.41233772 | 0.81556555 |
| CLNK | 2690.97471 | 0.379516961 | 0.29099836 | 1.30418933 | 0.19216904 | 0.6421543 |
| AC084048.1 | 50.7371784 | -1.026502653 | 0.61852459 | -1.6595988 | 0.0969952 | 0.48257296 |
| HS3ST1 | 18.1793569 | 0.428815617 | 0.59319614 | 0.7228901 | 0.46974741 | 0.84396182 |
| AC005699.1 | 27.2439196 | -0.101053444 | 0.65034585 | -0.1553842 | 0.87651847 | 0.97532685 |
| RAB28 | 168.019734 | 0.004448453 | 0.18866006 | 0.0235792 | 0.98118826 | 0.99714727 |
| BOD1L1 | 118.609669 | -0.24026318 | 0.19459533 | -1.2346811 | 0.2169492 | 0.67156502 |
| CPEB2 | 180.262743 | 0.031790499 | 0.37211609 | 0.08543167 | 0.93191822 | 0.9875946 |
| AC098829.1 | 51.421615 | -0.294666932 | 0.44702941 | -0.6591668 | 0.50978868 | 0.86281145 |
| CC2D2A | 27.4309717 | -0.851087937 | 0.37827389 | -2.2499251 | 0.0244537 | 0.24529563 |
| FBXL5 | 114.564785 | -0.093893879 | 0.17080551 | -0.5497122 | 0.58251676 | 0.89036744 |
| FAM200B | 37.034419 | 0.118496163 | 0.36335838 | 0.32611375 | 0.74433829 | 0.94546379 |
| TAPT1 | 89.6061617 | 0.141743927 | 0.2024719 | 0.70006717 | 0.48388536 | 0.85235034 |
| TAPT1-AS1 | 168.669978 | 0.392995246 | 0.5097259 | 0.77099328 | 0.44071091 | 0.82793978 |
| AC097515.1 | 24.3472294 | 0.225936127 | 0.64025324 | 0.35288556 | 0.72417423 | 0.93910295 |
| QDPR | 18.4848871 | 0.521121501 | 0.39006197 | 1.3359967 | 0.18155035 | 0.62984206 |
| LAP3 | 51.5497618 | -0.071590122 | 0.62127518 | -0.1152309 | 0.90826208 | 0.98250721 |
| MED28 | 102.768301 | -0.151764943 | 0.29897105 | -0.5076242 | 0.61171691 | 0.90011356 |
| FAM184B | 156.926562 | 1.034297825 | 0.57618527 | 1.79507857 | 0.07264118 | 0.42472542 |
| DCAF16 | 77.2459523 | 0.81809335 | 0.34967844 | 2.33955904 | 0.01930652 | 0.21826718 |
| LCORL | 150.566636 | 0.187288505 | 0.1613731 | 1.1605931 | 0.24580741 | 0.69938066 |
| AC024230.1 | 34.5676993 | 0.692759264 | 0.56228172 | 1.23205014 | 0.21793036 | 0.67202699 |
| SLIT2 | 1740.64149 | 0.330415561 | 0.50126766 | 0.65915994 | 0.50979307 | 0.86281145 |
| PACRGL | 55.416413 | -0.01698411 | 0.25003227 | -0.0679277 | 0.94584321 | 0.99134285 |
| KCNIP4 | 2618.39823 | -0.845717526 | 0.57055027 | -1.482284 | 0.13826475 | 0.57086516 |
| ADGRA3 | 26.3963687 | 1.105063342 | 0.3745768 | 2.95016495 | 0.00317604 | 0.07049798 |
| PPARGC1A | 698.387475 | 0.431208723 | 0.38227226 | 1.12801469 | 0.25931371 | 0.71066775 |
| DHX15 | 123.471432 | 0.225292395 | 0.2085061 | 1.08050744 | 0.27991627 | 0.73179753 |
| CCDC149 | 31.4671256 | 0.294656659 | 0.28361372 | 1.03893654 | 0.29883425 | 0.74296125 |
| SEPSECS | 31.6370157 | 0.003958388 | 0.44156229 | 0.00896451 | 0.99284746 | 0.99923337 |
| ZCCHC4 | 21.899696 | -0.25835194 | 0.3511337 | -0.7357652 | 0.46187363 | 0.83851286 |
| ANAPC4 | 87.0168927 | -1.242469733 | 0.55911964 | -2.2221894 | 0.02627051 | 0.25456937 |
| SEL1L3 | 14.9300659 | -1.307368839 | 0.65667174 | -1.9909016 | 0.0464917 | 0.34368082 |
| SMIM20 | 23.8501719 | 0.020733012 | 0.4454636 | 0.04654255 | 0.96287782 | 0.99306991 |
| RBPJ | 197.767958 | -0.330853771 | 0.23122801 | -1.4308551 | 0.15247175 | 0.59064536 |
| TBC1D19 | 106.219974 | -0.27197051 | 0.3837366 | -0.7087427 | 0.47848416 | 0.84833462 |
| STIM2 | 632.966343 | -2.888745538 | 0.51707607 | -5.5866935 | 2.31E-08 | 8.46E-06 |
| PCDH7 | 114.243253 | 0.8413547 | 1.04128453 | 0.80799693 | 0.41909235 | 0.8189355 |
| LINC02506 | 42.805576 | -1.137571929 | 1.22951775 | -0.925218 | 0.35485253 | 0.7810615 |
| ARAP2 | 114.926848 | -0.248953983 | 0.37168302 | -0.6698019 | 0.50298409 | 0.86010152 |
| C4orf19 | 195.785574 | -0.024841722 | 0.19659067 | -0.1263627 | 0.89944485 | 0.97936951 |
| RELL1 | 27.1236723 | -0.979377107 | 0.37462522 | -2.6142851 | 0.00894144 | 0.13745162 |
| PGM2 | 26.4756552 | -0.795816973 | 0.31997715 | -2.4871057 | 0.01287871 | 0.16919599 |
| TBC1D1 | 943.347652 | -0.159281577 | 0.30420625 | -0.5235973 | 0.60055865 | 0.89490801 |
| KLF3 | 62.4113747 | -0.372927512 | 0.2524486 | -1.4772414 | 0.13961095 | 0.57327809 |
| TLR1 | 13.6286716 | 0.009143955 | 0.47634898 | 0.01919592 | 0.98468482 | 0.99773864 |
| FAM114A1 | 53.0611927 | -0.234962788 | 0.38285778 | -0.6137078 | 0.53940846 | 0.86972442 |
| KLHL5 | 101.352593 | -0.797602839 | 0.31317352 | -2.5468399 | 0.01087033 | 0.15419892 |
| WDR19 | 36.0448281 | -0.259081029 | 0.25916047 | -0.9996935 | 0.31745887 | 0.75958338 |
| RFC1 | 74.1394863 | -0.094688637 | 0.20089191 | -0.4713412 | 0.63739709 | 0.91003118 |
| RPL9 | 18.9997232 | 0.092793648 | 0.31850786 | 0.29133865 | 0.77079233 | 0.95244403 |
| LIAS | 26.6471065 | 0.520837852 | 0.34958666 | 1.48986764 | 0.13625904 | 0.5673498 |
| UGDH | 23.2748723 | -0.5760571 | 0.37812446 | -1.523459 | 0.12764392 | 0.55102181 |
| UGDH-AS1 | 29.7287358 | -0.065909115 | 0.40687926 | -0.1619869 | 0.87131616 | 0.97434023 |
| SMIM14 | 94.8125953 | 0.042276721 | 0.25580607 | 0.16526863 | 0.86873255 | 0.97434023 |
| UBE2K | 282.346749 | 0.216148978 | 0.17111837 | 1.26315476 | 0.20653357 | 0.65836247 |
| PDS5A | 215.939985 | -0.075889486 | 0.19031661 | -0.3987539 | 0.69007456 | 0.92736537 |
| N4BP2 | 48.0455592 | 0.247062146 | 0.32939778 | 0.75004194 | 0.45322945 | 0.83364149 |
| RBM47 | 876.96316 | -0.086776152 | 0.2038866 | -0.4256099 | 0.67039214 | 0.92146857 |
| NSUN7 | 151.042512 | -0.27009382 | 0.35113671 | -0.7691985 | 0.4417755 | 0.8281114 |
| APBB2 | 291.868601 | 0.085160009 | 0.31493373 | 0.27040612 | 0.78684783 | 0.95577371 |
| LIMCH1 | 781.205451 | 0.299660701 | 0.28620616 | 1.04700995 | 0.29509499 | 0.74111836 |
| TMEM33 | 117.208368 | 0.362183063 | 0.28571058 | 1.26765717 | 0.20492041 | 0.65805651 |
| SLC30A9 | 210.46458 | 0.161381772 | 0.17372392 | 0.92895538 | 0.35291221 | 0.78010119 |
| ATP8A1 | 97.6172415 | 0.709372751 | 0.50619489 | 1.40138269 | 0.16109966 | 0.60060549 |
| GUF1 | 34.7134293 | -0.007854433 | 0.26086482 | -0.0301092 | 0.97597996 | 0.99556048 |
| GNPDA2 | 30.0645218 | 0.015538086 | 0.27281421 | 0.05695483 | 0.95458118 | 0.99228059 |
| COMMD8 | 21.2044416 | 0.181850164 | 0.32969352 | 0.55157336 | 0.58124069 | 0.8901472 |
| ATP10D | 214.30463 | 0.493177981 | 0.33874984 | 1.45587664 | 0.14542672 | 0.58098815 |
| AC107068.2 | 20.0836063 | -0.768901988 | 0.69549333 | -1.105549 | 0.26892175 | 0.72010324 |
| NFXL1 | 52.1481027 | -0.107823735 | 0.45732568 | -0.2357701 | 0.81361105 | 0.96264114 |
| NIPAL1 | 55.2346162 | 0.19251746 | 0.40932226 | 0.47033225 | 0.63811766 | 0.91025608 |
| CNGA1 | 31.2466862 | -0.188268173 | 0.62376383 | -0.3018261 | 0.76278467 | 0.95054102 |
| TXK | 19.9572889 | -0.739033377 | 0.53180609 | -1.389667 | 0.16463002 | 0.60511982 |
| TEC | 74.6147244 | -0.431571129 | 0.25194456 | -1.7129607 | 0.08671977 | 0.46422617 |
| SLAIN2 | 110.518013 | 0.27481265 | 0.20253472 | 1.35686687 | 0.17482352 | 0.62071528 |
| FRYL | 275.662964 | -0.061706775 | 0.19560315 | -0.3154692 | 0.75240541 | 0.94728927 |
| OCIAD1 | 166.406534 | 0.021238136 | 0.1999375 | 0.10622388 | 0.91540473 | 0.98430616 |
| OCIAD2 | 16.2429602 | -0.422780324 | 0.44629406 | -0.9473134 | 0.34347913 | 0.77370523 |
| CWH43 | 29.9796111 | 0.783803426 | 0.76300152 | 1.02726326 | 0.30429652 | 0.74820946 |
| DCUN1D4 | 97.2495364 | 0.245573692 | 0.27505314 | 0.89282273 | 0.37195211 | 0.7900012 |
| SGCB | 11.3681691 | -0.555017811 | 0.52254074 | -1.0621522 | 0.28816658 | NA |
| SCFD2 | 154.927325 | 0.357227946 | 0.28709954 | 1.24426511 | 0.21340201 | 0.66676543 |
| AC058822.1 | 75.2144334 | -0.709604303 | 0.46140869 | -1.5379084 | 0.12407101 | 0.5426009 |
| LNX1 | 149.776057 | -0.067630203 | 0.31808415 | -0.2126173 | 0.83162545 | 0.96701848 |
| CHIC2 | 82.1959131 | -0.427418874 | 0.19242159 | -2.2212625 | 0.02633319 | 0.2546679 |
| KIT | 833.481199 | -0.257066356 | 0.34477818 | -0.7455993 | 0.45590953 | 0.83506875 |
| TMEM165 | 196.540312 | -0.194041474 | 0.22097805 | -0.878103 | 0.37988785 | 0.79587736 |
| CLOCK | 117.024069 | 0.448552226 | 0.2739326 | 1.63745473 | 0.10153549 | 0.49220103 |
| EXOC1 | 110.06974 | -0.145205068 | 0.27776339 | -0.5227653 | 0.60113758 | 0.89490801 |
| CEP135 | 22.5833419 | 0.212504414 | 0.30530065 | 0.69604965 | 0.48639773 | 0.85363814 |
| AASDH | 54.5252608 | 0.151434403 | 0.22333351 | 0.67806395 | 0.49773115 | 0.85818031 |
| PAICS | 20.2359108 | 0.054105853 | 0.34214426 | 0.15813755 | 0.87434842 | 0.97434023 |
| SRP72 | 50.272963 | 0.055230863 | 0.22005 | 0.25099233 | 0.80182004 | 0.95890203 |
| REST | 107.00646 | 0.01277813 | 0.25249748 | 0.05060696 | 0.95963872 | 0.99238848 |
| NOA1 | 15.2909996 | -0.104657391 | 0.35295945 | -0.2965139 | 0.76683764 | 0.95205353 |
| POLR2B | 80.0237044 | -0.173775059 | 0.2178382 | -0.7977254 | 0.42502988 | 0.82241967 |
| IGFBP7 | 1029.13987 | -0.80820014 | 0.44667057 | -1.8093875 | 0.07039083 | 0.41992067 |
| AC104806.2 | 22.6237332 | 0.897879499 | 0.62035139 | 1.44737242 | 0.14779265 | 0.58466095 |
| CENPC | 102.687881 | 0.567870747 | 0.24883905 | 2.2820805 | 0.02248459 | 0.23562337 |
| STAP1 | 245.577412 | 0.864558282 | 0.41484769 | 2.08403781 | 0.03715673 | 0.30857899 |
| UBA6 | 179.757565 | -0.171608142 | 0.25766914 | -0.6660019 | 0.50540987 | 0.86197912 |
| UBA6-AS1 | 137.972824 | 0.484134245 | 0.38315239 | 1.26355533 | 0.20638967 | 0.65836247 |
| YTHDC1 | 135.597871 | -0.004697866 | 0.22260717 | -0.0211038 | 0.98316282 | 0.99745627 |
| UGT2B7 | 50.8155375 | -0.068900437 | 0.46257026 | -0.1489513 | 0.88159207 | 0.97630358 |
| ENAM | 35.839263 | -0.78778169 | 0.58160575 | -1.3544943 | 0.17557872 | 0.62231623 |
| UTP3 | 37.9549439 | -1.156549078 | 0.29381691 | -3.9362918 | 8.28E-05 | 0.00564171 |
| RUFY3 | 714.732523 | -0.749122396 | 0.32701605 | -2.2907817 | 0.02197604 | 0.23220872 |
| GRSF1 | 90.3941541 | -0.105596216 | 0.20275758 | -0.5208003 | 0.60250586 | 0.8956362 |
| MOB1B | 889.377554 | 0.342821273 | 0.36255134 | 0.94557994 | 0.34436288 | 0.77393882 |
| DCK | 28.2914137 | 0.275414075 | 0.66887584 | 0.41175665 | 0.6805178 | 0.92375164 |
| SLC4A4 | 13.5224984 | 0.959447957 | 0.45529348 | 2.10731759 | 0.03509006 | 0.2993611 |

| | | | | | | |
|---|---|---|---|---|---|---|
| ADAMTS3 | 109.877108 | 0.673719153 | 1.15298017 | 0.58432848 | 0.55899933 | 0.88004847 |
| COX18 | 19.3896542 | 0.576785703 | 0.4315558 | 1.33652638 | 0.18137729 | 0.62946679 |
| ANKRD17 | 682.412347 | 0.199017816 | 0.17903035 | 1.11164289 | 0.26629172 | 0.71747544 |
| AC053527.1 | 77.7920972 | 0.071188062 | 0.48824216 | 0.14580482 | 0.88407547 | 0.97662748 |
| RASSF6 | 48.7959219 | -0.648652522 | 0.42895259 | -1.5121776 | 0.13048868 | 0.55685884 |
| MTHFD2L | 33.8209253 | 0.212566014 | 0.26347852 | 0.80676792 | 0.4198002 | 0.81899879 |
| BTC | 13.4023326 | -0.331649178 | 0.6661458 | -0.4978627 | 0.61858079 | 0.90267467 |
| RCHY1 | 50.5710351 | 0.504949061 | 0.27830077 | 1.81440053 | 0.06961608 | 0.41838662 |
| THAP6 | 28.7001142 | 0.164326696 | 0.28670724 | 0.57315153 | 0.5665421 | 0.88382043 |
| CDKL2 | 30.0745331 | -0.072024956 | 0.39285395 | -0.1833377 | 0.85453302 | 0.97140165 |
| G3BP2 | 91.6150972 | 0.085663285 | 0.26252742 | 0.32630225 | 0.74419569 | 0.94546379 |
| USO1 | 184.788279 | 0.003095508 | 0.18911227 | 0.01636863 | 0.98694031 | 0.9983649 |
| SDAD1 | 41.7036802 | -0.162752408 | 0.26527716 | -0.6135184 | 0.53953365 | 0.86978169 |
| NUP54 | 58.5582369 | 0.286363015 | 0.24693243 | 1.15968169 | 0.24617843 | 0.69965741 |
| SCARB2 | 131.605 | -0.022290718 | 0.27930378 | -0.0798082 | 0.93638984 | 0.98871562 |
| SHROOM3 | 1270.09832 | -0.793318527 | 0.27448216 | -2.8902371 | 0.00384951 | 0.08135138 |
| AC107072.1 | 16.0505524 | -2.4891209 | 0.75038325 | -3.3171328 | 0.00090946 | 0.03073797 |
| 11-Sep | 280.298213 | -0.243632014 | 0.31249334 | -0.7796391 | 0.43560335 | 0.82590941 |
| CCNI | 317.18141 | 0.041568164 | 0.23921191 | 0.1737713 | 0.8620452 | 0.97302617 |
| CCNG2 | 76.0622407 | -0.480191858 | 0.2923609 | -1.6424627 | 0.10049416 | 0.48886326 |
| CNOT6L | 307.057328 | -0.358547507 | 0.37951164 | -0.9447602 | 0.3447813 | 0.7746997 |
| MRPL1 | 121.157286 | 0.213134421 | 0.27440134 | 0.77672514 | 0.43732095 | 0.82687287 |
| FRAS1 | 48.0423156 | 2.630284364 | 0.57132508 | 4.60383142 | 4.15E-06 | 0.00059752 |
| ANXA3 | 55.5989673 | 1.018795911 | 0.58078673 | 1.75416527 | 0.07940219 | 0.44494585 |
| BMP2K | 47.2964981 | 0.429518642 | 0.32758794 | 1.31115524 | 0.18980532 | 0.64181696 |
| PAQR3 | 21.9529031 | 0.164739043 | 0.43690776 | 0.3770568 | 0.70613139 | 0.93407096 |
| RASGEF1B | 455.189422 | 0.601587069 | 0.51251649 | 1.17379066 | 0.24047889 | 0.69470237 |
| HNRNPD | 171.654729 | 0.11506588 | 0.17124116 | 0.67195222 | 0.50161412 | 0.85942996 |
| HNRNPDL | 149.928463 | 0.29155165 | 0.22486946 | 1.29653733 | 0.19479043 | 0.6459894 |
| ENOPH1 | 29.9603641 | -0.232164709 | 0.33962334 | -0.6835947 | 0.49423115 | 0.85774549 |
| TMEM150C | 86.8066566 | 0.207451463 | 0.28258407 | 0.73412299 | 0.46287378 | 0.8392291 |
| SCD5 | 37.4854135 | 0.573014212 | 0.49744363 | 1.15191788 | 0.24935482 | 0.70373632 |
| SEC31A | 170.261564 | 0.086040151 | 0.17427056 | 0.49371593 | 0.62150681 | 0.90380166 |
| THAP9 | 12.5800665 | 0.384673056 | 0.43736623 | 0.87952162 | 0.37911852 | 0.79535699 |
| LIN54 | 36.2522418 | -0.026259446 | 0.30539698 | -0.0859846 | 0.93147864 | 0.98757533 |
| COPS4 | 37.2125187 | 0.304646577 | 0.24749581 | 1.2309161 | 0.21835425 | 0.67239244 |
| COQ2 | 12.5028426 | 0.633508765 | 0.39253492 | 1.6138915 | 0.10655096 | NA |
| HELQ | 27.3835832 | 0.224561425 | 0.38611561 | 0.58159115 | 0.5608421 | 0.88071368 |
| MRPS18C | 15.5717603 | 0.13834289 | 0.41486425 | 0.33346544 | 0.73878297 | 0.94376279 |
| ABRAXAS1 | 16.8893405 | 0.065983578 | 0.35107708 | 0.18794613 | 0.85091888 | 0.97027687 |
| GPAT3 | 365.767108 | 1.456486642 | 0.39329212 | 3.70332022 | 0.0002128 | 0.01141021 |
| CDS1 | 93.8217891 | 0.192453475 | 0.28548714 | 0.67412309 | 0.50023306 | 0.85942996 |
| WDFY3 | 377.16992 | -0.193294422 | 0.17883748 | -1.0808384 | 0.27976899 | 0.7316687 |
| ARHGAP24 | 261.847249 | 1.020114278 | 0.35856045 | 2.8450273 | 0.00444076 | 0.08992774 |
| MAPK10 | 481.949278 | 0.282342145 | 0.42207797 | 0.66893363 | 0.50353782 | 0.86067257 |
| AC104827.1 | 12.2541894 | 0.322773592 | 0.65308591 | 0.49422838 | 0.6211449 | NA |
| PTPN13 | 323.551274 | 0.635862932 | 0.33921905 | 1.87449061 | 0.06086284 | 0.39227209 |
| AC093827.5 | 13.7271047 | 0.11001223 | 0.47981559 | 0.22928023 | 0.81865112 | 0.9635539 |
| AFF1 | 457.996273 | -0.527280379 | 0.22594006 | -2.333718 | 0.01961049 | 0.21965565 |
| KLHL8 | 64.5986485 | -0.370230137 | 0.38019731 | -0.9737842 | 0.3301637 | 0.76685572 |
| HSD17B11 | 26.0070886 | -0.374378628 | 0.38726084 | -0.9667351 | 0.3336765 | 0.76777176 |
| NUDT9 | 65.1477701 | -0.19629308 | 0.30995801 | -0.6332893 | 0.52654477 | 0.86750588 |
| SPARCL1 | 737.149745 | -0.094150257 | 0.49236052 | -0.1912222 | 0.84835152 | 0.97027687 |
| SPP1 | 3666.48238 | -5.084494104 | 0.6070927 | -8.3751528 | 5.52E-17 | 1.07E-13 |
| PKD2 | 142.460784 | -0.16282176 | 0.29874565 | -0.545018 | 0.58574114 | 0.89059199 |
| PPM1K | 41.2406995 | -0.617147692 | 0.51897367 | -1.1891696 | 0.23437295 | 0.68933219 |
| HERC6 | 26.5782608 | -1.448362193 | 0.88211398 | -1.6419218 | 0.10060621 | 0.48916301 |
| HERC5 | 15.2784462 | -1.03620813 | 0.86403029 | -1.1992729 | 0.23042184 | 0.68523674 |
| PYURF | 24.0090553 | -0.079744498 | 0.41186632 | -0.1936174 | 0.84647545 | 0.97027687 |
| HERC3 | 92.1227087 | -0.15341786 | 0.29767586 | -0.5153856 | 0.60628357 | 0.89643487 |
| FAM13A-AS1 | 35.3778487 | -0.089415598 | 0.31421832 | -0.2845652 | 0.77597728 | 0.95495808 |
| FAM13A | 233.422959 | 0.437966146 | 0.36521751 | 1.19919262 | 0.23045306 | 0.68523674 |
| GPRIN3 | 174.730967 | 0.051154781 | 0.27859171 | 0.18361918 | 0.85431221 | 0.97137832 |
| SNCA | 31.0653533 | 1.44470131 | 1.14557679 | 1.26111259 | 0.20726828 | 0.65905456 |
| CCSER1 | 1178.97329 | 1.062565117 | 0.96705153 | 1.09876783 | 0.27186935 | 0.7232847 |
| SMARCAD1 | 87.0156467 | 0.002551069 | 0.20636314 | 0.01236204 | 0.99013677 | 0.99923337 |
| PDLIM5 | 458.677693 | -0.158688251 | 0.19602157 | -0.8095448 | 0.41820183 | 0.8188001 |
| BMPR1B | 489.270388 | 0.464541785 | 0.58918612 | 0.78844659 | 0.43043553 | 0.82469033 |
| UNC5C | 35.0065206 | 0.069318767 | 0.7150324 | 0.09694493 | 0.92277013 | 0.98672316 |
| STPG2 | 484.389812 | 1.379101574 | 0.31819224 | 4.33417731 | 1.46E-05 | 0.00155953 |
| STPG2-AS1 | 12.9445675 | -0.049318351 | 0.63351464 | -0.0778488 | 0.93794833 | 0.98871562 |
| RAP1GDS1 | 147.290675 | 0.011404608 | 0.18207662 | 0.06263631 | 0.95005611 | 0.99192022 |
| TSPAN5 | 514.494125 | -0.911868731 | 0.35397877 | -2.5760549 | 0.00999348 | 0.14754137 |
| EIF4E | 101.293304 | 0.220133986 | 0.22138926 | 0.99433 | 0.32006224 | 0.76106474 |
| METAP1 | 61.9085297 | 0.251792993 | 0.2197643 | 1.14574112 | 0.25190228 | 0.70579207 |
| ADH5 | 52.0314734 | -0.008919501 | 0.31403286 | -0.0284031 | 0.97734067 | 0.9957152 |
| TRMT10A | 10.4805417 | 0.14836372 | 0.38980787 | 0.38060729 | 0.70349467 | NA |
| LAMTOR3 | 32.1242263 | 0.024442231 | 0.36261723 | 0.06740505 | 0.94625925 | 0.99134285 |
| DNAJB14 | 128.846331 | -0.101269613 | 0.19836891 | -0.5105115 | 0.60969314 | 0.89919773 |
| H2AFZ | 76.8030828 | 0.311649948 | 0.57997311 | 0.53735241 | 0.59102421 | 0.89149106 |
| PPP3CA | 503.162976 | 0.132160817 | 0.2864662 | 0.46134873 | 0.64454843 | 0.9119294 |
| BANK1 | 35.7955933 | -0.034577441 | 0.41720899 | -0.082878 | 0.93394856 | 0.9881638 |
| AP002075.1 | 15.4874172 | -0.748608055 | 0.60929476 | -1.2286468 | 0.21920426 | 0.67305276 |
| NFKB1 | 148.206455 | -0.034649837 | 0.27722363 | -0.1249888 | 0.90053245 | 0.97940229 |
| MANBA | 83.5545451 | 0.472773375 | 0.31528765 | 1.49949855 | 0.13374435 | 0.56229288 |
| UBE2D3 | 314.321388 | -0.278893724 | 0.16630745 | -1.6769767 | 0.09354704 | 0.47792014 |
| CISD2 | 39.5269835 | -0.072626386 | 0.31879531 | -0.2278151 | 0.81978998 | 0.96389705 |
| SLC9B1 | 13.9074952 | 0.123039672 | 0.48355515 | 0.25444807 | 0.79914942 | 0.95848876 |
| TET2 | 227.683902 | 0.382246728 | 0.21151581 | 1.80717803 | 0.07073452 | 0.42119392 |
| PPA2 | 182.985033 | -0.073202759 | 0.26659114 | -0.2745881 | 0.7836327 | 0.95544332 |
| ARHGEF38 | 299.238303 | -0.749142194 | 0.30208009 | -2.4799456 | 0.01314024 | 0.17172 |
| INTS12 | 76.7853777 | -0.410660989 | 0.29708186 | -1.382316 | 0.16687471 | 0.60807088 |
| GSTCD | 71.2291556 | -0.030868735 | 0.22983636 | -0.1343074 | 0.89315946 | 0.97831181 |
| NPNT | 181.846279 | 0.988635772 | 0.38464086 | 2.57028274 | 0.01016155 | 0.14844189 |
| TBCK | 272.591854 | -0.156189274 | 0.29366643 | -0.5318595 | 0.59482332 | 0.89301752 |
| AIMP1 | 49.1200713 | -0.053065142 | 0.26959739 | -0.1968311 | 0.84395974 | 0.96983882 |
| PAPSS1 | 46.2977778 | -0.331651521 | 0.2635539 | -1.2583821 | 0.20825359 | 0.65973983 |
| SGMS2 | 122.379366 | 0.13620732 | 0.32460973 | 0.41960332 | 0.67477526 | 0.92193843 |
| AC096564.1 | 16.0415763 | -0.799401909 | 0.56267652 | -1.4207131 | 0.15540018 | 0.59533243 |
| HADH | 25.5579581 | 0.464488795 | 0.46212862 | 1.00510718 | 0.31484524 | 0.75742745 |
| LEF1 | 454.990184 | -0.500242184 | 0.61993184 | -0.8069309 | 0.41970627 | 0.81899879 |
| RPL34-AS1 | 25.1077759 | 1.103626081 | 0.43488457 | 2.53774485 | 0.01115693 | 0.15593979 |
| RPL34 | 139.378979 | 0.082983329 | 0.37011612 | 0.2242089 | 0.82259475 | 0.96521156 |
| OSTC | 33.0132295 | -0.154528044 | 0.28420636 | -0.5437178 | 0.58663571 | 0.89059199 |
| SEC24B | 168.805109 | -0.103686368 | 0.15210706 | -0.681667 | 0.49544952 | 0.85814183 |
| CASP6 | 26.0152245 | -0.128431564 | 0.34971649 | -0.3672448 | 0.71343643 | 0.93616842 |
| PLA2G12A | 38.5539686 | 0.155301716 | 0.36333308 | 0.42743621 | 0.66906164 | 0.92049996 |
| GAR1 | 16.2905783 | 0.284629974 | 0.43317321 | 0.6570812 | 0.51112871 | 0.86312853 |
| EGF | 159.345597 | 0.399979665 | 0.34629446 | 1.15502763 | 0.24807911 | 0.70279421 |
| ELOVL6 | 108.28174 | -0.48747505 | 0.53873352 | -0.9048538 | 0.36554287 | 0.78793182 |
| FAM241A | 39.4715362 | 0.049023806 | 0.378876 | 0.12939275 | 0.89704689 | 0.97889563 |
| AP1AR | 37.5285404 | 0.070420767 | 0.3311415 | 0.21266065 | 0.83159166 | 0.96701848 |
| ALPK1 | 162.494497 | -1.331794918 | 0.29679603 | -4.4872397 | 7.22E-06 | 0.00092089 |

| | | | | | | |
|---|---|---|---|---|---|---|
| ZGRF1 | 12.0945713 | -0.136897388 | 0.56509082 | -0.2422573 | 0.80858078 | NA |
| LARP7 | 67.643109 | -0.135931429 | 0.28298586 | -0.4803471 | 0.63098062 | 0.90768383 |
| ANK2 | 34.3332949 | 1.348524326 | 0.63353718 | 2.12856382 | 0.03329036 | 0.28961124 |
| CAMK2D | 53.4430449 | -0.797482404 | 0.36352661 | -2.1937387 | 0.0282542 | 0.26479779 |
| ARSJ | 188.0291 | -0.121551097 | 0.36059508 | -0.3370847 | 0.73605303 | 0.94340834 |
| UGT8 | 334.945216 | 0.105087646 | 0.2579152 | 0.40745038 | 0.68367723 | 0.92517705 |
| SNHG8 | 31.3400552 | 0.139194019 | 0.4576245 | 0.30416645 | 0.76100108 | 0.94990484 |
| METTL14 | 44.680822 | 0.192943426 | 0.24821911 | 0.77731093 | 0.43697535 | 0.82687287 |
| SEC24D | 71.8644265 | 0.072889448 | 0.35168961 | 0.20725505 | 0.83581068 | 0.96748214 |
| USP53 | 255.415986 | 0.022352483 | 0.23481598 | 0.09519149 | 0.92416273 | 0.98672316 |
| C4orf3 | 75.6704058 | -0.659621646 | 0.37741355 | -1.7477423 | 0.08050865 | 0.4486083 |
| PDE5A | 63.9955295 | -0.7730726 | 0.51828847 | -1.4915875 | 0.13580732 | 0.56635038 |
| AC108866.1 | 28.6734714 | 0.659045334 | 0.3523772 | 1.87028372 | 0.06144443 | 0.39288791 |
| PRDM5 | 19.1863331 | 1.057304035 | 0.58258009 | 1.81486468 | 0.06954471 | 0.41827125 |
| ANXA5 | 93.2929677 | -0.256930851 | 0.32903394 | -0.7808643 | 0.43488232 | 0.82565889 |
| EXOSC9 | 13.305979 | 0.267880602 | 0.39572834 | 0.67693056 | 0.49845002 | 0.858786 |
| BBS7 | 15.99187 | -0.056391847 | 0.35817672 | -0.1574414 | 0.87489698 | 0.97467563 |
| KIAA1109 | 282.516354 | 0.239882414 | 0.18752847 | 1.27917861 | 0.20083417 | 0.65358156 |
| SPATA5 | 96.8005301 | 0.313786229 | 0.21138372 | 1.48443893 | 0.13769251 | 0.56931687 |
| SPRY1 | 28.9480404 | 0.106252895 | 0.57524081 | 0.18471029 | 0.85345628 | 0.97131147 |
| LINC01091 | 183.869358 | 1.442228697 | 0.4730741 | 3.04863172 | 0.00229886 | 0.05602752 |
| ANKRD50 | 30.3814403 | 0.195757408 | 0.38303701 | 0.51106657 | 0.60930444 | 0.89906009 |
| INTU | 64.6276316 | 0.065453287 | 0.36174807 | 0.18093611 | 0.85641773 | 0.97166553 |
| HSPA4L | 105.693749 | -0.239838665 | 0.33679285 | -0.7121252 | 0.47638727 | 0.84663915 |
| MFSD8 | 60.0074498 | 0.080345884 | 0.2205593 | 0.36428246 | 0.7156471 | 0.93689626 |
| ABHD18 | 150.953087 | 0.054211091 | 0.24364252 | 0.22250258 | 0.82392266 | 0.96539788 |
| LARP1B | 135.912431 | 0.30116672 | 0.25715803 | 1.17113482 | 0.24154459 | 0.69556484 |
| PGRMC2 | 68.4880636 | -0.252727376 | 0.41103902 | -0.6148501 | 0.53865373 | 0.86972442 |
| AC108062.1 | 35.0554364 | -0.193753685 | 0.35521275 | -0.5454581 | 0.58543846 | 0.89050542 |
| AC078850.2 | 48.8199687 | 0.457583466 | 0.5354521 | 0.85457404 | 0.39278701 | 0.80400536 |
| JADE1 | 430.089017 | -0.565417584 | 0.26289131 | -2.1507656 | 0.03149471 | 0.28099688 |
| SCLT1 | 160.744442 | 0.001058978 | 0.26464529 | 0.0040015 | 0.99680727 | 0.99975875 |
| C4orf33 | 20.3700548 | -0.026500032 | 0.35980843 | -0.0736504 | 0.94128857 | 0.98985996 |
| AC096711.2 | 63.1306361 | 0.686420689 | 0.27798605 | 2.46926312 | 0.01353916 | 0.17531992 |
| AC109927.1 | 32.0188749 | -1.22886571 | 0.42133334 | -2.9166116 | 0.00353856 | 0.07661617 |
| NOCT | 21.3490975 | -1.015477695 | 0.39973596 | -2.5403711 | 0.01107349 | 0.15593979 |
| ELF2 | 475.755863 | -0.509604354 | 0.18759477 | -2.7165169 | 0.00659728 | 0.11656402 |
| NDUFC1 | 81.7285377 | -0.096660474 | 0.32342274 | -0.2988673 | 0.76504132 | 0.95205353 |
| NAA15 | 111.599412 | 0.174959836 | 0.22779217 | 0.7680678 | 0.4424469 | 0.82820049 |
| AC097376.2 | 153.577117 | -0.337537897 | 0.26209056 | -1.2878674 | 0.19779211 | 0.64914661 |
| MGST2 | 64.8638172 | -0.149533795 | 0.37239522 | -0.401546 | 0.68801821 | 0.9267526 |
| MAML3 | 308.97824 | -0.191558486 | 0.45670466 | -0.4194362 | 0.67489735 | 0.92193843 |
| SCOC | 96.1055634 | -0.263333513 | 0.28154188 | -0.9353262 | 0.34962019 | 0.77868979 |
| ELMOD2 | 33.9120036 | 0.09385693 | 0.29830925 | 0.31462963 | 0.75304288 | 0.94771649 |
| TBC1D9 | 163.80588 | 0.085772356 | 0.26857482 | 0.31936112 | 0.74945269 | 0.94684415 |
| RNF150 | 41.411101 | 0.915219939 | 0.75586619 | 1.21082269 | 0.22596337 | 0.6800635 |
| ZNF330 | 30.2177551 | 0.072872585 | 0.3454041 | 0.21097776 | 0.83290462 | 0.96701848 |
| IL15 | 55.1060712 | -0.621504374 | 0.39135708 | -1.5880749 | 0.11226939 | 0.5156312 |
| INPP4B | 63.6085935 | 1.029798093 | 0.57946187 | 1.77716284 | 0.07554145 | 0.43306861 |
| USP38 | 28.6513078 | -0.074960033 | 0.28748301 | -0.260746 | 0.79428842 | 0.95785825 |
| GAB1 | 60.522818 | -0.069904014 | 0.36373931 | -0.1921816 | 0.84759993 | 0.97027687 |
| SMARCA5 | 100.166269 | -0.126463521 | 0.18778716 | -0.6734407 | 0.50066694 | 0.85942996 |
| ANAPC10 | 99.2754725 | 0.101161109 | 0.25871515 | 0.39101347 | 0.69578728 | 0.92900737 |
| ABCE1 | 37.5984343 | 0.289946012 | 0.32199733 | 0.90046091 | 0.36787502 | 0.78877653 |
| OTUD4 | 116.687404 | 0.158767974 | 0.1796973 | 0.88353009 | 0.37694993 | 0.79432848 |
| SMAD1 | 111.592101 | -0.439486492 | 0.23494046 | -1.8706292 | 0.06139649 | 0.39288791 |
| MMAA | 18.4038133 | -0.185097214 | 0.3404149 | -0.5437401 | 0.58662037 | 0.89059199 |
| ZNF827 | 167.447856 | -0.139320468 | 0.35214909 | -0.3956292 | 0.69237858 | 0.928133 |
| LSM6 | 18.6277379 | 0.206264519 | 0.38660597 | 0.53352647 | 0.5936692 | 0.89260988 |
| SLC10A7 | 108.248023 | -0.134869046 | 0.37941 | -0.3554705 | 0.72223717 | 0.93855548 |
| TMEM184C | 46.7612821 | 0.300663888 | 0.3023462 | 0.99443582 | 0.32001074 | 0.76106474 |
| PRMT9 | 38.4694078 | 0.305182332 | 0.26971468 | 1.13150065 | 0.25784443 | 0.71022118 |
| NR3C2 | 1206.28796 | 0.268872028 | 0.31693446 | 0.84835214 | 0.39624189 | 0.80674902 |
| AC069272.1 | 13.6395043 | -0.261554411 | 0.70148459 | -0.3728584 | 0.70925385 | 0.93485232 |
| DCLK2 | 87.8434261 | 0.739939494 | 0.62045552 | 1.19257461 | 0.23303602 | 0.68763965 |
| LRBA | 1075.42828 | 0.190981105 | 0.22683307 | 0.84194559 | 0.39981841 | 0.80830282 |
| RPS3A | 109.724061 | 0.678167936 | 0.400429 | 1.69360343 | 0.09034067 | 0.47236403 |
| SH3D19 | 59.7711313 | -0.980807125 | 0.49001028 | -2.0016052 | 0.04532721 | 0.33821763 |
| FAM160A1 | 336.775051 | -0.136780975 | 0.20756458 | -0.6589803 | 0.50990841 | 0.86281145 |
| GATB | 22.5007864 | 0.150300521 | 0.3480103 | 0.43188526 | 0.66582481 | 0.91958319 |
| FBXW7 | 228.363223 | 0.224555102 | 0.18394075 | 1.2208013 | 0.22216126 | 0.67557645 |
| ARFIP1 | 178.526856 | -0.270470842 | 0.17989796 | -1.5034681 | 0.13271838 | 0.56045639 |
| FHDC1 | 40.8204929 | -0.548502708 | 0.32242671 | -1.7011702 | 0.08891103 | 0.4687332 |
| TRIM2 | 649.24508 | 0.072685848 | 0.3110021 | 0.23371498 | 0.81520626 | 0.96300833 |
| MND1 | 20.8544186 | 1.474933458 | 1.53031388 | 0.96381107 | 0.33514066 | 0.76777176 |
| TMEM131L | 17.352297 | -0.170261796 | 0.450135 | -0.3782461 | 0.7052478 | 0.93403928 |
| TLR2 | 26.0184876 | -2.060596015 | 0.60905881 | -3.3832464 | 0.00071634 | 0.02602446 |
| AC079298.3 | 34.7157257 | -0.843963466 | 0.60041437 | -1.405635 | 0.15983253 | 0.60024417 |
| PLRG1 | 21.5851522 | 0.333303535 | 0.38756572 | 0.85999231 | 0.38979328 | 0.80191835 |
| MAP9 | 20.028336 | -0.143970758 | 0.31216024 | -0.4612079 | 0.64464948 | 0.9119294 |
| CTSO | 24.8642798 | -0.466005152 | 0.43255937 | -1.0773207 | 0.28133703 | 0.73311066 |
| PDGFC | 130.111023 | -0.00042995 | 0.25889674 | -0.0016607 | 0.99867495 | 0.99982181 |
| TMEM144 | 12.9946841 | 0.084233034 | 0.38183474 | 0.22060076 | 0.82540331 | 0.96539788 |
| ETFDH | 132.02684 | 0.351351962 | 0.36631886 | 0.95914242 | 0.337487 | 0.76966704 |
| PPID | 45.850265 | -0.155602396 | 0.2873629 | -0.5414839 | 0.58817407 | 0.89073775 |
| FNIP2 | 971.271805 | -1.872840209 | 0.40698115 | -4.6017861 | 4.19E-06 | 0.00059752 |
| C4orf45 | 43.795054 | -2.02371972 | 0.74704363 | -2.7089712 | 0.00674922 | 0.11774717 |
| RAPGEF2 | 563.403196 | 0.09517953 | 0.22914047 | 0.41537635 | 0.67786642 | 0.9226233 |
| NAF1 | 30.3219339 | 0.203405815 | 0.28425849 | 0.71556635 | 0.47425915 | 0.84593347 |
| TMA16 | 16.4652974 | 0.067091921 | 0.32320536 | 0.20758294 | 0.83555462 | 0.96748214 |
| 1-Mar | 971.922809 | 0.253163473 | 1.02875259 | 0.24608781 | 0.80561426 | 0.96005568 |
| TMEM192 | 42.6946689 | -0.283518743 | 0.42049639 | -0.6742478 | 0.50015382 | 0.85942996 |
| KLHL2 | 139.210658 | -0.640511024 | 0.38122309 | -1.6801475 | 0.09292862 | 0.47759124 |
| MSMO1 | 107.320534 | -1.116769522 | 0.56830624 | -1.965084 | 0.04940451 | 0.35550725 |
| DDX60 | 53.2456168 | -1.632720502 | 1.15288369 | -1.4162057 | 0.15671527 | 0.59731328 |
| DDX60L | 62.2106495 | -1.864643275 | 1.19388742 | -1.561825 | 0.11832921 | 0.52869338 |
| PALLD | 158.086308 | 0.560225688 | 0.49531657 | 1.13104572 | 0.25803585 | 0.71022118 |
| CBR4 | 153.43222 | -0.023398538 | 0.26568038 | -0.0880703 | 0.92982084 | 0.98733028 |
| SH3RF1 | 375.578247 | -0.385092708 | 0.35926634 | -1.0718864 | 0.28377105 | 0.73550473 |
| NEK1 | 129.842233 | 0.113289661 | 0.17202954 | 0.65854773 | 0.51018624 | 0.86281145 |
| CLCN3 | 159.980156 | -0.025665372 | 0.18348089 | -0.1398804 | 0.88875452 | 0.97765744 |
| HPF1 | 21.2488301 | 0.019579814 | 0.35048803 | 0.05586443 | 0.95544981 | 0.99228059 |
| MFAP3L | 24.0439678 | 0.984022443 | 0.6285301 | 1.56559319 | 0.11744388 | 0.52594907 |
| AADAT | 9.59066862 | 0.311044458 | 0.52375041 | 0.59387916 | 0.55259294 | NA |
| GALNT7 | 200.48493 | 0.067472032 | 0.32181634 | 0.20966006 | 0.833933 | 0.96701848 |
| HMGB2 | 19.923321 | 0.419879867 | 0.3264952 | 1.28602154 | 0.19843554 | 0.6504984 |
| FBXO8 | 30.035864 | 0.071044326 | 0.27015115 | 0.26297991 | 0.79256606 | 0.95746995 |
| CEP44 | 42.9087223 | 0.303936937 | 0.30547273 | 0.9949724 | 0.31974969 | 0.76106474 |
| WDR17 | 14.6333877 | 0.738905294 | 0.42241788 | 1.74922825 | 0.08025157 | 0.44789428 |
| SPCS3 | 129.841722 | -0.019105526 | 0.23098516 | -0.0827132 | 0.93407957 | 0.9881638 |
| AC078881.1 | 26.4622307 | -0.151022768 | 0.36650826 | -0.4120583 | 0.6802967 | 0.92375164 |
| TENM3 | 171.15324 | -0.222051987 | 0.52394696 | -0.4238062 | 0.67170713 | 0.92170876 |
| DCTD | 33.0097919 | 0.042128056 | 0.27501923 | 0.15318222 | 0.87825458 | 0.97542218 |

| | | | | | | |
|---|---|---|---|---|---|---|
| WWC2 | 171.086142 | 0.540673527 | 0.19367939 | 2.79159036 | 0.00524497 | 0.09995328 |
| CDKN2AIP | 15.7275462 | 0.528077316 | 0.36687297 | 1.439401 | 0.15003694 | 0.58673121 |
| RWDD4 | 39.7955355 | 0.686340735 | 0.25235027 | 2.71979389 | 0.00653226 | 0.11604934 |
| TRAPPC11 | 72.6543507 | 0.038695943 | 0.31539292 | 0.12269122 | 0.90235161 | 0.98004822 |
| STOX2 | 417.01908 | -0.679832112 | 0.35248652 | -1.9286755 | 0.05377116 | 0.36782809 |
| IRF2 | 80.2619092 | -0.989393116 | 0.2677495 | -3.6952193 | 0.0002197 | 0.01158302 |
| CASP3 | 25.153243 | -0.2062192 | 0.27297668 | -0.7554462 | 0.44998128 | 0.83118301 |
| PRIMPOL | 29.4773789 | -0.076696115 | 0.48471686 | -0.1582287 | 0.8742766 | 0.97434023 |
| ACSL1 | 416.514392 | -1.314919542 | 0.24048486 | -5.4677852 | 4.56E-08 | 1.47E-05 |
| SLC25A4 | 117.519764 | 0.012196426 | 0.34212852 | 0.03564867 | 0.9715625 | 0.99526459 |
| CFAP97 | 63.5660853 | -0.269437828 | 0.214192 | -1.2579266 | 0.20841829 | 0.65973983 |
| SNX25 | 102.94155 | -0.265066469 | 0.22440507 | -1.1811964 | 0.2375247 | 0.69207518 |
| AC112722.1 | 12.5581135 | -0.892459315 | 0.46237811 | -1.9301504 | 0.0535882 | 0.36684184 |
| UFSP2 | 40.0876038 | -0.142616494 | 0.2693064 | -0.5295696 | 0.59641035 | 0.89360216 |
| SORBS2 | 188.678161 | 0.698468241 | 0.47308919 | 1.47639865 | 0.13983691 | 0.57374456 |
| FAM149A | 31.4101501 | -0.069196329 | 0.43097877 | -0.1605562 | 0.87244293 | 0.97434023 |
| CYP4V2 | 14.1125805 | 0.83012158 | 0.3928417 | 2.11311982 | 0.03459051 | 0.29719042 |
| FAT1 | 19.7900933 | 0.84893766 | 0.52821078 | 1.60719489 | 0.10801162 | 0.50589702 |
| FRG1-DT | 20.156837 | 1.077640815 | 0.45324727 | 2.37760021 | 0.01742571 | 0.20451748 |
| FRG1 | 21.0712227 | 0.170647819 | 0.29848702 | 0.57170935 | 0.5675189 | 0.88446872 |
| CCDC127 | 10.490375 | 0.496633452 | 0.42591472 | 1.16603965 | 0.24359841 | NA |
| SDHA | 162.006676 | 0.02688894 | 0.24133756 | 0.11141631 | 0.91128623 | 0.9835359 |
| PDCD6 | 59.905699 | -0.027723494 | 0.30019268 | -0.0923523 | 0.92641811 | 0.98672316 |
| EXOC3 | 44.5102272 | -0.062369459 | 0.26348122 | -0.2367131 | 0.81287936 | 0.96262567 |
| ZDHHC11B | 21.5605928 | -0.083278481 | 0.56316009 | -0.1478771 | 0.88243976 | 0.97662748 |
| BRD9 | 29.2329852 | -0.135736145 | 0.37388996 | -0.3630377 | 0.71657674 | 0.93689626 |
| SLC12A7 | 83.3660242 | -0.417399851 | 0.30952675 | -1.3485098 | 0.17749447 | 0.6251709 |
| CLPTM1L | 39.4666626 | -0.008962286 | 0.32511795 | -0.0275663 | 0.97800809 | 0.99609121 |
| LPCAT1 | 13.0945039 | -0.197563322 | 0.46122013 | -0.4283493 | 0.66839682 | 0.92049996 |
| MRPL36 | 13.8866962 | -0.796288448 | 0.47468759 | -1.6775 | 0.09344474 | 0.47792014 |
| NDUFS6 | 70.0657801 | 0.083133982 | 0.31657787 | 0.262602 | 0.79285736 | 0.95746995 |
| ADAMTS16 | 15.8711151 | 2.114125753 | 0.59744496 | 3.53861172 | 0.00040224 | 0.01726416 |
| ICE1 | 90.5669859 | 0.091903621 | 0.19979389 | 0.45999214 | 0.64552186 | 0.91225913 |
| MED10 | 36.211662 | -0.431736093 | 0.26393841 | -1.6357456 | 0.10189284 | 0.49220184 |
| UBE2QL1 | 107.424685 | 1.346549566 | 0.47235369 | 2.8507231 | 0.00436199 | 0.08870302 |
| NSUN2 | 39.7374771 | 0.287214888 | 0.33213929 | 0.86474228 | 0.38718026 | 0.80009555 |
| TENT4A | 60.8012619 | 0.045867665 | 0.26569895 | 0.17263021 | 0.86294211 | 0.97359204 |
| MTRR | 38.4745973 | 0.258783837 | 0.27744465 | 0.9327404 | 0.35095402 | 0.77900549 |
| MIR4458HG | 12.3930996 | 0.052200063 | 0.44337645 | 0.11773305 | 0.90627918 | NA |
| FAM173B | 24.8389611 | 0.728865849 | 0.37881952 | 1.92404514 | 0.05434893 | 0.37099548 |
| CCT5 | 57.4998496 | -0.25680935 | 0.25710763 | -0.9988399 | 0.31787227 | 0.75982282 |
| CMBL | 40.6304297 | -0.347914911 | 0.40297249 | -0.8633714 | 0.38793332 | 0.80079872 |
| 6-Mar | 279.755284 | -0.158987254 | 0.18244089 | -0.8714453 | 0.38351106 | 0.7980695 |
| ANKRD33B | 70.7069788 | -0.643201265 | 0.68098085 | -0.9445218 | 0.34490306 | 0.77479382 |
| DAP | 187.864988 | 0.020406439 | 0.32239873 | 0.06329565 | 0.94953108 | 0.99192022 |
| DNAH5 | 140.788094 | -3.822415582 | 0.81543836 | -4.6875592 | 2.76E-06 | 0.0004257 |
| TRIO | 403.57464 | -1.699014908 | 0.41673331 | -4.0769837 | 4.56E-05 | 0.00379781 |
| OTULIN | 45.2459261 | -0.410848607 | 0.3125097 | -1.3146747 | 0.18861924 | 0.64021224 |
| ANKH | 129.869039 | 0.363060797 | 0.38630316 | 0.93983388 | 0.34730278 | 0.77726831 |
| ZNF622 | 13.4889666 | -0.147848611 | 0.47407835 | -0.3118653 | 0.75514286 | 0.94795939 |
| RETREG1 | 303.49355 | -0.994040834 | 0.26570384 | -3.7411609 | 0.00018317 | 0.01033006 |
| MYO10 | 556.253988 | 0.031800808 | 0.280692 | 0.11329432 | 0.90979722 | 0.98329594 |
| LINC02241 | 28.9897424 | 0.640727796 | 0.35926918 | 1.78341988 | 0.07451798 | 0.43076544 |
| CDH12 | 18.565626 | -0.179487325 | 0.52969188 | -0.3388523 | 0.73472098 | 0.94282226 |
| C5orf17 | 60.1031334 | -0.985366179 | 1.06906992 | -0.9217042 | 0.35668292 | 0.78241164 |
| DROSHA | 73.3585366 | 0.166615036 | 0.19575796 | 0.85112775 | 0.39469839 | 0.80516812 |
| C5orf22 | 23.9466677 | 0.065458211 | 0.28794784 | 0.22732663 | 0.82016977 | 0.96389705 |
| PDZD2 | 173.506115 | 1.229131552 | 0.54474151 | 2.25635742 | 0.02404826 | 0.24298759 |
| GOLPH3 | 227.939289 | -0.289030832 | 0.24076124 | -1.2004874 | 0.22995012 | 0.68523674 |
| MTMR12 | 95.4090981 | 0.192648 | 0.24778272 | 0.77748764 | 0.43687113 | 0.82687287 |
| ZFR | 265.575716 | -0.025777053 | 0.13992208 | -0.1842243 | 0.85383747 | 0.97135895 |
| SUB1 | 122.440741 | 0.087778869 | 0.26818133 | 0.32731163 | 0.7434322 | 0.94546379 |
| TARS | 38.5688 | -0.322899966 | 0.35416795 | -0.9117142 | 0.36191917 | 0.78713361 |
| C1QTNF3-AN | 87.7179301 | -0.189266153 | 0.34297406 | -0.5518381 | 0.58105928 | 0.89012555 |
| RAI14 | 45.4753517 | -0.130355621 | 0.33158542 | -0.3931283 | 0.6942247 | 0.92841398 |
| RAD1 | 14.0154589 | 0.425484374 | 0.36894012 | 1.15326134 | 0.24880313 | 0.70361235 |
| BRIX1 | 19.5876686 | 0.636575755 | 0.39514232 | 1.61100374 | 0.10717891 | 0.50321172 |
| DNAJC21 | 58.4251922 | -0.144064122 | 0.19262151 | -0.747913 | 0.45451266 | 0.83385406 |
| PRLR | 288.396221 | 0.287816265 | 0.29720371 | 0.96841411 | 0.3328376 | 0.7674968 |
| SPEF2 | 18.5719727 | 0.522553252 | 0.46410271 | 1.12594312 | 0.2601896 | 0.71091822 |
| LMBRD2 | 41.8628226 | 0.255448607 | 0.2693326 | 0.94845039 | 0.34290022 | 0.7729798 |
| NADK2 | 50.8372617 | 0.348866858 | 0.40865371 | 0.853698 | 0.39327234 | 0.80400536 |
| RANBP3L | 89.1376082 | 0.846848964 | 0.37360496 | 2.26669627 | 0.02340879 | 0.23883233 |
| NIPBL | 482.607684 | -0.173762348 | 0.15403822 | -1.128047 | 0.25930008 | 0.71066775 |
| CPLANE1 | 147.299487 | 0.296040153 | 0.18772372 | 1.57699919 | 0.11479571 | 0.51928773 |
| NUP155 | 24.120242 | 0.610178508 | 0.354633 | 1.72059147 | 0.08532498 | 0.46032833 |
| WDR70 | 205.639169 | 0.113334637 | 0.18283921 | 0.61985958 | 0.53535024 | 0.86946459 |
| LIFR | 233.261244 | 0.547103144 | 0.32967892 | 1.65950294 | 0.09701449 | 0.48257296 |
| OSMR-AS1 | 121.852879 | -1.081313049 | 0.36358561 | -2.974026 | 0.0029392 | 0.06702971 |
| OSMR | 138.894814 | -1.90549587 | 0.39610635 | -4.8105663 | 1.51E-06 | 0.00027035 |
| RICTOR | 320.758927 | 0.239761424 | 0.2393495 | 1.00172102 | 0.31647835 | 0.75864788 |
| FYB1 | 80.5793684 | -1.643208736 | 0.81992319 | -2.0041008 | 0.04505926 | 0.33698907 |
| C9 | 24.92864 | 1.229565591 | 0.8934001 | 1.37627653 | 0.16873604 | 0.61072372 |
| TTC33 | 64.3348328 | -0.144083535 | 0.30255456 | -0.4762233 | 0.6339153 | 0.90805737 |
| PRKAA1 | 108.465332 | -0.167206002 | 0.23176328 | -0.7214517 | 0.47063167 | 0.84424147 |
| RPL37 | 123.207133 | 0.214873297 | 0.27774699 | 0.77362961 | 0.43914986 | 0.82745796 |
| CARD6 | 17.0203137 | 0.039339106 | 0.67349016 | 0.05841081 | 0.9534214 | 0.99217529 |
| C6 | 50.073041 | -1.262724132 | 1.20852783 | -1.0448449 | 0.29609466 | 0.74123516 |
| PLCXD3 | 25.2750044 | 1.84536517 | 0.70459333 | 2.61905 | 0.0088175 | 0.13641112 |
| OXCT1 | 126.102219 | 0.868970497 | 0.45814368 | 1.89672045 | 0.05786484 | 0.38201831 |
| C5orf51 | 19.5576414 | -0.347868074 | 0.34932075 | -0.9958414 | 0.3193272 | 0.76086314 |
| FBXO4 | 15.1397059 | -0.460598308 | 0.39475317 | -1.1668008 | 0.24329083 | 0.69729164 |
| SELENOP | 125.033984 | 0.295340238 | 0.35796357 | 0.82505668 | 0.40933941 | 0.81413754 |
| ZNF131 | 128.165839 | 0.046883333 | 0.1744614 | 0.26873183 | 0.78813605 | 0.95669124 |
| HMGCS1 | 172.233105 | -1.19812577 | 0.62725791 | -1.9101007 | 0.05612025 | 0.37694988 |
| CCL28 | 14.0156576 | -1.023538303 | 0.69376898 | -1.4753302 | 0.1401238 | 0.57374456 |
| TMEM267 | 36.3097994 | -0.017087657 | 0.27919137 | -0.0612041 | 0.95119666 | 0.99217308 |
| C5orf34 | 11.8932949 | -0.076036568 | 0.5618634 | -0.1353293 | 0.89235154 | NA |
| PAIP1 | 54.9649343 | 0.043459587 | 0.22135787 | 0.19633179 | 0.84435048 | 0.97005799 |
| NNT-AS1 | 119.226427 | 0.231109314 | 0.28370046 | 0.8146244 | 0.41528734 | 0.81726257 |
| NNT | 396.921318 | 0.386225336 | 0.25855098 | 1.4938073 | 0.13522599 | 0.56489756 |
| MRPS30-DT | 13.06176 | 0.153777419 | 0.49874618 | 0.30832801 | 0.75783275 | 0.94918855 |
| MRPS30 | 28.0187645 | 0.213731649 | 0.2876021 | 0.74315051 | 0.45739056 | 0.83569193 |
| PARP8 | 191.152943 | -0.988614106 | 0.38114351 | -2.5938107 | 0.00949187 | 0.14341304 |
| ITGA1 | 95.7466479 | 0.587602641 | 0.3717932 | 1.58045557 | 0.11400257 | 0.51864031 |
| ITGA2 | 171.409333 | -0.837866738 | 0.41582906 | -2.0149307 | 0.04391191 | 0.33287289 |
| MOCS2 | 31.9486492 | 0.273919364 | 0.26335201 | 1.04012635 | 0.2982812 | 0.74296125 |
| NDUFS4 | 153.450909 | 0.018093492 | 0.17967163 | 0.10070311 | 0.91978614 | 0.98584813 |
| ARL15 | 2430.49289 | -0.552354618 | 0.16989786 | -3.2510982 | 0.0011496 | 0.03605702 |
| SNX18 | 49.8209512 | 0.030945988 | 0.35261966 | 0.08776024 | 0.93006724 | 0.98748382 |
| AC026704.1 | 13.663009 | -1.385060282 | 0.65623765 | -2.1106078 | 0.03480604 | 0.29824962 |
| DHX29 | 76.9640649 | -0.074141499 | 0.20550672 | -0.3607741 | 0.71826833 | 0.93733389 |
| MTREX | 143.575402 | -0.090486551 | 0.19927748 | -0.4540731 | 0.64977618 | 0.91438111 |

| | | | | | | |
|---|---|---|---|---|---|---|
| PLPP1 | 138.993824 | 0.168685986 | 0.37576532 | 0.44891314 | 0.65349432 | 0.9158679 |
| SLC38A9 | 77.718935 | -0.117501382 | 0.22113416 | -0.5313579 | 0.59517079 | 0.893081 |
| IL6ST | 168.477816 | -0.285522218 | 0.20243562 | -1.4104347 | 0.15841137 | 0.59881912 |
| LINC01948 | 10.628009 | -0.002322406 | 0.68799409 | -0.0033756 | 0.99730665 | NA |
| MAP3K1 | 156.987714 | -0.254616864 | 0.20355756 | -1.2508347 | 0.21099479 | 0.6627751 |
| MIER3 | 30.5215434 | 0.023236097 | 0.26876443 | 0.08645525 | 0.93110453 | 0.98757533 |
| GPBP1 | 278.964933 | 0.002934799 | 0.23931596 | 0.01226328 | 0.99021556 | 0.99923337 |
| PDE4D | 4382.79138 | 0.509506647 | 0.43516295 | 1.17084105 | 0.24166267 | 0.69556484 |
| PART1 | 45.556407 | 0.810933626 | 0.82811427 | 0.97925329 | 0.32745484 | 0.76463938 |
| ELOVL7 | 258.58933 | -0.244305577 | 0.45436405 | -0.5376868 | 0.59079326 | 0.89149106 |
| ERCC8 | 60.9881694 | 0.499577432 | 0.21442029 | 2.32989814 | 0.01981154 | 0.22088723 |
| NDUFAF2 | 139.150917 | -0.418824081 | 0.32053608 | -1.3066363 | 0.19133627 | 0.6421543 |
| SMIM15 | 26.0674047 | 0.020111518 | 0.2803053 | 0.07174862 | 0.94280196 | 0.9903811 |
| ZSWIM6 | 980.702103 | 0.373709933 | 0.27240921 | 1.37186969 | 0.17010401 | 0.61252734 |
| KIF2A | 63.9956081 | -0.11242443 | 0.2683397 | -0.4189631 | 0.6752431 | 0.92193843 |
| IPO11 | 44.8737448 | -0.030531819 | 0.28692516 | -0.1064104 | 0.91525675 | 0.98425615 |
| RNF180 | 118.429564 | -0.035150955 | 0.19416703 | -0.1810346 | 0.8563404 | 0.97166553 |
| SREK1IP1 | 43.8912865 | 0.231268035 | 0.23832301 | 0.97039743 | 0.33184843 | 0.76685572 |
| CWC27 | 97.0651157 | 0.141827969 | 0.19395634 | 0.73123658 | 0.46463466 | 0.84133578 |
| PPWD1 | 65.5214661 | 0.146707062 | 0.30347936 | 0.48341694 | 0.62879972 | 0.90659442 |
| TRIM23 | 30.9444876 | 0.16588799 | 0.27570147 | 0.60169425 | 0.54737768 | 0.8729963 |
| TRAPPC13 | 56.281182 | 0.143107818 | 0.2276228 | 0.62870599 | 0.52954156 | 0.86758926 |
| NLN | 37.5324169 | -0.453952326 | 0.4434804 | -1.0236131 | 0.30601808 | 0.74897897 |
| ERBIN | 480.186971 | -0.310915405 | 0.20650186 | -1.50563 | 0.13216217 | 0.55956923 |
| SREK1 | 182.865638 | 0.173732326 | 0.2222916 | 0.78155145 | 0.43447823 | 0.82565889 |
| AC113378.1 | 16.7561304 | 1.335619476 | 0.77696405 | 1.71902352 | 0.08561009 | 0.46118484 |
| MAST4 | 725.533984 | -0.351612495 | 0.52800417 | -0.6659275 | 0.50545745 | 0.86197912 |
| PIK3R1 | 110.042457 | -0.81653585 | 0.4213647 | -1.9378364 | 0.05264319 | 0.36370293 |
| SLC30A5 | 54.1423928 | 0.507323315 | 0.22800414 | 2.22506188 | 0.02607708 | 0.25383383 |
| MRPS36 | 64.016107 | -0.203097584 | 0.35258053 | -0.5760318 | 0.56459372 | 0.88191823 |
| CDK7 | 29.6895149 | -0.120984285 | 0.32953194 | -0.3671398 | 0.71351476 | 0.93616842 |
| CCDC125 | 42.7897737 | 0.038402794 | 0.40981397 | 0.09370787 | 0.92534122 | 0.98672316 |
| AK6 | 24.4160288 | -0.176565275 | 0.36737731 | -0.4806102 | 0.63079358 | 0.90768383 |
| RAD17 | 50.0449988 | 0.085625492 | 0.22003117 | 0.38915165 | 0.69716397 | 0.92989421 |
| MARVELD2 | 39.5729943 | 0.382008868 | 0.25547109 | 1.49531153 | 0.13483316 | 0.56408911 |
| OCLN | 100.332207 | 0.032237487 | 0.29982593 | 0.10752068 | 0.91437592 | 0.98407488 |
| GTF2H2C | 62.9535363 | 0.169307742 | 0.3419626 | 0.49510603 | 0.62052528 | 0.90380166 |
| NAIP | 103.912175 | 0.824255025 | 0.30757144 | 2.6798815 | 0.00736482 | 0.12396174 |
| GTF2H2 | 53.1108517 | 0.421725583 | 0.48765066 | 0.86481085 | 0.38714262 | 0.80009555 |
| BDP1 | 211.34537 | 0.040606689 | 0.17624444 | 0.23039982 | 0.8177811 | 0.9635539 |
| MCCC2 | 86.2922928 | 0.177712213 | 0.20638445 | 0.86107364 | 0.38919748 | 0.80191835 |
| MRPS27 | 72.1235238 | -0.037123888 | 0.22839475 | -0.1625427 | 0.87087854 | 0.97434023 |
| PTCD2 | 11.3769836 | 0.196181934 | 0.56279407 | 0.34858565 | 0.7274004 | NA |
| TNPO1 | 168.692589 | 0.018306546 | 0.17908894 | 0.10222041 | 0.91858172 | 0.98521039 |
| FCHO2 | 202.407457 | -0.567008257 | 0.25493923 | -2.2240918 | 0.02614227 | 0.25383383 |
| TMEM171 | 29.3732907 | -1.256474613 | 0.5077052 | -2.4748114 | 0.01333066 | 0.17367489 |
| AC116345.3 | 21.3632928 | 0.502975786 | 0.68824276 | 0.73081159 | 0.46489425 | 0.84143009 |
| AC116345.1 | 16.6707129 | -1.427645039 | 0.70040539 | -2.0383125 | 0.04151869 | 0.32715785 |
| BTF3 | 77.3252368 | -0.193562324 | 0.43339838 | -0.4466152 | 0.65515289 | 0.9158679 |
| ANKRA2 | 25.5385147 | 0.014893327 | 0.4435617 | 0.03357668 | 0.97321472 | 0.99526459 |
| ARHGEF28 | 916.042728 | -0.19961668 | 0.22697296 | -0.8794734 | 0.37914466 | 0.79535699 |
| ENC1 | 15.9183004 | 0.67597348 | 0.72455388 | 0.93295129 | 0.35084512 | 0.77894201 |
| HEXB | 36.9600801 | -0.700226628 | 0.3402316 | -2.0580881 | 0.03958167 | 0.31915397 |
| GFM2 | 42.4823855 | 0.116751992 | 0.26504855 | 0.44049286 | 0.65958018 | 0.91732025 |
| NSA2 | 37.6107793 | -0.067881908 | 0.34765859 | -0.1952545 | 0.84519371 | 0.97027687 |
| FAM169A | 100.639557 | 0.048519357 | 0.28436509 | 0.17062346 | 0.86451985 | 0.97407858 |
| GCNT4 | 11.9438446 | -0.075544616 | 0.44583288 | -0.169446 | 0.86544581 | NA |
| HMGCR | 128.817577 | -0.746295555 | 0.46022541 | -1.621587 | 0.1048918 | 0.49923967 |
| COL4A3BP | 224.40394 | -0.125222523 | 0.20733952 | -0.6039491 | 0.54587747 | 0.87247631 |
| POLK | 142.288269 | -0.138750527 | 0.19552886 | -0.7096166 | 0.47794191 | 0.84771464 |
| ANKDD1B | 76.7633375 | 0.351267047 | 0.39559538 | 0.88794527 | 0.37457019 | 0.79261144 |
| POC5 | 26.812309 | -0.096737911 | 0.28091527 | -0.3443669 | 0.7305704 | 0.94155624 |
| IQGAP2 | 191.450213 | 0.090639906 | 0.45204335 | 0.20051153 | 0.84108054 | 0.96767658 |
| F2RL1 | 21.5668774 | -0.220048157 | 0.3507829 | -0.6273058 | 0.53045879 | 0.86784453 |
| ZBED3 | 21.492917 | 0.175042468 | 0.31613623 | 0.55369316 | 0.57978886 | 0.88972503 |
| WDR41 | 97.6152805 | 0.225329346 | 0.31291913 | 0.72008811 | 0.47147074 | 0.84517949 |
| TBCA | 179.440374 | -0.151084584 | 0.23569573 | -0.6410154 | 0.5215127 | 0.86703098 |
| AP3B1 | 295.558313 | -0.02576469 | 0.22275372 | -0.1156645 | 0.90791847 | 0.98250721 |
| SCAMP1-AS1 | 10.8638167 | -0.040846282 | 0.5139923 | -0.0794687 | 0.93665985 | NA |
| SCAMP1 | 152.151145 | 0.059633436 | 0.21426486 | 0.27831646 | 0.78076944 | 0.95537097 |
| ARSB | 24.0231902 | -0.127311071 | 0.45201195 | -0.2816542 | 0.77820866 | 0.95535696 |
| DMGDH | 80.6909481 | -0.924408967 | 0.50085633 | -1.845657 | 0.06494203 | 0.40484427 |
| BHMT2 | 42.1665386 | -0.843342614 | 0.40673244 | -2.073458 | 0.03812968 | 0.31326583 |
| JMY | 112.362154 | -0.833266377 | 0.35065896 | -2.3762872 | 0.01748785 | 0.20486971 |
| HOMER1 | 48.3468413 | 0.416491284 | 0.41712374 | 0.99848378 | 0.31804483 | 0.75986079 |
| TENT2 | 281.991854 | 0.002704937 | 0.21983553 | 0.01230437 | 0.99018278 | 0.99923337 |
| CMYA5 | 139.820912 | -0.230507056 | 0.38579948 | -0.5974789 | 0.55018768 | 0.87445431 |
| MTX3 | 25.2397863 | 0.233376894 | 0.32960807 | 0.70804362 | 0.47891817 | 0.84833462 |
| SERINC5 | 116.298324 | -0.232137951 | 0.41058646 | -0.5653814 | 0.57181437 | 0.88578992 |
| ZFYVE16 | 178.917693 | 0.132688984 | 0.27078654 | 0.49001322 | 0.62412454 | 0.90447745 |
| FAM151B | 26.9048612 | 0.608567588 | 0.62292425 | 0.97695279 | 0.32859253 | 0.76649357 |
| DHFR | 483.818527 | 0.062060834 | 0.42675383 | 0.14542537 | 0.88437504 | 0.97662748 |
| MSH3 | 107.966946 | 0.021375424 | 0.21310678 | 0.10030382 | 0.92010313 | 0.98586108 |
| RASGRF2 | 193.370972 | -0.665171561 | 0.33340634 | -1.9950777 | 0.0460344 | 0.34112583 |
| CKMT2-AS1 | 67.7548086 | 0.169097442 | 0.27834579 | 0.60750853 | 0.54351349 | 0.87233405 |
| CKMT2 | 119.53857 | -0.187774403 | 0.33214058 | -0.5653462 | 0.57183833 | 0.88578992 |
| ZCCHC9 | 33.3579163 | -0.152209123 | 0.33857802 | -0.4495541 | 0.65303202 | 0.9158679 |
| SSBP2 | 47.7539067 | 0.821685321 | 0.61306488 | 1.34029096 | 0.18015077 | 0.62768048 |
| ATG10 | 117.958188 | 0.047254603 | 0.33475176 | 0.14116312 | 0.88774108 | 0.9776239 |
| RPS23 | 268.613554 | 0.129572105 | 0.42689228 | 0.30352412 | 0.76149047 | 0.9499444 |
| ATP6AP1L | 30.9351952 | 0.671740339 | 0.37972489 | 1.76901847 | 0.07689079 | 0.43590923 |
| TMEM167A | 52.3723428 | 0.048080967 | 0.32856103 | 0.14633801 | 0.88365456 | 0.97662748 |
| XRCC4 | 53.670484 | -0.206222565 | 0.29961519 | -0.6882914 | 0.49126928 | 0.85654883 |
| COX7C | 203.519432 | -0.274686292 | 0.35674637 | -0.7699764 | 0.44131389 | 0.8281114 |
| RASA1 | 189.887054 | -0.130280188 | 0.23279509 | -0.5596346 | 0.5757287 | 0.88746465 |
| CCNH | 202.357219 | 0.161568741 | 0.28454209 | 0.56782018 | 0.57015708 | 0.88563341 |
| TMEM161B | 91.2268678 | 0.149760432 | 0.21981291 | 0.68130862 | 0.49567623 | 0.85814183 |
| TMEM161B-. | 186.133639 | 0.667767729 | 0.42393734 | 1.57515666 | 0.11522029 | 0.51983107 |
| MEF2C | 27.9774587 | -0.365771808 | 0.63423057 | -0.5767174 | 0.56413037 | 0.88191823 |
| CETN3 | 22.2184667 | 0.4255688 | 0.44642705 | 0.95327736 | 0.34044956 | 0.77175993 |
| LYSMD3 | 38.7436211 | 0.530132866 | 0.34867444 | 1.52042366 | 0.12840453 | 0.5528291 |
| ADGRV1 | 59.0557951 | -0.076478716 | 0.4985181 | -0.1534121 | 0.87807329 | 0.97542218 |
| ARRDC3 | 48.7780022 | -1.152411782 | 0.42229601 | -2.7289194 | 0.00635422 | 0.11309352 |
| ARRDC3-AS1 | 40.5080455 | -0.116750114 | 0.34401619 | -0.3393739 | 0.73432808 | 0.94277981 |
| FAM172A | 253.19501 | 0.13011046 | 0.22088547 | 0.58904037 | 0.55583419 | 0.87787661 |
| KIAA0825 | 49.0394106 | 0.923213713 | 0.25767368 | 3.58287944 | 0.00033983 | 0.01569679 |
| SLF1 | 12.5680113 | 0.493595409 | 0.36726633 | 1.34397132 | 0.17895766 | 0.62639418 |
| TTC37 | 161.576449 | -0.342776659 | 0.17992116 | -1.9051492 | 0.05676068 | 0.37858396 |
| ARSK | 38.0847365 | -0.364571926 | 0.38307833 | -0.9516903 | 0.34125408 | 0.77193691 |
| RHOBTB3 | 252.743988 | -0.099234979 | 0.28621823 | -0.3467109 | 0.72880853 | 0.94097305 |
| GLRX | 79.3902113 | -0.142520386 | 0.4467548 | -0.3190125 | 0.749717 | 0.94684415 |
| ELL2 | 162.072636 | -0.325763439 | 0.38552549 | -0.8449855 | 0.39811895 | 0.80714564 |
| AC104123.1 | 56.1730941 | -0.229757648 | 0.44433151 | -0.5170861 | 0.60509606 | 0.89600432 |

| | | | | | | |
|---|---|---|---|---|---|---|
| CAST | 256.163604 | -0.636436048 | 0.18325081 | -3.4730327 | 0.00051461 | 0.02047878 |
| ERAP1 | 70.8283018 | -0.408834333 | 0.28279257 | -1.4457039 | 0.14826026 | 0.58517098 |
| AC009126.1 | 27.7195328 | 0.294999784 | 0.49057541 | 0.60133422 | 0.5476174 | 0.87309151 |
| ERAP2 | 32.5529022 | 0.260526195 | 0.56832023 | 0.45841443 | 0.64665473 | 0.91276933 |
| LNPEP | 275.267031 | 0.576795342 | 0.21437161 | 2.69063311 | 0.00713166 | 0.12146099 |
| LIX1-AS1 | 42.835524 | 0.942494931 | 0.56710122 | 1.66195188 | 0.09652242 | 0.48257296 |
| RIOK2 | 13.2603142 | 0.367147237 | 0.46547009 | 0.78876655 | 0.43024846 | 0.82469033 |
| RGMB | 12.9504337 | 0.323653417 | 0.45262189 | 0.71506355 | 0.47456977 | 0.84593347 |
| CHD1 | 136.386058 | 0.098924311 | 0.25457364 | 0.38858819 | 0.6975808 | 0.9299799 |
| LINC02062 | 28.8230572 | -0.220478755 | 0.42056938 | -0.5242387 | 0.60011251 | 0.89486416 |
| FAM174A | 47.9351365 | 0.200297551 | 0.26474852 | 0.75655779 | 0.44931484 | 0.83118301 |
| SLCO4C1 | 33.8187867 | 0.639730498 | 0.91280136 | 0.70084306 | 0.48340096 | 0.85207874 |
| PAM | 212.488001 | -0.793429471 | 0.34117099 | -2.3256065 | 0.02003955 | 0.22256173 |
| GIN1 | 21.8431134 | 0.34508998 | 0.33309975 | 1.0359959 | 0.30020405 | 0.74437099 |
| PPIP5K2 | 78.272822 | 0.447458388 | 0.2118366 | 2.11228083 | 0.03466237 | 0.29754423 |
| C5orf30 | 12.5234446 | 0.031967055 | 0.57512866 | 0.05558244 | 0.95567445 | NA |
| NUDT12 | 28.2278207 | 0.121223399 | 0.31165364 | 0.38896834 | 0.69729956 | 0.92989421 |
| EFNA5 | 1350.41241 | -0.92219594 | 0.3974553 | -2.3202507 | 0.02032732 | 0.22348867 |
| FBXL17 | 638.854086 | 0.272099389 | 0.19956442 | 1.36346647 | 0.17273554 | 0.61746168 |
| FER | 407.956483 | 0.239517392 | 0.20672992 | 1.15860054 | 0.24661905 | 0.70024501 |
| AC008467.1 | 20.3730589 | -0.205312106 | 0.65748246 | -0.3122701 | 0.75483527 | 0.94795939 |
| PJA2 | 173.402963 | -0.205877281 | 0.15792979 | -1.3036 | 0.19236999 | 0.64233696 |
| MAN2A1 | 336.150671 | -0.746418765 | 0.38452211 | -1.9411595 | 0.05223893 | 0.36271841 |
| TMEM232 | 49.3103466 | 0.753667389 | 0.46996916 | 1.60365286 | 0.10879059 | 0.50661006 |
| SLC25A46 | 66.9049566 | 0.21039054 | 0.32159623 | 0.65420711 | 0.51297838 | 0.86357 |
| WDR36 | 41.5090266 | 0.255554942 | 0.23526921 | 1.08622351 | 0.27738012 | 0.72678432 |
| STARD4 | 25.2586508 | -0.648193049 | 0.44839343 | -1.44559 | 0.14829224 | 0.58517098 |
| NREP | 30.6173253 | 0.449364739 | 0.41139592 | 1.09229265 | 0.27470449 | 0.72554623 |
| EPB41L4A | 329.67599 | -0.396971587 | 0.22861754 | -1.7364004 | 0.08249306 | 0.45473406 |
| EPB41L4A-AS | 12.0166411 | 0.555546391 | 0.3578254 | 1.55256275 | 0.12052763 | NA |
| APC | 61.8328605 | 0.326027722 | 0.21930543 | 1.48663768 | 0.13711053 | 0.56856307 |
| SRP19 | 36.0737254 | -0.163200472 | 0.39141584 | -0.4169491 | 0.67671566 | 0.92237057 |
| REEP5 | 188.814698 | -0.063434159 | 0.23954697 | -0.2648089 | 0.79115671 | 0.95746995 |
| DCP2 | 107.509752 | 0.583181337 | 0.39679965 | 1.46971232 | 0.14163969 | 0.57654425 |
| MCC | 188.391619 | 0.420911099 | 0.35621675 | 1.18161513 | 0.23735845 | 0.69207518 |
| YTHDC2 | 128.912279 | 0.165960292 | 0.19541706 | 0.84926203 | 0.3957355 | 0.80624334 |
| PGGT1B | 52.2517215 | 0.116107068 | 0.30901187 | 0.37573659 | 0.70711273 | 0.93432317 |
| CCDC112 | 13.8206701 | -0.148957944 | 0.50071036 | -0.2974932 | 0.76608998 | 0.95205353 |
| FEM1C | 20.8628336 | -0.339238971 | 0.34108992 | -0.9945734 | 0.31994377 | 0.76106474 |
| TMED7-TICAI | 17.1129493 | -0.090898623 | 0.41655932 | -0.2182129 | 0.82726322 | 0.96561348 |
| ATG12 | 37.5105173 | -0.028511691 | 0.34670631 | -0.0822359 | 0.93445916 | 0.9881638 |
| AP3S1 | 54.1564513 | -0.119500716 | 0.27828897 | -0.4294123 | 0.66762318 | 0.92049996 |
| COMMD10 | 245.729132 | 0.218569173 | 0.18971099 | 1.15211656 | 0.24927318 | 0.70371066 |
| SEMA6A | 13.4515325 | 0.08419436 | 0.44210675 | 0.19043899 | 0.84896515 | 0.97027687 |
| SEMA6A-AS1 | 34.2109682 | 0.027704713 | 0.35830375 | 0.07732186 | 0.9383675 | 0.98871562 |
| DTWD2 | 59.751306 | 0.350293777 | 0.31869139 | 1.09916299 | 0.27169698 | 0.72303448 |
| DMXL1 | 278.769481 | -0.041705768 | 0.19569346 | -0.2131178 | 0.83123505 | 0.96701848 |
| TNFAIP8 | 52.3324953 | -0.39401876 | 0.38239659 | -1.030393 | 0.30282557 | 0.74712557 |
| HSD17B4 | 193.610308 | -0.331698434 | 0.36291073 | -0.9139946 | 0.3607197 | 0.78611122 |
| SRFBP1 | 47.8135634 | -0.166669516 | 0.26281035 | -0.6341817 | 0.52596225 | 0.86750588 |
| AC010255.1 | 17.9801771 | 0.813098792 | 1.73801285 | 0.46783244 | 0.63990442 | 0.91096398 |
| SNX2 | 102.995115 | -0.303668792 | 0.21604387 | -1.4055886 | 0.15984633 | 0.60024417 |
| SNX24 | 250.816953 | 0.300700364 | 0.24966052 | 1.20443698 | 0.22842073 | 0.68364117 |
| PPIC | 55.5213616 | -0.029716006 | 0.25926646 | -0.1146157 | 0.90874974 | 0.98281552 |
| CEP120 | 32.230279 | 0.111014334 | 0.3304387 | 0.33596044 | 0.7369007 | 0.94369536 |
| CSNK1G3 | 111.410322 | -0.273136553 | 0.31929452 | -0.8554376 | 0.39230891 | 0.80367402 |
| ZNF608 | 123.883236 | -0.441108112 | 0.47545582 | -0.9277584 | 0.35353293 | 0.78028099 |
| GRAMD2B | 415.529507 | -0.041134832 | 0.15528286 | -0.2649026 | 0.7910845 | 0.95746995 |
| AC093535.1 | 18.4278706 | -0.68532428 | 0.65967114 | -1.0388878 | 0.29885693 | 0.74296125 |
| ALDH7A1 | 28.1058611 | 0.494396358 | 0.34791037 | 1.4210452 | 0.15530362 | 0.59533243 |
| PHAX | 28.4297497 | 0.336851459 | 0.34633475 | 0.97261815 | 0.33074312 | 0.76685572 |
| C5orf63 | 27.0822654 | 0.262653796 | 0.34751225 | 0.75581162 | 0.44976215 | 0.83118301 |
| PRRC1 | 30.6359498 | 0.343576877 | 0.26595698 | 1.29185134 | 0.19640863 | 0.64685325 |
| LINC01184 | 57.9133818 | -0.001095298 | 0.39720088 | -0.0027575 | 0.9977998 | 0.99982181 |
| SLC12A2 | 44.0785036 | 0.671836761 | 0.45776654 | 1.4676406 | 0.14220187 | 0.57673923 |
| SLC27A6 | 17.730567 | 2.52548808 | 1.39127931 | 1.81522723 | 0.069489 | 0.41827125 |
| ISOC1 | 20.0044263 | 0.504660864 | 0.41462444 | 1.21715177 | 0.22354648 | 0.67606266 |
| HINT1 | 197.531238 | -0.377049742 | 0.37956587 | -0.993371 | 0.32052917 | 0.76131909 |
| LYRM7 | 31.964509 | 0.078566568 | 0.30416333 | 0.25830388 | 0.79617239 | 0.95848876 |
| CDC42SE2 | 448.716684 | -0.707296358 | 0.33241292 | -2.1277644 | 0.03335662 | 0.28966809 |
| P4HA2 | 14.296748 | -0.003295131 | 0.42697019 | -0.0077175 | 0.99384241 | 0.99923337 |
| SLC22A4 | 13.8521868 | -0.236157081 | 0.61709516 | -0.3826915 | 0.70194851 | 0.93157758 |
| SLC22A5 | 90.633083 | 0.706463641 | 0.40946359 | 1.72533933 | 0.08446634 | 0.4585327 |
| AC116366.3 | 86.1363805 | -0.346629778 | 0.57335505 | -0.6045639 | 0.54546879 | 0.87247631 |
| IRF1 | 21.5209047 | -1.448365568 | 0.49079745 | -2.9510454 | 0.003167 | 0.07045856 |
| KIF3A | 65.448321 | 0.33216563 | 0.23767129 | 1.39758417 | 0.16223797 | 0.60094226 |
| CCNI2 | 28.358317 | 0.9540106 | 0.66191049 | 1.44129851 | 0.14950037 | 0.58659144 |
| 8-Sep | 44.0615826 | 0.313021635 | 0.30864162 | 1.01419127 | 0.31049149 | 0.75369557 |
| SOWAHA | 15.1414701 | 0.349612688 | 0.41254506 | 0.84745335 | 0.39674248 | 0.80674902 |
| SHROOM1 | 12.8365271 | 0.313920665 | 0.39941361 | 0.78595385 | 0.43189452 | 0.8248269 |
| UQCRQ | 98.2260355 | 0.331525022 | 0.36695778 | 0.90344187 | 0.36629143 | 0.78793182 |
| AFF4 | 322.917331 | -0.404438451 | 0.22037322 | -1.8352432 | 0.06646966 | 0.40936869 |
| ZCCHC10 | 47.8726134 | -0.314144793 | 0.28427945 | -1.1050563 | 0.26913521 | 0.72031329 |
| HSPA4 | 103.567337 | 0.032995668 | 0.23126156 | 0.14267683 | 0.88654541 | 0.9771038 |
| FSTL4 | 94.3025209 | 0.107192277 | 0.80295917 | 0.13349655 | 0.89380069 | 0.97833263 |
| C5orf15 | 57.7616886 | 0.348257851 | 0.21421858 | 1.62571266 | 0.1040108 | 0.49650823 |
| VDAC1 | 178.270066 | -0.063694895 | 0.34045101 | -0.1870897 | 0.85159026 | 0.97041406 |
| TCF7 | 15.0684582 | -0.458315636 | 0.41479097 | -1.1049316 | 0.26918924 | 0.72031329 |
| SKP1 | 155.79046 | -0.110869895 | 0.3773695 | -0.2937967 | 0.76891329 | 0.95205353 |
| CDKL3 | 20.4224261 | -0.083048898 | 0.30730729 | -0.2702471 | 0.78697017 | 0.95577371 |
| UBE2B | 121.768367 | -0.408858338 | 0.25138443 | -1.6264267 | 0.10385892 | 0.4962717 |
| JADE2 | 31.3094827 | -0.316686235 | 0.29420921 | -1.0763981 | 0.28174926 | 0.73311066 |
| SAR1B | 55.7198488 | 0.237012403 | 0.23456126 | 1.0104499 | 0.31227979 | 0.75484355 |
| SEC24A | 59.5144956 | 0.187388311 | 0.25545364 | 0.73355116 | 0.46322234 | 0.83970411 |
| CAMLG | 27.7769471 | -0.515089165 | 0.37218517 | -1.3839594 | 0.16637088 | 0.60746969 |
| DDX46 | 105.173529 | -0.008810251 | 0.22921635 | -0.0384364 | 0.96933974 | 0.99486579 |
| C5orf24 | 39.87465 | 0.587485457 | 0.44698456 | 1.31433053 | 0.188735 | 0.64021224 |
| TXNDC15 | 19.9064271 | 0.444890515 | 0.32060238 | 1.38767065 | 0.16523736 | 0.60578861 |
| PCBD2 | 95.5549506 | 0.327959284 | 0.32400436 | 1.01220638 | 0.31143939 | 0.75450562 |
| H2AFY | 96.8276854 | -0.386569948 | 0.28309056 | -1.3655346 | 0.17208509 | 0.61617768 |
| CXCL14 | 15.5047917 | -1.222378134 | 0.77105606 | -1.5853298 | 0.11289143 | 0.51571293 |
| SMAD5 | 102.991751 | 0.266721368 | 0.17141008 | 1.55604248 | 0.11969799 | 0.53109067 |
| KLHL3 | 76.5487234 | 0.498932301 | 0.5065665 | 0.98492953 | 0.32465869 | 0.7632548 |
| HNRNPA0 | 70.5230849 | 0.14822976 | 0.34899058 | 0.42473856 | 0.67102727 | 0.92170876 |
| AC106791.1 | 29.7810553 | 0.31843141 | 0.59796605 | 0.53252423 | 0.59436296 | 0.89289841 |
| FAM13B | 265.733528 | -0.10573632 | 0.17449376 | -0.6059605 | 0.544541 | 0.87247631 |
| NME5 | 14.2409984 | 0.481761047 | 0.55142359 | 0.87366782 | 0.38229918 | 0.79731722 |
| BRD8 | 41.4407132 | 0.075451203 | 0.23613716 | 0.31952279 | 0.74933011 | 0.94684415 |
| CDC23 | 24.4460781 | 0.083844699 | 0.33595022 | 0.24957477 | 0.80291621 | 0.95926681 |
| FAM53C | 23.2436053 | 0.08940468 | 0.45866852 | 0.1949222 | 0.84545385 | 0.97027687 |
| KDM3B | 113.42111 | -0.14473304 | 0.16020562 | -0.9034205 | 0.36630278 | 0.78793182 |
| EGR1 | 43.0237813 | 1.110800463 | 0.80797147 | 1.3748016 | 0.16919296 | 0.61168883 |
| ETF1 | 70.2799483 | 0.246706529 | 0.31463437 | 0.78410545 | 0.43297823 | 0.8248269 |

| | | | | | | |
|---|---|---|---|---|---|---|
| HSPA9 | 180.872904 | 0.071408577 | 0.22856246 | 0.31242479 | 0.75471771 | 0.94795939 |
| CTNNA1 | 345.159927 | -0.585462824 | 0.18039404 | -3.2454666 | 0.00117258 | 0.03637703 |
| SIL1 | 142.728451 | 0.246394868 | 0.25669764 | 0.95986416 | 0.33712359 | 0.76966704 |
| PAIP2 | 76.905895 | -0.0531806 | 0.24775705 | -0.2146482 | 0.83004163 | 0.966913 |
| TMEM173 | 25.6679144 | -1.588056155 | 0.4968354 | -3.1963426 | 0.00139182 | 0.04030231 |
| UBE2D2 | 171.431138 | 0.103853408 | 0.23772589 | 0.436862 | 0.66221143 | 0.91812398 |
| CXXC5 | 80.129448 | 0.280040712 | 0.36882232 | 0.75928352 | 0.44768297 | 0.83066268 |
| NRG2 | 21.6582185 | 1.208161118 | 0.79328489 | 1.52298516 | 0.12776242 | 0.55107954 |
| PURA | 122.008744 | 0.161089126 | 0.18724412 | 0.86031606 | 0.38961484 | 0.80191835 |
| IGIP | 18.0196681 | 0.607239947 | 0.3675674 | 1.6520506 | 0.09852424 | 0.48476221 |
| CYSTM1 | 412.842866 | -0.281549873 | 0.3321922 | -0.8475511 | 0.39668801 | 0.80674902 |
| PFDN1 | 46.2727404 | 0.032734852 | 0.26872674 | 0.12181464 | 0.90304582 | 0.98016176 |
| SLC4A9 | 242.023824 | 1.327712968 | 0.76500251 | 1.73556682 | 0.08264046 | 0.45473406 |
| AC008438.1 | 75.989833 | 0.886696444 | 0.77721055 | 1.14087032 | 0.25392389 | 0.70701839 |
| ANKHD1-EIF4 | 12.3844752 | -0.221541604 | 0.44625653 | -0.4964445 | 0.61958083 | NA |
| SRA1 | 41.4387972 | -0.379731807 | 0.37853621 | -1.0031585 | 0.3157844 | 0.75793414 |
| SLC35A4 | 27.2364689 | 0.27778051 | 0.28891033 | 0.96147655 | 0.33631261 | 0.76884797 |
| NDUFA2 | 49.2827376 | 0.38570587 | 0.39082481 | 0.98690222 | 0.32369058 | 0.76304922 |
| IK | 21.9980648 | -0.495850255 | 0.35924993 | -1.3802376 | 0.1675135 | 0.60902585 |
| WDR55 | 16.8083469 | -0.369629427 | 0.39515935 | -0.9353933 | 0.34958563 | 0.77868979 |
| HARS | 28.4606844 | 0.049111699 | 0.29895036 | 0.16428045 | 0.86951037 | 0.97434023 |
| ZMAT2 | 24.0275439 | -0.437911072 | 0.29826852 | -1.4681773 | 0.14205606 | 0.57673923 |
| TAF7 | 51.8474773 | 0.211976936 | 0.27371309 | 0.7744494 | 0.43866508 | 0.82745796 |
| DIAPH1 | 352.014012 | -0.346199394 | 0.21907753 | -1.5802597 | 0.1140474 | 0.51864031 |
| HDAC3 | 27.2239067 | 0.612605874 | 0.3235051 | 1.89365135 | 0.0582713 | 0.38243048 |
| PCDH1 | 40.9982318 | -0.118500557 | 0.37395225 | -0.3168869 | 0.75132945 | 0.94719044 |
| DELE1 | 49.3072018 | 0.257855436 | 0.22676602 | 1.13709907 | 0.25549687 | 0.70849619 |
| RNF14 | 57.0928168 | 0.087755507 | 0.269062 | 0.32615348 | 0.74430824 | 0.94546379 |
| GNPDA1 | 23.3703933 | -0.948179132 | 0.50306619 | -1.8847999 | 0.05945685 | 0.38732806 |
| NDFIP1 | 204.563489 | 0.088742437 | 0.20371625 | 0.43561786 | 0.66311401 | 0.918408 |
| SPRY4-AS1 | 14.7668685 | -0.012371014 | 0.64183193 | -0.0192745 | 0.9846221 | 0.99773864 |
| ARHGAP26 | 30.3265876 | -0.478666934 | 0.33879449 | -1.4128534 | 0.15769884 | 0.59847934 |
| NR3C1 | 87.7357556 | -0.112662686 | 0.32533109 | -0.3463016 | 0.72911604 | 0.94106333 |
| YIPF5 | 23.973633 | -0.179934729 | 0.33053886 | -0.5443679 | 0.58618836 | 0.89059199 |
| PRELID2 | 51.6218859 | -1.505399589 | 0.48459646 | -3.1065014 | 0.00189315 | 0.04963134 |
| SH3RF2 | 16.5395312 | 1.536731706 | 0.67689244 | 2.2702746 | 0.02319093 | 0.23879078 |
| LARS | 74.4482205 | 0.271076749 | 0.24229139 | 1.11880472 | 0.26322346 | 0.71348562 |
| TCERG1 | 112.608206 | 0.094308179 | 0.18079347 | 0.52163488 | 0.60192458 | 0.89522668 |
| STK32A | 51.0774124 | 0.98879331 | 0.75083239 | 1.31692947 | 0.18786227 | 0.63942753 |
| SPINK1 | 484.331816 | -1.275697719 | 0.73763865 | -1.7294345 | 0.08373136 | 0.45665431 |
| AC091948.1 | 24.2652981 | 0.666196818 | 0.47201782 | 1.41138066 | 0.1581324 | 0.59870582 |
| FBXO38 | 58.2511143 | -0.101060436 | 0.19959023 | -0.5063396 | 0.61261827 | 0.90011356 |
| HTR4 | 18.0916314 | -0.81020293 | 1.26213286 | -0.6419316 | 0.52091762 | 0.8669958 |
| AFAP1L1 | 15.795608 | -0.544665711 | 0.84514861 | -0.6444615 | 0.51927624 | 0.86618562 |
| GRPEL2 | 21.4181623 | 0.666293432 | 0.48577093 | 1.37162064 | 0.17018157 | 0.61252734 |
| CSNK1A1 | 423.985116 | -0.239082468 | 0.19166042 | -1.2474275 | 0.2122408 | 0.66453706 |
| ARHGEF37 | 32.8719526 | -0.334535065 | 0.44217618 | -0.7565651 | 0.44931046 | 0.83118301 |
| SLC26A2 | 16.8383552 | 0.352883825 | 0.42136368 | 0.83748041 | 0.40232259 | 0.80890307 |
| HMGXB3 | 49.6147593 | -0.497415981 | 0.30140773 | -1.6503093 | 0.09887969 | 0.48485489 |
| TCOF1 | 16.1086565 | 0.424361619 | 0.5065522 | 0.83774509 | 0.40217388 | 0.80890307 |
| CD74 | 388.895084 | -2.519099453 | 0.59758757 | -4.2154482 | 2.49E-05 | 0.00244247 |
| RPS14 | 260.994346 | 0.397300822 | 0.36194412 | 1.09768552 | 0.27234184 | 0.72345643 |
| NDST1 | 56.4166165 | 0.111825483 | 0.29792915 | 0.37534254 | 0.70740573 | 0.93432317 |
| RBM22 | 18.1655678 | -0.2163027 | 0.31567629 | -0.6852041 | 0.49321511 | 0.85714521 |
| DCTN4 | 91.5694494 | 0.050504792 | 0.17802851 | 0.28368934 | 0.77664846 | 0.95512504 |
| GPX3 | 314.331175 | -1.394340064 | 0.55734413 | -2.5017579 | 0.01235784 | 0.16567149 |
| TNIP1 | 76.119994 | -0.783435117 | 0.33472731 | -2.3405175 | 0.01925704 | 0.21796181 |
| ANXA6 | 47.0881457 | 0.376226749 | 0.43778696 | 0.85938318 | 0.39012915 | 0.80191835 |
| ATOX1 | 67.8468978 | -0.01639636 | 0.31835258 | -0.0515038 | 0.95892409 | 0.99238848 |
| G3BP1 | 115.489644 | 0.340733188 | 0.21542442 | 1.58168324 | 0.1137219 | 0.51813172 |
| FAM114A2 | 27.3124537 | -0.751317181 | 0.35786843 | -2.0994229 | 0.03577963 | 0.3024108 |
| MFAP3 | 41.3488 | -0.371786481 | 0.26833096 | -1.3855519 | 0.16588377 | 0.60628206 |
| GALNT10 | 188.604554 | 0.462515136 | 0.37282172 | 1.24057992 | 0.21476097 | 0.66839951 |
| SAP30L-AS1 | 25.6751194 | 0.094524206 | 0.44529936 | 0.21227115 | 0.8318955 | 0.96701848 |
| SAP30L | 14.829472 | 0.61225393 | 0.40067638 | 1.52805096 | 0.12649989 | 0.54817868 |
| LARP1 | 174.070141 | 0.151166714 | 0.31559747 | 0.47898583 | 0.63194871 | 0.90805737 |
| FAXDC2 | 21.5816356 | 0.692322321 | 0.46554714 | 1.48711541 | 0.13698433 | 0.56856307 |
| CNOT8 | 32.9534011 | 0.149648078 | 0.25018831 | 0.59814178 | 0.54974534 | 0.87445431 |
| GEMIN5 | 16.1729459 | 0.550792287 | 0.32944081 | 1.67190061 | 0.09454392 | 0.47996672 |
| MRPL22 | 28.4931511 | 0.420761022 | 0.2801602 | 1.50185863 | 0.13313362 | 0.56148254 |
| TIMD4 | 41.2891833 | 0.431042517 | 0.56947448 | 0.75691279 | 0.44910211 | 0.83118301 |
| CYFIP2 | 235.207999 | 0.314749962 | 0.3995093 | 0.78784139 | 0.43078949 | 0.8248269 |
| CLINT1 | 215.740965 | -0.476296833 | 0.22678618 | -2.1002022 | 0.03571106 | 0.30226639 |
| RNF145 | 73.1609049 | -0.609591835 | 0.20937102 | -2.9115388 | 0.00359653 | 0.07769793 |
| UBLCP1 | 27.1751918 | 0.572180367 | 0.30567015 | 1.87188825 | 0.06122207 | 0.39273006 |
| AC008691.1 | 61.9566656 | -0.687311753 | 0.45030906 | -1.5263112 | 0.12693238 | 0.54952841 |
| TTC1 | 117.270664 | 0.097255142 | 0.30443623 | 0.31945982 | 0.74937786 | 0.94684415 |
| PWWP2A | 120.121686 | 0.60830403 | 0.2547085 | 2.38823606 | 0.01692946 | 0.20205205 |
| CCNJL | 84.9157976 | -0.20784284 | 0.34794273 | -0.5973478 | 0.55027517 | 0.87445431 |
| SLU7 | 19.2810389 | -0.130291277 | 0.44279292 | -0.2942488 | 0.76856781 | 0.95205353 |
| AC113414.1 | 449.694888 | -2.061201299 | 0.63137079 | -3.2646447 | 0.00109601 | 0.03485686 |
| CCNG1 | 67.4745017 | -0.906123342 | 0.35851363 | -2.5274446 | 0.01148959 | 0.15875935 |
| NUDCD2 | 18.1108993 | -0.439386375 | 0.44811102 | -0.9805302 | 0.32682449 | 0.76371899 |
| MAT2B | 45.3541064 | -0.118023295 | 0.26116301 | -0.4519143 | 0.65133074 | 0.91563886 |
| TENM2 | 11.7456499 | 0.316244332 | 0.65908747 | 0.47982149 | 0.63135433 | NA |
| WWC1 | 411.239877 | -0.887656097 | 0.35532243 | -2.4981708 | 0.0124836 | 0.16647826 |
| RARS | 51.9477594 | 0.036703916 | 0.28395662 | 0.12925889 | 0.8971528 | 0.97889563 |
| PANK3 | 129.028858 | -0.4677448 | 0.23222292 | -2.014206 | 0.04398791 | 0.33308568 |
| SLIT3 | 101.293398 | 0.125777924 | 1.52989427 | 0.08221347 | 0.93447696 | 0.9881638 |
| FOXI1 | 318.311598 | 0.013520702 | 0.22352006 | 0.06048988 | 0.95176547 | 0.99217529 |
| LINC01187 | 611.152988 | 0.425720211 | 0.35023295 | 1.21553444 | 0.22416232 | 0.6771276 |
| RANBP17 | 69.2556967 | -0.212266305 | 0.37586553 | -0.56474 | 0.57225062 | 0.88578992 |
| NPM1 | 80.9868682 | 0.063781658 | 0.30896038 | 0.2064396 | 0.83644754 | 0.96748214 |
| FBXW11 | 440.305063 | -0.251797218 | 0.2485311 | -1.0131417 | 0.3109925 | 0.75416279 |
| STK10 | 23.1710509 | -0.699308941 | 0.39703905 | -1.7613102 | 0.07818591 | 0.44093218 |
| UBTD2 | 71.8499243 | -0.014296352 | 0.20008613 | -0.071451 | 0.94303883 | 0.9904154 |
| NEURL1B | 18.059383 | 0.903559165 | 0.72121288 | 1.25283282 | 0.21026655 | 0.66220311 |
| DUSP1 | 141.787467 | -1.333154308 | 0.47380905 | -2.8136953 | 0.00489756 | 0.0961667 |
| ERGIC1 | 138.268084 | -0.529316846 | 0.2712382 | -1.9514834 | 0.05099957 | 0.35974306 |
| RPL26L1 | 13.9969994 | 0.005553658 | 0.40240036 | 0.01380133 | 0.98898849 | 0.99918637 |
| ATP6V0E1 | 343.293946 | -0.386065211 | 0.20533788 | -1.8801461 | 0.06008817 | 0.38930355 |
| CREBRF | 159.861182 | -0.458801334 | 0.2993944 | -1.5324313 | 0.12541605 | 0.54505746 |
| BNIP1 | 15.0451307 | 0.263547339 | 0.42650415 | 0.61792444 | 0.53662515 | 0.86963427 |
| BOD1 | 18.8003907 | 0.166760238 | 0.39634064 | 0.42074978 | 0.6739378 | 0.92189158 |
| CPEB4 | 779.562628 | 0.018548769 | 0.17137719 | 0.10823359 | 0.9138104 | 0.98400986 |
| SFXN1 | 97.0906965 | -0.107653999 | 0.21561754 | -0.4992822 | 0.61758059 | 0.90214869 |
| THOC3 | 60.3062071 | -1.24696657 | 0.50603249 | -2.4642026 | 0.01373185 | 0.17578162 |
| FAM153B | 9.89785218 | 0.586050945 | 0.89145645 | 0.65740839 | 0.51091836 | NA |
| SIMC1 | 71.774587 | -0.110617887 | 0.26941369 | -0.4105875 | 0.68137505 | 0.92386609 |
| KIAA1191 | 52.4932961 | -0.417417272 | 0.29316213 | -1.4238444 | 0.15449151 | 0.59443382 |
| ARL10 | 12.032242 | -0.464855424 | 0.51980198 | -0.8942933 | 0.37116499 | NA |
| CLTB | 42.5152603 | 0.008258819 | 0.29234598 | 0.02825015 | 0.97746264 | 0.99573488 |
| FAF2 | 82.5130179 | 0.057947145 | 0.24740091 | 0.23422365 | 0.81481135 | 0.96300833 |

| | | | | | | |
|---|---|---|---|---|---|---|
| RNF44 | 23.0695228 | 0.145271761 | 0.34765755 | 0.41785879 | 0.67605037 | 0.9220597 |
| TSPAN17 | 20.4889594 | -0.445943801 | 0.40537727 | -1.100071 | 0.27130117 | 0.72217929 |
| UIMC1 | 103.432314 | -0.250923828 | 0.20015776 | -1.2536302 | 0.20997643 | 0.6617191 |
| ZNF346 | 51.1736679 | 0.300268227 | 0.21453645 | 1.39961401 | 0.16162894 | 0.60060549 |
| NSD1 | 202.780197 | 0.142235404 | 0.29076057 | 0.48918395 | 0.62471147 | 0.90447745 |
| PRELID1 | 47.11257 | -0.013267815 | 0.39338461 | -0.0337273 | 0.97309458 | 0.99526459 |
| LMAN2 | 61.1563137 | -0.01896423 | 0.26181869 | -0.0724327 | 0.94225758 | 0.98991882 |
| RGS14 | 19.9892596 | -0.861660382 | 0.47162932 | -1.8269864 | 0.06770181 | 0.41458809 |
| DBN1 | 33.1934492 | 0.006450087 | 0.36452375 | 0.01769456 | 0.98588252 | 0.99804172 |
| DDX41 | 13.3596613 | -0.023854993 | 0.3849032 | -0.0619766 | 0.95058146 | 0.99209822 |
| FAM193B | 53.571021 | 0.209375955 | 0.2240725 | 0.93441167 | 0.35009158 | 0.77868979 |
| TMED9 | 65.5918319 | -0.347161152 | 0.29571961 | -1.1739538 | 0.24041353 | 0.69470237 |
| B4GALT7 | 22.2228598 | -0.204412447 | 0.47463549 | -0.4306725 | 0.66670653 | 0.92031498 |
| FAM153A | 25.8312892 | -0.722356758 | 0.72148007 | -1.0012151 | 0.31672281 | 0.75894548 |
| AC106795.3 | 17.3632112 | -1.323127196 | 0.61614374 | -2.1474327 | 0.03175885 | 0.28215622 |
| N4BP3 | 27.1861153 | 1.087195849 | 0.38943772 | 2.79170656 | 0.00524309 | 0.09995328 |
| RMND5B | 67.076435 | 0.527048379 | 0.2437592 | 2.16216819 | 0.03060521 | 0.27849022 |
| NHP2 | 30.6646262 | 0.028467672 | 0.33205977 | 0.08573057 | 0.9316806 | 0.98757533 |
| HNRNPAB | 39.7921745 | -0.095904712 | 0.34669619 | -0.2766247 | 0.78206833 | 0.95544332 |
| PHYKPL | 104.58676 | -0.440133841 | 0.26153476 | -1.6828885 | 0.09239668 | 0.4764741 |
| CLK4 | 46.1189284 | 0.359823636 | 0.32469741 | 1.1081814 | 0.26778348 | 0.71922155 |
| ZNF354A | 17.6034828 | -0.146688722 | 0.40521176 | -0.3620051 | 0.71734822 | 0.93714179 |
| ZNF354B | 13.1355227 | 0.299185819 | 0.39974381 | 0.7484439 | 0.45419245 | 0.83364149 |
| RUFY1 | 84.8068451 | 0.029832775 | 0.20632073 | 0.14459418 | 0.8850313 | 0.97662748 |
| HNRNPH1 | 213.502388 | 0.770360983 | 0.28083243 | 2.74313396 | 0.00608559 | 0.10972152 |
| CANX | 218.177078 | 0.086205258 | 0.21571799 | 0.39962016 | 0.6894363 | 0.92734996 |
| MAML1 | 39.3269882 | 0.288884493 | 0.24762789 | 1.16660724 | 0.24336902 | 0.69729164 |
| MGAT4B | 64.0508617 | -0.906945703 | 0.46094145 | -1.9675942 | 0.04911475 | 0.3550023 |
| SQSTM1 | 226.325404 | -1.337012385 | 0.41388942 | -3.2303613 | 0.00123634 | 0.03747651 |
| MRNIP | 28.3122395 | 0.015715752 | 0.35586306 | 0.04416236 | 0.96477499 | 0.99334616 |
| TBC1D9B | 39.8350025 | -0.193307765 | 0.32828005 | -0.5888502 | 0.55596179 | 0.87788204 |
| RNF130 | 199.126675 | -0.218214437 | 0.21323564 | -1.0233488 | 0.30614295 | 0.74897897 |
| MAPK9 | 57.6283099 | 0.122583045 | 0.21074837 | 0.58165595 | 0.56079845 | 0.88071368 |
| CNOT6 | 65.5751266 | 0.03519746 | 0.19906148 | 0.17681704 | 0.85965211 | 0.97287712 |
| MGAT1 | 42.5351413 | -0.337465797 | 0.41187178 | -0.8193467 | 0.41258861 | 0.81556555 |
| ZFP62 | 18.1085497 | 0.418722822 | 0.41803976 | 1.00163396 | 0.31652041 | 0.75864788 |
| BTNL9 | 57.6522804 | -0.380214947 | 0.40468366 | -0.9395362 | 0.34745552 | 0.77726831 |
| TRIM41 | 17.5035663 | 0.184416653 | 0.32806781 | 0.56212968 | 0.57402767 | 0.88637538 |
| RACK1 | 152.819751 | 0.099752143 | 0.37485959 | 0.26610536 | 0.79015807 | 0.95746824 |
| TRIM52 | 23.2735025 | 0.385460853 | 0.31724729 | 1.21501702 | 0.2243596 | 0.6771276 |
| AC138035.1 | 11.0424084 | 0.048311877 | 0.50968442 | 0.09478783 | 0.92448336 | NA |
| DUSP22 | 35.5091391 | -0.499498583 | 0.27376794 | -1.8245328 | 0.06807156 | 0.4151827 |
| EXOC2 | 135.172982 | -0.158420324 | 0.18400535 | -0.860955 | 0.38926284 | 0.80191835 |
| GMDS | 309.300123 | -0.694456514 | 0.24955279 | -2.7828041 | 0.00538913 | 0.10170156 |
| GMDS-DT | 487.446522 | -0.495145162 | 0.33364401 | -1.4840523 | 0.13779505 | 0.56949806 |
| WRNIP1 | 54.498953 | 0.288603969 | 0.28544774 | 1.01105711 | 0.31198909 | 0.75484355 |
| SERPINB1 | 144.743315 | -1.810676497 | 0.54248672 | -3.3377342 | 0.00084464 | 0.02926092 |
| SERPINB6 | 108.676665 | -0.309333984 | 0.2454979 | -1.260027 | 0.20765961 | 0.65934476 |
| NQO2 | 42.0292002 | 0.076304225 | 0.27634265 | 0.27612178 | 0.78245453 | 0.95544332 |
| RIPK1 | 70.5649775 | -0.217427933 | 0.27200347 | -0.7993572 | 0.42408332 | 0.82222831 |
| BPHL | 19.5008839 | 0.226113496 | 0.33761609 | 0.66973555 | 0.50302638 | 0.86010152 |
| TUBB2A | 26.1427624 | -0.761074484 | 0.46941039 | -1.6213414 | 0.10494445 | 0.49924529 |
| LINC02525 | 42.1917429 | -2.091287269 | 0.4637815 | -4.509208 | 6.51E-06 | 0.00084157 |
| PSMG4 | 168.22518 | -1.553398883 | 0.41221055 | -3.7684598 | 0.00016426 | 0.0098352 |
| SLC22A23 | 384.383813 | -0.015392684 | 0.28707845 | -0.0536184 | 0.95723921 | 0.99236316 |
| PRPF4B | 152.159496 | -0.153381803 | 0.19125454 | -0.8019773 | 0.42256608 | 0.82138304 |
| ECI2 | 51.4562343 | -0.116642715 | 0.33643814 | -0.3466988 | 0.72881758 | 0.94097305 |
| CDYL | 203.057265 | -0.375681503 | 0.32225292 | -1.165797 | 0.24369652 | 0.69755218 |
| LYRM4 | 66.0173115 | -0.173890524 | 0.21479997 | -0.8095463 | 0.41820098 | 0.8188001 |
| FARS2 | 224.018471 | 0.582951287 | 0.20665983 | 2.82082534 | 0.00479003 | 0.09501691 |
| LY86-AS1 | 280.943486 | -0.038297336 | 0.75908795 | -0.0504518 | 0.95976238 | 0.99238848 |
| LY86 | 34.5992203 | -2.008121771 | 0.79447615 | -2.5276049 | 0.01148435 | 0.15875935 |
| AL031123.1 | 72.2325399 | -0.134320235 | 0.59522159 | -0.2256643 | 0.82146256 | 0.96443801 |
| RREB1 | 234.37139 | -0.100358346 | 0.33096976 | -0.3032251 | 0.7617183 | 0.9499444 |
| SSR1 | 74.7780572 | 0.436787152 | 0.230401 | 1.89576938 | 0.05799055 | 0.38213879 |
| AL139095.4 | 14.7123619 | -0.263247106 | 0.77633945 | -0.3390876 | 0.73454372 | 0.94277981 |
| RIOK1 | 13.7370631 | 0.348973869 | 0.37089481 | 0.94089715 | 0.34675757 | 0.7765713 |
| DSP | 299.785489 | 0.178329219 | 0.28268003 | 0.63085185 | 0.5281374 | 0.86750588 |
| SNRNP48 | 22.163312 | 0.563377067 | 0.39038373 | 1.44313666 | 0.14898198 | 0.58629428 |
| BLOC1S5-TXI | 12.4926369 | -1.204129732 | 0.46840442 | -2.5707053 | 0.01014916 | NA |
| SLC35B3 | 63.7532725 | -0.152870156 | 0.35469531 | -0.4309901 | 0.66647559 | 0.92012714 |
| TFAP2A | 38.6265886 | 0.391092734 | 0.34089272 | 1.14726044 | 0.251274 | 0.70545786 |
| GCNT2 | 264.198022 | 0.408283129 | 0.28366386 | 1.43932022 | 0.15005982 | 0.58673121 |
| PAK1IP1 | 11.6274625 | -0.394430849 | 0.40922986 | -0.9638369 | 0.33512769 | NA |
| TMEM14C | 41.4850944 | 0.224651615 | 0.47461011 | 0.47333929 | 0.63597113 | 0.90881415 |
| MAK | 39.2979779 | -0.45431117 | 0.62705018 | -0.7245212 | 0.4687458 | 0.84326495 |
| SMIM13 | 14.2159393 | 0.034264014 | 0.36372313 | 0.09420356 | 0.92494746 | 0.98672316 |
| AL139807.1 | 16.4854766 | -0.493784787 | 0.5465266 | -0.9034963 | 0.36626253 | 0.78793182 |
| NEDD9 | 41.2692935 | 0.203187173 | 0.5959068 | 0.3409714 | 0.73312512 | 0.94269288 |
| TMEM170B | 21.4345696 | 0.391445559 | 0.37211439 | 1.05194953 | 0.29282271 | 0.74013384 |
| ADTRP | 199.017884 | 1.279458803 | 0.40237511 | 3.1797663 | 0.00147394 | 0.04156164 |
| HIVEP1 | 167.734368 | -0.656294485 | 0.31970699 | -2.0527999 | 0.04009199 | 0.32113318 |
| PHACTR1 | 20.786843 | -0.133037667 | 0.62329752 | -0.2134417 | 0.83098248 | 0.96701848 |
| TBC1D7 | 11.4682578 | 0.537962734 | 0.4184913 | 1.28548129 | 0.19862415 | NA |
| SIRT5 | 50.2038827 | 0.068047503 | 0.3508516 | 0.19394953 | 0.8462154 | 0.97027687 |
| NOL7 | 43.0684873 | -0.438712888 | 0.31347395 | -1.3995195 | 0.16165727 | 0.60060549 |
| RANBP9 | 169.602528 | -0.260072207 | 0.23787198 | -1.0933285 | 0.27424961 | 0.72525116 |
| MCUR1 | 29.9058217 | 0.00529853 | 0.39984103 | 0.01325159 | 0.98942707 | 0.99923337 |
| CD83 | 17.638919 | -0.118991035 | 0.51877265 | -0.2293703 | 0.81858113 | 0.9635539 |
| JARID2 | 183.326695 | 0.455932256 | 0.25319937 | 1.80068479 | 0.07175258 | 0.42284325 |
| DTNBP1 | 124.496886 | -0.66151447 | 0.39065052 | -1.6933664 | 0.09038574 | 0.47236403 |
| MYLIP | 20.4467141 | 0.253120753 | 0.45816034 | 0.55247199 | 0.58062502 | 0.89012555 |
| GMPR | 267.508145 | -0.126054942 | 0.41095403 | -0.3067373 | 0.75904331 | 0.94924107 |
| ATXN1 | 482.463722 | -0.423258842 | 0.35727793 | -1.1846767 | 0.23614529 | 0.69077483 |
| CAP2 | 17.093558 | 0.482245585 | 0.63777366 | 0.75613907 | 0.44956582 | 0.83118301 |
| FAM8A1 | 15.9606253 | 0.027700502 | 0.35252485 | 0.07857744 | 0.93736873 | 0.98871562 |
| NUP153 | 114.02506 | 0.380653102 | 0.21991026 | 1.73094745 | 0.08346114 | 0.4560197 |
| KIF13A | 378.053004 | -0.02526273 | 0.31626507 | -0.0798783 | 0.93633401 | 0.98871562 |
| TPMT | 25.5410826 | 0.051536348 | 0.31133258 | 0.16553471 | 0.86852313 | 0.97434023 |
| KDM1B | 18.7453654 | -0.088862397 | 0.4245826 | -0.2092935 | 0.83421909 | 0.96701848 |
| DEK | 86.4026313 | -0.336235523 | 0.29542166 | -1.1381546 | 0.25505593 | 0.70849619 |
| RNF144B | 111.170138 | -0.359062997 | 0.34658485 | -1.0360031 | 0.30020068 | 0.74437099 |
| AL022068.1 | 35.9982004 | 1.315022875 | 0.61760139 | 2.1292421 | 0.03323424 | 0.28938248 |
| ID4 | 16.217886 | 0.241303189 | 0.48864463 | 0.49382143 | 0.6214323 | 0.90380166 |
| MBOAT1 | 140.124075 | -0.738131511 | 0.42515912 | -1.73613 | 0.08254085 | 0.45473406 |
| AL158198.1 | 18.1114381 | -0.557988813 | 0.62117144 | -0.8982847 | 0.36903378 | 0.78889478 |
| E2F3 | 55.2933525 | -0.261654546 | 0.30415267 | -0.8602737 | 0.38963818 | 0.80191835 |
| CDKAL1 | 330.96406 | 0.02189177 | 0.21807616 | 0.10038589 | 0.92003797 | 0.98586108 |
| SOX4 | 71.9318852 | -0.152930422 | 0.49122022 | -0.3113276 | 0.75555157 | 0.94795939 |
| CASC15 | 164.394639 | 0.143356151 | 0.31816956 | 0.45056526 | 0.65230291 | 0.9158679 |
| NBAT1 | 12.0976709 | -0.31788526 | 0.64331115 | -0.4941392 | 0.62120787 | NA |
| DCDC2 | 170.409558 | -0.275522632 | 0.36610736 | -0.7525733 | 0.45170635 | 0.83236163 |
| MRS2 | 41.5233122 | 0.437669985 | 0.27278397 | 1.60445637 | 0.10861349 | 0.50659799 |

| | | | | | | |
|---|---|---|---|---|---|---|
| ALDH5A1 | 18.4569064 | 0.542910679 | 0.41010744 | 1.32382549 | 0.18556101 | 0.63613695 |
| TDP2 | 37.3788309 | -0.167494015 | 0.30542852 | -0.5483902 | 0.58342399 | 0.89036744 |
| ACOT13 | 33.7192213 | -0.375358402 | 0.34083864 | -1.1012789 | 0.27077529 | 0.7217698 |
| C6orf62 | 105.57554 | 0.183718723 | 0.35482326 | 0.51777531 | 0.60461505 | 0.89593125 |
| GMNN | 21.1654007 | 1.011174503 | 0.44066774 | 2.29464153 | 0.02175368 | 0.23061274 |
| CARMIL1 | 380.275744 | -0.350241948 | 0.37693465 | -0.9291848 | 0.35279332 | 0.78005362 |
| TRIM38 | 84.3973557 | -0.195227071 | 0.28527958 | -0.684336 | 0.49376305 | 0.85756519 |
| HIST1H1C | 24.1598759 | -1.185911824 | 0.38865632 | -3.0513123 | 0.00227843 | 0.05595143 |
| HFE | 11.4249185 | -0.124337758 | 0.52929268 | -0.234913 | 0.81427622 | NA |
| HIST1H4C | 32.0534364 | -1.805722455 | 0.6547879 | -2.7577212 | 0.00582058 | 0.10592804 |
| HIST1H2AC | 36.9769788 | 0.157833489 | 0.64519529 | 0.24462902 | 0.80674369 | 0.96005568 |
| HIST1H1E | 18.4142396 | -2.177990109 | 0.68940361 | -3.159238 | 0.00158182 | 0.04396469 |
| BTN3A2 | 28.632423 | -1.08909119 | 0.77221685 | -1.4103437 | 0.15843821 | 0.59881912 |
| BTN3A1 | 17.4151215 | -1.127162057 | 0.64607145 | -1.7446399 | 0.08104756 | 0.45026424 |
| BTN3A3 | 12.3273742 | -0.250835389 | 0.67137623 | -0.3736137 | 0.70869171 | NA |
| BTN2A1 | 26.4792662 | -0.149753956 | 0.28657672 | -0.5225615 | 0.60127945 | 0.89490801 |
| HCG11 | 11.8530276 | 0.609465037 | 0.44624982 | 1.36574855 | 0.1720179 | NA |
| HMGN4 | 21.9358684 | -0.223585415 | 0.40759021 | -0.5485544 | 0.58331127 | 0.89036744 |
| ZNF322 | 22.589503 | -0.115594538 | 0.33079243 | -0.3494474 | 0.72675349 | 0.94009907 |
| AL121944.1 | 11.4543003 | -0.860669035 | 0.54920846 | -1.5671081 | 0.11708942 | NA |
| ZSCAN16-AS | 20.4653613 | 0.084730333 | 0.4047929 | 0.20931774 | 0.83420021 | 0.96701848 |
| ZKSCAN8 | 22.1342733 | 0.316666117 | 0.41053136 | 0.77135671 | 0.44049553 | 0.82793978 |
| ZSCAN9 | 11.190364 | 0.093242295 | 0.3926578 | 0.23746452 | 0.81229644 | NA |
| ZSCAN31 | 31.6245115 | 0.426644678 | 0.54260281 | 0.78629279 | 0.43169598 | 0.8248269 |
| ZSCAN12 | 10.5439583 | -0.392693369 | 0.56441761 | -0.6957497 | 0.48658561 | NA |
| TRIM27 | 62.5866426 | -0.972162507 | 0.21793055 | -4.4608821 | 8.16E-06 | 0.00098271 |
| GABBR1 | 22.3578918 | -0.711864199 | 0.53850396 | -1.3219294 | 0.18619167 | 0.63705791 |
| HLA-F | 58.387795 | -2.361040806 | 0.60807354 | -3.8828212 | 0.00010325 | 0.00663271 |
| HLA-A | 790.269179 | -1.976334488 | 0.41651629 | -4.7449152 | 2.09E-06 | 0.00033723 |
| ZNRD1 | 11.3587952 | -0.220287077 | 0.61786639 | -0.3565287 | 0.72144469 | NA |
| PPP1R11 | 17.2554759 | -0.361886402 | 0.35713247 | -1.0133114 | 0.31091145 | 0.75416279 |
| TRIM26 | 61.134166 | 0.146415164 | 0.26059332 | 0.5618531 | 0.57421611 | 0.88644064 |
| HCG17 | 75.3370087 | 0.580704881 | 0.39062252 | 1.48661393 | 0.13711681 | 0.56856307 |
| HLA-E | 264.632424 | -2.273965012 | 0.44243346 | -5.1396768 | 2.75E-07 | 6.85E-05 |
| GNL1 | 36.8990029 | -0.289304044 | 0.40101143 | -0.7214359 | 0.47064136 | 0.84424147 |
| ABCF1 | 43.5683993 | -0.502454547 | 0.40155362 | -1.2512763 | 0.21083367 | 0.6625727 |
| PPP1R10 | 39.4544997 | 0.29948195 | 0.30455025 | 0.9833581 | 0.32543123 | 0.76334591 |
| MRPS18B | 41.3291257 | 0.167709336 | 0.34543955 | 0.48549547 | 0.62732493 | 0.90589724 |
| ATAT1 | 13.5335833 | -0.050494661 | 0.40278761 | -0.125363 | 0.90023618 | 0.97940229 |
| C6orf136 | 32.7548817 | 0.538043518 | 0.25780991 | 2.0869776 | 0.03689016 | 0.30715411 |
| DHX16 | 16.3012496 | -0.13958798 | 0.3356764 | -0.4158409 | 0.67752642 | 0.92259972 |
| MDC1 | 24.8509845 | 0.182158899 | 0.31017523 | 0.5872774 | 0.55701742 | 0.87869068 |
| TUBB | 71.1660421 | 0.152323963 | 0.44672309 | 0.34098072 | 0.73311809 | 0.94269288 |
| FLOT1 | 124.935517 | -0.451476116 | 0.20196867 | -2.235377 | 0.02539261 | 0.25137888 |
| IER3 | 126.139153 | -1.339855236 | 1.02350307 | -1.3090877 | 0.19050466 | 0.6421543 |
| DDR1 | 165.531172 | -0.346303122 | 0.22774786 | -1.5205549 | 0.12837157 | 0.5528291 |
| CCHCR1 | 15.4291535 | -0.422163775 | 0.42076008 | -1.0033361 | 0.31569873 | 0.75793414 |
| POU5F1 | 97.8455147 | -0.918951572 | 0.53367137 | -1.7219428 | 0.08507988 | 0.45976313 |
| HLA-C | 509.791514 | -1.475351419 | 0.53144553 | -2.7761103 | 0.00550135 | 0.10201977 |
| HLA-B | 809.05533 | -2.456093103 | 0.44460814 | -5.5241749 | 3.31E-08 | 1.11E-05 |
| MICA | 28.6692312 | -0.247453969 | 0.31329344 | -0.7898473 | 0.42961696 | 0.82469033 |
| MCCD1 | 15.2545326 | 1.050820638 | 0.51392723 | 2.04468759 | 0.04088567 | 0.32473787 |
| NFKBIL1 | 12.2371651 | -0.207801019 | 0.39453364 | -0.5267004 | 0.59840167 | NA |
| PRRC2A | 61.7679096 | 0.207191637 | 0.25094756 | 0.82563719 | 0.40900993 | 0.81399186 |
| BAG6 | 36.3874316 | -0.197614204 | 0.36637758 | -0.5393731 | 0.58962945 | 0.89149106 |
| C6orf47 | 12.0151424 | 0.024824561 | 0.49082137 | 0.05057759 | 0.95966212 | NA |
| DDAH2 | 30.4178292 | 0.287052888 | 0.37641541 | 0.762596 | 0.44570437 | 0.83035712 |
| CLIC1 | 119.764747 | -0.835527145 | 0.33174105 | -2.5186125 | 0.01178183 | 0.16141766 |
| VARS | 14.656009 | 0.283818608 | 0.39103101 | 0.72582123 | 0.46794837 | 0.84326495 |
| LSM2 | 19.4088872 | 0.370856198 | 0.36195895 | 1.02458084 | 0.30556101 | 0.74897897 |
| HSPA1A | 137.134996 | 0.559144594 | 0.39791883 | 1.4051725 | 0.15996999 | 0.60024417 |
| HSPA1B | 56.0925947 | 0.619448173 | 0.3726792 | 1.66214848 | 0.09648301 | 0.48257296 |
| C6orf48 | 35.0553143 | 0.534549482 | 0.33011877 | 1.61926412 | 0.10539045 | 0.50058098 |
| NEU1 | 34.0580102 | -0.820478195 | 0.47145013 | -1.7403287 | 0.08180132 | 0.45315408 |
| SLC44A4 | 25.6198674 | -1.431031932 | 0.70448629 | -2.0313127 | 0.04222328 | 0.32888549 |
| EHMT2 | 12.0392611 | 0.758926827 | 0.49864429 | 1.52198037 | 0.128014 | NA |
| NELFE | 13.3928137 | -0.084822941 | 0.45650531 | -0.1858093 | 0.8525943 | 0.97090452 |
| STK19 | 16.6265562 | 0.395061359 | 0.38693131 | 1.0210116 | 0.30724894 | 0.75038069 |
| ATF6B | 31.4137843 | 0.057143298 | 0.27896196 | 0.20484262 | 0.83769509 | 0.96748214 |
| AGPAT1 | 18.1580849 | 0.253033441 | 0.41010311 | 0.61699956 | 0.53723501 | 0.86963427 |
| RNF5 | 21.6709846 | -0.348165875 | 0.54205717 | -0.6423047 | 0.52067535 | 0.86689854 |
| PBX2 | 15.504393 | 0.191253014 | 0.37005929 | 0.51681722 | 0.60528376 | 0.89600432 |
| AL662796.1 | 42.0431834 | 0.482007435 | 0.73206057 | 0.65842562 | 0.51026468 | 0.86281145 |
| HLA-DRA | 137.96498 | -2.31527711 | 0.7331538 | -3.1579692 | 0.00158872 | 0.04403034 |
| HLA-DRB5 | 16.9691447 | -1.74810441 | 0.7517612 | -2.3253454 | 0.0200535 | 0.22256173 |
| HLA-DRB1 | 83.8864186 | -1.105298684 | 1.23350114 | -0.8960662 | 0.37021741 | 0.7900012 |
| HLA-DQB1 | 27.8799654 | -1.021793036 | 1.15967001 | -0.8811067 | 0.37826006 | 0.79504281 |
| PSMB8 | 69.7306667 | -1.407386824 | 0.46147222 | -3.0497758 | 0.00229012 | 0.05598343 |
| PSMB9 | 29.0871113 | -2.410625741 | 0.74614194 | -3.2307871 | 0.0012345 | 0.03747651 |
| TAP1 | 48.7373161 | -1.975201689 | 0.63341267 | -3.1183489 | 0.00181867 | 0.04819983 |
| HLA-DMB | 19.8191239 | -0.189976229 | 0.57538002 | -0.3301752 | 0.74126757 | 0.94497245 |
| BRD2 | 95.1157483 | 0.166546758 | 0.24297343 | 0.68545255 | 0.49305839 | 0.85714521 |
| HLA-DPA1 | 42.6416638 | -2.848073371 | 0.91666665 | -3.1069892 | 0.00189003 | 0.04963134 |
| HLA-DPB1 | 34.2463693 | -1.514240903 | 0.93447876 | -1.6204123 | 0.10514373 | 0.49994814 |
| RXRB | 16.1269424 | 0.116656374 | 0.33930697 | 0.34380777 | 0.73099086 | 0.94155624 |
| SLC39A7 | 37.6664908 | 0.047674253 | 0.50117245 | 0.09512545 | 0.92421519 | 0.98672316 |
| RING1 | 24.2757043 | 0.235210725 | 0.40330695 | 0.58320523 | 0.55975515 | 0.88037687 |
| VPS52 | 27.1861332 | 0.363815961 | 0.31869463 | 1.14158173 | 0.25362792 | 0.70701839 |
| RPS18 | 151.40008 | 0.534365182 | 0.37094256 | 1.44056045 | 0.1497089 | 0.58659144 |
| WDR46 | 11.4914182 | 0.480863886 | 0.5143171 | 0.93495606 | 0.34981095 | NA |
| PFDN6 | 19.3627623 | -0.053322081 | 0.32489054 | -0.1641232 | 0.86963414 | 0.97434023 |
| RGL2 | 21.7764061 | -0.024124028 | 0.34146694 | -0.0706482 | 0.94367774 | 0.99076459 |
| TAPBP | 156.393667 | -0.835601656 | 0.30841921 | -2.7093048 | 0.00674244 | 0.11774717 |
| DAXX | 11.1486848 | -0.114040212 | 0.55493424 | -0.2055022 | 0.83717977 | NA |
| PHF1 | 42.0952198 | 0.020368182 | 0.42422552 | 0.04801263 | 0.96170618 | 0.99282141 |
| CUTA | 45.8818729 | 0.323708093 | 0.43431337 | 0.74533301 | 0.45607043 | 0.83506875 |
| SYNGAP1 | 11.6122545 | 0.212593859 | 0.47801583 | 0.4447423 | 0.656506 | NA |
| ITPR3 | 28.9863981 | -1.279058558 | 0.49961379 | -2.5600946 | 0.01046437 | 0.15015439 |
| UQCC2 | 31.3613918 | -0.029480029 | 0.30893149 | -0.0954258 | 0.92397664 | 0.98672316 |
| LEMD2 | 31.2900533 | -0.162193871 | 0.28772137 | -0.5637185 | 0.5729457 | 0.88578992 |
| HMGA1 | 20.7199536 | -0.511887322 | 0.41600869 | -1.2304727 | 0.21852016 | 0.67250923 |
| SMIM29 | 11.4508422 | -1.009709431 | 0.4669542 | -2.1623308 | 0.03059269 | NA |
| NUDT3 | 62.6499498 | -0.237751001 | 0.50239597 | -0.4732343 | 0.63604603 | 0.90881415 |
| C6orf106 | 166.568348 | -0.060074598 | 0.16516514 | -0.3637244 | 0.71606379 | 0.93689626 |
| SNRPC | 22.1029345 | 0.065682838 | 0.31704467 | 0.20717219 | 0.83587539 | 0.96748214 |
| UHRF1BP1 | 76.4227042 | -0.001382245 | 0.22583592 | -0.0061206 | 0.99511652 | 0.99923337 |
| TAF11 | 42.4905154 | 0.036226001 | 0.28291288 | 0.12804649 | 0.89811218 | 0.97892709 |
| ANKS1A | 391.87204 | 0.381638455 | 0.25337422 | 1.50622451 | 0.13200955 | 0.55916709 |
| ZNF76 | 51.3328295 | 0.144450497 | 0.25456208 | 0.56744702 | 0.57041051 | 0.88563341 |
| PPARD | 39.3148542 | 0.49473825 | 0.34725125 | 1.42472703 | 0.15423612 | 0.59439427 |
| RPL10A | 176.818023 | 0.265941573 | 0.40616812 | 0.65475738 | 0.51262397 | 0.86346419 |
| TEAD3 | 31.3402164 | -1.339551555 | 0.3544864 | -3.7788517 | 0.00015755 | 0.00961174 |
| FKBP5 | 142.42043 | -2.398468928 | 0.79685765 | -3.0099089 | 0.00261326 | 0.06201986 |

| | | | | | | |
|---|---|---|---|---|---|---|
| SRPK1 | 66.3170806 | -0.24364759 | 0.30467981 | -0.7996841 | 0.42389386 | 0.82218966 |
| MAPK14 | 121.822682 | -0.809217635 | 0.30118322 | -2.6867953 | 0.00721412 | 0.12212377 |
| MAPK13 | 33.5035198 | -0.2871973 | 0.31538511 | -0.9106242 | 0.36249344 | 0.78714716 |
| BRPF3 | 30.3058352 | 0.191151493 | 0.29480321 | 0.64840371 | 0.51672387 | 0.86500999 |
| KCTD20 | 105.448609 | -0.140641621 | 0.20998346 | -0.6697748 | 0.50300139 | 0.86010152 |
| STK38 | 101.172473 | 0.232240208 | 0.21463985 | 1.08199949 | 0.27925276 | 0.73051557 |
| SRSF3 | 93.0676727 | 0.359535625 | 0.25689742 | 1.39952993 | 0.16165413 | 0.60060549 |
| CDKN1A | 37.2602866 | -3.314625881 | 0.55398996 | -5.9831876 | 2.19E-09 | 1.12E-06 |
| C6orf89 | 61.3153365 | 0.089822062 | 0.27122958 | 0.33116618 | 0.74051897 | 0.94414222 |
| MTCH1 | 71.5645273 | -0.176634401 | 0.22830045 | -0.7736927 | 0.43911252 | 0.82745796 |
| PIM1 | 28.7250396 | -0.673558243 | 0.39483063 | -1.7059422 | 0.08801884 | 0.46731404 |
| TBC1D22B | 56.3529122 | -0.17723814 | 0.22730599 | -0.7797337 | 0.43554765 | 0.82590941 |
| RNF8 | 37.8076856 | 0.022297745 | 0.24016831 | 0.09284216 | 0.92602896 | 0.98672316 |
| CMTR1 | 28.5525762 | 0.27454438 | 0.31895359 | 0.86076591 | 0.38936698 | 0.80191835 |
| ZFAND3 | 727.957361 | -0.016709514 | 0.18809613 | -0.088835 | 0.92921306 | 0.98733028 |
| BTBD9 | 546.766135 | -0.207476219 | 0.23688857 | -0.8758389 | 0.38111763 | 0.79688435 |
| GLO1 | 80.5730573 | -0.453500959 | 0.26389962 | -1.7184601 | 0.08571273 | 0.46138374 |
| SAYSD1 | 28.2549168 | -0.195910466 | 0.31811889 | -0.6158404 | 0.53799985 | 0.86972442 |
| DAAM2 | 14.9714079 | 1.25623903 | 0.609714 | 2.06037426 | 0.03936278 | 0.31818244 |
| MOCS1 | 12.3530494 | -0.024485758 | 0.4259355 | -0.057487 | 0.95415725 | NA |
| OARD1 | 43.9061039 | -0.228533292 | 0.2777207 | -0.8228889 | 0.41057116 | 0.81427553 |
| NFYA | 23.4661296 | 0.08711097 | 0.27464416 | 0.31717758 | 0.75110886 | 0.94718617 |
| FOXP4 | 21.3107611 | -0.280652838 | 0.36270482 | -0.7737775 | 0.43906237 | 0.82745796 |
| TFEB | 100.139367 | 0.36906027 | 0.33339253 | 1.10698421 | 0.26830076 | 0.71992181 |
| USP49 | 44.7931861 | 0.047149217 | 0.26341974 | 0.17898893 | 0.8579464 | 0.97243282 |
| CCND3 | 64.7544721 | -1.497387732 | 0.52909292 | -2.8301035 | 0.00465329 | 0.09325818 |
| TAF8 | 33.4929522 | 0.098003922 | 0.23877467 | 0.41044521 | 0.68147939 | 0.92387841 |
| C6orf132 | 18.8440563 | 0.159551012 | 0.46135227 | 0.34583337 | 0.72946794 | 0.94118636 |
| MRPS10 | 20.0573988 | 0.322212492 | 0.3671126 | 0.87769392 | 0.38010985 | 0.7959986 |
| TRERF1 | 25.8936833 | 0.919416502 | 0.73556095 | 1.24995285 | 0.21131677 | 0.66321402 |
| UBR2 | 244.641199 | -0.158305718 | 0.23501365 | -0.6736022 | 0.50056424 | 0.85942996 |
| BICRAL | 81.4088522 | 0.23505746 | 0.35086133 | 0.66994405 | 0.50289346 | 0.86010152 |
| RPL7L1 | 62.0039253 | 0.168764705 | 0.27474824 | 0.61425219 | 0.53904869 | 0.86972442 |
| CNPY3 | 13.6402668 | -0.120315862 | 0.40545222 | -0.2967449 | 0.7666613 | 0.95205353 |
| PEX6 | 25.4696027 | -0.256770337 | 0.43219802 | -0.5941034 | 0.55244292 | 0.87576439 |
| PPP2R5D | 12.4285538 | 0.783125276 | 0.34736301 | 2.25448665 | 0.02416557 | NA |
| MEA1 | 12.4990452 | -0.395684476 | 0.80101987 | -0.4939759 | 0.62132323 | NA |
| KLHDC3 | 20.5516049 | -0.102253102 | 0.30085652 | -0.3398733 | 0.73395194 | 0.94277981 |
| RRP36 | 33.9247927 | -0.207893002 | 0.2806369 | -0.74079 | 0.4588208 | 0.83645518 |
| CUL7 | 18.2188598 | 0.025776158 | 0.36040767 | 0.07151945 | 0.94298435 | 0.9904154 |
| KLC4 | 21.2574336 | 0.182308558 | 0.35685953 | 0.51086924 | 0.60944262 | 0.89906009 |
| MRPL2 | 19.3517138 | 0.412026352 | 0.39420194 | 1.04521645 | 0.29592294 | 0.74123516 |
| PTK7 | 30.7971077 | 0.719685897 | 0.50845529 | 1.41543596 | 0.15694071 | 0.59772062 |
| SRF | 10.7147711 | 0.724582984 | 0.43247429 | 1.67543596 | 0.09384873 | NA |
| CUL9 | 23.7420202 | 0.466327557 | 0.32676218 | 1.42711606 | 0.15354644 | 0.59291422 |
| DNPH1 | 26.7770363 | 0.485243565 | 0.44431387 | 1.09211889 | 0.27478085 | 0.72554623 |
| ZNF318 | 10.2385129 | 0.944779491 | 0.4004035 | 2.35956853 | 0.0182962 | NA |
| ABCC10 | 17.0335782 | 0.617733095 | 0.30333174 | 2.03649345 | 0.04170083 | 0.32806006 |
| TJAP1 | 19.8803241 | -0.026414514 | 0.30556904 | -0.0864437 | 0.93111372 | 0.98757533 |
| POLR1C | 24.9081748 | 0.238925394 | 0.31494111 | 0.75863514 | 0.44807084 | 0.83071238 |
| YIPF3 | 40.890274 | -0.190342967 | 0.34469832 | -0.5522016 | 0.58081023 | 0.89012555 |
| XPO5 | 124.432487 | 0.443419832 | 0.23203766 | 1.91098221 | 0.05600687 | 0.37674523 |
| POLH | 20.0964919 | 0.409497679 | 0.2904621 | 1.40981449 | 0.15859447 | 0.59881912 |
| GTPBP2 | 24.8495239 | -0.712979619 | 0.35505126 | -2.0081033 | 0.04463231 | 0.33560731 |
| MRPS18A | 24.4401356 | -0.528933919 | 0.34117004 | -1.5503528 | 0.12105687 | 0.53496658 |
| VEGFA | 208.280867 | -0.212408395 | 0.31483603 | -0.6746636 | 0.49988955 | 0.85942996 |
| C6orf223 | 12.9740269 | -2.803234025 | 0.70118359 | -3.9978603 | 6.39E-05 | 0.00492064 |
| MRPL14 | 50.6318978 | -0.117166571 | 0.333074 | -0.3517734 | 0.72500821 | 0.93940528 |
| TMEM63B | 35.2494635 | 0.357210111 | 0.28518324 | 1.25256346 | 0.21036462 | 0.66226229 |
| SLC29A1 | 38.2856072 | -0.130944058 | 0.40602619 | -0.3225015 | 0.7470728 | 0.9462156 |
| HSP90AB1 | 326.408821 | -0.109060539 | 0.22365254 | -0.4876338 | 0.62580925 | 0.90467209 |
| SLC35B2 | 16.9426343 | -0.035518965 | 0.46086628 | -0.07707 | 0.93856786 | 0.98871562 |
| CDC5L | 41.7470305 | 0.396205289 | 0.23776252 | 1.66639087 | 0.09563559 | 0.48081904 |
| SUPT3H | 267.478463 | -0.024941773 | 0.3306878 | -0.0754239 | 0.93987742 | 0.98891539 |
| RUNX2 | 15.6694082 | -0.489033823 | 0.53779702 | -0.9093279 | 0.36317708 | 0.78718296 |
| ENPP4 | 49.4310969 | -0.140494992 | 0.2652199 | -0.5297302 | 0.596299 | 0.89360216 |
| ENPP5 | 15.6410648 | -0.463843652 | 0.44582461 | -1.0404173 | 0.29814605 | 0.74290325 |
| RCAN2 | 1550.1846 | 0.511261081 | 0.33769838 | 1.51395773 | 0.13003656 | 0.55639816 |
| CYP39A1 | 37.5551801 | -0.075459527 | 0.50552035 | -0.149271 | 0.8813398 | 0.97630358 |
| SLC25A27 | 77.5040262 | 0.169401058 | 0.39551135 | 0.42830897 | 0.66842619 | 0.92049996 |
| ADGRF5 | 2399.6126 | 0.915459305 | 0.38293306 | 2.39065101 | 0.01681853 | 0.20115876 |
| ADGRF1 | 1225.5367 | -0.453798576 | 0.30532751 | -1.4862682 | 0.13720819 | 0.56856307 |
| TNFRSF21 | 198.631649 | -0.409625401 | 0.46731063 | -0.8765591 | 0.38072614 | 0.79677315 |
| CD2AP | 330.149102 | -0.31671642 | 0.19193652 | -1.6501102 | 0.09892039 | 0.48485489 |
| MUT | 46.7013198 | 0.103790551 | 0.32251072 | 0.32182047 | 0.74758871 | 0.9462156 |
| PKHD1 | 470.176534 | 1.066757092 | 0.72276032 | 1.47594861 | 0.13995769 | 0.57374456 |
| MCM3 | 15.0938585 | 0.269572817 | 0.39198027 | 0.68772037 | 0.49162889 | 0.85654883 |
| EFHC1 | 24.2610476 | 0.017816333 | 0.33414018 | 0.05331994 | 0.957477 | 0.99236316 |
| GSTA4 | 12.2365669 | -0.125504945 | 0.60508856 | -0.2074158 | 0.83568512 | NA |
| ICK | 16.5811399 | -0.443743134 | 0.3622486 | -1.2249685 | 0.22058707 | 0.67434435 |
| FBXO9 | 57.4067159 | 0.155006797 | 0.21682508 | 0.71489329 | 0.47467498 | 0.84593347 |
| ELOVL5 | 62.3018773 | -0.6293579 | 0.41393005 | -1.520445 | 0.12839916 | 0.5528291 |
| GCLC | 84.1656844 | -1.423764409 | 0.33997182 | -4.1878895 | 2.82E-05 | 0.0026346 |
| LRRC1 | 37.5477714 | 0.035363175 | 0.24729786 | 0.1429983 | 0.88629152 | 0.97693497 |
| FAM83B | 113.224113 | -0.642015389 | 0.20535499 | -3.1263686 | 0.0017698 | 0.04768616 |
| DST | 402.484933 | -1.096212903 | 0.33691109 | -3.2537157 | 0.00113906 | 0.03587305 |
| KIAA1586 | 10.9586124 | -0.368213941 | 0.39881791 | -0.9232633 | 0.35587001 | NA |
| ZNF451 | 118.767894 | 0.093575563 | 0.2317322 | 0.40380907 | 0.68635313 | 0.92597882 |
| ZNF451-AS1 | 16.7232597 | -0.77292144 | 0.66049171 | -1.1702213 | 0.24191194 | 0.69556484 |
| PRIM2 | 98.6725339 | 0.226715292 | 0.26617257 | 0.85176055 | 0.394347 | 0.80495916 |
| AL021368.2 | 11.883462 | 0.85861685 | 0.4436312 | 1.93542934 | 0.05293763 | NA |
| PTP4A1 | 53.744079 | -0.285203037 | 0.52474583 | -0.543507 | 0.58678077 | 0.89059199 |
| PHF3 | 168.275426 | -0.207855526 | 0.19341091 | -1.0746835 | 0.28251643 | 0.73410378 |
| EYS | 15.2022015 | -0.247023959 | 0.53176755 | -0.4645337 | 0.64226541 | 0.91146968 |
| LMBRD1 | 154.506914 | 0.183950668 | 0.26132198 | 0.70392345 | 0.48148045 | 0.85020971 |
| COL19A1 | 34.6201707 | 0.201461382 | 0.2781268 | 0.72435085 | 0.46885036 | 0.84326495 |
| FAM135A | 393.60886 | -1.027539426 | 0.2864954 | -3.5865826 | 0.00033504 | 0.0156695 |
| SDHAF4 | 22.1057652 | 0.114143839 | 0.51687243 | 0.22083561 | 0.82522044 | 0.96539788 |
| SMAP1 | 221.877787 | 0.03741051 | 0.19924697 | 0.18775949 | 0.85106519 | 0.97027687 |
| OGFRL1 | 40.97823 | 0.779722544 | 0.57771919 | 1.34965664 | 0.17712615 | 0.6249995 |
| AL136164.2 | 85.655106 | 0.723242474 | 0.47409391 | 1.52552575 | 0.12712801 | 0.54952841 |
| LINC00472 | 51.4556014 | -0.011163824 | 0.47910017 | -0.0233016 | 0.98140966 | 0.99714727 |
| OOEP | 173.16663 | 0.362826381 | 0.38330519 | 0.94657309 | 0.34385636 | 0.77370523 |
| MTO1 | 33.9170732 | 0.417112 | 0.24052678 | 1.73416036 | 0.08288963 | 0.45502516 |
| EEF1A1 | 844.608995 | 0.172176332 | 0.40598477 | 0.42409554 | 0.67149614 | 0.92170876 |
| SLC17A5 | 50.5683554 | -0.255547804 | 0.44123663 | -0.5791627 | 0.56247938 | 0.88191823 |
| COX7A2 | 136.087764 | -0.002169639 | 0.33798111 | -0.0064194 | 0.99487809 | 0.99923337 |
| TMEM30A | 101.419924 | -0.107480582 | 0.21035622 | -0.5109456 | 0.60938915 | 0.89906009 |
| FILIP1 | 76.5808904 | -4.803752527 | 0.78589044 | -6.1124964 | 9.81E-10 | 6.34E-07 |
| SENP6 | 387.966711 | 0.127395885 | 0.15245614 | 0.83562316 | 0.40336694 | 0.80927646 |
| MYO6 | 740.960731 | -0.432012424 | 0.27752283 | -1.5566735 | 0.11954803 | 0.53108939 |
| IRAK1BP1 | 25.8663571 | -0.00953606 | 0.41691388 | -0.022873 | 0.9817516 | 0.99714727 |
| PHIP | 221.521274 | 0.210655878 | 0.21379704 | 0.98530773 | 0.32447294 | 0.76304922 |

| | | | | | | |
|---|---|---|---|---|---|---|
| HMGN3 | 108.716634 | 0.161739586 | 0.35879085 | 0.45079072 | 0.65214039 | 0.9158679 |
| LCA5 | 15.1236132 | -0.14624605 | 0.39993904 | -0.3656709 | 0.7146107 | 0.93684603 |
| SH3BGRL2 | 16.3826416 | 0.599157476 | 0.4727062 | 1.26750501 | 0.20497477 | 0.65805651 |
| BCKDHB | 110.868729 | 0.184775669 | 0.39667332 | 0.4658132 | 0.64134922 | 0.91146968 |
| TENT5A | 65.6172414 | -0.569623641 | 0.58007404 | -0.9819844 | 0.32610755 | 0.76353952 |
| IBTK | 161.595547 | -0.09552195 | 0.17816938 | -0.53613 | 0.59186871 | 0.89161773 |
| TPBG | 13.3161566 | -1.472029439 | 0.50191294 | -2.9328382 | 0.00335879 | 0.07371098 |
| UBE3D | 65.9870435 | 0.509685638 | 0.26625083 | 1.91430632 | 0.05558103 | 0.37507329 |
| DOPEY1 | 140.415527 | -0.035186101 | 0.22034758 | -0.1596845 | 0.87312959 | 0.97434023 |
| PGM3 | 31.6146998 | -0.397707304 | 0.35890825 | -1.108103 | 0.26781734 | 0.71922155 |
| CYB5R4 | 45.4194548 | -0.154370552 | 0.33277456 | -0.4638893 | 0.64272707 | 0.91146968 |
| CEP162 | 54.1064475 | -0.167769952 | 0.23448123 | -0.7154942 | 0.47430375 | 0.84593347 |
| SNX14 | 141.091609 | 0.121178046 | 0.19082053 | 0.63503675 | 0.52540445 | 0.86747074 |
| SYNCRIP | 52.3579783 | 0.207282439 | 0.32792381 | 0.63210549 | 0.52731795 | 0.86750588 |
| ZNF292 | 455.767146 | -1.120163627 | 0.32586552 | -3.4375028 | 0.0005871 | 0.02282458 |
| SMIM8 | 71.1981007 | 0.790351476 | 0.34799726 | 2.2711428 | 0.02313833 | 0.23879078 |
| RARS2 | 89.8270507 | -0.0379603 | 0.19912735 | -0.1906333 | 0.84881291 | 0.97027687 |
| ORC3 | 48.6778137 | -0.146666837 | 0.33723966 | -0.4349039 | 0.6636322 | 0.91855484 |
| AKIRIN2 | 26.2151492 | -0.268111643 | 0.28216801 | -0.9501844 | 0.34201855 | 0.7722486 |
| RNGTT | 181.488509 | 0.191082492 | 0.18217792 | 1.04887845 | 0.29423406 | 0.7405942 |
| PNRC1 | 89.8957266 | -0.342940167 | 0.38381226 | -0.8935102 | 0.37158402 | 0.7900012 |
| PM20D2 | 38.0744946 | -0.24122805 | 0.26775997 | -0.9009115 | 0.36763535 | 0.78877653 |
| UBE2J1 | 43.9220479 | -0.127418702 | 0.31942583 | -0.3988992 | 0.68996749 | 0.92734996 |
| RRAGD | 80.2889287 | -0.741445303 | 0.38833785 | -1.909279 | 0.05622611 | 0.37709488 |
| ANKRD6 | 51.6251868 | -0.350048178 | 0.2662173 | -1.3148965 | 0.1885447 | 0.64021224 |
| LYRM2 | 44.1433039 | -0.390091247 | 0.34619608 | -1.1267928 | 0.25983009 | 0.71091822 |
| MDN1 | 72.543856 | 0.019460059 | 0.23079835 | 0.08431628 | 0.93280497 | 0.98809895 |
| CASP8AP2 | 46.2650316 | 0.36008076 | 0.30749135 | 1.17102727 | 0.24158782 | 0.69556484 |
| BACH2 | 98.474061 | -1.119474564 | 0.61539238 | -1.8191232 | 0.06889263 | 0.41687995 |
| MAP3K7 | 94.7603619 | -0.065971813 | 0.25196013 | -0.2618343 | 0.79344917 | 0.95762809 |
| MANEA | 46.0579092 | 0.23014622 | 0.28292435 | 0.81345497 | 0.41595726 | 0.81763864 |
| UFL1 | 63.3424141 | 0.008950161 | 0.1993568 | 0.04489519 | 0.96419085 | 0.99334616 |
| NDUFAF4 | 34.5700647 | 0.33393917 | 0.41836005 | 0.79820998 | 0.42474864 | 0.82241967 |
| MMS22L | 25.8047461 | 0.210163833 | 0.40449504 | 0.51957086 | 0.60336271 | 0.8956362 |
| FBXL4 | 121.12491 | -0.25044465 | 0.28657712 | -0.8739171 | 0.38216339 | 0.79731722 |
| FAXC | 20.2403242 | -0.726185577 | 0.56825614 | -1.2779195 | 0.20127783 | 0.65374585 |
| COQ3 | 13.9955999 | 0.680782333 | 0.35757857 | 1.90386782 | 0.0569274 | 0.37925535 |
| PNISR | 376.192778 | 0.183680275 | 0.14866009 | 1.23557223 | 0.21661761 | 0.67130697 |
| AL513550.1 | 19.00728 | -0.362896111 | 0.42206888 | -0.8598031 | 0.38989761 | 0.80191835 |
| USP45 | 60.6524265 | -0.209600136 | 0.34354423 | -0.6101111 | 0.54178823 | 0.87175518 |
| TSTD3 | 15.8361042 | -0.57068947 | 0.40778587 | -1.3994832 | 0.16166814 | 0.60060549 |
| CCNC | 126.155552 | -0.567546291 | 0.25897641 | -2.191498 | 0.02841577 | 0.26605501 |
| SIM1 | 323.260102 | 0.423392982 | 0.28610789 | 1.47983677 | 0.13891681 | 0.57182884 |
| ASCC3 | 216.075409 | -0.499174482 | 0.26185187 | -1.9063239 | 0.0566082 | 0.37816771 |
| HACE1 | 37.8665098 | 0.123370522 | 0.26005289 | 0.47440551 | 0.63521077 | 0.90881415 |
| PREP | 52.4538281 | 0.114617527 | 0.26527257 | 0.43207455 | 0.66568723 | 0.91958319 |
| ATG5 | 128.612029 | -0.044593916 | 0.22112544 | -0.201668 | 0.84017631 | 0.96748214 |
| CRYBG1 | 109.968746 | -0.670306994 | 0.76251221 | -0.8790771 | 0.3793595 | 0.79535699 |
| RTN4IP1 | 54.6139521 | -1.024364467 | 0.40125847 | -2.5528794 | 0.01068365 | 0.15239912 |
| QRSL1 | 69.4664218 | -0.320511188 | 0.3049871 | -1.0509008 | 0.29330415 | 0.74013384 |
| LINC02532 | 108.91333 | -2.107365454 | 0.59491082 | -3.5423216 | 0.00039662 | 0.01717513 |
| CD24 | 1021.30196 | -1.088334712 | 0.29758723 | -3.6571956 | 0.00025499 | 0.01282319 |
| C6orf203 | 26.0771368 | 0.372238757 | 0.25546038 | 1.45712913 | 0.14508074 | 0.58008375 |
| PDSS2 | 183.136009 | 0.238974699 | 0.19535332 | 1.2232948 | 0.22121837 | 0.67492938 |
| SEC63 | 93.654056 | 0.129249448 | 0.19218209 | 0.6725364 | 0.50124228 | 0.85942996 |
| OSTM1 | 75.7419203 | -0.462666006 | 0.38492659 | -1.2019591 | 0.22937939 | 0.68503082 |
| SNX3 | 119.546927 | -0.042755496 | 0.2383272 | -0.1793983 | 0.85762496 | 0.97243282 |
| AFG1L | 86.4229602 | 0.206078945 | 0.28410178 | 0.72537012 | 0.468225 | 0.84326495 |
| FOXO3 | 313.34487 | -0.277117359 | 0.24664855 | -1.1235313 | 0.26121192 | 0.71232939 |
| ARMC2 | 23.2602134 | 0.445916741 | 0.43725012 | 1.01982074 | 0.30781348 | 0.7512664 |
| SESN1 | 112.978051 | 0.419163751 | 0.25544421 | 1.64092093 | 0.10081383 | 0.4896816 |
| CEP57L1 | 33.8613647 | 0.256284136 | 0.29932389 | 0.85621009 | 0.39188158 | 0.80360124 |
| CD164 | 163.9972 | 0.051387557 | 0.2230522 | 0.23038355 | 0.81779374 | 0.9635539 |
| PPIL6 | 12.5918163 | 0.242601708 | 0.57236858 | 0.42385574 | 0.67167103 | 0.92170876 |
| ZBTB24 | 10.7491909 | -0.067916696 | 0.38372231 | -0.1769944 | 0.85951279 | NA |
| AK9 | 42.0264774 | 0.348896348 | 0.46835297 | 0.74494317 | 0.45630608 | 0.83506875 |
| FIG4 | 51.7909443 | -0.262695209 | 0.29998184 | -0.8757037 | 0.38119113 | 0.79688435 |
| CDC40 | 26.9523598 | 0.140484258 | 0.27768246 | 0.50591693 | 0.61291497 | 0.90011356 |
| CDK19 | 76.5592864 | 0.150853254 | 0.33049852 | 0.45644154 | 0.6480725 | 0.91397255 |
| AMD1 | 69.4247476 | 0.285444154 | 0.29588627 | 0.96470903 | 0.33469057 | 0.76777176 |
| GTF3C6 | 54.2640522 | -0.349989016 | 0.38696315 | -0.9044505 | 0.36575657 | 0.78793182 |
| RPF2 | 18.3714356 | -0.524567014 | 0.36713605 | -1.4288082 | 0.15305936 | 0.59174003 |
| SLC16A10 | 212.972674 | 0.421767082 | 0.23759972 | 1.77511605 | 0.07587873 | 0.43397621 |
| MFSD4B | 16.9175247 | 0.211343422 | 0.34312536 | 0.61593646 | 0.53793645 | 0.86972442 |
| REV3L | 196.458165 | 0.053082824 | 0.19772658 | 0.2684658 | 0.78834079 | 0.95682003 |
| TRAF3IP2-AS | 66.0866359 | -0.3349713 | 0.22024428 | -1.5209081 | 0.12828291 | 0.5528291 |
| TRAF3IP2 | 19.0967412 | -0.542739608 | 0.32502369 | -1.6698463 | 0.09494977 | 0.48035393 |
| TUBE1 | 29.6943756 | -0.209276394 | 0.38642212 | -0.5415746 | 0.58811162 | 0.89073775 |
| LAMA4 | 21.564648 | -0.993802977 | 0.85212879 | -1.1662591 | 0.2435097 | 0.69738531 |
| MARCKS | 129.828601 | 0.368677634 | 0.26784926 | 1.376437 | 0.16868638 | 0.61072372 |
| HDAC2 | 45.0208543 | 0.030258905 | 0.22200114 | 0.13630067 | 0.89158359 | 0.97793768 |
| FRK | 315.886195 | -0.019464131 | 0.25638092 | -0.0759188 | 0.9394837 | 0.98871562 |
| NT5DC1 | 256.393489 | 0.171745939 | 0.22302061 | 0.77009 | 0.44124651 | 0.8281114 |
| COL10A1 | 14.0441999 | -1.419653321 | 0.68563619 | -2.0705636 | 0.0383996 | 0.3141788 |
| TSPYL4 | 16.5922688 | 0.648420894 | 0.3676011 | 1.76392533 | 0.07774455 | 0.4397149 |
| DSE | 57.0123155 | 0.223649739 | 0.32170979 | 0.69519096 | 0.48693564 | 0.85365949 |
| TSPYL1 | 76.436559 | 0.160356532 | 0.21289785 | 0.75320878 | 0.45132447 | 0.83230402 |
| RWDD1 | 34.4078557 | 0.357108755 | 0.3057493 | 1.16797896 | 0.24281525 | 0.69621873 |
| ZUP1 | 26.2912738 | 0.14293759 | 0.30009225 | 0.47631217 | 0.633852 | 0.90805737 |
| KPNA5 | 78.2819353 | 0.010847966 | 0.21799434 | 0.0497626 | 0.96031157 | 0.99238848 |
| DCBLD1 | 29.6946733 | -0.169377688 | 0.34531558 | -0.4905012 | 0.62377931 | 0.90447745 |
| GOPC | 47.6885957 | 0.054162387 | 0.2395068 | 0.22614133 | 0.8210915 | 0.96423578 |
| NUS1 | 63.282559 | -0.0808587 | 0.22887578 | -0.3532864 | 0.72387374 | 0.93896433 |
| SLC35F1 | 12.6551197 | -0.40366446 | 0.45638817 | -0.8844762 | 0.37643922 | 0.79422885 |
| CEP85L | 51.8459905 | -0.330692127 | 0.32226611 | -1.0261462 | 0.3048227 | 0.74832335 |
| MCM9 | 109.85032 | 0.141282086 | 0.23887235 | 0.59145434 | 0.55421604 | 0.87712443 |
| AL365275.1 | 26.7218965 | 1.147556698 | 0.46594635 | 2.46285156 | 0.0137837 | 0.17615528 |
| FAM184A | 64.0302839 | 1.326139966 | 0.57956331 | 2.28817101 | 0.02212756 | 0.23355536 |
| MAN1A1 | 354.693834 | 0.283362181 | 0.27730185 | 1.02185462 | 0.30684971 | 0.74973356 |
| TBC1D32 | 52.8413536 | 0.518397337 | 0.32981008 | 1.57180563 | 0.11599564 | 0.52163082 |
| HSF2 | 20.3587629 | 0.455191474 | 0.38884454 | 1.17062585 | 0.2417492 | 0.69556484 |
| SERINC1 | 153.247533 | -0.137071777 | 0.20979335 | -0.6533657 | 0.51352055 | 0.86388299 |
| PKIB | 23.4479416 | 0.477494185 | 0.49127019 | 0.9719584 | 0.33107125 | 0.76685572 |
| SMPDL3A | 19.6075634 | -0.12705025 | 0.35531526 | -0.3575705 | 0.72066477 | 0.93806337 |
| NKAIN2 | 12.359824 | -0.369667762 | 0.48912874 | -0.7557678 | 0.44978841 | NA |
| RNF217-AS1 | 16.927139 | -2.001351688 | 0.65312124 | -3.0642882 | 0.00218189 | 0.05412864 |
| RNF217 | 247.751208 | -1.746288916 | 0.45303077 | -3.8546806 | 0.00011588 | 0.00720541 |
| TPD52L1 | 102.356131 | 0.034792225 | 0.50918961 | 0.06832862 | 0.94552404 | 0.99134285 |
| HDDC2 | 44.1471636 | 0.226714898 | 0.33591339 | 0.67492069 | 0.49972615 | 0.85942996 |
| NCOA7 | 875.971333 | -0.519412982 | 0.21976868 | -2.3634531 | 0.01810552 | 0.20906729 |
| NCOA7-AS1 | 37.7047867 | -0.89128183 | 0.55782243 | -1.5977877 | 0.11009023 | 0.5105769 |
| HINT3 | 36.9340461 | 0.001401821 | 0.25267377 | 0.00554795 | 0.9955734 | 0.99923337 |
| TRMT11 | 60.3291446 | 0.071507495 | 0.25065704 | 0.28528022 | 0.77542946 | 0.95493992 |

| | | | | | | |
|---|---|---|---|---|---|---|
| RNF146 | 46.8840786 | -0.132070208 | 0.34034814 | -0.3880445 | 0.69798314 | 0.93015138 |
| ECHDC1 | 69.9431809 | -0.295174962 | 0.23963989 | -1.2317438 | 0.21804479 | 0.67207958 |
| PTPRK | 641.213301 | 0.382432912 | 0.3661478 | 1.04447687 | 0.29626481 | 0.74123516 |
| LAMA2 | 249.264159 | 0.140284762 | 0.53657646 | 0.26144412 | 0.79375003 | 0.95770265 |
| AL356124.1 | 218.063138 | 0.168206832 | 0.40897556 | 0.41128823 | 0.6808612 | 0.92375164 |
| ARHGAP18 | 1327.98531 | 0.523578142 | 0.17237494 | 3.03743778 | 0.00238599 | 0.05737663 |
| L3MBTL3 | 22.9158868 | -1.388770539 | 0.35885065 | -3.8700516 | 0.00010881 | 0.00689856 |
| EPB41L2 | 16.7234765 | -0.270632497 | 0.36347589 | -0.744568 | 0.45653295 | 0.83506875 |
| AKAP7 | 44.9048811 | 0.269245826 | 0.53445615 | 0.50377533 | 0.61441926 | 0.90054121 |
| MED23 | 58.8541769 | 0.415135965 | 0.23001744 | 1.80480212 | 0.07110565 | 0.42236671 |
| STX7 | 102.05109 | -0.508403775 | 0.26143787 | -1.9446447 | 0.05181775 | 0.36134594 |
| SLC18B1 | 56.9734475 | -0.210242666 | 0.41063419 | -0.511995 | 0.60865449 | 0.89834883 |
| RPS12 | 179.368217 | 0.69571702 | 0.3673519 | 1.89387076 | 0.05824217 | 0.38243048 |
| EYA4 | 149.198908 | -0.217477716 | 0.77913859 | -0.2791258 | 0.78014825 | 0.95537097 |
| TBPL1 | 32.8150178 | 0.218167745 | 0.36943732 | 0.59054063 | 0.55482826 | 0.87754295 |
| SLC2A12 | 27.6201955 | 0.6885445 | 0.60209599 | 1.14357928 | 0.25279816 | 0.7066692 |
| HBS1L | 61.4609874 | 0.301556014 | 0.23759837 | 1.26918386 | 0.2043755 | 0.65771647 |
| AHI1 | 265.952341 | -0.312523296 | 0.19697772 | -1.5865921 | 0.11260506 | 0.51571293 |
| LINC00271 | 66.6715975 | 0.304900939 | 0.49249689 | 0.61909211 | 0.53585568 | 0.86946459 |
| PDE7B | 188.040919 | 0.954786148 | 0.356182 | 2.68061313 | 0.00734874 | 0.12396174 |
| AL138828.1 | 38.3990718 | -1.987710095 | 0.61995923 | -3.2061949 | 0.00134503 | 0.03953569 |
| BCLAF1 | 152.533603 | -0.030073207 | 0.18160583 | -0.165596 | 0.86847486 | 0.97434023 |
| MAP7 | 277.750109 | 0.18149345 | 0.22897336 | 0.79264002 | 0.42798759 | 0.82407405 |
| MAP3K5 | 289.928692 | -0.800354755 | 0.36871771 | -2.1706436 | 0.02995812 | 0.27570566 |
| PEX7 | 58.5473766 | 0.195338621 | 0.26234673 | 0.74458188 | 0.45652453 | 0.83506875 |
| IFNGR1 | 61.7379047 | -0.526845474 | 0.29219871 | -1.8030384 | 0.07138219 | 0.42281025 |
| TNFAIP3 | 15.1840276 | -1.624153226 | 0.61120397 | -2.6573015 | 0.0078769 | 0.12884636 |
| PERP | 69.0745785 | -1.119917037 | 0.28255521 | -3.9635334 | 7.38E-05 | 0.00526714 |
| ARFGEF3 | 63.3833865 | -1.425440744 | 0.42547406 | -3.3502413 | 0.00080741 | 0.02847962 |
| HEBP2 | 52.2270597 | -0.426468056 | 0.27368561 | -1.5582407 | 0.1191762 | 0.53056597 |
| NHSL1 | 50.1353687 | 0.606653069 | 0.60299799 | 1.00606152 | 0.31438598 | 0.75700972 |
| CCDC28A | 26.8854612 | 0.183701025 | 0.38397523 | 0.47841895 | 0.63235205 | 0.90805737 |
| REPS1 | 136.558683 | -0.146580856 | 0.25123553 | -0.58344 | 0.55959713 | 0.88037687 |
| ABRACL | 13.825136 | -1.746448831 | 0.60039122 | -2.9088514 | 0.00362759 | 0.07819478 |
| HECA | 29.6552654 | -0.518422074 | 0.41527002 | -1.2483975 | 0.21188551 | 0.66386489 |
| CITED2 | 55.1407906 | -0.422285283 | 0.50215789 | -0.8409412 | 0.40038085 | 0.80830282 |
| VTA1 | 49.8644419 | -0.097028115 | 0.2328973 | -0.4166133 | 0.67696127 | 0.92237262 |
| ADGRG6 | 13.1002402 | -1.549687328 | 0.69675464 | -2.2241507 | 0.02613831 | 0.25383383 |
| HIVEP2 | 213.757847 | -0.42460128 | 0.33443151 | -1.269621 | 0.20421965 | 0.65771647 |
| AIG1 | 189.177865 | 0.382046192 | 0.32353334 | 1.18085572 | 0.23766005 | 0.69207518 |
| ADAT2 | 16.9746826 | 0.542968371 | 0.41953844 | 1.29420409 | 0.19559493 | 0.6459894 |
| PEX3 | 52.0180333 | -0.048350027 | 0.31548145 | -0.1532579 | 0.87819489 | 0.97542218 |
| FUCA2 | 41.2058122 | -0.079927786 | 0.26142225 | -0.3057421 | 0.75980102 | 0.94952011 |
| PHACTR2 | 268.336239 | -0.34800372 | 0.30497214 | -1.1411 | 0.2538283 | 0.70701839 |
| PLAGL1 | 44.4541802 | -1.377459678 | 0.60363826 | -2.281929 | 0.02249353 | 0.23562337 |
| SF3B5 | 53.4895467 | -0.43656258 | 0.30779121 | -1.4183725 | 0.15608205 | 0.59606138 |
| UTRN | 58.2423007 | 0.662636583 | 0.53369995 | 1.24159011 | 0.21438783 | 0.66839951 |
| EPM2A | 32.6111556 | 0.290741979 | 0.43513756 | 0.66816108 | 0.50403078 | 0.86090835 |
| FBXO30 | 25.6472247 | -0.189455774 | 0.42289095 | -0.4480015 | 0.65415212 | 0.9158679 |
| AL356599.1 | 44.8189738 | 0.04192214 | 0.46141778 | 0.09085506 | 0.92760776 | 0.98690525 |
| SHPRH | 157.768198 | 0.197998324 | 0.24079025 | 0.82228547 | 0.41091444 | 0.81444015 |
| STXBP5 | 167.295943 | -0.356668439 | 0.33888592 | -1.0524735 | 0.29258237 | 0.74013384 |
| SAMD5 | 774.422459 | -0.575846078 | 0.31634165 | -1.8203296 | 0.06870882 | 0.41651909 |
| AL033504.1 | 425.306454 | -0.788209196 | 0.37565991 | -2.0981989 | 0.03588758 | 0.30296738 |
| AL513164.1 | 28.1456595 | -1.194877541 | 0.47130431 | -2.535257 | 0.01123649 | 0.15682578 |
| SASH1 | 315.346489 | -0.465288962 | 0.16572058 | -2.8076716 | 0.00499011 | 0.09739245 |
| UST | 731.579933 | -0.717326492 | 0.32137404 | -2.2320611 | 0.02561092 | 0.25205612 |
| TAB2 | 156.625583 | 0.190337313 | 0.16246377 | 1.17156778 | 0.24137063 | 0.69556484 |
| PPIL4 | 64.3596324 | 0.272293522 | 0.22834002 | 1.19249147 | 0.23306859 | 0.68763965 |
| GINM1 | 99.2957734 | -0.127209397 | 0.21388022 | -0.5947693 | 0.55199767 | 0.87576439 |
| KATNA1 | 44.6058469 | 0.058632758 | 0.24502453 | 0.23929342 | 0.81087807 | 0.96243713 |
| LATS1 | 86.3680948 | 0.182439562 | 0.21282032 | 0.85724692 | 0.39130843 | 0.80332153 |
| NUP43 | 24.4680634 | 0.335506482 | 0.28483877 | 1.17788207 | 0.23884362 | 0.69356592 |
| PCMT1 | 77.4221366 | 0.431896541 | 0.30849168 | 1.40002655 | 0.16150537 | 0.60060549 |
| LRP11 | 75.4431846 | -0.642736182 | 0.44307276 | -1.4506335 | 0.14688194 | 0.58364362 |
| PLEKHG1 | 39.6160872 | 0.047185109 | 0.67739394 | 0.06965682 | 0.94446681 | 0.99105669 |
| MTHFD1L | 248.212791 | -0.356097697 | 0.3088901 | -1.1528297 | 0.24898027 | 0.70371066 |
| RMND1 | 25.2024801 | 0.215485061 | 0.3352227 | 0.64281166 | 0.52034632 | 0.86680671 |
| ARMT1 | 25.5625815 | 0.32514219 | 0.384261 | 0.84614934 | 0.39746944 | 0.80674902 |
| ESR1 | 71.9682454 | -0.246127131 | 1.21871222 | -0.2019567 | 0.83995055 | 0.96748214 |
| SYNE1 | 122.040745 | 0.132910347 | 0.60106348 | 0.22112531 | 0.82499486 | 0.96539788 |
| MTRF1L | 23.395046 | 7.62E-05 | 0.39653228 | 0.00019218 | 0.99984666 | 0.99994975 |
| OPRM1 | 27.8869686 | 0.535571663 | 0.76393107 | 0.70107328 | 0.48325728 | 0.85205504 |
| IPCEF1 | 238.801486 | 1.352147112 | 0.6048069 | 2.23566746 | 0.02537356 | 0.25137888 |
| CNKSR3 | 107.673542 | -0.509890746 | 0.30410806 | -1.6766762 | 0.09360582 | 0.47792014 |
| SCAF8 | 264.795409 | 0.212899315 | 0.18516333 | 1.14979198 | 0.25022956 | 0.70453262 |
| TIAM2 | 159.079838 | -0.175534802 | 0.32379965 | -0.5421093 | 0.58774321 | 0.89073775 |
| TFB1M | 49.6375503 | 0.092150989 | 0.32151975 | 0.28661067 | 0.77441045 | 0.95470568 |
| ARID1B | 735.38085 | -0.028324067 | 0.2164759 | -0.1308417 | 0.89590056 | 0.97885533 |
| TMEM242 | 30.6741569 | 0.093065845 | 0.27766747 | 0.33517013 | 0.73749675 | 0.94376279 |
| ZDHHC14 | 112.638516 | -1.889756762 | 0.50581758 | -3.736044 | 0.00018694 | 0.0103617 |
| SNX9 | 249.745129 | -0.523089475 | 0.21306826 | -2.4550324 | 0.01408719 | 0.17815613 |
| SYNJ2 | 32.2790785 | -0.81113631 | 0.44061366 | -1.8409241 | 0.06563268 | 0.40715521 |
| SERAC1 | 25.2860828 | -0.644224966 | 0.45455229 | -1.4172736 | 0.15640295 | 0.59681693 |
| GTF2H5 | 33.1449559 | -0.167798448 | 0.42338644 | -0.3963246 | 0.69186561 | 0.928133 |
| TULP4 | 325.485206 | -0.016894285 | 0.26922274 | -0.0627521 | 0.94996393 | 0.99192022 |
| TMEM181 | 124.827252 | -0.394069343 | 0.28934903 | -1.3619169 | 0.1732241 | 0.61802607 |
| DYNLT1 | 128.281091 | -0.692481373 | 0.36132996 | -1.9164793 | 0.05530412 | 0.37383273 |
| SYTL3 | 47.1675909 | -1.277140741 | 0.45553745 | -2.8035911 | 0.0050537 | 0.09823817 |
| EZR | 479.497246 | -0.621741688 | 0.28319001 | -2.195493 | 0.02812825 | 0.26404526 |
| RSPH3 | 37.592712 | -0.117282547 | 0.35906874 | -0.3266298 | 0.74394791 | 0.94546379 |
| AL035530.2 | 21.2640618 | 0.444916104 | 0.38021424 | 1.17017211 | 0.24193171 | 0.69556484 |
| SOD2.1 | 27.3140782 | -0.328374473 | 0.37097603 | -0.8851636 | 0.3760684 | 0.79422241 |
| WTAP | 143.357009 | -0.379770199 | 0.33986885 | -1.1174022 | 0.26382239 | 0.71382907 |
| ACAT2 | 26.3087048 | -1.432230102 | 0.61840455 | -2.3160084 | 0.02055781 | 0.22428714 |
| TCP1 | 48.0480916 | 0.032287515 | 0.31524325 | 0.10242096 | 0.91842255 | 0.98521039 |
| MRPL18 | 29.6347207 | -0.240685972 | 0.40064049 | -0.600753 | 0.54800452 | 0.87341177 |
| IGF2R | 245.316921 | -0.277921317 | 0.22661126 | -1.2264233 | 0.22003944 | 0.67350139 |
| MAP3K4 | 89.5761075 | 0.440200912 | 0.17900127 | 2.4592055 | 0.01392449 | 0.17681258 |
| PRKN | 1052.19368 | 0.361511447 | 0.28865351 | 1.25240619 | 0.21042189 | 0.66226229 |
| PACRG | 1892.35692 | 0.14347844 | 0.4318903 | 0.33221038 | 0.73973041 | 0.94376279 |
| PACRG-AS3 | 32.0958934 | -0.318383135 | 0.30120671 | -1.0570254 | 0.2905 | 0.73979353 |
| QKI | 427.851018 | -0.12813654 | 0.24039539 | -0.5330241 | 0.59401689 | 0.89288778 |
| PDE10A | 249.74119 | -1.011496431 | 0.45970051 | -2.2003379 | 0.02778293 | 0.26203825 |
| SFT2D1 | 58.3702648 | -0.199457636 | 0.2370694 | -0.841347 | 0.40015355 | 0.80830282 |
| MPC1 | 275.069466 | -0.165006742 | 0.2998977 | -0.5502101 | 0.58217528 | 0.89036744 |
| RPS6KA2 | 57.9231699 | -0.362305398 | 0.30903752 | -1.172367 | 0.24104974 | 0.69556484 |
| Z94721.1 | 23.8890918 | 0.294776323 | 0.37981402 | 0.77610701 | 0.43768581 | 0.8270818 |
| FGFR1OP | 95.0961885 | 0.342211284 | 0.31680133 | 1.08020785 | 0.28004963 | 0.73179753 |
| AFDN | 351.199367 | -0.101288411 | 0.20879515 | -0.485109 | 0.62759902 | 0.90603677 |
| WDR27 | 65.3942815 | 0.238346069 | 0.32978902 | 0.72272288 | 0.46985016 | 0.8439901 |
| C6orf120 | 12.589484 | -0.66078029 | 0.39225751 | -1.6845574 | 0.092074 | 0.4764741 |
| PHF10 | 50.6018849 | -0.109051862 | 0.28888561 | -0.3774915 | 0.70580838 | 0.93407096 |

| | | | | | | |
|---|---|---|---|---|---|---|
| ERMARD | 21.7276281 | 0.039683221 | 0.4339209 | 0.09145266 | 0.92713292 | 0.98690525 |
| DLL1 | 70.4456591 | 0.17076802 | 0.26170946 | 0.65250993 | 0.51407227 | 0.86416832 |
| FAM120B | 265.474733 | -0.217918617 | 0.24032144 | -0.9067798 | 0.36452327 | 0.7878511 |
| PSMB1 | 91.9766838 | -0.412149886 | 0.25852768 | -1.5942195 | 0.11088687 | 0.51243576 |
| TBP | 17.7441038 | -0.062527753 | 0.39484716 | -0.1583594 | 0.87417362 | 0.97434023 |
| PDCD2 | 30.8669966 | -0.10946421 | 0.32646997 | -0.3352964 | 0.7374015 | 0.94376279 |
| FAM20C | 83.354337 | -1.460443047 | 0.38879027 | -3.7563776 | 0.00017239 | 0.01007342 |
| PRKAR1B | 34.2960382 | 0.72034745 | 0.40067486 | 1.7978354 | 0.0722031 | 0.42395281 |
| DNAAF5 | 23.702499 | 0.266839879 | 0.33931258 | 0.78641316 | 0.43162548 | 0.8248269 |
| SUN1 | 109.928648 | 0.06085866 | 0.16083525 | 0.37839132 | 0.70513992 | 0.93403928 |
| COX19 | 47.5892429 | 0.079061901 | 0.31840888 | 0.24830306 | 0.80389993 | 0.95961473 |
| C7orf50 | 70.739406 | 0.248655261 | 0.26100961 | 0.95266707 | 0.34075878 | 0.77193691 |
| MICALL2 | 15.2527278 | -0.646432789 | 0.39148272 | -1.6512422 | 0.09868913 | 0.48476221 |
| INTS1 | 34.2734455 | 0.178487756 | 0.38177524 | 0.46752051 | 0.64012752 | 0.91096398 |
| MAFK | 28.8520159 | -0.026926415 | 0.43639747 | -0.0617016 | 0.95080048 | 0.99212184 |
| PSMG3 | 20.6362252 | 0.067326871 | 0.41661766 | 0.1616035 | 0.8716181 | 0.97434023 |
| MAD1L1 | 48.3769589 | 0.199039505 | 0.24598766 | 0.80914426 | 0.41843217 | 0.8188001 |
| SNX8 | 15.4025773 | -0.882312536 | 0.55496217 | -1.5898607 | 0.11186621 | 0.51426646 |
| EIF3B | 50.0084648 | -0.179039378 | 0.32305794 | -0.5542021 | 0.57944056 | 0.88957613 |
| CHST12 | 19.6239698 | -0.282959308 | 0.46026951 | -0.6147687 | 0.53870745 | 0.86972442 |
| BRAT1 | 11.7970403 | 0.536149336 | 0.39704428 | 1.35035149 | 0.17690326 | NA |
| IQCE | 23.9101493 | 0.137032526 | 0.2841673 | 0.48222483 | 0.62964624 | 0.90724429 |
| GNA12 | 54.0890065 | 0.150126804 | 0.25107492 | 0.59793628 | 0.54988245 | 0.87445431 |
| SDK1 | 81.1407811 | 0.027499535 | 0.32514278 | 0.0845768 | 0.93259784 | 0.98800776 |
| FOXK1 | 121.437459 | -0.349923039 | 0.33173542 | -1.0548257 | 0.29150505 | 0.73994767 |
| AP5Z1 | 23.8887295 | -0.577798871 | 0.54171707 | -1.0666064 | 0.28614962 | 0.73672385 |
| RBAK-RBAKI | 50.0982879 | -0.244045815 | 0.22410939 | -1.0889585 | 0.27617221 | 0.72575599 |
| WIPI2 | 77.0451196 | -0.067851433 | 0.29055327 | -0.2335249 | 0.81535381 | 0.96300833 |
| SLC29A4 | 30.0231874 | 0.87748296 | 0.45405709 | 1.93253884 | 0.05329303 | 0.36627428 |
| TNRC18 | 167.975229 | 0.014103661 | 0.26885061 | 0.0524591 | 0.95816288 | 0.99238848 |
| FBXL18 | 23.0451654 | -0.299771816 | 0.46166196 | -0.6493318 | 0.5161239 | 0.8648129 |
| ACTB | 526.593247 | 0.19481896 | 0.4064362 | 0.47933467 | 0.63170056 | 0.90804615 |
| RNF216 | 118.055685 | 0.023967295 | 0.21581429 | 0.11105518 | 0.91157259 | 0.9835359 |
| CCZ1 | 22.2217179 | 0.117065977 | 0.38695925 | 0.30252792 | 0.76224965 | 0.95024054 |
| PMS2 | 30.1595253 | -0.060708905 | 0.27708187 | -0.219101 | 0.8265714 | 0.96542185 |
| AIMP2 | 27.2932979 | -0.065076344 | 0.29141651 | -0.2233104 | 0.82329391 | 0.96539788 |
| EIF2AK1 | 67.2633826 | -0.084265432 | 0.22335005 | -0.3772797 | 0.70596578 | 0.93407096 |
| USP42 | 73.9725658 | 0.204000151 | 0.25344591 | 0.80490606 | 0.42087389 | 0.81954914 |
| CYTH3 | 69.0679426 | 0.218667686 | 0.30173879 | 0.72469199 | 0.46864101 | 0.84326495 |
| RAC1 | 162.307549 | -0.347071682 | 0.2195219 | -1.5810344 | 0.11387017 | 0.51856369 |
| DAGLB | 15.1471928 | -0.115337154 | 0.42440353 | -0.2717629 | 0.7858043 | 0.95577371 |
| ZDHHC4 | 18.7852048 | 0.081403159 | 0.35370056 | 0.2301471 | 0.81797747 | 0.9635539 |
| C7orf26 | 16.5666175 | 0.347383305 | 0.33857475 | 1.02601658 | 0.30488377 | 0.74832335 |
| ZNF316 | 26.6363002 | 0.005075833 | 0.48654889 | 0.01043232 | 0.99167636 | 0.99923337 |
| AC073343.2 | 15.8390772 | -0.899041175 | 0.52347216 | -1.7174575 | 0.08589562 | 0.46168448 |
| ZNF12 | 32.0987297 | 0.290987686 | 0.29557276 | 0.98448749 | 0.32487588 | 0.7633033 |
| CCZ1B | 27.323585 | -0.052894264 | 0.33450687 | -0.1581261 | 0.87435744 | 0.97434023 |
| C1GALT1 | 176.197669 | 0.08309187 | 0.38512781 | 0.21575142 | 0.82918152 | 0.96669991 |
| COL28A1 | 94.1428486 | -0.685997197 | 0.44986027 | -1.5249117 | 0.12728112 | 0.54994515 |
| AC004982.2 | 19.9358745 | -0.585588398 | 0.50556581 | -1.1582832 | 0.24674847 | 0.70025166 |
| MIOS | 67.8597827 | 0.088919725 | 0.30979663 | 0.28702612 | 0.77409232 | 0.95457609 |
| RPA3 | 49.0498281 | -0.913811733 | 0.53924927 | -1.6945998 | 0.09015136 | 0.47226726 |
| UMAD1 | 475.810971 | -0.338770125 | 0.23016948 | -1.471829 | 0.14106706 | 0.57566279 |
| GLCCI1 | 542.995483 | -0.328704723 | 0.2926191 | -1.1233194 | 0.26130186 | 0.71237438 |
| ICA1 | 251.158854 | -0.916316434 | 0.37235486 | -2.4608688 | 0.01386011 | 0.17643441 |
| NDUFA4 | 235.757176 | 0.253830177 | 0.45701474 | 0.55540917 | 0.57861482 | 0.88932306 |
| PHF14 | 117.139393 | 0.175493435 | 0.19864134 | 0.88346885 | 0.37698301 | 0.79432848 |
| THSD7A | 1869.5426 | -0.370782715 | 0.54077145 | -0.6856551 | 0.49293061 | 0.85714521 |
| TMEM106B | 97.1688922 | 0.476091144 | 0.23566208 | 2.02022801 | 0.04335974 | 0.33179556 |
| VWDE | 24.0368116 | -0.438911389 | 0.44035124 | -0.9967302 | 0.31889548 | 0.76052272 |
| SCIN | 1092.41234 | 0.368022818 | 0.23960252 | 1.53597224 | 0.12454519 | 0.54320519 |
| ARL4A | 19.4626668 | 0.30067756 | 0.49071812 | 0.61272969 | 0.54005509 | 0.87014896 |
| AC011287.1 | 37.6368008 | -1.587026242 | 0.52000743 | -3.0519299 | 0.00227375 | 0.05595143 |
| ISPD | 43.4207313 | -0.043511984 | 0.28411872 | -0.1531472 | 0.8782822 | 0.97542218 |
| ANKMY2 | 42.4945599 | 0.67431613 | 0.28202275 | 2.39099909 | 0.01680259 | 0.20115876 |
| BZW2 | 34.8441454 | 0.384168163 | 0.47354446 | 0.81126102 | 0.41721579 | 0.8188001 |
| AC073332.1 | 17.196486 | -0.423552353 | 0.40537462 | -1.0448418 | 0.29609608 | 0.74123516 |
| SNX13 | 219.087034 | 0.020101042 | 0.21598095 | 0.09306859 | 0.92584907 | 0.98672316 |
| HDAC9 | 29.3558884 | -1.0388566 | 0.60597761 | -1.7143481 | 0.08646481 | 0.46337493 |
| TWISTNB | 16.2170458 | 0.196772441 | 0.36970252 | 0.53224534 | 0.59455608 | 0.89289841 |
| MACC1 | 209.723735 | -1.509470705 | 0.4212798 | -3.5830598 | 0.00033959 | 0.01569679 |
| ITGB8 | 62.9088978 | -2.14130658 | 0.48412388 | -4.4230551 | 9.73E-06 | 0.00111401 |
| SP4 | 48.0989676 | 0.131740298 | 0.23938017 | 0.55033924 | 0.58208671 | 0.89036744 |
| RAPGEF5 | 50.5731041 | -2.433470799 | 0.53601725 | -4.5399114 | 5.63E-06 | 0.0007478 |
| STEAP1B | 149.309155 | -0.000783485 | 0.36287844 | -0.0021591 | 0.9982773 | 0.99982181 |
| TOMM7 | 76.1677336 | 0.540030064 | 0.37240087 | 1.45013104 | 0.14702198 | 0.58364362 |
| FAM126A | 13.7781802 | 1.093589817 | 0.3814763 | 2.86673069 | 0.00414736 | 0.08574324 |
| KLHL7 | 42.1759568 | -0.146973046 | 0.29834896 | -0.4926213 | 0.62228021 | 0.90415189 |
| NUPL2 | 53.5199921 | 0.015990627 | 0.27311857 | 0.0585483 | 0.95331189 | 0.99217529 |
| MALSU1 | 36.4575748 | 0.223733986 | 0.28087972 | 0.79654731 | 0.42571399 | 0.82308728 |
| TRA2A | 235.382962 | 0.16628182 | 0.23420656 | 0.70997934 | 0.47771695 | 0.84771464 |
| CCDC126 | 54.7301118 | 0.338211402 | 0.20547573 | 1.64599199 | 0.09976541 | 0.48678293 |
| FAM221A | 31.1277394 | 0.57648592 | 0.29319613 | 1.96621259 | 0.04927406 | 0.35509537 |
| MPP6 | 13.9913774 | 0.28262615 | 0.50793555 | 0.55642128 | 0.5779229 | 0.88877358 |
| OSBPL3 | 540.786819 | -0.828097999 | 0.40962815 | -2.0215847 | 0.04321927 | 0.33179556 |
| CYCS | 163.50565 | 0.244149705 | 0.36799173 | 0.66346519 | 0.50703267 | 0.86269371 |
| HNRNPA2B1 | 370.297006 | 0.121131192 | 0.23380143 | 0.51809432 | 0.60439246 | 0.89593125 |
| CBX3 | 71.9146312 | -0.353738292 | 0.32884909 | -1.0756858 | 0.2820678 | 0.73352753 |
| SNX10 | 84.5077468 | -1.370480912 | 0.40609951 | -3.3747416 | 0.00073885 | 0.02625219 |
| AC004540.1 | 27.5850315 | 1.3527926 | 0.64291281 | 2.10416185 | 0.03536433 | 0.30090703 |
| SKAP2 | 151.849579 | -0.687988961 | 0.40688982 | -1.6908483 | 0.09086578 | 0.47310687 |
| HOTAIRM1 | 10.9791943 | -0.177204396 | 0.60308418 | -0.2938303 | 0.76888759 | NA |
| HOXA3 | 19.8371625 | -0.111358505 | 0.70017911 | -0.1590429 | 0.87363509 | 0.97434023 |
| HOXA7 | 12.5447618 | 0.539001758 | 0.70906867 | 0.76015452 | 0.44716222 | 0.83066268 |
| HOXA9 | 16.2701228 | 0.225028609 | 0.62670958 | 0.35906362 | 0.7195475 | 0.93786761 |
| HIBADH | 253.282006 | 0.290146761 | 0.27933753 | 1.03869596 | 0.29894616 | 0.74296125 |
| TAX1BP1 | 329.51071 | -0.226205255 | 0.14224858 | -1.590211 | 0.11178725 | 0.51426646 |
| JAZF1 | 75.9456631 | -0.844052153 | 0.35629486 | -2.3689709 | 0.01783766 | 0.20721591 |
| CPVL | 28.1813636 | 1.271140782 | 0.61308931 | 2.07333705 | 0.03814092 | 0.31326583 |
| AC007255.1 | 14.3727776 | -0.610393607 | 0.67370649 | -0.906023 | 0.36492366 | 0.78793182 |
| PRR15 | 11.3954313 | -0.513191894 | 0.73108954 | -0.7019549 | 0.48270728 | NA |
| WIPF3 | 15.1793338 | -0.626447282 | 0.55330725 | -1.132187 | 0.25755581 | 0.71022118 |
| SCRN1 | 116.018014 | -0.17542921 | 0.20885308 | -0.8399647 | 0.40092819 | 0.80835657 |
| FKBP14 | 32.8243719 | 0.148733458 | 0.35335739 | 0.42091509 | 0.67381709 | 0.92189158 |
| PLEKHA8 | 164.516939 | -0.439485794 | 0.2347372 | -1.8722461 | 0.06117257 | 0.39270279 |
| MTURN | 84.6345016 | 0.879021968 | 0.41827077 | 2.10156202 | 0.03559166 | 0.30204646 |
| ZNRF2 | 78.856805 | -0.309217474 | 0.21081011 | -1.4668057 | 0.14242892 | 0.57673923 |
| AC006978.2 | 64.4213833 | -0.310879486 | 0.25060445 | -1.2405186 | 0.21478364 | 0.66839951 |
| NOD1 | 27.5613658 | -0.31860732 | 0.39566002 | -0.8052553 | 0.42067238 | 0.81938194 |
| GGCT | 16.0442207 | 0.253744015 | 0.48243795 | 0.52596197 | 0.59891463 | 0.89476262 |
| AC005154.6 | 30.0766236 | 0.214628801 | 0.34290638 | 0.62591078 | 0.53137347 | 0.86840264 |
| GARS | 50.3617727 | -0.555670129 | 0.37957893 | -1.4639119 | 0.14321801 | 0.57698886 |
| AQP1 | 14.6960426 | -0.278974937 | 0.64447858 | -0.4328692 | 0.66510979 | 0.91953619 |

| | | | | | | |
|---|---|---|---|---|---|---|
| CCDC129 | 20.0698016 | -0.61287141 | 0.56744245 | -1.0800592 | 0.28011584 | 0.73179753 |
| PDE1C | 1727.28115 | 0.328440113 | 0.37182037 | 0.88333006 | 0.37705797 | 0.79432848 |
| LSM5 | 18.8357752 | 0.100333065 | 0.39878119 | 0.25159929 | 0.8013508 | 0.95877735 |
| AVL9 | 219.700614 | -0.455044692 | 0.18576204 | -2.4496107 | 0.01430107 | 0.17992271 |
| KBTBD2 | 63.1835758 | -0.084913213 | 0.30785778 | -0.2758196 | 0.78268662 | 0.95544332 |
| FKBP9 | 31.492304 | 0.299056444 | 0.30664377 | 0.97525687 | 0.32943286 | 0.76685572 |
| NT5C3A | 61.0618627 | -0.073065611 | 0.35833316 | -0.2039041 | 0.83842842 | 0.96748214 |
| RP9 | 13.8902815 | 0.325220109 | 0.40656769 | 0.79991626 | 0.42375932 | 0.82218966 |
| BBS9 | 116.201357 | 0.185308704 | 0.22744933 | 0.81472519 | 0.41522963 | 0.81726257 |
| DPY19L1 | 74.7445113 | -0.922113705 | 0.35544271 | -2.5942681 | 0.00947925 | 0.14341304 |
| HERPUD2 | 50.5771387 | 0.098408925 | 0.28916489 | 0.34032114 | 0.7336147 | 0.94277981 |
| 7-Sep | 145.133826 | 0.022083581 | 0.17791618 | 0.12412351 | 0.90121748 | 0.979694 |
| EEPD1 | 24.8439154 | 0.043244833 | 0.45821574 | 0.09437658 | 0.92481002 | 0.98672316 |
| KIAA0895 | 30.8329722 | 0.043183697 | 0.33560666 | 0.12867354 | 0.89761597 | 0.97892709 |
| EPDR1 | 32.7947789 | 0.593777792 | 0.35346462 | 1.67987898 | 0.09298086 | 0.47759124 |
| STARD3NL | 38.9057949 | -0.123847738 | 0.25038841 | -0.4946225 | 0.62086662 | 0.90380166 |
| VPS41 | 135.678993 | -0.248161114 | 0.21685924 | -1.1443419 | 0.25248188 | 0.70626085 |
| YAE1D1 | 15.6361917 | 0.5130117 | 0.35164443 | 1.45889329 | 0.14459448 | 0.57889578 |
| RALA | 140.473589 | -0.699331101 | 0.45646787 | -1.5320489 | 0.12551037 | 0.54505746 |
| LINC00265 | 24.9543458 | 0.061990806 | 0.44626176 | 0.13891131 | 0.88952023 | 0.97782578 |
| CDK13 | 295.348759 | -0.003307339 | 0.16899819 | -0.0195703 | 0.98438618 | 0.99773864 |
| MPLKIP | 28.2291745 | 0.409681889 | 0.40255578 | 1.01770217 | 0.30881952 | 0.75132915 |
| SUGCT | 12.4882145 | -0.044153585 | 0.40191549 | -0.1098579 | 0.91252209 | NA |
| MRPL32 | 27.8143934 | -0.038257321 | 0.36653685 | -0.1043751 | 0.91687168 | 0.98495314 |
| STK17A | 146.045651 | 0.35766258 | 0.29642858 | 1.20657251 | 0.22759682 | 0.6826497 |
| COA1 | 194.430446 | 0.026838431 | 0.1883598 | 0.14248492 | 0.88669699 | 0.97714523 |
| BLVRA | 36.7456583 | -0.605498142 | 0.25722017 | -2.3540072 | 0.01857225 | 0.21194213 |
| URGCP | 25.4672845 | 0.044434288 | 0.45110222 | 0.09850159 | 0.92153401 | 0.98619592 |
| UBE2D4 | 15.2871775 | -0.085704943 | 0.41904296 | -0.2045254 | 0.83794292 | 0.96748214 |
| DBNL | 44.3305423 | 0.040534646 | 0.28288549 | 0.14328995 | 0.8860612 | 0.97693497 |
| POLM | 21.8071283 | -0.109751763 | 0.31427005 | -0.3492276 | 0.72691848 | 0.94009907 |
| POLD2 | 20.6784525 | -0.073529292 | 0.37857527 | -0.1942263 | 0.84599865 | 0.97027687 |
| YKT6 | 20.3220309 | -0.039115291 | 0.41265421 | -0.0947895 | 0.92448202 | 0.98672316 |
| NUDCD3 | 112.209178 | -0.172027046 | 0.23300602 | -0.7382944 | 0.46033556 | 0.83713065 |
| DDX56 | 26.7907306 | 0.502711246 | 0.29983284 | 1.67663835 | 0.09361322 | 0.47792014 |
| TMED4 | 73.7650218 | 0.096034992 | 0.29913579 | 0.32104147 | 0.74817897 | 0.94656788 |
| OGDH | 301.637473 | 0.340252997 | 0.16737447 | 2.03288465 | 0.04206418 | 0.32888549 |
| ZMIZ2 | 57.862328 | -0.14538522 | 0.28753421 | -0.5056275 | 0.61311814 | 0.90011356 |
| PPIA | 251.440936 | -0.423707304 | 0.38306027 | -1.1061113 | 0.26867833 | 0.72009952 |
| H2AFV | 89.139817 | 0.411771258 | 0.31272957 | 1.31670073 | 0.18793896 | 0.63942753 |
| PURB | 50.8650044 | 0.381018889 | 0.27434368 | 1.38883786 | 0.16488206 | 0.60558727 |
| SNHG15 | 11.7921116 | 0.003588188 | 0.45423129 | 0.00789947 | 0.9936972 | NA |
| CCM2 | 63.8863399 | -0.057015987 | 0.25875351 | -0.2203487 | 0.82559963 | 0.96539788 |
| TBRG4 | 23.3539222 | 0.112620913 | 0.35797385 | 0.31460654 | 0.75306041 | 0.94771649 |
| ADCY1 | 62.2136773 | 1.154231039 | 0.57851203 | 1.99517204 | 0.04602412 | 0.34112583 |
| AC004870.4 | 52.1029688 | 0.621353625 | 0.70257159 | 0.88439902 | 0.37648085 | 0.79422885 |
| TNS3 | 695.280854 | -0.820491281 | 0.26258256 | -3.1246983 | 0.00177988 | 0.04771768 |
| HUS1 | 28.4578286 | 0.415231718 | 0.29500034 | 1.4075635 | 0.15926036 | 0.60024133 |
| UPP1 | 51.3751434 | -0.84753615 | 0.44404345 | -1.9086784 | 0.05630359 | 0.37709488 |
| GRB10 | 244.068199 | 0.347366325 | 0.30169841 | 1.15136944 | 0.24958028 | 0.70382428 |
| COBL | 120.963051 | -0.138909292 | 0.42720459 | -0.3251587 | 0.74506096 | 0.94595436 |
| SEC61G | 32.2962202 | 0.200172068 | 0.45102594 | 0.44381497 | 0.65717636 | 0.9158679 |
| EGFR | 464.200393 | -0.541548335 | 0.30106426 | -1.7987799 | 0.07205351 | 0.4235873 |
| LANCL2 | 43.2268006 | 0.301878188 | 0.38823717 | 0.77756127 | 0.4368277 | 0.82687287 |
| VOPP1 | 88.8933646 | -0.125799541 | 0.25298571 | -0.4972595 | 0.61900609 | 0.90295415 |
| NIPSNAP2 | 166.703917 | 0.01567165 | 0.19167507 | 0.08176155 | 0.93483633 | 0.98832833 |
| CCT6A | 59.7188385 | -0.073504867 | 0.26640384 | -0.2759152 | 0.7826132 | 0.95544332 |
| SUMF2 | 44.0010887 | -0.081037717 | 0.2222994 | -0.3645431 | 0.71545248 | 0.93689626 |
| PHKG1 | 13.3666012 | -0.034020795 | 0.44110527 | -0.0771262 | 0.93852311 | 0.98871562 |
| CHCHD2 | 209.262959 | -0.003478908 | 0.41431967 | -0.0083967 | 0.9933005 | 0.99923337 |
| NUPR2 | 30.5083243 | 0.507773191 | 0.48434487 | 1.04837115 | 0.29446764 | 0.74075106 |
| ZNF736 | 49.9676572 | -0.069826585 | 0.26785232 | -0.2606906 | 0.79433111 | 0.95785825 |
| ZNF680 | 72.8766313 | 0.186651077 | 0.32389722 | 0.57626638 | 0.56443515 | 0.88191823 |
| ZNF107 | 41.0363929 | -0.017250335 | 0.43657675 | -0.0395127 | 0.96848161 | 0.99441717 |
| ZNF138 | 48.1539933 | -0.054107098 | 0.30776208 | -0.1758082 | 0.86044462 | 0.97287712 |
| ZNF273 | 13.2233832 | 0.081066835 | 0.35547401 | 0.22805277 | 0.81960521 | 0.96389705 |
| ZNF117 | 20.5383719 | 0.007327856 | 0.35992839 | 0.02035921 | 0.98375683 | 0.99762255 |
| ZNF92 | 14.7622563 | 0.114085751 | 0.45412235 | 0.2512225 | 0.80164209 | 0.95890203 |
| VKORC1L1 | 43.0669385 | 0.158851649 | 0.28529642 | 0.5567951 | 0.57766743 | 0.88877358 |
| GUSB | 20.7980433 | 0.423898988 | 0.4528744 | 0.93601888 | 0.34926347 | 0.77868979 |
| ASL | 14.6241353 | 0.139395963 | 0.41473962 | 0.33610477 | 0.73679186 | 0.94369536 |
| AC068533.4 | 14.7199798 | 0.304206438 | 0.49451159 | 0.61516543 | 0.53844547 | 0.86972442 |
| CRCP | 20.1917478 | 0.304051301 | 0.31519539 | 0.96464387 | 0.33472322 | 0.76777176 |
| AC068533.3 | 15.7325818 | -0.55616973 | 0.49252412 | -1.1292233 | 0.25880363 | 0.71022613 |
| TPST1 | 238.271929 | -0.440162505 | 0.29197256 | -1.5075475 | 0.13167038 | 0.55870633 |
| AC027644.4 | 62.1938361 | 0.521177945 | 0.30781941 | 1.69312893 | 0.09043093 | 0.47236403 |
| AC027644.3 | 13.7377154 | 0.243612968 | 0.50072778 | 0.48651778 | 0.62660011 | 0.90541055 |
| TMEM248 | 76.4809411 | -0.027132838 | 0.19675918 | -0.1378987 | 0.89032046 | 0.97782578 |
| SBDS | 61.2815366 | -0.247810171 | 0.4151322 | -0.5969428 | 0.55054558 | 0.87459747 |
| TYW1 | 135.456159 | 0.105863102 | 0.23167891 | 0.45693887 | 0.64771498 | 0.91360118 |
| AC006480.2 | 12.8523291 | 0.279852754 | 0.36288336 | 0.77119203 | 0.44059312 | 0.82793978 |
| AUTS2 | 126.996603 | 0.616667425 | 0.3649579 | 1.68969471 | 0.09108638 | 0.47374684 |
| GALNT17 | 1230.00205 | 1.353026026 | 0.52461034 | 2.57910667 | 0.00990562 | 0.1471432 |
| CALN1 | 27.985131 | 2.030331895 | 0.53802922 | 3.77364613 | 0.00016088 | 0.00975328 |
| TYW1B | 85.2984696 | 0.491322967 | 0.24368697 | 2.01620535 | 0.04377851 | 0.33268382 |
| POM121 | 91.8282691 | 0.21342964 | 0.27782149 | 0.7682258 | 0.44235305 | 0.82820049 |
| NSUN5 | 33.0749717 | 0.612821588 | 0.31382258 | 1.95276447 | 0.05084751 | 0.35962249 |
| BAZ1B | 101.427936 | 0.239771344 | 0.20390366 | 1.17590504 | 0.23963283 | 0.69419674 |
| BCL7B | 27.3396926 | 0.215060455 | 0.37501138 | 0.57347715 | 0.56632167 | 0.88373877 |
| MLXIPL | 20.9064138 | 0.240670393 | 0.57177793 | 0.42091585 | 0.67381653 | 0.92189158 |
| VPS37D | 15.4557422 | -0.298854143 | 0.42255752 | -0.7072508 | 0.47941063 | 0.84833462 |
| DNAJC30 | 9.94661871 | -0.219665157 | 0.44119682 | -0.4978847 | 0.61856529 | NA |
| BUD23 | 43.5071503 | 0.26081584 | 0.30690972 | 0.8498129 | 0.39542911 | 0.80598075 |
| ABHD11 | 22.6113791 | -0.752848424 | 0.34732515 | -2.1675609 | 0.03019211 | 0.27654719 |
| CLDN3 | 25.4788487 | -1.425192969 | 0.48763851 | -2.9226424 | 0.00347075 | 0.0754849 |
| CLDN4 | 184.155138 | -0.972159878 | 0.36776968 | -2.6433932 | 0.00820797 | 0.13111608 |
| LIMK1 | 15.4053445 | -0.220869184 | 0.38665952 | -0.571224 | 0.56784783 | 0.88483919 |
| EIF4H | 70.8854251 | 0.157351203 | 0.25481474 | 0.61751216 | 0.53689696 | 0.86963427 |
| RFC2 | 10.5250492 | -0.231858531 | 0.40685324 | -0.5698825 | 0.56875741 | NA |
| CLIP2 | 18.8428055 | 0.328986539 | 0.50968072 | 0.64547574 | 0.51861893 | 0.86555466 |
| GTF2IRD1 | 200.545418 | -1.457315135 | 0.552963 | -2.6354659 | 0.00840219 | 0.13244159 |
| GTF2I | 297.60378 | 0.303515976 | 0.15833413 | 1.91693332 | 0.0552464 | 0.37383273 |
| GTF2IRD2 | 47.9025886 | 0.051785197 | 0.33799405 | 0.15321334 | 0.87823004 | 0.97542218 |
| CASTOR2 | 23.6255821 | 0.290861144 | 0.46255221 | 0.62881798 | 0.52946823 | 0.86758926 |
| RCC1L | 42.3865233 | -0.209320586 | 0.25945055 | -0.8067841 | 0.41979086 | 0.81899879 |
| GTF2IRD2B | 70.742572 | 0.20460812 | 0.26200327 | 0.78093727 | 0.43483939 | 0.82565889 |
| TRIM73 | 56.9785051 | 0.311468969 | 0.33073785 | 0.94173972 | 0.34632592 | 0.77642952 |
| POM121C | 81.614518 | 0.286815923 | 0.2198354 | 1.30468488 | 0.19200017 | 0.6421543 |
| SPDYE5 | 19.0481206 | -0.402947823 | 0.57394125 | -0.7020715 | 0.48263455 | 0.85134662 |
| HIP1 | 155.401406 | 0.582854495 | 0.64645 | 0.90162347 | 0.36725692 | 0.78856614 |
| RHBDD2 | 23.9105067 | -0.300739469 | 0.43916738 | -0.6847946 | 0.49347351 | 0.85721581 |
| POR | 49.8536916 | -0.669973343 | 0.32594576 | -2.055475 | 0.03983315 | 0.31990103 |
| TMEM120A | 11.0052887 | 0.296901737 | 0.40579064 | 0.73166236 | 0.46437468 | NA |

| | | | | | | |
|---|---|---|---|---|---|---|
| STYXL1 | 44.2664163 | 0.301012701 | 0.26049615 | 1.15553608 | 0.24787097 | 0.70240969 |
| MDH2 | 81.6701251 | 0.378758394 | 0.35181783 | 1.07657533 | 0.28167003 | 0.73311066 |
| HSPB1 | 149.947937 | -0.051926037 | 0.37791815 | -0.1374002 | 0.89071446 | 0.9779208 |
| YWHAG | 79.6269556 | -0.176907826 | 0.24722285 | -0.7155804 | 0.47425048 | 0.84593347 |
| DTX2 | 34.3791496 | 0.233882596 | 0.30805105 | 0.75923325 | 0.44771304 | 0.83066268 |
| POMZP3 | 16.2205525 | -0.170496075 | 0.40165995 | -0.4244787 | 0.67121677 | 0.92170876 |
| CCDC146 | 55.2967849 | -0.239944805 | 0.33233841 | -0.7219894 | 0.47030101 | 0.84423815 |
| GSAP | 65.5724037 | 0.202236636 | 0.30094567 | 0.67200381 | 0.50158127 | 0.85942996 |
| PTPN12 | 131.778915 | -0.932502864 | 0.23767888 | -3.9233729 | 8.73E-05 | 0.00580125 |
| APTR | 34.3945333 | 0.19947537 | 0.34313964 | 0.58132418 | 0.56102199 | 0.88072236 |
| RSBN1L | 116.365868 | 0.267254825 | 0.22699491 | 1.17736041 | 0.23905168 | 0.69356592 |
| TMEM60 | 12.754228 | 0.173151019 | 0.414159 | 0.41807861 | 0.67588965 | 0.92197013 |
| PHTF2 | 67.044745 | 0.280881999 | 0.30071944 | 0.9340334 | 0.35028667 | 0.77868979 |
| MAGI2 | 129.867429 | -0.697723395 | 0.70056825 | -0.9959392 | 0.31927968 | 0.76086314 |
| GNAI1 | 12.8784096 | -0.457016923 | 0.60112384 | -0.7602708 | 0.44709271 | 0.83066268 |
| SEMA3C | 891.183309 | 1.112933284 | 0.34473898 | 3.22833607 | 0.00124513 | 0.0376253 |
| CACNA2D1 | 83.4866085 | -0.910584767 | 0.71241715 | -1.2781623 | 0.20119219 | 0.65374585 |
| PCLO | 127.211144 | -0.030783908 | 0.56479704 | -0.0545044 | 0.95653333 | 0.99236316 |
| KIAA1324L | 38.216706 | -0.700370982 | 0.48572422 | -1.4419108 | 0.14932755 | 0.58659144 |
| DMTF1 | 89.7582661 | -0.187987903 | 0.33470671 | -0.5616496 | 0.57435475 | 0.88644064 |
| TMEM243 | 47.8449898 | -0.456109968 | 0.27907547 | -1.6343606 | 0.10218315 | 0.49238776 |
| CROT | 16.5869449 | 0.332179652 | 0.40823216 | 0.8137028 | 0.41581523 | 0.81763864 |
| SLC25A40 | 64.3848915 | -0.015498233 | 0.33243365 | -0.0466205 | 0.96281567 | 0.99306991 |
| DBF4 | 35.989231 | -0.697441811 | 0.38707478 | -1.801827 | 0.07157262 | 0.42281025 |
| ADAM22 | 221.322743 | -0.387741402 | 0.4614431 | -0.84028 | 0.40075142 | 0.80830282 |
| SRI | 74.2648357 | -0.050249732 | 0.40649998 | -0.1236156 | 0.90161965 | 0.9797999 |
| AC003991.1 | 27.0345941 | -0.710947132 | 0.6600193 | -1.0771611 | 0.28140829 | 0.73311066 |
| CFAP69 | 21.83375 | 0.013966948 | 0.43958252 | 0.03177321 | 0.97465291 | 0.99543096 |
| GTPBP10 | 53.2267859 | -0.279174559 | 0.31055709 | -0.8989476 | 0.36868056 | 0.78877653 |
| CLDN12 | 46.150891 | -0.290533624 | 0.27523319 | -1.0555908 | 0.2911552 | 0.73994767 |
| CDK14 | 229.196025 | 0.180446582 | 0.4096499 | 0.44048975 | 0.65958244 | 0.91732025 |
| AC079760.1 | 19.1763252 | 1.017962897 | 0.44748428 | 2.27485733 | 0.02291449 | 0.23772251 |
| MTERF1 | 114.579004 | -0.131081995 | 0.34163304 | -0.3836924 | 0.70120646 | 0.93127962 |
| AKAP9 | 445.506272 | 0.064623043 | 0.17094981 | 0.37802349 | 0.70541315 | 0.93407096 |
| KRIT1 | 145.107939 | 0.218780472 | 0.15936698 | 1.37280927 | 0.16981165 | 0.61252734 |
| ANKIB1 | 230.408856 | 0.125397011 | 0.16584004 | 0.7561323 | 0.44956988 | 0.83118301 |
| GATAD1 | 41.2910082 | -0.09943579 | 0.29833395 | -0.3333036 | 0.7389051 | 0.94376279 |
| AC007566.1 | 14.231572 | -0.56040762 | 0.42589575 | -1.3158329 | 0.18823015 | 0.63996932 |
| PEX1 | 46.7719007 | 0.211900868 | 0.25125065 | 0.84338434 | 0.39901353 | 0.80768598 |
| RBM48 | 10.8508241 | 0.178733525 | 0.49287191 | 0.36263686 | 0.71687617 | NA |
| FAM133B | 57.4609751 | 0.183033235 | 0.25260909 | 0.72457107 | 0.46871521 | 0.84326495 |
| CDK6 | 60.1506761 | 0.419183818 | 0.4539589 | 0.92339597 | 0.35580089 | 0.78153728 |
| SAMD9 | 15.1244375 | -1.319681542 | 0.66130671 | -1.9955665 | 0.04598112 | 0.34112583 |
| SAMD9L | 38.0990927 | -1.777440325 | 0.83713197 | -2.1232498 | 0.03373292 | 0.2921512 |
| HEPACAM2 | 134.528822 | 0.689401851 | 0.59866177 | 1.15157153 | 0.24949719 | 0.70382428 |
| VPS50 | 107.081312 | -0.008166964 | 0.24357854 | -0.0335291 | 0.97325268 | 0.99526459 |
| CALCR | 302.294023 | -0.183907719 | 0.62164604 | -0.2958399 | 0.76735233 | 0.95205353 |
| GNGT1 | 113.270772 | -0.526973501 | 0.48161899 | -1.0941709 | 0.27388002 | 0.72510729 |
| BET1 | 25.7624533 | -0.113305948 | 0.46612943 | -0.2430783 | 0.80794474 | 0.96054677 |
| CASD1 | 65.9193463 | -0.026983121 | 0.22193813 | -0.1215795 | 0.90323207 | 0.98016176 |
| SGCE | 30.9123024 | -0.575530484 | 0.40444482 | -1.4230136 | 0.15473221 | 0.59498864 |
| PPP1R9A | 328.839859 | -0.008670084 | 0.32552018 | -0.0266346 | 0.97875122 | 0.99610605 |
| AC002429.2 | 27.5455784 | -1.091501927 | 0.68454368 | -1.5944957 | 0.11082506 | 0.51243576 |
| PON2 | 35.5974489 | -0.271755432 | 0.29071879 | -0.9347708 | 0.34990643 | 0.77868979 |
| PDK4 | 85.7700949 | -0.725800863 | 0.99730557 | -0.7277618 | 0.46675943 | 0.84249469 |
| SLC25A13 | 168.801353 | -0.323460513 | 0.26795049 | -1.2071652 | 0.22736852 | 0.68217589 |
| SEM1 | 83.8095405 | 0.213745648 | 0.18899861 | 1.13093764 | 0.25808134 | 0.71022118 |
| SDHAF3 | 20.9780918 | 0.497600996 | 0.2953453 | 1.68481093 | 0.09202506 | 0.4764741 |
| ASNS | 18.8124544 | -0.445168768 | 0.43320176 | -1.0276246 | 0.30412646 | 0.74820946 |
| LMTK2 | 92.647331 | -0.105864756 | 0.32361833 | -0.3271284 | 0.74357075 | 0.94546379 |
| TECPR1 | 18.1503394 | -0.512546805 | 0.41746331 | -1.2277649 | 0.21953523 | 0.67305276 |
| BRI3 | 132.888116 | -0.573594978 | 0.33337586 | -1.7205654 | 0.08532971 | 0.46032833 |
| BAIAP2L1 | 491.59021 | 0.380252408 | 0.30907298 | 1.23029975 | 0.21858488 | 0.67250923 |
| TRRAP | 114.645832 | 0.168076008 | 0.18808769 | 0.8936045 | 0.37153354 | 0.7900012 |
| SMURF1 | 419.570481 | 0.022152755 | 0.22596302 | 0.09803708 | 0.92190285 | 0.986393 |
| ARPC1B | 31.6019294 | -0.834755303 | 0.3996421 | -2.0887572 | 0.03672959 | 0.3068708 |
| PDAP1 | 32.2551992 | -0.415333525 | 0.29575557 | -1.4043135 | 0.16022553 | 0.60024417 |
| BUD31 | 26.9273696 | 0.128151501 | 0.38886889 | 0.32954938 | 0.74174048 | 0.9450785 |
| CPSF4 | 14.6947877 | 0.047924628 | 0.40351916 | 0.11876667 | 0.90546021 | 0.98111752 |
| ZNF789 | 33.4405355 | 0.133294545 | 0.33897676 | 0.39322621 | 0.69415241 | 0.92841398 |
| ZNF394 | 24.0391231 | -0.65241396 | 0.29887259 | -2.1829167 | 0.02904194 | 0.26979454 |
| ZKSCAN5 | 20.198447 | 0.135929405 | 0.30652874 | 0.4434475 | 0.65744208 | 0.9158679 |
| FAM200A | 20.184424 | 0.107395868 | 0.34155143 | 0.31443543 | 0.75319035 | 0.94771649 |
| ZSCAN25 | 14.6568413 | 0.346589247 | 0.43476308 | 0.79719107 | 0.42534007 | 0.82269166 |
| TRIM4 | 45.1889598 | 0.123956383 | 0.218573 | 0.56711664 | 0.57063494 | 0.88563341 |
| ZKSCAN1 | 182.35651 | 0.216668744 | 0.22010076 | 0.98440709 | 0.32491539 | 0.7633033 |
| ZSCAN21 | 11.9253234 | 0.205451084 | 0.41651409 | 0.49326323 | 0.62182661 | NA |
| ZNF3 | 14.8140412 | -0.05958791 | 0.39376499 | -0.1513286 | 0.8797165 | 0.97581687 |
| COPS6 | 30.4715771 | -0.038314235 | 0.33187914 | -0.1154463 | 0.90809134 | 0.98250721 |
| AP4M1 | 16.4471071 | -0.143073839 | 0.35565252 | -0.4022855 | 0.68747395 | 0.92643753 |
| TAF6 | 13.7512068 | -0.217443519 | 0.35451057 | -0.6133626 | 0.53963662 | 0.86980312 |
| LAMTOR4 | 88.9737565 | 0.021543526 | 0.27856956 | 0.07733625 | 0.93835605 | 0.98871562 |
| CASTOR3 | 37.8891735 | -0.731648662 | 0.29275925 | -2.4991479 | 0.01244924 | 0.16647826 |
| SPDYE3 | 26.4901366 | -0.245429226 | 0.45765863 | -0.5362714 | 0.591771 | 0.891609 |
| PILRB | 65.3599379 | -0.183430573 | 0.29821698 | -0.615091 | 0.53849465 | 0.86972442 |
| PILRA | 12.4383063 | -0.384241522 | 0.45706155 | -0.8406779 | 0.40052843 | NA |
| MEPCE | 18.8570509 | 0.367073119 | 0.62828964 | 0.58424187 | 0.55905759 | 0.88004847 |
| PPP1R35 | 28.0997931 | -0.781337945 | 0.32395902 | -2.4118419 | 0.01587216 | 0.19341706 |
| TSC22D4 | 38.1134916 | 0.032545193 | 0.29331189 | 0.11095763 | 0.91164994 | 0.9835359 |
| AGFG2 | 17.84671 | 0.097633553 | 0.51622148 | 0.18913113 | 0.84999004 | 0.97027687 |
| SAP25 | 14.7830789 | 0.643365952 | 0.44227463 | 1.45467523 | 0.14575919 | 0.58207665 |
| LRCH4 | 25.0845614 | 0.486786921 | 0.33991783 | 1.43207235 | 0.15212312 | 0.59013416 |
| GNB2 | 81.1214605 | -0.489736784 | 0.33862851 | -1.4462361 | 0.14811098 | 0.58515729 |
| GIGYF1 | 41.4586706 | 0.163443039 | 0.35345232 | 0.46241892 | 0.64378093 | 0.9119294 |
| SLC12A9 | 12.0661757 | 1.009683968 | 0.43055157 | 2.34509414 | 0.01902228 | NA |
| TRIP6 | 12.8892057 | -0.251738925 | 0.45431799 | -0.5541029 | 0.57950841 | 0.88957613 |
| SRRT | 19.5433107 | 0.290493662 | 0.37516264 | 0.77431394 | 0.43874516 | 0.82745796 |
| TRIM56 | 80.1114146 | 0.122884183 | 0.34132283 | 0.36002334 | 0.71882968 | 0.93767415 |
| AP1S1 | 20.2847389 | -0.023133788 | 0.41137638 | -0.0562351 | 0.95515453 | 0.99228059 |
| PLOD3 | 18.5660186 | -0.296182991 | 0.41213916 | -0.718648 | 0.47235782 | 0.84570599 |
| ZNHIT1 | 38.7763649 | 0.133737353 | 0.3565994 | 0.37503527 | 0.70763423 | 0.93432317 |
| FIS1 | 91.4177709 | -0.112219747 | 0.33005155 | -0.3400067 | 0.73385146 | 0.94277981 |
| IFT22 | 11.5849835 | -0.382084614 | 0.5009738 | -0.7626838 | 0.44565198 | NA |
| CUX1 | 444.56853 | 0.194631896 | 0.27217122 | 0.7151083 | 0.47454213 | 0.84593347 |
| PRKRIP1 | 102.598753 | -0.092580506 | 0.21593167 | -0.4287491 | 0.66810583 | 0.92049996 |
| ORAI2 | 12.212752 | -0.094835207 | 0.75953608 | -0.1248594 | 0.90063487 | NA |
| ALKBH4 | 11.7101726 | 0.799313655 | 0.42287359 | 1.89019527 | 0.05873185 | NA |
| POLR2J | 21.6717529 | -0.061936857 | 0.4648381 | -0.1332439 | 0.89400047 | 0.97833263 |
| SPDYE2 | 14.7403021 | 0.191503876 | 0.5522114 | 0.3467945 | 0.72874571 | 0.94097305 |
| AC105052.3 | 29.1855179 | 0.576798102 | 0.40335772 | 1.42999148 | 0.15271946 | 0.59136879 |
| FAM185A | 54.6668057 | 0.348243111 | 0.34345858 | 1.01393044 | 0.31061595 | 0.75380903 |
| FBXL13 | 30.6258696 | -0.367291126 | 0.46532703 | -0.7893183 | 0.42992601 | 0.82469033 |
| ARMC10 | 38.6667721 | 0.01984523 | 0.3015528 | 0.06581013 | 0.94752899 | 0.99149867 |

| | | | | | | |
|---|---|---|---|---|---|---|
| NAPEPLD | 66.4352459 | 1.453785671 | 0.49161106 | 2.95718666 | 0.0031046 | 0.06938855 |
| PMPCB | 71.0834498 | 0.358175436 | 0.27741982 | 1.29109534 | 0.19667062 | 0.64685325 |
| DNAJC2 | 41.4275574 | 0.123935615 | 0.33310625 | 0.37206031 | 0.70984794 | 0.93489817 |
| PSMC2 | 41.5508609 | 0.240317595 | 0.2572177 | 0.93429651 | 0.35015097 | 0.77868979 |
| SLC26A5 | 50.3464384 | -0.50036269 | 0.35263828 | -1.418912 | 0.15592468 | 0.59597065 |
| ORC5 | 42.847881 | -0.002333848 | 0.25200355 | -0.0092612 | 0.99261076 | 0.99923337 |
| LHFPL3 | 50.0053947 | 0.382530911 | 0.46588795 | 0.82107921 | 0.41160115 | 0.8149686 |
| LINC01004 | 180.490732 | 0.78819948 | 0.25953499 | 3.03696813 | 0.00238971 | 0.05737663 |
| KMT2E-AS1 | 12.1937603 | -0.322252746 | 0.46510076 | -0.6928665 | 0.48839332 | NA |
| KMT2E | 422.185333 | 0.260846588 | 0.23813664 | 1.09536518 | 0.27335668 | 0.7244699 |
| SRPK2 | 507.244528 | 0.117519016 | 0.17449583 | 0.6734775 | 0.50064356 | 0.85942996 |
| PUS7 | 26.7430412 | 0.042225324 | 0.38529084 | 0.10959338 | 0.91273186 | 0.98376271 |
| RINT1 | 32.3482634 | -0.040397076 | 0.35744731 | -0.1130155 | 0.91001829 | 0.98331039 |
| ATXN7L1 | 180.884607 | 0.018257421 | 0.23327161 | 0.07826679 | 0.93761583 | 0.98871562 |
| CDHR3 | 15.941753 | 1.320905265 | 0.57452178 | 2.2991387 | 0.02149706 | 0.22951242 |
| SYPL1 | 140.130314 | 0.271157731 | 0.2208382 | 1.22785702 | 0.21950065 | 0.67305276 |
| NAMPT | 220.782267 | -2.421296379 | 0.43364085 | -5.5836446 | 2.36E-08 | 8.46E-06 |
| HBP1 | 75.3534409 | -0.62188083 | 0.2561055 | -2.4282213 | 0.01517308 | 0.18701259 |
| COG5 | 355.442357 | 0.066892108 | 0.24177752 | 0.27666802 | 0.78203503 | 0.95544332 |
| DUS4L | 12.1597868 | 0.008717002 | 0.35200314 | 0.02476399 | 0.98024321 | NA |
| BCAP29 | 68.1218767 | -0.112731295 | 0.28605831 | -0.394085 | 0.69351827 | 0.92826395 |
| CBLL1 | 18.8097791 | -0.241954315 | 0.36917542 | -0.6553912 | 0.51221592 | 0.86346419 |
| SLC26A3 | 385.281009 | 0.73152067 | 0.57189557 | 1.27911581 | 0.20085628 | 0.65358156 |
| DLD | 121.760528 | 0.341156323 | 0.23207206 | 1.47004479 | 0.14154962 | 0.57641954 |
| LAMB1 | 352.548738 | 0.182584668 | 0.33964664 | 0.53757243 | 0.59087227 | 0.89149106 |
| NRCAM | 114.33096 | 0.427880054 | 0.81559629 | 0.52462236 | 0.59984574 | 0.89476262 |
| PNPLA8 | 304.823869 | -0.075810494 | 0.30569247 | -0.2479959 | 0.80413754 | 0.95966218 |
| DNAJB9 | 20.7744741 | -0.537301944 | 0.38494125 | -1.3958024 | 0.16277399 | 0.60133611 |
| IMMP2L | 814.103771 | -0.030047005 | 0.29713829 | -0.1011213 | 0.91945418 | 0.98560124 |
| DOCK4 | 264.52098 | 0.782228665 | 0.35737266 | 2.18883189 | 0.02860906 | 0.26683451 |
| ZNF277 | 115.954136 | -0.023392268 | 0.21862547 | -0.106997 | 0.91479136 | 0.98407488 |
| IFRD1 | 96.9815269 | 0.596010172 | 0.49721455 | 1.19869817 | 0.23064533 | 0.68523674 |
| TMEM168 | 88.4506953 | 0.263618421 | 0.25359871 | 1.03951011 | 0.29856756 | 0.74296125 |
| BMT2 | 64.0550539 | 0.028318742 | 0.25855208 | 0.10952819 | 0.91278356 | 0.98376271 |
| SMIM30 | 13.5266645 | -0.216137658 | 0.39896374 | -0.5417476 | 0.58799238 | 0.89073775 |
| FOXP2 | 29.7503771 | -0.80857104 | 0.71804483 | -1.1260732 | 0.26013454 | 0.71091822 |
| MDFIC | 80.075138 | -0.184649138 | 0.50918999 | -0.3626331 | 0.71687899 | 0.93689626 |
| TES | 69.3539194 | -0.351866777 | 0.26900126 | -1.3080488 | 0.19085675 | 0.6421543 |
| CAV2 | 24.6675687 | 0.005108659 | 0.56813564 | 0.00899197 | 0.99282554 | 0.99923337 |
| MET | 193.300305 | -0.358796227 | 0.21315262 | -1.6832832 | 0.09232028 | 0.4764741 |
| CAPZA2 | 172.005573 | -0.380856496 | 0.17597911 | -2.1642142 | 0.03044791 | 0.27757967 |
| ST7 | 290.964382 | 0.391290546 | 0.18118269 | 2.15964635 | 0.03080006 | 0.27936455 |
| ST7-AS2 | 19.8633716 | -0.209240374 | 0.61064779 | -0.3426531 | 0.73185943 | 0.94189153 |
| CFTR | 496.497161 | -3.397076356 | 0.62262005 | -5.4560985 | 4.87E-08 | 1.48E-05 |
| AC000061.1 | 30.2557858 | -3.995735648 | 0.8490295 | -4.7062389 | 2.52E-06 | 0.00039477 |
| CTTNBP2 | 262.972956 | -0.977946099 | 0.28688855 | -3.4088014 | 0.00065249 | 0.02443687 |
| LSM8 | 52.3642269 | -0.127736834 | 0.23539982 | -0.5426378 | 0.58737923 | 0.89073775 |
| ANKRD7 | 17.1605695 | 0.388853735 | 0.37855463 | 1.0272064 | 0.30432329 | 0.74820946 |
| TSPAN12 | 281.166993 | -0.883061995 | 0.6119191 | -1.4431025 | 0.14899159 | 0.58629428 |
| ING3 | 42.4698137 | 0.32498588 | 0.27546216 | 1.1797841 | 0.2380861 | 0.69296239 |
| FAM3C | 171.355346 | -0.264763393 | 0.30111958 | -0.8792633 | 0.37925854 | 0.79535699 |
| AASS | 34.1523708 | -0.212588476 | 0.29575336 | -0.7188032 | 0.47226216 | 0.84570599 |
| CADPS2 | 221.39756 | 1.040042345 | 0.45784292 | 2.27161393 | 0.02310984 | 0.23879078 |
| AC004594.1 | 14.0343345 | 0.000790148 | 0.88615361 | 0.00089166 | 0.99928856 | 0.99982181 |
| NDUFA5 | 137.147134 | 0.284451789 | 0.31223953 | 0.91100506 | 0.36229271 | 0.78714716 |
| WASL | 172.95697 | 0.329735756 | 0.21706665 | 1.51905302 | 0.12874915 | 0.55308726 |
| HYAL4 | 52.9321404 | 0.50324359 | 1.28156169 | 0.39267996 | 0.69455587 | 0.92841398 |
| POT1 | 53.6783085 | 0.011502732 | 0.25814367 | 0.04455942 | 0.96445849 | 0.99334616 |
| POT1-AS1 | 29.0077051 | 0.302510873 | 0.28575546 | 1.0586355 | 0.28976581 | 0.73966536 |
| ZNF800 | 94.9650482 | 0.083071181 | 0.30425976 | 0.27302717 | 0.78483233 | 0.95555482 |
| ARF5 | 30.6918747 | -0.054827292 | 0.32537233 | -0.1685063 | 0.86618498 | 0.97434023 |
| SND1 | 219.749578 | -0.075260203 | 0.20084087 | -0.3747255 | 0.7078646 | 0.93432317 |
| LRRC4 | 14.8155135 | 0.212244305 | 0.39492313 | 0.53743194 | 0.59096929 | 0.89149106 |
| RBM28 | 46.8773852 | 0.204314968 | 0.31444153 | 0.64977094 | 0.5158402 | 0.86476217 |
| HILPDA | 36.3149376 | -3.960001447 | 1.28226851 | -3.0882779 | 0.0020132 | 0.05152847 |
| METTL2B | 43.7481342 | -0.17583686 | 0.24565033 | -0.7158014 | 0.47411396 | 0.84593347 |
| AC090114.2 | 12.0481679 | 0.999885442 | 0.51073213 | 1.95774924 | 0.05025944 | NA |
| CALU | 65.1052076 | -0.147748963 | 0.24521799 | -0.6025209 | 0.54682746 | 0.87269272 |
| KCP | 31.348898 | -0.135613631 | 1.13766122 | -0.1192039 | 0.90511383 | 0.98107098 |
| ATP6V1F | 154.165501 | -0.278874186 | 0.4046459 | -0.6891808 | 0.4907095 | 0.85652518 |
| TNPO3 | 123.785468 | -0.226092252 | 0.17726609 | -1.27544 | 0.20215356 | 0.6545025 |
| TSPAN33 | 34.8408013 | 0.711460706 | 0.4083182 | 1.74241733 | 0.08143545 | 0.45164312 |
| AHCYL2 | 705.501431 | 0.558912727 | 0.31568754 | 1.7704618 | 0.07665024 | 0.43573154 |
| NRF1 | 112.487086 | 0.08935304 | 0.22925632 | 0.3897517 | 0.69672016 | 0.92961401 |
| UBE2H | 390.922017 | -0.522052795 | 0.28035466 | -1.8621156 | 0.0625868 | 0.39783219 |
| ZC3HC1 | 22.997985 | 0.170939092 | 0.27059385 | 0.63171832 | 0.52757095 | 0.86750588 |
| KLHDC10 | 111.755997 | 0.321522456 | 0.24241551 | 1.32632788 | 0.18473112 | 0.6349723 |
| TMEM209 | 23.836976 | -0.179280248 | 0.36233625 | -0.4947897 | 0.62074857 | 0.90380166 |
| COPG2 | 32.2723521 | 0.223577408 | 0.30801341 | 0.72586908 | 0.46791903 | 0.84326495 |
| AC016831.7 | 1665.25808 | -1.00565153 | 0.27778667 | -3.6202296 | 0.00029434 | 0.01427558 |
| LINC00513 | 159.314274 | -1.637423744 | 0.51417982 | -3.1845352 | 0.00144987 | 0.04136384 |
| AC058791.1 | 44.8929452 | -1.681657322 | 0.59010735 | -2.8497481 | 0.00437539 | 0.08878921 |
| MKLN1 | 755.688807 | -0.231828789 | 0.18713894 | -1.2388057 | 0.21541745 | 0.66933563 |
| PODXL | 11.9722107 | -0.105473584 | 0.4089556 | -0.2579096 | 0.79647666 | NA |
| CHCHD3 | 446.475072 | 0.276263262 | 0.15919104 | 1.73541971 | 0.08266649 | 0.45473406 |
| EXOC4 | 553.863769 | -0.114520191 | 0.17429109 | -0.6570628 | 0.51114054 | 0.86312853 |
| SLC35B4 | 35.168753 | 0.229105588 | 0.28628369 | 0.80027469 | 0.42355167 | 0.82204213 |
| AKR1B1 | 44.4108263 | -0.944465406 | 0.39505712 | -2.3907059 | 0.01681602 | 0.20115876 |
| BPGM | 43.0278895 | -0.571744665 | 0.62381097 | -0.9165351 | 0.35938629 | 0.78473041 |
| AGBL3 | 13.5487381 | -0.538759327 | 0.40700264 | -1.3237244 | 0.1855946 | 0.63613695 |
| CYREN | 32.5163689 | -0.427261677 | 0.30093591 | -1.4197763 | 0.15567281 | 0.59590618 |
| TMEM140 | 47.0311255 | -2.801535866 | 0.62619365 | -4.4739129 | 7.68E-06 | 0.00095509 |
| WDR91 | 29.4806749 | -1.17592211 | 0.50616015 | -2.3232214 | 0.02016726 | 0.22331327 |
| CNOT4 | 200.54342 | -0.110148624 | 0.17036881 | -0.6465305 | 0.51793588 | 0.86500999 |
| NUP205 | 53.3620936 | 0.234674099 | 0.23857533 | 0.98364779 | 0.32528872 | 0.76334591 |
| STMP1 | 51.5213757 | -0.027856885 | 0.28107826 | -0.0991072 | 0.92105314 | 0.98614962 |
| MTPN | 89.6765708 | 0.179451057 | 0.21181745 | 0.84719677 | 0.39688545 | 0.80674902 |
| PTN | 20.7135689 | -1.255966769 | 0.70506853 | -1.78134 | 0.07485692 | 0.4319525 |
| DGKI | 256.178157 | 0.601452426 | 0.71076388 | 0.84620567 | 0.39743802 | 0.80674902 |
| CREB3L2 | 64.9584622 | 1.198532427 | 0.29058681 | 4.12452458 | 3.72E-05 | 0.00330602 |
| TRIM24 | 70.2482673 | 0.473123266 | 0.27257304 | 1.73576691 | 0.08260506 | 0.45473406 |
| SVOPL | 128.615798 | 0.596370812 | 0.6452321 | 0.92427332 | 0.35534403 | 0.78124141 |
| ATP6V0A4 | 1316.9121 | 0.527710974 | 0.22610147 | 2.33395636 | 0.01959801 | 0.21965565 |
| TMEM213 | 1459.58914 | 0.925482505 | 0.32165921 | 2.87721436 | 0.00401203 | 0.08381056 |
| KIAA1549 | 43.0533266 | -0.193634369 | 0.33415808 | -0.5794694 | 0.56227251 | 0.88191823 |
| ZC3HAV1 | 88.8856834 | -0.372012276 | 0.36454199 | -1.0204923 | 0.30749506 | 0.75074303 |
| TTC26 | 16.044346 | -0.077823148 | 0.3705039 | -0.2100468 | 0.83363117 | 0.96701848 |
| UBN2 | 150.284996 | -0.243598963 | 0.21759084 | -1.1195277 | 0.26291509 | 0.71348562 |
| LUC7L2 | 36.1511801 | 0.537659069 | 0.25237723 | 2.13037871 | 0.03314036 | 0.28919158 |
| HIPK2 | 1119.51419 | -0.465128697 | 0.33312532 | -1.3962574 | 0.16263699 | 0.60133611 |
| TBXAS1 | 269.526425 | -2.858470764 | 0.48928508 | -5.8421376 | 5.15E-09 | 2.38E-06 |
| PARP12 | 54.3082348 | -0.900444382 | 0.65405972 | -1.3767006 | 0.16860485 | 0.61070463 |
| KDM7A | 69.1335857 | -0.073700299 | 0.36460023 | -0.20214 | 0.83980726 | 0.96748214 |

| | | | | | | |
|---|---|---|---|---|---|---|
| SLC37A3 | 67.6026069 | -0.26845986 | 0.27801877 | -0.9656178 | 0.33423549 | 0.76777176 |
| MKRN1 | 63.4756879 | -0.063238747 | 0.25936868 | -0.243818 | 0.80737179 | 0.96021412 |
| NDUFB2 | 151.831677 | 0.018580925 | 0.33746417 | 0.05506044 | 0.95609031 | 0.99234582 |
| BRAF | 492.263668 | -0.081880915 | 0.16308997 | -0.5020598 | 0.61562547 | 0.90123257 |
| MRPS33 | 29.2412922 | 0.193145361 | 0.46003646 | 0.41984795 | 0.67459654 | 0.92189158 |
| AGK | 36.9489362 | 0.089725215 | 0.25226558 | 0.3556776 | 0.72208202 | 0.93852279 |
| KIAA1147 | 43.4186636 | -0.162976289 | 0.40830425 | -0.399154 | 0.68977971 | 0.92734996 |
| SSBP1 | 58.211889 | -0.254026318 | 0.23001759 | -1.1043778 | 0.26942929 | 0.72046011 |
| MGAM | 37.1965901 | -0.904352714 | 0.76082603 | -1.1886459 | 0.23457905 | 0.68936745 |
| GSTK1 | 71.9700677 | -0.05828318 | 0.33069603 | -0.176244 | 0.86010228 | 0.97287712 |
| CASP2 | 13.4990446 | 0.413980716 | 0.35030689 | 1.18176583 | 0.23729863 | 0.69207518 |
| TCAF2 | 26.2447155 | -0.943322239 | 0.79704487 | -1.1835246 | 0.23660129 | 0.69148312 |
| TCAF1 | 65.282981 | -0.148373119 | 0.24378149 | -0.6086316 | 0.54276865 | 0.87210113 |
| AC004889.1 | 22.7875296 | -0.010409172 | 0.41521552 | -0.0250693 | 0.97999967 | 0.99664466 |
| AC074386.1 | 65.5521813 | -0.094528231 | 0.44111089 | -0.2142958 | 0.83031636 | 0.966913 |
| ARHGEF5 | 19.743511 | -0.50468324 | 0.4792746 | -1.0530148 | 0.29233422 | 0.74013384 |
| TPK1 | 36.4979302 | -0.25892782 | 0.33260808 | -0.7784772 | 0.43628773 | 0.82656073 |
| CNTNAP2 | 14.3728905 | 0.127096245 | 0.43206981 | 0.29415673 | 0.76863814 | 0.95205353 |
| CUL1 | 135.381106 | -0.214144114 | 0.23702343 | -0.9034723 | 0.36627527 | 0.78793182 |
| EZH2 | 13.3490413 | -0.083558423 | 0.40363083 | -0.207017 | 0.83599662 | 0.96748214 |
| PDIA4 | 26.4837104 | 0.205953115 | 0.37682772 | 0.5465445 | 0.58469171 | 0.89036744 |
| ZNF786 | 12.7989086 | 0.420737175 | 0.41280243 | 1.01922165 | 0.30809775 | 0.7512664 |
| ZNF398 | 32.9376921 | 0.415675976 | 0.26601393 | 1.56260982 | 0.1181444 | 0.5281109 |
| ZNF282 | 21.9493377 | -0.025362492 | 0.43263861 | -0.0586228 | 0.95325254 | 0.99217529 |
| ZNF212 | 13.1760489 | 0.483045692 | 0.46163261 | 1.04638555 | 0.29538306 | 0.74111836 |
| ZNF783 | 23.0377407 | 0.308613349 | 0.32299366 | 0.95547805 | 0.339336 | 0.77045469 |
| ZNF777 | 10.2971329 | -0.516388594 | 0.47696276 | -1.0826602 | 0.27895928 | NA |
| ZNF746 | 16.7948007 | 0.181691049 | 0.35861796 | 0.50664236 | 0.61240578 | 0.90011356 |
| ZNF862 | 42.1039825 | 0.119264436 | 0.38081291 | 0.3131838 | 0.75414102 | 0.94795939 |
| ATP6V0E2 | 10.648488 | 0.86098694 | 0.53787723 | 1.60071274 | 0.10944056 | NA |
| ACTR3C | 187.536105 | -0.319995525 | 0.26149267 | -1.2237265 | 0.2210554 | 0.67492938 |
| LRRC61 | 19.631093 | -0.759553027 | 0.40325876 | -1.8835376 | 0.05962755 | 0.38765901 |
| RARRES2 | 30.8567302 | 0.611046043 | 0.62018275 | 0.98526773 | 0.32449258 | 0.76304922 |
| ABCB8 | 19.8210529 | 0.253135129 | 0.34939686 | 0.72449171 | 0.46876392 | 0.84326495 |
| SLC4A2 | 21.9739856 | 0.376669457 | 0.62272258 | 0.60487522 | 0.54526191 | 0.87247631 |
| FASTK | 24.0855525 | 1.100889559 | 0.30946595 | 3.55738511 | 0.00037456 | 0.01644018 |
| TMUB1 | 32.1284049 | -0.255542667 | 0.37060527 | -0.6895279 | 0.49049113 | 0.856329 |
| AGAP3 | 30.7830399 | -0.015080092 | 0.45838976 | -0.032898 | 0.97375595 | 0.99530377 |
| CHPF2 | 18.8272212 | -0.3130223 | 0.49673407 | -0.6301607 | 0.52858943 | 0.86750588 |
| SMARCD3 | 18.6528161 | -0.385006127 | 0.40799619 | -0.9436513 | 0.34534787 | 0.77507505 |
| NUB1 | 63.1532944 | -0.242606807 | 0.35658285 | -0.6803659 | 0.49627281 | 0.85814183 |
| RHEB | 127.012761 | -0.330265155 | 0.22924328 | -1.4406754 | 0.14967641 | 0.58659144 |
| PRKAG2 | 271.896117 | -0.688075887 | 0.36723591 | -1.8736618 | 0.06097706 | 0.39246665 |
| GALNT11 | 111.661696 | 0.107518527 | 0.19729452 | 0.54496459 | 0.58577787 | 0.89059199 |
| KMT2C | 498.02938 | 0.076992241 | 0.18195379 | 0.42314172 | 0.67219187 | 0.92176197 |
| ACTR3B | 53.2222002 | -0.078067863 | 0.23058606 | -0.3385628 | 0.73493911 | 0.94293405 |
| DPP6 | 105.611213 | -0.233644441 | 0.28952693 | -0.8069869 | 0.41967402 | 0.81899879 |
| PAXIP1-AS2 | 22.0757248 | 0.247654248 | 0.42495445 | 0.58277834 | 0.56004253 | 0.88037687 |
| PAXIP1 | 28.6010007 | 0.320112066 | 0.30647166 | 1.04450789 | 0.29625047 | 0.74123516 |
| INSIG1 | 105.847189 | -2.15406547 | 0.60055766 | -3.5867754 | 0.00033479 | 0.0156695 |
| AC009403.1 | 12.7415891 | -0.073462179 | 0.48380715 | -0.1518419 | 0.87931166 | 0.97567183 |
| RBM33 | 175.220678 | -0.193706802 | 0.13505878 | -1.4342407 | 0.15150359 | 0.58913705 |
| LINC01006 | 35.3737844 | 0.208322713 | 0.45793766 | 0.454915 | 0.64917039 | 0.91417939 |
| LMBR1 | 180.781376 | -0.1107409 | 0.21103242 | -0.5247578 | 0.5997516 | 0.89476262 |
| NOM1 | 20.1837649 | 0.2191306 | 0.38492384 | 0.56928301 | 0.5691641 | 0.88562534 |
| UBE3C | 201.257913 | 0.035995028 | 0.19858613 | 0.1812565 | 0.85616625 | 0.97166553 |
| DNAJB6 | 205.536316 | -0.674426582 | 0.22032706 | -3.0610248 | 0.00220581 | 0.0545825 |
| PTPRN2 | 12.056168 | -1.103092877 | 0.57442561 | -1.9203407 | 0.05481488 | NA |
| NCAPG2 | 15.067197 | 0.335784932 | 0.40457481 | 0.82996995 | 0.40655577 | 0.81177254 |
| ESYT2 | 225.397248 | -0.189042162 | 0.35518227 | -0.5322399 | 0.59455988 | 0.89289841 |
| WDR60 | 116.737935 | -0.166409467 | 0.24772298 | -0.6717563 | 0.50173887 | 0.85942996 |
| GTPBP6 | 39.2534743 | 0.084546215 | 0.28603409 | 0.2955809 | 0.76755016 | 0.95205353 |
| CSF2RA | 46.0183918 | -1.199747201 | 0.74381226 | -1.6129705 | 0.10675093 | 0.50217458 |
| IL3RA | 29.4267304 | 0.182268324 | 0.57008349 | 0.31972216 | 0.74917896 | 0.94684415 |
| SLC25A6 | 336.307811 | -0.258498778 | 0.4042377 | -0.6394722 | 0.52251578 | 0.86728831 |
| LINC00106 | 36.1475838 | 0.275104449 | 0.4396964 | 0.6256691 | 0.531532 | 0.86840264 |
| ASMTL-AS1 | 18.2715237 | 0.234434009 | 0.38322203 | 0.6117446 | 0.54070675 | 0.87051543 |
| ASMTL | 27.7570937 | -0.099531995 | 0.27867118 | -0.3571664 | 0.7209672 | 0.93818343 |
| AKAP17A | 23.9065876 | 0.218244419 | 0.33566536 | 0.65018451 | 0.51557304 | 0.86463667 |
| DHRSX | 156.409233 | -0.080236327 | 0.1780467 | -0.4506476 | 0.65224352 | 0.9158679 |
| CD99 | 23.020008 | -0.194434957 | 0.3602459 | -0.5397284 | 0.58938432 | 0.8914748 |
| ARSD | 80.3710924 | 0.134001453 | 0.33732747 | 0.39724441 | 0.69118723 | 0.92786587 |
| ARSE | 26.6883061 | 0.369171602 | 0.4995648 | 0.73898641 | 0.45991525 | 0.83690755 |
| PRKX | 92.5854418 | -0.323639734 | 0.64747309 | -0.4998505 | 0.61718036 | 0.90173965 |
| NLGN4X | 16.8726143 | 0.044853975 | 0.87809612 | 0.05108094 | 0.95926102 | 0.99238848 |
| PUDP | 38.5617498 | -0.386885291 | 0.37511386 | -1.031381 | 0.3023622 | 0.74690278 |
| ANOS1 | 19.7135127 | -1.531353119 | 1.03674634 | -1.477076 | 0.13965527 | 0.57327809 |
| TBL1X | 331.665108 | 0.26251714 | 0.3343226 | 0.78522103 | 0.43232399 | 0.8248269 |
| GPR143 | 15.5198578 | -0.155422619 | 0.44582822 | -0.3486155 | 0.72737801 | 0.94011548 |
| SHROOM2 | 126.954838 | 0.252055551 | 0.36642079 | 0.6878855 | 0.49152489 | 0.85654883 |
| WWC3 | 34.6631728 | -0.203773614 | 0.33273163 | -0.6124263 | 0.54025573 | 0.87022261 |
| CLCN4 | 14.0262171 | -0.165327866 | 0.44838498 | -0.3687186 | 0.71233751 | 0.9357596 |
| MID1 | 15.2930473 | 0.226904452 | 0.43936377 | 0.5164387 | 0.60554804 | 0.8962185 |
| HCCS | 12.3970283 | -0.223982439 | 0.36907756 | -0.6068709 | 0.54393658 | NA |
| ARHGAP6 | 536.164844 | -0.65728955 | 0.47513888 | -1.383363 | 0.16655359 | 0.60758548 |
| PRPS2 | 22.4572784 | 1.141540031 | 0.32662426 | 3.49496402 | 0.00047413 | 0.01945996 |
| TMSB4X | 508.652312 | 0.068128367 | 0.50651455 | 0.13450427 | 0.89300384 | 0.97831181 |
| RAB9A | 79.8317139 | -0.345493119 | 0.32516399 | -1.0625196 | 0.28799986 | 0.73803612 |
| TRAPPC2 | 25.2015239 | -0.047130449 | 0.41292377 | -0.1141384 | 0.9091281 | 0.98300553 |
| OFD1 | 119.621521 | -0.278463823 | 0.27886363 | -0.9985663 | 0.31800483 | 0.75986079 |
| GEMIN8 | 22.8492854 | -0.183617413 | 0.40994085 | -0.447912 | 0.65421673 | 0.9158679 |
| MOSPD2 | 33.535731 | -0.50184877 | 0.32463777 | -1.545873 | 0.12213524 | 0.53752806 |
| ASB9 | 38.3857767 | 1.388871196 | 0.53649726 | 2.58877594 | 0.00963177 | 0.14462571 |
| PIGA | 29.7393743 | 0.159007826 | 0.38512621 | 0.41287199 | 0.6797004 | 0.92366123 |
| PIR | 35.7098346 | -0.530025695 | 0.40415393 | -1.3114451 | 0.18970742 | 0.64181696 |
| CLTRN | 12.9269973 | -0.883483654 | 0.60615667 | -1.457517 | 0.14497372 | 0.58002047 |
| CA5B | 88.7562894 | 0.596263412 | 0.287993 | 2.07040936 | 0.03841403 | 0.3141788 |
| ZRSR2 | 43.7184732 | 0.413399945 | 0.28157943 | 1.4681468 | 0.14206435 | 0.57673923 |
| AP1S2 | 72.7320197 | -1.217802034 | 0.36640971 | -3.3236075 | 0.00088861 | 0.03024398 |
| CTPS2 | 46.7166649 | -0.072962281 | 0.25552102 | -0.2855432 | 0.77522804 | 0.95493992 |
| SYAP1 | 86.8624169 | -0.272311414 | 0.27074024 | -1.0058033 | 0.31451022 | 0.75700972 |
| TXLNG | 40.6355049 | -0.442477066 | 0.31043325 | -1.4253533 | 0.1540551 | 0.59416877 |
| RBBP7 | 54.7354553 | 0.179162004 | 0.24640086 | 0.72711598 | 0.46715491 | 0.84289484 |
| REPS2 | 41.4191914 | 0.105730277 | 0.72691635 | 0.14545041 | 0.88435527 | 0.97662748 |
| NHS | 80.4139293 | 1.054756122 | 0.5788878 | 1.82203895 | 0.06844908 | 0.41572605 |
| SCML1 | 127.962966 | -0.544380724 | 0.23614401 | -2.3052912 | 0.02115027 | 0.22694426 |
| RAI2 | 30.9386266 | -0.571103576 | 0.28975032 | -1.9710196 | 0.04872163 | 0.35268642 |
| CDKL5 | 54.8003257 | 0.445140007 | 0.27565807 | 1.61482669 | 0.10634823 | 0.50162992 |
| PHKA2 | 31.549734 | 0.487850069 | 0.24607504 | 1.98252558 | 0.04742045 | 0.34767826 |
| PDHA1 | 105.213794 | 0.151136301 | 0.30550499 | 0.49470976 | 0.62080501 | 0.90380166 |
| MAP3K15 | 21.6304028 | 0.661374496 | 0.56785366 | 1.16469179 | 0.24414377 | 0.69773111 |
| BCLAF3 | 51.5871593 | -0.039704282 | 0.30995814 | -0.1280956 | 0.8980733 | 0.97892709 |
| EIF1AX | 92.6157861 | 0.214502625 | 0.24692225 | 0.86870513 | 0.38500844 | 0.79869171 |

| | | | | | | |
|---|---|---|---|---|---|---|
| RPS6KA3 | 112.193756 | -0.748938131 | 0.32454067 | -2.3076865 | 0.02101658 | 0.22580589 |
| MBTPS2 | 27.584926 | -0.117577099 | 0.34802664 | -0.3378394 | 0.73548425 | 0.94318131 |
| SMS | 72.743779 | 0.001865063 | 0.26629397 | 0.00700377 | 0.99441184 | 0.99923337 |
| PRDX4 | 12.8448658 | -0.54159961 | 0.49060653 | -1.1039389 | 0.26961966 | 0.72046011 |
| ACOT9 | 32.0072861 | 0.503230851 | 0.41495628 | 1.21273223 | 0.22523221 | 0.67849456 |
| SAT1 | 417.30502 | -0.786261937 | 0.32591848 | -2.4124497 | 0.01584572 | 0.19341706 |
| APOO | 487.680229 | 0.297294238 | 0.37961629 | 0.78314404 | 0.43354253 | 0.82504445 |
| KLHL15 | 11.8473233 | 0.717637439 | 0.36131939 | 1.98615812 | 0.04701577 | NA |
| EIF2S3 | 58.8083236 | -0.025304989 | 0.30049047 | -0.0842123 | 0.93288765 | 0.98809895 |
| ZFX | 90.42616 | 0.256663122 | 0.32656202 | 0.78595523 | 0.43189372 | 0.8248269 |
| PDK3 | 23.975453 | -0.929564546 | 0.51009278 | -1.822344 | 0.0684028 | 0.41572605 |
| POLA1 | 83.6145112 | -0.012797546 | 0.26251523 | -0.0487497 | 0.96111875 | 0.99238848 |
| GK | 59.7480441 | 0.538613747 | 0.2607927 | 2.06529458 | 0.03889514 | 0.31669347 |
| TAB3 | 69.0666897 | -0.044130991 | 0.27048415 | -0.1631556 | 0.87039596 | 0.97434023 |
| DMD | 118.754388 | 0.918420755 | 0.36523368 | 2.51461132 | 0.01191637 | 0.1625721 |
| AF241726.2 | 68.9590482 | -0.401431219 | 0.26650315 | -1.5062907 | 0.13199256 | 0.55916709 |
| DYNLT3 | 133.738512 | -0.911874901 | 0.32436878 | -2.811229 | 0.00493527 | 0.09671126 |
| RPGR | 35.1939243 | -0.027997426 | 0.39311641 | -0.0712192 | 0.94322332 | 0.99050192 |
| TSPAN7 | 52.4227979 | 1.204951251 | 0.60022399 | 2.00750265 | 0.04469617 | 0.33582714 |
| MID1IP1 | 28.1828301 | -0.879814353 | 0.39402846 | -2.2328701 | 0.02555751 | 0.25205612 |
| BCOR | 94.6830033 | -0.126733098 | 0.40632999 | -0.311897 | 0.75511882 | 0.94795939 |
| ATP6AP2 | 452.144033 | 0.379265824 | 0.26170436 | 1.44921479 | 0.14727761 | 0.58405266 |
| CXorf38 | 55.4781716 | -0.23902139 | 0.26328911 | -0.9078286 | 0.36396877 | 0.78770574 |
| MED14 | 115.881919 | -0.226169402 | 0.23323602 | -0.9697019 | 0.33219512 | 0.76721253 |
| USP9X | 277.330787 | -0.014472938 | 0.15454167 | -0.0936507 | 0.92538662 | 0.98672316 |
| DDX3X | 129.135256 | -0.101692306 | 0.32190964 | -0.3159033 | 0.75207593 | 0.94728927 |
| CASK | 61.4689407 | 0.230140476 | 0.21929316 | 1.0494649 | 0.2939642 | 0.74044476 |
| MAOA | 134.700663 | -1.933171583 | 0.53840984 | -3.5905205 | 0.00033002 | 0.0156155 |
| FUNDC1 | 17.2370799 | -0.284128275 | 0.45231219 | -0.6281685 | 0.52989355 | 0.86758926 |
| KDM6A | 253.983021 | -0.155491418 | 0.33436196 | -0.4650392 | 0.64190339 | 0.91146968 |
| MIR222HG | 45.3337404 | -0.695722653 | 0.46701714 | -1.4897155 | 0.13629907 | 0.5673498 |
| KRBOX4 | 31.1072912 | -0.275435491 | 0.25589004 | -1.0763822 | 0.28175634 | 0.73311066 |
| ZNF674 | 15.5971269 | 0.631427105 | 0.32156142 | 1.96362827 | 0.04957322 | 0.35590543 |
| SLC9A7 | 31.5329117 | -0.350510668 | 0.35567048 | -0.9854927 | 0.32438211 | 0.76304922 |
| RP2 | 17.6974925 | -0.155279809 | 0.44147347 | -0.3517308 | 0.72504017 | 0.93940528 |
| JADE3 | 72.9712677 | 0.17975212 | 0.29017753 | 0.61945569 | 0.5356162 | 0.86946459 |
| RGN | 16.8921804 | -1.176339212 | 0.61344689 | -1.9175893 | 0.0551631 | 0.37383273 |
| NDUFB11 | 89.6140886 | 0.119962629 | 0.42472701 | 0.28244643 | 0.77760122 | 0.9551993 |
| RBM10 | 47.0897214 | 0.412055103 | 0.24362191 | 1.69137129 | 0.09076591 | 0.47310687 |
| UBA1 | 75.8446357 | -0.161193816 | 0.22841248 | -0.7057137 | 0.4803662 | 0.84944638 |
| CDK16 | 30.9263701 | -0.075189579 | 0.36213289 | -0.2076298 | 0.83551803 | 0.96748214 |
| USP11 | 30.8566001 | 0.180892202 | 0.28147198 | 0.64266504 | 0.52044147 | 0.86680671 |
| ZNF41 | 25.2359286 | 0.310768006 | 0.27036387 | 1.14944357 | 0.25037312 | 0.70453262 |
| ARAF | 27.298382 | 0.469450124 | 0.32612164 | 1.43949395 | 0.15001062 | 0.58673121 |
| TIMP1 | 26.5248353 | -3.292124683 | 0.53042559 | -6.2065721 | 5.42E-10 | 4.04E-07 |
| UXT | 22.408521 | 0.037121757 | 0.29585323 | 0.12547355 | 0.90014866 | 0.97940229 |
| ZNF81 | 46.8182623 | 0.035692967 | 0.28933371 | 0.12336263 | 0.90181995 | 0.9797999 |
| ZNF630 | 19.4258651 | -0.121975265 | 0.48168663 | -0.2532253 | 0.80009408 | 0.95861074 |
| EBP | 35.3582039 | 0.467262717 | 0.35777866 | 1.3060106 | 0.19154895 | 0.6421543 |
| TBC1D25 | 13.7832087 | 0.424142186 | 0.43335157 | 0.97874847 | 0.32770428 | 0.76503766 |
| RBM3 | 54.4859679 | -0.296622439 | 0.47041951 | -0.6305488 | 0.52833561 | 0.86750588 |
| WDR13 | 50.5266576 | -0.181757907 | 0.24286973 | -0.7483761 | 0.45423332 | 0.83364149 |
| HDAC6 | 24.2873292 | 0.076866312 | 0.33053687 | 0.23254989 | 0.81611095 | 0.9631678 |
| PCSK1N | 31.5239437 | -1.192015096 | 0.65457561 | -1.8210503 | 0.06859922 | 0.41614283 |
| TIMM17B | 20.8784256 | -0.595122121 | 0.37542579 | -1.5851924 | 0.11292262 | 0.51571293 |
| SLC35A2 | 9.48862791 | 0.354607655 | 0.49714978 | 0.71328133 | 0.47567169 | NA |
| OTUD5 | 48.0146113 | -0.028538498 | 0.32791747 | -0.0870295 | 0.93064805 | 0.98756002 |
| GRIPAP1 | 38.4823631 | -0.301919919 | 0.25079062 | -1.2038725 | 0.22863888 | 0.68387206 |
| TFE3 | 18.5189233 | -0.476864611 | 0.35996948 | -1.3247362 | 0.18525866 | 0.63578845 |
| CCDC120 | 13.4922565 | 0.147425387 | 0.39313599 | 0.37499845 | 0.70766162 | 0.93432317 |
| WDR45 | 30.7331131 | -0.193786898 | 0.29359183 | -0.6600555 | 0.50921822 | 0.86281145 |
| PLP2 | 34.2521766 | -1.339732767 | 0.4502435 | -2.9755738 | 0.00292441 | 0.06690277 |
| PPP1R3F | 9.99170977 | -0.129294639 | 0.40422713 | -0.3198564 | 0.74907718 | NA |
| CLCN5 | 116.298233 | 0.417584248 | 0.35858349 | 1.16453841 | 0.24420589 | 0.69773111 |
| MAGED1 | 149.759846 | -0.077272193 | 0.27662195 | -0.2793422 | 0.7799822 | 0.95537097 |
| FAM156A | 15.8983433 | 0.017267509 | 0.38610618 | 0.04472218 | 0.96432876 | 0.99334616 |
| TSPYL2 | 12.8448645 | -0.56202465 | 0.6897617 | -0.8148099 | 0.41518115 | 0.81726257 |
| KANTR | 55.2432209 | 0.518131987 | 0.20654194 | 2.50860423 | 0.01212092 | 0.16352285 |
| KDM5C | 46.4024659 | -0.077358118 | 0.28006247 | -0.2762174 | 0.78238112 | 0.95544332 |
| IQSEC2 | 19.0383917 | 0.070504251 | 0.40048173 | 0.17604861 | 0.86025576 | 0.97287712 |
| SMC1A | 62.6898171 | 0.046854126 | 0.26836842 | 0.17458882 | 0.86140273 | 0.97302617 |
| HSD17B10 | 23.4384171 | 0.593599916 | 0.37452887 | 1.58492434 | 0.11298352 | 0.51573656 |
| HUWE1 | 204.906645 | 0.283653907 | 0.17418616 | 1.62845262 | 0.10342895 | 0.495277 |
| PHF8 | 89.7957485 | -0.043490852 | 0.361766 | -0.1202182 | 0.90431031 | 0.98041914 |
| FAM120C | 25.5221322 | 0.372227574 | 0.2924068 | 1.2729785 | 0.20302567 | 0.65623093 |
| WNK3 | 101.925828 | 1.015454029 | 0.29657511 | 3.42393545 | 0.00061721 | 0.02349088 |
| TSR2 | 20.9576127 | -0.145636923 | 0.43635648 | -0.3337568 | 0.73856312 | 0.94376279 |
| GNL3L | 11.573193 | 0.343255693 | 0.47691519 | 0.71974159 | 0.47168411 | NA |
| MAGED2 | 54.668713 | -0.225496323 | 0.31171328 | -0.7234094 | 0.46942839 | 0.8435449 |
| USP51 | 11.2870909 | -1.38639104 | 0.64269381 | -2.1571564 | 0.03099348 | NA |
| RRAGB | 53.2538556 | -0.464157771 | 0.23772144 | -1.952528 | 0.05087555 | 0.35962249 |
| AL445472.1 | 80.1357234 | -0.581572721 | 0.33233658 | -1.749951 | 0.08012677 | 0.44789428 |
| UBQLN2 | 29.3506539 | -0.18532896 | 0.32584758 | -0.5687597 | 0.56951925 | 0.88562534 |
| NBDY | 96.8719093 | 0.190439676 | 0.23773521 | 0.80105792 | 0.42309812 | 0.82180534 |
| SPIN3 | 17.68986 | 0.172558047 | 0.41758658 | 0.413227 | 0.67944031 | 0.92356656 |
| FAAH2 | 256.881596 | -0.849506299 | 0.28625117 | -2.9676954 | 0.00300041 | 0.06784993 |
| ARHGEF9 | 69.654993 | -0.583325802 | 0.32444607 | -1.797913 | 0.0721908 | 0.42395281 |
| MTMR8 | 17.011964 | -0.138099405 | 0.38583814 | -0.3579206 | 0.72040276 | 0.93806337 |
| ZC4H2 | 35.1482058 | 0.317617267 | 0.31036853 | 1.02335525 | 0.30613991 | 0.74897897 |
| LAS1L | 18.2440384 | 0.476606812 | 0.30529587 | 1.56113088 | 0.11849287 | 0.52918088 |
| MSN | 165.091527 | -0.579862799 | 0.26930028 | -2.1532202 | 0.03130138 | 0.28061311 |
| OPHN1 | 197.586612 | -0.081133216 | 0.27288942 | -0.2973117 | 0.76622855 | 0.95205353 |
| YIPF6 | 69.2550106 | -0.070144561 | 0.21439941 | -0.3271677 | 0.74354105 | 0.94546379 |
| EDA | 49.5614307 | -0.106180453 | 0.41530321 | -0.2556697 | 0.79820588 | 0.95848876 |
| IGBP1 | 39.1862824 | -0.053209515 | 0.28705623 | -0.1853627 | 0.85294457 | 0.97107539 |
| DLG3 | 56.4514202 | 0.136657137 | 0.29436183 | 0.46424883 | 0.64246948 | 0.91146968 |
| SNX12 | 19.6278481 | -0.307846583 | 0.38079616 | -0.8084288 | 0.41884378 | 0.8189355 |
| MED12 | 11.8263872 | 0.11539984 | 0.38513942 | 0.29963134 | 0.76445838 | NA |
| ZMYM3 | 15.8082753 | 0.423315716 | 0.36968879 | 1.14505964 | 0.25218445 | 0.70617471 |
| NONO | 53.9833462 | -0.108901758 | 0.27275635 | -0.3992639 | 0.68969878 | 0.92734996 |
| TAF1 | 91.8550334 | 0.250715021 | 0.23458152 | 1.06877565 | 0.28517077 | 0.73659018 |
| OGT | 179.995589 | 0.589241019 | 0.28266075 | 2.08462271 | 0.03710356 | 0.30857899 |
| NHSL2 | 24.2600457 | 0.200099635 | 0.28018078 | 0.71418044 | 0.47511561 | 0.84594051 |
| PIN4 | 36.6410951 | 0.175172736 | 0.32808973 | 0.53391715 | 0.59339886 | 0.89239828 |
| RPS4X | 258.092405 | 0.272282849 | 0.40945343 | 0.66499101 | 0.50605624 | 0.86214551 |
| HDAC8 | 219.714898 | -0.184772725 | 0.26532763 | -0.6963946 | 0.48618175 | 0.85356796 |
| PHKA1 | 30.1431576 | 0.013890762 | 0.31349506 | 0.04430935 | 0.96465782 | 0.99334616 |
| CHIC1 | 100.396507 | -0.172491805 | 0.23010853 | -0.7496107 | 0.45348923 | 0.83364149 |
| XIST | 340.542559 | -0.870859759 | 3.42690227 | -0.2541245 | 0.79939939 | 0.95848876 |
| JPX | 234.802382 | -0.122923392 | 0.35205933 | -0.3491553 | 0.7269727 | 0.94009907 |
| FTX | 1189.56871 | -0.141614331 | 0.24136228 | -0.5867293 | 0.55738551 | 0.87898544 |
| RLIM | 43.147967 | -0.052061023 | 0.23667776 | -0.2199658 | 0.82589776 | 0.96539788 |
| ABCB7 | 94.5318127 | 0.136331118 | 0.2137256 | 0.63787923 | 0.5235523 | 0.86747074 |

| | | | | | | |
|---|---|---|---|---|---|---|
| UPRT | 20.0574996 | -0.05149173 | 0.31519133 | -0.1633666 | 0.87022981 | 0.97434023 |
| ZDHHC15 | 15.693532 | -0.092478814 | 0.44621711 | -0.2072507 | 0.83581406 | 0.96748214 |
| PBDC1 | 29.4652059 | -0.427735763 | 0.38866568 | -1.1005236 | 0.27110405 | 0.72194341 |
| ATRX | 373.64627 | -0.024663415 | 0.15191547 | -0.1623496 | 0.87103056 | 0.97434023 |
| MAGT1 | 75.0821362 | 0.263624978 | 0.2352911 | 1.12042054 | 0.2625346 | 0.71333232 |
| COX7B | 214.00464 | -0.051782468 | 0.37356419 | -0.1386173 | 0.88975255 | 0.97782578 |
| PGK1 | 191.371499 | -0.714633136 | 0.31681242 | -2.255698 | 0.02408955 | 0.24315158 |
| BRWD3 | 149.288742 | -0.675422406 | 0.25583831 | -2.6400363 | 0.00828972 | 0.13160433 |
| HMGN5 | 15.264439 | 0.132710739 | 0.41617414 | 0.31888271 | 0.74981545 | 0.94684415 |
| SH3BGRL | 98.3200997 | -0.122011503 | 0.19675757 | -0.6201109 | 0.53518481 | 0.86946459 |
| RPS6KA6 | 101.721223 | 0.02949949 | 0.26680373 | 0.11056626 | 0.91196031 | 0.98354625 |
| HDX | 34.9263227 | -0.652137442 | 0.38775222 | -1.6818407 | 0.09259974 | 0.47676089 |
| APOOL | 64.756768 | 0.233316114 | 0.20723009 | 1.12587951 | 0.26021652 | 0.71091822 |
| POF1B | 19.2549099 | -1.062309364 | 0.70733238 | -1.5018532 | 0.13313504 | 0.56148254 |
| CHM | 105.294597 | -0.315108601 | 0.20248064 | -1.5562406 | 0.11965088 | 0.53109067 |
| DIAPH2 | 107.672033 | 0.090518344 | 0.79853521 | 0.11335548 | 0.90974873 | 0.98329594 |
| TSPAN6 | 10.8559952 | 0.994446138 | 0.48398304 | 2.05471278 | 0.03990676 | NA |
| TRMT2B | 44.8216605 | -0.352355302 | 0.28646616 | -1.2300067 | 0.21869459 | 0.67250923 |
| HNRNPH2 | 34.4089536 | -0.361934305 | 0.35238564 | -1.0270972 | 0.30437469 | 0.74820946 |
| ARMCX4 | 150.7084 | 0.183256453 | 0.28472948 | 0.64361602 | 0.51982446 | 0.86667193 |
| ARMCX6 | 10.9500992 | 0.624221998 | 0.39779027 | 1.56922388 | 0.11659578 | NA |
| ARMCX3 | 58.673823 | 0.083674608 | 0.29870065 | 0.28012864 | 0.77937881 | 0.95537097 |
| ARMCX2 | 18.566169 | 0.350075603 | 0.35186598 | 0.99491177 | 0.31977918 | 0.76106474 |
| ZMAT1 | 74.9894002 | 0.345647831 | 0.40217506 | 0.85944622 | 0.39009438 | 0.80191835 |
| ARMCX5 | 11.0089146 | 0.096363652 | 0.38132054 | 0.25271037 | 0.80049204 | NA |
| GPRASP1 | 15.448013 | -0.268751574 | 0.38387095 | -0.7001092 | 0.48385912 | 0.85235034 |
| BHLHB9 | 13.5035356 | 0.463813642 | 0.46696336 | 0.9932549 | 0.32058575 | 0.76131909 |
| LINC00630 | 51.7980608 | -0.005980062 | 0.24503656 | -0.0244048 | 0.98052974 | 0.99687019 |
| BEX4 | 30.2387085 | -0.313739329 | 0.30081509 | -1.0429641 | 0.29696493 | 0.74160729 |
| TCEAL8 | 18.8460782 | -0.470736503 | 0.36559553 | -1.2875882 | 0.19788933 | 0.64914661 |
| TCEAL9 | 30.6057461 | -0.805630234 | 0.41995214 | -1.9183858 | 0.0550621 | 0.37349819 |
| BEX3 | 48.3852332 | 0.467453315 | 0.32886827 | 1.42139987 | 0.15520055 | 0.59533243 |
| TCEAL4 | 56.2424371 | -0.581117572 | 0.2966809 | -1.9587293 | 0.05014449 | 0.35738545 |
| TCEAL3 | 30.0283181 | -0.817110712 | 0.35545731 | -2.298759 | 0.02151863 | 0.22951242 |
| MORF4L2 | 92.9025067 | -0.031992587 | 0.30695349 | -0.1042262 | 0.91698987 | 0.98495314 |
| SLC25A53 | 41.9224696 | 0.42341383 | 0.29820931 | 1.41985449 | 0.15565005 | 0.59590618 |
| FAM199X | 59.0314297 | 0.319152406 | 0.20073452 | 1.58992286 | 0.11185219 | 0.51426646 |
| NRK | 23.6206122 | 2.040633584 | 0.56655714 | 3.60181424 | 0.000316 | 0.01510953 |
| RNF128 | 17.9018837 | 0.351510501 | 0.61982874 | 0.56710907 | 0.57064008 | 0.88563341 |
| TBC1D8B | 25.8095255 | -0.180626811 | 0.41602899 | -0.4341688 | 0.66416586 | 0.91876908 |
| MORC4 | 27.4548114 | -0.246241611 | 0.36316531 | -0.6780428 | 0.49774458 | 0.85818031 |
| RBM41 | 52.5421577 | 0.349097699 | 0.20582046 | 1.69612727 | 0.08986178 | 0.47218812 |
| MID2 | 22.3237301 | -0.409660376 | 0.40624396 | -1.0084098 | 0.31325779 | 0.75618668 |
| PSMD10 | 11.1929849 | 0.156750013 | 0.40062932 | 0.39125946 | 0.69560546 | NA |
| ATG4A | 14.8675266 | -0.284341269 | 0.41793132 | -0.6803541 | 0.4962803 | 0.85814183 |
| COL4A5 | 25.9302847 | 1.07912302 | 0.44988539 | 2.39866205 | 0.01645509 | 0.19852538 |
| ACSL4 | 119.953676 | -1.33912871 | 0.32630375 | -4.1039329 | 4.06E-05 | 0.00358182 |
| TMEM164 | 119.596498 | 0.89853778 | 0.32859412 | 2.73449138 | 0.00624767 | 0.11201928 |
| AMMECR1 | 100.394187 | -0.084271133 | 0.32292644 | -0.2609608 | 0.79412276 | 0.95784516 |
| ALG13 | 93.7340589 | -0.190354 | 0.39124495 | -0.4865341 | 0.62658855 | 0.90541055 |
| AMOT | 13.9255336 | 0.828514366 | 0.42199218 | 1.96334058 | 0.04960661 | 0.35590543 |
| PLS3 | 187.685971 | 0.290492452 | 0.22153113 | 1.31129402 | 0.18975845 | 0.64181696 |
| DANT2 | 223.723698 | -0.346832822 | 0.51168144 | -0.6778296 | 0.49787975 | 0.85826081 |
| KLHL13 | 50.649305 | 0.025822999 | 0.42875411 | 0.06022799 | 0.95197405 | 0.99217529 |
| WDR44 | 46.3414031 | 0.103644824 | 0.29906801 | 0.34655938 | 0.72892237 | 0.94098309 |
| IL13RA1 | 60.1473949 | -0.637998718 | 0.28531972 | -2.2360835 | 0.0253463 | 0.25137888 |
| PGRMC1 | 87.9959022 | 0.008757343 | 0.28405664 | 0.03082957 | 0.97540546 | 0.99552114 |
| SLC25A43 | 38.3337679 | -0.330367409 | 0.38918822 | -0.8488628 | 0.39595763 | 0.80638022 |
| SLC25A5 | 200.28457 | 0.428452538 | 0.42165984 | 1.01610943 | 0.30957728 | 0.75241784 |
| UBE2A | 37.4941107 | -0.117424543 | 0.34136124 | -0.3439891 | 0.73085449 | 0.94155624 |
| NKRF | 12.3504128 | -0.064191893 | 0.37534598 | -0.1710206 | 0.86420757 | NA |
| 6-Sep | 132.846853 | 0.387971128 | 0.40487688 | 0.95824471 | 0.33793937 | 0.76966704 |
| UPF3B | 32.0372089 | -0.181854629 | 0.38990634 | -0.4664059 | 0.64092497 | 0.91146968 |
| NDUFA1 | 107.316655 | -0.1606277 | 0.35998415 | -0.4462077 | 0.65544722 | 0.9158679 |
| NKAP | 19.5822894 | -0.29576919 | 0.33983789 | -0.8703244 | 0.38412317 | 0.79837042 |
| RHOXF1-AS1 | 23.3476297 | -0.112471382 | 0.32225488 | -0.3490137 | 0.72707899 | 0.9401048 |
| ZBTB33 | 12.2518288 | -0.253124105 | 0.42936456 | -0.5895319 | 0.55550452 | NA |
| TMEM255A | 53.5030762 | -3.832703646 | 1.17727446 | -3.2555736 | 0.00113164 | 0.03575526 |
| LAMP2 | 104.266263 | -0.443653306 | 0.22791582 | -1.9465665 | 0.05158672 | 0.36077232 |
| CUL4B | 43.0399138 | -0.133550979 | 0.24199688 | -0.5518707 | 0.58103697 | 0.89012555 |
| MCTS1 | 30.9984616 | 0.149552844 | 0.28265034 | 0.52910902 | 0.59672982 | 0.89363331 |
| C1GALT1C1 | 15.2634286 | -0.348672824 | 0.40294455 | -0.8653122 | 0.38686747 | 0.80009555 |
| THOC2 | 144.136144 | -0.163719215 | 0.18298586 | -0.8947097 | 0.3709423 | 0.7900012 |
| XIAP | 150.280056 | 0.060052943 | 0.17665639 | 0.33994209 | 0.73390014 | 0.94277981 |
| STAG2 | 174.548928 | -0.10682691 | 0.20671712 | -0.5167783 | 0.60531096 | 0.89600432 |
| AL442647.1 | 18.8811068 | -0.236992502 | 0.69904085 | -0.3390253 | 0.7345907 | 0.94277981 |
| SMARCA1 | 141.433563 | 0.23551649 | 0.31875544 | 0.73886266 | 0.4599904 | 0.83690755 |
| OCRL | 31.4954247 | 0.006847091 | 0.25552557 | 0.02679611 | 0.97862236 | 0.99610605 |
| ZDHHC9 | 83.5615067 | -0.861886872 | 0.28014327 | -3.0765931 | 0.00209381 | 0.05234522 |
| UTP14A | 15.0945764 | -0.130474176 | 0.39977163 | -0.3263718 | 0.74414309 | 0.94546379 |
| BCORL1 | 15.2653902 | 0.209321532 | 0.38861194 | 0.53863896 | 0.590136 | 0.89149106 |
| ELF4 | 17.205281 | 0.240713921 | 0.36663463 | 0.65654987 | 0.51147039 | 0.86312853 |
| AIFM1 | 62.0839554 | 0.511345888 | 0.29089684 | 1.75782549 | 0.07877721 | 0.44254372 |
| ZNF280C | 17.0703442 | -0.208856989 | 0.36147335 | -0.5777936 | 0.56340347 | 0.88191823 |
| RBMX2 | 12.5516849 | 0.517806358 | 0.36900292 | 1.40325814 | 0.16053988 | 0.60024417 |
| ENOX2 | 122.086632 | -0.297872288 | 0.34433595 | -0.865063 | 0.38700422 | 0.80009555 |
| FRMD7 | 15.3277983 | 2.380653342 | 0.79241383 | 3.00430564 | 0.00266188 | 0.0628229 |
| RAP2C | 77.7066965 | 0.697857516 | 0.32271296 | 2.16247129 | 0.03058187 | 0.27849022 |
| RAP2C-AS1 | 73.4715108 | 0.329662042 | 0.3626291 | 0.90908876 | 0.36330327 | 0.78718296 |
| MBNL3 | 54.3267585 | -0.049855722 | 0.29849638 | -0.1670229 | 0.86735205 | 0.97434023 |
| GPC4 | 16.5460887 | -0.062007867 | 0.42564195 | -0.1456808 | 0.88417337 | 0.97662748 |
| GPC3 | 20.4284728 | 2.59675359 | 0.51436113 | 5.04850277 | 4.45E-07 | 0.00010484 |
| PHF6 | 38.9455509 | 0.23635488 | 0.28856305 | 0.81907534 | 0.41274343 | 0.81556555 |
| HPRT1 | 127.427273 | -0.337933473 | 0.20368762 | -1.6590771 | 0.09710026 | 0.48257296 |
| FAM122B | 19.1216081 | 0.579113993 | 0.32891921 | 1.76065724 | 0.07829643 | 0.4412989 |
| FAM122C | 20.244199 | 0.106048227 | 0.54087868 | 0.19606657 | 0.84455806 | 0.97006669 |
| MOSPD1 | 11.5244297 | -0.237251616 | 0.42709992 | -0.5554944 | 0.57855654 | NA |
| RTL8C | 37.4303936 | -0.306771756 | 0.28235105 | -1.0864906 | 0.277262 | 0.72678432 |
| RTL8A | 23.5654269 | -0.64105865 | 0.31837786 | -2.0135152 | 0.04406046 | 0.33311495 |
| ZNF75D | 66.4272805 | -0.127514237 | 0.21338776 | -0.5975705 | 0.55012653 | 0.87445431 |
| INTS6L | 13.851907 | -0.672445546 | 0.47616838 | -1.4122012 | 0.15789073 | 0.59849163 |
| MMGT1 | 26.8789889 | 0.038899775 | 0.27648285 | 0.14069507 | 0.88811084 | 0.97765744 |
| SLC9A6 | 19.6737172 | 0.0193325 | 0.31790429 | 0.06081233 | 0.95150867 | 0.99217308 |
| HTATSF1 | 40.7416337 | 0.695609011 | 0.40361645 | 1.72344067 | 0.08480887 | 0.45956155 |
| ARHGEF6 | 34.4137755 | -0.162095944 | 0.33774722 | -0.4799327 | 0.63127523 | 0.90770379 |
| AL683813.1 | 15.2191631 | 0.110144094 | 0.52952879 | 0.20800397 | 0.83522586 | 0.96748214 |
| RBMX | 124.36706 | 0.405155004 | 0.2221916 | 1.82344883 | 0.06823544 | 0.41547228 |
| FGF13 | 338.873609 | -0.397498296 | 0.59529299 | -0.6677356 | 0.50430241 | 0.86122066 |
| ATP11C | 53.1200872 | 0.78805182 | 0.49136844 | 1.60379004 | 0.10876034 | 0.50661006 |
| FMR1 | 71.2062604 | -0.5238959 | 0.28008256 | -1.8705053 | 0.06141369 | 0.39288791 |
| IDS | 48.5226177 | -0.237448632 | 0.26102393 | -0.9096815 | 0.36299053 | 0.78718296 |
| TMEM185A | 19.5394879 | 0.330023628 | 0.35490686 | 0.92988799 | 0.35242908 | 0.77964174 |
| LINC00894 | 117.625694 | -0.243391673 | 0.4474946 | -0.5438986 | 0.58651128 | 0.89059199 |

| | | | | | | |
|---|---|---|---|---|---|---|
| MTM1 | 68.5366552 | 0.039094055 | 0.3522445 | 0.11098556 | 0.91162779 | 0.9835359 |
| MTMR1 | 73.6820337 | -0.066602435 | 0.20734014 | -0.3212231 | 0.74804136 | 0.94656788 |
| CD99L2 | 53.6321063 | 0.098112582 | 0.26511862 | 0.37007051 | 0.71132995 | 0.93570661 |
| HMGB3 | 16.3854624 | -0.444521088 | 0.361413 | -1.2299532 | 0.21871461 | 0.67250923 |
| VMA21 | 30.7766123 | 0.573432634 | 0.32040816 | 1.78969421 | 0.07350308 | 0.42788861 |
| CETN2 | 12.0523198 | -0.030773213 | 0.55534882 | -0.0554124 | 0.95580991 | NA |
| CCNQ | 12.416022 | 0.302585546 | 0.42024864 | 0.72001553 | 0.47151543 | NA |
| SLC6A8 | 31.8121411 | -1.073527409 | 0.63268311 | -1.6967853 | 0.08973726 | 0.47198427 |
| BCAP31 | 66.7418526 | -0.036743426 | 0.36570393 | -0.1004732 | 0.9199687 | 0.98586108 |
| IDH3G | 21.5420619 | -0.16312467 | 0.39580247 | -0.4121366 | 0.68023933 | 0.92375164 |
| SSR4 | 46.7411497 | -0.208830231 | 0.42375612 | -0.4928076 | 0.62214855 | 0.90415189 |
| HCFC1 | 21.6147615 | 0.28596916 | 0.35388832 | 0.80807741 | 0.41904602 | 0.8189355 |
| IRAK1 | 18.8402453 | -0.329473737 | 0.64044087 | -0.5144483 | 0.60693858 | 0.89690801 |
| MECP2 | 142.765539 | 0.125484815 | 0.22030487 | 0.5695962 | 0.56895161 | 0.88562534 |
| EMD | 35.9791299 | -0.186859295 | 0.27768475 | -0.6729188 | 0.50099893 | 0.85942996 |
| RPL10 | 379.240722 | 0.262103774 | 0.47167418 | 0.55568819 | 0.57842403 | 0.88917798 |
| DNASE1L1 | 17.4006089 | 0.001945452 | 0.34200459 | 0.00568838 | 0.99546135 | 0.99923337 |
| TAZ | 25.6691545 | -0.22919479 | 0.37073351 | -0.6182198 | 0.53643047 | 0.86963427 |
| ATP6AP1 | 128.533398 | 0.518984085 | 0.34351428 | 1.51080788 | 0.13083741 | 0.55687707 |
| GDI1 | 37.2628309 | -0.318953769 | 0.32998228 | -0.9665785 | 0.3337548 | 0.76777176 |
| FAM50A | 36.8091032 | -0.394156482 | 0.44703712 | -0.8817086 | 0.3779344 | 0.79470272 |
| PLXNA3 | 17.5026517 | 0.333292031 | 0.43681426 | 0.7630063 | 0.44545964 | 0.83035712 |
| UBL4A | 13.0760619 | 0.185555429 | 0.47417489 | 0.39132277 | 0.69555867 | 0.92900737 |
| FAM3A | 26.9643231 | 0.060004671 | 0.31195739 | 0.19234893 | 0.84746889 | 0.97027687 |
| IKBKG | 13.5693305 | -0.465979282 | 0.44705882 | -1.0423221 | 0.29726237 | 0.74160729 |
| DKC1 | 18.278295 | 0.245508636 | 0.34156413 | 0.71877756 | 0.47227799 | 0.84570599 |
| MPP1 | 24.6077002 | 0.535075846 | 0.36322748 | 1.47311501 | 0.14072003 | 0.57497231 |
| FUNDC2 | 51.0414185 | -0.075380389 | 0.35875708 | -0.2101154 | 0.8335776 | 0.96701848 |
| BRCC3 | 35.9886685 | -0.309005067 | 0.28288438 | -1.092337 | 0.27468502 | 0.72554623 |
| VBP1 | 28.3432064 | 0.357471665 | 0.33148277 | 1.07840195 | 0.28085442 | 0.73272938 |
| TMLHE-AS1 | 62.6139731 | -0.850586449 | 0.61714204 | -1.3782669 | 0.1681209 | 0.60986266 |
| TMLHE | 149.861964 | -0.358856034 | 0.27696956 | -1.2956515 | 0.19509556 | 0.6459894 |
| VAMP7 | 43.7692549 | 0.117067352 | 0.3054093 | 0.383313 | 0.70148771 | 0.93127962 |
| ZNF596 | 17.1717515 | -0.23440521 | 0.36241621 | -0.6467846 | 0.51777138 | 0.86500999 |
| FBXO25 | 60.4572094 | -0.099756152 | 0.23688224 | -0.4211213 | 0.67366652 | 0.92189158 |
| TDRP | 61.9402108 | 0.246767243 | 0.34617772 | 0.71283398 | 0.4759485 | 0.84662643 |
| ERICH1 | 102.851393 | -0.313971542 | 0.37149528 | -0.8451562 | 0.39802363 | 0.80714564 |
| AC100810.1 | 16.3160404 | -0.077971668 | 0.47957427 | -0.1625852 | 0.87084506 | 0.97434023 |
| AC246817.1 | 17.5199628 | 0.182074108 | 0.54521262 | 0.33395065 | 0.7384168 | 0.94376279 |
| CSMD1 | 52.6483055 | 0.515639561 | 0.83238366 | 0.61947343 | 0.53560452 | 0.86946459 |
| MCPH1 | 180.245427 | -0.150363327 | 0.27617301 | -0.5444534 | 0.58612953 | 0.89059199 |
| ANGPT2 | 35.2076326 | -0.593690295 | 0.65744851 | -0.9030217 | 0.36651436 | 0.78802412 |
| MCPH1-AS1 | 90.1671238 | -0.935099366 | 0.50372118 | -1.8563828 | 0.06339901 | 0.4001801 |
| AGPAT5 | 121.229813 | -1.062759633 | 0.28528648 | -3.7252365 | 0.00019513 | 0.01063361 |
| AF233439.1 | 101.486476 | -3.722113774 | 0.73780782 | -5.0448283 | 4.54E-07 | 0.00010484 |
| DEFB1 | 416.205538 | -1.344312985 | 0.62043229 | -2.166736 | 0.03025499 | 0.27663612 |
| FAM85B | 58.6693394 | 0.163930163 | 0.43447027 | 0.37731043 | 0.70594292 | 0.93407096 |
| MFHAS1 | 46.724534 | -2.312305347 | 0.56729177 | -4.0760425 | 4.58E-05 | 0.00379781 |
| ERI1 | 31.8106898 | -0.688677779 | 0.42195084 | -1.632128 | 0.10265253 | 0.49342395 |
| TNKS | 258.020332 | -0.376803652 | 0.22647837 | -1.6637512 | 0.09616217 | 0.48180425 |
| MSRA | 108.827452 | -0.263226168 | 0.31673136 | -0.8310707 | 0.40593367 | 0.8115705 |
| XKR6 | 209.397289 | -0.686370665 | 0.39826655 | -1.7233952 | 0.08481708 | 0.45956155 |
| MTMR9 | 49.5350221 | -0.006776402 | 0.24806016 | -0.0273176 | 0.97820644 | 0.99609121 |
| FDFT1 | 168.335217 | -1.396787344 | 0.55427692 | -2.5200172 | 0.01173491 | 0.16123037 |
| CTSB | 123.907279 | -1.110682166 | 0.47954245 | -2.316129 | 0.02055123 | 0.22428714 |
| FAM66D | 15.8966353 | 0.91377084 | 0.63994944 | 1.42787974 | 0.15332648 | 0.59253658 |
| LONRF1 | 47.2391096 | -0.835118756 | 0.36412318 | -2.2935062 | 0.02181888 | 0.23105147 |
| TRMT9B | 156.316901 | -0.573391469 | 0.44297007 | -1.2944249 | 0.19551871 | 0.6459894 |
| DLC1 | 136.552229 | 1.920518868 | 0.48828338 | 3.9332055 | 8.38E-05 | 0.00564208 |
| TUSC3 | 64.1549003 | -0.868315347 | 0.46259096 | -1.8770694 | 0.06050858 | 0.39102814 |
| MICU3 | 62.7438644 | 0.010403702 | 0.26477403 | 0.03929276 | 0.96865698 | 0.99441717 |
| ZDHHC2 | 87.4268652 | -0.340616314 | 0.26232627 | -1.2984453 | 0.19413436 | 0.6455015 |
| CNOT7 | 30.792243 | 0.175879402 | 0.34013305 | 0.51709001 | 0.60509333 | 0.89600432 |
| VPS37A | 57.0634228 | 0.062678902 | 0.28169095 | 0.22250946 | 0.82391731 | 0.96539788 |
| MTUS1 | 320.61795 | 0.172410806 | 0.23613261 | 0.730144 | 0.46530217 | 0.84143009 |
| AC027117.2 | 26.3226687 | 0.000796106 | 0.81279548 | 0.00097947 | 0.9992185 | 0.99982181 |
| PCM1 | 264.794277 | 0.12459344 | 0.20928851 | 0.59531908 | 0.55163021 | 0.87546024 |
| ASAH1 | 72.0656973 | -0.598105027 | 0.3233806 | -1.849539 | 0.06438002 | 0.40281695 |
| PSD3 | 1463.62212 | -1.420305416 | 0.45387192 | -3.1293089 | 0.00175218 | 0.04734303 |
| AC100800.1 | 18.8824355 | -1.834795222 | 0.58963634 | -3.1117404 | 0.00185988 | 0.04915759 |
| SH2D4A | 193.488195 | 0.049295918 | 0.37852131 | 0.13023287 | 0.89638219 | 0.97885533 |
| CSGALNACT1 | 128.644598 | -1.147971586 | 0.54891637 | -2.0913415 | 0.03649746 | 0.30603532 |
| INTS10 | 96.171854 | -0.149854103 | 0.23953662 | -0.6256 | 0.53157736 | 0.86840264 |
| ATP6V1B2 | 27.2715302 | -0.474009773 | 0.44943702 | -1.0546745 | 0.29157421 | 0.73994767 |
| XPO7 | 66.2445583 | -0.032296373 | 0.19746505 | -0.1635549 | 0.87008156 | 0.97434023 |
| DMTN | 75.0172378 | 0.275324272 | 0.20929755 | 1.31546822 | 0.18835259 | 0.64016122 |
| FAM160B2 | 32.5907157 | -0.676242263 | 0.32264064 | -2.0959612 | 0.03608563 | 0.30384602 |
| SLC39A14 | 28.8408887 | -0.77497327 | 0.35066339 | -2.2100205 | 0.02710374 | 0.25953238 |
| PPP3CC | 141.723442 | -0.086747636 | 0.22603467 | -0.3837802 | 0.70114135 | 0.93127962 |
| CCAR2 | 18.6317657 | 0.251162762 | 0.36855984 | 0.68147079 | 0.49557364 | 0.85814183 |
| BIN3 | 40.8139704 | -0.167358623 | 0.32975469 | -0.5075246 | 0.61178676 | 0.90011356 |
| TNFRSF10B | 77.205804 | -0.568455805 | 0.21133917 | -2.6897797 | 0.00714992 | 0.12146099 |
| TNFRSF10D | 22.950076 | -0.669785505 | 0.37441511 | -1.7888848 | 0.07363336 | 0.42788861 |
| TNFRSF10A | 18.7651318 | -0.948600175 | 0.54277466 | -1.7476869 | 0.08051825 | 0.4486083 |
| CHMP7 | 19.4260557 | 0.138587285 | 0.36027891 | 0.38466666 | 0.7004844 | 0.93127962 |
| R3HCC1 | 10.5958227 | -0.137549071 | 0.40437672 | -0.3401508 | 0.73374296 | NA |
| ENTPD4 | 52.3392062 | -0.353209702 | 0.25945865 | -1.3613333 | 0.17340839 | 0.61802607 |
| SLC25A37 | 79.9473569 | -0.918681445 | 0.46120895 | -1.9918986 | 0.04638219 | 0.34344063 |
| DOCK5 | 268.986903 | -1.57758152 | 0.38963659 | -4.0488537 | 5.15E-05 | 0.00412604 |
| KCTD9 | 61.6748635 | -0.044213671 | 0.2872264 | -0.1539332 | 0.87766243 | 0.97542218 |
| EBF2 | 115.558457 | 0.155074352 | 0.48030105 | 0.32286907 | 0.7467944 | 0.94618979 |
| PPP2R2A | 197.168296 | 0.180854112 | 0.23923257 | 0.75597613 | 0.44966351 | 0.83118301 |
| BNIP3L | 78.9544341 | -1.59536859 | 0.47198484 | -3.3801268 | 0.00072452 | 0.02606817 |
| TRIM35 | 16.6705852 | 0.581665117 | 0.40061955 | 1.45191395 | 0.14652553 | 0.58349206 |
| PTK2B | 123.06902 | -0.459210568 | 0.25300305 | -1.8150397 | 0.06951781 | 0.41827125 |
| EPHX2 | 37.3858368 | 0.585804655 | 0.4609833 | 1.27077197 | 0.20380978 | 0.65771647 |
| CLU | 115.117442 | -1.041151193 | 0.39386317 | -2.6434337 | 0.00820698 | 0.13111608 |
| CCDC25 | 33.9089478 | -0.015003722 | 0.28012985 | -0.0535599 | 0.95728582 | 0.99236316 |
| ELP3 | 276.891191 | 0.641203697 | 0.60856286 | 1.05363594 | 0.29204963 | 0.73999615 |
| ZNF395 | 26.1420173 | -0.681012983 | 0.39155135 | -1.7392686 | 0.08198752 | 0.45345012 |
| FBXO16 | 22.183318 | 0.486551094 | 0.37109282 | 1.31113045 | 0.18981369 | 0.64181696 |
| FZD3 | 34.6360319 | 0.488209691 | 0.41753635 | 1.16926274 | 0.24229779 | 0.69596937 |
| EXTL3 | 26.511417 | 0.026520502 | 0.46858892 | 0.05659652 | 0.95486661 | 0.99228059 |
| INTS9 | 115.113419 | 0.197524881 | 0.21563671 | 0.91600769 | 0.35966286 | 0.7850252 |
| HMBOX1 | 323.99639 | 0.373816464 | 0.19282905 | 1.93858996 | 0.05255129 | 0.36358594 |
| KIF13B | 412.522896 | -0.94964235 | 0.43844386 | -2.1659383 | 0.0303159 | 0.27689666 |
| AC108449.1 | 24.2596582 | -0.917088047 | 0.73415194 | -1.2491802 | 0.21159919 | 0.66359913 |
| SARAF | 123.782852 | 0.002737707 | 0.25810965 | 0.01060676 | 0.99153719 | 0.99923337 |
| LEPROTL1 | 38.539807 | -0.210448275 | 0.34497538 | -0.6100385 | 0.54183629 | 0.87175518 |
| DCTN6 | 22.3000937 | -0.170167132 | 0.32343918 | -0.5261179 | 0.59880632 | 0.89476262 |
| RBPMS | 1589.54732 | -1.026948975 | 0.27477078 | -3.7374752 | 0.00018588 | 0.0103617 |
| GTF2E2 | 73.5676695 | -0.270229473 | 0.23509791 | -1.1494337 | 0.25037717 | 0.70453262 |
| GSR | 102.640321 | -0.056989318 | 0.26947508 | -0.2114827 | 0.83251063 | 0.96701848 |

| | | | | | | |
|---|---|---|---|---|---|---|
| UBXN8 | 12.7815777 | 0.074119842 | 0.41714215 | 0.17768485 | 0.85897048 | 0.97287712 |
| PPP2CB | 139.713098 | -0.234023272 | 0.19628424 | -1.1922672 | 0.23315648 | 0.68763965 |
| WRN | 106.414396 | 0.09365442 | 0.22221775 | 0.42145337 | 0.67342406 | 0.92189158 |
| NRG1 | 141.184682 | -0.31950887 | 0.95833346 | -0.3334005 | 0.73883197 | 0.94376279 |
| FUT10 | 51.1163393 | 0.289796289 | 0.30430037 | 0.9523363 | 0.34092645 | 0.77193691 |
| TTI2 | 22.0465787 | -0.064581569 | 0.35015725 | -0.1844359 | 0.85367151 | 0.97133298 |
| MAK16 | 11.4164446 | 0.580061901 | 0.42438444 | 1.36683123 | 0.17167821 | NA |
| RNF122 | 15.6537424 | -0.13339393 | 0.5153615 | -0.2588357 | 0.79576205 | 0.95848876 |
| UNC5D | 35.0356178 | 0.678423474 | 0.29253936 | 2.31908439 | 0.02039046 | 0.22348867 |
| ERLIN2 | 37.6503684 | 0.346780481 | 0.28124741 | 1.23300863 | 0.21757255 | 0.67202699 |
| PLPBP | 32.6982612 | 0.071755285 | 0.31527209 | 0.22759796 | 0.81995881 | 0.96389705 |
| RAB11FIP1 | 81.3328717 | -0.650405049 | 0.24326075 | -2.673695 | 0.00750207 | 0.1252511 |
| ASH2L | 42.1255112 | 0.126734873 | 0.26782971 | 0.47319199 | 0.63607621 | 0.90881415 |
| LSM1 | 27.7935489 | -0.109717344 | 0.37556096 | -0.2921426 | 0.77017762 | 0.9522529 |
| BAG4 | 52.0861274 | 0.273054943 | 0.21296127 | 1.28218125 | 0.19977907 | 0.65269687 |
| DDHD2 | 64.9071687 | 0.261804687 | 0.2250278 | 1.16343266 | 0.24465401 | 0.69798349 |
| PLPP5 | 46.6047955 | -0.052195813 | 0.35619911 | -0.1465355 | 0.88349867 | 0.97662748 |
| NSD3 | 192.978571 | -0.263885361 | 0.19783648 | -1.3338559 | 0.18225108 | 0.63068306 |
| FGFR1 | 77.5738881 | -0.051810039 | 0.56225457 | -0.0921469 | 0.92658129 | 0.98672316 |
| TACC1 | 124.530463 | 0.231603132 | 0.5873769 | 0.39430071 | 0.69335904 | 0.92826395 |
| PLEKHA2 | 59.7883774 | -0.236043716 | 0.31842475 | -0.7412857 | 0.45852023 | 0.83645518 |
| TM2D2 | 18.8707405 | 0.078454394 | 0.38977351 | 0.20128201 | 0.84047807 | 0.96748214 |
| ADAM9 | 93.2817002 | -0.196192268 | 0.2249329 | -0.8722257 | 0.38308524 | 0.7980695 |
| TCIM | 162.691719 | -2.463298241 | 0.63597518 | -3.8732616 | 0.00010739 | 0.00685308 |
| GOLGA7 | 23.5076691 | 0.355451156 | 0.33839044 | 1.05041726 | 0.29352631 | 0.74013384 |
| GPAT4 | 54.1106307 | -0.442993284 | 0.41565765 | -1.0657648 | 0.28652998 | 0.7371955 |
| ANK1 | 18.153455 | -3.706478458 | 0.75396075 | -4.9160098 | 8.83E-07 | 0.00017849 |
| KAT6A | 223.107455 | -0.213400169 | 0.20451581 | -1.0434409 | 0.29674412 | 0.74160729 |
| AP3M2 | 18.6116754 | 0.545191632 | 0.4445992 | 1.2262542 | 0.22010303 | 0.67350139 |
| IKBKB | 79.4704052 | -0.433241694 | 0.24073187 | -1.7996857 | 0.07191029 | 0.42325835 |
| POLB | 21.6855565 | 0.356490716 | 0.44661909 | 0.79819855 | 0.42475528 | 0.82241967 |
| VDAC3 | 79.7086365 | -0.381634638 | 0.37027527 | -1.0306782 | 0.30269177 | 0.74712557 |
| SLC20A2 | 174.414094 | -0.387740676 | 0.25774349 | -1.5043665 | 0.13248703 | 0.55997449 |
| SMIM19 | 36.2020341 | -0.34028733 | 0.38527911 | -0.8832229 | 0.37711585 | 0.79432848 |
| RNF170 | 47.935323 | -0.188543878 | 0.32121885 | -0.5869639 | 0.55722793 | 0.87887982 |
| HOOK3 | 93.3886122 | -0.009881235 | 0.20096084 | -0.04917 | 0.96078386 | 0.99238848 |
| FNTA | 33.7630173 | -0.494527178 | 0.37901155 | -1.3047813 | 0.19196733 | 0.6421543 |
| HGSNAT | 114.44584 | -0.242702725 | 0.21063518 | -1.1522421 | 0.2492216 | 0.70371066 |
| SPIDR | 375.067507 | -0.252336416 | 0.26248924 | -0.961321 | 0.3363908 | 0.76884797 |
| CEBPD | 285.414235 | -1.805933569 | 0.48950861 | -3.6892784 | 0.00022489 | 0.01172819 |
| PRKDC | 109.260993 | 0.097719848 | 0.27981415 | 0.34923126 | 0.7269157 | 0.94009907 |
| UBE2V2 | 51.5702277 | -0.037926847 | 0.29795792 | -0.1272893 | 0.89871145 | 0.97908342 |
| PCMTD1 | 216.957022 | -0.107245445 | 0.18584678 | -0.5770638 | 0.56389637 | 0.88191823 |
| AC064807.1 | 12.3411628 | 0.091805164 | 0.47879541 | 0.19174195 | 0.84794435 | NA |
| RB1CC1 | 175.138298 | -0.456267889 | 0.29360778 | -1.5540047 | 0.12018329 | 0.53231866 |
| ATP6V1H | 333.750914 | 0.686448557 | 0.25846634 | 2.65585283 | 0.00791081 | 0.12896619 |
| TCEA1 | 66.717932 | 0.24296001 | 0.24628522 | 0.98649853 | 0.32388854 | 0.76304922 |
| LYPLA1 | 173.281532 | -0.127528347 | 0.17262009 | -0.7387805 | 0.46004032 | 0.83690755 |
| MRPL15 | 44.6624382 | 0.172414889 | 0.28078447 | 0.61404709 | 0.53918421 | 0.86972442 |
| TMEM68 | 36.4627052 | 0.472945659 | 0.32383875 | 1.46043565 | 0.14417038 | 0.57846905 |
| TGS1 | 55.4234299 | 0.343572864 | 0.38400147 | 0.89471758 | 0.37093808 | 0.7900012 |
| LYN | 46.5111154 | -0.986061539 | 0.37230159 | -2.6485558 | 0.00808365 | 0.13068568 |
| RPS20 | 130.252492 | 0.336334382 | 0.42095228 | 0.79898458 | 0.42429935 | 0.82231588 |
| PLAG1 | 84.6608958 | -0.536443869 | 0.380671 | -1.409206 | 0.15877425 | 0.59926468 |
| CHCHD7 | 28.6409143 | 0.313439773 | 0.36326984 | 0.86282907 | 0.38823145 | 0.80109289 |
| IMPAD1 | 64.1030212 | 0.469038604 | 0.2637089 | 1.77862259 | 0.07530166 | 0.43246068 |
| LINC01606 | 77.8502364 | 0.960015727 | 0.46669919 | 2.05703321 | 0.03968303 | 0.31970549 |
| FAM110B | 191.484712 | 0.68308693 | 0.26882688 | 2.54099195 | 0.01105385 | 0.15593979 |
| AC027698.1 | 15.5854209 | 0.419008991 | 0.56049231 | 0.74757313 | 0.45471767 | 0.83385406 |
| UBXN2B | 55.5014372 | -0.001707707 | 0.25325819 | -0.0067429 | 0.99461995 | 0.99923337 |
| SDCBP | 71.690378 | -0.404564611 | 0.27186816 | -1.4880912 | 0.13672685 | 0.5681351 |
| NSMAF | 43.8726448 | 0.014409542 | 0.29112627 | 0.04949585 | 0.96052415 | 0.99238848 |
| TOX | 102.109945 | 0.144960508 | 0.72427854 | 0.2001447 | 0.84136742 | 0.9677342 |
| CA8 | 521.223365 | 0.264256377 | 0.58834862 | 0.44914931 | 0.65332395 | 0.9158679 |
| RAB2A | 192.191984 | -0.061462572 | 0.19701275 | -0.3119726 | 0.75506138 | 0.94795939 |
| CHD7 | 77.6352476 | -0.893854212 | 0.36662058 | -2.4380906 | 0.01476507 | 0.18456339 |
| ASPH | 150.871161 | 0.100825246 | 0.34974083 | 0.2882856 | 0.77312813 | 0.95447666 |
| YTHDF3 | 126.125044 | 0.231196659 | 0.2733702 | 0.84572736 | 0.39770485 | 0.80705797 |
| CYP7B1 | 37.5336454 | 0.684652921 | 0.40948058 | 1.6720034 | 0.09452365 | 0.47996672 |
| ARMC1 | 34.4235176 | 0.651838241 | 0.27605333 | 2.36127654 | 0.01821214 | 0.20980736 |
| MTFR1 | 63.1171419 | -0.46151688 | 0.33634829 | -1.3721398 | 0.17001992 | 0.61252734 |
| PDE7A | 87.5886414 | 0.173027543 | 0.29344005 | 0.58965211 | 0.55542391 | 0.87787661 |
| ADHFE1 | 40.4390442 | 0.343890637 | 0.44685836 | 0.76957413 | 0.44155255 | 0.8281114 |
| VCPIP1 | 41.3834934 | -0.241059135 | 0.26696878 | -0.9029488 | 0.36655308 | 0.78802412 |
| SGK3 | 151.491702 | 0.091278154 | 0.25729159 | 0.3547654 | 0.72276535 | 0.93873366 |
| CSPP1 | 112.559631 | 0.339104334 | 0.26980819 | 1.25683486 | 0.20881344 | 0.66041421 |
| ARFGEF1 | 410.861207 | -0.134159963 | 0.19619448 | -0.6838111 | 0.49409446 | 0.85774549 |
| NCOA2 | 668.03685 | -0.144481536 | 0.18967167 | -0.7617455 | 0.44621192 | 0.83035712 |
| TRAM1 | 128.208884 | -0.260755445 | 0.15716951 | -1.6590715 | 0.09710139 | 0.48257296 |
| LACTB2 | 38.8414801 | -1.17380283 | 0.31709109 | -3.7017843 | 0.00021409 | 0.01141021 |
| TERF1 | 35.4644502 | 0.147054168 | 0.26331353 | 0.55847555 | 0.57651969 | 0.88809976 |
| RPL7 | 219.280522 | 0.624495351 | 0.46457411 | 1.34423191 | 0.1788734 | 0.62639418 |
| RDH10 | 62.0398879 | 0.044842241 | 0.58781374 | 0.07628648 | 0.93919118 | 0.98871562 |
| RDH10-AS1 | 25.6438193 | 0.109750398 | 0.64292428 | 0.17070501 | 0.86445572 | 0.97407858 |
| STAU2 | 172.247499 | 0.182730347 | 0.23182293 | 0.78823242 | 0.43056077 | 0.82469033 |
| AC027018.1 | 30.2107824 | -0.585473479 | 0.67365257 | -0.869103 | 0.38479079 | 0.79869171 |
| ELOC | 52.7635197 | 0.132045951 | 0.2867355 | 0.46051483 | 0.64514673 | 0.9122579 |
| TMEM70 | 22.8120569 | -0.97042351 | 0.57928993 | -1.6751949 | 0.09389601 | 0.4781144 |
| JPH1 | 20.3767174 | -0.075776245 | 0.49451438 | -0.1532337 | 0.87821402 | 0.97542218 |
| GDAP1 | 39.3097846 | 0.123004231 | 0.48220739 | 0.25508574 | 0.79865688 | 0.95848876 |
| HNF4G | 51.4832233 | -0.377945973 | 0.69975325 | -0.5401132 | 0.58911896 | 0.89133386 |
| PEX2 | 50.2376886 | -0.189468753 | 0.32143689 | -0.5894431 | 0.55556408 | 0.87787661 |
| PKIA | 41.8764715 | 0.244272255 | 0.30579936 | 0.79879911 | 0.42440691 | 0.82231588 |
| ZC2HC1A | 79.1566676 | -0.337595044 | 0.29096122 | -1.160275 | 0.24593685 | 0.69938066 |
| IL7 | 48.0280682 | -1.443028171 | 0.44991289 | -3.2073502 | 0.00133964 | 0.03949693 |
| TPD52 | 164.773605 | -0.261316963 | 0.26283734 | -0.9942155 | 0.32011795 | 0.76106474 |
| ZBTB10 | 52.4492497 | 0.236625425 | 0.27122448 | 0.87243387 | 0.38297173 | 0.7980695 |
| ZNF704 | 153.583537 | 0.557630043 | 0.36975482 | 1.50810758 | 0.131527 | 0.55834217 |
| PAG1 | 25.4527523 | 0.056384839 | 0.63507846 | 0.08878405 | 0.92925353 | 0.98733028 |
| IMPA1 | 43.5272487 | 0.171173493 | 0.28141609 | 0.60825766 | 0.5430166 | 0.87220748 |
| ZFAND1 | 49.812994 | -0.20687631 | 0.26885315 | -0.769477 | 0.44161021 | 0.8281114 |
| CHMP4C | 18.4681124 | -0.098544956 | 0.39253323 | -0.2510487 | 0.80177646 | 0.95890203 |
| SNX16 | 28.8376361 | -0.317597056 | 0.28918027 | -1.0982667 | 0.27208806 | 0.7232847 |
| LRRCC1 | 11.0027921 | 0.339285706 | 0.43882022 | 0.77317701 | 0.43941763 | NA |
| E2F5 | 29.0019321 | 0.390483361 | 0.33142347 | 1.1782007 | 0.23871661 | 0.69356592 |
| C8orf59 | 34.1452419 | -0.523067083 | 0.35626294 | -1.4682052 | 0.1420485 | 0.57673923 |
| CA13 | 44.306518 | -0.139979763 | 0.28654386 | -0.4885108 | 0.6251881 | 0.90447745 |
| CA1 | 15.8808036 | -0.67487594 | 0.66235893 | -1.0188976 | 0.30825157 | 0.7512664 |
| CA3 | 46.7992828 | -0.538375571 | 0.4022801 | -1.3383102 | 0.18079533 | 0.6286114 |
| CA2 | 214.618002 | 0.875040853 | 0.46406946 | 1.88558166 | 0.05935135 | 0.3871608 |
| ATP6V0D2 | 1534.99309 | 0.761762098 | 0.32922035 | 2.31383662 | 0.02067669 | 0.22444738 |
| AC023194.3 | 255.238385 | -0.064571574 | 0.51336513 | -0.125781 | 0.89990529 | 0.97940229 |
| WWP1 | 534.305048 | -0.114783358 | 0.18515594 | -0.619928 | 0.53530517 | 0.86946459 |

| | | | | | | |
|---|---|---|---|---|---|---|
| RMDN1 | 78.6165776 | -0.09487361 | 0.26928052 | -0.3523226 | 0.72459635 | 0.93935288 |
| CPNE3 | 71.3846297 | 0.178692191 | 0.2317994 | 0.77089151 | 0.44077124 | 0.82793978 |
| CNGB3 | 13.8967165 | -0.45131689 | 0.65588192 | -0.6881069 | 0.49138545 | 0.85654883 |
| AF117829.1 | 133.028192 | -0.151106276 | 0.2705782 | -0.558457 | 0.57653239 | 0.88809976 |
| RIPK2 | 21.1835381 | -0.895405226 | 0.32260686 | -2.7755306 | 0.00551117 | 0.10201977 |
| OSGIN2 | 30.88626 | -0.40893449 | 0.40563386 | -1.008137 | 0.31338871 | 0.75618668 |
| NBN | 40.8424286 | -0.320142 | 0.33441173 | -0.9573289 | 0.33840129 | 0.76998177 |
| DECR1 | 140.440173 | -0.189757148 | 0.33159282 | -0.5722595 | 0.56714617 | 0.88417195 |
| CALB1 | 35.8704926 | 3.554039377 | 1.3208385 | 2.69074484 | 0.00712927 | 0.12146099 |
| PIP4P2 | 39.4481393 | 0.092803636 | 0.39235182 | 0.23653168 | 0.81302013 | 0.96262567 |
| OTUD6B-AS1 | 51.8666044 | 0.22806525 | 0.35155236 | 0.64873764 | 0.51650797 | 0.86500817 |
| OTUD6B | 38.3467799 | 0.103021117 | 0.35304287 | 0.29180908 | 0.7704326 | 0.9522529 |
| LRRC69 | 49.2387987 | -0.062523374 | 0.56408967 | -0.1108394 | 0.91174368 | 0.9835359 |
| SLC26A7 | 3843.14886 | 1.849340924 | 0.70990999 | 2.60503578 | 0.00918647 | 0.13988811 |
| TRIQK | 56.1634694 | -0.305465011 | 0.32248346 | -0.9472269 | 0.34352316 | 0.77370523 |
| FAM92A | 15.379261 | -0.191223519 | 0.4362424 | -0.4383424 | 0.66113813 | 0.91798452 |
| RBM12B | 42.7317961 | -0.307605364 | 0.24323348 | -1.2646506 | 0.2059966 | 0.65836247 |
| TMEM67 | 18.8552844 | -0.173856631 | 0.33364109 | -0.5210888 | 0.60230492 | 0.89551774 |
| PDP1 | 73.2083823 | -0.165097221 | 0.34253787 | -0.4819824 | 0.62981845 | 0.90735764 |
| VIRMA | 57.129305 | 0.181980835 | 0.21070252 | 0.8636861 | 0.38776035 | 0.80079761 |
| ESRP1 | 70.4507571 | -0.297982145 | 0.25668901 | -1.1608684 | 0.24569544 | 0.69938066 |
| DPY19L4 | 55.3830656 | 0.549704966 | 0.2761454 | 1.990636 | 0.04652092 | 0.34368082 |
| INTS8 | 56.376031 | -0.073734211 | 0.33658817 | -0.2190636 | 0.82660051 | 0.96542185 |
| NDUFAF6 | 53.6898747 | 0.135077291 | 0.22429155 | 0.60223977 | 0.54701455 | 0.87284769 |
| TP53INP1 | 41.903961 | -0.064189172 | 0.29347211 | -0.2187232 | 0.82686564 | 0.96542185 |
| PLEKHF2 | 26.3912814 | -0.046264026 | 0.28550437 | -0.1620431 | 0.87127188 | 0.97434023 |
| C8orf37-AS1 | 35.7845263 | 0.135857142 | 0.48199982 | 0.2818614 | 0.77804979 | 0.95532696 |
| UQCRB | 172.169511 | 0.197578332 | 0.36291465 | 0.54442094 | 0.58615184 | 0.89059199 |
| MTERF3 | 18.1555192 | 0.107983888 | 0.33364957 | 0.32364462 | 0.7462071 | 0.94614106 |
| PTDSS1 | 18.9109825 | 0.304637316 | 0.33195798 | 0.91769843 | 0.35887677 | 0.78434408 |
| SDC2 | 220.834727 | 0.761258589 | 0.27301645 | 2.78832502 | 0.00529814 | 0.10037484 |
| CPQ | 339.329067 | 0.50589664 | 0.30553472 | 1.65577465 | 0.09776746 | 0.48351985 |
| TSPYL5 | 24.2101505 | 0.301108356 | 0.27022537 | 1.11428604 | 0.2651565 | 0.71643955 |
| MTDH | 209.604298 | -0.059487319 | 0.24159134 | -0.2462312 | 0.80550329 | 0.96005568 |
| LAPTM4B | 165.344159 | -0.483000389 | 0.33340401 | -1.448694 | 0.14742305 | 0.58421969 |
| RPL30 | 191.618218 | 0.258746396 | 0.45723153 | 0.56589797 | 0.57146314 | 0.88578992 |
| RIDA | 12.8504679 | 0.476538793 | 0.53552013 | 0.88986159 | 0.37354021 | 0.79181382 |
| POP1 | 12.1572705 | 0.691396652 | 0.42079781 | 1.64306144 | 0.10037022 | NA |
| NIPAL2 | 47.1145605 | 0.130573877 | 0.38455813 | 0.33954263 | 0.73420099 | 0.94277981 |
| STK3 | 820.504968 | -0.272103478 | 0.19382614 | -1.4038534 | 0.16036252 | 0.60024417 |
| AC104986.2 | 22.5449885 | -0.022535192 | 0.28612083 | -0.0787611 | 0.93722264 | 0.98871562 |
| VPS13B | 406.66571 | -0.191393179 | 0.18396223 | -1.0403939 | 0.29815694 | 0.74290325 |
| COX6C | 196.018179 | 0.189657739 | 0.26712065 | 0.71000777 | 0.47769932 | 0.84771464 |
| POLR2K | 32.3513768 | 0.18805657 | 0.435525 | 0.43179283 | 0.66589199 | 0.91958319 |
| SPAG1 | 42.2791041 | -0.566368579 | 0.28816767 | -1.9654134 | 0.04936642 | 0.35549683 |
| RNF19A | 148.965634 | -0.283651069 | 0.26701717 | -1.0622953 | 0.28810167 | 0.73803612 |
| ANKRD46 | 30.458312 | 0.597290047 | 0.32728809 | 1.82496726 | 0.06800597 | 0.4151827 |
| PABPC1 | 148.103861 | -0.12022378 | 0.31571285 | -0.380801 | 0.70335089 | 0.93304742 |
| YWHAZ | 254.9845 | 0.082973424 | 0.25777432 | 0.32188397 | 0.74754059 | 0.9462156 |
| ZNF706 | 63.6765538 | -0.203649692 | 0.31372036 | -0.649144 | 0.5162453 | 0.86486692 |
| GRHL2 | 201.76053 | -0.040705569 | 0.33257989 | -0.1223934 | 0.9025875 | 0.98016176 |
| NCALD | 34.4905141 | 1.279024381 | 0.52248576 | 2.44796026 | 0.01436675 | 0.18028134 |
| RRM2B | 65.4856765 | -0.278493744 | 0.31046678 | -0.8970162 | 0.36971026 | 0.78990958 |
| UBR5-AS1 | 22.217675 | -0.926205654 | 0.56593622 | -1.6365902 | 0.10171613 | 0.49220184 |
| UBR5 | 306.883874 | 0.064648493 | 0.18805197 | 0.34377992 | 0.7310118 | 0.94155624 |
| KLF10 | 48.698619 | -0.373628968 | 0.40171193 | -0.9300918 | 0.35232356 | 0.77964174 |
| AZIN1 | 85.4308792 | -0.187892584 | 0.27936569 | -0.6725686 | 0.5012218 | 0.85942996 |
| AZIN1-AS1 | 89.669206 | -0.452272317 | 0.35034257 | -1.2909431 | 0.19672341 | 0.64685325 |
| AP003354.2 | 14.4785235 | -0.015782578 | 0.48853885 | -0.0323057 | 0.97422828 | 0.9953169 |
| ATP6V1C1 | 79.5055918 | -0.448684244 | 0.258655 | -1.7346823 | 0.0827971 | 0.45496557 |
| BAALC | 89.0119144 | -1.754362181 | 0.60334115 | -2.907745 | 0.00364045 | 0.07820893 |
| BAALC-AS1 | 32.9336919 | -0.559381609 | 0.33542451 | -1.6676826 | 0.09537876 | 0.48081904 |
| FZD6 | 107.206952 | -0.096227431 | 0.22368432 | -0.430193 | 0.66705524 | 0.92032206 |
| SLC25A32 | 41.1328668 | 0.146882569 | 0.28324178 | 0.51857663 | 0.60405601 | 0.8958512 |
| DCAF13 | 21.2239604 | 0.114619507 | 0.38646865 | 0.29658164 | 0.76678593 | 0.95205353 |
| RIMS2 | 62.5353393 | -1.062705825 | 0.5743507 | -1.8502734 | 0.06427415 | 0.40275147 |
| LRP12 | 35.2613786 | 0.393302328 | 0.31745938 | 1.238906 | 0.21538031 | 0.66933563 |
| ZFPM2 | 21.7553141 | -0.659381156 | 0.57272123 | -1.1513126 | 0.24960366 | 0.70382428 |
| AC027031.2 | 20.5261229 | -0.210822282 | 0.42303795 | -0.4983531 | 0.61823517 | 0.90246519 |
| OXR1 | 1151.32614 | 0.35740648 | 0.29724568 | 1.20239418 | 0.22921084 | 0.68503082 |
| AC090579.1 | 110.758783 | -0.740325992 | 0.52980062 | -1.3973672 | 0.16230318 | 0.60094226 |
| ANGPT1 | 51.9406086 | 1.269950596 | 1.05463657 | 1.20415945 | 0.22852795 | 0.68375113 |
| EIF3E | 94.710159 | 0.055485383 | 0.19317739 | 0.28722504 | 0.77394001 | 0.95457609 |
| EMC2 | 79.9538052 | -0.014212623 | 0.30313478 | -0.0468855 | 0.96260449 | 0.99306991 |
| NUDCD1 | 33.4525578 | 0.325334034 | 0.30303543 | 1.07358415 | 0.28300911 | 0.7344003 |
| ENY2 | 50.6176141 | -0.090112896 | 0.34380264 | -0.2621065 | 0.79323936 | 0.95761317 |
| EBAG9 | 31.4649181 | -0.011095606 | 0.29452002 | -0.0376735 | 0.96994799 | 0.99486579 |
| TRPS1 | 330.106986 | 0.058597549 | 0.23786968 | 0.24634308 | 0.80541667 | 0.96005568 |
| EIF3H | 149.150389 | -0.105960402 | 0.18640222 | -0.5684503 | 0.56972922 | 0.88562534 |
| UTP23 | 22.9860488 | 0.45228664 | 0.34919392 | 1.29523058 | 0.1952407 | 0.6459894 |
| RAD21 | 67.8684995 | -0.063870171 | 0.22933475 | -0.2785019 | 0.78062708 | 0.95537097 |
| MED30 | 15.5186991 | 0.323516981 | 0.40988629 | 0.7892847 | 0.42994562 | 0.82469033 |
| EXT1 | 456.825949 | 0.345745735 | 0.23513494 | 1.47041414 | 0.14144962 | 0.57625424 |
| SAMD12 | 695.659465 | -0.657532839 | 0.26320395 | -2.4981876 | 0.01248301 | 0.16647826 |
| AC023590.1 | 75.0127356 | -1.178669219 | 0.5743625 | -2.0521347 | 0.04015657 | 0.32138511 |
| SAMD12-AS1 | 49.6139028 | -1.025827376 | 0.45970772 | -2.2314774 | 0.02564952 | 0.25207739 |
| MAL2 | 71.8112962 | 0.159405113 | 0.39028873 | 0.40842868 | 0.68295898 | 0.92474072 |
| TAF2 | 89.4397792 | 0.116662057 | 0.23382032 | 0.49893892 | 0.61782242 | 0.90214869 |
| DEPTOR | 135.927088 | -1.50096044 | 0.47721462 | -3.1452524 | 0.00165944 | 0.04508831 |
| MRPL13 | 46.7596481 | 0.013848354 | 0.29476528 | 0.04698095 | 0.96252841 | 0.99306991 |
| MTBP | 32.3584228 | -0.562723958 | 0.43895085 | -1.2819749 | 0.19985146 | 0.65271352 |
| SNTB1 | 1704.03353 | -0.845344016 | 0.2705495 | -3.1245448 | 0.0017808 | 0.04771768 |
| AC104958.1 | 21.7704771 | -1.507428434 | 0.6678027 | -2.2572961 | 0.02398958 | 0.24264747 |
| AC104958.2 | 18.0739148 | -1.836732039 | 0.45008024 | -4.080899 | 4.49E-05 | 0.003784 |
| ZHX2 | 215.843717 | -0.476623676 | 0.22307864 | -2.1365725 | 0.03263278 | 0.28645971 |
| AC104316.1 | 14.6571504 | -1.383386364 | 0.58173726 | -2.3780261 | 0.0174056 | 0.20451748 |
| DERL1 | 47.3922001 | -0.365155052 | 0.2241663 | -1.6289471 | 0.10332421 | 0.495277 |
| TBC1D31 | 19.9857625 | 0.229468617 | 0.31785358 | 0.72193183 | 0.47033639 | 0.84423815 |
| C8orf76 | 15.928308 | -0.412262481 | 0.47899056 | -0.8606902 | 0.38940869 | 0.80191835 |
| ATAD2 | 20.7628068 | -0.580497729 | 0.43268258 | -1.3416249 | 0.17971764 | 0.62729799 |
| WDYHV1 | 24.9794673 | -0.146199771 | 0.39382326 | -0.3712319 | 0.71046479 | 0.93540281 |
| FBXO32 | 45.112587 | -2.341162797 | 0.63669085 | -3.6770794 | 0.00023592 | 0.01217245 |
| FAM91A1 | 99.8430361 | -0.063972148 | 0.24919747 | -0.2567127 | 0.79740059 | 0.95848876 |
| FER1L6 | 93.8631865 | -0.961702106 | 0.58192453 | -1.6526234 | 0.09840753 | 0.48476221 |
| TMEM65 | 317.22679 | 0.701978569 | 0.28920831 | 2.427242 | 0.0152141 | 0.18728021 |
| RNF139 | 18.4774239 | 0.179082056 | 0.41013914 | 0.43663732 | 0.66237439 | 0.91812398 |
| TATDN1 | 60.6993893 | -0.274858171 | 0.25960438 | -1.0587578 | 0.28971008 | 0.73966536 |
| NDUFB9 | 92.4484396 | -0.227245922 | 0.33384092 | -0.6807012 | 0.49606057 | 0.85814183 |
| MTSS1 | 667.449224 | -1.421600523 | 0.30425736 | -4.672362 | 2.98E-06 | 0.00045129 |
| SQLE | 66.8102785 | -1.340328868 | 0.62345434 | -2.1498429 | 0.03156764 | 0.28099688 |
| WASHC5 | 49.4153706 | -0.357056611 | 0.2472563 | -1.4440749 | 0.14871791 | 0.58569376 |
| NSMCE2 | 348.498493 | -0.081809055 | 0.24787166 | -0.330046 | 0.74136519 | 0.94497272 |
| TRIB1 | 54.4713853 | -0.332057492 | 0.40979896 | -0.8102936 | 0.41777143 | 0.8188001 |

| | | | | | | |
|---|---|---|---|---|---|---|
| FAM84B | 23.0240238 | 0.45165802 | 0.4139634 | 1.09105787 | 0.27524742 | 0.72554623 |
| PCAT1 | 205.238116 | -0.346923837 | 0.41436944 | -0.8372332 | 0.40246152 | 0.80890307 |
| FAM49B | 375.103583 | -0.069339072 | 0.21026585 | -0.3297686 | 0.74157482 | 0.94501909 |
| ASAP1 | 68.6255863 | 0.013732031 | 0.2504368 | 0.05483232 | 0.95627205 | 0.99234582 |
| EFR3A | 128.294721 | -0.205603241 | 0.26087688 | -0.7881236 | 0.43062438 | 0.82469033 |
| LRRC6 | 20.0974171 | 0.539339546 | 0.32672678 | 1.65073567 | 0.09879256 | 0.48476221 |
| PHF20L1 | 93.6463961 | 0.345881405 | 0.25540235 | 1.35426085 | 0.17565317 | 0.62231623 |
| NDRG1 | 286.704753 | -1.520469635 | 0.56297435 | -2.7007796 | 0.00691772 | 0.11939831 |
| ST3GAL1 | 332.96877 | -1.008577312 | 0.50214541 | -2.0085363 | 0.04458633 | 0.33552163 |
| ZFAT | 44.5608164 | -0.209214285 | 0.23912217 | -0.8749263 | 0.381614 | 0.79725517 |
| AC083843.3 | 34.3365826 | -0.161263236 | 0.37995101 | -0.4244317 | 0.67125104 | 0.92170876 |
| KHDRBS3 | 36.4531633 | 1.35850104 | 0.33154472 | 4.09748964 | 4.18E-05 | 0.00364978 |
| TRAPPC9 | 416.762054 | -0.337233954 | 0.20251619 | -1.6652198 | 0.09586892 | 0.48108046 |
| CHRAC1 | 23.5783433 | 0.180137601 | 0.36775969 | 0.48982422 | 0.62425829 | 0.90447745 |
| AGO2 | 105.831875 | -0.224609005 | 0.29743721 | -0.7551476 | 0.4501604 | 0.83118301 |
| PTK2 | 672.32205 | 0.129499447 | 0.19567371 | 0.66181323 | 0.50809093 | 0.86281145 |
| DENND3 | 38.5033463 | -0.779228177 | 0.42766082 | -1.8220705 | 0.06844429 | 0.41572605 |
| SLC45A4 | 62.6433011 | 0.015475648 | 0.31301871 | 0.04944001 | 0.96056864 | 0.99238848 |
| PTP4A3 | 78.7393298 | -1.379738373 | 0.6784785 | -2.0335772 | 0.04199425 | 0.32888549 |
| TSNARE1 | 30.6661984 | -0.059724976 | 0.3250011 | -0.1837685 | 0.85419504 | 0.97135895 |
| JRK | 14.3447094 | 0.882287375 | 0.46808882 | 1.88487173 | 0.05944716 | 0.38732806 |
| THEM6 | 12.9084087 | 0.782791863 | 0.39571828 | 1.9781544 | 0.04791129 | 0.34992596 |
| LY6E | 51.5500053 | -2.503566618 | 1.33680051 | -1.872805 | 0.06109532 | 0.39246665 |
| GLI4 | 13.3152103 | 0.45950444 | 0.4007636 | 1.14657228 | 0.25155844 | 0.70564397 |
| TOP1MT | 11.3958415 | -0.283331768 | 0.39868101 | -0.7106729 | 0.47728699 | NA |
| RHPN1 | 22.2045139 | 0.32961642 | 0.42234544 | 0.78044271 | 0.43513034 | 0.82565889 |
| ZC3H3 | 31.9372714 | 0.276989606 | 0.3925428 | 0.70562906 | 0.48041885 | 0.84944638 |
| GSDMD | 30.8362658 | -0.204969549 | 0.36654977 | -0.5591861 | 0.5760347 | 0.88761502 |
| NAPRT | 11.9178965 | 0.116421332 | 0.59445342 | 0.19584601 | 0.84473069 | NA |
| EEF1D | 160.76854 | 0.086520754 | 0.35027375 | 0.24700896 | 0.80490129 | 0.95997099 |
| TSTA3 | 12.4785426 | -0.428390346 | 0.58285964 | -0.7349803 | 0.4623515 | NA |
| ZNF623 | 16.5676022 | -0.045893355 | 0.33670772 | -0.1363003 | 0.89158391 | 0.97793768 |
| FAM83H | 21.6730713 | -0.045750749 | 0.44075005 | -0.103802 | 0.91732646 | 0.98495314 |
| SCRIB | 27.1836548 | 0.28343113 | 0.29314314 | 0.96686938 | 0.33360934 | 0.76777176 |
| PUF60 | 34.0330992 | -0.305236308 | 0.40731712 | -0.7493825 | 0.45362672 | 0.83364149 |
| NRBP2 | 19.7377696 | -0.300363406 | 0.34125699 | -0.8801678 | 0.37876843 | 0.79535699 |
| PLEC | 37.9846558 | -0.020084697 | 0.2506701 | -0.080124 | 0.93613861 | 0.98871562 |
| PARP10 | 21.9424017 | -0.119273971 | 0.61264204 | -0.1946879 | 0.84563731 | 0.97027687 |
| GRINA | 39.7098343 | 0.22659615 | 0.40480916 | 0.55976043 | 0.57564286 | 0.88746465 |
| OPLAH | 14.2480101 | 0.221220838 | 0.46788007 | 0.47281526 | 0.63634499 | 0.9089304 |
| EXOSC4 | 13.2384886 | -0.159936765 | 0.45550364 | -0.3511207 | 0.72549778 | 0.93968502 |
| GPAA1 | 44.4430537 | 0.415983966 | 0.45448501 | 0.91528644 | 0.36004128 | 0.78516196 |
| CYC1 | 87.7506035 | -0.072671212 | 0.35336123 | -0.205657 | 0.83705885 | 0.96748214 |
| SHARPIN | 27.2951732 | 0.036465262 | 0.36928661 | 0.09874515 | 0.92134062 | 0.98614962 |
| MAF1 | 42.6132107 | -0.548931883 | 0.39551812 | -1.3878805 | 0.16517344 | 0.60578861 |
| MROH1 | 117.319934 | -0.285371212 | 0.33607364 | -0.849133 | 0.3958073 | 0.80624334 |
| HSF1 | 70.816608 | -0.070788611 | 0.26169376 | -0.2705017 | 0.7867743 | 0.95577371 |
| DGAT1 | 18.1020633 | 0.686406947 | 0.43050263 | 1.59443148 | 0.11083943 | 0.51243576 |
| SLC52A2 | 16.5351962 | -0.122524627 | 0.47878426 | -0.2559078 | 0.79802203 | 0.95848876 |
| ADCK5 | 9.75106023 | 0.51251266 | 0.5907068 | 0.86762614 | 0.38559903 | NA |
| CPSF1 | 14.3847509 | 0.20808397 | 0.3987786 | 0.52180325 | 0.60180732 | 0.89522668 |
| SLC39A4 | 17.4298301 | 0.040277152 | 0.51798789 | 0.07775694 | 0.9380214 | 0.98871562 |
| VPS28 | 77.1336148 | -0.224178567 | 0.25806623 | -0.8686862 | 0.38501881 | 0.79869171 |
| CYHR1 | 44.1506798 | 0.757806454 | 0.25541182 | 2.96699839 | 0.00300723 | 0.06784993 |
| KIFC2 | 17.091023 | 0.439764811 | 0.41926568 | 1.04889293 | 0.2942274 | 0.7405942 |
| PPP1R16A | 32.4530398 | 0.121924367 | 0.27985946 | 0.43566283 | 0.66308138 | 0.918408 |
| MFSD3 | 9.14575894 | 1.327878832 | 0.58793455 | 2.25854874 | 0.02391147 | NA |
| ZNF251 | 50.254187 | 0.677274181 | 0.2974799 | 2.2767057 | 0.02280381 | 0.23733575 |
| RPL8 | 372.596068 | 0.350907863 | 0.39921146 | 0.87900247 | 0.37939994 | 0.79535699 |
| ZNF7 | 19.9922721 | 0.266296381 | 0.33276476 | 0.80025415 | 0.42356356 | 0.82204213 |
| COMMD5 | 12.5454506 | 0.054758122 | 0.39710775 | 0.13789235 | 0.89032549 | 0.97782578 |
| ZNF250 | 18.8871522 | 0.726656682 | 0.35781016 | 2.03084417 | 0.0422708 | 0.32888549 |
| ZNF16 | 14.4764011 | 0.581430602 | 0.43230499 | 1.34495465 | 0.17863988 | 0.62639418 |
| C8orf33 | 16.1912616 | 0.05874808 | 0.45438335 | 0.12929188 | 0.89712669 | 0.97889563 |
| WASHC1 | 93.7380184 | 0.31477183 | 0.21294881 | 1.47815726 | 0.13936569 | 0.57281662 |
| CBWD1 | 29.1482295 | 0.387203483 | 0.34042909 | 1.13739833 | 0.25537181 | 0.70849619 |
| DOCK8 | 419.05929 | 0.055415092 | 0.28449048 | 0.19478716 | 0.84555958 | 0.97027687 |
| KANK1 | 224.795562 | 0.09196682 | 0.30215593 | 0.30436874 | 0.76084698 | 0.94990484 |
| LINC01230 | 135.932601 | -0.086009929 | 0.49562661 | -0.1735378 | 0.86222875 | 0.97307493 |
| DMRT2 | 393.336778 | 0.533634691 | 0.25947064 | 2.05662843 | 0.03972198 | 0.31975371 |
| SMARCA2 | 289.692451 | -0.10514444 | 0.25760771 | -0.4081572 | 0.68315827 | 0.92474072 |
| VLDLR-AS1 | 15.9010618 | -0.926252884 | 0.5412239 | -1.7114043 | 0.08700651 | 0.46422617 |
| VLDLR | 24.8775717 | -1.167248142 | 0.5150718 | -2.2661853 | 0.02344004 | 0.23883233 |
| PUM3 | 193.939263 | -1.080052387 | 0.31424557 | -3.4369693 | 0.00058826 | 0.02282458 |
| RFX3 | 147.847739 | -0.009237766 | 0.26449908 | -0.0349255 | 0.97213914 | 0.99526459 |
| RFX3-AS1 | 96.2418221 | -0.336071431 | 0.40280576 | -0.8343263 | 0.40409715 | 0.81027913 |
| GLIS3 | 1055.89435 | -0.531713184 | 0.31699102 | -1.6773762 | 0.09346894 | 0.47792014 |
| SLC1A1 | 17.8580085 | -1.267612707 | 0.6059884 | -2.0918102 | 0.0364555 | 0.30603532 |
| SPATA6L | 135.376232 | -1.519131036 | 0.53456365 | -2.8418151 | 0.00448575 | 0.09064954 |
| CDC37L1 | 52.4247405 | -0.085836169 | 0.25481062 | -0.3368626 | 0.73622048 | 0.9434983 |
| AK3 | 232.578282 | 0.563980225 | 0.26896777 | 2.09683199 | 0.03600845 | 0.30345958 |
| RCL1 | 49.1826411 | -0.108200528 | 0.25512357 | -0.4241103 | 0.67148539 | 0.92170876 |
| JAK2 | 120.86007 | 0.031735557 | 0.25357075 | 0.12515465 | 0.90040112 | 0.97940229 |
| PLGRKT | 62.4059174 | 0.039063591 | 0.26914741 | 0.14513828 | 0.8846017 | 0.97662748 |
| RIC1 | 202.032381 | 0.289243036 | 0.22884941 | 1.26390117 | 0.2062655 | 0.65836247 |
| ERMP1 | 148.373521 | 0.628949262 | 0.36307054 | 1.73230596 | 0.0832191 | 0.4560197 |
| KIAA2026 | 243.102332 | 0.530225349 | 0.27935896 | 1.8980073 | 0.05769512 | 0.38148784 |
| RANBP6 | 21.9188703 | 0.324747497 | 0.32352839 | 1.00376817 | 0.31549037 | 0.75786443 |
| UHRF2 | 79.218704 | 0.066042601 | 0.20362446 | 0.32433531 | 0.74568419 | 0.94614106 |
| KDM4C | 350.162487 | -0.311529666 | 0.19840772 | -1.5701489 | 0.11638047 | 0.52287658 |
| DMAC1 | 31.2365056 | 0.130051292 | 0.40131362 | 0.32406399 | 0.74588959 | 0.94614106 |
| PTPRD | 14.8475305 | 0.136002572 | 0.39606863 | 0.34338133 | 0.7313116 | 0.94155624 |
| LURAP1L | 14.0594892 | -3.149399241 | 0.71877744 | -4.3816056 | 1.18E-05 | 0.00129856 |
| MPDZ | 73.695267 | -0.007375436 | 0.32479139 | -0.0227082 | 0.98188301 | 0.99714727 |
| NFIB | 291.719558 | 0.406875642 | 0.33778038 | 1.20455675 | 0.22837446 | 0.68364117 |
| ZDHHC21 | 218.006719 | 0.252995577 | 0.29178948 | 0.86704831 | 0.38591554 | 0.7994129 |
| FREM1 | 422.343172 | 1.199460519 | 0.47621848 | 2.51871897 | 0.01177826 | 0.16141766 |
| TTC39B | 352.526737 | 0.197267695 | 0.26206161 | 0.75275312 | 0.45159828 | 0.8323206 |
| SNAPC3 | 93.8258284 | -0.130675806 | 0.24945089 | -0.5238538 | 0.6003802 | 0.89490801 |
| PSIP1 | 46.3394777 | 0.018870694 | 0.28681823 | 0.06579322 | 0.94754246 | 0.99149867 |
| CCDC171 | 65.839903 | 0.318652218 | 0.23471134 | 1.35763455 | 0.17457968 | 0.62052871 |
| BNC2 | 102.054828 | -0.234905848 | 0.57141892 | -0.4110922 | 0.68100495 | 0.92375164 |
| CNTLN | 49.0813988 | -0.058215268 | 0.27585367 | -0.2110368 | 0.83285858 | 0.96701848 |
| SH3GL2 | 54.1625589 | -0.981443187 | 0.76986795 | -1.2748202 | 0.2023729 | 0.65499405 |
| ADAMTSL1 | 243.096461 | 2.849963342 | 0.46798205 | 6.08989884 | 1.13E-09 | 6.48E-07 |
| RRAGA | 28.4282606 | -0.363771124 | 0.47506066 | -0.7657362 | 0.44383331 | 0.82999481 |
| HAUS6 | 17.6206495 | 0.51465211 | 0.35081171 | 1.46703231 | 0.14236727 | 0.57673923 |
| PLIN2 | 44.8915011 | -0.920221049 | 0.47168999 | -1.9509022 | 0.05106868 | 0.35974306 |
| DENND4C | 219.190144 | -0.160235394 | 0.19370859 | -0.8271982 | 0.40812475 | 0.81361336 |
| RPS6 | 346.819882 | 0.418548143 | 0.44812897 | 0.93399038 | 0.35030885 | 0.77868979 |
| SLC24A2 | 44.0330348 | 0.491552907 | 0.60268898 | 0.81559963 | 0.41472916 | 0.81726257 |
| MLLT3 | 156.67063 | -0.113483773 | 0.46259636 | -0.2453192 | 0.80620928 | 0.96005568 |
| FOCAD | 245.754205 | -0.134115896 | 0.3764907 | -0.3562263 | 0.72167108 | 0.93824012 |

| | | | | | | |
|---|---|---|---|---|---|---|
| KLHL9 | 36.5520889 | 0.13162618 | 0.26930966 | 0.48875401 | 0.62501586 | 0.90447745 |
| LINC01239 | 15.9893957 | -0.321032043 | 0.44911457 | -0.714811 | 0.47472583 | 0.84593347 |
| AL391117.1 | 61.243846 | 1.666358525 | 0.81517324 | 2.04417718 | 0.04093605 | 0.32473787 |
| ELAVL2 | 40.4719593 | 0.386350449 | 0.8124962 | 0.47551047 | 0.63442318 | 0.90805737 |
| TUSC1 | 28.7789738 | -0.203538365 | 0.3711118 | -0.5484557 | 0.58337907 | 0.89036744 |
| CAAP1 | 145.441385 | -0.195686433 | 0.24029803 | -0.8143489 | 0.41544511 | 0.81740722 |
| PLAA | 51.2420951 | 0.051714414 | 0.27254429 | 0.18974683 | 0.84950752 | 0.97027687 |
| IFT74 | 65.555477 | 0.008890506 | 0.28002316 | 0.03174918 | 0.97467208 | 0.99543096 |
| MOB3B | 168.647029 | 0.313178159 | 0.43747604 | 0.715875 | 0.47406854 | 0.84593347 |
| C9orf72 | 29.6765885 | -1.007625552 | 0.37259796 | -2.7043239 | 0.00684435 | 0.11876608 |
| ACO1 | 51.7865789 | -0.252865984 | 0.2901966 | -0.8713609 | 0.3835571 | 0.7980695 |
| DDX58 | 80.0233415 | -1.234202491 | 1.01440618 | -1.2166748 | 0.22372795 | 0.67606266 |
| TOPORS | 36.4569917 | 0.229685285 | 0.29748974 | 0.77207801 | 0.44006823 | 0.82793978 |
| NDUFB6 | 45.492505 | 0.158593796 | 0.33142373 | 0.47852276 | 0.63227818 | 0.90805737 |
| APTX | 69.9365457 | -0.402400192 | 0.26793322 | -1.5018675 | 0.13313133 | 0.56148254 |
| DNAJA1 | 97.7230186 | 0.002549297 | 0.35316944 | 0.00721834 | 0.99424065 | 0.99923337 |
| SMU1 | 41.9134975 | -0.005709162 | 0.21389808 | -0.026691 | 0.97870616 | 0.99610605 |
| B4GALT1 | 118.889335 | -0.700146936 | 0.22215804 | -3.1515714 | 0.00162394 | 0.04456355 |
| BAG1 | 60.7739558 | 0.516660686 | 0.47191434 | 1.09481879 | 0.27359603 | 0.72490618 |
| CHMP5 | 59.5385754 | -0.164213552 | 0.45474062 | -0.3611148 | 0.71801366 | 0.93733389 |
| NFX1 | 142.512608 | -0.218169327 | 0.26953365 | -0.8094326 | 0.41826635 | 0.8188001 |
| AQP7 | 17.3021 | -0.735161108 | 0.64312432 | -1.1431089 | 0.2529934 | 0.70680759 |
| AQP3 | 95.2028163 | -0.070104266 | 0.73867203 | -0.0949058 | 0.92438964 | 0.98672316 |
| UBE2R2 | 179.060961 | -0.070868778 | 0.21188239 | -0.3344722 | 0.73802323 | 0.94376279 |
| UBAP2 | 102.495016 | 0.01450372 | 0.26859254 | 0.05399897 | 0.95693599 | 0.99236316 |
| DCAF12 | 27.2956211 | 0.315854311 | 0.26592913 | 1.18773866 | 0.23493639 | 0.68936745 |
| UBAP1 | 120.325539 | -0.104133403 | 0.27497391 | -0.3787028 | 0.70490854 | 0.93397252 |
| NUDT2 | 13.6710149 | 0.356762082 | 0.38752619 | 0.92061412 | 0.35725193 | 0.78277472 |
| FAM219A | 45.4747306 | -0.558253845 | 0.32340502 | -1.7261756 | 0.08431583 | 0.4585327 |
| RPP25L | 12.3530217 | -0.263481144 | 0.48188364 | -0.5467734 | 0.58453444 | NA |
| DCTN3 | 21.8345804 | 0.359312798 | 0.51582047 | 0.69658499 | 0.48606255 | 0.85356796 |
| VCP | 107.99708 | -0.458056046 | 0.26484622 | -1.729517 | 0.0837166 | 0.45665431 |
| PIGO | 16.1739664 | -0.28268432 | 0.36076766 | -0.7835634 | 0.43329636 | 0.82501678 |
| STOML2 | 29.9755872 | -0.275110011 | 0.29147339 | -0.9438598 | 0.3452413 | 0.7750152 |
| FAM214B | 10.5821385 | -0.214822821 | 0.43507505 | -0.4937604 | 0.62147542 | NA |
| UNC13B | 132.838774 | 0.164344144 | 0.22029829 | 0.74600735 | 0.45566297 | 0.83489438 |
| RUSC2 | 57.7427817 | -0.315428165 | 0.31649801 | -0.9966197 | 0.31894912 | 0.76052272 |
| TESK1 | 15.5560022 | -0.007239722 | 0.43863465 | -0.0165051 | 0.98683141 | 0.99835885 |
| CCDC107 | 33.1858249 | 0.206249657 | 0.39231343 | 0.52572673 | 0.59907809 | 0.89476262 |
| TLN1 | 82.7021726 | -0.426366508 | 0.28471263 | -1.4975328 | 0.13425467 | 0.56229288 |
| CREB3 | 15.0132723 | -0.068135804 | 0.38050364 | -0.1790674 | 0.85788477 | 0.97243282 |
| GBA2 | 77.0415168 | 0.146493893 | 0.26701976 | 0.54862567 | 0.58326237 | 0.89036744 |
| RGP1 | 34.7442808 | -0.334929141 | 0.23817703 | -1.4062193 | 0.15965901 | 0.60024417 |
| AL133410.1 | 40.8156301 | -0.023909894 | 0.28022643 | -0.0853235 | 0.93200422 | 0.9875946 |
| HINT2 | 14.3858856 | 0.893863159 | 0.45565637 | 1.9617045 | 0.0497969 | 0.35621674 |
| TMEM8B | 25.0286764 | 0.26266874 | 0.46458545 | 0.56538305 | 0.57181325 | 0.88578992 |
| RECK | 29.6540401 | -0.426920461 | 0.34868967 | -1.2243565 | 0.22081777 | 0.67462437 |
| CLTA | 172.4023 | -0.590347894 | 0.3014517 | -1.9583499 | 0.05018897 | 0.35743981 |
| GNE | 64.3727053 | 0.299930327 | 0.36162587 | 0.82939401 | 0.40688149 | 0.8122557 |
| RNF38 | 291.156257 | 0.200826311 | 0.19791114 | 1.01472967 | 0.3102347 | 0.75363506 |
| EBLN3P | 64.6069156 | 0.102989913 | 0.26172499 | 0.39350432 | 0.69394704 | 0.92841398 |
| ZCCHC7 | 415.434818 | 0.21486254 | 0.19020722 | 1.12962348 | 0.25863491 | 0.71022613 |
| GRHPR | 34.4633667 | 0.455202365 | 0.36887032 | 1.23404444 | 0.21718635 | 0.6715676 |
| ZBTB5 | 26.17336 | 0.023045406 | 0.32853051 | 0.07014693 | 0.94407672 | 0.99086173 |
| AL138752.2 | 49.1183532 | -0.25871163 | 0.268652 | -0.9629991 | 0.33554799 | 0.76777176 |
| TRMT10B | 55.8529316 | 0.106405771 | 0.31238384 | 0.34062508 | 0.73338585 | 0.94273028 |
| DCAF10 | 135.251325 | -0.079598179 | 0.2399264 | -0.3317608 | 0.74006987 | 0.94376279 |
| GLIDR | 31.8101388 | 1.224540251 | 0.35138091 | 3.4849368 | 0.00049225 | 0.01973081 |
| CBWD6 | 12.2910381 | 0.040720368 | 0.56414787 | 0.07218031 | 0.94245841 | NA |
| FAM27C | 14.2199746 | 0.046579408 | 0.47523441 | 0.09801354 | 0.92192154 | 0.986393 |
| AL391987.2 | 11.4130228 | 0.108654675 | 0.46066393 | 0.23586539 | 0.81353713 | NA |
| AL512625.3 | 11.695102 | 0.658950633 | 0.48032033 | 1.37189828 | 0.17009511 | NA |
| CBWD5 | 36.4522452 | 0.040017855 | 0.29261396 | 0.13675989 | 0.89122059 | 0.97793768 |
| ZNF658 | 11.7738594 | 0.219993723 | 0.34804311 | 0.63208757 | 0.52732965 | NA |
| PIP5K1B | 34.5919698 | 0.559029825 | 0.60553783 | 0.92319554 | 0.35590531 | 0.78158966 |
| FAM189A2 | 151.349345 | -0.645253612 | 0.3743849 | -1.7235033 | 0.08479755 | 0.45956155 |
| APBA1 | 16.1793032 | -1.425540788 | 0.53892413 | -2.6451604 | 0.00816522 | 0.13111608 |
| PTAR1 | 112.274786 | 0.007534135 | 0.20447308 | 0.03684659 | 0.97060733 | 0.99502125 |
| MAMDC2-AS | 41.1958593 | 0.681526033 | 0.30586179 | 2.22821568 | 0.02586614 | 0.25292492 |
| MAMDC2 | 151.640742 | 0.275250672 | 0.46567255 | 0.59108201 | 0.55446547 | 0.87737603 |
| SMC5-AS1 | 62.9042563 | 0.383121443 | 0.20180645 | 1.89845983 | 0.05763553 | 0.38148784 |
| SMC5 | 186.499948 | -0.350935909 | 0.25235493 | -1.3906442 | 0.16433336 | 0.60496776 |
| KLF9 | 60.0610529 | -0.318238143 | 0.31530401 | -1.0093057 | 0.31282803 | 0.75558564 |
| TRPM3 | 36.560066 | 0.240978304 | 0.67584186 | 0.35656019 | 0.72142108 | 0.93824012 |
| CEMIP2 | 180.821725 | -1.770651079 | 0.58290948 | -3.037609 | 0.00238463 | 0.05737663 |
| ABHD17B | 15.7080716 | -0.250742154 | 0.4717031 | -0.5315677 | 0.59502541 | 0.893081 |
| C9orf85 | 41.5771026 | -0.027818853 | 0.26794737 | -0.1038221 | 0.91731054 | 0.98495314 |
| LINC01504 | 20.7909861 | -0.369376422 | 0.54183448 | -0.6817145 | 0.4954195 | 0.85814183 |
| ZFAND5 | 159.339943 | -0.204247107 | 0.2941034 | -0.6944738 | 0.48738513 | 0.85382621 |
| ALDH1A1 | 193.78627 | 0.412482028 | 0.36252588 | 1.13780024 | 0.25520391 | 0.70849619 |
| CARNMT1 | 71.7486276 | 0.311607973 | 0.24701395 | 1.26149952 | 0.20712893 | 0.65895397 |
| NMRK1 | 55.5643226 | -0.565635516 | 0.37983503 | -1.489161 | 0.13644498 | 0.56754558 |
| OSTF1 | 40.8878755 | -0.424193118 | 0.25214177 | -1.6823596 | 0.09249913 | 0.4764958 |
| RFK | 42.6784591 | -0.453733596 | 0.33265857 | -1.3639618 | 0.17257958 | 0.61726473 |
| GCNT1 | 21.3445154 | -1.97479119 | 0.48305771 | -4.0881061 | 4.35E-05 | 0.00370054 |
| VPS13A | 188.913792 | -0.074658334 | 0.16654299 | -0.4482827 | 0.65394921 | 0.9158679 |
| GNAQ | 745.795764 | 0.131398222 | 0.16726735 | 0.78555811 | 0.43212642 | 0.8248269 |
| CEP78 | 86.2267438 | 1.825987131 | 0.49999641 | 3.65200049 | 0.00026021 | 0.01294355 |
| TLE4 | 55.7124496 | -0.772380925 | 0.48700851 | -1.5859701 | 0.1127461 | 0.51571293 |
| TLE1 | 123.190146 | -0.003313392 | 0.34178289 | -0.0096944 | 0.99226508 | 0.99923337 |
| AL591368.1 | 9.63874622 | 1.806673141 | 0.87857468 | 2.05636832 | 0.03974703 | NA |
| RASEF | 44.1604689 | -0.748073068 | 0.44822296 | -1.6689754 | 0.09512226 | 0.48035393 |
| FRMD3 | 51.078926 | 0.316436442 | 0.26471019 | 1.19540711 | 0.23192801 | 0.68630314 |
| IDNK | 13.130595 | -0.666101036 | 0.39192969 | -1.6995421 | 0.08921708 | 0.4698185 |
| UBQLN1 | 133.786899 | 0.129322895 | 0.22646594 | 0.57104788 | 0.56796718 | 0.88488302 |
| GKAP1 | 49.3647705 | 0.62054564 | 0.30012891 | 2.06759703 | 0.03867793 | 0.31580466 |
| KIF27 | 38.5518354 | 0.212265241 | 0.33313992 | 0.63716544 | 0.52401709 | 0.86747074 |
| C9orf64 | 14.9171463 | -0.156183974 | 0.36840213 | -0.4239497 | 0.67160249 | 0.92170876 |
| HNRNPK | 195.779013 | 0.099349323 | 0.18969999 | 0.52371812 | 0.60047461 | 0.89490801 |
| RMI1 | 13.8403349 | 0.414183041 | 0.37263457 | 1.11149925 | 0.26635351 | 0.71747544 |
| AGTPBP1 | 68.718696 | -0.090284158 | 0.24520505 | -0.3681986 | 0.71272514 | 0.9357596 |
| NAA35 | 118.864122 | -0.092687517 | 0.18538793 | -0.4999652 | 0.61709957 | 0.90173965 |
| GOLM1 | 73.6689601 | -0.962304531 | 0.68544037 | -1.4039216 | 0.1603422 | 0.60024417 |
| ISCA1 | 47.9259527 | 0.136799557 | 0.37321806 | 0.36654056 | 0.71396175 | 0.93650155 |
| TUT7 | 102.127408 | -0.450451535 | 0.187337 | -2.4044986 | 0.01619467 | 0.1962825 |
| DAPK1 | 190.361301 | 0.10736837 | 0.43158171 | 0.24877878 | 0.80353191 | 0.95938063 |
| CTSL | 54.1367266 | -0.182058456 | 0.34801355 | -0.5231361 | 0.60087953 | 0.89490801 |
| SPIN1 | 219.281085 | 0.138624109 | 0.21166985 | 0.65490719 | 0.51252751 | 0.86346419 |
| SECISBP2 | 158.166055 | -0.173313814 | 0.18472968 | -0.9382023 | 0.34814046 | 0.77774355 |
| SEMA4D | 139.219576 | -0.575477158 | 0.24179444 | -2.3800264 | 0.0173114 | 0.20451748 |
| GADD45G | 84.0340994 | -1.754635405 | 0.38579775 | -4.5480707 | 5.41E-06 | 0.00072939 |
| SYK | 157.759655 | 0.221419643 | 0.27317213 | 0.8105499 | 0.4176242 | 0.8188001 |
| AUH | 266.659527 | 0.129036411 | 0.19714566 | 0.6545232 | 0.51277478 | 0.86346419 |

| | | | | | | |
|---|---|---|---|---|---|---|
| SPTLC1 | 83.8343679 | 0.24686223 | 0.20776279 | 1.18819267 | 0.23475751 | 0.68936745 |
| IARS | 73.4002012 | -0.1662547 | 0.28173287 | -0.5901147 | 0.55511377 | 0.87754295 |
| NOL8 | 25.6257267 | -0.061667449 | 0.3012582 | -0.2046997 | 0.83780679 | 0.96748214 |
| CENPP | 22.8267436 | 1.091419656 | 0.44857374 | 2.43308862 | 0.01497064 | 0.18522344 |
| IPPK | 19.7688253 | 0.29906135 | 0.39269564 | 0.76156015 | 0.44632256 | 0.83035712 |
| BICD2 | 23.6899704 | 0.542634493 | 0.33677485 | 1.61126785 | 0.10712135 | 0.50318505 |
| ZNF484 | 11.7358544 | -0.396875396 | 0.39277999 | -1.0104267 | 0.31229089 | NA |
| SUSD3 | 12.5888764 | -0.240521922 | 0.49832214 | -0.4826635 | 0.62933467 | 0.90706482 |
| CARD19 | 24.6920719 | -0.367202615 | 0.29253078 | -1.2552615 | 0.20938385 | 0.66142564 |
| NINJ1 | 16.0500654 | -0.13478462 | 0.40162662 | -0.3355968 | 0.73717492 | 0.94376279 |
| FAM120AOS | 21.7280922 | -2.55E-06 | 0.3287612 | -7.74E-06 | 0.99999382 | 0.99999382 |
| FAM120A | 202.237621 | 0.130630512 | 0.22203904 | 0.58832226 | 0.55631601 | 0.87807756 |
| PHF2 | 44.7542555 | 0.489588692 | 0.29321461 | 1.66972814 | 0.09497316 | 0.48035393 |
| PTPDC1 | 76.8904197 | 0.100890008 | 0.19923054 | 0.50639831 | 0.61257706 | 0.90011356 |
| AL158152.2 | 57.7660757 | -0.146889094 | 0.24720371 | -0.5942026 | 0.5523766 | 0.87576439 |
| ZNF169 | 19.4277482 | 0.048279375 | 0.31964955 | 0.15103846 | 0.87994538 | 0.97581687 |
| MFSD14B | 57.2116727 | -0.057922349 | 0.24482691 | -0.2365849 | 0.81297884 | 0.96262567 |
| FBP1 | 38.9339175 | 0.369161227 | 0.77595336 | 0.47575183 | 0.63425119 | 0.90805737 |
| C9orf3 | 438.607768 | -0.245700411 | 0.214933 | -1.1431488 | 0.25297681 | 0.70680759 |
| FANCC | 163.464863 | 0.258778784 | 0.23073353 | 1.12154826 | 0.26205456 | 0.71302363 |
| PTCH1 | 30.5010855 | -1.158056523 | 0.4628617 | -2.5019493 | 0.01235116 | 0.16567149 |
| LINC00476 | 35.7295466 | 0.404065419 | 0.3568855 | 1.13219904 | 0.25755076 | 0.71022118 |
| ERCC6L2 | 129.538282 | 0.114855338 | 0.25892749 | 0.4435811 | 0.65734548 | 0.9158679 |
| SLC35D2 | 18.5098676 | -0.093448676 | 0.36507654 | -0.2559701 | 0.79797393 | 0.95848876 |
| HABP4 | 28.5962912 | -0.580520893 | 0.47712446 | -1.2167075 | 0.22371554 | 0.67606266 |
| CDC14B | 159.835636 | 0.31270611 | 0.36439403 | 0.85815376 | 0.39080757 | 0.80263252 |
| AAED1 | 16.0955327 | -0.366426053 | 0.52836095 | -0.6935146 | 0.48798665 | 0.85426285 |
| AL589843.1 | 24.5991824 | -0.328365045 | 0.56958716 | -0.5764966 | 0.56427959 | 0.88191823 |
| ZNF510 | 13.1302657 | 0.385643024 | 0.36466142 | 1.05753723 | 0.29026647 | 0.73979353 |
| ZNF782 | 46.8539345 | -0.073538209 | 0.26980283 | -0.2725628 | 0.78518933 | 0.95577371 |
| MFSD14C | 110.106142 | 0.241549721 | 0.24376332 | 0.99091908 | 0.32172509 | 0.76227001 |
| CCDC180 | 20.5376509 | 0.812279968 | 0.5101833 | 1.59213358 | 0.11135469 | 0.51337478 |
| TDRD7 | 41.4290516 | -0.829738437 | 0.42871073 | -1.9354273 | 0.05293788 | 0.36495912 |
| TSTD2 | 46.3178842 | -0.189487575 | 0.22637933 | -0.8370357 | 0.40257251 | 0.80890307 |
| NCBP1 | 33.5076478 | 0.026143688 | 0.2556121 | 0.10227876 | 0.91853542 | 0.98521039 |
| XPA | 38.1265382 | 0.344354988 | 0.29676521 | 1.16036173 | 0.24590156 | 0.69938066 |
| TRMO | 12.6421438 | 0.354333471 | 0.47059087 | 0.75295441 | 0.45147731 | 0.8323206 |
| ANP32B | 58.1209528 | 0.349719289 | 0.24939988 | 1.40224321 | 0.16084263 | 0.60052869 |
| NANS | 22.1692921 | 0.499906878 | 0.34058504 | 1.4677887 | 0.14216163 | 0.57673923 |
| TRIM14 | 25.966337 | -1.748604016 | 0.68563979 | -2.5503246 | 0.01076227 | 0.15307037 |
| TBC1D2 | 50.9229145 | -0.805334938 | 0.39084579 | -2.0604928 | 0.03935145 | 0.31818244 |
| ANKS6 | 28.2370393 | 0.103057372 | 0.38594938 | 0.26702303 | 0.78945143 | 0.95719943 |
| TGFBR1 | 81.9225427 | -0.196956212 | 0.29152512 | -0.6756063 | 0.49929063 | 0.85942996 |
| ALG2 | 26.9412237 | 0.532603847 | 0.32169624 | 1.65561105 | 0.09780061 | 0.48351985 |
| SEC61B | 74.3090406 | -0.064442238 | 0.29346767 | -0.2195889 | 0.82619135 | 0.96539788 |
| AL359710.1 | 65.2902188 | 1.091881463 | 0.35126029 | 3.10846826 | 0.0018806 | 0.04957012 |
| STX17 | 91.1847258 | 0.227756161 | 0.28709802 | 0.79330453 | 0.42760042 | 0.82407405 |
| ERP44 | 107.132246 | 0.119336444 | 0.20661806 | 0.57757025 | 0.56355429 | 0.88191823 |
| INVS | 173.590995 | -0.155908651 | 0.26157031 | -0.5960487 | 0.55114268 | 0.87511606 |
| TEX10 | 37.0292671 | -0.333313424 | 0.292169 | -1.140824 | 0.25394315 | 0.70701839 |
| PLPPR1 | 127.421967 | 1.387938207 | 0.50383091 | 2.75476986 | 0.00587334 | 0.10648865 |
| MRPL50 | 26.0457947 | -0.401231237 | 0.42166338 | -0.9515439 | 0.34132833 | 0.77193691 |
| ZNF189 | 23.7832662 | 0.198436446 | 0.30036272 | 0.66065604 | 0.50883292 | 0.86281145 |
| ALDOB | 34.3281389 | -0.739693148 | 1.19521559 | -0.6188784 | 0.53599645 | 0.86946459 |
| RNF20 | 20.0373187 | 0.204121799 | 0.34811784 | 0.58635835 | 0.55763473 | 0.87923551 |
| SMC2 | 11.1446741 | 0.230002782 | 0.47464363 | 0.48457994 | 0.62797434 | NA |
| NIPSNAP3A | 21.2020874 | 0.253586552 | 0.31023737 | 0.81739525 | 0.41370259 | 0.81662904 |
| ABCA1 | 20.2109315 | -0.168214518 | 0.53262235 | -0.3158232 | 0.75213667 | 0.94728927 |
| SLC44A1 | 219.576217 | -0.424741816 | 0.30108984 | -1.4106813 | 0.1583386 | 0.59881912 |
| FSD1L | 97.9712672 | -0.217482435 | 0.18597872 | -1.1693942 | 0.24224485 | 0.69596937 |
| FKTN | 53.15796 | 0.423714376 | 0.23055446 | 1.83780599 | 0.066091 | 0.40885376 |
| TMEM38B | 29.194689 | 0.089729624 | 0.43499262 | 0.2062785 | 0.83657337 | 0.96748214 |
| ZNF462 | 20.6771231 | 1.473827588 | 0.57882709 | 2.54623117 | 0.0108893 | 0.15419892 |
| RAD23B | 149.814404 | -0.152682001 | 0.19690699 | -0.7754016 | 0.43810237 | 0.82709089 |
| ELP1 | 49.6843115 | 0.404147971 | 0.29346681 | 1.37715053 | 0.16846571 | 0.61056834 |
| FAM206A | 9.96661435 | 0.00670949 | 0.39048985 | 0.01718224 | 0.98629123 | NA |
| CTNNAL1 | 58.683025 | -0.166192699 | 0.25188778 | -0.6597887 | 0.50938946 | 0.86281145 |
| TMEM245 | 216.823187 | 0.428120606 | 0.23647628 | 1.81041671 | 0.07023119 | 0.41948432 |
| EPB41L4B | 59.0100556 | 0.306617675 | 0.42881861 | 0.71502884 | 0.47459122 | 0.84593347 |
| PTPN3 | 259.329171 | -0.507109596 | 0.21975543 | -2.3076089 | 0.0210209 | 0.22580589 |
| PALM2 | 15.2663533 | -0.8172959 | 0.64031526 | -1.2763961 | 0.20181553 | 0.6545025 |
| TXN | 91.3400346 | -0.874762258 | 0.37534821 | -2.3305353 | 0.01977787 | 0.22076569 |
| SVEP1 | 61.8722841 | 1.245130943 | 0.59935657 | 2.07744605 | 0.03776041 | 0.31172423 |
| LPAR1 | 57.343815 | -0.811446868 | 0.32905342 | -2.4660035 | 0.013663 | 0.1757707 |
| ECPAS | 241.489327 | -0.08084149 | 0.17791172 | -0.454391 | 0.6495474 | 0.91432445 |
| ZNF483 | 83.4752665 | -0.279100497 | 0.36947228 | -0.7554031 | 0.45000717 | 0.83118301 |
| PTGR1 | 376.107047 | 0.815935472 | 0.63782893 | 1.27923874 | 0.200813 | 0.65358156 |
| C9orf84 | 299.317969 | 1.591242932 | 0.55330958 | 2.87586369 | 0.00402924 | 0.08381056 |
| UGCG | 226.011681 | -1.219935729 | 0.32950843 | -3.7022899 | 0.00021366 | 0.01141021 |
| SUSD1 | 259.07698 | 0.205431043 | 0.34195761 | 0.60075003 | 0.54800649 | 0.87341177 |
| PTBP3 | 777.119894 | 0.348071072 | 0.27775302 | 1.25316757 | 0.21014473 | 0.66203439 |
| HSDL2 | 116.293862 | 0.148919285 | 0.34412091 | 0.43275279 | 0.66519437 | 0.91953619 |
| C9orf147 | 16.9421647 | 0.531360947 | 0.46686603 | 1.13814438 | 0.2550602 | 0.70849619 |
| KIAA1958 | 203.379387 | -0.091481662 | 0.20909308 | -0.4375165 | 0.66173684 | 0.91808148 |
| INIP | 37.7899862 | 0.304850935 | 0.27060137 | 1.12656833 | 0.25992504 | 0.71091822 |
| SNX30 | 33.3182681 | -0.121492 | 0.59282369 | -0.2049378 | 0.8376207 | 0.96748214 |
| FKBP15 | 43.4961552 | -0.396617845 | 0.2716352 | -1.4601121 | 0.14425927 | 0.57846905 |
| SLC31A1 | 51.2202356 | 0.084680248 | 0.25017899 | 0.33847866 | 0.73500251 | 0.94293405 |
| CDC26 | 23.1361574 | -0.259668753 | 0.44151889 | -0.588126 | 0.55644771 | 0.87807756 |
| PRPF4 | 15.6310924 | -0.220851337 | 0.49445003 | -0.4466606 | 0.65512015 | 0.9158679 |
| BSPRY | 26.4517725 | -0.396697952 | 0.34146277 | -1.1617605 | 0.24533278 | 0.6988922 |
| HDHD3 | 42.2168757 | -0.243417656 | 0.34544415 | -0.7046513 | 0.48102729 | 0.84974772 |
| ALAD | 11.7266907 | 0.443642071 | 0.56503501 | 0.78515855 | 0.43236061 | NA |
| RGS3 | 59.6064264 | 0.480847355 | 0.40034533 | 1.20108147 | 0.2297196 | 0.68523674 |
| ZNF618 | 172.548724 | 0.395849321 | 0.33045837 | 1.19787954 | 0.23096391 | 0.6853319 |
| KIF12 | 54.7750353 | 1.067875283 | 0.33388081 | 3.19837278 | 0.00138206 | 0.04025806 |
| AKNA | 24.6928541 | -0.543566547 | 0.5189598 | -1.0474155 | 0.29490798 | 0.74111836 |
| WHRN | 29.7512365 | -1.509939029 | 0.42607908 | -3.5437999 | 0.0003944 | 0.01717513 |
| ATP6V1G1 | 135.423485 | -0.468281636 | 0.28131722 | -1.6646035 | 0.09599189 | 0.48135478 |
| TMEM268 | 20.422512 | -0.471565816 | 0.45853521 | -1.0284179 | 0.3037533 | 0.74820946 |
| PAPPA | 12.5592231 | -0.966868174 | 0.53327088 | -1.8130901 | 0.06981792 | 0.41879921 |
| ASTN2 | 312.089181 | -0.329242339 | 0.42064565 | -0.7827071 | 0.43379913 | 0.82504445 |
| CDK5RAP2 | 102.787589 | 0.129538286 | 0.21441645 | 0.60414341 | 0.5457483 | 0.87247631 |
| MEGF9 | 232.23197 | 0.018452512 | 0.20361989 | 0.09062234 | 0.92779268 | 0.98690525 |
| FBXW2 | 57.3515478 | 0.143407908 | 0.25377807 | 0.5650918 | 0.57201133 | 0.88578992 |
| PSMD5 | 52.4070265 | 0.019825364 | 0.24852524 | 0.07977203 | 0.93641857 | 0.98871562 |
| CNTRL | 42.6627579 | -0.015588401 | 0.38960165 | -0.0400111 | 0.96808426 | 0.99411574 |
| RAB14 | 120.334464 | 0.025649393 | 0.19450577 | 0.13186957 | 0.89508746 | 0.97862357 |
| GSN | 110.536162 | -0.313985773 | 0.2067805 | -1.5184496 | 0.1289011 | 0.55324807 |
| STOM | 44.277604 | -1.185431321 | 0.4121562 | -2.87617 | 0.00402533 | 0.08381056 |
| DAB2IP | 194.904486 | -0.370558018 | 0.27976551 | -1.3245307 | 0.18532684 | 0.63578845 |
| TTLL11 | 24.4954297 | 0.322432006 | 0.32598211 | 0.98910951 | 0.32260957 | 0.76292283 |
| NDUFA8 | 50.2584383 | 0.138683559 | 0.40067105 | 0.34612822 | 0.72924636 | 0.94106333 |

| | | | | | | |
|---|---|---|---|---|---|---|
| RBM18 | 23.9092719 | 0.165760795 | 0.31576166 | 0.52495543 | 0.59961418 | 0.89476262 |
| MRRF | 97.4553808 | 0.052683871 | 0.28112275 | 0.18740522 | 0.85134293 | 0.97027687 |
| PDCL | 31.4605 | -0.093178469 | 0.26389736 | -0.353086 | 0.72402395 | 0.9390336 |
| RC3H2 | 94.2549235 | -0.161377954 | 0.25391003 | -0.6355714 | 0.52505582 | 0.86747074 |
| ZBTB26 | 11.1848373 | 0.682473375 | 0.37182399 | 1.83547431 | 0.06643544 | NA |
| RABGAP1 | 198.703065 | -0.022795659 | 0.229305 | -0.099412 | 0.92081119 | 0.98614962 |
| STRBP | 224.585819 | 0.314272429 | 0.22324691 | 1.40773472 | 0.15920964 | 0.60024133 |
| DENND1A | 296.648028 | -0.104217229 | 0.24039903 | -0.4335177 | 0.66463873 | 0.91903003 |
| PSMB7 | 65.9396599 | -0.084168815 | 0.23076543 | -0.3647375 | 0.71530733 | 0.93689626 |
| NR6A1 | 149.433315 | 0.027768181 | 0.3652884 | 0.07601714 | 0.93940546 | 0.98871562 |
| MIR181A2HC | 13.2480272 | -0.777153328 | 0.52473138 | -1.4810499 | 0.13859329 | 0.57119278 |
| RPL35 | 125.94597 | 0.125422286 | 0.3364848 | 0.37274279 | 0.70933988 | 0.93485232 |
| ARPC5L | 48.0357336 | -0.071150862 | 0.285936 | -0.2488349 | 0.80348848 | 0.95938063 |
| GOLGA1 | 45.0776565 | 0.142248345 | 0.3164074 | 0.44957338 | 0.65301809 | 0.9158679 |
| SCAI | 101.127183 | 0.588852825 | 0.38429791 | 1.53228215 | 0.12545282 | 0.54505746 |
| PPP6C | 70.2838382 | 0.046480315 | 0.22853276 | 0.20338578 | 0.83883351 | 0.96748214 |
| RABEPK | 14.8607256 | 0.172363009 | 0.4561724 | 0.3778462 | 0.70554485 | 0.93407096 |
| HSPA5 | 159.748147 | -0.175981887 | 0.3330978 | -0.528319 | 0.59727797 | 0.89363331 |
| GAPVD1 | 120.898694 | -0.098805322 | 0.18228007 | -0.5420523 | 0.58778251 | 0.89073775 |
| MAPKAP1 | 243.26161 | -0.224997091 | 0.29260311 | -0.7689498 | 0.44192313 | 0.8281114 |
| AL162584.1 | 19.8213705 | 0.653956398 | 0.62802448 | 1.04129125 | 0.2977404 | 0.74243749 |
| PBX3 | 235.598468 | -0.459730267 | 0.27590981 | -1.6662339 | 0.09566685 | 0.48081904 |
| MVB12B | 170.770225 | 0.384480886 | 0.4841989 | 0.79405567 | 0.42716302 | 0.82375285 |
| ZBTB43 | 85.2672278 | -0.41560495 | 0.31554594 | -1.3170981 | 0.18780576 | 0.63942753 |
| ZBTB34 | 14.019431 | -0.41404515 | 0.37359685 | -1.1082672 | 0.26774642 | 0.71922155 |
| RALGPS1 | 264.973641 | -0.026699901 | 0.24466024 | -0.1091305 | 0.91309896 | 0.98376271 |
| GARNL3 | 55.5768227 | 0.618158232 | 0.43110189 | 1.43390285 | 0.15159999 | 0.58915059 |
| SLC2A8 | 13.005945 | 0.401402325 | 0.38272656 | 1.04879664 | 0.29427172 | 0.7405942 |
| RPL12 | 53.9465597 | 0.082276775 | 0.32279281 | 0.25489036 | 0.79880778 | 0.95848876 |
| LRSAM1 | 79.7263832 | -0.528956713 | 0.31518554 | -1.6782391 | 0.09330042 | 0.47790456 |
| FAM129B | 58.5527852 | -0.671752769 | 0.4031579 | -1.6662275 | 0.09566812 | 0.48081904 |
| STXBP1 | 31.6764632 | -0.584609077 | 0.30729787 | -1.9024183 | 0.05711649 | 0.37973269 |
| PTRH1 | 13.1608932 | -0.010051126 | 0.37233917 | -0.0269945 | 0.97846409 | 0.99610605 |
| CDK9 | 24.5029758 | -0.09162396 | 0.31008906 | -0.2954763 | 0.76763007 | 0.95205353 |
| FPGS | 28.1424699 | 0.124321339 | 0.4230351 | 0.29387949 | 0.76884999 | 0.95205353 |
| DPM2 | 29.7407199 | -0.821000251 | 0.43819001 | -1.873617 | 0.06098323 | 0.39246665 |
| FAM102A | 97.4660342 | -0.969688996 | 0.48259652 | -2.0093162 | 0.04450361 | 0.33541961 |
| SLC25A25 | 80.1909831 | -0.943587869 | 0.50037715 | -1.8857533 | 0.0593282 | 0.3871608 |
| SLC25A25-AS | 28.4625662 | 0.573160749 | 0.3698773 | 1.549597 | 0.12123827 | 0.53552425 |
| PTGES2 | 47.4345745 | -0.130816186 | 0.27349253 | -0.4783172 | 0.63242445 | 0.90805737 |
| C9orf16 | 87.6718761 | 0.340088618 | 0.46616254 | 0.72954943 | 0.46566564 | 0.8416191 |
| CIZ1 | 32.4370522 | 0.247668694 | 0.27230012 | 0.9095431 | 0.36306352 | 0.78718296 |
| DNM1 | 15.694408 | 0.625083047 | 0.63576923 | 0.98319173 | 0.32551308 | 0.76334591 |
| GOLGA2 | 84.4163799 | 0.073214179 | 0.31392999 | 0.23321817 | 0.815592 | 0.96300833 |
| SWI5 | 25.7662656 | 0.453119882 | 0.46615997 | 0.97202658 | 0.33103733 | 0.76685572 |
| TRUB2 | 21.768954 | 0.284059546 | 0.39202967 | 0.72458685 | 0.46870553 | 0.84326495 |
| COQ4 | 46.6982894 | 0.264331405 | 0.22084115 | 1.19693002 | 0.23133383 | 0.68580018 |
| SLC27A4 | 55.3983393 | 0.198948732 | 0.26860829 | 0.7406649 | 0.45889665 | 0.83645518 |
| URM1 | 23.9531231 | -0.356282788 | 0.33518798 | -1.0629343 | 0.28781177 | 0.73803612 |
| ODF2 | 20.1486019 | 0.023604277 | 0.48501405 | 0.0486672 | 0.96118451 | 0.99238848 |
| GLE1 | 29.5035794 | 0.297964055 | 0.34778003 | 0.85676011 | 0.39157747 | 0.80332153 |
| SPTAN1 | 230.076485 | -0.242880503 | 0.37262659 | -0.6518067 | 0.51452588 | 0.86452469 |
| WDR34 | 24.0170716 | 0.281157603 | 0.35856746 | 0.78411355 | 0.43297349 | 0.8248269 |
| SET | 113.830761 | -0.293740809 | 0.26028636 | -1.1285294 | 0.25909642 | 0.71055562 |
| ZDHHC12 | 19.1499937 | -0.239892475 | 0.45678698 | -0.5251736 | 0.59946249 | 0.89476262 |
| ZER1 | 87.2455568 | 0.155622724 | 0.27423143 | 0.56748683 | 0.57038347 | 0.88563341 |
| TBC1D13 | 19.8484573 | 0.413793334 | 0.37716037 | 1.09712836 | 0.27258529 | 0.72345643 |
| ENDOG | 29.3740861 | 0.157219055 | 0.39027579 | 0.40284091 | 0.68706526 | 0.92614411 |
| KYAT1 | 18.1095311 | 1.081270732 | 0.48750131 | 2.2179853 | 0.02655583 | 0.25580099 |
| LRRC8A | 14.3658122 | 0.228766218 | 0.40262119 | 0.56819219 | 0.56990447 | 0.88562534 |
| PHYHD1 | 14.3548603 | 0.359288365 | 0.47045609 | 0.76370223 | 0.44504471 | 0.83035712 |
| NUP188 | 22.5121251 | -0.028628904 | 0.2962513 | -0.0966372 | 0.92301449 | 0.98672316 |
| SH3GLB2 | 54.6906509 | -0.676089094 | 0.28916071 | -2.3381084 | 0.01938162 | 0.21848783 |
| MIGA2 | 26.8424809 | 0.50418924 | 0.3452534 | 1.46034548 | 0.14419515 | 0.57846905 |
| CRAT | 13.9506495 | -0.05038859 | 0.37634947 | -0.1338878 | 0.89349132 | 0.97833263 |
| PTPA | 53.6321506 | -0.315367892 | 0.3268085 | -0.9649929 | 0.33454835 | 0.76777176 |
| IER5L | 22.2602665 | -2.443953121 | 0.55665514 | -4.3904258 | 1.13E-05 | 0.00126132 |
| LINC01503 | 18.0519845 | -1.87611206 | 0.50504251 | -3.7147607 | 0.0002034 | 0.01102202 |
| NTMT1 | 22.7716548 | -0.601191348 | 0.43855743 | -1.3708384 | 0.17042535 | 0.61294989 |
| PTGES | 29.5046336 | 0.345942801 | 0.5397222 | 0.64096456 | 0.52154571 | 0.86703098 |
| TOR1B | 24.6633315 | -0.017043135 | 0.36155642 | -0.0471382 | 0.96240305 | 0.99306991 |
| C9orf78 | 30.4337715 | 0.079072817 | 0.33373386 | 0.23693376 | 0.81270818 | 0.96262567 |
| USP20 | 24.4913142 | 0.124062629 | 0.31340698 | 0.39585151 | 0.69221457 | 0.928133 |
| FNBP1 | 185.716745 | -0.944215991 | 0.24435911 | -3.8640508 | 0.00011152 | 0.00699705 |
| GPR107 | 93.2490622 | -0.076593367 | 0.19916582 | -0.3845708 | 0.7005554 | 0.93127962 |
| FUBP3 | 114.307872 | -0.002080138 | 0.25600379 | -0.0081254 | 0.99351692 | 0.99923337 |
| ABL1 | 161.078652 | 0.293787559 | 0.2101997 | 1.39765924 | 0.16221542 | 0.60094226 |
| AIF1L | 51.9900862 | 1.365021535 | 0.56098145 | 2.43327394 | 0.01496298 | 0.18522344 |
| NUP214 | 120.71821 | 0.440598176 | 0.21668955 | 2.03331533 | 0.04202068 | 0.32888549 |
| AL157938.2 | 36.1147542 | -0.274535987 | 0.6469619 | -0.4243465 | 0.67131317 | 0.92170876 |
| PRRC2B | 258.873647 | -0.147758061 | 0.23222015 | -0.6362844 | 0.52459106 | 0.86747074 |
| POMT1 | 19.6667221 | 0.529542328 | 0.32995888 | 1.6048737 | 0.1085216 | 0.50659799 |
| UCK1 | 10.507366 | -0.068229949 | 0.52463293 | -0.1300527 | 0.8965247 | NA |
| RAPGEF1 | 154.010026 | -0.642083829 | 0.39422301 | -1.6287325 | 0.10336966 | 0.495277 |
| MED27 | 124.298816 | 0.073404544 | 0.20108156 | 0.3650486 | 0.71507512 | 0.93689626 |
| SETX | 193.122011 | -0.16356236 | 0.27768181 | -0.589028 | 0.5558425 | 0.87787661 |
| TTF1 | 23.9389834 | -0.08063226 | 0.27282582 | -0.2955448 | 0.76757772 | 0.95205353 |
| DDX31 | 27.5293911 | 0.10814787 | 0.29676794 | 0.36441898 | 0.71554517 | 0.93689626 |
| GTF3C4 | 13.7969118 | 0.588831342 | 0.4418805 | 1.33255788 | 0.18267693 | 0.6314919 |
| TSC1 | 78.7173832 | -0.323111058 | 0.24831772 | -1.3012002 | 0.19318994 | 0.64401245 |
| GTF3C5 | 22.3818727 | 0.235115474 | 0.42936839 | 0.5475845 | 0.58397724 | 0.89036744 |
| RALGDS | 10.1058102 | -0.253170975 | 0.48000353 | -0.5274357 | 0.59789109 | NA |
| ABO | 85.6714368 | -0.113209986 | 0.43125406 | -0.2625134 | 0.79292562 | 0.95746995 |
| SURF6 | 16.3098628 | 0.197646152 | 0.37269974 | 0.53030933 | 0.59589748 | 0.89345937 |
| MED22 | 12.4574338 | 0.386136745 | 0.46765241 | 0.82569177 | 0.40897896 | NA |
| RPL7A | 201.040036 | 0.364887589 | 0.36377446 | 1.00305995 | 0.31583194 | 0.75793414 |
| SURF1 | 17.9483713 | 0.248220962 | 0.39120226 | 0.634508 | 0.52574935 | 0.86749236 |
| SURF4 | 64.0823634 | -0.093593989 | 0.26460084 | -0.3537177 | 0.72355049 | 0.93896433 |
| REXO4 | 14.1040955 | 0.224701289 | 0.37129839 | 0.60517712 | 0.54506132 | 0.87247631 |
| CACFD1 | 16.9987233 | -0.381679137 | 0.50963773 | -0.7489224 | 0.45390395 | 0.83364149 |
| VAV2 | 103.429082 | -0.062658921 | 0.26710639 | -0.2345841 | 0.81453153 | 0.96294404 |
| BRD3 | 35.3043123 | 0.412589098 | 0.25084958 | 1.64476693 | 0.10001788 | 0.48752436 |
| WDR5 | 23.0718605 | 0.30056015 | 0.31206906 | 0.96312064 | 0.33548699 | 0.76777176 |
| RXRA | 35.1897161 | 0.153357398 | 0.40659721 | 0.37717277 | 0.70604521 | 0.93407096 |
| PPP1R26-AS: | 14.0948614 | 0.498466968 | 0.4246375 | 1.17386468 | 0.24044923 | 0.69470237 |
| PPP1R26 | 20.6301426 | 0.252163926 | 0.34770724 | 0.72521908 | 0.46831764 | 0.84326495 |
| MRPS2 | 15.5402585 | 0.126831581 | 0.50754103 | 0.24989424 | 0.80266914 | 0.95908975 |
| CAMSAP1 | 24.5734047 | -0.269846808 | 0.35710788 | -0.7556451 | 0.44986203 | 0.83118301 |
| UBAC1 | 61.4584645 | -0.166034438 | 0.2228005 | -0.7452157 | 0.45614132 | 0.83506875 |
| NACC2 | 31.1965565 | -0.169492344 | 0.43395129 | -0.3905792 | 0.69610831 | 0.92929405 |
| TMEM250 | 12.2866542 | -0.035726686 | 0.58032274 | -0.0615635 | 0.95091046 | NA |
| SNAPC4 | 14.1855098 | 0.098342301 | 0.34622416 | 0.28404228 | 0.77637798 | 0.95512504 |
| ENTR1 | 17.1337891 | -0.111476082 | 0.39153851 | -0.2847129 | 0.77586407 | 0.95493992 |

| | | | | | | |
|---|---|---|---|---|---|---|
| PMPCA | 35.3265925 | 0.20411783 | 0.35957851 | 0.56765859 | 0.57026682 | 0.88563341 |
| INPP5E | 12.3194354 | 0.216591182 | 0.40537596 | 0.53429706 | 0.59313603 | NA |
| SEC16A | 51.406114 | 0.033110173 | 0.29431674 | 0.11249844 | 0.91042821 | 0.98334456 |
| NOTCH1 | 26.0679679 | -0.756476965 | 0.41307521 | -1.8313299 | 0.06705132 | 0.4116442 |
| EGFL7 | 15.9318074 | -0.613343598 | 0.6769723 | -0.9060099 | 0.36493062 | 0.78793182 |
| AGPAT2 | 53.5969455 | -0.553300029 | 0.3357346 | -1.648028 | 0.09934692 | 0.48596326 |
| SNHG7 | 38.8271593 | 0.32989681 | 0.41715681 | 0.79082207 | 0.42904783 | 0.82469033 |
| RABL6 | 97.1900406 | 0.121574428 | 0.24280261 | 0.50071301 | 0.61657311 | 0.90160403 |
| PHPT1 | 84.8182158 | 0.036501285 | 0.37908735 | 0.09628727 | 0.92329242 | 0.98672316 |
| EDF1 | 83.7381876 | -0.193262553 | 0.42458014 | -0.4551851 | 0.64897608 | 0.91417939 |
| TRAF2 | 19.9176037 | -0.746313109 | 0.34801369 | -2.1444935 | 0.03199336 | 0.28341152 |
| FBXW5 | 57.7900575 | -0.639198411 | 0.39897828 | -1.6020882 | 0.1091361 | 0.5077315 |
| LCN12 | 10.1821381 | -0.241408125 | 0.51199388 | -0.4715059 | 0.63727952 | NA |
| PAXX | 22.844774 | 0.540461324 | 0.3727279 | 1.45001576 | 0.14705412 | 0.58364362 |
| ABCA2 | 10.9425363 | 0.891499788 | 0.56788369 | 1.56986333 | 0.11644691 | NA |
| UAP1L1 | 14.778807 | -0.420037539 | 0.67627233 | -0.6211071 | 0.53452916 | 0.86909171 |
| MAN1B1 | 29.2896859 | 0.19420538 | 0.29388994 | 0.66080989 | 0.50873424 | 0.86281145 |
| DPP7 | 42.3635733 | 0.111351643 | 0.38642337 | 0.2881597 | 0.7732245 | 0.95447666 |
| SSNA1 | 35.9953248 | -0.446682984 | 0.33340505 | -1.3397607 | 0.18032316 | 0.62805552 |
| TPRN | 17.8951748 | -0.256220273 | 0.35666978 | -0.7183683 | 0.4725302 | 0.84582819 |
| TMEM203 | 19.4049718 | 0.429331805 | 0.3611874 | 1.18866772 | 0.23457045 | 0.68936745 |
| TUBB4B | 132.697965 | -0.346487319 | 0.43254141 | -0.8010501 | 0.42310267 | 0.82180534 |
| NELFB | 38.0550734 | 0.504841819 | 0.30066897 | 1.67906193 | 0.09313998 | 0.47776719 |
| EXD3 | 43.3369931 | 0.208577784 | 0.24495485 | 0.85149482 | 0.39449454 | 0.80508012 |
| NOXA1 | 16.7950317 | -0.50546432 | 0.38633116 | -1.3083706 | 0.19074765 | 0.6421543 |
| NSMF | 25.2412435 | -0.658143712 | 0.3101597 | -2.1219511 | 0.03384184 | 0.29283309 |
| PNPLA7 | 14.641029 | -0.374989559 | 0.58408435 | -0.6420127 | 0.52086495 | 0.8669958 |
| MRPL41 | 111.451836 | -0.351360568 | 0.38394685 | -0.9151281 | 0.36012436 | 0.78516662 |
| DPH7 | 20.7069441 | 0.299378251 | 0.29411589 | 1.01789215 | 0.30872921 | 0.75132915 |
| ARRDC1 | 30.5691774 | 0.209587148 | 0.30588764 | 0.6851769 | 0.4932323 | 0.85714521 |
| ARRDC1-AS1 | 11.9560546 | 0.124508335 | 0.39466795 | 0.31547618 | 0.75240013 | NA |
| EHMT1 | 219.059082 | -0.102740245 | 0.1720893 | -0.597017 | 0.550496 | 0.87459747 |
| BET1L | 20.3688646 | -0.501307494 | 0.36197257 | -1.3849323 | 0.16607318 | 0.6067457 |
| RIC8A | 16.6435066 | -0.374758941 | 0.42994583 | -0.8716422 | 0.38340359 | 0.7980695 |
| SIRT3 | 16.9806048 | -0.06389746 | 0.35321268 | -0.1809036 | 0.85644321 | 0.97166553 |
| PSMD13 | 40.3965241 | -0.492267103 | 0.32825818 | -1.4996339 | 0.13370926 | 0.56229288 |
| PGGHG | 19.9757088 | -1.064123939 | 0.64384058 | -1.6527755 | 0.09837656 | 0.48476221 |
| IFITM2 | 207.662065 | -2.822061186 | 0.47338304 | -5.961475 | 2.50E-09 | 1.21E-06 |
| IFITM3 | 291.95425 | -2.620680893 | 0.54096527 | -4.8444531 | 1.27E-06 | 0.0002463 |
| SIGIRR | 38.070853 | -0.258938877 | 0.37121364 | -0.6975468 | 0.48546068 | 0.85356796 |
| ANO9 | 22.3724863 | -0.559452667 | 0.32927132 | -1.6990628 | 0.08930735 | 0.47003867 |
| PTDSS2 | 45.3491818 | -0.432293706 | 0.35525783 | -1.216845 | 0.22366321 | 0.67606266 |
| RNH1 | 88.5313085 | -0.848819551 | 0.30686559 | -2.7660956 | 0.00567319 | 0.10402636 |
| HRAS | 20.9014158 | 0.031589392 | 0.54084099 | 0.05840791 | 0.95342371 | 0.99217529 |
| RASSF7 | 39.409009 | -0.670083727 | 0.3393663 | -1.9745147 | 0.04832325 | 0.35179202 |
| MIR210HG | 13.0002066 | -2.291488503 | 0.72659811 | -3.1537221 | 0.00161203 | 0.04442228 |
| PHRF1 | 27.6968083 | 0.234347965 | 0.277666 | 0.8439923 | 0.39867371 | 0.80724646 |
| IRF7 | 22.5083435 | -1.260351192 | 0.80852231 | -1.5588329 | 0.11903593 | 0.53056597 |
| DEAF1 | 92.9711087 | -0.325804323 | 0.22332855 | -1.4588566 | 0.14460459 | 0.57889578 |
| AC131934.1 | 16.5791249 | -0.344167941 | 0.47027573 | -0.7318429 | 0.46426448 | 0.84112168 |
| EPS8L2 | 86.3581426 | -0.36067653 | 0.34335405 | -1.0504508 | 0.2935109 | 0.74013384 |
| AP006621.4 | 13.1856512 | -1.177315066 | 0.70879952 | -1.6609987 | 0.09671371 | 0.48257296 |
| TALDO1 | 93.147711 | -1.064448838 | 0.26743659 | -3.9801915 | 6.89E-05 | 0.005138 |
| GATD1 | 20.2651468 | -0.211615307 | 0.50201235 | -0.4215341 | 0.67336514 | 0.92189158 |
| AP006621.3 | 24.7363883 | 0.048413605 | 0.38714179 | 0.12505394 | 0.90048085 | 0.97940229 |
| RPLP2 | 137.457309 | 0.398139873 | 0.32255152 | 1.23434504 | 0.21707436 | 0.67156502 |
| PNPLA2 | 25.9416037 | -0.14844833 | 0.30024674 | -0.4944211 | 0.6210088 | 0.90380166 |
| CRACR2B | 13.3293659 | -0.207156665 | 0.65085296 | -0.3182849 | 0.75026884 | 0.94686544 |
| CD151 | 84.6580494 | -0.36882584 | 0.29254954 | -1.2607295 | 0.20740631 | 0.65905456 |
| POLR2L | 83.0854186 | 0.003937977 | 0.42923211 | 0.00917447 | 0.99267993 | 0.99923337 |
| CHID1 | 22.0300107 | 0.747842086 | 0.32206123 | 2.32204939 | 0.02023028 | 0.22348867 |
| AP2A2 | 102.63137 | -0.106506266 | 0.25761214 | -0.4134365 | 0.67928683 | 0.92348735 |
| TOLLIP | 41.430001 | -0.393140776 | 0.4051192 | -0.9704323 | 0.33183103 | 0.76685572 |
| MOB2 | 23.0040611 | -0.162730773 | 0.3441464 | -0.4728533 | 0.63631782 | 0.9089304 |
| DUSP8 | 12.7408758 | 0.012860506 | 0.70718413 | 0.01818551 | 0.98549086 | 0.99801117 |
| MRPL23 | 16.3121801 | 0.238820656 | 0.63764326 | 0.37453647 | 0.70800523 | 0.93432317 |
| CD81 | 283.838814 | -0.272568883 | 0.2843016 | -0.9587314 | 0.33769405 | 0.76966704 |
| TSSC4 | 22.1776807 | -0.209064279 | 0.40982384 | -0.5101321 | 0.60995895 | 0.89945299 |
| KCNQ1 | 239.422168 | -0.033263247 | 0.2736849 | -0.1215385 | 0.90326454 | 0.98016176 |
| KCNQ1OT1 | 253.416363 | -0.230984847 | 0.41228096 | -0.5602608 | 0.57530158 | 0.88744956 |
| CDKN1C | 69.9432063 | -1.586662614 | 0.70867279 | -2.2389213 | 0.02516104 | 0.25057706 |
| SLC22A18 | 33.8880388 | -0.220845005 | 0.66508589 | -0.3320549 | 0.73984783 | 0.94376279 |
| NAP1L4 | 250.369438 | -0.041256122 | 0.24949564 | -0.1653581 | 0.86866214 | 0.97434023 |
| CARS | 78.4004486 | -0.860449802 | 0.42554824 | -2.0219794 | 0.04317848 | 0.33179556 |
| OSBPL5 | 18.9766479 | -0.408192085 | 0.45166049 | -0.9037587 | 0.36612337 | 0.78793182 |
| ZNF195 | 25.5544115 | 0.098031106 | 0.34478012 | 0.28432935 | 0.776158 | 0.95505931 |
| AC127526.1 | 30.2613277 | -0.005642303 | 0.55311947 | -0.0102009 | 0.99186102 | 0.99923337 |
| NUP98 | 149.251815 | 0.238175132 | 0.24013795 | 0.99182628 | 0.32128227 | 0.76163052 |
| PGAP2 | 20.6107964 | 0.408001588 | 0.45344919 | 0.89977354 | 0.36824078 | 0.78877653 |
| RHOG | 15.2777489 | -0.876760011 | 0.45947801 | -1.9081653 | 0.05636985 | 0.37709488 |
| STIM1 | 367.824772 | 0.254804484 | 0.28206457 | 0.90335517 | 0.36633743 | 0.78793182 |
| RRM1 | 17.789032 | 0.375204484 | 0.31771887 | 1.18093234 | 0.2376296 | 0.69207518 |
| AC104389.4 | 32.918653 | 0.389852073 | 0.7607418 | 0.51246306 | 0.60832696 | 0.8980479 |
| TRIM5 | 14.4417602 | -0.710902054 | 0.5341933 | -1.3307955 | 0.1832563 | 0.63214299 |
| TRIM22 | 40.052544 | -2.421447141 | 0.81813044 | -2.9597324 | 0.00307906 | 0.06902196 |
| FAM160A2 | 19.7593955 | 0.381766739 | 0.29563386 | 1.29134985 | 0.19658239 | 0.64685325 |
| SMPD1 | 18.3818365 | 0.2785757 | 0.39416261 | 0.70675323 | 0.47971985 | 0.84857898 |
| APBB1 | 20.2095014 | -0.359107064 | 0.37239507 | -0.9643174 | 0.3348868 | 0.76777176 |
| ARFIP2 | 16.9858564 | -0.300330943 | 0.35587537 | -0.8439217 | 0.39871317 | 0.80724646 |
| AC084337.2 | 32.4359739 | -0.421062341 | 0.44592579 | -0.9442431 | 0.34504545 | 0.77492029 |
| DNHD1 | 47.8366252 | -0.825016191 | 0.43540961 | -1.8948047 | 0.05811828 | 0.38243048 |
| RRP8 | 12.0706747 | 0.283335554 | 0.39185984 | 0.72305331 | 0.46964713 | NA |
| ILK | 33.9162288 | -0.315448457 | 0.26494535 | -1.1906171 | 0.23380395 | 0.68840359 |
| TAF10 | 30.5827382 | -0.081613443 | 0.38979869 | -0.2093733 | 0.83415683 | 0.96701848 |
| TPP1 | 36.0979205 | -0.523719759 | 0.35712928 | -1.4664711 | 0.14251999 | 0.57673923 |
| MRPL17 | 22.5580798 | -0.385665573 | 0.44624547 | -0.8642454 | 0.38745312 | 0.80048887 |
| AC087280.2 | 22.052608 | 0.085283521 | 0.51912907 | 0.16428192 | 0.86950921 | 0.97434023 |
| ZNF215 | 11.5207666 | -0.154291762 | 0.41580901 | -0.371064 | 0.71058985 | NA |
| AC107884.1 | 24.8494327 | -0.018034227 | 0.41706178 | -0.0432411 | 0.96550931 | 0.99334616 |
| PPFIBP2 | 198.218755 | 0.291760441 | 0.2550115 | 1.14410699 | 0.25257927 | 0.70626085 |
| EIF3F | 50.4689104 | 0.255235942 | 0.36650069 | 0.69641327 | 0.48617006 | 0.85356796 |
| RIC3 | 47.7147465 | -0.267650231 | 0.41708489 | -0.6417164 | 0.52105732 | 0.8669958 |
| STK33 | 355.270925 | -0.050860175 | 0.28278473 | -0.1798547 | 0.85726661 | 0.97223034 |
| TRIM66 | 40.7541284 | 0.37553695 | 0.41405628 | 0.90697079 | 0.36442224 | 0.78780828 |
| RPL27A | 190.287962 | 0.190498423 | 0.33657098 | 0.56599777 | 0.5713953 | 0.88578992 |
| ST5 | 233.549034 | -0.650652895 | 0.39347436 | -1.6536094 | 0.0982069 | 0.48476221 |
| AC026894.1 | 16.6596267 | -0.84449615 | 0.63130219 | -1.3377051 | 0.18099259 | 0.62880664 |
| AKIP1 | 25.6829554 | -0.824721073 | 0.34957764 | -2.3591929 | 0.01831473 | 0.21024014 |
| TMEM9B | 86.688912 | 0.100872453 | 0.21033537 | 0.47957912 | 0.6315267 | 0.90793078 |
| TMEM9B-AS | 26.1893728 | -0.311033875 | 0.53877736 | -0.5772957 | 0.56373969 | 0.88191823 |
| DENND5A | 86.4889162 | -0.269660213 | 0.2767751 | -0.9742936 | 0.32991077 | 0.76685572 |
| TMEM41B | 102.789541 | 0.039480181 | 0.25297222 | 0.15606528 | 0.87598156 | 0.97532382 |
| IPO7 | 131.76304 | -0.035498179 | 0.21182982 | -0.1675788 | 0.86691468 | 0.97434023 |

| | | | | | | |
|---|---|---|---|---|---|---|
| ZNF143 | 33.0527032 | -0.784684644 | 0.28240227 | -2.778606 | 0.00545927 | 0.10201977 |
| SWAP70 | 67.3342144 | -0.418561602 | 0.35922197 | -1.1651893 | 0.24394236 | 0.69773111 |
| SBF2-AS1 | 88.1183859 | -0.25725099 | 0.60142226 | -0.4277377 | 0.66884208 | 0.92049996 |
| SBF2 | 879.026515 | -0.214971159 | 0.25094483 | -0.8566471 | 0.39163995 | 0.80332153 |
| AC011092.2 | 36.1134884 | -1.066282521 | 0.40988963 | -2.6013894 | 0.0092847 | 0.14114208 |
| AC100763.1 | 12.3703107 | -1.15680811 | 0.7169531 | -1.613506 | 0.10663463 | NA |
| ADM | 40.151964 | -0.752667029 | 0.73487089 | -1.0242167 | 0.30573294 | 0.74897897 |
| AMPD3 | 112.198224 | -0.13011244 | 0.33610021 | -0.3871239 | 0.69866445 | 0.9304129 |
| MTRNR2L8 | 157.925378 | 0.623925415 | 0.79853667 | 0.78133596 | 0.43460492 | 0.82565889 |
| RNF141 | 40.9260682 | 0.153082629 | 0.23800471 | 0.6431916 | 0.52009978 | 0.86677556 |
| MRVI1 | 10.2464147 | 0.975480309 | 0.68916255 | 1.41545752 | 0.15693439 | NA |
| CTR9 | 52.9821369 | 0.290751915 | 0.2507095 | 1.15971639 | 0.2461643 | 0.69965741 |
| EIF4G2 | 162.243561 | 0.047466898 | 0.26966103 | 0.17602432 | 0.86027484 | 0.97287712 |
| ZBED5 | 59.781115 | -0.080096042 | 0.27184245 | -0.2946414 | 0.76826783 | 0.95205353 |
| ZBED5-AS1 | 15.1790563 | 0.548971013 | 0.51604554 | 1.06380343 | 0.28741776 | 0.73803612 |
| GALNT18 | 31.2089462 | -1.400899156 | 0.59083313 | -2.3710572 | 0.01773729 | 0.20666126 |
| USP47 | 307.135888 | 0.144776617 | 0.2470559 | 0.58600754 | 0.55787045 | 0.87946422 |
| PARVA | 98.6185812 | 0.03394057 | 0.2718493 | 0.12485068 | 0.90064176 | 0.97940229 |
| TEAD1 | 295.817576 | 0.317980041 | 0.38630457 | 0.823133 | 0.41043237 | 0.81427553 |
| LINC00958 | 58.5939778 | -0.583966401 | 0.23655583 | -2.4686198 | 0.01356352 | 0.17531992 |
| ARNTL | 25.4534213 | -0.096385009 | 0.38242016 | -0.2520396 | 0.80101049 | 0.95877735 |
| BTBD10 | 68.1130872 | -0.056452307 | 0.21598613 | -0.2613701 | 0.79380715 | 0.95770265 |
| FAR1 | 168.629803 | 0.104083393 | 0.22396263 | 0.46473554 | 0.64212085 | 0.91146968 |
| RRAS2 | 52.1814795 | -0.841072648 | 0.45471005 | -1.84969 | 0.06435824 | 0.40281695 |
| COPB1 | 87.8850552 | -0.050965426 | 0.19755099 | -0.2579862 | 0.79641757 | 0.95848876 |
| PSMA1 | 21.4089332 | 0.319750527 | 0.58038724 | 0.55092619 | 0.58168428 | 0.89036744 |
| PDE3B | 35.9183455 | 0.074370873 | 0.52670129 | 0.14120124 | 0.88771097 | 0.9776239 |
| CYP2R1 | 17.6328685 | 0.714375873 | 0.35112976 | 2.03450678 | 0.04190053 | 0.32888549 |
| CALCB | 34.4261616 | 2.122567959 | 1.22059599 | 1.73896029 | 0.08204175 | 0.45345012 |
| CALCA | 98.0121794 | 2.555698432 | 1.2666883 | 2.0176222 | 0.04363062 | 0.33245643 |
| SOX6 | 379.961396 | 0.235013931 | 0.20518517 | 1.14537484 | 0.25205392 | 0.70601299 |
| C11orf58 | 112.334144 | -0.274997545 | 0.21269723 | -1.2929061 | 0.19604353 | 0.64659037 |
| PLEKHA7 | 660.325878 | 0.18579291 | 0.21205747 | 0.87614416 | 0.38095167 | 0.79688435 |
| RPS13 | 147.899908 | 0.196516116 | 0.46446542 | 0.42310172 | 0.67222105 | 0.92176197 |
| PIK3C2A | 279.456924 | -0.341650971 | 0.13830431 | -2.4702844 | 0.01350057 | 0.1750742 |
| NUCB2 | 69.650004 | 0.461875462 | 0.34396587 | 1.34279444 | 0.17933854 | 0.62668718 |
| NCR3LG1 | 28.3243734 | -0.918145953 | 0.47478288 | -1.9338228 | 0.05313492 | 0.36579755 |
| AC124798.1 | 11.0191251 | -0.194160254 | 0.60864659 | -0.3190033 | 0.74972403 | NA |
| KCNC1 | 13.3143171 | -0.858312618 | 0.67102344 | -1.2791097 | 0.20085842 | 0.65358156 |
| SERGEF | 157.341559 | 0.223774173 | 0.2378961 | 0.94063827 | 0.34689026 | 0.7765713 |
| HPS5 | 32.1481666 | -0.015479531 | 0.26302098 | -0.0588528 | 0.95306932 | 0.99217529 |
| GTF2H1 | 45.3555762 | 0.002767933 | 0.27817669 | 0.00995027 | 0.99206096 | 0.99923337 |
| LDHA | 345.967792 | -1.246257381 | 0.37546327 | -3.3192525 | 0.00090259 | 0.03061224 |
| TSG101 | 92.9145732 | -0.208249145 | 0.17041376 | -1.2220207 | 0.2216998 | 0.67499434 |
| UEVLD | 72.1521735 | 0.017778965 | 0.19353999 | 0.09186197 | 0.9268077 | 0.98672316 |
| SPTY2D1OS | 25.0508171 | 0.111337219 | 0.35863536 | 0.31044686 | 0.75622117 | 0.94796399 |
| SPTY2D1 | 42.0886789 | -0.186930153 | 0.32368306 | -0.5775098 | 0.5635951 | 0.88191823 |
| IGSF22 | 12.1937068 | -0.486629519 | 0.67278207 | -0.7233093 | 0.46948989 | NA |
| ZDHHC13 | 85.6724131 | -0.584269118 | 0.26178069 | -2.231903 | 0.02562137 | 0.25205612 |
| NAV2 | 192.037261 | 0.748338266 | 0.42774547 | 1.7494943 | 0.08020561 | 0.44789428 |
| HTATIP2 | 36.5926228 | -1.488018203 | 0.37737793 | -3.9430451 | 8.05E-05 | 0.00561438 |
| PRMT3 | 51.992937 | 0.235362604 | 0.25613697 | 0.91889353 | 0.35815126 | 0.78368312 |
| NELL1 | 51.740734 | 3.429713195 | 0.59030682 | 5.81005181 | 6.25E-09 | 2.75E-06 |
| ANO5 | 54.5341399 | -0.300237794 | 0.36773045 | -0.8164616 | 0.41423618 | 0.81718343 |
| FANCF | 10.3259684 | -0.094037836 | 0.48528494 | -0.1937786 | 0.84634924 | NA |
| GAS2 | 79.5708854 | 0.723386147 | 0.49884433 | 1.45012401 | 0.14702394 | 0.58364362 |
| SVIP | 83.0414974 | -0.108332845 | 0.29344268 | -0.3691789 | 0.71199439 | 0.9357596 |
| MUC15 | 16.7914362 | -0.496416591 | 0.710925 | -0.6982686 | 0.48500924 | 0.85356796 |
| CCDC34 | 24.1289404 | 0.263264641 | 0.63548141 | 0.41427591 | 0.67867205 | 0.92291026 |
| LGR4 | 799.475863 | -0.271675041 | 0.29905768 | -0.9084369 | 0.36364742 | 0.7873616 |
| LIN7C | 42.3880041 | 0.380278663 | 0.33535401 | 1.13396189 | 0.25681053 | 0.70972499 |
| BDNF-AS | 122.515631 | 0.166576428 | 0.24796755 | 0.67176705 | 0.50173201 | 0.85942996 |
| METTL15 | 226.426886 | 0.150898657 | 0.2557042 | 0.59012975 | 0.55510366 | 0.87754295 |
| ARL14EP | 40.0934748 | 0.003074077 | 0.28298972 | 0.01086286 | 0.99133286 | 0.99923337 |
| MPPED2 | 1229.64138 | 0.281650489 | 0.35541301 | 0.79245971 | 0.42809268 | 0.82407405 |
| AL136088.1 | 36.6328303 | 0.64100874 | 0.57283331 | 1.11901444 | 0.26313398 | 0.71348562 |
| DCDC1 | 14.2992501 | 0.122227121 | 0.55684525 | 0.21949926 | 0.82626116 | 0.96539788 |
| DNAJC24 | 35.3199595 | 0.00798908 | 0.29701423 | 0.02689797 | 0.97854111 | 0.99610605 |
| IMMP1L | 96.7110599 | 0.301989249 | 0.21605567 | 1.39773814 | 0.16219172 | 0.60094226 |
| ELP4 | 116.154762 | 0.253459808 | 0.22984816 | 1.10272718 | 0.27014566 | 0.72068562 |
| AC131571.1 | 16.4203078 | -0.400493089 | 0.73871112 | -0.5421512 | 0.58771438 | 0.89073775 |
| RCN1 | 27.877219 | -0.906634989 | 0.40214363 | -2.2545054 | 0.02416439 | 0.24365344 |
| EIF3M | 50.2025706 | 0.049866391 | 0.24686918 | 0.20199521 | 0.83992047 | 0.96748214 |
| PRRG4 | 67.1074237 | 0.226411374 | 0.3197066 | 0.70818486 | 0.47883047 | 0.84833462 |
| QSER1 | 64.1926041 | -0.101147798 | 0.21919188 | -0.4614578 | 0.6444702 | 0.9119294 |
| TCP11L1 | 15.566413 | -0.781298611 | 0.44299769 | -1.763663 | 0.07778874 | 0.4397149 |
| CSTF3 | 68.9630763 | 0.332967938 | 0.34853378 | 0.95533907 | 0.33940625 | 0.77045469 |
| HIPK3 | 244.507364 | -0.317748164 | 0.26264 | -1.209824 | 0.22634645 | 0.68070982 |
| AL049629.1 | 49.4296785 | -0.898082894 | 0.59504314 | -1.5092736 | 0.13122888 | 0.55812058 |
| CD59 | 11.2741662 | -0.638997386 | 0.38602225 | -1.6553382 | 0.09785591 | NA |
| FBXO3 | 61.1764584 | 0.359405672 | 0.21967827 | 1.63605476 | 0.10182813 | 0.49220184 |
| CAPRIN1 | 112.954181 | 0.114591865 | 0.17637992 | 0.64968772 | 0.51589396 | 0.86476217 |
| NAT10 | 30.1535436 | 0.334902768 | 0.28979646 | 1.15564823 | 0.24782507 | 0.70240969 |
| ABTB2 | 255.730636 | -0.139299849 | 0.45347602 | -0.3071824 | 0.75870454 | 0.94924107 |
| CAT | 189.899674 | 0.079056027 | 0.47644497 | 0.16592898 | 0.86821284 | 0.97434023 |
| ELF5 | 22.2511979 | 4.14704511 | 1.96735505 | 2.10792917 | 0.03503712 | 0.29932858 |
| EHF | 61.3962483 | -0.012799047 | 0.84639572 | -0.0151218 | 0.98793499 | 0.99905853 |
| APIP | 66.1527488 | 0.216052844 | 0.29356508 | 0.73596233 | 0.46175362 | 0.83846194 |
| PDHX | 152.131555 | -0.08618309 | 0.24166653 | -0.3566199 | 0.72137639 | 0.93824012 |
| CD44 | 97.2192669 | -1.684861225 | 0.5117129 | -3.2925908 | 0.00099269 | 0.03257351 |
| TRIM44 | 185.85242 | -0.140169855 | 0.21391717 | -0.6552529 | 0.51230492 | 0.86346419 |
| COMMD9 | 14.9459292 | 0.308374884 | 0.35489594 | 0.86891634 | 0.38489289 | 0.79869171 |
| PRR5L | 16.9831053 | 0.245519364 | 0.84880118 | 0.28925427 | 0.7723868 | 0.95391163 |
| TRAF6 | 31.6398485 | -0.398374566 | 0.28200956 | -1.4126279 | 0.15776516 | 0.59848342 |
| C11orf74 | 16.2504106 | 0.563546851 | 0.40078672 | 1.40610159 | 0.15969396 | 0.60024417 |
| API5 | 65.7195883 | 0.141957401 | 0.26340922 | 0.53892343 | 0.58993969 | 0.89149106 |
| TTC17 | 339.757628 | -0.086356807 | 0.12800389 | -0.674642 | 0.49990322 | 0.85942996 |
| HSD17B12 | 277.271548 | -0.301799246 | 0.36032523 | -0.8375746 | 0.40226968 | 0.80890307 |
| ALKBH3 | 22.3114672 | 0.420024542 | 0.33216761 | 1.26449579 | 0.20605213 | 0.65836247 |
| ACCS | 14.2580926 | -0.351240146 | 0.50700517 | -0.6927743 | 0.48845121 | 0.85461341 |
| EXT2 | 64.3754991 | 0.212841276 | 0.23000118 | 0.92539211 | 0.35476198 | 0.7810615 |
| CD82 | 40.5932479 | -0.196099406 | 0.65760786 | -0.2982011 | 0.76554966 | 0.95205353 |
| TP53I11 | 41.7875384 | -0.19329683 | 0.31485296 | -0.6139273 | 0.53926336 | 0.86972442 |
| PRDM11 | 22.6585401 | 0.540494118 | 0.33062063 | 1.63478644 | 0.10209382 | 0.49220184 |
| SYT13 | 21.036526 | 0.961665387 | 0.56448417 | 1.703618 | 0.08845248 | 0.46851772 |
| CRY2 | 84.5794053 | -0.427291688 | 0.32900396 | -1.2987433 | 0.19403203 | 0.64544264 |
| PEX16 | 10.5119817 | -0.211923581 | 0.41907327 | -0.5056958 | 0.61307024 | NA |
| LARGE2 | 14.2927887 | 0.625745388 | 0.44840029 | 1.3955062 | 0.16286325 | 0.6013603 |
| PHF21A | 209.355928 | -0.144258224 | 0.23165966 | -0.6227162 | 0.53347103 | 0.86909171 |
| DGKZ | 28.6139279 | 0.398325179 | 0.45730941 | 0.87101899 | 0.38374379 | 0.7980695 |
| AMBRA1 | 452.654991 | 0.13662071 | 0.24075394 | 0.56747031 | 0.57039469 | 0.88563341 |
| ATG13 | 41.3296348 | -0.174589759 | 0.28099972 | -0.6213165 | 0.53439141 | 0.86909171 |
| ARHGAP1 | 37.0593887 | -0.33328305 | 0.31336367 | -1.0635663 | 0.2875252 | 0.73803612 |

| | | | | | | |
|---|---|---|---|---|---|---|
| CKAP5 | 76.6201673 | 0.106775758 | 0.19473523 | 0.54831247 | 0.58347736 | 0.89036744 |
| LRP4 | 49.9213362 | -0.798034777 | 0.4345778 | -1.836345 | 0.06630664 | 0.40888394 |
| C11orf49 | 164.192513 | -0.068161983 | 0.20549966 | -0.331689 | 0.74012408 | 0.94376279 |
| ARFGAP2 | 28.7702609 | 0.075205535 | 0.33949268 | 0.22152328 | 0.82468501 | 0.96539788 |
| DDB2 | 12.848043 | -1.089225712 | 0.41878777 | -2.6009014 | 0.00929792 | 0.14114208 |
| ACP2 | 13.136441 | 0.189576179 | 0.48484204 | 0.39100606 | 0.69579275 | 0.92900737 |
| NR1H3 | 19.9386946 | 0.110878626 | 0.33406965 | 0.33190273 | 0.73996271 | 0.94376279 |
| MADD | 21.3682505 | 0.203713253 | 0.3529848 | 0.57711622 | 0.56386095 | 0.88191823 |
| SLC39A13 | 33.072595 | -0.443781236 | 0.36912028 | -1.2022673 | 0.22925998 | 0.68503082 |
| PSMC3 | 17.1575629 | 0.145873548 | 0.48211742 | 0.30256851 | 0.76221872 | 0.95024054 |
| CELF1 | 306.809718 | 0.14184824 | 0.22769472 | 0.62297554 | 0.53330059 | 0.86909171 |
| NDUFS3 | 46.2782308 | 0.51307696 | 0.41622445 | 1.23269298 | 0.21769033 | 0.67202699 |
| MTCH2 | 71.5584532 | -0.140020795 | 0.26669298 | -0.5250262 | 0.59956499 | 0.89476262 |
| FNBP4 | 181.125433 | -0.401340394 | 0.19890317 | -2.0177677 | 0.04361546 | 0.33245643 |
| NUP160 | 130.9093 | -0.017181805 | 0.21710872 | -0.0791392 | 0.93692193 | 0.98871562 |
| PTPRJ | 1440.21174 | 0.139792488 | 0.25843817 | 0.5409127 | 0.58856775 | 0.89093433 |
| TNKS1BP1 | 50.7213586 | -1.598135368 | 0.42618459 | -3.7498666 | 0.00017693 | 0.01015508 |
| SSRP1 | 25.0769768 | -0.027479245 | 0.31200244 | -0.0880738 | 0.92981801 | 0.98733028 |
| TIMM10 | 18.9981441 | -0.368170535 | 0.45135355 | -0.8157032 | 0.41466991 | 0.81726257 |
| UBE2L6 | 31.4427284 | -1.31468631 | 0.70798972 | -1.8569285 | 0.06332132 | 0.4001801 |
| SERPING1 | 284.926482 | -3.044374079 | 0.46676968 | -6.522219 | 6.93E-11 | 8.40E-08 |
| ZDHHC5 | 85.5222303 | -0.368512637 | 0.26823306 | -1.3738524 | 0.1694875 | 0.61207324 |
| MED19 | 14.5544915 | -0.105443139 | 0.40185901 | -0.2623884 | 0.79302202 | 0.95746995 |
| CTNND1 | 183.297195 | 0.117454562 | 0.17618811 | 0.66664297 | 0.50500022 | 0.86193811 |
| LPXN | 24.0765198 | 0.032783206 | 0.38327573 | 0.08553426 | 0.93183666 | 0.9875946 |
| FAM111A | 38.221231 | 0.069496223 | 0.26266319 | 0.26458303 | 0.7913307 | 0.95746995 |
| OSBP | 85.818634 | 0.106765553 | 0.20420559 | 0.52283364 | 0.60109003 | 0.89490801 |
| PATL1 | 49.4445622 | -0.625100858 | 0.30977941 | -2.0178903 | 0.04360269 | 0.33245643 |
| STX3 | 53.8648521 | -0.956083677 | 0.46851354 | -2.0406746 | 0.04128318 | 0.32628324 |
| MRPL16 | 32.420809 | 0.379921043 | 0.34088054 | 1.1145284 | 0.26505257 | 0.71635829 |
| PRPF19 | 21.7582536 | 0.075705397 | 0.40898573 | 0.18510523 | 0.8531465 | 0.9711913 |
| TMEM109 | 22.4204799 | 0.395652766 | 0.35774218 | 1.10597181 | 0.26873872 | 0.72009952 |
| VPS37C | 31.6781162 | -0.164582353 | 0.35740624 | -0.460491 | 0.64516384 | 0.9122579 |
| DDB1 | 110.707135 | 0.216998056 | 0.23385745 | 0.92790739 | 0.35345561 | 0.78028099 |
| TMEM138 | 12.2459362 | 0.020650279 | 0.39379635 | 0.05243898 | 0.95817892 | NA |
| CPSF7 | 56.5947738 | 0.259537087 | 0.24749111 | 1.04867238 | 0.29432893 | 0.7405942 |
| AP003108.2 | 30.6645995 | -0.122495204 | 0.43790437 | -0.2797305 | 0.77968429 | 0.95537097 |
| SYT7 | 18.702871 | 0.804508097 | 0.87446512 | 0.92000021 | 0.35757265 | 0.78312366 |
| TMEM258 | 29.9935514 | 0.227270941 | 0.36652896 | 0.62006272 | 0.5352165 | 0.86946459 |
| FADS1 | 17.2156131 | -1.055788311 | 0.89615959 | -1.1781253 | 0.23874664 | 0.69356592 |
| FTH1 | 985.280448 | -1.158347289 | 0.44139748 | -2.6242725 | 0.00868343 | 0.13512263 |
| ASRGL1 | 14.1547184 | 0.395633073 | 0.42525498 | 0.93034318 | 0.35219343 | 0.77964174 |
| AHNAK | 179.3537 | 0.013206533 | 0.2691907 | 0.04906014 | 0.96087137 | 0.99238848 |
| EML3 | 13.3995778 | -0.39649916 | 0.50906397 | -0.7788788 | 0.43605108 | 0.82643523 |
| B3GAT3 | 33.2117855 | -0.186079114 | 0.30775088 | -0.604642 | 0.54541688 | 0.87247631 |
| GANAB | 82.8694282 | 0.26106841 | 0.2336919 | 1.11714787 | 0.26393111 | 0.71392408 |
| UQCC3 | 27.2801999 | -0.077468491 | 0.40768241 | -0.1900217 | 0.84929215 | 0.97027687 |
| UBXN1 | 31.9818497 | -0.47326052 | 0.50670817 | -0.9339903 | 0.35030889 | 0.77868979 |
| TTC9C | 24.6181747 | -0.311049497 | 0.31676287 | -0.9819632 | 0.32611796 | 0.76353952 |
| POLR2G | 12.6748237 | 0.478639086 | 0.45199166 | 1.05895556 | 0.28962002 | 0.73966536 |
| TAF6L | 11.7350458 | 0.032025976 | 0.59535409 | 0.05379316 | 0.95709996 | NA |
| TMEM223 | 25.8056075 | -0.471151725 | 0.30769692 | -1.5312202 | 0.12571499 | 0.54560868 |
| TMEM179B | 39.1398813 | -0.261103617 | 0.39459478 | -0.6617006 | 0.5081631 | 0.86281145 |
| NXF1 | 15.594178 | 0.770144918 | 0.33764397 | 2.28093785 | 0.02255212 | 0.23595493 |
| STX5 | 50.2290329 | -0.064289313 | 0.22729404 | -0.2828465 | 0.77729454 | 0.95512504 |
| SLC3A2 | 67.0466658 | -0.686367776 | 0.49844221 | -1.3770258 | 0.16850427 | 0.61056834 |
| RARRES3 | 333.518447 | -1.670808179 | 0.46757057 | -3.5733819 | 0.0003524 | 0.01589899 |
| PLA2G16 | 55.3803066 | -0.94500345 | 0.38196102 | -2.4740835 | 0.01335785 | 0.17368784 |
| ATL3 | 79.4771334 | 0.064393188 | 0.21695832 | 0.29679981 | 0.76661935 | 0.95205353 |
| RTN3 | 161.147546 | 0.596601456 | 0.36233719 | 1.64653661 | 0.09965333 | 0.48648077 |
| SPINDOC | 11.2260755 | -0.282669692 | 0.52581246 | -0.5375865 | 0.59086253 | NA |
| MARK2 | 92.2784791 | 0.124670511 | 0.20592332 | 0.605422 | 0.54489864 | 0.87247631 |
| NAA40 | 27.9560086 | 0.260632293 | 0.31501641 | 0.827361 | 0.40803248 | 0.81361336 |
| OTUB1 | 36.0103129 | 0.125293606 | 0.27982984 | 0.44774926 | 0.65433417 | 0.9158679 |
| MACROD1 | 47.1643723 | 0.509009618 | 0.29910111 | 1.70179783 | 0.08879327 | 0.4687332 |
| STIP1 | 46.2233917 | 0.309737689 | 0.26367991 | 1.17467307 | 0.24012554 | 0.69470237 |
| NUDT22 | 32.8865756 | 0.370106176 | 0.32709488 | 1.13149487 | 0.25784686 | 0.71022118 |
| DNAJC4 | 26.7731751 | 0.568005198 | 0.33846769 | 1.6781667 | 0.09331456 | 0.47790456 |
| VEGFB | 73.6222251 | -0.452539346 | 0.33319951 | -1.3581633 | 0.17441187 | 0.62052871 |
| FKBP2 | 68.0565283 | 0.300356123 | 0.32979404 | 0.91073849 | 0.36243318 | 0.78714716 |
| PPP1R14B | 47.9503096 | 0.003332442 | 0.36587391 | 0.00910817 | 0.99273283 | 0.99923337 |
| BAD | 27.2019453 | -0.395224269 | 0.35618044 | -1.1096181 | 0.26716362 | 0.71846053 |
| GPR137 | 19.2764133 | 0.064253154 | 0.34176242 | 0.18800533 | 0.85087248 | 0.97027687 |
| ESRRA | 119.418281 | 0.213938428 | 0.3222465 | 0.66389683 | 0.50675634 | 0.86237483 |
| TRMT112 | 46.1073939 | -0.08679017 | 0.49209141 | -0.17637 | 0.86000326 | 0.97287712 |
| PRDX5 | 98.8887732 | 0.330974255 | 0.43330543 | 0.76383592 | 0.44496503 | 0.83035712 |
| SF1 | 158.907672 | 0.157090067 | 0.23142353 | 0.67879903 | 0.49726521 | 0.85814183 |
| MAP4K2 | 12.5081496 | 0.818249066 | 0.39473106 | 2.07292798 | 0.03817898 | NA |
| MEN1 | 11.6763014 | 0.42157952 | 0.43148436 | 0.97704472 | 0.32854701 | NA |
| EHD1 | 13.2347075 | 0.704626971 | 0.43879028 | 1.60583997 | 0.10830908 | 0.50631231 |
| ATG2A | 23.0886107 | -0.722053491 | 0.58104567 | -1.2426794 | 0.21398599 | 0.66784559 |
| PPP2R5B | 12.4981186 | -0.105806448 | 0.39748058 | -0.2661928 | 0.79009077 | NA |
| ZFPL1 | 13.0821953 | -0.081020197 | 0.43045614 | -0.1882194 | 0.85070466 | 0.97027687 |
| VPS51 | 64.4435989 | 0.081680056 | 0.26594447 | 0.307132 | 0.7587429 | 0.94924107 |
| TM7SF2 | 18.3373053 | 0.592915343 | 0.34859951 | 1.7008496 | 0.08897123 | 0.46877836 |
| FAU | 166.675477 | 0.364907529 | 0.34708866 | 1.05133809 | 0.29310334 | 0.74013384 |
| SYVN1 | 21.0146011 | 0.67234452 | 0.3316542 | 2.02724562 | 0.0426373 | 0.33083096 |
| MRPL49 | 25.3915673 | -0.481348532 | 0.39090316 | -1.2313754 | 0.2181825 | 0.67229041 |
| CAPN1 | 80.4503871 | -0.447143939 | 0.24138951 | -1.8523752 | 0.06397196 | 0.40111702 |
| DPF2 | 32.1144792 | 0.138707973 | 0.24121806 | 0.57503146 | 0.56527002 | 0.88252361 |
| SLC25A45 | 15.0146194 | -1.538233215 | 0.54640877 | -2.8151693 | 0.00487515 | 0.09611584 |
| FRMD8 | 25.032029 | -0.631313998 | 0.37566026 | -1.6805451 | 0.09285132 | 0.47759124 |
| NEAT1 | 5140.88093 | -1.280605336 | 0.43162471 | -2.9669417 | 0.00300778 | 0.06784993 |
| MALAT1 | 90522.2756 | -1.038528923 | 0.30433084 | -3.4124998 | 0.0006437 | 0.02420111 |
| SCYL1 | 31.5370091 | -0.221654282 | 0.35099622 | -0.6315005 | 0.52771333 | 0.86750588 |
| LTBP3 | 44.9969002 | 0.474087281 | 0.31905846 | 1.48589472 | 0.13730697 | 0.56857145 |
| SSSCA1 | 14.0317257 | -0.35164164 | 0.36972756 | -0.9510831 | 0.34156219 | 0.77193691 |
| FAM89B | 10.9292043 | -0.284475017 | 0.521355 | -0.5456455 | 0.58530963 | NA |
| EHBP1L1 | 18.2943254 | -0.887337684 | 0.55492175 | -1.5990321 | 0.10981347 | 0.50994182 |
| MAP3K11 | 25.3722333 | -0.048186555 | 0.2868346 | -0.1679942 | 0.86658783 | 0.97434023 |
| PCNX3 | 21.2074235 | 0.460638908 | 0.4644859 | 0.99171773 | 0.32133524 | 0.76163052 |
| SIPA1 | 19.1279229 | -0.236929613 | 0.52882874 | -0.4480271 | 0.65413363 | 0.9158679 |
| RELA | 32.2751341 | -0.481957871 | 0.28510841 | -1.6904372 | 0.09094434 | 0.47326187 |
| KAT5 | 28.0825008 | 0.057285952 | 0.42457236 | 0.13492624 | 0.89267019 | 0.97823448 |
| RNASEH2C | 18.5983883 | 0.332414226 | 0.52768024 | 0.6299539 | 0.52872474 | 0.86750588 |
| AP5B1 | 10.3595638 | -0.188181181 | 0.57354268 | -0.3281032 | 0.74283365 | NA |
| CFL1 | 169.049482 | -0.486088985 | 0.35031711 | -1.3875685 | 0.16526848 | 0.60578861 |
| MUS81 | 16.7456435 | -0.261003188 | 0.33429997 | -0.7807455 | 0.4349522 | 0.82565889 |
| EFEMP2 | 14.8071665 | -0.27654287 | 0.36648683 | -0.7545779 | 0.45050228 | 0.83156463 |
| CCDC85B | 12.2416601 | 0.343567107 | 0.34630697 | 0.99208834 | 0.32115443 | NA |
| C11orf68 | 10.9976783 | 0.395050313 | 0.4119775 | 0.95891235 | 0.3376029 | NA |
| DRAP1 | 26.0278204 | 0.314541268 | 0.48342856 | 0.65064684 | 0.51527449 | 0.86463667 |
| SART1 | 22.0092637 | -0.130788379 | 0.39223489 | -0.333444 | 0.73879914 | 0.94376279 |

| | | | | | | |
|---|---|---|---|---|---|---|
| BANF1 | 35.0218736 | 0.208959205 | 0.35686595 | 0.58553976 | 0.55818484 | 0.87981686 |
| SF3B2 | 53.7083941 | -0.006588755 | 0.28063341 | -0.0234782 | 0.98126886 | 0.99714727 |
| PACS1 | 414.489219 | -0.036870564 | 0.20696544 | -0.1781484 | 0.85860643 | 0.97287712 |
| KLC2 | 22.5608751 | -0.161742231 | 0.33665639 | -0.4804371 | 0.6309166 | 0.90768383 |
| RAB1B | 51.335261 | -0.0556764 | 0.294466 | -0.1890758 | 0.85003339 | 0.97027687 |
| YIF1A | 18.2290645 | -0.088925763 | 0.37087283 | -0.2397743 | 0.81050525 | 0.9623221 |
| BRMS1 | 15.8667696 | 0.10906588 | 0.34374168 | 0.31729024 | 0.75102338 | 0.94718617 |
| SLC29A2 | 41.3286933 | 0.265325961 | 0.31746408 | 0.83576688 | 0.40328606 | 0.80927646 |
| MRPL11 | 15.5072995 | 0.367309513 | 0.40079884 | 0.91644356 | 0.3594343 | 0.78473041 |
| ZDHHC24 | 15.4791279 | -0.263306369 | 0.43635109 | -0.6034278 | 0.54622415 | 0.87259128 |
| CCS | 36.3275002 | 0.474401533 | 0.33424836 | 1.4193085 | 0.15580909 | 0.59597065 |
| RBM14-RBM | 26.7717027 | 0.418757202 | 0.3073085 | 1.36266065 | 0.17298949 | 0.61781959 |
| RBM4 | 24.8422593 | 0.342774357 | 0.291117 | 1.17744533 | 0.2390178 | 0.69356592 |
| RBM4B | 30.8866097 | 0.101228949 | 0.35473224 | 0.28536721 | 0.77536283 | 0.95493992 |
| AP001157.1 | 11.9676492 | -0.004719077 | 0.53270833 | -0.0088586 | 0.99293191 | NA |
| SPTBN2 | 302.365386 | -0.089589878 | 0.39918689 | -0.2244309 | 0.82242202 | 0.96521156 |
| C11orf80 | 541.19099 | -0.161997452 | 0.424471 | -0.3816455 | 0.70272432 | 0.93247961 |
| RCE1 | 20.6304298 | 0.025960294 | 0.39674168 | 0.06543375 | 0.94782866 | 0.99149867 |
| PC | 12.5818994 | 1.117373393 | 0.45399951 | 2.46117751 | 0.01384818 | 0.17643441 |
| LRFN4 | 12.3812659 | 0.342312005 | 0.54256453 | 0.63091483 | 0.52809622 | NA |
| RHOD | 25.8819266 | -0.112922336 | 0.34888819 | -0.3236634 | 0.74619289 | 0.94614106 |
| KDM2A | 302.17532 | 0.049153934 | 0.18617706 | 0.26401713 | 0.79176672 | 0.95746995 |
| GRK2 | 20.3443233 | 0.284003193 | 0.46274596 | 0.61373457 | 0.53939074 | 0.86972442 |
| ANKRD13D | 21.2633662 | -0.115986055 | 0.31101621 | -0.3729261 | 0.70920346 | 0.93485232 |
| SSH3 | 25.6018003 | -0.358347087 | 0.4202865 | -0.8526257 | 0.39386689 | 0.80438874 |
| RAD9A | 94.900514 | 0.311046826 | 0.313332 | 0.99270687 | 0.32085283 | 0.76131909 |
| PPP1CA | 61.4944628 | -0.522449761 | 0.35881073 | -1.4560595 | 0.14537618 | 0.58098815 |
| RPS6KB2 | 26.5872931 | 0.499084895 | 0.34513347 | 1.44606342 | 0.14815941 | 0.58515729 |
| CORO1B | 30.0701652 | -0.768085989 | 0.33685607 | -2.2801607 | 0.02259816 | 0.23595493 |
| TMEM134 | 31.9024164 | -0.369697186 | 0.32592447 | -1.1343033 | 0.25666736 | 0.70972499 |
| AIP | 14.7401118 | -0.17104871 | 0.43706803 | -0.3913549 | 0.69553493 | 0.92900737 |
| PITPNM1 | 19.3649784 | -0.468820101 | 0.44360262 | -1.056847 | 0.29058142 | 0.73979353 |
| CDK2AP2 | 29.1959196 | -0.39503714 | 0.32584292 | -1.2123545 | 0.22537669 | 0.67852788 |
| GSTP1 | 275.698176 | -0.723568481 | 0.38076447 | -1.9003046 | 0.05739315 | 0.38078905 |
| NDUFV1 | 93.2959059 | 0.404434342 | 0.30096355 | 1.34379842 | 0.17901358 | 0.62639418 |
| NUDT8 | 14.9309916 | -0.501955619 | 0.52695252 | -0.9525633 | 0.34081139 | 0.77193691 |
| UNC93B1 | 11.8619975 | -0.816577802 | 0.74275636 | -1.0993885 | 0.27159864 | NA |
| NDUFS8 | 92.8765637 | -0.01443547 | 0.36771541 | -0.0392572 | 0.96868535 | 0.99441717 |
| TCIRG1 | 23.7213998 | 0.13862389 | 0.36039578 | 0.38464348 | 0.70050157 | 0.93127962 |
| CHKA | 140.515745 | -0.27820803 | 0.29977758 | -0.9280481 | 0.35338259 | 0.78028099 |
| KMT5B | 164.007311 | 0.168300193 | 0.26746709 | 0.62923702 | 0.5291939 | 0.86758926 |
| LRP5 | 43.1793425 | 0.404026553 | 0.36632926 | 1.10290551 | 0.2700682 | 0.72068562 |
| PPP6R3 | 393.981943 | 0.105869235 | 0.15811057 | 0.66958985 | 0.50311929 | 0.86010876 |
| TESMIN | 37.576221 | 0.055208016 | 0.38142852 | 0.14474014 | 0.88491605 | 0.97662748 |
| CPT1A | 277.792945 | 0.12444664 | 0.38903021 | 0.3198894 | 0.74905217 | 0.94684415 |
| MRPL21 | 15.7167852 | -0.398946941 | 0.36648289 | -1.0885827 | 0.27633794 | 0.72575599 |
| IGHMBP2 | 16.2163945 | -0.330579465 | 0.34055167 | -0.9707175 | 0.33168898 | 0.76685572 |
| TPCN2 | 20.3665804 | 0.050670942 | 0.43474359 | 0.11655363 | 0.90721379 | 0.98224955 |
| CCND1 | 31.9292114 | 0.607539825 | 0.32553862 | 1.86626036 | 0.06200495 | 0.39551484 |
| ORAOV1 | 18.4681504 | 0.358021715 | 0.38285876 | 0.93512739 | 0.34972265 | 0.77868979 |
| PPFIA1 | 141.518875 | 0.266215883 | 0.3452969 | 0.77097675 | 0.44072071 | 0.82793978 |
| AP002336.2 | 48.7948427 | -0.254870912 | 0.45030913 | -0.565991 | 0.57139991 | 0.88578992 |
| CTTN | 211.326704 | -0.835334767 | 0.2393902 | -3.4894275 | 0.00048406 | 0.01956394 |
| SHANK2 | 404.908599 | 0.515625669 | 0.28658059 | 1.79923446 | 0.0719816 | 0.42342118 |
| DHCR7 | 26.424654 | -0.637261278 | 0.48716505 | -1.3081014 | 0.19083893 | 0.6421543 |
| NADSYN1 | 48.5344245 | -0.483008778 | 0.23130672 | -2.0881744 | 0.0367821 | 0.30695956 |
| KRTAP5-8 | 31.5144785 | -1.879075126 | 0.91408289 | -2.0556945 | 0.03981198 | 0.31990103 |
| AP002495.2 | 45.356211 | 0.535395002 | 0.31328779 | 1.70895582 | 0.08745913 | 0.46528247 |
| RNF121 | 83.3284999 | 0.296894094 | 0.21585883 | 1.37540861 | 0.1690048 | 0.61124033 |
| NUMA1 | 122.640605 | 0.008811393 | 0.28529599 | 0.03088509 | 0.97536118 | 0.99552114 |
| LAMTOR1 | 73.4056845 | -0.951373618 | 0.28982869 | -3.2825378 | 0.00102877 | 0.03326363 |
| ANAPC15 | 17.1784731 | -0.003642776 | 0.43825613 | -0.008312 | 0.99336808 | 0.99923337 |
| FOLR1 | 20.5532677 | 0.619622091 | 0.83367664 | 0.74324032 | 0.4573362 | 0.83569193 |
| INPPL1 | 43.1471261 | 0.560942295 | 0.26929833 | 2.08297728 | 0.03725329 | 0.3091163 |
| CLPB | 36.6155699 | 0.746959395 | 0.28225412 | 2.64640737 | 0.00813518 | 0.13108178 |
| ARAP1 | 142.91559 | 0.076678104 | 0.2163435 | 0.35442758 | 0.72301847 | 0.93873366 |
| ATG16L2 | 13.3439681 | 0.170429481 | 0.37604631 | 0.45321408 | 0.6503946 | 0.91485319 |
| FCHSD2 | 88.8977327 | -0.457669803 | 0.20537203 | -2.2284914 | 0.02584777 | 0.25292492 |
| ARHGEF17 | 52.7142829 | 0.056823176 | 0.25854476 | 0.21978081 | 0.82604187 | 0.96539788 |
| FAM168A | 130.494137 | 0.480692206 | 0.23802873 | 2.01947137 | 0.04343825 | 0.3320339 |
| PLEKHB1 | 15.3237813 | -0.508637671 | 0.56291794 | -0.9035734 | 0.36622166 | 0.78793182 |
| RAB6A | 164.619548 | -0.236520357 | 0.19854264 | -1.1912824 | 0.23354274 | 0.68826429 |
| MRPL48 | 40.2720127 | 0.378848166 | 0.30478446 | 1.24300356 | 0.21386652 | 0.66782265 |
| COA4 | 31.5813579 | -0.376900899 | 0.30005376 | -1.2561113 | 0.20907564 | 0.66102793 |
| PAAF1 | 33.0188431 | 0.218159854 | 0.3978644 | 0.54832715 | 0.58346729 | 0.89036744 |
| UCP2 | 32.2343531 | -0.612293617 | 0.67355009 | -0.9090543 | 0.36332146 | 0.78718296 |
| C2CD3 | 75.7432509 | 0.323437243 | 0.22986449 | 1.40707789 | 0.15940429 | 0.60024133 |
| PPME1 | 99.0855084 | -0.272999008 | 0.27494176 | -0.992934 | 0.32074213 | 0.76131909 |
| P4HA3 | 19.1093123 | -1.091054283 | 0.59512409 | -1.8333223 | 0.06675465 | 0.41086302 |
| POLD3 | 30.7525778 | -0.092129825 | 0.27091866 | -0.3400645 | 0.73380793 | 0.94277981 |
| RNF169 | 182.002308 | 0.307496943 | 0.25922497 | 1.18621653 | 0.2355368 | 0.6902438 |
| XRRA1 | 103.225371 | 0.094306686 | 0.42305225 | 0.22291971 | 0.82359799 | 0.96539788 |
| SPCS2 | 118.382707 | 0.141031298 | 0.27278871 | 0.51699829 | 0.60515735 | 0.89600432 |
| NEU3 | 14.7445951 | 0.280498326 | 0.4041381 | 0.69406553 | 0.48764112 | 0.85396621 |
| ARRB1 | 42.8907797 | 0.419192193 | 0.43279077 | 0.96857933 | 0.33275512 | 0.7674968 |
| RPS3 | 297.895663 | 0.279598958 | 0.34610458 | 0.8078453 | 0.41917964 | 0.8189411 |
| SERPINH1 | 15.1393812 | 0.471215393 | 0.41887746 | 1.12494807 | 0.26061104 | 0.71118349 |
| UVRAG | 198.093415 | -0.531083675 | 0.20049924 | -2.6488064 | 0.00807766 | 0.13068568 |
| THAP12 | 28.9319559 | 0.733852406 | 0.29394436 | 2.49656911 | 0.01254012 | 0.16675032 |
| EMSY | 108.596223 | -0.163476034 | 0.20731389 | -0.7885436 | 0.43037882 | 0.82469033 |
| ACER3 | 81.2196797 | 0.054307057 | 0.22586929 | 0.24043577 | 0.80999245 | 0.96215121 |
| CAPN5 | 13.091867 | 0.85332687 | 0.56907472 | 1.4994988 | 0.13374428 | 0.56229288 |
| PAK1 | 100.165034 | -0.226627818 | 0.33234674 | -0.6819017 | 0.49530109 | 0.85814183 |
| CLNS1A | 55.3412099 | 0.043701057 | 0.23026446 | 0.18978637 | 0.84947654 | 0.97027687 |
| RSF1 | 204.527526 | -0.099892402 | 0.18676554 | -0.5348546 | 0.59275042 | 0.89198431 |
| AAMDC | 30.2977525 | 0.077115789 | 0.30756404 | 0.25073084 | 0.80202222 | 0.95890737 |
| AP002812.2 | 15.4515981 | 0.645093936 | 0.53729044 | 1.20064287 | 0.22988976 | 0.68523674 |
| INTS4 | 51.1537023 | 0.307745435 | 0.20765309 | 1.48201712 | 0.13833574 | 0.57086516 |
| ALG8 | 29.4206098 | 0.078120791 | 0.30528006 | 0.25589877 | 0.79802901 | 0.95848876 |
| GAB2 | 155.140492 | -0.414605077 | 0.34412862 | -1.2047969 | 0.22828174 | 0.68364117 |
| AP003086.2 | 39.9057479 | -0.888299371 | 0.44689932 | -1.9876946 | 0.04684548 | 0.34525746 |
| NARS2 | 61.7782917 | 0.653832003 | 0.21880229 | 2.98823198 | 0.00280596 | 0.0651145 |
| TENM4 | 26.177515 | -1.370304971 | 0.67833472 | -2.0201015 | 0.04337286 | 0.33179556 |
| PRCP | 39.1910916 | 0.547239631 | 0.34054658 | 1.60694504 | 0.10806643 | 0.50590942 |
| RAB30 | 21.9525368 | 0.267345255 | 0.53602667 | 0.49875365 | 0.61795295 | 0.90218898 |
| RAB30-AS1 | 47.5010168 | -0.009686443 | 0.33352513 | -0.0290426 | 0.9768306 | 0.9957152 |
| PCF11 | 62.1633882 | 0.213792498 | 0.34240267 | 0.62438912 | 0.53237207 | 0.86892295 |
| ANKRD42 | 29.1864524 | -0.09414814 | 0.301975 | -0.3117746 | 0.75521182 | 0.94795939 |
| CCDC90B | 31.1943809 | -0.116229531 | 0.30024745 | -0.3871125 | 0.69867295 | 0.9304129 |
| AP000446.1 | 26.3208328 | 0.687436378 | 0.28558409 | 2.40712419 | 0.0160787 | 0.19519825 |
| DLG2 | 63.4145482 | 0.656535858 | 0.44961903 | 1.46020478 | 0.1442338 | 0.57846905 |
| TMEM126B | 25.7965884 | -0.36128758 | 0.43706379 | -0.8266244 | 0.40845 | 0.81362627 |
| TMEM126A | 14.1812345 | 0.216889361 | 0.45397593 | 0.4777552 | 0.63282445 | 0.90805737 |

| | | | | | | |
|---|---|---|---|---|---|---|
| CREBZF | 55.0448051 | 0.059400466 | 0.21139547 | 0.28099215 | 0.77871642 | 0.95537097 |
| SYTL2 | 15.8525321 | -0.418533693 | 0.63741617 | -0.6566098 | 0.51143186 | 0.86312853 |
| PICALM | 1001.18224 | 0.174984833 | 0.31845134 | 0.54948688 | 0.58267137 | 0.89036744 |
| EED | 35.2182996 | 0.162830193 | 0.2863964 | 0.56854832 | 0.5696627 | 0.88562534 |
| HIKESHI | 37.2563338 | 0.385088263 | 0.28448252 | 1.35364472 | 0.17584975 | 0.6225246 |
| AP001831.1 | 48.186534 | -1.178085808 | 0.46404241 | -2.538746 | 0.01112506 | 0.15593979 |
| ME3 | 125.995295 | -0.008165046 | 0.27329534 | -0.0298763 | 0.97616573 | 0.99556048 |
| PRSS23 | 32.6052704 | -0.091995306 | 0.50650259 | -0.1816285 | 0.85587428 | 0.97166553 |
| TMEM135 | 143.260373 | -0.158735566 | 0.24520797 | -0.6473508 | 0.51740495 | 0.86500999 |
| CTSC | 23.5389127 | 0.709823362 | 0.41261124 | 1.72031997 | 0.08537429 | 0.46032833 |
| NOX4 | 53.773197 | 1.366354885 | 0.65982834 | 2.07077328 | 0.03837999 | 0.3141788 |
| CHORDC1 | 42.9590189 | 0.32429363 | 0.35110235 | 0.92364413 | 0.35567163 | 0.78153728 |
| SLC36A4 | 77.1910595 | -0.111865506 | 0.37035036 | -0.3020532 | 0.76261152 | 0.95044735 |
| CEP295 | 40.0755432 | 0.128876745 | 0.28800529 | 0.44748047 | 0.65452819 | 0.9158679 |
| TAF1D | 150.033282 | -0.564609028 | 0.2955289 | -1.9105036 | 0.0560684 | 0.37689779 |
| PANX1 | 15.2496833 | -0.675715905 | 0.34617852 | -1.951929 | 0.05094663 | 0.3596669 |
| MRE11 | 35.3426007 | 0.187970172 | 0.29243741 | 0.6427706 | 0.52037296 | 0.86680671 |
| ANKRD49 | 14.1132366 | 0.258114053 | 0.39094418 | 0.66023249 | 0.50910464 | 0.86281145 |
| CWC15 | 24.5088325 | 0.110609424 | 0.35615796 | 0.31056283 | 0.75613299 | 0.94796399 |
| SRSF8 | 27.0309061 | -0.122352025 | 0.34509335 | -0.3545476 | 0.72292852 | 0.93873366 |
| ENDOD1 | 68.7337763 | -1.617347147 | 0.45784598 | -3.5325136 | 0.00041163 | 0.01743582 |
| AP000787.1 | 36.6059148 | 0.060863614 | 0.35576963 | 0.17107591 | 0.86416407 | 0.97401714 |
| SESN3 | 22.4814297 | 0.046453083 | 0.39404579 | 0.11788753 | 0.90615678 | 0.98165297 |
| FAM76B | 29.3873249 | 0.133739309 | 0.28484045 | 0.46952359 | 0.63869543 | 0.91063717 |
| CEP57 | 105.373235 | -0.249929755 | 0.20938015 | -1.193665 | 0.23260905 | 0.68763908 |
| MTMR2 | 65.4774743 | -0.132613573 | 0.27691878 | -0.4788898 | 0.63201704 | 0.90805737 |
| MAML2 | 36.745391 | -0.038233097 | 0.33995815 | -0.1124641 | 0.91045541 | 0.98334456 |
| CCDC82 | 37.1441079 | 0.078423886 | 0.3170198 | 0.24737851 | 0.8046153 | 0.95987805 |
| CNTN5 | 15.1422248 | 0.217539802 | 0.69505408 | 0.31298255 | 0.75429392 | 0.94795939 |
| ARHGAP42 | 26.244288 | -1.014933352 | 0.41599314 | -2.4397839 | 0.01469605 | 0.18393768 |
| YAP1 | 323.205424 | -0.486256177 | 0.18009853 | -2.6999453 | 0.00693509 | 0.11948554 |
| BIRC3 | 92.5408533 | -1.255580991 | 0.61573252 | -2.0391663 | 0.04143343 | 0.32675147 |
| BIRC2 | 74.2928353 | 0.122445243 | 0.23816204 | 0.51412576 | 0.60716407 | 0.89696793 |
| TMEM123 | 214.09475 | -0.428672838 | 0.21432937 | -2.0000658 | 0.04549316 | 0.33918806 |
| MMP7 | 45.4807473 | -3.676201753 | 0.63400091 | -5.7984172 | 6.69E-09 | 2.82E-06 |
| DCUN1D5 | 33.6157994 | 0.305224218 | 0.30307059 | 1.00710604 | 0.31388382 | 0.75646164 |
| DYNC2H1 | 63.3825483 | -0.069495567 | 0.45439801 | -0.1529399 | 0.8784457 | 0.97548444 |
| CASP4 | 45.3661186 | -1.381454643 | 0.36704118 | -3.7637592 | 0.00016738 | 0.00994467 |
| MSANTD4 | 16.1065803 | -0.050830498 | 0.34015254 | -0.1494344 | 0.88121085 | 0.97630358 |
| KBTBD3 | 18.2721863 | -0.219088818 | 0.35410813 | -0.618706 | 0.53611008 | 0.86946459 |
| AASDHPPT | 34.8838732 | -0.012055899 | 0.33422317 | -0.0360714 | 0.97122542 | 0.99526459 |
| CWF19L2 | 93.3916555 | 0.017934689 | 0.19510188 | 0.09192474 | 0.92675783 | 0.98672316 |
| ALKBH8 | 17.0883543 | 0.126591955 | 0.32006217 | 0.39552302 | 0.69245694 | 0.928133 |
| SLC35F2 | 14.9653744 | 0.652395214 | 0.36952958 | 1.76547495 | 0.07748398 | 0.43864187 |
| CUL5 | 361.727845 | 0.290428358 | 0.34933594 | 0.83137269 | 0.40576312 | 0.8115705 |
| AP002433.1 | 15.3871825 | 1.012968208 | 0.80296584 | 1.26153337 | 0.20711674 | 0.65895397 |
| ACAT1 | 295.961025 | 0.516750396 | 0.32264443 | 1.60160956 | 0.10924198 | 0.50798042 |
| NPAT | 48.5650525 | 0.022605831 | 0.24944747 | 0.09062361 | 0.92779167 | 0.98690525 |
| ATM | 130.151105 | 0.099432042 | 0.2586508 | 0.38442581 | 0.70066288 | 0.93127962 |
| C11orf65 | 32.6782893 | 0.559900204 | 0.55999627 | 0.99982846 | 0.31739353 | 0.75958338 |
| KDELC2 | 32.1949587 | 0.868512437 | 0.54943136 | 1.58074783 | 0.11393571 | 0.51861866 |
| EXPH5 | 86.8032032 | 0.121869488 | 0.28383426 | 0.42936849 | 0.66765508 | 0.92049996 |
| DDX10 | 111.45444 | 0.468445829 | 0.21029594 | 2.2275553 | 0.02591018 | 0.25310049 |
| RDX | 148.251069 | -0.059311694 | 0.17569788 | -0.3375777 | 0.73568142 | 0.94318131 |
| ZC3H12C | 110.694021 | -0.72890586 | 0.35618222 | -2.0464409 | 0.04071301 | 0.32449976 |
| FDX1 | 54.2182391 | 0.357878663 | 0.26038929 | 1.37439853 | 0.16931799 | 0.61168883 |
| COLCA1 | 19.47675 | 1.246405677 | 0.49420339 | 2.52205003 | 0.01166731 | 0.16075699 |
| COLCA2 | 13.8442382 | 1.109409054 | 0.74061629 | 1.4979539 | 0.13414523 | 0.56229288 |
| SIK2 | 353.712142 | 0.04459032 | 0.25475847 | 0.17502978 | 0.86105623 | 0.97302617 |
| PPP2R1B | 106.470637 | -0.536076645 | 0.21783753 | -2.4609013 | 0.01385885 | 0.17643441 |
| CRYAB | 200.442112 | -2.189926307 | 0.64068883 | -3.418081 | 0.00063064 | 0.02386134 |
| DIXDC1 | 22.6386593 | -0.002826936 | 0.52834574 | -0.0053505 | 0.99573091 | 0.99923337 |
| DLAT | 56.5097798 | 0.522293528 | 0.28306752 | 1.84511996 | 0.06502009 | 0.40487342 |
| NKAPD1 | 13.9843313 | 0.251751278 | 0.34087287 | 0.73854888 | 0.46018098 | 0.83700647 |
| TIMM8B | 23.8207705 | -0.193439378 | 0.40762553 | -0.4745517 | 0.63510656 | 0.90881415 |
| AP002884.3 | 35.3929025 | -0.339532216 | 0.37744849 | -0.8995458 | 0.36836199 | 0.78877653 |
| IL18 | 740.201625 | -0.037790391 | 0.25784662 | -0.1465615 | 0.88347813 | 0.97662748 |
| BCO2 | 78.1668585 | 0.178423309 | 0.47927109 | 0.37228056 | 0.70968397 | 0.93485232 |
| PTS | 72.5425575 | -0.172104656 | 0.31966793 | -0.5383857 | 0.59031077 | 0.89149106 |
| AP002884.1 | 32.4883117 | -0.585704162 | 0.5377665 | -1.0891422 | 0.2760912 | 0.72575599 |
| TTC12 | 35.0284913 | -0.040344728 | 0.33383673 | -0.1208517 | 0.90380852 | 0.98029547 |
| DRD2 | 15.3631253 | 1.586243063 | 0.98317951 | 1.61338091 | 0.10666178 | 0.50199866 |
| ZW10 | 25.8282109 | 0.566404085 | 0.34817998 | 1.62675662 | 0.1037888 | 0.49618104 |
| USP28 | 81.5484895 | -0.335185196 | 0.2821986 | -1.1877635 | 0.2349266 | 0.68936745 |
| ZBTB16 | 247.643194 | -2.208291133 | 0.41578964 | -5.3110778 | 1.09E-07 | 3.02E-05 |
| AP002518.2 | 20.4430067 | -2.471404903 | 0.80789024 | -3.059085 | 0.00222014 | 0.05479739 |
| NNMT | 43.6853351 | -5.438138025 | 0.83928067 | -6.4795225 | 9.20E-11 | 9.92E-08 |
| REXO2 | 21.6842664 | -0.750366824 | 0.39168541 | -1.9157385 | 0.05539839 | 0.37420918 |
| CADM1 | 31.8089638 | -0.445686823 | 0.42401115 | -1.0511205 | 0.29320324 | 0.74013384 |
| BUD13 | 12.6481933 | -0.084622527 | 0.45379388 | -0.1864779 | 0.85207001 | 0.97053536 |
| ZPR1 | 11.1307945 | -0.170823508 | 0.58070447 | -0.294166 | 0.76863106 | NA |
| SIK3 | 681.630758 | 0.239943613 | 0.28346924 | 0.84645378 | 0.39729965 | 0.80674902 |
| PAFAH1B2 | 131.618905 | -0.131375256 | 0.23825083 | -0.5514157 | 0.58134871 | 0.8901472 |
| SIDT2 | 39.5650923 | 0.203143176 | 0.27813422 | 0.73037823 | 0.46515902 | 0.84143009 |
| TAGLN | 14.3452241 | 0.165621975 | 0.48091577 | 0.34438874 | 0.73055396 | 0.94155624 |
| PCSK7 | 96.2270312 | 0.143042045 | 0.27726269 | 0.51590802 | 0.60591865 | 0.89643487 |
| RNF214 | 30.8206804 | 0.18566278 | 0.29459611 | 0.63022822 | 0.52854528 | 0.86750588 |
| BACE1 | 20.9071946 | 0.232469918 | 0.40725524 | 0.57082119 | 0.56812086 | 0.8849803 |
| CEP164 | 19.7333964 | -0.188956128 | 0.42462493 | -0.4449954 | 0.6563231 | 0.9158679 |
| DSCAML1 | 404.239525 | -0.645259688 | 0.92630845 | -0.6965927 | 0.48605774 | 0.85356796 |
| FXYD2 | 75.7836495 | 0.675836478 | 0.53371465 | 1.26628803 | 0.20540998 | 0.65836247 |
| AP000757.1 | 29.7018452 | 1.785693565 | 0.6190833 | 2.88441567 | 0.00392141 | 0.08233259 |
| MPZL3 | 18.4458835 | -0.639355438 | 0.32583479 | -1.9622074 | 0.04973834 | 0.35621674 |
| UBE4A | 56.1221759 | 0.264780883 | 0.24266995 | 1.09111527 | 0.27522217 | 0.72554623 |
| ATP5MG | 180.707424 | 0.330634103 | 0.39803534 | 0.83066519 | 0.4061628 | 0.8115705 |
| KMT2A | 178.84555 | -0.208356945 | 0.2059155 | -1.0118565 | 0.31160666 | 0.75460103 |
| IFT46 | 12.3547004 | 0.107886659 | 0.52348575 | 0.20609283 | 0.8367184 | NA |
| ARCN1 | 59.7826614 | -0.323752127 | 0.24651217 | -1.3133312 | 0.18907137 | 0.64036044 |
| PHLDB1 | 12.6750527 | -0.629041288 | 0.4755198 | -1.3228498 | 0.18588532 | 0.63645873 |
| AP002954.1 | 28.9162819 | -0.429911804 | 0.44523566 | -0.9655826 | 0.33425309 | 0.76777176 |
| DDX6 | 170.273532 | -0.026006037 | 0.16402253 | -0.1585516 | 0.87402215 | 0.97434023 |
| BCL9L | 10.4190219 | -0.094979555 | 0.50084988 | -0.1896368 | 0.84959377 | NA |
| CCDC84 | 45.6787423 | 0.01394489 | 0.48246275 | 0.02890356 | 0.97694151 | 0.9957152 |
| RPS25 | 179.721377 | 0.223976686 | 0.3754347 | 0.5965796 | 0.55078809 | 0.87469622 |
| TRAPPC4 | 48.6082815 | -0.456958912 | 0.42554788 | -1.0738132 | 0.28290643 | 0.7344003 |
| SLC37A4 | 17.032117 | -0.353599619 | 0.51915175 | -0.6811103 | 0.49580168 | 0.85814183 |
| HYOU1 | 24.2989001 | 0.184846684 | 0.35889864 | 0.51503869 | 0.60652599 | 0.89643487 |
| VPS11 | 24.1445415 | -0.377265687 | 0.31477986 | -1.1985064 | 0.23071992 | 0.68523674 |
| HINFP | 13.899138 | -0.092010016 | 0.36446591 | -0.2524516 | 0.80069199 | 0.95873501 |
| NLRX1 | 16.102041 | -0.984523723 | 0.45560184 | -2.1609301 | 0.03070074 | 0.27909763 |
| CBL | 66.2120093 | -0.486648696 | 0.24168647 | -2.0135537 | 0.04405641 | 0.33311495 |
| RNF26 | 11.0877558 | -0.386310785 | 0.46325415 | -0.8339068 | 0.40433351 | NA |
| OAF | 47.274018 | 0.744248868 | 0.49628354 | 1.49964448 | 0.13370652 | 0.56229288 |

| | | | | | | |
|---|---|---|---|---|---|---|
| ARHGEF12 | 582.468682 | -0.851352993 | 0.26664279 | -3.1928597 | 0.00140871 | 0.04066823 |
| SC5D | 57.2223423 | -0.746697834 | 0.38269096 | -1.9511771 | 0.05103598 | 0.35974306 |
| SORL1 | 217.31142 | 0.440717835 | 0.44506786 | 0.99022614 | 0.3220636 | 0.76248108 |
| UBASH3B | 189.028154 | -2.034916385 | 0.61579903 | -3.3045138 | 0.00095141 | 0.03192078 |
| JHY | 18.809223 | -0.741631062 | 0.38920702 | -1.9054925 | 0.05671608 | 0.37858396 |
| HSPA8 | 185.428585 | 0.487474311 | 0.38011405 | 1.28244223 | 0.19968756 | 0.65269687 |
| VWA5A | 13.8145057 | -0.821017313 | 0.54190107 | -1.5150686 | 0.12975502 | 0.55568377 |
| TBRG1 | 53.1439006 | -0.060047423 | 0.31653674 | -0.1897013 | 0.84954322 | 0.97027687 |
| SIAE | 33.983967 | 0.228782335 | 0.36029691 | 0.63498278 | 0.52543965 | 0.86747074 |
| MSANTD2 | 59.6491099 | 0.640001611 | 0.23145901 | 2.76507538 | 0.00569096 | 0.10415539 |
| EI24 | 47.6048212 | 0.017760022 | 0.27629527 | 0.06427914 | 0.94874797 | 0.99179534 |
| STT3A | 27.4472285 | 0.327122883 | 0.3064309 | 1.06752576 | 0.28573449 | 0.73672385 |
| CDON | 18.1873754 | 0.092288071 | 0.84940069 | 0.10865081 | 0.91347946 | 0.98376271 |
| RPUSD4 | 12.9158769 | 0.27575212 | 0.54431715 | 0.50660193 | 0.61243416 | 0.90011356 |
| FAM118B | 30.3039014 | -0.555000354 | 0.28877339 | -1.9219235 | 0.05461539 | 0.37229041 |
| SRPRA | 52.1852857 | -0.364100341 | 0.24490327 | -1.4867108 | 0.1370912 | 0.56856307 |
| FOXRED1 | 27.9659579 | 0.053632637 | 0.33468268 | 0.16024921 | 0.87268476 | 0.97434023 |
| TIRAP | 16.4129249 | 0.541829398 | 0.3729491 | 1.452824 | 0.14627263 | 0.58340645 |
| DCPS | 67.6485764 | -1.660348532 | 0.44932652 | -3.6951937 | 0.00021972 | 0.01158302 |
| ST3GAL4 | 56.4187198 | -1.389001264 | 0.49989768 | -2.7785711 | 0.00545986 | 0.10201977 |
| ARHGAP32 | 124.640838 | -0.086489869 | 0.27159707 | -0.3184492 | 0.75014422 | 0.94684415 |
| BARX2 | 51.5381704 | -0.328652957 | 0.48537647 | -0.6771094 | 0.49833656 | 0.85874305 |
| TMEM45B | 18.548612 | -1.506248675 | 0.83133802 | -1.8118366 | 0.07001144 | 0.41946325 |
| NFRKB | 26.012414 | -0.125689043 | 0.2668365 | -0.4710339 | 0.63761651 | 0.91007654 |
| PRDM10 | 30.152226 | 0.412668043 | 0.28831877 | 1.43129092 | 0.15234685 | 0.59039732 |
| APLP2 | 277.373277 | -0.44793614 | 0.2399529 | -1.866767 | 0.06193414 | 0.39549779 |
| ST14 | 216.185486 | 0.041783893 | 0.24800904 | 0.1684773 | 0.8662078 | 0.97434023 |
| ZBTB44 | 112.359031 | 0.280100699 | 0.26343469 | 1.06326428 | 0.28766212 | 0.73803612 |
| SNX19 | 73.2730372 | 0.499520465 | 0.38188851 | 1.30802694 | 0.19086417 | 0.6421543 |
| NCAPD3 | 16.3472916 | 0.929328129 | 0.34249639 | 2.71339543 | 0.00665976 | 0.11681674 |
| VPS26B | 11.7730909 | 0.314010264 | 0.4322089 | 0.72652429 | 0.46751742 | NA |
| ACAD8 | 25.673301 | 0.514712998 | 0.35107813 | 1.46609248 | 0.14262309 | 0.57691577 |
| AP004550.1 | 14.030086 | -1.128201338 | 0.60159236 | -1.8753585 | 0.06074342 | 0.39176277 |
| ZMYND11 | 191.669167 | 0.302935611 | 0.13192378 | 2.29629272 | 0.02165915 | 0.23011364 |
| DIP2C | 765.983642 | -0.999555253 | 0.62511429 | -1.598996 | 0.10982149 | 0.50994182 |
| LARP4B | 228.615267 | -0.152482797 | 0.20571955 | -0.7412169 | 0.45856196 | 0.83645518 |
| GTPBP4 | 28.9487194 | -0.402552159 | 0.37007076 | -1.0877708 | 0.27669632 | 0.72584803 |
| IDI1 | 91.4321415 | -0.802123851 | 0.45936243 | -1.7461677 | 0.08078181 | 0.44978727 |
| WDR37 | 90.2500865 | -0.211408894 | 0.27735473 | -0.7622329 | 0.44592103 | 0.83035712 |
| PFKP | 30.0598701 | -2.376172744 | 0.39269217 | -6.0509807 | 1.44E-09 | 7.76E-07 |
| PITRM1 | 43.9471132 | -0.648016001 | 0.3807053 | -1.702146 | 0.088728 | 0.4687332 |
| KLF6 | 199.25586 | -0.683762494 | 0.479259 | -1.4267077 | 0.15366417 | 0.59313269 |
| AKR1C1 | 119.782746 | -0.721937602 | 0.74491791 | -0.9691506 | 0.33247008 | 0.76729949 |
| AKR1C2 | 67.9198001 | 1.160906912 | 0.7694462 | 1.50875644 | 0.13136104 | 0.55812617 |
| AKR1C3 | 59.0541199 | -1.639590451 | 0.74681354 | -2.1954482 | 0.02813146 | 0.26404526 |
| NET1 | 20.360671 | 0.258242203 | 0.31396314 | 0.82252397 | 0.41077875 | 0.8143376 |
| ASB13 | 32.2757636 | 0.289579027 | 0.28193792 | 1.02710211 | 0.30437239 | 0.74820946 |
| FAM208B | 156.268081 | -0.098533387 | 0.2174981 | -0.453031 | 0.65052641 | 0.91487459 |
| GDI2 | 97.1521341 | -0.418301819 | 0.18945071 | -2.2079718 | 0.02724624 | 0.25997792 |
| FBH1 | 100.284148 | 0.023927282 | 0.18359405 | 0.13032711 | 0.89630763 | 0.97885533 |
| IL15RA | 16.8582153 | -0.950294761 | 0.54079796 | -1.7572085 | 0.07888228 | 0.44254372 |
| RBM17 | 64.590219 | 0.388153728 | 0.22017392 | 1.76294142 | 0.07791037 | 0.44014595 |
| PFKFB3 | 161.24535 | -0.035586863 | 0.55890196 | -0.0636728 | 0.94923075 | 0.99185581 |
| AL137145.2 | 49.1994498 | -0.650683974 | 0.76996249 | -0.8450853 | 0.39806322 | 0.80714564 |
| PRKCQ | 47.0867289 | -0.568379875 | 0.42002069 | -1.3532188 | 0.17598575 | 0.6225246 |
| SFMBT2 | 61.2697178 | 0.14175942 | 0.40490211 | 0.35010788 | 0.72625774 | 0.94004538 |
| ITIH5 | 396.420244 | -0.334669114 | 0.40670752 | -0.8228742 | 0.41057955 | 0.81427553 |
| KIN | 23.229342 | 0.651945008 | 0.32089074 | 2.03167289 | 0.04218678 | 0.32888549 |
| ATP5F1C | 157.545903 | 0.17092275 | 0.24532634 | 0.69671584 | 0.48598064 | 0.85356796 |
| TAF3 | 158.353525 | 0.466476335 | 0.25565287 | 1.82464733 | 0.06805427 | 0.4151827 |
| GATA3 | 64.3853992 | 0.920910325 | 0.62846882 | 1.46532381 | 0.14283259 | 0.57698886 |
| CELF2 | 675.484237 | -1.331843617 | 0.44344578 | -3.0033968 | 0.00266984 | 0.0628579 |
| USP6NL | 118.274299 | -0.187761874 | 0.21279021 | -0.8823802 | 0.37757122 | 0.79432848 |
| ECHDC3 | 35.8249943 | -0.126640183 | 0.32876777 | -0.3851965 | 0.70009186 | 0.93127962 |
| PROSER2 | 43.4330648 | -0.598430208 | 0.34750422 | -1.7220804 | 0.08505495 | 0.45976313 |
| UPF2 | 92.6119566 | -0.137894415 | 0.19942465 | -0.6914612 | 0.48927574 | 0.85513057 |
| DHTKD1 | 43.6422382 | -0.02047748 | 0.25267713 | -0.0810421 | 0.93540849 | 0.98871562 |
| SEC61A2 | 14.3734103 | -0.747987895 | 0.46282658 | -1.6161299 | 0.10606622 | 0.50162992 |
| NUDT5 | 36.8118747 | 0.055430685 | 0.25774325 | 0.21506164 | 0.82971926 | 0.96675998 |
| CDC123 | 41.992804 | -0.057580814 | 0.24955452 | -0.2307344 | 0.81752114 | 0.9635539 |
| CAMK1D | 71.3029202 | -1.870486026 | 0.56114799 | -3.3333204 | 0.00085816 | 0.0296168 |
| OPTN | 72.9521846 | -0.620271104 | 0.36932094 | -1.6794908 | 0.09305644 | 0.47759124 |
| PHYH | 38.9830188 | 0.437341647 | 0.37957701 | 1.15218159 | 0.24924646 | 0.70371066 |
| SEPHS1 | 28.5533674 | 0.392890672 | 0.29232645 | 1.34401341 | 0.17894405 | 0.62639418 |
| BEND7 | 125.111637 | -0.143382577 | 0.23527236 | -0.6094323 | 0.54223793 | 0.87177844 |
| PRPF18 | 16.0558458 | 0.08535361 | 0.42159151 | 0.20245571 | 0.83956048 | 0.96748214 |
| AL157392.5 | 38.2335024 | 0.060256654 | 0.25134347 | 0.23973829 | 0.81053315 | 0.9623221 |
| FRMD4A | 29.14053 | -0.196107078 | 0.38028861 | -0.5156796 | 0.60607821 | 0.89643487 |
| FAM107B | 567.518275 | 1.105833793 | 0.32765413 | 3.37500332 | 0.00073815 | 0.02625219 |
| DCLRE1C | 32.7960553 | -0.027797797 | 0.3337246 | -0.0832956 | 0.93361649 | 0.9881638 |
| RPP38 | 32.3210644 | 0.064794851 | 0.25280542 | 0.25630325 | 0.79771669 | 0.95848876 |
| NMT2 | 24.154773 | 0.323736321 | 0.52118975 | 0.62114867 | 0.53450181 | 0.86909171 |
| FAM171A1 | 42.5620936 | 0.641200921 | 0.48300665 | 1.32751986 | 0.18433678 | 0.63442801 |
| MINDY3 | 52.4462199 | -0.046813457 | 0.23011381 | -0.2034361 | 0.83879418 | 0.96748214 |
| PTER | 125.253375 | -0.002271597 | 0.17613979 | -0.0128966 | 0.98971032 | 0.99923337 |
| RSU1 | 63.0413919 | 0.450037159 | 0.28376499 | 1.58595025 | 0.1127506 | 0.51571293 |
| CUBN | 32.7271645 | 1.239104253 | 0.51126512 | 2.42360411 | 0.01536735 | 0.18844914 |
| TRDMT1 | 18.6200741 | 0.3856134 | 0.55115195 | 0.69964988 | 0.48414598 | 0.85261729 |
| VIM | 20.0075362 | -0.76100935 | 0.51571268 | -1.475646 | 0.14003895 | 0.57374456 |
| ST8SIA6 | 123.893802 | 1.755482651 | 0.73299085 | 2.39495849 | 0.01662225 | 0.20004441 |
| ST8SIA6-AS1 | 17.7721471 | 1.326842844 | 0.66134326 | 2.00628466 | 0.04482589 | 0.33628081 |
| HACD1 | 13.8356856 | -0.112532781 | 0.47593496 | -0.2364457 | 0.81308683 | 0.96262567 |
| STAM | 49.6402633 | -0.071226334 | 0.24173465 | -0.2946468 | 0.76826373 | 0.95205353 |
| CACNB2 | 665.090921 | 0.867721412 | 0.67190383 | 1.29143691 | 0.19655222 | 0.64685325 |
| AL450384.2 | 42.7774299 | 0.296226522 | 0.34483088 | 0.85904872 | 0.39031364 | 0.8021276 |
| NSUN6 | 96.1800893 | 0.320889389 | 0.25995893 | 1.23438495 | 0.2170595 | 0.67156502 |
| ARL5B | 43.8202263 | -0.569595829 | 0.43255707 | -1.3168108 | 0.18790204 | 0.63942753 |
| MALRD1 | 67.6042588 | 0.841789553 | 0.45350752 | 1.85617553 | 0.06342854 | 0.4001801 |
| PLXDC2 | 114.504698 | 0.73679068 | 0.35793806 | 2.05843067 | 0.03954881 | 0.31915397 |
| NEBL | 463.529382 | 0.597955628 | 0.63569788 | 0.94062864 | 0.3468952 | 0.7765713 |
| MLLT10 | 235.297565 | 0.131941825 | 0.20917271 | 0.63077935 | 0.52818481 | 0.86750588 |
| DNAJC1 | 174.015119 | 0.112703974 | 0.19321893 | 0.58329675 | 0.55969355 | 0.88037687 |
| BMI1 | 21.2014926 | 0.125300651 | 0.30509172 | 0.41069831 | 0.68129377 | 0.92386609 |
| PIP4K2A | 83.0523894 | -0.154232788 | 0.29712925 | -0.5190764 | 0.60370746 | 0.8956362 |
| MSRB2 | 31.2992634 | -0.112687685 | 0.36487212 | -0.3088416 | 0.75744203 | 0.94912643 |
| OTUD1 | 20.8915302 | 0.108464498 | 0.39162146 | 0.2769626 | 0.78180882 | 0.95544332 |
| KIAA1217 | 664.131694 | -0.212401863 | 0.33063007 | -0.6424154 | 0.52060348 | 0.86689854 |
| ARHGAP21 | 219.224058 | -0.270690063 | 0.15365537 | -1.76167 | 0.07812507 | 0.44084536 |
| PDSS1 | 20.660109 | 0.449082764 | 0.36071288 | 1.24498678 | 0.21313661 | 0.66648134 |
| ABI1 | 146.804845 | -0.223059693 | 0.17789575 | -1.2538787 | 0.2098861 | 0.66164938 |
| ANKRD26 | 178.42428 | 0.180723549 | 0.28673982 | 0.63027016 | 0.52851784 | 0.86750588 |
| YME1L1 | 127.358095 | -0.244257621 | 0.18871155 | -1.2943438 | 0.1955467 | 0.6459894 |
| MASTL | 21.1139754 | -0.342966883 | 0.35115251 | -0.9766893 | 0.32872301 | 0.76649357 |

| | | | | | | |
|---|---|---|---|---|---|---|
| ACBD5 | 115.138219 | 0.622205305 | 0.32066394 | 1.94036565 | 0.05233527 | 0.36312739 |
| RAB18 | 124.764918 | -0.610123201 | 0.16694032 | -3.6547385 | 0.00025744 | 0.01287221 |
| MPP7 | 642.887747 | -0.212061935 | 0.291503 | -0.7274777 | 0.46693337 | 0.84265185 |
| WAC-AS1 | 21.0914045 | -0.350545182 | 0.41951921 | -0.8355879 | 0.40338677 | 0.80927646 |
| WAC | 333.044352 | -0.167953499 | 0.17878636 | -0.9394089 | 0.34752085 | 0.77726831 |
| SVIL | 236.37189 | -0.966755688 | 0.37784638 | -2.5585946 | 0.01050962 | 0.15058101 |
| MTPAP | 26.0051749 | 0.416595208 | 0.26471784 | 1.57373305 | 0.11554918 | 0.52043834 |
| MAP3K8 | 46.1078441 | -0.841334529 | 0.27262083 | -3.0860978 | 0.00202802 | 0.05174581 |
| ZNF438 | 91.0141804 | -0.234802313 | 0.26876998 | -0.8736181 | 0.38232628 | 0.79731722 |
| ZEB1 | 62.0097711 | -1.276317865 | 0.62420356 | -2.0447142 | 0.04088304 | 0.32473787 |
| ARHGAP12 | 218.959537 | -0.430429812 | 0.13528951 | -3.1815461 | 0.00146491 | 0.04146468 |
| KIF5B | 156.008493 | 0.126622966 | 0.18399763 | 0.68817717 | 0.49134122 | 0.85654883 |
| EPC1 | 121.43963 | 0.139714892 | 0.23510429 | 0.59426774 | 0.55233305 | 0.87576439 |
| CCDC7 | 83.1895843 | 0.17473998 | 0.43166504 | 0.40480456 | 0.68562117 | 0.92593602 |
| ITGB1 | 146.03383 | 0.026410623 | 0.19067056 | 0.13851443 | 0.88983387 | 0.97782578 |
| NRP1 | 26.5987255 | -1.702269106 | 0.72141847 | -2.359614 | 0.01829396 | 0.21024014 |
| PARD3 | 453.193479 | -0.329262354 | 0.26926314 | -1.2228274 | 0.22139488 | 0.67492938 |
| CUL2 | 67.0764605 | 0.074701501 | 0.21697186 | 0.34429119 | 0.73062731 | 0.94155624 |
| CREM | 141.793465 | 0.010682757 | 0.3313664 | 0.0322385 | 0.97428185 | 0.9953169 |
| CCNY | 274.869394 | 0.139575604 | 0.22211449 | 0.62839485 | 0.52974531 | 0.86758926 |
| AL117336.3 | 23.3770123 | 0.267299868 | 0.3613341 | 0.73975821 | 0.45944672 | 0.83645518 |
| ANKRD30A | 19.4504359 | 1.930963429 | 0.58665346 | 3.29148904 | 0.00099659 | 0.03257351 |
| ZNF248 | 96.5306465 | 0.190251773 | 0.24734427 | 0.769178 | 0.44178765 | 0.8281114 |
| ZNF33A | 181.542976 | -0.216345143 | 0.19289646 | -1.121561 | 0.26204914 | 0.71302363 |
| AL117339.5 | 11.2292894 | -0.586459688 | 0.49947193 | -1.1741595 | 0.24033116 | NA |
| ZNF33B | 103.854794 | -0.11054103 | 0.21839275 | -0.506157 | 0.61274641 | 0.90011356 |
| BMS1 | 48.2528291 | 0.07868375 | 0.24190897 | 0.32526181 | 0.74498293 | 0.94595436 |
| CSGALNACT2 | 25.6060307 | -0.338419493 | 0.35371582 | -0.9567553 | 0.33869077 | 0.77011262 |
| FXYD4 | 30.016711 | 0.917942024 | 0.66238185 | 1.38582002 | 0.16580187 | 0.60621113 |
| HNRNPF | 147.771568 | -0.480917305 | 0.29447523 | -1.6331333 | 0.10244096 | 0.49289549 |
| ZNF487 | 16.0977731 | 0.30165008 | 0.37798552 | 0.79804666 | 0.42484341 | 0.82241967 |
| ZNF32 | 19.235028 | -0.033123663 | 0.3860864 | -0.0857934 | 0.93163066 | 0.98757533 |
| TMEM72-AS | 37.0035954 | 0.183967205 | 0.45202898 | 0.406981 | 0.68402195 | 0.92538534 |
| TMEM72 | 82.0497825 | 1.281546829 | 0.60126828 | 2.13140602 | 0.03305571 | 0.28919158 |
| AL353801.1 | 17.8935929 | 1.084939122 | 0.71810483 | 1.51083669 | 0.13083007 | 0.55687707 |
| RASSF4 | 31.6515091 | 0.039119508 | 0.43106076 | 0.09075173 | 0.92768987 | 0.98690525 |
| DEPP1 | 32.3606578 | -2.300793675 | 0.44194089 | -5.2061118 | 1.93E-07 | 4.92E-05 |
| ZNF22 | 26.4215569 | -0.001948496 | 0.34529406 | -0.005643 | 0.99549756 | 0.99923337 |
| 8-Mar | 43.8779717 | -0.459096746 | 0.2747267 | -1.6711035 | 0.09470124 | 0.47996672 |
| ZFAND4 | 19.1938797 | -0.416163623 | 0.38181016 | -1.0899752 | 0.27572405 | 0.72554623 |
| WASHC2C | 25.5606111 | -0.421095345 | 0.43035772 | -0.9784775 | 0.32783822 | 0.7651662 |
| AGAP4 | 35.6350631 | 0.247577826 | 0.27129678 | 0.91257193 | 0.36146773 | 0.78668094 |
| TIMM23 | 39.5999532 | 0.158206641 | 0.29786172 | 0.53114123 | 0.59532091 | 0.893081 |
| NCOA4 | 75.9807143 | 0.179868039 | 0.20736141 | 0.86741328 | 0.38571561 | 0.7994129 |
| PTPN20 | 70.2969463 | 0.105907998 | 0.41077486 | 0.25782493 | 0.79654203 | 0.95848876 |
| AGAP9 | 29.5416952 | 0.225902089 | 0.40032583 | 0.56429556 | 0.572553 | 0.88578992 |
| MAPK8 | 312.039362 | 0.07971777 | 0.1563863 | 0.50974907 | 0.61022727 | 0.89964773 |
| ERCC6 | 56.2750077 | 0.196755422 | 0.23498672 | 0.83730443 | 0.40242147 | 0.80890307 |
| OGDHL | 196.535891 | 1.019325717 | 0.3944594 | 2.58410807 | 0.00976312 | 0.14592028 |
| PARG | 118.402705 | -0.118028942 | 0.21969728 | -0.5372344 | 0.59110569 | 0.89149106 |
| TIMM23B | 100.183539 | 0.065467568 | 0.4167855 | 0.15707736 | 0.87518388 | 0.97477134 |
| LINC00843 | 24.1320878 | 0.039699883 | 0.32043518 | 0.12389365 | 0.90139948 | 0.97978205 |
| AGAP6 | 36.7399785 | 0.288805926 | 0.27050381 | 1.06765935 | 0.2856742 | 0.73672385 |
| WASHC2A | 32.8546047 | 0.062069535 | 0.42038731 | 0.14764845 | 0.88262022 | 0.97662748 |
| SGMS1 | 496.209527 | 0.177938987 | 0.24130883 | 0.73739113 | 0.46088453 | 0.83769303 |
| ASAH2B | 12.0835096 | 0.502484568 | 0.43735026 | 1.14892939 | 0.2505851 | NA |
| PRKG1 | 1092.93782 | 0.60590742 | 0.32916365 | 1.84074829 | 0.06565845 | 0.40715521 |
| CSTF2T | 10.5782903 | -0.221417224 | 0.39613684 | -0.5589413 | 0.57620181 | NA |
| IPMK | 43.1039497 | -0.577087141 | 0.3047962 | -1.8933541 | 0.05831079 | 0.38243048 |
| CISD1 | 123.965778 | -0.000520157 | 0.25857063 | -0.0020117 | 0.99839493 | 0.99982181 |
| UBE2D1 | 36.0994757 | -0.128256945 | 0.29540291 | -0.4341763 | 0.66416041 | 0.91876908 |
| TFAM | 33.6379896 | 0.11175468 | 0.31611532 | 0.35352504 | 0.72369486 | 0.93896433 |
| BICC1 | 370.692868 | 0.121056627 | 0.41257558 | 0.29341685 | 0.76920355 | 0.95205353 |
| PHYHIPL | 12.1331821 | -1.380728799 | 0.68999766 | -2.001063 | 0.0453856 | NA |
| FAM13C | 180.029989 | -0.244135119 | 0.49392079 | -0.4942799 | 0.62110852 | 0.90380166 |
| CCDC6 | 60.5565659 | -0.448401891 | 0.29649372 | -1.5123487 | 0.13044518 | 0.55685884 |
| ANK3 | 671.54532 | -0.0806369 | 0.3133383 | -0.2573477 | 0.79691035 | 0.95848876 |
| RHOBTB1 | 21.9151725 | -0.658958926 | 0.42592572 | -1.5471217 | 0.1218339 | 0.53742103 |
| ARID5B | 84.0011884 | -0.420961914 | 0.58119161 | -0.7243083 | 0.46887647 | 0.84326495 |
| NRBF2 | 19.6988467 | -0.571261264 | 0.44331225 | -1.2886205 | 0.19753004 | 0.64884571 |
| JMJD1C | 837.348977 | 0.188260042 | 0.16485026 | 1.14200636 | 0.25345137 | 0.70701839 |
| REEP3 | 178.546697 | 0.09036516 | 0.16302584 | 0.55429962 | 0.57937381 | 0.88957613 |
| LINC01515 | 24.7678742 | 0.179659178 | 0.31346984 | 0.57313066 | 0.56655623 | 0.88382043 |
| CTNNA3 | 17.863425 | 0.745098284 | 0.4910338 | 1.51740731 | 0.12916388 | 0.55413075 |
| SIRT1 | 39.4146238 | 0.187402464 | 0.3051432 | 0.61414597 | 0.53911888 | 0.86972442 |
| HERC4 | 244.511817 | -0.097213307 | 0.15594302 | -0.6233899 | 0.53302832 | 0.86909171 |
| PBLD | 52.3568474 | 0.274255054 | 0.4158975 | 0.65942943 | 0.50962005 | 0.86281145 |
| HNRNPH3 | 109.056669 | 0.296923803 | 0.28087958 | 1.05712136 | 0.29045621 | 0.73979353 |
| RUFY2 | 63.0749086 | 0.001262113 | 0.2761306 | 0.00457071 | 0.99635311 | 0.99965093 |
| SLC25A16 | 71.2633211 | 0.29096433 | 0.28207721 | 1.031506 | 0.3023036 | 0.74690278 |
| TET1 | 14.2230357 | 0.819229098 | 0.375346 | 2.1825971 | 0.02906549 | 0.26979454 |
| CCAR1 | 150.561731 | 0.147887141 | 0.23341896 | 0.63356954 | 0.52636178 | 0.86750588 |
| STOX1 | 24.89805 | 1.012418613 | 0.48852445 | 2.07240109 | 0.03822805 | 0.31371582 |
| DDX50 | 62.5850352 | -0.209265946 | 0.19221836 | -1.0886887 | 0.2762912 | 0.72575599 |
| DDX21 | 69.2875055 | -0.309646039 | 0.26071169 | -1.1876953 | 0.23495349 | 0.68936745 |
| KIF1BP | 41.6978608 | -0.315780528 | 0.31673757 | -0.9969784 | 0.31877497 | 0.76052272 |
| VPS26A | 75.4971082 | -0.17345208 | 0.24981249 | -0.6943291 | 0.48747586 | 0.85383095 |
| SUPV3L1 | 31.8252109 | 0.409587922 | 0.25153251 | 1.62836971 | 0.10344652 | 0.495277 |
| HKDC1 | 39.9965535 | -2.506854503 | 0.54589478 | -4.5921936 | 4.39E-06 | 0.0006166 |
| HK1 | 128.426921 | -0.397011485 | 0.24243984 | -1.637567 | 0.10151204 | 0.49220103 |
| TSPAN15 | 37.825186 | -0.907737004 | 0.48286377 | -1.8799029 | 0.06012131 | 0.38930355 |
| H2AFY2 | 24.0719314 | -0.114573687 | 0.46064469 | -0.2487246 | 0.80357378 | 0.95938063 |
| AIFM2 | 31.4576696 | -0.453130002 | 0.3413667 | -1.3273995 | 0.18437655 | 0.63442801 |
| TYSND1 | 10.5341857 | 0.651556246 | 0.46025443 | 1.41564363 | 0.15687987 | NA |
| SAR1A | 62.3638637 | 0.05318384 | 0.28441092 | 0.18699648 | 0.85166339 | 0.97041406 |
| PPA1 | 85.4608557 | -0.942236908 | 0.38183591 | -2.4676487 | 0.01360037 | 0.17531992 |
| EIF4EBP2 | 78.1342513 | -0.066022651 | 0.25299572 | -0.2609635 | 0.79412065 | 0.95784516 |
| SGPL1 | 97.7729295 | -0.236737925 | 0.22476964 | -1.0532469 | 0.29222785 | 0.74010708 |
| PCBD1 | 61.9774468 | -0.13157221 | 0.57240622 | -0.2298581 | 0.81820203 | 0.9635539 |
| PSAP | 251.9037 | 0.006045377 | 0.32239846 | 0.01875126 | 0.98503954 | 0.99777861 |
| CHST3 | 24.7680092 | -0.102017393 | 0.50552121 | -0.2018064 | 0.84006811 | 0.96748214 |
| ASCC1 | 62.252424 | -0.03961831 | 0.30758699 | -0.1288036 | 0.89751306 | 0.97892709 |
| ANAPC16 | 183.389236 | -0.029930282 | 0.23353933 | -0.1281595 | 0.89802276 | 0.97892709 |
| DDIT4 | 22.035714 | -1.314987845 | 0.70200843 | -1.8731796 | 0.06104359 | 0.39246665 |
| DNAJB12 | 29.3075037 | -0.238269136 | 0.37377516 | -0.6374665 | 0.52382103 | 0.86747074 |
| MICU1 | 193.664687 | -0.098601001 | 0.18052044 | -0.5462041 | 0.58492565 | 0.89036744 |
| MCU | 348.718006 | 0.694205774 | 0.26409592 | 2.62861224 | 0.00857341 | 0.13434901 |
| P4HA1 | 158.165871 | -1.478084117 | 0.45619796 | -3.240006 | 0.00119527 | 0.03680678 |
| ECD | 36.08677 | 0.137886434 | 0.24304699 | 0.56732418 | 0.57049395 | 0.88563341 |
| FAM149B1 | 53.6658264 | -0.180783135 | 0.3207291 | -0.563663 | 0.57298349 | 0.88578992 |
| DNAJC9 | 52.8660984 | -0.082665082 | 0.31349463 | -0.263689 | 0.79201958 | 0.95746995 |
| MRPS16 | 45.7103418 | -0.460139305 | 0.36588343 | -1.2576118 | 0.2085322 | 0.65973983 |
| CFAP70 | 34.7621251 | 0.729040841 | 0.38036728 | 1.91667603 | 0.0552791 | 0.37383273 |

| | | | | | | |
|---|---|---|---|---|---|---|
| ANXA7 | 134.015583 | -0.418366575 | 0.22795088 | -1.8353365 | 0.06645584 | 0.40936869 |
| PPP3CB | 136.623664 | 0.097639517 | 0.15353937 | 0.63592495 | 0.52482534 | 0.86747074 |
| USP54 | 79.4039567 | 0.074813389 | 0.28166387 | 0.2656123 | 0.79053781 | 0.95746995 |
| AGAP5 | 14.7869353 | 0.308146225 | 0.43909153 | 0.70178129 | 0.48281557 | 0.85151109 |
| SEC24C | 33.3446136 | 0.295594538 | 0.31102429 | 0.95039052 | 0.34191386 | 0.77219195 |
| FUT11 | 31.5629901 | -0.914809552 | 0.44094898 | -2.0746381 | 0.03802009 | 0.31295081 |
| CHCHD1 | 31.9508261 | -0.083015086 | 0.53384805 | -0.1555032 | 0.87642462 | 0.97532685 |
| ZSWIM8 | 53.2975733 | -0.423770965 | 0.30111542 | -1.4073373 | 0.15932739 | 0.60024133 |
| CAMK2G | 26.999859 | 0.592234789 | 0.40592103 | 1.45899016 | 0.14456782 | 0.57889578 |
| VCL | 198.907726 | 0.666550378 | 0.32947387 | 2.02307507 | 0.04306541 | 0.33179556 |
| AP3M1 | 31.5977586 | 0.192706929 | 0.27926587 | 0.69004826 | 0.49016384 | 0.85591165 |
| ADK | 508.023121 | -0.006498894 | 0.182477 | -0.0356149 | 0.97158946 | 0.99526459 |
| KAT6B | 176.310993 | 0.152109187 | 0.30272624 | 0.5024645 | 0.61534082 | 0.90108785 |
| SAMD8 | 75.27482 | 0.27281161 | 0.28525823 | 0.95636719 | 0.33888675 | 0.77015605 |
| VDAC2 | 86.385153 | -0.240082274 | 0.3454369 | -0.6950105 | 0.48704872 | 0.85365949 |
| COMTD1 | 49.7268958 | 0.075049914 | 0.41025503 | 0.18293478 | 0.85484919 | 0.97152899 |
| AC010997.3 | 15.388158 | -0.281601018 | 0.65212431 | -0.4318211 | 0.66587147 | 0.91958319 |
| ZNF503-AS2 | 13.3585081 | 0.172073885 | 0.40895694 | 0.42076284 | 0.67392827 | 0.92189158 |
| LRMDA | 1768.49609 | 0.270344945 | 0.26943499 | 1.00337728 | 0.31567886 | 0.75793414 |
| KCNMA1 | 79.4469988 | -1.276694821 | 0.85233023 | -1.4978875 | 0.13416247 | 0.56229288 |
| DLG5 | 160.316822 | -1.440297924 | 0.42503965 | -3.3886202 | 0.00070245 | 0.02571241 |
| POLR3A | 27.9999715 | 0.280446191 | 0.34483223 | 0.81328301 | 0.41605581 | 0.81763864 |
| RPS24 | 404.23133 | 0.499211339 | 0.3907474 | 1.2775807 | 0.20139731 | 0.65374585 |
| ZMIZ1 | 223.541767 | 0.030639217 | 0.42105455 | 0.07276781 | 0.94199089 | 0.98991882 |
| PPIF | 32.0609475 | 0.11823545 | 0.49367567 | 0.23950026 | 0.81071769 | 0.9624234 |
| NUTM2B-AS | 168.440381 | -0.484966363 | 0.24055606 | -2.0160222 | 0.04379765 | 0.33268382 |
| ANXA11 | 270.488565 | -0.324184372 | 0.20801212 | -1.5584879 | 0.11911764 | 0.53056597 |
| FAM213A | 39.8249648 | -0.787703443 | 0.57568415 | -1.368291 | 0.17122101 | 0.61479659 |
| TSPAN14 | 169.59784 | -0.165672626 | 0.25921644 | -0.6391285 | 0.52273931 | 0.86740339 |
| GHITM | 115.406799 | 0.150188996 | 0.33855015 | 0.44362407 | 0.6573144 | 0.9158679 |
| CCSER2 | 216.964151 | -0.040976634 | 0.16166975 | -0.2534589 | 0.79991364 | 0.95861074 |
| WAPL | 165.892399 | -0.072158506 | 0.20849676 | -0.3460893 | 0.72927558 | 0.94106333 |
| BMPR1A | 169.493555 | 0.288773795 | 0.18313268 | 1.5768556 | 0.11482875 | 0.51928773 |
| GLUD1 | 240.446155 | -0.025195028 | 0.28082496 | -0.0897179 | 0.92851138 | 0.9872367 |
| SHLD2 | 207.826891 | 0.333044611 | 0.27919004 | 1.19289576 | 0.2329102 | 0.68763965 |
| NUTM2A-AS | 192.351564 | 0.22978539 | 0.188287 | 1.22039966 | 0.22231341 | 0.67557645 |
| MINPP1 | 15.2644905 | 0.064105708 | 0.41248659 | 0.15541283 | 0.87649587 | 0.97532685 |
| PAPSS2 | 56.4397047 | 0.34405875 | 1.14483111 | 0.30053232 | 0.76377115 | 0.9513483 |
| ATAD1 | 133.633586 | 0.378100117 | 0.18696457 | 2.02230889 | 0.04314445 | 0.33179556 |
| PTEN | 211.228604 | -0.028206424 | 0.23253651 | -0.1212989 | 0.90345429 | 0.98025107 |
| RNLS | 65.6034548 | 0.240922165 | 0.30245829 | 0.79654674 | 0.42571432 | 0.82308728 |
| ANKRD22 | 118.933348 | -0.418211785 | 0.3485456 | -1.1998768 | 0.23018718 | 0.68523674 |
| STAMBPL1 | 18.8768378 | 1.061401714 | 0.42219807 | 2.51398998 | 0.01193738 | 0.16263007 |
| LIPA | 35.2357254 | 0.257020583 | 0.39403801 | 0.65227359 | 0.5142247 | 0.86416832 |
| IFIT2 | 22.2202995 | -0.973729456 | 1.01070491 | -0.9634162 | 0.33533871 | 0.76777176 |
| IFIT3 | 47.2259311 | -1.756950716 | 1.07440219 | -1.6352821 | 0.10198991 | 0.49220184 |
| IFIT1 | 29.6360344 | -0.965119831 | 1.67631371 | -0.5757394 | 0.56479135 | 0.88191823 |
| IFIT5 | 13.6935034 | -0.733716017 | 0.69243146 | -1.0596226 | 0.28931633 | 0.73917583 |
| SLC16A12 | 17.7710346 | 1.664940258 | 0.42263006 | 3.93947431 | 8.17E-05 | 0.00561777 |
| PANK1 | 25.8538172 | 1.10719676 | 0.50261225 | 2.20288455 | 0.02760289 | 0.26121756 |
| RPP30 | 16.4462149 | -0.505416725 | 0.32055364 | -1.5766994 | 0.11486471 | 0.51928773 |
| PCGF5 | 229.914719 | -0.297654314 | 0.22530817 | -1.3210986 | 0.18646848 | 0.63778007 |
| HECTD2 | 73.2761661 | 0.036534201 | 0.26572441 | 0.13748907 | 0.89064424 | 0.9779208 |
| PPP1R3C | 36.5038392 | -0.487442519 | 0.8027246 | -0.6072351 | 0.54369493 | 0.87243024 |
| TNKS2 | 165.435949 | -0.141753763 | 0.234307 | -0.6049916 | 0.54518459 | 0.87247631 |
| BTAF1 | 158.722029 | 0.109760385 | 0.27100252 | 0.4050161 | 0.68546567 | 0.92591798 |
| CPEB3 | 415.265125 | 0.133327686 | 0.29958814 | 0.44503659 | 0.65629331 | 0.9158679 |
| 5-Mar | 91.8247415 | -0.187871961 | 0.22708394 | -0.8273239 | 0.40805352 | 0.81361336 |
| IDE | 82.6403882 | -0.023020558 | 0.23127632 | -0.099537 | 0.92071189 | 0.98614962 |
| EXOC6 | 181.640221 | 0.140283301 | 0.21249331 | 0.66017751 | 0.50913993 | 0.86281145 |
| MYOF | 169.633763 | -1.334315239 | 0.30477565 | -4.3780244 | 1.20E-05 | 0.00130525 |
| FRA10AC1 | 51.0311954 | -0.158860235 | 0.2513948 | -0.6319154 | 0.52744219 | 0.86750588 |
| PLCE1 | 15.5615295 | -0.283678038 | 0.54626651 | -0.5193034 | 0.60354921 | 0.8956362 |
| NOC3L | 18.3645774 | 0.297678878 | 0.3211743 | 0.92684527 | 0.35400688 | 0.78042425 |
| TBC1D12 | 76.4268694 | -0.464065065 | 0.22105706 | -2.0992999 | 0.03579047 | 0.3024108 |
| SORBS1 | 212.448423 | -0.835678592 | 0.36088733 | -2.3156219 | 0.02057892 | 0.22428714 |
| ALDH18A1 | 25.598745 | -0.105675088 | 0.35631437 | -0.2965782 | 0.76678853 | 0.95205353 |
| TCTN3 | 42.0459979 | 0.336273684 | 0.25332907 | 1.32741844 | 0.1843703 | 0.63442801 |
| ENTPD1-AS1 | 147.219307 | -0.074975908 | 0.35878074 | -0.2089742 | 0.8344684 | 0.96703798 |
| ZNF518A | 170.390929 | -0.005242384 | 0.23113436 | -0.0226811 | 0.98190464 | 0.99714727 |
| BLNK | 345.717857 | -1.437186411 | 0.32879696 | -4.3710453 | 1.24E-05 | 0.00133271 |
| TM9SF3 | 310.455878 | -0.001048736 | 0.18810607 | -0.0055752 | 0.99555163 | 0.99923337 |
| LCOR | 328.693364 | -0.09483115 | 0.20880508 | -0.4541611 | 0.64971286 | 0.91438111 |
| RRP12 | 21.4675982 | -0.04107505 | 0.32277571 | -0.1272557 | 0.89873802 | 0.97908342 |
| PGAM1 | 88.3012813 | -0.211957576 | 0.32070129 | -0.660919 | 0.50866426 | 0.86281145 |
| EXOSC1 | 27.5537237 | 0.04687915 | 0.27371696 | 0.17126871 | 0.86401248 | 0.97395945 |
| ZDHHC16 | 21.9989909 | -0.115263639 | 0.36702855 | -0.3140454 | 0.75348653 | 0.94791062 |
| MMS19 | 54.2793928 | -0.014448159 | 0.21193327 | -0.0681732 | 0.9456478 | 0.99134285 |
| UBTD1 | 14.2536545 | -0.259229901 | 0.37127797 | -0.6982097 | 0.48504603 | 0.85356796 |
| MORN4 | 14.8174931 | -0.162369017 | 0.40283977 | -0.403061 | 0.68690332 | 0.92610697 |
| ZFYVE27 | 34.8494375 | 0.181583199 | 0.27576917 | 0.65846084 | 0.51024205 | 0.86281145 |
| R3HCC1L | 115.067456 | -0.559322862 | 0.22959614 | -2.4361161 | 0.01484592 | 0.18484189 |
| HPS1 | 27.0407537 | -0.47064904 | 0.37053995 | -1.2701708 | 0.20402378 | 0.65771647 |
| GOT1 | 250.060532 | 0.012894963 | 0.26166975 | 0.04927953 | 0.96069653 | 0.99238848 |
| SLC25A28 | 30.0780813 | -0.520219207 | 0.4607838 | -1.1289876 | 0.25890305 | 0.71022613 |
| CUTC | 33.0578512 | -0.218282043 | 0.274503 | -0.79519 | 0.42650298 | 0.82329929 |
| COX15 | 36.2292822 | 0.204556895 | 0.28631037 | 0.71445855 | 0.47494368 | 0.84593347 |
| DNMBP | 104.213951 | 0.186255657 | 0.2348615 | 0.79304465 | 0.42775181 | 0.82407405 |
| ERLIN1 | 26.5407812 | -0.016448509 | 0.30488576 | -0.0539497 | 0.9569752 | 0.99236316 |
| CHUK | 37.2575256 | 0.64070269 | 0.33283249 | 1.92500042 | 0.05422931 | 0.37043962 |
| CWF19L1 | 26.5520182 | 0.191127012 | 0.33927748 | 0.56333539 | 0.57320653 | 0.88578992 |
| BLOC1S2 | 26.0564489 | -0.293271826 | 0.31629445 | -0.9272114 | 0.35381677 | 0.78035986 |
| SCD | 73.8196964 | -1.249151462 | 0.73098845 | -1.7088525 | 0.08747827 | 0.46528247 |
| OLMALINC | 61.7697669 | -0.470053424 | 0.56415985 | -0.8331919 | 0.4047365 | 0.81081042 |
| SEC31B | 18.2311799 | 0.53548977 | 0.33969963 | 1.57636252 | 0.11494228 | 0.51928773 |
| HIF1AN | 35.0358329 | -0.133232109 | 0.3067118 | -0.4343886 | 0.66400628 | 0.91876908 |
| PAX2 | 329.787518 | -0.33152619 | 0.44735954 | -0.7410733 | 0.45864902 | 0.83645518 |
| SLF2 | 63.0305567 | -0.158481339 | 0.24947007 | -0.6352719 | 0.52525107 | 0.86747074 |
| MRPL43 | 26.0138307 | -0.193595919 | 0.43196853 | -0.4481713 | 0.65402954 | 0.9158679 |
| LZTS2 | 34.3348206 | -0.223894462 | 0.33703067 | -0.6643148 | 0.50648885 | 0.86235197 |
| BTRC | 168.456963 | -0.002878488 | 0.22352366 | -0.0128778 | 0.9897253 | 0.99923337 |
| DPCD | 14.1437831 | -0.00171211 | 0.52201754 | -0.0032798 | 0.99738311 | 0.99982181 |
| POLL | 22.9232749 | 0.103317709 | 0.38170696 | 0.27067285 | 0.78664266 | 0.95577371 |
| FBXW4 | 156.108146 | -0.134838699 | 0.14778539 | -0.9123953 | 0.36156065 | 0.78670667 |
| OGA | 239.536054 | -0.327079596 | 0.19139378 | -1.7089354 | 0.08746292 | 0.46528247 |
| ARMH3 | 169.530717 | -0.04712452 | 0.18532633 | -0.2542786 | 0.79928033 | 0.95848876 |
| LDB1 | 23.6645979 | -0.084539743 | 0.29938048 | -0.2823823 | 0.7776504 | 0.9551993 |
| PPRC1 | 15.6534738 | -0.011128179 | 0.51895451 | -0.0214435 | 0.98289191 | 0.99728572 |
| NOLC1 | 19.8825952 | -0.1046963 | 0.37620254 | -0.2782977 | 0.78078385 | 0.95537097 |
| GBF1 | 200.620577 | 0.121497128 | 0.20321406 | 0.59787756 | 0.54992164 | 0.87445431 |
| NFKB2 | 14.291756 | -1.039526981 | 0.52341963 | -1.9860298 | 0.04703002 | 0.34586138 |
| FBXL15 | 36.7966076 | -0.014302676 | 0.32513908 | -0.0439894 | 0.96491285 | 0.99334616 |
| CUEDC2 | 19.2277949 | 0.092614494 | 0.30850439 | 0.30020478 | 0.76402096 | 0.9513483 |

| | | | | | | |
|---|---|---|---|---|---|---|
| ACTR1A | 33.2607974 | 0.381373493 | 0.28840955 | 1.32233309 | 0.18605726 | 0.63682266 |
| SUFU | 46.2512526 | 0.757698704 | 0.41121193 | 1.84259904 | 0.06538757 | 0.40631613 |
| TRIM8 | 135.657053 | 0.089727859 | 0.22477327 | 0.39919275 | 0.68975118 | 0.92734996 |
| ARL3 | 35.5123494 | -0.240704487 | 0.44230673 | -0.5442026 | 0.58630204 | 0.89059199 |
| SFXN2 | 10.4802063 | 1.063576077 | 0.49301265 | 2.15729979 | 0.03098231 | NA |
| WBP1L | 153.055967 | -0.32151543 | 0.20636831 | -1.557969 | 0.1192406 | 0.53056597 |
| CNNM2 | 80.7933157 | 0.608490876 | 0.2325726 | 2.61634814 | 0.00888759 | 0.13705821 |
| NT5C2 | 255.856836 | -0.256701773 | 0.23618829 | -1.0868522 | 0.27710212 | 0.72665331 |
| PCGF6 | 20.1130557 | 0.24968352 | 0.34498173 | 0.72375868 | 0.4692139 | 0.8433157 |
| ATP5MD | 64.4398751 | 0.1537303 | 0.35448716 | 0.43366958 | 0.6645284 | 0.91900848 |
| PDCD11 | 23.6862892 | 0.158355215 | 0.30185785 | 0.52460195 | 0.59985993 | 0.89476262 |
| NEURL1 | 99.8287157 | -0.687971653 | 0.58316023 | -1.1797301 | 0.23810759 | 0.69296239 |
| SH3PXD2A | 55.6827336 | -0.122379213 | 0.52772434 | -0.2318999 | 0.81661577 | 0.96340618 |
| STN1 | 21.3206664 | -0.356246293 | 0.40076631 | -0.8889128 | 0.37404995 | 0.7923487 |
| SLK | 98.9595181 | 0.114978189 | 0.20616662 | 0.55769545 | 0.57705236 | 0.88833644 |
| CFAP43 | 13.5491977 | 0.244565774 | 0.39048791 | 0.62630818 | 0.53111282 | 0.8681824 |
| GSTO1 | 76.6465611 | -0.896543072 | 0.25124176 | -3.5684477 | 0.0003591 | 0.01600474 |
| GSTO2 | 72.5771217 | 0.397786193 | 0.3718695 | 1.06969297 | 0.28475753 | 0.73638801 |
| LINC01435 | 42.7426514 | 0.634432747 | 0.48058616 | 1.32012279 | 0.18679402 | 0.63791877 |
| XPNPEP1 | 62.464004 | -0.513384677 | 0.25602225 | -2.0052346 | 0.04493797 | 0.336601 |
| ADD3 | 199.426224 | 0.377369281 | 0.26646661 | 1.41619724 | 0.15671776 | 0.59731328 |
| MXI1 | 55.4416742 | -1.298256414 | 0.32742202 | -3.9650859 | 7.34E-05 | 0.00526714 |
| SMNDC1 | 28.7119064 | 0.077163579 | 0.36602831 | 0.21081315 | 0.83303308 | 0.96701848 |
| SMC3 | 50.0687316 | 0.045851355 | 0.2656864 | 0.17257697 | 0.86298395 | 0.97359204 |
| PDCD4 | 84.5202016 | -0.576334336 | 0.35253525 | -1.6348275 | 0.10208521 | 0.49220184 |
| BBIP1 | 55.3600761 | -0.354633115 | 0.33614809 | -1.0549907 | 0.29142957 | 0.73994767 |
| SHOC2 | 137.690282 | -0.093331692 | 0.18978978 | -0.4917635 | 0.62288653 | 0.90430717 |
| GPAM | 20.2204158 | 1.103726626 | 0.43028616 | 2.56509905 | 0.01031464 | 0.14935229 |
| ACSL5 | 36.7199356 | -0.292120591 | 0.57430775 | -0.5086482 | 0.61099885 | 0.90002868 |
| ZDHHC6 | 34.3308205 | -0.189721372 | 0.27661842 | -0.6858595 | 0.49280172 | 0.85714521 |
| VTI1A | 387.032296 | -0.196537127 | 0.17558661 | -1.1193173 | 0.26300481 | 0.71348562 |
| TCF7L2 | 330.517142 | -0.493597296 | 0.22986206 | -2.147363 | 0.03176439 | 0.28215622 |
| HABP2 | 36.2939854 | -5.206185156 | 0.83670863 | -6.22222 | 4.90E-10 | 3.96E-07 |
| CASP7 | 50.6432879 | -0.322691087 | 0.24687253 | -1.3071162 | 0.19117325 | 0.6421543 |
| PLEKHS1 | 94.2983566 | -1.842676361 | 1.22904623 | -1.4992734 | 0.13380271 | 0.56229288 |
| NHLRC2 | 185.123894 | 0.049634046 | 0.15682586 | 0.31649147 | 0.7516295 | 0.94719044 |
| ADRB1 | 18.9601241 | 0.410730009 | 0.79324569 | 0.5177841 | 0.60460891 | 0.89593125 |
| CCDC186 | 103.675967 | -0.082283801 | 0.2966503 | -0.2773764 | 0.78149108 | 0.95540382 |
| AFAP1L2 | 17.5262998 | 1.015902995 | 0.8397974 | 1.2097001 | 0.22639399 | 0.68070982 |
| ABLIM1 | 122.93992 | 0.213112785 | 0.47987148 | 0.44410388 | 0.65696748 | 0.9158679 |
| FAM160B1 | 42.0976783 | -0.274077239 | 0.30680758 | -0.8933196 | 0.37168602 | 0.7900012 |
| TRUB1 | 28.0803237 | 0.043888471 | 0.31098755 | 0.14112614 | 0.8877703 | 0.9776239 |
| HSPA12A | 76.6372667 | 0.217190566 | 0.32132186 | 0.67592839 | 0.49908611 | 0.85942996 |
| ENO4 | 13.9425424 | -0.499983645 | 0.54171189 | -0.9229697 | 0.35602301 | 0.78167116 |
| SHTN1 | 199.665808 | -0.557006313 | 0.22341886 | -2.4931034 | 0.0126632 | 0.16734742 |
| PDZD8 | 201.724433 | -0.69238034 | 0.27272842 | -2.5387172 | 0.01112597 | 0.15593979 |
| EMX2OS | 156.699358 | -0.08071639 | 0.28892061 | -0.2793722 | 0.77995919 | 0.95537097 |
| EMX2 | 59.1544678 | 0.409052384 | 0.25385335 | 1.61137282 | 0.10709849 | 0.50318505 |
| RAB11FIP2 | 99.5114482 | -0.23191823 | 0.26660837 | -0.8698835 | 0.38436406 | 0.79852888 |
| CASC2 | 43.4627843 | -0.034649565 | 0.39594728 | -0.0875106 | 0.9302657 | 0.98756002 |
| FAM204A | 56.3330546 | 0.182132823 | 0.25924305 | 0.70255625 | 0.48233233 | 0.85112309 |
| CACUL1 | 154.13724 | -0.367156689 | 0.19912078 | -1.8438894 | 0.06519926 | 0.40540566 |
| EIF3A | 177.850513 | -0.157136596 | 0.32175849 | -0.4883681 | 0.6252891 | 0.90447745 |
| FAM45A | 39.5421956 | -0.673861801 | 0.24429635 | -2.7583786 | 0.00580889 | 0.10591394 |
| SFXN4 | 38.4542556 | 0.031299236 | 0.36324736 | 0.08616508 | 0.93133519 | 0.98757533 |
| PRDX3 | 71.9807652 | 0.357997142 | 0.27125218 | 1.31979454 | 0.18690362 | 0.63791877 |
| TIAL1 | 91.1116375 | 0.374288318 | 0.20646586 | 1.81283396 | 0.06985743 | 0.41879921 |
| BAG3 | 36.33018 | -0.76588997 | 0.33175846 | -2.308577 | 0.02096706 | 0.22580589 |
| INPP5F | 57.7871729 | 0.173819306 | 0.37168485 | 0.46765238 | 0.6400332 | 0.91096398 |
| MCMBP | 60.2320253 | 0.315716811 | 0.23191812 | 1.36132875 | 0.17340982 | 0.61802607 |
| SEC23IP | 87.2204725 | 0.33888792 | 0.19642298 | 1.72529672 | 0.08447401 | 0.4585327 |
| PLPP4 | 96.3847358 | 0.721880055 | 0.45165457 | 1.59830123 | 0.10997595 | 0.51041471 |
| WDR11-AS1 | 13.7719618 | 0.837359383 | 0.4226209 | 1.98134871 | 0.04755218 | 0.3482087 |
| WDR11 | 124.921446 | -0.064150967 | 0.16738531 | -0.3832533 | 0.70153198 | 0.93127962 |
| FGFR2 | 72.7812235 | 1.224576449 | 0.4606023 | 2.6586416 | 0.00784564 | 0.12855183 |
| ATE1 | 231.787956 | -0.380241544 | 0.17231367 | -2.2066824 | 0.02733625 | 0.26047315 |
| NSMCE4A | 19.3538863 | 0.088144643 | 0.37795202 | 0.23321649 | 0.8155933 | 0.96300833 |
| TACC2 | 188.946638 | -0.788012005 | 0.41876814 | -1.8817382 | 0.05987157 | 0.38898476 |
| PLEKHA1 | 390.14239 | -1.412735417 | 0.33135453 | -4.2635162 | 2.01E-05 | 0.00201235 |
| FAM24B | 16.5571314 | -0.003129232 | 0.52391241 | -0.0059728 | 0.99523441 | 0.99923337 |
| C10orf88 | 19.018626 | -0.09067755 | 0.36159636 | -0.2507701 | 0.80199187 | 0.95890737 |
| IKZF5 | 25.495469 | 0.081952489 | 0.2907208 | 0.28189414 | 0.77802469 | 0.95532696 |
| ACADSB | 27.9939773 | 0.32269012 | 0.39802349 | 0.81073135 | 0.41751996 | 0.8188001 |
| BUB3 | 31.5977437 | 0.171120063 | 0.26608336 | 0.64310697 | 0.52015469 | 0.86677556 |
| CPXM2 | 33.9032507 | 0.188140072 | 0.67786522 | 0.2775479 | 0.78135943 | 0.95540382 |
| CHST15 | 46.2258068 | -2.209640764 | 0.56872167 | -3.8852762 | 0.00010221 | 0.00660981 |
| OAT | 41.3302365 | 0.088845665 | 0.32412285 | 0.27411109 | 0.78399926 | 0.95544332 |
| LHPP | 33.5057193 | 0.034202727 | 0.3128968 | 0.10930993 | 0.91295667 | 0.98376271 |
| ABRAXAS2 | 32.6136931 | -0.124197886 | 0.29559053 | -0.4201687 | 0.67436223 | 0.92189158 |
| ZRANB1 | 76.7823197 | 0.069137463 | 0.29253669 | 0.23633775 | 0.81317059 | 0.96262567 |
| CTBP2 | 403.361109 | -0.600730302 | 0.25893958 | -2.3199633 | 0.02034286 | 0.22348867 |
| EDRF1 | 38.3739226 | 0.089188498 | 0.35687143 | 0.24991773 | 0.80265097 | 0.95908975 |
| UROS | 23.1883407 | 0.123798425 | 0.33258 | 0.37223653 | 0.70971675 | 0.93485232 |
| BCCIP | 42.5397281 | 0.182683923 | 0.29601599 | 0.61714207 | 0.53714102 | 0.86963427 |
| DHX32 | 140.136766 | -0.948252881 | 0.25892909 | -3.6622107 | 0.00025005 | 0.01276561 |
| FANK1 | 37.3518605 | -0.541066371 | 0.3872537 | -1.3971884 | 0.16235693 | 0.60094226 |
| DOCK1 | 597.271276 | -0.573599004 | 0.31515077 | -1.8200781 | 0.06874712 | 0.41651909 |
| FAM196A | 25.1243818 | -1.542610075 | 0.55080071 | -2.8006683 | 0.00509969 | 0.09853985 |
| FOXI2 | 11.7624106 | 0.417461664 | 0.67195937 | 0.62126027 | 0.53442839 | NA |
| MGMT | 125.736513 | 0.126361913 | 0.25854046 | 0.48875102 | 0.62501798 | 0.90447745 |
| C10orf143 | 12.7291421 | 1.193406679 | 0.47706954 | 2.50153611 | 0.01236558 | 0.16567149 |
| GLRX3 | 65.4359843 | -0.21575139 | 0.28543014 | -0.7558816 | 0.44972018 | 0.83118301 |
| PPP2R2D | 118.126365 | 0.021478838 | 0.22933191 | 0.0936583 | 0.92538059 | 0.98672316 |
| BNIP3 | 103.423752 | -1.208886497 | 0.53283213 | -2.2687943 | 0.02328084 | 0.23879078 |
| LRRC27 | 13.4822912 | -0.06199997 | 0.41695924 | -0.1486955 | 0.8817939 | 0.97638622 |
| PWWP2B | 16.2559421 | 0.252812967 | 0.39591125 | 0.63855969 | 0.52310941 | 0.86740339 |
| INPP5A | 82.109123 | -0.086668735 | 0.27823163 | -0.3114985 | 0.75542168 | 0.94795939 |
| KNDC1 | 14.1233389 | -0.788517074 | 0.59185833 | -1.3322733 | 0.18277038 | 0.63154788 |
| TUBGCP2 | 58.5644169 | -0.206855611 | 0.2215447 | -0.9336969 | 0.35046024 | 0.77868979 |
| ECHS1 | 78.1795596 | -0.161273997 | 0.37753271 | -0.4271789 | 0.66924905 | 0.92049996 |
| SCART1 | 22.8720737 | 0.408093474 | 0.40782662 | 1.00065432 | 0.31699396 | 0.7592201 |
| SLC6A12 | 43.4667645 | 1.554365609 | 0.83291994 | 1.86616447 | 0.06201836 | 0.39551484 |
| KDM5A | 158.133638 | -0.009051954 | 0.15622547 | -0.0579416 | 0.95379515 | 0.99217529 |
| CCDC77 | 12.7577332 | 0.320309821 | 0.43867358 | 0.73017806 | 0.46528135 | 0.84143009 |
| B4GALNT3 | 104.34988 | 0.031794798 | 0.53477944 | 0.05945404 | 0.95259047 | 0.99217529 |
| WNK1 | 287.92667 | 0.512125337 | 0.21604972 | 2.37040505 | 0.01776861 | 0.20666126 |
| RAD52 | 28.1337821 | 0.003771984 | 0.38127689 | 0.00989303 | 0.99210663 | 0.99923337 |
| ERC1 | 438.100978 | -0.217482727 | 0.13784251 | -1.5777624 | 0.11462022 | 0.51928773 |
| FBXL14 | 14.8332271 | 0.198221795 | 0.47384935 | 0.41832239 | 0.67571142 | 0.92193843 |
| ADIPOR2 | 347.692237 | -0.138600598 | 0.28158234 | -0.4922205 | 0.62256347 | 0.90429256 |
| DCP1B | 50.9799702 | -0.215207341 | 0.38317404 | -0.5616438 | 0.5743587 | 0.88644064 |
| CACNA1C | 17.6009073 | -0.531196015 | 0.5510284 | -0.9640084 | 0.3350417 | 0.76777176 |
| FKBP4 | 32.1027859 | 0.463940898 | 0.36313326 | 1.2776051 | 0.2013887 | 0.65374585 |

| | | | | | | |
|---|---|---|---|---|---|---|
| ITFG2 | 71.2061424 | 0.481766823 | 0.26295401 | 1.83213338 | 0.06693155 | 0.41142968 |
| TULP3 | 58.4262032 | -0.278687905 | 0.27956845 | -0.9968504 | 0.31883715 | 0.76052272 |
| TEAD4 | 36.1931391 | -1.636343876 | 0.45818938 | -3.5713265 | 0.00035518 | 0.01595011 |
| TSPAN9 | 360.115845 | -0.261627094 | 0.35785094 | -0.7311063 | 0.4647142 | 0.84133578 |
| PRMT8 | 23.6120633 | 2.020275762 | 0.80516199 | 2.50915441 | 0.01210206 | 0.16352285 |
| PARP11 | 31.1602685 | -0.477424951 | 0.29521507 | -1.6172107 | 0.10583282 | 0.50150383 |
| TIGAR | 20.5123926 | -0.188012041 | 0.3560706 | -0.528019 | 0.59748615 | 0.89363331 |
| C12orf4 | 34.7114678 | -0.14617735 | 0.31479495 | -0.4643573 | 0.64239174 | 0.91146968 |
| ANO2 | 14.2033889 | 0.816428087 | 0.55749708 | 1.46445268 | 0.1430703 | 0.57698886 |
| VWF | 28.1727508 | -0.440055195 | 0.69442906 | -0.6336935 | 0.52628086 | 0.86750588 |
| CD9 | 446.423052 | 0.18627338 | 0.46127724 | 0.40382088 | 0.68634444 | 0.92597882 |
| TNFRSF1A | 81.7673495 | -0.559511715 | 0.28260902 | -1.9798084 | 0.04772506 | 0.34885689 |
| SCNN1A | 359.096858 | 0.146005954 | 0.30064735 | 0.48563859 | 0.62722343 | 0.90589724 |
| LTBR | 43.6993241 | -0.764274562 | 0.30822633 | -2.4795888 | 0.0131534 | 0.17172 |
| TAPBPL | 16.3657032 | -0.187161248 | 0.37988234 | -0.4926821 | 0.6222372 | 0.90415189 |
| MRPL51 | 40.5243537 | 0.217513375 | 0.47341365 | 0.45945734 | 0.64590578 | 0.91242041 |
| NCAPD2 | 17.4785149 | 0.027801165 | 0.40913877 | 0.06795045 | 0.94582508 | 0.99134285 |
| GAPDH | 477.160033 | -0.799986401 | 0.44516087 | -1.7970726 | 0.0723241 | 0.42440638 |
| CHD4 | 29.7330598 | -0.131586256 | 0.37172974 | -0.3539837 | 0.72335113 | 0.93891422 |
| ING4 | 17.6224275 | -0.059931862 | 0.37898301 | -0.1581387 | 0.87434754 | 0.97434023 |
| ZNF384 | 18.2884636 | 0.061929848 | 0.32850959 | 0.18851762 | 0.8504709 | 0.97027687 |
| COPS7A | 28.5882942 | 0.121522574 | 0.34110637 | 0.35626005 | 0.72164582 | 0.93824012 |
| MLF2 | 54.3454966 | -0.005267945 | 0.23086706 | -0.0228181 | 0.98179538 | 0.99714727 |
| PTMS | 84.3218905 | -0.571602549 | 0.37676125 | -1.5171479 | 0.12922934 | 0.55416651 |
| USP5 | 21.0287632 | 0.048635945 | 0.34667433 | 0.1402929 | 0.88842858 | 0.97765744 |
| TPI1 | 148.526863 | -0.329388216 | 0.44236304 | -0.7446106 | 0.45650715 | 0.83506875 |
| LRRC23 | 26.3422823 | -0.418813343 | 0.29855166 | -1.402817 | 0.16067142 | 0.60035161 |
| ATN1 | 57.1243504 | -0.301771104 | 0.31299188 | -0.96415 | 0.33497075 | 0.76777176 |
| C12orf57 | 25.8533003 | -0.085168537 | 0.38635233 | -0.2204427 | 0.82552642 | 0.96539788 |
| PTPN6 | 21.3085845 | 0.150985914 | 0.32546093 | 0.46391409 | 0.6427093 | 0.91146968 |
| PHB2 | 56.7568896 | -0.126656927 | 0.3466344 | -0.3653905 | 0.71481991 | 0.93689626 |
| EMG1.1 | 12.4101921 | -0.679170499 | 0.39816291 | -1.7057603 | 0.08805271 | NA |
| LPCAT3 | 128.538237 | -0.064534922 | 0.220969 | -0.2920542 | 0.7702452 | 0.9522529 |
| C1S | 29.1602566 | -0.37148134 | 0.38324553 | -0.9693038 | 0.33239364 | 0.76729949 |
| C1RL | 39.3533428 | -1.25825023 | 0.30754121 | -4.0913223 | 4.29E-05 | 0.00368188 |
| CLSTN3 | 11.1465717 | 0.171084767 | 0.40421738 | 0.4232494 | 0.6721133 | NA |
| PEX5 | 10.6485065 | 1.03215073 | 0.45835425 | 2.25186245 | 0.02433097 | NA |
| FOXJ2 | 12.980673 | 0.510407126 | 0.38672886 | 1.31980614 | 0.18689975 | 0.63791877 |
| NECAP1 | 23.1209549 | 0.560953879 | 0.33657198 | 1.6666684 | 0.09558036 | 0.48081904 |
| LINC00937 | 11.699787 | -0.442344521 | 0.39614499 | -1.1166228 | 0.26415565 | NA |
| RIMKLB | 113.239684 | -0.624657235 | 0.39822264 | -1.5686131 | 0.11673813 | 0.52399807 |
| A2ML1-AS1 | 17.059863 | -0.633554032 | 0.59372883 | -1.0670764 | 0.28593733 | 0.73672385 |
| PHC1 | 25.4194887 | -0.04958239 | 0.30315606 | -0.163554 | 0.87008225 | 0.97434023 |
| M6PR | 57.4490766 | -0.249444194 | 0.30570447 | -0.8159651 | 0.41452007 | 0.81725499 |
| CLEC2D | 20.983133 | -0.300025687 | 0.37029354 | -0.8102374 | 0.41780373 | 0.8188001 |
| TMEM52B | 48.8354216 | 0.304827804 | 1.05584787 | 0.28870428 | 0.77280768 | 0.95420501 |
| GABARAPL1 | 251.892475 | -0.730980841 | 0.25504784 | -2.8660539 | 0.00415624 | 0.08574324 |
| KLRD1 | 146.624958 | -1.123703718 | 0.64979409 | -1.7293228 | 0.08375134 | 0.45665431 |
| MAGOHB | 12.5096939 | 0.368744027 | 0.37552316 | 0.98194749 | 0.32612572 | NA |
| YBX3 | 193.491782 | -1.244172311 | 0.28133544 | -4.4223802 | 9.76E-06 | 0.00111401 |
| SMIM10L1 | 18.3680785 | -0.04896006 | 0.47978346 | -0.1020462 | 0.91872004 | 0.98524979 |
| AC244131.2 | 24.0271813 | -0.942743305 | 0.68439201 | -1.3774902 | 0.16836073 | 0.61050434 |
| ETV6 | 260.029766 | -1.086129199 | 0.36659253 | -2.9627696 | 0.00304885 | 0.06861674 |
| BCL2L14 | 43.0457309 | -0.087816643 | 0.37282914 | -0.2355413 | 0.81378866 | 0.96265244 |
| LRP6 | 159.035961 | 0.897932825 | 0.22930913 | 3.91581805 | 9.01E-05 | 0.00594525 |
| MANSC1 | 12.9850183 | -0.434345696 | 0.39792595 | -1.0915239 | 0.27504241 | 0.72554623 |
| BORCS5 | 36.2493985 | 0.186430917 | 0.29731375 | 0.62705112 | 0.53062573 | 0.86786739 |
| DUSP16 | 69.0509127 | 0.030340831 | 0.37028462 | 0.08193921 | 0.93469505 | 0.98828668 |
| CREBL2 | 45.7252749 | 0.145160389 | 0.26593374 | 0.54585172 | 0.58516787 | 0.89036744 |
| CDKN1B | 27.3030494 | -0.093272889 | 0.34855728 | -0.267597 | 0.78900953 | 0.95715263 |
| GPRC5A | 53.4173218 | -1.072174451 | 0.53065366 | -2.0204788 | 0.04333375 | 0.33179556 |
| GPRC5D-AS1 | 44.2427954 | 0.189676134 | 0.29015858 | 0.65369817 | 0.51330628 | 0.86382217 |
| HEBP1 | 39.8965512 | 0.416504958 | 0.31366567 | 1.32786274 | 0.18422346 | 0.63442801 |
| FAM234B | 34.3913452 | 1.505246983 | 0.40005196 | 3.76262867 | 0.00016814 | 0.00994467 |
| ATF7IP | 203.5798 | 0.110616637 | 0.24885966 | 0.44449405 | 0.65668543 | 0.9158679 |
| PLBD1 | 37.3400103 | -0.047592791 | 0.41873723 | -0.1136579 | 0.90950899 | 0.98329594 |
| PLBD1-AS1 | 16.065019 | 0.534080452 | 0.51328356 | 1.04051736 | 0.2980996 | 0.74290325 |
| AC010168.1 | 24.8590472 | 0.200686284 | 0.57547086 | 0.34873405 | 0.72728898 | 0.94011548 |
| H2AFJ | 90.7457294 | -0.510340361 | 0.38233057 | -1.3348144 | 0.18193709 | 0.63058858 |
| WBP11 | 45.1992278 | 0.136300781 | 0.24913256 | 0.54710143 | 0.58430905 | 0.89036744 |
| C12orf60 | 18.7657624 | -0.523023052 | 0.51757508 | -1.0105259 | 0.31224338 | 0.75484355 |
| MGP | 23.4332803 | -1.053279499 | 0.50386022 | -2.09042 | 0.03658008 | 0.30641349 |
| ERP27 | 214.523296 | -1.122097597 | 0.45730939 | -2.4536946 | 0.0141397 | 0.17828198 |
| RERG | 139.814269 | -1.902558865 | 0.47513765 | -4.0042267 | 6.22E-05 | 0.00482832 |
| EPS8 | 1823.87269 | 0.373321728 | 0.20444739 | 1.82600391 | 0.06784967 | 0.4151827 |
| STRAP | 73.8518425 | 0.176979716 | 0.32312278 | 0.54771662 | 0.5838865 | 0.89036744 |
| DERA | 135.982007 | -0.398625809 | 0.32117089 | -1.2411642 | 0.2145451 | 0.66839951 |
| MGST1 | 19.43716 | -1.511690308 | 0.57023261 | -2.6510064 | 0.00802523 | 0.13039321 |
| RERGL | 22.8708619 | -0.211201256 | 0.54644291 | -0.386502 | 0.69912495 | 0.93075927 |
| PIK3C2G | 282.136029 | 1.092672599 | 0.48161939 | 2.2687471 | 0.02328371 | 0.23879078 |
| PLEKHA5 | 239.987487 | -0.249711229 | 0.2577891 | -0.9686648 | 0.33271245 | 0.7674968 |
| AEBP2 | 199.283407 | 0.256448831 | 0.32157098 | 0.79748747 | 0.42516798 | 0.82252281 |
| PYROXD1 | 39.9817379 | -0.398163705 | 0.27557693 | -1.4448369 | 0.14850371 | 0.58556342 |
| RECQL | 28.1126415 | -0.199352867 | 0.26873721 | -0.7418134 | 0.4582004 | 0.83645518 |
| GOLT1B | 44.55157 | -0.19269455 | 0.26462456 | -0.7281809 | 0.46650286 | 0.84234507 |
| SPX | 16.0358434 | 0.4422643 | 0.47459405 | 0.93187915 | 0.35139898 | 0.77910174 |
| LDHB | 286.240154 | -0.096924445 | 0.38301183 | -0.2530586 | 0.80022291 | 0.95864669 |
| AC010197.1 | 42.6912494 | -1.993494425 | 0.58069837 | -3.4329258 | 0.00059711 | 0.02307539 |
| ABCC9 | 31.4223697 | -3.569731012 | 0.90793119 | -3.9317198 | 8.43E-05 | 0.00564208 |
| AC008250.1 | 45.5507183 | -2.633040099 | 0.76028467 | -3.4632292 | 0.00053373 | 0.02101475 |
| CMAS | 31.5303903 | -0.062157996 | 0.36277537 | -0.1713402 | 0.86395629 | 0.97395945 |
| C2CD5 | 118.243653 | 0.107970984 | 0.23578359 | 0.45792407 | 0.64700699 | 0.91313368 |
| ETNK1 | 232.125807 | 0.016270407 | 0.25901526 | 0.0628164 | 0.9499127 | 0.99192022 |
| SOX5 | 175.40036 | 2.131639393 | 0.77570911 | 2.74798807 | 0.00599622 | 0.10851367 |
| LRMP | 23.5654012 | -0.558620814 | 0.65476875 | -0.8531574 | 0.39357201 | 0.80438874 |
| ETFRF1 | 43.7426919 | 0.048025783 | 0.29218582 | 0.16436726 | 0.86944203 | 0.97434023 |
| KRAS | 86.2909412 | 0.035056279 | 0.2266882 | 0.15464536 | 0.87710091 | 0.97541592 |
| RASSF8-AS1 | 11.9891348 | -0.344032232 | 0.53137544 | -0.6474372 | 0.51734902 | NA |
| RASSF8 | 77.8830039 | 0.094823492 | 0.38737281 | 0.24478613 | 0.80662203 | 0.96005568 |
| AC022509.2 | 16.6469833 | 1.286258123 | 0.55599935 | 2.31341661 | 0.02069974 | 0.22444738 |
| AC024145.1 | 43.1768662 | -0.325161741 | 0.57704478 | -0.5634948 | 0.57309799 | 0.88578992 |
| ITPR2 | 1600.97011 | 0.541015422 | 0.31503576 | 1.71731435 | 0.08592175 | 0.46168448 |
| AC023051.1 | 50.2005681 | -0.761532302 | 0.68393559 | -1.1134562 | 0.26551256 | 0.71660319 |
| INTS13 | 18.4512867 | 0.109967322 | 0.30817936 | 0.35682897 | 0.72121985 | 0.93824012 |
| FGFR1OP2 | 68.1082756 | 0.408612746 | 0.27002927 | 1.51321652 | 0.13022467 | 0.55685884 |
| TM7SF3 | 71.5001921 | -0.269134574 | 0.38534423 | -0.6984264 | 0.48491058 | 0.85356796 |
| MED21 | 32.4659302 | -0.203520954 | 0.3001388 | -0.6780894 | 0.49771498 | 0.85818031 |
| STK38L | 37.9505977 | 0.05910408 | 0.2506091 | 0.23584171 | 0.8135555 | 0.96264114 |
| PPFIBP1 | 147.209581 | 0.124005434 | 0.2267851 | 0.54679709 | 0.58451814 | 0.89036744 |
| MRPS35 | 88.6615585 | 0.210292269 | 0.24622982 | 0.85404873 | 0.39307799 | 0.80400536 |
| KLHL42 | 51.3531997 | -0.116151956 | 0.39457037 | -0.2943758 | 0.76847078 | 0.95205353 |
| CCDC91 | 324.308534 | -0.236875059 | 0.20895018 | -1.1336437 | 0.25694403 | 0.70987101 |
| ERGIC2 | 64.4060099 | 0.207490101 | 0.22293978 | 0.93070022 | 0.35200866 | 0.77964174 |

| | | | | | | |
|---|---|---|---|---|---|---|
| TMTC1 | 127.208517 | -1.048283395 | 1.37066636 | -0.7647984 | 0.44439162 | 0.83035712 |
| IPO8 | 67.6104425 | 0.32682502 | 0.21183676 | 1.54281542 | 0.12287557 | 0.53921501 |
| CAPRIN2 | 39.0868704 | -0.049870957 | 0.28138438 | -0.1772343 | 0.85932438 | 0.97287712 |
| DDX11 | 13.553753 | 0.811698344 | 0.45814441 | 1.77170849 | 0.07644296 | 0.43515065 |
| SINHCAF | 155.50856 | -0.209089159 | 0.31626605 | -0.6611179 | 0.50853668 | 0.86281145 |
| DENND5B | 180.9846 | -0.16186155 | 0.26410936 | -0.6128581 | 0.5399702 | 0.87014896 |
| AMN1 | 29.4195876 | 0.101290605 | 0.2746231 | 0.36883498 | 0.71225073 | 0.9357596 |
| KIAA1551 | 32.3818642 | -0.176474623 | 0.32155117 | -0.5488228 | 0.58312704 | 0.89036744 |
| BICD1 | 34.4319205 | -0.622194355 | 0.4781844 | -1.3011599 | 0.19320374 | 0.64401245 |
| FGD4 | 848.182164 | 0.01774218 | 0.19238474 | 0.09222239 | 0.92652135 | 0.98672316 |
| DNM1L | 163.648005 | -0.046253366 | 0.13479899 | -0.3431284 | 0.73150184 | 0.94155624 |
| YARS2 | 33.41035 | 0.220112034 | 0.23944831 | 0.91924656 | 0.35796662 | 0.78363265 |
| PKP2 | 53.9213868 | 0.032358437 | 0.52935149 | 0.06112845 | 0.95125691 | 0.99217308 |
| SYT10 | 103.379121 | 0.899665111 | 0.42736664 | 2.10513648 | 0.03527943 | 0.30071219 |
| ALG10B | 11.045397 | 0.775199962 | 0.42750974 | 1.81329191 | 0.06978681 | NA |
| CPNE8 | 27.7618984 | -1.509665098 | 0.52904604 | -2.8535609 | 0.00432322 | 0.08811382 |
| KIF21A | 398.10529 | 0.137644966 | 0.34430524 | 0.39977598 | 0.68932152 | 0.92734996 |
| SLC2A13 | 12.75886 | -0.73323737 | 0.3878544 | -1.8904965 | 0.05869159 | 0.38425985 |
| LRRK2 | 107.358278 | 0.947303399 | 0.97470423 | 0.97188805 | 0.33110624 | 0.76685572 |
| GXYLT1 | 58.2480121 | 0.140855264 | 0.25758709 | 0.54682579 | 0.58449842 | 0.89036744 |
| YAF2 | 229.769947 | -0.024896698 | 0.17182254 | -0.1448977 | 0.88479161 | 0.97662748 |
| PPHLN1 | 126.621944 | -0.028425643 | 0.22880036 | -0.1242378 | 0.90112703 | 0.979694 |
| ZCRB1 | 29.6313337 | -0.196754448 | 0.30354853 | -0.6481812 | 0.51686777 | 0.86500999 |
| PUS7L | 48.2529533 | 0.541795422 | 0.30276871 | 1.78946965 | 0.07353921 | 0.42788861 |
| IRAK4 | 83.2419384 | 0.274143574 | 0.37312996 | 0.73471338 | 0.46251407 | 0.8390474 |
| TWF1 | 77.0712964 | 0.432070174 | 0.39586745 | 1.09145163 | 0.2750742 | 0.72554623 |
| TMEM117 | 1662.88394 | 0.167515501 | 0.32886887 | 0.50936867 | 0.61049383 | 0.89983135 |
| ANO6 | 99.4865337 | -0.53636216 | 0.28597345 | -1.8755663 | 0.06071486 | 0.39176277 |
| AC008124.1 | 12.3587351 | 0.249601597 | 0.49685219 | 0.5023659 | 0.61541016 | NA |
| ARID2 | 244.864554 | -0.035094515 | 0.21391116 | -0.1640612 | 0.86968299 | 0.97434023 |
| SCAF11 | 313.932925 | -0.098326488 | 0.14568284 | -0.6749353 | 0.49971689 | 0.85942996 |
| SLC38A1 | 279.710096 | -0.402771071 | 0.30026268 | -1.3413957 | 0.17979201 | 0.62733185 |
| SLC38A2 | 137.735366 | 0.527006618 | 0.57205433 | 0.92125275 | 0.35691849 | 0.7825111 |
| AC008014.1 | 340.365547 | 0.079991278 | 0.40785308 | 0.19612768 | 0.84451023 | 0.97006669 |
| SLC38A4 | 35.3004194 | 0.808342199 | 0.46937706 | 1.72215957 | 0.08504062 | 0.45976313 |
| AMIGO2 | 18.8770782 | 0.417268773 | 0.68586006 | 0.60838763 | 0.54293042 | 0.87220748 |
| PCED1B | 218.978631 | 0.287597246 | 0.24921632 | 1.15400648 | 0.24849751 | 0.70336326 |
| RPAP3 | 36.5330856 | 0.416727903 | 0.29692928 | 1.40345844 | 0.16048018 | 0.60024417 |
| RAPGEF3 | 78.4991056 | -0.101896971 | 0.46618011 | -0.2185785 | 0.82697837 | 0.96542185 |
| SLC48A1 | 44.9268425 | -0.888876737 | 0.34487096 | -2.5774184 | 0.00995414 | 0.14718772 |
| HDAC7 | 104.526687 | -1.148354537 | 0.39600971 | -2.8998141 | 0.00373384 | 0.07960055 |
| VDR | 38.5812084 | 0.542735597 | 0.37656121 | 1.44129449 | 0.1495015 | 0.58659144 |
| TMEM106C | 109.739741 | -0.236364414 | 0.41589482 | -0.5683274 | 0.56981269 | 0.88562534 |
| SENP1 | 33.4626511 | -0.021431003 | 0.30454067 | -0.0703716 | 0.94389792 | 0.99086173 |
| PFKM | 30.50071 | -0.09901813 | 0.32301298 | -0.3065454 | 0.75918945 | 0.94924107 |
| ASB8 | 29.8335684 | -0.076420799 | 0.32379722 | -0.2360144 | 0.81342152 | 0.96264114 |
| ZNF641 | 13.384439 | 0.009191713 | 0.34990103 | 0.02626947 | 0.97904241 | 0.99623022 |
| KANSL2 | 31.7306891 | 0.519082472 | 0.28669119 | 1.81059791 | 0.07020312 | 0.41948432 |
| CCNT1 | 49.4366078 | 0.14358883 | 0.2147597 | 0.66860229 | 0.50374921 | 0.86073055 |
| ADCY6 | 46.6746213 | 0.625072379 | 0.27513686 | 2.2718598 | 0.02309498 | 0.23879078 |
| CACNB3 | 14.6173965 | 0.386705112 | 0.33741035 | 1.14609737 | 0.25175486 | 0.70574787 |
| DDX23 | 28.8908558 | 0.051731629 | 0.30193887 | 0.17133147 | 0.86396314 | 0.97395945 |
| CCDC65 | 22.8010505 | -0.650076879 | 0.51226756 | -1.2690183 | 0.20443455 | 0.65771647 |
| PRKAG1 | 39.661683 | 0.093971837 | 0.30032212 | 0.31290348 | 0.75435399 | 0.94795939 |
| KMT2D | 37.0637162 | 0.190172512 | 0.35425382 | 0.53682558 | 0.5913881 | 0.89151328 |
| LMBR1L | 31.3745424 | -0.495617994 | 0.31926903 | -1.5523522 | 0.12057797 | 0.53341392 |
| TUBA1B | 109.233014 | 0.028656907 | 0.42858909 | 0.06686336 | 0.94669048 | 0.99135244 |
| TUBA1A | 12.4279063 | -1.27819411 | 0.52344296 | -2.4418976 | 0.01461029 | NA |
| TUBA1C | 63.4306516 | -0.588539523 | 0.34101402 | -1.7258514 | 0.08437415 | 0.4585327 |
| SPATS2 | 113.687939 | 0.198977008 | 0.24145071 | 0.82408954 | 0.40988868 | 0.8142372 |
| MCRS1 | 20.1213892 | 0.155296421 | 0.30629955 | 0.50700833 | 0.61214898 | 0.90011356 |
| PRPF40B | 68.471461 | 0.16826509 | 0.22393356 | 0.75140631 | 0.45240817 | 0.83316488 |
| FMNL3 | 35.7643212 | 0.079121152 | 0.58328894 | 0.13564658 | 0.89210068 | 0.97794037 |
| TMBIM6 | 383.854264 | 0.034934086 | 0.32724915 | 0.10675073 | 0.91498673 | 0.98407488 |
| NCKAP5L | 22.5815154 | 0.366958161 | 0.37297641 | 0.98386427 | 0.32518225 | 0.76334591 |
| BCDIN3D | 11.7793395 | -0.454208168 | 0.38437722 | -1.181673 | 0.23733549 | NA |
| AQP2 | 25.4946826 | 1.832300445 | 0.69295128 | 2.64419809 | 0.00818847 | 0.13111608 |
| AC025154.2 | 14.9438414 | -0.419443377 | 0.50316037 | -0.8336177 | 0.40449646 | 0.81056858 |
| AQP6 | 526.113274 | 0.673916588 | 0.55185104 | 1.22119293 | 0.22201298 | 0.67551 |
| RACGAP1 | 70.0692513 | 0.732622076 | 0.36261799 | 2.02036883 | 0.04334514 | 0.33179556 |
| SMARCD1 | 27.0744161 | -0.080937558 | 0.34121481 | -0.2372041 | 0.81249844 | 0.96262567 |
| AC074032.1 | 68.5413621 | -0.135096499 | 0.27259531 | -0.4955936 | 0.62018115 | 0.90380166 |
| CERS5 | 48.1436479 | -0.447648093 | 0.22799623 | -1.9634013 | 0.04959956 | 0.35590543 |
| LIMA1 | 892.477061 | -0.639152411 | 0.28913318 | -2.2105814 | 0.02706484 | 0.25953238 |
| LARP4 | 225.771429 | 0.350834354 | 0.17412853 | 2.01480109 | 0.04392549 | 0.33287289 |
| DIP2B | 142.340744 | -0.24401465 | 0.20969512 | -1.1636639 | 0.24456023 | 0.69792121 |
| ATF1 | 49.3634849 | 0.061979166 | 0.28599302 | 0.21671566 | 0.82842995 | 0.96638336 |
| METTL7A | 74.4807964 | 1.372329204 | 0.42438369 | 3.23369921 | 0.00122198 | 0.03740723 |
| SLC11A2 | 73.5309184 | -0.453588375 | 0.23828777 | -1.9035319 | 0.05697117 | 0.37928648 |
| LETMD1 | 67.0230535 | 0.340389263 | 0.21331506 | 1.59571132 | 0.11055326 | 0.51186951 |
| CSRNP2 | 18.825499 | 0.245015731 | 0.37270354 | 0.65740115 | 0.51092302 | 0.86312853 |
| TFCP2 | 136.652042 | 0.536137994 | 0.29506087 | 1.817042 | 0.06921068 | 0.41776205 |
| POU6F1 | 23.1549006 | 0.24658874 | 0.2878442 | 0.85667433 | 0.39162489 | 0.80332153 |
| DAZAP2 | 151.11033 | -0.235673196 | 0.25030788 | -0.9415333 | 0.34643164 | 0.77642952 |
| SMAGP | 15.2317935 | 0.393417715 | 0.42370021 | 0.92852848 | 0.3531335 | 0.7802722 |
| ACVR1B | 146.719988 | -0.52908043 | 0.29153557 | -1.8148058 | 0.06955377 | 0.41827125 |
| NR4A1 | 83.0885763 | 1.85740042 | 0.49584793 | 3.74590736 | 0.00017974 | 0.01020367 |
| ATG101 | 33.0045855 | 0.477758877 | 0.40325784 | 1.18474789 | 0.23611714 | 0.69077483 |
| AC078864.2 | 31.9062334 | -1.705898831 | 0.63816908 | -2.6731142 | 0.00751507 | 0.1252511 |
| AC078864.1 | 67.7925272 | -1.199510157 | 0.27985006 | -4.2862601 | 1.82E-05 | 0.00183599 |
| KRT7 | 31.2874952 | -1.881936588 | 0.7130711 | -2.6391991 | 0.00831022 | 0.13171422 |
| KRT8 | 182.540439 | -0.914829592 | 0.32384283 | -2.8249185 | 0.00472926 | 0.09401676 |
| KRT18 | 144.915628 | -1.227868416 | 0.40456226 | -3.0350543 | 0.00240492 | 0.05759942 |
| EIF4B | 139.095703 | 0.379688322 | 0.2644905 | 1.43554614 | 0.15113153 | 0.58850897 |
| AC068888.1 | 58.5982041 | -0.411669565 | 0.38755412 | -1.0622247 | 0.28813368 | 0.73803612 |
| TNS2 | 44.352451 | -0.741565276 | 0.51887422 | -1.4291812 | 0.15295216 | 0.59174003 |
| SPRYD3 | 14.4106107 | -0.183205232 | 0.45138205 | -0.4058762 | 0.68483357 | 0.92581857 |
| CSAD | 55.2312589 | 0.636071761 | 0.34224467 | 1.85852936 | 0.06309388 | 0.40000694 |
| ZNF740 | 11.1646737 | 0.241002886 | 0.41710821 | 0.57779463 | 0.56340278 | NA |
| MFSD5 | 11.315014 | 0.351680081 | 0.51912013 | 0.67745413 | 0.49811786 | NA |
| PFDN5 | 122.227796 | 0.299021451 | 0.40821218 | 0.73251476 | 0.46385444 | 0.84068563 |
| C12orf10 | 13.0344612 | 0.045727472 | 0.42030774 | 0.10879522 | 0.91336492 | 0.98376271 |
| SP1 | 61.1785844 | -0.205745447 | 0.2608299 | -0.7888108 | 0.43022259 | 0.82469033 |
| PRR13 | 89.5324492 | -0.096563952 | 0.26173541 | -0.3689373 | 0.71217447 | 0.9357596 |
| PCBP2 | 183.162353 | 0.14168987 | 0.22924817 | 0.61806326 | 0.53653364 | 0.86963427 |
| MAP3K12 | 11.9601286 | -1.429801968 | 0.56138142 | -2.546935 | 0.01086737 | NA |
| ATP5MC2 | 144.509772 | 0.410275044 | 0.30013545 | 1.36696631 | 0.17163587 | 0.61538348 |
| CALCOCO1 | 33.1126328 | -0.312856016 | 0.33523831 | -0.9332347 | 0.35069881 | 0.77894201 |
| SMUG1 | 26.528951 | -0.003158193 | 0.32192427 | -0.0098104 | 0.99217259 | 0.99923337 |
| LINC02381 | 30.9811566 | 0.014446924 | 0.3559394 | 0.04058816 | 0.96762423 | 0.99393148 |
| CBX5 | 99.252065 | 0.617894857 | 0.28723978 | 2.15114657 | 0.03146463 | 0.28099688 |
| HNRNPA1 | 73.5155134 | 0.675143293 | 0.34674132 | 1.94710943 | 0.05152162 | 0.36077232 |
| COPZ1 | 56.6254791 | -0.441543187 | 0.22118112 | -1.9962969 | 0.04590162 | 0.3409232 |

| | | | | | | | |
|---|---|---|---|---|---|---|---|
| CD63 | 186.823862 | | -0.527118069 | 0.47648279 | -1.1062689 | 0.26861017 | 0.72009952 |
| ORMDL2 | 25.6924845 | | -0.816650488 | 0.38801855 | -2.1046687 | 0.03532016 | 0.30079505 |
| PYM1 | 14.7247407 | | 0.350777097 | 0.35478279 | 0.98870944 | 0.32280532 | 0.76296579 |
| RAB5B | 57.1288426 | | 0.040095699 | 0.27475461 | 0.14593276 | 0.88397447 | 0.97662748 |
| SUOX | 20.1646137 | | 0.238706005 | 0.47196746 | 0.50576793 | 0.61301958 | 0.90011356 |
| IKZF4 | 13.9403949 | | -0.382745187 | 0.43124618 | -0.8875329 | 0.37479205 | 0.7926388 |
| RPS26 | 32.7176325 | | -0.552079932 | 0.32641818 | -1.6913272 | 0.09077434 | 0.47310687 |
| ERBB3 | 28.6830384 | | -0.76741684 | 0.35381314 | -2.1689891 | 0.03008351 | 0.27617753 |
| PA2G4 | 23.7859856 | | -0.202031455 | 0.35777089 | -0.5646951 | 0.5722812 | 0.88578992 |
| RPL41 | 521.605867 | | 0.119555559 | 0.37385675 | 0.31978975 | 0.74912772 | 0.94684415 |
| ESYT1 | 23.4522761 | | -0.362787689 | 0.3414613 | -1.0624562 | 0.28802862 | 0.73803612 |
| MYL6 | 181.625951 | | 0.509101812 | 0.37531969 | 1.35644845 | 0.17495653 | 0.62095803 |
| SMARCC2 | 77.948499 | | -0.414677811 | 0.21301552 | -1.9467024 | 0.05157043 | 0.36077232 |
| RNF41 | 20.6127837 | | 0.229714979 | 0.32469708 | 0.70747474 | 0.47927151 | 0.84833462 |
| ANKRD52 | 21.3697273 | | 0.263194515 | 0.29594585 | 0.88933335 | 0.37382395 | 0.79224216 |
| CS | 65.2958312 | | 0.363997843 | 0.24642257 | 1.47712865 | 0.13964116 | 0.57327809 |
| PAN2 | 20.9298701 | | 0.691978284 | 0.36232273 | 1.90983956 | 0.05615387 | 0.37694988 |
| STAT2 | 43.3963465 | | -0.244207828 | 0.45537019 | -0.5362842 | 0.59176217 | 0.891609 |
| SPRYD4 | 10.8736429 | | 0.155709232 | 0.4192455 | 0.37140347 | 0.71033704 | NA |
| RBMS2 | 36.0070331 | | 0.06975448 | 0.31901485 | 0.2186559 | 0.8269181 | 0.96542185 |
| BAZ2A | 77.3900839 | | -0.120977089 | 0.23882928 | -0.5065421 | 0.61247614 | 0.90011356 |
| ATP5F1B | 298.597675 | | 0.087068739 | 0.38536886 | 0.2259361 | 0.82125112 | 0.96430648 |
| PTGES3 | 185.121141 | | -0.013533888 | 0.19686219 | -0.068748 | 0.94519018 | 0.99127958 |
| NACA | 122.339865 | | 0.132537592 | 0.43477395 | 0.30484253 | 0.76048609 | 0.94962861 |
| NEMP1 | 15.2848038 | | -0.071467736 | 0.44655416 | -0.1600427 | 0.87284743 | 0.97434023 |
| STAT6 | 113.638322 | | -0.47880138 | 0.2477884 | -1.9322994 | 0.05332256 | 0.36627428 |
| R3HDM2 | 200.429085 | | 0.197054323 | 0.22097217 | 0.89176082 | 0.37252115 | 0.79086345 |
| MARS | 28.5169175 | | 0.19474687 | 0.38361134 | 0.50766714 | 0.6116868 | 0.90011356 |
| DDIT3 | 44.6852765 | | -1.108513734 | 0.64228541 | -1.7258896 | 0.08436727 | 0.4585327 |
| MBD6 | 20.7593408 | | 0.250527539 | 0.39648776 | 0.63186702 | 0.52747377 | 0.86750588 |
| DCTN2 | 47.1605092 | | -0.093911796 | 0.25369472 | -0.3701764 | 0.71125107 | 0.93570661 |
| PIP4K2C | 39.7248378 | | -0.25941005 | 0.22602197 | -1.1477205 | 0.25108396 | 0.70545786 |
| DTX3 | 15.6889051 | | 0.204864897 | 0.32829481 | 0.62402722 | 0.53260971 | 0.86909171 |
| OS9 | 77.2415536 | | -0.132155241 | 0.33266838 | -0.3972582 | 0.69117707 | 0.92786587 |
| TSPAN31 | 26.5587043 | | -0.256670848 | 0.33684537 | -0.7619842 | 0.44606944 | 0.83035712 |
| CDK4 | 14.9479945 | | -0.857824971 | 0.44279829 | -1.9372816 | 0.05271093 | 0.36391177 |
| 9-Mar | 25.3053053 | | -0.696915644 | 0.36866637 | -1.8903694 | 0.05870857 | 0.38425985 |
| TSFM | 26.2112492 | | -0.072410512 | 0.41655646 | -0.1738312 | 0.86199812 | 0.97302617 |
| AVIL | 14.2053396 | | 0.256214446 | 0.632236 | 0.40525128 | 0.68529282 | 0.92591798 |
| CTDSP2 | 57.5605551 | | -0.452972599 | 0.40412676 | -1.1208676 | 0.26234422 | 0.71321158 |
| LRIG3 | 16.6170576 | | 0.183441763 | 0.38571901 | 0.47558393 | 0.63437082 | 0.90805737 |
| SLC16A7 | 456.688104 | | -0.088926345 | 0.40469999 | -0.219734 | 0.82607833 | 0.96539788 |
| FAM19A2 | 30.2849618 | | 0.422179729 | 0.5170838 | 0.81646288 | 0.41423544 | 0.81718343 |
| USP15 | 187.58201 | | 0.075889544 | 0.24787144 | 0.30616494 | 0.75947906 | 0.94934883 |
| MON2 | 359.679524 | | 0.294268622 | 0.29356302 | 1.00240357 | 0.31614872 | 0.75850669 |
| AC048341.1 | 25.0241871 | | 0.524478697 | 0.50666514 | 1.03515845 | 0.30059492 | 0.74458947 |
| PPM1H | 64.3667032 | | -0.649389949 | 0.36236175 | -1.792104 | 0.0731163 | 0.42673173 |
| AVPR1A | 49.5110954 | | -0.791338341 | 0.58619364 | -1.3499606 | 0.17702861 | 0.62488264 |
| DPY19L2 | 457.675913 | | 0.225746267 | 0.35916226 | 0.62853561 | 0.52965313 | 0.86758926 |
| RXYLT1 | 37.1254102 | | 0.219105033 | 0.33584283 | 0.65240349 | 0.51414092 | 0.86416832 |
| SRGAP1 | 319.779224 | | -0.199158459 | 0.35786518 | -0.5565181 | 0.5778567 | 0.88877358 |
| AC020611.2 | 17.3335114 | | -0.867504099 | 0.67179177 | -1.2913289 | 0.19658967 | 0.64685325 |
| XPOT | 32.8600154 | | -0.057959365 | 0.32750319 | -0.1769734 | 0.85952925 | 0.97287712 |
| TBK1 | 131.271899 | | -0.391920226 | 0.26584497 | -1.4742435 | 0.14041604 | 0.57469854 |
| RASSF3 | 14.08382 | | -0.700403657 | 0.47969191 | -1.4601115 | 0.14425945 | 0.57846905 |
| GNS | 124.540067 | | -0.315939544 | 0.28383799 | -1.1130982 | 0.26566628 | 0.7168186 |
| TBC1D30 | 45.4814896 | | -0.522899457 | 0.46737697 | -1.1187959 | 0.2632272 | 0.71348562 |
| LEMD3 | 108.957992 | | -0.016575525 | 0.19859787 | -0.0834628 | 0.93348359 | 0.9881638 |
| LLPH | 12.8577789 | | -0.632693166 | 0.41857954 | -1.5115244 | 0.13065491 | 0.55687707 |
| HELB | 25.0577071 | | -0.701720556 | 0.40231605 | -1.7442023 | 0.08112383 | 0.45042996 |
| GRIP1 | 1236.73487 | | 1.07462608 | 0.22205928 | 4.83936574 | 1.30E-06 | 0.00024774 |
| CAND1 | 114.004573 | | 0.025357207 | 0.22511724 | 0.11264 | 0.91031597 | 0.98334456 |
| DYRK2 | 16.7708283 | | -0.484947849 | 0.40530525 | -1.1965003 | 0.23150138 | 0.68587765 |
| MDM1 | 10.960816 | | 0.61410751 | 0.38114569 | 1.61121461 | 0.10713295 | NA |
| RAP1B | 264.948658 | | 0.028719062 | 0.28380058 | 0.10119451 | 0.91939605 | 0.98560124 |
| NUP107 | 46.3875407 | | -0.145441365 | 0.23664735 | -0.6145912 | 0.53882476 | 0.86972442 |
| SLC35E3 | 24.5669528 | | 0.110937654 | 0.49894524 | 0.22234435 | 0.82404583 | 0.96539788 |
| MDM2 | 76.0734438 | | -0.421099016 | 0.30893646 | -1.3630603 | 0.17286352 | 0.61759711 |
| CPM | 119.343081 | | -1.285333084 | 0.29047876 | -4.424878 | 9.65E-06 | 0.00111401 |
| CPSF6 | 85.3570201 | | 0.430373265 | 0.24271595 | 1.77315611 | 0.07620284 | 0.43480446 |
| YEATS4 | 14.2945141 | | 0.156167526 | 0.33965542 | 0.45978223 | 0.64567254 | 0.91231226 |
| FRS2 | 107.725832 | | 0.146175729 | 0.21034061 | 0.69494773 | 0.48708806 | 0.85365949 |
| CCT2 | 47.2679428 | | -0.067089924 | 0.31278937 | -0.2144891 | 0.83016563 | 0.966913 |
| RAB3IP | 86.2331835 | | -0.123513154 | 0.24453254 | -0.505099 | 0.61348927 | 0.9002782 |
| MYRFL | 19.5016714 | | 1.055932614 | 0.797899 | 1.32339132 | 0.18570528 | 0.63629148 |
| AC025159.1 | 18.3188157 | | 0.469167185 | 0.35980333 | 1.30395454 | 0.19224908 | 0.6421543 |
| CNOT2 | 167.267449 | | 0.007037385 | 0.21546114 | 0.03266197 | 0.97394415 | 0.9953169 |
| ZFC3H1 | 136.764273 | | -0.074770144 | 0.21269222 | -0.3515415 | 0.72518214 | 0.93940528 |
| TMEM19 | 18.3718213 | | 0.536997653 | 0.41332495 | 1.29921421 | 0.19387043 | 0.64534767 |
| RAB21 | 126.63245 | | -0.379904684 | 0.18731748 | -2.0281326 | 0.04254671 | 0.3306916 |
| TBC1D15 | 111.760388 | | -0.616441765 | 0.24020698 | -2.5662941 | 0.01027916 | 0.14935229 |
| TRHDE | 41.8611504 | | -0.139816702 | 0.48224701 | -0.2899276 | 0.77187165 | 0.95341335 |
| ATXN7L3B | 43.2212211 | | 0.302308039 | 0.26577197 | 1.13747148 | 0.25534124 | 0.70849619 |
| CAPS2 | 107.330349 | | 0.042151595 | 0.45235334 | 0.0931829 | 0.92575826 | 0.98672316 |
| KRR1 | 33.9714507 | | 0.187373724 | 0.36531218 | 0.51291398 | 0.60801149 | 0.89780963 |
| AC078923.1 | 128.416922 | | -0.019706581 | 0.62987679 | -0.0312864 | 0.97504113 | 0.99543096 |
| PHLDA1 | 112.650975 | | -0.339867672 | 0.34556963 | -0.9834998 | 0.32536151 | 0.76334591 |
| NAP1L1 | 111.863463 | | -0.115400373 | 0.18704857 | -0.6169541 | 0.53726502 | 0.86963427 |
| OSBPL8 | 186.553453 | | -0.265392339 | 0.25444739 | -1.0430146 | 0.29694153 | 0.74160729 |
| ZDHHC17 | 112.907221 | | 0.114562251 | 0.19011039 | 0.60260911 | 0.54676876 | 0.87269272 |
| PAWR | 181.424347 | | -0.147021576 | 0.19309798 | -0.7613833 | 0.44642817 | 0.83035712 |
| PPP1R12A | 180.220194 | | -0.030960869 | 0.16935904 | -0.182812 | 0.85494552 | 0.97152899 |
| LIN7A | 316.813543 | | -0.017626951 | 0.39462124 | -0.044668 | 0.96437192 | 0.99334616 |
| ACSS3 | 113.347609 | | 0.123278086 | 0.4503612 | 0.27373158 | 0.7842909 | 0.95544332 |
| CCDC59 | 20.9138033 | | -0.025860164 | 0.44481476 | -0.0581369 | 0.95363956 | 0.99217529 |
| METTL25 | 61.4552522 | | 0.432889643 | 0.24722157 | 1.7510189 | 0.07994265 | 0.44745744 |
| TMTC2 | 118.284864 | | -0.512217655 | 0.5223378 | -0.9806253 | 0.32677756 | 0.76371899 |
| LRRIQ1 | 16.0964086 | | 0.866771366 | 0.48438667 | 1.78942035 | 0.07354714 | 0.42788861 |
| C12orf29 | 14.3663077 | | -0.303099348 | 0.37984854 | -0.7979479 | 0.42490072 | 0.82241967 |
| CEP290 | 78.0796158 | | -0.002523822 | 0.27011103 | -0.0093436 | 0.99254496 | 0.99923337 |
| TMTC3 | 56.3680784 | | 0.075385264 | 0.20795066 | 0.36251515 | 0.7169671 | 0.93689626 |
| KITLG | 28.5391935 | | -0.665793778 | 0.42596375 | -1.5630292 | 0.11804572 | 0.5279131 |
| DUSP6 | 59.9793625 | | 0.220183077 | 0.45102391 | 0.48818494 | 0.62541885 | 0.90447745 |
| POC1B | 53.4359796 | | -0.107264477 | 0.29806102 | -0.3598742 | 0.7189412 | 0.93767415 |
| POC1B-AS1 | 22.405176 | | 0.815148585 | 0.44577828 | 1.82859645 | 0.06746008 | 0.4136301 |
| ATP2B1 | 117.011496 | | 0.541204603 | 0.25079116 | 2.15798911 | 0.03092868 | 0.27936455 |
| BTG1 | 206.613147 | | -0.631804182 | 0.31192887 | -2.0254752 | 0.04281859 | 0.33147675 |
| EEA1 | 126.409621 | | -0.140948176 | 0.3144273 | -0.4482695 | 0.65395869 | 0.9158679 |
| AC138123.1 | 16.9513614 | | 0.630457458 | 0.49672323 | 1.26923288 | 0.20435802 | 0.65771647 |
| NUDT4 | 11.8360958 | | 1.030843519 | 0.54675523 | 1.88538391 | 0.05937803 | NA |
| UBE2N | 71.3151406 | | 0.332273359 | 0.21680135 | 1.53261663 | 0.12537034 | 0.54505746 |
| MRPL42 | 55.5170466 | | 0.464590667 | 0.20939056 | 2.21877565 | 0.02650199 | 0.25574536 |
| CRADD | 347.509674 | | -0.079559114 | 0.19982505 | -0.3981438 | 0.69052416 | 0.92771252 |

| | | | | | | |
|---|---|---|---|---|---|---|
| AC012085.2 | 28.7324567 | -1.26700333 | 0.64754012 | -1.9566407 | 0.05038973 | 0.35799473 |
| CEP83 | 42.4070466 | 0.221876578 | 0.25138234 | 0.88262594 | 0.37743841 | 0.79432848 |
| TMCC3 | 51.1276565 | 0.439055704 | 0.41799019 | 1.05039715 | 0.29353555 | 0.74013384 |
| NDUFA12 | 82.9647158 | 0.070326257 | 0.34048311 | 0.20654844 | 0.83636253 | 0.96748214 |
| NR2C1 | 50.7029333 | 0.243744763 | 0.23752888 | 1.02616896 | 0.30481195 | 0.74832335 |
| FGD6 | 27.9741122 | -0.017247083 | 0.42440924 | -0.0406379 | 0.9675846 | 0.99393148 |
| VEZT | 152.406093 | -0.229940064 | 0.20362809 | -1.1292158 | 0.2588068 | 0.71022613 |
| METAP2 | 79.3181405 | 0.272174453 | 0.19389989 | 1.40368544 | 0.16041254 | 0.60024417 |
| USP44 | 11.2014661 | -0.15045249 | 0.57215586 | -0.2629572 | 0.79258359 | NA |
| NTN4 | 44.6185226 | 0.562242465 | 0.56635079 | 0.99274596 | 0.32083377 | 0.76131909 |
| SNRPF | 17.7964402 | 0.078839747 | 0.42898388 | 0.18378254 | 0.85418406 | 0.97135895 |
| LTA4H | 31.9630753 | -0.107921735 | 0.30212517 | -0.3572087 | 0.72093558 | 0.93818343 |
| ELK3 | 14.1684749 | -1.518854822 | 0.69188684 | -2.1952359 | 0.02814668 | 0.26404526 |
| CDK17 | 74.5937494 | -0.021630995 | 0.28478851 | -0.0759546 | 0.93945522 | 0.98871562 |
| NEDD1 | 31.5523025 | 0.219742618 | 0.27280073 | 0.80550598 | 0.42052776 | 0.81938194 |
| TMPO | 76.2919812 | 0.034554122 | 0.23870966 | 0.14475376 | 0.88490529 | 0.97662748 |
| SLC25A3 | 254.25624 | -0.059431811 | 0.41311821 | -0.1438615 | 0.88560983 | 0.97662748 |
| APAF1 | 26.9851952 | -0.203063707 | 0.33593396 | -0.6044751 | 0.54552783 | 0.87247631 |
| UHRF1BP1L | 101.493807 | 0.052671859 | 0.17086721 | 0.30826195 | 0.75788302 | 0.94918855 |
| ACTR6 | 30.4214916 | -0.353123377 | 0.29421434 | -1.2002249 | 0.23005201 | 0.68523674 |
| SCYL2 | 102.131813 | -0.032217129 | 0.25477964 | -0.126451 | 0.89937497 | 0.97936951 |
| ANO4 | 47.8124584 | -4.311259425 | 0.79283873 | -5.4377508 | 5.40E-08 | 1.59E-05 |
| UTP20 | 59.5279354 | -0.237455797 | 0.47036244 | -0.5048358 | 0.61367417 | 0.9002782 |
| ARL1 | 66.218199 | -0.156996149 | 0.33821443 | -0.4641912 | 0.64251079 | 0.91146968 |
| CHPT1 | 93.3645192 | -0.883035648 | 0.2852561 | -3.095589 | 0.00196422 | 0.05060915 |
| GNPTAB | 106.364006 | 0.102721698 | 0.34632763 | 0.29660266 | 0.76676988 | 0.95205353 |
| DRAM1 | 143.879026 | -0.726662788 | 0.37350187 | -1.9455399 | 0.05171003 | 0.36111397 |
| WASHC3 | 119.113729 | 0.316665036 | 0.25119041 | 1.26065734 | 0.20743233 | 0.65905456 |
| NUP37 | 31.2233416 | 0.282371053 | 0.29307287 | 0.96348412 | 0.33530463 | 0.76777176 |
| HSP90B1 | 181.072644 | 0.247229689 | 0.28563132 | 0.86555524 | 0.38673411 | 0.80009555 |
| C12orf73 | 12.6354278 | 0.428508809 | 0.38560445 | 1.11126521 | 0.26645421 | 0.71754743 |
| TDG | 53.164876 | 0.287141901 | 0.27496233 | 1.04429544 | 0.29634872 | 0.7412539 |
| HCFC2 | 46.8572203 | 0.547512899 | 0.24843821 | 2.20381919 | 0.02753706 | 0.26098352 |
| NFYB | 19.0065823 | 0.34691874 | 0.35240854 | 0.98442205 | 0.32490804 | 0.7633033 |
| TXNRD1 | 341.156309 | -0.449682642 | 0.3093283 | -1.4537391 | 0.14601865 | 0.5828728 |
| SLC41A2 | 14.1104669 | -1.552814692 | 0.43828648 | -3.5429217 | 0.00039572 | 0.01717513 |
| C12orf45 | 14.2831288 | 1.04049083 | 0.43643958 | 2.38404325 | 0.01712359 | 0.20380223 |
| WASHC4 | 180.023631 | -0.402610821 | 0.18839958 | -2.1370049 | 0.03259759 | 0.28641003 |
| APPL2 | 67.4370002 | -0.275530686 | 0.24285868 | -1.134531 | 0.25657189 | 0.70972499 |
| C12orf75 | 685.515187 | -0.05144665 | 0.25603105 | -0.2009391 | 0.84074619 | 0.96748214 |
| CKAP4 | 19.918265 | 0.035561434 | 0.34444845 | 0.10324167 | 0.91777117 | 0.98519419 |
| TCP11L2 | 55.1988816 | -0.394776783 | 0.29897646 | -1.3204276 | 0.18669228 | 0.63791877 |
| POLR3B | 38.2718595 | 0.424410189 | 0.2398182 | 1.76971634 | 0.07677441 | 0.43580177 |
| RIC8B | 76.8541605 | 0.06076626 | 0.21901326 | 0.27745471 | 0.78143098 | 0.95540382 |
| TMEM263 | 72.1182497 | 0.328942116 | 0.26413614 | 1.24535066 | 0.21300288 | 0.66628439 |
| MTERF2 | 17.4695137 | 0.541776999 | 0.35124349 | 1.5424542 | 0.12296326 | 0.53921501 |
| CRY1 | 66.2927514 | 0.097302447 | 0.42649732 | 0.22814316 | 0.81953495 | 0.96389705 |
| BTBD11 | 23.8937801 | 2.386943446 | 0.9297946 | 2.56717286 | 0.01025315 | 0.14933265 |
| PWP1 | 34.5496103 | 0.090075948 | 0.24996216 | 0.36035833 | 0.71857918 | 0.93761341 |
| PRDM4 | 105.08113 | 0.601937025 | 0.22203413 | 2.71101126 | 0.00670783 | 0.11744765 |
| SART3 | 35.4217064 | 0.029282715 | 0.24251712 | 0.12074494 | 0.90389306 | 0.98029547 |
| ISCU | 143.375587 | -0.083724722 | 0.35108402 | -0.2384749 | 0.81151279 | 0.96256453 |
| CORO1C | 15.1766307 | -0.320457406 | 0.41627916 | -0.7698137 | 0.44141041 | 0.8281114 |
| SSH1 | 59.6359983 | 0.197656251 | 0.22923569 | 0.86224032 | 0.38855528 | 0.80123007 |
| USP30 | 21.4114614 | 0.193142036 | 0.31895407 | 0.60554811 | 0.54481487 | 0.87247631 |
| UNG | 17.6874027 | 0.420856568 | 0.33336415 | 1.26245298 | 0.20678583 | 0.65851037 |
| ACACB | 275.662553 | 0.402931932 | 0.29365888 | 1.37210878 | 0.17002958 | 0.61252734 |
| KCTD10 | 35.9047085 | 0.023325365 | 0.26268079 | 0.08879737 | 0.92924295 | 0.98733028 |
| UBE3B | 39.9765512 | 0.088819936 | 0.30280171 | 0.29332706 | 0.76927217 | 0.95205353 |
| MMAB | 46.8818231 | -0.535466181 | 0.32873772 | -1.6288553 | 0.10334365 | 0.495277 |
| MVK | 18.2968242 | -0.723843827 | 0.45475016 | -1.5917396 | 0.11144323 | 0.51353887 |
| FAM222A | 23.2642066 | 1.331263744 | 0.42632897 | 3.12262087 | 0.00179248 | 0.04789836 |
| TRPV4 | 50.2847138 | 0.669556714 | 0.54069263 | 1.2383315 | 0.21559317 | 0.66962975 |
| GLTP | 120.781012 | 0.683659295 | 0.3601159 | 1.89844241 | 0.05763783 | 0.38148784 |
| TCHP | 40.0546309 | -0.435445934 | 0.36386746 | -1.1967158 | 0.23141735 | 0.68583815 |
| GIT2 | 55.0632116 | -0.293620475 | 0.26093614 | -1.125258 | 0.26047973 | 0.7111324 |
| ANKRD13A | 142.179008 | -0.553716063 | 0.26494232 | -2.0899495 | 0.03662234 | 0.30650274 |
| C12orf76 | 18.7261526 | 0.142613032 | 0.42726439 | 0.33378169 | 0.7385443 | 0.94376279 |
| IFT81 | 34.1785873 | -0.113327385 | 0.30500605 | -0.3715578 | 0.71022209 | 0.93526396 |
| ATP2A2 | 171.752872 | 0.170019783 | 0.2378637 | 0.71477818 | 0.47474612 | 0.84593347 |
| ANAPC7 | 43.831215 | 0.112102345 | 0.29934024 | 0.37449808 | 0.70803379 | 0.93432317 |
| ARPC3 | 158.400128 | -0.578866459 | 0.26652532 | -2.1719005 | 0.02986317 | 0.27509283 |
| GPN3 | 18.2865457 | 0.30203882 | 0.3668733 | 0.82327829 | 0.41034976 | 0.81427553 |
| VPS29 | 64.8503687 | -0.103146886 | 0.28346043 | -0.3638846 | 0.71594418 | 0.93689626 |
| PPTC7 | 95.2675132 | 0.204461522 | 0.20089624 | 1.01774691 | 0.30879825 | 0.75132915 |
| TCTN1 | 19.900433 | 0.099150269 | 0.38683741 | 0.25630993 | 0.79771153 | 0.95848876 |
| PPP1CC | 39.674439 | 0.282632291 | 0.30945304 | 0.91332852 | 0.36106979 | 0.78616766 |
| SH2B3 | 15.3981237 | -1.218803835 | 0.48172163 | -2.5300999 | 0.01140301 | 0.1580708 |
| ATXN2 | 282.510089 | 0.119250947 | 0.16203603 | 0.73595326 | 0.46175914 | 0.83846194 |
| BRAP | 36.8644894 | -0.007906052 | 0.24446452 | -0.0323403 | 0.97420068 | 0.9953169 |
| ACAD10 | 47.1465195 | 0.184199271 | 0.26507061 | 0.69490642 | 0.48711395 | 0.85365949 |
| ALDH2 | 60.7360199 | -0.570874491 | 0.39377064 | -1.449764 | 0.14712435 | 0.5836835 |
| MAPKAPK5-A | 14.6551041 | -0.362669776 | 0.47457391 | -0.7642008 | 0.44474757 | 0.83035712 |
| MAPKAPK5 | 47.51313 | 0.069598037 | 0.23180048 | 0.30024976 | 0.76398665 | 0.9513483 |
| TMEM116 | 297.137027 | 0.523760926 | 0.32424131 | 1.61534297 | 0.10623644 | 0.50162992 |
| ERP29 | 122.049889 | -0.10186487 | 0.24845635 | -0.409991 | 0.68181255 | 0.92420091 |
| NAA25 | 82.5523866 | 0.181262613 | 0.18759954 | 0.966221 | 0.33393361 | 0.76777176 |
| TRAFD1 | 36.7010968 | -0.797037949 | 0.34625616 | -2.3018737 | 0.0213423 | 0.2284992 |
| HECTD4 | 210.48952 | -0.027609259 | 0.15304991 | -0.1803938 | 0.8568434 | 0.97197766 |
| RPL6 | 269.778538 | 0.205773643 | 0.40780973 | 0.50458248 | 0.61385212 | 0.90040308 |
| PTPN11 | 91.8298264 | 0.091454488 | 0.27865906 | 0.32819492 | 0.74276429 | 0.94546379 |
| OAS1 | 18.725557 | -2.016188053 | 1.31229332 | -1.5363852 | 0.12444393 | 0.54320519 |
| OAS3 | 16.8415363 | -3.408567872 | 1.64032466 | -2.0779837 | 0.03771086 | 0.31172423 |
| DDX54 | 28.904015 | -0.006395361 | 0.3658757 | -0.0174796 | 0.98605401 | 0.99804172 |
| TPCN1 | 101.056312 | -0.536670512 | 0.33855469 | -1.5851811 | 0.11292518 | 0.51571293 |
| SLC8B1 | 36.2169001 | 0.10533281 | 0.41084923 | 0.25637826 | 0.79765878 | 0.95848876 |
| PLBD2 | 13.7263572 | -0.287955581 | 0.42966483 | -0.6701865 | 0.50273889 | 0.86010152 |
| SDSL | 10.1147556 | 0.478357685 | 0.63708898 | 0.7508491 | 0.45274348 | NA |
| RBM19 | 37.4705121 | -0.35484131 | 0.33561886 | -1.0572746 | 0.29038626 | 0.73979353 |
| TBX3 | 14.3511012 | 0.654453178 | 1.02514797 | 0.63839874 | 0.52321415 | 0.86740339 |
| MED13L | 581.109222 | -0.019154608 | 0.20286044 | -0.0944226 | 0.92477347 | 0.98672316 |
| C12orf49 | 27.784177 | -1.262821761 | 0.72355793 | -1.7452946 | 0.0809336 | 0.45014674 |
| FBXW8 | 69.882033 | 0.016595253 | 0.22155948 | 0.07490202 | 0.94029267 | 0.98913772 |
| FBXO21 | 177.547614 | -0.272604398 | 0.2492673 | -1.0936228 | 0.27412046 | 0.72510729 |
| WSB2 | 48.3940294 | -0.140675189 | 0.34716819 | -0.4052076 | 0.68532492 | 0.92591798 |
| VSIG10 | 32.4280411 | 0.45072133 | 0.45276066 | 0.99549579 | 0.31949519 | 0.76106474 |
| PEBP1 | 229.304875 | 0.154938429 | 0.39030634 | 0.39696621 | 0.69139238 | 0.92797926 |
| TAOK3 | 283.502962 | 0.453234337 | 0.26184374 | 1.73093442 | 0.08346346 | 0.4560197 |
| SUDS3 | 45.4065012 | 0.019363889 | 0.24551657 | 0.07886999 | 0.93713603 | 0.98871562 |
| CCDC60 | 14.4283294 | -0.766945877 | 0.78297526 | -0.9795276 | 0.32731936 | 0.76450704 |
| AC002070.1 | 74.9927651 | -1.030639088 | 0.54355163 | -1.89612 | 0.05794419 | 0.38209286 |
| TMEM233 | 16.5266901 | -1.308974007 | 0.61815852 | -2.1175378 | 0.03421423 | 0.29474074 |
| PRKAB1 | 68.6485675 | -0.347010731 | 0.23627813 | -1.4686536 | 0.14192677 | 0.57673923 |

| | | | | | | |
|---|---|---|---|---|---|---|
| CIT | 50.5747116 | -0.374504452 | 0.45595902 | -0.8213555 | 0.4114438 | 0.81482336 |
| BICDL1 | 141.971882 | 0.289806818 | 0.32341611 | 0.89608033 | 0.37020986 | 0.7900012 |
| RAB35 | 31.2513525 | -0.320748724 | 0.54881762 | -0.5844359 | 0.55892707 | 0.88004847 |
| GCN1 | 45.0259534 | 0.1114569 | 0.23531225 | 0.47365532 | 0.63574572 | 0.90881415 |
| RPLP0 | 262.429687 | 0.177406098 | 0.38635398 | 0.4591802 | 0.64610477 | 0.91252421 |
| PXN | 88.3494999 | -0.264246552 | 0.2556483 | -1.0336331 | 0.30130771 | 0.74591425 |
| GATC | 22.4165513 | -0.173437723 | 0.3513026 | -0.4936989 | 0.62151881 | 0.90380166 |
| SRSF9 | 69.923637 | 0.275012449 | 0.3507442 | 0.78408267 | 0.4329916 | 0.8248269 |
| DYNLL1 | 118.307492 | -0.301058632 | 0.49006931 | -0.6143185 | 0.53900489 | 0.86972442 |
| COQ5 | 21.2577822 | 0.23986253 | 0.32402485 | 0.74025967 | 0.45914245 | 0.83645518 |
| RNF10 | 125.967213 | 0.074356422 | 0.2055253 | 0.3617872 | 0.71751106 | 0.93717562 |
| POP5 | 24.9524103 | -0.060544742 | 0.34666041 | -0.1746514 | 0.86135352 | 0.97302617 |
| MLEC | 130.45381 | 0.326892856 | 0.2240738 | 1.45886248 | 0.14460296 | 0.57889578 |
| UNC119B | 20.3586705 | 0.074347751 | 0.31215672 | 0.23817443 | 0.81174581 | 0.96256453 |
| ACADS | 64.6134644 | -0.116942897 | 0.34073749 | -0.3432053 | 0.73144405 | 0.94155624 |
| AC069234.1 | 32.0449356 | -0.091995314 | 0.86990057 | -0.1057538 | 0.91577767 | 0.98437981 |
| SPPL3 | 359.933593 | -0.129372778 | 0.23690581 | -0.5460937 | 0.5850015 | 0.89036744 |
| C12orf43 | 10.7361195 | 0.168934075 | 0.49378168 | 0.34212301 | 0.73225832 | NA |
| P2RX4 | 53.5271326 | -1.021431955 | 0.36736949 | -2.7803941 | 0.0054293 | 0.10201977 |
| CAMKK2 | 40.7162584 | 0.135797752 | 0.33222743 | 0.40874937 | 0.6827236 | 0.92474072 |
| ANAPC5 | 87.3922845 | 0.556832717 | 0.18554161 | 3.00112049 | 0.00268988 | 0.06304501 |
| RNF34 | 59.5913049 | -0.117817056 | 0.2828707 | -0.416505 | 0.67704052 | 0.92237262 |
| KDM2B | 65.5969727 | 0.296144911 | 0.31011563 | 0.95494995 | 0.33960301 | 0.77056121 |
| ORAI1 | 27.4414288 | -0.020066306 | 0.31569348 | -0.0635626 | 0.94931849 | 0.99185581 |
| TMEM120B | 110.489564 | 0.496121211 | 0.35618455 | 1.39287684 | 0.16365704 | 0.60337258 |
| RHOF | 25.3408897 | -0.150714187 | 0.44571302 | -0.3381418 | 0.73525636 | 0.94313498 |
| SETD1B | 23.690589 | 0.323691566 | 0.38904234 | 0.83202143 | 0.40539685 | 0.8115705 |
| WDR66 | 12.71635 | -0.674214891 | 0.53608744 | -1.2576584 | 0.20851531 | 0.65973983 |
| BCL7A | 70.6490761 | -0.122793235 | 0.33529236 | -0.3662274 | 0.71419543 | 0.93655477 |
| MLXIP | 141.068482 | -1.131608739 | 0.30943272 | -3.6570429 | 0.00025514 | 0.01282319 |
| CLIP1 | 424.080628 | 0.241476175 | 0.2533734 | 0.9530447 | 0.34056742 | 0.77184673 |
| ZCCHC8 | 66.7621661 | 0.07288821 | 0.23028822 | 0.31650863 | 0.75161647 | 0.94719044 |
| RSRC2 | 115.491601 | -0.331380894 | 0.18728418 | -1.7694014 | 0.07682691 | 0.43580177 |
| KNTC1 | 19.7295565 | 0.700325014 | 0.42037786 | 1.66594173 | 0.09572502 | 0.48085588 |
| AC026333.3 | 177.082189 | 0.678807216 | 0.64653759 | 1.04991144 | 0.29375883 | 0.74041796 |
| HCAR1 | 18.0096238 | 0.171375758 | 0.58365137 | 0.29362693 | 0.769043 | 0.95205353 |
| DENR | 112.460537 | -0.315506205 | 0.20066623 | -1.5722935 | 0.11588251 | 0.5213638 |
| HIP1R | 100.57204 | 0.02019938 | 0.27580913 | 0.0732368 | 0.94161768 | 0.98988745 |
| VPS37B | 197.910161 | -0.115745954 | 0.23095589 | -0.5011604 | 0.61625822 | 0.90160403 |
| ABCB9 | 13.9860484 | 0.834932749 | 0.50346572 | 1.65837059 | 0.09724269 | 0.48257296 |
| ARL6IP4 | 75.1572227 | 0.076978724 | 0.28952577 | 0.26587866 | 0.79033266 | 0.95746995 |
| PITPNM2 | 86.4046784 | 0.274462802 | 0.23272185 | 1.17935984 | 0.23825492 | 0.69303797 |
| MPHOSPH9 | 51.0089056 | 0.266205024 | 0.27521861 | 0.96724935 | 0.3334194 | 0.76777176 |
| C12orf65 | 41.2352988 | -0.258386572 | 0.30220579 | -0.8550021 | 0.39255001 | 0.80399812 |
| SBNO1 | 94.4426519 | 0.127657268 | 0.25507303 | 0.50047341 | 0.61674177 | 0.90164208 |
| KMT5A | 17.7354978 | 0.222301082 | 0.4483531 | 0.49581699 | 0.62002354 | 0.90380166 |
| RILPL2 | 15.0859581 | 0.170004048 | 0.52775694 | 0.32212565 | 0.7473575 | 0.9462156 |
| SNRNP35 | 40.2437988 | 0.234900794 | 0.22233037 | 1.05653941 | 0.29072184 | 0.73979353 |
| RILPL1 | 46.7825947 | -0.472264523 | 0.29255991 | -1.6142489 | 0.10647344 | 0.50162992 |
| TMED2 | 52.1505869 | 0.353306248 | 0.28624406 | 1.23428323 | 0.21709738 | 0.67156502 |
| DDX55 | 19.0067586 | 0.272258259 | 0.34380154 | 0.79190528 | 0.42841591 | 0.82420355 |
| EIF2B1 | 21.3853747 | -0.23316088 | 0.30325267 | -0.7688667 | 0.44197244 | 0.8281114 |
| GTF2H3 | 25.7939144 | -0.056263682 | 0.31401115 | -0.1791773 | 0.85779846 | 0.97243282 |
| TCTN2 | 12.7247499 | 0.525117965 | 0.36360537 | 1.44419748 | 0.14868343 | 0.58569376 |
| ATP6V0A2 | 27.2177191 | 0.027410583 | 0.34431552 | 0.07960891 | 0.93654831 | 0.98871562 |
| DNAH10 | 13.5927845 | -0.331954006 | 0.52878774 | -0.6277642 | 0.53015843 | 0.86764582 |
| CCDC92 | 74.7554219 | -0.102814641 | 0.23639725 | -0.4349232 | 0.66361819 | 0.91855484 |
| RFLNA | 12.0800957 | -0.379518345 | 0.47024936 | -0.8070577 | 0.41963327 | NA |
| NCOR2 | 114.420145 | 0.246295691 | 0.37332957 | 0.65972726 | 0.50942887 | 0.86281145 |
| SCARB1 | 15.3052297 | -0.230294883 | 0.62446819 | -0.3687856 | 0.71228753 | 0.9357596 |
| UBC | 362.954877 | -0.862640396 | 0.33466162 | -2.5776496 | 0.00994748 | 0.14718772 |
| DHX37 | 12.7243636 | -0.475937254 | 0.4942806 | -0.9628888 | 0.33560333 | 0.76777176 |
| BRI3BP | 44.6502478 | 0.643116079 | 0.35251709 | 1.82435432 | 0.06809852 | 0.4151827 |
| AACS | 40.4465386 | -0.293323181 | 0.44289149 | -0.6622913 | 0.50778455 | 0.86281145 |
| SLC15A4 | 34.229272 | -0.114285531 | 0.34416779 | -0.3320634 | 0.73984138 | 0.94376279 |
| RIMBP2 | 40.9254959 | -1.298138133 | 1.08305118 | -1.1985935 | 0.23068604 | 0.68523674 |
| STX2 | 33.8812092 | 0.228658178 | 0.35478717 | 0.64449393 | 0.51925519 | 0.86618562 |
| RAN | 65.2178859 | -0.268960913 | 0.37750503 | -0.7124697 | 0.47617394 | 0.84663915 |
| SFSWAP | 79.6112365 | 0.154830619 | 0.22725436 | 0.6813098 | 0.49567548 | 0.85814183 |
| ULK1 | 94.5033967 | -0.011113258 | 0.30800525 | -0.0360814 | 0.97121746 | 0.99526459 |
| PUS1 | 15.0969983 | 0.114842487 | 0.34428243 | 0.33357057 | 0.73870363 | 0.94376279 |
| EP400 | 82.8432081 | -0.12646839 | 0.2081918 | -0.6074609 | 0.54354505 | 0.87233405 |
| DDX51 | 10.9844243 | -0.082562177 | 0.46480594 | -0.1776272 | 0.85901576 | NA |
| FBRSL1 | 57.2198911 | -0.042743325 | 0.37411377 | -0.1142522 | 0.90903786 | 0.98300553 |
| ANKLE2 | 43.6748232 | -0.257016957 | 0.42936973 | -0.5985912 | 0.5494455 | 0.87413832 |
| GOLGA3 | 68.2514009 | 0.169242873 | 0.30057018 | 0.56307274 | 0.57338536 | 0.88592513 |
| CHFR | 76.797545 | -0.24299945 | 0.23003647 | -1.0563518 | 0.2908075 | 0.73979353 |
| ZNF26 | 31.9623079 | -0.139226954 | 0.31021881 | -0.4488024 | 0.65357419 | 0.9158679 |
| ZNF84 | 54.0232899 | 0.291639833 | 0.23720741 | 1.22947183 | 0.21889495 | 0.67256289 |
| ZNF140 | 35.1236612 | -0.134619073 | 0.2840329 | -0.4739559 | 0.63553135 | 0.90881415 |
| ZNF891 | 18.4075574 | 0.788973896 | 0.34892932 | 2.2611281 | 0.02375133 | 0.24175011 |
| ZNF268 | 13.8398629 | 0.657231554 | 0.41258973 | 1.59294212 | 0.11117318 | 0.5130256 |
| MPHOSPH8 | 153.907623 | -0.014698442 | 0.226866 | -0.0647891 | 0.94834193 | 0.99179534 |
| PSPC1 | 179.044509 | 0.134027319 | 0.20659214 | 0.64875324 | 0.51649789 | 0.86500817 |
| ZMYM5 | 62.5135963 | 0.268665758 | 0.22234244 | 1.20834221 | 0.22691566 | 0.68135521 |
| ZMYM2 | 406.887154 | 0.203215756 | 0.21942159 | 0.92614293 | 0.3543717 | 0.78087359 |
| CRYL1 | 213.346151 | 0.24340489 | 0.34011224 | 0.71566049 | 0.47420102 | 0.84593347 |
| IFT88 | 106.698492 | 0.046722072 | 0.28816926 | 0.16213413 | 0.87120023 | 0.97434023 |
| EEF1AKMT1 | 14.7023519 | 0.291550766 | 0.35570323 | 0.8196461 | 0.41241789 | 0.81556555 |
| XPO4 | 113.349626 | 0.272457819 | 0.21469355 | 1.2690545 | 0.20442163 | 0.65771647 |
| LATS2 | 48.6156559 | 0.159168709 | 0.43763231 | 0.3637042 | 0.7160789 | 0.93689626 |
| SAP18 | 78.6770553 | -0.210614944 | 0.38355386 | -0.5491144 | 0.58292695 | 0.89036744 |
| MRPL57 | 56.058074 | -0.29169807 | 0.40827791 | -0.7144596 | 0.47494303 | 0.84593347 |
| ZDHHC20 | 116.902774 | -0.140496254 | 0.25843559 | -0.5436413 | 0.58668836 | 0.89059199 |
| MICU2 | 223.207324 | 0.17953557 | 0.21317354 | 0.84220384 | 0.39967387 | 0.80830282 |
| FGF9 | 209.078865 | 0.177373345 | 0.5891572 | 0.30106285 | 0.76336657 | 0.9510218 |
| AL136962.1 | 41.069153 | -0.399605991 | 1.11369953 | -0.3588095 | 0.71973759 | 0.93798934 |
| SACS | 20.180174 | -0.943602831 | 0.44508261 | -2.1200622 | 0.0340008 | 0.29368455 |
| MIPEP | 83.2381048 | -0.218847698 | 0.2874858 | -0.761247 | 0.44650956 | 0.83035712 |
| PARP4 | 230.269519 | -0.780299081 | 0.27846258 | -2.8021685 | 0.00507604 | 0.09827853 |
| MTMR6 | 35.5796995 | -0.109388338 | 0.2710448 | -0.4035803 | 0.68652139 | 0.92597882 |
| NUP58 | 78.3505363 | -0.084839884 | 0.32223411 | -0.2632865 | 0.79232978 | 0.95746995 |
| RNF6 | 30.9672444 | -0.161646519 | 0.46927817 | -0.3444579 | 0.73050197 | 0.94155624 |
| CDK8 | 203.98826 | 0.039773519 | 0.24305295 | 0.16364138 | 0.87001347 | 0.97434023 |
| WASF3 | 24.0389271 | 0.006383893 | 0.41474646 | 0.01539228 | 0.98771922 | 0.99904864 |
| USP12 | 136.157441 | -0.098084896 | 0.26567548 | -0.3691906 | 0.71198565 | 0.9357596 |
| LINC02340 | 13.6449857 | -0.189265224 | 0.40504183 | -0.4672733 | 0.64030437 | 0.91096398 |
| RPL21 | 124.568613 | 0.689367673 | 0.42195198 | 1.6337586 | 0.10230955 | 0.49275205 |
| GTF3A | 21.2886085 | -0.363738089 | 0.37990453 | -0.957446 | 0.33834217 | 0.76998177 |
| MTIF3 | 66.7133791 | -0.302872255 | 0.25599269 | -1.1831285 | 0.2367582 | 0.69173331 |
| LNX2 | 84.3687477 | -0.33266681 | 0.23834547 | -1.3957337 | 0.1627947 | 0.60133611 |
| POLR1D | 58.726514 | -0.22976283 | 0.30171933 | -0.7615118 | 0.44635144 | 0.83035712 |
| PAN3 | 347.134412 | -0.100913502 | 0.19407792 | -0.5199639 | 0.60308877 | 0.8956362 |

| | | | | | | |
|---|---|---|---|---|---|---|
| POMP | 46.5011467 | -0.051641435 | 0.32304534 | -0.1598582 | 0.8729928 | 0.97434023 |
| SLC46A3 | 18.4085547 | -0.107010141 | 0.34899558 | -0.3066232 | 0.7591302 | 0.94924107 |
| SLC7A1 | 46.5306367 | -0.718847301 | 0.31905047 | -2.2530834 | 0.02425389 | 0.24374605 |
| UBL3 | 308.532761 | 0.283420439 | 0.25681991 | 1.10357659 | 0.26977685 | 0.72049434 |
| KATNAL1 | 21.3984472 | 0.154925659 | 0.2909761 | 0.53243432 | 0.59442522 | 0.89289841 |
| HMGB1 | 194.221074 | 0.35406103 | 0.17942417 | 1.97331843 | 0.04845929 | 0.35179202 |
| USPL1 | 52.8530874 | -0.028759093 | 0.32244682 | -0.0891902 | 0.92893076 | 0.98733028 |
| HSPH1 | 105.318215 | 0.223640678 | 0.35174128 | 0.63581016 | 0.52490016 | 0.86747074 |
| B3GLCT | 55.7570626 | -0.135328525 | 0.23411584 | -0.5780409 | 0.56323653 | 0.88191823 |
| FRY | 94.1681755 | -0.064617385 | 0.37927439 | -0.1703711 | 0.86471832 | 0.97418905 |
| N4BP2L1 | 35.4711 | -0.90799965 | 0.57317337 | -1.5841623 | 0.11315679 | 0.51614362 |
| N4BP2L2 | 573.449437 | -0.181171413 | 0.24040282 | -0.753616 | 0.45107983 | 0.83215563 |
| PDS5B | 154.012612 | 0.160095849 | 0.2406552 | 0.66524992 | 0.50589066 | 0.86213981 |
| STARD13 | 87.939839 | 1.997241814 | 0.77561828 | 2.57503191 | 0.01002308 | 0.14754137 |
| AL139383.1 | 25.1940051 | 0.911448867 | 0.62963314 | 1.4475872 | 0.14773253 | 0.58466095 |
| RFC3 | 18.6569936 | 0.223725719 | 0.36256197 | 0.6170689 | 0.53718928 | 0.86963427 |
| LINC02343 | 380.851539 | 0.346493816 | 0.42837955 | 0.80884771 | 0.41860275 | 0.81880353 |
| LINC00457 | 24.8833567 | -0.481135074 | 0.69552753 | -0.6917556 | 0.48909081 | 0.85496141 |
| NBEA | 147.968918 | -0.389148626 | 0.24087695 | -1.6155494 | 0.10619176 | 0.50162992 |
| SPART | 94.8795505 | -0.698088546 | 0.29509215 | -2.3656629 | 0.01799782 | 0.20807972 |
| RFXAP | 10.7720623 | 0.117151762 | 0.41264075 | 0.2839074 | 0.77648135 | NA |
| SMAD9 | 50.5598586 | -0.233593164 | 1.13878753 | -0.2051244 | 0.8374749 | 0.96748214 |
| ALG5 | 41.5096783 | -0.064995508 | 0.25565314 | -0.2542332 | 0.79931542 | 0.95848876 |
| EXOSC8 | 20.2834346 | -0.093108098 | 0.33890143 | -0.274735 | 0.78351981 | 0.95544332 |
| SUPT20H | 95.0971915 | 0.08790573 | 0.20113659 | 0.43704494 | 0.66207876 | 0.91810778 |
| LINC00571 | 49.3204387 | -0.835018518 | 0.33254929 | -2.5109616 | 0.01204028 | 0.1634359 |
| UFM1 | 71.8069215 | -0.283618981 | 0.28336704 | -1.0008891 | 0.31688044 | 0.75913564 |
| FREM2 | 72.2993143 | 0.35118822 | 0.4122441 | 0.85189387 | 0.394273 | 0.80495916 |
| PROSER1 | 32.9504489 | 0.088044511 | 0.32017218 | 0.27499114 | 0.78332305 | 0.95544332 |
| NHLRC3 | 47.4367503 | 0.404075913 | 0.28571776 | 1.41424849 | 0.15728895 | 0.59804205 |
| COG6 | 79.3144168 | 0.725565214 | 0.22796991 | 3.18272365 | 0.00145897 | 0.04146468 |
| LINC00598 | 40.8078978 | 0.1451271 | 0.36665593 | 0.39581277 | 0.69224315 | 0.928133 |
| FOXO1 | 817.920906 | -0.121559377 | 0.36143443 | -0.3363248 | 0.73662592 | 0.94365644 |
| MRPS31 | 74.5269413 | 0.005848376 | 0.26759726 | 0.02185514 | 0.98256351 | 0.99719767 |
| SLC25A15 | 17.1227484 | 0.398504124 | 0.45205115 | 0.88154654 | 0.37802207 | 0.79471481 |
| ELF1 | 359.234052 | 0.117832549 | 0.2795905 | 0.42144689 | 0.67342878 | 0.92189158 |
| WBP4 | 50.8194917 | 0.239970421 | 0.24401478 | 0.98342577 | 0.32539793 | 0.76334591 |
| AL354696.2 | 13.8690793 | -0.077776959 | 0.54388686 | -0.1430021 | 0.88628853 | 0.97693497 |
| MTRF1 | 22.6006939 | 0.089588349 | 0.32522358 | 0.27546696 | 0.78295751 | 0.95544332 |
| NAA16 | 75.9002796 | 0.093071595 | 0.24168031 | 0.3851021 | 0.70016177 | 0.93127962 |
| RGCC | 82.2674265 | 2.037745591 | 0.79001241 | 2.57938427 | 0.00989766 | 0.1471432 |
| VWA8 | 412.803701 | 0.213311816 | 0.23518955 | 0.9069783 | 0.36441827 | 0.78780828 |
| DGKH | 172.052554 | -1.174517022 | 0.48766548 | -2.4084482 | 0.0160205 | 0.19473542 |
| AKAP11 | 66.5497445 | 0.274050826 | 0.2394964 | 1.14427951 | 0.25250774 | 0.70626085 |
| EPSTI1 | 65.6162357 | -2.442783186 | 1.1719849 | -2.0843128 | 0.03713172 | 0.30857899 |
| DNAJC15 | 66.3703436 | -0.334820655 | 0.24276253 | -1.3792106 | 0.16782984 | 0.60954339 |
| ENOX1 | 17.0748632 | 0.169370069 | 0.65903259 | 0.25699802 | 0.79718031 | 0.95848876 |
| CCDC122 | 64.1109957 | -0.66560494 | 0.42451835 | -1.5679062 | 0.11690304 | 0.52425309 |
| TSC22D1 | 121.81452 | -0.616317865 | 0.44535645 | -1.3838755 | 0.16639659 | 0.60746969 |
| NUFIP1 | 12.1014123 | 0.471483186 | 0.41152153 | 1.14570721 | 0.25191632 | NA |
| AL359706.1 | 16.526806 | -0.794269 | 0.62256494 | -1.275801 | 0.20202586 | 0.6545025 |
| GPALPP1 | 43.838628 | 0.210421028 | 0.26603723 | 0.79094578 | 0.42897563 | 0.82469033 |
| GTF2F2 | 105.637713 | 0.241865049 | 0.2294262 | 1.0542172 | 0.29178349 | 0.73994767 |
| TPT1 | 517.608317 | -0.312931002 | 0.46563739 | -0.6720487 | 0.5015527 | 0.85942996 |
| SLC25A30 | 44.1590126 | -0.31883263 | 0.36277847 | -0.8788632 | 0.37947548 | 0.79535699 |
| COG3 | 62.6772868 | 0.112646727 | 0.19502607 | 0.5775983 | 0.56353535 | 0.88191823 |
| ZC3H13 | 113.45999 | 0.199173053 | 0.21206211 | 0.93922038 | 0.34761761 | 0.77726831 |
| AL138686.1 | 43.9686975 | 0.000548997 | 0.44191498 | 0.00124231 | 0.99900878 | 0.99982181 |
| LRCH1 | 231.829225 | -0.720644529 | 0.22229005 | -3.2419109 | 0.00118731 | 0.03667809 |
| ESD | 68.8582879 | -0.181210911 | 0.21972846 | -0.8247039 | 0.40953972 | 0.81413754 |
| SUCLA2 | 112.660107 | 0.120497509 | 0.27038345 | 0.44565415 | 0.65584709 | 0.9158679 |
| NUDT15 | 11.8960297 | -0.336903524 | 0.38856097 | -0.8670545 | 0.38591217 | NA |
| MED4 | 72.6534599 | 0.101640529 | 0.23532837 | 0.43190938 | 0.66580728 | 0.91958319 |
| ITM2B | 587.531928 | -0.028282832 | 0.27633926 | -0.1023482 | 0.91848028 | 0.98521039 |
| RB1 | 238.123 | -0.432892648 | 0.23850521 | -1.8150239 | 0.06952024 | 0.41827125 |
| LPAR6 | 12.7993341 | -0.889249006 | 0.50231984 | -1.7702845 | 0.07667977 | 0.43573154 |
| RCBTB2 | 55.1649641 | 0.101395475 | 0.32627985 | 0.31076229 | 0.75598134 | 0.94796399 |
| FNDC3A | 786.430493 | 0.269335777 | 0.3037141 | 0.88680696 | 0.37518284 | 0.79287005 |
| CDADC1 | 34.048428 | 0.745400156 | 0.33706109 | 2.211469 | 0.02700338 | 0.25933936 |
| CAB39L | 122.931903 | 0.45581511 | 0.36881969 | 1.23587521 | 0.21650495 | 0.67117227 |
| SETDB2 | 130.894533 | 0.522783165 | 0.35434385 | 1.47535554 | 0.14011698 | 0.57374456 |
| PHF11 | 70.0787402 | 0.008061321 | 0.38672278 | 0.02084522 | 0.98336913 | 0.99749828 |
| RCBTB1 | 37.9573576 | 0.352142853 | 0.30561461 | 1.15224482 | 0.24922049 | 0.70371066 |
| EBPL | 31.8497797 | -0.000201071 | 0.41742316 | -0.0004817 | 0.99961566 | 0.99982181 |
| KPNA3 | 101.203943 | 0.045606885 | 0.19167442 | 0.23793934 | 0.81192814 | 0.96256453 |
| SPRYD7 | 18.9000725 | 0.384648428 | 0.2934413 | 1.31081897 | 0.18991893 | 0.64181696 |
| DLEU2 | 195.636483 | -0.948512464 | 0.27312931 | -3.4727598 | 0.00051514 | 0.02047878 |
| TRIM13 | 49.7618346 | -0.113076731 | 0.25307976 | -0.4468027 | 0.65501749 | 0.9158679 |
| RNASEH2B | 40.8836068 | -0.151896422 | 0.27117187 | -0.5601482 | 0.57537838 | 0.88744956 |
| INTS6 | 219.985749 | 0.733587042 | 0.31247412 | 2.34767296 | 0.0188911 | 0.21532746 |
| INTS6-AS1 | 12.1412334 | -0.111767135 | 0.39379951 | -0.2838174 | 0.77655036 | NA |
| WDFY2 | 48.1435429 | 0.601293306 | 0.3555686 | 1.69107539 | 0.09082241 | 0.47310687 |
| ATP7B | 22.3213762 | 0.221739437 | 0.40632189 | 0.54572358 | 0.58525596 | 0.89036744 |
| ALG11 | 17.4906289 | 0.097201465 | 0.40414663 | 0.24051039 | 0.80993461 | 0.96215121 |
| UTP14C | 23.6147749 | 0.655024931 | 0.46166467 | 1.41883271 | 0.15594779 | 0.59597065 |
| NEK3 | 38.3802389 | -0.232374363 | 0.36300591 | -0.6401393 | 0.52208202 | 0.86728831 |
| VPS36 | 78.5336002 | -0.215595189 | 0.20799203 | -1.036555 | 0.29994328 | 0.74437099 |
| SUGT1 | 55.6889592 | 0.654890603 | 0.25885293 | 2.52997174 | 0.01140717 | 0.1580708 |
| DIAPH3 | 13.7799834 | 2.09351865 | 0.59156683 | 3.53893854 | 0.00040174 | 0.01726416 |
| TDRD3 | 153.126768 | 0.296450752 | 0.20494427 | 1.44649448 | 0.14803856 | 0.58515649 |
| PCDH9 | 19.5394409 | 1.118930007 | 0.65373832 | 1.711587 | 0.08697281 | 0.46422617 |
| DACH1 | 73.177432 | 0.92423012 | 0.70352329 | 1.31371645 | 0.18894164 | 0.64036044 |
| MZT1 | 14.9804414 | -0.191032147 | 0.45138449 | -0.4232138 | 0.67213928 | 0.92176197 |
| DIS3 | 41.0817871 | 0.172974923 | 0.24298991 | 0.71186053 | 0.47655115 | 0.84677526 |
| PIBF1 | 175.691071 | 0.085508419 | 0.18333014 | 0.46641767 | 0.64091656 | 0.91146968 |
| KLF5 | 37.1205631 | 0.151083752 | 0.58960121 | 0.25624736 | 0.79775984 | 0.95848876 |
| KLF12 | 153.697691 | -0.00811183 | 0.24236326 | -0.0334697 | 0.97330002 | 0.99526459 |
| TBC1D4 | 836.041623 | -0.142684168 | 0.23057486 | -0.6188193 | 0.53603542 | 0.86946459 |
| COMMD6 | 78.9462099 | 0.297456389 | 0.43821283 | 0.67879434 | 0.49726818 | 0.85814183 |
| UCHL3 | 67.0729752 | -0.12958775 | 0.27802515 | -0.4661008 | 0.64114334 | 0.91146968 |
| LMO7-AS1 | 26.5380728 | -1.32419316 | 0.44581561 | -2.970271 | 0.00297537 | 0.06759041 |
| LMO7 | 645.910842 | 0.255151675 | 0.38232636 | 0.66736616 | 0.50453828 | 0.8614718 |
| AL365394.1 | 34.2106665 | 2.316471222 | 0.6502832 | 3.56224985 | 0.00036769 | 0.01625927 |
| KCTD12 | 178.491762 | 1.68419158 | 0.36771099 | 4.58020465 | 4.65E-06 | 0.00063463 |
| AC001226.2 | 30.1154121 | -0.675024596 | 0.56225298 | -1.200571 | 0.22991767 | 0.68523674 |
| FBXL3 | 89.754666 | 0.233611884 | 0.2612901 | 0.89407095 | 0.37128393 | 0.7900012 |
| MYCBP2 | 273.293924 | 0.089967713 | 0.24255728 | 0.37091327 | 0.71070213 | 0.93551509 |
| SLAIN1 | 13.4644966 | -0.336941847 | 0.41600599 | -0.8099447 | 0.41797196 | 0.8188001 |
| RNF219 | 21.3951821 | -0.020576078 | 0.37108437 | -0.0554485 | 0.95578114 | 0.99234582 |
| RBM26 | 218.346716 | 0.271639844 | 0.22208214 | 1.22315033 | 0.22127292 | 0.67492938 |
| NDFIP2 | 97.7084628 | 0.160755154 | 0.23872617 | 0.67338724 | 0.50070096 | 0.85942996 |
| GPC5 | 20.5662929 | 0.729781002 | 0.6129429 | 1.19061825 | 0.23380348 | 0.68840359 |
| GPC6 | 64.918006 | 0.4401404 | 1.17786312 | 0.37367704 | 0.70864461 | 0.93470937 |

| | | | | | | |
|---|---|---|---|---|---|---|
| TGDS | 21.3161612 | -0.032908447 | 0.31421277 | -0.104733 | 0.91658767 | 0.98495314 |
| GPR180 | 16.7695303 | 0.068149716 | 0.51120633 | 0.13331157 | 0.89394698 | 0.97833263 |
| ABCC4 | 999.375464 | -0.007997559 | 0.30576064 | -0.0261563 | 0.97913269 | 0.99623022 |
| DNAJC3-DT | 19.9584328 | -0.113606628 | 0.42544925 | -0.2670274 | 0.78944803 | 0.95719943 |
| DNAJC3 | 146.557892 | 0.306597069 | 0.25375486 | 1.20824119 | 0.2269545 | 0.68135521 |
| UGGT2 | 195.659197 | 0.035245163 | 0.18929416 | 0.18619256 | 0.85229376 | 0.97067623 |
| HS6ST3 | 804.207606 | -0.281943488 | 0.5911342 | -0.4769534 | 0.63339528 | 0.90805737 |
| OXGR1 | 30.3514311 | 1.693694092 | 0.64039542 | 2.64476298 | 0.00817482 | 0.13111608 |
| MBNL2 | 216.964338 | 0.524050104 | 0.26088895 | 2.00870946 | 0.04456796 | 0.33552163 |
| RAP2A | 13.5492103 | -0.106490522 | 0.39407882 | -0.2702265 | 0.78698604 | 0.95577371 |
| IPO5 | 60.6947058 | 0.200127477 | 0.19824001 | 1.0095211 | 0.31272479 | 0.7555244 |
| FARP1 | 417.770058 | 0.047026919 | 0.18236985 | 0.25786565 | 0.79651059 | 0.95848876 |
| STK24 | 263.244135 | -0.21702386 | 0.18484544 | -1.1740829 | 0.24036183 | 0.69470237 |
| DOCK9 | 92.6544424 | 0.031593467 | 0.22695248 | 0.13920741 | 0.88928626 | 0.97778408 |
| UBAC2 | 279.0205 | 0.331841614 | 0.17734774 | 1.87113531 | 0.06132633 | 0.39288791 |
| TM9SF2 | 233.757176 | -0.195302463 | 0.19384249 | -1.0075317 | 0.31367931 | 0.75632348 |
| CLYBL | 347.044151 | 0.010750182 | 0.39382879 | 0.02729659 | 0.97822318 | 0.99609121 |
| PCCA | 670.866256 | 0.014163069 | 0.2586701 | 0.0547534 | 0.95633492 | 0.99234582 |
| GGACT | 41.236274 | -0.469463117 | 0.25724455 | -1.8249681 | 0.06800584 | 0.4151827 |
| TMTC4 | 40.6388952 | 0.27087859 | 0.33559036 | 0.80717035 | 0.41956835 | 0.81899879 |
| FGF14 | 38.2094105 | 1.372601333 | 0.63288257 | 2.16880887 | 0.0300972 | 0.27617753 |
| TPP2 | 115.925627 | -0.129621038 | 0.25789144 | -0.5026186 | 0.61523244 | 0.90106518 |
| EFNB2 | 35.6999519 | 0.638370458 | 0.43622091 | 1.46341095 | 0.14335497 | 0.57698886 |
| ARGLU1 | 197.442636 | -0.033366105 | 0.21267696 | -0.1568863 | 0.87533444 | 0.9748271 |
| LIG4 | 24.2548555 | -0.152258773 | 0.34331364 | -0.4434976 | 0.65740586 | 0.9158679 |
| ABHD13 | 35.1756731 | 0.265663474 | 0.32825761 | 0.80931397 | 0.41833457 | 0.8188001 |
| TNFSF13B | 35.6325435 | 0.090458779 | 0.54673923 | 0.16545141 | 0.86858869 | 0.97434023 |
| IRS2 | 66.1001519 | -0.394386672 | 0.50425237 | -0.7821216 | 0.43414312 | 0.82539949 |
| AL355974.2 | 14.2686609 | 0.368029156 | 0.70416859 | 0.52264353 | 0.60122234 | 0.89490801 |
| COL4A2 | 11.8530271 | -0.285254451 | 0.58700416 | -0.4859496 | 0.62700288 | NA |
| RAB20 | 49.506173 | -0.417738195 | 0.33201258 | -1.2581999 | 0.20831947 | 0.65973983 |
| NAXD | 42.3940051 | 0.39190758 | 0.26810774 | 1.46175408 | 0.14380861 | 0.57830611 |
| CARS2 | 74.4190468 | 0.373014306 | 0.22742448 | 1.64016776 | 0.10097029 | 0.49019611 |
| ANKRD10 | 259.487127 | -0.146106209 | 0.20956954 | -0.6971729 | 0.48569458 | 0.85356796 |
| ARHGEF7 | 228.557185 | -0.416865474 | 0.18946776 | -2.2001922 | 0.02779326 | 0.26203825 |
| TUBGCP3 | 55.8230266 | 0.236993341 | 0.24261905 | 0.97681259 | 0.32866194 | 0.76649357 |
| ATP11A | 384.588535 | -1.33184117 | 0.42267075 | -3.1510134 | 0.00162705 | 0.04456355 |
| MCF2L | 67.6737653 | 0.340779473 | 0.32285683 | 1.05551266 | 0.29119092 | 0.73994767 |
| PCID2 | 55.1869863 | 0.181983778 | 0.23105754 | 0.78761236 | 0.43092348 | 0.8248269 |
| CUL4A | 122.234004 | 0.062630393 | 0.23723399 | 0.26400261 | 0.79177791 | 0.95746995 |
| LAMP1 | 117.281628 | 0.008147613 | 0.2862115 | 0.0284671 | 0.9772896 | 0.9957152 |
| GRTP1 | 31.7697821 | 0.736715464 | 0.37358733 | 1.97200335 | 0.04860922 | 0.35213553 |
| DCUN1D2 | 46.3485367 | 0.189372303 | 0.22987827 | 0.82379383 | 0.41005671 | 0.81427553 |
| TMCO3 | 45.7098406 | -0.388055166 | 0.31693807 | -1.224388 | 0.2208059 | 0.67462437 |
| TFDP1 | 41.2573588 | 0.313864414 | 0.32006373 | 0.98063099 | 0.32677475 | 0.76371899 |
| GAS6 | 195.309998 | 0.517021207 | 0.41173787 | 1.25570476 | 0.20922303 | 0.6612784 |
| CDC16 | 65.0726763 | -0.000742293 | 0.22759252 | -0.0032615 | 0.9973977 | 0.99982181 |
| UPF3A | 77.4021017 | 0.279859544 | 0.22296004 | 1.25520047 | 0.20940599 | 0.66142564 |
| CHAMP1 | 20.2436634 | 0.218993385 | 0.34950642 | 0.626579 | 0.53093524 | 0.86803839 |
| TTC5 | 15.032078 | 0.316270251 | 0.36313603 | 0.87094153 | 0.38378608 | 0.7980695 |
| CCNB1IP1 | 40.7633896 | 0.56184147 | 0.27476637 | 2.04479706 | 0.04087487 | 0.32473787 |
| TEP1 | 36.3424625 | -1.168348961 | 0.335207 | -3.4854551 | 0.0004913 | 0.01973081 |
| OSGEP | 16.6874475 | 0.255350245 | 0.40600227 | 0.62893797 | 0.52938967 | 0.86758926 |
| APEX1 | 19.8221998 | -0.029863528 | 0.43599777 | -0.0684947 | 0.94539185 | 0.99134285 |
| PIP4P1 | 18.7115512 | -0.395663232 | 0.44540218 | -0.888328 | 0.37436434 | 0.7923487 |
| PNP | 19.355112 | -0.678579639 | 0.3251183 | -2.0871776 | 0.03687208 | 0.30715411 |
| METTL17 | 20.3126477 | -0.055471067 | 0.3722649 | -0.1490097 | 0.88154601 | 0.97630358 |
| NDRG2 | 40.9683146 | 0.58399847 | 0.39867201 | 1.46485946 | 0.14295927 | 0.57698886 |
| ZNF219 | 12.4026555 | 0.087555936 | 0.60377087 | 0.14501517 | 0.88469889 | NA |
| HNRNPC | 330.314356 | 0.283410067 | 0.2099353 | 1.34998766 | 0.17701994 | 0.62488264 |
| SUPT16H | 62.4788048 | -0.072424665 | 0.24670636 | -0.2935663 | 0.76908935 | 0.95205353 |
| CHD8 | 87.5555471 | 0.004666824 | 0.26999299 | 0.01728498 | 0.98620927 | 0.99804172 |
| RAB2B | 39.9732453 | 0.267423012 | 0.27224015 | 0.98230555 | 0.32594934 | 0.76353952 |
| TOX4 | 56.7694361 | -0.300524249 | 0.22457859 | -1.3381696 | 0.18084114 | 0.6286114 |
| METTL3 | 19.2444822 | -0.217934763 | 0.34509612 | -0.631519 | 0.52770121 | 0.86750588 |
| DAD1 | 73.9897805 | -0.097459342 | 0.36607993 | -0.2662242 | 0.79006653 | 0.95746824 |
| ABHD4 | 16.6562812 | -0.889359498 | 0.37559072 | -2.3678953 | 0.0178896 | 0.20757069 |
| OXA1L | 25.6305627 | 0.181417325 | 0.3391949 | 0.53484686 | 0.59275576 | 0.89198431 |
| MRPL52 | 18.8313774 | 0.442345877 | 0.41187505 | 1.07398076 | 0.2828313 | 0.7344003 |
| MMP14 | 23.4162372 | -1.031808534 | 0.55378676 | -1.8631874 | 0.0624359 | 0.39739386 |
| LRP10 | 140.777046 | -1.36300336 | 0.41852317 | -3.2566975 | 0.00112716 | 0.03573039 |
| RBM23 | 63.175821 | 0.092335222 | 0.19501282 | 0.47348283 | 0.63586874 | 0.90881415 |
| PRMT5 | 11.0208053 | -0.229715005 | 0.42579772 | -0.5394933 | 0.58954654 | NA |
| AL132780.1 | 25.6831989 | -0.231416949 | 0.54882561 | -0.4216584 | 0.67327434 | 0.92189158 |
| PSMB5 | 62.8518561 | -0.20299145 | 0.29874649 | -0.6794773 | 0.4968355 | 0.85814183 |
| ACIN1 | 107.126899 | 0.070265759 | 0.18952737 | 0.370742 | 0.7108297 | 0.93555605 |
| C14orf119 | 25.7700308 | -0.306249182 | 0.35853006 | -0.8541799 | 0.3930053 | 0.80400536 |
| HOMEZ | 13.2007714 | -0.2725795 | 0.39940544 | -0.6824632 | 0.49494613 | 0.85814183 |
| BCL2L2-PABI | 21.6194598 | 0.210384466 | 0.31311184 | 0.67191476 | 0.50163796 | 0.85942996 |
| PABPN1 | 63.3530095 | 0.746106963 | 0.2460794 | 3.03197649 | 0.00242958 | 0.05804665 |
| NGDN | 19.6955609 | 0.259663538 | 0.36894038 | 0.70380893 | 0.48155177 | 0.85020971 |
| ZFHX2 | 10.8552793 | 0.245181881 | 0.55773615 | 0.43960192 | 0.66022545 | NA |
| AP1G2 | 27.0663999 | -0.401729786 | 0.35022329 | -1.1470676 | 0.25135369 | 0.70545786 |
| DHRS4-AS1 | 21.5247921 | 0.205489035 | 0.34731804 | 0.59164515 | 0.55408824 | 0.87706525 |
| DHRS4L2 | 26.1820254 | 0.237663301 | 0.45281366 | 0.52485895 | 0.59968125 | 0.89476262 |
| PSME1 | 60.5031662 | -0.408079839 | 0.40082315 | -1.0181045 | 0.30862832 | 0.75132915 |
| EMC9 | 11.0950717 | 0.43100647 | 0.63001097 | 0.68412534 | 0.49389602 | NA |
| PSME2 | 15.0548625 | -0.060879117 | 0.46412682 | -0.1311691 | 0.89564151 | 0.97885533 |
| AL136295.1 | 16.453609 | -0.010094225 | 0.33893157 | -0.0297825 | 0.97624053 | 0.99556048 |
| GMPR2 | 16.6950026 | -0.487205703 | 0.34904741 | -1.3958153 | 0.16277013 | 0.60133611 |
| TINF2 | 14.6364018 | -0.43675493 | 0.49227549 | -0.8872165 | 0.37496236 | 0.79274953 |
| DHRS1 | 12.7351915 | 0.597704922 | 0.53720786 | 1.11261387 | 0.2658743 | 0.71698101 |
| KHNYN | 19.4659048 | -0.254496488 | 0.3423404 | -0.7434018 | 0.45723843 | 0.83569193 |
| SDR39U1 | 24.6725131 | 0.117474491 | 0.2782204 | 0.42223536 | 0.67285323 | 0.92189158 |
| STXBP6 | 48.2630243 | 3.444969598 | 1.56332462 | 2.20361757 | 0.02755125 | 0.26098352 |
| NOVA1 | 92.1304304 | 0.402121715 | 0.32983516 | 1.21915963 | 0.22278361 | 0.67606266 |
| AL110292.1 | 40.7387716 | 1.899169236 | 0.68016202 | 2.79223066 | 0.0052346 | 0.09995328 |
| PRKD1 | 439.415038 | -2.026249584 | 0.41993128 | -4.8251933 | 1.40E-06 | 0.00025804 |
| G2E3 | 44.8768303 | 0.351571017 | 0.22293517 | 1.57701008 | 0.11479321 | 0.51928773 |
| SCFD1 | 145.254173 | 0.206903206 | 0.18349557 | 1.12756512 | 0.25950362 | 0.71066775 |
| COCH | 13.6373639 | -0.473707865 | 0.48766404 | -0.9713816 | 0.3313583 | 0.76685572 |
| STRN3 | 279.301909 | -0.004306638 | 0.24213723 | -0.0177859 | 0.98580962 | 0.99804172 |
| AP4S1 | 13.7480363 | 0.298046928 | 0.43875522 | 0.67930115 | 0.49694707 | 0.85814183 |
| HECTD1 | 390.199864 | 0.052729069 | 0.2294238 | 0.2298326 | 0.81822185 | 0.9635539 |
| AL136418.1 | 58.9241053 | -0.761672494 | 0.80677198 | -0.9440988 | 0.34511914 | 0.77492029 |
| HEATR5A | 60.9401983 | -0.13269936 | 0.33658883 | -0.3942477 | 0.6933982 | 0.92826395 |
| NUBPL | 137.955487 | -0.040670722 | 0.37318193 | -0.1089836 | 0.91321547 | 0.98376271 |
| ARHGAP5 | 280.733311 | -0.097883319 | 0.15042895 | -0.6506947 | 0.5152436 | 0.86463667 |
| AKAP6 | 147.666417 | -0.409914846 | 0.34534043 | -1.1869877 | 0.23523247 | 0.68976873 |
| NPAS3 | 60.133948 | -0.560540099 | 0.68077287 | -0.8233878 | 0.41028748 | 0.81427553 |
| EGLN3 | 19.087463 | 0.175962083 | 0.65926603 | 0.26690604 | 0.78954151 | 0.95719943 |
| EAPP | 48.1095909 | -0.19161744 | 0.23272402 | -0.8233677 | 0.41029892 | 0.81427553 |
| SNX6 | 135.52009 | -0.184145455 | 0.18203044 | -1.011619 | 0.31172024 | 0.75460103 |

| | | | | | | |
|---|---|---|---|---|---|---|
| CFL2 | 21.9748395 | 0.061325391 | 0.3342913 | 0.18344896 | 0.85444577 | 0.97140165 |
| BAZ1A | 57.5861348 | -0.926198181 | 0.45416449 | -2.0393452 | 0.04141559 | 0.32675147 |
| SRP54 | 93.4155431 | -0.11165317 | 0.19852938 | -0.5624012 | 0.57384268 | 0.88637538 |
| FAM177A1 | 71.6477728 | -0.263564196 | 0.25059044 | -1.0517727 | 0.29290383 | 0.74013384 |
| PPP2R3C | 57.2370342 | -0.044109067 | 0.31270018 | -0.1410587 | 0.8878236 | 0.9776239 |
| NFKBIA | 51.7391066 | 0.101842369 | 0.5352789 | 0.19026038 | 0.8491051 | 0.97027687 |
| RALGAPA1 | 453.614006 | -0.03979703 | 0.26207529 | -0.1518534 | 0.87930255 | 0.97567183 |
| BRMS1L | 23.3481708 | 0.551035125 | 0.36900234 | 1.49331063 | 0.13535589 | 0.5651968 |
| MBIP | 49.0115129 | 0.077464091 | 0.29076287 | 0.26641672 | 0.78991829 | 0.95741689 |
| SLC25A21 | 21.5900354 | 0.262155546 | 0.59273826 | 0.44227877 | 0.65828749 | 0.91665069 |
| MIPOL1 | 112.520187 | 0.435769997 | 0.28238456 | 1.54317924 | 0.12278729 | 0.53921501 |
| SEC23A | 146.148197 | -0.104649784 | 0.26231575 | -0.3989459 | 0.6899331 | 0.92734996 |
| GEMIN2 | 11.2023801 | 0.282454877 | 0.42179228 | 0.66965398 | 0.5030784 | NA |
| TRAPPC6B | 100.511009 | 0.621465379 | 0.20871778 | 2.97753921 | 0.00290572 | 0.06679036 |
| PNN | 123.668304 | 0.392305891 | 0.18340833 | 2.13897533 | 0.03243767 | 0.28526325 |
| MIA2 | 171.817875 | -0.013135863 | 0.19571502 | -0.0671173 | 0.94648832 | 0.99134285 |
| FBXO33 | 70.5214003 | 0.364254232 | 0.35259045 | 1.03308027 | 0.30156634 | 0.74603251 |
| AL049875.1 | 33.8151589 | 0.775545268 | 0.44734758 | 1.73365254 | 0.08297975 | 0.45526221 |
| KLHL28 | 57.5190909 | 0.111837974 | 0.25560577 | 0.43754088 | 0.66171914 | 0.91808148 |
| TOGARAM1 | 78.6947201 | 0.344472153 | 0.19094503 | 1.80403829 | 0.0712253 | 0.42274211 |
| PRPF39 | 46.1271802 | -0.347456561 | 0.2673748 | -1.2995112 | 0.19376854 | 0.64522994 |
| FKBP3 | 41.9722384 | 0.119930697 | 0.38025305 | 0.31539707 | 0.75246019 | 0.94728927 |
| FANCM | 21.3484108 | 0.541925444 | 0.47764702 | 1.13457306 | 0.25655425 | 0.70972499 |
| MIS18BP1 | 33.889155 | -0.299464817 | 0.28623184 | -1.0462317 | 0.29545409 | 0.74111836 |
| RPS29 | 101.7233 | 0.406248163 | 0.2780687 | 1.46096332 | 0.14402551 | 0.57846905 |
| RPL36AL | 119.156423 | -0.385509132 | 0.40230355 | -0.9582544 | 0.33793451 | 0.76966704 |
| DNAAF2 | 14.4295716 | 0.588389035 | 0.40196526 | 1.46378081 | 0.14325385 | 0.57698886 |
| KLHDC1 | 38.5911463 | 0.287511158 | 0.36105601 | 0.79630626 | 0.42585405 | 0.82319336 |
| KLHDC2 | 79.3274968 | 0.453643801 | 0.27068448 | 1.67591359 | 0.09375512 | 0.4781144 |
| NEMF | 247.484174 | -0.040596308 | 0.21190026 | -0.1915822 | 0.84806952 | 0.97027687 |
| ARF6 | 46.8784759 | 0.070967226 | 0.30460706 | 0.23297958 | 0.81577726 | 0.96300833 |
| LINC01588 | 19.0754294 | -0.629378489 | 0.46435249 | -1.3553895 | 0.17529349 | 0.62192644 |
| VCPKMT | 24.1378185 | 0.370116554 | 0.33604608 | 1.10138633 | 0.27072855 | 0.7217698 |
| SOS2 | 252.100102 | 0.163569805 | 0.15069134 | 1.08546253 | 0.27771685 | 0.72708595 |
| L2HGDH | 70.4446485 | 0.308840739 | 0.34627782 | 0.89188716 | 0.37245342 | 0.79086345 |
| ATP5S | 54.5671724 | 0.250362689 | 0.21761701 | 1.15047391 | 0.24994874 | 0.70438778 |
| CDKL1 | 121.329844 | 0.395856457 | 0.30327831 | 1.30525805 | 0.19180499 | 0.6421543 |
| MAP4K5 | 174.249797 | -0.248990483 | 0.26285153 | -0.9472666 | 0.34350292 | 0.77370523 |
| ATL1 | 20.7643496 | 0.326109611 | 0.4656299 | 0.70036227 | 0.48370109 | 0.85235034 |
| SAV1 | 113.077092 | 0.723526077 | 0.31088513 | 2.32731005 | 0.01994877 | 0.22190718 |
| NIN | 19.7736071 | 0.3120168 | 0.46872407 | 0.66567267 | 0.50562035 | 0.86210535 |
| TMX1 | 25.5189161 | 0.176221236 | 0.35699644 | 0.49362183 | 0.62157328 | 0.90380166 |
| RTRAF | 52.3021284 | -0.002460049 | 0.32049869 | -0.0076757 | 0.99387575 | 0.99923337 |
| TXNDC16 | 23.1897391 | 0.108851267 | 0.43104312 | 0.25252988 | 0.80063152 | 0.95873501 |
| ERO1A | 367.490504 | -1.269823489 | 0.37668754 | -3.371026 | 0.00074889 | 0.02651173 |
| AL133453.1 | 54.3565928 | -2.646825312 | 0.59051806 | -4.4822089 | 7.39E-06 | 0.00093063 |
| PSMC6 | 91.645845 | -0.096426977 | 0.1925072 | -0.5009006 | 0.61644107 | 0.90160403 |
| STYX | 48.4168 | 0.122106577 | 0.29369684 | 0.41575721 | 0.67758767 | 0.92259972 |
| GNPNAT1 | 92.7948523 | -0.510617079 | 0.48506231 | -1.0526835 | 0.29248609 | 0.74013384 |
| FERMT2 | 118.286643 | -0.015171321 | 0.26684284 | -0.0568549 | 0.95466079 | 0.99228059 |
| DDHD1 | 25.227115 | 0.123925936 | 0.34061745 | 0.36382733 | 0.71598696 | 0.93689626 |
| CNIH1 | 29.7118177 | 0.258434829 | 0.28404377 | 0.90984156 | 0.36290607 | 0.78718296 |
| GMFB | 34.0741689 | -0.223272853 | 0.28474872 | -0.7841048 | 0.43297859 | 0.8248269 |
| CGRRF1 | 15.782636 | 0.254777298 | 0.34958415 | 0.72880105 | 0.46612337 | 0.84226007 |
| SAMD4A | 165.261271 | -0.183577873 | 0.47797244 | -0.3840763 | 0.70092192 | 0.93127962 |
| GCH1 | 18.887184 | -0.118825051 | 0.42003033 | -0.2828964 | 0.77725629 | 0.95512504 |
| SOCS4 | 38.9134395 | 0.092193343 | 0.26087368 | 0.35340223 | 0.72378691 | 0.93896433 |
| MAPK1IP1L | 132.684188 | -0.030994927 | 0.20133415 | -0.1539477 | 0.87765098 | 0.97542218 |
| LGALS3 | 358.220484 | -0.192622167 | 0.31561105 | -0.610315 | 0.54165315 | 0.87174971 |
| FBXO34 | 288.796313 | 0.38648353 | 0.26443172 | 1.46156265 | 0.1438611 | 0.57830611 |
| AL158801.3 | 14.5184863 | -0.110163568 | 0.52975516 | -0.2079519 | 0.83526656 | 0.96748214 |
| ATG14 | 38.2902471 | -0.1683031 | 0.28615272 | -0.5881583 | 0.55642605 | 0.87807756 |
| KTN1-AS1 | 12.6722584 | 0.020700006 | 0.47971962 | 0.04315022 | 0.96558179 | 0.99334616 |
| KTN1 | 341.373004 | 0.130248509 | 0.2690399 | 0.48412339 | 0.6282983 | 0.90624439 |
| PELI2 | 141.113953 | -0.991081972 | 0.29740846 | -3.3323934 | 0.00086102 | 0.0296168 |
| TMEM260 | 100.62292 | -0.059674835 | 0.25735336 | -0.231879 | 0.81663201 | 0.96340618 |
| EXOC5 | 90.9755607 | -0.244198123 | 0.19818366 | -1.2321809 | 0.2178815 | 0.67202699 |
| AP5M1 | 69.6232744 | -0.015458412 | 0.22325999 | -0.0692395 | 0.94479898 | 0.99121467 |
| NAA30 | 48.1191502 | -0.072253015 | 0.23981771 | -0.3012831 | 0.76319865 | 0.95093474 |
| CCDC198 | 190.757079 | 0.651300553 | 0.56897117 | 1.14469869 | 0.25233399 | 0.70621866 |
| ARMH4 | 62.0310204 | 0.306998301 | 0.27466844 | 1.11770503 | 0.26369299 | 0.71380385 |
| ACTR10 | 75.3625762 | 0.0523308 | 0.20587091 | 0.2541923 | 0.799347 | 0.95848876 |
| PSMA3 | 67.1156402 | -0.199576228 | 0.2831448 | -0.7048557 | 0.48090003 | 0.84967765 |
| ARID4A | 99.1975326 | -0.185560056 | 0.22741425 | -0.8159562 | 0.41452521 | 0.81725499 |
| TIMM9 | 24.2203606 | 0.159276811 | 0.28857172 | 0.5519488 | 0.58098343 | 0.89012555 |
| KIAA0586 | 72.5169077 | -0.205806278 | 0.20721162 | -0.9932178 | 0.32060381 | 0.76131909 |
| DAAM1 | 107.790471 | 0.160621031 | 0.26463365 | 0.60695619 | 0.54387999 | 0.87243854 |
| L3HYPDH | 13.9302281 | -0.362383331 | 0.41748674 | -0.8680116 | 0.38538798 | 0.7991545 |
| JKAMP | 48.3523861 | 0.177765253 | 0.23142705 | 0.7681265 | 0.44241203 | 0.82820049 |
| RTN1 | 54.2734847 | 1.528442442 | 0.48263091 | 3.1668971 | 0.00154075 | 0.04294614 |
| LRRC9 | 50.7126895 | 0.438479353 | 0.55262635 | 0.79344633 | 0.42751782 | 0.82407405 |
| AL157911.1 | 134.421196 | 1.574452839 | 0.2357644 | 6.6780772 | 2.42E-11 | 3.35E-08 |
| PCNX4 | 600.325536 | 1.271347019 | 0.3400937 | 3.73822576 | 0.00018532 | 0.0103617 |
| DHRS7 | 164.912868 | 0.580697171 | 0.32868746 | 1.7667153 | 0.07727592 | 0.4378367 |
| PPM1A | 131.564123 | 0.29905941 | 0.24345335 | 1.22840541 | 0.21929482 | 0.67305276 |
| MNAT1 | 252.528366 | 0.073529633 | 0.22388691 | 0.3284231 | 0.74259178 | 0.94546379 |
| SLC38A6 | 31.7513394 | 0.228050288 | 0.3852637 | 0.59193297 | 0.55389548 | 0.87691576 |
| PRKCH | 28.9783992 | -0.225706227 | 0.31568885 | -0.7149642 | 0.47463116 | 0.84593347 |
| TMEM30B | 38.3915602 | 0.119963609 | 0.29295235 | 0.40949871 | 0.68217371 | 0.92456127 |
| HIF1A | 922.718296 | -1.621545528 | 0.33806008 | -4.7966193 | 1.61E-06 | 0.00027951 |
| HIF1A-AS2 | 148.23172 | -1.894727865 | 0.56875933 | -3.331335 | 0.00086431 | 0.0296246 |
| SNAPC1 | 74.353814 | -0.582916346 | 0.38010909 | -1.5335501 | 0.12514037 | 0.54482117 |
| PPP2R5E | 332.320077 | -0.75642274 | 0.28821231 | -2.6245331 | 0.00867679 | 0.13512263 |
| WDR89 | 24.4288978 | -0.317653483 | 0.3599039 | -0.8826064 | 0.37744898 | 0.79432848 |
| SGPP1 | 33.6810247 | -0.224566043 | 0.29815069 | -0.7531965 | 0.45133187 | 0.83230402 |
| SYNE2 | 1355.75065 | -0.720121949 | 0.31879311 | -2.2589006 | 0.02388957 | 0.24214086 |
| ESR2 | 112.690501 | -0.100754993 | 0.77504536 | -0.1299988 | 0.89656735 | 0.97885533 |
| MTHFD1 | 15.5400926 | -0.022071142 | 0.34525828 | -0.0639265 | 0.94902878 | 0.99185581 |
| AL122035.1 | 11.9896663 | -0.901101715 | 0.47016905 | -1.9165483 | 0.05529534 | NA |
| ZBTB25 | 32.9075064 | -0.477037329 | 0.44507752 | -1.0718073 | 0.2838066 | 0.73550473 |
| ZBTB1 | 85.0764273 | 0.003925414 | 0.28107414 | 0.01396576 | 0.9888573 | 0.99915789 |
| PLEKHG3 | 61.660703 | -0.675845592 | 0.26925115 | -2.5100937 | 0.01206991 | 0.163517 |
| CHURC1-FNT | 34.4526845 | -0.864734827 | 0.37649233 | -2.2968193 | 0.02162908 | 0.23004616 |
| GPX2 | 15.3847634 | -1.709275363 | 0.83385086 | -2.0498574 | 0.04037835 | 0.32236211 |
| RAB15 | 45.2774229 | 0.061245141 | 0.55708991 | 0.10993762 | 0.91245884 | 0.98376271 |
| MAX | 84.7948783 | -0.614550289 | 0.2095244 | -2.9330727 | 0.00335625 | 0.07371098 |
| FUT8 | 63.1253982 | -0.186813895 | 0.39845253 | -0.4688486 | 0.63917789 | 0.91069705 |
| AL359232.1 | 49.9198999 | -0.10776275 | 0.3663029 | -0.2941903 | 0.76861251 | 0.95205353 |
| GPHN | 1404.37506 | -0.57946791 | 0.27740679 | -2.0888743 | 0.03671904 | 0.3068708 |
| MPP5 | 146.559005 | 0.328945045 | 0.18264017 | 1.8010553 | 0.07169416 | 0.42284325 |
| ATP6V1D | 129.178684 | 0.835860337 | 0.27168212 | 3.07661151 | 0.00209368 | 0.05234522 |
| EIF2S1 | 49.9467498 | 0.03447051 | 0.23622828 | 0.14592034 | 0.88398428 | 0.97662748 |
| PLEK2 | 46.5398733 | -0.961415751 | 0.48284784 | -1.9911361 | 0.04646593 | 0.34368082 |

| | | | | | | |
|---|---|---|---|---|---|---|
| PLEKHH1 | 83.2151732 | -0.33333259 | 0.39774039 | -0.8380657 | 0.40199379 | 0.80890307 |
| PIGH | 38.6093216 | -0.311550102 | 0.4140971 | -0.75236 | 0.45183458 | 0.83243978 |
| ARG2 | 79.4202035 | 0.028550412 | 0.7062844 | 0.04042339 | 0.96775558 | 0.99393148 |
| VTI1B | 44.5598591 | 0.356560849 | 0.37592887 | 0.94847956 | 0.34288537 | 0.7729798 |
| RDH11 | 11.9363085 | -1.587044055 | 0.50671649 | -3.1320158 | 0.00173611 | NA |
| ZFYVE26 | 31.0742703 | -0.330084464 | 0.35411624 | -0.9321359 | 0.35126629 | 0.77910174 |
| RAD51B | 254.031696 | 0.163021953 | 0.43367776 | 0.37590572 | 0.70698699 | 0.93432317 |
| ZFP36L1 | 127.609187 | -0.672581874 | 0.2518199 | -2.6708845 | 0.00756517 | 0.12565432 |
| ACTN1 | 17.7877627 | -0.398562365 | 0.429716 | -0.9275018 | 0.35366605 | 0.78035986 |
| DCAF5 | 179.629551 | -0.157667211 | 0.22681398 | -0.6951389 | 0.48696829 | 0.85365949 |
| AL359317.2 | 12.8153788 | -0.079553416 | 0.55188334 | -0.144149 | 0.88538283 | 0.97662748 |
| EXD2 | 41.6654589 | 0.19859447 | 0.30648953 | 0.64796494 | 0.51700763 | 0.86500999 |
| GALNT16 | 36.0948353 | -0.787340243 | 0.62136406 | -1.2671158 | 0.20511388 | 0.65805651 |
| ERH | 51.9406018 | 0.010778837 | 0.32463081 | 0.03320337 | 0.97351241 | 0.99526459 |
| SLC39A9 | 71.0630667 | 0.273360435 | 0.25323508 | 1.079473 | 0.28037692 | 0.73207433 |
| PLEKHD1 | 10.9911445 | 0.535489572 | 0.6706073 | 0.79851438 | 0.42457205 | NA |
| SUSD6 | 148.739734 | -0.469802301 | 0.20236891 | -2.3215142 | 0.02025911 | 0.22348867 |
| SRSF5 | 273.037579 | 0.275217396 | 0.21776045 | 1.26385388 | 0.20628248 | 0.65836247 |
| SMOC1 | 75.574909 | 0.348366851 | 0.66001885 | 0.52781349 | 0.59762879 | 0.89363331 |
| AL357153.2 | 33.6792591 | -0.14401537 | 0.4595449 | -0.3133869 | 0.75398671 | 0.94795939 |
| MED6 | 32.9363851 | -0.300095906 | 0.32446491 | -0.9248948 | 0.35502063 | 0.7810615 |
| TTC9 | 73.8684052 | -2.347378667 | 0.56553686 | -4.1507085 | 3.31E-05 | 0.00299743 |
| MAP3K9 | 44.0379398 | -0.178845479 | 0.29680551 | -0.6025679 | 0.54679617 | 0.87269272 |
| PCNX1 | 321.792602 | -0.165874684 | 0.24164522 | -0.6864389 | 0.49243637 | 0.85714521 |
| SIPA1L1 | 189.361495 | -0.45132901 | 0.26801667 | -1.6839587 | 0.09218965 | 0.4764741 |
| DPF3 | 16.6910739 | 1.256796033 | 0.69324233 | 1.81292454 | 0.06984346 | 0.41879921 |
| DCAF4 | 25.1326469 | 0.157072947 | 0.35276847 | 0.4452579 | 0.65613339 | 0.9158679 |
| ZFYVE1 | 77.1842228 | -0.450292939 | 0.25061456 | -1.7967549 | 0.07237454 | 0.4244456 |
| RBM25 | 190.189506 | 0.122763044 | 0.22476705 | 0.54617902 | 0.58494289 | 0.89036744 |
| PSEN1 | 200.891135 | -0.407807284 | 0.17438076 | -2.3386025 | 0.01935602 | 0.21848783 |
| PAPLN | 18.2084582 | -0.610193725 | 0.48731509 | -1.2521544 | 0.21051362 | 0.66233607 |
| NUMB | 488.072252 | -0.194629787 | 0.19527358 | -0.9967031 | 0.31890863 | 0.76052272 |
| HEATR4 | 23.6285126 | -1.862732443 | 0.4726561 | -3.9409889 | 8.11E-05 | 0.00561777 |
| ACOT2 | 13.0633033 | -1.2073068 | 0.67679434 | -1.7838607 | 0.07444631 | 0.43060774 |
| DNAL1 | 31.0682911 | 0.486767468 | 0.25881999 | 1.88071822 | 0.06001026 | 0.38930355 |
| PNMA1 | 13.8559956 | 0.286175277 | 0.37707522 | 0.75893419 | 0.44789193 | 0.83066268 |
| ELMSAN1 | 61.9786657 | -0.706082392 | 0.40160981 | -1.7581303 | 0.07872534 | 0.44254372 |
| AC005480.1 | 18.8016095 | -0.700690865 | 0.56317527 | -1.244179 | 0.21343368 | 0.66676543 |
| COQ6 | 15.5490852 | -0.106986418 | 0.38860104 | -0.2753117 | 0.78307677 | 0.95544332 |
| ENTPD5 | 36.4579459 | 0.292974054 | 0.3081712 | 0.95068604 | 0.34176378 | 0.77203275 |
| BBOF1 | 26.569382 | 0.12615588 | 0.40535532 | 0.31122295 | 0.75563113 | 0.94795939 |
| ALDH6A1 | 45.0980254 | 0.568353993 | 0.35391819 | 1.60589089 | 0.10829789 | 0.50631231 |
| LIN52 | 62.3998291 | 0.495593307 | 0.24797578 | 1.99855532 | 0.04565649 | 0.3398833 |
| ABCD4 | 58.2795209 | 0.060600149 | 0.23280985 | 0.2602989 | 0.79463322 | 0.95786532 |
| NPC2 | 103.661256 | -0.417742023 | 0.43027266 | -0.9708775 | 0.33160931 | 0.76685572 |
| ISCA2 | 25.7215953 | -0.21419276 | 0.43290011 | -0.4947856 | 0.62075144 | 0.90380166 |
| AREL1 | 73.8528105 | 0.337463808 | 0.24100939 | 1.4002102 | 0.16145038 | 0.60060549 |
| FCF1 | 23.5641754 | -0.023061137 | 0.29295581 | -0.0787188 | 0.93725627 | 0.98871562 |
| YLPM1 | 124.613716 | 0.399844207 | 0.17749057 | 2.25276309 | 0.02427409 | 0.24374605 |
| DLST | 71.4093695 | -0.028085003 | 0.24501724 | -0.1146246 | 0.90874268 | 0.98281552 |
| EIF2B2 | 15.7148721 | -0.356860982 | 0.37592908 | -0.9492774 | 0.34247955 | 0.77274985 |
| MLH3 | 53.2757134 | -0.514791944 | 0.33489988 | -1.5371518 | 0.12425615 | 0.54316569 |
| ACYP1 | 39.1846353 | 0.685186823 | 0.40635091 | 1.68619487 | 0.09175827 | 0.47622003 |
| ZC2HC1C | 14.582453 | -0.33254898 | 0.50511391 | -0.6583643 | 0.51030405 | 0.86281145 |
| NEK9 | 66.1979551 | -0.12120892 | 0.19496275 | -0.621703 | 0.5341372 | 0.86909171 |
| AL049780.2 | 13.4633289 | -0.020661912 | 0.58246563 | -0.0354732 | 0.97170243 | 0.99526459 |
| TMED10 | 141.77981 | 0.029949257 | 0.1792433 | 0.16708718 | 0.86730145 | 0.97434023 |
| AL691403.1 | 88.799825 | 0.445513951 | 0.54045872 | 0.82432559 | 0.40975458 | 0.81413754 |
| AF111167.1 | 12.9224262 | -0.978034683 | 0.73736252 | -1.326396 | 0.18470857 | 0.6349723 |
| FOS | 43.8965323 | 0.848623068 | 0.64074814 | 1.32442533 | 0.18536183 | 0.63578845 |
| JDP2 | 14.8242118 | -0.600258475 | 0.5010572 | -1.1979839 | 0.23092327 | 0.6853319 |
| FLVCR2 | 51.307656 | 0.796002151 | 0.49182532 | 1.61846516 | 0.10556239 | 0.50071158 |
| TTLL5 | 129.892255 | -0.190692254 | 0.25182469 | -0.7572421 | 0.44890483 | 0.83118301 |
| ERG28 | 33.1635742 | -0.413374015 | 0.3341788 | -1.2369845 | 0.21609282 | 0.67053754 |
| IFT43 | 82.5447478 | -0.027802859 | 0.26765501 | -0.1038757 | 0.91726797 | 0.98495314 |
| GPATCH2L | 140.844125 | -0.141354148 | 0.23225021 | -0.6086287 | 0.54277057 | 0.87210113 |
| ANGEL1 | 13.6799934 | 0.856460668 | 0.35697852 | 2.39919382 | 0.01643121 | 0.19848417 |
| IRF2BPL | 46.1977443 | 0.054733043 | 0.28458975 | 0.19232261 | 0.84748951 | 0.97027687 |
| POMT2 | 14.8401188 | 0.554027639 | 0.3591442 | 1.54263285 | 0.12291988 | 0.53921501 |
| GSTZ1 | 19.4325116 | -0.15452002 | 0.42328148 | -0.3650526 | 0.71507211 | 0.93689626 |
| TMED8 | 48.0136785 | 0.37210386 | 0.26191037 | 1.4207298 | 0.15539533 | 0.59533243 |
| VIPAS39 | 19.9701437 | -0.23616726 | 0.37186114 | -0.6350953 | 0.52536626 | 0.86747074 |
| AHSA1 | 40.5209846 | 0.491911643 | 0.28297141 | 1.7383793 | 0.082144 | 0.45375673 |
| SPTLC2 | 383.820081 | 0.704073551 | 0.32829371 | 2.14464529 | 0.03198122 | 0.28341152 |
| ALKBH1 | 36.1504064 | 0.116669208 | 0.23464414 | 0.49721765 | 0.61903558 | 0.90295415 |
| SLIRP | 60.4017347 | -0.130839115 | 0.29892324 | -0.4377014 | 0.66160277 | 0.91808148 |
| SNW1 | 55.8565338 | 0.02846083 | 0.20551847 | 0.13848308 | 0.88985864 | 0.97782578 |
| ADCK1 | 27.3080865 | 0.799008575 | 0.28769098 | 2.77731537 | 0.005481 | 0.10201977 |
| NRXN3 | 891.214485 | 0.406417689 | 0.56797283 | 0.71555833 | 0.47426411 | 0.84593347 |
| AC008056.1 | 38.6121598 | -1.028902588 | 0.67434665 | -1.525777 | 0.12706542 | 0.54952841 |
| CEP128 | 36.1076955 | 0.487496187 | 0.32230222 | 1.51254367 | 0.13039561 | 0.55685884 |
| GTF2A1 | 54.0725927 | 0.174331325 | 0.20371628 | 0.85575547 | 0.39213305 | 0.8036532 |
| STON2 | 139.834067 | 0.222460796 | 0.36892714 | 0.60299385 | 0.54651278 | 0.87269272 |
| SEL1L | 134.264132 | 0.049021403 | 0.23259949 | 0.21075456 | 0.8330788 | 0.96701848 |
| GALC | 12.025854 | -0.062959561 | 0.3934052 | -0.1600374 | 0.87285158 | NA |
| SPATA7 | 53.4668134 | 0.328812989 | 0.29789795 | 1.10377729 | 0.26968976 | 0.72046011 |
| PTPN21 | 113.753564 | 0.462429467 | 0.22914198 | 2.01809141 | 0.04358174 | 0.33245643 |
| ZC3H14 | 111.050206 | 0.426014736 | 0.21595111 | 1.97273697 | 0.04852554 | 0.35179202 |
| EML5 | 31.801596 | 0.044237736 | 0.46586233 | 0.09495882 | 0.92434754 | 0.98672316 |
| TTC8 | 43.6908195 | 0.110556436 | 0.29963786 | 0.36896685 | 0.71215243 | 0.9357596 |
| FOXN3 | 441.131026 | 0.22595204 | 0.4056272 | 0.55704361 | 0.57749764 | 0.88873982 |
| EFCAB11 | 35.4385827 | 0.620000458 | 0.33020177 | 1.87764121 | 0.06043026 | 0.39078238 |
| TDP1 | 19.2418717 | 0.117490043 | 0.35401429 | 0.33187938 | 0.73998034 | 0.94376279 |
| PSMC1 | 34.9541864 | -0.047685916 | 0.32571959 | -0.1464017 | 0.88360425 | 0.97662748 |
| NRDE2 | 34.2545002 | -0.152298827 | 0.32970562 | -0.4619237 | 0.64413607 | 0.9119294 |
| CALM1 | 96.9448848 | 0.16234267 | 0.36023602 | 0.45065641 | 0.6522372 | 0.9158679 |
| TTC7B | 126.087213 | -0.328484217 | 0.28204254 | -1.164662 | 0.24415585 | 0.69773111 |
| RPS6KA5 | 39.1741597 | -0.509132329 | 0.54278687 | -0.9379968 | 0.34824607 | 0.77780034 |
| DGLUCY | 202.362745 | -0.251869911 | 0.23503606 | -1.0716224 | 0.28388966 | 0.73550473 |
| PPP4R3A | 136.483325 | -0.003766444 | 0.19434562 | -0.0193801 | 0.98453786 | 0.99773864 |
| CATSPERB | 32.0203306 | 0.312125989 | 0.38794806 | 0.80455613 | 0.42107586 | 0.81961398 |
| TC2N | 53.6159363 | 0.119753702 | 0.43848377 | 0.27310863 | 0.78476971 | 0.95555482 |
| TRIP11 | 150.231114 | 0.125834211 | 0.18972411 | 0.66324841 | 0.50717147 | 0.86277855 |
| ATXN3 | 54.2714878 | 0.061780249 | 0.22069416 | 0.27993604 | 0.77952657 | 0.95537097 |
| NDUFB1 | 63.0881963 | 0.298817712 | 0.30326182 | 0.98534564 | 0.32445432 | 0.76304922 |
| CPSF2 | 60.9936248 | 0.019841723 | 0.21059805 | 0.09421608 | 0.92493751 | 0.98672316 |
| LGMN | 72.9059523 | -0.75568846 | 0.42248542 | -1.7886735 | 0.07366742 | 0.42788861 |
| GOLGA5 | 30.4428704 | 0.188100707 | 0.25202609 | 0.7463541 | 0.45545354 | 0.8346683 |
| ITPK1 | 94.0153058 | 0.202235793 | 0.36351044 | 0.55634109 | 0.5779777 | 0.88877358 |
| MOAP1 | 27.1026284 | -0.786559613 | 0.34897145 | -2.2539368 | 0.02420014 | 0.24374605 |
| UBR7 | 17.4061504 | -0.465421606 | 0.33942755 | -1.3711957 | 0.17031394 | 0.61277643 |
| BTBD7 | 205.564447 | 0.090659031 | 0.13797096 | 0.65708776 | 0.51112449 | 0.86312853 |
| DDX24 | 155.295987 | -0.645627429 | 0.2363907 | -2.7311879 | 0.00631065 | 0.11273164 |

| | | | | | | |
|---|---|---|---|---|---|---|
| IFI27L1 | 13.9739615 | 0.278328945 | 0.40847472 | 0.68138597 | 0.4956273 | 0.85814183 |
| IFI27 | 67.1230633 | -1.646141433 | 1.57863113 | -1.0427651 | 0.2970571 | 0.74160729 |
| IFI27L2 | 16.0991067 | -1.253350313 | 0.51661046 | -2.4261032 | 0.01526193 | 0.1876308 |
| PPP4R4 | 18.0579945 | -0.274891464 | 0.46528077 | -0.5908077 | 0.55464927 | 0.87752372 |
| SERPINA1 | 51.436666 | -1.081106337 | 0.66862523 | -1.6169093 | 0.10589786 | 0.50156703 |
| DICER1 | 243.82189 | -0.159201361 | 0.19154036 | -0.8311635 | 0.40588125 | 0.8115705 |
| DICER1-AS1 | 18.7754583 | 0.244056531 | 0.44148768 | 0.55280485 | 0.58039705 | 0.89012555 |
| CLMN | 451.893373 | -0.302358913 | 0.46290585 | -0.6531758 | 0.51364293 | 0.86393903 |
| GLRX5 | 48.5769261 | 0.102049187 | 0.37094858 | 0.27510333 | 0.78323686 | 0.95544332 |
| ATG2B | 92.0169636 | -0.132937644 | 0.2448906 | -0.542845 | 0.58723651 | 0.89073775 |
| GSKIP | 36.6382491 | -0.011587667 | 0.32847267 | -0.0352774 | 0.97185853 | 0.99526459 |
| AK7 | 19.1605031 | -1.938121586 | 0.48160316 | -4.0243124 | 5.71E-05 | 0.00450632 |
| PAPOLA | 239.308983 | -0.00460413 | 0.21235664 | -0.0216811 | 0.98270232 | 0.99719767 |
| VRK1 | 16.7377783 | 0.351905837 | 0.39324571 | 0.89487521 | 0.3708538 | 0.7900012 |
| SETD3 | 153.58981 | 0.473476644 | 0.22774532 | 2.07897418 | 0.03761972 | 0.31147067 |
| CCNK | 43.596994 | -0.116846136 | 0.25973039 | -0.4498747 | 0.65280078 | 0.9158679 |
| CCDC85C | 41.7580095 | -0.242622034 | 0.32743296 | -0.7409823 | 0.45870416 | 0.83645518 |
| EVL | 78.915863 | -0.385516056 | 0.38675146 | -0.9968057 | 0.31885883 | 0.76052272 |
| YY1 | 133.724515 | 0.06746917 | 0.29402083 | 0.22947072 | 0.81850308 | 0.9635539 |
| SLC25A29 | 157.280906 | -0.383234783 | 0.34815934 | -1.1007454 | 0.27100749 | 0.72194341 |
| WARS | 50.4009891 | -1.246607841 | 0.44596553 | -2.7953009 | 0.00518514 | 0.09959581 |
| WDR25 | 26.7695972 | 0.348209025 | 0.43424909 | 0.80186473 | 0.42263121 | 0.82138304 |
| PPP2R5C | 169.1322 | 0.029700798 | 0.21353902 | 0.13908839 | 0.8893803 | 0.97778408 |
| DYNC1H1 | 132.168087 | -0.293560898 | 0.28880735 | -1.0164592 | 0.30941076 | 0.75220159 |
| HSP90AA1 | 478.738885 | 0.134028555 | 0.2921336 | 0.45879199 | 0.64638354 | 0.91265217 |
| WDR20 | 136.121182 | -0.000325392 | 0.1779785 | -0.0018283 | 0.99854126 | 0.99982181 |
| MOK | 29.7950919 | 0.619204374 | 0.32975605 | 1.87776503 | 0.06041332 | 0.39078238 |
| ZNF839 | 23.6262917 | 0.020764006 | 0.30171015 | 0.06882104 | 0.94513207 | 0.99127958 |
| CINP | 20.9820065 | 0.326506175 | 0.34708384 | 0.94071268 | 0.34685212 | 0.7765713 |
| TECPR2 | 47.1800931 | 0.465835742 | 0.34398328 | 1.35423949 | 0.17565998 | 0.62231623 |
| ANKRD9 | 84.4504265 | -0.045659512 | 0.26329568 | -0.1734153 | 0.86232496 | 0.97307493 |
| RCOR1 | 107.360455 | 0.166676103 | 0.19449008 | 0.85699024 | 0.39145028 | 0.80332153 |
| TRAF3 | 93.4571791 | -0.272999535 | 0.32576584 | -0.8380238 | 0.40201733 | 0.80890307 |
| AMN | 48.4012983 | 0.343865888 | 0.54170042 | 0.63478977 | 0.52556553 | 0.86747074 |
| CDC42BPB | 281.097769 | -0.052770879 | 0.23994738 | -0.2199269 | 0.8259281 | 0.96539788 |
| EIF5 | 102.317889 | -0.388947824 | 0.37482057 | -1.0376907 | 0.29941407 | 0.74336229 |
| MARK3 | 372.520879 | -0.066267569 | 0.22785767 | -0.2908288 | 0.77118227 | 0.95268314 |
| CKB | 602.7937 | -0.201131873 | 0.45994187 | -0.4372985 | 0.66189491 | 0.91810778 |
| BAG5 | 63.6803304 | -0.740130138 | 0.26176263 | -2.8274859 | 0.00469151 | 0.0936371 |
| KLC1 | 107.865414 | -0.2264655 | 0.1819319 | -1.2447817 | 0.213212 | 0.66650224 |
| AL049840.1 | 12.3282968 | -1.665173127 | 0.60058365 | -2.7725915 | 0.00556119 | NA |
| ZFYVE21 | 55.6300164 | -0.353481365 | 0.31204989 | -1.132772 | 0.25731003 | 0.71022118 |
| PPP1R13B | 160.760915 | 0.119673301 | 0.19481146 | 0.61430318 | 0.539015 | 0.86972442 |
| ATP5MPL | 83.9983037 | 0.600675829 | 0.332341 | 1.80740813 | 0.07069867 | 0.42119392 |
| TDRD9 | 15.767528 | -1.278310282 | 0.59439432 | -2.1506098 | 0.03150701 | 0.28099688 |
| INF2 | 14.2926931 | -0.791226225 | 0.46992871 | -1.6837154 | 0.09223668 | 0.4764741 |
| SIVA1 | 33.3545936 | 0.372110417 | 0.33258433 | 1.11884531 | 0.26320614 | 0.71348562 |
| AKT1 | 45.6929703 | -0.314557603 | 0.30853124 | -1.0195324 | 0.30795027 | 0.7512664 |
| CEP170B | 21.466557 | 0.066728541 | 0.36741749 | 0.18161504 | 0.85588485 | 0.97166553 |
| NUDT14 | 23.6900131 | -0.573515768 | 0.36150043 | -1.5864871 | 0.11262886 | 0.51571293 |
| BRF1 | 30.4766587 | 0.076114696 | 0.3314311 | 0.22965465 | 0.81836013 | 0.9635539 |
| BTBD6 | 16.6700846 | 0.059597633 | 0.36184746 | 0.16470375 | 0.86917717 | 0.97434023 |
| PACS2 | 48.5159962 | 0.108642109 | 0.33303845 | 0.32621491 | 0.74426176 | 0.94546379 |
| MTA1 | 29.336592 | 0.329327867 | 0.33317707 | 0.98844696 | 0.3229338 | 0.76304922 |
| NIPA1 | 70.0821195 | 0.031660296 | 0.23901223 | 0.13246308 | 0.89461802 | 0.97850297 |
| NIPA2 | 52.3449674 | 0.061180122 | 0.30169814 | 0.20278588 | 0.8393024 | 0.96748214 |
| CYFIP1 | 84.3774026 | -0.37173846 | 0.23158515 | -1.6051913 | 0.10845171 | 0.50659799 |
| TUBGCP5 | 23.5202852 | 0.481010935 | 0.34678498 | 1.38705816 | 0.16542403 | 0.60594576 |
| PWRN1 | 94.0604768 | 1.120215655 | 1.38420516 | 0.80928441 | 0.41835157 | 0.8188001 |
| AC124312.1 | 208.088048 | 0.677044043 | 0.14943978 | 4.53054766 | 5.88E-06 | 0.00077116 |
| PWAR6 | 45.6026532 | 0.686207238 | 0.27848354 | 2.46408543 | 0.01373634 | 0.17578162 |
| UBE3A | 185.485897 | 0.177768247 | 0.15990578 | 1.1117062 | 0.26626449 | 0.71747544 |
| HERC2 | 194.854574 | 0.031547504 | 0.21659865 | 0.14564959 | 0.88419802 | 0.97662748 |
| TJP1 | 188.157209 | -0.579975736 | 0.18797161 | -3.0854433 | 0.00203249 | 0.05174581 |
| FAN1 | 42.4465674 | 0.028010184 | 0.32048398 | 0.08739964 | 0.93035386 | 0.98756002 |
| MTMR10 | 168.303097 | -0.450015159 | 0.22376268 | -2.011127 | 0.04431205 | 0.33449561 |
| KLF13 | 64.6977382 | 0.106882563 | 0.42950229 | 0.24885214 | 0.80347516 | 0.95938063 |
| OTUD7A | 49.4716498 | 0.081875255 | 0.26615015 | 0.30762806 | 0.75836537 | 0.94924107 |
| LINC02256 | 151.638756 | -0.045873765 | 0.30555038 | -0.1501349 | 0.88065821 | 0.97616097 |
| AC123768.3 | 20.1031552 | 0.443559481 | 0.4925957 | 0.90045342 | 0.367879 | 0.78877653 |
| GOLGA8N | 16.034122 | 0.399657309 | 0.35892178 | 1.11349416 | 0.26549626 | 0.71660319 |
| FMN1 | 141.15539 | -0.448880114 | 0.41619457 | -1.0785343 | 0.28079539 | 0.73272938 |
| AVEN | 58.3637924 | 0.627113314 | 0.32392705 | 1.93597082 | 0.05287127 | 0.36475915 |
| EMC7 | 47.2080414 | -0.081402467 | 0.27857624 | -0.2922089 | 0.77012688 | 0.9522529 |
| KATNBL1 | 143.102747 | -0.241679228 | 0.16811183 | -1.4376099 | 0.15054478 | 0.58759787 |
| EMC4 | 18.2860319 | -0.04758517 | 0.40645102 | -0.1170748 | 0.90680078 | 0.98202161 |
| SLC12A6 | 81.3545973 | -0.124740038 | 0.24312861 | -0.513062 | 0.60790798 | 0.89779346 |
| NOP10 | 56.3626004 | -0.080297266 | 0.48667959 | -0.16499 | 0.86895185 | 0.97434023 |
| LPCAT4 | 16.3284842 | 1.103799927 | 0.43108403 | 2.56052151 | 0.01045152 | 0.15015439 |
| GOLGA8A | 41.4084376 | 0.297136608 | 0.28914196 | 1.02764956 | 0.3041147 | 0.74820946 |
| GOLGA8B | 56.9146658 | 0.392827765 | 0.3607955 | 1.08878233 | 0.27624988 | 0.72575599 |
| AQR | 83.1032084 | 0.022861086 | 0.18774985 | 0.12176354 | 0.90308629 | 0.98016176 |
| ZNF770 | 43.9790676 | 0.168360245 | 0.25711118 | 0.65481496 | 0.5125869 | 0.86346419 |
| AC021231.1 | 72.5860239 | 0.065898043 | 0.36082604 | 0.18263106 | 0.85508751 | 0.97157653 |
| DPH6 | 263.957518 | 0.163552302 | 0.25113114 | 0.65126254 | 0.51487703 | 0.86463667 |
| DPH6-DT | 34.633775 | 0.665945123 | 0.47565704 | 1.40005312 | 0.16149741 | 0.60060549 |
| C15orf41 | 44.5577888 | 0.736817384 | 0.31452014 | 2.34267152 | 0.01914623 | 0.21721455 |
| SPRED1 | 73.8263829 | -0.276898927 | 0.37548498 | -0.7374434 | 0.46085274 | 0.83769303 |
| FAM98B | 36.7405538 | 0.127901284 | 0.23951804 | 0.53399436 | 0.59334544 | 0.89239828 |
| RASGRP1 | 24.2157084 | -1.04364079 | 0.47860538 | -2.1805872 | 0.02921396 | 0.2709134 |
| C15orf53 | 38.2977741 | -1.647435057 | 0.74592116 | -2.2085914 | 0.02720307 | 0.25997792 |
| AC013652.1 | 56.5400224 | 0.038172509 | 0.64898107 | 0.05881914 | 0.95309616 | 0.99217529 |
| THBS1 | 17.3586275 | -1.500582222 | 0.57952337 | -2.5893386 | 0.00961605 | 0.14462571 |
| EIF2AK4 | 54.9059777 | 0.046503316 | 0.25597919 | 0.18166835 | 0.85584301 | 0.97166553 |
| SRP14 | 269.374736 | -0.430219366 | 0.39531154 | -1.0883046 | 0.27646066 | 0.72575599 |
| SRP14-AS1 | 19.8388632 | 0.320592438 | 0.53247192 | 0.60208327 | 0.54711871 | 0.87287031 |
| IVD | 73.6394053 | 0.125412573 | 0.28226616 | 0.44430609 | 0.6568213 | 0.9158679 |
| BAHD1 | 28.8752305 | 0.260130453 | 0.33478328 | 0.77701149 | 0.43715199 | 0.82687287 |
| CCDC32 | 44.8840137 | -0.24207748 | 0.29826724 | -0.8116127 | 0.41701391 | 0.8188001 |
| RMDN3 | 29.5916947 | 0.465785926 | 0.38158677 | 1.22065535 | 0.22221654 | 0.67557645 |
| GCHFR | 18.8766537 | -0.222432754 | 0.44020937 | -0.5052885 | 0.61335619 | 0.90022016 |
| DNAJC17 | 37.9125929 | 0.096378915 | 0.26919586 | 0.35802524 | 0.72032442 | 0.93806337 |
| ZFYVE19 | 15.8441588 | 0.423794307 | 0.34518479 | 1.22773171 | 0.2195477 | 0.67305276 |
| SPINT1-AS1 | 44.343316 | 0.14152999 | 0.34367419 | 0.41181443 | 0.68047545 | 0.92375164 |
| SPINT1 | 65.020755 | -0.075123643 | 0.24517623 | -0.3064067 | 0.75929499 | 0.94924107 |
| VPS18 | 13.0154743 | -0.219025166 | 0.52347554 | -0.4184057 | 0.6756505 | 0.92193843 |
| INO80 | 110.19159 | 0.066653549 | 0.29352859 | 0.22707685 | 0.82036398 | 0.96389705 |
| CHP1 | 132.325914 | 0.269705893 | 0.29310035 | 0.92018278 | 0.35747725 | 0.78309154 |
| NDUFAF1 | 16.2825715 | 0.060182309 | 0.33057275 | 0.18205466 | 0.85553983 | 0.97166553 |
| RTF1 | 73.5298935 | 0.073410392 | 0.34489518 | 0.21284841 | 0.8314452 | 0.96701848 |
| RPAP1 | 18.3077629 | 0.025431128 | 0.34529024 | 0.07365145 | 0.94128773 | 0.98985996 |
| MGA | 152.667498 | 0.07301792 | 0.21738997 | 0.33588449 | 0.73695797 | 0.94369536 |
| MAPKBP1 | 62.7035742 | 0.215430634 | 0.25844825 | 0.83355424 | 0.40453222 | 0.81056858 |

| | | | | | | |
|---|---|---|---|---|---|---|
| EHD4 | 51.6000819 | -0.456213471 | 0.32994561 | -1.3826929 | 0.16675904 | 0.60787774 |
| PLA2G4F | 29.7464053 | 0.156952932 | 0.62154871 | 0.25251912 | 0.80063984 | 0.95873501 |
| VPS39 | 58.4775476 | -0.127348579 | 0.2418429 | -0.5265756 | 0.59848832 | 0.89450488 |
| TMEM87A | 196.201679 | -0.216150356 | 0.20517181 | -1.053509 | 0.29210776 | 0.73999615 |
| GANC | 36.2257746 | -0.126805432 | 0.3602069 | -0.352035 | 0.72481202 | 0.93940528 |
| ZNF106 | 92.2337992 | 0.152001955 | 0.18292281 | 0.83096226 | 0.40599495 | 0.8115705 |
| SNAP23 | 60.4816641 | -0.064913087 | 0.22894042 | -0.283537 | 0.7767652 | 0.95512504 |
| LRRC57 | 19.9683259 | -0.069652233 | 0.32204175 | -0.2162832 | 0.82876697 | 0.96657925 |
| HAUS2 | 17.6258132 | 0.213429551 | 0.38250222 | 0.55798252 | 0.57685632 | 0.88821715 |
| STARD9 | 27.0737757 | -0.396061772 | 0.55176117 | -0.7178138 | 0.47287211 | 0.84593347 |
| TTBK2 | 123.974806 | -0.195789414 | 0.20702726 | -0.945718 | 0.34429243 | 0.77393882 |
| UBR1 | 159.807763 | -0.126916572 | 0.20245913 | -0.626875 | 0.53074117 | 0.86786739 |
| AC068724.3 | 35.3957023 | 0.019648455 | 0.43052697 | 0.04563815 | 0.96359866 | 0.99329511 |
| TMEM62 | 49.4030476 | -0.484334866 | 0.28209301 | -1.7169332 | 0.08599137 | 0.46168448 |
| CCNDBP1 | 49.118813 | -0.411122449 | 0.2957037 | -1.3903189 | 0.16443206 | 0.60496776 |
| ADAL | 30.7326042 | 0.451426 | 0.335066 | 1.34727486 | 0.17789173 | 0.62542578 |
| ZSCAN29 | 13.7397354 | -0.024104315 | 0.36854458 | -0.0654041 | 0.94785228 | 0.99149867 |
| TUBGCP4 | 29.1069865 | -0.094439728 | 0.32287386 | -0.2924973 | 0.76990643 | 0.95219845 |
| TP53BP1 | 130.525372 | 0.265949858 | 0.1845286 | 1.44123921 | 0.14951712 | 0.58659144 |
| PPIP5K1 | 58.0991518 | 0.041470142 | 0.23537036 | 0.17619102 | 0.86014388 | 0.97287712 |
| CATSPER2 | 60.0873366 | 0.38027308 | 0.44704282 | 0.8506413 | 0.39496864 | 0.8052114 |
| PDIA3 | 170.743851 | 0.216069254 | 0.26303988 | 0.82143153 | 0.41140051 | 0.81482336 |
| SERF2 | 356.173959 | -0.185338237 | 0.31716564 | -0.5843579 | 0.55897957 | 0.88004847 |
| MFAP1 | 23.6710973 | 0.051744704 | 0.31323708 | 0.16519342 | 0.86879174 | 0.97434023 |
| CASC4 | 259.45835 | -0.214835361 | 0.22216196 | -0.9670214 | 0.33353336 | 0.76777176 |
| AC025043.1 | 13.6386342 | -1.104107924 | 0.69883132 | -1.5799348 | 0.1141218 | 0.51869892 |
| CTDSPL2 | 120.406924 | -0.033411915 | 0.21395892 | -0.1561604 | 0.87590657 | 0.97532382 |
| EIF3J-DT | 38.3289374 | -0.595207573 | 0.29715849 | -2.0029971 | 0.0451776 | 0.33761383 |
| EIF3J | 61.7483433 | 0.140816449 | 0.23309484 | 0.60411654 | 0.54576616 | 0.87247631 |
| SPG11 | 222.661712 | -0.216547999 | 0.18423334 | -1.1754007 | 0.23983443 | 0.6944034 |
| B2M | 1841.47693 | -1.286089925 | 0.37793335 | -3.4029543 | 0.00066661 | 0.02477424 |
| TRIM69 | 17.0766691 | -0.639574037 | 0.46628496 | -1.3716377 | 0.17017625 | 0.61252734 |
| SORD | 27.2018945 | 0.447698703 | 0.37584567 | 1.19117695 | 0.23358413 | 0.68826429 |
| DUOX1 | 21.1619356 | -0.907923978 | 0.44703362 | -2.0309971 | 0.04225529 | 0.32888549 |
| SPATA5L1 | 18.6218491 | 0.098392044 | 0.35439631 | 0.27763281 | 0.78129424 | 0.95540382 |
| SLC30A4 | 11.6334581 | 0.644089699 | 0.41978956 | 1.53431567 | 0.12495201 | NA |
| SQOR | 29.5543162 | -0.821784392 | 0.31951368 | -2.571985 | 0.01011173 | 0.1479393 |
| SEMA6D | 43.1274127 | -0.151210172 | 0.61528096 | -0.2457579 | 0.80586963 | 0.96005568 |
| MYEF2 | 121.771111 | -0.106204484 | 0.29268491 | -0.3628629 | 0.71670733 | 0.93689626 |
| SLC12A1 | 28.6379066 | 2.260417062 | 0.73019499 | 3.09563486 | 0.00196392 | 0.05060915 |
| DUT | 44.2123248 | 0.450282663 | 0.32472189 | 1.38667173 | 0.16554188 | 0.60594576 |
| FBN1 | 15.5856859 | 0.400392275 | 0.63321681 | 0.63231466 | 0.52718128 | 0.86750588 |
| CEP152 | 111.332439 | 0.650718286 | 0.44090596 | 1.47586639 | 0.13997976 | 0.57374456 |
| EID1 | 74.2748145 | -0.033394934 | 0.3662186 | -0.0911885 | 0.92734279 | 0.98690525 |
| SECISBP2L | 219.622583 | -0.15483565 | 0.23291708 | -0.6647673 | 0.50619937 | 0.86214551 |
| COPS2 | 43.4208776 | 0.132016211 | 0.238913 | 0.55257022 | 0.58055774 | 0.89012555 |
| GALK2 | 79.3443472 | 0.115379953 | 0.2346857 | 0.49163605 | 0.62297665 | 0.90430717 |
| FAM227B | 41.3653581 | 0.513192501 | 0.35691575 | 1.43785335 | 0.15047567 | 0.58759787 |
| FGF7 | 46.5206862 | 1.910575424 | 0.62277341 | 3.06785002 | 0.00215605 | 0.05376264 |
| DTWD1 | 47.66709 | 0.513430832 | 0.26346041 | 1.94879693 | 0.05131968 | 0.36077232 |
| GABPB1 | 62.2006573 | -0.028116939 | 0.27730727 | -0.1013927 | 0.91923872 | 0.9855881 |
| GABPB1-AS1 | 58.6678782 | 1.02034707 | 0.4326754 | 2.3582276 | 0.01836243 | 0.21053851 |
| USP8 | 119.496227 | -0.057849161 | 0.20714029 | -0.2792753 | 0.78003358 | 0.95537097 |
| TRPM7 | 196.564632 | -0.037470665 | 0.17506194 | -0.2140423 | 0.83051405 | 0.96694276 |
| SPPL2A | 182.326662 | -0.177499143 | 0.1886912 | -0.9406859 | 0.34686587 | 0.7765713 |
| AC021752.1 | 21.1245502 | 0.676478079 | 0.59576008 | 1.13548743 | 0.25617114 | 0.70955457 |
| AP4E1 | 59.6657831 | -0.040485265 | 0.24830469 | -0.1630467 | 0.87048165 | 0.97434023 |
| MIR4713HG | 23.7085059 | -0.529152437 | 0.81328065 | -0.6506394 | 0.51527929 | 0.86463667 |
| GLDN | 23.813368 | -2.257210872 | 0.75594441 | -2.9859482 | 0.002827 | 0.06539018 |
| DMXL2 | 76.0323104 | 0.193183049 | 0.30201742 | 0.63964208 | 0.52240532 | 0.86728831 |
| LYSMD2 | 13.5593434 | -0.180421783 | 0.46375207 | -0.3890479 | 0.69724069 | 0.92989421 |
| TMOD3 | 155.67113 | -0.160055666 | 0.19726794 | -0.8113618 | 0.41715795 | 0.8188001 |
| LEO1 | 18.272373 | 0.271986117 | 0.35674056 | 0.76241993 | 0.44580941 | 0.83035712 |
| MAPK6 | 88.8496817 | -0.474920262 | 0.23492343 | -2.021596 | 0.04321811 | 0.33179556 |
| GNB5 | 63.0126398 | -0.69420847 | 0.24574844 | -2.8248744 | 0.00472992 | 0.09401676 |
| MYO5C | 86.0630894 | -0.236891374 | 0.33095475 | -0.7157818 | 0.47412612 | 0.84593347 |
| MYO5A | 105.489213 | 0.152539771 | 0.20599074 | 0.74051761 | 0.45898598 | 0.83645518 |
| ARPP19 | 133.025563 | 0.000280043 | 0.22349964 | 0.00125299 | 0.99900026 | 0.99982181 |
| FAM214A | 154.457113 | -0.197151445 | 0.22681563 | -0.8692146 | 0.38472979 | 0.79869171 |
| WDR72 | 718.451508 | 0.354139424 | 0.33207619 | 1.06644027 | 0.28622466 | 0.73672385 |
| UNC13C | 25.9091533 | 2.315193963 | 1.37567713 | 1.68294865 | 0.09238503 | 0.4764741 |
| RSL24D1 | 32.9590691 | 0.017429425 | 0.32922724 | 0.05294041 | 0.95777939 | 0.99238848 |
| RAB27A | 111.467942 | -0.436989278 | 0.2614444 | -1.6714425 | 0.09463431 | 0.47996672 |
| PIGB | 51.7020516 | -0.04383044 | 0.30129569 | -0.1454732 | 0.8843373 | 0.97662748 |
| CCPG1 | 296.11298 | 0.425831906 | 0.33511957 | 1.27068649 | 0.2038402 | 0.65771647 |
| C15orf65 | 17.9163263 | 0.955021216 | 0.66570297 | 1.43460562 | 0.15139951 | 0.58907953 |
| DNAAF4 | 19.7740082 | 0.775357053 | 0.43367655 | 1.78786946 | 0.07379707 | 0.42813679 |
| RFX7 | 109.786801 | -0.097082484 | 0.22646069 | -0.4286946 | 0.66814546 | 0.92049996 |
| TEX9 | 39.4853315 | -0.345164611 | 0.32350424 | -1.0669555 | 0.28599195 | 0.73672385 |
| ZNF280D | 71.9508807 | 0.043065355 | 0.33085644 | 0.13016327 | 0.89643726 | 0.97885533 |
| AC090517.4 | 13.203916 | -0.344947881 | 0.42259849 | -0.8162544 | 0.41435464 | 0.81725092 |
| TCF12 | 667.446376 | -0.413578255 | 0.23822593 | -1.7360758 | 0.08255044 | 0.45473406 |
| CGNL1 | 885.857757 | -0.445177835 | 0.37393741 | -1.1905143 | 0.23384431 | 0.68840359 |
| POLR2M | 23.309976 | 0.353501965 | 0.3016561 | 1.17187075 | 0.24124895 | 0.69556484 |
| ALDH1A2 | 91.4341154 | -2.580955102 | 0.52985789 | -4.8710327 | 1.11E-06 | 0.00021977 |
| LIPC | 25.4059797 | 0.090264539 | 0.50931644 | 0.17722683 | 0.85933022 | 0.97287712 |
| ADAM10 | 207.355836 | -0.535953999 | 0.17354568 | -3.088259 | 0.00201333 | 0.05152847 |
| MINDY2 | 223.990757 | -0.164346803 | 0.16570969 | -0.9917755 | 0.32130707 | 0.76163052 |
| RNF111 | 189.622358 | 0.02567273 | 0.21511939 | 0.11934177 | 0.90500459 | 0.9810622 |
| SLTM | 232.216946 | 0.002492072 | 0.20711607 | 0.01203225 | 0.99039989 | 0.99923337 |
| MYO1E | 185.506826 | -0.460724066 | 0.26406694 | -1.7447245 | 0.08103283 | 0.45026424 |
| GTF2A2 | 47.3314526 | -0.366134552 | 0.37772631 | -0.9693117 | 0.33238967 | 0.76729949 |
| BNIP2 | 66.1171195 | 0.102763614 | 0.24050367 | 0.42728501 | 0.66917175 | 0.92049996 |
| ANXA2 | 64.8446153 | -0.795814387 | 0.6082468 | -1.3083741 | 0.19074644 | 0.6421543 |
| ICE2 | 81.4642101 | 0.248386097 | 0.29057576 | 0.85480666 | 0.39265819 | 0.80400536 |
| RORA-AS1 | 59.2194672 | -0.382983665 | 0.50132803 | -0.7639383 | 0.44490403 | 0.83035712 |
| RORA | 1190.23624 | -0.268127041 | 0.4995122 | -0.5367778 | 0.59142113 | 0.89151328 |
| AC012404.1 | 43.3849886 | -1.227736461 | 0.66277897 | -1.8524071 | 0.06396739 | 0.40111702 |
| AC009554.1 | 45.1447924 | -0.693814954 | 0.49090386 | -1.4133418 | 0.15755525 | 0.59838915 |
| VPS13C | 315.731452 | -0.480622009 | 0.18100655 | -2.6552742 | 0.0079244 | 0.12897089 |
| TLN2 | 64.6642396 | 0.340414888 | 0.4719053 | 0.72136272 | 0.47068638 | 0.84424147 |
| TPM1 | 200.864428 | -0.682069054 | 0.32188829 | -2.118962 | 0.03409368 | 0.29400113 |
| LACTB | 52.4929707 | -0.210132381 | 0.37382904 | -0.5621082 | 0.57404228 | 0.88637538 |
| RPS27L | 66.9710273 | -0.63026267 | 0.34412212 | -1.8315087 | 0.06702465 | 0.4116442 |
| RAB8B | 63.6412517 | -0.5422161 | 0.27773276 | -1.9522944 | 0.05090327 | 0.35962249 |
| APH1B | 60.1339028 | -1.209382219 | 0.25412478 | -4.7590094 | 1.95E-06 | 0.00033107 |
| CA12 | 1926.50663 | -0.033040039 | 0.32035653 | -0.1031352 | 0.91785566 | 0.98519419 |
| LINC02568 | 45.1268972 | -1.884281591 | 0.43502536 | -4.3314293 | 1.48E-05 | 0.00156196 |
| USP3 | 224.848526 | -0.140084518 | 0.20778461 | -0.6741814 | 0.50019599 | 0.85942996 |
| USP3-AS1 | 44.5188783 | -0.517994923 | 0.45572025 | -1.1366511 | 0.25568417 | 0.70861042 |
| HERC1 | 231.201901 | -0.049985457 | 0.16772159 | -0.2980264 | 0.76568303 | 0.95205353 |
| DAPK2 | 22.0647318 | -0.762919657 | 0.89440199 | -0.8529941 | 0.39366255 | 0.80438874 |
| FAM96A | 51.1380198 | -0.211519706 | 0.33058427 | -0.639836 | 0.52227924 | 0.86728831 |

| | | | | | | |
|---|---|---|---|---|---|---|
| SNX1 | 101.761357 | -0.02453799 | 0.1750823 | -0.1401512 | 0.88854055 | 0.97765744 |
| PPIB | 126.904071 | -0.620392894 | 0.31485008 | -1.970439 | 0.04878807 | 0.35290405 |
| CSNK1G1 | 27.489487 | 0.075094759 | 0.32959563 | 0.22783906 | 0.81977136 | 0.96389705 |
| TRIP4 | 54.2170342 | 0.274861091 | 0.25724055 | 1.06849828 | 0.28529581 | 0.73659018 |
| ZNF609 | 271.575229 | -0.151070141 | 0.23950842 | -0.6307509 | 0.52820343 | 0.86750588 |
| OAZ2 | 61.1367442 | -0.268263734 | 0.28915665 | -0.9277453 | 0.35353968 | 0.78028099 |
| RBPMS2 | 20.8868443 | 0.505359871 | 0.37208512 | 1.35818351 | 0.17440546 | 0.62052871 |
| SPG21 | 52.4753023 | -0.23650268 | 0.32168904 | -0.7351904 | 0.46222357 | 0.83894774 |
| MTFMT | 33.7882297 | 0.202145648 | 0.25628382 | 0.78875698 | 0.43025406 | 0.82469033 |
| UBAP1L | 13.2543215 | 0.513684323 | 0.53203916 | 0.96550096 | 0.33429395 | 0.76777176 |
| PDCD7 | 30.3778328 | -0.219166155 | 0.27949811 | -0.7841418 | 0.4329569 | 0.8248269 |
| CLPX | 66.0179759 | -0.292219241 | 0.23589404 | -1.2387733 | 0.21542947 | 0.66933563 |
| PARP16 | 16.6180444 | 0.143379496 | 0.46756771 | 0.3066497 | 0.75911002 | 0.94924107 |
| DPP8 | 101.902063 | 0.107806544 | 0.19226364 | 0.56072246 | 0.57498675 | 0.88712765 |
| HACD3 | 124.99066 | 0.304510803 | 0.26833443 | 1.13481822 | 0.25645149 | 0.70972499 |
| INTS14 | 33.4259952 | -0.367548897 | 0.25628739 | -1.4341279 | 0.15153577 | 0.58913705 |
| SLC24A1 | 14.0704425 | 0.355255782 | 0.40160916 | 0.88458087 | 0.37638273 | 0.79422885 |
| DENND4A | 362.399381 | -0.536000663 | 0.27289902 | -1.9640989 | 0.04951862 | 0.35590543 |
| RAB11A | 258.345643 | -0.602484435 | 0.39023629 | -1.5438965 | 0.12261342 | 0.53899578 |
| DIS3L | 27.1879265 | 0.232625739 | 0.39074519 | 0.59533871 | 0.55161708 | 0.87546024 |
| TIPIN | 23.4927232 | 0.576570896 | 0.40468264 | 1.4247483 | 0.15422997 | 0.59439427 |
| MAP2K1 | 142.400345 | 0.146998327 | 0.23126946 | 0.63561495 | 0.52502743 | 0.86747074 |
| SNAPC5 | 18.3766666 | -1.007576211 | 0.4547899 | -2.2154762 | 0.02672739 | 0.25719813 |
| RPL4 | 193.125417 | -0.089431884 | 0.40525817 | -0.2206788 | 0.82534254 | 0.96539788 |
| ZWILCH | 14.1710837 | 0.249353104 | 0.41265179 | 0.60427001 | 0.54566414 | 0.87247631 |
| SMAD6 | 19.3984932 | 1.067319925 | 0.42543751 | 2.50875838 | 0.01211563 | 0.16352285 |
| SMAD3 | 386.660979 | -1.728593053 | 0.34980122 | -4.9416439 | 7.75E-07 | 0.00015988 |
| AAGAB | 113.695175 | -0.282701383 | 0.21060734 | -1.342315 | 0.17949389 | 0.62696821 |
| IQCH | 72.4329935 | 0.129487003 | 0.42468239 | 0.30490316 | 0.76043991 | 0.94962861 |
| IQCH-AS1 | 56.614528 | 0.620272504 | 0.33762393 | 1.83716985 | 0.06618483 | 0.40888394 |
| C15orf61 | 18.0139305 | -0.292441262 | 0.40387538 | -0.7240879 | 0.4690118 | 0.84326495 |
| MAP2K5 | 194.662755 | -0.405875168 | 0.39574502 | -1.0255977 | 0.30508127 | 0.74842902 |
| PIAS1 | 295.846678 | 0.276466907 | 0.16228358 | 1.7036037 | 0.08845515 | 0.46851772 |
| FEM1B | 45.6728279 | 0.084628813 | 0.25784492 | 0.32821594 | 0.74274839 | 0.94546379 |
| CORO2B | 26.8189018 | 0.180895525 | 0.3830245 | 0.47228186 | 0.63672562 | 0.90920631 |
| ANP32A | 133.026482 | 0.675568152 | 0.26973865 | 2.50452858 | 0.01226147 | 0.16518924 |
| SPESP1 | 29.9431709 | 1.052857033 | 0.42670736 | 2.46739832 | 0.01360989 | 0.17531992 |
| GLCE | 113.919986 | 0.293354344 | 0.34485241 | 0.85066636 | 0.39495471 | 0.8052114 |
| AC026992.1 | 15.1288441 | 2.010771358 | 0.59225808 | 3.39509316 | 0.00068605 | 0.02520723 |
| PAQR5 | 205.667865 | 0.071056093 | 0.27651061 | 0.2569742 | 0.79719869 | 0.95848876 |
| KIF23 | 22.5644413 | 1.061192894 | 0.71595177 | 1.48221282 | 0.13828368 | 0.57086516 |
| RPLP1 | 362.420418 | 0.529270617 | 0.39254265 | 1.34831365 | 0.17755752 | 0.6251709 |
| TLE3 | 61.5758628 | -0.740613269 | 0.29686553 | -2.4947769 | 0.01260363 | 0.16711862 |
| UACA | 128.147296 | 0.330055182 | 0.31199541 | 1.05788472 | 0.290108 | 0.73979353 |
| LRRC49 | 34.3349958 | -0.553888586 | 0.44004569 | -1.258707 | 0.20813619 | 0.65973983 |
| THSD4 | 211.176288 | 1.359743219 | 0.54820763 | 2.48034348 | 0.01312559 | 0.17172 |
| THSD4-AS1 | 66.961276 | 0.114037648 | 0.26654799 | 0.42783159 | 0.66877374 | 0.92049996 |
| MYO9A | 536.851474 | -0.372654266 | 0.16972703 | -2.1956094 | 0.02811991 | 0.26404526 |
| AC022872.1 | 16.8202253 | -0.567686491 | 0.58221345 | -0.9750487 | 0.32953608 | 0.76685572 |
| SENP8 | 14.9624094 | 0.089222099 | 0.38638376 | 0.23091576 | 0.81738025 | 0.9635539 |
| PKM | 463.72558 | -0.50380952 | 0.19785495 | -2.5463579 | 0.01088535 | 0.15419892 |
| PARP6 | 37.7694336 | 0.226248056 | 0.29957391 | 0.75523285 | 0.45010928 | 0.83118301 |
| AC009690.2 | 28.1763779 | -0.745443556 | 0.59578359 | -1.2511985 | 0.21086205 | 0.6625727 |
| AC009690.1 | 99.2058552 | -0.12512546 | 0.39250125 | -0.31879 | 0.74988579 | 0.94684415 |
| ARIH1 | 370.297567 | 0.175598521 | 0.17427656 | 1.00758545 | 0.31365352 | 0.75632348 |
| BBS4 | 49.9582292 | -0.363227304 | 0.38391927 | -0.9461034 | 0.34409587 | 0.77387665 |
| ADPGK | 46.7618283 | -0.039222265 | 0.26974329 | -0.1454059 | 0.88439041 | 0.97662748 |
| NEO1 | 130.481245 | -0.402267671 | 0.33045843 | -1.2173019 | 0.22348938 | 0.67606266 |
| NPTN | 225.129825 | 0.146148767 | 0.18542311 | 0.78819068 | 0.43058518 | 0.82469033 |
| CD276 | 16.592804 | 0.149945655 | 0.41552471 | 0.36085858 | 0.71820517 | 0.93733389 |
| C15orf59 | 23.2893895 | 1.865125311 | 0.49675031 | 3.75465352 | 0.00017358 | 0.01008227 |
| C15orf59-AS | 15.896818 | 1.037396796 | 0.69271862 | 1.49757313 | 0.13424419 | 0.56229288 |
| LOXL1 | 16.8134453 | 0.593167772 | 0.38492928 | 1.54097856 | 0.123322 | 0.54029966 |
| PML | 35.5336102 | -0.831434853 | 0.34137815 | -2.4355245 | 0.01487022 | 0.18484189 |
| CLK3 | 27.579024 | 0.35169508 | 0.28649829 | 1.22756434 | 0.21961056 | 0.67305276 |
| EDC3 | 40.1130028 | 0.234556049 | 0.25930071 | 0.90457155 | 0.36569241 | 0.78793182 |
| CSK | 14.1940044 | 0.595695099 | 0.42189776 | 1.41194184 | 0.15796709 | 0.59854717 |
| ULK3 | 16.5068386 | 1.055421299 | 0.40877474 | 2.58191417 | 0.0098254 | 0.14662523 |
| SCAMP2 | 56.3817055 | -0.081526588 | 0.21627902 | -0.376951 | 0.70621003 | 0.93407096 |
| FAM219B | 26.5904038 | 0.233354077 | 0.27600356 | 0.84547488 | 0.39784575 | 0.80714564 |
| COX5A | 130.827301 | 0.119103989 | 0.36262505 | 0.32844942 | 0.74257188 | 0.94546379 |
| SCAMP5 | 28.154374 | 0.71015208 | 0.6885704 | 1.03134274 | 0.30238012 | 0.74690278 |
| PPCDC | 31.6782256 | 0.662009483 | 0.41683349 | 1.58818687 | 0.11224409 | 0.5156312 |
| COMMD4 | 28.401538 | -0.159616974 | 0.32176013 | -0.4960744 | 0.6198419 | 0.90380166 |
| NEIL1 | 12.7194115 | 0.535722789 | 0.36984451 | 1.44850817 | 0.14747498 | 0.58421969 |
| MAN2C1 | 43.768949 | 0.065758054 | 0.28547385 | 0.23034703 | 0.81782212 | 0.9635539 |
| SIN3A | 81.4439718 | -0.087995573 | 0.22353173 | -0.3936603 | 0.69383184 | 0.92841398 |
| PTPN9 | 131.339028 | -0.095687102 | 0.28706337 | -0.3333309 | 0.73888449 | 0.94376279 |
| SNUPN | 17.1750679 | -0.106737323 | 0.33734274 | -0.3164062 | 0.75169424 | 0.94719044 |
| IMP3 | 23.3354187 | 0.020032758 | 0.45906014 | 0.04363863 | 0.96519245 | 0.99334616 |
| UBE2Q2 | 45.678353 | -0.848230159 | 0.37077735 | -2.2877076 | 0.02215456 | 0.23358607 |
| FBXO22 | 73.9151545 | 0.107855676 | 0.28417965 | 0.37953342 | 0.70429179 | 0.93328284 |
| NRG4 | 19.1345042 | -0.218970339 | 0.65936074 | -0.3320949 | 0.7398176 | 0.94376279 |
| ETFA | 102.897627 | 0.200924164 | 0.19548659 | 1.02781557 | 0.30403659 | 0.74820946 |
| SCAPER | 357.566701 | -0.344487418 | 0.1668172 | -2.0650594 | 0.03891738 | 0.31669347 |
| RCN2 | 76.250579 | 0.320696572 | 0.25006257 | 1.2824653 | 0.19967947 | 0.65269687 |
| TSPAN3 | 108.963289 | -0.2794946 | 0.21432665 | -1.304059 | 0.19221347 | 0.6421543 |
| PEAK1 | 86.5200889 | 0.39109312 | 0.31863997 | 1.22738247 | 0.21967887 | 0.67305276 |
| HMG20A | 91.9020704 | 0.142931224 | 0.22580497 | 0.6329853 | 0.52674325 | 0.86750588 |
| LINGO1 | 663.789751 | 0.70809783 | 0.5233346 | 1.35304991 | 0.17603969 | 0.6225246 |
| TBC1D2B | 26.3262925 | -0.007025987 | 0.40456385 | -0.0173668 | 0.98614398 | 0.99804172 |
| CIB2 | 14.5979612 | -0.631615013 | 0.46194189 | -1.3673041 | 0.17153002 | 0.61532589 |
| IDH3A | 124.541918 | -0.207514436 | 0.18417612 | -1.1267174 | 0.25986199 | 0.71091822 |
| ACSBG1 | 17.0158503 | -0.033085247 | 0.78729968 | -0.0420237 | 0.9664798 | 0.99351999 |
| DNAJA4 | 51.3150122 | -0.383794147 | 0.40330347 | -0.9516262 | 0.34128659 | 0.77193691 |
| WDR61 | 48.0843056 | 0.057629559 | 0.26421967 | 0.2181123 | 0.82734161 | 0.96561348 |
| IREB2 | 114.290834 | -0.179881054 | 0.24630944 | -0.7303052 | 0.46520367 | 0.84143009 |
| PSMA4 | 59.2916308 | -0.155295804 | 0.27983641 | -0.5549521 | 0.57892742 | 0.88938803 |
| MORF4L1 | 148.671022 | 0.019076948 | 0.1932807 | 0.09870074 | 0.92137588 | 0.98614962 |
| CTSH | 68.2946793 | 0.10915006 | 0.49901452 | 0.21873123 | 0.82685942 | 0.96542185 |
| TMED3 | 29.0029288 | -0.543639877 | 0.3329342 | -1.6328749 | 0.10249532 | 0.49291254 |
| ZFAND6 | 287.602896 | -0.309207259 | 0.19361415 | -1.5970282 | 0.11025942 | 0.5107528 |
| FAH | 18.6655182 | 0.286672673 | 0.45443372 | 0.63083495 | 0.52814845 | 0.86750588 |
| CTXND1 | 15.092294 | 1.177671377 | 0.72804329 | 1.61758428 | 0.10575222 | 0.50136683 |
| AC016705.2 | 258.570565 | -1.107600848 | 0.27227934 | -4.0678843 | 4.74E-05 | 0.00389989 |
| ARNT2 | 309.925997 | -0.468913707 | 0.41467564 | -1.1307964 | 0.25814081 | 0.71022118 |
| ABHD17C | 52.2851911 | -0.299371489 | 0.26124191 | -1.1459551 | 0.25181375 | 0.70574787 |
| AC027808.1 | 20.9000664 | 0.291299667 | 0.35433635 | 0.82209931 | 0.41102038 | 0.81448369 |
| MESD | 80.5522617 | -0.120000142 | 0.23727654 | -0.5057396 | 0.61303946 | 0.90011356 |
| AC104041.1 | 29.6515804 | -0.583626999 | 0.82389064 | -0.7083792 | 0.4787098 | 0.84833462 |
| EFL1 | 49.4625281 | -0.264409363 | 0.27871221 | -0.9486824 | 0.34278218 | 0.7729798 |
| RPS17 | 20.6542967 | 0.499275996 | 0.58487274 | 0.85364894 | 0.39329953 | 0.80400536 |
| WHAMM | 46.7472641 | -0.444111382 | 0.28719324 | -1.5463852 | 0.12201157 | 0.53747904 |

| | | | | | | |
|---|---|---|---|---|---|---|
| HOMER2 | 16.4105499 | 0.557511874 | 0.462204 | 1.20620305 | 0.22773921 | 0.68265461 |
| C15orf40 | 22.4615953 | 0.066887202 | 0.36188199 | 0.18483153 | 0.85336117 | 0.97131147 |
| BTBD1 | 97.4829568 | -0.315702518 | 0.21902838 | -1.4413772 | 0.14947815 | 0.58659144 |
| HDGFL3 | 79.5930504 | -0.944420911 | 0.29547051 | -3.1963288 | 0.00139188 | 0.04030231 |
| UBE2Q2L | 21.1760114 | -0.212644987 | 0.45176856 | -0.4706945 | 0.6378589 | 0.91025608 |
| GOLGA6L4 | 12.3746431 | 0.036909183 | 0.487951 | 0.07564117 | 0.93970458 | NA |
| WDR73 | 46.6247982 | -0.057252914 | 0.23346303 | -0.2452333 | 0.80627578 | 0.96005568 |
| SEC11A | 132.993822 | 0.350661766 | 0.21240014 | 1.65094885 | 0.09874902 | 0.48476221 |
| ZNF592 | 59.5951129 | -0.002040667 | 0.2776562 | -0.0073496 | 0.9941359 | 0.99923337 |
| ALPK3 | 114.833259 | -0.943562056 | 0.56253572 | -1.6773371 | 0.09347657 | 0.47792014 |
| PDE8A | 451.173041 | -0.433075654 | 0.27822782 | -1.5565505 | 0.11957724 | 0.53108939 |
| AKAP13 | 412.946967 | -0.307199468 | 0.22768721 | -1.3492171 | 0.17726725 | 0.6251709 |
| AC087286.2 | 12.7155354 | -0.650722683 | 0.50213832 | -1.2959033 | 0.19500881 | 0.6459894 |
| KLHL25 | 20.8436005 | 0.085039703 | 0.38738553 | 0.21952215 | 0.82624333 | 0.96539788 |
| MRPL46 | 16.8316027 | -0.358889107 | 0.38464769 | -0.9330333 | 0.35080277 | 0.77894201 |
| MRPS11 | 22.5732843 | 0.175803948 | 0.33376405 | 0.52673123 | 0.59838025 | 0.89448118 |
| DET1 | 28.5478354 | -0.119664413 | 0.39533083 | -0.3026944 | 0.7621228 | 0.95024054 |
| AEN | 12.2536286 | -0.425135511 | 0.46665045 | -0.9110363 | 0.36227623 | NA |
| ISG20 | 23.6978756 | -2.205309922 | 1.0303418 | -2.1403673 | 0.03232509 | 0.28504855 |
| ABHD2 | 240.922141 | 0.58427897 | 0.34777963 | 1.68002642 | 0.09295217 | 0.47759124 |
| POLG | 28.2445267 | 0.306562136 | 0.36563346 | 0.83844114 | 0.40178299 | 0.80890307 |
| MIR9-3HG | 10.6078501 | 0.106834806 | 0.51988971 | 0.20549513 | 0.8371853 | NA |
| RHCG | 633.886486 | 0.866656986 | 0.60496364 | 1.43257698 | 0.15197876 | 0.59013416 |
| PEX11A | 12.8557823 | -0.428735935 | 0.55269064 | -0.775725 | 0.43791139 | 0.8270818 |
| ANPEP | 13.0175001 | -0.777012299 | 0.61749507 | -1.2583296 | 0.2082726 | 0.65973983 |
| ZNF710 | 62.9699327 | -0.592350989 | 0.30689493 | -1.9301427 | 0.05358916 | 0.36684184 |
| ZNF710-AS1 | 65.5854826 | 0.600686144 | 0.54472462 | 1.1027336 | 0.27014287 | 0.72068562 |
| IDH2 | 361.17026 | -0.262625014 | 0.28364682 | -0.9258874 | 0.35450451 | 0.78097851 |
| IDH2-DT | 20.8699496 | 0.136643302 | 0.40284071 | 0.33919933 | 0.73445957 | 0.94277981 |
| SEMA4B | 163.430482 | -1.645628811 | 0.37000654 | -4.4475669 | 8.68E-06 | 0.00102735 |
| CIB1 | 168.844472 | -0.673746708 | 0.34342761 | -1.9618304 | 0.04978224 | 0.35621674 |
| IQGAP1 | 215.948598 | -0.323620745 | 0.23641027 | -1.3688946 | 0.17103223 | 0.61444911 |
| CRTC3 | 176.774503 | -0.209353752 | 0.25770633 | -0.8123733 | 0.41657745 | 0.818473 |
| CRTC3-AS1 | 14.4331872 | 0.132033063 | 0.48226193 | 0.27377874 | 0.78425467 | 0.95544332 |
| FURIN | 40.8838065 | -0.799854487 | 0.40440212 | -1.9778692 | 0.04794346 | 0.34992596 |
| MAN2A2 | 16.5783609 | 0.102211591 | 0.35770908 | 0.28573944 | 0.77507769 | 0.95493992 |
| HDDC3 | 13.0365197 | 0.174555173 | 0.41698473 | 0.41861287 | 0.67549908 | 0.92193843 |
| UNC45A | 33.0368692 | -0.142834149 | 0.28254665 | -0.5055241 | 0.61319076 | 0.90011356 |
| RCCD1 | 13.7447053 | -0.08216077 | 0.51662037 | -0.1590351 | 0.87364122 | 0.97434023 |
| SLCO3A1 | 99.0373731 | -0.129648457 | 0.60881002 | -0.2129539 | 0.83136293 | 0.96701848 |
| FAM174B | 70.9408173 | -0.390527106 | 0.5347412 | -0.7303105 | 0.46520042 | 0.84143009 |
| AC013394.1 | 121.630945 | 0.077152534 | 0.18061375 | 0.42716866 | 0.66925649 | 0.92049996 |
| CHD2 | 219.300211 | -0.043480188 | 0.25838372 | -0.1682776 | 0.86636491 | 0.97434023 |
| MCTP2 | 30.5698076 | -0.906994679 | 0.46630008 | -1.945088 | 0.05176439 | 0.36123352 |
| AC012409.2 | 27.7617364 | 0.135791319 | 0.55632988 | 0.24408418 | 0.80716563 | 0.96016552 |
| NR2F2-AS1 | 146.775457 | 0.726659269 | 0.41305834 | 1.75921705 | 0.07854065 | 0.4421357 |
| NR2F2 | 64.7731268 | 0.422840226 | 0.30825601 | 1.37171768 | 0.17015134 | 0.61252734 |
| LINC00923 | 31.3623349 | -0.051118827 | 0.8763978 | -0.0583283 | 0.9534871 | 0.99217529 |
| ARRDC4 | 82.6032396 | 0.145269364 | 0.36765441 | 0.39512477 | 0.69275081 | 0.928133 |
| FAM169B | 17.9954433 | -4.051870935 | 0.93698264 | -4.3243821 | 1.53E-05 | 0.00159539 |
| IGF1R | 562.746645 | -0.268430812 | 0.21784571 | -1.2322061 | 0.2178721 | 0.67202699 |
| AC036108.1 | 16.1133809 | -0.310347206 | 0.45746386 | -0.6784081 | 0.49751295 | 0.85818031 |
| SYNM | 13.0845504 | -0.963457082 | 0.856158 | -1.1253263 | 0.2604508 | 0.7111324 |
| TTC23 | 63.6358285 | -0.107977802 | 0.26549735 | -0.4067001 | 0.68422825 | 0.92548169 |
| LRRC28 | 85.1951665 | 0.416954369 | 0.26400407 | 1.57934825 | 0.1142562 | 0.51869892 |
| MEF2A | 323.006563 | 0.304440542 | 0.21259158 | 1.43204419 | 0.15213118 | 0.59013416 |
| AC090825.1 | 33.4401094 | -0.181229559 | 0.35629229 | -0.5086542 | 0.61099466 | 0.90002868 |
| LINS1 | 23.3209772 | 0.400893346 | 0.32682797 | 1.22661883 | 0.21996589 | 0.67350139 |
| ASB7 | 32.7795729 | -0.222361573 | 0.26840491 | -0.8284557 | 0.40741248 | 0.81294693 |
| AC015712.1 | 10.9033513 | -0.132816979 | 0.48176651 | -0.2756874 | 0.78278814 | NA |
| ALDH1A3 | 26.0199113 | -1.286125415 | 0.55175049 | -2.330991 | 0.01975384 | 0.22075139 |
| CHSY1 | 86.3803853 | -0.729746861 | 0.31142921 | -2.343219 | 0.01911816 | 0.21715001 |
| SELENOS | 64.1735533 | -0.367982948 | 0.38174925 | -0.9639389 | 0.33507656 | 0.76777176 |
| SNRPA1 | 47.1183103 | -0.476689837 | 0.24359834 | -1.9568682 | 0.05036297 | 0.35799473 |
| PCSK6 | 45.552674 | -0.583772573 | 0.60622054 | -0.9629706 | 0.33556227 | 0.76777176 |
| TM2D3 | 62.391094 | -0.225736961 | 0.27629309 | -0.8170199 | 0.41391703 | 0.81688611 |
| TARSL2 | 76.7296877 | -0.272673965 | 0.19285745 | -1.4138628 | 0.1574022 | 0.59804205 |
| SNRNP25 | 16.404729 | 0.799321043 | 0.35684707 | 2.23995405 | 0.02509391 | 0.25042272 |
| MPG | 36.3289777 | -0.858128376 | 0.28858732 | -2.9735485 | 0.00294378 | 0.06702971 |
| NPRL3 | 58.0112938 | 0.155172956 | 0.23667291 | 0.65564309 | 0.51205379 | 0.86346419 |
| LUC7L | 121.10685 | 0.288549209 | 0.17104213 | 1.68700662 | 0.09160208 | 0.4756639 |
| FAM234A | 17.5191385 | 1.001140943 | 0.46566614 | 2.14991141 | 0.03156222 | 0.28099688 |
| AXIN1 | 28.3933993 | -0.466541475 | 0.33265676 | -1.402471 | 0.16077463 | 0.60050595 |
| MRPL28 | 28.100281 | 0.111536432 | 0.33323084 | 0.33471221 | 0.73784219 | 0.94376279 |
| TMEM8A | 90.2467907 | -0.434830033 | 0.30789195 | -1.4122813 | 0.15786716 | 0.59849163 |
| NME4 | 14.4138015 | 0.411634323 | 0.53529109 | 0.76899155 | 0.44189833 | 0.8281114 |
| RAB11FIP3 | 121.764921 | 0.016976252 | 0.31790488 | 0.05340041 | 0.95741288 | 0.99236316 |
| CAPN15 | 32.2465721 | 0.439096877 | 0.25790937 | 1.70252391 | 0.0886572 | 0.4687332 |
| PIGQ | 44.8501545 | 1.077148314 | 0.38411409 | 2.80424056 | 0.00504353 | 0.09823735 |
| RAB40C | 79.8758905 | -0.018951028 | 0.28260513 | -0.0670583 | 0.94653527 | 0.99134285 |
| METTL26 | 11.8264788 | 0.53325363 | 0.50021253 | 1.06605413 | 0.28639917 | NA |
| MCRIP2 | 97.7831157 | 0.320485069 | 0.33694098 | 0.95116084 | 0.34152274 | 0.77193691 |
| RHOT2 | 40.1697338 | 0.194676777 | 0.2436632 | 0.79895846 | 0.4243145 | 0.82231588 |
| STUB1 | 61.6774709 | 0.238361115 | 0.38217221 | 0.6237008 | 0.5328241 | 0.86909171 |
| JMJD8 | 18.2007931 | 0.598807294 | 0.37064109 | 1.61559878 | 0.10618109 | 0.50162992 |
| METRN | 35.0787198 | 0.260413403 | 0.36211696 | 0.71914169 | 0.47205362 | 0.84570599 |
| FAM173A | 46.0648275 | 0.151197527 | 0.23990003 | 0.63025223 | 0.52852957 | 0.86750588 |
| HAGHL | 20.2315621 | -0.122839492 | 0.53527313 | -0.2294894 | 0.81848858 | 0.9635539 |
| NARFL | 16.3567081 | -0.004419129 | 0.34019631 | -0.0129899 | 0.98963582 | 0.99923337 |
| LMF1 | 62.2868475 | 0.27743729 | 0.32128551 | 0.86352256 | 0.38785022 | 0.80079761 |
| CACNA1H | 13.292812 | -0.066569026 | 0.74112384 | -0.0898217 | 0.92842888 | 0.9872367 |
| UBE2I | 90.9585701 | -0.528149068 | 0.28378552 | -1.8610853 | 0.06273213 | 0.39823408 |
| TSR3 | 25.1575027 | -0.145993627 | 0.37014277 | -0.3944252 | 0.69326716 | 0.92826395 |
| GNPTG | 25.4470196 | 0.104457248 | 0.34390635 | 0.30373748 | 0.7613279 | 0.9499444 |
| UNKL | 22.4784151 | -0.046587709 | 0.43395119 | -0.107357 | 0.91450575 | 0.98407488 |
| CLCN7 | 44.5923929 | -0.599253741 | 0.44807012 | -1.3374106 | 0.18108863 | 0.62891505 |
| IFT140 | 21.0896183 | 0.413305149 | 0.31963013 | 1.29307317 | 0.19598575 | 0.64659037 |
| JPT2 | 50.0988748 | -0.762892247 | 0.29768726 | -2.5627306 | 0.01038526 | 0.14952531 |
| MAPK8IP3 | 77.6870909 | -0.160541729 | 0.22382844 | -0.7172535 | 0.4732177 | 0.84593347 |
| NME3 | 56.6884006 | 0.133527408 | 0.35658751 | 0.37445901 | 0.70806285 | 0.93432317 |
| MRPS34 | 35.9724762 | 0.207383263 | 0.4601703 | 0.45066634 | 0.65223005 | 0.9158679 |
| SPSB3 | 36.0436586 | 0.415315916 | 0.3555133 | 1.16821485 | 0.24272011 | 0.69621873 |
| NUBP2 | 15.6901608 | 0.588879102 | 0.34442588 | 1.70974115 | 0.08731375 | 0.46528247 |
| HAGH | 74.2565036 | -0.060127989 | 0.23432264 | -0.2566034 | 0.79748494 | 0.95848876 |
| FAHD1 | 20.9886732 | -0.391594824 | 0.37019523 | -1.0578062 | 0.29014379 | 0.73979353 |
| NDUFB10 | 61.9585326 | 0.318154895 | 0.41765363 | 0.76176735 | 0.44619887 | 0.83035712 |
| RPS2 | 243.371699 | 0.341832369 | 0.37525617 | 0.91093072 | 0.36233188 | 0.78714716 |
| GFER | 12.7389232 | 0.341124471 | 0.38553484 | 0.88480843 | 0.37625996 | 0.79422885 |
| ZNF598 | 12.5346561 | 0.04349135 | 0.42364882 | 0.10265897 | 0.91823364 | NA |
| TSC2 | 41.4742228 | 0.381629279 | 0.27282228 | 1.39882008 | 0.16186694 | 0.60088379 |
| PKD1 | 39.4383305 | 0.405134104 | 0.27902536 | 1.45196159 | 0.14651229 | 0.58349206 |
| SNHG19 | 14.5140688 | 0.441407326 | 0.35931332 | 1.22847471 | 0.21926882 | 0.67305276 |
| TRAF7 | 17.6099101 | -0.435578492 | 0.36542212 | -1.1919872 | 0.23326626 | 0.6877455 |

| | | | | | | |
|---|---|---|---|---|---|---|
| MLST8 | 11.3578041 | 0.006933649 | 0.60505886 | 0.01145946 | 0.99085687 | NA |
| PGP | 61.8906752 | -0.163122007 | 0.29582145 | -0.5514205 | 0.58134546 | 0.8901472 |
| E4F1 | 19.2006943 | 0.375217256 | 0.36568196 | 1.02607538 | 0.30485605 | 0.74832335 |
| ECI1 | 161.888873 | 0.146548304 | 0.2667848 | 0.5493128 | 0.58279081 | 0.89036744 |
| RNPS1 | 60.8436358 | 0.097963769 | 0.22383464 | 0.43766135 | 0.66163179 | 0.91808148 |
| ABCA3 | 18.816568 | 0.397917541 | 0.44565295 | 0.89288659 | 0.37191791 | 0.7900012 |
| TBC1D24 | 12.0202268 | 0.925767975 | 0.55132122 | 1.67918074 | 0.09311682 | NA |
| PDPK1 | 106.308105 | 0.104338563 | 0.21392957 | 0.48772389 | 0.62574543 | 0.90467209 |
| KCTD5 | 10.5351546 | -0.17454935 | 0.56369947 | -0.3096497 | 0.75682739 | NA |
| SRRM2 | 273.395639 | -0.02065591 | 0.22292165 | -0.09266 | 0.92617371 | 0.98672316 |
| ELOB | 73.5095626 | -0.019891913 | 0.3488587 | -0.05702 | 0.95452929 | 0.99228059 |
| PRSS22 | 13.3092289 | -0.651561477 | 0.52423263 | -1.2428862 | 0.21390979 | 0.66782265 |
| FLYWCH2 | 11.1020487 | -0.576376922 | 0.44698723 | -1.2894707 | 0.19723451 | NA |
| FLYWCH1 | 30.9233004 | 0.263608756 | 0.31310763 | 0.841911 | 0.39983777 | 0.80830282 |
| TNFRSF12A | 60.7773319 | -1.624494311 | 0.70663266 | -2.2989234 | 0.02150929 | 0.22951242 |
| HCFC1R1 | 12.8669103 | 0.24511659 | 0.43434991 | 0.56432978 | 0.57252972 | 0.88578992 |
| BICDL2 | 30.9608608 | -0.185969169 | 0.34038208 | -0.5463542 | 0.5848225 | 0.89036744 |
| IL32 | 15.1659457 | -2.747769638 | 0.56965179 | -4.8235952 | 1.41E-06 | 0.00025804 |
| ZNF213-AS1 | 14.884823 | -0.02248423 | 0.43644746 | -0.0515165 | 0.95891398 | 0.99238848 |
| ZNF213 | 22.4244455 | -0.035308489 | 0.37036986 | -0.0953331 | 0.92405029 | 0.98672316 |
| ZNF263 | 16.2861876 | 0.176402921 | 0.42491943 | 0.4151444 | 0.6780362 | 0.9226233 |
| TIGD7 | 14.4442643 | 0.149419823 | 0.33616697 | 0.44448097 | 0.65669488 | 0.9158679 |
| ZNF75A | 53.0817567 | 0.032618476 | 0.51971513 | 0.06276222 | 0.94995585 | 0.99192022 |
| AC025283.3 | 17.0924097 | 0.102168236 | 0.43652892 | 0.23404689 | 0.81494857 | 0.96300833 |
| NAA60 | 27.7432437 | 0.090686383 | 0.28509663 | 0.31808998 | 0.75041668 | 0.94692882 |
| CLUAP1 | 36.1742301 | -0.073666027 | 0.29148903 | -0.2527232 | 0.80048215 | 0.95873501 |
| SLX4 | 11.448347 | -0.683495329 | 0.46991445 | -1.4545101 | 0.14580493 | NA |
| DNASE1 | 108.198228 | -0.133483969 | 0.22489053 | -0.5935509 | 0.55281256 | 0.87587522 |
| TRAP1 | 50.5176795 | 0.683516312 | 0.31276877 | 2.18537261 | 0.02886154 | 0.26841507 |
| CREBBP | 201.505327 | 0.298945262 | 0.23882939 | 1.25171055 | 0.21067535 | 0.66241521 |
| ADCY9 | 121.687359 | 0.408366378 | 0.37617345 | 1.08558001 | 0.27766485 | 0.72708595 |
| SRL | 22.7831384 | 0.454567074 | 1.86875693 | 0.24324569 | 0.80781507 | 0.96054677 |
| GLIS2 | 26.5332474 | 0.003759393 | 0.40964827 | 0.00917712 | 0.99267782 | 0.99923337 |
| VASN | 28.9245421 | -0.893267144 | 0.33944423 | -2.6315579 | 0.00849944 | 0.13356532 |
| DNAJA3 | 46.6756189 | -0.325825698 | 0.38982588 | -0.8358237 | 0.40325411 | 0.80927646 |
| NMRAL1 | 29.6028786 | -0.380303991 | 0.32400818 | -1.1737481 | 0.24049593 | 0.69470237 |
| HMOX2 | 47.420567 | -0.005677928 | 0.25801587 | -0.0220061 | 0.98244307 | 0.99719767 |
| CDIP1 | 38.0185284 | 0.407827699 | 0.24232504 | 1.68297798 | 0.09237935 | 0.4764741 |
| UBALD1 | 17.7878211 | 0.671171133 | 0.3317291 | 2.0232507 | 0.04304731 | 0.33179556 |
| MGRN1 | 75.0021551 | 0.029588759 | 0.25673782 | 0.11524893 | 0.90824782 | 0.98250721 |
| NUDT16L1 | 25.6359774 | 0.31471452 | 0.38408325 | 0.81939143 | 0.41256312 | 0.81556555 |
| ANKS3 | 44.238164 | 0.174249023 | 0.30165759 | 0.57763844 | 0.56350825 | 0.88191823 |
| ZNF500 | 13.4945837 | 0.597987174 | 0.46699434 | 1.28050197 | 0.20036865 | 0.65330283 |
| SMIM22 | 18.2319134 | -1.45794559 | 0.61784921 | -2.359711 | 0.01828918 | 0.21024014 |
| ROGDI | 15.7697931 | 0.608855818 | 0.5810638 | 1.04782955 | 0.29471715 | 0.74099438 |
| GLYR1 | 121.703462 | 0.258161132 | 0.19696674 | 1.31068388 | 0.18996459 | 0.64181696 |
| UBN1 | 56.4982686 | 0.234482676 | 0.24143689 | 0.97119654 | 0.33145042 | 0.76685572 |
| PPL | 32.5776981 | 0.647573098 | 0.36846407 | 1.75749319 | 0.07883378 | 0.44254372 |
| NAGPA | 12.2363786 | -0.336922714 | 0.47524371 | -0.7089472 | 0.47835722 | NA |
| ALG1 | 13.3146314 | 0.472592908 | 0.46050088 | 1.02625842 | 0.30476979 | 0.74832335 |
| C16orf89 | 10.2275278 | 1.248367729 | 0.6411192 | 1.94716947 | 0.05151442 | NA |
| RBFOX1 | 46.0047992 | 1.56058576 | 0.56607242 | 2.7568659 | 0.00583583 | 0.10600661 |
| METTL22 | 15.0259194 | -0.228193945 | 0.38235779 | -0.5968074 | 0.550636 | 0.87459787 |
| ABAT | 20.043642 | 0.257875733 | 0.32211694 | 0.80056557 | 0.42338319 | 0.822021 |
| PMM2 | 69.9903447 | -0.11215981 | 0.26546615 | -0.4225014 | 0.67265911 | 0.92189158 |
| CARHSP1 | 40.444312 | -0.133463077 | 0.47777077 | -0.2793454 | 0.77997975 | 0.95537097 |
| USP7 | 97.9837721 | 0.283454636 | 0.19906764 | 1.42391118 | 0.15447219 | 0.59443382 |
| C16orf72 | 76.0274468 | 0.108098156 | 0.21997535 | 0.49141032 | 0.62313627 | 0.90431205 |
| AC087190.1 | 12.5999277 | -0.260724943 | 0.50170097 | -0.519682 | 0.60328526 | 0.8956362 |
| ATF7IP2 | 108.342412 | 0.203525992 | 0.35813334 | 0.56829669 | 0.56983353 | 0.88562534 |
| EMP2 | 21.5041816 | 0.255077295 | 0.86753312 | 0.294026 | 0.76873803 | 0.95205353 |
| NUBP1 | 24.9047178 | -0.114704621 | 0.30799331 | -0.3724257 | 0.70957593 | 0.93485232 |
| CIITA | 27.2121312 | -1.612270974 | 0.48484414 | -3.3253387 | 0.00088311 | 0.03016262 |
| DEXI | 29.9943019 | -0.069151549 | 0.34195949 | -0.2022215 | 0.83974359 | 0.96748214 |
| CLEC16A | 398.864176 | -0.898701289 | 0.27336421 | -3.2875601 | 0.0010106 | 0.03289524 |
| AC007014.1 | 24.8512608 | -1.413036594 | 0.58024945 | -2.4352226 | 0.01488263 | 0.18484189 |
| AC007220.1 | 15.0309403 | -0.752188103 | 0.55097052 | -1.3652057 | 0.1721884 | 0.6163201 |
| SOCS1 | 81.8873988 | -4.292035302 | 0.50339416 | -8.526192 | 1.51E-17 | 3.67E-14 |
| AC099489.1 | 80.9627784 | -0.024243477 | 0.64560934 | -0.0375513 | 0.97004543 | 0.99486579 |
| LITAF | 772.273161 | -0.844679739 | 0.28586004 | -2.9548717 | 0.00312799 | 0.0697506 |
| SNN | 14.8872602 | -0.917823687 | 0.66084828 | -1.3888569 | 0.16487627 | 0.60558727 |
| TXNDC11 | 121.386061 | 0.38571933 | 0.27256184 | 1.41516261 | 0.15702082 | 0.59776373 |
| ZC3H7A | 90.9169054 | 0.087998565 | 0.23625848 | 0.37246732 | 0.70954494 | 0.93485232 |
| RSL1D1 | 24.511903 | 0.208351189 | 0.29534811 | 0.70544278 | 0.48053473 | 0.84949642 |
| AC007216.4 | 42.511846 | 0.286474499 | 0.31078569 | 0.92177506 | 0.35664592 | 0.78241164 |
| GSPT1 | 173.438304 | -0.184842669 | 0.24010821 | -0.7698307 | 0.44140033 | 0.8281114 |
| SNX29 | 526.99669 | -0.389786111 | 0.30182824 | -1.291417 | 0.19655913 | 0.64685325 |
| AC010333.2 | 69.5089485 | 0.768418269 | 0.59247905 | 1.29695434 | 0.1946469 | 0.6459894 |
| CPPED1 | 289.161741 | -0.265649873 | 0.3156717 | -0.8415384 | 0.40004636 | 0.80830282 |
| AC109597.1 | 16.322704 | -0.39676129 | 0.52540693 | -0.7551505 | 0.4501587 | 0.83118301 |
| ERCC4 | 21.5626753 | -0.013044563 | 0.32316744 | -0.0403647 | 0.96780236 | 0.99393148 |
| MKL2 | 317.693604 | -0.199385161 | 0.21336156 | -0.9344943 | 0.35004897 | 0.77868979 |
| MIR193BHG | 19.1814329 | -1.538201974 | 0.96508132 | -1.5938574 | 0.11096798 | 0.5125664 |
| PARN | 124.998734 | 0.099272022 | 0.28377343 | 0.34982846 | 0.72646744 | 0.94009907 |
| BFAR | 60.3516295 | -0.625061543 | 0.32783274 | -1.9066477 | 0.05656623 | 0.37814775 |
| NOMO1 | 16.2579591 | 0.603131255 | 0.39415446 | 1.53019009 | 0.12596968 | 0.54646955 |
| PDXDC1 | 201.07191 | -0.128419548 | 0.18786401 | -0.6835772 | 0.49424221 | 0.85774549 |
| NTAN1 | 16.3359635 | 0.025086541 | 0.48732217 | 0.05147835 | 0.95894435 | 0.99238848 |
| RRN3 | 35.4749642 | -0.08787449 | 0.23548176 | -0.373169 | 0.70902268 | 0.93482669 |
| C16orf45 | 66.5346374 | -0.719470241 | 0.36689283 | -1.960982 | 0.04988112 | 0.35629373 |
| MARF1 | 45.3635261 | 0.181026205 | 0.32032737 | 0.56512874 | 0.5719862 | 0.88578992 |
| NDE1 | 17.9645232 | 0.514077664 | 0.30219374 | 1.70115258 | 0.08891434 | 0.4687332 |
| FOPNL | 28.3602031 | 0.182577848 | 0.34554862 | 0.52837094 | 0.5972419 | 0.89363331 |
| ABCC1 | 125.502931 | -0.288476461 | 0.40915283 | -0.705058 | 0.48077417 | 0.84961003 |
| RPS15A | 45.8515345 | 0.468556077 | 0.38605335 | 1.21370809 | 0.22485921 | 0.6780026 |
| SMG1 | 240.569967 | 0.016746901 | 0.20263268 | 0.08264659 | 0.93413255 | 0.9881638 |
| COQ7 | 24.1437385 | 0.120688427 | 0.3258344 | 0.37039805 | 0.71108592 | 0.93563937 |
| AC099518.5 | 11.8084704 | 0.388526085 | 0.53386226 | 0.72776465 | 0.46675767 | NA |
| SYT17 | 287.453977 | 0.163173555 | 0.46138926 | 0.35365702 | 0.72359593 | 0.93896433 |
| GDE1 | 40.2650446 | -0.000356182 | 0.33184941 | -0.0010733 | 0.99914361 | 0.99982181 |
| CCP110 | 18.1652265 | 0.340491091 | 0.4138761 | 0.82268846 | 0.41068518 | 0.81431853 |
| VPS35L | 97.687789 | -0.110609339 | 0.26549235 | -0.4166197 | 0.67695661 | 0.92237262 |
| KNOP1 | 30.5306861 | -0.223747784 | 0.35213764 | -0.6353987 | 0.52516844 | 0.86747074 |
| IQCK | 123.05953 | 0.210894157 | 0.32868934 | 0.64162153 | 0.52111895 | 0.8669958 |
| GPRC5B | 125.761185 | 0.056981894 | 0.39500784 | 0.14425509 | 0.88529904 | 0.97662748 |
| UMOD | 60.9489349 | 0.933433518 | 0.618583 | 1.5089867 | 0.13130218 | 0.55812058 |
| ACSM3 | 25.0236467 | 0.311629243 | 0.38082011 | 0.81831088 | 0.41317969 | 0.8160951 |
| THUMPD1 | 59.425832 | 0.272343834 | 0.27513135 | 0.9898684 | 0.32223845 | 0.762555 |
| ERI2 | 20.4619467 | 0.079638233 | 0.41699779 | 0.19097999 | 0.84854127 | 0.97027687 |
| LYRM1 | 15.8950641 | -0.00920113 | 0.44248756 | -0.0207941 | 0.98340991 | 0.99749828 |
| TMEM159 | 16.9099205 | -0.910095517 | 0.42188545 | -2.15721 | 0.03098931 | 0.27936455 |
| METTL9 | 67.1992278 | -0.096606606 | 0.22138031 | -0.436383 | 0.66255886 | 0.91824845 |
| NPIPB4 | 18.3478988 | 0.100434196 | 0.47606339 | 0.21096812 | 0.83291215 | 0.96701848 |

| | | | | | | |
|---|---|---|---|---|---|---|
| UQCRC2 | 149.60235 | 0.020251051 | 0.21774539 | 0.09300335 | 0.9259009 | 0.98672316 |
| MOSMO | 70.5960581 | -0.140184248 | 0.24848105 | -0.5641648 | 0.57264201 | 0.88578992 |
| EEF2K | 15.327878 | 0.083827686 | 0.49241063 | 0.17023939 | 0.86482187 | 0.97419257 |
| POLR3E | 41.9189191 | -0.034182884 | 0.24317759 | -0.1405676 | 0.88821157 | 0.97765744 |
| CDR2 | 142.827621 | 0.076599247 | 0.22776257 | 0.33631183 | 0.73663573 | 0.94365644 |
| NPIPB5 | 60.4990629 | -0.271329652 | 0.35129619 | -0.7723672 | 0.43989698 | 0.82793978 |
| USP31 | 24.1072185 | -0.578552649 | 0.38607457 | -1.4985516 | 0.13399001 | 0.56229288 |
| COG7 | 70.2595912 | -0.141358636 | 0.23337301 | -0.6057197 | 0.54470087 | 0.87247631 |
| GGA2 | 124.273622 | -0.06148562 | 0.2500526 | -0.2458907 | 0.80576681 | 0.96005568 |
| EARS2 | 15.3551727 | 0.167751446 | 0.3493728 | 0.48015028 | 0.63112054 | 0.90770379 |
| UBFD1 | 21.1428698 | 0.292894447 | 0.37426572 | 0.78258422 | 0.43387131 | 0.82504445 |
| NDUFAB1 | 54.3069097 | 0.259415621 | 0.35709825 | 0.72645447 | 0.46756021 | 0.84315561 |
| PALB2 | 12.4233703 | 0.010389304 | 0.46050719 | 0.02256057 | 0.9820008 | NA |
| DCTN5 | 47.8134629 | 0.169197644 | 0.26166999 | 0.64660698 | 0.51788634 | 0.86500999 |
| RBBP6 | 106.520396 | 0.056147242 | 0.26055637 | 0.21548981 | 0.82938546 | 0.96669991 |
| TNRC6A | 579.428837 | -0.209830739 | 0.29368439 | -0.714477 | 0.47493227 | 0.84593347 |
| AC008731.1 | 18.7808206 | -1.02370629 | 0.50360545 | -2.0327546 | 0.04207732 | 0.32888549 |
| ARHGAP17 | 74.06717 | -0.272413261 | 0.27779557 | -0.9806249 | 0.32677773 | 0.76371899 |
| LCMT1 | 49.4093256 | 0.38913944 | 0.23379482 | 1.6644485 | 0.09602283 | 0.48135478 |
| NSMCE1 | 21.1914841 | 0.042097702 | 0.37982634 | 0.11083408 | 0.91174792 | 0.9835359 |
| GTF3C1 | 40.9074685 | 0.278831768 | 0.27657556 | 1.00815765 | 0.31337878 | 0.75618668 |
| KIAA0556 | 84.0409126 | -0.120194874 | 0.22015737 | -0.5459498 | 0.58510045 | 0.89036744 |
| XPO6 | 75.2267825 | -0.283370332 | 0.39432539 | -0.7186205 | 0.47237475 | 0.84570599 |
| NUPR1 | 34.4144381 | -0.912215508 | 0.87037559 | -1.0480711 | 0.29460585 | 0.74090659 |
| SGF29 | 13.7375088 | 0.742106377 | 0.34950956 | 2.12327917 | 0.03373047 | 0.2921512 |
| SULT1A1 | 10.6472744 | 0.992637878 | 0.6419865 | 1.54619744 | 0.12205689 | NA |
| ATXN2L | 52.2405942 | 0.174858207 | 0.23340206 | 0.74917165 | 0.45375375 | 0.83364149 |
| TUFM | 112.795254 | 0.155552772 | 0.37098848 | 0.41929272 | 0.67500221 | 0.92193843 |
| SH2B1 | 53.3996768 | 0.062898577 | 0.29451094 | 0.21356958 | 0.83088273 | 0.96701848 |
| RABEP2 | 39.86709 | 0.159627654 | 0.29056128 | 0.5493769 | 0.58274682 | 0.89036744 |
| NFATC2IP | 81.8566893 | 0.070005657 | 0.29050057 | 0.24098286 | 0.8095684 | 0.96200092 |
| SPNS1 | 16.5927967 | 0.015071933 | 0.47784269 | 0.03154162 | 0.9748376 | 0.99543096 |
| QPRT | 43.8728575 | -0.410302064 | 0.51775881 | -0.7924579 | 0.42809372 | 0.82407405 |
| MAZ | 19.9634817 | -0.313583514 | 0.49143179 | -0.6381018 | 0.52340741 | 0.86747074 |
| AC009133.1 | 19.4146033 | -1.989877706 | 0.61666048 | -3.2268611 | 0.00125156 | 0.03770234 |
| MVP | 97.0153846 | -1.632666587 | 0.3299416 | -4.9483502 | 7.48E-07 | 0.00015783 |
| CDIPT | 25.0842437 | 0.0506902 | 0.28909303 | 0.17534217 | 0.86081078 | 0.97302617 |
| KCTD13 | 23.3765015 | 0.265856152 | 0.32621153 | 0.81498086 | 0.41508327 | 0.81726257 |
| TMEM219 | 70.0793451 | 0.159691631 | 0.33515745 | 0.4764675 | 0.63374135 | 0.90805737 |
| TAOK2 | 17.1533181 | 0.541797022 | 0.35507195 | 1.52587954 | 0.12703987 | 0.54952841 |
| INO80E | 26.3164343 | 0.878556868 | 0.27559329 | 3.18787468 | 0.00143323 | 0.0412531 |
| PPP4C | 41.7514261 | -0.506635865 | 0.3032221 | -1.6708408 | 0.09475312 | 0.47996672 |
| YPEL3 | 34.8610196 | -0.291863026 | 0.34738269 | -0.8401772 | 0.40080902 | 0.80830282 |
| MAPK3 | 28.8660423 | -0.662617302 | 0.29235597 | -2.2664743 | 0.02342236 | 0.23883233 |
| NPIPB13 | 16.0386577 | -0.090450038 | 0.50232722 | -0.180062 | 0.85710391 | 0.97215948 |
| CD2BP2 | 23.2307689 | 0.106000739 | 0.33703332 | 0.31451116 | 0.75313284 | 0.94771649 |
| TBC1D10B | 16.6164773 | 0.834794345 | 0.41302955 | 2.0211492 | 0.04326432 | 0.33179556 |
| DCTPP1 | 14.4063585 | -0.491667873 | 0.4100782 | -1.1989613 | 0.23054301 | 0.68523674 |
| SEPHS2 | 66.4427089 | -0.975235776 | 0.45529052 | -2.1420076 | 0.03219287 | 0.28439965 |
| ZNF768 | 19.4130433 | 0.26434174 | 0.45386175 | 0.58242789 | 0.5602785 | 0.88054139 |
| ZNF785 | 19.3310037 | 0.379339784 | 0.42732455 | 0.88770886 | 0.37469738 | 0.7926388 |
| PRR14 | 10.9973649 | 0.18709436 | 0.42185367 | 0.44350535 | 0.65740025 | NA |
| FBRS | 26.0765759 | -0.230294013 | 0.32246385 | -0.71417 | 0.47512205 | 0.84594051 |
| PHKG2 | 19.2465389 | 0.191304912 | 0.33154006 | 0.57701899 | 0.56392662 | 0.88191823 |
| RNF40 | 31.7325512 | 0.469667506 | 0.28078684 | 1.67268346 | 0.09438963 | 0.4798634 |
| ZNF629 | 18.2335998 | 0.058931847 | 0.47980989 | 0.12282333 | 0.902247 | 0.98004434 |
| BCL7C | 43.1462051 | 0.378009551 | 0.23003431 | 1.64327462 | 0.10032613 | 0.48878123 |
| FBXL19 | 22.0657104 | 0.391002424 | 0.3399186 | 1.15028251 | 0.25002753 | 0.70440519 |
| SETD1A | 18.6060393 | 0.158085871 | 0.43541641 | 0.36306825 | 0.7165539 | 0.93689626 |
| STX4 | 15.6285213 | 0.029901454 | 0.38629673 | 0.0774054 | 0.93830104 | 0.98871562 |
| BCKDK | 22.2663075 | -0.439877341 | 0.31880848 | -1.3797542 | 0.16766234 | 0.60933859 |
| KAT8 | 36.3831798 | 0.152284261 | 0.3308007 | 0.46035047 | 0.64526468 | 0.91225913 |
| PRSS8 | 61.6010947 | -0.658271299 | 0.32479028 | -2.026758 | 0.04268717 | 0.33083096 |
| FUS | 136.900541 | 0.376060525 | 0.1996046 | 1.88402739 | 0.05956127 | 0.38765901 |
| AC026471.4 | 14.7310866 | 0.027208588 | 0.42311871 | 0.06430486 | 0.94872749 | 0.99179534 |
| C16orf58 | 25.7167297 | -0.232092766 | 0.31185826 | -0.7442252 | 0.45674026 | 0.83529044 |
| ZNF720 | 63.7317602 | 0.034831153 | 0.33544356 | 0.10383611 | 0.91729941 | 0.98495314 |
| ZNF267 | 19.2847347 | -0.42743413 | 0.39707043 | -1.0764693 | 0.28171742 | 0.73311066 |
| VPS35 | 60.8386337 | -0.474978138 | 0.27681734 | -1.715854 | 0.08618877 | 0.46240657 |
| C16orf87 | 16.6658367 | 0.432504497 | 0.3358297 | 1.2878685 | 0.19779174 | 0.64914661 |
| GPT2 | 32.2588716 | -0.398543528 | 0.78571735 | -0.5072352 | 0.61198977 | 0.90011356 |
| DNAJA2 | 42.6895779 | 0.116528743 | 0.22225583 | 0.52430005 | 0.60006987 | 0.89486416 |
| ITFG1 | 388.192591 | -0.087177547 | 0.20651429 | -0.4221381 | 0.67292424 | 0.92189158 |
| PHKB | 348.023002 | -0.142791443 | 0.17939593 | -0.795957 | 0.42605704 | 0.82328415 |
| LONP2 | 205.453935 | 0.141087608 | 0.16191252 | 0.87138172 | 0.38354576 | 0.7980695 |
| SIAH1 | 50.4082292 | -0.037417626 | 0.27613007 | -0.1355073 | 0.89221083 | 0.97794037 |
| N4BP1 | 135.384422 | -0.012745314 | 0.29601464 | -0.0430564 | 0.9656566 | 0.99334616 |
| CNEP1R1 | 11.8480378 | 0.953189985 | 0.43875391 | 2.17249343 | 0.02981846 | NA |
| TENT4B | 44.5422647 | 0.539345919 | 0.30552146 | 1.7653291 | 0.07750847 | 0.43864187 |
| BRD7 | 56.3147448 | 0.146352111 | 0.23773532 | 0.61560946 | 0.53815231 | 0.86972442 |
| CYLD | 48.2385915 | -0.076747609 | 0.23667864 | -0.3242693 | 0.74573418 | 0.94614106 |
| SALL1 | 13.5001652 | -1.183250693 | 0.68534336 | -1.7265079 | 0.08425609 | 0.4585327 |
| AC087564.1 | 16.5610232 | -0.365803443 | 0.55895396 | -0.6544429 | 0.51282652 | 0.86346419 |
| HNRNPA1P4! | 12.9735201 | -0.11959454 | 0.52485394 | -0.2278625 | 0.81975312 | 0.96389705 |
| TOX3 | 455.838807 | -0.074915689 | 0.32466926 | -0.2307446 | 0.8175132 | 0.9635539 |
| CASC16 | 149.387499 | -2.449815349 | 0.68786395 | -3.5614824 | 0.00036877 | 0.01625927 |
| CHD9 | 532.026395 | 0.152339702 | 0.20045131 | 0.75998359 | 0.44726439 | 0.83066268 |
| RBL2 | 142.481712 | -0.318024658 | 0.31082581 | -1.0231604 | 0.30623202 | 0.74897897 |
| AKTIP | 17.4631897 | 0.623012737 | 0.37544052 | 1.65941794 | 0.0970316 | 0.48257296 |
| FTO | 225.364244 | -0.026375055 | 0.21365023 | -0.1234497 | 0.90175101 | 0.9797999 |
| LPCAT2 | 26.9101818 | 0.170658971 | 0.60223332 | 0.28337683 | 0.77688799 | 0.95512504 |
| AMFR | 284.219399 | 0.578492679 | 0.24766746 | 2.33576379 | 0.01950356 | 0.21947157 |
| AC092140.1 | 13.6098782 | 0.977228858 | 0.75141726 | 1.30051426 | 0.19342477 | 0.6443384 |
| NUDT21 | 64.9728304 | 0.531868332 | 0.20792235 | 2.55801427 | 0.01052718 | 0.15061006 |
| OGFOD1 | 37.6221291 | 0.192124635 | 0.30911135 | 0.6215386 | 0.53424531 | 0.86909171 |
| BBS2 | 47.2611077 | 0.406834651 | 0.24179944 | 1.68252936 | 0.09246624 | 0.4764958 |
| MT2A | 287.069174 | -2.874865314 | 0.98520989 | -2.9180232 | 0.00352258 | 0.0764408 |
| MT1E | 69.2119825 | -2.10148093 | 1.02577298 | -2.0486803 | 0.04049338 | 0.32301466 |
| MT1G | 104.413583 | -1.516872378 | 0.99828783 | -1.519474 | 0.12864324 | 0.55308726 |
| MT1H | 34.4190991 | -1.409225577 | 1.21040517 | -1.1642594 | 0.24431891 | 0.69784848 |
| MT1X | 135.576261 | -2.079642931 | 0.68193422 | -3.049624 | 0.00229128 | 0.05598343 |
| NUP93 | 39.6037209 | -0.352549786 | 0.32342107 | -1.0900644 | 0.27568478 | 0.72554623 |
| SLC12A3 | 99.876341 | 1.003677259 | 0.73731153 | 1.36126619 | 0.17342958 | 0.61802607 |
| HERPUD1 | 126.418599 | -0.516512322 | 0.40258269 | -1.2829968 | 0.19949318 | 0.65266113 |
| NLRC5 | 89.7445105 | -2.625672116 | 0.7236333 | -3.6284567 | 0.00028512 | 0.01390829 |
| CPNE2 | 36.5679582 | -2.433656687 | 0.6489555 | -3.7501133 | 0.00017675 | 0.01015508 |
| FAM192A | 75.142331 | 0.051977091 | 0.20775999 | 0.25017854 | 0.80244928 | 0.95908975 |
| RSPRY1 | 74.2935336 | 0.124603672 | 0.18852742 | 0.66093129 | 0.50865638 | 0.86281145 |
| ARL2BP | 99.6970559 | -0.308245092 | 0.20065963 | -1.536159 | 0.12449939 | 0.54320519 |
| PLLP | 121.68262 | 0.334084353 | 0.42288667 | 0.79000918 | 0.4295224 | 0.82469033 |
| CX3CL1 | 17.2557811 | 0.706609325 | 0.49922536 | 1.41541151 | 0.15694788 | 0.59772062 |
| CIAPIN1 | 46.4318826 | -0.05544797 | 0.23754069 | -0.2334251 | 0.81543129 | 0.96300833 |
| COQ9 | 81.9136924 | 0.110074579 | 0.25063254 | 0.43918711 | 0.66052597 | 0.91752855 |

| | | | | | | |
|---|---|---|---|---|---|---|
| POLR2C | 17.2818575 | 0.260606924 | 0.42575069 | 0.61211158 | 0.54046394 | 0.87026896 |
| ADGRG1 | 157.281525 | -0.371202304 | 0.42001074 | -0.8837924 | 0.37680828 | 0.79432848 |
| KIFC3 | 74.2884993 | 0.263871437 | 0.27564509 | 0.95728693 | 0.33842245 | 0.76998177 |
| USB1 | 13.3220175 | -0.513563047 | 0.41100654 | -1.2495252 | 0.21147303 | 0.66341798 |
| MMP15 | 14.7559817 | 0.522380124 | 0.46401232 | 1.12578933 | 0.2602547 | 0.71091822 |
| CFAP20 | 10.7110139 | -0.277946969 | 0.42537376 | -0.6534182 | 0.51348668 | NA |
| CSNK2A2 | 45.229223 | -0.085385653 | 0.30175371 | -0.2829647 | 0.77720388 | 0.95512504 |
| CNOT1 | 149.212583 | 0.265208272 | 0.14956192 | 1.77323391 | 0.07618996 | 0.43480446 |
| SLC38A7 | 19.6678364 | -0.536099684 | 0.54398811 | -0.9854989 | 0.32437907 | 0.76304922 |
| GOT2 | 76.5114412 | 0.441986227 | 0.23805884 | 1.85662601 | 0.06336438 | 0.4001801 |
| AC138305.1 | 686.393014 | 0.350372518 | 0.26843556 | 1.30523882 | 0.19181154 | 0.6421543 |
| AC010542.3 | 13.6337868 | -0.391236195 | 0.53223072 | -0.7350876 | 0.46228615 | 0.83894774 |
| TK2 | 16.757857 | -0.365403378 | 0.44511666 | -0.820916 | 0.41169413 | 0.81498633 |
| CMTM4 | 294.024447 | 0.26169249 | 0.2019928 | 1.29555356 | 0.19512934 | 0.6459894 |
| DYNC1LI2 | 101.301985 | 0.076413242 | 0.19332667 | 0.39525452 | 0.69265506 | 0.928133 |
| AC018557.1 | 15.7617748 | 0.059083776 | 0.5424248 | 0.10892529 | 0.91326175 | 0.98376271 |
| NAE1 | 37.9968807 | -0.094095604 | 0.23068123 | -0.4079032 | 0.68334477 | 0.92485618 |
| PDP2 | 26.1988403 | -0.041870242 | 0.47342825 | -0.0884405 | 0.92952655 | 0.98733028 |
| CDH16 | 267.86842 | 0.389669556 | 0.39078823 | 0.99713738 | 0.31869783 | 0.76052272 |
| RRAD | 45.0950512 | -1.115360232 | 0.7381546 | -1.5110117 | 0.13078547 | 0.55687707 |
| FAM96B | 81.6857792 | -0.282712208 | 0.48493589 | -0.5829888 | 0.55990082 | 0.88037687 |
| CES2 | 37.9003611 | -0.545589098 | 0.26130532 | -2.0879372 | 0.0368035 | 0.30695956 |
| CBFB | 161.265017 | 0.038523582 | 0.26775275 | 0.14387745 | 0.88559725 | 0.97662748 |
| C16orf70 | 16.5738649 | 0.061052974 | 0.42081966 | 0.14508109 | 0.88464685 | 0.97662748 |
| TRADD | 15.9884311 | -0.273950326 | 0.41124825 | -0.6661434 | 0.50531942 | 0.86197912 |
| NOL3 | 22.3628073 | -1.52073245 | 0.54394345 | -2.7957547 | 0.00517787 | 0.09959581 |
| KIAA0895L | 24.0305754 | -0.05322835 | 0.41601839 | -0.1279471 | 0.89819083 | 0.97892709 |
| E2F4 | 23.269917 | 0.297346563 | 0.33294769 | 0.89307291 | 0.37181813 | 0.7900012 |
| ELMO3 | 13.9904481 | 0.242408883 | 0.45567403 | 0.53197871 | 0.59474074 | 0.89301752 |
| TMEM208 | 34.2756288 | -0.065533579 | 0.32334519 | -0.2026737 | 0.83939005 | 0.96748214 |
| ZDHHC1 | 30.6503544 | 0.279008609 | 0.33589205 | 0.83064964 | 0.40617158 | 0.8115705 |
| HSD11B2 | 173.463102 | 1.361256424 | 0.75218376 | 1.80973919 | 0.07033624 | 0.41985326 |
| ATP6V0D1 | 51.1920065 | 0.067681557 | 0.44797301 | 0.15108401 | 0.87990945 | 0.97581687 |
| AC009061.2 | 13.0747385 | 0.954016228 | 0.55737972 | 1.71160916 | 0.08696872 | 0.46422617 |
| RIPOR1 | 29.4581463 | -0.414472157 | 0.3596458 | -1.1524454 | 0.24913808 | 0.70371066 |
| CTCF | 71.4078302 | 0.022671255 | 0.28787153 | 0.07875477 | 0.93722768 | 0.98871562 |
| ENKD1 | 10.9876799 | -0.029578381 | 0.44365298 | -0.0666701 | 0.94684435 | NA |
| GFOD2 | 27.0228213 | -0.255788928 | 0.32629111 | -0.7839286 | 0.43308202 | 0.82483715 |
| RANBP10 | 77.7584608 | 0.010594258 | 0.33872785 | 0.03127661 | 0.97504895 | 0.99543096 |
| NUTF2 | 15.9814715 | 0.054876322 | 0.39126391 | 0.14025399 | 0.88845932 | 0.97765744 |
| PSKH1 | 28.581252 | 0.095360823 | 0.38139817 | 0.25002958 | 0.80256448 | 0.95908975 |
| DUS2 | 18.4871619 | -0.253773418 | 0.33560505 | -0.7561669 | 0.44954916 | 0.83118301 |
| NFATC3 | 116.897242 | 0.265470596 | 0.17554556 | 1.51226036 | 0.13046764 | 0.55685884 |
| ESRP2 | 15.3511866 | 0.081759278 | 0.34230288 | 0.23885068 | 0.81122137 | 0.96247289 |
| PLA2G15 | 23.2924402 | 0.238538934 | 0.38521004 | 0.61924381 | 0.53575575 | 0.86946459 |
| SLC7A6 | 61.531682 | -0.242076796 | 0.38964059 | -0.6212823 | 0.53441391 | 0.86909171 |
| SLC7A6OS | 30.983434 | -0.296095054 | 0.32141608 | -0.9212204 | 0.35693537 | 0.7825111 |
| PRMT7 | 25.4174913 | 0.65171954 | 0.3783472 | 1.7225436 | 0.08497109 | 0.45976313 |
| ZFP90 | 21.8151684 | 0.182185931 | 0.28570647 | 0.63766821 | 0.52368969 | 0.86747074 |
| CDH1 | 379.296131 | 0.354605016 | 0.17292063 | 2.05068081 | 0.04029804 | 0.32205065 |
| TANGO6 | 59.5962553 | 0.254683466 | 0.26241504 | 0.97053686 | 0.33177896 | 0.76685572 |
| CHTF8 | 41.3109081 | 0.075038671 | 0.28591643 | 0.26244967 | 0.79297479 | 0.95746995 |
| UTP4 | 17.376583 | 0.34290317 | 0.37309161 | 0.91908573 | 0.35805073 | 0.78363992 |
| SNTB2 | 48.5687117 | 0.234450139 | 0.25919546 | 0.90453026 | 0.3657143 | 0.78793182 |
| VPS4A | 31.9868838 | -0.026244423 | 0.30485849 | -0.0860872 | 0.93139707 | 0.98757533 |
| COG8 | 12.3633303 | 0.840355763 | 0.40892008 | 2.05506114 | 0.0398731 | NA |
| TERF2 | 36.0106346 | -0.102792852 | 0.24621379 | -0.4174943 | 0.67631689 | 0.92229234 |
| CYB5B | 103.379197 | -0.340229917 | 0.22340172 | -1.5229512 | 0.12777092 | 0.55107954 |
| NFAT5 | 1228.56154 | 0.212010127 | 0.23435203 | 0.90466519 | 0.36564279 | 0.78793182 |
| NQO1 | 39.1084282 | -1.900065325 | 0.45551393 | -4.1712563 | 3.03E-05 | 0.00277205 |
| WWP2 | 182.769807 | 0.053859281 | 0.35353451 | 0.15234519 | 0.87891469 | 0.97556614 |
| PDPR | 58.1824931 | 0.100802699 | 0.35186575 | 0.28648056 | 0.77451008 | 0.95470568 |
| AC009060.1 | 12.5657954 | -0.857057605 | 0.50685319 | -1.6909386 | 0.09084853 | 0.47310687 |
| EXOSC6 | 26.3384221 | 0.083131433 | 0.43464408 | 0.19126324 | 0.84831936 | 0.97027687 |
| AARS | 53.2923221 | -0.473422956 | 0.26946495 | -1.7569 | 0.07893486 | 0.44258275 |
| DDX19A | 10.1743283 | -0.03981273 | 0.48364878 | -0.0823174 | 0.93439429 | NA |
| ST3GAL2 | 29.5086517 | 0.268010384 | 0.30444454 | 0.88032579 | 0.37868285 | 0.79535699 |
| FUK | 16.7167375 | -0.19461739 | 0.38651231 | -0.5035218 | 0.61459743 | 0.90054121 |
| COG4 | 33.872379 | 0.300363326 | 0.26367094 | 1.13915977 | 0.25463652 | 0.70793186 |
| SF3B3 | 71.4312152 | 0.096389537 | 0.22778058 | 0.42316836 | 0.67217243 | 0.92176197 |
| VAC14 | 40.1573561 | -0.365844587 | 0.48015299 | -0.7619334 | 0.44609977 | 0.83035712 |
| HYDIN | 48.9579916 | 0.257823296 | 0.44158612 | 0.58385734 | 0.5593163 | 0.88031285 |
| CMTR2 | 31.2807117 | 0.284873232 | 0.29075411 | 0.97977369 | 0.32719784 | 0.76440729 |
| ZNF19 | 21.1589403 | -0.231857291 | 0.50510215 | -0.4590305 | 0.64621227 | 0.91254316 |
| MARVELD3 | 13.837895 | -0.080785531 | 0.39814723 | -0.2029037 | 0.83921033 | 0.96748214 |
| PHLPP2 | 37.3887304 | -0.709708878 | 0.283583 | -2.5026496 | 0.01232675 | 0.16567149 |
| AP1G1 | 181.10527 | -0.021815449 | 0.1976846 | -0.1103548 | 0.91212798 | 0.98361772 |
| IST1 | 119.493764 | -0.075856662 | 0.19181559 | -0.3954666 | 0.69249855 | 0.928133 |
| ZNF821 | 23.968741 | -0.18253878 | 0.39443202 | -0.4627889 | 0.64351565 | 0.91192138 |
| TXNL4B | 20.1354728 | -0.015045049 | 0.30317221 | -0.0496254 | 0.96042089 | 0.99238848 |
| DHX38 | 21.9039866 | -0.042357655 | 0.39524148 | -0.1071691 | 0.91465487 | 0.98407488 |
| LINC01572 | 87.8317791 | -0.189740243 | 0.41089676 | -0.4617711 | 0.64424548 | 0.9119294 |
| AC004943.2 | 40.5128476 | 0.357496728 | 0.3325054 | 1.07516067 | 0.2823028 | 0.73374523 |
| ZFHX3 | 427.804645 | 0.368219244 | 0.31586216 | 1.16575928 | 0.24371179 | 0.69755218 |
| AC138627.1 | 299.487231 | 1.934148048 | 0.67218964 | 2.87738449 | 0.00400987 | 0.08381056 |
| PSMD7 | 60.1742571 | -0.492478203 | 0.35027465 | -1.4059773 | 0.15973087 | 0.60024417 |
| NPIPB15 | 12.7962982 | -1.790946495 | 0.84164291 | -2.1279173 | 0.03334394 | 0.28966809 |
| GLG1 | 346.618599 | 0.1083643 | 0.22438183 | 0.48294597 | 0.6291341 | 0.9069105 |
| RFWD3 | 34.1291627 | -0.117454674 | 0.30039616 | -0.3909993 | 0.69579779 | 0.92900737 |
| FA2H | 73.3652205 | 0.23875176 | 0.43337781 | 0.55090906 | 0.58169602 | 0.89036744 |
| WDR59 | 97.5137675 | 0.051284844 | 0.20159316 | 0.25439773 | 0.7991883 | 0.95848876 |
| ZNRF1 | 51.25271 | -0.209160939 | 0.4210765 | -0.4967291 | 0.61938012 | 0.90332088 |
| LDHD | 10.4310725 | 0.289509608 | 0.50677805 | 0.57127495 | 0.56781328 | NA |
| BCAR1 | 64.9335822 | 0.000473403 | 0.28297845 | 0.00167293 | 0.9986652 | 0.99982181 |
| CFDP1 | 140.865822 | -0.043143685 | 0.20522516 | -0.2102261 | 0.83349121 | 0.96701848 |
| GABARAPL2 | 68.5655493 | -0.160247548 | 0.40774264 | -0.3930115 | 0.69431099 | 0.92841398 |
| ADAT1 | 21.5156598 | -0.143984688 | 0.3283555 | -0.4385024 | 0.66102211 | 0.91795482 |
| KARS | 28.4181036 | -0.02547597 | 0.36877585 | -0.0690825 | 0.94492392 | 0.99121467 |
| TERF2IP | 108.852585 | -0.381958457 | 0.32696357 | -1.1681988 | 0.24272657 | 0.69621873 |
| MON1B | 10.8736083 | -0.654860962 | 0.59530018 | -1.1000517 | 0.2713096 | NA |
| ADAMTS18 | 49.7449339 | -0.829011668 | 0.31817062 | -2.6055569 | 0.0091725 | 0.13988811 |
| NUDT7 | 16.4319211 | 0.597540377 | 0.42863483 | 1.39405464 | 0.16330111 | 0.60228925 |
| VAT1L | 118.772702 | -0.658799508 | 0.42609069 | -1.5461486 | 0.12206869 | 0.53747904 |
| WWOX | 1404.75181 | 0.118889823 | 0.34737224 | 0.34225482 | 0.73215913 | 0.94194042 |
| AC046158.1 | 12.2065533 | -0.763413715 | 0.62927025 | -1.2131731 | 0.22506366 | NA |
| MAF | 32.1255021 | -2.839672174 | 0.64557577 | -4.398666 | 1.09E-05 | 0.0012285 |
| CDYL2 | 206.071501 | 0.23038703 | 0.37344996 | 0.61691539 | 0.53729053 | 0.86963427 |
| CMC2 | 96.9910174 | 0.288457963 | 0.26502804 | 1.08840543 | 0.27641617 | 0.72575599 |
| ATMIN | 26.3194746 | 0.164384837 | 0.510753 | 0.32184801 | 0.74756784 | 0.9462156 |
| GAN | 44.6122221 | 0.134058575 | 0.27317312 | 0.49074585 | 0.62360621 | 0.90447745 |
| CMIP | 293.312617 | -0.907874125 | 0.38768673 | -2.3417725 | 0.01919241 | 0.21748409 |
| PLCG2 | 1675.5602 | -0.931728338 | 0.54998957 | -1.6940836 | 0.09024939 | 0.47226726 |
| AC092142.1 | 16.8162735 | 0.566711148 | 0.46577167 | 1.2167145 | 0.22371286 | 0.67606266 |

| | | | | | | |
|---|---|---|---|---|---|---|
| MPHOSPH6 | 62.2197888 | 0.50477348 | 0.25421579 | 1.98561028 | 0.04707661 | 0.34594179 |
| HSBP1 | 66.6131542 | -0.020726777 | 0.28435095 | -0.0728915 | 0.94189243 | 0.98991882 |
| MBTPS1 | 85.9290035 | 0.000280199 | 0.23617775 | 0.00118639 | 0.9990534 | 0.99982181 |
| HSDL1 | 19.3877541 | -0.081204524 | 0.46330813 | -0.1752711 | 0.86086662 | 0.97302617 |
| TAF1C | 12.2844008 | 0.29646637 | 0.37065618 | 0.79984198 | 0.42380236 | NA |
| ATP2C2 | 47.6980794 | 0.125376994 | 0.73990998 | 0.16944898 | 0.8654435 | 0.97434023 |
| TLDC1 | 42.9260136 | -0.544880043 | 0.36160666 | -1.5068308 | 0.13185404 | 0.55916709 |
| KLHL36 | 62.3334158 | 0.222455078 | 0.2622568 | 0.84823379 | 0.39630778 | 0.80674902 |
| USP10 | 115.964479 | -0.098775853 | 0.22026222 | -0.4484466 | 0.65383089 | 0.9158679 |
| ZDHHC7 | 59.5344553 | 0.134485803 | 0.21157968 | 0.63562722 | 0.52501942 | 0.86747074 |
| KIAA0513 | 27.8433082 | -0.30639678 | 0.31516258 | -0.9721864 | 0.33095781 | 0.76685572 |
| GSE1 | 341.267092 | -0.234080493 | 0.35348465 | -0.6622084 | 0.50783769 | 0.86281145 |
| C16orf74 | 23.3158995 | 1.733688723 | 0.64855703 | 2.67314767 | 0.00751432 | 0.1252511 |
| EMC8 | 23.5132904 | 0.075634096 | 0.33257077 | 0.22742256 | 0.82009518 | 0.96389705 |
| COX4I1 | 422.665222 | 0.201990659 | 0.39545753 | 0.51077712 | 0.60950713 | 0.89906009 |
| MTHFSD | 15.1003289 | -0.21003668 | 0.37150413 | -0.5653684 | 0.57182324 | 0.88578992 |
| FBXO31 | 21.1850465 | -0.521661573 | 0.45469371 | -1.1472813 | 0.25126539 | 0.70545786 |
| MAP1LC3B | 98.3406303 | -1.010165999 | 0.353494 | -2.857661 | 0.00426776 | 0.08752068 |
| ZCCHC14 | 184.907506 | -1.033765791 | 0.49209081 | -2.1007622 | 0.03566184 | 0.30211342 |
| KLHDC4 | 57.8492401 | -0.103239263 | 0.26932724 | -0.3833228 | 0.70148046 | 0.93127962 |
| BANP | 59.1170412 | -0.018527323 | 0.35770372 | -0.0517952 | 0.95869191 | 0.99238848 |
| ZFPM1 | 19.7381956 | 0.271972047 | 0.35343042 | 0.76952077 | 0.44158422 | 0.8281114 |
| ZC3H18 | 46.7179826 | 0.041282366 | 0.33066885 | 0.12484504 | 0.90064623 | 0.97940229 |
| CYBA | 111.331754 | -1.327609423 | 0.47627073 | -2.78751 | 0.00531148 | 0.10043153 |
| MVD | 39.304187 | -1.072166537 | 0.52989376 | -2.0233613 | 0.04303591 | 0.33179556 |
| SNAI3-AS1 | 11.671512 | -0.836794671 | 0.53067195 | -1.5768587 | 0.11482804 | NA |
| PIEZO1 | 25.0439314 | -0.323833269 | 0.35211351 | -0.9196843 | 0.35773776 | 0.78330841 |
| APRT | 63.8043827 | -0.404110272 | 0.38104281 | -1.0605377 | 0.28890004 | 0.73881634 |
| GALNS | 25.3346511 | 0.023710559 | 0.42544531 | 0.05573116 | 0.95555598 | 0.99228059 |
| TRAPPC2L | 16.5537709 | 0.21287896 | 0.37771331 | 0.5635993 | 0.57302686 | 0.88578992 |
| ACSF3 | 27.2013344 | -0.138496207 | 0.30002445 | -0.4616164 | 0.64435643 | 0.9119294 |
| ANKRD11 | 280.199291 | -0.124941008 | 0.28442486 | -0.439276 | 0.66046158 | 0.91752855 |
| SPG7 | 112.753253 | 0.39148856 | 0.33691348 | 1.16198546 | 0.24524137 | 0.69883704 |
| RPL13 | 508.140839 | 0.264856543 | 0.34703477 | 0.76319888 | 0.4453448 | 0.83035712 |
| CHMP1A | 32.0888631 | 0.139944613 | 0.35010704 | 0.3997195 | 0.68936312 | 0.92734996 |
| CDK10 | 36.7840181 | 0.334982895 | 0.25864051 | 1.29516791 | 0.19526231 | 0.6459894 |
| VPS9D1 | 23.9587637 | 0.426198487 | 0.29440147 | 1.44767787 | 0.14770716 | 0.58466095 |
| ZNF276 | 19.3614287 | 0.507126383 | 0.43290705 | 1.17144403 | 0.24142034 | 0.69556484 |
| SPIRE2 | 29.8369536 | 0.443448328 | 0.36969922 | 1.19948407 | 0.23033977 | 0.68523674 |
| TCF25 | 257.050176 | -0.330454799 | 0.20845641 | -1.5852465 | 0.11291033 | 0.51571293 |
| DEF8 | 20.4725942 | -0.905815502 | 0.32255095 | -2.8082866 | 0.00498059 | 0.09739245 |
| GAS8 | 29.6716136 | -0.31885172 | 0.3004031 | -1.0614129 | 0.2885023 | 0.73815415 |
| FAM157C | 47.4649535 | 0.232388534 | 0.43029915 | 0.54006273 | 0.58915377 | 0.89133386 |
| LINC02193 | 10.7205119 | 0.097676248 | 0.55609554 | 0.17564652 | 0.86057165 | NA |
| RPH3AL | 85.4949192 | 0.310673486 | 0.41714879 | 0.74475461 | 0.45642008 | 0.83506875 |
| VPS53 | 182.730517 | -0.10605199 | 0.18890872 | -0.5613928 | 0.57452981 | 0.88656365 |
| GLOD4 | 43.1459993 | -0.149427249 | 0.28181141 | -0.5302385 | 0.59594661 | 0.89345937 |
| NXN | 13.1096895 | -0.02128752 | 0.38832762 | -0.0548185 | 0.9562831 | 0.99234582 |
| TIMM22 | 14.401458 | -0.205600245 | 0.40810264 | -0.5037954 | 0.61440514 | 0.90054121 |
| ABR | 178.38351 | -0.799520034 | 0.40923078 | -1.9537143 | 0.05073501 | 0.35921868 |
| YWHAE | 260.132726 | -0.040226725 | 0.16175813 | -0.2486844 | 0.80360491 | 0.95938063 |
| CRK | 90.4061932 | -0.038155798 | 0.33828343 | -0.1127924 | 0.91019515 | 0.98334456 |
| MYO1C | 91.8160467 | -0.493945494 | 0.27785481 | -1.7777108 | 0.07545137 | 0.43299712 |
| INPP5K | 15.1181032 | -0.523846531 | 0.49327087 | -1.0619855 | 0.28824225 | 0.73805799 |
| PITPNA | 113.849958 | -0.067425118 | 0.19575204 | -0.3444415 | 0.73051431 | 0.94155624 |
| SLC43A2 | 407.370376 | 0.145283011 | 0.53992349 | 0.26908074 | 0.78786755 | 0.95655322 |
| SCARF1 | 16.5642111 | -0.187211338 | 0.43654209 | -0.4288506 | 0.66803195 | 0.92049996 |
| RILP | 19.5473416 | 0.312799904 | 0.39616028 | 0.78957916 | 0.42977358 | 0.82469033 |
| PRPF8 | 113.574924 | 0.32191113 | 0.22942508 | 1.40312092 | 0.16058079 | 0.60024417 |
| MIR22HG | 49.6172177 | 0.035829234 | 0.57322601 | 0.06250455 | 0.95016104 | 0.99192022 |
| WDR81 | 26.2170685 | -0.826121322 | 0.61402842 | -1.3454122 | 0.17849216 | 0.62617502 |
| SERPINF2 | 33.4668434 | -0.035462666 | 0.45922889 | -0.0772222 | 0.93844678 | 0.98871562 |
| SMYD4 | 28.788982 | -0.096668511 | 0.34403077 | -0.280988 | 0.77871962 | 0.95537097 |
| RPA1 | 40.9619902 | -0.027456821 | 0.28241794 | -0.0972205 | 0.92255127 | 0.98672316 |
| SMG6 | 189.260592 | -0.09903678 | 0.27204463 | -0.3640461 | 0.71582361 | 0.93689626 |
| TSR1 | 20.9740534 | -0.435814928 | 0.40858967 | -1.0666323 | 0.28613792 | 0.73672385 |
| SGSM2 | 60.4051102 | 0.288030256 | 0.21153602 | 1.36161332 | 0.17331995 | 0.61802607 |
| MNT | 24.7672742 | -0.20198941 | 0.29116359 | -0.6937317 | 0.48785049 | 0.85417865 |
| METTL16 | 95.5476179 | 0.12166895 | 0.23067011 | 0.52745867 | 0.59787511 | 0.89386384 |
| PAFAH1B1 | 354.794055 | 0.129932885 | 0.19884417 | 0.65344075 | 0.51347218 | 0.86388299 |
| CLUH | 55.286945 | 0.240715833 | 0.38594292 | 0.62370838 | 0.53281912 | 0.86909171 |
| RAP1GAP2 | 62.6038143 | 0.419685157 | 0.39499143 | 1.06251713 | 0.28800099 | 0.73803612 |
| CTNS | 26.621264 | 0.186751376 | 0.57275901 | 0.32605577 | 0.74438216 | 0.94546379 |
| TAX1BP3 | 16.1247047 | -0.212982474 | 0.49474268 | -0.4304914 | 0.66683822 | 0.92032206 |
| EMC6 | 34.52138 | 0.039943701 | 0.46493387 | 0.08591265 | 0.93153586 | 0.98757533 |
| ITGAE | 17.9641446 | 0.202672612 | 0.37276334 | 0.54370317 | 0.58664575 | 0.89059199 |
| NCBP3 | 80.1786546 | 0.448376234 | 0.22157941 | 2.02354652 | 0.04301683 | 0.33179556 |
| CAMKK1 | 11.5015698 | 0.209263035 | 0.51731932 | 0.40451425 | 0.6858346 | NA |
| ZZEF1 | 122.689981 | 0.045556063 | 0.23930511 | 0.19036811 | 0.84902068 | 0.97027687 |
| CYB5D2 | 37.6560673 | -0.07122754 | 0.23892909 | -0.2981116 | 0.76561796 | 0.95205353 |
| ANKFY1 | 87.299837 | -0.061877391 | 0.24984139 | -0.2476667 | 0.80439231 | 0.95973006 |
| UBE2G1 | 113.19521 | 0.193325292 | 0.23850884 | 0.81055817 | 0.41761945 | 0.8188001 |
| GGT6 | 18.9151275 | 0.682962631 | 0.59548126 | 1.14690868 | 0.25141936 | 0.70545786 |
| SMTNL2 | 16.6707708 | 1.446295809 | 0.54374772 | 2.65986549 | 0.00781719 | 0.12855183 |
| PELP1 | 27.1866936 | 0.073059776 | 0.31856586 | 0.22933963 | 0.81860495 | 0.9635539 |
| MED11 | 18.5063735 | 0.176120434 | 0.39691486 | 0.44372345 | 0.65724254 | 0.9158679 |
| PSMB6 | 46.448581 | -0.110432935 | 0.42064286 | -0.2625337 | 0.79290999 | 0.95746995 |
| MINK1 | 72.7322401 | -0.317665014 | 0.34597629 | -0.9181699 | 0.35852992 | 0.78398112 |
| SLC25A11 | 41.65353 | -0.088672254 | 0.3915375 | -0.2264719 | 0.82083439 | 0.96405056 |
| RNF167 | 23.4515378 | -0.081973276 | 0.31082858 | -0.263725 | 0.79199181 | 0.95746995 |
| PFN1 | 121.05493 | -0.805344599 | 0.36139891 | -2.2284091 | 0.02585325 | 0.25292492 |
| SPAG7 | 45.3249382 | 0.262260605 | 0.26700136 | 0.98224447 | 0.32597942 | 0.76353952 |
| CAMTA2 | 26.9504418 | 0.246274957 | 0.35945248 | 0.68513911 | 0.49325614 | 0.85714521 |
| KIF1C | 32.9390739 | 0.334091666 | 0.45878781 | 0.72820519 | 0.46648799 | 0.84234507 |
| ZNF594 | 16.4830772 | 0.299323496 | 0.44428433 | 0.67372058 | 0.50048897 | 0.85942996 |
| AC087500.1 | 24.4636268 | 0.546913031 | 0.37311357 | 1.46580849 | 0.14270047 | 0.57698814 |
| RABEP1 | 192.211549 | -0.01483273 | 0.25234038 | -0.0587806 | 0.95312682 | 0.99217529 |
| NUP88 | 102.467158 | -0.054133877 | 0.25266217 | -0.214254 | 0.830349 | 0.966913 |
| RPAIN | 45.2114988 | -0.013449461 | 0.26942104 | -0.0499199 | 0.96018625 | 0.99238848 |
| C1QBP | 44.242673 | -0.174535948 | 0.36002435 | -0.4847893 | 0.62782582 | 0.90603677 |
| DERL2 | 35.1331786 | -0.175872872 | 0.33792363 | -0.5204515 | 0.6027489 | 0.8956362 |
| MIS12 | 13.896469 | 0.337483055 | 0.39822982 | 0.84745802 | 0.39673988 | 0.80674902 |
| PITPNM3 | 18.0362408 | 0.0508946 | 0.47478559 | 0.10719491 | 0.91463435 | 0.98407488 |
| KIAA0753 | 43.8878197 | 0.104872476 | 0.35463808 | 0.2957169 | 0.76744629 | 0.95205353 |
| TXNDC17 | 20.2627668 | -0.018904329 | 0.41022209 | -0.0460832 | 0.96324397 | 0.99314929 |
| MED31 | 15.1798309 | -0.08384009 | 0.4372711 | -0.1917348 | 0.84794994 | 0.97027687 |
| XAF1 | 46.4677716 | -2.573704758 | 1.50384165 | -1.7114201 | 0.0870036 | 0.46422617 |
| ALOX12-AS1 | 39.9148152 | -0.861963177 | 0.45448519 | -1.8965705 | 0.05788465 | 0.38201831 |
| SLC16A11 | 14.3472006 | 1.167167375 | 0.47361634 | 2.46437311 | 0.01372532 | 0.17578162 |
| ACADVL | 204.365364 | -0.073253674 | 0.27741181 | -0.2640611 | 0.79173282 | 0.95746995 |
| DVL2 | 17.1206605 | -0.462047327 | 0.38713678 | -1.1934989 | 0.23267403 | 0.68763908 |
| PHF23 | 15.5604038 | -0.010468006 | 0.46225584 | -0.0226455 | 0.98193306 | 0.99714727 |
| ELP5 | 30.488491 | -0.414388252 | 0.38857491 | -1.0664308 | 0.28622895 | 0.73672385 |

| | | | | | | |
|---|---|---|---|---|---|---|
| EIF5A | 35.4255199 | -0.293531643 | 0.36575033 | -0.8025465 | 0.42223691 | 0.82112114 |
| GPS2 | 27.8525511 | -0.185203158 | 0.32024783 | -0.578312 | 0.5630535 | 0.88191823 |
| ZBTB4 | 105.321385 | 0.304616798 | 0.23333889 | 1.30546948 | 0.19173303 | 0.6421543 |
| POLR2A | 129.723006 | 0.019783866 | 0.32570929 | 0.06074087 | 0.95156558 | 0.99217308 |
| EIF4A1 | 39.2401029 | -0.595371384 | 0.37276564 | -1.5971735 | 0.11022705 | 0.5107528 |
| MPDU1 | 33.4141984 | 0.03883497 | 0.33453625 | 0.11608599 | 0.9075844 | 0.98250721 |
| FXR2 | 24.1813969 | 0.456951702 | 0.34133968 | 1.33870082 | 0.18066809 | 0.6286114 |
| SAT2 | 33.721901 | -0.743304323 | 0.46923878 | -1.5840642 | 0.11317912 | 0.51614362 |
| TP53 | 13.8932345 | 0.199338725 | 0.36815375 | 0.5414551 | 0.58819394 | 0.89073775 |
| KDM6B | 31.866168 | -0.06203708 | 0.49124235 | -0.1262861 | 0.89950546 | 0.97936951 |
| NAA38 | 86.8007022 | -0.028538657 | 0.33543044 | -0.0850807 | 0.93219723 | 0.98769122 |
| CHD3 | 31.3223441 | -0.086485115 | 0.34140752 | -0.2533193 | 0.80002148 | 0.95861074 |
| TRAPPC1 | 17.9099522 | -0.964947794 | 0.55268318 | -1.745933 | 0.0808226 | 0.44978727 |
| PER1 | 50.1057909 | 0.028824639 | 0.48327296 | 0.05964464 | 0.95243867 | 0.99217529 |
| TMEM107 | 13.7092448 | 0.231821353 | 0.42347247 | 0.54742957 | 0.58408365 | 0.89036744 |
| CTC1 | 28.2406814 | -0.039317066 | 0.30164424 | -0.1303425 | 0.89629545 | 0.97885533 |
| RANGRF | 14.5312312 | 0.251298673 | 0.50486579 | 0.49775342 | 0.61865785 | 0.90267467 |
| NDEL1 | 61.6531319 | 0.039648901 | 0.30571554 | 0.12969213 | 0.89681001 | 0.97885533 |
| MYH10 | 87.7936174 | 0.138054626 | 0.38602253 | 0.35763359 | 0.72061753 | 0.93806337 |
| NTN1 | 250.496839 | -0.392661101 | 0.34870859 | -1.1260437 | 0.26014704 | 0.71091822 |
| STX8 | 209.464848 | 0.188397138 | 0.2385442 | 0.78977875 | 0.42965699 | 0.82469033 |
| USP43 | 64.0718297 | 0.263631767 | 0.56374802 | 0.46764114 | 0.64004124 | 0.91096398 |
| SCO1 | 33.9582317 | 0.064855859 | 0.34473218 | 0.18813404 | 0.85077157 | 0.97027687 |
| TMEM220 | 34.5846283 | -0.003049043 | 0.27456481 | -0.011105 | 0.99113967 | 0.99923337 |
| TMEM220-A! | 35.9306373 | 0.114731617 | 0.26105223 | 0.43949679 | 0.66030161 | 0.91752855 |
| MAP2K4 | 219.560696 | -0.100768887 | 0.18750826 | -0.5374104 | 0.59098417 | 0.89149106 |
| ARHGAP44 | 21.291691 | -0.337952765 | 0.36876902 | -0.9164348 | 0.35943889 | 0.78473041 |
| ELAC2 | 29.7566189 | -0.148780588 | 0.30396427 | -0.4894674 | 0.62451086 | 0.90447745 |
| COX10 | 68.8410269 | 0.457864427 | 0.29079905 | 1.57450455 | 0.11537085 | 0.52026836 |
| TRIM16 | 11.8815129 | 0.516227812 | 0.40274149 | 1.28178452 | 0.19991825 | NA |
| ZSWIM7 | 49.9678915 | 0.019803658 | 0.25870012 | 0.07655063 | 0.93898103 | 0.98871562 |
| TTC19 | 156.135873 | 0.056867134 | 0.3007813 | 0.18906473 | 0.85004209 | 0.97027687 |
| NCOR1 | 575.91848 | 0.082753673 | 0.29874972 | 0.277 | 0.7817801 | 0.95544332 |
| PIGL | 99.5832643 | 0.271028267 | 0.24885335 | 1.08910836 | 0.27610611 | 0.72575599 |
| CENPV | 11.6724479 | 0.979274834 | 0.42540163 | 2.30200067 | 0.02133513 | NA |
| UBB | 230.388954 | -0.166052088 | 0.43238311 | -0.3840393 | 0.70094935 | 0.93127962 |
| LRRC75A | 25.3167651 | 0.330750564 | 0.45673359 | 0.72416519 | 0.46896433 | 0.84326495 |
| AC098850.3 | 67.3821272 | -0.257165921 | 0.55680145 | -0.4618629 | 0.64417966 | 0.9119294 |
| MPRIP | 188.289024 | -0.098282534 | 0.20318105 | -0.483719 | 0.6285853 | 0.90652355 |
| COPS3 | 27.3010335 | 0.177421102 | 0.27539794 | 0.64423541 | 0.5194228 | 0.86618562 |
| MED9 | 53.502139 | -0.046598926 | 0.28843925 | -0.1615554 | 0.87165596 | 0.97434023 |
| RASD1 | 182.51818 | -1.920869191 | 0.45779182 | -4.1959448 | 2.72E-05 | 0.00262501 |
| PEMT | 12.5673066 | 0.437526378 | 0.52169266 | 0.83866693 | 0.40165624 | 0.80890307 |
| RAI1 | 89.8527637 | -0.397863566 | 0.35690457 | -1.1147618 | 0.26495252 | 0.71628748 |
| SREBF1 | 167.363253 | -0.774882242 | 0.37699606 | -2.0554121 | 0.03983922 | 0.31990103 |
| TOM1L2 | 125.756233 | 0.459925286 | 0.20353339 | 2.25970432 | 0.02383961 | 0.24214052 |
| ATPAF2 | 17.5371918 | 0.287360348 | 0.3181102 | 0.90333584 | 0.36634768 | 0.78793182 |
| GID4 | 11.3469183 | -0.348464214 | 0.43733683 | -0.7967868 | 0.42557487 | NA |
| DRG2 | 14.4268427 | 0.586451469 | 0.43811291 | 1.33858523 | 0.18070574 | 0.6286114 |
| ALKBH5 | 61.9858089 | 0.039411479 | 0.25440898 | 0.15491387 | 0.87688922 | 0.97532685 |
| FLII | 52.98968 | -0.643692138 | 0.25410767 | -2.5331472 | 0.01130435 | 0.15732016 |
| MIEF2 | 11.706851 | 0.361467579 | 0.36876682 | 0.98020636 | 0.32698427 | NA |
| TOP3A | 31.6740415 | -0.249793521 | 0.27273945 | -0.9158687 | 0.35973578 | 0.7850252 |
| SMCR8 | 19.6150519 | -0.372833334 | 0.40139979 | -0.9288329 | 0.35297568 | 0.78010119 |
| SHMT1 | 27.4804583 | 0.270070287 | 0.36128966 | 0.74751735 | 0.45475134 | 0.83385406 |
| TVP23B | 23.3551445 | 0.017030734 | 0.3483062 | 0.04889587 | 0.96100228 | 0.99238848 |
| PRPSAP2 | 63.5798515 | 0.348431461 | 0.23999467 | 1.45183002 | 0.14654887 | 0.58349206 |
| FAM83G | 10.4907816 | -1.259105241 | 0.78299564 | -1.6080616 | 0.10782168 | NA |
| EPN2 | 141.073717 | -0.536696681 | 0.26884053 | -1.9963384 | 0.0458971 | 0.3409232 |
| B9D1 | 16.430026 | -0.382604435 | 0.43066038 | -0.8884134 | 0.37431844 | 0.7923487 |
| AC004448.2 | 23.654942 | 0.289657606 | 0.39148759 | 0.73988962 | 0.45936698 | 0.83645518 |
| SLC47A1 | 29.5526908 | 0.540877683 | 0.391771 | 1.38059652 | 0.16740305 | 0.60898823 |
| ALDH3A2 | 40.1107712 | 0.360866719 | 0.34003134 | 1.06127486 | 0.28856501 | 0.73815415 |
| ULK2 | 36.7472969 | 0.207275742 | 0.32413804 | 0.6394675 | 0.52251885 | 0.86728831 |
| AKAP10 | 123.83025 | -0.016055323 | 0.17339498 | -0.0925939 | 0.92622616 | 0.98672316 |
| SPECC1 | 35.2478808 | -0.273023809 | 0.35400651 | -0.7712395 | 0.44056498 | 0.82793978 |
| CCDC144NL-/ | 36.465478 | -0.682448858 | 0.50159945 | -1.3605455 | 0.17365738 | 0.61861056 |
| USP22 | 147.60874 | 0.279415412 | 0.21263693 | 1.31404931 | 0.18882961 | 0.64021224 |
| DHRS7B | 95.3374772 | -0.045420716 | 0.2144255 | -0.2118252 | 0.83224344 | 0.96701848 |
| TMEM11 | 18.1161841 | -0.219087037 | 0.38515311 | -0.568831 | 0.56947082 | 0.88562534 |
| AC087294.1 | 17.1384687 | -0.462804508 | 0.39736759 | -1.164676 | 0.24415015 | 0.69773111 |
| NATD1 | 10.6116636 | -0.277719468 | 0.4797476 | -0.5788866 | 0.56266568 | NA |
| MAP2K3 | 21.0244496 | 0.310876017 | 0.43654364 | 0.71213044 | 0.476384 | 0.84663915 |
| WSB1 | 598.218415 | -0.883808088 | 0.3308635 | -2.6712167 | 0.00755768 | 0.12565432 |
| KSR1 | 47.4038791 | 0.037334087 | 0.24794917 | 0.15057154 | 0.88031372 | 0.97600218 |
| NLK | 211.321572 | 0.120511623 | 0.17916104 | 0.67264412 | 0.50117372 | 0.85942996 |
| IFT20 | 14.369875 | 0.119054299 | 0.35663621 | 0.3338256 | 0.73851116 | 0.94376279 |
| TNFAIP1 | 32.5547252 | -0.890778726 | 0.36847622 | -2.417466 | 0.01562899 | 0.19117434 |
| POLDIP2 | 34.1389765 | -0.315959231 | 0.3802454 | -0.830935 | 0.40601033 | 0.8115705 |
| TMEM199 | 11.0059143 | -0.243665023 | 0.4712476 | -0.5170637 | 0.6051117 | NA |
| SARM1 | 12.7336147 | -0.01293842 | 0.51436375 | -0.0251542 | 0.97993195 | 0.99664466 |
| AC005726.1 | 15.8812188 | -0.441606543 | 0.37306233 | -1.1837339 | 0.2365184 | 0.69144921 |
| KIAA0100 | 64.2897382 | 0.374889496 | 0.30324033 | 1.23627849 | 0.21635506 | 0.67099191 |
| SDF2 | 27.0854415 | -0.098875387 | 0.30665488 | -0.3224321 | 0.74712534 | 0.9462156 |
| SUPT6H | 75.6433174 | -0.450288109 | 0.30461019 | -1.4782437 | 0.13934255 | 0.57281662 |
| RAB34 | 34.5266132 | -0.459780333 | 0.34294664 | -1.340676 | 0.18002567 | 0.62768048 |
| RPL23A | 169.068971 | 0.381900808 | 0.38411606 | 0.99423285 | 0.32010952 | 0.76106474 |
| NEK8 | 17.003977 | -0.071315299 | 0.40793589 | -0.1748199 | 0.86122117 | 0.97302617 |
| TRAF4 | 43.9161864 | 0.161700994 | 0.30132714 | 0.53662936 | 0.59152366 | 0.89151328 |
| FAM222B | 133.784342 | 0.044256163 | 0.28635331 | 0.15455091 | 0.87717538 | 0.97541592 |
| ERAL1 | 12.630577 | 0.054727779 | 0.40400064 | 0.13546458 | 0.89224456 | 0.97794037 |
| FLOT2 | 103.647477 | -0.635440219 | 0.22393943 | -2.8375539 | 0.00454607 | 0.09167744 |
| PHF12 | 55.723238 | -0.009377072 | 0.30975061 | -0.030273 | 0.97584935 | 0.99556048 |
| MYO18A | 49.889543 | -0.501704357 | 0.39104605 | -1.2829803 | 0.19949899 | 0.65266113 |
| NUFIP2 | 170.528988 | -0.190817397 | 0.23994681 | -0.7952487 | 0.42646882 | 0.82329929 |
| TAOK1 | 245.61275 | -0.133057123 | 0.17476734 | -0.7613386 | 0.44645485 | 0.83035712 |
| ABHD15-AS1 | 23.31532 | -0.533647788 | 0.53380859 | -0.9996988 | 0.31745631 | 0.75958338 |
| GIT1 | 15.2789437 | 0.025959315 | 0.43408993 | 0.0598017 | 0.95231357 | 0.99217529 |
| SSH2 | 353.616359 | -0.138638025 | 0.25576541 | -0.5420515 | 0.58778303 | 0.89073775 |
| AC023389.1 | 15.61205 | -0.084841364 | 0.41612701 | -0.2038833 | 0.83844467 | 0.96748214 |
| NSRP1 | 155.961531 | 0.025357317 | 0.26150252 | 0.09696777 | 0.92275199 | 0.98672316 |
| AC104984.4 | 18.9897141 | -0.247558949 | 0.45766346 | -0.5409192 | 0.58856328 | 0.89093433 |
| BLMH | 31.2080751 | 0.182561828 | 0.28958226 | 0.63043167 | 0.52841219 | 0.86750588 |
| CPD | 151.75137 | -0.373612955 | 0.22376959 | -1.6696324 | 0.09499212 | 0.48035393 |
| GOSR1 | 99.2165608 | -0.075367736 | 0.17999863 | -0.4187128 | 0.67542602 | 0.92193843 |
| CRLF3 | 68.5484012 | 0.038233047 | 0.29436156 | 0.12988465 | 0.89665769 | 0.97885533 |
| TEFM | 24.9900177 | 0.217165003 | 0.37267589 | 0.58271814 | 0.56008306 | 0.88037687 |
| RNF135 | 17.0606924 | 0.13867283 | 0.32968482 | 0.42062243 | 0.67403081 | 0.92189158 |
| NF1 | 606.804151 | -0.265999586 | 0.22199345 | -1.1982317 | 0.23082684 | 0.6853319 |
| OMG | 16.6422253 | -0.596801863 | 0.75956132 | -0.7857191 | 0.43203205 | 0.8248269 |
| RAB11FIP4 | 42.2131428 | 0.027010357 | 0.30577507 | 0.08833407 | 0.92961116 | 0.98733028 |
| COPRS | 49.8120149 | 0.137123809 | 0.39318257 | 0.34875353 | 0.72727436 | 0.94011548 |
| UTP6 | 30.0407188 | 0.322324957 | 0.30542009 | 1.05534957 | 0.29126548 | 0.73994767 |

| | | | | | | |
|---|---|---|---|---|---|---|
| SUZ12 | 164.158936 | 0.256340713 | 0.19047115 | 1.34582438 | 0.17835916 | 0.62598987 |
| LRRC37B | 45.4495422 | -0.344651222 | 0.24702725 | -1.3951951 | 0.16295701 | 0.60147753 |
| RHOT1 | 82.0148441 | 0.419635729 | 0.1988833 | 2.10995962 | 0.03486183 | 0.29846407 |
| C17orf75 | 16.1121727 | 0.537875244 | 0.43085606 | 1.24838733 | 0.21188925 | 0.66386489 |
| ZNF207 | 183.895938 | 0.113220477 | 0.19544295 | 0.57930192 | 0.56238547 | 0.88191823 |
| PSMD11 | 70.5267943 | 0.000788679 | 0.20751843 | 0.00380052 | 0.99696763 | 0.99975875 |
| MYO1D | 374.217954 | -0.153190973 | 0.23370895 | -0.6554776 | 0.51216033 | 0.86346419 |
| TMEM98 | 13.4952098 | -0.179583149 | 0.48979476 | -0.3666498 | 0.71388026 | 0.93650155 |
| ASIC2 | 37.857297 | -1.251121164 | 0.6361422 | -1.9667319 | 0.04921413 | 0.35506449 |
| ZNF830 | 10.8831151 | 0.083633923 | 0.51277196 | 0.16310159 | 0.87043845 | NA |
| LIG3 | 16.4149154 | 0.093158533 | 0.33091979 | 0.28151393 | 0.77831624 | 0.95535696 |
| RFFL | 13.7836436 | -0.685400444 | 0.43554955 | -1.5736452 | 0.1155695 | 0.52043834 |
| SLFN5 | 33.7258996 | -1.900714328 | 0.45381259 | -4.1883243 | 2.81E-05 | 0.0026346 |
| SLFN13 | 16.5063444 | -0.274624265 | 0.49504378 | -0.5547474 | 0.57906743 | 0.88946224 |
| LINC02001 | 13.9988047 | 0.499649571 | 0.33968398 | 1.47092475 | 0.14131147 | 0.57617539 |
| AP2B1 | 160.382889 | -0.216373104 | 0.21240903 | -1.0186625 | 0.30836323 | 0.75132915 |
| TAF15 | 255.18003 | -0.209600653 | 0.1794522 | -1.1680027 | 0.24280569 | 0.69621873 |
| AC015849.3 | 17.8730222 | -0.697632983 | 0.49974645 | -1.3959739 | 0.16272236 | 0.60133611 |
| ZNHIT3 | 18.3137194 | -0.05856973 | 0.36577627 | -0.1601245 | 0.87278303 | 0.97434023 |
| MYO19 | 171.214477 | -0.048287898 | 0.28463816 | -0.1696466 | 0.86528807 | 0.97434023 |
| GGNBP2 | 111.659615 | 0.031906264 | 0.247164 | 0.12908945 | 0.89728687 | 0.97892709 |
| AATF | 60.9816972 | 0.150404296 | 0.27086109 | 0.55528203 | 0.57870177 | 0.88932306 |
| ACACA | 295.671659 | 0.059831312 | 0.286442 | 0.20887758 | 0.83454381 | 0.96703798 |
| TADA2A | 40.9890635 | 0.2900733 | 0.3228344 | 0.89852041 | 0.36890817 | 0.78889478 |
| SYNRG | 152.311057 | -0.201193404 | 0.16524204 | -1.2175679 | 0.22338821 | 0.67606266 |
| DDX52 | 151.442345 | -1.120987839 | 0.3055319 | -3.6689715 | 0.00024353 | 0.01249854 |
| AC243571.2 | 81.7764815 | -4.26334534 | 0.73766136 | -5.7795427 | 7.49E-09 | 3.03E-06 |
| HNF1B | 154.221172 | -0.909156732 | 0.21762497 | -4.1776306 | 2.95E-05 | 0.00272119 |
| MRPL45 | 28.1214663 | 0.380301456 | 0.31238372 | 1.21741767 | 0.22344534 | 0.67606266 |
| SOCS7 | 26.0842504 | 0.535369185 | 0.27136768 | 1.97285537 | 0.04851204 | 0.35179202 |
| MLLT6 | 201.787266 | -0.410973012 | 0.27306475 | -1.5050387 | 0.13231413 | 0.55993651 |
| CISD3 | 26.5486906 | 0.269257959 | 0.35041795 | 0.76839088 | 0.442255 | 0.82820049 |
| PCGF2 | 22.8420345 | -0.37094181 | 0.40204994 | -0.9226262 | 0.35620203 | 0.78188723 |
| PSMB3 | 53.8918668 | -0.557247556 | 0.26978968 | -2.0654888 | 0.03887678 | 0.31669347 |
| PIP4K2B | 26.9133784 | 0.571293439 | 0.28043889 | 2.0371406 | 0.04163595 | 0.32781555 |
| CWC25 | 28.4142664 | -0.481331871 | 0.30520453 | -1.5770797 | 0.11477719 | 0.51928773 |
| RPL23 | 146.882736 | 0.38750829 | 0.36911881 | 1.04981996 | 0.2938009 | 0.74041796 |
| LASP1 | 43.2650678 | -0.039389136 | 0.32405485 | -0.1215508 | 0.90325476 | 0.98016176 |
| RPL19 | 250.255385 | 0.006759917 | 0.45162905 | 0.01496785 | 0.98805783 | 0.99907859 |
| STAC2 | 297.181868 | -0.635813629 | 0.39363878 | -1.615221 | 0.10626284 | 0.50162992 |
| FBXL20 | 265.874995 | -0.504702032 | 0.22352238 | -2.2579485 | 0.02394887 | 0.24248855 |
| MED1 | 49.9775158 | 0.36500116 | 0.26681874 | 1.36797424 | 0.17132014 | 0.61480036 |
| CDK12 | 104.75598 | 0.089003172 | 0.1879966 | 0.47342968 | 0.63590666 | 0.90881415 |
| STARD3 | 39.9430433 | 0.036957832 | 0.25669293 | 0.14397682 | 0.88551877 | 0.97662748 |
| PGAP3 | 28.0693788 | 0.347405321 | 0.47723428 | 0.7279555 | 0.46664083 | 0.84243738 |
| ERBB2 | 150.798708 | 0.593342505 | 0.30598811 | 1.93910317 | 0.05248878 | 0.36358594 |
| MIEN1 | 44.3741564 | -1.118933877 | 0.28771222 | -3.8890731 | 0.00010063 | 0.00655094 |
| GRB7 | 67.4091266 | -0.925769766 | 0.40848648 | -2.2663413 | 0.0234305 | 0.23883233 |
| ORMDL3 | 14.1543582 | 0.458533624 | 0.34735049 | 1.3200892 | 0.18680524 | 0.63791877 |
| PSMD3 | 29.4520961 | 0.035505039 | 0.29965437 | 0.11848664 | 0.90568208 | 0.98124832 |
| MED24 | 37.9493872 | 0.154355713 | 0.25425854 | 0.60708171 | 0.54379669 | 0.87243854 |
| THRA | 26.000858 | 0.455201526 | 0.30389651 | 1.49788337 | 0.13416355 | 0.56229288 |
| MSL1 | 53.8454122 | -0.290954623 | 0.27556708 | -1.0558395 | 0.29104153 | 0.73994767 |
| CASC3 | 50.0357989 | 0.055306133 | 0.19875446 | 0.27826361 | 0.78081001 | 0.95537097 |
| WIPF2 | 108.346456 | -0.043894807 | 0.22273671 | -0.1970704 | 0.84377247 | 0.9697385 |
| RARA | 11.2126398 | 0.205693444 | 0.41678589 | 0.49352305 | 0.62164306 | NA |
| IGFBP4 | 11.9813163 | -0.724029484 | 0.43704767 | -1.6566373 | 0.09759282 | NA |
| TNS4 | 15.9644128 | 1.748811653 | 0.61676128 | 2.83547573 | 0.00457575 | 0.09208458 |
| KRT10 | 43.5780928 | -0.160883287 | 0.49659994 | -0.3239696 | 0.74596105 | 0.94614106 |
| KRT19 | 19.5412711 | -1.130690157 | 0.69849655 | -1.6187484 | 0.10550141 | 0.50066717 |
| EIF1 | 361.486412 | -0.581350696 | 0.3649234 | -1.593076 | 0.11114315 | 0.5130256 |
| JUP | 143.5596 | -0.262458694 | 0.31178594 | -0.8417913 | 0.39990478 | 0.80830282 |
| NT5C3B | 24.9130751 | 0.037941198 | 0.34289344 | 0.11065011 | 0.91189381 | 0.98354625 |
| ACLY | 42.3254175 | -0.423140254 | 0.32361606 | -1.3075379 | 0.1910301 | 0.6421543 |
| CNP | 30.8571451 | -0.347683413 | 0.35715458 | -0.9734816 | 0.330314 | 0.76685572 |
| DNAJC7 | 83.9050376 | -0.075999037 | 0.21384947 | -0.3553857 | 0.72230068 | 0.93855548 |
| NKIRAS2 | 12.1817743 | 0.274456159 | 0.48874729 | 0.56155024 | 0.57442249 | NA |
| GHDC | 10.3887804 | -0.288665139 | 0.43432633 | -0.6646273 | 0.5062889 | NA |
| STAT5B | 196.069394 | -1.114942436 | 0.40659603 | -2.742138 | 0.00610407 | 0.10985059 |
| STAT5A | 21.4367294 | -1.562330934 | 0.61560643 | -2.537873 | 0.01115285 | 0.15593979 |
| STAT3 | 272.4694 | -0.818642656 | 0.27138419 | -3.0165452 | 0.00255673 | 0.06093438 |
| ATP6V0A1 | 66.641234 | -0.511722216 | 0.27611648 | -1.8532839 | 0.06384168 | 0.40107793 |
| NAGLU | 27.1198363 | -1.350910769 | 0.63407164 | -2.1305333 | 0.03312761 | 0.28919158 |
| COASY | 25.765068 | -0.03768916 | 0.39126709 | -0.0963259 | 0.92326173 | 0.98672316 |
| MLX | 24.790947 | -0.088017894 | 0.43779867 | -0.2010465 | 0.84066221 | 0.96748214 |
| RETREG3 | 76.7421307 | 0.203438906 | 0.2412839 | 0.8431516 | 0.39914366 | 0.80778083 |
| TUBG1 | 10.5330652 | 0.082968909 | 0.54750476 | 0.15154007 | 0.8795497 | NA |
| TUBG2 | 15.8549163 | -0.074101934 | 0.39725582 | -0.1865345 | 0.85202559 | 0.97053536 |
| PLEKHH3 | 17.2071063 | 0.542551816 | 0.41116819 | 1.31953744 | 0.1869895 | 0.6379874 |
| EZH1 | 51.403562 | -0.206711332 | 0.2563984 | -0.8062115 | 0.42012092 | 0.81913024 |
| VPS25 | 25.6001417 | -0.210916177 | 0.29712537 | -0.7098558 | 0.47779354 | 0.84771464 |
| COA3 | 103.490922 | -0.155222061 | 0.40318963 | -0.3849853 | 0.70024834 | 0.93127962 |
| BECN1 | 40.9268387 | -0.463828683 | 0.33309213 | -1.3924937 | 0.16377296 | 0.60357056 |
| PSME3 | 27.5601843 | 0.048723461 | 0.32666657 | 0.14915349 | 0.88143252 | 0.97630358 |
| LINC00671 | 32.2641761 | 1.464575821 | 0.65805615 | 2.22560919 | 0.02604037 | 0.25383383 |
| RUNDC1 | 32.365746 | 0.204841029 | 0.31726667 | 0.64564307 | 0.51851054 | 0.86552266 |
| RPL27 | 105.353607 | -0.131778577 | 0.31193686 | -0.4224527 | 0.6726946 | 0.92189158 |
| IFI35 | 26.2937053 | -1.522403506 | 0.73121258 | -2.0820259 | 0.03734011 | 0.30957182 |
| VAT1 | 16.6334603 | -1.374448557 | 0.56706032 | -2.4238137 | 0.01535848 | 0.18844914 |
| BRCA1 | 21.2905778 | -0.472650805 | 0.60383925 | -0.7827428 | 0.43377818 | 0.82504445 |
| NBR1 | 96.4782554 | -0.16037384 | 0.25371701 | -0.6320973 | 0.52732329 | 0.86750588 |
| LINC00854 | 106.444489 | 0.315236119 | 0.59620043 | 0.52874185 | 0.59698454 | 0.89363331 |
| ARL4D | 105.77811 | -0.560496939 | 0.36108008 | -1.5522788 | 0.12059554 | 0.53341392 |
| DHX8 | 33.198638 | -0.062397998 | 0.29527295 | -0.2113231 | 0.83263515 | 0.96701848 |
| DUSP3 | 66.541481 | -0.130340149 | 0.34278948 | -0.3802338 | 0.70377187 | 0.93306088 |
| PYY | 125.756166 | -2.380074036 | 0.66527305 | -3.5775897 | 0.00034678 | 0.0157922 |
| TMEM101 | 980.802584 | -1.80452638 | 0.51640817 | -3.49438 | 0.00047516 | 0.01945996 |
| LSM12 | 59.8344361 | -0.140580837 | 0.28499295 | -0.4932783 | 0.62181596 | 0.90401901 |
| G6PC3 | 18.8667549 | 0.107931894 | 0.37698293 | 0.28630446 | 0.77464494 | 0.95470568 |
| HDAC5 | 66.5529104 | 0.264742261 | 0.34002572 | 0.77859482 | 0.43621843 | 0.82656073 |
| ASB16-AS1 | 14.3269447 | -0.153024873 | 0.36582849 | -0.4182968 | 0.67573016 | 0.92193843 |
| TMUB2 | 26.338265 | 0.0081768 | 0.28603335 | 0.02858688 | 0.97719408 | 0.9957152 |
| ATXN7L3 | 20.386037 | 0.153062177 | 0.52187536 | 0.29329259 | 0.76929851 | 0.95205353 |
| UBTF | 60.6881955 | 0.468844294 | 0.34553031 | 1.35688327 | 0.1748183 | 0.62071528 |
| AC003102.1 | 12.3971552 | -0.802444581 | 0.49411994 | -1.6239874 | 0.10437849 | NA |
| SLC4A1 | 876.582799 | 3.52381078 | 0.73393201 | 4.80127689 | 1.58E-06 | 0.00027805 |
| RUNDC3A-AS | 14.2501449 | 0.540194332 | 0.4775738 | 1.13112222 | 0.25800365 | 0.71022118 |
| SLC25A39 | 202.604264 | 0.320323312 | 0.27255563 | 1.17525848 | 0.23989132 | 0.6944034 |
| GRN | 85.3615813 | -0.895514606 | 0.44401325 | -2.0168644 | 0.04370967 | 0.33268382 |
| GPATCH8 | 413.626582 | 0.070320331 | 0.15796484 | 0.44516446 | 0.65620091 | 0.9158679 |
| MEIOC | 21.2537582 | 0.311345212 | 0.41995058 | 0.74138536 | 0.45845982 | 0.83645518 |
| CCDC43 | 15.9969613 | 0.743167533 | 0.38266383 | 1.94208981 | 0.05212623 | 0.36245478 |
| DBF4B | 13.1994927 | -0.007441964 | 0.37217481 | -0.0199959 | 0.98404666 | 0.99762255 |

| | | | | | | |
|---|---|---|---|---|---|---|
| EFTUD2 | 40.1619547 | -0.042230046 | 0.27703907 | -0.1524335 | 0.878845 | 0.97556614 |
| DCAKD | 41.1702782 | 0.300311294 | 0.31707745 | 0.94712283 | 0.34357619 | 0.77370523 |
| NMT1 | 78.1567043 | -0.288028706 | 0.18238094 | -1.5792698 | 0.11427418 | 0.51869892 |
| HEXIM1 | 18.4423306 | 0.510436685 | 0.63823143 | 0.79976738 | 0.42384558 | 0.82218966 |
| AC138150.1 | 36.6292241 | 0.432419814 | 0.37980193 | 1.13854032 | 0.25489493 | 0.70844723 |
| MAP3K14 | 36.3182993 | -0.636709244 | 0.27405857 | -2.3232597 | 0.0201652 | 0.22331327 |
| ARHGAP27 | 26.5527479 | 0.20993252 | 0.43674478 | 0.48067551 | 0.63074714 | 0.90768383 |
| PLEKHM1 | 40.5619763 | -0.366283376 | 0.3540033 | -1.0346892 | 0.3008141 | 0.74493152 |
| LINC02210-C | 24.6662324 | 0.410805202 | 0.39860478 | 1.03060781 | 0.30272477 | 0.74712557 |
| KANSL1 | 535.251765 | 0.478336337 | 0.31543479 | 1.51643495 | 0.12940941 | 0.55444843 |
| ARL17B | 115.422529 | 1.897483447 | 1.00212841 | 1.8934534 | 0.0582976 | 0.38243048 |
| LRRC37A2 | 33.104959 | 0.327066433 | 0.37787783 | 0.86553485 | 0.3867453 | 0.80009555 |
| ARL17A | 33.3539046 | -0.097554376 | 0.33934608 | -0.2874775 | 0.77374669 | 0.95457609 |
| NSF | 141.835614 | 0.077784501 | 0.35134345 | 0.22139164 | 0.8247875 | 0.96539788 |
| AC005670.2 | 30.2512535 | -0.06677203 | 0.32960257 | -0.2025835 | 0.83946061 | 0.96748214 |
| CDC27 | 181.433167 | -0.309288529 | 0.21492625 | -1.4390449 | 0.1501378 | 0.58675934 |
| AC068234.1 | 176.278585 | -1.724196008 | 0.71716387 | -2.4041869 | 0.01620848 | 0.1962825 |
| AC068234.2 | 23.1921073 | -1.679136063 | 0.51276948 | -3.2746412 | 0.00105796 | 0.03398092 |
| EFCAB13 | 145.171847 | -1.097240224 | 0.49738517 | -2.2060172 | 0.0273828 | 0.26050089 |
| NPEPPS | 316.649183 | -0.033673376 | 0.17430238 | -0.1931894 | 0.84681062 | 0.97027687 |
| AC025682.1 | 29.2703306 | -0.201926794 | 0.33496895 | -0.6028224 | 0.54662683 | 0.87269272 |
| KPNB1 | 135.954777 | -0.176740103 | 0.23546466 | -0.7506014 | 0.45289258 | 0.83343921 |
| MRPL10 | 11.6115088 | -0.095195426 | 0.4576303 | -0.2080182 | 0.83521477 | NA |
| SCRN2 | 19.4271514 | 0.086157202 | 0.42226452 | 0.20403609 | 0.8383253 | 0.96748214 |
| SP2 | 18.0964496 | 0.204650959 | 0.37360238 | 0.54777745 | 0.58384472 | 0.89036744 |
| PNPO | 23.5457136 | 0.218367403 | 0.3219956 | 0.6781689 | 0.49766461 | 0.85818031 |
| PRR15L | 12.0871341 | -0.734546213 | 0.73504107 | -0.9993268 | 0.31763643 | NA |
| CDK5RAP3 | 33.9196012 | 0.19534141 | 0.26271513 | 0.74354837 | 0.45714975 | 0.83569193 |
| NFE2L1 | 148.15875 | -0.172560323 | 0.33069895 | -0.5218049 | 0.6018062 | 0.89522668 |
| CBX1 | 33.0171752 | 0.305238011 | 0.29829614 | 1.02327174 | 0.30617938 | 0.74897897 |
| SNX11 | 16.2403889 | -0.195974252 | 0.39361298 | -0.4978856 | 0.61856465 | 0.90267467 |
| SKAP1 | 568.465237 | -0.341761914 | 0.2580286 | -1.3245118 | 0.18533314 | 0.63578845 |
| SKAP1-AS1 | 12.7123396 | -1.071200337 | 0.68059409 | -1.5739195 | 0.11550605 | 0.52043834 |
| HOXB2 | 33.175338 | -0.692292815 | 0.35531931 | -1.9483681 | 0.05137093 | 0.36077232 |
| HOXB-AS1 | 15.0917525 | -0.107171814 | 0.47252475 | -0.2268068 | 0.820574 | 0.96389705 |
| HOXB3 | 199.583885 | -0.271058598 | 0.20313019 | -1.3344082 | 0.1820701 | 0.63058858 |
| HOXB-AS3 | 61.764788 | 0.021973254 | 0.43259502 | 0.05079405 | 0.95948963 | 0.99238848 |
| HOXB7 | 54.1240031 | -0.342191691 | 0.3563125 | -0.9603696 | 0.33686924 | 0.76939761 |
| AC103702.2 | 29.8520773 | -0.509721387 | 0.34090144 | -1.4952163 | 0.134858 | 0.56408911 |
| LINC02086 | 64.2999572 | -0.606268085 | 0.27681146 | -2.1901842 | 0.02851088 | 0.26643114 |
| CALCOCO2 | 77.7889067 | -0.132142842 | 0.25461672 | -0.5189873 | 0.6037696 | 0.8956362 |
| ATP5MC1 | 98.0098365 | 0.480259063 | 0.39275595 | 1.22279257 | 0.22140805 | 0.67492938 |
| UBE2Z | 104.71963 | -0.236536537 | 0.26206671 | -0.9025814 | 0.36674811 | 0.78809407 |
| SNF8 | 32.8180145 | 0.295070264 | 0.25087754 | 1.17615259 | 0.23953391 | 0.69419674 |
| B4GALNT2 | 27.1610337 | -0.649253229 | 0.646416 | -1.0043892 | 0.31519107 | 0.75786443 |
| ZNF652 | 189.440768 | -0.14760428 | 0.20932019 | -0.7051603 | 0.48071051 | 0.84961003 |
| AC091180.5 | 15.0668682 | -0.686253149 | 0.58086635 | -1.1814304 | 0.2374318 | 0.69207518 |
| PHB | 105.462578 | -0.405641893 | 0.3059794 | -1.3257163 | 0.18493367 | 0.63544336 |
| SPOP | 103.954283 | -0.260735115 | 0.25112161 | -1.0382823 | 0.29913867 | 0.74324925 |
| SLC35B1 | 16.7776722 | -0.425334971 | 0.3493864 | -1.217377 | 0.2234608 | 0.67606266 |
| FAM117A | 16.4072457 | -0.035851329 | 0.41255802 | -0.0869001 | 0.93075093 | 0.98756115 |
| KAT7 | 37.5792777 | -0.193487767 | 0.24913302 | -0.7766444 | 0.4373686 | 0.82687287 |
| ITGA3 | 123.539379 | -1.766666908 | 0.44895408 | -3.9350726 | 8.32E-05 | 0.00564171 |
| PDK2 | 13.5181482 | 0.892593599 | 0.38945383 | 2.2919112 | 0.02191077 | 0.23177148 |
| XYLT2 | 18.4527509 | 0.092534453 | 0.40633385 | 0.22773011 | 0.81985606 | 0.96389705 |
| MRPL27 | 54.5824511 | -0.404260579 | 0.28519162 | -1.4175051 | 0.15633531 | 0.59679357 |
| LRRC59 | 20.5515529 | -0.14091806 | 0.48314328 | -0.2916693 | 0.77053949 | 0.9522529 |
| ACSF2 | 16.5625269 | 0.820970404 | 0.45520298 | 1.80352597 | 0.07130565 | 0.422778 |
| RSAD1 | 22.4751411 | 0.884194777 | 0.34926343 | 2.53159847 | 0.01135439 | 0.15779028 |
| EPN3 | 13.95875 | 1.529345918 | 0.56256912 | 2.71850314 | 0.0065578 | 0.11607791 |
| SPATA20 | 22.4020382 | 0.09629384 | 0.35795938 | 0.26900773 | 0.78792373 | 0.95655322 |
| ABCC3 | 108.21528 | -0.3152425 | 0.5962447 | -0.5287133 | 0.59700435 | 0.89363331 |
| ANKRD40 | 39.3813055 | 0.328400894 | 0.3431471 | 0.95702658 | 0.33855384 | 0.76998177 |
| AC005921.2 | 11.516044 | 0.15327707 | 0.45974358 | 0.33339687 | 0.73883472 | NA |
| LUC7L3 | 247.553516 | 0.333156825 | 0.15753604 | 2.11479746 | 0.03444721 | 0.29622161 |
| TOB1 | 54.206948 | 0.316572468 | 0.2849644 | 1.11091934 | 0.26660307 | 0.71754988 |
| SPAG9 | 466.010608 | -0.310344016 | 0.24793415 | -1.2517195 | 0.21067209 | 0.66241521 |
| AC005920.2 | 36.2851454 | -0.505924965 | 0.55817951 | -0.906384 | 0.36473265 | 0.78793182 |
| MBTD1 | 99.4758545 | 0.06614255 | 0.23988523 | 0.27572581 | 0.78275867 | 0.95544332 |
| UTP18 | 32.3359075 | -0.059589087 | 0.27460981 | -0.2169955 | 0.82821187 | 0.96636016 |
| TOM1L1 | 195.193246 | -0.204093959 | 0.22679042 | -0.8999232 | 0.36816112 | 0.78877653 |
| COX11 | 33.5377959 | 0.677152983 | 0.40926148 | 1.65457297 | 0.09801115 | 0.48431389 |
| STXBP4 | 132.451903 | 0.387144885 | 0.39773855 | 0.97336525 | 0.33037181 | 0.76685572 |
| MMD | 15.9533836 | -0.448102823 | 0.45363785 | -0.9877986 | 0.32325131 | 0.76304922 |
| PCTP | 21.753346 | -0.299555462 | 0.42031032 | -0.7127007 | 0.47603098 | 0.84662643 |
| DGKE | 11.4056924 | -0.166090346 | 0.45267063 | -0.3669121 | 0.71368456 | NA |
| TRIM25 | 25.9571518 | 0.066420671 | 0.41999256 | 0.15814726 | 0.87434076 | 0.97434023 |
| COIL | 20.1218447 | -0.012841284 | 0.43161518 | -0.0297517 | 0.97626508 | 0.99556048 |
| SCPEP1 | 63.8057463 | 0.274101192 | 0.32579127 | 0.84133989 | 0.40015755 | 0.80830282 |
| AKAP1 | 60.2075021 | 0.797852382 | 0.29395533 | 2.71419599 | 0.00664368 | 0.11674591 |
| MSI2 | 1245.3173 | 0.279442945 | 0.2629362 | 1.06277854 | 0.2878824 | 0.73803612 |
| MRPS23 | 31.6713026 | -0.172760786 | 0.27180444 | -0.6356069 | 0.52503265 | 0.86747074 |
| CUEDC1 | 150.603079 | 0.003404118 | 0.35368554 | 0.0096247 | 0.99232072 | 0.99923337 |
| VEZF1 | 67.6104055 | -0.034797939 | 0.2189811 | -0.1589084 | 0.87374103 | 0.97434023 |
| AC015813.2 | 37.6911231 | 0.144974485 | 0.34449936 | 0.42082657 | 0.67388173 | 0.92189158 |
| SRSF1 | 18.9083489 | -0.112356105 | 0.36629172 | -0.3067394 | 0.75904173 | 0.94924107 |
| DYNLL2 | 49.2320629 | 0.546274403 | 0.3242398 | 1.68478515 | 0.09203003 | 0.4764741 |
| MKS1 | 10.3543674 | 0.450853657 | 0.56291479 | 0.800927 | 0.42317391 | NA |
| TSPOAP1 | 12.1575953 | 0.813064813 | 0.58182056 | 1.39744943 | 0.16227846 | NA |
| SUPT4H1 | 16.2056941 | 0.11875536 | 0.44783379 | 0.26517731 | 0.79087287 | 0.95746995 |
| RNF43 | 34.2206763 | 0.473705697 | 0.35158996 | 1.34732431 | 0.17787581 | 0.62542578 |
| MTMR4 | 24.5356041 | 0.050686438 | 0.41131678 | 0.12322969 | 0.90192522 | 0.97980453 |
| RAD51C | 16.7839947 | 0.128723685 | 0.39787871 | 0.32352494 | 0.74629773 | 0.94614106 |
| PPM1E | 113.307433 | 0.67272478 | 1.25544112 | 0.53584734 | 0.59206406 | 0.89177351 |
| TRIM37 | 58.0248003 | 0.103143861 | 0.300374 | 0.34338478 | 0.731309 | 0.94155624 |
| SKA2 | 29.3533583 | 0.737176013 | 0.31777942 | 2.31977266 | 0.02035318 | 0.22348867 |
| YPEL2 | 102.780025 | -0.288664373 | 0.36180279 | -0.79785 | 0.42495751 | 0.82241967 |
| DHX40 | 36.4927761 | -0.334053306 | 0.28326268 | -1.1793057 | 0.23827646 | 0.69303797 |
| CLTC | 316.491289 | -0.025192409 | 0.15953265 | -0.1579138 | 0.87452471 | 0.97437281 |
| PTRH2 | 42.3310462 | -0.620397602 | 0.39847156 | -1.5569432 | 0.11948397 | 0.53108939 |
| VMP1 | 226.26569 | -0.260113518 | 0.23548985 | -1.1045636 | 0.26934872 | 0.72046011 |
| TUBD1 | 19.736067 | 0.171078228 | 0.32576807 | 0.52515345 | 0.59947652 | 0.89476262 |
| RPS6KB1 | 51.5435041 | 0.310711426 | 0.23794671 | 1.30580255 | 0.19161971 | 0.6421543 |
| RNFT1 | 12.5332657 | 0.031907899 | 0.35733322 | 0.08929452 | 0.92884785 | NA |
| HEATR6 | 15.5167227 | 0.129287592 | 0.43555043 | 0.29683725 | 0.76659076 | 0.95205353 |
| USP32 | 140.623127 | -0.705693177 | 0.31118009 | -2.2677967 | 0.0233416 | 0.23883233 |
| APPBP2 | 125.1666 | 0.266613289 | 0.21460159 | 1.24236397 | 0.2141023 | 0.66787972 |
| PPM1D | 41.0395308 | 0.918449557 | 0.29794582 | 3.08260591 | 0.00205197 | 0.05196888 |
| BCAS3 | 403.812354 | 0.025358913 | 0.21958057 | 0.11548796 | 0.90805836 | 0.98250721 |
| TBX2 | 56.4605495 | 0.341486013 | 0.34563666 | 0.9879913 | 0.32315691 | 0.76304922 |
| NACA2 | 18.8983351 | 0.105798993 | 0.49036276 | 0.21575658 | 0.82917749 | 0.96669991 |
| INTS2 | 34.4056096 | 0.391929687 | 0.27316822 | 1.43475577 | 0.15135671 | 0.58907953 |
| MED13 | 292.464259 | 0.032966026 | 0.187417 | 0.17589667 | 0.86037512 | 0.97287712 |

| | | | | | | |
|---|---|---|---|---|---|---|
| METTL2A | 20.0018602 | 0.165856462 | 0.35288921 | 0.46999585 | 0.63835798 | 0.910465 |
| TLK2 | 180.655365 | 0.11244382 | 0.26298913 | 0.42756072 | 0.66897097 | 0.92049996 |
| AC005821.1 | 12.1659833 | 1.346798025 | 0.82799294 | 1.62658153 | 0.10382601 | NA |
| 10-Mar | 29.7856182 | 0.949855188 | 0.53470246 | 1.77641823 | 0.07566401 | 0.43325908 |
| TANC2 | 394.519836 | -0.159406218 | 0.20861852 | -0.7641039 | 0.44480535 | 0.83035712 |
| AC015923.1 | 31.5969317 | -0.841146261 | 0.62554751 | -1.3446561 | 0.17873633 | 0.62639418 |
| CYB561 | 20.302756 | 0.07992994 | 0.32076661 | 0.2491841 | 0.80321838 | 0.95938063 |
| DCAF7 | 117.318204 | 0.186976822 | 0.22493086 | 0.83126355 | 0.40582476 | 0.8115705 |
| TACO1 | 12.2890347 | 0.238162537 | 0.4622958 | 0.51517348 | 0.6064318 | NA |
| MAP3K3 | 123.719487 | 0.130238093 | 0.25725857 | 0.50625366 | 0.61267859 | 0.90011356 |
| CCDC47 | 34.3223679 | 0.24876893 | 0.32508801 | 0.76523563 | 0.44413124 | 0.83007188 |
| DDX42 | 67.8518889 | 0.149191976 | 0.21710392 | 0.68719153 | 0.49196204 | 0.85689205 |
| FTSJ3 | 13.3387972 | 0.217651063 | 0.37218182 | 0.58479768 | 0.55868377 | 0.88004847 |
| PSMC5 | 23.5282653 | 0.39733735 | 0.33403783 | 1.18949805 | 0.23424374 | 0.68933219 |
| SMARCD2 | 18.5085625 | 0.324569826 | 0.34853129 | 0.93125017 | 0.35172417 | 0.77928836 |
| ERN1 | 24.4742411 | -0.542703464 | 0.4288374 | -1.2655227 | 0.20568401 | 0.65836247 |
| SNHG25 | 14.4857628 | -0.135018813 | 0.42330788 | -0.3189613 | 0.74975589 | 0.94684415 |
| TEX2 | 41.2655167 | 0.454734955 | 0.27457943 | 1.65611442 | 0.09769865 | 0.48350863 |
| POLG2 | 20.0815469 | 0.62643822 | 0.34601068 | 1.81045919 | 0.07022461 | 0.41948432 |
| DDX5 | 484.849612 | -0.517951069 | 0.21890792 | -2.3660682 | 0.01797813 | 0.20807972 |
| CEP95 | 101.451562 | -0.243252329 | 0.2024185 | -1.2017297 | 0.22946826 | 0.68508528 |
| SMURF2 | 121.267314 | 0.175719792 | 0.23049238 | 0.76236705 | 0.44584097 | 0.83035712 |
| LRRC37A3 | 120.941103 | 0.40536787 | 0.28845674 | 1.40529865 | 0.15993249 | 0.60024417 |
| GNA13 | 87.2510204 | -0.332515849 | 0.24267189 | -1.3702281 | 0.17061571 | 0.61319058 |
| CEP112 | 90.5503278 | -0.576713999 | 0.45799057 | -1.2592268 | 0.20794843 | 0.65973983 |
| PRKCA | 879.560335 | 0.329974909 | 0.3420901 | 0.96458478 | 0.33475283 | 0.76777176 |
| PRKCA-AS1 | 41.182393 | -0.298113055 | 0.51207486 | -0.5821669 | 0.56045424 | 0.88067489 |
| HELZ | 259.339192 | -0.127460276 | 0.19436213 | -0.6557876 | 0.5119608 | 0.86346419 |
| PSMD12 | 45.6535973 | -0.094496611 | 0.28452712 | -0.3321181 | 0.73980007 | 0.94376279 |
| PITPNC1 | 22.9654335 | -0.651963891 | 0.51615409 | -1.2631187 | 0.20654652 | 0.65836247 |
| NOL11 | 25.2591574 | 0.29278379 | 0.28544054 | 1.02572601 | 0.30502075 | 0.74842902 |
| BPTF | 330.520191 | 0.132577426 | 0.22888973 | 0.57921963 | 0.56244099 | 0.88191823 |
| AC005332.7 | 15.7043892 | -0.191574449 | 0.36077976 | -0.5310011 | 0.59541802 | 0.893081 |
| AMZ2 | 16.7115703 | -0.204536047 | 0.32564767 | -0.62809 | 0.52994498 | 0.86758926 |
| ARSG | 36.5520209 | 0.735354536 | 0.35373552 | 2.07882582 | 0.03763336 | 0.31147067 |
| WIPI1 | 38.7821724 | -0.793284352 | 0.37759621 | -2.1008801 | 0.0356515 | 0.30211342 |
| PRKAR1A | 232.57454 | 0.075886752 | 0.1720691 | 0.44102487 | 0.65919499 | 0.91725598 |
| ABCA6 | 14.7538235 | 1.206132256 | 0.50878525 | 2.37061166 | 0.01775868 | 0.20666126 |
| ABCA10 | 87.3387078 | 0.89392758 | 0.54821467 | 1.63061594 | 0.10297138 | 0.49444882 |
| ABCA5 | 488.016245 | 0.599528544 | 0.29469545 | 2.03440043 | 0.04191124 | 0.32888549 |
| MAP2K6 | 60.7627862 | 1.659714925 | 0.77339497 | 2.14601206 | 0.03187202 | 0.28285321 |
| KCNJ16 | 19.7812153 | 1.021038621 | 0.42888981 | 2.38065488 | 0.01728189 | 0.20451748 |
| SOX9 | 20.5110951 | -2.402678673 | 0.74535592 | -3.223532 | 0.0012662 | 0.03790788 |
| SLC39A11 | 108.910185 | -0.135802877 | 0.26977583 | -0.5033916 | 0.614689 | 0.90054121 |
| COG1 | 21.2008965 | -0.090223523 | 0.31877932 | -0.2830282 | 0.77715525 | 0.95512504 |
| FAM104A | 23.5469472 | -0.165312122 | 0.31552636 | -0.5239249 | 0.60033076 | 0.89490801 |
| C17orf80 | 19.4593736 | 0.026919478 | 0.34601151 | 0.07779937 | 0.93798765 | 0.98871562 |
| CDC42EP4 | 39.3256156 | -1.022142049 | 0.34866437 | -2.9315931 | 0.00337228 | 0.07384005 |
| RPL38 | 83.7867455 | 0.313076116 | 0.34924853 | 0.89642787 | 0.37002429 | 0.7900012 |
| GPRC5C | 145.330748 | -0.350069501 | 0.26579099 | -1.3170856 | 0.18780992 | 0.63942753 |
| SLC9A3R1 | 88.0835667 | -0.067742685 | 0.27664585 | -0.2448715 | 0.80655593 | 0.96005568 |
| NAT9 | 16.8679622 | 0.174768318 | 0.33072003 | 0.52844794 | 0.59718847 | 0.89363331 |
| TMEM104 | 17.4089249 | -0.123693263 | 0.53039391 | -0.2332102 | 0.81559819 | 0.96300833 |
| KCTD2 | 28.256191 | 0.252353175 | 0.42074266 | 0.59978034 | 0.54865264 | 0.87371493 |
| ATP5PD | 65.5399008 | 0.200596729 | 0.40802202 | 0.49163211 | 0.62297944 | 0.90430717 |
| SLC16A5 | 223.934716 | -1.062648723 | 0.52263943 | -2.0332349 | 0.0420288 | 0.32888549 |
| ARMC7 | 126.41979 | -1.070107674 | 0.51725682 | -2.0688131 | 0.03856363 | 0.31513667 |
| NT5C | 18.8515832 | 0.019009332 | 0.45633745 | 0.0416563 | 0.96677269 | 0.9937157 |
| JPT1 | 62.3043398 | -0.522687832 | 0.36239109 | -1.4423308 | 0.14920908 | 0.58659144 |
| SUMO2 | 28.5757626 | -0.115476855 | 0.40749263 | -0.2833839 | 0.77688256 | 0.95512504 |
| NUP85 | 18.9347328 | 0.277443627 | 0.33810406 | 0.82058649 | 0.41188185 | 0.81519158 |
| GGA3 | 37.4141986 | 0.248698031 | 0.28684976 | 0.86699753 | 0.38594336 | 0.7994129 |
| MRPS7 | 39.0651841 | -0.078823075 | 0.33583629 | -0.2347068 | 0.81443628 | 0.96294404 |
| GRB2 | 101.998386 | -0.061559645 | 0.19015439 | -0.3237351 | 0.74613861 | 0.94614106 |
| AC011933.4 | 12.9769796 | -0.553771633 | 0.45569598 | -1.2152217 | 0.22428156 | 0.6771276 |
| TMEM94 | 30.7303929 | 0.412428492 | 0.27518041 | 1.49875671 | 0.13393676 | 0.56229288 |
| CASKIN2 | 16.4001165 | 0.155430027 | 0.38402606 | 0.40473822 | 0.68566994 | 0.92593602 |
| TSEN54 | 22.4109551 | 0.099752479 | 0.31309028 | 0.31860611 | 0.75002522 | 0.94684415 |
| LLGL2 | 306.798947 | -0.381332943 | 0.19579546 | -1.9476088 | 0.05146179 | 0.36077232 |
| MYO15B | 38.8028342 | 0.148731281 | 0.61340507 | 0.24246829 | 0.80841732 | 0.96098628 |
| RECQL5 | 51.1365934 | -0.008165554 | 0.40546524 | -0.0201387 | 0.98393271 | 0.99762255 |
| SMIM5 | 66.3012112 | 0.466520444 | 0.49663161 | 0.93936921 | 0.34754122 | 0.77726831 |
| SMIM6 | 66.7324899 | -0.017124237 | 0.44463805 | -0.0385128 | 0.96927886 | 0.99486579 |
| SAP30BP | 45.8229007 | 0.140959027 | 0.30044337 | 0.46917004 | 0.6389481 | 0.91063717 |
| ITGB4 | 12.4894193 | -0.13233584 | 0.66976901 | -0.1975843 | 0.84337032 | NA |
| GALK1 | 11.4232182 | 0.00982213 | 0.48603389 | 0.02020873 | 0.98387686 | NA |
| H3F3B | 218.712086 | -0.629974728 | 0.27149263 | -2.3204119 | 0.02031861 | 0.22348867 |
| UNK | 73.1604273 | 0.315175031 | 0.21898045 | 1.43928385 | 0.15007012 | 0.58673121 |
| WBP2 | 68.0369573 | -0.910701053 | 0.25819907 | -3.5271275 | 0.00042009 | 0.01771702 |
| TRIM47 | 16.7814871 | -1.67397184 | 0.48345308 | -3.4625322 | 0.00053512 | 0.02101475 |
| TRIM65 | 10.7037771 | 0.239670053 | 0.47845786 | 0.50092197 | 0.61642604 | NA |
| ACOX1 | 69.1540455 | 0.183874637 | 0.21916305 | 0.83898555 | 0.40147742 | 0.80890307 |
| TEN1 | 17.4904724 | -0.549715606 | 0.4287224 | -1.2822181 | 0.19976616 | 0.65269687 |
| SRP68 | 29.6844381 | -0.272782885 | 0.29975603 | -0.9100164 | 0.36281389 | 0.78718296 |
| EXOC7 | 56.6492928 | 0.057736092 | 0.30431203 | 0.18972662 | 0.84952336 | 0.97027687 |
| RNF157 | 26.9914805 | -0.545769968 | 0.32698503 | -1.6690977 | 0.09509801 | 0.48035393 |
| UBALD2 | 37.3500696 | -0.424393586 | 0.40356366 | -1.051615 | 0.29297623 | 0.74013384 |
| PRPSAP1 | 41.1842708 | 0.150272001 | 0.25513498 | 0.5889902 | 0.55586785 | 0.87787661 |
| UBE2O | 56.4560776 | 0.087626018 | 0.24401496 | 0.35910101 | 0.71951953 | 0.93786761 |
| RHBDF2 | 26.0023791 | -1.251502815 | 0.54070416 | -2.3145796 | 0.02063595 | 0.22444738 |
| MXRA7 | 41.060379 | 0.380081551 | 0.35793935 | 1.06186022 | 0.28829915 | 0.73805799 |
| JMJD6 | 27.8712124 | -0.434104736 | 0.38514412 | -1.1271228 | 0.25969055 | 0.71091822 |
| AC005837.2 | 12.6038365 | 0.356203044 | 0.78485685 | 0.4538446 | 0.64994068 | 0.91447992 |
| SRSF2 | 97.1269821 | 0.891657985 | 0.33828988 | 2.63578082 | 0.00839439 | 0.13244159 |
| MFSD11 | 54.0940587 | -0.035561299 | 0.21935409 | -0.1621182 | 0.87121274 | 0.97434023 |
| SEC14L1 | 226.485053 | -1.111239601 | 0.29778203 | -3.7317214 | 0.00019018 | 0.01042206 |
| 9-Sep | 370.7602 | -0.183971042 | 0.34732477 | -0.5296802 | 0.5963337 | 0.89360216 |
| TNRC6C | 83.3608183 | 0.520983717 | 0.21409837 | 2.43338473 | 0.0149584 | 0.18522344 |
| TMC6 | 11.7812254 | -0.002749525 | 0.41748571 | -0.0065859 | 0.99474524 | NA |
| SYNGR2 | 37.9126758 | -0.21354201 | 0.35626226 | -0.5993955 | 0.54890915 | 0.87371493 |
| AFMID | 29.5881571 | 0.997377541 | 0.54297987 | 1.83685916 | 0.06623069 | 0.40888394 |
| SOCS3 | 12.6068504 | -2.431430992 | 0.69606942 | -3.4930869 | 0.00047747 | 0.01945996 |
| PGS1 | 33.563348 | -0.189109519 | 0.26632527 | -0.7100697 | 0.47766089 | 0.84771464 |
| CYTH1 | 357.982491 | -1.692013383 | 0.42370494 | -3.9933766 | 6.51E-05 | 0.00493631 |
| USP36 | 83.3839226 | -0.817525647 | 0.51992419 | -1.5723939 | 0.11585922 | 0.5213638 |
| TIMP2 | 24.5605773 | 0.026824018 | 0.40033909 | 0.06700324 | 0.94657912 | 0.99134285 |
| LGALS3BP | 93.9841366 | -0.596525863 | 0.63347485 | -0.9416725 | 0.34636033 | 0.77642952 |
| CANT1 | 19.0896814 | 0.249902341 | 0.3644062 | 0.6857796 | 0.4928521 | 0.85714521 |
| ENGASE | 16.5317954 | 0.129770414 | 0.36405624 | 0.35645705 | 0.72149831 | 0.93824012 |
| TBC1D16 | 127.425893 | -0.11751329 | 0.29662504 | -0.3961678 | 0.69198124 | 0.928133 |
| CCDC40 | 28.9496017 | 0.35375322 | 0.37958016 | 0.93195919 | 0.35135761 | 0.77910174 |
| GAA | 29.1367655 | 0.37062571 | 0.3191721 | 1.16120961 | 0.24555667 | 0.69932463 |
| EIF4A3 | 25.797564 | -0.178692173 | 0.3580805 | -0.4990279 | 0.6177597 | 0.90214869 |

| | | | | | | |
|---|---|---|---|---|---|---|
| AC087741.1 | 14.9542016 | 0.647636877 | 0.43427052 | 1.49132131 | 0.13587716 | 0.56639812 |
| SGSH | 13.9302214 | 0.231656589 | 0.55659624 | 0.41620222 | 0.67726204 | 0.92254483 |
| SLC26A11 | 37.7356232 | -0.161218511 | 0.39675259 | -0.4063452 | 0.68448898 | 0.92562987 |
| RNF213 | 556.128911 | -1.628071439 | 0.44103947 | -3.6914416 | 0.00022299 | 0.01169173 |
| AC124319.2 | 150.758984 | 0.372985025 | 0.27046506 | 1.37905069 | 0.16787913 | 0.60954339 |
| ENDOV | 32.4849962 | 0.336528645 | 0.36757838 | 0.91552894 | 0.35991402 | 0.78506093 |
| RPTOR | 146.911061 | -0.085186092 | 0.24770135 | -0.3439065 | 0.73091664 | 0.94155624 |
| TEPSIN | 11.2508634 | -0.323026715 | 0.55024607 | -0.5870587 | 0.55716432 | NA |
| NDUFAF8 | 30.654312 | 0.021636123 | 0.43726252 | 0.04948085 | 0.9605361 | 0.99238848 |
| SLC38A10 | 66.0586257 | 0.207998652 | 0.20071638 | 1.03628138 | 0.30007089 | 0.74437099 |
| LINC00482 | 31.718025 | -0.705673334 | 0.32826014 | -2.1497381 | 0.03157594 | 0.28099688 |
| BAHCC1 | 13.8886343 | -0.290376225 | 0.50958409 | -0.5698299 | 0.5687931 | 0.88562534 |
| ACTG1 | 448.817749 | 0.187172757 | 0.40440721 | 0.46283239 | 0.64348451 | 0.91192138 |
| FAAP100 | 11.4866168 | 0.224261614 | 0.54605019 | 0.41069781 | 0.68129414 | NA |
| NPLOC4 | 139.637704 | -0.052490929 | 0.27629101 | -0.1899842 | 0.84932151 | 0.97027687 |
| OXLD1 | 29.4523657 | -0.590768136 | 0.32803983 | -1.8009037 | 0.07171806 | 0.42284325 |
| CCDC137 | 32.0313733 | -0.288143937 | 0.34703721 | -0.8302969 | 0.40637092 | 0.8115705 |
| ARL16 | 17.832483 | 0.55657704 | 0.36048875 | 1.54395121 | 0.12260015 | 0.53899578 |
| HGS | 41.2716525 | -0.33413952 | 0.38496337 | -0.8679775 | 0.38540668 | 0.7991545 |
| GCGR | 36.9890836 | 0.345570864 | 0.61936589 | 0.557943 | 0.5768833 | 0.88821715 |
| MCRIP1 | 30.8674491 | 0.096194347 | 0.41804809 | 0.23010354 | 0.81801131 | 0.9635539 |
| P4HB | 121.434826 | -0.448347954 | 0.30531231 | -1.4684896 | 0.14197128 | 0.57673923 |
| ARHGDIA | 96.9364076 | -0.886071723 | 0.50650636 | -1.7493792 | 0.08022549 | 0.44789428 |
| ANAPC11 | 63.3661062 | -0.120351348 | 0.47851373 | -0.2515108 | 0.80141925 | 0.95877735 |
| PCYT2 | 15.3281564 | 0.124691378 | 0.48815214 | 0.25543549 | 0.79838677 | 0.95848876 |
| SIRT7 | 18.1521101 | 0.194308818 | 0.36304719 | 0.53521642 | 0.59250021 | 0.89198431 |
| MAFG | 17.3845438 | -1.367496305 | 0.61641779 | -2.2184569 | 0.02652369 | 0.25574536 |
| ASPSCR1 | 19.0972077 | -0.00262098 | 0.46866332 | -0.0055925 | 0.99553789 | 0.99923337 |
| CENPX | 15.7386348 | 0.079340261 | 0.49261197 | 0.16106036 | 0.87204586 | 0.97434023 |
| DCXR | 29.374995 | 0.288589283 | 0.41682567 | 0.69235007 | 0.48871751 | 0.85461689 |
| RFNG | 14.1481339 | 0.244052512 | 0.38447533 | 0.63476767 | 0.52557995 | 0.86747074 |
| GPS1 | 23.163891 | -0.128315597 | 0.37264317 | -0.3443391 | 0.73059131 | 0.94155624 |
| DUS1L | 35.700971 | -0.306565735 | 0.30214925 | -1.0146169 | 0.31028848 | 0.75363506 |
| FASN | 23.4683679 | -1.539297041 | 0.51144545 | -3.0096993 | 0.00261506 | 0.06201986 |
| CCDC57 | 77.2222308 | 0.09364589 | 0.23869009 | 0.39233255 | 0.69481252 | 0.92858659 |
| SLC16A3 | 99.8152106 | -1.495930735 | 0.45204941 | -3.3092195 | 0.00093556 | 0.03151033 |
| CSNK1D | 94.4957835 | -0.173222345 | 0.22546935 | -0.7682745 | 0.44232414 | 0.82820049 |
| OGFOD3 | 26.4957758 | 0.338760609 | 0.32227935 | 1.05113966 | 0.29319445 | 0.74013384 |
| HEXDC | 63.1181074 | 0.005964109 | 0.20724293 | 0.02877834 | 0.97704137 | 0.9957152 |
| CYBC1 | 24.5027934 | -0.933996229 | 0.32698526 | -2.8563863 | 0.00428493 | 0.08768748 |
| NARF | 39.8697229 | -0.56177078 | 0.39938742 | -1.406581 | 0.15955166 | 0.60024417 |
| FOXK2 | 169.669264 | -0.583383643 | 0.24500097 | -2.3811483 | 0.01725876 | 0.20451748 |
| WDR45B | 166.446564 | -0.992696484 | 0.30048433 | -3.3036548 | 0.00095433 | 0.03192078 |
| RAB40B | 159.415845 | -0.60292038 | 0.40591041 | -1.4853533 | 0.13745025 | 0.56880008 |
| FN3KRP | 11.9978456 | -0.016784016 | 0.38834616 | -0.0432192 | 0.96552679 | NA |
| FN3K | 43.8682142 | 0.382340219 | 0.39322432 | 0.97232088 | 0.33089094 | 0.76685572 |
| TBCD | 68.3800096 | -0.082090313 | 0.32579649 | -0.2519681 | 0.80106574 | 0.95877735 |
| METRNL | 83.5772849 | -0.146376181 | 0.35543146 | -0.4118267 | 0.68046641 | 0.92375164 |
| USP14 | 101.5274 | 0.127678565 | 0.2116657 | 0.60320857 | 0.54636994 | 0.87268046 |
| THOC1 | 73.5866451 | 0.018275487 | 0.24687847 | 0.07402625 | 0.9409895 | 0.98976341 |
| ENOSF1 | 233.68116 | -0.633657967 | 0.35602007 | -1.7798378 | 0.07510251 | 0.43243667 |
| YES1 | 148.450229 | -0.64025806 | 0.24069754 | -2.6600109 | 0.00781381 | 0.12855183 |
| METTL4 | 23.9224634 | 0.161690912 | 0.38213501 | 0.42312509 | 0.67220399 | 0.92176197 |
| SMCHD1 | 217.280293 | -0.160413939 | 0.28471171 | -0.5634259 | 0.57314493 | 0.88578992 |
| AP001011.1 | 34.3681679 | -0.41793253 | 0.51386441 | -0.8133129 | 0.41603871 | 0.81763864 |
| LPIN2 | 99.9064495 | 0.404486234 | 0.37690405 | 1.07318092 | 0.28318995 | 0.73467304 |
| MYL12A | 112.202966 | -0.741953658 | 0.38948882 | -1.9049421 | 0.05678759 | 0.37858396 |
| MYL12B | 245.547865 | -0.310926538 | 0.39607172 | -0.7850258 | 0.43243842 | 0.8248269 |
| TGIF1 | 26.2704408 | -1.584848755 | 0.49765047 | -3.1846624 | 0.00144923 | 0.04136384 |
| DLGAP1 | 18.1359476 | -0.772701231 | 0.47708887 | -1.619617 | 0.10531458 | 0.50051517 |
| DLGAP1-AS1 | 12.0185461 | -0.759680069 | 0.49497093 | -1.5347973 | 0.12483361 | NA |
| LINC00667 | 20.4920517 | 0.301540036 | 0.31253961 | 0.96480584 | 0.33464207 | 0.76777176 |
| EPB41L3 | 20.0720336 | 0.539069853 | 0.52708436 | 1.02273923 | 0.30643116 | 0.74908826 |
| L3MBTL4 | 427.793923 | -0.238026254 | 0.28108898 | -0.8468004 | 0.39710641 | 0.80674902 |
| AP001021.2 | 32.5483818 | -0.803962872 | 0.73745119 | -1.0901913 | 0.27562888 | 0.72554623 |
| ARHGAP28 | 76.8488904 | 0.958750891 | 0.6028164 | 1.59045256 | 0.11173283 | 0.51426646 |
| LAMA1 | 36.5222485 | 0.356155893 | 0.52785857 | 0.6747184 | 0.49985469 | 0.85942996 |
| PTPRM | 137.492253 | 0.286025272 | 0.46166948 | 0.61954554 | 0.53555703 | 0.86946459 |
| RAB12 | 60.4661053 | 0.048684739 | 0.2394523 | 0.20331707 | 0.83888722 | 0.96748214 |
| NDUFV2 | 107.546785 | 0.108806 | 0.28118397 | 0.38695663 | 0.69878832 | 0.9304388 |
| ANKRD12 | 480.927853 | -0.510016516 | 0.20269613 | -2.5161631 | 0.01186403 | 0.162086 |
| TWSG1 | 60.3187415 | 0.261034763 | 0.24979012 | 1.04501638 | 0.2960154 | 0.74123516 |
| RALBP1 | 109.536014 | 0.293523627 | 0.28859112 | 1.0170917 | 0.30910981 | 0.75165835 |
| PPP4R1 | 118.939521 | 0.046523215 | 0.26772262 | 0.17377395 | 0.86204312 | 0.97302617 |
| RAB31 | 98.7293321 | 0.038866334 | 0.51806693 | 0.07502184 | 0.94019734 | 0.98913772 |
| VAPA | 321.613511 | -0.026812087 | 0.25398041 | -0.1055675 | 0.91592548 | 0.9844296 |
| NAPG | 84.7463374 | -0.254917654 | 0.2170992 | -1.174199 | 0.24031534 | 0.69470237 |
| GNAL | 22.4671458 | -1.316491789 | 0.50246462 | -2.6200686 | 0.00879121 | 0.13622159 |
| MPPE1 | 52.8281548 | 0.167639411 | 0.30898011 | 0.54255728 | 0.58743466 | 0.89073775 |
| IMPA2 | 189.346148 | -0.197155625 | 0.34326125 | -0.5743603 | 0.56572404 | 0.88294821 |
| AFG3L2 | 142.800361 | 0.293217831 | 0.25229026 | 1.16222415 | 0.24514443 | 0.69883704 |
| SPIRE1 | 446.820595 | -0.132501089 | 0.22481195 | -0.5893863 | 0.55560215 | 0.87787661 |
| PSMG2 | 65.4637497 | 0.092814793 | 0.24172943 | 0.3839615 | 0.70100698 | 0.93127962 |
| CEP76 | 15.1636589 | 0.004781366 | 0.4398883 | 0.0108695 | 0.99132756 | 0.99923337 |
| PTPN2 | 153.991537 | -0.129679782 | 0.22954799 | -0.5649354 | 0.57211772 | 0.88578992 |
| SEH1L | 32.2717791 | 0.145780458 | 0.26163171 | 0.5571972 | 0.5773927 | 0.88871933 |
| CEP192 | 152.534523 | 0.259363346 | 0.29691982 | 0.87351309 | 0.38238347 | 0.79731722 |
| LDLRAD4 | 106.50048 | 0.253841865 | 1.009088 | 0.25155573 | 0.80138448 | 0.95877735 |
| FAM210A | 51.6373043 | -0.48714315 | 0.31138074 | -1.5644614 | 0.11770925 | 0.52689421 |
| RNMT | 89.9471865 | -0.384391667 | 0.2357664 | -1.630392 | 0.10301867 | 0.49444882 |
| ROCK1 | 246.98667 | -0.059296743 | 0.16393364 | -0.3617119 | 0.71756735 | 0.93717562 |
| ESCO1 | 79.7223269 | -0.478139186 | 0.22443085 | -2.1304521 | 0.03313431 | 0.28919158 |
| SNRPD1 | 26.9094678 | 0.245624085 | 0.31284968 | 0.78511853 | 0.43238407 | 0.8248269 |
| ABHD3 | 224.291028 | -1.519950611 | 0.51898425 | -2.9287028 | 0.0034038 | 0.07436223 |
| MIB1 | 249.903277 | 0.098229683 | 0.18249705 | 0.53825355 | 0.59040201 | 0.89149106 |
| AC090912.1 | 16.4387792 | -0.597764565 | 0.48916761 | -1.2220036 | 0.22170628 | 0.67499434 |
| RBBP8 | 280.093045 | 0.107530641 | 0.27715917 | 0.38797432 | 0.69803504 | 0.93015138 |
| CABLES1 | 30.0700874 | -1.466328491 | 0.45756165 | -3.2046577 | 0.00135223 | 0.03962733 |
| TMEM241 | 24.4071156 | -0.779859195 | 0.31240466 | -2.496311 | 0.01254925 | 0.16675032 |
| RIOK3 | 106.370903 | -0.415324665 | 0.227801 | -1.8231907 | 0.06827452 | 0.41547228 |
| RMC1 | 43.3201161 | -0.513834282 | 0.27614487 | -1.8607417 | 0.06278066 | 0.39828148 |
| NPC1 | 267.012977 | -0.386180966 | 0.33025117 | -1.1693553 | 0.2422605 | 0.69596937 |
| ANKRD29 | 53.2197237 | 0.008736219 | 0.51554709 | 0.01694553 | 0.98648007 | 0.99810751 |
| LAMA3 | 128.163995 | -0.825948454 | 0.50900475 | -1.6226734 | 0.10465925 | 0.49862215 |
| TTC39C | 15.0230347 | -1.268346643 | 0.54087563 | -2.3449876 | 0.01902771 | 0.2163761 |
| OSBPL1A | 81.6107049 | -0.515270796 | 0.51581123 | -0.9989523 | 0.31781782 | 0.75982282 |
| IMPACT | 16.6442815 | 0.931696976 | 0.34552888 | 2.69643738 | 0.00700856 | 0.12011132 |
| SS18 | 177.194276 | -0.602432344 | 0.19122912 | -3.150317 | 0.00163093 | 0.04456355 |
| TAF4B | 19.7127951 | -0.551463092 | 0.54864885 | -1.0051294 | 0.31483455 | 0.75742745 |
| KCTD1 | 262.014847 | -0.197622548 | 0.39252824 | -0.5034607 | 0.6146404 | 0.90054121 |
| DSC2 | 51.4367447 | -0.184916956 | 0.4137318 | -0.4469489 | 0.65491199 | 0.9158679 |
| DSG2 | 382.744989 | 0.441345989 | 0.39485101 | 1.11775321 | 0.26367241 | 0.71380385 |
| B4GALT6 | 63.3272715 | -0.572466218 | 0.37679875 | -1.5192891 | 0.12868975 | 0.55308726 |

| | | | | | | |
|---|---|---|---|---|---|---|
| TRAPPC8 | 154.215512 | 0.124538578 | 0.20432682 | 0.60950675 | 0.5421886 | 0.87177844 |
| RNF138 | 51.9779611 | 0.325451016 | 0.22896148 | 1.42142258 | 0.15519395 | 0.59533243 |
| GAREM1 | 390.843529 | -0.575673181 | 0.33358057 | -1.7257396 | 0.08439428 | 0.4585327 |
| CCDC178 | 32.903584 | -0.135011532 | 0.41837357 | -0.3227057 | 0.74691814 | 0.94620686 |
| ASXL3 | 102.018836 | 0.183523296 | 0.37743648 | 0.48623624 | 0.62679968 | 0.90542917 |
| MAPRE2 | 190.084879 | 1.28615064 | 0.36810539 | 3.4939739 | 0.00047589 | 0.01945996 |
| ZNF397 | 43.2354531 | -0.072071962 | 0.26665769 | -0.270279 | 0.78694565 | 0.95577371 |
| ZSCAN30 | 35.2694854 | -0.00650057 | 0.35921408 | -0.0180966 | 0.98556175 | 0.99801117 |
| ZNF24 | 64.9216833 | -0.12648635 | 0.25684377 | -0.4924641 | 0.62239126 | 0.90417781 |
| GALNT1 | 54.2729137 | 0.015436589 | 0.268005 | 0.05759814 | 0.95406873 | 0.99228059 |
| RPRD1A | 139.310178 | 0.2398883 | 0.16275631 | 1.47391087 | 0.14050559 | 0.57482254 |
| SLC39A6 | 22.900679 | 0.293740253 | 0.39904091 | 0.73611563 | 0.46166033 | 0.83846194 |
| ELP2 | 84.3516733 | -0.240939575 | 0.22021201 | -1.0941255 | 0.27389993 | 0.72510729 |
| MOCOS | 28.7344917 | -1.406179652 | 0.52479702 | -2.6794734 | 0.00737381 | 0.12396174 |
| FHOD3 | 106.270424 | 0.754221365 | 0.61329903 | 1.22977752 | 0.21878043 | 0.67250923 |
| TPGS2 | 47.6164009 | -0.934562659 | 0.2890178 | -3.2335817 | 0.00122248 | 0.03740723 |
| KIAA1328 | 274.626926 | 0.488028349 | 0.24648159 | 1.97997893 | 0.0477059 | 0.34885689 |
| AC016205.1 | 40.3604139 | 0.372993573 | 0.34802305 | 1.07174961 | 0.28383251 | 0.73550473 |
| PIK3C3 | 120.746473 | -0.271523676 | 0.22439375 | -1.2100323 | 0.22626651 | 0.68070982 |
| LINC00907 | 22.4436148 | -0.391972962 | 0.41195073 | -0.9515045 | 0.34134835 | 0.77193691 |
| SETBP1 | 687.659324 | 0.413876426 | 0.23263261 | 1.77909893 | 0.07522354 | 0.43245501 |
| SLC14A2 | 86.5031088 | 3.846783514 | 1.09749658 | 3.50505285 | 0.00045652 | 0.0190052 |
| EPG5 | 124.538191 | 0.768317728 | 0.46986432 | 1.63519061 | 0.1020091 | 0.49220184 |
| ATP5F1A | 220.920078 | 0.357233913 | 0.34616309 | 1.03198151 | 0.30208078 | 0.74690278 |
| HAUS1 | 19.7244726 | 0.590215835 | 0.4011682 | 1.47124283 | 0.14122546 | 0.57606684 |
| C18orf25 | 64.2489902 | -0.068380484 | 0.23043809 | -0.2967412 | 0.76666407 | 0.95205353 |
| LOXHD1 | 45.5941992 | 1.76494749 | 0.83207045 | 2.1211515 | 0.03390906 | 0.29315315 |
| ST8SIA5 | 69.3068694 | 0.791126653 | 0.5150479 | 1.53602539 | 0.12453215 | 0.54320519 |
| PIAS2 | 204.640091 | -0.211289389 | 0.26736615 | -0.7902623 | 0.4293746 | 0.82469033 |
| KATNAL2 | 88.5572543 | 0.635090242 | 0.38341066 | 1.656423 | 0.09763619 | 0.48344615 |
| HDHD2 | 11.5034534 | 0.597287181 | 0.39799852 | 1.50072713 | 0.13342615 | NA |
| SMAD2 | 234.095631 | -0.105663665 | 0.17012442 | -0.6210964 | 0.5345362 | 0.86909171 |
| CTIF | 42.6074696 | -0.175916079 | 0.36894292 | -0.4768111 | 0.63349664 | 0.90805737 |
| SMAD7 | 21.6146406 | 0.755807057 | 0.5436624 | 1.39021396 | 0.16446392 | 0.60496776 |
| DYM | 264.077883 | -0.136443361 | 0.17649617 | -0.773067 | 0.43948274 | 0.82760292 |
| LIPG | 45.149696 | -1.113019868 | 0.64673219 | -1.7209904 | 0.08525257 | 0.46032833 |
| ACAA2 | 52.6381079 | 0.005145453 | 0.36187658 | 0.01421881 | 0.98865542 | 0.99915789 |
| SNHG22 | 21.8776842 | -0.523538898 | 0.46351938 | -1.1294865 | 0.25869265 | 0.71022613 |
| MYO5B | 160.273693 | -0.424745043 | 0.27299725 | -1.5558583 | 0.11974178 | 0.53109067 |
| MBD1 | 25.7177134 | -0.071299643 | 0.32085045 | -0.2222208 | 0.82414201 | 0.96539788 |
| CXXC1 | 12.2080079 | 0.371517501 | 0.45049449 | 0.82468822 | 0.40954862 | NA |
| ME2 | 39.4936939 | -0.022085671 | 0.24903521 | -0.0886849 | 0.92933231 | 0.98733028 |
| AC091551.1 | 12.5096021 | -0.56468227 | 0.42120413 | -1.340638 | 0.18003801 | NA |
| SMAD4 | 64.4028848 | 0.283086 | 0.20165476 | 1.40381509 | 0.16037392 | 0.60024417 |
| MEX3C | 20.8224105 | -0.174095935 | 0.34634646 | -0.5026641 | 0.61520045 | 0.90106518 |
| MBD2 | 117.498966 | -0.537136945 | 0.28916616 | -1.8575373 | 0.06323474 | 0.4001801 |
| POLI | 42.6488215 | 0.01594731 | 0.28561322 | 0.05583534 | 0.95547298 | 0.99228059 |
| CCDC68 | 46.2533538 | -1.482686942 | 0.48817628 | -3.0371958 | 0.0023879 | 0.05737663 |
| TXNL1 | 55.9617185 | 0.125976175 | 0.28135788 | 0.44774354 | 0.65433829 | 0.9158679 |
| WDR7 | 132.763786 | 0.344314673 | 0.22374643 | 1.53886107 | 0.12383822 | 0.54182711 |
| FECH | 24.7199021 | 0.542166112 | 0.34797475 | 1.55806165 | 0.11921864 | 0.53056597 |
| NARS | 88.6679974 | -0.362725873 | 0.24405697 | -1.4862344 | 0.13721713 | 0.56856307 |
| AC027097.2 | 36.1574782 | -0.369006146 | 0.3501604 | -1.0538203 | 0.29196518 | 0.73999615 |
| ATP8B1 | 125.395326 | -0.048670389 | 0.29569998 | -0.1645938 | 0.8692637 | 0.97434023 |
| NEDD4L | 2302.02208 | 0.459530307 | 0.23660717 | 1.94216561 | 0.05211705 | 0.36245478 |
| MALT1 | 42.5875671 | -0.077108623 | 0.28420815 | -0.2713104 | 0.78615232 | 0.95577371 |
| ZNF532 | 247.299882 | -1.2192479 | 0.36416006 | -3.3481099 | 0.00081365 | 0.02859558 |
| SEC11C | 29.5333683 | 0.481622559 | 0.33633964 | 1.43195301 | 0.15215727 | 0.59013416 |
| LMAN1 | 127.048269 | 0.481649989 | 0.2401802 | 2.00536926 | 0.04492359 | 0.336601 |
| CCBE1 | 115.844069 | 0.207535192 | 0.5723343 | 0.36261183 | 0.71689487 | 0.93689626 |
| AC105094.2 | 28.2930503 | -0.609277338 | 0.63574773 | -0.9583634 | 0.33787956 | 0.76966704 |
| RNF152 | 918.346152 | 0.250722557 | 0.37385454 | 0.670642 | 0.50244862 | 0.86010152 |
| PIGN | 102.489164 | 0.022009174 | 0.16545349 | 0.13302332 | 0.89417493 | 0.97839784 |
| KIAA1468 | 125.825806 | -0.037636391 | 0.18168656 | -0.2071501 | 0.83589263 | 0.96748214 |
| ZCCHC2 | 93.6515847 | -0.249211212 | 0.24324633 | -1.024522 | 0.3055888 | 0.74897897 |
| PHLPP1 | 218.958846 | 0.0637129 | 0.23336099 | 0.27302293 | 0.78483559 | 0.95555482 |
| BCL2 | 345.871677 | -0.631758655 | 0.34991819 | -1.8054467 | 0.0710048 | 0.42202609 |
| KDSR | 68.7696403 | -0.877347371 | 0.3517046 | -2.4945576 | 0.01261143 | 0.16711862 |
| VPS4B | 66.7825295 | 0.051625207 | 0.2090985 | 0.2468942 | 0.80499011 | 0.95997099 |
| TMX3 | 63.5736013 | 0.152764666 | 0.19722503 | 0.77457036 | 0.43859358 | 0.82745796 |
| RTTN | 40.1675487 | 0.558662865 | 0.30296838 | 1.84396426 | 0.06518835 | 0.40540566 |
| SOCS6 | 27.5059008 | -0.24057084 | 0.35615503 | -0.6754666 | 0.49937934 | 0.85942996 |
| LINC01541 | 20.1743034 | -0.554160926 | 0.86515877 | -0.6405309 | 0.52182751 | 0.86703098 |
| TIMM21 | 14.7989404 | 0.267702228 | 0.48848915 | 0.54802083 | 0.5836776 | 0.89036744 |
| CYB5A | 164.548471 | 0.842975232 | 0.62497673 | 1.34881059 | 0.17739781 | 0.6251709 |
| CNDP2 | 157.406594 | -0.846912877 | 0.24895197 | -3.4019128 | 0.00066916 | 0.02477424 |
| LINC00909 | 13.6178133 | -0.468038839 | 0.40219001 | -1.1637257 | 0.2445352 | 0.69792121 |
| ZNF407 | 245.425213 | -0.043583262 | 0.17778619 | -0.2451443 | 0.80634474 | 0.96005568 |
| ZADH2 | 22.7104986 | 0.130454657 | 0.31377199 | 0.41576259 | 0.67758373 | 0.92259972 |
| TSHZ1 | 23.3684818 | 0.471305652 | 0.40277475 | 1.17014696 | 0.24194183 | 0.69556484 |
| ZNF516 | 94.0692493 | -0.196817976 | 0.28976616 | -0.6792304 | 0.4969919 | 0.85814183 |
| ZNF236-DT | 10.9122567 | 0.45027262 | 0.46164267 | 0.97537045 | 0.32937654 | NA |
| ZNF236 | 84.1937232 | 0.65989848 | 0.33002581 | 1.99953596 | 0.0455504 | 0.33935394 |
| MBP | 283.270526 | -0.850749918 | 0.24373506 | -3.49047 | 0.00048217 | 0.01956394 |
| ATP9B | 215.903771 | -0.052072191 | 0.19790036 | -0.2631233 | 0.79245556 | 0.95746995 |
| NFATC1 | 14.8549467 | -0.230709823 | 0.50303902 | -0.4586321 | 0.64649841 | 0.91268149 |
| CTDP1 | 43.2209457 | -0.359353902 | 0.26683896 | -1.346707 | 0.17807461 | 0.62584192 |
| PQLC1 | 27.1656616 | 0.363525143 | 0.28457388 | 1.27743681 | 0.20144808 | 0.65374585 |
| HSBP1L1 | 40.6445909 | -0.685376999 | 0.4115624 | -1.6653052 | 0.09585188 | 0.48108046 |
| TXNL4A | 60.2171263 | -0.090960524 | 0.19378773 | -0.4693823 | 0.63879642 | 0.91063717 |
| ADNP2 | 89.9083317 | 0.175776796 | 0.36590771 | 0.4803856 | 0.63095323 | 0.90768383 |
| PARD6G-AS1 | 47.1011056 | 0.666769959 | 0.67608099 | 0.98622793 | 0.32402128 | 0.76304922 |
| C20orf96 | 11.0742764 | 0.285858726 | 0.47923814 | 0.59648576 | 0.55085076 | NA |
| ZCCHC3 | 17.5005902 | 0.135527642 | 0.37328668 | 0.36306583 | 0.71655571 | 0.93689626 |
| NRSN2-AS1 | 63.6659378 | 0.167535012 | 0.25054334 | 0.66868676 | 0.50369532 | 0.86073055 |
| SOX12 | 14.9949703 | 0.674675462 | 0.34920461 | 1.93203479 | 0.05335521 | 0.36627428 |
| NRSN2 | 18.1973952 | -0.224979276 | 0.373833 | -0.6018176 | 0.54729556 | 0.8729963 |
| RBCK1 | 68.5970259 | -0.533605003 | 0.281517 | -1.8954628 | 0.05803111 | 0.38214651 |
| TBC1D20 | 28.2982542 | 0.010318337 | 0.29747924 | 0.03468591 | 0.9723302 | 0.99526459 |
| CSNK2A1 | 209.756514 | 0.142176006 | 0.23676589 | 0.60049193 | 0.54817844 | 0.87341177 |
| SLC52A3 | 23.2619366 | 0.195559043 | 0.48930841 | 0.39966418 | 0.68940388 | 0.92734996 |
| PSMF1 | 63.5961463 | 0.178012951 | 0.1962466 | 0.90708809 | 0.36436022 | 0.78780828 |
| SDCBP2 | 69.3129818 | -3.082866269 | 0.62108555 | -4.9636741 | 6.92E-07 | 0.0001491 |
| AL136531.2 | 25.9884183 | -0.476579014 | 0.41787595 | -1.1404797 | 0.25408652 | 0.7071191 |
| SDCBP2-AS1 | 57.0822394 | 0.457265379 | 0.1977237 | 2.31264831 | 0.02074198 | 0.22444738 |
| NSFL1C | 44.7887595 | 0.024683223 | 0.25234949 | 0.09781364 | 0.92208027 | 0.98645403 |
| STK35 | 34.6123857 | -0.173883144 | 0.33280812 | -0.5224727 | 0.60134128 | 0.89490801 |
| SNRPB | 43.5102632 | -0.813966134 | 0.37514531 | -2.1697356 | 0.03002688 | 0.27607655 |
| NOP56 | 14.8197993 | 0.473619552 | 0.44258158 | 1.07012937 | 0.28456108 | 0.73638801 |
| IDH3B | 22.4466988 | 0.127999334 | 0.32481026 | 0.39407417 | 0.69352628 | 0.92826395 |
| EBF4 | 14.9345028 | -1.258995673 | 0.49347475 | -2.5512869 | 0.01073259 | 0.15287248 |
| PCED1A | 15.9868642 | 0.436816032 | 0.43061246 | 1.01440639 | 0.31038887 | 0.75363506 |
| PTPRA | 148.677416 | 0.063039466 | 0.20212108 | 0.31188961 | 0.75512442 | 0.94795939 |

| | | | | | | |
|---|---|---|---|---|---|---|
| MRPS26 | 44.0420597 | -0.14673221 | 0.44792516 | -0.327582 | 0.74322775 | 0.94546379 |
| UBOX5-AS1 | 17.0276468 | -0.311025534 | 0.51499257 | -0.6039418 | 0.54588234 | 0.87247631 |
| UBOX5 | 18.0582309 | 0.2533822 | 0.34795004 | 0.72821432 | 0.4664824 | 0.84234507 |
| LZTS3 | 18.7573748 | 1.105417586 | 0.61400745 | 1.80033252 | 0.07180815 | 0.42291381 |
| DDRGK1 | 46.6931586 | -0.251675645 | 0.36524033 | -0.6890686 | 0.4907801 | 0.85652518 |
| ITPA | 35.7145445 | -0.283516285 | 0.29980564 | -0.9456669 | 0.34431849 | 0.77393882 |
| C20orf194 | 365.40099 | 0.39367984 | 0.15353326 | 2.56413394 | 0.01034336 | 0.14940917 |
| ATRN | 102.151883 | 0.535687129 | 0.21335637 | 2.51076227 | 0.01204708 | 0.1634359 |
| CENPB | 31.8754845 | -0.234083755 | 0.4544439 | -0.5150993 | 0.6064836 | 0.89643487 |
| CDC25B | 15.8577809 | -0.352517108 | 0.4539662 | -0.7765272 | 0.43743776 | 0.82687287 |
| MAVS | 81.7229734 | 0.256133 | 0.3422427 | 0.7483958 | 0.45422145 | 0.83364149 |
| PANK2 | 74.5738328 | 0.173770701 | 0.2981177 | 0.58289294 | 0.55996537 | 0.88037687 |
| RNF24 | 193.689607 | 0.006464299 | 0.60898412 | 0.01061489 | 0.9915307 | 0.99923337 |
| PRNP | 48.4667359 | -1.124125938 | 0.3628204 | -3.0982986 | 0.00194635 | 0.05060915 |
| SLC23A2 | 34.7058366 | 0.461676829 | 0.23205495 | 1.98951509 | 0.04664438 | 0.34406879 |
| TMEM230 | 53.1272706 | 0.248728903 | 0.28936931 | 0.85955524 | 0.39003426 | 0.80191835 |
| PCNA | 16.8924658 | 0.56773415 | 0.46282106 | 1.22668176 | 0.21994223 | 0.67350139 |
| CDS2 | 69.6907713 | -0.273096527 | 0.22436398 | -1.2172031 | 0.22352696 | 0.67606266 |
| GPCPD1 | 105.49157 | -0.105141575 | 0.45606216 | -0.2305422 | 0.81767047 | 0.9635539 |
| SHLD1 | 32.3145986 | 0.639232674 | 0.32846091 | 1.94614538 | 0.05163728 | 0.36086571 |
| TRMT6 | 11.1223841 | -0.279201812 | 0.46905544 | -0.5952427 | 0.55168127 | NA |
| MCM8 | 18.1775657 | -0.035490156 | 0.34497654 | -0.102877 | 0.91806058 | 0.98521039 |
| CRLS1 | 62.3708696 | 0.283473947 | 0.2445103 | 1.15935382 | 0.24631199 | 0.69972061 |
| TMX4 | 97.8160632 | 0.181142601 | 0.29090897 | 0.62267795 | 0.53349618 | 0.86909171 |
| PLCB1 | 24.2234407 | -0.114639981 | 0.57597438 | -0.1990366 | 0.84223412 | 0.96831468 |
| PLCB4 | 223.758143 | -0.174260834 | 0.44443499 | -0.3920952 | 0.69498786 | 0.92863367 |
| PAK5 | 44.7404785 | 1.218378267 | 0.46410334 | 2.62523052 | 0.00865903 | 0.13512263 |
| MKKS | 34.9052851 | 0.570975004 | 0.30068342 | 1.89892416 | 0.05757445 | 0.38148784 |
| SLX4IP | 163.752676 | 0.010786583 | 0.37364367 | 0.02886864 | 0.97696936 | 0.9957152 |
| JAG1 | 79.2796269 | 0.020313901 | 0.41637352 | 0.04878769 | 0.96108849 | 0.99238848 |
| BTBD3 | 38.2261431 | 0.014486132 | 0.27162563 | 0.05333124 | 0.95746799 | 0.99236316 |
| SPTLC3 | 150.99101 | -0.481482712 | 0.30005733 | -1.6046358 | 0.10857399 | 0.50659799 |
| TASP1 | 145.123873 | -0.012451161 | 0.22128471 | -0.0562676 | 0.95512862 | 0.99228059 |
| ESF1 | 59.7863383 | 0.129078923 | 0.28343205 | 0.455414 | 0.64881142 | 0.91417939 |
| NDUFAF5 | 49.8090253 | 0.271916918 | 0.24789108 | 1.09692095 | 0.27267595 | 0.72345643 |
| MACROD2 | 676.233789 | 0.911709768 | 0.42044005 | 2.16846558 | 0.03012328 | 0.27617753 |
| KIF16B | 287.520813 | 0.042132786 | 0.26549525 | 0.15869506 | 0.87390913 | 0.97434023 |
| SNRPB2 | 29.1126445 | -0.156294916 | 0.35854358 | -0.4359161 | 0.66289761 | 0.918408 |
| DSTN | 166.831806 | -0.150023107 | 0.23614056 | -0.6353128 | 0.52522446 | 0.86747074 |
| RRBP1 | 74.5809289 | -0.45716403 | 0.34276781 | -1.3337426 | 0.18228823 | 0.63068306 |
| SNX5 | 42.3787802 | -0.026618754 | 0.3441778 | -0.0773401 | 0.93835297 | 0.98871562 |
| OVOL2 | 24.4228178 | -0.138345273 | 0.34101546 | -0.4056862 | 0.68497317 | 0.92581857 |
| MGME1 | 30.8193532 | -0.248338373 | 0.33565485 | -0.7398623 | 0.45938355 | 0.83645518 |
| ZNF133 | 31.5284 | 0.390466322 | 0.31681993 | 1.23245503 | 0.21777915 | 0.67202699 |
| DZANK1 | 22.0682078 | 0.166947727 | 0.40814626 | 0.40903897 | 0.68251106 | 0.92474072 |
| POLR3F | 13.0745215 | 0.237582365 | 0.37083793 | 0.64066361 | 0.52174126 | 0.86703098 |
| SEC23B | 113.882644 | -0.252395498 | 0.28500583 | -0.8855801 | 0.37584382 | 0.79392096 |
| DTD1 | 13.1291326 | -0.749607115 | 0.42120873 | -1.7796571 | 0.07513209 | 0.43243667 |
| AL121761.1 | 27.2408797 | -0.034289877 | 0.39284812 | -0.0872853 | 0.93044472 | 0.98756002 |
| RIN2 | 228.454472 | -0.133096968 | 0.33898229 | -0.3926369 | 0.69458765 | 0.92841398 |
| NAA20 | 40.6505512 | -0.013881276 | 0.37638168 | -0.0368808 | 0.97058001 | 0.99502125 |
| CRNKL1 | 17.8288412 | 0.262068967 | 0.33959128 | 0.77171877 | 0.44028101 | 0.82793978 |
| RALGAPA2 | 759.098426 | -1.757345898 | 0.2750235 | -6.3898028 | 1.66E-10 | 1.60E-07 |
| KIZ | 50.5994091 | 0.44782248 | 0.40209257 | 1.11372982 | 0.26539511 | 0.71660319 |
| KIZ-AS1 | 20.6041169 | 0.073274408 | 0.48753667 | 0.15029517 | 0.88053174 | 0.97613233 |
| XRN2 | 139.991414 | -0.609166538 | 0.18019724 | -3.3805541 | 0.0007234 | 0.02606817 |
| GZF1 | 15.1123956 | -0.249913797 | 0.43044418 | -0.5805951 | 0.56151335 | 0.88133972 |
| NAPB | 64.0758839 | -0.139738152 | 0.31809071 | -0.4393028 | 0.66044211 | 0.91752855 |
| CST3 | 154.865849 | -0.384768076 | 0.33704179 | -1.1416035 | 0.25361888 | 0.70701839 |
| AL035661.1 | 20.7740128 | 0.008937683 | 0.41184538 | 0.02170155 | 0.98268603 | 0.99719767 |
| APMAP | 54.0416317 | -0.21853049 | 0.36155639 | -0.6044161 | 0.54556706 | 0.87247631 |
| ACSS1 | 95.6820137 | 0.395783962 | 0.35637212 | 1.11059182 | 0.26674409 | 0.71773028 |
| ENTPD6 | 50.4795961 | -0.368871344 | 0.25120073 | -1.4684326 | 0.14198675 | 0.57673923 |
| PYGB | 153.02641 | -0.094211874 | 0.45073565 | -0.209018 | 0.83443417 | 0.96703798 |
| ABHD12 | 118.00221 | -0.254522893 | 0.29472018 | -0.8636086 | 0.38780292 | 0.80079761 |
| NINL | 196.789524 | 0.26689192 | 0.39942698 | 0.66818702 | 0.50401422 | 0.86090835 |
| ZNF337-AS1 | 21.4648652 | 0.198378633 | 0.3375253 | 0.58774448 | 0.55670382 | 0.87833881 |
| ZNF337 | 30.5245185 | 0.092276938 | 0.33092848 | 0.27884254 | 0.78036567 | 0.95537097 |
| FAM182B | 23.2839556 | -0.074868256 | 0.48314585 | -0.1549599 | 0.87685289 | 0.97532685 |
| HM13 | 60.5610977 | -0.010108745 | 0.26429204 | -0.0382484 | 0.96948964 | 0.99486579 |
| ID1 | 28.7141528 | -0.118460981 | 0.57571667 | -0.2057626 | 0.83697633 | 0.96748214 |
| BCL2L1 | 109.458385 | 0.196780437 | 0.40466285 | 0.48628243 | 0.62676693 | 0.90542917 |
| AL117381.1 | 22.5649285 | -0.377642296 | 0.46892067 | -0.8053437 | 0.42062138 | 0.81938194 |
| DUSP15 | 69.5813178 | 1.049516032 | 0.40757728 | 2.5750111 | 0.01002369 | 0.14754137 |
| PDRG1 | 11.2938359 | 0.532915521 | 0.44213095 | 1.20533411 | 0.22807435 | NA |
| TM9SF4 | 76.7411015 | 0.179727341 | 0.25286709 | 0.71075813 | 0.47723413 | 0.84767828 |
| PLAGL2 | 30.9319883 | -0.023921954 | 0.30764756 | -0.0777577 | 0.93802083 | 0.98871562 |
| POFUT1 | 41.1203899 | 0.717246637 | 0.38650441 | 1.85572691 | 0.06349249 | 0.4001801 |
| KIF3B | 160.095045 | 0.408079959 | 0.22070167 | 1.84901168 | 0.06445613 | 0.40285079 |
| ASXL1 | 457.010899 | 0.258341483 | 0.19399144 | 1.3317159 | 0.18295356 | 0.63154788 |
| NOL4L | 27.6290759 | 0.331241554 | 0.34468466 | 0.96099883 | 0.33655275 | 0.76903692 |
| MAPRE1 | 45.2148305 | 0.526975851 | 0.27193441 | 1.93787848 | 0.05263805 | 0.36370293 |
| AL035071.2 | 11.4515603 | 0.40102619 | 0.60580268 | 0.66197494 | 0.50798728 | NA |
| CDK5RAP1 | 46.1746264 | 0.092142529 | 0.2615894 | 0.35224106 | 0.72465748 | 0.93935288 |
| SNTA1 | 18.0842771 | -0.152850626 | 0.37059029 | -0.4124518 | 0.68000831 | 0.92375164 |
| CBFA2T2 | 256.368767 | 0.179962576 | 0.18214247 | 0.98803192 | 0.32313702 | 0.76304922 |
| NECAB3 | 14.0020891 | 0.014638171 | 0.45134384 | 0.03243241 | 0.97412721 | 0.9953169 |
| PXMP4 | 13.6804903 | 1.311355673 | 0.43780812 | 2.9952749 | 0.00274198 | 0.0639355 |
| CHMP4B | 58.4571875 | -0.336515443 | 0.32897031 | -1.0229356 | 0.3063383 | 0.74905004 |
| RALY | 103.471255 | -0.026847991 | 0.28473741 | -0.0942904 | 0.92487851 | 0.98672316 |
| EIF2S2 | 78.6424185 | -0.63862852 | 0.32766456 | -1.9490314 | 0.05129168 | 0.36077232 |
| AHCY | 18.5199739 | -0.142709371 | 0.33980737 | -0.4199714 | 0.67450636 | 0.92189158 |
| ITCH | 211.423719 | -0.035089453 | 0.19249275 | -0.1822897 | 0.85535535 | 0.97166553 |
| DYNLRB1 | 127.650718 | -0.358465722 | 0.36194171 | -0.9903963 | 0.32198047 | 0.76248108 |
| MAP1LC3A | 60.6272047 | -0.942412442 | 0.32084787 | -2.9372563 | 0.0033113 | 0.07299917 |
| PIGU | 72.2679208 | 0.273507472 | 0.29895517 | 0.91487788 | 0.36025574 | 0.78527656 |
| NCOA6 | 197.657178 | 0.060425159 | 0.14581545 | 0.41439477 | 0.67858502 | 0.92291026 |
| TP53INP2 | 55.7499719 | 0.08143909 | 0.40504592 | 0.20106137 | 0.84065058 | 0.96748214 |
| GGT7 | 66.7768037 | -0.370381 | 0.24840655 | -1.4910275 | 0.13595428 | 0.56647616 |
| ACSS2 | 78.0863133 | -0.593141884 | 0.41510455 | -1.4288976 | 0.15303367 | 0.59174003 |
| GSS | 33.1010612 | -0.137268522 | 0.38565858 | -0.3559328 | 0.72189092 | 0.93840016 |
| TRPC4AP | 143.131804 | -0.197517347 | 0.22981877 | -0.8594483 | 0.39009324 | 0.80191835 |
| EDEM2 | 25.1725178 | -0.256001672 | 0.29017658 | -0.8822272 | 0.37765395 | 0.79432848 |
| PROCR | 13.8044422 | -0.588050992 | 0.55261555 | -1.0641231 | 0.28727292 | 0.73803612 |
| MMP24OS | 37.2806931 | 0.212280745 | 0.33573935 | 0.63227841 | 0.52720497 | 0.86750588 |
| MMP24 | 32.6840359 | -0.375361276 | 0.40277256 | -0.9319435 | 0.35136572 | 0.77910174 |
| UQCC1 | 93.9587914 | 0.145286605 | 0.2194021 | 0.66219333 | 0.50784733 | 0.86281145 |
| CEP250 | 13.7625792 | -1.286920386 | 0.51702911 | -2.4890676 | 0.01280786 | 0.16857023 |
| ERGIC3 | 36.9888452 | 0.28633063 | 0.35381613 | 0.80926393 | 0.41836335 | 0.8188001 |
| CPNE1 | 43.8885584 | -0.164194039 | 0.33320746 | -0.4927682 | 0.62217638 | 0.90415189 |
| ROMO1 | 47.9891386 | -0.459196567 | 0.32867432 | -1.3971172 | 0.16237832 | 0.60094226 |
| RBM39 | 333.981625 | -0.045769649 | 0.16342647 | -0.2800626 | 0.77942944 | 0.95537097 |
| PHF20 | 130.198999 | 0.099727704 | 0.29750368 | 0.33521503 | 0.73746289 | 0.94376279 |

| | | | | | | |
|---|---|---|---|---|---|---|
| SCAND1 | 55.400476 | -0.435118176 | 0.37712929 | -1.153764 | 0.24859696 | 0.70343946 |
| NORAD | 239.73802 | 0.148971048 | 0.46717413 | 0.31887692 | 0.74981984 | 0.94684415 |
| EPB41L1 | 243.890859 | 0.098788656 | 0.37913887 | 0.26056061 | 0.79443137 | 0.95786007 |
| AAR2 | 18.4801862 | 0.05424389 | 0.46434625 | 0.11681776 | 0.90700447 | 0.98213254 |
| DLGAP4 | 85.4034067 | -0.006963841 | 0.21145584 | -0.0329328 | 0.97372814 | 0.99530377 |
| DLGAP4-AS1 | 22.4538086 | 0.313025422 | 0.44933696 | 0.6966385 | 0.48602906 | 0.85356796 |
| RAB5IF | 16.5513958 | -0.03156549 | 0.43021701 | -0.0733711 | 0.94151082 | 0.9898824 |
| NDRG3 | 23.9801778 | 0.321723597 | 0.42154113 | 0.76320809 | 0.4453393 | 0.83035712 |
| SAMHD1 | 15.6560713 | -0.966883615 | 0.86892228 | -1.1127389 | 0.26582058 | 0.71698101 |
| RBL1 | 19.0328124 | 1.015968794 | 0.3926007 | 2.58779158 | 0.00965934 | 0.14481546 |
| MROH8 | 13.5380185 | 0.030518525 | 0.43776298 | 0.06971472 | 0.94442072 | 0.99105669 |
| RPN2 | 106.681436 | 0.378112641 | 0.27381863 | 1.38088719 | 0.16731365 | 0.60898402 |
| MANBAL | 26.8035216 | -0.027909396 | 0.30644382 | -0.0910751 | 0.92743293 | 0.98690525 |
| SRC | 28.3571527 | 0.35515692 | 0.29385463 | 1.20861434 | 0.22681105 | 0.68135521 |
| BLCAP | 90.3857915 | 0.150954193 | 0.22180264 | 0.6805789 | 0.49613798 | 0.85814183 |
| CTNNBL1 | 43.0760631 | 0.117584584 | 0.26279455 | 0.44743921 | 0.65455797 | 0.9158679 |
| TTI1 | 32.5960977 | -0.179739125 | 0.28968702 | -0.6204597 | 0.53495517 | 0.86946459 |
| RPRD1B | 76.9930658 | -0.051914379 | 0.29452265 | -0.1762662 | 0.86008484 | 0.97287712 |
| TGM2 | 15.9483357 | -3.863119356 | 0.76988859 | -5.0177641 | 5.23E-07 | 0.00011793 |
| RALGAPB | 125.206453 | 0.099371348 | 0.18197802 | 0.54606237 | 0.58502307 | 0.89036744 |
| ACTR5 | 10.9150549 | 0.191207888 | 0.44546348 | 0.42923358 | 0.66775325 | NA |
| DHX35 | 30.8278903 | 0.051994975 | 0.26009287 | 0.19990927 | 0.84155154 | 0.9677342 |
| TOP1 | 156.158847 | -0.387398583 | 0.17814674 | -2.1746038 | 0.02965982 | 0.2737395 |
| PLCG1 | 19.7960541 | 0.095459812 | 0.31404114 | 0.30397232 | 0.76114898 | 0.9499444 |
| ZHX3 | 82.2122419 | 0.526279129 | 0.37017505 | 1.42170339 | 0.15511237 | 0.59533243 |
| LPIN3 | 28.8790329 | -0.127108739 | 0.2747942 | -0.4625598 | 0.64367996 | 0.9119294 |
| CHD6 | 184.559407 | 0.022107403 | 0.20690569 | 0.10684773 | 0.91490978 | 0.98407488 |
| PTPRT | 21.6082126 | 0.052916128 | 0.96385989 | 0.05490023 | 0.95621795 | 0.99234582 |
| SRSF6 | 61.4483722 | 0.994486683 | 0.30453113 | 3.26563226 | 0.0010922 | 0.0348498 |
| L3MBTL1 | 26.8567513 | -0.584302289 | 0.28828861 | -2.0267963 | 0.04268325 | 0.33083096 |
| SGK2 | 123.61859 | -0.336212638 | 0.31936933 | -1.0527393 | 0.29246049 | 0.74013384 |
| IFT52 | 32.5361752 | -0.183947522 | 0.27076871 | -0.6793529 | 0.49691426 | 0.85814183 |
| OSER1 | 38.6495144 | -0.288951366 | 0.40841431 | -0.7074957 | 0.47925849 | 0.84833462 |
| FITM2 | 17.1118997 | 0.71579515 | 0.38545439 | 1.85701649 | 0.06330881 | 0.4001801 |
| TTPAL | 14.0456049 | -0.286318144 | 0.42532469 | -0.6731755 | 0.50083567 | 0.85942996 |
| SERINC3 | 76.4100883 | -0.107340359 | 0.2139256 | -0.5017649 | 0.6158329 | 0.9014002 |
| PKIG | 51.0568963 | 0.083767143 | 0.24472185 | 0.34229531 | 0.73212866 | 0.94194042 |
| YWHAB | 152.608282 | -0.331426254 | 0.31668282 | -1.0465558 | 0.29530449 | 0.74111836 |
| PABPC1L | 12.544188 | 0.209512585 | 0.39275176 | 0.53344785 | 0.5937236 | 0.89260988 |
| TOMM34 | 14.6977271 | 0.021280707 | 0.40151991 | 0.05300038 | 0.95773161 | 0.99238848 |
| STK4 | 55.3814912 | 0.021118132 | 0.25295125 | 0.08348696 | 0.93346434 | 0.9881638 |
| SLPI | 49.7189 | -5.394430189 | 0.87319721 | -6.1777914 | 6.50E-10 | 4.50E-07 |
| SDC4 | 219.840978 | 0.124589248 | 0.31463599 | 0.39597901 | 0.69212051 | 0.928133 |
| PIGT | 21.7063219 | 0.805926789 | 0.36418349 | 2.21296903 | 0.02689978 | 0.25860047 |
| WFDC2 | 712.705177 | -3.619570145 | 0.39099405 | -9.2573536 | 2.10E-20 | 1.02E-16 |
| DNTTIP1 | 15.0847738 | 0.071038752 | 0.38654032 | 0.18378096 | 0.85418529 | 0.97135895 |
| SNX21 | 15.3830619 | 0.001540348 | 0.4064091 | 0.00379014 | 0.99697591 | 0.99975875 |
| ACOT8 | 18.2107725 | 0.134036642 | 0.34691364 | 0.3863689 | 0.69922348 | 0.9307627 |
| CTSA | 60.4982792 | -0.996753881 | 0.62154389 | -1.6036742 | 0.10878589 | 0.50661006 |
| ZNF335 | 23.7698293 | 0.53125618 | 0.30983469 | 1.71464398 | 0.08641052 | 0.46333999 |
| NCOA5 | 42.8474939 | 0.27005172 | 0.34741077 | 0.77732686 | 0.43696595 | 0.82687287 |
| CD40 | 12.913166 | -0.84079757 | 0.57929494 | -1.4514154 | 0.14666423 | 0.58349206 |
| SLC35C2 | 40.0762619 | -0.017848456 | 0.27178879 | -0.0656703 | 0.9476403 | 0.99149867 |
| ELMO2 | 37.328537 | 0.194390353 | 0.32721108 | 0.59408243 | 0.55245698 | 0.87576439 |
| SLC13A3 | 19.2191041 | -0.54227319 | 0.60252886 | -0.8999954 | 0.36812271 | 0.78877653 |
| ZMYND8 | 202.468892 | 0.114573275 | 0.36497181 | 0.31392363 | 0.75357905 | 0.94791062 |
| NCOA3 | 159.63209 | -0.03281973 | 0.2409343 | -0.1362186 | 0.89164848 | 0.97793768 |
| ARFGEF2 | 279.253773 | -0.154338433 | 0.20266709 | -0.7615367 | 0.44633654 | 0.83035712 |
| CSE1L | 44.2222772 | 0.314286153 | 0.26470892 | 1.18728963 | 0.2351134 | 0.68962805 |
| STAU1 | 141.386441 | -0.002362926 | 0.16894784 | -0.0139861 | 0.98884105 | 0.99915789 |
| DDX27 | 24.0951343 | 0.113820052 | 0.38659736 | 0.29441498 | 0.76844082 | 0.95205353 |
| ZNFX1 | 49.8234867 | -1.107174798 | 0.45731325 | -2.4210425 | 0.01547607 | 0.18954277 |
| ZFAS1 | 46.8871476 | -0.439968482 | 0.26390727 | -1.6671328 | 0.09548799 | 0.48081904 |
| B4GALT5 | 173.354026 | -0.208035934 | 0.26819174 | -0.7756985 | 0.43792702 | 0.8270818 |
| SLC9A8 | 64.135892 | -0.40685949 | 0.29925036 | -1.3595957 | 0.17395791 | 0.61945363 |
| RNF114 | 69.3690121 | -0.523300462 | 0.2984315 | -1.7535027 | 0.07951575 | 0.44532492 |
| CEBPB | 26.4620784 | -2.683306803 | 0.57745608 | -4.6467721 | 3.37E-06 | 0.00049554 |
| PTPN1 | 117.305388 | -0.819023564 | 0.493173 | -1.6607226 | 0.09676917 | 0.48257296 |
| RIPOR3 | 45.446329 | 0.676568887 | 0.62780874 | 1.07766719 | 0.2811823 | 0.73311066 |
| PARD6B | 173.04551 | -0.116129464 | 0.32133849 | -0.3613929 | 0.71780572 | 0.93723455 |
| BCAS4 | 34.5178846 | -0.858345616 | 0.43483259 | -1.973968 | 0.04838538 | 0.35179202 |
| ADNP | 221.02045 | 0.262114086 | 0.18702614 | 1.40148372 | 0.16106947 | 0.60060549 |
| DPM1 | 58.1795577 | -0.036470484 | 0.19757813 | -0.1845877 | 0.85355247 | 0.97131147 |
| NFATC2 | 30.8609984 | -1.054902899 | 0.39948042 | -2.6406874 | 0.0082738 | 0.13156712 |
| ATP9A | 220.797653 | 0.056834208 | 0.23292847 | 0.24399855 | 0.80723194 | 0.96016552 |
| ZFP64 | 61.4909887 | 0.062255537 | 0.41894256 | 0.1486016 | 0.88186801 | 0.97638622 |
| ZNF217 | 110.005134 | -0.366914332 | 0.30522346 | -1.2021171 | 0.22931817 | 0.68503082 |
| PFDN4 | 15.293013 | -0.236188547 | 0.37756529 | -0.6255568 | 0.53160566 | 0.86840264 |
| FAM210B | 43.3511662 | 0.018075336 | 0.31513384 | 0.05735765 | 0.9542603 | 0.99228059 |
| CSTF1 | 13.3951458 | 0.325364859 | 0.42787338 | 0.76042323 | 0.44700164 | 0.83066268 |
| RTF2 | 36.287837 | -0.342309071 | 0.29344519 | -1.1665179 | 0.2434051 | 0.69729164 |
| RAE1 | 21.6612978 | 0.519282368 | 0.29263466 | 1.7745074 | 0.07597926 | 0.43403934 |
| RBM38 | 11.4486028 | 0.03789516 | 0.50479376 | 0.07507058 | 0.94015856 | NA |
| PCK1 | 88.4638656 | -0.226320282 | 0.7375547 | -0.3068522 | 0.75895587 | 0.94924107 |
| PMEPA1 | 24.0507608 | -0.444532315 | 0.44898031 | -0.9900931 | 0.3221286 | 0.76248108 |
| RAB22A | 68.0226881 | 0.107254592 | 0.27656274 | 0.38781287 | 0.69815452 | 0.93015138 |
| VAPB | 114.266834 | -0.118185038 | 0.19547612 | -0.6046009 | 0.54544421 | 0.87247631 |
| NPEPL1 | 20.6663342 | 0.298621358 | 0.35362641 | 0.84445434 | 0.39841557 | 0.80714564 |
| GNAS | 385.149301 | -0.468450454 | 0.21858948 | -2.1430604 | 0.03210825 | 0.28394452 |
| NELFCD | 37.7896887 | -0.048285549 | 0.26695983 | -0.180872 | 0.85646807 | 0.97166553 |
| CTSZ | 59.700892 | -0.247005744 | 0.3436651 | -0.7187397 | 0.47230132 | 0.84570599 |
| ATP5F1E | 270.187488 | -0.301384904 | 0.27743634 | -1.0863209 | 0.27733703 | 0.72678432 |
| PRELID3B | 22.0673543 | -0.05427562 | 0.3337335 | -0.1626316 | 0.87080849 | 0.97434023 |
| SYCP2 | 18.9120812 | -0.430349473 | 0.6327945 | -0.6800778 | 0.49645522 | 0.85814183 |
| FAM217B | 14.1117497 | 0.291753991 | 0.43843385 | 0.66544586 | 0.50576536 | 0.86213981 |
| MIR646HG | 40.6810792 | 0.670992294 | 0.6697069 | 1.00191934 | 0.31638255 | 0.75864788 |
| TAF4 | 62.6763851 | 0.247628961 | 0.24183755 | 1.02394751 | 0.30586007 | 0.74897897 |
| LSM14B | 14.9759428 | 0.131479804 | 0.35720924 | 0.36807504 | 0.71281728 | 0.9357596 |
| PSMA7 | 113.882045 | -0.413760518 | 0.32374374 | -1.2780495 | 0.20123198 | 0.65374585 |
| SS18L1 | 48.442751 | 0.458128311 | 0.28020894 | 1.63495252 | 0.102059 | 0.49220184 |
| MTG2 | 35.2951163 | 0.435588235 | 0.32660937 | 1.33366729 | 0.18231292 | 0.63068306 |
| OSBPL2 | 83.0938805 | -0.436041033 | 0.23969234 | -1.8191696 | 0.06888555 | 0.41687995 |
| ADRM1 | 16.863718 | -0.088333135 | 0.52238916 | -0.1690945 | 0.86572231 | 0.97434023 |
| LAMA5 | 46.2896923 | -0.290093383 | 0.31008929 | -0.9355156 | 0.34952262 | 0.77868979 |
| RPS21 | 14.7500271 | 0.405855032 | 0.44684659 | 0.9082648 | 0.36373833 | 0.78738269 |
| SLCO4A1 | 45.9872273 | 0.013629181 | 0.6789756 | 0.02007315 | 0.98398502 | 0.99762255 |
| MRGBP | 11.3671155 | -0.508870377 | 0.45779744 | -1.1115623 | 0.26632639 | NA |
| OGFR | 41.6176957 | -0.886793241 | 0.35233137 | -2.5169296 | 0.01183825 | 0.16196195 |
| TCFL5 | 20.5550678 | 0.655407614 | 0.41280946 | 1.58767586 | 0.11235965 | 0.51571293 |
| DIDO1 | 102.200831 | 0.025894621 | 0.1923959 | 0.13459029 | 0.89293582 | 0.97831181 |
| GID8 | 34.6998039 | 0.360171788 | 0.28368737 | 1.26960812 | 0.20422426 | 0.65771647 |
| YTHDF1 | 40.0124909 | 0.222928291 | 0.28806176 | 0.7738906 | 0.43899549 | 0.82745796 |
| ARFGAP1 | 30.84544 | 0.188959303 | 0.34505905 | 0.5476144 | 0.5839567 | 0.89036744 |

| | | | | | | |
|---|---|---|---|---|---|---|
| KCNQ2 | 14.7875414 | -2.435356003 | 1.05685252 | -2.304348 | 0.02120312 | 0.22725991 |
| EEF1A2 | 52.5894787 | -5.277318018 | 0.96662852 | -5.4595099 | 4.77E-08 | 1.48E-05 |
| PPDPF | 193.795142 | -0.49416363 | 0.39820081 | -1.240991 | 0.21460907 | 0.66839951 |
| HELZ2 | 29.5043296 | -1.348605208 | 0.8521853 | -1.5825258 | 0.11352959 | 0.5174986 |
| GMEB2 | 23.83517 | -0.706120395 | 0.32237973 | -2.1903374 | 0.02849977 | 0.26643114 |
| MHENCR | 11.2413218 | 0.201968846 | 0.39114043 | 0.51635891 | 0.60560376 | NA |
| ARFRP1 | 20.979633 | 0.229122762 | 0.36666288 | 0.62488671 | 0.53204542 | 0.86853594 |
| SLC2A4RG | 20.9811498 | 0.388002593 | 0.33956697 | 1.14263938 | 0.25318835 | 0.70701839 |
| ZBTB46 | 56.4112788 | -0.416134306 | 0.24545226 | -1.6953778 | 0.09000378 | 0.47226726 |
| TPD52L2 | 35.2483532 | 0.176083136 | 0.33751443 | 0.52170551 | 0.60187539 | 0.89522668 |
| DNAJC5 | 106.241838 | -0.119149907 | 0.23647013 | -0.5038687 | 0.61435361 | 0.90054121 |
| UCKL1 | 24.5640735 | 0.636886518 | 0.35527002 | 1.79268296 | 0.07302363 | 0.42644747 |
| ZNF512B | 12.5176045 | 0.199262468 | 0.52624016 | 0.3786531 | 0.70494548 | NA |
| PRPF6 | 49.1104525 | 0.075585159 | 0.27638423 | 0.27347855 | 0.78448538 | 0.95548822 |
| PCMTD2 | 110.652527 | -0.072972749 | 0.3268948 | -0.2232301 | 0.82335645 | 0.96539788 |
| MIER2 | 16.4347057 | 0.320301293 | 0.33376119 | 0.95967209 | 0.33722028 | 0.76966704 |
| SHC2 | 16.4426957 | -2.001872393 | 0.7202266 | -2.7795036 | 0.00544421 | 0.10201977 |
| TPGS1 | 22.9182091 | -0.348010411 | 0.5059775 | -0.6877982 | 0.49157988 | 0.85654883 |
| CDC34 | 22.6701911 | 0.164566407 | 0.34592308 | 0.47573122 | 0.63426588 | 0.90805737 |
| BSG | 350.708824 | -0.116684841 | 0.419989 | -0.2778283 | 0.78114414 | 0.95540382 |
| HCN2 | 11.2993309 | 1.040376392 | 0.66532514 | 1.56371122 | 0.1178854 | NA |
| RNF126 | 20.1919747 | -0.022019066 | 0.32499139 | -0.0677528 | 0.94598244 | 0.99134285 |
| PALM | 37.6037844 | 0.428853957 | 0.49448747 | 0.86726962 | 0.3857943 | 0.7994129 |
| PTBP1 | 73.2492926 | 0.135869633 | 0.28709908 | 0.47324998 | 0.63603484 | 0.90881415 |
| MED16 | 29.7698532 | -0.164129135 | 0.36638229 | -0.4479723 | 0.65417315 | 0.9158679 |
| R3HDM4 | 31.9596979 | -0.451844 | 0.43478124 | -1.0392445 | 0.29869105 | 0.74296125 |
| ARID3A | 70.4306024 | -0.520757181 | 0.35140677 | -1.4819213 | 0.13836124 | 0.57086516 |
| WDR18 | 13.4198725 | 0.187108588 | 0.43955761 | 0.42567478 | 0.67034484 | 0.92146857 |
| TMEM259 | 82.4132318 | 0.107122996 | 0.21429724 | 0.49988043 | 0.61715927 | 0.90173965 |
| CNN2 | 20.2121345 | 0.340018235 | 0.39421884 | 0.86251138 | 0.38840617 | 0.80109289 |
| ABCA7 | 12.99605 | 0.604011382 | 0.49291012 | 1.22539863 | 0.22042506 | 0.67406151 |
| POLR2E | 32.6030531 | -0.113965086 | 0.26806869 | -0.4251339 | 0.67073907 | 0.92170876 |
| GPX4 | 144.41569 | -0.113113058 | 0.31941249 | -0.3541285 | 0.7232426 | 0.93889899 |
| SBNO2 | 77.1820328 | -1.845917899 | 0.44006768 | -4.1946228 | 2.73E-05 | 0.00262501 |
| STK11 | 55.9593655 | 0.163305648 | 0.25027686 | 0.65249998 | 0.51407869 | 0.86416832 |
| ATP5F1D | 176.398073 | 0.045898508 | 0.36531193 | 0.12564196 | 0.90001534 | 0.97940229 |
| MIDN | 84.1415664 | -0.933110523 | 0.48622116 | -1.9191072 | 0.05497077 | 0.37340089 |
| CIRBP | 130.864169 | 0.267286862 | 0.20475412 | 1.30540406 | 0.1917553 | 0.6421543 |
| C19orf24 | 32.2158193 | -0.274442047 | 0.33049864 | -0.8303878 | 0.40631955 | 0.8115705 |
| MUM1 | 41.6522078 | -0.155948077 | 0.27247577 | -0.5723374 | 0.56709341 | 0.88417195 |
| NDUFS7 | 56.336012 | 0.222540398 | 0.40416522 | 0.55061739 | 0.58189598 | 0.89036744 |
| DAZAP1 | 69.1113169 | -0.077894515 | 0.26316976 | -0.2959858 | 0.76724092 | 0.95205353 |
| RPS15 | 353.382758 | -0.002445492 | 0.33934799 | -0.0072064 | 0.99425014 | 0.99923337 |
| C19orf25 | 40.7032941 | -0.044545144 | 0.32712087 | -0.1361733 | 0.89168424 | 0.97793768 |
| MBD3 | 31.5616025 | -0.054487618 | 0.4997899 | -0.109021 | 0.9131858 | 0.98376271 |
| TCF3 | 52.5141045 | 0.448387111 | 0.34276458 | 1.30814892 | 0.1908228 | 0.6421543 |
| REXO1 | 23.1278453 | 0.177310911 | 0.46553679 | 0.38087411 | 0.70329666 | 0.93304742 |
| ABHD17A | 38.0888871 | -0.046672971 | 0.24876229 | -0.1876208 | 0.85117395 | 0.97027687 |
| SCAMP4 | 36.1337603 | 0.144653647 | 0.31037886 | 0.46605508 | 0.64117607 | 0.91146968 |
| CSNK1G2 | 47.3962423 | -0.456803082 | 0.37661277 | -1.2129251 | 0.22515845 | 0.67848307 |
| BTBD2 | 37.7447442 | 0.32215451 | 0.29509438 | 1.09169991 | 0.27496502 | 0.72554623 |
| MKNK2 | 48.7950354 | 0.073358699 | 0.22691336 | 0.32328946 | 0.74647604 | 0.94614106 |
| MOB3A | 20.0842698 | -0.395190321 | 0.49314126 | -0.8013735 | 0.42291547 | 0.82177084 |
| AP3D1 | 68.0379949 | -0.154841631 | 0.33261761 | -0.4655245 | 0.64155593 | 0.91146968 |
| DOT1L | 60.611023 | 0.094204164 | 0.34107282 | 0.27619956 | 0.78239479 | 0.95544332 |
| PLEKHJ1 | 83.6364309 | 0.358089628 | 0.2420655 | 1.47930879 | 0.1390578 | 0.57203592 |
| SF3A2 | 12.2242223 | 0.327496956 | 0.38617309 | 0.84805742 | 0.39640599 | NA |
| LSM7 | 49.6900553 | -0.069090927 | 0.32718371 | -0.2111686 | 0.83275571 | 0.96701848 |
| SPPL2B | 33.6023217 | 0.24341689 | 0.30190325 | 0.80627448 | 0.4200846 | 0.81913024 |
| TIMM13 | 47.88331 | 0.323617201 | 0.26351572 | 1.22807551 | 0.21941862 | 0.67305276 |
| LMNB2 | 14.2975626 | 0.952584274 | 0.35858753 | 2.65649022 | 0.00789587 | 0.12893936 |
| GADD45B | 61.7771144 | -2.125411309 | 0.53532712 | -3.9703038 | 7.18E-05 | 0.00526714 |
| GNG7 | 58.7096731 | 1.016345206 | 0.35110617 | 2.89469478 | 0.00379527 | 0.08055614 |
| DIRAS1 | 11.4302742 | 0.419160289 | 0.61359568 | 0.68312132 | 0.49453018 | NA |
| SGTA | 40.0596403 | -0.032752801 | 0.36940948 | -0.0886626 | 0.92935006 | 0.98733028 |
| THOP1 | 15.7117382 | 0.43567458 | 0.41916682 | 1.0393823 | 0.29862697 | 0.74296125 |
| ZNF555 | 19.2055058 | -0.378957184 | 0.34217266 | -1.1075028 | 0.2680766 | 0.71965004 |
| TLE2 | 70.4048967 | 0.554043994 | 0.38947932 | 1.42252481 | 0.15487396 | 0.59498864 |
| AES | 230.832264 | -0.168280294 | 0.28392516 | -0.5926924 | 0.55338701 | 0.87638432 |
| GNA11 | 228.320037 | -0.091158598 | 0.3722073 | -0.2449135 | 0.8065234 | 0.96005568 |
| NCLN | 23.2785444 | 0.751030194 | 0.43376123 | 1.73143688 | 0.08337387 | 0.4560197 |
| NFIC | 211.490793 | -0.079364207 | 0.27276712 | -0.2909596 | 0.77108223 | 0.95268089 |
| SMIM24 | 91.4756691 | 0.517933241 | 0.40749564 | 1.27101542 | 0.20372316 | 0.65771647 |
| DOHH | 13.7671988 | 0.005065161 | 0.42997871 | 0.01178003 | 0.99060112 | 0.99923337 |
| FZR1 | 23.9588495 | 0.448137081 | 0.31306629 | 1.43144468 | 0.15230281 | 0.59039732 |
| MFSD12 | 35.4632911 | 0.104641986 | 0.26313427 | 0.39767525 | 0.69086958 | 0.92786587 |
| HMG20B | 24.5337852 | -0.037758207 | 0.34732067 | -0.1087128 | 0.91343028 | 0.98376271 |
| PIP5K1C | 26.0516345 | -0.219610814 | 0.33950686 | -0.6468524 | 0.51772751 | 0.86500999 |
| TJP3 | 41.1574838 | 0.374052237 | 0.3018472 | 1.23921057 | 0.21526753 | 0.66933563 |
| MRPL54 | 39.615762 | -0.375670211 | 0.40481646 | -0.9280013 | 0.35340688 | 0.78028099 |
| DAPK3 | 22.6562626 | -0.323820039 | 0.33965831 | -0.95337 | 0.34040264 | 0.77175993 |
| EEF2 | 307.786547 | -0.097589145 | 0.38485798 | -0.2535718 | 0.79982636 | 0.95861074 |
| PIAS4 | 20.2889508 | 0.086509915 | 0.3929932 | 0.22013082 | 0.82576927 | 0.96539788 |
| ZBTB7A | 78.8958569 | 0.168225991 | 0.41914769 | 0.40135255 | 0.68816058 | 0.9267526 |
| MAP2K2 | 81.0714601 | -0.204920059 | 0.21118615 | -0.970329 | 0.3318825 | 0.76685572 |
| SIRT6 | 12.6595326 | -0.307432782 | 0.40514422 | -0.7588231 | 0.4479584 | 0.83066268 |
| YJU2 | 10.7106296 | 0.536728975 | 0.47472014 | 1.13062189 | 0.25821427 | NA |
| STAP2 | 22.4333809 | -0.670261707 | 0.35123219 | -1.9083152 | 0.0563505 | 0.37709488 |
| SH3GL1 | 24.2803346 | -0.310391166 | 0.41231591 | -0.7527994 | 0.45157048 | 0.8323206 |
| UBXN6 | 35.4799416 | -0.503234546 | 0.34935229 | -1.4404787 | 0.149732 | 0.58659144 |
| HDGFL2 | 20.1342184 | 0.588625325 | 0.32955756 | 1.78610778 | 0.07408181 | 0.42901107 |
| PLIN5 | 46.8418025 | 0.173536484 | 0.28702357 | 0.60460709 | 0.5454401 | 0.87247631 |
| MYDGF | 55.1735222 | -0.60314425 | 0.30727291 | -1.9628943 | 0.04965846 | 0.35601407 |
| DPP9 | 27.851247 | 0.04940104 | 0.35920993 | 0.13752693 | 0.89061431 | 0.9779208 |
| TICAM1 | 17.9892157 | -0.726779014 | 0.32953401 | -2.205475 | 0.02742078 | 0.26051084 |
| PLIN3 | 63.6819976 | -1.233289809 | 0.38910523 | -3.1695534 | 0.00152673 | 0.04267815 |
| KDM4B | 142.678361 | -0.273592649 | 0.28759855 | -0.9513005 | 0.34145185 | 0.77193691 |
| PTPRS | 45.6224856 | 0.834229338 | 0.38317629 | 2.17714239 | 0.02946995 | 0.27224616 |
| TINCR | 16.8980236 | -0.014034357 | 0.39025773 | -0.0359618 | 0.97131285 | 0.99526459 |
| SAFB2 | 77.0489008 | -0.049584959 | 0.2026886 | -0.2446362 | 0.80673817 | 0.96005568 |
| SAFB | 92.3522144 | 0.194817515 | 0.21325713 | 0.91353341 | 0.36096207 | 0.78616766 |
| RPL36 | 218.770255 | -0.115545555 | 0.35357872 | -0.3267888 | 0.74382765 | 0.94546379 |
| C19orf70 | 70.2934819 | -0.035023963 | 0.30988763 | -0.1130215 | 0.91001352 | 0.98331039 |
| LONP1 | 61.2190389 | -0.527804792 | 0.36651312 | -1.4400707 | 0.14984741 | 0.58673121 |
| CAPS | 43.079149 | 0.727412271 | 0.61433876 | 1.18405726 | 0.2363904 | 0.69128335 |
| RANBP3 | 33.4332406 | 0.245681414 | 0.26421483 | 0.92985475 | 0.35244629 | 0.77964174 |
| RFX2 | 19.5694962 | 0.369116766 | 0.36559526 | 1.00963226 | 0.31267151 | 0.7555244 |
| MLLT1 | 29.4945178 | 0.23078673 | 0.41639688 | 0.55424702 | 0.5794098 | 0.88957613 |
| CLPP | 21.1569512 | -0.646018558 | 0.38969779 | -1.6577424 | 0.09736947 | 0.4826182 |
| ALKBH7 | 30.5395796 | 0.395556517 | 0.42659857 | 0.92723357 | 0.35380528 | 0.78035986 |
| GTF2F1 | 22.9364421 | 0.198293157 | 0.4085283 | 0.48538414 | 0.62740388 | 0.90589724 |
| KHSRP | 33.3305382 | 0.126548928 | 0.43244467 | 0.29263612 | 0.7698003 | 0.95218859 |
| SLC25A23 | 24.1091955 | -0.051184349 | 0.34535443 | -0.1482082 | 0.88217848 | 0.9766185 |

| | | | | | | |
|---|---|---|---|---|---|---|
| CRB3 | 15.9597483 | -0.327183507 | 0.54751055 | -0.5975839 | 0.55011759 | 0.87445431 |
| AC010503.4 | 16.7906538 | -0.037376363 | 0.64363866 | -0.0580704 | 0.95369254 | 0.99217529 |
| TRIP10 | 20.8652537 | -0.2447188 | 0.52853713 | -0.4630116 | 0.64335608 | 0.91192138 |
| SH2D3A | 31.5557477 | -1.071606509 | 0.51887031 | -2.0652685 | 0.0388976 | 0.31669347 |
| ZNF557 | 11.3776341 | -0.068856907 | 0.41571292 | -0.1656357 | 0.86844364 | NA |
| INSR | 453.577579 | -0.534236714 | 0.2387254 | -2.2378713 | 0.02522945 | 0.25100069 |
| ZNF358 | 13.9495751 | 0.504735433 | 0.47218985 | 1.06892478 | 0.28510356 | 0.73659018 |
| PNPLA6 | 23.4719932 | 1.020424201 | 0.43807177 | 2.32935393 | 0.01984032 | 0.22095423 |
| CAMSAP3 | 17.3574117 | 0.213141046 | 0.45444603 | 0.46901289 | 0.63906042 | 0.91066345 |
| XAB2 | 13.1227421 | -0.086568978 | 0.42698228 | -0.2027461 | 0.83933351 | 0.96748214 |
| STXBP2 | 23.7918227 | -0.032082022 | 0.39930064 | -0.0803455 | 0.93596245 | 0.98871562 |
| EVI5L | 54.5953195 | 0.374620411 | 0.42454946 | 0.88239522 | 0.37756312 | 0.79432848 |
| MAP2K7 | 44.0462571 | -0.273393615 | 0.397407 | -0.6879436 | 0.49148828 | 0.85654883 |
| TIMM44 | 24.3668759 | 0.361491738 | 0.31996673 | 1.12977914 | 0.2585693 | 0.71022613 |
| ELAVL1 | 69.4090354 | 0.052738942 | 0.23070462 | 0.22859942 | 0.81918028 | 0.9638584 |
| FBN3 | 74.8897743 | 0.968814939 | 0.42153199 | 2.29831887 | 0.02154365 | 0.22951242 |
| CERS4 | 162.193609 | -0.240874954 | 0.46654066 | -0.5163 | 0.60564488 | 0.89622507 |
| AC022146.2 | 40.5355511 | -0.579728984 | 0.43027875 | -1.3473335 | 0.17787284 | 0.62542578 |
| CD320 | 20.7862536 | 0.730839532 | 0.38430877 | 1.90169879 | 0.05721054 | 0.38009745 |
| RPS28 | 61.5211701 | 0.081316878 | 0.37489154 | 0.21690774 | 0.82828024 | 0.96636016 |
| ANGPTL4 | 16.6087511 | -2.519234269 | 0.95746478 | -2.6311509 | 0.00850963 | 0.13356532 |
| RAB11B | 48.9969333 | -0.048141751 | 0.31824365 | -0.1512732 | 0.87976017 | 0.97581687 |
| 2-Mar | 22.4927829 | 0.375734575 | 0.38801724 | 0.96834506 | 0.33287207 | 0.7674968 |
| HNRNPM | 156.147909 | 0.282941857 | 0.21104648 | 1.34066135 | 0.18003043 | 0.62768048 |
| ZNF558 | 11.6076977 | 0.093407789 | 0.43230112 | 0.21607113 | 0.82893231 | NA |
| ZNF317 | 18.7555724 | 0.135362737 | 0.41672222 | 0.32482726 | 0.74531181 | 0.94614106 |
| ZNF559-ZNF | 15.9466157 | 0.305419972 | 0.36341201 | 0.84042343 | 0.40067101 | 0.80830282 |
| ZNF266 | 30.4162137 | -0.32276693 | 0.33924059 | -0.9514396 | 0.34138126 | 0.77193691 |
| ZNF426 | 18.0337278 | 0.069497559 | 0.32856938 | 0.21151563 | 0.83248494 | 0.96701848 |
| ZNF121 | 37.735363 | 0.046751704 | 0.31831949 | 0.14687038 | 0.88323433 | 0.97662748 |
| ZNF561 | 20.9996754 | 0.146272192 | 0.34002833 | 0.43017648 | 0.66706727 | 0.92032206 |
| ZNF562 | 51.0744965 | -0.082498589 | 0.27821612 | -0.296527 | 0.76682765 | 0.95205353 |
| ZNF846 | 44.2499506 | 0.332014581 | 0.26230484 | 1.26575851 | 0.20559955 | 0.65836247 |
| FBXL12 | 19.5090784 | 0.229068896 | 0.33494775 | 0.68389441 | 0.49404185 | 0.85774549 |
| UBL5 | 48.1520379 | -0.157911289 | 0.35458348 | -0.445343 | 0.65607186 | 0.9158679 |
| PIN1 | 32.4221415 | -0.077181664 | 0.32800655 | -0.2353053 | 0.81397182 | 0.96275169 |
| C19orf66 | 25.6988585 | -0.822434153 | 0.62981 | -1.3058449 | 0.19160532 | 0.6421543 |
| EIF3G | 23.0366271 | 0.324741706 | 0.36810893 | 0.88218915 | 0.37767453 | 0.79432848 |
| DNMT1 | 23.2750494 | 0.43905021 | 0.3444534 | 1.27462876 | 0.20244067 | 0.65499484 |
| MRPL4 | 31.4684062 | 0.040012811 | 0.30518896 | 0.13110832 | 0.89568962 | 0.97885533 |
| AC011511.2 | 29.3955416 | 0.331923811 | 0.68530683 | 0.48434335 | 0.62814221 | 0.906154 |
| ICAM1 | 14.8535771 | -2.502341609 | 0.69477737 | -3.6016453 | 0.00031621 | 0.01510953 |
| TYK2 | 26.4803966 | 0.220523943 | 0.36384039 | 0.60610078 | 0.54444783 | 0.87247631 |
| CDC37 | 54.8124846 | -0.231380249 | 0.26309799 | -0.8794451 | 0.37915997 | 0.79535699 |
| PDE4A | 43.2502211 | -0.017694446 | 0.39699583 | -0.0445709 | 0.96444937 | 0.99334616 |
| KEAP1 | 23.5231422 | -0.045756121 | 0.34109943 | -0.1341431 | 0.89328945 | 0.97831181 |
| ATG4D | 28.8670443 | 0.038512638 | 0.34687108 | 0.11102868 | 0.9115936 | 0.9835359 |
| KRI1 | 14.4216843 | 0.05035326 | 0.42892475 | 0.11739416 | 0.90654771 | 0.98196682 |
| AP1M2 | 35.5372496 | -0.132563756 | 0.36000456 | -0.368228 | 0.71270324 | 0.9357596 |
| SLC44A2 | 32.9457673 | 0.420111363 | 0.39130395 | 1.07361902 | 0.28299347 | 0.7344003 |
| ILF3-DT | 25.820468 | -0.013342964 | 0.39326458 | -0.0339287 | 0.97293399 | 0.99526459 |
| ILF3 | 102.646434 | 0.219853623 | 0.31835472 | 0.69059325 | 0.48982119 | 0.85562139 |
| QTRT1 | 25.0933187 | 0.904443922 | 0.28141309 | 3.21393695 | 0.00130928 | 0.03883809 |
| DNM2 | 225.680589 | -0.563224609 | 0.30365698 | -1.8548054 | 0.06362402 | 0.40032348 |
| TMED1 | 27.0157734 | -0.532809829 | 0.44799686 | -1.189316 | 0.23431535 | 0.68933219 |
| CARM1 | 21.0108323 | 0.081125082 | 0.36626748 | 0.22149136 | 0.82470987 | 0.96539788 |
| YIPF2 | 21.6477796 | 0.232943464 | 0.34250731 | 0.68011239 | 0.4964333 | 0.85814183 |
| TIMM29 | 15.9129647 | -0.230188103 | 0.34777059 | -0.6618964 | 0.50803761 | 0.86281145 |
| SMARCA4 | 121.245431 | 0.237293678 | 0.29710194 | 0.79869448 | 0.42446759 | 0.82231588 |
| LDLR | 179.192167 | -2.158471755 | 0.5643489 | -3.8247116 | 0.00013093 | 0.00803782 |
| KANK2 | 46.4027529 | -0.435062383 | 0.34509339 | -1.2607091 | 0.20741366 | 0.65905456 |
| DOCK6 | 62.187497 | -0.339626767 | 0.37048672 | -0.9167043 | 0.35929761 | 0.78473041 |
| CCDC159 | 12.3105648 | -0.125270633 | 0.51322097 | -0.2440871 | 0.80716334 | NA |
| EPOR | 28.4890398 | 0.070208616 | 0.41099724 | 0.17082503 | 0.86436135 | 0.97407858 |
| RGL3 | 262.62207 | -0.309672923 | 0.2597085 | -1.1923865 | 0.23310971 | 0.68763965 |
| PRKCSH | 50.997982 | 0.175553928 | 0.30697986 | 0.57187442 | 0.56740705 | 0.88443651 |
| ECSIT | 37.7749439 | 0.279362004 | 0.24987023 | 1.11802837 | 0.26355488 | 0.71380385 |
| ELOF1 | 14.4382769 | -0.189558214 | 0.5147039 | -0.3682859 | 0.71266003 | 0.9357596 |
| ZNF627 | 26.69977 | 0.402774533 | 0.32581194 | 1.23621785 | 0.2163776 | 0.67099191 |
| AC008543.1 | 23.8745057 | 0.116266284 | 0.40832768 | 0.2847377 | 0.7758451 | 0.95493992 |
| ZNF823 | 22.7469485 | 0.236950742 | 0.31556337 | 0.75088164 | 0.45272389 | 0.83328686 |
| ZNF441 | 15.4047666 | -0.801799 | 0.42839607 | -1.8716301 | 0.06125779 | 0.39273006 |
| ZNF440 | 41.0508245 | -0.830141736 | 0.31777109 | -2.6123891 | 0.00899119 | 0.13780744 |
| ZNF439 | 21.7844272 | -0.33679787 | 0.32036587 | -1.0512914 | 0.29312479 | 0.74013384 |
| ZNF69 | 38.138675 | 0.445309544 | 0.45850915 | 0.9712119 | 0.33144277 | 0.76685572 |
| ZNF433 | 10.6782032 | -0.283710823 | 0.45958047 | -0.6173257 | 0.53701993 | NA |
| ZNF844 | 27.8072624 | -0.113304306 | 0.31226316 | -0.3628488 | 0.71671785 | 0.93689626 |
| ZNF136 | 19.0458028 | 0.170697063 | 0.38820517 | 0.43970837 | 0.66014834 | 0.91752855 |
| ZNF44 | 124.844214 | 0.163757625 | 0.21178853 | 0.7732129 | 0.43939639 | 0.82760292 |
| AC008758.5 | 17.0544262 | -0.424303574 | 0.45036728 | -0.9421279 | 0.34612717 | 0.77642952 |
| ZNF490 | 30.8499346 | -0.393610305 | 0.41207115 | -0.9551999 | 0.33947663 | 0.77045469 |
| ZNF791 | 53.2466556 | -0.064846075 | 0.25488916 | -0.2544089 | 0.79917966 | 0.95848876 |
| MAN2B1 | 11.2317711 | 0.317445803 | 0.57575112 | 0.55135942 | 0.58138731 | NA |
| WDR83 | 12.18743 | 0.104295829 | 0.57832738 | 0.18034047 | 0.85688529 | NA |
| DHPS | 18.9705451 | -0.016965856 | 0.41775789 | -0.0406117 | 0.96760546 | 0.99393148 |
| TNPO2 | 34.7472086 | 0.420589439 | 0.35229927 | 1.19384136 | 0.23254003 | 0.68763908 |
| TRIR | 77.038096 | 0.016560094 | 0.3808516 | 0.04348175 | 0.96531751 | 0.99334616 |
| ASNA1 | 22.0622306 | 0.03980085 | 0.31042815 | 0.12821276 | 0.8979806 | 0.97892709 |
| HOOK2 | 130.368556 | -0.693372898 | 0.23904802 | -2.9005591 | 0.00372498 | 0.07958649 |
| JUNB | 66.606618 | -1.729885022 | 0.52559769 | -3.2912721 | 0.00099735 | 0.03257351 |
| DNASE2 | 20.6628234 | -0.268890622 | 0.35495022 | -0.7575446 | 0.44872366 | 0.83118301 |
| GCDH | 19.9215182 | 0.201217081 | 0.30770939 | 0.65391922 | 0.51316385 | 0.86373232 |
| FARSA | 23.4547366 | -0.14118222 | 0.3635558 | -0.3883371 | 0.69776656 | 0.93009971 |
| CALR | 179.273808 | -0.045725103 | 0.3426927 | -0.1334289 | 0.89385421 | 0.97833263 |
| AC092069.1 | 29.7285631 | 0.032974289 | 0.39490259 | 0.08349981 | 0.93345413 | 0.9881638 |
| RAD23A | 38.7550072 | -0.258795937 | 0.37338434 | -0.6931087 | 0.48824134 | 0.85440032 |
| GADD45GIP1 | 67.572305 | -0.665211019 | 0.33916475 | -1.9613212 | 0.04984156 | 0.35627353 |
| NFIX | 25.5386902 | -0.219288106 | 0.47543052 | -0.4612411 | 0.64462562 | 0.9119294 |
| NACC1 | 18.9383093 | 0.346861366 | 0.41660678 | 0.83258694 | 0.40507772 | 0.81132644 |
| STX10 | 12.9879923 | -0.353235991 | 0.38849518 | -0.9092416 | 0.3632226 | 0.78718296 |
| IER2 | 56.7721618 | 0.294415121 | 0.50934112 | 0.57803133 | 0.56324297 | 0.88191823 |
| CCDC130 | 47.2733706 | -0.063158235 | 0.2757277 | -0.2290602 | 0.81882215 | 0.9635539 |
| MRI1 | 19.0285144 | -0.053944742 | 0.42763749 | -0.126146 | 0.89961639 | 0.97938035 |
| C19orf53 | 57.2450099 | -0.888296031 | 0.36849499 | -2.4106055 | 0.01592607 | 0.19383042 |
| ZSWIM4 | 16.9598778 | -0.785090931 | 0.42359234 | -1.8534115 | 0.0638234 | 0.40107793 |
| CC2D1A | 36.5091303 | -0.340111605 | 0.24511534 | -1.3875574 | 0.16527185 | 0.60578861 |
| DCAF15 | 10.880062 | -0.100219007 | 0.46659498 | -0.214788 | 0.8299326 | NA |
| RFX1 | 25.4422979 | 0.665781805 | 0.31677309 | 2.10176252 | 0.03557408 | 0.30204646 |
| PALM3 | 25.3156736 | 1.138014001 | 0.52815171 | 2.15471042 | 0.0311845 | 0.28008302 |
| SAMD1 | 12.7106111 | 0.124485071 | 0.43316802 | 0.28738287 | 0.77381917 | 0.95457609 |
| PRKACA | 30.3080271 | 0.038775915 | 0.3051112 | 0.12708781 | 0.8988709 | 0.97911821 |
| ADGRL1 | 25.5580006 | 0.816097903 | 0.40315651 | 2.02427067 | 0.04294231 | 0.33179556 |
| DDX39A | 13.3077906 | -0.445927283 | 0.63612536 | -0.7010054 | 0.48329967 | 0.85205504 |

| | | | | | | |
|---|---|---|---|---|---|---|
| PKN1 | 46.4802555 | -0.431014404 | 0.47270689 | -0.9118006 | 0.36187372 | 0.78713361 |
| GIPC1 | 66.7968499 | -0.396736635 | 0.43325562 | -0.9157103 | 0.35981886 | 0.7850299 |
| DNAJB1 | 41.0789394 | -0.012873696 | 0.36712566 | -0.0350662 | 0.97202697 | 0.99526459 |
| TECR | 46.1973036 | 0.553121911 | 0.29252007 | 1.89088533 | 0.05863965 | 0.38425985 |
| NDUFB7 | 88.0326244 | 0.542269966 | 0.32000462 | 1.69456915 | 0.09015718 | 0.47226726 |
| ZNF333 | 11.6004136 | 0.196232902 | 0.40809674 | 0.48084898 | 0.63062383 | NA |
| ILVBL | 24.4993316 | 0.297403761 | 0.33069736 | 0.89932305 | 0.36848061 | 0.78877653 |
| BRD4 | 196.355625 | -0.008011995 | 0.23290118 | -0.0344008 | 0.97255752 | 0.99526459 |
| AKAP8 | 42.7110649 | -0.080089117 | 0.22866212 | -0.3502509 | 0.7261504 | 0.94003188 |
| AKAP8L | 116.313564 | -0.031512419 | 0.21821026 | -0.1444131 | 0.88517428 | 0.97662748 |
| WIZ | 11.711013 | -0.241926498 | 0.44081787 | -0.5488128 | 0.58313391 | NA |
| TPM4 | 20.134245 | 1.571496384 | 0.82970996 | 1.89403101 | 0.05822089 | 0.38243048 |
| RAB8A | 23.3386115 | 0.151424507 | 0.31688152 | 0.47785843 | 0.63275097 | 0.90805737 |
| FAM32A | 13.6786888 | -0.232620333 | 0.54324742 | -0.4282033 | 0.66850312 | 0.92049996 |
| AP1M1 | 17.3945995 | -0.063668125 | 0.31391652 | -0.2028187 | 0.83927678 | 0.96748214 |
| EPS15L1 | 84.5824057 | 0.239441562 | 0.24215 | 0.98881503 | 0.32275365 | 0.76296579 |
| C19orf44 | 12.7981879 | 0.165530074 | 0.37583006 | 0.44043862 | 0.65961946 | 0.91732025 |
| TMEM38A | 96.4083062 | 0.632664512 | 0.35703118 | 1.77201475 | 0.07639211 | 0.43515065 |
| SIN3B | 47.9788171 | 0.014799632 | 0.3472368 | 0.04262115 | 0.96600353 | 0.99334616 |
| CPAMD8 | 225.126757 | -0.969405 | 0.49572766 | -1.9555193 | 0.05052181 | 0.35823214 |
| MYO9B | 49.150495 | -0.18954057 | 0.39419065 | -0.4808348 | 0.63063394 | 0.90768383 |
| OCEL1 | 13.9742052 | 0.437463366 | 0.45867721 | 0.95374995 | 0.34021023 | 0.77157803 |
| NR2F6 | 52.066762 | -0.188812715 | 0.27144726 | -0.6955779 | 0.48669321 | 0.85365949 |
| ABHD8 | 11.6025384 | -0.007810371 | 0.46158506 | -0.0169208 | 0.98649983 | NA |
| MRPL34 | 52.4992563 | -0.129624731 | 0.41990418 | -0.3087007 | 0.75754918 | 0.94913809 |
| DDA1 | 18.4705279 | -0.239218191 | 0.4346228 | -0.5504041 | 0.58204221 | 0.89036744 |
| BST2 | 46.2302923 | -1.603558635 | 1.23573851 | -1.2976521 | 0.19440692 | 0.6459894 |
| MVB12A | 15.015027 | 0.568922796 | 0.44298713 | 1.28428742 | 0.19904141 | 0.65182365 |
| SLC27A1 | 18.0487294 | 0.903455408 | 0.39149793 | 2.30768887 | 0.02101645 | 0.22580589 |
| AC010618.3 | 18.7917889 | 0.675802747 | 0.41236247 | 1.63885609 | 0.10124323 | 0.49127532 |
| PGLS | 57.9473571 | -0.470759349 | 0.25366988 | -1.8557952 | 0.06348275 | 0.4001801 |
| COLGALT1 | 16.8622763 | 0.888556661 | 0.38425673 | 2.31240365 | 0.02075545 | 0.22444738 |
| RPL18A | 18.568793 | -0.583370351 | 0.39295099 | -1.4845881 | 0.13765298 | 0.56931687 |
| CCDC124 | 22.5089413 | -0.169647292 | 0.45183581 | -0.3754623 | 0.70731671 | 0.93432317 |
| ARRDC2 | 12.334949 | -0.722532481 | 0.51854816 | -1.3933759 | 0.16350616 | NA |
| MAST3 | 11.9191636 | 0.686348415 | 0.41136229 | 1.66847675 | 0.09522112 | NA |
| JUND | 163.339654 | -0.371891263 | 0.49111588 | -0.7572373 | 0.44890771 | 0.83118301 |
| LSM4 | 65.077601 | 0.069002567 | 0.26282126 | 0.26254561 | 0.79290083 | 0.95746995 |
| PGPEP1 | 35.7611533 | 0.187634912 | 0.47442752 | 0.39549753 | 0.69247574 | 0.928133 |
| GDF15 | 133.152067 | -2.554590401 | 0.64614551 | -3.9535837 | 7.70E-05 | 0.00545107 |
| SSBP4 | 27.2282011 | -0.341344927 | 0.28780776 | -1.1860171 | 0.23561554 | 0.690266 |
| ISYNA1 | 12.2144223 | 0.654972302 | 0.6603717 | 0.99182371 | 0.32128353 | NA |
| ELL | 50.3005105 | -0.212263767 | 0.25253808 | -0.8405218 | 0.40061588 | 0.80830282 |
| FKBP8 | 48.563003 | -0.22804372 | 0.2746276 | -0.8303744 | 0.40632715 | 0.8115705 |
| KXD1 | 23.4181329 | -0.257092245 | 0.28595736 | -0.899058 | 0.36862179 | 0.78877653 |
| UBA52 | 107.183817 | 0.333755934 | 0.38020483 | 0.87783192 | 0.38003494 | 0.7959986 |
| REX1BD | 29.005349 | 0.973249055 | 0.45150539 | 2.15556463 | 0.03111768 | 0.27974186 |
| KLHL26 | 14.002872 | 0.60510201 | 0.57559671 | 1.05126037 | 0.29313902 | 0.74013384 |
| CRTC1 | 114.860037 | 0.506783234 | 0.31655751 | 1.60091993 | 0.10939466 | 0.50844665 |
| UPF1 | 46.9866907 | 0.136167187 | 0.25479492 | 0.53441876 | 0.59305184 | 0.89215305 |
| COPE | 64.4935212 | -0.034710242 | 0.29189097 | -0.1189151 | 0.90534262 | 0.9810997 |
| DDX49 | 11.7293169 | -0.252297724 | 0.48463733 | -0.5205908 | 0.60265188 | NA |
| SUGP2 | 97.513652 | 0.168196893 | 0.19298899 | 0.87153619 | 0.38346145 | 0.7980695 |
| ARMC6 | 10.6905514 | -0.165446111 | 0.51000756 | -0.3243993 | 0.74563573 | NA |
| SLC25A42 | 17.4690383 | 0.333073686 | 0.53272019 | 0.62523195 | 0.53181884 | 0.86846407 |
| TMEM161A | 15.1560776 | -0.055994017 | 0.34664765 | -0.1615301 | 0.87167593 | 0.97434023 |
| RFXANK | 10.6049126 | 0.41850158 | 0.40109089 | 1.04340834 | 0.29675921 | NA |
| SUGP1 | 20.442616 | 0.356472724 | 0.37652194 | 0.94675152 | 0.34376541 | 0.77370523 |
| MAU2 | 66.3349777 | 0.105703823 | 0.2019171 | 0.52350109 | 0.60062559 | 0.89490801 |
| GATAD2A | 124.756784 | -0.204455392 | 0.2553387 | -0.8007223 | 0.42329242 | 0.82200931 |
| ATP13A1 | 21.6396301 | 0.03178784 | 0.32969237 | 0.09641667 | 0.92318965 | 0.98672316 |
| ZNF101 | 11.0487856 | -0.35085348 | 0.40465934 | -0.8670342 | 0.38592328 | NA |
| ZNF14 | 18.4314991 | 0.089124395 | 0.37771678 | 0.23595562 | 0.81346711 | 0.96264114 |
| ZNF253 | 30.9542279 | 0.195187044 | 0.35342059 | 0.55227978 | 0.58075668 | 0.89012555 |
| ZNF93 | 12.2069202 | -0.026149851 | 0.38630677 | -0.0676919 | 0.94603087 | NA |
| ZNF90 | 135.580214 | 0.132894098 | 0.37150181 | 0.35772126 | 0.72055192 | 0.93806337 |
| AC011447.3 | 23.9521432 | -0.129882867 | 0.44919899 | -0.2891433 | 0.77247174 | 0.95391163 |
| ZNF486 | 17.7145414 | -0.133293468 | 0.475571 | -0.2802809 | 0.779262 | 0.95537097 |
| ZNF737 | 33.6146293 | -0.292262252 | 0.33641069 | -0.8687662 | 0.384975 | 0.79869171 |
| AC010636.2 | 17.7313458 | -0.77253431 | 0.54548854 | -1.4162247 | 0.15670974 | 0.59731328 |
| AC010329.1 | 15.2774886 | -0.860465531 | 0.69425232 | -1.2394133 | 0.21519249 | 0.66933563 |
| ZNF66 | 35.2901563 | -0.679761496 | 0.335416 | -2.0266222 | 0.04270107 | 0.33083096 |
| ZNF85 | 21.6107381 | -0.676146819 | 0.3991342 | -1.6940338 | 0.09025887 | 0.47226726 |
| ZNF430 | 42.9577217 | -0.215273207 | 0.25705763 | -0.8374512 | 0.40233902 | 0.80890307 |
| ZNF714 | 26.1869223 | -0.042590231 | 0.31201118 | -0.1365023 | 0.89142424 | 0.97793768 |
| ZNF431 | 99.3060532 | 0.540461355 | 0.20048903 | 2.69571532 | 0.00702377 | 0.12015968 |
| ZNF708 | 59.5773345 | 0.111443417 | 0.25908421 | 0.43014362 | 0.66709117 | 0.92032206 |
| ZNF738 | 12.2672349 | 0.184809337 | 0.34580346 | 0.5344346 | 0.59304088 | NA |
| ZNF493 | 34.0609725 | 0.278560975 | 0.33682989 | 0.82700788 | 0.4082326 | 0.81361336 |
| AC010615.4 | 46.6813261 | -0.810403951 | 0.65660558 | -1.2342325 | 0.21711628 | 0.67156502 |
| ZNF429 | 62.862767 | -0.185425015 | 0.23483911 | -0.7895832 | 0.42977122 | 0.82469033 |
| AC123912.1 | 24.7886308 | -0.186251214 | 0.43081495 | -0.432323 | 0.66550667 | 0.91958319 |
| AC123912.4 | 34.5665331 | 0.050681198 | 0.46282406 | 0.10950424 | 0.91280256 | 0.98376271 |
| ZNF100 | 38.6186632 | -0.13558832 | 0.42821381 | -0.316637 | 0.75151909 | 0.94719044 |
| ZNF43 | 44.8012871 | 0.044743449 | 0.29872338 | 0.14978221 | 0.88093644 | 0.97624626 |
| ZNF208 | 115.363864 | -0.423776743 | 0.30476335 | -1.3905108 | 0.16437381 | 0.60496776 |
| ZNF257 | 16.549018 | 0.080590387 | 0.37439392 | 0.2152556 | 0.82956805 | 0.96669991 |
| ZNF676 | 17.288943 | 0.120945127 | 0.56363535 | 0.21458045 | 0.83009444 | 0.966913 |
| ZNF98 | 65.9096365 | 0.147086377 | 0.32428489 | 0.45357148 | 0.65013729 | 0.91462388 |
| ZNF99 | 13.7620279 | -0.262297332 | 0.57401971 | -0.4569483 | 0.6477082 | 0.91360118 |
| ZNF91 | 264.654813 | -0.282330822 | 0.23093546 | -1.2225529 | 0.2214986 | 0.67492938 |
| ZNF675 | 24.0675635 | 0.182629982 | 0.29638795 | 0.61618558 | 0.53777205 | 0.86972442 |
| ZNF726 | 13.0315242 | 1.100011926 | 0.53872666 | 2.04187393 | 0.04116404 | 0.32568611 |
| ZNF254 | 165.487217 | -0.199749168 | 0.19989766 | -0.9992572 | 0.31767014 | 0.75971409 |
| LINC00662 | 49.1755557 | -0.169080823 | 0.26605531 | -0.63551 | 0.52509582 | 0.86747074 |
| AC006504.5 | 77.7172112 | 0.18821659 | 0.32299351 | 0.58272561 | 0.56007803 | 0.88037687 |
| UQCRFS1 | 125.878317 | -0.046806742 | 0.43543821 | -0.1074934 | 0.91439755 | 0.98407488 |
| POP4 | 12.3951766 | -0.041505469 | 0.38237236 | -0.1085473 | 0.9135616 | NA |
| C19orf12 | 50.6040792 | -0.519976353 | 0.23352373 | -2.2266532 | 0.02597046 | 0.25343407 |
| URI1 | 140.48933 | 0.002565323 | 0.17726362 | 0.0144718 | 0.98845358 | 0.99915789 |
| ZNF507 | 42.9075006 | -0.014682104 | 0.23507213 | -0.0624579 | 0.95019821 | 0.99192022 |
| DPY19L3 | 37.0103318 | -0.018245454 | 0.3402827 | -0.0536185 | 0.9572391 | 0.99236316 |
| PDCD5 | 22.9747132 | -0.180358597 | 0.36519303 | -0.493872 | 0.62139661 | 0.90380166 |
| ANKRD27 | 44.3230883 | -0.195594913 | 0.3192703 | -0.6126311 | 0.5401203 | 0.87014896 |
| NUDT19 | 15.5126729 | 0.252499837 | 0.41836653 | 0.60353737 | 0.54615126 | 0.87259128 |
| CEP89 | 30.9626818 | 0.245519966 | 0.29830606 | 0.8230472 | 0.41048115 | 0.81427553 |
| RHPN2 | 126.187647 | -0.842040448 | 0.3019543 | -2.7886354 | 0.00529306 | 0.10037484 |
| GPATCH1 | 14.5106961 | 0.479787997 | 0.39397894 | 1.21780114 | 0.22329955 | 0.67606266 |
| CEBPG | 15.1079414 | -0.282678179 | 0.44553196 | -0.6344734 | 0.52577192 | 0.86749236 |
| PEPD | 43.9111491 | -0.3247884 | 0.37135854 | -0.8745952 | 0.38179419 | 0.79731722 |
| LSM14A | 187.342685 | 0.056588495 | 0.17812108 | 0.31769678 | 0.75071495 | 0.94705879 |
| KIAA0355 | 103.713405 | 0.139559822 | 0.1666072 | 0.83765782 | 0.40222291 | 0.80890307 |
| GPI | 135.860737 | -0.213609512 | 0.39101399 | -0.5462963 | 0.58486225 | 0.89036744 |

| | | | | | | |
|---|---|---|---|---|---|---|
| UBA2 | 90.2218174 | 0.04089149 | 0.21568531 | 0.18958867 | 0.84963147 | 0.97027687 |
| SCGB2B2 | 40.7027464 | 0.06238416 | 0.39265756 | 0.15887676 | 0.87376597 | 0.97434023 |
| ZNF302 | 60.996124 | -0.518456234 | 0.34303309 | -1.5113884 | 0.13068952 | 0.55687707 |
| GRAMD1A | 39.5852516 | -0.980445734 | 0.37444954 | -2.6183654 | 0.00883521 | 0.13646747 |
| HPN | 118.745567 | -0.313791001 | 0.43217024 | -0.7260819 | 0.46778854 | 0.84326495 |
| LSR | 53.3163126 | -1.059646587 | 0.40036181 | -2.6467224 | 0.0081276 | 0.13108178 |
| USF2 | 44.2423053 | 0.129162058 | 0.41555095 | 0.31082123 | 0.75593653 | 0.94796399 |
| DMKN | 88.6495643 | -0.413924528 | 0.31497105 | -1.3141669 | 0.18879005 | 0.64021224 |
| TMEM147 | 29.0907819 | 0.272791692 | 0.37795596 | 0.72175523 | 0.47044498 | 0.84424147 |
| RBM42 | 16.6545305 | 0.351500975 | 0.43798735 | 0.80253682 | 0.4222425 | 0.82112114 |
| COX6B1 | 146.624485 | 0.06977696 | 0.29167384 | 0.23922941 | 0.8109277 | 0.96243713 |
| KMT2B | 28.4621199 | -0.213109562 | 0.33431213 | -0.6374569 | 0.5238273 | 0.86747074 |
| PROSER3 | 11.252323 | 0.00999718 | 0.42407681 | 0.02357398 | 0.98119242 | NA |
| SDHAF1 | 13.774967 | -0.522799252 | 0.43995087 | -1.1883128 | 0.2347102 | 0.68936745 |
| TBCB | 24.739272 | 0.038300849 | 0.35757955 | 0.10711141 | 0.9147006 | 0.98407488 |
| CAPNS1 | 114.415808 | -0.291293098 | 0.31622643 | -0.9211535 | 0.35697027 | 0.7825111 |
| COX7A1 | 24.5748168 | 0.963239073 | 0.86111505 | 1.11859509 | 0.26331292 | 0.71348562 |
| ZNF565 | 52.6999164 | -0.394026704 | 0.43591642 | -0.9039043 | 0.36604617 | 0.78793182 |
| ZNF146 | 64.8946754 | 0.020977341 | 0.27238694 | 0.07701302 | 0.93861319 | 0.98871562 |
| LINC00665 | 20.5779856 | 0.116782645 | 0.43975109 | 0.26556533 | 0.79057399 | 0.95746995 |
| ZFP14 | 44.0770109 | -0.090798076 | 0.32131002 | -0.2825871 | 0.77749335 | 0.9551993 |
| ZFP82 | 19.3519599 | -0.268794374 | 0.34435642 | -0.7805702 | 0.4350553 | 0.82565889 |
| ZNF566 | 27.138215 | 0.409040122 | 0.32523993 | 1.25765654 | 0.208516 | 0.65973983 |
| ZNF260 | 22.926552 | 0.324070504 | 0.30530588 | 1.06146173 | 0.28848012 | 0.73815415 |
| ZNF529 | 55.2763014 | 0.129946624 | 0.30496998 | 0.42609645 | 0.67003757 | 0.9212423 |
| ZNF529-AS1 | 32.3330118 | -0.241749594 | 0.32452472 | -0.7449343 | 0.45631144 | 0.83506875 |
| ZNF382 | 16.219139 | -0.427336723 | 0.4564401 | -0.9362383 | 0.34915048 | 0.77868979 |
| ZNF461 | 16.3745785 | 0.086198013 | 0.3397051 | 0.25374365 | 0.79969361 | 0.95861074 |
| ZNF567 | 25.8324568 | -0.087283381 | 0.28022357 | -0.3114777 | 0.75543752 | 0.94795939 |
| AC020928.1 | 37.2848096 | 0.135629683 | 0.2825722 | 0.47998241 | 0.6312399 | 0.90770379 |
| ZNF790 | 37.8051685 | 0.222145169 | 0.3241602 | 0.68529441 | 0.49315816 | 0.85714521 |
| ZNF345 | 36.9267515 | 0.468459103 | 0.32284896 | 1.4510163 | 0.14677532 | 0.58349206 |
| ZNF568 | 26.2246535 | 0.247646689 | 0.33874037 | 0.73108113 | 0.4647296 | 0.84133578 |
| ZNF420 | 74.7952236 | 0.300043967 | 0.30764014 | 0.97530826 | 0.32940738 | 0.76685572 |
| ZNF383 | 16.2709241 | 0.414005942 | 0.35331573 | 1.17177331 | 0.24128808 | 0.69556484 |
| HKR1 | 80.7297287 | 0.317923638 | 0.2362382 | 1.34577576 | 0.17837485 | 0.62598987 |
| ZNF569 | 24.5856241 | 0.102049021 | 0.35543103 | 0.28711342 | 0.77402547 | 0.95457609 |
| ZNF793 | 18.0681102 | 0.267705651 | 0.43019641 | 0.62228705 | 0.53375314 | 0.86909171 |
| ZNF540 | 31.8508421 | 0.106511924 | 0.37037054 | 0.28758207 | 0.77366668 | 0.95457609 |
| ZNF571 | 18.7804124 | -0.442388928 | 0.35033189 | -1.2627709 | 0.20667152 | 0.65836247 |
| ZFP30 | 24.2436589 | -0.144195573 | 0.33761696 | -0.4270981 | 0.66930786 | 0.92049996 |
| ZNF573 | 36.004167 | 0.134331991 | 0.25860435 | 0.51944984 | 0.60344708 | 0.8956362 |
| SIPA1L3 | 277.55523 | 0.22085065 | 0.26387345 | 0.83695669 | 0.40261691 | 0.80890307 |
| SPINT2 | 25.0983744 | -0.15201902 | 0.31850671 | -0.4772867 | 0.63315798 | 0.90805737 |
| CATSPERG | 11.4343381 | -0.299792936 | 0.44756826 | -0.6698262 | 0.5029686 | NA |
| PSMD8 | 67.4608131 | -0.506919126 | 0.23563314 | -2.1513065 | 0.03145201 | 0.28099688 |
| EIF3K | 89.2116615 | 0.566634994 | 0.26441047 | 2.14301274 | 0.03211208 | 0.28394452 |
| ACTN4 | 278.81953 | -0.218989126 | 0.25026425 | -0.8750316 | 0.38155672 | 0.79725517 |
| ECH1 | 53.9647234 | -0.317796878 | 0.42906178 | -0.7406786 | 0.45888833 | 0.83645518 |
| HNRNPL | 52.4333872 | -0.053697656 | 0.34607415 | -0.1551623 | 0.87669338 | 0.97532685 |
| SIRT2 | 39.0431621 | -0.234865659 | 0.27910705 | -0.8414895 | 0.40007375 | 0.80830282 |
| NFKBIB | 16.4077914 | -0.527782788 | 0.41769708 | -1.2635539 | 0.20639018 | 0.65836247 |
| MRPS12 | 52.4935954 | 0.149809226 | 0.37083946 | 0.40397326 | 0.68623238 | 0.92597882 |
| FBXO17 | 45.9277991 | 0.030412565 | 0.39628452 | 0.07674427 | 0.93882699 | 0.98871562 |
| PAK4 | 73.6288655 | 0.592080814 | 0.26673446 | 2.21973872 | 0.02643651 | 0.25541249 |
| SAMD4B | 108.962318 | 0.037319118 | 0.2403724 | 0.15525542 | 0.87661996 | 0.97532685 |
| PAF1 | 19.1749118 | 0.150809976 | 0.3332521 | 0.45254021 | 0.65087986 | 0.91513765 |
| MED29 | 22.8007144 | 0.239488107 | 0.32414382 | 0.73883285 | 0.4600085 | 0.83690755 |
| ZFP36 | 39.1832375 | -1.805524582 | 0.49362691 | -3.6576705 | 0.00025452 | 0.01282319 |
| RPS16 | 158.55535 | 0.315864108 | 0.38255191 | 0.82567646 | 0.40898765 | 0.81399186 |
| SUPT5H | 63.1614418 | -0.250003799 | 0.27368781 | -0.9134634 | 0.36099886 | 0.78616766 |
| TIMM50 | 16.9640341 | 0.245629831 | 0.31495267 | 0.77989443 | 0.43545302 | 0.82590941 |
| FBL | 15.3067122 | 0.339343186 | 0.39112069 | 0.86761757 | 0.38560372 | 0.7993922 |
| PSMC4 | 23.6042152 | -0.845283594 | 0.31663739 | -2.6695634 | 0.00759499 | 0.12593407 |
| ZNF546 | 14.041969 | -0.026246045 | 0.47616719 | -0.0551194 | 0.95604335 | 0.99234582 |
| ZNF780B | 64.3425712 | 0.178941358 | 0.34238316 | 0.52263481 | 0.60122841 | 0.89490801 |
| ZNF780A | 18.4869055 | 0.590225618 | 0.29656732 | 1.99019103 | 0.0465699 | 0.34378081 |
| MAP3K10 | 15.9086194 | -0.225365754 | 0.3852082 | -0.5850492 | 0.55851462 | 0.88004847 |
| AKT2 | 79.0348462 | -0.117090637 | 0.18776675 | -0.6235962 | 0.53289279 | 0.86909171 |
| PLD3 | 102.745944 | -0.106837374 | 0.22353742 | -0.4779396 | 0.63269323 | 0.90805737 |
| SERTAD1 | 14.8733927 | -0.225478201 | 0.56972686 | -0.3957654 | 0.69227807 | 0.928133 |
| SERTAD3 | 12.4452367 | -0.291729453 | 0.47133909 | -0.6189375 | 0.53595751 | NA |
| BLVRB | 72.4611568 | -0.683136592 | 0.26187499 | -2.6086363 | 0.00909038 | 0.13886093 |
| SPTBN4 | 17.6647667 | 0.602724709 | 0.43987488 | 1.37021852 | 0.1706187 | 0.61319058 |
| SHKBP1 | 16.8748655 | -0.244912251 | 0.38659357 | -0.6335135 | 0.52639835 | 0.86750588 |
| LTBP4 | 13.0704614 | -0.526924794 | 0.51752853 | -1.018156 | 0.30860381 | 0.75132915 |
| COQ8B | 19.4858867 | -0.455886747 | 0.41812532 | -1.0903113 | 0.27557605 | 0.72554623 |
| ITPKC | 31.4997692 | -0.933778433 | 0.34549379 | -2.7027358 | 0.00687714 | 0.11912187 |
| SNRPA | 20.2961522 | 0.147010965 | 0.34995093 | 0.42009023 | 0.67441954 | 0.92189158 |
| AC008537.1 | 21.5819505 | -0.673213817 | 0.69175704 | -0.973194 | 0.33045688 | 0.76685572 |
| HNRNPUL1 | 50.8505635 | -0.163224448 | 0.25713294 | -0.6347862 | 0.52556784 | 0.86747074 |
| TGFB1 | 60.6299088 | -0.709675404 | 0.28237521 | -2.5132355 | 0.01196294 | 0.16274972 |
| CCDC97 | 13.2829791 | -0.160367404 | 0.42511959 | -0.3772289 | 0.70600349 | 0.93407096 |
| B3GNT8 | 10.0739495 | 0.06524495 | 0.50790254 | 0.12845959 | 0.89778528 | NA |
| DMAC2 | 18.3407777 | 0.381906821 | 0.32235522 | 1.18473905 | 0.23612064 | 0.69077483 |
| RPS19 | 394.131246 | -0.197686099 | 0.26408225 | -0.7485777 | 0.45411175 | 0.83364149 |
| ARHGEF1 | 23.5485781 | 0.317703578 | 0.32715738 | 0.9711032 | 0.33149689 | 0.76685572 |
| RABAC1 | 65.5270494 | -0.26916213 | 0.37643247 | -0.7150343 | 0.47458784 | 0.84593347 |
| GSK3A | 19.2452193 | 0.20759339 | 0.32099633 | 0.64671577 | 0.51781592 | 0.86500999 |
| CIC | 22.9230689 | -0.356362591 | 0.39707245 | -0.897475 | 0.36946552 | 0.7895606 |
| MEGF8 | 30.3482321 | 1.041077287 | 0.39507757 | 2.63512123 | 0.00841072 | 0.13244159 |
| LIPE-AS1 | 29.573978 | -0.032573206 | 0.392433 | -0.0830032 | 0.93384897 | 0.9881638 |
| ETHE1 | 11.8951434 | 0.114710806 | 0.36955346 | 0.31040382 | 0.75625389 | NA |
| ZNF428 | 38.8926988 | 0.166426805 | 0.42664892 | 0.39007905 | 0.69647809 | 0.9295666 |
| SMG9 | 15.2720945 | 0.377847858 | 0.35418926 | 1.06679649 | 0.28606375 | 0.73672385 |
| ZNF283 | 14.806869 | 0.195469908 | 0.37909261 | 0.51562575 | 0.60611583 | 0.89643487 |
| ZNF45 | 11.7512036 | 0.538993364 | 0.49801213 | 1.08228963 | 0.27912385 | NA |
| ZNF155 | 18.5342166 | 0.183752719 | 0.51810756 | 0.35466133 | 0.72284332 | 0.93873366 |
| ZNF224 | 25.7947625 | -0.236482901 | 0.33376446 | -0.7085323 | 0.47861475 | 0.84833462 |
| ZNF234 | 14.2176171 | -0.355159469 | 0.52036122 | -0.6825249 | 0.49490713 | 0.85814183 |
| ZNF226 | 27.4393752 | -0.323370893 | 0.28421181 | -1.1377813 | 0.25521181 | 0.70849619 |
| ZNF227 | 16.9683304 | 0.308303598 | 0.42234239 | 0.72998497 | 0.46539937 | 0.841449 |
| ZNF235 | 67.4860141 | -0.112540133 | 0.3540944 | -0.3178252 | 0.75061751 | 0.94705879 |
| ZNF229 | 14.3394641 | -0.215768166 | 0.52177872 | -0.4135243 | 0.67922254 | 0.92348735 |
| ZNF180 | 15.2441075 | 0.46974856 | 0.32083284 | 1.46415359 | 0.14315199 | 0.57698886 |
| AC243964.2 | 34.4180651 | -0.647794127 | 0.41046378 | -1.5782005 | 0.11451956 | 0.51928773 |
| PVR | 15.9313665 | -0.335362607 | 0.42729507 | -0.7848502 | 0.43254142 | 0.8248269 |
| CEACAM19 | 19.8239962 | 0.295233784 | 0.59048285 | 0.49998706 | 0.61708419 | 0.90173965 |
| BCL3 | 17.8027664 | -2.330906317 | 0.44323052 | -5.2589031 | 1.45E-07 | 3.90E-05 |
| CBLC | 27.865875 | -0.710419887 | 0.33180393 | -2.1410834 | 0.03226731 | 0.28479793 |
| BCAM | 109.274875 | 0.276172255 | 0.42056996 | 0.65666187 | 0.51139836 | 0.86312853 |
| NECTIN2 | 111.934122 | -0.218083886 | 0.21583187 | -1.0104341 | 0.31228734 | 0.75484355 |
| TOMM40 | 26.6898959 | -0.171221116 | 0.39855686 | -0.4296027 | 0.66748465 | 0.92049996 |

| | | | | | | |
|---|---|---|---|---|---|---|
| APOE | 594.469789 | -0.115311059 | 0.57677065 | -0.1999253 | 0.84153899 | 0.9677342 |
| APOC1 | 13.8706514 | 0.45373104 | 0.57833053 | 0.78455315 | 0.43271561 | 0.8248269 |
| CLPTM1 | 52.5288593 | 0.271918311 | 0.26502339 | 1.02601629 | 0.30488391 | 0.74832335 |
| RELB | 43.8555419 | -1.267675558 | 0.46986092 | -2.6979804 | 0.00697615 | 0.11997996 |
| CLASRP | 63.7070491 | -0.11505604 | 0.31333203 | -0.3672017 | 0.7134686 | 0.93616842 |
| MARK4 | 51.1667746 | 0.413402177 | 0.24443214 | 1.69127588 | 0.09078412 | 0.47310687 |
| TRAPPC6A | 25.8833153 | 0.413240686 | 0.43520508 | 0.94953094 | 0.34235064 | 0.77263872 |
| PPP1R13L | 23.2408664 | -0.129212698 | 0.30894361 | -0.4182404 | 0.67577137 | 0.92193843 |
| ERCC1 | 13.3734941 | -0.011205814 | 0.48855262 | -0.0229368 | 0.98170072 | 0.99714727 |
| VASP | 31.6794301 | -0.037850715 | 0.43720711 | -0.0865739 | 0.93101023 | 0.98757533 |
| OPA3 | 80.753456 | 0.341682132 | 0.345932 | 0.98771472 | 0.32329238 | 0.76304922 |
| EML2 | 121.75208 | -0.176869615 | 0.3056131 | -0.578737 | 0.56276664 | 0.88191823 |
| GIPR | 32.8672229 | -0.140031756 | 0.45689694 | -0.3064843 | 0.7592359 | 0.94924107 |
| SNRPD2 | 61.2290771 | -0.266814586 | 0.38752482 | -0.6885097 | 0.49113188 | 0.85654883 |
| FBXO46 | 18.2056725 | 0.664638582 | 0.34609045 | 1.92041875 | 0.05480503 | 0.37279717 |
| DMPK | 19.0519455 | 0.51967614 | 0.49285464 | 1.05442071 | 0.29169034 | 0.73994767 |
| SYMPK | 83.1292026 | -0.1558686 | 0.24195553 | -0.6442035 | 0.51944348 | 0.86618562 |
| MYPOP | 21.3034193 | 0.882055745 | 0.41514496 | 2.12469338 | 0.03361221 | 0.29162648 |
| PPP5C | 42.0757058 | 0.058025148 | 0.23022054 | 0.25204158 | 0.80100893 | 0.95877735 |
| CCDC8 | 11.2238108 | 0.98870705 | 0.45886952 | 2.15465835 | 0.03118858 | NA |
| CALM3 | 69.4925932 | 0.491780169 | 0.37372135 | 1.31590067 | 0.18820739 | 0.63996932 |
| PRKD2 | 47.0867907 | -0.362363226 | 0.27203478 | -1.3320474 | 0.18284461 | 0.63154788 |
| STRN4 | 22.4166071 | 0.087667663 | 0.33476869 | 0.26187533 | 0.79341756 | 0.95762809 |
| FKRP | 12.03426 | 0.013494136 | 0.4260707 | 0.03167112 | 0.97473433 | NA |
| AP2S1 | 34.7824717 | -0.173614085 | 0.31443329 | -0.5521492 | 0.58084614 | 0.89012555 |
| ARHGAP35 | 171.658788 | 0.032399212 | 0.18259418 | 0.17743836 | 0.85916408 | 0.97287712 |
| TMEM160 | 38.9586599 | -0.137889952 | 0.3807815 | -0.3621236 | 0.7172597 | 0.93714179 |
| ZC3H4 | 33.7093033 | 0.299251812 | 0.34365534 | 0.87079052 | 0.38386855 | 0.7980695 |
| SAE1 | 39.9687596 | 0.126273748 | 0.31306577 | 0.40334574 | 0.68669389 | 0.92602958 |
| CCDC9 | 13.8141174 | 0.275311764 | 0.40826293 | 0.67434916 | 0.50008936 | 0.85942996 |
| INAFM1 | 12.8278323 | -0.393207382 | 0.44101638 | -0.8915936 | 0.3726108 | 0.7908807 |
| C5AR2 | 10.9656865 | 2.012133843 | 0.85042177 | 2.36604226 | 0.01797939 | NA |
| DHX34 | 24.5152174 | -0.254585266 | 0.36843292 | -0.6909949 | 0.48956875 | 0.85533446 |
| NAPA-AS1 | 14.0915592 | -0.910328987 | 0.56635609 | -1.6073439 | 0.10797896 | 0.50589702 |
| NAPA | 77.6779176 | -0.574165415 | 0.27832797 | -2.0629095 | 0.03912123 | 0.31755308 |
| NOP53 | 33.8767193 | 0.40204187 | 0.47929752 | 0.83881483 | 0.40157323 | 0.80890307 |
| SELENOW | 132.937827 | -0.202428172 | 0.25554865 | -0.7921316 | 0.42828393 | 0.82411309 |
| CARD8 | 30.3032204 | 0.40128334 | 0.35441752 | 1.13223338 | 0.25753633 | 0.71022118 |
| EMP3 | 15.3126195 | -0.126885833 | 0.53470947 | -0.2372986 | 0.81242511 | 0.96262567 |
| TMEM143 | 13.4355468 | 0.331910858 | 0.53323707 | 0.6224452 | 0.53364917 | 0.86909171 |
| KDELR1 | 60.2128557 | -0.377475429 | 0.24069189 | -1.5682931 | 0.11681275 | 0.52409049 |
| CYTH2 | 30.0383449 | -0.00554488 | 0.42283524 | -0.0131136 | 0.98953718 | 0.99923337 |
| LMTK3 | 15.2251921 | -0.359734895 | 0.38513116 | -0.9340581 | 0.3502739 | 0.77868979 |
| RPL18 | 196.84567 | 0.332426573 | 0.25682378 | 1.29437614 | 0.19553553 | 0.6459894 |
| BCAT2 | 25.8470089 | 0.021718824 | 0.29919606 | 0.07259061 | 0.9421319 | 0.98991882 |
| PLEKHA4 | 15.4573929 | -1.725837144 | 0.78966617 | -2.1855275 | 0.02885019 | 0.26841507 |
| PPP1R15A | 17.8932841 | -1.295373549 | 0.52416546 | -2.4713066 | 0.01346204 | 0.17480823 |
| NUCB1 | 70.1436419 | -0.383930001 | 0.2891591 | -1.3277466 | 0.18426185 | 0.63442801 |
| BAX | 26.6385116 | -0.230598051 | 0.37532581 | -0.6143943 | 0.53895478 | 0.86972442 |
| FTL | 1175.76333 | -1.360963639 | 0.51538239 | -2.6406871 | 0.00827381 | 0.13156712 |
| GYS1 | 38.4421192 | 0.446388422 | 0.32478745 | 1.3744017 | 0.16931701 | 0.61168883 |
| RUVBL2 | 11.4747479 | 0.121505519 | 0.4334099 | 0.28034782 | 0.77921067 | NA |
| SNRNP70 | 173.553209 | 0.373400409 | 0.2950787 | 1.26542652 | 0.20571847 | 0.65836247 |
| TRPM4 | 47.4559953 | 0.340497697 | 0.40200988 | 0.84698839 | 0.39700159 | 0.80674902 |
| SLC6A16 | 43.726152 | -0.111712312 | 0.3583943 | -0.3117023 | 0.75526681 | 0.94795939 |
| TEAD2 | 20.2602267 | -0.412011445 | 0.45497972 | -0.90556 | 0.36516878 | 0.78793182 |
| PIH1D1 | 17.2530758 | -0.356818238 | 0.52982734 | -0.6734614 | 0.50065383 | 0.85942996 |
| RPL13A | 240.428114 | 0.065445753 | 0.31965961 | 0.20473576 | 0.83777858 | 0.96748214 |
| RPS11 | 215.30327 | 0.318154052 | 0.40621207 | 0.78322157 | 0.43349701 | 0.82504445 |
| FCGRT | 50.1494654 | 0.344788279 | 0.28394142 | 1.21429369 | 0.2246356 | 0.67753895 |
| NOSIP | 33.4417843 | -0.01810215 | 0.28400995 | -0.0637377 | 0.94917906 | 0.99185581 |
| PRRG2 | 14.9305561 | -0.416126854 | 0.39266328 | -1.059755 | 0.2892561 | 0.73917583 |
| PRR12 | 21.1155453 | 0.511305512 | 0.40764885 | 1.25427929 | 0.20974051 | 0.66164938 |
| RRAS | 13.4231153 | -1.276887179 | 0.54162386 | -2.3575165 | 0.01839764 | 0.21069315 |
| SCAF1 | 22.3372218 | -0.163955032 | 0.363772 | -0.4507082 | 0.65219986 | 0.9158679 |
| IRF3 | 60.9183906 | -0.244963781 | 0.2552979 | -0.9595213 | 0.33729618 | 0.76966704 |
| PRMT1 | 44.093828 | 0.275150568 | 0.26677787 | 1.03138451 | 0.30236054 | 0.74690278 |
| AP2A1 | 26.9150877 | -0.248133808 | 0.40706146 | -0.6095733 | 0.54214449 | 0.87177844 |
| MED25 | 31.3264416 | 0.21083844 | 0.27178106 | 0.77576575 | 0.43788731 | 0.8270818 |
| PTOV1 | 39.8754351 | -0.148574009 | 0.29233021 | -0.5082403 | 0.6112848 | 0.90011356 |
| PNKP | 50.3792313 | -0.786890598 | 0.37908213 | -2.0757787 | 0.03791443 | 0.31246383 |
| TBC1D17 | 20.0882237 | -0.172496891 | 0.39215856 | -0.4398652 | 0.66003477 | 0.91752855 |
| ATF5 | 18.2019844 | 0.544125295 | 0.43979 | 1.23723889 | 0.21599841 | 0.67045905 |
| VRK3 | 34.8272132 | -0.293789235 | 0.34061394 | -0.8625285 | 0.38839675 | 0.80109289 |
| MYH14 | 96.9537189 | -0.229099279 | 0.20688217 | -1.1073901 | 0.26812528 | 0.71965004 |
| KCNC3 | 55.5585054 | -1.408050675 | 0.4170491 | -3.3762228 | 0.00073488 | 0.02625219 |
| NR1H2 | 54.110407 | -1.07568925 | 0.2713611 | -3.964051 | 7.37E-05 | 0.00526714 |
| EMC10 | 115.17066 | -0.057188085 | 0.30201405 | -0.1893557 | 0.84981403 | 0.97027687 |
| JOSD2 | 13.5104656 | 0.384090989 | 0.42837838 | 0.89661619 | 0.36992376 | 0.7900012 |
| CLEC11A | 11.4156211 | 0.022650017 | 0.4390814 | 0.05158501 | 0.95885937 | NA |
| ETFB | 87.570492 | 0.44796308 | 0.3168147 | 1.41395926 | 0.15737386 | 0.59804205 |
| ZNF175 | 13.5921838 | -0.260600744 | 0.38128989 | -0.6834714 | 0.494309 | 0.85774549 |
| SPACA6 | 15.5736472 | 0.0987481 | 0.46975948 | 0.21020991 | 0.83350384 | 0.96701848 |
| ZNF577 | 27.3769065 | 0.354017599 | 0.40208615 | 0.88045211 | 0.37861444 | 0.79535699 |
| ZNF350-AS1 | 21.7950858 | 0.083855471 | 0.71565737 | 0.11717265 | 0.90672324 | 0.98202161 |
| ZNF350 | 16.4531476 | -0.449105969 | 0.51656287 | -0.869412 | 0.38462181 | 0.79869171 |
| ZNF615 | 14.0084524 | 0.358895913 | 0.38241151 | 0.93850708 | 0.34798388 | 0.77757282 |
| ZNF432 | 27.1118752 | 0.044486341 | 0.33697352 | 0.13201732 | 0.89497059 | 0.97862357 |
| AC011468.3 | 18.5993325 | 0.102077935 | 0.50621294 | 0.20165019 | 0.84019021 | 0.96748214 |
| PPP2R1A | 125.807152 | -0.40740579 | 0.26861304 | -1.5167015 | 0.12934207 | 0.55440483 |
| ZNF766 | 35.5267377 | -0.152248854 | 0.25796029 | -0.5902027 | 0.55505478 | 0.87754295 |
| ZNF480 | 26.474456 | -0.341290671 | 0.31284481 | -1.0909264 | 0.27530525 | 0.72554623 |
| ZNF880 | 11.4605336 | 0.331308646 | 0.41054489 | 0.80699736 | 0.419668 | NA |
| ZNF528 | 19.5581478 | -0.315941444 | 0.3883987 | -0.8134462 | 0.41596229 | 0.81763864 |
| ZNF808 | 24.0679139 | 0.295451042 | 0.31593048 | 0.9351774 | 0.34969689 | 0.77868979 |
| ZNF701 | 36.5869203 | 0.320708593 | 0.26604163 | 1.20548272 | 0.22801701 | 0.68327617 |
| ZNF83 | 99.5534892 | 0.038610272 | 0.22144209 | 0.17435832 | 0.86158386 | 0.97302617 |
| ZNF611 | 61.456328 | -0.03639945 | 0.26793802 | -0.1358503 | 0.89193965 | 0.97793768 |
| ZNF600 | 15.6256855 | -0.13161577 | 0.42905493 | -0.3067574 | 0.75902804 | 0.94924107 |
| ZNF28 | 21.4782055 | -0.01426555 | 0.30478063 | -0.046806 | 0.96266788 | 0.99306991 |
| AC010487.3 | 18.8666765 | 0.855743554 | 0.52331088 | 1.63524894 | 0.10199687 | 0.49220184 |
| ZNF160 | 67.9582315 | -0.651867949 | 0.33901071 | -1.9228535 | 0.05449844 | 0.37175451 |
| ZNF347 | 22.7626258 | -0.62231063 | 0.31318396 | -1.987045 | 0.04691741 | 0.34529505 |
| ZNF677 | 20.521243 | -0.235111383 | 0.30173842 | -0.7791894 | 0.43586815 | 0.82624996 |
| ZNF525 | 20.9043913 | -0.577458412 | 0.38401384 | -1.5037438 | 0.13264735 | 0.56040041 |
| ZNF765 | 36.4311138 | -0.717808398 | 0.31929958 | -2.2480718 | 0.02457161 | 0.24622377 |
| ZNF761 | 37.870338 | -0.251885704 | 0.37791097 | -0.6665213 | 0.50507796 | 0.86193811 |
| ZNF813 | 13.9476924 | 0.40240319 | 0.47012235 | 0.85595418 | 0.39202312 | 0.80360124 |
| ZNF331 | 48.6039637 | 0.169031361 | 0.3130229 | 0.53999679 | 0.58919925 | 0.89133386 |
| NDUFA3 | 29.0961076 | -0.018709783 | 0.49621412 | -0.0377051 | 0.96992284 | 0.99486579 |
| PRPF31 | 30.1527358 | -0.078594744 | 0.3463142 | -0.2269464 | 0.82046546 | 0.96389705 |
| CNOT3 | 20.5402247 | 0.715551781 | 0.36813287 | 1.94373239 | 0.05192773 | 0.36159296 |
| LENG1 | 10.3138149 | 0.025698109 | 0.40242072 | 0.06385881 | 0.94908265 | NA |

| | | | | | | |
|---|---|---|---|---|---|---|
| TMC4 | 71.7856601 | 0.309003591 | 0.22952321 | 1.34628475 | 0.1782107 | 0.62598987 |
| MBOAT7 | 35.6758636 | -0.434087721 | 0.38409083 | -1.1301695 | 0.25840483 | 0.71022613 |
| TSEN34 | 20.4725897 | -0.348361498 | 0.48576454 | -0.7171406 | 0.47328732 | 0.84593347 |
| RPS9 | 377.0241 | 0.154364222 | 0.37536979 | 0.4112324 | 0.68090213 | 0.92375164 |
| LENG8-AS1 | 11.9572198 | -0.376024185 | 0.43810777 | -0.8582915 | 0.39073151 | NA |
| LENG8 | 114.02577 | 0.515372724 | 0.28961927 | 1.77948354 | 0.07516052 | 0.43243667 |
| AC011476.3 | 13.9760587 | -1.00698126 | 0.65863234 | -1.5288974 | 0.12628989 | 0.54761372 |
| RDH13 | 41.342733 | 0.222098453 | 0.30443327 | 0.72954724 | 0.46566698 | 0.8416191 |
| EPS8L1 | 20.439259 | -0.043857251 | 0.44403871 | -0.098769 | 0.9213217 | 0.98614962 |
| PPP1R12C | 30.780447 | 0.25094997 | 0.37952075 | 0.6612286 | 0.50846573 | 0.86281145 |
| PPP6R1 | 40.2161944 | 0.217856774 | 0.34424001 | 0.63286302 | 0.5268231 | 0.86750588 |
| HSPBP1 | 13.185448 | 0.09893465 | 0.50329128 | 0.19657533 | 0.84415988 | 0.9699539 |
| TMEM238 | 23.0893177 | 0.478133779 | 0.49178446 | 0.97224255 | 0.3309299 | 0.76685572 |
| RPL28 | 301.121319 | -0.213551557 | 0.37492318 | -0.5695875 | 0.56895751 | 0.88562534 |
| UBE2S | 13.9146474 | 0.198383836 | 0.51733579 | 0.38347209 | 0.70136976 | 0.93127962 |
| ISOC2 | 26.7593846 | -0.068515432 | 0.51752081 | -0.1323916 | 0.89467452 | 0.97850297 |
| NAT14 | 10.8041447 | 0.160132283 | 0.51883963 | 0.30863541 | 0.75759888 | NA |
| CCDC106 | 13.7769097 | 0.847760629 | 0.5148206 | 1.64671076 | 0.09961751 | 0.48648077 |
| U2AF2 | 25.7260205 | -0.038615139 | 0.31866094 | -0.1211794 | 0.90354895 | 0.98025107 |
| EPN1 | 85.1784548 | 0.459543145 | 0.22858678 | 2.0103662 | 0.04439245 | 0.33484193 |
| ZNF787 | 24.7442704 | -0.434050958 | 0.36746172 | -1.1812141 | 0.23751768 | 0.69207518 |
| ZNF444 | 22.4664457 | 0.444414771 | 0.30346873 | 1.46444999 | 0.14307104 | 0.57698886 |
| ZSCAN5A | 74.8741231 | 0.212003834 | 0.2368635 | 0.89504646 | 0.37076225 | 0.7900012 |
| ZNF582 | 12.1839871 | -0.267943632 | 0.44827763 | -0.5977181 | 0.55002808 | NA |
| ZNF667-AS1 | 21.5439715 | 0.397626758 | 0.33531356 | 1.18583559 | 0.23568723 | 0.69026756 |
| ZNF471 | 18.532272 | 0.073458286 | 0.35756126 | 0.20544252 | 0.8372264 | 0.96748214 |
| ZNF470 | 12.9451442 | -0.513250094 | 0.51071897 | -1.004956 | 0.31491803 | 0.75742745 |
| ZNF264 | 38.9176337 | 0.076580144 | 0.24513696 | 0.31239738 | 0.75473854 | 0.94795939 |
| ZNF460 | 15.8664694 | -0.537735773 | 0.42386296 | -1.2686548 | 0.20456421 | 0.65790847 |
| AC005261.1 | 57.5303588 | -0.124827077 | 0.35837041 | -0.3483186 | 0.72760093 | 0.94027832 |
| AC004076.1 | 24.1534015 | -0.201568764 | 0.42538168 | -0.4738539 | 0.63560411 | 0.90881415 |
| ZNF549 | 23.1493058 | -0.251167979 | 0.37003083 | -0.6787758 | 0.49727992 | 0.85814183 |
| ZNF134 | 11.4910897 | 0.076291882 | 0.41730475 | 0.18282055 | 0.85493883 | NA |
| AC003006.1 | 14.5891152 | 0.102264429 | 0.36825675 | 0.27769872 | 0.78124364 | 0.95540382 |
| ZNF154 | 12.9711717 | 0.226892763 | 0.36119753 | 0.6281681 | 0.52989383 | 0.86758926 |
| ZNF586 | 108.822062 | 0.202072982 | 0.22602743 | 0.89401975 | 0.37131133 | 0.7900012 |
| ZNF552 | 32.9544004 | 0.029607108 | 0.38984942 | 0.07594498 | 0.93946287 | 0.98871562 |
| ZNF814 | 55.8269281 | -0.32343642 | 0.26547252 | -1.2183424 | 0.2230939 | 0.67606266 |
| ZNF587 | 30.2856297 | 0.317633091 | 0.26150549 | 1.21463261 | 0.22450625 | 0.67735943 |
| ZNF606 | 20.9513223 | 0.202900337 | 0.29314782 | 0.69214342 | 0.48884726 | 0.8546897 |
| ZNF135 | 33.995441 | 0.00950681 | 0.25640407 | 0.03707745 | 0.97042325 | 0.99502125 |
| ZSCAN18 | 84.0978393 | -0.644851663 | 0.28060024 | -2.2981151 | 0.02155524 | 0.22951242 |
| ZNF329 | 21.4040039 | -0.086137772 | 0.34865417 | -0.2470579 | 0.8048634 | 0.95997099 |
| ZNF274 | 48.4384576 | -0.155544721 | 0.3485636 | -0.4462449 | 0.65542037 | 0.9158679 |
| AC020915.3 | 21.0756712 | -0.085294723 | 0.36190682 | -0.2356814 | 0.81367987 | 0.96264114 |
| RPS5 | 126.179532 | 0.183557034 | 0.3843197 | 0.47761547 | 0.63292392 | 0.90805737 |
| TRIM28 | 43.1132249 | 0.140508145 | 0.34811422 | 0.40362656 | 0.68648735 | 0.92597882 |
| CHMP2A | 43.9089416 | 0.130709702 | 0.32885305 | 0.39747146 | 0.69101983 | 0.92786587 |
| UBE2M | 39.2340114 | -0.244317444 | 0.33819547 | -0.7224149 | 0.47003943 | 0.84415485 |
| MZF1-AS1 | 12.6688677 | 0.109877381 | 0.56200489 | 0.19550965 | 0.84499398 | 0.97027687 |
| MZF1 | 44.5839061 | -0.56504541 | 0.31672106 | -1.7840475 | 0.07441595 | 0.43060774 |
| RPS4Y1 | 63.2189704 | 0.232587217 | 1.69626787 | 0.13711703 | 0.89093829 | 0.97793768 |
| ZFY | 34.7491431 | 0.586100268 | 1.92029491 | 0.30521367 | 0.76020342 | 0.94952011 |
| LINC00278 | 116.73689 | 0.267269256 | 1.96677647 | 0.13589203 | 0.89190663 | 0.97793768 |
| TTTY15 | 29.0828069 | 0.060875212 | 1.59860772 | 0.03808014 | 0.96962378 | 0.99486579 |
| USP9Y | 122.189568 | -0.033466606 | 2.20589778 | -0.0151714 | 0.98789542 | 0.99905853 |
| DDX3Y | 49.0179642 | 0.408453969 | 2.02929835 | 0.20127842 | 0.84048087 | 0.96748214 |
| UTY | 344.568093 | -0.054509634 | 2.16477513 | -0.0251803 | 0.97991117 | 0.99664466 |
| TTTY14 | 335.688655 | 0.706821389 | 0.75639233 | 0.93446398 | 0.35006461 | 0.77868979 |
| KDM5D | 35.8807267 | 0.174643333 | 1.92988622 | 0.09049411 | 0.92789458 | 0.98690541 |
| TTTY10 | 12.2948063 | 0.675523514 | 1.04185014 | 0.64838837 | 0.51673379 | NA |
| EIF1AY | 22.1556927 | 0.0678786 | 1.85512283 | 0.03658981 | 0.97081206 | 0.99512597 |
| IL17RA | 29.4987977 | -0.10420889 | 0.33290376 | -0.3130301 | 0.75425781 | 0.94795939 |
| HDHD5 | 16.5815407 | 0.296280733 | 0.47087316 | 0.62921558 | 0.52920793 | 0.86758926 |
| ADA2 | 12.4535881 | 1.26186049 | 0.67319235 | 1.87444271 | 0.06086943 | NA |
| CECR2 | 41.4526019 | 0.527929155 | 0.6897563 | 0.76538504 | 0.4440423 | 0.83006558 |
| ATP6V1E1 | 155.181285 | -0.189363562 | 0.22910221 | -0.8265462 | 0.40849432 | 0.81362627 |
| BCL2L13 | 180.795232 | 0.131222452 | 0.27664654 | 0.47433252 | 0.6352628 | 0.90881415 |
| MICAL3 | 98.1406666 | -0.440866674 | 0.26577721 | -1.6587828 | 0.09715957 | 0.48257296 |
| PEX26 | 41.8230466 | -0.269828543 | 0.30724554 | -0.8782179 | 0.37982547 | 0.79587736 |
| DGCR2 | 66.2472146 | 0.498406196 | 0.32461154 | 1.53539271 | 0.12468739 | 0.543581 |
| SLC25A1 | 35.8214625 | -0.199680933 | 0.29977751 | -0.6660971 | 0.50534903 | 0.86197912 |
| CLTCL1 | 22.009636 | -0.05957246 | 0.33801917 | -0.1762399 | 0.86010549 | 0.97287712 |
| HIRA | 22.0330784 | 0.080191221 | 0.36090516 | 0.22219472 | 0.82416231 | 0.96539788 |
| MRPL40 | 27.1033691 | -0.469944554 | 0.35258666 | -1.3328483 | 0.18258158 | 0.63138727 |
| UFD1 | 20.4107787 | -0.546412735 | 0.41819122 | -1.3066098 | 0.19134527 | 0.6421543 |
| TXNRD2 | 37.2856496 | 0.028029769 | 0.29926617 | 0.09366167 | 0.92537792 | 0.98672316 |
| COMT | 59.7943659 | -0.105732292 | 0.24109378 | -0.4385525 | 0.66098579 | 0.91795482 |
| TANGO2 | 65.4453761 | -0.644301993 | 0.4843717 | -1.3301809 | 0.18345867 | 0.63239131 |
| DGCR8 | 26.2271698 | 0.449229528 | 0.35150291 | 1.27802505 | 0.2012406 | 0.65374585 |
| RANBP1 | 19.8484379 | 0.057979533 | 0.37694032 | 0.15381622 | 0.87775465 | 0.97542218 |
| ZDHHC8 | 15.7628551 | 1.049132397 | 0.42576902 | 2.46408817 | 0.01373623 | 0.17578162 |
| DGCR6L | 11.4104637 | 0.895687246 | 0.57188586 | 1.56619933 | 0.11730196 | NA |
| KLHL22 | 62.6867181 | 0.503766639 | 0.26784398 | 1.88082122 | 0.05999624 | 0.38930355 |
| MED15 | 82.4224286 | -0.506406869 | 0.30064983 | -1.6843744 | 0.09210934 | 0.4764741 |
| PI4KA | 130.280403 | -0.071327504 | 0.27567175 | -0.2587407 | 0.79583531 | 0.95848876 |
| SNAP29 | 24.6828329 | -0.053451603 | 0.36996382 | -0.1444779 | 0.88512309 | 0.97662748 |
| CRKL | 34.9829326 | 0.185399982 | 0.25611519 | 0.72389295 | 0.46913146 | 0.8433157 |
| LZTR1 | 10.3636564 | -0.213694322 | 0.43798419 | -0.4879042 | 0.62561771 | NA |
| THAP7 | 12.8977609 | 0.192320998 | 0.40881232 | 0.47043836 | 0.63804186 | 0.91025608 |
| HIC2 | 35.0123799 | 0.992048669 | 0.35443742 | 2.79893886 | 0.00512708 | 0.0988722 |
| UBE2L3 | 94.7784105 | -0.210460183 | 0.17276618 | -1.2181793 | 0.22315584 | 0.67606266 |
| SDF2L1 | 33.9491295 | 0.113003647 | 0.33300141 | 0.33934886 | 0.73434694 | 0.94277981 |
| PPIL2 | 92.3081104 | 0.216595049 | 0.24455135 | 0.8856833 | 0.37578821 | 0.79392096 |
| MAPK1 | 174.455215 | 0.181788159 | 0.16661958 | 1.09103722 | 0.2752565 | 0.72554623 |
| PPM1F | 14.9353139 | 0.421798701 | 0.42806009 | 0.98537265 | 0.32444106 | 0.76304922 |
| TOP3B | 20.7357056 | 0.541306027 | 0.36551029 | 1.48095975 | 0.1386173 | 0.57119278 |
| AC245060.5 | 21.7932783 | 0.908923825 | 0.38946938 | 2.33374914 | 0.01960886 | 0.21965565 |
| BCR | 114.30137 | -0.221709658 | 0.38409119 | -0.5772318 | 0.56378287 | 0.88191823 |
| CHCHD10 | 277.499977 | -0.063385355 | 0.32574161 | -0.1945878 | 0.84571563 | 0.97027687 |
| SMARCB1 | 48.8105379 | 0.304406569 | 0.27211063 | 1.11868681 | 0.26327377 | 0.71348562 |
| DDT | 90.73461 | 0.117681043 | 0.43183674 | 0.27251281 | 0.78522774 | 0.95577371 |
| CABIN1 | 141.883501 | 0.012025538 | 0.26101307 | 0.04607255 | 0.96325243 | 0.99314929 |
| GUCD1 | 31.6664807 | -0.432156731 | 0.31993433 | -1.350767 | 0.17677009 | 0.62442456 |
| SNRPD3 | 52.1963786 | 0.263113301 | 0.23909711 | 1.10044535 | 0.27113813 | 0.72194341 |
| SGSM1 | 100.347651 | -0.179223137 | 0.28814825 | -0.6219824 | 0.53395345 | 0.86909171 |
| KIAA1671 | 90.6731438 | -0.190363991 | 0.32933615 | -0.5780234 | 0.56324834 | 0.88191823 |
| LRP5L | 57.9422628 | 0.028913857 | 0.49882455 | 0.05796398 | 0.95377732 | 0.99217529 |
| GRK3 | 35.9218898 | -0.264026834 | 0.40470012 | -0.6524012 | 0.51414241 | 0.86416832 |
| HPS4 | 65.1680467 | -0.208189226 | 0.4113996 | -0.5060511 | 0.61282077 | 0.90011356 |
| SRRD | 13.0738055 | 0.019284254 | 0.39350361 | 0.04900655 | 0.96091408 | 0.99238848 |
| TFIP11 | 26.2049725 | -0.223970516 | 0.29500134 | -0.7592186 | 0.44772179 | 0.83066268 |
| TPST2 | 74.1886789 | -0.502948331 | 0.26026852 | -1.9324209 | 0.05330757 | 0.36627428 |

| | | | | | | |
|---|---|---|---|---|---|---|
| MIATNB | 53.9410147 | -0.498773953 | 0.27033078 | -1.8450506 | 0.06503018 | 0.40487342 |
| PITPNB | 163.608454 | -0.042613829 | 0.25376873 | -0.1679239 | 0.86664317 | 0.97434023 |
| TTC28 | 221.655262 | -0.00123162 | 0.29499493 | -0.0041751 | 0.9966688 | 0.99975875 |
| HSCB | 35.8744592 | -0.136633768 | 0.27907275 | -0.4895991 | 0.62441761 | 0.90447745 |
| CCDC117 | 25.604731 | 0.245330604 | 0.39662781 | 0.6185411 | 0.53621872 | 0.86949541 |
| XBP1 | 64.9606537 | -0.550588611 | 0.2380695 | -2.3127222 | 0.02073792 | 0.22444738 |
| Z93930.2 | 74.01662 | 0.168499595 | 0.36967595 | 0.45580351 | 0.64853128 | 0.91417939 |
| ZNRF3 | 296.840401 | 1.063631862 | 0.30088094 | 3.53505892 | 0.00040768 | 0.01742087 |
| KREMEN1 | 108.268771 | -0.047905904 | 0.29528551 | -0.1622359 | 0.87112011 | 0.97434023 |
| RHBDD3 | 19.6468968 | -0.122649674 | 0.43978522 | -0.2788854 | 0.78033278 | 0.95537097 |
| EWSR1 | 122.198289 | 0.210149642 | 0.22733694 | 0.92439726 | 0.35527952 | 0.78124141 |
| GAS2L1 | 20.0428818 | 0.140288701 | 0.42785728 | 0.32788667 | 0.74299735 | 0.94546379 |
| AP1B1 | 85.726026 | 0.204332682 | 0.29422391 | 0.69448021 | 0.48738111 | 0.85382621 |
| THOC5 | 13.3550229 | 0.255330752 | 0.37532034 | 0.68030086 | 0.49631398 | 0.85814183 |
| NIPSNAP1 | 17.4990027 | 0.391161127 | 0.43548873 | 0.89821182 | 0.36907263 | 0.78889478 |
| NF2 | 103.614322 | 1.490323973 | 0.23373664 | 6.37608209 | 1.82E-10 | 1.60E-07 |
| UQCR10 | 202.566909 | -0.38912785 | 0.39975297 | -0.9734208 | 0.33034421 | 0.76685572 |
| ASCC2 | 37.3747267 | 0.355267442 | 0.31000953 | 1.14598878 | 0.25179979 | 0.70574787 |
| MTMR3 | 154.66944 | 0.071133086 | 0.19368201 | 0.36726739 | 0.71341957 | 0.93616842 |
| TBC1D10A | 14.7386668 | -0.1698378 | 0.44721657 | -0.3797663 | 0.70411887 | 0.93318118 |
| SF3A1 | 28.1718795 | 0.400490172 | 0.31701663 | 1.26330967 | 0.20647791 | 0.65836247 |
| RNF215 | 15.4611713 | -0.291205973 | 0.36760473 | -0.7921715 | 0.42826071 | 0.82411309 |
| PES1 | 15.1273692 | 0.272942193 | 0.40518214 | 0.6736284 | 0.50054759 | 0.85942996 |
| DUSP18 | 13.2857826 | 0.300200937 | 0.43064009 | 0.69710402 | 0.4857377 | 0.85356796 |
| OSBP2 | 39.8694218 | 0.355578646 | 0.34957333 | 1.01717898 | 0.30906829 | 0.75165835 |
| MORC2 | 21.2772882 | -0.06107573 | 0.35462064 | -0.1722284 | 0.86325801 | 0.97368057 |
| SMTN | 12.6165843 | 1.031864031 | 0.56421844 | 1.82883785 | 0.0674239 | 0.4136301 |
| SELENOM | 107.025176 | -0.455756535 | 0.42335461 | -1.0765361 | 0.28168755 | 0.73311066 |
| INPP5J | 66.6264045 | 0.785371298 | 0.57174263 | 1.37364482 | 0.16955198 | 0.61207822 |
| RNF185 | 50.8206911 | 0.097958365 | 0.30848295 | 0.31754872 | 0.75082728 | 0.94707732 |
| LIMK2 | 57.7273069 | -0.394026859 | 0.24245783 | -1.6251356 | 0.10413367 | 0.49685025 |
| PIK3IP1 | 31.5116007 | -1.182499179 | 0.45717723 | -2.5865224 | 0.00969499 | 0.14512556 |
| PIK3IP1-AS1 | 20.5163257 | -1.525324958 | 0.42393004 | -3.5980582 | 0.0003206 | 0.0152443 |
| PATZ1 | 26.2315593 | 0.32286159 | 0.4205677 | 0.76768042 | 0.44267707 | 0.82847146 |
| DRG1 | 48.1285899 | 0.135644661 | 0.24002428 | 0.56512891 | 0.57198609 | 0.88578992 |
| EIF4ENIF1 | 51.4144322 | 0.017893884 | 0.25233517 | 0.07091316 | 0.94346687 | 0.99065042 |
| SFI1 | 40.601204 | -0.030515826 | 0.3632506 | -0.0840076 | 0.93305036 | 0.98816339 |
| PISD | 149.06194 | 0.303747453 | 0.28590565 | 1.0624045 | 0.28805209 | 0.73803612 |
| PRR14L | 124.168555 | 0.069264607 | 0.2428682 | 0.28519422 | 0.77549535 | 0.95493992 |
| YWHAH | 71.0548719 | -1.026351549 | 0.23898948 | -4.294547 | 1.75E-05 | 0.00180637 |
| RTCB | 59.0364185 | -0.109198923 | 0.22865961 | -0.4775611 | 0.63296265 | 0.90805737 |
| FBXO7 | 83.3509552 | -0.265014667 | 0.2717609 | -0.9751759 | 0.32947304 | 0.76685572 |
| TIMP3 | 50.1713358 | 1.498009509 | 0.48625442 | 3.08071137 | 0.00206507 | 0.05202897 |
| LARGE1 | 123.893979 | -0.382386205 | 0.3709389 | -1.0308603 | 0.30260632 | 0.74712557 |
| HMGXB4 | 34.9399319 | 0.153440009 | 0.29560647 | 0.51906851 | 0.60371297 | 0.8956362 |
| TOM1 | 86.8395412 | -0.580872155 | 0.29530476 | -1.967026 | 0.04918021 | 0.35506449 |
| APOL6 | 44.8219324 | -1.080187971 | 0.51991236 | -2.0776347 | 0.03774302 | 0.31172423 |
| RBFOX2 | 360.106212 | 0.071807048 | 0.22177202 | 0.32378768 | 0.74609878 | 0.94614106 |
| APOL2 | 42.8222274 | -1.115046583 | 0.41186056 | -2.70734 | 0.00678248 | 0.11790324 |
| APOL1 | 27.7611801 | -1.809336591 | 0.4911343 | -3.6839956 | 0.00022961 | 0.01191006 |
| MYH9 | 191.935839 | -0.25323159 | 0.24400301 | -1.0378216 | 0.29935313 | 0.74336229 |
| TXN2 | 46.1191041 | 0.011953236 | 0.27514776 | 0.04344297 | 0.96534842 | 0.99334616 |
| EIF3D | 54.2921267 | -0.046812299 | 0.287832 | -0.1626376 | 0.8708038 | 0.97434023 |
| IFT27 | 24.8234347 | 0.019251919 | 0.31122541 | 0.06185844 | 0.95067556 | 0.99209822 |
| PVALB | 15.5819905 | 1.205576455 | 0.67770879 | 1.77890042 | 0.07525609 | 0.43245501 |
| TST | 24.9107223 | -0.659450102 | 0.3536536 | -1.864678 | 0.06222655 | 0.3965818 |
| MPST | 21.1892263 | -0.052645246 | 0.37423623 | -0.1406738 | 0.88812761 | 0.97765744 |
| CARD10 | 11.0385748 | 0.078021027 | 0.53141697 | 0.14681697 | 0.88327649 | NA |
| GGA1 | 47.3412228 | 0.244942938 | 0.25791483 | 0.94970475 | 0.3422623 | 0.7726191 |
| LGALS1 | 17.2549514 | -1.683964816 | 0.57919797 | -2.907408 | 0.00364438 | 0.07820893 |
| TRIOBP | 95.2271274 | -0.265570968 | 0.27686138 | -0.95922 | 0.33744794 | 0.76966704 |
| H1F0 | 129.915094 | -1.145445876 | 0.6170651 | -1.8562804 | 0.06341359 | 0.4001801 |
| GCAT | 12.9106743 | -0.045649306 | 0.56873061 | -0.0802653 | 0.93602629 | 0.98871562 |
| ANKRD54 | 16.4978043 | 0.070333729 | 0.32075633 | 0.21927464 | 0.82643611 | 0.96542185 |
| EIF3L | 88.8435944 | 0.314253567 | 0.23972207 | 1.31090797 | 0.18988886 | 0.64181696 |
| MICALL1 | 22.1431522 | -0.147672877 | 0.33756681 | -0.4374627 | 0.66177585 | 0.91808148 |
| POLR2F | 25.5700869 | 0.87154477 | 0.3981395 | 2.18904373 | 0.02859366 | 0.26683451 |
| PICK1 | 23.0969846 | 0.410249856 | 0.34194925 | 1.19973901 | 0.23024072 | 0.68523674 |
| AL031587.2 | 10.9644621 | -0.611798066 | 0.51898474 | -1.1788363 | 0.23846335 | NA |
| PLA2G6 | 26.2924987 | 0.20547532 | 0.35167439 | 0.58427718 | 0.55903384 | 0.88004847 |
| MAFF | 18.6138365 | -2.054446025 | 0.62788217 | -3.2720248 | 0.0010678 | 0.03418376 |
| TMEM184B | 35.6342 | 0.129011452 | 0.34098278 | 0.37835181 | 0.70516926 | 0.93403928 |
| CSNK1E | 19.697759 | 0.59714342 | 0.36009793 | 1.65828062 | 0.09726084 | 0.48257296 |
| TPTEP2-CSNI | 85.1333226 | -0.381930527 | 0.3148671 | -1.2129896 | 0.22513378 | 0.67848307 |
| DDX17 | 663.874236 | 0.141618363 | 0.18906255 | 0.74905561 | 0.45382369 | 0.83364149 |
| CBY1 | 26.1168417 | 0.358028577 | 0.31191898 | 1.14782557 | 0.25104058 | 0.70545786 |
| TOMM22 | 14.7953829 | 0.611648717 | 0.53589187 | 1.14136592 | 0.25371768 | 0.70701839 |
| JOSD1 | 35.8972162 | -0.458145182 | 0.38275662 | -1.1969621 | 0.23132131 | 0.68580018 |
| GTPBP1 | 37.7912862 | -0.661408545 | 0.35515607 | -1.8623039 | 0.06256026 | 0.39783219 |
| SUN2 | 40.4101066 | 0.092766192 | 0.31799274 | 0.29172425 | 0.77049747 | 0.9522529 |
| AL021707.1 | 13.3883967 | -0.56761192 | 0.41031028 | -1.3833724 | 0.16655071 | 0.60758548 |
| CBX6 | 29.8555592 | 0.363000399 | 0.55269829 | 0.65677858 | 0.5113233 | 0.86312853 |
| CBX7 | 21.228078 | 0.074952217 | 0.28510305 | 0.26289518 | 0.79263137 | 0.95746995 |
| RPL3 | 219.086279 | 0.654353484 | 0.39163504 | 1.67082465 | 0.09475632 | 0.47996672 |
| SYNGR1 | 16.4225789 | 0.381485233 | 0.42924692 | 0.88873143 | 0.37414744 | 0.7923487 |
| TAB1 | 19.5131652 | 0.457751111 | 0.37882881 | 1.20833235 | 0.22691945 | 0.68135521 |
| MIEF1 | 20.2512245 | 0.375202048 | 0.32387836 | 1.15846595 | 0.24667394 | 0.70024501 |
| ATF4 | 63.7234088 | -0.22670112 | 0.45834029 | -0.4946131 | 0.62087324 | 0.90380166 |
| RPS19BP1 | 49.2040889 | 0.438739564 | 0.52467916 | 0.83620543 | 0.40303934 | 0.80924894 |
| FAM83F | 55.9357601 | 0.271116441 | 0.37409439 | 0.72472736 | 0.46861931 | 0.84326495 |
| TNRC6B | 664.969241 | 0.001193569 | 0.22203224 | 0.00537566 | 0.99571087 | 0.99923337 |
| MKL1 | 196.097663 | 0.255077792 | 0.19614033 | 1.30048623 | 0.19343437 | 0.6443384 |
| SLC25A17 | 45.6889318 | 0.341904518 | 0.27554679 | 1.24082198 | 0.21467152 | 0.66839951 |
| ST13 | 130.505229 | -0.029950505 | 0.25890777 | -0.1156802 | 0.90790599 | 0.98250721 |
| XPNPEP3 | 58.0075323 | 0.054627743 | 0.21142565 | 0.25837803 | 0.79611517 | 0.95848876 |
| RBX1 | 64.9725009 | -0.339979322 | 0.22848744 | -1.4879563 | 0.13676242 | 0.5681351 |
| EP300 | 171.349876 | 0.245580326 | 0.20210959 | 1.21508499 | 0.22433368 | 0.6771276 |
| L3MBTL2 | 19.0343149 | 0.048068098 | 0.31895702 | 0.150704 | 0.88020922 | 0.97599788 |
| RANGAP1 | 23.5663499 | -0.16416782 | 0.35895915 | -0.457344 | 0.64742379 | 0.913589 |
| ZC3H7B | 46.1198407 | 0.45097886 | 0.26454534 | 1.7047318 | 0.08824446 | 0.46799957 |
| TEF | 14.6919202 | -0.137869838 | 0.54119225 | -0.2547521 | 0.79891461 | 0.95848876 |
| TOB2 | 47.2284718 | 0.677859556 | 0.26425499 | 2.5651722 | 0.01031246 | 0.14935229 |
| PHF5A | 11.4550143 | -0.447904251 | 0.51441562 | -0.870705 | 0.38391525 | NA |
| ACO2 | 162.898128 | 0.237665909 | 0.23626589 | 1.0059256 | 0.31445136 | 0.75700972 |
| POLR3H | 12.4880492 | -0.266773031 | 0.47734171 | -0.5588722 | 0.57624892 | NA |
| PMM1 | 17.2458438 | 0.606822848 | 0.36128158 | 1.67963962 | 0.09302745 | 0.47759124 |
| DESI1 | 65.5688733 | -0.637446028 | 0.31240657 | -2.0404373 | 0.04130679 | 0.32628324 |
| XRCC6 | 131.165908 | -0.150497391 | 0.20478486 | -0.7349049 | 0.46239742 | 0.83899271 |
| SNU13 | 100.464421 | -0.369120151 | 0.31709844 | -1.1640554 | 0.24440156 | 0.69787904 |
| CCDC134 | 33.2398788 | 0.232129759 | 0.56456432 | 0.41116619 | 0.68095068 | 0.92375164 |
| SREBF2 | 508.850334 | -0.551712296 | 0.30370477 | -1.8166073 | 0.06927727 | 0.41790391 |
| WBP2NL | 24.7712457 | 0.823087412 | 0.4288064 | 1.91948489 | 0.054923 | 0.37333782 |
| NAGA | 12.8744773 | -0.104947453 | 0.40388079 | -0.2598476 | 0.79498134 | 0.95804683 |

| | | | | | | |
|---|---|---|---|---|---|---|
| SMDT1 | 64.7436667 | 0.012359353 | 0.41779014 | 0.02958268 | 0.97639988 | 0.99559327 |
| NDUFA6 | 75.4449413 | -0.002646907 | 0.43033646 | -0.0061508 | 0.99509242 | 0.99923337 |
| TCF20 | 147.381463 | -0.363374787 | 0.20893768 | -1.7391539 | 0.08200769 | 0.45345012 |
| RRP7A | 24.4730128 | -0.049164175 | 0.39834286 | -0.1234218 | 0.90177313 | 0.9797999 |
| SERHL2 | 16.8096034 | -0.262012129 | 0.36790208 | -0.712179 | 0.47635394 | 0.84663915 |
| POLDIP3 | 37.4480134 | -0.018444291 | 0.24088055 | -0.0765703 | 0.9389654 | 0.98871562 |
| CYB5R3 | 95.9266999 | -0.289916673 | 0.30220034 | -0.9593526 | 0.33738116 | 0.76966704 |
| A4GALT | 64.0423131 | -1.851444479 | 0.35532185 | -5.2106125 | 1.88E-07 | 4.92E-05 |
| ARFGAP3 | 134.209085 | -0.854801474 | 0.21315563 | -4.0102224 | 6.07E-05 | 0.0047453 |
| PACSIN2 | 194.101379 | 0.092813033 | 0.28147338 | 0.32974 | 0.74159643 | 0.94501909 |
| MCAT | 13.6781281 | 0.099100946 | 0.41605802 | 0.23819021 | 0.81173357 | 0.96256453 |
| TSPO | 161.110777 | -0.466686756 | 0.35414317 | -1.3177912 | 0.18757355 | 0.63942735 |
| EFCAB6 | 51.3051515 | 0.06234998 | 0.3672493 | 0.16977563 | 0.8651866 | 0.97434023 |
| SAMM50 | 17.0480566 | 0.665995877 | 0.36636059 | 1.81786987 | 0.06908402 | 0.41751716 |
| PARVB | 176.124387 | 0.084529916 | 0.31172074 | 0.27117194 | 0.7862588 | 0.95577371 |
| PHF21B | 18.3212401 | 0.76944233 | 0.80792175 | 0.95237235 | 0.34090818 | 0.77193691 |
| NUP50-DT | 117.973645 | -0.800715765 | 0.28453132 | -2.8141569 | 0.00489054 | 0.0961667 |
| NUP50 | 49.3647898 | -0.024577965 | 0.26718728 | -0.0919878 | 0.92670775 | 0.98672316 |
| KIAA0930 | 15.48707 | -1.020970953 | 0.49537291 | -2.0610149 | 0.03930162 | 0.31818244 |
| FAM118A | 18.2397748 | 0.353513793 | 0.55670571 | 0.63501018 | 0.52542178 | 0.86747074 |
| ATXN10 | 187.410572 | -0.114876445 | 0.14467236 | -0.7940455 | 0.42716892 | 0.82375285 |
| MIRLET7BHG | 92.6860207 | 0.062876905 | 0.30511368 | 0.20607697 | 0.83673078 | 0.96748214 |
| PPARA | 132.233312 | 0.602339191 | 0.24730456 | 2.43561703 | 0.01486641 | 0.18484189 |
| TTC38 | 34.5172981 | -0.06048006 | 0.43717836 | -0.1383418 | 0.88997026 | 0.97782578 |
| TRMU | 34.5605321 | 0.121139831 | 0.32453205 | 0.3732754 | 0.70894349 | 0.93482669 |
| CELSR1 | 48.7583874 | 0.146607889 | 0.36286893 | 0.40402437 | 0.6861948 | 0.92597882 |
| GRAMD4 | 44.6205304 | 0.496599105 | 0.34463419 | 1.44094555 | 0.14960007 | 0.58659144 |
| CERK | 26.1017951 | -0.364181673 | 0.47029867 | -0.7743625 | 0.43871642 | 0.82745796 |
| TBC1D22A | 240.383679 | -0.326058691 | 0.31216086 | -1.0445214 | 0.29624424 | 0.74123516 |
| BRD1 | 55.4894259 | 0.068340127 | 0.277326 | 0.24642525 | 0.80535306 | 0.96005568 |
| ZBED4 | 35.3299177 | 0.300181983 | 0.33552009 | 0.89467663 | 0.37095998 | 0.7900012 |
| ALG12 | 19.229803 | 0.18209801 | 0.39849186 | 0.45696795 | 0.64769408 | 0.91360118 |
| CRELD2 | 17.6592863 | 0.491904817 | 0.36632575 | 1.34280711 | 0.17933444 | 0.62668718 |
| PIM3 | 118.070244 | -1.243685924 | 0.41579896 | -2.9910751 | 0.00277997 | 0.06466599 |
| TRABD | 31.9421235 | 0.161809537 | 0.30512539 | 0.53030506 | 0.59590044 | 0.89345937 |
| SELENOO | 48.2880431 | 0.126395185 | 0.29790123 | 0.42428554 | 0.67135758 | 0.92170876 |
| TUBGCP6 | 15.09219 | 0.348502633 | 0.37845101 | 0.92086592 | 0.35712044 | 0.7826634 |
| PLXNB2 | 47.426049 | -0.475795756 | 0.35040104 | -1.3578606 | 0.17450793 | 0.62052871 |
| PPP6R2 | 200.777344 | 0.502846771 | 0.2209789 | 2.275542 | 0.02287344 | 0.23759542 |
| SBF1 | 16.166574 | -0.136559768 | 0.4256782 | -0.3208052 | 0.74835804 | 0.94667097 |
| LMF2 | 17.3618539 | -0.485382245 | 0.44433198 | -1.0923865 | 0.27466326 | 0.72554623 |
| NCAPH2 | 10.4748449 | 0.115487437 | 0.50299905 | 0.22959772 | 0.81840438 | NA |
| ARSA | 13.2630439 | -0.314485424 | 0.39003884 | -0.8062926 | 0.42007416 | 0.81913024 |
| RABL2B | 14.2117558 | 0.388085447 | 0.36942179 | 1.05052126 | 0.29347852 | 0.74013384 |
| SMIM34B | 143.702697 | 0.783552119 | 0.55228092 | 1.41875645 | 0.15597003 | 0.59597065 |
| FP671120.1 | 412.713352 | -0.554606565 | 0.43004313 | -1.2896534 | 0.19717104 | 0.64788587 |
| FP236383.1 | 271.648849 | -0.514861398 | 0.52354392 | -0.9834159 | 0.3254028 | 0.76334591 |
| CR381653.1 | 65.8590546 | -0.131522488 | 0.75413157 | -0.1744026 | 0.86154909 | 0.97302617 |
| RBM11 | 17.0984946 | -1.529388582 | 0.47639565 | -3.2103328 | 0.00132581 | 0.03920851 |
| HSPA13 | 20.1569468 | -0.266127664 | 0.37823979 | -0.7035951 | 0.48168496 | 0.8502901 |
| NRIP1 | 39.1667608 | -0.381624843 | 0.275035 | -1.3875501 | 0.16527408 | 0.60578861 |
| USP25 | 262.0101 | -0.04931464 | 0.25988047 | -0.1897589 | 0.84949804 | 0.97027687 |
| MIR99AHG | 543.578903 | 1.462900127 | 0.32802407 | 4.45973408 | 8.21E-06 | 0.00098271 |
| CXADR | 93.7089023 | 0.447728556 | 0.34511198 | 1.29734285 | 0.19451325 | 0.6459894 |
| BTG3 | 72.1939755 | 0.000962759 | 0.32254862 | 0.00298485 | 0.99761844 | 0.99982181 |
| C21orf91 | 20.8399427 | 0.103558426 | 0.31078679 | 0.33321373 | 0.73897295 | 0.94376279 |
| LINC01687 | 321.77421 | 1.265368554 | 0.94566254 | 1.33807622 | 0.18087159 | 0.6286114 |
| MRPL39 | 19.4603919 | -0.060189853 | 0.37600335 | -0.160078 | 0.87281966 | 0.97434023 |
| ATP5PF | 126.030096 | -0.005541343 | 0.39052907 | -0.0141893 | 0.98867894 | 0.99915789 |
| GABPA | 34.6990902 | -0.024470607 | 0.25127122 | -0.0973872 | 0.92241889 | 0.98670746 |
| APP | 540.126293 | -0.049562831 | 0.31946648 | -0.1551425 | 0.87670897 | 0.97532685 |
| ADAMTS1 | 262.874437 | -4.671736709 | 0.67005637 | -6.9721547 | 3.12E-12 | 5.05E-09 |
| ADAMTS5 | 50.3529133 | 3.223074695 | 0.64605235 | 4.98887546 | 6.07E-07 | 0.00013389 |
| LTN1 | 83.0519414 | 0.227561643 | 0.23378572 | 0.973377 | 0.33036597 | 0.76685572 |
| USP16 | 64.6370088 | 0.094413868 | 0.23275532 | 0.4056357 | 0.6850103 | 0.92581857 |
| CCT8 | 51.7960386 | -0.106092906 | 0.26091141 | -0.4066242 | 0.68428398 | 0.92548169 |
| MAP3K7CL | 77.0327597 | -0.781173272 | 0.29285797 | -2.6674134 | 0.00764376 | 0.12652637 |
| BACH1 | 417.148631 | -1.341218961 | 0.42831406 | -3.1313914 | 0.0017398 | 0.04713986 |
| GRIK1 | 33.0432068 | -2.528061487 | 0.59655698 | -4.2377536 | 2.26E-05 | 0.00223464 |
| CLDN8 | 112.787245 | 1.274893737 | 0.36866064 | 3.45817696 | 0.00054384 | 0.02127132 |
| SOD1 | 94.3265561 | -0.150875936 | 0.3603436 | -0.4187002 | 0.67543525 | 0.92193843 |
| SCAF4 | 56.4658621 | 0.138167975 | 0.24278837 | 0.56908811 | 0.56929635 | 0.88562534 |
| HUNK | 23.8590527 | -1.586978663 | 0.71151195 | -2.2304315 | 0.02571881 | 0.25224718 |
| URB1 | 18.188331 | 0.162790718 | 0.42837611 | 0.3800182 | 0.70393191 | 0.93306088 |
| SYNJ1 | 72.5874759 | 0.201800286 | 0.26495961 | 0.76162661 | 0.44628289 | 0.83035712 |
| PAXBP1 | 92.5747539 | 0.29661157 | 0.26093558 | 1.13672337 | 0.25565395 | 0.70861042 |
| C21orf62-AS | 20.2750835 | 0.220227598 | 0.44979587 | 0.48961676 | 0.62440511 | 0.90447745 |
| AP000282.1 | 142.889763 | 1.012519131 | 0.54977525 | 1.84169645 | 0.06551956 | 0.40687565 |
| OLIG1 | 16.245218 | 0.763252434 | 0.57019274 | 1.33858672 | 0.18070525 | 0.6286114 |
| IFNAR1 | 124.632037 | -0.453017638 | 0.21662266 | -2.0912754 | 0.03650339 | 0.30603532 |
| IFNGR2 | 167.152836 | -0.630990978 | 0.29779036 | -2.11891 | 0.03409807 | 0.29400113 |
| TMEM50B | 114.941872 | 0.372481774 | 0.33941245 | 1.0974311 | 0.27245299 | 0.72345643 |
| GART | 37.4962529 | 0.279835707 | 0.2616107 | 1.06966459 | 0.28477031 | 0.73638801 |
| SON | 277.206581 | -0.254925779 | 0.21487331 | -1.1864004 | 0.23546421 | 0.69023959 |
| AP000311.1 | 56.3203078 | -0.622102968 | 0.43400952 | -1.4333855 | 0.15174769 | 0.58948844 |
| ITSN1 | 195.167682 | 0.125068024 | 0.16690901 | 0.7493186 | 0.4536652 | 0.83364149 |
| LINC00649 | 15.1393111 | -0.396833541 | 0.46221621 | -0.8585453 | 0.39059144 | 0.80235853 |
| MRPS6 | 491.420922 | 1.158726847 | 0.62154911 | 1.86425629 | 0.06228572 | 0.39669829 |
| LINC00310 | 41.5941162 | 1.956688388 | 0.79662069 | 2.45623596 | 0.01404009 | 0.17779231 |
| AP000317.1 | 36.471452 | 1.076796734 | 0.81136563 | 1.32714117 | 0.18446199 | 0.63449691 |
| RCAN1 | 43.8815926 | 0.65665964 | 0.31375726 | 2.09289063 | 0.03635892 | 0.30561656 |
| CLIC6 | 20.5851618 | -0.045419577 | 0.52138066 | -0.087114 | 0.93058086 | 0.98756002 |
| RUNX1 | 52.0218915 | -1.044319713 | 0.43889613 | -2.3794234 | 0.01733975 | 0.20451748 |
| SETD4 | 47.688416 | 0.054956831 | 0.23542279 | 0.23343887 | 0.81542063 | 0.96300833 |
| CBR1 | 114.930805 | -1.255842899 | 0.26444381 | -4.7489972 | 2.04E-06 | 0.00033609 |
| CBR3 | 53.1868709 | -0.182317464 | 0.41747336 | -0.4367164 | 0.66231703 | 0.91812398 |
| DOPEY2 | 93.5276184 | 0.428420659 | 0.52772498 | 0.81182562 | 0.4168917 | 0.81875875 |
| MORC3 | 110.596998 | -0.219962063 | 0.23423031 | -0.9390845 | 0.34768734 | 0.77726831 |
| HLCS | 165.447755 | -0.328216758 | 0.25959425 | -1.2643452 | 0.20610614 | 0.65836247 |
| PIGP | 17.0535002 | -0.195338337 | 0.48287165 | -0.4045347 | 0.68581957 | 0.92597882 |
| TTC3 | 304.873992 | 0.446899971 | 0.18378914 | 2.43159068 | 0.01503269 | 0.1857542 |
| VPS26C | 66.9773205 | -0.398798973 | 0.24698787 | -1.61465 | 0.1063865 | 0.50162992 |
| DYRK1A | 412.161281 | 0.101867297 | 0.21270492 | 0.47891369 | 0.63200003 | 0.90805737 |
| KCNJ15 | 49.1640608 | 0.079405085 | 0.46140509 | 0.17209408 | 0.86336357 | 0.97368057 |
| ETS2 | 182.519055 | -2.973386198 | 0.33581104 | -8.8543431 | 8.42E-19 | 2.72E-15 |
| AP001043.1 | 34.2033433 | -0.458876501 | 0.86939031 | -0.5278141 | 0.59762834 | 0.89363331 |
| BRWD1 | 312.986169 | -0.00670049 | 0.1587996 | -0.0421946 | 0.96634355 | 0.99351999 |
| HMGN1 | 108.054171 | -0.06589053 | 0.18772479 | -0.3509954 | 0.72559184 | 0.93968502 |
| WRB | 26.3177948 | 0.478804908 | 0.42893316 | 1.11626927 | 0.26430689 | 0.71474125 |
| DSCAM | 108.77815 | -1.565060219 | 0.46307454 | -3.3797155 | 0.00072561 | 0.02606817 |
| BACE2 | 719.829641 | -1.654326834 | 0.65472335 | -2.526757 | 0.01151211 | 0.15884421 |
| FAM3B | 53.0408345 | -0.702953419 | 0.25082781 | -2.8025338 | 0.00507029 | 0.09827853 |
| MX1 | 126.426088 | -1.978081459 | 1.11283835 | -1.7775101 | 0.07548434 | 0.43299712 |

| | | | | | | | |
|---|---|---|---|---|---|---|---|
| TMPRSS2 | 590.835615 | | 0.132877889 | 0.24148315 | 0.55025739 | 0.58214284 | 0.89036744 |
| RIPK4 | 13.9860164 | | -0.312168117 | 0.4339612 | -0.7193457 | 0.47192795 | 0.84570599 |
| PRDM15 | 16.2642783 | | 0.344381927 | 0.39784233 | 0.86562416 | 0.3866963 | 0.80009555 |
| C2CD2 | 53.381983 | | 0.147646064 | 0.28968453 | 0.50967881 | 0.6102765 | 0.89964773 |
| ZBTB21 | 41.6932948 | | 0.191738852 | 0.29471818 | 0.65058373 | 0.51531523 | 0.86463667 |
| UMODL1 | 13.5816582 | | 0.803768523 | 0.9163792 | 0.87711345 | 0.38042502 | 0.79631478 |
| ABCG1 | 42.0242426 | | -0.596164905 | 0.48036226 | -1.2410736 | 0.21457857 | 0.66839951 |
| SLC37A1 | 93.0767343 | | -1.25199529 | 0.39121974 | -3.2002355 | 0.00137315 | 0.04011925 |
| PDE9A | 85.4527869 | | -0.245071959 | 0.34527196 | -0.709794 | 0.4778319 | 0.84771464 |
| WDR4 | 15.8542126 | | 0.233570295 | 0.49772418 | 0.46927657 | 0.63887196 | 0.91063717 |
| NDUFV3 | 73.6126333 | | -0.001868282 | 0.25631087 | -0.0072891 | 0.99418417 | 0.99923337 |
| PKNOX1 | 49.3855201 | | -0.209861912 | 0.35550605 | -0.5903188 | 0.55497692 | 0.87754295 |
| RRP1B | 25.9276805 | | 0.296932258 | 0.35293709 | 0.84131781 | 0.40016991 | 0.80830282 |
| PDXK | 99.5697581 | | 0.168633015 | 0.28669581 | 0.58819491 | 0.55640147 | 0.87807756 |
| CSTB | 152.123517 | | -0.614401064 | 0.41036842 | -1.4971938 | 0.13434284 | 0.5624193 |
| AGPAT3 | 131.161213 | | 0.065997421 | 0.23449555 | 0.28144423 | 0.7783697 | 0.95535696 |
| TRAPPC10 | 135.331568 | | -0.623782588 | 0.2520752 | -2.4745893 | 0.01333895 | 0.17367489 |
| PFKL | 134.437496 | | -1.003791221 | 0.18744311 | -5.3551781 | 8.55E-08 | 2.44E-05 |
| C21orf2 | 23.6047396 | | 0.409224388 | 0.35884723 | 1.1403861 | 0.25412548 | 0.7071191 |
| UBE2G2 | 83.6715719 | | 0.21723156 | 0.20904544 | 1.03915955 | 0.29873054 | 0.74296125 |
| SUMO3 | 48.9765547 | | -0.155644508 | 0.38812833 | -0.401013 | 0.68841055 | 0.9267526 |
| PTTG1IP | 148.259716 | | -0.238345658 | 0.2836121 | -0.8403931 | 0.40068801 | 0.80830282 |
| FAM207A | 13.98914 | | 0.061860628 | 0.42539039 | 0.14542084 | 0.88437861 | 0.97662748 |
| ADARB1 | 47.4526222 | | 0.083773202 | 0.46067462 | 0.18184896 | 0.85570126 | 0.97166553 |
| POFUT2 | 24.5326383 | | 0.303760417 | 0.33871105 | 0.89681283 | 0.3698188 | 0.78996749 |
| COL18A1 | 149.870879 | | -0.367082717 | 0.22348162 | -1.6425633 | 0.10047331 | 0.48886326 |
| PCBP3 | 87.8479624 | | -0.657690288 | 0.36415711 | -1.8060619 | 0.07090867 | 0.42197185 |
| COL6A1 | 121.096639 | | 0.890396911 | 0.58637079 | 1.51848784 | 0.12889146 | 0.55324807 |
| LSS | 45.422251 | | -0.78618454 | 0.47423018 | -1.6578121 | 0.09735539 | 0.4826182 |
| MCM3AP-AS | 11.9791499 | | -0.070067886 | 0.44286316 | -0.1582157 | 0.87428687 | NA |
| MCM3AP | 79.4148322 | | 0.116323958 | 0.19560643 | 0.59468371 | 0.55205492 | 0.87576439 |
| YBEY | 15.792101 | | 0.533452575 | 0.39910555 | 1.33662029 | 0.18134661 | 0.62946679 |
| PCNT | 31.5644228 | | 0.228441089 | 0.35281732 | 0.64747697 | 0.51732329 | 0.86500999 |
| DIP2A | 129.664807 | | 0.219214826 | 0.16839503 | 1.30178917 | 0.19298847 | 0.64373732 |
| PRMT2 | 106.981506 | | -0.375996676 | 0.28625205 | -1.3135161 | 0.18900909 | 0.64036044 |
| AL592183.1 | 74.0766296 | | -0.148304919 | 0.44363548 | -0.3342945 | 0.73815731 | 0.94376279 |
| AL354822.1 | 13.7926501 | | -0.565507254 | 0.64714546 | -0.8738488 | 0.38220062 | 0.79731722 |

**Collecting duct intercalated cells type B**

*This means that positive fold changes imply higher mean value in controls than in AKI and vice versa.

| | baseMean | log2 fold change control/AKI* | lfcSE | stat | pvalue | padj |
|---|---|---|---|---|---|---|
| AL669831.5 | 6.85036901 | -0.014335148 | 0.63622379 | -0.0225316 | 0.9820239 | 0.99777713 |
| NOC2L | 4.32635912 | 0.514956812 | 0.71158268 | 0.72367811 | 0.46926337 | 0.99617422 |
| ISG15 | 73.1567311 | -1.421805822 | 2.17457692 | -0.653831 | 0.51322069 | 0.99665574 |
| AGRN | 12.2711978 | -0.44270921 | 0.61873982 | -0.7155014 | 0.47429927 | 0.99617422 |
| C1orf159 | 7.38299122 | -0.931007222 | 0.58473285 | -1.5921925 | 0.11134147 | 0.8083966 |
| AL390719.2 | 2.79983962 | -0.904078292 | 0.75916217 | -1.1908895 | 0.23369695 | 0.92960234 |
| SDF4 | 13.2809507 | -0.092980106 | 0.43377243 | -0.2143523 | 0.83027236 | 0.99665574 |
| B3GALT6 | 3.13529671 | -0.351682406 | 0.90777552 | -0.3874112 | 0.69845181 | 0.99665574 |
| UBE2J2 | 10.8502805 | -0.395018915 | 0.47832369 | -0.8258402 | 0.40889477 | 0.99279547 |
| ACAP3 | 4.77179459 | 0.119222174 | 0.85206932 | 0.13992075 | 0.88872261 | 0.99665574 |
| INTS11 | 7.89135668 | 0.763510273 | 0.5259927 | 1.45156058 | 0.14662383 | 0.85169374 |
| CPTP | 5.45870049 | -0.079122012 | 0.62635479 | -0.1263214 | 0.89947752 | 0.99777713 |
| DVL1 | 5.66247424 | 0.190825368 | 0.6195729 | 0.30799502 | 0.75808612 | 0.99665574 |
| AURKAIP1 | 25.010472 | 0.420678022 | 0.51323463 | 0.81966025 | 0.41240981 | 0.99279547 |
| CCNL2 | 24.2245333 | -0.571422051 | 0.35702 | -1.6005323 | 0.10948055 | 0.80453999 |
| MRPL20 | 21.1558157 | 0.148161844 | 0.33847271 | 0.43773645 | 0.66157734 | 0.99665574 |
| VWA1 | 6.50836665 | -1.561422057 | 0.63434908 | -2.4614555 | 0.01383745 | 0.39608255 |
| ATAD3C | 4.62484246 | -1.23550648 | 0.85391098 | -1.4468797 | 0.14793061 | 0.85308336 |
| ATAD3B | 8.57115054 | -0.694288134 | 0.66421947 | -1.0452692 | 0.29589858 | 0.97286866 |
| ATAD3A | 5.02987711 | 0.751785167 | 0.79621196 | 0.9442023 | 0.34506628 | 0.97788427 |
| SSU72 | 22.5773763 | 0.152074607 | 0.43433765 | 0.35012992 | 0.72624119 | 0.99665574 |
| MIB2 | 4.68663638 | 0.639086502 | 0.66973256 | 0.95424134 | 0.33996149 | 0.97788427 |
| CDK11B | 6.1679364 | -0.052104765 | 0.70293206 | -0.0741249 | 0.94091101 | 0.99777713 |
| SLC35E2B | 7.00258062 | -0.460972125 | 0.5134232 | -0.8978405 | 0.36927061 | 0.98532838 |
| CDK11A | 4.23278121 | -1.458603912 | 0.8381085 | -1.7403521 | 0.08179721 | 0.75124322 |
| SLC35E2A | 7.25853404 | -0.342318624 | 0.58459774 | -0.5855627 | 0.55816943 | 0.99665574 |
| NADK | 12.2247804 | 0.702412092 | 0.46854964 | 1.49911992 | 0.13384253 | 0.84351899 |
| GNB1 | 97.9028032 | -0.724018189 | 0.427765 | -1.6925606 | 0.09053914 | 0.76326474 |
| PRKCZ | 16.0370395 | 1.135915896 | 0.43194428 | 2.62977416 | 0.00854416 | 0.31119582 |
| FAAP20 | 13.1922234 | 0.146381688 | 0.45750599 | 0.31995578 | 0.74900185 | 0.99665574 |
| SKI | 14.4139254 | -0.074894513 | 0.4557948 | -0.1643163 | 0.86948215 | 0.99665574 |
| MORN1 | 4.86781624 | 0.131671812 | 0.61532602 | 0.21398707 | 0.83055714 | 0.99665574 |
| RER1 | 22.6157582 | 0.100024878 | 0.51369719 | 0.19471564 | 0.84561557 | 0.99665574 |
| PEX10 | 4.90471173 | -0.407153927 | 0.58423732 | -0.6968982 | 0.48586651 | 0.99640678 |
| PANK4 | 2.87252796 | 0.59804242 | 0.81925678 | 0.72998165 | 0.4654014 | 0.99617422 |
| TNFRSF14-A | 4.69784715 | -0.26327262 | 0.72068733 | -0.3653077 | 0.71488174 | 0.99665574 |
| TNFRSF14 | 10.5317113 | 0.228910906 | 0.68307793 | 0.33511682 | 0.73753696 | 0.99665574 |
| LINC00982 | 7.19576218 | 0.689855651 | 0.53566449 | 1.28785027 | 0.19779809 | 0.90381759 |
| PRDM16 | 61.0618915 | -0.445344509 | 0.43125734 | -1.0326653 | 0.30176054 | 0.97358745 |
| ARHGEF16 | 5.9831691 | -0.013694586 | 0.64107906 | -0.0213618 | 0.98295707 | 0.99777713 |
| TPRG1L | 8.20611174 | -0.042907121 | 0.53385802 | -0.0803718 | 0.93594157 | 0.99777713 |
| WRAP73 | 3.37239962 | 0.825990031 | 0.8859325 | 0.93233969 | 0.351161 | 0.97838306 |
| LRRC47 | 6.5651721 | 0.678790091 | 0.64573076 | 1.05119678 | 0.29316822 | 0.97072515 |
| CEP104 | 12.1647757 | -0.31120461 | 0.53664075 | -0.5799124 | 0.56197371 | 0.99665574 |
| C1orf174 | 3.27842003 | 0.347465319 | 0.83987224 | 0.41371211 | 0.67908496 | 0.99665574 |
| NPHP4 | 5.47554662 | -0.276515601 | 0.65275377 | -0.4236139 | 0.67184738 | 0.99665574 |
| RPL22 | 15.7182922 | 0.79010276 | 0.51887273 | 1.52272939 | 0.12782642 | 0.83778831 |
| RNF207 | 7.4570565 | -0.286574189 | 0.61716876 | -0.4643368 | 0.64240643 | 0.99665574 |
| ICMT | 8.33840892 | 0.294513392 | 0.50646518 | 0.58150768 | 0.56089834 | 0.99665574 |
| ACOT7 | 4.44653687 | 0.704927696 | 0.71882985 | 0.98066002 | 0.32676043 | 0.97788427 |
| NOL9 | 10.7585306 | 0.76186961 | 0.56445203 | 1.34975086 | 0.17709591 | 0.88584149 |
| ZBTB48 | 2.54596463 | 0.230354135 | 0.81351492 | 0.28315908 | 0.7770549 | 0.99665574 |
| KLHL21 | 8.05456258 | -0.579729129 | 0.591593 | -0.9799459 | 0.32711283 | 0.97788427 |
| PHF13 | 3.21983802 | -0.395941706 | 0.85417355 | -0.4635378 | 0.64297895 | 0.99665574 |
| THAP3 | 4.18308121 | -0.723703928 | 0.72046836 | -1.0044909 | 0.31514204 | 0.97652251 |
| DNAJC11 | 18.8523018 | 0.240611296 | 0.42074474 | 0.57187 | 0.56741005 | 0.99665574 |
| CAMTA1 | 85.4418843 | 0.148721 | 0.3286923 | 0.45246269 | 0.6509357 | 0.99665574 |
| VAMP3 | 10.6551454 | -0.442789101 | 0.57107341 | -0.7753628 | 0.43812529 | 0.99617422 |
| PER3 | 22.5019506 | -0.287785289 | 0.6969028 | -0.412949 | 0.679644 | 0.99665574 |
| PARK7 | 35.5177207 | 0.188539694 | 0.65931089 | 0.28596478 | 0.77490509 | 0.99665574 |
| ERRFI1 | 15.78837 | -1.205533406 | 0.75458158 | -1.5976184 | 0.11012793 | 0.80612054 |
| RERE | 253.714551 | -0.040435976 | 0.27042221 | -0.149529 | 0.88113619 | 0.99665574 |
| ENO1 | 75.7334702 | -0.581511607 | 0.35451737 | -1.6402909 | 0.10094471 | 0.78386461 |
| GPR157 | 3.78943433 | -0.525450904 | 0.79645475 | -0.6597373 | 0.50942243 | 0.99665574 |
| H6PD | 10.4436061 | -0.155021802 | 0.4459544 | -0.3476181 | 0.72812704 | 0.99665574 |
| SPSB1 | 4.16658612 | -1.666546706 | 0.99255891 | -1.6790406 | 0.09314413 | 0.76709603 |
| SLC25A33 | 12.1107131 | -0.477222981 | 0.52088881 | -0.9161705 | 0.35957746 | 0.9819332 |
| TMEM201 | 2.88818283 | 1.014988758 | 0.81764611 | 1.2413546 | 0.21447478 | 0.91280851 |
| CLSTN1 | 19.3910964 | -0.443237193 | 0.35927068 | -1.2337138 | 0.21730958 | 0.91584128 |
| CTNNBIP1 | 5.70345399 | -0.60516589 | 0.58339016 | -1.0373262 | 0.29958387 | 0.97358745 |
| LZIC | 8.73015985 | -0.012732503 | 0.51902846 | -0.0245314 | 0.98042872 | 0.99777713 |
| NMNAT1 | 12.1840167 | 0.118375308 | 0.43416658 | 0.27264952 | 0.78512264 | 0.99665574 |
| UBE4B | 38.6399112 | -0.299340819 | 0.31953533 | -0.9368004 | 0.34886125 | 0.97838306 |
| KIF1B | 84.617566 | -0.070787645 | 0.31730847 | -0.2230878 | 0.82346718 | 0.99665574 |
| PGD | 9.0915598 | 0.330235663 | 0.45624446 | 0.72381297 | 0.46918056 | 0.99617422 |
| DFFA | 9.88745539 | -0.086908443 | 0.50985666 | -0.1704566 | 0.86465105 | 0.99665574 |
| PEX14 | 30.3576594 | 0.008379807 | 0.40333618 | 0.02077623 | 0.98342416 | 0.99777713 |
| CASZ1 | 52.9189872 | -0.064477425 | 0.4087693 | -0.1577355 | 0.87466524 | 0.99665574 |
| SRM | 4.60227055 | -0.123787754 | 0.89204735 | -0.1387681 | 0.88963337 | 0.99665574 |
| EXOSC10 | 9.10569245 | -0.846979528 | 0.51039985 | -1.6594431 | 0.09702653 | 0.77606444 |
| MTOR | 24.6550816 | 0.062527335 | 0.36093388 | 0.17323764 | 0.86246463 | 0.99665574 |
| UBIAD1 | 2.8119876 | 0.202595797 | 0.89061496 | 0.22747855 | 0.82005165 | 0.99665574 |
| FBXO2 | 4.11938176 | -1.810111557 | 0.71763271 | -2.522337 | 0.0116578 | 0.37335809 |
| FBXO44 | 3.06206633 | -0.684910128 | 0.93763298 | -0.7304672 | 0.46510466 | 0.99617422 |
| MAD2L2 | 4.57970565 | -0.933285225 | 0.71347083 | -1.3080916 | 0.19084223 | 0.89964233 |
| AGTRAP | 6.30213816 | -1.286181289 | 0.75256503 | -1.7090633 | 0.08743922 | 0.75597266 |
| MTHFR | 6.17998716 | -0.907806968 | 0.58904002 | -1.5411635 | 0.12327701 | 0.82915695 |
| CLCN6 | 8.52144691 | -0.780887565 | 0.79762162 | -0.9790201 | 0.32757007 | 0.97788427 |
| KIAA2013 | 9.18547811 | 0.436116526 | 0.59241402 | 0.73616847 | 0.46162818 | 0.99617422 |
| PLOD1 | 5.05902495 | -0.492913799 | 0.61697515 | -0.79892 | 0.42433679 | 0.99518711 |
| MFN2 | 14.1338735 | 0.526070006 | 0.6474815 | 0.81248654 | 0.41651251 | 0.992853 |
| VPS13D | 89.0462139 | -0.095510309 | 0.25276305 | -0.377865 | 0.70553089 | 0.99665574 |
| DHRS3 | 21.9980751 | -0.199859385 | 0.56096894 | -0.3562753 | 0.7216344 | 0.99665574 |
| PRDM2 | 47.7095684 | -0.472886047 | 0.27714122 | -1.7062999 | 0.08795225 | 0.75650018 |
| KAZN | 313.956202 | -0.931897612 | 0.30731343 | -3.0324012 | 0.00242617 | 0.15277306 |
| TMEM51-AS | 20.9511453 | -0.495751519 | 0.5355826 | -0.9256304 | 0.3546381 | 0.98022903 |
| TMEM51 | 143.893451 | -0.097476293 | 0.29708312 | -0.3281112 | 0.7428276 | 0.99665574 |
| C1orf195 | 5.59535445 | -0.917578366 | 0.74524434 | -1.231245 | 0.21823125 | 0.91662582 |
| EFHD2 | 4.37377097 | -0.222628765 | 0.74473253 | -0.2989379 | 0.76498744 | 0.99665574 |
| CASP9 | 5.45713541 | 0.373331302 | 0.62011318 | 0.60203736 | 0.54714927 | 0.99665574 |
| DNAJC16 | 12.5599405 | -0.024315925 | 0.4588938 | -0.0529881 | 0.95774136 | 0.99777713 |
| DDI2 | 17.671688 | -0.305986889 | 0.46772036 | -0.654209 | 0.51297714 | 0.99665574 |
| PLEKHM2 | 11.8762434 | -0.425393838 | 0.57882197 | -0.7349304 | 0.46238191 | 0.99617422 |
| UQCRHL | 5.97888307 | 0.130411664 | 0.65713867 | 0.1984538 | 0.84269003 | 0.99665574 |
| AL450998.2 | 2.21857323 | -1.572361782 | 0.95089231 | -1.6535645 | 0.09821603 | 0.78203824 |
| SPEN | 46.1713753 | -0.183928922 | 0.34659373 | -0.5306758 | 0.59564342 | 0.99665574 |
| ZBTB17 | 6.99493963 | 0.201566039 | 0.5475459 | 0.36812629 | 0.71277907 | 0.99665574 |
| SRARP | 2.61171047 | 0.561015848 | 1.02576256 | 0.54692564 | 0.58442982 | 0.99665574 |
| CLCNKA | 25.0518976 | 2.034667039 | 0.5118978 | 3.97475245 | 7.05E-05 | 0.01742067 |
| CLCNKB | 54.5990382 | 0.903226764 | 0.64312083 | 1.40444333 | 0.16018688 | 0.86797143 |
| FAM131C | 4.06623725 | 0.690552806 | 0.82723942 | 0.83476777 | 0.40384847 | 0.99194293 |

| | | | | | | |
|---|---|---|---|---|---|---|
| EPHA2 | 5.46042377 | 0.792598158 | 0.73272793 | 1.08170867 | 0.279382 | 0.95980166 |
| FBXO42 | 37.9346354 | 0.248035628 | 0.32482242 | 0.76360378 | 0.44510339 | 0.99617422 |
| SZRD1 | 18.3115678 | -0.237350226 | 0.45454595 | -0.5221699 | 0.60155202 | 0.99665574 |
| NECAP2 | 6.49706839 | -0.653561621 | 0.56648475 | -1.1537144 | 0.24861727 | 0.93575175 |
| NBPF1 | 6.26894141 | -1.039359745 | 0.60289602 | -1.7239453 | 0.08471772 | 0.75373899 |
| CROCC | 7.09500754 | 0.619218994 | 0.55327142 | 1.1191957 | 0.26305666 | 0.94557817 |
| ATP13A2 | 4.48983179 | -0.294291838 | 0.75099922 | -0.391867 | 0.69515646 | 0.99665574 |
| SDHB | 21.6753873 | -0.184983863 | 0.36219205 | -0.5107342 | 0.60953719 | 0.99665574 |
| RCC2 | 9.40539299 | 0.289425213 | 0.53801452 | 0.53795056 | 0.59061118 | 0.99665574 |
| ARHGEF10L | 30.9529386 | -0.636065253 | 0.53956905 | -1.1788394 | 0.23846213 | 0.93121101 |
| KLHDC7A | 3.89769691 | -0.400609825 | 0.99616218 | -0.4021532 | 0.68757127 | 0.99665574 |
| UBR4 | 31.2606269 | 0.000819428 | 0.33204806 | 0.0024678 | 0.99803098 | 0.99919905 |
| AL035413.1 | 5.7538753 | -2.144609629 | 0.63683672 | -3.3675973 | 0.00075826 | 0.07295564 |
| EMC1 | 7.48650494 | 0.333309614 | 0.59672698 | 0.558563 | 0.57646 | 0.99665574 |
| MRTO4 | 5.1293632 | -0.725906067 | 0.66132275 | -1.0976578 | 0.27235395 | 0.9504731 |
| AKR7A2 | 6.45397624 | 0.369250875 | 0.70454242 | 0.52410028 | 0.6002088 | 0.99665574 |
| PQLC2 | 2.6267441 | 0.006459656 | 0.88036634 | 0.00733746 | 0.9941456 | 0.9983802 |
| CAPZB | 79.3037392 | -0.290430423 | 0.34894648 | -0.8323065 | 0.40523597 | 0.99194293 |
| NBL1 | 4.68583487 | -1.327379967 | 0.79401452 | -1.6717326 | 0.09457706 | 0.77075743 |
| TMCO4 | 46.4746345 | -0.090607424 | 0.45343068 | -0.1998264 | 0.84161635 | 0.99665574 |
| OTUD3 | 5.8162413 | 0.359183591 | 0.58529908 | 0.6136753 | 0.53942991 | 0.99665574 |
| VWA5B1 | 24.9615817 | -0.036764449 | 0.78691764 | -0.0467196 | 0.96273674 | 0.99777713 |
| CAMK2N1 | 65.2270819 | 0.799145523 | 0.3675773 | 2.17408832 | 0.0296985 | 0.55812946 |
| MUL1 | 5.75346037 | 0.256345525 | 0.75761339 | 0.33835928 | 0.73509245 | 0.99665574 |
| CDA | 55.6758934 | -0.393823574 | 0.37624823 | -1.0467121 | 0.29523238 | 0.97209026 |
| PINK1 | 19.9223089 | 0.26705075 | 0.39466707 | 0.67664817 | 0.49862921 | 0.99640678 |
| DDOST | 9.24342279 | -0.143093703 | 0.50956441 | -0.2808157 | 0.77885174 | 0.99665574 |
| HP1BP3 | 58.9668425 | 0.333941913 | 0.30618721 | 1.0906462 | 0.27542859 | 0.95593806 |
| EIF4G3 | 135.272825 | -0.349878271 | 0.37058992 | -0.9441117 | 0.34511258 | 0.97788427 |
| ECE1 | 32.5737795 | 0.369761455 | 0.50491742 | 0.73232065 | 0.46397288 | 0.99617422 |
| RAP1GAP | 16.1827203 | 0.089006302 | 0.57794202 | 0.15400559 | 0.87760533 | 0.99665574 |
| USP48 | 33.8460518 | 0.236611563 | 0.35174004 | 0.67268873 | 0.50114534 | 0.99640678 |
| HSPG2 | 14.1575107 | 0.656578474 | 0.57481147 | 1.14225013 | 0.25335006 | 0.94076968 |
| CDC42 | 50.9330829 | -0.485456445 | 0.35004566 | -1.3868375 | 0.16549132 | 0.87495175 |
| ZBTB40 | 14.0736287 | -0.429768298 | 0.47331314 | -0.9079999 | 0.36387826 | 0.98362663 |
| EPHB2 | 4.03958242 | 0.213214606 | 0.85273007 | 0.25003763 | 0.80255825 | 0.99665574 |
| KDM1A | 27.6699488 | 0.22174823 | 0.34753874 | 0.63805326 | 0.52343901 | 0.99665574 |
| AL031428.1 | 6.25894223 | -0.34850354 | 0.7676283 | -0.4540004 | 0.64982855 | 0.99665574 |
| LUZP1 | 26.5856457 | -0.021274964 | 0.43068904 | -0.0493975 | 0.96060252 | 0.99777713 |
| LINC01355 | 4.45605524 | 0.653659444 | 0.82225434 | 0.79496016 | 0.42663668 | 0.99525676 |
| HNRNPR | 35.4900755 | -0.017360151 | 0.37114058 | -0.0467751 | 0.96269245 | 0.99777713 |
| TCEA3 | 12.3431527 | 0.467436489 | 0.59422507 | 0.78663206 | 0.43149729 | 0.99617422 |
| ASAP3 | 13.4292487 | -0.357296553 | 0.52474628 | -0.6808939 | 0.49593861 | 0.99640678 |
| ID3 | 12.636544 | 1.081605027 | 0.80455522 | 1.34435151 | 0.17883475 | 0.88813816 |
| RPL11 | 107.029845 | 0.551020714 | 0.45258592 | 1.21749415 | 0.22341626 | 0.92191316 |
| ELOA | 9.28260422 | 0.102753767 | 0.55514015 | 0.18509518 | 0.85315438 | 0.99665574 |
| ELOA-AS1 | 4.18110488 | -0.274473944 | 0.72605829 | -0.3780329 | 0.70540613 | 0.99665574 |
| PITHD1 | 11.2677958 | -0.083103096 | 0.55083353 | -0.1508679 | 0.88007993 | 0.99665574 |
| LYPLA2 | 10.3358579 | 0.517423016 | 0.6566318 | 0.78799567 | 0.43069924 | 0.99617422 |
| GALE | 10.283336 | -0.305641154 | 0.46979188 | -0.6505884 | 0.51531221 | 0.99665574 |
| HMGCL | 10.1685789 | 0.033291297 | 0.58812738 | 0.05660559 | 0.95485938 | 0.99777713 |
| FUCA1 | 9.80450664 | -0.078755936 | 0.60489994 | -0.1301966 | 0.89641086 | 0.99777713 |
| PNRC2 | 14.347247 | -0.277343277 | 0.39364106 | -0.7045588 | 0.48108483 | 0.99640678 |
| SRSF10 | 25.940578 | -0.123373113 | 0.37592359 | -0.3281867 | 0.74277052 | 0.99665574 |
| IFNLR1 | 5.99046654 | -0.508542355 | 0.56895789 | -0.8938137 | 0.37142158 | 0.98532838 |
| NIPAL3 | 13.6266315 | -0.403576099 | 0.58474539 | -0.6901741 | 0.49008473 | 0.99640678 |
| RCAN3 | 9.14998408 | -0.229184416 | 0.55484543 | -0.4130599 | 0.6795627 | 0.99665574 |
| SRRM1 | 26.4910997 | 0.093539947 | 0.35275875 | 0.26516691 | 0.79088089 | 0.99665574 |
| CLIC4 | 16.0541102 | -0.131711965 | 0.63943826 | -0.2059807 | 0.83680596 | 0.99665574 |
| SYF2 | 15.4665116 | -0.085223312 | 0.50424336 | -0.1690123 | 0.865787 | 0.99665574 |
| RSRP1 | 74.5985639 | -0.044166188 | 0.30815195 | -0.143326 | 0.88603272 | 0.99665574 |
| TMEM50A | 19.2350681 | -0.175826439 | 0.43235629 | -0.4066702 | 0.6842502 | 0.99665574 |
| MACO1 | 47.1838774 | -0.272356053 | 0.32735035 | -0.8320017 | 0.40540797 | 0.99194293 |
| LDLRAP1 | 6.01737455 | 0.34097303 | 0.62956399 | 0.54160187 | 0.58809281 | 0.99665574 |
| MAN1C1 | 55.3639537 | -0.568789015 | 0.49687992 | -1.1447213 | 0.25232464 | 0.94076968 |
| AL031280.1 | 5.60664667 | 0.049330133 | 0.84568359 | 0.05833167 | 0.95348444 | 0.99777713 |
| MTFR1L | 10.8288222 | 0.857499037 | 0.76917454 | 1.11483024 | 0.26492318 | 0.94557817 |
| PAQR7 | 2.40908922 | 1.157729947 | 1.05195897 | 1.10054667 | 0.27109401 | 0.94865166 |
| STMN1 | 6.68934662 | -0.793252698 | 0.64064505 | -1.2382094 | 0.21563844 | 0.913434 |
| PAFAH2 | 5.90649871 | 0.053012342 | 0.77311121 | 0.06857014 | 0.94533179 | 0.99777713 |
| ZNF593 | 7.69005033 | -0.072530303 | 0.70553791 | -0.1028014 | 0.91812057 | 0.99777713 |
| CNKSR1 | 6.50859207 | -0.794714298 | 0.56727578 | -1.4009311 | 0.16123467 | 0.86946655 |
| CEP85 | 5.48836983 | 0.580937415 | 0.68409644 | 0.84920397 | 0.3957678 | 0.99179694 |
| SH3BGRL3 | 14.6459695 | -1.016758574 | 0.44137884 | -2.3035961 | 0.02124533 | 0.4733368 |
| DHDDS | 14.3585653 | -0.800880544 | 0.57403122 | -1.3951864 | 0.16295964 | 0.87135275 |
| HMGN2 | 17.958651 | 0.408076281 | 0.40532605 | 1.00678522 | 0.314038 | 0.9755712 |
| RPS6KA1 | 9.76154699 | -0.817747723 | 0.48549105 | -1.6843724 | 0.09210972 | 0.76658136 |
| ARID1A | 52.144073 | 0.155716881 | 0.28861916 | 0.53952371 | 0.58952554 | 0.99665574 |
| GPN2 | 7.03862649 | 0.022860657 | 0.71707833 | 0.03188028 | 0.97456753 | 0.99777713 |
| GPATCH3 | 2.44773403 | -0.314770016 | 0.89873483 | -0.3502368 | 0.72616099 | 0.99665574 |
| NUDC | 14.7042765 | 0.025551166 | 0.46288399 | 0.05519993 | 0.95597918 | 0.99777713 |
| NR0B2 | 3.1924327 | 0.103899843 | 0.82184701 | 0.12642237 | 0.8993976 | 0.99777713 |
| KDF1 | 2.56169403 | 0.279824291 | 0.92291871 | 0.30319495 | 0.7617413 | 0.99665574 |
| SLC9A1 | 4.67542995 | 0.066892587 | 0.68616588 | 0.09748749 | 0.92233927 | 0.99777713 |
| WDTC1 | 18.7656072 | 0.169776513 | 0.44574817 | 0.38087989 | 0.70329238 | 0.99665574 |
| TMEM222 | 5.472443 | -0.413116678 | 0.71941672 | -0.5742384 | 0.56580651 | 0.99665574 |
| WASF2 | 35.3087054 | -0.160342867 | 0.33865846 | -0.4734648 | 0.6358816 | 0.99665574 |
| AHDC1 | 6.14897579 | 0.46391988 | 0.79325253 | 0.58483252 | 0.55866033 | 0.99665574 |
| IFI6 | 48.2886808 | -0.053869706 | 1.53235419 | -0.0351549 | 0.97195625 | 0.99777713 |
| FAM76A | 7.66251816 | 0.255919304 | 0.50723161 | 0.50454132 | 0.61388104 | 0.99665574 |
| STX12 | 19.0796232 | -0.48323276 | 0.40854964 | -1.1828006 | 0.23688818 | 0.9305595 |
| PPP1R8 | 7.51713294 | 0.394750111 | 0.60185142 | 0.65589296 | 0.511893 | 0.99665574 |
| RPA2 | 8.02246441 | -0.419818021 | 0.5568113 | -0.7539682 | 0.45086831 | 0.99617422 |
| EYA3 | 52.0547484 | 0.189900123 | 0.33817588 | 0.56154249 | 0.57442778 | 0.99665574 |
| DNAJC8 | 14.137935 | 0.039230009 | 0.49353857 | 0.07948722 | 0.9366451 | 0.99777713 |
| ATP5IF1 | 36.0509816 | 0.013116433 | 0.41415122 | 0.03167064 | 0.97473471 | 0.99777713 |
| SESN2 | 6.5148651 | -0.344652942 | 0.65734896 | -0.5243074 | 0.60006473 | 0.99665574 |
| PHACTR4 | 81.4157253 | -0.158371601 | 0.26352474 | -0.6009743 | 0.54785709 | 0.99665574 |
| RCC1 | 10.8169052 | -0.572567312 | 0.53112076 | -1.078036 | 0.28101768 | 0.96059387 |
| TRNAU1AP | 21.0974157 | -0.658593056 | 0.54003862 | -1.2195296 | 0.22264326 | 0.92191316 |
| TAF12 | 8.46723695 | 0.059328977 | 0.54998081 | 0.10787463 | 0.91409514 | 0.99777713 |
| GMEB1 | 8.46857843 | -0.345857379 | 0.55092923 | -0.627771 | 0.53015398 | 0.99665574 |
| YTHDF2 | 28.8453806 | -0.23525294 | 0.34910527 | -0.6738739 | 0.50039147 | 0.99640678 |
| EPB41 | 61.0226404 | -0.488657878 | 0.33377618 | -1.4640286 | 0.14318613 | 0.84969914 |
| SRSF4 | 61.2717634 | -0.437642878 | 0.34111345 | -1.2829834 | 0.1994979 | 0.90381759 |
| MECR | 6.85342826 | 0.63925069 | 0.58786301 | 1.08741438 | 0.27685372 | 0.95828005 |
| PTPRU | 3.29503731 | -0.281700165 | 0.71303352 | -0.3950728 | 0.69278915 | 0.99665574 |
| PUM1 | 92.3824291 | 0.133043946 | 0.29846394 | 0.44576221 | 0.65576902 | 0.99665574 |
| SNRNP40 | 11.470275 | 0.183614198 | 0.43097773 | 0.42604103 | 0.67007795 | 0.99665574 |
| ZCCHC17 | 9.74272804 | -0.407487723 | 0.4798332 | -0.8492279 | 0.39575452 | 0.99179694 |
| SERINC2 | 24.5885458 | -0.137702633 | 0.39947815 | -0.3447063 | 0.73031519 | 0.99665574 |
| TINAGL1 | 11.5441827 | -0.020891059 | 0.60347577 | -0.0346179 | 0.97238443 | 0.99777713 |
| PEF1 | 6.96155571 | -0.232261861 | 0.59278678 | -0.3918135 | 0.69519602 | 0.99665574 |

| | | | | | | |
|---|---|---|---|---|---|---|
| PTP4A2 | 9.23277907 | -0.752837221 | 0.62500733 | -1.2045254 | 0.22838656 | 0.92646258 |
| KHDRBS1 | 43.6448539 | -0.442035223 | 0.40334695 | -1.0959181 | 0.27311462 | 0.95186117 |
| TMEM39B | 2.9139718 | -0.366433368 | 0.75768621 | -0.4836215 | 0.62865449 | 0.99665574 |
| KPNA6 | 35.1021168 | -0.040967827 | 0.3327881 | -0.1231048 | 0.90202408 | 0.99777713 |
| TXLNA | 9.22204458 | -0.061501371 | 0.56123942 | -0.1095813 | 0.9127414 | 0.99777713 |
| EIF3I | 11.5738568 | -0.054229128 | 0.48316955 | -0.1122362 | 0.9106361 | 0.99777713 |
| HDAC1 | 17.4784454 | 0.385780965 | 0.39094303 | 0.98679586 | 0.32374272 | 0.97700345 |
| MARCKSL1 | 9.39307142 | -0.500670751 | 0.92387259 | -0.5419262 | 0.58786936 | 0.99665574 |
| FAM229A | 3.76699897 | 1.498424952 | 0.9136013 | 1.64013005 | 0.10097813 | 0.78386461 |
| BSDC1 | 15.1652612 | -0.318036895 | 0.39666873 | -0.8017695 | 0.4226863 | 0.99518711 |
| ZBTB8A | 3.49237052 | 0.046252319 | 0.81742273 | 0.05658311 | 0.95487729 | 0.99777713 |
| ZBTB8OS | 19.4889727 | -0.068182826 | 0.50967788 | -0.1337763 | 0.89357946 | 0.99777713 |
| RBBP4 | 26.4060954 | 0.015661458 | 0.33841592 | 0.04627873 | 0.96308809 | 0.99777713 |
| SYNC | 6.17316367 | -0.222565744 | 0.62273166 | -0.3574023 | 0.72079064 | 0.99665574 |
| KIAA1522 | 41.2289984 | -0.51090688 | 0.32672393 | -1.5637265 | 0.11788181 | 0.82599629 |
| YARS | 14.6932101 | -0.878285554 | 0.48219566 | -1.8214298 | 0.06854155 | 0.70708655 |
| S100PBP | 18.2653753 | 0.095137703 | 0.37667002 | 0.25257572 | 0.80059609 | 0.99665574 |
| RNF19B | 9.96867973 | -0.751713432 | 0.68916741 | -1.0907559 | 0.2753803 | 0.95593806 |
| AL020995.1 | 3.771367 | -1.325575488 | 0.77344082 | -1.713868 | 0.08655297 | 0.75427976 |
| AK2 | 22.9732257 | -0.004819294 | 0.35994094 | -0.0133891 | 0.98931734 | 0.99777713 |
| ZNF362 | 4.85150279 | -0.436622632 | 0.76352764 | -0.5718492 | 0.56742417 | 0.99665574 |
| PHC2 | 24.1852312 | -0.623484794 | 0.40776569 | -1.5290271 | 0.12625772 | 0.83633607 |
| SMIM12 | 9.3364724 | -0.220373283 | 0.54333692 | -0.4055923 | 0.68504217 | 0.99665574 |
| ZMYM1 | 14.0674153 | 0.171603929 | 0.46608365 | 0.36818269 | 0.71273702 | 0.99665574 |
| SFPQ | 56.0665702 | 0.305009997 | 0.32942581 | 0.92588373 | 0.35450641 | 0.98022903 |
| ZMYM4 | 115.30853 | 0.254761138 | 0.22118385 | 1.15180715 | 0.24940033 | 0.9369778 |
| KIAA0319L | 48.7196138 | 0.160387106 | 0.27766098 | 0.57763645 | 0.56350959 | 0.99665574 |
| PSMB2 | 12.5585109 | -0.018520032 | 0.44297213 | -0.0418086 | 0.9666513 | 0.99777713 |
| AGO4 | 17.839783 | -0.674202477 | 0.43730701 | -1.5417143 | 0.12314303 | 0.82915695 |
| AGO1 | 9.80264311 | 0.590311742 | 0.50523619 | 1.16838769 | 0.24265042 | 0.9342309 |
| AGO3 | 64.2740511 | -0.025893779 | 0.27942624 | -0.0926677 | 0.92616758 | 0.99777713 |
| ADPRHL2 | 4.53729227 | 0.134086039 | 0.77833474 | 0.17227297 | 0.86322293 | 0.99665574 |
| TRAPPC3 | 7.19395527 | -0.014903903 | 0.60024344 | -0.0248298 | 0.98019075 | 0.99777713 |
| MAP7D1 | 5.87146961 | -0.462547346 | 0.78669622 | -0.5879618 | 0.55655791 | 0.99665574 |
| THRAP3 | 57.4210348 | 0.005315492 | 0.35505529 | 0.01497089 | 0.98805541 | 0.99777713 |
| EVA1B | 3.14363573 | 0.386462608 | 0.86674893 | 0.44587607 | 0.65568677 | 0.99665574 |
| STK40 | 16.4990237 | -0.487406552 | 0.46911226 | -1.0389977 | 0.29880581 | 0.97358745 |
| LSM10 | 2.70539649 | 0.52415407 | 0.91153812 | 0.57502156 | 0.56527672 | 0.99665574 |
| MRPS15 | 8.07875656 | -0.310570196 | 0.6409198 | -0.4845695 | 0.62798174 | 0.99665574 |
| MEAF6 | 9.55036795 | 0.128556968 | 0.50322655 | 0.25546539 | 0.79836367 | 0.99665574 |
| SNIP1 | 4.86176722 | 0.419838885 | 0.69831856 | 0.60121399 | 0.54769747 | 0.99665574 |
| GNL2 | 4.02720987 | -0.776717981 | 0.84012533 | -0.9245263 | 0.35521235 | 0.98022903 |
| C1orf109 | 3.53357493 | -0.435934528 | 0.7046663 | -0.6186397 | 0.53615376 | 0.99665574 |
| C1orf122 | 13.558538 | -0.419268492 | 0.49121123 | -0.8535401 | 0.39335985 | 0.99179694 |
| MTF1 | 7.1585961 | 0.313955887 | 0.55385396 | 0.56685681 | 0.57081147 | 0.99665574 |
| INPP5B | 19.7218088 | -0.007917072 | 0.43134293 | -0.0183545 | 0.98535607 | 0.99777713 |
| SF3A3 | 8.04758801 | -0.006723669 | 0.55702633 | -0.0120706 | 0.99036925 | 0.99777713 |
| UTP11 | 3.19223624 | -0.452906914 | 0.75912483 | -0.5966172 | 0.550763 | 0.99665574 |
| RRAGC | 15.2947427 | -0.921337755 | 0.75122558 | -1.2264462 | 0.22003081 | 0.91768188 |
| AL139260.1 | 26.0458812 | -1.802583169 | 0.79523787 | -2.266722 | 0.02340721 | 0.49335204 |
| RHBDL2 | 3.81277201 | -2.500207373 | 0.75045646 | -3.3315822 | 0.00086354 | 0.08031696 |
| AKIRIN1 | 26.4557049 | -0.160133373 | 0.48178234 | -0.332377 | 0.7396046 | 0.99665574 |
| NDUFS5 | 33.3998731 | -0.396502202 | 0.36192344 | -1.0955416 | 0.27327941 | 0.95186117 |
| MACF1 | 258.897882 | -0.496679002 | 0.26159946 | -1.898624 | 0.05761393 | 0.67545594 |
| PABPC4 | 9.29673258 | -0.372280811 | 0.49407919 | -0.7534841 | 0.45115906 | 0.99617422 |
| PPIE | 5.83769735 | 0.240854684 | 0.77770342 | 0.30969992 | 0.75678917 | 0.99665574 |
| TRIT1 | 10.7647745 | 0.618502842 | 0.49006605 | 1.26208057 | 0.20691979 | 0.90890506 |
| CAP1 | 23.3950179 | -0.308741827 | 0.41426917 | -0.7452686 | 0.45610933 | 0.99617422 |
| PPT1 | 6.17122388 | -0.53804073 | 0.67474413 | -0.7973997 | 0.42521896 | 0.99518711 |
| RLF | 34.9546505 | -0.314638381 | 0.38016744 | -0.8276311 | 0.40787947 | 0.99279547 |
| ZMPSTE24 | 9.39744817 | -0.57578628 | 0.52482585 | -1.0970997 | 0.27259781 | 0.95099677 |
| SMAP2 | 10.6429986 | 0.432547383 | 0.51046905 | 0.84735281 | 0.3967985 | 0.99179694 |
| AL603839.3 | 5.05573676 | 0.423210618 | 0.67185066 | 0.62991769 | 0.52874844 | 0.99665574 |
| ZNF684 | 5.41986736 | -0.371487125 | 0.63798797 | -0.5822792 | 0.56037863 | 0.99665574 |
| NFYC | 32.268393 | 0.642117356 | 0.3709083 | 1.73120244 | 0.08341567 | 0.75242737 |
| CITED4 | 16.6882429 | -0.748506404 | 0.5317671 | -1.4075831 | 0.15925454 | 0.86757292 |
| CTPS1 | 3.07754397 | -0.477954541 | 0.83650588 | -0.5713702 | 0.56774872 | 0.99665574 |
| SCMH1 | 79.3407558 | -0.165351154 | 0.27067538 | -0.6108836 | 0.54127663 | 0.99665574 |
| HIVEP3 | 10.0725839 | -0.600889497 | 0.68439189 | -0.8779904 | 0.37994894 | 0.98679664 |
| FOXJ3 | 51.4524629 | 0.130281939 | 0.28875303 | 0.45118813 | 0.65185396 | 0.99665574 |
| PPCS | 10.1239298 | -0.090756721 | 0.54489117 | -0.1665594 | 0.86771678 | 0.99665574 |
| CCDC30 | 14.7803869 | -0.236171993 | 0.46422347 | -0.5087463 | 0.61093003 | 0.99665574 |
| PPIH | 3.31090598 | 0.156848351 | 0.81657989 | 0.19207962 | 0.84767984 | 0.99665574 |
| YBX1 | 37.7204312 | -0.511012114 | 0.39202761 | -1.3035105 | 0.19240052 | 0.89964233 |
| P3H1 | 3.04492614 | -1.285301482 | 0.82364726 | -1.5604999 | 0.11864179 | 0.82619588 |
| SVBP | 5.4788846 | 0.439880836 | 0.83452114 | 0.52710568 | 0.5981202 | 0.99665574 |
| ERMAP | 4.60307587 | -0.046903923 | 0.68240334 | -0.0687334 | 0.94520181 | 0.99777713 |
| SLC2A1 | 13.6959017 | -0.619211055 | 0.63265672 | -0.9787473 | 0.32770486 | 0.97788427 |
| EBNA1BP2 | 5.02497954 | 0.564775038 | 0.71362779 | 0.79141402 | 0.42870243 | 0.99589919 |
| ELOVL1 | 5.01742035 | -0.217358376 | 0.82568293 | -0.2632468 | 0.79236037 | 0.99665574 |
| MED8 | 4.93076083 | 0.14428376 | 0.75856272 | 0.19020676 | 0.84914711 | 0.99665574 |
| SZT2 | 9.427282 | -0.088672224 | 0.48600414 | -0.1824516 | 0.85522835 | 0.99665574 |
| PTPRF | 27.3911765 | -0.039610582 | 0.42839333 | -0.0924631 | 0.9263301 | 0.99777713 |
| IPO13 | 3.71804205 | 0.665231316 | 0.89956406 | 0.7395041 | 0.45960095 | 0.99617422 |
| ATP6V0B | 108.586252 | 0.186859416 | 0.41757392 | 0.44748824 | 0.65452258 | 0.99665574 |
| B4GALT2 | 6.69414262 | -0.289119516 | 0.73813651 | -0.3916884 | 0.69528845 | 0.99665574 |
| CCDC24 | 5.76970099 | -0.073883039 | 0.66844057 | -0.1105305 | 0.9119887 | 0.99777713 |
| DMAP1 | 4.68069512 | 0.453022646 | 0.71245787 | 0.63585886 | 0.52486842 | 0.99665574 |
| ERI3 | 19.4882505 | 0.173079409 | 0.40846647 | 0.42372978 | 0.67176289 | 0.99665574 |
| RNF220 | 6.40961935 | 0.920275952 | 0.59616009 | 1.54367253 | 0.12266768 | 0.82915695 |
| TMEM53 | 4.93704109 | 0.778032483 | 0.7544994 | 1.03119032 | 0.30245158 | 0.97358745 |
| RPS8 | 62.6047898 | -0.385934522 | 0.37311891 | -1.0343472 | 0.30097386 | 0.97358745 |
| EIF2B3 | 14.7043506 | 0.383141814 | 0.45346801 | 0.84491475 | 0.39815844 | 0.99179694 |
| HECTD3 | 2.28804727 | -0.285463957 | 0.91679627 | -0.3113712 | 0.75551844 | 0.99665574 |
| UROD | 4.75956948 | 0.148185544 | 0.83147107 | 0.17822093 | 0.85854948 | 0.99665574 |
| ZSWIM5 | 28.7541548 | 0.319788975 | 0.52324845 | 0.61116087 | 0.54109309 | 0.99665574 |
| TESK2 | 4.96865244 | -0.281934733 | 0.67312528 | -0.4188444 | 0.67532988 | 0.99665574 |
| PRDX1 | 55.6242414 | -0.4666826 | 0.4289081 | -1.0880713 | 0.27656363 | 0.95823721 |
| AKR1A1 | 5.24263394 | -0.551826905 | 0.64833546 | -0.8511441 | 0.39468931 | 0.99179694 |
| NASP | 15.0896314 | 0.53102807 | 0.3918687 | 1.35511732 | 0.17538018 | 0.88428483 |
| GPBP1L1 | 46.2432361 | 0.056791616 | 0.27574123 | 0.20595982 | 0.8368223 | 0.99665574 |
| TMEM69 | 3.91722755 | 0.066974354 | 0.69342974 | 0.0965842 | 0.92305661 | 0.99777713 |
| IPP | 11.3933915 | 0.24345276 | 0.5042438 | 0.48280764 | 0.62923232 | 0.99665574 |
| MAST2 | 21.5446686 | -0.50835792 | 0.43928276 | -1.1572453 | 0.24717215 | 0.93441937 |
| TSPAN1 | 24.4385514 | -1.83756044 | 0.74660964 | -2.4612064 | 0.01384707 | 0.39608255 |
| POMGNT1 | 8.65592541 | -0.26771442 | 0.58355745 | -0.4587627 | 0.64640455 | 0.99665574 |
| LURAP1 | 4.76653955 | -0.379965552 | 0.71067189 | -0.5346568 | 0.5928872 | 0.99665574 |
| LRRC41 | 11.2498643 | -0.473716918 | 0.51600528 | -0.9180467 | 0.35859444 | 0.9819332 |
| UQCRH | 47.2532246 | 0.534061915 | 0.35205637 | 1.51697841 | 0.12927213 | 0.83778831 |
| NSUN4 | 6.94205163 | -0.364417193 | 0.58219465 | -0.625937 | 0.53135624 | 0.99665574 |
| FAAH | 5.43970691 | 0.353554159 | 0.70386754 | 0.50230212 | 0.61545502 | 0.99665574 |
| MKNK1 | 12.3972286 | -0.637430554 | 0.53625705 | -1.1886661 | 0.2345711 | 0.92960234 |
| ATPAF1 | 10.0355094 | 0.185776089 | 0.52461958 | 0.35411581 | 0.72325209 | 0.99665574 |

| | | | | | | |
|---|---|---|---|---|---|---|
| EFCAB14 | 23.1052024 | -0.004490711 | 0.3580132 | -0.0125434 | 0.98999206 | 0.99777713 |
| PDZK1IP1 | 15.5774118 | -0.674606678 | 0.74444529 | -0.906187 | 0.36483686 | 0.98362663 |
| CMPK1 | 20.0627571 | -0.189578651 | 0.38225219 | -0.4959518 | 0.61992844 | 0.99665574 |
| TRABD2B | 19.8863943 | -1.651868241 | 0.7383565 | -2.2372231 | 0.02527176 | 0.50908112 |
| SPATA6 | 24.1576135 | -0.668593461 | 0.55064077 | -1.2142099 | 0.22466759 | 0.92318867 |
| AGBL4 | 22.8197694 | 1.357873009 | 0.65903897 | 2.06038347 | 0.0393619 | 0.60188673 |
| FAF1 | 167.89764 | 0.03025425 | 0.20820375 | 0.14531078 | 0.8844655 | 0.99665574 |
| RNF11 | 29.3967569 | 0.285416522 | 0.39617329 | 0.72043353 | 0.47125811 | 0.99617422 |
| TTC39A | 7.52631926 | -1.098158413 | 0.61465308 | -1.7866313 | 0.0739971 | 0.72810353 |
| EPS15 | 55.9329579 | -0.440847102 | 0.35332404 | -1.2477133 | 0.21213607 | 0.91043197 |
| OSBPL9 | 55.7311724 | -0.058577469 | 0.28275932 | -0.2071637 | 0.83588201 | 0.99665574 |
| NRDC | 52.7921187 | 0.232797401 | 0.31538885 | 0.7381282 | 0.46043655 | 0.99617422 |
| TXNDC12 | 8.96231281 | 0.107006968 | 0.47137356 | 0.22701097 | 0.82041521 | 0.99665574 |
| BTF3L4 | 16.1960269 | -0.013751186 | 0.38197993 | -0.0359998 | 0.97128255 | 0.99777713 |
| ZFYVE9 | 37.6530139 | -0.349789743 | 0.34218941 | -1.0222109 | 0.30668109 | 0.97376371 |
| CC2D1B | 11.2708462 | -0.150631567 | 0.69202673 | -0.2176673 | 0.82768837 | 0.99665574 |
| PRPF38A | 8.48545128 | 0.352070617 | 0.52912235 | 0.66538602 | 0.50580362 | 0.99665574 |
| TUT4 | 52.5002935 | 0.302509686 | 0.24580341 | 1.23069767 | 0.21843596 | 0.91662582 |
| ZYG11B | 46.9006515 | 0.79886937 | 0.36864203 | 2.16705995 | 0.03023028 | 0.56130882 |
| ECHDC2 | 46.7762741 | 0.185124265 | 0.3724556 | 0.49703713 | 0.61916287 | 0.99665574 |
| SCP2 | 31.8374649 | 0.356268538 | 0.39575129 | 0.90023342 | 0.36799605 | 0.98532838 |
| CPT2 | 7.76521391 | -0.30078221 | 0.55846821 | -0.5385843 | 0.59017372 | 0.99665574 |
| C1orf123 | 3.13587944 | 0.214430031 | 0.79776923 | 0.26878704 | 0.78809357 | 0.99665574 |
| MAGOH | 8.59224102 | 0.08150085 | 0.58339958 | 0.13969988 | 0.88889712 | 0.99665574 |
| NDC1 | 5.38692003 | 0.301020495 | 0.79959177 | 0.37646773 | 0.7065692 | 0.99665574 |
| YIPF1 | 9.46835503 | -0.688916287 | 0.53823772 | -1.279948 | 0.20056343 | 0.90530367 |
| HSPB11 | 11.1165385 | 0.21316259 | 0.60770608 | 0.35076593 | 0.72576396 | 0.99665574 |
| LRRC42 | 5.41645016 | -0.647709117 | 0.68453038 | -0.9462095 | 0.34404176 | 0.97788427 |
| TMEM59 | 75.4172964 | -0.388292842 | 0.30754109 | -1.2625722 | 0.20674296 | 0.90890506 |
| TCEANC2 | 7.64755205 | -0.793792405 | 0.52086048 | -1.524002 | 0.12750822 | 0.83778831 |
| MRPL37 | 7.84722761 | 0.120270611 | 0.59039366 | 0.20371257 | 0.83857812 | 0.99665574 |
| SSBP3 | 91.0302597 | -0.673367036 | 0.48131902 | -1.3990036 | 0.16181191 | 0.87026479 |
| ACOT11 | 68.855004 | 0.459123315 | 0.34838427 | 1.31786465 | 0.18754896 | 0.89729653 |
| MROH7-TTC4 | 3.64452128 | 2.175079704 | 0.9594421 | 2.26702549 | 0.02338867 | 0.49335204 |
| DHCR24 | 9.64106386 | -1.447722325 | 0.96440243 | -1.50116 | 0.13331419 | 0.84208054 |
| TMEM61 | 6.93808135 | 0.629964945 | 0.86349485 | 0.72955264 | 0.46566368 | 0.99617422 |
| BSND | 8.41197729 | 0.800040361 | 0.57478486 | 1.39189534 | 0.16395409 | 0.8734454 |
| USP24 | 59.1055506 | -0.023456765 | 0.32207091 | -0.0728311 | 0.94194056 | 0.99777713 |
| PLPP3 | 132.867475 | 0.237767208 | 0.46192229 | 0.51473421 | 0.60673876 | 0.99665574 |
| PRKAA2 | 134.128712 | 0.288637185 | 0.28731253 | 1.00461049 | 0.31508444 | 0.97652251 |
| FYB2 | 89.3999666 | -0.762615083 | 0.47882735 | -1.5926723 | 0.11123371 | 0.8083966 |
| DAB1 | 22.7819941 | 1.04670477 | 0.70590542 | 1.48278331 | 0.13813199 | 0.84656101 |
| TACSTD2 | 24.597894 | -1.272194256 | 0.52933672 | -2.4033743 | 0.01624455 | 0.42281551 |
| MYSM1 | 28.5146126 | -0.059986042 | 0.34123772 | -0.1757896 | 0.86045924 | 0.99665574 |
| JUN | 23.0007283 | -0.360456585 | 0.66995226 | -0.5380332 | 0.5905541 | 0.99665574 |
| LINC01135 | 7.96236431 | -0.35767205 | 0.87457023 | -0.4089689 | 0.68256246 | 0.99665574 |
| FGGY | 33.4228056 | -0.135817335 | 0.41358292 | -0.328392 | 0.74261527 | 0.99665574 |
| HOOK1 | 21.9322266 | 0.354357083 | 0.40585343 | 0.8731159 | 0.38259991 | 0.98914365 |
| NFIA | 27.0628149 | 0.516897303 | 0.59136632 | 0.87407295 | 0.38207853 | 0.98914365 |
| TM2D1 | 24.4951747 | -0.059446669 | 0.34069059 | -0.1744887 | 0.86148138 | 0.99665574 |
| PATJ | 229.782002 | -0.061270166 | 0.18802778 | -0.325857 | 0.74453256 | 0.99665574 |
| USP1 | 8.95606277 | -0.058066263 | 0.54332945 | -0.1068712 | 0.91489117 | 0.99777713 |
| DOCK7 | 57.2903182 | 0.466379188 | 0.32768294 | 1.42326356 | 0.15465977 | 0.86196782 |
| ATG4C | 8.49109622 | 0.625181546 | 0.60692573 | 1.03007916 | 0.30297285 | 0.97376371 |
| ALG6 | 4.07100022 | 0.412408061 | 0.84356715 | 0.48888587 | 0.6249225 | 0.99665574 |
| ITGB3BP | 14.9937834 | -0.362175152 | 0.67775085 | -0.534378 | 0.59308003 | 0.99665574 |
| EFCAB7 | 6.6612814 | -0.075272567 | 0.71796078 | -0.1048422 | 0.91650105 | 0.99777713 |
| PGM1 | 19.6972989 | -0.852484428 | 0.42522715 | -2.0047742 | 0.04498719 | 0.6313648 |
| RAVER2 | 15.8796361 | -0.163771426 | 0.60310501 | -0.2715471 | 0.78597027 | 0.99665574 |
| JAK1 | 62.5995345 | -0.177459392 | 0.34287794 | -0.5175585 | 0.60476635 | 0.99665574 |
| AK4 | 7.49937502 | 0.991086518 | 0.9879358 | 1.00318919 | 0.31576959 | 0.97652251 |
| DNAJC6 | 8.4923085 | -0.571718553 | 0.66391812 | -0.8611281 | 0.38916749 | 0.99176253 |
| LEPR | 8.84306017 | -0.968795694 | 0.63400406 | -1.5280591 | 0.12649785 | 0.83655267 |
| SGIP1 | 14.8304965 | -0.280372365 | 0.89741682 | -0.3124216 | 0.75472016 | 0.99665574 |
| WDR78 | 5.96539376 | -0.760603295 | 0.68426708 | -1.1115591 | 0.26632777 | 0.94557817 |
| MIER1 | 40.2575967 | 0.354310016 | 0.31700391 | 1.11768343 | 0.26370222 | 0.94557817 |
| SLC35D1 | 11.9163777 | 0.283571339 | 0.43210905 | 0.65624948 | 0.51166362 | 0.99665574 |
| SERBP1 | 35.6841033 | -0.207942672 | 0.31263721 | -0.6651245 | 0.50597086 | 0.99665574 |
| GADD45A | 39.9071725 | 0.393558191 | 0.57482874 | 0.68465295 | 0.49356295 | 0.99640678 |
| GNG12 | 44.1988607 | -0.466408226 | 0.29552784 | -1.578221 | 0.11451486 | 0.81638817 |
| GNG12-AS1 | 10.5410184 | -1.309342321 | 0.70966474 | -1.8450153 | 0.06503531 | 0.69039578 |
| WLS | 14.7198618 | -0.115948827 | 0.62863442 | -0.1844456 | 0.85366393 | 0.99665574 |
| LRRC40 | 11.1902936 | 0.215500781 | 0.50995694 | 0.42258623 | 0.67259717 | 0.99665574 |
| SRSF11 | 65.6380516 | 0.115378763 | 0.22947335 | 0.50279811 | 0.61510623 | 0.99665574 |
| ANKRD13C | 27.064331 | 0.025794202 | 0.39324084 | 0.06559391 | 0.94770114 | 0.99777713 |
| PTGER3 | 236.048233 | -0.219756067 | 0.26871751 | -0.8177959 | 0.41347376 | 0.99279547 |
| ZRANB2-AS1 | 4.43175958 | -0.768817638 | 0.81501862 | -0.943313 | 0.34552084 | 0.97788427 |
| ZRANB2 | 46.7867459 | -0.057739728 | 0.32198795 | -0.1793226 | 0.85768437 | 0.99665574 |
| NEGR1 | 6.26473127 | -0.824436271 | 0.69607698 | -1.1844039 | 0.23625323 | 0.9305595 |
| CRYZ | 25.9344482 | 0.570859245 | 0.53289588 | 1.07123975 | 0.28406164 | 0.96059387 |
| TYW3 | 7.48738391 | 0.413479851 | 0.55097687 | 0.75044866 | 0.45298454 | 0.99617422 |
| ACADM | 33.9908064 | 0.518138267 | 0.29866558 | 1.73484425 | 0.0827684 | 0.75242737 |
| RABGGTB | 8.88349186 | 0.305637374 | 0.52494157 | 0.58223123 | 0.56041094 | 0.99665574 |
| PIGK | 10.9990447 | 0.453581282 | 0.49240647 | 0.92115216 | 0.35697099 | 0.9819332 |
| AC118549.1 | 73.9329983 | 0.288707113 | 0.27862654 | 1.03617952 | 0.3001184 | 0.97358745 |
| USP33 | 46.8659719 | 0.120528239 | 0.2915066 | 0.41346659 | 0.67926479 | 0.99665574 |
| MIGA1 | 27.4872931 | 0.116231392 | 0.3968943 | 0.29285226 | 0.76963508 | 0.99665574 |
| NEXN | 12.9637261 | 0.320405258 | 0.60216535 | 0.5320885 | 0.5946647 | 0.99665574 |
| FUBP1 | 30.2455406 | 0.253414633 | 0.33059808 | 0.7665339 | 0.44335869 | 0.99617422 |
| GIPC2 | 8.74157856 | 0.300424458 | 0.63163249 | 0.47563174 | 0.63433676 | 0.99665574 |
| PTGFR | 41.0742135 | 1.11034875 | 0.57352041 | 1.93602307 | 0.05286488 | 0.66411295 |
| IFI44L | 17.0126481 | -1.95959779 | 2.40161387 | -0.8159504 | 0.41452851 | 0.99279547 |
| IFI44 | 7.2172313 | -0.025857704 | 1.71388699 | -0.0150872 | 0.98796264 | 0.99777713 |
| ADGRL2 | 53.6847914 | 0.4371512 | 0.36514792 | 1.19718935 | 0.23123276 | 0.92774608 |
| LINC01725 | 40.6256221 | 0.608096616 | 0.50779971 | 1.19751273 | 0.23110677 | 0.92774608 |
| PRKACB | 30.0769293 | 0.818749117 | 0.31117196 | 2.63117898 | 0.00850892 | 0.31119582 |
| RPF1 | 5.68840153 | -0.437630436 | 0.60266199 | -0.7261623 | 0.46773926 | 0.99617422 |
| GNG5 | 20.6430323 | -0.146436629 | 0.39388844 | -0.3717718 | 0.71006274 | 0.99665574 |
| SPATA1 | 3.5941299 | -1.405982953 | 0.98434113 | -1.4283493 | 0.15319135 | 0.8618391 |
| CTBS | 9.22464345 | 0.075909095 | 0.58856748 | 0.12897263 | 0.89737931 | 0.99777713 |
| SSX2IP | 6.84472614 | 1.435265315 | 0.67984132 | 2.11117692 | 0.03475711 | 0.58178209 |
| SYDE2 | 7.40900548 | 0.155639307 | 0.55135035 | 0.28228749 | 0.77772308 | 0.99665574 |
| C1orf52 | 7.68927261 | -0.41910751 | 0.52536489 | -0.7977456 | 0.42501816 | 0.99518711 |
| BCL10 | 7.87357043 | 0.053824837 | 0.52965905 | 0.10162167 | 0.91905698 | 0.99777713 |
| AL078459.1 | 10.5789255 | -0.925912261 | 0.68673302 | -1.3482856 | 0.17756653 | 0.886783 |
| DDAH1 | 55.8990778 | -0.132873385 | 0.38391586 | -0.3461003 | 0.72926736 | 0.99665574 |
| ZNHIT6 | 14.6329125 | -0.384594505 | 0.39881103 | -0.9643527 | 0.33486912 | 0.97788427 |
| COL24A1 | 10.1004571 | -0.876552021 | 0.65112912 | -1.3462031 | 0.17823704 | 0.88811213 |
| ODF2L | 21.363046 | -0.383810596 | 0.3594458 | -1.0677843 | 0.28561781 | 0.96391256 |
| SH3GLB1 | 36.0972986 | -0.016089604 | 0.38639377 | -0.0416404 | 0.96678534 | 0.99777713 |
| SELENOF | 34.0056688 | 0.014646365 | 0.28976651 | 0.0505454 | 0.95968777 | 0.99777713 |
| HS2ST1 | 20.2711187 | -0.504180253 | 0.50793623 | -0.9926054 | 0.32090228 | 0.97700345 |
| LMO4 | 8.19754697 | -1.017788797 | 0.63010623 | -1.6152654 | 0.10625323 | 0.79596216 |

| | | | | | | |
|---|---|---|---|---|---|---|
| PKN2-AS1 | 3.86404893 | 0.299046077 | 0.84430713 | 0.35419111 | 0.72319567 | 0.99665574 |
| PKN2 | 97.2211809 | 0.262578551 | 0.20782699 | 1.26344781 | 0.20642829 | 0.90890506 |
| GTF2B | 11.7586814 | -0.060511458 | 0.45998242 | -0.1315517 | 0.89533891 | 0.99777713 |
| KYAT3 | 15.325972 | -0.227568328 | 0.40223458 | -0.5657602 | 0.57155679 | 0.99665574 |
| GBP3 | 4.17345648 | -2.237351069 | 0.83014715 | -2.6951259 | 0.0070362 | 0.27541714 |
| GBP2 | 13.3997314 | -3.095779362 | 0.59797653 | -5.1770918 | 2.25E-07 | 0.00022848 |
| LRRC8B | 11.0657585 | -0.089150587 | 0.55214867 | -0.1614612 | 0.87173017 | 0.99665574 |
| LRRC8D | 2.45160779 | 0.873381436 | 0.9485236 | 0.92077987 | 0.35716537 | 0.9819332 |
| ZNF326 | 19.8085011 | -0.372685072 | 0.39445703 | -0.9448053 | 0.34475831 | 0.97788427 |
| ZNF644 | 92.9605603 | 0.067100354 | 0.25501966 | 0.26311836 | 0.79245936 | 0.99665574 |
| HFM1 | 9.15923891 | -0.156169535 | 0.49396971 | -0.316152 | 0.7518871 | 0.99665574 |
| TGFBR3 | 20.7587341 | -1.055279791 | 0.55401639 | -1.9047808 | 0.05680857 | 0.67545594 |
| BTBD8 | 8.78816636 | -0.649397122 | 0.71934722 | -0.9027589 | 0.36665385 | 0.98365837 |
| GLMN | 8.68900754 | 0.34277079 | 0.5601851 | 0.61188844 | 0.54061157 | 0.99665574 |
| RPAP2 | 33.6579149 | 0.257046518 | 0.37159743 | 0.69173385 | 0.48910449 | 0.99640678 |
| EVI5 | 60.4932717 | 0.085545685 | 0.27151632 | 0.31506646 | 0.75271119 | 0.99665574 |
| RPL5 | 50.0698162 | 0.063148493 | 0.40160528 | 0.1572402 | 0.87505555 | 0.99665574 |
| FAM69A | 6.32626887 | -1.458660038 | 0.5923629 | -2.4624433 | 0.0137994 | 0.39608255 |
| MTF2 | 38.0261081 | 0.292366749 | 0.3110473 | 0.93994305 | 0.34724678 | 0.97804112 |
| TMED5 | 25.6423882 | 0.23306085 | 0.41770253 | 0.55795891 | 0.57687244 | 0.99665574 |
| CCDC18 | 6.62138975 | 0.608606478 | 0.59055852 | 1.03056082 | 0.30274682 | 0.97374595 |
| DR1 | 7.20991955 | -0.034831925 | 0.5802098 | -0.0600333 | 0.95212909 | 0.99777713 |
| FNBP1L | 125.517438 | 0.123546394 | 0.25558981 | 0.48337761 | 0.62882764 | 0.99665574 |
| BCAR3 | 15.5989042 | -0.100170913 | 0.43053315 | -0.2326671 | 0.81601989 | 0.99665574 |
| AL109613.1 | 2.35584139 | -1.104535263 | 0.98965736 | -1.1160785 | 0.26438855 | 0.94557817 |
| DNTTIP2 | 5.81735967 | -0.969480086 | 0.66383084 | -1.4604324 | 0.14417127 | 0.84969914 |
| GCLM | 14.5895356 | -1.325019244 | 0.57256244 | -2.3141917 | 0.02065721 | 0.47061299 |
| ARHGAP29 | 31.694049 | 0.179466712 | 0.67541436 | 0.2657135 | 0.79045987 | 0.99665574 |
| ABCD3 | 69.614244 | 1.010830722 | 0.28214585 | 3.58265319 | 0.00034012 | 0.04789107 |
| SLC44A3 | 30.0249934 | 0.140193933 | 0.57872287 | 0.24224709 | 0.80858871 | 0.99665574 |
| CNN3 | 62.1015379 | -0.330290889 | 0.40973891 | -0.8061009 | 0.42018469 | 0.99518711 |
| AC105942.1 | 4.11739231 | -0.838860242 | 0.82896809 | -1.0119331 | 0.31157004 | 0.97430509 |
| ALG14 | 9.34033901 | 0.572160933 | 0.60577403 | 0.94451216 | 0.34490799 | 0.97788427 |
| PTBP2 | 61.1345907 | -0.060196989 | 0.31762309 | -0.1895233 | 0.84968266 | 0.99665574 |
| DPYD | 33.5495378 | -0.456590265 | 0.46370673 | -0.9846531 | 0.3247945 | 0.97700345 |
| SNX7 | 16.1631899 | 0.05397326 | 0.41023591 | 0.13156639 | 0.89532727 | 0.99777713 |
| FRRS1 | 14.3614561 | 0.537615757 | 0.52939756 | 1.01552368 | 0.30985627 | 0.97376371 |
| AGL | 29.0857452 | 0.604997464 | 0.29383103 | 2.05899784 | 0.03949445 | 0.60209727 |
| SASS6 | 3.99497719 | -0.213290953 | 0.7470577 | -0.285508 | 0.77525497 | 0.99665574 |
| TRMT13 | 14.1676924 | 0.277386722 | 0.39966706 | 0.69404449 | 0.48765431 | 0.99640678 |
| DBT | 13.8295283 | 0.087803274 | 0.47327034 | 0.18552457 | 0.85281761 | 0.99665574 |
| RTCA | 13.6043605 | 0.004320269 | 0.48925774 | 0.00883025 | 0.99295457 | 0.99777713 |
| CDC14A | 41.7238601 | 0.428289794 | 0.48086704 | 0.89066157 | 0.37311076 | 0.98532838 |
| EXTL2 | 11.671544 | 0.275212233 | 0.5756856 | 0.47805995 | 0.63260753 | 0.99665574 |
| SLC30A7 | 22.221015 | -0.243276736 | 0.36340424 | -0.6694384 | 0.50321589 | 0.99640678 |
| DPH5 | 5.46454508 | -0.055667923 | 0.82942751 | -0.0671161 | 0.94648929 | 0.99777713 |
| AC093157.1 | 6.67353513 | 0.318281527 | 0.55715697 | 0.57126006 | 0.56782337 | 0.99665574 |
| RNPC3 | 28.5654681 | 0.028798764 | 0.42242055 | 0.06817558 | 0.94564587 | 0.99777713 |
| AMY2B | 5.80507652 | -0.625225578 | 0.6544467 | -0.9553499 | 0.33940079 | 0.97788427 |
| VAV3 | 72.8549891 | -0.334124182 | 0.40299768 | -0.829097 | 0.4070495 | 0.99250037 |
| SLC25A24 | 14.5018186 | -0.645160757 | 0.39766741 | -1.6223627 | 0.10472572 | 0.79232041 |
| FAM102B | 3.85162111 | -0.073642388 | 0.75245604 | -0.0978694 | 0.92203603 | 0.99777713 |
| PRPF38B | 23.2401802 | -0.028637302 | 0.45995736 | -0.0622608 | 0.95035516 | 0.99777713 |
| STXBP3 | 37.1233591 | 0.66354673 | 0.31419877 | 2.11186925 | 0.03469767 | 0.58178209 |
| AL449266.1 | 9.5792412 | -0.006963856 | 0.78475196 | -0.008874 | 0.9929197 | 0.99777713 |
| GPSM2 | 81.1584985 | -0.068138632 | 0.64061727 | -0.106364 | 0.91529353 | 0.99777713 |
| WDR47 | 5.32513354 | -0.422332794 | 0.63364081 | -0.6665177 | 0.50508027 | 0.99665574 |
| TAF13 | 6.89022615 | -0.310483465 | 0.59968951 | -0.5177404 | 0.60463943 | 0.99665574 |
| TMEM167B | 6.30541706 | -0.3897183 | 0.60515314 | -0.6439995 | 0.51957578 | 0.99665574 |
| SARS | 16.8933823 | 0.453576352 | 0.47653206 | 0.95182757 | 0.34118444 | 0.97788427 |
| CELSR2 | 3.05521975 | 0.576696763 | 0.85213629 | 0.67676588 | 0.49855451 | 0.99640678 |
| SORT1 | 36.1619947 | 0.397133782 | 0.27460789 | 1.44618492 | 0.14812534 | 0.85372071 |
| PSMA5 | 16.5863685 | -0.279737714 | 0.45537622 | -0.6143002 | 0.53901695 | 0.99665574 |
| SYPL2 | 3.50609574 | -0.96471921 | 0.75802312 | -1.2726778 | 0.20313239 | 0.9066717 |
| CYB561D1 | 2.45460171 | -0.894892822 | 0.85419088 | -1.0476497 | 0.29480003 | 0.97161335 |
| GNAI3 | 24.8052938 | -0.262276664 | 0.39391038 | -0.6658283 | 0.50552087 | 0.99665574 |
| GSTM4 | 3.96434353 | -0.354599025 | 0.79797315 | -0.4443746 | 0.65677175 | 0.99665574 |
| GSTM3 | 7.48362844 | -0.422383594 | 0.8682508 | -0.4864765 | 0.62662938 | 0.99665574 |
| LINC01768 | 4.4641826 | 4.086649955 | 1.17889708 | 3.46650273 | 0.00052728 | 0.06288856 |
| AHCYL1 | 29.5204647 | -0.325746138 | 0.3561266 | -0.914692 | 0.36035337 | 0.9819332 |
| STRIP1 | 9.3420854 | -0.268044217 | 0.55864157 | -0.4798143 | 0.63135944 | 0.99665574 |
| KCNC4 | 9.23078464 | 1.202513607 | 0.55855836 | 2.15288802 | 0.03132748 | 0.56612833 |
| RBM15 | 8.09188769 | 0.773541548 | 0.57180865 | 1.3527979 | 0.1761202 | 0.88428483 |
| SLC16A4 | 6.84435346 | 0.057912426 | 0.78073079 | 0.0741772 | 0.94086939 | 0.99777713 |
| LAMTOR5 | 29.2076422 | 0.135887211 | 0.36876053 | 0.36849717 | 0.71250256 | 0.99665574 |
| LRIF1 | 5.56658965 | -1.009270549 | 0.61452645 | -1.6423549 | 0.10051647 | 0.78386461 |
| DRAM2 | 13.1416122 | -0.286413244 | 0.60789536 | -0.4711555 | 0.63752969 | 0.99665574 |
| CEPT1 | 20.6400686 | -0.043516884 | 0.36503131 | -0.1192141 | 0.90510572 | 0.99777713 |
| DENND2D | 8.25127771 | 0.030693249 | 0.68972066 | 0.04450099 | 0.96450507 | 0.99777713 |
| WDR77 | 3.50352803 | 0.117825025 | 0.77940275 | 0.15117348 | 0.87983887 | 0.99665574 |
| ATP5PB | 28.3279379 | 0.203908658 | 0.38969671 | 0.52324962 | 0.60080055 | 0.99665574 |
| RAP1A | 59.7319056 | -0.222928126 | 0.36718 | -0.6071358 | 0.54376079 | 0.99665574 |
| INKA2 | 3.90313405 | -0.075119284 | 0.8642743 | -0.086916 | 0.93073826 | 0.99777713 |
| DDX20 | 4.34419992 | 0.520160778 | 0.74544626 | 0.69778441 | 0.48531203 | 0.99640678 |
| CTTNBP2NL | 16.3709111 | -0.734108284 | 0.41577594 | -1.7656343 | 0.07745722 | 0.74141484 |
| ST7L | 17.7789456 | 0.076592267 | 0.38655153 | 0.19814245 | 0.84293361 | 0.99665574 |
| CAPZA1 | 28.9279334 | -0.390733333 | 0.35751457 | -1.0929158 | 0.27443076 | 0.95443536 |
| MOV10 | 10.4091948 | 0.508493626 | 0.50975864 | 0.9975184 | 0.31851295 | 0.97700345 |
| RHOC | 12.531206 | -0.386364774 | 0.53395384 | -0.7235921 | 0.4693162 | 0.99617422 |
| SLC16A1 | 8.27262134 | -0.04468075 | 0.87218772 | -0.0512284 | 0.95914355 | 0.99777713 |
| SLC16A1-AS1 | 5.61710748 | -0.163855462 | 0.77129665 | -0.2124416 | 0.83176257 | 0.99665574 |
| LRIG2 | 16.9929007 | -0.025006692 | 0.40999389 | -0.0609928 | 0.95136491 | 0.99777713 |
| MAGI3 | 445.880806 | 0.775886497 | 0.27071198 | 2.86609588 | 0.00415568 | 0.20883915 |
| PHTF1 | 6.58338611 | -0.090522853 | 0.63476863 | -0.1426076 | 0.88660007 | 0.99665574 |
| RSBN1 | 18.2644526 | -0.244170881 | 0.39653091 | -0.6157676 | 0.53804792 | 0.99665574 |
| HIPK1 | 17.0460867 | 0.01735138 | 0.43999225 | 0.03943565 | 0.96854306 | 0.99777713 |
| TRIM33 | 54.9414603 | 0.150378442 | 0.29363252 | 0.51213144 | 0.60855902 | 0.99665574 |
| BCAS2 | 6.18956574 | -0.109502156 | 0.76311783 | -0.1434931 | 0.88590075 | 0.99665574 |
| NRAS | 7.12043502 | -0.880316216 | 0.68823863 | -1.2790857 | 0.20086687 | 0.90586667 |
| CSDE1 | 47.2450461 | -0.00259668 | 0.3052246 | -0.0085074 | 0.99321213 | 0.99783813 |
| SIKE1 | 6.95636153 | -0.357756497 | 0.51799843 | -0.6906517 | 0.48978445 | 0.99640678 |
| VANGL1 | 2.75072902 | 0.028194027 | 0.99863541 | 0.02823255 | 0.97747667 | 0.99777713 |
| SLC22A15 | 25.9527806 | 0.013986247 | 0.43979833 | 0.0318015 | 0.97463035 | 0.99777713 |
| MAB21L3 | 4.39376195 | -0.746439924 | 1.04794547 | -0.7122889 | 0.4762859 | 0.99617422 |
| ATP1A1 | 160.94428 | -0.788446237 | 0.39832963 | -1.9793813 | 0.04777309 | 0.6410909 |
| LINC01762 | 14.5453304 | 1.194644711 | 0.8868871 | 1.34700878 | 0.1779774 | 0.8879601 |
| CD58 | 9.01037489 | -0.893993948 | 0.48228186 | -1.8536753 | 0.06378563 | 0.68630818 |
| IGSF3 | 7.87457371 | -0.247338807 | 0.54760498 | -0.4516738 | 0.65150401 | 0.99665574 |
| PTGFRN | 3.88597181 | -0.836444663 | 0.75373541 | -1.1097325 | 0.26711433 | 0.94557817 |
| TTF2 | 5.06040552 | -1.556888928 | 0.66130681 | -2.3542611 | 0.01855957 | 0.45138305 |
| MAN1A2 | 47.449435 | 0.273637197 | 0.30704922 | 0.8911835 | 0.37283074 | 0.98532838 |
| GDAP2 | 15.877594 | -0.621546417 | 0.40599898 | -1.5309064 | 0.12579254 | 0.83633607 |
| WDR3 | 3.55456351 | -0.380987628 | 0.72417062 | -0.526102 | 0.59881732 | 0.99665574 |

| | | | | | | |
|---|---|---|---|---|---|---|
| SPAG17 | 7.1944319 | -1.171640715 | 0.5891655 | -1.9886445 | 0.04674046 | 0.64060657 |
| WARS2 | 14.7556258 | 0.049451911 | 0.47819832 | 0.10341298 | 0.91763521 | 0.99777713 |
| AL359915.2 | 28.1552496 | 0.232836714 | 0.46567312 | 0.50000033 | 0.61707484 | 0.99665574 |
| HMGCS2 | 4.20145821 | 0.877521174 | 1.39148024 | 0.63063862 | 0.52827684 | 0.99665574 |
| NOTCH2 | 4.23651761 | -1.377694729 | 0.82051038 | -1.6790704 | 0.09313832 | 0.76709603 |
| SEC22B | 16.815891 | -0.403147492 | 0.51999194 | -0.7752957 | 0.43816498 | 0.99617422 |
| LINC00623 | 21.0299182 | -0.330465698 | 0.36976464 | -0.893719 | 0.37147224 | 0.98532838 |
| AC239800.3 | 3.38071852 | -0.130911257 | 0.74407798 | -0.1759376 | 0.860343 | 0.99665574 |
| NBPF15 | 2.85540574 | -1.113903898 | 0.81872639 | -1.3605325 | 0.17366146 | 0.88205405 |
| FP700111.1 | 5.60450312 | 0.114889278 | 0.62306375 | 0.1843941 | 0.8537043 | 0.99665574 |
| SRGAP2B | 5.28998495 | -0.337908511 | 0.68773076 | -0.4913384 | 0.62318715 | 0.99665574 |
| GPR89A | 6.33983934 | -0.242173971 | 0.6199165 | -0.3906558 | 0.69605168 | 0.99665574 |
| PDZK1 | 13.2097149 | 1.414393778 | 0.62002231 | 2.2811982 | 0.02253672 | 0.48612187 |
| RNF115 | 53.2239185 | -0.204402115 | 0.32366898 | -0.6315159 | 0.52770324 | 0.99665574 |
| POLR3C | 6.73854414 | -0.027729545 | 0.59971069 | -0.0462382 | 0.96312039 | 0.99777713 |
| PEX11B | 3.63152885 | 0.164185153 | 0.6925161 | 0.23708496 | 0.81259088 | 0.99665574 |
| RBM8A | 12.6342093 | -0.008257631 | 0.42336548 | -0.0195047 | 0.98443846 | 0.99777713 |
| TXNIP | 131.513475 | -0.09008723 | 0.66180163 | -0.1361242 | 0.89172308 | 0.99760413 |
| NBPF12 | 8.32364658 | -0.0211482 | 0.58786412 | -0.0359746 | 0.97130258 | 0.99777713 |
| PRKAB2 | 6.66260746 | 0.056454392 | 0.61902305 | 0.09119917 | 0.92733433 | 0.99777713 |
| AC242426.2 | 11.1382645 | -0.53214084 | 0.43991763 | -1.2096374 | 0.22641804 | 0.92318867 |
| FMO5 | 5.24743552 | -1.181800116 | 0.7086341 | -1.6677156 | 0.09537221 | 0.77178333 |
| CHD1L | 15.9397841 | 0.430453028 | 0.45688319 | 0.94215116 | 0.34611526 | 0.97788427 |
| BCL9 | 18.8351334 | 0.210487059 | 0.48898717 | 0.43045518 | 0.66686456 | 0.99665574 |
| ACP6 | 8.40793189 | 1.047318247 | 0.63982555 | 1.63688094 | 0.10165535 | 0.7843028 |
| GPR89B | 4.68173911 | -0.177808728 | 0.61344766 | -0.2898515 | 0.77192984 | 0.99665574 |
| NBPF11 | 10.7249121 | -0.551374627 | 0.49223905 | -1.1201359 | 0.26265586 | 0.94557817 |
| AC239809.3 | 4.47934969 | 0.843911399 | 0.91268261 | 0.92464937 | 0.35514832 | 0.98022903 |
| PDE4DIP | 12.7015051 | -0.111482628 | 0.58406402 | -0.190874 | 0.84862432 | 0.99665574 |
| NBPF9 | 2.62099155 | -0.719508308 | 0.90448037 | -0.7954936 | 0.42632646 | 0.99518711 |
| AC245297.3 | 25.5164622 | -0.500152705 | 0.53673271 | -0.9318469 | 0.35141566 | 0.97838306 |
| NBPF19 | 3.51621651 | -0.667876598 | 0.83851411 | -0.7965001 | 0.4257414 | 0.99518711 |
| SF3B4 | 4.85083205 | 0.334319856 | 0.72532774 | 0.46092247 | 0.64485423 | 0.99665574 |
| OTUD7B | 27.589641 | -0.196475227 | 0.34223889 | -0.574088 | 0.56590828 | 0.99665574 |
| VPS45 | 25.7434821 | 0.023459671 | 0.32354307 | 0.07250865 | 0.94219712 | 0.99777713 |
| ANP32E | 6.44006026 | 0.518971459 | 0.79608791 | 0.6519022 | 0.51446427 | 0.99665574 |
| APH1A | 10.7888951 | 0.665130819 | 0.4651695 | 1.42986766 | 0.15275501 | 0.86116133 |
| MRPS21 | 17.0624974 | -0.110789192 | 0.41406564 | -0.2675643 | 0.7890347 | 0.99665574 |
| PRPF3 | 13.5210641 | -0.000392299 | 0.57519559 | -0.000682 | 0.99945582 | 0.99975166 |
| RPRD2 | 47.4625939 | 0.002421563 | 0.30149962 | 0.00803173 | 0.99359168 | 0.99802164 |
| TARS2 | 3.18079107 | -0.789861989 | 0.83061166 | -0.9509402 | 0.34163475 | 0.97788427 |
| MCL1 | 22.5026234 | -0.617773231 | 0.3496823 | -1.76667 | 0.07728351 | 0.74124906 |
| ENSA | 33.1624806 | -0.660560124 | 0.33078804 | -1.9969287 | 0.04583293 | 0.63477358 |
| GOLPH3L | 13.4643177 | 0.238547948 | 0.50952019 | 0.46818154 | 0.63965477 | 0.99665574 |
| ARNT | 20.2286633 | 0.41352524 | 0.38320988 | 1.07910902 | 0.28053913 | 0.96059387 |
| SETDB1 | 13.6285938 | -0.37114086 | 0.43545807 | -0.8522999 | 0.39404768 | 0.99179694 |
| CERS2 | 18.7856992 | -0.345207797 | 0.53830879 | -0.6412821 | 0.52133941 | 0.99665574 |
| MINDY1 | 39.5490357 | 0.431202328 | 0.2824514 | 1.52664256 | 0.12684992 | 0.83669777 |
| PRUNE1 | 18.6015937 | 0.370683908 | 0.47644777 | 0.77801584 | 0.43655968 | 0.99617422 |
| BNIPL | 2.4141187 | 0.439510437 | 0.99306577 | 0.44257939 | 0.65807 | 0.99665574 |
| C1orf56 | 5.69528546 | 0.936884409 | 0.86436748 | 1.08389595 | 0.27841094 | 0.95969433 |
| CDC42SE1 | 10.1055133 | -0.12802522 | 0.57838753 | -0.2213485 | 0.82482108 | 0.99665574 |
| GABPB2 | 18.6016177 | -0.118327998 | 0.46730874 | -0.2532116 | 0.80010469 | 0.99665574 |
| SCNM1 | 4.47663032 | -0.959190356 | 0.66482714 | -1.4427665 | 0.14908625 | 0.85485259 |
| VPS72 | 5.2012367 | 0.519824577 | 0.66257189 | 0.78455573 | 0.43271409 | 0.99617422 |
| PIP5K1A | 62.9660544 | -0.057929089 | 0.33093591 | -0.1750462 | 0.86104329 | 0.99665574 |
| PSMD4 | 13.7138631 | 0.018520818 | 0.4427634 | 0.04183006 | 0.96663417 | 0.99777713 |
| PI4KB | 16.8385287 | -0.753373131 | 0.46167473 | -1.6318267 | 0.10271601 | 0.78770791 |
| SELENBP1 | 2.99604798 | 3.024871853 | 1.34958984 | 2.24132678 | 0.02500492 | 0.50908112 |
| PSMB4 | 9.32701413 | 0.309220613 | 0.52039169 | 0.59420744 | 0.55237338 | 0.99665574 |
| POGZ | 71.0572376 | 0.512670677 | 0.28311115 | 1.81084597 | 0.0701647 | 0.71132971 |
| CGN | 9.94790389 | -0.17912334 | 0.60962715 | -0.2938244 | 0.76889208 | 0.99665574 |
| TUFT1 | 10.8987798 | -0.884650434 | 0.50380196 | -1.7559488 | 0.07909716 | 0.74455616 |
| SNX27 | 19.9808189 | 0.069722294 | 0.38132676 | 0.18284134 | 0.85492251 | 0.99665574 |
| MRPL9 | 3.41160135 | -0.470786611 | 0.70538287 | -0.66742 | 0.50450391 | 0.99665574 |
| RORC | 7.64986997 | 0.4711125 | 0.72606837 | 0.64885418 | 0.51643263 | 0.99665574 |
| THEM4 | 24.2732821 | 0.366117625 | 0.33351227 | 1.09776359 | 0.27230774 | 0.9504731 |
| S100A10 | 35.9419145 | -1.626301809 | 0.6460752 | -2.517202 | 0.0118291 | 0.37359313 |
| S100A11 | 105.722708 | -1.604601982 | 0.40103121 | -4.0011898 | 6.30E-05 | 0.01672481 |
| FLG-AS1 | 16.3034379 | 1.651489221 | 0.5655501 | 2.92014661 | 0.00349867 | 0.19069619 |
| S100A6 | 239.089826 | -1.285444689 | 0.4984788 | -2.5787349 | 0.00991628 | 0.34338499 |
| S100A4 | 14.9068041 | -2.175425597 | 0.74172462 | -2.9329289 | 0.00335781 | 0.18400786 |
| S100A2 | 46.8092088 | -0.627177976 | 0.56058258 | -1.1187968 | 0.26322685 | 0.94557817 |
| S100A16 | 7.65932643 | -1.150886037 | 0.87424037 | -1.3164412 | 0.188026 | 0.89739428 |
| S100A14 | 4.92898366 | -2.044148762 | 0.8477878 | -2.4111562 | 0.01590204 | 0.41983038 |
| S100A13 | 29.0839781 | -0.352989534 | 0.46156282 | -0.7647703 | 0.44440834 | 0.99617422 |
| CHTOP | 14.9404301 | 0.715227205 | 0.42530339 | 1.68168704 | 0.09262954 | 0.76709603 |
| ILF2 | 12.3498175 | -0.119333244 | 0.47296569 | -0.2523085 | 0.80080265 | 0.99665574 |
| INTS3 | 10.1387958 | 0.433060476 | 0.47796221 | 0.90605589 | 0.36490627 | 0.98362663 |
| GATAD2B | 64.2470282 | 0.054953252 | 0.29761468 | 0.18464563 | 0.85350699 | 0.99665574 |
| DENND4B | 7.80859874 | -0.150320603 | 0.54615561 | -0.275234 | 0.78313645 | 0.99665574 |
| CRTC2 | 5.29398269 | 1.263246047 | 0.67281732 | 1.87754686 | 0.06044318 | 0.6834742 |
| SLC39A1 | 11.7234454 | -0.490223182 | 0.6382802 | -0.7680376 | 0.44246486 | 0.99617422 |
| JTB | 14.7237906 | 0.445968895 | 0.47671299 | 0.93550817 | 0.34952647 | 0.97838306 |
| RAB13 | 17.34601 | -0.357884621 | 0.40885927 | -0.8753247 | 0.38139726 | 0.98839606 |
| RPS27 | 24.9706504 | 0.591881854 | 0.45002464 | 1.31522099 | 0.18843564 | 0.89739428 |
| TPM3 | 24.7897559 | -0.422291669 | 0.36527513 | -1.156092 | 0.24764352 | 0.93441937 |
| C1orf43 | 19.6766307 | -0.363624969 | 0.43139553 | -0.8429039 | 0.3992822 | 0.99179694 |
| UBAP2L | 22.3872437 | 0.237314715 | 0.42185268 | 0.56255353 | 0.57373894 | 0.99665574 |
| HAX1 | 5.85475079 | 0.255755137 | 0.73253117 | 0.34913891 | 0.72698502 | 0.99665574 |
| IL6R | 13.3639151 | -0.627603755 | 0.4619198 | -1.3586856 | 0.17424625 | 0.88281284 |
| SHE | 7.36639744 | 0.985395576 | 0.59520341 | 1.65556104 | 0.09781075 | 0.7801773 |
| TDRD10 | 8.14584227 | 1.140329593 | 0.66021133 | 1.72721905 | 0.08412833 | 0.75343906 |
| UBE2Q1 | 8.87860808 | 0.253530959 | 0.51349652 | 0.49373452 | 0.62149368 | 0.99665574 |
| ADAR | 35.198776 | -0.257097817 | 0.44970759 | -0.5717 | 0.56752525 | 0.99665574 |
| PMVK | 12.1062766 | 0.339148487 | 0.47750522 | 0.71025085 | 0.47754859 | 0.99617422 |
| PBXIP1 | 7.4592048 | 0.2304852 | 0.58228319 | 0.39583008 | 0.69223039 | 0.99665574 |
| PYGO2 | 4.29827375 | 0.691749568 | 0.69280566 | 0.99847563 | 0.31804878 | 0.97700345 |
| FLAD1 | 3.47016839 | 0.622865286 | 0.7260628 | 0.85786696 | 0.39096593 | 0.99179694 |
| ZBTB7B | 3.12279648 | -0.136383828 | 0.80941059 | -0.1684977 | 0.86619175 | 0.99665574 |
| EFNA1 | 27.0915782 | -1.332486927 | 0.5235529 | -2.5450856 | 0.0109251 | 0.36314303 |
| SLC50A1 | 3.35122054 | -0.271478179 | 0.75198415 | -0.3610158 | 0.71808762 | 0.99665574 |
| DPM3 | 10.0190122 | 0.510645022 | 0.57497852 | 0.88811148 | 0.37448079 | 0.98535336 |
| MUC1 | 4.02760093 | -0.213739093 | 0.66524864 | -0.321292 | 0.74798909 | 0.99665574 |
| MTX1 | 3.31210814 | 0.295443338 | 0.81844829 | 0.36097985 | 0.71811451 | 0.99665574 |
| GBA | 4.22222248 | 0.142802499 | 0.73385682 | 0.19459177 | 0.84571255 | 0.99665574 |
| SCAMP3 | 7.70511781 | -0.157431622 | 0.55492349 | -0.2836997 | 0.77664053 | 0.99665574 |
| CLK2 | 4.76002072 | -0.24561607 | 0.65086861 | -0.3773666 | 0.70590119 | 0.99665574 |
| FDPS | 16.3900668 | -0.731177946 | 0.44418249 | -1.6461206 | 0.09973894 | 0.78288816 |
| RUSC1 | 3.20588292 | -0.077712059 | 0.75755223 | -0.1025831 | 0.91829385 | 0.99777713 |
| ASH1L | 122.943015 | -0.075713557 | 0.18454144 | -0.4102794 | 0.68160098 | 0.99665574 |
| YY1AP1 | 19.4796845 | -0.226334329 | 0.36640511 | -0.6177161 | 0.53676251 | 0.99665574 |
| DAP3 | 24.9075879 | -0.487505592 | 0.36252593 | -1.3447468 | 0.178707 | 0.88813816 |

| | | | | | | |
|---|---|---|---|---|---|---|
| GON4L | 25.3049105 | 0.233608894 | 0.40842768 | 0.57197125 | 0.56734145 | 0.99665574 |
| RIT1 | 8.29650591 | -1.062007086 | 0.7319655 | -1.4508977 | 0.14680834 | 0.85169374 |
| KHDC4 | 21.4595552 | -0.604302336 | 0.34814426 | -1.7357814 | 0.08260249 | 0.75228386 |
| ARHGEF2 | 5.86822543 | -0.882160567 | 0.62119335 | -1.4201063 | 0.15557675 | 0.8636413 |
| SSR2 | 15.5974131 | -0.620076071 | 0.46614847 | -1.3302115 | 0.18344859 | 0.89532394 |
| UBQLN4 | 4.58924327 | 0.082671795 | 0.68067345 | 0.12145588 | 0.90332996 | 0.99777713 |
| LAMTOR2 | 6.85681877 | 0.91044436 | 0.65829173 | 1.38304086 | 0.16665234 | 0.87495175 |
| RAB25 | 9.66027833 | -0.09470082 | 0.59517526 | -0.1591142 | 0.87357892 | 0.99665574 |
| LMNA | 21.4475951 | -0.718527719 | 0.46608255 | -1.5416319 | 0.12316307 | 0.82915695 |
| SEMA4A | 2.49621127 | 0.25199323 | 0.93591212 | 0.26924882 | 0.78773822 | 0.99665574 |
| SLC25A44 | 5.15546077 | 0.048064683 | 0.66404274 | 0.07238191 | 0.94229798 | 0.99777713 |
| SMG5 | 17.5988977 | -0.406939215 | 0.44532505 | -0.9138027 | 0.36082055 | 0.9819332 |
| GLMP | 3.94927491 | -0.205877983 | 0.89327571 | -0.2304753 | 0.81772246 | 0.99665574 |
| CCT3 | 19.5298208 | 0.185127939 | 0.47743175 | 0.38775791 | 0.69819519 | 0.99665574 |
| RHBG | 48.085235 | 2.062031769 | 0.67675729 | 3.04692952 | 0.00231192 | 0.1502451 |
| MEF2D | 9.39028637 | -0.15331273 | 0.54666969 | -0.2804486 | 0.77913338 | 0.99665574 |
| NAXE | 8.99289688 | 1.008931108 | 0.65018437 | 1.55176155 | 0.12071929 | 0.82659025 |
| GPATCH4 | 4.81940012 | -0.23809947 | 0.66175982 | -0.3597974 | 0.71899864 | 0.99665574 |
| ISG20L2 | 2.5243468 | -0.564438533 | 0.9239534 | -0.610895 | 0.54126908 | 0.99665574 |
| RRNAD1 | 5.75196286 | -0.018731286 | 0.60673228 | -0.0308724 | 0.9753713 | 0.99777713 |
| MRPL24 | 4.19755521 | 1.285940206 | 0.90333064 | 1.42355428 | 0.15457554 | 0.86196782 |
| HDGF | 14.1698549 | -0.192416489 | 0.4366681 | -0.440647 | 0.65946857 | 0.99665574 |
| PRCC | 11.2798011 | 0.508672962 | 0.42815063 | 1.18807009 | 0.2348058 | 0.92960234 |
| NTRK1 | 18.8134141 | 0.416131247 | 0.57608649 | 0.72234162 | 0.47008447 | 0.99617422 |
| INSRR | 68.2878016 | 0.713559105 | 0.32295251 | 2.20948616 | 0.02714084 | 0.53118509 |
| ARHGEF11 | 28.626412 | 0.158925773 | 0.34978485 | 0.45435294 | 0.64957482 | 0.99665574 |
| ETV3 | 4.65682331 | 0.029218822 | 0.78290361 | 0.0373211 | 0.97022898 | 0.99777713 |
| DUSP23 | 14.8859632 | 0.106437391 | 0.61441271 | 0.17323436 | 0.86246722 | 0.99665574 |
| TAGLN2 | 12.9974865 | -1.147905323 | 0.55045688 | -2.0853683 | 0.03703588 | 0.58591936 |
| IGSF8 | 4.23609656 | 1.128320727 | 0.80824629 | 1.39601102 | 0.16271118 | 0.87135275 |
| PEA15 | 3.64434615 | -0.544636841 | 0.76982274 | -0.7074834 | 0.4792661 | 0.99617422 |
| DCAF8 | 9.03499114 | 0.704773731 | 0.62994435 | 1.11878728 | 0.2632309 | 0.94557817 |
| COPA | 39.0344746 | -0.41885832 | 0.39478787 | -1.0609706 | 0.28870326 | 0.96676791 |
| NCSTN | 7.84862254 | 1.103914133 | 0.58262329 | 1.89473052 | 0.05812811 | 0.67545594 |
| F11R | 12.2833918 | -0.4126657 | 0.47826503 | -0.862839 | 0.38822601 | 0.99168679 |
| AL591806.3 | 3.64802206 | -0.018355886 | 1.06102781 | -0.0173001 | 0.98619721 | 0.99777713 |
| USF1 | 2.68908378 | -0.255759568 | 0.8402682 | -0.3043785 | 0.76083955 | 0.99665574 |
| PFDN2 | 19.6255529 | -0.494926932 | 0.5148614 | -0.9612819 | 0.33641046 | 0.97788427 |
| NIT1 | 8.70578389 | -0.187945083 | 0.51288079 | -0.3664498 | 0.71402943 | 0.99665574 |
| DEDD | 3.85901109 | 0.412385907 | 0.8298599 | 0.49693437 | 0.61923533 | 0.99665574 |
| UFC1 | 9.75643798 | -0.15058092 | 0.59739872 | -0.252061 | 0.80099391 | 0.99665574 |
| B4GALT3 | 3.78554433 | 0.559715836 | 0.96295019 | 0.58125108 | 0.56107125 | 0.99665574 |
| NDUFS2 | 18.8273944 | 0.601448522 | 0.38619598 | 1.5573661 | 0.11938359 | 0.82625998 |
| SDHC | 23.2293493 | 0.1463747 | 0.35448702 | 0.41291977 | 0.67966539 | 0.99665574 |
| AL592295.4 | 2.88899525 | -1.196652437 | 0.84309987 | -1.4193484 | 0.15579747 | 0.8636413 |
| DUSP12 | 4.23964665 | 0.188731874 | 0.72269843 | 0.26114886 | 0.79397771 | 0.99665574 |
| ATF6 | 41.1585002 | 0.241008776 | 0.27209543 | 0.88575091 | 0.37575177 | 0.98637037 |
| NOS1AP | 51.4778773 | -0.830236699 | 0.46987653 | -1.7669252 | 0.07724075 | 0.74124906 |
| UHMK1 | 51.1894664 | 0.618502437 | 0.30119367 | 2.05350413 | 0.04002371 | 0.60713077 |
| UAP1 | 22.794493 | 1.531248265 | 0.62212054 | 2.46133695 | 0.01384203 | 0.39608255 |
| DDR2 | 5.13081505 | 0.16141619 | 0.96411622 | 0.167424 | 0.86703645 | 0.99665574 |
| HSD17B7 | 4.25455204 | -1.080344618 | 0.80029012 | -1.3499412 | 0.17703484 | 0.88584149 |
| PBX1 | 26.2150417 | 0.734914806 | 0.64500506 | 1.13939387 | 0.25453891 | 0.94099699 |
| MGST3 | 53.5037117 | 0.058145685 | 0.35776009 | 0.16252703 | 0.87089085 | 0.99665574 |
| ALDH9A1 | 9.60432557 | 1.229388644 | 0.59174238 | 2.0775741 | 0.0377486 | 0.58998295 |
| TMCO1 | 24.4433741 | -0.654176515 | 0.37322404 | -1.7527717 | 0.07964121 | 0.7446402 |
| UCK2 | 5.5834257 | -0.645035586 | 0.67801997 | -0.9513519 | 0.34142577 | 0.97788427 |
| FAM78B | 2.99699178 | 0.027480295 | 0.76267491 | 0.03603147 | 0.97125727 | 0.99777713 |
| POGK | 9.08220879 | 0.298813071 | 0.51879253 | 0.57597798 | 0.56463007 | 0.99665574 |
| TADA1 | 4.32577723 | -1.080619849 | 0.75103802 | -1.4388351 | 0.15019726 | 0.85593018 |
| POU2F1 | 60.2027021 | -0.127032983 | 0.33222324 | -0.3823724 | 0.70218519 | 0.99665574 |
| CREG1 | 6.59851737 | 0.822120648 | 0.70610345 | 1.16430624 | 0.24429993 | 0.93426912 |
| MPZL1 | 19.0860008 | -1.022052786 | 0.47372899 | -2.157463 | 0.03096961 | 0.56271575 |
| MPC2 | 14.2102308 | 0.363639276 | 0.47093085 | 0.77217128 | 0.44001299 | 0.99617422 |
| DCAF6 | 71.5673557 | 0.058248501 | 0.37894886 | 0.15371072 | 0.87783784 | 0.99665574 |
| TIPRL | 8.430217 | -0.186304086 | 0.61029472 | -0.305269 | 0.76016125 | 0.99665574 |
| SFT2D2 | 13.0549361 | -0.346628809 | 0.40348001 | -0.8590979 | 0.39028653 | 0.99179694 |
| ATP1B1 | 175.597651 | 0.39753757 | 0.27176892 | 1.46277788 | 0.14352817 | 0.84969914 |
| NME7 | 29.5450413 | -0.602297627 | 0.321122 | -1.8756037 | 0.06070971 | 0.6834742 |
| BLZF1 | 10.3477498 | 0.331470657 | 0.53468699 | 0.61993402 | 0.53530123 | 0.99665574 |
| SLC19A2 | 29.3819946 | 1.354842871 | 0.57966257 | 2.33729575 | 0.01942381 | 0.46057537 |
| SCYL3 | 9.13392536 | 0.744805926 | 0.59567591 | 1.25035428 | 0.21117016 | 0.91043197 |
| KIFAP3 | 18.0405041 | 0.275501782 | 0.43014399 | 0.64048734 | 0.52185581 | 0.99665574 |
| GORAB | 2.975757 | -0.743929007 | 0.74527707 | -0.9981912 | 0.31818666 | 0.97700345 |
| FMO4 | 6.909784 | 0.393968084 | 0.6160273 | 0.63953023 | 0.52247805 | 0.99665574 |
| PRRC2C | 77.7721962 | -0.464661218 | 0.27898041 | -1.6655694 | 0.09579922 | 0.77210482 |
| VAMP4 | 5.50356578 | -0.122276707 | 0.6830617 | -0.1790127 | 0.85792774 | 0.99665574 |
| METTL13 | 3.31222458 | 0.018359005 | 0.77377773 | 0.02372646 | 0.9810708 | 0.99777713 |
| DNM3 | 15.2251574 | -0.20073338 | 0.61439859 | -0.3267152 | 0.74388327 | 0.99665574 |
| PIGC | 7.66482119 | 0.349077237 | 0.61067163 | 0.57162838 | 0.56757376 | 0.99665574 |
| SUCO | 26.1847294 | -0.062761302 | 0.39342905 | -0.1595238 | 0.8732562 | 0.99665574 |
| AL645568.1 | 4.66906156 | 0.634606275 | 1.03766496 | 0.61157146 | 0.54082132 | 0.99665574 |
| PRDX6 | 36.1499168 | -0.066223913 | 0.52273475 | -0.1266874 | 0.89918781 | 0.99777713 |
| KLHL20 | 16.2716691 | -0.676536239 | 0.46193462 | -1.4645714 | 0.14303789 | 0.84969914 |
| DARS2 | 4.27012847 | -0.467761243 | 0.71951025 | -0.6501106 | 0.51562078 | 0.99665574 |
| ZBTB37 | 6.84742586 | 0.311374677 | 0.56646899 | 0.54967647 | 0.5825413 | 0.99665574 |
| RC3H1 | 50.0832738 | 0.019723598 | 0.2853527 | 0.06912007 | 0.94489405 | 0.99777713 |
| RABGAP1L | 156.432306 | -0.128361191 | 0.20054911 | -0.6400487 | 0.52214096 | 0.99665574 |
| CACYBP | 6.82727828 | -0.053203805 | 0.56068489 | -0.0948907 | 0.92440161 | 0.99777713 |
| MRPS14 | 7.90661895 | 0.474134395 | 0.58190753 | 0.81479337 | 0.4151906 | 0.99279547 |
| KIAA0040 | 27.2341806 | -1.491785872 | 0.47152578 | -3.1637419 | 0.00155755 | 0.1152586 |
| COP1 | 87.7459395 | 0.205418469 | 0.25115648 | 0.81789038 | 0.41341979 | 0.99279547 |
| PAPPA2 | 7.26792431 | 0.1194467 | 0.60619381 | 0.19704375 | 0.8437933 | 0.99665574 |
| RASAL2 | 54.7131972 | 0.125858976 | 0.41726479 | 0.30162855 | 0.76293524 | 0.99665574 |
| RALGPS2 | 92.8250881 | -0.061879561 | 0.34552083 | -0.1790907 | 0.8578665 | 0.99665574 |
| ANGPTL1 | 17.9525979 | 1.236290089 | 0.52902956 | 2.33690173 | 0.01944429 | 0.46057537 |
| FAM20B | 12.878876 | 0.180535003 | 0.42134654 | 0.42847155 | 0.66830784 | 0.99665574 |
| TOR3A | 4.14772674 | 0.134688806 | 0.71536748 | 0.18827918 | 0.8506578 | 0.99665574 |
| ABL2 | 36.7512426 | -0.143005002 | 0.36766041 | -0.3889595 | 0.69730612 | 0.99665574 |
| SOAT1 | 46.9557978 | 0.771729771 | 0.40166293 | 1.92133682 | 0.05468926 | 0.66961316 |
| TOR1AIP2 | 29.5217335 | -0.0923813 | 0.38235614 | -0.2416106 | 0.8090819 | 0.99665574 |
| TOR1AIP1 | 13.5082506 | 0.255643308 | 0.41972425 | 0.60907443 | 0.5424751 | 0.99665574 |
| CEP350 | 53.572098 | 0.235067645 | 0.27993585 | 0.8397197 | 0.40106557 | 0.99194293 |
| QSOX1 | 11.4385706 | 0.597417402 | 0.55737252 | 1.07184581 | 0.28378929 | 0.96059387 |
| ACBD6 | 65.50893 | -0.356793182 | 0.26709037 | -1.3358519 | 0.18159768 | 0.8928406 |
| XPR1 | 43.8713628 | 0.156160898 | 0.3435169 | 0.45459451 | 0.64940098 | 0.99665574 |
| KIAA1614 | 2.68142692 | -1.021975878 | 0.81691117 | -1.2510245 | 0.21092555 | 0.91043197 |
| STX6 | 18.4577376 | 0.228397086 | 0.43012866 | 0.53099713 | 0.59542077 | 0.99665574 |
| MR1 | 12.009339 | 0.2912729 | 0.51914125 | 0.56106676 | 0.57475203 | 0.99665574 |
| GLUL | 24.2484431 | -0.845273472 | 0.44465321 | -1.9009724 | 0.05730563 | 0.67545594 |
| RNASEL | 2.93176458 | -0.171019173 | 1.00966107 | -0.1693828 | 0.86549559 | 0.99665574 |
| DHX9 | 36.1697897 | -0.075232547 | 0.32375991 | -0.2323714 | 0.81624955 | 0.99665574 |
| LAMC1 | 40.2420487 | 0.18871848 | 0.33707648 | 0.55986843 | 0.57556918 | 0.99665574 |

| | | | | | | |
|---|---|---|---|---|---|---|
| SMG7 | 60.8390076 | 0.198539172 | 0.27335192 | 0.72631344 | 0.46764664 | 0.99617422 |
| ARPC5 | 18.489907 | -0.687358247 | 0.50668497 | -1.3565791 | 0.17491498 | 0.88415395 |
| RGL1 | 5.81366886 | 0.16060129 | 0.56179332 | 0.28587255 | 0.77497574 | 0.99665574 |
| COLGALT2 | 5.24671005 | 2.348564686 | 0.84908254 | 2.7660028 | 0.0056748 | 0.23482111 |
| TSEN15 | 16.0387764 | 0.676059206 | 0.49280589 | 1.37185701 | 0.17010796 | 0.87918701 |
| C1orf21 | 16.2199437 | 0.369930307 | 0.47622183 | 0.7768025 | 0.4372753 | 0.99617422 |
| EDEM3 | 23.4846133 | 1.127509938 | 0.4054 | 2.78122824 | 0.00541536 | 0.22875404 |
| RNF2 | 13.7245123 | 0.329712191 | 0.40849537 | 0.80713814 | 0.4195869 | 0.99518711 |
| TRMT1L | 9.96694013 | -0.156805722 | 0.52074428 | -0.3011185 | 0.76332415 | 0.99665574 |
| SWT1 | 22.7272737 | -0.178705593 | 0.42605681 | -0.4194408 | 0.67489404 | 0.99665574 |
| IVNS1ABP | 47.5941697 | 0.154899272 | 0.51659642 | 0.29984581 | 0.76429477 | 0.99665574 |
| TPR | 34.2447127 | -0.244846926 | 0.28505402 | -0.8589492 | 0.39036853 | 0.99179694 |
| ODR4 | 21.2059168 | 0.033089452 | 0.35755985 | 0.09254241 | 0.92626709 | 0.99777713 |
| PLA2G4A | 7.65468738 | -0.026379704 | 0.75120683 | -0.0351164 | 0.9719869 | 0.99777713 |
| BRINP3 | 11.4737759 | 1.052576005 | 1.01728541 | 1.03469095 | 0.30081326 | 0.97358745 |
| AL390957.1 | 113.762878 | 0.724752222 | 0.79172507 | 0.91540896 | 0.35997698 | 0.9819332 |
| AL136987.1 | 5.33263188 | -1.054834142 | 1.17787014 | -0.8955437 | 0.37049653 | 0.98532838 |
| AL136454.1 | 23.3436948 | 0.380584429 | 0.4655581 | 0.81747999 | 0.41365418 | 0.99279547 |
| UCHL5 | 10.3898498 | 0.205361371 | 0.49507928 | 0.41480502 | 0.67828464 | 0.99665574 |
| TROVE2 | 27.8320877 | -0.012973608 | 0.31332725 | -0.0414059 | 0.96697228 | 0.99777713 |
| GLRX2 | 4.09997547 | -1.455574763 | 0.68940532 | -2.1113483 | 0.03474238 | 0.58178209 |
| CDC73 | 59.4480319 | 0.12480886 | 0.32693091 | 0.38175914 | 0.70264003 | 0.99665574 |
| ZBTB41 | 9.68163735 | 0.259686375 | 0.46610584 | 0.55714036 | 0.57743154 | 0.99665574 |
| CRB1 | 5.29757166 | -2.425568143 | 1.30987159 | -1.8517603 | 0.06406026 | 0.68724116 |
| DENND1B | 74.45172 | 0.331453381 | 0.33638029 | 0.98535316 | 0.32445063 | 0.97700345 |
| NEK7 | 98.9272968 | -0.101689745 | 0.29950658 | -0.3395242 | 0.73421484 | 0.99665574 |
| ATP6V1G3 | 99.458827 | 0.576585846 | 0.64316481 | 0.89648227 | 0.36999525 | 0.98532838 |
| PTPRC | 27.9358387 | 1.503072835 | 0.4235604 | 3.54866235 | 0.00038719 | 0.0523382 |
| ZNF281 | 5.03271555 | 0.353682672 | 0.61127395 | 0.57859928 | 0.56285959 | 0.99665574 |
| DDX59 | 7.9299269 | -0.944303694 | 0.49329795 | -1.9142664 | 0.05558612 | 0.67167119 |
| CAMSAP2 | 49.5414123 | -0.058061789 | 0.43130567 | -0.1346187 | 0.89291339 | 0.99773598 |
| TMEM9 | 5.9211023 | 0.425922781 | 0.64054709 | 0.66493594 | 0.50609146 | 0.99665574 |
| CSRP1 | 7.13877056 | -0.020999855 | 0.57691179 | -0.0364005 | 0.97096305 | 0.99777713 |
| AC096677.1 | 7.30399366 | -0.218478987 | 0.64394057 | -0.3392844 | 0.7343955 | 0.99665574 |
| IPO9 | 15.9275084 | -0.383673528 | 0.49066946 | -0.7819389 | 0.43425049 | 0.99617422 |
| TIMM17A | 10.5563237 | -0.371804557 | 0.62119962 | -0.5985267 | 0.54948855 | 0.99665574 |
| RNPEP | 12.2695721 | -0.280774264 | 0.45902206 | -0.6116792 | 0.54075 | 0.99665574 |
| ELF3 | 54.7295139 | -1.402285374 | 0.47500544 | -2.952146 | 0.00315574 | 0.17694235 |
| ARL8A | 8.90533751 | 0.000363197 | 0.7513348 | 0.0004834 | 0.9996143 | 0.99981154 |
| PPP1R12B | 64.8841334 | 0.004407986 | 0.36756478 | 0.01199241 | 0.99043167 | 0.99777713 |
| KDM5B | 39.4958994 | -0.411996589 | 0.27584287 | -1.4935916 | 0.1352824 | 0.84373478 |
| KLHL12 | 8.44384298 | 0.439051743 | 0.52156161 | 0.84180226 | 0.39989865 | 0.99194293 |
| ADIPOR1 | 6.07463232 | -0.339782106 | 0.60235136 | -0.5640929 | 0.57269094 | 0.99665574 |
| CYB5R1 | 6.50890239 | 0.940723881 | 0.75649299 | 1.24353285 | 0.21367154 | 0.91243155 |
| TMEM183A | 2.70028481 | 0.122529776 | 0.78775013 | 0.15554396 | 0.8763925 | 0.99665574 |
| BTG2 | 10.7165707 | -0.167883752 | 0.62915534 | -0.2668399 | 0.78959243 | 0.99665574 |
| ATP2B4 | 12.3319522 | -2.074034856 | 0.60070795 | -3.4526509 | 0.00055511 | 0.06395083 |
| SNRPE | 10.020086 | 0.758006693 | 0.64553571 | 1.1742289 | 0.24030335 | 0.93216487 |
| SOX13 | 10.0382865 | -0.749740717 | 0.48983739 | -1.530591 | 0.1258705 | 0.83633607 |
| ETNK2 | 2.64232172 | 0.469800189 | 1.09624827 | 0.42855273 | 0.66824875 | 0.99665574 |
| GOLT1A | 7.3140579 | -1.921159015 | 0.68564103 | -2.8019896 | 0.00507885 | 0.22386693 |
| PLEKHA6 | 43.7769611 | 0.38875707 | 0.28038407 | 1.38651627 | 0.16558932 | 0.87495175 |
| PPP1R15B | 9.98000545 | 0.568466954 | 0.60778786 | 0.93530489 | 0.3496312 | 0.97838306 |
| MDM4 | 37.7497504 | 0.167944805 | 0.35751243 | 0.46975934 | 0.63852696 | 0.99665574 |
| LRRN2 | 11.7722505 | 0.465386458 | 0.41942902 | 1.10957143 | 0.26718375 | 0.94557817 |
| RBBP5 | 8.13167726 | -0.462691583 | 0.55873184 | -0.8281103 | 0.40760804 | 0.99279547 |
| DSTYK | 13.8483438 | -0.258883444 | 0.47210707 | -0.5483575 | 0.58344647 | 0.99665574 |
| NUAK2 | 13.9526046 | -1.889097637 | 0.74468234 | -2.5367832 | 0.01118762 | 0.37065396 |
| CDK18 | 5.40196566 | 0.277669835 | 0.65358925 | 0.42483844 | 0.67095446 | 0.99665574 |
| MFSD4A | 4.42359098 | 1.204227871 | 0.83592229 | 1.44059788 | 0.14969832 | 0.85525834 |
| ELK4 | 20.8906866 | 0.029476264 | 0.3514925 | 0.08386029 | 0.93316751 | 0.99777713 |
| NUCKS1 | 53.8036885 | 0.715980367 | 0.30338428 | 2.35997843 | 0.018276 | 0.44944531 |
| RAB29 | 5.92846703 | 0.053209086 | 0.7177056 | 0.07413776 | 0.94090077 | 0.99777713 |
| SLC41A1 | 3.95809727 | -0.684394758 | 0.74480444 | -0.9188919 | 0.35815212 | 0.9819332 |
| RHEX | 12.7891271 | -0.204864588 | 0.78161981 | -0.2621026 | 0.79324234 | 0.99665574 |
| SRGAP2 | 5.38624109 | -0.764071475 | 0.66775159 | -1.1442451 | 0.25252201 | 0.94076968 |
| EIF2D | 4.50247387 | 0.999306985 | 0.68993825 | 1.4484006 | 0.14750505 | 0.85284617 |
| MAPKAPK2 | 27.5163207 | -0.262963883 | 0.4089357 | -0.6430446 | 0.52019518 | 0.99665574 |
| PIGR | 20.7716985 | -0.430898759 | 0.87634014 | -0.4917026 | 0.62292957 | 0.99665574 |
| C1orf116 | 5.99873583 | -1.368960752 | 0.65100966 | -2.1028271 | 0.03548089 | 0.58349146 |
| PFKFB2 | 13.2070257 | 0.571780012 | 0.4444318 | 1.28654162 | 0.1982541 | 0.90381759 |
| YOD1 | 4.69730853 | -0.354123657 | 0.69846767 | -0.5070008 | 0.61215427 | 0.99665574 |
| CD55 | 12.5787465 | -0.572284445 | 0.51779758 | -1.1052281 | 0.26906075 | 0.94673212 |
| CD46 | 71.0151493 | -0.065154366 | 0.29648611 | -0.2197552 | 0.82606181 | 0.99665574 |
| MIR29B2CHG | 36.3465473 | -0.031937077 | 0.35996224 | -0.0887234 | 0.92930173 | 0.99777713 |
| PLXNA2 | 5.03613886 | 0.545348292 | 0.73239363 | 0.74461092 | 0.45650697 | 0.99617422 |
| G0S2 | 8.89563516 | -2.181848664 | 1.23985858 | -1.7597561 | 0.07844917 | 0.74431421 |
| TRAF3IP3 | 6.98734 | 2.342778027 | 0.83282143 | 2.81306164 | 0.00490722 | 0.22110862 |
| IRF6 | 10.8204606 | 0.119857245 | 0.5556961 | 0.21568848 | 0.82923058 | 0.99665574 |
| UTP25 | 2.7127952 | 1.278504814 | 0.96174264 | 1.32936273 | 0.18372833 | 0.89532394 |
| HHAT | 6.23237531 | -0.566926055 | 0.58076812 | -0.9761659 | 0.32898224 | 0.97788427 |
| RCOR3 | 44.5949802 | -0.016494413 | 0.39064945 | -0.0422231 | 0.96632088 | 0.99777713 |
| TRAF5 | 3.54820241 | -1.35155949 | 0.83964091 | -1.6096875 | 0.10746609 | 0.79992016 |
| LINC00467 | 6.29221735 | -0.033840989 | 0.55733955 | -0.0607188 | 0.95158316 | 0.99777713 |
| SLC30A1 | 4.22092544 | -0.009033939 | 0.68008437 | -0.0132836 | 0.98940157 | 0.99777713 |
| LPGAT1 | 59.6200523 | -0.603993754 | 0.32006672 | -1.8870871 | 0.05914862 | 0.6784101 |
| INTS7 | 15.0044705 | 0.201002848 | 0.44072201 | 0.45607627 | 0.64833513 | 0.99665574 |
| PPP2R5A | 153.443712 | 0.163038421 | 0.34408776 | 0.47382802 | 0.63562255 | 0.99665574 |
| AL360091.3 | 21.4464079 | -0.124251296 | 0.62527735 | -0.1987139 | 0.84248656 | 0.99665574 |
| TMEM206 | 3.00403824 | -0.594095836 | 0.84323227 | -0.7045459 | 0.48109288 | 0.99640678 |
| NENF | 12.5292487 | 0.605080272 | 0.64255534 | 0.94167807 | 0.34635749 | 0.97788427 |
| ATF3 | 4.04742316 | 0.668262435 | 0.90575924 | 0.73779256 | 0.46064051 | 0.99617422 |
| NSL1 | 16.8750889 | -0.071349884 | 0.35678788 | -0.1999784 | 0.84149744 | 0.99665574 |
| TATDN3 | 7.29229494 | 0.045422991 | 0.78830418 | 0.05762115 | 0.9540504 | 0.99777713 |
| FLVCR1 | 5.48830823 | -0.878317662 | 0.7206069 | -1.2188582 | 0.222898 | 0.92191316 |
| ANGEL2 | 13.4355168 | 0.315293931 | 0.4519044 | 0.69770052 | 0.4853645 | 0.99640678 |
| RPS6KC1 | 40.3736909 | -0.008925283 | 0.34739804 | -0.0256918 | 0.97950316 | 0.99777713 |
| PROX1 | 5.09539062 | 0.409535936 | 0.70571267 | 0.58031541 | 0.56170194 | 0.99665574 |
| SMYD2 | 14.151949 | -0.46747142 | 0.49277432 | -0.9486521 | 0.34279756 | 0.97788427 |
| PTPN14 | 41.773983 | -0.620741427 | 0.35847592 | -1.7316126 | 0.08334256 | 0.75242737 |
| KCTD3 | 27.2614139 | 0.316040989 | 0.3677038 | 0.85949884 | 0.39006536 | 0.99179694 |
| ESRRG | 631.253721 | 0.094152561 | 0.32110517 | 0.29321409 | 0.76935852 | 0.99665574 |
| GPATCH2 | 50.2333412 | -0.300327211 | 0.32211603 | -0.9323572 | 0.35115194 | 0.97838306 |
| SPATA17 | 9.19738109 | -0.3175849 | 0.76387178 | -0.4157568 | 0.67758796 | 0.99665574 |
| RRP15 | 5.89572146 | -0.44220157 | 0.53690775 | -0.8236081 | 0.41016228 | 0.99279547 |
| LYPLAL1 | 43.1047843 | 0.252686927 | 0.47233889 | 0.53496955 | 0.59267091 | 0.99665574 |
| ZC3H11B | 2.69010633 | 0.333574765 | 0.90327887 | 0.36929322 | 0.71190918 | 0.99665574 |
| EPRS | 18.0270097 | 0.053547383 | 0.35481718 | 0.15091542 | 0.88004244 | 0.99665574 |
| BPNT1 | 5.27970728 | -0.604693002 | 0.85934518 | -0.7036672 | 0.48164006 | 0.99640678 |
| IARS2 | 16.8435448 | -0.146160337 | 0.39864472 | -0.3666431 | 0.71388526 | 0.99665574 |
| RAB3GAP2 | 53.4509546 | 0.016581037 | 0.29259677 | 0.05666856 | 0.95480922 | 0.99777713 |
| MARK1 | 15.79579 | -0.336907633 | 0.4572941 | -0.7367417 | 0.46127944 | 0.99617422 |
| 2-Mar | 11.3433282 | 1.285149122 | 0.52861695 | 2.43115385 | 0.01505082 | 0.41291622 |

| | | | | | | |
|---|---|---|---|---|---|---|
| AL513314.2 | 4.71982446 | 0.050960067 | 0.59761169 | 0.08527287 | 0.93204446 | 0.99777713 |
| TAF1A | 3.69441964 | 1.913707343 | 0.89052569 | 2.14896365 | 0.03163728 | 0.56814511 |
| MIA3 | 12.8895489 | -0.446281318 | 0.41971133 | -1.0633054 | 0.28764349 | 0.96559334 |
| AIDA | 5.80793758 | -0.267581262 | 0.6516593 | -0.4106153 | 0.68135466 | 0.99665574 |
| BROX | 13.2846427 | 0.450694525 | 0.49036101 | 0.9191076 | 0.3580393 | 0.9819332 |
| FAM177B | 5.28006714 | -0.406843404 | 0.96876856 | -0.4199593 | 0.67451516 | 0.99665574 |
| AC093152.1 | 7.76636455 | -0.258156793 | 0.61951468 | -0.4167081 | 0.67689192 | 0.99665574 |
| SUSD4 | 8.72856126 | -0.217302795 | 0.5701704 | -0.381119 | 0.70311492 | 0.99665574 |
| CAPN2 | 50.0517239 | -0.406563998 | 0.3139003 | -1.295201 | 0.19525089 | 0.90381759 |
| TP53BP2 | 12.3414126 | -0.315316458 | 0.53438828 | -0.5900512 | 0.55515631 | 0.99665574 |
| FBXO28 | 22.0804063 | -0.056097583 | 0.38454375 | -0.1458809 | 0.88401542 | 0.99665574 |
| DEGS1 | 6.14334548 | -1.106973553 | 0.73644207 | -1.5031373 | 0.13280364 | 0.84089786 |
| NVL | 18.3799882 | -0.001383204 | 0.42568848 | -0.0032493 | 0.99740741 | 0.99888534 |
| CNIH4 | 15.5574294 | -0.402516047 | 0.4901246 | -0.8212525 | 0.41150246 | 0.99279547 |
| WDR26 | 42.2211408 | 0.214402745 | 0.29675034 | 0.72250211 | 0.46998583 | 0.99617422 |
| CNIH3 | 9.93419414 | -0.068101046 | 0.55979486 | -0.1216536 | 0.90317338 | 0.99777713 |
| DNAH14 | 55.2301474 | -0.111171516 | 0.28560688 | -0.3892466 | 0.69709372 | 0.99665574 |
| LBR | 12.6108997 | -0.55126792 | 0.5289045 | -1.0422825 | 0.29728071 | 0.97345988 |
| ENAH | 6.94156885 | -0.466577367 | 0.72931151 | -0.6397505 | 0.52233485 | 0.99665574 |
| SRP9 | 23.9535979 | 0.435397085 | 0.36551899 | 1.191175 | 0.23358489 | 0.92960234 |
| EPHX1 | 3.93714883 | -0.541309905 | 0.67656644 | -0.8000839 | 0.4236622 | 0.99518711 |
| TMEM63A | 11.5580745 | -0.935081556 | 0.54523655 | -1.7150016 | 0.08634493 | 0.75397496 |
| SDE2 | 5.20507913 | 0.435110898 | 0.78523536 | 0.55411527 | 0.57949996 | 0.99665574 |
| H3F3A | 6.01566596 | -0.669437626 | 0.60439349 | -1.1076189 | 0.26802645 | 0.94660092 |
| ACBD3 | 24.2420935 | -0.744789841 | 0.43645107 | -1.7064681 | 0.08792096 | 0.75650018 |
| PARP1 | 13.4665184 | -0.023931887 | 0.46678362 | -0.0512698 | 0.95911056 | 0.99777713 |
| STUM | 12.1752373 | 0.940623665 | 0.70106439 | 1.34170795 | 0.17969071 | 0.88820302 |
| ITPKB | 11.6523406 | -0.048189092 | 0.67612141 | -0.0712728 | 0.94318061 | 0.99777713 |
| PSEN2 | 16.7292493 | 1.357276603 | 0.47285911 | 2.87036153 | 0.00410003 | 0.20783038 |
| COQ8A | 21.6327562 | 0.785145272 | 0.42203983 | 1.86035822 | 0.06283486 | 0.68628329 |
| CDC42BPA | 136.245692 | 0.470856867 | 0.26527892 | 1.77495021 | 0.07590611 | 0.7342902 |
| ZNF678 | 15.2043368 | 0.001113375 | 0.37894294 | 0.00293811 | 0.99765573 | 0.99903534 |
| SNAP47 | 5.19270268 | 0.856290378 | 0.66012001 | 1.2971738 | 0.1945714 | 0.90333151 |
| JMJD4 | 2.53987284 | -0.630466121 | 0.92458739 | -0.6818892 | 0.49530904 | 0.99640678 |
| ARF1 | 38.4922868 | -0.564590188 | 0.48180232 | -1.1718295 | 0.2412655 | 0.93253689 |
| C1orf35 | 3.26802404 | 0.542255182 | 0.74995991 | 0.72304556 | 0.46965189 | 0.99617422 |
| MRPL55 | 5.4310157 | 0.851380527 | 0.67918077 | 1.25354039 | 0.21000911 | 0.91043197 |
| GUK1 | 35.0817122 | -0.053878171 | 0.36802959 | -0.1463963 | 0.88360855 | 0.99665574 |
| IBA57 | 3.43959542 | 0.278082314 | 0.75651734 | 0.36758221 | 0.71318478 | 0.99665574 |
| OBSCN | 10.1357769 | 0.179345924 | 0.54503964 | 0.32905116 | 0.74211702 | 0.99665574 |
| TRIM11 | 2.56775847 | 0.065344888 | 0.82581529 | 0.07912773 | 0.93693103 | 0.99777713 |
| RNF187 | 6.94475981 | 0.931477107 | 0.60512074 | 1.53932437 | 0.12372513 | 0.82915695 |
| RHOU | 8.01391918 | -0.087771684 | 0.5577975 | -0.157354 | 0.87496584 | 0.99665574 |
| RAB4A | 10.169556 | 0.036699002 | 0.67789467 | 0.05413673 | 0.95682623 | 0.99777713 |
| NUP133 | 12.8124187 | 0.135616196 | 0.45694607 | 0.29678819 | 0.76662822 | 0.99665574 |
| ABCB10 | 6.05841618 | 0.20841588 | 0.57753858 | 0.36086919 | 0.71819724 | 0.99665574 |
| TAF5L | 5.65331042 | 0.131261315 | 0.7085325 | 0.185258 | 0.85302668 | 0.99665574 |
| GALNT2 | 18.2577996 | 0.162906345 | 0.41349683 | 0.39397241 | 0.69360141 | 0.99665574 |
| COG2 | 9.84816921 | 0.680788922 | 0.45515616 | 1.49572603 | 0.13472507 | 0.84373478 |
| C1orf198 | 4.84830499 | 0.490493269 | 0.78312473 | 0.62632841 | 0.53109955 | 0.99665574 |
| TTC13 | 6.99974798 | -0.04627288 | 0.56623827 | -0.0817198 | 0.93486953 | 0.99777713 |
| ARV1 | 7.48358013 | 0.339608111 | 0.55499954 | 0.61190701 | 0.54059928 | 0.99665574 |
| C1orf131 | 4.7628208 | -0.244789024 | 0.61814295 | -0.3960071 | 0.69209976 | 0.99665574 |
| GNPAT | 14.8011625 | 0.524592147 | 0.45752755 | 1.14658046 | 0.25155506 | 0.94071013 |
| EXOC8 | 2.92284317 | -0.223055355 | 0.89568806 | -0.2490324 | 0.80333571 | 0.99665574 |
| SPRTN | 3.16597043 | -0.274725348 | 0.94427374 | -0.2909382 | 0.77109855 | 0.99665574 |
| EGLN1 | 75.4174783 | -0.099440464 | 0.34218829 | -0.2906016 | 0.77135604 | 0.99665574 |
| DISC1 | 18.2935781 | -2.061528589 | 0.97387278 | -2.1168356 | 0.0342738 | 0.58178209 |
| SIPA1L2 | 31.5406503 | -1.324637043 | 0.69914001 | -1.8946663 | 0.05813662 | 0.67545594 |
| NTPCR | 9.69881547 | 0.686865977 | 0.48472472 | 1.41702281 | 0.15647626 | 0.86532131 |
| PCNX2 | 23.0208306 | -0.820627584 | 0.40664486 | -2.0180449 | 0.04358659 | 0.62204719 |
| MAP3K21 | 24.8385487 | 0.285547493 | 0.43130625 | 0.66205276 | 0.50793741 | 0.99665574 |
| SLC35F3 | 279.460988 | 0.190827182 | 0.44975568 | 0.42429077 | 0.67135377 | 0.99665574 |
| AL355472.3 | 20.61704 | 1.281143822 | 0.36280409 | 3.53122758 | 0.00041364 | 0.05517682 |
| COA6 | 9.47314286 | 1.041781735 | 0.59483832 | 1.75136958 | 0.07988227 | 0.7446402 |
| TARBP1 | 14.7860284 | 0.133325615 | 0.48794103 | 0.27324124 | 0.78466778 | 0.99665574 |
| IRF2BP2 | 19.6086798 | -0.314629963 | 0.53520047 | -0.5878731 | 0.55661747 | 0.99665574 |
| TOMM20 | 26.7452297 | -0.028146475 | 0.43660104 | -0.0644673 | 0.94859817 | 0.99777713 |
| ARID4B | 105.6244 | -0.087640062 | 0.24366657 | -0.3596721 | 0.71909237 | 0.99665574 |
| B3GALNT2 | 16.2524057 | -0.124834215 | 0.45504564 | -0.2743334 | 0.78382842 | 0.99665574 |
| LYST | 48.9757001 | 0.618839909 | 0.31803371 | 1.94583119 | 0.05167502 | 0.65584036 |
| GPR137B | 9.53160357 | 0.046009792 | 0.54766588 | 0.0840107 | 0.93304793 | 0.99777713 |
| ERO1B | 10.323631 | 0.220834556 | 0.66970975 | 0.32974666 | 0.74159139 | 0.99665574 |
| EDARADD | 10.1204252 | 0.091436412 | 0.60842762 | 0.15028314 | 0.88054124 | 0.99665574 |
| LGALS8 | 9.41801581 | -0.45946452 | 0.50427316 | -0.9111421 | 0.3622205 | 0.98265758 |
| HEATR1 | 9.75750702 | -0.32651209 | 0.4944844 | -0.6603082 | 0.50905609 | 0.99665574 |
| MTR | 61.927002 | 0.643142218 | 0.3505951 | 1.83443014 | 0.06659017 | 0.69668848 |
| RGS7 | 48.2927443 | 0.402202249 | 0.69786059 | 0.57633609 | 0.56438803 | 0.99665574 |
| FH | 8.2938509 | 0.01048006 | 0.57016063 | 0.01838089 | 0.985335 | 0.99777713 |
| OPN3 | 10.1513102 | -0.731424474 | 0.63266353 | -1.1561034 | 0.24763886 | 0.93441937 |
| CEP170 | 13.9383514 | -0.933267001 | 0.49124263 | -1.8998086 | 0.05745824 | 0.67545594 |
| SDCCAG8 | 60.4891449 | -0.007536553 | 0.35793664 | -0.0210555 | 0.98320134 | 0.99777713 |
| AKT3 | 43.7551363 | -0.52392546 | 0.5388248 | -0.9723485 | 0.33087722 | 0.97788427 |
| ZBTB18 | 6.26083821 | 0.336657364 | 0.54736475 | 0.61505123 | 0.53852089 | 0.99665574 |
| AL645465.1 | 3.54100162 | -1.748299241 | 0.90328394 | -1.9354925 | 0.05292989 | 0.66411295 |
| ADSS | 15.2097897 | -0.350630647 | 0.49587073 | -0.7071009 | 0.47950377 | 0.99617422 |
| CATSPERE | 14.0332706 | -0.312466355 | 0.60828109 | -0.5136874 | 0.60747054 | 0.99665574 |
| DESI2 | 17.0623671 | 0.116536792 | 0.4897898 | 0.23793226 | 0.81193364 | 0.99665574 |
| COX20 | 19.0195696 | 0.011684237 | 0.39189594 | 0.02981464 | 0.97621488 | 0.99777713 |
| HNRNPU | 68.6953704 | 0.032756592 | 0.27268531 | 0.12012599 | 0.90438335 | 0.99777713 |
| EFCAB2 | 26.0472909 | 0.448079562 | 0.43051388 | 1.04080167 | 0.2979676 | 0.97358745 |
| KIF26B | 8.24819099 | -1.133690852 | 0.70043509 | -1.6185523 | 0.10554362 | 0.79494888 |
| SMYD3 | 120.622007 | -0.04911314 | 0.27697966 | -0.1773168 | 0.85925957 | 0.99665574 |
| TFB2M | 4.00592952 | 1.35800545 | 0.71898049 | 1.88879319 | 0.05891955 | 0.67724076 |
| CNST | 16.4071922 | -0.099645928 | 0.3894838 | -0.255841 | 0.79807361 | 0.99665574 |
| SCCPDH | 15.7706082 | -0.293380815 | 0.53440075 | -0.5489903 | 0.58301213 | 0.99665574 |
| AHCTF1 | 28.6175279 | 0.169189677 | 0.29384178 | 0.57578496 | 0.56476054 | 0.99665574 |
| ZNF124 | 8.70527925 | -0.670102991 | 0.47603653 | -1.4076714 | 0.15922841 | 0.86757292 |
| AL390728.6 | 3.5327868 | -0.411654356 | 0.74572064 | -0.5520222 | 0.58093314 | 0.99665574 |
| ZNF496 | 6.22548601 | -0.731591404 | 0.59668987 | -1.2260832 | 0.22016738 | 0.91768188 |
| SH3BP5L | 3.10590079 | 0.869747562 | 0.89046048 | 0.97673909 | 0.32869834 | 0.97788427 |
| ZNF672 | 6.90931008 | 0.249280913 | 0.60155391 | 0.41439497 | 0.67858487 | 0.99665574 |
| ZNF692 | 3.42244753 | -0.454986835 | 0.69865882 | -0.6512289 | 0.51489872 | 0.99665574 |
| FAM110C | 7.35283958 | -0.971067541 | 0.62925835 | -1.5431937 | 0.12278378 | 0.82915695 |
| SH3YL1 | 29.7798241 | 0.446712034 | 0.36150738 | 1.23569272 | 0.2165728 | 0.91522096 |
| ACP1 | 10.87694 | 0.187074189 | 0.50067251 | 0.37364582 | 0.70866784 | 0.99665574 |
| TMEM18 | 2.49457768 | -0.323334595 | 0.8611439 | -0.375471 | 0.70731018 | 0.99665574 |
| LINC01250 | 3.41377804 | 0.71830736 | 0.78726268 | 0.91241129 | 0.36155225 | 0.9819332 |
| EIPR1 | 43.7023588 | 0.043048711 | 0.38408118 | 0.11208232 | 0.91075813 | 0.99777713 |
| TRAPPC12 | 45.1269912 | 0.120276104 | 0.2973503 | 0.40449296 | 0.68585025 | 0.99665574 |
| ADI1 | 21.7295676 | -0.203218727 | 0.48124023 | -0.4222812 | 0.67281974 | 0.99665574 |
| AC108488.2 | 2.71598776 | 0.771713227 | 0.85087822 | 0.90696084 | 0.36442751 | 0.98362663 |
| RPS7 | 47.1965551 | 0.716546182 | 0.43554259 | 1.64518051 | 0.09993259 | 0.78293401 |

| | | | | | | |
|---|---|---|---|---|---|---|
| RNF144A | 6.44062508 | -0.760961185 | 0.57609639 | -1.3208921 | 0.18653733 | 0.89532394 |
| ID2 | 6.0494217 | -0.547381432 | 0.77055137 | -0.7103763 | 0.47747081 | 0.99617422 |
| KIDINS220 | 62.1547309 | 0.087398946 | 0.26370572 | 0.33142605 | 0.74032269 | 0.99665574 |
| MBOAT2 | 109.533054 | 0.202934667 | 0.30475989 | 0.66588379 | 0.50548539 | 0.99665574 |
| AC093904.4 | 2.8763073 | 1.949090663 | 1.05118312 | 1.85418755 | 0.06371233 | 0.68630818 |
| ASAP2 | 208.58298 | -0.050108262 | 0.35218926 | -0.1422765 | 0.88686159 | 0.99665574 |
| ITGB1BP1 | 15.7239926 | -0.739805957 | 0.42708604 | -1.7322176 | 0.08323482 | 0.75242737 |
| CPSF3 | 8.17728874 | 0.59902924 | 0.67031678 | 0.89365097 | 0.37150867 | 0.98532838 |
| IAH1 | 12.4878853 | -0.571550123 | 0.4496026 | -1.271234 | 0.20364541 | 0.90707783 |
| ADAM17 | 27.2246521 | -0.302417387 | 0.42988928 | -0.7034774 | 0.48175829 | 0.99640678 |
| YWHAQ | 31.871898 | -0.457631096 | 0.4292705 | -1.066067 | 0.28639338 | 0.96460334 |
| TAF1B | 22.1135625 | -0.152784395 | 0.41190975 | -0.3709172 | 0.71069923 | 0.99665574 |
| KLF11 | 5.10609135 | 1.208900228 | 0.80811656 | 1.49594785 | 0.13466725 | 0.84373478 |
| CYS1 | 9.83517619 | -0.048118512 | 0.52687815 | -0.0913276 | 0.92723229 | 0.99777713 |
| HPCAL1 | 18.9598265 | 0.521933747 | 0.4054065 | 1.2874331 | 0.19794337 | 0.90381759 |
| ODC1 | 6.71004647 | 0.134749468 | 0.57137038 | 0.23583559 | 0.81356025 | 0.99665574 |
| NOL10 | 26.6182098 | 0.746134333 | 0.36628394 | 2.0370381 | 0.04164622 | 0.61367648 |
| ATP6V1C2 | 95.1638313 | 1.269571512 | 0.42346065 | 2.99808614 | 0.00271681 | 0.16107021 |
| PDIA6 | 27.7406368 | -0.185775416 | 0.40053432 | -0.463819 | 0.64277745 | 0.99665574 |
| PQLC3 | 12.0391421 | 0.818585045 | 0.58831028 | 1.39141721 | 0.16409895 | 0.87375796 |
| ROCK2 | 76.1387716 | 0.330537515 | 0.25734849 | 1.28439654 | 0.19900324 | 0.90381759 |
| E2F6 | 5.07799225 | -0.355091956 | 0.79350601 | -0.4474975 | 0.65451589 | 0.99665574 |
| LPIN1 | 22.6216236 | -0.535495045 | 0.49963545 | -1.0717715 | 0.28382267 | 0.96059387 |
| MIR3681HG | 5.5529001 | 0.36747694 | 0.79991631 | 0.45939423 | 0.64595109 | 0.99665574 |
| TRIB2 | 12.7248104 | 0.038000118 | 0.48965655 | 0.07760566 | 0.93814174 | 0.99777713 |
| NBAS | 60.7531914 | -0.37021537 | 0.35439703 | -1.0446345 | 0.29619192 | 0.97286866 |
| DDX1 | 19.4283609 | 0.282679967 | 0.38494878 | 0.73433138 | 0.4627468 | 0.99617422 |
| SMC6 | 11.0629855 | 0.308430864 | 0.53652155 | 0.57487135 | 0.56537831 | 0.99665574 |
| GEN1 | 3.94904703 | -0.148508902 | 0.6870371 | -0.2161585 | 0.82886421 | 0.99665574 |
| RDH14 | 6.45828767 | 0.444782852 | 0.60769367 | 0.73191951 | 0.4642177 | 0.99617422 |
| LINC01376 | 11.3874983 | 0.062235102 | 0.62522215 | 0.09954078 | 0.92070891 | 0.99777713 |
| TTC32 | 3.39384461 | 1.049183497 | 0.76504357 | 1.37140359 | 0.17024918 | 0.87918701 |
| WDR35 | 10.8193241 | 0.886885649 | 0.61446251 | 1.44335193 | 0.14892136 | 0.85485259 |
| LAPTM4A | 35.9681495 | -0.000569344 | 0.35276317 | -0.001614 | 0.99871225 | 0.99940231 |
| SDC1 | 5.49041273 | 0.431926323 | 0.67151117 | 0.64321539 | 0.52008435 | 0.99665574 |
| PUM2 | 88.3926458 | 0.109837223 | 0.23280103 | 0.47180728 | 0.63706435 | 0.99665574 |
| RHOB | 7.69931787 | 0.133460617 | 0.77277283 | 0.17270356 | 0.86288445 | 0.99665574 |
| HS1BP3 | 3.68807098 | -0.749018593 | 0.78518182 | -0.9539429 | 0.34011257 | 0.97788427 |
| LDAH | 26.272256 | -0.329324854 | 0.40702782 | -0.8090967 | 0.41845955 | 0.99443758 |
| KLHL29 | 8.63626111 | -4.709297208 | 1.55174532 | -3.034839 | 0.00240664 | 0.15249081 |
| ATAD2B | 58.0164345 | 0.251351006 | 0.31493407 | 0.79810675 | 0.42480854 | 0.99518711 |
| UBXN2A | 16.04723 | -0.442860463 | 0.49005765 | -0.9036905 | 0.36615952 | 0.98365837 |
| SF3B6 | 16.4976622 | 0.569962681 | 0.4474532 | 1.27379283 | 0.20273685 | 0.9066717 |
| FAM228B | 3.48202276 | -0.93949098 | 0.78613863 | -1.1950704 | 0.23205954 | 0.92842131 |
| TP53I3 | 13.4194139 | 0.191914966 | 0.57133188 | 0.33590803 | 0.73694022 | 0.99665574 |
| ITSN2 | 81.3087233 | -0.129329101 | 0.29880077 | -0.4328272 | 0.66514031 | 0.99665574 |
| NCOA1 | 137.102143 | 0.007310623 | 0.21591618 | 0.03385862 | 0.97298989 | 0.99777713 |
| PTRHD1 | 3.12381219 | 0.951904858 | 0.99950061 | 0.95238046 | 0.34090406 | 0.97788427 |
| DNAJC27 | 9.79128685 | -0.253146572 | 0.49410796 | -0.5123305 | 0.60841973 | 0.99665574 |
| DNMT3A | 28.2146975 | 0.489987137 | 0.36514152 | 1.34191023 | 0.1796251 | 0.88820302 |
| DTNB | 87.7202784 | 0.198547186 | 0.26974412 | 0.73605752 | 0.46169569 | 0.99617422 |
| ASXL2 | 53.8607566 | -0.213780493 | 0.33215752 | -0.6436118 | 0.51982722 | 0.99665574 |
| RAB10 | 90.1542874 | -0.158236085 | 0.30470322 | -0.5193121 | 0.60354308 | 0.99665574 |
| HADHA | 43.2169187 | 0.006779204 | 0.33180443 | 0.02043133 | 0.9836993 | 0.99777713 |
| HADHB | 36.848272 | 0.038459282 | 0.34278063 | 0.11219794 | 0.91066646 | 0.99777713 |
| SELENOI | 40.6180005 | 0.713814869 | 0.37960973 | 1.88039139 | 0.06005476 | 0.68195946 |
| AC013472.2 | 20.2540332 | 0.824952323 | 0.64089463 | 1.28718869 | 0.19802853 | 0.90381759 |
| MAPRE3 | 14.2321103 | -0.088869351 | 0.55154018 | -0.1611294 | 0.87199147 | 0.99665574 |
| TMEM214 | 4.53767662 | 0.059827263 | 0.81781407 | 0.07315509 | 0.9416827 | 0.99777713 |
| AGBL5 | 5.31848983 | -0.20889144 | 0.75387909 | -0.2770888 | 0.78171194 | 0.99665574 |
| OST4 | 21.8293443 | -0.282828956 | 0.49600381 | -0.5702153 | 0.56853169 | 0.99665574 |
| PREB | 2.53740238 | 0.341858048 | 0.90611975 | 0.3772769 | 0.70596784 | 0.99665574 |
| ATRAID | 12.0815703 | 0.344364289 | 0.50366248 | 0.68372036 | 0.49415177 | 0.99640678 |
| MPV17 | 4.13198777 | 0.268886446 | 0.66402767 | 0.40493259 | 0.68552706 | 0.99665574 |
| GTF3C2 | 13.2373384 | 0.245332013 | 0.52611802 | 0.46630604 | 0.64099645 | 0.99665574 |
| EIF2B4 | 3.10173247 | 0.158447655 | 0.77269378 | 0.2050588 | 0.83752619 | 0.99665574 |
| SNX17 | 7.12631126 | 0.762431972 | 0.57585038 | 1.32401054 | 0.18549954 | 0.89532394 |
| ZNF513 | 2.58462865 | -0.942721595 | 0.95298203 | -0.9892333 | 0.32254899 | 0.97700345 |
| PPM1G | 24.9460461 | -0.039823453 | 0.389475 | -0.1022491 | 0.91855899 | 0.99777713 |
| NRBP1 | 9.50462364 | -0.107752557 | 0.53597793 | -0.2010392 | 0.84066795 | 0.99665574 |
| IFT172 | 7.76549145 | 0.294328111 | 0.55185363 | 0.53334453 | 0.59379511 | 0.99665574 |
| ZNF512 | 15.494838 | -0.185267881 | 0.44471643 | -0.4165978 | 0.67697263 | 0.99665574 |
| GPN1 | 3.97137935 | -0.263123579 | 0.76233352 | -0.3451555 | 0.7299775 | 0.99665574 |
| SUPT7L | 7.16664472 | -0.07695224 | 0.58892611 | -0.1306654 | 0.89604004 | 0.99777713 |
| SLC4A1AP | 6.54571062 | 0.042072208 | 0.57269532 | 0.07346351 | 0.94143728 | 0.99777713 |
| MRPL33 | 23.0816776 | -0.050463636 | 0.44454608 | -0.1135172 | 0.90962051 | 0.99777713 |
| RBKS | 12.4725411 | 0.164579561 | 0.57575768 | 0.28584866 | 0.77499404 | 0.99665574 |
| BABAM2 | 112.866936 | 0.130965847 | 0.29552874 | 0.44315773 | 0.65765165 | 0.99665574 |
| FOSL2 | 39.5293063 | 0.064808634 | 0.35964855 | 0.1801999 | 0.85699563 | 0.99665574 |
| PPP1CB | 27.535168 | 0.12430746 | 0.46888008 | 0.26511567 | 0.79092035 | 0.99665574 |
| TRMT61B | 11.2278784 | 0.6853841 | 0.44995451 | 1.52322976 | 0.12770123 | 0.83778831 |
| WDR43 | 18.4523254 | -0.475182678 | 0.48565341 | -0.9784399 | 0.32785679 | 0.97788427 |
| CLIP4 | 13.1773332 | -1.706237263 | 0.60159747 | -2.8361776 | 0.00456571 | 0.21492549 |
| YPEL5 | 14.2870753 | 0.051080507 | 0.46593975 | 0.109629 | 0.91270361 | 0.99777713 |
| LCLAT1 | 17.3703729 | -0.472675452 | 0.39638566 | -1.1924636 | 0.23307953 | 0.92924083 |
| CAPN13 | 4.52919202 | -0.135732641 | 0.69633646 | -0.1949239 | 0.8454525 | 0.99665574 |
| GALNT14 | 18.6134221 | -0.302856903 | 0.57749905 | -0.5244284 | 0.59998061 | 0.99665574 |
| MEMO1 | 2.84922819 | 0.209884857 | 0.8160292 | 0.25720263 | 0.79702235 | 0.99665574 |
| DPY30 | 11.8274571 | 0.324638871 | 0.57122912 | 0.56831639 | 0.56982015 | 0.99665574 |
| SPAST | 19.1638603 | 0.884821044 | 0.43853581 | 2.01767114 | 0.04362552 | 0.62204719 |
| SLC30A6 | 16.4196458 | 0.09479907 | 0.35690287 | 0.26561588 | 0.79053505 | 0.99665574 |
| YIPF4 | 23.8194606 | -0.36653656 | 0.42659436 | -0.8592157 | 0.39022154 | 0.99179694 |
| BIRC6 | 141.286709 | 0.064525674 | 0.2690509 | 0.23982701 | 0.81046436 | 0.99665574 |
| TTC27 | 21.289371 | 0.543253205 | 0.3888919 | 1.39692599 | 0.16243583 | 0.87130922 |
| LINC00486 | 23.6509981 | 0.198465609 | 0.38046555 | 0.52163885 | 0.60192181 | 0.99665574 |
| LTBP1 | 4.93784485 | 0.497774925 | 0.67889053 | 0.73321825 | 0.46342532 | 0.99617422 |
| RASGRP3 | 8.73149839 | 0.427407664 | 0.65482156 | 0.65270861 | 0.51394415 | 0.99665574 |
| FAM98A | 8.09935676 | 0.621592616 | 0.53366218 | 1.16476798 | 0.24411293 | 0.93426912 |
| LINC01320 | 50.4062656 | -1.457552578 | 0.60863674 | -2.3947825 | 0.01663023 | 0.42361121 |
| AC012593.1 | 19.0529731 | -0.450918407 | 0.92178497 | -0.4891796 | 0.62471455 | 0.99665574 |
| CRIM1 | 41.1257423 | -0.836796006 | 0.37390726 | -2.2379774 | 0.02522253 | 0.50908112 |
| FEZ2 | 25.1768636 | 0.133283295 | 0.4013113 | 0.33211946 | 0.73979905 | 0.99665574 |
| STRN | 62.906182 | -0.063467725 | 0.31788733 | -0.1996548 | 0.84175058 | 0.99665574 |
| HEATR5B | 21.6436436 | -0.03934061 | 0.32936885 | -0.1194424 | 0.90492486 | 0.99777713 |
| GPATCH11 | 3.15933818 | -0.021432001 | 0.73368146 | -0.0292116 | 0.97669584 | 0.99777713 |
| EIF2AK2 | 24.5509811 | 0.225901275 | 0.66431167 | 0.34005315 | 0.7338165 | 0.99665574 |
| CEBPZOS | 11.263341 | -0.341955585 | 0.58900805 | -0.5805618 | 0.56153581 | 0.99665574 |
| CEBPZ | 15.3883463 | -0.353073877 | 0.43208423 | -0.8171413 | 0.41384767 | 0.99279547 |
| NDUFAF7 | 9.49556859 | 0.485795475 | 0.49536661 | 0.98067868 | 0.32675122 | 0.97788427 |
| PRKD3 | 20.4295162 | 0.171095958 | 0.39169107 | 0.43681353 | 0.66224659 | 0.99665574 |
| CDC42EP3 | 4.61349272 | -1.332940132 | 0.79181329 | -1.6834021 | 0.09229728 | 0.76670359 |
| RMDN2 | 14.4103572 | 0.816795157 | 0.45761887 | 1.78488083 | 0.07428065 | 0.72810353 |
| ATL2 | 138.496659 | 0.614176374 | 0.26956445 | 2.27840271 | 0.0227026 | 0.48762481 |

| | | | | | | |
|---|---|---|---|---|---|---|
| HNRNPLL | 14.4982787 | -0.062838154 | 0.43516433 | -0.144401 | 0.88518385 | 0.99665574 |
| GALM | 8.66192125 | 0.387954002 | 0.54038463 | 0.71792198 | 0.47280539 | 0.99617422 |
| SRSF7 | 19.4314038 | -0.048735896 | 0.57241917 | -0.0851402 | 0.93214991 | 0.99777713 |
| DHX57 | 11.7052977 | 0.082946457 | 0.43526518 | 0.19056534 | 0.84886615 | 0.99665574 |
| SOS1 | 89.8391478 | -0.059879005 | 0.24467578 | -0.244728 | 0.80666708 | 0.99665574 |
| MAP4K3 | 99.9350473 | 0.118120474 | 0.262007 | 0.45082946 | 0.65211246 | 0.99665574 |
| AC007388.1 | 42.8448043 | -0.204089585 | 0.39171228 | -0.5210191 | 0.60235346 | 0.99665574 |
| THUMPD2 | 11.8100106 | -0.520380869 | 0.52839414 | -0.9848347 | 0.32470529 | 0.97700345 |
| SLC8A1 | 24.3686941 | 0.862307197 | 1.28607476 | 0.67049539 | 0.50254204 | 0.99640678 |
| PKDCC | 2.79233371 | 0.513688481 | 1.17241716 | 0.4381448 | 0.66128132 | 0.99665574 |
| EML4 | 51.4470543 | -0.147046986 | 0.44142917 | -0.3331157 | 0.73904695 | 0.99665574 |
| COX7A2L | 24.2883405 | -0.193277713 | 0.47121716 | -0.410167 | 0.68168347 | 0.99665574 |
| MTA3 | 27.5177553 | 0.154716058 | 0.37015349 | 0.41797811 | 0.67596312 | 0.99665574 |
| ZFP36L2 | 26.4851215 | -0.530544079 | 0.35660494 | -1.4877643 | 0.13681307 | 0.84475183 |
| THADA | 44.5379002 | 0.113840213 | 0.28474461 | 0.3997976 | 0.6893056 | 0.99665574 |
| PLEKHH2 | 21.6481872 | -1.381911803 | 0.61447317 | -2.2489376 | 0.02451647 | 0.50422081 |
| DYNC2LI1 | 13.3794566 | -0.448882962 | 0.47419077 | -0.9466295 | 0.34382761 | 0.97788427 |
| LRPPRC | 41.8485037 | 0.447310136 | 0.30985641 | 1.44360458 | 0.14885024 | 0.85485259 |
| PPM1B | 39.8686055 | 0.08029411 | 0.28098773 | 0.28575665 | 0.77506452 | 0.99665574 |
| SLC3A1 | 14.7218188 | 0.561507633 | 0.66823167 | 0.84028887 | 0.40074644 | 0.99194293 |
| PREPL | 22.4572462 | -0.340091563 | 0.3984828 | -0.8534661 | 0.39340088 | 0.99179694 |
| CAMKMT | 93.7921652 | -0.466468491 | 0.34707485 | -1.3439997 | 0.17894849 | 0.88813816 |
| SRBD1 | 29.8214833 | 0.340128874 | 0.33500377 | 1.01529866 | 0.30996348 | 0.97376371 |
| PRKCE | 172.060674 | 0.318354343 | 0.3731769 | 0.85309232 | 0.39360811 | 0.99179694 |
| EPAS1 | 28.8164049 | -0.495640776 | 0.44255728 | -1.1199472 | 0.26273627 | 0.94557817 |
| RHOQ | 22.2939732 | -0.53643805 | 0.35130788 | -1.5269742 | 0.12676744 | 0.83669777 |
| AC018682.1 | 3.38731245 | -0.906754588 | 0.91135095 | -0.9949565 | 0.3197574 | 0.97700345 |
| PIGF | 11.4906324 | 0.218663267 | 0.48551512 | 0.45037375 | 0.65244097 | 0.99665574 |
| CRIPT | 3.92514704 | -0.320474885 | 0.80132633 | -0.3999306 | 0.68920767 | 0.99665574 |
| SOCS5 | 13.1524715 | 0.248776229 | 0.42031625 | 0.59187869 | 0.55393183 | 0.99665574 |
| MCFD2 | 17.216504 | 0.120542402 | 0.4600472 | 0.26202182 | 0.79330462 | 0.99665574 |
| TTC7A | 11.2756178 | -0.540085514 | 0.46889934 | -1.1518155 | 0.24939691 | 0.9369778 |
| AC106869.1 | 5.38420807 | -0.125457186 | 0.91406264 | -0.1372523 | 0.89083138 | 0.99712197 |
| EPCAM | 65.9074673 | -0.265735761 | 0.34459549 | -0.7711528 | 0.44061639 | 0.99617422 |
| MSH2 | 13.4437369 | -0.019388342 | 0.4443782 | -0.0436303 | 0.96519912 | 0.99777713 |
| MSH6 | 9.22205187 | 0.455190672 | 0.54221491 | 0.83950232 | 0.40118749 | 0.99194293 |
| FBXO11 | 73.2016001 | -0.15296331 | 0.25454314 | -0.6009327 | 0.54788478 | 0.99665574 |
| FOXN2 | 8.39971628 | -0.788373592 | 0.51656947 | -1.5261715 | 0.12696716 | 0.8369266 |
| PPP1R21 | 31.7322292 | -0.159025922 | 0.33429137 | -0.4757105 | 0.63428063 | 0.99665574 |
| AC010967.1 | 9.45720343 | 0.883426801 | 1.71491059 | 0.51514452 | 0.60645203 | 0.99665574 |
| GPR75-ASB3 | 10.7052969 | -0.965310479 | 0.4546378 | -2.1232517 | 0.03373277 | 0.57865105 |
| ERLEC1 | 10.1916881 | -0.078567847 | 0.47384062 | -0.1658107 | 0.86830592 | 0.99665574 |
| PSME4 | 53.0070432 | -0.066413762 | 0.34497071 | -0.19252 | 0.8473349 | 0.99665574 |
| ACYP2 | 49.2721692 | 0.345479005 | 0.32157601 | 1.07433077 | 0.28267446 | 0.96059387 |
| SPTBN1 | 91.9689937 | -1.173129696 | 0.31247107 | -3.7543625 | 0.00017378 | 0.02986125 |
| AC093110.1 | 4.02281741 | -1.972933675 | 0.92000215 | -2.1444881 | 0.0319938 | 0.56904056 |
| EML6 | 6.07728613 | 0.502826143 | 0.75591881 | 0.66518539 | 0.50593193 | 0.99665574 |
| RTN4 | 129.790736 | -0.090039424 | 0.30989096 | -0.290552 | 0.77139401 | 0.99665574 |
| CLHC1 | 8.44024529 | 0.771597602 | 0.61370834 | 1.25727084 | 0.20865558 | 0.91043197 |
| RPS27A | 99.4475817 | 0.990815897 | 0.44973731 | 2.20309919 | 0.02758776 | 0.53563234 |
| MTIF2 | 11.0255766 | 0.047466818 | 0.4314657 | 0.11001296 | 0.9123991 | 0.99777713 |
| AC012358.3 | 4.59054671 | -0.385765361 | 0.92255279 | -0.4181499 | 0.67583752 | 0.99665574 |
| CCDC88A | 7.71202526 | -0.827146581 | 0.55916978 | -1.4792405 | 0.13907605 | 0.84668189 |
| CFAP36 | 9.7573421 | -1.000407869 | 0.5153889 | -1.9410738 | 0.05224933 | 0.65962952 |
| PPP4R3B | 36.9571054 | -0.075987013 | 0.29540239 | -0.2572322 | 0.79699951 | 0.99665574 |
| PNPT1 | 13.6170181 | 0.311716 | 0.51921365 | 0.60036172 | 0.5482652 | 0.99665574 |
| EFEMP1 | 21.0334972 | -0.109170323 | 0.58774572 | -0.1857441 | 0.85264541 | 0.99665574 |
| VRK2 | 27.8726792 | -0.051566644 | 0.41021691 | -0.1257058 | 0.89996481 | 0.99777713 |
| FANCL | 16.9976126 | 1.167876042 | 0.51108482 | 2.28509242 | 0.02230741 | 0.48357743 |
| LINC01122 | 4.16599753 | 0.129141702 | 1.24893442 | 0.10340151 | 0.91764432 | 0.99777713 |
| PAPOLG | 9.70937443 | -0.162093832 | 0.47707145 | -0.3397685 | 0.7340309 | 0.99665574 |
| REL | 11.5402907 | 0.421331644 | 0.49746826 | 0.84695181 | 0.39702198 | 0.99179694 |
| PUS10 | 4.10064563 | -0.510964084 | 0.91969247 | -0.5555815 | 0.57849701 | 0.99665574 |
| PEX13 | 10.2034382 | 0.14734962 | 0.5757179 | 0.25594066 | 0.79799666 | 0.99665574 |
| KIAA1841 | 3.93177349 | -0.230319641 | 0.77743119 | -0.2962573 | 0.76703362 | 0.99665574 |
| C2orf74 | 2.92529108 | 0.22594368 | 0.92071315 | 0.24540073 | 0.80614616 | 0.99665574 |
| USP34 | 183.08853 | -0.282312246 | 0.37630033 | -0.7502312 | 0.45311547 | 0.99617422 |
| AC016727.1 | 5.7875468 | -0.16712891 | 0.54823255 | -0.3048504 | 0.7604801 | 0.99665574 |
| XPO1 | 60.3594853 | -0.031027062 | 0.41270384 | -0.07518 | 0.94007152 | 0.99777713 |
| FAM161A | 4.32641533 | -0.064264201 | 0.83794385 | -0.0766927 | 0.93886799 | 0.99777713 |
| CCT4 | 16.2285138 | 0.630386488 | 0.43641021 | 1.44448151 | 0.14860358 | 0.85485259 |
| COMMD1 | 64.4255007 | 0.053530829 | 0.32746959 | 0.16346809 | 0.8701499 | 0.99665574 |
| AC018462.1 | 3.15494764 | -1.232214461 | 0.82603367 | -1.4917243 | 0.13577145 | 0.84475183 |
| B3GNT2 | 29.2733864 | -0.665607719 | 0.3975463 | -1.6742898 | 0.09407366 | 0.76974878 |
| EHBP1 | 35.9034303 | 0.239711014 | 0.42633435 | 0.56226062 | 0.57393846 | 0.99665574 |
| WDPCP | 46.756977 | 0.276343697 | 0.37419335 | 0.7385051 | 0.46020757 | 0.99617422 |
| MDH1 | 20.7734031 | 0.353016606 | 0.41308571 | 0.8545844 | 0.39278126 | 0.99179694 |
| UGP2 | 48.0802715 | -0.305061089 | 0.34826158 | -0.8759539 | 0.38105509 | 0.98820924 |
| VPS54 | 42.2709241 | 0.294305558 | 0.33495945 | 0.87863042 | 0.3796017 | 0.98679664 |
| PELI1 | 6.84647192 | -0.393793419 | 0.68408204 | -0.5756523 | 0.5648502 | 0.99665574 |
| AC008074.1 | 3.48770498 | 0.108917251 | 0.75035286 | 0.14515471 | 0.88458872 | 0.99665574 |
| AFTPH | 40.2411484 | -0.2497105 | 0.27896009 | -0.8951478 | 0.3707081 | 0.98532838 |
| SERTAD2 | 5.84792849 | -0.200252523 | 0.66901568 | -0.2993241 | 0.76469276 | 0.99665574 |
| CEP68 | 9.20619253 | -0.352104525 | 0.52947474 | -0.6650072 | 0.50604588 | 0.99665574 |
| RAB1A | 59.0547247 | -0.408743728 | 0.40786084 | -1.0021647 | 0.31626407 | 0.97663269 |
| ACTR2 | 44.9485843 | -0.04786077 | 0.25546699 | -0.1873462 | 0.8513892 | 0.99665574 |
| SPRED2 | 41.2177891 | 0.020371927 | 0.33036392 | 0.06166511 | 0.95082953 | 0.99777713 |
| ETAA1 | 5.02440928 | 0.731340934 | 0.68782657 | 1.06326357 | 0.28766244 | 0.96559334 |
| C1D | 9.22962212 | 0.956203351 | 0.56583207 | 1.68990659 | 0.09104583 | 0.76412898 |
| PNO1 | 3.3777738 | -1.300119534 | 0.85435658 | -1.5217528 | 0.12807103 | 0.83778831 |
| APLF | 8.54120192 | 0.564024151 | 0.61453954 | 0.91779961 | 0.35872379 | 0.9819332 |
| GFPT1 | 31.9010949 | -1.010850327 | 0.53471379 | -1.8904512 | 0.05869764 | 0.67545594 |
| NFU1 | 8.99171051 | 0.060251795 | 0.48406805 | 0.12446968 | 0.90094341 | 0.99777713 |
| AAK1 | 70.4907383 | -0.259683467 | 0.26448534 | -0.9818445 | 0.32617648 | 0.97788427 |
| ANXA4 | 133.62282 | 0.136066185 | 0.35374139 | 0.38464875 | 0.70049767 | 0.99665574 |
| GMCL1 | 23.5450718 | 0.136276196 | 0.40000792 | 0.34068374 | 0.73334168 | 0.99665574 |
| MXD1 | 12.1259511 | -0.974225235 | 0.57688083 | -1.6887807 | 0.09126145 | 0.76526768 |
| PCBP1 | 28.2025279 | 0.145086514 | 0.53918015 | 0.26908727 | 0.78786253 | 0.99665574 |
| C2orf42 | 5.74843138 | -0.293391108 | 0.63380155 | -0.4629069 | 0.6434311 | 0.99665574 |
| PCYOX1 | 17.1664251 | -0.21241539 | 0.47864853 | -0.4437816 | 0.65720053 | 0.99665574 |
| SNRPG | 11.6912489 | 0.410815485 | 0.52166752 | 0.78750443 | 0.43098663 | 0.99617422 |
| FAM136A | 6.4274476 | 0.102001582 | 0.57441506 | 0.1775747 | 0.85905699 | 0.99665574 |
| TGFA | 7.00682713 | -1.011487633 | 0.91927573 | -1.1003093 | 0.27119738 | 0.94865166 |
| ATP6V1B1 | 15.3249335 | 2.940717216 | 0.74181122 | 3.96423931 | 7.36E-05 | 0.01777297 |
| AC007040.2 | 33.3343524 | -0.101183911 | 0.87988931 | -0.1149962 | 0.90844815 | 0.99777713 |
| NAGK | 6.4621381 | -0.691988001 | 0.67197219 | -1.0297867 | 0.30311016 | 0.97376371 |
| MCEE | 8.55095918 | -0.357503582 | 0.5260565 | -0.6795916 | 0.49676309 | 0.99640678 |
| MPHOSPH10 | 6.98611653 | -0.311033709 | 0.61079676 | -0.5092262 | 0.61059369 | 0.99665574 |
| PAIP2B | 2.56332095 | -0.158622051 | 1.02125002 | -0.1553215 | 0.87656789 | 0.99665574 |
| ZNF638 | 106.624354 | 0.052737375 | 0.1971898 | 0.26744474 | 0.78912676 | 0.99665574 |
| EXOC6B | 137.114945 | -0.058630494 | 0.31763308 | -0.1845856 | 0.85355408 | 0.99665574 |
| SPR | 4.06458201 | -0.172572771 | 0.8342704 | -0.2068547 | 0.83612332 | 0.99665574 |
| EMX1 | 5.72504953 | 1.327493866 | 1.08917114 | 1.21881109 | 0.2229159 | 0.92191316 |

| | | | | | | |
|---|---|---|---|---|---|---|
| SFXN5 | 10.9593184 | -0.616962938 | 0.45579109 | -1.353609 | 0.17586114 | 0.88428483 |
| RAB11FIP5 | 4.00415281 | 0.191998381 | 0.6874734 | 0.27928118 | 0.78002905 | 0.99665574 |
| PRADC1 | 2.96370861 | 0.676829626 | 0.81428781 | 0.83119214 | 0.4058651 | 0.99194293 |
| CCT7 | 15.4645583 | 0.181484068 | 0.44528488 | 0.40756845 | 0.68359053 | 0.99665574 |
| ALMS1 | 22.1383782 | 0.25438555 | 0.34694183 | 0.73322249 | 0.46342274 | 0.99617422 |
| NAT8 | 5.10509671 | 2.366392902 | 1.25025026 | 1.89273538 | 0.05839307 | 0.67545594 |
| TPRKB | 2.77707687 | -0.601278429 | 0.80942453 | -0.7428468 | 0.45757443 | 0.99617422 |
| DUSP11 | 4.18924783 | -0.439441127 | 0.74372886 | -0.590862 | 0.55461288 | 0.99665574 |
| STAMBP | 14.0588705 | -0.39946787 | 0.45146634 | -0.8848231 | 0.37625202 | 0.98666399 |
| DGUOK | 14.5361525 | -0.6431367 | 0.42466948 | -1.5144406 | 0.12991413 | 0.83778831 |
| TET3 | 25.0166859 | -0.691228237 | 0.50016962 | -1.3819877 | 0.16697549 | 0.87495175 |
| BOLA3 | 3.60542658 | 0.307541556 | 0.86255268 | 0.35654814 | 0.72143011 | 0.99665574 |
| MOB1A | 25.7116948 | 0.546339355 | 0.34788023 | 1.5704812 | 0.11630321 | 0.82007861 |
| DCTN1 | 2.63169616 | -0.2689851 | 0.93404604 | -0.2879784 | 0.77336326 | 0.99665574 |
| RTKN | 9.67086103 | -0.403863976 | 0.4771184 | -0.8464649 | 0.39729345 | 0.99179694 |
| MOGS | 5.29471995 | -0.664327081 | 0.6554307 | -1.0135733 | 0.31078638 | 0.97395745 |
| TTC31 | 3.49078791 | 0.306483001 | 0.77801921 | 0.3939273 | 0.69363472 | 0.99665574 |
| AUP1 | 5.13169287 | 0.404955986 | 0.7363067 | 0.54998275 | 0.5823312 | 0.99665574 |
| HTRA2 | 3.89941303 | 0.279081161 | 0.76053155 | 0.3669554 | 0.71365229 | 0.99665574 |
| POLE4 | 7.67312451 | 1.131379305 | 0.5842226 | 1.93655519 | 0.05279974 | 0.66411295 |
| TACR1 | 3.8037896 | -1.298476573 | 0.86025168 | -1.5094148 | 0.13119283 | 0.83966722 |
| AC007681.1 | 2.63843344 | -0.335207911 | 0.89966687 | -0.3725911 | 0.70945276 | 0.99665574 |
| EVA1A | 4.00564488 | -0.1515874 | 0.79084855 | -0.1916769 | 0.8479953 | 0.99665574 |
| MRPL19 | 23.1816479 | 0.497455285 | 0.51483067 | 0.96625028 | 0.33391896 | 0.97788427 |
| GCFC2 | 24.6916109 | 0.246381568 | 0.41236067 | 0.59749046 | 0.55017997 | 0.99665574 |
| CTNNA2 | 3.21689425 | 0.496801685 | 0.86651703 | 0.5733317 | 0.56642012 | 0.99665574 |
| SUCLG1 | 38.5681729 | 0.676317759 | 0.34339405 | 1.96950926 | 0.04889464 | 0.64375827 |
| DNAH6 | 4.04851555 | 0.591067006 | 0.87942382 | 0.67210711 | 0.50151551 | 0.99640678 |
| TMSB10 | 72.71865 | -1.385219087 | 0.48800208 | -2.8385516 | 0.00453188 | 0.21492549 |
| KCMF1 | 32.1734175 | -0.310065489 | 0.3143164 | -0.9864757 | 0.32389974 | 0.97700345 |
| TCF7L1 | 40.4847246 | 0.964553188 | 0.5217238 | 1.84878127 | 0.0644894 | 0.68846743 |
| TGOLN2 | 39.2524672 | -0.197428286 | 0.31804071 | -0.6207642 | 0.53475478 | 0.99665574 |
| RETSAT | 6.16184847 | -0.495161734 | 0.57125208 | -0.8668008 | 0.38605119 | 0.99051797 |
| ELMOD3 | 12.6487761 | -0.655639908 | 0.50522457 | -1.2977198 | 0.19438365 | 0.90314457 |
| CAPG | 14.5018259 | 1.160267509 | 0.56589078 | 2.05033823 | 0.04033144 | 0.60754844 |
| MAT2A | 14.8452564 | 0.68782173 | 0.41978131 | 1.63852397 | 0.10131244 | 0.78386461 |
| GGCX | 6.01695057 | 0.003179957 | 0.59663965 | 0.00532978 | 0.99574747 | 0.99869666 |
| VAMP8 | 32.0456729 | -0.119434742 | 0.4263325 | -0.2801446 | 0.77936658 | 0.99665574 |
| RNF181 | 5.49371733 | -0.035318944 | 0.80035493 | -0.0441291 | 0.9648015 | 0.99777713 |
| USP39 | 14.6306593 | -0.544199441 | 0.55565497 | -0.9793837 | 0.32739041 | 0.97788427 |
| C2orf68 | 6.96625037 | 0.150405343 | 0.5677751 | 0.26490302 | 0.79108417 | 0.99665574 |
| ST3GAL5 | 7.45106015 | -0.569591115 | 0.57590083 | -0.9890438 | 0.32264174 | 0.97700345 |
| POLR1A | 11.3004194 | -0.412098512 | 0.55841466 | -0.7379794 | 0.46052697 | 0.99617422 |
| PTCD3 | 21.857349 | 0.307352218 | 0.39699559 | 0.77419554 | 0.43881517 | 0.99617422 |
| IMMT | 34.9299275 | 0.154686152 | 0.280628 | 0.55121424 | 0.58148682 | 0.99665574 |
| MRPL35 | 6.27441711 | 0.243035162 | 0.5938063 | 0.40928357 | 0.68233157 | 0.99665574 |
| REEP1 | 2.95636991 | 0.279341546 | 0.83448216 | 0.33474837 | 0.73781491 | 0.99665574 |
| KDM3A | 32.2096116 | 0.407660782 | 0.47774038 | 0.8533103 | 0.39348725 | 0.99179694 |
| CHMP3 | 12.8857914 | 0.054054706 | 0.41888331 | 0.12904478 | 0.89732221 | 0.99777713 |
| RNF103-CHM | 7.56890108 | -3.976552623 | 1.21068111 | -3.2845583 | 0.00102142 | 0.08775589 |
| RMND5A | 25.0037743 | 0.062132631 | 0.39596569 | 0.15691418 | 0.87531248 | 0.99665574 |
| PLGLB1 | 4.18305321 | 0.831179061 | 0.81331745 | 1.02196143 | 0.30679916 | 0.97376371 |
| CYTOR | 4.99676291 | -2.941804963 | 0.6981631 | -4.2136357 | 2.51E-05 | 0.00821807 |
| KRCC1 | 12.8196331 | -0.40860647 | 0.48390965 | -0.8443859 | 0.39845382 | 0.99179694 |
| THNSL2 | 4.21804908 | 0.398049259 | 0.74667689 | 0.53309439 | 0.59396824 | 0.99665574 |
| EIF2AK3 | 19.6462102 | -0.687191838 | 0.49913443 | -1.3767671 | 0.16858428 | 0.87566218 |
| RPIA | 10.9118593 | 1.143307813 | 0.57556602 | 1.98640602 | 0.04698826 | 0.64060657 |
| MAL | 130.327323 | 0.278104418 | 0.52144058 | 0.53333866 | 0.59379917 | 0.99665574 |
| MRPS5 | 42.0564297 | 0.305697204 | 0.26636828 | 1.14764867 | 0.25111363 | 0.93975266 |
| AC092835.1 | 3.21510293 | -0.122194029 | 0.82693536 | -0.1477673 | 0.88252639 | 0.99665574 |
| FAHD2A | 3.5489945 | 0.049622091 | 0.75354344 | 0.06585167 | 0.94749592 | 0.99777713 |
| ANKRD36C | 12.3623297 | -2.477728965 | 0.61031448 | -4.0597578 | 4.91E-05 | 0.01383377 |
| STARD7 | 46.6152165 | 0.072273724 | 0.29113274 | 0.24825007 | 0.80394093 | 0.99665574 |
| TMEM127 | 16.5920843 | 0.332484969 | 0.39691063 | 0.83768219 | 0.40220922 | 0.99194293 |
| CIAO1 | 10.9595338 | -0.411017395 | 0.47339797 | -0.8682281 | 0.3852695 | 0.99051797 |
| SNRNP200 | 18.3498523 | -0.34321126 | 0.44203884 | -0.7764278 | 0.43749641 | 0.99617422 |
| KANSL3 | 26.6720667 | 0.473597423 | 0.37267202 | 1.27081562 | 0.20379425 | 0.90707783 |
| LMAN2L | 7.83305681 | 0.032595403 | 0.5316455 | 0.06131041 | 0.951112 | 0.99777713 |
| CNNM4 | 16.2711849 | 0.565789067 | 0.70276406 | 0.80509107 | 0.42076713 | 0.99518711 |
| ANKRD23 | 4.43945367 | 0.117410819 | 0.65065578 | 0.18044998 | 0.85679932 | 0.99665574 |
| ANKRD36 | 17.2441867 | -0.217167833 | 0.64537732 | -0.3364975 | 0.73649576 | 0.99665574 |
| AC092683.1 | 12.691125 | -1.236340461 | 0.54298943 | -2.2769144 | 0.02279134 | 0.4879643 |
| ANKRD36B | 7.51010349 | -0.762936855 | 0.95602843 | -0.7980274 | 0.42485461 | 0.99518711 |
| COX5B | 47.1582342 | 0.257902152 | 0.45416317 | 0.56786232 | 0.57012846 | 0.99665574 |
| ACTR1B | 5.54239347 | 0.221140158 | 0.60708927 | 0.364263 | 0.71566163 | 0.99665574 |
| TMEM131 | 53.8150993 | 0.483250414 | 0.30912727 | 1.56327331 | 0.11798832 | 0.82599629 |
| INPP4A | 84.6349178 | -0.58452831 | 0.61670717 | -0.9478215 | 0.34322034 | 0.97788427 |
| COA5 | 7.24588719 | -0.084415072 | 0.5436438 | -0.1552764 | 0.87660339 | 0.99665574 |
| UNC50 | 9.6798729 | -0.120203192 | 0.4523174 | -0.2657497 | 0.79043202 | 0.99665574 |
| MGAT4A | 24.131515 | 0.290553222 | 0.39571653 | 0.73424585 | 0.46279892 | 0.99617422 |
| TSGA10 | 29.0817249 | 0.172106514 | 0.42430128 | 0.40562336 | 0.68501936 | 0.99665574 |
| AC079447.1 | 9.37795427 | 0.11138308 | 0.58014742 | 0.19199099 | 0.84774926 | 0.99665574 |
| MITD1 | 7.95486783 | -0.177725975 | 0.69184573 | -0.2568867 | 0.79726623 | 0.99665574 |
| TXNDC9 | 7.18443638 | -0.197513023 | 0.58032793 | -0.3403473 | 0.73359503 | 0.99665574 |
| EIF5B | 31.7026288 | -0.629547643 | 0.36332422 | -1.7327434 | 0.08314129 | 0.75242737 |
| REV1 | 59.7996609 | 0.105325057 | 0.27467705 | 0.38345052 | 0.70138575 | 0.99665574 |
| LONRF2 | 13.8010721 | -0.213526548 | 0.58496903 | -0.365022 | 0.71509501 | 0.99665574 |
| CHST10 | 5.45348757 | 0.056942839 | 0.6615786 | 0.08607116 | 0.93140985 | 0.99777713 |
| PDCL3 | 5.04955884 | -0.228794061 | 0.70441336 | -0.3248009 | 0.7453318 | 0.99665574 |
| NPAS2 | 11.920122 | -1.398817365 | 0.61006319 | -2.2929057 | 0.02185343 | 0.47850999 |
| RPL31 | 29.053153 | 0.372062282 | 0.3488373 | 1.06657826 | 0.28616232 | 0.96449041 |
| TBC1D8 | 28.3654402 | -1.047853566 | 0.36337365 | -2.8836806 | 0.00393057 | 0.20347549 |
| CNOT11 | 10.3377522 | 0.046495933 | 0.44340052 | 0.10486215 | 0.91648519 | 0.99777713 |
| RNF149 | 82.8173247 | -0.612606957 | 0.37433966 | -1.6365003 | 0.10173492 | 0.7843028 |
| MAP4K4 | 51.6209107 | -0.677014972 | 0.37038763 | -1.8278552 | 0.06757128 | 0.70477126 |
| IL1R1 | 19.6819678 | 0.043481893 | 0.5483262 | 0.07929932 | 0.93679455 | 0.99777713 |
| SLC9A2 | 12.0061393 | -0.025880719 | 0.67366755 | -0.0384176 | 0.9693547 | 0.99777713 |
| MFSD9 | 3.34196095 | -1.150746576 | 0.78924139 | -1.4580413 | 0.14482915 | 0.85018987 |
| TMEM182 | 3.01021212 | 1.274671126 | 0.8735082 | 1.4592549 | 0.14449496 | 0.85018987 |
| POU3F3 | 17.3593197 | 0.622314517 | 0.42223768 | 1.47384885 | 0.14052229 | 0.84969914 |
| MRPS9 | 15.6782201 | 0.148873118 | 0.46521286 | 0.32001075 | 0.74896018 | 0.99665574 |
| TGFBRAP1 | 7.79895801 | 0.404315002 | 0.57582702 | 0.70214663 | 0.48258772 | 0.99640678 |
| C2orf49 | 6.06484502 | 0.237004001 | 0.61639587 | 0.38449966 | 0.70060815 | 0.99665574 |
| NCK2 | 16.9060861 | -0.78103125 | 0.39808828 | -1.9619549 | 0.04976774 | 0.64444758 |
| UXS1 | 12.3078569 | -0.155096666 | 0.52236624 | -0.2969117 | 0.7665339 | 0.99665574 |
| SULT1C2 | 63.9884909 | -0.109902 | 0.38425952 | -0.2860098 | 0.77487059 | 0.99665574 |
| GCC2 | 58.0195783 | -0.060542895 | 0.26493896 | -0.2285164 | 0.81924482 | 0.99665574 |
| GCC2-AS1 | 13.3067112 | 0.236270001 | 0.64602804 | 0.36572716 | 0.71456868 | 0.99665574 |
| LIMS1 | 256.12961 | -0.422411808 | 0.40927316 | -1.0321024 | 0.30202415 | 0.97358745 |
| RANBP2 | 48.1173855 | 0.065834896 | 0.27411887 | 0.24016915 | 0.81019913 | 0.99665574 |
| CCDC138 | 9.7190123 | 0.41038967 | 0.57094789 | 0.71878657 | 0.47227244 | 0.99617422 |
| 10-Sep | 47.5618252 | -0.914944287 | 0.30386359 | -3.0110363 | 0.00260358 | 0.1571135 |
| SOWAHC | 10.9952325 | -0.561397097 | 0.44592139 | -1.2589598 | 0.20804486 | 0.91026892 |

| | | | | | | |
|---|---|---|---|---|---|---|
| NPHP1 | 3.77786877 | -0.101819139 | 0.86970587 | -0.1170731 | 0.90680215 | 0.99777713 |
| SMIM37 | 11.7930766 | 0.042237068 | 0.66111117 | 0.063888 | 0.94905941 | 0.99777713 |
| LINC01106 | 3.88922758 | 1.068398732 | 0.70317539 | 1.51939153 | 0.12866397 | 0.83778831 |
| AC112229.3 | 1.87199246 | -0.969910717 | 1.06261321 | -0.9127599 | 0.36136884 | 0.9819332 |
| MIR4435-2H | 19.5401289 | -2.021058724 | 0.40542241 | -4.9850691 | 6.19E-07 | 0.00048303 |
| BCL2L11 | 4.32412299 | -0.912466961 | 0.67403161 | -1.3537451 | 0.17581773 | 0.88428483 |
| ANAPC1 | 19.1040216 | 0.220443596 | 0.41757824 | 0.52790968 | 0.59756203 | 0.99665574 |
| MERTK | 5.39861479 | -0.282354193 | 0.64513333 | -0.437668 | 0.661627 | 0.99665574 |
| TMEM87B | 18.9158681 | -0.909300106 | 0.39457417 | -2.30451 | 0.02119403 | 0.4733368 |
| ZC3H8 | 13.5485394 | 0.139903293 | 0.40759159 | 0.34324382 | 0.73141504 | 0.99665574 |
| ZC3H6 | 14.6924194 | -0.293774604 | 0.58737883 | -0.500145 | 0.61697296 | 0.99665574 |
| RGPD8 | 5.4498961 | 0.029294735 | 0.69485368 | 0.04215957 | 0.96637149 | 0.99777713 |
| TTL | 11.6152095 | -0.696386883 | 0.46204728 | -1.5071767 | 0.13176538 | 0.84074439 |
| POLR1B | 5.59798217 | -0.385003458 | 0.6263732 | -0.6146551 | 0.53878255 | 0.99665574 |
| CHCHD5 | 6.21575989 | -0.111457505 | 0.64060652 | -0.1739875 | 0.86187531 | 0.99665574 |
| SLC20A1 | 13.4433444 | -0.245730326 | 0.58831856 | -0.4176824 | 0.67617932 | 0.99665574 |
| PSD4 | 6.38935476 | -0.785236332 | 0.64561285 | -1.216265 | 0.22388398 | 0.92265683 |
| PAX8 | 265.918252 | -0.816459281 | 0.28277034 | -2.8873582 | 0.00388492 | 0.20301691 |
| CBWD2 | 8.67675286 | 0.758314992 | 0.54032845 | 1.40343339 | 0.16048764 | 0.86820903 |
| RABL2A | 2.679361 | -0.273904066 | 0.92521542 | -0.2960436 | 0.76719682 | 0.99665574 |
| SLC35F5 | 56.051436 | -0.575660763 | 0.33298167 | -1.7288062 | 0.08384379 | 0.75242737 |
| ACTR3 | 52.1354613 | -0.798316731 | 0.39302246 | -2.0312242 | 0.04223226 | 0.61533334 |
| DDX18 | 14.8151539 | -0.197048345 | 0.54794455 | -0.3596137 | 0.71913607 | 0.99665574 |
| CCDC93 | 28.1110636 | -0.622865688 | 0.35834326 | -1.7381817 | 0.08217881 | 0.75124322 |
| INSIG2 | 25.1676925 | 0.277662133 | 0.45685174 | 0.60777296 | 0.54333806 | 0.99665574 |
| C2orf76 | 3.56082327 | 0.558317503 | 0.78988588 | 0.70683313 | 0.47967019 | 0.99617422 |
| DBI | 60.1888562 | -0.301450406 | 0.45826103 | -0.6578137 | 0.51065782 | 0.99665574 |
| TMEM37 | 4.34819732 | -1.168350686 | 0.97038106 | -1.2040123 | 0.22858484 | 0.92646258 |
| CFAP221 | 14.1657765 | -0.660976846 | 0.58008932 | -1.1394398 | 0.25451977 | 0.94099699 |
| PTPN4 | 43.1628667 | 0.624007119 | 0.38018742 | 1.64131448 | 0.10073215 | 0.78386461 |
| EPB41L5 | 66.2571064 | 0.333119839 | 0.25518934 | 1.30538307 | 0.19176244 | 0.89964233 |
| RALB | 7.2664089 | -0.259425386 | 0.56433715 | -0.4596993 | 0.64573208 | 0.99665574 |
| TFCP2L1 | 485.64959 | 0.20094743 | 0.31405128 | 0.63985546 | 0.52226657 | 0.99665574 |
| CLASP1 | 98.7336587 | 0.065606062 | 0.29858905 | 0.21972025 | 0.82608903 | 0.99665574 |
| NIFK-AS1 | 5.03231761 | -0.083615372 | 0.72943797 | -0.1146299 | 0.90873851 | 0.99777713 |
| NIFK | 5.99481759 | 1.123312255 | 0.59551914 | 1.88627398 | 0.05925804 | 0.67882265 |
| TSN | 11.3955737 | 0.367428672 | 0.49908777 | 0.73620051 | 0.46160868 | 0.99617422 |
| BIN1 | 5.96665765 | 0.80889072 | 0.72263736 | 1.11935912 | 0.26298696 | 0.94557817 |
| ERCC3 | 9.79335034 | -0.711142241 | 0.51609844 | -1.3779198 | 0.16822806 | 0.87566218 |
| MAP3K2 | 57.3852212 | 0.255271382 | 0.2677311 | 0.95346181 | 0.34035614 | 0.97788427 |
| IWS1 | 22.889875 | -0.332165313 | 0.38494685 | -0.8628862 | 0.38820006 | 0.99168679 |
| WDR33 | 46.5289706 | 0.21539622 | 0.32852031 | 0.65565571 | 0.51204567 | 0.99665574 |
| SFT2D3 | 2.53778291 | 1.040619615 | 0.9831152 | 1.05849204 | 0.28983118 | 0.96676791 |
| POLR2D | 4.33304489 | -0.135578058 | 0.65640036 | -0.2065478 | 0.83636302 | 0.99665574 |
| AMMECR1L | 8.3218622 | 0.139291727 | 0.57165227 | 0.24366514 | 0.80749018 | 0.99665574 |
| SAP130 | 11.3692651 | -0.069038012 | 0.44909866 | -0.1537257 | 0.87782601 | 0.99665574 |
| UGGT1 | 22.1092372 | 0.333009031 | 0.41565555 | 0.80116585 | 0.42303564 | 0.99518711 |
| HS6ST1 | 21.8418128 | -0.663906153 | 0.48689002 | -1.3635649 | 0.17270454 | 0.88157681 |
| SMPD4 | 11.7284439 | -0.57873306 | 0.45036063 | -1.2850436 | 0.19877704 | 0.90381759 |
| MZT2B | 17.2114239 | 0.403826595 | 0.51155979 | 0.78940253 | 0.42987678 | 0.99617422 |
| CCDC115 | 5.45095597 | -0.321571021 | 0.66352859 | -0.4846378 | 0.62793331 | 0.99665574 |
| IMP4 | 5.00125135 | -0.170936007 | 0.63718354 | -0.2682681 | 0.78849297 | 0.99665574 |
| PTPN18 | 9.43905451 | 0.696949673 | 0.49927952 | 1.39591081 | 0.16274136 | 0.87135275 |
| FAM168B | 38.5267415 | -0.183498857 | 0.36958202 | -0.4965037 | 0.61953904 | 0.99665574 |
| PLEKHB2 | 62.7334764 | -0.031706429 | 0.33464013 | -0.0947478 | 0.92451512 | 0.99777713 |
| MZT2A | 8.46950706 | -0.2707816 | 0.50728606 | -0.5337848 | 0.59349042 | 0.99665574 |
| GPR39 | 13.340715 | 0.028269787 | 0.5633814 | 0.05017877 | 0.95997993 | 0.99777713 |
| NCKAP5 | 28.216703 | -0.510376879 | 0.66660336 | -0.765638 | 0.44389172 | 0.99617422 |
| MGAT5 | 52.0394505 | -0.141025585 | 0.41490634 | -0.3398974 | 0.7339338 | 0.99665574 |
| TMEM163 | 55.4546006 | -0.2676961 | 0.56014822 | -0.4779023 | 0.63271977 | 0.99665574 |
| CCNT2-AS1 | 9.71748158 | -0.746743695 | 0.91859007 | -0.8129238 | 0.41626177 | 0.992853 |
| ACMSD | 3.23238462 | -0.182509963 | 1.13088163 | -0.1613873 | 0.87178835 | 0.99665574 |
| CCNT2 | 48.2576996 | -0.338770141 | 0.45156797 | -0.7502085 | 0.45312913 | 0.99617422 |
| RAB3GAP1 | 48.1346018 | -0.1714702 | 0.27797309 | -0.616859 | 0.53732773 | 0.99665574 |
| ZRANB3 | 19.4963659 | -0.18354471 | 0.50123902 | -0.366182 | 0.71422927 | 0.99665574 |
| R3HDM1 | 42.6040939 | 0.238691759 | 0.27724145 | 0.86095265 | 0.38926412 | 0.99176253 |
| UBXN4 | 44.606394 | -0.073915868 | 0.34229328 | -0.2159431 | 0.82903211 | 0.99665574 |
| DARS | 36.4811433 | -0.296733795 | 0.33382889 | -0.8888799 | 0.37406762 | 0.98535336 |
| HNMT | 19.830209 | 0.05811093 | 0.52915369 | 0.10981862 | 0.91255322 | 0.99777713 |
| AC114763.1 | 7.31172946 | -0.450349773 | 0.72432473 | -0.6217512 | 0.53410548 | 0.99665574 |
| SPOPL | 31.8561359 | 0.372422132 | 0.30869562 | 1.20643803 | 0.22764864 | 0.92389989 |
| NXPH2 | 16.7566745 | 1.023053792 | 0.97522393 | 1.04904501 | 0.29415741 | 0.97161335 |
| LRP1B | 56.5610513 | -3.079602887 | 0.67478829 | -4.563806 | 5.02E-06 | 0.00268041 |
| GTDC1 | 19.9825802 | 0.514989471 | 0.42685336 | 1.20647868 | 0.22763298 | 0.92389989 |
| TEX41 | 21.5587696 | 1.319723962 | 1.19944139 | 1.10028216 | 0.2712092 | 0.94865166 |
| ACVR2A | 21.501587 | 0.640633678 | 0.36655265 | 1.74772622 | 0.08051145 | 0.74618467 |
| ORC4 | 37.4302214 | 0.543791882 | 0.33185354 | 1.63865024 | 0.10128612 | 0.78386461 |
| MBD5 | 131.865994 | 0.05110474 | 0.21317616 | 0.23973009 | 0.81053951 | 0.99665574 |
| EPC2 | 54.2461444 | -0.056825055 | 0.35588548 | -0.1596723 | 0.87313922 | 0.99665574 |
| AC105402.3 | 5.90041171 | 2.115045774 | 1.21629182 | 1.73892955 | 0.08204716 | 0.75124322 |
| MMADHC | 8.31216359 | 0.87833321 | 0.54964532 | 1.59799997 | 0.11004298 | 0.80608076 |
| AC007364.1 | 3.72954515 | -0.632822566 | 0.80222552 | -0.7888338 | 0.43020918 | 0.99617422 |
| NMI | 4.59703017 | -0.646209889 | 0.81402912 | -0.7938412 | 0.42728786 | 0.99582628 |
| RIF1 | 37.5986537 | 0.24829667 | 0.38912331 | 0.63809251 | 0.52341346 | 0.99665574 |
| ARL5A | 9.82597061 | 0.178917636 | 0.54166439 | 0.33031087 | 0.74116508 | 0.99665574 |
| CACNB4 | 6.90846237 | 0.08088337 | 0.64501027 | 0.12539858 | 0.90020801 | 0.99777713 |
| STAM2 | 26.1471849 | 0.219921958 | 0.32367635 | 0.67945018 | 0.49685266 | 0.99640678 |
| FMNL2 | 66.5343282 | -0.531901373 | 0.33548385 | -1.5854754 | 0.11285837 | 0.81261232 |
| PRPF40A | 37.3458944 | 0.296779488 | 0.31029803 | 0.95643368 | 0.33885317 | 0.97788427 |
| ARL6IP6 | 14.0802476 | -0.119903515 | 0.48439372 | -0.2475332 | 0.80449562 | 0.99665574 |
| KCNJ3 | 27.460158 | 0.726374673 | 0.76757786 | 0.94632051 | 0.34398513 | 0.97788427 |
| LINC01876 | 12.6053055 | -1.24208558 | 0.80209271 | -1.5485561 | 0.12148846 | 0.82659025 |
| GPD2 | 26.4762542 | 0.07927464 | 0.35850199 | 0.22112748 | 0.82499318 | 0.99665574 |
| CYTIP | 22.1840577 | 1.329003994 | 0.70243026 | 1.89200846 | 0.05848985 | 0.67545594 |
| ACVR1 | 39.7425658 | -0.127641117 | 0.34412406 | -0.370916 | 0.7107001 | 0.99665574 |
| UPP2 | 6.71775705 | 0.665622327 | 0.72283621 | 0.92084807 | 0.35712976 | 0.9819332 |
| CCDC148 | 17.508009 | 0.477759139 | 0.51808972 | 0.92215522 | 0.35644762 | 0.9819332 |
| PKP4 | 103.647004 | 0.495557205 | 0.31917462 | 1.55262095 | 0.12051372 | 0.82659025 |
| TANC1 | 57.8804184 | -0.791759097 | 0.307422 | -2.5754796 | 0.01001012 | 0.34338499 |
| WDSUB1 | 31.4101259 | -0.130000174 | 0.36461502 | -0.3565409 | 0.72143551 | 0.99665574 |
| BAZ2B | 162.005709 | 0.106490228 | 0.2062071 | 0.51642367 | 0.60555854 | 0.99665574 |
| AC009961.1 | 3.68096952 | -0.487521181 | 0.72529931 | -0.6721655 | 0.50147833 | 0.99640678 |
| 7-Mar | 55.5476316 | 0.031939943 | 0.30976171 | 0.10311133 | 0.91787461 | 0.99777713 |
| PLA2R1 | 4.01645379 | -0.352285076 | 0.66623827 | -0.5287674 | 0.59696682 | 0.99665574 |
| ITGB6 | 43.9169118 | -1.730324011 | 1.25384105 | -1.3800186 | 0.16758091 | 0.87566218 |
| RBMS1 | 50.410286 | -0.0128098 | 0.38482666 | -0.0332872 | 0.97344556 | 0.99777713 |
| TANK | 30.4142671 | -0.102414845 | 0.41525138 | -0.2466334 | 0.80519198 | 0.99665574 |
| PSMD14 | 21.8056794 | 0.104564441 | 0.44589468 | 0.2345048 | 0.81459311 | 0.99665574 |
| IFIH1 | 5.45809799 | -0.632401045 | 0.90337856 | -0.7000399 | 0.48390238 | 0.99640678 |
| GCA | 8.06576156 | -0.875669262 | 0.64707183 | -1.3532798 | 0.17596624 | 0.88428483 |
| FIGN | 8.39744152 | 0.695622233 | 0.74362761 | 0.93544433 | 0.34955936 | 0.97838306 |
| GRB14 | 46.8693298 | 0.954979614 | 0.48454722 | 1.97087007 | 0.04873874 | 0.64337675 |
| COBLL1 | 336.021199 | 0.16495611 | 0.25585914 | 0.64471454 | 0.51911219 | 0.99665574 |

| | | | | | | |
|---|---|---|---|---|---|---|
| AC019197.1 | 170.813349 | -0.348622451 | 0.73108802 | -0.4768543 | 0.63346589 | 0.99665574 |
| SLC38A11 | 39.7848262 | 0.6150517 | 0.7482958 | 0.8219366 | 0.41111298 | 0.99279547 |
| SCN3A | 13.0396352 | -2.250257596 | 0.71559674 | -3.1445889 | 0.0016632 | 0.11874336 |
| SCN2A | 18.1980573 | 0.065949821 | 0.88154936 | 0.07481126 | 0.94036488 | 0.99777713 |
| CSRNP3 | 10.6260035 | 0.660773764 | 0.76995147 | 0.85820183 | 0.39078103 | 0.99179694 |
| GALNT3 | 11.5136473 | -0.669210166 | 0.46169959 | -1.4494494 | 0.14721213 | 0.85282092 |
| TTC21B | 18.8953846 | -0.041097814 | 0.37589705 | -0.1093326 | 0.91293866 | 0.99777713 |
| AC010127.1 | 3.66027211 | -0.895876355 | 1.03916805 | -0.8621092 | 0.38862742 | 0.99176253 |
| SCN9A | 17.1228207 | -1.07725697 | 0.55850339 | -1.928828 | 0.05375222 | 0.6653339 |
| STK39 | 17.7923555 | -1.480716637 | 0.81126087 | -1.8252041 | 0.06797024 | 0.7053043 |
| CERS6 | 35.1657441 | -0.396707219 | 0.56991764 | -0.6960782 | 0.48637989 | 0.99640678 |
| LRP2 | 20.3578145 | 1.010990866 | 0.59026092 | 1.71278639 | 0.08675184 | 0.75427976 |
| FASTKD1 | 38.2445923 | 0.150106211 | 0.37469753 | 0.40060636 | 0.68870997 | 0.99665574 |
| PPIG | 32.905227 | 0.101216002 | 0.33150747 | 0.30532042 | 0.76012212 | 0.99665574 |
| KLHL23 | 44.5679376 | 0.725815134 | 0.33586479 | 2.16103371 | 0.03069274 | 0.56271575 |
| SSB | 15.6004853 | 0.928266552 | 0.4679849 | 1.98353956 | 0.04730719 | 0.64060657 |
| METTL5 | 3.90211859 | -0.459986239 | 0.67755422 | -0.6788921 | 0.4972062 | 0.99640678 |
| UBR3 | 100.857516 | 0.171358819 | 0.26682185 | 0.64222184 | 0.52072915 | 0.99665574 |
| MYO3B | 19.5557502 | 0.40416163 | 0.74295047 | 0.54399539 | 0.58644464 | 0.99665574 |
| GORASP2 | 14.9744299 | -0.262582376 | 0.445794 | -0.5890218 | 0.55584666 | 0.99665574 |
| TLK1 | 62.0173038 | 0.107819758 | 0.26655519 | 0.40449319 | 0.68585008 | 0.99665574 |
| METTL8 | 6.47817153 | -0.054922534 | 0.66425542 | -0.0826829 | 0.93410372 | 0.99777713 |
| DCAF17 | 7.87634727 | 0.358410875 | 0.5941221 | 0.60326131 | 0.54633487 | 0.99665574 |
| DYNC1I2 | 35.7945703 | -0.413957124 | 0.37438972 | -1.1056851 | 0.26886285 | 0.94673212 |
| SLC25A12 | 39.3793776 | -0.62139933 | 0.41503793 | -1.4972109 | 0.13433838 | 0.84373478 |
| HAT1 | 12.8362971 | -0.012490246 | 0.42005263 | -0.029735 | 0.97627844 | 0.99777713 |
| METAP1D | 4.73247303 | 1.484777301 | 0.81468243 | 1.8225228 | 0.0683757 | 0.70708655 |
| ITGA6 | 205.994785 | 0.216349834 | 0.47665287 | 0.45389391 | 0.64990518 | 0.99665574 |
| AC078883.1 | 7.66664842 | -1.326678352 | 0.72980142 | -1.8178621 | 0.06908521 | 0.7094165 |
| PDK1 | 14.3576165 | -0.511156253 | 0.67918086 | -0.752607 | 0.45168612 | 0.99617422 |
| MAP3K20 | 5.48708393 | -0.422282419 | 0.64270771 | -0.6570365 | 0.51115745 | 0.99665574 |
| SP3 | 28.0918094 | 0.085248935 | 0.35326952 | 0.24131416 | 0.80931164 | 0.99665574 |
| OLA1 | 52.8780643 | 0.012469919 | 0.37312027 | 0.03342064 | 0.97333915 | 0.99777713 |
| CIR1 | 14.3906256 | 0.143262501 | 0.41257246 | 0.34724204 | 0.72840949 | 0.99665574 |
| SCRN3 | 6.09081114 | -0.301502712 | 0.58672079 | -0.5138777 | 0.60733752 | 0.99665574 |
| GPR155 | 6.21840687 | 0.324812948 | 0.91944372 | 0.35327116 | 0.72388517 | 0.99665574 |
| AC010894.3 | 37.9637737 | -0.755832383 | 0.37548116 | -2.0129702 | 0.04411776 | 0.6273014 |
| WIPF1 | 4.50413574 | -0.275275911 | 0.70283482 | -0.3916652 | 0.69530564 | 0.99665574 |
| CHN1 | 5.40632256 | -1.02643063 | 1.0667786 | -0.9621777 | 0.33596033 | 0.97788427 |
| ATF2 | 37.0285089 | 0.199335616 | 0.26996372 | 0.73837927 | 0.46028401 | 0.99617422 |
| ATP5MC3 | 48.0104235 | 0.210584961 | 0.44483943 | 0.47339545 | 0.63593108 | 0.99665574 |
| LNPK | 14.2327887 | 0.622864497 | 0.48313759 | 1.28920726 | 0.19732604 | 0.90381759 |
| HOXD10 | 7.25171037 | 0.851536204 | 0.67074635 | 1.26953536 | 0.20425019 | 0.90819667 |
| HOXD-AS2 | 2.51881713 | 0.405537277 | 1.03391907 | 0.3922331 | 0.69488599 | 0.99665574 |
| HOXD9 | 2.83607785 | 0.831927663 | 0.87608154 | 0.94960072 | 0.34231517 | 0.97788427 |
| HOXD8 | 7.68824016 | 0.189893026 | 0.59011069 | 0.32179222 | 0.74761011 | 0.99665574 |
| HOXD3 | 7.59087556 | -0.606766611 | 0.48898226 | -1.2408765 | 0.21465136 | 0.91280851 |
| HAGLR | 3.43217869 | -0.480298606 | 0.78603592 | -0.611039 | 0.54117376 | 0.99665574 |
| MTX2 | 54.3169617 | 0.44036971 | 0.27133074 | 1.6229997 | 0.10458947 | 0.79188053 |
| AC074286.1 | 9.647254 | -0.104131952 | 0.65254059 | -0.1595793 | 0.87321251 | 0.99665574 |
| HNRNPA3 | 54.8942856 | 0.191108863 | 0.26998062 | 0.7078614 | 0.47903133 | 0.99617422 |
| NFE2L2 | 127.020912 | -0.649692593 | 0.24024718 | -2.7042673 | 0.00684552 | 0.26899182 |
| AGPS | 29.5651931 | -0.003794521 | 0.34975583 | -0.0108491 | 0.99134387 | 0.99777713 |
| RBM45 | 5.55952657 | -0.096623261 | 0.59210083 | -0.1631872 | 0.87037106 | 0.99665574 |
| AC009948.1 | 8.71224867 | -0.567485373 | 0.79424732 | -0.7144945 | 0.47492143 | 0.99617422 |
| PRKRA | 22.7226096 | 0.334668962 | 0.43159945 | 0.77541564 | 0.4380941 | 0.99617422 |
| PJVK | 2.89157031 | -0.477587276 | 0.92101248 | -0.5185459 | 0.60407742 | 0.99665574 |
| PLEKHA3 | 12.2547667 | 0.512557461 | 0.49711568 | 1.03106275 | 0.30251139 | 0.97358745 |
| TTN-AS1 | 29.3521911 | -0.267258971 | 0.29770535 | -0.8977298 | 0.3693296 | 0.98532838 |
| SESTD1 | 99.7411239 | 0.011093289 | 0.29995853 | 0.03698274 | 0.97049877 | 0.99777713 |
| ZNF385B | 9.25800288 | 0.922111694 | 0.94982501 | 0.97082272 | 0.33163657 | 0.97788427 |
| CWC22 | 13.6542633 | -0.023914316 | 0.47302152 | -0.0505565 | 0.95967892 | 0.99777713 |
| UBE2E3 | 39.8184476 | -0.096256482 | 0.37313819 | -0.2579647 | 0.79643415 | 0.99665574 |
| SSFA2 | 29.8754767 | -0.398587059 | 0.47605082 | -0.8372784 | 0.40243611 | 0.99194293 |
| PDE1A | 53.3397916 | 0.3039351 | 0.7968414 | 0.38142484 | 0.70288804 | 0.99665574 |
| DNAJC10 | 14.6987113 | -0.221929749 | 0.56298546 | -0.3942016 | 0.69343223 | 0.99665574 |
| NCKAP1 | 63.3944437 | -0.00031932 | 0.23913946 | -0.0013353 | 0.9989346 | 0.99940262 |
| NUP35 | 8.38799507 | 1.086056764 | 0.66292986 | 1.63826799 | 0.1013658 | 0.78386461 |
| LINC01473 | 6.20062129 | 0.484102333 | 0.68001413 | 0.7119004 | 0.47652646 | 0.99617422 |
| ZC3H15 | 20.8131929 | -0.099121695 | 0.37096066 | -0.2672027 | 0.78931309 | 0.99665574 |
| ITGAV | 55.3521553 | -0.178170517 | 0.47502966 | -0.3750724 | 0.70760662 | 0.99665574 |
| AC017101.1 | 4.6442429 | 0.027500726 | 0.83910473 | 0.03277389 | 0.9738549 | 0.99777713 |
| AC007319.1 | 12.7616617 | 0.604446074 | 0.77558855 | 0.77933857 | 0.4357803 | 0.99617422 |
| TFPI | 19.5556041 | -1.562980356 | 0.74838666 | -2.0884664 | 0.03675579 | 0.58591936 |
| LINC01090 | 6.89788486 | -1.81363595 | 0.72023448 | -2.5181188 | 0.01179835 | 0.37359313 |
| GULP1 | 134.622286 | -0.778910648 | 0.42118435 | -1.8493343 | 0.06440956 | 0.68846743 |
| WDR75 | 7.45398245 | -0.244747401 | 0.51176247 | -0.4782441 | 0.63247645 | 0.99665574 |
| SLC40A1 | 5.94282678 | 0.236823564 | 0.68451953 | 0.3459705 | 0.72936489 | 0.99665574 |
| ASNSD1 | 4.10888024 | 0.38555462 | 0.78754377 | 0.48956595 | 0.62444107 | 0.99665574 |
| ANKAR | 16.7919472 | 0.005023966 | 0.43738263 | 0.01148643 | 0.99083536 | 0.99777713 |
| ORMDL1 | 12.9854903 | 0.016293495 | 0.55642067 | 0.02928269 | 0.97663913 | 0.99777713 |
| PMS1 | 12.9158347 | -0.184192441 | 0.41855643 | -0.440066 | 0.65988933 | 0.99665574 |
| C2orf88 | 4.47993979 | 0.075808964 | 0.77478244 | 0.09784549 | 0.92205499 | 0.99777713 |
| HIBCH | 27.8289447 | 0.593486337 | 0.39280541 | 1.51089146 | 0.13081611 | 0.83831459 |
| INPP1 | 5.3346801 | -0.574096922 | 0.74330169 | -0.7723606 | 0.43990089 | 0.99617422 |
| MFSD6 | 13.0519591 | -0.321638686 | 0.44761157 | -0.7185665 | 0.47240805 | 0.99617422 |
| NAB1 | 6.35411669 | -0.243922393 | 0.73496164 | -0.3318845 | 0.73997646 | 0.99665574 |
| GLS | 157.035745 | 0.533741385 | 0.37832182 | 1.4108131 | 0.15829973 | 0.86661434 |
| STAT1 | 33.436856 | -0.813168626 | 0.79807435 | -1.0189134 | 0.3082441 | 0.97376371 |
| MYO1B | 23.6243122 | -0.251483977 | 0.45718234 | -0.5500737 | 0.58226883 | 0.99665574 |
| SLC39A10 | 7.52523634 | -0.980836083 | 0.65509681 | -1.4972384 | 0.13433124 | 0.84373478 |
| DNAH7 | 7.02746829 | -0.767949043 | 0.55836227 | -1.3753598 | 0.16901991 | 0.87601071 |
| GTF3C3 | 17.9152001 | -0.132589746 | 0.43360398 | -0.3057854 | 0.75976808 | 0.99665574 |
| PGAP1 | 26.4966829 | -0.294870975 | 0.51903725 | -0.5681114 | 0.56995933 | 0.99665574 |
| ANKRD44 | 40.9130345 | 0.03644001 | 0.45937745 | 0.07932477 | 0.9367743 | 0.99777713 |
| SF3B1 | 50.2145621 | 0.467879067 | 0.2800689 | 1.67058557 | 0.09480357 | 0.77178333 |
| COQ10B | 9.27044929 | -0.660668144 | 0.53507466 | -1.2347214 | 0.2169342 | 0.9156801 |
| HSPD1 | 42.1490233 | 0.140846545 | 0.31921909 | 0.44122219 | 0.65905215 | 0.99665574 |
| MOB4 | 5.01876175 | 0.782014694 | 0.74619486 | 1.04800332 | 0.29463708 | 0.97161335 |
| AC011997.1 | 2.52611161 | -0.525434562 | 0.92009497 | -0.5710656 | 0.56795519 | 0.99665574 |
| PLCL1 | 351.462353 | 0.693583341 | 0.35866944 | 1.93376761 | 0.05314171 | 0.66476617 |
| FTCDNL1 | 9.12016768 | -0.653687896 | 0.53714028 | -1.216978 | 0.22361259 | 0.92191316 |
| C2orf69 | 6.35985303 | -0.2356426 | 0.6010828 | -0.3920302 | 0.69503591 | 0.99665574 |
| TYW5 | 8.59667124 | -0.295311722 | 0.52503799 | -0.5624578 | 0.57380414 | 0.99665574 |
| MAIP1 | 3.89355923 | -0.291636129 | 0.81314021 | -0.3586542 | 0.71985381 | 0.99665574 |
| SPATS2L | 31.6094297 | -0.14121817 | 0.40639532 | -0.3474897 | 0.72822349 | 0.99665574 |
| KCTD18 | 3.8938726 | -0.478664624 | 0.66346107 | -0.721466 | 0.47062284 | 0.99617422 |
| BZW1 | 9.42668114 | 0.095902333 | 0.52037856 | 0.1842934 | 0.85378329 | 0.99665574 |
| CLK1 | 27.5565771 | 0.423425528 | 0.33878848 | 1.24982269 | 0.21136433 | 0.91043197 |
| PPIL3 | 6.2165138 | -0.557165466 | 0.54582826 | -1.0207706 | 0.30736312 | 0.97376371 |
| NIF3L1 | 4.68767676 | -0.25039644 | 0.69823534 | -0.3586132 | 0.71988444 | 0.99665574 |
| ORC2 | 25.3265636 | -0.10891799 | 0.40920792 | -0.2661678 | 0.79010994 | 0.99665574 |
| AC005037.1 | 6.16919092 | -0.450260015 | 0.83245998 | -0.5408789 | 0.58859108 | 0.99665574 |

| | | | | | | |
|---|---|---|---|---|---|---|
| FAM126B | 33.9852877 | 0.134876905 | 0.31647881 | 0.42617989 | 0.66997677 | 0.99665574 |
| NDUFB3 | 19.0482071 | 0.693189675 | 0.44953528 | 1.54201394 | 0.12307021 | 0.82915695 |
| CFLAR | 45.3724916 | -0.44767001 | 0.29973519 | -1.4935517 | 0.13529283 | 0.84373478 |
| CFLAR-AS1 | 5.03513677 | -1.806389407 | 0.86432715 | -2.0899371 | 0.03662345 | 0.58591936 |
| CASP10 | 5.40605035 | -0.760638314 | 0.82062523 | -0.926901 | 0.35397796 | 0.98022903 |
| CASP8 | 10.3058641 | -1.112840332 | 0.5863504 | -1.8979101 | 0.05770793 | 0.67545594 |
| ALS2CR12 | 3.03116838 | -0.669591946 | 0.83888949 | -0.7981885 | 0.42476111 | 0.99518711 |
| TRAK2 | 24.1186757 | -0.49833641 | 0.47177854 | -1.0562931 | 0.29083433 | 0.96703132 |
| STRADB | 9.04081326 | -0.358781206 | 0.71638799 | -0.5008197 | 0.61649803 | 0.99665574 |
| TMEM237 | 2.79153275 | 1.269175777 | 0.89399474 | 1.41966806 | 0.15570434 | 0.8636413 |
| ALS2 | 17.9684174 | -0.61582419 | 0.38020955 | -1.6196968 | 0.10529743 | 0.79427477 |
| FZD7 | 5.1384523 | 0.29897466 | 1.17744448 | 0.25391826 | 0.7995587 | 0.99665574 |
| SUMO1 | 21.1371907 | -0.291780462 | 0.39096991 | -0.746299 | 0.4554868 | 0.99617422 |
| NOP58 | 22.1567168 | -0.046121926 | 0.3769636 | -0.1223511 | 0.90262093 | 0.99777713 |
| BMPR2 | 90.0453637 | -0.467760127 | 0.26168368 | -1.7875021 | 0.07385637 | 0.72810353 |
| FAM117B | 50.3752566 | 0.766876163 | 0.41262623 | 1.85852502 | 0.06309449 | 0.68630818 |
| ICA1L | 12.739692 | -0.528315235 | 0.49009296 | -1.0779899 | 0.28103829 | 0.96059387 |
| WDR12 | 10.2280762 | 0.021815453 | 0.57061998 | 0.03823114 | 0.96950339 | 0.99777713 |
| CARF | 15.155418 | 0.333955619 | 0.52891213 | 0.63140094 | 0.5277784 | 0.99665574 |
| NBEAL1 | 143.375719 | 0.331967114 | 0.25179574 | 1.31839844 | 0.1873703 | 0.89728868 |
| CYP20A1 | 31.5929733 | 0.026621662 | 0.28378647 | 0.09380878 | 0.92526106 | 0.99777713 |
| ABI2 | 37.9811529 | -0.129325184 | 0.29103196 | -0.4443676 | 0.6567768 | 0.99665574 |
| RAPH1 | 48.1573421 | -0.676288839 | 0.31292874 | -2.1611593 | 0.03068304 | 0.56271575 |
| PARD3B | 213.043473 | 0.580142629 | 0.27624239 | 2.10012167 | 0.03571814 | 0.58404919 |
| NRP2 | 12.8614222 | 0.34119351 | 0.54786006 | 0.62277493 | 0.53343243 | 0.99665574 |
| INO80D | 72.1354065 | 0.706249351 | 0.53505355 | 1.31996014 | 0.18684833 | 0.89605882 |
| NDUFS1 | 43.5758155 | 0.273500497 | 0.28843306 | 0.94822866 | 0.34301306 | 0.97788427 |
| EEF1B2 | 21.9101746 | 0.55188447 | 0.49229778 | 1.1210379 | 0.26227173 | 0.94557817 |
| ZDBF2 | 11.2823088 | 0.215104849 | 0.46992625 | 0.45774171 | 0.64713802 | 0.99665574 |
| FASTKD2 | 10.4779538 | 0.252419421 | 0.45243075 | 0.55791836 | 0.57690013 | 0.99665574 |
| KLF7 | 25.8374549 | -0.105184299 | 0.44578458 | -0.2359532 | 0.81346899 | 0.99665574 |
| LINC01802 | 5.42125432 | 1.38901009 | 0.78071391 | 1.77915376 | 0.07521456 | 0.73097508 |
| CREB1 | 23.5286781 | 0.265914486 | 0.36003357 | 0.73858247 | 0.46016057 | 0.99617422 |
| METTL21A | 6.17221456 | -0.257704291 | 0.58203326 | -0.4427656 | 0.65793531 | 0.99665574 |
| CCNYL1 | 8.35551351 | -0.223133893 | 0.72524014 | -0.307669 | 0.75833423 | 0.99665574 |
| PLEKHM3 | 14.5044593 | -0.16779399 | 0.48373952 | -0.3468685 | 0.72869013 | 0.99665574 |
| IDH1 | 9.54137856 | 1.226281962 | 0.61703807 | 1.98736841 | 0.04688159 | 0.64060657 |
| PIKFYVE | 28.2816229 | 0.282895604 | 0.41471434 | 0.6821457 | 0.49514683 | 0.99640678 |
| PTH2R | 60.0523088 | 1.178282509 | 0.59401268 | 1.98359825 | 0.04730064 | 0.64060657 |
| RPE | 12.7021402 | -0.115300965 | 0.46325822 | -0.2488914 | 0.80344482 | 0.99665574 |
| KANSL1L | 98.1674042 | -0.048146218 | 0.46030252 | -0.1045969 | 0.91669567 | 0.99777713 |
| AC007038.1 | 10.7779615 | -0.471696232 | 0.60658685 | -0.7776236 | 0.43679096 | 0.99617422 |
| ACADL | 9.02796869 | 0.010630039 | 0.64430195 | 0.01649854 | 0.98683667 | 0.99777713 |
| LANCL1 | 14.8144272 | -0.024839779 | 0.43354669 | -0.0572944 | 0.95431071 | 0.99777713 |
| ERBB4 | 330.897468 | -0.288466839 | 0.36256112 | -0.7956364 | 0.42624338 | 0.99518711 |
| IKZF2 | 11.6747699 | 0.20434991 | 0.78596252 | 0.25999956 | 0.79486411 | 0.99665574 |
| SPAG16 | 36.7027162 | -0.632702871 | 0.52302122 | -1.2097078 | 0.22639102 | 0.92318867 |
| AC068051.1 | 6.2304035 | -0.897708478 | 0.84939144 | -1.0568843 | 0.29056439 | 0.96676791 |
| BARD1 | 6.0898378 | 1.125092572 | 0.62340783 | 1.80474567 | 0.07111449 | 0.71523681 |
| AC016708.1 | 6.53396127 | -0.586877515 | 0.76779984 | -0.7643626 | 0.44465117 | 0.99617422 |
| ATIC | 9.00746936 | 0.132334805 | 0.5394078 | 0.2453335 | 0.80619821 | 0.99665574 |
| MREG | 11.4863034 | 0.712746534 | 0.47978977 | 1.48553925 | 0.13740104 | 0.84598921 |
| PECR | 17.3352496 | 0.293417338 | 0.45691488 | 0.64217067 | 0.52076237 | 0.99665574 |
| XRCC5 | 67.6843046 | -0.161574408 | 0.24647171 | -0.6555495 | 0.51211402 | 0.99665574 |
| SMARCAL1 | 6.74502125 | -0.178765503 | 0.58406518 | -0.3060712 | 0.75955047 | 0.99665574 |
| RPL37A | 55.7084537 | 0.631339609 | 0.42540479 | 1.48409143 | 0.13778467 | 0.84656101 |
| IGFBP2 | 2.93062488 | -0.656095748 | 1.00498406 | -0.6528419 | 0.51385818 | 0.99665574 |
| IGFBP5 | 935.612562 | -0.364606736 | 0.43196856 | -0.8440585 | 0.39863671 | 0.99179694 |
| AC007563.2 | 60.619955 | 0.552387253 | 0.59351697 | 0.9307017 | 0.35200789 | 0.97838306 |
| DIRC3-AS1 | 3.43314256 | 2.634554982 | 1.14293499 | 2.30507859 | 0.02116217 | 0.4733368 |
| DIRC3 | 16.7285147 | 0.113513468 | 0.71033192 | 0.15980342 | 0.87303593 | 0.99665574 |
| TNS1 | 35.458503 | 0.204772124 | 0.48017404 | 0.42645397 | 0.66977709 | 0.99665574 |
| ARPC2 | 32.104906 | -0.427914012 | 0.3421591 | -1.2506287 | 0.21106996 | 0.91043197 |
| AAMP | 8.15044831 | 0.396641636 | 0.55027669 | 0.72080399 | 0.47103012 | 0.99617422 |
| PNKD | 22.0980747 | -0.444720305 | 0.38264981 | -1.1622123 | 0.24514925 | 0.93426912 |
| TMBIM1 | 17.9560626 | -0.908359038 | 0.44751706 | -2.0297752 | 0.04237939 | 0.61641649 |
| CTDSP1 | 13.9411165 | 0.574466027 | 0.42696604 | 1.34546069 | 0.17847651 | 0.88813816 |
| USP37 | 24.4411901 | 0.006293842 | 0.35168384 | 0.01789631 | 0.98572157 | 0.99777713 |
| CNOT9 | 11.0106413 | -0.240554491 | 0.45882321 | -0.5242858 | 0.60007978 | 0.99665574 |
| PLCD4 | 6.40091424 | 1.277941619 | 0.79430946 | 1.60887121 | 0.10764451 | 0.80026315 |
| ZNF142 | 4.84395179 | 0.236139524 | 0.60613456 | 0.38958268 | 0.69684516 | 0.99665574 |
| STK36 | 2.55837048 | 0.080210013 | 0.82704582 | 0.09698376 | 0.92273929 | 0.99777713 |
| TTLL4 | 4.79427816 | -1.472941858 | 0.70478295 | -2.0899227 | 0.03662475 | 0.58591936 |
| CYP27A1 | 8.03983746 | 0.462342611 | 0.64413415 | 0.71777379 | 0.47289678 | 0.99617422 |
| CNPPD1 | 10.8184771 | -0.238675221 | 0.52503959 | -0.4545852 | 0.64940769 | 0.99665574 |
| RETREG2 | 13.5888533 | -0.150284714 | 0.49001909 | -0.3066915 | 0.75907817 | 0.99665574 |
| ZFAND2B | 5.18278666 | -0.010726264 | 0.61528048 | -0.0174331 | 0.98609108 | 0.99777713 |
| ANKZF1 | 5.68393971 | -0.331060483 | 0.58852781 | -0.5625231 | 0.57375968 | 0.99665574 |
| STK16 | 3.43948 | -1.028680765 | 0.70448009 | -1.4601985 | 0.14423553 | 0.84969914 |
| TUBA4A | 9.897005 | 0.340792104 | 0.67723566 | 0.50321051 | 0.61481627 | 0.99665574 |
| DNAJB2 | 10.5840471 | -0.033978558 | 0.49702004 | -0.0683646 | 0.94549543 | 0.99777713 |
| DNPEP | 5.14960525 | -0.363870798 | 0.6823337 | -0.533274 | 0.59384395 | 0.99665574 |
| GMPPA | 2.82648666 | -0.002951639 | 0.83736287 | -0.0035249 | 0.99718752 | 0.99884397 |
| CHPF | 6.05571018 | -0.78672921 | 0.6399399 | -1.2293798 | 0.21892943 | 0.91662582 |
| OBSL1 | 5.12474772 | -0.009917536 | 0.77350922 | -0.0128215 | 0.98977022 | 0.99777713 |
| SGPP2 | 63.5451785 | -0.269728186 | 0.38930037 | -0.6928537 | 0.4884014 | 0.99640678 |
| FARSB | 7.87320048 | 0.411142939 | 0.49580584 | 0.82924182 | 0.40696759 | 0.99250037 |
| ACSL3 | 68.4536075 | 0.296059076 | 0.35854548 | 0.82572251 | 0.40896152 | 0.99279547 |
| AP1S3 | 27.9116379 | 0.257772233 | 0.56840248 | 0.453503 | 0.65018658 | 0.99665574 |
| WDFY1 | 18.9559077 | -0.678691086 | 0.44816857 | -1.5143656 | 0.12993313 | 0.83778831 |
| MRPL44 | 4.91071705 | 0.01321967 | 0.61071759 | 0.02164613 | 0.98273024 | 0.99777713 |
| SERPINE2 | 5.996786 | -3.67149136 | 1.11222486 | -3.3010333 | 0.00096329 | 0.08492068 |
| CUL3 | 66.981088 | 0.505291699 | 0.25316047 | 1.99593444 | 0.04594106 | 0.63540308 |
| DOCK10 | 32.3784091 | 0.172027874 | 0.83911999 | 0.20500986 | 0.83756442 | 0.99665574 |
| IRS1 | 33.4459992 | 0.522624521 | 0.59865142 | 0.87300305 | 0.38266141 | 0.98914365 |
| RHBDD1 | 22.8928617 | 0.836008672 | 0.4034922 | 2.07193266 | 0.03827172 | 0.5945274 |
| COL4A4 | 67.8321164 | 0.542889021 | 0.53958609 | 1.00612122 | 0.31435726 | 0.97596515 |
| COL4A3 | 57.78214 | 0.572248577 | 0.47424592 | 1.20664944 | 0.22756718 | 0.92389989 |
| AC097662.1 | 9.56313206 | -0.536255684 | 0.88447646 | -0.6062973 | 0.54431736 | 0.99665574 |
| MFF | 29.2194317 | 0.403996028 | 0.40044357 | 1.00887131 | 0.31303635 | 0.97520189 |
| AGFG1 | 46.4773982 | 0.494140384 | 0.3576113 | 1.38178068 | 0.16703905 | 0.87495175 |
| DNER | 2.30512363 | 1.040460354 | 0.99592988 | 1.04471246 | 0.29615589 | 0.97286866 |
| TRIP12 | 74.6188068 | 0.101224682 | 0.22135409 | 0.45729754 | 0.6474572 | 0.99665574 |
| FBXO36 | 4.14623237 | -0.368595849 | 0.7322609 | -0.5033668 | 0.61470639 | 0.99665574 |
| SP110 | 18.2284137 | -1.031244501 | 0.80527935 | -1.2806047 | 0.20033256 | 0.90466435 |
| SP140L | 11.1312793 | -1.120406856 | 0.63508042 | -1.7641968 | 0.07769885 | 0.74141484 |
| SP100 | 24.7735812 | -0.075036853 | 0.46311104 | -0.1620278 | 0.87128398 | 0.99665574 |
| CAB39 | 76.1678946 | 0.195003829 | 0.34558078 | 0.56427857 | 0.57256457 | 0.99665574 |
| ITM2C | 20.1686352 | -0.268604488 | 0.48077304 | -0.5586929 | 0.57637132 | 0.99665574 |
| C2orf72 | 9.39992975 | 1.00699842 | 0.72007021 | 1.39847255 | 0.16197121 | 0.87065966 |
| PSMD1 | 36.6587654 | 0.240812027 | 0.34412007 | 0.6997907 | 0.48405802 | 0.99640678 |
| ARMC9 | 34.4381484 | -0.164848231 | 0.34648723 | -0.47577 | 0.63423824 | 0.99665574 |
| NCL | 34.9253766 | -0.204121391 | 0.40797616 | -0.5003268 | 0.61684501 | 0.99665574 |

| | | | | | | |
|---|---|---|---|---|---|---|
| PTMA | 118.95973 | 0.173648747 | 0.34386032 | 0.50499793 | 0.61356029 | 0.99665574 |
| PDE6D | 11.8464925 | -0.503256547 | 0.49376941 | -1.0192137 | 0.30810152 | 0.97376371 |
| COPS7B | 6.94115656 | 0.507886175 | 0.58487654 | 0.86836476 | 0.38519468 | 0.99051797 |
| DIS3L2 | 53.2271181 | 0.036386328 | 0.27525225 | 0.13219266 | 0.89483191 | 0.99777713 |
| EIF4E2 | 10.6386824 | -0.274105506 | 0.42969402 | -0.6379086 | 0.52353318 | 0.99665574 |
| EFHD1 | 83.7358522 | 0.376722677 | 0.28852119 | 1.30570195 | 0.19165393 | 0.89964233 |
| GIGYF2 | 109.714234 | 0.348177106 | 0.22917144 | 1.51928667 | 0.12869035 | 0.83778831 |
| ATG16L1 | 5.48076227 | -0.01439949 | 0.60402561 | -0.0238392 | 0.98098087 | 0.99777713 |
| DGKD | 8.60416092 | 0.591530908 | 0.50770053 | 1.16511777 | 0.24397133 | 0.93426912 |
| USP40 | 34.1637445 | 0.273507666 | 0.41281512 | 0.66254275 | 0.50762345 | 0.99665574 |
| SH3BP4 | 25.8034879 | -0.924167517 | 0.427691 | -2.1608299 | 0.03070848 | 0.56271575 |
| AGAP1 | 112.679846 | -0.520712614 | 0.25345567 | -2.0544524 | 0.03993193 | 0.60713077 |
| IQCA1 | 4.33655865 | -0.312250879 | 0.772602 | -0.4041549 | 0.68609882 | 0.99665574 |
| COPS8 | 6.09346577 | 0.093681849 | 0.58410008 | 0.16038664 | 0.87257651 | 0.99665574 |
| RAB17 | 20.9758654 | -0.184498298 | 0.34558218 | -0.5338768 | 0.59342679 | 0.99665574 |
| LRRFIP1 | 45.0613109 | -1.084047645 | 0.3994614 | -2.7137732 | 0.00665217 | 0.26446936 |
| ILKAP | 18.9391942 | 0.030881466 | 0.36244052 | 0.08520423 | 0.93209903 | 0.99777713 |
| PER2 | 7.70412387 | 0.015534019 | 0.62408142 | 0.02489101 | 0.9801419 | 0.99777713 |
| TRAF3IP1 | 13.137449 | 0.360122735 | 0.44412949 | 0.81085076 | 0.41745138 | 0.99392253 |
| ASB1 | 4.26317416 | -0.022095789 | 0.7085947 | -0.0311825 | 0.97512396 | 0.99777713 |
| HDAC4 | 37.7422466 | -0.564583955 | 0.4310022 | -1.3099329 | 0.19021854 | 0.8994569 |
| NDUFA10 | 34.4049918 | -0.017000642 | 0.30822582 | -0.0551564 | 0.95601382 | 0.99777713 |
| COPS9 | 9.49912801 | -0.08113549 | 0.5364769 | -0.1512376 | 0.87978827 | 0.99665574 |
| GPC1 | 3.61551261 | 0.353495812 | 0.80295232 | 0.44024508 | 0.65975961 | 0.99665574 |
| ANKMY1 | 7.04496628 | 0.885981424 | 0.6938519 | 1.27690278 | 0.20163657 | 0.90614933 |
| RNPEPL1 | 8.85588051 | -0.120529101 | 0.49760802 | -0.242217 | 0.80861206 | 0.99665574 |
| CAPN10 | 3.91009594 | -0.235666686 | 0.77564749 | -0.3038322 | 0.76125573 | 0.99665574 |
| SNED1 | 50.1961944 | 0.038882996 | 0.75501575 | 0.05149958 | 0.95892743 | 0.99777713 |
| AC005237.1 | 11.8049438 | -1.534840722 | 0.84937693 | -1.8070196 | 0.07075923 | 0.71378813 |
| MTERF4 | 11.8749602 | -0.007132927 | 0.48234467 | -0.014788 | 0.98820129 | 0.99777713 |
| PPP1R7 | 10.3147966 | -0.364350691 | 0.65569482 | -0.5556711 | 0.57843568 | 0.99665574 |
| HDLBP | 53.7990361 | -0.057882843 | 0.36823054 | -0.1571919 | 0.87509364 | 0.99665574 |
| 2-Sep | 52.691216 | -0.191363511 | 0.27320714 | -0.7004338 | 0.48365645 | 0.99640678 |
| FARP2 | 18.1992505 | -0.772214763 | 0.36344356 | -2.1247171 | 0.03361022 | 0.57831228 |
| STK25 | 10.9298211 | -0.008234626 | 0.46581468 | -0.0176779 | 0.98589581 | 0.99777713 |
| BOK | 8.1220277 | 0.806068311 | 0.55925032 | 1.44133725 | 0.14948943 | 0.85525834 |
| THAP4 | 15.4088201 | 0.095273447 | 0.42668733 | 0.22328633 | 0.82331267 | 0.99665574 |
| ATG4B | 19.6259397 | 0.331164462 | 0.39225248 | 0.84426352 | 0.39852217 | 0.99179694 |
| DTYMK | 2.42340743 | 0.473109618 | 1.03229867 | 0.45830691 | 0.64673196 | 0.99665574 |
| ING5 | 13.1845402 | 0.817747678 | 0.55475833 | 1.47406112 | 0.14046514 | 0.84969914 |
| D2HGDH | 13.4256018 | 0.405920327 | 0.46207739 | 0.87846828 | 0.37968964 | 0.98679664 |
| CHL1 | 25.900206 | 2.186603902 | 1.2955767 | 1.68774563 | 0.09146007 | 0.76595534 |
| LINC01266 | 10.659606 | -0.45106691 | 0.71237694 | -0.6331857 | 0.52661237 | 0.99665574 |
| CNTN4 | 92.7005374 | -0.338479459 | 0.44756642 | -0.7562664 | 0.44948947 | 0.99617422 |
| TRNT1 | 15.4612886 | -0.294599188 | 0.37954182 | -0.7761969 | 0.43763273 | 0.99617422 |
| CRBN | 14.1205319 | 0.48464722 | 0.52963281 | 0.91506269 | 0.36015872 | 0.9819332 |
| AC034195.1 | 15.2359178 | 0.737750459 | 0.66275974 | 1.11314917 | 0.26564437 | 0.94557817 |
| SUMF1 | 83.6862511 | 0.470457941 | 0.40608135 | 1.15853126 | 0.2466473 | 0.93441937 |
| SETMAR | 6.4463612 | 1.596382841 | 0.72745668 | 2.19447136 | 0.02820154 | 0.54458518 |
| ITPR1 | 51.4265556 | 1.507420012 | 0.43131153 | 3.49496807 | 0.00047412 | 0.05861723 |
| BHLHE40 | 54.2208152 | -0.853745921 | 0.70345594 | -1.2136452 | 0.22488324 | 0.92318867 |
| ARL8B | 31.1371642 | 0.38245447 | 0.40292609 | 0.94919261 | 0.34252266 | 0.97788427 |
| AC026202.2 | 4.37442725 | -0.755238138 | 0.93484091 | -0.8078788 | 0.41916037 | 0.99476033 |
| EDEM1 | 6.3104695 | 0.315668203 | 0.5893523 | 0.53561886 | 0.59222199 | 0.99665574 |
| LMCD1-AS1 | 12.3504135 | -0.579823931 | 0.6623704 | -0.8753772 | 0.38136872 | 0.98839606 |
| RAD18 | 3.89327376 | -0.876480687 | 0.83079533 | -1.0549899 | 0.29142994 | 0.96837651 |
| SRGAP3 | 36.2052248 | 0.430729159 | 0.5264192 | 0.81822463 | 0.41322893 | 0.99279547 |
| THUMPD3-A! | 23.129722 | -0.516849483 | 0.40762153 | -1.2679641 | 0.20481076 | 0.90834797 |
| THUMPD3 | 7.29254377 | -0.101203012 | 0.63402462 | -0.15962 | 0.87318042 | 0.99665574 |
| SETD5 | 67.8133328 | -0.464696015 | 0.24483734 | -1.8979785 | 0.05769891 | 0.67545594 |
| MTMR14 | 14.257654 | -0.124922244 | 0.39848935 | -0.3134895 | 0.75390877 | 0.99665574 |
| BRPF1 | 2.02946275 | -0.804854203 | 0.97125048 | -0.8286783 | 0.40728646 | 0.99279547 |
| OGG1 | 6.16777298 | -0.445926666 | 0.67150239 | -0.6640731 | 0.50664353 | 0.99665574 |
| CAMK1 | 11.2754002 | 0.002701475 | 0.45415943 | 0.0059483 | 0.99525397 | 0.99860301 |
| TADA3 | 11.3547829 | -0.511197309 | 0.54086257 | -0.9451519 | 0.34458131 | 0.97788427 |
| RPUSD3 | 3.5174179 | -0.254654508 | 0.90003293 | -0.2829391 | 0.77722352 | 0.99665574 |
| JAGN1 | 4.50094583 | -0.323227907 | 0.80329061 | -0.4023798 | 0.68740454 | 0.99665574 |
| IL17RC | 3.28948515 | -0.3533167 | 0.89337861 | -0.3954837 | 0.69248593 | 0.99665574 |
| CRELD1 | 5.00787117 | 0.219806531 | 0.66183666 | 0.33211598 | 0.73980168 | 0.99665574 |
| EMC3 | 11.5462658 | -0.086841545 | 0.51203219 | -0.1696017 | 0.86532336 | 0.99665574 |
| BRK1 | 13.4711583 | 0.282278094 | 0.47429934 | 0.59514756 | 0.55174484 | 0.99665574 |
| VHL | 7.12658493 | -0.694787778 | 0.5662259 | -1.2270505 | 0.21980361 | 0.91768188 |
| SEC13 | 10.5937082 | -1.133174282 | 0.55496694 | -2.0418771 | 0.04116372 | 0.61100706 |
| ATP2B2 | 5.40202025 | 1.399468691 | 0.92243847 | 1.51714042 | 0.12923124 | 0.83778831 |
| ATG7 | 58.8692767 | -0.378409136 | 0.42889843 | -0.8822815 | 0.37762462 | 0.98679664 |
| VGLL4 | 76.2348186 | 0.229041495 | 0.29128738 | 0.78630765 | 0.43168727 | 0.99617422 |
| TAMM41 | 17.03386 | -0.317933199 | 0.3847672 | -0.8263002 | 0.40863385 | 0.99279547 |
| PPARG | 93.4021919 | 0.91979886 | 0.43637733 | 2.10780623 | 0.03504775 | 0.58349146 |
| TSEN2 | 13.8194455 | -0.141191755 | 0.53257724 | -0.2651104 | 0.79092443 | 0.99665574 |
| MKRN2 | 4.04293067 | -0.034448617 | 0.8032813 | -0.0428849 | 0.96579331 | 0.99777713 |
| RAF1 | 40.3860775 | 0.156067279 | 0.26931279 | 0.57950193 | 0.56225055 | 0.99665574 |
| RPL32 | 75.686411 | 0.434366818 | 0.49341562 | 0.88032645 | 0.37868249 | 0.98679664 |
| IQSEC1 | 6.21639171 | -0.043114663 | 0.61117249 | -0.0705442 | 0.94376054 | 0.99777713 |
| NUP210 | 10.3323162 | 0.204917826 | 0.54767334 | 0.37416068 | 0.70828478 | 0.99665574 |
| HDAC11 | 4.1600137 | 0.192315339 | 0.71610726 | 0.26855661 | 0.7882709 | 0.99665574 |
| CHCHD4 | 2.16937254 | -0.462000821 | 0.9368883 | -0.4931226 | 0.62192594 | 0.99665574 |
| TMEM43 | 3.32746846 | -1.094335636 | 0.80309168 | -1.3626534 | 0.17299176 | 0.88157681 |
| XPC | 14.904172 | -0.420882841 | 0.37701579 | -1.1163533 | 0.26427092 | 0.94557817 |
| LSM3 | 11.4683372 | -0.119229583 | 0.51681394 | -0.2307012 | 0.81754696 | 0.99665574 |
| SLC6A6 | 10.7763627 | -1.141166236 | 0.73351118 | -1.5557585 | 0.11976553 | 0.82659025 |
| CCDC174 | 7.6495081 | 0.118279799 | 0.57352203 | 0.2062341 | 0.83660805 | 0.99665574 |
| FGD5-AS1 | 16.8359183 | -0.224368355 | 0.43527297 | -0.5154659 | 0.60622752 | 0.99665574 |
| NR2C2 | 27.2965179 | -0.178129098 | 0.35426329 | -0.5028156 | 0.61509395 | 0.99665574 |
| MRPS25 | 26.0425568 | 0.271774584 | 0.32483266 | 0.83666028 | 0.40278355 | 0.99194293 |
| RBSN | 7.08486891 | 0.40591693 | 0.62758717 | 0.64678972 | 0.51776805 | 0.99665574 |
| CAPN7 | 33.2916344 | -0.472269827 | 0.2984819 | -1.5822394 | 0.11359492 | 0.81582066 |
| METTL6 | 8.23615536 | -0.416994911 | 0.52311588 | -0.7971368 | 0.42537159 | 0.99518711 |
| EAF1 | 4.0534804 | 0.032291546 | 0.81405525 | 0.03966751 | 0.9683582 | 0.99777713 |
| COLQ | 3.14866166 | -0.700559933 | 0.78733327 | -0.8897883 | 0.37357957 | 0.98532838 |
| HACL1 | 12.9710071 | -0.20850364 | 0.49069951 | -0.424911 | 0.67090153 | 0.99665574 |
| BTD | 3.30375832 | 0.547176781 | 0.72954864 | 0.75002097 | 0.45324208 | 0.99617422 |
| ANKRD28 | 94.3276509 | 0.025588085 | 0.2622267 | 0.09758001 | 0.9222658 | 0.99777713 |
| DPH3 | 3.57133352 | -1.239769524 | 0.82473238 | -1.5032386 | 0.13277754 | 0.84089786 |
| OXNAD1 | 25.9717186 | 0.250640844 | 0.39805043 | 0.62967108 | 0.52890981 | 0.99665574 |
| RFTN1 | 38.9474913 | -0.050387293 | 0.47096858 | -0.1069865 | 0.91479967 | 0.99777713 |
| PLCL2 | 295.699916 | -1.190360979 | 0.60966026 | -1.9524989 | 0.050879 | 0.65127694 |
| TBC1D5 | 157.292843 | 0.345830243 | 0.33172491 | 1.04252117 | 0.29717011 | 0.97341216 |
| KCNH8 | 7.19823712 | -1.628139826 | 0.72397597 | -2.2488866 | 0.02451971 | 0.50422081 |
| RAB5A | 27.7548073 | -0.144757498 | 0.35399341 | -0.4089271 | 0.68259316 | 0.99665574 |
| KAT2B | 22.3974708 | 0.22905799 | 0.41698994 | 0.54931299 | 0.58279067 | 0.99665574 |
| ZNF385D | 8.65269947 | -1.16617784 | 0.90022701 | -1.2954264 | 0.19517317 | 0.90381759 |
| UBE2E2 | 176.459581 | 0.275895503 | 0.33436423 | 0.82513462 | 0.40929517 | 0.99279547 |
| UBE2E1 | 79.5196021 | 0.06468774 | 0.28764952 | 0.22488388 | 0.82206961 | 0.99665574 |

| | | | | | | |
|---|---|---|---|---|---|---|
| NKIRAS1 | 7.49967113 | 0.436105317 | 0.56416575 | 0.7730092 | 0.43951693 | 0.99617422 |
| RPL15 | 125.53268 | -0.35287325 | 0.41238099 | -0.8556972 | 0.3921653 | 0.99179694 |
| NR1D2 | 34.7102426 | -0.191069221 | 0.37540896 | -0.5089629 | 0.61077825 | 0.99665574 |
| THRB | 35.2330113 | -0.09946418 | 0.45076919 | -0.2206543 | 0.82536158 | 0.99665574 |
| RARB | 21.27223 | 0.813066339 | 0.3766517 | 2.15866897 | 0.03087586 | 0.56271575 |
| TOP2B | 36.8062998 | 0.205616106 | 0.29799438 | 0.68999994 | 0.49019423 | 0.99640678 |
| NGLY1 | 24.0789532 | -0.125297514 | 0.37855074 | -0.3309927 | 0.74065004 | 0.99665574 |
| OXSM | 3.07260752 | 0.434262033 | 0.88566165 | 0.49032498 | 0.62390395 | 0.99665574 |
| NEK10 | 8.58737629 | 0.096118043 | 0.78819536 | 0.12194698 | 0.90294101 | 0.99777713 |
| SLC4A7 | 16.1756131 | -1.126679679 | 0.62025088 | -1.8164903 | 0.0692952 | 0.7094165 |
| CMC1 | 41.0295386 | 0.484197142 | 0.37442027 | 1.29319157 | 0.19594481 | 0.90381759 |
| AZI2 | 17.2963103 | -0.074264023 | 0.43042221 | -0.1725376 | 0.86301489 | 0.99665574 |
| TGFBR2 | 12.5158387 | -0.550800445 | 0.89659358 | -0.6143257 | 0.53900015 | 0.99665574 |
| STT3B | 57.5386636 | 0.822285045 | 0.38312603 | 2.1462521 | 0.03185287 | 0.56852889 |
| OSBPL10 | 41.7110694 | -0.898297166 | 0.5512298 | -1.6296237 | 0.10318104 | 0.78828139 |
| GPD1L | 75.6130123 | 0.301405493 | 0.29469218 | 1.02278077 | 0.30641151 | 0.97376371 |
| CMTM8 | 9.89341497 | -0.057037326 | 0.49192738 | -0.1159466 | 0.90769483 | 0.99777713 |
| CMTM7 | 7.34465468 | -1.832381657 | 0.5333779 | -3.4354285 | 0.00059162 | 0.06631714 |
| CMTM6 | 21.6758518 | -0.338347082 | 0.40572284 | -0.8339365 | 0.40431677 | 0.99194293 |
| DYNC1LI1 | 10.9022053 | -0.068815203 | 0.58626814 | -0.1173784 | 0.90656022 | 0.99777713 |
| CNOT10 | 18.4116552 | 0.240213555 | 0.43988505 | 0.54608256 | 0.58500919 | 0.99665574 |
| GLB1 | 20.2167497 | 0.223198057 | 0.35423001 | 0.63009359 | 0.52863335 | 0.99665574 |
| CRTAP | 8.77732525 | 0.135217114 | 0.49344431 | 0.2740271 | 0.78406379 | 0.99665574 |
| FBXL2 | 32.2832478 | -0.699700348 | 0.37597666 | -1.8610207 | 0.06274125 | 0.68628329 |
| UBP1 | 20.8298092 | 0.466047741 | 0.45708766 | 1.01960255 | 0.30791699 | 0.97376371 |
| CLASP2 | 69.0669764 | 0.137631044 | 0.33488367 | 0.41098165 | 0.68108599 | 0.99665574 |
| PDCD6IP | 51.8376037 | -0.006770294 | 0.30047592 | -0.0225319 | 0.98202366 | 0.99777713 |
| STAC | 19.1564695 | 0.332433071 | 0.46551009 | 0.71412645 | 0.47514899 | 0.99617422 |
| TRANK1 | 13.3861525 | -1.604560416 | 0.63740472 | -2.5173337 | 0.01182468 | 0.37359313 |
| EPM2AIP1 | 16.7218622 | 0.367428317 | 0.42640765 | 0.86168323 | 0.38886186 | 0.99176253 |
| MLH1 | 6.71013022 | 0.546751743 | 0.63861006 | 0.85615899 | 0.39190984 | 0.99179694 |
| LRRFIP2 | 106.450803 | -0.782819294 | 0.29109152 | -2.6892549 | 0.00716117 | 0.27640407 |
| GOLGA4 | 121.254779 | -0.285025526 | 0.37785763 | -0.7543199 | 0.45065716 | 0.99617422 |
| ITGA9 | 18.0958843 | -0.922949441 | 0.70774835 | -1.3040644 | 0.19221162 | 0.89964233 |
| CTDSPL | 131.287647 | 0.124669308 | 0.37843318 | 0.32943546 | 0.74182657 | 0.99665574 |
| PLCD1 | 3.3206326 | 0.507204786 | 0.80469903 | 0.63030371 | 0.52849589 | 0.99665574 |
| ACAA1 | 13.2126733 | 0.29626244 | 0.48027846 | 0.61685557 | 0.53733 | 0.99665574 |
| OXSR1 | 32.7721271 | -0.129495629 | 0.30660439 | -0.4223541 | 0.67276656 | 0.99665574 |
| XYLB | 2.60698622 | 0.593335277 | 0.93463322 | 0.63483222 | 0.52553785 | 0.99665574 |
| ACVR2B | 6.23455801 | 0.100439373 | 0.62058176 | 0.16184713 | 0.87142624 | 0.99665574 |
| EXOG | 6.57234071 | -0.201633267 | 0.578998 | -0.3482452 | 0.72765606 | 0.99665574 |
| WDR48 | 13.7210033 | -0.277742287 | 0.4786501 | -0.5802616 | 0.5617382 | 0.99665574 |
| GORASP1 | 5.9312802 | 0.247702832 | 0.57404666 | 0.43150296 | 0.6661027 | 0.99665574 |
| TTC21A | 3.16772745 | 0.735735392 | 0.88339767 | 0.83284734 | 0.40493083 | 0.99194293 |
| SLC25A38 | 6.18631051 | 0.352509553 | 0.59710103 | 0.59036836 | 0.55494372 | 0.99665574 |
| RPSA | 53.8500153 | 0.733274552 | 0.36923752 | 1.9859156 | 0.0470427 | 0.64060657 |
| MYRIP | 166.634022 | -0.901205372 | 0.52791953 | -1.7070885 | 0.0878056 | 0.75650018 |
| EIF1B | 15.2238042 | -0.835694011 | 0.44015951 | -1.8986163 | 0.05761495 | 0.67545594 |
| ENTPD3-AS1 | 4.86841452 | 0.29681003 | 0.70842151 | 0.41897377 | 0.6752353 | 0.99665574 |
| RPL14 | 102.506063 | 0.203543536 | 0.48537259 | 0.41935524 | 0.67495653 | 0.99665574 |
| ZNF621 | 13.3638834 | -0.395641812 | 0.45215833 | -0.8750072 | 0.38156997 | 0.98844371 |
| CTNNB1 | 33.8478319 | 0.330240877 | 0.32033401 | 1.03092669 | 0.3025752 | 0.97358745 |
| ULK4 | 34.2495429 | -0.7188179 | 0.36248166 | -1.9830463 | 0.04736226 | 0.64060657 |
| TRAK1 | 48.488137 | -0.783189541 | 0.27524554 | -2.8454213 | 0.00443527 | 0.21310323 |
| SEC22C | 8.39754619 | 0.277518935 | 0.66193457 | 0.41925433 | 0.67503027 | 0.99665574 |
| SS18L2 | 4.20500716 | 1.042152394 | 0.96619831 | 1.07861127 | 0.28076105 | 0.96059387 |
| NKTR | 72.7977809 | -0.241679469 | 0.25238033 | -0.9576003 | 0.33826437 | 0.97788427 |
| AC006059.1 | 7.33009782 | -1.623770977 | 0.82698563 | -1.9634815 | 0.04959025 | 0.64444758 |
| AC099329.3 | 4.77241533 | -0.832294913 | 0.68313007 | -1.218355 | 0.22308911 | 0.92191316 |
| SNRK | 25.0590822 | 0.138949536 | 0.48462096 | 0.28671797 | 0.77432828 | 0.99665574 |
| ANO10 | 38.3821448 | -0.113140602 | 0.30826639 | -0.3670222 | 0.71360248 | 0.99665574 |
| ABHD5 | 9.61596574 | -0.79729336 | 0.47084266 | -1.6933329 | 0.09039212 | 0.76326341 |
| TCAIM | 46.4133115 | 0.762617647 | 0.45678516 | 1.66953243 | 0.09501191 | 0.77178333 |
| ZNF445 | 12.9898197 | 0.921143065 | 0.44153717 | 2.08621862 | 0.03695882 | 0.58591936 |
| ZNF852 | 4.09305827 | 0.189029323 | 0.6641757 | 0.28460741 | 0.77594494 | 0.99665574 |
| AC099669.1 | 13.1045776 | -0.252453875 | 0.47062428 | -0.5364234 | 0.59166597 | 0.99665574 |
| ZNF197 | 8.05089861 | -0.250973923 | 0.50385456 | -0.4981079 | 0.61840801 | 0.99665574 |
| ZNF35 | 2.97400479 | -0.189842699 | 1.02542902 | -0.1851349 | 0.85312323 | 0.99665574 |
| ZNF502 | 3.38270398 | 0.65719532 | 0.82867016 | 0.79307227 | 0.42773572 | 0.99589919 |
| KIAA1143 | 4.83748819 | -0.256072358 | 0.63822949 | -0.401223 | 0.68825594 | 0.99665574 |
| ZDHHC3 | 9.6150677 | -0.188973146 | 0.4560442 | -0.4143746 | 0.67859977 | 0.99665574 |
| EXOSC7 | 4.03575682 | -0.690156313 | 0.64309542 | -1.0731787 | 0.28319094 | 0.96059387 |
| CDCP1 | 13.2119851 | -0.810917992 | 0.80186573 | -1.011289 | 0.31187813 | 0.97442101 |
| LARS2 | 18.7132722 | -0.268856408 | 0.43989793 | -0.6111791 | 0.54108104 | 0.99665574 |
| LIMD1 | 14.1117582 | 0.112405227 | 0.46154906 | 0.24353907 | 0.80758783 | 0.99665574 |
| SACM1L | 20.7052953 | 0.215824574 | 0.42339933 | 0.50974236 | 0.61023197 | 0.99665574 |
| LZTFL1 | 4.68520545 | 0.683678714 | 0.94156223 | 0.72611103 | 0.46777071 | 0.99617422 |
| FYCO1 | 11.1088113 | -0.446473603 | 0.48876205 | -0.9134785 | 0.36099095 | 0.9819332 |
| LTF | 24.219742 | -1.272000555 | 1.40515198 | -0.9052406 | 0.36533796 | 0.98365837 |
| ALS2CL | 7.82166931 | 0.213384085 | 0.56401078 | 0.37833335 | 0.70518298 | 0.99665574 |
| PTH1R | 14.5605108 | 1.251979135 | 0.79917161 | 1.56659611 | 0.11720913 | 0.82404032 |
| CCDC12 | 27.8454679 | -0.239558325 | 0.29354873 | -0.8160768 | 0.41445618 | 0.99279547 |
| NBEAL2 | 6.52208724 | 0.545484378 | 0.56717795 | 0.96175174 | 0.33617433 | 0.97788427 |
| SETD2 | 107.192405 | 0.316936041 | 0.25769508 | 1.22988783 | 0.21873911 | 0.91662582 |
| KIF9-AS1 | 24.2944559 | 0.03375446 | 0.43085407 | 0.07834314 | 0.9375551 | 0.99777713 |
| KLHL18 | 12.8028154 | -0.243410116 | 0.46913366 | -0.5188503 | 0.60386517 | 0.99665574 |
| PTPN23 | 6.78198645 | 0.080620771 | 0.65769914 | 0.12258002 | 0.90243968 | 0.99777713 |
| SCAP | 19.0608615 | 0.408445149 | 0.37919737 | 1.07713075 | 0.28142186 | 0.96059387 |
| ELP6 | 6.12332281 | 0.110549105 | 0.62255039 | 0.17757455 | 0.85905711 | 0.99665574 |
| SMARCC1 | 85.5943321 | -0.248948222 | 0.27396141 | -0.9086981 | 0.36350949 | 0.98362663 |
| DHX30 | 12.5518465 | 0.999230414 | 0.43248433 | 2.31044304 | 0.02086364 | 0.47318919 |
| MAP4 | 112.233413 | -0.339081333 | 0.28082923 | -1.2074289 | 0.227267 | 0.92389989 |
| ZNF589 | 3.17501795 | 0.177647328 | 0.80042675 | 0.22194077 | 0.82435999 | 0.99665574 |
| NME6 | 4.65299219 | -0.342936445 | 0.65589732 | -0.5228508 | 0.60107808 | 0.99665574 |
| PLXNB1 | 16.9201649 | 0.421433728 | 0.49312956 | 0.85461056 | 0.39276678 | 0.99179694 |
| TMA7 | 26.8426717 | 0.353099735 | 0.52808613 | 0.66864043 | 0.50372488 | 0.99665574 |
| SHISA5 | 12.8995092 | 0.035992476 | 0.53841084 | 0.06684947 | 0.94670154 | 0.99777713 |
| UQCRC1 | 17.8059115 | 0.65682981 | 0.42438579 | 1.54771865 | 0.12169005 | 0.82687246 |
| NCKIPSD | 2.34200416 | -0.352059867 | 1.03039134 | -0.3416759 | 0.73259483 | 0.99665574 |
| IP6K2 | 22.9246814 | -0.23131481 | 0.47617859 | -0.4857732 | 0.62712796 | 0.99665574 |
| PRKAR2A | 34.5578978 | -0.152207318 | 0.34251637 | -0.4443797 | 0.65676809 | 0.99665574 |
| SLC25A20 | 5.38487772 | 0.212521945 | 0.74758949 | 0.28427626 | 0.77619868 | 0.99665574 |
| ARIH2 | 32.5206388 | 0.194877272 | 0.33692715 | 0.57839587 | 0.56299688 | 0.99665574 |
| P4HTM | 6.97135475 | 0.221910979 | 0.61424007 | 0.36127727 | 0.71789218 | 0.99665574 |
| WDR6 | 19.7022966 | 0.526099159 | 0.50988691 | 1.03179577 | 0.3021678 | 0.97358745 |
| DALRD3 | 4.5367909 | 1.400655634 | 0.82113321 | 1.70575934 | 0.0880529 | 0.75650018 |
| NDUFAF3 | 7.93084329 | 0.174475785 | 0.56951203 | 0.30636014 | 0.75933045 | 0.99665574 |
| IMPDH2 | 6.93852457 | 1.087587527 | 0.59064337 | 1.84136077 | 0.06556871 | 0.69221233 |
| QRICH1 | 19.4490574 | -0.300704275 | 0.37713502 | -0.7973385 | 0.42525446 | 0.99518711 |
| QARS | 7.56838828 | 0.490427618 | 0.58821354 | 0.83375779 | 0.40441749 | 0.99194293 |
| USP19 | 4.33157458 | -0.222781081 | 0.83561144 | -0.2666085 | 0.78977064 | 0.99665574 |
| LAMB2 | 8.29157911 | -0.008042403 | 0.65416281 | -0.0122942 | 0.9901909 | 0.99777713 |
| KLHDC8B | 2.9673965 | 0.386211856 | 0.90234799 | 0.42800766 | 0.66864554 | 0.99665574 |

| | | | | | | |
|---|---|---|---|---|---|---|
| C3orf62 | 2.32967766 | -0.470575496 | 1.03497885 | -0.4546716 | 0.64934551 | 0.99665574 |
| USP4 | 25.1057186 | -0.587705954 | 0.46099733 | -1.2748576 | 0.20235965 | 0.90614933 |
| RHOA | 46.0707009 | -0.3297692 | 0.36797792 | -0.8961657 | 0.37016426 | 0.98532838 |
| TCTA | 6.25569143 | -0.042353861 | 0.56575007 | -0.0748632 | 0.94032356 | 0.99777713 |
| DAG1 | 18.1830813 | -0.584327706 | 0.42240296 | -1.3833419 | 0.16656006 | 0.87495175 |
| APEH | 7.79831145 | 0.404121426 | 0.64319685 | 0.62830132 | 0.52980657 | 0.99665574 |
| MST1 | 4.2570648 | 1.550208615 | 0.81570055 | 1.90046289 | 0.0573724 | 0.67545594 |
| RNF123 | 3.89120138 | 0.277297468 | 0.72167111 | 0.38424355 | 0.70079794 | 0.99665574 |
| IP6K1 | 28.566503 | 0.245057379 | 0.48927298 | 0.50086023 | 0.61646949 | 0.99665574 |
| RBM6 | 119.882389 | 0.091376742 | 0.24782394 | 0.36871636 | 0.71233915 | 0.99665574 |
| RBM5 | 36.2179791 | 0.076504052 | 0.29168338 | 0.26228457 | 0.79310205 | 0.99665574 |
| SEMA3F | 14.4631449 | 0.659479442 | 0.46413071 | 1.42089162 | 0.15534827 | 0.86297028 |
| GNAI2 | 15.0598401 | -0.718770159 | 0.47468712 | -1.5141977 | 0.1299757 | 0.83778831 |
| SEMA3B | 8.82751526 | -0.290137181 | 0.54710051 | -0.5303179 | 0.59589155 | 0.99665574 |
| IFRD2 | 3.76602035 | 0.090998502 | 0.71842602 | 0.12666371 | 0.89920657 | 0.99777713 |
| HYAL2 | 3.7771728 | 0.451806702 | 0.9149804 | 0.49378839 | 0.62145563 | 0.99665574 |
| TUSC2 | 4.5863413 | -0.016481054 | 0.72294288 | -0.0227972 | 0.98181206 | 0.99777713 |
| RASSF1 | 3.03951675 | -1.215463432 | 1.1300705 | -1.0755642 | 0.28212218 | 0.96059387 |
| TMEM115 | 3.9474349 | -0.424955149 | 0.68008529 | -0.6248557 | 0.53206579 | 0.99665574 |
| HEMK1 | 4.74127544 | 0.04337873 | 0.66121575 | 0.0656045 | 0.9476927 | 0.99777713 |
| CISH | 6.55446415 | 0.270869092 | 0.6784103 | 0.39927031 | 0.68969404 | 0.99665574 |
| MAPKAPK3 | 8.23404292 | -0.298930625 | 0.57419316 | -0.5206099 | 0.60263857 | 0.99665574 |
| DOCK3 | 38.2987246 | -0.864246737 | 0.50383134 | -1.7153493 | 0.0862812 | 0.75397496 |
| MANF | 9.24591285 | -0.866634651 | 0.62962304 | -1.3764341 | 0.16868727 | 0.87566218 |
| RBM15B | 6.20646954 | -0.082086755 | 0.90420507 | -0.0907833 | 0.92766475 | 0.99777713 |
| DCAF1 | 20.2215276 | 0.066159825 | 0.40277849 | 0.16425858 | 0.86952758 | 0.99665574 |
| RAD54L2 | 25.9650953 | 0.209988978 | 0.37022717 | 0.56718953 | 0.57058542 | 0.99665574 |
| TEX264 | 19.7042625 | 0.15378845 | 0.39854844 | 0.38587141 | 0.6995919 | 0.99665574 |
| PCBP4 | 3.01919942 | 0.896754784 | 0.87895095 | 1.02025578 | 0.30760717 | 0.97376371 |
| RPL29 | 94.4147903 | -0.304224921 | 0.43040689 | -0.706831 | 0.47967153 | 0.99617422 |
| ALAS1 | 6.29542137 | 0.505160603 | 0.64758253 | 0.78007139 | 0.43534885 | 0.99617422 |
| WDR82 | 18.5248421 | 0.140633047 | 0.36568124 | 0.38457823 | 0.70054992 | 0.99665574 |
| BAP1 | 4.67673182 | -0.0140634 | 0.77654598 | -0.0181102 | 0.98555094 | 0.99777713 |
| PHF7 | 2.89847143 | 0.084588739 | 0.89729694 | 0.09427062 | 0.92489419 | 0.99777713 |
| NISCH | 14.5277432 | -0.18676987 | 0.45291149 | -0.4123761 | 0.68006378 | 0.99665574 |
| SMIM4 | 11.1852911 | -0.677907789 | 0.51192821 | -1.3242243 | 0.18542855 | 0.89532394 |
| PBRM1 | 52.0868428 | -0.403862057 | 0.39612401 | -1.0195344 | 0.30794932 | 0.97376371 |
| GNL3 | 9.74125452 | -0.359421923 | 0.70778261 | -0.507814 | 0.61158378 | 0.99665574 |
| GLT8D1 | 3.24176497 | 0.038677885 | 0.9605811 | 0.04026509 | 0.96788178 | 0.99777713 |
| SPCS1 | 20.6902537 | 0.112731576 | 0.45325329 | 0.24871651 | 0.80358008 | 0.99665574 |
| NEK4 | 12.5899499 | 0.11694174 | 0.44437578 | 0.26315958 | 0.79242759 | 0.99665574 |
| SFMBT1 | 13.2233819 | 0.287972348 | 0.42548964 | 0.67680225 | 0.49853143 | 0.99640678 |
| RFT1 | 2.51492394 | 0.341167969 | 0.8860682 | 0.38503579 | 0.7002109 | 0.99665574 |
| PRKCD | 19.9935688 | 0.949865466 | 0.53251092 | 1.78374832 | 0.07446457 | 0.72810353 |
| TKT | 20.0629583 | -0.980580291 | 0.43665333 | -2.2456723 | 0.024725 | 0.50638808 |
| DCP1A | 27.5351627 | -0.341405433 | 0.3467563 | -0.9845688 | 0.32483593 | 0.97700345 |
| CACNA1D | 58.5484305 | -0.517267696 | 0.43095104 | -1.2002934 | 0.2300254 | 0.92774608 |
| CHDH | 8.70727468 | -0.786945135 | 0.75458807 | -1.0428804 | 0.29700367 | 0.97341216 |
| IL17RB | 3.7973049 | -0.752908597 | 1.27919605 | -0.5885795 | 0.55614338 | 0.99665574 |
| ACTR8 | 5.23238277 | 1.041771048 | 0.68157909 | 1.52846685 | 0.12639667 | 0.83642913 |
| SELENOK | 10.2147877 | -0.451330863 | 0.54309615 | -0.8310331 | 0.40595495 | 0.99194293 |
| CACNA2D3 | 31.327614 | 0.506987191 | 0.63246858 | 0.8016006 | 0.42278403 | 0.99518711 |
| ERC2 | 17.5811573 | -1.092461745 | 0.41524414 | -2.6308902 | 0.00851615 | 0.31119582 |
| CCDC66 | 24.6674227 | 0.163342852 | 0.41126173 | 0.39717494 | 0.69123846 | 0.99665574 |
| FAM208A | 30.4920404 | 0.026273597 | 0.39057227 | 0.06726949 | 0.94636717 | 0.99777713 |
| ARHGEF3 | 92.9681272 | -1.997703306 | 0.52066732 | -3.8368133 | 0.00012464 | 0.02485289 |
| APPL1 | 21.1931965 | 1.007484399 | 0.39899358 | 2.52506417 | 0.01156772 | 0.3725572 |
| PDE12 | 7.59123618 | -0.083485755 | 0.69609231 | -0.1199349 | 0.90453472 | 0.99777713 |
| ARF4 | 21.5346851 | -0.607501937 | 0.50680639 | -1.1986864 | 0.2306499 | 0.92774608 |
| DENND6A | 25.5335179 | -0.480083837 | 0.35771211 | -1.3420956 | 0.17956501 | 0.88820302 |
| DENND6A-AS | 5.56146305 | -0.932874571 | 0.81597018 | -1.1432704 | 0.25292633 | 0.94076968 |
| SLMAP | 75.3612918 | -0.161523649 | 0.30754729 | -0.5251994 | 0.59944459 | 0.99665574 |
| FLNB | 71.596024 | -0.085828348 | 0.30931718 | -0.2774768 | 0.781414 | 0.99665574 |
| ABHD6 | 3.74039217 | 0.999596946 | 0.89672256 | 1.11472265 | 0.26496929 | 0.94557817 |
| RPP14 | 3.22095231 | 0.509982533 | 0.78485327 | 0.64978074 | 0.51583386 | 0.99665574 |
| PXK | 82.9051715 | 1.074932062 | 0.45286841 | 2.37360795 | 0.01761524 | 0.43770425 |
| PDHB | 10.899904 | 0.855221844 | 0.59989705 | 1.42561436 | 0.15397969 | 0.86196782 |
| KCTD6 | 3.10627809 | 0.016697732 | 0.91331823 | 0.01828249 | 0.9854135 | 0.99777713 |
| FAM107A | 11.9082325 | 1.209162718 | 0.68153534 | 1.77417464 | 0.07603427 | 0.73482879 |
| FHIT | 632.05105 | 0.419905092 | 0.46187957 | 0.90912245 | 0.36328549 | 0.98362663 |
| PTPRG | 179.210956 | -0.527716914 | 0.44419169 | -1.1880387 | 0.23481817 | 0.92960234 |
| THOC7 | 10.2477371 | 0.351815805 | 0.56976185 | 0.6174787 | 0.53691903 | 0.99665574 |
| ATXN7 | 7.64514238 | 1.511228258 | 0.66313334 | 2.27892063 | 0.02267178 | 0.48762481 |
| PSMD6-AS2 | 3.02533327 | -0.328742594 | 0.78796343 | -0.4172054 | 0.67652818 | 0.99665574 |
| PSMD6 | 5.2721314 | 1.005038635 | 0.61584184 | 1.63197524 | 0.1026847 | 0.78770791 |
| PRICKLE2 | 18.0354656 | 0.152533222 | 0.47532445 | 0.32090338 | 0.74828361 | 0.99665574 |
| ADAMTS9 | 15.3381594 | -1.000336806 | 0.69956538 | -1.4299404 | 0.15273412 | 0.86116133 |
| ADAMTS9-AS | 14.6584681 | 0.732095969 | 0.76237978 | 0.96027727 | 0.33691569 | 0.97788427 |
| ADAMTS9-AS | 55.8902703 | 0.218725295 | 0.76790781 | 0.28483275 | 0.77577228 | 0.99665574 |
| MAGI1 | 2078.5482 | -0.165348224 | 0.23618281 | -0.7000858 | 0.48387375 | 0.99640678 |
| MAGI1-AS1 | 33.2969552 | -0.7640514 | 0.63467714 | -1.2038426 | 0.22865041 | 0.92646258 |
| SLC25A26 | 24.2564391 | -0.639551709 | 0.43808483 | -1.459881 | 0.14432278 | 0.84969914 |
| LRIG1 | 5.09781459 | 0.026782199 | 0.70654021 | 0.03790612 | 0.96976253 | 0.99777713 |
| SUCLG2 | 52.6991243 | -0.084084656 | 0.3102207 | -0.2710479 | 0.78635422 | 0.99665574 |
| SUCLG2-AS1 | 30.6173572 | 0.851976392 | 0.62160201 | 1.37061396 | 0.17049533 | 0.87918701 |
| FAM19A1 | 7.41803797 | 0.138221719 | 0.65021536 | 0.21257837 | 0.83165585 | 0.99665574 |
| EOGT | 3.97297907 | -0.219888466 | 0.76380103 | -0.2878871 | 0.77343316 | 0.99665574 |
| TMF1 | 26.1712129 | -0.521898885 | 0.46756375 | -1.1162091 | 0.26433266 | 0.94557817 |
| UBA3 | 17.884959 | -0.135236233 | 0.38513038 | -0.351144 | 0.7254803 | 0.99665574 |
| ARL6IP5 | 19.7497489 | -0.663487803 | 0.45415288 | -1.4609349 | 0.1440333 | 0.84969914 |
| FRMD4B | 12.4767405 | -0.143429431 | 0.57508977 | -0.2494036 | 0.80304864 | 0.99665574 |
| MITF | 77.4857832 | 0.785712964 | 0.39840534 | 1.97214466 | 0.0485931 | 0.64337675 |
| FOXP1 | 638.096548 | -0.140127005 | 0.28280122 | -0.4954965 | 0.62024971 | 0.99665574 |
| EIF4E3 | 6.11385733 | -1.101124712 | 0.56225608 | -1.9584043 | 0.05018259 | 0.64726607 |
| RYBP | 32.4118197 | 0.367470212 | 0.40757447 | 0.90160263 | 0.36726799 | 0.98449574 |
| SHQ1 | 11.7408747 | -0.026280413 | 0.43492961 | -0.0604245 | 0.95181753 | 0.99777713 |
| GXYLT2 | 4.86045927 | -1.133628121 | 0.74545066 | -1.5207286 | 0.12832797 | 0.83778831 |
| PPP4R2 | 33.4480316 | 0.262953762 | 0.36988005 | 0.71091632 | 0.47713609 | 0.99617422 |
| PDZRN3 | 55.3120206 | -1.285791062 | 0.56624833 | -2.2707194 | 0.02316397 | 0.49335204 |
| CNTN3 | 14.4727327 | -1.093843731 | 0.63656829 | -1.7183447 | 0.08573377 | 0.75373899 |
| ZNF717 | 10.7849994 | 0.040704207 | 0.64170793 | 0.06343105 | 0.94942326 | 0.99777713 |
| ROBO2 | 5.18462022 | -1.333409041 | 1.07157911 | -1.2443403 | 0.21337435 | 0.91189898 |
| GBE1 | 67.2926978 | 0.066159069 | 0.38768405 | 0.17065202 | 0.86449739 | 0.99665574 |
| CHMP2B | 13.5622987 | 0.300493221 | 0.48910454 | 0.61437422 | 0.53896807 | 0.99665574 |
| CGGBP1 | 43.8360642 | 0.096019095 | 0.34816744 | 0.27578425 | 0.78271378 | 0.99665574 |
| ZNF654 | 56.9405593 | -0.221123974 | 0.31233856 | -0.7079624 | 0.47896858 | 0.99617422 |
| C3orf38 | 5.53612145 | 0.626245046 | 0.6324012 | 0.99026543 | 0.3220444 | 0.97700345 |
| ARL13B | 6.52389682 | -0.708057017 | 0.56212794 | -1.2596012 | 0.20781327 | 0.91026892 |
| NSUN3 | 12.6636602 | -0.156812617 | 0.40790237 | -0.3844366 | 0.70065485 | 0.99665574 |
| MTRNR2L12 | 115.697749 | -0.254756492 | 0.53348292 | -0.4775345 | 0.63298157 | 0.99665574 |
| ARL6 | 11.9049377 | -0.047098103 | 0.53712836 | -0.087685 | 0.93012704 | 0.99777713 |
| CRYBG3 | 47.5134353 | -0.290058943 | 0.351944 | -0.8241622 | 0.40984741 | 0.99279547 |
| RIOX2 | 10.89346 | -0.172856008 | 0.58213223 | -0.296936 | 0.76651538 | 0.99665574 |

| | | | | | | |
|---|---|---|---|---|---|---|
| CLDND1 | 8.56853531 | -0.172344455 | 0.48564912 | -0.3548744 | 0.72268367 | 0.99665574 |
| CPOX | 2.26823678 | -0.232244599 | 0.84405138 | -0.2751546 | 0.78319749 | 0.99665574 |
| ST3GAL6 | 41.1523703 | 0.703604495 | 0.47113219 | 1.49343328 | 0.13532381 | 0.84373478 |
| DCBLD2 | 15.1269854 | 0.039787174 | 0.40026967 | 0.09940092 | 0.92081995 | 0.99777713 |
| CMSS1 | 30.9927132 | -0.765731894 | 0.35881895 | -2.1340341 | 0.03283999 | 0.5740203 |
| FILIP1L | 7.16871612 | -1.653094503 | 0.65923026 | -2.5076132 | 0.01215496 | 0.37684104 |
| TBC1D23 | 17.5425026 | -0.479127021 | 0.35191211 | -1.3614962 | 0.17335692 | 0.88157681 |
| NIT2 | 5.71915604 | -0.647407836 | 0.58794929 | -1.1011287 | 0.27084065 | 0.94865166 |
| TOMM70 | 11.5080742 | -0.740070154 | 0.46083472 | -1.605934 | 0.1082884 | 0.80237722 |
| TFG | 37.0006305 | -0.107750302 | 0.41610183 | -0.2589518 | 0.79567246 | 0.99665574 |
| SENP7 | 29.0451829 | -0.462749695 | 0.35570368 | -1.3009415 | 0.19327847 | 0.90153775 |
| TRMT10C | 7.97462149 | -0.553466008 | 0.52126295 | -1.0617789 | 0.28833607 | 0.96676791 |
| PCNP | 29.2182214 | 0.022008828 | 0.36696352 | 0.05997552 | 0.95217513 | 0.99777713 |
| ZBTB11 | 11.454951 | -0.276575118 | 0.49086178 | -0.5634481 | 0.57312982 | 0.99665574 |
| RPL24 | 37.4824435 | 0.044801602 | 0.50684924 | 0.08839236 | 0.92956483 | 0.99777713 |
| CEP97 | 5.1959975 | 1.347733549 | 0.85586604 | 1.57470152 | 0.11532536 | 0.81721642 |
| NFKBIZ | 19.9543533 | -2.40631398 | 0.59675832 | -4.0323091 | 5.52E-05 | 0.01513343 |
| ZPLD1 | 13.6173806 | -2.007358755 | 0.98255523 | -2.0429984 | 0.0410526 | 0.61100706 |
| ALCAM | 25.3260367 | -1.380641232 | 0.72859981 | -1.894924 | 0.05810248 | 0.67545594 |
| CBLB | 54.2248545 | -0.062885336 | 0.42589482 | -0.1476546 | 0.88261535 | 0.99665574 |
| BBX | 102.432354 | 0.030076485 | 0.36631359 | 0.08210584 | 0.93456255 | 0.99777713 |
| CD47 | 9.77524104 | -1.985782835 | 0.56706565 | -3.501857 | 0.00046203 | 0.05855044 |
| IFT57 | 6.52727882 | 0.035899952 | 0.67537952 | 0.05315523 | 0.95760823 | 0.99777713 |
| DZIP3 | 10.9108758 | -0.264546293 | 0.63956353 | -0.4136357 | 0.67914094 | 0.99665574 |
| NECTIN3 | 9.64667077 | -0.31925047 | 0.53580991 | -0.5958279 | 0.55129023 | 0.99665574 |
| PHLDB2 | 196.118417 | 0.238923735 | 0.29979984 | 0.79694418 | 0.42548345 | 0.99518711 |
| ABHD10 | 28.0982217 | 0.48373792 | 0.5018333 | 0.96394145 | 0.33507528 | 0.97788427 |
| ATG3 | 7.74634157 | 0.494563149 | 0.58620121 | 0.84367473 | 0.39885119 | 0.99179694 |
| SLC35A5 | 6.78537434 | -0.62731369 | 0.560264 | -1.1196752 | 0.26285221 | 0.94557817 |
| GTPBP8 | 6.4830807 | 0.453584494 | 0.59806661 | 0.75841802 | 0.44820077 | 0.99617422 |
| NEPRO | 10.3072833 | 0.173576938 | 0.47601939 | 0.36464258 | 0.71537823 | 0.99665574 |
| CFAP44 | 9.16917493 | -0.049781053 | 0.49978572 | -0.0996048 | 0.92065809 | 0.99777713 |
| SPICE1 | 9.43532536 | 0.019973441 | 0.54975057 | 0.03633182 | 0.97101777 | 0.99777713 |
| USF3 | 16.9847466 | -0.234231354 | 0.4609618 | -0.5081362 | 0.61135786 | 0.99665574 |
| NAA50 | 38.6497702 | 0.002587725 | 0.41653283 | 0.00621253 | 0.99504315 | 0.9985287 |
| ATP6V1A | 202.985651 | 0.368978789 | 0.34025674 | 1.08441287 | 0.27818178 | 0.95958042 |
| GRAMD1C | 47.7209122 | -0.43585081 | 0.4298738 | -1.0139041 | 0.31062851 | 0.97376371 |
| ZDHHC23 | 12.4401071 | 0.276298647 | 0.47041058 | 0.58735636 | 0.5569644 | 0.99665574 |
| CCDC191 | 9.45870694 | -0.530983982 | 0.71586406 | -0.7417386 | 0.45824575 | 0.99617422 |
| QTRT2 | 6.26689449 | 0.064805044 | 0.61142129 | 0.10599082 | 0.91558964 | 0.99777713 |
| ZBTB20 | 1513.39964 | 0.032683752 | 0.23402227 | 0.13966086 | 0.88892795 | 0.99665574 |
| ZBTB20-AS1 | 5.50773034 | -0.048092699 | 0.8715781 | -0.0551789 | 0.95599596 | 0.99777713 |
| ZBTB20-AS5 | 25.2965974 | -0.508832571 | 0.66245098 | -0.768106 | 0.44242421 | 0.99617422 |
| ZBTB20-AS2 | 5.66199081 | -0.668648056 | 0.7257679 | -0.9212974 | 0.35689519 | 0.9819332 |
| LSAMP | 9.62315048 | 0.290210238 | 0.7706726 | 0.37656748 | 0.70649506 | 0.99665574 |
| B4GALT4 | 7.07929052 | -0.5542936 | 0.51391968 | -1.0785607 | 0.28078359 | 0.96059387 |
| TMEM39A | 8.55345813 | -0.472847467 | 0.66904853 | -0.7067461 | 0.47972425 | 0.99617422 |
| POGLUT1 | 4.95467961 | -0.945985635 | 0.73985294 | -1.2786131 | 0.20103334 | 0.90591604 |
| TIMMDC1 | 10.4297039 | -0.420179815 | 0.5122787 | -0.8202172 | 0.41209228 | 0.99279547 |
| COX17 | 12.1626223 | 0.413287994 | 0.48597508 | 0.85043043 | 0.39508582 | 0.99179694 |
| GSK3B | 62.6297665 | -0.192899549 | 0.39614464 | -0.4869422 | 0.62629928 | 0.99665574 |
| LRRC58 | 8.46718071 | 0.019103438 | 0.53860185 | 0.03546857 | 0.97170611 | 0.99777713 |
| NDUFB4 | 15.9819738 | -0.08655488 | 0.39916476 | -0.21684 | 0.82833305 | 0.99665574 |
| HGD | 4.38134088 | 0.130826475 | 1.03942096 | 0.12586476 | 0.89983897 | 0.99777713 |
| RABL3 | 7.61477929 | -0.488994226 | 0.60293994 | -0.8110165 | 0.41735621 | 0.99392253 |
| GTF2E1 | 2.50095713 | 1.279993242 | 0.95044381 | 1.34673216 | 0.17806651 | 0.88796768 |
| GOLGB1 | 65.6000206 | -0.441098025 | 0.37381932 | -1.1799765 | 0.23800955 | 0.9305595 |
| IQCB1 | 29.4455518 | -0.214004008 | 0.39394782 | -0.5432293 | 0.58697192 | 0.99665574 |
| ILDR1 | 4.15389134 | -0.596211509 | 0.68729336 | -0.8674775 | 0.38568045 | 0.99051797 |
| CASR | 16.1298942 | 0.893435263 | 0.76423527 | 1.16905788 | 0.24238031 | 0.9342309 |
| CCDC58 | 5.50049777 | 0.88304427 | 0.74730547 | 1.18163763 | 0.23734951 | 0.9305595 |
| FAM162A | 25.5378834 | 0.107135933 | 0.41255369 | 0.25968967 | 0.79510316 | 0.99665574 |
| AC083798.2 | 6.13869862 | 0.223196327 | 0.65980249 | 0.33827748 | 0.73515409 | 0.99665574 |
| KPNA1 | 33.5442019 | -0.013052085 | 0.37150479 | -0.035133 | 0.97197367 | 0.99777713 |
| PARP9 | 19.9177347 | -0.96955916 | 0.7815338 | -1.2405851 | 0.21475907 | 0.91288363 |
| DTX3L | 7.56143561 | -0.612316355 | 0.8042408 | -0.7613595 | 0.44644238 | 0.99617422 |
| PARP14 | 37.6765096 | -0.547560341 | 0.72672921 | -0.7534586 | 0.45117441 | 0.99617422 |
| HSPBAP1 | 8.97530053 | -0.626879876 | 0.53324267 | -1.1755996 | 0.23975491 | 0.93157178 |
| DIRC2 | 12.9782655 | 0.088518357 | 0.63877789 | 0.13857455 | 0.88978636 | 0.99665574 |
| PDIA5 | 5.21500748 | -1.890433195 | 0.73414421 | -2.5750162 | 0.01002354 | 0.34338499 |
| SEC22A | 24.5401288 | 0.213793301 | 0.45654459 | 0.46828569 | 0.6395803 | 0.99665574 |
| HACD2 | 15.4747255 | 0.723854021 | 0.42133644 | 1.71799529 | 0.08579747 | 0.75373899 |
| MYLK | 7.66921015 | -0.79348316 | 0.5158165 | -1.5383051 | 0.12397403 | 0.82915695 |
| CCDC14 | 19.5191526 | 0.371380751 | 0.43720693 | 0.8494393 | 0.39563689 | 0.99179694 |
| KALRN | 11.0691095 | -0.989078705 | 0.54830111 | -1.8038969 | 0.07124746 | 0.71586402 |
| UMPS | 5.81115787 | 1.149185956 | 0.62280264 | 1.84518479 | 0.06501066 | 0.69039578 |
| ITGB5 | 23.9378005 | 0.333587779 | 0.37782096 | 0.88292555 | 0.3772765 | 0.98679664 |
| ZNF148 | 74.7240311 | -0.033900807 | 0.23118282 | -0.1466407 | 0.88341564 | 0.99665574 |
| SNX4 | 35.159155 | -0.09473645 | 0.27505933 | -0.3444219 | 0.73052904 | 0.99665574 |
| OSBPL11 | 11.5031684 | 0.009581128 | 0.44242991 | 0.0216557 | 0.9827226 | 0.99777713 |
| SLC41A3 | 6.82282785 | -0.13437393 | 0.6171281 | -0.2177407 | 0.82763112 | 0.99665574 |
| ALDH1L1 | 22.6311418 | 0.25421945 | 0.61683694 | 0.41213396 | 0.68024124 | 0.99665574 |
| KLF15 | 5.69606855 | 0.356633389 | 0.62306944 | 0.57238145 | 0.56706358 | 0.99665574 |
| ZXDC | 40.7851671 | -0.219343146 | 0.26556425 | -0.8259513 | 0.40883169 | 0.99279547 |
| TXNRD3NB | 3.12477374 | 0.130856256 | 0.82893328 | 0.15786102 | 0.87456631 | 0.99665574 |
| TXNRD3 | 4.26347842 | 0.710346253 | 0.69127989 | 1.02758125 | 0.30414685 | 0.97376371 |
| CHCHD6 | 21.7040392 | 1.356533208 | 0.39803555 | 3.4080705 | 0.00065424 | 0.06805261 |
| PLXNA1 | 5.21594427 | 0.252193184 | 0.60182577 | 0.41904684 | 0.6751819 | 0.99665574 |
| TPRA1 | 5.23470491 | 0.504441299 | 0.67777412 | 0.74426167 | 0.45671818 | 0.99617422 |
| PODXL2 | 4.83584752 | 0.109758479 | 0.63874512 | 0.17183455 | 0.86356761 | 0.99665574 |
| MGLL | 15.2026204 | -1.651838034 | 0.65830919 | -2.5092131 | 0.01210005 | 0.37648636 |
| KBTBD12 | 8.99025423 | 0.202629538 | 0.56161381 | 0.36079871 | 0.71824993 | 0.99665574 |
| SEC61A1 | 19.944167 | -0.621597143 | 0.56205504 | -1.1059364 | 0.26875404 | 0.94673212 |
| RUVBL1 | 6.74745309 | -0.117682626 | 0.62325186 | -0.1888203 | 0.85023363 | 0.99665574 |
| EEFSEC | 16.7971268 | 0.164779191 | 0.41798621 | 0.3942216 | 0.69341744 | 0.99665574 |
| GATA2 | 13.3108096 | 1.22007565 | 0.52547655 | 2.32184607 | 0.02024123 | 0.46637624 |
| GATA2-AS1 | 5.95832048 | -0.176568919 | 0.68281838 | -0.2585884 | 0.79595283 | 0.99665574 |
| RPN1 | 16.485846 | -0.612336055 | 0.48319495 | -1.267265 | 0.20506055 | 0.90834797 |
| RAB7A | 66.2238076 | -0.205742913 | 0.34925866 | -0.5890846 | 0.5558045 | 0.99665574 |
| ACAD9 | 10.9786283 | 0.113323867 | 0.43848264 | 0.2584455 | 0.7960631 | 0.99665574 |
| CNBP | 30.305603 | -0.840146176 | 0.57704279 | -1.4559513 | 0.14540609 | 0.85062144 |
| COPG1 | 11.6894783 | -0.108950704 | 0.5975706 | -0.1823227 | 0.85532946 | 0.99665574 |
| HMCES | 4.68956522 | 0.346724853 | 0.71679425 | 0.48371601 | 0.62858743 | 0.99665574 |
| H1FX | 10.8791139 | -1.492787377 | 0.78286977 | -1.9068144 | 0.05654462 | 0.67545594 |
| MBD4 | 16.1634398 | -0.359034568 | 0.40969874 | -0.876338 | 0.38084633 | 0.98820924 |
| IFT122 | 13.7494256 | -0.262346923 | 0.42762559 | -0.6134968 | 0.53954791 | 0.99665574 |
| TMCC1 | 135.085682 | -0.110741767 | 0.31044268 | -0.3567221 | 0.72129985 | 0.99665574 |
| PIK3R4 | 9.02410594 | 0.137328177 | 0.50581462 | 0.27149903 | 0.78600725 | 0.99665574 |
| ATP2C1 | 27.5987337 | 0.359603179 | 0.34197575 | 1.05154583 | 0.29300797 | 0.97072515 |
| NEK11 | 9.54228166 | -1.257798738 | 0.63565246 | -1.9787522 | 0.04784391 | 0.6410909 |
| NUDT16 | 9.5663884 | 0.11385626 | 0.47942474 | 0.23748515 | 0.81228043 | 0.99665574 |
| MRPL3 | 15.9271605 | 0.07171757 | 0.41196667 | 0.17408586 | 0.86179799 | 0.99665574 |
| ACPP | 5.07985728 | -2.336796075 | 0.68503308 | -3.4112164 | 0.00064674 | 0.06805261 |

| | | | | | | |
|---|---|---|---|---|---|---|
| DNAJC13 | 26.9401187 | 0.324930394 | 0.36931018 | 0.87983058 | 0.37895109 | 0.98679664 |
| UBA5 | 12.5275331 | 0.34604392 | 0.5298899 | 0.65304872 | 0.51372487 | 0.99665574 |
| CDV3 | 21.3478176 | -0.92985179 | 0.47333884 | -1.9644528 | 0.04947761 | 0.64444758 |
| TOPBP1 | 9.27995787 | -0.212066826 | 0.52740268 | -0.4020966 | 0.68761294 | 0.99665574 |
| SRPRB | 5.54245477 | -0.431836237 | 0.62213581 | -0.6941189 | 0.48760764 | 0.99640678 |
| RYK | 46.7919846 | 0.136157644 | 0.26286789 | 0.51796985 | 0.6044793 | 0.99665574 |
| AMOTL2 | 21.3716695 | 0.411896135 | 0.52685485 | 0.78180192 | 0.43433099 | 0.99617422 |
| ANAPC13 | 7.36319677 | -0.121030024 | 0.60512493 | -0.2000083 | 0.84147407 | 0.99665574 |
| CEP63 | 12.896294 | -0.037829265 | 0.58229172 | -0.0649662 | 0.94820093 | 0.99777713 |
| EPHB1 | 3.95167498 | -1.190318672 | 0.72044614 | -1.6521966 | 0.09849448 | 0.78203824 |
| PPP2R3A | 103.765599 | -0.748033917 | 0.37849854 | -1.9763191 | 0.04811864 | 0.64187738 |
| MSL2 | 17.0595906 | 0.504821714 | 0.38358718 | 1.31605472 | 0.18815568 | 0.89739428 |
| PCCB | 9.85604186 | -0.090277433 | 0.47159254 | -0.191431 | 0.84818793 | 0.99665574 |
| STAG1 | 160.23565 | -0.268195427 | 0.32303125 | -0.8302461 | 0.40639967 | 0.99250037 |
| NCK1 | 13.1432943 | -0.331605933 | 0.41383978 | -0.8012906 | 0.42296342 | 0.99518711 |
| DBR1 | 3.50919475 | 0.341427052 | 0.74442473 | 0.4586455 | 0.64648876 | 0.99665574 |
| ARMC8 | 32.8047621 | 0.101065447 | 0.34191741 | 0.29558438 | 0.7675475 | 0.99665574 |
| MRAS | 14.233544 | -0.10506684 | 0.48485125 | -0.2166991 | 0.82844283 | 0.99665574 |
| ESYT3 | 2.17983611 | -0.527889138 | 0.88476534 | -0.5966431 | 0.55074569 | 0.99665574 |
| CEP70 | 29.6213353 | 0.445086922 | 0.40932179 | 1.08737657 | 0.27687042 | 0.95828005 |
| FAIM | 6.17445537 | 0.195259372 | 0.57046604 | 0.34228045 | 0.73213985 | 0.99665574 |
| PIK3CB | 42.8214471 | 0.434459619 | 0.30061043 | 1.44525797 | 0.14838544 | 0.85424848 |
| MRPS22 | 3.8116666 | 0.187183621 | 0.74407278 | 0.25156628 | 0.80137633 | 0.99665574 |
| COPB2 | 23.1484012 | -0.493807712 | 0.43339761 | -1.1393872 | 0.25454167 | 0.94099699 |
| AC097103.2 | 13.0486561 | 0.136543968 | 0.51857636 | 0.26330542 | 0.79231518 | 0.99665574 |
| NMNAT3 | 13.3830273 | 0.102026063 | 0.51368004 | 0.19861792 | 0.84256163 | 0.99665574 |
| SLC25A36 | 45.4422256 | 0.246043309 | 0.32676988 | 0.75295591 | 0.45147641 | 0.99617422 |
| PXYLP1 | 3.01263476 | -0.137567963 | 0.76504402 | -0.1798171 | 0.85729619 | 0.99665574 |
| ZBTB38 | 26.9828137 | -0.453312968 | 0.34489352 | -1.3143563 | 0.18872633 | 0.89739428 |
| RASA2 | 42.3391788 | -0.521456266 | 0.38859482 | -1.3419023 | 0.17962769 | 0.88820302 |
| RNF7 | 10.9305343 | -0.414714758 | 0.65774006 | -0.6305147 | 0.52835791 | 0.99665574 |
| ATP1B3 | 25.8643011 | -0.588913799 | 0.5849273 | -1.0068154 | 0.3140235 | 0.9755712 |
| TFDP2 | 58.9638891 | 0.057212803 | 0.41140554 | 0.13906668 | 0.88939746 | 0.99665574 |
| GK5 | 28.5068879 | 0.242513613 | 0.39481697 | 0.61424313 | 0.53905468 | 0.99665574 |
| XRN1 | 82.597662 | -0.219989821 | 0.35071625 | -0.6272587 | 0.53048967 | 0.99665574 |
| ATR | 41.3017047 | 0.078758721 | 0.37759288 | 0.20858105 | 0.8347753 | 0.99665574 |
| PLS1 | 27.7197315 | 0.481580868 | 0.35473963 | 1.35756152 | 0.17460286 | 0.88366595 |
| TRPC1 | 133.871614 | 0.704443663 | 0.34676517 | 2.03147179 | 0.04220716 | 0.61533334 |
| PCOLCE2 | 6.57205261 | 0.442155966 | 0.74234381 | 0.59562154 | 0.55142808 | 0.99665574 |
| U2SURP | 62.5937749 | 0.523238171 | 0.34350929 | 1.52321402 | 0.12770517 | 0.83778831 |
| CHST2 | 30.9717814 | -0.499003595 | 0.66225787 | -0.7534884 | 0.4511565 | 0.99617422 |
| SLC9A9 | 12.9469002 | -1.499253499 | 0.80895979 | -1.8533103 | 0.0638379 | 0.68630818 |
| C3orf58 | 4.64178396 | -0.499241339 | 0.67134398 | -0.7436446 | 0.4570915 | 0.99617422 |
| AC107021.1 | 7.37324153 | -3.236110911 | 0.85838546 | -3.7699973 | 0.00016325 | 0.02986125 |
| PLOD2 | 56.2113461 | -1.360669298 | 0.47610285 | -2.8579314 | 0.00426413 | 0.20883915 |
| PLSCR4 | 5.64463782 | -3.082639762 | 0.92942381 | -3.3167213 | 0.0009108 | 0.08160886 |
| PLSCR2 | 5.96942339 | 0.102159141 | 0.70592114 | 0.1447175 | 0.88493392 | 0.99665574 |
| PLSCR1 | 44.8620658 | -1.234553348 | 0.51170825 | -2.4126118 | 0.01583868 | 0.41983038 |
| GYG1 | 13.9325043 | -0.654934483 | 0.55061641 | -1.1894569 | 0.23425993 | 0.92960234 |
| HLTF | 9.89134249 | -0.004638556 | 0.51401087 | -0.0090242 | 0.9927998 | 0.99777713 |
| HPS3 | 9.82560314 | -0.416754893 | 0.56002394 | -0.7441734 | 0.4567716 | 0.99617422 |
| WWTR1 | 44.8165114 | -0.363267815 | 0.46655911 | -0.7786105 | 0.4362092 | 0.99617422 |
| COMMD2 | 8.05748054 | -0.865244596 | 0.50781714 | -1.7038507 | 0.08840898 | 0.75650018 |
| ANKUB1 | 16.0529094 | -1.051996644 | 0.58145982 | -1.8092336 | 0.07041472 | 0.71264656 |
| RNF13 | 36.9409974 | 0.19888479 | 0.35044376 | 0.5675227 | 0.57035911 | 0.99665574 |
| AC117386.2 | 10.767642 | 1.337103063 | 0.7306391 | 1.83004586 | 0.06724308 | 0.70207041 |
| TSC22D2 | 56.7870713 | -0.230403802 | 0.33039199 | -0.697365 | 0.48557441 | 0.99640678 |
| SERP1 | 30.0183286 | -0.148157167 | 0.38520886 | -0.3846152 | 0.70052256 | 0.99665574 |
| EIF2A | 13.6736315 | -0.098648934 | 0.42329503 | -0.2330501 | 0.81572253 | 0.99665574 |
| SELENOT | 20.0282933 | 0.20711008 | 0.40493967 | 0.5114591 | 0.60902962 | 0.99665574 |
| SIAH2 | 2.6811043 | -0.328964303 | 1.05743628 | -0.3110961 | 0.75572757 | 0.99665574 |
| MBNL1 | 139.194284 | 0.180383492 | 0.271463 | 0.66448647 | 0.506379 | 0.99665574 |
| RAP2B | 30.0978693 | 0.71884124 | 0.47396677 | 1.51664903 | 0.12935532 | 0.83778831 |
| DHX36 | 43.6519354 | 0.003224737 | 0.27464504 | 0.01174147 | 0.99063188 | 0.99777713 |
| MME | 4.51458217 | 1.130890621 | 0.76511269 | 1.47807067 | 0.13938886 | 0.84714487 |
| PLCH1 | 30.1948234 | 0.198078564 | 0.66119453 | 0.29957683 | 0.76449996 | 0.99665574 |
| GMPS | 18.0453884 | 0.585006329 | 0.4359558 | 1.34189367 | 0.17963048 | 0.88820302 |
| SSR3 | 11.7479499 | -0.091170679 | 0.53763471 | -0.1695774 | 0.86534252 | 0.99665574 |
| TIPARP | 4.20260037 | -1.137919706 | 0.66919471 | -1.7004314 | 0.08904981 | 0.75864448 |
| LINC00886 | 20.9610454 | 0.888634652 | 0.60586951 | 1.46670964 | 0.14245506 | 0.84969914 |
| CCNL1 | 30.1825745 | -0.596230126 | 0.38212506 | -1.560301 | 0.11868876 | 0.82619588 |
| VEPH1 | 8.98605185 | 1.708331019 | 0.60253594 | 2.83523507 | 0.0045792 | 0.21492549 |
| RSRC1 | 58.472155 | -0.078584079 | 0.27245441 | -0.2884302 | 0.77301745 | 0.99665574 |
| GFM1 | 17.3756762 | -0.512856473 | 0.39616264 | -1.2945604 | 0.19547191 | 0.90381759 |
| RARRES1 | 3.25868072 | -1.892476944 | 0.87477969 | -2.1633755 | 0.03051231 | 0.56271575 |
| MFSD1 | 9.36477911 | -0.388513997 | 0.63848373 | -0.6084948 | 0.54285939 | 0.99665574 |
| IQCJ-SCHIP1 | 13.890743 | -0.902168956 | 0.65914989 | -1.3686856 | 0.17109757 | 0.87968373 |
| SMC4 | 8.12253063 | -0.319544013 | 0.50957088 | -0.6270845 | 0.53060383 | 0.99665574 |
| KPNA4 | 34.0853523 | -0.274455139 | 0.36815525 | -0.7454875 | 0.45597707 | 0.99617422 |
| PPM1L | 71.93444 | 0.144256553 | 0.38296907 | 0.37667938 | 0.70641189 | 0.99665574 |
| NMD3 | 16.4839608 | 0.412732008 | 0.41627178 | 0.99149649 | 0.3214432 | 0.97700345 |
| LINC01322 | 4.47189819 | -0.081808749 | 0.8351246 | -0.0979599 | 0.92196411 | 0.99777713 |
| PDCD10 | 26.3078066 | -0.559170536 | 0.32735369 | -1.7081541 | 0.08760776 | 0.75650018 |
| GOLIM4 | 28.6579648 | 0.794684225 | 0.34160104 | 2.32635188 | 0.01999979 | 0.46397674 |
| MECOM | 586.717818 | 0.206807189 | 0.25416666 | 0.81366765 | 0.41583537 | 0.992853 |
| MYNN | 5.94001291 | -0.147063993 | 0.55615126 | -0.2644316 | 0.79144733 | 0.99665574 |
| SEC62 | 37.7290119 | -0.0397771 | 0.33169524 | -0.1199206 | 0.90454603 | 0.99777713 |
| GPR160 | 17.4616892 | -0.317791989 | 0.63360512 | -0.5015616 | 0.61597594 | 0.99665574 |
| PHC3 | 62.904835 | 0.261165953 | 0.27051553 | 0.96543792 | 0.33432552 | 0.97788427 |
| PRKCI | 46.3450675 | -1.216403142 | 0.52430283 | -2.3200393 | 0.02033875 | 0.46756073 |
| SKIL | 17.5276676 | -0.172070886 | 0.41653071 | -0.4131049 | 0.67952975 | 0.99665574 |
| RPL22L1 | 2.83201312 | -0.249495075 | 0.9076881 | -0.2748687 | 0.78341709 | 0.99665574 |
| TNIK | 15.9943758 | -0.005866515 | 0.56180571 | -0.0104422 | 0.99166844 | 0.99777713 |
| PLD1 | 14.4618027 | -2.01211911 | 0.66997802 | -3.0032614 | 0.00267103 | 0.16023015 |
| FNDC3B | 165.579863 | -0.477126057 | 0.28353282 | -1.6827895 | 0.09241584 | 0.76670359 |
| TNFSF10 | 15.2956454 | -1.210628688 | 0.52616877 | -2.3008372 | 0.02140083 | 0.4733368 |
| NCEH1 | 24.2883264 | -0.02433794 | 0.51215204 | -0.0475209 | 0.96209805 | 0.99777713 |
| ECT2 | 5.43579581 | -1.093234887 | 0.78004054 | -1.4015103 | 0.16106152 | 0.86928072 |
| NLGN1 | 65.527955 | 0.104231967 | 0.42962289 | 0.24261269 | 0.80830545 | 0.99665574 |
| NAALADL2 | 136.007185 | -0.075644699 | 0.44766299 | -0.1689769 | 0.86581482 | 0.99665574 |
| TBL1XR1 | 137.412078 | -0.355099746 | 0.48046527 | -0.7390747 | 0.45986162 | 0.99617422 |
| KCNMB2-AS1 | 9.55843729 | -0.065426341 | 0.96872562 | -0.0675386 | 0.94615296 | 0.99777713 |
| ZMAT3 | 7.79687279 | -0.384066427 | 0.52545678 | -0.7309192 | 0.46482854 | 0.99617422 |
| PIK3CA | 48.2027182 | -0.079808902 | 0.28736949 | -0.2777223 | 0.78122558 | 0.99665574 |
| KCNMB3 | 9.17550526 | -0.765842212 | 0.60195476 | -1.2722588 | 0.2032812 | 0.9066717 |
| ZNF639 | 5.88560518 | 0.069999884 | 0.81388677 | 0.08600691 | 0.93146093 | 0.99777713 |
| MFN1 | 29.4390766 | -0.779047977 | 0.42059016 | -1.8522734 | 0.06398657 | 0.68717782 |
| GNB4 | 5.18380336 | -2.356511594 | 0.93281002 | -2.5262503 | 0.01152873 | 0.3725572 |
| ACTL6A | 6.32282448 | 0.308188651 | 0.64644813 | 0.4767415 | 0.6335462 | 0.99665574 |
| MRPL47 | 6.93874511 | 0.624064672 | 0.662226 | 0.94237416 | 0.34600111 | 0.97788427 |
| NDUFB5 | 16.1349658 | 0.513837492 | 0.53953331 | 0.95237399 | 0.34090734 | 0.97788427 |
| USP13 | 3.00278154 | 0.451044265 | 0.88199039 | 0.51139363 | 0.60907546 | 0.99665574 |
| TTC14 | 32.9557975 | 0.576738096 | 0.4174263 | 1.38165251 | 0.16707842 | 0.87495175 |

| | | | | | | |
|---|---|---|---|---|---|---|
| FXR1 | 37.0998864 | 0.562169243 | 0.28533854 | 1.97018336 | 0.04881736 | 0.64357659 |
| DNAJC19 | 6.01439492 | -0.2385522 | 0.7515198 | -0.3174264 | 0.7509201 | 0.99665574 |
| ATP11B | 40.3551598 | 0.018222041 | 0.29732187 | 0.06128725 | 0.95113044 | 0.99777713 |
| DCUN1D1 | 20.161931 | 0.157798859 | 0.3687412 | 0.42793933 | 0.6686953 | 0.99665574 |
| MCCC1 | 29.4158015 | -0.106968431 | 0.30366142 | -0.3522622 | 0.72464165 | 0.99665574 |
| B3GNT5 | 5.11888328 | -0.980508774 | 0.71913521 | -1.3634554 | 0.17273904 | 0.88157681 |
| KLHL24 | 52.6163512 | 0.002968885 | 0.30915501 | 0.00960322 | 0.99233785 | 0.99777713 |
| YEATS2 | 21.2483201 | -0.452043507 | 0.50858801 | -0.8888206 | 0.3740995 | 0.98535336 |
| ABCC5 | 66.8100176 | -0.351306349 | 0.3164357 | -1.1101982 | 0.26691362 | 0.94557817 |
| EIF2B5 | 5.73971895 | -0.107833769 | 0.59919368 | -0.1799648 | 0.8571802 | 0.99665574 |
| DVL3 | 12.1443831 | 0.078165267 | 0.53139725 | 0.14709385 | 0.88305795 | 0.99665574 |
| AP2M1 | 17.3053256 | -0.072375781 | 0.46466782 | -0.1557581 | 0.87622369 | 0.99665574 |
| ABCF3 | 4.2038037 | 0.198571208 | 0.65641342 | 0.30250937 | 0.76226379 | 0.99665574 |
| ALG3 | 3.10795554 | -0.210878146 | 0.75420743 | -0.2796023 | 0.77978263 | 0.99665574 |
| PSMD2 | 8.72670268 | 0.319172695 | 0.66749785 | 0.47816288 | 0.63253428 | 0.99665574 |
| EIF4G1 | 19.7760384 | -0.201489478 | 0.47620253 | -0.4231172 | 0.67220976 | 0.99665574 |
| FAM131A | 3.06244019 | -0.114144474 | 0.81467148 | -0.140111 | 0.88857225 | 0.99665574 |
| POLR2H | 13.840132 | 0.028956545 | 0.52980133 | 0.05465548 | 0.95641294 | 0.99777713 |
| MAGEF1 | 8.07331507 | -0.389347624 | 0.61185127 | -0.6363436 | 0.52455251 | 0.99665574 |
| VPS8 | 40.7683177 | 0.311094721 | 0.27157129 | 1.14553612 | 0.25198714 | 0.94076968 |
| EHHADH | 9.62555057 | 0.229659439 | 0.60087519 | 0.38220822 | 0.70230692 | 0.99665574 |
| MAP3K13 | 35.746362 | -0.076215557 | 0.37012488 | -0.2059185 | 0.83685458 | 0.99665574 |
| TMEM41A | 6.23546896 | -0.120864938 | 0.74788629 | -0.1616087 | 0.871614 | 0.99665574 |
| LIPH | 4.20529773 | -1.670783294 | 0.85401982 | -1.9563753 | 0.05042096 | 0.64929786 |
| SENP2 | 15.36202 | 0.354281938 | 0.40366763 | 0.87765754 | 0.38012959 | 0.98679664 |
| IGF2BP2 | 17.5059295 | 1.075112835 | 0.92178481 | 1.1663382 | 0.24347773 | 0.93426912 |
| TRA2B | 26.8957772 | -0.062281577 | 0.33985161 | -0.1832611 | 0.85459316 | 0.99665574 |
| ETV5 | 4.53306148 | -2.104808859 | 0.84508416 | -2.49065 | 0.01275097 | 0.38590337 |
| AC068631.2 | 4.16206027 | -0.827992745 | 0.77003299 | -1.0752692 | 0.28225423 | 0.96059387 |
| EIF4A2 | 39.618816 | -0.179249443 | 0.39278188 | -0.4563587 | 0.64813204 | 0.99665574 |
| ST6GAL1 | 137.640486 | -0.355291094 | 0.2991009 | -1.1878637 | 0.23488713 | 0.92960234 |
| BCL6 | 47.3935922 | -1.71549819 | 0.52844336 | -3.2463237 | 0.00116906 | 0.09557995 |
| LPP | 389.230029 | -0.469715926 | 0.35334216 | -1.3293515 | 0.18373204 | 0.89532394 |
| TPRG1 | 24.2533572 | 2.152609152 | 0.74300749 | 2.89715672 | 0.00376562 | 0.20198844 |
| P3H2 | 69.6038528 | -0.097353796 | 0.50466481 | -0.1929078 | 0.84703115 | 0.99665574 |
| CLDN1 | 49.4286107 | -1.052464909 | 0.3758612 | -2.8001424 | 0.00510801 | 0.22417735 |
| CLDN16 | 3.61012392 | -0.675578186 | 0.70502206 | -0.9582369 | 0.33794329 | 0.97788427 |
| CCDC50 | 39.1320664 | -0.032421801 | 0.4176024 | -0.077638 | 0.93811604 | 0.99777713 |
| FGF12 | 16.0629754 | -3.467466673 | 1.51867265 | -2.2832219 | 0.0224173 | 0.48457695 |
| MB21D2 | 3.78921991 | -1.300198403 | 0.69503539 | -1.8706938 | 0.06138753 | 0.6834742 |
| OPA1 | 35.1531546 | 0.184202707 | 0.28301618 | 0.65085574 | 0.51513961 | 0.99665574 |
| HES1 | 6.33076885 | -0.242269824 | 0.9494629 | -0.2551651 | 0.79859556 | 0.99665574 |
| ATP13A3 | 38.9177787 | 0.523043673 | 0.32669398 | 1.60102022 | 0.10937244 | 0.80453999 |
| LSG1 | 7.93107694 | -0.001641812 | 0.58920787 | -0.0027865 | 0.99777672 | 0.99905781 |
| XXYLT1 | 7.25139099 | 0.247589702 | 0.56175115 | 0.44074623 | 0.65939672 | 0.99665574 |
| ACAP2 | 59.7281405 | -0.121316972 | 0.30774259 | -0.3942157 | 0.69342177 | 0.99665574 |
| PPP1R2 | 14.0775801 | -0.398678509 | 0.47251285 | -0.8437411 | 0.3988141 | 0.99179694 |
| MUC20-OT1 | 229.957019 | 0.533827771 | 0.35990315 | 1.48325396 | 0.13800695 | 0.84656101 |
| TNK2 | 7.71772683 | -0.169105016 | 0.52995882 | -0.3190909 | 0.74965762 | 0.99665574 |
| TFRC | 31.5658073 | -1.440071767 | 0.54610043 | -2.637009 | 0.00836406 | 0.31119582 |
| LINC00885 | 2.20047379 | 0.168358026 | 1.18460488 | 0.14212167 | 0.8869839 | 0.99665574 |
| PCYT1A | 8.38532677 | 0.083298454 | 0.55302699 | 0.15062276 | 0.8802733 | 0.99665574 |
| UBXN7 | 18.7792726 | 0.351857808 | 0.4086978 | 0.86092415 | 0.38927982 | 0.99176253 |
| RNF168 | 12.8250888 | 0.144067103 | 0.42705203 | 0.33735257 | 0.73585113 | 0.99665574 |
| WDR53 | 2.52145457 | -0.834683081 | 0.86467178 | -0.9653178 | 0.33438565 | 0.97788427 |
| PIGX | 4.69095056 | -0.272010265 | 0.76910414 | -0.3536716 | 0.72358504 | 0.99665574 |
| PAK2 | 41.6198325 | -0.366206987 | 0.28030495 | -1.3064592 | 0.19139643 | 0.89964233 |
| SENP5 | 42.9359204 | -0.471346267 | 0.37741063 | -1.248895 | 0.21170347 | 0.91043197 |
| NCBP2 | 9.4529888 | -0.160910882 | 0.4800908 | -0.3351676 | 0.73749866 | 0.99665574 |
| NCBP2-AS2 | 5.49185739 | -0.305256738 | 0.64170968 | -0.4756929 | 0.63429318 | 0.99665574 |
| DLG1 | 114.254557 | -0.036901201 | 0.30635443 | -0.1204526 | 0.90412459 | 0.99777713 |
| BDH1 | 18.8060483 | 0.636533015 | 0.53602098 | 1.18751512 | 0.2350245 | 0.92960234 |
| AC024560.1 | 5.83912502 | 4.252340302 | 1.30884387 | 3.24892861 | 0.00115841 | 0.09547898 |
| RUBCN | 9.21009138 | -0.605156599 | 0.49240118 | -1.228991 | 0.2190752 | 0.91662582 |
| FYTTD1 | 11.674937 | 0.714490031 | 0.46718641 | 1.52934678 | 0.1261785 | 0.83633607 |
| LRCH3 | 38.8061241 | -0.335386261 | 0.31444741 | -1.0665893 | 0.28615731 | 0.96449041 |
| IQCG | 4.87765517 | -0.306374377 | 0.61550774 | -0.4977588 | 0.61865408 | 0.99665574 |
| RPL35A | 44.081038 | 0.392030242 | 0.42083447 | 0.9315545 | 0.35156681 | 0.97838306 |
| LMLN | 5.61350447 | -0.176223422 | 0.68680098 | -0.2565859 | 0.79749849 | 0.99665574 |
| ZNF595 | 9.60542197 | -0.240276292 | 0.60417869 | -0.3976908 | 0.69085814 | 0.99665574 |
| ZNF718 | 18.399809 | -0.206034111 | 0.42058095 | -0.4898798 | 0.62421896 | 0.99665574 |
| ZNF141 | 21.319134 | 0.199137699 | 0.33929623 | 0.58691398 | 0.55726148 | 0.99665574 |
| AC092574.2 | 3.39145815 | 0.035552491 | 0.94459796 | 0.0376377 | 0.96997655 | 0.99777713 |
| ZNF721 | 45.7920945 | -0.181983654 | 0.27440515 | -0.6631933 | 0.50720677 | 0.99665574 |
| PIGG | 11.7677452 | -0.596789281 | 0.41920299 | -1.4236284 | 0.15455407 | 0.86196782 |
| ATP5ME | 13.7314514 | 1.055625039 | 0.45224139 | 2.33420703 | 0.01958488 | 0.4617478 |
| PCGF3 | 17.4486177 | -0.155958078 | 0.45237661 | -0.3447527 | 0.73028026 | 0.99665574 |
| AC139887.2 | 22.9898598 | -0.107651272 | 0.39868478 | -0.270016 | 0.78714794 | 0.99665574 |
| GAK | 28.6127301 | -0.38499982 | 0.40812902 | -0.9433287 | 0.34551279 | 0.97788427 |
| TMEM175 | 4.12592143 | 0.554350422 | 0.67558544 | 0.8205482 | 0.41190366 | 0.99279547 |
| IDUA | 4.91855918 | 1.383989214 | 0.77571452 | 1.78414761 | 0.07439968 | 0.72810353 |
| FGFRL1 | 4.23374271 | 0.245025168 | 0.69379582 | 0.3531661 | 0.72396392 | 0.99665574 |
| RNF212 | 8.109092 | 0.871677085 | 0.54415145 | 1.6019016 | 0.10917737 | 0.80453999 |
| AC092535.3 | 18.092059 | 0.7351885 | 0.59169307 | 1.24251666 | 0.214046 | 0.9124923 |
| CTBP1-AS | 3.91861502 | 0.994794496 | 0.88971121 | 1.11810943 | 0.26352026 | 0.94557817 |
| CTBP1 | 12.9933394 | 0.42265823 | 0.42616721 | 0.99176619 | 0.32131159 | 0.97700345 |
| CTBP1-DT | 2.17032024 | 0.36216166 | 0.86998424 | 0.41628531 | 0.67720124 | 0.99665574 |
| MAEA | 16.891596 | -0.256450488 | 0.41625795 | -0.6160855 | 0.53783809 | 0.99665574 |
| UVSSA | 12.2089053 | -0.584387458 | 0.44735161 | -1.306327 | 0.1914414 | 0.89964233 |
| SLBP | 18.1445079 | 0.230109473 | 0.38885043 | 0.59176859 | 0.55400556 | 0.99665574 |
| TMEM129 | 8.06435409 | 0.62828643 | 0.55870034 | 1.12454993 | 0.2607798 | 0.94557817 |
| LETM1 | 8.13478672 | 0.419716165 | 0.59860532 | 0.70115676 | 0.48320519 | 0.99640678 |
| NSD2 | 31.7326786 | -0.238347357 | 0.35295655 | -0.6752881 | 0.49949275 | 0.99640678 |
| NELFA | 12.7903209 | 0.643328893 | 0.46701509 | 1.37753343 | 0.16834739 | 0.87566218 |
| NAT8L | 1.77289568 | 1.222436046 | 1.10729789 | 1.1039812 | 0.26960129 | 0.94771773 |
| POLN | 4.62067978 | 0.217945155 | 0.80959092 | 0.26920405 | 0.78777266 | 0.99665574 |
| MXD4 | 14.7826506 | -0.340841111 | 0.47070196 | -0.7241124 | 0.46899675 | 0.99617422 |
| ZFYVE28 | 11.3331754 | -0.483017837 | 0.49771151 | -0.9704775 | 0.33180852 | 0.97788427 |
| RNF4 | 16.450253 | -0.142762477 | 0.4277784 | -0.33373 | 0.73858332 | 0.99665574 |
| FAM193A | 47.5065356 | -0.159588884 | 0.36894099 | -0.4325594 | 0.66533491 | 0.99665574 |
| TNIP2 | 7.51198765 | -0.702899817 | 0.66107173 | -1.0632731 | 0.2876581 | 0.96559334 |
| SH3BP2 | 8.92967283 | -1.561819386 | 0.56386388 | -2.7698518 | 0.00560818 | 0.23322054 |
| ADD1 | 42.3283407 | 0.06272487 | 0.28450648 | 0.22046903 | 0.82550589 | 0.99665574 |
| MFSD10 | 14.7642757 | 0.157585687 | 0.46388085 | 0.33971156 | 0.73407376 | 0.99665574 |
| NOP14 | 5.96191224 | -0.527289419 | 0.60060943 | -0.877924 | 0.37998497 | 0.98679664 |
| GRK4 | 4.36248461 | 0.116798258 | 0.69494509 | 0.16806833 | 0.86652953 | 0.99665574 |
| HTT | 31.7785167 | 0.074009584 | 0.29741433 | 0.24884337 | 0.80348194 | 0.99665574 |
| RGS12 | 11.2729853 | -0.904408745 | 0.57877265 | -1.5626322 | 0.11813914 | 0.82599629 |
| LRPAP1 | 18.3124186 | -0.411550771 | 0.38017567 | -1.0825279 | 0.27901803 | 0.95980166 |
| TMEM128 | 4.21838349 | -0.456810271 | 0.68027934 | -0.671504 | 0.50189954 | 0.99640678 |
| ZBTB49 | 3.51824521 | -0.113490672 | 0.71596845 | -0.1585135 | 0.87405218 | 0.99665574 |
| STX18 | 57.0499564 | 0.61775725 | 0.31321753 | 1.97229464 | 0.04857598 | 0.64337675 |
| STX18-AS1 | 26.1078499 | -0.225592976 | 0.45408664 | -0.496806 | 0.61932585 | 0.99665574 |

| | | | | | | |
|---|---|---|---|---|---|---|
| EVC2 | 14.0241525 | 0.035610305 | 0.46904573 | 0.07592075 | 0.93948215 | 0.99777713 |
| EVC | 15.2875809 | -0.537704352 | 0.52889984 | -1.0166468 | 0.30932146 | 0.97376371 |
| CRMP1 | 3.65155353 | -1.28098622 | 1.10200502 | -1.1624141 | 0.24506728 | 0.93426912 |
| MAN2B2 | 8.16316013 | 0.438041054 | 0.58695713 | 0.74629139 | 0.45549141 | 0.99617422 |
| MRFAP1 | 16.3774854 | -0.266873716 | 0.49653911 | -0.5374677 | 0.59094461 | 0.99665574 |
| MRFAP1L1 | 4.55949657 | 0.609836177 | 0.68102472 | 0.89546848 | 0.37053671 | 0.98532838 |
| BLOC1S4 | 4.8985172 | -0.751912944 | 0.71923258 | -1.0454378 | 0.29582066 | 0.97286866 |
| KIAA0232 | 50.5366768 | 0.028994261 | 0.27864216 | 0.10405554 | 0.91712527 | 0.99777713 |
| TBC1D14 | 77.5679212 | 0.808980667 | 0.39069852 | 2.07060078 | 0.03839612 | 0.59519856 |
| TADA2B | 4.68644613 | 0.479337215 | 0.65528975 | 0.73148896 | 0.46448055 | 0.99617422 |
| GRPEL1 | 7.95108876 | 0.093420091 | 0.57283552 | 0.16308362 | 0.8704526 | 0.99665574 |
| SORCS2 | 6.62938111 | 1.397646716 | 0.74608831 | 1.87329931 | 0.06102706 | 0.6834742 |
| AFAP1 | 5.28379317 | -1.585635645 | 0.75067184 | -2.1122887 | 0.03466169 | 0.58178209 |
| ABLIM2 | 3.52280451 | -0.120567439 | 0.75093031 | -0.1605574 | 0.87244199 | 0.99665574 |
| SH3TC1 | 38.0110152 | -0.16745867 | 0.41071661 | -0.4077231 | 0.68347694 | 0.99665574 |
| ACOX3 | 2.88079957 | -0.92106528 | 0.82990586 | -1.1098431 | 0.26706665 | 0.94557817 |
| TRMT44 | 4.69704382 | 0.34702553 | 0.95194262 | 0.36454459 | 0.71545138 | 0.99665574 |
| SLC2A9 | 10.9948955 | 0.024133953 | 0.55913014 | 0.04316339 | 0.96557129 | 0.99777713 |
| WDR1 | 37.788814 | 0.447481009 | 0.36635109 | 1.221454 | 0.22191417 | 0.92191316 |
| ZNF518B | 4.28835129 | -0.886533229 | 0.82486826 | -1.0747574 | 0.28248338 | 0.96059387 |
| CLNK | 412.289879 | 0.491030372 | 0.43639826 | 1.12518865 | 0.2605091 | 0.94557817 |
| AC084048.1 | 4.37621725 | -1.289315322 | 1.15008978 | -1.1210562 | 0.26226393 | 0.94557817 |
| HS3ST1 | 4.2956502 | -1.26300166 | 0.76599307 | -1.6488422 | 0.09917998 | 0.78261212 |
| AC005699.1 | 4.29855344 | 0.900194589 | 2.25448897 | 0.39928986 | 0.68967964 | 0.99665574 |
| RAB28 | 45.5465664 | 0.141230489 | 0.3188592 | 0.4429243 | 0.65782049 | 0.99665574 |
| BOD1L1 | 42.0059345 | -0.734838412 | 0.46322186 | -1.5863638 | 0.1126568 | 0.81249398 |
| LINC01182 | 15.7980242 | 1.21848379 | 0.56167403 | 2.16937889 | 0.03005393 | 0.55923879 |
| CPEB2 | 51.2563569 | -0.197284641 | 0.52877415 | -0.3730981 | 0.70907542 | 0.99665574 |
| AC098829.1 | 13.3470697 | 0.44710089 | 0.5443392 | 0.82136449 | 0.41143869 | 0.99279547 |
| CC2D2A | 3.73436675 | -0.969879388 | 1.22637722 | -0.7908492 | 0.42903202 | 0.99599877 |
| FBXL5 | 30.0434678 | 0.539659523 | 0.28850151 | 1.87056051 | 0.06140602 | 0.6834742 |
| FAM200B | 8.34571612 | 0.108650246 | 0.6989084 | 0.15545706 | 0.876461 | 0.99665574 |
| PROM1 | 8.36096223 | -0.498780523 | 1.08386683 | -0.4601862 | 0.6453826 | 0.99665574 |
| TAPT1 | 20.3647518 | 0.192556199 | 0.46762246 | 0.41177706 | 0.68050284 | 0.99665574 |
| TAPT1-AS1 | 31.4069151 | 0.73230379 | 0.61036995 | 1.19977038 | 0.23022853 | 0.92774608 |
| LDB2 | 5.06346087 | -0.560579837 | 0.88772109 | -0.6314819 | 0.52772546 | 0.99665574 |
| QDPR | 6.87389531 | 0.223357534 | 0.67881237 | 0.32904164 | 0.74212422 | 0.99665574 |
| LAP3 | 11.4374679 | -0.503332688 | 0.8571961 | -0.587185 | 0.55707947 | 0.99665574 |
| MED28 | 16.7805404 | 0.211560156 | 0.41663577 | 0.50778203 | 0.61160622 | 0.99665574 |
| FAM184B | 3.55205328 | -0.779161199 | 0.71653648 | -1.0873992 | 0.27686042 | 0.95828005 |
| DCAF16 | 15.7674198 | 0.275384556 | 0.40952741 | 0.67244474 | 0.50130061 | 0.99640678 |
| LCORL | 51.6558913 | -0.411364936 | 0.32722398 | -1.2571357 | 0.20870452 | 0.91043197 |
| AC024230.1 | 7.69712691 | 1.366262019 | 0.62033221 | 2.20246829 | 0.02763225 | 0.53563234 |
| SLIT2 | 8.26111826 | 0.195153314 | 0.86687745 | 0.22512215 | 0.82188425 | 0.99665574 |
| PACRGL | 12.1162795 | 0.576867692 | 0.51762599 | 1.11444886 | 0.26508667 | 0.94557817 |
| KCNIP4 | 700.444243 | -3.540735552 | 0.65126304 | -5.4367212 | 5.43E-08 | 6.11E-05 |
| ADGRA3 | 19.8240105 | 0.331762413 | 0.41237379 | 0.80451867 | 0.42109749 | 0.99518711 |
| PPARGC1A | 224.947469 | 0.224726553 | 0.27344099 | 0.82184661 | 0.4111642 | 0.99279547 |
| DHX15 | 39.2993684 | 0.533130783 | 0.3442853 | 1.54851451 | 0.12149847 | 0.82659025 |
| CCDC149 | 7.2551221 | -0.294151485 | 0.50955268 | -0.5772739 | 0.56375442 | 0.99665574 |
| SEPSECS | 4.9162142 | -0.8849121 | 0.58882323 | -1.5028485 | 0.13287812 | 0.84089786 |
| ZCCHC4 | 4.60543183 | -0.575324112 | 0.72982876 | -0.7883001 | 0.43052117 | 0.99617422 |
| ANAPC4 | 17.0839503 | -0.849907092 | 0.39099812 | -2.1736859 | 0.02972873 | 0.55812946 |
| SEL1L3 | 5.29784202 | -0.904996345 | 0.79196225 | -1.1427266 | 0.25315211 | 0.94076968 |
| SMIM20 | 4.29874617 | -0.151884993 | 0.66002444 | -0.2301203 | 0.81799832 | 0.99665574 |
| RBPJ | 52.8400893 | 0.415029866 | 0.26125687 | 1.58858931 | 0.11215314 | 0.81214897 |
| TBC1D19 | 19.5619905 | 0.431061341 | 0.48030208 | 0.89747964 | 0.36946303 | 0.98532838 |
| STIM2 | 19.7007103 | -0.805502815 | 0.49695193 | -1.6208868 | 0.10504192 | 0.7935283 |
| ARAP2 | 19.6042296 | 0.055611173 | 0.63606859 | 0.08742952 | 0.9303301 | 0.99777713 |
| C4orf19 | 62.2623588 | -0.345637247 | 0.3210193 | -1.0766868 | 0.2816202 | 0.96059387 |
| RELL1 | 5.41944131 | -2.036878781 | 0.78222787 | -2.6039455 | 0.00921574 | 0.32667543 |
| PGM2 | 5.14250943 | 0.157765331 | 0.64606207 | 0.24419532 | 0.80707956 | 0.99665574 |
| TBC1D1 | 235.305258 | 0.034048379 | 0.29869683 | 0.11398976 | 0.90924591 | 0.99777713 |
| KLF3 | 22.0782041 | 0.518090884 | 0.46709613 | 1.10917401 | 0.26735512 | 0.94557817 |
| FAM114A1 | 11.7636015 | -0.998527114 | 0.52459401 | -1.9034283 | 0.05698467 | 0.67545594 |
| KLHL5 | 33.3469435 | -0.556746017 | 0.41256274 | -1.3494821 | 0.17718218 | 0.88584149 |
| WDR19 | 11.4601385 | -0.523169766 | 0.50692295 | -1.0320499 | 0.30204876 | 0.97358745 |
| RFC1 | 17.299784 | 0.227772146 | 0.41575023 | 0.54785814 | 0.58378931 | 0.99665574 |
| RPL9 | 4.72425792 | 0.579525199 | 0.67144845 | 0.86309709 | 0.38808409 | 0.99168679 |
| LIAS | 7.30764743 | 1.449088273 | 0.65080071 | 2.22662368 | 0.02597244 | 0.51788242 |
| UGDH | 8.30000876 | 0.023299571 | 0.5420825 | 0.0429816 | 0.9657162 | 0.99777713 |
| UGDH-AS1 | 8.78931681 | -0.331386135 | 0.57677194 | -0.5745531 | 0.56559355 | 0.99665574 |
| SMIM14 | 29.6100394 | -0.005367388 | 0.38229058 | -0.0140401 | 0.98879801 | 0.99777713 |
| UBE2K | 83.2069161 | 0.137583247 | 0.31460064 | 0.43732666 | 0.66187446 | 0.99665574 |
| PDS5A | 62.3080506 | 0.005453711 | 0.26311378 | 0.02072757 | 0.98346297 | 0.99777713 |
| N4BP2 | 13.0771523 | 0.061551799 | 0.43421839 | 0.14175309 | 0.88727504 | 0.99665574 |
| RBM47 | 240.017838 | -0.089179256 | 0.25342344 | -0.3518982 | 0.7249146 | 0.99665574 |
| NSUN7 | 30.7170548 | 0.233446883 | 0.43758757 | 0.53348609 | 0.59369714 | 0.99665574 |
| APBB2 | 102.446014 | 0.504911753 | 0.36268587 | 1.39214618 | 0.16387814 | 0.8734454 |
| UCHL1 | 3.38040975 | -0.707914495 | 0.73619787 | -0.9615818 | 0.3362597 | 0.97788427 |
| LIMCH1 | 82.4368777 | 0.454301301 | 0.44571095 | 1.01927336 | 0.30807321 | 0.97376371 |
| TMEM33 | 30.5243782 | -0.025177812 | 0.28904055 | -0.0871082 | 0.93058548 | 0.99777713 |
| SLC30A9 | 47.2515038 | -0.098303823 | 0.27853269 | -0.3529346 | 0.72413747 | 0.99665574 |
| ATP8A1 | 26.0661051 | 0.091889905 | 0.42609356 | 0.21565664 | 0.8292554 | 0.99665574 |
| KCTD8 | 14.4271157 | -2.480000013 | 0.80385652 | -3.0851277 | 0.00203465 | 0.13751507 |
| GUF1 | 9.27551127 | 0.134919086 | 0.53348004 | 0.25290372 | 0.80034262 | 0.99665574 |
| GNPDA2 | 5.49702031 | 0.299586961 | 0.67508825 | 0.44377451 | 0.65720562 | 0.99665574 |
| COMMD8 | 4.58887782 | -0.011104601 | 0.6576013 | -0.0168865 | 0.98652714 | 0.99777713 |
| ATP10D | 65.3005437 | 1.813395607 | 0.38062964 | 4.76419969 | 1.90E-06 | 0.00120138 |
| AC107068.2 | 3.23575266 | -1.143591006 | 0.97912719 | -1.1679698 | 0.24281894 | 0.9342309 |
| NFXL1 | 15.0991189 | -0.522633816 | 0.44089505 | -1.1853928 | 0.23586217 | 0.9305595 |
| NIPAL1 | 5.98664627 | -0.333280637 | 0.72097069 | -0.4622666 | 0.64389018 | 0.99665574 |
| TXK | 3.68674475 | -0.158589875 | 0.84229277 | -0.1882836 | 0.85065438 | 0.99665574 |
| TEC | 13.2314193 | -0.139333052 | 0.43482772 | -0.3204328 | 0.74864028 | 0.99665574 |
| SLAIN2 | 32.8547328 | 0.047690034 | 0.34614543 | 0.13777456 | 0.89041859 | 0.99702493 |
| FRYL | 70.5897294 | 0.190477322 | 0.30709701 | 0.6202513 | 0.53509235 | 0.99665574 |
| OCIAD1 | 41.1052045 | 0.329145069 | 0.33017306 | 0.99688651 | 0.3188196 | 0.97700345 |
| OCIAD2 | 5.83381067 | 0.484002052 | 0.72363222 | 0.66885089 | 0.5035906 | 0.99665574 |
| DCUN1D4 | 20.3158791 | -0.094049072 | 0.46642347 | -0.2016388 | 0.84019911 | 0.99665574 |
| SGCB | 3.88528349 | -0.929974376 | 0.83605731 | -1.1123333 | 0.26599488 | 0.94557817 |
| USP46 | 6.58690924 | -0.061876131 | 0.5824008 | -0.1062432 | 0.91538939 | 0.99777713 |
| SCFD2 | 50.0288746 | -0.424326573 | 0.34298188 | -1.2371691 | 0.21602432 | 0.91441171 |
| AC058822.1 | 23.2235188 | -0.162940177 | 0.61235997 | -0.2660856 | 0.79017328 | 0.99665574 |
| LNX1 | 49.2405344 | 0.284298968 | 0.43463013 | 0.65411702 | 0.51303641 | 0.99665574 |
| CHIC2 | 22.4716495 | 0.034462449 | 0.37696334 | 0.09142122 | 0.9271579 | 0.99777713 |
| KIT | 19.4893412 | 0.557758737 | 0.59934799 | 0.93060917 | 0.35205577 | 0.97838306 |
| TMEM165 | 40.9603116 | -0.200316179 | 0.30741102 | -0.6516233 | 0.51464422 | 0.99665574 |
| CLOCK | 26.9202942 | 0.483372903 | 0.35663651 | 1.35536571 | 0.17530107 | 0.88428483 |
| EXOC1 | 30.5106264 | 0.097326129 | 0.39792359 | 0.24458497 | 0.8067778 | 0.99665574 |
| CEP135 | 9.3338659 | -0.408443683 | 0.51129606 | -0.7988399 | 0.42438327 | 0.99518711 |
| AASDH | 13.703312 | 0.126496479 | 0.41437814 | 0.30526823 | 0.76016187 | 0.99665574 |
| PPAT | 3.55651943 | 1.078492032 | 0.94156971 | 1.14541921 | 0.25203555 | 0.94076968 |
| PAICS | 7.00066912 | -0.444929466 | 0.57571615 | -0.7728278 | 0.43962428 | 0.99617422 |

| | | | | | | |
|---|---|---|---|---|---|---|
| SRP72 | 16.2518025 | 0.098564027 | 0.41591758 | 0.23697971 | 0.81267253 | 0.99665574 |
| REST | 25.7728353 | 0.089606295 | 0.37274658 | 0.24039468 | 0.8100243 | 0.99665574 |
| NOA1 | 3.89116283 | 0.372350616 | 0.77414272 | 0.48098446 | 0.63052755 | 0.99665574 |
| POLR2B | 22.3440372 | 0.226235663 | 0.45070535 | 0.50195912 | 0.61569628 | 0.99665574 |
| IGFBP7 | 117.172223 | -1.438618915 | 0.38053371 | -3.7805295 | 0.0001565 | 0.02938052 |
| AC104806.2 | 3.58280978 | -0.629416556 | 0.9107918 | -0.6910652 | 0.48952454 | 0.99640678 |
| CENPC | 20.3438581 | -0.18820443 | 0.46300799 | -0.406482 | 0.68438845 | 0.99665574 |
| UBA6 | 31.4417857 | -0.373666624 | 0.44107707 | -0.8471685 | 0.39690118 | 0.99179694 |
| UBA6-AS1 | 30.5626951 | -0.481841799 | 0.51745744 | -0.9311719 | 0.35176468 | 0.97838306 |
| YTHDC1 | 44.3018824 | 0.163148872 | 0.29429884 | 0.55436465 | 0.57932931 | 0.99665574 |
| UGT2B7 | 29.8184251 | 0.368762712 | 0.74712312 | 0.49357689 | 0.62160502 | 0.99665574 |
| UTP3 | 5.28155349 | -0.158533368 | 0.60220263 | -0.2632559 | 0.79235338 | 0.99665574 |
| RUFY3 | 66.6905695 | -0.980192902 | 0.33078232 | -2.9632566 | 0.00304403 | 0.17337273 |
| GRSF1 | 16.0271298 | 0.296514511 | 0.43891564 | 0.67556151 | 0.49931909 | 0.99640678 |
| MOB1B | 78.3955515 | 0.502064194 | 0.44767081 | 1.12150309 | 0.26207378 | 0.94557817 |
| SLC4A4 | 15.2741701 | 1.323331146 | 1.00134004 | 1.3215602 | 0.18631464 | 0.89532394 |
| COX18 | 3.87190164 | -0.08232582 | 0.75210416 | -0.1094607 | 0.91283712 | 0.99777713 |
| ANKRD17 | 163.16303 | 0.229023201 | 0.25460608 | 0.89951978 | 0.36837586 | 0.98532838 |
| AC053527.1 | 11.9256953 | 0.066192976 | 0.51326396 | 0.12896479 | 0.89738551 | 0.99777713 |
| MTHFD2L | 13.6336762 | 0.140122185 | 0.50754794 | 0.27607675 | 0.78248912 | 0.99665574 |
| PARM1 | 3.30728585 | -1.115233661 | 1.12516061 | -0.9911773 | 0.32159901 | 0.97700345 |
| RCHY1 | 15.4452441 | 0.029110774 | 0.40752151 | 0.07143371 | 0.94305258 | 0.99777713 |
| THAP6 | 5.90548583 | -0.266660706 | 0.63021332 | -0.4231277 | 0.6722021 | 0.99665574 |
| CDKL2 | 3.52936308 | -1.453940037 | 0.81391477 | -1.7863542 | 0.07404193 | 0.72810353 |
| G3BP2 | 20.9870139 | -0.302991724 | 0.37889304 | -0.7996761 | 0.42389846 | 0.99518711 |
| USO1 | 42.769858 | 0.150120065 | 0.28361244 | 0.52931411 | 0.59658757 | 0.99665574 |
| SDAD1 | 10.4463188 | -0.582539114 | 0.60491384 | -0.9630117 | 0.33554164 | 0.97788427 |
| NUP54 | 22.7142924 | 0.071238809 | 0.34882113 | 0.20422733 | 0.83817585 | 0.99665574 |
| SCARB2 | 38.4276686 | 0.432134546 | 0.29346929 | 1.47250345 | 0.14088498 | 0.84969914 |
| SHROOM3 | 102.244591 | -0.614416688 | 0.50110147 | -1.2261323 | 0.2201489 | 0.91768188 |
| 11-Sep | 25.3667013 | -0.288088817 | 0.51756659 | -0.5566217 | 0.57778589 | 0.99665574 |
| CCNI | 87.4253375 | -0.015708783 | 0.32388681 | -0.0485008 | 0.96131709 | 0.99777713 |
| CCNG2 | 22.8488172 | -0.158751295 | 0.42501248 | -0.3735215 | 0.70876035 | 0.99665574 |
| CNOT6L | 51.8051989 | 0.370368978 | 0.44337992 | 0.83533097 | 0.40353139 | 0.99194293 |
| MRPL1 | 32.4248574 | -0.063344094 | 0.45714376 | -0.1385649 | 0.88979396 | 0.99665574 |
| FRAS1 | 53.0402273 | -0.26074807 | 0.41652502 | -0.6260082 | 0.53130958 | 0.99665574 |
| BMP2K | 21.5021861 | 0.666802046 | 0.4576915 | 1.45688098 | 0.14514924 | 0.85059132 |
| PAQR3 | 3.62702346 | -0.601223249 | 0.72092083 | -0.8339657 | 0.40430031 | 0.99194293 |
| RASGEF1B | 112.381873 | 1.659474973 | 0.55345134 | 2.99841172 | 0.00271391 | 0.16107021 |
| HNRNPD | 45.416445 | -0.137544412 | 0.31875269 | -0.4315082 | 0.66609887 | 0.99665574 |
| HNRNPDL | 46.5741964 | 0.440409889 | 0.32347469 | 1.3614972 | 0.17335662 | 0.88157681 |
| ENOPH1 | 7.98146935 | 0.04824422 | 0.52981093 | 0.09105931 | 0.92744546 | 0.99777713 |
| TMEM150C | 15.7011602 | -0.071476678 | 0.42936925 | -0.166469 | 0.86778786 | 0.99665574 |
| SEC31A | 65.7454507 | -0.826271475 | 0.33298894 | -2.4813781 | 0.01308755 | 0.38795772 |
| THAP9 | 3.4357489 | 0.717615921 | 0.75695383 | 0.9480313 | 0.34311352 | 0.97788427 |
| LIN54 | 7.17612543 | -0.300098068 | 0.61668613 | -0.4866302 | 0.62652045 | 0.99665574 |
| COPS4 | 10.3417074 | -0.103948835 | 0.4523547 | -0.229795 | 0.81825109 | 0.99665574 |
| COQ2 | 4.39352055 | 0.655485239 | 0.83450344 | 0.78547937 | 0.43217256 | 0.99617422 |
| HPSE | 12.5423047 | -0.825994708 | 0.98087474 | -0.8421001 | 0.39973193 | 0.99179694 |
| HELQ | 9.94177868 | -0.604833631 | 0.47777975 | -1.2659256 | 0.20553971 | 0.90874907 |
| MRPS18C | 3.966493 | 0.121294538 | 0.7385268 | 0.16423851 | 0.86954339 | 0.99665574 |
| ABRAXAS1 | 5.64730902 | 0.044616093 | 0.72762441 | 0.06131748 | 0.95110637 | 0.99777713 |
| GPAT3 | 54.6301951 | 1.966667585 | 0.50573364 | 3.88874189 | 0.00010077 | 0.02321722 |
| CDS1 | 16.1018804 | -0.114053394 | 0.40361415 | -0.2825803 | 0.77749861 | 0.99665574 |
| WDFY3 | 99.1569864 | -0.301672753 | 0.27248785 | -1.1071053 | 0.26824839 | 0.94660092 |
| ARHGAP24 | 59.9185345 | 1.390412682 | 0.50691883 | 2.74287047 | 0.00609047 | 0.24797265 |
| MAPK10 | 46.3638617 | 0.015519837 | 0.61852785 | 0.02509157 | 0.97998192 | 0.99777713 |
| PTPN13 | 37.5071552 | 0.54026429 | 0.49058468 | 1.10126612 | 0.27078085 | 0.94865166 |
| AC093827.5 | 5.22104222 | 0.40650911 | 0.75658478 | 0.53729486 | 0.59106396 | 0.99665574 |
| AFF1 | 166.335894 | -0.238858343 | 0.29216354 | -0.8175501 | 0.41361411 | 0.99279547 |
| KLHL8 | 24.6946929 | -0.126673118 | 0.43854149 | -0.2888509 | 0.77269546 | 0.99665574 |
| HSD17B11 | 8.88343458 | -0.813808901 | 0.64578813 | -1.2601794 | 0.20760465 | 0.90994205 |
| NUDT9 | 20.8036004 | -0.477603951 | 0.46907106 | -1.0181911 | 0.30858717 | 0.97376371 |
| SPP1 | 781.961217 | -4.145334471 | 0.59005086 | -7.025385 | 2.13E-12 | 1.08E-08 |
| PKD2 | 54.800204 | -0.913794117 | 0.3242573 | -2.8181142 | 0.00483066 | 0.22060024 |
| PPM1K | 9.68606026 | -0.139702633 | 0.52270853 | -0.2672668 | 0.78926375 | 0.99665574 |
| HERC6 | 5.45948339 | -0.927680976 | 1.18881911 | -0.7803382 | 0.43519183 | 0.99617422 |
| HERC5 | 9.04321253 | -0.030217073 | 0.95719831 | -0.0315682 | 0.97481637 | 0.99777713 |
| PYURF | 6.46222323 | -0.189851941 | 0.64504561 | -0.2943233 | 0.76851088 | 0.99665574 |
| HERC3 | 18.8127484 | 0.232801739 | 0.49420423 | 0.47106383 | 0.63759515 | 0.99665574 |
| FAM13A-AS1 | 7.04812352 | 1.013293336 | 0.76940489 | 1.31698322 | 0.18784425 | 0.89739428 |
| FAM13A | 45.5004277 | 0.113000573 | 0.44735035 | 0.25259972 | 0.80057755 | 0.99665574 |
| GPRIN3 | 38.6425232 | 1.563926554 | 0.6444411 | 2.42679517 | 0.01523285 | 0.41291622 |
| CCSER1 | 94.9106069 | 0.342909097 | 0.52330648 | 0.65527393 | 0.5122914 | 0.99665574 |
| SMARCAD1 | 25.6950817 | 0.118079184 | 0.35002217 | 0.33734772 | 0.73585479 | 0.99665574 |
| PDLIM5 | 95.927584 | 0.292233696 | 0.28721436 | 1.01747593 | 0.30892708 | 0.97376371 |
| BMPR1B | 356.933607 | 0.045149069 | 0.336662 | 0.13410801 | 0.89331717 | 0.99773598 |
| UNC5C | 3.45885602 | -1.506682726 | 1.29714325 | -1.1615392 | 0.2454227 | 0.93426912 |
| STPG2 | 76.2937985 | 0.647941848 | 0.55663632 | 1.16403085 | 0.24441151 | 0.93426912 |
| RAP1GDS1 | 27.7971412 | 0.200614734 | 0.40638662 | 0.49365487 | 0.62154995 | 0.99665574 |
| TSPAN5 | 70.8912601 | 0.691258525 | 0.5490643 | 1.25897555 | 0.20803916 | 0.91026892 |
| EIF4E | 24.2161973 | 0.043150682 | 0.46118662 | 0.09356447 | 0.92545513 | 0.99777713 |
| METAP1 | 17.765081 | 0.697371537 | 0.3760427 | 1.85450091 | 0.06366753 | 0.68630818 |
| ADH5 | 15.218344 | 0.393345544 | 0.39306616 | 1.00071079 | 0.31696665 | 0.97700345 |
| AP002026.1 | 6.16550051 | 1.691599263 | 0.72603961 | 2.32989942 | 0.01981147 | 0.46397674 |
| TRMT10A | 3.12504584 | -1.084372142 | 0.71125028 | -1.5245999 | 0.12735891 | 0.83778831 |
| LAMTOR3 | 8.89819818 | 0.093062733 | 0.69149586 | 0.13458176 | 0.89294256 | 0.99773598 |
| DNAJB14 | 36.2070918 | 0.109305944 | 0.3060963 | 0.35709659 | 0.7210195 | 0.99665574 |
| H2AFZ | 14.5532424 | -0.267750837 | 0.60906256 | -0.4396114 | 0.66021859 | 0.99665574 |
| DDIT4L | 4.44509085 | 2.886116455 | 1.18494245 | 2.43565961 | 0.01486467 | 0.40950537 |
| PPP3CA | 263.10667 | 0.346974947 | 0.30632495 | 1.13270221 | 0.25733933 | 0.94557817 |
| AP002075.1 | 2.48738383 | -1.285088995 | 1.31950372 | -0.9739184 | 0.33009704 | 0.97788427 |
| NFKB1 | 28.9117247 | -0.601890425 | 0.43706891 | -1.3771065 | 0.16847933 | 0.87566218 |
| MANBA | 15.1436507 | 0.3502539 | 0.57041784 | 0.6140304 | 0.53919524 | 0.99665574 |
| UBE2D3 | 74.364418 | 0.18426134 | 0.21343302 | 0.86332162 | 0.38796066 | 0.99168679 |
| CISD2 | 8.24729666 | -0.070781508 | 0.52680249 | -0.1343606 | 0.89311742 | 0.99773598 |
| TET2 | 43.5501585 | 0.467919428 | 0.32689307 | 1.43141432 | 0.15231151 | 0.85976284 |
| PPA2 | 42.7551884 | 0.197046887 | 0.29524593 | 0.66739916 | 0.5045172 | 0.99665574 |
| ARHGEF38 | 44.7594263 | -0.365389231 | 0.47938803 | -0.7621993 | 0.44594105 | 0.99617422 |
| INTS12 | 13.3465517 | -0.120301068 | 0.42729761 | -0.2815393 | 0.77829679 | 0.99665574 |
| GSTCD | 15.6715533 | -0.766458901 | 0.41210679 | -1.8598551 | 0.06290603 | 0.68628329 |
| NPNT | 46.8698526 | 0.977509177 | 0.3034203 | 3.22163403 | 0.00127462 | 0.10255619 |
| TBCK | 71.4640392 | -0.101758249 | 0.39036269 | -0.2606762 | 0.79434225 | 0.99665574 |
| AIMP1 | 11.308367 | -0.611161862 | 0.5463243 | -1.1186796 | 0.26327684 | 0.94557817 |
| PAPSS1 | 20.7366088 | 0.131894057 | 0.4308928 | 0.30609483 | 0.75953244 | 0.99665574 |
| SGMS2 | 42.0183237 | 0.181605037 | 0.42764236 | 0.42466569 | 0.6710804 | 0.99665574 |
| AC096564.1 | 7.77939389 | 0.824514446 | 0.92998964 | 0.88658455 | 0.37530261 | 0.98569969 |
| HADH | 5.75359155 | 0.015189146 | 0.69004108 | 0.02201195 | 0.98243843 | 0.99777713 |
| RPL34-AS1 | 4.72674462 | 0.721956518 | 0.76955681 | 0.93814584 | 0.34816947 | 0.97838306 |
| RPL34 | 50.6146291 | 0.527141656 | 0.40938252 | 1.2876506 | 0.19786762 | 0.90381759 |
| OSTC | 8.04946522 | 0.162560188 | 0.59585203 | 0.27281973 | 0.7849918 | 0.99665574 |
| SEC24B | 55.3628873 | -0.135827687 | 0.32745552 | -0.4147974 | 0.67829024 | 0.99665574 |
| CASP6 | 6.76139941 | -1.289523818 | 0.64746962 | -1.991636 | 0.04641101 | 0.64011028 |

| | | | | | | |
|---|---|---|---|---|---|---|
| PLA2G12A | 8.79109515 | -0.096975176 | 0.5462482 | -0.1775295 | 0.85909249 | 0.99665574 |
| GAR1 | 3.10136084 | -0.368633361 | 0.86881016 | -0.4242968 | 0.67134938 | 0.99665574 |
| EGF | 26.4022971 | 3.570742467 | 1.18393779 | 3.01598826 | 0.00256143 | 0.1564325 |
| ELOVL6 | 22.8816109 | 0.093729786 | 0.46949673 | 0.19963885 | 0.84176304 | 0.99665574 |
| ENPEP | 7.37219881 | 1.389659587 | 0.84754004 | 1.63963887 | 0.10108028 | 0.78386461 |
| FAM241A | 10.6628795 | -0.569967452 | 0.43283796 | -1.3168149 | 0.18790069 | 0.89739428 |
| AP1AR | 15.5149861 | -0.192967031 | 0.39634081 | -0.4868715 | 0.62634942 | 0.99665574 |
| ALPK1 | 29.2744396 | -0.810589888 | 0.49834663 | -1.6265584 | 0.10383093 | 0.79062881 |
| ZGRF1 | 4.34417616 | -0.579843982 | 0.724345 | -0.800508 | 0.42341652 | 0.99518711 |
| LARP7 | 20.3675808 | 0.145875273 | 0.37546974 | 0.38851406 | 0.69763565 | 0.99665574 |
| ANK2 | 11.0250468 | 1.192897322 | 0.65496838 | 1.82130522 | 0.06856047 | 0.70708655 |
| CAMK2D | 77.2633418 | -0.936222956 | 0.40666037 | -2.3022233 | 0.02132258 | 0.4733368 |
| ARSJ | 107.368185 | 0.614248711 | 0.34134181 | 1.79951209 | 0.07193771 | 0.71852664 |
| UGT8 | 59.6862161 | 0.352124502 | 0.39783279 | 0.88510679 | 0.37609904 | 0.98655788 |
| SNHG8 | 6.44554311 | 0.658548536 | 0.66481043 | 0.99058093 | 0.32189025 | 0.97700345 |
| METTL14 | 11.0846245 | 0.095616478 | 0.42889144 | 0.22293865 | 0.82358326 | 0.99665574 |
| SEC24D | 15.2887397 | -2.018803897 | 0.80461592 | -2.509028 | 0.01210639 | 0.37648636 |
| USP53 | 70.8006424 | 0.071191412 | 0.32439947 | 0.21945601 | 0.82629485 | 0.99665574 |
| C4orf3 | 20.4527063 | 0.129147049 | 0.4157659 | 0.31062444 | 0.75608615 | 0.99665574 |
| PDE5A | 20.1568417 | -1.277889787 | 0.5212577 | -2.4515509 | 0.01422421 | 0.40056949 |
| AC108866.1 | 14.9512547 | -0.84059065 | 0.48421002 | -1.7360042 | 0.08256309 | 0.75228386 |
| PRDM5 | 10.6424242 | -0.690706245 | 0.49968288 | -1.3822892 | 0.16688292 | 0.87495175 |
| ANXA5 | 55.9478032 | -0.123293801 | 0.32325523 | -0.3814132 | 0.70289669 | 0.99665574 |
| EXOSC9 | 2.45289421 | 0.886566471 | 1.01227511 | 0.87581573 | 0.38113022 | 0.98820924 |
| BBS7 | 5.15067158 | -0.224767505 | 0.77287049 | -0.2908217 | 0.77118769 | 0.99665574 |
| KIAA1109 | 70.4961838 | 0.149477222 | 0.28505848 | 0.52437388 | 0.60001852 | 0.99665574 |
| SPATA5 | 38.4366996 | 0.339602246 | 0.34390789 | 0.98748023 | 0.32340727 | 0.97700345 |
| SPRY1 | 5.88542689 | -0.967045222 | 0.61124 | -1.582104 | 0.11362584 | 0.81582066 |
| LINC01091 | 67.0690424 | 0.834265423 | 0.34834338 | 2.39495129 | 0.01662257 | 0.42361121 |
| ANKRD50 | 5.24161611 | -0.045932631 | 0.65533304 | -0.0700905 | 0.94412162 | 0.99777713 |
| INTU | 8.55652271 | 0.273679532 | 0.69854602 | 0.39178454 | 0.69521742 | 0.99665574 |
| HSPA4L | 19.6651072 | 0.244640718 | 0.42586706 | 0.57445325 | 0.56566113 | 0.99665574 |
| MFSD8 | 14.0343585 | 0.117430542 | 0.47616861 | 0.24661547 | 0.80520583 | 0.99665574 |
| ABHD18 | 43.4897215 | -0.262771922 | 0.29486508 | -0.8911599 | 0.37284341 | 0.98532838 |
| LARP1B | 48.9071866 | -0.099418312 | 0.33774865 | -0.2943559 | 0.76848595 | 0.99665574 |
| PGRMC2 | 25.6702686 | -0.244585944 | 0.68590284 | -0.3565898 | 0.72139893 | 0.99665574 |
| AC108062.1 | 13.4263706 | -0.514738558 | 0.44775893 | -1.1495886 | 0.25031336 | 0.93814301 |
| AC078850.2 | 8.68608633 | -0.797597426 | 0.91750199 | -0.8693141 | 0.38467534 | 0.99051797 |
| JADE1 | 104.90353 | -0.519978706 | 0.32157338 | -1.616983 | 0.10588194 | 0.79596216 |
| SCLT1 | 32.1464841 | -0.174991387 | 0.29859306 | -0.5860531 | 0.55783983 | 0.99665574 |
| C4orf33 | 3.55291623 | 0.096505267 | 0.92299873 | 0.10455623 | 0.91672795 | 0.99777713 |
| AC096711.2 | 12.8482753 | 0.258582936 | 0.59048013 | 0.43791979 | 0.66144443 | 0.99665574 |
| AC109927.1 | 6.5121095 | 0.971116107 | 0.67272337 | 1.44355935 | 0.14886297 | 0.85485259 |
| NOCT | 3.03178248 | -1.715288846 | 1.15117552 | -1.4900324 | 0.13621571 | 0.84475183 |
| ELF2 | 152.763006 | -0.26770652 | 0.29617007 | -0.9038946 | 0.36605131 | 0.98365837 |
| NDUFC1 | 17.4780254 | 0.835931974 | 0.53670079 | 1.55753819 | 0.11934276 | 0.82625998 |
| NAA15 | 30.55138 | 0.208173839 | 0.31424121 | 0.66246511 | 0.50767319 | 0.99665574 |
| AC097376.2 | 36.4141996 | -1.355597397 | 0.38662023 | -3.5062764 | 0.00045442 | 0.0583157 |
| RAB33B | 3.341692 | 0.40022032 | 0.87875869 | 0.45543825 | 0.64879398 | 0.99665574 |
| SETD7 | 7.84411024 | -1.014927912 | 0.54855124 | -1.8501971 | 0.06428515 | 0.68846743 |
| MGST2 | 21.6516334 | 0.788173923 | 0.61490033 | 1.28179134 | 0.19991585 | 0.90381759 |
| MAML3 | 73.105005 | -0.197465766 | 0.47601962 | -0.4148269 | 0.67826859 | 0.99665574 |
| SCOC | 27.2259815 | -0.076654136 | 0.43350752 | -0.1768231 | 0.85964736 | 0.99665574 |
| ELMOD2 | 8.04496019 | -0.223186911 | 0.48355126 | -0.4615579 | 0.64439838 | 0.99665574 |
| TBC1D9 | 48.14767 | -0.650418832 | 0.34013501 | -1.9122372 | 0.05584577 | 0.67320387 |
| RNF150 | 142.632342 | -0.385417067 | 0.39368973 | -0.9789868 | 0.32758648 | 0.97788427 |
| ZNF330 | 9.32063245 | 0.59461007 | 0.55203307 | 1.07712763 | 0.28142325 | 0.96059387 |
| IL15 | 7.0299353 | -1.974650066 | 0.94701066 | -2.0851403 | 0.03705657 | 0.58591936 |
| INPP4B | 181.767335 | 0.253222972 | 0.33576355 | 0.75417052 | 0.45074684 | 0.99617422 |
| AC104596.1 | 3.51715005 | 1.463268848 | 1.01061122 | 1.44790481 | 0.14764368 | 0.85288409 |
| USP38 | 10.849153 | 0.14510877 | 0.46979651 | 0.30887579 | 0.75741601 | 0.99665574 |
| GAB1 | 51.7909467 | 0.5492973 | 0.37324554 | 1.47167812 | 0.14110782 | 0.84969914 |
| SMARCA5 | 29.0053036 | -0.025610476 | 0.30099319 | -0.0850866 | 0.93219257 | 0.99777713 |
| ANAPC10 | 27.0702884 | 0.501299486 | 0.33485479 | 1.49706529 | 0.13437627 | 0.84373478 |
| ABCE1 | 9.14308877 | -0.119416881 | 0.53803285 | -0.2219509 | 0.8243521 | 0.99665574 |
| OTUD4 | 33.3575747 | 0.13880546 | 0.3751569 | 0.36999308 | 0.71138764 | 0.99665574 |
| SMAD1 | 42.0936583 | 0.189730033 | 0.47566856 | 0.39887024 | 0.68998882 | 0.99665574 |
| MMAA | 4.33989964 | 0.349987526 | 0.71205975 | 0.49151427 | 0.62306276 | 0.99665574 |
| ZNF827 | 46.0767254 | 0.289650746 | 0.26795577 | 1.08096478 | 0.27971278 | 0.95996214 |
| LSM6 | 5.61452662 | 0.387566168 | 0.62197739 | 0.62311938 | 0.53320607 | 0.99665574 |
| SLC10A7 | 26.0302769 | -0.324911539 | 0.39018271 | -0.8327164 | 0.4050047 | 0.99194293 |
| TMEM184C | 13.4356935 | 0.328717209 | 0.45203166 | 0.72719953 | 0.46710373 | 0.99617422 |
| PRMT9 | 8.93290087 | -0.582796475 | 0.45870017 | -1.270539 | 0.20389268 | 0.90707783 |
| ARHGAP10 | 6.28072845 | 0.314352923 | 0.78226175 | 0.40185133 | 0.68779344 | 0.99665574 |
| NR3C2 | 363.117911 | 0.152313498 | 0.23528401 | 0.64736017 | 0.51739887 | 0.99665574 |
| AC069272.1 | 4.18962527 | -1.469107815 | 0.86935461 | -1.6898833 | 0.09105029 | 0.76412898 |
| DCLK2 | 69.5647768 | 0.602454004 | 0.41412636 | 1.45475888 | 0.14573603 | 0.85107825 |
| LRBA | 223.028941 | 0.060137067 | 0.23265104 | 0.25848613 | 0.79603175 | 0.99665574 |
| RPS3A | 32.6531325 | 0.535050391 | 0.40637538 | 1.31664076 | 0.18795907 | 0.89739428 |
| SH3D19 | 62.0144245 | -0.416684069 | 0.40192335 | -1.0367252 | 0.29986394 | 0.97358745 |
| FAM160A1 | 85.2833229 | -0.705954973 | 0.37939909 | -1.8607187 | 0.06278391 | 0.68628329 |
| GATB | 6.59009588 | 0.671740365 | 0.60040804 | 1.1188064 | 0.26322274 | 0.94557817 |
| FBXW7 | 68.6507407 | 0.270261476 | 0.2702408 | 1.00007651 | 0.31727348 | 0.97700345 |
| ARFIP1 | 44.7639508 | 0.393825198 | 0.36293312 | 1.08511783 | 0.27786947 | 0.95948252 |
| FHDC1 | 3.51549904 | -1.019229856 | 0.76549929 | -1.3314576 | 0.18303848 | 0.89500374 |
| TRIM2 | 107.176113 | 0.005198739 | 0.30106669 | 0.01726773 | 0.98622303 | 0.99777713 |
| TMEM131L | 17.3648842 | -0.125508804 | 0.42703124 | -0.2939101 | 0.76882658 | 0.99665574 |
| PLRG1 | 7.08132548 | 0.970562362 | 0.57946182 | 1.67493756 | 0.09394648 | 0.76974878 |
| MAP9 | 5.25389407 | 0.149365907 | 0.70574909 | 0.21164166 | 0.83238661 | 0.99665574 |
| GUCY1A1 | 7.76619936 | -3.409232861 | 0.76587286 | -4.4514345 | 8.53E-06 | 0.00344882 |
| CTSO | 4.04504279 | 0.211615813 | 0.71624145 | 0.29545318 | 0.76764771 | 0.99665574 |
| PDGFC | 23.5347298 | -0.098800829 | 0.60628835 | -0.1629601 | 0.87054982 | 0.99665574 |
| ETFDH | 16.1133768 | 0.148400603 | 0.43745852 | 0.33923354 | 0.7344338 | 0.99665574 |
| PPID | 9.8140036 | 0.4925318 | 0.5049286 | 0.9754484 | 0.32933788 | 0.97788427 |
| FNIP2 | 140.607668 | -0.651977304 | 0.37994198 | -1.7159917 | 0.08616355 | 0.75397496 |
| C4orf45 | 5.28122915 | -1.567158951 | 0.90224334 | -1.7369582 | 0.08239455 | 0.75228386 |
| RAPGEF2 | 124.110168 | 0.205210625 | 0.29638635 | 0.69237542 | 0.4887016 | 0.99640678 |
| NAF1 | 7.71593806 | -0.331950793 | 0.49780905 | -0.6668235 | 0.50488485 | 0.99665574 |
| TMA16 | 5.26483547 | 0.47248607 | 0.63500625 | 0.74406523 | 0.45683702 | 0.99617422 |
| 1-Mar | 3.29845148 | -0.570058639 | 0.82264582 | -0.6929576 | 0.48833619 | 0.99640678 |
| TMEM192 | 12.5627184 | -0.165346413 | 0.45790621 | -0.3610923 | 0.71803044 | 0.99665574 |
| KLHL2 | 24.0473476 | -0.300628392 | 0.45045548 | -0.6673876 | 0.5045246 | 0.99665574 |
| MSMO1 | 17.1214875 | -1.410304576 | 0.65441987 | -2.1550455 | 0.03115828 | 0.56407608 |
| DDX60 | 13.2669899 | -0.989547149 | 1.88039422 | -0.5262445 | 0.59871833 | 0.99665574 |
| DDX60L | 10.6550426 | 0.081608528 | 1.03469768 | 0.07887186 | 0.93713455 | 0.99777713 |
| PALLD | 60.9393312 | -0.850266764 | 0.72220074 | -1.1773275 | 0.23906483 | 0.93157178 |
| CBR4 | 63.1170978 | -0.454568335 | 0.34668627 | -1.3111807 | 0.18979673 | 0.8994569 |
| SH3RF1 | 109.034634 | -0.550239262 | 0.34333137 | -1.6026478 | 0.10901244 | 0.80434362 |
| NEK1 | 35.4048111 | 0.097963882 | 0.31166894 | 0.31432033 | 0.75327776 | 0.99665574 |
| CLCN3 | 41.8463998 | -0.137336185 | 0.31363446 | -0.4378861 | 0.66146882 | 0.99665574 |
| HPF1 | 6.75167448 | 0.497828786 | 0.54433367 | 0.91456547 | 0.36041979 | 0.9819332 |
| MFAP3L | 4.68615995 | -0.422014663 | 0.74502218 | -0.5664458 | 0.57109079 | 0.99665574 |
| AADAT | 4.58192795 | 0.141406646 | 0.9080228 | 0.15573028 | 0.87624563 | 0.99665574 |

| | | | | | | |
|---|---|---|---|---|---|---|
| AC105285.1 | 3.01960753 | 0.309008394 | 1.17857712 | 0.26218767 | 0.79317676 | 0.99665574 |
| GALNT7 | 68.2530429 | 0.34204438 | 0.23333283 | 1.46590764 | 0.14267345 | 0.84969914 |
| HMGB2 | 9.79845234 | 0.287727452 | 0.50382548 | 0.57108556 | 0.56794165 | 0.99665574 |
| FBXO8 | 8.25545509 | 0.068008268 | 0.64455625 | 0.10551177 | 0.91596973 | 0.99777713 |
| CEP44 | 11.4392139 | 0.176233762 | 0.50497666 | 0.34899387 | 0.72709391 | 0.99665574 |
| WDR17 | 4.94646783 | 0.17608393 | 0.65620748 | 0.26833575 | 0.78844089 | 0.99665574 |
| ASB5 | 3.55404234 | -0.409196033 | 1.1297344 | -0.3622055 | 0.71719846 | 0.99665574 |
| SPCS3 | 26.953585 | -0.106515073 | 0.42676122 | -0.2495894 | 0.8029049 | 0.99665574 |
| AC078881.1 | 8.37066724 | -0.576585683 | 0.90109946 | -0.6398691 | 0.52225772 | 0.99665574 |
| TENM3 | 5.40604372 | 0.965565966 | 0.95598505 | 1.01002203 | 0.31248473 | 0.97476007 |
| DCTD | 9.10737289 | 0.566577989 | 0.51500477 | 1.10014125 | 0.27127058 | 0.94865166 |
| WWC2 | 54.0032145 | 0.351440111 | 0.32401486 | 1.08464195 | 0.27808027 | 0.95958042 |
| CDKN2AIP | 4.74563705 | -0.655041578 | 0.80820925 | -0.8104851 | 0.41766141 | 0.9941891 |
| RWDD4 | 9.66688323 | 0.242766665 | 0.50937912 | 0.47659328 | 0.63365177 | 0.99665574 |
| TRAPPC11 | 18.6320231 | -0.153915607 | 0.38525131 | -0.39952 | 0.68951009 | 0.99665574 |
| STOX2 | 97.7193069 | -0.074106266 | 0.49739435 | -0.148989 | 0.88156235 | 0.99665574 |
| IRF2 | 15.7103892 | 0.408715787 | 0.4994251 | 0.81837254 | 0.4131445 | 0.99279547 |
| CASP3 | 7.05659983 | 0.396811834 | 0.54832078 | 0.72368557 | 0.4692588 | 0.99617422 |
| PRIMPOL | 7.50630155 | -0.113416072 | 0.72909915 | -0.1555564 | 0.87638266 | 0.99665574 |
| ACSL1 | 48.0402187 | 0.752173458 | 0.47575884 | 1.58099732 | 0.11387865 | 0.81638817 |
| SLC25A4 | 17.740016 | -0.176348181 | 0.55204585 | -0.3194448 | 0.74938924 | 0.99665574 |
| CFAP97 | 15.2419949 | 0.242008471 | 0.41851841 | 0.57825048 | 0.56309502 | 0.99665574 |
| SNX25 | 25.1110857 | -0.644105467 | 0.33622837 | -1.9156785 | 0.05540603 | 0.67128562 |
| AC112722.1 | 2.75057077 | -0.714116332 | 0.87412345 | -0.8169514 | 0.41395623 | 0.99279547 |
| UFSP2 | 7.35584256 | -0.292704147 | 0.5291559 | -0.5531529 | 0.58015868 | 0.99665574 |
| SORBS2 | 54.7689361 | 0.231546172 | 0.5875591 | 0.3940815 | 0.69352087 | 0.99665574 |
| TLR3 | 4.159327 | 0.012670777 | 0.91772329 | 0.01380675 | 0.98898416 | 0.99777713 |
| FAM149A | 7.0117632 | -0.013594942 | 0.60479354 | -0.0224786 | 0.98206614 | 0.99777713 |
| CYP4V2 | 5.90381339 | 0.514401837 | 0.58440249 | 0.88021842 | 0.378741 | 0.98679664 |
| KLKB1 | 4.47117974 | 2.04937534 | 0.81563095 | 2.51262576 | 0.01198364 | 0.37496952 |
| FAT1 | 9.52034184 | -0.462209881 | 0.5948412 | -0.7770307 | 0.43714065 | 0.99617422 |
| FRG1-DT | 10.9167557 | 1.007800134 | 0.71678164 | 1.40600717 | 0.15972199 | 0.86757292 |
| FRG1 | 5.90888731 | -0.207089549 | 0.61635791 | -0.3359891 | 0.73687907 | 0.99665574 |
| CCDC127 | 2.21621376 | 0.390713529 | 1.06522405 | 0.36679 | 0.71377567 | 0.99665574 |
| SDHA | 37.066027 | -0.11358493 | 0.30600402 | -0.3711877 | 0.71049773 | 0.99665574 |
| PDCD6 | 10.9597997 | -0.293527592 | 0.56991747 | -0.5150353 | 0.60652838 | 0.99665574 |
| EXOC3 | 13.153037 | -0.267699725 | 0.57328906 | -0.4669542 | 0.64053266 | 0.99665574 |
| SLC9A3 | 7.57619902 | 0.00511715 | 0.68780205 | 0.00743986 | 0.99406391 | 0.9983802 |
| TPPP | 3.27937555 | 0.638271346 | 0.89935462 | 0.70969931 | 0.47789062 | 0.99617422 |
| ZDHHC11B | 8.66254007 | 0.278352516 | 0.72496532 | 0.38395287 | 0.70101338 | 0.99665574 |
| BRD9 | 9.63066253 | -0.714721982 | 0.47532771 | -1.5036405 | 0.13267397 | 0.84089786 |
| SLC12A7 | 10.1568527 | -0.48230235 | 0.5671669 | -0.8503711 | 0.39511879 | 0.99179694 |
| CLPTM1L | 11.7690245 | 0.315079939 | 0.47852043 | 0.65844616 | 0.51025148 | 0.99665574 |
| LPCAT1 | 3.80684188 | -0.472692074 | 0.69299428 | -0.682101 | 0.49517511 | 0.99640678 |
| MRPL36 | 4.91458292 | -0.38189284 | 0.74870413 | -0.5100718 | 0.61000118 | 0.99665574 |
| NDUFS6 | 21.5336635 | 0.179628871 | 0.4676458 | 0.38411309 | 0.70089463 | 0.99665574 |
| ADAMTS16 | 3.98683968 | 0.502887334 | 0.88767391 | 0.5665226 | 0.57103857 | 0.99665574 |
| ICE1 | 26.9850508 | -0.166084982 | 0.36454302 | -0.4555978 | 0.64867926 | 0.99665574 |
| MED10 | 8.4711884 | 0.289740607 | 0.71892609 | 0.40301863 | 0.68693452 | 0.99665574 |
| UBE2QL1 | 12.6523699 | 1.693477589 | 0.61929737 | 2.73451442 | 0.00624724 | 0.25132729 |
| NSUN2 | 14.5661716 | 0.185004443 | 0.39465428 | 0.46877597 | 0.63922978 | 0.99665574 |
| SRD5A1 | 4.20975509 | 0.429712855 | 0.70385714 | 0.61051147 | 0.54152305 | 0.99665574 |
| TENT4A | 14.6152368 | 0.199539604 | 0.48018063 | 0.41555113 | 0.67773849 | 0.99665574 |
| MTRR | 10.1209505 | 0.413070745 | 0.53349554 | 0.77427216 | 0.43876986 | 0.99617422 |
| MIR4458HG | 3.64133452 | -0.319382808 | 0.83379912 | -0.3830453 | 0.7016862 | 0.99665574 |
| SEMA5A | 77.6703274 | 0.346483801 | 0.60465786 | 0.57302455 | 0.56662807 | 0.99665574 |
| FAM173B | 6.18274287 | 0.49327703 | 0.60702797 | 0.81261005 | 0.41644167 | 0.992853 |
| CCT5 | 14.9361122 | 0.568023902 | 0.48095553 | 1.18103206 | 0.23758999 | 0.9305595 |
| CMBL | 5.80830453 | 0.182719241 | 0.98182118 | 0.18610236 | 0.85236449 | 0.99665574 |
| 6-Mar | 61.7336322 | -0.304582657 | 0.22887139 | -1.3308027 | 0.18325394 | 0.89532394 |
| ANKRD33B | 6.38394842 | -0.579722958 | 1.02389383 | -0.5661944 | 0.57126163 | 0.99665574 |
| DAP | 58.4780759 | -0.116319755 | 0.26220413 | -0.4436229 | 0.65731525 | 0.99665574 |
| DNAH5 | 22.5955575 | -4.348861429 | 0.91235335 | -4.7666416 | 1.87E-06 | 0.00120138 |
| TRIO | 119.192313 | -1.19015101 | 0.31025358 | -3.8360589 | 0.00012502 | 0.02485289 |
| OTULIN | 10.2571835 | 0.072676599 | 0.50946227 | 0.14265354 | 0.88656381 | 0.99665574 |
| ANKH | 15.735109 | -0.382393553 | 0.46568686 | -0.8211388 | 0.41156721 | 0.99279547 |
| ZNF622 | 4.62885813 | -0.669848569 | 0.71106727 | -0.9420326 | 0.34617594 | 0.97788427 |
| RETREG1 | 72.4539881 | -0.934156909 | 0.44458996 | -2.1011651 | 0.03562648 | 0.58349146 |
| MYO10 | 167.153898 | -0.197819203 | 0.40526209 | -0.4881266 | 0.62546018 | 0.99665574 |
| LINC02241 | 8.33965876 | 0.546225372 | 0.54625219 | 0.9999509 | 0.31733427 | 0.97700345 |
| CDH12 | 63.8602918 | 1.913807947 | 0.56665189 | 3.37739619 | 0.00073176 | 0.07202465 |
| C5orf17 | 13.9746167 | -0.764763182 | 1.00495141 | -0.7609952 | 0.44665995 | 0.99617422 |
| CDH6 | 9.22702094 | 1.061551067 | 0.79074392 | 1.34247136 | 0.17944321 | 0.88820302 |
| DROSHA | 20.4553642 | 0.276801673 | 0.33368442 | 0.82953131 | 0.40680383 | 0.99250037 |
| C5orf22 | 5.84406884 | 0.110715071 | 0.62771136 | 0.17637895 | 0.85999624 | 0.99665574 |
| PDZD2 | 347.446794 | 0.665626015 | 0.42951699 | 1.54970824 | 0.12121156 | 0.82659025 |
| GOLPH3 | 38.653279 | 0.473565408 | 0.37964226 | 1.24739909 | 0.2122512 | 0.91043197 |
| MTMR12 | 18.3086567 | -0.460651509 | 0.36763983 | -1.2529967 | 0.21020689 | 0.91043197 |
| ZFR | 58.4071041 | -0.141186104 | 0.27554811 | -0.5123828 | 0.60838315 | 0.99665574 |
| SUB1 | 40.5656186 | 0.347166442 | 0.31435091 | 1.10439139 | 0.26942339 | 0.94742087 |
| NPR3 | 3.81882464 | 0.828173333 | 0.79724253 | 1.03879723 | 0.29889905 | 0.97358745 |
| TARS | 8.6047267 | -0.11348851 | 0.61138484 | -0.1856253 | 0.85273859 | 0.99665574 |
| C1QTNF3-AN | 22.4668007 | -0.182289059 | 0.5195117 | -0.3508854 | 0.72567434 | 0.99665574 |
| RAI14 | 35.1640242 | 0.203161038 | 0.37386627 | 0.54340563 | 0.58685055 | 0.99665574 |
| RAD1 | 7.36678019 | 0.473126842 | 0.62154877 | 0.76120631 | 0.44653385 | 0.99617422 |
| BRIX1 | 5.31418946 | -0.091982804 | 0.62647374 | -0.1468263 | 0.88326915 | 0.99665574 |
| DNAJC21 | 14.9035499 | 0.014587176 | 0.41786714 | 0.03490865 | 0.97215259 | 0.99777713 |
| PRLR | 41.6499222 | 0.94199172 | 0.55831364 | 1.68720885 | 0.0915632 | 0.76595534 |
| SPEF2 | 6.4004384 | 1.111809133 | 0.83016857 | 1.33925708 | 0.180487 | 0.89040251 |
| LMBRD2 | 10.3996455 | 0.846173451 | 0.48293956 | 1.75213116 | 0.07975126 | 0.7446402 |
| NADK2 | 8.11670155 | 0.399633726 | 0.57066756 | 0.70029165 | 0.48374519 | 0.99640678 |
| RANBP3L | 12.3253906 | 0.749840368 | 1.05354431 | 0.71173121 | 0.47663124 | 0.99617422 |
| NIPBL | 137.256249 | 0.098705012 | 0.24435215 | 0.40394575 | 0.68625262 | 0.99665574 |
| CPLANE1 | 38.0441505 | -0.32328103 | 0.3287592 | -0.9833368 | 0.32544168 | 0.97788427 |
| NUP155 | 7.96498025 | 0.529002699 | 0.52248977 | 1.01246518 | 0.31131569 | 0.97430509 |
| WDR70 | 68.5051484 | 0.182418967 | 0.29306912 | 0.62244349 | 0.53365029 | 0.99665574 |
| LIFR | 31.3925027 | 0.368358168 | 0.45130535 | 0.81620608 | 0.41438228 | 0.99279547 |
| OSMR-AS1 | 19.7538403 | -1.668020508 | 0.48188553 | -3.4614455 | 0.00053728 | 0.0633369 |
| OSMR | 27.0382146 | -3.249996442 | 0.5080479 | -6.3970276 | 1.58E-10 | 3.08E-07 |
| RICTOR | 89.4786795 | 0.391083574 | 0.28219228 | 1.38587625 | 0.16578469 | 0.87495175 |
| FYB1 | 17.7473964 | -4.61687321 | 1.02178227 | -4.5184511 | 6.23E-06 | 0.0030073 |
| TTC33 | 14.4012397 | -0.062905984 | 0.40739274 | -0.1544112 | 0.87728556 | 0.99665574 |
| PRKAA1 | 20.2877469 | 0.19823796 | 0.40531012 | 0.48910193 | 0.62476953 | 0.99665574 |
| RPL37 | 31.8705117 | 0.077281931 | 0.40960166 | 0.18867582 | 0.8503469 | 0.99665574 |
| PLCXD3 | 7.34903558 | -0.367443325 | 0.85109192 | -0.4317317 | 0.66593646 | 0.99665574 |
| OXCT1 | 105.617592 | 0.501742906 | 0.48508372 | 1.0343429 | 0.30097589 | 0.97358745 |
| C5orf51 | 5.05057235 | 0.294770884 | 0.74251556 | 0.3969895 | 0.6913752 | 0.99665574 |
| FBXO4 | 1.93308748 | -0.68029068 | 0.99554133 | -0.6833375 | 0.49439363 | 0.99640678 |
| GHR | 5.28245868 | -0.761727496 | 0.65172108 | -1.1687937 | 0.24248676 | 0.9342309 |
| SELENOP | 50.6548828 | 0.629798802 | 0.51191978 | 1.23026854 | 0.21859656 | 0.91662582 |
| ZNF131 | 27.3823351 | -0.045053394 | 0.40444452 | -0.1113957 | 0.91130254 | 0.99777713 |
| HMGCS1 | 21.67067 | -1.364824597 | 0.69938008 | -1.9514777 | 0.05100025 | 0.65200573 |
| TMEM267 | 11.5267389 | -0.884198671 | 0.4256365 | -2.0773563 | 0.03776869 | 0.58998295 |

| | | | | | | |
|---|---|---|---|---|---|---|
| PAIP1 | 14.7378331 | -0.686147458 | 0.51194716 | -1.3402701 | 0.18015756 | 0.88920999 |
| NNT-AS1 | 21.4867674 | 0.075063262 | 0.4194645 | 0.17895021 | 0.85797679 | 0.99665574 |
| NNT | 69.771428 | 0.048277573 | 0.30589234 | 0.15782538 | 0.8745944 | 0.99665574 |
| MRPS30 | 7.56590008 | 0.961606735 | 0.5929402 | 1.62176005 | 0.10485474 | 0.79270493 |
| PARP8 | 19.3322707 | -1.748291786 | 0.4704007 | -3.7166011 | 0.00020192 | 0.03411789 |
| ITGA1 | 8.30232803 | 0.174909842 | 0.63012318 | 0.2775804 | 0.78133448 | 0.99665574 |
| ITGA2 | 32.2488516 | -1.103026792 | 0.47191339 | -2.3373501 | 0.01942099 | 0.46057537 |
| MOCS2 | 5.78008747 | 0.713608937 | 0.73291832 | 0.97365411 | 0.33022831 | 0.97788427 |
| NDUFS4 | 41.1575223 | 0.110089428 | 0.32828535 | 0.33534675 | 0.73736353 | 0.99665574 |
| ARL15 | 949.017502 | -0.871172718 | 0.27638527 | -3.1520229 | 0.00162144 | 0.11753244 |
| SNX18 | 24.8153419 | 0.676456633 | 0.51104359 | 1.32367697 | 0.18561035 | 0.89532394 |
| DHX29 | 16.8785606 | 0.052781303 | 0.40918938 | 0.12898991 | 0.89736563 | 0.99777713 |
| MTREX | 34.5152668 | 0.118318896 | 0.30229631 | 0.39140039 | 0.6955013 | 0.99665574 |
| PLPP1 | 80.6349793 | 0.475252498 | 0.46842127 | 1.01458352 | 0.3103044 | 0.97376371 |
| SLC38A9 | 21.353535 | -0.639424658 | 0.40609041 | -1.574587 | 0.11535182 | 0.81721642 |
| IL6ST | 63.899324 | 0.79786392 | 0.28258998 | 2.82339776 | 0.00475176 | 0.21929238 |
| AC008892.1 | 3.92961671 | -1.042118835 | 0.79922942 | -1.3039045 | 0.19226615 | 0.89964233 |
| MAP3K1 | 44.003413 | 0.046551418 | 0.41665182 | 0.11172738 | 0.91103957 | 0.99777713 |
| MIER3 | 8.95504611 | 0.481541748 | 0.57855758 | 0.8323143 | 0.40523156 | 0.99194293 |
| GPBP1 | 80.964494 | 0.197643612 | 0.2580987 | 0.76576755 | 0.44381463 | 0.99617422 |
| PDE4D | 844.122062 | 1.069086869 | 0.3707705 | 2.88341943 | 0.00393383 | 0.20347549 |
| ELOVL7 | 41.4273906 | -0.121651854 | 0.44074668 | -0.2760131 | 0.78253801 | 0.99665574 |
| ERCC8 | 12.4083559 | 0.265578672 | 0.49161164 | 0.54022047 | 0.589045 | 0.99665574 |
| NDUFAF2 | 29.2762336 | -0.163527336 | 0.42687183 | -0.383083 | 0.70165823 | 0.99665574 |
| SMIM15 | 7.65917244 | 0.391597178 | 0.56377651 | 0.69459647 | 0.48730822 | 0.99640678 |
| ZSWIM6 | 181.50795 | 0.31405964 | 0.48299583 | 0.65023261 | 0.51554198 | 0.99665574 |
| KIF2A | 20.3067473 | 0.024647579 | 0.32527805 | 0.07577388 | 0.939599 | 0.99777713 |
| DIMT1 | 1.99590837 | 0.377734392 | 1.11805638 | 0.33784915 | 0.73547687 | 0.99665574 |
| IPO11 | 10.1218369 | -0.166850189 | 0.5278392 | -0.3161004 | 0.75192629 | 0.99665574 |
| RNF180 | 21.9409288 | -0.269743313 | 0.36833729 | -0.7323269 | 0.46396905 | 0.99617422 |
| SREK1IP1 | 13.2723014 | -0.287126827 | 0.42684254 | -0.6726762 | 0.5011533 | 0.99640678 |
| CWC27 | 36.0202754 | -0.288464727 | 0.31453801 | -0.9171061 | 0.35908704 | 0.9819332 |
| PPWD1 | 10.6515199 | -0.108943608 | 0.46419581 | -0.2346932 | 0.81444685 | 0.99665574 |
| TRIM23 | 6.97183105 | 0.392735361 | 0.55607708 | 0.70626065 | 0.48002606 | 0.99617422 |
| TRAPPC13 | 11.49998 | 0.536077359 | 0.52952238 | 1.01237905 | 0.31135685 | 0.97430509 |
| NLN | 7.848645 | -0.639773144 | 0.55015794 | -1.16289 | 0.24487415 | 0.93426912 |
| ERBIN | 114.768311 | 0.124610485 | 0.22122879 | 0.56326524 | 0.57325428 | 0.99665574 |
| SREK1 | 45.0639617 | 0.182760954 | 0.36066722 | 0.50673015 | 0.61234417 | 0.99665574 |
| MAST4 | 96.3731986 | -0.666613826 | 0.40590988 | -1.6422705 | 0.10053396 | 0.78386461 |
| PIK3R1 | 54.0011904 | 0.366690359 | 0.42530948 | 0.86217302 | 0.38859231 | 0.99176253 |
| SLC30A5 | 13.9990865 | -0.163484199 | 0.42720634 | -0.3826821 | 0.70195552 | 0.99665574 |
| MRPS36 | 13.1256889 | -0.162522258 | 0.45631594 | -0.3561617 | 0.72171948 | 0.99665574 |
| CDK7 | 6.82072005 | 0.219693464 | 0.58550117 | 0.37522293 | 0.70749468 | 0.99665574 |
| CCDC125 | 9.81509001 | -0.147522901 | 0.46066083 | -0.3202419 | 0.74878496 | 0.99665574 |
| AK6 | 6.75454493 | -0.03239068 | 0.61577127 | -0.0526018 | 0.95804918 | 0.99777713 |
| RAD17 | 15.2292702 | -0.019410295 | 0.48763798 | -0.0398047 | 0.96824881 | 0.99777713 |
| MARVELD2 | 10.3074547 | 0.437694073 | 0.46068582 | 0.95009236 | 0.34206533 | 0.97788427 |
| OCLN | 33.8202514 | 0.375160253 | 0.4304271 | 0.87159999 | 0.38342664 | 0.98955375 |
| GTF2H2C | 12.2798277 | 0.516653503 | 0.47587125 | 1.08570018 | 0.27761166 | 0.95946581 |
| NAIP | 28.3291821 | 0.214550821 | 0.42064077 | 0.51005713 | 0.61001144 | 0.99665574 |
| GTF2H2 | 9.17521276 | -0.259453345 | 0.55957172 | -0.4636641 | 0.64288839 | 0.99665574 |
| BDP1 | 51.5224537 | -0.143112085 | 0.30991114 | -0.4617843 | 0.64423605 | 0.99665574 |
| MCCC2 | 21.96181 | 0.138940606 | 0.34066908 | 0.40784624 | 0.68338656 | 0.99665574 |
| MRPS27 | 18.0471466 | 0.194280038 | 0.42457527 | 0.4575868 | 0.64724933 | 0.99665574 |
| PTCD2 | 2.90900399 | 0.157251825 | 0.78881702 | 0.19935146 | 0.84198783 | 0.99665574 |
| TNPO1 | 46.3858191 | -0.049392164 | 0.26685427 | -0.1850904 | 0.85315813 | 0.99665574 |
| FCHO2 | 41.7425721 | -0.261244992 | 0.39161179 | -0.667102 | 0.504707 | 0.99665574 |
| BTF3 | 23.5013068 | -0.38630177 | 0.4458092 | -0.8665182 | 0.38620607 | 0.99051797 |
| ANKRA2 | 6.36407205 | -0.125387417 | 0.63145197 | -0.19857 | 0.84259912 | 0.99665574 |
| ARHGEF28 | 108.92263 | -0.074988793 | 0.39759374 | -0.1886066 | 0.85040118 | 0.99665574 |
| ENC1 | 3.73283485 | 0.863571933 | 0.92438982 | 0.93420754 | 0.35019685 | 0.97838306 |
| HEXB | 9.0331481 | -0.023662717 | 0.50127042 | -0.0472055 | 0.96234945 | 0.99777713 |
| GFM2 | 9.06447631 | 0.642469157 | 0.50568494 | 1.27049295 | 0.20390909 | 0.90707783 |
| NSA2 | 10.9145991 | -0.213111062 | 0.5165538 | -0.4125631 | 0.67992671 | 0.99665574 |
| FAM169A | 25.5919495 | 0.240290429 | 0.3740123 | 0.64246664 | 0.52057024 | 0.99665574 |
| AC116337.3 | 2.99248225 | 0.004557102 | 0.87401498 | 0.00521399 | 0.99583986 | 0.99869666 |
| GCNT4 | 3.02516155 | 1.606909748 | 0.89212619 | 1.80121351 | 0.07166923 | 0.71725834 |
| HMGCR | 13.0530767 | -1.168657798 | 0.62082422 | -1.8824294 | 0.05977773 | 0.68092881 |
| COL4A3BP | 54.1429657 | 0.328062962 | 0.33964218 | 0.9659076 | 0.33409042 | 0.97788427 |
| POLK | 33.3488179 | 0.054233698 | 0.34061087 | 0.1592248 | 0.87349176 | 0.99665574 |
| ANKDD1B | 10.1450326 | 1.238658676 | 0.75753561 | 1.6351161 | 0.10202471 | 0.78476974 |
| POC5 | 4.65303665 | -0.80071113 | 0.65511838 | -1.2222388 | 0.22161732 | 0.92117932 |
| IQGAP2 | 235.855055 | 0.323432684 | 0.25805535 | 1.25334618 | 0.21007975 | 0.91043197 |
| F2RL1 | 7.27420143 | -0.639846439 | 0.70718035 | -0.9047854 | 0.3655791 | 0.98365837 |
| ZBED3 | 4.92424315 | -0.858906958 | 0.59829378 | -1.435594 | 0.1511179 | 0.85612189 |
| WDR41 | 17.5353674 | -0.040047937 | 0.47734476 | -0.0838973 | 0.93313809 | 0.99777713 |
| TBCA | 60.1060705 | 0.137465842 | 0.24830324 | 0.55362081 | 0.57983838 | 0.99665574 |
| AP3B1 | 79.082809 | 0.364803026 | 0.34961427 | 1.04344431 | 0.29674256 | 0.97341216 |
| SCAMP1 | 30.7211399 | 0.887197639 | 0.48291474 | 1.83717242 | 0.06618445 | 0.69459411 |
| LHFPL2 | 2.43522919 | -0.117466374 | 0.87583757 | -0.1341189 | 0.89330855 | 0.99773598 |
| ARSB | 8.66134535 | -0.366587878 | 0.6132211 | -0.597807 | 0.54996871 | 0.99665574 |
| DMGDH | 17.184892 | 0.838179917 | 0.57978376 | 1.44567678 | 0.14826788 | 0.85405669 |
| BHMT2 | 12.9889179 | -0.135360104 | 0.52592995 | -0.2573729 | 0.79689094 | 0.99665574 |
| BHMT | 4.36271413 | 2.135247647 | 1.00354047 | 2.12771453 | 0.03336076 | 0.5761693 |
| JMY | 33.0979383 | -0.085001893 | 0.44703229 | -0.1901471 | 0.84919386 | 0.99665574 |
| HOMER1 | 20.9819967 | 1.42598124 | 0.58995459 | 2.41710337 | 0.01564457 | 0.41847166 |
| TENT2 | 75.7687546 | 0.161443452 | 0.27796832 | 0.58079803 | 0.56137658 | 0.99665574 |
| CMYA5 | 11.7103813 | -0.628997593 | 0.60919125 | -1.0325125 | 0.30183208 | 0.97358745 |
| MTX3 | 7.12676807 | 0.237986374 | 0.53636406 | 0.44370306 | 0.65725728 | 0.99665574 |
| THBS4 | 3.8074506 | -0.700172731 | 0.75071145 | -0.9326789 | 0.35098578 | 0.97838306 |
| SERINC5 | 46.1578874 | -0.893510215 | 0.5400293 | -1.6545588 | 0.09801403 | 0.78118418 |
| AC010260.1 | 3.28745223 | -1.657041174 | 0.949383 | -1.7453875 | 0.08091745 | 0.74735643 |
| ZFYVE16 | 67.6492528 | 0.564721551 | 0.30228067 | 1.86820263 | 0.06173383 | 0.68358351 |
| FAM151B | 8.53857633 | 0.327039702 | 0.4951306 | 0.660512 | 0.50892532 | 0.99665574 |
| DHFR | 181.916753 | -0.414147552 | 0.49460465 | -0.8373305 | 0.40240683 | 0.99194293 |
| MSH3 | 28.8825424 | -0.306178208 | 0.38777058 | -0.789586 | 0.4297696 | 0.99617422 |
| RASGRF2 | 5.56001823 | 0.024371484 | 1.01501895 | 0.02401087 | 0.98084394 | 0.99777713 |
| CKMT2-AS1 | 13.5513321 | 0.322156321 | 0.41225504 | 0.78144907 | 0.43453842 | 0.99617422 |
| CKMT2 | 9.33257578 | -0.266342838 | 0.71047613 | -0.3748794 | 0.70775019 | 0.99665574 |
| ZCCHC9 | 3.91187017 | -0.839523242 | 0.71592996 | -1.1726332 | 0.24094293 | 0.93216487 |
| SSBP2 | 105.280835 | 0.16355176 | 0.30962006 | 0.52823373 | 0.59733712 | 0.99665574 |
| ATG10 | 34.5423852 | 0.100697744 | 0.36346622 | 0.27704843 | 0.78174292 | 0.99665574 |
| RPS23 | 89.374662 | 0.43866288 | 0.39487283 | 1.11089658 | 0.26661287 | 0.94557817 |
| ATP6AP1L | 6.72330191 | -0.421245189 | 0.64367067 | -0.6544421 | 0.51282701 | 0.99665574 |
| TMEM167A | 14.1693729 | -0.1219348 | 0.51035963 | -0.2389194 | 0.81116811 | 0.99665574 |
| XRCC4 | 20.3667553 | -0.456155589 | 0.56245608 | -0.8110066 | 0.41736188 | 0.99392253 |
| COX7C | 58.8956681 | -0.198692928 | 0.39893652 | -0.4980565 | 0.61844422 | 0.99665574 |
| RASA1 | 51.1583342 | -0.296659702 | 0.28020058 | -1.0587405 | 0.28971798 | 0.96676791 |
| CCNH | 41.0310638 | 0.262571858 | 0.4513339 | 0.58176852 | 0.56072261 | 0.99665574 |
| TMEM161B | 24.9101901 | -0.079226141 | 0.45104442 | -0.1756504 | 0.86056859 | 0.99665574 |
| TMEM161B-. | 27.4091513 | 0.144144908 | 0.37175638 | 0.38774024 | 0.69820827 | 0.99665574 |
| CETN3 | 4.48193507 | 0.896154115 | 0.88504115 | 1.01255644 | 0.31127208 | 0.97430509 |
| LYSMD3 | 7.86118622 | -0.479359045 | 0.70738568 | -0.6776488 | 0.49799441 | 0.99640678 |

| | | | | | | |
|---|---|---|---|---|---|---|
| ADGRV1 | 21.4831415 | 0.646572984 | 0.3752018 | 1.72326727 | 0.0848402 | 0.75373899 |
| ARRDC3 | 37.6757843 | 0.505721688 | 0.44160375 | 1.14519338 | 0.25212906 | 0.94076968 |
| ARRDC3-AS1 | 2.60485553 | 0.074068043 | 0.84295671 | 0.08786696 | 0.92998242 | 0.99777713 |
| FAM172A | 82.9866475 | -0.044927053 | 0.21221498 | -0.2117054 | 0.83233689 | 0.99665574 |
| KIAA0825 | 16.5875331 | -0.298551336 | 0.4630029 | -0.6448153 | 0.51904692 | 0.99665574 |
| SLF1 | 5.49487192 | -0.157105603 | 0.69528778 | -0.2259577 | 0.82123435 | 0.99665574 |
| MCTP1 | 4.58978196 | 0.055594646 | 0.83061721 | 0.06693173 | 0.94663605 | 0.99777713 |
| TTC37 | 37.6142057 | -0.332759169 | 0.36681959 | -0.9071467 | 0.36432924 | 0.98362663 |
| ARSK | 5.05263042 | -0.015222546 | 0.6749059 | -0.0225551 | 0.98200519 | 0.99777713 |
| RHOBTB3 | 108.885642 | -0.338785797 | 0.29001731 | -1.1681571 | 0.24274338 | 0.9342309 |
| GLRX | 19.0670689 | -0.373703821 | 0.49862324 | -0.7494713 | 0.45357318 | 0.99617422 |
| ELL2 | 38.38724 | -0.080270059 | 0.32599145 | -0.2462336 | 0.80550138 | 0.99665574 |
| AC104123.1 | 8.04152098 | 0.290100443 | 0.88625553 | 0.32733273 | 0.74341624 | 0.99665574 |
| CAST | 53.6909003 | -0.946959999 | 0.38969223 | -2.4300202 | 0.01509798 | 0.41291622 |
| ERAP1 | 13.2945813 | -0.587131197 | 0.63114168 | -0.9302685 | 0.35223211 | 0.97838483 |
| AC009126.1 | 2.62507377 | 0.930439894 | 0.96235223 | 0.96683923 | 0.33362441 | 0.97788427 |
| ERAP2 | 7.15941245 | -1.361366992 | 0.77278004 | -1.7616488 | 0.07812866 | 0.74233211 |
| LNPEP | 71.2218035 | 0.651264719 | 0.3339264 | 1.95032415 | 0.0511375 | 0.65232219 |
| LIX1-AS1 | 8.61447577 | 0.04310744 | 0.61250357 | 0.07037908 | 0.94389194 | 0.99777713 |
| RIOK2 | 3.24582692 | -0.671405867 | 0.8852603 | -0.7584276 | 0.44819503 | 0.99617422 |
| CHD1 | 40.9106583 | 0.154760719 | 0.30150087 | 0.51330106 | 0.60774074 | 0.99665574 |
| LINC02062 | 7.99391436 | 0.144496405 | 0.84682357 | 0.17063342 | 0.86451202 | 0.99665574 |
| FAM174A | 7.05212581 | -0.254564106 | 0.5772042 | -0.4410295 | 0.65919161 | 0.99665574 |
| SLCO4C1 | 9.10775382 | 0.710897219 | 0.72951665 | 0.97447702 | 0.32981974 | 0.97788427 |
| PAM | 38.9666487 | -1.156157764 | 0.46398277 | -2.4918118 | 0.01270934 | 0.38590337 |
| GIN1 | 4.69876642 | 1.220254182 | 0.74832967 | 1.63063717 | 0.1029669 | 0.78770791 |
| PPIP5K2 | 23.8285442 | 0.310523431 | 0.3462077 | 0.89692815 | 0.36975726 | 0.98532838 |
| NUDT12 | 5.22137149 | 0.610749596 | 0.63937514 | 0.95522887 | 0.33946197 | 0.97788427 |
| EFNA5 | 158.161542 | -2.351032734 | 0.55713203 | -4.2198843 | 2.44E-05 | 0.00821807 |
| FBXL17 | 161.549161 | 0.161301568 | 0.24594392 | 0.65584693 | 0.51192262 | 0.99665574 |
| FER | 84.4455866 | 0.226580777 | 0.29880992 | 0.75827731 | 0.44828499 | 0.99617422 |
| AC008467.1 | 4.81437652 | -0.33352756 | 0.79706366 | -0.4184453 | 0.67562155 | 0.99665574 |
| PJA2 | 38.9164804 | -0.204598412 | 0.33987006 | -0.6019901 | 0.54718072 | 0.99665574 |
| MAN2A1 | 202.137862 | 0.775459159 | 0.46260575 | 1.67628515 | 0.09368235 | 0.7690297 |
| TMEM232 | 22.6827675 | 0.030625462 | 0.5543543 | 0.05524529 | 0.95594305 | 0.99777713 |
| SLC25A46 | 18.3323007 | -0.060244891 | 0.40466268 | -0.1488768 | 0.88165084 | 0.99665574 |
| WDR36 | 10.5699507 | -0.170380652 | 0.47454187 | -0.3590424 | 0.71956337 | 0.99665574 |
| NREP | 6.21941396 | -0.526645204 | 0.76805208 | -0.6856894 | 0.49290896 | 0.99640678 |
| EPB41L4A | 85.6242638 | -0.961661975 | 0.31810944 | -3.0230538 | 0.00250238 | 0.15473119 |
| EPB41L4A-AS | 3.29751404 | -0.62902264 | 0.72210491 | -0.8710959 | 0.38370179 | 0.98956212 |
| APC | 17.0295879 | -0.000480252 | 0.37229206 | -0.00129 | 0.99897074 | 0.99940262 |
| SRP19 | 13.1505568 | -0.321605197 | 0.4431429 | -0.725737 | 0.46800002 | 0.99617422 |
| REEP5 | 64.5743173 | 0.487636146 | 0.33500366 | 1.4556144 | 0.14549924 | 0.85067551 |
| DCP2 | 26.3188968 | 0.008636037 | 0.37634361 | 0.02294721 | 0.98169238 | 0.99777713 |
| MCC | 19.7495505 | 0.460519399 | 0.49687054 | 0.92683981 | 0.35400971 | 0.98022903 |
| YTHDC2 | 34.4181729 | 0.688977172 | 0.33289786 | 2.06963536 | 0.0384865 | 0.59541251 |
| PGGT1B | 13.3530821 | 0.112646806 | 0.40684869 | 0.27687641 | 0.78187501 | 0.99665574 |
| CCDC112 | 3.17139343 | -0.097391295 | 0.78716952 | -0.1237234 | 0.90153428 | 0.99777713 |
| FEM1C | 4.26792275 | -0.151644804 | 0.70191296 | -0.216045 | 0.82895265 | 0.99665574 |
| TMED7-TICAI | 3.39919036 | 0.385226549 | 0.83314831 | 0.46237452 | 0.64381277 | 0.99665574 |
| ATG12 | 11.7256696 | -0.096839372 | 0.49506993 | -0.1956075 | 0.84491742 | 0.99665574 |
| AP3S1 | 16.1386114 | -0.28484488 | 0.46305323 | -0.615145 | 0.53845896 | 0.99665574 |
| COMMD10 | 65.4225354 | 0.196636615 | 0.22599044 | 0.87011033 | 0.38424012 | 0.99033485 |
| SEMA6A | 6.85351651 | -1.289942699 | 0.56076802 | -2.3003143 | 0.02143042 | 0.4733368 |
| SEMA6A-AS1 | 8.46069059 | -0.736157799 | 0.5199197 | -1.4159067 | 0.15680282 | 0.86532131 |
| DTWD2 | 21.9131749 | 0.621899519 | 0.43987187 | 1.41381971 | 0.15741484 | 0.86532131 |
| AC008629.1 | 3.96449304 | -0.566178982 | 0.82879167 | -0.6831379 | 0.49451972 | 0.99640678 |
| DMXL1 | 145.465009 | -0.036761988 | 0.26244782 | -0.1400735 | 0.88860191 | 0.99665574 |
| TNFAIP8 | 70.055818 | -0.567376373 | 0.39975619 | -1.419306 | 0.15580982 | 0.8636413 |
| HSD17B4 | 39.8955738 | 0.013265454 | 0.40294826 | 0.03292098 | 0.9737376 | 0.99777713 |
| SRFBP1 | 12.7044546 | 0.193548665 | 0.52320759 | 0.3699271 | 0.71143681 | 0.99665574 |
| SNX2 | 24.0600895 | 0.10026215 | 0.38415465 | 0.26099424 | 0.79409695 | 0.99665574 |
| SNX24 | 49.6879166 | 0.363143379 | 0.26703866 | 1.35989067 | 0.17386452 | 0.88264323 |
| PPIC | 9.24788418 | -0.326703975 | 0.52469161 | -0.622659 | 0.5335086 | 0.99665574 |
| CEP120 | 8.83936427 | 0.185576144 | 0.49040518 | 0.37841391 | 0.70512313 | 0.99665574 |
| CSNK1G3 | 26.6821597 | 0.247667858 | 0.34753843 | 0.71263446 | 0.47607198 | 0.99617422 |
| ZNF608 | 36.6435329 | 0.249156717 | 0.46642887 | 0.53417945 | 0.59321738 | 0.99665574 |
| GRAMD2B | 69.1405215 | -0.028791197 | 0.45146917 | -0.0637722 | 0.94915159 | 0.99777713 |
| ALDH7A1 | 10.7649176 | -0.059000012 | 0.45152969 | -0.130667 | 0.89603877 | 0.99777713 |
| PHAX | 6.44124723 | -0.330413308 | 0.69788155 | -0.4734518 | 0.63589085 | 0.99665574 |
| 3-Mar | 6.81513933 | -1.867395919 | 0.89707447 | -2.0816509 | 0.03737436 | 0.58653453 |
| C5orf63 | 6.44253459 | -0.556758077 | 0.61044476 | -0.9120532 | 0.36174073 | 0.9819332 |
| PRRC1 | 14.5766464 | 0.24706982 | 0.67587375 | 0.36555617 | 0.71469628 | 0.99665574 |
| LINC01184 | 13.7572288 | 0.354969171 | 0.49620276 | 0.71537122 | 0.47437969 | 0.99617422 |
| SLC12A2 | 9.06539775 | 0.643061598 | 0.66714917 | 0.96389477 | 0.33509869 | 0.97788427 |
| SLC27A6 | 3.46096958 | 4.930058831 | 1.48319702 | 3.32394063 | 0.00088755 | 0.08137272 |
| ISOC1 | 4.46283812 | 0.701755698 | 0.7092864 | 0.9893827 | 0.32247594 | 0.97700345 |
| HINT1 | 59.0263708 | -0.379273892 | 0.41089129 | -0.9230517 | 0.35598028 | 0.98175411 |
| LYRM7 | 7.86532406 | 0.01250975 | 0.63571177 | 0.01967834 | 0.98429997 | 0.99777713 |
| CDC42SE2 | 138.049561 | -0.558115763 | 0.32859962 | -1.6984675 | 0.08941956 | 0.75955113 |
| P4HA2 | 2.78002964 | -1.250492492 | 0.91707139 | -1.3635716 | 0.17270244 | 0.88157681 |
| SLC22A4 | 2.57041471 | -1.003250286 | 1.06428005 | -0.9426563 | 0.34585674 | 0.97788427 |
| SLC22A5 | 14.6725259 | 0.893589941 | 0.65634556 | 1.36146262 | 0.17336754 | 0.88157681 |
| AC116366.3 | 19.4659664 | 0.658908106 | 0.5911086 | 1.11469891 | 0.26497947 | 0.94557817 |
| IRF1 | 5.15753636 | -0.987246275 | 0.6853905 | -1.4404143 | 0.14975022 | 0.85525834 |
| KIF3A | 13.9801456 | 0.123864183 | 0.44712973 | 0.27702069 | 0.78176422 | 0.99665574 |
| 8-Sep | 8.95539948 | 0.284721818 | 0.49894496 | 0.57064775 | 0.56823844 | 0.99665574 |
| SOWAHA | 3.27781726 | 0.272665174 | 0.89557202 | 0.30445924 | 0.76077804 | 0.99665574 |
| UQCRQ | 20.6590011 | 0.059452983 | 0.46595346 | 0.12759425 | 0.89847008 | 0.99777713 |
| AFF4 | 87.9803472 | -0.535837679 | 0.36259382 | -1.4777904 | 0.13946388 | 0.84714487 |
| ZCCHC10 | 10.656666 | -0.114754715 | 0.51781574 | -0.221613 | 0.82461514 | 0.99665574 |
| HSPA4 | 22.9287933 | 0.280810124 | 0.4235386 | 0.66300953 | 0.50732445 | 0.99665574 |
| FSTL4 | 14.7836927 | 0.815343283 | 0.69965283 | 1.16535408 | 0.2438757 | 0.93426912 |
| C5orf15 | 14.6731254 | 1.032660139 | 0.45979752 | 2.24590194 | 0.02471029 | 0.50638808 |
| VDAC1 | 43.3137207 | 0.436466596 | 0.33663257 | 1.29656675 | 0.1947803 | 0.90333151 |
| TCF7 | 4.91787231 | 0.373628958 | 0.80165268 | 0.46607336 | 0.64116299 | 0.99665574 |
| SKP1 | 41.6746039 | 0.192310473 | 0.4304197 | 0.44679756 | 0.65502123 | 0.99665574 |
| CDKL3 | 5.6314746 | -0.746420768 | 0.77359307 | -0.9648752 | 0.33460733 | 0.97788427 |
| UBE2B | 44.1664854 | -0.355035585 | 0.31070123 | -1.1426913 | 0.25316679 | 0.94076968 |
| CDKN2AIPNL | 2.36386282 | 0.663579197 | 0.8583495 | 0.77308742 | 0.43947065 | 0.99617422 |
| JADE2 | 10.7614651 | -0.411275057 | 0.70450889 | -0.5837755 | 0.55937134 | 0.99665574 |
| SAR1B | 15.0501338 | 0.316648359 | 0.40224812 | 0.78719662 | 0.43116678 | 0.99617422 |
| SEC24A | 19.1033258 | -0.367512455 | 0.44452788 | -0.8267478 | 0.40838001 | 0.99279547 |
| CAMLG | 9.27533872 | -0.108180911 | 0.51673215 | -0.2093559 | 0.83417044 | 0.99665574 |
| DDX46 | 35.0728708 | 0.043186866 | 0.28723079 | 0.15035598 | 0.88048377 | 0.99665574 |
| C5orf24 | 21.2889163 | 0.732611846 | 0.42121193 | 1.7392951 | 0.08198287 | 0.75124322 |
| TXNDC15 | 6.55649533 | -0.63138079 | 0.56812795 | -1.1113356 | 0.26642394 | 0.94557817 |
| PCBD2 | 29.965888 | -0.100982722 | 0.34821735 | -0.2899991 | 0.77181691 | 0.99665574 |
| H2AFY | 42.9101341 | -0.507492549 | 0.48855199 | -1.0387688 | 0.29891229 | 0.97358745 |
| CXCL14 | 8.21437463 | 0.117124165 | 0.80380981 | 0.14571129 | 0.88414931 | 0.99665574 |
| SMAD5 | 24.2375734 | 0.523156014 | 0.34077624 | 1.53518925 | 0.12473735 | 0.83196528 |
| KLHL3 | 40.590304 | 0.273019283 | 0.61402654 | 0.44463759 | 0.65658168 | 0.99665574 |
| HNRNPA0 | 22.1849809 | -0.220614404 | 0.52481149 | -0.4203688 | 0.67421602 | 0.99665574 |

| | | | | | | |
|---|---|---|---|---|---|---|
| AC106791.1 | 4.03014793 | -0.031110355 | 0.78504367 | -0.0396288 | 0.96838905 | 0.99777713 |
| FAM13B | 59.5851295 | 0.081143042 | 0.34396672 | 0.23590376 | 0.81350735 | 0.99665574 |
| BRD8 | 12.4976731 | 0.546942782 | 0.50455316 | 1.08401419 | 0.27835851 | 0.95969433 |
| CDC23 | 4.67158274 | 0.418127138 | 0.68829078 | 0.60748618 | 0.54352831 | 0.99665574 |
| FAM53C | 4.24519429 | 0.306227982 | 0.70262211 | 0.43583596 | 0.66295575 | 0.99665574 |
| KDM3B | 30.1624743 | 0.031580401 | 0.34518784 | 0.09148758 | 0.92710517 | 0.99777713 |
| EGR1 | 12.3166515 | 1.429615114 | 0.97736278 | 1.46272719 | 0.14354204 | 0.84969914 |
| ETF1 | 23.3633913 | -0.330570745 | 0.44524526 | -0.7424464 | 0.45781691 | 0.99617422 |
| HSPA9 | 42.1425044 | 0.191864848 | 0.36049314 | 0.53222885 | 0.5945675 | 0.99665574 |
| CTNNA1 | 96.8397133 | -0.35042437 | 0.26887703 | -1.3032886 | 0.19247625 | 0.89964233 |
| SIL1 | 35.8010302 | 0.42714404 | 0.38704921 | 1.10359103 | 0.26977058 | 0.94798412 |
| PAIP2 | 20.0873201 | -0.271454925 | 0.38265797 | -0.7093931 | 0.47808057 | 0.99617422 |
| TMEM173 | 3.49541144 | -1.317356265 | 0.74766518 | -1.7619602 | 0.07807603 | 0.74233211 |
| UBE2D2 | 70.5840941 | -0.591517679 | 0.33405971 | -1.7706945 | 0.07661152 | 0.73943494 |
| CXXC5 | 27.377135 | 0.85735992 | 0.51339449 | 1.66998271 | 0.09492278 | 0.77178333 |
| AC008667.1 | 2.451077 | 1.265698639 | 1.0306537 | 1.22805424 | 0.21942661 | 0.91727062 |
| PURA | 28.2240056 | 0.390230719 | 0.31237024 | 1.24925704 | 0.21157108 | 0.91043197 |
| IGIP | 3.75716651 | 0.808937099 | 0.79023768 | 1.02366303 | 0.30599446 | 0.97376371 |
| CYSTM1 | 94.3978818 | 0.050504108 | 0.33136208 | 0.15241366 | 0.87886068 | 0.99665574 |
| PFDN1 | 13.2013072 | 0.275161652 | 0.41851785 | 0.65746694 | 0.51088072 | 0.99665574 |
| SLC4A9 | 530.304392 | 0.930221942 | 0.57651563 | 1.61352424 | 0.10663066 | 0.79764531 |
| AC008438.1 | 108.293441 | 0.930892901 | 0.72205123 | 1.28923387 | 0.19731679 | 0.90381759 |
| ANKHD1-EIF4 | 1.93661999 | 0.41057801 | 1.2282292 | 0.33428452 | 0.73816487 | 0.99665574 |
| SRA1 | 7.28658795 | -0.426821234 | 0.5323118 | -0.8018256 | 0.42265384 | 0.99518711 |
| SLC35A4 | 9.77911317 | 1.074456359 | 0.58151926 | 1.84767115 | 0.06464993 | 0.68846743 |
| NDUFA2 | 16.8353314 | 0.425841898 | 0.54704932 | 0.77843419 | 0.43631309 | 0.99617422 |
| IK | 5.83867229 | 0.010439984 | 0.66536277 | 0.01569066 | 0.98748118 | 0.99777713 |
| WDR55 | 4.43659883 | -0.56985661 | 0.75236381 | -0.7574216 | 0.44879731 | 0.99617422 |
| HARS | 9.63202074 | 0.066753946 | 0.46885451 | 0.14237668 | 0.88678248 | 0.99665574 |
| ZMAT2 | 7.55379363 | 0.465925521 | 0.53989948 | 0.86298568 | 0.38814534 | 0.99168679 |
| TAF7 | 9.79874796 | -0.070134233 | 0.5791037 | -0.1211082 | 0.9036053 | 0.99777713 |
| DIAPH1 | 90.8519927 | -0.47524087 | 0.33700456 | -1.4101912 | 0.15848323 | 0.86661434 |
| HDAC3 | 7.42994474 | -0.063345937 | 0.56479778 | -0.1121568 | 0.91069905 | 0.99777713 |
| PCDH1 | 13.6250476 | 0.24022589 | 0.58872702 | 0.40804292 | 0.68324217 | 0.99665574 |
| DELE1 | 10.156883 | 0.282529678 | 0.52856999 | 0.53451706 | 0.59298385 | 0.99665574 |
| RNF14 | 12.5661875 | 0.486720691 | 0.46312854 | 1.05094082 | 0.29328577 | 0.97072515 |
| GNPDA1 | 6.06501017 | 0.304099143 | 0.63562016 | 0.47842904 | 0.63234487 | 0.99665574 |
| NDFIP1 | 60.564503 | 0.051245866 | 0.25462336 | 0.20126145 | 0.84049415 | 0.99665574 |
| ARHGAP26 | 15.8990889 | -1.158416659 | 0.61388069 | -1.8870388 | 0.05915511 | 0.6784101 |
| NR3C1 | 19.0573467 | 0.047696337 | 0.45450522 | 0.10494123 | 0.91642244 | 0.99777713 |
| YIPF5 | 6.90680924 | -0.35250957 | 0.64486439 | -0.5466414 | 0.58462512 | 0.99665574 |
| PRELID2 | 11.6972 | -0.571768859 | 0.79919619 | -0.7154299 | 0.47434344 | 0.99617422 |
| SH3RF2 | 4.32060667 | 1.229329048 | 0.80756135 | 1.52227326 | 0.12794063 | 0.83778831 |
| LARS | 30.7635918 | 0.93729958 | 0.36335547 | 2.57956648 | 0.00989244 | 0.34338499 |
| TCERG1 | 28.6754115 | 0.063817304 | 0.34281223 | 0.18615819 | 0.85232071 | 0.99665574 |
| STK32A | 6.22830936 | -0.473157543 | 0.80620345 | -0.586896 | 0.55727359 | 0.99665574 |
| SPINK5 | 6.98038359 | -1.041475198 | 0.65670887 | -1.5859009 | 0.11276179 | 0.81249398 |
| AC091948.1 | 5.53049098 | 0.291361408 | 0.72095491 | 0.40413264 | 0.68611519 | 0.99665574 |
| FBXO38 | 18.553786 | 0.003144331 | 0.36314837 | 0.00865853 | 0.99309158 | 0.9978159 |
| AFAP1L1 | 2.66026903 | 1.38035787 | 1.17114969 | 1.17863488 | 0.2385436 | 0.93121101 |
| GRPEL2 | 3.1018926 | -0.405510019 | 0.87676972 | -0.4625046 | 0.64371951 | 0.99665574 |
| CSNK1A1 | 124.798138 | -0.135893589 | 0.23942148 | -0.5675915 | 0.5703124 | 0.99665574 |
| ARHGEF37 | 6.8426162 | -0.52962193 | 0.5966054 | -0.8877257 | 0.37468834 | 0.98562282 |
| SLC26A2 | 4.16649866 | -0.474377405 | 0.91972605 | -0.5157812 | 0.60600723 | 0.99665574 |
| HMGXB3 | 13.1674758 | -0.663138852 | 0.48408576 | -1.3698789 | 0.17072472 | 0.87951271 |
| TCOF1 | 4.44128821 | 0.347756334 | 0.66774297 | 0.5207937 | 0.60251049 | 0.99665574 |
| CD74 | 30.1475806 | -2.411350151 | 0.82097951 | -2.9371624 | 0.00331231 | 0.18250085 |
| RPS14 | 90.7128806 | 0.885826368 | 0.38134943 | 2.3228732 | 0.02018597 | 0.46616245 |
| NDST1 | 16.2891529 | 0.013808071 | 0.49768302 | 0.02774471 | 0.97786576 | 0.99777713 |
| RBM22 | 7.38368259 | -0.013760499 | 0.56231172 | -0.0244713 | 0.98047668 | 0.99777713 |
| DCTN4 | 19.349337 | 0.164378408 | 0.3569582 | 0.46049764 | 0.64515906 | 0.99665574 |
| GPX3 | 124.529119 | 0.113874675 | 0.59911945 | 0.19007007 | 0.84925422 | 0.99665574 |
| TNIP1 | 21.1007148 | -0.381096699 | 0.51327958 | -0.7424739 | 0.45780025 | 0.99617422 |
| ANXA6 | 5.67331604 | 0.866795932 | 0.68963885 | 1.25688385 | 0.2087957 | 0.91043197 |
| SLC36A1 | 4.382967 | -1.246851153 | 1.09798577 | -1.1355804 | 0.2561322 | 0.94314473 |
| SPARC | 6.15528575 | -0.163013896 | 0.79441335 | -0.2052003 | 0.8374156 | 0.99665574 |
| ATOX1 | 17.7698495 | 0.809455282 | 0.40530933 | 1.99712966 | 0.0458111 | 0.63477358 |
| G3BP1 | 35.5225458 | 0.061257627 | 0.32648929 | 0.18762523 | 0.85117045 | 0.99665574 |
| FAM114A2 | 8.34915309 | -0.735156924 | 0.49221926 | -1.4935558 | 0.13529176 | 0.84373478 |
| MFAP3 | 12.8725604 | -0.795074079 | 0.46224685 | -1.7200205 | 0.08542871 | 0.75373899 |
| GALNT10 | 42.9658385 | -1.292473468 | 0.39741996 | -3.2521604 | 0.00114531 | 0.09547898 |
| SAP30L-AS1 | 5.47566858 | -0.644819517 | 0.84124323 | -0.7665078 | 0.4433742 | 0.99617422 |
| SAP30L | 4.80387412 | -0.179891645 | 0.63716627 | -0.2823308 | 0.77768989 | 0.99665574 |
| LARP1 | 49.9401651 | 0.273800533 | 0.32502414 | 0.84240061 | 0.39956375 | 0.99179694 |
| FAXDC2 | 8.62467008 | -0.162363635 | 0.84308714 | -0.1925823 | 0.84728613 | 0.99665574 |
| CNOT8 | 9.2048558 | 0.201050797 | 0.4560651 | 0.44083793 | 0.65933033 | 0.99665574 |
| GEMIN5 | 4.93709859 | 0.701746875 | 0.71574255 | 0.98044593 | 0.32686604 | 0.97788427 |
| MRPL22 | 7.03620348 | 0.286246192 | 0.53055223 | 0.539525 | 0.58952465 | 0.99665574 |
| SGCD | 31.3677885 | -1.989526003 | 1.26689116 | -1.5704001 | 0.11632206 | 0.82007861 |
| HAVCR2 | 3.41103414 | -0.874048402 | 0.77596743 | -1.1263983 | 0.25999696 | 0.94557817 |
| CYFIP2 | 46.1364581 | 0.780279321 | 0.33085071 | 2.35840304 | 0.01835375 | 0.44944531 |
| CLINT1 | 37.3715757 | -0.098507622 | 0.41387135 | -0.2380151 | 0.8118694 | 0.99665574 |
| RNF145 | 11.7805095 | -0.426807014 | 0.47270588 | -0.9029018 | 0.36657799 | 0.98365837 |
| UBLCP1 | 9.03792015 | 0.178039144 | 0.46025503 | 0.38682715 | 0.69888417 | 0.99665574 |
| AC008691.1 | 4.1911962 | -1.509267271 | 0.85818121 | -1.7586813 | 0.07863165 | 0.74431421 |
| TTC1 | 20.9413414 | 0.134836445 | 0.33989849 | 0.39669622 | 0.69159149 | 0.99665574 |
| PWWP2A | 22.5117895 | 0.385762917 | 0.41778794 | 0.92334623 | 0.35582681 | 0.98159786 |
| CCNJL | 13.6526153 | 1.098323669 | 0.51951326 | 2.11413982 | 0.03450333 | 0.58178209 |
| SLU7 | 6.55004394 | 0.522036082 | 0.55660198 | 0.93789835 | 0.34829665 | 0.97838306 |
| AC113414.1 | 34.3479276 | -2.489343913 | 0.82615918 | -3.0131529 | 0.00258549 | 0.15695605 |
| CCNG1 | 16.3052269 | -1.05305264 | 0.39339502 | -2.6768327 | 0.00743217 | 0.28540676 |
| NUDCD2 | 3.98853752 | 0.406073264 | 0.76923651 | 0.5278913 | 0.59757478 | 0.99665574 |
| MAT2B | 13.5988317 | 0.272698075 | 0.47949484 | 0.56871952 | 0.5695465 | 0.99665574 |
| WWC1 | 108.997042 | -0.982613828 | 0.41064859 | -2.3928338 | 0.01671881 | 0.4237383 |
| RARS | 17.5335238 | -0.163332486 | 0.40929523 | -0.3990579 | 0.68985055 | 0.99665574 |
| PANK3 | 22.7066284 | -0.268914616 | 0.44748269 | -0.6009498 | 0.54787345 | 0.99665574 |
| FOXI1 | 99.4034142 | -0.230508846 | 0.41192969 | -0.559583 | 0.57576391 | 0.99665574 |
| LINC01187 | 219.887426 | 0.819069941 | 0.33103605 | 2.47426204 | 0.01335117 | 0.39134822 |
| RANBP17 | 39.2022257 | -0.078044379 | 0.36186974 | -0.2156698 | 0.82924514 | 0.99665574 |
| NPM1 | 28.5740387 | -0.090624932 | 0.37644731 | -0.2407374 | 0.80975868 | 0.99665574 |
| FBXW11 | 78.2427478 | -0.500705474 | 0.35222713 | -1.4215415 | 0.15515941 | 0.86239367 |
| STK10 | 4.5714208 | -1.000354769 | 0.62790989 | -1.5931502 | 0.1111265 | 0.8083966 |
| UBTD2 | 22.0561968 | -0.289679678 | 0.44663682 | -0.6485799 | 0.51660993 | 0.99665574 |
| DUSP1 | 19.4913742 | -0.475180124 | 0.68172889 | -0.6970221 | 0.48578894 | 0.99640678 |
| ERGIC1 | 25.6236978 | -0.428256094 | 0.41259655 | -1.0379536 | 0.29929164 | 0.97358745 |
| RPL26L1 | 4.07278159 | -0.59897754 | 0.72396763 | -0.827354 | 0.40803642 | 0.99279547 |
| ATP6V0E1 | 121.89605 | -0.161565378 | 0.25056968 | -0.6447922 | 0.51906185 | 0.99665574 |
| CREBRF | 59.7042895 | -0.011968312 | 0.42395508 | -0.0282301 | 0.9774786 | 0.99777713 |
| BNIP1 | 3.10546781 | 0.307120167 | 0.81902051 | 0.37498471 | 0.70767184 | 0.99665574 |
| BOD1 | 6.31131361 | -0.191470127 | 0.56876714 | -0.3366406 | 0.73638783 | 0.99665574 |
| CPEB4 | 162.803965 | 0.197135726 | 0.22670905 | 0.86955383 | 0.38454428 | 0.99047507 |
| SFXN1 | 29.8692786 | -0.104138042 | 0.28482356 | -0.365623 | 0.71464641 | 0.99665574 |
| THOC3 | 6.92232296 | 0.071218775 | 0.72647405 | 0.09803347 | 0.92190571 | 0.99777713 |

| | | | | | | |
|---|---|---|---|---|---|---|
| SIMC1 | 27.1654757 | 0.529477776 | 0.35744679 | 1.4812772 | 0.13853272 | 0.84656101 |
| KIAA1191 | 16.1647258 | -0.050655834 | 0.46649293 | -0.1085886 | 0.91352877 | 0.99777713 |
| ARL10 | 4.07988342 | -0.318150438 | 0.72681461 | -0.4377326 | 0.66158015 | 0.99665574 |
| CLTB | 13.4258666 | -0.582628203 | 0.49025471 | -1.1884194 | 0.23466822 | 0.92960234 |
| FAF2 | 20.4913006 | -0.355351037 | 0.3686728 | -0.9638656 | 0.3351133 | 0.97788427 |
| RNF44 | 5.74636507 | 0.058645584 | 0.61181879 | 0.0958545 | 0.92363613 | 0.99777713 |
| TSPAN17 | 4.69585371 | -0.648101257 | 0.67008988 | -0.9671856 | 0.33345128 | 0.97788427 |
| UIMC1 | 29.9800948 | -0.050669492 | 0.34171059 | -0.1482819 | 0.88212031 | 0.99665574 |
| ZNF346 | 14.5062882 | 0.382467115 | 0.38495765 | 0.99353038 | 0.32045156 | 0.97700345 |
| NSD1 | 69.5156307 | 0.196392823 | 0.27604356 | 0.71145592 | 0.47680176 | 0.99617422 |
| PRELID1 | 13.8161208 | -0.246438445 | 0.53900417 | -0.4572107 | 0.64751964 | 0.99665574 |
| LMAN2 | 19.5658076 | -0.304653594 | 0.44148331 | -0.6900682 | 0.4901513 | 0.99640678 |
| RGS14 | 4.01939146 | 0.656449581 | 0.87184685 | 0.75294139 | 0.45148513 | 0.99617422 |
| DBN1 | 11.6234125 | 0.04610266 | 0.49746232 | 0.09267568 | 0.92616122 | 0.99777713 |
| DDX41 | 2.84848285 | 0.222577965 | 0.83854965 | 0.26543206 | 0.79067664 | 0.99665574 |
| FAM193B | 12.8677479 | -0.23627998 | 0.44373496 | -0.53248 | 0.59439361 | 0.99665574 |
| TMED9 | 17.9752078 | -1.100197697 | 0.51513707 | -2.1357378 | 0.03270079 | 0.57356513 |
| B4GALT7 | 5.6016878 | 0.447273355 | 0.61913804 | 0.72241297 | 0.47004061 | 0.99617422 |
| RMND5B | 11.1500598 | -0.616647296 | 0.51242841 | -1.2033823 | 0.2288284 | 0.92646258 |
| NHP2 | 6.84198225 | 0.215485164 | 0.73673507 | 0.29248664 | 0.76991458 | 0.99665574 |
| HNRNPAB | 13.1011327 | -0.835361517 | 0.56703634 | -1.4732063 | 0.14069541 | 0.84969914 |
| PHYKPL | 26.3115655 | -0.558690858 | 0.35937573 | -1.5546149 | 0.12003782 | 0.82659025 |
| CLK4 | 12.1593816 | 0.189393187 | 0.54802379 | 0.345593 | 0.72964861 | 0.99665574 |
| ZNF354A | 5.66130367 | 0.139955234 | 0.64267808 | 0.2177688 | 0.82760926 | 0.99665574 |
| ZNF354B | 4.47826031 | -0.26001087 | 0.66687228 | -0.3898961 | 0.69661341 | 0.99665574 |
| ZNF354C | 3.99102605 | -1.496342033 | 0.70443626 | -2.1241695 | 0.03365597 | 0.57831228 |
| RUFY1 | 21.5322352 | -0.288480117 | 0.43052348 | -0.6700683 | 0.50281427 | 0.99640678 |
| HNRNPH1 | 69.7874446 | 0.562505019 | 0.28079674 | 2.00324623 | 0.04515087 | 0.6313648 |
| CANX | 66.468962 | -0.077665542 | 0.34510432 | -0.2250495 | 0.8219408 | 0.99665574 |
| MAML1 | 9.72694096 | 0.318342975 | 0.57309507 | 0.55548022 | 0.57856624 | 0.99665574 |
| MGAT4B | 14.4418072 | -0.479944266 | 0.41359121 | -1.1604315 | 0.24587317 | 0.93426912 |
| SQSTM1 | 63.1738283 | -1.172616362 | 0.51971111 | -2.256285 | 0.02405279 | 0.50069513 |
| MRNIP | 8.75397809 | 0.082051941 | 0.6227664 | 0.13175396 | 0.8951789 | 0.99777713 |
| TBC1D9B | 14.1844183 | -0.161146598 | 0.39407648 | -0.4089221 | 0.6825968 | 0.99665574 |
| RNF130 | 53.9971632 | -0.832052089 | 0.4153411 | -2.0032982 | 0.04514529 | 0.6313648 |
| MAPK9 | 15.6514531 | 0.196376818 | 0.468553 | 0.41911335 | 0.67513329 | 0.99665574 |
| CNOT6 | 20.1060881 | 0.112099816 | 0.42403068 | 0.26436723 | 0.79149696 | 0.99665574 |
| MGAT1 | 16.8444441 | 0.227876716 | 0.6166361 | 0.36954813 | 0.71171921 | 0.99665574 |
| LINC00847 | 2.27685964 | 0.538909084 | 1.05305784 | 0.51175639 | 0.60882151 | 0.99665574 |
| ZFP62 | 5.30697008 | -0.280468254 | 0.79480108 | -0.3528786 | 0.72417949 | 0.99665574 |
| BTNL9 | 4.14577808 | -0.144686557 | 0.69434475 | -0.2083786 | 0.8349334 | 0.99665574 |
| TRIM41 | 5.58803633 | -0.058202131 | 0.59829991 | -0.0972792 | 0.92250468 | 0.99777713 |
| RACK1 | 54.4641596 | 0.056149162 | 0.41594057 | 0.13499323 | 0.89261722 | 0.99773598 |
| TRIM52 | 7.34440479 | 0.393138184 | 0.58694831 | 0.66980035 | 0.50298507 | 0.99640678 |
| DUSP22 | 6.60592446 | -0.597359171 | 0.56138622 | -1.0640788 | 0.28729301 | 0.96559334 |
| EXOC2 | 27.0908408 | -0.522773798 | 0.43160765 | -1.2112246 | 0.22580934 | 0.92318867 |
| GMDS | 70.1765849 | -0.350472377 | 0.40066089 | -0.8747357 | 0.38171774 | 0.98846857 |
| GMDS-DT | 92.3106643 | -0.303180191 | 0.39339438 | -0.7706775 | 0.44089812 | 0.99617422 |
| WRNIP1 | 12.2253293 | 0.057577432 | 0.41763237 | 0.1378663 | 0.89034608 | 0.99702493 |
| SERPINB1 | 29.5423349 | -0.616372666 | 0.53287485 | -1.1566931 | 0.24739776 | 0.93441937 |
| SERPINB6 | 24.4779826 | 0.356793153 | 0.41624672 | 0.85716749 | 0.39135233 | 0.99179694 |
| NQO2 | 9.41520579 | -0.665847754 | 0.62039791 | -1.0732592 | 0.28315484 | 0.96059387 |
| RIPK1 | 13.906559 | -0.400295381 | 0.54491436 | -0.7346024 | 0.46258169 | 0.99617422 |
| BPHL | 6.41358217 | 0.154658816 | 0.70595296 | 0.21907808 | 0.82658923 | 0.99665574 |
| PSMG4 | 12.2257078 | -0.857193754 | 0.61460887 | -1.394698 | 0.16310693 | 0.8716636 |
| SLC22A23 | 68.1724668 | 0.506838688 | 0.48801945 | 1.03856248 | 0.29900826 | 0.97358745 |
| PRPF4B | 35.6915665 | -0.170795606 | 0.38014861 | -0.4492864 | 0.65322506 | 0.99665574 |
| ECI2 | 7.91391495 | 0.455562645 | 0.67200588 | 0.67791468 | 0.49782579 | 0.99640678 |
| CDYL | 54.4406587 | -0.600122776 | 0.38985347 | -1.5393547 | 0.12371772 | 0.82915695 |
| LYRM4 | 16.7370232 | -0.196179047 | 0.43009916 | -0.4561252 | 0.64829998 | 0.99665574 |
| FARS2 | 77.0014579 | 0.366342875 | 0.37688199 | 0.97203605 | 0.33103262 | 0.97788427 |
| LY86-AS1 | 22.4615122 | -0.367568678 | 1.23245793 | -0.2982403 | 0.76551973 | 0.99665574 |
| AL031123.1 | 13.585481 | -0.036817565 | 0.74146877 | -0.0496549 | 0.96039739 | 0.99777713 |
| RREB1 | 67.2818686 | -0.04670915 | 0.35826507 | -0.130376 | 0.89626899 | 0.99777713 |
| SSR1 | 20.7848252 | -0.075981019 | 0.36701678 | -0.2070233 | 0.83599168 | 0.99665574 |
| AL139095.4 | 2.3439713 | -1.607750458 | 0.95695715 | -1.6800653 | 0.09294462 | 0.76709603 |
| RIOK1 | 4.55222468 | 0.155242586 | 0.71457253 | 0.21725238 | 0.82801166 | 0.99665574 |
| DSP | 76.6979019 | -0.156898907 | 0.24937338 | -0.6291726 | 0.52923605 | 0.99665574 |
| SNRNP48 | 7.43300555 | 0.142132009 | 0.52675291 | 0.26982672 | 0.78729357 | 0.99665574 |
| BLOC1S5-TXI | 4.50539702 | 1.051348265 | 0.80740463 | 1.30213306 | 0.1928709 | 0.90024181 |
| SLC35B3 | 7.88382461 | -0.189171753 | 0.50770606 | -0.3726009 | 0.70944548 | 0.99665574 |
| TFAP2A | 7.94909204 | 0.514946455 | 0.50139599 | 1.02702548 | 0.30440846 | 0.97376371 |
| GCNT2 | 18.982728 | -0.096150616 | 0.59726014 | -0.1609862 | 0.8721043 | 0.99665574 |
| PAK1IP1 | 2.8458626 | -0.53171239 | 0.79296893 | -0.6705337 | 0.50251763 | 0.99640678 |
| TMEM14C | 10.8379006 | -0.21719572 | 0.56087215 | -0.3872464 | 0.6985738 | 0.99665574 |
| MAK | 8.55902224 | -0.828602755 | 0.92126226 | -0.8994211 | 0.36842838 | 0.98532838 |
| NEDD9 | 10.7793451 | -0.043688442 | 0.92847448 | -0.047054 | 0.96247019 | 0.99777713 |
| TMEM170B | 3.32906947 | -0.107088086 | 0.74301674 | -0.1441261 | 0.88540092 | 0.99665574 |
| HIVEP1 | 32.7626685 | -1.088806642 | 0.3893236 | -2.7966623 | 0.00516335 | 0.22466099 |
| PHACTR1 | 13.1043432 | -0.533652869 | 0.60596424 | -0.8806673 | 0.37849794 | 0.98679664 |
| TBC1D7 | 3.79712169 | 0.448731628 | 0.88513987 | 0.50696126 | 0.612182 | 0.99665574 |
| SIRT5 | 8.87598288 | 0.983438225 | 0.77292679 | 1.27235624 | 0.20324658 | 0.9066717 |
| NOL7 | 12.9114249 | -0.438315921 | 0.60390186 | -0.7258065 | 0.46795737 | 0.99617422 |
| RANBP9 | 61.9905086 | 0.215157938 | 0.30889169 | 0.69654817 | 0.4860856 | 0.99640678 |
| MCUR1 | 9.06129605 | 0.159983307 | 0.48627582 | 0.32899704 | 0.74215792 | 0.99665574 |
| JARID2 | 45.5695547 | 0.616325521 | 0.36575872 | 1.68506037 | 0.09197692 | 0.76658136 |
| DTNBP1 | 35.1598043 | -1.071993958 | 0.40729502 | -2.6319839 | 0.00848879 | 0.31119582 |
| MYLIP | 12.495244 | 0.488323226 | 0.4805833 | 1.01610528 | 0.30957925 | 0.97376371 |
| GMPR | 22.7533089 | 0.29624075 | 0.66895512 | 0.44284099 | 0.65788075 | 0.99665574 |
| ATXN1 | 78.4287805 | 0.005933137 | 0.50817934 | 0.01167528 | 0.99068468 | 0.99777713 |
| CAP2 | 10.1113468 | 0.199144216 | 0.55940009 | 0.35599604 | 0.72184353 | 0.99665574 |
| FAM8A1 | 4.37015665 | -0.08125472 | 0.68127064 | -0.1192694 | 0.90506195 | 0.99777713 |
| NUP153 | 27.5903028 | 0.178032966 | 0.34939654 | 0.50954416 | 0.61037086 | 0.99665574 |
| KIF13A | 38.0069292 | -0.283843412 | 0.32503401 | -0.873273 | 0.38251432 | 0.98914365 |
| TPMT | 6.77711003 | -0.675709686 | 0.60239365 | -1.1217079 | 0.26198668 | 0.94557817 |
| KDM1B | 11.6982602 | -1.194581794 | 0.54645859 | -2.1860427 | 0.02881249 | 0.55075144 |
| DEK | 23.8156134 | -0.407037823 | 0.38011394 | -1.0708311 | 0.28424539 | 0.96059387 |
| RNF144B | 10.2965377 | -2.212515174 | 0.5279213 | -4.1909943 | 2.78E-05 | 0.00872941 |
| AL022068.1 | 4.86436714 | -0.92197725 | 0.69479345 | -1.3269803 | 0.18451519 | 0.89532394 |
| ID4 | 4.49390759 | -0.637633336 | 0.84990527 | -0.7502405 | 0.45310988 | 0.99617422 |
| MBOAT1 | 36.4357075 | 0.546467807 | 0.55235808 | 0.98933613 | 0.32249871 | 0.97700345 |
| E2F3 | 8.72729628 | -0.787932965 | 0.60297953 | -1.3067325 | 0.19130356 | 0.89964233 |
| CDKAL1 | 118.876794 | -0.002785336 | 0.30087607 | -0.0092574 | 0.99261375 | 0.99777713 |
| SOX4 | 40.782087 | -1.403683327 | 0.76459799 | -1.8358449 | 0.0663806 | 0.69521333 |
| CASC15 | 30.9087557 | -0.753761605 | 0.40937476 | -1.8412508 | 0.06558481 | 0.69221233 |
| DCDC2 | 115.215762 | -0.489088302 | 0.43673357 | -1.119878 | 0.26276577 | 0.94557817 |
| MRS2 | 12.7345027 | 0.549569397 | 0.4537061 | 1.21128943 | 0.2257845 | 0.92318867 |
| ALDH5A1 | 5.49376865 | 0.113549913 | 0.63082315 | 0.18000277 | 0.8571504 | 0.99665574 |
| TDP2 | 9.54543237 | 0.027620316 | 0.53371659 | 0.0517509 | 0.95872718 | 0.99777713 |
| ACOT13 | 5.11349652 | -0.276277072 | 0.80080307 | -0.345 | 0.73009436 | 0.99665574 |
| C6orf62 | 38.119186 | 0.44984291 | 0.31611315 | 1.42304397 | 0.15472341 | 0.86196782 |
| GMNN | 15.7632433 | 0.382997843 | 0.44389667 | 0.86280855 | 0.38824274 | 0.99168679 |
| RIPOR2 | 32.5840575 | 0.531383775 | 0.38462349 | 1.38156869 | 0.16710417 | 0.87495175 |

| | | | | | | |
|---|---|---|---|---|---|---|
| CARMIL1 | 207.493532 | 0.476252145 | 0.26599449 | 1.79045868 | 0.0733802 | 0.72792151 |
| SLC17A1 | 7.91069181 | 1.306624723 | 1.04558502 | 1.24965899 | 0.21142415 | 0.91043197 |
| TRIM38 | 16.3591293 | -0.494548233 | 0.48314471 | -1.0236027 | 0.30602297 | 0.97376371 |
| HIST1H1C | 7.23957367 | -2.15087116 | 0.68244117 | -3.1517312 | 0.00162306 | 0.11753244 |
| HIST1H4C | 12.350524 | -1.229108498 | 0.61863262 | -1.9868149 | 0.04694292 | 0.64060657 |
| HIST1H2AC | 6.95962493 | -1.811462863 | 0.60471816 | -2.995549 | 0.00273951 | 0.1614166 |
| HIST1H1E | 2.6755165 | -2.029247491 | 0.89968448 | -2.2555102 | 0.02410132 | 0.50069513 |
| BTN3A2 | 5.58654907 | -1.059549505 | 1.05322092 | -1.0060088 | 0.31441134 | 0.97596515 |
| BTN3A1 | 4.99923905 | -0.869307598 | 0.99490858 | -0.8737563 | 0.382251 | 0.98914365 |
| BTN2A1 | 5.63203488 | -0.104186337 | 0.64553223 | -0.161396 | 0.8717815 | 0.99665574 |
| HCG11 | 3.43180387 | -1.036261393 | 0.75928342 | -1.3647887 | 0.17231949 | 0.88157681 |
| HMGN4 | 7.90159194 | -0.876906618 | 0.64945621 | -1.3502167 | 0.17694648 | 0.88584149 |
| ABT1 | 3.93172784 | 0.01702035 | 0.73297756 | 0.02322083 | 0.98147412 | 0.99777713 |
| ZNF322 | 4.25531689 | 0.69525342 | 0.78369832 | 0.88714421 | 0.37500127 | 0.98569969 |
| AL121944.1 | 4.18560094 | -0.157010082 | 0.84558907 | -0.1856813 | 0.85269469 | 0.99665574 |
| ZNF165 | 4.71252991 | -0.707006211 | 0.92433415 | -0.7648816 | 0.44434203 | 0.99617422 |
| ZSCAN16-AS | 6.66127291 | -0.742067856 | 0.69285525 | -1.0710287 | 0.28415653 | 0.96059387 |
| ZKSCAN8 | 6.07943716 | -0.139570775 | 0.79293443 | -0.1760181 | 0.86027976 | 0.99665574 |
| ZSCAN9 | 3.76570468 | 0.077410855 | 0.76527404 | 0.10115442 | 0.91942788 | 0.99777713 |
| ZSCAN31 | 4.92294115 | -0.017465753 | 1.00542706 | -0.0173715 | 0.98614026 | 0.99777713 |
| ZSCAN12 | 2.72709465 | -1.157404527 | 0.89972528 | -1.2863977 | 0.1983043 | 0.90381759 |
| ZSCAN23 | 1.8769922 | 0.531040052 | 0.96912893 | 0.54795604 | 0.58372209 | 0.99665574 |
| TRIM27 | 16.0644136 | -1.41363579 | 0.44575049 | -3.1713611 | 0.00151726 | 0.1139409 |
| GABBR1 | 3.56185135 | -0.019226161 | 0.89738904 | -0.0214246 | 0.98290699 | 0.99777713 |
| HLA-F | 8.61054891 | -1.915230485 | 0.83223896 | -2.3012988 | 0.02137475 | 0.4733368 |
| HLA-A | 130.577946 | -0.74861524 | 0.43213351 | -1.7323703 | 0.08320766 | 0.75242737 |
| PPP1R11 | 5.82880274 | -0.492693959 | 0.79005461 | -0.6236201 | 0.53287709 | 0.99665574 |
| TRIM26 | 13.318693 | -0.433537069 | 0.48011266 | -0.9029903 | 0.36653105 | 0.98365837 |
| HCG17 | 17.0109329 | 0.019982706 | 0.45310085 | 0.04410212 | 0.96482301 | 0.99777713 |
| HLA-E | 63.5127499 | -1.380145456 | 0.47429438 | -2.9098921 | 0.00361554 | 0.19601232 |
| GNL1 | 10.742229 | -0.804307073 | 0.48291587 | -1.6655222 | 0.09580863 | 0.77210482 |
| PRR3 | 4.07037984 | -0.654285632 | 0.73274038 | -0.8929297 | 0.37189483 | 0.98532838 |
| ABCF1 | 13.5950492 | -1.083881463 | 0.54984792 | -1.9712386 | 0.04869659 | 0.64337675 |
| PPP1R10 | 13.2294212 | 0.045117154 | 0.42815237 | 0.1053764 | 0.91607714 | 0.99777713 |
| MRPS18B | 11.7995627 | -0.592154846 | 0.5270703 | -1.1234836 | 0.26123216 | 0.94557817 |
| ATAT1 | 4.59014735 | 0.389493648 | 0.78492126 | 0.49622002 | 0.61973918 | 0.99665574 |
| C6orf136 | 8.74062006 | 0.961944043 | 0.53762064 | 1.78926175 | 0.07357267 | 0.72792151 |
| DHX16 | 5.71805922 | -0.092245643 | 0.57635999 | -0.1600487 | 0.87284275 | 0.99665574 |
| MDC1 | 6.06606834 | -0.531980928 | 0.52938761 | -1.0048987 | 0.31494562 | 0.97652251 |
| TUBB | 26.6849164 | -1.106246325 | 0.75693447 | -1.4614823 | 0.14388313 | 0.84969914 |
| FLOT1 | 27.3278021 | -1.190767926 | 0.35924717 | -3.3146202 | 0.00091768 | 0.08160886 |
| IER3 | 12.2847643 | -2.003511965 | 0.83087306 | -2.4113334 | 0.01589431 | 0.41983038 |
| DDR1 | 25.2684865 | 0.229219702 | 0.38803974 | 0.59071193 | 0.55471345 | 0.99665574 |
| VARS2 | 2.9927628 | -0.394088659 | 0.81766244 | -0.4819699 | 0.62982735 | 0.99665574 |
| CCHCR1 | 1.8388055 | 0.582921606 | 1.1428131 | 0.51007606 | 0.60999817 | 0.99665574 |
| POU5F1 | 16.7208854 | 0.576796564 | 0.52901974 | 1.090312 | 0.27557573 | 0.95612141 |
| HLA-C | 115.852775 | -0.737562207 | 0.62132786 | -1.1870741 | 0.23519839 | 0.92960234 |
| HLA-B | 183.385456 | -0.852817468 | 0.57647435 | -1.4793676 | 0.1390421 | 0.84668189 |
| MICA | 5.5708125 | -1.201776859 | 0.60560244 | -1.984432 | 0.0472077 | 0.64060657 |
| NFKBIL1 | 4.06159487 | 0.726941782 | 0.79196535 | 0.91789594 | 0.35867335 | 0.9819332 |
| PRRC2A | 12.8922466 | -0.079773328 | 0.55259917 | -0.1443602 | 0.88521604 | 0.99665574 |
| BAG6 | 8.18200323 | 0.288730924 | 0.5800609 | 0.49775967 | 0.61865344 | 0.99665574 |
| C6orf47 | 3.69613386 | -0.325266074 | 0.78918649 | -0.4121536 | 0.68022682 | 0.99665574 |
| GPANK1 | 3.72608578 | -0.149591279 | 0.80835385 | -0.1850567 | 0.85318458 | 0.99665574 |
| DDAH2 | 7.41601762 | 0.932695561 | 0.60446205 | 1.54301757 | 0.12282651 | 0.82915695 |
| CLIC1 | 31.7137656 | -0.642457779 | 0.33538992 | -1.9155548 | 0.05542179 | 0.67128562 |
| VARS | 2.98249333 | 0.012586263 | 0.84026121 | 0.01497899 | 0.98804894 | 0.99777713 |
| LSM2 | 4.8007908 | 0.042022206 | 0.69521608 | 0.06044481 | 0.95180137 | 0.99777713 |
| HSPA1A | 36.3620479 | 0.828719469 | 0.47003896 | 1.76308677 | 0.07788586 | 0.74141484 |
| HSPA1B | 15.0762923 | 1.097496145 | 0.65054312 | 1.68704597 | 0.09159451 | 0.76595534 |
| C6orf48 | 10.8245326 | 0.004818708 | 0.45672972 | 0.01055046 | 0.99158211 | 0.99777713 |
| NEU1 | 8.68419042 | -1.065269243 | 0.69575436 | -1.5310996 | 0.12574477 | 0.83633607 |
| EHMT2 | 4.4234105 | 0.295977147 | 0.73749638 | 0.40132691 | 0.68817945 | 0.99665574 |
| NELFE | 3.00214661 | 0.204432669 | 0.88207893 | 0.23176233 | 0.81672262 | 0.99665574 |
| SKIV2L | 2.31128576 | 0.54661735 | 0.98847676 | 0.55298958 | 0.58027054 | 0.99665574 |
| STK19 | 3.86188309 | 0.122200763 | 0.68683221 | 0.17791938 | 0.85878629 | 0.99665574 |
| ATF6B | 7.82595727 | 0.304012697 | 0.68648853 | 0.44285182 | 0.65787292 | 0.99665574 |
| AGPAT1 | 7.15293517 | 0.714879457 | 0.57255578 | 1.24857609 | 0.21182016 | 0.91043197 |
| RNF5 | 8.02917737 | 0.146016902 | 0.63547872 | 0.22977465 | 0.81826688 | 0.99665574 |
| PBX2 | 5.73119203 | 0.086196886 | 0.59495169 | 0.14488048 | 0.88480524 | 0.99665574 |
| HLA-DRA | 10.1668553 | -0.748287491 | 0.81905491 | -0.9135987 | 0.36092776 | 0.9819332 |
| HLA-DRB1 | 5.20186436 | -0.630324956 | 1.23501092 | -0.5103801 | 0.60978522 | 0.99665574 |
| PSMB8 | 16.3956902 | -0.864589367 | 0.5447664 | -1.5870828 | 0.1124939 | 0.81249398 |
| PSMB9 | 6.86911474 | -0.821656945 | 1.28224731 | -0.6407944 | 0.52165625 | 0.99665574 |
| TAP1 | 8.37821905 | -0.632537289 | 0.80651197 | -0.7842875 | 0.43287141 | 0.99617422 |
| BRD2 | 23.0987334 | -0.347506156 | 0.49070079 | -0.7081834 | 0.47883137 | 0.99617422 |
| RXRB | 3.68759797 | 0.688364156 | 0.86404191 | 0.79667913 | 0.42563741 | 0.99518711 |
| SLC39A7 | 12.0881816 | -0.648177715 | 0.70777499 | -0.9157963 | 0.35977374 | 0.9819332 |
| RING1 | 6.85483408 | -0.538115501 | 0.6617921 | -0.8131186 | 0.41615004 | 0.992853 |
| VPS52 | 6.82264866 | -0.305680161 | 0.62100069 | -0.492238 | 0.62255108 | 0.99665574 |
| RPS18 | 47.6825532 | 0.463588361 | 0.37964475 | 1.22111094 | 0.22204402 | 0.92191316 |
| WDR46 | 3.99289675 | 0.252665391 | 0.69380785 | 0.364172 | 0.71572958 | 0.99665574 |
| PFDN6 | 4.85575092 | 0.645450866 | 0.74188828 | 0.87001087 | 0.38429447 | 0.99033485 |
| RGL2 | 3.69624141 | -0.09762073 | 0.76319054 | -0.1279113 | 0.89821913 | 0.99777713 |
| TAPBP | 26.9007699 | -0.748515714 | 0.47026431 | -1.5916915 | 0.11145403 | 0.8083966 |
| DAXX | 3.3317432 | -0.348601765 | 0.78425738 | -0.4444992 | 0.65668172 | 0.99665574 |
| PHF1 | 7.58093369 | 0.287483671 | 0.56475036 | 0.50904557 | 0.61072028 | 0.99665574 |
| CUTA | 18.4616746 | 0.533968492 | 0.46203306 | 1.15569326 | 0.24780664 | 0.93441937 |
| SYNGAP1 | 3.00393163 | 0.15142455 | 0.91094549 | 0.16622789 | 0.86797761 | 0.99665574 |
| ITPR3 | 7.32556663 | -1.050025063 | 0.6130466 | -1.7127981 | 0.08674969 | 0.75427976 |
| UQCC2 | 8.04020842 | 0.095540152 | 0.57008157 | 0.16759032 | 0.86690559 | 0.99665574 |
| LEMD2 | 6.83405529 | -0.483932298 | 0.55947433 | -0.8649768 | 0.38705154 | 0.99164228 |
| HMGA1 | 4.8943099 | 0.097928631 | 0.69776394 | 0.14034636 | 0.88838633 | 0.99665574 |
| SMIM29 | 4.43027529 | 0.873430316 | 0.99086413 | 0.88148343 | 0.37805622 | 0.98679664 |
| NUDT3 | 17.4629695 | -0.538016816 | 0.47567773 | -1.1310532 | 0.2580327 | 0.94557817 |
| C6orf106 | 31.944803 | -0.100484702 | 0.35550245 | -0.2826554 | 0.77744097 | 0.99665574 |
| SNRPC | 6.33389845 | -0.580525689 | 0.57864017 | -1.0032585 | 0.31573614 | 0.97652251 |
| UHRF1BP1 | 16.267402 | 0.372779731 | 0.44875761 | 0.83069284 | 0.40614717 | 0.9921735 |
| TAF11 | 10.0165999 | 0.577327623 | 0.52214242 | 1.10568994 | 0.26886075 | 0.94673212 |
| ANKS1A | 77.7382694 | -0.157426185 | 0.26612648 | -0.5915465 | 0.55415432 | 0.99665574 |
| ZNF76 | 10.1983814 | -0.374453018 | 0.46541658 | -0.8045545 | 0.42107679 | 0.99518711 |
| PPARD | 11.8870895 | 0.281787361 | 0.4479595 | 0.62904651 | 0.52931861 | 0.99665574 |
| RPL10A | 49.7804985 | 1.101878265 | 0.54516566 | 2.02118061 | 0.04326107 | 0.62038248 |
| TEAD3 | 6.25526819 | -1.604315382 | 0.77827823 | -2.0613648 | 0.03926825 | 0.60188673 |
| FKBP5 | 19.6630342 | -0.771299188 | 1.36224161 | -0.5661985 | 0.57125883 | 0.99665574 |
| SRPK1 | 20.9421272 | -0.204289931 | 0.34796266 | -0.587103 | 0.55713452 | 0.99665574 |
| MAPK14 | 32.5192991 | -0.647529527 | 0.33313765 | -1.9437296 | 0.05192806 | 0.65723681 |
| MAPK13 | 7.5623654 | -0.60652315 | 0.54225944 | -1.118511 | 0.26334881 | 0.94557817 |
| BRPF3 | 10.3951999 | 0.235109592 | 0.53373875 | 0.44049564 | 0.65957817 | 0.99665574 |
| KCTD20 | 26.7352277 | -0.044725943 | 0.35657822 | -0.1254309 | 0.90018239 | 0.99777713 |
| STK38 | 39.2958116 | -0.266594019 | 0.40459522 | -0.6589154 | 0.5099501 | 0.99665574 |
| SRSF3 | 31.9453412 | 0.295881251 | 0.43791434 | 0.67566012 | 0.49925646 | 0.99640678 |
| CDKN1A | 8.63738482 | -1.654840103 | 0.71776618 | -2.305542 | 0.02113623 | 0.4733368 |

| | | | | | | |
|---|---|---|---|---|---|---|
| PPIL1 | 3.2857277 | 0.016585432 | 0.77554009 | 0.02138565 | 0.98293802 | 0.99777713 |
| C6orf89 | 18.8828555 | -0.265074889 | 0.37561437 | -0.7057102 | 0.48036838 | 0.99617422 |
| MTCH1 | 19.1920426 | -0.514165055 | 0.43383506 | -1.1851625 | 0.23595319 | 0.9305595 |
| PIM1 | 16.0347067 | 0.027759161 | 0.56131186 | 0.04945408 | 0.96055743 | 0.99777713 |
| TBC1D22B | 14.8611675 | 0.174752537 | 0.43808798 | 0.39889827 | 0.68996817 | 0.99665574 |
| RNF8 | 9.66433871 | -0.434113726 | 0.46543716 | -0.9327011 | 0.35097434 | 0.97838306 |
| CMTR1 | 5.48956436 | 0.623557903 | 0.68962868 | 0.9041937 | 0.3658927 | 0.98365837 |
| ZFAND3 | 326.015955 | 0.167606413 | 0.30879142 | 0.54278196 | 0.58727994 | 0.99665574 |
| BTBD9 | 177.610914 | -0.177099484 | 0.30672261 | -0.577393 | 0.56367401 | 0.99665574 |
| GLO1 | 18.8385051 | -0.190027741 | 0.435293 | -0.4365513 | 0.66243676 | 0.99665574 |
| SAYSD1 | 5.42941943 | -0.05580571 | 0.66102532 | -0.084423 | 0.93272016 | 0.99777713 |
| DAAM2 | 2.50936694 | 2.135407234 | 1.16019157 | 1.84056435 | 0.06568543 | 0.69222334 |
| MOCS1 | 5.76780851 | 0.441786421 | 0.61849475 | 0.71429292 | 0.47504607 | 0.99617422 |
| OARD1 | 7.18352794 | -0.119836919 | 0.63054461 | -0.190053 | 0.84926757 | 0.99665574 |
| NFYA | 6.47725396 | 0.208043735 | 0.72915128 | 0.28532314 | 0.77539658 | 0.99665574 |
| FOXP4 | 5.5358663 | 0.127159054 | 0.56103042 | 0.22665269 | 0.82069382 | 0.99665574 |
| TFEB | 27.176797 | 0.156098485 | 0.45870598 | 0.34030183 | 0.73362924 | 0.99665574 |
| USP49 | 8.33998435 | 0.849252722 | 0.63539009 | 1.33658478 | 0.18135821 | 0.89279728 |
| CCND3 | 15.2312515 | -0.867293139 | 0.65942889 | -1.3152186 | 0.18843645 | 0.89739428 |
| TAF8 | 8.6415666 | -0.34044838 | 0.5050086 | -0.6741437 | 0.50021994 | 0.99640678 |
| MRPS10 | 5.17824144 | 0.75365913 | 0.64020548 | 1.17721443 | 0.23910993 | 0.93157178 |
| TRERF1 | 5.57935609 | -0.812296672 | 0.64223633 | -1.264794 | 0.20594517 | 0.90890506 |
| UBR2 | 72.0057919 | 0.145355366 | 0.30954865 | 0.46957196 | 0.63866086 | 0.99665574 |
| TBCC | 2.42566365 | -0.592399469 | 1.01098517 | -0.5859626 | 0.55790067 | 0.99665574 |
| BICRAL | 31.6384219 | -0.075587649 | 0.34979899 | -0.2160888 | 0.82891852 | 0.99665574 |
| RPL7L1 | 18.3225519 | -0.122816771 | 0.38992685 | -0.3149739 | 0.75278149 | 0.99665574 |
| C6orf226 | 2.98852351 | -0.642095002 | 0.87797096 | -0.7313397 | 0.4645717 | 0.99617422 |
| CNPY3 | 4.4684993 | -0.602728408 | 0.79194548 | -0.7610731 | 0.4466134 | 0.99617422 |
| PEX6 | 7.47596012 | -0.245897052 | 0.53764919 | -0.4573559 | 0.64741522 | 0.99665574 |
| PPP2R5D | 2.33044691 | -0.092991652 | 0.91571576 | -0.1015508 | 0.91911325 | 0.99777713 |
| MEA1 | 5.72076986 | 0.037762007 | 0.67008384 | 0.05635415 | 0.95505968 | 0.99777713 |
| KLHDC3 | 5.27912653 | 0.257833739 | 0.63412913 | 0.40659501 | 0.68430546 | 0.99665574 |
| RRP36 | 8.78452415 | 0.223323973 | 0.5634728 | 0.39633496 | 0.69185794 | 0.99665574 |
| CUL7 | 6.65268003 | -0.168139003 | 0.64832472 | -0.2593438 | 0.79536998 | 0.99665574 |
| KLC4 | 7.56571304 | -0.085691364 | 0.56532289 | -0.1515795 | 0.8795186 | 0.99665574 |
| MRPL2 | 4.68944506 | 0.219056796 | 0.65286303 | 0.33553255 | 0.7372234 | 0.99665574 |
| PTK7 | 10.9437675 | 0.22511403 | 0.57547929 | 0.3911766 | 0.6956667 | 0.99665574 |
| SRF | 3.49243679 | 1.075258359 | 1.07018365 | 1.0047419 | 0.31502114 | 0.97652251 |
| CUL9 | 4.31089141 | 0.161299273 | 0.74065895 | 0.21777807 | 0.82760204 | 0.99665574 |
| DNPH1 | 6.5462222 | -0.320461286 | 0.67136311 | -0.4773293 | 0.63312765 | 0.99665574 |
| ZNF318 | 5.51537626 | -0.270432166 | 0.61526667 | -0.4395365 | 0.66027283 | 0.99665574 |
| ABCC10 | 6.61701127 | -0.368651644 | 0.51916987 | -0.710079 | 0.47765512 | 0.99617422 |
| TJAP1 | 5.70160671 | -0.335387912 | 0.59663169 | -0.5621356 | 0.57402364 | 0.99665574 |
| POLR1C | 8.26188109 | -0.324322844 | 0.57341846 | -0.5655954 | 0.57166886 | 0.99665574 |
| YIPF3 | 10.6659198 | -0.062144417 | 0.59216785 | -0.1049439 | 0.91642031 | 0.99777713 |
| XPO5 | 39.7536668 | 0.401986061 | 0.39598299 | 1.01515991 | 0.31002961 | 0.97376371 |
| POLH | 7.25516953 | -0.197649876 | 0.56353996 | -0.3507291 | 0.72579158 | 0.99665574 |
| GTPBP2 | 5.11250372 | 0.179542017 | 0.70070237 | 0.25623149 | 0.79777209 | 0.99665574 |
| MAD2L1BP | 2.43242709 | -0.33611487 | 0.99796598 | -0.3367999 | 0.73626773 | 0.99665574 |
| MRPS18A | 6.05954663 | 0.498636662 | 0.59078758 | 0.84402022 | 0.39865811 | 0.99179694 |
| VEGFA | 57.5528579 | 0.800635472 | 0.32847115 | 2.43746052 | 0.01479083 | 0.40950537 |
| MRPL14 | 11.8056411 | -0.183859543 | 0.52293677 | -0.3515904 | 0.72514547 | 0.99665574 |
| TMEM63B | 6.7394847 | -0.12759399 | 0.53534551 | -0.2383395 | 0.81161777 | 0.99665574 |
| SLC29A1 | 7.1286705 | 0.72295675 | 0.60432497 | 1.19630461 | 0.23157771 | 0.92795844 |
| HSP90AB1 | 80.1163624 | 0.112640596 | 0.26882962 | 0.41900367 | 0.67521345 | 0.99665574 |
| SLC35B2 | 4.53588268 | -0.658993361 | 0.76890154 | -0.8570582 | 0.39141273 | 0.99179694 |
| CDC5L | 9.78021201 | -0.016249458 | 0.50415582 | -0.032231 | 0.97428782 | 0.99777713 |
| SUPT3H | 113.065956 | 0.164539881 | 0.35300843 | 0.46610752 | 0.64113854 | 0.99665574 |
| RUNX2 | 31.9911309 | -1.203195346 | 0.58883116 | -2.0433622 | 0.04101659 | 0.61100706 |
| ENPP4 | 11.7698501 | 0.504689013 | 0.49349402 | 1.02268516 | 0.30645673 | 0.97376371 |
| ENPP5 | 3.1891637 | -0.379694406 | 0.85949926 | -0.4417623 | 0.65866119 | 0.99665574 |
| RCAN2 | 382.001007 | 0.681421168 | 0.35243482 | 1.93346719 | 0.05317867 | 0.66476617 |
| CYP39A1 | 9.69125395 | 0.163895684 | 0.6632004 | 0.24712844 | 0.80480882 | 0.99665574 |
| SLC25A27 | 27.9522072 | 0.371064446 | 0.44845579 | 0.82742704 | 0.40799506 | 0.99279547 |
| ADGRF1 | 60.2350819 | -0.772161166 | 0.63040663 | -1.2248621 | 0.22062719 | 0.91857862 |
| TNFRSF21 | 5.86556309 | -1.16331439 | 0.74581107 | -1.5597977 | 0.1188077 | 0.82619588 |
| CD2AP | 86.5471131 | -0.50734334 | 0.26541348 | -1.9115206 | 0.05593771 | 0.67351132 |
| PTCHD4 | 8.37576218 | -2.732067756 | 1.15652834 | -2.3623008 | 0.0181619 | 0.44799357 |
| MUT | 12.5105051 | 0.839956856 | 0.48386085 | 1.73594713 | 0.08257318 | 0.75228386 |
| PKHD1 | 81.7042543 | 1.376594407 | 0.62110018 | 2.21638062 | 0.02666545 | 0.52453943 |
| MCM3 | 4.78918393 | -0.003678986 | 0.65214163 | -0.0056414 | 0.99549884 | 0.99865103 |
| EFHC1 | 8.39100263 | -0.002899436 | 0.6799938 | -0.0042639 | 0.9965979 | 0.998802 |
| TRAM2 | 4.15412244 | 0.238559904 | 0.7573796 | 0.31498063 | 0.75277636 | 0.99665574 |
| TMEM14A | 2.88908243 | 0.846087634 | 0.85806253 | 0.98604426 | 0.32411139 | 0.97700345 |
| GSTA4 | 4.29177377 | -0.821103215 | 0.82431784 | -0.9961003 | 0.31920143 | 0.97700345 |
| ICK | 5.84396699 | 0.044964462 | 0.6112119 | 0.07356608 | 0.94135566 | 0.99777713 |
| FBXO9 | 17.923693 | 0.301523837 | 0.39717051 | 0.75917982 | 0.447745 | 0.99617422 |
| ELOVL5 | 18.8248498 | -0.487601576 | 0.63262734 | -0.7707564 | 0.44085133 | 0.99617422 |
| GCLC | 18.1087798 | -0.904174222 | 0.4201572 | -2.1519903 | 0.03139812 | 0.56639521 |
| LRRC1 | 20.6071626 | -0.291517401 | 0.45027107 | -0.6474265 | 0.51735598 | 0.99665574 |
| TINAG | 6.78488454 | 2.419836519 | 0.97315782 | 2.48658179 | 0.01289769 | 0.38590337 |
| FAM83B | 47.0540055 | -0.508342587 | 0.34779763 | -1.4616045 | 0.14384961 | 0.84969914 |
| DST | 72.3136447 | -0.679716826 | 0.40516631 | -1.6776242 | 0.09342048 | 0.76763189 |
| KIAA1586 | 3.02741771 | 0.371516887 | 0.80965028 | 0.45886094 | 0.64633403 | 0.99665574 |
| ZNF451 | 33.0983524 | 0.132326127 | 0.3027379 | 0.43709799 | 0.66204029 | 0.99665574 |
| ZNF451-AS1 | 3.30042665 | -0.477191843 | 0.96270303 | -0.4956792 | 0.62012078 | 0.99665574 |
| PRIM2 | 25.1584445 | 0.267523637 | 0.35629566 | 0.75084731 | 0.45274455 | 0.99617422 |
| AL021368.2 | 3.3035651 | 0.957085038 | 0.84959362 | 1.12652097 | 0.25994507 | 0.94557817 |
| PTP4A1 | 17.4183168 | -0.761201079 | 0.74653341 | -1.0196477 | 0.30789557 | 0.97376371 |
| PHF3 | 61.4389528 | 0.231737479 | 0.27928228 | 0.82976076 | 0.40667406 | 0.99250037 |
| EYS | 3.55604569 | 0.153980242 | 0.96130456 | 0.16017842 | 0.87274053 | 0.99665574 |
| LMBRD1 | 46.9624033 | 0.200915928 | 0.29818697 | 0.67379177 | 0.50044371 | 0.99640678 |
| COL19A1 | 10.8922322 | -0.170868529 | 0.49288587 | -0.3466696 | 0.72883958 | 0.99665574 |
| FAM135A | 181.511192 | -0.864343761 | 0.40262474 | -2.1467726 | 0.03181139 | 0.56852889 |
| SDHAF4 | 2.64741149 | -0.035937836 | 0.8136466 | -0.0441689 | 0.96476981 | 0.99777713 |
| SMAP1 | 80.8178915 | -0.038686853 | 0.27545192 | -0.1404487 | 0.88830551 | 0.99665574 |
| OGFRL1 | 43.8664529 | 0.441946325 | 0.43391443 | 1.01851033 | 0.30843549 | 0.97376371 |
| AL136164.2 | 26.9147895 | 0.932760364 | 0.45261539 | 2.06082333 | 0.0393199 | 0.60188673 |
| LINC00472 | 11.5273823 | -0.345917112 | 0.52408377 | -0.6600416 | 0.5092271 | 0.99665574 |
| OOEP | 49.3866618 | 0.39064199 | 0.35422898 | 1.10279513 | 0.27011615 | 0.94821243 |
| MTO1 | 9.71418712 | 0.183250294 | 0.4608833 | 0.39760671 | 0.69092011 | 0.99665574 |
| EEF1A1 | 285.746871 | 0.181282886 | 0.41826965 | 0.43341152 | 0.66471583 | 0.99665574 |
| SLC17A5 | 8.23112582 | -1.671178336 | 0.5504198 | -3.0361886 | 0.00239589 | 0.15249081 |
| COX7A2 | 31.4717751 | 0.706373223 | 0.38675694 | 1.82640092 | 0.06778989 | 0.70516795 |
| TMEM30A | 28.7558261 | 0.043375131 | 0.36383811 | 0.11921547 | 0.90510465 | 0.99777713 |
| SENP6 | 96.2410211 | 0.080140113 | 0.24376986 | 0.32875317 | 0.74234226 | 0.99665574 |
| MYO6 | 225.311865 | -0.385714846 | 0.37183908 | -1.0373166 | 0.29958834 | 0.97358745 |
| MEI4 | 4.61659412 | 0.105084024 | 0.6374716 | 0.16484503 | 0.86906596 | 0.99665574 |
| IRAK1BP1 | 14.5462076 | 0.297431743 | 0.5684071 | 0.52327239 | 0.60078471 | 0.99665574 |
| PHIP | 72.4008062 | -0.00616233 | 0.32756205 | -0.0188127 | 0.98499051 | 0.99777713 |
| HMGN3 | 33.7554395 | 0.381550882 | 0.4284845 | 0.89046601 | 0.37321571 | 0.98532838 |
| LCA5 | 3.17183753 | 0.073799844 | 0.95945799 | 0.07691826 | 0.93868857 | 0.99777713 |
| SH3BGRL2 | 5.87386297 | -0.636057913 | 0.67121253 | -0.9476252 | 0.34332029 | 0.97788427 |

| | | | | | | |
|---|---|---|---|---|---|---|
| BCKDHB | 46.7983952 | 0.466270773 | 0.38363695 | 1.2153959 | 0.22421513 | 0.92318867 |
| TENT5A | 8.53086034 | 0.0168257 | 0.57657505 | 0.02918215 | 0.97671932 | 0.99777713 |
| IBTK | 42.5313159 | -0.003891596 | 0.38044137 | -0.0102292 | 0.99183845 | 0.99777713 |
| UBE3D | 25.9989756 | -0.028947093 | 0.32528998 | -0.0889886 | 0.92909099 | 0.99777713 |
| DOPEY1 | 38.6588889 | -0.112571531 | 0.29219522 | -0.3852614 | 0.70004377 | 0.99665574 |
| PGM3 | 9.254339 | -0.834778073 | 0.65879125 | -1.2671359 | 0.20510668 | 0.90834797 |
| CYB5R4 | 14.0175899 | 0.077683905 | 0.40543753 | 0.19160512 | 0.84805153 | 0.99665574 |
| CEP162 | 13.7381017 | -0.065130823 | 0.40822763 | -0.1595454 | 0.87323923 | 0.99665574 |
| SNX14 | 42.9141395 | 0.365388517 | 0.30536618 | 1.19655855 | 0.23147866 | 0.92792831 |
| SYNCRIP | 17.2942331 | -0.230861512 | 0.43556191 | -0.5300315 | 0.59609012 | 0.99665574 |
| ZNF292 | 128.572173 | -0.617620758 | 0.29555409 | -2.0897047 | 0.03664434 | 0.58591936 |
| SMIM8 | 19.6404794 | 0.325621683 | 0.48012446 | 0.67820265 | 0.49764321 | 0.99640678 |
| RARS2 | 25.0277255 | 0.392395212 | 0.36825759 | 1.06554549 | 0.28662915 | 0.96477573 |
| ORC3 | 14.4255636 | 0.698943537 | 0.45921506 | 1.52203965 | 0.12799915 | 0.83778831 |
| AKIRIN2 | 8.0565604 | 0.135197332 | 0.58513054 | 0.23105499 | 0.81727208 | 0.99665574 |
| RNGTT | 58.2802115 | 0.278568275 | 0.2908173 | 0.95788067 | 0.33812293 | 0.97788427 |
| PNRC1 | 16.204682 | -0.728993823 | 0.46387017 | -1.5715471 | 0.11605562 | 0.81933976 |
| PM20D2 | 12.6546913 | -0.576936875 | 0.50304874 | -1.1468807 | 0.25143095 | 0.94059297 |
| UBE2J1 | 14.6553787 | -0.545715317 | 0.40979626 | -1.3316747 | 0.1829671 | 0.89500374 |
| RRAGD | 29.7587191 | 0.666232531 | 0.52665433 | 1.2650281 | 0.20586125 | 0.90890506 |
| ANKRD6 | 24.4637415 | -0.303310721 | 0.38067678 | -0.7967671 | 0.42558633 | 0.99518711 |
| LYRM2 | 11.2005972 | -0.461562146 | 0.54787512 | -0.8424587 | 0.39953126 | 0.99179694 |
| MDN1 | 27.1283513 | 0.138710907 | 0.4012874 | 0.34566475 | 0.72959468 | 0.99665574 |
| CASP8AP2 | 15.4670073 | -0.018170968 | 0.42299466 | -0.0429579 | 0.96573508 | 0.99777713 |
| BACH2 | 60.4434678 | -1.066260622 | 0.45543553 | -2.3411889 | 0.01922244 | 0.45961574 |
| MAP3K7 | 24.5856783 | 0.218706221 | 0.37390544 | 0.58492388 | 0.5585989 | 0.99665574 |
| MANEA | 17.4983552 | 1.128815354 | 0.40037935 | 2.81936454 | 0.00481188 | 0.22060024 |
| UFL1 | 16.4787745 | -0.257045924 | 0.43630583 | -0.5891416 | 0.55576627 | 0.99665574 |
| NDUFAF4 | 10.3373374 | 0.094985257 | 0.64741603 | 0.1467144 | 0.88335745 | 0.99665574 |
| KLHL32 | 12.3270608 | -0.237592851 | 0.49430007 | -0.4806652 | 0.63075445 | 0.99665574 |
| MMS22L | 11.9792333 | -0.963259694 | 0.55412142 | -1.7383549 | 0.0821483 | 0.75124322 |
| FBXL4 | 26.1168291 | 0.271563779 | 0.40295846 | 0.67392499 | 0.500359 | 0.99640678 |
| COQ3 | 3.49944656 | 0.373245384 | 0.90400556 | 0.41287952 | 0.67969488 | 0.99665574 |
| PNISR | 79.2502254 | 0.4411427 | 0.22953723 | 1.92187863 | 0.05462103 | 0.66958648 |
| AL513550.1 | 3.20040558 | -0.250786423 | 0.80856538 | -0.3101622 | 0.75643761 | 0.99665574 |
| USP45 | 16.8130141 | -0.105085166 | 0.46567559 | -0.2256617 | 0.82146451 | 0.99665574 |
| CCNC | 28.046753 | 0.086599538 | 0.44610137 | 0.19412524 | 0.84607782 | 0.99665574 |
| SIM1 | 55.7951023 | 0.194593017 | 0.40553461 | 0.47984318 | 0.6313389 | 0.99665574 |
| ASCC3 | 59.5003321 | -0.423806111 | 0.33767978 | -1.2550533 | 0.20945943 | 0.91043197 |
| HACE1 | 30.449026 | -0.601740888 | 0.70743402 | -0.8505965 | 0.39499355 | 0.99179694 |
| PREP | 11.1369622 | 0.239386806 | 0.55623592 | 0.4303692 | 0.6669271 | 0.99665574 |
| ATG5 | 37.3164671 | 0.086013728 | 0.3019612 | 0.28485026 | 0.77575886 | 0.99665574 |
| CRYBG1 | 19.2227916 | -0.77349689 | 0.55257848 | -1.3997955 | 0.16157455 | 0.87011714 |
| RTN4IP1 | 10.8190261 | -0.627225827 | 0.53419998 | -1.1741405 | 0.24033876 | 0.93216487 |
| QRSL1 | 15.5334561 | -0.331586002 | 0.46980807 | -0.7057904 | 0.48031852 | 0.99617422 |
| LINC02532 | 54.6253314 | -0.537433622 | 0.55117425 | -0.9750703 | 0.3295254 | 0.97788427 |
| CD24 | 92.9713945 | -2.302426551 | 0.33649151 | -6.8424507 | 7.78E-12 | 2.10E-08 |
| C6orf203 | 10.4114318 | 0.458767191 | 0.4830045 | 0.94981971 | 0.34220387 | 0.97788427 |
| PDSS2 | 59.92082 | 0.205518346 | 0.266103 | 0.7723263 | 0.43992119 | 0.99617422 |
| SEC63 | 33.0527959 | 0.01624176 | 0.35230665 | 0.0461012 | 0.96322959 | 0.99777713 |
| OSTM1 | 17.0700299 | -0.054139892 | 0.40190033 | -0.1347097 | 0.89284137 | 0.99773598 |
| SNX3 | 30.255748 | -0.570195912 | 0.29202108 | -1.9525848 | 0.05086882 | 0.65127694 |
| AFG1L | 23.6387262 | 0.573527134 | 0.43483812 | 1.31894402 | 0.18718783 | 0.89683847 |
| FOXO3 | 102.100264 | 0.193679808 | 0.2989574 | 0.64785085 | 0.51708142 | 0.99665574 |
| ARMC2 | 6.6891018 | 0.492539062 | 0.64229092 | 0.7668473 | 0.44317231 | 0.99617422 |
| SESN1 | 45.6379913 | 0.491945401 | 0.33448022 | 1.47077577 | 0.14135177 | 0.84969914 |
| CEP57L1 | 6.74477289 | 0.766056954 | 0.60924834 | 1.25738045 | 0.2086159 | 0.91043197 |
| CD164 | 41.4001905 | 0.215246587 | 0.37276958 | 0.5774253 | 0.56365219 | 0.99665574 |
| AK9 | 10.6666772 | -0.188740433 | 0.65008076 | -0.2903338 | 0.77156086 | 0.99665574 |
| FIG4 | 14.9753091 | -0.614097566 | 0.40551604 | -1.5143607 | 0.12993437 | 0.83778831 |
| WASF1 | 8.08629822 | -0.646439908 | 0.55398395 | -1.1668928 | 0.24325365 | 0.93426912 |
| CDC40 | 9.18191523 | 0.515609326 | 0.57218775 | 0.90111913 | 0.36752498 | 0.9849242 |
| CDK19 | 29.8218431 | 0.423480395 | 0.3825925 | 1.10687061 | 0.26834988 | 0.94660092 |
| AMD1 | 19.5361284 | -0.507898889 | 0.39502899 | -1.2857256 | 0.19853884 | 0.90381759 |
| GTF3C6 | 15.9708184 | -0.261631892 | 0.4473725 | -0.5848189 | 0.5586695 | 0.99665574 |
| RPF2 | 5.06587983 | -0.810907346 | 0.73700657 | -1.1002715 | 0.27121384 | 0.94865166 |
| SLC16A10 | 57.9831761 | -0.243443829 | 0.38394799 | -0.6340542 | 0.52604546 | 0.99665574 |
| MFSD4B | 6.62301235 | 0.228341436 | 0.5282077 | 0.43229479 | 0.66552718 | 0.99665574 |
| REV3L | 74.9016102 | -0.108551817 | 0.28017156 | -0.3874477 | 0.69842482 | 0.99665574 |
| TRAF3IP2-AS | 23.5791567 | -0.515273448 | 0.4080379 | -1.2628078 | 0.20665826 | 0.90890506 |
| TRAF3IP2 | 4.35529698 | -0.602044154 | 0.73464789 | -0.8195003 | 0.41250104 | 0.99279547 |
| TUBE1 | 6.42751363 | 1.082286521 | 0.64267333 | 1.68403832 | 0.09217426 | 0.76658136 |
| MARCKS | 71.6427305 | -0.673460445 | 0.39113544 | -1.7218088 | 0.08510415 | 0.75373899 |
| HDAC2 | 15.2998715 | -0.602855346 | 0.4000006 | -1.5071361 | 0.13177578 | 0.84074439 |
| FRK | 63.670624 | -0.214497038 | 0.37544893 | -0.5713082 | 0.56779077 | 0.99665574 |
| NT5DC1 | 57.5365063 | 0.00785635 | 0.33033497 | 0.02378298 | 0.98102572 | 0.99777713 |
| COL10A1 | 2.54939249 | -1.917593955 | 1.38354069 | -1.3860047 | 0.16574546 | 0.87495175 |
| TSPYL4 | 4.03154923 | 0.575411646 | 0.76365226 | 0.75349957 | 0.45114977 | 0.99617422 |
| DSE | 12.1356155 | 0.457887093 | 0.48300804 | 0.94799063 | 0.34313422 | 0.97788427 |
| TSPYL1 | 14.7063138 | 0.623242089 | 0.4419355 | 1.41025578 | 0.15846417 | 0.86661434 |
| RWDD1 | 10.4375094 | -0.218796063 | 0.6139651 | -0.3563656 | 0.72156676 | 0.99665574 |
| ZUP1 | 8.00346757 | -0.23937008 | 0.54544513 | -0.4388527 | 0.66076826 | 0.99665574 |
| KPNA5 | 16.2676763 | 0.538967791 | 0.41156428 | 1.30955921 | 0.19034499 | 0.89963522 |
| GPRC6A | 20.3889357 | 0.999922669 | 0.58104757 | 1.72089639 | 0.08526963 | 0.75373899 |
| DCBLD1 | 8.44770659 | -0.563262201 | 0.50659211 | -1.1118653 | 0.26619606 | 0.94557817 |
| GOPC | 15.2155054 | -0.169044079 | 0.44731829 | -0.3779056 | 0.70550074 | 0.99665574 |
| NUS1 | 21.9366308 | -0.076427151 | 0.40452448 | -0.1889308 | 0.85014702 | 0.99665574 |
| SLC35F1 | 3.79982874 | -1.048368266 | 0.75430138 | -1.3898533 | 0.16457343 | 0.87444728 |
| CEP85L | 10.4714584 | -0.190924064 | 0.59437823 | -0.3212164 | 0.74804637 | 0.99665574 |
| MCM9 | 32.9752322 | 0.256065395 | 0.3137631 | 0.81611062 | 0.41443687 | 0.99279547 |
| AL365275.1 | 10.7365211 | 0.02693843 | 0.80215171 | 0.03358271 | 0.97320991 | 0.99777713 |
| FAM184A | 11.5438479 | 0.062899446 | 0.50813399 | 0.12378516 | 0.90148538 | 0.99777713 |
| MAN1A1 | 50.8037983 | -1.327676285 | 0.53256593 | -2.4929801 | 0.01266759 | 0.38590337 |
| TBC1D32 | 21.0313445 | -0.200883377 | 0.41646121 | -0.482358 | 0.62955169 | 0.99665574 |
| HSF2 | 6.24018392 | -0.380945589 | 0.56086045 | -0.6792163 | 0.49700079 | 0.99640678 |
| SERINC1 | 35.3060551 | -0.603564889 | 0.32037227 | -1.8839486 | 0.05957193 | 0.68011288 |
| PKIB | 9.58943115 | -0.296331372 | 0.69134583 | -0.4286297 | 0.66819271 | 0.99665574 |
| SMPDL3A | 7.52862617 | 0.575755738 | 0.6887544 | 0.83593765 | 0.40318998 | 0.99194293 |
| NKAIN2 | 5.35253769 | 0.368600567 | 0.74472448 | 0.49494891 | 0.62063619 | 0.99665574 |
| RNF217 | 65.0828329 | -1.662935356 | 0.44246191 | -3.7583696 | 0.00017102 | 0.02986125 |
| TPD52L1 | 13.0064046 | -1.045374162 | 0.6183285 | -1.6906453 | 0.09090457 | 0.76412898 |
| HDDC2 | 12.7383013 | -0.256039218 | 0.54186685 | -0.4725132 | 0.63656055 | 0.99665574 |
| NCOA7 | 268.46308 | 0.480076831 | 0.28322554 | 1.69503366 | 0.09006904 | 0.76220358 |
| NCOA7-AS1 | 8.03839537 | -0.290987575 | 0.5778028 | -0.5036105 | 0.61453509 | 0.99665574 |
| HINT3 | 9.68795396 | 0.276601406 | 0.48347276 | 0.57211374 | 0.56724492 | 0.99665574 |
| TRMT11 | 13.0911018 | -0.226977628 | 0.45135544 | -0.50288 | 0.61504863 | 0.99665574 |
| AC020559.1 | 3.35286212 | 0.52865271 | 0.82958925 | 0.63724634 | 0.5239644 | 0.99665574 |
| RNF146 | 11.7527304 | 0.149363642 | 0.62357527 | 0.23952785 | 0.8106963 | 0.99665574 |
| ECHDC1 | 21.0551315 | -0.296468306 | 0.408751 | -0.725303 | 0.46826619 | 0.99617422 |
| PTPRK | 122.802764 | -0.142249949 | 0.28949472 | -0.4913732 | 0.62316251 | 0.99665574 |
| LAMA2 | 112.47931 | -1.246450738 | 1.226666 | -1.0161289 | 0.30956802 | 0.97376371 |
| AL356124.1 | 21.1479176 | -0.207507302 | 0.48282617 | -0.4297764 | 0.66735829 | 0.99665574 |
| ARHGAP18 | 157.031596 | -0.138266604 | 0.3470353 | -0.3984223 | 0.69031892 | 0.99665574 |

| | | | | | | |
|---|---|---|---|---|---|---|
| L3MBTL3 | 8.9793608 | -0.853954069 | 0.55123089 | -1.5491767 | 0.12133924 | 0.82659025 |
| EPB41L2 | 10.9561145 | 0.452330709 | 0.62524809 | 0.72344197 | 0.4694084 | 0.99617422 |
| AKAP7 | 14.4360131 | 0.034409815 | 0.59813902 | 0.05752812 | 0.9541245 | 0.99777713 |
| MED23 | 17.3367432 | -0.01555489 | 0.4170201 | -0.0373001 | 0.97024573 | 0.99777713 |
| ENPP1 | 12.9922962 | 0.926629598 | 1.02205463 | 0.90663412 | 0.36460031 | 0.98362663 |
| CTGF | 9.94811402 | -0.564992666 | 0.77063916 | -0.7331481 | 0.46346808 | 0.99617422 |
| STX7 | 32.8121667 | -0.277196095 | 0.31140911 | -0.8901348 | 0.37339349 | 0.98532838 |
| SLC18B1 | 8.70057332 | 0.675972082 | 0.53894671 | 1.25424659 | 0.20975239 | 0.91043197 |
| RPS12 | 46.1769054 | 0.852241339 | 0.35885818 | 2.37486948 | 0.01755516 | 0.43770425 |
| EYA4 | 43.1988568 | 0.607498528 | 0.45506825 | 1.33496135 | 0.181889 | 0.89342536 |
| TBPL1 | 7.01940247 | 1.167456903 | 0.62498134 | 1.86798683 | 0.06176391 | 0.68358351 |
| SGK1 | 5.15938242 | -0.277567371 | 0.76538777 | -0.3626493 | 0.71686685 | 0.99665574 |
| HBS1L | 16.2380766 | 0.457771254 | 0.3784265 | 1.20967018 | 0.22640548 | 0.92318867 |
| AHI1 | 57.1382358 | -0.267445259 | 0.33043022 | -0.809385 | 0.41829373 | 0.99432434 |
| LINC00271 | 14.3998121 | 0.503481209 | 0.61802313 | 0.81466402 | 0.41526465 | 0.99279547 |
| PDE7B | 50.2039284 | -0.10319902 | 0.46593406 | -0.2214885 | 0.82471212 | 0.99665574 |
| AL138828.1 | 7.92041207 | -4.361034951 | 0.97384988 | -4.4781388 | 7.53E-06 | 0.00344882 |
| BCLAF1 | 38.8051848 | 0.11912845 | 0.32689066 | 0.36442904 | 0.71553765 | 0.99665574 |
| MAP7 | 92.4188782 | 0.301756342 | 0.20689841 | 1.4584759 | 0.14470941 | 0.85018987 |
| MAP3K5 | 133.312829 | -0.364869592 | 0.47898153 | -0.7617613 | 0.44620248 | 0.99617422 |
| PEX7 | 14.6061377 | 0.844088476 | 0.40534084 | 2.08241657 | 0.03730444 | 0.58653453 |
| IFNGR1 | 11.0591441 | -0.366272229 | 0.43833118 | -0.8356061 | 0.40337651 | 0.99194293 |
| AL357060.1 | 5.95905159 | 1.782531568 | 1.25369572 | 1.42182153 | 0.15507807 | 0.86239367 |
| TNFAIP3 | 8.06395055 | 0.974754965 | 1.07427646 | 0.90735951 | 0.36421671 | 0.98362663 |
| PERP | 29.1573655 | -0.315419452 | 0.30730239 | -1.0264139 | 0.30469651 | 0.97376371 |
| ARFGEF3 | 9.04319338 | -1.532234043 | 0.60926439 | -2.5148919 | 0.01190689 | 0.37488215 |
| HEBP2 | 16.4154428 | -0.551996472 | 0.49771846 | -1.1090536 | 0.26740704 | 0.94557817 |
| NHSL1 | 48.087804 | 1.11650898 | 0.42490836 | 2.62764656 | 0.00859778 | 0.31119582 |
| CCDC28A | 6.57883781 | 0.182274102 | 0.57891722 | 0.31485348 | 0.7528729 | 0.99665574 |
| REPS1 | 68.7813254 | 0.457113416 | 0.35530894 | 1.28652381 | 0.19826031 | 0.90381759 |
| ABRACL | 3.92779659 | -1.314226768 | 0.78575411 | -1.6725675 | 0.09441248 | 0.77065513 |
| HECA | 8.88692597 | 0.081067622 | 0.50575097 | 0.16029158 | 0.87265139 | 0.99665574 |
| CITED2 | 22.2937431 | -0.813107404 | 0.77542617 | -1.0485942 | 0.29436492 | 0.97161335 |
| VTA1 | 14.5049668 | 0.046830581 | 0.38704205 | 0.1209961 | 0.90369412 | 0.99777713 |
| HIVEP2 | 70.2180397 | -0.203697142 | 0.37253053 | -0.5467932 | 0.58452085 | 0.99665574 |
| AIG1 | 36.9782596 | 0.704584494 | 0.50571901 | 1.39323316 | 0.16354929 | 0.8726646 |
| ADAT2 | 4.26132928 | 1.056436389 | 0.75493097 | 1.39938144 | 0.16169863 | 0.87011714 |
| PEX3 | 10.6502131 | -0.117486121 | 0.45984572 | -0.2554903 | 0.79834443 | 0.99665574 |
| FUCA2 | 7.98069063 | -0.516367976 | 0.60775294 | -0.8496347 | 0.39552822 | 0.99179694 |
| PHACTR2 | 50.7658069 | 0.198919438 | 0.39295495 | 0.50621435 | 0.61270618 | 0.99665574 |
| PLAGL1 | 19.859442 | -0.72546039 | 0.42395921 | -1.7111561 | 0.0870523 | 0.75584737 |
| SF3B5 | 16.1325494 | -0.0116016 | 0.39659612 | -0.0292529 | 0.97666286 | 0.99777713 |
| UTRN | 51.6910581 | -0.189021435 | 0.32473299 | -0.5820826 | 0.56051102 | 0.99665574 |
| EPM2A | 11.8828683 | 0.919281308 | 0.57745659 | 1.59194876 | 0.11139622 | 0.8083966 |
| FBXO30 | 7.34163624 | 1.201531844 | 0.64117358 | 1.8739572 | 0.06093632 | 0.6834742 |
| AL356599.1 | 7.62758321 | -1.069557382 | 0.6790361 | -1.5751112 | 0.11523077 | 0.81721642 |
| SHPRH | 46.0443853 | -0.038024879 | 0.31224323 | -0.1217797 | 0.90307351 | 0.99777713 |
| RAB32 | 3.87958809 | -1.054778568 | 0.74443945 | -1.4168762 | 0.15651912 | 0.86532131 |
| STXBP5-AS1 | 3.1843462 | -1.093016357 | 0.93299933 | -1.1715082 | 0.24139457 | 0.93253689 |
| STXBP5 | 35.7161351 | -0.117899154 | 0.38581892 | -0.3055816 | 0.75992325 | 0.99665574 |
| SAMD5 | 15.8764771 | -0.890684031 | 0.58877577 | -1.5127729 | 0.13033735 | 0.83789479 |
| AL033504.1 | 8.03038043 | -1.617549478 | 0.74345126 | -2.1757304 | 0.02957543 | 0.5576677 |
| SASH1 | 50.1886277 | -1.15997705 | 0.44281216 | -2.6195691 | 0.00880409 | 0.31539185 |
| UST | 29.1021074 | -0.201450508 | 0.63129328 | -0.3191076 | 0.74964489 | 0.99665574 |
| TAB2 | 39.9113693 | 0.301174643 | 0.29480942 | 1.02159097 | 0.30697453 | 0.97376371 |
| PPIL4 | 17.662048 | 0.409622629 | 0.39477866 | 1.03760074 | 0.29945598 | 0.97358745 |
| GINM1 | 23.1937255 | -0.332478653 | 0.34070087 | -0.9758668 | 0.3291305 | 0.97788427 |
| KATNA1 | 12.2130455 | 0.290307514 | 0.50514112 | 0.57470576 | 0.56549031 | 0.99665574 |
| LATS1 | 29.4300106 | -0.070600594 | 0.38798757 | -0.1819661 | 0.85560931 | 0.99665574 |
| NUP43 | 8.12401012 | 0.775765128 | 0.54984019 | 1.41089201 | 0.15827646 | 0.86661434 |
| PCMT1 | 19.6303127 | -0.660410906 | 0.41820873 | -1.5791419 | 0.11430352 | 0.81638817 |
| LRP11 | 6.40265673 | -1.322891534 | 0.9131813 | -1.4486625 | 0.14743185 | 0.85284617 |
| PLEKHG1 | 39.2806069 | -0.5260503 | 0.40941824 | -1.2848727 | 0.19883679 | 0.90381759 |
| MTHFD1L | 36.2615099 | -1.994639974 | 0.44813985 | -4.450932 | 8.55E-06 | 0.00344882 |
| AKAP12 | 6.45418904 | -1.195816404 | 0.8127026 | -1.4714071 | 0.14118105 | 0.84969914 |
| RMND1 | 10.524325 | -0.125144887 | 0.50654119 | -0.2470577 | 0.80486359 | 0.99665574 |
| ARMT1 | 7.66102243 | 1.670215533 | 0.65456377 | 2.55164677 | 0.01072152 | 0.35991631 |
| ESR1 | 33.2088984 | -1.385546209 | 0.65538391 | -2.1140986 | 0.03450685 | 0.58178209 |
| SYNE1 | 37.2373324 | -0.890786954 | 0.41705459 | -2.1359001 | 0.03268755 | 0.57356513 |
| MTRF1L | 5.71073166 | -0.061119517 | 0.6882355 | -0.0888061 | 0.929236 | 0.99777713 |
| IPCEF1 | 17.3160793 | 1.886582779 | 0.54973046 | 3.43183234 | 0.00059952 | 0.06631714 |
| CNKSR3 | 111.742064 | 0.098584432 | 0.33075577 | 0.29805808 | 0.76565883 | 0.99665574 |
| SCAF8 | 76.3575128 | 0.230979158 | 0.31384155 | 0.73597379 | 0.46174665 | 0.99617422 |
| TIAM2 | 37.593853 | 0.275937885 | 0.41009671 | 0.67286052 | 0.50103603 | 0.99640678 |
| TFB1M | 17.1123065 | -0.034918129 | 0.41645472 | -0.0838462 | 0.93317875 | 0.99777713 |
| ARID1B | 252.108804 | 0.047795951 | 0.23535293 | 0.20308203 | 0.83907091 | 0.99665574 |
| TMEM242 | 6.79479253 | -0.095007484 | 0.60593987 | -0.1567936 | 0.87540753 | 0.99665574 |
| ZDHHC14 | 22.8452901 | -0.52302279 | 0.86224524 | -0.6065824 | 0.54412808 | 0.99665574 |
| SNX9 | 41.7317521 | -0.502469832 | 0.3631303 | -1.3837177 | 0.16644491 | 0.87495175 |
| SYNJ2 | 4.75519765 | -1.350888283 | 0.67385824 | -2.0047069 | 0.04499439 | 0.6313648 |
| SERAC1 | 5.53651677 | -1.033890453 | 0.71785807 | -1.4402436 | 0.1497985 | 0.85525834 |
| GTF2H5 | 8.30089863 | -0.462157676 | 0.75730718 | -0.6102645 | 0.54168662 | 0.99665574 |
| TULP4 | 62.7636874 | -0.570439215 | 0.35066357 | -1.6267422 | 0.10379187 | 0.79062881 |
| TMEM181 | 23.1369199 | -0.646478842 | 0.58994951 | -1.0958206 | 0.27315728 | 0.95186117 |
| DYNLT1 | 21.6697884 | -0.333897003 | 0.40907608 | -0.8162223 | 0.41437303 | 0.99279547 |
| SYTL3 | 5.54549038 | -0.781743306 | 0.72893085 | -1.0724519 | 0.28351709 | 0.96059387 |
| EZR | 100.206775 | -0.790625888 | 0.4740731 | -1.6677299 | 0.09536935 | 0.77178333 |
| RSPH3 | 7.26717509 | -0.680106429 | 0.57098973 | -1.191101 | 0.23361394 | 0.92960234 |
| AL035530.2 | 5.63078846 | 0.380893831 | 0.70669767 | 0.53897706 | 0.58990268 | 0.99665574 |
| SOD2.1 | 8.16366399 | -0.424621437 | 0.54980153 | -0.7723177 | 0.4399263 | 0.99617422 |
| WTAP | 34.4995386 | -0.426533548 | 0.34411632 | -1.239504 | 0.21515892 | 0.913434 |
| ACAT2 | 7.1501408 | -1.460074169 | 0.64612959 | -2.2597234 | 0.02383842 | 0.49867035 |
| TCP1 | 13.7243432 | 0.465083417 | 0.47451593 | 0.98012181 | 0.32702599 | 0.97788427 |
| MRPL18 | 6.63417993 | -0.206172781 | 0.54810441 | -0.376156 | 0.7068009 | 0.99665574 |
| IGF2R | 79.487731 | -0.349040205 | 0.37930247 | -0.920216 | 0.35745989 | 0.9819332 |
| SLC22A1 | 2.60462278 | -0.957611602 | 1.03557543 | -0.9247145 | 0.35511444 | 0.98022903 |
| MAP3K4 | 26.7269925 | 0.730138471 | 0.33981483 | 2.14863625 | 0.03166325 | 0.56814511 |
| PRKN | 262.805888 | 0.315639625 | 0.37904806 | 0.83271663 | 0.40500456 | 0.99194293 |
| PACRG | 567.225742 | 0.029315296 | 0.41904589 | 0.06995724 | 0.94422769 | 0.99777713 |
| PACRG-AS2 | 3.77347173 | -0.828705365 | 1.0813701 | -0.7663476 | 0.44346951 | 0.99617422 |
| PACRG-AS3 | 13.4636694 | -0.527157741 | 0.72887401 | -0.7232495 | 0.46952663 | 0.99617422 |
| QKI | 162.856049 | -0.034000031 | 0.37084497 | -0.0916826 | 0.92695022 | 0.99777713 |
| PDE10A | 50.95315 | -0.980652159 | 0.60708831 | -1.6153369 | 0.10623775 | 0.79596216 |
| SFT2D1 | 16.1089853 | 0.112922587 | 0.40169057 | 0.28111834 | 0.77861963 | 0.99665574 |
| MPC1 | 51.403114 | 0.041602653 | 0.34917681 | 0.11914495 | 0.90516052 | 0.99777713 |
| RPS6KA2 | 26.8485552 | -0.571142125 | 0.39035133 | -1.4631489 | 0.14342664 | 0.84969914 |
| Z94721.1 | 10.7917264 | -0.446062848 | 0.5939269 | -0.75104 | 0.4526286 | 0.99617422 |
| FGFR1OP | 23.4699097 | 0.644798427 | 0.39705351 | 1.62395852 | 0.10438466 | 0.79169635 |
| AFDN | 114.082426 | 0.333834282 | 0.24076877 | 1.38653482 | 0.16558366 | 0.87495175 |
| WDR27 | 25.8619243 | 0.094073922 | 0.45378855 | 0.20730784 | 0.83576945 | 0.99665574 |
| C6orf120 | 2.21393596 | 0.144542008 | 0.903073 | 0.16005573 | 0.87283717 | 0.99665574 |
| PHF10 | 16.1701995 | 0.348217355 | 0.43947096 | 0.79235579 | 0.42815325 | 0.99589919 |
| ERMARD | 5.7874688 | 0.212346716 | 0.60968758 | 0.34828775 | 0.72762409 | 0.99665574 |

| | | | | | | |
|---|---|---|---|---|---|---|
| DLL1 | 41.1534865 | 0.085420136 | 0.63099967 | 0.13537271 | 0.8923172 | 0.99773598 |
| FAM120B | 93.6887502 | -0.601075251 | 0.33437968 | -1.7975831 | 0.07224311 | 0.71945049 |
| PSMB1 | 24.830695 | -0.17883095 | 0.46573247 | -0.3839778 | 0.70099487 | 0.99665574 |
| TBP | 6.17792749 | 0.381107111 | 0.57845951 | 0.6588311 | 0.51000424 | 0.99665574 |
| PDCD2 | 7.9238921 | -0.98365391 | 0.58447905 | -1.6829584 | 0.09238315 | 0.76670359 |
| FAM20C | 24.1656503 | -0.793564965 | 0.51587537 | -1.5382881 | 0.12397818 | 0.82915695 |
| PRKAR1B | 8.27318794 | -0.063160269 | 0.54662005 | -0.1155469 | 0.90801162 | 0.99777713 |
| DNAAF5 | 6.99218785 | -0.912699497 | 0.78395222 | -1.1642285 | 0.24433143 | 0.93426912 |
| SUN1 | 24.7742502 | 0.111011284 | 0.31599725 | 0.35130458 | 0.72535986 | 0.99665574 |
| ADAP1 | 3.81760917 | 0.892669551 | 0.83359224 | 1.07087075 | 0.28422755 | 0.96059387 |
| COX19 | 13.8747322 | -0.409346046 | 0.39754457 | -1.0296859 | 0.30315747 | 0.97376371 |
| C7orf50 | 22.1759451 | 0.456964788 | 0.39119333 | 1.16813031 | 0.24275421 | 0.9342309 |
| GPR146 | 3.70236993 | 0.228408892 | 0.89406369 | 0.25547273 | 0.798358 | 0.99665574 |
| MICALL2 | 3.53199403 | -1.541373055 | 0.75256819 | -2.0481507 | 0.04054524 | 0.60895944 |
| INTS1 | 11.7149133 | -0.006826614 | 0.62974403 | -0.0108403 | 0.99135086 | 0.99777713 |
| MAFK | 5.66085938 | 0.480726803 | 0.71399088 | 0.67329544 | 0.50075935 | 0.99640678 |
| PSMG3 | 4.32332161 | 0.783978642 | 0.86464112 | 0.90670988 | 0.36456024 | 0.98362663 |
| MAD1L1 | 15.3916276 | 0.110079802 | 0.46658464 | 0.23592676 | 0.8134895 | 0.99665574 |
| MRM2 | 3.4779031 | -0.668529206 | 0.93341246 | -0.7162206 | 0.47385516 | 0.99617422 |
| SNX8 | 6.25552463 | -1.409117798 | 0.83301057 | -1.6915965 | 0.09072293 | 0.76391115 |
| EIF3B | 16.5139004 | -0.057301456 | 0.54217006 | -0.1056891 | 0.91582904 | 0.99777713 |
| CHST12 | 9.91055613 | -0.407086015 | 0.4938495 | -0.8243119 | 0.40976236 | 0.99279547 |
| BRAT1 | 3.31140173 | 0.399983192 | 0.8407983 | 0.47571836 | 0.63427504 | 0.99665574 |
| IQCE | 5.40800487 | 1.176075475 | 0.85405926 | 1.377042 | 0.16849926 | 0.87566218 |
| GNA12 | 16.25417 | -0.315760813 | 0.37658017 | -0.8384956 | 0.40175242 | 0.99194293 |
| SDK1 | 71.995973 | -0.439340606 | 0.36097613 | -1.2170905 | 0.2235698 | 0.92191316 |
| FOXK1 | 37.6540284 | -0.326360406 | 0.39047748 | -0.8357983 | 0.40326839 | 0.99194293 |
| AP5Z1 | 6.44324254 | -0.588328199 | 0.715418 | -0.8223559 | 0.41087438 | 0.99279547 |
| RBAK-RBAKD | 15.7921745 | -0.505371979 | 0.49796179 | -1.014881 | 0.31016254 | 0.97376371 |
| WIPI2 | 18.6559604 | 0.210704183 | 0.39254258 | 0.53676771 | 0.59142808 | 0.99665574 |
| SLC29A4 | 2.76157528 | 0.835473388 | 1.04051479 | 0.80294235 | 0.42200806 | 0.99518711 |
| TNRC18 | 49.8825806 | -0.39732144 | 0.38665473 | -1.0275872 | 0.30414406 | 0.97376371 |
| FBXL18 | 4.52109203 | -0.447242456 | 0.76539044 | -0.5843324 | 0.55899667 | 0.99665574 |
| ACTB | 224.783335 | 0.210957951 | 0.42180558 | 0.50013078 | 0.616983 | 0.99665574 |
| RNF216 | 33.0097144 | -0.19846562 | 0.28884715 | -0.6870956 | 0.49202246 | 0.99640678 |
| CCZ1 | 7.12610826 | -0.748092235 | 0.65889812 | -1.1353686 | 0.25622091 | 0.94314473 |
| PMS2 | 11.9156751 | 0.43024937 | 0.60768074 | 0.70801878 | 0.4789336 | 0.99617422 |
| AIMP2 | 11.8272665 | -0.504011025 | 0.53101444 | -0.9491475 | 0.3425456 | 0.97788427 |
| EIF2AK1 | 22.3353633 | 0.305264119 | 0.39586283 | 0.7711361 | 0.44062627 | 0.99617422 |
| USP42 | 21.9858645 | 0.12660368 | 0.37757501 | 0.33530736 | 0.73739324 | 0.99665574 |
| CYTH3 | 20.6091749 | -0.386055812 | 0.38632603 | -0.9993005 | 0.31764912 | 0.97700345 |
| RAC1 | 44.936901 | -0.49207745 | 0.28518026 | -1.7254962 | 0.08443809 | 0.75373899 |
| DAGLB | 4.27217602 | -0.701641167 | 0.70544154 | -0.9946128 | 0.31992463 | 0.97700345 |
| ZDHHC4 | 10.3649949 | 0.018725172 | 0.45779593 | 0.04090288 | 0.96737332 | 0.99777713 |
| C7orf26 | 6.84669346 | -0.111194853 | 0.55866173 | -0.1990379 | 0.8422331 | 0.99665574 |
| ZNF316 | 7.21088167 | -0.874171486 | 0.72362563 | -1.2080438 | 0.2270304 | 0.92360922 |
| AC073343.2 | 4.79745604 | -0.958001681 | 0.63129309 | -1.5175228 | 0.12913473 | 0.83778831 |
| ZNF12 | 11.1049425 | -0.099318279 | 0.57273621 | -0.1734102 | 0.86232904 | 0.99665574 |
| CCZ1B | 8.11094785 | -0.797120586 | 0.49850129 | -1.5990341 | 0.10981302 | 0.80522301 |
| C1GALT1 | 31.513374 | 0.398211797 | 0.3529396 | 1.12827178 | 0.25920516 | 0.94557817 |
| COL28A1 | 13.3640923 | 0.013202919 | 0.59101701 | 0.02233932 | 0.98217728 | 0.99777713 |
| AC004982.2 | 3.64657196 | 1.151814174 | 0.97879313 | 1.17676977 | 0.23928741 | 0.93157178 |
| MIOS | 13.5498676 | 0.498416397 | 0.48234429 | 1.03332082 | 0.30145379 | 0.97358745 |
| RPA3 | 9.62828861 | -1.149930717 | 0.67915218 | -1.6931856 | 0.09042014 | 0.76326341 |
| UMAD1 | 81.929818 | -0.126187195 | 0.24898051 | -0.5068155 | 0.61228425 | 0.99665574 |
| GLCCI1 | 100.771587 | 0.396508817 | 0.5261332 | 0.7536282 | 0.4510725 | 0.99617422 |
| ICA1 | 49.3107097 | -0.492884586 | 0.35645222 | -1.3827508 | 0.1667413 | 0.87495175 |
| NDUFA4 | 57.2141979 | 0.06918177 | 0.47830647 | 0.144639 | 0.8849959 | 0.99665574 |
| PHF14 | 41.4762437 | -0.155110036 | 0.34942395 | -0.4439021 | 0.65711334 | 0.99665574 |
| THSD7A | 174.071389 | -0.674386478 | 0.50563775 | -1.3337344 | 0.1822909 | 0.89451364 |
| TMEM106B | 18.6332674 | 0.59141286 | 0.45478788 | 1.30041474 | 0.19345886 | 0.90153775 |
| SCIN | 249.02282 | -0.060455923 | 0.33158659 | -0.1823232 | 0.8553291 | 0.99665574 |
| ARL4A | 4.93838858 | -0.571142935 | 0.93403773 | -0.6114774 | 0.54088359 | 0.99665574 |
| AC011287.1 | 4.99454636 | -0.340354034 | 0.88419833 | -0.3849295 | 0.70028964 | 0.99665574 |
| ETV1 | 10.2419803 | -1.352098587 | 0.82933203 | -1.6303465 | 0.10302828 | 0.78770791 |
| DGKB | 56.1284886 | -1.240679647 | 1.28435812 | -0.965992 | 0.33404819 | 0.97788427 |
| AGMO | 5.31014616 | 0.195081509 | 1.04471963 | 0.18673097 | 0.85187157 | 0.99665574 |
| ISPD | 34.380763 | -0.478027535 | 0.48533333 | -0.9849468 | 0.32465018 | 0.97700345 |
| ANKMY2 | 9.15289028 | -0.003494963 | 0.55407036 | -0.0063078 | 0.99496714 | 0.9985287 |
| BZW2 | 8.71156139 | -0.929331988 | 0.5375089 | -1.7289611 | 0.08381606 | 0.75242737 |
| AC073332.1 | 4.66132768 | -1.227910075 | 0.79225291 | -1.5498966 | 0.12116634 | 0.82659025 |
| SNX13 | 48.4160473 | 0.352879297 | 0.28138302 | 1.25408883 | 0.20980972 | 0.91043197 |
| HDAC9 | 15.9334676 | -3.289261454 | 0.84318782 | -3.9009831 | 9.58E-05 | 0.02258719 |
| TWISTNB | 4.93748035 | -0.389383264 | 0.65219145 | -0.5970383 | 0.55048182 | 0.99665574 |
| MACC1 | 32.5024725 | -1.536612781 | 0.46930126 | -3.2742567 | 0.0010594 | 0.08950192 |
| ITGB8 | 29.2920092 | -1.682888925 | 0.54480611 | -3.0889685 | 0.00200853 | 0.13751507 |
| SP4 | 16.2026163 | 0.575011462 | 0.49913058 | 1.15202612 | 0.24931034 | 0.9369778 |
| DNAH11 | 39.2162998 | -0.675122159 | 1.38233819 | -0.4883915 | 0.62527259 | 0.99665574 |
| RAPGEF5 | 6.85084651 | -3.753291886 | 0.74726737 | -5.0226894 | 5.10E-07 | 0.00043047 |
| STEAP1B | 49.9005444 | 0.059909472 | 0.4011846 | 0.14933143 | 0.88129211 | 0.99665574 |
| TOMM7 | 24.40134 | 0.444565748 | 0.50575229 | 0.87901875 | 0.37939111 | 0.98679664 |
| FAM126A | 3.65910599 | -0.39781795 | 0.80466727 | -0.4943881 | 0.62103209 | 0.99665574 |
| KLHL7 | 11.0290219 | -0.42273682 | 0.44950871 | -0.9404419 | 0.34699095 | 0.97804112 |
| NUPL2 | 13.73743 | -0.424305224 | 0.4425902 | -0.9586864 | 0.33771673 | 0.97788427 |
| MALSU1 | 9.05111196 | 0.377545455 | 0.59695641 | 0.63245062 | 0.52709246 | 0.99665574 |
| TRA2A | 68.1186568 | 0.06279027 | 0.3716005 | 0.16897251 | 0.86581826 | 0.99665574 |
| CCDC126 | 13.2100434 | -0.052411551 | 0.46063992 | -0.1137799 | 0.9094123 | 0.99777713 |
| FAM221A | 9.69155142 | 1.147118202 | 0.55008708 | 2.08533931 | 0.03703851 | 0.58591936 |
| MPP6 | 10.6441708 | 1.165036072 | 0.85810727 | 1.35768116 | 0.17456488 | 0.88366595 |
| OSBPL3 | 112.807628 | 0.240343419 | 0.4545741 | 0.5287222 | 0.59699817 | 0.99665574 |
| CYCS | 33.5761795 | 0.554896994 | 0.48067287 | 1.15441713 | 0.2483292 | 0.93520111 |
| HNRNPA2B1 | 108.139148 | 0.484134384 | 0.22925695 | 2.11175448 | 0.03470751 | 0.58178209 |
| CBX3 | 18.3441745 | -0.330661236 | 0.45173706 | -0.7319772 | 0.46418247 | 0.99617422 |
| SNX10 | 14.6317385 | -2.355072659 | 0.62598802 | -3.7621689 | 0.00016845 | 0.02986125 |
| AC004540.1 | 10.785956 | 0.643083453 | 0.67176396 | 0.95730568 | 0.33841299 | 0.97788427 |
| SKAP2 | 29.329983 | -0.908838558 | 0.41853748 | -2.1714628 | 0.02989621 | 0.55886856 |
| HOXA3 | 4.29027207 | -0.368434388 | 0.71888733 | -0.5125064 | 0.60829661 | 0.99665574 |
| HOXA9 | 3.33200574 | 0.532679658 | 0.82463344 | 0.64595932 | 0.5183057 | 0.99665574 |
| HIBADH | 53.0579708 | 0.58305957 | 0.38998381 | 1.49508662 | 0.13489184 | 0.84373478 |
| TAX1BP1 | 79.2601418 | -0.001326661 | 0.27215712 | -0.0048746 | 0.99611064 | 0.99877061 |
| JAZF1 | 11.3816969 | -1.001269657 | 0.65007182 | -1.5402447 | 0.12350073 | 0.82915695 |
| CPVL | 7.89414612 | -0.013056312 | 0.82715071 | -0.0157847 | 0.98740617 | 0.99777713 |
| SCRN1 | 24.4672098 | -0.232809014 | 0.35755031 | -0.6511224 | 0.51496748 | 0.99665574 |
| FKBP14 | 4.68784286 | -1.109704859 | 0.74620598 | -1.4871294 | 0.13698064 | 0.84475183 |
| PLEKHA8 | 28.5616358 | -0.216017876 | 0.39539323 | -0.5463368 | 0.58483443 | 0.99665574 |
| MTURN | 31.5753173 | 0.214746338 | 0.36445446 | 0.5892268 | 0.55570914 | 0.99665574 |
| ZNRF2 | 16.8230698 | -0.668507333 | 0.45990122 | -1.453589 | 0.1460603 | 0.85169374 |
| AC006978.2 | 15.6862882 | -0.175703235 | 0.50662373 | -0.3468121 | 0.72873249 | 0.99665574 |
| AC005154.6 | 8.23804453 | 0.169461879 | 0.53666053 | 0.31577109 | 0.75217626 | 0.99665574 |
| GARS | 16.9628003 | -0.093250558 | 0.53237906 | -0.1751582 | 0.86095533 | 0.99665574 |
| AQP1 | 7.87406643 | -0.123625475 | 0.6566289 | -0.188273 | 0.85066267 | 0.99665574 |
| CCDC129 | 4.13460426 | -0.534140247 | 0.74406149 | -0.7178711 | 0.47283677 | 0.99617422 |
| PDE1C | 538.087515 | 0.40370092 | 0.30563988 | 1.32083849 | 0.18655522 | 0.89532394 |

| | | | | | | |
|---|---|---|---|---|---|---|
| LSM5 | 5.93310699 | 0.49158027 | 0.63106379 | 0.7789708 | 0.43599691 | 0.99617422 |
| AVL9 | 41.8931843 | 0.065413443 | 0.267109 | 0.24489419 | 0.80653836 | 0.99665574 |
| KBTBD2 | 17.9294202 | -0.610335137 | 0.6253576 | -0.9759778 | 0.32907547 | 0.97788427 |
| FKBP9 | 5.25047863 | -0.32931178 | 0.87087944 | -0.378137 | 0.70532879 | 0.99665574 |
| NT5C3A | 16.1005974 | 0.827091469 | 0.53304213 | 1.55164371 | 0.1207475 | 0.82659025 |
| RP9 | 2.99321974 | -0.473920878 | 0.85930019 | -0.5515196 | 0.58127755 | 0.99665574 |
| BBS9 | 36.6113668 | 0.092496146 | 0.36284653 | 0.25491809 | 0.79878636 | 0.99665574 |
| DPY19L1 | 20.5953931 | -1.093604172 | 0.43444528 | -2.5172426 | 0.01182773 | 0.37359313 |
| HERPUD2 | 15.18731 | -0.057905105 | 0.3914313 | -0.1479317 | 0.88239666 | 0.99665574 |
| SEPT7-AS1 | 3.47466409 | -0.722119536 | 0.82806615 | -0.8720554 | 0.38317817 | 0.98947027 |
| 7-Sep | 48.5582275 | -0.061585832 | 0.29437313 | -0.2092101 | 0.83428423 | 0.99665574 |
| EEPD1 | 16.9232199 | -1.834778912 | 0.64470698 | -2.8459114 | 0.00442845 | 0.21310323 |
| KIAA0895 | 7.70777632 | -0.461338826 | 0.61714137 | -0.7475416 | 0.45473672 | 0.99617422 |
| EPDR1 | 4.8466083 | 0.678973965 | 0.75505495 | 0.89923781 | 0.368526 | 0.98532838 |
| STARD3NL | 10.8184502 | -0.054911149 | 0.47334874 | -0.1160057 | 0.90764803 | 0.99777713 |
| TRGC2 | 2.57162322 | 1.245647677 | 0.95296782 | 1.30712459 | 0.1911704 | 0.89964233 |
| TRGC1 | 21.9607106 | 1.72520199 | 0.65700282 | 2.62586696 | 0.00864286 | 0.31119582 |
| VPS41 | 46.2755875 | 0.392675035 | 0.34832854 | 1.12731226 | 0.25961048 | 0.94557817 |
| YAE1D1 | 3.85571541 | 0.059579004 | 0.69092024 | 0.08623138 | 0.93128249 | 0.99777713 |
| RALA | 38.0171244 | -0.630204401 | 0.39712288 | -1.5869255 | 0.11252953 | 0.81249398 |
| LINC00265 | 6.09741933 | 1.1349434 | 0.57287776 | 1.98112665 | 0.04757707 | 0.64060657 |
| CDK13 | 103.805609 | 0.253541947 | 0.19773152 | 1.28225354 | 0.19975372 | 0.90381759 |
| MPLKIP | 9.14599223 | -0.052760338 | 0.55320905 | -0.0953714 | 0.92401981 | 0.99777713 |
| SUGCT | 19.3666333 | 0.717025133 | 0.64095622 | 1.11868036 | 0.26327653 | 0.94557817 |
| MRPL32 | 9.44051035 | -0.268850539 | 0.52874524 | -0.508469 | 0.61112451 | 0.99665574 |
| STK17A | 6.73984932 | -1.595438852 | 0.51924581 | -3.072608 | 0.00212197 | 0.13969177 |
| COA1 | 41.3985491 | 0.142391569 | 0.31849376 | 0.44707805 | 0.6548187 | 0.99665574 |
| BLVRA | 5.74523249 | -1.880957419 | 0.65697685 | -2.8630498 | 0.00419585 | 0.20883915 |
| URGCP | 5.65956178 | -0.310961975 | 0.84500908 | -0.3679984 | 0.71287444 | 0.99665574 |
| UBE2D4 | 4.74409808 | -0.70587482 | 0.75805584 | -0.9311647 | 0.35176839 | 0.97838306 |
| AC004951.1 | 2.37043292 | 0.474794386 | 0.90084903 | 0.52705211 | 0.5981574 | 0.99665574 |
| DBNL | 9.50035005 | 0.666721449 | 0.51764698 | 1.28798481 | 0.19775125 | 0.90381759 |
| POLM | 6.34543146 | -0.557268299 | 0.58394112 | -0.9543227 | 0.3399203 | 0.97788427 |
| POLD2 | 3.06430749 | -0.419302979 | 0.84481156 | -0.4963272 | 0.61966356 | 0.99665574 |
| YKT6 | 6.13244477 | -0.537905387 | 0.80157145 | -0.6710636 | 0.50218004 | 0.99640678 |
| NUDCD3 | 35.6282478 | 0.143555135 | 0.31277557 | 0.4589717 | 0.64625449 | 0.99665574 |
| DDX56 | 8.35158451 | 1.121323328 | 0.53221648 | 2.10689328 | 0.03512683 | 0.58349146 |
| TMED4 | 21.0465321 | 0.526873761 | 0.36083749 | 1.4601414 | 0.14425122 | 0.84969914 |
| OGDH | 73.0051501 | 0.241093312 | 0.27239763 | 0.88507859 | 0.37611424 | 0.98655788 |
| ZMIZ2 | 14.1380644 | 0.54269508 | 0.40928492 | 1.32595914 | 0.18485323 | 0.89532394 |
| PPIA | 83.4956581 | 0.086741183 | 0.44942345 | 0.19300547 | 0.84695469 | 0.99665574 |
| H2AFV | 29.5629062 | 0.502174752 | 0.31524638 | 1.59295963 | 0.11116925 | 0.8083966 |
| PURB | 14.5314846 | 0.231986855 | 0.45501871 | 0.50984025 | 0.61016338 | 0.99665574 |
| SNHG15 | 4.11633826 | -1.610249017 | 0.97576288 | -1.6502462 | 0.09889258 | 0.78203824 |
| CCM2 | 24.4435754 | -0.702591841 | 0.34567898 | -2.032498 | 0.04210326 | 0.61533334 |
| TBRG4 | 5.68613428 | 0.306303095 | 0.69184205 | 0.44273559 | 0.657957 | 0.99665574 |
| ADCY1 | 33.162242 | -0.564650749 | 1.21069813 | -0.4663844 | 0.64094036 | 0.99665574 |
| TNS3 | 156.15424 | -1.327269517 | 0.47451767 | -2.7970919 | 0.00515649 | 0.22466099 |
| HUS1 | 7.63486686 | 0.477519504 | 0.56129973 | 0.85073888 | 0.39491442 | 0.99179694 |
| UPP1 | 17.6122365 | -0.031232553 | 0.7023473 | -0.0444688 | 0.96453071 | 0.99777713 |
| GRB10 | 115.598542 | 0.207469621 | 0.35760506 | 0.58016411 | 0.56180396 | 0.99665574 |
| COBL | 39.2665031 | -0.584558436 | 0.42299333 | -1.3819566 | 0.16698502 | 0.87495175 |
| SEC61G | 8.83525694 | 0.367623959 | 0.59496811 | 0.61788851 | 0.53664883 | 0.99665574 |
| EGFR | 225.597299 | -0.278533141 | 0.24047415 | -1.1582665 | 0.24675531 | 0.93441937 |
| LANCL2 | 9.49545436 | 0.283318415 | 0.57460455 | 0.49306678 | 0.6219654 | 0.99665574 |
| VOPP1 | 20.672961 | 0.039620721 | 0.43090234 | 0.09194826 | 0.92673915 | 0.99777713 |
| NIPSNAP2 | 51.4204773 | -0.171836264 | 0.28207012 | -0.609197 | 0.54239388 | 0.99665574 |
| PSPH | 4.86453713 | -0.034785695 | 0.64640975 | -0.0538137 | 0.9570836 | 0.99777713 |
| CCT6A | 18.5526041 | 0.048446813 | 0.56134294 | 0.0863052 | 0.93122381 | 0.99777713 |
| SUMF2 | 12.6327952 | -0.396949646 | 0.43969836 | -0.9027772 | 0.36664417 | 0.98365837 |
| PHKG1 | 4.13024192 | -0.093944796 | 0.74036716 | -0.1268895 | 0.89902788 | 0.99777713 |
| CHCHD2 | 52.3603531 | 0.337398658 | 0.50601571 | 0.66677506 | 0.50491583 | 0.99665574 |
| NUPR2 | 5.884761 | 0.502379124 | 0.70550204 | 0.71208742 | 0.47641064 | 0.99617422 |
| ZNF736 | 10.638851 | -0.16346321 | 0.62433448 | -0.2618199 | 0.79346027 | 0.99665574 |
| ZNF680 | 17.2080418 | -0.019232895 | 0.43162763 | -0.044559 | 0.96445882 | 0.99777713 |
| ZNF107 | 3.56356887 | 0.015718921 | 0.87370464 | 0.01799112 | 0.98564594 | 0.99777713 |
| ZNF138 | 9.33369696 | -0.311751952 | 0.48879469 | -0.6377973 | 0.52360561 | 0.99665574 |
| ZNF117 | 5.67124753 | 0.019069618 | 0.62228374 | 0.03064457 | 0.975553 | 0.99777713 |
| VKORC1L1 | 28.0583089 | -0.163667664 | 0.33808435 | -0.484103 | 0.62831274 | 0.99665574 |
| GUSB | 5.12419162 | 0.371668969 | 0.6599882 | 0.56314487 | 0.57333624 | 0.99665574 |
| AC068533.4 | 3.25007083 | 0.188949272 | 0.80340597 | 0.2351853 | 0.81406492 | 0.99665574 |
| CRCP | 4.95202864 | -0.739038609 | 0.63211349 | -1.1691549 | 0.24234121 | 0.9342309 |
| AC068533.3 | 4.2938841 | 0.586323449 | 0.77752217 | 0.75409226 | 0.45079383 | 0.99617422 |
| TPST1 | 73.5395276 | 0.013839209 | 0.24531031 | 0.05641512 | 0.95501112 | 0.99777713 |
| AC027644.4 | 16.7735051 | 0.079420188 | 0.39885369 | 0.19912111 | 0.84216801 | 0.99665574 |
| AC027644.3 | 6.3767741 | -0.640899165 | 0.68564538 | -0.9347385 | 0.34992306 | 0.97838306 |
| TMEM248 | 21.1163996 | 0.253940976 | 0.39198616 | 0.64783149 | 0.51709395 | 0.99665574 |
| SBDS | 16.4779762 | -0.268791674 | 0.46842161 | -0.5738242 | 0.56608674 | 0.99665574 |
| TYW1 | 33.1059831 | 0.201878335 | 0.29435879 | 0.68582405 | 0.49282407 | 0.99640678 |
| AC006480.2 | 3.13207723 | -0.546306065 | 0.77608528 | -0.7039253 | 0.4814793 | 0.99640678 |
| AUTS2 | 56.5596024 | -0.713384871 | 0.45938416 | -1.5529157 | 0.12044328 | 0.82659025 |
| GALNT17 | 91.1930712 | -0.055084166 | 1.20019456 | -0.045896 | 0.96339312 | 0.99777713 |
| CALN1 | 5.23965104 | 1.92676538 | 0.74784669 | 2.57641762 | 0.009983 | 0.34338499 |
| TYW1B | 25.1200212 | 0.155783838 | 0.39312619 | 0.3962693 | 0.69190637 | 0.99665574 |
| POM121 | 25.8915893 | 0.23867283 | 0.41617111 | 0.57349687 | 0.56630832 | 0.99665574 |
| NSUN5 | 7.41067699 | 0.755463839 | 0.6674214 | 1.13191431 | 0.25767046 | 0.94557817 |
| BAZ1B | 32.3531145 | -0.042013878 | 0.38722748 | -0.1084992 | 0.9135997 | 0.99777713 |
| BCL7B | 8.43204388 | -0.111400302 | 0.57595035 | -0.19342 | 0.84663007 | 0.99665574 |
| TBL2 | 3.15989706 | -0.159959504 | 0.75623296 | -0.2115215 | 0.83248038 | 0.99665574 |
| MLXIPL | 7.32533441 | 1.731342508 | 0.74767042 | 2.31564933 | 0.02057742 | 0.46985116 |
| VPS37D | 6.03819362 | 0.424074352 | 0.59028739 | 0.71842015 | 0.47249827 | 0.99617422 |
| DNAJC30 | 2.92123296 | -0.055691904 | 0.93433528 | -0.0596059 | 0.95246951 | 0.99777713 |
| BUD23 | 12.5243552 | 0.208546339 | 0.47086054 | 0.44290468 | 0.65783468 | 0.99665574 |
| ABHD11 | 5.18647942 | -0.484485109 | 0.78486058 | -0.6172881 | 0.5370447 | 0.99665574 |
| CLDN3 | 7.07316301 | -0.560080974 | 0.6295287 | -0.889683 | 0.37363613 | 0.98532838 |
| CLDN4 | 21.1147484 | -1.395024283 | 0.50905909 | -2.7403976 | 0.00613649 | 0.24884698 |
| LIMK1 | 3.09548554 | -0.66769934 | 0.83195643 | -0.8025653 | 0.42222605 | 0.99518711 |
| EIF4H | 17.7963575 | -0.124033003 | 0.41993474 | -0.2953626 | 0.76771692 | 0.99665574 |
| RFC2 | 3.54141843 | 0.733844252 | 0.94923871 | 0.77308715 | 0.4394708 | 0.99617422 |
| CLIP2 | 5.56041059 | -1.217550145 | 0.61409301 | -1.9826803 | 0.04740315 | 0.64060657 |
| GTF2IRD1 | 41.4072697 | -1.50897993 | 0.49638054 | -3.0399659 | 0.00236605 | 0.15249081 |
| GTF2I | 87.3722714 | 0.041789673 | 0.26128601 | 0.15993843 | 0.87292958 | 0.99665574 |
| GTF2IRD2 | 10.8190396 | -0.770869413 | 0.54338547 | -1.4186419 | 0.15600343 | 0.86424197 |
| CASTOR2 | 16.9261601 | 0.067681131 | 0.46606932 | 0.14521688 | 0.88453964 | 0.99665574 |
| RCC1L | 13.7984331 | -0.253821402 | 0.40900136 | -0.6205882 | 0.53487063 | 0.99665574 |
| GTF2IRD2B | 17.8555259 | -0.383758813 | 0.42401269 | -0.9050644 | 0.36543125 | 0.98365837 |
| TRIM73 | 9.2870213 | -0.795598267 | 0.49099758 | -1.6203711 | 0.10515259 | 0.79377288 |
| POM121C | 22.0503503 | -0.240291071 | 0.45063955 | -0.5332223 | 0.59387969 | 0.99665574 |
| SPDYE5 | 5.63802073 | -1.339819863 | 0.64564544 | -2.0751635 | 0.03797138 | 0.59132698 |
| HIP1 | 29.0725642 | 0.762466842 | 0.47166096 | 1.61655702 | 0.10597394 | 0.79596216 |
| RHBDD2 | 6.94326857 | 0.480182391 | 0.85157079 | 0.56387842 | 0.57283688 | 0.99665574 |
| POR | 11.1388162 | -1.035975673 | 0.53121831 | -1.9501882 | 0.05115369 | 0.65232219 |
| TMEM120A | 3.97078582 | 0.584110225 | 0.77599545 | 0.75272377 | 0.45161592 | 0.99617422 |

| | | | | | | |
|---|---|---|---|---|---|---|
| STYXL1 | 10.0278254 | 0.529564866 | 0.48538335 | 1.09102397 | 0.27526234 | 0.95593806 |
| MDH2 | 16.8812272 | 0.228491125 | 0.56889535 | 0.40163999 | 0.68794899 | 0.99665574 |
| HSPB1 | 35.8682262 | 0.697593836 | 0.47031055 | 1.48326216 | 0.13800477 | 0.84656101 |
| YWHAG | 17.5423385 | -0.231001303 | 0.44698079 | -0.5168036 | 0.60529323 | 0.99665574 |
| DTX2 | 8.92985897 | -0.351856389 | 0.49286581 | -0.713899 | 0.47528966 | 0.99617422 |
| POMZP3 | 5.66888332 | -0.76196955 | 0.68167798 | -1.1177852 | 0.26365875 | 0.94557817 |
| CCDC146 | 14.5817021 | 0.23499189 | 0.63360685 | 0.37087966 | 0.71072717 | 0.99665574 |
| FGL2 | 7.74082743 | 0.541992856 | 0.94277888 | 0.57488863 | 0.56536662 | 0.99665574 |
| GSAP | 12.3008201 | 0.940083629 | 0.59930177 | 1.56863149 | 0.11673383 | 0.8218386 |
| PTPN12 | 36.5529505 | -0.677832237 | 0.3326215 | -2.0378485 | 0.04156508 | 0.61348464 |
| APTR | 3.60320836 | -0.328269931 | 0.66501463 | -0.4936281 | 0.62156883 | 0.99665574 |
| RSBN1L | 17.4558178 | -0.126359674 | 0.39492755 | -0.3199566 | 0.74900123 | 0.99665574 |
| PHTF2 | 12.2396176 | -0.020436753 | 0.4730096 | -0.0432058 | 0.96553749 | 0.99777713 |
| MAGI2 | 203.929654 | -0.484628684 | 0.49129983 | -0.9864214 | 0.32392635 | 0.97700345 |
| GNAI1 | 11.2124725 | -1.18863212 | 0.63548778 | -1.8704248 | 0.06142485 | 0.6834742 |
| SEMA3C | 182.577094 | 0.679997975 | 0.51456024 | 1.32151285 | 0.18633042 | 0.89532394 |
| CACNA2D1 | 29.0626073 | -1.615141694 | 0.42663564 | -3.7857637 | 0.00015324 | 0.02931169 |
| PCLO | 55.4653029 | 0.657980231 | 0.32336792 | 2.03477277 | 0.04187374 | 0.61480373 |
| KIAA1324L | 7.31852984 | 0.007262575 | 0.79848052 | 0.00909549 | 0.99274295 | 0.99777713 |
| DMTF1 | 31.9841767 | 0.245310781 | 0.38126184 | 0.64341812 | 0.51995283 | 0.99665574 |
| TMEM243 | 12.1136772 | -0.154839532 | 0.499233 | -0.3101548 | 0.75644321 | 0.99665574 |
| CROT | 5.82475123 | -0.402579281 | 0.66372212 | -0.6065479 | 0.54415096 | 0.99665574 |
| ABCB1 | 6.8851692 | 0.687004529 | 0.87697584 | 0.78337908 | 0.43340454 | 0.99617422 |
| SLC25A40 | 12.299083 | 0.120799418 | 0.51873712 | 0.23287213 | 0.8158607 | 0.99665574 |
| DBF4 | 3.88925416 | -0.873365502 | 0.76661939 | -1.1392426 | 0.25460197 | 0.94099699 |
| ADAM22 | 28.8095902 | 0.202581444 | 0.47717248 | 0.42454553 | 0.67116801 | 0.99665574 |
| SRI | 20.3682795 | 0.133482586 | 0.43900696 | 0.30405574 | 0.76108542 | 0.99665574 |
| CFAP69 | 4.89164669 | 0.519375455 | 0.78270431 | 0.66356535 | 0.50696854 | 0.99665574 |
| GTPBP10 | 11.3874829 | -0.025935886 | 0.48631145 | -0.0533318 | 0.95746751 | 0.99777713 |
| CLDN12 | 11.0736526 | 0.075046295 | 0.50157024 | 0.14962271 | 0.88106229 | 0.99665574 |
| CDK14 | 53.4060999 | 0.504224402 | 0.34463429 | 1.46307091 | 0.14344798 | 0.84969914 |
| AC079760.1 | 8.92010278 | 0.190102596 | 0.7708163 | 0.24662503 | 0.80519843 | 0.99665574 |
| MTERF1 | 37.3647316 | -0.194237601 | 0.36764175 | -0.5283339 | 0.5972676 | 0.99665574 |
| AKAP9 | 117.397599 | -0.055038696 | 0.20237264 | -0.2719671 | 0.78564734 | 0.99665574 |
| KRIT1 | 35.162733 | 0.000678773 | 0.28729492 | 0.00236263 | 0.99811489 | 0.99919905 |
| ANKIB1 | 54.8111808 | -0.051206075 | 0.24797594 | -0.2064961 | 0.83640337 | 0.99665574 |
| GATAD1 | 10.6613257 | -0.669354798 | 0.48158448 | -1.3899011 | 0.16455891 | 0.87444728 |
| AC007566.1 | 5.37423244 | -0.141151713 | 0.63476194 | -0.2223695 | 0.82402623 | 0.99665574 |
| PEX1 | 11.6662793 | 0.334577509 | 0.46551809 | 0.71872075 | 0.472313 | 0.99617422 |
| RBM48 | 2.44791085 | -0.673814748 | 0.92252769 | -0.7304006 | 0.46514537 | 0.99617422 |
| FAM133B | 17.7847987 | 0.206509725 | 0.34997721 | 0.5900662 | 0.55514627 | 0.99665574 |
| CDK6 | 31.0155683 | 0.233192773 | 0.46032698 | 0.50658072 | 0.61244904 | 0.99665574 |
| SAMD9 | 4.27367674 | -0.702417724 | 1.13628427 | -0.6181708 | 0.53646277 | 0.99665574 |
| HEPACAM2 | 62.7371143 | 1.080614766 | 0.47809737 | 2.26023995 | 0.02380636 | 0.49867035 |
| VPS50 | 33.880983 | 0.322610666 | 0.28809578 | 1.11980351 | 0.2627975 | 0.94557817 |
| GNGT1 | 6.25092056 | -1.186295602 | 0.9701668 | -1.2227749 | 0.22141473 | 0.92071475 |
| BET1 | 6.98900009 | 0.400628437 | 0.63922879 | 0.62673716 | 0.53083154 | 0.99665574 |
| CASD1 | 23.1617969 | 0.139827767 | 0.39468151 | 0.35428 | 0.72312905 | 0.99665574 |
| SGCE | 7.64329771 | -1.291664137 | 0.73431206 | -1.7590126 | 0.07857538 | 0.74431421 |
| PPP1R9A | 107.730098 | -0.066680373 | 0.23958963 | -0.2783108 | 0.78077382 | 0.99665574 |
| AC002429.2 | 9.946002 | -1.295739264 | 0.72594257 | -1.784906 | 0.07427656 | 0.72810353 |
| PON2 | 15.317229 | -1.148331802 | 0.5133848 | -2.2367857 | 0.02530035 | 0.50908112 |
| PDK4 | 34.9243126 | 0.746249332 | 0.82418659 | 0.90543736 | 0.36523373 | 0.98365837 |
| SLC25A13 | 39.0815714 | -0.301075901 | 0.36146671 | -0.8329284 | 0.40488509 | 0.99194293 |
| SEM1 | 27.1332556 | -0.088377548 | 0.35200042 | -0.2510723 | 0.80175823 | 0.99665574 |
| SDHAF3 | 5.54325341 | -0.00616344 | 0.60786527 | -0.0101395 | 0.99191 | 0.99777713 |
| TAC1 | 20.8963187 | -7.201529253 | 2.97558765 | -2.420204 | 0.0155118 | 0.41713165 |
| ASNS | 5.51595917 | -2.644116642 | 0.94286998 | -2.8043279 | 0.00504216 | 0.22322013 |
| LMTK2 | 30.6768546 | 0.187446477 | 0.35160138 | 0.53312213 | 0.59394904 | 0.99665574 |
| TECPR1 | 3.82927815 | -0.760722623 | 0.75568001 | -1.0066729 | 0.31409197 | 0.9755712 |
| BRI3 | 38.8639117 | 0.108330873 | 0.39958871 | 0.27110594 | 0.78630955 | 0.99665574 |
| BAIAP2L1 | 103.005559 | 0.224462085 | 0.35266082 | 0.6364815 | 0.52446264 | 0.99665574 |
| TRRAP | 35.2420744 | 0.166151942 | 0.28760167 | 0.5777155 | 0.56345621 | 0.99665574 |
| SMURF1 | 85.153288 | 0.123149668 | 0.38852385 | 0.3169681 | 0.75126781 | 0.99665574 |
| ARPC1B | 7.25222191 | -0.805982536 | 0.5597695 | -1.4398472 | 0.14991064 | 0.85525834 |
| PDAP1 | 9.33025946 | -0.11733912 | 0.48290387 | -0.2429865 | 0.80801586 | 0.99665574 |
| BUD31 | 8.057839 | -0.135499366 | 0.50529122 | -0.2681609 | 0.78857544 | 0.99665574 |
| CPSF4 | 3.49880444 | 0.240129784 | 0.75021631 | 0.32008073 | 0.74890713 | 0.99665574 |
| ZNF789 | 7.40465696 | -0.218028355 | 0.60102295 | -0.3627621 | 0.7167826 | 0.99665574 |
| ZNF394 | 4.18680279 | 0.459505025 | 0.77965408 | 0.58937039 | 0.55561284 | 0.99665574 |
| ZKSCAN5 | 4.44786523 | -0.855902947 | 0.77587 | -1.1031525 | 0.26996093 | 0.94821243 |
| FAM200A | 5.20949179 | -0.05195199 | 0.67619366 | -0.07683 | 0.93875875 | 0.99777713 |
| ZSCAN25 | 4.12156329 | -0.189767591 | 0.78428878 | -0.2419614 | 0.80881009 | 0.99665574 |
| TRIM4 | 13.3738781 | -0.064748234 | 0.42459013 | -0.1524959 | 0.87879586 | 0.99665574 |
| AZGP1 | 12.5879076 | 1.254075675 | 0.65404143 | 1.91742543 | 0.05518391 | 0.67080867 |
| ZKSCAN1 | 35.5089738 | -0.12637878 | 0.312814 | -0.4040061 | 0.6862082 | 0.99665574 |
| ZNF3 | 7.52396372 | 0.229551698 | 0.65138576 | 0.35240515 | 0.72453444 | 0.99665574 |
| COPS6 | 7.56901865 | -0.177574018 | 0.5539048 | -0.3205858 | 0.74852429 | 0.99665574 |
| MCM7 | 2.90548058 | 0.156518234 | 0.9447216 | 0.16567657 | 0.86841149 | 0.99665574 |
| AP4M1 | 4.14756283 | -0.114255099 | 0.66148835 | -0.1727243 | 0.86286816 | 0.99665574 |
| TAF6 | 3.92898504 | 0.001742268 | 0.78939949 | 0.00220708 | 0.99823901 | 0.99922463 |
| LAMTOR4 | 26.5856004 | 0.296413641 | 0.4195802 | 0.70645289 | 0.47990654 | 0.99617422 |
| STAG3 | 2.74414849 | -1.541192465 | 0.84385664 | -1.8263676 | 0.0677949 | 0.70516795 |
| CASTOR3 | 18.879825 | -0.483896874 | 0.373149 | -1.2967926 | 0.19470255 | 0.90333151 |
| SPDYE3 | 5.37444715 | -1.467681625 | 0.85196472 | -1.7227023 | 0.08494237 | 0.75373899 |
| PILRB | 15.3534333 | -0.808837817 | 0.52500692 | -1.5406232 | 0.12340853 | 0.82915695 |
| MEPCE | 6.20301848 | -0.322200218 | 0.56510203 | -0.5701629 | 0.56856721 | 0.99665574 |
| PPP1R35 | 6.40659757 | -0.469184192 | 0.55340763 | -0.8478094 | 0.39654413 | 0.99179694 |
| TSC22D4 | 11.8151213 | 0.493191894 | 0.50281761 | 0.98085644 | 0.32666354 | 0.97788427 |
| AGFG2 | 5.16482526 | -0.547622307 | 0.63375567 | -0.8640906 | 0.38753813 | 0.99168679 |
| SAP25 | 2.22366189 | 0.470247362 | 0.91075146 | 0.51632897 | 0.60562467 | 0.99665574 |
| LRCH4 | 9.14694088 | 0.476718686 | 0.61022786 | 0.78121422 | 0.43467651 | 0.99617422 |
| PCOLCE | 3.64381667 | 0.543071712 | 0.78918922 | 0.68813879 | 0.49136539 | 0.99640678 |
| GNB2 | 22.9193244 | -0.18248499 | 0.47951651 | -0.3805604 | 0.70352948 | 0.99665574 |
| GIGYF1 | 14.3507401 | 0.087604614 | 0.43905642 | 0.19952929 | 0.84184874 | 0.99665574 |
| POP7 | 3.68060209 | 1.128105219 | 0.95755204 | 1.17811375 | 0.23875126 | 0.93157178 |
| SLC12A9 | 3.55729561 | -0.012382145 | 0.80697301 | -0.0153439 | 0.98775779 | 0.99777713 |
| TRIP6 | 5.00790527 | 0.060403279 | 0.65647959 | 0.0920109 | 0.92668938 | 0.99777713 |
| SRRT | 6.83214086 | 0.523837063 | 0.64271863 | 0.81503326 | 0.41505327 | 0.99279547 |
| TRIM56 | 21.8017653 | -0.609647881 | 0.41839791 | -1.4571007 | 0.14508859 | 0.85059132 |
| AP1S1 | 4.70599522 | 0.214955998 | 0.80313226 | 0.26764707 | 0.78897099 | 0.99665574 |
| PLOD3 | 3.97246637 | -0.07525689 | 0.70916614 | -0.1061203 | 0.91548694 | 0.99777713 |
| ZNHIT1 | 12.4889779 | -0.115391791 | 0.41952579 | -0.2750529 | 0.78327559 | 0.99665574 |
| FIS1 | 24.4828952 | 0.506510176 | 0.38725768 | 1.30794094 | 0.19089334 | 0.89964233 |
| CUX1 | 129.981378 | 0.008698607 | 0.22436767 | 0.03876943 | 0.96907422 | 0.99777713 |
| PRKRIP1 | 30.0390315 | -0.081429928 | 0.37575949 | -0.2167076 | 0.82843624 | 0.99665574 |
| ORAI2 | 3.28017386 | -1.556991471 | 1.22287972 | -1.2732172 | 0.20294099 | 0.9066717 |
| ALKBH4 | 2.75300581 | -0.609953959 | 0.8402375 | -0.7259304 | 0.46788143 | 0.99617422 |
| POLR2J | 8.26270164 | 0.344375517 | 0.68673884 | 0.50146504 | 0.61604388 | 0.99665574 |
| SPDYE2 | 3.52809254 | -0.068573087 | 0.83198124 | -0.0824214 | 0.93431159 | 0.99777713 |
| AC105052.3 | 8.84606616 | 0.206294149 | 0.63721752 | 0.32374212 | 0.74613328 | 0.99665574 |
| FAM185A | 18.0987721 | -0.018954812 | 0.42891772 | -0.0441922 | 0.96475121 | 0.99777713 |
| FBXL13 | 6.69105345 | -0.786236609 | 0.68038195 | -1.1555812 | 0.24785249 | 0.93441937 |

| | | | | | | |
|---|---|---|---|---|---|---|
| ARMC10 | 11.0896747 | -0.340582747 | 0.60627517 | -0.5617627 | 0.57427774 | 0.99665574 |
| NAPEPLD | 18.0978652 | -0.145060899 | 0.36767303 | -0.3945378 | 0.69318405 | 0.99665574 |
| PMPCB | 19.8621755 | 0.129731526 | 0.3688028 | 0.35176394 | 0.7250153 | 0.99665574 |
| DNAJC2 | 11.4349588 | -0.666401552 | 0.50347545 | -1.3236029 | 0.18563497 | 0.89532394 |
| PSMC2 | 8.96989632 | 0.15517361 | 0.50747255 | 0.30577735 | 0.75977418 | 0.99665574 |
| ORC5 | 11.0508802 | 0.256512762 | 0.46979079 | 0.54601488 | 0.58505571 | 0.99665574 |
| LHFPL3 | 14.577094 | -0.381238636 | 0.46676887 | -0.8167611 | 0.41406499 | 0.99279547 |
| LINC01004 | 29.6481508 | 0.032816458 | 0.39146173 | 0.08383056 | 0.93319115 | 0.99777713 |
| KMT2E-AS1 | 2.72644541 | -1.430399636 | 0.81489226 | -1.7553236 | 0.07920397 | 0.7446402 |
| KMT2E | 103.836912 | 0.002644529 | 0.2385916 | 0.01108391 | 0.9911565 | 0.99777713 |
| SRPK2 | 123.509001 | -0.088163259 | 0.26251721 | -0.335838 | 0.73699302 | 0.99665574 |
| PUS7 | 6.85776538 | -0.026426632 | 0.5898136 | -0.0448051 | 0.96426269 | 0.99777713 |
| RINT1 | 7.71516021 | -0.057030044 | 0.5223614 | -0.1091774 | 0.91306181 | 0.99777713 |
| EFCAB10 | 3.71563324 | -1.422320587 | 0.83440594 | -1.7045907 | 0.08827079 | 0.75650018 |
| ATXN7L1 | 43.6809026 | -0.108480862 | 0.39452843 | -0.2749634 | 0.78334439 | 0.99665574 |
| CDHR3 | 5.80792573 | -0.852083731 | 0.84289936 | -1.0108962 | 0.31206613 | 0.97442101 |
| SYPL1 | 21.871227 | 0.203623103 | 0.45230884 | 0.450186 | 0.65257633 | 0.99665574 |
| NAMPT | 43.313012 | -1.63361774 | 0.46407731 | -3.5201414 | 0.00043132 | 0.05678819 |
| HBP1 | 21.3738304 | -0.158234017 | 0.40481887 | -0.3908761 | 0.69588883 | 0.99665574 |
| COG5 | 123.469227 | 0.396302247 | 0.27177279 | 1.45821163 | 0.14478222 | 0.85018987 |
| DUS4L | 3.3399702 | 0.065935056 | 0.85392475 | 0.07721413 | 0.9384532 | 0.99777713 |
| BCAP29 | 68.0441403 | 1.371095381 | 0.51752336 | 2.64934007 | 0.00806491 | 0.30282256 |
| SLC26A4-AS1 | 2.65329429 | 2.322038241 | 1.08231632 | 2.145434 | 0.03191816 | 0.56869303 |
| SLC26A4 | 467.772246 | 1.621108272 | 0.65561611 | 2.47264863 | 0.01341159 | 0.39134822 |
| CBLL1 | 5.2074889 | 0.656691104 | 0.79586124 | 0.82513266 | 0.40929628 | 0.99279547 |
| SLC26A3 | 89.6380812 | 1.60293506 | 0.56636772 | 2.83020203 | 0.00465186 | 0.21732984 |
| DLD | 24.9656221 | 0.227161677 | 0.37411198 | 0.60720235 | 0.54371663 | 0.99665574 |
| LAMB1 | 65.5556871 | -0.326857062 | 0.45366639 | -0.7204789 | 0.47123019 | 0.99617422 |
| NRCAM | 12.9964402 | -0.673888648 | 0.81000613 | -0.831955 | 0.40543435 | 0.99194293 |
| PNPLA8 | 63.9190496 | -0.384669501 | 0.52761211 | -0.7290763 | 0.46595497 | 0.99617422 |
| DNAJB9 | 4.91276876 | -0.729352498 | 0.7012541 | -1.0400688 | 0.29830794 | 0.97358745 |
| IMMP2L | 305.963235 | 0.255683705 | 0.39514962 | 0.64705542 | 0.51759608 | 0.99665574 |
| DOCK4 | 42.1420899 | -0.03402268 | 0.43608603 | -0.0780183 | 0.93781351 | 0.99777713 |
| ZNF277 | 28.1987372 | -0.076940727 | 0.41998754 | -0.1831976 | 0.85464294 | 0.99665574 |
| IFRD1 | 24.9242405 | -0.114489305 | 0.45828344 | -0.249822 | 0.80272497 | 0.99665574 |
| LSMEM1 | 3.03690664 | 0.937743919 | 1.05187732 | 0.89149552 | 0.37266339 | 0.98532838 |
| TMEM168 | 19.0367599 | -0.047533186 | 0.43550825 | -0.1091442 | 0.91308814 | 0.99777713 |
| BMT2 | 22.955786 | 0.278447005 | 0.41302495 | 0.67416509 | 0.50020636 | 0.99640678 |
| SMIM30 | 4.28885714 | -0.078300115 | 0.82005391 | -0.0954817 | 0.92393225 | 0.99777713 |
| FOXP2 | 18.9475501 | -0.311523161 | 1.31537299 | -0.2368326 | 0.81278669 | 0.99665574 |
| MDFIC | 49.1783424 | 0.358406722 | 0.40541162 | 0.88405637 | 0.37666578 | 0.98679664 |
| TES | 21.6785047 | -1.251157849 | 0.51550258 | -2.4270642 | 0.01522156 | 0.41291622 |
| CAV2 | 5.03548589 | 0.354389084 | 0.79023917 | 0.44845801 | 0.65382269 | 0.99665574 |
| MET | 21.8253179 | -0.773368003 | 0.41221827 | -1.8761129 | 0.06063978 | 0.6834742 |
| CAPZA2 | 43.0957251 | -0.12801593 | 0.28454357 | -0.4498992 | 0.65278311 | 0.99665574 |
| ST7 | 85.7064955 | 0.152331622 | 0.34392034 | 0.44292705 | 0.6578185 | 0.99665574 |
| ST7-AS2 | 4.35359074 | -0.600007508 | 0.77672888 | -0.77248 | 0.4398302 | 0.99617422 |
| CFTR | 103.731803 | -1.297296648 | 0.54612604 | -2.3754528 | 0.01752743 | 0.43770425 |
| AC000061.1 | 5.09002193 | -2.247427589 | 1.07119403 | -2.0980584 | 0.03589999 | 0.58508511 |
| CTTNBP2 | 31.0331942 | -0.508438298 | 0.44199557 | -1.1503244 | 0.25001027 | 0.93755294 |
| LSM8 | 12.9756308 | -0.158233795 | 0.42001894 | -0.3767301 | 0.70637416 | 0.99665574 |
| TSPAN12 | 7.81764602 | 0.265614448 | 0.68164323 | 0.38966785 | 0.69678218 | 0.99665574 |
| ING3 | 7.62158783 | 0.059559692 | 0.5781447 | 0.10301866 | 0.91794816 | 0.99777713 |
| FAM3C | 28.9793707 | -0.71330596 | 0.39233275 | -1.8181148 | 0.0690466 | 0.7094165 |
| AASS | 14.5305359 | 0.384442759 | 0.42003096 | 0.91527243 | 0.36004863 | 0.9819332 |
| CADPS2 | 80.6867672 | -0.087935327 | 0.46010803 | -0.1911189 | 0.84843246 | 0.99665574 |
| AC004594.1 | 3.62894826 | -1.751080433 | 2.30958735 | -0.7581789 | 0.44834388 | 0.99617422 |
| SLC13A1 | 5.65393038 | 2.632194382 | 1.08454168 | 2.42701081 | 0.0152238 | 0.41291622 |
| IQUB | 4.78019645 | 0.864492107 | 1.14303006 | 0.75631616 | 0.44945966 | 0.99617422 |
| NDUFA5 | 31.5188245 | 0.385954238 | 0.39332277 | 0.98126595 | 0.32646161 | 0.97788427 |
| WASL | 39.8515117 | 0.487923044 | 0.26397202 | 1.84838922 | 0.06454606 | 0.68846743 |
| POT1 | 9.86160769 | -0.320604385 | 0.48544434 | -0.6604349 | 0.50897478 | 0.99665574 |
| POT1-AS1 | 6.53087178 | 0.318068373 | 0.72949587 | 0.43601121 | 0.6628286 | 0.99665574 |
| GRM8 | 97.4913287 | -2.345750861 | 0.81503497 | -2.8780984 | 0.0040008 | 0.20588904 |
| ZNF800 | 21.3899559 | 0.247580255 | 0.34920812 | 0.70897622 | 0.47833924 | 0.99617422 |
| ARF5 | 6.53601983 | -0.042356228 | 0.56973049 | -0.0743443 | 0.94073641 | 0.99777713 |
| SND1 | 78.9381462 | -0.672939893 | 0.46724159 | -1.4402397 | 0.1497996 | 0.85525834 |
| RBM28 | 11.2026964 | 0.156959635 | 0.45942004 | 0.34164734 | 0.73261631 | 0.99665574 |
| METTL2B | 9.96746091 | -0.286981769 | 0.51489032 | -0.5573649 | 0.57727817 | 0.99665574 |
| AC090114.2 | 2.80402362 | -1.303109699 | 0.82696371 | -1.5757762 | 0.1150774 | 0.81721642 |
| FAM71F2 | 5.0020709 | -0.382313467 | 0.77580685 | -0.4927947 | 0.62215769 | 0.99665574 |
| CALU | 20.2107 | -1.324597058 | 0.72668235 | -1.8228006 | 0.06833359 | 0.70708655 |
| ATP6V1F | 40.3894366 | 0.018227517 | 0.55259265 | 0.03298545 | 0.97368619 | 0.99777713 |
| TNPO3 | 38.2757687 | -0.140176931 | 0.29508582 | -0.4750378 | 0.63476001 | 0.99665574 |
| TSPAN33 | 5.36011364 | 2.053631317 | 0.69437972 | 2.95750474 | 0.0031014 | 0.17565358 |
| AHCYL2 | 75.5618817 | 0.221292943 | 0.32867986 | 0.67327807 | 0.5007704 | 0.99640678 |
| NRF1 | 37.0079364 | 0.200478776 | 0.28067335 | 0.71427791 | 0.47505535 | 0.99617422 |
| UBE2H | 182.206297 | -0.125736181 | 0.27648268 | -0.4547705 | 0.64927432 | 0.99665574 |
| ZC3HC1 | 6.78996351 | 0.740128795 | 0.61183652 | 1.2096839 | 0.22640021 | 0.92318867 |
| KLHDC10 | 29.8334516 | 0.421545166 | 0.33357486 | 1.26371982 | 0.20633061 | 0.90890506 |
| TMEM209 | 7.2738442 | 0.360678747 | 0.5417533 | 0.66576197 | 0.50556325 | 0.99665574 |
| CEP41 | 4.50036835 | -0.404751847 | 0.80065638 | -0.505525 | 0.61319012 | 0.99665574 |
| MEST | 4.03185524 | 0.026886552 | 1.01041333 | 0.02660946 | 0.97877123 | 0.99777713 |
| COPG2 | 10.8532734 | -0.052181374 | 0.44659163 | -0.1168436 | 0.90698399 | 0.99777713 |
| AC016831.7 | 362.59672 | -0.710558142 | 0.3282818 | -2.1644762 | 0.03042782 | 0.56271575 |
| LINC00513 | 30.642022 | -1.206669543 | 0.60997075 | -1.9782417 | 0.04790145 | 0.6410909 |
| AC058791.1 | 7.81953539 | -1.129896091 | 0.75492945 | -1.496691 | 0.13447369 | 0.84373478 |
| MKLN1 | 127.435021 | 0.076319901 | 0.30142023 | 0.25320099 | 0.8001129 | 0.99665574 |
| PODXL | 3.49257549 | -1.648451473 | 0.84693983 | -1.946362 | 0.05161128 | 0.65584036 |
| CHCHD3 | 149.033938 | 0.348185209 | 0.19178364 | 1.81551044 | 0.0694455 | 0.7094165 |
| EXOC4 | 169.683379 | -0.18045381 | 0.34707981 | -0.5199202 | 0.60311919 | 0.99665574 |
| LRGUK | 4.16073375 | 0.276235702 | 0.85046075 | 0.32480712 | 0.74532706 | 0.99665574 |
| SLC35B4 | 11.441287 | -0.104395008 | 0.44488723 | -0.234655 | 0.81447653 | 0.99665574 |
| AKR1B1 | 16.7110524 | -0.187089724 | 0.66024948 | -0.2833622 | 0.77689923 | 0.99665574 |
| BPGM | 22.2940931 | -0.148347378 | 0.59356353 | -0.2499267 | 0.80264403 | 0.99665574 |
| CALD1 | 9.74370672 | -1.042112863 | 0.70038978 | -1.4879042 | 0.13677617 | 0.84475183 |
| AGBL3 | 3.55638396 | -0.582670598 | 0.88357469 | -0.6594469 | 0.50960883 | 0.99665574 |
| CYREN | 7.79053455 | -0.413904708 | 0.58896086 | -0.7027712 | 0.48219837 | 0.99640678 |
| TMEM140 | 10.7132871 | -1.423024421 | 0.68241285 | -2.0852837 | 0.03704355 | 0.58591936 |
| WDR91 | 7.94423678 | -1.07557209 | 0.6165564 | -1.7444829 | 0.08107492 | 0.74789587 |
| CNOT4 | 59.1126395 | -0.141234395 | 0.31938614 | -0.4422058 | 0.65834031 | 0.99665574 |
| NUP205 | 14.3566299 | 0.479373582 | 0.41305987 | 1.16054262 | 0.24582795 | 0.93426912 |
| STMP1 | 16.8652375 | 0.181683617 | 0.50406369 | 0.36043782 | 0.71851975 | 0.99665574 |
| MTPN | 27.5472161 | 0.466839735 | 0.38685988 | 1.20674115 | 0.22753185 | 0.92389989 |
| PTN | 18.3396737 | 0.062643472 | 0.80787058 | 0.07754147 | 0.9381928 | 0.99777713 |
| DGKI | 346.143669 | 1.325527185 | 0.48916005 | 2.7098026 | 0.00673233 | 0.26557324 |
| CREB3L2 | 26.1845678 | 0.59631869 | 0.39400266 | 1.51348898 | 0.1301555 | 0.83778831 |
| TRIM24 | 28.2319071 | -0.28835882 | 0.33049628 | -0.8725025 | 0.38293432 | 0.98934457 |
| SVOPL | 56.5317697 | 0.183740755 | 0.53876872 | 0.34103827 | 0.73307477 | 0.99665574 |
| ATP6V0A4 | 540.773191 | 0.132308755 | 0.29188511 | 0.45329052 | 0.65033956 | 0.99665574 |
| TMEM213 | 388.89685 | 0.1182814 | 0.37872652 | 0.31231349 | 0.75480229 | 0.99665574 |
| KIAA1549 | 12.0601471 | 0.522401781 | 0.56045467 | 0.93210354 | 0.35128302 | 0.97838306 |
| ZC3HAV1 | 19.0186863 | -0.407202779 | 0.50311549 | -0.8093624 | 0.4183067 | 0.99432434 |

| | | | | | | |
|---|---|---|---|---|---|---|
| TTC26 | 2.76566751 | -1.017426231 | 0.88853915 | -1.145055 | 0.25218636 | 0.94076968 |
| UBN2 | 25.2014435 | 0.379232185 | 0.41356565 | 0.91698183 | 0.35915216 | 0.9819332 |
| LUC7L2 | 9.79428011 | 0.489344684 | 0.47845893 | 1.0227517 | 0.30642526 | 0.97376371 |
| HIPK2 | 289.608825 | -0.902538682 | 0.2833883 | -3.1848128 | 0.00144848 | 0.11041095 |
| TBXAS1 | 41.5522576 | -2.644997024 | 0.51435135 | -5.1423935 | 2.71E-07 | 0.00025 |
| PARP12 | 13.1404283 | -0.629848849 | 0.71771204 | -0.8775788 | 0.38017236 | 0.98679664 |
| KDM7A | 46.7716052 | -0.008041389 | 0.52356509 | -0.0153589 | 0.98774584 | 0.99777713 |
| SLC37A3 | 13.8031886 | -0.317430138 | 0.51742129 | -0.6134849 | 0.53955578 | 0.99665574 |
| MKRN1 | 15.9721165 | -0.269468889 | 0.43131078 | -0.6247673 | 0.53212376 | 0.99665574 |
| DENND2A | 6.81109603 | -0.038773522 | 1.08559441 | -0.0357164 | 0.97150849 | 0.99777713 |
| NDUFB2 | 35.5773002 | 0.260138809 | 0.44510985 | 0.58443733 | 0.55892612 | 0.99665574 |
| BRAF | 107.218573 | 0.219777205 | 0.22851711 | 0.96175385 | 0.33617327 | 0.97788427 |
| MRPS33 | 7.41418672 | -0.049813864 | 0.660421 | -0.0754274 | 0.93987463 | 0.99777713 |
| TMEM178B | 6.65043646 | -1.239985473 | 0.8370859 | -1.4813121 | 0.13852342 | 0.84656101 |
| AGK | 11.2569243 | 0.390504285 | 0.50344506 | 0.77566415 | 0.43794731 | 0.99617422 |
| AC004918.1 | 3.39776236 | 1.869016293 | 1.03337673 | 1.80864949 | 0.07050547 | 0.71264656 |
| KIAA1147 | 8.9937742 | 0.068762119 | 0.50923901 | 0.13502917 | 0.89258881 | 0.99773598 |
| SSBP1 | 13.9424985 | -0.294848426 | 0.41635964 | -0.708158 | 0.47884712 | 0.99617422 |
| GSTK1 | 19.4842181 | 0.093039581 | 0.56647573 | 0.16424284 | 0.86953998 | 0.99665574 |
| TMEM139 | 2.43502501 | 0.221477338 | 0.90811358 | 0.24388726 | 0.80731813 | 0.99665574 |
| CASP2 | 2.76959946 | -0.56100891 | 0.77886109 | -0.7202939 | 0.47134404 | 0.99617422 |
| TCAF2 | 6.70923884 | -1.32169079 | 1.18168304 | -1.1184816 | 0.26336134 | 0.94557817 |
| TCAF1 | 12.4012411 | -0.264583988 | 0.56588183 | -0.4675605 | 0.64009892 | 0.99665574 |
| AC004889.1 | 5.34091999 | 0.446244955 | 0.75688783 | 0.58957872 | 0.55547312 | 0.99665574 |
| AC074386.1 | 13.3111383 | -0.199058746 | 0.3862404 | -0.5153753 | 0.60629082 | 0.99665574 |
| ARHGEF5 | 4.00650739 | -0.26954577 | 0.65986063 | -0.4084889 | 0.68291475 | 0.99665574 |
| TPK1 | 9.4960449 | -0.754853495 | 0.48305679 | -1.5626599 | 0.1181326 | 0.82599629 |
| CNTNAP2 | 3.02189548 | 0.31838139 | 0.7977678 | 0.3990903 | 0.68982667 | 0.99665574 |
| CUL1 | 38.7060205 | 0.322862439 | 0.30807646 | 1.04799451 | 0.29464113 | 0.97161335 |
| EZH2 | 2.98155314 | 0.083054943 | 0.77927426 | 0.10657986 | 0.9151223 | 0.99777713 |
| PDIA4 | 5.65500371 | 0.385358021 | 0.71453651 | 0.53931187 | 0.58967168 | 0.99665574 |
| ZNF786 | 4.0150839 | -0.858015769 | 0.76971573 | -1.1147177 | 0.2649714 | 0.94557817 |
| ZNF425 | 2.53048181 | -0.27420278 | 1.09246608 | -0.2509943 | 0.8018185 | 0.99665574 |
| ZNF398 | 9.71535989 | -0.285261768 | 0.46978355 | -0.6072196 | 0.5437052 | 0.99665574 |
| ZNF282 | 6.66112361 | 0.016719311 | 0.60291099 | 0.02773098 | 0.97787672 | 0.99777713 |
| ZNF212 | 3.39260049 | -0.966785982 | 0.77296214 | -1.2507546 | 0.21102401 | 0.91043197 |
| ZNF783 | 4.26865001 | 0.144549658 | 0.81783824 | 0.17674602 | 0.85970789 | 0.99665574 |
| ZNF777 | 4.75254513 | -0.470634085 | 0.64019061 | -0.7351468 | 0.46225009 | 0.99617422 |
| ZNF746 | 6.45410634 | 0.624675493 | 0.65012257 | 0.96085803 | 0.33662356 | 0.97788427 |
| KRBA1 | 2.82836234 | -0.071195779 | 0.81524452 | -0.0873306 | 0.93040875 | 0.99777713 |
| ZNF467 | 2.50664285 | 1.778730629 | 1.04186182 | 1.70726156 | 0.08777345 | 0.75650018 |
| ZNF862 | 9.02774658 | -0.055525398 | 0.64625989 | -0.0859181 | 0.93153155 | 0.99777713 |
| ATP6V0E2-A | 2.14742242 | 0.563806548 | 1.01373486 | 0.55616766 | 0.57809624 | 0.99665574 |
| ATP6V0E2 | 6.71339469 | 0.389047397 | 0.58923561 | 0.66025778 | 0.50908842 | 0.99665574 |
| ACTR3C | 67.2155843 | -1.055714183 | 0.44872764 | -2.3526837 | 0.01863848 | 0.45192386 |
| LRRC61 | 9.72827446 | -1.111263993 | 0.52412456 | -2.1202288 | 0.03398676 | 0.58178209 |
| RARRES2 | 36.2103822 | 0.048369197 | 0.53779854 | 0.08993925 | 0.92833549 | 0.99777713 |
| TMEM176B | 8.0125566 | -0.840260505 | 0.69271411 | -1.2129975 | 0.22513076 | 0.92318867 |
| TMEM176A | 5.61704119 | -0.532330499 | 0.96214759 | -0.5532732 | 0.58007634 | 0.99665574 |
| AOC1 | 2.81190174 | 1.660673526 | 1.07959096 | 1.53824327 | 0.12398914 | 0.82915695 |
| ABCB8 | 4.31740279 | -0.513729745 | 0.6569821 | -0.7819539 | 0.43424164 | 0.99617422 |
| SLC4A2 | 8.15924364 | 0.365543221 | 0.79891797 | 0.45754788 | 0.6472773 | 0.99665574 |
| FASTK | 6.20423006 | 0.286741055 | 0.61069097 | 0.46953544 | 0.63868696 | 0.99665574 |
| TMUB1 | 9.29939908 | 0.17571814 | 0.55583767 | 0.31613212 | 0.75190223 | 0.99665574 |
| AGAP3 | 8.17989735 | 0.019881558 | 0.52760235 | 0.03768285 | 0.96994055 | 0.99777713 |
| CHPF2 | 4.30740479 | -0.120151635 | 0.76047889 | -0.1579947 | 0.87446097 | 0.99665574 |
| SMARCD3 | 7.21926081 | -1.835907563 | 0.72714437 | -2.5248185 | 0.01157581 | 0.3725572 |
| NUB1 | 13.5711159 | -0.07332864 | 0.50927699 | -0.1439858 | 0.8855117 | 0.99665574 |
| RHEB | 47.157584 | -0.443925161 | 0.31375782 | -1.4148656 | 0.15710789 | 0.86532131 |
| PRKAG2 | 93.636748 | 0.006609844 | 0.55824472 | 0.01184041 | 0.99055294 | 0.99777713 |
| GALNT11 | 35.9730697 | -0.179925888 | 0.34854103 | -0.5162258 | 0.60569668 | 0.99665574 |
| KMT2C | 156.446738 | 0.017080439 | 0.28242398 | 0.060478 | 0.95177493 | 0.99777713 |
| ACTR3B | 12.3240096 | 0.534962185 | 0.46868819 | 1.14140316 | 0.25370219 | 0.94099699 |
| DPP6 | 34.0453868 | -0.032448217 | 0.46119021 | -0.0703576 | 0.94390907 | 0.99777713 |
| PAXIP1-AS2 | 4.66151855 | 0.929762232 | 0.75742368 | 1.22753256 | 0.21962249 | 0.91740124 |
| PAXIP1 | 9.44097353 | 0.157373694 | 0.48036754 | 0.32761101 | 0.7432058 | 0.99665574 |
| PAXIP1-AS1 | 3.49165536 | -0.259050719 | 0.81026994 | -0.3197092 | 0.74918882 | 0.99665574 |
| INSIG1 | 10.9349148 | -2.329915896 | 1.08378134 | -2.1498026 | 0.03157084 | 0.56814511 |
| AC009403.1 | 2.19259584 | -0.746770291 | 0.95945979 | -0.7783237 | 0.43637821 | 0.99617422 |
| RBM33 | 49.6912979 | -0.011714579 | 0.2798322 | -0.0418629 | 0.96660801 | 0.99777713 |
| LINC01006 | 12.2244708 | 0.409712549 | 0.54873939 | 0.74664323 | 0.45527895 | 0.99617422 |
| LMBR1 | 48.2894846 | 0.306496621 | 0.29271508 | 1.04708176 | 0.29506187 | 0.97184446 |
| NOM1 | 5.1187153 | -0.659684365 | 0.65032334 | -1.0143944 | 0.31039459 | 0.97376371 |
| UBE3C | 67.871117 | -0.289499734 | 0.32362084 | -0.8945646 | 0.37101991 | 0.98532838 |
| DNAJB6 | 44.7038185 | -0.006784914 | 0.29670477 | -0.0228676 | 0.98175592 | 0.99777713 |
| ESYT2 | 51.3612639 | -0.095483978 | 0.29679996 | -0.3217116 | 0.74767122 | 0.99665574 |
| WDR60 | 22.07636 | -0.93563366 | 0.34396343 | -2.7201545 | 0.00652514 | 0.26146993 |
| GTPBP6 | 13.2862409 | -0.17192788 | 0.57625533 | -0.2983536 | 0.76543326 | 0.99665574 |
| LINC00685 | 3.38995491 | -0.990585746 | 0.9459953 | -1.047136 | 0.29503685 | 0.97184446 |
| PPP2R3B | 3.42061711 | -0.263079422 | 0.71687799 | -0.3669794 | 0.71363442 | 0.99665574 |
| IL3RA | 5.07211493 | -1.825281268 | 1.00747432 | -1.8117397 | 0.07002642 | 0.71124502 |
| SLC25A6 | 112.566787 | 0.107243984 | 0.36266144 | 0.29571378 | 0.76744867 | 0.99665574 |
| LINC00106 | 17.6716163 | -0.479942682 | 0.63431921 | -0.7566264 | 0.4492737 | 0.99617422 |
| ASMTL-AS1 | 8.82073203 | 0.087228158 | 0.64312274 | 0.13563221 | 0.89211204 | 0.99773598 |
| ASMTL | 8.08355684 | -0.874159204 | 0.49529847 | -1.764914 | 0.07757822 | 0.74141484 |
| AKAP17A | 7.54615934 | -0.75721952 | 0.50515121 | -1.4989958 | 0.13387473 | 0.84351899 |
| DHRSX | 50.3862273 | -0.194564259 | 0.29601301 | -0.6572828 | 0.5109991 | 0.99665574 |
| CD99 | 6.12449136 | -0.782830076 | 0.65294116 | -1.198929 | 0.23055556 | 0.92774608 |
| ARSD | 15.2450857 | 0.078926116 | 0.52825129 | 0.14941017 | 0.88122999 | 0.99665574 |
| ARSE | 9.01340553 | 0.327327172 | 0.54867529 | 0.59657721 | 0.55078969 | 0.99665574 |
| PRKX | 35.7499545 | -0.952380843 | 0.38200222 | -2.4931291 | 0.01266228 | 0.38590337 |
| BX890604.1 | 2.67984718 | 1.251066854 | 0.91510183 | 1.36713403 | 0.1715833 | 0.8807653 |
| NLGN4X | 18.3942632 | 0.748250007 | 0.84994929 | 0.88034664 | 0.37867155 | 0.98679664 |
| PUDP | 6.84887774 | -0.04166657 | 0.74189021 | -0.0561627 | 0.95521218 | 0.99777713 |
| STS | 2.92334152 | 0.214946481 | 1.04081473 | 0.20651752 | 0.83638668 | 0.99665574 |
| PNPLA4 | 3.58677891 | 0.131483002 | 0.8019373 | 0.16395671 | 0.86976522 | 0.99665574 |
| ANOS1 | 4.77014746 | 0.481767038 | 1.31169401 | 0.36728615 | 0.71340559 | 0.99665574 |
| TBL1X | 212.35792 | -0.074992609 | 0.43173596 | -0.1737002 | 0.86210111 | 0.99665574 |
| SHROOM2 | 20.436383 | -0.374303314 | 0.47429939 | -0.789171 | 0.43001208 | 0.99617422 |
| WWC3 | 8.06306504 | -0.445248298 | 0.62181107 | -0.7160508 | 0.47396 | 0.99617422 |
| MID1 | 15.3013586 | -0.765180103 | 0.57635512 | -1.3276192 | 0.18430395 | 0.89532394 |
| AC073529.1 | 11.0675595 | 0.331021904 | 0.57925634 | 0.57146013 | 0.56768778 | 0.99665574 |
| HCCS | 3.95047748 | -0.336025566 | 0.83084478 | -0.4044384 | 0.68589034 | 0.99665574 |
| ARHGAP6 | 126.991461 | 0.565575787 | 0.3675554 | 1.53874978 | 0.12386539 | 0.82915695 |
| MSL3 | 3.35208443 | -0.309488372 | 0.74833641 | -0.4135685 | 0.67919014 | 0.99665574 |
| FRMPD4 | 21.7373837 | 1.634051656 | 0.58123009 | 2.81136797 | 0.00493313 | 0.2212925 |
| PRPS2 | 4.6144599 | 0.406255499 | 0.73583985 | 0.55209772 | 0.58088141 | 0.99665574 |
| TMSB4X | 147.264306 | -0.472860537 | 0.46456535 | -1.0178558 | 0.30874649 | 0.97376371 |
| RAB9A | 15.2670815 | -0.431225805 | 0.47604131 | -0.905858 | 0.36501104 | 0.98364751 |
| TRAPPC2 | 5.78453862 | -0.596131696 | 0.65734869 | -0.9068729 | 0.36447403 | 0.98362663 |
| OFD1 | 30.7926597 | -0.386248186 | 0.33537474 | -1.1516914 | 0.24944793 | 0.9369778 |
| GEMIN8 | 6.0430808 | -0.20930051 | 0.62352796 | -0.3356714 | 0.73711867 | 0.99665574 |
| MOSPD2 | 10.1443183 | -0.267424547 | 0.49853217 | -0.5364239 | 0.59166565 | 0.99665574 |

| | | | | | | |
|---|---|---|---|---|---|---|
| PIGA | 2.35569479 | -0.098970628 | 0.82659061 | -0.1197336 | 0.90469422 | 0.99777713 |
| PIR | 3.82782713 | -0.759080494 | 0.90200832 | -0.8415449 | 0.40004275 | 0.99194293 |
| BMX | 2.69470343 | -0.567478629 | 0.81814091 | -0.6936197 | 0.48792075 | 0.99640678 |
| CLTRN | 9.68135465 | -1.573637468 | 0.55744915 | -2.8229256 | 0.00475876 | 0.21929238 |
| CA5B | 16.0162349 | -0.204352276 | 0.38413226 | -0.5319842 | 0.59473697 | 0.99665574 |
| ZRSR2 | 8.81626739 | 0.172798436 | 0.478649 | 0.36101284 | 0.71808985 | 0.99665574 |
| AP1S2 | 13.5656722 | -0.942481496 | 0.53782187 | -1.7524046 | 0.07970427 | 0.7446402 |
| CTPS2 | 13.2671672 | -0.233152834 | 0.48066679 | -0.4850612 | 0.6276329 | 0.99665574 |
| SYAP1 | 17.3425095 | -0.121348883 | 0.49959365 | -0.2428952 | 0.80808661 | 0.99665574 |
| TXLNG | 8.74241841 | 0.23810433 | 0.60109346 | 0.39611865 | 0.6920175 | 0.99665574 |
| RBBP7 | 15.1422134 | 0.329385317 | 0.38111518 | 0.8642671 | 0.38744118 | 0.99168679 |
| REPS2 | 6.50228544 | -1.345275491 | 0.76788325 | -1.7519271 | 0.07978634 | 0.7446402 |
| NHS | 143.721872 | -0.67023969 | 0.44189509 | -1.5167394 | 0.12933249 | 0.83778831 |
| SCML1 | 52.8558806 | -0.123220499 | 0.47756209 | -0.2580199 | 0.79639159 | 0.99665574 |
| RAI2 | 8.45885929 | -0.518459439 | 0.59917121 | -0.8652943 | 0.38687727 | 0.99144635 |
| SCML2 | 3.77606735 | 0.451647448 | 0.74456853 | 0.60658949 | 0.54412338 | 0.99665574 |
| CDKL5 | 14.1087641 | 0.281706898 | 0.43229834 | 0.65164927 | 0.51462745 | 0.99665574 |
| PHKA2 | 10.4745948 | -0.239809101 | 0.5030702 | -0.4766911 | 0.63358208 | 0.99665574 |
| ADGRG2 | 2.82517732 | -1.497762438 | 1.00066499 | -1.4967671 | 0.13445387 | 0.84373478 |
| PDHA1 | 24.6827637 | 0.116255895 | 0.38714862 | 0.30028751 | 0.76395786 | 0.99665574 |
| SH3KBP1 | 2.8768945 | -0.855490787 | 0.85890538 | -0.9960245 | 0.31923825 | 0.97700345 |
| BCLAF3 | 20.5991859 | -0.007269515 | 0.42017995 | -0.017301 | 0.98619652 | 0.99777713 |
| EIF1AX | 34.8632528 | 0.580051276 | 0.48960771 | 1.18472659 | 0.23612557 | 0.9305595 |
| RPS6KA3 | 83.0895407 | -0.389906917 | 0.34667201 | -1.1247142 | 0.26071018 | 0.94557817 |
| MBTPS2 | 7.79554973 | 0.612728762 | 0.63400184 | 0.96644635 | 0.33382088 | 0.97788427 |
| SMS | 21.2117819 | -0.263812319 | 0.45521872 | -0.5795287 | 0.56223247 | 0.99665574 |
| PRDX4 | 3.88728006 | -1.274738548 | 0.87794166 | -1.4519627 | 0.14651198 | 0.85169374 |
| ACOT9 | 9.77349477 | -0.817628873 | 0.55577662 | -1.4711466 | 0.14125148 | 0.84969914 |
| SAT1 | 134.492299 | -0.258174821 | 0.33646939 | -0.7673055 | 0.44289988 | 0.99617422 |
| APOO | 130.380266 | 0.470777917 | 0.42042103 | 1.11977728 | 0.26280868 | 0.94557817 |
| KLHL15 | 4.0357457 | 1.055548023 | 0.73564198 | 1.43486649 | 0.15132515 | 0.85657975 |
| EIF2S3 | 17.5283079 | -0.259314755 | 0.3998794 | -0.6484824 | 0.51667299 | 0.99665574 |
| ZFX | 27.4057445 | 0.059793056 | 0.42766708 | 0.13981215 | 0.88880841 | 0.99665574 |
| PDK3 | 6.0649096 | 1.19252259 | 0.8799954 | 1.35514639 | 0.17537092 | 0.88428483 |
| POLA1 | 17.7838654 | 0.513334122 | 0.43477597 | 1.1806865 | 0.23772728 | 0.9305595 |
| GK | 28.8318018 | 1.991086014 | 0.40848846 | 4.87427723 | 1.09E-06 | 0.00079082 |
| TAB3 | 15.7856534 | 0.134705144 | 0.43486406 | 0.3097638 | 0.75674058 | 0.99665574 |
| DMD | 8.51896119 | 0.202620057 | 0.66516192 | 0.30461765 | 0.76065738 | 0.99665574 |
| AF241726.2 | 11.8454733 | 0.444715376 | 0.53388944 | 0.83297279 | 0.40486008 | 0.99194293 |
| DYNLT3 | 16.4349186 | -0.825109544 | 0.50292346 | -1.6406265 | 0.10087498 | 0.78386461 |
| RPGR | 7.21482972 | -0.199409657 | 0.60600542 | -0.3290559 | 0.74211344 | 0.99665574 |
| TSPAN7 | 8.68927003 | 0.90821183 | 0.69438819 | 1.30793098 | 0.19089672 | 0.89964233 |
| MID1IP1 | 4.17619309 | -1.096099206 | 0.76767734 | -1.4278124 | 0.15334587 | 0.8618391 |
| BCOR | 23.154072 | -0.276494018 | 0.40455815 | -0.6834469 | 0.49432449 | 0.99640678 |
| ATP6AP2 | 141.496758 | 0.351461705 | 0.29797779 | 1.1794896 | 0.23820328 | 0.93059916 |
| CXorf38 | 16.0114171 | -0.048222522 | 0.43034091 | -0.1120566 | 0.91077856 | 0.99777713 |
| MED14 | 29.1551081 | -0.253058583 | 0.40200535 | -0.6294906 | 0.52902793 | 0.99665574 |
| USP9X | 88.0812127 | 0.119059713 | 0.32867407 | 0.36224249 | 0.71717083 | 0.99665574 |
| DDX3X | 28.3216373 | -0.236476606 | 0.34504062 | -0.6853587 | 0.49311757 | 0.99640678 |
| CASK | 23.6424716 | -0.020752429 | 0.3485397 | -0.0595411 | 0.95252115 | 0.99777713 |
| MAOA | 12.1182545 | 0.18381764 | 0.67177777 | 0.27362864 | 0.78437002 | 0.99665574 |
| FUNDC1 | 3.8006174 | 0.997303058 | 0.98086357 | 1.01676022 | 0.30926751 | 0.97376371 |
| KDM6A | 75.9667106 | 0.53416691 | 0.39069224 | 1.36723195 | 0.17155261 | 0.8807653 |
| MIR222HG | 23.8760701 | 0.694832745 | 0.54483628 | 1.27530557 | 0.20220111 | 0.90614933 |
| KRBOX4 | 9.9587584 | 0.262318876 | 0.50600414 | 0.51841251 | 0.60417049 | 0.99665574 |
| ZNF674 | 9.11155278 | 0.983894532 | 0.49648468 | 1.98172182 | 0.04751038 | 0.64060657 |
| SLC9A7 | 18.2486961 | -0.393852934 | 0.46029957 | -0.8556448 | 0.39219428 | 0.99179694 |
| RP2 | 2.97706362 | -0.557799094 | 0.7942934 | -0.7022583 | 0.48251812 | 0.99640678 |
| JADE3 | 22.1777167 | 0.979389809 | 0.44454537 | 2.20312676 | 0.02758582 | 0.53563234 |
| RGN | 4.92312931 | -1.900966584 | 0.95063673 | -1.9996772 | 0.04553513 | 0.63477358 |
| NDUFB11 | 19.9173712 | 0.730135343 | 0.43914648 | 1.6626237 | 0.09638778 | 0.7740159 |
| RBM10 | 18.8080453 | 0.063113517 | 0.35242658 | 0.17908274 | 0.85787274 | 0.99665574 |
| UBA1 | 24.8432585 | -0.110517819 | 0.34771781 | -0.3178377 | 0.75060807 | 0.99665574 |
| CDK16 | 10.2465504 | 0.533317922 | 0.48497643 | 1.09967801 | 0.27147244 | 0.94870306 |
| USP11 | 8.82049126 | 0.227008603 | 0.56790248 | 0.39973167 | 0.68935417 | 0.99665574 |
| ZNF41 | 6.15281322 | -0.015346216 | 0.60055445 | -0.0255534 | 0.97961354 | 0.99777713 |
| ARAF | 7.68337996 | 0.240785476 | 0.50956849 | 0.47252819 | 0.63654983 | 0.99665574 |
| TIMP1 | 4.30334751 | -3.195998831 | 0.92492923 | -3.4553982 | 0.00054948 | 0.06395083 |
| ELK1 | 3.13265103 | 0.252222919 | 0.84195854 | 0.29956691 | 0.76450753 | 0.99665574 |
| UXT | 3.80897377 | 0.500012175 | 0.87550507 | 0.57111283 | 0.56792317 | 0.99665574 |
| ZNF81 | 9.23074937 | 0.170250654 | 0.53666148 | 0.31724031 | 0.75106127 | 0.99665574 |
| ZNF182 | 2.4810766 | 0.060947332 | 0.92507888 | 0.06588339 | 0.94747067 | 0.99777713 |
| EBP | 8.1427132 | -0.375019428 | 0.59764101 | -0.6274995 | 0.53033187 | 0.99665574 |
| TBC1D25 | 3.92755753 | -0.840665734 | 0.68893147 | -1.2202458 | 0.22237171 | 0.92191316 |
| RBM3 | 16.8631576 | 0.056173246 | 0.5356527 | 0.10486878 | 0.91647993 | 0.99777713 |
| WDR13 | 9.4354174 | -0.071419131 | 0.50500873 | -0.1414216 | 0.88753691 | 0.99665574 |
| HDAC6 | 5.74359192 | 0.117602721 | 0.61560616 | 0.19103565 | 0.84849767 | 0.99665574 |
| PCSK1N | 18.8118458 | -0.563464329 | 0.59526856 | -0.9465716 | 0.34385711 | 0.97788427 |
| TIMM17B | 5.96300996 | 0.132528573 | 0.81407682 | 0.16279615 | 0.87067894 | 0.99665574 |
| PQBP1 | 4.08805413 | 0.951810016 | 0.75178081 | 1.26607384 | 0.20548665 | 0.90874907 |
| SLC35A2 | 3.34111568 | -0.309631931 | 0.84791425 | -0.3651689 | 0.71498532 | 0.99665574 |
| OTUD5 | 12.8328306 | -0.085240368 | 0.43437254 | -0.1962379 | 0.84442394 | 0.99665574 |
| GRIPAP1 | 9.42604259 | -0.441894782 | 0.48723927 | -0.9069359 | 0.3644407 | 0.98362663 |
| TFE3 | 6.46028038 | 0.318208395 | 0.67341579 | 0.47252886 | 0.63654935 | 0.99665574 |
| CCDC120 | 4.20174169 | 0.183576832 | 0.72404423 | 0.25354367 | 0.79984812 | 0.99665574 |
| WDR45 | 13.0404782 | -0.025065337 | 0.47628013 | -0.0526273 | 0.95802886 | 0.99777713 |
| PLP2 | 5.44677333 | -2.65752802 | 0.73586893 | -3.6114149 | 0.00030453 | 0.0434836 |
| CLCN5 | 76.1547429 | 1.023336687 | 0.29143724 | 3.51134499 | 0.00044585 | 0.05794847 |
| SHROOM4 | 5.04293551 | -1.16840573 | 0.75345134 | -1.5507381 | 0.12096445 | 0.82659025 |
| MAGED1 | 44.4149507 | 0.257480891 | 0.32705644 | 0.78726746 | 0.43112531 | 0.99617422 |
| FAM156A | 2.3585203 | -0.677210893 | 0.99284205 | -0.6820933 | 0.49517997 | 0.99640678 |
| KANTR | 18.6927554 | -0.35398372 | 0.49751401 | -0.711505 | 0.47677134 | 0.99617422 |
| KDM5C | 9.06383717 | 0.174232664 | 0.77694718 | 0.22425291 | 0.82256052 | 0.99665574 |
| IQSEC2 | 4.12392105 | -0.694395334 | 0.79281994 | -0.875855 | 0.38110885 | 0.98820924 |
| SMC1A | 12.5934322 | 0.017680522 | 0.48227535 | 0.03666064 | 0.97075559 | 0.99777713 |
| HSD17B10 | 6.31937969 | 0.339539333 | 0.66404041 | 0.5113233 | 0.60912469 | 0.99665574 |
| HUWE1 | 50.5528209 | 0.338621724 | 0.26595251 | 1.27324132 | 0.20293242 | 0.9066717 |
| PHF8 | 21.2174717 | -0.172957314 | 0.4101945 | -0.4216471 | 0.67328264 | 0.99665574 |
| FAM120C | 8.77885448 | 0.109859781 | 0.49031548 | 0.22405938 | 0.8227111 | 0.99665574 |
| WNK3 | 30.3579716 | 1.264823085 | 0.34765516 | 3.63815422 | 0.0002746 | 0.04093948 |
| TSR2 | 5.8434503 | -0.91354628 | 0.74031462 | -1.2339974 | 0.21720389 | 0.91584128 |
| FGD1 | 2.65127286 | 0.020437583 | 0.96702669 | 0.02113446 | 0.9831384 | 0.99777713 |
| GNL3L | 3.58972673 | 0.179924273 | 0.70393856 | 0.25559656 | 0.79826238 | 0.99665574 |
| MAGED2 | 15.0155696 | -0.401438725 | 0.49287096 | -0.8144905 | 0.415364 | 0.99279547 |
| RRAGB | 15.3990444 | -0.076251578 | 0.42728901 | -0.1784543 | 0.85836618 | 0.99665574 |
| AL445472.1 | 8.98259797 | -0.501414018 | 0.58704242 | -0.8541359 | 0.39302968 | 0.99179694 |
| KLF8 | 9.5361268 | 0.452688833 | 0.63080508 | 0.71763663 | 0.47298137 | 0.99617422 |
| UBQLN2 | 6.51911326 | -0.458518388 | 0.57923584 | -0.7915919 | 0.42859869 | 0.99589919 |
| NBDY | 28.415572 | 0.110398501 | 0.33294454 | 0.33158225 | 0.74020472 | 0.99665574 |
| SPIN3 | 5.86768235 | 0.529983942 | 0.59744835 | 0.8870791 | 0.37503631 | 0.98569969 |
| FAAH2 | 66.4970004 | -1.039036067 | 0.41732158 | -2.4897732 | 0.01278246 | 0.38590337 |
| ARHGEF9 | 7.80658042 | -0.531240169 | 0.53064718 | -1.0011175 | 0.31677002 | 0.97700345 |
| MTMR8 | 2.98839975 | 0.064486064 | 0.79132967 | 0.08149077 | 0.93505167 | 0.99777713 |

| | | | | | | | |
|---|---|---|---|---|---|---|---|
| ZC4H2 | 5.79052276 | | -0.820267508 | 0.70699742 | -1.1602129 | 0.24596215 | 0.93426912 |
| LAS1L | 4.12746455 | | 0.167509469 | 0.7401745 | 0.22631078 | 0.82095971 | 0.99665574 |
| MSN | 28.6700436 | | -1.032207986 | 0.45354888 | -2.2758473 | 0.02285515 | 0.4879643 |
| HEPH | 4.01091719 | | 0.314803466 | 1.17091421 | 0.26885272 | 0.78804303 | 0.99665574 |
| OPHN1 | 32.3304357 | | 0.048778904 | 0.48855662 | 0.09984289 | 0.92046906 | 0.99777713 |
| YIPF6 | 16.1841027 | | -0.397150204 | 0.43767971 | -0.9073992 | 0.36419575 | 0.98362663 |
| EDA | 8.01623328 | | -1.476431823 | 0.697398 | -2.1170577 | 0.03425495 | 0.58178209 |
| IGBP1 | 11.0788469 | | 0.035830326 | 0.51821235 | 0.06914217 | 0.94487645 | 0.99777713 |
| PDZD11 | 3.22272869 | | 0.283958509 | 0.9034803 | 0.31429408 | 0.75329769 | 0.99665574 |
| DLG3 | 9.74375911 | | -0.396330669 | 0.47365334 | -0.8367526 | 0.40273164 | 0.99194293 |
| SNX12 | 5.02551889 | | 0.139524539 | 0.684933 | 0.20370538 | 0.83858373 | 0.99665574 |
| MED12 | 1.80946218 | | 0.297490088 | 1.04807695 | 0.28384375 | 0.77653013 | 0.99665574 |
| ZMYM3 | 2.23064149 | | 1.834069202 | 1.03754543 | 1.76770014 | 0.07711105 | 0.74099697 |
| NONO | 19.7123222 | | 0.242001271 | 0.36117441 | 0.67003992 | 0.50283234 | 0.99640678 |
| TAF1 | 22.1736055 | | 0.115692652 | 0.3759521 | 0.30773243 | 0.75828594 | 0.99665574 |
| OGT | 38.6259058 | | -0.068141338 | 0.42414266 | -0.1606566 | 0.87236384 | 0.99665574 |
| GCNA | 3.75268855 | | 0.353048708 | 0.78397667 | 0.45033063 | 0.65247206 | 0.99665574 |
| NHSL2 | 5.91912451 | | 0.357118989 | 0.63415204 | 0.56314412 | 0.57333675 | 0.99665574 |
| PIN4 | 9.43985173 | | -0.112970909 | 0.58144202 | -0.1942944 | 0.84594539 | 0.99665574 |
| RPS4X | 84.3574197 | | 0.126889558 | 0.40863989 | 0.31051682 | 0.75616797 | 0.99665574 |
| HDAC8 | 56.3408565 | | -0.227478514 | 0.29502581 | -0.7710462 | 0.44067957 | 0.99617422 |
| PHKA1 | 5.20946341 | | 0.450170695 | 0.69349664 | 0.64913176 | 0.51625321 | 0.99665574 |
| CHIC1 | 19.1998606 | | -0.39572944 | 0.40047572 | -0.9881484 | 0.32307997 | 0.97700345 |
| XIST | 81.6449425 | | 1.282009364 | 4.03413454 | 0.31779043 | 0.75064391 | 0.99665574 |
| JPX | 61.6920453 | | -0.053614463 | 0.39674151 | -0.135137 | 0.89250354 | 0.99773598 |
| FTX | 472.67041 | | -0.228816558 | 0.28083937 | -0.8147595 | 0.41520996 | 0.99279547 |
| RLIM | 11.664305 | | 0.093379178 | 0.495228 | 0.18855796 | 0.85043929 | 0.99665574 |
| NEXMIF | 5.17697335 | | -1.859761745 | 0.64390849 | -2.8882392 | 0.00387405 | 0.20301691 |
| ABCB7 | 25.0373082 | | 0.118691817 | 0.33755934 | 0.35161764 | 0.72512504 | 0.99665574 |
| UPRT | 5.45817182 | | 1.049292365 | 0.7080917 | 1.48185944 | 0.1383777 | 0.84656101 |
| ZDHHC15 | 7.64273616 | | -0.178849168 | 0.54198055 | -0.3299919 | 0.74140611 | 0.99665574 |
| PBDC1 | 11.3004125 | | -1.209679532 | 0.53706416 | -2.252393 | 0.02429745 | 0.50168544 |
| ATRX | 101.470692 | | 0.098838599 | 0.30670387 | 0.32226069 | 0.74725521 | 0.99665574 |
| MAGT1 | 17.0506519 | | -0.210334625 | 0.46153071 | -0.4557327 | 0.64858222 | 0.99665574 |
| COX7B | 41.230003 | | 0.323216942 | 0.38851928 | 0.83191996 | 0.40545412 | 0.99194293 |
| PGK1 | 39.8868417 | | -0.049914881 | 0.4652047 | -0.1072966 | 0.91455369 | 0.99777713 |
| BRWD3 | 30.9463947 | | 0.254639946 | 0.43672276 | 0.58307002 | 0.55984617 | 0.99665574 |
| HMGN5 | 4.95894352 | | 0.163200541 | 0.65410521 | 0.24950197 | 0.80297251 | 0.99665574 |
| SH3BGRL | 32.7411171 | | -0.084359314 | 0.44505238 | -0.1895492 | 0.84966241 | 0.99665574 |
| POU3F4 | 3.69047672 | | -0.344918386 | 0.78949698 | -0.4368837 | 0.66219567 | 0.99665574 |
| RPS6KA6 | 29.4684106 | | 0.02846888 | 0.47300882 | 0.06018678 | 0.95200687 | 0.99777713 |
| HDX | 9.78677406 | | -1.788246706 | 0.73412808 | -2.4358784 | 0.01485568 | 0.40950537 |
| APOOL | 31.3858106 | | 0.26878403 | 0.31523937 | 0.8526347 | 0.39386191 | 0.99179694 |
| ZNF711 | 2.71958165 | | 0.922492975 | 0.90727132 | 1.0167774 | 0.30925933 | 0.97376371 |
| CHM | 25.5676624 | | 0.062108849 | 0.319401 | 0.19445414 | 0.8458203 | 0.99665574 |
| PCDH11X | 2.51505359 | | 0.551850327 | 0.95400158 | 0.5784585 | 0.56295461 | 0.99665574 |
| DIAPH2 | 85.5768082 | | 0.137640669 | 0.27392469 | 0.50247631 | 0.61533251 | 0.99665574 |
| DIAPH2-AS1 | 3.92668261 | | -2.73347043 | 0.80652119 | -3.3892109 | 0.00070094 | 0.07140296 |
| TSPAN6 | 6.44955432 | | 1.14506717 | 0.59688902 | 1.9183921 | 0.05506131 | 0.67068154 |
| TRMT2B | 12.5723764 | | -0.087422626 | 0.4350477 | -0.2009495 | 0.84073805 | 0.99665574 |
| HNRNPH2 | 10.0934893 | | 0.189760991 | 0.50520402 | 0.37561259 | 0.70720493 | 0.99665574 |
| ARMCX4 | 24.1271817 | | 0.227637342 | 0.52263944 | 0.43555331 | 0.66316085 | 0.99665574 |
| ARMCX6 | 3.09294748 | | -0.047252412 | 0.78225905 | -0.0604051 | 0.95183302 | 0.99777713 |
| ARMCX3 | 12.2731223 | | -0.736144651 | 0.46132211 | -1.5957281 | 0.11054952 | 0.80757694 |
| ZMAT1 | 21.6920704 | | 0.795828135 | 0.44265023 | 1.79787127 | 0.07219741 | 0.71945049 |
| TCEAL2 | 2.71501245 | | -0.377488809 | 0.79192785 | -0.4766707 | 0.63359663 | 0.99665574 |
| BEX5 | 20.2106732 | | 1.95234614 | 1.23308843 | 1.58329776 | 0.11335361 | 0.81502052 |
| ARMCX5 | 2.7716263 | | 0.596355692 | 0.8166038 | 0.73028767 | 0.46521436 | 0.99617422 |
| LINC00630 | 11.9588757 | | 0.780476207 | 0.4940571 | 1.57972875 | 0.114169 | 0.81638817 |
| BEX4 | 14.1725283 | | 0.599547201 | 0.62169131 | 0.96438086 | 0.33485502 | 0.97788427 |
| TCEAL8 | 5.9643829 | | 0.169160173 | 0.62144842 | 0.27220308 | 0.78546587 | 0.99665574 |
| BEX2 | 7.49649505 | | -0.773511498 | 0.79438296 | -0.9737262 | 0.33019251 | 0.97788427 |
| TCEAL9 | 11.1189481 | | -0.85688633 | 0.61301402 | -1.397825 | 0.16216562 | 0.87124274 |
| BEX3 | 17.468101 | | 1.365433115 | 0.63253172 | 2.15867927 | 0.03087506 | 0.56271575 |
| TCEAL4 | 17.328037 | | -0.248035038 | 0.39623478 | -0.62598 | 0.53132807 | 0.99665574 |
| TCEAL3 | 5.55857469 | | -0.478438838 | 0.63482717 | -0.7536521 | 0.45105814 | 0.99617422 |
| MORF4L2 | 28.3973791 | | -0.02383728 | 0.42773212 | -0.0557295 | 0.95555733 | 0.99777713 |
| SLC25A53 | 13.4815131 | | 0.278212009 | 0.43835915 | 0.63466682 | 0.52564574 | 0.99665574 |
| FAM199X | 20.0299038 | | 0.690857533 | 0.38624111 | 1.78866907 | 0.07366813 | 0.72792151 |
| IL1RAPL2 | 2.84532738 | | 0.956446817 | 0.81950193 | 1.16710746 | 0.24316697 | 0.93426912 |
| NRK | 16.4207101 | | 0.430459358 | 0.68325703 | 0.63001087 | 0.52868747 | 0.99665574 |
| RNF128 | 4.70308503 | | 0.899543685 | 0.98529322 | 0.91297055 | 0.36125803 | 0.9819332 |
| TBC1D8B | 6.97608714 | | 0.125854698 | 0.70815283 | 0.17772251 | 0.85894091 | 0.99665574 |
| MORC4 | 8.36812891 | | -0.215535271 | 0.55354485 | -0.3893727 | 0.69700044 | 0.99665574 |
| RBM41 | 16.2757377 | | -0.026072673 | 0.44047934 | -0.0591916 | 0.95279951 | 0.99777713 |
| NUP62CL | 3.28261607 | | 0.658177125 | 0.84914462 | 0.77510604 | 0.438277 | 0.99617422 |
| MID2 | 6.28415183 | | -0.065805166 | 0.73838129 | -0.0891208 | 0.92898587 | 0.99777713 |
| ATG4A | 2.33997936 | | 0.437352455 | 0.91850968 | 0.47615443 | 0.63396436 | 0.99665574 |
| COL4A5 | 14.4981072 | | 0.415355939 | 0.65190152 | 0.63714522 | 0.52403026 | 0.99665574 |
| ACSL4 | 168.867085 | | -1.119630647 | 0.51088965 | -2.1915313 | 0.02841337 | 0.54555819 |
| TMEM164 | 13.9308669 | | 0.362859718 | 0.45752157 | 0.7930986 | 0.42772038 | 0.99589919 |
| AMMECR1 | 17.1363627 | | -0.121740348 | 0.38002574 | -0.3203476 | 0.74870481 | 0.99665574 |
| ALG13 | 15.1134416 | | 0.115071539 | 0.48451904 | 0.23749642 | 0.81227169 | 0.99665574 |
| PLS3 | 48.7606853 | | 0.58433343 | 0.32759815 | 1.78368966 | 0.07447411 | 0.72810353 |
| DANT2 | 123.687522 | | -0.039573571 | 0.45189982 | -0.0875716 | 0.93021721 | 0.99777713 |
| KLHL13 | 16.7027297 | | -1.096270912 | 0.71038917 | -1.5431977 | 0.12278282 | 0.82915695 |
| WDR44 | 17.055227 | | 0.145847612 | 0.42586542 | 0.34247348 | 0.7319946 | 0.99665574 |
| IL13RA1 | 14.9376195 | | -0.490741781 | 0.40605848 | -1.2085495 | 0.22683596 | 0.92318867 |
| PGRMC1 | 22.0989971 | | 0.158937775 | 0.35537444 | 0.44724031 | 0.65470156 | 0.99665574 |
| SLC25A43 | 7.38980572 | | -1.198939905 | 0.58696841 | -2.042597 | 0.04109235 | 0.61100706 |
| SLC25A5 | 38.5131657 | | -0.058276654 | 0.51523724 | -0.1131064 | 0.90994616 | 0.99777713 |
| CXorf56 | 3.52449732 | | -0.007388549 | 0.71230243 | -0.0103728 | 0.99172388 | 0.99777713 |
| UBE2A | 10.1933523 | | 0.298613999 | 0.51499067 | 0.57984352 | 0.56202015 | 0.99665574 |
| NKRF | 3.2449761 | | -0.301992192 | 0.70760406 | -0.4267813 | 0.66953862 | 0.99665574 |
| 6-Sep | 12.810441 | | -0.512603904 | 0.47811119 | -1.0721437 | 0.28365549 | 0.96059387 |
| RPL39 | 3.43681025 | | 1.867789664 | 1.11490618 | 1.67528865 | 0.09387761 | 0.76974878 |
| UPF3B | 10.4164847 | | -0.69309998 | 0.51653204 | -1.3418335 | 0.17964999 | 0.88820302 |
| NDUFA1 | 24.3946586 | | 0.00914704 | 0.35961495 | 0.02543565 | 0.97970748 | 0.99777713 |
| NKAP | 4.1334398 | | -0.184210175 | 0.69819373 | -0.2638382 | 0.7919046 | 0.99665574 |
| RHOXF1-AS1 | 5.16472311 | | 0.04113393 | 0.65873748 | 0.06244358 | 0.95020959 | 0.99777713 |
| ZBTB33 | 2.31709984 | | -0.77268091 | 0.9214549 | -0.8385445 | 0.40172498 | 0.99194293 |
| TMEM255A | 15.5884786 | | -0.467185012 | 0.59032822 | -0.7913987 | 0.42871134 | 0.99589919 |
| LAMP2 | 28.3703242 | | -0.279341086 | 0.35995148 | -0.7760521 | 0.43771825 | 0.99617422 |
| CUL4B | 8.81286098 | | 0.28365501 | 0.53874429 | 0.5265114 | 0.59853293 | 0.99665574 |
| MCTS1 | 6.76534044 | | -0.305518631 | 0.59847275 | -0.5104972 | 0.60970321 | 0.99665574 |
| C1GALT1C1 | 3.77772584 | | -0.003014785 | 0.88033251 | -0.0034246 | 0.99726757 | 0.99884397 |
| THOC2 | 39.4835376 | | 0.460279437 | 0.29535959 | 1.55836973 | 0.11914563 | 0.82619588 |
| XIAP | 38.173629 | | -0.19649527 | 0.37555991 | -0.5232062 | 0.60083078 | 0.99665574 |
| STAG2 | 55.9348642 | | 0.151893781 | 0.29770997 | 0.51020724 | 0.60990628 | 0.99665574 |
| SMARCA1 | 13.0123122 | | 0.290247943 | 0.6143005 | 0.47248528 | 0.63658045 | 0.99665574 |
| OCRL | 5.00039257 | | -0.484875459 | 0.63645063 | -0.761843 | 0.44615371 | 0.99617422 |
| ZDHHC9 | 19.2933905 | | -0.789109191 | 0.41412731 | -1.9054749 | 0.05671837 | 0.67545594 |
| UTP14A | 3.87822547 | | -0.685899402 | 0.75135407 | -0.9128844 | 0.36130334 | 0.9819332 |

| | | | | | | |
|---|---|---|---|---|---|---|
| BCORL1 | 3.95620132 | 0.458688257 | 0.82107742 | 0.55864191 | 0.57640613 | 0.99665574 |
| ELF4 | 4.90966461 | -0.022386778 | 0.66023634 | -0.0339072 | 0.97295114 | 0.99777713 |
| AIFM1 | 12.8568585 | 0.492571722 | 0.45824421 | 1.07491095 | 0.2824146 | 0.96059387 |
| ZNF280C | 4.25398459 | 1.644503965 | 0.81173885 | 2.02590275 | 0.04277475 | 0.61874363 |
| SLC25A14 | 3.34387959 | 0.558320429 | 0.82004921 | 0.68083771 | 0.49597418 | 0.99640678 |
| RBMX2 | 3.13150143 | -0.346275914 | 0.78671946 | -0.4401517 | 0.65982723 | 0.99665574 |
| ENOX2 | 24.4144107 | 0.057620593 | 0.34577269 | 0.16664298 | 0.86765097 | 0.99665574 |
| STK26 | 4.56203222 | -0.933649354 | 0.8816888 | -1.058933 | 0.2896303 | 0.96676791 |
| FRMD7 | 7.65587697 | 1.561591592 | 0.60941514 | 2.56244307 | 0.01039386 | 0.35124327 |
| RAP2C | 13.665579 | 0.886029792 | 0.51649755 | 1.71545789 | 0.08626131 | 0.75397496 |
| RAP2C-AS1 | 18.1888808 | 1.144510191 | 0.50091911 | 2.2848204 | 0.02232336 | 0.48357743 |
| MBNL3 | 10.7092972 | 0.475781764 | 0.50853546 | 0.93559211 | 0.34948324 | 0.97838306 |
| GPC4 | 6.94679152 | -0.902282298 | 0.64212291 | -1.4051551 | 0.15997515 | 0.86757484 |
| GPC3 | 20.6805142 | 2.476449731 | 0.66863785 | 3.70372349 | 0.00021246 | 0.03474028 |
| PHF6 | 9.71633773 | -0.175942743 | 0.49823887 | -0.3531293 | 0.72399151 | 0.99665574 |
| HPRT1 | 31.1331437 | -0.051949367 | 0.51912478 | -0.1000711 | 0.92028791 | 0.99777713 |
| FAM122B | 5.72134357 | -0.407341886 | 0.65551625 | -0.6214062 | 0.53433237 | 0.99665574 |
| MOSPD1 | 3.20349584 | -0.069437295 | 0.80389728 | -0.0863758 | 0.93116766 | 0.99777713 |
| RTL8C | 9.89295346 | -0.114506193 | 0.50870258 | -0.2250946 | 0.8219057 | 0.99665574 |
| RTL8A | 5.79898796 | -0.471390763 | 0.66018908 | -0.7140239 | 0.47521242 | 0.99617422 |
| ZNF75D | 11.2437984 | -0.358837928 | 0.55016833 | -0.652233 | 0.51425089 | 0.99665574 |
| INTS6L | 4.70810222 | 0.514862825 | 0.74419177 | 0.6918416 | 0.48903681 | 0.99640678 |
| MMGT1 | 6.355309 | 0.020625348 | 0.56885394 | 0.03625772 | 0.97107686 | 0.99777713 |
| SLC9A6 | 4.57552827 | -0.283496949 | 0.66688673 | -0.4251051 | 0.67076006 | 0.99665574 |
| HTATSF1 | 8.26753827 | -0.212791641 | 0.50679175 | -0.4198798 | 0.67457323 | 0.99665574 |
| AL683813.1 | 4.43515397 | -0.211970827 | 0.61903955 | -0.3424189 | 0.73203569 | 0.99665574 |
| RBMX | 34.5355477 | 0.239760341 | 0.32496871 | 0.73779516 | 0.46063894 | 0.99617422 |
| FGF13 | 17.3988809 | -1.588726894 | 1.2924623 | -1.2292249 | 0.21898748 | 0.91662582 |
| ATP11C | 16.1697436 | -0.867265742 | 0.56665152 | -1.5305099 | 0.12589057 | 0.83633607 |
| FMR1 | 20.1568489 | 0.096770064 | 0.44206499 | 0.21890461 | 0.82672436 | 0.99665574 |
| IDS | 12.704669 | -0.253621299 | 0.49500294 | -0.5123632 | 0.60839683 | 0.99665574 |
| TMEM185A | 5.05847549 | 0.165145786 | 0.64766886 | 0.25498491 | 0.79873475 | 0.99665574 |
| LINC00894 | 25.1161346 | 0.339629821 | 0.40331227 | 0.84210138 | 0.39973121 | 0.99179694 |
| MTM1 | 11.3840869 | -0.24053354 | 0.55875267 | -0.430483 | 0.66684431 | 0.99665574 |
| MTMR1 | 15.6300225 | 0.213971343 | 0.4461927 | 0.47954918 | 0.63154799 | 0.99665574 |
| CD99L2 | 9.29904558 | -0.071427583 | 0.52274943 | -0.1366383 | 0.89131672 | 0.99736963 |
| HMGB3 | 2.58595473 | -0.407079761 | 0.82839028 | -0.4914106 | 0.62313607 | 0.99665574 |
| VMA21 | 11.1351598 | 0.617804623 | 0.51625873 | 1.19669574 | 0.23142517 | 0.92792831 |
| CETN2 | 2.8028599 | 0.286500274 | 0.81828667 | 0.35012213 | 0.72624704 | 0.99665574 |
| SLC6A8 | 7.24757897 | -0.627747198 | 0.53195609 | -1.1800733 | 0.23797105 | 0.9305595 |
| BCAP31 | 15.5667054 | 0.168259045 | 0.44570757 | 0.37750995 | 0.70579467 | 0.99665574 |
| IDH3G | 6.10132208 | -0.251825215 | 0.61989781 | -0.4062367 | 0.68456872 | 0.99665574 |
| SSR4 | 12.8397685 | -0.068642265 | 0.46457461 | -0.1477529 | 0.88253775 | 0.99665574 |
| RENBP | 2.52404057 | -1.431660797 | 1.16134778 | -1.232758 | 0.21766605 | 0.91640302 |
| HCFC1 | 6.71014534 | 0.446187909 | 0.5589692 | 0.79823344 | 0.42473504 | 0.99518711 |
| IRAK1 | 7.18359858 | -1.297625078 | 0.82173057 | -1.5791369 | 0.11430466 | 0.81638817 |
| MECP2 | 42.5201563 | 0.036943192 | 0.38445378 | 0.09609268 | 0.92344697 | 0.99777713 |
| FLNA | 6.03610242 | -0.176411291 | 1.08612126 | -0.1624232 | 0.8709726 | 0.99665574 |
| EMD | 9.4954116 | 0.2872248 | 0.48875889 | 0.58766153 | 0.5567595 | 0.99665574 |
| RPL10 | 102.124372 | 0.638540098 | 0.6002366 | 1.063814 | 0.28741297 | 0.96559334 |
| DNASE1L1 | 2.60894634 | -0.509588807 | 0.78913876 | -0.6457531 | 0.51843926 | 0.99665574 |
| TAZ | 6.3330182 | -0.57122039 | 0.62245687 | -0.9176867 | 0.35878291 | 0.9819332 |
| AC244090.1 | 3.29242327 | 1.748636104 | 0.9811881 | 1.78216195 | 0.07472283 | 0.72883798 |
| ATP6AP1 | 20.7979175 | -0.043096263 | 0.5268927 | -0.0817932 | 0.93481113 | 0.99777713 |
| GDI1 | 9.53748854 | -0.430359585 | 0.49321696 | -0.8725563 | 0.38290494 | 0.98934457 |
| FAM50A | 8.54154101 | 0.084992783 | 0.67630724 | 0.12567185 | 0.89999169 | 0.99777713 |
| PLXNA3 | 5.56414218 | -1.00095558 | 0.68737269 | -1.456205 | 0.14533595 | 0.85062144 |
| UBL4A | 3.62930342 | 0.101997641 | 0.82085805 | 0.12425734 | 0.90111152 | 0.99777713 |
| FAM3A | 3.97730246 | -0.375531541 | 0.66497885 | -0.564727 | 0.57225944 | 0.99665574 |
| G6PD | 2.6838592 | 0.220364913 | 0.86924285 | 0.25351363 | 0.79987133 | 0.99665574 |
| IKBKG | 3.1368312 | 0.373370053 | 0.80932818 | 0.46133332 | 0.64455949 | 0.99665574 |
| DKC1 | 6.77016996 | -0.164763103 | 0.53528346 | -0.3078053 | 0.75823046 | 0.99665574 |
| MPP1 | 4.46929157 | -0.023722098 | 0.71204426 | -0.0333155 | 0.97342301 | 0.99777713 |
| F8 | 2.59381281 | -0.93449722 | 0.93953764 | -0.9946352 | 0.31991372 | 0.97700345 |
| F8A1 | 2.1187921 | 1.040699691 | 1.05610036 | 0.98541742 | 0.32441908 | 0.97700345 |
| FUNDC2 | 14.9019483 | -0.169891844 | 0.55725792 | -0.3048711 | 0.76046432 | 0.99665574 |
| BRCC3 | 8.53662222 | 0.282154673 | 0.58739239 | 0.48035125 | 0.63097765 | 0.99665574 |
| VBP1 | 6.91843123 | -0.225330732 | 0.57655511 | -0.3908225 | 0.69592842 | 0.99665574 |
| TMLHE-AS1 | 8.6021096 | -1.275684966 | 0.7907506 | -1.6132583 | 0.1066884 | 0.79764531 |
| TMLHE | 24.830004 | -0.403008281 | 0.47091423 | -0.8557998 | 0.39210855 | 0.99179694 |
| VAMP7 | 8.65929868 | 0.121792427 | 0.48247539 | 0.25243241 | 0.80070685 | 0.99665574 |
| ZNF596 | 4.94558798 | -0.1075053 | 0.62591663 | -0.1717566 | 0.8636289 | 0.99665574 |
| FBXO25 | 22.1101285 | -0.040688274 | 0.41728628 | -0.0975069 | 0.92232389 | 0.99777713 |
| TDRP | 12.7542451 | 0.074549578 | 0.51388291 | 0.14507114 | 0.88465471 | 0.99665574 |
| ERICH1 | 28.8124169 | -0.346894223 | 0.42685921 | -0.8126666 | 0.41640924 | 0.992853 |
| AC100810.1 | 2.92353916 | 0.468655194 | 0.94073736 | 0.49817857 | 0.61835819 | 0.99665574 |
| AC019257.8 | 4.05792082 | -0.050616336 | 0.81025046 | -0.06247 | 0.95018856 | 0.99777713 |
| CSMD1 | 13.8381127 | 2.765700811 | 0.83985851 | 3.29305566 | 0.00099105 | 0.08661422 |
| MCPH1 | 35.2787915 | -0.172154378 | 0.41562926 | -0.4142018 | 0.67872634 | 0.99665574 |
| ANGPT2 | 5.50695471 | -0.941529538 | 0.84531344 | -1.113823 | 0.26535513 | 0.94557817 |
| MCPH1-AS1 | 9.73904123 | -0.887805394 | 0.53815528 | -1.6497198 | 0.09900027 | 0.78227959 |
| AGPAT5 | 18.4835767 | -0.708965419 | 0.58534854 | -1.2111851 | 0.22582448 | 0.92318867 |
| AF233439.1 | 58.3722638 | -0.406051528 | 0.85616139 | -0.4742698 | 0.63530749 | 0.99665574 |
| DEFB1 | 144.827611 | 1.50442719 | 0.52550549 | 2.86281915 | 0.0041989 | 0.20883915 |
| FAM85B | 17.3464976 | -0.102181222 | 0.51647163 | -0.1978448 | 0.8431665 | 0.99665574 |
| PRAG1 | 10.7682268 | 0.624541631 | 0.56014322 | 1.11496775 | 0.26486424 | 0.94557817 |
| MFHAS1 | 198.092413 | -0.773986291 | 0.37275132 | -2.0764146 | 0.03785562 | 0.59043117 |
| ERI1 | 12.7129053 | -1.167199583 | 0.5953429 | -1.9605501 | 0.04993153 | 0.64567071 |
| TNKS | 63.2761964 | -0.211671358 | 0.2874067 | -0.7364872 | 0.46143425 | 0.99617422 |
| MSRA | 32.4723085 | 0.013584112 | 0.32529085 | 0.0417599 | 0.9666901 | 0.99777713 |
| XKR6 | 109.247216 | -0.329496204 | 0.37981943 | -0.8675075 | 0.38566401 | 0.99051797 |
| MTMR9 | 15.6151591 | 0.037423117 | 0.3872401 | 0.0966406 | 0.92301181 | 0.99777713 |
| FDFT1 | 25.0115938 | -1.342274323 | 0.46422803 | -2.8914116 | 0.00383515 | 0.20301691 |
| CTSB | 44.0118807 | 0.382905685 | 0.34460838 | 1.11113282 | 0.26651118 | 0.94557817 |
| FAM66D | 2.9877214 | -0.230788287 | 0.87057966 | -0.2650973 | 0.79093453 | 0.99665574 |
| LONRF1 | 10.59987 | -0.054097311 | 0.43790468 | -0.1235367 | 0.90168209 | 0.99777713 |
| TRMT9B | 81.3232339 | 0.130445636 | 0.47628854 | 0.27387943 | 0.78417728 | 0.99665574 |
| DLC1 | 15.3409465 | -0.195580153 | 0.59908606 | -0.3264642 | 0.74407317 | 0.99665574 |
| TUSC3 | 18.6253338 | -0.621459091 | 0.4403475 | -1.4112924 | 0.15815841 | 0.86661434 |
| MSR1 | 5.86610784 | -0.966084909 | 0.77053085 | -1.2537913 | 0.20991785 | 0.91043197 |
| MICU3 | 20.0006278 | -0.330320001 | 0.40161117 | -0.8224871 | 0.41079973 | 0.99279547 |
| ZDHHC2 | 17.8923311 | -0.463644688 | 0.40685348 | -1.1395864 | 0.25445866 | 0.94099699 |
| CNOT7 | 8.87674706 | -0.244149722 | 0.48372857 | -0.5047246 | 0.61375226 | 0.99665574 |
| VPS37A | 14.9188817 | -0.308922832 | 0.42849212 | -0.7209534 | 0.47093821 | 0.99617422 |
| SLC7A2 | 5.69762287 | 1.749917704 | 0.83393312 | 2.09839095 | 0.03587062 | 0.58508511 |
| MTUS1 | 61.7434644 | -0.39127785 | 0.4210363 | -0.9293209 | 0.35272278 | 0.97912036 |
| AC027117.2 | 5.73393418 | -0.157210524 | 0.6313025 | -0.2490257 | 0.80334093 | 0.99665574 |
| PCM1 | 93.43895 | 0.161254447 | 0.23551601 | 0.68468572 | 0.49354226 | 0.99640678 |
| ASAH1 | 22.9777049 | -0.106887815 | 0.36157625 | -0.2956162 | 0.76752316 | 0.99665574 |
| PSD3 | 250.02537 | -0.873499972 | 0.49350707 | -1.7699847 | 0.07672969 | 0.73943494 |
| SH2D4A | 42.6416567 | -0.463409397 | 0.50876962 | -0.9108433 | 0.36237795 | 0.98282173 |
| CSGALNACT1 | 31.3299902 | -1.289607831 | 0.61881901 | -2.0839822 | 0.03716178 | 0.58591936 |
| INTS10 | 25.7505246 | 0.046283044 | 0.32836583 | 0.14094964 | 0.88790973 | 0.99665574 |

| | | | | | | |
|---|---|---|---|---|---|---|
| ATP6V1B2 | 8.09360977 | -0.07928981 | 0.53858041 | -0.14722 | 0.88295837 | 0.99665574 |
| XPO7 | 20.1647174 | 0.051496327 | 0.36520418 | 0.14100695 | 0.88786445 | 0.99665574 |
| DMTN | 19.5567156 | 0.237984563 | 0.45679109 | 0.52099213 | 0.60237225 | 0.99665574 |
| FAM160B2 | 6.77841627 | -0.849685828 | 0.64607951 | -1.3151413 | 0.18846243 | 0.89739428 |
| SLC39A14 | 8.15276511 | -1.393801378 | 0.78056455 | -1.7856324 | 0.07415879 | 0.72810353 |
| PPP3CC | 11.4932513 | -0.653293783 | 0.52287486 | -1.2494266 | 0.21150907 | 0.91043197 |
| CCAR2 | 4.29692621 | 0.247000712 | 0.701743 | 0.35198173 | 0.72485197 | 0.99665574 |
| BIN3 | 9.9106287 | -0.251241711 | 0.57816398 | -0.434551 | 0.6638884 | 0.99665574 |
| TNFRSF10B | 11.402443 | -0.747507088 | 0.47507337 | -1.5734561 | 0.11561325 | 0.81792539 |
| TNFRSF10D | 6.18462906 | -1.419478095 | 0.6139468 | -2.3120539 | 0.02077471 | 0.47222879 |
| CHMP7 | 5.33886829 | 0.631634344 | 0.6923712 | 0.91227704 | 0.3616229 | 0.9819332 |
| R3HCC1 | 4.77220331 | 0.61984142 | 0.70604635 | 0.87790471 | 0.37999543 | 0.98679664 |
| ENTPD4 | 11.2489311 | -0.482172542 | 0.45754253 | -1.0538311 | 0.29196025 | 0.96918567 |
| SLC25A37 | 13.8650789 | -1.758969171 | 0.55030974 | -3.1963257 | 0.0013919 | 0.10938814 |
| DOCK5 | 100.496633 | -0.584322373 | 0.57624595 | -1.0140156 | 0.31057532 | 0.97376371 |
| KCTD9 | 12.9646696 | -0.03613159 | 0.44474284 | -0.0812415 | 0.93524987 | 0.99777713 |
| PPP2R2A | 47.8997072 | 0.13628436 | 0.30920746 | 0.4407538 | 0.65939125 | 0.99665574 |
| BNIP3L | 20.4471156 | -0.884037274 | 0.66744708 | -1.3245054 | 0.18533525 | 0.89532394 |
| DPYSL2 | 3.67535659 | -0.738887331 | 0.88703693 | -0.8329837 | 0.4048539 | 0.99194293 |
| TRIM35 | 2.91635366 | -0.371996081 | 1.1858413 | -0.313698 | 0.7537504 | 0.99665574 |
| PTK2B | 23.7522474 | -0.312047416 | 0.4577806 | -0.6816528 | 0.49545853 | 0.99640678 |
| EPHX2 | 7.80078585 | 0.867292813 | 0.69222266 | 1.25291017 | 0.2102384 | 0.91043197 |
| CLU | 33.0463566 | -0.763162215 | 0.64115322 | -1.1902962 | 0.23393001 | 0.92960234 |
| CCDC25 | 6.79460724 | -0.240725859 | 0.57634261 | -0.4176784 | 0.67618227 | 0.99665574 |
| ELP3 | 24.3226116 | 0.203699011 | 0.38052905 | 0.53530476 | 0.59243913 | 0.99665574 |
| ZNF395 | 5.95898556 | 0.547107372 | 0.64338526 | 0.85035733 | 0.39512645 | 0.99179694 |
| FZD3 | 7.5911875 | -0.037714407 | 0.61123831 | -0.0617016 | 0.95080043 | 0.99777713 |
| EXTL3 | 7.92849893 | -0.367956963 | 0.64154555 | -0.5735477 | 0.56627392 | 0.99665574 |
| INTS9 | 23.0940418 | -0.171079815 | 0.40423014 | -0.4232238 | 0.67213199 | 0.99665574 |
| HMBOX1 | 158.640977 | 0.160434931 | 0.33428714 | 0.4799315 | 0.6312761 | 0.99665574 |
| KIF13B | 293.075221 | -0.679388566 | 0.28293059 | -2.4012552 | 0.01633894 | 0.42281551 |
| AC108449.1 | 12.674678 | -0.775850306 | 0.73746952 | -1.0520439 | 0.29277941 | 0.97063365 |
| SARAF | 31.5155056 | -0.199686416 | 0.28826511 | -0.6927179 | 0.48848658 | 0.99640678 |
| LEPROTL1 | 10.7990129 | -0.431950505 | 0.45324446 | -0.9530188 | 0.34058053 | 0.97788427 |
| DCTN6 | 6.00957322 | 0.259943625 | 0.62534007 | 0.41568362 | 0.67764153 | 0.99665574 |
| RBPMS | 266.571224 | -0.836021039 | 0.37654573 | -2.2202377 | 0.02640264 | 0.52177372 |
| GTF2E2 | 16.2593137 | -0.193874785 | 0.43976436 | -0.4408606 | 0.65931391 | 0.99665574 |
| GSR | 16.3794744 | -0.077810434 | 0.41664055 | -0.1867567 | 0.85185136 | 0.99665574 |
| PPP2CB | 35.0454034 | -0.105472015 | 0.34079195 | -0.3094909 | 0.75694812 | 0.99665574 |
| WRN | 37.3407623 | 0.673261652 | 0.30847899 | 2.18252026 | 0.02907116 | 0.55281655 |
| NRG1 | 12.0336017 | 1.171679004 | 1.4523735 | 0.80673394 | 0.41981978 | 0.99518711 |
| FUT10 | 11.3042275 | -0.002034716 | 0.51239088 | -0.003971 | 0.99683159 | 0.998802 |
| TTI2 | 5.10309159 | -0.546199929 | 0.58508524 | -0.9335391 | 0.35054171 | 0.97838306 |
| UNC5D | 24.6396927 | 0.91243365 | 0.61629389 | 1.48051711 | 0.1387353 | 0.84668189 |
| ERLIN2 | 10.8653653 | 0.233345553 | 0.48521471 | 0.48091195 | 0.63057908 | 0.99665574 |
| PLPBP | 7.96225556 | 0.270926941 | 0.55654946 | 0.4867976 | 0.62640177 | 0.99665574 |
| RAB11FIP1 | 19.8512332 | -1.276836444 | 0.40311086 | -3.1674573 | 0.00153778 | 0.11463266 |
| ASH2L | 11.1012372 | 0.08373838 | 0.51217734 | 0.16349489 | 0.87012879 | 0.99665574 |
| LSM1 | 7.86900534 | -0.667556922 | 0.55674486 | -1.1990356 | 0.2305141 | 0.92774608 |
| BAG4 | 16.6128024 | 0.104882037 | 0.43671569 | 0.24016091 | 0.81020552 | 0.99665574 |
| DDHD2 | 16.2817623 | -0.219490255 | 0.36792352 | -0.5965649 | 0.55079791 | 0.99665574 |
| PLPP5 | 4.81307136 | -1.240133669 | 0.73781381 | -1.680822 | 0.0927975 | 0.76709603 |
| NSD3 | 47.1436696 | -0.355176408 | 0.31449005 | -1.1293725 | 0.25874074 | 0.94557817 |
| FGFR1 | 7.24512401 | 0.394798703 | 0.56830297 | 0.69469759 | 0.48724484 | 0.99640678 |
| TACC1 | 29.2224412 | -1.239419288 | 0.43357743 | -2.8585881 | 0.00425531 | 0.20883915 |
| PLEKHA2 | 13.5326555 | -0.46972991 | 0.44253597 | -1.0614502 | 0.28848534 | 0.96676791 |
| TM2D2 | 3.88084078 | -0.42250861 | 0.78046295 | -0.5413564 | 0.58826196 | 0.99665574 |
| ADAM9 | 25.9215069 | 0.134381126 | 0.36206079 | 0.37115625 | 0.71052116 | 0.99665574 |
| TCIM | 37.3362048 | -0.979893825 | 0.58855636 | -1.6649108 | 0.09593056 | 0.7724734 |
| GOLGA7 | 10.0690112 | 0.281460351 | 0.48099901 | 0.58515785 | 0.55844158 | 0.99665574 |
| GPAT4 | 13.9160684 | 0.085056703 | 0.40690785 | 0.20903186 | 0.83442337 | 0.99665574 |
| KAT6A | 68.6137931 | -0.013134445 | 0.25786437 | -0.0509355 | 0.95937694 | 0.99777713 |
| AP3M2 | 4.52461959 | -0.420022636 | 0.65081425 | -0.6453802 | 0.51868083 | 0.99665574 |
| IKBKB | 22.2549784 | -0.60489983 | 0.35523537 | -1.7028142 | 0.08860285 | 0.75650018 |
| POLB | 6.42844098 | -1.028031979 | 0.59237688 | -1.7354357 | 0.08266366 | 0.75228386 |
| VDAC3 | 16.7952454 | -0.523807316 | 0.41481582 | -1.2627467 | 0.20668022 | 0.90890506 |
| SLC20A2 | 38.5110541 | 0.055143954 | 0.30295984 | 0.18201737 | 0.85556909 | 0.99665574 |
| SMIM19 | 8.92363252 | -0.237988654 | 0.48901459 | -0.4866698 | 0.62649232 | 0.99665574 |
| THAP1 | 3.35837874 | -0.058601527 | 0.75334804 | -0.0777881 | 0.9379966 | 0.99777713 |
| RNF170 | 10.9271134 | -0.085034667 | 0.53796939 | -0.158066 | 0.8744048 | 0.99665574 |
| HOOK3 | 27.1509573 | -0.192009253 | 0.30458408 | -0.6303982 | 0.52843409 | 0.99665574 |
| FNTA | 11.0423797 | -0.024849408 | 0.60636553 | -0.0409809 | 0.96731112 | 0.99777713 |
| HGSNAT | 26.2343804 | -0.173138049 | 0.41462065 | -0.4175818 | 0.67625288 | 0.99665574 |
| SPIDR | 146.162399 | -0.544303333 | 0.26651397 | -2.042307 | 0.04112109 | 0.61100706 |
| CEBPD | 80.5605211 | -1.211873867 | 0.33449748 | -3.6229686 | 0.00029124 | 0.04218004 |
| PRKDC | 89.9382829 | 0.654921721 | 0.27046257 | 2.42148745 | 0.01545713 | 0.41676711 |
| MCM4 | 10.3976695 | -0.2267864 | 0.73481347 | -0.3086313 | 0.75760205 | 0.99665574 |
| UBE2V2 | 15.1500253 | -0.02695281 | 0.42724735 | -0.0630848 | 0.94969898 | 0.99777713 |
| PXDNL | 10.7999621 | 0.553092046 | 0.50338998 | 1.09873472 | 0.27188379 | 0.94948601 |
| PCMTD1 | 68.4266619 | 0.347048208 | 0.40589591 | 0.85501774 | 0.39254132 | 0.99179694 |
| AC064807.1 | 3.22794889 | -0.722532066 | 1.00742582 | -0.7172062 | 0.47324686 | 0.99617422 |
| RB1CC1 | 47.4445946 | -0.339475873 | 0.26236842 | -1.29389 | 0.19570342 | 0.90381759 |
| ATP6V1H | 96.7457878 | 0.344293906 | 0.28598244 | 1.20389878 | 0.2286287 | 0.92646258 |
| TCEA1 | 20.7691703 | 0.12028844 | 0.43064288 | 0.27932295 | 0.779997 | 0.99665574 |
| LYPLA1 | 43.796725 | -0.512844657 | 0.30185161 | -1.6989959 | 0.08931995 | 0.75955113 |
| MRPL15 | 17.2188439 | -0.742628984 | 0.47816912 | -1.5530676 | 0.12040698 | 0.82659025 |
| TMEM68 | 8.29989415 | -0.090770093 | 0.56452664 | -0.1607897 | 0.872259 | 0.99665574 |
| TGS1 | 12.7663547 | 0.18956873 | 0.46032023 | 0.41181925 | 0.68047191 | 0.99665574 |
| LYN | 15.5810894 | -1.027601818 | 0.44748574 | -2.2963901 | 0.02165359 | 0.47619103 |
| RPS20 | 35.2289401 | 0.602834752 | 0.44007177 | 1.36985554 | 0.170732 | 0.87951271 |
| PLAG1 | 18.7653305 | 0.22024247 | 0.4698659 | 0.46873474 | 0.63925925 | 0.99665574 |
| CHCHD7 | 7.60464698 | 0.597113991 | 0.57682424 | 1.03517492 | 0.30058723 | 0.97358745 |
| IMPAD1 | 16.4366452 | 0.286253552 | 0.39970468 | 0.71616262 | 0.47389094 | 0.99617422 |
| LINC01606 | 9.63052951 | 0.242858595 | 0.90571842 | 0.26813918 | 0.78859219 | 0.99665574 |
| FAM110B | 62.4678472 | 0.451466596 | 0.44994128 | 1.00339004 | 0.31567271 | 0.97652251 |
| AC027698.1 | 4.89832754 | -0.634124644 | 0.91872987 | -0.6902188 | 0.49005659 | 0.99640678 |
| UBXN2B | 11.0371447 | -0.28980543 | 0.46843574 | -0.6186664 | 0.53613618 | 0.99665574 |
| SDCBP | 15.9822998 | -0.546562317 | 0.4622601 | -1.1823696 | 0.23705906 | 0.9305595 |
| NSMAF | 12.8639428 | -0.452345693 | 0.50905833 | -0.888593 | 0.37422183 | 0.98535336 |
| TOX | 17.4395625 | -0.103983609 | 0.69135106 | -0.1504064 | 0.88044401 | 0.99665574 |
| CA8 | 408.332064 | -0.503445229 | 0.74602604 | -0.674836 | 0.49977997 | 0.99640678 |
| LINC01301 | 5.31442582 | 0.438615717 | 0.67920197 | 0.64578099 | 0.5184212 | 0.99665574 |
| RAB2A | 68.4776133 | -0.182565069 | 0.23979814 | -0.7613281 | 0.44646111 | 0.99617422 |
| CHD7 | 58.2612991 | 0.073972045 | 0.28690232 | 0.25783007 | 0.79653806 | 0.99665574 |
| CLVS1 | 2.34640996 | -0.485727441 | 1.00715381 | -0.4822773 | 0.62960896 | 0.99665574 |
| ASPH | 19.1598784 | -0.359785289 | 0.42475159 | -0.8470487 | 0.39696796 | 0.99179694 |
| YTHDF3 | 29.4789155 | 0.106002349 | 0.33802929 | 0.31358924 | 0.75383303 | 0.99665574 |
| CYP7B1 | 27.4441189 | 0.684243129 | 0.46488881 | 1.47184255 | 0.1410634 | 0.84969914 |
| ARMC1 | 12.4939701 | 0.392078787 | 0.51112762 | 0.76708589 | 0.44303045 | 0.99617422 |
| MTFR1 | 16.5578229 | -0.779676961 | 0.36638641 | -2.1280182 | 0.03333557 | 0.5761693 |
| PDE7A | 56.4694725 | 0.063930999 | 0.40498788 | 0.15785904 | 0.87456787 | 0.99665574 |
| ADHFE1 | 9.88605581 | 0.586729642 | 0.64974823 | 0.90301076 | 0.36652019 | 0.98365837 |
| VCPIP1 | 8.70233204 | -0.137315526 | 0.55468605 | -0.2475554 | 0.80447843 | 0.99665574 |

| | | | | | | |
|---|---|---|---|---|---|---|
| SGK3 | 17.7333625 | 0.136112252 | 0.42022662 | 0.32390202 | 0.74601222 | 0.99665574 |
| CSPP1 | 52.7085755 | 0.635114838 | 0.42725444 | 1.48650262 | 0.13714623 | 0.84522093 |
| ARFGEF1 | 136.568056 | 0.138881288 | 0.26812548 | 0.51797125 | 0.60447833 | 0.99665574 |
| NCOA2 | 149.640458 | 0.008849211 | 0.27135658 | 0.03261101 | 0.97398479 | 0.99777713 |
| TRAM1 | 28.7000134 | 0.079620272 | 0.37675435 | 0.21133206 | 0.83262817 | 0.99665574 |
| LACTB2 | 8.43543189 | -1.36185515 | 0.5268466 | -2.5849178 | 0.00974022 | 0.34168293 |
| TERF1 | 10.5903962 | 0.290788762 | 0.46485436 | 0.6255481 | 0.53161139 | 0.99665574 |
| RPL7 | 78.8503582 | 0.643949185 | 0.46400225 | 1.38781478 | 0.16519345 | 0.87495175 |
| RDH10 | 26.8197055 | 0.253428149 | 0.51347137 | 0.49355848 | 0.62161803 | 0.99665574 |
| RDH10-AS1 | 6.23235856 | -0.721329319 | 0.76755452 | -0.939776 | 0.34733247 | 0.97804112 |
| STAU2 | 45.0037883 | -0.117486181 | 0.30931563 | -0.3798262 | 0.70407443 | 0.99665574 |
| AC027018.1 | 5.45329067 | -0.888688966 | 0.93686329 | -0.9485791 | 0.34283471 | 0.97788427 |
| ELOC | 10.5080063 | -1.054524243 | 0.54402195 | -1.9383855 | 0.05257621 | 0.66213369 |
| TMEM70 | 4.81959223 | -0.311091587 | 0.71435295 | -0.4354872 | 0.66320881 | 0.99665574 |
| GDAP1 | 20.635907 | 0.386924708 | 0.4702017 | 0.82289092 | 0.41057003 | 0.99279547 |
| MIR2052HG | 4.60132975 | 1.192116703 | 0.79991054 | 1.49031253 | 0.13614208 | 0.84475183 |
| HNF4G | 38.4790907 | 0.434798554 | 0.49418144 | 0.87983587 | 0.37894823 | 0.98679664 |
| PEX2 | 12.531078 | -0.315020581 | 0.51352923 | -0.6134424 | 0.53958388 | 0.99665574 |
| PKIA | 5.26986376 | -0.407204198 | 0.86313956 | -0.471771 | 0.63709027 | 0.99665574 |
| ZC2HC1A | 19.0772654 | 0.359546279 | 0.56387111 | 0.63763912 | 0.52370862 | 0.99665574 |
| IL7 | 10.2318917 | -1.368489247 | 0.76567373 | -1.7873008 | 0.07388889 | 0.72810353 |
| TPD52 | 89.5416593 | 0.066267771 | 0.35341299 | 0.18750802 | 0.85126233 | 0.99665574 |
| AC034114.2 | 8.14959101 | 1.302817822 | 1.12506558 | 1.15799278 | 0.24686698 | 0.93441937 |
| ZBTB10 | 18.5509618 | -0.275646547 | 0.54268691 | -0.5079292 | 0.61150297 | 0.99665574 |
| ZNF704 | 58.1869318 | 0.256197096 | 0.48553882 | 0.52765522 | 0.59773866 | 0.99665574 |
| IMPA1 | 12.3702919 | 0.237647035 | 0.4538808 | 0.52358909 | 0.60056437 | 0.99665574 |
| ZFAND1 | 10.3478169 | 0.485664799 | 0.48819502 | 0.99481719 | 0.31982518 | 0.97700345 |
| CHMP4C | 6.85355448 | -0.436582091 | 0.54261909 | -0.804583 | 0.42106036 | 0.99518711 |
| SNX16 | 10.0904418 | -0.123838346 | 0.49415122 | -0.2506082 | 0.80211704 | 0.99665574 |
| RALYL | 17.8027056 | 0.895291922 | 0.91438504 | 0.97911917 | 0.3275211 | 0.97788427 |
| LRRCC1 | 2.63109077 | 0.245254044 | 0.93717128 | 0.26169607 | 0.79355577 | 0.99665574 |
| E2F5 | 7.56081231 | -0.15582792 | 0.62525765 | -0.2492219 | 0.80318911 | 0.99665574 |
| C8orf59 | 8.38943188 | 0.653644975 | 0.56137794 | 1.16435814 | 0.2442789 | 0.93426912 |
| CA13 | 10.1823256 | -0.771599754 | 0.49569143 | -1.5566131 | 0.11956238 | 0.82625998 |
| CA3 | 12.0590773 | -0.670377563 | 0.56181238 | -1.193241 | 0.23277501 | 0.92924083 |
| CA2 | 55.7920683 | 0.684043346 | 0.52030841 | 1.31468824 | 0.1886147 | 0.89739428 |
| ATP6V0D2 | 374.919506 | 0.561803504 | 0.30493603 | 1.8423651 | 0.06542176 | 0.69221233 |
| AC023194.3 | 55.6402328 | 0.481344748 | 0.50945409 | 0.94482458 | 0.34474844 | 0.97788427 |
| WWP1 | 93.8630662 | 0.050654998 | 0.28595794 | 0.17714143 | 0.8593973 | 0.99665574 |
| RMDN1 | 19.7042889 | -0.082274355 | 0.38718061 | -0.2124961 | 0.83172005 | 0.99665574 |
| CPNE3 | 23.0450489 | 0.301787179 | 0.38683839 | 0.78013761 | 0.43530988 | 0.99617422 |
| CNGB3 | 3.77256541 | -1.026816918 | 1.20555322 | -0.8517392 | 0.39435886 | 0.99179694 |
| AF117829.1 | 37.833204 | 0.384395256 | 0.38023218 | 1.01094878 | 0.31204095 | 0.97442101 |
| RIPK2 | 5.43220396 | -1.35966366 | 0.63808527 | -2.1308495 | 0.03310155 | 0.5761693 |
| OSGIN2 | 10.1785072 | 0.481107871 | 0.7127192 | 0.67503144 | 0.49965579 | 0.99640678 |
| NBN | 12.2636077 | -0.133575513 | 0.45312102 | -0.2947899 | 0.76815436 | 0.99665574 |
| DECR1 | 26.2856384 | 0.332408978 | 0.41576694 | 0.79950797 | 0.42399593 | 0.99518711 |
| CALB1 | 14.3971414 | 0.832139079 | 1.94735959 | 0.4273166 | 0.66914874 | 0.99665574 |
| TMEM64 | 3.35462679 | 0.51317942 | 0.88009143 | 0.58309785 | 0.55982743 | 0.99665574 |
| OTUD6B-AS1 | 7.51477976 | 0.272891814 | 0.65429433 | 0.41707807 | 0.67662131 | 0.99665574 |
| OTUD6B | 7.04155247 | 0.980856811 | 0.62185606 | 1.57730522 | 0.11472531 | 0.81638817 |
| LRRC69 | 6.38338408 | 0.106018362 | 0.76300584 | 0.1389483 | 0.88949101 | 0.99665574 |
| SLC26A7 | 4.68897019 | 2.172007885 | 1.18096496 | 1.83918064 | 0.06588862 | 0.69292411 |
| TRIQK | 11.1939931 | -0.528063074 | 0.53147241 | -0.9935851 | 0.3204249 | 0.97700345 |
| LINC00535 | 4.25279127 | -0.795078452 | 0.71615213 | -1.1102089 | 0.26690903 | 0.94557817 |
| FAM92A | 30.238866 | -0.121455682 | 0.45972903 | -0.2641897 | 0.79163373 | 0.99665574 |
| RBM12B | 14.6140196 | 0.530639073 | 0.53489664 | 0.9920404 | 0.32117782 | 0.97700345 |
| TMEM67 | 14.6156938 | -0.455564114 | 0.44775026 | -1.0174514 | 0.30893876 | 0.97376371 |
| PDP1 | 22.1292173 | 0.203804509 | 0.61183892 | 0.33310158 | 0.73905761 | 0.99665574 |
| VIRMA | 15.4717709 | 0.887052167 | 0.48691639 | 1.82177513 | 0.06848911 | 0.70708655 |
| ESRP1 | 19.8487976 | -0.300529353 | 0.42843192 | -0.7014635 | 0.48301381 | 0.99640678 |
| DPY19L4 | 17.027159 | 0.167603041 | 0.44345968 | 0.37794426 | 0.705472 | 0.99665574 |
| INTS8 | 10.7705613 | 0.052768331 | 0.50065021 | 0.1053996 | 0.91605874 | 0.99777713 |
| NDUFAF6 | 11.944513 | 0.206072078 | 0.51550717 | 0.39974629 | 0.68934339 | 0.99665574 |
| TP53INP1 | 7.17728125 | 0.085233133 | 0.53910011 | 0.15810261 | 0.87437595 | 0.99665574 |
| PLEKHF2 | 9.03411911 | 0.904750456 | 0.63926112 | 1.41530655 | 0.15697864 | 0.86532131 |
| C8orf37-AS1 | 21.8789841 | 0.139866483 | 0.75854699 | 0.18438737 | 0.85370958 | 0.99665574 |
| UQCRB | 49.0683111 | 0.699960644 | 0.4170641 | 1.67830473 | 0.09328763 | 0.76763189 |
| MTERF3 | 5.54255167 | 0.755323945 | 0.65786018 | 1.1481527 | 0.25090553 | 0.93973926 |
| PTDSS1 | 5.11434283 | -0.807315401 | 0.65267336 | -1.2369363 | 0.21611072 | 0.91441171 |
| SDC2 | 97.2790444 | 0.467082133 | 0.36211213 | 1.28988261 | 0.19709142 | 0.90381759 |
| CPQ | 71.5295716 | 0.461658883 | 0.34819167 | 1.32587572 | 0.18488087 | 0.89532394 |
| TSPYL5 | 4.38080381 | -0.202209754 | 0.66861441 | -0.302431 | 0.76232349 | 0.99665574 |
| MTDH | 90.406439 | -0.266999393 | 0.34765379 | -0.7680037 | 0.44248499 | 0.99617422 |
| LAPTM4B | 22.9629062 | -0.385147815 | 0.37186278 | -1.0357256 | 0.30033016 | 0.97358745 |
| RPL30 | 54.9045089 | 0.211348433 | 0.40932908 | 0.51632889 | 0.60562472 | 0.99665574 |
| RIDA | 5.08329795 | 0.462186545 | 0.76652908 | 0.60296022 | 0.54653515 | 0.99665574 |
| POP1 | 3.76217609 | -1.191405137 | 0.68086463 | -1.7498414 | 0.08014568 | 0.74542836 |
| NIPAL2 | 7.39429989 | -1.241790831 | 0.6493558 | -1.9123427 | 0.05583225 | 0.67320387 |
| STK3 | 180.113093 | -0.112080398 | 0.27466817 | -0.4080575 | 0.68323149 | 0.99665574 |
| AC104986.2 | 3.89206721 | -0.470861015 | 0.7911242 | -0.5951796 | 0.5517234 | 0.99665574 |
| VPS13B | 112.716103 | -0.086431382 | 0.23026285 | -0.3753596 | 0.70739301 | 0.99665574 |
| COX6C | 51.4623515 | -0.016160071 | 0.29438747 | -0.0548939 | 0.95622301 | 0.99777713 |
| POLR2K | 9.90995211 | 0.59047452 | 0.72286268 | 0.81685573 | 0.41401088 | 0.99279547 |
| SPAG1 | 8.77723055 | -0.018173839 | 0.54475677 | -0.0333614 | 0.9733864 | 0.99777713 |
| RNF19A | 47.8480317 | -0.054927852 | 0.42228772 | -0.1300721 | 0.89650938 | 0.99777713 |
| ANKRD46 | 8.49584883 | 0.044045777 | 0.59180491 | 0.07442618 | 0.94067128 | 0.99777713 |
| PABPC1 | 48.3032377 | -0.150463885 | 0.33049348 | -0.4552704 | 0.64891475 | 0.99665574 |
| YWHAZ | 68.1492721 | 0.10645566 | 0.36267792 | 0.29352672 | 0.76911958 | 0.99665574 |
| ZNF706 | 18.84545 | 0.102275028 | 0.36332896 | 0.28149429 | 0.77833131 | 0.99665574 |
| GRHL2 | 48.7090705 | -0.087328718 | 0.29720375 | -0.2938345 | 0.76888436 | 0.99665574 |
| RRM2B | 15.7291912 | -0.197295759 | 0.38911614 | -0.5070357 | 0.6121298 | 0.99665574 |
| UBR5-AS1 | 4.46381329 | -0.745640645 | 0.83321961 | -0.8948909 | 0.37084541 | 0.98532838 |
| UBR5 | 88.3771466 | -0.136464498 | 0.26515789 | -0.5146537 | 0.60679501 | 0.99665574 |
| KLF10 | 11.4540989 | -1.119489151 | 0.6131946 | -1.825667 | 0.06790043 | 0.70530184 |
| AZIN1 | 18.875176 | 0.209649837 | 0.42033624 | 0.49876699 | 0.61794355 | 0.99665574 |
| AZIN1-AS1 | 16.2382282 | 0.044923381 | 0.51354157 | 0.08747759 | 0.93029189 | 0.99777713 |
| ATP6V1C1 | 22.0283067 | -0.535643332 | 0.551171 | -0.9718279 | 0.33113619 | 0.97788427 |
| BAALC | 7.14040251 | 0.352516838 | 0.74137331 | 0.47549168 | 0.63443657 | 0.99665574 |
| BAALC-AS1 | 6.38434569 | 0.279042002 | 0.74794419 | 0.37307864 | 0.70908991 | 0.99665574 |
| FZD6 | 25.8213982 | 0.228539997 | 0.31074452 | 0.73545945 | 0.46205973 | 0.99617422 |
| SLC25A32 | 11.0431511 | 0.364473713 | 0.50460525 | 0.72229472 | 0.4701133 | 0.99617422 |
| DCAF13 | 7.06679897 | 0.387069647 | 0.60845238 | 0.63615438 | 0.5246758 | 0.99665574 |
| DPYS | 7.51989332 | 1.521556246 | 1.03745999 | 1.46661679 | 0.14248033 | 0.84969914 |
| LRP12 | 18.7635827 | 0.091529226 | 0.44361658 | 0.20632508 | 0.83653699 | 0.99665574 |
| ZFPM2 | 9.5964489 | -1.793055193 | 0.78200083 | -2.292907 | 0.02185336 | 0.47850999 |
| AC027031.2 | 2.56653081 | -0.100070994 | 0.87680555 | -0.1141313 | 0.90913368 | 0.99777713 |
| OXR1 | 241.333849 | 0.487326151 | 0.31476345 | 1.54822979 | 0.12156698 | 0.82659025 |
| AC090579.1 | 24.1285178 | -0.01063386 | 0.7697853 | -0.0138141 | 0.98897833 | 0.99777713 |
| ANGPT1 | 13.2450556 | 2.303053796 | 0.78140589 | 2.94732077 | 0.00320541 | 0.17855163 |
| EIF3E | 26.6995501 | 0.486840234 | 0.3971258 | 1.22590935 | 0.22023279 | 0.91768188 |
| EMC2 | 17.7969142 | 0.134703435 | 0.53434318 | 0.25209161 | 0.80097025 | 0.99665574 |
| NUDCD1 | 8.03280471 | 0.764497343 | 0.56512914 | 1.35278345 | 0.17612482 | 0.88428483 |

| | | | | | | |
|---|---|---|---|---|---|---|
| ENY2 | 14.4377409 | 0.02742806 | 0.49249694 | 0.05569184 | 0.9555873 | 0.99777713 |
| EBAG9 | 8.54438112 | 0.153755333 | 0.50309769 | 0.30561725 | 0.75989609 | 0.99665574 |
| CSMD3 | 134.322404 | 0.287701852 | 0.62914204 | 0.45729237 | 0.64746092 | 0.99665574 |
| TRPS1 | 83.3898233 | 0.186518402 | 0.34745414 | 0.53681445 | 0.59139579 | 0.99665574 |
| EIF3H | 43.9624356 | 0.327510383 | 0.28311711 | 1.15680179 | 0.24735335 | 0.93441937 |
| UTP23 | 4.35538545 | -0.188919799 | 0.69030889 | -0.2736743 | 0.78433493 | 0.99665574 |
| RAD21 | 25.5693227 | 0.063693216 | 0.35171781 | 0.18109181 | 0.85629551 | 0.99665574 |
| MED30 | 3.16972207 | -0.673730924 | 0.85051188 | -0.7921476 | 0.42827463 | 0.99589919 |
| EXT1 | 179.009045 | 0.07621724 | 0.3205632 | 0.23776042 | 0.81206692 | 0.99665574 |
| SAMD12 | 190.219964 | -0.171034463 | 0.34417812 | -0.4969359 | 0.61923426 | 0.99665574 |
| AC023590.1 | 19.1667879 | -1.147999029 | 0.79269159 | -1.4482291 | 0.14755299 | 0.85284617 |
| SAMD12-AS1 | 16.5147657 | -0.930569114 | 0.44649359 | -2.0841713 | 0.03714459 | 0.58591936 |
| MAL2 | 41.4442293 | 0.583547154 | 0.33994475 | 1.71659412 | 0.08605336 | 0.75397496 |
| TAF2 | 25.9004864 | 0.016465766 | 0.40662377 | 0.04049386 | 0.9676994 | 0.99777713 |
| DEPTOR | 49.516553 | -0.844775571 | 0.5111331 | -1.6527507 | 0.09838162 | 0.78203824 |
| MRPL13 | 9.432395 | 1.024896392 | 0.66169641 | 1.54889218 | 0.12140764 | 0.82659025 |
| MTBP | 6.02544231 | -0.612080662 | 0.62597419 | -0.9778049 | 0.32817081 | 0.97788427 |
| SNTB1 | 402.455127 | -0.318057987 | 0.24409481 | -1.30301 | 0.19257134 | 0.89967199 |
| AC104958.1 | 5.82413777 | -1.260105396 | 1.01350659 | -1.2433125 | 0.21375271 | 0.91243155 |
| ZHX2 | 69.5224441 | -0.763468026 | 0.27777912 | -2.7484716 | 0.00598738 | 0.24620771 |
| DERL1 | 7.66085767 | 0.665603152 | 0.57364965 | 1.16029559 | 0.24592848 | 0.93426912 |
| TBC1D31 | 3.23911158 | 0.34129995 | 0.83394973 | 0.40925722 | 0.6823509 | 0.99665574 |
| C8orf76 | 3.148523 | 0.501910533 | 0.76128382 | 0.65929489 | 0.50970642 | 0.99665574 |
| WDYHV1 | 6.74171511 | -0.097132611 | 0.60750446 | -0.1598879 | 0.87296938 | 0.99665574 |
| FBXO32 | 13.6912163 | -0.560091216 | 0.60715981 | -0.9224774 | 0.3562796 | 0.9819332 |
| FAM91A1 | 23.242862 | -0.180415559 | 0.35509741 | -0.5080734 | 0.61140185 | 0.99665574 |
| TMEM65 | 47.1351628 | 0.73263942 | 0.32334251 | 2.2658308 | 0.02346174 | 0.49347545 |
| RNF139 | 2.77352728 | 0.064226614 | 0.90471514 | 0.07099098 | 0.94340494 | 0.99777713 |
| TATDN1 | 16.2741564 | 0.173219428 | 0.44664243 | 0.38782573 | 0.698145 | 0.99665574 |
| NDUFB9 | 22.3854586 | 0.509220961 | 0.53487244 | 0.95204188 | 0.34107574 | 0.97788427 |
| MTSS1 | 55.8047635 | -0.590980392 | 0.36402097 | -1.623479 | 0.10448705 | 0.79169635 |
| SQLE | 8.33725996 | -2.323765856 | 0.8360555 | -2.7794397 | 0.00544528 | 0.2290631 |
| WASHC5 | 16.6944056 | -0.486644923 | 0.40941677 | -1.1886297 | 0.23458542 | 0.92960234 |
| NSMCE2 | 103.28128 | -0.166613745 | 0.29353338 | -0.5676143 | 0.57029689 | 0.99665574 |
| TRIB1 | 14.1175927 | -1.392431756 | 0.58467792 | -2.3815364 | 0.01724059 | 0.43370986 |
| FAM84B | 9.9118566 | 0.500304105 | 0.58960448 | 0.8485419 | 0.39613625 | 0.99179694 |
| PCAT1 | 58.3337865 | -0.474228138 | 0.28466799 | -1.6658991 | 0.09573352 | 0.77210482 |
| PVT1 | 12.4401336 | 0.030189837 | 0.51949713 | 0.05811358 | 0.95365816 | 0.99777713 |
| FAM49B | 75.1975136 | -0.101257393 | 0.22715833 | -0.4457569 | 0.65577286 | 0.99665574 |
| ASAP1 | 33.2590863 | -0.890156606 | 0.35716553 | -2.4922803 | 0.01269258 | 0.38590337 |
| EFR3A | 34.2804238 | 0.112249024 | 0.32937131 | 0.34079782 | 0.73325579 | 0.99665574 |
| KCNQ3 | 9.78022493 | -1.429254169 | 1.61932413 | -0.8826239 | 0.37743951 | 0.98679664 |
| LRRC6 | 6.02525161 | -0.566990515 | 0.77059418 | -0.7357835 | 0.46186244 | 0.99617422 |
| PHF20L1 | 32.8420186 | 0.378405116 | 0.38504296 | 0.98276076 | 0.3257252 | 0.97788427 |
| TG | 6.36919204 | -1.331556268 | 1.07896111 | -1.2341096 | 0.21716206 | 0.91584128 |
| NDRG1 | 136.102736 | -0.370565122 | 0.54850972 | -0.6755853 | 0.49930396 | 0.99640678 |
| ST3GAL1 | 68.0472203 | -0.603018665 | 0.41181612 | -1.4642911 | 0.14311444 | 0.84969914 |
| ZFAT | 12.1878124 | -0.477396709 | 0.54400009 | -0.8775673 | 0.38017857 | 0.98679664 |
| AC083843.3 | 6.70784668 | 0.847604544 | 0.56112131 | 1.5105549 | 0.13090189 | 0.83833441 |
| KHDRBS3 | 27.667053 | 0.637210313 | 0.51864984 | 1.22859444 | 0.2192239 | 0.9168696 |
| TRAPPC9 | 119.955277 | -0.233933279 | 0.31255564 | -0.7484532 | 0.45418682 | 0.99617422 |
| CHRAC1 | 6.764939 | 0.409120172 | 0.58411918 | 0.7004053 | 0.48367423 | 0.99640678 |
| AGO2 | 28.3695448 | -0.303868497 | 0.41187983 | -0.7377601 | 0.46066025 | 0.99617422 |
| PTK2 | 174.475961 | -0.169566036 | 0.26096791 | -0.6497582 | 0.51584843 | 0.99665574 |
| DENND3 | 4.14989237 | -1.832335398 | 0.80778905 | -2.268334 | 0.02330885 | 0.49335204 |
| SLC45A4 | 13.575012 | 0.363478351 | 0.52533117 | 0.69190326 | 0.48899808 | 0.99640678 |
| PTP4A3 | 5.13606415 | -2.220245723 | 0.79048866 | -2.8087003 | 0.00497419 | 0.22209534 |
| TSNARE1 | 11.2023375 | 0.471205826 | 0.47817271 | 0.98543019 | 0.3244128 | 0.97700345 |
| JRK | 3.21207204 | 0.126738952 | 0.76337376 | 0.16602477 | 0.86813746 | 0.99665574 |
| THEM6 | 4.08465301 | 0.543305349 | 0.81075391 | 0.67012363 | 0.50277899 | 0.99640678 |
| LY6E | 8.78405015 | -0.478900596 | 1.16016651 | -0.4127861 | 0.67976334 | 0.99665574 |
| GLI4 | 2.21672264 | -0.204137929 | 0.89935396 | -0.2269829 | 0.82043707 | 0.99665574 |
| TOP1MT | 4.22178685 | 0.852459895 | 0.7268244 | 1.17285537 | 0.24085381 | 0.93216487 |
| RHPN1 | 2.745751 | 1.69998204 | 0.97295053 | 1.74724407 | 0.08059501 | 0.74618467 |
| ZC3H3 | 6.64250302 | -0.461413433 | 0.63565942 | -0.7258815 | 0.4679114 | 0.99617422 |
| GSDMD | 7.67586823 | -0.333352436 | 0.65432546 | -0.5094597 | 0.61043006 | 0.99665574 |
| NAPRT | 2.91034659 | 0.792599863 | 0.88998546 | 0.89057619 | 0.37315658 | 0.98532838 |
| EEF1D | 51.3618834 | 0.621366773 | 0.40964667 | 1.51683589 | 0.12930812 | 0.83778831 |
| TSTA3 | 3.91349562 | 0.053364284 | 0.80377944 | 0.0663917 | 0.94706598 | 0.99777713 |
| ZNF623 | 6.24318294 | 0.25282525 | 0.62477623 | 0.40466528 | 0.68572357 | 0.99665574 |
| FAM83H | 7.18030987 | -0.163309365 | 0.57276926 | -0.2851224 | 0.77555034 | 0.99665574 |
| IQANK1 | 3.10022322 | -0.490375855 | 0.75225396 | -0.6518754 | 0.51448155 | 0.99665574 |
| SCRIB | 7.23821124 | 0.400113702 | 0.5575674 | 0.71760598 | 0.47300027 | 0.99617422 |
| PUF60 | 7.99153067 | -0.077689959 | 0.68582451 | -0.1132796 | 0.90980885 | 0.99777713 |
| NRBP2 | 4.30658206 | -0.406771271 | 0.8130648 | -0.5002938 | 0.61686823 | 0.99665574 |
| PLEC | 8.33311005 | -0.315427993 | 0.55898801 | -0.564284 | 0.57256087 | 0.99665574 |
| PARP10 | 5.60687037 | 0.756549748 | 0.79922502 | 0.94660418 | 0.34384051 | 0.97788427 |
| GRINA | 10.562003 | 1.023510812 | 0.51021717 | 2.00602972 | 0.04485308 | 0.6313648 |
| OPLAH | 1.74625789 | 2.336251831 | 1.33698791 | 1.74739937 | 0.08056809 | 0.74618467 |
| EXOSC4 | 2.53326977 | 0.387548047 | 0.96308072 | 0.40240453 | 0.68738633 | 0.99665574 |
| GPAA1 | 10.4239679 | 0.039065873 | 0.5345071 | 0.07308766 | 0.94173636 | 0.99777713 |
| CYC1 | 17.188766 | 0.439500702 | 0.55640249 | 0.78989708 | 0.42958788 | 0.99617422 |
| SHARPIN | 6.80570579 | -0.558233311 | 0.5478115 | -1.0190245 | 0.30819136 | 0.97376371 |
| MAF1 | 9.48259454 | -0.144629828 | 0.54345342 | -0.266131 | 0.79013829 | 0.99665574 |
| MROH1 | 28.5521495 | -0.573147213 | 0.44795913 | -1.2794632 | 0.200734 | 0.90567036 |
| BOP1 | 1.97306213 | 1.72513126 | 1.25628858 | 1.37319664 | 0.16969122 | 0.87861574 |
| HSF1 | 14.2596202 | -0.191836407 | 0.44525947 | -0.4308418 | 0.66658338 | 0.99665574 |
| DGAT1 | 4.77000723 | 1.226006148 | 0.68837767 | 1.781008 | 0.07491114 | 0.72883798 |
| SLC52A2 | 5.84901457 | -0.473001276 | 0.61068503 | -0.7745421 | 0.43861027 | 0.99617422 |
| ADCK5 | 2.20500091 | 0.2922257 | 0.9483585 | 0.30813843 | 0.757977 | 0.99665574 |
| CPSF1 | 3.54356598 | 0.141077145 | 0.90211554 | 0.15638478 | 0.87572973 | 0.99665574 |
| VPS28 | 21.5861803 | -0.514419851 | 0.40613979 | -1.2666079 | 0.20529554 | 0.90846188 |
| CYHR1 | 11.292155 | 0.651261036 | 0.50665489 | 1.2854135 | 0.19864782 | 0.90381759 |
| KIFC2 | 4.13832829 | -0.066716285 | 0.77504338 | -0.0860807 | 0.93140226 | 0.99777713 |
| PPP1R16A | 7.1870758 | 1.131763706 | 0.60527491 | 1.86983417 | 0.06150685 | 0.6834742 |
| LRRC14 | 2.33003109 | 0.496731757 | 0.91313529 | 0.54398484 | 0.5864519 | 0.99665574 |
| ARHGAP39 | 4.46798326 | 0.315620732 | 0.65884065 | 0.47905474 | 0.63189969 | 0.99665574 |
| ZNF251 | 17.8852825 | -0.254626203 | 0.34124498 | -0.7461683 | 0.45556573 | 0.99617422 |
| RPL8 | 104.014972 | 0.478317729 | 0.39441812 | 1.21271744 | 0.22523787 | 0.92318867 |
| ZNF7 | 5.60112166 | -1.223507667 | 0.70379063 | -1.7384541 | 0.08213084 | 0.75124322 |
| COMMD5 | 3.66877641 | 0.146834695 | 0.75765958 | 0.19380035 | 0.84633221 | 0.99665574 |
| ZNF250 | 5.42445795 | 0.270502984 | 0.67629957 | 0.39997509 | 0.68917486 | 0.99665574 |
| ZNF16 | 4.07713424 | -0.815133917 | 0.63888532 | -1.2758689 | 0.20200186 | 0.90614933 |
| C8orf33 | 5.97225327 | -0.249025496 | 0.58976956 | -0.422242 | 0.67284837 | 0.99665574 |
| WASHC1 | 25.524864 | 0.621301196 | 0.3589275 | 1.73099357 | 0.08345291 | 0.75242737 |
| CBWD1 | 7.44346695 | -0.124008399 | 0.65843901 | -0.188337 | 0.85061252 | 0.99665574 |
| DOCK8 | 84.2992821 | -0.472924478 | 0.47777946 | -0.9898384 | 0.32225309 | 0.97700345 |
| KANK1 | 91.0244506 | 0.350947421 | 0.30754767 | 1.14111552 | 0.25382185 | 0.94099699 |
| SMARCA2 | 44.9535164 | -0.106611724 | 0.44863211 | -0.2376373 | 0.81216242 | 0.99665574 |
| PUM3 | 35.6745677 | -0.41839205 | 0.41859093 | -0.9995249 | 0.3175405 | 0.97700345 |
| RFX3 | 31.7629159 | 0.377983924 | 0.41463693 | 0.91160217 | 0.36197818 | 0.98226306 |
| RFX3-AS1 | 22.6750096 | -0.330383629 | 0.35860953 | -0.9212907 | 0.35689867 | 0.9819332 |
| GLIS3 | 115.712653 | -1.686358705 | 0.39111613 | -4.3116573 | 1.62E-05 | 0.00586684 |

| | | | | | | |
|---|---|---|---|---|---|---|
| SLC1A1 | 16.9528258 | -2.068387952 | 0.48496648 | -4.2650122 | 2.00E-05 | 0.00698792 |
| SPATA6L | 16.5339466 | -1.39088096 | 0.49243453 | -2.8244992 | 0.00473546 | 0.21929238 |
| CDC37L1 | 13.3453998 | 0.053154317 | 0.48666633 | 0.10922128 | 0.91302699 | 0.99777713 |
| AK3 | 52.1686704 | 0.297414501 | 0.3656203 | 0.81345184 | 0.41595905 | 0.992853 |
| RCL1 | 11.5664473 | 0.681330346 | 0.50138901 | 1.35888569 | 0.17418282 | 0.88281284 |
| JAK2 | 18.0224986 | 0.446761184 | 0.42191258 | 1.05889516 | 0.28964753 | 0.96676791 |
| PLGRKT | 10.947386 | -0.289368722 | 0.46588336 | -0.6211184 | 0.53452173 | 0.99665574 |
| RIC1 | 39.2814005 | -0.165031906 | 0.3965917 | -0.4161255 | 0.6773182 | 0.99665574 |
| ERMP1 | 33.4867433 | 0.924752099 | 0.37743784 | 2.45007787 | 0.01428253 | 0.40109782 |
| KIAA2026 | 49.1224719 | 0.278374851 | 0.28448767 | 0.97851289 | 0.32782072 | 0.97788427 |
| RANBP6 | 5.36096015 | 0.220406678 | 0.70543577 | 0.31244046 | 0.75470581 | 0.99665574 |
| UHRF2 | 24.7206195 | 0.004963686 | 0.38461976 | 0.01290544 | 0.98970324 | 0.99777713 |
| GLDC | 4.30733988 | 0.454831471 | 0.73087262 | 0.62231292 | 0.53373613 | 0.99665574 |
| KDM4C | 92.723693 | -0.467378996 | 0.23848905 | -1.9597503 | 0.05002498 | 0.64605506 |
| DMAC1 | 7.11166655 | 0.313783774 | 0.72068998 | 0.43539356 | 0.66327679 | 0.99665574 |
| PTPRD | 11.9946394 | 0.767602587 | 0.602685 | 1.27363811 | 0.2027917 | 0.9066717 |
| MPDZ | 17.169851 | 0.281182824 | 0.40637779 | 0.69192468 | 0.48898463 | 0.99640678 |
| NFIB | 37.5883803 | 0.099778053 | 0.58095823 | 0.17174738 | 0.86363614 | 0.99665574 |
| ZDHHC21 | 58.2613401 | 0.589214932 | 0.30234543 | 1.94881377 | 0.05131767 | 0.65359112 |
| FREM1 | 111.051678 | 1.183615232 | 0.38972578 | 3.03704627 | 0.00238909 | 0.15249081 |
| TTC39B | 78.0503206 | 0.320683707 | 0.40066874 | 0.80037116 | 0.42349578 | 0.99518711 |
| SNAPC3 | 21.4429732 | 0.007692435 | 0.34325161 | 0.02241049 | 0.98212052 | 0.99777713 |
| PSIP1 | 10.9357376 | 0.125956388 | 0.6247868 | 0.20159899 | 0.84023024 | 0.99665574 |
| CCDC171 | 21.6647377 | -0.836577441 | 0.39730368 | -2.1056373 | 0.03523587 | 0.58349146 |
| BNC2 | 28.0996785 | 0.36059546 | 0.58186447 | 0.61972414 | 0.53543942 | 0.99665574 |
| CNTLN | 16.7020566 | -0.595967198 | 0.64450281 | -0.9246929 | 0.35512566 | 0.98022903 |
| SH3GL2 | 37.2370406 | -1.690287359 | 1.30074873 | -1.2994726 | 0.19378179 | 0.90235789 |
| ADAMTSL1 | 101.779811 | 0.647340134 | 0.6261628 | 1.0338208 | 0.30121994 | 0.97358745 |
| RRAGA | 5.30077502 | -1.042932181 | 0.75268987 | -1.3856068 | 0.16586701 | 0.87495175 |
| HAUS6 | 4.06381388 | 0.117767232 | 0.9717333 | 0.12119296 | 0.9035382 | 0.99777713 |
| PLIN2 | 13.3538691 | -1.220394649 | 0.70554878 | -1.7297098 | 0.08368213 | 0.75242737 |
| DENND4C | 60.982452 | 0.018232123 | 0.25420785 | 0.07172132 | 0.94282369 | 0.99777713 |
| RPS6 | 103.450224 | 0.276867351 | 0.44874537 | 0.61698097 | 0.53724728 | 0.99665574 |
| SLC24A2 | 11.848799 | 1.9302943 | 0.63854179 | 3.02297255 | 0.00250305 | 0.15473119 |
| MLLT3 | 100.819256 | -0.194624436 | 0.46637459 | -0.4173136 | 0.67644908 | 0.99665574 |
| FOCAD | 46.9231389 | 0.07820232 | 0.50210862 | 0.15574781 | 0.87623181 | 0.99665574 |
| KLHL9 | 9.49213922 | 0.71185883 | 0.51693118 | 1.37708628 | 0.16848558 | 0.87566218 |
| LINC01239 | 3.53708794 | -0.621860866 | 1.15545672 | -0.5381949 | 0.59044253 | 0.99665574 |
| AL391117.1 | 5.14790129 | 1.814308585 | 1.1158747 | 1.62590708 | 0.10396942 | 0.79072919 |
| TUSC1 | 6.93542662 | 0.576928801 | 0.66330336 | 0.86978122 | 0.38441998 | 0.99040654 |
| CAAP1 | 30.8650719 | 0.728514989 | 0.36250709 | 2.00965723 | 0.04446748 | 0.63050533 |
| PLAA | 9.79670651 | 0.096299997 | 0.6166865 | 0.15615713 | 0.87590916 | 0.99665574 |
| IFT74 | 16.6013293 | 0.173056085 | 0.47161267 | 0.36694537 | 0.71365977 | 0.99665574 |
| MOB3B | 48.3377141 | -0.175830811 | 0.436378 | -0.4029323 | 0.68699799 | 0.99665574 |
| C9orf72 | 6.64666759 | -0.276979916 | 0.61378002 | -0.451269 | 0.65179566 | 0.99665574 |
| ACO1 | 11.421579 | 0.411721115 | 0.52831394 | 0.77931148 | 0.43579625 | 0.99617422 |
| DDX58 | 21.4638507 | -0.800410991 | 0.96184644 | -0.8321609 | 0.40531814 | 0.99194293 |
| TOPORS | 6.88170171 | -0.203585421 | 0.59645159 | -0.3413277 | 0.73285693 | 0.99665574 |
| NDUFB6 | 12.9372124 | -0.356819398 | 0.40590107 | -0.8790797 | 0.37935805 | 0.98679664 |
| APTX | 15.3336383 | -0.44380319 | 0.39999852 | -1.1095121 | 0.26720934 | 0.94557817 |
| DNAJA1 | 23.9350165 | -0.437520814 | 0.37723264 | -1.159817 | 0.24612334 | 0.93441937 |
| SMU1 | 11.6488952 | -0.483622044 | 0.45352961 | -1.0663516 | 0.28626471 | 0.96449041 |
| B4GALT1 | 26.3341275 | -0.515391213 | 0.35196196 | -1.4643378 | 0.14310167 | 0.84969914 |
| BAG1 | 17.2393841 | 0.121242331 | 0.45030604 | 0.26924429 | 0.7877417 | 0.99665574 |
| CHMP5 | 14.0067998 | 0.083660972 | 0.59274733 | 0.14114104 | 0.88775853 | 0.99665574 |
| NFX1 | 38.3183726 | -0.894335611 | 0.36360792 | -2.4596153 | 0.0139086 | 0.39608255 |
| AQP7 | 7.08061582 | 0.190257204 | 0.78533218 | 0.24226335 | 0.80857611 | 0.99665574 |
| AQP3 | 16.5973842 | 0.147262011 | 0.92776564 | 0.1587276 | 0.87388349 | 0.99665574 |
| UBE2R2 | 49.3625678 | 0.092247468 | 0.3066779 | 0.30079594 | 0.7635701 | 0.99665574 |
| UBAP2 | 40.0317598 | -0.028545821 | 0.42267602 | -0.0675359 | 0.94615505 | 0.99777713 |
| DCAF12 | 8.45346678 | 0.252154503 | 0.48329053 | 0.52174517 | 0.60184777 | 0.99665574 |
| UBAP1 | 51.6585388 | -0.592655261 | 0.38073091 | -1.556625 | 0.11955954 | 0.82625998 |
| NUDT2 | 4.61368506 | -1.077204553 | 0.80481774 | -1.3384453 | 0.18075131 | 0.89083946 |
| FAM219A | 14.2958154 | -0.50094164 | 0.47394241 | -1.0569673 | 0.2905265 | 0.96676791 |
| RPP25L | 4.94037524 | 0.107402251 | 0.74315554 | 0.1445219 | 0.88508836 | 0.99665574 |
| DCTN3 | 5.56080655 | 0.765840121 | 0.78901783 | 0.97062461 | 0.33173525 | 0.97788427 |
| VCP | 21.5236301 | -0.387587324 | 0.47859818 | -0.8098387 | 0.41803289 | 0.99432434 |
| PIGO | 3.32371669 | 1.239623141 | 0.83178931 | 1.49030906 | 0.13614299 | 0.84475183 |
| STOML2 | 7.43799265 | -0.234806547 | 0.58838124 | -0.3990721 | 0.68984007 | 0.99665574 |
| UNC13B | 58.6911068 | 0.451850867 | 0.26600797 | 1.69863657 | 0.08938768 | 0.75955113 |
| RUSC2 | 11.424567 | -1.101488443 | 0.62224747 | -1.7701775 | 0.07669758 | 0.73943494 |
| TESK1 | 3.53538573 | 1.063609587 | 0.90062983 | 1.18096198 | 0.23761783 | 0.9305595 |
| CCDC107 | 5.55297263 | 0.232676308 | 0.72198326 | 0.32227382 | 0.74724526 | 0.99665574 |
| TLN1 | 26.0249864 | -0.091851135 | 0.41287182 | -0.2224689 | 0.82394889 | 0.99665574 |
| CREB3 | 5.15319022 | -0.483459229 | 0.6722337 | -0.7191833 | 0.47202801 | 0.99617422 |
| GBA2 | 20.0437286 | -0.089462605 | 0.45256708 | -0.1976781 | 0.84329692 | 0.99665574 |
| RGP1 | 8.80204953 | -0.198082259 | 0.53134568 | -0.3727936 | 0.70930208 | 0.99665574 |
| AL133410.1 | 7.03922093 | -0.778180473 | 0.68481977 | -1.1363289 | 0.25581895 | 0.94314473 |
| HINT2 | 4.90705683 | 0.788053682 | 0.65827775 | 1.19714464 | 0.23125018 | 0.92774608 |
| TMEM8B | 3.79323663 | 1.531459204 | 0.77678759 | 1.97152892 | 0.04866341 | 0.64337675 |
| RECK | 5.17948951 | -0.228275148 | 0.66172981 | -0.3449673 | 0.73011895 | 0.99665574 |
| CLTA | 55.2310845 | -0.383196571 | 0.32242546 | -1.1884811 | 0.23464392 | 0.92960234 |
| GNE | 13.3816434 | 0.227639361 | 0.5144288 | 0.44250897 | 0.65812094 | 0.99665574 |
| RNF38 | 79.2562824 | -0.096350826 | 0.27025226 | -0.3565218 | 0.72144982 | 0.99665574 |
| EBLN3P | 13.9295906 | 0.305213034 | 0.45225892 | 0.6748635 | 0.49976249 | 0.99640678 |
| ZCCHC7 | 105.9755 | -0.142952035 | 0.32402871 | -0.4411709 | 0.65908928 | 0.99665574 |
| GRHPR | 10.986443 | -0.109549483 | 0.51048223 | -0.2146 | 0.8300792 | 0.99665574 |
| ZBTB5 | 5.85644502 | -0.163930336 | 0.63605907 | -0.2577282 | 0.79661671 | 0.99665574 |
| AL138752.2 | 14.7683059 | -0.793492663 | 0.4132847 | -1.9199662 | 0.05486217 | 0.67040291 |
| FRMPD1 | 2.96103464 | 0.820080147 | 0.94721709 | 0.86577845 | 0.38661167 | 0.99126684 |
| TRMT10B | 12.3273625 | 0.015888731 | 0.47725142 | 0.03329216 | 0.9734416 | 0.99777713 |
| DCAF10 | 36.4799551 | 0.259946839 | 0.31687102 | 0.82035535 | 0.41201356 | 0.99279547 |
| ANKRD18A | 2.89624372 | -1.258093191 | 0.81911383 | -1.5359198 | 0.12455804 | 0.83196528 |
| GLIDR | 9.63245436 | 1.259162328 | 0.52752619 | 2.38691908 | 0.01699023 | 0.42847502 |
| BMS1P14 | 4.46319766 | 1.345026571 | 0.86097532 | 1.56221269 | 0.11823789 | 0.82611699 |
| CNTNAP3B | 20.6714684 | 0.058984737 | 0.662758 | 0.08899891 | 0.92908278 | 0.99777713 |
| FAM27C | 3.75583178 | 0.678386256 | 0.80525292 | 0.84245116 | 0.39953547 | 0.99179694 |
| CBWD5 | 7.95080965 | -0.562014852 | 0.53111628 | -1.0581767 | 0.28997491 | 0.96676791 |
| ZNF658 | 3.75061404 | -0.127135064 | 0.71406414 | -0.1780443 | 0.85868817 | 0.99665574 |
| PGM5 | 5.25218697 | -0.179705014 | 0.77968033 | -0.2304855 | 0.81771453 | 0.99665574 |
| PIP5K1B | 56.8601987 | -0.513782621 | 0.72494613 | -0.7087183 | 0.47849928 | 0.99617422 |
| FAM189A2 | 11.7096044 | -1.00921364 | 0.71384484 | -1.4137717 | 0.15742894 | 0.86532131 |
| PTAR1 | 34.3161874 | 0.463854935 | 0.31272124 | 1.48328566 | 0.13799853 | 0.84656101 |
| MAMDC2-AS | 9.48677894 | 0.14078585 | 0.53356434 | 0.26385918 | 0.79188843 | 0.99665574 |
| MAMDC2 | 52.8472638 | 1.436108319 | 0.5624435 | 2.55333792 | 0.01066959 | 0.35936322 |
| SMC5-AS1 | 15.9566946 | -0.323383522 | 0.38325898 | -0.8437728 | 0.39879635 | 0.99179694 |
| SMC5 | 43.9561922 | -0.484182691 | 0.31900884 | -1.517772 | 0.12907188 | 0.83778831 |
| KLF9 | 10.772364 | 0.133334776 | 0.6135525 | 0.217316 | 0.82796209 | 0.99665574 |
| TRPM3 | 21.8266657 | 1.626852023 | 1.29245595 | 1.25872918 | 0.20812817 | 0.91026892 |
| CEMIP2 | 62.7325083 | -1.41166189 | 0.47405483 | -2.9778452 | 0.00290283 | 0.16913131 |
| ABHD17B | 9.92982432 | 0.490810333 | 0.50215411 | 0.97740977 | 0.32836633 | 0.97788427 |
| C9orf85 | 19.8199791 | 0.265652176 | 0.40154718 | 0.66157152 | 0.50824587 | 0.99665574 |
| GDA | 8.9617016 | -0.811540387 | 0.90681762 | -0.8949323 | 0.37082328 | 0.98532838 |

| | | | | | | |
|---|---|---|---|---|---|---|
| ZFAND5 | 65.919427 | 0.184336737 | 0.32544703 | 0.56641088 | 0.5711145 | 0.99665574 |
| ALDH1A1 | 28.3635622 | -0.485746687 | 0.5092665 | -0.9538163 | 0.34017663 | 0.97788427 |
| C9orf41-AS1 | 3.41839295 | 0.186668919 | 0.98278712 | 0.1899383 | 0.84935748 | 0.99665574 |
| CARNMT1 | 24.3700157 | 0.541477826 | 0.36207355 | 1.49549123 | 0.13478629 | 0.84373478 |
| NMRK1 | 7.81150561 | -0.356966619 | 0.73613881 | -0.4849175 | 0.62773484 | 0.99665574 |
| OSTF1 | 10.6484544 | -0.370401493 | 0.4921893 | -0.752559 | 0.45171496 | 0.99617422 |
| PCSK5 | 5.79638786 | -1.210125471 | 0.91321979 | -1.3251196 | 0.18513148 | 0.89532394 |
| RFK | 10.3979515 | 0.129272553 | 0.49189729 | 0.26280395 | 0.79270169 | 0.99665574 |
| GCNT1 | 12.0356239 | -0.909963732 | 0.63035315 | -1.4435777 | 0.14885781 | 0.85485259 |
| PRUNE2 | 9.44495752 | -0.642055175 | 0.85082146 | -0.7546297 | 0.45047117 | 0.99617422 |
| VPS13A | 45.8163534 | 0.239341454 | 0.30499675 | 0.78473445 | 0.43260928 | 0.99617422 |
| GNAQ | 289.300598 | 0.207317056 | 0.2798338 | 0.74085782 | 0.45877966 | 0.99617422 |
| CEP78 | 34.6175617 | 1.142562809 | 0.38707051 | 2.95182085 | 0.00315906 | 0.17694235 |
| TLE4 | 18.6517067 | 0.300360115 | 0.47237992 | 0.63584438 | 0.52487786 | 0.99665574 |
| TLE1 | 46.6628894 | 0.166791725 | 0.42941598 | 0.38841527 | 0.69770875 | 0.99665574 |
| AL591368.1 | 2.21255684 | 2.944700874 | 1.45573317 | 2.0228301 | 0.04309067 | 0.61964991 |
| RASEF | 3.52432971 | -1.61335306 | 0.71156301 | -2.2673369 | 0.02336965 | 0.49335204 |
| FRMD3 | 10.4297376 | 0.084575637 | 0.57309752 | 0.14757635 | 0.88267712 | 0.99665574 |
| IDNK | 2.65384454 | 0.062705622 | 1.06003933 | 0.05915405 | 0.95282941 | 0.99777713 |
| UBQLN1 | 29.3640526 | 0.142204444 | 0.32378305 | 0.43919668 | 0.66051903 | 0.99665574 |
| GKAP1 | 15.7673358 | 1.288090548 | 0.46250828 | 2.78501076 | 0.00535259 | 0.22875404 |
| KIF27 | 6.79068459 | 0.778510884 | 0.59691183 | 1.30423094 | 0.19215485 | 0.89964233 |
| C9orf64 | 2.76959765 | -0.301870221 | 0.78104331 | -0.3864961 | 0.69912926 | 0.99665574 |
| HNRNPK | 53.0236387 | 0.34339615 | 0.35704966 | 0.96176019 | 0.33617008 | 0.97788427 |
| RMI1 | 3.74464884 | 0.293018358 | 0.78368522 | 0.37389803 | 0.70848018 | 0.99665574 |
| AGTPBP1 | 31.2312981 | -0.119646802 | 0.32468184 | -0.3685048 | 0.7124969 | 0.99665574 |
| NAA35 | 31.8244969 | 0.310097493 | 0.32441548 | 0.95586528 | 0.3391403 | 0.97788427 |
| GOLM1 | 60.440048 | 0.574326893 | 0.45505653 | 1.26210009 | 0.20691277 | 0.90890506 |
| ISCA1 | 14.1571082 | 0.125836398 | 0.46877506 | 0.26843663 | 0.78836325 | 0.99665574 |
| TUT7 | 26.8390754 | -0.491468396 | 0.30068078 | -1.6345189 | 0.10214995 | 0.78513738 |
| DAPK1 | 61.2588104 | -0.416535889 | 0.39829994 | -1.0457845 | 0.29566056 | 0.97286866 |
| CTSL | 17.0151329 | -0.065378973 | 0.53069994 | -0.1231939 | 0.90195359 | 0.99777713 |
| SPIN1 | 69.0910552 | 0.028905636 | 0.24505079 | 0.11795774 | 0.90610115 | 0.99777713 |
| SECISBP2 | 38.1856149 | 0.253498852 | 0.35404968 | 0.71599798 | 0.4739926 | 0.99617422 |
| SEMA4D | 32.65419 | -0.929287269 | 0.39172693 | -2.3722834 | 0.01767853 | 0.4382027 |
| GADD45G | 17.0169163 | -1.749919672 | 0.83794674 | -2.0883424 | 0.03676696 | 0.58591936 |
| SYK | 46.8571545 | -0.326810688 | 0.35038228 | -0.9327261 | 0.35096142 | 0.97838306 |
| AUH | 53.4010111 | -0.014827869 | 0.45863935 | -0.0323301 | 0.97420878 | 0.99777713 |
| NFIL3 | 3.02563466 | -1.552724258 | 0.96568643 | -1.6078969 | 0.10785775 | 0.80026315 |
| SPTLC1 | 21.1628432 | 0.139306189 | 0.37778655 | 0.36874312 | 0.71231921 | 0.99665574 |
| IARS | 19.7861146 | 0.125841599 | 0.44529999 | 0.2825996 | 0.77748379 | 0.99665574 |
| NOL8 | 7.05453403 | -0.104219701 | 0.5620147 | -0.1854395 | 0.85288436 | 0.99665574 |
| CENPP | 14.6217129 | 0.129626937 | 0.61561242 | 0.21056583 | 0.83322608 | 0.99665574 |
| IPPK | 6.91347307 | 0.631932156 | 0.65808485 | 0.96025939 | 0.33692469 | 0.97788427 |
| BICD2 | 8.40262706 | 0.302195845 | 0.57815149 | 0.52269319 | 0.60118778 | 0.99665574 |
| ZNF484 | 3.72740213 | -0.921109126 | 0.66067572 | -1.3941925 | 0.16325949 | 0.8716636 |
| FGD3 | 4.61154997 | 1.799343113 | 0.89317087 | 2.01455643 | 0.04395115 | 0.62581001 |
| SUSD3 | 4.68413533 | 1.442647443 | 0.80127461 | 1.80044072 | 0.07179108 | 0.71776916 |
| CARD19 | 7.32057544 | 0.140056131 | 0.56896256 | 0.24616054 | 0.80555796 | 0.99665574 |
| NINJ1 | 5.16753312 | -0.849666661 | 0.71976048 | -1.1804853 | 0.23780726 | 0.9305595 |
| FAM120AOS | 5.43553616 | -0.375376533 | 0.58807497 | -0.6383141 | 0.52326925 | 0.99665574 |
| FAM120A | 41.6164091 | 0.211477122 | 0.33868122 | 0.62441349 | 0.53235607 | 0.99665574 |
| PHF2 | 9.27842898 | 0.894609249 | 0.54180304 | 1.65117059 | 0.09870375 | 0.78203824 |
| PTPDC1 | 12.6060652 | -0.684801061 | 0.44149227 | -1.5511054 | 0.12087642 | 0.82659025 |
| AL158152.2 | 12.952409 | 0.375332988 | 0.43131068 | 0.87021491 | 0.38418297 | 0.99033485 |
| ZNF169 | 5.69235198 | 0.393593264 | 0.61900979 | 0.63584335 | 0.52487853 | 0.99665574 |
| MFSD14B | 15.1020917 | -0.020349891 | 0.45701062 | -0.0445283 | 0.96448332 | 0.99777713 |
| FBP1 | 5.59722221 | 0.981712174 | 1.05807075 | 0.92783226 | 0.35349459 | 0.98022903 |
| C9orf3 | 84.3719345 | -0.009485259 | 0.24237261 | -0.039135 | 0.96878273 | 0.99777713 |
| FANCC | 41.9791033 | 0.337734039 | 0.29510247 | 1.14446361 | 0.25243142 | 0.94076968 |
| PTCH1 | 11.3434952 | 0.205016976 | 0.72029412 | 0.28462953 | 0.77592799 | 0.99665574 |
| LINC00476 | 15.0624441 | -0.273990255 | 0.38134023 | -0.7184929 | 0.47245342 | 0.99617422 |
| ERCC6L2 | 27.3234683 | -0.184781564 | 0.34391981 | -0.537281 | 0.59107356 | 0.99665574 |
| SLC35D2 | 3.85352667 | -0.149748037 | 0.75757482 | -0.1976677 | 0.8433051 | 0.99665574 |
| HABP4 | 9.07701362 | -0.293412035 | 0.51377189 | -0.571094 | 0.56793594 | 0.99665574 |
| CDC14B | 32.5877692 | -0.094858595 | 0.42528298 | -0.2230482 | 0.82349801 | 0.99665574 |
| AAED1 | 3.57694974 | -0.793023179 | 0.75414635 | -1.0515508 | 0.2930057 | 0.97072515 |
| AL589843.1 | 3.59028258 | -0.369214227 | 0.85364902 | -0.4325129 | 0.66536867 | 0.99665574 |
| ZNF510 | 3.46506506 | 0.443293851 | 0.78148924 | 0.56724242 | 0.57054949 | 0.99665574 |
| ZNF782 | 10.8275235 | 0.48356366 | 0.47190901 | 1.02469682 | 0.30550626 | 0.97376371 |
| MFSD14C | 35.2108432 | -0.299196212 | 0.31602477 | -0.9467492 | 0.34376658 | 0.97788427 |
| TDRD7 | 11.8430431 | 0.20044899 | 0.57711382 | 0.34733008 | 0.72834335 | 0.99665574 |
| TSTD2 | 9.93043106 | -0.295220494 | 0.50318887 | -0.5866992 | 0.55740577 | 0.99665574 |
| NCBP1 | 8.1287087 | 0.419166002 | 0.57772618 | 0.72554441 | 0.46811811 | 0.99617422 |
| XPA | 12.0616853 | 0.87544579 | 0.53133558 | 1.64763253 | 0.0994281 | 0.78261212 |
| TRMO | 3.64718695 | -0.541764746 | 0.72263943 | -0.7497027 | 0.45343377 | 0.99617422 |
| ANP32B | 14.1214504 | -0.312830527 | 0.4467597 | -0.700221 | 0.48378929 | 0.99640678 |
| NANS | 3.52317855 | -0.912828774 | 0.76162849 | -1.1985224 | 0.23071373 | 0.92774608 |
| TRIM14 | 5.13111016 | -1.872927449 | 1.08917981 | -1.719576 | 0.08550954 | 0.75373899 |
| TBC1D2 | 6.14064997 | 0.09268932 | 0.81984472 | 0.11305717 | 0.90998523 | 0.99777713 |
| ANKS6 | 7.78222599 | 0.267498962 | 0.58710421 | 0.45562433 | 0.64866014 | 0.99665574 |
| TGFBR1 | 17.3073296 | 0.344968066 | 0.46612993 | 0.74006848 | 0.45925845 | 0.99617422 |
| ALG2 | 5.92837402 | -0.468770821 | 0.84382271 | -0.5555324 | 0.57853058 | 0.99665574 |
| SEC61B | 22.9398321 | 0.228251292 | 0.44823183 | 0.50922597 | 0.61059384 | 0.99665574 |
| AL359710.1 | 20.6918124 | 1.502327636 | 0.488061 | 3.07815549 | 0.00208286 | 0.13801343 |
| STX17 | 20.3312665 | 0.601277158 | 0.38720596 | 1.55286131 | 0.12045627 | 0.82659025 |
| ERP44 | 27.2416893 | 0.535518298 | 0.38510895 | 1.39056309 | 0.16435795 | 0.87444728 |
| INVS | 57.9089385 | -0.320892799 | 0.36736031 | -0.8735097 | 0.38238532 | 0.98914365 |
| TEX10 | 6.70904799 | -0.29386066 | 0.61776865 | -0.4756808 | 0.63430183 | 0.99665574 |
| PLPPR1 | 8.81384181 | 1.857299069 | 0.71238597 | 2.60715279 | 0.00912986 | 0.32476678 |
| MRPL50 | 5.27558956 | 0.162379273 | 0.67550636 | 0.24038156 | 0.81003447 | 0.99665574 |
| ZNF189 | 9.85348889 | -0.537438597 | 0.63166972 | -0.8508222 | 0.39486815 | 0.99179694 |
| ALDOB | 22.6262863 | 2.357676738 | 1.23577138 | 1.90785834 | 0.05640953 | 0.67545594 |
| RNF20 | 7.34181492 | 0.628302369 | 0.53211711 | 1.18075956 | 0.23769825 | 0.9305595 |
| SMC2 | 2.80611418 | 0.063051411 | 0.80176093 | 0.07864116 | 0.93731805 | 0.99777713 |
| NIPSNAP3A | 4.41065721 | -0.630357794 | 0.86979299 | -0.7247216 | 0.46862282 | 0.99617422 |
| ABCA1 | 20.4635177 | -0.200660757 | 0.64299198 | -0.3120735 | 0.75498467 | 0.99665574 |
| SLC44A1 | 29.97825 | -0.269965378 | 0.34560596 | -0.7811364 | 0.4347223 | 0.99617422 |
| FSD1L | 19.4277001 | -0.71743924 | 0.36528674 | -1.964044 | 0.04952499 | 0.64444758 |
| FKTN | 11.7548161 | -0.629131526 | 0.5277011 | -1.1922119 | 0.23317817 | 0.92924083 |
| TMEM38B | 4.18399769 | -0.155388948 | 0.85353827 | -0.1820527 | 0.85554137 | 0.99665574 |
| LINC01505 | 5.17719804 | 0.499172858 | 0.87988077 | 0.56731875 | 0.57049764 | 0.99665574 |
| ZNF462 | 6.61882738 | 0.547761055 | 0.73261011 | 0.74768427 | 0.45465062 | 0.99617422 |
| AL807761.4 | 3.14485604 | 1.44845967 | 1.05335169 | 1.37509597 | 0.16910169 | 0.87601071 |
| RAD23B | 53.4840107 | 0.05195107 | 0.27323239 | 0.19013511 | 0.84920326 | 0.99665574 |
| ELP1 | 10.694347 | 1.040434641 | 0.54879304 | 1.89585975 | 0.05797859 | 0.67545594 |
| FAM206A | 2.77591239 | -0.48031002 | 0.82629677 | -0.5812803 | 0.56105157 | 0.99665574 |
| CTNNAL1 | 11.3785809 | 0.011442447 | 0.53155936 | 0.02152619 | 0.98282591 | 0.99777713 |
| TMEM245 | 38.2037214 | 0.386917559 | 0.34079963 | 1.1353227 | 0.25624014 | 0.94314473 |
| EPB41L4B | 11.6611254 | 0.533839581 | 0.47084512 | 1.13379019 | 0.25688257 | 0.94460482 |
| PTPN3 | 68.45147 | -0.392534242 | 0.31144405 | -1.2603684 | 0.2075365 | 0.90994205 |
| PALM2 | 8.8687688 | -0.750107205 | 0.51505496 | -1.4563634 | 0.14529218 | 0.85062144 |
| TXN | 22.7948407 | -0.875166618 | 0.46035727 | -1.9010596 | 0.05729421 | 0.67545594 |

| | | | | | | |
|---|---|---|---|---|---|---|
| SVEP1 | 5.08383138 | 0.530523165 | 0.92459363 | 0.57379063 | 0.56610949 | 0.99665574 |
| LPAR1 | 22.7894645 | -1.214464488 | 0.57325868 | -2.1185278 | 0.03413039 | 0.58178209 |
| ECPAS | 62.0153158 | 0.152450357 | 0.35880937 | 0.42487842 | 0.67092531 | 0.99665574 |
| ZNF483 | 11.2292949 | -0.227093577 | 0.51053171 | -0.4448178 | 0.65645145 | 0.99665574 |
| PTGR1 | 32.3754008 | -0.612126349 | 0.40014039 | -1.5297789 | 0.12607146 | 0.83633607 |
| C9orf84 | 8.04637488 | 0.93999801 | 0.64215361 | 1.46382111 | 0.14324283 | 0.84969914 |
| UGCG | 42.7970335 | -1.013632845 | 0.3011432 | -3.3659496 | 0.00076281 | 0.07295564 |
| SUSD1 | 61.4812159 | 0.786556465 | 0.28937623 | 2.71811012 | 0.0065656 | 0.26205526 |
| PTBP3 | 226.566242 | 0.953503625 | 0.27792489 | 3.43079614 | 0.00060181 | 0.06631714 |
| HSDL2 | 28.6331338 | 0.147042463 | 0.40528154 | 0.36281559 | 0.71674264 | 0.99665574 |
| C9orf147 | 6.71448136 | -0.031369873 | 0.63805737 | -0.0491647 | 0.96078807 | 0.99777713 |
| KIAA1958 | 71.5396801 | 0.2889203 | 0.30519484 | 0.94667492 | 0.34380446 | 0.97788427 |
| INIP | 9.58998525 | 0.170629328 | 0.50652075 | 0.33686542 | 0.73621835 | 0.99665574 |
| SNX30 | 17.5964901 | 0.029100331 | 0.4695495 | 0.061975 | 0.95058274 | 0.99777713 |
| FKBP15 | 8.56424947 | 0.348374446 | 0.52545831 | 0.6629916 | 0.50733593 | 0.99665574 |
| SLC31A1 | 7.39668957 | 0.092419304 | 0.513992 | 0.17980689 | 0.85730417 | 0.99665574 |
| CDC26 | 6.75581786 | -1.415282896 | 0.71080753 | -1.9910916 | 0.04647082 | 0.64011028 |
| PRPF4 | 4.86909904 | 0.168287737 | 0.84763369 | 0.19853828 | 0.84262394 | 0.99665574 |
| BSPRY | 5.95803912 | -0.777383116 | 0.59785722 | -1.3002822 | 0.19350426 | 0.90153775 |
| HDHD3 | 9.59792599 | -0.041549676 | 0.55510576 | -0.07485 | 0.94033404 | 0.99777713 |
| ALAD | 3.52784819 | -0.01566778 | 0.77290451 | -0.0202713 | 0.98382695 | 0.99777713 |
| POLE3 | 2.26821318 | 0.368216492 | 1.07188607 | 0.34352204 | 0.73120576 | 0.99665574 |
| RGS3 | 7.6041921 | 0.516886578 | 0.58902097 | 0.8775351 | 0.38019606 | 0.98679664 |
| ZNF618 | 72.2859236 | 0.795656407 | 0.31455541 | 2.52946345 | 0.01142371 | 0.3725572 |
| KIF12 | 19.0893727 | 0.812089185 | 0.60008001 | 1.35330152 | 0.17595932 | 0.88428483 |
| COL27A1 | 7.35410749 | -0.440243143 | 0.74675354 | -0.5895428 | 0.55549724 | 0.99665574 |
| WHRN | 6.76783152 | -0.352401347 | 0.59109967 | -0.5961792 | 0.5510555 | 0.99665574 |
| ATP6V1G1 | 33.9662897 | 0.188246014 | 0.40028181 | 0.47028371 | 0.63815233 | 0.99665574 |
| TMEM268 | 5.80494508 | -0.471380705 | 0.62317293 | -0.7564204 | 0.44939719 | 0.99617422 |
| PAPPA | 15.689915 | -0.051370788 | 1.32341372 | -0.0388169 | 0.96903639 | 0.99777713 |
| ASTN2 | 120.587607 | -0.404692886 | 0.33704252 | -1.2007176 | 0.22986077 | 0.92774608 |
| CDK5RAP2 | 15.5621883 | 0.584399977 | 0.47536788 | 1.22936361 | 0.21893551 | 0.91662582 |
| MEGF9 | 71.6054471 | -0.017558441 | 0.31461406 | -0.0558095 | 0.9554936 | 0.99777713 |
| FBXW2 | 13.4403624 | 0.099980777 | 0.54374345 | 0.18387491 | 0.85411159 | 0.99665574 |
| PSMD5 | 7.75387873 | 0.470231035 | 0.55401275 | 0.84877295 | 0.39600765 | 0.99179694 |
| CNTRL | 5.38421783 | -0.518766784 | 0.61710873 | -0.8406408 | 0.40054919 | 0.99194293 |
| RAB14 | 31.9242214 | 0.020733873 | 0.30509814 | 0.06795804 | 0.94581903 | 0.99777713 |
| GSN | 19.9628258 | -0.303730796 | 0.38161082 | -0.7959177 | 0.42607989 | 0.99518711 |
| STOM | 9.85352287 | -0.844113273 | 0.5595659 | -1.5085145 | 0.1314229 | 0.84006367 |
| DAB2IP | 40.1566345 | -0.253375065 | 0.36831833 | -0.6879241 | 0.49150057 | 0.99640678 |
| TTLL11 | 5.57259612 | 0.178916263 | 0.60769046 | 0.29442007 | 0.76843694 | 0.99665574 |
| NDUFA8 | 12.2738694 | 0.530391585 | 0.47760718 | 1.11051844 | 0.26677569 | 0.94557817 |
| RBM18 | 4.75312729 | -0.134113072 | 0.72317007 | -0.1854516 | 0.85287481 | 0.99665574 |
| MRRF | 20.2899634 | 0.300756791 | 0.4326196 | 0.69519918 | 0.48693049 | 0.99640678 |
| PDCL | 7.33301951 | -0.300290321 | 0.55766729 | -0.5384758 | 0.59024863 | 0.99665574 |
| RC3H2 | 24.5436616 | -0.32665355 | 0.41005552 | -0.7966081 | 0.42567867 | 0.99518711 |
| ZBTB26 | 3.54200131 | -1.612841895 | 0.74091136 | -2.1768352 | 0.02949287 | 0.5576677 |
| RABGAP1 | 64.3804228 | 0.21521472 | 0.32777647 | 0.6565899 | 0.51144464 | 0.99665574 |
| STRBP | 61.9461934 | 0.432464536 | 0.28670264 | 1.50840792 | 0.13145016 | 0.84006367 |
| DENND1A | 65.5197621 | 0.150169927 | 0.26715927 | 0.56209888 | 0.57404865 | 0.99665574 |
| NEK6 | 12.3006628 | -0.913412072 | 0.51236592 | -1.7827339 | 0.07462964 | 0.72883798 |
| PSMB7 | 19.8785051 | 0.318970607 | 0.40841103 | 0.7810039 | 0.4348002 | 0.99617422 |
| NR6A1 | 95.7024735 | -0.304017591 | 0.4190598 | -0.7254754 | 0.46816041 | 0.99617422 |
| MIR181A2H( | 6.76701736 | -0.991793614 | 0.70182391 | -1.4131659 | 0.15760695 | 0.86532131 |
| RPL35 | 36.3424112 | 0.93498836 | 0.43101899 | 2.16925096 | 0.03006364 | 0.55923879 |
| ARPC5L | 11.4934083 | -0.172563957 | 0.51491418 | -0.3351315 | 0.7375259 | 0.99665574 |
| GOLGA1 | 14.5479991 | -0.133282054 | 0.44246903 | -0.3012235 | 0.7632441 | 0.99665574 |
| SCAI | 48.9667721 | 0.925393645 | 0.37577951 | 2.46259737 | 0.01379347 | 0.39608255 |
| PPP6C | 25.8835605 | 0.403724494 | 0.42919814 | 0.94064829 | 0.34688513 | 0.97804112 |
| RABEPK | 5.04396202 | -0.044496965 | 0.61670857 | -0.0721523 | 0.94248067 | 0.99777713 |
| HSPA5 | 37.00532 | 0.414459463 | 0.60625623 | 0.68363745 | 0.49420413 | 0.99640678 |
| GAPVD1 | 34.488218 | 0.079384434 | 0.33210506 | 0.2390341 | 0.81107914 | 0.99665574 |
| MAPKAP1 | 75.9367256 | 0.109809868 | 0.41606987 | 0.2639217 | 0.79184025 | 0.99665574 |
| PBX3 | 56.3433531 | -0.481666641 | 0.38602162 | -1.2477712 | 0.21211487 | 0.91043197 |
| MVB12B | 179.582295 | -0.046391306 | 0.35054221 | -0.1323416 | 0.89471413 | 0.99777713 |
| ZBTB43 | 25.2146239 | -0.683514968 | 0.3625438 | -1.8853307 | 0.0593852 | 0.6795115 |
| ZBTB34 | 4.45664422 | 0.836418233 | 0.81337981 | 1.02832431 | 0.3037973 | 0.97376371 |
| RALGPS1 | 64.0119806 | 0.054477169 | 0.2832354 | 0.19233884 | 0.8474768 | 0.99665574 |
| GARNL3 | 8.26048436 | -0.557834027 | 0.58218469 | -0.9581736 | 0.33797521 | 0.97788427 |
| SLC2A8 | 5.77723047 | 0.447898816 | 0.59897556 | 0.74777478 | 0.45459602 | 0.99617422 |
| RPL12 | 16.4604193 | 0.361540484 | 0.49192372 | 0.73495232 | 0.46236853 | 0.99617422 |
| LRSAM1 | 13.2976699 | -0.053942047 | 0.52183173 | -0.1033706 | 0.91766886 | 0.99777713 |
| FAM129B | 11.9045204 | -1.025029238 | 0.50601325 | -2.0256964 | 0.0427959 | 0.61874363 |
| STXBP1 | 7.28054436 | -1.527066406 | 0.56793149 | -2.6888215 | 0.00717047 | 0.27640407 |
| PTRH1 | 3.05220643 | 0.663584993 | 0.81228854 | 0.81693261 | 0.41396694 | 0.99279547 |
| CDK9 | 5.48218537 | 0.004578729 | 0.74275017 | 0.00616456 | 0.99508142 | 0.9985287 |
| FPGS | 6.24859201 | 0.68748371 | 0.72297025 | 0.95091562 | 0.34164721 | 0.97788427 |
| DPM2 | 7.79357899 | 0.423447297 | 0.56075821 | 0.75513348 | 0.4501689 | 0.99617422 |
| FAM102A | 17.8955519 | -0.209592618 | 0.39566145 | -0.5297272 | 0.59630111 | 0.99665574 |
| SLC25A25 | 9.64678799 | -1.396857353 | 0.50359245 | -2.7737853 | 0.00554082 | 0.23211926 |
| SLC25A25-AS | 13.4991841 | 0.598243359 | 0.48013922 | 1.24597893 | 0.21277213 | 0.91073943 |
| PTGES2 | 9.67485028 | -0.272521871 | 0.54913572 | -0.4962742 | 0.61970099 | 0.99665574 |
| C9orf16 | 24.9088642 | 0.069015296 | 0.55027279 | 0.12542015 | 0.90019094 | 0.99777713 |
| CIZ1 | 7.45592855 | -0.290754184 | 0.59554131 | -0.4882183 | 0.6253952 | 0.99665574 |
| DNM1 | 6.85146768 | -0.668179487 | 0.75037617 | -0.8904594 | 0.37321928 | 0.98532838 |
| GOLGA2 | 19.1971751 | -0.750933506 | 0.47495914 | -1.5810487 | 0.11386691 | 0.81638817 |
| SWI5 | 9.88719037 | -0.187945602 | 0.46438713 | -0.4047175 | 0.68568517 | 0.99665574 |
| TRUB2 | 5.2469951 | 0.210389564 | 0.64475214 | 0.32631076 | 0.74418925 | 0.99665574 |
| COQ4 | 13.5631271 | 0.565621718 | 0.44967728 | 1.2578392 | 0.20844991 | 0.91043197 |
| SLC27A4 | 12.573024 | -0.1983049 | 0.46982552 | -0.422082 | 0.67296516 | 0.99665574 |
| URM1 | 7.59800033 | -0.363572953 | 0.55467552 | -0.6554696 | 0.51216544 | 0.99665574 |
| ODF2 | 5.5275861 | -0.344839178 | 0.63961201 | -0.5391381 | 0.58979159 | 0.99665574 |
| GLE1 | 9.01041036 | 0.095254646 | 0.4975084 | 0.19146339 | 0.84816256 | 0.99665574 |
| SPTAN1 | 30.9080833 | 0.049306583 | 0.41108496 | 0.11994256 | 0.90452865 | 0.99777713 |
| WDR34 | 7.48767867 | 0.243944265 | 0.56252582 | 0.43365878 | 0.66453624 | 0.99665574 |
| SET | 39.5258295 | -0.020969016 | 0.32193993 | -0.0651333 | 0.94806785 | 0.99777713 |
| ZDHHC12 | 5.37683363 | -0.087316368 | 0.77257239 | -0.1130203 | 0.91001446 | 0.99777713 |
| ZER1 | 21.2386449 | -0.357812989 | 0.35839808 | -0.9983675 | 0.31810119 | 0.97700345 |
| TBC1D13 | 6.64571494 | -0.195371816 | 0.56859581 | -0.343604 | 0.73114409 | 0.99665574 |
| ENDOG | 6.61325894 | 0.182728619 | 0.64538511 | 0.28313114 | 0.77707631 | 0.99665574 |
| SPOUT1 | 3.77464839 | 0.165547928 | 0.75255627 | 0.2199808 | 0.82588611 | 0.99665574 |
| KYAT1 | 7.1757405 | -1.146834781 | 0.61724506 | -1.8579894 | 0.06317052 | 0.68630818 |
| LRRC8A | 4.10836925 | -0.546473489 | 0.76842701 | -0.7111586 | 0.47698595 | 0.99617422 |
| PHYHD1 | 3.27451843 | 0.158534132 | 0.78323885 | 0.20240841 | 0.83959745 | 0.99665574 |
| NUP188 | 6.4953346 | 1.127369718 | 0.64334717 | 1.75235047 | 0.07971356 | 0.7446402 |
| SH3GLB2 | 12.8706629 | 0.267564671 | 0.47969848 | 0.55777677 | 0.57699683 | 0.99665574 |
| MIGA2 | 8.13299167 | -0.001996299 | 0.54723664 | -0.003648 | 0.99708935 | 0.99884397 |
| CRAT | 4.26753384 | -0.263211161 | 0.72391699 | -0.363593 | 0.71616194 | 0.99665574 |
| PTPA | 11.2520858 | -0.518111266 | 0.52951192 | -0.9784695 | 0.32784216 | 0.97788427 |
| IER5L | 4.44202795 | -1.823786314 | 0.7603685 | -2.3985559 | 0.01645987 | 0.42361121 |
| LINC01503 | 7.30128267 | -2.01119174 | 1.15956303 | -1.7344393 | 0.08284016 | 0.75242737 |
| NTMT1 | 5.00013345 | -0.62216247 | 0.62837122 | -0.9901193 | 0.32211581 | 0.97700345 |
| ASB6 | 3.19495861 | 0.197210216 | 0.81417007 | 0.24222239 | 0.80860785 | 0.99665574 |

| | | | | | | |
|---|---|---|---|---|---|---|
| TOR1B | 5.2166384 | -0.334683387 | 0.6620449 | -0.5055297 | 0.61318681 | 0.99665574 |
| TOR1A | 3.35221852 | 0.285370826 | 0.79870661 | 0.35729118 | 0.72087383 | 0.99665574 |
| C9orf78 | 10.3994671 | -0.073988072 | 0.50660582 | -0.1460466 | 0.88388458 | 0.99665574 |
| USP20 | 6.73043448 | 0.407030436 | 0.58415927 | 0.6967799 | 0.48594055 | 0.99640678 |
| FNBP1 | 32.1044952 | 0.170553909 | 0.41181921 | 0.41414753 | 0.67876606 | 0.99665574 |
| GPR107 | 21.9926151 | -0.453873071 | 0.46947802 | -0.9667611 | 0.33366349 | 0.97788427 |
| FUBP3 | 26.0696244 | 0.02504047 | 0.4113378 | 0.06087568 | 0.95145821 | 0.99777713 |
| EXOSC2 | 3.04738797 | 0.509992568 | 0.84134451 | 0.60616378 | 0.544406 | 0.99665574 |
| ABL1 | 43.8474872 | 0.234114476 | 0.29813537 | 0.78526233 | 0.43229978 | 0.99617422 |
| AIF1L | 26.8837313 | 0.635362845 | 0.53748778 | 1.18209729 | 0.2371671 | 0.9305595 |
| NUP214 | 34.3989603 | 0.185808038 | 0.29185131 | 0.6366531 | 0.52435083 | 0.99665574 |
| AL157938.2 | 9.55205815 | -0.317245821 | 0.90243597 | -0.3515439 | 0.72518038 | 0.99665574 |
| PRRC2B | 80.15218 | -0.22645057 | 0.25624327 | -0.8837327 | 0.3768405 | 0.98679664 |
| POMT1 | 4.64334053 | -0.545570918 | 0.60119341 | -0.9074799 | 0.36415309 | 0.98362663 |
| UCK1 | 3.87835036 | 0.149935355 | 0.71549034 | 0.20955609 | 0.83401416 | 0.99665574 |
| RAPGEF1 | 61.1261887 | 0.031563787 | 0.36798678 | 0.08577424 | 0.93164588 | 0.99777713 |
| MED27 | 29.8764643 | -0.182201055 | 0.39194681 | -0.4648617 | 0.64203051 | 0.99665574 |
| SETX | 43.4551576 | -0.033431265 | 0.3453473 | -0.0968048 | 0.92288144 | 0.99777713 |
| TTF1 | 8.20566235 | -0.634848685 | 0.5025177 | -1.263336 | 0.20646846 | 0.90890506 |
| DDX31 | 8.32021058 | 0.209336269 | 0.55644255 | 0.37620464 | 0.70676477 | 0.99665574 |
| GTF3C4 | 3.42687145 | -0.348094919 | 0.75873505 | -0.4587832 | 0.64638984 | 0.99665574 |
| TSC1 | 21.0906614 | -0.343538926 | 0.38087692 | -0.9019683 | 0.36707368 | 0.98423512 |
| GTF3C5 | 5.94403031 | -0.260328406 | 0.57834285 | -0.4501282 | 0.65261803 | 0.99665574 |
| RALGDS | 3.33633882 | -0.57695893 | 0.7482487 | -0.7710791 | 0.44066005 | 0.99617422 |
| ABO | 12.2741494 | -0.942211938 | 0.62246107 | -1.5136881 | 0.13010496 | 0.83778831 |
| SURF6 | 3.54322745 | 0.329663259 | 0.75161542 | 0.4386063 | 0.66094683 | 0.99665574 |
| MED22 | 2.80511964 | -0.26180165 | 0.90999869 | -0.2876945 | 0.77358057 | 0.99665574 |
| RPL7A | 61.2747418 | 0.521541091 | 0.42968906 | 1.21376394 | 0.22483788 | 0.92318867 |
| SURF1 | 5.34347103 | 0.083207881 | 0.6680972 | 0.12454457 | 0.90088411 | 0.99777713 |
| SURF2 | 2.8861624 | -0.08462299 | 0.98528387 | -0.0858869 | 0.93155632 | 0.99777713 |
| SURF4 | 19.476496 | -0.155006529 | 0.45023097 | -0.3442822 | 0.73063403 | 0.99665574 |
| REXO4 | 4.70340911 | -0.946359635 | 0.68327634 | -1.3850321 | 0.16604266 | 0.87495175 |
| CACFD1 | 3.82958892 | 0.430727155 | 0.72513153 | 0.59399866 | 0.55251301 | 0.99665574 |
| VAV2 | 45.5155057 | 0.140725455 | 0.29637256 | 0.47482619 | 0.63491087 | 0.99665574 |
| BRD3 | 12.2513381 | 0.055407949 | 0.47303066 | 0.11713395 | 0.90675391 | 0.99777713 |
| WDR5 | 4.30433623 | -0.748377117 | 0.81530676 | -0.9179086 | 0.3586667 | 0.9819332 |
| RXRA | 11.8178083 | 0.199622033 | 0.46436218 | 0.42988435 | 0.66727977 | 0.99665574 |
| PPP1R26-AS1 | 4.42341608 | 1.07003011 | 0.79344036 | 1.34859552 | 0.17746692 | 0.88672235 |
| PPP1R26 | 6.1704083 | 0.428851792 | 0.63233478 | 0.67820371 | 0.49764254 | 0.99640678 |
| MRPS2 | 5.6415249 | 0.536002476 | 0.69535781 | 0.77082973 | 0.44080786 | 0.99617422 |
| CAMSAP1 | 11.8932766 | 0.065225753 | 0.49180498 | 0.13262524 | 0.89448977 | 0.99777713 |
| UBAC1 | 11.9194296 | -0.40389421 | 0.50684965 | -0.7968718 | 0.42552547 | 0.99518711 |
| NACC2 | 12.4505666 | -0.306471401 | 0.49285006 | -0.621835 | 0.53405039 | 0.99665574 |
| TMEM250 | 3.29304072 | -0.615506147 | 0.96105345 | -0.6404494 | 0.52188045 | 0.99665574 |
| QSOX2 | 2.71802208 | 0.706757888 | 0.94261225 | 0.74978645 | 0.45338333 | 0.99617422 |
| GPSM1 | 4.55647441 | 1.609483727 | 0.72247379 | 2.22773996 | 0.02589786 | 0.51788242 |
| SNAPC4 | 3.77598286 | -0.998840177 | 0.697567 | -1.4318914 | 0.15217491 | 0.85976284 |
| ENTR1 | 5.85718841 | -0.411549832 | 0.63904375 | -0.6440088 | 0.5195697 | 0.99665574 |
| PMPCA | 8.14346099 | -0.092611014 | 0.55967547 | -0.1654727 | 0.86857193 | 0.99665574 |
| INPP5E | 3.16522276 | 0.37304952 | 0.8566644 | 0.43546752 | 0.66322311 | 0.99665574 |
| SEC16A | 13.8920226 | -0.320006177 | 0.4988949 | -0.64143 | 0.52124331 | 0.99665574 |
| NOTCH1 | 5.2015178 | -0.914165049 | 0.74951614 | -1.2196736 | 0.22258863 | 0.92191316 |
| EGFL7 | 5.64216522 | -0.283534993 | 0.64293722 | -0.4409995 | 0.65921336 | 0.99665574 |
| AGPAT2 | 19.1864538 | -0.451280382 | 0.46115792 | -0.978581 | 0.32778704 | 0.97788427 |
| FAM69B | 2.41220185 | 0.792365971 | 0.9426548 | 0.84056854 | 0.40058969 | 0.99194293 |
| SNHG7 | 18.431759 | 0.341812541 | 0.46104439 | 0.74138748 | 0.45845853 | 0.99617422 |
| RABL6 | 19.0884281 | 0.150813848 | 0.42737131 | 0.35288716 | 0.72417304 | 0.99665574 |
| PHPT1 | 16.0793242 | 0.232826772 | 0.55155285 | 0.42212958 | 0.67293044 | 0.99665574 |
| EDF1 | 27.9315301 | 0.331906516 | 0.44497771 | 0.74589471 | 0.45573102 | 0.99617422 |
| TRAF2 | 3.06769537 | 0.563344955 | 0.84862243 | 0.66383463 | 0.50679616 | 0.99665574 |
| FBXW5 | 12.2257064 | 0.041398703 | 0.48706952 | 0.08499547 | 0.93226499 | 0.99777713 |
| PAXX | 5.81123602 | 0.906625953 | 0.75814025 | 1.19585519 | 0.23175307 | 0.92810069 |
| UAP1L1 | 2.799142 | 0.187093944 | 1.1242432 | 0.16641768 | 0.86782826 | 0.99665574 |
| MAN1B1 | 6.22505668 | 0.242443375 | 0.64055154 | 0.3784916 | 0.70506544 | 0.99665574 |
| DPP7 | 9.04541994 | 0.103122651 | 0.52039697 | 0.19816151 | 0.8429187 | 0.99665574 |
| SSNA1 | 8.53528081 | -0.334938107 | 0.51474756 | -0.6506842 | 0.51525036 | 0.99665574 |
| TPRN | 5.21654267 | 0.611515159 | 0.63942231 | 0.95635568 | 0.33889257 | 0.97788427 |
| TMEM203 | 4.20028969 | 0.097893011 | 0.85353855 | 0.11469079 | 0.90869021 | 0.99777713 |
| NDOR1 | 3.55630303 | 0.630291028 | 0.85444925 | 0.73765765 | 0.46072251 | 0.99617422 |
| RNF208 | 2.61244614 | 1.040769769 | 0.92802709 | 1.12148641 | 0.26208087 | 0.94557817 |
| TUBB4B | 26.223648 | -0.539254154 | 0.56060896 | -0.9619078 | 0.3360959 | 0.97788427 |
| NELFB | 9.95175322 | 0.412276514 | 0.49675855 | 0.82993339 | 0.40657644 | 0.99250037 |
| EXD3 | 11.9223514 | 0.58116675 | 0.4693243 | 1.23830526 | 0.2156029 | 0.913434 |
| NOXA1 | 3.56566135 | -1.395408383 | 0.74412537 | -1.8752329 | 0.06076069 | 0.6834742 |
| NSMF | 4.16059505 | 0.273826618 | 0.71306099 | 0.3840157 | 0.70096681 | 0.99665574 |
| PNPLA7 | 4.58606954 | -1.558272577 | 0.69545325 | -2.2406575 | 0.02504827 | 0.50908112 |
| MRPL41 | 29.8627457 | -0.105749177 | 0.36774707 | -0.2875595 | 0.77368392 | 0.99665574 |
| DPH7 | 5.61116401 | 0.099844394 | 0.64672123 | 0.15438552 | 0.87730577 | 0.99665574 |
| ZMYND19 | 2.28728889 | 0.450083929 | 0.93968989 | 0.47897071 | 0.63195947 | 0.99665574 |
| ARRDC1 | 9.80476151 | 0.203929193 | 0.59015042 | 0.3455546 | 0.72967747 | 0.99665574 |
| ARRDC1-AS1 | 4.01295003 | 0.568585933 | 0.71269361 | 0.79779856 | 0.4249874 | 0.99518711 |
| EHMT1 | 69.1865233 | -0.478498484 | 0.33583856 | -1.4247872 | 0.15421872 | 0.86196782 |
| BET1L | 6.38634397 | -0.804434011 | 0.58663001 | -1.37128 | 0.17028768 | 0.87918701 |
| RIC8A | 5.37647547 | 0.232176628 | 0.64419106 | 0.36041578 | 0.71853622 | 0.99665574 |
| SIRT3 | 4.09750458 | 0.588922542 | 0.67433217 | 0.87334191 | 0.38247675 | 0.98914365 |
| PSMD13 | 10.4086608 | 0.61953143 | 0.50818185 | 1.21911365 | 0.22280106 | 0.92191316 |
| IFITM2 | 32.9241287 | -1.47629178 | 0.65893439 | -2.240423 | 0.02506347 | 0.50908112 |
| IFITM3 | 65.6280452 | -1.366757282 | 0.65183378 | -2.0967881 | 0.03601234 | 0.58508511 |
| PKP3 | 3.78448488 | -0.176078361 | 0.88879127 | -0.1981099 | 0.84295907 | 0.99665574 |
| SIGIRR | 8.78123957 | 0.037026095 | 0.62152144 | 0.05957332 | 0.95249547 | 0.99777713 |
| ANO9 | 3.02815147 | 0.78460184 | 0.92182584 | 0.85113891 | 0.39469219 | 0.99179694 |
| PTDSS2 | 10.4815031 | 0.342109993 | 0.5020471 | 0.68143008 | 0.4955994 | 0.99640678 |
| RNH1 | 14.6437316 | -0.663873891 | 0.47751838 | -1.3902583 | 0.16445046 | 0.87444728 |
| HRAS | 12.1651457 | -0.596263629 | 0.51310593 | -1.1620673 | 0.24520813 | 0.93426912 |
| RASSF7 | 11.8444842 | -0.198487355 | 0.51348383 | -0.3865504 | 0.69908912 | 0.99665574 |
| PHRF1 | 9.76180125 | -0.071521947 | 0.4674907 | -0.1529912 | 0.87840524 | 0.99665574 |
| IRF7 | 5.18266617 | -0.241734084 | 1.02207831 | -0.2365123 | 0.81303517 | 0.99665574 |
| DEAF1 | 28.1833851 | -0.90433517 | 0.36375734 | -2.4860946 | 0.01291536 | 0.38590337 |
| AC131934.1 | 4.77195299 | -0.917774828 | 0.79088832 | -1.1604354 | 0.24587157 | 0.93426912 |
| EPS8L2 | 25.3033596 | -0.191542756 | 0.532286 | -0.3598493 | 0.71895982 | 0.99665574 |
| AP006621.4 | 2.85271855 | -1.685308092 | 1.31447901 | -1.2821111 | 0.19980368 | 0.90381759 |
| TALDO1 | 20.3713765 | -0.086641074 | 0.4150981 | -0.2087243 | 0.83466344 | 0.99665574 |
| GATD1 | 6.54462408 | 0.109424269 | 0.66525995 | 0.16448348 | 0.86935055 | 0.99665574 |
| AP006621.3 | 4.86819301 | -1.111002137 | 0.79076178 | -1.404977 | 0.16002811 | 0.86757484 |
| RPLP2 | 45.1469354 | 0.231047578 | 0.34377319 | 0.67209306 | 0.50152445 | 0.99640678 |
| PNPLA2 | 6.39809301 | -0.117373698 | 0.59857002 | -0.1960902 | 0.84453959 | 0.99665574 |
| CD151 | 16.7973615 | -0.801704437 | 0.40746938 | -1.9675207 | 0.04912321 | 0.64444758 |
| POLR2L | 25.1821001 | 0.482007677 | 0.38910088 | 1.23877303 | 0.21542956 | 0.913434 |
| TSPAN4 | 3.25499622 | -1.567655885 | 0.74267949 | -2.1108108 | 0.03478858 | 0.58178209 |
| CHID1 | 6.43972145 | 1.033869073 | 0.59299831 | 1.7434604 | 0.08125323 | 0.74885928 |
| AP2A2 | 38.7691712 | 0.078627876 | 0.35171927 | 0.22355294 | 0.82310519 | 0.99665574 |
| TOLLIP | 11.9594417 | 0.245218843 | 0.5529931 | 0.44343925 | 0.65744805 | 0.99665574 |
| MOB2 | 8.46802422 | 0.35886864 | 0.58016541 | 0.61856263 | 0.53620453 | 0.99665574 |

| | | | | | | |
|---|---|---|---|---|---|---|
| MRPL23 | 3.56892697 | 0.276561683 | 0.91654473 | 0.30174379 | 0.76284738 | 0.99665574 |
| CD81 | 39.5034382 | -0.344944514 | 0.48882588 | -0.7056593 | 0.48040006 | 0.99617422 |
| TSSC4 | 7.23696215 | -0.503466479 | 0.56337294 | -0.8936646 | 0.37150135 | 0.98532838 |
| KCNQ1 | 53.4963136 | -0.118889623 | 0.32855372 | -0.3618575 | 0.71745853 | 0.99665574 |
| KCNQ1OT1 | 78.9321426 | 0.392616961 | 0.40653827 | 0.96575646 | 0.33416606 | 0.97788427 |
| CDKN1C | 38.1070166 | 1.353467265 | 0.58165451 | 2.32692644 | 0.01996918 | 0.46397674 |
| SLC22A18 | 6.6967403 | -0.49252154 | 0.64588754 | -0.76255 | 0.44573181 | 0.99617422 |
| NAP1L4 | 57.7971555 | 0.271810418 | 0.32056772 | 0.84790327 | 0.39649184 | 0.99179694 |
| CARS | 20.78288 | -0.968084452 | 0.46369376 | -2.0877669 | 0.03681887 | 0.58591936 |
| OSBPL5 | 5.05550246 | -0.472208664 | 0.80875249 | -0.5838729 | 0.55930582 | 0.99665574 |
| ZNF195 | 8.69059486 | -0.439566169 | 0.5382204 | -0.8167029 | 0.41409822 | 0.99279547 |
| AC127526.1 | 2.76107024 | -0.514136036 | 0.96487005 | -0.5328552 | 0.59413381 | 0.99665574 |
| NUP98 | 29.7731966 | 0.014755786 | 0.36341036 | 0.04060365 | 0.96761188 | 0.99777713 |
| PGAP2 | 5.08147513 | 0.107294186 | 0.65408801 | 0.16403631 | 0.86970256 | 0.99665574 |
| RHOG | 6.6802855 | -0.840391052 | 0.69979323 | -1.2009134 | 0.2297848 | 0.92774608 |
| STIM1 | 82.7690926 | 0.340671368 | 0.29889254 | 1.13977875 | 0.25437849 | 0.94099699 |
| RRM1 | 7.16752684 | 0.425479363 | 0.62422648 | 0.68161057 | 0.49548523 | 0.99640678 |
| FAM160A2 | 6.6336256 | 0.302883383 | 0.57230224 | 0.52923676 | 0.59664122 | 0.99665574 |
| SMPD1 | 5.62792314 | 0.309474753 | 0.69473629 | 0.44545643 | 0.65598994 | 0.99665574 |
| APBB1 | 6.27279428 | -0.220131411 | 0.64536805 | -0.3410944 | 0.73303253 | 0.99665574 |
| ARFIP2 | 5.61744244 | -0.543240314 | 0.59771419 | -0.908863 | 0.36342244 | 0.98362663 |
| AC084337.2 | 4.93642943 | -0.321120046 | 0.71666305 | -0.4480767 | 0.65409781 | 0.99665574 |
| DNHD1 | 5.47938031 | -1.230680393 | 0.57662839 | -2.1342695 | 0.03282073 | 0.5740203 |
| RRP8 | 3.55455597 | -0.286397267 | 0.71685162 | -0.399521 | 0.68950936 | 0.99665574 |
| ILK | 10.7881649 | 0.523228879 | 0.52867051 | 0.98970694 | 0.32231738 | 0.97700345 |
| TAF10 | 9.86301833 | 0.748458684 | 0.65889312 | 1.13593337 | 0.25598445 | 0.94314473 |
| TPP1 | 8.47473168 | -0.010713336 | 0.6295548 | -0.0170173 | 0.9864228 | 0.99777713 |
| MRPL17 | 7.23009028 | 0.686026479 | 0.68637205 | 0.99949652 | 0.31755422 | 0.97700345 |
| SYT9 | 3.20227166 | -0.673891763 | 0.84652523 | -0.7960681 | 0.42599243 | 0.99518711 |
| AC107884.1 | 3.52173581 | 0.432020055 | 0.88766397 | 0.48669324 | 0.62647573 | 0.99665574 |
| PPFIBP2 | 26.8181688 | 0.428517756 | 0.34847952 | 1.22967846 | 0.21881754 | 0.91662582 |
| EIF3F | 16.5383864 | 0.61594933 | 0.45576551 | 1.35146102 | 0.1765478 | 0.88518376 |
| TUB | 8.83683261 | 1.261794319 | 0.54219948 | 2.32717728 | 0.01995583 | 0.46397674 |
| RIC3 | 20.5163245 | -0.252977435 | 0.47248604 | -0.5354178 | 0.59236099 | 0.99665574 |
| STK33 | 64.4239847 | -0.056638779 | 0.36558705 | -0.1549256 | 0.87688 | 0.99665574 |
| TRIM66 | 18.6040208 | -0.145289773 | 0.45040919 | -0.3225728 | 0.74701877 | 0.99665574 |
| RPL27A | 56.8142999 | 0.52105048 | 0.52513949 | 0.99221348 | 0.3210934 | 0.97700345 |
| ST5 | 23.2495545 | -0.76259446 | 0.52045853 | -1.4652358 | 0.1428566 | 0.84969914 |
| AKIP1 | 3.41996237 | 0.687875582 | 0.88234657 | 0.77959796 | 0.43562756 | 0.99617422 |
| TMEM9B | 18.1642523 | 1.021000509 | 0.42574955 | 2.39812471 | 0.01647925 | 0.42361121 |
| TMEM9B-AS | 4.10794066 | -0.053600833 | 0.82975751 | -0.0645982 | 0.94849393 | 0.99777713 |
| DENND5A | 19.5589217 | -0.205632189 | 0.44555848 | -0.4615156 | 0.64442873 | 0.99665574 |
| TMEM41B | 23.9503531 | -0.383805493 | 0.41837435 | -0.9173734 | 0.35894702 | 0.9819332 |
| IPO7 | 33.4868508 | 0.056689216 | 0.32207444 | 0.17601278 | 0.8602839 | 0.99665574 |
| ZNF143 | 7.13158035 | -0.327874551 | 0.66521494 | -0.4928851 | 0.62209376 | 0.99665574 |
| SWAP70 | 23.6633302 | -1.464291223 | 0.43256118 | -3.3851656 | 0.00071135 | 0.07140296 |
| SBF2-AS1 | 19.4296576 | -0.885969864 | 0.63223356 | -1.4013332 | 0.16111447 | 0.86928072 |
| SBF2 | 185.075322 | 0.148098051 | 0.19304079 | 0.76718527 | 0.44297137 | 0.99617422 |
| AC011092.2 | 6.67231544 | -0.714247942 | 0.750264 | -0.9519955 | 0.34109927 | 0.97788427 |
| ADM | 12.4185329 | -1.020034246 | 0.50822424 | -2.0070555 | 0.04474375 | 0.6313648 |
| AMPD3 | 16.0428654 | 0.524564022 | 0.48083998 | 1.09093263 | 0.27530253 | 0.95593806 |
| MTRNR2L8 | 70.7128147 | -0.288790235 | 0.56083532 | -0.5149288 | 0.6066028 | 0.99665574 |
| RNF141 | 8.91986411 | 0.274431754 | 0.56745749 | 0.48361641 | 0.62865812 | 0.99665574 |
| MRVI1 | 3.41328687 | 0.078136017 | 0.75225031 | 0.1038697 | 0.91727275 | 0.99777713 |
| CTR9 | 14.0710075 | 0.221852869 | 0.41993661 | 0.52830085 | 0.59729054 | 0.99665574 |
| EIF4G2 | 47.5544864 | -0.203752088 | 0.28035597 | -0.7267621 | 0.46737172 | 0.99617422 |
| ZBED5 | 16.9758388 | 0.267330463 | 0.37131036 | 0.719965 | 0.47154654 | 0.99617422 |
| GALNT18 | 13.3324856 | -0.49057236 | 0.5694749 | -0.8614468 | 0.38899198 | 0.99176253 |
| USP47 | 62.6335218 | -0.008441974 | 0.24167412 | -0.0349312 | 0.97213458 | 0.99777713 |
| PARVA | 34.0582712 | 0.215689656 | 0.31522259 | 0.68424555 | 0.49382012 | 0.99640678 |
| TEAD1 | 83.043864 | -0.028577407 | 0.39016779 | -0.0732439 | 0.94161204 | 0.99777713 |
| LINC00958 | 6.33361269 | -0.480100388 | 0.58857207 | -0.8157037 | 0.41466965 | 0.99279547 |
| ARNTL | 8.21367939 | 0.203206645 | 0.54325664 | 0.37405276 | 0.70836506 | 0.99665574 |
| BTBD10 | 14.4012436 | -0.033132307 | 0.43360431 | -0.0764114 | 0.93909181 | 0.99777713 |
| FAR1 | 41.5415422 | -0.203567606 | 0.43640507 | -0.4664648 | 0.64088284 | 0.99665574 |
| RRAS2 | 25.8725152 | -0.85904164 | 0.52410594 | -1.6390611 | 0.10120054 | 0.78386461 |
| COPB1 | 23.6050188 | -0.208307678 | 0.44140034 | -0.4719246 | 0.63698061 | 0.99665574 |
| CYP2R1 | 3.53121846 | -0.09527401 | 0.86029643 | -0.1107456 | 0.91181812 | 0.99777713 |
| SOX6 | 256.620929 | 0.705600978 | 0.2356807 | 2.99388525 | 0.0027545 | 0.1614166 |
| C11orf58 | 29.1773202 | -0.250222283 | 0.32622279 | -0.7670288 | 0.44306438 | 0.99617422 |
| PLEKHA7 | 137.462286 | -0.026103386 | 0.29051229 | -0.0898529 | 0.92840407 | 0.99777713 |
| RPS13 | 37.564421 | 0.077155389 | 0.43527838 | 0.17725528 | 0.85930788 | 0.99665574 |
| PIK3C2A | 56.8898431 | -0.185383569 | 0.29654053 | -0.6251542 | 0.53186984 | 0.99665574 |
| NUCB2 | 5.17210642 | -0.209862181 | 0.72199375 | -0.2906704 | 0.77130344 | 0.99665574 |
| KCNC1 | 3.62446041 | -2.66761122 | 0.97105466 | -2.7471278 | 0.00601197 | 0.24620771 |
| SERGEF | 41.2870855 | 0.308870121 | 0.37126798 | 0.83193311 | 0.4054467 | 0.99194293 |
| SAAL1 | 3.00360157 | 0.281536774 | 0.81769078 | 0.34430714 | 0.73061531 | 0.99665574 |
| HPS5 | 8.3343948 | 0.293892191 | 0.49854642 | 0.58949815 | 0.55552715 | 0.99665574 |
| GTF2H1 | 13.8348834 | -0.277997341 | 0.51409235 | -0.5407537 | 0.58867736 | 0.99665574 |
| LDHA | 54.5752959 | -1.536678399 | 0.38452482 | -3.9963048 | 6.43E-05 | 0.01672481 |
| TSG101 | 26.6734415 | 0.130989834 | 0.30527053 | 0.42909427 | 0.66785462 | 0.99665574 |
| UEVLD | 19.2861256 | -0.329423343 | 0.39197347 | -0.8404225 | 0.40067152 | 0.99194293 |
| SPTY2D1OS | 5.25075915 | -1.230364157 | 0.70712867 | -1.7399438 | 0.08186888 | 0.75124322 |
| SPTY2D1 | 17.9361707 | -0.815816867 | 0.72907354 | -1.1189775 | 0.26314975 | 0.94557817 |
| ZDHHC13 | 20.5794592 | 0.104551873 | 0.43033219 | 0.2429562 | 0.80803933 | 0.99665574 |
| NAV2 | 69.8504115 | -2.036526272 | 0.59601131 | -3.4169256 | 0.00063333 | 0.06758588 |
| HTATIP2 | 5.39427496 | -2.471313461 | 0.7752579 | -3.1877307 | 0.00143394 | 0.11024264 |
| PRMT3 | 18.743943 | -0.034866609 | 0.35657608 | -0.0977817 | 0.92210565 | 0.99777713 |
| NELL1 | 27.406892 | 2.746227707 | 0.86157851 | 3.18743758 | 0.00143539 | 0.11024264 |
| ANO5 | 5.88195995 | -0.037301301 | 0.85617472 | -0.0435674 | 0.96524924 | 0.99777713 |
| SVIP | 12.3154692 | -0.086085806 | 0.55897946 | -0.1540053 | 0.87760555 | 0.99665574 |
| SLC5A12 | 5.97201654 | 2.051606632 | 1.03611717 | 1.98009134 | 0.04769327 | 0.6410909 |
| BBOX1 | 7.03609246 | -0.722155706 | 0.90322894 | -0.7995268 | 0.42398504 | 0.99518711 |
| BBOX1-AS1 | 5.62529154 | -1.304314212 | 1.11309847 | -1.1717869 | 0.24128262 | 0.93253689 |
| CCDC34 | 2.58877727 | -0.187007762 | 1.16092674 | -0.1610849 | 0.87202654 | 0.99665574 |
| LGR4 | 208.136383 | -0.031850122 | 0.40186328 | -0.0792561 | 0.93682891 | 0.99777713 |
| LIN7C | 12.9338651 | 0.522657428 | 0.44550336 | 1.17318403 | 0.24072202 | 0.93216487 |
| BDNF-AS | 31.08712 | 0.600711574 | 0.40589472 | 1.47996894 | 0.13888154 | 0.84668189 |
| METTL15 | 50.1766014 | 0.138455932 | 0.47194991 | 0.29336997 | 0.76923937 | 0.99665574 |
| ARL14EP | 7.26396187 | -0.140028044 | 0.58248507 | -0.2403977 | 0.810022 | 0.99665574 |
| MPPED2 | 14.0261618 | 0.006341467 | 0.77086428 | 0.00822644 | 0.99343633 | 0.99796447 |
| DCDC1 | 5.8971136 | -0.004419155 | 0.77864277 | -0.0056755 | 0.99547166 | 0.99865103 |
| DNAJC24 | 11.9919263 | -0.628652653 | 0.57838297 | -1.0869142 | 0.27707473 | 0.95828005 |
| IMMP1L | 27.154998 | 0.446636255 | 0.35206026 | 1.26863582 | 0.20457098 | 0.90834797 |
| ELP4 | 41.2522055 | 0.121528763 | 0.37169666 | 0.32695683 | 0.74370053 | 0.99665574 |
| AC131571.1 | 4.66768067 | -1.141161715 | 0.86750629 | -1.3154506 | 0.18835849 | 0.89739428 |
| RCN1 | 8.4451141 | -1.052288426 | 0.65941282 | -1.5957961 | 0.11053432 | 0.80757694 |
| EIF3M | 17.6794496 | -0.270369475 | 0.442783 | -0.6106139 | 0.54145519 | 0.99665574 |
| CCDC73 | 3.22825204 | -2.583193003 | 0.92014827 | -2.807366 | 0.00499484 | 0.22209534 |
| PRRG4 | 10.78709 | -1.212393875 | 0.55293733 | -2.1926425 | 0.02833315 | 0.5450502 |
| QSER1 | 22.6730379 | 0.012155631 | 0.4214731 | 0.02884082 | 0.97699154 | 0.99777713 |
| TCP11L1 | 4.54241166 | -0.576167359 | 0.650616 | -0.8855721 | 0.37584817 | 0.98637037 |
| CSTF3 | 17.892358 | 0.436923517 | 0.3805812 | 1.14804282 | 0.25095089 | 0.93973926 |

| | | | | | | |
|---|---|---|---|---|---|---|
| HIPK3 | 98.7263287 | -0.032665292 | 0.24561853 | -0.132992 | 0.89419973 | 0.99777713 |
| FBXO3 | 13.5638108 | 0.332944799 | 0.44085304 | 0.75522855 | 0.45011186 | 0.99617422 |
| CAPRIN1 | 32.7344246 | 0.142347895 | 0.32008003 | 0.44472594 | 0.65651782 | 0.99665574 |
| NAT10 | 8.18231381 | 0.268632126 | 0.52892594 | 0.5078823 | 0.61153589 | 0.99665574 |
| ABTB2 | 74.4312082 | -0.121646514 | 0.46808385 | -0.2598819 | 0.79495489 | 0.99665574 |
| CAT | 26.1069764 | 0.067846001 | 0.44755701 | 0.15159186 | 0.87950885 | 0.99665574 |
| EHF | 6.43210886 | -2.037090387 | 0.95736987 | -2.1277987 | 0.03335378 | 0.5761693 |
| APIP | 16.3023599 | 0.634690114 | 0.49680898 | 1.2775335 | 0.20141396 | 0.90591604 |
| PDHX | 32.0391876 | -0.124548679 | 0.31859953 | -0.3909255 | 0.69585231 | 0.99665574 |
| TRIM44 | 60.9018969 | -0.091167372 | 0.24657356 | -0.369737 | 0.71157845 | 0.99665574 |
| COMMD9 | 3.59314575 | -0.620096438 | 0.83992464 | -0.7382763 | 0.46034658 | 0.99617422 |
| TRAF6 | 9.11459084 | 0.42985907 | 0.56894896 | 0.75553187 | 0.44992992 | 0.99617422 |
| C11orf74 | 3.74288776 | -0.200711145 | 0.89083141 | -0.2253077 | 0.82173994 | 0.99665574 |
| API5 | 22.5200883 | 0.064612413 | 0.36619583 | 0.17644224 | 0.85994652 | 0.99665574 |
| TTC17 | 80.7505807 | -0.10672507 | 0.24176152 | -0.4414477 | 0.65888891 | 0.99665574 |
| HSD17B12 | 92.9920943 | 0.332889096 | 0.23657072 | 1.40714408 | 0.15938467 | 0.86757292 |
| ALKBH3 | 4.78491789 | -0.354737077 | 0.59326641 | -0.5979389 | 0.54988069 | 0.99665574 |
| ACCS | 5.6870328 | -0.444043126 | 0.77407078 | -0.5736467 | 0.56620692 | 0.99665574 |
| EXT2 | 20.7640059 | 0.141386658 | 0.37039792 | 0.38171558 | 0.70267234 | 0.99665574 |
| CD82 | 23.7304847 | -0.288981475 | 0.57379147 | -0.503635 | 0.61451788 | 0.99665574 |
| PRDM11 | 6.59644801 | 0.008865465 | 0.52758901 | 0.01680373 | 0.98659319 | 0.99777713 |
| CRY2 | 11.3003572 | 0.128049221 | 0.42480839 | 0.30142818 | 0.76308801 | 0.99665574 |
| PEX16 | 3.94302416 | 0.334903056 | 0.66609141 | 0.50278843 | 0.61511303 | 0.99665574 |
| LARGE2 | 5.42009651 | 1.016526289 | 0.64419928 | 1.57796867 | 0.11457281 | 0.81638817 |
| PHF21A | 67.4799119 | -0.141345657 | 0.34240483 | -0.4128028 | 0.67975113 | 0.99665574 |
| DGKZ | 6.14445631 | -0.128124164 | 0.65912121 | -0.1943863 | 0.84587338 | 0.99665574 |
| AMBRA1 | 132.61006 | 0.220230379 | 0.31289489 | 0.70384779 | 0.48152757 | 0.99640678 |
| ATG13 | 13.0478659 | 0.223457467 | 0.40961794 | 0.54552656 | 0.58539142 | 0.99665574 |
| ARHGAP1 | 9.88530137 | -0.403024058 | 0.58557376 | -0.688255 | 0.49129223 | 0.99640678 |
| CKAP5 | 18.5726017 | -0.22603509 | 0.35668148 | -0.6337169 | 0.52626558 | 0.99665574 |
| LRP4 | 16.9364635 | -1.159129847 | 0.55676883 | -2.0818871 | 0.03735278 | 0.58653453 |
| C11orf49 | 50.2337953 | -0.137023902 | 0.28928332 | -0.4736668 | 0.63573754 | 0.99665574 |
| ARFGAP2 | 9.36222091 | 0.645191866 | 0.55013867 | 1.17278042 | 0.24088387 | 0.93216487 |
| DDB2 | 2.65423102 | 0.977455647 | 0.97321468 | 1.00435769 | 0.31520624 | 0.97652251 |
| ACP2 | 3.962772 | 0.561125091 | 0.72005618 | 0.7792796 | 0.43581503 | 0.99617422 |
| NR1H3 | 4.24606984 | -0.493933899 | 0.68836809 | -0.7175433 | 0.47303894 | 0.99617422 |
| MADD | 7.84914213 | 0.344695706 | 0.54091106 | 0.63725024 | 0.52396186 | 0.99665574 |
| SLC39A13 | 8.15610521 | -0.199083896 | 0.62167806 | -0.3202363 | 0.74878919 | 0.99665574 |
| PSMC3 | 6.11591173 | 0.483620722 | 0.71971591 | 0.67196058 | 0.50160879 | 0.99640678 |
| CELF1 | 87.7471162 | 0.128720833 | 0.33564537 | 0.38350248 | 0.70134723 | 0.99665574 |
| NDUFS3 | 10.9975236 | -0.595389813 | 0.53136925 | -1.1204822 | 0.26250832 | 0.94557817 |
| MTCH2 | 26.5693693 | 0.053729412 | 0.35552048 | 0.15112888 | 0.87987405 | 0.99665574 |
| FNBP4 | 53.3469115 | -0.253247039 | 0.25096654 | -1.0090868 | 0.31293298 | 0.97520189 |
| NUP160 | 53.5414806 | 0.227154601 | 0.29642928 | 0.76630286 | 0.44349612 | 0.99617422 |
| PTPRJ | 243.907144 | -0.388041052 | 0.36472694 | -1.0639221 | 0.28736399 | 0.96559334 |
| TNKS1BP1 | 5.21009095 | -0.865194322 | 0.67525565 | -1.2812841 | 0.2000939 | 0.90398928 |
| SSRP1 | 6.46583469 | -0.028313139 | 0.62627332 | -0.0452089 | 0.96394079 | 0.99777713 |
| TIMM10 | 7.39839565 | 0.012329878 | 0.7088033 | 0.01739534 | 0.98612122 | 0.99777713 |
| UBE2L6 | 4.93554521 | -1.139674506 | 1.27139094 | -0.8963997 | 0.37003931 | 0.98532838 |
| SERPING1 | 48.6537593 | -2.021029714 | 0.55217947 | -3.6600957 | 0.00025212 | 0.03872732 |
| ZDHHC5 | 19.5553017 | -0.093958927 | 0.42647417 | -0.2203156 | 0.82562535 | 0.99665574 |
| MED19 | 3.72431575 | -0.197770585 | 0.87586015 | -0.2258016 | 0.82135577 | 0.99665574 |
| CTNND1 | 39.6584378 | 0.043997142 | 0.30660852 | 0.14349615 | 0.88589835 | 0.99665574 |
| LPXN | 8.47422194 | 0.722063439 | 0.58966721 | 1.22452702 | 0.22075348 | 0.9187187 |
| GLYAT | 5.82535424 | 2.65077328 | 1.1761266 | 2.25381628 | 0.02420773 | 0.50168544 |
| FAM111A | 10.9796323 | 0.245092206 | 0.57716909 | 0.42464541 | 0.67109519 | 0.99665574 |
| OSBP | 32.5154325 | -0.451923471 | 0.44330155 | -1.0194493 | 0.3079897 | 0.97376371 |
| PATL1 | 14.2861773 | -1.044272975 | 0.52691452 | -1.9818641 | 0.04749445 | 0.64060657 |
| STX3 | 19.6794517 | -1.030771038 | 0.49007824 | -2.1032785 | 0.03544143 | 0.58349146 |
| MRPL16 | 6.46020079 | -0.233177792 | 0.71056798 | -0.3281569 | 0.74279303 | 0.99665574 |
| PRPF19 | 7.91416814 | -0.040386586 | 0.63631693 | -0.0634693 | 0.94939281 | 0.99777713 |
| TMEM109 | 5.62539017 | 1.125673203 | 0.64074515 | 1.75681892 | 0.07894868 | 0.74454113 |
| VPS37C | 6.04273123 | -0.338542232 | 0.57439711 | -0.5893871 | 0.55560165 | 0.99665574 |
| DDB1 | 30.7538494 | 0.167293917 | 0.34409805 | 0.48618095 | 0.62683888 | 0.99665574 |
| TKFC | 4.85623183 | 0.159490763 | 0.69367379 | 0.22992185 | 0.81815249 | 0.99665574 |
| CYB561A3 | 3.8074361 | -0.554519762 | 0.77536389 | -0.7151736 | 0.4745018 | 0.99617422 |
| TMEM138 | 3.14910345 | -0.534541095 | 0.86149998 | -0.6204772 | 0.53494366 | 0.99665574 |
| CPSF7 | 18.736424 | 0.021668732 | 0.39371156 | 0.05503707 | 0.95610893 | 0.99777713 |
| AP003108.2 | 7.38359319 | -0.755527635 | 0.50661352 | -1.4913294 | 0.13587504 | 0.84475183 |
| TMEM258 | 7.80500403 | 0.346396541 | 0.71967922 | 0.48132075 | 0.63028855 | 0.99665574 |
| FADS1 | 3.72442797 | -2.555405505 | 1.03112102 | -2.4782789 | 0.01320179 | 0.39020332 |
| FADS3 | 2.7379584 | 0.050287401 | 0.99701319 | 0.05043805 | 0.95977332 | 0.99777713 |
| FTH1 | 205.661021 | -0.495736282 | 0.43678781 | -1.1349591 | 0.25639247 | 0.94314473 |
| ASRGL1 | 5.44477835 | 0.275284624 | 0.71575784 | 0.38460581 | 0.70052949 | 0.99665574 |
| AHNAK | 50.0600216 | -0.63794339 | 0.37175304 | -1.7160408 | 0.08615457 | 0.75397496 |
| B3GAT3 | 7.21904142 | -0.270589231 | 0.58256641 | -0.4644779 | 0.64230538 | 0.99665574 |
| GANAB | 19.1168281 | 0.199545065 | 0.36997861 | 0.53934216 | 0.58965078 | 0.99665574 |
| INTS5 | 2.76823732 | 0.886998892 | 0.8273233 | 1.07213092 | 0.28366123 | 0.96059387 |
| UQCC3 | 6.845737 | 0.24033129 | 0.66777167 | 0.3599004 | 0.71892162 | 0.99665574 |
| UBXN1 | 8.71668646 | 0.067151805 | 0.563467 | 0.11917611 | 0.90513583 | 0.99777713 |
| TTC9C | 7.47980222 | -0.399684575 | 0.62858847 | -0.6358446 | 0.52487772 | 0.99665574 |
| POLR2G | 4.35158846 | 0.604346538 | 0.78072254 | 0.77408619 | 0.43887982 | 0.99617422 |
| TAF6L | 4.51903419 | 0.721226809 | 0.64903278 | 1.11123325 | 0.26646796 | 0.94557817 |
| TMEM223 | 5.24940815 | -0.510255918 | 0.79599221 | -0.6410313 | 0.52150235 | 0.99665574 |
| TMEM179B | 11.8711749 | -0.332223365 | 0.62560863 | -0.5310403 | 0.59539089 | 0.99665574 |
| NXF1 | 6.00980777 | 0.342276554 | 0.63376267 | 0.54007055 | 0.58914838 | 0.99665574 |
| STX5 | 12.1565256 | -0.661771525 | 0.4965109 | -1.3328439 | 0.18258304 | 0.89461655 |
| SLC3A2 | 40.8528948 | -0.877169963 | 0.85180848 | -1.0297737 | 0.30311626 | 0.97376371 |
| SLC22A6 | 5.55648423 | 1.098563139 | 1.04625196 | 1.04999865 | 0.29371874 | 0.9713044 |
| RARRES3 | 96.0584394 | -1.239706562 | 0.72997997 | -1.6982748 | 0.08945591 | 0.75955113 |
| PLA2G16 | 21.3861899 | 0.075272321 | 0.4576527 | 0.16447477 | 0.86935741 | 0.99665574 |
| ATL3 | 20.2364226 | 0.090777166 | 0.38540708 | 0.2355358 | 0.81379289 | 0.99665574 |
| RTN3 | 54.5288572 | -0.304201759 | 0.2419652 | -1.2572128 | 0.20867658 | 0.91043197 |
| C11orf95 | 2.24690133 | 0.703385765 | 0.98989033 | 0.71056939 | 0.47735112 | 0.99617422 |
| MARK2 | 29.0059974 | 0.144907658 | 0.36233807 | 0.39992391 | 0.68921256 | 0.99665574 |
| NAA40 | 7.5568157 | 0.403922915 | 0.53558132 | 0.75417663 | 0.45074317 | 0.99617422 |
| OTUB1 | 10.5820702 | 0.566129498 | 0.55823344 | 1.01414472 | 0.3105137 | 0.97376371 |
| MACROD1 | 13.8851965 | 0.341644169 | 0.61466347 | 0.55582313 | 0.57833178 | 0.99665574 |
| STIP1 | 13.0675295 | -0.082753008 | 0.49406908 | -0.1674928 | 0.86698232 | 0.99665574 |
| NUDT22 | 8.99983114 | 0.058729039 | 0.50124367 | 0.11716664 | 0.906728 | 0.99777713 |
| AP001453.1 | 1.93286182 | 0.527516091 | 1.24146724 | 0.42491342 | 0.67089979 | 0.99665574 |
| DNAJC4 | 4.04529295 | 0.982881314 | 0.72483421 | 1.35600846 | 0.17509648 | 0.88428483 |
| VEGFB | 18.2974465 | 0.531368941 | 0.53603859 | 0.9912886 | 0.32154468 | 0.97700345 |
| FKBP2 | 19.2069287 | 0.662889847 | 0.45808524 | 1.44708841 | 0.14787216 | 0.85308336 |
| PPP1R14B | 13.3760954 | -0.171171345 | 0.53682899 | -0.3188564 | 0.74983543 | 0.99665574 |
| BAD | 8.04365508 | 0.212418193 | 0.6930418 | 0.30650127 | 0.75922301 | 0.99665574 |
| GPR137 | 5.11371724 | -0.048195318 | 0.59716782 | -0.0807065 | 0.93567538 | 0.99777713 |
| ESRRA | 32.3600959 | -0.326124927 | 0.41224704 | -0.791091 | 0.42889089 | 0.99589919 |
| TRMT112 | 13.0919249 | 0.695295405 | 0.55968529 | 1.2422971 | 0.21412697 | 0.9124923 |
| PRDX5 | 27.8747742 | 0.23045711 | 0.41095028 | 0.56079073 | 0.57494021 | 0.99665574 |
| SF1 | 54.2475239 | 0.098762583 | 0.29292988 | 0.33715435 | 0.73600055 | 0.99665574 |
| MAP4K2 | 2.34931867 | 0.256945715 | 1.07369662 | 0.23930942 | 0.81086566 | 0.99665574 |
| MEN1 | 3.16475914 | 1.409708521 | 0.83059309 | 1.69723121 | 0.08965296 | 0.76058717 |

| | | | | | | |
|---|---|---|---|---|---|---|
| EHD1 | 3.76898045 | -0.166423463 | 0.78502769 | -0.2119969 | 0.83210943 | 0.99665574 |
| ATG2A | 8.37997029 | 0.379028424 | 0.56291535 | 0.67333112 | 0.50073666 | 0.99640678 |
| PPP2R5B | 3.08520213 | 0.720892693 | 0.87871064 | 0.82039828 | 0.4119891 | 0.99279547 |
| ZFPL1 | 3.41100677 | -0.437852648 | 0.74047076 | -0.5913166 | 0.55430831 | 0.99665574 |
| VPS51 | 19.6754156 | 0.441611137 | 0.34436432 | 1.28239517 | 0.19970406 | 0.90381759 |
| TM7SF2 | 3.94874893 | 0.58997263 | 0.71950836 | 0.81996633 | 0.4122353 | 0.99279547 |
| FAU | 49.7207279 | 0.227416411 | 0.33205254 | 0.6848808 | 0.49341914 | 0.99640678 |
| SYVN1 | 3.88774431 | -0.133055173 | 0.76898576 | -0.1730268 | 0.86263033 | 0.99665574 |
| MRPL49 | 5.5917214 | -0.67857696 | 0.62428625 | -1.0869644 | 0.27705252 | 0.95828005 |
| CAPN1 | 16.4243589 | -0.770066213 | 0.4128388 | -1.8652951 | 0.06214005 | 0.6847563 |
| DPF2 | 8.9845358 | -0.048814791 | 0.5051385 | -0.0966364 | 0.92301511 | 0.99777713 |
| SLC25A45 | 3.02802899 | 0.584744138 | 0.82738454 | 0.70673805 | 0.47972928 | 0.99617422 |
| FRMD8 | 6.63109022 | -0.248725378 | 0.63459595 | -0.3919429 | 0.6951004 | 0.99665574 |
| NEAT1 | 1338.02 | -0.769960789 | 0.33688077 | -2.2855587 | 0.02228009 | 0.48357743 |
| MALAT1 | 18227.4228 | -0.408743536 | 0.33630988 | -1.2153777 | 0.22422206 | 0.92318867 |
| SCYL1 | 8.84354129 | -0.782503063 | 0.71056905 | -1.1012344 | 0.27079467 | 0.94865166 |
| LTBP3 | 11.0847569 | 0.910509554 | 0.58032205 | 1.56897287 | 0.11665426 | 0.8218386 |
| SSSCA1 | 3.89178113 | -0.264467217 | 0.9632605 | -0.2745542 | 0.78365876 | 0.99665574 |
| FAM89B | 4.87267901 | -0.289576072 | 0.74229927 | -0.3901069 | 0.69645749 | 0.99665574 |
| MAP3K11 | 4.80054889 | 0.268957198 | 0.86211354 | 0.31197422 | 0.75506012 | 0.99665574 |
| PCNX3 | 5.16134791 | 0.356326903 | 0.70151708 | 0.5079376 | 0.6114971 | 0.99665574 |
| RELA | 8.18857051 | -0.246628747 | 0.53767779 | -0.4586925 | 0.64645503 | 0.99665574 |
| KAT5 | 7.08234224 | 0.672006728 | 0.73556898 | 0.91358763 | 0.36093357 | 0.9819332 |
| RNASEH2C | 6.94324238 | 0.455936214 | 0.59345542 | 0.76827374 | 0.44232457 | 0.99617422 |
| AP5B1 | 2.27733748 | -0.560472605 | 1.25138049 | -0.4478834 | 0.65423732 | 0.99665574 |
| CFL1 | 43.7504283 | -0.604707402 | 0.377059 | -1.6037474 | 0.10876974 | 0.80313739 |
| MUS81 | 3.22944639 | 0.635947727 | 1.01915095 | 0.62399758 | 0.53262917 | 0.99665574 |
| FIBP | 3.15798683 | 1.604330738 | 0.95815323 | 1.67439892 | 0.09405222 | 0.76974878 |
| CCDC85B | 6.0156036 | 0.632363441 | 0.75577104 | 0.83671298 | 0.40275392 | 0.99194293 |
| C11orf68 | 3.26546937 | -0.34181447 | 0.84507863 | -0.4044765 | 0.68586233 | 0.99665574 |
| DRAP1 | 6.6184788 | 1.208789967 | 0.65658095 | 1.84103723 | 0.0656161 | 0.69221233 |
| SART1 | 5.79853625 | -0.023244497 | 0.65326782 | -0.0355819 | 0.97161576 | 0.99777713 |
| EIF1AD | 3.62495526 | -0.076495338 | 0.79938812 | -0.0956924 | 0.92376491 | 0.99777713 |
| BANF1 | 12.7323592 | 0.378124608 | 0.52588452 | 0.71902594 | 0.47212494 | 0.99617422 |
| SF3B2 | 15.6331802 | -0.285299364 | 0.41247288 | -0.6916803 | 0.48913812 | 0.99640678 |
| PACS1 | 106.25205 | -0.51861677 | 0.31422713 | -1.6504519 | 0.09885054 | 0.78203824 |
| KLC2 | 5.77125857 | 0.025111574 | 0.68608969 | 0.03660101 | 0.97080314 | 0.99777713 |
| RAB1B | 9.84588335 | -0.326770056 | 0.61696877 | -0.5296379 | 0.596363 | 0.99665574 |
| YIF1A | 5.09561705 | 0.815347279 | 0.70883788 | 1.15025919 | 0.25003714 | 0.93755294 |
| BRMS1 | 4.50189347 | -0.273551933 | 0.92389474 | -0.2960856 | 0.76716471 | 0.99665574 |
| SLC29A2 | 5.89104587 | 0.282799332 | 0.62218448 | 0.45452649 | 0.64944993 | 0.99665574 |
| MRPL11 | 6.01911162 | 0.935204646 | 0.68331017 | 1.36863857 | 0.17111228 | 0.87968373 |
| ZDHHC24 | 4.02918058 | -0.212237698 | 0.65919555 | -0.3219647 | 0.74747944 | 0.99665574 |
| CTSF | 4.33825688 | -0.175933401 | 0.77331902 | -0.2275043 | 0.82003163 | 0.99665574 |
| CCS | 8.11859866 | 0.767658092 | 0.55425937 | 1.38501599 | 0.16604758 | 0.87495175 |
| RBM14-RBM | 7.3228199 | 0.215633317 | 0.50747814 | 0.42491153 | 0.67090117 | 0.99665574 |
| RBM4 | 3.67555274 | -0.314134459 | 0.76153192 | -0.4125033 | 0.67997053 | 0.99665574 |
| RBM4B | 5.85486031 | 0.05334393 | 0.60167349 | 0.08865927 | 0.92935271 | 0.99777713 |
| SPTBN2 | 18.4375051 | 0.187961325 | 0.52998374 | 0.35465489 | 0.72284815 | 0.99665574 |
| C11orf80 | 61.0069274 | -0.63698276 | 0.37577557 | -1.6951149 | 0.09005363 | 0.76220358 |
| RCE1 | 2.33844344 | 1.742040103 | 1.07846669 | 1.61529339 | 0.10624718 | 0.79596216 |
| PC | 3.22854433 | 0.456168433 | 0.75766567 | 0.60207088 | 0.54712696 | 0.99665574 |
| LRFN4 | 2.72334365 | 0.696667103 | 0.94874907 | 0.7343007 | 0.46276549 | 0.99617422 |
| RHOD | 3.82329484 | 0.779779783 | 0.8358479 | 0.93292067 | 0.35086093 | 0.97838306 |
| KDM2A | 90.3218058 | 0.176919871 | 0.26346059 | 0.6715231 | 0.50188735 | 0.99640678 |
| GRK2 | 7.55672335 | 0.260828248 | 0.63633048 | 0.40989432 | 0.68188347 | 0.99665574 |
| ANKRD13D | 7.85578168 | -1.012299221 | 0.66546902 | -1.5211816 | 0.12821427 | 0.83778831 |
| SSH3 | 7.85805544 | -0.239337665 | 0.53932578 | -0.443772 | 0.65720746 | 0.99665574 |
| RAD9A | 25.6094874 | 0.067701984 | 0.35897187 | 0.18859969 | 0.85040658 | 0.99665574 |
| PPP1CA | 15.6229234 | 0.00704658 | 0.49576215 | 0.01421363 | 0.98865954 | 0.99777713 |
| RPS6KB2 | 4.90892702 | 0.637013653 | 0.64097472 | 0.99382025 | 0.32031038 | 0.97700345 |
| CORO1B | 9.03956387 | -0.200186303 | 0.49200802 | -0.4068761 | 0.68409899 | 0.99665574 |
| TMEM134 | 6.33473067 | 0.793519021 | 0.69890542 | 1.13537397 | 0.25621866 | 0.94314473 |
| AIP | 4.6459098 | 0.431925803 | 0.779221 | 0.55430463 | 0.57937039 | 0.99665574 |
| PITPNM1 | 2.55483144 | -0.94970137 | 0.85612561 | -1.1093014 | 0.26730016 | 0.94557817 |
| CDK2AP2 | 12.7968236 | -0.470928041 | 0.66729649 | -0.7057253 | 0.48035896 | 0.99617422 |
| GSTP1 | 100.870143 | -0.592240153 | 0.34259736 | -1.728677 | 0.08386693 | 0.75242737 |
| NDUFV1 | 18.0129232 | 0.665845987 | 0.45610028 | 1.45986752 | 0.14432649 | 0.84969914 |
| NUDT8 | 2.91353133 | 1.134741417 | 0.93229553 | 1.21714776 | 0.223548 | 0.92191316 |
| UNC93B1 | 4.5921468 | -0.772871111 | 0.9070945 | -0.8520293 | 0.39419782 | 0.99179694 |
| NDUFS8 | 21.015534 | -0.369773073 | 0.46098179 | -0.8021425 | 0.42247055 | 0.99518711 |
| TCIRG1 | 3.80037185 | 0.568225257 | 0.94191301 | 0.60326724 | 0.54633093 | 0.99665574 |
| CHKA | 35.4358039 | 0.272393602 | 0.38858204 | 0.70099381 | 0.48330688 | 0.99640678 |
| KMT5B | 37.6085283 | 0.249051466 | 0.33853084 | 0.73568325 | 0.46192348 | 0.99617422 |
| LRP5 | 26.0156648 | 0.102073861 | 0.54708943 | 0.18657619 | 0.85199294 | 0.99665574 |
| PPP6R3 | 122.088856 | 0.07223682 | 0.27640739 | 0.26134185 | 0.79382889 | 0.99665574 |
| TESMIN | 8.78389654 | 0.674248796 | 0.54975742 | 1.22644784 | 0.22003019 | 0.91768188 |
| CPT1A | 65.819299 | 0.124287006 | 0.34014103 | 0.36539845 | 0.71481399 | 0.99665574 |
| MRPL21 | 4.38018394 | -0.872621806 | 0.73460013 | -1.1878868 | 0.23487801 | 0.92960234 |
| IGHMBP2 | 7.06239923 | 0.781626242 | 0.58123217 | 1.34477458 | 0.17869804 | 0.88813816 |
| TPCN2 | 6.65810474 | 0.297013347 | 0.67228934 | 0.44179392 | 0.65863834 | 0.99665574 |
| CCND1 | 3.13026917 | -1.016827321 | 0.86343688 | -1.177651 | 0.23893576 | 0.93157178 |
| ORAOV1 | 5.97839687 | 0.195505134 | 0.73640306 | 0.26548658 | 0.79063464 | 0.99665574 |
| PPFIA1 | 64.3055708 | 0.350324426 | 0.27986037 | 1.25178291 | 0.21064898 | 0.91043197 |
| AP002336.2 | 22.3204465 | -0.475897601 | 0.47462295 | -1.0026856 | 0.31601257 | 0.97652251 |
| AP000487.1 | 2.66673573 | -0.266463381 | 0.82897046 | -0.3214389 | 0.74787778 | 0.99665574 |
| CTTN | 45.3047789 | -0.690063562 | 0.319704 | -2.1584452 | 0.03089324 | 0.56271575 |
| SHANK2 | 139.388321 | 0.438799757 | 0.31136555 | 1.40927524 | 0.1587538 | 0.8671584 |
| DHCR7 | 4.27653777 | -0.441174653 | 0.95173886 | -0.4635459 | 0.64297312 | 0.99665574 |
| NADSYN1 | 12.3456448 | 0.667679172 | 0.43294389 | 1.54218409 | 0.12302886 | 0.82915695 |
| KRTAP5-7 | 6.7393015 | 0.221289472 | 1.0651649 | 0.20775137 | 0.8354231 | 0.99665574 |
| KRTAP5-8 | 19.9541095 | 0.721688241 | 0.80326881 | 0.89843926 | 0.36895141 | 0.98532838 |
| AP002495.2 | 17.6568928 | 0.194059349 | 0.39643034 | 0.4895169 | 0.62447579 | 0.99665574 |
| RNF121 | 24.2804654 | -0.342183332 | 0.4324209 | -0.7913201 | 0.42875724 | 0.99589919 |
| IL18BP | 3.18512549 | -0.025541048 | 0.99930331 | -0.0255589 | 0.9796092 | 0.99777713 |
| NUMA1 | 34.8417645 | 0.260012086 | 0.36539333 | 0.71159505 | 0.47671557 | 0.99617422 |
| LAMTOR1 | 21.1867318 | -0.81172715 | 0.46004871 | -1.7644374 | 0.07765837 | 0.74141484 |
| ANAPC15 | 5.50668837 | -0.243399191 | 0.66855167 | -0.3640694 | 0.7158062 | 0.99665574 |
| FOLR1 | 5.29423616 | -0.570407832 | 0.88590882 | -0.6438674 | 0.51966141 | 0.99665574 |
| INPPL1 | 11.0534317 | 0.174902259 | 0.52088717 | 0.33577763 | 0.73703856 | 0.99665574 |
| CLPB | 14.0404258 | 0.10781177 | 0.42809935 | 0.2518382 | 0.80116613 | 0.99665574 |
| ARAP1 | 18.8286936 | -0.055705075 | 0.42731764 | -0.1303599 | 0.89628171 | 0.99777713 |
| ATG16L2 | 4.48420157 | 0.335318329 | 0.85379027 | 0.39274086 | 0.69451089 | 0.99665574 |
| FCHSD2 | 34.3115389 | -0.654356509 | 0.35105773 | -1.863957 | 0.06232775 | 0.68533482 |
| ARHGEF17 | 7.07592168 | 0.091856975 | 0.51018595 | 0.18004607 | 0.8571164 | 0.99665574 |
| FAM168A | 60.1378044 | -0.473444311 | 0.43992818 | -1.0761855 | 0.28184432 | 0.96059387 |
| PLEKHB1 | 2.5922083 | -0.345478104 | 1.10418195 | -0.3128815 | 0.75437069 | 0.99665574 |
| RAB6A | 56.7310527 | -0.519802259 | 0.31173628 | -1.6674423 | 0.09542649 | 0.77178333 |
| MRPL48 | 12.8934076 | 0.085200571 | 0.40630781 | 0.20969464 | 0.83390601 | 0.99665574 |
| COA4 | 8.21078034 | -0.458038893 | 0.54748755 | -0.8366197 | 0.40280635 | 0.99194293 |
| PAAF1 | 8.66516172 | 0.719417699 | 0.59431979 | 1.21048923 | 0.22609122 | 0.92318867 |
| UCP2 | 6.82948508 | 0.766365603 | 0.82479024 | 0.92916425 | 0.35280397 | 0.97912036 |
| C2CD3 | 16.6369307 | -0.004893617 | 0.43189491 | -0.0113306 | 0.9909597 | 0.99777713 |

| | | | | | | |
|---|---|---|---|---|---|---|
| PPME1 | 21.2482095 | -0.085311673 | 0.34822182 | -0.2449923 | 0.80646237 | 0.99665574 |
| POLD3 | 7.39244696 | -0.104465649 | 0.59296399 | -0.1761754 | 0.86015617 | 0.99665574 |
| RNF169 | 31.3763094 | -0.114127316 | 0.32687006 | -0.3491519 | 0.72697525 | 0.99665574 |
| XRRA1 | 25.3470482 | 0.458440194 | 0.7362866 | 0.62263824 | 0.53352228 | 0.99665574 |
| SPCS2 | 28.8160518 | 0.373164877 | 0.34245551 | 1.08967404 | 0.27585675 | 0.95644178 |
| NEU3 | 6.09722805 | 0.353218151 | 0.62153346 | 0.5683011 | 0.56983053 | 0.99665574 |
| ARRB1 | 3.67130045 | -0.883534721 | 0.86309364 | -1.0236835 | 0.30598479 | 0.97376371 |
| RPS3 | 97.0786625 | 0.395343897 | 0.3247423 | 1.21740805 | 0.223449 | 0.92191316 |
| SERPINH1 | 3.01363063 | 0.721109807 | 0.95718016 | 0.75336895 | 0.45122824 | 0.99617422 |
| UVRAG | 38.5988067 | -0.345391902 | 0.30980281 | -1.1148766 | 0.26490331 | 0.94557817 |
| THAP12 | 9.36841048 | -0.048463705 | 0.63187441 | -0.0766983 | 0.93886354 | 0.99777713 |
| EMSY | 27.362797 | -0.276735217 | 0.35690413 | -0.7753769 | 0.43811699 | 0.99617422 |
| ACER3 | 31.449391 | 0.062550875 | 0.32097501 | 0.19487771 | 0.84548868 | 0.99665574 |
| CAPN5 | 3.2229411 | -0.295530008 | 0.99934369 | -0.2957241 | 0.76744079 | 0.99665574 |
| PAK1 | 15.8852221 | 0.128932451 | 0.42279921 | 0.30494961 | 0.76040453 | 0.99665574 |
| CLNS1A | 14.6058886 | -0.111845893 | 0.49013972 | -0.2281919 | 0.81949709 | 0.99665574 |
| RSF1 | 58.1553673 | 0.079137448 | 0.33907964 | 0.23338897 | 0.81545938 | 0.99665574 |
| AAMDC | 11.8401912 | -0.851741987 | 0.48155691 | -1.7687255 | 0.0769397 | 0.74075464 |
| AP002812.2 | 6.09073311 | -0.403947304 | 0.64726319 | -0.6240851 | 0.53257171 | 0.99665574 |
| INTS4 | 17.9735565 | -0.149172511 | 0.55801724 | -0.267326 | 0.78921818 | 0.99665574 |
| ALG8 | 6.66631823 | -0.481603182 | 0.59613816 | -0.8078718 | 0.41916441 | 0.99476033 |
| GAB2 | 34.0868276 | -0.675249097 | 0.46992366 | -1.4369336 | 0.15073686 | 0.85612189 |
| AP003086.2 | 9.29018851 | -1.730010574 | 0.71636854 | -2.4149728 | 0.01573638 | 0.41872822 |
| NARS2 | 19.9672822 | 0.532233483 | 0.3538517 | 1.50411452 | 0.13255188 | 0.84089786 |
| TENM4 | 9.99126217 | -0.678225935 | 0.86756546 | -0.7817577 | 0.43435701 | 0.99617422 |
| MIR4300HG | 2.97760261 | 3.880570002 | 1.23882007 | 3.13247267 | 0.00173341 | 0.12036484 |
| PRCP | 9.11306598 | 0.718394687 | 0.54135047 | 1.32704176 | 0.18449487 | 0.89532394 |
| RAB30 | 9.04639676 | -0.653895724 | 0.6480613 | -1.0090029 | 0.31297325 | 0.97520189 |
| RAB30-AS1 | 15.0869386 | -0.286898264 | 0.51569957 | -0.5563283 | 0.57798645 | 0.99665574 |
| PCF11 | 15.7580478 | -0.244019663 | 0.39002047 | -0.6256586 | 0.53153889 | 0.99665574 |
| ANKRD42 | 6.07640063 | -0.234217163 | 0.6194502 | -0.3781049 | 0.70535266 | 0.99665574 |
| CCDC90B | 9.12408558 | -0.400223613 | 0.48571067 | -0.8239959 | 0.40994187 | 0.99279547 |
| AP000446.1 | 6.19387354 | 0.110706076 | 0.66969245 | 0.16530883 | 0.86870091 | 0.99665574 |
| DLG2 | 39.1578621 | -0.370514932 | 0.45439278 | -0.8154067 | 0.41483953 | 0.99279547 |
| TMEM126B | 6.85635111 | -0.419727145 | 0.55903316 | -0.750809 | 0.4527676 | 0.99617422 |
| TMEM126A | 4.85755417 | 0.226354709 | 0.68191825 | 0.33193819 | 0.73993593 | 0.99665574 |
| CREBZF | 21.1125628 | 0.215637989 | 0.35265973 | 0.6114619 | 0.54089383 | 0.99665574 |
| SYTL2 | 14.0347452 | -0.987570102 | 0.53250517 | -1.8545737 | 0.06365712 | 0.68630818 |
| PICALM | 439.349492 | 0.676483347 | 0.36474487 | 1.85467543 | 0.06364259 | 0.68630818 |
| EED | 11.5532877 | 0.449182488 | 0.57232691 | 0.78483551 | 0.43255001 | 0.99617422 |
| HIKESHI | 9.08979666 | 0.352548193 | 0.51717605 | 0.68167928 | 0.49544177 | 0.99640678 |
| AP001831.1 | 7.0531376 | -1.3322981 | 0.74805486 | -1.7810166 | 0.07490974 | 0.72883798 |
| ME3 | 25.319177 | 0.20635742 | 0.40534427 | 0.50909173 | 0.61068793 | 0.99665574 |
| PRSS23 | 19.4301078 | -1.237175746 | 0.52008662 | -2.3787879 | 0.01736967 | 0.43587549 |
| FZD4 | 5.48180337 | 0.624506439 | 0.72633969 | 0.85979942 | 0.38989962 | 0.99179694 |
| TMEM135 | 44.746112 | 0.327347678 | 0.39881008 | 0.82081094 | 0.41175396 | 0.99279547 |
| CTSC | 5.20279837 | 0.2481415 | 0.79359456 | 0.31268045 | 0.75452345 | 0.99665574 |
| NOX4 | 15.3471516 | 2.418469808 | 0.82335471 | 2.93733648 | 0.00331045 | 0.18250085 |
| CHORDC1 | 12.0104952 | 0.037926943 | 0.49146217 | 0.07717164 | 0.938487 | 0.99777713 |
| SLC36A4 | 24.4794118 | -0.607616714 | 0.37975615 | -1.6000181 | 0.10959457 | 0.80453999 |
| CEP295 | 11.6535525 | -0.699888641 | 0.43862157 | -1.5956549 | 0.11056587 | 0.80757694 |
| TAF1D | 27.2314305 | -0.482485269 | 0.37755011 | -1.277937 | 0.20127163 | 0.90591604 |
| PANX1 | 5.19161583 | -0.655729461 | 0.66332017 | -0.9885565 | 0.32288018 | 0.97700345 |
| MRE11 | 14.4622764 | 0.186031881 | 0.45420773 | 0.40957445 | 0.68211814 | 0.99665574 |
| ANKRD49 | 4.78382874 | -0.472005671 | 0.65536129 | -0.7202221 | 0.47138827 | 0.99617422 |
| AMOTL1 | 30.2789922 | -0.776378997 | 0.49033057 | -1.5833787 | 0.11333517 | 0.81502052 |
| CWC15 | 7.99926991 | 0.39624995 | 0.54286663 | 0.72992137 | 0.46543825 | 0.99617422 |
| SRSF8 | 7.64994669 | 0.232791348 | 0.52259213 | 0.44545514 | 0.65599087 | 0.99665574 |
| ENDOD1 | 4.84666546 | -1.976071221 | 0.71002398 | -2.7831049 | 0.00538414 | 0.22875404 |
| AP000787.1 | 4.38422243 | -1.48581154 | 0.79924133 | -1.8590274 | 0.06302325 | 0.68628329 |
| SESN3 | 5.26021544 | -0.690513134 | 0.81866091 | -0.8434666 | 0.39896754 | 0.99179694 |
| FAM76B | 7.30790267 | 0.166322256 | 0.55918628 | 0.29743623 | 0.76613349 | 0.99665574 |
| CEP57 | 21.558398 | -0.164312048 | 0.46041258 | -0.35688 | 0.72118163 | 0.99665574 |
| MTMR2 | 25.5327149 | -0.218972061 | 0.43392666 | -0.5046292 | 0.6138193 | 0.99665574 |
| MAML2 | 65.3649315 | 0.414377866 | 0.65340199 | 0.63418519 | 0.52595997 | 0.99665574 |
| CCDC82 | 13.3033303 | -0.298992037 | 0.43092867 | -0.6938318 | 0.48778767 | 0.99640678 |
| JRKL | 3.58081739 | -0.047621536 | 0.88306533 | -0.0539275 | 0.9569929 | 0.99777713 |
| CNTN5 | 5.28710183 | 0.172747441 | 1.21321106 | 0.14238861 | 0.88677306 | 0.99665574 |
| ARHGAP42 | 29.406591 | -0.331650722 | 0.54939319 | -0.6036673 | 0.54606484 | 0.99665574 |
| PGR | 19.9079552 | 0.323405124 | 0.57856138 | 0.55898153 | 0.57617432 | 0.99665574 |
| YAP1 | 101.00714 | -0.363201721 | 0.34790495 | -1.0439682 | 0.29650008 | 0.97341216 |
| BIRC3 | 7.95815369 | -1.021492584 | 1.09166749 | -0.9357177 | 0.34941856 | 0.97838306 |
| BIRC2 | 18.660606 | 0.401346352 | 0.35072084 | 1.14434704 | 0.25247974 | 0.94076968 |
| TMEM123 | 45.8363199 | -0.043082282 | 0.29905143 | -0.1440631 | 0.88545063 | 0.99665574 |
| MMP7 | 13.7224126 | -5.406980361 | 0.69992305 | -7.7251069 | 1.12E-14 | 1.13E-10 |
| DCUN1D5 | 9.74527922 | -0.187339611 | 0.53892057 | -0.3476201 | 0.72812553 | 0.99665574 |
| DYNC2H1 | 26.0091907 | 0.809506831 | 0.41167278 | 1.96638414 | 0.04925425 | 0.64444758 |
| CASP4 | 10.3586148 | -2.331458773 | 0.63780554 | -3.6554382 | 0.00025674 | 0.03884867 |
| MSANTD4 | 6.01816354 | 0.009845078 | 0.55892528 | 0.0176143 | 0.98594655 | 0.99777713 |
| KBTBD3 | 3.54195731 | 0.046826346 | 0.82495412 | 0.05676236 | 0.95473449 | 0.99777713 |
| AASDHPPT | 9.3978223 | -0.249650039 | 0.51672523 | -0.4831389 | 0.62899714 | 0.99665574 |
| CWF19L2 | 29.9489591 | 0.358319137 | 0.34606443 | 1.03541162 | 0.30047672 | 0.97358745 |
| ALKBH8 | 4.46138418 | 0.321713655 | 0.65762784 | 0.48920322 | 0.62469783 | 0.99665574 |
| CUL5 | 102.945379 | 0.60678523 | 0.30554888 | 1.98588597 | 0.04704599 | 0.64060657 |
| ACAT1 | 81.0117939 | 1.110633507 | 0.26533292 | 4.18581125 | 2.84E-05 | 0.00872941 |
| NPAT | 10.2538331 | 0.60172861 | 0.47318105 | 1.27166675 | 0.20349155 | 0.90702762 |
| ATM | 43.3094393 | -0.088561874 | 0.37289693 | -0.2374969 | 0.81227131 | 0.99665574 |
| C11orf65 | 7.8737181 | -0.225854083 | 0.75714113 | -0.2982985 | 0.76547533 | 0.99665574 |
| KDELC2 | 4.33937091 | 0.535765959 | 0.78919937 | 0.67887277 | 0.49721848 | 0.99640678 |
| EXPH5 | 17.8897171 | -0.781263549 | 0.39808765 | -1.9625415 | 0.04969947 | 0.64444758 |
| DDX10 | 29.8335824 | 0.521514106 | 0.36331134 | 1.43544682 | 0.15115981 | 0.85612189 |
| RDX | 41.2127408 | -0.131181196 | 0.27959498 | -0.469183 | 0.63893887 | 0.99665574 |
| ZC3H12C | 19.635542 | -0.215784599 | 0.41805501 | -0.5161632 | 0.60574045 | 0.99665574 |
| FDX1 | 17.709375 | -0.026306394 | 0.36106705 | -0.0728574 | 0.94191962 | 0.99777713 |
| COLCA1 | 2.62664455 | 1.22613349 | 0.95360308 | 1.2857902 | 0.19851629 | 0.90381759 |
| COLCA2 | 2.43949219 | 0.610281234 | 0.92909555 | 0.65685519 | 0.51127403 | 0.99665574 |
| SIK2 | 66.1939873 | 0.173387874 | 0.27403877 | 0.63271293 | 0.52692112 | 0.99665574 |
| PPP2R1B | 19.3361703 | -0.111598248 | 0.41278004 | -0.2703577 | 0.78688511 | 0.99665574 |
| CRYAB | 63.0184807 | -1.795412147 | 0.73573309 | -2.4403036 | 0.01467492 | 0.40872075 |
| DIXDC1 | 4.85637076 | -0.452209629 | 0.68595067 | -0.6592451 | 0.50973838 | 0.99665574 |
| DLAT | 13.0227142 | 0.015518072 | 0.44176778 | 0.03512722 | 0.9719783 | 0.99777713 |
| NKAPD1 | 4.41907467 | -0.996174432 | 0.72764104 | -1.3690465 | 0.17098473 | 0.87968373 |
| TIMM8B | 11.5902741 | -0.28743958 | 0.54381641 | -0.52856 | 0.59711074 | 0.99665574 |
| AP002884.3 | 14.2414761 | -0.494066876 | 0.48434484 | -1.0200726 | 0.30769405 | 0.97376371 |
| IL18 | 322.334784 | -0.557973682 | 0.24865094 | -2.2440039 | 0.02483215 | 0.50755712 |
| BCO2 | 94.407806 | -0.628232672 | 0.38056704 | -1.6507806 | 0.09878339 | 0.78203824 |
| PTS | 24.4884068 | -0.730812925 | 0.36403375 | -2.0075417 | 0.04469202 | 0.6313648 |
| AP002884.1 | 5.66855854 | -1.019541006 | 0.7297459 | -1.3971178 | 0.16237814 | 0.87130922 |
| TTC12 | 7.26571478 | -0.226516792 | 0.57348095 | -0.3949857 | 0.69285342 | 0.99665574 |
| ZW10 | 5.25631621 | 0.180356211 | 0.67495219 | 0.26721331 | 0.78930493 | 0.99665574 |
| USP28 | 13.4205344 | 0.709880894 | 0.49923199 | 1.42194593 | 0.15504195 | 0.86239367 |
| ZBTB16 | 76.0057178 | -0.3841267 | 0.56174664 | -0.6838077 | 0.49409658 | 0.99640678 |
| AP002518.2 | 4.08938828 | -1.800932992 | 1.23225957 | -1.4614883 | 0.14388147 | 0.84969914 |

| | | | | | | | |
|---|---|---|---|---|---|---|---|
| NNMT | 12.3310464 | | -2.504245277 | 0.73239926 | -3.4192351 | 0.00062797 | 0.06758588 |
| C11orf71 | 2.27043767 | | 0.368382978 | 0.9215849 | 0.39972766 | 0.68935711 | 0.99665574 |
| REXO2 | 9.02182294 | | -0.620393925 | 0.52200079 | -1.1884923 | 0.23463951 | 0.92960234 |
| CADM1 | 24.2880999 | | 0.702936909 | 0.60406031 | 1.16368663 | 0.24455103 | 0.93426912 |
| BUD13 | 3.12485331 | | -0.300187803 | 0.83898151 | -0.3578003 | 0.72049279 | 0.99665574 |
| ZPR1 | 3.87143003 | | -0.505625667 | 0.74526271 | -0.6784529 | 0.49748456 | 0.99640678 |
| SIK3 | 253.772072 | | 0.494578798 | 0.25806835 | 1.91646437 | 0.05530601 | 0.67128562 |
| PAFAH1B2 | 43.4401821 | | -0.023173419 | 0.33297512 | -0.069595 | 0.94451598 | 0.99777713 |
| SIDT2 | 6.98757145 | | -0.31297507 | 0.59131807 | -0.5292838 | 0.5966086 | 0.99665574 |
| TAGLN | 4.69318801 | | 0.693086767 | 0.81571252 | 0.84967038 | 0.39550837 | 0.99179694 |
| PCSK7 | 20.0037869 | | -0.172227991 | 0.35632579 | -0.4833442 | 0.62885138 | 0.99665574 |
| RNF214 | 11.9825406 | | 0.498147646 | 0.45072575 | 1.1052123 | 0.26906761 | 0.94673212 |
| BACE1 | 8.17958536 | | -0.392655418 | 0.55305657 | -0.7099733 | 0.47772067 | 0.99617422 |
| CEP164 | 8.72770016 | | -0.254590147 | 0.65230589 | -0.3902926 | 0.69632021 | 0.99665574 |
| DSCAML1 | 19.1937992 | | -0.036675945 | 0.97305272 | -0.0376916 | 0.96993355 | 0.99777713 |
| FXYD2 | 16.4610543 | | 0.787560589 | 0.64698273 | 1.21728224 | 0.22349685 | 0.92191316 |
| AP000757.1 | 14.6486581 | | 1.818453569 | 0.67996221 | 2.67434505 | 0.00748754 | 0.28644773 |
| MPZL3 | 3.62093794 | | -0.30760485 | 0.8807507 | -0.349253 | 0.72689936 | 0.99665574 |
| UBE4A | 19.2133154 | | 0.335467727 | 0.41834382 | 0.80189478 | 0.42261382 | 0.99518711 |
| ATP5MG | 48.7097198 | | 0.591978607 | 0.40289668 | 1.46930623 | 0.14174975 | 0.84969914 |
| KMT2A | 54.6771385 | | 0.096612614 | 0.28107702 | 0.34372292 | 0.73105467 | 0.99665574 |
| AP001267.3 | 2.14005143 | | 0.378127667 | 1.27010438 | 0.29771385 | 0.76592157 | 0.99665574 |
| IFT46 | 5.03114026 | | 0.80949467 | 0.6885088 | 1.1757216 | 0.23970614 | 0.93157178 |
| ARCN1 | 13.4379534 | | -0.403214003 | 0.42574954 | -0.9470685 | 0.34360385 | 0.97788427 |
| AP002954.1 | 6.76903377 | | 0.004586741 | 0.69353225 | 0.00661359 | 0.99472315 | 0.9985287 |
| DDX6 | 41.44284 | | 0.235301338 | 0.30047342 | 0.783102 | 0.43356722 | 0.99617422 |
| BCL9L | 4.17272122 | | -0.861752945 | 0.75455569 | -1.1420667 | 0.25342629 | 0.94076968 |
| CCDC84 | 25.6302317 | | 0.016486959 | 0.3948955 | 0.04175018 | 0.96669785 | 0.99777713 |
| RPS25 | 60.6196218 | | 0.203180079 | 0.46533966 | 0.43662747 | 0.66238154 | 0.99665574 |
| TRAPPC4 | 11.3250572 | | 0.168596623 | 0.48069423 | 0.3507357 | 0.72578664 | 0.99665574 |
| SLC37A4 | 4.0119761 | | -0.919517837 | 0.82750264 | -1.1111963 | 0.26648388 | 0.94557817 |
| HYOU1 | 7.40406646 | | 0.103780922 | 0.70149809 | 0.14794185 | 0.88238866 | 0.99665574 |
| VPS11 | 5.55990689 | | -0.082003373 | 0.61497418 | -0.1333444 | 0.893921 | 0.99777713 |
| H2AFX | 2.87761805 | | -0.420970961 | 1.01757916 | -0.4136985 | 0.67909493 | 0.99665574 |
| HINFP | 3.53196077 | | -0.636740207 | 0.75020738 | -0.8487523 | 0.39601916 | 0.99179694 |
| NLRX1 | 4.3657275 | | -0.597513147 | 0.65026279 | -0.9188795 | 0.3581586 | 0.9819332 |
| CBL | 17.3876763 | | -0.016585547 | 0.38995783 | -0.0425316 | 0.96607489 | 0.99777713 |
| NECTIN1 | 3.85965103 | | 0.626580613 | 0.81424819 | 0.76952042 | 0.44158443 | 0.99617422 |
| OAF | 10.4134091 | | 0.863381681 | 0.54735509 | 1.57737032 | 0.11471034 | 0.81638817 |
| POU2F3 | 1.86179511 | | 0.997680664 | 1.34233511 | 0.74324262 | 0.45733481 | 0.99617422 |
| ARHGEF12 | 163.960779 | | -0.389544877 | 0.33137978 | -1.175524 | 0.23978514 | 0.93157178 |
| SC5D | 9.6035985 | | -0.02860532 | 0.60767483 | -0.0470734 | 0.96245473 | 0.99777713 |
| SORL1 | 28.9249573 | | 1.00358042 | 0.63002333 | 1.59292579 | 0.11117684 | 0.8083966 |
| UBASH3B | 35.5951649 | | -2.434615445 | 0.66005916 | -3.6884806 | 0.0002256 | 0.03630324 |
| HSPA8 | 36.5162612 | | 0.293985257 | 0.30302125 | 0.97018034 | 0.33195661 | 0.97788427 |
| GRAMD1B | 59.1675299 | | 0.83517001 | 0.41806764 | 1.99769111 | 0.04575016 | 0.63477358 |
| ZNF202 | 2.76774294 | | 0.078974439 | 0.81228562 | 0.09722496 | 0.92254775 | 0.99777713 |
| TBRG1 | 13.4348277 | | -0.134767058 | 0.44777318 | -0.3009717 | 0.76343607 | 0.99665574 |
| SIAE | 7.41641462 | | 0.154521607 | 0.62000191 | 0.24922763 | 0.80318471 | 0.99665574 |
| MSANTD2 | 15.1720117 | | 0.581081179 | 0.40274157 | 1.44281401 | 0.14907287 | 0.85485259 |
| EI24 | 14.5412523 | | 0.443876608 | 0.4110309 | 1.07991055 | 0.28018202 | 0.96059387 |
| STT3A | 8.00899065 | | -0.211107507 | 0.65495113 | -0.3223256 | 0.74720605 | 0.99665574 |
| CDON | 5.55109954 | | 0.16586242 | 0.5955888 | 0.27848478 | 0.78064024 | 0.99665574 |
| RPUSD4 | 4.19988549 | | 0.243042489 | 0.6988364 | 0.34778167 | 0.72800416 | 0.99665574 |
| FAM118B | 5.93753412 | | 0.038384355 | 0.55204754 | 0.06953089 | 0.94456705 | 0.99777713 |
| SRPRA | 12.4392418 | | -0.656021472 | 0.42613501 | -1.5394686 | 0.12368994 | 0.82915695 |
| FOXRED1 | 7.19759636 | | -0.191234565 | 0.55809717 | -0.3426546 | 0.73185832 | 0.99665574 |
| TIRAP | 3.08697451 | | -0.352694509 | 0.87764619 | -0.4018641 | 0.68778403 | 0.99665574 |
| DCPS | 10.7586146 | | 0.451748347 | 0.57558585 | 0.78484964 | 0.43254173 | 0.99617422 |
| ST3GAL4 | 4.73821899 | | -0.258544372 | 0.80026047 | -0.3230753 | 0.74663824 | 0.99665574 |
| KCNJ1 | 9.21946629 | | 1.852456816 | 0.96040908 | 1.9288206 | 0.05375314 | 0.6653339 |
| ARHGAP32 | 53.1944775 | | 0.460973166 | 0.33867571 | 1.3611049 | 0.17348054 | 0.88157681 |
| TMEM45B | 11.7451019 | | -1.280492408 | 0.45984789 | -2.7846 | 0.00535938 | 0.22875404 |
| NFRKB | 8.48078638 | | 0.327598081 | 0.50667243 | 0.6465678 | 0.5179117 | 0.99665574 |
| PRDM10 | 9.2151448 | | -0.449857443 | 0.47165011 | -0.9537948 | 0.3401875 | 0.97788427 |
| APLP2 | 85.9874125 | | -0.230599521 | 0.23151556 | -0.9960433 | 0.31922912 | 0.97700345 |
| ST14 | 43.0994283 | | -0.182815965 | 0.34043942 | -0.537 | 0.59126762 | 0.99665574 |
| ZBTB44 | 37.8814454 | | 0.033672272 | 0.41824803 | 0.0805079 | 0.93583331 | 0.99777713 |
| AP002986.1 | 4.67392065 | | 0.59736966 | 0.80124344 | 0.74555326 | 0.45593733 | 0.99617422 |
| SNX19 | 22.2131451 | | 0.12154718 | 0.41095425 | 0.29576816 | 0.76740714 | 0.99665574 |
| NCAPD3 | 4.30611706 | | -0.166206844 | 0.69574207 | -0.2388915 | 0.81118974 | 0.99665574 |
| VPS26B | 5.25784164 | | 0.294725663 | 0.59078787 | 0.49886885 | 0.61787178 | 0.99665574 |
| ACAD8 | 4.54456805 | | -0.193951311 | 0.87034403 | -0.2228444 | 0.8236566 | 0.99665574 |
| ZMYND11 | 55.6525867 | | 0.280338079 | 0.27319345 | 1.02615227 | 0.30481982 | 0.97376371 |
| DIP2C | 162.602354 | | 0.217194756 | 0.52304285 | 0.41525232 | 0.6779572 | 0.99665574 |
| LARP4B | 71.0489259 | | 0.141767588 | 0.28185492 | 0.5029807 | 0.61497784 | 0.99665574 |
| GTPBP4 | 8.97151151 | | -0.23245288 | 0.57366825 | -0.4052044 | 0.68532729 | 0.99665574 |
| IDI1 | 13.0843932 | | -1.303921747 | 0.70221794 | -1.8568619 | 0.0633308 | 0.68630818 |
| WDR37 | 25.0211913 | | 0.082664439 | 0.36471402 | 0.2266555 | 0.82069163 | 0.99665574 |
| ADARB2 | 104.704842 | | 0.163209778 | 1.45476471 | 0.11218981 | 0.91067291 | 0.99777713 |
| PFKP | 17.8902644 | | -0.706640905 | 0.54763816 | -1.2903427 | 0.19693169 | 0.90381759 |
| PITRM1 | 8.79817906 | | -0.143977095 | 0.48783846 | -0.2951327 | 0.76789249 | 0.99665574 |
| KLF6 | 84.733578 | | -0.50801131 | 0.65038896 | -0.7810885 | 0.43475047 | 0.99617422 |
| AKR1C1 | 18.0430047 | | -3.05821977 | 0.65803312 | -4.6475165 | 3.36E-06 | 0.00189217 |
| AKR1C2 | 7.14156565 | | -0.85834004 | 0.83912803 | -1.0228952 | 0.3063574 | 0.97376371 |
| AKR1C3 | 4.8410308 | | 0.295801217 | 0.96406585 | 0.30682678 | 0.75897523 | 0.99665574 |
| NET1 | 4.72037538 | | -0.043868268 | 0.74728032 | -0.0587039 | 0.95318795 | 0.99777713 |
| ASB13 | 8.08278571 | | 1.174533927 | 0.6305321 | 1.86276628 | 0.06249516 | 0.6856882 |
| FAM208B | 46.5486505 | | -0.198150669 | 0.37628175 | -0.5266019 | 0.5984701 | 0.99665574 |
| AL365356.1 | 3.5594157 | | 0.872797556 | 0.98017727 | 0.89044868 | 0.37322501 | 0.98532838 |
| GDI2 | 29.9158548 | | -0.651125874 | 0.33706722 | -1.931739 | 0.05339173 | 0.66496977 |
| ANKRD16 | 3.00470537 | | -0.577808146 | 0.84548605 | -0.6834035 | 0.49435189 | 0.99640678 |
| FBH1 | 25.9815995 | | 0.26314748 | 0.31657606 | 0.83122988 | 0.40584378 | 0.99194293 |
| RBM17 | 16.3000392 | | 0.833374213 | 0.42025129 | 1.98303786 | 0.0473632 | 0.64060657 |
| PFKFB3 | 17.7871155 | | -0.961563279 | 0.77880027 | -1.2346725 | 0.21695242 | 0.9156801 |
| PRKCQ | 15.7995437 | | -0.373142398 | 0.53775693 | -0.6938867 | 0.48775325 | 0.99640678 |
| SFMBT2 | 18.2868833 | | 0.217503114 | 0.49696592 | 0.43766203 | 0.6616313 | 0.99665574 |
| ITIH5 | 96.752174 | | -0.774109292 | 0.52967664 | -1.4614752 | 0.14388507 | 0.84969914 |
| KIN | 7.91246519 | | 0.085519969 | 0.55317482 | 0.15459845 | 0.87713789 | 0.99665574 |
| ATP5F1C | 29.2361235 | | 0.270244805 | 0.38837399 | 0.69583652 | 0.48653121 | 0.99640678 |
| TAF3 | 43.1073097 | | 0.311195478 | 0.32999313 | 0.94303623 | 0.34566237 | 0.97788427 |
| GATA3 | 12.9542632 | | -0.623416103 | 0.64888124 | -0.9607553 | 0.33667521 | 0.97788427 |
| SFTA1P | 3.67934924 | | -0.666818316 | 1.11754598 | -0.5966809 | 0.55072046 | 0.99665574 |
| CELF2 | 669.189927 | | -0.110743706 | 0.30247816 | -0.3661213 | 0.71427455 | 0.99665574 |
| CELF2-AS2 | 5.45327876 | | -1.00147885 | 0.82529534 | -1.2134794 | 0.2249466 | 0.92318867 |
| USP6NL | 37.3864288 | | 0.006800367 | 0.26924409 | 0.02525726 | 0.97984977 | 0.99777713 |
| ECHDC3 | 8.16122267 | | 0.302226568 | 0.71893258 | 0.42038235 | 0.67420616 | 0.99665574 |
| PROSER2 | 16.375558 | | -0.097571186 | 0.4521775 | -0.2157807 | 0.82915868 | 0.99665574 |
| UPF2 | 30.1775828 | | 0.204308193 | 0.33904581 | 0.60259761 | 0.54677641 | 0.99665574 |
| DHTKD1 | 15.8018364 | | 0.612433922 | 0.44682791 | 1.37062594 | 0.17049159 | 0.87918701 |
| SEC61A2 | 4.66405159 | | -0.040204285 | 0.71161407 | -0.0564973 | 0.95494563 | 0.99777713 |
| NUDT5 | 15.3895557 | | 0.017352697 | 0.4827392 | 0.03594632 | 0.97132516 | 0.99777713 |
| CDC123 | 14.0790931 | | -0.189158386 | 0.48976421 | -0.3862234 | 0.69933124 | 0.99665574 |

| | | | | | | |
|---|---|---|---|---|---|---|
| CAMK1D | 54.2967721 | 0.60244912 | 1.13674678 | 0.52997653 | 0.5961282 | 0.99665574 |
| OPTN | 11.3411112 | -1.465292809 | 0.63683512 | -2.3008982 | 0.02139738 | 0.4733368 |
| PHYH | 12.9174346 | 0.619539331 | 0.60243227 | 1.02839666 | 0.30376328 | 0.97376371 |
| SEPHS1 | 9.00473687 | -0.092203381 | 0.47512069 | -0.1940631 | 0.84612649 | 0.99665574 |
| BEND7 | 31.6510378 | 0.0803785 | 0.34926813 | 0.23013408 | 0.81798758 | 0.99665574 |
| PRPF18 | 3.73090197 | -0.178836289 | 0.68728637 | -0.2602064 | 0.79470459 | 0.99665574 |
| AL157392.5 | 9.23678105 | -0.713610054 | 0.49666927 | -1.4367912 | 0.15077733 | 0.85612189 |
| FRMD4A | 12.3822939 | -0.897646664 | 0.45439247 | -1.9754875 | 0.04821285 | 0.6422889 |
| FAM107B | 106.317833 | 0.809810131 | 0.32477566 | 2.49344467 | 0.01265103 | 0.38590337 |
| HSPA14.1 | 2.90544506 | 0.409487238 | 0.90524286 | 0.4523507 | 0.65101636 | 0.99665574 |
| DCLRE1C | 8.20246427 | 0.40948605 | 0.52284726 | 0.78318485 | 0.43351857 | 0.99617422 |
| RPP38 | 11.2281219 | -0.140319768 | 0.46611182 | -0.3010431 | 0.76338159 | 0.99665574 |
| NMT2 | 4.50761791 | -1.030654558 | 0.72200932 | -1.427481 | 0.15344131 | 0.8618391 |
| FAM171A1 | 13.0528052 | -0.553314079 | 0.74897395 | -0.7387628 | 0.46005105 | 0.99617422 |
| MINDY3 | 19.3852333 | -0.265970993 | 0.34130046 | -0.7792869 | 0.43581071 | 0.99617422 |
| PTER | 28.5777585 | 0.859034976 | 0.4199469 | 2.04558 | 0.04079771 | 0.61100706 |
| RSU1 | 22.4351582 | -0.415544039 | 0.43150087 | -0.9630202 | 0.33553741 | 0.97788427 |
| CUBN | 25.4604903 | 2.137662534 | 1.24936498 | 1.71099925 | 0.08708125 | 0.75584737 |
| TRDMT1 | 3.15623842 | -0.240254164 | 0.8485838 | -0.2831237 | 0.77708203 | 0.99665574 |
| VIM | 5.87776385 | -1.485202193 | 0.73599362 | -2.0179553 | 0.04359592 | 0.62204719 |
| STAM | 14.6541172 | -0.141464692 | 0.44508563 | -0.317837 | 0.75060856 | 0.99665574 |
| CACNB2 | 36.7805163 | 1.669623036 | 0.57766266 | 2.89030807 | 0.00384864 | 0.20301691 |
| AL450384.2 | 6.58907025 | -0.03972046 | 0.63768583 | -0.0622884 | 0.95033313 | 0.99777713 |
| NSUN6 | 23.5078707 | 0.474082815 | 0.48365924 | 0.98020005 | 0.32698738 | 0.97788427 |
| ARL5B | 8.53704515 | -0.640218171 | 0.61291987 | -1.0445381 | 0.29623649 | 0.97286866 |
| MALRD1 | 5.1486625 | 1.081541242 | 0.88691014 | 1.21944851 | 0.22267401 | 0.92191316 |
| PLXDC2 | 128.454697 | -0.109513007 | 0.40831568 | -0.2682067 | 0.78854021 | 0.99665574 |
| NEBL | 356.938448 | 0.054806442 | 0.2733609 | 0.20049115 | 0.84109648 | 0.99665574 |
| MLLT10 | 78.9599012 | -0.033848931 | 0.33689143 | -0.1004743 | 0.91996779 | 0.99777713 |
| DNAJC1 | 73.5155243 | -0.407340758 | 0.39767969 | -1.0242936 | 0.30569663 | 0.97376371 |
| BMI1 | 8.19179035 | 0.738711014 | 0.57344441 | 1.28819988 | 0.1976764 | 0.90381759 |
| PIP4K2A | 31.931323 | -0.953706889 | 0.48611231 | -1.9619065 | 0.04977337 | 0.64444758 |
| MSRB2 | 10.6117713 | 0.397081853 | 0.55477914 | 0.71574763 | 0.47414719 | 0.99617422 |
| OTUD1 | 6.08879286 | 0.561591396 | 0.80493877 | 0.69768213 | 0.48537601 | 0.99640678 |
| KIAA1217 | 172.296873 | -0.315147889 | 0.35398483 | -0.8902864 | 0.37331211 | 0.98532838 |
| ARHGAP21 | 33.5976519 | -0.049540108 | 0.31580782 | -0.1568679 | 0.87534896 | 0.99665574 |
| PDSS1 | 7.60699461 | -1.164614087 | 0.59212738 | -1.9668303 | 0.04920277 | 0.64444758 |
| ABI1 | 47.9613113 | 0.066420443 | 0.28150839 | 0.23594481 | 0.8134755 | 0.99665574 |
| ANKRD26 | 29.3942043 | -0.279470241 | 0.35785484 | -0.7809598 | 0.43482614 | 0.99617422 |
| YME1L1 | 29.6584171 | -0.315405089 | 0.36672286 | -0.8600639 | 0.38975383 | 0.99179694 |
| MASTL | 7.06742132 | -0.048248624 | 0.55729777 | -0.086576 | 0.93100852 | 0.99777713 |
| ACBD5 | 20.9171247 | -0.355286778 | 0.37634776 | -0.9440385 | 0.34514998 | 0.97788427 |
| RAB18 | 29.2855116 | -0.23466615 | 0.43967785 | -0.5337229 | 0.59353324 | 0.99665574 |
| MPP7 | 45.9084919 | 0.156651485 | 0.35642351 | 0.43950941 | 0.66029247 | 0.99665574 |
| WAC-AS1 | 7.21495971 | -0.979520533 | 0.59642653 | -1.6423155 | 0.10052464 | 0.78386461 |
| WAC | 102.642578 | 0.037605158 | 0.2565479 | 0.14658143 | 0.88346241 | 0.99665574 |
| BAMBI | 4.0414672 | 0.819075811 | 0.70434983 | 1.1628821 | 0.24487734 | 0.93426912 |
| SVIL | 37.2012402 | -0.809437964 | 0.45380361 | -1.7836746 | 0.07447656 | 0.72810353 |
| MTPAP | 6.69283401 | 0.483888212 | 0.6115552 | 0.79124209 | 0.42880273 | 0.99589919 |
| MAP3K8 | 37.6615811 | -0.025166563 | 0.40696169 | -0.0618401 | 0.95069015 | 0.99777713 |
| ZNF438 | 19.0056273 | -0.507236692 | 0.44798635 | -1.1322593 | 0.25752545 | 0.94557817 |
| ZEB1 | 14.1778038 | -1.338183996 | 0.70188595 | -1.9065548 | 0.05657827 | 0.67545594 |
| ARHGAP12 | 50.1960809 | -0.076005345 | 0.30242393 | -0.2513205 | 0.8015663 | 0.99665574 |
| KIF5B | 37.7802662 | 0.098513229 | 0.34094623 | 0.28894066 | 0.77262679 | 0.99665574 |
| EPC1 | 57.4747904 | 0.110147826 | 0.26457698 | 0.41631673 | 0.67717825 | 0.99665574 |
| CCDC7 | 29.5832056 | 0.248758364 | 0.45630115 | 0.54516269 | 0.58564163 | 0.99665574 |
| ITGB1 | 39.9280967 | -0.386564456 | 0.28764731 | -1.3438834 | 0.17898609 | 0.88813816 |
| NRP1 | 31.3856941 | -1.850003973 | 0.60694867 | -3.0480402 | 0.00230339 | 0.1502451 |
| PARD3 | 241.611356 | 0.253729035 | 0.25279775 | 1.00368393 | 0.31553098 | 0.97652251 |
| CUL2 | 15.1218122 | -0.142404984 | 0.42081578 | -0.3384022 | 0.73506012 | 0.99665574 |
| CREM | 37.1820083 | -0.053268582 | 0.38871328 | -0.1370382 | 0.89100057 | 0.99712593 |
| CCNY | 78.4658304 | -0.086700095 | 0.30995265 | -0.2797205 | 0.77969199 | 0.99665574 |
| AL117336.3 | 4.94046425 | -0.576873955 | 0.65588224 | -0.8795389 | 0.37910914 | 0.98679664 |
| ANKRD30A | 6.51290635 | 1.447706861 | 0.89783899 | 1.61243483 | 0.10686735 | 0.79791531 |
| ZNF248 | 18.929814 | -0.080686122 | 0.47941435 | -0.1683014 | 0.86634615 | 0.99665574 |
| ZNF33A | 52.9429352 | -0.119933196 | 0.26224527 | -0.4573322 | 0.64743231 | 0.99665574 |
| ZNF33B | 32.53421 | -0.019304349 | 0.39391694 | -0.0490061 | 0.9609144 | 0.99777713 |
| BMS1 | 12.3355659 | -0.292605513 | 0.42320057 | -0.691411 | 0.48930732 | 0.99640678 |
| CSGALNACT2 | 6.74120425 | -0.090711607 | 0.6495928 | -0.1396438 | 0.88894143 | 0.99665574 |
| FXYD4 | 10.8517166 | 1.04575441 | 0.80956823 | 1.29174339 | 0.19644603 | 0.90381759 |
| HNRNPF | 31.7278238 | -0.595133808 | 0.33658708 | -1.7681422 | 0.07703714 | 0.74098911 |
| ZNF487 | 3.50408541 | -1.861086372 | 0.76638952 | -2.4283818 | 0.01516637 | 0.41291622 |
| ZNF32 | 8.37530878 | 0.587857982 | 0.57949777 | 1.01442665 | 0.31037921 | 0.97376371 |
| TMEM72-AS | 3.84293218 | 0.027511818 | 1.16356325 | 0.02364445 | 0.98113621 | 0.99777713 |
| TMEM72 | 27.1775277 | 0.808219825 | 0.45447848 | 1.77834563 | 0.07534711 | 0.73097508 |
| RASSF4 | 13.4023876 | 1.266065913 | 0.6572715 | 1.92624497 | 0.05407379 | 0.6653339 |
| DEPP1 | 21.0844538 | -1.884032236 | 0.80940438 | -2.3276774 | 0.01992924 | 0.46397674 |
| ZNF22 | 5.59010251 | -0.010524581 | 0.77216764 | -0.0136299 | 0.98912524 | 0.99777713 |
| 8-Mar | 23.1289562 | -0.664523994 | 0.45015179 | -1.476222 | 0.1398843 | 0.84918982 |
| ZFAND4 | 10.7344101 | 0.529615819 | 0.44291723 | 1.19574444 | 0.2317963 | 0.92810069 |
| WASHC2C | 13.3082255 | -0.336153083 | 0.53196094 | -0.6319131 | 0.52744367 | 0.99665574 |
| AGAP4 | 7.89616524 | 0.763142679 | 0.54268515 | 1.40623469 | 0.15965445 | 0.86757292 |
| TIMM23 | 12.0249929 | -0.051493253 | 0.4819667 | -0.1068399 | 0.91491603 | 0.99777713 |
| NCOA4 | 20.3109916 | 0.260375719 | 0.39850464 | 0.65338189 | 0.51351011 | 0.99665574 |
| PTPN20 | 16.5732402 | 0.025360414 | 0.51498864 | 0.04924461 | 0.96072436 | 0.99777713 |
| AGAP9 | 6.37254349 | -0.493655289 | 0.57633756 | -0.8565385 | 0.39170001 | 0.99179694 |
| MAPK8 | 88.1455261 | 0.076386892 | 0.24518543 | 0.31154743 | 0.75538449 | 0.99665574 |
| VSTM4 | 10.0264063 | 1.412861089 | 0.53697983 | 2.63112508 | 0.00851027 | 0.31119582 |
| ERCC6 | 15.093375 | 0.688641765 | 0.41681992 | 1.65213258 | 0.09850753 | 0.78203824 |
| OGDHL | 65.8172002 | 0.331099081 | 0.44820328 | 0.73872525 | 0.46007385 | 0.99617422 |
| PARG | 31.3384501 | 0.344387618 | 0.3300864 | 1.04332569 | 0.29679748 | 0.97341216 |
| TIMM23B | 27.0225128 | -0.172825444 | 0.55198713 | -0.3130969 | 0.75420707 | 0.99665574 |
| LINC00843 | 6.5806337 | 0.408470685 | 0.53891187 | 0.75795452 | 0.44847821 | 0.99617422 |
| AGAP6 | 7.97750156 | 0.064035801 | 0.52545654 | 0.12186698 | 0.90300437 | 0.99777713 |
| WASHC2A | 11.0944208 | -0.900888341 | 0.83838963 | -1.0745461 | 0.28257798 | 0.96059387 |
| SGMS1 | 158.434929 | -0.214720963 | 0.34569237 | -0.6211331 | 0.53451207 | 0.99665574 |
| PRKG1 | 27.4114338 | -1.013744744 | 0.63335664 | -1.6005907 | 0.1094676 | 0.80453999 |
| IPMK | 11.2410294 | -0.460026805 | 0.45526941 | -1.0104496 | 0.31227992 | 0.97442101 |
| CISD1 | 32.3660066 | 0.789208474 | 0.37550585 | 2.10172084 | 0.03557774 | 0.58349146 |
| UBE2D1 | 13.158781 | 0.58291345 | 0.44054866 | 1.32315338 | 0.18578438 | 0.89532394 |
| TFAM | 9.58713133 | 0.394352099 | 0.48704058 | 0.80969044 | 0.41811812 | 0.99432434 |
| BICC1 | 153.070651 | 0.105888034 | 0.51147941 | 0.20702306 | 0.83599185 | 0.99665574 |
| FAM13C | 14.4869272 | 0.222692549 | 0.93260373 | 0.23878582 | 0.81127166 | 0.99665574 |
| CCDC6 | 19.5314348 | -0.27347137 | 0.43889283 | -0.6230937 | 0.53322293 | 0.99665574 |
| ANK3 | 183.585199 | -0.162424242 | 0.37638082 | -0.4315423 | 0.66607411 | 0.99665574 |
| RHOBTB1 | 10.7498365 | -0.221320815 | 0.7990481 | -0.2769806 | 0.78179501 | 0.99665574 |
| ARID5B | 26.5311302 | -1.307102062 | 0.5455012 | -2.3961488 | 0.01656836 | 0.42361121 |
| ADO | 3.33422024 | -0.161190149 | 0.81839576 | -0.1969587 | 0.84385987 | 0.99665574 |
| NRBF2 | 4.09874387 | -0.69750909 | 0.70055433 | -0.9956531 | 0.31941873 | 0.97700345 |
| JMJD1C | 215.671771 | 0.171751377 | 0.20796825 | 0.82585383 | 0.40888701 | 0.99279547 |
| REEP3 | 38.8696266 | 0.205585012 | 0.32282672 | 0.63682774 | 0.52423705 | 0.99665574 |
| SIRT1 | 13.2905965 | 0.236337958 | 0.4669922 | 0.50608545 | 0.61279666 | 0.99665574 |
| HERC4 | 72.777215 | -0.380954657 | 0.32941094 | -1.1564724 | 0.24748799 | 0.93441937 |

| | | | | | | |
|---|---|---|---|---|---|---|
| PBLD | 7.64590178 | 0.609239898 | 0.52218181 | 1.16671987 | 0.24332351 | 0.93426912 |
| HNRNPH3 | 25.0345764 | 0.392564695 | 0.3771846 | 1.04077604 | 0.2979795 | 0.97358745 |
| RUFY2 | 18.8821678 | 0.155081094 | 0.46495945 | 0.33353681 | 0.73872911 | 0.99665574 |
| SLC25A16 | 24.5912874 | 0.334257656 | 0.35914921 | 0.930693 | 0.35201239 | 0.97838306 |
| TET1 | 4.08048504 | 0.620383057 | 0.82708419 | 0.75008453 | 0.4532038 | 0.99617422 |
| CCAR1 | 47.4303556 | 0.036130731 | 0.3587613 | 0.10070967 | 0.91978094 | 0.99777713 |
| STOX1 | 4.62561527 | 1.199844281 | 0.84343632 | 1.42256654 | 0.15486185 | 0.86215785 |
| DDX50 | 17.6944444 | -0.013322544 | 0.40871703 | -0.032596 | 0.97399675 | 0.99777713 |
| DDX21 | 20.8781173 | -0.126833371 | 0.48566101 | -0.2611562 | 0.79397207 | 0.99665574 |
| KIF1BP | 8.19172369 | -0.688775899 | 0.48709955 | -1.4140352 | 0.15735157 | 0.86532131 |
| VPS26A | 19.8766744 | -0.139255226 | 0.39242372 | -0.3548593 | 0.72269497 | 0.99665574 |
| SUPV3L1 | 7.02594315 | 0.452858584 | 0.60514613 | 0.74834583 | 0.45425159 | 0.99617422 |
| HKDC1 | 6.08730372 | -2.705304595 | 0.87822321 | -3.0804294 | 0.00206702 | 0.13786503 |
| HK1 | 29.1456343 | -0.520166614 | 0.36391419 | -1.4293661 | 0.15289903 | 0.86116133 |
| TSPAN15 | 8.47925877 | -2.230479496 | 0.56055964 | -3.9790226 | 6.92E-05 | 0.01742067 |
| H2AFY2 | 8.50222128 | 0.192842831 | 0.53122618 | 0.36301454 | 0.71659402 | 0.99665574 |
| AIFM2 | 8.59607096 | -0.727777025 | 0.55684337 | -1.306969 | 0.19122324 | 0.89964233 |
| TYSND1 | 3.51480055 | -0.281798174 | 0.74210332 | -0.379729 | 0.70414657 | 0.99665574 |
| SAR1A | 16.0908909 | -0.080870958 | 0.50203115 | -0.1610875 | 0.87202446 | 0.99665574 |
| PPA1 | 29.8087676 | -1.337200677 | 0.36806651 | -3.6330409 | 0.0002801 | 0.04115449 |
| EIF4EBP2 | 25.2641935 | 0.774954455 | 0.4772064 | 1.62393977 | 0.10438866 | 0.79169635 |
| SGPL1 | 25.693729 | -0.130092835 | 0.33730638 | -0.3856815 | 0.69973256 | 0.99665574 |
| PCBD1 | 14.6627877 | -0.09634436 | 0.51509771 | -0.1870409 | 0.85162853 | 0.99665574 |
| PSAP | 68.0583027 | 0.05414851 | 0.36772901 | 0.14725112 | 0.88293381 | 0.99665574 |
| CHST3 | 14.0931605 | -0.373870046 | 0.41963122 | -0.8909491 | 0.37295649 | 0.98532838 |
| ASCC1 | 14.4662152 | -0.865676979 | 0.4127958 | -2.0971071 | 0.0359841 | 0.58508511 |
| ANAPC16 | 45.4724884 | -0.070145976 | 0.34925117 | -0.2008468 | 0.84081838 | 0.99665574 |
| DDIT4 | 7.68912757 | -0.571757808 | 0.84360016 | -0.6777592 | 0.49792435 | 0.99640678 |
| DNAJB12 | 7.39838491 | -2.81E-05 | 0.53589004 | -5.24E-05 | 0.99995819 | 0.99995819 |
| MICU1 | 51.5635964 | -0.679482215 | 0.33902588 | -2.0042193 | 0.04504657 | 0.6313648 |
| MCU | 75.317111 | 0.142051109 | 0.58728111 | 0.24187924 | 0.80887374 | 0.99665574 |
| P4HA1 | 29.6342746 | -0.045764101 | 0.78831637 | -0.058053 | 0.95370644 | 0.99777713 |
| ECD | 7.94334957 | 0.084992119 | 0.52689845 | 0.16130645 | 0.87185204 | 0.99665574 |
| FAM149B1 | 14.4748063 | -0.040374616 | 0.459569 | -0.0878532 | 0.92999334 | 0.99777713 |
| DNAJC9 | 12.2675918 | 0.225881226 | 0.54283951 | 0.41611051 | 0.67732914 | 0.99665574 |
| MRPS16 | 12.0623879 | 0.049681254 | 0.44088418 | 0.1126855 | 0.9102799 | 0.99777713 |
| CFAP70 | 9.95806421 | -0.048788152 | 0.61220772 | -0.0796922 | 0.9364821 | 0.99777713 |
| ANXA7 | 27.9951149 | -0.154884943 | 0.32205556 | -0.4809262 | 0.63056898 | 0.99665574 |
| PPP3CB | 33.7443391 | 0.250328848 | 0.33424708 | 0.74893353 | 0.45389727 | 0.99617422 |
| USP54 | 14.0268908 | 0.033441923 | 0.44334315 | 0.07543124 | 0.93987161 | 0.99777713 |
| AGAP5 | 3.12728134 | 0.456774243 | 0.74205005 | 0.61555719 | 0.53818681 | 0.99665574 |
| SEC24C | 10.4652882 | 0.077967483 | 0.46235531 | 0.1686311 | 0.86608682 | 0.99665574 |
| FUT11 | 6.93149933 | 0.092989463 | 0.66140122 | 0.14059464 | 0.88819019 | 0.99665574 |
| CHCHD1 | 9.53228633 | 0.803162989 | 0.64361841 | 1.24788691 | 0.21207248 | 0.91043197 |
| ZSWIM8 | 13.7370017 | 0.062101874 | 0.48376687 | 0.12837149 | 0.89785499 | 0.99777713 |
| CAMK2G | 6.53027067 | -0.673414982 | 0.56704827 | -1.1875796 | 0.23499906 | 0.92960234 |
| VCL | 47.1036965 | 0.066347635 | 0.39839696 | 0.1665365 | 0.86773476 | 0.99665574 |
| AP3M1 | 7.88185351 | 0.690578815 | 0.59651461 | 1.15768969 | 0.24699069 | 0.93441937 |
| ADK | 118.147956 | -0.289585908 | 0.2704755 | -1.0706549 | 0.28432465 | 0.96059387 |
| KAT6B | 62.570272 | 0.245059822 | 0.2733244 | 0.89658963 | 0.36993793 | 0.98532838 |
| SAMD8 | 15.1555457 | -0.312100826 | 0.46003291 | -0.6784315 | 0.49749813 | 0.99640678 |
| VDAC2 | 24.0827285 | -0.26195588 | 0.38387226 | -0.6824038 | 0.49498367 | 0.99640678 |
| COMTD1 | 25.9622208 | 1.10495295 | 0.50642298 | 2.1818776 | 0.02911857 | 0.55281655 |
| AC010997.3 | 2.47456278 | -0.077814838 | 0.95034691 | -0.0818805 | 0.93474178 | 0.99777713 |
| ZNF503-AS2 | 4.08511746 | 0.540694193 | 0.8731986 | 0.61921102 | 0.53577735 | 0.99665574 |
| LRMDA | 460.53504 | 0.426861889 | 0.30488352 | 1.40008188 | 0.1614888 | 0.87011714 |
| KCNMA1 | 11.4662808 | 1.018888728 | 0.98466531 | 1.0347564 | 0.30078269 | 0.97358745 |
| DLG5 | 89.7904544 | -0.813445299 | 0.44542708 | -1.8262143 | 0.06781799 | 0.70516795 |
| POLR3A | 10.1135045 | -0.319846779 | 0.44615224 | -0.7169005 | 0.47343548 | 0.99617422 |
| RPS24 | 107.230911 | 0.480684179 | 0.3631894 | 1.32350829 | 0.1856664 | 0.89532394 |
| ZMIZ1 | 61.2296255 | -0.120254236 | 0.43636062 | -0.2755845 | 0.78286719 | 0.99665574 |
| PPIF | 13.0090856 | 0.045661704 | 0.5057363 | 0.09028757 | 0.92805869 | 0.99777713 |
| NUTM2B-AS | 38.2281278 | -0.255085096 | 0.29754955 | -0.8572861 | 0.39128679 | 0.99179694 |
| TMEM254 | 3.16617642 | 0.091541243 | 0.82623604 | 0.11079309 | 0.91178043 | 0.99777713 |
| ANXA11 | 84.8577265 | 0.124001435 | 0.28082945 | 0.44155424 | 0.6588118 | 0.99665574 |
| FAM213A | 23.3337151 | -0.320863011 | 0.43464529 | -0.7382181 | 0.46038194 | 0.99617422 |
| TSPAN14 | 59.3547109 | -0.120125879 | 0.43386228 | -0.2768756 | 0.78187563 | 0.99665574 |
| NRG3 | 57.291499 | 0.597729865 | 1.05845942 | 0.56471685 | 0.57226637 | 0.99665574 |
| GHITM | 30.4448485 | 0.237804395 | 0.41416938 | 0.57417184 | 0.56585153 | 0.99665574 |
| CCSER2 | 52.7983301 | 0.063465793 | 0.27730552 | 0.22886596 | 0.8189731 | 0.99665574 |
| WAPL | 45.6351181 | 0.134087488 | 0.26850411 | 0.4993871 | 0.6175067 | 0.99665574 |
| BMPR1A | 52.0313835 | 0.102196024 | 0.28704119 | 0.35603261 | 0.72181614 | 0.99665574 |
| MMRN2 | 3.01368244 | 1.21401993 | 0.85241571 | 1.42421111 | 0.15438537 | 0.86196782 |
| ADIRF | 11.126579 | 0.821427386 | 0.64285348 | 1.27778321 | 0.20132588 | 0.90591604 |
| GLUD1 | 88.120003 | 0.259160944 | 0.34581112 | 0.74942918 | 0.45359857 | 0.99617422 |
| SHLD2 | 52.3934384 | 0.073687535 | 0.31001052 | 0.23769366 | 0.8121187 | 0.99665574 |
| NUTM2A-AS | 48.4989403 | 0.083600173 | 0.31628708 | 0.26431738 | 0.79153537 | 0.99665574 |
| MINPP1 | 6.15049743 | 0.45552038 | 0.62695994 | 0.7265542 | 0.46749909 | 0.99617422 |
| ATAD1 | 29.9735318 | 0.10275056 | 0.30083819 | 0.34154759 | 0.73269138 | 0.99665574 |
| PTEN | 49.8986711 | 0.107857487 | 0.27671147 | 0.38978322 | 0.69669685 | 0.99665574 |
| RNLS | 17.5740699 | -0.055881974 | 0.51957539 | -0.1075532 | 0.91435016 | 0.99777713 |
| ANKRD22 | 5.50551181 | -1.316984072 | 0.87628433 | -1.5029187 | 0.13286002 | 0.84089786 |
| LIPA | 10.0320982 | 1.319264949 | 0.64315563 | 2.05123751 | 0.04024382 | 0.60713077 |
| IFIT2 | 5.11703632 | 0.205760897 | 1.19002991 | 0.17290397 | 0.86272691 | 0.99665574 |
| IFIT3 | 14.8773978 | -0.191906869 | 1.36948531 | -0.1401307 | 0.88855676 | 0.99665574 |
| IFIT1 | 14.2978113 | 1.918101557 | 1.88825246 | 1.01580779 | 0.30972092 | 0.97376371 |
| IFIT5 | 3.05344804 | -0.138028974 | 1.0655348 | -0.1295396 | 0.89693068 | 0.99777713 |
| SLC16A12 | 17.6409543 | 1.774512072 | 0.89352874 | 1.9859597 | 0.0470378 | 0.64060657 |
| PANK1 | 6.06812054 | 2.277337511 | 0.96737273 | 2.3541469 | 0.01856527 | 0.45138305 |
| KIF20B | 2.79713894 | -0.288678807 | 0.78851908 | -0.3661025 | 0.71428859 | 0.99665574 |
| RPP30 | 3.36958079 | -0.231625844 | 0.79710088 | -0.2905854 | 0.77136846 | 0.99665574 |
| PCGF5 | 60.2141768 | 0.1417554 | 0.2802364 | 0.50584221 | 0.61296742 | 0.99665574 |
| HECTD2 | 20.9650787 | -0.076488717 | 0.38706675 | -0.1976112 | 0.84334929 | 0.99665574 |
| PPP1R3C | 6.51169929 | -0.335530959 | 0.98604588 | -0.3402793 | 0.73364624 | 0.99665574 |
| TNKS2 | 48.346239 | -0.302190268 | 0.35463063 | -0.8521268 | 0.39414371 | 0.99179694 |
| BTAF1 | 38.3347452 | -0.169401058 | 0.43680721 | -0.3878165 | 0.69815181 | 0.99665574 |
| CPEB3 | 61.8499352 | 0.564380598 | 0.40764535 | 1.38448924 | 0.1662087 | 0.87495175 |
| 5-Mar | 18.42728 | -0.668827285 | 0.44191092 | -1.5134889 | 0.13015551 | 0.83778831 |
| IDE | 17.6274257 | 0.578325616 | 0.42439172 | 1.36271654 | 0.17297186 | 0.88157681 |
| EXOC6 | 38.5337936 | -0.244288923 | 0.47302645 | -0.5164382 | 0.6055484 | 0.99665574 |
| MYOF | 21.255875 | -1.227766455 | 0.55234519 | -2.2228246 | 0.02622763 | 0.52034391 |
| FRA10AC1 | 12.4730906 | 0.364146031 | 0.44982093 | 0.80953554 | 0.41820717 | 0.99432434 |
| PLCE1 | 25.0278138 | -0.607114435 | 0.58778362 | -1.0328876 | 0.30165648 | 0.97358745 |
| NOC3L | 4.89723988 | 0.050627727 | 0.59945908 | 0.08445569 | 0.93269413 | 0.99777713 |
| TBC1D12 | 17.8000751 | -0.128105509 | 0.40695239 | -0.3147924 | 0.7529193 | 0.99665574 |
| HELLS | 4.06496715 | -1.033765908 | 0.72004926 | -1.4356878 | 0.15109121 | 0.85612189 |
| SORBS1 | 9.5949535 | -1.155435809 | 0.657311 | -1.7578221 | 0.07877778 | 0.74431421 |
| ALDH18A1 | 9.87322716 | 0.122964074 | 0.50943992 | 0.2413711 | 0.80926751 | 0.99665574 |
| TCTN3 | 11.1115583 | 0.329911957 | 0.43475153 | 0.75885174 | 0.44794125 | 0.99617422 |
| ENTPD1-AS1 | 30.6962807 | -0.116373884 | 0.48230116 | -0.2412888 | 0.80933126 | 0.99665574 |
| ZNF518A | 46.1099895 | -0.052597531 | 0.30660089 | -0.1715505 | 0.86379094 | 0.99665574 |
| BLNK | 26.4611252 | -0.913334546 | 0.41858322 | -2.1819665 | 0.02911201 | 0.55281655 |
| TM9SF3 | 50.7852503 | -0.156315105 | 0.25104671 | -0.6226535 | 0.53351226 | 0.99665574 |

| | | | | | | |
|---|---|---|---|---|---|---|
| LCOR | 99.1122241 | 0.27057327 | 0.30318701 | 0.89243027 | 0.37216235 | 0.98532838 |
| RRP12 | 3.824478 | -0.031759848 | 0.83273906 | -0.038139 | 0.96957684 | 0.99777713 |
| PGAM1 | 23.7149268 | -0.322435609 | 0.39607907 | -0.8140688 | 0.41560555 | 0.99279547 |
| EXOSC1 | 6.0525978 | -0.669454983 | 0.57673085 | -1.1607754 | 0.24573325 | 0.93426912 |
| ZDHHC16 | 6.71213368 | -1.268739477 | 0.68040788 | -1.8646749 | 0.06222699 | 0.68496987 |
| MMS19 | 13.976146 | -0.09790999 | 0.4696415 | -0.2084781 | 0.83485564 | 0.99665574 |
| UBTD1 | 6.5301397 | -0.285460969 | 0.64559635 | -0.4421663 | 0.65836885 | 0.99665574 |
| ZFYVE27 | 9.34963966 | -0.294719708 | 0.55872248 | -0.5274885 | 0.59785437 | 0.99665574 |
| R3HCC1L | 24.3284886 | -0.022484424 | 0.3799627 | -0.0591753 | 0.95281245 | 0.99777713 |
| HPS1 | 7.83245121 | -0.078163055 | 0.56291529 | -0.138854 | 0.8895655 | 0.99665574 |
| GOT1 | 43.0569254 | 0.209627188 | 0.34102164 | 0.61470347 | 0.53875056 | 0.99665574 |
| AL391684.1 | 2.72514436 | -0.051322471 | 1.06796519 | -0.0480563 | 0.96167136 | 0.99777713 |
| SLC25A28 | 7.26776196 | -0.732074963 | 0.57758985 | -1.2674651 | 0.20498904 | 0.90834797 |
| CUTC | 8.3706273 | -0.277605577 | 0.51471363 | -0.5393399 | 0.58965237 | 0.99665574 |
| COX15 | 10.1068782 | -0.198772321 | 0.45871201 | -0.433327 | 0.66477719 | 0.99665574 |
| ABCC2 | 5.13475231 | -1.853673191 | 0.90831693 | -2.040778 | 0.0412729 | 0.61108334 |
| DNMBP | 22.4672385 | 0.583357097 | 0.36955037 | 1.5785591 | 0.11443722 | 0.81638817 |
| ERLIN1 | 6.22744898 | -0.011739236 | 0.56816332 | -0.0206617 | 0.9835155 | 0.99777713 |
| CHUK | 12.133668 | 0.158717812 | 0.45186592 | 0.35124979 | 0.72540096 | 0.99665574 |
| CWF19L1 | 9.29130902 | 0.228172335 | 0.47503507 | 0.48032735 | 0.63099465 | 0.99665574 |
| BLOC1S2 | 7.6167134 | 0.579696748 | 0.56700091 | 1.02239121 | 0.30659578 | 0.97376371 |
| SCD | 4.65292675 | -2.232104243 | 2.12594608 | -1.0499346 | 0.2937482 | 0.9713044 |
| OLMALINC | 4.31687916 | 0.941221885 | 0.97311945 | 0.96722132 | 0.33343341 | 0.97788427 |
| SEC31B | 3.68777109 | 0.691188566 | 0.84821736 | 0.81487199 | 0.41514559 | 0.99279547 |
| HIF1AN | 7.7972471 | 0.418202878 | 0.50297107 | 0.83146507 | 0.40571095 | 0.99194293 |
| PAX2 | 42.9513733 | 0.201896917 | 0.38404208 | 0.52571561 | 0.59908582 | 0.99665574 |
| SLF2 | 16.5343611 | -0.460822855 | 0.42120703 | -1.0940531 | 0.27393169 | 0.95331759 |
| MRPL43 | 8.72396927 | 0.015049741 | 0.6464393 | 0.02328098 | 0.98142614 | 0.99777713 |
| LZTS2 | 6.60882257 | -0.061071844 | 0.61850977 | -0.0987403 | 0.92134447 | 0.99777713 |
| BTRC | 47.1254233 | 0.353449899 | 0.35437688 | 0.99738418 | 0.31857807 | 0.97700345 |
| POLL | 3.14292293 | -0.629449486 | 0.93506999 | -0.6731576 | 0.50084702 | 0.99640678 |
| FBXW4 | 41.9464652 | -0.066322799 | 0.34536394 | -0.1920374 | 0.84771289 | 0.99665574 |
| OGA | 68.9757929 | -0.184711808 | 0.2715319 | -0.6802582 | 0.49634097 | 0.99640678 |
| ARMH3 | 49.5764023 | 0.141564247 | 0.3012917 | 0.46985778 | 0.63845663 | 0.99665574 |
| LDB1 | 5.80067348 | 0.979030691 | 0.63153065 | 1.55025048 | 0.12108141 | 0.82659025 |
| PPRC1 | 5.97363109 | -1.141466976 | 0.74349578 | -1.5352703 | 0.12471745 | 0.83196528 |
| NOLC1 | 5.67228446 | -0.467745202 | 0.54578375 | -0.8570156 | 0.39143625 | 0.99179694 |
| GBF1 | 55.4157657 | -0.477250909 | 0.33854238 | -1.4097228 | 0.15862155 | 0.86690309 |
| NFKB2 | 3.91742219 | -0.742878274 | 0.82146711 | -0.9043311 | 0.36581985 | 0.98365837 |
| FBXL15 | 7.67500972 | 0.578001567 | 0.64401074 | 0.897503 | 0.36945057 | 0.98532838 |
| CUEDC2 | 6.91441968 | 0.324158237 | 0.61258178 | 0.52916728 | 0.59668941 | 0.99665574 |
| ACTR1A | 10.0588456 | -0.723827764 | 0.51383687 | -1.4086723 | 0.15893209 | 0.86757292 |
| SUFU | 14.0184975 | 0.506527796 | 0.40812983 | 1.24109476 | 0.21457075 | 0.91280851 |
| TRIM8 | 31.4518437 | 0.568396078 | 0.31533583 | 1.80251031 | 0.07146515 | 0.71725834 |
| ARL3 | 8.14475508 | 0.476093962 | 0.62343529 | 0.76366219 | 0.44506858 | 0.99617422 |
| SFXN2 | 6.07485285 | 1.117766229 | 0.58906991 | 1.89751032 | 0.05776062 | 0.67545594 |
| WBP1L | 32.0015124 | 0.001564247 | 0.34456971 | 0.00453971 | 0.99637785 | 0.998802 |
| CNNM2 | 70.7417885 | 0.411207283 | 0.42367675 | 0.97056844 | 0.33176323 | 0.97788427 |
| NT5C2 | 80.5966341 | -0.619479485 | 0.3243002 | -1.9102038 | 0.05610698 | 0.67401304 |
| PCGF6 | 8.7239558 | 0.190217948 | 0.52916372 | 0.35946899 | 0.71924427 | 0.99665574 |
| ATP5MD | 20.078821 | 0.69336109 | 0.40724419 | 1.70256841 | 0.08864887 | 0.75650018 |
| PDCD11 | 7.76143572 | 0.626816955 | 0.6849817 | 0.91508569 | 0.36014665 | 0.9819332 |
| NEURL1 | 58.158877 | -0.513052329 | 0.67141285 | -0.7641384 | 0.44478478 | 0.99617422 |
| SH3PXD2A | 8.72685887 | 0.318073147 | 0.5523472 | 0.57585727 | 0.56471166 | 0.99665574 |
| AL121929.2 | 2.02649192 | 0.953314132 | 1.28464898 | 0.74208142 | 0.45803801 | 0.99617422 |
| STN1 | 4.03787565 | -0.903689614 | 0.75406435 | -1.1984251 | 0.23075158 | 0.92774608 |
| SLK | 26.299512 | -0.206110683 | 0.30706016 | -0.6712388 | 0.50206844 | 0.99640678 |
| CFAP43 | 3.62554708 | -0.623718218 | 0.90096366 | -0.692279 | 0.48876214 | 0.99640678 |
| GSTO1 | 13.8931284 | -0.025986051 | 0.44582694 | -0.0582873 | 0.95351978 | 0.99777713 |
| GSTO2 | 15.5437325 | 0.224509008 | 0.46263046 | 0.48528799 | 0.62747207 | 0.99665574 |
| XPNPEP1 | 10.1041124 | -0.101412498 | 0.45573606 | -0.2225246 | 0.8239055 | 0.99665574 |
| ADD3 | 41.9368446 | 0.932411158 | 0.37531957 | 2.48431266 | 0.01298018 | 0.38590337 |
| MXI1 | 18.41326 | -0.572594085 | 0.42694304 | -1.3411487 | 0.17987219 | 0.88823393 |
| SMNDC1 | 9.06090259 | -0.375053184 | 0.56197527 | -0.6673838 | 0.50452702 | 0.99665574 |
| SMC3 | 12.0561418 | 0.114980237 | 0.51094301 | 0.22503534 | 0.82195178 | 0.99665574 |
| PDCD4 | 14.5685055 | -0.394788293 | 0.55829107 | -0.707137 | 0.47948132 | 0.99617422 |
| BBIP1 | 10.9788535 | 0.039824456 | 0.52321951 | 0.07611424 | 0.93932821 | 0.99777713 |
| SHOC2 | 33.1422555 | 0.132401892 | 0.39618769 | 0.33418982 | 0.73823633 | 0.99665574 |
| GPAM | 3.70581682 | -0.220644515 | 0.75721486 | -0.2913896 | 0.77075339 | 0.99665574 |
| ZDHHC6 | 7.81586478 | 0.420269996 | 0.52799726 | 0.79597004 | 0.42604945 | 0.99518711 |
| VTI1A | 126.712968 | -0.154410801 | 0.26248579 | -0.5882635 | 0.55635546 | 0.99665574 |
| TCF7L2 | 96.194562 | 0.246745976 | 0.36552324 | 0.67504866 | 0.49964485 | 0.99640678 |
| CASP7 | 3.88784222 | -0.855431473 | 0.85950901 | -0.995256 | 0.31961179 | 0.97700345 |
| NHLRC2 | 22.6779915 | 0.028068905 | 0.37689254 | 0.07447456 | 0.94063278 | 0.99777713 |
| CCDC186 | 26.8533417 | -0.341170146 | 0.36490822 | -0.9349478 | 0.34981519 | 0.97838306 |
| ABLIM1 | 83.9548658 | -0.988777722 | 0.45241108 | -2.1855736 | 0.02884682 | 0.55075144 |
| FAM160B1 | 10.3878066 | -0.10119835 | 0.47161064 | -0.2145803 | 0.83009456 | 0.99665574 |
| TRUB1 | 7.98403695 | 0.205575514 | 0.62493344 | 0.32895586 | 0.74218905 | 0.99665574 |
| HSPA12A | 9.44178277 | 0.023828146 | 0.58476657 | 0.04074813 | 0.96749669 | 0.99777713 |
| SHTN1 | 51.5722094 | -0.665835158 | 0.34948756 | -1.9051755 | 0.05675726 | 0.67545594 |
| PDZD8 | 30.9660614 | 0.514955904 | 0.43316211 | 1.18882951 | 0.23450676 | 0.92960234 |
| EMX2OS | 16.2188519 | -0.160839655 | 0.52797339 | -0.3046359 | 0.76064347 | 0.99665574 |
| EMX2 | 8.33205279 | 0.024855774 | 0.57395305 | 0.04330628 | 0.96545738 | 0.99777713 |
| RAB11FIP2 | 19.3218361 | 0.104835881 | 0.38441066 | 0.27271845 | 0.78506965 | 0.99665574 |
| CASC2 | 8.61351734 | -0.532735453 | 0.70496293 | -0.7556929 | 0.44983336 | 0.99617422 |
| FAM204A | 15.2930144 | -0.04835353 | 0.40267901 | -0.1200796 | 0.9044201 | 0.99777713 |
| CACUL1 | 36.7796197 | -0.17022054 | 0.29927632 | -0.5687738 | 0.56950963 | 0.99665574 |
| EIF3A | 50.3060699 | -0.218152185 | 0.41351997 | -0.5275493 | 0.59781216 | 0.99665574 |
| FAM45A | 9.74068961 | -0.709859862 | 0.48862959 | -1.4527566 | 0.14629134 | 0.85169374 |
| SFXN4 | 19.8341454 | 0.541041812 | 0.39173769 | 1.38113289 | 0.1672381 | 0.87495175 |
| PRDX3 | 21.9004886 | 0.713702652 | 0.36111073 | 1.97640942 | 0.04810843 | 0.64187738 |
| GRK5 | 6.29308519 | -1.246430101 | 0.90412028 | -1.3786109 | 0.16801474 | 0.87566218 |
| TIAL1 | 17.8671603 | 0.477688536 | 0.39035032 | 1.22374318 | 0.22104913 | 0.91957162 |
| BAG3 | 9.83302488 | -0.169328698 | 0.50691405 | -0.3340383 | 0.73835067 | 0.99665574 |
| INPP5F | 13.3255873 | -0.285723837 | 0.39630109 | -0.7209767 | 0.47092387 | 0.99617422 |
| MCMBP | 17.1679939 | -0.168138094 | 0.42802506 | -0.392823 | 0.6944502 | 0.99665574 |
| SEC23IP | 25.3482889 | -0.582286414 | 0.40833141 | -1.4260143 | 0.15386422 | 0.86196782 |
| PLPP4 | 5.08433646 | 1.040732924 | 0.72367555 | 1.43812089 | 0.15039976 | 0.85612189 |
| WDR11-AS1 | 3.3635449 | -0.879006584 | 0.8070695 | -1.0891337 | 0.27609494 | 0.95694033 |
| WDR11 | 29.0712941 | 0.281147675 | 0.31527213 | 0.89176191 | 0.37252056 | 0.98532838 |
| FGFR2 | 26.7413939 | -0.110759304 | 0.65046857 | -0.1702762 | 0.86479294 | 0.99665574 |
| ATE1 | 57.259622 | 0.063573431 | 0.39383661 | 0.16142082 | 0.87176197 | 0.99665574 |
| NSMCE4A | 5.79627852 | 0.944645481 | 0.60578463 | 1.55937513 | 0.11890762 | 0.82619588 |
| TACC2 | 62.5415952 | -0.166143634 | 0.58380225 | -0.2845889 | 0.77595912 | 0.99665574 |
| PLEKHA1 | 64.33415 | -0.886038428 | 0.32922005 | -2.6913258 | 0.00711686 | 0.27640407 |
| C10orf88 | 4.83491653 | 0.110069901 | 0.71353668 | 0.15425963 | 0.87740503 | 0.99665574 |
| PSTK | 2.68934523 | -0.641509745 | 0.76337742 | -0.8403572 | 0.40070811 | 0.99194293 |
| IKZF5 | 6.85642228 | 0.356501862 | 0.60734366 | 0.5869854 | 0.55721351 | 0.99665574 |
| ACADSB | 11.9037495 | -0.002284782 | 0.54240569 | -0.0042123 | 0.99663907 | 0.998802 |
| HMX2 | 42.6757148 | -0.555046171 | 0.84846886 | -0.6541739 | 0.51299978 | 0.99665574 |
| BUB3 | 17.2307912 | 0.342217263 | 0.38119103 | 0.89775791 | 0.36931463 | 0.98532838 |
| CPXM2 | 23.8600208 | 0.01662724 | 0.65214648 | 0.02549617 | 0.9796592 | 0.99777713 |
| OAT | 9.09967648 | -0.435768627 | 0.56825141 | -0.7668589 | 0.44316543 | 0.99617422 |

| | | | | | | |
|---|---|---|---|---|---|---|
| LHPP | 6.08102381 | 0.500457654 | 0.64982839 | 0.77013818 | 0.44121793 | 0.99617422 |
| ABRAXAS2 | 6.89670686 | 0.034813937 | 0.55737555 | 0.06246047 | 0.95019614 | 0.99777713 |
| ZRANB1 | 24.884206 | -0.01555451 | 0.31553148 | -0.0492962 | 0.96068323 | 0.99777713 |
| CTBP2 | 112.764801 | -0.415141928 | 0.2796322 | -1.4845999 | 0.13764985 | 0.84656101 |
| EDRF1 | 11.7982104 | -0.707907391 | 0.4653077 | -1.5213748 | 0.12816582 | 0.83778831 |
| UROS | 6.78348073 | -0.589821836 | 0.69437338 | -0.8494304 | 0.39564186 | 0.99179694 |
| BCCIP | 12.9274018 | 0.556573442 | 0.50053296 | 1.11196161 | 0.26615466 | 0.94557817 |
| DHX32 | 26.2196172 | -0.165249759 | 0.3248246 | -0.5087354 | 0.61093773 | 0.99665574 |
| FANK1 | 5.40951701 | -0.273186841 | 0.72809892 | -0.3752057 | 0.70750752 | 0.99665574 |
| DOCK1 | 139.797484 | -0.343601092 | 0.21407029 | -1.6050854 | 0.10847501 | 0.80237722 |
| FAM196A | 6.73541877 | -1.746620399 | 0.72604437 | -2.4056662 | 0.01614301 | 0.42281551 |
| MGMT | 31.6994464 | 0.179114429 | 0.39962984 | 0.44820083 | 0.65400826 | 0.99665574 |
| AL157832.3 | 2.19703019 | 0.502926849 | 0.92372974 | 0.54445237 | 0.58613021 | 0.99665574 |
| C10orf143 | 3.27792526 | -0.383607803 | 0.73010335 | -0.5254158 | 0.59929421 | 0.99665574 |
| GLRX3 | 18.4298276 | -0.017587654 | 0.48319869 | -0.0363984 | 0.9709647 | 0.99777713 |
| PPP2R2D | 36.7604261 | 0.156242177 | 0.43375104 | 0.36021165 | 0.71868887 | 0.99665574 |
| BNIP3 | 31.1436967 | -0.665919402 | 0.5346925 | -1.245425 | 0.21297558 | 0.91073943 |
| STK32C | 5.19673277 | 0.544132218 | 0.62772536 | 0.86683167 | 0.38603425 | 0.99051797 |
| LRRC27 | 3.22522124 | 0.019723144 | 0.7778355 | 0.02535645 | 0.97977065 | 0.99777713 |
| PWWP2B | 4.4304292 | 0.687508513 | 0.73835482 | 0.93113567 | 0.35178339 | 0.97838306 |
| INPP5A | 21.3860279 | 0.47316464 | 0.42243075 | 1.12009989 | 0.2626712 | 0.94557817 |
| ADGRA1 | 4.46942298 | 2.013253479 | 0.80970159 | 2.48641415 | 0.01290377 | 0.38590337 |
| KNDC1 | 2.01309487 | -0.987800427 | 1.18062583 | -0.8366753 | 0.40277513 | 0.99194293 |
| TUBGCP2 | 14.7423308 | -0.232924277 | 0.39897043 | -0.5838134 | 0.55934588 | 0.99665574 |
| ECHS1 | 19.5074344 | 0.357553626 | 0.49033667 | 0.72920025 | 0.46587918 | 0.99617422 |
| SCART1 | 4.18749259 | 0.61880208 | 0.86439313 | 0.71588038 | 0.47406522 | 0.99617422 |
| SLC6A12 | 14.8040568 | -1.618161899 | 0.63891526 | -2.5326706 | 0.01131973 | 0.3725572 |
| SLC6A13 | 4.68639177 | 1.129308271 | 1.10566482 | 1.02138392 | 0.30707258 | 0.97376371 |
| KDM5A | 40.4014748 | 0.253201602 | 0.30540396 | 0.82907111 | 0.40706417 | 0.99250037 |
| CCDC77 | 3.33960452 | 0.284254222 | 0.99047951 | 0.28698647 | 0.77412268 | 0.99665574 |
| B4GALNT3 | 14.9695568 | -0.358007243 | 0.56421872 | -0.6345186 | 0.52574246 | 0.99665574 |
| WNK1 | 53.5968911 | 0.528641954 | 0.28699665 | 1.84197954 | 0.06547814 | 0.69221233 |
| RAD52 | 9.19430652 | 0.203160931 | 0.5341738 | 0.3803274 | 0.7037024 | 0.99665574 |
| ERC1 | 128.289402 | -0.264637861 | 0.27835689 | -0.9507143 | 0.34174945 | 0.97788427 |
| FBXL14 | 3.21424767 | -0.506670745 | 0.8093199 | -0.6260451 | 0.53128537 | 0.99665574 |
| ADIPOR2 | 50.8156798 | 0.162296495 | 0.37774635 | 0.42964411 | 0.66745454 | 0.99665574 |
| DCP1B | 9.97722813 | 0.097784989 | 0.59002291 | 0.16573083 | 0.86836878 | 0.99665574 |
| CACNA1C | 6.31149943 | -1.389582515 | 1.02238025 | -1.3591641 | 0.1740946 | 0.88281284 |
| FKBP4 | 11.1951953 | 0.856971903 | 0.60085127 | 1.42626296 | 0.15379245 | 0.86196782 |
| ITFG2 | 17.0048273 | -0.31719207 | 0.53921503 | -0.5882478 | 0.55636596 | 0.99665574 |
| TULP3 | 14.6997757 | 0.053954615 | 0.45923223 | 0.11748874 | 0.90647277 | 0.99777713 |
| TEAD4 | 9.74022444 | -2.334640064 | 0.69949997 | -3.3375842 | 0.0008451 | 0.07932995 |
| TSPAN9 | 53.7767173 | -0.350893005 | 0.45690676 | -0.7679751 | 0.44250199 | 0.99617422 |
| PARP11 | 7.10823102 | -0.081365081 | 0.62610848 | -0.1299536 | 0.8966031 | 0.99777713 |
| TIGAR | 4.81062449 | -0.802625058 | 0.7552787 | -1.0626873 | 0.2879238 | 0.96590717 |
| C12orf4 | 8.72093014 | 0.057154293 | 0.52357761 | 0.10916107 | 0.91307474 | 0.99777713 |
| ANO2 | 8.8521474 | 1.211616973 | 0.59653061 | 2.03110613 | 0.04224423 | 0.61533334 |
| VWF | 12.4265572 | -0.090852806 | 0.48591006 | -0.1869745 | 0.8516806 | 0.99665574 |
| CD9 | 276.847718 | 0.203676604 | 0.3717505 | 0.54788522 | 0.58377072 | 0.99665574 |
| TNFRSF1A | 14.169121 | -1.496842128 | 0.46884346 | -3.1926266 | 0.00140985 | 0.1099467 |
| SCNN1A | 111.036143 | 0.204320197 | 0.30141081 | 0.67787945 | 0.49784813 | 0.99640678 |
| LTBR | 10.8849141 | -1.14756456 | 0.47495523 | -2.4161531 | 0.01568546 | 0.41847166 |
| TAPBPL | 4.58747386 | 0.213033646 | 0.73612602 | 0.28939834 | 0.77227657 | 0.99665574 |
| MRPL51 | 13.4221346 | 0.111350569 | 0.54045662 | 0.20603054 | 0.83676706 | 0.99665574 |
| NCAPD2 | 5.61043882 | 0.107319249 | 0.70032357 | 0.15324238 | 0.87820714 | 0.99665574 |
| GAPDH | 128.830084 | -0.538662173 | 0.4519088 | -1.191971 | 0.23327263 | 0.92924083 |
| CHD4 | 8.33655477 | -0.614461318 | 0.51537274 | -1.1922658 | 0.23315702 | 0.92924083 |
| ING4 | 4.46688172 | 0.570018409 | 0.84065682 | 0.67806315 | 0.49773166 | 0.99640678 |
| ZNF384 | 5.74972997 | 0.40444943 | 0.58048809 | 0.69674028 | 0.48596535 | 0.99640678 |
| COPS7A | 7.10880355 | -0.051577469 | 0.57639908 | -0.0894822 | 0.92869868 | 0.99777713 |
| MLF2 | 12.033203 | 0.681481079 | 0.51049203 | 1.33494951 | 0.18189287 | 0.89342536 |
| PTMS | 15.8974457 | -0.85063851 | 0.65743974 | -1.2938654 | 0.19571193 | 0.90381759 |
| USP5 | 5.65719972 | -0.229725513 | 0.67102644 | -0.3423494 | 0.73208795 | 0.99665574 |
| TPI1 | 43.5992043 | 0.225931544 | 0.36501586 | 0.61896363 | 0.53594032 | 0.99665574 |
| LRRC23 | 5.40874054 | -0.411592445 | 0.68113413 | -0.6042752 | 0.54566071 | 0.99665574 |
| ATN1 | 13.3496821 | -0.307371774 | 0.51377872 | -0.5982571 | 0.54966839 | 0.99665574 |
| C12orf57 | 11.459442 | 0.736997618 | 0.5204239 | 1.41614869 | 0.15673198 | 0.86532131 |
| PTPN6 | 4.07580921 | -0.379165049 | 0.68797253 | -0.551134 | 0.58154182 | 0.99665574 |
| PHB2 | 17.4576868 | 0.474989172 | 0.46051192 | 1.03143729 | 0.3023358 | 0.97358745 |
| LPCAT3 | 39.8535925 | -0.430085551 | 0.3825537 | -1.1242488 | 0.26090747 | 0.94557817 |
| C1S | 8.73632812 | -0.070771453 | 0.57649507 | -0.1227616 | 0.90229589 | 0.99777713 |
| C1RL | 9.12321008 | -1.966008703 | 0.51574283 | -3.8119942 | 0.00013785 | 0.02687547 |
| PEX5 | 4.16788523 | 0.500975047 | 0.75378678 | 0.66461108 | 0.50629928 | 0.99665574 |
| CD163L1 | 3.40590637 | -0.813154578 | 0.75896556 | -1.0713985 | 0.28399028 | 0.96059387 |
| FOXJ2 | 3.77905189 | 1.242804061 | 0.91873958 | 1.35272724 | 0.17614279 | 0.88428483 |
| NECAP1 | 6.28034076 | 0.948114483 | 0.68905314 | 1.37596715 | 0.1688318 | 0.87595538 |
| LINC00937 | 2.68662037 | 0.398514339 | 0.88326921 | 0.45118106 | 0.65185906 | 0.99665574 |
| RIMKLB | 55.711983 | 0.109874783 | 0.66429296 | 0.1654011 | 0.86862829 | 0.99665574 |
| A2ML1-AS1 | 7.46349036 | 0.844858741 | 0.95312715 | 0.88640717 | 0.37539816 | 0.98569969 |
| PHC1 | 4.55131643 | 0.407497939 | 0.62970302 | 0.64712718 | 0.51754963 | 0.99665574 |
| M6PR | 18.1875331 | -0.357040608 | 0.42147305 | -0.8471256 | 0.39692512 | 0.99179694 |
| CLEC2D | 9.23465971 | -0.430863412 | 0.55216942 | -0.7803102 | 0.43520833 | 0.99617422 |
| TMEM52B | 2.53134997 | 2.020951825 | 1.27256329 | 1.58809533 | 0.11226478 | 0.81237714 |
| GABARAPL1 | 46.1390458 | -0.679348356 | 0.36087265 | -1.8825155 | 0.05976606 | 0.68092881 |
| KLRD1 | 7.18102817 | -1.015453162 | 0.89418097 | -1.1356238 | 0.25611406 | 0.94314473 |
| MAGOHB | 2.85954936 | -0.050734698 | 0.92864554 | -0.054633 | 0.95643084 | 0.99777713 |
| YBX3 | 46.6303882 | -1.214696154 | 0.40902092 | -2.9697653 | 0.00298027 | 0.17070061 |
| SMIM10L1 | 5.23072931 | -0.587827917 | 0.73969482 | -0.7946898 | 0.42679397 | 0.99525676 |
| AC244131.2 | 6.83289706 | -0.632623518 | 0.74589771 | -0.8481371 | 0.39636163 | 0.99179694 |
| ETV6 | 83.060516 | -0.829117539 | 0.35219802 | -2.3541232 | 0.01856646 | 0.45138305 |
| BCL2L14 | 9.20242182 | -0.520924718 | 0.51652248 | -1.0085228 | 0.31320353 | 0.97520189 |
| LRP6 | 29.0689348 | 0.35559127 | 0.37124304 | 0.95783957 | 0.33814366 | 0.97788427 |
| MANSC1 | 3.34196907 | 0.135479754 | 0.82894165 | 0.16343702 | 0.87017436 | 0.99665574 |
| BORCS5 | 11.9927575 | -0.19064448 | 0.51712459 | -0.3686626 | 0.71237925 | 0.99665574 |
| DUSP16 | 22.896749 | -0.412282159 | 0.43904974 | -0.9390329 | 0.34771384 | 0.97811401 |
| CREBL2 | 16.0049531 | -0.225773763 | 0.42848616 | -0.5269103 | 0.59825589 | 0.99665574 |
| CDKN1B | 6.29545996 | -0.189814626 | 0.58741924 | -0.3231331 | 0.74659441 | 0.99665574 |
| APOLD1 | 2.35714234 | 0.383778761 | 0.93830155 | 0.40901431 | 0.68252916 | 0.99665574 |
| GPRC5A | 4.87064621 | -1.532138069 | 0.88104676 | -1.7389974 | 0.08203521 | 0.75124322 |
| GPRC5D-AS1 | 9.83177713 | 1.019295231 | 0.54536172 | 1.869026 | 0.0616192 | 0.6834742 |
| HEBP1 | 9.23678026 | 0.438912959 | 0.58478909 | 0.75054916 | 0.45292403 | 0.99617422 |
| FAM234B | 6.71546204 | 1.299582462 | 0.66151914 | 1.96454251 | 0.04946721 | 0.64444758 |
| ATF7IP | 53.5845643 | 0.540871194 | 0.34508959 | 1.56733557 | 0.11703627 | 0.82339608 |
| PLBD1 | 5.60180171 | 0.374571957 | 0.76082721 | 0.49232198 | 0.62249175 | 0.99665574 |
| PLBD1-AS1 | 2.86396682 | -0.686902444 | 1.04054776 | -0.6601354 | 0.50916692 | 0.99665574 |
| AC010168.1 | 2.33279808 | -1.095264414 | 0.8787471 | -1.2463932 | 0.21262007 | 0.91066426 |
| H2AFJ | 18.3938054 | -0.685633191 | 0.45711482 | -1.4999146 | 0.13363653 | 0.84329092 |
| WBP11 | 11.5678559 | -0.606594182 | 0.5245833 | -1.1563353 | 0.24754404 | 0.93441937 |
| MGP | 11.7594687 | 0.166081242 | 0.83400373 | 0.19913729 | 0.84215535 | 0.99665574 |
| ERP27 | 6.39594283 | -1.036266649 | 0.88508467 | -1.1708108 | 0.24167486 | 0.93266071 |
| RERG | 19.5686039 | -0.713531157 | 0.82263363 | -0.8673742 | 0.38573703 | 0.99051797 |
| EPS8 | 543.218437 | 0.898614461 | 0.27382465 | 3.28171501 | 0.00103178 | 0.08790058 |
| STRAP | 17.181566 | 0.241180727 | 0.46224265 | 0.52176217 | 0.60183593 | 0.99665574 |

| | | | | | | |
|---|---|---|---|---|---|---|
| DERA | 49.0740503 | -0.612349014 | 0.34427254 | -1.7786752 | 0.07529303 | 0.73097508 |
| MGST1 | 17.61895 | -1.031986003 | 0.63252694 | -1.6315289 | 0.10277876 | 0.78770791 |
| PIK3C2G | 18.6614766 | -0.58028937 | 0.71943991 | -0.8065849 | 0.41990568 | 0.99518711 |
| PLEKHA5 | 233.714645 | 0.196170187 | 0.1945974 | 1.00808228 | 0.31341497 | 0.97556062 |
| AEBP2 | 49.9782059 | 0.199465472 | 0.30660091 | 0.65057038 | 0.51532385 | 0.99665574 |
| PYROXD1 | 9.19057245 | -0.297567289 | 0.45966019 | -0.6473636 | 0.51739663 | 0.99665574 |
| RECQL | 8.63078716 | -0.55732346 | 0.48799617 | -1.1420652 | 0.25342691 | 0.94076968 |
| GOLT1B | 9.91597344 | 0.010951636 | 0.54936215 | 0.01993518 | 0.98409508 | 0.99777713 |
| LDHB | 62.5668559 | -0.274135092 | 0.43745013 | -0.6266659 | 0.53087824 | 0.99665574 |
| CMAS | 6.72495921 | -0.464449131 | 0.65523649 | -0.7088267 | 0.47843201 | 0.99617422 |
| C2CD5 | 29.0191548 | 0.295852686 | 0.37960026 | 0.77937956 | 0.43575616 | 0.99617422 |
| ETNK1 | 53.6178272 | 0.440446298 | 0.31943545 | 1.37882723 | 0.16794803 | 0.87566218 |
| SOX5 | 11.6036552 | -0.836043692 | 0.54070646 | -1.5462062 | 0.12205477 | 0.82879523 |
| ETFRF1 | 7.79798436 | -0.089897243 | 0.63554892 | -0.1414482 | 0.88751589 | 0.99665574 |
| KRAS | 25.8584224 | -0.553093656 | 0.36151484 | -1.5299335 | 0.12603319 | 0.83633607 |
| RASSF8-AS1 | 5.83636359 | -0.763460923 | 0.77547286 | -0.9845102 | 0.32486473 | 0.97700345 |
| RASSF8 | 36.3498732 | -0.154133933 | 0.30430822 | -0.506506 | 0.61250149 | 0.99665574 |
| ITPR2 | 122.33643 | 0.346030158 | 0.32992902 | 1.04880181 | 0.29426934 | 0.97161335 |
| INTS13 | 5.07262218 | -0.035171671 | 0.61955299 | -0.0567694 | 0.95472887 | 0.99777713 |
| FGFR1OP2 | 15.108329 | 0.296440425 | 0.40263471 | 0.73625155 | 0.46157763 | 0.99617422 |
| TM7SF3 | 14.3342633 | -0.551219682 | 0.45454521 | -1.212684 | 0.22525067 | 0.92318867 |
| MED21 | 8.85482902 | 0.310325823 | 0.49848208 | 0.62254159 | 0.53358581 | 0.99665574 |
| STK38L | 11.9579871 | 0.035169737 | 0.46880473 | 0.07502001 | 0.94019879 | 0.99777713 |
| PPFIBP1 | 32.478279 | -0.703120683 | 0.40136802 | -1.7518104 | 0.07980641 | 0.7446402 |
| MRPS35 | 18.4365605 | 0.203165245 | 0.41924319 | 0.48459998 | 0.62796013 | 0.99665574 |
| KLHL42 | 19.3227956 | 0.241955649 | 0.41206746 | 0.58717485 | 0.55708628 | 0.99665574 |
| CCDC91 | 83.1984159 | -0.072117018 | 0.23139294 | -0.3116647 | 0.75529533 | 0.99665574 |
| ERGIC2 | 14.0221686 | 0.132670609 | 0.47814638 | 0.2774686 | 0.78142031 | 0.99665574 |
| IPO8 | 18.4629844 | 0.378971829 | 0.3775551 | 1.00375239 | 0.31549798 | 0.97652251 |
| CAPRIN2 | 11.5239533 | -0.808531882 | 0.48969026 | -1.6511088 | 0.09871637 | 0.78203824 |
| DDX11 | 2.8904914 | 0.558126955 | 0.91530573 | 0.60977107 | 0.54201347 | 0.99665574 |
| SINHCAF | 43.2758009 | -0.183954598 | 0.27065304 | -0.6796694 | 0.49671379 | 0.99640678 |
| DENND5B | 40.1242134 | -0.115843836 | 0.36014746 | -0.3216567 | 0.7477128 | 0.99665574 |
| AMN1 | 8.37545468 | 0.100569703 | 0.55632295 | 0.18077576 | 0.85654359 | 0.99665574 |
| KIAA1551 | 10.4916484 | -0.352397581 | 0.56983543 | -0.6184199 | 0.53629857 | 0.99665574 |
| BICD1 | 41.2479817 | -0.310592426 | 0.36406523 | -0.853123 | 0.39359109 | 0.99179694 |
| FGD4 | 229.295293 | -0.392968622 | 0.22446123 | -1.7507194 | 0.07999426 | 0.74470325 |
| DNM1L | 37.3267813 | 0.171023294 | 0.30768472 | 0.5558394 | 0.57832065 | 0.99665574 |
| YARS2 | 9.47944512 | -1.043530768 | 0.50870292 | -2.051356 | 0.04023229 | 0.60713077 |
| PKP2 | 9.10557105 | -0.054048346 | 0.52224094 | -0.1034931 | 0.91757161 | 0.99777713 |
| SYT10 | 3.73238061 | 2.562122996 | 0.97535322 | 2.62686681 | 0.00861751 | 0.31119582 |
| ALG10B | 3.46017187 | 1.024818278 | 0.86661279 | 1.18255614 | 0.2369851 | 0.9305595 |
| CPNE8 | 39.9126055 | -1.052066522 | 0.52633407 | -1.998857 | 0.04562383 | 0.63477358 |
| KIF21A | 53.1777564 | 0.295196605 | 0.44484086 | 0.66360048 | 0.50694605 | 0.99665574 |
| SLC2A13 | 16.1205263 | -0.417040899 | 0.67141908 | -0.6211335 | 0.53451177 | 0.99665574 |
| LRRK2 | 20.7646063 | 1.040490092 | 0.6315932 | 1.64740546 | 0.09947473 | 0.78261212 |
| GXYLT1 | 10.4314308 | 0.378165034 | 0.52739027 | 0.7170497 | 0.47334343 | 0.99617422 |
| YAF2 | 41.3608151 | 0.429636897 | 0.29315775 | 1.46554851 | 0.14277133 | 0.84969914 |
| PPHLN1 | 35.3730421 | -0.087908673 | 0.30184448 | -0.2912383 | 0.77086907 | 0.99665574 |
| ZCRB1 | 8.85354081 | -0.098878663 | 0.6560344 | -0.1507218 | 0.8801952 | 0.99665574 |
| PRICKLE1 | 14.8874162 | -1.190784906 | 0.71428002 | -1.6671122 | 0.09549211 | 0.77178333 |
| PUS7L | 8.65438019 | 0.117642831 | 0.50308185 | 0.23384431 | 0.81510585 | 0.99665574 |
| IRAK4 | 12.3724675 | -0.098026043 | 0.52475425 | -0.1868037 | 0.85181453 | 0.99665574 |
| TWF1 | 12.5600116 | 0.680790236 | 0.46030419 | 1.47900074 | 0.13914012 | 0.84668189 |
| TMEM117 | 258.605797 | 0.397012523 | 0.27327576 | 1.45279087 | 0.14628183 | 0.85169374 |
| ANO6 | 27.7129818 | -0.409485914 | 0.3426414 | -1.1950859 | 0.23205346 | 0.92842131 |
| AC008124.1 | 3.67535949 | 0.099438336 | 0.76723757 | 0.12960567 | 0.89687842 | 0.99777713 |
| ARID2 | 80.0864424 | -0.007884978 | 0.3240227 | -0.0243346 | 0.98058568 | 0.99777713 |
| SCAF11 | 80.5155362 | -0.063313689 | 0.21019588 | -0.3012128 | 0.76325223 | 0.99665574 |
| SLC38A1 | 51.8017899 | -0.572141675 | 0.44461871 | -1.2868142 | 0.19815905 | 0.90381759 |
| SLC38A2 | 35.2794024 | -0.299574229 | 0.43775112 | -0.6843483 | 0.49375527 | 0.99640678 |
| AC008014.1 | 45.0547929 | 0.625239012 | 0.56114422 | 1.1142216 | 0.26518413 | 0.94557817 |
| PCED1B | 7.02608559 | -1.331445232 | 0.61658078 | -2.1594011 | 0.03081906 | 0.56271575 |
| RPAP3 | 10.2409943 | 0.924947605 | 0.54078554 | 1.71037786 | 0.08719602 | 0.75595306 |
| RAPGEF3 | 10.4821454 | 0.276000828 | 0.62100162 | 0.44444462 | 0.65672116 | 0.99665574 |
| SLC48A1 | 6.02320219 | 0.359654348 | 0.75291936 | 0.47767977 | 0.63287814 | 0.99665574 |
| HDAC7 | 21.6412385 | -0.8594742 | 0.40042215 | -2.1464202 | 0.03183947 | 0.56852889 |
| VDR | 24.3382794 | 0.113143556 | 0.33220512 | 0.34058342 | 0.73341722 | 0.99665574 |
| TMEM106C | 9.90097518 | -0.564280031 | 0.50335526 | -1.1210373 | 0.26227198 | 0.94557817 |
| SENP1 | 12.5051703 | -0.654191571 | 0.43793346 | -1.493815 | 0.13522398 | 0.84373478 |
| PFKM | 6.31045703 | 0.341785728 | 0.65415665 | 0.522483 | 0.60133408 | 0.99665574 |
| ASB8 | 7.82943268 | 0.336049363 | 0.53824832 | 0.6243389 | 0.53240504 | 0.99665574 |
| ZNF641 | 4.34390062 | -0.912118877 | 0.67884097 | -1.3436415 | 0.17906435 | 0.88813816 |
| KANSL2 | 5.23601389 | 0.215447992 | 0.7576997 | 0.28434483 | 0.77614613 | 0.99665574 |
| CCNT1 | 14.554552 | -0.080904544 | 0.47826043 | -0.1691642 | 0.86566749 | 0.99665574 |
| ADCY6 | 6.48923393 | 0.251830228 | 0.73256606 | 0.34376453 | 0.73102338 | 0.99665574 |
| DDX23 | 6.49794615 | 0.037713586 | 0.62064282 | 0.06076536 | 0.95154608 | 0.99777713 |
| CCDC65 | 4.65947358 | -0.06161136 | 0.8876068 | -0.0694129 | 0.94466096 | 0.99777713 |
| PRKAG1 | 10.1214465 | -0.56861237 | 0.58779461 | -0.9673657 | 0.33336123 | 0.97788427 |
| KMT2D | 9.28522963 | -0.214964812 | 0.48306905 | -0.4449981 | 0.65632112 | 0.99665574 |
| LMBR1L | 7.87317755 | 0.590048001 | 0.59102105 | 0.99835361 | 0.31810792 | 0.97700345 |
| TUBA1B | 25.6249557 | -0.590839308 | 0.54682636 | -1.080488 | 0.27992494 | 0.96036515 |
| TUBA1A | 4.32010216 | -1.559051345 | 1.05382506 | -1.4794214 | 0.13902772 | 0.84668189 |
| TUBA1C | 21.4010206 | -0.830997338 | 0.67844453 | -1.2248567 | 0.22062921 | 0.91857862 |
| SPATS2 | 17.2274165 | -0.290641123 | 0.44612479 | -0.6514794 | 0.51473705 | 0.99665574 |
| MCRS1 | 4.62107067 | 0.226555084 | 0.69814577 | 0.32450972 | 0.74555217 | 0.99665574 |
| PRPF40B | 15.3350809 | 0.348572036 | 0.38587264 | 0.90333442 | 0.36634843 | 0.98365837 |
| FMNL3 | 3.54208592 | -0.191642946 | 0.79113097 | -0.2422392 | 0.80859481 | 0.99665574 |
| TMBIM6 | 93.6894021 | -0.116856893 | 0.30649646 | -0.3812667 | 0.70300536 | 0.99665574 |
| NCKAP5L | 4.17466335 | -0.084404738 | 0.69155701 | -0.1220503 | 0.90285919 | 0.99777713 |
| BCDIN3D | 2.84755082 | -0.559117782 | 0.86542504 | -0.6460615 | 0.51823954 | 0.99665574 |
| AQP2 | 62.7490284 | 2.070408861 | 1.26916473 | 1.63131611 | 0.10282364 | 0.78770791 |
| SMARCD1 | 7.56597935 | -0.287994378 | 0.59693296 | -0.4824568 | 0.62948147 | 0.99665574 |
| AC074032.1 | 16.96288 | -0.536964316 | 0.49761421 | -1.0790775 | 0.28055317 | 0.96059387 |
| CERS5 | 13.3498041 | -0.278474263 | 0.51481061 | -0.5409257 | 0.58855883 | 0.99665574 |
| LIMA1 | 57.8876052 | -0.516832876 | 0.34460648 | -1.499777 | 0.13367218 | 0.84329092 |
| LARP4 | 65.2029972 | 0.110972471 | 0.26866791 | 0.41304699 | 0.67957219 | 0.99665574 |
| DIP2B | 44.582847 | -0.626614666 | 0.30962109 | -2.0238113 | 0.04298957 | 0.61964991 |
| ATF1 | 14.7020516 | -0.092026382 | 0.45516373 | -0.202183 | 0.83977364 | 0.99665574 |
| METTL7A | 17.4284244 | 0.75643441 | 0.41833636 | 1.80819668 | 0.0705759 | 0.71264784 |
| SLC11A2 | 30.8257479 | -0.057297742 | 0.36164471 | -0.1584366 | 0.87411281 | 0.99665574 |
| LETMD1 | 17.8184185 | -0.021225769 | 0.38014612 | -0.0558358 | 0.9554726 | 0.99777713 |
| CSRNP2 | 5.13367659 | -0.415647198 | 0.72179982 | -0.5758483 | 0.56471772 | 0.99665574 |
| TFCP2 | 38.9634567 | 0.137243852 | 0.33344258 | 0.41159665 | 0.68063508 | 0.99665574 |
| POU6F1 | 8.75297922 | -1.043217406 | 0.51882842 | -2.0107175 | 0.0443553 | 0.62979556 |
| DAZAP2 | 38.0974661 | 0.172850891 | 0.36771929 | 0.47006207 | 0.63831067 | 0.99665574 |
| ACVR1B | 31.0848524 | -0.357190672 | 0.32433433 | -1.1013039 | 0.2707644 | 0.94865166 |
| NR4A1 | 17.597824 | 1.485276081 | 0.82746757 | 1.79496592 | 0.07265913 | 0.72288348 |
| ATG101 | 6.07899898 | 0.055698377 | 0.64487545 | 0.08637075 | 0.9311717 | 0.99777713 |
| KRT7 | 288.604698 | 0.944017748 | 0.49907986 | 1.89151642 | 0.05855544 | 0.67545594 |
| KRT8 | 49.4566776 | -0.588740614 | 0.42229701 | -1.3941387 | 0.16327571 | 0.8716636 |
| KRT18 | 42.4549261 | -0.634732772 | 0.41516841 | -1.5288561 | 0.12630011 | 0.83633607 |
| EIF4B | 40.7704343 | 0.03928636 | 0.33520608 | 0.11720062 | 0.90670108 | 0.99777713 |

| | | | | | | |
|---|---|---|---|---|---|---|
| AC068888.1 | 8.54748973 | -0.184664234 | 0.58786718 | -0.3141258 | 0.75342551 | 0.99665574 |
| TNS2 | 11.395631 | -0.301173603 | 0.66134323 | -0.4553968 | 0.64882378 | 0.99665574 |
| SPRYD3 | 4.22213278 | 0.43520892 | 0.77002125 | 0.5651908 | 0.571944 | 0.99665574 |
| CSAD | 15.1829706 | 0.499262146 | 0.50536561 | 0.98792269 | 0.32319051 | 0.97700345 |
| ZNF740 | 2.33111843 | 0.057164689 | 0.8723869 | 0.06552676 | 0.9477546 | 0.99777713 |
| MFSD5 | 3.92196883 | -0.392604645 | 0.91974195 | -0.4268639 | 0.66947845 | 0.99665574 |
| PFDN5 | 31.7648451 | 0.563431357 | 0.45725096 | 1.23221471 | 0.21786889 | 0.91662582 |
| C12orf10 | 3.53812974 | 0.397619296 | 0.82046523 | 0.48462663 | 0.62794122 | 0.99665574 |
| AAAS | 2.43848267 | 1.174873633 | 1.01328283 | 1.15947256 | 0.24626362 | 0.93441937 |
| SP1 | 18.7820204 | -0.574344163 | 0.38574955 | -1.4889043 | 0.13651257 | 0.84475183 |
| PRR13 | 22.0063835 | 0.13744882 | 0.34064903 | 0.403491 | 0.68658705 | 0.99665574 |
| PCBP2 | 62.0835647 | 0.244381853 | 0.32073758 | 0.76193707 | 0.44609756 | 0.99617422 |
| MAP3K12 | 4.67319911 | -1.488254453 | 0.83834104 | -1.7752375 | 0.07585868 | 0.7342902 |
| ATP5MC2 | 45.85568 | 0.846149313 | 0.45645962 | 1.85372218 | 0.06377892 | 0.68630818 |
| CALCOCO1 | 8.33962057 | 1.313171815 | 0.61559291 | 2.13318216 | 0.03290979 | 0.57425028 |
| SMUG1 | 4.82304057 | -0.016314478 | 0.64956379 | -0.0251161 | 0.9799624 | 0.99777713 |
| LINC02381 | 6.79035261 | 0.060389524 | 0.69184974 | 0.08728705 | 0.93044335 | 0.99777713 |
| CBX5 | 18.1883089 | 0.152271028 | 0.52927114 | 0.28769947 | 0.77357679 | 0.99665574 |
| HNRNPA1 | 22.9865549 | 0.429149359 | 0.42470747 | 1.0104587 | 0.31227558 | 0.97442101 |
| COPZ1 | 14.8426096 | -0.212700455 | 0.40624771 | -0.5235733 | 0.60057535 | 0.99665574 |
| PPP1R1A | 4.93529987 | -0.569904833 | 0.83326753 | -0.6839398 | 0.49401318 | 0.99640678 |
| CD63 | 41.6051847 | -1.117432587 | 0.44310244 | -2.521838 | 0.01167435 | 0.37335809 |
| ORMDL2 | 6.08629587 | -0.318253568 | 0.68985678 | -0.4613328 | 0.64455985 | 0.99665574 |
| PYM1 | 3.87513409 | -0.000236606 | 0.84682369 | -0.0002794 | 0.99977707 | 0.99987569 |
| RAB5B | 15.6660643 | 0.307004921 | 0.41230079 | 0.74461395 | 0.45650513 | 0.99617422 |
| SUOX | 4.21511228 | 0.104921564 | 0.75572449 | 0.13883573 | 0.88957996 | 0.99665574 |
| IKZF4 | 2.70144531 | -0.037072756 | 0.81138789 | -0.0456905 | 0.9635569 | 0.99777713 |
| RPS26 | 7.78680646 | -0.324874331 | 0.54400542 | -0.5971895 | 0.55038086 | 0.99665574 |
| ERBB3 | 13.2633637 | -0.578241322 | 0.59493063 | -0.9719475 | 0.33107668 | 0.97788427 |
| PA2G4 | 7.05142566 | 0.298544521 | 0.61300479 | 0.48701826 | 0.62624539 | 0.99665574 |
| RPL41 | 143.600885 | 0.261550718 | 0.36326883 | 0.71999218 | 0.47152981 | 0.99617422 |
| ESYT1 | 5.85040246 | -0.879406593 | 0.60234713 | -1.4599664 | 0.1442993 | 0.84969914 |
| MYL6B | 4.5131951 | 0.443314702 | 0.68504012 | 0.64713685 | 0.51754338 | 0.99665574 |
| MYL6 | 65.6686955 | 0.263184014 | 0.40862058 | 0.6440792 | 0.51952408 | 0.99665574 |
| SMARCC2 | 16.6023668 | -0.013515888 | 0.4311077 | -0.0313515 | 0.97498919 | 0.99777713 |
| RNF41 | 4.26847555 | -0.516929325 | 0.68760799 | -0.7517791 | 0.45218391 | 0.99617422 |
| NABP2 | 1.78724355 | 0.119481116 | 1.00554631 | 0.11882209 | 0.90541631 | 0.99777713 |
| ANKRD52 | 6.25759113 | -0.092698805 | 0.60190952 | -0.1540079 | 0.87760353 | 0.99665574 |
| COQ10A | 2.50953901 | 0.079061272 | 1.15688833 | 0.06833959 | 0.94551531 | 0.99777713 |
| CS | 14.2487915 | 0.180787877 | 0.44117738 | 0.40978501 | 0.68196366 | 0.99665574 |
| AC073896.2 | 3.67610871 | 1.004796065 | 0.88502201 | 1.13533454 | 0.25623517 | 0.94314473 |
| PAN2 | 7.3426853 | 1.047094486 | 0.70314851 | 1.48915125 | 0.13644755 | 0.84475183 |
| STAT2 | 12.7901892 | -0.194812563 | 0.55312482 | -0.3522036 | 0.72468556 | 0.99665574 |
| TIMELESS | 2.49587771 | 1.55188532 | 0.95867177 | 1.61878692 | 0.10549312 | 0.79494888 |
| SPRYD4 | 3.50715566 | 0.848686944 | 0.83402304 | 1.01758213 | 0.30887658 | 0.97376371 |
| RBMS2 | 14.1194244 | -0.600796641 | 0.4816922 | -1.2472625 | 0.21230125 | 0.91043197 |
| BAZ2A | 18.9677339 | 0.271503552 | 0.38785481 | 0.70001337 | 0.48391896 | 0.99640678 |
| ATP5F1B | 69.2431586 | 0.709226296 | 0.41098588 | 1.72567071 | 0.08440667 | 0.75373899 |
| PTGES3 | 54.9462399 | 0.222248094 | 0.28533677 | 0.77889749 | 0.43604011 | 0.99617422 |
| NACA | 35.6286807 | 0.162653928 | 0.37183893 | 0.43743114 | 0.66179871 | 0.99665574 |
| NEMP1 | 3.63154811 | 0.525201636 | 0.7436531 | 0.70624548 | 0.48003549 | 0.99617422 |
| STAT6 | 29.3288713 | -0.639172024 | 0.33695625 | -1.8968991 | 0.05784125 | 0.67545594 |
| SHMT2 | 3.04192586 | 0.21551685 | 0.81675627 | 0.26386923 | 0.79188068 | 0.99665574 |
| R3HDM2 | 66.7448478 | 0.151559709 | 0.24286218 | 0.62405644 | 0.53259052 | 0.99665574 |
| MARS | 8.18785923 | -0.253776753 | 0.7067321 | -0.3590848 | 0.71953165 | 0.99665574 |
| DDIT3 | 12.2752084 | 0.72796413 | 0.96209249 | 0.75664672 | 0.44926154 | 0.99617422 |
| MBD6 | 7.81188059 | 0.420399482 | 0.61413256 | 0.68454192 | 0.49363303 | 0.99640678 |
| DCTN2 | 16.0158972 | -0.267467885 | 0.37901935 | -0.705684 | 0.48038468 | 0.99617422 |
| PIP4K2C | 9.93069417 | 0.491341765 | 0.47088761 | 1.04343745 | 0.29674574 | 0.97341216 |
| DTX3 | 2.4366263 | 1.02769401 | 1.0085356 | 1.01899627 | 0.30820474 | 0.97376371 |
| OS9 | 22.285791 | -0.218547893 | 0.3993898 | -0.5472045 | 0.58423825 | 0.99665574 |
| TSPAN31 | 5.12405667 | -0.096553406 | 0.75547534 | -0.1278048 | 0.89830342 | 0.99777713 |
| CDK4 | 5.632804 | -0.815525421 | 0.78816745 | -1.0347109 | 0.30080396 | 0.97358745 |
| 9-Mar | 5.50522663 | -0.636329467 | 0.69761414 | -0.912151 | 0.36168921 | 0.9819332 |
| TSFM | 6.88060453 | 0.830983434 | 0.62072346 | 1.33873374 | 0.18065737 | 0.89080954 |
| AVIL | 13.6792721 | -0.572989505 | 0.6144638 | -0.9325033 | 0.35107649 | 0.97838306 |
| CTDSP2 | 21.8700462 | 0.157197024 | 0.40942714 | 0.38394383 | 0.70102008 | 0.99665574 |
| LRIG3 | 7.56257133 | 0.149269551 | 0.73034008 | 0.20438362 | 0.83805373 | 0.99665574 |
| SLC16A7 | 24.3966873 | -0.795994599 | 0.58475602 | -1.3612423 | 0.17343715 | 0.88157681 |
| FAM19A2 | 8.1431922 | 0.263353292 | 0.74422149 | 0.35386413 | 0.72344071 | 0.99665574 |
| USP15 | 51.8488629 | 0.018372055 | 0.28277446 | 0.0649707 | 0.94819733 | 0.99777713 |
| MON2 | 86.3421176 | 0.062959526 | 0.2625459 | 0.23980388 | 0.8104823 | 0.99665574 |
| AC048341.1 | 5.5383813 | -1.11358171 | 0.66745579 | -1.6683977 | 0.09523681 | 0.77178333 |
| PPM1H | 28.9816517 | -1.277422585 | 0.51981021 | -2.4574788 | 0.01399161 | 0.39733028 |
| AVPR1A | 52.3741926 | 0.773056981 | 0.44921477 | 1.72090733 | 0.08526764 | 0.75373899 |
| DPY19L2 | 110.009158 | 0.59880352 | 0.49916391 | 1.19961301 | 0.23028967 | 0.92774608 |
| RXYLT1 | 9.41719055 | -0.390156751 | 0.5031705 | -0.7753967 | 0.43810528 | 0.99617422 |
| SRGAP1 | 29.714577 | -0.744058489 | 0.51973618 | -1.431608 | 0.15225603 | 0.85976284 |
| XPOT | 6.06685585 | -0.168385097 | 0.6743168 | -0.2497121 | 0.80280996 | 0.99665574 |
| TBK1 | 25.8184514 | -0.245468317 | 0.35723664 | -0.6871309 | 0.49200027 | 0.99640678 |
| RASSF3 | 4.24023614 | -1.628516064 | 0.69425148 | -2.3457149 | 0.01899063 | 0.45540143 |
| GNS | 29.0987704 | -0.378151051 | 0.40685345 | -0.9294527 | 0.35265451 | 0.97912036 |
| TBC1D30 | 10.2862091 | -0.456874347 | 0.53039684 | -0.8613821 | 0.38902763 | 0.99176253 |
| LEMD3 | 30.5565029 | 0.120341682 | 0.38720123 | 0.31079881 | 0.75595357 | 0.99665574 |
| LLPH | 4.98770057 | -1.192408176 | 0.90254742 | -1.3211585 | 0.18644853 | 0.89532394 |
| HELB | 9.63103433 | -1.272609633 | 0.65087188 | -1.9552383 | 0.05055495 | 0.64929786 |
| GRIP1 | 47.5496102 | -0.363006097 | 0.59818126 | -0.6068497 | 0.54395069 | 0.99665574 |
| CAND1 | 30.3713059 | 0.416536383 | 0.35871431 | 1.16119255 | 0.2455636 | 0.93426912 |
| DYRK2 | 2.05444739 | 1.116894004 | 1.01560818 | 1.09972924 | 0.27145011 | 0.94870306 |
| MDM1 | 4.201345 | 0.801201068 | 0.68706987 | 1.16611294 | 0.24356879 | 0.93426912 |
| RAP1B | 52.7178071 | -0.728450888 | 0.33764741 | -2.1574307 | 0.03097212 | 0.56271575 |
| NUP107 | 14.4974165 | -0.209267041 | 0.44339539 | -0.4719649 | 0.63695188 | 0.99665574 |
| SLC35E3 | 8.76791171 | -0.666037053 | 0.49357414 | -1.3494164 | 0.17720325 | 0.88584149 |
| MDM2 | 16.348 | -0.625825511 | 0.48355179 | -1.2942264 | 0.19558722 | 0.90381759 |
| CPM | 9.99973017 | 1.348561383 | 0.86418191 | 1.56050638 | 0.11864026 | 0.82619588 |
| CPSF6 | 20.6369853 | 0.642647306 | 0.41580436 | 1.54555213 | 0.12221277 | 0.82915695 |
| YEATS4 | 3.46798615 | 0.134375179 | 0.84909755 | 0.15825647 | 0.87425471 | 0.99665574 |
| FRS2 | 28.5678654 | -0.415935966 | 0.32411529 | -1.2832963 | 0.19938827 | 0.90381759 |
| CCT2 | 12.8364221 | -0.467639314 | 0.48327219 | -0.967652 | 0.33321819 | 0.97788427 |
| RAB3IP | 25.1020958 | -0.076405071 | 0.39694132 | -0.1924845 | 0.84736267 | 0.99665574 |
| MYRFL | 5.22761864 | 0.107610299 | 1.06727545 | 0.10082711 | 0.91968771 | 0.99777713 |
| AC025159.1 | 5.21915137 | 1.175236006 | 1.08479015 | 1.08337636 | 0.2786414 | 0.95969433 |
| CNOT2 | 50.900375 | 0.330984203 | 0.26366329 | 1.25532911 | 0.20935931 | 0.91043197 |
| TSPAN8 | 64.8735935 | 0.952536819 | 0.42573498 | 2.23739382 | 0.02526061 | 0.50908112 |
| ZFC3H1 | 41.4419759 | 0.010602611 | 0.41253495 | 0.02570112 | 0.97949573 | 0.99777713 |
| TMEM19 | 6.06537622 | -0.109838339 | 0.60492359 | -0.1815739 | 0.85591713 | 0.99665574 |
| RAB21 | 34.7349538 | 0.24584787 | 0.32188702 | 0.76377068 | 0.44500391 | 0.99617422 |
| TBC1D15 | 27.4367722 | -0.507029499 | 0.39717739 | -1.276582 | 0.20174987 | 0.90614933 |
| TRHDE | 105.240422 | 0.711257665 | 0.37829378 | 1.88017278 | 0.06008453 | 0.68195946 |
| AC090502.1 | 6.35785004 | 0.081105432 | 0.71706314 | 0.1131078 | 0.90994509 | 0.99777713 |
| ATXN7L3B | 12.1339701 | -0.117425416 | 0.43897699 | -0.2674979 | 0.78908585 | 0.99665574 |
| CAPS2 | 16.1754495 | 0.30207904 | 0.50071127 | 0.60329987 | 0.54630922 | 0.99665574 |
| KRR1 | 10.6610043 | 0.66368325 | 0.4627219 | 1.43430264 | 0.15148592 | 0.85701128 |

| | | | | | | |
|---|---|---|---|---|---|---|
| PHLDA1 | 14.313171 | -0.404471489 | 0.65432563 | -0.6181502 | 0.53647636 | 0.99665574 |
| NAP1L1 | 30.5201085 | -0.423685075 | 0.45733214 | -0.9264275 | 0.35422386 | 0.98022903 |
| OSBPL8 | 52.274961 | -0.340411335 | 0.30641412 | -1.1109519 | 0.26658907 | 0.94557817 |
| ZDHHC17 | 24.8788535 | 0.07731022 | 0.37304969 | 0.2072384 | 0.83582368 | 0.99665574 |
| AC027288.3 | 4.56993566 | -0.231924703 | 1.09923955 | -0.2109865 | 0.83289781 | 0.99665574 |
| PAWR | 50.3815273 | 0.016932322 | 0.28048848 | 0.06036726 | 0.95186313 | 0.99777713 |
| AC073569.2 | 5.21591144 | -1.388775355 | 0.98627918 | -1.4080956 | 0.15910277 | 0.86757292 |
| PPP1R12A | 58.8854594 | 0.188716486 | 0.29904749 | 0.63105859 | 0.52800221 | 0.99665574 |
| ACSS3 | 17.3709768 | -0.722187 | 0.62176492 | -1.1615113 | 0.24543402 | 0.93426912 |
| CCDC59 | 4.04254373 | 0.776956204 | 0.80526318 | 0.96484755 | 0.33462118 | 0.97788427 |
| METTL25 | 29.4341588 | 0.126701552 | 0.30744863 | 0.41210641 | 0.68026143 | 0.99665574 |
| TMTC2 | 45.7036272 | -0.775394684 | 0.49238869 | -1.5747613 | 0.11531154 | 0.81721642 |
| C12orf29 | 2.23222205 | -1.326158276 | 0.958113 | -1.3841356 | 0.16631695 | 0.87495175 |
| CEP290 | 17.771431 | 0.282456608 | 0.37415574 | 0.75491721 | 0.45029866 | 0.99617422 |
| TMTC3 | 22.9511305 | -0.035691076 | 0.37575997 | -0.0949837 | 0.92432777 | 0.99777713 |
| KITLG | 16.9747003 | -1.815936618 | 0.99991992 | -1.816082 | 0.06935779 | 0.7094165 |
| POC1B | 13.8603399 | -0.36954091 | 0.52062725 | -0.7097994 | 0.47782854 | 0.99617422 |
| POC1B-AS1 | 2.5773461 | 0.663406867 | 0.95007419 | 0.69826849 | 0.4850093 | 0.99640678 |
| ATP2B1 | 44.9498507 | 0.347154816 | 0.59120689 | 0.58719684 | 0.55707152 | 0.99665574 |
| BTG1 | 99.3417196 | -0.409094759 | 0.32450367 | -1.2606784 | 0.20742474 | 0.90994205 |
| AC025164.1 | 2.82439282 | -1.103204476 | 0.9401238 | -1.1734672 | 0.24060849 | 0.93216487 |
| EEA1 | 50.5374829 | -0.003318176 | 0.3598547 | -0.0092209 | 0.99264291 | 0.99777713 |
| AC138123.1 | 10.9619408 | 1.197528883 | 0.75933102 | 1.57708411 | 0.11477617 | 0.81638817 |
| NUDT4 | 39.1492804 | 0.351164093 | 0.35439672 | 0.9908785 | 0.32174491 | 0.97700345 |
| UBE2N | 20.8284188 | 0.219408069 | 0.4525429 | 0.48483375 | 0.62779428 | 0.99665574 |
| MRPL42 | 14.3698543 | -0.091372786 | 0.39177012 | -0.2332306 | 0.81558233 | 0.99665574 |
| CRADD | 86.9148188 | -0.343272629 | 0.33686992 | -1.0190065 | 0.30819989 | 0.97376371 |
| AC012085.2 | 8.34954275 | -0.903356888 | 0.82884093 | -1.0899038 | 0.27575552 | 0.95641787 |
| CEP83 | 13.1563247 | 0.472102522 | 0.43641288 | 1.08177952 | 0.27935051 | 0.95980166 |
| TMCC3 | 8.63235916 | 0.87932492 | 0.66715877 | 1.31801448 | 0.1874988 | 0.89729653 |
| NDUFA12 | 15.3570944 | 0.174575512 | 0.4975894 | 0.35084251 | 0.7257065 | 0.99665574 |
| NR2C1 | 17.2305171 | -0.113406922 | 0.43097704 | -0.2631391 | 0.79244335 | 0.99665574 |
| FGD6 | 8.93463077 | -0.881925879 | 0.56762931 | -1.5537004 | 0.1202559 | 0.82659025 |
| VEZT | 38.1393158 | -0.247712678 | 0.28041242 | -0.883387 | 0.37702723 | 0.98679664 |
| METAP2 | 23.7200174 | -0.079222361 | 0.37267719 | -0.2125764 | 0.83165741 | 0.99665574 |
| NTN4 | 50.9393765 | -0.086033476 | 0.52192457 | -0.1648389 | 0.86907078 | 0.99665574 |
| SNRPF | 6.15007984 | 0.26686961 | 0.60313723 | 0.44246914 | 0.65814976 | 0.99665574 |
| LTA4H | 9.62488503 | 0.258737131 | 0.58726215 | 0.440582 | 0.65951564 | 0.99665574 |
| CDK17 | 22.4166329 | -0.045270439 | 0.4414053 | -0.1025598 | 0.91831235 | 0.99777713 |
| NEDD1 | 8.56341494 | -0.391328257 | 0.52095912 | -0.7511688 | 0.45255105 | 0.99617422 |
| TMPO | 22.5788767 | 0.142522717 | 0.43697592 | 0.32615691 | 0.74430564 | 0.99665574 |
| SLC25A3 | 57.1271746 | 0.411637422 | 0.39714876 | 1.03648169 | 0.29997748 | 0.97358745 |
| APAF1 | 7.92509735 | -0.523837878 | 0.55515068 | -0.9435959 | 0.34537621 | 0.97788427 |
| ANKS1B | 5.1142502 | 1.736576405 | 0.81562645 | 2.12913203 | 0.03324334 | 0.5761693 |
| UHRF1BP1L | 20.0986314 | -0.178812978 | 0.39472059 | -0.4530115 | 0.65054044 | 0.99665574 |
| ACTR6 | 7.10132858 | 0.153707236 | 0.5749051 | 0.26736106 | 0.78919118 | 0.99665574 |
| SCYL2 | 26.4739536 | 0.145337853 | 0.34198946 | 0.42497758 | 0.67085302 | 0.99665574 |
| NR1H4 | 8.37885569 | -1.182767996 | 0.93217196 | -1.2688303 | 0.20450161 | 0.90834797 |
| ANO4 | 18.395731 | -1.324617574 | 1.57831691 | -0.8392596 | 0.40132367 | 0.99194293 |
| UTP20 | 16.9322231 | 0.422023188 | 0.44163994 | 0.95558202 | 0.33928345 | 0.97788427 |
| ARL1 | 16.8483926 | 0.1792931 | 0.37174774 | 0.48229776 | 0.62959444 | 0.99665574 |
| CHPT1 | 35.5230475 | -0.222945191 | 0.41067809 | -0.5428709 | 0.58721869 | 0.99665574 |
| GNPTAB | 26.4261594 | -0.396556586 | 0.34155491 | -1.1610332 | 0.2456284 | 0.93426912 |
| DRAM1 | 21.0504964 | -1.339889429 | 0.39556073 | -3.3873166 | 0.0007058 | 0.07140296 |
| WASHC3 | 18.3384959 | 0.1760047 | 0.56647465 | 0.31070181 | 0.75602732 | 0.99665574 |
| NUP37 | 5.51886327 | 0.043880991 | 0.66803055 | 0.0656871 | 0.94762694 | 0.99777713 |
| HSP90B1 | 49.6868655 | 0.306181155 | 0.44982724 | 0.68066388 | 0.4960842 | 0.99640678 |
| C12orf73 | 3.07358506 | -0.598556584 | 0.94301774 | -0.6347246 | 0.52560803 | 0.99665574 |
| TDG | 11.3582661 | -0.174015174 | 0.52714719 | -0.3301074 | 0.74131883 | 0.99665574 |
| HCFC2 | 10.9560964 | 0.20822439 | 0.45721223 | 0.45542174 | 0.64880586 | 0.99665574 |
| NFYB | 5.74995073 | 0.026670782 | 0.71358196 | 0.03737592 | 0.97018527 | 0.99777713 |
| TXNRD1 | 89.6685967 | -0.083557205 | 0.41451144 | -0.20158 | 0.8402451 | 0.99665574 |
| SLC41A2 | 6.87720244 | -1.672438634 | 0.70451167 | -2.3738977 | 0.01760143 | 0.43770425 |
| C12orf45 | 3.74242787 | -0.414854483 | 0.73130982 | -0.567276 | 0.5705267 | 0.99665574 |
| WASHC4 | 42.6748122 | 0.093121366 | 0.29907446 | 0.31136515 | 0.75552304 | 0.99665574 |
| APPL2 | 19.0450489 | -0.436716734 | 0.37203146 | -1.1738704 | 0.24044693 | 0.93216487 |
| C12orf75 | 44.296052 | -0.096157517 | 0.40177817 | -0.2393299 | 0.81084981 | 0.99665574 |
| CKAP4 | 3.79545781 | -0.585862424 | 0.69229915 | -0.8462562 | 0.39740986 | 0.99179694 |
| TCP11L2 | 7.74483407 | -0.175635093 | 0.66459385 | -0.2642743 | 0.79156854 | 0.99665574 |
| POLR3B | 9.86585615 | 0.252090716 | 0.57871178 | 0.43560668 | 0.66312213 | 0.99665574 |
| RIC8B | 15.2044796 | 0.11483979 | 0.47226774 | 0.2431667 | 0.80787626 | 0.99665574 |
| TMEM263 | 13.1414351 | -0.944489005 | 0.44179997 | -2.1378205 | 0.03253132 | 0.57257379 |
| MTERF2 | 3.63712329 | -0.303989553 | 0.82093447 | -0.370297 | 0.71116123 | 0.99665574 |
| CRY1 | 15.0058385 | 0.123303757 | 0.58520531 | 0.21070171 | 0.83312004 | 0.99665574 |
| BTBD11 | 15.1005185 | 0.918425131 | 0.65343264 | 1.40553912 | 0.15986103 | 0.86757484 |
| PWP1 | 8.82911556 | -0.123856606 | 0.47941013 | -0.2583521 | 0.79613519 | 0.99665574 |
| PRDM4 | 34.9209062 | -0.028004748 | 0.398309 | -0.0703091 | 0.94394764 | 0.99777713 |
| AC126177.4 | 3.95021784 | 0.59729459 | 0.82536882 | 0.72366992 | 0.46926841 | 0.99617422 |
| SART3 | 10.1432661 | -0.086209084 | 0.46861016 | -0.1839676 | 0.85403888 | 0.99665574 |
| ISCU | 31.1875274 | -0.297695922 | 0.3988394 | -0.7464055 | 0.4554225 | 0.99617422 |
| CORO1C | 6.76984503 | -1.507810936 | 0.60163353 | -2.506195 | 0.01220383 | 0.37720254 |
| SSH1 | 16.7150684 | 0.058103987 | 0.35436356 | 0.16396716 | 0.86975699 | 0.99665574 |
| USP30 | 5.45164222 | 0.181152181 | 0.72122904 | 0.2511715 | 0.80168152 | 0.99665574 |
| UNG | 4.65766744 | -0.028630343 | 0.8566132 | -0.0334227 | 0.97333749 | 0.99777713 |
| ACACB | 33.6884177 | 0.362558065 | 0.38590914 | 0.93949073 | 0.34747885 | 0.97804112 |
| KCTD10 | 6.73837344 | -1.106317448 | 0.71262222 | -1.5524599 | 0.12055221 | 0.82659025 |
| UBE3B | 8.40073812 | -0.420974986 | 0.54768646 | -0.7686423 | 0.44210568 | 0.99617422 |
| MMAB | 7.6611948 | -0.87969142 | 0.5805877 | -1.5151741 | 0.12972833 | 0.83778831 |
| MVK | 3.17591975 | -0.992751065 | 0.79929983 | -1.2420259 | 0.21422702 | 0.91253508 |
| GLTP | 13.9251364 | 0.509455081 | 0.47597388 | 1.07034251 | 0.28446517 | 0.96066218 |
| TCHP | 8.62797311 | -1.104776927 | 0.5421493 | -2.0377725 | 0.04157269 | 0.61348464 |
| GIT2 | 18.1604606 | -0.374093459 | 0.38290667 | -0.9769834 | 0.32857737 | 0.97788427 |
| ANKRD13A | 36.5823746 | 0.292332493 | 0.3637032 | 0.80376662 | 0.42153177 | 0.99518711 |
| C12orf76 | 5.62713233 | 0.01630815 | 0.65249245 | 0.02499362 | 0.98006005 | 0.99777713 |
| IFT81 | 7.17744411 | -0.75441536 | 0.52082871 | -1.4484904 | 0.14747996 | 0.85284617 |
| ATP2A2 | 62.6981079 | -0.100337626 | 0.3363318 | -0.2983293 | 0.76545186 | 0.99665574 |
| ANAPC7 | 12.2815288 | 0.177571589 | 0.45166299 | 0.39315063 | 0.69420823 | 0.99665574 |
| ARPC3 | 41.6512138 | -0.727773495 | 0.30403681 | -2.3937019 | 0.0166793 | 0.4237383 |
| GPN3 | 3.33498182 | -0.725456903 | 0.85873129 | -0.8448008 | 0.39822206 | 0.99179694 |
| VPS29 | 18.7372131 | -0.636796068 | 0.38865302 | -1.6384694 | 0.10132381 | 0.78386461 |
| PPTC7 | 28.612661 | 0.125635935 | 0.40765999 | 0.30818805 | 0.75793925 | 0.99665574 |
| TCTN1 | 6.10412935 | -0.341387815 | 0.68415319 | -0.4989932 | 0.61778415 | 0.99665574 |
| PPP1CC | 15.1191672 | -0.126171506 | 0.45077898 | -0.2798966 | 0.77955683 | 0.99665574 |
| SH2B3 | 2.72044928 | -0.868458537 | 0.89194366 | -0.9736697 | 0.33022055 | 0.97788427 |
| ATXN2 | 78.0840055 | -0.111308723 | 0.29224936 | -0.380869 | 0.70330045 | 0.99665574 |
| BRAP | 10.51848 | 0.113024761 | 0.5144346 | 0.21970677 | 0.82609954 | 0.99665574 |
| ACAD10 | 11.8776681 | 0.04109368 | 0.44756162 | 0.09181681 | 0.92684359 | 0.99777713 |
| ALDH2 | 18.2867752 | 0.308840956 | 0.54774406 | 0.56384173 | 0.57286185 | 0.99665574 |
| MAPKAPK5-A | 2.94658439 | -0.196744107 | 0.82437597 | -0.2386582 | 0.81137061 | 0.99665574 |
| MAPKAPK5 | 14.2037417 | -0.310482135 | 0.46949345 | -0.661313 | 0.50841159 | 0.99665574 |
| TMEM116 | 59.465816 | 0.346791013 | 0.35501056 | 0.97684705 | 0.32864488 | 0.97788427 |
| ERP29 | 33.9857438 | -0.182576904 | 0.41006189 | -0.4452423 | 0.65614465 | 0.99665574 |
| NAA25 | 18.6584896 | -0.112204141 | 0.39835057 | -0.2816719 | 0.77819514 | 0.99665574 |

| | | | | | | |
|---|---|---|---|---|---|---|
| TRAFD1 | 7.29358057 | -0.467836649 | 0.63317679 | -0.7388721 | 0.45998468 | 0.99617422 |
| HECTD4 | 61.6256266 | -0.173224946 | 0.28855912 | -0.6003101 | 0.5482996 | 0.99665574 |
| RPL6 | 78.5350165 | 0.191728515 | 0.43768291 | 0.43805347 | 0.66134753 | 0.99665574 |
| PTPN11 | 24.0301818 | -0.236747831 | 0.39845023 | -0.5941717 | 0.55239731 | 0.99665574 |
| OAS1 | 6.60880611 | 1.131555504 | 1.50544599 | 0.75164138 | 0.45226675 | 0.99617422 |
| DDX54 | 9.89642282 | 0.551632462 | 0.51409844 | 1.07300941 | 0.28326689 | 0.96059387 |
| TPCN1 | 26.6893814 | 0.260729071 | 0.46575921 | 0.5597937 | 0.57562016 | 0.99665574 |
| SLC8B1 | 11.3676816 | -1.273634825 | 0.46372414 | -2.7465355 | 0.00602284 | 0.24620771 |
| PLBD2 | 4.04144561 | -0.894666695 | 0.74938613 | -1.1938661 | 0.23253035 | 0.92920486 |
| RBM19 | 11.5469266 | -0.103676665 | 0.51826796 | -0.2000445 | 0.84144576 | 0.99665574 |
| TBX3 | 10.1799956 | 0.859842685 | 0.61120639 | 1.40679598 | 0.1594879 | 0.86757292 |
| MED13L | 217.61346 | -0.176989848 | 0.23531411 | -0.752143 | 0.45196509 | 0.99617422 |
| MAP1LC3B2 | 2.96131895 | -1.68467772 | 0.76808575 | -2.1933459 | 0.02828247 | 0.5450502 |
| C12orf49 | 8.42551244 | -1.201018085 | 0.61116237 | -1.9651375 | 0.04939833 | 0.64444758 |
| FBXW8 | 16.7914997 | -0.429042358 | 0.40075017 | -1.0705981 | 0.28435019 | 0.96059387 |
| FBXO21 | 39.8938937 | 0.370590895 | 0.31782082 | 1.16603717 | 0.24359942 | 0.93426912 |
| NOS1 | 6.03339232 | 1.35355613 | 1.05516156 | 1.28279515 | 0.19956385 | 0.90381759 |
| KSR2 | 13.497063 | 0.3149014 | 1.05534983 | 0.2983858 | 0.76540873 | 0.99665574 |
| WSB2 | 10.1994777 | -0.548216328 | 0.55603807 | -0.9859331 | 0.32416596 | 0.97700345 |
| VSIG10 | 10.9030231 | -0.033105957 | 0.77603384 | -0.0426605 | 0.9659722 | 0.99777713 |
| PEBP1 | 105.736098 | 0.226796095 | 0.37638178 | 0.60256928 | 0.54679526 | 0.99665574 |
| TAOK3 | 55.2841215 | 0.040207348 | 0.29230081 | 0.13755469 | 0.89059236 | 0.99710937 |
| SUDS3 | 11.9889427 | 0.303838025 | 0.43534772 | 0.69792033 | 0.48522702 | 0.99640678 |
| AC002070.1 | 8.67590726 | -1.78918048 | 0.77218318 | -2.3170415 | 0.02050147 | 0.46985116 |
| PRKAB1 | 15.1513055 | -0.544799809 | 0.38521897 | -1.4142601 | 0.15728556 | 0.86532131 |
| CIT | 14.6485617 | -0.231422584 | 0.73371823 | -0.3154107 | 0.75244984 | 0.99665574 |
| BICDL1 | 44.5559129 | 0.835762436 | 0.52074592 | 1.60493323 | 0.1085085 | 0.80237722 |
| RAB35 | 9.00238236 | -0.056421715 | 0.57558144 | -0.0980256 | 0.92191197 | 0.99777713 |
| GCN1 | 13.2568634 | 0.057924882 | 0.43472911 | 0.13324362 | 0.89400071 | 0.99777713 |
| RPLP0 | 89.7095543 | 0.120490321 | 0.45657721 | 0.26389911 | 0.79185766 | 0.99665574 |
| PXN | 27.7184429 | -0.035015142 | 0.34865431 | -0.1004294 | 0.92000343 | 0.99777713 |
| GATC | 5.95068588 | 0.311658367 | 0.63084638 | 0.4940321 | 0.62128351 | 0.99665574 |
| SRSF9 | 18.2589076 | 0.451581577 | 0.49042854 | 0.92078976 | 0.3571602 | 0.9819332 |
| DYNLL1 | 29.0850703 | -0.078672993 | 0.41625244 | -0.1890031 | 0.8500904 | 0.99665574 |
| COQ5 | 6.06324374 | -0.110221194 | 0.60537584 | -0.1820707 | 0.85552725 | 0.99665574 |
| RNF10 | 31.980025 | 0.003685237 | 0.34630705 | 0.01064153 | 0.99150945 | 0.99777713 |
| POP5 | 8.0063257 | -0.395875799 | 0.557907 | -0.7095731 | 0.47796891 | 0.99617422 |
| MLEC | 21.2902842 | -0.312108867 | 0.35409709 | -0.8814217 | 0.3780896 | 0.98679664 |
| UNC119B | 5.26991844 | 0.87929327 | 0.76363169 | 1.15146252 | 0.24954201 | 0.93698403 |
| ACADS | 6.77696706 | 1.308873356 | 0.65153632 | 2.00890315 | 0.04454741 | 0.63075643 |
| SPPL3 | 74.0146461 | 0.097187342 | 0.29801536 | 0.32611521 | 0.74433719 | 0.99665574 |
| C12orf43 | 3.10636606 | 0.259451812 | 0.82920695 | 0.31289151 | 0.75436309 | 0.99665574 |
| P2RX4 | 9.05023799 | -0.036393082 | 0.52427917 | -0.0694155 | 0.94465892 | 0.99777713 |
| CAMKK2 | 5.20170902 | -0.184439501 | 0.62540688 | -0.2949112 | 0.76806171 | 0.99665574 |
| ANAPC5 | 23.7763486 | 0.189866078 | 0.38987522 | 0.48699191 | 0.62626406 | 0.99665574 |
| RNF34 | 13.1717787 | -0.364506403 | 0.45416401 | -0.8025876 | 0.42221314 | 0.99518711 |
| KDM2B | 13.6127311 | -0.140838634 | 0.41811065 | -0.3368454 | 0.73623347 | 0.99665574 |
| ORAI1 | 3.88666312 | -0.247181739 | 0.73547494 | -0.3360845 | 0.73680713 | 0.99665574 |
| TMEM120B | 12.9228257 | 0.276979471 | 0.43671657 | 0.63423164 | 0.52592966 | 0.99665574 |
| RHOF | 3.50617723 | -0.769358134 | 0.75012031 | -1.0256463 | 0.30505832 | 0.97376371 |
| SETD1B | 9.00983402 | -0.089448798 | 0.46601598 | -0.1919436 | 0.84778636 | 0.99665574 |
| BCL7A | 14.1588342 | 0.048242668 | 0.52541824 | 0.09181765 | 0.92684292 | 0.99777713 |
| MLXIP | 19.7407004 | -1.398369434 | 0.37232958 | -3.7557302 | 0.00017284 | 0.02986125 |
| CLIP1 | 62.3159855 | -0.570140584 | 0.33073982 | -1.7238341 | 0.08473779 | 0.75373899 |
| ZCCHC8 | 15.7902961 | 0.015287636 | 0.39315478 | 0.03888452 | 0.96898246 | 0.99777713 |
| RSRC2 | 25.9808543 | -0.491355526 | 0.38507066 | -1.2760139 | 0.2019506 | 0.90614933 |
| KNTC1 | 5.60486179 | 1.293135649 | 0.67368589 | 1.91949343 | 0.05492192 | 0.67040291 |
| DENR | 27.8809324 | -0.543940802 | 0.35416609 | -1.5358354 | 0.12457876 | 0.83196528 |
| HIP1R | 23.0252338 | 0.053297858 | 0.32941981 | 0.16179312 | 0.87146877 | 0.99665574 |
| VPS37B | 28.5815213 | 0.118849001 | 0.34647908 | 0.34301927 | 0.73158397 | 0.99665574 |
| ABCB9 | 3.79632213 | -0.209065143 | 0.86510337 | -0.2416649 | 0.80903981 | 0.99665574 |
| ARL6IP4 | 22.5212022 | 0.729125443 | 0.39210968 | 1.8594936 | 0.0629572 | 0.68628329 |
| PITPNM2 | 15.7119183 | -1.358441495 | 0.45720293 | -2.9712003 | 0.00296638 | 0.17070061 |
| MPHOSPH9 | 14.8046159 | 0.255498601 | 0.48492496 | 0.52688276 | 0.59827501 | 0.99665574 |
| C12orf65 | 10.1845011 | -0.431867705 | 0.54205533 | -0.7967225 | 0.42561218 | 0.99518711 |
| SBNO1 | 28.0177786 | -0.28197113 | 0.36382013 | -0.7750289 | 0.43832256 | 0.99617422 |
| KMT5A | 5.93069356 | 0.005776381 | 0.61142848 | 0.00944735 | 0.99246221 | 0.99777713 |
| RILPL2 | 5.11520545 | -0.65096813 | 0.92212903 | -0.7059404 | 0.48022521 | 0.99617422 |
| SNRNP35 | 9.72154191 | -0.275396045 | 0.47638632 | -0.5780939 | 0.5632007 | 0.99665574 |
| RILPL1 | 15.4137143 | -0.290612241 | 0.51124051 | -0.5684453 | 0.56973267 | 0.99665574 |
| TMED2 | 13.9161332 | 0.058221147 | 0.40411752 | 0.14406984 | 0.88544532 | 0.99665574 |
| DDX55 | 6.20000217 | -0.039236943 | 0.59010055 | -0.066492 | 0.94698616 | 0.99777713 |
| EIF2B1 | 7.03288766 | 0.007345435 | 0.56527721 | 0.01299439 | 0.98963226 | 0.99777713 |
| GTF2H3 | 5.64970584 | -0.036184656 | 0.59657238 | -0.0606543 | 0.95163456 | 0.99777713 |
| ATP6V0A2 | 5.57310244 | 0.209152699 | 0.62675893 | 0.33370517 | 0.73860204 | 0.99665574 |
| CCDC92 | 20.7167971 | -0.097715542 | 0.36402789 | -0.2684287 | 0.78836933 | 0.99665574 |
| RFLNA | 2.79829042 | -0.238260216 | 0.91001525 | -0.26182 | 0.7934602 | 0.99665574 |
| NCOR2 | 31.3566898 | 0.199606274 | 0.30646359 | 0.65132134 | 0.51483907 | 0.99665574 |
| UBC | 85.1137002 | -0.817835525 | 0.33721416 | -2.4252704 | 0.01529699 | 0.41354899 |
| DHX37 | 3.79749472 | -0.384461155 | 0.72750104 | -0.5284682 | 0.59717442 | 0.99665574 |
| BRI3BP | 8.93700438 | 0.623486436 | 0.51208519 | 1.21754436 | 0.22339717 | 0.92191316 |
| AACS | 7.72443156 | -0.83302986 | 0.49609234 | -1.6791831 | 0.09311637 | 0.76709603 |
| SLC15A4 | 9.44396454 | -0.383801245 | 0.65012588 | -0.5903491 | 0.55495661 | 0.99665574 |
| RIMBP2 | 14.9616605 | -0.558435552 | 0.62000086 | -0.9007013 | 0.36774718 | 0.98525922 |
| STX2 | 13.0350648 | -0.273527272 | 0.43154409 | -0.6338339 | 0.52618923 | 0.99665574 |
| RAN | 21.0228146 | -0.057460364 | 0.49537111 | -0.1159946 | 0.90765684 | 0.99777713 |
| SFSWAP | 23.0818915 | -0.124601868 | 0.39341828 | -0.316716 | 0.7514591 | 0.99665574 |
| ULK1 | 14.3638079 | 0.202798838 | 0.41681424 | 0.48654489 | 0.62658088 | 0.99665574 |
| PUS1 | 4.36201892 | -0.256513225 | 0.68844928 | -0.3725957 | 0.70944939 | 0.99665574 |
| EP400 | 25.7193198 | 0.234401436 | 0.41192053 | 0.56904528 | 0.56932541 | 0.99665574 |
| DDX51 | 2.56467906 | 0.49775958 | 0.89353053 | 0.55707059 | 0.5774792 | 0.99665574 |
| FBRSL1 | 19.2177915 | -0.367781917 | 0.42500839 | -0.8653521 | 0.38684555 | 0.99144635 |
| POLE | 3.16323385 | -0.321545978 | 0.79487162 | -0.4045257 | 0.6858262 | 0.99665574 |
| PGAM5 | 3.28178959 | 0.17814013 | 0.89798405 | 0.19837783 | 0.84274946 | 0.99665574 |
| ANKLE2 | 21.058298 | -0.424691557 | 0.42011559 | -1.0108922 | 0.31206804 | 0.97442101 |
| GOLGA3 | 19.2608963 | -0.642081738 | 0.62416418 | -1.0287065 | 0.30361762 | 0.97376371 |
| CHFR | 17.1271315 | -0.429404021 | 0.46336199 | -0.9267139 | 0.35407507 | 0.98022903 |
| ZNF26 | 11.3053187 | 0.240385207 | 0.49489155 | 0.4857331 | 0.62715641 | 0.99665574 |
| ZNF84 | 16.1848981 | -0.151757013 | 0.39497385 | -0.3842204 | 0.70081509 | 0.99665574 |
| ZNF140 | 5.35643371 | 0.142987911 | 0.73483055 | 0.19458624 | 0.84571688 | 0.99665574 |
| ZNF891 | 4.70836059 | 0.172146073 | 0.65619644 | 0.26233923 | 0.79305992 | 0.99665574 |
| ZNF268 | 5.13784159 | 0.535895686 | 0.67601309 | 0.79272974 | 0.4279353 | 0.99589919 |
| MPHOSPH8 | 50.9362061 | -0.197551888 | 0.28358449 | -0.6966244 | 0.48603786 | 0.99640678 |
| PSPC1 | 48.4627769 | 0.265089141 | 0.3095306 | 0.85642304 | 0.39176383 | 0.99179694 |
| ZMYM5 | 16.7485567 | -0.606649012 | 0.42314653 | -1.4336618 | 0.1516688 | 0.85756735 |
| ZMYM2 | 98.3513077 | 0.309515528 | 0.24149839 | 1.28164632 | 0.19996674 | 0.90381759 |
| CRYL1 | 22.2879343 | 0.936574087 | 0.58576751 | 1.59888363 | 0.10984646 | 0.80522301 |
| IFT88 | 23.7805966 | 0.429564463 | 0.39725591 | 1.08132932 | 0.27955065 | 0.95996214 |
| EEF1AKMT1 | 3.01478256 | 0.308356914 | 0.99663721 | 0.30939735 | 0.75701928 | 0.99665574 |
| XPO4 | 33.2937513 | 0.467478001 | 0.29205094 | 1.60067281 | 0.10944941 | 0.80453999 |
| LATS2 | 24.0427232 | -0.597046623 | 0.4520192 | -1.3208435 | 0.18655355 | 0.89532394 |
| SAP18 | 21.9739409 | -0.231247253 | 0.3521463 | -0.6566795 | 0.51138702 | 0.99665574 |
| MRPL57 | 16.9798977 | -0.142138483 | 0.47961425 | -0.29636 | 0.76695516 | 0.99665574 |

| | | | | | | |
|---|---|---|---|---|---|---|
| ZDHHC20 | 30.489063 | 0.098246469 | 0.42075731 | 0.23349914 | 0.81537384 | 0.99665574 |
| MICU2 | 60.1191666 | 0.124591622 | 0.34978392 | 0.35619597 | 0.72169381 | 0.99665574 |
| FGF9 | 65.333794 | 0.507165771 | 0.4568836 | 1.11005467 | 0.26697547 | 0.94557817 |
| AL136962.1 | 26.4458824 | 0.145095643 | 1.41932905 | 0.10222833 | 0.91857544 | 0.99777713 |
| SACS | 48.083379 | -0.76210256 | 0.75504108 | -1.0093525 | 0.31280563 | 0.97520189 |
| MIPEP | 22.1505685 | -0.659026529 | 0.36985704 | -1.7818412 | 0.07477512 | 0.72883798 |
| PARP4 | 39.4631345 | -0.24327873 | 0.38489036 | -0.6320728 | 0.52733931 | 0.99665574 |
| MTMR6 | 9.52415278 | 0.162157363 | 0.46058924 | 0.35206503 | 0.72478949 | 0.99665574 |
| NUP58 | 25.9619988 | 0.182607345 | 0.34408784 | 0.53069979 | 0.59562683 | 0.99665574 |
| SHISA2 | 5.6880854 | 0.186491898 | 0.89469032 | 0.20844296 | 0.83488312 | 0.99665574 |
| RNF6 | 5.59881508 | 0.047221695 | 0.67129075 | 0.07034462 | 0.94391937 | 0.99777713 |
| CDK8 | 55.9238993 | 0.100572372 | 0.25756724 | 0.39047035 | 0.69618878 | 0.99665574 |
| WASF3 | 8.63365443 | -0.106103204 | 0.48396879 | -0.2192356 | 0.8264665 | 0.99665574 |
| USP12 | 37.4261837 | -0.157084163 | 0.29480357 | -0.5328435 | 0.59414193 | 0.99665574 |
| RPL21 | 30.6704978 | 0.879489465 | 0.53108167 | 1.65603429 | 0.09771487 | 0.78002629 |
| GTF3A | 8.75048538 | -0.469721202 | 0.52443727 | -0.8956671 | 0.37043059 | 0.98532838 |
| MTIF3 | 15.9401799 | -0.211355795 | 0.48529273 | -0.4355223 | 0.66318337 | 0.99665574 |
| LNX2 | 16.4638026 | -0.925625629 | 0.44019103 | -2.1027817 | 0.03548486 | 0.58349146 |
| POLR1D | 14.6382587 | 0.606845883 | 0.49336974 | 1.23000223 | 0.21869627 | 0.91662582 |
| PAN3 | 93.5191326 | 0.212321669 | 0.26806295 | 0.792059 | 0.42832628 | 0.99589919 |
| POMP | 12.3616386 | -0.309832714 | 0.50356113 | -0.6152832 | 0.5383677 | 0.99665574 |
| SLC46A3 | 2.88822971 | -1.128064336 | 0.88646512 | -1.2725423 | 0.20318051 | 0.9066717 |
| SLC7A1 | 9.25263363 | -1.2504447 | 0.6428419 | -1.9451823 | 0.05175304 | 0.65584036 |
| UBL3 | 86.6771127 | 0.797412509 | 0.31551612 | 2.52732732 | 0.01149343 | 0.3725572 |
| KATNAL1 | 5.63964023 | -0.199723718 | 0.67915377 | -0.2940773 | 0.76869882 | 0.99665574 |
| HMGB1 | 49.2669663 | 0.075399245 | 0.26590625 | 0.28355575 | 0.77675086 | 0.99665574 |
| USPL1 | 6.99802275 | -0.320457127 | 0.60906817 | -0.5261433 | 0.59878866 | 0.99665574 |
| HSPH1 | 22.6369949 | 0.701890458 | 0.43563342 | 1.61119518 | 0.10713719 | 0.79864472 |
| B3GLCT | 16.3345742 | 0.478470489 | 0.50940604 | 0.93927133 | 0.34759145 | 0.97804112 |
| FRY | 14.8988288 | -1.259724646 | 0.65325671 | -1.9283761 | 0.05380836 | 0.6653339 |
| N4BP2L1 | 10.7177965 | 0.014440012 | 0.51498719 | 0.02803955 | 0.9776306 | 0.99777713 |
| N4BP2L2 | 176.547073 | 0.068505172 | 0.19807719 | 0.34585089 | 0.72945478 | 0.99665574 |
| PDS5B | 46.1400771 | 0.221751769 | 0.30995626 | 0.71542922 | 0.47434386 | 0.99617422 |
| KL | 4.1611019 | 2.069627741 | 0.90049473 | 2.29832299 | 0.02154341 | 0.47479808 |
| STARD13 | 17.5102494 | -0.720652505 | 0.73166947 | -0.9849427 | 0.32465222 | 0.97700345 |
| AL139383.1 | 4.6404683 | -1.606477727 | 0.61612655 | -2.6073827 | 0.00912373 | 0.32476678 |
| RFC3 | 4.74372377 | -0.046266805 | 0.65272534 | -0.0708825 | 0.94349127 | 0.99777713 |
| LINC02343 | 24.9893305 | 0.023899718 | 0.66333886 | 0.03602943 | 0.9712589 | 0.99777713 |
| NBEA | 54.8902959 | 0.440020385 | 0.31346716 | 1.40372085 | 0.16040199 | 0.86820896 |
| SPART | 16.695779 | -0.269260534 | 0.44932497 | -0.5992557 | 0.54900241 | 0.99665574 |
| RFXAP | 3.24462299 | -0.605831223 | 0.93635583 | -0.6470096 | 0.51762572 | 0.99665574 |
| ALG5 | 8.69913657 | 0.277489263 | 0.55950501 | 0.49595492 | 0.61992622 | 0.99665574 |
| EXOSC8 | 8.01460644 | -0.740247743 | 0.57674849 | -1.2834845 | 0.19932239 | 0.90381759 |
| SUPT20H | 23.0975218 | 0.404757872 | 0.34426486 | 1.17571649 | 0.23970819 | 0.93157178 |
| LINC00571 | 8.37106295 | -1.037927425 | 0.75048775 | -1.3830038 | 0.16666369 | 0.87495175 |
| UFM1 | 16.4081965 | -0.069526926 | 0.44989752 | -0.1545395 | 0.87718439 | 0.99665574 |
| FREM2 | 11.6294886 | -0.211019913 | 0.61589007 | -0.3426259 | 0.73187989 | 0.99665574 |
| PROSER1 | 7.47938256 | -0.248487563 | 0.5119357 | -0.4853882 | 0.62740098 | 0.99665574 |
| NHLRC3 | 12.1231609 | 0.218418261 | 0.50494099 | 0.43256195 | 0.66533303 | 0.99665574 |
| COG6 | 22.936321 | -0.35884505 | 0.38204559 | -0.9392728 | 0.34759068 | 0.97804112 |
| LINC00598 | 8.31486086 | 0.98286006 | 0.57935648 | 1.69646857 | 0.08979718 | 0.76117374 |
| FOXO1 | 260.00454 | 0.360273187 | 0.32994067 | 1.09193323 | 0.27486245 | 0.95528129 |
| MRPS31 | 16.7885057 | 0.33350457 | 0.45641185 | 0.7307097 | 0.46495649 | 0.99617422 |
| SLC25A15 | 3.15641266 | 0.04817552 | 0.81607839 | 0.05903296 | 0.95292586 | 0.99777713 |
| ELF1 | 86.0826473 | -0.61743876 | 0.34269807 | -1.801699 | 0.07159278 | 0.71725834 |
| WBP4 | 12.2200658 | -0.049759639 | 0.47072334 | -0.1057089 | 0.91581333 | 0.99777713 |
| MTRF1 | 6.68428515 | -0.121431149 | 0.58825391 | -0.2064264 | 0.83645784 | 0.99665574 |
| NAA16 | 20.593091 | 0.608008879 | 0.37996161 | 1.60018502 | 0.10955754 | 0.80453999 |
| VWA8 | 83.922158 | 0.640495029 | 0.28830765 | 2.2215679 | 0.02631252 | 0.52100847 |
| DGKH | 22.9806727 | -1.39073741 | 0.52531825 | -2.6474188 | 0.00811088 | 0.30342484 |
| AKAP11 | 18.4659664 | -0.07441617 | 0.46667967 | -0.1594588 | 0.87330743 | 0.99665574 |
| EPSTI1 | 14.2550844 | -0.296645949 | 1.12617628 | -0.2634099 | 0.79223468 | 0.99665574 |
| DNAJC15 | 18.0379266 | -0.456647157 | 0.47607339 | -0.9591949 | 0.33746058 | 0.97788427 |
| ENOX1 | 4.86061124 | -0.762418473 | 0.71552792 | -1.0655328 | 0.28663489 | 0.96477573 |
| CCDC122 | 7.69826662 | -0.151223691 | 0.69438474 | -0.2177808 | 0.82759987 | 0.99665574 |
| SMIM2-AS1 | 7.27067553 | -0.842247517 | 1.1016147 | -0.7645573 | 0.44453523 | 0.99617422 |
| TSC22D1 | 38.6390503 | -0.028682916 | 0.53081669 | -0.0540354 | 0.95690692 | 0.99777713 |
| NUFIP1 | 4.40546274 | -0.330575875 | 0.69231984 | -0.4774901 | 0.63301316 | 0.99665574 |
| GPALPP1 | 11.9922183 | 0.144855668 | 0.49447853 | 0.29294633 | 0.76956318 | 0.99665574 |
| GTF2F2 | 32.9185944 | -0.023407234 | 0.35690607 | -0.0655837 | 0.94770924 | 0.99777713 |
| TPT1 | 196.031722 | -0.257449958 | 0.44017811 | -0.5848768 | 0.55863057 | 0.99665574 |
| SLC25A30 | 16.084154 | 0.660986992 | 0.70941353 | 0.93173722 | 0.35147235 | 0.97838306 |
| COG3 | 19.4337379 | -0.376129313 | 0.41190479 | -0.9131462 | 0.36116563 | 0.9819332 |
| ZC3H13 | 32.0755573 | 0.042991998 | 0.32269269 | 0.13322892 | 0.89401234 | 0.99777713 |
| AL138686.1 | 12.7780751 | 0.378710415 | 0.60020447 | 0.630969 | 0.5280608 | 0.99665574 |
| LRCH1 | 48.6859925 | -0.596127127 | 0.39126452 | -1.5235911 | 0.12761089 | 0.83778831 |
| ESD | 14.7733514 | -0.218350923 | 0.44649375 | -0.4890347 | 0.62481715 | 0.99665574 |
| SUCLA2 | 33.810675 | -0.342461629 | 0.42791784 | -0.8002976 | 0.42353838 | 0.99518711 |
| NUDT15 | 2.87864223 | 1.034636839 | 0.9921641 | 1.04280818 | 0.29703714 | 0.97341216 |
| MED4 | 17.2944163 | -0.195463086 | 0.35487628 | -0.5507922 | 0.58177613 | 0.99665574 |
| ITM2B | 114.984152 | -0.209547221 | 0.42121766 | -0.4974797 | 0.61885084 | 0.99665574 |
| RB1 | 54.7608869 | -0.066829216 | 0.31045276 | -0.2152637 | 0.82956172 | 0.99665574 |
| RCBTB2 | 7.40451985 | -0.438822217 | 0.71484186 | -0.6138731 | 0.53929916 | 0.99665574 |
| FNDC3A | 110.215003 | 0.196615256 | 0.4466372 | 0.44021245 | 0.65978324 | 0.99665574 |
| CDADC1 | 5.06616561 | 0.300449697 | 0.62994883 | 0.47694302 | 0.63340269 | 0.99665574 |
| CAB39L | 30.2641916 | 0.991575748 | 0.37102052 | 2.67256306 | 0.00752742 | 0.28689101 |
| SETDB2 | 22.915262 | 0.245502177 | 0.40670543 | 0.60363634 | 0.54608545 | 0.99665574 |
| PHF11 | 16.0416914 | -0.424085958 | 0.47173752 | -0.8989871 | 0.36865952 | 0.98532838 |
| RCBTB1 | 5.92769848 | -0.316015968 | 0.65926185 | -0.4793482 | 0.63169095 | 0.99665574 |
| EBPL | 7.42134903 | -0.583245516 | 0.76205982 | -0.765354 | 0.44406079 | 0.99617422 |
| KPNA3 | 31.4610031 | -0.136260255 | 0.29409076 | -0.4633272 | 0.64312984 | 0.99665574 |
| SPRYD7 | 4.91469075 | -0.473854994 | 0.71205272 | -0.6654774 | 0.50574519 | 0.99665574 |
| DLEU2 | 35.8750128 | -1.082598859 | 0.37840475 | -2.8609547 | 0.00422367 | 0.20883915 |
| TRIM13 | 8.80051474 | -0.386751567 | 0.50427105 | -0.7669518 | 0.4431102 | 0.99617422 |
| RNASEH2B | 8.7137853 | -0.072342037 | 0.50504136 | -0.1432398 | 0.88610078 | 0.99665574 |
| INTS6 | 49.054767 | 0.948220588 | 0.45063361 | 2.10419409 | 0.03536152 | 0.58349146 |
| INTS6-AS1 | 2.97324443 | -0.73019969 | 0.86911237 | -0.8401672 | 0.40081466 | 0.99194293 |
| WDFY2 | 55.1136476 | 0.238724428 | 0.37858391 | 0.63057204 | 0.52832039 | 0.99665574 |
| ATP7B | 4.01762435 | -0.512754488 | 0.81213064 | -0.6313695 | 0.52779898 | 0.99665574 |
| ALG11 | 3.90642538 | -0.074657173 | 0.7239747 | -0.1031212 | 0.91786674 | 0.99777713 |
| UTP14C | 6.05696328 | -0.091060607 | 0.63481607 | -0.1434441 | 0.88593947 | 0.99665574 |
| NEK3 | 7.87024933 | -0.175203656 | 0.56640925 | -0.3093234 | 0.75707551 | 0.99665574 |
| VPS36 | 17.2148956 | -0.287102714 | 0.42422706 | -0.6767666 | 0.49855404 | 0.99640678 |
| CKAP2 | 1.79829961 | -0.284345981 | 1.19622938 | -0.2377019 | 0.81211232 | 0.99665574 |
| SUGT1 | 14.1841633 | 0.99241157 | 0.42841268 | 2.31648506 | 0.0205318 | 0.46985116 |
| TDRD3 | 43.6668921 | 0.007720263 | 0.330064 | 0.0233902 | 0.98133902 | 0.99777713 |
| PCDH9 | 39.4434836 | 0.283740545 | 0.75366038 | 0.3764833 | 0.70655763 | 0.99665574 |
| DACH1 | 304.869227 | 0.52080713 | 0.38145539 | 1.36531596 | 0.17215377 | 0.88157681 |
| MZT1 | 4.23540553 | 0.127018727 | 0.6717801 | 0.18907783 | 0.85003181 | 0.99665574 |
| BORA | 2.97790719 | -0.64801099 | 0.79523916 | -0.814863 | 0.41515072 | 0.99279547 |
| DIS3 | 10.6986703 | 0.493138693 | 0.47749883 | 1.03275371 | 0.30171917 | 0.97358745 |
| PIBF1 | 45.71161 | -0.28869411 | 0.28953142 | -0.9971081 | 0.31871206 | 0.97700345 |
| KLF5 | 10.993026 | -0.715058137 | 0.54450648 | -1.3132225 | 0.189108 | 0.89839592 |

| | | | | | | |
|---|---|---|---|---|---|---|
| KLF12 | 41.1793221 | 0.663348468 | 0.48448192 | 1.36919138 | 0.17093946 | 0.87968373 |
| TBC1D4 | 255.962385 | -0.068657179 | 0.25781623 | -0.2663028 | 0.79000603 | 0.99665574 |
| COMMD6 | 20.002491 | 0.676647287 | 0.52494412 | 1.28898917 | 0.19740185 | 0.90381759 |
| UCHL3 | 17.8597828 | -0.46885565 | 0.40317899 | -1.162897 | 0.24487129 | 0.93426912 |
| LMO7 | 86.8327296 | 0.588956577 | 0.34161439 | 1.72403912 | 0.08470078 | 0.75373899 |
| KCTD12 | 13.6475959 | 0.473983397 | 0.47295586 | 1.00217258 | 0.31626025 | 0.97663269 |
| AC001226.2 | 5.6663149 | -0.617349359 | 0.86188602 | -0.7162773 | 0.47382016 | 0.99617422 |
| FBXL3 | 28.4777421 | 0.241576131 | 0.35594932 | 0.67868126 | 0.49733984 | 0.99640678 |
| MYCBP2 | 53.4764369 | -0.184621888 | 0.27214469 | -0.6783961 | 0.4975206 | 0.99640678 |
| SLAIN1 | 4.50348642 | -0.752083083 | 0.78135189 | -0.9625408 | 0.33577801 | 0.97788427 |
| RNF219 | 4.89795038 | -0.204566051 | 0.61043025 | -0.3351178 | 0.73753621 | 0.99665574 |
| RBM26 | 57.598031 | -0.01268127 | 0.26099095 | -0.0485889 | 0.9612469 | 0.99777713 |
| NDFIP2 | 24.7287744 | 0.286664868 | 0.36035718 | 0.79550204 | 0.42632153 | 0.99518711 |
| GPC5 | 41.039523 | 2.236066093 | 1.79727853 | 1.24413999 | 0.21344804 | 0.91189898 |
| GPC6 | 604.696264 | -2.051787404 | 1.23353923 | -1.6633337 | 0.09624565 | 0.77378145 |
| TGDS | 7.37018908 | -0.117976091 | 0.55449874 | -0.2127617 | 0.83151284 | 0.99665574 |
| GPR180 | 7.63182556 | -0.885400816 | 0.67375633 | -1.3141261 | 0.18880376 | 0.89739428 |
| ABCC4 | 59.2972216 | -0.308959378 | 0.35312368 | -0.8749325 | 0.38161064 | 0.98844371 |
| CLDN10 | 15.6350676 | -0.250642796 | 0.83182305 | -0.3013174 | 0.76317244 | 0.99665574 |
| CLDN10-AS1 | 9.8576897 | 0.2205299 | 0.7733159 | 0.2851744 | 0.77551053 | 0.99665574 |
| DNAJC3-DT | 3.18716046 | -1.326701647 | 0.79351667 | -1.6719266 | 0.09453879 | 0.77075743 |
| DNAJC3 | 37.1103727 | -0.09940405 | 0.36497867 | -0.2723558 | 0.78534847 | 0.99665574 |
| UGGT2 | 58.1950843 | -0.005199875 | 0.30374091 | -0.0171194 | 0.98634133 | 0.99777713 |
| MBNL2 | 81.707732 | 0.507259743 | 0.2954805 | 1.71672831 | 0.08602883 | 0.75397496 |
| RAP2A | 3.6352229 | -0.221555589 | 0.71100928 | -0.3116072 | 0.75533909 | 0.99665574 |
| IPO5 | 21.1619174 | -0.004062338 | 0.35645963 | -0.0113963 | 0.99090723 | 0.99777713 |
| FARP1 | 163.407873 | -0.033267089 | 0.27602513 | -0.120522 | 0.90406969 | 0.99777713 |
| STK24 | 58.7926481 | 0.174361694 | 0.33938786 | 0.51375348 | 0.60742436 | 0.99665574 |
| DOCK9 | 28.4313815 | -0.065931416 | 0.32813395 | -0.2009284 | 0.84075459 | 0.99665574 |
| UBAC2 | 67.7904186 | 0.036966304 | 0.26478697 | 0.13960771 | 0.88896995 | 0.99665574 |
| TM9SF2 | 36.8493843 | -0.361631461 | 0.33577125 | -1.0770174 | 0.28147251 | 0.96059387 |
| CLYBL | 49.7168103 | 0.208442514 | 0.36297511 | 0.57426118 | 0.56579108 | 0.99665574 |
| PCCA | 162.68237 | 0.196091367 | 0.23357835 | 0.83951004 | 0.40118316 | 0.99194293 |
| GGACT | 11.8327878 | 0.005769647 | 0.63126978 | 0.00913975 | 0.99270764 | 0.99777713 |
| TMTC4 | 8.81597645 | -0.307072455 | 0.49203542 | -0.6240861 | 0.53257106 | 0.99665574 |
| FGF14 | 10.1778725 | 1.358366128 | 0.86295869 | 1.57408014 | 0.11546892 | 0.81747482 |
| TPP2 | 27.6827247 | 0.342390841 | 0.33109881 | 1.0341047 | 0.30108722 | 0.97358745 |
| TEX30 | 3.30433666 | -0.315032861 | 0.89617413 | -0.3515309 | 0.72519013 | 0.99665574 |
| EFNB2 | 27.9418122 | -1.765592944 | 0.75635383 | -2.3343478 | 0.01957752 | 0.4617478 |
| ARGLU1 | 57.2630076 | -0.424288845 | 0.28993628 | -1.4633865 | 0.14336164 | 0.84969914 |
| LIG4 | 5.02867859 | -0.494985357 | 0.71383069 | -0.6934212 | 0.48804524 | 0.99640678 |
| ABHD13 | 7.53707691 | 0.232251499 | 0.61575767 | 0.37718003 | 0.70603982 | 0.99665574 |
| TNFSF13B | 4.38531235 | 0.245279232 | 0.85148944 | 0.28805904 | 0.77330154 | 0.99665574 |
| IRS2 | 19.4537903 | 0.21336586 | 0.50605644 | 0.42162463 | 0.67329902 | 0.99665574 |
| COL4A1 | 9.98717402 | -0.488662798 | 0.85990582 | -0.5682748 | 0.5698484 | 0.99665574 |
| COL4A2 | 8.36656473 | -0.797460804 | 0.62072403 | -1.2847268 | 0.19888776 | 0.90381759 |
| RAB20 | 5.60543129 | -0.77147389 | 0.7125396 | -1.0827102 | 0.27893708 | 0.95980166 |
| NAXD | 7.72440645 | 0.579503019 | 0.54510497 | 1.06310352 | 0.28773501 | 0.96559334 |
| CARS2 | 14.6269627 | 0.259245909 | 0.3980792 | 0.65124204 | 0.51489026 | 0.99665574 |
| ANKRD10 | 81.9419441 | 0.001116841 | 0.27316195 | 0.00408857 | 0.9967378 | 0.998802 |
| ARHGEF7 | 87.7400313 | -0.046395785 | 0.35882383 | -0.1292996 | 0.89712057 | 0.99777713 |
| TUBGCP3 | 13.3998512 | -0.040530747 | 0.41174419 | -0.0984367 | 0.92158552 | 0.99777713 |
| ATP11A | 38.516382 | -0.877214211 | 0.30723406 | -2.8551985 | 0.00430099 | 0.20963212 |
| MCF2L | 26.8851183 | 0.793228408 | 0.4499037 | 1.76310708 | 0.07788243 | 0.74141484 |
| PCID2 | 12.1947552 | -0.193004453 | 0.41616507 | -0.463769 | 0.64281327 | 0.99665574 |
| CUL4A | 36.202065 | -0.20114401 | 0.33879713 | -0.5937005 | 0.55271248 | 0.99665574 |
| LAMP1 | 34.3621565 | -0.423187316 | 0.55928979 | -0.7566512 | 0.44925883 | 0.99617422 |
| GRTP1 | 9.89068805 | 0.386679977 | 0.6480791 | 0.59665552 | 0.55073739 | 0.99665574 |
| DCUN1D2 | 12.8050442 | -0.058581511 | 0.4516045 | -0.1297186 | 0.89678905 | 0.99777713 |
| TMCO3 | 13.7702124 | -0.447861367 | 0.43549124 | -1.028405 | 0.30375936 | 0.97376371 |
| TFDP1 | 20.7941847 | -0.25423212 | 0.57042878 | -0.445686 | 0.65582408 | 0.99665574 |
| GAS6 | 25.4599288 | 0.10293977 | 0.43741416 | 0.23533708 | 0.81394712 | 0.99665574 |
| CDC16 | 18.9101266 | 0.320209569 | 0.42616053 | 0.75138251 | 0.45242248 | 0.99617422 |
| UPF3A | 23.0720708 | -0.109672678 | 0.37579899 | -0.2918387 | 0.77040998 | 0.99665574 |
| CHAMP1 | 5.1232738 | -0.181472961 | 0.69530631 | -0.2609971 | 0.79409471 | 0.99665574 |
| TTC5 | 4.30286213 | 0.695843352 | 0.73330709 | 0.94891126 | 0.34266575 | 0.97788427 |
| CCNB1IP1 | 9.99937408 | -0.039243412 | 0.48713735 | -0.0805592 | 0.93579249 | 0.99777713 |
| PARP2 | 2.53470017 | -0.068154907 | 0.87801111 | -0.0776242 | 0.93812699 | 0.99777713 |
| TEP1 | 7.40117479 | -0.118274349 | 0.60426484 | -0.1957326 | 0.84481944 | 0.99665574 |
| OSGEP | 5.95471609 | 0.173960763 | 0.65748969 | 0.26458326 | 0.79133052 | 0.99665574 |
| APEX1 | 8.98015176 | -0.263005674 | 0.52182008 | -0.504016 | 0.61425013 | 0.99665574 |
| PIP4P1 | 3.80905084 | -0.762819407 | 0.82917596 | -0.9199729 | 0.35758692 | 0.9819332 |
| PNP | 7.4574816 | 0.27478431 | 0.60217149 | 0.45632235 | 0.64815819 | 0.99665574 |
| METTL17 | 7.01733088 | 0.821312228 | 0.56596792 | 1.45116392 | 0.14673422 | 0.85169374 |
| NDRG2 | 12.2612943 | 1.222385438 | 0.63244227 | 1.93280161 | 0.05326064 | 0.66496977 |
| ARHGEF40 | 3.67684521 | -0.109013506 | 0.69455381 | -0.1569547 | 0.87528053 | 0.99665574 |
| ZNF219 | 4.634041 | 0.180272437 | 0.90120972 | 0.20003384 | 0.84145412 | 0.99665574 |
| HNRNPC | 84.525718 | 0.279829236 | 0.25370779 | 1.10295877 | 0.27004507 | 0.94821243 |
| SUPT16H | 15.7500207 | 0.24514083 | 0.37818188 | 0.64820883 | 0.51684989 | 0.99665574 |
| CHD8 | 21.8612604 | -0.363634758 | 0.38661792 | -0.9405533 | 0.34693383 | 0.97804112 |
| RAB2B | 11.1975728 | -0.676781546 | 0.43050491 | -1.5720646 | 0.11593556 | 0.81906252 |
| TOX4 | 13.1826263 | 0.053721471 | 0.46901168 | 0.11454186 | 0.90880826 | 0.99777713 |
| METTL3 | 5.47476902 | -0.012035477 | 0.58354335 | -0.0206248 | 0.98354494 | 0.99777713 |
| DAD1 | 18.8849061 | -0.375010488 | 0.39481511 | -0.9498382 | 0.34219445 | 0.97788427 |
| ABHD4 | 4.42924437 | 0.491039042 | 0.73167289 | 0.67111826 | 0.50214519 | 0.99640678 |
| OXA1L | 8.66133578 | 1.233306591 | 0.52956748 | 2.32889414 | 0.01986467 | 0.46397674 |
| SLC7A7 | 3.50597967 | 1.368416803 | 1.03500209 | 1.32213917 | 0.18612181 | 0.89532394 |
| MRPL52 | 5.96381565 | -0.027337895 | 0.68908682 | -0.0396726 | 0.96835411 | 0.99777713 |
| MMP14 | 4.32979888 | -0.584107177 | 0.88155706 | -0.6625858 | 0.50759588 | 0.99665574 |
| LRP10 | 20.3863371 | -1.401879112 | 0.55447654 | -2.5282929 | 0.01146187 | 0.3725572 |
| RBM23 | 13.4942125 | -0.164025187 | 0.53741775 | -0.3052098 | 0.76020634 | 0.99665574 |
| PRMT5 | 2.60989922 | -0.133446709 | 0.86774542 | -0.1537856 | 0.87777883 | 0.99665574 |
| C14orf93 | 3.08036855 | 0.110332569 | 0.97126761 | 0.11359647 | 0.90955769 | 0.99777713 |
| PSMB5 | 11.9871578 | -0.542946679 | 0.55008903 | -0.987016 | 0.32363479 | 0.97700345 |
| ACIN1 | 29.5402588 | 0.080127775 | 0.31113901 | 0.25753047 | 0.7967693 | 0.99665574 |
| C14orf119 | 6.61834489 | -0.989510846 | 0.58872399 | -1.6807721 | 0.0928072 | 0.76709603 |
| SLC7A8 | 5.89652515 | 1.693259419 | 0.82136525 | 2.06151821 | 0.03925363 | 0.60188673 |
| HOMEZ | 2.36381364 | -0.298784614 | 1.01360288 | -0.2947748 | 0.7681659 | 0.99665574 |
| BCL2L2-PABI | 5.75335964 | -0.63956546 | 0.54412099 | -1.1754104 | 0.23983057 | 0.93157178 |
| PABPN1 | 20.6992298 | 0.951768949 | 0.50519467 | 1.88396474 | 0.05956974 | 0.68011288 |
| NGDN | 5.66774125 | -0.594883552 | 0.63632851 | -0.9348686 | 0.34985601 | 0.97838306 |
| ZFHX2 | 2.88179707 | 0.07745978 | 0.8355607 | 0.09270395 | 0.92613876 | 0.99777713 |
| AP1G2 | 6.67423436 | 0.777949533 | 0.59480593 | 1.3079048 | 0.1909056 | 0.89964233 |
| DHRS4-AS1 | 4.72993524 | -0.261415448 | 0.86165738 | -0.3033868 | 0.76159513 | 0.99665574 |
| DHRS4L2 | 5.79780366 | -0.375459389 | 0.8326177 | -0.4509385 | 0.65203386 | 0.99665574 |
| NRL | 2.54930964 | 0.37997367 | 0.87036537 | 0.436568 | 0.66242467 | 0.99665574 |
| PSME1 | 14.9331502 | 0.130446407 | 0.53033804 | 0.24596842 | 0.80570668 | 0.99665574 |
| EMC9 | 3.68651531 | 0.688162917 | 0.83779635 | 0.82139641 | 0.41142051 | 0.99279547 |
| PSME2 | 3.26593422 | 0.369126171 | 0.92342558 | 0.3997357 | 0.68935119 | 0.99665574 |
| AL136295.1 | 3.04453303 | -0.892067121 | 0.94142349 | -0.9475726 | 0.34334707 | 0.97788427 |
| GMPR2 | 4.15757155 | 0.402243276 | 0.76877082 | 0.52322912 | 0.60081482 | 0.99665574 |
| TINF2 | 3.15053421 | -0.348904561 | 0.86696923 | -0.4024417 | 0.687359 | 0.99665574 |
| DHRS1 | 4.49502241 | -0.084569493 | 0.79596121 | -0.1062483 | 0.91538538 | 0.99777713 |

| | | | | | | |
|---|---|---|---|---|---|---|
| KHNYN | 3.16358017 | 0.295016637 | 0.81848199 | 0.36044365 | 0.71851539 | 0.99665574 |
| SDR39U1 | 6.03072003 | 1.011894198 | 0.60706263 | 1.66686953 | 0.09554035 | 0.77178333 |
| STXBP6 | 7.2830154 | 0.129143614 | 0.84372209 | 0.15306416 | 0.87834768 | 0.99665574 |
| NOVA1 | 9.93221604 | -0.355515781 | 0.8975132 | -0.396112 | 0.69202238 | 0.99665574 |
| PRKD1 | 75.8367465 | -1.322489797 | 0.42156139 | -3.1371226 | 0.00170615 | 0.12036484 |
| G2E3 | 10.3355275 | 1.172127235 | 0.59165741 | 1.98109112 | 0.04758106 | 0.64060657 |
| SCFD1 | 39.9588728 | -0.342525017 | 0.38872651 | -0.8811465 | 0.37823851 | 0.98679664 |
| COCH | 5.41187235 | -1.42514043 | 0.79111498 | -1.8014327 | 0.0716347 | 0.71725834 |
| STRN3 | 79.1860755 | 0.226980279 | 0.30347279 | 0.74794278 | 0.45449467 | 0.99617422 |
| AP4S1 | 6.09365615 | -0.079169263 | 0.5382273 | -0.1470926 | 0.88305892 | 0.99665574 |
| HECTD1 | 96.4898645 | 0.531715816 | 0.2581782 | 2.05949152 | 0.03944718 | 0.60209727 |
| AL136418.1 | 4.00764949 | -0.099652897 | 1.08671065 | -0.0917014 | 0.92693528 | 0.99777713 |
| HEATR5A | 12.5664045 | 0.187080544 | 0.44218275 | 0.42308422 | 0.67223381 | 0.99665574 |
| AL139353.1 | 2.75732144 | -0.948297727 | 0.95356238 | -0.994479 | 0.31998974 | 0.97700345 |
| NUBPL | 29.9410681 | 0.580415376 | 0.38314889 | 1.51485595 | 0.12980889 | 0.83778831 |
| ARHGAP5 | 95.4451727 | 0.142124513 | 0.24861704 | 0.57166039 | 0.56755208 | 0.99665574 |
| AKAP6 | 46.7962497 | 0.034545738 | 0.4184158 | 0.08256318 | 0.93419888 | 0.99777713 |
| NPAS3 | 26.4683177 | 1.432077938 | 0.80047959 | 1.78902493 | 0.0736108 | 0.72792151 |
| EAPP | 13.3610101 | -0.068797731 | 0.38006866 | -0.181014 | 0.85635662 | 0.99665574 |
| SNX6 | 39.2767144 | -0.073422041 | 0.28676022 | -0.2560398 | 0.79792007 | 0.99665574 |
| CFL2 | 9.15624815 | -0.091010525 | 0.58120533 | -0.1565893 | 0.87556855 | 0.99665574 |
| BAZ1A | 14.9102543 | -1.11226934 | 0.44993544 | -2.4720643 | 0.01343354 | 0.39134822 |
| SRP54 | 26.4202661 | -0.115539922 | 0.4611536 | -0.2505454 | 0.80216559 | 0.99665574 |
| FAM177A1 | 17.8169628 | -0.184876776 | 0.633996 | -0.2916056 | 0.7705882 | 0.99665574 |
| PPP2R3C | 17.3383032 | -0.359394918 | 0.45758921 | -0.7854095 | 0.4322135 | 0.99617422 |
| NFKBIA | 11.6623636 | -0.908566419 | 0.81082399 | -1.120547 | 0.26248072 | 0.94557817 |
| RALGAPA1 | 86.9970392 | 0.245180682 | 0.27433907 | 0.89371405 | 0.3714749 | 0.98532838 |
| BRMS1L | 7.39844317 | 0.28719104 | 0.61560006 | 0.46652211 | 0.64084182 | 0.99665574 |
| MBIP | 7.97810011 | 0.271617154 | 0.62711208 | 0.43312378 | 0.66492485 | 0.99665574 |
| SLC25A21 | 3.48630725 | 0.357872749 | 1.0998818 | 0.32537382 | 0.74489816 | 0.99665574 |
| MIPOL1 | 25.346126 | -0.237965211 | 0.41593468 | -0.5721216 | 0.5672396 | 0.99665574 |
| SEC23A | 31.1080066 | -0.670213763 | 0.41498777 | -1.6150205 | 0.10630625 | 0.79596216 |
| GEMIN2 | 2.53710716 | -0.377229175 | 0.90025371 | -0.4190254 | 0.67519757 | 0.99665574 |
| TRAPPC6B | 16.9883781 | -0.127729187 | 0.47744872 | -0.2675244 | 0.78906542 | 0.99665574 |
| PNN | 32.2410943 | 0.197529686 | 0.42808776 | 0.46142335 | 0.6444949 | 0.99665574 |
| MIA2 | 50.6088332 | 0.355087261 | 0.36312603 | 0.97786231 | 0.32814243 | 0.97788427 |
| FBXO33 | 22.1681396 | -0.011172079 | 0.35130471 | -0.0318017 | 0.97463021 | 0.99777713 |
| AL049875.1 | 6.46552056 | 0.029427666 | 0.63238089 | 0.04653472 | 0.96288406 | 0.99777713 |
| KLHL28 | 15.1474299 | -0.25476174 | 0.54572538 | -0.4668314 | 0.64062051 | 0.99665574 |
| TOGARAM1 | 23.617293 | -0.226655449 | 0.30726573 | -0.7376529 | 0.46072542 | 0.99617422 |
| PRPF39 | 12.4785611 | 0.164290516 | 0.4589768 | 0.3579495 | 0.72038111 | 0.99665574 |
| FKBP3 | 12.4730758 | -0.017947599 | 0.46205257 | -0.0388432 | 0.9690154 | 0.99777713 |
| FANCM | 5.68580948 | -0.26437577 | 0.5931808 | -0.4456917 | 0.65581995 | 0.99665574 |
| MIS18BP1 | 10.633288 | 0.646608054 | 0.64963386 | 0.99534228 | 0.31956982 | 0.97700345 |
| RPS29 | 31.4203273 | 0.585623718 | 0.35562885 | 1.64672727 | 0.09961412 | 0.78285884 |
| RPL36AL | 38.2087348 | -0.157244589 | 0.45574406 | -0.3450283 | 0.73007311 | 0.99665574 |
| DNAAF2 | 3.86380626 | 0.227216657 | 0.7217041 | 0.31483354 | 0.75288804 | 0.99665574 |
| POLE2 | 2.67464454 | -1.090743965 | 0.86344227 | -1.2632506 | 0.20649913 | 0.90890506 |
| KLHDC1 | 9.6612635 | -0.089201309 | 0.75409813 | -0.1182887 | 0.90583889 | 0.99777713 |
| KLHDC2 | 13.5840688 | -0.090617929 | 0.40681369 | -0.2227504 | 0.82372974 | 0.99665574 |
| NEMF | 51.7109818 | 0.163114408 | 0.29424548 | 0.55434805 | 0.57934067 | 0.99665574 |
| ARF6 | 10.3759281 | 0.547610726 | 0.48616749 | 1.12638287 | 0.2600035 | 0.94557817 |
| VCPKMT | 5.60345554 | -0.325561098 | 0.64441353 | -0.5052053 | 0.61341468 | 0.99665574 |
| SOS2 | 63.0853042 | 0.020688517 | 0.27373205 | 0.07557945 | 0.93975369 | 0.99777713 |
| L2HGDH | 19.6255685 | -0.086567305 | 0.43644601 | -0.198346 | 0.84277439 | 0.99665574 |
| ATP5S | 15.1389704 | 0.149450232 | 0.43362962 | 0.3446495 | 0.73035788 | 0.99665574 |
| CDKL1 | 15.5823194 | 0.409832949 | 0.46665413 | 0.87823706 | 0.37981508 | 0.98679664 |
| MAP4K5 | 51.9110878 | 0.168349205 | 0.27856091 | 0.6043533 | 0.54560877 | 0.99665574 |
| ATL1 | 7.39228767 | 0.4701604 | 0.63131959 | 0.74472645 | 0.4564371 | 0.99617422 |
| SAV1 | 16.2737743 | 0.156230069 | 0.47093294 | 0.33174589 | 0.74008115 | 0.99665574 |
| NIN | 38.7473257 | 0.69524392 | 0.3677513 | 1.89052741 | 0.05868746 | 0.67545594 |
| TMX1 | 7.39983784 | 0.329695152 | 0.59259664 | 0.55635677 | 0.57796698 | 0.99665574 |
| FRMD6 | 4.78124699 | -0.120917125 | 1.09250497 | -0.1106788 | 0.91187107 | 0.99777713 |
| RTRAF | 15.3687381 | -0.156000932 | 0.42463417 | -0.3673772 | 0.71333764 | 0.99665574 |
| TXNDC16 | 5.41475883 | -0.164397151 | 0.74490491 | -0.2206955 | 0.82532955 | 0.99665574 |
| ERO1A | 56.0457208 | -0.490032254 | 0.55265572 | -0.8866863 | 0.37524782 | 0.98569969 |
| AL133453.1 | 6.75254957 | -1.609844352 | 0.9648337 | -1.66852 | 0.09521254 | 0.77178333 |
| PSMC6 | 23.6869441 | -0.402066602 | 0.39908329 | -1.0074754 | 0.31370637 | 0.9755712 |
| STYX | 10.4320541 | 1.428281533 | 0.57481742 | 2.48475687 | 0.01296399 | 0.38590337 |
| GNPNAT1 | 23.8523585 | -0.352989 | 0.61500013 | -0.5739657 | 0.56599099 | 0.99665574 |
| FERMT2 | 21.1057476 | -0.358363765 | 0.42051889 | -0.8521942 | 0.39410631 | 0.99179694 |
| DDHD1 | 9.47333372 | -0.106554599 | 0.57874473 | -0.1841133 | 0.85392458 | 0.99665574 |
| BMP4 | 3.127495 | 1.324822962 | 0.80805348 | 1.63952387 | 0.1011042 | 0.78386461 |
| CNIH1 | 8.47008918 | 0.309471665 | 0.61300958 | 0.50483985 | 0.61367133 | 0.99665574 |
| GMFB | 10.2306215 | -0.329700473 | 0.49520876 | -0.6657808 | 0.50555123 | 0.99665574 |
| CGRRF1 | 4.53759538 | -0.050866654 | 0.72799614 | -0.0698721 | 0.94429542 | 0.99777713 |
| SAMD4A | 75.6444581 | -0.239567886 | 0.40788524 | -0.5873414 | 0.55697444 | 0.99665574 |
| GCH1 | 5.42817599 | 0.272735728 | 0.87271929 | 0.31251255 | 0.75465103 | 0.99665574 |
| WDHD1 | 2.5520715 | -0.539488797 | 1.11978235 | -0.4817801 | 0.6299622 | 0.99665574 |
| SOCS4 | 8.88410847 | -0.785711664 | 0.59790329 | -1.3141116 | 0.18880864 | 0.89739428 |
| MAPK1IP1L | 30.6968238 | 0.144628572 | 0.3197824 | 0.45227183 | 0.65107317 | 0.99665574 |
| LGALS3 | 69.7178736 | 0.077895088 | 0.51122479 | 0.15236954 | 0.87889548 | 0.99665574 |
| FBXO34 | 70.7610658 | 0.243387893 | 0.36315903 | 0.67019646 | 0.50273256 | 0.99640678 |
| AL158801.3 | 3.85764975 | 0.186114259 | 0.88885641 | 0.20938619 | 0.83414677 | 0.99665574 |
| ATG14 | 9.23382424 | 0.405747746 | 0.66672169 | 0.60857139 | 0.54280857 | 0.99665574 |
| KTN1 | 81.9217461 | -0.078100203 | 0.23069505 | -0.338543 | 0.73495401 | 0.99665574 |
| PELI2 | 70.4698126 | -0.019989637 | 0.33313649 | -0.0600043 | 0.95215217 | 0.99777713 |
| TMEM260 | 38.2011194 | -0.184369232 | 0.3185069 | -0.5788547 | 0.56268719 | 0.99665574 |
| EXOC5 | 23.8653655 | -0.119217364 | 0.36103548 | -0.3302095 | 0.74124163 | 0.99665574 |
| AP5M1 | 18.5303748 | 0.830168759 | 0.41970956 | 1.97796007 | 0.04793321 | 0.6410909 |
| NAA30 | 14.0921348 | 0.457765219 | 0.43378738 | 1.05527555 | 0.29129932 | 0.96825983 |
| CCDC198 | 24.9614706 | 2.210325857 | 0.57209873 | 3.86353918 | 0.00011176 | 0.024106 |
| ARMH4 | 13.1031111 | 0.191231093 | 0.58734671 | 0.32558469 | 0.7447386 | 0.99665574 |
| ACTR10 | 19.7748884 | 0.474264434 | 0.48340429 | 0.98109272 | 0.32654702 | 0.97788427 |
| PSMA3 | 15.4668318 | 0.129063875 | 0.49073269 | 0.2630024 | 0.79254873 | 0.99665574 |
| ARID4A | 26.9266423 | -0.528574212 | 0.4604779 | -1.1478818 | 0.25101735 | 0.93973926 |
| TIMM9 | 9.48834027 | 0.090876106 | 0.51659566 | 0.17591341 | 0.86036197 | 0.99665574 |
| KIAA0586 | 20.4410617 | -0.077771526 | 0.39950921 | -0.1946677 | 0.84565313 | 0.99665574 |
| DACT1 | 2.30042976 | 0.632394932 | 1.11313959 | 0.56811827 | 0.56995466 | 0.99665574 |
| DAAM1 | 29.6167773 | 0.023683564 | 0.33386272 | 0.07093803 | 0.94344708 | 0.99777713 |
| L3HYPDH | 4.33317411 | -0.036073099 | 0.84763827 | -0.0425572 | 0.96605453 | 0.99777713 |
| JKAMP | 9.49740872 | 0.142450706 | 0.54773264 | 0.26007343 | 0.79480713 | 0.99665574 |
| RTN1 | 51.4372205 | 1.462092868 | 0.56781698 | 2.57493685 | 0.01002584 | 0.34338499 |
| LRRC9 | 22.5775221 | -0.055451129 | 0.84845252 | -0.0653556 | 0.94789087 | 0.99777713 |
| AL157911.1 | 66.6154937 | 0.995562764 | 0.42041013 | 2.36807509 | 0.01788091 | 0.44213812 |
| PCNX4 | 130.836913 | 0.796509672 | 0.32656646 | 2.4390431 | 0.01472621 | 0.40902558 |
| DHRS7 | 32.8870749 | 0.672836612 | 0.38345432 | 1.75467216 | 0.0793154 | 0.7446402 |
| PPM1A | 37.6093706 | 0.344064051 | 0.36101179 | 0.9530549 | 0.34056226 | 0.97788427 |
| SIX4 | 3.80877564 | -0.395559362 | 0.7897877 | -0.5008426 | 0.61648187 | 0.99665574 |
| MNAT1 | 79.6440945 | 0.218395025 | 0.29071917 | 0.75122334 | 0.45251825 | 0.99617422 |
| TRMT5 | 2.62315189 | 1.37505068 | 1.11812546 | 1.22978211 | 0.21877871 | 0.91662582 |
| SLC38A6 | 5.86827936 | 0.44568073 | 0.63309964 | 0.70396617 | 0.48145384 | 0.99640678 |
| PRKCH | 10.5857368 | -1.346500559 | 0.67413956 | -1.9973617 | 0.0457859 | 0.63477358 |

| | | | | | | | |
|---|---|---|---|---|---|---|---|
| TMEM30B | 9.74929718 | | -0.218392997 | 0.48026737 | -0.4547321 | 0.64930197 | 0.99665574 |
| HIF1A | 130.734445 | | -2.179325801 | 0.45977977 | -4.7399341 | 2.14E-06 | 0.00127493 |
| HIF1A-AS2 | 24.4405875 | | -2.817967886 | 0.63454627 | -4.4409179 | 8.96E-06 | 0.00344882 |
| SNAPC1 | 12.6667516 | | -0.809702641 | 0.59371663 | -1.3637864 | 0.17263481 | 0.88157681 |
| KCNH5 | 14.3841425 | | -0.514910144 | 1.01051623 | -0.5095516 | 0.61036565 | 0.99665574 |
| PPP2R5E | 80.7958975 | | -0.687405813 | 0.41422366 | -1.659504 | 0.09701428 | 0.77606444 |
| WDR89 | 5.59752884 | | 0.124648858 | 0.6322086 | 0.19716413 | 0.8436991 | 0.99665574 |
| SGPP1 | 10.1083335 | | -0.926015128 | 0.71301394 | -1.2987336 | 0.19403538 | 0.90235789 |
| SYNE2 | 338.262959 | | -1.001496326 | 0.38912699 | -2.5737005 | 0.01006173 | 0.34345402 |
| ESR2 | 20.4402571 | | -1.292480269 | 0.66531095 | -1.942671 | 0.05205593 | 0.65803363 |
| MTHFD1 | 4.76976955 | | -0.55314319 | 0.67808757 | -0.8157401 | 0.41464883 | 0.99279547 |
| ZBTB25 | 6.7155197 | | -0.154457736 | 0.66183263 | -0.2333788 | 0.81546724 | 0.99665574 |
| ZBTB1 | 12.04464 | | -0.101394794 | 0.51987096 | -0.1950384 | 0.8453629 | 0.99665574 |
| PLEKHG3 | 9.166103 | | -0.52967764 | 0.45219482 | -1.1713483 | 0.2414588 | 0.93253689 |
| CHURC1-FNT | 6.28420703 | | -0.80224421 | 0.58246783 | -1.3773193 | 0.16841354 | 0.87566218 |
| RAB15 | 3.20949497 | | -0.664302 | 0.83617199 | -0.7944562 | 0.42692991 | 0.99525676 |
| MAX | 19.9523976 | | -0.542187866 | 0.3923777 | -1.3818009 | 0.16703284 | 0.87495175 |
| FUT8 | 16.0079549 | | -0.446527213 | 0.43921793 | -1.0166416 | 0.30932396 | 0.97376371 |
| AL359232.1 | 40.4548085 | | 0.651127897 | 0.50722964 | 1.28369449 | 0.19924887 | 0.90381759 |
| GPHN | 808.001176 | | -0.036365193 | 0.46517631 | -0.0781751 | 0.93768879 | 0.99777713 |
| MPP5 | 28.5228965 | | -0.021960247 | 0.35806864 | -0.0613297 | 0.95109663 | 0.99777713 |
| ATP6V1D | 24.3437323 | | 0.202480928 | 0.49359875 | 0.41021362 | 0.68164925 | 0.99665574 |
| EIF2S1 | 13.663363 | | 0.18771622 | 0.39982546 | 0.46949541 | 0.63871556 | 0.99665574 |
| PLEK2 | 14.2055129 | | -0.077223565 | 0.49804627 | -0.155053 | 0.87677954 | 0.99665574 |
| PLEKHH1 | 15.4347306 | | 0.078383884 | 0.4627853 | 0.16937419 | 0.86550232 | 0.99665574 |
| PIGH | 8.49653037 | | 0.169386482 | 0.59029836 | 0.28695062 | 0.77415013 | 0.99665574 |
| ARG2 | 22.985297 | | 0.943480393 | 0.68482194 | 1.37770177 | 0.16829538 | 0.87566218 |
| VTI1B | 11.4623898 | | -0.193028108 | 0.50547502 | -0.3818747 | 0.70255433 | 0.99665574 |
| ZFYVE26 | 8.54456954 | | -0.112755292 | 0.50014223 | -0.2254465 | 0.82163198 | 0.99665574 |
| RAD51B | 68.9790562 | | 0.25941339 | 0.43671112 | 0.59401599 | 0.55250142 | 0.99665574 |
| ZFP36L1 | 42.5052717 | | -0.161752675 | 0.40380381 | -0.4005724 | 0.68873494 | 0.99665574 |
| ACTN1 | 11.7464739 | | -1.422173571 | 0.63125061 | -2.2529461 | 0.02426255 | 0.50168544 |
| DCAF5 | 44.8491947 | | 0.103638606 | 0.293769 | 0.35278945 | 0.72424629 | 0.99665574 |
| AL359317.2 | 3.61835443 | | -0.902658509 | 1.08520649 | -0.831785 | 0.4055303 | 0.99194293 |
| EXD2 | 8.95890873 | | 0.395450961 | 0.53686358 | 0.73659488 | 0.46136875 | 0.99617422 |
| ERH | 17.1550033 | | -0.028990069 | 0.43016613 | -0.0673927 | 0.94626906 | 0.99777713 |
| SLC39A9 | 22.373373 | | -0.335379223 | 0.46129082 | -0.7270451 | 0.46719833 | 0.99617422 |
| SUSD6 | 37.0264885 | | -0.901302498 | 0.37538076 | -2.4010354 | 0.01634876 | 0.42281551 |
| SRSF5 | 55.8237104 | | 0.340622052 | 0.31383744 | 1.08534551 | 0.27776866 | 0.95946581 |
| AL357153.2 | 9.62142447 | | -0.63609215 | 0.48087253 | -1.3227875 | 0.18590607 | 0.89532394 |
| MED6 | 9.41722708 | | -0.277955964 | 0.48173907 | -0.5769845 | 0.56394994 | 0.99665574 |
| TTC9 | 9.64821608 | | 0.167169952 | 0.57639475 | 0.29002685 | 0.7717957 | 0.99665574 |
| MAP3K9 | 14.3768257 | | 0.293010089 | 0.5119123 | 0.57238338 | 0.56706228 | 0.99665574 |
| PCNX1 | 77.1382044 | | 0.272065547 | 0.29493476 | 0.92246009 | 0.35628864 | 0.9819332 |
| SIPA1L1 | 58.2537058 | | -0.451320087 | 0.3378191 | -1.3359816 | 0.1815553 | 0.8928406 |
| DCAF4 | 3.48645392 | | -0.613688087 | 0.71105334 | -0.863069 | 0.38809954 | 0.99168679 |
| ZFYVE1 | 23.2896752 | | -0.520544163 | 0.37691372 | -1.3810698 | 0.1672575 | 0.87495175 |
| RBM25 | 41.6805989 | | 0.107738934 | 0.28673458 | 0.37574447 | 0.70710687 | 0.99665574 |
| PSEN1 | 39.9941398 | | -0.145450169 | 0.28025221 | -0.5189974 | 0.60376255 | 0.99665574 |
| PAPLN | 3.68359436 | | -0.815878414 | 0.75863692 | -1.0754531 | 0.28217193 | 0.96059387 |
| NUMB | 131.891843 | | -0.257897304 | 0.25201614 | -1.0233364 | 0.3061488 | 0.97376371 |
| HEATR4 | 7.06199428 | | -1.173273289 | 0.54833609 | -2.1396974 | 0.03237923 | 0.5724009 |
| AC005225.2 | 4.09480742 | | -0.218871639 | 0.68755282 | -0.3183343 | 0.75023138 | 0.99665574 |
| DNAL1 | 6.83044472 | | 0.52167737 | 0.63228878 | 0.82506189 | 0.40933646 | 0.99279547 |
| PNMA1 | 3.51101565 | | -0.326689365 | 0.82953411 | -0.3938227 | 0.69371194 | 0.99665574 |
| ELMSAN1 | 13.30604 | | -0.609833391 | 0.40952143 | -1.4891367 | 0.13645137 | 0.84475183 |
| AC005480.1 | 5.95336861 | | -0.377479921 | 0.77699195 | -0.4858222 | 0.62709325 | 0.99665574 |
| COQ6 | 3.62234629 | | -0.286364153 | 0.80059927 | -0.3576873 | 0.72057737 | 0.99665574 |
| ENTPD5 | 13.3782968 | | 0.091890171 | 0.50196524 | 0.18306083 | 0.85475029 | 0.99665574 |
| BBOF1 | 7.72680296 | | 0.851104623 | 0.75170805 | 1.13222762 | 0.25753875 | 0.94557817 |
| ALDH6A1 | 15.7940285 | | 1.036380143 | 0.58947213 | 1.75814952 | 0.07872207 | 0.74431421 |
| LIN52 | 22.4155582 | | 0.641297452 | 0.46247357 | 1.38666832 | 0.16554292 | 0.87495175 |
| ABCD4 | 15.9431275 | | -0.235522435 | 0.40843982 | -0.5766393 | 0.56418317 | 0.99665574 |
| NPC2 | 23.7490418 | | -0.356343354 | 0.50983537 | -0.6989381 | 0.48459073 | 0.99640678 |
| ISCA2 | 8.19076603 | | -0.488153304 | 0.6344359 | -0.7694289 | 0.44163875 | 0.99617422 |
| LTBP2 | 2.98189457 | | 2.531108665 | 0.96410725 | 2.62533931 | 0.00865627 | 0.31119582 |
| AREL1 | 21.3449058 | | 0.193971567 | 0.39821577 | 0.48710167 | 0.62618629 | 0.99665574 |
| FCF1 | 6.56578246 | | -0.387907982 | 0.54864296 | -0.7070317 | 0.47954676 | 0.99617422 |
| YLPM1 | 37.9036594 | | 0.409762082 | 0.36243515 | 1.1305804 | 0.25823174 | 0.94557817 |
| DLST | 17.6868106 | | -0.322144961 | 0.38391759 | -0.8390993 | 0.40141362 | 0.99194293 |
| EIF2B2 | 3.35051231 | | -0.650584529 | 0.86824294 | -0.7493116 | 0.4536694 | 0.99617422 |
| MLH3 | 22.6496572 | | 0.366451195 | 0.45435991 | 0.80652184 | 0.41994202 | 0.99518711 |
| ACYP1 | 13.7561875 | | 0.692330721 | 0.48435273 | 1.42939366 | 0.15289112 | 0.86116133 |
| ZC2HC1C | 5.16922853 | | -0.692120571 | 0.69997724 | -0.9887758 | 0.32277284 | 0.97700345 |
| NEK9 | 19.0862725 | | -0.057535109 | 0.37873735 | -0.151913 | 0.87925559 | 0.99665574 |
| TMED10 | 28.1593209 | | -0.431673144 | 0.37748368 | -1.1435545 | 0.25280846 | 0.94076968 |
| AL691403.1 | 23.7412433 | | -1.413977896 | 0.60284552 | -2.3455062 | 0.01900126 | 0.45540143 |
| AF111167.1 | 4.56088464 | | -1.38828111 | 0.82030608 | -1.692394 | 0.09057087 | 0.76326474 |
| FOS | 10.7068889 | | 0.606727783 | 0.90695568 | 0.66897181 | 0.50351346 | 0.99665574 |
| TTLL5 | 24.0926385 | | -0.283414765 | 0.34377912 | -0.8244095 | 0.40970693 | 0.99279547 |
| ERG28 | 6.96698158 | | -0.621348402 | 0.58783598 | -1.0570098 | 0.29050711 | 0.96676791 |
| IFT43 | 17.7140521 | | 0.298004888 | 0.40820146 | 0.73004366 | 0.4653635 | 0.99617422 |
| GPATCH2L | 37.8720961 | | -0.09032801 | 0.32160554 | -0.2808658 | 0.7788133 | 0.99665574 |
| ESRRB | 15.1077435 | | 1.353311349 | 0.71333108 | 1.89717143 | 0.05780532 | 0.67545594 |
| ANGEL1 | 5.43194939 | | 0.428653646 | 0.63627408 | 0.67369339 | 0.50050626 | 0.99640678 |
| IRF2BPL | 7.50465321 | | 1.116242881 | 0.76535433 | 1.4584655 | 0.14471228 | 0.85018987 |
| POMT2 | 5.16134205 | | 0.132169814 | 0.61889224 | 0.21355869 | 0.83089122 | 0.99665574 |
| GSTZ1 | 3.42453224 | | -0.132865592 | 0.78742196 | -0.1687349 | 0.86600514 | 0.99665574 |
| TMED8 | 14.1366579 | | 0.205097967 | 0.46569915 | 0.44040873 | 0.65964111 | 0.99665574 |
| VIPAS39 | 4.99592857 | | 0.306128398 | 0.63802676 | 0.47980495 | 0.63136609 | 0.99665574 |
| AHSA1 | 11.6932605 | | 0.64074091 | 0.45695748 | 1.40218936 | 0.16085871 | 0.86928072 |
| SPTLC2 | 130.406797 | | 0.517340192 | 0.24182294 | 2.13933468 | 0.03240857 | 0.5724009 |
| ALKBH1 | 8.06344398 | | 0.092342615 | 0.53555308 | 0.17242477 | 0.8631036 | 0.99665574 |
| SLIRP | 12.5475086 | | -0.543281627 | 0.53862945 | -1.0086371 | 0.31314872 | 0.97520189 |
| SNW1 | 14.134443 | | -0.02809837 | 0.38300782 | -0.0733624 | 0.94151774 | 0.99777713 |
| ADCK1 | 6.09649972 | | 0.967279206 | 0.72313939 | 1.337611 | 0.18102328 | 0.89174635 |
| NRXN3 | 27.7574751 | | -1.42796356 | 0.7875173 | -1.8132472 | 0.0697937 | 0.71112415 |
| DIO2 | 8.78218922 | | 1.278124862 | 1.78853412 | 0.71462146 | 0.47484298 | 0.99617422 |
| CEP128 | 8.15012901 | | -0.801468658 | 0.52322618 | -1.5317824 | 0.12557614 | 0.83633607 |
| GTF2A1 | 13.7120114 | | 0.537799224 | 0.46097882 | 1.16664628 | 0.24335325 | 0.93426912 |
| STON2 | 36.9265877 | | 0.194727369 | 0.47756487 | 0.40775061 | 0.68345677 | 0.99665574 |
| SEL1L | 44.3092387 | | -0.064468446 | 0.39731544 | -0.1622601 | 0.87110103 | 0.99665574 |
| GALC | 4.07109085 | | 0.372578272 | 0.73650302 | 0.50587474 | 0.61294459 | 0.99665574 |
| SPATA7 | 7.69332625 | | -0.012292409 | 0.52280962 | -0.0235122 | 0.9812417 | 0.99777713 |
| PTPN21 | 44.5380258 | | 0.282515005 | 0.37280561 | 0.75780781 | 0.44856604 | 0.99617422 |
| ZC3H14 | 34.0129321 | | 0.495036664 | 0.28952494 | 1.7098239 | 0.08729844 | 0.75595306 |
| EML5 | 18.4953036 | | -0.124632668 | 0.4681409 | -0.266229 | 0.79006287 | 0.99665574 |
| TTC8 | 14.9430328 | | 0.148923514 | 0.47623331 | 0.31271125 | 0.75450005 | 0.99665574 |
| FOXN3 | 303.9611 | | -0.361898815 | 0.4104714 | -0.8816663 | 0.37795727 | 0.98679664 |
| AL356805.1 | 5.19483122 | | -0.701363278 | 0.62827905 | -1.1163245 | 0.26428327 | 0.94557817 |
| EFCAB11 | 7.8907836 | | -0.174943111 | 0.70287478 | -0.2488966 | 0.8034408 | 0.99665574 |
| TDP1 | 5.82820334 | | 1.450454541 | 0.72744686 | 1.9938976 | 0.04616325 | 0.63760626 |
| PSMC1 | 8.89015378 | | -0.192475346 | 0.54102966 | -0.3557575 | 0.7220222 | 0.99665574 |

| | | | | | | |
|---|---|---|---|---|---|---|
| NRDE2 | 10.4086039 | -0.441670159 | 0.45750874 | -0.9653808 | 0.3343541 | 0.97788427 |
| CALM1 | 30.8477197 | -0.162762904 | 0.34026008 | -0.4783485 | 0.63240217 | 0.99665574 |
| TTC7B | 22.6097209 | 0.003683192 | 0.54744284 | 0.00672799 | 0.99463188 | 0.9985287 |
| RPS6KA5 | 13.1402985 | -2.384010613 | 0.61583949 | -3.8711558 | 0.00010832 | 0.02396939 |
| DGLUCY | 47.9299398 | -0.58454279 | 0.3928072 | -1.4881163 | 0.13672023 | 0.84475183 |
| CCDC88C | 2.92622604 | -0.527180769 | 0.88127463 | -0.5982026 | 0.54970476 | 0.99665574 |
| PPP4R3A | 37.4277807 | 0.011620629 | 0.3058155 | 0.03799882 | 0.96968862 | 0.99777713 |
| CATSPERB | 3.41827753 | -0.248036371 | 0.71398224 | -0.3473985 | 0.72829195 | 0.99665574 |
| TC2N | 9.71061849 | -0.600821466 | 0.49513458 | -1.2134508 | 0.2249575 | 0.92318867 |
| TRIP11 | 39.8208333 | -0.043731214 | 0.27319881 | -0.160071 | 0.87282512 | 0.99665574 |
| ATXN3 | 19.7646078 | -0.063238136 | 0.35860841 | -0.1763431 | 0.86002437 | 0.99665574 |
| NDUFB1 | 15.0651619 | 0.463194841 | 0.42711284 | 1.08447884 | 0.27815254 | 0.95958042 |
| CPSF2 | 13.5957667 | 0.1232155 | 0.51071425 | 0.24126114 | 0.80935273 | 0.99665574 |
| LGMN | 13.3846461 | -0.073194239 | 0.49667039 | -0.1473698 | 0.8828401 | 0.99665574 |
| GOLGA5 | 10.8987121 | -0.22022004 | 0.52652968 | -0.4182481 | 0.67576573 | 0.99665574 |
| ITPK1 | 64.0572563 | 0.312742085 | 0.30083244 | 1.03958897 | 0.29853091 | 0.97358745 |
| MOAP1 | 5.15763348 | 0.028139644 | 0.65224625 | 0.04314267 | 0.96558781 | 0.99777713 |
| GON7 | 3.87591092 | -1.038461813 | 0.79260362 | -1.3101906 | 0.19013137 | 0.8994569 |
| UBR7 | 8.11208443 | -0.087468514 | 0.60182101 | -0.1453397 | 0.88444263 | 0.99665574 |
| BTBD7 | 54.2347225 | -0.159137596 | 0.23925195 | -0.6651465 | 0.50595679 | 0.99665574 |
| DDX24 | 32.3135926 | -0.273209438 | 0.31133311 | -0.877547 | 0.3801896 | 0.98679664 |
| IFI27L1 | 2.90897043 | 0.111485354 | 0.85023347 | 0.13112322 | 0.89567783 | 0.99777713 |
| IFI27 | 16.701678 | 1.564528753 | 1.64998023 | 0.9482106 | 0.34302225 | 0.97788427 |
| SERPINA1 | 16.2996176 | -1.515713162 | 1.11115112 | -1.3640927 | 0.17253838 | 0.88157681 |
| DICER1 | 50.920548 | -0.046582779 | 0.2871723 | -0.162212 | 0.87113895 | 0.99665574 |
| CLMN | 119.200904 | -0.706811496 | 0.34558292 | -2.0452732 | 0.04082793 | 0.61100706 |
| GLRX5 | 12.6865488 | 0.223861873 | 0.4635296 | 0.48295054 | 0.62913086 | 0.99665574 |
| C14orf132 | 2.00717745 | 0.767800127 | 0.92310989 | 0.83175376 | 0.40554795 | 0.99194293 |
| ATG2B | 30.0867264 | 0.185432978 | 0.33609522 | 0.5517275 | 0.58113506 | 0.99665574 |
| GSKIP | 6.21227491 | 0.470231353 | 0.61321269 | 0.76683239 | 0.44318118 | 0.99617422 |
| PAPOLA | 63.1771968 | 0.701441911 | 0.24716733 | 2.83792329 | 0.00454081 | 0.21492549 |
| VRK1 | 4.99126115 | -0.071179678 | 0.73116358 | -0.0973512 | 0.92244747 | 0.99777713 |
| SETD3 | 31.1100191 | 0.096569902 | 0.30720747 | 0.31434751 | 0.75325712 | 0.99665574 |
| CCNK | 8.30040654 | 0.053638458 | 0.55050007 | 0.09743588 | 0.92238025 | 0.99777713 |
| CCDC85C | 11.5798022 | -0.008717198 | 0.5010211 | -0.0173989 | 0.98611842 | 0.99777713 |
| EVL | 23.3291027 | -0.98461817 | 0.45353847 | -2.1709695 | 0.02993348 | 0.55886856 |
| DEGS2 | 3.85192002 | -1.770909446 | 1.12101789 | -1.5797334 | 0.11416793 | 0.81638817 |
| YY1 | 34.5301311 | -0.344078577 | 0.33376885 | -1.0308888 | 0.30259296 | 0.97358745 |
| SLC25A29 | 19.4308052 | -0.543407184 | 0.56928989 | -0.9545351 | 0.33981285 | 0.97788427 |
| WARS | 6.75005558 | -1.091951753 | 1.01697572 | -1.0737245 | 0.28294617 | 0.96059387 |
| WDR25 | 5.84552773 | -0.216471165 | 0.62197442 | -0.3480387 | 0.72781112 | 0.99665574 |
| PPP2R5C | 39.653392 | 0.27772953 | 0.32929123 | 0.84341612 | 0.39899576 | 0.99179694 |
| DYNC1H1 | 36.4735963 | -0.465573034 | 0.35534458 | -1.3102016 | 0.19012764 | 0.8994569 |
| HSP90AA1 | 143.680814 | 0.428583778 | 0.32545061 | 1.31689346 | 0.18787434 | 0.89739428 |
| WDR20 | 38.4781458 | 0.021998407 | 0.33566996 | 0.06553582 | 0.94774739 | 0.99777713 |
| MOK | 6.28741709 | 0.159529143 | 0.80847439 | 0.19732121 | 0.84357619 | 0.99665574 |
| ZNF839 | 5.33267702 | 0.176582697 | 0.64814917 | 0.27244145 | 0.78528261 | 0.99665574 |
| CINP | 4.3573248 | 0.079622842 | 0.6709083 | 0.11867917 | 0.90552954 | 0.99777713 |
| TECPR2 | 15.7741717 | -0.434246966 | 0.5123195 | -0.8476097 | 0.39665539 | 0.99179694 |
| ANKRD9 | 14.2678495 | 0.06374123 | 0.42184706 | 0.15110033 | 0.87989657 | 0.99665574 |
| RCOR1 | 42.1720161 | 0.487349298 | 0.46078013 | 1.05766128 | 0.2902099 | 0.96676791 |
| TRAF3 | 17.3236765 | 0.07676722 | 0.41776744 | 0.18375587 | 0.85420498 | 0.99665574 |
| CDC42BPB | 80.6166325 | -0.08864979 | 0.29894357 | -0.2965436 | 0.76681501 | 0.99665574 |
| EIF5 | 26.6383403 | -0.848456959 | 0.46071033 | -1.8416278 | 0.06552961 | 0.69221233 |
| MARK3 | 88.4463563 | -0.073764464 | 0.32477501 | -0.2271248 | 0.82032669 | 0.99665574 |
| CKB | 53.6064877 | 0.590373325 | 0.41479612 | 1.42328554 | 0.1546534 | 0.86196782 |
| TRMT61A | 2.94803937 | 0.830219444 | 0.86976521 | 0.95453282 | 0.339814 | 0.97788427 |
| BAG5 | 10.9698575 | 0.008626751 | 0.48113146 | 0.01793013 | 0.98569459 | 0.99777713 |
| KLC1 | 22.5916749 | -1.010779922 | 0.48601652 | -2.0797234 | 0.03755091 | 0.58839433 |
| ZFYVE21 | 8.29856488 | -0.397447478 | 0.52544653 | -0.7563995 | 0.44940972 | 0.99617422 |
| PPP1R13B | 70.2257078 | -0.544991743 | 0.50213803 | -1.0853425 | 0.27776999 | 0.95946581 |
| ATP5MPL | 20.7708945 | 0.334523379 | 0.42243575 | 0.79189174 | 0.4284238 | 0.99589919 |
| SIVA1 | 10.894765 | 0.667639412 | 0.49668457 | 1.34419197 | 0.17888632 | 0.88813816 |
| AKT1 | 11.5105025 | -0.075649547 | 0.58998894 | -0.128222 | 0.89797331 | 0.99777713 |
| CEP170B | 7.35328613 | -1.025945798 | 0.64460905 | -1.5915783 | 0.11147948 | 0.8083966 |
| AHNAK2 | 2.6556429 | -1.508893312 | 0.92181947 | -1.6368642 | 0.10165884 | 0.7843028 |
| NUDT14 | 6.57861437 | -0.323370124 | 0.61973743 | -0.5217857 | 0.60181955 | 0.99665574 |
| BRF1 | 9.54050355 | 0.167769526 | 0.50139951 | 0.3346025 | 0.73792496 | 0.99665574 |
| BTBD6 | 5.31580585 | 0.248891867 | 0.63935401 | 0.38928647 | 0.69706424 | 0.99665574 |
| PACS2 | 19.608448 | -0.361222111 | 0.48808591 | -0.740079 | 0.45925208 | 0.99617422 |
| MTA1 | 7.2454381 | 0.402168018 | 0.57063037 | 0.7047785 | 0.48094809 | 0.99640678 |
| NIPA1 | 11.6055861 | -0.218797989 | 0.42526112 | -0.5145027 | 0.60690058 | 0.99665574 |
| NIPA2 | 15.9949209 | -0.476683271 | 0.46066183 | -1.0347792 | 0.30077204 | 0.97358745 |
| CYFIP1 | 18.4730145 | -0.308222909 | 0.41345516 | -0.7454809 | 0.45598108 | 0.99617422 |
| TUBGCP5 | 10.522401 | -0.328461997 | 0.47175948 | -0.6962489 | 0.48627299 | 0.99640678 |
| PWRN3 | 6.3437326 | -0.570170029 | 1.67264936 | -0.3408784 | 0.73319513 | 0.99665574 |
| PWRN1 | 48.5422807 | -1.290988844 | 0.84538764 | -1.5270969 | 0.12673692 | 0.83669777 |
| AC124312.1 | 64.9202908 | 0.352602756 | 0.29145983 | 1.20978167 | 0.22636268 | 0.92318867 |
| PWAR6 | 13.2048844 | 0.936441239 | 0.48470434 | 1.93198443 | 0.05336143 | 0.66496977 |
| UBE3A | 59.4338177 | 0.467156072 | 0.29010088 | 1.61032283 | 0.1073274 | 0.79947476 |
| GABRB3 | 3.9655965 | -0.013709096 | 0.94641636 | -0.0144853 | 0.98844283 | 0.99777713 |
| HERC2 | 51.557238 | 0.082209079 | 0.28001478 | 0.29358835 | 0.76907248 | 0.99665574 |
| FAM189A1 | 24.9206733 | -0.157827925 | 0.54382135 | -0.2902202 | 0.77164781 | 0.99665574 |
| AC022613.1 | 4.60064699 | -1.220235509 | 0.74514299 | -1.6375857 | 0.10150815 | 0.7843028 |
| TJP1 | 58.0529137 | -0.100093328 | 0.28236145 | -0.3544865 | 0.7229743 | 0.99665574 |
| FAN1 | 11.350198 | 0.099929899 | 0.52550356 | 0.19016027 | 0.84918354 | 0.99665574 |
| MTMR10 | 38.0056515 | -0.319739166 | 0.4472023 | -0.7149766 | 0.47462352 | 0.99617422 |
| KLF13 | 21.5874256 | -0.850217584 | 0.40462349 | -2.1012561 | 0.03561849 | 0.58349146 |
| OTUD7A | 20.8342301 | 0.224820357 | 0.45624865 | 0.4927584 | 0.62218331 | 0.99665574 |
| LINC02256 | 27.9358036 | -0.293876682 | 0.51464942 | -0.571023 | 0.56798402 | 0.99665574 |
| AC123768.3 | 4.14807131 | -0.192886873 | 0.77473666 | -0.2489709 | 0.8033833 | 0.99665574 |
| GOLGA8N | 4.16808594 | 0.282138583 | 0.80234399 | 0.35164292 | 0.72510607 | 0.99665574 |
| FMN1 | 14.5377779 | -1.908760345 | 0.60900869 | -3.1342087 | 0.00172318 | 0.12036484 |
| RYR3 | 7.96134672 | 0.17950658 | 0.67993246 | 0.26400649 | 0.79177492 | 0.99665574 |
| AVEN | 21.35757 | 0.25037078 | 0.39617234 | 0.63197441 | 0.5274036 | 0.99665574 |
| EMC7 | 10.2865008 | -0.131356516 | 0.5470612 | -0.240113 | 0.81024264 | 0.99665574 |
| KATNBL1 | 35.2092338 | -0.243410309 | 0.33746067 | -0.7212998 | 0.47072509 | 0.99617422 |
| EMC4 | 5.31571698 | 0.450993038 | 0.60992666 | 0.73942175 | 0.45965094 | 0.99617422 |
| SLC12A6 | 14.0422792 | -0.43952225 | 0.59557613 | -0.7379783 | 0.46052765 | 0.99617422 |
| NOP10 | 18.0780675 | -0.591241062 | 0.48771748 | -1.2122614 | 0.22541235 | 0.92318867 |
| GOLGA8A | 12.6069827 | -0.2902241 | 0.55637484 | -0.5216341 | 0.6019251 | 0.99665574 |
| GOLGA8B | 15.6356689 | -0.141851481 | 0.41561295 | -0.3413067 | 0.73287271 | 0.99665574 |
| AQR | 21.4909354 | 0.413005032 | 0.33163325 | 1.24536678 | 0.21299696 | 0.91073943 |
| ZNF770 | 12.8206753 | 0.105901736 | 0.43018689 | 0.24617611 | 0.80554591 | 0.99665574 |
| AC021231.1 | 23.629322 | 0.284311097 | 0.56150701 | 0.50633579 | 0.61262094 | 0.99665574 |
| DPH6 | 84.1717017 | 0.588877973 | 0.38921256 | 1.51299839 | 0.13028007 | 0.83789479 |
| DPH6-DT | 10.6137932 | 1.006125611 | 0.97188644 | 1.0352296 | 0.3005617 | 0.97358745 |
| C15orf41 | 14.58915 | -0.939639724 | 0.62915278 | -1.4935001 | 0.13530633 | 0.84373478 |
| SPRED1 | 27.67775 | -0.46003475 | 0.38374764 | -1.198795 | 0.23060767 | 0.92774608 |
| FAM98B | 10.203208 | -0.288610029 | 0.53856955 | -0.5358826 | 0.59203972 | 0.99665574 |
| RASGRP1 | 4.41567155 | -1.401545151 | 0.93018974 | -1.5067304 | 0.13187978 | 0.84087875 |
| C15orf53 | 90.4305776 | -2.407200283 | 1.08507572 | -2.218463 | 0.02652328 | 0.52313821 |
| AC013652.1 | 18.6635051 | -1.322356395 | 0.77596056 | -1.7041541 | 0.0883523 | 0.75650018 |

| | | | | | | |
|---|---|---|---|---|---|---|
| THBS1 | 15.5289698 | -1.111182626 | 0.82153622 | -1.3525668 | 0.17619406 | 0.88428483 |
| FSIP1 | 3.53402502 | -1.06300085 | 0.82480557 | -1.2887896 | 0.19747124 | 0.90381759 |
| EIF2AK4 | 14.7459523 | -0.222110181 | 0.44210041 | -0.5023976 | 0.61538787 | 0.99665574 |
| SRP14 | 81.4011333 | -0.256423572 | 0.32754741 | -0.7828594 | 0.43370968 | 0.99617422 |
| SRP14-AS1 | 4.72460497 | 0.060520799 | 0.96822502 | 0.06250696 | 0.95015912 | 0.99777713 |
| BMF | 6.66708734 | 0.369833225 | 0.52880785 | 0.69937166 | 0.48431979 | 0.99640678 |
| IVD | 15.0460136 | -0.136970979 | 0.40657793 | -0.3368874 | 0.73620179 | 0.99665574 |
| BAHD1 | 8.20895454 | -0.661012064 | 0.5590362 | -1.1824137 | 0.23704159 | 0.9305595 |
| CCDC32 | 14.180633 | -0.301316418 | 0.50467003 | -0.5970563 | 0.5504698 | 0.99665574 |
| RMDN3 | 7.68913874 | -0.813160385 | 0.53016308 | -1.5337929 | 0.1250806 | 0.83370619 |
| GCHFR | 6.15704431 | -0.482052349 | 0.71640107 | -0.6728806 | 0.50102328 | 0.99640678 |
| DNAJC17 | 10.0311194 | -0.304133452 | 0.5096314 | -0.5967714 | 0.55066001 | 0.99665574 |
| ZFYVE19 | 2.95416185 | 0.358031255 | 0.96183475 | 0.37223781 | 0.7097158 | 0.99665574 |
| SPINT1-AS1 | 5.96943586 | 0.144372509 | 0.66285169 | 0.21780514 | 0.82758094 | 0.99665574 |
| SPINT1 | 16.5060469 | -0.074412808 | 0.43996245 | -0.1691345 | 0.86569089 | 0.99665574 |
| VPS18 | 4.22063066 | -1.191164971 | 0.89343954 | -1.3332351 | 0.18245466 | 0.89461655 |
| INO80 | 32.882702 | -0.142503938 | 0.35399666 | -0.4025573 | 0.68727393 | 0.99665574 |
| CHP1 | 22.2298993 | -0.633587231 | 0.50797066 | -1.247291 | 0.21229082 | 0.91043197 |
| NDUFAF1 | 2.44314614 | -0.813552707 | 0.9235454 | -0.8809017 | 0.37837103 | 0.98679664 |
| RTF1 | 21.3702161 | 0.038001151 | 0.39148046 | 0.09707036 | 0.92267052 | 0.99777713 |
| RPAP1 | 4.90410305 | -0.150902694 | 0.64297996 | -0.2346927 | 0.81444727 | 0.99665574 |
| MGA | 31.9015595 | 0.010367257 | 0.30744027 | 0.03372121 | 0.97309947 | 0.99777713 |
| MAPKBP1 | 21.9812657 | -0.011723186 | 0.50246665 | -0.0233313 | 0.98138603 | 0.99777713 |
| EHD4 | 9.67884634 | -0.387223918 | 0.59086803 | -0.6553476 | 0.51224401 | 0.99665574 |
| PLA2G4F | 8.70259573 | 1.2367129 | 0.81823841 | 1.51143345 | 0.13067805 | 0.83795959 |
| VPS39 | 18.7558707 | -0.81450007 | 0.49323198 | -1.6513529 | 0.09866653 | 0.78203824 |
| TMEM87A | 45.350585 | -0.241071088 | 0.29121663 | -0.8278067 | 0.40777996 | 0.99279547 |
| GANC | 10.1306928 | -0.942721804 | 0.54841929 | -1.71898 | 0.08561801 | 0.75373899 |
| ZNF106 | 20.943003 | 0.051875747 | 0.45045285 | 0.11516355 | 0.90831549 | 0.99777713 |
| SNAP23 | 18.2390886 | 0.155166691 | 0.39634498 | 0.39149402 | 0.69543211 | 0.99665574 |
| LRRC57 | 3.31826923 | 0.258330776 | 0.812965 | 0.31776371 | 0.75066418 | 0.99665574 |
| HAUS2 | 4.45957541 | -1.170805729 | 0.72116213 | -1.6234986 | 0.10448285 | 0.79169635 |
| TTBK2 | 29.4138359 | -0.284117064 | 0.41112012 | -0.6910804 | 0.48951501 | 0.99640678 |
| UBR1 | 44.3834109 | -0.245667982 | 0.29325819 | -0.8377191 | 0.40218849 | 0.99194293 |
| AC068724.3 | 10.1230653 | 0.316510463 | 0.55358265 | 0.57174925 | 0.56749187 | 0.99665574 |
| TMEM62 | 10.5018722 | 0.020614476 | 0.51161946 | 0.0402926 | 0.96785986 | 0.99777713 |
| CCNDBP1 | 11.5880909 | 0.108813883 | 0.54465975 | 0.19978323 | 0.84165012 | 0.99665574 |
| ADAL | 9.36698977 | 0.448475897 | 0.48962745 | 0.91595334 | 0.35969137 | 0.9819332 |
| ZSCAN29 | 3.11555652 | 0.383061858 | 0.76292973 | 0.50209323 | 0.61560194 | 0.99665574 |
| TUBGCP4 | 8.00869671 | -0.237154465 | 0.48427583 | -0.4897095 | 0.62433949 | 0.99665574 |
| TP53BP1 | 26.9121812 | -0.146523065 | 0.34077845 | -0.4299658 | 0.66722055 | 0.99665574 |
| PPIP5K1 | 13.8907086 | 0.108578607 | 0.42090816 | 0.2579627 | 0.79643569 | 0.99665574 |
| CATSPER2 | 11.3119518 | -0.093042736 | 0.59199395 | -0.1571684 | 0.87511214 | 0.99665574 |
| PDIA3 | 30.6758794 | 0.465423926 | 0.37807159 | 1.23104707 | 0.21830526 | 0.91662582 |
| SERF2 | 140.531471 | 0.050324177 | 0.40199439 | 0.12518627 | 0.90037609 | 0.99777713 |
| MFAP1 | 6.78439025 | -0.15366757 | 0.54175886 | -0.2836457 | 0.77668191 | 0.99665574 |
| FRMD5 | 3.77074026 | 0.178338057 | 0.92303467 | 0.1932084 | 0.84679576 | 0.99665574 |
| CASC4 | 59.1013123 | -0.309870479 | 0.24176203 | -1.2817169 | 0.19994198 | 0.90381759 |
| AC025043.1 | 4.81222101 | 0.795567833 | 0.98493752 | 0.80773431 | 0.41924355 | 0.99476033 |
| CTDSPL2 | 34.5848945 | 0.208126484 | 0.32267735 | 0.64499875 | 0.518928 | 0.99665574 |
| EIF3J-DT | 6.88850355 | -0.427066437 | 0.62864993 | -0.679339 | 0.49692307 | 0.99640678 |
| EIF3J | 19.9808276 | -0.533626732 | 0.41540675 | -1.2845885 | 0.19893612 | 0.90381759 |
| SPG11 | 59.4305067 | -0.494039475 | 0.26340465 | -1.8755913 | 0.06071142 | 0.6834742 |
| B2M | 366.816083 | -0.515493497 | 0.48768339 | -1.0570249 | 0.29050022 | 0.96676791 |
| TRIM69 | 5.16727282 | -0.525124446 | 0.77923433 | -0.673898 | 0.50037617 | 0.99640678 |
| SORD | 7.24370278 | -0.65666596 | 0.55784792 | -1.1771415 | 0.23913902 | 0.93157178 |
| GATM | 11.3053507 | 2.662268742 | 0.98243067 | 2.7098795 | 0.00673077 | 0.26557324 |
| SPATA5L1 | 4.71907534 | 0.106493946 | 0.70043127 | 0.15204054 | 0.87915496 | 0.99665574 |
| SQOR | 4.19937027 | -0.335931811 | 1.04719237 | -0.3207928 | 0.74836739 | 0.99665574 |
| MYEF2 | 23.0990278 | 0.017128356 | 0.3866404 | 0.04430048 | 0.96466489 | 0.99777713 |
| SLC12A1 | 51.9288626 | 3.007109781 | 0.89407912 | 3.36335981 | 0.00077 | 0.07295564 |
| DUT | 10.774279 | 0.517699775 | 0.57373477 | 0.90233294 | 0.36688003 | 0.98397614 |
| CEP152 | 10.3217749 | 0.233579679 | 0.6692638 | 0.34900988 | 0.72708189 | 0.99665574 |
| EID1 | 32.1821161 | -0.317047777 | 0.38138732 | -0.8313013 | 0.40580344 | 0.99194293 |
| SECISBP2L | 48.4540718 | -0.116825496 | 0.28233396 | -0.4137848 | 0.67903173 | 0.99665574 |
| COPS2 | 11.1271936 | 0.245460327 | 0.43943185 | 0.55858566 | 0.57644453 | 0.99665574 |
| GALK2 | 22.5714751 | 0.072341264 | 0.41486972 | 0.17437104 | 0.86157387 | 0.99665574 |
| FAM227B | 10.776999 | -0.255410359 | 0.67348941 | -0.3792344 | 0.7045138 | 0.99665574 |
| DTWD1 | 11.9353975 | 0.80977198 | 0.74731452 | 1.08357587 | 0.2785529 | 0.95969433 |
| ATP8B4 | 28.6961927 | 1.602876577 | 0.56299537 | 2.84705109 | 0.00441263 | 0.21310323 |
| GABPB1 | 15.6829303 | 0.17018704 | 0.52753035 | 0.3226109 | 0.74698994 | 0.99665574 |
| GABPB1-AS1 | 12.9052776 | 0.316887508 | 0.63432901 | 0.49956332 | 0.61738259 | 0.99665574 |
| USP8 | 31.7287548 | -0.143803881 | 0.29903981 | -0.4808854 | 0.63059794 | 0.99665574 |
| USP50 | 4.35256161 | -0.288073093 | 0.66326408 | -0.4343264 | 0.66405144 | 0.99665574 |
| TRPM7 | 64.7573531 | -0.125486978 | 0.367227 | -0.341715 | 0.73256537 | 0.99665574 |
| SPPL2A | 45.5062731 | 0.271301273 | 0.34905346 | 0.77724848 | 0.43701218 | 0.99617422 |
| AP4E1 | 16.2266523 | -0.048733792 | 0.52268605 | -0.0932372 | 0.92571511 | 0.99777713 |
| DMXL2 | 16.6064662 | -0.04282847 | 0.43096096 | -0.099379 | 0.92083735 | 0.99777713 |
| LYSMD2 | 4.13867256 | 0.512685704 | 0.89763932 | 0.57114889 | 0.56789872 | 0.99665574 |
| TMOD2 | 4.43366931 | -0.490530245 | 0.71350662 | -0.6874922 | 0.49177261 | 0.99640678 |
| TMOD3 | 37.8745938 | -0.476139709 | 0.32893321 | -1.447527 | 0.14774938 | 0.85300864 |
| LEO1 | 6.61350003 | 0.677980464 | 0.64087962 | 1.05789051 | 0.29010536 | 0.96676791 |
| MAPK6 | 19.6227041 | -0.721156928 | 0.38041264 | -1.8957228 | 0.0579967 | 0.67545594 |
| GNB5 | 9.98397158 | -0.799431042 | 0.47654877 | -1.677543 | 0.09343635 | 0.76763189 |
| MYO5C | 18.8756452 | -0.859945964 | 0.56432399 | -1.5238515 | 0.12754581 | 0.83778831 |
| MYO5A | 26.2952772 | -0.227681211 | 0.42126201 | -0.5404741 | 0.58887011 | 0.99665574 |
| ARPP19 | 33.2078957 | -0.066942058 | 0.35116023 | -0.1906311 | 0.84881463 | 0.99665574 |
| FAM214A | 33.4667221 | -0.02909679 | 0.39291735 | -0.0740532 | 0.94096805 | 0.99777713 |
| WDR72 | 806.917911 | 0.547902085 | 0.32405647 | 1.69076114 | 0.09088243 | 0.76412898 |
| UNC13C | 83.8270698 | 0.225050516 | 0.48411701 | 0.46486802 | 0.64202597 | 0.99665574 |
| RSL24D1 | 10.7867183 | -0.578150553 | 0.4576779 | -1.2632258 | 0.20650803 | 0.90890506 |
| RAB27A | 16.7840447 | -1.210645344 | 0.48174614 | -2.5130359 | 0.01196972 | 0.37496952 |
| PIGB | 8.06754444 | -0.492165648 | 0.4737918 | -1.0387804 | 0.29890686 | 0.97358745 |
| CCPG1 | 30.5077548 | 0.051192989 | 0.37310076 | 0.13720956 | 0.89086516 | 0.99712197 |
| DNAAF4 | 4.69737815 | 0.03301372 | 0.64721303 | 0.05100905 | 0.95931831 | 0.99777713 |
| PRTG | 4.12708143 | -1.422961387 | 0.69682887 | -2.0420528 | 0.04114629 | 0.61100706 |
| NEDD4 | 4.70393167 | -0.572221731 | 0.79424972 | -0.7204557 | 0.47124447 | 0.99617422 |
| RFX7 | 31.5911365 | -0.711119949 | 0.37953271 | -1.8736724 | 0.0609756 | 0.6834742 |
| TEX9 | 9.27926559 | -0.544875601 | 0.59201718 | -0.9203713 | 0.35737878 | 0.9819332 |
| ZNF280D | 14.722977 | 0.860469104 | 0.41685092 | 2.06421306 | 0.03899752 | 0.60084629 |
| AC090517.4 | 5.52814836 | 0.414884924 | 0.62933937 | 0.65923879 | 0.50974244 | 0.99665574 |
| TCF12 | 183.561628 | 0.006963919 | 0.22787195 | 0.03056067 | 0.97561991 | 0.99777713 |
| CGNL1 | 214.331801 | 0.243463211 | 0.35300228 | 0.68969302 | 0.49038725 | 0.99640678 |
| POLR2M | 5.79062923 | 0.015503754 | 0.63636463 | 0.024363 | 0.98056306 | 0.99777713 |
| ALDH1A2 | 16.7765304 | -1.380255391 | 0.73675061 | -1.8734364 | 0.06100815 | 0.6834742 |
| ADAM10 | 63.3265639 | 0.048276418 | 0.28125121 | 0.17164875 | 0.86371368 | 0.99665574 |
| MINDY2 | 57.8664458 | 0.045403324 | 0.30589166 | 0.14842943 | 0.88200388 | 0.99665574 |
| RNF111 | 50.7172925 | -0.215826244 | 0.24774666 | -0.871157 | 0.38366843 | 0.98956212 |
| SLTM | 59.9545856 | 0.003128573 | 0.27385582 | 0.01142416 | 0.99088504 | 0.99777713 |
| MYO1E | 48.8294021 | -0.610993437 | 0.37120828 | -1.6459585 | 0.09977229 | 0.78288816 |
| FAM81A | 3.05778304 | 0.431789994 | 1.05517711 | 0.40921092 | 0.68238488 | 0.99665574 |
| GTF2A2 | 14.3249541 | 0.170229341 | 0.45383279 | 0.37509264 | 0.70759157 | 0.99665574 |
| BNIP2 | 15.7843272 | -0.635832331 | 0.43827842 | -1.4507498 | 0.14684954 | 0.85169374 |

| | | | | | | |
|---|---|---|---|---|---|---|
| ANXA2 | 29.5542285 | -3.434730939 | 0.50262366 | -6.8336038 | 8.28E-12 | 2.10E-08 |
| ICE2 | 21.4966611 | 0.408295003 | 0.36098983 | 1.13104294 | 0.25803702 | 0.94557817 |
| RORA-AS1 | 12.1075006 | 0.114178607 | 0.79058648 | 0.14442266 | 0.88516672 | 0.99665574 |
| RORA | 655.045979 | 0.703187397 | 0.584683 | 1.20268144 | 0.22909962 | 0.92719039 |
| AC012404.1 | 20.8067299 | -0.483146879 | 0.86005419 | -0.5617633 | 0.5742773 | 0.99665574 |
| AC009554.1 | 9.3282761 | -2.405358488 | 0.65296822 | -3.6837298 | 0.00022985 | 0.03640903 |
| VPS13C | 72.7228439 | -0.642367712 | 0.30723245 | -2.0908199 | 0.03654421 | 0.58591936 |
| TLN2 | 13.5537871 | 0.062037196 | 0.59508139 | 0.10424993 | 0.91697101 | 0.99777713 |
| TPM1 | 48.6683799 | -0.382301835 | 0.34161851 | -1.1190899 | 0.26310178 | 0.94557817 |
| LACTB | 8.69749949 | -0.510534594 | 0.54420592 | -0.9381276 | 0.34817884 | 0.97838306 |
| RPS27L | 12.1292527 | -0.391070578 | 0.53393869 | -0.732426 | 0.4639086 | 0.99617422 |
| RAB8B | 11.8505683 | 0.154904639 | 0.48349741 | 0.32038359 | 0.74867756 | 0.99665574 |
| APH1B | 8.69868533 | -0.198011163 | 0.5058367 | -0.3914527 | 0.69546261 | 0.99665574 |
| CA12 | 1258.06054 | 0.224947421 | 0.35519101 | 0.63331395 | 0.52652865 | 0.99665574 |
| LINC02568 | 25.9665656 | 0.326578997 | 0.79137124 | 0.41267484 | 0.67984486 | 0.99665574 |
| USP3 | 112.222001 | -0.0172733 | 0.35472822 | -0.0486945 | 0.96116279 | 0.99777713 |
| USP3-AS1 | 27.9391791 | -0.373740908 | 0.46683563 | -0.8005835 | 0.4233728 | 0.99518711 |
| HERC1 | 93.275961 | 0.133533318 | 0.24827896 | 0.53783582 | 0.5906904 | 0.99665574 |
| DAPK2 | 8.63911317 | -0.436789381 | 0.9109697 | -0.4794774 | 0.63159904 | 0.99665574 |
| FAM96A | 13.715605 | -0.267616673 | 0.42836827 | -0.624735 | 0.53214497 | 0.99665574 |
| SNX1 | 23.520729 | -0.145845712 | 0.34583415 | -0.4217215 | 0.67322828 | 0.99665574 |
| PPIB | 30.816745 | -0.479177217 | 0.44276453 | -1.0822394 | 0.27914616 | 0.95980166 |
| CSNK1G1 | 7.01242885 | 0.09269802 | 0.57082795 | 0.16239222 | 0.87099699 | 0.99665574 |
| TRIP4 | 16.3275722 | -0.468228698 | 0.39446374 | -1.1870006 | 0.23522738 | 0.92960234 |
| ZNF609 | 84.8913856 | 0.163344972 | 0.26607601 | 0.61390341 | 0.53927916 | 0.99665574 |
| OAZ2 | 22.4620538 | -0.411666815 | 0.50002979 | -0.8232846 | 0.41034619 | 0.99279547 |
| RBPMS2 | 5.85215496 | -0.028601406 | 0.74300454 | -0.0384943 | 0.96929361 | 0.99777713 |
| SPG21 | 12.8522525 | 0.185296501 | 0.55096659 | 0.33631168 | 0.73663584 | 0.99665574 |
| MTFMT | 8.85518456 | 0.310251036 | 0.48296322 | 0.64239061 | 0.5206196 | 0.99665574 |
| UBAP1L | 6.18873259 | 0.06117096 | 0.64701699 | 0.09454305 | 0.92467779 | 0.99777713 |
| PDCD7 | 6.43493193 | -0.133458945 | 0.56257578 | -0.2372284 | 0.81247962 | 0.99665574 |
| CLPX | 20.3085327 | 0.226166529 | 0.42660751 | 0.5301513 | 0.59600703 | 0.99665574 |
| PARP16 | 4.17714425 | -0.465695557 | 0.65110156 | -0.7152426 | 0.47445916 | 0.99617422 |
| DPP8 | 26.4859201 | -0.094423427 | 0.33158066 | -0.2847676 | 0.7758222 | 0.99665574 |
| HACD3 | 31.0689346 | 0.050740996 | 0.30476239 | 0.16649363 | 0.8677685 | 0.99665574 |
| INTS14 | 6.89297326 | 0.106945901 | 0.54409377 | 0.19655785 | 0.84417356 | 0.99665574 |
| SLC24A1 | 5.41447475 | -0.842387852 | 0.74545361 | -1.1300339 | 0.25846194 | 0.94557817 |
| DENND4A | 78.0402124 | -0.117945837 | 0.29618068 | -0.3982226 | 0.69046611 | 0.99665574 |
| RAB11A | 41.9601214 | -0.386387816 | 0.40615181 | -0.9513384 | 0.34143261 | 0.97788427 |
| DIS3L | 6.26178252 | -0.156721437 | 0.60153631 | -0.2605353 | 0.7944509 | 0.99665574 |
| TIPIN | 5.2835358 | 0.053315439 | 0.66551678 | 0.08011134 | 0.93614871 | 0.99777713 |
| MAP2K1 | 19.8919176 | -0.007042004 | 0.44004461 | -0.0160029 | 0.98723205 | 0.99777713 |
| SNAPC5 | 4.31644902 | 0.301687203 | 0.733109 | 0.41151753 | 0.68069309 | 0.99665574 |
| RPL4 | 63.9849929 | 0.138717784 | 0.34841287 | 0.39814196 | 0.69052554 | 0.99665574 |
| ZWILCH | 2.57921858 | -1.452086135 | 0.90322243 | -1.6076728 | 0.10790686 | 0.80026315 |
| SMAD6 | 7.24026862 | 0.178945939 | 0.70315137 | 0.25449135 | 0.79911599 | 0.99665574 |
| SMAD3 | 57.6271984 | -0.404579164 | 0.32374262 | -1.2496939 | 0.2114114 | 0.91043197 |
| AAGAB | 24.0760671 | 0.189621093 | 0.39760259 | 0.47691111 | 0.63342542 | 0.99665574 |
| IQCH | 9.68095092 | -0.165212813 | 0.60601348 | -0.2726223 | 0.78514354 | 0.99665574 |
| IQCH-AS1 | 16.7380585 | -0.070730304 | 0.64192027 | -0.1101855 | 0.91226226 | 0.99777713 |
| C15orf61 | 4.0355904 | 0.588203272 | 0.80973153 | 0.72641764 | 0.46758278 | 0.99617422 |
| MAP2K5 | 40.821765 | 0.079497232 | 0.35552129 | 0.22360751 | 0.82306272 | 0.99665574 |
| PIAS1 | 80.1349871 | 0.315786942 | 0.24760279 | 1.27537716 | 0.20217578 | 0.90614933 |
| FEM1B | 10.6471225 | 0.172829138 | 0.55206851 | 0.31305741 | 0.75423705 | 0.99665574 |
| CORO2B | 18.951075 | 0.028946941 | 0.67668273 | 0.04277772 | 0.96587873 | 0.99777713 |
| ANP32A | 20.4976043 | 0.187667635 | 0.43026662 | 0.43616592 | 0.66271635 | 0.99665574 |
| SPESP1 | 3.00278681 | -0.608706836 | 0.88652038 | -0.6866248 | 0.49231923 | 0.99640678 |
| GLCE | 11.321494 | 0.513473663 | 0.50474903 | 1.01728509 | 0.30901783 | 0.97376371 |
| PAQR5 | 22.5999779 | -1.144469956 | 0.46891228 | -2.440691 | 0.01465919 | 0.40872075 |
| RPLP1 | 114.631332 | 0.736678279 | 0.38212614 | 1.92784059 | 0.05387496 | 0.6653339 |
| TLE3 | 11.0111744 | -1.379765891 | 0.4476222 | -3.082434 | 0.00205315 | 0.13784675 |
| UACA | 47.2498648 | 0.086075924 | 0.46974884 | 0.18323818 | 0.85461114 | 0.99665574 |
| LRRC49 | 4.98990628 | -0.363995348 | 0.673601 | -0.5403723 | 0.58894028 | 0.99665574 |
| THSD4 | 15.6397554 | -0.870274802 | 0.60038423 | -1.4495298 | 0.1471897 | 0.85282092 |
| THSD4-AS1 | 18.0242755 | 0.157965007 | 0.41596286 | 0.37975747 | 0.70412545 | 0.99665574 |
| MYO9A | 136.146796 | -0.42290137 | 0.31934669 | -1.3242704 | 0.18541325 | 0.89532394 |
| AC022872.1 | 5.08934817 | -0.65739962 | 0.97318828 | -0.6755112 | 0.49935102 | 0.99640678 |
| SENP8 | 2.81594653 | -1.197864886 | 1.00481561 | -1.1921241 | 0.23321259 | 0.92924083 |
| PKM | 74.4400537 | -0.650837751 | 0.36331046 | -1.7914093 | 0.07322763 | 0.72782524 |
| PARP6 | 12.5328157 | -0.664995799 | 0.45640585 | -1.4570273 | 0.14510884 | 0.85059132 |
| AC009690.2 | 7.52174855 | -1.516067458 | 0.69300648 | -2.1876671 | 0.02869386 | 0.54990241 |
| AC009690.1 | 29.4344183 | -0.526172106 | 0.60959608 | -0.8631488 | 0.38805568 | 0.99168679 |
| ARIH1 | 111.605843 | -0.243508558 | 0.33074615 | -0.73624 | 0.46158463 | 0.99617422 |
| BBS4 | 10.7858756 | 0.5999612 | 0.61878309 | 0.96958242 | 0.33225468 | 0.97788427 |
| ADPGK | 13.6728672 | -0.233852835 | 0.4005453 | -0.5838362 | 0.55933054 | 0.99665574 |
| NEO1 | 42.5491785 | -0.604573929 | 0.45682166 | -1.3234353 | 0.18569064 | 0.89532394 |
| NPTN | 49.7478566 | 0.382931474 | 0.28994937 | 1.32068393 | 0.18660678 | 0.89532394 |
| CD276 | 3.99404397 | -0.427151751 | 0.81492016 | -0.5241639 | 0.60016453 | 0.99665574 |
| PML | 4.9632279 | -0.398827252 | 0.8644042 | -0.4613898 | 0.64451899 | 0.99665574 |
| ARID3B | 3.49209303 | 0.161417248 | 0.75246983 | 0.21451657 | 0.83014425 | 0.99665574 |
| CLK3 | 5.52011678 | 0.476236924 | 0.57769236 | 0.82437809 | 0.40972476 | 0.99279547 |
| EDC3 | 11.2177361 | 0.227911046 | 0.47898146 | 0.47582436 | 0.63419951 | 0.99665574 |
| CSK | 3.23886882 | 0.609760775 | 1.04344062 | 0.58437515 | 0.55896794 | 0.99665574 |
| ULK3 | 3.79829667 | 0.988041691 | 0.82331945 | 1.20007088 | 0.23011181 | 0.92774608 |
| SCAMP2 | 14.0912088 | 0.578146029 | 0.49015776 | 1.17951009 | 0.23819512 | 0.93059916 |
| FAM219B | 4.72818068 | -0.313935544 | 0.6229742 | -0.5039302 | 0.6143104 | 0.99665574 |
| COX5A | 31.7910884 | 0.141776144 | 0.43192672 | 0.3282412 | 0.7427293 | 0.99665574 |
| SCAMP5 | 4.20465295 | -0.906515541 | 0.72229676 | -1.2550458 | 0.20946212 | 0.91043197 |
| PPCDC | 7.85925112 | 0.131356184 | 0.56415891 | 0.23283543 | 0.8158892 | 0.99665574 |
| COMMD4 | 9.50758701 | 0.164149422 | 0.47411897 | 0.3462199 | 0.72917746 | 0.99665574 |
| NEIL1 | 4.33628022 | 0.285448658 | 0.77840434 | 0.36671 | 0.71383535 | 0.99665574 |
| MAN2C1 | 8.93122431 | -0.006212973 | 0.55136718 | -0.0112683 | 0.99100939 | 0.99777713 |
| SIN3A | 20.5634374 | 0.126112583 | 0.38159582 | 0.33048733 | 0.74103176 | 0.99665574 |
| PTPN9 | 35.8107336 | -0.132644146 | 0.29572446 | -0.4485397 | 0.65376377 | 0.99665574 |
| SNUPN | 3.75507319 | -0.182724709 | 0.72744816 | -0.2511859 | 0.8016704 | 0.99665574 |
| IMP3 | 6.93094789 | 0.078048161 | 0.5948359 | 0.13120957 | 0.89560953 | 0.99777713 |
| SNX33 | 3.38426969 | 0.164789196 | 0.76689928 | 0.21487724 | 0.82986303 | 0.99665574 |
| UBE2Q2 | 13.3439226 | -0.941716585 | 0.51878187 | -1.8152458 | 0.06948615 | 0.7094165 |
| FBXO22 | 13.9810569 | 0.299246937 | 0.43044874 | 0.69519762 | 0.48693147 | 0.99640678 |
| NRG4 | 9.88729605 | 0.610576056 | 0.722311 | 0.8453091 | 0.39793828 | 0.99179694 |
| ETFA | 29.4500384 | -0.07538863 | 0.29489932 | -0.2556419 | 0.79822734 | 0.99665574 |
| SCAPER | 81.6098589 | -0.012435354 | 0.34296172 | -0.0362587 | 0.97107606 | 0.99777713 |
| RCN2 | 18.6641184 | 0.380733705 | 0.39935176 | 0.95337932 | 0.34039792 | 0.97788427 |
| TSPAN3 | 21.2368939 | 0.572013454 | 0.44425949 | 1.28756608 | 0.19789705 | 0.90381759 |
| PEAK1 | 36.6579046 | 0.062165755 | 0.32206575 | 0.19302194 | 0.84694179 | 0.99665574 |
| HMG20A | 23.5442234 | 0.189694602 | 0.35831457 | 0.52940801 | 0.59652244 | 0.99665574 |
| LINGO1 | 165.352523 | 1.132147066 | 0.53822808 | 2.10347083 | 0.03542463 | 0.58349146 |
| TBC1D2B | 5.93226592 | -0.570507303 | 0.60347904 | -0.9453639 | 0.34447312 | 0.97788427 |
| IDH3A | 30.4086152 | 0.351498178 | 0.36287491 | 0.96864832 | 0.33272068 | 0.97788427 |
| DNAJA4 | 2.4893282 | 0.640413599 | 1.08622499 | 0.5895773 | 0.55547407 | 0.99665574 |
| WDR61 | 8.62182731 | 0.081166576 | 0.49792653 | 0.16300914 | 0.87051124 | 0.99665574 |
| IREB2 | 28.0972928 | -0.048879375 | 0.3470467 | -0.1408438 | 0.88799334 | 0.99665574 |
| PSMA4 | 15.1036874 | 0.499816062 | 0.42172153 | 1.18518034 | 0.23594615 | 0.9305595 |

| | | | | | | |
|---|---|---|---|---|---|---|
| MORF4L1 | 36.7784509 | -0.054311864 | 0.27701046 | -0.1960643 | 0.84455983 | 0.99665574 |
| CTSH | 14.7033489 | 0.195323385 | 0.58638774 | 0.33309596 | 0.73906185 | 0.99665574 |
| TMED3 | 6.44668773 | -1.453107235 | 0.75290202 | -1.9300084 | 0.05360579 | 0.6653339 |
| MTHFS | 1.96900976 | -0.671579992 | 0.96105109 | -0.6987974 | 0.48467866 | 0.99640678 |
| ZFAND6 | 72.4599877 | 0.438437961 | 0.30128625 | 1.45522062 | 0.14560819 | 0.85082182 |
| FAH | 3.1592045 | 0.460457469 | 0.8469096 | 0.5436914 | 0.58665385 | 0.99665574 |
| ARNT2 | 22.8003105 | -1.475279217 | 0.42485169 | -3.4724569 | 0.00051572 | 0.06224222 |
| ABHD17C | 10.0120857 | 0.188335042 | 0.54774746 | 0.34383554 | 0.73096997 | 0.99665574 |
| CEMIP | 7.08572387 | -1.834892811 | 0.78943722 | -2.3243049 | 0.02010916 | 0.46544896 |
| AC027808.1 | 5.0920007 | 0.296286021 | 0.62429206 | 0.47459521 | 0.63507552 | 0.99665574 |
| MESD | 15.6654685 | -0.01151708 | 0.49847863 | -0.0231045 | 0.98156695 | 0.99777713 |
| EFL1 | 15.9200706 | -0.373727462 | 0.3968107 | -0.9418281 | 0.34628066 | 0.97788427 |
| RPS17 | 5.86760742 | 0.993661033 | 0.74918807 | 1.32631722 | 0.18473465 | 0.89532394 |
| WHAMM | 8.59427164 | -0.085588135 | 0.55649143 | -0.1537996 | 0.87776778 | 0.99665574 |
| HOMER2 | 5.00158414 | 0.11353619 | 0.78078372 | 0.14541311 | 0.88438471 | 0.99665574 |
| C15orf40 | 7.15538724 | -0.38717608 | 0.60400722 | -0.6410123 | 0.52151467 | 0.99665574 |
| BTBD1 | 24.0118104 | 0.243653585 | 0.3379412 | 0.72099403 | 0.47091319 | 0.99617422 |
| HDGFL3 | 19.4222329 | -0.518911471 | 0.48414336 | -1.0718137 | 0.28380373 | 0.96059387 |
| ADAMTSL3 | 225.799816 | 0.583218212 | 0.63506679 | 0.91835729 | 0.35843184 | 0.9819332 |
| UBE2Q2L | 3.72126447 | 0.308632171 | 0.92800801 | 0.3325749 | 0.73945519 | 0.99665574 |
| AC048382.5 | 2.28743747 | -0.216029307 | 0.85968109 | -0.2512901 | 0.80158986 | 0.99665574 |
| ZSCAN2 | 2.90368861 | 0.370251941 | 0.8401026 | 0.44072229 | 0.65941406 | 0.99665574 |
| WDR73 | 11.5160689 | -0.348892737 | 0.48586609 | -0.7180841 | 0.47270541 | 0.99617422 |
| SEC11A | 47.2162506 | 0.018474571 | 0.28829643 | 0.06408186 | 0.94890505 | 0.99777713 |
| ZNF592 | 19.3030963 | -0.639218268 | 0.34384395 | -1.859036 | 0.06302204 | 0.68628329 |
| ALPK3 | 20.7083672 | -0.267963061 | 0.56603088 | -0.4734071 | 0.63592273 | 0.99665574 |
| SLC28A1 | 4.32154579 | -0.355943486 | 1.07805419 | -0.3301722 | 0.74126988 | 0.99665574 |
| PDE8A | 125.902068 | -0.17515493 | 0.42054438 | -0.4164957 | 0.67704731 | 0.99665574 |
| AKAP13 | 102.459404 | -0.02363151 | 0.28789796 | -0.0820829 | 0.93458076 | 0.99777713 |
| AC087286.2 | 4.3527347 | -0.192973101 | 1.00092763 | -0.1927943 | 0.8471201 | 0.99665574 |
| KLHL25 | 4.87150782 | -0.356836509 | 0.64950399 | -0.5493985 | 0.58273201 | 0.99665574 |
| MRPL46 | 4.7007058 | 0.970303187 | 0.89571595 | 1.08327109 | 0.27868811 | 0.95969433 |
| MRPS11 | 5.30077763 | 0.055082005 | 0.65245464 | 0.08442273 | 0.93272033 | 0.99777713 |
| DET1 | 5.09062156 | 0.872080128 | 0.69406604 | 1.25648004 | 0.20894198 | 0.91043197 |
| ABHD2 | 23.4824825 | 0.135815155 | 0.4400343 | 0.30864675 | 0.75759026 | 0.99665574 |
| POLG | 5.64396157 | 0.127234993 | 0.57746588 | 0.22033335 | 0.82561155 | 0.99665574 |
| RHCG | 3.81108466 | 3.80889239 | 1.19085154 | 3.19846115 | 0.00138163 | 0.10938814 |
| ANPEP | 6.49042969 | -0.200840285 | 0.89918963 | -0.223357 | 0.82325768 | 0.99665574 |
| ZNF710 | 7.26123422 | -0.828433604 | 0.62692324 | -1.3214275 | 0.18635887 | 0.89532394 |
| ZNF710-AS1 | 3.94429281 | 0.007829539 | 0.7202112 | 0.01087117 | 0.99132623 | 0.99777713 |
| IDH2 | 42.9854153 | -0.229253306 | 0.43114223 | -0.5317348 | 0.59490972 | 0.99665574 |
| SEMA4B | 26.5230844 | -1.355877494 | 0.39420407 | -3.439532 | 0.00058272 | 0.06631714 |
| CIB1 | 28.469699 | 0.142598141 | 0.47173693 | 0.30228318 | 0.7624362 | 0.99665574 |
| AC091167.6 | 2.55554503 | -0.084032648 | 0.88881528 | -0.0945446 | 0.92467659 | 0.99777713 |
| IQGAP1 | 68.0905195 | -0.809313503 | 0.28504887 | -2.8392096 | 0.00452254 | 0.21492549 |
| CRTC3 | 43.7862833 | -0.346426772 | 0.33328032 | -1.0394456 | 0.29859753 | 0.97358745 |
| CRTC3-AS1 | 3.64281042 | -0.283180143 | 0.78758614 | -0.3595545 | 0.71918031 | 0.99665574 |
| FURIN | 7.32149359 | -0.333707785 | 0.54492362 | -0.6123937 | 0.54027732 | 0.99665574 |
| MAN2A2 | 4.06403378 | -0.073653608 | 0.78325315 | -0.0940355 | 0.92508095 | 0.99777713 |
| HDDC3 | 3.47753345 | -0.362284409 | 0.9903305 | -0.3658217 | 0.71449811 | 0.99665574 |
| UNC45A | 7.25855591 | 0.14814109 | 0.61789965 | 0.23974943 | 0.81052451 | 0.99665574 |
| RCCD1 | 2.46295379 | 0.201443048 | 0.872461 | 0.2308906 | 0.81739979 | 0.99665574 |
| AC104035.1 | 10.568872 | 0.547011442 | 0.56769135 | 0.96357191 | 0.3352606 | 0.97788427 |
| SLCO3A1 | 11.7904897 | -0.049138257 | 0.70458195 | -0.069741 | 0.9443998 | 0.99777713 |
| FAM174B | 5.56990968 | -0.865704962 | 0.68870518 | -1.2570037 | 0.20875229 | 0.91043197 |
| AC013394.1 | 37.116916 | 0.012569062 | 0.28457332 | 0.0441681 | 0.96477041 | 0.99777713 |
| CHD2 | 74.9788777 | -0.523694372 | 0.37106712 | -1.4113198 | 0.15815034 | 0.86661434 |
| MCTP2 | 5.06098951 | -2.89691645 | 0.74852385 | -3.8701726 | 0.00010876 | 0.02396939 |
| NR2F2-AS1 | 20.9586555 | -0.149219986 | 0.65726178 | -0.2270328 | 0.82039823 | 0.99665574 |
| NR2F2 | 9.9258957 | 0.103113063 | 0.69375222 | 0.14863097 | 0.88184484 | 0.99665574 |
| ARRDC4 | 7.3159417 | 0.283722959 | 0.86352359 | 0.32856422 | 0.74248509 | 0.99665574 |
| IGF1R | 91.6147817 | 0.088507256 | 0.34650174 | 0.25543092 | 0.79839029 | 0.99665574 |
| TTC23 | 11.8005879 | -0.810929017 | 0.57389699 | -1.4130219 | 0.15764928 | 0.86532131 |
| LRRC28 | 22.386505 | 0.1047696 | 0.45540367 | 0.23005875 | 0.81804612 | 0.99665574 |
| MEF2A | 129.992172 | 0.250469363 | 0.25476006 | 0.9831579 | 0.32552973 | 0.97788427 |
| AC090825.1 | 5.62299074 | 0.33194552 | 0.76828758 | 0.43205894 | 0.66569857 | 0.99665574 |
| ADAMTS17 | 8.20088192 | -1.254710245 | 0.93553191 | -1.3411731 | 0.17986425 | 0.88823393 |
| LINS1 | 5.91171043 | 0.607131721 | 0.56885935 | 1.06727915 | 0.2858458 | 0.96436098 |
| ASB7 | 9.40699717 | 0.123349434 | 0.51910995 | 0.23761716 | 0.81217804 | 0.99665574 |
| ALDH1A3 | 9.52950813 | 0.286599935 | 0.61158188 | 0.46862071 | 0.63934077 | 0.99665574 |
| CHSY1 | 17.657665 | -0.642293915 | 0.44713206 | -1.4364747 | 0.1508673 | 0.85612189 |
| SELENOS | 19.4157857 | -0.686048891 | 0.52503907 | -1.3066626 | 0.19132734 | 0.89964233 |
| SNRPA1 | 9.49873666 | 0.571772146 | 0.56767746 | 1.00721306 | 0.3138324 | 0.9755712 |
| TM2D3 | 12.7708974 | 0.281966944 | 0.47987259 | 0.5875871 | 0.55680947 | 0.99665574 |
| TARSL2 | 17.9957888 | -0.412327726 | 0.3848974 | -1.0712666 | 0.28404958 | 0.96059387 |
| SNRNP25 | 3.97947151 | 0.385282709 | 0.71887004 | 0.535956 | 0.59198896 | 0.99665574 |
| MPG | 8.27594994 | -0.88693904 | 0.5071651 | -1.7488172 | 0.08032263 | 0.74618467 |
| NPRL3 | 16.4391091 | -0.809586955 | 0.42545872 | -1.9028566 | 0.05705925 | 0.67545594 |
| LUC7L | 33.5918226 | 0.136413965 | 0.31605125 | 0.43161977 | 0.66601779 | 0.99665574 |
| FAM234A | 5.34528079 | 0.168484492 | 0.67378146 | 0.25005807 | 0.80254244 | 0.99665574 |
| AXIN1 | 7.6384459 | -0.153734466 | 0.56647517 | -0.2713878 | 0.78609276 | 0.99665574 |
| MRPL28 | 5.48581508 | 0.233739467 | 0.77530837 | 0.30147935 | 0.76304899 | 0.99665574 |
| TMEM8A | 16.3773932 | -0.143455634 | 0.38476065 | -0.3728438 | 0.70926468 | 0.99665574 |
| NME4 | 3.84881077 | -0.241445557 | 0.82155115 | -0.2938899 | 0.76884206 | 0.99665574 |
| DECR2 | 2.53098619 | 0.675748992 | 0.83616615 | 0.80815157 | 0.41900333 | 0.99476033 |
| RAB11FIP3 | 36.3337215 | 0.181707592 | 0.35308478 | 0.51462878 | 0.60681245 | 0.99665574 |
| CAPN15 | 9.05281203 | -0.690873917 | 0.54656294 | -1.2640336 | 0.20621797 | 0.90890506 |
| PIGQ | 8.21656678 | 0.855176911 | 0.60830226 | 1.40584208 | 0.15977102 | 0.86757292 |
| RAB40C | 18.040226 | -0.162443236 | 0.39818123 | -0.4079631 | 0.68330079 | 0.99665574 |
| METTL26 | 4.16072353 | 0.568791752 | 0.77359439 | 0.73525837 | 0.46218216 | 0.99617422 |
| MCRIP2 | 20.5864358 | 0.766625476 | 0.54694003 | 1.40166276 | 0.16101598 | 0.86928072 |
| RHOT2 | 9.54311912 | -0.081087253 | 0.55347832 | -0.1465048 | 0.88352287 | 0.99665574 |
| STUB1 | 14.3563084 | -0.032754862 | 0.51516092 | -0.0635818 | 0.94930322 | 0.99777713 |
| JMJD8 | 4.83823485 | -0.130171587 | 0.78826543 | -0.1651367 | 0.86883635 | 0.99665574 |
| METRN | 9.51562253 | -0.440769701 | 0.60296878 | -0.7309992 | 0.46477963 | 0.99617422 |
| FAM173A | 10.1584312 | -0.149879998 | 0.64329473 | -0.2329881 | 0.81577067 | 0.99665574 |
| HAGHL | 5.45836933 | 0.672999832 | 0.69285942 | 0.97133678 | 0.3313806 | 0.97788427 |
| NARFL | 4.51912796 | 0.090300206 | 0.63277848 | 0.14270429 | 0.88652373 | 0.99665574 |
| LMF1 | 13.9439589 | -0.44218918 | 0.45218538 | -0.9778936 | 0.32812696 | 0.97788427 |
| UBE2I | 24.5070759 | -0.636471523 | 0.37012762 | -1.7196002 | 0.08550515 | 0.75373899 |
| TSR3 | 8.88152243 | -0.273934934 | 0.54158598 | -0.5058014 | 0.61299609 | 0.99665574 |
| GNPTG | 8.6405264 | -0.253427151 | 0.66876429 | -0.3789484 | 0.70472619 | 0.99665574 |
| UNKL | 4.68053366 | -0.532437394 | 0.61877223 | -0.860474 | 0.38952783 | 0.99179694 |
| CLCN7 | 9.91095469 | -0.660041535 | 0.64600236 | -1.0217324 | 0.30690758 | 0.97376371 |
| TELO2 | 3.71253128 | 0.706583971 | 0.74794289 | 0.9447031 | 0.34481047 | 0.97788427 |
| IFT140 | 6.38809688 | 0.170618165 | 0.56458362 | 0.30220176 | 0.76249826 | 0.99665574 |
| JPT2 | 10.0656811 | -0.439289018 | 0.47186753 | -0.9309584 | 0.35187511 | 0.97838306 |
| MAPK8IP3 | 19.5693426 | -0.513510862 | 0.34570534 | -1.4854004 | 0.13743778 | 0.84598921 |
| NME3 | 12.5222407 | 0.34440925 | 0.56681862 | 0.6076181 | 0.5434408 | 0.99665574 |
| MRPS34 | 10.4957271 | -0.389639558 | 0.46232263 | -0.8427871 | 0.39934751 | 0.99179694 |
| EME2 | 3.00548455 | 1.12882291 | 0.95531136 | 1.18162826 | 0.23735323 | 0.9305595 |
| SPSB3 | 12.6878763 | 0.22709859 | 0.44608255 | 0.50909544 | 0.61068533 | 0.99665574 |
| NUBP2 | 4.69281092 | 0.663368913 | 0.70615077 | 0.93941541 | 0.34751751 | 0.97804112 |

| | | | | | | |
|---|---|---|---|---|---|---|
| HAGH | 15.9512595 | -0.120753378 | 0.4445168 | -0.2716509 | 0.78589048 | 0.99665574 |
| FAHD1 | 5.13791037 | -1.193905495 | 0.70842752 | -1.6852895 | 0.09193272 | 0.76658136 |
| MSRB1 | 2.67882267 | -0.36531433 | 1.12051269 | -0.3260243 | 0.74440599 | 0.99665574 |
| NDUFB10 | 20.274287 | 0.447547299 | 0.42686049 | 1.04846268 | 0.29442549 | 0.97161335 |
| RPS2 | 65.9958499 | 0.248471579 | 0.43828557 | 0.56691709 | 0.57077051 | 0.99665574 |
| SNHG9 | 2.96200699 | 0.025091462 | 1.15586069 | 0.02170803 | 0.98268086 | 0.99777713 |
| GFER | 2.9544614 | 0.400059393 | 0.87018685 | 0.45973964 | 0.64570311 | 0.99665574 |
| ZNF598 | 4.49142962 | 0.320134105 | 0.77551998 | 0.4127993 | 0.67975367 | 0.99665574 |
| TSC2 | 12.2169007 | -0.728473791 | 0.46743832 | -1.5584383 | 0.11912937 | 0.82619588 |
| PKD1 | 7.90792417 | 0.156732138 | 0.49361356 | 0.31751992 | 0.75084913 | 0.99665574 |
| SNHG19 | 3.85413649 | -0.381158191 | 0.66223757 | -0.5755611 | 0.56491187 | 0.99665574 |
| TRAF7 | 6.07754668 | -0.255984096 | 0.69272273 | -0.3695333 | 0.71173028 | 0.99665574 |
| MLST8 | 2.33588567 | 0.668422902 | 1.10437437 | 0.60525028 | 0.54501271 | 0.99665574 |
| PGP | 10.3535462 | -0.268525781 | 0.4970183 | -0.5402734 | 0.58900848 | 0.99665574 |
| E4F1 | 6.11670169 | 0.093786763 | 0.72566749 | 0.12924206 | 0.89716611 | 0.99777713 |
| ECI1 | 34.4825545 | 0.091245452 | 0.48683717 | 0.18742499 | 0.85132743 | 0.99665574 |
| RNPS1 | 17.6423173 | -0.19171007 | 0.42700606 | -0.4489633 | 0.6534581 | 0.99665574 |
| ABCA3 | 4.44090674 | -2.049537198 | 0.72814654 | -2.8147318 | 0.0048818 | 0.22094485 |
| AMDHD2 | 4.01356865 | -0.356540818 | 1.18107642 | -0.3018779 | 0.76274518 | 0.99665574 |
| PDPK1 | 32.6652275 | -0.783664211 | 0.40526698 | -1.9336986 | 0.05315019 | 0.66476617 |
| AC093525.4 | 2.78606012 | -0.753230695 | 0.8043911 | -0.9363986 | 0.349068 | 0.97838306 |
| KCTD5 | 3.04630776 | -0.759262364 | 0.87948051 | -0.8633078 | 0.38796827 | 0.99168679 |
| SRRM2 | 75.5759075 | -0.134772943 | 0.33409617 | -0.4033957 | 0.68665718 | 0.99665574 |
| ELOB | 18.433625 | 0.018929696 | 0.48230401 | 0.03924847 | 0.96869229 | 0.99777713 |
| FLYWCH2 | 5.32635429 | -0.559996499 | 0.70165038 | -0.7981133 | 0.42480474 | 0.99518711 |
| FLYWCH1 | 9.46879343 | 0.236046355 | 0.46364656 | 0.50910839 | 0.61067625 | 0.99665574 |
| TNFRSF12A | 8.16585685 | -0.558885781 | 0.94763549 | -0.5897687 | 0.55534571 | 0.99665574 |
| HCFC1R1 | 6.72174578 | 0.259149312 | 0.83318664 | 0.31103393 | 0.75577483 | 0.99665574 |
| BICDL2 | 3.05430239 | 0.36761964 | 0.91627724 | 0.40121005 | 0.68826548 | 0.99665574 |
| IL32 | 5.36377844 | -1.872190982 | 0.79814879 | -2.3456666 | 0.01899309 | 0.45540143 |
| ZNF213-AS1 | 3.63482965 | -0.006403575 | 0.69993508 | -0.0091488 | 0.9927004 | 0.99777713 |
| ZNF205 | 2.87385342 | 0.101441679 | 0.80905379 | 0.12538311 | 0.90022026 | 0.99777713 |
| ZNF213 | 8.1203409 | -0.605619816 | 0.5437665 | -1.1137498 | 0.26538655 | 0.94557817 |
| ZNF263 | 5.61397062 | -0.068240846 | 0.62247135 | -0.1096289 | 0.91270369 | 0.99777713 |
| TIGD7 | 15.4007224 | -2.603104462 | 1.43912272 | -1.8088134 | 0.07048 | 0.71264656 |
| ZNF75A | 25.927475 | -0.584177992 | 0.66056846 | -0.8843565 | 0.37650381 | 0.98679664 |
| AC025283.3 | 5.53190208 | -0.122977303 | 0.64803459 | -0.1897697 | 0.84948963 | 0.99665574 |
| ZSCAN32 | 3.83579169 | -0.631347636 | 0.7243975 | -0.8715486 | 0.38345467 | 0.98955375 |
| NAA60 | 6.53728989 | 0.196903779 | 0.55874742 | 0.35240213 | 0.72453671 | 0.99665574 |
| CLUAP1 | 8.80279773 | 0.576920314 | 0.56203506 | 1.02648457 | 0.30466323 | 0.97376371 |
| SLX4 | 2.51921408 | 0.557298341 | 0.86841512 | 0.64174187 | 0.5210408 | 0.99665574 |
| DNASE1 | 23.0135445 | 0.170331537 | 0.47149235 | 0.36126045 | 0.71790476 | 0.99665574 |
| TRAP1 | 14.9533372 | 0.125859549 | 0.41754196 | 0.3014297 | 0.76308685 | 0.99665574 |
| CREBBP | 58.7822851 | -0.148858589 | 0.34245974 | -0.4346747 | 0.66379856 | 0.99665574 |
| ADCY9 | 23.7986715 | -0.440647437 | 0.40083674 | -1.099319 | 0.27162895 | 0.94892291 |
| GLIS2 | 9.98313983 | 0.414395024 | 0.57634467 | 0.71900556 | 0.47213749 | 0.99617422 |
| VASN | 11.5838651 | -0.226095284 | 0.48487176 | -0.4662991 | 0.64100139 | 0.99665574 |
| DNAJA3 | 8.23597213 | -0.100386639 | 0.51299679 | -0.1956867 | 0.84485541 | 0.99665574 |
| NMRAL1 | 8.23515279 | 0.386795186 | 0.55625984 | 0.69534983 | 0.48683609 | 0.99640678 |
| HMOX2 | 12.1219509 | -0.331097907 | 0.45217495 | -0.7322341 | 0.4640257 | 0.99617422 |
| CDIP1 | 8.25865132 | 0.152467055 | 0.69485289 | 0.2194235 | 0.82632016 | 0.99665574 |
| UBALD1 | 4.16255738 | 0.61656883 | 0.78447954 | 0.7859591 | 0.43189145 | 0.99617422 |
| MGRN1 | 20.0971769 | -0.313982409 | 0.4430298 | -0.7087162 | 0.4785006 | 0.99617422 |
| NUDT16L1 | 7.47529387 | 0.531304383 | 0.66111561 | 0.80364822 | 0.42160016 | 0.99518711 |
| ANKS3 | 14.4889292 | -0.26227678 | 0.43087714 | -0.6087043 | 0.54272043 | 0.99665574 |
| ZNF500 | 4.23941936 | 0.600070579 | 0.66256813 | 0.90567378 | 0.36510855 | 0.9836488 |
| SMIM22 | 4.76733574 | -0.102118422 | 0.70522447 | -0.1448027 | 0.88486663 | 0.99665574 |
| ROGDI | 3.44942827 | 0.796788226 | 0.8949785 | 0.89028756 | 0.3733115 | 0.98532838 |
| GLYR1 | 34.3010049 | -0.279022331 | 0.37012352 | -0.7538627 | 0.45093165 | 0.99617422 |
| UBN1 | 12.5637259 | -0.606785336 | 0.6486607 | -0.9354433 | 0.34955987 | 0.97838306 |
| PPL | 11.0374652 | -0.552628183 | 0.53220973 | -1.0383654 | 0.29909996 | 0.97358745 |
| NAGPA | 2.11785513 | 0.326521235 | 1.05887777 | 0.30836537 | 0.75780433 | 0.99665574 |
| ALG1 | 2.28315435 | -0.221770303 | 1.02590161 | -0.2161711 | 0.82885436 | 0.99665574 |
| RBFOX1 | 16.6657027 | -0.025286079 | 0.73909759 | -0.0342121 | 0.97270802 | 0.99777713 |
| METTL22 | 3.08531502 | -1.587253407 | 0.87906411 | -1.8056174 | 0.07097813 | 0.71510751 |
| ABAT | 5.57713857 | 0.432067894 | 0.67951678 | 0.63584581 | 0.52487693 | 0.99665574 |
| PMM2 | 18.3718378 | -1.403348916 | 0.59668763 | -2.3518988 | 0.01867786 | 0.45192386 |
| CARHSP1 | 10.7758588 | -1.156254727 | 0.52432283 | -2.2052344 | 0.02743765 | 0.53492871 |
| USP7 | 26.7184363 | 0.187435777 | 0.37148473 | 0.50455849 | 0.61386897 | 0.99665574 |
| C16orf72 | 23.3484474 | -0.058146428 | 0.39238615 | -0.1481867 | 0.88219539 | 0.99665574 |
| AC087190.1 | 2.66295453 | -0.910232382 | 1.18098016 | -0.7707432 | 0.44085918 | 0.99617422 |
| GRIN2A | 13.3481967 | 0.960641192 | 0.6896679 | 1.39290402 | 0.16364882 | 0.87273629 |
| ATF7IP2 | 35.0948975 | 0.070856588 | 0.49314435 | 0.14368326 | 0.88575059 | 0.99665574 |
| NUBP1 | 5.29935464 | -0.764150954 | 0.57590585 | -1.3268679 | 0.1845524 | 0.89532394 |
| DEXI | 6.8571782 | 0.46972116 | 0.65042877 | 0.72217156 | 0.47018901 | 0.99617422 |
| CLEC16A | 74.151106 | -0.769653788 | 0.26907599 | -2.8603585 | 0.00423162 | 0.20883915 |
| AC007014.1 | 5.06410491 | -2.379812331 | 0.7617851 | -3.1239943 | 0.00178414 | 0.1230449 |
| SOCS1 | 30.4452268 | -2.247469479 | 0.65974482 | -3.4065739 | 0.00065784 | 0.06805261 |
| AC099489.1 | 8.94343138 | -1.811955364 | 0.5573244 | -3.2511682 | 0.00114932 | 0.09547898 |
| LITAF | 161.188242 | -1.461273808 | 0.32796508 | -4.4555774 | 8.37E-06 | 0.00344882 |
| SNN | 4.8560686 | -1.117438379 | 0.81956301 | -1.3634563 | 0.17273874 | 0.88157681 |
| TXNDC11 | 32.6207569 | -0.284378255 | 0.40821603 | -0.6966367 | 0.4860302 | 0.99640678 |
| ZC3H7A | 27.2344355 | -0.307970819 | 0.31200611 | -0.9870666 | 0.32360997 | 0.97700345 |
| RSL1D1 | 9.06078886 | -0.528542305 | 0.51928261 | -1.0178317 | 0.30875794 | 0.97376371 |
| AC007216.4 | 14.8462949 | 0.154574152 | 0.57285354 | 0.26983189 | 0.78728959 | 0.99665574 |
| GSPT1 | 42.1632297 | 0.097717215 | 0.28931768 | 0.33775058 | 0.73555115 | 0.99665574 |
| SNX29 | 232.197837 | -0.636358458 | 0.38278918 | -1.6624254 | 0.09642751 | 0.7740159 |
| AC010333.2 | 9.90721199 | 0.933834048 | 0.6159985 | 1.51596806 | 0.12952743 | 0.83778831 |
| CPPED1 | 32.8171996 | -0.074358783 | 0.42803283 | -0.1737221 | 0.86208383 | 0.99665574 |
| ERCC4 | 3.51811447 | 0.300544756 | 0.76400965 | 0.39337822 | 0.69404015 | 0.99665574 |
| MKL2 | 59.2786134 | 0.14761527 | 0.2988223 | 0.49399014 | 0.62131315 | 0.99665574 |
| MIR193BHG | 7.70923084 | 0.16184918 | 0.7601854 | 0.21290751 | 0.83139911 | 0.99665574 |
| PARN | 33.714236 | 0.225996965 | 0.29336715 | 0.77035538 | 0.44108911 | 0.99617422 |
| BFAR | 15.2941177 | -0.448357077 | 0.44905876 | -0.9984374 | 0.31806729 | 0.97700345 |
| NOMO1 | 4.13513805 | 0.204917267 | 0.82995685 | 0.24690111 | 0.80498476 | 0.99665574 |
| PDXDC1 | 58.0755244 | -0.273260586 | 0.2848871 | -0.959189 | 0.33746352 | 0.97788427 |
| NTAN1 | 6.81293943 | -1.269935153 | 0.58482555 | -2.1714769 | 0.02989514 | 0.55886856 |
| RRN3 | 8.88178124 | -0.265843768 | 0.50090219 | -0.5307299 | 0.59560597 | 0.99665574 |
| C16orf45 | 16.602736 | -1.006804259 | 0.48731835 | -2.0660094 | 0.0388276 | 0.59913877 |
| MARF1 | 14.2959317 | 0.640550141 | 0.42643048 | 1.50212092 | 0.13306588 | 0.84156077 |
| NDE1 | 5.0176221 | -0.515615263 | 0.68481833 | -0.7529227 | 0.45149637 | 0.99617422 |
| FOPNL | 5.53278312 | -0.396827569 | 0.68867662 | -0.5762176 | 0.56446813 | 0.99665574 |
| ABCC1 | 16.0673145 | 0.373319619 | 0.43074269 | 0.86668823 | 0.38611285 | 0.99051797 |
| RPS15A | 11.070955 | 1.19786672 | 0.62902962 | 1.90430891 | 0.05686996 | 0.67545594 |
| SMG1 | 69.2138672 | 0.268309254 | 0.26632179 | 1.00746263 | 0.3137125 | 0.9755712 |
| COQ7 | 5.40327845 | -0.408467833 | 0.62738209 | -0.6510671 | 0.51500318 | 0.99665574 |
| GDE1 | 7.04959746 | -0.106319791 | 0.63223569 | -0.1681648 | 0.86645363 | 0.99665574 |
| CCP110 | 4.18103533 | -0.617774871 | 0.6773721 | -0.912017 | 0.36175977 | 0.9819332 |
| VPS35L | 30.2037909 | 0.135452917 | 0.39931246 | 0.33921535 | 0.73444751 | 0.99665574 |
| KNOP1 | 6.72842575 | -0.73654478 | 0.70504734 | -1.0446742 | 0.29617356 | 0.97286866 |
| IQCK | 30.2258047 | -0.192570053 | 0.41255519 | -0.466774 | 0.64066155 | 0.99665574 |
| GPRC5B | 38.5860943 | 0.011699513 | 0.41649025 | 0.02809073 | 0.97758979 | 0.99777713 |
| UMOD | 41.4180609 | 2.264379999 | 0.87425546 | 2.59006674 | 0.00959573 | 0.33778312 |

| | | | | | | |
|---|---|---|---|---|---|---|
| ACSM2A | 8.09830714 | 1.722354539 | 0.95086618 | 1.81135325 | 0.07008619 | 0.71124502 |
| ACSM2B | 10.8241269 | 1.374919932 | 1.42324024 | 0.96604909 | 0.33401962 | 0.97788427 |
| ACSM3 | 9.63548862 | -0.695447175 | 0.77264009 | -0.900092 | 0.36807128 | 0.98532838 |
| THUMPD1 | 18.5784889 | -0.047405411 | 0.40177985 | -0.1179885 | 0.90607675 | 0.99777713 |
| ERI2 | 4.74574314 | -0.607957585 | 0.70114835 | -0.8670884 | 0.38589359 | 0.99051797 |
| LYRM1 | 3.64023288 | -0.492061624 | 0.88536418 | -0.5557731 | 0.57836595 | 0.99665574 |
| METTL9 | 19.5619784 | 0.109021814 | 0.39706411 | 0.2745698 | 0.78364677 | 0.99665574 |
| NPIPB4 | 7.35064674 | 0.237125438 | 0.67011591 | 0.35385735 | 0.72344579 | 0.99665574 |
| UQCRC2 | 34.5015319 | 0.151770599 | 0.31560219 | 0.4808921 | 0.63059319 | 0.99665574 |
| MOSMO | 24.6060651 | -0.001389321 | 0.40567277 | -0.0034247 | 0.99726746 | 0.99884397 |
| EEF2K | 4.10376265 | -1.008997155 | 0.88551745 | -1.1394436 | 0.2545182 | 0.94099699 |
| POLR3E | 9.15778397 | 0.004803 | 0.50719867 | 0.00946966 | 0.99244442 | 0.99777713 |
| CDR2 | 36.3825828 | -0.040480333 | 0.33086224 | -0.122348 | 0.90262342 | 0.99777713 |
| NPIPB5 | 17.8121579 | -0.36617535 | 0.45927679 | -0.7972869 | 0.42528445 | 0.99518711 |
| USP31 | 11.4652639 | -0.844229803 | 0.57404509 | -1.4706681 | 0.1413809 | 0.84969914 |
| SCNN1G | 7.71755338 | 1.540943503 | 1.26575509 | 1.21741047 | 0.22344808 | 0.92191316 |
| SCNN1B | 4.64859425 | 1.739925508 | 1.08181225 | 1.60834332 | 0.10776001 | 0.80026315 |
| COG7 | 14.6134975 | -0.206516831 | 0.46182752 | -0.4471731 | 0.6547501 | 0.99665574 |
| GGA2 | 23.5742773 | -0.433757716 | 0.48812605 | -0.8886183 | 0.37420828 | 0.98535336 |
| EARS2 | 4.94012473 | -0.34385352 | 0.65978818 | -0.5211574 | 0.6022571 | 0.99665574 |
| UBFD1 | 7.12316996 | 0.353786764 | 0.61128714 | 0.57875709 | 0.56275309 | 0.99665574 |
| NDUFAB1 | 16.034836 | 0.901975689 | 0.49020064 | 1.84001327 | 0.06576629 | 0.69235581 |
| PALB2 | 4.09989327 | -0.680159679 | 0.67794425 | -1.0032679 | 0.31573164 | 0.97652251 |
| DCTN5 | 9.77921055 | 0.031673321 | 0.72581799 | 0.0436381 | 0.96519288 | 0.99777713 |
| RBBP6 | 19.1413577 | 0.123811707 | 0.51350931 | 0.24110898 | 0.80947065 | 0.99665574 |
| TNRC6A | 78.8450755 | 0.110827785 | 0.31774781 | 0.34879166 | 0.72724572 | 0.99665574 |
| ARHGAP17 | 16.8179696 | -0.356183461 | 0.40840601 | -0.8721308 | 0.38313702 | 0.98947027 |
| LCMT1 | 14.1731254 | 0.423083715 | 0.46279184 | 0.91419874 | 0.36061243 | 0.9819332 |
| NSMCE1 | 7.96193978 | 0.287604285 | 0.53368349 | 0.53890422 | 0.58995295 | 0.99665574 |
| GTF3C1 | 10.3087737 | -0.287353202 | 0.54238502 | -0.5297956 | 0.59625365 | 0.99665574 |
| KIAA0556 | 13.5038367 | -0.591117916 | 0.53219573 | -1.1107153 | 0.26669093 | 0.94557817 |
| XPO6 | 21.4123408 | -0.764470316 | 0.46013377 | -1.6614089 | 0.09663136 | 0.77503855 |
| NUPR1 | 7.45049633 | -2.24309192 | 1.13635984 | -1.9739275 | 0.04838999 | 0.64337675 |
| SGF29 | 3.98675509 | 0.116358052 | 0.67769694 | 0.17169629 | 0.8636763 | 0.99665574 |
| SULT1A1 | 3.70408641 | -0.207478374 | 0.82101387 | -0.25271 | 0.80049236 | 0.99665574 |
| ATXN2L | 12.0584097 | -0.288402433 | 0.44453341 | -0.6487756 | 0.51648342 | 0.99665574 |
| TUFM | 31.8992505 | 0.143503934 | 0.46748442 | 0.30697052 | 0.75886581 | 0.99665574 |
| SH2B1 | 19.758015 | 0.688415505 | 0.4208734 | 1.63568308 | 0.10190593 | 0.78445129 |
| RABEP2 | 11.9292764 | -0.216383909 | 0.42153324 | -0.5133258 | 0.6077234 | 0.99665574 |
| NFATC2IP | 20.9006524 | 0.169791232 | 0.52062326 | 0.3261307 | 0.74432547 | 0.99665574 |
| SPNS1 | 5.99241705 | -0.678273682 | 0.97749376 | -0.6938905 | 0.48775085 | 0.99640678 |
| QPRT | 2.39070576 | 0.085120477 | 1.2695677 | 0.06704682 | 0.94654443 | 0.99777713 |
| KIF22 | 2.72701587 | 0.936830845 | 0.89256302 | 1.0495963 | 0.29390376 | 0.97150189 |
| MAZ | 5.60010488 | -0.122020848 | 0.61571663 | -0.198177 | 0.84290661 | 0.99665574 |
| AC009133.1 | 6.14765438 | -1.318025562 | 0.70322375 | -1.874262 | 0.06089432 | 0.6834742 |
| MVP | 22.9094968 | -1.397776738 | 0.43987901 | -3.1776391 | 0.00148479 | 0.11233469 |
| CDIPT | 7.92624097 | 0.467307596 | 0.54546119 | 0.85672015 | 0.39159957 | 0.99179694 |
| KCTD13 | 9.34219564 | -0.526025201 | 0.52559766 | -1.0008134 | 0.31691701 | 0.97700345 |
| TMEM219 | 11.386779 | 0.721438396 | 0.50103233 | 1.43990388 | 0.1498946 | 0.85525834 |
| TAOK2 | 4.86817902 | 0.655931783 | 0.67807249 | 0.96734758 | 0.3333703 | 0.97788427 |
| INO80E | 6.63242128 | 0.321144909 | 0.64707828 | 0.49629994 | 0.61968281 | 0.99665574 |
| PPP4C | 13.6935247 | -0.307155395 | 0.457062 | -0.6720213 | 0.50157014 | 0.99640678 |
| YPEL3 | 11.685099 | 0.22683644 | 0.52960427 | 0.42831309 | 0.66842319 | 0.99665574 |
| MAPK3 | 2.81659356 | 0.247711446 | 0.97574996 | 0.25386775 | 0.79959773 | 0.99665574 |
| NPIPB13 | 6.14737399 | -0.309109622 | 0.80487928 | -0.3840447 | 0.70094532 | 0.99665574 |
| CD2BP2 | 6.41094338 | -0.417101022 | 0.59961576 | -0.6956138 | 0.4866707 | 0.99640678 |
| TBC1D10B | 4.77622597 | -0.166544034 | 0.63668332 | -0.2615806 | 0.79364477 | 0.99665574 |
| ZNF771 | 2.63737146 | -0.319834591 | 0.80326847 | -0.3981665 | 0.69050746 | 0.99665574 |
| DCTPP1 | 3.76949042 | -0.280902447 | 0.83714684 | -0.3355474 | 0.73721219 | 0.99665574 |
| SEPHS2 | 16.8285881 | -0.195559691 | 0.40983458 | -0.4771674 | 0.63324294 | 0.99665574 |
| ZNF768 | 7.14217234 | 0.689245108 | 0.64452036 | 1.0693923 | 0.28489294 | 0.96178641 |
| ZNF688 | 2.13075363 | 0.603164072 | 1.07883448 | 0.55908861 | 0.57610125 | 0.99665574 |
| ZNF785 | 4.97893197 | 0.378737661 | 0.75654193 | 0.50061688 | 0.61664078 | 0.99665574 |
| PRR14 | 4.71398313 | 0.079623913 | 0.65669819 | 0.12124887 | 0.90349392 | 0.99777713 |
| FBRS | 5.18024182 | 0.088206331 | 0.6084688 | 0.14496443 | 0.88473896 | 0.99665574 |
| PHKG2 | 4.73844464 | 0.493168221 | 0.70718231 | 0.6973707 | 0.48557083 | 0.99640678 |
| RNF40 | 8.61960688 | 1.117016566 | 0.65015399 | 1.71807999 | 0.08578203 | 0.75373899 |
| ZNF629 | 3.90131627 | -0.499691213 | 0.71744653 | -0.6964857 | 0.48612474 | 0.99640678 |
| BCL7C | 17.4452134 | 0.789066201 | 0.40964548 | 1.92621729 | 0.05407725 | 0.6653339 |
| MIR762HG | 2.70874897 | -0.97986389 | 0.95021322 | -1.0312042 | 0.30244506 | 0.97358745 |
| FBXL19 | 6.35006829 | 0.187227848 | 0.62033096 | 0.30181929 | 0.76278983 | 0.99665574 |
| SETD1A | 3.42075845 | -1.159290162 | 0.84281749 | -1.3754937 | 0.16897843 | 0.87601071 |
| STX4 | 3.60254524 | -0.308738999 | 0.679124 | -0.4546136 | 0.64938726 | 0.99665574 |
| BCKDK | 4.59341416 | 1.208782736 | 0.85943927 | 1.40647836 | 0.15958212 | 0.86757292 |
| KAT8 | 11.9394433 | 0.285455007 | 0.58138012 | 0.49099547 | 0.62342965 | 0.99665574 |
| PRSS8 | 17.6483182 | 0.008932726 | 0.53178375 | 0.01679767 | 0.98659803 | 0.99777713 |
| FUS | 44.205055 | 0.69587832 | 0.29121076 | 2.38960376 | 0.01686656 | 0.4264169 |
| AC026471.4 | 3.15912079 | -0.909172437 | 0.87702986 | -1.0366493 | 0.29989931 | 0.97358745 |
| C16orf58 | 7.64519023 | -0.310336116 | 0.57038812 | -0.5440789 | 0.58638721 | 0.99665574 |
| ZNF720 | 12.4646869 | 0.036448662 | 0.43895373 | 0.08303532 | 0.93382346 | 0.99777713 |
| ZNF267 | 3.5298491 | -0.496344713 | 0.7311732 | -0.6788333 | 0.4972435 | 0.99640678 |
| VPS35 | 16.9291007 | -0.508015891 | 0.42919676 | -1.1836433 | 0.23655428 | 0.9305595 |
| C16orf87 | 5.68216476 | -1.067735098 | 0.5807009 | -1.8387006 | 0.06595923 | 0.69294788 |
| GPT2 | 3.56579405 | 0.584525714 | 0.99333808 | 0.58844589 | 0.55623304 | 0.99665574 |
| DNAJA2 | 10.7094756 | 0.660008062 | 0.55870389 | 1.18131996 | 0.23747564 | 0.9305595 |
| NETO2 | 3.53280254 | -1.090785533 | 0.84251211 | -1.2946823 | 0.19542984 | 0.90381759 |
| ITFG1 | 93.8152446 | -0.188892407 | 0.22770205 | -0.8295595 | 0.40678787 | 0.99250037 |
| PHKB | 73.9036583 | -0.110403341 | 0.26291549 | -0.4199195 | 0.67454426 | 0.99665574 |
| LONP2 | 48.400599 | 0.153189841 | 0.28055299 | 0.54602819 | 0.58504656 | 0.99665574 |
| SIAH1 | 16.3703921 | 0.412087187 | 0.3973781 | 1.03701534 | 0.29972871 | 0.97358745 |
| N4BP1 | 41.8875058 | 0.497514563 | 0.28205648 | 1.76388277 | 0.07775172 | 0.74141484 |
| CNEP1R1 | 2.70762626 | 0.883732508 | 0.8339092 | 1.05974669 | 0.28925986 | 0.96676791 |
| HEATR3 | 4.47622035 | 1.295200956 | 0.80745778 | 1.60404789 | 0.1087035 | 0.80313739 |
| TENT4B | 15.1552589 | 0.891478177 | 0.46259848 | 1.92711004 | 0.05396592 | 0.6653339 |
| BRD7 | 15.9243193 | 0.214197556 | 0.40871758 | 0.52407229 | 0.60022826 | 0.99665574 |
| CYLD | 12.1269588 | 0.004841521 | 0.46982619 | 0.01030492 | 0.99177801 | 0.99777713 |
| SALL1 | 8.35705382 | -0.866130203 | 0.53717905 | -1.6123678 | 0.10688193 | 0.79791531 |
| AC087564.1 | 18.0779647 | 0.854277024 | 0.58070229 | 1.47111013 | 0.14126134 | 0.84969914 |
| HNRNPA1P4 | 4.24558919 | -0.298067976 | 0.70255829 | -0.4242608 | 0.67137559 | 0.99665574 |
| TOX3 | 110.627517 | -0.007221096 | 0.42263303 | -0.017086 | 0.98636803 | 0.99777713 |
| CHD9 | 139.942908 | -0.125882353 | 0.2239052 | -0.5622127 | 0.57397109 | 0.99665574 |
| RBL2 | 37.8752538 | 0.149775443 | 0.38077899 | 0.39333957 | 0.6940687 | 0.99665574 |
| AKTIP | 5.97610697 | 0.169652214 | 0.60803007 | 0.27901945 | 0.7802299 | 0.99665574 |
| FTO | 75.5017678 | 0.005900153 | 0.23553447 | 0.02505006 | 0.98001503 | 0.99777713 |
| LPCAT2 | 4.22860869 | -0.249900757 | 0.8312482 | -0.3006331 | 0.76369426 | 0.99665574 |
| AMFR | 52.2288123 | 0.490261254 | 0.35301405 | 1.38878681 | 0.16489759 | 0.87495175 |
| NUDT21 | 17.1279374 | 0.66218989 | 0.40199584 | 1.64725557 | 0.09950553 | 0.78261212 |
| OGFOD1 | 7.44560153 | 0.512907512 | 0.61574728 | 0.8329838 | 0.40485386 | 0.99194293 |
| BBS2 | 10.0387358 | 0.075504719 | 0.49465641 | 0.15264074 | 0.8786816 | 0.99665574 |
| MT2A | 84.2858745 | -0.459839666 | 0.68501611 | -0.671283 | 0.50204026 | 0.99640678 |
| MT1E | 26.0565597 | -1.250361445 | 0.77372564 | -1.6160269 | 0.1060885 | 0.79596216 |
| MT1G | 48.0617984 | 0.677804576 | 1.25705549 | 0.5392002 | 0.58974872 | 0.99665574 |
| MT1H | 14.1358311 | 1.123838286 | 1.18036184 | 0.95211337 | 0.34103949 | 0.97788427 |

| | | | | | | |
|---|---|---|---|---|---|---|
| MT1X | 17.6845424 | -0.735529429 | 0.80976245 | -0.9083274 | 0.36370526 | 0.98362663 |
| NUP93 | 11.5562196 | 0.455344696 | 0.49069886 | 0.9279514 | 0.35343278 | 0.98022903 |
| SLC12A3 | 9.82869136 | 1.803011384 | 0.87995286 | 2.04898634 | 0.04046345 | 0.60863273 |
| HERPUD1 | 28.379563 | 0.305179486 | 0.35825281 | 0.8518551 | 0.39429452 | 0.99179694 |
| NLRC5 | 27.8405477 | -0.925122169 | 0.79475328 | -1.1640369 | 0.24440904 | 0.93426912 |
| CPNE2 | 10.4487397 | -0.850688638 | 0.79761802 | -1.0665364 | 0.28618124 | 0.96449041 |
| FAM192A | 13.4402672 | 0.418300611 | 0.52612314 | 0.79506218 | 0.42657733 | 0.99525676 |
| RSPRY1 | 20.5294239 | -0.010021572 | 0.37656912 | -0.0266128 | 0.97876854 | 0.99777713 |
| ARL2BP | 15.782485 | 0.591083754 | 0.47945187 | 1.2328323 | 0.21763834 | 0.91640302 |
| PLLP | 21.6809922 | 0.74337614 | 0.57891764 | 1.28407928 | 0.19911422 | 0.90381759 |
| CX3CL1 | 3.70083477 | -0.290447798 | 0.92487767 | -0.3140391 | 0.7534913 | 0.99665574 |
| CIAPIN1 | 12.7749081 | -0.246518276 | 0.5279433 | -0.4669408 | 0.64054222 | 0.99665574 |
| COQ9 | 24.8851459 | 0.362017062 | 0.37201801 | 0.97311704 | 0.33049514 | 0.97788427 |
| POLR2C | 5.25957687 | -0.034314709 | 0.70724422 | -0.0485189 | 0.9613027 | 0.99777713 |
| ADGRG1 | 36.5548627 | 0.149185397 | 0.39225587 | 0.38032674 | 0.70370289 | 0.99665574 |
| KIFC3 | 20.7004676 | 0.253558306 | 0.50118772 | 0.50591484 | 0.61291643 | 0.99665574 |
| USB1 | 2.70707914 | -0.768178603 | 0.8094771 | -0.9489813 | 0.34263014 | 0.97788427 |
| MMP15 | 2.56012821 | 0.578284218 | 1.00661571 | 0.5744836 | 0.56564059 | 0.99665574 |
| CFAP20 | 3.39781967 | -0.117976138 | 0.87632814 | -0.1346255 | 0.89290796 | 0.99773598 |
| CSNK2A2 | 11.541127 | -0.144160707 | 0.45782431 | -0.3148822 | 0.75285113 | 0.99665574 |
| SETD6 | 4.40947988 | 1.446874263 | 0.75049428 | 1.92789514 | 0.05386817 | 0.6653339 |
| CNOT1 | 43.6388649 | 0.020511448 | 0.28672351 | 0.07153738 | 0.94297008 | 0.99777713 |
| SLC38A7 | 7.27777798 | -1.052859999 | 0.56262848 | -1.8713237 | 0.06130023 | 0.6834742 |
| GOT2 | 14.6636791 | 0.145789725 | 0.39433807 | 0.36970746 | 0.71160047 | 0.99665574 |
| AC138305.1 | 92.6620244 | -0.169551932 | 0.31942692 | -0.5308004 | 0.59555711 | 0.99665574 |
| AC092138.2 | 3.24507846 | 3.813803086 | 1.35419654 | 2.81628477 | 0.00485826 | 0.22086557 |
| TK2 | 2.7849733 | -1.02851123 | 0.91389745 | -1.1254121 | 0.26041445 | 0.94557817 |
| CMTM4 | 65.5192267 | 0.049107068 | 0.29711207 | 0.1652813 | 0.86872258 | 0.99665574 |
| DYNC1LI2 | 35.0189224 | -0.390008561 | 0.34813042 | -1.1202944 | 0.26258832 | 0.94557817 |
| AC018557.1 | 5.96162611 | 0.625916874 | 0.79999212 | 0.7824038 | 0.4339773 | 0.99617422 |
| NAE1 | 10.0293499 | 0.436193332 | 0.47812541 | 0.91229901 | 0.36161134 | 0.9819332 |
| PDP2 | 7.46064501 | 0.401528972 | 0.57503504 | 0.6982687 | 0.48500917 | 0.99640678 |
| CDH16 | 36.0034812 | 0.719733189 | 0.6027211 | 1.1941397 | 0.23242333 | 0.92916801 |
| RRAD | 7.50785309 | -1.610727296 | 1.01224561 | -1.5912416 | 0.11155522 | 0.8083966 |
| FAM96B | 15.4101354 | -0.30528121 | 0.46787572 | -0.6524836 | 0.51408928 | 0.99665574 |
| CES2 | 8.07836366 | -0.034942297 | 0.67791458 | -0.0515438 | 0.9588922 | 0.99777713 |
| CBFB | 49.6348472 | 0.297533968 | 0.30466519 | 0.97659326 | 0.32877056 | 0.97788427 |
| C16orf70 | 5.23929674 | -0.420384043 | 0.64119524 | -0.6556256 | 0.51206502 | 0.99665574 |
| TRADD | 2.11156341 | -0.218300716 | 1.01755893 | -0.2145337 | 0.83013086 | 0.99665574 |
| KIAA0895L | 3.45289354 | -1.408463797 | 0.71680451 | -1.9649204 | 0.04942345 | 0.64444758 |
| E2F4 | 6.77740897 | 0.847542491 | 0.65159873 | 1.30071232 | 0.19335694 | 0.90153775 |
| LRRC29 | 2.46083268 | -0.479721523 | 1.0234314 | -0.4687383 | 0.63925669 | 0.99665574 |
| TMEM208 | 11.1097962 | -0.394808061 | 0.51528806 | -0.766189 | 0.44356383 | 0.99617422 |
| ZDHHC1 | 3.22642605 | -0.26147906 | 0.82355766 | -0.3174994 | 0.7508647 | 0.99665574 |
| HSD11B2 | 51.7790523 | 1.209772326 | 0.58027233 | 2.08483544 | 0.03708424 | 0.58591936 |
| ATP6V0D1 | 19.7896834 | -0.321893663 | 0.53141389 | -0.6057306 | 0.54469365 | 0.99665574 |
| AC009061.2 | 5.29194534 | 0.689930223 | 0.68529931 | 1.0067575 | 0.31405132 | 0.9755712 |
| AC027682.6 | 3.41733875 | 1.411660683 | 0.90245075 | 1.56425233 | 0.11775832 | 0.82599629 |
| RIPOR1 | 15.1325403 | 0.453046208 | 0.54808341 | 0.82660085 | 0.40846334 | 0.99279547 |
| CTCF | 24.3800262 | -0.062847733 | 0.41869196 | -0.1501049 | 0.88068182 | 0.99665574 |
| GFOD2 | 6.6002686 | -0.951351724 | 0.63178271 | -1.5058211 | 0.13211311 | 0.84089786 |
| RANBP10 | 26.4558137 | -0.299205837 | 0.37535835 | -0.7971205 | 0.42538105 | 0.99518711 |
| CENPT | 2.73032145 | 0.782490895 | 0.86086694 | 0.90895684 | 0.36337291 | 0.98362663 |
| NUTF2 | 5.0521768 | 0.27606187 | 0.64889963 | 0.42543077 | 0.67052268 | 0.99665574 |
| PSKH1 | 6.81970023 | -0.130073963 | 0.66706607 | -0.1949941 | 0.84539755 | 0.99665574 |
| DUS2 | 4.66441037 | 0.441335275 | 0.7032392 | 0.6275749 | 0.53028245 | 0.99665574 |
| NFATC3 | 36.9273543 | 0.301905074 | 0.32609957 | 0.92580642 | 0.35454659 | 0.98022903 |
| AC020978.5 | 2.67615488 | 0.514762001 | 1.11967327 | 0.45974305 | 0.64570067 | 0.99665574 |
| ESRP2 | 4.87834847 | 0.947740231 | 0.69379768 | 1.36601815 | 0.17193327 | 0.88157681 |
| PLA2G15 | 11.2548315 | 0.540813398 | 0.56667785 | 0.95435775 | 0.33990259 | 0.97788427 |
| SLC7A6 | 19.7437254 | 0.558027145 | 0.39210757 | 1.4231481 | 0.15469323 | 0.86196782 |
| SLC7A6OS | 8.45198055 | -0.364293217 | 0.57932335 | -0.6288254 | 0.52946339 | 0.99665574 |
| PRMT7 | 6.68729369 | 0.357133839 | 0.73230096 | 0.48768724 | 0.62577139 | 0.99665574 |
| ZFP90 | 7.60951072 | -0.060583152 | 0.70372493 | -0.0860893 | 0.93139547 | 0.99777713 |
| CDH1 | 64.3121406 | -0.504980674 | 0.24012146 | -2.1030218 | 0.03546386 | 0.58349146 |
| TANGO6 | 20.133682 | -0.079549692 | 0.39756523 | -0.2000922 | 0.84140849 | 0.99665574 |
| CHTF8 | 9.98277894 | 0.140653546 | 0.53829267 | 0.26129567 | 0.7938645 | 0.99665574 |
| UTP4 | 5.98586531 | 0.491611777 | 0.6776594 | 0.72545556 | 0.4681726 | 0.99617422 |
| SNTB2 | 10.998939 | -0.195184716 | 0.49861467 | -0.391454 | 0.69546167 | 0.99665574 |
| VPS4A | 6.39638468 | 0.243730876 | 0.60412893 | 0.40344182 | 0.68662322 | 0.99665574 |
| COG8 | 3.95341591 | 0.119955761 | 0.71118122 | 0.16867116 | 0.8660553 | 0.99665574 |
| NIP7 | 4.15072205 | 0.351576833 | 0.7833708 | 0.44880002 | 0.65357592 | 0.99665574 |
| TERF2 | 8.79728986 | -0.059652857 | 0.48544413 | -0.1228831 | 0.90219971 | 0.99777713 |
| CYB5B | 21.3784768 | 0.07527103 | 0.52470469 | 0.14345408 | 0.88593157 | 0.99665574 |
| NFAT5 | 157.258464 | -0.156773479 | 0.25090121 | -0.6248415 | 0.53207512 | 0.99665574 |
| NQO1 | 5.75389665 | -1.383650661 | 0.82910777 | -1.668843 | 0.09514851 | 0.77178333 |
| NOB1 | 2.4001733 | 0.250304695 | 0.90648648 | 0.27612623 | 0.78245111 | 0.99665574 |
| WWP2 | 57.3024237 | -0.177995085 | 0.32054011 | -0.5552974 | 0.57869126 | 0.99665574 |
| PDPR | 18.0979581 | -0.150069644 | 0.48741 | -0.307892 | 0.75816451 | 0.99665574 |
| AC009060.1 | 4.48795104 | -0.280243946 | 0.72387516 | -0.387144 | 0.69864958 | 0.99665574 |
| EXOSC6 | 9.60291918 | 0.939077334 | 0.55743129 | 1.68465128 | 0.09205587 | 0.76658136 |
| AARS | 12.3600143 | 0.222412367 | 0.48482308 | 0.45874954 | 0.64641403 | 0.99665574 |
| DDX19A | 2.81164423 | -0.316715026 | 0.81476446 | -0.3887197 | 0.69748348 | 0.99665574 |
| ST3GAL2 | 10.1029596 | 0.031042916 | 0.6139799 | 0.05056015 | 0.95967602 | 0.99777713 |
| FUK | 4.80961059 | -0.019829545 | 0.62759111 | -0.0315963 | 0.97479401 | 0.99777713 |
| COG4 | 10.6434091 | -0.099608006 | 0.55527557 | -0.1793848 | 0.85763555 | 0.99665574 |
| SF3B3 | 16.0963495 | 0.636483486 | 0.40809464 | 1.55964675 | 0.11884338 | 0.82619588 |
| MTSS1L | 3.06194556 | 0.854380029 | 0.78955605 | 1.0821018 | 0.2792073 | 0.95980166 |
| VAC14 | 12.683583 | -0.538837757 | 0.63646174 | -0.8466145 | 0.39721001 | 0.99179694 |
| HYDIN | 8.57452499 | 0.97099661 | 0.64544728 | 1.50437789 | 0.1324841 | 0.84089786 |
| CMTR2 | 13.4052447 | 0.413538723 | 0.61743746 | 0.66976618 | 0.50300686 | 0.99640678 |
| ZNF19 | 3.81784003 | -0.957893468 | 0.73449336 | -1.3041554 | 0.19218061 | 0.89964233 |
| MARVELD3 | 4.90223399 | -0.467438452 | 0.70990821 | -0.6584491 | 0.51024957 | 0.99665574 |
| PHLPP2 | 9.69606088 | -0.883926881 | 0.49413507 | -1.7888365 | 0.07364114 | 0.72792151 |
| AP1G1 | 49.1026003 | 0.156660565 | 0.35948897 | 0.43578685 | 0.66299138 | 0.99665574 |
| IST1 | 32.2208747 | 0.242056967 | 0.30248858 | 0.80021852 | 0.4235842 | 0.99518711 |
| ZNF821 | 16.0453386 | -0.311532864 | 0.51747174 | -0.6020288 | 0.547155 | 0.99665574 |
| TXNL4B | 4.74924757 | -0.276639234 | 0.64270276 | -0.4304311 | 0.6668821 | 0.99665574 |
| DHX38 | 5.02937594 | 0.315162877 | 0.63727825 | 0.49454517 | 0.62092122 | 0.99665574 |
| LINC01572 | 7.43853253 | 0.124505861 | 0.79800765 | 0.15602089 | 0.87601656 | 0.99665574 |
| AC004943.2 | 6.92800171 | 0.009048833 | 0.65890986 | 0.01373304 | 0.98904297 | 0.99777713 |
| ZFHX3 | 73.3245527 | 0.088750427 | 0.31858637 | 0.27857572 | 0.78057045 | 0.99665574 |
| AC138627.1 | 11.0538143 | 0.377572805 | 0.67793188 | 0.556948 | 0.57756296 | 0.99665574 |
| PSMD7 | 13.2207043 | -0.319751156 | 0.47752299 | -0.6696037 | 0.50311046 | 0.99640678 |
| NPIPB15 | 3.51565651 | -0.384378373 | 1.01967402 | -0.376962 | 0.70620183 | 0.99665574 |
| GLG1 | 83.6444728 | 0.020620895 | 0.24980083 | 0.08254935 | 0.93420988 | 0.99777713 |
| RFWD3 | 10.1439627 | -0.958288252 | 0.49020113 | -1.9548879 | 0.0505963 | 0.64929786 |
| FA2H | 7.81441827 | 1.750140571 | 0.7033504 | 2.48829114 | 0.01283586 | 0.38590337 |
| WDR59 | 29.237481 | 0.245011115 | 0.35576858 | 0.68868114 | 0.49102394 | 0.99640678 |
| ZNRF1 | 16.1376666 | -0.541524508 | 0.53170107 | -1.0184755 | 0.30845204 | 0.97376371 |
| LDHD | 3.36534549 | 0.704640371 | 0.98040282 | 0.71872536 | 0.47231015 | 0.99617422 |
| BCAR1 | 14.4444353 | -0.126260967 | 0.49854402 | -0.2532594 | 0.80006775 | 0.99665574 |
| CFDP1 | 45.4560034 | -0.373398642 | 0.3337913 | -1.118659 | 0.26328565 | 0.94557817 |

| | | | | | | |
|---|---|---|---|---|---|---|
| GABARAPL2 | 18.1108165 | -0.203013357 | 0.58378124 | -0.3477559 | 0.72802353 | 0.99665574 |
| ADAT1 | 5.82832923 | -0.479710648 | 0.69874743 | -0.6865294 | 0.49237934 | 0.99640678 |
| KARS | 9.18877338 | -0.355601866 | 0.46899197 | -0.7582259 | 0.44831575 | 0.99617422 |
| TERF2IP | 26.5605383 | 0.171811224 | 0.36567281 | 0.4698496 | 0.63846247 | 0.99665574 |
| MON1B | 2.27229084 | -0.86936099 | 1.25456461 | -0.6929583 | 0.4883357 | 0.99640678 |
| ADAMTS18 | 27.68408 | -1.0730874 | 0.48174146 | -2.2275172 | 0.02591273 | 0.51788242 |
| NUDT7 | 3.43367746 | 0.022721498 | 0.95224686 | 0.02386093 | 0.98096354 | 0.99777713 |
| VAT1L | 7.5655178 | -1.51174418 | 1.00921664 | -1.4979382 | 0.1341493 | 0.84373478 |
| WWOX | 722.598657 | -1.119455443 | 0.40240491 | -2.7819129 | 0.00540395 | 0.22875404 |
| AC046158.1 | 4.39306895 | -1.661471012 | 0.86980631 | -1.9101621 | 0.05611235 | 0.67401304 |
| MAF | 13.5380294 | -1.540644387 | 0.73856137 | -2.0860073 | 0.03697797 | 0.58591936 |
| CDYL2 | 13.7400967 | -0.159038423 | 0.46784727 | -0.3399366 | 0.73390425 | 0.99665574 |
| CMC2 | 26.2682141 | 0.774713599 | 0.47004457 | 1.6481705 | 0.09931769 | 0.78261212 |
| ATMIN | 8.54666451 | 0.105415028 | 0.53155489 | 0.19831447 | 0.84279903 | 0.99665574 |
| GAN | 10.3360298 | -0.361385197 | 0.55564741 | -0.6503858 | 0.51544303 | 0.99665574 |
| CMIP | 81.8220987 | -0.803567011 | 0.3914929 | -2.0525711 | 0.04011419 | 0.60713077 |
| PLCG2 | 710.594706 | 0.461137242 | 0.40368425 | 1.14232162 | 0.25332036 | 0.94076968 |
| MPHOSPH6 | 12.4611124 | -0.262949737 | 0.44618675 | -0.5893266 | 0.55564218 | 0.99665574 |
| HSBP1 | 21.723829 | 0.374517533 | 0.44224001 | 0.84686488 | 0.39707044 | 0.99179694 |
| AC009119.2 | 3.01107794 | -0.68233804 | 0.72242916 | -0.9445051 | 0.34491158 | 0.97788427 |
| MBTPS1 | 23.4378959 | -0.181490463 | 0.37285547 | -0.4867582 | 0.62642968 | 0.99665574 |
| HSDL1 | 5.030834 | -0.689965826 | 0.72706751 | -0.9489708 | 0.34263547 | 0.97788427 |
| TAF1C | 3.73821052 | 1.134654162 | 0.87020388 | 1.30389463 | 0.19226951 | 0.89964233 |
| ATP2C2 | 11.1206966 | 0.836420514 | 1.0189412 | 0.82087221 | 0.41171906 | 0.99279547 |
| TLDC1 | 8.18951483 | -0.695970401 | 0.60116753 | -1.1576979 | 0.24698733 | 0.93441937 |
| KLHL36 | 11.8700836 | 0.269252441 | 0.44011297 | 0.61178029 | 0.54068313 | 0.99665574 |
| USP10 | 30.4854044 | 0.383180974 | 0.30985495 | 1.23664628 | 0.21621843 | 0.91448579 |
| ZDHHC7 | 18.3391467 | -0.26403851 | 0.37253041 | -0.7087704 | 0.478467 | 0.99617422 |
| KIAA0513 | 2.2421201 | -0.878153557 | 1.13035763 | -0.7768812 | 0.43722888 | 0.99617422 |
| GSE1 | 91.0613178 | -0.386375327 | 0.40992998 | -0.9425398 | 0.34591634 | 0.97788427 |
| EMC8 | 6.44107517 | -0.39956759 | 0.56739769 | -0.7042108 | 0.4813015 | 0.99640678 |
| COX4I1 | 90.7730149 | -0.054802235 | 0.33023323 | -0.1659501 | 0.86819623 | 0.99665574 |
| MTHFSD | 3.1737344 | 0.414394434 | 0.7601747 | 0.54513053 | 0.58566375 | 0.99665574 |
| FBXO31 | 8.54780415 | -0.170753118 | 0.56740586 | -0.3009365 | 0.76346294 | 0.99665574 |
| MAP1LC3B | 31.6438231 | -0.718227697 | 0.36480147 | -1.968818 | 0.048974 | 0.64396678 |
| ZCCHC14 | 69.8062649 | -1.215677536 | 0.40849326 | -2.9760039 | 0.00292031 | 0.16917783 |
| KLHDC4 | 15.2460073 | 0.008563556 | 0.43959792 | 0.01948043 | 0.98445785 | 0.99777713 |
| BANP | 22.914369 | 0.441096127 | 0.40803028 | 1.08103772 | 0.27968034 | 0.95996214 |
| ZFPM1 | 8.26430333 | -0.0948668 | 0.52185025 | -0.1817893 | 0.85574807 | 0.99665574 |
| ZC3H18 | 12.1831531 | -0.613272037 | 0.46847675 | -1.3090768 | 0.19050835 | 0.89964233 |
| CYBA | 11.944561 | -1.698154243 | 0.69400404 | -2.4468939 | 0.01440933 | 0.40354073 |
| MVD | 7.09518544 | -0.921908863 | 0.65575396 | -1.4058762 | 0.1597609 | 0.86757292 |
| SNAI3-AS1 | 3.16301188 | -1.390329866 | 0.8463654 | -1.6427064 | 0.10044369 | 0.78386461 |
| PIEZO1 | 5.58966751 | -0.462509593 | 0.79422577 | -0.5823402 | 0.56033755 | 0.99665574 |
| APRT | 15.9343513 | -0.49421797 | 0.48738088 | -1.0140282 | 0.31056928 | 0.97376371 |
| GALNS | 6.39516135 | 0.347893565 | 0.53507979 | 0.65017138 | 0.51558152 | 0.99665574 |
| TRAPPC2L | 6.03033301 | 0.272717485 | 0.62127968 | 0.4389609 | 0.66068987 | 0.99665574 |
| ACSF3 | 7.97237827 | -0.175460891 | 0.56720582 | -0.3093425 | 0.75706097 | 0.99665574 |
| ZNF778 | 3.06446366 | -0.246447537 | 0.82683016 | -0.2980631 | 0.76565503 | 0.99665574 |
| ANKRD11 | 98.7614696 | -0.458943687 | 0.37965108 | -1.2088565 | 0.22671797 | 0.92318867 |
| SPG7 | 33.3220451 | -0.112867255 | 0.38355214 | -0.2942683 | 0.76855286 | 0.99665574 |
| RPL13 | 160.873152 | 0.522683889 | 0.46644215 | 1.12057602 | 0.26246838 | 0.94557817 |
| CHMP1A | 10.3187908 | 0.298824219 | 0.65227851 | 0.45812366 | 0.6468636 | 0.99665574 |
| SPATA33 | 2.7804614 | -1.469128138 | 0.87123269 | -1.6862638 | 0.091745 | 0.76615391 |
| CDK10 | 10.2183187 | -0.167103266 | 0.49282323 | -0.3390734 | 0.73455441 | 0.99665574 |
| VPS9D1 | 8.03447437 | -0.118327135 | 0.6040512 | -0.1958892 | 0.84469685 | 0.99665574 |
| ZNF276 | 4.27829632 | 0.642196104 | 0.69914711 | 0.91854217 | 0.35833509 | 0.9819332 |
| SPIRE2 | 11.2246703 | 1.065353968 | 0.55950018 | 1.90411728 | 0.05689491 | 0.67545594 |
| TCF25 | 53.4827915 | -0.491404207 | 0.33132534 | -1.4831471 | 0.13803534 | 0.84656101 |
| DEF8 | 5.39269843 | -0.556007764 | 0.61334393 | -0.9065187 | 0.36466135 | 0.98362663 |
| GAS8 | 5.85200033 | -0.384267742 | 0.65950014 | -0.5826651 | 0.56011874 | 0.99665574 |
| FAM157C | 8.91593368 | 0.109482601 | 0.46985433 | 0.23301392 | 0.81575059 | 0.99665574 |
| LINC02193 | 3.86869344 | -0.710305767 | 0.71680799 | -0.9909289 | 0.32172029 | 0.97700345 |
| RPH3AL | 19.320613 | -0.033720997 | 0.37082708 | -0.0909346 | 0.92754459 | 0.99777713 |
| AC129507.2 | 2.92144379 | -0.430516006 | 0.91364914 | -0.471205 | 0.63749438 | 0.99665574 |
| C17orf97 | 1.40124129 | 2.221080444 | 1.32207538 | 1.67999532 | 0.09295823 | 0.76709603 |
| VPS53 | 41.9189646 | -0.688236454 | 0.4017651 | -1.713032 | 0.08670666 | 0.75427976 |
| GLOD4 | 9.46023854 | 0.057329043 | 0.60260146 | 0.09513592 | 0.92420687 | 0.99777713 |
| TIMM22 | 3.37332678 | 0.261985914 | 0.79379572 | 0.33004198 | 0.74136824 | 0.99665574 |
| ABR | 20.6723284 | 0.189089768 | 0.47250057 | 0.4001895 | 0.68901695 | 0.99665574 |
| YWHAE | 78.2399631 | -0.133857133 | 0.21919658 | -0.6106716 | 0.541417 | 0.99665574 |
| CRK | 26.1375818 | -0.435817753 | 0.51630377 | -0.8441111 | 0.39860731 | 0.99179694 |
| MYO1C | 23.8736318 | -0.502978742 | 0.46855522 | -1.0734674 | 0.28306147 | 0.96059387 |
| INPP5K | 5.96418477 | -0.780734594 | 0.65065884 | -1.1999139 | 0.23017278 | 0.92774608 |
| PITPNA | 27.2941073 | -0.333681229 | 0.34374702 | -0.9707174 | 0.331689 | 0.97788427 |
| SLC43A2 | 99.2955112 | 0.298483442 | 0.66176528 | 0.45104126 | 0.65195981 | 0.99665574 |
| SCARF1 | 7.26141111 | -0.392619631 | 0.62609372 | -0.627094 | 0.53059761 | 0.99665574 |
| RILP | 5.70136009 | 0.717875802 | 0.63246027 | 1.13505281 | 0.25635319 | 0.94314473 |
| PRPF8 | 31.1067442 | 0.119999201 | 0.34846496 | 0.34436518 | 0.73057167 | 0.99665574 |
| MIR22HG | 10.9522777 | -0.982478521 | 0.72296645 | -1.3589545 | 0.174161 | 0.88281284 |
| WDR81 | 9.6291616 | -0.46338869 | 0.77316397 | -0.5993408 | 0.54894566 | 0.99665574 |
| SERPINF2 | 13.7248798 | 1.768988399 | 0.49433927 | 3.57849052 | 0.00034558 | 0.04799363 |
| SMYD4 | 7.00844241 | -0.241059387 | 0.64904736 | -0.3714049 | 0.71033596 | 0.99665574 |
| RPA1 | 11.2184331 | -0.425579726 | 0.50253481 | -0.8468662 | 0.39706972 | 0.99179694 |
| SMG6 | 59.780459 | -0.263274843 | 0.34041742 | -0.7733883 | 0.43929262 | 0.99617422 |
| SRR | 3.20661673 | -0.168746103 | 0.75206689 | -0.2243765 | 0.82246439 | 0.99665574 |
| TSR1 | 4.80843077 | -0.890093854 | 0.78762075 | -1.1301046 | 0.25843214 | 0.94557817 |
| SGSM2 | 9.62991511 | -0.158346581 | 0.47736179 | -0.3317119 | 0.74010682 | 0.99665574 |
| MNT | 7.26937462 | -0.956228394 | 0.55301907 | -1.7291056 | 0.08379019 | 0.75242737 |
| METTL16 | 23.2564303 | -0.027988102 | 0.33992972 | -0.082335 | 0.93438035 | 0.99777713 |
| PAFAH1B1 | 79.3505965 | -0.234375865 | 0.31622681 | -0.7411638 | 0.45859412 | 0.99617422 |
| CLUH | 16.727273 | 0.011079329 | 0.42477338 | 0.02608292 | 0.9791912 | 0.99777713 |
| RAP1GAP2 | 25.7030508 | 0.169668777 | 0.42872331 | 0.39575356 | 0.69228684 | 0.99665574 |
| TRPV1 | 2.33024171 | -0.126585163 | 0.86151634 | -0.146933 | 0.88318492 | 0.99665574 |
| CTNS | 9.026306 | -1.872409956 | 0.87500222 | -2.1398917 | 0.03236352 | 0.5724009 |
| TAX1BP3 | 6.35080572 | -0.338079731 | 0.79626799 | -0.4245803 | 0.67114263 | 0.99665574 |
| EMC6 | 8.99493288 | 0.665312601 | 0.6864288 | 0.9692376 | 0.33242666 | 0.97788427 |
| ITGAE | 4.10730869 | 0.099753751 | 0.71512258 | 0.13949182 | 0.88906152 | 0.99665574 |
| NCBP3 | 17.9254429 | 0.210002785 | 0.36510133 | 0.57519042 | 0.56516252 | 0.99665574 |
| ZZEF1 | 35.2197671 | 0.000658597 | 0.33404578 | 0.00197158 | 0.99842691 | 0.99925101 |
| CYB5D2 | 9.32399016 | -0.438885891 | 0.55067096 | -0.7970021 | 0.42544983 | 0.99518711 |
| ANKFY1 | 22.500611 | 0.35378431 | 0.41099502 | 0.86079951 | 0.38934848 | 0.99176253 |
| UBE2G1 | 36.959745 | 0.288910942 | 0.3226165 | 0.89552439 | 0.37050683 | 0.98532838 |
| MYBBP1A | 2.75616807 | -0.582785055 | 0.78371329 | -0.7436202 | 0.45710626 | 0.99617422 |
| GGT6 | 3.84381613 | 1.528400092 | 0.82713134 | 1.8478324 | 0.06462659 | 0.68846743 |
| SMTNL2 | 4.54122 | 2.913694102 | 1.00327119 | 2.90419392 | 0.003682 | 0.19855382 |
| PELP1 | 7.35897446 | 0.288637356 | 0.57790761 | 0.49945242 | 0.6174607 | 0.99665574 |
| ARRB2 | 4.03422514 | 0.243797368 | 0.80996378 | 0.30099787 | 0.76341612 | 0.99665574 |
| MED11 | 4.0603918 | -0.893558651 | 0.94393589 | -0.9466307 | 0.34382702 | 0.97788427 |
| PSMB6 | 10.8708268 | 0.380155575 | 0.49710133 | 0.76474464 | 0.44442362 | 0.99617422 |
| MINK1 | 12.6664957 | -0.61116975 | 0.48004488 | -1.2731513 | 0.20296436 | 0.9066717 |
| SLC25A11 | 11.8548676 | 0.759370751 | 0.50458536 | 1.50494011 | 0.13233947 | 0.84089786 |
| RNF167 | 10.1904304 | 0.317898754 | 0.63257994 | 0.50254321 | 0.61528547 | 0.99665574 |

| | | | | | | |
|---|---|---|---|---|---|---|
| PFN1 | 36.1624203 | -0.416939338 | 0.50834946 | -0.8201825 | 0.41211207 | 0.99279547 |
| SPAG7 | 12.1643468 | 0.392924487 | 0.58797583 | 0.66826639 | 0.50396356 | 0.99665574 |
| CAMTA2 | 6.4381682 | 0.539465259 | 0.5677164 | 0.95023723 | 0.34199172 | 0.97788427 |
| KIF1C | 10.2202203 | -0.26453009 | 0.48497075 | -0.5454558 | 0.58544009 | 0.99665574 |
| ZFP3 | 3.60255103 | 0.415975578 | 1.00991866 | 0.41189018 | 0.68041992 | 0.99665574 |
| ZNF594 | 5.89019633 | 0.175276342 | 0.61649849 | 0.28430944 | 0.77617325 | 0.99665574 |
| AC087500.1 | 6.84287488 | 0.381793332 | 0.68501834 | 0.55734761 | 0.57728996 | 0.99665574 |
| RABEP1 | 52.4818093 | -0.465915606 | 0.32701475 | -1.4247541 | 0.15422829 | 0.86196782 |
| NUP88 | 29.7501166 | 0.481703858 | 0.4078779 | 1.18100015 | 0.23760266 | 0.9305595 |
| RPAIN | 14.3523214 | -0.289656076 | 0.43377021 | -0.6677639 | 0.50428433 | 0.99665574 |
| C1QBP | 14.1608344 | -0.696582516 | 0.52377582 | -1.3299249 | 0.18354301 | 0.89532394 |
| DERL2 | 7.65534103 | 0.186146388 | 0.54819097 | 0.33956486 | 0.73418425 | 0.99665574 |
| MIS12 | 4.50234905 | 0.565199185 | 0.83424215 | 0.67750015 | 0.49808867 | 0.99640678 |
| KIAA0753 | 11.2468568 | -0.139666662 | 0.56042666 | -0.2492149 | 0.80319457 | 0.99665574 |
| TXNDC17 | 6.13181713 | 1.058051608 | 0.69878739 | 1.51412521 | 0.12999409 | 0.83778831 |
| MED31 | 3.62899247 | 0.236519243 | 0.7447297 | 0.31759072 | 0.75079541 | 0.99665574 |
| XAF1 | 7.92174431 | 0.991005983 | 2.16393613 | 0.45796453 | 0.64697793 | 0.99665574 |
| ALOX12-AS1 | 5.90094265 | -1.090337065 | 0.79159219 | -1.3773974 | 0.1683894 | 0.87566218 |
| ACADVL | 42.2307386 | 0.469684115 | 0.412501 | 1.13862539 | 0.25485943 | 0.94126227 |
| DVL2 | 5.55342348 | -0.094427033 | 0.61671699 | -0.1531124 | 0.87830962 | 0.99665574 |
| PHF23 | 4.58907277 | -0.15259034 | 0.61228204 | -0.2492158 | 0.80319388 | 0.99665574 |
| ELP5 | 4.98243376 | -0.332905942 | 0.67630639 | -0.4922413 | 0.62254877 | 0.99665574 |
| EIF5A | 13.8844163 | -0.12789017 | 0.65815948 | -0.1943149 | 0.84592935 | 0.99665574 |
| GPS2 | 9.99351049 | 0.424435998 | 0.56309709 | 0.75375279 | 0.45099768 | 0.99617422 |
| ZBTB4 | 23.9176281 | -0.086084993 | 0.4767487 | -0.1805668 | 0.85670761 | 0.99665574 |
| POLR2A | 28.2545117 | -0.423151691 | 0.39961106 | -1.0589089 | 0.28964128 | 0.96676791 |
| EIF4A1 | 12.8457203 | -0.538674852 | 0.54390654 | -0.9903813 | 0.3219878 | 0.97700345 |
| MPDU1 | 7.3836095 | 0.552873363 | 0.57591195 | 0.95999633 | 0.33705706 | 0.97788427 |
| FXR2 | 5.5407635 | 0.006931798 | 0.58526821 | 0.0118438 | 0.99055024 | 0.99777713 |
| SAT2 | 9.46276545 | -0.115071493 | 0.51973671 | -0.2214034 | 0.82477832 | 0.99665574 |
| TP53 | 5.34734846 | 0.378625813 | 0.68877601 | 0.54970819 | 0.58251954 | 0.99665574 |
| KDM6B | 10.2029596 | -0.812620614 | 0.72373686 | -1.1228122 | 0.26151724 | 0.94557817 |
| NAA38 | 16.2457505 | -0.221204786 | 0.37749388 | -0.5859824 | 0.55788731 | 0.99665574 |
| CHD3 | 9.86444655 | 0.423974594 | 0.63507246 | 0.66760035 | 0.50438874 | 0.99665574 |
| TRAPPC1 | 4.05792604 | -0.209539869 | 0.74503452 | -0.2812485 | 0.77851978 | 0.99665574 |
| CNTROB | 2.66083919 | 0.022106651 | 0.83127697 | 0.0265936 | 0.97878387 | 0.99777713 |
| PER1 | 6.13163604 | 0.805788432 | 0.7161042 | 1.12523909 | 0.26048773 | 0.94557817 |
| BORCS6 | 2.23065914 | 0.626146363 | 0.91008723 | 0.68800698 | 0.49144839 | 0.99640678 |
| CTC1 | 6.73018109 | 1.243576505 | 0.65754033 | 1.89125512 | 0.0585903 | 0.67545594 |
| SLC25A35 | 4.47085332 | 3.097041469 | 0.92038846 | 3.3649286 | 0.00076563 | 0.07295564 |
| RANGRF | 4.06449326 | 0.919362575 | 0.79594161 | 1.15506284 | 0.24806469 | 0.93455216 |
| NDEL1 | 18.3233652 | -0.470430548 | 0.56495222 | -0.8326909 | 0.4050191 | 0.99194293 |
| MYH10 | 35.6181577 | 0.212920711 | 0.29922484 | 0.71157433 | 0.47672841 | 0.99617422 |
| NTN1 | 68.4354368 | -0.255305242 | 0.3765827 | -0.6779527 | 0.49780171 | 0.99640678 |
| STX8 | 59.7044477 | -0.110802665 | 0.30905386 | -0.3585222 | 0.71995257 | 0.99665574 |
| USP43 | 7.69176823 | -1.047692214 | 0.58276218 | -1.7978041 | 0.07220806 | 0.71945049 |
| GAS7 | 4.40273703 | 0.623086157 | 0.7003683 | 0.889655 | 0.37365117 | 0.98532838 |
| SCO1 | 10.4391557 | -0.055891905 | 0.46059819 | -0.1213463 | 0.90341672 | 0.99777713 |
| TMEM220 | 6.50601593 | -0.97962636 | 0.60606399 | -1.6163745 | 0.10601338 | 0.79596216 |
| TMEM220-A! | 5.18401342 | 0.128698002 | 0.60746262 | 0.2118616 | 0.83221501 | 0.99665574 |
| MAP2K4 | 57.1175922 | -0.153209508 | 0.27633046 | -0.5544431 | 0.57927566 | 0.99665574 |
| ARHGAP44 | 5.80831808 | 0.400423204 | 0.86232159 | 0.46435484 | 0.64239354 | 0.99665574 |
| ELAC2 | 7.0777287 | -0.253721243 | 0.51539492 | -0.4922851 | 0.62251781 | 0.99665574 |
| COX10 | 21.5776736 | 0.351725362 | 0.42318239 | 0.83114366 | 0.40589248 | 0.99194293 |
| TRIM16 | 2.50880233 | 0.855405915 | 0.96451233 | 0.88687919 | 0.37514395 | 0.98569969 |
| AC015922.4 | 2.58409548 | 0.394468875 | 0.86783878 | 0.45454166 | 0.64943902 | 0.99665574 |
| ZSWIM7 | 13.7403851 | -0.580800636 | 0.38559783 | -1.5062342 | 0.13200707 | 0.84089786 |
| TTC19 | 31.2202311 | -0.261487201 | 0.3811941 | -0.6859686 | 0.49273289 | 0.99640678 |
| NCOR1 | 94.4852824 | 0.129801338 | 0.21138308 | 0.61405738 | 0.53917741 | 0.99665574 |
| PIGL | 19.5004215 | 0.157287601 | 0.38871857 | 0.40463105 | 0.68574873 | 0.99665574 |
| UBB | 61.2832299 | -0.328452896 | 0.2854266 | -1.1507438 | 0.24983764 | 0.93755294 |
| LRRC75A | 4.43104178 | 1.038374084 | 0.76660495 | 1.35451003 | 0.17557372 | 0.88428483 |
| AC098850.3 | 15.2072432 | 1.32264114 | 0.81125026 | 1.63037376 | 0.10302253 | 0.78770791 |
| MPRIP | 40.8357475 | 0.051690608 | 0.28922597 | 0.17872049 | 0.85815718 | 0.99665574 |
| COPS3 | 6.81715155 | 0.328585797 | 0.637619 | 0.5153325 | 0.60632069 | 0.99665574 |
| NT5M | 3.73591861 | 0.547691087 | 0.79115542 | 0.69226737 | 0.48876944 | 0.99640678 |
| MED9 | 8.89735555 | 0.266923432 | 0.51724456 | 0.51604879 | 0.60582033 | 0.99665574 |
| RASD1 | 22.3893988 | -0.786011468 | 1.22178284 | -0.6433316 | 0.52000898 | 0.99665574 |
| PEMT | 3.62886714 | -0.230575794 | 0.7367213 | -0.3129756 | 0.7542992 | 0.99665574 |
| RAI1 | 24.152323 | -0.711389057 | 0.38093279 | -1.8674923 | 0.06183287 | 0.68360044 |
| SREBF1 | 16.7289506 | -0.33518736 | 0.63350685 | -0.5290982 | 0.5967373 | 0.99665574 |
| TOM1L2 | 35.317069 | 0.262496754 | 0.30396219 | 0.86358359 | 0.38781668 | 0.99168679 |
| ATPAF2 | 4.89437683 | 0.349041094 | 0.67242068 | 0.51908144 | 0.60370395 | 0.99665574 |
| GID4 | 5.49025705 | -1.252487406 | 0.73607178 | -1.7015832 | 0.08883352 | 0.75743838 |
| DRG2 | 5.14537848 | 0.207504153 | 0.61686628 | 0.33638433 | 0.73658106 | 0.99665574 |
| ALKBH5 | 14.1319972 | -0.509851023 | 0.52735559 | -0.9668069 | 0.33364058 | 0.97788427 |
| FLII | 11.3992589 | -0.395816965 | 0.49726102 | -0.7959944 | 0.42603532 | 0.99518711 |
| MIEF2 | 3.31361804 | -0.001064191 | 0.85621011 | -0.0012429 | 0.9990083 | 0.99940262 |
| TOP3A | 10.5728036 | -0.361439878 | 0.43910747 | -0.823124 | 0.4104375 | 0.99279547 |
| SMCR8 | 4.72054236 | -0.287295582 | 0.78596947 | -0.3655302 | 0.71471566 | 0.99665574 |
| SHMT1 | 11.9202051 | -0.057593127 | 0.47339609 | -0.1216595 | 0.9031687 | 0.99777713 |
| TVP23B | 6.96125571 | -0.529817708 | 0.56353169 | -0.9401738 | 0.34712844 | 0.97804112 |
| PRPSAP2 | 17.6367632 | 0.591456838 | 0.47490735 | 1.2454152 | 0.21297917 | 0.91073943 |
| SLC5A10 | 3.15334688 | -0.498310396 | 1.09439787 | -0.4553284 | 0.64887302 | 0.99665574 |
| FAM83G | 2.24368683 | -1.905202679 | 1.2963707 | -1.4696434 | 0.14165835 | 0.84969914 |
| EPN2 | 30.8496478 | -0.320109328 | 0.38436123 | -0.8328346 | 0.40493803 | 0.99194293 |
| B9D1 | 2.89107856 | -0.79029696 | 0.88725781 | -0.8907185 | 0.3730802 | 0.98532838 |
| MAPK7 | 1.96056533 | -0.077208033 | 1.2986465 | -0.0594527 | 0.95259154 | 0.99777713 |
| AC004448.2 | 6.47598947 | -0.823299506 | 0.6624578 | -1.2427954 | 0.21394323 | 0.9124923 |
| SLC47A1 | 11.0646908 | 2.420404533 | 0.69187867 | 3.49830776 | 0.00046822 | 0.05860271 |
| ALDH3A2 | 8.13972839 | 0.353509328 | 0.60160766 | 0.58760776 | 0.5567956 | 0.99665574 |
| ULK2 | 11.2882769 | -0.577742341 | 0.58662377 | -0.9848601 | 0.3246928 | 0.97700345 |
| AKAP10 | 39.7026132 | -0.145832956 | 0.3367004 | -0.4331238 | 0.66492483 | 0.99665574 |
| SPECC1 | 4.68074695 | -1.044297856 | 0.66862381 | -1.5618616 | 0.11832059 | 0.82612547 |
| CCDC144NL-/ | 8.30022576 | -0.551219929 | 0.70872178 | -0.7777663 | 0.43670679 | 0.99617422 |
| USP22 | 37.8949807 | 0.008197393 | 0.36269325 | 0.02260145 | 0.98196819 | 0.99777713 |
| DHRS7B | 23.575961 | -0.959273559 | 0.37374485 | -2.5666536 | 0.01026851 | 0.3493362 |
| TMEM11 | 5.17589859 | -0.155810935 | 0.62386776 | -0.2497499 | 0.80278074 | 0.99665574 |
| AC087294.1 | 2.94320251 | 0.170393786 | 0.7930932 | 0.21484712 | 0.82988652 | 0.99665574 |
| MAP2K3 | 5.66842955 | -0.785110834 | 0.76159979 | -1.0308706 | 0.3026015 | 0.97358745 |
| WSB1 | 184.845056 | -0.585695538 | 0.4114941 | -1.4233388 | 0.15463795 | 0.86196782 |
| KSR1 | 17.5794992 | -0.785067268 | 0.42886876 | -1.8305536 | 0.06716719 | 0.70200104 |
| NLK | 56.2171006 | -0.257288328 | 0.28025588 | -0.9180479 | 0.35859377 | 0.9819332 |
| IFT20 | 4.29132982 | -0.37409009 | 0.68697188 | -0.5445493 | 0.5860635 | 0.99665574 |
| TNFAIP1 | 5.68904086 | -0.220143517 | 0.71127786 | -0.3095042 | 0.75693798 | 0.99665574 |
| POLDIP2 | 9.27468004 | -0.277063966 | 0.46441495 | -0.5965871 | 0.55078309 | 0.99665574 |
| TMEM199 | 3.42771249 | 0.34485507 | 0.81353116 | 0.42389903 | 0.67163945 | 0.99665574 |
| SARM1 | 2.85432248 | -0.44668728 | 0.77459728 | -0.5766703 | 0.56416217 | 0.99665574 |
| AC005726.1 | 4.08501712 | -0.954679226 | 0.79327028 | -1.2034728 | 0.2287934 | 0.92646258 |
| KIAA0100 | 18.6397054 | 0.158999515 | 0.39626831 | 0.40124207 | 0.68824191 | 0.99665574 |
| SDF2 | 7.90443462 | -0.294408699 | 0.58224421 | -0.5056447 | 0.6131061 | 0.99665574 |
| SUPT6H | 20.6841344 | 0.056073278 | 0.43759262 | 0.12814037 | 0.89803789 | 0.99777713 |
| RAB34 | 9.37586777 | -0.155548456 | 0.5969805 | -0.2605587 | 0.79443285 | 0.99665574 |

| | | | | | | |
|---|---|---|---|---|---|---|
| RPL23A | 63.2349854 | 0.45228184 | 0.40817505 | 1.10805852 | 0.26783654 | 0.94643669 |
| TLCD1 | 3.52425111 | 0.685734295 | 0.86909206 | 0.78902377 | 0.43009811 | 0.99617422 |
| NEK8 | 6.61579401 | -0.256693558 | 0.61039367 | -0.4205377 | 0.67409269 | 0.99665574 |
| TRAF4 | 13.0901122 | 0.017995716 | 0.47964723 | 0.03751865 | 0.97007147 | 0.99777713 |
| FAM222B | 59.3410818 | -0.299889439 | 0.41601637 | -0.7208597 | 0.47099583 | 0.99617422 |
| ERAL1 | 4.04657782 | 0.830698909 | 0.75941811 | 1.09386239 | 0.27401533 | 0.95331759 |
| FLOT2 | 24.0586662 | -0.577957414 | 0.40260308 | -1.4355514 | 0.15113003 | 0.85612189 |
| DHRS13 | 3.37058593 | -1.171588438 | 0.85467845 | -1.3707944 | 0.17043906 | 0.87918701 |
| PHF12 | 14.2700437 | 0.264182043 | 0.4024038 | 0.65650981 | 0.51149616 | 0.99665574 |
| MYO18A | 10.5111954 | -0.501292846 | 0.52594266 | -0.9531321 | 0.34052313 | 0.97788427 |
| NUFIP2 | 43.6044813 | -0.023600568 | 0.33332925 | -0.0708026 | 0.94355489 | 0.99777713 |
| TAOK1 | 77.9837997 | -0.124424182 | 0.35433904 | -0.3511444 | 0.72547999 | 0.99665574 |
| ABHD15-AS1 | 5.41715393 | -0.869182446 | 0.65894489 | -1.3190518 | 0.18715179 | 0.89683847 |
| ABHD15 | 3.3833845 | 0.116293533 | 0.73146112 | 0.15898799 | 0.87367834 | 0.99665574 |
| GIT1 | 4.37648058 | -0.375200497 | 0.62353471 | -0.6017315 | 0.54735285 | 0.99665574 |
| SSH2 | 102.244533 | -0.65270497 | 0.29198145 | -2.235433 | 0.02538894 | 0.50968916 |
| AC023389.1 | 3.36482755 | -1.133083344 | 0.78550663 | -1.4424873 | 0.14916495 | 0.85485259 |
| NSRP1 | 33.9626246 | -0.116267187 | 0.32519158 | -0.3575344 | 0.72069175 | 0.99665574 |
| AC104984.4 | 9.80475379 | -0.728848341 | 0.58870918 | -1.2380448 | 0.21569945 | 0.913434 |
| BLMH | 9.42627695 | 0.060890627 | 0.530609 | 0.11475611 | 0.90863844 | 0.99777713 |
| CPD | 29.1598552 | 0.122882376 | 0.39064671 | 0.3145614 | 0.75309469 | 0.99665574 |
| GOSR1 | 26.7467541 | -0.240529677 | 0.35257706 | -0.6822045 | 0.49510962 | 0.99640678 |
| CRLF3 | 17.3190629 | -0.315750197 | 0.43365961 | -0.7281061 | 0.46654866 | 0.99617422 |
| ATAD5 | 3.54151763 | 0.047407593 | 0.81882673 | 0.05789698 | 0.95383069 | 0.99777713 |
| TEFM | 8.2148962 | 0.236606237 | 0.54042358 | 0.43781627 | 0.66151947 | 0.99665574 |
| RNF135 | 5.62596697 | 0.879478845 | 0.79454489 | 1.10689636 | 0.26833874 | 0.94660092 |
| NF1 | 171.19909 | -0.043268779 | 0.26694837 | -0.1620867 | 0.87123758 | 0.99665574 |
| OMG | 4.47693287 | -2.353576798 | 0.76273671 | -3.0857002 | 0.00203073 | 0.13751507 |
| RAB11FIP4 | 10.5575825 | 0.208776736 | 0.47921955 | 0.43565989 | 0.66308351 | 0.99665574 |
| COPRS | 14.6936926 | 1.542136722 | 0.58119415 | 2.65339341 | 0.00796869 | 0.30144259 |
| UTP6 | 8.90001298 | 0.287085519 | 0.55713729 | 0.51528685 | 0.60635259 | 0.99665574 |
| AC004253.1 | 3.2434351 | -0.569692416 | 0.91082143 | -0.625471 | 0.53166196 | 0.99665574 |
| SUZ12 | 37.9344676 | 0.202368239 | 0.30803386 | 0.65696752 | 0.5112018 | 0.99665574 |
| LRRC37B | 10.3575159 | -0.842396131 | 0.63150891 | -1.3339418 | 0.18222293 | 0.89451364 |
| RHOT1 | 22.0712619 | 0.427105058 | 0.39796841 | 1.07321347 | 0.28317535 | 0.96059387 |
| C17orf75 | 3.52754126 | 0.542800164 | 0.81509099 | 0.66593812 | 0.50545065 | 0.99665574 |
| ZNF207 | 50.5782103 | -0.04419381 | 0.26929559 | -0.1641089 | 0.8696454 | 0.99665574 |
| PSMD11 | 17.1220567 | 0.632715126 | 0.44850306 | 1.41072644 | 0.15832529 | 0.86661434 |
| MYO1D | 58.6034782 | -0.82042009 | 0.38045295 | -2.1564298 | 0.03105012 | 0.56312363 |
| ZNF830 | 2.68497622 | -0.239431267 | 0.77864868 | -0.3074959 | 0.75846595 | 0.99665574 |
| LIG3 | 5.76825913 | -0.668457651 | 0.63234937 | -1.0571018 | 0.29046513 | 0.96676791 |
| SLFN5 | 4.65476468 | -1.029075458 | 0.88491613 | -1.1629073 | 0.2448671 | 0.93426912 |
| LINC02001 | 4.02286842 | -0.814846287 | 0.65751401 | -1.2392835 | 0.21524052 | 0.913434 |
| AP2B1 | 34.613228 | -0.615936096 | 0.38834134 | -1.5860688 | 0.1127237 | 0.81249398 |
| TAF15 | 61.8151731 | 0.099738221 | 0.29128211 | 0.34241107 | 0.73204156 | 0.99665574 |
| ZNHIT3 | 6.30378247 | 0.84806624 | 0.63458166 | 1.33641782 | 0.18141274 | 0.89279728 |
| MYO19 | 46.5404484 | 0.115752549 | 0.3976575 | 0.29108604 | 0.77098551 | 0.99665574 |
| GGNBP2 | 31.2463785 | -0.166802986 | 0.44312094 | -0.3764277 | 0.70659898 | 0.99665574 |
| DHRS11 | 3.54962139 | 1.027365363 | 0.90147576 | 1.13964835 | 0.25443283 | 0.94099699 |
| LHX1-DT | 6.65049897 | 0.298184811 | 0.60723954 | 0.49104972 | 0.62339128 | 0.99665574 |
| LHX1 | 13.1177051 | 0.49607285 | 0.68336623 | 0.72592533 | 0.46788455 | 0.99617422 |
| AATF | 29.6028828 | 0.212341692 | 0.40188548 | 0.52836368 | 0.59724694 | 0.99665574 |
| ACACA | 44.8908922 | -0.383744888 | 0.38715169 | -0.9912003 | 0.32158776 | 0.97700345 |
| TADA2A | 15.256667 | -0.15772275 | 0.45265981 | -0.3484355 | 0.72751314 | 0.99665574 |
| SYNRG | 35.2817341 | 0.00718105 | 0.31097689 | 0.02309191 | 0.98157696 | 0.99777713 |
| DDX52 | 31.6788079 | -0.254576779 | 0.36372968 | -0.6999065 | 0.48398568 | 0.99640678 |
| AC243571.2 | 11.0127025 | -2.452860884 | 0.99697723 | -2.4602978 | 0.01388218 | 0.39608255 |
| HNF1B | 41.7573834 | -0.372603542 | 0.49466539 | -0.7532436 | 0.45130354 | 0.99617422 |
| MRPL45 | 7.5095991 | 0.330979159 | 0.55445911 | 0.59694061 | 0.55054704 | 0.99665574 |
| SOCS7 | 13.0529571 | -0.561772897 | 0.43540278 | -1.2902373 | 0.19696829 | 0.90381759 |
| MLLT6 | 31.6866515 | -0.086287211 | 0.34801597 | -0.2479404 | 0.80418054 | 0.99665574 |
| CISD3 | 9.51260859 | 0.126517563 | 0.61510611 | 0.20568413 | 0.83703766 | 0.99665574 |
| PCGF2 | 7.94195557 | -0.462819242 | 0.65515896 | -0.7064228 | 0.47992523 | 0.99617422 |
| PSMB3 | 9.74555648 | 0.256010791 | 0.59027778 | 0.4337124 | 0.6644973 | 0.99665574 |
| PIP4K2B | 9.60043822 | 0.442615719 | 0.54747115 | 0.80847314 | 0.41881826 | 0.99476033 |
| CWC25 | 6.42185673 | -0.43454694 | 0.60095307 | -0.7230963 | 0.46962072 | 0.99617422 |
| RPL23 | 41.3762646 | 0.449965546 | 0.36589218 | 1.22977633 | 0.21878087 | 0.91662582 |
| LASP1 | 12.3488505 | -0.44304475 | 0.45740708 | -0.9686005 | 0.33274453 | 0.97788427 |
| RPL19 | 77.2780581 | 0.229150719 | 0.47252816 | 0.48494616 | 0.62771453 | 0.99665574 |
| STAC2 | 52.3106345 | -0.783676458 | 0.77108563 | -1.0163287 | 0.30947289 | 0.97376371 |
| FBXL20 | 60.6214882 | 0.091926333 | 0.31029506 | 0.29625459 | 0.76703567 | 0.99665574 |
| MED1 | 13.0026874 | 0.093506869 | 0.53482416 | 0.17483666 | 0.86120798 | 0.99665574 |
| CDK12 | 26.5710408 | -0.27318333 | 0.35479058 | -0.7699847 | 0.44130895 | 0.99617422 |
| STARD3 | 6.32815025 | 0.016340086 | 0.74598153 | 0.02190414 | 0.98252442 | 0.99777713 |
| PGAP3 | 4.33699655 | 0.106257433 | 0.87822962 | 0.12099049 | 0.90369857 | 0.99777713 |
| ERBB2 | 30.7494781 | 0.282966934 | 0.4110694 | 0.68836779 | 0.4912212 | 0.99640678 |
| MIEN1 | 11.8351749 | -0.61515061 | 0.47576485 | -1.2929719 | 0.19602076 | 0.90381759 |
| GRB7 | 16.6099287 | -0.309832353 | 0.47755984 | -0.6487823 | 0.51647913 | 0.99665574 |
| ORMDL3 | 3.32098758 | 0.934430835 | 0.79667053 | 1.17292004 | 0.24082787 | 0.93216487 |
| PSMD3 | 8.97332616 | -0.198526165 | 0.51307328 | -0.3869353 | 0.6988041 | 0.99665574 |
| MED24 | 8.95194094 | -0.146090954 | 0.52491081 | -0.2783158 | 0.78076997 | 0.99665574 |
| THRA | 12.1209381 | 0.43601969 | 0.634026 | 0.68770002 | 0.49164171 | 0.99640678 |
| MSL1 | 13.559805 | 0.520692485 | 0.5434211 | 0.95817496 | 0.33797454 | 0.97788427 |
| CASC3 | 16.6058728 | -0.150959194 | 0.43516929 | -0.3468976 | 0.72866823 | 0.99665574 |
| WIPF2 | 28.6712636 | -0.348555923 | 0.43805093 | -0.7956973 | 0.42620803 | 0.99518711 |
| RARA | 4.0287034 | -0.279379764 | 0.85600741 | -0.3263754 | 0.74414034 | 0.99665574 |
| IGFBP4 | 6.64984898 | -1.251535709 | 0.58528132 | -2.138349 | 0.03248843 | 0.57257379 |
| KRT10 | 12.1681446 | -0.581443233 | 0.65790212 | -0.8837838 | 0.37681294 | 0.98679664 |
| TMEM99 | 3.41984768 | 0.60217561 | 0.822504 | 0.73212484 | 0.46409238 | 0.99617422 |
| KRT19 | 25.5317759 | -2.111384511 | 0.76235239 | -2.769565 | 0.00561312 | 0.23322054 |
| EIF1 | 96.8566663 | -0.521877925 | 0.422203 | -1.2360829 | 0.21642773 | 0.91498931 |
| JUP | 66.6312919 | 0.174090814 | 0.45706358 | 0.38088971 | 0.70328509 | 0.99665574 |
| NT5C3B | 8.09140838 | -0.08144317 | 0.54408927 | -0.1496871 | 0.88101145 | 0.99665574 |
| ACLY | 10.7983896 | -0.580087845 | 0.50634707 | -1.1456329 | 0.2519471 | 0.94076968 |
| CNP | 8.31004658 | 0.192180156 | 0.59465158 | 0.32318111 | 0.74655809 | 0.99665574 |
| DNAJC7 | 26.4018227 | -0.04677992 | 0.34334486 | -0.1362476 | 0.89162553 | 0.99760413 |
| NKIRAS2 | 3.32819008 | -0.308944025 | 0.77585581 | -0.3981977 | 0.69048444 | 0.99665574 |
| KAT2A | 3.9952003 | 1.222263868 | 0.75203236 | 1.62528095 | 0.10410271 | 0.79114938 |
| GHDC | 2.78097692 | -0.049426427 | 1.02364453 | -0.0482848 | 0.9614893 | 0.99777713 |
| STAT5B | 29.2778319 | -0.282509586 | 0.32935022 | -0.8577787 | 0.3910147 | 0.99179694 |
| STAT5A | 4.90735309 | -0.666623295 | 0.7233734 | -0.921548 | 0.3567644 | 0.9819332 |
| STAT3 | 76.7048121 | -1.442534928 | 0.35127454 | -4.1065741 | 4.02E-05 | 0.01163178 |
| ATP6V0A1 | 26.3574085 | 0.008320376 | 0.32453923 | 0.0256375 | 0.97954647 | 0.99777713 |
| NAGLU | 8.17783525 | -0.514066358 | 0.64890718 | -0.7922032 | 0.42824219 | 0.99589919 |
| COASY | 4.13013453 | -0.292321316 | 0.86190156 | -0.3391586 | 0.73449026 | 0.99665574 |
| MLX | 5.93734636 | 0.124041844 | 0.59897866 | 0.20708892 | 0.83594041 | 0.99665574 |
| RETREG3 | 16.1359022 | 0.203188463 | 0.42950539 | 0.47307547 | 0.63615934 | 0.99665574 |
| TUBG1 | 2.23279775 | 0.603827048 | 0.94000858 | 0.64236333 | 0.5206373 | 0.99665574 |
| TUBG2 | 5.22978559 | -0.94182808 | 0.64300443 | -1.4647303 | 0.14299451 | 0.84969914 |
| EZH1 | 16.216994 | 0.152874817 | 0.40741889 | 0.37522761 | 0.7074912 | 0.99665574 |
| VPS25 | 7.1285611 | -0.271549274 | 0.53203847 | -0.5103941 | 0.60977542 | 0.99665574 |
| COA3 | 19.5888662 | 0.606849812 | 0.41205844 | 1.47272752 | 0.14082452 | 0.84969914 |
| BECN1 | 8.56045006 | -0.270563031 | 0.62150401 | -0.4353359 | 0.6633186 | 0.99665574 |

| | | | | | | |
|---|---|---|---|---|---|---|
| PSME3 | 8.09805429 | -0.00910529 | 0.54625553 | -0.0166686 | 0.98670103 | 0.99777713 |
| LINC00671 | 6.55423796 | 0.010991197 | 0.77177788 | 0.0142414 | 0.98863739 | 0.99777713 |
| RUNDC1 | 7.08901089 | 0.740365169 | 0.64368648 | 1.1501953 | 0.25006344 | 0.93755294 |
| RPL27 | 24.2359839 | 0.354879653 | 0.41491457 | 0.85530777 | 0.39238079 | 0.99179694 |
| IFI35 | 5.10839408 | -0.798141541 | 1.386272 | -0.5757467 | 0.5647864 | 0.99665574 |
| VAT1 | 6.19654192 | -1.867166693 | 0.57861573 | -3.2269546 | 0.00125115 | 0.10147353 |
| BRCA1 | 6.04257769 | -0.173574026 | 0.68251156 | -0.2543166 | 0.79925097 | 0.99665574 |
| NBR1 | 22.3374352 | 0.217155687 | 0.51854153 | 0.41878167 | 0.6753757 | 0.99665574 |
| LINC00854 | 34.0294596 | 0.400766595 | 0.78618235 | 0.5097629 | 0.61021758 | 0.99665574 |
| ARL4D | 36.3056934 | -0.466149619 | 0.50110952 | -0.930235 | 0.35224942 | 0.97838483 |
| DHX8 | 9.40251437 | -0.452784894 | 0.4755476 | -0.9521337 | 0.34102918 | 0.97788427 |
| DUSP3 | 10.7482699 | -0.387332322 | 0.57403106 | -0.6747585 | 0.49982923 | 0.99640678 |
| TMEM101 | 17.3372772 | -1.711204356 | 0.59598604 | -2.8712155 | 0.00408897 | 0.20783038 |
| LSM12 | 13.7954425 | -0.035566633 | 0.48945033 | -0.0726665 | 0.94207152 | 0.99777713 |
| G6PC3 | 4.52541541 | -0.347399281 | 0.79965149 | -0.4344384 | 0.66397015 | 0.99665574 |
| HDAC5 | 21.2179121 | -0.192627908 | 0.43241847 | -0.4454664 | 0.65598272 | 0.99665574 |
| TMUB2 | 7.50555684 | 0.368822706 | 0.62591792 | 0.58925091 | 0.55569297 | 0.99665574 |
| ATXN7L3 | 5.64947058 | -0.044686016 | 0.69587436 | -0.0642156 | 0.94879853 | 0.99777713 |
| UBTF | 13.4319569 | 0.159446812 | 0.48059933 | 0.33176662 | 0.74006549 | 0.99665574 |
| SLC25A39 | 14.5262336 | 0.058037176 | 0.52241816 | 0.11109333 | 0.91154233 | 0.99777713 |
| GRN | 14.2029814 | 0.338408189 | 0.60876555 | 0.55589248 | 0.57828436 | 0.99665574 |
| GPATCH8 | 101.794534 | 0.144750424 | 0.22438091 | 0.64511023 | 0.51885575 | 0.99665574 |
| MEIOC | 5.49862169 | -0.213452947 | 0.63997879 | -0.3335313 | 0.73873328 | 0.99665574 |
| CCDC43 | 4.66313354 | 0.177154315 | 0.63647657 | 0.27833596 | 0.78075448 | 0.99665574 |
| EFTUD2 | 9.4674816 | -0.34546836 | 0.51313492 | -0.6732505 | 0.50078791 | 0.99640678 |
| DCAKD | 9.19389342 | 0.230577229 | 0.57173146 | 0.40329638 | 0.68673021 | 0.99665574 |
| NMT1 | 14.5565231 | 0.06806332 | 0.42845115 | 0.158859 | 0.87377997 | 0.99665574 |
| ACBD4 | 5.15179538 | 0.410105387 | 0.71909648 | 0.57030648 | 0.56846984 | 0.99665574 |
| HEXIM1 | 7.65742005 | 0.717374998 | 0.81407966 | 0.88120983 | 0.37820426 | 0.98679664 |
| AC138150.1 | 6.96294202 | 0.650966684 | 0.7511997 | 0.86656942 | 0.38617797 | 0.99051797 |
| MAP3K14 | 7.2261802 | -0.578135278 | 0.55011189 | -1.0509412 | 0.29328557 | 0.97072515 |
| ARHGAP27 | 5.1268741 | 0.145109326 | 0.67488759 | 0.21501259 | 0.82975751 | 0.99665574 |
| PLEKHM1 | 9.78646623 | -0.805332265 | 0.44313439 | -1.8173545 | 0.06916286 | 0.7094165 |
| LINC02210-C | 5.5129192 | 0.647157533 | 0.71217989 | 0.90869954 | 0.36350875 | 0.98362663 |
| KANSL1 | 174.047416 | 0.451757238 | 0.27540745 | 1.64032324 | 0.10093798 | 0.78386461 |
| ARL17B | 36.9711174 | 2.454265593 | 1.17367183 | 2.09110036 | 0.03651907 | 0.58591936 |
| LRRC37A2 | 8.82952727 | 0.784492706 | 0.61099034 | 1.28396909 | 0.19915277 | 0.90381759 |
| ARL17A | 11.7249578 | -0.756594299 | 0.62366626 | -1.2131397 | 0.22507642 | 0.92318867 |
| NSF | 44.3791354 | 0.462600217 | 0.38171343 | 1.21190449 | 0.22554894 | 0.92318867 |
| AC005670.2 | 5.77858472 | -0.285506979 | 0.65794457 | -0.4339377 | 0.66433366 | 0.99665574 |
| CDC27 | 48.5498191 | -0.166427671 | 0.29165644 | -0.5706292 | 0.56825104 | 0.99665574 |
| AC068234.1 | 29.9858634 | -2.061869365 | 0.62050729 | -3.3228769 | 0.00089094 | 0.08137272 |
| AC068234.2 | 6.61045936 | -1.451732893 | 0.94181268 | -1.5414242 | 0.12321357 | 0.82915695 |
| EFCAB13 | 15.8446867 | -1.189954149 | 0.55991061 | -2.1252574 | 0.03356514 | 0.57831228 |
| NPEPPS | 89.9811991 | -0.127808524 | 0.28297818 | -0.451655 | 0.65151751 | 0.99665574 |
| AC025682.1 | 10.4175915 | 0.450377513 | 0.59142344 | 0.76151448 | 0.44634983 | 0.99617422 |
| KPNB1 | 44.6268483 | -0.481465434 | 0.36432521 | -1.3215266 | 0.18632585 | 0.89532394 |
| MRPL10 | 4.6091294 | 0.137904934 | 0.74282384 | 0.18564958 | 0.85271957 | 0.99665574 |
| SCRN2 | 3.23016222 | -0.046935696 | 0.9371231 | -0.0500851 | 0.96005454 | 0.99777713 |
| SP2 | 6.3704646 | 0.386296696 | 0.58933241 | 0.65548185 | 0.51215757 | 0.99665574 |
| PNPO | 5.00318256 | 1.206633685 | 0.88576372 | 1.36225232 | 0.17311827 | 0.88157681 |
| CDK5RAP3 | 8.28557019 | 0.209333228 | 0.48706967 | 0.42978088 | 0.66735505 | 0.99665574 |
| NFE2L1 | 34.6603935 | -0.03747454 | 0.33443999 | -0.1120516 | 0.91078248 | 0.99777713 |
| CBX1 | 12.9462596 | -0.293772799 | 0.49050089 | -0.5989241 | 0.54922351 | 0.99665574 |
| SNX11 | 5.0913479 | 0.125909522 | 0.81641191 | 0.15422303 | 0.87743388 | 0.99665574 |
| SKAP1 | 132.881387 | 0.034290242 | 0.28056077 | 0.12222038 | 0.90272449 | 0.99777713 |
| HOXB2 | 6.76142251 | 0.134557288 | 0.65117701 | 0.20663704 | 0.83629333 | 0.99665574 |
| HOXB-AS1 | 3.35427568 | -0.309553086 | 0.87375275 | -0.35428 | 0.72312909 | 0.99665574 |
| HOXB3 | 75.0541021 | -0.072901538 | 0.29689134 | -0.2455496 | 0.80603094 | 0.99665574 |
| HOXB-AS3 | 23.7304178 | -0.116700353 | 0.4964606 | -0.2350647 | 0.81415853 | 0.99665574 |
| HOXB7 | 12.9716996 | -0.433763219 | 0.517448 | -0.838274 | 0.40187682 | 0.99194293 |
| AC103702.2 | 5.43751828 | -0.289864341 | 0.68264483 | -0.4246196 | 0.67111404 | 0.99665574 |
| LINC02086 | 10.5970449 | 0.519330933 | 0.72203766 | 0.71925741 | 0.47198233 | 0.99617422 |
| TTLL6 | 5.04840437 | 1.409045094 | 0.75857831 | 1.85748138 | 0.0632427 | 0.68630818 |
| CALCOCO2 | 23.3679428 | 0.226866298 | 0.37693819 | 0.60186604 | 0.54726331 | 0.99665574 |
| ATP5MC1 | 22.2297399 | 0.524386645 | 0.47304985 | 1.10852302 | 0.267636 | 0.94605781 |
| UBE2Z | 19.8735192 | -0.186065372 | 0.44788672 | -0.4154295 | 0.67782749 | 0.99665574 |
| SNF8 | 10.5722932 | -0.371017873 | 0.47393753 | -0.7828413 | 0.43372031 | 0.99617422 |
| B4GALNT2 | 47.5396968 | -0.654926282 | 0.69702987 | -0.9395957 | 0.34742497 | 0.97804112 |
| ZNF652 | 38.7209851 | 0.08956653 | 0.28769135 | 0.31132855 | 0.75555087 | 0.99665574 |
| PHB | 25.5035335 | 0.191801751 | 0.41360729 | 0.46372913 | 0.64284182 | 0.99665574 |
| SPOP | 26.9321855 | -0.060977582 | 0.31021689 | -0.1965643 | 0.84416847 | 0.99665574 |
| SLC35B1 | 4.11236828 | -0.179457431 | 0.70470633 | -0.2546556 | 0.79898909 | 0.99665574 |
| KAT7 | 11.8839898 | 0.140221447 | 0.47598326 | 0.29459324 | 0.76830463 | 0.99665574 |
| ITGA3 | 20.930275 | -2.160452435 | 0.4750464 | -4.5478767 | 5.42E-06 | 0.00274688 |
| PDK2 | 4.23936309 | -0.092376223 | 0.86369704 | -0.1069544 | 0.91482513 | 0.99777713 |
| XYLT2 | 3.94936885 | 0.4585703 | 0.79665189 | 0.57562193 | 0.56487076 | 0.99665574 |
| MRPL27 | 12.3137018 | -0.127696322 | 0.49564367 | -0.2576373 | 0.7966868 | 0.99665574 |
| LRRC59 | 7.256738 | -0.908034789 | 0.75788694 | -1.1981138 | 0.23087272 | 0.92774608 |
| ACSF2 | 4.97716091 | 1.690996615 | 0.79438065 | 2.12869814 | 0.03327924 | 0.5761693 |
| RSAD1 | 4.7383057 | 0.375936748 | 0.7576187 | 0.49620838 | 0.6197474 | 0.99665574 |
| SPATA20 | 3.57918982 | 0.933330649 | 0.8441807 | 1.10560529 | 0.2688974 | 0.94673212 |
| ABCC3 | 9.71386556 | -1.059718813 | 0.62232204 | -1.7028463 | 0.08859683 | 0.75650018 |
| ANKRD40 | 9.71473348 | -0.490278147 | 0.64084221 | -0.7650528 | 0.44424008 | 0.99617422 |
| LUC7L3 | 58.3817975 | 0.258629003 | 0.24458701 | 1.05741102 | 0.29032404 | 0.96676791 |
| TOB1 | 9.69889211 | 0.391177751 | 0.57994988 | 0.67450269 | 0.49999178 | 0.99640678 |
| SPAG9 | 156.21643 | -0.0246718 | 0.35002956 | -0.0704849 | 0.94380771 | 0.99777713 |
| AC005920.2 | 12.0575331 | -0.302131808 | 0.59234692 | -0.5100589 | 0.61001021 | 0.99665574 |
| MBTD1 | 34.0051167 | -0.14913948 | 0.33346951 | -0.4472357 | 0.65470487 | 0.99665574 |
| UTP18 | 9.85986891 | 0.161608644 | 0.52456315 | 0.30808234 | 0.75801968 | 0.99665574 |
| TOM1L1 | 37.624121 | 0.313902552 | 0.32302638 | 0.97175516 | 0.33117237 | 0.97788427 |
| COX11 | 7.9104384 | -0.009254204 | 0.53614236 | -0.0172607 | 0.98622862 | 0.99777713 |
| STXBP4 | 25.5778713 | -0.076837359 | 0.45690175 | -0.1681704 | 0.86644921 | 0.99665574 |
| MMD | 21.9702152 | 0.819025837 | 0.33398193 | 2.45230583 | 0.0141944 | 0.40056949 |
| PCTP | 2.76167877 | -0.349660853 | 1.08453803 | -0.3224053 | 0.74714564 | 0.99665574 |
| DGKE | 3.10232578 | -2.065732111 | 0.97888589 | -2.110289 | 0.03483347 | 0.58178209 |
| TRIM25 | 5.06549107 | -0.503111612 | 0.71592065 | -0.7027477 | 0.48221298 | 0.99640678 |
| COIL | 5.58598061 | 0.123912479 | 0.6196445 | 0.1999735 | 0.84150131 | 0.99665574 |
| SCPEP1 | 12.5904774 | -0.363277545 | 0.5073302 | -0.7160574 | 0.4739559 | 0.99617422 |
| AKAP1 | 22.0178654 | -0.150544597 | 0.39101106 | -0.3850137 | 0.7002273 | 0.99665574 |
| MSI2 | 407.266424 | 0.158843642 | 0.2917197 | 0.54450776 | 0.58609211 | 0.99665574 |
| MRPS23 | 6.3283105 | -0.286263959 | 0.54241688 | -0.5277564 | 0.59766845 | 0.99665574 |
| CUEDC1 | 23.014712 | -0.008744293 | 0.3357424 | -0.0260447 | 0.97922172 | 0.99777713 |
| VEZF1 | 20.3287778 | 0.31216584 | 0.39895436 | 0.78246003 | 0.43394427 | 0.99617422 |
| AC015813.2 | 6.09131554 | 0.680559764 | 0.61792786 | 1.10135796 | 0.2707409 | 0.94865166 |
| SRSF1 | 3.98311419 | 0.916844421 | 0.7653324 | 1.19796891 | 0.23092912 | 0.92774608 |
| DYNLL2 | 18.0685138 | 0.719340101 | 0.40882659 | 1.75952375 | 0.07848859 | 0.74431421 |
| TSPOAP1 | 4.53856365 | 0.034461946 | 0.89021251 | 0.03871204 | 0.96911997 | 0.99777713 |
| SUPT4H1 | 7.27579756 | 0.374168522 | 0.62533598 | 0.59834798 | 0.54960777 | 0.99665574 |
| RNF43 | 11.1743838 | -0.214919405 | 0.57824111 | -0.3716778 | 0.71013273 | 0.99665574 |
| MTMR4 | 7.04520762 | -0.100843695 | 0.54905439 | -0.183668 | 0.85427395 | 0.99665574 |
| RAD51C | 3.98414257 | 0.510340783 | 0.69022528 | 0.73938291 | 0.45967451 | 0.99617422 |
| PPM1E | 265.620918 | 1.417393034 | 0.52642894 | 2.69246791 | 0.00709254 | 0.27640407 |

| | | | | | | |
|---|---|---|---|---|---|---|
| TRIM37 | 13.1266268 | -0.568231917 | 0.43359167 | -1.3105231 | 0.19001892 | 0.8994569 |
| SKA2 | 8.11841311 | -0.013888935 | 0.54945222 | -0.0252778 | 0.97983339 | 0.99777713 |
| YPEL2 | 48.5583995 | -0.26585394 | 0.5069201 | -0.5244494 | 0.59996601 | 0.99665574 |
| DHX40 | 9.37123055 | -0.454247746 | 0.57179099 | -0.7944297 | 0.42694532 | 0.99525676 |
| CLTC | 60.487988 | -0.302478347 | 0.28790218 | -1.0506289 | 0.29342907 | 0.97088247 |
| PTRH2 | 6.29215762 | -0.307230586 | 0.6496876 | -0.4728897 | 0.63629186 | 0.99665574 |
| VMP1 | 74.0123356 | -0.437686173 | 0.29703603 | -1.4735121 | 0.14061301 | 0.84969914 |
| TUBD1 | 3.83406223 | -0.177011131 | 0.95882659 | -0.1846122 | 0.85353318 | 0.99665574 |
| RPS6KB1 | 14.8243355 | -0.200885359 | 0.46395331 | -0.4329862 | 0.66502483 | 0.99665574 |
| RNFT1 | 4.22572896 | -0.986491834 | 0.75676211 | -1.3035693 | 0.19238048 | 0.89964233 |
| HEATR6 | 4.37597384 | 0.402933056 | 0.74504953 | 0.54081378 | 0.58863594 | 0.99665574 |
| USP32 | 50.4705377 | -0.341871031 | 0.34853128 | -0.9808905 | 0.32664672 | 0.97788427 |
| APPBP2 | 33.1134508 | -0.168055466 | 0.35572059 | -0.4724367 | 0.63661511 | 0.99665574 |
| PPM1D | 11.8799294 | 0.448872572 | 0.5360274 | 0.83740602 | 0.40236438 | 0.99194293 |
| BCAS3 | 126.439611 | -0.052668987 | 0.33101603 | -0.1591131 | 0.87357977 | 0.99665574 |
| AC005746.2 | 3.3505272 | 0.597386264 | 0.93895522 | 0.63622445 | 0.52463014 | 0.99665574 |
| TBX2 | 15.5064623 | 0.434117485 | 0.42886581 | 1.0122455 | 0.31142069 | 0.97430509 |
| NACA2 | 5.51177188 | -0.551686585 | 0.7451342 | -0.7403855 | 0.45906609 | 0.99617422 |
| INTS2 | 8.93717792 | 0.091241546 | 0.50319247 | 0.18132534 | 0.85611222 | 0.99665574 |
| MED13 | 97.2767581 | -0.110393907 | 0.28264775 | -0.3905706 | 0.69611464 | 0.99665574 |
| METTL2A | 7.09714919 | -0.087087941 | 0.73309865 | -0.1187943 | 0.90543832 | 0.99777713 |
| TLK2 | 52.7748688 | -0.055516691 | 0.27434275 | -0.2023625 | 0.83963332 | 0.99665574 |
| 10-Mar | 5.02787117 | -0.355960291 | 1.04666785 | -0.3400891 | 0.73378946 | 0.99665574 |
| TANC2 | 68.0716232 | -0.484350689 | 0.34687591 | -1.3963226 | 0.16261736 | 0.87135275 |
| AC015923.1 | 5.49400913 | -1.611513559 | 0.83043961 | -1.9405548 | 0.0523123 | 0.65962952 |
| CYB561 | 3.30382196 | -0.850984258 | 0.73579636 | -1.1565486 | 0.24745683 | 0.93441937 |
| DCAF7 | 38.6874186 | -0.157811369 | 0.33755839 | -0.4675084 | 0.64013621 | 0.99665574 |
| TACO1 | 2.61673395 | -0.376091013 | 0.89042357 | -0.4223732 | 0.67275267 | 0.99665574 |
| MAP3K3 | 37.7379001 | -0.116387527 | 0.34320317 | -0.3391214 | 0.7345183 | 0.99665574 |
| CCDC47 | 9.77354611 | 0.125801753 | 0.59526344 | 0.21133795 | 0.83262357 | 0.99665574 |
| DDX42 | 23.4028957 | -0.203991527 | 0.40676838 | -0.5014931 | 0.61602415 | 0.99665574 |
| FTSJ3 | 4.47571474 | -0.042614295 | 0.67967283 | -0.0626982 | 0.95000679 | 0.99777713 |
| PSMC5 | 7.85456089 | 1.127476998 | 0.65339074 | 1.72557847 | 0.08442328 | 0.75373899 |
| SMARCD2 | 5.87092096 | -0.271366628 | 0.76628492 | -0.3541328 | 0.72323936 | 0.99665574 |
| ERN1 | 7.16690985 | -0.85699119 | 0.59984029 | -1.428699 | 0.15309078 | 0.86176253 |
| SNHG25 | 3.00946578 | 0.321064485 | 0.88591247 | 0.36241107 | 0.71704487 | 0.99665574 |
| TEX2 | 9.88495201 | 0.182087813 | 0.57313747 | 0.31770356 | 0.75070981 | 0.99665574 |
| POLG2 | 7.41962802 | 0.218593173 | 0.69200203 | 0.31588516 | 0.75208967 | 0.99665574 |
| DDX5 | 89.95171 | -0.270238647 | 0.29143474 | -0.9272698 | 0.35378645 | 0.98022903 |
| CEP95 | 28.8807896 | -0.486133882 | 0.43055958 | -1.1290746 | 0.25886637 | 0.94557817 |
| SMURF2 | 30.9090733 | -0.089364742 | 0.30855291 | -0.2896253 | 0.77210287 | 0.99665574 |
| LRRC37A3 | 28.8046351 | 0.4755634 | 0.45015894 | 1.05643443 | 0.29076978 | 0.96703132 |
| GNA13 | 20.6174663 | -0.4444151 | 0.35070736 | -1.2671964 | 0.20508508 | 0.90834797 |
| CEP112 | 22.6733544 | -1.110923059 | 0.71725999 | -1.5488429 | 0.12141948 | 0.82659025 |
| PRKCA | 139.998106 | -0.345723684 | 0.35005749 | -0.9876197 | 0.32333892 | 0.97700345 |
| PRKCA-AS1 | 4.74848265 | 0.031025682 | 0.90026723 | 0.03446275 | 0.97250815 | 0.99777713 |
| HELZ | 68.469922 | 0.01983151 | 0.25203221 | 0.07868641 | 0.93728206 | 0.99777713 |
| PSMD12 | 13.2575088 | -0.033764361 | 0.44089418 | -0.0765816 | 0.93895644 | 0.99777713 |
| PITPNC1 | 15.7847951 | -0.624111832 | 0.67316567 | -0.9271296 | 0.35385924 | 0.98022903 |
| NOL11 | 7.27883838 | -0.059437121 | 0.60478956 | -0.0982774 | 0.92171206 | 0.99777713 |
| BPTF | 78.6091768 | -0.325272647 | 0.27670086 | -1.175539 | 0.23977916 | 0.93157178 |
| AC005332.7 | 2.41442585 | -0.61933502 | 0.83620565 | -0.7406492 | 0.45890619 | 0.99617422 |
| AMZ2 | 4.35330262 | -0.032943643 | 0.66831808 | -0.0492934 | 0.96068551 | 0.99777713 |
| ARSG | 8.43192294 | 0.366678018 | 0.54444435 | 0.67349036 | 0.50063538 | 0.99640678 |
| WIPI1 | 5.62979748 | -0.917115247 | 0.69973724 | -1.3106566 | 0.1899738 | 0.8994569 |
| PRKAR1A | 47.220773 | 0.050862029 | 0.34798375 | 0.14616208 | 0.88379344 | 0.99665574 |
| FAM20A | 8.24964091 | 0.007536101 | 1.17541388 | 0.00641144 | 0.99488444 | 0.9985287 |
| ABCA9 | 3.32785102 | 0.985420494 | 0.9704163 | 1.0154616 | 0.30988584 | 0.97376371 |
| ABCA6 | 3.5932527 | -0.710538919 | 0.78045162 | -0.9104202 | 0.36260095 | 0.98316352 |
| ABCA10 | 16.8252069 | -0.38759549 | 0.56980375 | -0.6802263 | 0.49636119 | 0.99640678 |
| ABCA5 | 86.8115601 | 0.092648487 | 0.43416115 | 0.21339654 | 0.83101768 | 0.99665574 |
| AC005208.1 | 8.90650897 | -0.976973104 | 0.77281092 | -1.2641813 | 0.20616496 | 0.90890506 |
| KCNJ16 | 27.6565595 | -0.400056979 | 0.58922668 | -0.6789526 | 0.4971679 | 0.99640678 |
| SOX9 | 12.8139145 | -0.075723039 | 0.63236975 | -0.1197449 | 0.90468525 | 0.99777713 |
| SLC39A11 | 35.4425625 | 0.115819352 | 0.36498849 | 0.3173233 | 0.7509983 | 0.99665574 |
| COG1 | 5.09867576 | -0.298843094 | 0.71385704 | -0.4186316 | 0.67548541 | 0.99665574 |
| FAM104A | 7.37764297 | 0.669425409 | 0.56563072 | 1.18350257 | 0.23661003 | 0.9305595 |
| C17orf80 | 5.56147485 | -0.38791351 | 0.62568846 | -0.6199787 | 0.53527182 | 0.99665574 |
| CDC42EP4 | 6.52919242 | -0.401282584 | 0.58131885 | -0.6902969 | 0.49000751 | 0.99640678 |
| SDK2 | 14.5068132 | -0.213203943 | 0.83889127 | -0.2541497 | 0.79937993 | 0.99665574 |
| RPL38 | 19.893163 | 0.329127872 | 0.57254001 | 0.57485568 | 0.56538891 | 0.99665574 |
| GPRC5C | 28.9793042 | -0.202504501 | 0.37636265 | -0.5380569 | 0.5905378 | 0.99665574 |
| SLC9A3R1 | 9.66463957 | 1.150405314 | 0.61278011 | 1.87735419 | 0.06046957 | 0.6834742 |
| NAT9 | 4.74863471 | -0.897040056 | 0.61809355 | -1.4513014 | 0.14669594 | 0.85169374 |
| TMEM104 | 4.64714445 | 0.711966842 | 0.76002307 | 0.93677004 | 0.34887686 | 0.97838306 |
| MRPL58 | 4.03958189 | 0.44497533 | 0.79576888 | 0.55917659 | 0.57604121 | 0.99665574 |
| KCTD2 | 8.90640634 | -0.08937022 | 0.59059704 | -0.1513218 | 0.87972186 | 0.99665574 |
| ATP5PD | 15.5135394 | 0.469750788 | 0.59746443 | 0.7862406 | 0.43172655 | 0.99617422 |
| SLC16A5 | 40.3507708 | -0.807162578 | 0.65431462 | -1.2336001 | 0.21735196 | 0.91584128 |
| ARMC7 | 16.5615812 | -0.307156989 | 0.57932805 | -0.5301953 | 0.59597656 | 0.99665574 |
| NT5C | 13.3963929 | 0.340508605 | 0.46962371 | 0.72506689 | 0.46841099 | 0.99617422 |
| JPT1 | 20.6022672 | -0.355374189 | 0.44221844 | -0.8036169 | 0.42161829 | 0.99518711 |
| SUMO2 | 6.90037469 | -0.756750017 | 0.53411633 | -1.4168262 | 0.15653374 | 0.86532131 |
| NUP85 | 6.1937155 | -0.842916731 | 0.65940507 | -1.2782988 | 0.20114408 | 0.90591604 |
| GGA3 | 8.77528914 | -0.013414131 | 0.48305033 | -0.0277696 | 0.97784588 | 0.99777713 |
| MRPS7 | 9.92947359 | -0.046146141 | 0.55520418 | -0.0831156 | 0.9337596 | 0.99777713 |
| GRB2 | 31.9338224 | -0.159158117 | 0.34960092 | -0.4552566 | 0.64892466 | 0.99665574 |
| AC011933.4 | 3.46605735 | -1.520006442 | 0.75053809 | -2.0252223 | 0.04284455 | 0.61874363 |
| TMEM94 | 10.1118276 | 0.006022803 | 0.50005198 | 0.01204435 | 0.99039023 | 0.99777713 |
| CASKIN2 | 4.5309188 | -0.227117878 | 0.67302814 | -0.3374567 | 0.73577262 | 0.99665574 |
| TSEN54 | 7.07706194 | 0.48081171 | 0.58174472 | 0.82649948 | 0.40852081 | 0.99279547 |
| LLGL2 | 69.6615272 | -0.559883841 | 0.32066121 | -1.7460292 | 0.08080588 | 0.74735643 |
| MYO15B | 11.8419252 | 0.279542153 | 0.45155331 | 0.61906789 | 0.53587164 | 0.99665574 |
| RECQL5 | 8.26470771 | -0.119474296 | 0.53846393 | -0.2218798 | 0.82440742 | 0.99665574 |
| SMIM5 | 13.1882368 | 1.341196835 | 0.55846216 | 2.40158947 | 0.01632402 | 0.42281551 |
| SMIM6 | 8.27565671 | 1.238502765 | 0.71835178 | 1.72408952 | 0.08469168 | 0.75373899 |
| SAP30BP | 15.5907535 | -0.431917688 | 0.40063661 | -1.0780784 | 0.28099876 | 0.96059387 |
| ITGB4 | 7.87651589 | -0.531018083 | 0.61104979 | -0.8690259 | 0.38483297 | 0.99051797 |
| H3F3B | 58.9637203 | -0.395823584 | 0.2916191 | -1.3573308 | 0.17467613 | 0.88366595 |
| UNK | 17.1006819 | -0.066163897 | 0.45932253 | -0.1440467 | 0.88546359 | 0.99665574 |
| WBP2 | 13.1978001 | 0.734408581 | 0.41041256 | 1.78943984 | 0.07354401 | 0.72792151 |
| TRIM47 | 5.49713193 | -0.059221446 | 0.63845292 | -0.0927577 | 0.92609603 | 0.99777713 |
| TRIM65 | 1.43279446 | -0.358801529 | 1.19143882 | -0.3011498 | 0.76330029 | 0.99665574 |
| ACOX1 | 18.1252442 | 0.541069914 | 0.43818378 | 1.23480132 | 0.21690446 | 0.9156801 |
| TEN1 | 2.68404377 | 0.634611748 | 0.95813422 | 0.66234118 | 0.50775259 | 0.99665574 |
| EVPL | 2.90778345 | -1.233361116 | 0.87085779 | -1.4162601 | 0.15669937 | 0.86532131 |
| SRP68 | 8.24867068 | -0.070935337 | 0.64992626 | -0.1091437 | 0.91308854 | 0.99777713 |
| EXOC7 | 15.6093281 | -0.302802531 | 0.4445173 | -0.681194 | 0.49574872 | 0.99640678 |
| RNF157 | 7.88837725 | -0.128818158 | 0.66843033 | -0.1927174 | 0.84718029 | 0.99665574 |
| UBALD2 | 9.13602375 | 0.036179241 | 0.56276424 | 0.06428845 | 0.94874055 | 0.99777713 |
| PRPSAP1 | 8.83137413 | -0.195514013 | 0.51620479 | -0.3787528 | 0.70487144 | 0.99665574 |
| UBE2O | 13.2041483 | -0.421755796 | 0.4600572 | -0.9167464 | 0.35927553 | 0.9819332 |
| RHBDF2 | 4.05317633 | -1.158599491 | 0.69239097 | -1.6733313 | 0.09426209 | 0.7700476 |

| | | | | | | |
|---|---|---|---|---|---|---|
| MXRA7 | 14.7455005 | -0.205168366 | 0.53758717 | -0.3816467 | 0.70272345 | 0.99665574 |
| JMJD6 | 7.26992269 | -1.115675513 | 0.61516517 | -1.8136195 | 0.06973633 | 0.71112415 |
| METTL23 | 2.79068979 | -0.750692235 | 0.76044912 | -0.9871696 | 0.32355951 | 0.97700345 |
| SRSF2 | 27.2403709 | 0.200500178 | 0.3856112 | 0.51995424 | 0.60309547 | 0.99665574 |
| MFSD11 | 13.4436205 | -0.367906565 | 0.50898103 | -0.7228296 | 0.46978456 | 0.99617422 |
| SEC14L1 | 36.7407164 | -0.045504739 | 0.35129841 | -0.129533 | 0.89693591 | 0.99777713 |
| 9-Sep | 72.472422 | 0.123525259 | 0.24824087 | 0.49760242 | 0.6187643 | 0.99665574 |
| TNRC6C | 27.7536185 | -0.259538464 | 0.48114116 | -0.5394227 | 0.58959522 | 0.99665574 |
| TMC6 | 2.83039646 | 0.296396625 | 0.84905001 | 0.34909207 | 0.72702019 | 0.99665574 |
| SYNGR2 | 8.4475843 | -0.230803382 | 0.57081724 | -0.4043385 | 0.68596382 | 0.99665574 |
| AFMID | 6.09662191 | 1.089059568 | 0.75665571 | 1.43930661 | 0.15006367 | 0.85564988 |
| PGS1 | 7.35113868 | -0.25592176 | 0.54149699 | -0.472619 | 0.63648503 | 0.99665574 |
| CYTH1 | 131.109378 | -1.929933803 | 0.43510887 | -4.4355193 | 9.19E-06 | 0.00344882 |
| USP36 | 17.0276569 | -0.810455556 | 0.60014347 | -1.3504364 | 0.17687606 | 0.88584149 |
| TIMP2 | 20.9885994 | 0.066468572 | 0.43064966 | 0.15434488 | 0.87733781 | 0.99665574 |
| LGALS3BP | 29.1457663 | -0.409869245 | 0.63707578 | -0.6433603 | 0.51999036 | 0.99665574 |
| CANT1 | 5.01155552 | 0.980385827 | 0.7253161 | 1.35166699 | 0.17648187 | 0.88518376 |
| ENGASE | 3.80314348 | -0.321831469 | 0.73069431 | -0.4404461 | 0.65961403 | 0.99665574 |
| TBC1D16 | 22.6732078 | -0.417109136 | 0.41891156 | -0.9956974 | 0.31939721 | 0.97700345 |
| GAA | 6.26076454 | 0.186833332 | 0.75608315 | 0.24710686 | 0.80482553 | 0.99665574 |
| EIF4A3 | 5.1701853 | -0.717553552 | 0.76097293 | -0.9429423 | 0.34571043 | 0.97788427 |
| AC087741.1 | 3.80418375 | 0.309087786 | 0.72403882 | 0.42689394 | 0.66945658 | 0.99665574 |
| SGSH | 3.99436563 | -0.120361996 | 0.83939664 | -0.1433911 | 0.88598132 | 0.99665574 |
| SLC26A11 | 8.95923707 | -0.727425938 | 0.5398571 | -1.3474416 | 0.17783804 | 0.88770163 |
| RNF213 | 91.359848 | -1.726574511 | 0.61998488 | -2.7848655 | 0.00535499 | 0.22875404 |
| AC124319.2 | 21.5629887 | -0.156559181 | 0.45606085 | -0.3432857 | 0.73138352 | 0.99665574 |
| ENDOV | 5.65183701 | -0.788465344 | 0.66433754 | -1.1868445 | 0.23528897 | 0.92960234 |
| RPTOR | 49.3291539 | 0.278546393 | 0.32561713 | 0.85544145 | 0.3923068 | 0.99179694 |
| CHMP6 | 4.17737088 | 0.760544734 | 0.73347951 | 1.03689977 | 0.29978257 | 0.97358745 |
| TEPSIN | 2.34989475 | -0.028316774 | 0.93705445 | -0.0302189 | 0.97589246 | 0.99777713 |
| NDUFAF8 | 9.11802014 | 0.791240624 | 0.70778153 | 1.11791646 | 0.26360267 | 0.94557817 |
| SLC38A10 | 19.1964595 | 0.209885067 | 0.34810129 | 0.60294251 | 0.54654693 | 0.99665574 |
| BAHCC1 | 5.07794358 | 0.728264331 | 0.74935426 | 0.97185586 | 0.33112226 | 0.97788427 |
| ACTG1 | 148.061782 | 0.102469996 | 0.43141911 | 0.23751844 | 0.81225461 | 0.99665574 |
| FAAP100 | 2.6659641 | 0.284690716 | 0.89262537 | 0.31893639 | 0.74977475 | 0.99665574 |
| NPLOC4 | 26.4302013 | -0.181959061 | 0.40774546 | -0.4462565 | 0.65541197 | 0.99665574 |
| OXLD1 | 5.239444 | -0.813699713 | 0.63993716 | -1.2715306 | 0.20353993 | 0.90702762 |
| CCDC137 | 6.80557164 | -0.334072551 | 0.59845049 | -0.5582292 | 0.57668787 | 0.99665574 |
| ARL16 | 5.06037434 | 0.423482118 | 0.61547486 | 0.68805754 | 0.49141654 | 0.99640678 |
| HGS | 10.757639 | -0.815007402 | 0.61219853 | -1.3312796 | 0.18309704 | 0.89500374 |
| GCGR | 11.0474143 | 0.486023204 | 0.64519002 | 0.75330242 | 0.45126821 | 0.99617422 |
| MCRIP1 | 12.1395256 | -0.052381075 | 0.45792293 | -0.1143884 | 0.9089299 | 0.99777713 |
| P4HB | 31.8359482 | -0.740842703 | 0.42754577 | -1.7327799 | 0.0831348 | 0.75242737 |
| ARHGDIA | 31.4622068 | -0.302964312 | 0.51676115 | -0.5862753 | 0.55769051 | 0.99665574 |
| ALYREF | 2.96055739 | -0.106222141 | 0.99381634 | -0.1068831 | 0.91488174 | 0.99777713 |
| ANAPC11 | 13.2832336 | 0.347789612 | 0.44869001 | 0.77512226 | 0.43826742 | 0.99617422 |
| PCYT2 | 5.5160208 | 0.083911909 | 0.6869086 | 0.12215877 | 0.90277328 | 0.99777713 |
| SIRT7 | 5.83044516 | -0.682435286 | 0.64165465 | -1.0635554 | 0.28753014 | 0.96559334 |
| MAFG | 5.28320737 | -1.527887984 | 0.81582584 | -1.8728115 | 0.06109443 | 0.6834742 |
| ASPSCR1 | 9.073099 | -0.693141159 | 0.49415071 | -1.4026918 | 0.16070876 | 0.86894153 |
| CENPX | 5.28174399 | 1.232001738 | 0.92676471 | 1.32935762 | 0.18373001 | 0.89532394 |
| DCXR | 9.98613518 | 0.832510519 | 0.5293077 | 1.57282904 | 0.11575841 | 0.81838128 |
| RFNG | 3.98554514 | 0.139918464 | 0.8730786 | 0.16025873 | 0.87267727 | 0.99665574 |
| GPS1 | 6.58008193 | -0.003878462 | 0.72874312 | -0.0053221 | 0.99575358 | 0.99869666 |
| DUS1L | 6.45944586 | 0.214337717 | 0.74098977 | 0.28925867 | 0.77238344 | 0.99665574 |
| FASN | 4.13008072 | -1.218829938 | 0.76583434 | -1.591506 | 0.11149575 | 0.8083966 |
| CCDC57 | 21.0211419 | -0.035464568 | 0.34520545 | -0.1027347 | 0.91817356 | 0.99777713 |
| SLC16A3 | 26.6749936 | -0.598234551 | 0.49901071 | -1.1988411 | 0.23058973 | 0.92774608 |
| CSNK1D | 21.5247251 | -0.601712048 | 0.4337763 | -1.3871483 | 0.16539655 | 0.87495175 |
| OGFOD3 | 5.94784252 | -0.044465089 | 0.59365445 | -0.0749006 | 0.94029378 | 0.99777713 |
| HEXDC | 12.7210888 | 0.004626077 | 0.49196287 | 0.00940331 | 0.99249736 | 0.99777713 |
| CYBC1 | 7.09448937 | -1.002262218 | 0.69041538 | -1.45168 | 0.14659059 | 0.85169374 |
| NARF | 9.91447549 | -1.480156611 | 0.70684067 | -2.0940456 | 0.03625592 | 0.58591936 |
| FOXK2 | 38.793588 | -0.414599332 | 0.32932064 | -1.2589534 | 0.20804717 | 0.91026892 |
| WDR45B | 27.3712251 | -0.192816095 | 0.47985295 | -0.4018233 | 0.68781407 | 0.99665574 |
| RAB40B | 40.5394046 | -0.119823685 | 0.46879256 | -0.2556007 | 0.79825922 | 0.99665574 |
| FN3KRP | 5.53609296 | -0.225500538 | 0.68755672 | -0.3279737 | 0.74293153 | 0.99665574 |
| FN3K | 9.61183043 | -0.006389919 | 0.53754333 | -0.0118873 | 0.99051556 | 0.99777713 |
| TBCD | 19.1131735 | 0.048921842 | 0.47557856 | 0.10286806 | 0.91806769 | 0.99777713 |
| METRNL | 29.1867801 | 0.345158029 | 0.49188871 | 0.70169943 | 0.48286663 | 0.99640678 |
| AC144831.1 | 3.37342164 | 0.993217257 | 1.04527933 | 0.95019315 | 0.34201412 | 0.97788427 |
| USP14 | 26.2972876 | -0.070721118 | 0.390533 | -0.1810887 | 0.85629795 | 0.99665574 |
| THOC1 | 19.7316042 | -0.158012626 | 0.40349611 | -0.3916088 | 0.69534729 | 0.99665574 |
| ENOSF1 | 43.3596921 | -0.210360609 | 0.34967434 | -0.6015901 | 0.54744704 | 0.99665574 |
| YES1 | 32.1977099 | 0.187532092 | 0.32867105 | 0.57057685 | 0.56828652 | 0.99665574 |
| METTL4 | 6.94472823 | 0.20904238 | 0.56002113 | 0.37327588 | 0.70894313 | 0.99665574 |
| SMCHD1 | 49.4554493 | 0.349467098 | 0.27407147 | 1.27509475 | 0.20227571 | 0.90614933 |
| AP001011.1 | 7.87012293 | -0.389892315 | 0.84710594 | -0.4602639 | 0.64532678 | 0.99665574 |
| LPIN2 | 14.6614653 | -0.987653954 | 0.56672706 | -1.742733 | 0.08138027 | 0.74934889 |
| MYL12A | 27.6529759 | -0.656342686 | 0.32367634 | -2.0277747 | 0.04258325 | 0.61849421 |
| MYL12B | 67.072505 | -0.286686141 | 0.3779342 | -0.758561 | 0.44811522 | 0.99617422 |
| TGIF1 | 5.51380422 | -1.646271201 | 0.71553804 | -2.3007459 | 0.021406 | 0.4733368 |
| DLGAP1 | 5.21167831 | -0.788029532 | 0.73095272 | -1.0780855 | 0.2809956 | 0.96059387 |
| LINC00667 | 4.15425248 | 0.475075951 | 0.68643672 | 0.69208995 | 0.48888084 | 0.99640678 |
| EPB41L3 | 46.8855403 | -0.021542732 | 0.57291851 | -0.0376017 | 0.97000522 | 0.99777713 |
| L3MBTL4 | 93.0123602 | 0.20354652 | 0.36395458 | 0.55926352 | 0.57598189 | 0.99665574 |
| AP001021.2 | 5.0055986 | -1.586656361 | 0.94417894 | -1.6804615 | 0.09286756 | 0.76709603 |
| ARHGAP28 | 4.85895648 | -0.509643162 | 0.70958422 | -0.7182279 | 0.4726168 | 0.99617422 |
| LAMA1 | 6.72119283 | -1.401932777 | 0.75972087 | -1.8453261 | 0.06499011 | 0.69039578 |
| PTPRM | 89.8469545 | -0.126049396 | 0.32609095 | -0.3865467 | 0.69909179 | 0.99665574 |
| RAB12 | 12.4817625 | 0.064971463 | 0.49006887 | 0.13257619 | 0.89452856 | 0.99777713 |
| MTCL1 | 4.89816036 | -2.230003913 | 0.99702315 | -2.2366621 | 0.02530843 | 0.50908112 |
| NDUFV2 | 22.4344856 | 0.296188624 | 0.46715854 | 0.63402164 | 0.5260667 | 0.99665574 |
| ANKRD12 | 112.137555 | -0.632886036 | 0.36119674 | -1.752192 | 0.0797408 | 0.7446402 |
| TWSG1 | 16.1263349 | 0.266297073 | 0.50576962 | 0.52651852 | 0.59852799 | 0.99665574 |
| RALBP1 | 36.0879287 | -0.231331941 | 0.34553825 | -0.6694829 | 0.5031875 | 0.99640678 |
| PPP4R1 | 26.3417508 | -0.340675211 | 0.40277583 | -0.8458184 | 0.39765405 | 0.99179694 |
| RAB31 | 5.72552792 | -1.155863705 | 0.80972881 | -1.4274701 | 0.15344442 | 0.8618391 |
| VAPA | 73.5539847 | -0.063498519 | 0.28336592 | -0.2240866 | 0.82268988 | 0.99665574 |
| NAPG | 21.9217073 | 0.08061086 | 0.39237349 | 0.20544421 | 0.83722508 | 0.99665574 |
| GNAL | 5.17138166 | -3.050081494 | 0.89086379 | -3.4237349 | 0.00061767 | 0.06733249 |
| MPPE1 | 13.2397545 | 0.097757588 | 0.4151723 | 0.23546269 | 0.81384963 | 0.99665574 |
| IMPA2 | 45.4562697 | -0.290662498 | 0.34177397 | -0.8504524 | 0.3950736 | 0.99179694 |
| AFG3L2 | 36.3583111 | 0.144283162 | 0.37233586 | 0.3875081 | 0.69838009 | 0.99665574 |
| SPIRE1 | 127.828509 | 0.415371844 | 0.35131633 | 1.18233003 | 0.23707477 | 0.9305595 |
| PSMG2 | 21.0290639 | 0.516112788 | 0.38742468 | 1.3321629 | 0.18280666 | 0.89487878 |
| CEP76 | 6.54794125 | -0.127855125 | 0.63950503 | -0.1999283 | 0.84153669 | 0.99665574 |
| PTPN2 | 42.3142345 | 0.196909479 | 0.33601729 | 0.58600997 | 0.55786882 | 0.99665574 |
| SEH1L | 8.80542301 | -0.15209997 | 0.65168602 | -0.2333946 | 0.81545504 | 0.99665574 |
| CEP192 | 36.6052417 | 0.732020615 | 0.38941704 | 1.87978576 | 0.06013728 | 0.68195946 |
| LDLRAD4 | 10.1387467 | 0.366104488 | 0.77372049 | 0.47317409 | 0.63608898 | 0.99665574 |
| FAM210A | 12.4292368 | -0.569766934 | 0.5482496 | -1.0392473 | 0.29868972 | 0.97358745 |
| RNMT | 22.2795738 | -0.269556678 | 0.42176329 | -0.6391184 | 0.52274591 | 0.99665574 |

| | | | | | | |
|---|---|---|---|---|---|---|
| ROCK1 | 65.4492549 | 0.088436084 | 0.26216053 | 0.33733562 | 0.73586391 | 0.99665574 |
| ESCO1 | 19.8888295 | -0.091671358 | 0.42874712 | -0.2138122 | 0.83069352 | 0.99665574 |
| SNRPD1 | 7.54812208 | 0.101450761 | 0.57270923 | 0.17714183 | 0.85939699 | 0.99665574 |
| ABHD3 | 66.5066261 | -0.365155273 | 0.52640114 | -0.6936825 | 0.48788133 | 0.99640678 |
| MIB1 | 50.0461901 | 0.192989757 | 0.33237887 | 0.58063185 | 0.5614886 | 0.99665574 |
| RBBP8 | 52.8416125 | -0.034366906 | 0.30037587 | -0.114413 | 0.9089104 | 0.99777713 |
| CABLES1 | 8.63518894 | -0.55269935 | 0.63717312 | -0.8674241 | 0.38570966 | 0.99051797 |
| TMEM241 | 4.90237087 | -1.295589231 | 0.67343266 | -1.9238586 | 0.05437231 | 0.66734438 |
| RIOK3 | 29.0679767 | -0.72641713 | 0.38609309 | -1.8814559 | 0.05990993 | 0.68166875 |
| RMC1 | 9.26134853 | -0.203914539 | 0.47858173 | -0.4260809 | 0.67004889 | 0.99665574 |
| NPC1 | 24.7286915 | -1.211985201 | 0.53671085 | -2.2581716 | 0.02393496 | 0.49928529 |
| LAMA3 | 4.59072768 | -0.327678638 | 1.12292023 | -0.2918094 | 0.77043239 | 0.99665574 |
| TTC39C | 3.67633259 | -1.434486009 | 0.82855776 | -1.7313048 | 0.08339742 | 0.75242737 |
| OSBPL1A | 23.0448716 | -0.183225385 | 0.55156311 | -0.332193 | 0.73974355 | 0.99665574 |
| IMPACT | 4.51528781 | -0.12958239 | 0.70722505 | -0.1832265 | 0.85462028 | 0.99665574 |
| SS18 | 32.9970265 | -0.529270752 | 0.36751223 | -1.4401446 | 0.14982649 | 0.85525834 |
| TAF4B | 5.030808 | -0.724306971 | 0.71569519 | -1.0120327 | 0.3115224 | 0.97430509 |
| KCTD1 | 49.7703749 | -0.312319775 | 0.36279967 | -0.8608601 | 0.38931507 | 0.99176253 |
| CDH2 | 5.59772475 | 0.352259507 | 0.79817105 | 0.44133336 | 0.65897168 | 0.99665574 |
| DSC2 | 11.2830404 | -0.456067911 | 0.51882003 | -0.8790484 | 0.37937504 | 0.98679664 |
| DSG2 | 39.2247087 | 0.731606287 | 0.35658586 | 2.05169742 | 0.04019908 | 0.60713077 |
| B4GALT6 | 3.43120148 | -0.621472687 | 0.72608661 | -0.8559209 | 0.39204155 | 0.99179694 |
| TRAPPC8 | 34.5075005 | 0.188020486 | 0.37607461 | 0.49995527 | 0.61710657 | 0.99665574 |
| RNF138 | 14.5737093 | 0.411305984 | 0.46567356 | 0.8832496 | 0.37710143 | 0.98679664 |
| GAREM1 | 200.860735 | 0.092291222 | 0.32323273 | 0.28552561 | 0.77524149 | 0.99665574 |
| CCDC178 | 16.5916511 | -1.566582343 | 0.56242327 | -2.7854152 | 0.00534592 | 0.22875404 |
| ASXL3 | 18.0313503 | -0.448858478 | 0.46111241 | -0.9734253 | 0.33034198 | 0.97788427 |
| DTNA | 5.97088728 | -3.099690635 | 0.74526127 | -4.1592 | 3.19E-05 | 0.00952269 |
| MAPRE2 | 29.5281181 | 0.172902344 | 0.47190476 | 0.36639245 | 0.71407225 | 0.99665574 |
| ZNF397 | 13.016678 | -0.411909626 | 0.41778416 | -0.9859388 | 0.32416313 | 0.97700345 |
| ZSCAN30 | 8.25604791 | 0.18009866 | 0.59831883 | 0.30100784 | 0.76340852 | 0.99665574 |
| ZNF24 | 15.0657175 | 0.210611801 | 0.43530913 | 0.48382124 | 0.62851274 | 0.99665574 |
| GALNT1 | 16.9158746 | -0.222466278 | 0.4255561 | -0.522766 | 0.60113707 | 0.99665574 |
| RPRD1A | 30.481365 | 0.005161974 | 0.31898863 | 0.01618231 | 0.98708894 | 0.99777713 |
| SLC39A6 | 9.0855122 | -0.469843187 | 0.55514638 | -0.8463411 | 0.39736249 | 0.99179694 |
| ELP2 | 19.9326834 | 0.255520469 | 0.39118989 | 0.65318782 | 0.5136352 | 0.99665574 |
| MOCOS | 7.81985344 | -1.764252286 | 0.85262557 | -2.0691994 | 0.03852738 | 0.59541251 |
| FHOD3 | 60.3613981 | 0.846291883 | 0.33219383 | 2.54758456 | 0.01084716 | 0.36173838 |
| TPGS2 | 10.0060734 | -1.09684886 | 0.53750977 | -2.0406119 | 0.04128941 | 0.61108334 |
| KIAA1328 | 60.4787325 | -0.269999311 | 0.39827718 | -0.6779181 | 0.49782362 | 0.99640678 |
| AC016205.1 | 12.157597 | 0.644626277 | 0.5800217 | 1.11138304 | 0.26640351 | 0.94557817 |
| PIK3C3 | 31.8732431 | 0.239661788 | 0.29776968 | 0.80485626 | 0.42090263 | 0.99518711 |
| LINC00907 | 5.59235057 | -0.245784703 | 0.61428279 | -0.4001165 | 0.68907068 | 0.99665574 |
| SETBP1 | 127.48312 | -0.655557082 | 0.28664376 | -2.2870098 | 0.02219525 | 0.48357743 |
| EPG5 | 20.9853488 | 0.096289734 | 0.40415656 | 0.2382486 | 0.81168828 | 0.99665574 |
| ATP5F1A | 45.8521418 | 0.506509593 | 0.31134108 | 1.62686401 | 0.10376598 | 0.79062881 |
| HAUS1 | 5.47118616 | 1.535547863 | 0.67053558 | 2.29003191 | 0.02201947 | 0.48110637 |
| C18orf25 | 21.62144 | 0.423052867 | 0.39422893 | 1.07311471 | 0.28321965 | 0.96059387 |
| ST8SIA5 | 5.37345441 | -0.299727618 | 0.78813683 | -0.380299 | 0.70372351 | 0.99665574 |
| PIAS2 | 50.8315142 | 0.033829769 | 0.3564113 | 0.09491778 | 0.92438013 | 0.99777713 |
| KATNAL2 | 35.6526175 | 0.34628198 | 0.50686184 | 0.68318811 | 0.49448799 | 0.99640678 |
| HDHD2 | 3.29767375 | 1.331366691 | 0.84300489 | 1.57931076 | 0.11426479 | 0.81638817 |
| SMAD2 | 63.5296239 | -0.242103485 | 0.31559397 | -0.767136 | 0.44300068 | 0.99617422 |
| CTIF | 17.7961802 | -0.554079935 | 0.56266198 | -0.9847474 | 0.32474815 | 0.97700345 |
| SMAD7 | 5.38879611 | 0.186170921 | 0.95147216 | 0.19566618 | 0.84487146 | 0.99665574 |
| DYM | 72.2149348 | -0.030987255 | 0.27196192 | -0.1139397 | 0.90928561 | 0.99777713 |
| ACAA2 | 10.2747871 | -0.953429482 | 0.54444816 | -1.751185 | 0.07991404 | 0.7446402 |
| MYO5B | 45.1867325 | 0.149709289 | 0.38812371 | 0.3857257 | 0.69969983 | 0.99665574 |
| MBD1 | 7.75082796 | -0.347112263 | 0.59517864 | -0.5832069 | 0.55975405 | 0.99665574 |
| CXXC1 | 4.64749931 | 1.140903768 | 0.76719728 | 1.48710613 | 0.13698678 | 0.84475183 |
| MAPK4 | 3.14071704 | -0.563234552 | 0.79406928 | -0.7093015 | 0.47813738 | 0.99617422 |
| ME2 | 8.73333971 | -0.418825613 | 0.56026925 | -0.7475435 | 0.45473558 | 0.99617422 |
| SMAD4 | 17.4222701 | 0.576999061 | 0.39729982 | 1.45230133 | 0.14641784 | 0.85169374 |
| MEX3C | 4.34741411 | -1.036594793 | 0.73334524 | -1.4135154 | 0.15750424 | 0.86532131 |
| MBD2 | 39.5894958 | -0.938506492 | 0.32501388 | -2.8875889 | 0.00388207 | 0.20301691 |
| POLI | 12.8032799 | 0.266631616 | 0.48322117 | 0.55177966 | 0.58109932 | 0.99665574 |
| RAB27B | 3.46855139 | 0.438209131 | 0.9033092 | 0.48511532 | 0.62759455 | 0.99665574 |
| CCDC68 | 33.5506633 | 0.309590707 | 0.64527388 | 0.47978187 | 0.63138251 | 0.99665574 |
| TCF4 | 17.7156964 | -1.676034737 | 0.65730752 | -2.5498487 | 0.01077697 | 0.36058379 |
| TXNL1 | 13.1742468 | -0.190033483 | 0.46557103 | -0.4081729 | 0.68314673 | 0.99665574 |
| WDR7 | 38.4728547 | 0.091986286 | 0.35931317 | 0.25600589 | 0.79794629 | 0.99665574 |
| FECH | 4.9761335 | 0.05003445 | 0.62942574 | 0.07949222 | 0.93664112 | 0.99777713 |
| NARS | 25.728363 | 0.173611197 | 0.36034286 | 0.48179447 | 0.62995196 | 0.99665574 |
| AC027097.2 | 10.2198326 | -0.141958984 | 0.52109325 | -0.2724253 | 0.78529502 | 0.99665574 |
| ATP8B1 | 26.053421 | -0.582359251 | 0.38731951 | -1.5035629 | 0.13269395 | 0.84089786 |
| NEDD4L | 738.010891 | 0.363293786 | 0.31377176 | 1.15782817 | 0.24693416 | 0.93441937 |
| MALT1 | 12.4368918 | 0.140680379 | 0.47254525 | 0.29770774 | 0.76592623 | 0.99665574 |
| ZNF532 | 39.9232086 | -0.638291311 | 0.41886592 | -1.5238559 | 0.12754471 | 0.83778831 |
| SEC11C | 8.05253757 | -0.156506382 | 0.64214566 | -0.2437241 | 0.8074445 | 0.99665574 |
| LMAN1 | 27.8433047 | 0.138223005 | 0.44254828 | 0.3123343 | 0.75478648 | 0.99665574 |
| CCBE1 | 54.8947799 | 0.582443739 | 0.46110106 | 1.26315855 | 0.20653221 | 0.90890506 |
| AC010776.3 | 1.03712355 | 2.575026353 | 1.98807976 | 1.29523292 | 0.19523989 | 0.90381759 |
| CDH20 | 7.08007421 | 0.265903263 | 0.88509762 | 0.30042253 | 0.76385488 | 0.99665574 |
| AC105094.2 | 9.13760165 | -0.021470664 | 0.58879128 | -0.0364657 | 0.97091106 | 0.99777713 |
| LINC01544 | 4.85638391 | 0.433621891 | 0.75066722 | 0.57764863 | 0.56350137 | 0.99665574 |
| RNF152 | 655.886615 | 0.501051026 | 0.32049678 | 1.56335746 | 0.11796854 | 0.82599629 |
| PIGN | 37.3479087 | 0.38241687 | 0.30504027 | 1.25366027 | 0.20996551 | 0.91043197 |
| KIAA1468 | 37.4916883 | 0.188445682 | 0.34608768 | 0.54450273 | 0.58609557 | 0.99665574 |
| ZCCHC2 | 19.4486431 | -0.355233288 | 0.39972075 | -0.8887037 | 0.37416237 | 0.98535336 |
| PHLPP1 | 39.9409794 | 0.268148768 | 0.36151622 | 0.74173371 | 0.45824869 | 0.99617422 |
| BCL2 | 49.3700149 | 0.381906362 | 0.44707504 | 0.85423325 | 0.39297576 | 0.99179694 |
| KDSR | 20.0360928 | -0.505094756 | 0.38014239 | -1.3286988 | 0.18394735 | 0.89532394 |
| VPS4B | 21.7712116 | -0.251026177 | 0.32869116 | -0.7637144 | 0.44503745 | 0.99617422 |
| TMX3 | 18.7824226 | -0.161416311 | 0.38880286 | -0.4151624 | 0.67802306 | 0.99665574 |
| DOK6 | 6.25842732 | -0.989511545 | 0.79380449 | -1.2465432 | 0.21256506 | 0.91066426 |
| RTTN | 11.7809794 | 0.736665401 | 0.48844014 | 1.50819996 | 0.13150336 | 0.84006367 |
| SOCS6 | 9.1320839 | -0.089678934 | 0.51830637 | -0.173023 | 0.86263333 | 0.99665574 |
| TIMM21 | 4.45332614 | 0.043213486 | 0.68959111 | 0.06266538 | 0.95003297 | 0.99777713 |
| CYB5A | 34.3404175 | 0.2023578 | 0.52908139 | 0.38247008 | 0.70211272 | 0.99665574 |
| CNDP2 | 43.8416314 | -0.516422318 | 0.46302423 | -1.1153246 | 0.26471134 | 0.94557817 |
| LINC00909 | 2.52195214 | 0.64704606 | 0.90849003 | 0.71222142 | 0.47632767 | 0.99617422 |
| ZNF407 | 72.0568539 | 0.198374143 | 0.39531358 | 0.50181465 | 0.61579791 | 0.99665574 |
| ZADH2 | 7.99795133 | -0.771295194 | 0.7989689 | -0.9653632 | 0.33436292 | 0.97788427 |
| TSHZ1 | 9.80343967 | 0.05951676 | 0.55008944 | 0.1081947 | 0.91384125 | 0.99777713 |
| ZNF516 | 26.3386778 | -0.653257805 | 0.53928625 | -1.2113378 | 0.22576598 | 0.92318867 |
| ZNF236 | 16.4897275 | 0.484600428 | 0.43454816 | 1.11518233 | 0.26477229 | 0.94557817 |
| MBP | 38.6194973 | -1.136699606 | 0.32566386 | -3.4904075 | 0.00048228 | 0.05890843 |
| ATP9B | 64.8320816 | 0.046615392 | 0.34619465 | 0.13465082 | 0.89288796 | 0.99773598 |
| NFATC1 | 4.56994894 | -0.154607658 | 0.67747005 | -0.2282133 | 0.81948044 | 0.99665574 |
| CTDP1 | 8.24528216 | -0.169965721 | 0.52455411 | -0.3240194 | 0.74592333 | 0.99665574 |
| PQLC1 | 11.7587805 | -0.03178126 | 0.48788047 | -0.0651415 | 0.94806135 | 0.99777713 |
| HSBP1L1 | 9.86964251 | 0.044110819 | 0.54652394 | 0.08071159 | 0.93567132 | 0.99777713 |
| TXNL4A | 17.6626166 | -0.095929651 | 0.53960724 | -0.1777768 | 0.85889826 | 0.99665574 |

| | | | | | | |
|---|---|---|---|---|---|---|
| ADNP2 | 18.8359137 | -0.318476514 | 0.47195084 | -0.6748087 | 0.49979734 | 0.99640678 |
| PARD6G-AS1 | 3.6808848 | -1.43311599 | 0.88117253 | -1.6263739 | 0.10387014 | 0.79062881 |
| C20orf96 | 5.47103459 | 0.848335212 | 0.83641825 | 1.0142476 | 0.31046462 | 0.97376371 |
| ZCCHC3 | 3.43794038 | 0.048593501 | 0.69677259 | 0.06974083 | 0.94439994 | 0.99777713 |
| NRSN2-AS1 | 20.3153415 | 0.239792438 | 0.33399004 | 0.71796285 | 0.4727802 | 0.99617422 |
| SOX12 | 4.8518919 | 0.354544451 | 0.69530474 | 0.50991232 | 0.61011289 | 0.99665574 |
| NRSN2 | 4.35329135 | -0.855336998 | 0.74817262 | -1.1432348 | 0.2529411 | 0.94076968 |
| RBCK1 | 16.0643601 | 0.610971135 | 0.48224525 | 1.26693034 | 0.2051802 | 0.90834797 |
| TBC1D20 | 8.62449748 | 0.061495772 | 0.50222598 | 0.12244642 | 0.90254548 | 0.99777713 |
| CSNK2A1 | 53.9113409 | -0.279570579 | 0.30402727 | -0.9195576 | 0.35780401 | 0.9819332 |
| SLC52A3 | 7.44640442 | -1.181384813 | 0.78001708 | -1.5145627 | 0.12988317 | 0.83778831 |
| PSMF1 | 21.1335149 | -0.408598135 | 0.4142783 | -0.986289 | 0.32399131 | 0.97700345 |
| AL136531.2 | 8.33388055 | -1.678364153 | 0.69693445 | -2.4082095 | 0.01603098 | 0.42104164 |
| SDCBP2-AS1 | 7.5169275 | 0.868232328 | 0.56334924 | 1.54119729 | 0.12326878 | 0.82915695 |
| NSFL1C | 9.95669282 | -0.458250889 | 0.48024376 | -0.9542048 | 0.33998 | 0.97788427 |
| STK35 | 6.3771659 | -0.527483775 | 0.56577049 | -0.9323282 | 0.35116694 | 0.97838306 |
| SNRPB | 12.6350831 | -0.209367751 | 0.53100223 | -0.3942879 | 0.6933685 | 0.99665574 |
| NOP56 | 5.62139008 | 0.259152568 | 0.8466101 | 0.30610616 | 0.75952382 | 0.99665574 |
| IDH3B | 5.23862551 | 0.269614235 | 0.61237082 | 0.44027936 | 0.65973478 | 0.99665574 |
| PCED1A | 6.60939974 | 0.441380663 | 0.59708407 | 0.73922699 | 0.45976917 | 0.99617422 |
| PTPRA | 45.5142268 | -0.352834841 | 0.31538026 | -1.1187601 | 0.26324252 | 0.94557817 |
| MRPS26 | 13.1700738 | -0.060973243 | 0.58222008 | -0.1047254 | 0.91659369 | 0.99777713 |
| UBOX5-AS1 | 3.81288673 | -0.017247999 | 0.99638987 | -0.0173105 | 0.98618892 | 0.99777713 |
| UBOX5 | 4.93894793 | -0.119420703 | 0.64762807 | -0.184397 | 0.85370199 | 0.99665574 |
| LZTS3 | 5.52452282 | 0.829516661 | 0.71968789 | 1.15260612 | 0.24907209 | 0.93695468 |
| DDRGK1 | 10.5316148 | -0.546684211 | 0.46577269 | -1.1737146 | 0.24050935 | 0.93216487 |
| ITPA | 10.701871 | -0.031049061 | 0.4677161 | -0.0663844 | 0.94707178 | 0.99777713 |
| C20orf194 | 110.664626 | -0.069917942 | 0.24733615 | -0.2826839 | 0.77741918 | 0.99665574 |
| ATRN | 39.7132559 | 0.425049468 | 0.36016273 | 1.18015952 | 0.23793678 | 0.9305595 |
| CENPB | 8.09730643 | 0.493481101 | 0.70786264 | 0.69714246 | 0.48571365 | 0.99640678 |
| CDC25B | 3.11533158 | -0.472129643 | 0.75083233 | -0.6288084 | 0.52947453 | 0.99665574 |
| AP5S1 | 2.10404699 | 0.024727234 | 2.57598942 | 0.00959912 | 0.99234113 | 0.99777713 |
| MAVS | 29.2906318 | 0.091577992 | 0.35761047 | 0.25608308 | 0.79788668 | 0.99665574 |
| PANK2 | 15.0614686 | -0.390416328 | 0.46270074 | -0.8437772 | 0.39879392 | 0.99179694 |
| RNF24 | 22.9621875 | -1.384924708 | 0.43996871 | -3.14778 | 0.00164515 | 0.11828777 |
| PRNP | 9.47099169 | -0.604237708 | 0.54319214 | -1.112383 | 0.2659735 | 0.94557817 |
| SLC23A2 | 8.10740471 | 0.319083864 | 0.62497584 | 0.51055392 | 0.60966345 | 0.99665574 |
| TMEM230 | 20.4376928 | 0.334303872 | 0.37947737 | 0.88095865 | 0.3783402 | 0.98679664 |
| PCNA | 5.49095706 | 0.44343095 | 0.69125063 | 0.64149084 | 0.52120383 | 0.99665574 |
| CDS2 | 20.2736341 | -0.382801617 | 0.40717154 | -0.9401483 | 0.34714152 | 0.97804112 |
| GPCPD1 | 28.1121151 | 0.391646677 | 0.46725373 | 0.83818844 | 0.40192488 | 0.99194293 |
| SHLD1 | 14.3076982 | 0.381941867 | 0.41507973 | 0.92016506 | 0.35748651 | 0.9819332 |
| TRMT6 | 3.51136438 | -0.25283568 | 0.74811126 | -0.3379653 | 0.7353893 | 0.99665574 |
| MCM8 | 3.41454311 | -0.663742778 | 0.78961818 | -0.840587 | 0.40057934 | 0.99194293 |
| CRLS1 | 19.576238 | 0.519302324 | 0.40368624 | 1.28640086 | 0.1983032 | 0.90381759 |
| TMX4 | 11.7082832 | -0.164873743 | 0.52345484 | -0.3149722 | 0.75278273 | 0.99665574 |
| PLCB1 | 52.1140053 | 0.899720547 | 0.8004868 | 1.12396675 | 0.26102713 | 0.94557817 |
| PLCB4 | 19.3436084 | 0.13969502 | 0.61796282 | 0.22605732 | 0.82115684 | 0.99665574 |
| MKKS | 6.61545296 | 0.274376639 | 0.62887622 | 0.43629673 | 0.66262146 | 0.99665574 |
| SLX4IP | 65.5065306 | -0.032024239 | 0.37874985 | -0.0845525 | 0.93261718 | 0.99777713 |
| JAG1 | 21.1045233 | -1.034951154 | 0.6135662 | -1.6867799 | 0.09164567 | 0.76595534 |
| AL050403.2 | 3.31998022 | 0.598008417 | 0.8572819 | 0.69756333 | 0.48545032 | 0.99640678 |
| BTBD3 | 9.67630202 | -0.254530562 | 0.51764061 | -0.4917129 | 0.62292233 | 0.99665574 |
| SPTLC3 | 115.454476 | -0.186494115 | 0.30483975 | -0.6117775 | 0.54068495 | 0.99665574 |
| TASP1 | 39.7177706 | -0.046390085 | 0.33105745 | -0.140127 | 0.88855965 | 0.99665574 |
| ESF1 | 18.0752046 | 0.143794423 | 0.5145271 | 0.2794691 | 0.77988485 | 0.99665574 |
| NDUFAF5 | 13.2651987 | 0.141501577 | 0.43053374 | 0.32866548 | 0.74240855 | 0.99665574 |
| MACROD2 | 232.377434 | 0.282108294 | 0.46091798 | 0.61205748 | 0.54049973 | 0.99665574 |
| KIF16B | 54.7133476 | 0.511119847 | 0.33142433 | 1.54219168 | 0.12302702 | 0.82915695 |
| SNRPB2 | 10.8528236 | 0.802594148 | 0.53448681 | 1.50161637 | 0.13319621 | 0.84185986 |
| DSTN | 43.3545775 | -0.72189574 | 0.38157749 | -1.8918719 | 0.05850805 | 0.67545594 |
| RRBP1 | 22.916995 | -0.96458283 | 0.51695445 | -1.8658952 | 0.06205603 | 0.68457455 |
| SNX5 | 9.96214891 | 0.332313081 | 0.53220316 | 0.6244102 | 0.53235823 | 0.99665574 |
| OVOL2 | 7.70783422 | -0.416102067 | 0.55251812 | -0.7531012 | 0.4513891 | 0.99617422 |
| MGME1 | 7.29887779 | 0.410944538 | 0.58914118 | 0.69753151 | 0.48547023 | 0.99640678 |
| KAT14 | 3.06335981 | 1.27041218 | 0.87392021 | 1.45369356 | 0.14603128 | 0.85169374 |
| ZNF133 | 8.48358029 | -0.288344819 | 0.61142586 | -0.4715941 | 0.63721655 | 0.99665574 |
| DZANK1 | 3.11157743 | -0.065913936 | 0.84576325 | -0.0779343 | 0.93788034 | 0.99777713 |
| POLR3F | 3.56991075 | -0.205197483 | 0.71906254 | -0.2853681 | 0.77536218 | 0.99665574 |
| RBBP9 | 2.42728278 | 0.478289326 | 1.07626748 | 0.44439634 | 0.65675606 | 0.99665574 |
| SEC23B | 19.4567414 | 0.425335206 | 0.38345125 | 1.10922891 | 0.26733144 | 0.94557817 |
| DTD1 | 3.45658812 | -0.45994155 | 0.78383297 | -0.5867852 | 0.55734801 | 0.99665574 |
| AL121761.1 | 9.52374169 | -0.14108426 | 0.52357714 | -0.2694622 | 0.78757401 | 0.99665574 |
| RIN2 | 56.6974831 | -1.068381364 | 0.52490938 | -2.0353635 | 0.04181432 | 0.61480373 |
| NAA20 | 15.6057306 | -0.519689624 | 0.48488193 | -1.0717859 | 0.2838162 | 0.96059387 |
| CRNKL1 | 4.47802817 | -0.370805425 | 0.64039509 | -0.579026 | 0.56257161 | 0.99665574 |
| RALGAPA2 | 187.231384 | -0.482195814 | 0.38632152 | -1.2481723 | 0.21196798 | 0.91043197 |
| KIZ | 15.3671668 | 0.124457829 | 0.43116607 | 0.28865405 | 0.77284613 | 0.99665574 |
| KIZ-AS1 | 5.31323365 | -0.189868407 | 0.67792632 | -0.2800723 | 0.77942201 | 0.99665574 |
| XRN2 | 30.8654239 | 0.004284983 | 0.33655347 | 0.01273195 | 0.98984165 | 0.99777713 |
| NAPB | 14.3892085 | 0.526972823 | 0.44130495 | 1.19412397 | 0.23242948 | 0.92916801 |
| CST3 | 29.9989638 | -0.475449233 | 0.42383361 | -1.1217827 | 0.26195482 | 0.94557817 |
| AL035661.1 | 2.77553371 | 0.111187794 | 0.84538206 | 0.13152372 | 0.89536102 | 0.99777713 |
| APMAP | 10.1333553 | 0.188530317 | 0.50892549 | 0.37044778 | 0.71104888 | 0.99665574 |
| ACSS1 | 18.9625096 | 0.234317227 | 0.41489947 | 0.56475662 | 0.57223931 | 0.99665574 |
| ENTPD6 | 9.92191864 | -0.091387205 | 0.45874851 | -0.1992098 | 0.84209862 | 0.99665574 |
| PYGB | 18.7455762 | -0.586154549 | 0.36777897 | -1.5937685 | 0.11098788 | 0.8083966 |
| ABHD12 | 33.1197762 | 0.344488216 | 0.28498761 | 1.20878314 | 0.22674617 | 0.92318867 |
| NINL | 77.5059291 | -0.3067557 | 0.33404848 | -0.9182969 | 0.35846343 | 0.9819332 |
| ZNF337-AS1 | 5.17152438 | 0.670359032 | 0.66035149 | 1.01515487 | 0.31003201 | 0.97376371 |
| ZNF337 | 9.32524383 | -0.354923406 | 0.48954419 | -0.7250079 | 0.46844718 | 0.99617422 |
| FAM182B | 6.35044501 | -0.589179893 | 0.6250229 | -0.9426533 | 0.34585827 | 0.97788427 |
| HM13 | 22.0436811 | -0.459447689 | 0.52721635 | -0.8714595 | 0.38350332 | 0.98955375 |
| ID1 | 13.2540937 | 0.593324612 | 0.769993 | 0.77055844 | 0.4409687 | 0.99617422 |
| BCL2L1 | 44.3534666 | 0.771376582 | 0.31341284 | 2.46121566 | 0.01384671 | 0.39608255 |
| AL117381.1 | 5.6795633 | -0.267733606 | 0.61552756 | -0.4349661 | 0.66358705 | 0.99665574 |
| DUSP15 | 13.9672139 | 0.862776468 | 0.44770333 | 1.9271165 | 0.05396511 | 0.6653339 |
| PDRG1 | 3.59225401 | -0.680464063 | 0.72609771 | -0.9371522 | 0.34868027 | 0.97838306 |
| TM9SF4 | 24.6711029 | 0.177361721 | 0.39825811 | 0.44534365 | 0.65607143 | 0.99665574 |
| PLAGL2 | 18.7501971 | 0.603908395 | 0.44292343 | 1.36346003 | 0.17273757 | 0.88157681 |
| POFUT1 | 25.4661481 | 0.199799115 | 0.34682312 | 0.57608361 | 0.56455867 | 0.99665574 |
| KIF3B | 56.2700454 | -0.079455701 | 0.26136894 | -0.3039982 | 0.76112922 | 0.99665574 |
| ASXL1 | 267.445075 | -0.373443436 | 0.33295858 | -1.1215913 | 0.26203627 | 0.94557817 |
| NOL4L | 7.57156997 | 0.745874125 | 0.70475943 | 1.05833863 | 0.28990109 | 0.96676791 |
| MAPRE1 | 13.1708523 | 0.110711268 | 0.57214197 | 0.19350314 | 0.84656495 | 0.99665574 |
| AL035071.2 | 2.7208144 | -0.519787806 | 0.91634816 | -0.5672383 | 0.57055227 | 0.99665574 |
| AL035071.1 | 4.09915397 | 0.042507313 | 0.75907569 | 0.05599878 | 0.95534278 | 0.99777713 |
| BPIFA2 | 9.86766566 | -2.466500132 | 0.93017876 | -2.651641 | 0.00801017 | 0.30188501 |
| CDK5RAP1 | 11.2075624 | 0.445670493 | 0.45492225 | 0.979663 | 0.32725249 | 0.97788427 |
| SNTA1 | 4.82836736 | -0.512016717 | 0.66995604 | -0.7642542 | 0.44471578 | 0.99617422 |
| CBFA2T2 | 68.4959896 | 0.272860818 | 0.3422163 | 0.79733436 | 0.42525687 | 0.99518711 |
| NECAB3 | 4.13179167 | 0.430843663 | 0.80360451 | 0.53613893 | 0.59186254 | 0.99665574 |
| PXMP4 | 3.03349837 | -0.505085693 | 0.81670633 | -0.6184422 | 0.53628386 | 0.99665574 |

| | | | | | | |
|---|---|---|---|---|---|---|
| CHMP4B | 16.9619107 | 0.176278876 | 0.36828212 | 0.47865173 | 0.63218641 | 0.99665574 |
| RALY | 28.9083342 | -0.480773732 | 0.43506623 | -1.1050587 | 0.26913415 | 0.94673212 |
| EIF2S2 | 21.6837585 | -0.915850924 | 0.40403661 | -2.2667523 | 0.02340536 | 0.49335204 |
| AHCY | 6.53718371 | 0.423839805 | 0.57428258 | 0.73803355 | 0.46049406 | 0.99617422 |
| ITCH | 69.7758098 | -0.124924205 | 0.2882798 | -0.4333436 | 0.66476517 | 0.99665574 |
| DYNLRB1 | 37.0412277 | -0.471288995 | 0.37716556 | -1.2495547 | 0.21146226 | 0.91043197 |
| MAP1LC3A | 11.9364677 | 0.635817095 | 0.96289613 | 0.66031742 | 0.50905015 | 0.99665574 |
| PIGU | 15.1119574 | 0.025923229 | 0.51987445 | 0.0498644 | 0.96023044 | 0.99777713 |
| NCOA6 | 50.0050892 | -0.104632918 | 0.29021396 | -0.3605372 | 0.71844547 | 0.99665574 |
| TP53INP2 | 12.7260836 | -0.682133569 | 0.76183569 | -0.8953815 | 0.3705832 | 0.98532838 |
| GGT7 | 15.089727 | -0.626426602 | 0.40390504 | -1.5509255 | 0.12091955 | 0.82659025 |
| ACSS2 | 17.8847778 | 0.092283686 | 0.42917812 | 0.21502421 | 0.82974845 | 0.99665574 |
| GSS | 6.99749412 | 0.412968082 | 0.57342738 | 0.72017503 | 0.47141724 | 0.99617422 |
| TRPC4AP | 37.5933261 | -0.347736438 | 0.36447224 | -0.9540821 | 0.34004209 | 0.97788427 |
| EDEM2 | 7.83042887 | -0.400608462 | 0.5760357 | -0.6954577 | 0.48676851 | 0.99640678 |
| MMP24OS | 5.74569642 | 0.757664606 | 0.65094976 | 1.16393715 | 0.24444948 | 0.93426912 |
| MMP24 | 11.2106086 | 0.23890744 | 0.50367362 | 0.47432987 | 0.6352647 | 0.99665574 |
| UQCC1 | 23.5025912 | 0.377160296 | 0.34978491 | 1.07826349 | 0.28091618 | 0.96059387 |
| CEP250 | 3.8935892 | -0.866970792 | 0.65058851 | -1.3325947 | 0.18266485 | 0.89461655 |
| ERGIC3 | 12.9503969 | 0.181712703 | 0.43702306 | 0.4157966 | 0.67755885 | 0.99665574 |
| CPNE1 | 15.0568523 | 0.474037688 | 0.46361311 | 1.02248552 | 0.30655116 | 0.97376371 |
| ROMO1 | 12.7253503 | 0.817909476 | 0.5744795 | 1.42374005 | 0.15452174 | 0.86196782 |
| RBM39 | 95.8597154 | -0.060511626 | 0.23936132 | -0.2528045 | 0.80041926 | 0.99665574 |
| PHF20 | 38.9070914 | 0.090744037 | 0.34935897 | 0.25974441 | 0.79506094 | 0.99665574 |
| SCAND1 | 27.5312361 | -0.270704671 | 0.35036407 | -0.7726382 | 0.43973652 | 0.99617422 |
| NORAD | 76.6774963 | -0.361541061 | 0.44411751 | -0.8140662 | 0.41560702 | 0.99279547 |
| EPB41L1 | 56.0861113 | 0.185637189 | 0.40074131 | 0.46323447 | 0.64319631 | 0.99665574 |
| AAR2 | 5.36706769 | 0.546776227 | 0.61465853 | 0.88956096 | 0.37370168 | 0.98532838 |
| DLGAP4 | 22.9035858 | -0.011667173 | 0.53384055 | -0.0218552 | 0.98256349 | 0.99777713 |
| DLGAP4-AS1 | 10.013129 | -0.019338597 | 0.50817827 | -0.0380548 | 0.96964403 | 0.99777713 |
| MYL9 | 3.85715652 | -0.106717742 | 0.72622394 | -0.1469488 | 0.88317243 | 0.99665574 |
| RAB5IF | 4.56141332 | 0.081109923 | 0.68836823 | 0.11782927 | 0.90620294 | 0.99777713 |
| NDRG3 | 11.6611981 | 0.313149015 | 0.49014627 | 0.6388889 | 0.52289521 | 0.99665574 |
| SOGA1 | 40.4390134 | -0.319216584 | 0.42051245 | -0.7591133 | 0.4477848 | 0.99617422 |
| TLDC2 | 82.0095879 | 0.089820766 | 0.46410064 | 0.19353726 | 0.84653823 | 0.99665574 |
| SAMHD1 | 41.9899525 | -0.176043593 | 0.69006275 | -0.2551124 | 0.79863626 | 0.99665574 |
| RBL1 | 5.82927667 | -0.107598282 | 0.60355198 | -0.1782751 | 0.85850694 | 0.99665574 |
| RPN2 | 23.8309642 | 0.231331105 | 0.36941724 | 0.62620549 | 0.53118017 | 0.99665574 |
| MANBAL | 7.51056525 | 0.221174464 | 0.53308968 | 0.41489166 | 0.67822121 | 0.99665574 |
| SRC | 10.0338506 | 0.612012483 | 0.48800053 | 1.25412257 | 0.20979746 | 0.91043197 |
| BLCAP | 19.6295089 | -0.458577928 | 0.49161597 | -0.932797 | 0.35092477 | 0.97838306 |
| CTNNBL1 | 12.7867928 | -0.121540045 | 0.42682895 | -0.2847512 | 0.77583478 | 0.99665574 |
| TTI1 | 8.429966 | 0.244123382 | 0.52721107 | 0.46304677 | 0.64333084 | 0.99665574 |
| RPRD1B | 16.0469447 | 0.835480085 | 0.41230481 | 2.02636513 | 0.04272738 | 0.61874363 |
| TGM2 | 6.19506995 | -2.701356732 | 0.85689438 | -3.1524967 | 0.00161881 | 0.11753244 |
| RALGAPB | 35.2006997 | 0.1604779 | 0.33986054 | 0.47218751 | 0.63679296 | 0.99665574 |
| ACTR5 | 2.86588689 | 0.423429394 | 0.82446601 | 0.51358017 | 0.60754555 | 0.99665574 |
| DHX35 | 7.87849301 | 0.302777146 | 0.54655236 | 0.55397647 | 0.57959495 | 0.99665574 |
| TOP1 | 52.766565 | -0.172815562 | 0.34456463 | -0.5015476 | 0.61598579 | 0.99665574 |
| PLCG1 | 7.593705 | -0.745280072 | 0.665007 | -1.1207101 | 0.26241128 | 0.94557817 |
| ZHX3 | 50.3399757 | 0.061771173 | 0.46868255 | 0.13179747 | 0.89514449 | 0.99777713 |
| LPIN3 | 8.0339528 | -0.135518845 | 0.53163743 | -0.2549084 | 0.79879385 | 0.99665574 |
| CHD6 | 53.7080208 | 0.440070655 | 0.28996008 | 1.51769395 | 0.12909157 | 0.83778831 |
| PTPRT | 2.36200648 | 3.169160857 | 1.21961963 | 2.59848298 | 0.00936367 | 0.33076262 |
| SRSF6 | 19.7376566 | 0.654280955 | 0.37808714 | 1.73050308 | 0.08354043 | 0.75242737 |
| L3MBTL1 | 4.22335189 | -0.339936526 | 0.67972294 | -0.5001104 | 0.61699733 | 0.99665574 |
| SGK2 | 11.1447652 | 1.024162975 | 0.50743231 | 2.01832433 | 0.04355749 | 0.62204719 |
| IFT52 | 8.56356782 | -0.194147073 | 0.54215201 | -0.3581045 | 0.72026511 | 0.99665574 |
| OSER1 | 14.5528382 | 0.097448818 | 0.5235504 | 0.18613073 | 0.85234225 | 0.99665574 |
| FITM2 | 2.99989048 | 0.981746881 | 1.01265882 | 0.96947447 | 0.33230851 | 0.97788427 |
| TTPAL | 3.31494389 | 0.404405188 | 0.79863159 | 0.50637264 | 0.61259508 | 0.99665574 |
| SERINC3 | 18.0918121 | -0.106781451 | 0.35098036 | -0.3042377 | 0.76094682 | 0.99665574 |
| PKIG | 13.3026854 | -0.170767929 | 0.470445 | -0.3629923 | 0.71661061 | 0.99665574 |
| YWHAB | 41.9183584 | -0.371265367 | 0.37064945 | -1.0016617 | 0.316507 | 0.97700345 |
| PABPC1L | 6.32112684 | 0.257540337 | 0.62835598 | 0.40986375 | 0.6819059 | 0.99665574 |
| TOMM34 | 4.05448615 | 1.371483693 | 0.8005742 | 1.71312503 | 0.08668954 | 0.75427976 |
| STK4 | 18.0852764 | -0.50745332 | 0.43279446 | -1.1725042 | 0.24099468 | 0.93216487 |
| SLPI | 43.4216019 | -5.983260788 | 1.02245378 | -5.8518643 | 4.86E-09 | 7.04E-06 |
| SDC4 | 63.6883051 | 0.17254629 | 0.41521676 | 0.41555714 | 0.67773409 | 0.99665574 |
| PIGT | 6.87030352 | -0.133027554 | 0.71743315 | -0.1854215 | 0.85289842 | 0.99665574 |
| WFDC2 | 1006.65492 | -1.213864532 | 0.60495092 | -2.0065504 | 0.04479755 | 0.6313648 |
| DNTTIP1 | 5.12278705 | -0.273276542 | 0.78006803 | -0.350324 | 0.72609557 | 0.99665574 |
| SNX21 | 6.52671873 | 0.032668462 | 0.75547944 | 0.04324203 | 0.9655086 | 0.99777713 |
| ACOT8 | 6.98172669 | -0.039630228 | 0.54219958 | -0.0730916 | 0.94173324 | 0.99777713 |
| ZSWIM3 | 5.22258449 | -0.250237383 | 0.68062437 | -0.3676586 | 0.71312784 | 0.99665574 |
| CTSA | 22.3971337 | -0.406513166 | 0.61330365 | -0.6628253 | 0.50744246 | 0.99665574 |
| PCIF1 | 2.18727748 | -0.364247873 | 0.98499321 | -0.3697973 | 0.71153349 | 0.99665574 |
| ZNF335 | 6.60775097 | -0.50749022 | 0.62558991 | -0.8112187 | 0.4172401 | 0.99392253 |
| NCOA5 | 14.7494421 | -0.047718533 | 0.54207737 | -0.088029 | 0.92985362 | 0.99777713 |
| CDH22 | 8.20186341 | -1.804009185 | 0.7508843 | -2.4025129 | 0.01628286 | 0.42281551 |
| SLC35C2 | 11.0236408 | 0.3705192 | 0.49895277 | 0.74259373 | 0.45772769 | 0.99617422 |
| ELMO2 | 8.06595071 | 0.146340216 | 0.50909461 | 0.2874519 | 0.77376632 | 0.99665574 |
| ZNF334 | 7.47566554 | 0.199679517 | 0.73111418 | 0.27311673 | 0.78476348 | 0.99665574 |
| SLC13A3 | 14.4640635 | 0.575920062 | 0.64021151 | 0.8995778 | 0.36834498 | 0.98532838 |
| TP53RK | 3.03088581 | 0.571723755 | 0.80205171 | 0.71282655 | 0.47595309 | 0.99617422 |
| ZMYND8 | 42.9346581 | 0.117817774 | 0.47209502 | 0.24956369 | 0.80292478 | 0.99665574 |
| NCOA3 | 45.8071095 | -0.152961613 | 0.2830664 | -0.5403736 | 0.5889394 | 0.99665574 |
| SULF2 | 4.15911636 | -0.819825231 | 0.70888461 | -1.1565003 | 0.24747659 | 0.93441937 |
| ARFGEF2 | 63.7547387 | -0.126899511 | 0.22731857 | -0.5582452 | 0.57667693 | 0.99665574 |
| CSE1L | 14.7606884 | -0.271449245 | 0.39506874 | -0.6870937 | 0.49202368 | 0.99640678 |
| STAU1 | 44.3992562 | -0.237945077 | 0.25627751 | -0.9284665 | 0.35316565 | 0.97985587 |
| DDX27 | 7.35110095 | -0.099699167 | 0.51707245 | -0.1928147 | 0.8471041 | 0.99665574 |
| ZNFX1 | 13.9900969 | -1.24738895 | 0.60732081 | -2.053921 | 0.03998334 | 0.60713077 |
| ZFAS1 | 13.3974296 | -0.589583092 | 0.51107652 | -1.1536102 | 0.24866001 | 0.93575175 |
| B4GALT5 | 43.0429021 | 0.022746077 | 0.30311548 | 0.07504096 | 0.94018212 | 0.99777713 |
| SLC9A8 | 15.4668413 | 0.157674811 | 0.54112332 | 0.29138425 | 0.77075746 | 0.99665574 |
| RNF114 | 12.6271991 | -0.058890299 | 0.42692913 | -0.1379393 | 0.8902884 | 0.99702493 |
| CEBPB | 4.92618713 | -1.710540874 | 0.77704477 | -2.2013415 | 0.02771186 | 0.53615045 |
| PTPN1 | 28.1619734 | -0.486567594 | 0.40241449 | -1.2091205 | 0.22661657 | 0.92318867 |
| RIPOR3 | 8.31427301 | -0.334773912 | 0.58366519 | -0.5735718 | 0.56625758 | 0.99665574 |
| PARD6B | 41.0938151 | 0.273594112 | 0.36900079 | 0.74144588 | 0.45842314 | 0.99617422 |
| BCAS4 | 13.1910168 | 0.825084847 | 0.6646725 | 1.24134043 | 0.21448002 | 0.91280851 |
| ADNP | 59.3908192 | 0.12814221 | 0.36500269 | 0.35107196 | 0.72553436 | 0.99665574 |
| DPM1 | 15.2544191 | 0.385681803 | 0.41691179 | 0.92509209 | 0.35491801 | 0.98022903 |
| NFATC2 | 6.285982 | 0.056409066 | 0.6468733 | 0.08720265 | 0.93051043 | 0.99777713 |
| ATP9A | 47.1559552 | -0.794370095 | 0.32099109 | -2.4747419 | 0.01333325 | 0.39134822 |
| ZFP64 | 46.4202041 | -0.032842789 | 0.50020195 | -0.0656591 | 0.94764927 | 0.99777713 |
| AL157838.1 | 3.48519523 | -1.285446458 | 1.27969159 | -1.0044971 | 0.31513908 | 0.97652251 |
| ZNF217 | 27.5846282 | 0.248584517 | 0.41412601 | 0.60026299 | 0.54833098 | 0.99665574 |
| PFDN4 | 4.90288314 | -0.359191892 | 0.70240535 | -0.5113741 | 0.60908914 | 0.99665574 |
| FAM210B | 4.23018901 | 0.434054833 | 0.94238416 | 0.46059224 | 0.64509118 | 0.99665574 |
| CSTF1 | 3.0613611 | -1.364314405 | 0.7808071 | -1.747313 | 0.08058306 | 0.74618467 |
| RTF2 | 9.35805161 | -0.615760545 | 0.59135678 | -1.0412674 | 0.29775145 | 0.97358745 |

| | | | | | | |
|---|---|---|---|---|---|---|
| RAE1 | 6.32642414 | -0.122493788 | 0.62539385 | -0.1958666 | 0.84471455 | 0.99665574 |
| RBM38 | 2.03923752 | 1.273017696 | 0.98460962 | 1.29291617 | 0.19604005 | 0.90381759 |
| PCK1 | 25.2673565 | 1.600461111 | 0.93015733 | 1.72063485 | 0.08531711 | 0.75373899 |
| PMEPA1 | 3.77484202 | -1.68816753 | 1.02605071 | -1.6453061 | 0.09990669 | 0.78293401 |
| RAB22A | 20.5249637 | 0.073957627 | 0.44373115 | 0.16667215 | 0.86762802 | 0.99665574 |
| VAPB | 32.0438717 | 0.282055401 | 0.34457609 | 0.81855767 | 0.41303883 | 0.99279547 |
| NPEPL1 | 5.75298361 | 0.536682638 | 0.69403532 | 0.77327857 | 0.43935753 | 0.99617422 |
| GNAS | 123.119381 | -0.310802751 | 0.29509919 | -1.0532145 | 0.29224269 | 0.9698057 |
| NELFCD | 9.87027086 | -0.010515363 | 0.46648049 | -0.0225419 | 0.98201568 | 0.99777713 |
| CTSZ | 13.7728793 | -0.840800146 | 0.47847825 | -1.757238 | 0.07887725 | 0.74454113 |
| ATP5F1E | 59.8538072 | 0.194185296 | 0.40992396 | 0.47371053 | 0.63570634 | 0.99665574 |
| PRELID3B | 7.33289461 | 1.087110942 | 0.72769797 | 1.49390404 | 0.1352007 | 0.84373478 |
| SYCP2 | 8.33517252 | 0.729797633 | 0.52565997 | 1.38834546 | 0.16503188 | 0.87495175 |
| FAM217B | 4.17710828 | 0.676681486 | 0.73890112 | 0.91579436 | 0.35977476 | 0.9819332 |
| MIR646HG | 20.2721035 | 1.244317594 | 0.91731474 | 1.35647837 | 0.17494701 | 0.88415395 |
| TAF4 | 15.7889501 | 0.314525003 | 0.50022372 | 0.62876867 | 0.52950051 | 0.99665574 |
| LSM14B | 3.92195018 | 0.438696166 | 0.75078809 | 0.58431423 | 0.55900892 | 0.99665574 |
| PSMA7 | 36.0002319 | -0.357881116 | 0.43667695 | -0.8195558 | 0.41246939 | 0.99279547 |
| SS18L1 | 10.9183382 | -0.745561708 | 0.59134475 | -1.2607903 | 0.20738442 | 0.90994205 |
| MTG2 | 11.3801555 | -0.079773708 | 0.42288369 | -0.1886422 | 0.85037326 | 0.99665574 |
| OSBPL2 | 14.3752948 | 0.039681531 | 0.39145504 | 0.10136932 | 0.91925729 | 0.99777713 |
| ADRM1 | 5.10129521 | -0.129675075 | 0.63522259 | -0.2041412 | 0.83824319 | 0.99665574 |
| LAMA5 | 14.6755672 | -0.324405287 | 0.47247323 | -0.6866109 | 0.49232793 | 0.99640678 |
| RPS21 | 6.74515552 | 1.086688065 | 0.63767675 | 1.70413626 | 0.08835563 | 0.75650018 |
| SLCO4A1 | 9.64357211 | -0.63821831 | 0.5262989 | -1.2126537 | 0.22526225 | 0.92318867 |
| MRGBP | 3.06232577 | -0.191431397 | 0.89034709 | -0.2150076 | 0.82976139 | 0.99665574 |
| OGFR | 9.527744 | -1.065335304 | 0.7135239 | -1.4930618 | 0.135421 | 0.84382181 |
| TCFL5 | 6.94769649 | -0.884387207 | 0.58500203 | -1.5117678 | 0.13059295 | 0.83795959 |
| DIDO1 | 27.9284512 | 0.054827625 | 0.30038374 | 0.18252528 | 0.85517052 | 0.99665574 |
| GID8 | 11.2705964 | -0.332145624 | 0.46899167 | -0.7082122 | 0.47881349 | 0.99617422 |
| YTHDF1 | 10.7120287 | -0.34777398 | 0.58322779 | -0.5962919 | 0.55098027 | 0.99665574 |
| ARFGAP1 | 7.04214073 | -0.233572835 | 0.78332403 | -0.2981816 | 0.76556454 | 0.99665574 |
| PPDPF | 58.9603764 | 0.08408331 | 0.49784886 | 0.16889325 | 0.86588061 | 0.99665574 |
| GMEB2 | 5.2610904 | -0.37225693 | 0.62298484 | -0.5975377 | 0.55014844 | 0.99665574 |
| MHENCR | 3.15665282 | -0.712718311 | 0.82025087 | -0.8689028 | 0.38490028 | 0.99051797 |
| ARFRP1 | 5.75601232 | 0.808741314 | 0.61106436 | 1.32349613 | 0.18567044 | 0.89532394 |
| SLC2A4RG | 6.57892985 | 0.200454919 | 0.65198127 | 0.30745503 | 0.75849705 | 0.99665574 |
| ZBTB46 | 10.5747194 | -0.495758729 | 0.51494394 | -0.9627431 | 0.33567646 | 0.97788427 |
| TPD52L2 | 11.280739 | 0.062532806 | 0.62997981 | 0.0992616 | 0.92093056 | 0.99777713 |
| DNAJC5 | 30.0422703 | -0.347999229 | 0.39105106 | -0.8899074 | 0.37351561 | 0.98532838 |
| UCKL1 | 7.75707395 | 0.744883811 | 0.50285074 | 1.4813219 | 0.13852081 | 0.84656101 |
| ZNF512B | 4.38124157 | 0.359431676 | 0.76079597 | 0.47244161 | 0.63661161 | 0.99665574 |
| PRPF6 | 14.5375338 | -0.326880362 | 0.44985625 | -0.7266329 | 0.46745085 | 0.99617422 |
| TCEA2 | 3.94311054 | 0.165706894 | 0.71429821 | 0.23198559 | 0.8165492 | 0.99665574 |
| PCMTD2 | 48.5319978 | 0.539755644 | 0.56665639 | 0.95252723 | 0.34082966 | 0.97788427 |
| MIER2 | 4.24777123 | 0.422172188 | 0.69498872 | 0.60745185 | 0.54355109 | 0.99665574 |
| TPGS1 | 6.79348707 | -0.321067528 | 0.54482154 | -0.5893077 | 0.55565488 | 0.99665574 |
| CDC34 | 5.49773143 | -0.013963582 | 0.66541817 | -0.0209847 | 0.98325788 | 0.99777713 |
| BSG | 84.9557208 | 0.032111039 | 0.37863909 | 0.08480645 | 0.93241526 | 0.99777713 |
| HCN2 | 2.96877601 | 1.783445249 | 1.25331941 | 1.42297744 | 0.15474269 | 0.86196782 |
| RNF126 | 5.06022709 | -0.025282013 | 0.65907401 | -0.0383599 | 0.96940073 | 0.99777713 |
| PALM | 2.67984062 | 0.549198295 | 0.92893489 | 0.5912129 | 0.55437778 | 0.99665574 |
| PTBP1 | 19.1345707 | -0.085214387 | 0.50074549 | -0.170175 | 0.86487247 | 0.99665574 |
| MED16 | 7.28445417 | 0.71271359 | 0.55889286 | 1.27522401 | 0.20222997 | 0.90614933 |
| R3HDM4 | 5.75557732 | -0.554310887 | 0.68198744 | -0.8127875 | 0.41633988 | 0.992853 |
| ARID3A | 22.2076462 | 0.152119435 | 0.46830588 | 0.32482922 | 0.74531033 | 0.99665574 |
| WDR18 | 4.43000717 | 0.45149829 | 0.72623038 | 0.62170119 | 0.53413837 | 0.99665574 |
| TMEM259 | 16.5850474 | 0.118408569 | 0.44048621 | 0.26881334 | 0.78807333 | 0.99665574 |
| CNN2 | 7.20039788 | 0.570345818 | 0.70004014 | 0.81473302 | 0.41522515 | 0.99279547 |
| ABCA7 | 3.14208683 | -0.124167602 | 0.77664863 | -0.1598762 | 0.87297863 | 0.99665574 |
| POLR2E | 9.34177854 | -0.949540223 | 0.55564245 | -1.7089051 | 0.08746853 | 0.75597266 |
| GPX4 | 40.8263139 | 0.419979194 | 0.5823273 | 0.72120815 | 0.47078146 | 0.99617422 |
| SBNO2 | 16.5859971 | -1.560954759 | 0.5437795 | -2.8705656 | 0.00409738 | 0.20783038 |
| STK11 | 15.8459858 | 0.245024231 | 0.38029149 | 0.64430637 | 0.51937678 | 0.99665574 |
| ATP5F1D | 43.1862761 | 0.741054938 | 0.35032837 | 2.11531525 | 0.03440309 | 0.58178209 |
| MIDN | 23.6810953 | -0.749893953 | 0.57289847 | -1.3089474 | 0.19055218 | 0.89964233 |
| CIRBP | 38.2856829 | 0.508827474 | 0.40146131 | 1.26743839 | 0.20499858 | 0.90834797 |
| C19orf24 | 7.80003716 | 0.637584102 | 0.65456614 | 0.97405604 | 0.33002871 | 0.97788427 |
| MUM1 | 17.29302 | 0.355418685 | 0.36325074 | 0.97843899 | 0.32785725 | 0.97788427 |
| NDUFS7 | 11.4891583 | 1.203962055 | 0.49960937 | 2.4098068 | 0.01596097 | 0.42029172 |
| DAZAP1 | 21.1660044 | 0.042515485 | 0.3859363 | 0.11016192 | 0.91228096 | 0.99777713 |
| RPS15 | 97.349655 | 0.155463819 | 0.40362578 | 0.38516821 | 0.7001128 | 0.99665574 |
| C19orf25 | 12.8122679 | -0.467961748 | 0.45054947 | -1.0386468 | 0.29896905 | 0.97358745 |
| MBD3 | 9.96273791 | -0.212770368 | 0.75494001 | -0.2818375 | 0.77806815 | 0.99665574 |
| UQCR11 | 1.45908637 | -1.102869376 | 1.17122489 | -0.9416376 | 0.34637822 | 0.97788427 |
| TCF3 | 22.4990053 | 0.526595376 | 0.40929836 | 1.28658071 | 0.19824047 | 0.90381759 |
| REXO1 | 7.24456271 | -0.719575371 | 0.58084957 | -1.2388326 | 0.21540751 | 0.913434 |
| ABHD17A | 11.4045938 | -0.248469748 | 0.48513494 | -0.5121663 | 0.60853465 | 0.99665574 |
| SCAMP4 | 8.82024086 | -0.218314031 | 0.49916974 | -0.4373543 | 0.66185442 | 0.99665574 |
| CSNK1G2 | 11.8269449 | -0.654415833 | 0.5037896 | -1.2989864 | 0.1939486 | 0.90235789 |
| BTBD2 | 12.4760457 | 0.212194377 | 0.45919016 | 0.46210567 | 0.64400554 | 0.99665574 |
| MKNK2 | 10.0740478 | 0.465104348 | 0.46651074 | 0.9969853 | 0.31877164 | 0.97700345 |
| MOB3A | 7.77142962 | -0.27066679 | 0.72226855 | -0.3747454 | 0.70784985 | 0.99665574 |
| AP3D1 | 20.8642999 | -0.342175095 | 0.43233208 | -0.7914636 | 0.42867353 | 0.99589919 |
| DOT1L | 16.0908018 | 0.424697396 | 0.46183551 | 0.91958584 | 0.35778923 | 0.9819332 |
| PLEKHJ1 | 22.573216 | -0.088746666 | 0.49868792 | -0.1779603 | 0.85875413 | 0.99665574 |
| SF3A2 | 2.64639712 | -0.368986207 | 1.05815085 | -0.3487085 | 0.72730815 | 0.99665574 |
| LSM7 | 16.0689423 | -0.03257306 | 0.41553136 | -0.0783889 | 0.93751868 | 0.99777713 |
| SPPL2B | 9.12366811 | -0.024595817 | 0.45840431 | -0.0536553 | 0.9572098 | 0.99777713 |
| TIMM13 | 14.7264885 | 0.485896411 | 0.46954451 | 1.03482504 | 0.30075063 | 0.97358745 |
| LMNB2 | 4.76428519 | -0.967262603 | 0.62734273 | -1.5418408 | 0.12311229 | 0.82915695 |
| GADD45B | 31.2850696 | -1.417576072 | 0.7852395 | -1.8052786 | 0.0710311 | 0.71510751 |
| GNG7 | 16.3092936 | 0.678619905 | 0.51157436 | 1.32653228 | 0.18466345 | 0.89532394 |
| DIRAS1 | 2.41322646 | 0.536899128 | 0.91971952 | 0.58376398 | 0.55937912 | 0.99665574 |
| SGTA | 11.7162451 | -0.249638875 | 0.47342323 | -0.5273059 | 0.59798116 | 0.99665574 |
| THOP1 | 6.75100085 | 0.032526785 | 0.63313684 | 0.05137402 | 0.95902749 | 0.99777713 |
| ZNF554 | 2.33002206 | -0.806100825 | 0.91130833 | -0.8845533 | 0.37639758 | 0.98679046 |
| ZNF555 | 4.90132991 | 0.818718624 | 0.71676483 | 1.14224163 | 0.2533536 | 0.94076968 |
| TLE2 | 16.00113 | 0.546340476 | 0.55221933 | 0.98935412 | 0.32248991 | 0.97700345 |
| AES | 62.4611459 | 0.053075499 | 0.36696532 | 0.14463356 | 0.8850002 | 0.99665574 |
| GNA11 | 48.622099 | 0.066865655 | 0.36594795 | 0.18271903 | 0.85501848 | 0.99665574 |
| NCLN | 7.08670653 | -0.177127688 | 0.56259831 | -0.3148386 | 0.75288417 | 0.99665574 |
| NFIC | 55.3333517 | -0.115779134 | 0.28316855 | -0.40887 | 0.68263505 | 0.99665574 |
| SMIM24 | 34.5423673 | 1.092231848 | 0.46303868 | 2.35883503 | 0.0183324 | 0.44944531 |
| DOHH | 3.59857205 | -0.757411621 | 0.87183213 | -0.8687586 | 0.38497921 | 0.99051797 |
| FZR1 | 8.03385845 | -0.002682156 | 0.52729534 | -0.0050866 | 0.99594147 | 0.99869977 |
| MFSD12 | 8.77322223 | -0.504142505 | 0.64961656 | -0.7760617 | 0.43771258 | 0.99617422 |
| HMG20B | 6.67744584 | -0.407265501 | 0.64382074 | -0.6325759 | 0.52701061 | 0.99665574 |
| PIP5K1C | 7.82566479 | -0.376926214 | 0.5266894 | -0.7156518 | 0.47420639 | 0.99617422 |
| TJP3 | 8.32100622 | -0.094196096 | 0.55950576 | -0.1683559 | 0.8663033 | 0.99665574 |
| MRPL54 | 11.5766283 | 0.173952913 | 0.51354282 | 0.33873108 | 0.73481232 | 0.99665574 |
| DAPK3 | 6.65686015 | -0.521070417 | 0.55737323 | -0.934868 | 0.34985631 | 0.97838306 |
| EEF2 | 105.948299 | -0.11461506 | 0.38270746 | -0.2994848 | 0.76457018 | 0.99665574 |

| | | | | | | |
|---|---|---|---|---|---|---|
| PIAS4 | 7.7972653 | 0.244368078 | 0.55485777 | 0.44041571 | 0.65963605 | 0.99665574 |
| ZBTB7A | 21.695281 | -0.138456921 | 0.35583461 | -0.3891047 | 0.69719869 | 0.99665574 |
| MAP2K2 | 18.9779687 | 0.064609854 | 0.4710301 | 0.13716714 | 0.89089868 | 0.99712197 |
| SIRT6 | 3.75116721 | -0.615925782 | 0.69047971 | -0.8920259 | 0.37237905 | 0.98532838 |
| YJU2 | 4.63155316 | -0.185991348 | 0.67675807 | -0.2748269 | 0.7834492 | 0.99665574 |
| SH3GL1 | 10.2281622 | -0.311584851 | 0.48983322 | -0.636104 | 0.52470866 | 0.99665574 |
| UBXN6 | 8.85408966 | 0.247814143 | 0.5619739 | 0.44097091 | 0.65923406 | 0.99665574 |
| HDGFL2 | 5.85562947 | 0.614109945 | 0.6409138 | 0.95817869 | 0.33797266 | 0.97788427 |
| PLIN5 | 16.1281001 | 0.551827745 | 0.58262088 | 0.94714721 | 0.34356377 | 0.97788427 |
| MYDGF | 13.6076129 | -0.70443539 | 0.56822251 | -1.2397175 | 0.21507991 | 0.913434 |
| DPP9 | 9.80783264 | -0.20863799 | 0.56061764 | -0.3721574 | 0.70977568 | 0.99665574 |
| FEM1A | 3.29412026 | -0.230109517 | 0.79875437 | -0.2880855 | 0.77328132 | 0.99665574 |
| PLIN3 | 7.63476216 | -0.943052535 | 0.58633291 | -1.6083909 | 0.10774959 | 0.80026315 |
| KDM4B | 38.3209083 | -0.257985683 | 0.3142967 | -0.8208349 | 0.41174034 | 0.99279547 |
| PTPRS | 10.3842517 | 0.214945141 | 0.55155612 | 0.38970675 | 0.69675341 | 0.99665574 |
| TINCR | 8.04657457 | -0.387129262 | 0.5301208 | -0.7302661 | 0.46522753 | 0.99617422 |
| SAFB2 | 22.4146861 | -0.139172753 | 0.40316442 | -0.345201 | 0.72994328 | 0.99665574 |
| SAFB | 25.8333279 | 0.251062879 | 0.47216153 | 0.53173091 | 0.59491238 | 0.99665574 |
| RPL36 | 55.5259051 | 0.390633312 | 0.50007837 | 0.78114419 | 0.4347177 | 0.99617422 |
| C19orf70 | 20.3835455 | 0.354006444 | 0.56488045 | 0.62669269 | 0.5308607 | 0.99665574 |
| LONP1 | 14.8616045 | -0.300958323 | 0.51449921 | -0.5849539 | 0.55857871 | 0.99665574 |
| CAPS | 8.72470732 | 0.891404393 | 0.80623956 | 1.10563217 | 0.26888576 | 0.94673212 |
| RANBP3 | 8.87279761 | -0.660154862 | 0.61153483 | -1.0795049 | 0.2803627 | 0.96059387 |
| RFX2 | 3.97233264 | -0.520569068 | 0.85939151 | -0.6057415 | 0.54468645 | 0.99665574 |
| MLLT1 | 13.1747042 | 0.296293157 | 0.55712177 | 0.53182836 | 0.59484488 | 0.99665574 |
| CLPP | 5.65731217 | -0.141610799 | 0.6444883 | -0.2197259 | 0.8260846 | 0.99665574 |
| ALKBH7 | 13.0990017 | 0.93570447 | 0.58965973 | 1.58685498 | 0.11254549 | 0.81249398 |
| GTF2F1 | 8.96974126 | -0.449963113 | 0.48266316 | -0.9322508 | 0.35120693 | 0.97838306 |
| KHSRP | 13.6314027 | 0.118788178 | 0.49935724 | 0.23788216 | 0.81197249 | 0.99665574 |
| SLC25A23 | 12.2484205 | -0.790657285 | 0.60577699 | -1.3051953 | 0.19182635 | 0.89964233 |
| CRB3 | 5.39569288 | -0.060177904 | 0.8515453 | -0.0706691 | 0.94366115 | 0.99777713 |
| AC010503.4 | 5.76428299 | -0.016557311 | 0.69806508 | -0.0237189 | 0.98107686 | 0.99777713 |
| TRIP10 | 5.86334053 | 0.170518755 | 0.70564799 | 0.24164847 | 0.80905256 | 0.99665574 |
| SH2D3A | 10.0705846 | -1.754229375 | 0.52906753 | -3.3157003 | 0.00091414 | 0.08160886 |
| ZNF557 | 2.10810722 | -0.467048341 | 0.91274914 | -0.5116941 | 0.60886512 | 0.99665574 |
| INSR | 101.56425 | -0.379752334 | 0.22830667 | -1.6633431 | 0.09624377 | 0.77378145 |
| ZNF358 | 6.56926506 | -0.035028352 | 0.73623913 | -0.0475774 | 0.96205303 | 0.99777713 |
| MCOLN1 | 2.75757542 | -0.7416141 | 0.90833387 | -0.8164554 | 0.41423972 | 0.99279547 |
| PNPLA6 | 7.62189826 | 0.311859968 | 0.72664689 | 0.42917677 | 0.66779458 | 0.99665574 |
| CAMSAP3 | 7.38937485 | 0.34708158 | 0.52068924 | 0.66658104 | 0.50503978 | 0.99665574 |
| XAB2 | 3.36031855 | 1.411479234 | 0.90179733 | 1.56518454 | 0.11753965 | 0.82579136 |
| STXBP2 | 7.83431852 | 0.919727873 | 0.60812346 | 1.51240321 | 0.13043132 | 0.83795959 |
| EVI5L | 17.7641871 | 0.543959949 | 0.45284434 | 1.20120734 | 0.22967078 | 0.92774608 |
| MAP2K7 | 11.0838672 | -0.280322562 | 0.47854365 | -0.5857826 | 0.55802159 | 0.99665574 |
| TIMM44 | 7.11621884 | -0.153991698 | 0.69081876 | -0.2229119 | 0.8236041 | 0.99665574 |
| ELAVL1 | 25.1455713 | 0.126258623 | 0.41744317 | 0.30245704 | 0.76230368 | 0.99665574 |
| FBN3 | 19.4909392 | 0.083423042 | 0.71103873 | 0.1173256 | 0.90660204 | 0.99777713 |
| CERS4 | 132.716065 | 0.134393589 | 0.53364093 | 0.25184273 | 0.80116263 | 0.99665574 |
| AC022146.2 | 27.1750628 | -0.391757794 | 0.64099213 | -0.6111741 | 0.54108432 | 0.99665574 |
| CD320 | 4.54321239 | -0.393461805 | 0.73908721 | -0.5323618 | 0.59447543 | 0.99665574 |
| RPS28 | 23.5145738 | 0.748160952 | 0.47939055 | 1.5606502 | 0.11860631 | 0.82619588 |
| RAB11B | 8.98406255 | 0.32359292 | 0.49590395 | 0.65253145 | 0.51405839 | 0.99665574 |
| 2-Mar | 6.24048314 | -0.862236025 | 0.69389547 | -1.2426022 | 0.21401447 | 0.9124923 |
| HNRNPM | 41.5493015 | -0.013553044 | 0.36718855 | -0.0369103 | 0.97055652 | 0.99777713 |
| ZNF317 | 4.42247534 | -0.051959173 | 0.73915946 | -0.0702949 | 0.94395891 | 0.99777713 |
| ZNF559-ZNF | 3.72141231 | 0.795973486 | 0.7877402 | 1.01045177 | 0.31227889 | 0.97442101 |
| ZNF266 | 7.22109276 | -0.408462263 | 0.51392915 | -0.7947832 | 0.42673961 | 0.99525676 |
| ZNF426 | 4.49007904 | -0.355435659 | 0.60626755 | -0.5862686 | 0.55769499 | 0.99665574 |
| ZNF121 | 7.38317874 | 0.425057222 | 0.60027763 | 0.70810106 | 0.4788825 | 0.99617422 |
| ZNF561 | 4.59089236 | -0.118984897 | 0.73827846 | -0.1611653 | 0.87196318 | 0.99665574 |
| ZNF562 | 15.8492161 | -0.322210503 | 0.38491301 | -0.8370995 | 0.40253662 | 0.99194293 |
| ZNF846 | 8.30309944 | -0.224330374 | 0.59428517 | -0.3774793 | 0.70581741 | 0.99665574 |
| FBXL12 | 4.15354353 | -0.140065179 | 0.64938787 | -0.215688 | 0.82923095 | 0.99665574 |
| UBL5 | 12.9426296 | 0.302248697 | 0.54564968 | 0.55392445 | 0.57963055 | 0.99665574 |
| PIN1 | 7.65759354 | 0.082652771 | 0.71699483 | 0.11527666 | 0.90822584 | 0.99777713 |
| C19orf66 | 7.11204668 | -0.072101224 | 0.91912863 | -0.0784452 | 0.93747392 | 0.99777713 |
| EIF3G | 8.53604275 | 0.69034798 | 0.57603125 | 1.19845577 | 0.23073963 | 0.92774608 |
| DNMT1 | 5.68717634 | -0.013836651 | 0.66082723 | -0.0209384 | 0.98329481 | 0.99777713 |
| MRPL4 | 7.57871664 | 0.017633763 | 0.57546528 | 0.03064262 | 0.97555455 | 0.99777713 |
| TYK2 | 8.40615533 | -0.10844031 | 0.4868343 | -0.2227458 | 0.82373333 | 0.99665574 |
| CDC37 | 12.4633353 | -0.491803061 | 0.51409359 | -0.9566411 | 0.33874843 | 0.97788427 |
| KEAP1 | 6.93391112 | 0.077156071 | 0.57297033 | 0.1346598 | 0.89288086 | 0.99773598 |
| ATG4D | 6.84083583 | -0.382285394 | 0.69623902 | -0.5490721 | 0.582956 | 0.99665574 |
| KRI1 | 3.33050135 | -0.059280312 | 0.74312075 | -0.0797721 | 0.9364185 | 0.99777713 |
| AP1M2 | 10.8942587 | 0.170977873 | 0.51321293 | 0.33315192 | 0.73901961 | 0.99665574 |
| SLC44A2 | 11.3773073 | 0.633450153 | 0.51589906 | 1.22785676 | 0.21950074 | 0.91727062 |
| ILF3-DT | 7.48013512 | 0.560323227 | 0.62471774 | 0.89692222 | 0.36976042 | 0.98532838 |
| ILF3 | 30.8715243 | 0.201371422 | 0.43875822 | 0.45895761 | 0.64626461 | 0.99665574 |
| QTRT1 | 6.90957404 | 0.991410244 | 0.64010391 | 1.54882703 | 0.12142331 | 0.82659025 |
| DNM2 | 46.0042119 | -0.521449562 | 0.28220104 | -1.8477946 | 0.06463206 | 0.68846743 |
| TMED1 | 4.50703852 | 0.019942143 | 0.78823659 | 0.02529969 | 0.97981592 | 0.99777713 |
| CARM1 | 5.0326186 | -0.261032594 | 0.62979935 | -0.4144695 | 0.67853033 | 0.99665574 |
| YIPF2 | 6.21662093 | 0.162465427 | 0.60178201 | 0.26997389 | 0.78718034 | 0.99665574 |
| TIMM29 | 4.9702203 | -0.489671976 | 0.60497164 | -0.8094131 | 0.41827757 | 0.99432434 |
| SMARCA4 | 35.5301338 | -0.316865084 | 0.41687447 | -0.7600971 | 0.44719653 | 0.99617422 |
| LDLR | 21.3601006 | -2.910384745 | 0.75503746 | -3.854623 | 0.00011591 | 0.02448076 |
| KANK2 | 11.2579785 | 0.514298206 | 0.5234945 | 0.98243287 | 0.32588664 | 0.97788427 |
| DOCK6 | 10.8546424 | -0.074811083 | 0.55702363 | -0.134305 | 0.89316137 | 0.99773598 |
| EPOR | 8.2925095 | 0.842311751 | 0.80612551 | 1.04488909 | 0.29607423 | 0.97286866 |
| RGL3 | 61.4754332 | 0.033524756 | 0.40404697 | 0.08297242 | 0.93387347 | 0.99777713 |
| PRKCSH | 13.8515326 | -0.0699016 | 0.45239459 | -0.1545147 | 0.87720395 | 0.99665574 |
| ECSIT | 9.3249757 | 0.332969067 | 0.54217519 | 0.61413557 | 0.53912575 | 0.99665574 |
| ELOF1 | 3.95790366 | 0.298751101 | 0.75945889 | 0.39337363 | 0.69404354 | 0.99665574 |
| ZNF627 | 6.37180241 | -0.188534732 | 0.65879637 | -0.2861806 | 0.77473981 | 0.99665574 |
| AC008543.1 | 3.19968268 | 0.420386631 | 0.84825191 | 0.49559173 | 0.62018249 | 0.99665574 |
| ZNF823 | 8.792587 | -0.452136359 | 0.55533691 | -0.8141659 | 0.41554994 | 0.99279547 |
| ZNF441 | 5.0269646 | 0.323355691 | 0.62855554 | 0.51444251 | 0.60694265 | 0.99665574 |
| ZNF440 | 8.45858354 | -0.574204609 | 0.56731794 | -1.012139 | 0.3114716 | 0.97430509 |
| ZNF439 | 9.50053581 | -0.632400342 | 0.53006768 | -1.1930558 | 0.23284751 | 0.92924083 |
| ZNF69 | 10.922034 | -1.061515209 | 0.45391511 | -2.3385765 | 0.01935736 | 0.46057537 |
| ZNF844 | 11.4969185 | -0.239185396 | 0.42515208 | -0.5625878 | 0.57371557 | 0.99665574 |
| ZNF788P | 3.29824258 | -0.219501235 | 0.80116987 | -0.2739759 | 0.78410314 | 0.99665574 |
| ZNF136 | 5.0667888 | -0.185782207 | 0.77542658 | -0.2395871 | 0.81065037 | 0.99665574 |
| ZNF44 | 48.5225978 | -0.270635064 | 0.36037154 | -0.750989 | 0.45265927 | 0.99617422 |
| AC008758.5 | 2.38077861 | -0.177230399 | 0.94483901 | -0.1875774 | 0.85120798 | 0.99665574 |
| ZNF490 | 7.15997081 | -0.831860569 | 0.64825776 | -1.283225 | 0.19941326 | 0.90381759 |
| ZNF791 | 12.1617761 | -0.282299412 | 0.43056361 | -0.6556509 | 0.51204877 | 0.99665574 |
| WDR83 | 1.86478403 | -1.330081048 | 1.261891 | -1.054038 | 0.29186553 | 0.96918567 |
| DHPS | 4.85801603 | 0.644447094 | 0.62985812 | 1.02316232 | 0.30623111 | 0.97376371 |
| TNPO2 | 13.0438167 | -0.08671554 | 0.4128786 | -0.2100267 | 0.83364682 | 0.99665574 |
| TRIR | 24.2827477 | -0.097039838 | 0.39323055 | -0.2467759 | 0.80508163 | 0.99665574 |
| ASNA1 | 4.54862481 | 0.265468809 | 0.77805924 | 0.34119357 | 0.73295786 | 0.99665574 |
| HOOK2 | 21.6969217 | -0.678852014 | 0.39799856 | -1.7056645 | 0.08807056 | 0.75650018 |

| | | | | | | |
|---|---|---|---|---|---|---|
| JUNB | 15.8775115 | -1.512812118 | 0.56753212 | -2.6655974 | 0.00768517 | 0.2918061 |
| DNASE2 | 5.70615646 | -0.364442875 | 0.64729604 | -0.5630235 | 0.57341889 | 0.99665574 |
| GCDH | 6.01418336 | 0.073966793 | 0.56125838 | 0.13178742 | 0.89515244 | 0.99777713 |
| FARSA | 4.7563053 | 0.474322715 | 0.64243841 | 0.73831625 | 0.4603223 | 0.99617422 |
| CALR | 37.7555973 | -0.025764465 | 0.41378497 | -0.0622653 | 0.95035152 | 0.99777713 |
| AC092069.1 | 5.11848114 | 0.294440522 | 0.71726928 | 0.41050207 | 0.68143769 | 0.99665574 |
| RAD23A | 11.9286954 | 0.187089053 | 0.49475903 | 0.37814176 | 0.70532528 | 0.99665574 |
| GADD45GIP1 | 18.9197104 | -0.192655736 | 0.42006183 | -0.4586366 | 0.64649514 | 0.99665574 |
| NFIX | 3.87810454 | 0.519863405 | 0.78473989 | 0.66246588 | 0.50767269 | 0.99665574 |
| NACC1 | 4.97431553 | -0.335181719 | 0.64403574 | -0.5204396 | 0.6027572 | 0.99665574 |
| STX10 | 4.07101372 | -0.511529274 | 0.67532487 | -0.7574566 | 0.44887636 | 0.99617422 |
| IER2 | 11.3961412 | 0.984359188 | 0.62426053 | 1.57684034 | 0.11483227 | 0.81638817 |
| CCDC130 | 10.9612662 | -0.64988886 | 0.47503012 | -1.3681003 | 0.17128068 | 0.88010314 |
| MRI1 | 5.19995119 | -0.507946147 | 0.66800106 | -0.7603972 | 0.44701719 | 0.99617422 |
| C19orf53 | 20.7345341 | 0.265428753 | 0.53927295 | 0.49219742 | 0.62257979 | 0.99665574 |
| ZSWIM4 | 4.64111865 | -0.121737366 | 0.82082811 | -0.1483104 | 0.88209779 | 0.99665574 |
| CC2D1A | 8.59295607 | -0.450035452 | 0.50973393 | -0.8828831 | 0.37729945 | 0.98679664 |
| DCAF15 | 3.1686954 | -1.200509836 | 0.81827807 | -1.4671172 | 0.14234417 | 0.84969914 |
| RFX1 | 5.72153566 | -0.263851709 | 0.56090058 | -0.4704073 | 0.63806407 | 0.99665574 |
| SAMD1 | 4.70291558 | 0.144706552 | 0.67741214 | 0.21361671 | 0.83084597 | 0.99665574 |
| PRKACA | 9.73889011 | 0.015966964 | 0.60372128 | 0.02644758 | 0.97890035 | 0.99777713 |
| ADGRL1 | 2.50563619 | 1.164882295 | 0.98004057 | 1.18860619 | 0.23459467 | 0.92960234 |
| DDX39A | 6.40642315 | -1.085204843 | 0.83114125 | -1.3056804 | 0.19166126 | 0.89964233 |
| PKN1 | 10.7902924 | -0.41492035 | 0.6932509 | -0.598514 | 0.54949704 | 0.99665574 |
| GIPC1 | 17.3506897 | -0.30342181 | 0.47582286 | -0.6376781 | 0.52368326 | 0.99665574 |
| DNAJB1 | 9.71058217 | 0.303315505 | 0.55537694 | 0.5461435 | 0.5849673 | 0.99665574 |
| TECR | 7.76347469 | 0.540212267 | 0.57352732 | 0.94191201 | 0.34623769 | 0.97788427 |
| NDUFB7 | 25.1924061 | 0.431091688 | 0.40764017 | 1.05752994 | 0.2902698 | 0.96676791 |
| ZNF333 | 4.47526804 | 0.185609134 | 0.64447257 | 0.28800161 | 0.77334551 | 0.99665574 |
| ILVBL | 6.65114837 | 0.579024342 | 0.56532486 | 1.02423293 | 0.30572527 | 0.97376371 |
| BRD4 | 55.9749015 | -0.132206862 | 0.33361355 | -0.3962874 | 0.69189299 | 0.99665574 |
| AKAP8 | 10.6312045 | 0.112425393 | 0.48588077 | 0.23138473 | 0.81701592 | 0.99665574 |
| AKAP8L | 31.6343432 | -0.163320957 | 0.31507026 | -0.5183636 | 0.6042046 | 0.99665574 |
| WIZ | 3.85281373 | -0.51851705 | 0.69404834 | -0.7470907 | 0.45500882 | 0.99617422 |
| TPM4 | 7.27201732 | -1.29939369 | 0.66455828 | -1.9552742 | 0.05055071 | 0.64929786 |
| RAB8A | 7.90208816 | -0.195419642 | 0.60387728 | -0.3236082 | 0.74623468 | 0.99665574 |
| FAM32A | 6.51837497 | -0.522278341 | 0.60668331 | -0.8608747 | 0.38930703 | 0.99176253 |
| AP1M1 | 4.60952482 | 0.634964071 | 0.70524237 | 0.90034872 | 0.3679347 | 0.98532838 |
| EPS15L1 | 22.5771621 | -0.054516172 | 0.38564969 | -0.1413619 | 0.88758405 | 0.99665574 |
| C19orf44 | 2.48646345 | -0.064362052 | 1.01801559 | -0.0632231 | 0.94958889 | 0.99777713 |
| TMEM38A | 18.8276738 | -1.060626619 | 0.55785934 | -1.9012438 | 0.05727008 | 0.67545594 |
| SIN3B | 12.718928 | -0.151270657 | 0.44316126 | -0.3413445 | 0.73284425 | 0.99665574 |
| MYO9B | 16.0939679 | -0.297285761 | 0.48750675 | -0.6098085 | 0.54198867 | 0.99665574 |
| USE1 | 4.56224381 | -0.325779603 | 0.70724045 | -0.4606349 | 0.6450606 | 0.99665574 |
| NR2F6 | 11.0954611 | -0.275872688 | 0.56396567 | -0.4891658 | 0.62472435 | 0.99665574 |
| ABHD8 | 2.11515962 | 1.283649396 | 1.05158343 | 1.22068241 | 0.22220629 | 0.92191316 |
| MRPL34 | 18.9592935 | -0.592560742 | 0.41206523 | -1.4380266 | 0.15042652 | 0.85612189 |
| DDA1 | 5.76004009 | -0.409501014 | 0.57818696 | -0.7082502 | 0.47878992 | 0.99617422 |
| BST2 | 8.08452515 | 0.386448531 | 1.22568911 | 0.31529083 | 0.75254085 | 0.99665574 |
| MVB12A | 6.25547615 | -0.480412162 | 0.61049479 | -0.7869226 | 0.43132716 | 0.99617422 |
| SLC27A1 | 4.48908201 | 0.286020184 | 0.72284449 | 0.39568702 | 0.69233593 | 0.99665574 |
| AC010618.3 | 4.37505145 | -0.488606286 | 0.67152311 | -0.727609 | 0.46685294 | 0.99617422 |
| PGLS | 12.4422659 | -0.479767423 | 0.5850338 | -0.8200679 | 0.41217742 | 0.99279547 |
| COLGALT1 | 3.10329949 | -1.095872543 | 0.87603708 | -1.2509431 | 0.21095524 | 0.91043197 |
| MAP1S | 3.2505661 | 0.100136418 | 0.85932324 | 0.1165294 | 0.90723299 | 0.99777713 |
| RPL18A | 4.0902928 | 0.137442187 | 0.66894804 | 0.20546018 | 0.8372126 | 0.99665574 |
| CCDC124 | 6.8032626 | 0.103300612 | 0.67385785 | 0.15329734 | 0.8781638 | 0.99665574 |
| ARRDC2 | 3.57964292 | -0.6006462 | 0.72995414 | -0.8228547 | 0.41059062 | 0.99279547 |
| MAST3 | 3.45299648 | 0.460256906 | 0.71792959 | 0.6410892 | 0.52146473 | 0.99665574 |
| JUND | 50.3273697 | -0.055977692 | 0.58442834 | -0.095782 | 0.92369375 | 0.99777713 |
| LSM4 | 16.1689957 | 0.385273244 | 0.43028738 | 0.89538588 | 0.37058084 | 0.98532838 |
| PGPEP1 | 9.08565165 | -0.127224581 | 0.5476568 | -0.2323071 | 0.81629947 | 0.99665574 |
| GDF15 | 21.3967955 | -0.614341605 | 0.79676333 | -0.7710465 | 0.44067935 | 0.99617422 |
| SSBP4 | 5.92282472 | -1.663434656 | 0.78682085 | -2.1141212 | 0.03450492 | 0.58178209 |
| ELL | 14.5386795 | 0.032037699 | 0.40866923 | 0.07839518 | 0.9375137 | 0.99777713 |
| FKBP8 | 13.3815142 | 0.625273411 | 0.42585297 | 1.46828472 | 0.1420269 | 0.84969914 |
| KXD1 | 5.83055259 | -0.632290553 | 0.66093462 | -0.9566613 | 0.33873825 | 0.97788427 |
| UBA52 | 36.337744 | 0.739074338 | 0.54904914 | 1.34609871 | 0.17827068 | 0.88811213 |
| REX1BD | 8.19378785 | 0.305046537 | 0.70812001 | 0.43078367 | 0.66662567 | 0.99665574 |
| KLHL26 | 2.50548311 | 0.336246657 | 0.93794522 | 0.35849285 | 0.71997452 | 0.99665574 |
| CRTC1 | 27.0967237 | 0.600635681 | 0.41068124 | 1.46253497 | 0.14359467 | 0.84969914 |
| UPF1 | 14.4223464 | 0.053124146 | 0.40089246 | 0.1325147 | 0.89457719 | 0.99777713 |
| COPE | 12.7073124 | 0.177521414 | 0.6322472 | 0.28077849 | 0.7788803 | 0.99665574 |
| DDX49 | 3.19933663 | -0.55294328 | 0.88361221 | -0.625776 | 0.5314619 | 0.99665574 |
| SUGP2 | 25.9451512 | 0.179648689 | 0.35881847 | 0.50066734 | 0.61660526 | 0.99665574 |
| ARMC6 | 3.58766331 | 0.179331729 | 0.81455609 | 0.22015885 | 0.82574745 | 0.99665574 |
| SLC25A42 | 4.92489926 | -0.086469335 | 0.59437839 | -0.1454786 | 0.88433301 | 0.99665574 |
| TMEM161A | 4.8448293 | 0.446656288 | 0.7289823 | 0.61271212 | 0.54006671 | 0.99665574 |
| RFXANK | 3.45958407 | 0.746343572 | 0.77289091 | 0.9656519 | 0.3342184 | 0.97788427 |
| SUGP1 | 5.47164862 | 0.0356848 | 0.58071318 | 0.06144996 | 0.95100087 | 0.99777713 |
| MAU2 | 14.9191639 | -0.044308184 | 0.39763671 | -0.1114288 | 0.91127632 | 0.99777713 |
| GATAD2A | 34.755543 | -0.354803931 | 0.3121714 | -1.1365677 | 0.25571905 | 0.94314473 |
| ATP13A1 | 7.16486837 | 0.349809743 | 0.63806151 | 0.54823828 | 0.5835283 | 0.99665574 |
| ZNF101 | 3.48584921 | 0.214947453 | 0.73382973 | 0.29291189 | 0.7695895 | 0.99665574 |
| ZNF14 | 5.28585988 | 0.69158951 | 0.70028277 | 0.98758608 | 0.32335541 | 0.97700345 |
| AC011477.4 | 3.15868001 | 0.349611625 | 0.81539274 | 0.4287647 | 0.66809447 | 0.99665574 |
| ZNF253 | 7.24987604 | -0.811716182 | 0.5525435 | -1.4690539 | 0.14181817 | 0.84969914 |
| ZNF93 | 3.91664853 | -0.492807627 | 0.67945002 | -0.7253037 | 0.46826572 | 0.99617422 |
| ZNF682 | 2.07851788 | -0.41179348 | 1.0783964 | -0.3818572 | 0.70256725 | 0.99665574 |
| ZNF90 | 35.2215934 | 0.342118195 | 0.43604906 | 0.78458648 | 0.43269606 | 0.99617422 |
| AC011447.3 | 6.48950223 | -0.74628846 | 0.57296971 | -1.302492 | 0.19274825 | 0.90008372 |
| ZNF486 | 4.57434031 | -0.90836419 | 0.78620787 | -1.1553741 | 0.24793728 | 0.93441937 |
| AC008554.1 | 6.05476775 | -0.921571212 | 1.03966842 | -0.8864088 | 0.37539729 | 0.98569969 |
| ZNF737 | 10.2052333 | -0.026012176 | 0.59559722 | -0.0436741 | 0.96516418 | 0.99777713 |
| ZNF66 | 16.4570782 | 0.044256108 | 0.549531 | 0.08053432 | 0.9358123 | 0.99777713 |
| ZNF85 | 5.80257141 | 0.008992845 | 0.73562412 | 0.01222478 | 0.99024628 | 0.99777713 |
| ZNF430 | 14.6796782 | -0.167109045 | 0.51375503 | -0.3252699 | 0.74497681 | 0.99665574 |
| ZNF714 | 9.40022255 | 0.512376287 | 0.54210163 | 0.94516648 | 0.34457389 | 0.97788427 |
| ZNF431 | 57.9676973 | 0.945458487 | 0.36884011 | 2.56332879 | 0.01036738 | 0.35124327 |
| ZNF708 | 20.1918823 | -0.188065571 | 0.36889786 | -0.5098039 | 0.61018885 | 0.99665574 |
| ZNF738 | 4.05060623 | -0.422960937 | 0.66982306 | -0.6314517 | 0.52774519 | 0.99665574 |
| ZNF493 | 6.7505999 | 0.385452195 | 0.61728492 | 0.62443157 | 0.5323442 | 0.99665574 |
| AC010615.4 | 10.555125 | -0.710164075 | 0.79404571 | -0.8943617 | 0.37112839 | 0.98532838 |
| ZNF429 | 16.835126 | -0.481349532 | 0.43024546 | -1.1187789 | 0.26323448 | 0.94557817 |
| AC123912.1 | 4.23008984 | 0.130327322 | 0.79140297 | 0.16467884 | 0.86919677 | 0.99665574 |
| ZNF100 | 16.943642 | -0.265604941 | 0.48482123 | -0.547841 | 0.58380109 | 0.99665574 |
| ZNF43 | 16.1680721 | -0.453048672 | 0.49606353 | -0.9132876 | 0.3610913 | 0.9819332 |
| ZNF208 | 34.5459178 | -0.5987473 | 0.50654163 | -1.1820298 | 0.23719388 | 0.9305595 |
| ZNF257 | 2.31440613 | 0.470540082 | 0.95935251 | 0.49047673 | 0.62379659 | 0.99665574 |
| ZNF676 | 3.50194322 | -0.008546372 | 0.77459664 | -0.0110333 | 0.99119686 | 0.99777713 |
| ZNF98 | 16.8120407 | 0.597136597 | 0.45533208 | 1.31143097 | 0.1897122 | 0.8994569 |
| ZNF91 | 70.1561188 | 0.030382754 | 0.39998014 | 0.07596066 | 0.9394504 | 0.99777713 |
| ZNF675 | 5.95136237 | -0.275198801 | 0.56957034 | -0.4831691 | 0.62897566 | 0.99665574 |

| | | | | | | |
|---|---|---|---|---|---|---|
| ZNF726 | 3.5813299 | -0.096899926 | 0.85763053 | -0.1129856 | 0.91004195 | 0.99777713 |
| ZNF254 | 46.9890966 | -0.52709835 | 0.36036016 | -1.4626987 | 0.14354983 | 0.84969914 |
| LINC00662 | 13.2649991 | -0.84420063 | 0.38271821 | -2.2058021 | 0.02739786 | 0.53492871 |
| AC006504.5 | 7.35237943 | 0.216881804 | 0.62713747 | 0.34582817 | 0.72947185 | 0.99665574 |
| LINC01532 | 3.01283519 | 0.057923561 | 1.15673969 | 0.05007484 | 0.96006275 | 0.99777713 |
| UQCRFS1 | 27.6099183 | 0.664432756 | 0.48156424 | 1.37973857 | 0.16766715 | 0.87566218 |
| POP4 | 3.94124847 | -0.484468996 | 0.70605216 | -0.686166 | 0.49260843 | 0.99640678 |
| C19orf12 | 13.5262278 | -0.479714187 | 0.43112916 | -1.1126925 | 0.26584051 | 0.94557817 |
| URI1 | 40.9555027 | -0.056213357 | 0.29377561 | -0.1913479 | 0.84825301 | 0.99665574 |
| ZNF507 | 12.9755298 | 0.030369816 | 0.43830005 | 0.06929001 | 0.94475877 | 0.99777713 |
| DPY19L3 | 18.1553963 | -0.207393264 | 0.37917246 | -0.5469629 | 0.58440423 | 0.99665574 |
| PDCD5 | 5.74919213 | 0.241388367 | 0.60196967 | 0.40099755 | 0.68842192 | 0.99665574 |
| ANKRD27 | 15.4412773 | -0.584754494 | 0.3739334 | -1.5637932 | 0.11786615 | 0.82599629 |
| NUDT19 | 3.27771569 | 0.608131004 | 0.79753661 | 0.76251171 | 0.44575466 | 0.99617422 |
| CEP89 | 6.92990381 | -0.125019565 | 0.71189762 | -0.1756145 | 0.86059679 | 0.99665574 |
| RHPN2 | 32.3290385 | -0.749602348 | 0.36183099 | -2.0716919 | 0.03829418 | 0.5945274 |
| GPATCH1 | 3.74969139 | -0.207140132 | 0.75139025 | -0.2756758 | 0.78279706 | 0.99665574 |
| CEBPG | 4.39120507 | -1.926084562 | 0.7623453 | -2.5265251 | 0.01151971 | 0.3725572 |
| PEPD | 12.4406887 | -0.060393806 | 0.51928628 | -0.1163016 | 0.90741355 | 0.99777713 |
| LSM14A | 65.0097621 | -0.031367896 | 0.33344094 | -0.0940733 | 0.92505091 | 0.99777713 |
| KIAA0355 | 30.1504891 | -0.146169284 | 0.41143913 | -0.3552634 | 0.72239224 | 0.99665574 |
| GPI | 37.5029114 | -0.621439545 | 0.56428874 | -1.1012794 | 0.27077509 | 0.94865166 |
| UBA2 | 25.889366 | -0.19918265 | 0.37912507 | -0.5253745 | 0.59932287 | 0.99665574 |
| SCGB2B2 | 17.2210048 | -0.911141222 | 0.47473707 | -1.9192544 | 0.05495214 | 0.67040291 |
| ZNF302 | 15.8468122 | -0.986346067 | 0.38919855 | -2.5343005 | 0.01126721 | 0.37207476 |
| ZNF181 | 1.82342531 | 0.22645614 | 0.9816879 | 0.23068038 | 0.81756311 | 0.99665574 |
| GRAMD1A | 8.00289563 | -1.00861888 | 0.54370522 | -1.855084 | 0.06358423 | 0.68630818 |
| HPN | 27.9513319 | -0.569273775 | 0.47088425 | -1.2089463 | 0.22668347 | 0.92318867 |
| LSR | 14.4626803 | -0.573445999 | 0.45855618 | -1.2505469 | 0.21109985 | 0.91043197 |
| USF2 | 17.2632435 | 0.085765976 | 0.46653454 | 0.18383629 | 0.85414189 | 0.99665574 |
| MAG | 2.57062726 | 0.25862421 | 1.09238519 | 0.23675185 | 0.81284931 | 0.99665574 |
| CD22 | 5.66237016 | 1.159149653 | 0.88433176 | 1.31076334 | 0.18993773 | 0.8994569 |
| DMKN | 13.4802892 | -0.780176529 | 0.55833045 | -1.3973383 | 0.16231186 | 0.87130922 |
| TMEM147 | 6.79585894 | 0.692099781 | 0.55188242 | 1.25407108 | 0.20981617 | 0.91043197 |
| RBM42 | 4.43444583 | -0.024849776 | 0.68680734 | -0.0361816 | 0.97113757 | 0.99777713 |
| COX6B1 | 38.9465514 | 0.293295385 | 0.32226066 | 0.9101185 | 0.36276002 | 0.98333184 |
| KMT2B | 7.69486476 | -0.118263939 | 0.52886114 | -0.22362 | 0.82305298 | 0.99665574 |
| PROSER3 | 2.42525618 | -0.203658447 | 0.95266587 | -0.2137774 | 0.83072064 | 0.99665574 |
| SDHAF1 | 2.06950427 | 0.761655148 | 1.01974542 | 0.74690715 | 0.45511961 | 0.99617422 |
| TBCB | 6.42339323 | -0.895894212 | 0.60087902 | -1.4909727 | 0.13596867 | 0.84475183 |
| CAPNS1 | 28.3045463 | -0.525008257 | 0.43493584 | -1.2070936 | 0.22739611 | 0.92389989 |
| COX7A1 | 1.49215712 | 2.521630903 | 1.47487696 | 1.7097229 | 0.08731713 | 0.75595306 |
| ZNF565 | 13.0508358 | -0.999537162 | 0.4251988 | -2.3507526 | 0.01873549 | 0.45223898 |
| ZNF146 | 16.703133 | -0.591294213 | 0.42711092 | -1.3844043 | 0.16623468 | 0.87495175 |
| LINC00665 | 3.88958289 | 0.757324409 | 0.81816845 | 0.92563385 | 0.3546363 | 0.98022903 |
| ZFP14 | 10.9037079 | -0.589257183 | 0.53812829 | -1.0950125 | 0.27351117 | 0.95221713 |
| ZFP82 | 5.93431912 | 0.135978974 | 0.67501691 | 0.20144529 | 0.84035041 | 0.99665574 |
| ZNF566 | 7.52603883 | 0.351479949 | 0.62463056 | 0.56270053 | 0.57363882 | 0.99665574 |
| ZNF260 | 7.93222132 | 0.096991621 | 0.564482 | 0.17182411 | 0.86357581 | 0.99665574 |
| ZNF529 | 9.78295046 | 0.030087471 | 0.47981621 | 0.06270624 | 0.95000043 | 0.99777713 |
| ZNF529-AS1 | 5.37447748 | 0.137580625 | 0.69735329 | 0.19728971 | 0.84360083 | 0.99665574 |
| ZNF382 | 4.6742944 | -0.778918777 | 0.62461875 | -1.2470307 | 0.21238623 | 0.91043197 |
| ZNF461 | 4.32319302 | 0.072518178 | 0.78057542 | 0.09290349 | 0.92598024 | 0.99777713 |
| ZNF567 | 4.53103565 | -0.76544431 | 0.67977435 | -1.126027 | 0.26015407 | 0.94557817 |
| AC020928.1 | 7.74133298 | 0.18977664 | 0.55191587 | 0.34385067 | 0.73095859 | 0.99665574 |
| ZNF790 | 7.17582241 | 0.048391217 | 0.73523882 | 0.065817 | 0.94752352 | 0.99777713 |
| ZNF345 | 10.1197844 | -0.609361417 | 0.45727411 | -1.3325955 | 0.18266458 | 0.89461655 |
| ZNF568 | 9.52316037 | -0.313813108 | 0.53720824 | -0.5841554 | 0.55911575 | 0.99665574 |
| ZNF420 | 17.0514505 | -0.347983865 | 0.40943529 | -0.8499118 | 0.39537415 | 0.99179694 |
| ZNF383 | 4.85445679 | 0.736308212 | 0.75266941 | 0.97826244 | 0.32794454 | 0.97788427 |
| HKR1 | 17.7925664 | 0.631799202 | 0.41571682 | 1.51978263 | 0.12856562 | 0.83778831 |
| ZNF569 | 6.26067898 | -0.132656625 | 0.55865067 | -0.237459 | 0.81230073 | 0.99665574 |
| ZNF793 | 6.62388884 | -0.041633144 | 0.55198312 | -0.0754247 | 0.93987683 | 0.99777713 |
| ZNF540 | 5.80145818 | 0.142215544 | 0.69792668 | 0.2037686 | 0.83853433 | 0.99665574 |
| ZNF571 | 3.53647648 | 0.31910113 | 0.78068482 | 0.40874515 | 0.6827267 | 0.99665574 |
| ZFP30 | 8.36316976 | -0.737576359 | 0.58803751 | -1.2543016 | 0.20973242 | 0.91043197 |
| ZNF573 | 7.58029122 | 0.162319172 | 0.52467363 | 0.3093717 | 0.75703879 | 0.99665574 |
| SIPA1L3 | 96.0104671 | -0.061561559 | 0.33371416 | -0.1844739 | 0.85364169 | 0.99665574 |
| SPINT2 | 4.71625012 | -0.601363799 | 0.72757166 | -0.8265355 | 0.4085004 | 0.99279547 |
| CATSPERG | 3.35462075 | -0.824853933 | 0.92870242 | -0.8881789 | 0.37444451 | 0.98535336 |
| PSMD8 | 14.199317 | -0.010944314 | 0.40070513 | -0.0273126 | 0.97821038 | 0.99777713 |
| EIF3K | 23.6907905 | 0.736296196 | 0.35179274 | 2.09298291 | 0.03635068 | 0.58591936 |
| ACTN4 | 65.9280812 | 0.076434767 | 0.25800223 | 0.29625622 | 0.76703441 | 0.99665574 |
| ECH1 | 14.0344193 | -0.176016379 | 0.5484598 | -0.3209285 | 0.74826458 | 0.99665574 |
| HNRNPL | 14.1629538 | 0.153287956 | 0.51082834 | 0.30007724 | 0.76411824 | 0.99665574 |
| SIRT2 | 10.4318531 | -0.40006868 | 0.4662568 | -0.8580436 | 0.39086837 | 0.99179694 |
| NFKBIB | 4.43036282 | -0.159479696 | 0.69239257 | -0.2303313 | 0.81783432 | 0.99665574 |
| MRPS12 | 10.7294649 | 0.304851501 | 0.50627515 | 0.60214589 | 0.54707703 | 0.99665574 |
| FBXO17 | 19.2859687 | -0.074544307 | 0.57335481 | -0.1300143 | 0.89655514 | 0.99777713 |
| PAK4 | 17.849666 | 0.376239505 | 0.37591476 | 1.00086388 | 0.31689262 | 0.97700345 |
| NCCRP1 | 18.9936258 | 0.572090835 | 0.64417624 | 0.88809676 | 0.37448871 | 0.98535336 |
| SAMD4B | 24.1132177 | -0.065948856 | 0.33028421 | -0.1996731 | 0.84173629 | 0.99665574 |
| PAF1 | 5.4572688 | -0.196896611 | 0.59042294 | -0.333484 | 0.73876896 | 0.99665574 |
| MED29 | 8.02620971 | -1.123847241 | 0.55557899 | -2.0228397 | 0.04308968 | 0.61964991 |
| ZFP36 | 13.7493608 | -1.963945824 | 0.79405916 | -2.4732991 | 0.0133872 | 0.39134822 |
| RPS16 | 45.6906904 | 0.424436444 | 0.66460994 | 0.63862488 | 0.52306699 | 0.99665574 |
| SUPT5H | 18.8966471 | -0.249408542 | 0.44152869 | -0.5648751 | 0.57215875 | 0.99665574 |
| TIMM50 | 4.45519709 | 1.004257946 | 0.70947008 | 1.41550429 | 0.15692069 | 0.86532131 |
| EID2 | 2.70484962 | -0.623709724 | 0.89741082 | -0.6950103 | 0.48704888 | 0.99640678 |
| FBL | 5.09822958 | 0.593666982 | 0.67446048 | 0.88021019 | 0.37874546 | 0.98679664 |
| PSMC4 | 7.50825686 | 0.356631851 | 0.59222175 | 0.6021931 | 0.54704561 | 0.99665574 |
| ZNF546 | 3.28184226 | -0.70154523 | 0.75884986 | -0.9244849 | 0.35523392 | 0.98022903 |
| ZNF780B | 20.908197 | -0.147318067 | 0.46184055 | -0.3189804 | 0.7497414 | 0.99665574 |
| ZNF780A | 4.45592275 | 0.675784631 | 0.75317782 | 0.89724447 | 0.36958848 | 0.98532838 |
| MAP3K10 | 2.84201597 | -0.493532392 | 0.8848454 | -0.5577612 | 0.57700749 | 0.99665574 |
| AKT2 | 16.2819416 | 0.105760468 | 0.3755901 | 0.28158482 | 0.77826188 | 0.99665574 |
| PLD3 | 18.4857447 | 0.03008147 | 0.38882434 | 0.07736519 | 0.93833303 | 0.99777713 |
| SERTAD3 | 3.05155826 | -0.268487079 | 0.80898987 | -0.3318794 | 0.73998032 | 0.99665574 |
| BLVRB | 23.3645837 | -0.180278773 | 0.56119274 | -0.3212422 | 0.74802688 | 0.99665574 |
| SPTBN4 | 4.69467342 | 2.527527956 | 0.76811199 | 3.29057217 | 0.00099984 | 0.08663558 |
| SHKBP1 | 3.37527993 | 0.551439804 | 0.90371045 | 0.61019522 | 0.54173249 | 0.99665574 |
| COQ8B | 4.65677045 | -0.654052787 | 0.77632591 | -0.8424977 | 0.39950943 | 0.99179694 |
| ITPKC | 7.10662843 | -1.376270416 | 0.59679574 | -2.3060996 | 0.02110507 | 0.4733368 |
| SNRPA | 7.21748725 | -0.832175115 | 0.69757742 | -1.1929502 | 0.23288888 | 0.92924083 |
| AC008537.1 | 4.02324753 | -1.840025823 | 1.00185652 | -1.8366161 | 0.06626659 | 0.69473699 |
| HNRNPUL1 | 13.5317127 | -0.197000962 | 0.46038906 | -0.4279011 | 0.66872316 | 0.99665574 |
| TGFB1 | 10.133911 | -0.995379457 | 0.62000072 | -1.6054489 | 0.10839506 | 0.80237722 |
| CCDC97 | 3.77216914 | -0.680366604 | 0.72192192 | -0.9424379 | 0.34596847 | 0.97788427 |
| DMAC2 | 5.26776048 | 0.066548425 | 0.62894445 | 0.1058097 | 0.91573334 | 0.99777713 |
| RPS19 | 118.40472 | -0.022190548 | 0.34305117 | -0.0646858 | 0.94842415 | 0.99777713 |
| ARHGEF1 | 4.42652697 | 0.576105273 | 0.71217437 | 0.80893851 | 0.41855052 | 0.99443758 |
| RABAC1 | 20.0139781 | 0.258794136 | 0.52007203 | 0.4976121 | 0.61875747 | 0.99665574 |
| GRIK5 | 4.57201718 | 1.599567242 | 0.78623452 | 2.0344658 | 0.04190465 | 0.61480373 |

| | | | | | | |
|---|---|---|---|---|---|---|
| DEDD2 | 3.03298371 | 0.004251639 | 1.03372199 | 0.00411294 | 0.99671836 | 0.998802 |
| GSK3A | 5.26669089 | 0.2035415 | 0.68926299 | 0.2953031 | 0.76776235 | 0.99665574 |
| CIC | 6.36357296 | -0.312083472 | 0.67364849 | -0.4632735 | 0.64316836 | 0.99665574 |
| PAFAH1B3 | 3.52340774 | -0.30659535 | 0.84100267 | -0.3645593 | 0.7154404 | 0.99665574 |
| MEGF8 | 10.6943606 | 0.126659923 | 0.50357974 | 0.2515191 | 0.8014128 | 0.99665574 |
| LIPE-AS1 | 7.11069622 | 0.323156275 | 0.75384836 | 0.42867544 | 0.66815944 | 0.99665574 |
| ETHE1 | 3.89911553 | -0.872761496 | 0.82745887 | -1.0547491 | 0.29154009 | 0.96842511 |
| ZNF428 | 13.9131156 | -0.230895984 | 0.59445949 | -0.3884133 | 0.69771019 | 0.99665574 |
| SMG9 | 4.49241381 | -0.050649167 | 0.66470142 | -0.0761984 | 0.93926128 | 0.99777713 |
| ZNF283 | 4.9368458 | 0.433148535 | 0.66202683 | 0.6542764 | 0.51293375 | 0.99665574 |
| ZNF45 | 2.33583098 | -0.619733766 | 0.84064292 | -0.737214 | 0.46099219 | 0.99617422 |
| ZNF155 | 5.76034227 | -1.235613381 | 0.56718224 | -2.1785121 | 0.02936793 | 0.55650856 |
| ZNF224 | 7.83837587 | -0.968117863 | 0.51955474 | -1.8633607 | 0.06241154 | 0.68551266 |
| ZNF234 | 3.583593 | 0.298098511 | 0.84012837 | 0.35482496 | 0.72272073 | 0.99665574 |
| ZNF226 | 7.08430609 | 1.037441443 | 0.57267314 | 1.8115769 | 0.0700516 | 0.71124502 |
| ZNF227 | 3.54434814 | -0.727811916 | 0.69128311 | -1.052842 | 0.2924134 | 0.97005465 |
| ZNF235 | 9.85805763 | -0.604498431 | 0.54609636 | -1.1069446 | 0.26831787 | 0.94660092 |
| ZNF229 | 2.42964671 | -0.592909776 | 0.81003933 | -0.7319518 | 0.46419797 | 0.99617422 |
| ZNF180 | 2.88602526 | -0.251051857 | 0.89167518 | -0.2815508 | 0.77828797 | 0.99665574 |
| AC243964.2 | 7.44007455 | -0.792194309 | 0.59915559 | -1.3221846 | 0.18610668 | 0.89532394 |
| PVR | 4.39605263 | -0.266058814 | 0.68467426 | -0.3885918 | 0.69757812 | 0.99665574 |
| CEACAM19 | 3.15397043 | -0.336542617 | 0.80486921 | -0.4181333 | 0.67584966 | 0.99665574 |
| CBLC | 3.72428799 | -1.247092196 | 0.75160003 | -1.6592498 | 0.09706547 | 0.77606444 |
| BCAM | 28.5183934 | -0.241050146 | 0.48999514 | -0.491944 | 0.62275897 | 0.99665574 |
| NECTIN2 | 20.0531537 | -0.243098466 | 0.38804472 | -0.6264702 | 0.53100656 | 0.99665574 |
| TOMM40 | 8.32413669 | -0.325305675 | 0.66516322 | -0.4890614 | 0.62479821 | 0.99665574 |
| APOE | 31.5961269 | 1.21007727 | 0.63213415 | 1.91427288 | 0.0555853 | 0.67167119 |
| CLPTM1 | 12.304177 | -0.098971644 | 0.56008786 | -0.1767074 | 0.85973826 | 0.99665574 |
| RELB | 9.02963758 | -0.64306029 | 0.66958978 | -0.9603795 | 0.33686426 | 0.97788427 |
| CLASRP | 15.7279323 | -0.364304584 | 0.40190273 | -0.9064496 | 0.3646979 | 0.98362663 |
| GEMIN7-AS1 | 4.04611857 | -0.052096669 | 0.96187117 | -0.0541618 | 0.95680626 | 0.99777713 |
| MARK4 | 11.9716419 | -0.474241331 | 0.52495557 | -0.9033933 | 0.36631721 | 0.98365837 |
| TRAPPC6A | 6.51718006 | 0.315528577 | 0.68458228 | 0.46090673 | 0.64486552 | 0.99665574 |
| PPP1R13L | 5.31045073 | -0.627873685 | 0.6582028 | -0.9539213 | 0.34012347 | 0.97788427 |
| ERCC1 | 4.2283666 | 0.238336632 | 0.72005288 | 0.33099879 | 0.7406454 | 0.99665574 |
| VASP | 7.12635176 | 0.133030912 | 0.60976113 | 0.2181689 | 0.82729752 | 0.99665574 |
| OPA3 | 14.568404 | -1.12708265 | 0.52197172 | -2.1592792 | 0.03082852 | 0.56271575 |
| EML2 | 9.39821625 | -0.69366954 | 0.67987305 | -1.0202927 | 0.30758964 | 0.97376371 |
| SNRPD2 | 13.6586617 | -0.331224538 | 0.48650385 | -0.6808261 | 0.49598151 | 0.99640678 |
| FBXO46 | 6.63379052 | 0.266109186 | 0.54802062 | 0.48558243 | 0.62726325 | 0.99665574 |
| DMPK | 3.60129518 | -0.979125543 | 0.90498528 | -1.0819243 | 0.27928618 | 0.95980166 |
| SYMPK | 18.367565 | 0.495286373 | 0.3903977 | 1.26867133 | 0.20455831 | 0.90834797 |
| IRF2BP1 | 2.92796698 | 0.82589138 | 0.92443392 | 0.89340229 | 0.37164178 | 0.98532838 |
| MYPOP | 3.77957073 | -0.072468387 | 0.7478193 | -0.0969063 | 0.92280082 | 0.99777713 |
| PPP5C | 11.8241654 | 0.033805183 | 0.43408214 | 0.07787739 | 0.93792559 | 0.99777713 |
| CCDC8 | 2.48109956 | 0.999017362 | 0.93838346 | 1.06461528 | 0.28705006 | 0.96559334 |
| PPP5D1 | 3.27905094 | -0.126611964 | 0.72115212 | -0.175569 | 0.86063255 | 0.99665574 |
| CALM3 | 25.50054 | 0.186368514 | 0.41840235 | 0.44542894 | 0.6560098 | 0.99665574 |
| PRKD2 | 11.8035288 | -0.23837986 | 0.5409719 | -0.4406511 | 0.65946561 | 0.99665574 |
| STRN4 | 5.51651536 | 0.159719687 | 0.66883515 | 0.23880277 | 0.81125852 | 0.99665574 |
| FKRP | 4.32022151 | 0.205043731 | 0.70728388 | 0.28990302 | 0.77189043 | 0.99665574 |
| SLC1A5 | 3.13033258 | -0.464817043 | 0.92833651 | -0.5006989 | 0.61658307 | 0.99665574 |
| AP2S1 | 7.37716959 | -0.36731995 | 0.57460078 | -0.6392611 | 0.52265307 | 0.99665574 |
| ARHGAP35 | 30.9673781 | -0.005169379 | 0.41536393 | -0.0124454 | 0.99007025 | 0.99777713 |
| TMEM160 | 12.2795374 | -0.049036841 | 0.60115521 | -0.081571 | 0.93498785 | 0.99777713 |
| ZC3H4 | 10.2324436 | -0.44877673 | 0.4709427 | -0.9529328 | 0.34062414 | 0.97788427 |
| SAE1 | 11.2926349 | -0.616650848 | 0.56742166 | -1.0867594 | 0.27714314 | 0.95828005 |
| CCDC9 | 2.62587845 | -0.589055298 | 0.8216743 | -0.7168963 | 0.47343806 | 0.99617422 |
| INAFM1 | 2.13647526 | -0.369652594 | 1.05266262 | -0.3511596 | 0.72546861 | 0.99665574 |
| C5AR1 | 2.83256519 | -0.237493496 | 0.76956541 | -0.3086073 | 0.75762027 | 0.99665574 |
| DHX34 | 6.06961034 | -0.524033076 | 0.61533262 | -0.8516257 | 0.39442187 | 0.99179694 |
| NAPA-AS1 | 3.26926023 | -2.135416824 | 0.87035472 | -2.4535018 | 0.01414728 | 0.40056949 |
| NAPA | 23.7575922 | -0.534071731 | 0.45377637 | -1.1769492 | 0.23921579 | 0.93157178 |
| NOP53 | 12.7132376 | 0.472884977 | 0.46381445 | 1.01955638 | 0.3079389 | 0.97376371 |
| SELENOW | 18.850856 | 0.239250787 | 0.50954125 | 0.46954155 | 0.63868259 | 0.99665574 |
| LIG1 | 1.88352878 | 0.006294617 | 0.9684779 | 0.00649949 | 0.99481419 | 0.9985287 |
| ZSWIM9 | 3.64046096 | -0.258984382 | 0.72162444 | -0.3588908 | 0.71967677 | 0.99665574 |
| CARD8 | 7.98880038 | 0.912417424 | 0.63544056 | 1.43588163 | 0.15103603 | 0.85612189 |
| TMEM143 | 4.17713343 | -0.669524031 | 0.71908296 | -0.9310804 | 0.35181199 | 0.97838306 |
| KDELR1 | 13.024377 | -0.217771127 | 0.43925385 | -0.4957751 | 0.62005309 | 0.99665574 |
| GRWD1 | 2.22949936 | -0.044334323 | 0.99910472 | -0.044374 | 0.96460625 | 0.99777713 |
| CYTH2 | 6.15710644 | -1.094441723 | 0.74013798 | -1.4786996 | 0.13922063 | 0.84668189 |
| RPL18 | 57.3893953 | 0.58426117 | 0.38250834 | 1.5274469 | 0.12664992 | 0.83669777 |
| SPHK2 | 3.7427291 | -0.922895392 | 0.80955983 | -1.1399965 | 0.25428775 | 0.94099699 |
| BCAT2 | 8.09947168 | 0.06684674 | 0.61959669 | 0.1078875 | 0.91408493 | 0.99777713 |
| PPP1R15A | 6.96840723 | -1.735922914 | 1.03726299 | -1.673561 | 0.0942169 | 0.7700476 |
| NUCB1 | 17.7562083 | -0.034136139 | 0.44050088 | -0.0774939 | 0.93823063 | 0.99777713 |
| BAX | 5.80622975 | -0.60789657 | 0.64851783 | -0.9373629 | 0.34857191 | 0.97838306 |
| FTL | 280.124358 | -0.241213067 | 0.57750991 | -0.4176778 | 0.67618271 | 0.99665574 |
| GYS1 | 10.6166676 | -0.277427066 | 0.64441739 | -0.4305083 | 0.6668259 | 0.99665574 |
| RUVBL2 | 3.86151446 | 1.494694343 | 0.87755477 | 1.70324907 | 0.08852147 | 0.75650018 |
| SNRNP70 | 46.3620512 | 0.405978726 | 0.37829302 | 1.07318589 | 0.28318772 | 0.96059387 |
| TRPM4 | 6.58028118 | 0.158569897 | 0.7495572 | 0.21155143 | 0.83245701 | 0.99665574 |
| SLC6A16 | 11.3586583 | 0.523782191 | 0.49411683 | 1.06003713 | 0.28912771 | 0.96676791 |
| TEAD2 | 3.18331991 | -1.29012558 | 1.00294898 | -1.2863322 | 0.19832714 | 0.90381759 |
| PIH1D1 | 7.10444287 | -0.170200403 | 0.59325773 | -0.2868912 | 0.77419565 | 0.99665574 |
| RPL13A | 80.148994 | 0.419352453 | 0.39849993 | 1.05232755 | 0.2926493 | 0.97051965 |
| RPS11 | 66.9111292 | 0.421299185 | 0.39810756 | 1.05825468 | 0.28993935 | 0.96676791 |
| FCGRT | 13.1757434 | 0.371716416 | 0.57733467 | 0.64384912 | 0.51967328 | 0.99665574 |
| NOSIP | 9.34740406 | -0.529308684 | 0.51434943 | -1.0290838 | 0.30344028 | 0.97376371 |
| PRRG2 | 4.80213533 | -0.227847056 | 0.97049639 | -0.2347737 | 0.81438436 | 0.99665574 |
| PRR12 | 5.74316411 | -0.014558166 | 0.61089137 | -0.023831 | 0.9809874 | 0.99777713 |
| SCAF1 | 5.85183296 | 0.046492114 | 0.57281018 | 0.08116496 | 0.93531077 | 0.99777713 |
| IRF3 | 10.6007581 | -0.363880506 | 0.44852321 | -0.8112858 | 0.41720156 | 0.99392253 |
| PRMT1 | 10.5529229 | 0.527564333 | 0.54909344 | 0.96079154 | 0.336657 | 0.97788427 |
| AP2A1 | 7.11625559 | -0.281178544 | 0.63544646 | -0.4424897 | 0.65813485 | 0.99665574 |
| MED25 | 10.4590101 | 0.33644631 | 0.68239508 | 0.49303742 | 0.62198614 | 0.99665574 |
| PTOV1 | 13.5419106 | 0.701115881 | 0.49944498 | 1.40379003 | 0.16038138 | 0.86820896 |
| PNKP | 6.36879976 | -0.958517682 | 0.52748994 | -1.8171298 | 0.06919724 | 0.7094165 |
| TBC1D17 | 6.92464892 | 0.79733359 | 0.57133658 | 1.39555844 | 0.1628475 | 0.87135275 |
| ATF5 | 2.20951257 | 0.716660613 | 0.99334815 | 0.72145965 | 0.47062676 | 0.99617422 |
| VRK3 | 9.98516325 | 0.125678276 | 0.4908957 | 0.25601828 | 0.79793672 | 0.99665574 |
| MYH14 | 16.7004827 | -0.175512226 | 0.45841096 | -0.3828709 | 0.70181548 | 0.99665574 |
| KCNC3 | 10.2848327 | -0.093804441 | 0.73044402 | -0.1284211 | 0.89781572 | 0.99777713 |
| NR1H2 | 13.5632807 | -0.023013659 | 0.44007175 | -0.0522952 | 0.95829344 | 0.99777713 |
| EMC10 | 22.9284447 | -0.16723835 | 0.40495396 | -0.4129811 | 0.67962043 | 0.99665574 |
| JOSD2 | 5.72392429 | 0.226211605 | 0.76640351 | 0.29515993 | 0.76787171 | 0.99665574 |
| CLEC11A | 3.54434747 | -0.556304168 | 0.88248442 | -0.6303841 | 0.5284433 | 0.99665574 |
| C19orf48 | 3.38910737 | 0.852327677 | 0.85008548 | 1.00263761 | 0.31603574 | 0.97652251 |
| CTU1 | 2.90232914 | -1.093092485 | 1.05140205 | -1.0396522 | 0.2985015 | 0.97358745 |
| ETFB | 23.5704559 | 1.01976143 | 0.52412034 | 1.94566278 | 0.05169526 | 0.65584036 |
| ZNF175 | 5.19656673 | 0.066614383 | 0.66646604 | 0.09995165 | 0.92038271 | 0.99777713 |
| SPACA6 | 4.52898947 | 0.147177382 | 0.71347975 | 0.20628109 | 0.83657135 | 0.99665574 |

| | | | | | | |
|---|---|---|---|---|---|---|
| ZNF577 | 5.87072307 | 0.588799521 | 0.68728106 | 0.8567085 | 0.39160601 | 0.99179694 |
| ZNF350-AS1 | 5.03883917 | 1.51183088 | 0.73263999 | 2.06353857 | 0.03906149 | 0.6009186 |
| ZNF350 | 4.86268354 | -0.03299401 | 0.90448242 | -0.0364783 | 0.97090096 | 0.99777713 |
| ZNF615 | 2.38836754 | 1.15656893 | 0.89071561 | 1.29847161 | 0.19412533 | 0.90235789 |
| ZNF432 | 6.81850643 | -0.868492094 | 0.77143453 | -1.1258144 | 0.26024409 | 0.94557817 |
| ZNF616 | 3.09477024 | -0.643789573 | 0.75398807 | -0.8538458 | 0.39319042 | 0.99179694 |
| AC011468.3 | 6.12604453 | 0.643396926 | 0.80196007 | 0.8022805 | 0.42239072 | 0.99518711 |
| PPP2R1A | 32.1756482 | -0.142161404 | 0.38271665 | -0.3714534 | 0.71029985 | 0.99665574 |
| ZNF766 | 6.81469775 | 0.036886088 | 0.54829852 | 0.06727373 | 0.94636379 | 0.99777713 |
| ZNF480 | 7.179967 | -0.821697285 | 0.65537322 | -1.2537853 | 0.20992007 | 0.91043197 |
| ZNF528 | 3.370449 | -0.083385817 | 0.72059048 | -0.1157187 | 0.90787546 | 0.99777713 |
| ZNF808 | 6.86011314 | 0.482231148 | 0.60044608 | 0.80312149 | 0.42190452 | 0.99518711 |
| ZNF701 | 7.16104129 | 0.439341431 | 0.50703619 | 0.8664893 | 0.38622189 | 0.99051797 |
| ZNF83 | 26.2728624 | 0.135594835 | 0.42883456 | 0.31619381 | 0.7518554 | 0.99665574 |
| ZNF611 | 19.208673 | -0.847536301 | 0.42440074 | -1.9970189 | 0.04582313 | 0.63477358 |
| ZNF600 | 3.91250187 | -0.542323506 | 0.83541785 | -0.6491644 | 0.51623214 | 0.99665574 |
| ZNF28 | 4.3333294 | 0.443516141 | 0.7664717 | 0.57864647 | 0.56282774 | 0.99665574 |
| ZNF160 | 17.2637708 | -0.246332422 | 0.41384368 | -0.5952306 | 0.55168934 | 0.99665574 |
| ZNF347 | 4.86119977 | 0.128438128 | 0.62558548 | 0.20530868 | 0.83733096 | 0.99665574 |
| ZNF677 | 4.69858504 | -0.996787531 | 0.7038853 | -1.4161221 | 0.15673976 | 0.86532131 |
| ZNF525 | 3.77604883 | 0.118320211 | 0.679505 | 0.17412707 | 0.8617656 | 0.99665574 |
| ZNF765 | 8.21338072 | 0.164885267 | 0.59524573 | 0.2770037 | 0.78177727 | 0.99665574 |
| ZNF761 | 9.59746079 | -0.188755261 | 0.44743658 | -0.4218593 | 0.67312775 | 0.99665574 |
| ZNF813 | 4.44268555 | -0.258961093 | 0.69317309 | -0.3735879 | 0.70871092 | 0.99665574 |
| ZNF331 | 10.9397617 | -0.055169176 | 0.70022092 | -0.0787882 | 0.93720106 | 0.99777713 |
| NDUFA3 | 7.22741151 | 0.068882159 | 0.80000491 | 0.08610217 | 0.9313852 | 0.99777713 |
| PRPF31 | 10.7427669 | -0.12820494 | 0.48521144 | -0.2642249 | 0.79160663 | 0.99665574 |
| CNOT3 | 5.45739289 | -0.035188175 | 0.64497719 | -0.0545572 | 0.95649121 | 0.99777713 |
| TMC4 | 22.6736865 | -0.203897055 | 0.38582612 | -0.5284688 | 0.59717402 | 0.99665574 |
| MBOAT7 | 10.7865736 | -0.863260514 | 0.55389662 | -1.5585228 | 0.11910937 | 0.82619588 |
| TSEN34 | 5.47035835 | 0.038647333 | 0.64177079 | 0.06021984 | 0.95198055 | 0.99777713 |
| RPS9 | 127.324669 | 0.35758418 | 0.47654291 | 0.75037142 | 0.45303104 | 0.99617422 |
| LENG8-AS1 | 5.02852732 | -1.230431775 | 0.6593197 | -1.8662142 | 0.06201141 | 0.68457455 |
| LENG8 | 23.8357151 | 0.638601372 | 0.46733892 | 1.36646305 | 0.17179367 | 0.8813989 |
| AC011476.3 | 4.94476123 | -0.449200104 | 1.00077286 | -0.4488532 | 0.65353756 | 0.99665574 |
| RDH13 | 13.5006033 | -0.247848184 | 0.46598602 | -0.531879 | 0.59480979 | 0.99665574 |
| EPS8L1 | 4.10134532 | -0.466640707 | 0.73749391 | -0.6327384 | 0.5269045 | 0.99665574 |
| PPP1R12C | 8.60084417 | -0.387528526 | 0.55083698 | -0.7035267 | 0.48172757 | 0.99640678 |
| PPP6R1 | 7.67579567 | -0.444366039 | 0.63840446 | -0.6960572 | 0.48639299 | 0.99640678 |
| HSPBP1 | 4.37863757 | -0.322368743 | 0.70481119 | -0.4573831 | 0.64739569 | 0.99665574 |
| TMEM238 | 7.0807919 | 0.744302292 | 0.6010527 | 1.23833116 | 0.2155933 | 0.913434 |
| RPL28 | 81.0740617 | 0.455843543 | 0.35102098 | 1.29862194 | 0.19407371 | 0.90235789 |
| UBE2S | 5.06922181 | 0.123947791 | 0.65900955 | 0.18808193 | 0.85081242 | 0.99665574 |
| ISOC2 | 9.40467511 | 0.685505517 | 0.58540957 | 1.17098448 | 0.24160502 | 0.93266071 |
| FIZ1 | 2.47689964 | 0.110796144 | 0.87112567 | 0.12718732 | 0.89879214 | 0.99777713 |
| ZNF524 | 3.92681307 | 0.265645792 | 0.71722875 | 0.37037806 | 0.71110082 | 0.99665574 |
| CCDC106 | 3.65617083 | 0.495932768 | 0.82477756 | 0.60129275 | 0.54764501 | 0.99665574 |
| U2AF2 | 9.04927686 | 0.971674261 | 0.54667815 | 1.77741557 | 0.07549989 | 0.73175707 |
| EPN1 | 30.9208665 | 0.347025637 | 0.37521514 | 0.9248711 | 0.35503296 | 0.98022903 |
| ZNF787 | 7.14115612 | -0.679481463 | 0.68317289 | -0.9945966 | 0.31993249 | 0.97700345 |
| ZNF444 | 6.97995707 | -0.05919031 | 0.62885426 | -0.094124 | 0.92501062 | 0.99777713 |
| ZSCAN5A | 18.893461 | -0.210337887 | 0.43333489 | -0.4853934 | 0.62739733 | 0.99665574 |
| ZNF582 | 3.19927565 | 0.811199041 | 0.8952393 | 0.90612537 | 0.3648695 | 0.98362663 |
| ZNF667 | 2.63523137 | -0.112936304 | 0.91819883 | -0.1229977 | 0.90210895 | 0.99777713 |
| ZNF667-AS1 | 7.17751635 | 1.449890964 | 0.6417049 | 2.25943572 | 0.02385629 | 0.49867035 |
| ZNF471 | 4.60310455 | -0.141988076 | 0.82553955 | -0.1719943 | 0.86344203 | 0.99665574 |
| ZFP28 | 2.68953377 | 0.525871435 | 0.99180799 | 0.53021496 | 0.5959629 | 0.99665574 |
| ZNF470 | 1.85069449 | 0.800218381 | 1.16883839 | 0.68462705 | 0.49357929 | 0.99640678 |
| ZNF264 | 11.0674776 | 0.041680085 | 0.46401248 | 0.08982535 | 0.928426 | 0.99777713 |
| ZNF460 | 2.67194612 | -0.286799159 | 0.86567012 | -0.3313031 | 0.74041558 | 0.99665574 |
| AC005261.1 | 12.1800397 | -0.342872809 | 0.54581078 | -0.6281899 | 0.52987956 | 0.99665574 |
| ZNF304 | 2.34572841 | -0.295264168 | 0.89222033 | -0.3309319 | 0.74069593 | 0.99665574 |
| AC004076.1 | 7.52485874 | -0.062864105 | 0.52736263 | -0.1192047 | 0.90511318 | 0.99777713 |
| ZNF549 | 4.53213969 | -0.077821945 | 0.75789101 | -0.1026822 | 0.91821517 | 0.99777713 |
| ZNF550 | 4.45576511 | -0.154324854 | 0.73525854 | -0.209892 | 0.833752 | 0.99665574 |
| ZNF134 | 2.78754284 | 1.021394698 | 0.90769866 | 1.12525747 | 0.26047995 | 0.94557817 |
| AC003006.1 | 3.96090923 | -0.709232742 | 0.68356906 | -1.0375437 | 0.29948256 | 0.97358745 |
| ZNF154 | 4.41277531 | -0.49319725 | 0.67065391 | -0.7353976 | 0.46209741 | 0.99617422 |
| ZNF671 | 1.78508177 | -0.264574755 | 1.00864819 | -0.2623063 | 0.79308532 | 0.99665574 |
| ZNF586 | 30.0558889 | -0.040570477 | 0.33030002 | -0.1228292 | 0.90224237 | 0.99777713 |
| ZNF552 | 9.68868232 | -0.62393617 | 0.45516276 | -1.3707979 | 0.17043796 | 0.87918701 |
| ZNF814 | 18.4202447 | -0.562613174 | 0.33938645 | -1.6577361 | 0.09737074 | 0.77789171 |
| ZNF587 | 7.84382847 | -0.571897837 | 0.5908633 | -0.9679021 | 0.33309326 | 0.97788427 |
| ZNF418 | 3.7242661 | 0.502796488 | 0.85633926 | 0.58714637 | 0.55710541 | 0.99665574 |
| C19orf18 | 2.10912287 | -0.094339446 | 0.96584156 | -0.0976759 | 0.92218965 | 0.99777713 |
| ZNF606 | 5.59167269 | 0.413242241 | 0.72598357 | 0.56921707 | 0.56920884 | 0.99665574 |
| ZNF135 | 8.74345561 | -0.24197246 | 0.52856488 | -0.4577914 | 0.64710231 | 0.99665574 |
| ZSCAN18 | 10.9557758 | -0.356762917 | 0.64286361 | -0.554959 | 0.57892273 | 0.99665574 |
| ZNF329 | 5.36314484 | -0.419771229 | 0.71505133 | -0.5870505 | 0.5571698 | 0.99665574 |
| ZNF274 | 10.5621115 | -0.493045095 | 0.60464406 | -0.8154303 | 0.41482604 | 0.99279547 |
| AC020915.3 | 4.54295305 | -1.09212952 | 0.86654103 | -1.2603321 | 0.20754957 | 0.90994205 |
| AC010642.2 | 2.93031123 | 0.066099783 | 0.91000476 | 0.07263674 | 0.94209519 | 0.99777713 |
| RPS5 | 35.1993278 | 0.431633502 | 0.55117393 | 0.78311669 | 0.43355859 | 0.99617422 |
| SLC27A5 | 2.89387419 | -0.185862817 | 0.75759102 | -0.245334 | 0.80619785 | 0.99665574 |
| TRIM28 | 11.1366301 | 0.607689619 | 0.55473451 | 1.09546028 | 0.27331504 | 0.95186117 |
| CHMP2A | 11.105938 | -0.110865618 | 0.49480853 | -0.2240576 | 0.82271247 | 0.99665574 |
| UBE2M | 10.2590559 | -0.44849245 | 0.56946399 | -0.7875695 | 0.43094858 | 0.99617422 |
| MZF1-AS1 | 2.86136363 | -0.012985306 | 0.94041565 | -0.013808 | 0.98898312 | 0.99777713 |
| MZF1 | 13.38624 | -0.264654233 | 0.4282595 | -0.6179763 | 0.53659095 | 0.99665574 |
| RPS4Y1 | 21.2700444 | 0.358813141 | 2.21099637 | 0.16228572 | 0.87108086 | 0.99665574 |
| ZFY | 12.2112332 | 0.583730323 | 1.96454948 | 0.2971319 | 0.76636581 | 0.99665574 |
| LINC00278 | 45.8879744 | -0.044987155 | 2.45569497 | -0.0183195 | 0.98538395 | 0.99777713 |
| TBL1Y | 4.31208502 | 0.46824142 | 1.88110588 | 0.24891816 | 0.80342409 | 0.99665574 |
| TTTY15 | 8.0653839 | -0.547980676 | 1.871664 | -0.2927773 | 0.7696924 | 0.99665574 |
| USP9Y | 32.8952089 | 0.466719235 | 2.34239367 | 0.19924885 | 0.84206809 | 0.99665574 |
| DDX3Y | 7.62773495 | -0.147432741 | 1.85772441 | -0.079362 | 0.93674469 | 0.99777713 |
| UTY | 105.329538 | -0.661212159 | 2.13252369 | -0.3100609 | 0.75651467 | 0.99665574 |
| NLGN4Y | 42.593778 | -0.483802466 | 2.13028182 | -0.2271073 | 0.82034034 | 0.99665574 |
| TTTY14 | 72.1921421 | 0.271629771 | 1.25844034 | 0.21584636 | 0.82910751 | 0.99665574 |
| KDM5D | 8.3741756 | 0.213358199 | 1.89764544 | 0.11243312 | 0.91048 | 0.99777713 |
| TTTY10 | 5.18919814 | 0.170662113 | 1.11986286 | 0.15239555 | 0.87887497 | 0.99665574 |
| EIF1AY | 8.29005569 | 0.113368175 | 1.94474956 | 0.05829449 | 0.95351406 | 0.99777713 |
| IL17RA | 7.88947949 | -0.645383308 | 0.50012051 | -1.2904556 | 0.19689253 | 0.90381759 |
| HDHD5 | 8.86795384 | -0.423945678 | 0.46962163 | -0.9027388 | 0.36666453 | 0.98365837 |
| CECR2 | 8.62317153 | 0.351916616 | 0.71514255 | 0.49209296 | 0.62265363 | 0.99665574 |
| ATP6V1E1 | 36.9893565 | 0.279599488 | 0.37251295 | 0.75057656 | 0.45290754 | 0.99617422 |
| BCL2L13 | 38.6929427 | -0.200168364 | 0.33057814 | -0.60551 | 0.54484019 | 0.99665574 |
| MICAL3 | 43.8727152 | -0.455490731 | 0.37608829 | -1.2111271 | 0.22584669 | 0.92318867 |
| PEX26 | 11.2865139 | -0.10037759 | 0.45137585 | -0.2223814 | 0.824017 | 0.99665574 |
| DGCR2 | 22.0972187 | 0.095716648 | 0.44654556 | 0.21434912 | 0.83027482 | 0.99665574 |
| SLC25A1 | 10.190097 | 0.914069314 | 0.55867262 | 1.63614483 | 0.10180928 | 0.7843028 |
| CLTCL1 | 2.67496323 | 0.650569822 | 0.81444462 | 0.79878951 | 0.42441247 | 0.99518711 |
| HIRA | 8.03720751 | 0.282103403 | 0.60326605 | 0.46762685 | 0.64005146 | 0.99665574 |

| | | | | | | |
|---|---|---|---|---|---|---|
| MRPL40 | 7.4155935 | -0.69106594 | 0.66450839 | -1.0399657 | 0.29835583 | 0.97358745 |
| UFD1 | 8.19046014 | -0.773409822 | 0.56395486 | -1.3714038 | 0.17024912 | 0.87918701 |
| TXNRD2 | 10.4368651 | -0.000987217 | 0.51231274 | -0.001927 | 0.99846249 | 0.99925101 |
| COMT | 21.8868561 | 0.144491043 | 0.43442785 | 0.33260078 | 0.73943565 | 0.99665574 |
| TANGO2 | 10.7844054 | -0.136651665 | 0.56727316 | -0.2408922 | 0.80963869 | 0.99665574 |
| DGCR8 | 8.66255778 | -0.389733395 | 0.53563492 | -0.7276101 | 0.46685228 | 0.99617422 |
| TRMT2A | 2.25172317 | 1.595554646 | 0.97082223 | 1.64350856 | 0.10027776 | 0.78386461 |
| RANBP1 | 5.21198648 | 0.2665839 | 0.68480087 | 0.38928674 | 0.69706404 | 0.99665574 |
| ZDHHC8 | 4.34105681 | -0.055184467 | 0.73082284 | -0.07551 | 0.9398089 | 0.99777713 |
| DGCR6L | 4.65021557 | 0.812493796 | 0.71345284 | 1.1388192 | 0.25477857 | 0.94126227 |
| KLHL22 | 15.7349618 | -0.100483144 | 0.4364089 | -0.23025 | 0.81789752 | 0.99665574 |
| MED15 | 23.62176 | -0.294221239 | 0.39488361 | -0.7450835 | 0.45622127 | 0.99617422 |
| PI4KA | 37.238654 | -0.357661201 | 0.39161569 | -0.9132964 | 0.36108667 | 0.9819332 |
| SNAP29 | 7.17876338 | -0.139924746 | 0.55297215 | -0.2530412 | 0.80023636 | 0.99665574 |
| CRKL | 6.46926618 | 0.980903153 | 0.66246295 | 1.48069134 | 0.13868884 | 0.84668189 |
| LZTR1 | 3.0927816 | 0.331341504 | 0.86988359 | 0.38090327 | 0.70327503 | 0.99665574 |
| THAP7 | 3.0212434 | 0.694379281 | 0.82769264 | 0.83893374 | 0.4015065 | 0.99194293 |
| HIC2 | 10.4328672 | -0.024061685 | 0.49400229 | -0.0487076 | 0.96115229 | 0.99777713 |
| UBE2L3 | 26.8263654 | 0.03025998 | 0.35138553 | 0.08611618 | 0.93137406 | 0.99777713 |
| YDJC | 2.74388133 | -0.629018405 | 0.89503156 | -0.7027891 | 0.48218721 | 0.99640678 |
| SDF2L1 | 8.05066687 | 0.189761426 | 0.81593429 | 0.2325695 | 0.81609572 | 0.99665574 |
| PPIL2 | 19.9078101 | 0.35685108 | 0.33699261 | 1.05892849 | 0.28963235 | 0.96676791 |
| YPEL1 | 5.04625801 | 0.112790971 | 0.59998935 | 0.18798829 | 0.85088583 | 0.99665574 |
| MAPK1 | 47.853788 | 0.003821473 | 0.29147098 | 0.01311099 | 0.98953924 | 0.99777713 |
| PPM1F | 3.0458848 | 0.688112546 | 0.8818999 | 0.78026151 | 0.43523696 | 0.99617422 |
| TOP3B | 4.78996151 | 0.271060652 | 0.6849295 | 0.39574971 | 0.69228968 | 0.99665574 |
| AC245060.5 | 7.28875656 | -0.013484992 | 0.68972809 | -0.0195512 | 0.98440142 | 0.99777713 |
| BCR | 32.6586005 | 0.043625628 | 0.36353607 | 0.12000357 | 0.90448032 | 0.99777713 |
| CHCHD10 | 68.9714027 | 0.178584109 | 0.34223291 | 0.52182038 | 0.6017954 | 0.99665574 |
| SMARCB1 | 10.558654 | -0.060011873 | 0.42777328 | -0.140289 | 0.88843168 | 0.99665574 |
| DDT | 43.5802026 | 0.779987828 | 0.5235239 | 1.48988008 | 0.13625577 | 0.84475183 |
| CABIN1 | 35.1786385 | -0.04257317 | 0.34272127 | -0.124221 | 0.90114031 | 0.99777713 |
| ADORA2A-A! | 6.26999079 | 1.220647881 | 0.6364073 | 1.91802934 | 0.05510729 | 0.67068154 |
| GUCD1 | 6.90258762 | 0.573891906 | 0.61528887 | 0.93271947 | 0.35096483 | 0.97838306 |
| SNRPD3 | 12.8573106 | -0.191210397 | 0.46471539 | -0.411457 | 0.68073747 | 0.99665574 |
| SGSM1 | 16.3833355 | -0.212936567 | 0.6396432 | -0.332899 | 0.73921054 | 0.99665574 |
| KIAA1671 | 27.28812 | -0.473893677 | 0.31848506 | -1.487962 | 0.13676091 | 0.84475183 |
| LRP5L | 10.7551286 | -0.156140993 | 0.45944725 | -0.3398453 | 0.73397302 | 0.99665574 |
| GRK3 | 12.517561 | -0.422204028 | 0.53756164 | -0.7854058 | 0.43221569 | 0.99617422 |
| HPS4 | 12.7051335 | -0.467845026 | 0.42690486 | -1.0959 | 0.27312255 | 0.95186117 |
| SRRD | 3.23277686 | 0.131293805 | 0.88487867 | 0.14837492 | 0.88204689 | 0.99665574 |
| TFIP11 | 5.01001581 | -0.453426955 | 0.59672266 | -0.7598621 | 0.44733699 | 0.99617422 |
| TPST2 | 9.48534755 | -0.412074896 | 0.6687675 | -0.6161706 | 0.53778191 | 0.99665574 |
| MIATNB | 9.94221196 | -0.445386873 | 0.6335505 | -0.7030014 | 0.48205489 | 0.99640678 |
| PITPNB | 39.7925321 | -0.124069002 | 0.34225362 | -0.362506 | 0.71697392 | 0.99665574 |
| TTC28 | 60.3558721 | -0.00555873 | 0.31593387 | -0.0175946 | 0.98596227 | 0.99777713 |
| HSCB | 6.842409 | -0.172637441 | 0.52893602 | -0.3263862 | 0.74413214 | 0.99665574 |
| CCDC117 | 6.13833916 | 0.234674093 | 0.58016199 | 0.40449753 | 0.68584689 | 0.99665574 |
| XBP1 | 12.245882 | -0.980071143 | 0.51742045 | -1.8941485 | 0.05820531 | 0.67545594 |
| Z93930.2 | 9.64315485 | -0.703120475 | 0.55093132 | -1.2762398 | 0.20187077 | 0.90614933 |
| ZNRF3 | 93.6723285 | 0.660324385 | 0.2901607 | 2.2757196 | 0.0228628 | 0.4879643 |
| KREMEN1 | 15.3873264 | -0.065561551 | 0.56973505 | -0.1150738 | 0.90838666 | 0.99777713 |
| RHBDD3 | 3.35013642 | -0.784420907 | 0.98140191 | -0.7992861 | 0.42412453 | 0.99518711 |
| EWSR1 | 30.2452576 | 0.347933267 | 0.42376084 | 0.82106044 | 0.41161184 | 0.99279547 |
| GAS2L1 | 4.94447002 | -0.251864743 | 0.8599712 | -0.2928758 | 0.76961709 | 0.99665574 |
| AP1B1 | 15.2736769 | 0.277935955 | 0.37709777 | 0.7370395 | 0.46109833 | 0.99617422 |
| THOC5 | 3.81552848 | -0.319482962 | 0.73063355 | -0.4372684 | 0.66191671 | 0.99665574 |
| NIPSNAP1 | 4.26731132 | -0.663459353 | 0.83751337 | -0.7921776 | 0.42825711 | 0.99589919 |
| NF2 | 11.7826484 | 0.119333274 | 0.43833484 | 0.27224227 | 0.78543574 | 0.99665574 |
| ZMAT5 | 2.72936879 | 0.206128299 | 0.88287829 | 0.23347306 | 0.81539408 | 0.99665574 |
| UQCR10 | 41.7862939 | -0.303923117 | 0.42927062 | -0.7079989 | 0.47894595 | 0.99617422 |
| ASCC2 | 12.3792488 | -0.301443727 | 0.43437812 | -0.6939662 | 0.48770341 | 0.99640678 |
| MTMR3 | 48.774154 | 0.109529093 | 0.3716933 | 0.29467599 | 0.76824141 | 0.99665574 |
| TBC1D10A | 5.5653991 | -1.148268033 | 0.60423001 | -1.9003823 | 0.05738297 | 0.67545594 |
| SF3A1 | 8.06045294 | 0.760624685 | 0.725901 | 1.04783529 | 0.2947145 | 0.97161335 |
| RNF215 | 4.51489982 | -0.239033102 | 0.68914079 | -0.3468567 | 0.72869897 | 0.99665574 |
| PES1 | 3.29983212 | -0.251834048 | 0.87175161 | -0.2888828 | 0.77267106 | 0.99665574 |
| DUSP18 | 3.01735334 | -0.793612851 | 0.97567809 | -0.8133962 | 0.41599094 | 0.992853 |
| OSBP2 | 16.1578088 | -0.375333782 | 0.52905261 | -0.7094451 | 0.47804831 | 0.99617422 |
| MORC2 | 8.26004348 | -0.597532164 | 0.49898327 | -1.1974994 | 0.23111196 | 0.92774608 |
| SMTN | 3.27170194 | 1.441135147 | 0.85051727 | 1.69442196 | 0.09018513 | 0.76254949 |
| SELENOM | 16.5313641 | -0.065313073 | 0.55359025 | -0.1179809 | 0.9060828 | 0.99777713 |
| INPP5J | 6.35690278 | 1.433457758 | 0.6689456 | 2.14286147 | 0.03212422 | 0.57035969 |
| RNF185 | 18.5687012 | -0.587182011 | 0.55534983 | -1.0573191 | 0.29036596 | 0.96676791 |
| LIMK2 | 11.9374013 | -0.96597894 | 0.48981336 | -1.9721368 | 0.04859399 | 0.64337675 |
| PIK3IP1 | 4.75209434 | -1.914825913 | 0.79953725 | -2.3949177 | 0.0166241 | 0.42361121 |
| PATZ1 | 5.3007507 | 0.898136579 | 0.68515997 | 1.31084217 | 0.18991109 | 0.8994569 |
| DRG1 | 13.7044042 | -0.220059687 | 0.45666583 | -0.4818834 | 0.62988877 | 0.99665574 |
| EIF4ENIF1 | 13.7925958 | -0.433869533 | 0.5371803 | -0.8076795 | 0.41927509 | 0.99476033 |
| SFI1 | 10.0018492 | -0.177171464 | 0.48302305 | -0.3667971 | 0.71377035 | 0.99665574 |
| PISD | 41.1229516 | -0.463446367 | 0.51601062 | -0.8981334 | 0.36911444 | 0.98532838 |
| PRR14L | 31.6283921 | 0.024833188 | 0.39123564 | 0.06347374 | 0.94938927 | 0.99777713 |
| YWHAH | 10.9144235 | -1.348233024 | 0.60583073 | -2.2254286 | 0.02605248 | 0.51788242 |
| RTCB | 14.6645975 | -0.251713143 | 0.4031312 | -0.6243951 | 0.53236814 | 0.99665574 |
| FBXO7 | 15.5887281 | 0.156791074 | 0.41371379 | 0.37898441 | 0.70469944 | 0.99665574 |
| SYN3 | 3.00575507 | 0.964524122 | 0.96193529 | 1.00269128 | 0.31600984 | 0.97652251 |
| TIMP3 | 32.3885402 | 0.331397035 | 0.65409086 | 0.50665291 | 0.61239838 | 0.99665574 |
| LARGE1 | 43.344566 | -0.853892495 | 0.36634015 | -2.3308733 | 0.01976004 | 0.46397674 |
| HMGXB4 | 10.4840976 | -0.349816987 | 0.53393297 | -0.6551702 | 0.51235816 | 0.99665574 |
| TOM1 | 16.5720575 | -0.253457674 | 0.39763178 | -0.637418 | 0.52385258 | 0.99665574 |
| HMOX1 | 6.9138578 | -3.370399389 | 0.99786813 | -3.3776 | 0.00073121 | 0.07202465 |
| APOL6 | 7.44394464 | -1.52777379 | 1.03771434 | -1.4722489 | 0.14095368 | 0.84969914 |
| RBFOX2 | 91.2234986 | 0.324185665 | 0.24724163 | 1.31120989 | 0.18978686 | 0.8994569 |
| APOL4 | 8.99967386 | -2.664215329 | 0.97400143 | -2.73533 | 0.00623178 | 0.25132729 |
| APOL2 | 19.4975976 | -2.137302577 | 0.82938155 | -2.5769835 | 0.00996667 | 0.34338499 |
| APOL1 | 13.7730155 | -2.662964994 | 0.87952646 | -3.0277259 | 0.00246402 | 0.15419867 |
| MYH9 | 43.9034775 | -0.251279482 | 0.29299809 | -0.8576148 | 0.39110523 | 0.99179694 |
| TXN2 | 14.2787771 | -0.040094694 | 0.4476388 | -0.0895693 | 0.92862948 | 0.99777713 |
| EIF3D | 11.8523474 | -0.105877411 | 0.62865305 | -0.1684195 | 0.8662533 | 0.99665574 |
| IFT27 | 3.55441358 | 0.464088012 | 0.82466867 | 0.56275694 | 0.57360041 | 0.99665574 |
| PVALB | 4.31406478 | 0.537980183 | 0.9909232 | 0.54290805 | 0.58719311 | 0.99665574 |
| TST | 11.7456658 | 0.240469945 | 0.46719035 | 0.51471514 | 0.60675209 | 0.99665574 |
| MPST | 10.7047227 | 0.217224275 | 0.4953703 | 0.43850888 | 0.66101744 | 0.99665574 |
| CARD10 | 6.97135872 | 0.157248265 | 0.83932647 | 0.18735054 | 0.8513858 | 0.99665574 |
| CDC42EP1 | 5.53574518 | 0.796403325 | 0.71613264 | 1.11208913 | 0.26609983 | 0.94557817 |
| GGA1 | 14.6019163 | 0.354965375 | 0.424916 | 0.83537777 | 0.40350504 | 0.99194293 |
| TRIOBP | 20.4128702 | -0.27294012 | 0.37641517 | -0.7251039 | 0.46838828 | 0.99617422 |
| H1F0 | 34.6417146 | -0.625137979 | 0.88500413 | -0.7063673 | 0.47995975 | 0.99617422 |
| GCAT | 3.12316226 | 0.466190807 | 0.77695248 | 0.60002487 | 0.54848966 | 0.99665574 |
| ANKRD54 | 5.26777559 | 0.134560223 | 0.61888248 | 0.21742451 | 0.82787753 | 0.99665574 |
| EIF3L | 37.6588896 | 0.596277071 | 0.31492206 | 1.89341154 | 0.05830316 | 0.67545594 |
| MICALL1 | 6.33910586 | -0.004628437 | 0.71076377 | -0.0065119 | 0.99480428 | 0.9985287 |
| POLR2F | 5.69712184 | 0.388755008 | 0.65649523 | 0.5921673 | 0.55373856 | 0.99665574 |

| | | | | | | |
|---|---|---|---|---|---|---|
| PICK1 | 4.28293659 | 0.154238895 | 0.74409031 | 0.20728518 | 0.83578715 | 0.99665574 |
| PLA2G6 | 4.69311436 | 0.531944539 | 0.6876045 | 0.77361992 | 0.43915559 | 0.99617422 |
| MAFF | 5.7791125 | -2.733560034 | 0.74304501 | -3.678862 | 0.00023428 | 0.03654 |
| TMEM184B | 12.1429905 | 0.242041035 | 0.58066301 | 0.41683564 | 0.67679864 | 0.99665574 |
| CSNK1E | 3.25867515 | 1.194810568 | 0.86805112 | 1.3764288 | 0.16868892 | 0.87566218 |
| TPTEP2-CSNI | 14.7503593 | -0.206569962 | 0.58759325 | -0.3515526 | 0.72517378 | 0.99665574 |
| DDX17 | 161.301472 | 0.357524074 | 0.19127043 | 1.86920723 | 0.06159399 | 0.6834742 |
| CBY1 | 5.4787528 | -0.132847764 | 0.61893535 | -0.2146392 | 0.83004866 | 0.99665574 |
| TOMM22 | 5.67071748 | 0.796942717 | 0.7601707 | 1.04837337 | 0.29446662 | 0.97161335 |
| JOSD1 | 7.22692088 | -0.658124648 | 0.54287528 | -1.2122944 | 0.22539972 | 0.92318867 |
| GTPBP1 | 6.36599696 | -0.526095212 | 0.55141482 | -0.9540825 | 0.3400419 | 0.97788427 |
| SUN2 | 13.5889262 | 0.267196348 | 0.47301938 | 0.564874 | 0.57215947 | 0.99665574 |
| AL021707.1 | 3.28383556 | -0.907925227 | 0.79402028 | -1.1434535 | 0.25285037 | 0.94076968 |
| CBX6 | 9.19117663 | -0.323843099 | 0.60026481 | -0.5395004 | 0.58954163 | 0.99665574 |
| CBX7 | 7.20541632 | -0.431467166 | 0.70022497 | -0.6161836 | 0.53777333 | 0.99665574 |
| RPL3 | 67.2982895 | 0.679092837 | 0.45143159 | 1.50430952 | 0.13250169 | 0.84089786 |
| TAB1 | 5.25790469 | 0.289418423 | 0.62796999 | 0.46087938 | 0.64488515 | 0.99665574 |
| MIEF1 | 5.02224126 | -0.134752288 | 0.59508218 | -0.2264432 | 0.82085676 | 0.99665574 |
| ATF4 | 20.4995115 | -0.449669608 | 0.74594248 | -0.6028208 | 0.54662794 | 0.99665574 |
| RPS19BP1 | 16.247157 | 0.175716705 | 0.47298611 | 0.371505 | 0.71026143 | 0.99665574 |
| FAM83F | 10.443952 | 0.045176131 | 0.4976468 | 0.09077951 | 0.92766779 | 0.99777713 |
| TNRC6B | 190.318985 | -0.078499479 | 0.34042949 | -0.2305895 | 0.81763369 | 0.99665574 |
| MKL1 | 93.1164291 | 0.187343984 | 0.28198589 | 0.66437361 | 0.50645122 | 0.99665574 |
| SLC25A17 | 14.3419648 | -0.100351128 | 0.39775827 | -0.2522917 | 0.80081557 | 0.99665574 |
| ST13 | 45.9152281 | 0.408312981 | 0.35329591 | 1.1557252 | 0.24779358 | 0.93441937 |
| XPNPEP3 | 16.5534585 | -0.270072391 | 0.55376514 | -0.487702 | 0.6257609 | 0.99665574 |
| RBX1 | 19.7866752 | 0.83294974 | 0.43285482 | 1.92431665 | 0.05431491 | 0.66734438 |
| EP300 | 49.1329085 | -0.095084104 | 0.29539304 | -0.3218901 | 0.74753593 | 0.99665574 |
| L3MBTL2 | 3.80938888 | 0.601186662 | 0.70390213 | 0.85407706 | 0.39306229 | 0.99179694 |
| RANGAP1 | 5.14467901 | -0.577161335 | 0.59896519 | -0.9635975 | 0.33524778 | 0.97788427 |
| ZC3H7B | 14.8632427 | -0.395635903 | 0.4004963 | -0.9878641 | 0.32321922 | 0.97700345 |
| TEF | 4.1283622 | 1.126050835 | 0.8375931 | 1.34438886 | 0.17882267 | 0.88813816 |
| TOB2 | 13.0151728 | 0.233801468 | 0.48948245 | 0.47765036 | 0.63289908 | 0.99665574 |
| PHF5A | 3.3319089 | -0.545921532 | 0.90361381 | -0.6041536 | 0.54574153 | 0.99665574 |
| ACO2 | 43.75443 | -0.356788562 | 0.33705338 | -1.0585521 | 0.28980381 | 0.96676791 |
| POLR3H | 5.47449154 | -0.484811879 | 0.7159604 | -0.677149 | 0.49831141 | 0.99640678 |
| PMM1 | 6.16190982 | 0.241525628 | 0.63968445 | 0.37756995 | 0.70575009 | 0.99665574 |
| DESI1 | 10.4931573 | 0.020173175 | 0.57159732 | 0.03529263 | 0.9718464 | 0.99777713 |
| XRCC6 | 30.8928392 | 0.014523707 | 0.32656161 | 0.04447463 | 0.96452607 | 0.99777713 |
| SNU13 | 29.8622719 | -0.128517683 | 0.36812409 | -0.3491151 | 0.72700289 | 0.99665574 |
| C22orf46 | 2.41999335 | -0.6115491 | 0.87908309 | -0.695667 | 0.4866374 | 0.99640678 |
| CCDC134 | 3.40947113 | 0.153946472 | 0.85312129 | 0.18045086 | 0.85679863 | 0.99665574 |
| SREBF2 | 68.8944665 | -0.595628402 | 0.38423883 | -1.5501515 | 0.12110515 | 0.82659025 |
| WBP2NL | 5.14424863 | 0.618033893 | 0.76856362 | 0.80414149 | 0.42131526 | 0.99518711 |
| SMDT1 | 15.6329427 | -0.215858445 | 0.49793691 | -0.4335056 | 0.66464749 | 0.99665574 |
| NDUFA6 | 14.8829702 | -0.06505213 | 0.46658203 | -0.1394227 | 0.88911613 | 0.99665574 |
| TCF20 | 34.3091599 | -0.013931515 | 0.43331764 | -0.0321508 | 0.97435178 | 0.99777713 |
| RRP7A | 8.75861536 | 0.562212436 | 0.56226158 | 0.9999126 | 0.31735281 | 0.97700345 |
| POLDIP3 | 9.69783342 | 0.169429601 | 0.54259589 | 0.31225744 | 0.75484488 | 0.99665574 |
| CYB5R3 | 27.4561014 | -0.621819717 | 0.4180612 | -1.4873892 | 0.13691204 | 0.84475183 |
| A4GALT | 9.25638749 | -2.802424106 | 0.72879867 | -3.8452651 | 0.00012042 | 0.02485289 |
| ARFGAP3 | 30.3188919 | -1.001623953 | 0.31177539 | -3.2126459 | 0.00131518 | 0.10498682 |
| PACSIN2 | 41.0355056 | -0.444648449 | 0.30300069 | -1.4674833 | 0.14224464 | 0.84969914 |
| BIK | 3.17225882 | -0.08178166 | 0.78865937 | -0.1036971 | 0.91740976 | 0.99777713 |
| MCAT | 3.04389691 | 0.746378277 | 0.80612961 | 0.92587876 | 0.35450899 | 0.98022903 |
| TSPO | 26.5393453 | -0.374510419 | 0.50500268 | -0.7416009 | 0.4583292 | 0.99617422 |
| TTLL12 | 3.69500976 | 1.016683527 | 0.74594021 | 1.36295579 | 0.17289644 | 0.88157681 |
| EFCAB6 | 12.0285324 | 0.021898285 | 0.5120121 | 0.04276908 | 0.96588561 | 0.99777713 |
| SAMM50 | 4.9707052 | 0.137340195 | 0.68001316 | 0.20196697 | 0.83994255 | 0.99665574 |
| PARVB | 60.653012 | 0.609239639 | 0.36965606 | 1.64812568 | 0.09932688 | 0.78261212 |
| RTL6 | 2.29612169 | 0.067657192 | 0.98138615 | 0.06894044 | 0.94503703 | 0.99777713 |
| NUP50-DT | 15.8828676 | 0.159614546 | 0.54424281 | 0.29327819 | 0.76930952 | 0.99665574 |
| NUP50 | 10.9250155 | -0.105632033 | 0.45742241 | -0.2309289 | 0.81737007 | 0.99665574 |
| KIAA0930 | 4.9901044 | -1.680711257 | 0.79466509 | -2.1149932 | 0.03443053 | 0.58178209 |
| FAM118A | 3.49536439 | 0.324040147 | 1.09141567 | 0.29689893 | 0.76654367 | 0.99665574 |
| ATXN10 | 60.5641581 | -0.018135337 | 0.25399313 | -0.0714009 | 0.9430787 | 0.99777713 |
| MIRLET7BHG | 24.9471919 | -0.747926016 | 0.41198583 | -1.8154168 | 0.06945987 | 0.7094165 |
| PPARA | 34.9436412 | 0.143301466 | 0.32685671 | 0.4384229 | 0.66107975 | 0.99665574 |
| TTC38 | 8.87319297 | -0.213739727 | 0.6032627 | -0.3543062 | 0.72310941 | 0.99665574 |
| TRMU | 8.02783882 | -0.287799983 | 0.49050372 | -0.5867437 | 0.55737584 | 0.99665574 |
| CELSR1 | 16.5252584 | 0.587313619 | 0.45955391 | 1.27800809 | 0.20124657 | 0.90591604 |
| GRAMD4 | 8.43765825 | -0.852910922 | 0.48324845 | -1.7649532 | 0.07757162 | 0.74141484 |
| CERK | 8.73528838 | -0.398429965 | 0.51534066 | -0.773139 | 0.43944012 | 0.99617422 |
| TBC1D22A | 83.0097518 | -0.310666619 | 0.37911503 | -0.8194521 | 0.41252851 | 0.99279547 |
| BRD1 | 18.3336149 | -0.643172065 | 0.44004632 | -1.4616008 | 0.14385062 | 0.84969914 |
| ZBED4 | 9.42966675 | -0.42265711 | 0.61551093 | -0.6866769 | 0.49228638 | 0.99640678 |
| ALG12 | 2.58115955 | 0.463149354 | 0.8667999 | 0.53432096 | 0.59311949 | 0.99665574 |
| CRELD2 | 3.02229871 | 0.160249887 | 0.84703678 | 0.18918882 | 0.84994483 | 0.99665574 |
| PIM3 | 18.4958324 | -0.520152739 | 0.42627591 | -1.2202255 | 0.2223794 | 0.92191316 |
| TRABD | 10.9983598 | 0.678766468 | 0.5243261 | 1.29455023 | 0.19547543 | 0.90381759 |
| SELENOO | 14.3646779 | -0.211212028 | 0.40846385 | -0.5170887 | 0.60509427 | 0.99665574 |
| TUBGCP6 | 3.93083163 | 1.201475163 | 0.7717805 | 1.5567576 | 0.11952805 | 0.82625998 |
| PLXNB2 | 29.7627226 | 0.204362194 | 0.38576676 | 0.52975584 | 0.59628122 | 0.99665574 |
| PPP6R2 | 57.902488 | 0.292428023 | 0.29277231 | 0.99882404 | 0.31787994 | 0.97700345 |
| SBF1 | 5.141354 | -0.045902512 | 0.59510818 | -0.0771331 | 0.9385177 | 0.99777713 |
| MIOX | 5.2505424 | 2.494713531 | 0.95001723 | 2.62596661 | 0.00864033 | 0.31119582 |
| LMF2 | 3.47286322 | 0.113122072 | 0.78066667 | 0.14490445 | 0.88478631 | 0.99665574 |
| NCAPH2 | 2.36370951 | -0.15632495 | 0.91149061 | -0.1715047 | 0.86382691 | 0.99665574 |
| ARSA | 3.64064766 | 0.653028224 | 0.81556232 | 0.80070916 | 0.42330004 | 0.99518711 |
| RABL2B | 3.81580557 | -0.978078755 | 0.7219829 | -1.3547118 | 0.1755094 | 0.88428483 |
| FP671120.1 | 132.529757 | -0.962292948 | 0.55885858 | -1.7218899 | 0.08508946 | 0.75373899 |
| FP236383.1 | 77.8039301 | -0.778649769 | 0.52978946 | -1.4697343 | 0.14163372 | 0.84969914 |
| CR381653.1 | 7.70733801 | 0.865771331 | 0.73108155 | 1.18423359 | 0.23632061 | 0.9305595 |
| HSPA13 | 3.86399703 | 0.972366569 | 0.73485244 | 1.32321336 | 0.18576443 | 0.89532394 |
| AF127577.4 | 3.53818759 | 0.354659243 | 0.75046478 | 0.47258612 | 0.63650848 | 0.99665574 |
| NRIP1 | 37.8052047 | 0.174408901 | 0.33674923 | 0.51791922 | 0.60451463 | 0.99665574 |
| AJ009632.2 | 7.19465668 | 0.842154671 | 0.64931861 | 1.29698219 | 0.19463732 | 0.90333151 |
| USP25 | 88.0359966 | 0.427613006 | 0.28291625 | 1.51144732 | 0.13067452 | 0.83795959 |
| MIR99AHG | 117.900042 | 0.545432499 | 0.39092995 | 1.395218 | 0.16295011 | 0.87135275 |
| CXADR | 18.9428807 | -0.076876799 | 0.52326474 | -0.1469176 | 0.88319706 | 0.99665574 |
| BTG3 | 11.4913573 | 0.214149401 | 0.66405704 | 0.32248645 | 0.7470842 | 0.99665574 |
| C21orf91 | 2.95036646 | -0.61825246 | 0.8616738 | -0.7175018 | 0.47306455 | 0.99617422 |
| LINC01687 | 25.3962813 | 0.816906828 | 0.46510821 | 1.75638015 | 0.07902352 | 0.74455432 |
| MRPL39 | 4.45080428 | -0.711425408 | 0.62363319 | -1.1407754 | 0.2539634 | 0.94099699 |
| ATP5PF | 34.2936864 | 0.31366142 | 0.40625579 | 0.77207865 | 0.44006784 | 0.99617422 |
| GABPA | 9.69262552 | 0.207054185 | 0.50018 | 0.41395935 | 0.67890388 | 0.99665574 |
| APP | 160.257265 | -0.559058912 | 0.27660405 | -2.0211523 | 0.043264 | 0.62038248 |
| ADAMTS1 | 20.7902082 | -4.714719157 | 0.73952617 | -6.3753243 | 1.83E-10 | 3.08E-07 |
| N6AMT1 | 4.5153108 | -0.732880352 | 0.86749751 | -0.8448213 | 0.39821063 | 0.99179694 |
| LTN1 | 18.6441016 | 0.284619108 | 0.45200447 | 0.62968207 | 0.52890261 | 0.99665574 |
| USP16 | 15.5917362 | -0.019214304 | 0.52509537 | -0.036592 | 0.9708103 | 0.99777713 |
| CCT8 | 13.9881772 | 0.30909947 | 0.40585868 | 0.76159384 | 0.44630245 | 0.99617422 |
| MAP3K7CL | 10.5329704 | -0.076929097 | 0.69683425 | -0.110398 | 0.91209375 | 0.99777713 |

| | | | | | | |
|---|---|---|---|---|---|---|
| BACH1 | 212.361434 | -0.671617391 | 0.36856246 | -1.8222621 | 0.06841523 | 0.70708655 |
| GRIK1 | 8.35702443 | -1.158214601 | 0.67045605 | -1.7275027 | 0.08407743 | 0.75343906 |
| CLDN8 | 16.8486654 | 1.081937938 | 0.57840315 | 1.87056025 | 0.06140606 | 0.6834742 |
| TIAM1 | 56.2307435 | 0.579061914 | 0.4137437 | 1.39956671 | 0.16164311 | 0.87011714 |
| SOD1 | 28.2217336 | -0.197242643 | 0.41541536 | -0.4748083 | 0.63492364 | 0.99665574 |
| SCAF4 | 21.2164301 | 0.196399864 | 0.35924299 | 0.5467048 | 0.58458156 | 0.99665574 |
| URB1 | 7.86164085 | 0.169497604 | 0.55172201 | 0.30721559 | 0.75867928 | 0.99665574 |
| EVA1C | 48.6910341 | -0.584942163 | 0.5000627 | -1.1697376 | 0.24210657 | 0.93397124 |
| SYNJ1 | 14.2187334 | 0.275235507 | 0.38958362 | 0.70648635 | 0.47988574 | 0.99617422 |
| PAXBP1 | 29.230161 | -0.069611132 | 0.36731708 | -0.1895124 | 0.84969126 | 0.99665574 |
| C21orf62-AS | 3.65077521 | -0.153988059 | 0.80706736 | -0.1907995 | 0.84868267 | 0.99665574 |
| AP000282.1 | 6.05328351 | 0.757678551 | 0.93026607 | 0.8144751 | 0.41537283 | 0.99279547 |
| IFNAR1 | 29.6247109 | -1.054588396 | 0.33654722 | -3.1335525 | 0.00172704 | 0.12036484 |
| IFNGR2 | 31.9442059 | -0.578650112 | 0.3590468 | -1.6116287 | 0.10704277 | 0.79852803 |
| TMEM50B | 19.6215011 | 0.141960835 | 0.4742079 | 0.29936413 | 0.76466223 | 0.99665574 |
| GART | 11.0519035 | 0.036721005 | 0.46312931 | 0.07928888 | 0.93680285 | 0.99777713 |
| SON | 70.859842 | -0.393408181 | 0.27404701 | -1.43555 | 0.15113043 | 0.85612189 |
| AP000311.1 | 12.771732 | -0.943293004 | 0.63650252 | -1.4819941 | 0.13834185 | 0.84656101 |
| ITSN1 | 52.5496029 | -0.008235751 | 0.34177271 | -0.0240972 | 0.98077511 | 0.99777713 |
| MRPS6 | 59.9202126 | 0.963863526 | 0.44305775 | 2.17548056 | 0.02959412 | 0.5576677 |
| LINC00310 | 5.55052404 | -0.086138119 | 0.67520898 | -0.1275725 | 0.89848727 | 0.99777713 |
| AP000317.1 | 5.86621751 | -0.155406398 | 0.82238172 | -0.1889711 | 0.85011545 | 0.99665574 |
| RCAN1 | 11.0223765 | 0.135848414 | 0.5949905 | 0.22832031 | 0.81939724 | 0.99665574 |
| RUNX1 | 16.5686246 | -2.004702904 | 0.56347059 | -3.5577773 | 0.00037401 | 0.05123884 |
| SETD4 | 8.96756936 | 0.141805362 | 0.46760109 | 0.3032614 | 0.76169066 | 0.99665574 |
| CBR1 | 18.3231204 | -1.010191934 | 0.41803266 | -2.4165383 | 0.01566888 | 0.41847166 |
| CBR3 | 16.6624925 | 0.436899389 | 0.67437663 | 0.64785666 | 0.51707767 | 0.99665574 |
| DOPEY2 | 34.1870384 | 0.631358102 | 0.4317951 | 1.46217062 | 0.14369446 | 0.84969914 |
| MORC3 | 29.2309226 | 0.121729913 | 0.32162016 | 0.37848968 | 0.70506686 | 0.99665574 |
| SIM2 | 9.72613199 | -0.627937086 | 0.9606034 | -0.6536903 | 0.51331138 | 0.99665574 |
| HLCS | 41.9711541 | -0.09144207 | 0.30439785 | -0.3004031 | 0.76386967 | 0.99665574 |
| PIGP | 4.31539626 | -0.118696491 | 0.85090337 | -0.1394947 | 0.88905926 | 0.99665574 |
| TTC3 | 60.9961816 | 0.048774366 | 0.34872514 | 0.13986478 | 0.88876683 | 0.99665574 |
| VPS26C | 13.4838767 | 0.146547231 | 0.4522868 | 0.32401395 | 0.74592748 | 0.99665574 |
| DYRK1A | 120.880668 | 0.066786847 | 0.31760334 | 0.21028383 | 0.83344615 | 0.99665574 |
| KCNJ6 | 30.2507378 | 1.459504258 | 0.6586484 | 2.21590798 | 0.0266978 | 0.52453943 |
| KCNJ15 | 12.4602706 | 1.711004061 | 0.84219117 | 2.03161007 | 0.04219315 | 0.61533334 |
| ETS2 | 22.0669278 | -2.916361565 | 0.52384363 | -5.5672369 | 2.59E-08 | 3.28E-05 |
| PSMG1 | 4.63192106 | 0.014080922 | 0.65113799 | 0.0216251 | 0.98274701 | 0.99777713 |
| BRWD1 | 102.496358 | 0.352324197 | 0.32265628 | 1.09194897 | 0.27485553 | 0.95528129 |
| HMGN1 | 31.5530193 | 0.126491178 | 0.3426161 | 0.36919216 | 0.7119845 | 0.99665574 |
| WRB | 7.11979225 | -0.04819451 | 0.5610924 | -0.0858941 | 0.93155062 | 0.99777713 |
| PCP4 | 14.7932627 | 0.796436846 | 0.63830131 | 1.24774433 | 0.2121247 | 0.91043197 |
| DSCAM | 9.90327437 | -2.832367645 | 0.76362819 | -3.7090926 | 0.000208 | 0.03456948 |
| BACE2 | 174.161312 | -0.202210129 | 0.5961984 | -0.3391658 | 0.7344848 | 0.99665574 |
| FAM3B | 17.6076634 | -0.439453585 | 0.47017064 | -0.9346683 | 0.34995927 | 0.97838306 |
| MX1 | 32.3357665 | 0.099674514 | 1.25029031 | 0.0797211 | 0.93645908 | 0.99777713 |
| TMPRSS2 | 68.2943933 | 0.272687385 | 0.43491109 | 0.6269957 | 0.53066206 | 0.99665574 |
| RIPK4 | 4.95584797 | -1.116145276 | 0.68629447 | -1.6263358 | 0.10387824 | 0.79062881 |
| PRDM15 | 3.9710894 | -1.172957241 | 0.64440142 | -1.8202276 | 0.06872436 | 0.70805642 |
| C2CD2 | 11.4479404 | -0.761087456 | 0.43609275 | -1.7452422 | 0.08094273 | 0.74735643 |
| ZBTB21 | 11.0416722 | 0.484744973 | 0.46495918 | 1.04255384 | 0.29715498 | 0.97341216 |
| ABCG1 | 9.15816928 | -0.064898992 | 1.00810486 | -0.0643772 | 0.94866987 | 0.99777713 |
| SLC37A1 | 14.9023678 | -1.84907147 | 0.61249998 | -3.0188923 | 0.00253701 | 0.15587986 |
| PDE9A | 23.4773518 | -0.131558706 | 0.38349973 | -0.3430477 | 0.73156256 | 0.99665574 |
| WDR4 | 5.67631724 | -0.312460365 | 0.5823273 | -0.5365717 | 0.59156348 | 0.99665574 |
| NDUFV3 | 19.0815635 | 0.30520073 | 0.37895162 | 0.80538179 | 0.4205994 | 0.99518711 |
| PKNOX1 | 18.6876748 | -0.621393599 | 0.50436951 | -1.2320205 | 0.21794141 | 0.91662582 |
| RRP1B | 6.09767954 | -0.328371354 | 0.54719769 | -0.6000964 | 0.548442 | 0.99665574 |
| PDXK | 37.3457849 | 0.00437805 | 0.33519611 | 0.01306116 | 0.989579 | 0.99777713 |
| CSTB | 40.1854487 | -0.229387073 | 0.43180521 | -0.5312281 | 0.59526071 | 0.99665574 |
| RRP1 | 3.06823535 | -0.595846668 | 0.91452395 | -0.6515375 | 0.51469957 | 0.99665574 |
| AGPAT3 | 31.4293127 | -0.09934366 | 0.34146818 | -0.2909309 | 0.77110413 | 0.99665574 |
| TRAPPC10 | 34.9682806 | -0.147193857 | 0.28410815 | -0.5180909 | 0.60439482 | 0.99665574 |
| PFKL | 23.8723218 | -0.208014432 | 0.39752739 | -0.5232707 | 0.6007859 | 0.99665574 |
| C21orf2 | 5.10525843 | 0.612489743 | 0.71730497 | 0.85387633 | 0.39317351 | 0.99179694 |
| UBE2G2 | 23.8987031 | -0.005745216 | 0.40012984 | -0.0143584 | 0.98854406 | 0.99777713 |
| SUMO3 | 8.44981884 | -0.273164387 | 0.51480518 | -0.530617 | 0.5956842 | 0.99665574 |
| PTTG1IP | 39.3534623 | -0.659825958 | 0.28670955 | -2.3013742 | 0.02137049 | 0.4733368 |
| FAM207A | 3.98929766 | -0.003695275 | 0.82188942 | -0.0044961 | 0.99641266 | 0.998802 |
| ADARB1 | 4.42570304 | 0.887964283 | 0.86740047 | 1.0237074 | 0.3059735 | 0.97376371 |
| POFUT2 | 5.43508203 | -0.259336262 | 0.69588605 | -0.3726706 | 0.70939363 | 0.99665574 |
| COL18A1 | 64.1182075 | 0.195754702 | 0.35706935 | 0.54822601 | 0.58353673 | 0.99665574 |
| COL6A1 | 53.1271296 | 0.987217099 | 0.44599975 | 2.21349249 | 0.02686371 | 0.52677814 |
| LSS | 10.9902337 | -0.987052315 | 0.59441792 | -1.6605359 | 0.09680669 | 0.77583097 |
| MCM3AP-AS | 3.02510639 | -0.580720829 | 0.84074943 | -0.6907181 | 0.48974272 | 0.99640678 |
| MCM3AP | 20.6547386 | 0.148253338 | 0.37006537 | 0.40061392 | 0.6887044 | 0.99665574 |
| YBEY | 2.78742018 | 0.143876214 | 0.77088094 | 0.1866387 | 0.85194392 | 0.99665574 |
| PCNT | 10.5693713 | -0.241443709 | 0.48820317 | -0.4945558 | 0.62091371 | 0.99665574 |
| DIP2A | 32.2208749 | -0.673949429 | 0.37194794 | -1.8119456 | 0.0699946 | 0.71124502 |
| PRMT2 | 25.013152 | -0.587087307 | 0.34141445 | -1.7195737 | 0.08550996 | 0.75373899 |
| AC011043.1 | 3.7841956 | 2.262443621 | 1.01646246 | 2.22580145 | 0.02602748 | 0.51788242 |
| AL592183.1 | 17.8492647 | -0.250431206 | 0.46590977 | -0.5375101 | 0.59091531 | 0.99665574 |

**Endothelial cells**

*This means that positive fold changes imply higher mean value in controls than in AKI and vice versa.

| | baseMean | log2 fold change control/AKI* | lfcSE | stat | pvalue | padj |
|---|---|---|---|---|---|---|
| ANGPTL4 | 123.454957 | -5.076443763 | 0.73134595 | -6.9412345 | 3.89E-12 | 1.59E-08 |
| SERPINE1 | 816.722209 | -4.747771629 | 1.12746238 | -4.2110244 | 2.54E-05 | 0.00345277 |
| ADAMTS4 | 178.195367 | -4.055582278 | 1.26486117 | -3.2063458 | 0.00134432 | 0.04507706 |
| STC2 | 63.7775825 | -3.988356556 | 1.24524261 | -3.2028751 | 0.00136063 | 0.04507706 |
| INHBB | 123.708098 | -3.96487199 | 0.56703995 | -6.9922269 | 2.71E-12 | 1.59E-08 |
| HILPDA | 102.502663 | -3.723499096 | 1.22021201 | -3.0515182 | 0.00227687 | 0.06286489 |
| SLPI | 167.258874 | -3.583151567 | 0.74993631 | -4.7779412 | 1.77E-06 | 0.00042624 |
| GRAMD1B | 117.300221 | -3.544233637 | 0.72030513 | -4.9204615 | 8.63E-07 | 0.00023551 |
| STC1 | 346.606761 | -3.504180825 | 0.98764821 | -3.548005 | 0.00038816 | 0.02112867 |
| VCAN | 224.117247 | -3.466034121 | 0.54100917 | -6.4066088 | 1.49E-10 | 2.44E-07 |
| RAB3C | 150.960344 | -3.417371888 | 0.69002182 | -4.9525563 | 7.32E-07 | 0.00023083 |
| ACKR1 | 751.083754 | -3.235102967 | 1.25031153 | -2.5874375 | 0.00966927 | 0.13832807 |
| NNMT | 454.473825 | -3.181954706 | 0.48370617 | -6.5782802 | 4.76E-11 | 1.30E-07 |
| NAMPT | 1457.72557 | -3.159325899 | 0.76178941 | -4.1472432 | 3.37E-05 | 0.00417212 |
| ANKRD37 | 200.681445 | -3.087737934 | 1.52287927 | -2.0275658 | 0.04260458 | 0.29149941 |
| CP | 224.591414 | -3.029110478 | 0.9675007 | -3.1308613 | 0.00174294 | 0.05379909 |
| TCTEX1D1 | 69.4929894 | -3.012621769 | 0.76280392 | -3.9494052 | 7.83E-05 | 0.00766536 |
| SELE | 353.568081 | -2.971944358 | 1.80318336 | -1.6481654 | 0.09931874 | 0.42376187 |
| WFDC2 | 237.065251 | -2.925859447 | 0.58313064 | -5.0175025 | 5.23E-07 | 0.00018624 |
| CEBPD | 687.773079 | -2.842184706 | 0.66982544 | -4.2431722 | 2.20E-05 | 0.00324088 |
| IER3 | 124.812381 | -2.768844438 | 0.61558012 | -4.4979432 | 6.86E-06 | 0.00124771 |
| DEPP1 | 671.265787 | -2.768357511 | 0.51146684 | -5.4125846 | 6.21E-08 | 2.99E-05 |
| SPRY1 | 770.039111 | -2.706308893 | 0.58086688 | -4.6590863 | 3.18E-06 | 0.00063392 |
| ADM | 125.953362 | -2.661433768 | 1.01342933 | -2.6261661 | 0.00863527 | 0.13154285 |
| LITAF | 155.343468 | -2.647878274 | 0.53467676 | -4.9522973 | 7.33E-07 | 0.00023083 |
| SOCS3 | 379.092797 | -2.601230715 | 0.85173501 | -3.0540376 | 0.00225784 | 0.06286489 |
| SPP1 | 2239.94144 | -2.575849273 | 0.65631458 | -3.9247174 | 8.68E-05 | 0.00814512 |
| CDKN1A | 187.67161 | -2.557947659 | 0.96699941 | -2.6452422 | 0.00816325 | 0.12821467 |
| SLC39A14 | 273.495805 | -2.490028138 | 0.55806167 | -4.4619229 | 8.12E-06 | 0.00141327 |
| CXCL2 | 166.432677 | -2.48020122 | 1.05683637 | -2.3468167 | 0.01893457 | 0.19411645 |
| CCDC85C | 63.3184663 | -2.47008458 | 0.44325154 | -5.5726474 | 2.51E-08 | 1.58E-05 |
| TNFAIP3 | 368.302923 | -2.451156088 | 0.98164313 | -2.4969931 | 0.01252514 | 0.15866671 |
| GFRA1 | 99.1617139 | -2.44135826 | 1.27486835 | -1.9149885 | 0.05549397 | 0.32988018 |
| C11orf96 | 208.678329 | -2.43140111 | 0.72081786 | -3.3731144 | 0.00074323 | 0.03040928 |
| AP000695.2 | 63.4659322 | -2.417122291 | 0.64157123 | -3.7675042 | 0.00016489 | 0.01230754 |
| HIST1H4C | 58.7492086 | -2.405675145 | 0.70865907 | -3.3946862 | 0.00068707 | 0.02883241 |
| LYVE1 | 122.94912 | -2.392271477 | 0.51007082 | -4.6900771 | 2.73E-06 | 0.00057302 |
| LINC02147 | 1495.65786 | -2.380692433 | 0.59843891 | -3.9781712 | 6.94E-05 | 0.0071036 |
| F2RL3 | 188.363881 | -2.365900332 | 0.67574402 | -3.5011784 | 0.00046321 | 0.02326599 |
| ADAMTS9 | 3568.57614 | -2.348856452 | 0.71266795 | -3.2958637 | 0.0009812 | 0.03649603 |
| ITGB6 | 56.5787001 | -2.315195162 | 0.73483732 | -3.1506227 | 0.00162923 | 0.0514748 |
| C1R | 136.324947 | -2.311210806 | 0.62907773 | -3.673967 | 0.00023881 | 0.01538749 |
| ABCA1 | 147.505196 | -2.283243561 | 0.58563397 | -3.8987553 | 9.67E-05 | 0.00867938 |
| SEMA3A | 371.808046 | -2.250131802 | 0.68669817 | -3.2767406 | 0.00105013 | 0.03823551 |
| EGLN3 | 161.83769 | -2.227300644 | 0.93477173 | -2.3827214 | 0.01718519 | 0.18698165 |
| AC068234.1 | 95.1880609 | -2.22073573 | 0.71050648 | -3.1255672 | 0.00177463 | 0.05418567 |
| C4orf19 | 190.146557 | -2.215996187 | 0.53585011 | -4.1354777 | 3.54E-05 | 0.00432621 |
| PLAUR | 69.5955571 | -2.210438084 | 0.64336861 | -3.4357257 | 0.00059097 | 0.02686611 |
| DGKG | 245.730653 | -2.179595462 | 0.64484242 | -3.3800435 | 0.00072474 | 0.02983026 |
| SDCBP2 | 55.5907625 | -2.166596977 | 0.34050216 | -6.3629464 | 1.98E-10 | 2.70E-07 |
| PVT1 | 122.661639 | -2.151284183 | 0.40556865 | -5.3043651 | 1.13E-07 | 4.87E-05 |
| IGFBP3 | 883.470416 | -2.14810621 | 1.17785115 | -1.8237502 | 0.06818985 | 0.36142549 |
| PAPSS2 | 227.12745 | -2.138137112 | 0.87862861 | -2.4334936 | 0.0149539 | 0.17581575 |
| TIMP1 | 579.744294 | -2.124072984 | 0.71060893 | -2.9890885 | 0.00279811 | 0.07081266 |
| CEBPB | 158.802311 | -2.091613425 | 0.62576094 | -3.342512 | 0.00083024 | 0.03297047 |
| BCL6 | 128.996619 | -2.085154898 | 0.43099309 | -4.838024 | 1.31E-06 | 0.00033534 |
| ACKR3 | 78.2625044 | -2.074727296 | 0.56643196 | -3.6628006 | 0.00024947 | 0.01582508 |
| ABCC3 | 55.4102743 | -2.055770679 | 0.42662194 | -4.8187176 | 1.44E-06 | 0.00035828 |
| COL13A1 | 56.5807835 | -2.033437231 | 0.50365979 | -4.0373229 | 5.41E-05 | 0.00582119 |
| TIAM1 | 211.825301 | -2.021521982 | 0.70506372 | -2.8671479 | 0.00414189 | 0.08875226 |
| GDF15 | 74.9650315 | -2.018559673 | 0.9447539 | -2.1365984 | 0.03263067 | 0.25224169 |
| RELL1 | 470.598726 | -2.010037368 | 0.48448815 | -4.1487854 | 3.34E-05 | 0.00417212 |
| HIC1 | 87.1078451 | -1.995921162 | 0.56664154 | -3.52237 | 0.00042771 | 0.02229251 |
| MAFF | 71.6302435 | -1.970561121 | 0.73245595 | -2.6903476 | 0.00713776 | 0.12018171 |
| SERPINA1 | 95.3381615 | -1.952034265 | 0.74998335 | -2.6027701 | 0.00924739 | 0.13620125 |
| SAMD5 | 103.858019 | -1.951955616 | 0.59514018 | -3.279825 | 0.00103872 | 0.03823551 |
| CDC42EP3 | 196.080727 | -1.931220088 | 0.3275195 | -5.8965042 | 3.71E-09 | 3.04E-06 |
| CEMIP2 | 1019.97911 | -1.926709577 | 0.56068472 | -3.4363511 | 0.00058961 | 0.02686611 |
| DUSP23 | 162.313452 | -1.904049357 | 0.40395031 | -4.7135732 | 2.43E-06 | 0.00054188 |
| SLC16A7 | 264.636808 | -1.897342028 | 0.542378 | -3.4981914 | 0.00046842 | 0.02326599 |
| ABLIM2 | 156.761097 | -1.888746221 | 0.5538225 | -3.4103819 | 0.00064872 | 0.02779306 |
| PDE4B | 408.049202 | -1.886150503 | 1.05525313 | -1.7873915 | 0.07387423 | 0.37640899 |
| APCDD1 | 71.1381637 | -1.866624911 | 1.26283326 | -1.4781246 | 0.13937443 | 0.49091481 |
| CHST15 | 331.636769 | -1.863338811 | 0.65580347 | -2.8413067 | 0.00449291 | 0.09141064 |
| GALNT15 | 767.562782 | -1.848079637 | 0.47113447 | -3.9226161 | 8.76E-05 | 0.00814512 |
| TACR1 | 420.35239 | -1.845376846 | 0.39206617 | -4.7067995 | 2.52E-06 | 0.00054188 |
| KCNJ2 | 53.1677709 | -1.84479911 | 0.58350074 | -3.1616054 | 0.00156902 | 0.05054839 |
| TNFRSF1B | 408.818399 | -1.808565596 | 0.48907213 | -3.6979527 | 0.00021735 | 0.01457818 |
| DDIT4 | 628.81694 | -1.804562012 | 0.62549209 | -2.8850277 | 0.00391379 | 0.08541619 |
| CSGALNACT1 | 951.12489 | -1.789157639 | 0.53973601 | -3.3148754 | 0.00091684 | 0.0349812 |
| CCL2 | 234.615547 | -1.783558628 | 1.06699586 | -1.6715703 | 0.09460908 | 0.41378199 |
| OPCML | 196.504566 | -1.783242725 | 1.09883396 | -1.62285 | 0.10462147 | 0.43670609 |
| RDH10 | 85.2522641 | -1.778896288 | 0.65770601 | -2.7046983 | 0.00683665 | 0.11703823 |
| UGCG | 824.646172 | -1.777452934 | 0.63805348 | -2.7857429 | 0.00534052 | 0.10080241 |
| PNP | 116.530243 | -1.755538329 | 0.66485802 | -2.640471 | 0.00827909 | 0.12904337 |
| SEMA6B | 252.502286 | -1.746134015 | 0.47401992 | -3.6836722 | 0.0002299 | 0.01505003 |
| THBS1 | 136.735181 | -1.743215513 | 0.55765313 | -3.1259854 | 0.0017721 | 0.05418567 |
| TGM2 | 1051.11662 | -1.740616188 | 0.51369081 | -3.3884511 | 0.00070289 | 0.02932644 |
| TLL1 | 1773.08578 | -1.735577974 | 0.4752939 | -3.651589 | 0.00026062 | 0.01627997 |
| TBC1D2 | 85.7890014 | -1.730667785 | 0.50438318 | -3.4312559 | 0.00060079 | 0.02694332 |
| CPXM2 | 65.6871403 | -1.723733108 | 0.99504468 | -1.7323173 | 0.08321709 | 0.39543345 |
| BMP1 | 86.4833408 | -1.722020053 | 0.47856225 | -3.5983199 | 0.00032028 | 0.01872034 |
| 3-Mar | 789.302097 | -1.715416214 | 0.67206246 | -2.5524655 | 0.01069635 | 0.14539575 |
| RPL18A | 44.1876479 | -1.712341896 | 0.54935605 | -3.1169983 | 0.00182703 | 0.05451467 |
| EIF4EBP1 | 50.6910173 | -1.707161988 | 0.57905797 | -2.9481711 | 0.0031966 | 0.07516604 |
| GAPDH | 902.312833 | -1.704867879 | 0.52347969 | -3.2567985 | 0.00112676 | 0.0400883 |
| BMX | 56.9960576 | -1.699944321 | 0.43845519 | -3.8771221 | 0.0001057 | 0.00910461 |
| CLDN5 | 1031.62809 | -1.69419222 | 0.57748334 | -2.9337508 | 0.00334893 | 0.07718524 |
| PTCHD4 | 114.033258 | -1.677659267 | 0.94840802 | -1.7689214 | 0.07690699 | 0.38356763 |
| RPL22L1 | 50.3081748 | -1.677637385 | 0.62780306 | -2.6722351 | 0.00753478 | 0.12282297 |
| FTH1 | 1831.3912 | -1.674561669 | 0.48182097 | -3.4754853 | 0.00050993 | 0.02469087 |
| SLCO4A1 | 291.803523 | -1.662500694 | 0.60741719 | -2.7369997 | 0.00620023 | 0.11005752 |
| ABCG1 | 486.355302 | -1.66161127 | 0.38136175 | -4.3570475 | 1.32E-05 | 0.00207453 |
| ITPKC | 147.667367 | -1.660256221 | 0.39452658 | -4.2082239 | 2.57E-05 | 0.00345277 |
| FAM89A | 58.8011502 | -1.652044228 | 0.41974461 | -3.9358319 | 8.29E-05 | 0.00788889 |
| FBLN2 | 240.109929 | -1.646976559 | 0.55910552 | -2.9457347 | 0.00322189 | 0.07554353 |
| ZNF385D | 1011.71814 | -1.64674434 | 1.1211986 | -1.4687356 | 0.14190451 | 0.49378081 |
| MAN1C1 | 220.48915 | -1.62983566 | 0.53791688 | -3.0299024 | 0.00244633 | 0.06478413 |
| AP002518.2 | 128.224557 | -1.623908956 | 1.02333032 | -1.5868864 | 0.11253838 | 0.44956377 |
| PCDH17 | 714.099028 | -1.620294585 | 0.53042226 | -3.0547259 | 0.00225266 | 0.06286489 |
| ARSG | 76.9543628 | -1.620058165 | 0.42469116 | -3.8146737 | 0.00013636 | 0.01083358 |

| | | | | | | |
|---|---|---|---|---|---|---|
| YBX3 | 660.325936 | -1.616480886 | 0.42915464 | -3.7666629 | 0.00016544 | 0.01230754 |
| MGST1 | 83.2744738 | -1.613751782 | 0.75001922 | -2.1516139 | 0.03142778 | 0.24759838 |
| ADAM19 | 144.945774 | -1.610651298 | 0.54587258 | -2.9505994 | 0.00317158 | 0.07516604 |
| ELL2 | 276.932947 | -1.58938494 | 0.6293802 | -2.5253177 | 0.01155937 | 0.15149764 |
| TMEM70 | 53.6077866 | -1.586116732 | 0.53667821 | -2.9554334 | 0.0031223 | 0.07448917 |
| GALNT18 | 1600.77967 | -1.579934507 | 0.59506062 | -2.6550816 | 0.00792892 | 0.1259852 |
| TMEM273 | 50.9550445 | -1.579216401 | 0.58687846 | -2.6908747 | 0.0071265 | 0.12018171 |
| GADD45A | 79.996872 | -1.577692293 | 0.45092337 | -3.4988036 | 0.00046735 | 0.02326599 |
| GXYLT2 | 47.4510558 | -1.575703701 | 0.41897083 | -3.7608912 | 0.00016931 | 0.01248159 |
| IL33 | 134.81745 | -1.571114513 | 0.63793282 | -2.4628213 | 0.01378486 | 0.16892472 |
| DUSP6 | 449.605175 | -1.568922211 | 0.45116425 | -3.4774968 | 0.00050612 | 0.02465222 |
| MT2A | 1264.97669 | -1.563299672 | 0.66580231 | -2.3479938 | 0.01887484 | 0.19403618 |
| GADD45B | 124.453426 | -1.562331707 | 0.64922201 | -2.4064676 | 0.01610763 | 0.18205631 |
| RAB13 | 143.478227 | -1.558466856 | 0.41195095 | -3.783137 | 0.00015486 | 0.01206908 |
| VWF | 1223.01858 | -1.555429277 | 0.56584329 | -2.7488694 | 0.00598012 | 0.10663031 |
| MRC1 | 95.8055956 | -1.554911277 | 0.64292145 | -2.4185096 | 0.01558423 | 0.17910924 |
| SBNO2 | 238.704314 | -1.554170885 | 0.45625992 | -3.4063279 | 0.00065843 | 0.02802281 |
| ENO1 | 386.771348 | -1.551962522 | 0.33248098 | -4.6678234 | 3.04E-06 | 0.00062274 |
| ACTN1 | 406.339089 | -1.53957484 | 0.46203582 | -3.3321547 | 0.00086176 | 0.03348148 |
| AFF3 | 210.926945 | -1.538781625 | 0.97447746 | -1.5790839 | 0.11431682 | 0.45476644 |
| MTHFD1L | 405.714599 | -1.534074921 | 0.46439302 | -3.3033979 | 0.00095521 | 0.03585533 |
| HIST1H1E | 49.0153124 | -1.533802647 | 0.45041726 | -3.4052928 | 0.00066093 | 0.02802281 |
| NAV2 | 730.68557 | -1.528271812 | 0.54836844 | -2.7869434 | 0.00532078 | 0.10080241 |
| LGALS3 | 170.187784 | -1.525858124 | 0.47972661 | -3.1806827 | 0.00146928 | 0.04790102 |
| RASSF2 | 97.2579401 | -1.523233347 | 0.603977 | -2.5220055 | 0.01166879 | 0.15162005 |
| ACSM3 | 196.794682 | -1.521114247 | 0.51753868 | -2.9391315 | 0.00329133 | 0.07646344 |
| NEK6 | 107.019769 | -1.519595541 | 0.47440733 | -3.2031451 | 0.00135935 | 0.04507706 |
| HIST1H1C | 47.7994802 | -1.518284437 | 0.39344924 | -3.8589081 | 0.00011389 | 0.00960826 |
| NR2F1 | 183.150034 | -1.508607302 | 0.50866825 | -2.9657981 | 0.00301899 | 0.07244682 |
| INSR | 2902.70023 | -1.507864489 | 0.41448377 | -3.6379337 | 0.00027483 | 0.01678334 |
| FAM213A | 204.322113 | -1.502711461 | 0.5022272 | -2.9920949 | 0.0027707 | 0.07063129 |
| IL1RAP | 148.429247 | -1.502153454 | 0.5226011 | -2.8743787 | 0.00404823 | 0.08740551 |
| ARHGAP26-A | 67.7704188 | -1.496752796 | 0.65528826 | -2.2841136 | 0.02236486 | 0.2108594 |
| NHP2 | 71.567667 | -1.493194787 | 0.57589324 | -2.5928326 | 0.00951891 | 0.13774528 |
| P4HA1 | 396.700147 | -1.491988748 | 0.63894981 | -2.3350641 | 0.01954007 | 0.19632601 |
| SLC16A14 | 62.548911 | -1.490518626 | 0.615008 | -2.423576 | 0.01536854 | 0.17781377 |
| SPSB1 | 164.333763 | -1.490384372 | 0.64734015 | -2.3023203 | 0.02131712 | 0.20628732 |
| FTL | 2011.864 | -1.478953944 | 0.61781769 | -2.3938356 | 0.01667322 | 0.18464043 |
| PLCXD3 | 154.213224 | -1.478575718 | 1.10116683 | -1.3427354 | 0.17935766 | 0.54322277 |
| CPNE5 | 178.569661 | -1.473700882 | 0.56618489 | -2.6028615 | 0.00924493 | 0.13620125 |
| METRNL | 93.6002571 | -1.473226088 | 0.57583061 | -2.5584366 | 0.0105144 | 0.14485284 |
| COL8A1 | 355.286198 | -1.473186485 | 1.05674966 | -1.3940733 | 0.16329547 | 0.52357299 |
| IVNS1ABP | 366.270579 | -1.456264858 | 0.71550227 | -2.0353043 | 0.04182026 | 0.28963374 |
| NLGN1 | 150.442032 | -1.455900329 | 0.47694021 | -3.0525846 | 0.0022688 | 0.06286489 |
| XBP1 | 92.3163638 | -1.453961291 | 0.44913856 | -3.2372222 | 0.00120699 | 0.04185098 |
| CFH | 259.422703 | -1.445611122 | 0.63620061 | -2.2722567 | 0.02307101 | 0.21247788 |
| SDF2L1 | 63.1741628 | -1.441959814 | 0.55281605 | -2.6083899 | 0.00909693 | 0.13583969 |
| OAF | 82.2615427 | -1.438131712 | 0.53558628 | -2.6851541 | 0.00724964 | 0.12102126 |
| GRAMD1A | 256.524374 | -1.43301835 | 0.34387335 | -4.1672853 | 3.08E-05 | 0.00394125 |
| HIF1A-AS2 | 422.542105 | -1.432274938 | 0.91083735 | -1.5724816 | 0.11583891 | 0.45792745 |
| FBLIM1 | 88.574951 | -1.427224411 | 0.51159042 | -2.7897794 | 0.0052744 | 0.100373 |
| BCL3 | 117.043173 | -1.424284835 | 0.49055504 | -2.903415 | 0.00369117 | 0.08250421 |
| PKHD1L1 | 324.474603 | -1.416408465 | 1.19233488 | -1.1879284 | 0.23486162 | 0.59949266 |
| ARRDC3 | 910.124185 | -1.409608451 | 0.67256089 | -2.0958823 | 0.03609264 | 0.26728148 |
| MYO1B | 348.561089 | -1.407195303 | 0.56339887 | -2.4976892 | 0.01250057 | 0.15866671 |
| EFEMP1 | 115.893482 | -1.406265526 | 0.73841939 | -1.9044266 | 0.05685465 | 0.33347614 |
| SRM | 66.6895679 | -1.406041854 | 0.5530366 | -2.5424029 | 0.01100932 | 0.14749418 |
| KLHL29 | 70.1834512 | -1.405170792 | 0.55850927 | -2.515931 | 0.01187184 | 0.15189817 |
| S100A6 | 1663.94738 | -1.404400833 | 0.71106293 | -1.9750725 | 0.04825992 | 0.30798222 |
| STEAP1B | 305.457674 | -1.402845577 | 0.35362108 | -3.9670869 | 7.28E-05 | 0.00727039 |
| AC008050.1 | 93.9767144 | -1.400171926 | 0.55004169 | -2.5455742 | 0.01090982 | 0.14707585 |
| TFPI | 1415.08759 | -1.395076456 | 0.41593368 | -3.3540839 | 0.00079628 | 0.03194107 |
| GYPC | 341.023682 | -1.389448881 | 0.42494935 | -3.2696811 | 0.00107669 | 0.03881293 |
| RPL29 | 743.555554 | -1.374665889 | 0.54605621 | -2.5174439 | 0.01182098 | 0.15189817 |
| PLCB4 | 814.769682 | -1.373318785 | 0.5098865 | -2.6933813 | 0.00707313 | 0.1198332 |
| CD200 | 159.291272 | -1.372555985 | 0.57416508 | -2.390525 | 0.0168243 | 0.1855435 |
| RPGR | 535.857267 | -1.371110727 | 0.4244017 | -3.2306909 | 0.00123491 | 0.04245924 |
| CXCL14 | 77.0959592 | -1.369943527 | 1.00145976 | -1.3679466 | 0.17132878 | 0.5332763 |
| PXDN | 222.806516 | -1.369006203 | 0.56006813 | -2.4443565 | 0.01451108 | 0.17422157 |
| NFIL3 | 50.9641045 | -1.366261565 | 0.65933569 | -2.072179 | 0.03824875 | 0.27652057 |
| PDE3A | 745.11158 | -1.365220719 | 0.61190007 | -2.2311171 | 0.02567337 | 0.22302039 |
| GNAL | 85.988373 | -1.362161556 | 0.45824005 | -2.9725939 | 0.00295295 | 0.07188756 |
| HSPA5 | 365.316514 | -1.351113428 | 0.61305659 | -2.2038968 | 0.02753161 | 0.23083108 |
| SLC16A3 | 101.722115 | -1.35016118 | 0.42362531 | -3.1871589 | 0.00143678 | 0.0472175 |
| PLIN2 | 442.37024 | -1.349470577 | 0.61292723 | -2.2016816 | 0.02768781 | 0.23121895 |
| ERO1A | 354.474153 | -1.348040875 | 0.6214779 | -2.169089 | 0.03007593 | 0.24082201 |
| RPL15 | 704.093049 | -1.346541237 | 0.51400519 | -2.6197036 | 0.00880062 | 0.13311551 |
| ADAMTS9-A! | 157.843999 | -1.345568379 | 0.55788196 | -2.4119231 | 0.01586863 | 0.18072864 |
| MT1X | 194.397269 | -1.343457995 | 0.61677458 | -2.1781993 | 0.0293912 | 0.23789138 |
| MGLL | 230.031322 | -1.341984187 | 0.48347577 | -2.775701 | 0.00550828 | 0.10220927 |
| RETREG1 | 55.7748835 | -1.339635758 | 0.49384795 | -2.7126482 | 0.00667479 | 0.11547531 |
| SH3PXD2B | 110.814033 | -1.330860365 | 0.26826467 | -4.9609975 | 7.01E-07 | 0.00023083 |
| AF241726.2 | 113.305427 | -1.33028945 | 0.49308644 | -2.6978829 | 0.0069782 | 0.11847013 |
| ISOC2 | 43.2117194 | -1.323536424 | 0.47340784 | -2.7957636 | 0.00517773 | 0.09919793 |
| FAM43A | 115.848602 | -1.323518526 | 0.50052354 | -2.6442683 | 0.00818677 | 0.12833787 |
| RYR2 | 105.934276 | -1.320315736 | 1.09234143 | -1.2087024 | 0.22677719 | 0.59326015 |
| LDHA | 384.991555 | -1.319237761 | 0.44275606 | -2.9796041 | 0.00288621 | 0.07188756 |
| LDLRAD3 | 408.819856 | -1.318625479 | 0.37698813 | -3.4977904 | 0.00046913 | 0.02326599 |
| AC004448.2 | 40.5201207 | -1.317117083 | 0.43240984 | -3.0459924 | 0.00231914 | 0.06319319 |
| APRT | 119.508959 | -1.31575099 | 0.45538356 | -2.8893247 | 0.0038607 | 0.0851496 |
| SLC1A5 | 45.3012869 | -1.315590957 | 0.49010817 | -2.6842869 | 0.00726847 | 0.12102126 |
| RPS6 | 698.142419 | -1.313702566 | 0.58430987 | -2.2482978 | 0.02455721 | 0.21795189 |
| DLGAP1 | 305.52261 | -1.312472481 | 0.52118374 | -2.5182529 | 0.01179386 | 0.15189817 |
| EIF4A1 | 136.798694 | -1.308289773 | 0.52769127 | -2.4792712 | 0.01316512 | 0.16347517 |
| RPLP0 | 501.744343 | -1.304923723 | 0.5391396 | -2.4203819 | 0.01550422 | 0.17869154 |
| STX11 | 54.181729 | -1.302907424 | 0.51463594 | -2.531707 | 0.01135088 | 0.15054173 |
| SLC2A1 | 97.0239992 | -1.295753765 | 0.58518222 | -2.214274 | 0.02680994 | 0.22805167 |
| SOX7 | 142.51075 | -1.294279792 | 0.52916164 | -2.4459063 | 0.01444885 | 0.17387496 |
| FBLN5 | 238.627208 | -1.294224818 | 0.89362353 | -1.4482887 | 0.14753634 | 0.50136364 |
| PAICS | 51.314506 | -1.290389662 | 0.44313599 | -2.9119496 | 0.00359181 | 0.08141758 |
| RARRES1 | 52.8343316 | -1.289299899 | 0.31826713 | -4.0509992 | 5.10E-05 | 0.00556437 |
| MARCKS | 695.247144 | -1.287339097 | 0.51168169 | -2.5158983 | 0.01187294 | 0.15189817 |
| PROCR | 194.603667 | -1.286619594 | 0.46298258 | -2.7789806 | 0.00545298 | 0.10194119 |
| TSR1 | 40.5664335 | -1.282484803 | 0.47264094 | -2.7134442 | 0.00665878 | 0.11544233 |
| DDX21 | 217.401849 | -1.278254499 | 0.46000695 | -2.7787721 | 0.00545648 | 0.10194119 |
| MCTP2 | 193.358837 | -1.276511409 | 0.61184594 | -2.0863282 | 0.03694891 | 0.270683 |
| CD93 | 814.207435 | -1.275243494 | 0.45037869 | -2.8314917 | 0.00463314 | 0.09292407 |
| S100A11 | 698.126896 | -1.270862688 | 0.49585388 | -2.5629782 | 0.01037786 | 0.14395446 |
| H6PD | 85.7629277 | -1.267014035 | 0.31768012 | -3.9883328 | 6.65E-05 | 0.00689229 |
| PPP1R3B | 88.1147362 | -1.264351707 | 0.54149854 | -2.3349125 | 0.01954799 | 0.19632601 |
| TMTC2 | 1386.6744 | -1.260542115 | 0.42407945 | -2.9724197 | 0.00295462 | 0.07188756 |
| AL592295.4 | 45.7847901 | -1.259950118 | 0.33237016 | -3.790804 | 0.00015016 | 0.01181503 |
| TRABD2B | 106.251338 | -1.259676633 | 0.70935801 | -1.7757981 | 0.07576619 | 0.38160963 |

| | | | | | | |
|---|---|---|---|---|---|---|
| C1QBP | 75.276777 | -1.259203129 | 0.43333855 | -2.9058184 | 0.00366294 | 0.08246847 |
| ZMAT3 | 73.1500866 | -1.240974065 | 0.53014833 | -2.3408054 | 0.01924219 | 0.19499826 |
| C1RL | 137.093971 | -1.233361124 | 0.28072935 | -4.393417 | 1.12E-05 | 0.00179036 |
| RPS8 | 509.547433 | -1.233284845 | 0.51692598 | -2.3858055 | 0.01704176 | 0.18593697 |
| CYBA | 130.594058 | -1.224742076 | 0.5255043 | -2.3306033 | 0.01977429 | 0.19704872 |
| GALNT1 | 424.063921 | -1.223472883 | 0.20672879 | -5.9182512 | 3.25E-09 | 3.04E-06 |
| RPS19 | 829.442401 | -1.223309183 | 0.50435077 | -2.4255127 | 0.01528678 | 0.17738647 |
| TKT | 90.2607067 | -1.222254294 | 0.5678345 | -2.1524833 | 0.03135931 | 0.24759838 |
| ENPP2 | 1904.30382 | -1.21413848 | 0.87173854 | -1.3927782 | 0.16368686 | 0.52357299 |
| RPL27 | 205.455092 | -1.213872415 | 0.53966303 | -2.2493155 | 0.02449243 | 0.21761298 |
| PDK1 | 146.262506 | -1.212792206 | 0.53523648 | -2.2658997 | 0.02345752 | 0.21378373 |
| EPB41L3 | 467.83959 | -1.210552793 | 0.47305563 | -2.5590073 | 0.01049715 | 0.14485284 |
| GCLM | 93.1202317 | -1.206633248 | 0.42161314 | -2.8619441 | 0.00421051 | 0.08903002 |
| EIF2S2 | 229.034801 | -1.205744524 | 0.37288066 | -3.2335936 | 0.00122243 | 0.04220746 |
| IRAK1 | 45.4321909 | -1.203812482 | 0.49446812 | -2.4345604 | 0.01490989 | 0.17555058 |
| LAMB1 | 172.794531 | -1.200583247 | 0.38358842 | -3.1298735 | 0.00174882 | 0.05379909 |
| GPR146 | 52.2178123 | -1.2000052 | 0.5960387 | -2.0133008 | 0.04408299 | 0.29738759 |
| THSD7A | 2718.26802 | -1.198924788 | 0.5113609 | -2.3445766 | 0.0190487 | 0.19411645 |
| SNAI1 | 45.0757757 | -1.198235211 | 0.69305582 | -1.7289159 | 0.08382415 | 0.39616527 |
| ARHGAP26 | 4081.74788 | -1.197858696 | 0.51609609 | -2.3209994 | 0.02028688 | 0.20000907 |
| SLC25A6 | 188.344535 | -1.197598825 | 0.48938031 | -2.4471741 | 0.01439813 | 0.17370104 |
| DBI | 150.229512 | -1.195255061 | 0.49071882 | -2.4357229 | 0.01486206 | 0.17549246 |
| TNIP1 | 277.896056 | -1.194749995 | 0.60077102 | -1.9886945 | 0.04673494 | 0.30359341 |
| IL1R1 | 467.638066 | -1.193299812 | 0.55622332 | -2.145361 | 0.031924 | 0.24974576 |
| RHOU | 111.494189 | -1.191806667 | 0.51461431 | -2.3159221 | 0.02056252 | 0.20151271 |
| SV2B | 137.291325 | -1.186502144 | 0.61961786 | -1.9148934 | 0.0555061 | 0.32988018 |
| NCL | 241.543911 | -1.185837965 | 0.41069376 | -2.887402 | 0.00388438 | 0.08521676 |
| RPS26 | 65.0653586 | -1.183293751 | 0.50515556 | -2.3424344 | 0.0191584 | 0.1947132 |
| ASRGL1 | 491.817264 | -1.182177067 | 0.37691234 | -3.1364775 | 0.00170991 | 0.05338195 |
| PKM | 307.428195 | -1.177323657 | 0.26204678 | -4.4927995 | 7.03E-06 | 0.00125045 |
| AP2S1 | 149.574564 | -1.177200625 | 0.41659804 | -2.8257469 | 0.00471705 | 0.09368843 |
| C19orf53 | 130.743732 | -1.167916942 | 0.48017302 | -2.4322836 | 0.01500396 | 0.17589882 |
| PABPC1 | 422.59526 | -1.166947566 | 0.33995015 | -3.4327021 | 0.0005976 | 0.02694332 |
| RAPGEF5 | 3195.35857 | -1.165870128 | 0.40821887 | -2.8559928 | 0.00429025 | 0.08955895 |
| SLC7A1 | 86.2326319 | -1.162206046 | 0.5836472 | -1.9912818 | 0.04644991 | 0.30359341 |
| TNS1 | 856.171305 | -1.158716111 | 0.37378258 | -3.0999736 | 0.00193538 | 0.05676419 |
| C1QTNF1 | 95.0971696 | -1.156982675 | 0.69158641 | -1.6729401 | 0.09433908 | 0.41317143 |
| SMIM3 | 155.325242 | -1.156891432 | 0.57037928 | -2.0282845 | 0.04253122 | 0.291241 |
| EDN1 | 189.221354 | -1.156058842 | 0.60250149 | -1.9187651 | 0.05501407 | 0.32883866 |
| CNKSR3 | 465.040053 | -1.155337835 | 0.33678742 | -3.4304661 | 0.00060255 | 0.02694332 |
| APOLD1 | 457.162817 | -1.155208597 | 0.89028555 | -1.2975709 | 0.19443484 | 0.5592479 |
| TCAF2 | 391.356867 | -1.154402385 | 0.48457989 | -2.3822746 | 0.01720606 | 0.18698165 |
| FKBP5 | 839.03194 | -1.152066689 | 0.60557791 | -1.9024252 | 0.05711559 | 0.33407922 |
| PGS1 | 223.149668 | -1.147427044 | 0.30626026 | -3.746575 | 0.00017927 | 0.01298167 |
| IL18R1 | 209.863003 | -1.146475467 | 0.62831101 | -1.8246942 | 0.06804719 | 0.36134337 |
| ZDHHC9 | 66.9420124 | -1.146259635 | 0.3195817 | -3.5867499 | 0.00033483 | 0.01921832 |
| F11R | 136.363063 | -1.146052834 | 0.32027109 | -3.5783836 | 0.00034573 | 0.01951085 |
| MRPL36 | 48.8851496 | -1.145511781 | 0.50787873 | -2.2554829 | 0.02410304 | 0.21636008 |
| CD14 | 57.6657928 | -1.144465526 | 0.95940643 | -1.1928891 | 0.23291279 | 0.59910683 |
| SEMA4B | 42.2309678 | -1.142164871 | 0.45139601 | -2.5302946 | 0.01139668 | 0.15075694 |
| RPS5 | 203.166232 | -1.140196006 | 0.50499783 | -2.2578236 | 0.02395666 | 0.2161382 |
| SERPINE2 | 269.337982 | -1.139148987 | 0.43978803 | -2.5902228 | 0.00959138 | 0.13793724 |
| CMTM7 | 130.33951 | -1.137773026 | 0.44255566 | -2.5709151 | 0.01014302 | 0.14236887 |
| IL4R | 845.792644 | -1.136947051 | 0.39942661 | -2.8464479 | 0.004421 | 0.091042 |
| BGN | 81.4203561 | -1.134606401 | 1.02182244 | -1.1103753 | 0.26683733 | 0.63217426 |
| EEF1B2 | 134.213598 | -1.134093696 | 0.47355533 | -2.3948494 | 0.01662719 | 0.18464043 |
| CHST12 | 161.457025 | -1.13352206 | 0.66373635 | -1.7077896 | 0.08767538 | 0.40026508 |
| EIF5A | 106.62398 | -1.132691319 | 0.52742574 | -2.1475845 | 0.03174678 | 0.24859704 |
| BZW2 | 63.0874101 | -1.132281125 | 0.31437942 | -3.6016389 | 0.00031622 | 0.01861588 |
| CTPS1 | 43.4996043 | -1.131903255 | 0.48274364 | -2.3447295 | 0.01904089 | 0.19411645 |
| SLC45A4 | 166.724891 | -1.129644552 | 0.52714464 | -2.1429499 | 0.03211712 | 0.25029944 |
| FOSL2 | 505.923953 | -1.129281857 | 0.53117349 | -2.1260132 | 0.03350216 | 0.25525899 |
| NDUFB4 | 110.688789 | -1.125116132 | 0.32934517 | -3.4162217 | 0.00063497 | 0.02763334 |
| KLHL3 | 144.577561 | -1.124868601 | 0.42415613 | -2.6520154 | 0.00800129 | 0.12664322 |
| AL136531.2 | 442.833091 | -1.123394326 | 0.39507198 | -2.8435181 | 0.00446185 | 0.09127824 |
| TIMM8B | 63.2356492 | -1.121275484 | 0.46927222 | -2.3893924 | 0.01687627 | 0.18561624 |
| CD53 | 117.171064 | -1.118314549 | 0.67513473 | -1.6564317 | 0.09763443 | 0.42093918 |
| SH3BGRL3 | 327.132377 | -1.117921458 | 0.60049866 | -1.8616552 | 0.06265171 | 0.35114993 |
| PDZRN3 | 148.596394 | -1.114025661 | 0.70988273 | -1.5693094 | 0.11657585 | 0.4592091 |
| SLC39A1 | 96.8353502 | -1.113682223 | 0.43913705 | -2.5360698 | 0.01121044 | 0.14940555 |
| GRPEL1 | 51.3161609 | -1.110874035 | 0.4010015 | -2.7702491 | 0.00560134 | 0.10277085 |
| HIF1A | 1403.97161 | -1.109296343 | 0.58150111 | -1.9076427 | 0.05643741 | 0.3322499 |
| COL18A1 | 292.592272 | -1.108643471 | 0.43970777 | -2.5213188 | 0.01169159 | 0.15162005 |
| SH3BP5 | 1839.70889 | -1.107711986 | 0.43571464 | -2.5422877 | 0.01101295 | 0.14749418 |
| IL32 | 215.380197 | -1.107619285 | 0.60203994 | -1.8397771 | 0.06580097 | 0.35579065 |
| IFITM3 | 2508.56448 | -1.106501759 | 0.49181579 | -2.2498297 | 0.02445976 | 0.21755892 |
| SULF2 | 525.478785 | -1.104340754 | 0.43088949 | -2.5629327 | 0.01037922 | 0.14395446 |
| GADD45GIP1 | 164.183288 | -1.103701152 | 0.4842261 | -2.2793095 | 0.02264867 | 0.21131824 |
| MCUB | 87.9922875 | -1.102599671 | 0.40425089 | -2.7275133 | 0.00638137 | 0.11278347 |
| NOLC1 | 41.7989372 | -1.102164211 | 0.43623709 | -2.5265257 | 0.0115197 | 0.15149764 |
| AL138828.1 | 157.426576 | -1.101702923 | 0.63668295 | -1.7303792 | 0.08356255 | 0.3954843 |
| AKAP12 | 972.166167 | -1.09934114 | 0.76072664 | -1.4451198 | 0.14842423 | 0.50188244 |
| C19orf24 | 62.440175 | -1.096101852 | 0.41391799 | -2.6481136 | 0.00809423 | 0.12762062 |
| RPL6 | 341.08485 | -1.095956097 | 0.46436091 | -2.3601386 | 0.01826811 | 0.19288766 |
| SH3TC1 | 131.514723 | -1.090504118 | 0.27619121 | -3.9483665 | 7.87E-05 | 0.00766536 |
| ABLIM1 | 607.72115 | -1.090364345 | 0.34477668 | -3.1625235 | 0.00156408 | 0.05054839 |
| RPL36AL | 258.959952 | -1.086403842 | 0.39425565 | -2.7555822 | 0.00585878 | 0.10526109 |
| RPL4 | 309.252862 | -1.084134238 | 0.47056681 | -2.3038901 | 0.02122881 | 0.20628732 |
| SLC4A7 | 317.305866 | -1.083681579 | 0.46199094 | -2.3456771 | 0.01899255 | 0.19411645 |
| ZNF22 | 93.7044932 | -1.080844682 | 0.37712985 | -2.8659749 | 0.00415727 | 0.08875226 |
| ANO2 | 625.098504 | -1.08051012 | 0.67196997 | -1.6079738 | 0.10784091 | 0.44278082 |
| YBX1 | 283.615245 | -1.079455122 | 0.3715402 | -2.9053522 | 0.0036684 | 0.08246847 |
| CTSL | 183.739264 | -1.077059992 | 0.44977869 | -2.3946443 | 0.01663649 | 0.18464043 |
| PARK7 | 200.02518 | -1.075509111 | 0.40437702 | -2.6596692 | 0.00782174 | 0.12501039 |
| RPL19 | 520.468171 | -1.074741744 | 0.54933528 | -1.9564404 | 0.0504133 | 0.31490993 |
| SIPA1L2 | 636.720115 | -1.072899243 | 0.40595907 | -2.6428754 | 0.00822053 | 0.12859845 |
| CAVIN3 | 105.783601 | -1.072140673 | 0.62232602 | -1.7227958 | 0.08492545 | 0.39686932 |
| UCK2 | 133.763691 | -1.072021242 | 0.46775936 | -2.2918221 | 0.02191591 | 0.20902438 |
| S100A16 | 250.029191 | -1.070659251 | 0.43656529 | -2.4524608 | 0.01418828 | 0.17200405 |
| ADAMTS1 | 645.579713 | -1.069490895 | 0.47103888 | -2.2704939 | 0.02317763 | 0.21286486 |
| CCDC3 | 386.475478 | -1.068541728 | 0.44061803 | -2.4250976 | 0.01530427 | 0.17738647 |
| YARS | 125.863299 | -1.066172797 | 0.42841728 | -2.4886316 | 0.01282358 | 0.16143895 |
| SINHCAF | 164.890039 | -1.065213404 | 0.27345671 | -3.8953639 | 9.81E-05 | 0.00867938 |
| IFITM2 | 1388.06726 | -1.064644049 | 0.56926681 | -1.8702022 | 0.06145574 | 0.34778328 |
| RCC1 | 75.5105013 | -1.062404531 | 0.48139219 | -2.2069418 | 0.02731813 | 0.23022064 |
| NPC2 | 140.760942 | -1.062025829 | 0.40143495 | -2.6455739 | 0.00815525 | 0.12821467 |
| ABLIM3 | 137.752486 | -1.061970375 | 0.34582969 | -3.0707901 | 0.00213493 | 0.06003487 |
| PPIB | 258.957925 | -1.061958454 | 0.4791246 | -2.2164557 | 0.02666031 | 0.22752964 |
| MRPL40 | 50.8965781 | -1.058881061 | 0.50130644 | -2.1122431 | 0.0346656 | 0.26000791 |
| FAM129A | 102.948347 | -1.058330349 | 0.42934119 | -2.4650101 | 0.01370094 | 0.16834054 |
| MFAP3 | 239.768545 | -1.057827312 | 0.19386617 | -5.4564824 | 4.86E-08 | 2.48E-05 |
| TXNRD1 | 783.829832 | -1.055732456 | 0.39416416 | -2.678408 | 0.0073973 | 0.12155049 |
| TPI1 | 224.457701 | -1.055395078 | 0.40619504 | -2.5982471 | 0.0093701 | 0.13676431 |

| | | | | | | |
|---|---|---|---|---|---|---|
| IFT43 | 269.525364 | -1.055108498 | 0.27096379 | -3.8939096 | 9.86E-05 | 0.00867938 |
| VEGFB | 54.0313355 | -1.05466994 | 0.33793678 | -3.120909 | 0.00180294 | 0.05451467 |
| CSRP2 | 52.2750396 | -1.053109724 | 0.4641522 | -2.2688888 | 0.02327509 | 0.2129727 |
| RPL14 | 540.764389 | -1.050602061 | 0.55478436 | -1.8937125 | 0.05826318 | 0.33857513 |
| MPZL2 | 289.379527 | -1.049540322 | 0.48860357 | -2.1480406 | 0.03171053 | 0.24855101 |
| SPARCL1 | 985.254881 | -1.04823193 | 0.63237636 | -1.6576077 | 0.09739667 | 0.42080093 |
| RFK | 151.952187 | -1.047503572 | 0.47113399 | -2.2233666 | 0.02619109 | 0.22519168 |
| SULF1 | 364.96234 | -1.046461447 | 0.97817393 | -1.0698112 | 0.28470429 | 0.64764125 |
| RPL32 | 571.829328 | -1.045883252 | 0.51776044 | -2.0200138 | 0.04338195 | 0.29533655 |
| DPM2 | 41.3746151 | -1.042778931 | 0.42407976 | -2.4589217 | 0.0139355 | 0.16969375 |
| AC074286.1 | 228.076338 | -1.042308484 | 1.18977101 | -0.8760581 | 0.38099847 | 0.71543195 |
| BAG3 | 110.957199 | -1.042298143 | 0.36985175 | -2.8181512 | 0.00483011 | 0.09480113 |
| PLOD2 | 420.105145 | -1.039343577 | 0.57989858 | -1.7922851 | 0.07308731 | 0.37402969 |
| PHF10 | 185.715255 | -1.039261595 | 0.41169778 | -2.5243313 | 0.01159186 | 0.15149764 |
| PUS7 | 88.5035169 | -1.038935732 | 0.42804748 | -2.4271507 | 0.01521793 | 0.177259 |
| GNL3 | 69.8068649 | -1.037439456 | 0.39679413 | -2.6145534 | 0.00893442 | 0.13439403 |
| FAM107A | 234.948823 | -1.034880427 | 0.53198959 | -1.945302 | 0.05173864 | 0.31875077 |
| UBC | 1252.94254 | -1.034729473 | 0.45541694 | -2.2720487 | 0.02308357 | 0.21247788 |
| ZNRF3 | 117.887088 | -1.030176986 | 0.29117502 | -3.5379992 | 0.00040317 | 0.02142306 |
| SOX4 | 177.514368 | -1.02956121 | 0.45743533 | -2.2507252 | 0.02440295 | 0.21755892 |
| RPL28 | 515.471739 | -1.025952873 | 0.47702738 | -2.1507211 | 0.03149822 | 0.24759838 |
| TLE3 | 100.953556 | -1.023937294 | 0.44370467 | -2.3077 | 0.02101583 | 0.20497318 |
| LIMK2 | 666.48807 | -1.023817065 | 0.43528019 | -2.3520874 | 0.01866839 | 0.19394842 |
| MIR4435-2H( | 421.49763 | -1.022649729 | 0.37352761 | -2.7378157 | 0.00618487 | 0.11002352 |
| LRRC32 | 757.703761 | -1.022204726 | 0.44662078 | -2.2887532 | 0.0220937 | 0.2102241 |
| CD320 | 75.1424368 | -1.021078467 | 0.54231207 | -1.8828245 | 0.05972415 | 0.34296331 |
| EFNA5 | 185.131029 | -1.020638613 | 0.72513683 | -1.4075118 | 0.15927569 | 0.51679341 |
| INSIG1 | 69.8451997 | -1.019896427 | 0.5819087 | -1.7526743 | 0.07965793 | 0.38738328 |
| AC007681.1 | 252.690079 | -1.019130759 | 0.54997632 | -1.8530448 | 0.06387594 | 0.3533385 |
| EPHB1 | 83.5150044 | -1.017841569 | 0.46327619 | -2.1970513 | 0.02801678 | 0.23214303 |
| APEX1 | 48.5769863 | -1.01780174 | 0.43312484 | -2.3499039 | 0.01877826 | 0.19394842 |
| UFC1 | 76.1570252 | -1.013301138 | 0.53050976 | -1.9100518 | 0.05612654 | 0.33089589 |
| LAPTM5 | 49.5552613 | -1.012130221 | 0.48235224 | -2.0983218 | 0.03587673 | 0.26597323 |
| FLT1 | 7180.33784 | -1.010660383 | 0.58385926 | -1.731 | 0.08345177 | 0.3954379 |
| FUCA2 | 41.7438146 | -1.010405177 | 0.32814681 | -3.0791254 | 0.00207609 | 0.05884254 |
| SERPINB1 | 131.956905 | -1.009432869 | 0.35107248 | -2.8752834 | 0.00403665 | 0.087386 |
| S100A13 | 148.500977 | -1.007825021 | 0.44146371 | -2.282917 | 0.02243526 | 0.2108594 |
| PXN | 797.082779 | -1.007375371 | 0.26716886 | -3.7705568 | 0.00016288 | 0.01230754 |
| SYNPO | 446.878724 | -1.007220402 | 0.32643359 | -3.0855293 | 0.0020319 | 0.05828136 |
| OOEP | 334.54519 | -1.006199364 | 0.53352136 | -1.8859589 | 0.05930049 | 0.34172953 |
| CENPB | 52.4759762 | -1.005298242 | 0.55039674 | -1.8264975 | 0.06777536 | 0.36107148 |
| PMP22 | 468.361553 | -1.004971197 | 0.35269937 | -2.8493705 | 0.00438058 | 0.09052099 |
| MCAM | 388.074353 | -1.00348394 | 0.51476404 | -1.9494057 | 0.051247 | 0.31793341 |
| RPS18 | 318.232028 | -1.002895122 | 0.53306329 | -1.8813809 | 0.05992012 | 0.34384735 |
| RPS15 | 638.607307 | -1.00288619 | 0.52497722 | -1.9103423 | 0.05608915 | 0.33089589 |
| TWISTNB | 43.5633374 | -1.002868902 | 0.44213549 | -2.2682389 | 0.02331465 | 0.2129727 |
| DIXDC1 | 114.921299 | -1.002831227 | 0.4305931 | -2.3289533 | 0.01986154 | 0.19704872 |
| EID1 | 261.06319 | -1.002401599 | 0.52387981 | -1.913419 | 0.05569443 | 0.32988018 |
| MT1E | 229.896686 | -1.001835428 | 0.67590596 | -1.4822113 | 0.13828409 | 0.48991657 |
| NEURL1B | 353.862004 | -1.001299082 | 0.42675353 | -2.346317 | 0.01895997 | 0.19411645 |
| WFS1 | 64.9915426 | -1.000538946 | 0.28959543 | -3.4549542 | 0.00055039 | 0.02618495 |
| MANF | 74.5930604 | -0.999037843 | 0.47029005 | -2.1243015 | 0.03364494 | 0.25585395 |
| TEX264 | 75.8330116 | -0.996592009 | 0.38767379 | -2.5706974 | 0.0101494 | 0.14236887 |
| FGFR1 | 456.878323 | -0.994970103 | 0.65852193 | -1.5109142 | 0.13081032 | 0.47936448 |
| GGACT | 86.2159056 | -0.994089547 | 0.35304764 | -2.8157377 | 0.00486654 | 0.09504268 |
| MPG | 83.7364354 | -0.992122665 | 0.31797524 | -3.1201255 | 0.00180774 | 0.05451467 |
| CD63 | 309.317018 | -0.991108806 | 0.46335182 | -2.1389984 | 0.0324358 | 0.25182364 |
| LEPROTL1 | 201.696843 | -0.990863164 | 0.50174743 | -1.9748246 | 0.04828806 | 0.30798222 |
| MRPL57 | 99.3337395 | -0.989855037 | 0.39694285 | -2.4936966 | 0.01264206 | 0.15947378 |
| TSPAN1 | 51.0389101 | -0.98966335 | 0.33967448 | -2.9135641 | 0.00357328 | 0.08141758 |
| FOXC1 | 69.78504 | -0.988751335 | 0.29840901 | -3.3134098 | 0.00092166 | 0.0349812 |
| SSR2 | 151.940214 | -0.988299306 | 0.44061442 | -2.2430026 | 0.02489664 | 0.21937533 |
| COX7C | 176.853007 | -0.988183615 | 0.43350762 | -2.2795069 | 0.02263695 | 0.21131824 |
| GPI | 120.361147 | -0.988038374 | 0.44993515 | -2.1959573 | 0.028095 | 0.23222363 |
| CPVL | 58.0217057 | -0.98793288 | 0.74503065 | -1.3260299 | 0.18482981 | 0.5500631 |
| STAB1 | 170.532138 | -0.987780171 | 0.52168258 | -1.8934505 | 0.05829798 | 0.33857513 |
| RPL5 | 271.664485 | -0.986910819 | 0.31158486 | -3.1673902 | 0.00153814 | 0.04994676 |
| BACE2 | 403.559849 | -0.986327759 | 0.23461349 | -4.204054 | 2.62E-05 | 0.00346031 |
| PLD1 | 248.454185 | -0.98538065 | 0.34555399 | -2.8515968 | 0.00435002 | 0.0903458 |
| ALDH18A1 | 45.4204154 | -0.985001949 | 0.2873984 | -3.4273049 | 0.0006096 | 0.02711082 |
| CRYAB | 614.454502 | -0.983754349 | 1.13260443 | -0.8685772 | 0.38507844 | 0.71699718 |
| NDRG1 | 734.566898 | -0.983204659 | 0.50697888 | -1.9393405 | 0.0524599 | 0.32059696 |
| AC243964.2 | 94.8048776 | -0.982926066 | 0.24100093 | -4.0785156 | 4.53E-05 | 0.00522377 |
| EGLN1 | 377.034336 | -0.981219236 | 0.44624125 | -2.1988537 | 0.02788832 | 0.23190345 |
| UQCRQ | 99.4395359 | -0.980424339 | 0.45027752 | -2.1773779 | 0.02945238 | 0.23815102 |
| RPL23 | 265.115297 | -0.97938495 | 0.49794781 | -1.9668426 | 0.04920136 | 0.31155957 |
| VIM | 1530.06691 | -0.976255229 | 0.60035021 | -1.6261429 | 0.10391926 | 0.43532543 |
| NID1 | 309.236901 | -0.975875701 | 0.41899878 | -2.3290657 | 0.01985558 | 0.19704872 |
| RPS2 | 389.168688 | -0.975495999 | 0.57445 | -1.6981391 | 0.08948151 | 0.40581146 |
| TPT1 | 1240.24161 | -0.974969823 | 0.5163634 | -1.8881467 | 0.05900626 | 0.34075389 |
| EEF2 | 417.893284 | -0.973259195 | 0.42428465 | -2.2938826 | 0.02179724 | 0.20902438 |
| EIF3I | 119.451109 | -0.973000685 | 0.43733674 | -2.2248318 | 0.02609253 | 0.22518506 |
| AC105942.1 | 54.337145 | -0.971864395 | 0.39519497 | -2.4592023 | 0.01392461 | 0.16969375 |
| PLPP5 | 48.1475335 | -0.970828119 | 0.26209049 | -3.7041715 | 0.00021208 | 0.01434275 |
| AC011476.3 | 73.4669015 | -0.970726666 | 0.47705947 | -2.0348127 | 0.04186972 | 0.28963374 |
| KRT10 | 104.754885 | -0.970346069 | 0.54299177 | -1.7870364 | 0.0739316 | 0.3764669 |
| TNFRSF10D | 206.175994 | -0.967125158 | 0.3960674 | -2.4418197 | 0.01461344 | 0.17457199 |
| SNRPB | 107.12276 | -0.966766766 | 0.42174491 | -2.2923022 | 0.02188821 | 0.20902438 |
| FAM110B | 59.9372568 | -0.96642045 | 0.63114912 | -1.5312078 | 0.12571805 | 0.47103973 |
| GSTO1 | 228.622543 | -0.966258091 | 0.3122572 | -3.0944301 | 0.00197191 | 0.05722046 |
| BHLHE40 | 470.989684 | -0.965866965 | 0.64618916 | -1.4947124 | 0.13498951 | 0.484753 |
| C9orf84 | 172.585924 | -0.964060234 | 0.63707156 | -1.5132683 | 0.13021151 | 0.47888574 |
| SH3GL1 | 263.192914 | -0.96307008 | 0.31647255 | -3.0431394 | 0.00234124 | 0.0632289 |
| BNIP3 | 85.8041833 | -0.962320498 | 0.53874611 | -1.7862226 | 0.07406322 | 0.37666831 |
| RSL1D1 | 58.2852711 | -0.956595051 | 0.3515951 | -2.7207292 | 0.00651381 | 0.1136514 |
| JAK3 | 74.9082777 | -0.955792774 | 0.6304512 | -1.5160456 | 0.12950782 | 0.4777902 |
| EDNRB | 151.515871 | -0.954905972 | 0.48461242 | -1.970453 | 0.04878648 | 0.30995325 |
| NEAT1 | 11801.758 | -0.954854135 | 0.42619227 | -2.2404304 | 0.02506299 | 0.22005414 |
| CISD1 | 112.348942 | -0.954121472 | 0.34629899 | -2.7551957 | 0.0058657 | 0.10526109 |
| GTF3C6 | 72.9530663 | -0.953296566 | 0.40423214 | -2.3582899 | 0.01835935 | 0.19310354 |
| RPS9 | 667.789753 | -0.950752651 | 0.51874536 | -1.8327926 | 0.06683342 | 0.35877642 |
| PRNP | 88.5164585 | -0.950372714 | 0.42723087 | -2.2244945 | 0.02611519 | 0.22518506 |
| H2AFJ | 165.226393 | -0.950187646 | 0.59510312 | -1.5966773 | 0.11033766 | 0.44631216 |
| GRAMD4 | 49.6868308 | -0.946941384 | 0.36374432 | -2.6033159 | 0.00923268 | 0.13620125 |
| WIPI1 | 137.701539 | -0.945482999 | 0.35470522 | -2.6655458 | 0.00768635 | 0.12405 |
| ATP6V1F | 108.362428 | -0.943836928 | 0.43520019 | -2.168742 | 0.03010228 | 0.24082201 |
| ICAM2 | 423.389059 | -0.942698649 | 0.44472566 | -2.1197307 | 0.03402876 | 0.2573543 |
| RPL10A | 286.493859 | -0.941977842 | 0.55690516 | -1.6914511 | 0.09075068 | 0.40735756 |
| G6PC3 | 40.2333144 | -0.941742909 | 0.44253482 | -2.1280651 | 0.03333169 | 0.25490607 |
| ZNF90 | 240.399255 | -0.941443659 | 0.45864923 | -2.0526442 | 0.0401071 | 0.28442544 |
| NUP50-DT | 96.9873597 | -0.937460727 | 0.24467111 | -3.8315137 | 0.00012736 | 0.01043639 |
| PNKP | 69.2550881 | -0.937116462 | 0.46961255 | -1.9955098 | 0.0459873 | 0.30250329 |
| CD99 | 324.489273 | -0.936240064 | 0.40355876 | -2.3199597 | 0.02034306 | 0.20014099 |

| | | | | | | |
|---|---|---|---|---|---|---|
| TGIF1 | 44.5388251 | -0.935471696 | 0.29161923 | -3.2078533 | 0.0013373 | 0.04503334 |
| NUCB1 | 232.4808 | -0.935155898 | 0.32882308 | -2.8439485 | 0.00445582 | 0.09127824 |
| SPECC1 | 195.470619 | -0.934729033 | 1.03259432 | -0.9052239 | 0.3653468 | 0.70668224 |
| TAGLN2 | 548.023441 | -0.933840532 | 0.48187977 | -1.9379119 | 0.05263397 | 0.32094174 |
| BAX | 90.5313542 | -0.933721922 | 0.43100983 | -2.1663587 | 0.03028378 | 0.241768 |
| SLC46A3 | 57.3906982 | -0.931099477 | 0.41289818 | -2.2550341 | 0.02413119 | 0.21636008 |
| RPL23A | 357.478368 | -0.929561793 | 0.49658759 | -1.871899 | 0.06122058 | 0.34778328 |
| VEGFC | 1255.64297 | -0.925334659 | 0.36229035 | -2.5541245 | 0.01064552 | 0.14515134 |
| IER5L | 94.3823567 | -0.925267313 | 0.39396037 | -2.3486304 | 0.0188426 | 0.19394842 |
| NDUFS2 | 131.24935 | -0.925029788 | 0.43302799 | -2.1361893 | 0.03266398 | 0.25224169 |
| RASSF4 | 42.6720991 | -0.924111721 | 0.38318365 | -2.4116679 | 0.01587974 | 0.18072864 |
| ARPC5L | 127.250941 | -0.923966121 | 0.40781362 | -2.2656578 | 0.02347234 | 0.21378373 |
| ATP5F1E | 347.71966 | -0.923446654 | 0.47594023 | -1.9402576 | 0.05234839 | 0.32059696 |
| NDUFAF8 | 46.7446976 | -0.92256811 | 0.3983214 | -2.3161399 | 0.02055063 | 0.20151271 |
| LGALS1 | 255.376636 | -0.920161667 | 0.56820991 | -1.6194045 | 0.10536026 | 0.43797647 |
| KCNT2 | 77.5990179 | -0.920091473 | 0.53253066 | -1.7277718 | 0.08402914 | 0.39616527 |
| AC010894.3 | 314.466849 | -0.918602531 | 0.38545863 | -2.3831417 | 0.01716558 | 0.18698165 |
| SIRPA | 41.6443348 | -0.918108894 | 0.36171784 | -2.5381908 | 0.01114272 | 0.14874535 |
| TPK1 | 342.898291 | -0.918042223 | 0.29328822 | -3.1301707 | 0.00174705 | 0.05379909 |
| PGRMC1 | 105.694493 | -0.91630203 | 0.39558941 | -2.3162956 | 0.02054213 | 0.20151271 |
| RPL31 | 174.524561 | -0.914997884 | 0.3924761 | -2.3313468 | 0.01973508 | 0.19704872 |
| NPR3 | 91.3825635 | -0.914381968 | 0.61139599 | -1.4955642 | 0.13476725 | 0.48474744 |
| BIN3 | 180.821692 | -0.913646535 | 0.23483897 | -3.8905235 | 0.00010003 | 0.00870777 |
| PTGR1 | 134.730478 | -0.912902528 | 0.27433868 | -3.3276479 | 0.00087582 | 0.03373359 |
| CPE | 148.988479 | -0.9116339 | 0.91298764 | -0.9985172 | 0.31802861 | 0.67298374 |
| H1FX | 196.33975 | -0.911207973 | 0.62052886 | -1.4684377 | 0.14198537 | 0.49378081 |
| HCLS1 | 90.9671555 | -0.910072023 | 0.37261466 | -2.4423945 | 0.01459019 | 0.17454903 |
| COTL1 | 204.648579 | -0.910010594 | 0.36619989 | -2.4850105 | 0.01295476 | 0.16283994 |
| AL136454.1 | 143.082758 | -0.909963321 | 0.48626358 | -1.8713376 | 0.0612983 | 0.34778328 |
| PPA1 | 225.26773 | -0.909447907 | 0.45142028 | -2.0146368 | 0.04394272 | 0.29693087 |
| PDLIM1 | 567.656673 | -0.908961957 | 0.61150614 | -1.4864315 | 0.13716504 | 0.48888307 |
| C16orf45 | 366.312228 | -0.907220756 | 0.36348741 | -2.4958794 | 0.01256453 | 0.15891121 |
| GGT5 | 121.083032 | -0.90682549 | 0.54730879 | -1.6568809 | 0.09754355 | 0.42092574 |
| NANS | 52.7517287 | -0.906308078 | 0.31154402 | -2.9090851 | 0.00362488 | 0.08194035 |
| RPL41 | 800.407015 | -0.906031026 | 0.50751222 | -1.7852398 | 0.07422243 | 0.37724356 |
| CBX6 | 74.1790438 | -0.904842946 | 0.48733339 | -1.8567227 | 0.06335062 | 0.3533385 |
| AK4 | 117.479154 | -0.904797623 | 0.49495093 | -1.8280552 | 0.06754127 | 0.3605285 |
| AC018816.1 | 279.910331 | -0.904427242 | 0.42179181 | -2.1442504 | 0.03201283 | 0.24982585 |
| HNRNPAB | 116.357171 | -0.904013007 | 0.42257975 | -2.139272 | 0.03241365 | 0.25182364 |
| SGIP1 | 115.266807 | -0.903561608 | 0.68956335 | -1.3103388 | 0.19008125 | 0.55552647 |
| NEU1 | 42.5595169 | -0.903319167 | 0.30010222 | -3.0100383 | 0.00261215 | 0.06807389 |
| AGPAT5 | 122.410294 | -0.903176122 | 0.3022202 | -2.9884704 | 0.00280378 | 0.07081266 |
| LAPTM4B | 155.792242 | -0.90279553 | 0.46189587 | -1.9545434 | 0.05063698 | 0.3155845 |
| RPS4X | 412.253237 | -0.897840013 | 0.49076431 | -1.8294729 | 0.06732879 | 0.35986382 |
| RPS4Y1 | 146.633925 | -0.897439226 | 2.29423218 | -0.3911719 | 0.69567016 | 0.89184849 |
| CREM | 328.060365 | -0.897240757 | 0.48733055 | -1.8411338 | 0.06560195 | 0.35579065 |
| RACK1 | 335.320712 | -0.896725339 | 0.46477846 | -1.9293608 | 0.05368609 | 0.32491861 |
| GLIS3 | 804.437142 | -0.896361033 | 0.31302356 | -2.8635578 | 0.00418912 | 0.08880727 |
| STOML2 | 50.9159543 | -0.895328805 | 0.46810143 | -1.9126812 | 0.05578888 | 0.32988018 |
| AC068888.1 | 84.4629244 | -0.895322146 | 0.30154834 | -2.9690833 | 0.0029869 | 0.07188756 |
| C9orf16 | 106.457229 | -0.894982007 | 0.48975013 | -1.8274258 | 0.06763578 | 0.36079762 |
| ADGRG1 | 259.023199 | -0.894734488 | 0.35704219 | -2.5059629 | 0.01221184 | 0.15551732 |
| SQSTM1 | 469.965054 | -0.894558572 | 0.35452223 | -2.5232792 | 0.01162661 | 0.15149764 |
| PPRC1 | 47.1912055 | -0.894518451 | 0.5059847 | -1.7678765 | 0.07708155 | 0.38356763 |
| SEPHS2 | 71.3275022 | -0.892587427 | 0.39352945 | -2.2681592 | 0.02331951 | 0.2129727 |
| RPS13 | 307.0742 | -0.892116303 | 0.55082096 | -1.6196121 | 0.10531562 | 0.43797647 |
| IGFBP2 | 128.919575 | -0.890346662 | 0.74745026 | -1.1911785 | 0.23358353 | 0.59918934 |
| CDC42EP5 | 130.792411 | -0.889744019 | 0.3966737 | -2.2430124 | 0.02489602 | 0.21937533 |
| FURIN | 124.662947 | -0.889614404 | 0.5482311 | -1.6226996 | 0.10465363 | 0.43670609 |
| FAM13A | 359.892046 | -0.888623753 | 0.61433338 | -1.4464846 | 0.14804134 | 0.50152657 |
| PYGL | 82.8778283 | -0.888484735 | 0.620997 | -1.4307392 | 0.15250498 | 0.50874369 |
| ICAM1 | 300.74427 | -0.888391748 | 0.48888002 | -1.8171979 | 0.06918682 | 0.36386894 |
| FKBP4 | 54.2273389 | -0.888386055 | 0.36899999 | -2.4075503 | 0.01605995 | 0.1817684 |
| SLC25A5 | 116.358655 | -0.886980257 | 0.53000002 | -1.6735476 | 0.09421955 | 0.4129612 |
| SNRPD2 | 109.765026 | -0.885488861 | 0.45592493 | -1.9421813 | 0.05211516 | 0.32016393 |
| MT1M | 75.450367 | -0.884746261 | 0.63114294 | -1.401816 | 0.1609702 | 0.51944206 |
| TXN | 134.72473 | -0.88467423 | 0.39128051 | -2.2609719 | 0.023761 | 0.21508435 |
| C14orf119 | 47.5435731 | -0.883457378 | 0.29700366 | -2.9745673 | 0.00293402 | 0.07188756 |
| TNIP2 | 91.6965389 | -0.88311712 | 0.40460356 | -2.1826726 | 0.02905993 | 0.23661432 |
| TAF4B | 47.697237 | -0.88215124 | 0.36473529 | -2.4186067 | 0.01558007 | 0.17910924 |
| FBL | 41.6506941 | -0.880631593 | 0.36850828 | -2.38972 | 0.01686122 | 0.18561624 |
| NOP56 | 72.9232823 | -0.877722217 | 0.42921476 | -2.0449488 | 0.0408599 | 0.28670444 |
| RPS3 | 596.641595 | -0.876713741 | 0.52162399 | -1.6807389 | 0.09281364 | 0.41187311 |
| SLC20A1 | 165.976868 | -0.87554599 | 0.39886533 | -2.1950918 | 0.02815702 | 0.23250138 |
| SFXN1 | 79.3144585 | -0.8731698 | 0.3688622 | -2.3671979 | 0.01792335 | 0.19122132 |
| ROBO1 | 825.858802 | -0.872430938 | 0.63844083 | -1.3665024 | 0.17178132 | 0.53377758 |
| TOMM7 | 153.949504 | -0.872195991 | 0.50689536 | -1.7206628 | 0.08531203 | 0.39686932 |
| PMM2 | 165.836452 | -0.871307768 | 0.33658563 | -2.588666 | 0.00963485 | 0.13807701 |
| MRPS12 | 47.1091112 | -0.871191996 | 0.48111414 | -1.8107803 | 0.07017487 | 0.36582906 |
| POLR1A | 66.758874 | -0.87001221 | 0.49054532 | -1.7735613 | 0.07613574 | 0.38261461 |
| PRDX1 | 467.606121 | -0.869724231 | 0.51825179 | -1.6781886 | 0.09331029 | 0.4122884 |
| IARS | 157.087891 | -0.869020154 | 0.32811826 | -2.6484967 | 0.00808506 | 0.12762062 |
| NIT1 | 101.281215 | -0.868314483 | 0.3005289 | -2.8892878 | 0.00386116 | 0.0851496 |
| ST6GALNAC | 5633.99896 | -0.867839624 | 0.37226942 | -2.3312138 | 0.01974209 | 0.19704872 |
| RSPO3 | 65.6554853 | -0.86751018 | 0.64222069 | -1.3507976 | 0.17676027 | 0.54037749 |
| RPL36 | 383.648126 | -0.866281981 | 0.49451521 | -1.7517802 | 0.0798116 | 0.38738328 |
| BACH1 | 549.099952 | -0.865770756 | 0.388551 | -2.2282037 | 0.02586694 | 0.22446357 |
| SFMBT2 | 656.543164 | -0.864146768 | 0.37718818 | -2.2910229 | 0.02196209 | 0.20921508 |
| IRF2BP2 | 199.775837 | -0.863570046 | 0.53373278 | -1.617982 | 0.10566647 | 0.43832302 |
| XYLT1 | 182.936706 | -0.86319399 | 0.44239699 | -1.9511751 | 0.05103622 | 0.31710683 |
| CHSY1 | 639.023292 | -0.862866114 | 0.33830413 | -2.5505633 | 0.0107549 | 0.1457075 |
| SNHG8 | 47.7721465 | -0.861029693 | 0.66247939 | -1.2997079 | 0.1937011 | 0.55851167 |
| FPGS | 86.4453476 | -0.860977884 | 0.38268471 | -2.2498361 | 0.02445935 | 0.21755892 |
| TRIM5 | 86.5012964 | -0.85930335 | 0.42612051 | -2.0165735 | 0.04374004 | 0.29605023 |
| HYOU1 | 40.6632613 | -0.859193321 | 0.43938557 | -1.9554427 | 0.05053084 | 0.31516299 |
| RELB | 178.159439 | -0.858964373 | 0.62087119 | -1.3834824 | 0.16651699 | 0.52671378 |
| DSP | 58.0300106 | -0.858225729 | 0.51707092 | -1.6597834 | 0.09695804 | 0.41934863 |
| BTF3 | 203.619681 | -0.855786616 | 0.47468709 | -1.8028437 | 0.07141277 | 0.36926912 |
| EIF1 | 741.471209 | -0.855234333 | 0.35177618 | -2.4311889 | 0.01504937 | 0.1761788 |
| TRAF3IP2 | 57.144213 | -0.854361656 | 0.28742383 | -2.9724803 | 0.00295404 | 0.07188756 |
| ZFAS1 | 103.807814 | -0.854097988 | 0.23407223 | -3.6488652 | 0.0002634 | 0.01632889 |
| SF3B5 | 108.264023 | -0.854036971 | 0.44326614 | -1.9266912 | 0.05401813 | 0.32559425 |
| TIMM13 | 59.7860011 | -0.853993437 | 0.44620431 | -1.9139067 | 0.05563207 | 0.32988018 |
| CST3 | 343.214878 | -0.853296285 | 0.41686813 | -2.0469214 | 0.0406658 | 0.28612922 |
| RAN | 142.998314 | -0.852975948 | 0.57664074 | -1.4792155 | 0.13908272 | 0.49082099 |
| MCTP1 | 941.33812 | -0.852507924 | 0.63222914 | -1.3484161 | 0.1775246 | 0.54070521 |
| RPL35A | 308.267676 | -0.85168237 | 0.55048586 | -1.5471467 | 0.12182788 | 0.46672169 |
| GTF3A | 64.6954385 | -0.850905355 | 0.2547691 | -3.339908 | 0.00083806 | 0.03297047 |
| CNBP | 329.774329 | -0.848380851 | 0.38136427 | -2.2245945 | 0.02610847 | 0.22518506 |
| FXYD5 | 113.62507 | -0.847620536 | 0.40547147 | -2.0904567 | 0.0365768 | 0.26964679 |
| NUPR1 | 96.1535209 | -0.847620066 | 0.55506599 | -1.5270618 | 0.12674565 | 0.47358888 |
| NFKB1 | 588.744088 | -0.847043326 | 0.50888221 | -1.6645175 | 0.09600906 | 0.41722898 |
| GCNT1 | 53.7389894 | -0.847005714 | 0.46584519 | -1.8182129 | 0.06903161 | 0.36373833 |

| | | | | | | |
|---|---|---|---|---|---|---|
| TC2N | 75.8475879 | -0.846975114 | 0.35871144 | -2.36116 | 0.01821787 | 0.19288766 |
| DDX18 | 134.995617 | -0.845822904 | 0.20014809 | -4.2259854 | 2.38E-05 | 0.00335641 |
| ARHGDIA | 221.689102 | -0.845676262 | 0.46089767 | -1.834846 | 0.0665285 | 0.35767274 |
| NFKBIA | 968.669028 | -0.845594412 | 0.55989314 | -1.5102782 | 0.13097245 | 0.47974377 |
| SERBP1 | 286.336029 | -0.845587271 | 0.28193327 | -2.9992461 | 0.00270649 | 0.06964521 |
| PYURF | 69.8434151 | -0.842416828 | 0.48033779 | -1.7538009 | 0.07946463 | 0.38738328 |
| STX6 | 99.3640087 | -0.842313252 | 0.28556089 | -2.94968 | 0.00318103 | 0.07516604 |
| SERPING1 | 274.855438 | -0.841477003 | 0.34537717 | -2.4364002 | 0.01483426 | 0.17549246 |
| RPL13 | 926.990598 | -0.841231709 | 0.50627403 | -1.6616134 | 0.09659032 | 0.41820029 |
| NPM1 | 127.873736 | -0.840768694 | 0.35339909 | -2.3790912 | 0.01735538 | 0.1883542 |
| TUBA1C | 112.267186 | -0.840273571 | 0.34461627 | -2.4382876 | 0.01475703 | 0.17549246 |
| TSPO | 212.377666 | -0.839899735 | 0.50922458 | -1.64937 | 0.09907186 | 0.42376187 |
| CSTB | 188.57644 | -0.839857786 | 0.3716454 | -2.2598364 | 0.02383141 | 0.21548334 |
| RPS23 | 596.788555 | -0.83821239 | 0.59792359 | -1.4018721 | 0.16095345 | 0.51944206 |
| NTMT1 | 42.5392311 | -0.837662515 | 0.30844617 | -2.7157495 | 0.00661259 | 0.11488503 |
| RPL13A | 425.978762 | -0.836852919 | 0.48561132 | -1.7232978 | 0.08483469 | 0.39686932 |
| ATRAID | 85.0206338 | -0.836682858 | 0.51139219 | -1.6360885 | 0.10182107 | 0.43079198 |
| CD9 | 1374.34902 | -0.836035364 | 0.46130563 | -1.8123242 | 0.06993611 | 0.36567873 |
| AC007100.1 | 371.650795 | -0.835530596 | 0.52219219 | -1.6000442 | 0.10958877 | 0.44576178 |
| RPLP2 | 327.266416 | -0.832084217 | 0.52602274 | -1.5818408 | 0.11368592 | 0.45258538 |
| PPARD | 119.960015 | -0.83111832 | 0.28183817 | -2.9489203 | 0.00318886 | 0.07516604 |
| HSP90B1 | 492.609231 | -0.829907805 | 0.50732009 | -1.6358662 | 0.10186759 | 0.43079198 |
| UQCR10 | 130.736303 | -0.828577842 | 0.46756485 | -1.7721132 | 0.07637577 | 0.38287714 |
| IRX5 | 38.6478709 | -0.827570204 | 0.41372411 | -2.0002948 | 0.04546844 | 0.30053976 |
| NOP14 | 47.1867523 | -0.827354202 | 0.38494402 | -2.1492845 | 0.03161186 | 0.24825319 |
| CMIP | 1449.38957 | -0.826809687 | 0.42864254 | -1.9289026 | 0.05374296 | 0.32491861 |
| SLC9B2 | 76.5798588 | -0.825344264 | 0.25410265 | -3.2480742 | 0.00116189 | 0.04098165 |
| SDC2 | 75.8324747 | -0.825308547 | 0.37417999 | -2.2056459 | 0.0274088 | 0.23052871 |
| PSMB5 | 129.240016 | -0.8252953 | 0.33777817 | -2.4433056 | 0.01455341 | 0.17436388 |
| TRMT10C | 50.0793229 | -0.825286244 | 0.35696799 | -2.3119334 | 0.02078135 | 0.20341364 |
| TNFRSF1A | 532.051957 | -0.824908419 | 0.37687883 | -2.1887895 | 0.02861214 | 0.23392531 |
| RPL37A | 346.59733 | -0.824427087 | 0.41078588 | -2.0069509 | 0.04475489 | 0.29946125 |
| ANKDD1A | 78.4381951 | -0.823989776 | 0.32365335 | -2.5459022 | 0.01089957 | 0.14707585 |
| ISG20 | 153.964935 | -0.822767025 | 0.71635662 | -1.1485439 | 0.2507441 | 0.61583146 |
| NACA | 261.082048 | -0.822194528 | 0.43962915 | -1.8702002 | 0.06145602 | 0.34778328 |
| COA4 | 59.8286291 | -0.822009971 | 0.37389839 | -2.1984849 | 0.02791457 | 0.23190345 |
| BCAR3 | 325.199044 | -0.821461748 | 0.31706844 | -2.5908026 | 0.00957524 | 0.13793724 |
| VCAM1 | 819.720728 | -0.820585731 | 0.69248432 | -1.1849882 | 0.23602212 | 0.60018541 |
| FAM210A | 103.83855 | -0.820453334 | 0.46568694 | -1.7618131 | 0.07810088 | 0.38566641 |
| RPL27A | 351.905559 | -0.819990168 | 0.49862967 | -1.6444873 | 0.10007559 | 0.42607623 |
| HOOK2 | 83.4474436 | -0.819436684 | 0.27770253 | -2.9507714 | 0.00316981 | 0.07516604 |
| NAGLU | 52.4767783 | -0.819155218 | 0.40053473 | -2.045154 | 0.04083967 | 0.28670444 |
| TIMM10 | 42.3867225 | -0.818915279 | 0.35183881 | -2.3275297 | 0.01993709 | 0.19732169 |
| AL136985.3 | 103.887606 | -0.818794127 | 0.93132175 | -0.8791743 | 0.37930679 | 0.71402518 |
| SFRP1 | 85.7427241 | -0.816001986 | 0.34949418 | -2.3348085 | 0.01955343 | 0.19632601 |
| ILF3-DT | 50.853278 | -0.815733875 | 0.34068119 | -2.39442 | 0.01664667 | 0.18464043 |
| AC016205.1 | 70.25763 | -0.815168655 | 0.48681352 | -1.6744988 | 0.09403261 | 0.41280519 |
| SET | 234.331622 | -0.814691346 | 0.3509917 | -2.3211128 | 0.02028075 | 0.20000907 |
| RNF4 | 191.181063 | -0.812667726 | 0.3050276 | -2.6642433 | 0.00771617 | 0.12405 |
| UBE2E2 | 816.954991 | -0.812548015 | 0.47372305 | -1.7152385 | 0.08630151 | 0.39864674 |
| UPP1 | 202.596382 | -0.8113175 | 0.40481574 | -2.0041649 | 0.0450524 | 0.29976402 |
| RPL24 | 237.916318 | -0.810938957 | 0.45179311 | -1.7949343 | 0.07266416 | 0.37296554 |
| PSMD8 | 112.042666 | -0.810852132 | 0.35611966 | -2.2769092 | 0.02279165 | 0.21136949 |
| DGKI | 122.435482 | -0.810663826 | 0.63599997 | -1.2746287 | 0.20244069 | 0.56693091 |
| RPS16 | 286.259077 | -0.810608955 | 0.48205172 | -1.6815809 | 0.09265014 | 0.41185145 |
| PHB | 98.9920712 | -0.810553343 | 0.3558074 | -2.2780677 | 0.02272254 | 0.21131824 |
| HINT3 | 54.3922482 | -0.810403808 | 0.28741184 | -2.8196605 | 0.00480745 | 0.09480113 |
| DOCK5 | 123.098071 | -0.809851251 | 0.5014711 | -1.614951 | 0.10632131 | 0.43901385 |
| PDLIM3 | 82.3363897 | -0.808594449 | 0.72193089 | -1.1200441 | 0.26269496 | 0.6283639 |
| RPL7 | 351.809539 | -0.808570143 | 0.52578282 | -1.5378405 | 0.12408761 | 0.46936415 |
| RPS25 | 379.772847 | -0.808426321 | 0.50083626 | -1.6141529 | 0.10649426 | 0.43901385 |
| TCEAL4 | 129.443012 | -0.808054571 | 0.29681208 | -2.7224451 | 0.00648008 | 0.11362684 |
| MRPL27 | 57.496361 | -0.807716547 | 0.3315157 | -2.4364353 | 0.01483282 | 0.17549246 |
| NUDT14 | 51.7175712 | -0.807146706 | 0.4781253 | -1.6881489 | 0.09138264 | 0.40862523 |
| HEBP2 | 86.4032729 | -0.80595878 | 0.29672344 | -2.7161951 | 0.0066037 | 0.11488503 |
| EVL | 220.022234 | -0.801385669 | 0.29004353 | -2.7629841 | 0.00572756 | 0.10415242 |
| C4orf3 | 229.071003 | -0.801176818 | 0.35971659 | -2.2272446 | 0.02593093 | 0.22478053 |
| SLC29A1 | 38.7642291 | -0.800654689 | 0.5294888 | -1.5121277 | 0.13050138 | 0.47929621 |
| SLC6A16 | 47.2354912 | -0.800625937 | 0.51795487 | -1.5457446 | 0.12216626 | 0.46736163 |
| RPL8 | 548.761386 | -0.800354102 | 0.53475973 | -1.4966611 | 0.13448145 | 0.48474744 |
| CFL1 | 449.631846 | -0.799979861 | 0.49466923 | -1.6172016 | 0.10583477 | 0.43872641 |
| NOVA1 | 69.5962848 | -0.799464012 | 0.34189576 | -2.3383268 | 0.0193703 | 0.19568789 |
| IGFBP4 | 1367.72856 | -0.79926146 | 0.53934373 | -1.4819148 | 0.13836296 | 0.48991657 |
| FUNDC2 | 89.6535334 | -0.797180179 | 0.31657135 | -2.5181691 | 0.01179667 | 0.15189817 |
| PRSS23 | 617.19901 | -0.797172621 | 0.46189782 | -1.7258636 | 0.08437196 | 0.39616527 |
| RHOJ | 815.87291 | -0.796773949 | 0.4038757 | -1.9728197 | 0.04851611 | 0.30895509 |
| RFX2 | 200.293948 | -0.79670065 | 0.67023617 | -1.1886864 | 0.23456308 | 0.59949266 |
| PRUNE2 | 82.1919454 | -0.796370834 | 0.53964955 | -1.4757185 | 0.14001947 | 0.49118733 |
| ODC1 | 44.3537388 | -0.79624463 | 0.38440721 | -2.0713572 | 0.03832543 | 0.27652057 |
| F8 | 727.46561 | -0.794494919 | 0.57043918 | -1.3927776 | 0.16368705 | 0.52357299 |
| ITGB4 | 102.21261 | -0.793835938 | 0.54857419 | -1.4470895 | 0.14787187 | 0.50143858 |
| SELENOM | 232.025059 | -0.793389209 | 0.55593264 | -1.4271319 | 0.15354188 | 0.5097092 |
| PFDN2 | 129.989425 | -0.792958137 | 0.39502628 | -2.0073554 | 0.04471183 | 0.29946125 |
| PGK1 | 261.087455 | -0.792933965 | 0.33227997 | -2.3863429 | 0.01701687 | 0.18593697 |
| MTHFD1 | 43.2593029 | -0.792006539 | 0.26565133 | -2.9813761 | 0.00286956 | 0.07180923 |
| RUVBL1 | 108.560296 | -0.79179833 | 0.27952135 | -2.8326935 | 0.00461576 | 0.09280289 |
| PFKP | 55.8340364 | -0.791673514 | 0.52458588 | -1.50914 | 0.13126302 | 0.48037803 |
| ARMC9 | 214.699369 | -0.791557033 | 0.33822662 | -2.3403156 | 0.01926745 | 0.19499826 |
| ESR2 | 269.287939 | -0.790099582 | 0.61652845 | -1.2815298 | 0.20000764 | 0.56449294 |
| SREBF1 | 61.8450479 | -0.788086481 | 0.31955005 | -2.466238 | 0.01365406 | 0.16826979 |
| ABHD2 | 210.940615 | -0.786667158 | 0.2764173 | -2.8459404 | 0.00442805 | 0.091042 |
| B3GNT5 | 106.061445 | -0.785851343 | 0.38741621 | -2.0284421 | 0.04251515 | 0.291241 |
| PSMA7 | 242.380982 | -0.785138221 | 0.4010505 | -1.9577042 | 0.05026474 | 0.31422181 |
| SPCS1 | 148.156864 | -0.783699656 | 0.40930597 | -1.9147037 | 0.0555303 | 0.32988018 |
| EBPL | 71.2111735 | -0.783693356 | 0.45785832 | -1.7116503 | 0.08696112 | 0.39969788 |
| ZFP36L2 | 347.213761 | -0.783056533 | 0.26295306 | -2.9779328 | 0.002902 | 0.07188756 |
| SEC13 | 127.841002 | -0.782458432 | 0.2875365 | -2.721249 | 0.00650357 | 0.1136514 |
| TARS | 87.2073532 | -0.78177862 | 0.33948958 | -2.302806 | 0.02128976 | 0.20628732 |
| PRELID1 | 111.690658 | -0.781772968 | 0.50648041 | -1.5435404 | 0.12269972 | 0.46808475 |
| ACER2 | 395.488956 | -0.781295672 | 0.70202011 | -1.1129249 | 0.26574068 | 0.63060082 |
| NR2F2 | 666.836821 | -0.780577309 | 0.47605472 | -1.6396798 | 0.10107176 | 0.42853379 |
| SLC22A23 | 597.275613 | -0.779605454 | 0.58979354 | -1.3218277 | 0.18622553 | 0.55236793 |
| TCEAL9 | 164.789252 | -0.77859365 | 0.45294666 | -1.7189522 | 0.08562308 | 0.39787261 |
| TIPARP | 69.6135093 | -0.778069081 | 0.56022943 | -1.3888401 | 0.16488139 | 0.52480753 |
| RPL7A | 295.158124 | -0.777461267 | 0.50636322 | -1.5353826 | 0.12468988 | 0.47038289 |
| RPL38 | 168.263064 | -0.777309567 | 0.46568684 | -1.669168 | 0.0950841 | 0.41475115 |
| MTUS1 | 728.250922 | -0.774854434 | 0.60459611 | -1.2816067 | 0.19998064 | 0.56449294 |
| NSA2 | 59.2576727 | -0.773731796 | 0.33223555 | -2.3288652 | 0.01986621 | 0.19704872 |
| MAP2K7 | 57.4870556 | -0.773662981 | 0.37387018 | -2.0693359 | 0.03851458 | 0.27738487 |
| CLEC1A | 298.066703 | -0.77362445 | 0.33874874 | -2.2837707 | 0.02238501 | 0.2108594 |
| CLEC14A | 414.467592 | -0.773359187 | 0.45412221 | -1.7029759 | 0.08857258 | 0.40312938 |
| CCDC124 | 45.1065186 | -0.772899754 | 0.42759822 | -1.8075374 | 0.07067853 | 0.36751516 |
| PFKFB3 | 222.32861 | -0.771706269 | 0.38734796 | -1.9922818 | 0.04634015 | 0.30359341 |

| | | | | | | |
|---|---|---|---|---|---|---|
| RBM3 | 152.45696 | -0.771183176 | 0.43259459 | -1.7826926 | 0.07463636 | 0.37864186 |
| LIMS1 | 457.266496 | -0.771074898 | 0.33806877 | -2.2808226 | 0.02255894 | 0.21131824 |
| MRPL51 | 93.6062243 | -0.770468263 | 0.43427546 | -1.7741465 | 0.07603893 | 0.38243795 |
| MAP3K5 | 244.362671 | -0.770296698 | 0.59980714 | -1.2842406 | 0.19905778 | 0.56402001 |
| TNFRSF10B | 188.893405 | -0.77022322 | 0.45433152 | -1.695289 | 0.09002061 | 0.40630925 |
| MAP1B | 158.128915 | -0.769954726 | 0.442388 | -1.7404512 | 0.08177982 | 0.393187 |
| ALPK1 | 285.046118 | -0.768478888 | 0.31407538 | -2.4467976 | 0.01441317 | 0.17370104 |
| NQO1 | 112.174367 | -0.767922398 | 0.62147144 | -1.2356519 | 0.21658797 | 0.58218265 |
| H1F0 | 94.3751894 | -0.767011285 | 0.67981358 | -1.1282671 | 0.25920714 | 0.62403413 |
| VAC14 | 78.6266558 | -0.766919975 | 0.4035737 | -1.900322 | 0.05739088 | 0.33544968 |
| FES | 144.141757 | -0.766465484 | 0.38456907 | -1.9930503 | 0.04625594 | 0.30359341 |
| INTS10 | 202.149963 | -0.76416266 | 0.2289312 | -3.3379577 | 0.00084397 | 0.03304389 |
| GAK | 405.810252 | -0.764076939 | 0.41322403 | -1.8490622 | 0.06444883 | 0.35347506 |
| SRPRA | 87.912911 | -0.763689465 | 0.28993733 | -2.6339812 | 0.00843902 | 0.13029523 |
| ALDH1A1 | 105.556315 | -0.762889534 | 0.35372326 | -2.1567412 | 0.03102584 | 0.24529895 |
| BANF1 | 86.0262607 | -0.762288428 | 0.49505049 | -1.5398196 | 0.12360434 | 0.46913465 |
| SARS | 127.732941 | -0.760578492 | 0.27033297 | -2.8134877 | 0.00490073 | 0.09526334 |
| ADGRG6 | 110.45235 | -0.760263265 | 0.60814955 | -1.2501255 | 0.21125371 | 0.5768065 |
| IFI27L2 | 65.0999771 | -0.759770151 | 0.59821808 | -1.2700555 | 0.20406487 | 0.56856072 |
| AC069368.1 | 84.7341718 | -0.759230848 | 0.49998287 | -1.5185137 | 0.12888494 | 0.47679271 |
| WDR89 | 55.6334109 | -0.759169648 | 0.26282538 | -2.8884944 | 0.00387091 | 0.0851496 |
| RNH1 | 107.830592 | -0.758588476 | 0.39112614 | -1.9394983 | 0.0524407 | 0.32059696 |
| PGM2L1 | 90.7162442 | -0.758510559 | 0.37559865 | -2.019471 | 0.04343829 | 0.29547428 |
| PCDH7 | 123.774494 | -0.757763309 | 0.63559792 | -1.1922054 | 0.2331807 | 0.59910683 |
| MRPL41 | 120.021081 | -0.757362607 | 0.40582365 | -1.8662358 | 0.06200839 | 0.3492186 |
| TAF10 | 43.8420014 | -0.757183659 | 0.47305928 | -1.6006105 | 0.10946321 | 0.44576178 |
| KIF3C | 45.3201761 | -0.757022322 | 0.41134059 | -1.8403783 | 0.06571271 | 0.35579065 |
| RPL10 | 814.053377 | -0.756944723 | 0.57973822 | -1.3056664 | 0.19166603 | 0.55597416 |
| GNS | 218.187101 | -0.755807784 | 0.22133155 | -3.4148216 | 0.00063824 | 0.02763334 |
| REV3L | 753.640153 | -0.755667802 | 0.3409616 | -2.2162842 | 0.02667204 | 0.22752964 |
| SNRPF | 52.2074568 | -0.755471638 | 0.34524567 | -2.1882146 | 0.02865397 | 0.23400743 |
| NRP2 | 224.922994 | -0.754903866 | 0.49930917 | -1.5118966 | 0.13056016 | 0.47929621 |
| JUNB | 363.607595 | -0.754696197 | 0.62666432 | -1.204307 | 0.22847096 | 0.59497702 |
| SSR4 | 75.2891996 | -0.75461318 | 0.66555002 | -1.1338189 | 0.25687053 | 0.62114998 |
| CYP7B1 | 64.2156653 | -0.752994104 | 0.44540963 | -1.6905654 | 0.09091984 | 0.40744637 |
| CCT3 | 153.904987 | -0.752821141 | 0.33264945 | -2.2631065 | 0.02362912 | 0.21448172 |
| RAB32 | 43.3970962 | -0.751764405 | 0.40907727 | -1.8377076 | 0.0661055 | 0.35600847 |
| MRPL17 | 99.2078798 | -0.751557946 | 0.48364879 | -1.5539333 | 0.12020033 | 0.46382142 |
| TSEN2 | 99.7266299 | -0.751501222 | 0.33096761 | -2.2706186 | 0.02317008 | 0.21286486 |
| CARHSP1 | 407.594526 | -0.750735048 | 0.32127412 | -2.336743 | 0.01945255 | 0.19627649 |
| EEF1A1 | 1316.99409 | -0.749524473 | 0.49719968 | -1.5074919 | 0.13168463 | 0.48084574 |
| SDCBP2-AS1 | 229.762139 | -0.74826886 | 0.42914011 | -1.743647 | 0.08122067 | 0.39164922 |
| ROMO1 | 58.9069107 | -0.748020817 | 0.52197577 | -1.4330566 | 0.15184166 | 0.50784035 |
| CDK4 | 41.4669136 | -0.747784987 | 0.34272594 | -2.1818745 | 0.0291188 | 0.23685798 |
| HPRT1 | 124.961282 | -0.747670977 | 0.25035331 | -2.9864633 | 0.00282225 | 0.07105983 |
| WAC-AS1 | 52.621927 | -0.747032433 | 0.48115271 | -1.5525891 | 0.12052134 | 0.46432492 |
| ARL5B | 103.102367 | -0.746184399 | 0.53613784 | -1.3917772 | 0.16398989 | 0.52357299 |
| ITPA | 49.2921779 | -0.74586004 | 0.36193672 | -2.0607471 | 0.03932717 | 0.28057468 |
| CCT2 | 92.1484172 | -0.745804503 | 0.3178705 | -2.3462527 | 0.01896325 | 0.19411645 |
| CLPP | 46.7443502 | -0.745357529 | 0.33983271 | -2.1933072 | 0.02828526 | 0.23285538 |
| CYC1 | 58.581386 | -0.745319671 | 0.38870415 | -1.9174472 | 0.05518114 | 0.32935616 |
| ZFYVE28 | 300.282442 | -0.745152621 | 0.3070394 | -2.4268957 | 0.01522863 | 0.177259 |
| CEACAM19 | 87.9658651 | -0.744904094 | 0.46885703 | -1.588766 | 0.11211324 | 0.44883692 |
| CLU | 212.077691 | -0.744843748 | 0.46581483 | -1.5990125 | 0.10981782 | 0.44576178 |
| STAT5A | 101.275196 | -0.744511785 | 0.31348272 | -2.3749691 | 0.01755042 | 0.18854202 |
| SLIT3 | 221.402367 | -0.743402165 | 1.03846405 | -0.715867 | 0.47407347 | 0.77932979 |
| BSG | 175.972646 | -0.743345928 | 0.38434659 | -1.934051 | 0.05310685 | 0.3223838 |
| FAM126A | 250.63472 | -0.743047724 | 0.30333528 | -2.4495921 | 0.01430181 | 0.17286813 |
| MAMDC2 | 256.837914 | -0.741466677 | 0.54938547 | -1.3496292 | 0.17713495 | 0.54037749 |
| NRIP1 | 343.895573 | -0.741340567 | 0.37558436 | -1.9738324 | 0.04840081 | 0.30846092 |
| AZIN1-AS1 | 67.0122208 | -0.741117937 | 0.55905923 | -1.3256519 | 0.18495502 | 0.55006698 |
| P4HB | 253.016847 | -0.740985023 | 0.3769118 | -1.9659375 | 0.04930583 | 0.31155957 |
| PCDH1 | 106.157792 | -0.740652404 | 0.42870039 | -1.7276691 | 0.08404757 | 0.39616527 |
| FAM110D | 106.396757 | -0.739727848 | 0.66751216 | -1.1081863 | 0.26778136 | 0.63276201 |
| RBKS | 71.4856814 | -0.739467764 | 0.41721764 | -1.7723789 | 0.07633167 | 0.38287714 |
| ASAH1 | 176.827466 | -0.738579353 | 0.36380197 | -2.0301686 | 0.0423394 | 0.291241 |
| TMEM132A | 45.6900448 | -0.73765351 | 0.51715172 | -1.4263774 | 0.15375944 | 0.50978388 |
| POLR2L | 182.267209 | -0.737618001 | 0.4364493 | -1.6900428 | 0.09101976 | 0.40767089 |
| RBP5 | 124.598538 | -0.737467942 | 0.37178604 | -1.9835816 | 0.0473025 | 0.30550621 |
| RAB20 | 96.7344347 | -0.737434464 | 0.40883709 | -1.8037367 | 0.07127259 | 0.36912889 |
| DDX31 | 60.4840663 | -0.737022509 | 0.43832067 | -1.6814688 | 0.0926719 | 0.41185145 |
| PGAM1 | 166.568454 | -0.736741619 | 0.36589895 | -2.0135111 | 0.04406088 | 0.29738759 |
| OSMR | 733.930969 | -0.735285104 | 0.20504575 | -3.5859564 | 0.00033585 | 0.01921832 |
| LSM4 | 76.2297401 | -0.734845972 | 0.32388026 | -2.2688816 | 0.02327552 | 0.2129727 |
| ASPH | 335.136302 | -0.734811167 | 0.34408727 | -2.1355372 | 0.03271716 | 0.25233222 |
| LY96 | 78.0569905 | -0.734228786 | 0.34244338 | -2.1440881 | 0.03202582 | 0.24982585 |
| SLC2A3 | 1230.14677 | -0.733971276 | 0.63224512 | -1.1608967 | 0.24568391 | 0.61070214 |
| MCFD2 | 128.924348 | -0.732190217 | 0.30217465 | -2.4230697 | 0.01538998 | 0.17781377 |
| APOO | 733.842256 | -0.732039855 | 0.52982105 | -1.3816738 | 0.16707189 | 0.52764813 |
| AC027018.1 | 76.6103858 | -0.73180456 | 0.51493214 | -1.421167 | 0.15526822 | 0.51129169 |
| ASXL3 | 52.9044321 | -0.73176775 | 0.61069867 | -1.1982468 | 0.23082094 | 0.59732678 |
| RRP12 | 48.428625 | -0.731073351 | 0.39454586 | -1.852949 | 0.06388967 | 0.3533385 |
| ARF1 | 334.21819 | -0.730922883 | 0.2724832 | -2.6824512 | 0.00730848 | 0.12106337 |
| BTG1 | 427.984023 | -0.730507178 | 0.53624515 | -1.3622635 | 0.17311475 | 0.53517113 |
| FADS1 | 39.4740381 | -0.730051786 | 0.48173807 | -1.5154538 | 0.12965752 | 0.4777902 |
| NDUFA12 | 149.836088 | -0.729932151 | 0.47898804 | -1.5239048 | 0.12753251 | 0.47479008 |
| NFKB2 | 73.9977103 | -0.729685364 | 0.61823013 | -1.1802811 | 0.23788841 | 0.60096874 |
| AC078883.1 | 103.575659 | -0.728128647 | 0.50200079 | -1.4504532 | 0.14693218 | 0.50118636 |
| ITPRIP | 556.567078 | -0.728044766 | 0.35144875 | -2.0715531 | 0.03830713 | 0.27652057 |
| RPL35 | 247.436924 | -0.727475301 | 0.46360989 | -1.569154 | 0.11661206 | 0.4592091 |
| PRDX6 | 143.353931 | -0.726970584 | 0.29531986 | -2.461638 | 0.01383042 | 0.16916939 |
| CLIC1 | 408.404902 | -0.725593973 | 0.45728338 | -1.5867491 | 0.11256949 | 0.44956377 |
| MYDGF | 183.446535 | -0.725590392 | 0.35993638 | -2.0158851 | 0.04381199 | 0.29629216 |
| NR2F6 | 57.2151534 | -0.725407183 | 0.41690355 | -1.739988 | 0.08186112 | 0.39328747 |
| GABRE | 172.287942 | -0.725399099 | 0.52340368 | -1.3859266 | 0.16576931 | 0.5261375 |
| TUBB6 | 335.23906 | -0.725222698 | 0.34747043 | -2.0871494 | 0.03687463 | 0.27062343 |
| PTMA | 1252.78861 | -0.723599538 | 0.44418948 | -1.6290335 | 0.10330593 | 0.43424379 |
| P2RY8 | 240.383486 | -0.723591403 | 0.47021303 | -1.5388587 | 0.1238388 | 0.46936415 |
| HNRNPH2 | 62.3583596 | -0.722626542 | 0.42115989 | -1.7158009 | 0.08619848 | 0.39864674 |
| UBL5 | 72.2243996 | -0.721615372 | 0.51473157 | -1.4019256 | 0.16093746 | 0.51944206 |
| ACAA2 | 49.6632275 | -0.721307144 | 0.48688367 | -1.4814774 | 0.13847941 | 0.48991657 |
| CTSA | 93.4673781 | -0.720735635 | 0.3810642 | -1.8913759 | 0.05857418 | 0.33921624 |
| BCR | 401.185654 | -0.720599969 | 0.31647785 | -2.2769365 | 0.02279002 | 0.21136949 |
| LRP5L | 52.5601261 | -0.720433096 | 0.50104825 | -1.4378517 | 0.15047613 | 0.5057732 |
| RPS6KA2 | 1050.51125 | -0.720000717 | 0.24956841 | -2.8849834 | 0.00391434 | 0.08541619 |
| GUCY1A1 | 70.7934209 | -0.719661244 | 0.65287063 | -1.102303 | 0.27032997 | 0.63548122 |
| SCAND1 | 136.584777 | -0.717557142 | 0.41102856 | -1.7457598 | 0.0808527 | 0.39055112 |
| MRPL14 | 56.0095771 | -0.716559975 | 0.40315223 | -1.777393 | 0.0755036 | 0.38092632 |
| AC098617.1 | 200.63323 | -0.716539943 | 0.39671513 | -1.8061825 | 0.07088983 | 0.36807837 |
| MTHFD2L | 119.943905 | -0.716456657 | 0.23064892 | -3.106265 | 0.00189467 | 0.05597138 |
| ATP11A | 661.763937 | -0.715519655 | 0.39050267 | -1.8323041 | 0.06690613 | 0.35877642 |
| SYNJ2 | 438.962213 | -0.715034043 | 0.39258741 | -1.8213372 | 0.06855562 | 0.36169608 |
| PPIA | 446.68518 | -0.714366832 | 0.47434967 | -1.5059921 | 0.13206921 | 0.48092078 |

| | | | | | | |
|---|---|---|---|---|---|---|
| LRRK1 | 118.921286 | -0.713568575 | 0.24128162 | -2.9574096 | 0.00310236 | 0.07422979 |
| MRPL34 | 59.2428427 | -0.713500265 | 0.50782892 | -1.4050013 | 0.16002091 | 0.51818404 |
| NDUFS8 | 81.9369011 | -0.712876072 | 0.36948773 | -1.9293633 | 0.05368578 | 0.32491861 |
| PTP4A2 | 82.8850001 | -0.712590794 | 0.40635274 | -1.7536261 | 0.07949459 | 0.38738328 |
| HSP90AB1 | 564.492855 | -0.712068955 | 0.32834252 | -2.1686772 | 0.0301072 | 0.24082201 |
| TCEAL3 | 45.8759621 | -0.711461732 | 0.31519122 | -2.2572384 | 0.02399318 | 0.21622928 |
| AFAP1L1 | 534.376695 | -0.711186772 | 0.34996569 | -2.0321614 | 0.04213732 | 0.29097861 |
| CXorf36 | 433.821494 | -0.71106844 | 0.29740844 | -2.3908819 | 0.01680796 | 0.1855435 |
| RHBDF2 | 100.697045 | -0.710589131 | 0.4556549 | -1.5594897 | 0.11888052 | 0.46285439 |
| BNIP3L | 262.131941 | -0.710360024 | 0.43969227 | -1.6155845 | 0.10618417 | 0.43901385 |
| H2AFZ | 162.105547 | -0.710214943 | 0.73453648 | -0.9668886 | 0.33359973 | 0.6852158 |
| CNIH1 | 88.4093135 | -0.708924429 | 0.3534306 | -2.0058377 | 0.04487357 | 0.29946125 |
| RPF2 | 62.0827024 | -0.708716907 | 0.27435957 | -2.5831682 | 0.00978976 | 0.1397149 |
| TMSB10 | 2791.17544 | -0.708696909 | 0.62469782 | -1.1344636 | 0.25660015 | 0.62095131 |
| PITPNM2 | 655.598353 | -0.7086732 | 0.35325121 | -2.0061452 | 0.04484076 | 0.29946125 |
| CERK | 133.733693 | -0.7072968 | 0.3072504 | -2.3020207 | 0.021334 | 0.20628732 |
| XPO5 | 237.815921 | -0.707270066 | 0.36076759 | -1.960459 | 0.04994217 | 0.31316226 |
| FP236383.1 | 715.169669 | -0.707067003 | 0.42707806 | -1.655592 | 0.09780448 | 0.42128893 |
| S1PR1 | 281.661035 | -0.706933933 | 0.46954298 | -1.5055788 | 0.13217534 | 0.48092078 |
| HDGF | 121.505487 | -0.706899649 | 0.32656179 | -2.1646735 | 0.0304127 | 0.24194918 |
| LINC01515 | 113.001998 | -0.706715124 | 0.68312052 | -1.0345395 | 0.30088404 | 0.65926326 |
| TOM1 | 57.9278377 | -0.706649735 | 0.31338471 | -2.2548954 | 0.02413989 | 0.21636008 |
| DCDC2 | 45.2967621 | -0.706047988 | 0.5000824 | -1.4118633 | 0.15799022 | 0.51474819 |
| NARS | 106.738422 | -0.705305359 | 0.33410133 | -2.1110523 | 0.03476782 | 0.26053578 |
| APLP2 | 347.577869 | -0.705234637 | 0.26865922 | -2.6250156 | 0.0086645 | 0.13154285 |
| HM13 | 175.140312 | -0.705029199 | 0.2513502 | -2.8049678 | 0.00503216 | 0.09734793 |
| TMEM255B | 450.725092 | -0.704637948 | 0.45504588 | -1.5484987 | 0.12150227 | 0.46569232 |
| IFI27 | 1397.00415 | -0.70462858 | 0.53165894 | -1.3253395 | 0.18505858 | 0.55006698 |
| SLC1A1 | 154.059531 | -0.704517919 | 0.51953628 | -1.3560514 | 0.1750828 | 0.53840758 |
| DAP | 167.18944 | -0.704458897 | 0.29711313 | -2.3710124 | 0.01773944 | 0.18975401 |
| CNTNAP3B | 1514.45411 | -0.704253995 | 0.47441408 | -1.4844711 | 0.13768399 | 0.48962215 |
| PFN1 | 356.6825 | -0.704244481 | 0.4781249 | -1.4729299 | 0.14076993 | 0.49248412 |
| MSMO1 | 53.2138818 | -0.703791959 | 0.42481524 | -1.6567013 | 0.09757988 | 0.42092574 |
| GSTP1 | 329.648656 | -0.703775947 | 0.4327639 | -1.6262353 | 0.1038996 | 0.43532543 |
| HINT1 | 371.722247 | -0.703135683 | 0.46293726 | -1.5188574 | 0.1287984 | 0.47679271 |
| SELENOS | 148.440125 | -0.702652704 | 0.35101872 | -2.0017528 | 0.04531132 | 0.30043379 |
| LTF | 100.312237 | -0.70159694 | 0.55601018 | -1.2618419 | 0.20700569 | 0.57162978 |
| PER1 | 131.102616 | -0.700226071 | 0.68603615 | -1.0206839 | 0.30740421 | 0.66400081 |
| CIAO1 | 85.4785125 | -0.69999686 | 0.27277759 | -2.5661817 | 0.0102825 | 0.14331772 |
| FAM114A2 | 108.613561 | -0.699692609 | 0.21355407 | -3.276419 | 0.00105132 | 0.03823551 |
| SSH1 | 379.784853 | -0.699603525 | 0.38193919 | -1.8317144 | 0.06699398 | 0.35889508 |
| PLEKHA7 | 570.251289 | -0.698885512 | 0.44514818 | -1.5700064 | 0.11641361 | 0.4592091 |
| 2-Mar | 141.212721 | -0.6983982 | 0.37874012 | -1.8440037 | 0.0651826 | 0.35535588 |
| RCN1 | 89.136996 | -0.69831153 | 0.53654139 | -1.3015054 | 0.19308551 | 0.55752247 |
| RPL11 | 638.329873 | -0.698308354 | 0.53245872 | -1.3114789 | 0.18969603 | 0.55535814 |
| DPM3 | 57.6538605 | -0.69821538 | 0.45899204 | -1.5211928 | 0.12821147 | 0.47578036 |
| NR5A2 | 382.482339 | -0.698098131 | 0.63392337 | -1.1012343 | 0.27079471 | 0.63561102 |
| NAA38 | 227.437391 | -0.697183274 | 0.39746668 | -1.7540672 | 0.07941899 | 0.38738328 |
| SNRK | 1443.5095 | -0.69706103 | 0.35008583 | -1.9911147 | 0.04646828 | 0.30359341 |
| FARSB | 91.6003582 | -0.696763742 | 0.40337135 | -1.7273506 | 0.08410472 | 0.39616527 |
| NINJ1 | 54.3152825 | -0.696379528 | 0.3656858 | -1.9043111 | 0.05686968 | 0.33347614 |
| RPS7 | 399.91366 | -0.696287619 | 0.55767683 | -1.2485504 | 0.21182957 | 0.57690664 |
| LHFPL3 | 53.4464407 | -0.695409048 | 0.31529044 | -2.2056142 | 0.02741103 | 0.23052871 |
| SELENOK | 103.109303 | -0.694731245 | 0.49153093 | -1.4134029 | 0.15753729 | 0.51438736 |
| DIAPH2 | 770.111799 | -0.694645184 | 0.25671785 | -2.7058702 | 0.00681257 | 0.11693671 |
| PKP4-AS1 | 51.8647002 | -0.694613507 | 0.62296306 | -1.1150156 | 0.26484375 | 0.63018797 |
| CMSS1 | 453.500551 | -0.694580695 | 0.31060339 | -2.2362303 | 0.02533669 | 0.22138474 |
| ALCAM | 113.876131 | -0.694075369 | 0.39250173 | -1.7683371 | 0.07700457 | 0.38356763 |
| PA2G4 | 62.5306927 | -0.693989498 | 0.41277314 | -1.6812855 | 0.09270747 | 0.41185145 |
| HNRNPF | 223.604771 | -0.693718971 | 0.23085669 | -3.0049767 | 0.00265601 | 0.0688219 |
| GNG12-AS1 | 84.9617027 | -0.693289368 | 0.3267472 | -2.1217913 | 0.03385527 | 0.25638251 |
| MLXIP | 210.064147 | -0.692341301 | 0.26927742 | -2.5711079 | 0.01013737 | 0.14236887 |
| DIRC2 | 86.769184 | -0.692314129 | 0.27432957 | -2.5236584 | 0.01161407 | 0.15149764 |
| AP001831.1 | 47.7039238 | -0.691427037 | 0.44965322 | -1.5376895 | 0.12412455 | 0.46936415 |
| DDX24 | 241.639554 | -0.691362859 | 0.2204961 | -3.1354879 | 0.00171569 | 0.05338195 |
| CYB5D2 | 70.6392137 | -0.691162386 | 0.26493779 | -2.6087724 | 0.00908677 | 0.13583969 |
| FAM198B | 364.429804 | -0.690850391 | 0.60715398 | -1.1378504 | 0.25518296 | 0.6208774 |
| MYO5C | 175.175072 | -0.69036858 | 0.86516724 | -0.7979597 | 0.42489388 | 0.74186041 |
| HIVEP2 | 761.12527 | -0.68876538 | 0.48871729 | -1.4093329 | 0.15873674 | 0.51586289 |
| ZFP36 | 238.330419 | -0.688764897 | 1.00391307 | -0.6860802 | 0.49266252 | 0.79102628 |
| NKIRAS2 | 45.6148188 | -0.688648994 | 0.35822776 | -1.9223775 | 0.05455827 | 0.32727552 |
| DDB2 | 113.286044 | -0.687923899 | 0.52014374 | -1.322565 | 0.18598008 | 0.55200399 |
| IGFBP7 | 2567.15011 | -0.687307791 | 0.5058827 | -1.3586307 | 0.17426363 | 0.53666198 |
| KCTD12 | 601.090419 | -0.686758876 | 0.56522575 | -1.215017 | 0.22435962 | 0.59083342 |
| BLVRB | 81.6733408 | -0.686591425 | 0.34372441 | -1.9975056 | 0.04577029 | 0.30185088 |
| RGS7BP | 110.59646 | -0.686204082 | 0.66268494 | -1.0354907 | 0.30043982 | 0.65926326 |
| EIF4B | 218.833228 | -0.685892112 | 0.20098524 | -3.4126492 | 0.00064335 | 0.02770794 |
| CDC42EP2 | 49.6281604 | -0.685394033 | 0.44483464 | -1.5407838 | 0.1233694 | 0.46870024 |
| ELOVL1 | 63.0288875 | -0.68520913 | 0.38774539 | -1.7671625 | 0.07720101 | 0.38356763 |
| PRMT3 | 111.066533 | -0.685125263 | 0.23037622 | -2.9739409 | 0.00294002 | 0.07188756 |
| MGAT4B | 70.8767139 | -0.684572185 | 0.36435223 | -1.8788747 | 0.0602616 | 0.34478399 |
| APH1B | 62.2977772 | -0.683902141 | 0.46276943 | -1.4778464 | 0.1394489 | 0.49091481 |
| CASC15 | 641.788558 | -0.68343877 | 0.41823926 | -1.6340856 | 0.10224087 | 0.43132417 |
| PIK3R1 | 221.825892 | -0.682958136 | 0.28104363 | -2.4300787 | 0.01509555 | 0.17646692 |
| AC007563.2 | 219.08366 | -0.682525202 | 0.56998352 | -1.1974473 | 0.23113227 | 0.59758464 |
| KIZ-AS1 | 42.9362634 | -0.682214755 | 0.48272313 | -1.413263 | 0.1575784 | 0.51438736 |
| CKAP4 | 61.1743265 | -0.681323381 | 0.42994787 | -1.5846651 | 0.11304245 | 0.45079257 |
| HNRNPA0 | 137.202179 | -0.680311043 | 0.39811893 | -1.7088136 | 0.08748549 | 0.40016418 |
| HSCB | 60.5393233 | -0.680107489 | 0.2697151 | -2.5215773 | 0.011683 | 0.15162005 |
| NUDCD2 | 44.487795 | -0.679928648 | 0.36067235 | -1.8851699 | 0.05940691 | 0.34186127 |
| TIMM9 | 51.9080219 | -0.679071584 | 0.26467822 | -2.5656496 | 0.01029828 | 0.14331772 |
| LDHB | 169.440129 | -0.67813046 | 0.38325857 | -1.7693811 | 0.0768303 | 0.38356763 |
| NSMCE1 | 63.6881133 | -0.677581727 | 0.32406377 | -2.0908901 | 0.03653792 | 0.26964679 |
| C1orf122 | 59.989181 | -0.677536354 | 0.34495809 | -1.9641121 | 0.04951709 | 0.31166266 |
| TPM4 | 501.42088 | -0.676622842 | 0.30030286 | -2.2531348 | 0.02425065 | 0.21687765 |
| TMSB4X | 2749.07695 | -0.676006353 | 0.58937802 | -1.1469826 | 0.2513888 | 0.61661694 |
| SYTL3 | 121.030922 | -0.675266437 | 0.43800045 | -1.5417026 | 0.12314588 | 0.46855087 |
| ATP6V0E1 | 363.336201 | -0.675126368 | 0.30339552 | -2.2252351 | 0.02606546 | 0.22518506 |
| LHX6 | 46.4937382 | -0.674906808 | 0.68998306 | -0.9781498 | 0.32800023 | 0.68053395 |
| TTF2 | 45.1640183 | -0.674209476 | 0.38377518 | -1.7567824 | 0.07895491 | 0.38686531 |
| RPL30 | 456.868074 | -0.67345138 | 0.52640578 | -1.2793389 | 0.20077775 | 0.56536969 |
| GRN | 227.295597 | -0.672928602 | 0.42675325 | -1.5768564 | 0.11482856 | 0.45569454 |
| CYB5R3 | 272.80596 | -0.671427262 | 0.39325087 | -1.7073764 | 0.08775211 | 0.40026508 |
| SNAPC1 | 96.6591308 | -0.670972896 | 0.43052252 | -1.5585082 | 0.11911282 | 0.46290743 |
| GTPBP4 | 69.9061644 | -0.670448409 | 0.34865625 | -1.9229496 | 0.05448637 | 0.32727552 |
| NOP10 | 145.212003 | -0.670286427 | 0.53116691 | -1.261913 | 0.20698009 | 0.57162978 |
| PLIN3 | 49.7999512 | -0.670087844 | 0.32846868 | -2.0400357 | 0.04134677 | 0.28834707 |
| EIF3D | 116.153267 | -0.669810047 | 0.33007502 | -2.0292661 | 0.0424312 | 0.291241 |
| RPS3A | 194.754555 | -0.668979449 | 0.49375818 | -1.3548727 | 0.17545813 | 0.53854985 |
| AP002884.1 | 66.0055491 | -0.667879311 | 0.37910511 | -1.7617259 | 0.07811561 | 0.38566641 |
| RPS24 | 567.340486 | -0.667187082 | 0.5167689 | -1.2910744 | 0.19667789 | 0.56172669 |
| CLMN | 62.6756395 | -0.665746855 | 0.58435304 | -1.1392888 | 0.25458273 | 0.62019961 |
| PPDPF | 150.938971 | -0.665453436 | 0.4170287 | -1.5957018 | 0.11055539 | 0.44631216 |

| | | | | | | |
|---|---|---|---|---|---|---|
| RPL3 | 340.154832 | -0.664834521 | 0.53297188 | -1.2474101 | 0.21224716 | 0.57690664 |
| NDUFS5 | 163.862073 | -0.664186255 | 0.38311099 | -1.7336654 | 0.08297747 | 0.39515849 |
| NLGN4Y | 159.484895 | -0.664040301 | 1.86583529 | -0.3558944 | 0.72191966 | 0.90162538 |
| RPLP1 | 723.174537 | -0.663099118 | 0.54040242 | -1.2270469 | 0.21980496 | 0.58687522 |
| ZBTB7C | 207.817278 | -0.66265175 | 0.4883815 | -1.3568322 | 0.17483453 | 0.53784624 |
| GARS | 100.152389 | -0.66218271 | 0.3413904 | -1.9396641 | 0.05242052 | 0.32059696 |
| FARSA | 41.7243151 | -0.66201943 | 0.42821301 | -1.5460049 | 0.12210337 | 0.46733953 |
| TMTC4 | 61.2160581 | -0.661183484 | 0.38700493 | -1.7084627 | 0.08755053 | 0.40023797 |
| SSRP1 | 66.6484308 | -0.660372171 | 0.39016513 | -1.6925453 | 0.09054204 | 0.40709096 |
| RPS14 | 507.674723 | -0.659641585 | 0.48743694 | -1.353286 | 0.17596428 | 0.53949634 |
| ERRFI1 | 98.4131324 | -0.659518718 | 0.62006494 | -1.0636285 | 0.28749704 | 0.65078515 |
| IPMK | 127.242155 | -0.659444815 | 0.37267847 | -1.7694738 | 0.07681483 | 0.38356763 |
| DEDD | 63.7019262 | -0.658354899 | 0.27578064 | -2.3872412 | 0.01697535 | 0.18593697 |
| CYREN | 65.5087607 | -0.657778215 | 0.22127381 | -2.972689 | 0.00295203 | 0.07188756 |
| CPD | 319.295672 | -0.657511631 | 0.37317682 | -1.7619305 | 0.07808104 | 0.38566641 |
| GOLM1 | 145.386339 | -0.657358316 | 0.53869239 | -1.2202851 | 0.2223568 | 0.58808846 |
| PSMD7 | 119.519226 | -0.657116576 | 0.31887395 | -2.0607408 | 0.03932777 | 0.28057468 |
| UBE2M | 64.4405863 | -0.657030883 | 0.42945266 | -1.529925 | 0.12603528 | 0.47177251 |
| TMEM101 | 46.456959 | -0.656945595 | 0.33768722 | -1.9454263 | 0.05172369 | 0.31875077 |
| MSRB2 | 56.4218495 | -0.655962193 | 0.36873211 | -1.7789668 | 0.0752452 | 0.38047182 |
| TTC7A | 73.163466 | -0.654201256 | 0.29440525 | -2.2221114 | 0.02627578 | 0.22538227 |
| AFMID | 96.8151541 | -0.654194553 | 0.50983982 | -1.2831374 | 0.19944394 | 0.56449294 |
| KIAA1549L | 81.7151849 | -0.654146833 | 0.50995618 | -1.2827511 | 0.1995793 | 0.56449294 |
| TMED10 | 219.817376 | -0.653811368 | 0.32598122 | -2.0056719 | 0.04489127 | 0.29946125 |
| PPT1 | 69.1288421 | -0.653456678 | 0.40364938 | -1.618872 | 0.10547481 | 0.43812201 |
| SRP14 | 1020.35443 | -0.653341196 | 0.67566378 | -0.966962 | 0.33356303 | 0.6852158 |
| TSPAN5 | 119.558819 | -0.652607709 | 0.44911174 | -1.4531077 | 0.14619387 | 0.50011031 |
| RALGAPA2 | 2623.2909 | -0.652021686 | 0.2471545 | -2.6381138 | 0.00833686 | 0.12969682 |
| DDOST | 78.4078587 | -0.651698171 | 0.37548444 | -1.7356197 | 0.08263111 | 0.39477007 |
| FKBP10 | 59.078969 | -0.651237592 | 0.37980187 | -1.7146772 | 0.08640443 | 0.39864674 |
| DEGS1 | 106.171247 | -0.651215756 | 0.40050252 | -1.6259966 | 0.10395037 | 0.43532543 |
| LINC00513 | 151.983835 | -0.650091278 | 0.53544738 | -1.2141086 | 0.22470625 | 0.59105472 |
| CANX | 391.905026 | -0.649918227 | 0.36678651 | -1.7719251 | 0.07640699 | 0.38287714 |
| EIF1B | 87.7886119 | -0.649687443 | 0.37985315 | -1.7103648 | 0.08719844 | 0.39981956 |
| BNC2 | 530.01422 | -0.64947363 | 0.55295303 | -1.1745548 | 0.24017288 | 0.60471836 |
| IL15RA | 323.953129 | -0.649414399 | 0.33356534 | -1.9468881 | 0.05154815 | 0.31859402 |
| PCSK6 | 149.580197 | -0.649076331 | 0.6291232 | -1.0317158 | 0.30220529 | 0.65926326 |
| NDUFAF2 | 222.37951 | -0.64859877 | 0.27309966 | -2.3749527 | 0.0175512 | 0.18854202 |
| CCDC137 | 47.2671664 | -0.648203375 | 0.28516963 | -2.2730449 | 0.02302347 | 0.21247788 |
| TEAD2 | 117.214144 | -0.648194503 | 0.34967812 | -1.8536891 | 0.06378366 | 0.3533385 |
| GJA1 | 298.532243 | -0.648161021 | 0.5274598 | -1.2288349 | 0.21913371 | 0.58619521 |
| COX7B | 132.602786 | -0.648019163 | 0.38543151 | -1.6812823 | 0.0927081 | 0.41185145 |
| NCBP2 | 57.8866085 | -0.647945549 | 0.29764127 | -2.1769345 | 0.02948546 | 0.2381831 |
| PEA15 | 202.493271 | -0.647832347 | 0.37901362 | -1.7092587 | 0.08740303 | 0.40003816 |
| SLC39A7 | 60.0145712 | -0.647131091 | 0.44673223 | -1.4485883 | 0.14745258 | 0.50134527 |
| PHYKPL | 277.449326 | -0.647062455 | 0.3317519 | -1.9504408 | 0.0511236 | 0.31740851 |
| RPL18 | 360.734384 | -0.646843332 | 0.4754769 | -1.3604096 | 0.17370035 | 0.53536344 |
| FRMD8 | 149.668124 | -0.645222817 | 0.30959607 | -2.0840795 | 0.03715294 | 0.27144867 |
| RPL34 | 296.553753 | -0.644162695 | 0.5133739 | -1.2547632 | 0.20956472 | 0.57507314 |
| PPM1G | 191.007349 | -0.643959428 | 0.33986902 | -1.8947282 | 0.05812842 | 0.33831183 |
| ARID5A | 110.626513 | -0.643957353 | 0.45397168 | -1.4184968 | 0.15604579 | 0.51234409 |
| LAMTOR1 | 121.579483 | -0.643057436 | 0.31678659 | -2.029939 | 0.04236274 | 0.291241 |
| EIF3B | 173.431398 | -0.642449334 | 0.27315155 | -2.3519886 | 0.01867335 | 0.19394842 |
| EIF5B | 230.076902 | -0.642154455 | 0.33407901 | -1.9221634 | 0.0545852 | 0.32727552 |
| AC097515.1 | 44.5760007 | -0.641752763 | 0.62479594 | -1.0271398 | 0.30435465 | 0.66086613 |
| YIPF5 | 68.7092673 | -0.641624305 | 0.32066266 | -2.0009324 | 0.04539968 | 0.30053976 |
| SHROOM2 | 282.980439 | -0.641377341 | 0.42203802 | -1.5197146 | 0.12858272 | 0.47653642 |
| FNDC3B | 2135.61964 | -0.640839534 | 0.29065506 | -2.2048112 | 0.02746735 | 0.23076519 |
| CCNY | 2142.44503 | -0.640746351 | 0.40974404 | -1.5637722 | 0.11787107 | 0.46042456 |
| MRPS25 | 84.2275359 | -0.640710783 | 0.26525216 | -2.4154781 | 0.01571457 | 0.17984936 |
| UBB | 425.050317 | -0.64004915 | 0.45952003 | -1.3928645 | 0.16366076 | 0.52357299 |
| RPS11 | 405.162516 | -0.639520866 | 0.51020444 | -1.25346 | 0.21003834 | 0.57579356 |
| MRPS7 | 49.6864571 | -0.638731634 | 0.44901422 | -1.4225198 | 0.1548754 | 0.51082038 |
| BCAR1 | 191.732132 | -0.637768481 | 0.29462398 | -2.1646862 | 0.03041172 | 0.24194918 |
| OST4 | 107.327298 | -0.637413486 | 0.57135399 | -1.1156192 | 0.26458517 | 0.63018314 |
| RABGGTB | 58.085089 | -0.63718036 | 0.44910925 | -1.4187647 | 0.15596762 | 0.51234409 |
| PDE7B | 823.274393 | -0.636096026 | 0.47281586 | -1.3453356 | 0.17851687 | 0.54244469 |
| PSMC4 | 50.4702381 | -0.635845449 | 0.37084704 | -1.714576 | 0.08642299 | 0.39864674 |
| TPP1 | 102.351021 | -0.635034802 | 0.42597388 | -1.4907834 | 0.13601836 | 0.48583075 |
| XXYLT1 | 165.369456 | -0.634879659 | 0.26981944 | -2.3529797 | 0.01862365 | 0.19394842 |
| PECAM1 | 3023.74571 | -0.634836709 | 0.31185058 | -2.035708 | 0.04177969 | 0.28963374 |
| SLC35E4 | 62.9508213 | -0.634824415 | 0.34783052 | -1.8250969 | 0.0679864 | 0.36125501 |
| PCTP | 54.6277156 | -0.634406214 | 0.46485779 | -1.3647318 | 0.17233736 | 0.53438295 |
| MYRIP | 388.148182 | -0.634220765 | 0.62488668 | -1.0149372 | 0.31013574 | 0.6648266 |
| CTDSPL | 188.885893 | -0.632857052 | 0.40406402 | -1.5662297 | 0.11729486 | 0.46012647 |
| OSBPL10 | 484.425164 | -0.632023711 | 0.34272078 | -1.8441359 | 0.06516333 | 0.35535588 |
| CNN3 | 867.617674 | -0.631708514 | 0.49268094 | -1.2821858 | 0.19977747 | 0.56449294 |
| MKL2 | 550.099492 | -0.6315098 | 0.47295646 | -1.3352388 | 0.18179821 | 0.5467309 |
| FGF13 | 117.058575 | -0.630908195 | 0.66313563 | -0.9514014 | 0.34140063 | 0.69116603 |
| CLEC2B | 274.758415 | -0.630544531 | 0.60866633 | -1.0359445 | 0.30022804 | 0.65926326 |
| CBR1 | 59.146264 | -0.630404824 | 0.39556664 | -1.5936754 | 0.11100876 | 0.44762779 |
| PPP1R14B | 51.1212162 | -0.629562154 | 0.39534211 | -1.592449 | 0.11128385 | 0.44770684 |
| SEC61A1 | 182.721674 | -0.629346583 | 0.37349497 | -1.6850202 | 0.09198467 | 0.41019648 |
| NOP58 | 138.712398 | -0.628944766 | 0.19126391 | -3.2883608 | 0.00100773 | 0.03731322 |
| ANAPC11 | 105.177926 | -0.626467395 | 0.48889215 | -1.281402 | 0.20005249 | 0.56449294 |
| PTMS | 385.056695 | -0.626320161 | 0.5014584 | -1.2489972 | 0.21166608 | 0.57690664 |
| NR2F1-AS1 | 598.010625 | -0.626189021 | 0.30572465 | -2.0482124 | 0.04053919 | 0.28580837 |
| SSBP3 | 442.753356 | -0.625951786 | 0.49476639 | -1.2651461 | 0.20581895 | 0.57060151 |
| PGM2 | 67.4133571 | -0.62589293 | 0.25306315 | -2.4732678 | 0.01338838 | 0.16599561 |
| ABHD5 | 86.8563224 | -0.625379833 | 0.31309075 | -1.9974395 | 0.04577746 | 0.30185088 |
| FNBP1L | 465.822106 | -0.624715964 | 0.33869857 | -1.84446 | 0.06511613 | 0.35535588 |
| UBXN6 | 57.7941132 | -0.624630853 | 0.36871814 | -1.6940606 | 0.09025378 | 0.4068172 |
| GFOD1 | 745.120865 | -0.624510039 | 0.31088087 | -2.0088404 | 0.04455406 | 0.29933161 |
| RPSA | 297.788297 | -0.624367142 | 0.43591418 | -1.4323167 | 0.15205321 | 0.50814312 |
| NARF | 193.927267 | -0.623425867 | 0.22288122 | -2.7971216 | 0.00515601 | 0.09916296 |
| CCDC85B | 432.270754 | -0.623267644 | 0.55117684 | -1.1307943 | 0.25814167 | 0.62293521 |
| POLDIP2 | 46.4275165 | -0.62296286 | 0.29970283 | -2.0786018 | 0.03765396 | 0.27413022 |
| USP36 | 198.036483 | -0.621502916 | 0.40354158 | -1.5401211 | 0.12353083 | 0.4690732 |
| EI24 | 95.361619 | -0.620843446 | 0.29189944 | -2.1269086 | 0.03342768 | 0.25516667 |
| ANKRD29 | 91.312726 | -0.620676691 | 1.05481441 | -0.5884227 | 0.55624864 | 0.82504669 |
| SIK2 | 695.765679 | -0.620598698 | 0.39521384 | -1.5702858 | 0.11634863 | 0.4592091 |
| MTRNR2L8 | 97.0804134 | -0.619426262 | 0.65769598 | -0.9418124 | 0.34628867 | 0.69521104 |
| PUF60 | 70.3337233 | -0.619399164 | 0.46540985 | -1.3308682 | 0.18322238 | 0.54813352 |
| SPNS2 | 180.167729 | -0.61919006 | 0.49093497 | -1.2612466 | 0.20722002 | 0.5718993 |
| SLC25A3 | 221.823208 | -0.618234775 | 0.39407737 | -1.5688157 | 0.11669088 | 0.45929845 |
| RPS28 | 160.488686 | -0.617924392 | 0.48042917 | -1.2861925 | 0.19837589 | 0.56364928 |
| CHURC1-FNT | 43.5655948 | -0.617861065 | 0.33410322 | -1.8493119 | 0.06441278 | 0.35347506 |
| MET | 145.171873 | -0.61780799 | 0.37485265 | -1.6481356 | 0.09932485 | 0.42376187 |
| TNKS1BP1 | 106.625638 | -0.61768845 | 0.21085014 | -2.929514 | 0.00339492 | 0.077817 |
| B4GALT3 | 38.9893113 | -0.617245019 | 0.45051047 | -1.3701014 | 0.17065526 | 0.53193492 |
| RNF215 | 44.8892007 | -0.617067438 | 0.2814472 | -2.1924803 | 0.02834485 | 0.23303779 |
| EMC4 | 49.9567436 | -0.616400853 | 0.38342954 | -1.6075988 | 0.10792308 | 0.44289597 |
| TSR3 | 45.5849333 | -0.615430909 | 0.40370426 | -1.5244598 | 0.12739389 | 0.47449441 |

| | | | | | | |
|---|---|---|---|---|---|---|
| RIOX2 | 48.6813724 | -0.615104318 | 0.28368043 | -2.1683001 | 0.03013586 | 0.24082201 |
| RCL1 | 121.790685 | -0.615097759 | 0.2625671 | -2.3426307 | 0.01914833 | 0.1947132 |
| ADI1 | 66.3158706 | -0.61438674 | 0.38025923 | -1.615705 | 0.10615811 | 0.43901385 |
| PLOD1 | 100.763717 | -0.614161973 | 0.3505555 | -1.7519679 | 0.07977933 | 0.38738328 |
| LACTB2 | 54.1908293 | -0.613658019 | 0.35660093 | -1.7208537 | 0.08527738 | 0.39686932 |
| MYEF2 | 64.6948359 | -0.61283264 | 0.64416388 | -0.9513614 | 0.34142096 | 0.69116603 |
| IFNAR1 | 222.243256 | -0.612438251 | 0.24699856 | -2.4795216 | 0.01315588 | 0.16347517 |
| IFRD2 | 41.6697804 | -0.612335472 | 0.44253522 | -1.3836988 | 0.16645069 | 0.52671378 |
| ADAT1 | 48.0292559 | -0.612311257 | 0.26822978 | -2.2827862 | 0.02244296 | 0.2108594 |
| BASP1 | 62.1757946 | -0.611890222 | 0.74122132 | -0.8255162 | 0.40907858 | 0.73419295 |
| AGPAT2 | 41.6450062 | -0.611766419 | 0.29925297 | -2.0443119 | 0.04092274 | 0.28670444 |
| LSM7 | 118.305286 | -0.611545905 | 0.46567738 | -1.3132394 | 0.18910228 | 0.55483111 |
| FAM198B-AS | 48.3123027 | -0.611373899 | 0.64094987 | -0.953856 | 0.34015652 | 0.69059971 |
| LRRC59 | 48.854484 | -0.61109007 | 0.45090492 | -1.3552526 | 0.17533708 | 0.53847091 |
| URB1 | 75.0612661 | -0.610744711 | 0.44511297 | -1.3721117 | 0.17002867 | 0.53193492 |
| EPHA3 | 154.539081 | -0.610725123 | 1.0205566 | -0.5984236 | 0.54955734 | 0.82355262 |
| RPL12 | 92.7415098 | -0.61062761 | 0.4026048 | -1.5166923 | 0.12934438 | 0.47762865 |
| TIMM17A | 48.2904727 | -0.610479345 | 0.35871375 | -1.7018565 | 0.08878226 | 0.40361404 |
| HIPK3 | 884.92984 | -0.610322874 | 0.32259699 | -1.8919051 | 0.05850363 | 0.3390476 |
| SNRPD1 | 69.1390501 | -0.610184465 | 0.39663721 | -1.5383944 | 0.12395221 | 0.46936415 |
| FSTL1 | 175.677548 | -0.609380453 | 0.3342027 | -1.8233858 | 0.06824498 | 0.36142549 |
| HTRA1 | 195.794953 | -0.609216114 | 0.65949338 | -0.9237638 | 0.3556093 | 0.70018061 |
| REXO2 | 80.4509542 | -0.608908216 | 0.39091325 | -1.5576556 | 0.11931491 | 0.46290743 |
| STAT3 | 990.212112 | -0.6087677 | 0.27769593 | -2.1922097 | 0.02836437 | 0.23303779 |
| HSPB8 | 47.0554497 | -0.607335814 | 0.35577826 | -1.7070627 | 0.08781039 | 0.40030777 |
| SEC61G | 84.9679887 | -0.607011622 | 0.49253267 | -1.2324291 | 0.21778882 | 0.58393378 |
| PSMB1 | 221.14963 | -0.60668673 | 0.41374629 | -1.4663255 | 0.14255964 | 0.49489199 |
| STAU2 | 592.658271 | -0.606394393 | 0.2770511 | -2.1887457 | 0.02861533 | 0.23392531 |
| RBP7 | 390.061938 | -0.606233201 | 1.16000279 | -0.5226136 | 0.6012432 | 0.8499944 |
| SERPINH1 | 255.858293 | -0.606080567 | 0.52500405 | -1.1544303 | 0.24832381 | 0.61372207 |
| ELF4 | 53.2976415 | -0.60593793 | 0.21890346 | -2.76806 | 0.00563911 | 0.10300181 |
| RBM8A | 112.322663 | -0.605797309 | 0.28998946 | -2.0890321 | 0.03670483 | 0.2703471 |
| CD81 | 767.372064 | -0.60525452 | 0.42441858 | -1.4260792 | 0.15384548 | 0.50978388 |
| LGALS9 | 81.1013884 | -0.604706396 | 0.58491779 | -1.0338314 | 0.30121498 | 0.65926326 |
| DLL4 | 260.359999 | -0.604398241 | 0.32374659 | -1.8668868 | 0.06191739 | 0.3489463 |
| ALDH2 | 73.5174129 | -0.604105992 | 0.4928782 | -1.2256699 | 0.22032291 | 0.58688228 |
| PLEKHG2 | 65.6586025 | -0.604045854 | 0.34771849 | -1.7371692 | 0.08235731 | 0.39457252 |
| HSD17B10 | 41.8663368 | -0.60401567 | 0.49701044 | -1.2152977 | 0.22425255 | 0.59083342 |
| IDI1 | 66.8044944 | -0.603565182 | 0.36489567 | -1.654076 | 0.09811207 | 0.42166547 |
| ASAP3 | 72.0756754 | -0.603429108 | 0.35963599 | -1.6778886 | 0.09336886 | 0.41231599 |
| RIOK1 | 54.7407627 | -0.602877719 | 0.32543382 | -1.8525355 | 0.06394896 | 0.3533385 |
| NAT10 | 74.2553519 | -0.602827783 | 0.35126452 | -1.7161647 | 0.0861319 | 0.39864674 |
| DNAJC5 | 287.919818 | -0.602567317 | 0.33394005 | -1.8044176 | 0.07116586 | 0.36904326 |
| SNU13 | 225.539435 | -0.60194816 | 0.32516834 | -1.8511893 | 0.06414232 | 0.35347506 |
| ENOSF1 | 318.406646 | -0.601308911 | 0.33100829 | -1.8165978 | 0.06927873 | 0.36386894 |
| PPP1CB | 218.369942 | -0.601063236 | 0.39517904 | -1.5209897 | 0.12826243 | 0.47578036 |
| CNDP2 | 207.765556 | -0.599807447 | 0.25772265 | -2.3273369 | 0.01994734 | 0.19732169 |
| HS3ST1 | 113.8745 | -0.599536693 | 0.58195012 | -1.0302201 | 0.30290671 | 0.65940027 |
| MBTPS2 | 60.3066668 | -0.599451051 | 0.25213263 | -2.3775227 | 0.01742937 | 0.18853893 |
| SLC30A1 | 72.8026944 | -0.598553576 | 0.45283521 | -1.3217912 | 0.1862377 | 0.55236793 |
| RSL24D1 | 69.9326544 | -0.59848447 | 0.30101024 | -1.9882528 | 0.04678373 | 0.30359341 |
| HLA-B | 1953.62613 | -0.598323652 | 0.4666475 | -1.2821748 | 0.19978134 | 0.56449294 |
| ANKRD16 | 42.3179295 | -0.59817689 | 0.4093302 | -1.4613554 | 0.14391794 | 0.49679586 |
| MEDAG | 55.6959898 | -0.597677117 | 0.52549981 | -1.1373498 | 0.25539207 | 0.6208774 |
| ATP5MC1 | 73.5334736 | -0.597557812 | 0.38061777 | -1.5699682 | 0.1164225 | 0.4592091 |
| UGP2 | 194.017001 | -0.597398215 | 0.27039756 | -2.2093329 | 0.0271515 | 0.22976287 |
| PALMD | 634.550133 | -0.596928479 | 0.49717931 | -1.2006302 | 0.22989469 | 0.59648134 |
| GPS2 | 75.5445241 | -0.596686285 | 0.35704475 | -1.6711807 | 0.094686 | 0.41389718 |
| GMEB2 | 52.0338194 | -0.596682892 | 0.28117222 | -2.1221261 | 0.03382715 | 0.25638251 |
| ATP6V1B2 | 85.5997369 | -0.596559923 | 0.32190367 | -1.853225 | 0.06385012 | 0.3533385 |
| CALR | 320.305175 | -0.596499436 | 0.53676716 | -1.1112815 | 0.26644718 | 0.63198183 |
| HMOX2 | 79.5945796 | -0.596476319 | 0.50669483 | -1.1771905 | 0.2391195 | 0.60299379 |
| RRP15 | 59.9171148 | -0.595923477 | 0.29525125 | -2.0183605 | 0.04355373 | 0.29561649 |
| PSMB3 | 117.55927 | -0.595115463 | 0.37095775 | -1.6042675 | 0.1086551 | 0.44474428 |
| SMDT1 | 70.2526887 | -0.595073613 | 0.37477432 | -1.5878185 | 0.11232738 | 0.44910158 |
| SHROOM1 | 80.0442383 | -0.594893392 | 0.64828769 | -0.917638 | 0.35880844 | 0.70242891 |
| FAM96B | 96.7540465 | -0.594803841 | 0.43189401 | -1.3771986 | 0.16845084 | 0.53037061 |
| KCTD10 | 89.7431036 | -0.593931651 | 0.31330802 | -1.8956797 | 0.05800241 | 0.33805823 |
| ADGRE3 | 43.3556215 | -0.593857314 | 0.38456507 | -1.544231 | 0.12253237 | 0.46802258 |
| SND1 | 439.843072 | -0.593241628 | 0.30492437 | -1.9455369 | 0.05171039 | 0.31875077 |
| TMED9 | 145.974129 | -0.593035566 | 0.46692832 | -1.2700784 | 0.20405671 | 0.56856072 |
| MRPL43 | 41.1457683 | -0.592860956 | 0.38647605 | -1.5340173 | 0.12502539 | 0.47038289 |
| EIF2S1 | 114.471118 | -0.59246816 | 0.28022817 | -2.1142348 | 0.03449522 | 0.2592051 |
| MRPS15 | 41.6175952 | -0.591731754 | 0.3506056 | -1.6877419 | 0.09146079 | 0.40875132 |
| RNF144B | 712.559001 | -0.591563382 | 0.38965275 | -1.518181 | 0.12896879 | 0.47688729 |
| MICALL2 | 67.9804892 | -0.591057084 | 0.25160645 | -2.3491332 | 0.01881717 | 0.19394842 |
| BFAR | 105.394882 | -0.589944805 | 0.23877051 | -2.4707608 | 0.0134826 | 0.16640738 |
| PPME1 | 153.837592 | -0.589722088 | 0.22684747 | -2.5996414 | 0.00933212 | 0.13676431 |
| RPS29 | 194.838843 | -0.589715539 | 0.40596389 | -1.4526305 | 0.14632637 | 0.50016236 |
| ABCE1 | 118.236754 | -0.589176644 | 0.29044743 | -2.0285139 | 0.04250783 | 0.291241 |
| TMEM241 | 181.77657 | -0.589120333 | 0.2949156 | -1.9975896 | 0.04576117 | 0.30185088 |
| SLIRP | 106.491764 | -0.58790764 | 0.19782845 | -2.9718052 | 0.00296054 | 0.07188756 |
| SEMA3F | 389.914485 | -0.587740792 | 0.34863734 | -1.6858229 | 0.09182992 | 0.40973546 |
| GPX3 | 1024.89623 | -0.58771298 | 0.59035115 | -0.9955312 | 0.31947798 | 0.67395826 |
| PRKAR1A | 343.252879 | -0.587601879 | 0.30722673 | -1.9126001 | 0.05579927 | 0.32988018 |
| SPARC | 1167.51441 | -0.587134459 | 0.53454548 | -1.0983807 | 0.27203827 | 0.63693538 |
| GTF2H3 | 43.3255725 | -0.587024293 | 0.20670718 | -2.8398834 | 0.004513 | 0.09141064 |
| RPN1 | 141.951705 | -0.586944521 | 0.27741595 | -2.1157562 | 0.03436555 | 0.25870587 |
| ADAMTSL1 | 404.27455 | -0.586386734 | 0.64804272 | -0.9048582 | 0.36554054 | 0.70668224 |
| ELMO1-AS1 | 66.8424475 | -0.586304416 | 0.4493729 | -1.3047169 | 0.19198926 | 0.55646778 |
| PKP4 | 2188.83945 | -0.585936496 | 0.33230458 | -1.7632513 | 0.07785811 | 0.38566158 |
| COMT | 167.749829 | -0.585709472 | 0.28206219 | -2.076526 | 0.03784533 | 0.27527852 |
| ATP6V1G1 | 159.229487 | -0.585624556 | 0.39216747 | -1.4933022 | 0.1353581 | 0.48508022 |
| CMAS | 69.4331485 | -0.585182657 | 0.21806771 | -2.6834906 | 0.0072858 | 0.12102126 |
| KIAA1522 | 69.5605508 | -0.584557162 | 0.37536335 | -1.5573101 | 0.11939687 | 0.46290743 |
| ZDHHC24 | 53.5597107 | -0.58451009 | 0.33049979 | -1.7685642 | 0.07696663 | 0.38356763 |
| WDR43 | 156.91954 | -0.584369433 | 0.32909085 | -1.7757086 | 0.07578097 | 0.38160963 |
| C19orf12 | 42.5489306 | -0.58416136 | 0.26932551 | -2.1689789 | 0.03008428 | 0.24082201 |
| CCNG1 | 49.7790872 | -0.583629728 | 0.30136474 | -1.9366225 | 0.05279151 | 0.3215522 |
| TNFRSF10A | 121.855122 | -0.583146841 | 0.41645726 | -1.4002561 | 0.16143663 | 0.52009288 |
| ESYT2 | 673.202467 | -0.58270894 | 0.51810971 | -1.1246825 | 0.26072358 | 0.62580682 |
| PHPT1 | 95.9389883 | -0.582705523 | 0.4002985 | -1.4556775 | 0.14548179 | 0.49835302 |
| RPS27 | 109.176373 | -0.58243387 | 0.49931406 | -1.166468 | 0.24342528 | 0.608518 |
| TTTY14 | 398.89677 | -0.582312879 | 1.49657154 | -0.3890979 | 0.69720371 | 0.89283536 |
| ATP8A1 | 279.966583 | -0.581824278 | 0.69926503 | -0.8320512 | 0.40538007 | 0.73260271 |
| PDIA4 | 80.9731501 | -0.581634107 | 0.43133644 | -1.3484465 | 0.17751482 | 0.54070521 |
| AFAP1 | 578.212089 | -0.581364892 | 0.34159758 | -1.7018999 | 0.08877413 | 0.40361404 |
| PRDM1 | 85.9599724 | -0.580906857 | 0.68596929 | -0.8468409 | 0.3970838 | 0.72680823 |
| RELN | 166.168205 | -0.580411827 | 1.09004751 | -0.5324647 | 0.59440418 | 0.84547356 |
| CDC37 | 378.72677 | -0.58023801 | 0.2900646 | -2.0003751 | 0.04545977 | 0.30053976 |
| SAP18 | 147.997437 | -0.579920581 | 0.42341861 | -1.3696152 | 0.17080704 | 0.53193492 |
| AHSA1 | 66.3382249 | -0.579322942 | 0.27119952 | -2.1361503 | 0.03266716 | 0.25224169 |
| SLC35B4 | 74.8099467 | -0.579153828 | 0.2675604 | -2.1645723 | 0.03042046 | 0.24194918 |

| | | | | | | |
|---|---|---|---|---|---|---|
| PES1 | 87.5857398 | -0.578952742 | 0.24685216 | -2.345342 | 0.01900963 | 0.19411645 |
| OSBP2 | 75.7535472 | -0.578615939 | 0.41239421 | -1.4030652 | 0.16059741 | 0.51944206 |
| NDUFA2 | 71.6242469 | -0.577531026 | 0.43338685 | -1.3325993 | 0.18266333 | 0.54790102 |
| SELENOW | 339.704092 | -0.577393541 | 0.51534502 | -1.1204019 | 0.26254254 | 0.62818292 |
| ATP5MC2 | 218.375523 | -0.577147786 | 0.32786226 | -1.7603361 | 0.07835083 | 0.38566641 |
| UQCRB | 158.728232 | -0.576968614 | 0.4993058 | -1.1555416 | 0.24786871 | 0.61296757 |
| ANGEL1 | 126.718925 | -0.57641909 | 0.32037027 | -1.7992278 | 0.07198266 | 0.37116201 |
| LYRM1 | 67.4582008 | -0.575862574 | 0.38185969 | -1.5080476 | 0.13154235 | 0.48069696 |
| C8orf33 | 45.6566331 | -0.575654066 | 0.38783719 | -1.4842673 | 0.13773802 | 0.48962215 |
| RPS20 | 242.44601 | -0.575650703 | 0.45799542 | -1.2568918 | 0.2087928 | 0.57372449 |
| HSPA9 | 195.539434 | -0.575217042 | 0.26446849 | -2.1749927 | 0.02963066 | 0.23885783 |
| COPE | 73.8869884 | -0.574709704 | 0.40434199 | -1.4213456 | 0.15521632 | 0.51129169 |
| C8orf59 | 61.0047063 | -0.574401783 | 0.25836057 | -2.2232563 | 0.02619852 | 0.22519168 |
| TPM3 | 356.378003 | -0.57418334 | 0.2440584 | -2.3526474 | 0.0186403 | 0.19394842 |
| UMPS | 42.9271833 | -0.573813793 | 0.26043257 | -2.2033104 | 0.02757288 | 0.23094047 |
| CES2 | 48.0309753 | -0.573210662 | 0.34333703 | -1.6695276 | 0.09501286 | 0.41466145 |
| BEX4 | 49.3795984 | -0.573103602 | 0.42023299 | -1.3637758 | 0.17263813 | 0.53470772 |
| PSMB6 | 96.9580682 | -0.572959073 | 0.45982087 | -1.2460484 | 0.21274662 | 0.57730926 |
| BID | 52.9459687 | -0.572454519 | 0.39827765 | -1.4373252 | 0.1506256 | 0.5057732 |
| SCFD2 | 408.936376 | -0.572223167 | 0.4194461 | -1.3642353 | 0.17249353 | 0.53461168 |
| TXNL4B | 46.2441272 | -0.57133029 | 0.31984825 | -1.7862543 | 0.0740581 | 0.37666831 |
| PUM3 | 246.78991 | -0.571236374 | 0.29432772 | -1.9408174 | 0.05228043 | 0.32059696 |
| FLNA | 320.159772 | -0.570406176 | 0.48541979 | -1.1750781 | 0.23996346 | 0.60456313 |
| JTB | 94.4747553 | -0.57028526 | 0.42481113 | -1.3424442 | 0.179452 | 0.54326885 |
| ERP29 | 96.7837366 | -0.570188819 | 0.42169016 | -1.3521511 | 0.17632698 | 0.54007689 |
| LINC01006 | 54.1378974 | -0.569705762 | 0.60995078 | -0.9340192 | 0.35029397 | 0.69858573 |
| PCBP1 | 213.184483 | -0.569248448 | 0.43529252 | -1.3077377 | 0.19096228 | 0.55586607 |
| AC026471.4 | 42.5420268 | -0.568418044 | 0.31140756 | -1.8253187 | 0.06795295 | 0.36125501 |
| SRA1 | 64.6908539 | -0.568362374 | 0.28272938 | -2.0102699 | 0.04440263 | 0.29898844 |
| IKBIP | 103.486626 | -0.568340659 | 0.238981 | -2.3781835 | 0.01739817 | 0.18853893 |
| RPL37 | 234.647185 | -0.568160683 | 0.38962276 | -1.4582328 | 0.14477639 | 0.49729058 |
| KDELR1 | 97.444306 | -0.568089337 | 0.29260611 | -1.9414814 | 0.05219991 | 0.32044401 |
| HYAL2 | 716.527226 | -0.567510021 | 0.45999013 | -1.2337439 | 0.21729835 | 0.583192 |
| LACTB | 73.0987264 | -0.566975038 | 0.32954897 | -1.7204576 | 0.08534929 | 0.39686932 |
| PHACTR1 | 923.909746 | -0.566945528 | 0.3875584 | -1.4628648 | 0.14350438 | 0.49569285 |
| FOXK1 | 314.488644 | -0.566933537 | 0.27191441 | -2.0849706 | 0.03707197 | 0.27134164 |
| TMEM134 | 42.2692017 | -0.565358182 | 0.39658416 | -1.4255693 | 0.15399272 | 0.50978388 |
| ZBTB25 | 83.3481838 | -0.564593469 | 0.41812128 | -1.3503103 | 0.17691647 | 0.54037749 |
| CIB1 | 97.0660583 | -0.564417597 | 0.38179024 | -1.4783448 | 0.13931552 | 0.49091481 |
| GIMAP4 | 287.334611 | -0.564062064 | 0.45274227 | -1.2458789 | 0.21280884 | 0.57730926 |
| MAP3K8 | 118.723233 | -0.56378791 | 0.38051302 | -1.4816521 | 0.13843289 | 0.48991657 |
| MPV17 | 53.2034229 | -0.563454158 | 0.21722524 | -2.5938706 | 0.00949022 | 0.13769231 |
| POFUT1 | 54.6122334 | -0.562989778 | 0.26810194 | -2.0999094 | 0.03573682 | 0.26584942 |
| RASA3 | 72.2328792 | -0.562150662 | 0.5120476 | -1.0978484 | 0.27227067 | 0.63711493 |
| IL17RA | 102.573921 | -0.562140606 | 0.20950027 | -2.6832453 | 0.00729115 | 0.12102126 |
| ATP5MC3 | 152.857559 | -0.562136106 | 0.48837729 | -1.1510283 | 0.24972058 | 0.6149919 |
| YWHAG | 219.165395 | -0.561802445 | 0.24491025 | -2.2939115 | 0.02179558 | 0.20902438 |
| MDM2 | 240.144291 | -0.561381785 | 0.31879964 | -1.7609235 | 0.07825135 | 0.38566641 |
| SH3RF3 | 2133.14451 | -0.561298001 | 0.67070955 | -0.8368719 | 0.40266456 | 0.73037606 |
| TMEM258 | 63.9387509 | -0.560868443 | 0.4875193 | -1.1504538 | 0.24995701 | 0.61500658 |
| SCIN | 330.739059 | -0.56005244 | 1.02394162 | -0.5469574 | 0.584408 | 0.83895258 |
| TSPAN31 | 49.1473601 | -0.559870978 | 0.43387486 | -1.2903974 | 0.19691272 | 0.56183292 |
| VCP | 256.224487 | -0.559359079 | 0.32455874 | -1.7234448 | 0.08480811 | 0.39686932 |
| LRPAP1 | 106.06307 | -0.559291711 | 0.36095448 | -1.5494799 | 0.1212664 | 0.46542849 |
| SEC24D | 245.291815 | -0.559265369 | 0.22620505 | -2.4723823 | 0.01342159 | 0.1660879 |
| SLC25A39 | 49.6048759 | -0.558585422 | 0.40586368 | -1.3762883 | 0.16873241 | 0.53071749 |
| AL390860.1 | 80.0105144 | -0.55857863 | 0.58304667 | -0.9580342 | 0.33804553 | 0.68878364 |
| FZD6 | 159.54772 | -0.558192906 | 0.35675048 | -1.5646592 | 0.11766285 | 0.46042456 |
| DHX32 | 130.543252 | -0.557584697 | 0.20711308 | -2.6921752 | 0.00709877 | 0.120019 |
| COL6A2 | 68.9295198 | -0.556693287 | 0.68433826 | -0.8134768 | 0.41594475 | 0.73795362 |
| SERF2 | 840.622404 | -0.556448404 | 0.56739027 | -0.9807154 | 0.32673308 | 0.68007671 |
| VASH1 | 118.418637 | -0.555457359 | 0.43211861 | -1.285428 | 0.19864276 | 0.56401586 |
| NDUFA8 | 75.1414576 | -0.555402425 | 0.37814803 | -1.4687434 | 0.1419024 | 0.49378081 |
| NDUFA4 | 215.518463 | -0.555399366 | 0.52602741 | -1.0558373 | 0.29104255 | 0.65374724 |
| VPS37A | 214.034205 | -0.555332713 | 0.38054746 | -1.4592995 | 0.1444827 | 0.49679586 |
| CALU | 225.448325 | -0.555309559 | 0.30840782 | -1.800569 | 0.07177084 | 0.37030316 |
| PRR13 | 87.1089401 | -0.555199032 | 0.26081479 | -2.1287099 | 0.03327827 | 0.25490607 |
| MLF2 | 89.6868604 | -0.554920537 | 0.35041844 | -1.583594 | 0.11328615 | 0.45132451 |
| HEATR3 | 57.2372363 | -0.554888991 | 0.26620208 | -2.0844653 | 0.03711786 | 0.27143474 |
| AHI1 | 436.92457 | -0.554774464 | 0.15236159 | -3.6411701 | 0.0002714 | 0.01669835 |
| ZNF354A | 49.5669646 | -0.554386034 | 0.38441656 | -1.4421492 | 0.14926029 | 0.50283523 |
| SRSF4 | 654.63099 | -0.553412193 | 0.27842323 | -1.9876653 | 0.04684872 | 0.3037742 |
| UQCRFS1 | 73.8942768 | -0.553276994 | 0.4610474 | -1.2000436 | 0.2301224 | 0.59648134 |
| CPLX1 | 100.437238 | -0.552929157 | 0.53227872 | -1.0387963 | 0.29889949 | 0.65856711 |
| UROS | 40.972422 | -0.552373741 | 0.38151226 | -1.4478532 | 0.14765812 | 0.50136364 |
| TRAF3 | 167.289409 | -0.55205878 | 0.58372739 | -0.9457476 | 0.34427733 | 0.6938969 |
| GPAT4 | 135.56916 | -0.5509557 | 0.41356399 | -1.3322139 | 0.1827899 | 0.54790102 |
| AAED1 | 49.6944491 | -0.550676732 | 0.44705638 | -1.2317836 | 0.21802991 | 0.58419738 |
| ATP6V0B | 144.316349 | -0.550383394 | 0.35500484 | -1.5503546 | 0.12105643 | 0.46542849 |
| MRPS26 | 48.6057416 | -0.550376426 | 0.47668367 | -1.1545947 | 0.24825645 | 0.61372207 |
| G3BP2 | 177.393358 | -0.549861705 | 0.21695838 | -2.5344109 | 0.01126366 | 0.14987072 |
| LAMA3 | 122.810869 | -0.549557771 | 1.10851183 | -0.4957618 | 0.62006251 | 0.85868532 |
| GNA13 | 290.099208 | -0.549202774 | 0.41959271 | -1.3088949 | 0.19056994 | 0.55558405 |
| PDAP1 | 83.0793582 | -0.549010202 | 0.35328202 | -1.5540282 | 0.1201777 | 0.46382142 |
| CSNK1G2 | 88.8664611 | -0.548423406 | 0.273127 | -2.0079429 | 0.04464936 | 0.29946125 |
| KDSR | 120.858829 | -0.54732302 | 0.23388802 | -2.3401071 | 0.01927821 | 0.19499826 |
| PCNX2 | 84.9705452 | -0.547226645 | 0.49011149 | -1.116535 | 0.26419319 | 0.63018314 |
| NUDT4 | 134.652029 | -0.54628511 | 0.60475136 | -0.9033218 | 0.36635511 | 0.70695551 |
| C12orf57 | 90.8245984 | -0.546215765 | 0.63437967 | -0.8610234 | 0.38922514 | 0.72202943 |
| RPS12 | 319.660301 | -0.545642184 | 0.53184953 | -1.0259334 | 0.30492299 | 0.66115125 |
| MOB3B | 80.0922951 | -0.545215501 | 0.39073442 | -1.3953608 | 0.16290706 | 0.52335809 |
| RFX3-AS1 | 165.155551 | -0.5448389 | 0.42068422 | -1.2951256 | 0.1952769 | 0.56007526 |
| UTP20 | 112.686804 | -0.544792159 | 0.3836385 | -1.4200665 | 0.15558834 | 0.51180943 |
| PHB2 | 85.5149295 | -0.544677131 | 0.37571588 | -1.4497048 | 0.14714085 | 0.50127566 |
| PRR5L | 176.780927 | -0.543960126 | 0.30955488 | -1.7572332 | 0.07887806 | 0.38686531 |
| TUBA1B | 354.186085 | -0.543649221 | 0.59769856 | -0.9095709 | 0.36304884 | 0.70525625 |
| SLC3A2 | 270.932652 | -0.543517434 | 0.25181247 | -2.1584215 | 0.03089508 | 0.24450137 |
| ITGA5 | 456.81561 | -0.542626652 | 0.27278197 | -1.9892321 | 0.04667559 | 0.30359341 |
| PCAT19 | 399.104965 | -0.542263115 | 0.24652056 | -2.1996669 | 0.02783054 | 0.23190345 |
| BCAP31 | 151.560237 | -0.54126716 | 0.42840803 | -1.2634384 | 0.20643167 | 0.57126492 |
| BDNF-AS | 228.861409 | -0.541027665 | 0.40258278 | -1.3438917 | 0.17898341 | 0.54285442 |
| CTSS | 79.9608966 | -0.540999839 | 0.3704716 | -1.4603004 | 0.14420752 | 0.49679586 |
| TM4SF1 | 1279.95455 | -0.539663726 | 0.5617403 | -0.9606997 | 0.33670319 | 0.68748823 |
| BAALC-AS1 | 68.4974982 | -0.539556704 | 0.433111 | -1.24577 | 0.21284884 | 0.57730926 |
| SIAE | 53.9537037 | -0.539544659 | 0.41399305 | -1.3032698 | 0.19248266 | 0.55696095 |
| VDAC1 | 137.539421 | -0.539338344 | 0.28295441 | -1.9060963 | 0.05663772 | 0.33295004 |
| TMEM192 | 71.2970309 | -0.539014587 | 0.40596827 | -1.3277259 | 0.18426868 | 0.54912489 |
| TUBB | 286.346366 | -0.538603627 | 0.64484081 | -0.8352505 | 0.40357666 | 0.73065697 |
| GRAPL | 535.362413 | -0.538422493 | 0.48715821 | -1.1052313 | 0.26905939 | 0.63347822 |
| SEC61B | 168.248864 | -0.538113919 | 0.41430805 | -1.2988256 | 0.19400377 | 0.55913011 |
| NDST1 | 219.701342 | -0.538089866 | 0.29007551 | -1.8549993 | 0.06359633 | 0.3533385 |
| PLEKHO1 | 161.557309 | -0.537670131 | 0.46686691 | -1.1516561 | 0.24946242 | 0.6146796 |
| MLH3 | 123.858482 | -0.537135219 | 0.26425285 | -2.0326563 | 0.04208726 | 0.29087842 |

| | | | | | | |
|---|---|---|---|---|---|---|
| LONP1 | 71.2202476 | -0.536606459 | 0.27295164 | -1.9659397 | 0.04930558 | 0.31155957 |
| EXTL3 | 86.8877971 | -0.53658322 | 0.22709184 | -2.3628467 | 0.01813517 | 0.192636 |
| PTRH2 | 81.0964768 | -0.53639576 | 0.26194498 | -2.0477421 | 0.04058528 | 0.28580837 |
| IMP3 | 43.8381473 | -0.535772361 | 0.35688785 | -1.5012345 | 0.13329491 | 0.48236854 |
| SNX8 | 73.8921126 | -0.535210031 | 0.4391831 | -1.218649 | 0.22297745 | 0.58858855 |
| MCF2L | 328.870057 | -0.534658679 | 0.45103082 | -1.185415 | 0.23585342 | 0.60018541 |
| ADIPOR2 | 580.624475 | -0.534645402 | 0.38941686 | -1.3729385 | 0.16977146 | 0.53166471 |
| CLPB | 61.2150808 | -0.534572662 | 0.30644902 | -1.7444097 | 0.08108767 | 0.39123843 |
| YWHAB | 439.833705 | -0.534321629 | 0.36295844 | -1.472129 | 0.14098606 | 0.49255036 |
| RPS15A | 92.3771034 | -0.534320535 | 0.59698551 | -0.895031 | 0.37077052 | 0.70984233 |
| RRBP1 | 309.79897 | -0.533885747 | 0.29440716 | -1.8134265 | 0.06976606 | 0.3652226 |
| FP671120.1 | 1265.30278 | -0.533838953 | 0.3462828 | -1.5416271 | 0.12316423 | 0.46855087 |
| DIMT1 | 50.7093211 | -0.532637871 | 0.29784054 | -1.7883323 | 0.0737224 | 0.37609297 |
| TRMT112 | 116.8739 | -0.531910729 | 0.48344439 | -1.1002521 | 0.27122228 | 0.63561102 |
| PIK3C2B | 785.538875 | -0.531522516 | 0.37319374 | -1.4242536 | 0.15437308 | 0.51037454 |
| ST13 | 173.156136 | -0.531043664 | 0.20662622 | -2.5700691 | 0.01016782 | 0.14236887 |
| SDC4 | 71.2827935 | -0.530128933 | 0.34176114 | -1.551168 | 0.12086144 | 0.46513092 |
| PCSK5 | 793.797725 | -0.529052928 | 0.32149937 | -1.64558 | 0.09985027 | 0.42533822 |
| VPS25 | 40.1501357 | -0.528951543 | 0.30900901 | -1.7117674 | 0.08693955 | 0.39969788 |
| PSMD2 | 94.9045432 | -0.528942116 | 0.31973242 | -1.6543275 | 0.098061 | 0.42166547 |
| MLEC | 106.080637 | -0.528865699 | 0.31592536 | -1.6740211 | 0.09412646 | 0.41293638 |
| GATA6 | 100.620982 | -0.528787937 | 0.37008501 | -1.4288283 | 0.15305359 | 0.50905923 |
| PSAP | 450.625836 | -0.528648586 | 0.36080082 | -1.4652089 | 0.14286392 | 0.49500775 |
| CYBC1 | 74.6545956 | -0.527865322 | 0.35791506 | -1.4748341 | 0.14025716 | 0.49118733 |
| CFAP97 | 121.06424 | -0.527602037 | 0.2889696 | -1.8258047 | 0.06787969 | 0.36115704 |
| FDFT1 | 89.4811992 | -0.527374778 | 0.31526643 | -1.6727908 | 0.0943685 | 0.41317143 |
| MTR | 261.475065 | -0.527184985 | 0.18082201 | -2.9154913 | 0.00355129 | 0.08117377 |
| ARL4A | 76.7634358 | -0.527170778 | 0.60625804 | -0.8695485 | 0.38454718 | 0.71699718 |
| ANPEP | 46.5480041 | -0.526798721 | 0.68413112 | -0.770026 | 0.44128449 | 0.75607852 |
| KCTD20 | 184.56088 | -0.526767532 | 0.21652054 | -2.4328756 | 0.01497945 | 0.17586342 |
| PRPF19 | 65.5219216 | -0.52613127 | 0.3413795 | -1.5411918 | 0.12327012 | 0.46870024 |
| LRP10 | 116.216659 | -0.525403269 | 0.35071455 | -1.4980937 | 0.1341089 | 0.48408165 |
| A4GALT | 144.104605 | -0.52487642 | 0.24662678 | -2.1282216 | 0.03331872 | 0.25490607 |
| AP1S1 | 45.3464292 | -0.524574983 | 0.41873113 | -1.2527728 | 0.21028839 | 0.57593532 |
| ZHX2 | 447.958971 | -0.522949206 | 0.2958441 | -1.7676513 | 0.07711922 | 0.38356763 |
| EMC7 | 84.427017 | -0.52261846 | 0.29339324 | -1.7812901 | 0.07486507 | 0.37933183 |
| POLE4 | 107.332603 | -0.522204873 | 0.48302908 | -1.0811044 | 0.27965067 | 0.64286511 |
| SYAP1 | 103.636984 | -0.521499499 | 0.21336825 | -2.4441289 | 0.01452024 | 0.17422157 |
| GHITM | 103.327518 | -0.521331105 | 0.3158889 | -1.6503622 | 0.09886887 | 0.42358324 |
| UTP4 | 47.089142 | -0.521327197 | 0.33335344 | -1.5638872 | 0.11784405 | 0.46042456 |
| HSPD1 | 186.696717 | -0.521279306 | 0.3152832 | -1.6533685 | 0.09825589 | 0.42206191 |
| ALPK3 | 84.8068981 | -0.521137255 | 0.4122672 | -1.2640764 | 0.20620259 | 0.57121049 |
| TMEM147 | 48.6317901 | -0.520644756 | 0.52391738 | -0.9937536 | 0.32034286 | 0.67435802 |
| BCL6B | 109.600007 | -0.520502641 | 0.39396575 | -1.3211876 | 0.18643883 | 0.55256392 |
| RFLNB | 93.4018229 | -0.520290776 | 0.37296893 | -1.3949976 | 0.16301655 | 0.52335809 |
| RFC3 | 204.142206 | -0.519934891 | 0.57134375 | -0.9100211 | 0.36281136 | 0.70519842 |
| CEP68 | 186.005449 | -0.519856004 | 0.32576868 | -1.5957827 | 0.11053733 | 0.44631216 |
| HADHA | 192.188702 | -0.519213622 | 0.19062545 | -2.7237371 | 0.00645479 | 0.11362684 |
| NRCAM | 126.249065 | -0.518780299 | 0.62792785 | -0.8261782 | 0.40870302 | 0.73419295 |
| FIS1 | 197.534537 | -0.517710587 | 0.42152382 | -1.2281882 | 0.21937632 | 0.58626926 |
| TSPYL2 | 43.123247 | -0.517552344 | 0.39866096 | -1.2982268 | 0.19420941 | 0.55913011 |
| RAI14 | 870.586544 | -0.517417439 | 0.38889443 | -1.330483 | 0.18335917 | 0.54819669 |
| CRTAP | 91.0934482 | -0.516756188 | 0.34404254 | -1.5020125 | 0.13309389 | 0.48236854 |
| NORAD | 289.070504 | -0.516749829 | 0.31464202 | -1.6423421 | 0.10051913 | 0.4268542 |
| RAD23A | 112.229879 | -0.516618965 | 0.32002107 | -1.614328 | 0.1064563 | 0.43901385 |
| RIN3 | 404.075572 | -0.516547219 | 0.29643883 | -1.7425086 | 0.08141948 | 0.39205363 |
| AC093157.1 | 87.0461698 | -0.515572281 | 0.24374404 | -2.1152201 | 0.0344112 | 0.25881141 |
| WDR12 | 66.8980659 | -0.515148352 | 0.36800913 | -1.3998249 | 0.16156575 | 0.52030403 |
| CNPY3 | 39.4306956 | -0.514776975 | 0.43191448 | -1.1918493 | 0.23332033 | 0.59910683 |
| POLR3E | 45.6530497 | -0.513773803 | 0.26068872 | -1.9708325 | 0.04874303 | 0.30991784 |
| MARCKSL1 | 152.432752 | -0.513558486 | 0.45036863 | -1.140307 | 0.25415844 | 0.61966901 |
| AC010615.4 | 48.2669244 | -0.513074228 | 0.49957405 | -1.0270234 | 0.30440946 | 0.66086613 |
| RPS19BP1 | 127.467042 | -0.51264749 | 0.48051917 | -1.0668617 | 0.2860343 | 0.64873023 |
| C1S | 98.5673339 | -0.512333543 | 0.34980885 | -1.46461 | 0.14302735 | 0.49529954 |
| JOSD1 | 103.568216 | -0.512015912 | 0.28925249 | -1.7701349 | 0.07670468 | 0.38356763 |
| FMO2 | 169.146105 | -0.511743637 | 1.04718264 | -0.4886861 | 0.62506392 | 0.86016045 |
| MANSC1 | 80.1878561 | -0.511685445 | 0.39509272 | -1.2951022 | 0.19528498 | 0.56007526 |
| ENTR1 | 40.5649159 | -0.511331551 | 0.29456333 | -1.7358968 | 0.08258208 | 0.39477007 |
| CINP | 52.0164975 | -0.511272195 | 0.22256238 | -2.2972085 | 0.02160688 | 0.20825014 |
| AGTRAP | 70.9489876 | -0.511267803 | 0.35662281 | -1.4336374 | 0.15167576 | 0.50771689 |
| AC112722.1 | 46.6899163 | -0.510756035 | 0.44772056 | -1.140792 | 0.25395649 | 0.61961636 |
| E2F3 | 124.946668 | -0.509976943 | 0.3551997 | -1.4357471 | 0.15107431 | 0.50677567 |
| PLEKHB2 | 169.251561 | -0.508737812 | 0.23852428 | -2.1328554 | 0.03293659 | 0.25378541 |
| AC109927.1 | 81.7704522 | -0.508469763 | 0.39498592 | -1.2873111 | 0.19798588 | 0.56327864 |
| MZT2B | 124.803565 | -0.508313466 | 0.4629306 | -1.0980338 | 0.27218971 | 0.63710767 |
| RPS27L | 146.994991 | -0.507753625 | 0.3415981 | -1.4864065 | 0.13717164 | 0.48888307 |
| ARMC10 | 69.0627553 | -0.507399756 | 0.2692645 | -1.8843916 | 0.05951203 | 0.34222554 |
| CEBPZ | 119.413463 | -0.507013624 | 0.25595047 | -1.9809052 | 0.0476019 | 0.30623143 |
| FAM162A | 90.5290976 | -0.506630565 | 0.50079297 | -1.0116567 | 0.31170223 | 0.66561244 |
| ORAI1 | 63.9676525 | -0.506495074 | 0.54970055 | -0.9214018 | 0.35684069 | 0.70058911 |
| RPN2 | 167.971147 | -0.506367846 | 0.29930694 | -1.6918012 | 0.09068389 | 0.40729106 |
| MRPL54 | 50.3107117 | -0.505635702 | 0.40443677 | -1.2502219 | 0.2112185 | 0.5768065 |
| Z93930.2 | 112.715068 | -0.505351512 | 0.42740249 | -1.1823785 | 0.23705556 | 0.60074482 |
| CCT7 | 122.622877 | -0.504663994 | 0.37565669 | -1.3434181 | 0.17913663 | 0.54311783 |
| C22orf34 | 464.829102 | -0.504651099 | 0.5745271 | -0.8783765 | 0.37973944 | 0.71418244 |
| MAP2K3 | 126.7616 | -0.504409909 | 0.32339782 | -1.5597196 | 0.11882617 | 0.46285439 |
| OXA1L | 50.7167811 | -0.503991588 | 0.30089451 | -1.6749777 | 0.09393861 | 0.41280519 |
| TNFAIP8 | 83.7962487 | -0.503719641 | 0.38274772 | -1.3160618 | 0.18815331 | 0.55388137 |
| TCF4-AS1 | 62.9493874 | -0.502792819 | 0.50313699 | -0.999316 | 0.31764166 | 0.67251273 |
| CHMP2B | 86.4651385 | -0.502713935 | 0.27141697 | -1.8521832 | 0.06399953 | 0.3533793 |
| MBOAT7 | 64.723431 | -0.502549337 | 0.39079246 | -1.285975 | 0.19845178 | 0.56366919 |
| SEC23B | 93.0308286 | -0.502316347 | 0.2144781 | -2.3420403 | 0.01917864 | 0.1947132 |
| SLC8B1 | 112.91572 | -0.502259466 | 0.32764089 | -1.5329572 | 0.1252864 | 0.47093183 |
| COX4I1 | 293.681921 | -0.502159251 | 0.41714547 | -1.2037989 | 0.22866732 | 0.59497702 |
| CENPX | 41.6268147 | -0.501991249 | 0.48722051 | -1.0303163 | 0.30286153 | 0.65940027 |
| PDCD4 | 262.962137 | -0.501588302 | 0.49857881 | -1.0060361 | 0.31439819 | 0.66883106 |
| CPPED1 | 105.31093 | -0.500709421 | 0.37377416 | -1.3396042 | 0.18037407 | 0.5444489 |
| LSM2 | 72.5304492 | -0.500631847 | 0.42902869 | -1.166896 | 0.24325239 | 0.608518 |
| TMX4 | 100.259512 | -0.500211299 | 0.25192364 | -1.9855671 | 0.04708141 | 0.30480001 |
| DMAC1 | 46.0990163 | -0.49967437 | 0.26764624 | -1.8669209 | 0.06191263 | 0.3489463 |
| CYCS | 81.5283354 | -0.499505854 | 0.44189791 | -1.1303648 | 0.25832253 | 0.6230042 |
| ACAA1 | 62.3185248 | -0.499219836 | 0.34866563 | -1.4318011 | 0.15220075 | 0.50814312 |
| SLC23A2 | 255.028413 | -0.498751468 | 0.28734931 | -1.7356974 | 0.08261735 | 0.39477007 |
| CCT5 | 138.622878 | -0.498614839 | 0.34397216 | -1.4495791 | 0.14717594 | 0.50127566 |
| TOMM20 | 145.782432 | -0.498339124 | 0.37267941 | -1.3371791 | 0.18116417 | 0.54524118 |
| AARS | 86.4874961 | -0.497589358 | 0.3178139 | -1.5656627 | 0.11742761 | 0.46042456 |
| MRPL20 | 145.610027 | -0.496687815 | 0.3220485 | -1.5422765 | 0.12300643 | 0.46838604 |
| ITGB3BP | 100.951626 | -0.496629129 | 0.32049119 | -1.5495874 | 0.12124057 | 0.46542849 |
| TSFM | 40.2806907 | -0.496408944 | 0.3404532 | -1.4580828 | 0.14481773 | 0.49729058 |
| AL590226.1 | 81.9308293 | -0.496028913 | 0.55119388 | -0.8999173 | 0.36816426 | 0.70903462 |
| TTTY10 | 50.6347737 | -0.49594748 | 1.54903938 | -0.3201645 | 0.7488436 | 0.91065347 |
| NDUFAF3 | 61.1207682 | -0.495713778 | 0.48042985 | -1.031813 | 0.30215971 | 0.65926326 |
| CSNK1D | 188.599066 | -0.495658726 | 0.21010252 | -2.359128 | 0.01831794 | 0.19310354 |

| | | | | | | |
|---|---|---|---|---|---|---|
| HLA-A | 1680.91238 | -0.495495764 | 0.46005096 | -1.0770454 | 0.28145999 | 0.64471757 |
| POLR2J | 53.5075714 | -0.495238844 | 0.49813705 | -0.9941819 | 0.32013432 | 0.67434487 |
| ACTB | 1707.76835 | -0.495189788 | 0.47052558 | -1.0524184 | 0.29260762 | 0.65502684 |
| KDM6B | 70.2657526 | -0.494584333 | 0.41770468 | -1.1840526 | 0.23639222 | 0.60018541 |
| LAMTOR5 | 156.229774 | -0.494435808 | 0.37834896 | -1.3068248 | 0.19127221 | 0.55590015 |
| DARS | 267.480978 | -0.494303609 | 0.2386728 | -2.0710512 | 0.03835401 | 0.27652057 |
| GRINA | 80.8938594 | -0.494066593 | 0.37011582 | -1.3348973 | 0.18190997 | 0.54686603 |
| AC005699.1 | 130.571136 | -0.493969388 | 0.90437596 | -0.5461992 | 0.58492905 | 0.83895258 |
| LARP1 | 264.460423 | -0.493387683 | 0.30821333 | -1.6007993 | 0.10942138 | 0.44576178 |
| TP53 | 63.7259677 | -0.493128256 | 0.3278662 | -1.5040533 | 0.13256763 | 0.48191957 |
| PHC2 | 624.138549 | -0.492563196 | 0.40879399 | -1.2049179 | 0.228235 | 0.59497702 |
| MDK | 52.3518193 | -0.492460533 | 0.7384825 | -0.6668547 | 0.50486494 | 0.79785821 |
| KALRN | 2016.82762 | -0.492237531 | 0.34020179 | -1.4468987 | 0.1479253 | 0.50143858 |
| MPDU1 | 44.9367987 | -0.492174372 | 0.35363261 | -1.3917675 | 0.16399282 | 0.52357299 |
| SPATA6 | 293.36228 | -0.491338852 | 0.33839035 | -1.4519884 | 0.14650484 | 0.50035439 |
| ABCD4 | 201.064801 | -0.491098567 | 0.23513071 | -2.0886194 | 0.036742 | 0.27037642 |
| PTS | 170.797708 | -0.490606298 | 0.188229 | -2.6064331 | 0.00914907 | 0.13593083 |
| ZNF331 | 63.5879627 | -0.489595466 | 0.27998598 | -1.7486428 | 0.08035278 | 0.38871437 |
| SURF4 | 142.891379 | -0.488302903 | 0.33644528 | -1.4513591 | 0.1466799 | 0.50053446 |
| RNF169 | 202.783879 | -0.487977487 | 0.28178866 | -1.7317145 | 0.08332442 | 0.39543345 |
| TBC1D22B | 101.868141 | -0.487937574 | 0.35439415 | -1.3768218 | 0.16856736 | 0.53053336 |
| RPS27A | 618.078962 | -0.487272644 | 0.54519375 | -0.8937605 | 0.37145004 | 0.70991341 |
| CBX3 | 156.644721 | -0.487016993 | 0.26896719 | -1.810693 | 0.07018839 | 0.36582906 |
| ETFB | 88.9827846 | -0.486571384 | 0.36531574 | -1.3319201 | 0.18288644 | 0.5479055 |
| VAMP5 | 459.658441 | -0.485749404 | 0.54861902 | -0.8854039 | 0.37593885 | 0.71257833 |
| ARL1 | 81.8293389 | -0.485594336 | 0.33276584 | -1.4592674 | 0.14449153 | 0.49679586 |
| DCTD | 51.1511668 | -0.48556346 | 0.33674417 | -1.4419358 | 0.14932049 | 0.50283523 |
| M6PR | 94.6238783 | -0.485229875 | 0.27628833 | -1.7562446 | 0.07904666 | 0.38686531 |
| TSPAN3 | 132.123766 | -0.484763182 | 0.25938017 | -1.8689293 | 0.06163266 | 0.34830113 |
| DOK5 | 145.599861 | -0.484292535 | 0.37588463 | -1.2884074 | 0.19760419 | 0.56300834 |
| LPCAT1 | 80.620693 | -0.48416644 | 0.33492849 | -1.4455815 | 0.14829461 | 0.50185887 |
| INPP5D | 729.706665 | -0.484077871 | 0.38776885 | -1.2483671 | 0.21189666 | 0.57690664 |
| AHNAK | 553.38012 | -0.483956981 | 0.33031439 | -1.4651405 | 0.1428826 | 0.49500775 |
| AKAP8 | 96.4087995 | -0.483383616 | 0.23525141 | -2.0547533 | 0.03990284 | 0.28368803 |
| SARAF | 266.873963 | -0.483280215 | 0.36686647 | -1.3173191 | 0.18773168 | 0.55388137 |
| RPL7L1 | 136.714123 | -0.482806774 | 0.32679835 | -1.4773844 | 0.13957263 | 0.49091481 |
| ATXN7L3B | 74.4994904 | -0.482797325 | 0.25136313 | -1.9207166 | 0.05476745 | 0.32800002 |
| TACC1 | 2329.05286 | -0.482569693 | 0.45857702 | -1.0523198 | 0.29265283 | 0.65502684 |
| TSC22D4 | 72.1261554 | -0.482151569 | 0.24924499 | -1.9344484 | 0.05305803 | 0.32232652 |
| AMD1 | 97.4704678 | -0.481763387 | 0.28335967 | -1.7001833 | 0.08909645 | 0.4043684 |
| EDEM2 | 69.2596579 | -0.481540123 | 0.20166525 | -2.3878191 | 0.01694868 | 0.18591296 |
| MPHOSPH10 | 67.4295053 | -0.481523964 | 0.37281954 | -1.2915738 | 0.19650477 | 0.56164812 |
| PIM3 | 258.29277 | -0.48111639 | 0.40513045 | -1.1875592 | 0.23500712 | 0.59949266 |
| SH3RF1 | 203.299954 | -0.481103972 | 0.26098006 | -1.8434511 | 0.06526318 | 0.35545467 |
| ELL | 145.327413 | -0.480623631 | 0.38552897 | -1.2466602 | 0.21252211 | 0.57720441 |
| RBM42 | 44.150644 | -0.480598808 | 0.40071518 | -1.1993526 | 0.23039085 | 0.5967991 |
| DKC1 | 44.3502287 | -0.48036056 | 0.31691875 | -1.5157215 | 0.12958979 | 0.4777902 |
| NF2 | 89.5260906 | -0.479596576 | 0.27859729 | -1.7214689 | 0.08516577 | 0.39686932 |
| TBC1D30 | 123.075032 | -0.479536523 | 0.38608093 | -1.2420622 | 0.21421362 | 0.57870917 |
| S100A10 | 465.625157 | -0.479187907 | 0.51254673 | -0.9349156 | 0.34983182 | 0.69858573 |
| LAMTOR2 | 49.9646935 | -0.478859485 | 0.39199011 | -1.2216112 | 0.22185471 | 0.58750144 |
| NDUFA1 | 110.132395 | -0.478737289 | 0.39792172 | -1.2030941 | 0.2289399 | 0.59511283 |
| KLF9 | 491.52115 | -0.478668328 | 0.58659631 | -0.8160098 | 0.41449454 | 0.73671465 |
| SLC50A1 | 40.5237579 | -0.478387679 | 0.2371163 | -2.0175234 | 0.04364093 | 0.29562392 |
| LINC02360 | 173.153909 | -0.477830683 | 1.50919379 | -0.3166132 | 0.75153711 | 0.9112603 |
| TSHZ3 | 215.709635 | -0.476220406 | 0.30820659 | -1.5451337 | 0.12231392 | 0.46748939 |
| AC010737.1 | 699.831968 | -0.475916125 | 1.44674458 | -0.3289566 | 0.74218852 | 0.90850686 |
| SNRPB2 | 77.1865884 | -0.474634365 | 0.29829865 | -1.5911382 | 0.11157849 | 0.44839019 |
| GRAP | 71.681761 | -0.474625038 | 0.34094872 | -1.3920716 | 0.16390071 | 0.52357299 |
| GLRX | 141.666415 | -0.47444237 | 0.55414479 | -0.8561704 | 0.39190353 | 0.72247779 |
| AKT1 | 83.7294252 | -0.474416119 | 0.24390876 | -1.9450557 | 0.05176827 | 0.31875077 |
| RXYLT1 | 45.0029145 | -0.474338764 | 0.30382459 | -1.5612257 | 0.1184705 | 0.46185998 |
| ZDBF2 | 69.9513975 | -0.474324143 | 0.35443828 | -1.3382419 | 0.18081761 | 0.54518441 |
| AP2M1 | 190.17172 | -0.473277352 | 0.37332729 | -1.2677277 | 0.2048952 | 0.56960077 |
| ASMTL | 59.2833524 | -0.473273677 | 0.36276881 | -1.3046151 | 0.19202394 | 0.55646778 |
| LCP2 | 84.9217695 | -0.473032029 | 0.50347308 | -0.9395379 | 0.34745466 | 0.69635598 |
| ACTG1 | 1124.98306 | -0.472852705 | 0.46583688 | -1.0150607 | 0.3100769 | 0.6648266 |
| SRPRB | 44.6209842 | -0.472652081 | 0.42314307 | -1.117003 | 0.26399304 | 0.63018314 |
| XRN2 | 352.313247 | -0.472419675 | 0.16824177 | -2.8079809 | 0.00498532 | 0.09667028 |
| HSD17B12 | 563.58488 | -0.472011785 | 0.19453268 | -2.4263882 | 0.01524995 | 0.177259 |
| YIPF1 | 52.7783384 | -0.471601173 | 0.30151625 | -1.5640987 | 0.1177944 | 0.46042456 |
| SACS | 64.3692473 | -0.471310608 | 0.20977237 | -2.2467717 | 0.02465462 | 0.2185794 |
| WDYHV1 | 48.5077136 | -0.470944672 | 0.33283827 | -1.4149355 | 0.15708742 | 0.51417853 |
| VAMP8 | 132.588614 | -0.470307129 | 0.47808496 | -0.9837313 | 0.32524765 | 0.67912772 |
| FAU | 352.830067 | -0.470269148 | 0.52584502 | -0.8943113 | 0.37115536 | 0.70984233 |
| DPP7 | 68.8329727 | -0.47017667 | 0.41081126 | -1.1445077 | 0.25241313 | 0.61777104 |
| DTD1 | 43.6643196 | -0.470161145 | 0.33334649 | -1.4104278 | 0.15841342 | 0.51542623 |
| DAB2IP | 437.099646 | -0.469488277 | 0.2801222 | -1.6760124 | 0.09373577 | 0.41280519 |
| ATP6AP1 | 103.59893 | -0.468826731 | 0.38950146 | -1.2036585 | 0.22872161 | 0.59497702 |
| SLC25A43 | 73.6671223 | -0.467966519 | 0.39913865 | -1.172441 | 0.24102005 | 0.60559848 |
| UBTD1 | 97.3638035 | -0.467313083 | 0.37537713 | -1.2449162 | 0.21316255 | 0.57758664 |
| COX6B1 | 145.730262 | -0.466916322 | 0.33129474 | -1.4093683 | 0.15872628 | 0.51586289 |
| SOD1 | 164.071936 | -0.466615519 | 0.41997256 | -1.1110619 | 0.26654169 | 0.6320228 |
| SRSF9 | 128.047024 | -0.46654524 | 0.35144304 | -1.3275131 | 0.184339 | 0.54912489 |
| DYNLL1 | 339.479081 | -0.466538861 | 0.55391487 | -0.8422573 | 0.39964393 | 0.72825165 |
| IFI16 | 743.04559 | -0.466521597 | 0.19438482 | -2.3999899 | 0.01639553 | 0.18378711 |
| LEPR | 467.323943 | -0.46593581 | 0.52476176 | -0.8878997 | 0.37459471 | 0.71257833 |
| TRAF7 | 52.7466519 | -0.465924436 | 0.42343279 | -1.1003504 | 0.27117949 | 0.63561102 |
| ITPR1 | 350.115468 | -0.46590408 | 0.27516002 | -1.6932114 | 0.09041524 | 0.40693312 |
| EIF3J | 162.894774 | -0.465880973 | 0.24018801 | -1.9396512 | 0.05242209 | 0.32059696 |
| ADIRF | 345.899625 | -0.465800749 | 0.86612438 | -0.5377989 | 0.59071588 | 0.84242385 |
| EIF1AX | 187.413779 | -0.465715178 | 0.26158529 | -1.7803569 | 0.07501757 | 0.37986929 |
| FAN1 | 65.9633036 | -0.464822257 | 0.25663688 | -1.811206 | 0.07010897 | 0.36582906 |
| CDKL1 | 104.953015 | -0.464665736 | 0.31817152 | -1.4604253 | 0.14417322 | 0.49679586 |
| CAPNS1 | 257.145008 | -0.463055725 | 0.37768014 | -1.2260526 | 0.22017887 | 0.58688069 |
| SLC35E3 | 113.860694 | -0.462915689 | 0.30089846 | -1.5384449 | 0.12393988 | 0.46936415 |
| NUDCD1 | 205.19097 | -0.462850191 | 0.19250904 | -2.4043037 | 0.01620331 | 0.18288505 |
| SNRPA | 68.1585224 | -0.462109816 | 0.29552939 | -1.5636679 | 0.11789559 | 0.46042456 |
| CSRNP1 | 63.3887466 | -0.46193471 | 0.68639742 | -0.6729843 | 0.50095726 | 0.79642461 |
| UTY | 438.290434 | -0.461305848 | 2.0429803 | -0.2258004 | 0.82135664 | 0.93726124 |
| TMCO1 | 140.660215 | -0.461236554 | 0.20814123 | -2.2159788 | 0.02669296 | 0.22752964 |
| RPL21 | 185.453589 | -0.461005521 | 0.49988839 | -0.9222169 | 0.35641545 | 0.70046372 |
| GUCD1 | 48.1234175 | -0.460837178 | 0.31490329 | -1.4634245 | 0.14335127 | 0.49538989 |
| SIVA1 | 90.4160905 | -0.460807921 | 0.33897708 | -1.3594073 | 0.17401755 | 0.53613916 |
| NOL7 | 107.483796 | -0.460776044 | 0.32322857 | -1.4255425 | 0.15400046 | 0.50978388 |
| SLC24A2 | 78.8754534 | -0.460706758 | 1.44196259 | -0.3194998 | 0.74934754 | 0.91075944 |
| EPN1 | 98.8442883 | -0.460631872 | 0.26949731 | -1.7092263 | 0.08740905 | 0.40003816 |
| IRAK3 | 653.954692 | -0.460604233 | 0.44640048 | -1.0318184 | 0.3021572 | 0.65926326 |
| VPS28 | 164.161401 | -0.460593292 | 0.35319174 | -1.3040885 | 0.1922034 | 0.55659725 |
| TMA16 | 73.1039774 | -0.460580779 | 0.26930158 | -1.7102788 | 0.08721434 | 0.39981956 |
| KXD1 | 61.0822595 | -0.460495398 | 0.22737105 | -2.0253035 | 0.04283621 | 0.29235086 |
| VAT1 | 122.571008 | -0.460306082 | 0.44811377 | -1.0272081 | 0.3043225 | 0.66086613 |
| TSHZ2 | 376.384218 | -0.459819613 | 0.49929305 | -0.9209414 | 0.35708105 | 0.70058911 |

| | | | | | | |
|---|---|---|---|---|---|---|
| RAB29 | 65.9959634 | -0.459445791 | 0.34829351 | -1.3191339 | 0.18712434 | 0.55379329 |
| CLEC2D | 68.763056 | -0.459338302 | 0.47546874 | -0.9660747 | 0.33400682 | 0.6852158 |
| PTTG1IP | 564.126082 | -0.459247941 | 0.36876846 | -1.2453558 | 0.21300099 | 0.57753052 |
| ARL6IP5 | 188.233623 | -0.459236381 | 0.47532166 | -0.9661592 | 0.33396454 | 0.6852158 |
| ZNF589 | 78.7420262 | -0.458882799 | 0.31666193 | -1.4491252 | 0.14730261 | 0.50127566 |
| KMT2B | 56.4585751 | -0.45828365 | 0.24496157 | -1.8708389 | 0.06136741 | 0.34778328 |
| ATP6V0A1 | 180.641727 | -0.458072936 | 0.26290614 | -1.742344 | 0.08144827 | 0.39205363 |
| ICMT | 57.9689186 | -0.45791774 | 0.31517382 | -1.4529054 | 0.14625003 | 0.50011031 |
| FAM129B | 168.031102 | -0.457811415 | 0.41238955 | -1.1101431 | 0.26693737 | 0.63222575 |
| EBLN3P | 65.163833 | -0.457411806 | 0.27050542 | -1.6909525 | 0.09084588 | 0.40744637 |
| PCBP3 | 103.4763 | -0.456764991 | 0.45171271 | -1.0111847 | 0.31192803 | 0.66561244 |
| AL606489.1 | 79.5719786 | -0.456754976 | 0.52569203 | -0.8688642 | 0.38492142 | 0.71699718 |
| MAN1A1 | 930.602041 | -0.455070176 | 0.24475949 | -1.8592544 | 0.06299108 | 0.35232811 |
| SMIM37 | 73.0254334 | -0.454196016 | 0.48944231 | -0.9279868 | 0.3534144 | 0.69977303 |
| AURKAIP1 | 107.357535 | -0.45417068 | 0.48592059 | -0.9346603 | 0.3499634 | 0.69858573 |
| SLBP | 96.2281006 | -0.453950796 | 0.22650964 | -2.0041125 | 0.04505802 | 0.29976402 |
| TRIM8 | 136.247857 | -0.453567138 | 0.37760318 | -1.2011741 | 0.22968369 | 0.59648134 |
| IL18BP | 68.5706773 | -0.453381351 | 0.42123684 | -1.0763098 | 0.28178871 | 0.64471757 |
| SBDS | 147.623807 | -0.452959232 | 0.376272 | -1.203808 | 0.22866381 | 0.59497702 |
| TMEM223 | 60.069487 | -0.45292045 | 0.33421919 | -1.3551599 | 0.17536661 | 0.53847091 |
| CAPZB | 528.186978 | -0.452795825 | 0.17203945 | -2.6319303 | 0.00849013 | 0.13062917 |
| PPP1CA | 82.4931743 | -0.452699491 | 0.43354992 | -1.0441692 | 0.29640709 | 0.65753344 |
| OPA3 | 93.422825 | -0.452469565 | 0.32211186 | -1.404697 | 0.1601114 | 0.51827199 |
| CRTC3 | 335.156475 | -0.452139696 | 0.15931626 | -2.838001 | 0.0045397 | 0.09149853 |
| SLC26A5 | 98.8361665 | -0.451828365 | 0.39734323 | -1.1371236 | 0.25548661 | 0.62092277 |
| RRP36 | 83.6889923 | -0.451767485 | 0.2795502 | -1.6160514 | 0.10608321 | 0.43901385 |
| ELMSAN1 | 266.782908 | -0.4517319 | 0.31511423 | -1.4335497 | 0.15170082 | 0.50771689 |
| MIR181A1HG | 202.965614 | -0.451609041 | 0.55818922 | -0.8090608 | 0.41848015 | 0.73821752 |
| SLC25A11 | 46.9276428 | -0.451295479 | 0.38458894 | -1.1734489 | 0.24061583 | 0.6051624 |
| SLC25A32 | 117.609794 | -0.450392362 | 0.27866027 | -1.6162776 | 0.1060343 | 0.43901385 |
| EPRS | 148.3855 | -0.450284161 | 0.2198239 | -2.0483858 | 0.04052221 | 0.28580837 |
| ECHDC1 | 137.539632 | -0.449558205 | 0.21727792 | -2.0690469 | 0.03854168 | 0.27738487 |
| CHCHD2 | 239.176952 | -0.449377092 | 0.46778513 | -0.9606485 | 0.33672893 | 0.68748823 |
| USP3-AS1 | 67.4425274 | -0.448753345 | 0.46126378 | -0.9728779 | 0.33061398 | 0.68317773 |
| APMAP | 60.7143348 | -0.448736811 | 0.3125731 | -1.435622 | 0.15110994 | 0.50677567 |
| NQO2 | 91.4227596 | -0.448524276 | 0.23444575 | -1.9131261 | 0.05573191 | 0.32988018 |
| UBA52 | 234.6403 | -0.448509658 | 0.54968917 | -0.8159332 | 0.41453833 | 0.73671465 |
| COX5B | 166.293517 | -0.448473269 | 0.41321926 | -1.0853155 | 0.27778194 | 0.64148182 |
| COPS9 | 78.2256524 | -0.448226851 | 0.46141333 | -0.9714215 | 0.3313384 | 0.68433673 |
| COL3A1 | 225.838865 | -0.447938213 | 1.1663434 | -0.3840535 | 0.70093883 | 0.8944316 |
| C19orf70 | 68.5216568 | -0.447818231 | 0.44246977 | -1.0120877 | 0.3114961 | 0.66561244 |
| EPCAM | 60.8189839 | -0.447519743 | 0.42735335 | -1.047189 | 0.2950124 | 0.65671864 |
| GANAB | 166.235889 | -0.447329475 | 0.3404003 | -1.3141277 | 0.18880322 | 0.55432448 |
| KDM5B | 233.244688 | -0.447265555 | 0.28496145 | -1.5695651 | 0.11651631 | 0.4592091 |
| IVD | 92.1490275 | -0.446975182 | 0.28721765 | -1.5562247 | 0.11965468 | 0.46316663 |
| CDV3 | 171.179793 | -0.44658911 | 0.26152993 | -1.7076023 | 0.08771016 | 0.40026508 |
| CHMP3 | 74.6331751 | -0.446436628 | 0.46932457 | -0.9512322 | 0.34148653 | 0.69116603 |
| ABL2 | 415.020505 | -0.446148203 | 0.17980345 | -2.4813105 | 0.01309003 | 0.16344193 |
| ANXA5 | 328.382289 | -0.445704707 | 0.35698425 | -1.2485276 | 0.21183789 | 0.57690664 |
| ANP32E | 74.6515432 | -0.445296767 | 0.21847307 | -2.0382227 | 0.04152767 | 0.28896334 |
| PLEKHG1 | 2813.10499 | -0.44495274 | 0.39816724 | -1.1175021 | 0.26377969 | 0.63003771 |
| LIX1-AS1 | 61.9621141 | -0.44398236 | 0.44775732 | -0.9915692 | 0.32140773 | 0.67541844 |
| DDRGK1 | 53.2345753 | -0.443781225 | 0.29447309 | -1.5070349 | 0.13180171 | 0.48092078 |
| MYH9 | 1318.42092 | -0.443591199 | 0.339658 | -1.3059937 | 0.19155471 | 0.55590015 |
| BTG2 | 87.3634562 | -0.443237928 | 0.49608349 | -0.8934745 | 0.37160314 | 0.7099763 |
| DAD1 | 188.121361 | -0.442965253 | 0.44157724 | -1.0031433 | 0.31579173 | 0.6703304 |
| RBBP8 | 111.933533 | -0.442952483 | 0.2489335 | -1.7794009 | 0.07517406 | 0.38042632 |
| PPP1R15A | 76.808165 | -0.442788088 | 0.53538062 | -0.8270529 | 0.40820709 | 0.73419295 |
| CTSZ | 84.3315324 | -0.442675966 | 0.34760978 | -1.2734854 | 0.20284586 | 0.56730017 |
| SETBP1 | 766.952425 | -0.442557478 | 0.21840429 | -2.0263223 | 0.04273177 | 0.29212534 |
| SF3B6 | 98.334981 | -0.442440978 | 0.4539969 | -0.9745463 | 0.32978538 | 0.68216508 |
| ACER3 | 523.622915 | -0.442331513 | 0.35446541 | -1.2478834 | 0.21207375 | 0.57690664 |
| NFIC | 371.785021 | -0.44221991 | 0.2522483 | -1.7531136 | 0.07958251 | 0.38738328 |
| PSMC3 | 43.4232602 | -0.441962467 | 0.43446187 | -1.0172641 | 0.30902781 | 0.66424338 |
| SRGN | 683.293498 | -0.44194702 | 0.69398461 | -0.6368254 | 0.52423859 | 0.80889528 |
| MRPL32 | 59.8192931 | -0.441618937 | 0.35169599 | -1.2556838 | 0.20923065 | 0.57454176 |
| MCRIP1 | 87.7730122 | -0.441551985 | 0.36071259 | -1.2241103 | 0.22091063 | 0.58706366 |
| LAMP2 | 223.464462 | -0.441281641 | 0.27497355 | -1.6048148 | 0.10853456 | 0.44451366 |
| BAIAP2L1 | 504.798824 | -0.440549819 | 0.51687499 | -0.8523334 | 0.39402908 | 0.72408262 |
| DDX39A | 40.668158 | -0.440530964 | 0.35908338 | -1.2268208 | 0.21988995 | 0.58687522 |
| NDUFA6 | 55.4397701 | -0.440303348 | 0.50049749 | -0.8797314 | 0.37900484 | 0.7138888 |
| ITPR2 | 678.75888 | -0.440233811 | 0.34205241 | -1.2870361 | 0.1980817 | 0.56327864 |
| BCAS2 | 56.4232914 | -0.440015834 | 0.29140299 | -1.5099908 | 0.13104577 | 0.47979755 |
| GFM1 | 136.62832 | -0.439980571 | 0.25706106 | -1.71158 | 0.0869741 | 0.39969788 |
| MCF2L2 | 106.201988 | -0.439333228 | 0.40372592 | -1.0881967 | 0.27650826 | 0.64080066 |
| EMC6 | 49.5188093 | -0.43925328 | 0.57778569 | -0.7602356 | 0.44711374 | 0.76128417 |
| NKAP | 49.4583866 | -0.438705264 | 0.26130292 | -1.6789145 | 0.09316871 | 0.41205855 |
| VAMP3 | 89.7243 | -0.438673128 | 0.282314 | -1.5538483 | 0.12022061 | 0.46382142 |
| ELOB | 158.440401 | -0.438513601 | 0.51155564 | -0.8572159 | 0.3913256 | 0.72247779 |
| GNA14-AS1 | 433.952319 | -0.438385447 | 0.5358854 | -0.8180582 | 0.41332396 | 0.73622771 |
| FLII | 95.7246739 | -0.438236819 | 0.2930185 | -1.4955944 | 0.1347594 | 0.48474744 |
| GRK5 | 537.709559 | -0.438154063 | 0.46050328 | -0.9514678 | 0.34136693 | 0.69116603 |
| PMEPA1 | 59.6654878 | -0.438023428 | 0.54628119 | -0.8018278 | 0.42265259 | 0.74011688 |
| ALG8 | 59.7350831 | -0.437395831 | 0.33843733 | -1.2923983 | 0.19621924 | 0.56122406 |
| WDR73 | 80.5587094 | -0.436303987 | 0.23535205 | -1.8538355 | 0.06376269 | 0.3533385 |
| C1orf43 | 139.23385 | -0.43628315 | 0.2198459 | -1.9844952 | 0.04720066 | 0.30523972 |
| LYRM4 | 173.001733 | -0.435981531 | 0.34822459 | -1.2520125 | 0.21056532 | 0.5762729 |
| ELOC | 128.552443 | -0.435820401 | 0.30485464 | -1.4296007 | 0.15283166 | 0.50900346 |
| RHOC | 378.368932 | -0.435632404 | 0.45806316 | -0.9510313 | 0.34158849 | 0.69120143 |
| ATP5F1D | 135.94203 | -0.434689684 | 0.30523503 | -1.4241147 | 0.15441328 | 0.51037454 |
| BTG3 | 58.9489609 | -0.434640505 | 0.26677141 | -1.6292619 | 0.10325758 | 0.43424379 |
| AP1AR | 89.0962102 | -0.433620876 | 0.27999918 | -1.5486505 | 0.12146576 | 0.46569232 |
| RPL22 | 78.5907625 | -0.433036062 | 0.30749223 | -1.4082829 | 0.1590473 | 0.51646194 |
| CD24 | 141.633315 | -0.432818145 | 0.2902272 | -1.491308 | 0.13588066 | 0.48555083 |
| KRI1 | 45.8768392 | -0.432806918 | 0.27050542 | -1.5999935 | 0.10960002 | 0.44576178 |
| PDIA3 | 346.742987 | -0.43263754 | 0.34907232 | -1.2393923 | 0.21520026 | 0.58008061 |
| AL513327.1 | 104.885392 | -0.432342815 | 0.68869266 | -0.6277732 | 0.5301525 | 0.8131655 |
| MDN1 | 200.25603 | -0.432114885 | 0.23353708 | -1.8503053 | 0.06426957 | 0.35347506 |
| ZBTB16 | 1081.45135 | -0.432043838 | 0.76916129 | -0.5617077 | 0.57431517 | 0.83430769 |
| EEA1 | 565.238878 | -0.43091008 | 0.27379961 | -1.5738155 | 0.1155301 | 0.45692741 |
| LMAN2 | 129.392206 | -0.430706126 | 0.32955912 | -1.3069161 | 0.1912412 | 0.55590015 |
| JCAD | 336.174754 | -0.430409541 | 0.28573011 | -1.5063499 | 0.13197736 | 0.48092078 |
| PLSCR1 | 845.612338 | -0.430267715 | 0.37523125 | -1.1466735 | 0.2515166 | 0.61661694 |
| SPNS1 | 45.2249696 | -0.430174512 | 0.33789975 | -1.2730833 | 0.2029885 | 0.56730017 |
| PPP1R11 | 44.1490717 | -0.429861094 | 0.33222309 | -1.2938929 | 0.19570242 | 0.56052955 |
| RABEPK | 59.9392372 | -0.429613637 | 0.36224131 | -1.1859874 | 0.23562726 | 0.60010516 |
| IL13RA1 | 259.15006 | -0.429496927 | 0.41635575 | -1.0315624 | 0.30227718 | 0.65926326 |
| GTF3C1 | 163.25864 | -0.429287716 | 0.24943395 | -1.7210477 | 0.08524218 | 0.39686932 |
| RAB5IF | 44.2726197 | -0.428875028 | 0.4419854 | -0.9703375 | 0.33187827 | 0.68476044 |
| TLE1 | 150.467348 | -0.428783153 | 0.35697547 | -1.2011558 | 0.22969078 | 0.59648134 |
| CLPTM1L | 92.3430973 | -0.42815624 | 0.28779965 | -1.4876885 | 0.13683305 | 0.4881015 |
| ACBD3 | 169.71765 | -0.42780135 | 0.22221603 | -1.9251598 | 0.05420938 | 0.32641304 |
| RECQL | 124.080325 | -0.427435857 | 0.16724053 | -2.555815 | 0.01059394 | 0.14485284 |

| | | | | | | |
|---|---|---|---|---|---|---|
| URM1 | 69.217903 | -0.42740884 | 0.28626299 | -1.4930636 | 0.13542055 | 0.48508022 |
| NDUFB11 | 95.9174655 | -0.426829116 | 0.48909406 | -0.8726933 | 0.38283026 | 0.71637321 |
| TRAPPC3 | 60.7038328 | -0.426809001 | 0.31130986 | -1.3710102 | 0.17037177 | 0.53193492 |
| POMP | 250.777009 | -0.426736155 | 0.33806741 | -1.2622812 | 0.2068476 | 0.57162978 |
| MYL12A | 665.419418 | -0.42654702 | 0.52092765 | -0.818822 | 0.41288798 | 0.73622771 |
| EPB41L1 | 48.7286468 | -0.426544915 | 0.51623472 | -0.8262616 | 0.40865573 | 0.73419295 |
| CPAMD8 | 176.159379 | -0.425912749 | 0.65675507 | -0.6485108 | 0.51665463 | 0.80498569 |
| NXN | 1192.36664 | -0.425835554 | 0.42065155 | -1.0123237 | 0.31138328 | 0.66561244 |
| HIVEP1 | 253.179538 | -0.425208083 | 0.36462746 | -1.1661439 | 0.24355627 | 0.608518 |
| POLR1C | 70.1659684 | -0.425177192 | 0.37398819 | -1.1368733 | 0.25559125 | 0.62095131 |
| BCL2L14 | 123.816658 | -0.425054227 | 0.30070134 | -1.4135428 | 0.15749618 | 0.51438736 |
| ABCC4 | 326.835239 | -0.42491558 | 0.37248064 | -1.1407722 | 0.25396471 | 0.61961636 |
| TPM1 | 197.023578 | -0.424671088 | 0.30512905 | -1.3917753 | 0.16399044 | 0.52357299 |
| PIN4 | 88.395537 | -0.424095433 | 0.2671055 | -1.5877451 | 0.112344 | 0.44910158 |
| WNT5B | 79.8205049 | -0.42339808 | 0.54717642 | -0.7737871 | 0.43905667 | 0.75352365 |
| EIF3F | 101.491243 | -0.422742018 | 0.34676744 | -1.2190938 | 0.22280861 | 0.58858855 |
| LRRC1 | 93.8129762 | -0.422599733 | 0.48067582 | -0.8791783 | 0.37930463 | 0.71402518 |
| XPNPEP1 | 107.906666 | -0.422576243 | 0.2454826 | -1.7214102 | 0.08517642 | 0.39686932 |
| EEF1D | 284.254727 | -0.422555358 | 0.30642056 | -1.3790046 | 0.16789333 | 0.52916221 |
| MRPS21 | 125.263924 | -0.422080821 | 0.27710457 | -1.5231825 | 0.12771306 | 0.47495385 |
| OGFOD3 | 40.3543128 | -0.422065168 | 0.35425719 | -1.1914089 | 0.2334931 | 0.59914521 |
| KCNIP4 | 4946.8846 | -0.421866739 | 0.66037588 | -0.6388282 | 0.5229347 | 0.80854347 |
| HOXB7 | 60.9974139 | -0.421310794 | 0.40538717 | -1.0392801 | 0.29867451 | 0.65856711 |
| SLC9A1 | 60.2096635 | -0.421113001 | 0.33341943 | -1.2630128 | 0.20658459 | 0.57138701 |
| SMIM19 | 55.8163607 | -0.420942975 | 0.3435081 | -1.2254237 | 0.22041562 | 0.58693817 |
| DHX40 | 85.1211575 | -0.420468816 | 0.31434659 | -1.3375962 | 0.18102809 | 0.54524118 |
| GNG12 | 321.191719 | -0.420350494 | 0.18727927 | -2.2445116 | 0.0247995 | 0.21915154 |
| MCM3 | 59.8238118 | -0.419944633 | 0.34424491 | -1.2199008 | 0.22250249 | 0.58828363 |
| AC048341.1 | 82.0633214 | -0.419275472 | 0.35314618 | -1.1872576 | 0.23512604 | 0.59949266 |
| GPX4 | 184.351451 | -0.418944887 | 0.43795652 | -0.9565901 | 0.33887417 | 0.68942777 |
| DRAM2 | 102.334104 | -0.41888988 | 0.2512972 | -1.6669103 | 0.09553225 | 0.41575423 |
| MRFAP1 | 188.870391 | -0.418801996 | 0.41080107 | -1.0194764 | 0.30797685 | 0.66414532 |
| EIF5 | 188.854181 | -0.418522693 | 0.37440934 | -1.1178212 | 0.26364337 | 0.62989598 |
| ATP5MPL | 87.5426684 | -0.418369525 | 0.35750028 | -1.1702635 | 0.24189495 | 0.60662775 |
| TNXB | 295.896677 | -0.418163917 | 0.57259204 | -0.7302999 | 0.46520688 | 0.77477727 |
| DOCK9 | 1921.62386 | -0.41814333 | 0.26303454 | -1.5896898 | 0.11190474 | 0.44839019 |
| MRPS34 | 72.1966724 | -0.418030553 | 0.49558262 | -0.8435133 | 0.39894141 | 0.72771679 |
| CASP3 | 78.6130421 | -0.417990419 | 0.21099019 | -1.9810893 | 0.04758126 | 0.30623143 |
| ELOA | 65.5832505 | -0.417953053 | 0.2787971 | -1.4991298 | 0.13383996 | 0.48375105 |
| EEFSEC | 176.345095 | -0.417828242 | 0.28693577 | -1.4561734 | 0.14534469 | 0.49826376 |
| ATN1 | 178.169765 | -0.417267283 | 0.23962951 | -1.7413018 | 0.0816307 | 0.39270076 |
| VAMP7 | 71.4258962 | -0.416597964 | 0.23729359 | -1.7556225 | 0.07915289 | 0.3870932 |
| MPZL1 | 232.742953 | -0.416223766 | 0.31728566 | -1.3118266 | 0.18957866 | 0.55529769 |
| MIEF1 | 39.5320449 | -0.415831526 | 0.32866067 | -1.2652306 | 0.20578869 | 0.57060151 |
| ATP1B3 | 532.804569 | -0.415444383 | 0.42709667 | -0.9727174 | 0.33069375 | 0.68317773 |
| BMS1 | 119.248194 | -0.415364403 | 0.23199488 | -1.7904034 | 0.07338909 | 0.37510489 |
| EIF3A | 307.201987 | -0.415040394 | 0.29071773 | -1.4276405 | 0.15339537 | 0.50963635 |
| PPTC7 | 99.9559215 | -0.4150089 | 0.21678031 | -1.9144216 | 0.05556631 | 0.32988018 |
| GYG1 | 58.2264057 | -0.41495381 | 0.20847518 | -1.9904231 | 0.04654435 | 0.30359341 |
| CANT1 | 42.3422059 | -0.414722217 | 0.32531278 | -1.2748414 | 0.20236537 | 0.56693091 |
| MXI1 | 100.114863 | -0.414120405 | 0.40002478 | -1.0352369 | 0.3005583 | 0.65926326 |
| MRPS36 | 58.4801405 | -0.414023682 | 0.22283339 | -1.8579966 | 0.0631695 | 0.35284368 |
| GNAI2 | 688.904152 | -0.413913661 | 0.28417 | -1.4565706 | 0.14523496 | 0.49826376 |
| CBLB | 705.626363 | -0.41378694 | 0.36002362 | -1.1493327 | 0.2504188 | 0.61573829 |
| ZNF208 | 162.415593 | -0.413108916 | 0.26869595 | -1.5374587 | 0.12418103 | 0.46936415 |
| SLC30A7 | 371.204454 | -0.412621695 | 0.30213898 | -1.3656685 | 0.17204302 | 0.53377758 |
| MAP1LC3B | 147.695658 | -0.412122115 | 0.40379816 | -1.0206141 | 0.30743729 | 0.66400081 |
| USP37 | 210.601297 | -0.411741979 | 0.26576719 | -1.5492581 | 0.12131968 | 0.46542849 |
| CNTNAP3 | 114.080993 | -0.411505177 | 0.54301381 | -0.7578171 | 0.44856046 | 0.76192883 |
| GTF2H5 | 63.0104633 | -0.411419827 | 0.39875685 | -1.0317561 | 0.30218637 | 0.65926326 |
| FAM107B | 179.15354 | -0.411350477 | 0.5102504 | -0.8061737 | 0.42014268 | 0.73901389 |
| THBD | 214.055356 | -0.411205906 | 0.4927887 | -0.8344467 | 0.40402931 | 0.73096878 |
| HSPA13 | 48.468505 | -0.411120943 | 0.28644967 | -1.4352292 | 0.15122179 | 0.50694302 |
| LPCAT2 | 106.964926 | -0.410718037 | 0.67160922 | -0.6115432 | 0.54084004 | 0.81928059 |
| CASD1 | 198.517809 | -0.41064856 | 0.22383893 | -1.8345717 | 0.06656918 | 0.35767274 |
| KLRD1 | 88.2153527 | -0.410464568 | 0.72333838 | -0.5674586 | 0.57040266 | 0.83368112 |
| WDR27 | 208.567325 | -0.410424748 | 0.32888198 | -1.2479393 | 0.21205328 | 0.57690664 |
| ACSF3 | 63.3423358 | -0.410404657 | 0.27710624 | -1.4810372 | 0.13859666 | 0.49011948 |
| C3orf58 | 80.2726732 | -0.410079534 | 0.3624209 | -1.1315008 | 0.25784437 | 0.62258498 |
| PDCD11 | 57.1825175 | -0.409847689 | 0.48070466 | -0.8525977 | 0.39388243 | 0.72408262 |
| PSME1 | 135.060128 | -0.409844301 | 0.40625879 | -1.0088257 | 0.31305825 | 0.6674715 |
| PRMT1 | 138.322614 | -0.409774102 | 0.42996918 | -0.9530313 | 0.3405742 | 0.69116603 |
| OSTC | 94.6448538 | -0.409623129 | 0.39785383 | -1.029582 | 0.30320628 | 0.6597014 |
| TRIM28 | 83.2445248 | -0.408761123 | 0.36837104 | -1.1096451 | 0.26715199 | 0.63237047 |
| HGS | 86.1430997 | -0.408526514 | 0.31355808 | -1.3028735 | 0.19261795 | 0.55703491 |
| PDE10A | 1599.35131 | -0.40845144 | 0.44242005 | -0.9232209 | 0.35589209 | 0.70018061 |
| PROS1 | 280.412143 | -0.407950451 | 0.32899547 | -1.239988 | 0.21497982 | 0.57982196 |
| PFKL | 164.311423 | -0.40793415 | 0.23226557 | -1.7563264 | 0.07903269 | 0.38686531 |
| AGAP6 | 82.4758269 | -0.407439689 | 0.31424057 | -1.2965852 | 0.19477393 | 0.55995114 |
| RAB1B | 94.0604802 | -0.40733752 | 0.40861791 | -0.9968665 | 0.3188293 | 0.67380686 |
| CRYBG3 | 469.190968 | -0.407163335 | 0.24863891 | -1.6375688 | 0.10151167 | 0.42995342 |
| ZNHIT1 | 106.027742 | -0.407074551 | 0.42590093 | -0.9557963 | 0.33917513 | 0.68955779 |
| MEPCE | 51.9395713 | -0.406922412 | 0.38356357 | -1.0608995 | 0.28873556 | 0.65160592 |
| MARS | 70.5521297 | -0.406875777 | 0.3345422 | -1.2162166 | 0.22390242 | 0.59051502 |
| GMDS | 931.537063 | -0.406810841 | 0.21524159 | -1.8900197 | 0.05875533 | 0.34002466 |
| EFNA1 | 249.889383 | -0.406479432 | 0.30825975 | -1.3186264 | 0.18729405 | 0.55388137 |
| F2R | 162.981254 | -0.40634086 | 0.32398005 | -1.2542157 | 0.20976361 | 0.57542597 |
| GABPB1-AS1 | 167.901536 | -0.406154315 | 0.37687771 | -1.077682 | 0.28117571 | 0.64449883 |
| DTNA | 158.35734 | -0.406141563 | 1.19063414 | -0.3411137 | 0.73301802 | 0.90508002 |
| TNFAIP2 | 101.167901 | -0.406116485 | 0.48934437 | -0.8299196 | 0.40658424 | 0.73348299 |
| KDM4B | 242.354098 | -0.405756498 | 0.30137276 | -1.3463609 | 0.17818616 | 0.54184221 |
| PDIA6 | 216.32464 | -0.405668114 | 0.33419916 | -1.2138514 | 0.22480449 | 0.59112311 |
| ZNF292 | 730.8216 | -0.405182306 | 0.36609624 | -1.1067645 | 0.26839577 | 0.63347822 |
| PILRB | 133.89853 | -0.405020689 | 0.35360284 | -1.1454113 | 0.25203882 | 0.61768005 |
| TALDO1 | 93.5756392 | -0.404389624 | 0.27355938 | -1.4782517 | 0.13934041 | 0.49091481 |
| CCT6A | 127.444209 | -0.404181204 | 0.29425727 | -1.373564 | 0.16957707 | 0.5312985 |
| ERN1 | 82.9085533 | -0.404029017 | 0.28594516 | -1.4129598 | 0.15766754 | 0.51443122 |
| HTATSF1 | 40.6951221 | -0.403488038 | 0.39660859 | -1.0173457 | 0.30898901 | 0.66424338 |
| EDF1 | 180.575748 | -0.403463683 | 0.45265572 | -0.8913257 | 0.37275446 | 0.71084822 |
| PDCD6 | 92.2943948 | -0.403196942 | 0.34834454 | -1.1574659 | 0.24708204 | 0.61194684 |
| SLC39A11 | 244.137143 | -0.403008498 | 0.37952741 | -1.0618693 | 0.28829503 | 0.65138125 |
| NMRAL1 | 48.8090431 | -0.402345171 | 0.28470112 | -1.4132195 | 0.1575912 | 0.51438736 |
| BLVRA | 45.8538303 | -0.402268229 | 0.29368617 | -1.3697214 | 0.17077388 | 0.53193492 |
| MRPL3 | 89.7613001 | -0.40225059 | 0.29867375 | -1.3467892 | 0.17804812 | 0.54178558 |
| NDUFB10 | 98.6633856 | -0.402244913 | 0.49506177 | -0.8125146 | 0.41649641 | 0.73802299 |
| CREG1 | 50.4064424 | -0.402073456 | 0.31396225 | -1.2806427 | 0.2003192 | 0.56505068 |
| MYL12B | 760.796133 | -0.401646602 | 0.55750471 | -0.7204362 | 0.47125644 | 0.77841975 |
| EMC1 | 44.7002886 | -0.401356492 | 0.30619016 | -1.3108079 | 0.18992266 | 0.55544572 |
| RGL3 | 128.617499 | -0.401250862 | 0.54208721 | -0.7401961 | 0.45918098 | 0.76976328 |
| DYNLT3 | 68.4892216 | -0.401095657 | 0.41015861 | -0.9779038 | 0.32812192 | 0.68061385 |
| PRKDC | 375.18346 | -0.401073106 | 0.22067203 | -1.8175077 | 0.06913941 | 0.36386894 |
| TMEM167B | 55.1397254 | -0.400985734 | 0.25984973 | -1.5431447 | 0.12279567 | 0.46808475 |
| GLO1 | 109.403268 | -0.400619346 | 0.18077011 | -2.2161813 | 0.02667909 | 0.22752964 |

| | | | | | | |
|---|---|---|---|---|---|---|
| ACTR3B | 76.8761715 | -0.400520424 | 0.30929482 | -1.2949471 | 0.19533848 | 0.56007526 |
| UQCC3 | 50.0040677 | -0.400495236 | 0.39584023 | -1.0117598 | 0.31165291 | 0.66561244 |
| GNB2 | 212.262923 | -0.400461976 | 0.37744924 | -1.0609691 | 0.28870394 | 0.65160592 |
| GSTK1 | 114.56148 | -0.400393727 | 0.30472444 | -1.3139535 | 0.18886187 | 0.55432448 |
| FAM120C | 71.5434432 | -0.400086899 | 0.2319862 | -1.7246151 | 0.08459687 | 0.3964812 |
| CMTM6 | 165.785081 | -0.400021357 | 0.28775257 | -1.3901574 | 0.16448109 | 0.52418378 |
| ETF1 | 136.617864 | -0.399860602 | 0.22694418 | -1.7619337 | 0.0780805 | 0.38566641 |
| DCAF13 | 58.4331541 | -0.399780174 | 0.28709556 | -1.3924986 | 0.16377146 | 0.52357299 |
| MIDN | 291.939663 | -0.399663714 | 0.37319547 | -1.0709233 | 0.28420393 | 0.64726501 |
| CYYR1-AS1 | 60.0859846 | -0.399574539 | 0.6375022 | -0.6267814 | 0.53080252 | 0.81324789 |
| IRX3 | 190.51673 | -0.399446094 | 0.42765426 | -0.9340398 | 0.35028337 | 0.69858573 |
| THSD4-AS1 | 218.037236 | -0.399286618 | 0.41285745 | -0.9671295 | 0.3334793 | 0.6852158 |
| ERCC1 | 42.8970063 | -0.39905916 | 0.28131561 | -1.4185461 | 0.15603139 | 0.51234409 |
| DYNLT1 | 183.854183 | -0.398446319 | 0.27836362 | -1.4313879 | 0.15231908 | 0.50833076 |
| TLDC1 | 49.7601966 | -0.398199258 | 0.37249981 | -1.0689918 | 0.28507334 | 0.6478076 |
| RNF145 | 460.914869 | -0.398032285 | 0.36819235 | -1.0810444 | 0.27967735 | 0.64286511 |
| EIF4G1 | 168.47808 | -0.397941353 | 0.23505228 | -1.6929908 | 0.09045721 | 0.40693312 |
| RALB | 173.274376 | -0.397936002 | 0.26116815 | -1.5236774 | 0.12758932 | 0.47479008 |
| TXLNA | 92.2887611 | -0.397783966 | 0.28111326 | -1.415031 | 0.15705941 | 0.51417853 |
| AL356599.1 | 68.2950386 | -0.39747453 | 0.38539746 | -1.0313367 | 0.30238297 | 0.65926326 |
| TMEM59 | 299.495362 | -0.397472298 | 0.40716494 | -0.9761948 | 0.32896794 | 0.68172735 |
| TXNDC12 | 98.3850992 | -0.397393466 | 0.23741524 | -1.673833 | 0.09416342 | 0.41293638 |
| HLA-C | 1104.23925 | -0.39721269 | 0.58242843 | -0.681994 | 0.49524275 | 0.79337734 |
| MBD3 | 53.0180106 | -0.397042241 | 0.34600537 | -1.1475031 | 0.25117375 | 0.61648314 |
| XKR6 | 331.827554 | -0.39672322 | 0.43455937 | -0.9129322 | 0.36127823 | 0.70386371 |
| ELOVL7 | 176.798077 | -0.396580363 | 0.50696753 | -0.7822599 | 0.43406186 | 0.74808934 |
| ERI1 | 73.2860503 | -0.39584175 | 0.27743033 | -1.426815 | 0.15363323 | 0.50978388 |
| NASP | 207.116046 | -0.395686982 | 0.24018805 | -1.6474049 | 0.09947484 | 0.42413631 |
| PTP4A1 | 79.0226239 | -0.395465224 | 0.40918667 | -0.9664665 | 0.33381078 | 0.6852158 |
| WDR60 | 310.558711 | -0.395331481 | 0.25386985 | -1.5572211 | 0.11941801 | 0.46290743 |
| NCSTN | 68.5932408 | -0.395281141 | 0.26630498 | -1.4843175 | 0.13772471 | 0.48962215 |
| IPO7 | 238.124662 | -0.394846708 | 0.28142495 | -1.4030267 | 0.16060888 | 0.51944206 |
| CENPP | 177.120005 | -0.394771063 | 0.41866463 | -0.9429291 | 0.34571716 | 0.69497757 |
| HCFC1R1 | 52.2696342 | -0.394387918 | 0.47214338 | -0.8353139 | 0.403541 | 0.73065697 |
| TPGS1 | 48.5724262 | -0.394277531 | 0.32747508 | -1.2039925 | 0.22859249 | 0.59497702 |
| SYNGR2 | 97.3479739 | -0.394211833 | 0.47735277 | -0.8258291 | 0.40890102 | 0.73419295 |
| ARF6 | 88.5082811 | -0.394064898 | 0.28911287 | -1.363014 | 0.17287809 | 0.53502507 |
| EFCAB14 | 212.584859 | -0.393906983 | 0.26679613 | -1.4764344 | 0.13982732 | 0.49118733 |
| OAT | 48.2191762 | -0.393689664 | 0.49283349 | -0.798829 | 0.4243896 | 0.74157166 |
| ESRRA | 58.4628991 | -0.393650613 | 0.32901889 | -1.1964377 | 0.23152578 | 0.59795465 |
| CTSC | 111.300113 | -0.393641825 | 0.30073142 | -1.3089481 | 0.19055193 | 0.55558405 |
| EDIL3 | 428.976495 | -0.393397597 | 0.95423472 | -0.412265 | 0.68014519 | 0.88287247 |
| KARS | 69.996014 | -0.392991393 | 0.33550306 | -1.1713496 | 0.24145827 | 0.60620642 |
| COX15 | 59.9010715 | -0.392815334 | 0.20458622 | -1.9200478 | 0.05485186 | 0.32810875 |
| AP1B1 | 105.831999 | -0.392803423 | 0.26127445 | -1.5034131 | 0.13273256 | 0.48209077 |
| STX4 | 62.8489138 | -0.392338221 | 0.24229123 | -1.6192836 | 0.10538625 | 0.43797647 |
| PABPC4 | 54.1668733 | -0.392188282 | 0.25423363 | -1.5426294 | 0.12292072 | 0.46827757 |
| CTBS | 111.551888 | -0.392059534 | 0.34681012 | -1.1304732 | 0.2582769 | 0.6230042 |
| NDUFS6 | 112.835485 | -0.392025977 | 0.36004437 | -1.0888268 | 0.27623025 | 0.64068436 |
| LYRM2 | 65.737608 | -0.391995787 | 0.26302148 | -1.4903565 | 0.13613051 | 0.4860192 |
| TMEM167A | 98.2637018 | -0.391971412 | 0.28325098 | -1.3838308 | 0.16641028 | 0.52671378 |
| STOM | 1181.27495 | -0.391889414 | 0.47300071 | -0.8285176 | 0.40737743 | 0.73419295 |
| ELP6 | 49.0437328 | -0.391727993 | 0.35827454 | -1.0933738 | 0.27422971 | 0.63975272 |
| TPD52L2 | 104.279406 | -0.391401726 | 0.23064049 | -1.6970209 | 0.08969271 | 0.40583359 |
| LAMTOR4 | 180.025017 | -0.391371916 | 0.43760903 | -0.8943415 | 0.3711392 | 0.70984233 |
| CCDC107 | 73.2568225 | -0.391314474 | 0.4424163 | -0.8844938 | 0.37642971 | 0.71257833 |
| GPR137B | 69.9482048 | -0.39126929 | 0.40707156 | -0.9611806 | 0.33646136 | 0.68745652 |
| ATP1A1 | 677.745888 | -0.391110163 | 0.36056849 | -1.0847042 | 0.27805268 | 0.64148182 |
| MAGI2 | 935.654146 | -0.391085297 | 0.33057261 | -1.1830541 | 0.23678769 | 0.6006304 |
| ASNSD1 | 52.1453223 | -0.39067983 | 0.32580187 | -1.1991332 | 0.23047617 | 0.59683118 |
| RWDD1 | 75.6943902 | -0.390574012 | 0.4146761 | -0.9418773 | 0.34625545 | 0.69521104 |
| PRPF31 | 66.2063173 | -0.389485233 | 0.2978683 | -1.3075753 | 0.1910174 | 0.55586607 |
| METTL8 | 82.9831764 | -0.389374596 | 0.31460902 | -1.237646 | 0.21584736 | 0.58158674 |
| VWA1 | 217.813115 | -0.389011171 | 0.97755172 | -0.3979443 | 0.69067122 | 0.88832346 |
| MAGI1 | 2513.0468 | -0.388949702 | 0.24469893 | -1.5895031 | 0.11194686 | 0.44839019 |
| PTPRE | 657.294116 | -0.38820457 | 0.34324845 | -1.1309725 | 0.25806665 | 0.62293521 |
| MAP2K1 | 295.733118 | -0.387975645 | 0.34571457 | -1.1222427 | 0.26175927 | 0.62704219 |
| AAK1 | 692.854455 | -0.387434526 | 0.20259769 | -1.9123344 | 0.05583332 | 0.32988018 |
| RAB21 | 259.591166 | -0.387399363 | 0.27082238 | -1.4304555 | 0.15258633 | 0.50880764 |
| TMEM160 | 77.9066207 | -0.387271892 | 0.41786234 | -0.926793 | 0.35403402 | 0.69977303 |
| DOC2B | 163.425397 | -0.386837537 | 0.36893674 | -1.04852 | 0.29439912 | 0.65671864 |
| OCIAD2 | 62.9601964 | -0.38680892 | 0.4038158 | -0.9578846 | 0.33812097 | 0.68878364 |
| TSSC4 | 48.2311758 | -0.386580591 | 0.49047242 | -0.7881801 | 0.43059138 | 0.74477473 |
| HK1 | 326.553521 | -0.386250046 | 0.16275466 | -2.3732042 | 0.01763451 | 0.1890691 |
| SHISA3 | 52.7527836 | -0.386170415 | 0.9632997 | -0.4008829 | 0.68850631 | 0.88739126 |
| SUMO3 | 110.34774 | -0.38560813 | 0.42229522 | -0.9131245 | 0.36117704 | 0.70386371 |
| KIAA1551 | 389.124607 | -0.385207106 | 0.30035717 | -1.2824968 | 0.19966843 | 0.56449294 |
| TMEM179B | 84.7867894 | -0.384778561 | 0.31597974 | -1.2177318 | 0.22332592 | 0.5893183 |
| KIF26B | 569.059566 | -0.384335133 | 0.64864743 | -0.5925178 | 0.55350392 | 0.82372318 |
| H3F3B | 902.974797 | -0.384172285 | 0.35268214 | -1.0892876 | 0.27602707 | 0.64068436 |
| BRK1 | 144.940456 | -0.383706364 | 0.37599291 | -1.0205149 | 0.30748433 | 0.66400081 |
| CHMP2A | 93.346665 | -0.383361851 | 0.45370227 | -0.8449635 | 0.39813123 | 0.72753637 |
| PIK3R3 | 183.541871 | -0.383232558 | 0.53449956 | -0.7169932 | 0.47337828 | 0.77899012 |
| NPAS3 | 952.216602 | -0.383180098 | 0.53109169 | -0.7214952 | 0.4706049 | 0.77797169 |
| DPP9 | 93.5109058 | -0.383059124 | 0.28551691 | -1.3416338 | 0.17971477 | 0.5436621 |
| LSAMP | 76.0604823 | -0.382790725 | 0.40638085 | -0.9419507 | 0.34621789 | 0.69521104 |
| SPTLC2 | 657.659305 | -0.38227615 | 0.41347468 | -0.9245455 | 0.35520238 | 0.70018061 |
| HNRNPA1 | 131.013615 | -0.382160474 | 0.36916988 | -1.0351887 | 0.30058082 | 0.65926326 |
| SSU72 | 127.159893 | -0.381989194 | 0.29939055 | -1.2758893 | 0.20199466 | 0.56693091 |
| FCGRT | 201.456486 | -0.38195541 | 0.3856663 | -0.990378 | 0.3219894 | 0.6761919 |
| OS9 | 167.115383 | -0.38150246 | 0.25714057 | -1.4836339 | 0.13790607 | 0.48979401 |
| CDIP1 | 51.5442686 | -0.381372201 | 0.27813795 | -1.3711621 | 0.17032444 | 0.53193492 |
| TXNL4A | 105.351558 | -0.381219087 | 0.28361782 | -1.3441295 | 0.17890651 | 0.54282239 |
| CISD2 | 53.3306302 | -0.381029792 | 0.23828592 | -1.5990445 | 0.10981071 | 0.44576178 |
| NFE2L1 | 173.485165 | -0.380862753 | 0.25366382 | -1.5014469 | 0.13324001 | 0.48236854 |
| UPB1 | 53.1874246 | -0.380519424 | 0.54535021 | -0.6977524 | 0.48533204 | 0.78823909 |
| NDUFB7 | 118.091852 | -0.380059263 | 0.37854026 | -1.0040128 | 0.31537245 | 0.66961411 |
| DEXI | 68.5727872 | -0.378561746 | 0.28672109 | -1.3203136 | 0.18673035 | 0.55302731 |
| ECE1 | 800.433837 | -0.378479037 | 0.25913546 | -1.4605452 | 0.14414031 | 0.49679586 |
| COMMD6 | 97.9974606 | -0.378333925 | 0.49764283 | -0.7602519 | 0.447104 | 0.76128417 |
| PSMA4 | 141.830813 | -0.378186177 | 0.31211845 | -1.2116752 | 0.22563674 | 0.59165529 |
| TCP1 | 107.627215 | -0.378170952 | 0.29299621 | -1.2907025 | 0.19680685 | 0.56172669 |
| NPRL3 | 96.1783483 | -0.37785087 | 0.32318949 | -1.169131 | 0.24235084 | 0.6070269 |
| JMJD8 | 40.3716037 | -0.377038279 | 0.40471015 | -0.9316255 | 0.35153012 | 0.69889281 |
| SUPT6H | 152.474012 | -0.376943257 | 0.24509623 | -1.5379399 | 0.12406332 | 0.46936415 |
| RNF24 | 205.28728 | -0.376888327 | 0.4204271 | -0.8964416 | 0.37001697 | 0.70984233 |
| KIAA1191 | 66.9651857 | -0.3768473 | 0.26468174 | -1.4237752 | 0.15451156 | 0.51037454 |
| 10-Sep | 229.036231 | -0.376386051 | 0.17909204 | -2.1016347 | 0.03558529 | 0.26544614 |
| UQCRH | 118.573033 | -0.376034742 | 0.38730766 | -0.9708942 | 0.33160099 | 0.68453353 |
| EFCAB13 | 109.562675 | -0.375764752 | 0.31143868 | -1.2065449 | 0.22760745 | 0.59434823 |
| TTLL5 | 337.043482 | -0.375655407 | 0.2130905 | -1.7628914 | 0.07791881 | 0.38566641 |
| ACSS3 | 59.7011213 | -0.375388492 | 0.98057418 | -0.3828252 | 0.70184939 | 0.89443605 |
| TGFB1I1 | 48.8545469 | -0.375035711 | 0.35106487 | -1.0682804 | 0.28539406 | 0.64806045 |

| | | | | | | |
|---|---|---|---|---|---|---|
| NCAPD2 | 45.6540007 | -0.374969732 | 0.25614632 | -1.4638888 | 0.14322433 | 0.49538989 |
| GART | 90.0621104 | -0.374959897 | 0.32144982 | -1.1664648 | 0.24342658 | 0.608518 |
| ZCRB1 | 45.1879774 | -0.374831221 | 0.24710687 | -1.516879 | 0.12929724 | 0.47762865 |
| NDUFB3 | 96.9093087 | -0.374727686 | 0.54395068 | -0.6889001 | 0.49088613 | 0.79073252 |
| PURB | 124.530496 | -0.374599839 | 0.23019315 | -1.6273284 | 0.10366738 | 0.43503085 |
| MSL3 | 55.7559804 | -0.374525491 | 0.36604318 | -1.023173 | 0.30622606 | 0.66324139 |
| SNHG25 | 45.2379926 | -0.374002928 | 0.4208829 | -0.8886152 | 0.37420994 | 0.71229586 |
| PSMC1 | 102.86357 | -0.373947597 | 0.31160579 | -1.2000663 | 0.2301136 | 0.59648134 |
| TOP3A | 67.1066696 | -0.373697993 | 0.22460631 | -1.6637912 | 0.09615417 | 0.41727618 |
| CHMP4B | 134.800717 | -0.373503414 | 0.22025089 | -1.6958088 | 0.08992209 | 0.40608855 |
| TMEM263 | 86.5358947 | -0.373426579 | 0.25819108 | -1.4463187 | 0.14808784 | 0.50152657 |
| DHRS7B | 179.145039 | -0.37318666 | 0.30479174 | -1.2243989 | 0.22080179 | 0.58706366 |
| DNAAF5 | 53.3665578 | -0.372502713 | 0.27593433 | -1.3499687 | 0.17702602 | 0.54037749 |
| LTBR | 133.076637 | -0.37243267 | 0.23654155 | -1.5744915 | 0.11537385 | 0.4568354 |
| FLOT1 | 375.475821 | -0.372009672 | 0.27551153 | -1.3502508 | 0.17693554 | 0.54037749 |
| AC078881.1 | 89.4527126 | -0.371805905 | 0.42161758 | -0.8818558 | 0.37785479 | 0.71305115 |
| TMEM41A | 47.7542389 | -0.37150532 | 0.27875399 | -1.3327354 | 0.18261864 | 0.54790102 |
| RTL8C | 64.5265979 | -0.370683262 | 0.38315974 | -0.9674379 | 0.33332516 | 0.6852158 |
| BRI3 | 403.32007 | -0.370670997 | 0.3251355 | -1.1400508 | 0.25426513 | 0.61966901 |
| RUNDC1 | 39.6135274 | -0.369803086 | 0.26691648 | -1.3854637 | 0.16591073 | 0.5261375 |
| ZRANB3 | 142.518132 | -0.369668182 | 0.23577198 | -1.5679055 | 0.1169032 | 0.4596919 |
| GTF2F1 | 47.7712454 | -0.369259086 | 0.27468318 | -1.3443091 | 0.17884847 | 0.54282239 |
| AC114763.1 | 95.6672293 | -0.369038756 | 0.34516011 | -1.0691814 | 0.28498795 | 0.64779344 |
| SPCS2 | 191.377497 | -0.368934646 | 0.33536326 | -1.1001045 | 0.27128658 | 0.63561102 |
| PDZK1IP1 | 125.380197 | -0.368623964 | 0.68927025 | -0.5348032 | 0.59278592 | 0.84426894 |
| AAR2 | 42.7698254 | -0.368546171 | 0.22846084 | -1.61317 | 0.10670759 | 0.43932066 |
| ALDOB | 169.8956 | -0.368534198 | 1.1070456 | -0.3328988 | 0.73921064 | 0.908116 |
| ARPC5 | 195.085274 | -0.368472471 | 0.22369582 | -1.6472032 | 0.09951628 | 0.42413631 |
| TFDP1 | 72.1089781 | -0.368384582 | 0.35795203 | -1.0291451 | 0.30341149 | 0.65997242 |
| MTRF1L | 55.0109833 | -0.36795743 | 0.27750435 | -1.3259519 | 0.18485562 | 0.5500631 |
| MORC4 | 47.6905925 | -0.367701758 | 0.38616116 | -0.9521977 | 0.34099674 | 0.69116603 |
| MAP3K12 | 45.8054364 | -0.366995162 | 0.33988182 | -1.0797728 | 0.28024335 | 0.64362372 |
| MMP24 | 52.2816654 | -0.366782715 | 0.33579309 | -1.0922879 | 0.27470659 | 0.63988729 |
| COLGALT1 | 93.6342128 | -0.366662205 | 0.33817183 | -1.0842482 | 0.27825475 | 0.64157752 |
| ZBTB17 | 55.7748626 | -0.366503168 | 0.32927179 | -1.1130719 | 0.26567756 | 0.63060082 |
| KLHL9 | 64.9019616 | -0.366169597 | 0.34836767 | -1.051101 | 0.29321222 | 0.6556551 |
| SERINC5 | 206.695396 | -0.366074953 | 0.34188579 | -1.0707522 | 0.28428088 | 0.64726501 |
| TMEM181 | 231.924343 | -0.365941632 | 0.20241791 | -1.807852 | 0.07062954 | 0.36751516 |
| GFOD2 | 275.601091 | -0.365783074 | 0.22864431 | -1.5997909 | 0.10964498 | 0.44576178 |
| PPP1R18 | 84.8861254 | -0.365012134 | 0.30751641 | -1.186968 | 0.23524025 | 0.59949266 |
| CCDC130 | 80.128763 | -0.364850037 | 0.23286077 | -1.5668163 | 0.11715765 | 0.45991971 |
| ABHD17B | 112.618312 | -0.364761735 | 0.37577683 | -0.9706871 | 0.3317041 | 0.68457368 |
| MED29 | 48.9274751 | -0.363927834 | 0.29676308 | -1.2263245 | 0.22007659 | 0.58688069 |
| POLD3 | 88.4860598 | -0.36385745 | 0.21882699 | -1.6627631 | 0.09635986 | 0.41764444 |
| ATP5PF | 148.153829 | -0.363834273 | 0.43801766 | -0.8306384 | 0.40617796 | 0.73307328 |
| LRP4 | 138.169673 | -0.363829327 | 0.41528529 | -0.8760949 | 0.38097845 | 0.71543195 |
| SERPINB6 | 912.127472 | -0.363409195 | 0.42768351 | -0.8497152 | 0.39548342 | 0.7252893 |
| GATAD1 | 92.5293582 | -0.362705414 | 0.27976545 | -1.2964625 | 0.19481619 | 0.55995114 |
| ECH1 | 73.9232448 | -0.362151141 | 0.52726733 | -0.6868454 | 0.49218015 | 0.79094859 |
| SLC15A4 | 54.1533016 | -0.362112128 | 0.27498798 | -1.3168289 | 0.18789598 | 0.55388137 |
| CHD4 | 55.4289825 | -0.361518632 | 0.3372152 | -1.072071 | 0.28368815 | 0.64699557 |
| PDXK | 71.4948328 | -0.361294337 | 0.28891249 | -1.2505321 | 0.21110525 | 0.5768065 |
| RTF1 | 164.780975 | -0.361117309 | 0.31085852 | -1.1616774 | 0.24536654 | 0.6104646 |
| RRAGA | 79.7891811 | -0.360907209 | 0.55611379 | -0.6489809 | 0.51635075 | 0.80483629 |
| LAPTM4A | 317.993903 | -0.360762572 | 0.38819553 | -0.9293321 | 0.352717 | 0.69970503 |
| WDR35 | 61.6817685 | -0.360484208 | 0.36762328 | -0.9805805 | 0.32679967 | 0.68007671 |
| PELI2 | 333.649706 | -0.360083999 | 0.30329778 | -1.1872293 | 0.2351372 | 0.59949266 |
| SRGAP1 | 385.923427 | -0.360061854 | 0.31459063 | -1.144541 | 0.25239937 | 0.61777104 |
| TRAF3IP2-AS | 275.772117 | -0.35976331 | 0.2509095 | -1.4338369 | 0.1516188 | 0.50771689 |
| MTPN | 202.085282 | -0.359718323 | 0.25158488 | -1.429809 | 0.15277184 | 0.50900346 |
| TRIM27 | 158.563699 | -0.35952193 | 0.2141507 | -1.6788268 | 0.09318581 | 0.41205855 |
| GDI2 | 312.468052 | -0.359445327 | 0.22112541 | -1.625527 | 0.10405031 | 0.43552105 |
| OLA1 | 246.607904 | -0.35919059 | 0.26565897 | -1.352074 | 0.17635164 | 0.54007689 |
| ZNF333 | 64.9529178 | -0.35914881 | 0.32114253 | -1.1183471 | 0.26341878 | 0.62972711 |
| WDR45 | 52.1460375 | -0.359021684 | 0.21144182 | -1.6979691 | 0.08951359 | 0.40581146 |
| SAT2 | 88.689356 | -0.358801217 | 0.3880054 | -0.9247325 | 0.35510505 | 0.70018061 |
| COX6C | 201.024501 | -0.358483147 | 0.37897238 | -0.9459347 | 0.34418187 | 0.6938754 |
| CABLES1 | 271.253968 | -0.358312155 | 0.57328921 | -0.6250112 | 0.53196373 | 0.81427134 |
| TMED2 | 87.5366358 | -0.358057968 | 0.33950103 | -1.0546594 | 0.2915811 | 0.65416882 |
| RALGPS2 | 52.062635 | -0.357787514 | 0.34331309 | -1.042161 | 0.29733704 | 0.65795352 |
| ETV6 | 1704.06892 | -0.357774601 | 0.34613338 | -1.0336322 | 0.30130816 | 0.65926326 |
| TFAM | 48.0523258 | -0.357392616 | 0.25506541 | -1.4011803 | 0.16116018 | 0.51944206 |
| TGOLN2 | 289.62822 | -0.357385351 | 0.25218313 | -1.417166 | 0.15643441 | 0.5126563 |
| PREP | 144.973695 | -0.356681989 | 0.21896246 | -1.6289641 | 0.10332062 | 0.43424379 |
| PDCD5 | 56.0483247 | -0.356652469 | 0.25428315 | -1.40258 | 0.16074212 | 0.51944206 |
| MOGS | 46.6070602 | -0.356597906 | 0.37461664 | -0.9519009 | 0.34114725 | 0.69116603 |
| PARVB | 345.16568 | -0.356448769 | 0.23655778 | -1.5068148 | 0.13185813 | 0.48092078 |
| PSMD14 | 182.980359 | -0.356361679 | 0.36987559 | -0.9634636 | 0.33531491 | 0.68631363 |
| HDAC2 | 134.635965 | -0.356145081 | 0.24224113 | -1.4702089 | 0.14150517 | 0.49336891 |
| PIGK | 100.465604 | -0.356140651 | 0.22620799 | -1.5743947 | 0.11539623 | 0.4568354 |
| ABCA5 | 90.7875215 | -0.356005225 | 0.44433359 | -0.8012116 | 0.42300916 | 0.74026603 |
| RAI1 | 166.76864 | -0.355226358 | 0.34390413 | -1.0329226 | 0.3016401 | 0.65926326 |
| RCAN1 | 140.0604 | -0.355140024 | 0.3361202 | -1.0565864 | 0.2907004 | 0.65333738 |
| MYO5A | 121.04468 | -0.354914344 | 0.48291818 | -0.7349368 | 0.46237798 | 0.77248653 |
| RTN4 | 1270.01949 | -0.354640244 | 0.35718662 | -0.992871 | 0.32077282 | 0.67473727 |
| ENY2 | 159.281708 | -0.35377839 | 0.34669801 | -1.0204223 | 0.3075282 | 0.66400081 |
| HPS5 | 65.513085 | -0.353694149 | 0.25973815 | -1.3617336 | 0.17328199 | 0.53536344 |
| TCOF1 | 46.0391689 | -0.353414075 | 0.35506055 | -0.9953628 | 0.31955982 | 0.67395826 |
| FAAP20 | 68.7277317 | -0.352993661 | 0.42713194 | -0.8264277 | 0.40856152 | 0.73419295 |
| TBC1D16 | 138.510979 | -0.352462773 | 0.28491876 | -1.2370641 | 0.21606329 | 0.58197692 |
| LMAN1 | 180.353588 | -0.352042866 | 0.32420542 | -1.0858636 | 0.27753934 | 0.64148182 |
| SDF4 | 147.114366 | -0.352035398 | 0.40531225 | -0.8685536 | 0.38509136 | 0.71699718 |
| ANKRD50 | 201.502854 | -0.351955913 | 0.39668142 | -0.8872508 | 0.37494388 | 0.71257833 |
| UBALD2 | 80.3845137 | -0.35174353 | 0.30002261 | -1.1723901 | 0.24104048 | 0.60559848 |
| UBE2G2 | 203.301587 | -0.351729143 | 0.2575107 | -1.3658816 | 0.17197612 | 0.53377758 |
| PNPLA8 | 212.863574 | -0.351649254 | 0.33123077 | -1.0616443 | 0.2883972 | 0.65138125 |
| RBM19 | 71.8891299 | -0.351237037 | 0.38087938 | -0.9221739 | 0.35643785 | 0.70046372 |
| SNRPG | 81.9675461 | -0.349853367 | 0.36268617 | -0.9646173 | 0.33473652 | 0.68581595 |
| VPS37C | 53.1293252 | -0.349646956 | 0.33208021 | -1.0528991 | 0.29238724 | 0.65502684 |
| TTI2 | 43.5916049 | -0.349558963 | 0.34549724 | -1.0117562 | 0.31165465 | 0.66561244 |
| XRCC6 | 226.202518 | -0.349531911 | 0.23677481 | -1.4762208 | 0.13988462 | 0.49118733 |
| LETM1 | 55.6785319 | -0.349057229 | 0.35655353 | -0.9789757 | 0.32759201 | 0.68017808 |
| HOXD3 | 139.602241 | -0.348661729 | 0.43894242 | -0.7943223 | 0.42700785 | 0.74286176 |
| AKR1A1 | 58.9899372 | -0.348463044 | 0.37636299 | -0.9258696 | 0.35451376 | 0.70004491 |
| C19orf25 | 79.1462391 | -0.348332457 | 0.3251056 | -1.071444 | 0.28396982 | 0.64709691 |
| UTP23 | 49.2645867 | -0.348135411 | 0.28400023 | -1.2258279 | 0.22026344 | 0.58688228 |
| COPZ1 | 79.1677142 | -0.347883254 | 0.26015186 | -1.3372315 | 0.18114709 | 0.54524118 |
| AC024230.1 | 51.1460227 | -0.347865854 | 0.60324537 | -0.5766573 | 0.56417098 | 0.83092353 |
| CARD19 | 42.1627666 | -0.347600878 | 0.35186511 | -0.9878811 | 0.3232109 | 0.67664579 |
| ZNF428 | 69.8283677 | -0.34751995 | 0.34870802 | -0.9965929 | 0.31896213 | 0.67391353 |
| DNHD1 | 94.019106 | -0.347404686 | 0.27815719 | -1.248951 | 0.21168301 | 0.57690664 |
| PPCS | 48.7561114 | -0.347336325 | 0.30610913 | -1.1346813 | 0.25650886 | 0.62095131 |
| CHD7 | 408.932145 | -0.346863731 | 0.34617851 | -1.0019794 | 0.31635355 | 0.67134883 |

| | | | | | | |
|---|---|---|---|---|---|---|
| EFHD2 | 69.2302065 | -0.346838525 | 0.3427733 | -1.0118598 | 0.31160509 | 0.66561244 |
| RNF122 | 45.8543651 | -0.346676895 | 0.48838789 | -0.7098393 | 0.47780382 | 0.78129872 |
| BCCIP | 76.8253086 | -0.346422305 | 0.18905221 | -1.8324161 | 0.06688945 | 0.35877642 |
| LRMDA | 3962.6869 | -0.346234316 | 0.33049284 | -1.0476303 | 0.29480897 | 0.65671864 |
| CHCHD3 | 696.830379 | -0.345245587 | 0.18729921 | -1.8432838 | 0.06528759 | 0.35545467 |
| RPS6KA5 | 250.36315 | -0.344889718 | 0.25460266 | -1.3546194 | 0.17553883 | 0.53859553 |
| NUP107 | 146.35444 | -0.344375639 | 0.18601729 | -1.8513099 | 0.06412499 | 0.35347506 |
| GPS1 | 47.672117 | -0.344020525 | 0.39369901 | -0.8738161 | 0.38221843 | 0.71588105 |
| CLTA | 288.938608 | -0.343705996 | 0.29653733 | -1.1590649 | 0.24642975 | 0.61160108 |
| MRPS11 | 59.3698096 | -0.34368853 | 0.28088417 | -1.2235952 | 0.22110496 | 0.58706366 |
| IMPAD1 | 103.796058 | -0.343610946 | 0.24741257 | -1.3888177 | 0.1648882 | 0.52480753 |
| PSMB8 | 166.506011 | -0.343550405 | 0.37199455 | -0.9235361 | 0.35572788 | 0.70018061 |
| MLLT1 | 104.283587 | -0.343446785 | 0.41654035 | -0.8245222 | 0.40964288 | 0.73419295 |
| PSMA5 | 128.597689 | -0.342634525 | 0.30700297 | -1.1160626 | 0.26439535 | 0.63018314 |
| PNRC2 | 97.0790125 | -0.342353788 | 0.28259849 | -1.2114494 | 0.2257232 | 0.59165529 |
| HPS4 | 64.3486758 | -0.342065833 | 0.32180093 | -1.0629734 | 0.28779401 | 0.65121935 |
| RNF141 | 50.3094114 | -0.341739705 | 0.29023297 | -1.1774669 | 0.23900922 | 0.60290148 |
| PSMD3 | 83.6261416 | -0.341153251 | 0.27878267 | -1.2237248 | 0.22105608 | 0.58706366 |
| RTRAF | 115.318561 | -0.341103508 | 0.43150713 | -0.7904933 | 0.42923972 | 0.74408005 |
| YWHAE | 503.7195 | -0.340026486 | 0.15918402 | -2.1360592 | 0.03267459 | 0.25224169 |
| FCHSD2 | 1887.76667 | -0.339524615 | 0.26814139 | -1.2662149 | 0.20543616 | 0.57043912 |
| HLA-E | 1881.44902 | -0.339317803 | 0.37151278 | -0.9133409 | 0.3610633 | 0.70386371 |
| TM9SF4 | 144.183635 | -0.33910016 | 0.14400332 | -2.3548079 | 0.01853228 | 0.19394842 |
| OXNAD1 | 100.929221 | -0.339088676 | 0.23785357 | -1.4256195 | 0.15397821 | 0.50978388 |
| APP | 4121.71781 | -0.338918673 | 0.22193363 | -1.5271173 | 0.12673186 | 0.47358888 |
| AL121761.1 | 102.349568 | -0.338914981 | 0.43642785 | -0.7765659 | 0.43741494 | 0.75196773 |
| ZSWIM4 | 74.6503418 | -0.338870006 | 0.27575013 | -1.2289024 | 0.21910839 | 0.58619521 |
| IFNGR2 | 201.559479 | -0.338731319 | 0.23823201 | -1.4218548 | 0.15506842 | 0.51125095 |
| GUK1 | 300.087341 | -0.3387128 | 0.38891064 | -0.8709271 | 0.38379398 | 0.71699718 |
| RETREG2 | 53.8249619 | -0.338650453 | 0.35197332 | -0.9621481 | 0.33597522 | 0.68706501 |
| DZIP3 | 76.8892357 | -0.338135063 | 0.49929205 | -0.677229 | 0.49826066 | 0.79513036 |
| EIF2A | 123.490885 | -0.338046341 | 0.29148614 | -1.1597338 | 0.24615719 | 0.61132148 |
| AIP | 49.6057268 | -0.337819395 | 0.42609411 | -0.7928281 | 0.42787797 | 0.74338119 |
| RNF220 | 102.843758 | -0.337566855 | 0.38156714 | -0.8846853 | 0.37632636 | 0.71257833 |
| MRPL4 | 39.3466278 | -0.337257131 | 0.43267407 | -0.7794716 | 0.43570199 | 0.74971913 |
| HMGB1 | 510.883121 | -0.337168655 | 0.23456755 | -1.4374054 | 0.15060284 | 0.5057732 |
| CLUH | 40.8767271 | -0.337129109 | 0.40388682 | -0.8347118 | 0.40387998 | 0.73096878 |
| PEBP1 | 259.892325 | -0.3368197 | 0.3489823 | -0.9651484 | 0.3344705 | 0.68544254 |
| DMD | 80.4097844 | -0.336677115 | 0.46496653 | -0.7240889 | 0.46901115 | 0.77737861 |
| ATP5F1B | 206.38784 | -0.336466434 | 0.32290856 | -1.0419867 | 0.29741783 | 0.65795352 |
| BZW1 | 83.9424757 | -0.336446498 | 0.26294817 | -1.2795164 | 0.20071527 | 0.56536969 |
| ENDOD1 | 165.312253 | -0.336112804 | 0.31208006 | -1.0770083 | 0.28147657 | 0.64471757 |
| AKTIP | 48.6591711 | -0.335905555 | 0.25030248 | -1.3419985 | 0.17959648 | 0.54350518 |
| TMBIM6 | 441.958732 | -0.335814952 | 0.28155315 | -1.1927231 | 0.23297782 | 0.59910683 |
| METRN | 50.3628669 | -0.335797406 | 0.4629411 | -0.7253567 | 0.46823326 | 0.77664704 |
| SH2B2 | 84.5655132 | -0.335236575 | 0.60441061 | -0.5546504 | 0.57913382 | 0.83649569 |
| CHD1L | 113.243928 | -0.334718498 | 0.22546842 | -1.4845471 | 0.13766383 | 0.48962215 |
| AC097103.2 | 97.6735872 | -0.334654337 | 0.37422213 | -0.8942666 | 0.37117928 | 0.70984233 |
| PHAX | 57.9359233 | -0.334370798 | 0.29805273 | -1.1218512 | 0.26192573 | 0.6270738 |
| MB21D2 | 150.816945 | -0.334149586 | 0.71914985 | -0.4646453 | 0.6421855 | 0.86490071 |
| TWF1 | 51.7197904 | -0.333829233 | 0.35144668 | -0.9498716 | 0.34217749 | 0.692051 |
| BTF3L4 | 113.683238 | -0.333818114 | 0.17765213 | -1.879055 | 0.06023699 | 0.34478399 |
| RHOG | 53.9654526 | -0.333718553 | 0.34462626 | -0.9683492 | 0.33287002 | 0.6852158 |
| ZNF462 | 286.609855 | -0.333532978 | 0.31391237 | -1.0625035 | 0.28800719 | 0.65121935 |
| NOC2L | 51.4037925 | -0.333453956 | 0.33512391 | -0.9950169 | 0.31972804 | 0.67413928 |
| RRM2B | 71.909201 | -0.333387531 | 0.36002812 | -0.9260041 | 0.35444383 | 0.70004491 |
| SYNCRIP | 121.781414 | -0.333312345 | 0.35100768 | -0.949587 | 0.34232212 | 0.6921189 |
| WBP1L | 214.16102 | -0.332957109 | 0.17754826 | -1.8753049 | 0.06075079 | 0.34618645 |
| SNRNP200 | 145.963027 | -0.332939026 | 0.24064191 | -1.3835454 | 0.16649768 | 0.52671378 |
| RHOQ | 128.584934 | -0.332818223 | 0.45299111 | -0.7347125 | 0.46251461 | 0.77255707 |
| LINC00667 | 49.7955857 | -0.332549087 | 0.25890903 | -1.2844244 | 0.19899348 | 0.56402001 |
| SUSD6 | 574.249561 | -0.332541489 | 0.40365665 | -0.8238226 | 0.41004034 | 0.73419295 |
| BMT2 | 147.121888 | -0.332368396 | 0.22235234 | -1.4947825 | 0.1349712 | 0.484753 |
| EOGT | 118.412282 | -0.332078151 | 0.33328633 | -0.9963749 | 0.319068 | 0.67394237 |
| FGL2 | 265.662999 | -0.332041651 | 0.52698595 | -0.6300769 | 0.5286443 | 0.81207005 |
| NOL8 | 82.9512095 | -0.331624566 | 0.33549008 | -0.988478 | 0.3229186 | 0.67633553 |
| BCL2L11 | 69.1903001 | -0.331608051 | 0.35050052 | -0.9460986 | 0.3440983 | 0.6938754 |
| ATF6 | 543.90447 | -0.331513251 | 0.28748931 | -1.1531324 | 0.24885603 | 0.61434376 |
| GBE1 | 717.790416 | -0.331342386 | 0.48261593 | -0.686555 | 0.49236319 | 0.79099316 |
| TMEM120B | 58.4597061 | -0.330491173 | 0.30638221 | -1.0786892 | 0.28072631 | 0.64400993 |
| TNFAIP1 | 160.029802 | -0.330429377 | 0.27716651 | -1.1921692 | 0.2331949 | 0.59910683 |
| IGF2R | 490.974549 | -0.330082251 | 0.30001743 | -1.1002102 | 0.27124053 | 0.63561102 |
| MYO19 | 105.205851 | -0.32923177 | 0.3298661 | -0.998077 | 0.31824201 | 0.6730872 |
| LRRC41 | 78.6793953 | -0.328783151 | 0.25330833 | -1.2979563 | 0.19430235 | 0.55913011 |
| SSNA1 | 75.1981448 | -0.328700441 | 0.37585716 | -0.8745355 | 0.38182668 | 0.71553552 |
| PEX5 | 40.7904783 | -0.327880411 | 0.29455686 | -1.1131311 | 0.26565211 | 0.63060082 |
| ZSWIM5 | 65.6665274 | -0.327848697 | 0.34415632 | -0.9526157 | 0.34078484 | 0.69116603 |
| CDA | 107.324289 | -0.327445097 | 0.38991763 | -0.8397802 | 0.40103165 | 0.72892981 |
| ARCN1 | 111.66005 | -0.327415245 | 0.24889398 | -1.3154808 | 0.18834837 | 0.55388137 |
| ANXA2 | 567.5164 | -0.327364531 | 0.32691329 | -1.0013803 | 0.31664299 | 0.67161471 |
| SUN2 | 137.137532 | -0.327289615 | 0.3859312 | -0.8480517 | 0.39640917 | 0.7264986 |
| ADIPOR1 | 54.2809838 | -0.327227938 | 0.323618 | -1.0111549 | 0.31194228 | 0.66561244 |
| PARP1 | 85.7409282 | -0.326298444 | 0.29951163 | -1.089435 | 0.27596211 | 0.64068436 |
| ATP5F1A | 133.014938 | -0.326115989 | 0.226133 | -1.4421424 | 0.14926221 | 0.50283523 |
| SLC25A1 | 42.9876607 | -0.326088115 | 0.39237464 | -0.8310632 | 0.40593793 | 0.73307328 |
| CEP57 | 148.282479 | -0.325998787 | 0.19372397 | -1.6828005 | 0.09241372 | 0.41143716 |
| ZNF394 | 64.7101764 | -0.325964328 | 0.29592443 | -1.101512 | 0.27067387 | 0.63561102 |
| CD151 | 281.730513 | -0.32543284 | 0.51695482 | -0.6295189 | 0.52900939 | 0.81217332 |
| AMN1 | 99.0323648 | -0.325404161 | 0.35609224 | -0.9138199 | 0.36081151 | 0.70386371 |
| APH1A | 97.4163188 | -0.325003372 | 0.37863866 | -0.858347 | 0.39070086 | 0.72247779 |
| HELB | 67.8911589 | -0.325000589 | 0.39981942 | -0.8128685 | 0.41629348 | 0.73802299 |
| MAGED1 | 89.7206409 | -0.324603227 | 0.29078166 | -1.1163126 | 0.26428835 | 0.63018314 |
| TRIM24 | 241.213171 | -0.324594 | 0.29843525 | -1.087653 | 0.27674832 | 0.64098811 |
| TUFM | 121.099309 | -0.324324708 | 0.3850952 | -0.8421936 | 0.3996796 | 0.72825165 |
| INTS9 | 159.894243 | -0.324090698 | 0.21830706 | -1.4845635 | 0.13765948 | 0.48962215 |
| LPCAT3 | 140.181513 | -0.323644813 | 0.23749526 | -1.3627422 | 0.17296377 | 0.53502507 |
| ASCC3 | 593.5081 | -0.323600277 | 0.24374412 | -1.3276229 | 0.18430272 | 0.54912489 |
| SERINC1 | 307.647098 | -0.32335898 | 0.28480751 | -1.1353597 | 0.25622462 | 0.62095131 |
| SEL1L | 226.618692 | -0.322886369 | 0.2060081 | -1.5673479 | 0.11703339 | 0.45985048 |
| ZMIZ1 | 630.647829 | -0.322573397 | 0.38277652 | -0.8427199 | 0.39938511 | 0.72820151 |
| TSC22D3 | 262.212531 | -0.321066796 | 0.53513602 | -0.5999723 | 0.54852469 | 0.82355262 |
| IFT88 | 207.418173 | -0.320549185 | 0.28252661 | -1.1345805 | 0.25655112 | 0.62095131 |
| SLC11A2 | 83.8446766 | -0.32042155 | 0.23744615 | -1.3494493 | 0.17719268 | 0.54037749 |
| THEM4 | 65.1376925 | -0.319947826 | 0.40167166 | -0.7965407 | 0.42571782 | 0.74197066 |
| C7orf50 | 100.520244 | -0.319799235 | 0.31413137 | -1.018043 | 0.30865754 | 0.66414532 |
| HPS1 | 58.1554851 | -0.31913636 | 0.28822135 | -1.1072613 | 0.26818096 | 0.63336518 |
| IRAK4 | 100.751939 | -0.318891938 | 0.27483891 | -1.1602867 | 0.24593208 | 0.61094785 |
| PPM1F | 380.448925 | -0.318782447 | 0.32422735 | -0.9832065 | 0.3255058 | 0.67949336 |
| SNRPD3 | 121.404929 | -0.318764627 | 0.28069906 | -1.1356099 | 0.25611986 | 0.62095131 |
| CTSH | 76.3909183 | -0.318580335 | 0.43530602 | -0.7318537 | 0.46425784 | 0.77428988 |
| ABCF1 | 92.548502 | -0.317509733 | 0.28738487 | -1.1048241 | 0.26923582 | 0.63363725 |
| YES1 | 845.45638 | -0.317086711 | 0.29912171 | -1.0600592 | 0.28911769 | 0.65210861 |
| LRRC23 | 46.6180471 | -0.317081415 | 0.34404426 | -0.9216297 | 0.35672175 | 0.70058911 |

| | | | | | | |
|---|---|---|---|---|---|---|
| IPO5 | 113.144892 | -0.316760702 | 0.20254712 | -1.5638865 | 0.11784423 | 0.46042456 |
| NUDT3 | 75.9583185 | -0.316584359 | 0.42251835 | -0.7492795 | 0.45368874 | 0.76578691 |
| TLN1 | 188.20731 | -0.316523333 | 0.256145 | -1.2357194 | 0.2165629 | 0.58218265 |
| NGDN | 55.0187711 | -0.316333274 | 0.29207644 | -1.0830496 | 0.2787864 | 0.64226044 |
| DHRS7 | 69.5829426 | -0.316288325 | 0.24654262 | -1.2828951 | 0.19952882 | 0.56449294 |
| GOT2 | 79.0355198 | -0.316047675 | 0.24976524 | -1.2653789 | 0.20573553 | 0.57060151 |
| CAV2 | 153.706473 | -0.315600863 | 0.32874295 | -0.9600232 | 0.33704353 | 0.68778733 |
| ZC3H15 | 142.76821 | -0.315529138 | 0.25402407 | -1.242123 | 0.21419118 | 0.57870917 |
| MRPS18B | 67.0891131 | -0.315071256 | 0.217869 | -1.44615 | 0.14813512 | 0.50152657 |
| SIN3B | 210.301507 | -0.315008324 | 0.30309692 | -1.039299 | 0.2986657 | 0.65856711 |
| DHRS4L2 | 48.1737598 | -0.31467746 | 0.39374187 | -0.7991974 | 0.42417598 | 0.74152442 |
| VDAC2 | 94.9005991 | -0.3146108 | 0.37624482 | -0.8361864 | 0.40305005 | 0.73065099 |
| LDLR | 132.226871 | -0.314373589 | 0.4458359 | -0.705133 | 0.48072749 | 0.78473327 |
| FAM167B | 313.754673 | -0.313588276 | 0.70735812 | -0.4433232 | 0.65753197 | 0.87254693 |
| PPP1R15B | 89.2689464 | -0.313530501 | 0.33304259 | -0.9414126 | 0.34649344 | 0.69545151 |
| TATDN3 | 46.162127 | -0.313288964 | 0.44181078 | -0.7091021 | 0.4782611 | 0.78162783 |
| COPG1 | 71.2839022 | -0.313112113 | 0.24076992 | -1.3004619 | 0.19344271 | 0.55822662 |
| EMP2 | 316.443757 | -0.313041383 | 0.34269303 | -0.9134746 | 0.36099299 | 0.70386371 |
| RALBP1 | 135.116503 | -0.31258982 | 0.24577626 | -1.2718471 | 0.20342746 | 0.56736432 |
| ANXA4 | 211.027128 | -0.312543717 | 0.23328375 | -1.3397578 | 0.18032411 | 0.5444489 |
| TMEM14C | 79.8892061 | -0.312453183 | 0.43284896 | -0.7218527 | 0.47038506 | 0.77792258 |
| SPIRE1 | 375.995966 | -0.312443369 | 0.28352905 | -1.1019801 | 0.27047032 | 0.63561102 |
| CLTB | 106.400865 | -0.312070667 | 0.34191355 | -0.9127181 | 0.36139085 | 0.70386371 |
| REEP5 | 232.970703 | -0.311216242 | 0.30152703 | -1.0321338 | 0.30200944 | 0.65926326 |
| DLAT | 45.4532847 | -0.311124681 | 0.35531284 | -0.8756359 | 0.381228 | 0.71543195 |
| ANKRD28 | 611.660966 | -0.311063127 | 0.4807627 | -0.6470201 | 0.51761894 | 0.80546868 |
| DPYD-AS1 | 65.8746705 | -0.311014521 | 0.66926346 | -0.4647116 | 0.64213797 | 0.86490071 |
| HOXD8 | 48.5065846 | -0.310925859 | 0.36997305 | -0.8404014 | 0.40068338 | 0.72864885 |
| ATP5PB | 130.790241 | -0.310886212 | 0.32093774 | -0.9686808 | 0.3327045 | 0.6852158 |
| ILF3 | 176.0705 | -0.31084941 | 0.25159435 | -1.2355183 | 0.21663768 | 0.58218265 |
| TCAIM | 171.389009 | -0.310764863 | 0.27925007 | -1.1128551 | 0.26577067 | 0.63060082 |
| NPC1 | 155.212405 | -0.310651094 | 0.38366094 | -0.8097022 | 0.41811137 | 0.73821752 |
| JUN | 476.703581 | -0.310294912 | 0.87844392 | -0.3532325 | 0.72391417 | 0.90260394 |
| UBXN1 | 85.2490275 | -0.310026881 | 0.40934917 | -0.7573654 | 0.44883101 | 0.76209291 |
| MRPS23 | 47.6932318 | -0.309892457 | 0.30621439 | -1.0120114 | 0.3115326 | 0.66561244 |
| NCOA4 | 133.160575 | -0.309804039 | 0.27979918 | -1.1072371 | 0.26819142 | 0.63336518 |
| GINM1 | 168.54616 | -0.309689127 | 0.15645803 | -1.9793751 | 0.04777378 | 0.30661402 |
| CCDC25 | 68.4684193 | -0.309529787 | 0.234255 | -1.3213369 | 0.18638904 | 0.55256392 |
| SERP1 | 186.852393 | -0.309313025 | 0.29010748 | -1.0662015 | 0.28633258 | 0.64886721 |
| DDX10 | 307.688325 | -0.309068645 | 0.3137027 | -0.9852279 | 0.32451215 | 0.67845758 |
| PPP3CA | 1105.69932 | -0.309048263 | 0.3315123 | -0.9322377 | 0.3512137 | 0.69858573 |
| DNAJC18 | 126.404931 | -0.308467907 | 0.41803793 | -0.7378945 | 0.46057854 | 0.770424 |
| GLRX5 | 60.9109272 | -0.308441695 | 0.46166036 | -0.6681139 | 0.5040609 | 0.79776858 |
| PTP4A3 | 45.194029 | -0.308224209 | 0.76225367 | -0.4043591 | 0.68594867 | 0.88671976 |
| INPP5B | 145.867598 | -0.308193004 | 0.24495745 | -1.2581492 | 0.20833782 | 0.57338509 |
| PTPN23 | 48.5681989 | -0.308146179 | 0.3269112 | -0.942599 | 0.34588604 | 0.69497757 |
| CCDC59 | 59.555952 | -0.307973719 | 0.21137898 | -1.4569742 | 0.14512351 | 0.49813157 |
| SUMF2 | 100.533703 | -0.3079354 | 0.20100039 | -1.5320139 | 0.125519 | 0.47103973 |
| WWTR1 | 2594.82142 | -0.307899627 | 0.33364362 | -0.9228399 | 0.35609067 | 0.70018061 |
| ILF2 | 101.874054 | -0.307731395 | 0.33266802 | -0.9250405 | 0.35494484 | 0.70018061 |
| PLEKHA1 | 331.480142 | -0.3071453 | 0.1472147 | -2.0863766 | 0.03694452 | 0.270683 |
| NAA25 | 184.073048 | -0.306968062 | 0.2737966 | -1.1211537 | 0.26222245 | 0.62760056 |
| SLC9A8 | 146.270477 | -0.306852749 | 0.34176532 | -0.8978464 | 0.36926746 | 0.70965609 |
| ARHGEF7 | 725.347842 | -0.306514816 | 0.28960527 | -1.0583883 | 0.28987847 | 0.65303974 |
| DHFR | 351.455056 | -0.306292487 | 0.28987486 | -1.056637 | 0.2906773 | 0.65333738 |
| CAMLG | 67.9825396 | -0.306272046 | 0.21306437 | -1.4374625 | 0.15058661 | 0.5057732 |
| DCAKD | 56.1349386 | -0.306144209 | 0.28011179 | -1.0929358 | 0.27442197 | 0.63975272 |
| KRCC1 | 109.318789 | -0.306134382 | 0.33205418 | -0.9219411 | 0.35655932 | 0.7005342 |
| MVD | 39.5376755 | -0.306114434 | 0.33534308 | -0.9128396 | 0.36132692 | 0.70386371 |
| HES1 | 509.750424 | -0.305965811 | 0.89330366 | -0.3425104 | 0.7319668 | 0.9046495 |
| PTDSS2 | 56.2183097 | -0.305963773 | 0.27370548 | -1.1178577 | 0.26362778 | 0.62989598 |
| ZNF124 | 66.3653947 | -0.305865731 | 0.27476598 | -1.1131863 | 0.26562842 | 0.63060082 |
| CCDC60 | 52.6466134 | -0.305796486 | 0.51839664 | -0.589889 | 0.55526506 | 0.82487928 |
| LINC01376 | 80.58354 | -0.305432901 | 0.52587101 | -0.5808133 | 0.56136626 | 0.82940393 |
| TUBB4B | 159.140755 | -0.305300244 | 0.61481926 | -0.4965691 | 0.61949295 | 0.85855421 |
| ACO1 | 153.856035 | -0.305274012 | 0.2039292 | -1.4969608 | 0.13440346 | 0.48474744 |
| STARD3NL | 100.269128 | -0.305051854 | 0.16414895 | -1.8583845 | 0.06311443 | 0.35277692 |
| ARF5 | 46.5166283 | -0.30485871 | 0.37347581 | -0.8162743 | 0.41434327 | 0.73671465 |
| AC244131.2 | 74.4366439 | -0.304489935 | 0.56353425 | -0.540322 | 0.58897501 | 0.84140756 |
| SEC24A | 148.119412 | -0.304324762 | 0.23691034 | -1.2845567 | 0.19894723 | 0.56402001 |
| CD36 | 231.213079 | -0.304165771 | 0.87894819 | -0.3460565 | 0.72930022 | 0.90408479 |
| GABARAPL2 | 211.667766 | -0.303653546 | 0.58621011 | -0.5179944 | 0.60446219 | 0.85105198 |
| UGGT2 | 480.14659 | -0.303483602 | 0.1993975 | -1.522003 | 0.12800832 | 0.47548439 |
| NONO | 140.890205 | -0.30312902 | 0.2916811 | -1.0392481 | 0.29868938 | 0.65856711 |
| WWP1 | 629.075335 | -0.302036056 | 0.25940217 | -1.1643544 | 0.24428042 | 0.60883916 |
| NR1H2 | 72.8894972 | -0.302012997 | 0.29097756 | -1.0379254 | 0.29930479 | 0.65856711 |
| PCYOX1 | 93.0622716 | -0.301929656 | 0.43237522 | -0.6983047 | 0.48498665 | 0.78805515 |
| RALA | 280.885201 | -0.30169495 | 0.31418659 | -0.9602413 | 0.33693376 | 0.68773484 |
| SYNC | 49.8573453 | -0.301629381 | 0.49892627 | -0.604557 | 0.54547337 | 0.82355262 |
| MRPL18 | 55.0807458 | -0.301466856 | 0.36398653 | -0.8282363 | 0.40753669 | 0.73419295 |
| EXOC3 | 77.3535554 | -0.301459408 | 0.18555934 | -1.6245984 | 0.10424816 | 0.43590329 |
| C1orf115 | 49.6898421 | -0.300805557 | 0.44571112 | -0.6748891 | 0.49974625 | 0.79545295 |
| KIAA1143 | 56.8593026 | -0.300747495 | 0.25321284 | -1.1877261 | 0.23494135 | 0.59949266 |
| SPDYE3 | 64.7808704 | -0.300743748 | 0.35121037 | -0.8563066 | 0.39182823 | 0.72247779 |
| CD74 | 1919.09379 | -0.300652702 | 0.40961887 | -0.7339816 | 0.46295997 | 0.77282771 |
| DISC1 | 544.046043 | -0.300615482 | 0.32939427 | -0.9126312 | 0.36143655 | 0.70386371 |
| SMG9 | 52.5292058 | -0.299592604 | 0.286093 | -1.0471861 | 0.29501378 | 0.65671864 |
| GNG5 | 196.189055 | -0.299532221 | 0.45921884 | -0.6522647 | 0.51423045 | 0.80398873 |
| ATP6AP2 | 151.450652 | -0.299403904 | 0.28757581 | -1.0411303 | 0.29781506 | 0.6579563 |
| CHKA | 139.410039 | -0.299237734 | 0.22974359 | -1.3024857 | 0.19275041 | 0.55703491 |
| CCT8 | 85.2683638 | -0.299191407 | 0.25096202 | -1.192178 | 0.23319145 | 0.59910683 |
| DNAJC3 | 284.876967 | -0.299057132 | 0.24483894 | -1.2214443 | 0.22191783 | 0.58750144 |
| SLC36A4 | 115.445407 | -0.298622952 | 0.26324851 | -1.1343766 | 0.2566366 | 0.62095131 |
| MAP7D3 | 62.608798 | -0.298464106 | 0.20116051 | -1.4837112 | 0.13788556 | 0.48979401 |
| HAX1 | 42.0020727 | -0.298434406 | 0.37606061 | -0.7935806 | 0.42743963 | 0.74309294 |
| MDH1 | 91.736481 | -0.298423012 | 0.26925653 | -1.1083223 | 0.26772266 | 0.63276201 |
| GOLGB1 | 326.101235 | -0.298317443 | 0.22452963 | -1.3286329 | 0.18396913 | 0.54902239 |
| EHD2 | 82.2435843 | -0.298096833 | 0.39619633 | -0.7523967 | 0.4518125 | 0.76467047 |
| UBE2N | 135.961953 | -0.298071241 | 0.18285266 | -1.6301171 | 0.10307676 | 0.43388739 |
| MAP1S | 43.6202919 | -0.297944729 | 0.42442885 | -0.7019898 | 0.48268552 | 0.78697262 |
| ZNF330 | 63.738764 | -0.297853596 | 0.40706158 | -0.7317163 | 0.46434174 | 0.77428988 |
| SF3A3 | 75.2079814 | -0.297671375 | 0.28794759 | -1.0337693 | 0.30124403 | 0.65926326 |
| RMND5B | 64.9790397 | -0.297489507 | 0.27860224 | -1.0677929 | 0.28561393 | 0.64813609 |
| BMP2K | 267.386509 | -0.297156321 | 0.20633292 | -1.4401789 | 0.14981678 | 0.50388438 |
| POLR2K | 62.3106741 | -0.297023487 | 0.54970161 | -0.5403359 | 0.58896544 | 0.84140756 |
| ZFP82 | 44.7699217 | -0.296941094 | 0.29256554 | -1.0149558 | 0.3101269 | 0.6648266 |
| DNAJC8 | 103.123744 | -0.296442576 | 0.24902757 | -1.1904007 | 0.23388896 | 0.59949266 |
| RND3 | 67.8978231 | -0.29638042 | 0.49384821 | -0.6001448 | 0.54840976 | 0.82355262 |
| ATP13A1 | 41.3489367 | -0.295739267 | 0.26250521 | -1.1266034 | 0.25991019 | 0.6252109 |
| AC087286.2 | 68.6285517 | -0.295688978 | 0.47328453 | -0.6247594 | 0.53212896 | 0.8143653 |
| ARFGAP3 | 193.720625 | -0.295329622 | 0.14045261 | -2.1026995 | 0.03549205 | 0.26499217 |
| FGD4 | 278.292136 | -0.295146102 | 0.44913891 | -0.6571377 | 0.51109239 | 0.80177277 |
| MEGF8 | 46.6129211 | -0.295063479 | 0.28087995 | -1.0504968 | 0.29348978 | 0.6556551 |

| | | | | | | |
|---|---|---|---|---|---|---|
| DERL1 | 93.7734508 | -0.295003812 | 0.21647548 | -1.3627586 | 0.17295861 | 0.53502507 |
| HMGN1 | 170.389107 | -0.294465182 | 0.27253203 | -1.0804792 | 0.27992886 | 0.64326252 |
| C2orf69 | 46.5530841 | -0.294200545 | 0.35559347 | -0.8273508 | 0.40803826 | 0.73419295 |
| AGAP3 | 148.888945 | -0.293791438 | 0.24008197 | -1.223713 | 0.2210605 | 0.58706366 |
| TMED8 | 77.2104732 | -0.293499324 | 0.24604475 | -1.1928697 | 0.23292041 | 0.59910683 |
| WASF3 | 63.828765 | -0.293410086 | 0.21565547 | -1.3605501 | 0.1736559 | 0.53536344 |
| CCDC144NL-/ | 53.0420215 | -0.293256243 | 0.46456456 | -0.6312497 | 0.52787726 | 0.81165345 |
| EDEM3 | 243.01998 | -0.293182862 | 0.20766972 | -1.4117747 | 0.15801631 | 0.51474819 |
| WDR75 | 105.30173 | -0.292915601 | 0.1980399 | -1.4790737 | 0.13912063 | 0.49082099 |
| SOCS4 | 74.3315358 | -0.292717314 | 0.25398514 | -1.1524978 | 0.24911658 | 0.61434376 |
| RLF | 346.437978 | -0.292621211 | 0.3193769 | -0.9162253 | 0.35954872 | 0.70353591 |
| SCAMP3 | 77.0445875 | -0.292423413 | 0.29076195 | -1.0057142 | 0.31455309 | 0.66883106 |
| INTU | 62.1503334 | -0.29223354 | 0.28581888 | -1.0224431 | 0.30657124 | 0.663319 |
| CATSPERE | 60.0050796 | -0.291877893 | 0.36101612 | -0.8084899 | 0.41880861 | 0.73821752 |
| FAM196B | 60.7669118 | -0.291835143 | 1.34174598 | -0.217504 | 0.82781557 | 0.94018985 |
| PEX6 | 50.8413946 | -0.291324963 | 0.33082501 | -0.8806014 | 0.3785336 | 0.71322599 |
| SELP | 176.049028 | -0.291070599 | 1.00275104 | -0.290272 | 0.77160812 | 0.91702269 |
| GNA14 | 2765.57026 | -0.29103991 | 0.35023282 | -0.8309898 | 0.40597938 | 0.73307328 |
| SYF2 | 80.7970092 | -0.291033376 | 0.32947287 | -0.8833303 | 0.37705782 | 0.71257833 |
| BUD23 | 73.5865409 | -0.290902669 | 0.34176769 | -0.8511708 | 0.3946745 | 0.72461789 |
| CRCP | 42.1915874 | -0.290898242 | 0.27658048 | -1.0517671 | 0.29290643 | 0.655236 |
| TERF2IP | 277.492512 | -0.290895309 | 0.32353482 | -0.8991159 | 0.36859092 | 0.70919884 |
| COL4A2 | 1239.93122 | -0.290669466 | 0.38838887 | -0.748398 | 0.45422011 | 0.76620968 |
| MKL1 | 711.577869 | -0.29033399 | 0.33961138 | -0.8549007 | 0.39260615 | 0.72325442 |
| NUDT9 | 100.775058 | -0.290298869 | 0.23352498 | -1.243117 | 0.21382473 | 0.57825348 |
| CRYL1 | 128.969361 | -0.290214053 | 0.27776223 | -1.0448291 | 0.29610197 | 0.65753344 |
| TAX1BP3 | 44.7034223 | -0.290058076 | 0.47134328 | -0.615386 | 0.53829981 | 0.81807397 |
| SVIL | 1077.61306 | -0.289817582 | 0.41600023 | -0.6966765 | 0.48600527 | 0.78826504 |
| STAT5B | 396.435027 | -0.289785866 | 0.18227481 | -1.5898294 | 0.11187326 | 0.44839019 |
| SLC39A8 | 57.6970028 | -0.289774895 | 0.4196844 | -0.6904591 | 0.48990555 | 0.79046901 |
| NUP37 | 45.2626658 | -0.289687656 | 0.25989424 | -1.1146367 | 0.26500614 | 0.63020787 |
| TANK | 369.777987 | -0.289312609 | 0.25684999 | -1.1263875 | 0.26000155 | 0.6252109 |
| HNRNPM | 423.423378 | -0.289271437 | 0.27172625 | -1.0645693 | 0.28707086 | 0.65000024 |
| PDE4D | 3656.47386 | -0.288975464 | 0.30956842 | -0.9334785 | 0.35057297 | 0.69858573 |
| EIPR1 | 112.72943 | -0.288958582 | 0.34924842 | -0.8273726 | 0.40802589 | 0.73419295 |
| EVA1C | 967.528153 | -0.288761638 | 0.31593752 | -0.9139834 | 0.36072559 | 0.70386371 |
| HEATR1 | 114.440685 | -0.288648986 | 0.2870823 | -1.0054573 | 0.31467672 | 0.66883106 |
| MRPS16 | 63.6052249 | -0.288593118 | 0.27601621 | -1.0455658 | 0.29576153 | 0.65731032 |
| RHBDD2 | 47.9122158 | -0.288412007 | 0.26324458 | -1.0956047 | 0.27325179 | 0.63887536 |
| TMEM30A | 266.947052 | -0.287968613 | 0.40757335 | -0.7065443 | 0.47984973 | 0.78391103 |
| BSDC1 | 156.023642 | -0.28785771 | 0.32661176 | -0.8813452 | 0.37813101 | 0.71305115 |
| PLEKHJ1 | 43.0682668 | -0.287610093 | 0.29300849 | -0.981576 | 0.32630879 | 0.67997183 |
| LYN | 223.533035 | -0.287463731 | 0.32057508 | -0.8967126 | 0.36987227 | 0.70984233 |
| LIMD1 | 127.357772 | -0.287396069 | 0.33082859 | -0.8687159 | 0.38500252 | 0.71699718 |
| CCDC178 | 145.098139 | -0.287335362 | 1.10816008 | -0.2592905 | 0.79541112 | 0.92847059 |
| SDHC | 131.873871 | -0.287295157 | 0.18121571 | -1.5853766 | 0.11288079 | 0.45058708 |
| ARPC3 | 316.49293 | -0.287115795 | 0.26284829 | -1.0923251 | 0.27469022 | 0.63988729 |
| RELA | 100.105878 | -0.28698998 | 0.27130169 | -1.057826 | 0.29013477 | 0.65314246 |
| AFG3L2 | 103.638846 | -0.286872681 | 0.29796934 | -0.9627591 | 0.33566845 | 0.68669373 |
| PLPBP | 53.9289053 | -0.286612464 | 0.25906159 | -1.1063487 | 0.2685756 | 0.63347822 |
| PRKAG1 | 73.3468302 | -0.286598559 | 0.35088255 | -0.8167934 | 0.41404649 | 0.73671465 |
| CNOT2 | 672.601538 | -0.286360098 | 0.41981008 | -0.6821182 | 0.49516421 | 0.79337734 |
| B2M | 5671.49537 | -0.28617553 | 0.51949543 | -0.5508721 | 0.58172137 | 0.83789748 |
| HMGCS1 | 75.452341 | -0.286020891 | 0.24709114 | -1.1575522 | 0.24704684 | 0.61194684 |
| PRPF40B | 106.757245 | -0.286006289 | 0.28392756 | -1.0073213 | 0.31378039 | 0.66814075 |
| CORO1B | 59.1776708 | -0.285651338 | 0.42820116 | -0.6670961 | 0.50471073 | 0.79776858 |
| HSD17B11 | 145.914755 | -0.28522287 | 0.25304069 | -1.1271818 | 0.25966562 | 0.62495404 |
| GOLGA4 | 697.15413 | -0.284882969 | 0.20712995 | -1.3753828 | 0.1690128 | 0.53091431 |
| RNF11 | 175.655245 | -0.284737369 | 0.29197991 | -0.9751951 | 0.3294635 | 0.68216508 |
| PRKCA | 882.676923 | -0.28441946 | 0.40707058 | -0.6986982 | 0.48474069 | 0.78781193 |
| MAP2K2 | 137.005909 | -0.284386485 | 0.21687912 | -1.3112672 | 0.18976749 | 0.55535814 |
| CTSB | 239.312659 | -0.284330569 | 0.39914143 | -0.7123554 | 0.4762447 | 0.78051479 |
| SLC25A37 | 436.234249 | -0.284010112 | 0.49144481 | -0.5779085 | 0.56332593 | 0.83089872 |
| SSR1 | 166.64369 | -0.283992773 | 0.29668388 | -0.9572235 | 0.33845448 | 0.68911993 |
| PDE12 | 46.093282 | -0.283977869 | 0.24324195 | -1.1674708 | 0.2430203 | 0.60833132 |
| PLAT | 1445.30589 | -0.283817474 | 0.4314201 | -0.657868 | 0.51062296 | 0.80177277 |
| MAT2A | 130.965735 | -0.283549165 | 0.473667 | -0.5986255 | 0.54942262 | 0.82355262 |
| TAF1 | 336.964254 | -0.283240681 | 0.22193118 | -1.2762546 | 0.20186554 | 0.56693091 |
| HMGN2 | 97.6050961 | -0.283221632 | 0.34895083 | -0.8116377 | 0.41699958 | 0.73821752 |
| MAN1B1 | 57.4899923 | -0.283138007 | 0.28346858 | -0.9988338 | 0.31787519 | 0.67283307 |
| TXNDC9 | 68.6487518 | -0.283073274 | 0.30199575 | -0.9373419 | 0.34858271 | 0.69793304 |
| COX7A2 | 134.644331 | -0.282606904 | 0.50910299 | -0.5551075 | 0.57882111 | 0.83649569 |
| FKBP2 | 79.1521031 | -0.282335706 | 0.36922633 | -0.7646684 | 0.44446903 | 0.75968568 |
| SPIDR | 1202.54425 | -0.282223655 | 0.20783877 | -1.357897 | 0.17449636 | 0.53700779 |
| CCDC47 | 67.5191707 | -0.282146807 | 0.30558555 | -0.9232989 | 0.35585145 | 0.70018061 |
| ANAPC16 | 324.528639 | -0.282126784 | 0.24230876 | -1.1643276 | 0.24429127 | 0.60883916 |
| EMC10 | 72.7102396 | -0.282047885 | 0.33755015 | -0.8355733 | 0.40339501 | 0.73065697 |
| 6-Sep | 194.866048 | -0.282023571 | 0.29106248 | -0.9689451 | 0.33257256 | 0.6852158 |
| MKRN1 | 73.4662082 | -0.28168973 | 0.22571411 | -1.2479934 | 0.21203346 | 0.57690664 |
| DESI1 | 67.2100817 | -0.281686435 | 0.2681485 | -1.0504867 | 0.29349439 | 0.6556551 |
| PELP1 | 42.5633586 | -0.281484938 | 0.39242916 | -0.7172885 | 0.47319607 | 0.77899012 |
| AK2 | 141.004001 | -0.281300041 | 0.17423955 | -1.6144442 | 0.1064311 | 0.43901385 |
| DCTN3 | 54.662337 | -0.28085649 | 0.46787702 | -0.6002784 | 0.54832068 | 0.82355262 |
| CDIPT | 64.8886156 | -0.280565872 | 0.428629 | -0.6545658 | 0.51274737 | 0.80317989 |
| SDF2 | 66.3207718 | -0.280306498 | 0.3397871 | -0.8249474 | 0.40940143 | 0.73419295 |
| PIN1 | 60.1515915 | -0.280151484 | 0.26479066 | -1.0580112 | 0.29005033 | 0.65313205 |
| AC116049.2 | 54.6616669 | -0.280111062 | 0.51458372 | -0.544345 | 0.58620411 | 0.83950091 |
| EAPP | 104.971036 | -0.279881215 | 0.29267481 | -0.9562874 | 0.33892708 | 0.68955779 |
| ENPP4 | 64.9000025 | -0.279322307 | 0.5024748 | -0.5558932 | 0.57828389 | 0.83635509 |
| SRI | 87.6426084 | -0.279244889 | 0.34715188 | -0.8043882 | 0.42117279 | 0.73910721 |
| SMC1A | 114.834707 | -0.279108883 | 0.27474117 | -1.0158976 | 0.30967817 | 0.6648266 |
| TATDN1 | 80.1806406 | -0.279085786 | 0.26687375 | -1.0457596 | 0.29567205 | 0.65731032 |
| EIF3G | 61.2652133 | -0.278905098 | 0.46993824 | -0.5934931 | 0.55285121 | 0.8236419 |
| PARP4 | 275.273061 | -0.278795421 | 0.32798748 | -0.8500185 | 0.3953148 | 0.72514258 |
| AGFG2 | 72.1150499 | -0.278684442 | 0.34641655 | -0.8044779 | 0.42112105 | 0.73910721 |
| WLS | 114.027771 | -0.278447416 | 0.33276317 | -0.8367735 | 0.40271988 | 0.73037606 |
| NRN1 | 84.2499098 | -0.2777737 | 0.55606382 | -0.4995356 | 0.61740208 | 0.85790478 |
| TANC1 | 1474.41764 | -0.277703007 | 0.27554482 | -1.0078324 | 0.31353492 | 0.66807452 |
| BAG6 | 75.3954218 | -0.277558377 | 0.29735325 | -0.9334298 | 0.35059811 | 0.69858573 |
| ZNF358 | 50.7714107 | -0.276641835 | 0.54390417 | -0.5086224 | 0.61101694 | 0.85449687 |
| NFAT5 | 2155.91909 | -0.276468408 | 0.19364851 | -1.4276815 | 0.15338354 | 0.50963635 |
| GPATCH2L | 302.277917 | -0.276435075 | 0.26304218 | -1.0509154 | 0.29329745 | 0.6556551 |
| AP001439.1 | 69.4760556 | -0.276256681 | 0.42526193 | -0.6496154 | 0.51594071 | 0.80483629 |
| MUT | 50.6408295 | -0.276207159 | 0.25943432 | -1.0646516 | 0.28703364 | 0.65000024 |
| ACTR2 | 432.599943 | -0.275939953 | 0.21932686 | -1.258122 | 0.20834764 | 0.57338509 |
| FAM155A | 384.999634 | -0.275779428 | 0.58902411 | -0.4681972 | 0.63964359 | 0.86399909 |
| MRPL52 | 51.4964121 | -0.275750985 | 0.36064269 | -0.7646099 | 0.44450385 | 0.75968568 |
| FAM84B | 229.172117 | -0.275454689 | 0.52771455 | -0.5219767 | 0.60168657 | 0.85011818 |
| PRPF8 | 186.384335 | -0.275302838 | 0.23096655 | -1.1919598 | 0.23327701 | 0.59910683 |
| ALDH7A1 | 128.069223 | -0.275278413 | 0.42735109 | -0.6441505 | 0.51947786 | 0.80617483 |
| AL138752.2 | 92.9287312 | -0.274877432 | 0.19889856 | -1.3819981 | 0.16697228 | 0.52764813 |
| DUS1L | 53.1552044 | -0.274861699 | 0.29535403 | -0.9306177 | 0.35205134 | 0.69889281 |
| ERH | 131.48003 | -0.274539379 | 0.3246721 | -0.8455897 | 0.39778168 | 0.72733685 |

| | | | | | | |
|---|---|---|---|---|---|---|
| LRRC58 | 98.5055686 | -0.273503104 | 0.2415871 | -1.1321097 | 0.25758831 | 0.62233395 |
| AUTS2 | 1214.6212 | -0.273454126 | 0.20778684 | -1.316032 | 0.1881633 | 0.55388137 |
| MFNG | 75.662845 | -0.27295617 | 0.36075075 | -0.7566337 | 0.44926935 | 0.76241624 |
| POGLUT1 | 83.723222 | -0.272955326 | 0.28575569 | -0.9552052 | 0.33947393 | 0.68965122 |
| PPP2R3A | 150.685206 | -0.272922481 | 0.1909528 | -1.4292667 | 0.15292758 | 0.50905923 |
| MAP3K3 | 463.595373 | -0.27288306 | 0.24012093 | -1.1364401 | 0.25577242 | 0.62095131 |
| AC007566.1 | 64.235024 | -0.272759106 | 0.42546662 | -0.6410823 | 0.52146924 | 0.80772643 |
| DUS2 | 55.3947556 | -0.272724161 | 0.30130345 | -0.9051478 | 0.36538708 | 0.70668224 |
| SNX25 | 574.826629 | -0.272615023 | 0.53464666 | -0.5098976 | 0.61012324 | 0.85388036 |
| LY6E | 316.270573 | -0.27248404 | 0.41603839 | -0.6549493 | 0.51250042 | 0.80294676 |
| SLC12A7 | 151.511778 | -0.272351237 | 0.35745579 | -0.7619159 | 0.44611023 | 0.76084202 |
| CCDC91 | 338.278831 | -0.272201827 | 0.24396389 | -1.1157464 | 0.26453071 | 0.63018314 |
| AIMP1 | 78.5125208 | -0.272170501 | 0.24962904 | -1.0902998 | 0.27558108 | 0.64048591 |
| LEMD2 | 77.9371156 | -0.271958106 | 0.21690904 | -1.2537887 | 0.20991881 | 0.57565873 |
| GOLIM4 | 203.537584 | -0.271950431 | 0.26488668 | -1.0266671 | 0.30457726 | 0.66092151 |
| MTMR12 | 267.034598 | -0.271857219 | 0.28739124 | -0.9459482 | 0.34417503 | 0.6938754 |
| NUP98 | 296.15327 | -0.271555048 | 0.23226454 | -1.1691628 | 0.24233806 | 0.6070269 |
| SUPT4H1 | 44.4595018 | -0.271551618 | 0.40744507 | -0.6664742 | 0.50510806 | 0.7980883 |
| COL4A1 | 1108.89194 | -0.271026876 | 0.46964894 | -0.577084 | 0.56388273 | 0.83089872 |
| GFPT1 | 160.583215 | -0.270858964 | 0.25050292 | -1.0812607 | 0.27958117 | 0.64286511 |
| ZNF197 | 70.4841565 | -0.270514 | 0.31942691 | -0.8468729 | 0.39706596 | 0.72680823 |
| CCDC65 | 46.9120618 | -0.2704888 | 0.41675533 | -0.649035 | 0.51631574 | 0.80483629 |
| POLR2E | 68.9298101 | -0.270350664 | 0.26498842 | -1.0202358 | 0.30761665 | 0.66400081 |
| PRPF38B | 134.294929 | -0.270334273 | 0.29345307 | -0.9212181 | 0.35693658 | 0.70058911 |
| TRIM66 | 102.089919 | -0.270149263 | 0.32998076 | -0.8186819 | 0.41296795 | 0.73622771 |
| ATG3 | 78.871332 | -0.270004163 | 0.22066626 | -1.2235861 | 0.22110842 | 0.58706366 |
| ZNF143 | 80.2401246 | -0.269963168 | 0.26846968 | -1.005563 | 0.31462584 | 0.66883106 |
| NADSYN1 | 73.2007444 | -0.269655707 | 0.27985857 | -0.9635428 | 0.33527519 | 0.68631363 |
| INSIG2 | 60.8600393 | -0.269545878 | 0.39913748 | -0.6753209 | 0.49947191 | 0.79532568 |
| PDCD7 | 60.0394188 | -0.269448791 | 0.26602504 | -1.01287 | 0.31112224 | 0.66561244 |
| DDX54 | 53.5176562 | -0.269272984 | 0.3946532 | -0.6823028 | 0.4950475 | 0.79337519 |
| TCEAL8 | 56.5065389 | -0.269077796 | 0.44981918 | -0.598191 | 0.54971248 | 0.82355262 |
| LHFPL2 | 116.816861 | -0.268907132 | 0.26294195 | -1.0226863 | 0.30645618 | 0.66324542 |
| NFX1 | 236.63115 | -0.268483058 | 0.23527645 | -1.1411387 | 0.25381222 | 0.61961636 |
| ORAI2 | 98.2358262 | -0.268317878 | 0.36754471 | -0.7300279 | 0.46537316 | 0.77477727 |
| DNAJA2 | 58.8325322 | -0.268261099 | 0.26929812 | -0.9961492 | 0.31917768 | 0.67394237 |
| C6orf89 | 149.554538 | -0.268201631 | 0.20047918 | -1.3378029 | 0.18096069 | 0.54524118 |
| L3MBTL2 | 39.7347698 | -0.268147404 | 0.23619862 | -1.1352624 | 0.25626541 | 0.62095131 |
| DNAJC15 | 155.479508 | -0.267804495 | 0.17123602 | -1.5639495 | 0.11782943 | 0.46042456 |
| COPS6 | 88.0225621 | -0.267720411 | 0.30381402 | -0.8811983 | 0.37821048 | 0.71305115 |
| TMEM230 | 131.245848 | -0.266943209 | 0.20672958 | -1.2912676 | 0.1966109 | 0.56172669 |
| SOCS5 | 196.496385 | -0.266723865 | 0.23028752 | -1.1582211 | 0.24677382 | 0.61160108 |
| HSPA8 | 288.970448 | -0.266606537 | 0.41889844 | -0.6364467 | 0.5244853 | 0.80889528 |
| ZC3H4 | 67.9811371 | -0.266545222 | 0.23485258 | -1.134947 | 0.25639754 | 0.62095131 |
| KNOP1 | 81.7520817 | -0.266149695 | 0.38085345 | -0.6988244 | 0.48466176 | 0.78781193 |
| MAGED2 | 84.1734341 | -0.266082131 | 0.4658856 | -0.5711319 | 0.56791023 | 0.83343068 |
| ZNF565 | 95.4031523 | -0.266004064 | 0.25612844 | -1.0385573 | 0.29901067 | 0.65856711 |
| OTUB1 | 53.7061077 | -0.265980173 | 0.22719598 | -1.1707081 | 0.24171613 | 0.60636513 |
| LAMP1 | 270.0627 | -0.265682087 | 0.39714926 | -0.6689729 | 0.50351278 | 0.79776858 |
| TCF4 | 4500.04923 | -0.26562628 | 0.32067766 | -0.828328 | 0.40748477 | 0.73419295 |
| OARD1 | 62.9618333 | -0.265596066 | 0.29795652 | -0.891392 | 0.3727189 | 0.71084822 |
| GOLGA7 | 71.5709351 | -0.265446103 | 0.33359239 | -0.7957199 | 0.42619486 | 0.74203245 |
| WTAP | 395.451408 | -0.265412881 | 0.48158255 | -0.5511265 | 0.58154699 | 0.83789748 |
| PIK3IP1 | 37.6267728 | -0.26524115 | 0.71289123 | -0.372064 | 0.70984521 | 0.89727558 |
| ZCCHC17 | 100.362256 | -0.265009516 | 0.234591 | -1.1296662 | 0.25861692 | 0.62353042 |
| XRCC4 | 131.956545 | -0.264896502 | 0.18197957 | -1.4556387 | 0.14549253 | 0.49835302 |
| DBNL | 93.9372638 | -0.264771588 | 0.18340217 | -1.4436666 | 0.14883277 | 0.50222622 |
| ALKBH5 | 119.554328 | -0.264234027 | 0.25067209 | -1.0541023 | 0.29183609 | 0.65445183 |
| SRSF7 | 205.750407 | -0.264014968 | 0.32337278 | -0.8164415 | 0.41424766 | 0.73671465 |
| AC027097.2 | 176.751258 | -0.263832368 | 0.23091993 | -1.1425275 | 0.25323482 | 0.61894281 |
| UBE2B | 334.814092 | -0.26376721 | 0.27686949 | -0.952677 | 0.34075373 | 0.69116603 |
| SNX5 | 84.6130701 | -0.263691431 | 0.20191678 | -1.3059411 | 0.19157258 | 0.55590015 |
| FKBP8 | 79.096581 | -0.263680505 | 0.41329482 | -0.6379962 | 0.52347618 | 0.80861173 |
| HACD3 | 130.603687 | -0.263645247 | 0.24383064 | -1.0812638 | 0.27957979 | 0.64286511 |
| PTDSS1 | 68.3388334 | -0.263522639 | 0.30936584 | -0.8518156 | 0.39431647 | 0.72444806 |
| APAF1 | 99.9375028 | -0.26342697 | 0.27847219 | -0.9459723 | 0.34416273 | 0.6938754 |
| IFT57 | 66.5093109 | -0.263283179 | 0.22666702 | -1.1615416 | 0.24542171 | 0.6104646 |
| FAM69B | 63.3446499 | -0.263006754 | 0.57991968 | -0.4535227 | 0.65017238 | 0.86766991 |
| EFCC1 | 74.9668105 | -0.262857504 | 0.3113226 | -0.8443252 | 0.39848773 | 0.72770737 |
| GSDMD | 162.521375 | -0.262742567 | 0.40611427 | -0.6469671 | 0.51765324 | 0.80546868 |
| CPEB2 | 450.382501 | -0.262467469 | 0.25270816 | -1.0386189 | 0.29898201 | 0.65856711 |
| NNT-AS1 | 62.6301814 | -0.262459626 | 0.30624451 | -0.8570264 | 0.3914303 | 0.72247779 |
| THUMPD2 | 87.0658756 | -0.262366622 | 0.34800254 | -0.7539216 | 0.45089631 | 0.76390984 |
| RNF180 | 290.17249 | -0.262056258 | 0.16191997 | -1.6184308 | 0.1055698 | 0.4382941 |
| RALGDS | 96.1469046 | -0.261726554 | 0.4417644 | -0.5924573 | 0.55354439 | 0.82372318 |
| USP53 | 303.600309 | -0.261643611 | 0.3806178 | -0.6874182 | 0.49181921 | 0.79094859 |
| TMEM8A | 61.8522868 | -0.261518893 | 0.2951041 | -0.886192 | 0.37551407 | 0.71257833 |
| RRAGC | 57.8522203 | -0.261306535 | 0.35961363 | -0.7266313 | 0.46745182 | 0.77620906 |
| 2-Sep | 526.964191 | -0.261124873 | 0.15899338 | -1.6423632 | 0.10051475 | 0.4268542 |
| B3GAT3 | 56.6530377 | -0.260955274 | 0.30433322 | -0.8574656 | 0.39118761 | 0.72247779 |
| PJA2 | 376.401213 | -0.260952776 | 0.2174183 | -1.2002337 | 0.23004857 | 0.59648134 |
| TMEM65 | 377.651718 | -0.260847631 | 0.26986916 | -0.9665707 | 0.33375868 | 0.6852158 |
| HNRNPR | 351.117942 | -0.260767158 | 0.15457512 | -1.6869931 | 0.09160468 | 0.40917089 |
| SUPT16H | 132.540305 | -0.260764749 | 0.19416852 | -1.3429816 | 0.17927792 | 0.54322277 |
| MGMT | 271.29686 | -0.260494442 | 0.36222872 | -0.7191435 | 0.47205248 | 0.77858009 |
| ZNF117 | 64.5208119 | -0.260379089 | 0.20675878 | -1.2593375 | 0.20790844 | 0.57302618 |
| U2AF2 | 60.5671954 | -0.260268109 | 0.20573282 | -1.2650782 | 0.20584329 | 0.57060151 |
| SNX3 | 280.719821 | -0.260265377 | 0.22624203 | -1.1503848 | 0.24998544 | 0.61500658 |
| DTNBP1 | 229.472315 | -0.259856257 | 0.29642053 | -0.8766473 | 0.38067824 | 0.71529048 |
| UBE2L6 | 79.7808429 | -0.259783663 | 0.49233761 | -0.5276535 | 0.59773986 | 0.84800715 |
| SRP72 | 153.091873 | -0.259655759 | 0.18872359 | -1.3758521 | 0.16886742 | 0.53071749 |
| P3H1 | 45.6304081 | -0.259590198 | 0.28728689 | -0.9035922 | 0.36621167 | 0.70695551 |
| HRH1 | 164.571938 | -0.259203902 | 0.37830933 | -0.6851639 | 0.49324049 | 0.79156441 |
| CEP170 | 454.577953 | -0.259161271 | 0.14718988 | -1.7607275 | 0.07828454 | 0.38566641 |
| MCC | 751.430913 | -0.25889347 | 0.31390653 | -0.824747 | 0.40951524 | 0.73419295 |
| UFD1 | 66.6374295 | -0.258581216 | 0.24112888 | -1.0723776 | 0.28355044 | 0.6468618 |
| DCTN6 | 67.2045345 | -0.258146227 | 0.39877837 | -0.6473426 | 0.51741023 | 0.80546868 |
| TGFBRAP1 | 46.754389 | -0.258005701 | 0.35573636 | -0.7252722 | 0.46828504 | 0.77664704 |
| CYB5B | 290.486231 | -0.257963478 | 0.14089035 | -1.830952 | 0.0671077 | 0.35896806 |
| SLC35A4 | 50.4997032 | -0.257880889 | 0.37217358 | -0.6929049 | 0.48836924 | 0.78931968 |
| PEPD | 142.683745 | -0.257727758 | 0.25934701 | -0.9937564 | 0.32034147 | 0.67435802 |
| ENOPH1 | 73.2180311 | -0.257006636 | 0.22448691 | -1.1448625 | 0.25226613 | 0.61777104 |
| PRRC1 | 89.7625533 | -0.256886674 | 0.27714155 | -0.9269151 | 0.35397063 | 0.69977303 |
| ZNF121 | 67.4202034 | -0.256821299 | 0.24915538 | -1.0307676 | 0.30264981 | 0.65940027 |
| OSBPL3 | 236.906016 | -0.256813809 | 0.32421096 | -0.7921195 | 0.42829103 | 0.74378299 |
| CDH16 | 61.4651861 | -0.256741532 | 0.3043109 | -0.8436817 | 0.39884731 | 0.72770737 |
| MRPL42 | 123.180082 | -0.256739598 | 0.18724976 | -1.3711077 | 0.17034137 | 0.53193492 |
| GIMAP7 | 410.851642 | -0.256595631 | 0.34720016 | -0.7390424 | 0.45988127 | 0.7702023 |
| DHX38 | 66.6289114 | -0.256390723 | 0.24805681 | -1.0335968 | 0.30132469 | 0.65926326 |
| CISD3 | 48.2620734 | -0.256203143 | 0.33974973 | -0.7540937 | 0.45079293 | 0.76389285 |
| MBD4 | 82.9322731 | -0.256185024 | 0.24680598 | -1.0380017 | 0.29926927 | 0.65856711 |
| EFL1 | 92.3925299 | -0.255895099 | 0.36501531 | -0.7010531 | 0.4832699 | 0.78745472 |
| DHDDS | 57.8957963 | -0.255629191 | 0.3714925 | -0.688114 | 0.491381 | 0.79094859 |

| | | | | | | |
|---|---|---|---|---|---|---|
| LAMA5 | 306.630112 | -0.255374699 | 0.37710497 | -0.6771979 | 0.49828039 | 0.79513036 |
| IQCH | 53.1073333 | -0.255032912 | 0.35034706 | -0.7279436 | 0.46664809 | 0.77550392 |
| TMED4 | 67.705628 | -0.254830624 | 0.26841295 | -0.9493976 | 0.34241841 | 0.6921189 |
| ATXN10 | 426.761225 | -0.254743824 | 0.18267189 | -1.3945431 | 0.16315367 | 0.52335809 |
| HMGXB4 | 103.551355 | -0.253549507 | 0.24951654 | -1.0161631 | 0.30955171 | 0.6648266 |
| CLINT1 | 254.16006 | -0.253532962 | 0.16579361 | -1.5292083 | 0.12621282 | 0.47202901 |
| ANO10 | 431.803609 | -0.253520067 | 0.18963675 | -1.3368721 | 0.18126441 | 0.54532597 |
| KHDRBS1 | 378.348267 | -0.253459972 | 0.16359748 | -1.5492902 | 0.12131199 | 0.46542849 |
| FOXRED1 | 48.1689891 | -0.253386453 | 0.35043288 | -0.723067 | 0.46963872 | 0.77792258 |
| NFE2L3 | 46.7412394 | -0.253096998 | 0.3079857 | -0.8217817 | 0.41120118 | 0.73470224 |
| PGRMC2 | 151.66218 | -0.253096377 | 0.25725227 | -0.9838451 | 0.3251917 | 0.67912772 |
| RNF144A | 517.709111 | -0.252832461 | 0.42399685 | -0.5963074 | 0.55096988 | 0.8236419 |
| SDCCAG8 | 657.573522 | -0.252746037 | 0.22102059 | -1.1435407 | 0.25281417 | 0.61828402 |
| CACHD1 | 460.566996 | -0.252712716 | 0.42072017 | -0.600667 | 0.54806181 | 0.82355262 |
| ARHGEF28 | 289.296614 | -0.252617324 | 0.48679734 | -0.5189374 | 0.60380443 | 0.85085787 |
| FAM193A | 360.522727 | -0.252599825 | 0.25134147 | -1.0050065 | 0.3148937 | 0.66894474 |
| USP10 | 191.239802 | -0.25227263 | 0.29181799 | -0.8644862 | 0.38732085 | 0.72055795 |
| THAP6 | 53.1419584 | -0.25209156 | 0.34740885 | -0.7256337 | 0.46806337 | 0.77664704 |
| MYL6 | 623.118813 | -0.251753066 | 0.48173701 | -0.5225944 | 0.60125654 | 0.8499944 |
| ZSCAN18 | 81.135726 | -0.251388155 | 0.33920813 | -0.741103 | 0.458631 | 0.76968366 |
| CACNA1C | 1632.14218 | -0.250953501 | 0.43364612 | -0.5787057 | 0.56278774 | 0.83082829 |
| POLR2M | 41.2809911 | -0.250916458 | 0.39814914 | -0.6302072 | 0.52855902 | 0.81207005 |
| KLHL2 | 534.863148 | -0.250677476 | 0.19970531 | -1.2552369 | 0.20939278 | 0.57479406 |
| TFE3 | 63.1839509 | -0.250205579 | 0.23989708 | -1.0429705 | 0.29696194 | 0.65795352 |
| SPAG7 | 60.2072317 | -0.250185423 | 0.30581886 | -0.8180837 | 0.4133094 | 0.73622771 |
| TMEM208 | 55.5589607 | -0.250024554 | 0.49702197 | -0.5030453 | 0.61493244 | 0.85680098 |
| COPS7A | 47.9649241 | -0.249791007 | 0.29428535 | -0.8488054 | 0.39598956 | 0.7260548 |
| SLC26A2 | 91.9946529 | -0.249469353 | 0.34285656 | -0.7276202 | 0.46684613 | 0.7756755 |
| CLIP1 | 360.892004 | -0.249446339 | 0.20899146 | -1.1935719 | 0.23264546 | 0.59910683 |
| KCNMB4 | 337.751244 | -0.249172003 | 0.4494397 | -0.5544059 | 0.57930112 | 0.83649569 |
| ADGRL2 | 1720.90676 | -0.249058483 | 0.51218997 | -0.4862619 | 0.62678146 | 0.86052754 |
| RTL8A | 49.8844225 | -0.248888708 | 0.50540877 | -0.4924503 | 0.62240104 | 0.85940448 |
| AC093827.5 | 53.0813661 | -0.248580377 | 0.23735488 | -1.0472941 | 0.29496394 | 0.65671864 |
| TINF2 | 41.8312254 | -0.248559066 | 0.33287447 | -0.7467051 | 0.45524158 | 0.76710126 |
| HMGN5 | 49.1505859 | -0.248249873 | 0.30668332 | -0.8094665 | 0.41824687 | 0.73821752 |
| CERS6 | 245.628708 | -0.248218658 | 0.19417304 | -1.2783374 | 0.20113048 | 0.56597343 |
| MCCC1 | 88.497386 | -0.247903018 | 0.29254693 | -0.8473957 | 0.39677459 | 0.72680823 |
| CDK2AP2 | 46.5984512 | -0.247841663 | 0.47676659 | -0.5198386 | 0.6031761 | 0.85055833 |
| TMF1 | 175.973774 | -0.247629158 | 0.16964492 | -1.459691 | 0.14437502 | 0.49679586 |
| SLC16A1-AS1 | 140.590622 | -0.24740207 | 0.3510284 | -0.7047922 | 0.48093958 | 0.78475146 |
| G3BP1 | 330.215293 | -0.24729207 | 0.19997663 | -1.2366048 | 0.21623382 | 0.58205308 |
| RNPS1 | 138.076909 | -0.247019142 | 0.20770311 | -1.1892896 | 0.23432574 | 0.59949266 |
| ELAC2 | 49.7628358 | -0.246512049 | 0.30427731 | -0.8101559 | 0.4178506 | 0.73821752 |
| NEK9 | 159.878568 | -0.246407871 | 0.17354217 | -1.4198731 | 0.15564461 | 0.51180943 |
| CSDE1 | 366.63978 | -0.246267338 | 0.16322549 | -1.5087554 | 0.1313613 | 0.48052281 |
| HBP1 | 195.523873 | -0.246225772 | 0.39526512 | -0.6229383 | 0.53332509 | 0.81479963 |
| TAF7 | 100.009524 | -0.245987986 | 0.21263834 | -1.1568374 | 0.24733879 | 0.61239737 |
| MYOF | 324.741469 | -0.245708424 | 0.55571698 | -0.4421467 | 0.65838306 | 0.87304304 |
| KYAT3 | 100.892102 | -0.245566974 | 0.29813508 | -0.8236769 | 0.41012317 | 0.73419295 |
| CTSO | 65.7768404 | -0.245206051 | 0.2737056 | -0.8958752 | 0.37031943 | 0.70984233 |
| COX7A2L | 130.354017 | -0.245094563 | 0.2820422 | -0.8689996 | 0.38484735 | 0.71699718 |
| SIPA1L3 | 448.924136 | -0.244856308 | 0.31108103 | -0.7871142 | 0.43121499 | 0.74490865 |
| NAP1L1 | 305.44424 | -0.24485442 | 0.26688043 | -0.9174686 | 0.35889713 | 0.70242891 |
| NUDC | 97.0510762 | -0.244842269 | 0.26946504 | -0.9086235 | 0.3635489 | 0.70562634 |
| PSMC2 | 88.0911088 | -0.244540007 | 0.22377394 | -1.0927993 | 0.27448193 | 0.63975272 |
| NETO2 | 96.0723869 | -0.244352757 | 0.50297267 | -0.4858172 | 0.6270968 | 0.86052754 |
| TRAPPC11 | 150.236273 | -0.244291871 | 0.21473281 | -1.1376551 | 0.25526453 | 0.6208774 |
| PGLS | 79.7302172 | -0.243694585 | 0.42333942 | -0.5756482 | 0.56485298 | 0.83113895 |
| EFTUD2 | 96.1812124 | -0.243451983 | 0.23128661 | -1.0525987 | 0.29252497 | 0.65502684 |
| CWC25 | 67.1855535 | -0.243261685 | 0.23589231 | -1.0312404 | 0.30242809 | 0.65926326 |
| HDLBP | 315.808103 | -0.243232017 | 0.16563978 | -1.4684396 | 0.14198485 | 0.49378081 |
| LDAH | 173.829802 | -0.242890459 | 0.24559908 | -0.9889714 | 0.32267715 | 0.6761919 |
| DRG1 | 99.2765421 | -0.24271854 | 0.19076701 | -1.2723297 | 0.20325599 | 0.567308 |
| HS3ST3A1 | 127.88443 | -0.242654241 | 0.57707187 | -0.4204922 | 0.6741259 | 0.87910389 |
| BNIP2 | 251.402976 | -0.24219207 | 0.22911144 | -1.0570929 | 0.29046921 | 0.65333738 |
| LSM3 | 71.5076616 | -0.242050636 | 0.48235571 | -0.5018094 | 0.61580159 | 0.85728214 |
| LARS2 | 132.640085 | -0.241471767 | 0.31429444 | -0.7682979 | 0.44231021 | 0.75704339 |
| LAYN | 62.8146034 | -0.241465902 | 0.27238388 | -0.8864912 | 0.37535292 | 0.71257833 |
| DGUOK | 99.7882239 | -0.241463167 | 0.19747859 | -1.2227309 | 0.22143136 | 0.58750144 |
| IK | 55.5238585 | -0.241248083 | 0.29850987 | -0.8081746 | 0.4189901 | 0.7382851 |
| PCCB | 60.6816388 | -0.241120992 | 0.25981666 | -0.9280428 | 0.35338535 | 0.69977303 |
| ANP32B | 127.651944 | -0.241044436 | 0.22916586 | -1.051834 | 0.29287573 | 0.655236 |
| IRF2BPL | 112.798002 | -0.241014725 | 0.52174146 | -0.4619428 | 0.64412232 | 0.86523823 |
| MBP | 87.30989 | -0.240904606 | 0.42981885 | -0.5604794 | 0.57515249 | 0.834481 |
| RCC1L | 79.1013689 | -0.240619928 | 0.2889878 | -0.83263 | 0.40505341 | 0.73217407 |
| XRCC5 | 525.32262 | -0.240405405 | 0.17596088 | -1.3662435 | 0.17186256 | 0.53377758 |
| DNAJA3 | 63.5777189 | -0.240245913 | 0.22071529 | -1.0884879 | 0.27637979 | 0.64068436 |
| RANBP10 | 184.629335 | -0.240233898 | 0.25139127 | -0.9556175 | 0.33926551 | 0.68957022 |
| LARS | 216.682166 | -0.239556242 | 0.24775718 | -0.9668993 | 0.33359438 | 0.6852158 |
| ZYX | 71.5424238 | -0.239451973 | 0.29971349 | -0.7989362 | 0.42432738 | 0.74157166 |
| VIPAS39 | 43.6384161 | -0.239426209 | 0.28473204 | -0.8408826 | 0.40041372 | 0.72864885 |
| PPP1CC | 96.7307566 | -0.239298522 | 0.25267475 | -0.9470615 | 0.34360746 | 0.69374287 |
| DLG1 | 840.08528 | -0.239296786 | 0.22187685 | -1.0785117 | 0.28080544 | 0.64401091 |
| CERS2 | 81.0088689 | -0.239218696 | 0.31386565 | -0.7621691 | 0.4459591 | 0.76074282 |
| ECHS1 | 92.7990508 | -0.239210813 | 0.34720841 | -0.6889545 | 0.49085187 | 0.79073252 |
| HACL1 | 101.264374 | -0.239199636 | 0.22161126 | -1.0793659 | 0.28042466 | 0.64385822 |
| GNAS | 753.082372 | -0.239178618 | 0.3782666 | -0.6323017 | 0.52718975 | 0.81084479 |
| RPTOR | 274.295763 | -0.238938773 | 0.25729703 | -0.9286495 | 0.35307073 | 0.69977303 |
| ACP1 | 71.2415074 | -0.238834928 | 0.20621676 | -1.1581742 | 0.24679296 | 0.61160108 |
| MFAP1 | 52.1711373 | -0.238778827 | 0.22799611 | -1.0472934 | 0.29496427 | 0.65671864 |
| GNL2 | 44.0875071 | -0.237998711 | 0.43551906 | -0.5464714 | 0.58474194 | 0.83895258 |
| MFHAS1 | 263.035335 | -0.237980697 | 0.26298363 | -0.9049259 | 0.36550464 | 0.70668224 |
| TXNDC15 | 89.9995006 | -0.237868097 | 0.226021 | -1.0524159 | 0.29260878 | 0.65502684 |
| VBP1 | 44.589107 | -0.237737897 | 0.24029133 | -0.9893736 | 0.32248038 | 0.6761919 |
| BUD31 | 56.8451886 | -0.237401905 | 0.30332323 | -0.7826697 | 0.4338211 | 0.74798947 |
| FBXO28 | 185.917839 | -0.237252943 | 0.23589602 | -1.0057522 | 0.31453479 | 0.66883106 |
| PLSCR4 | 578.843799 | -0.237132049 | 0.35403275 | -0.6698026 | 0.50298365 | 0.79776858 |
| CCT4 | 146.361479 | -0.237122217 | 0.27117077 | -0.8744387 | 0.38187939 | 0.71553552 |
| GRIPAP1 | 97.1414637 | -0.236956672 | 0.23222354 | -1.0203818 | 0.30754743 | 0.66400081 |
| MZF1 | 68.7103467 | -0.23655984 | 0.32997287 | -0.7169069 | 0.47343153 | 0.77899012 |
| DAZAP1 | 134.984489 | -0.236545453 | 0.22582328 | -1.0474804 | 0.29487809 | 0.65671864 |
| NDUFC1 | 87.6588017 | -0.236106082 | 0.26978333 | -0.8751693 | 0.3814818 | 0.71543195 |
| SYMPK | 157.294186 | -0.235879367 | 0.25530002 | -0.9239301 | 0.35552272 | 0.70018061 |
| DEK | 202.52171 | -0.235872905 | 0.19311147 | -1.2214339 | 0.22192176 | 0.58750144 |
| CCDC28A | 45.3227351 | -0.235781506 | 0.4180406 | -0.5640158 | 0.57274337 | 0.8342245 |
| TACSTD2 | 47.5061765 | -0.235718671 | 0.76190621 | -0.3093802 | 0.75703236 | 0.91355195 |
| COPB2 | 151.56733 | -0.235524337 | 0.17698539 | -1.3307558 | 0.18326938 | 0.54813352 |
| EMC3 | 116.285439 | -0.235503676 | 0.19256229 | -1.223 | 0.22132969 | 0.58746055 |
| HDHD5 | 58.4906411 | -0.235180507 | 0.23492351 | -1.001094 | 0.31678138 | 0.67173413 |
| ALDH9A1 | 79.9833063 | -0.235065268 | 0.30663089 | -0.7666066 | 0.44331544 | 0.75844663 |
| ACOX3 | 57.2646807 | -0.234944273 | 0.21873074 | -1.0741255 | 0.28276642 | 0.64582577 |
| MRPL28 | 57.9906709 | -0.23487757 | 0.45634969 | -0.5146877 | 0.60677127 | 0.85196053 |
| SHISA5 | 255.251988 | -0.234789059 | 0.2846214 | -0.8249171 | 0.40941866 | 0.73419295 |

| | | | | | | |
|---|---|---|---|---|---|---|
| PRRC2C | 796.59864 | -0.234752282 | 0.23987792 | -0.9786323 | 0.3277617 | 0.6802115 |
| PEX13 | 74.3934425 | -0.234725636 | 0.2793743 | -0.8401833 | 0.4008056 | 0.72868078 |
| TRNT1 | 123.054892 | -0.234601752 | 0.22974126 | -1.0211564 | 0.30718035 | 0.66400081 |
| NACC2 | 214.950562 | -0.234508595 | 0.24283696 | -0.9657039 | 0.33419238 | 0.6852158 |
| MECP2 | 451.805692 | -0.234436263 | 0.32451906 | -0.7224114 | 0.47004159 | 0.77792258 |
| TUBG2 | 42.1816957 | -0.233914889 | 0.28338848 | -0.8254213 | 0.40913244 | 0.73419295 |
| ABL1 | 648.19369 | -0.233540016 | 0.28786422 | -0.8112853 | 0.41720184 | 0.73821752 |
| COQ10B | 73.3381909 | -0.233391524 | 0.24973906 | -0.9345415 | 0.35002463 | 0.69858573 |
| EMC8 | 50.9718142 | -0.233135392 | 0.25603269 | -0.9105689 | 0.36252258 | 0.70497203 |
| BRD9 | 69.7268442 | -0.233103175 | 0.29594586 | -0.7876548 | 0.43089866 | 0.74483391 |
| SWI5 | 47.549534 | -0.232986252 | 0.42998588 | -0.5418463 | 0.5879244 | 0.8409165 |
| ISCU | 120.697468 | -0.232655662 | 0.47348226 | -0.4913714 | 0.62316376 | 0.85940448 |
| OSER1 | 43.0439158 | -0.232029659 | 0.395753 | -0.5862992 | 0.55767448 | 0.82656227 |
| MRRF | 139.076257 | -0.231924654 | 0.28789639 | -0.8055837 | 0.4204829 | 0.73901389 |
| GPCPD1 | 295.984044 | -0.231756568 | 0.47166736 | -0.491356 | 0.62317471 | 0.85940448 |
| CYP26B1 | 234.83894 | -0.231715849 | 1.10303267 | -0.2100716 | 0.83361178 | 0.9431004 |
| SDK1 | 146.597785 | -0.231220825 | 0.38812635 | -0.595736 | 0.55135163 | 0.8236419 |
| TOPORS | 64.6565657 | -0.231195581 | 0.25597507 | -0.9031957 | 0.36642204 | 0.70695551 |
| PIGL | 185.296182 | -0.231184332 | 0.38189751 | -0.605357 | 0.54494181 | 0.82334912 |
| PEF1 | 76.6410551 | -0.230926272 | 0.24770199 | -0.9322746 | 0.35119463 | 0.69858573 |
| PAWR | 199.621465 | -0.230906871 | 0.22752126 | -1.0148804 | 0.31016283 | 0.6648266 |
| GRHPR | 66.2806878 | -0.230835253 | 0.28685992 | -0.8046968 | 0.42099467 | 0.73910721 |
| NCOR2 | 453.939242 | -0.230522306 | 0.36699025 | -0.6281429 | 0.52991036 | 0.8130989 |
| EPAS1 | 3405.33722 | -0.230409337 | 0.38317773 | -0.6013119 | 0.54763225 | 0.82355262 |
| FAM118B | 64.8151116 | -0.230344205 | 0.26828198 | -0.8585899 | 0.39056679 | 0.72247779 |
| GAA | 50.6713202 | -0.230340241 | 0.38788817 | -0.5938316 | 0.55262477 | 0.8236419 |
| BTBD2 | 59.2732189 | -0.230162527 | 0.33062153 | -0.696151 | 0.48633424 | 0.78826504 |
| SCO1 | 67.6892803 | -0.230113762 | 0.2669556 | -0.8619926 | 0.38869157 | 0.72189357 |
| TFPI2 | 58.0821127 | -0.230075382 | 0.61492028 | -0.3741548 | 0.70828914 | 0.89680567 |
| BUB3 | 84.2068092 | -0.229991826 | 0.27927789 | -0.8235232 | 0.41021053 | 0.73419295 |
| NCLN | 56.9369671 | -0.229971879 | 0.31499231 | -0.7300873 | 0.46533683 | 0.77477727 |
| PPP2R1B | 169.355388 | -0.229862261 | 0.28866394 | -0.7962971 | 0.42585937 | 0.74197066 |
| DDT | 90.6344664 | -0.229709563 | 0.40191729 | -0.5715344 | 0.56763743 | 0.83332923 |
| AC012409.2 | 75.0556226 | -0.229575398 | 0.72271046 | -0.3176589 | 0.7507437 | 0.91109327 |
| PSMB4 | 86.4290713 | -0.229429295 | 0.47453601 | -0.4834813 | 0.62875402 | 0.86052754 |
| TTC1 | 140.766653 | -0.229333119 | 0.25642565 | -0.8943455 | 0.37113708 | 0.70984233 |
| CUEDC2 | 46.7828147 | -0.229293086 | 0.318941 | -0.7189201 | 0.47219017 | 0.77858009 |
| EIF3H | 330.194594 | -0.22893052 | 0.21853071 | -1.0475897 | 0.29482769 | 0.65671864 |
| DDX3Y | 103.524944 | -0.228888318 | 2.43567746 | -0.0939732 | 0.92513047 | 0.97581112 |
| ITGA9 | 877.798296 | -0.228784946 | 0.27704366 | -0.8258083 | 0.40891285 | 0.73419295 |
| SYPL1 | 227.951011 | -0.228649882 | 0.30486734 | -0.749998 | 0.45325593 | 0.76568057 |
| SMIM4 | 53.6134447 | -0.228125678 | 0.28075609 | -0.8125404 | 0.4164816 | 0.73802299 |
| SPRTN | 46.7267736 | -0.228073365 | 0.34188898 | -0.6670977 | 0.5047097 | 0.79776858 |
| NR2F2-AS1 | 544.236282 | -0.227978154 | 0.4665563 | -0.4886402 | 0.62509646 | 0.86016045 |
| WBP2 | 133.496668 | -0.227959795 | 0.25574481 | -0.8913565 | 0.37273795 | 0.71084822 |
| MID1 | 550.563727 | -0.227030642 | 0.24449096 | -0.928585 | 0.35310419 | 0.69977303 |
| TP53BP2 | 103.581737 | -0.226981323 | 0.27674306 | -0.8201879 | 0.41210897 | 0.73582539 |
| C6orf48 | 51.60008 | -0.226764005 | 0.44438051 | -0.5102924 | 0.60984661 | 0.85375662 |
| ZNF480 | 44.2384508 | -0.226540516 | 0.29670789 | -0.7635136 | 0.44515715 | 0.76016714 |
| CA2 | 115.264143 | -0.226481631 | 0.71825096 | -0.3153238 | 0.75251581 | 0.91163598 |
| SRFBP1 | 129.393901 | -0.226333505 | 0.26523419 | -0.8533346 | 0.39347379 | 0.72360479 |
| NPDC1 | 78.2403903 | -0.22587563 | 0.43781198 | -0.5159193 | 0.60591081 | 0.85156118 |
| CLPTM1 | 101.352839 | -0.225683299 | 0.22268331 | -1.013472 | 0.31083477 | 0.66561244 |
| PIGT | 72.9401692 | -0.224669196 | 0.39589122 | -0.5675023 | 0.57037293 | 0.83368112 |
| MVP | 181.457081 | -0.224616478 | 0.33600259 | -0.6684963 | 0.50381687 | 0.79776858 |
| SLC3A1 | 69.9274741 | -0.22446521 | 0.6134635 | -0.3658982 | 0.71444102 | 0.89980314 |
| SMG6 | 532.186673 | -0.22438893 | 0.28967831 | -0.7746142 | 0.43856767 | 0.75300027 |
| IGFBP5 | 4716.48657 | -0.224327281 | 0.61630129 | -0.3639896 | 0.71586575 | 0.9001716 |
| RAC1 | 604.134285 | -0.223975123 | 0.26025624 | -0.8605946 | 0.38946134 | 0.72202943 |
| FAM228B | 84.3911959 | -0.22372315 | 0.32966539 | -0.6786371 | 0.49736786 | 0.7946635 |
| STX12 | 491.774278 | -0.223295321 | 0.3002644 | -0.7436623 | 0.45708079 | 0.76850053 |
| MYO10 | 589.791863 | -0.222926527 | 0.34193312 | -0.6519595 | 0.51442733 | 0.80411821 |
| USP22 | 240.537159 | -0.222758293 | 0.16233816 | -1.3721868 | 0.17000528 | 0.53193492 |
| ZDHHC13 | 145.981641 | -0.222350048 | 0.33063809 | -0.6724877 | 0.50127327 | 0.79642461 |
| ACTR1A | 89.8693761 | -0.222342275 | 0.26455968 | -0.8404239 | 0.40067077 | 0.72864885 |
| MALL | 95.2001042 | -0.22214644 | 0.53130075 | -0.4181181 | 0.6758608 | 0.88033151 |
| STK3 | 1068.97098 | -0.222097427 | 0.2514148 | -0.8833904 | 0.37702537 | 0.71257833 |
| NFIB | 2027.10349 | -0.222097397 | 0.23086366 | -0.9620284 | 0.33603533 | 0.68706501 |
| HYAL1 | 176.954544 | -0.222043176 | 0.47438422 | -0.4680661 | 0.63973732 | 0.86399909 |
| EIF2AK3 | 149.297518 | -0.221934172 | 0.29417438 | -0.7544307 | 0.45059066 | 0.76375998 |
| EDEM1 | 72.7370712 | -0.221785406 | 0.26620067 | -0.8331512 | 0.40475947 | 0.73196613 |
| ZCCHC9 | 43.6259968 | -0.221580043 | 0.24475629 | -0.9053089 | 0.36530176 | 0.70668224 |
| PFDN5 | 184.967275 | -0.221433347 | 0.41189957 | -0.5375906 | 0.5908597 | 0.84248213 |
| LAMC1 | 488.914642 | -0.221091233 | 0.34583569 | -0.6392956 | 0.52263065 | 0.80854347 |
| TAF1D | 222.876682 | -0.220828714 | 0.26092911 | -0.8463169 | 0.39737598 | 0.72680823 |
| ATR | 307.188855 | -0.220302048 | 0.28318128 | -0.7779541 | 0.43659605 | 0.75103332 |
| RAD23B | 302.551081 | -0.220146275 | 0.1602578 | -1.3737008 | 0.16953458 | 0.5312985 |
| NDUFS3 | 50.6885759 | -0.22007118 | 0.41767429 | -0.5268966 | 0.59826537 | 0.84839769 |
| MRPL22 | 72.6926524 | -0.219892875 | 0.2759911 | -0.796739 | 0.42560262 | 0.74197066 |
| TFG | 248.080243 | -0.219616997 | 0.20332518 | -1.0801269 | 0.28008568 | 0.64344221 |
| ZC3H18 | 110.006102 | -0.219172276 | 0.26629129 | -0.8230546 | 0.41047693 | 0.73419295 |
| TNPO2 | 86.4294928 | -0.219158792 | 0.23401256 | -0.9365258 | 0.34900256 | 0.69809042 |
| PLEKHG5 | 56.2592823 | -0.219099767 | 0.32660484 | -0.6708405 | 0.50232212 | 0.79691777 |
| EMD | 55.8392023 | -0.219070035 | 0.3612702 | -0.6063883 | 0.54425692 | 0.82246618 |
| CARS | 108.587061 | -0.218619761 | 0.24735038 | -0.8838465 | 0.37677909 | 0.71257833 |
| GTPBP6 | 82.7926059 | -0.218508311 | 0.35460349 | -0.6162046 | 0.53775947 | 0.81807397 |
| VCPIP1 | 72.5237292 | -0.218423511 | 0.19423751 | -1.1245176 | 0.26079349 | 0.62580682 |
| PSMC5 | 51.1512513 | -0.218291692 | 0.43231544 | -0.5049361 | 0.61360369 | 0.85573702 |
| SEPSECS | 51.1536159 | -0.218286839 | 0.26134986 | -0.8352285 | 0.40358909 | 0.73065697 |
| FAM98A | 55.8953296 | -0.218240203 | 0.28374837 | -0.7691329 | 0.44181443 | 0.75666963 |
| ZNF277 | 182.671772 | -0.218183996 | 0.20998083 | -1.0390663 | 0.29877392 | 0.65856711 |
| NHS | 118.854948 | -0.217986434 | 0.49967392 | -0.4362574 | 0.66265 | 0.87493354 |
| COIL | 44.9507882 | -0.217829776 | 0.27606477 | -0.7890532 | 0.43008093 | 0.74420645 |
| ACVRL1 | 261.819113 | -0.217281918 | 0.26383585 | -0.8235496 | 0.4101955 | 0.73419295 |
| IKZF2 | 49.3527476 | -0.217136792 | 0.36202385 | -0.5997859 | 0.54864895 | 0.82355262 |
| DDX55 | 53.2718419 | -0.217064476 | 0.30397149 | -0.7140949 | 0.47516853 | 0.78022377 |
| NAALADL2 | 338.64549 | -0.216991563 | 0.39311024 | -0.5519865 | 0.58095757 | 0.83774521 |
| HDAC8 | 613.955948 | -0.216621938 | 0.18563992 | -1.1668931 | 0.24325355 | 0.608518 |
| MGAT1 | 269.988928 | -0.216572541 | 0.28557246 | -0.7583804 | 0.44822331 | 0.76164496 |
| CUTA | 58.0054911 | -0.216518013 | 0.49930135 | -0.433642 | 0.66454846 | 0.8755434 |
| RTF2 | 85.0342974 | -0.2163098 | 0.26972007 | -0.8019789 | 0.42256517 | 0.74011688 |
| ALG5 | 82.4622826 | -0.216185401 | 0.25138164 | -0.8599888 | 0.38979521 | 0.72202943 |
| IFNGR1 | 245.580441 | -0.215831461 | 0.38028559 | -0.567551 | 0.5703399 | 0.83368112 |
| ZNF253 | 44.4935771 | -0.215684269 | 0.36304445 | -0.5940988 | 0.55244604 | 0.8236419 |
| LYRM7 | 44.3111216 | -0.215586445 | 0.31663087 | -0.6808763 | 0.49594978 | 0.79381653 |
| TSHZ1 | 230.501129 | -0.215297957 | 0.27294602 | -0.7887932 | 0.4302329 | 0.74431201 |
| UNC45A | 75.9524147 | -0.215168453 | 0.25242274 | -0.8524131 | 0.39398485 | 0.72408262 |
| FBXO31 | 50.6368489 | -0.21515301 | 0.3017896 | -0.7129239 | 0.47589287 | 0.78022377 |
| ZC3H7B | 122.610693 | -0.215057125 | 0.3024902 | -0.7109557 | 0.47711171 | 0.78098272 |
| PCNA | 43.5552263 | -0.215024163 | 0.47688739 | -0.4508909 | 0.65206821 | 0.8685756 |
| DGKD | 106.925813 | -0.214948164 | 0.21872842 | -0.9827171 | 0.32574669 | 0.67964945 |
| HMGB2 | 89.0899583 | -0.214788716 | 0.57840277 | -0.371348 | 0.71037834 | 0.89734887 |
| LYSMD3 | 55.6252068 | -0.214705413 | 0.27537356 | -0.7796878 | 0.43557464 | 0.74971913 |

| | | | | | | |
|---|---|---|---|---|---|---|
| ATP6V0D1 | 85.6811148 | -0.214572367 | 0.32686705 | -0.6564515 | 0.51153365 | 0.80205051 |
| MPC2 | 65.5931199 | -0.214465406 | 0.34533511 | -0.6210356 | 0.53457619 | 0.81566976 |
| PCCA | 786.318186 | -0.213767711 | 0.17622832 | -1.2130157 | 0.22512383 | 0.59122266 |
| NFXL1 | 83.8166459 | -0.21358997 | 0.25870275 | -0.8256193 | 0.4090201 | 0.73419295 |
| RNMT | 179.738326 | -0.213575305 | 0.40538246 | -0.5268489 | 0.59829852 | 0.84839769 |
| ADPGK | 91.5504039 | -0.213545328 | 0.29612146 | -0.721141 | 0.47082276 | 0.77817464 |
| ZNF440 | 46.905833 | -0.212632658 | 0.34138858 | -0.6228464 | 0.53338545 | 0.81479963 |
| GRAMD2B | 486.51016 | -0.212593543 | 0.49714396 | -0.4276297 | 0.66892071 | 0.87628452 |
| TECPR1 | 199.56256 | -0.212559006 | 0.29080539 | -0.7309321 | 0.4648206 | 0.77464921 |
| EXOSC1 | 65.0445449 | -0.212481881 | 0.25095718 | -0.8466858 | 0.39717027 | 0.72680823 |
| TMEM51 | 178.965586 | -0.21230607 | 0.4255885 | -0.4988529 | 0.61788299 | 0.85842725 |
| CHD2 | 329.09777 | -0.212248959 | 0.23092265 | -0.9191344 | 0.35802526 | 0.70156148 |
| RAB8B | 427.231107 | -0.211882457 | 0.37218976 | -0.5692861 | 0.569162 | 0.83368112 |
| HNRNPUL1 | 137.015087 | -0.211606502 | 0.20718914 | -1.0213205 | 0.30710264 | 0.66400081 |
| PCK1 | 131.124146 | -0.211563734 | 1.20451644 | -0.175642 | 0.86057517 | 0.95279212 |
| RIC8A | 41.2740958 | -0.211071979 | 0.348976 | -0.6048324 | 0.54529039 | 0.82355262 |
| SIL1 | 217.396153 | -0.210986495 | 0.21882112 | -0.9641962 | 0.33494756 | 0.68607656 |
| AL360091.3 | 67.8255893 | -0.210969757 | 0.49715176 | -0.4243569 | 0.67130558 | 0.87752293 |
| ATP5S | 47.3365673 | -0.210667292 | 0.26613662 | -0.7915757 | 0.42860809 | 0.74408005 |
| SLC16A2 | 51.7958228 | -0.210663578 | 0.29537481 | -0.7132077 | 0.47571727 | 0.78022377 |
| AMBRA1 | 875.107408 | -0.210563845 | 0.27076779 | -0.7776547 | 0.4367726 | 0.75117911 |
| CAVIN1 | 349.193092 | -0.210522762 | 0.42594223 | -0.4942519 | 0.62112828 | 0.85890999 |
| ALG14 | 78.5428407 | -0.210497594 | 0.20408156 | -1.0314386 | 0.3023352 | 0.65926326 |
| TIMP2 | 469.402071 | -0.210338891 | 0.45244186 | -0.4648971 | 0.64200517 | 0.86490071 |
| TMEM39B | 62.1833161 | -0.210228891 | 0.35258741 | -0.5962462 | 0.55101079 | 0.8236419 |
| GOLT1B | 64.1228923 | -0.209996347 | 0.20251712 | -1.0369313 | 0.29976787 | 0.65926326 |
| PIAS1 | 1046.47094 | -0.209904324 | 0.25654298 | -0.8182034 | 0.41324108 | 0.73622771 |
| NDUFB6 | 79.4351774 | -0.209833189 | 0.32754596 | -0.6406221 | 0.52176822 | 0.80772643 |
| ARHGAP28 | 148.253192 | -0.209609038 | 0.37164092 | -0.5640096 | 0.57274762 | 0.8342245 |
| CCDC69 | 220.616187 | -0.209522257 | 0.24457451 | -0.8566807 | 0.39162138 | 0.72247779 |
| SGCB | 48.4986948 | -0.209450624 | 0.26676057 | -0.7851634 | 0.4323578 | 0.7464101 |
| NDUFB2 | 124.246269 | -0.209207936 | 0.3151684 | -0.6637973 | 0.50682005 | 0.79909604 |
| CLSTN3 | 90.5002219 | -0.208971804 | 0.30066236 | -0.6950381 | 0.48703141 | 0.78826504 |
| ZNF100 | 88.1795072 | -0.208802673 | 0.37026798 | -0.5639231 | 0.57280646 | 0.8342245 |
| ZNF765 | 63.5458076 | -0.208798718 | 0.35295068 | -0.5915804 | 0.55413157 | 0.82401134 |
| GLTP | 86.9115231 | -0.208772412 | 0.23260436 | -0.897543 | 0.36942925 | 0.70980031 |
| SSB | 113.013228 | -0.208687229 | 0.26744735 | -0.7802927 | 0.43521858 | 0.74960928 |
| PPCDC | 192.884361 | -0.208593224 | 0.34670691 | -0.6016414 | 0.54741288 | 0.82355262 |
| PRDM4 | 66.9410448 | -0.208510861 | 0.32122658 | -0.6491084 | 0.51626834 | 0.80483629 |
| DFFA | 67.355302 | -0.208485701 | 0.29327696 | -0.7108833 | 0.47715654 | 0.78098272 |
| PGP | 83.7213385 | -0.208458967 | 0.30492455 | -0.6836411 | 0.4942018 | 0.7926408 |
| CBX1 | 65.8616476 | -0.208227657 | 0.27856703 | -0.7474957 | 0.4547644 | 0.76681168 |
| HMCN1 | 210.0906 | -0.208038607 | 0.65161572 | -0.3192658 | 0.74952497 | 0.91078628 |
| CCZ1B | 71.5913792 | -0.207818026 | 0.30042368 | -0.6917498 | 0.48909446 | 0.78955612 |
| DST | 662.084481 | -0.207695789 | 0.28091376 | -0.7393578 | 0.45968974 | 0.77003914 |
| NBR1 | 180.770458 | -0.207585974 | 0.21851452 | -0.9499871 | 0.3421188 | 0.692051 |
| ATP6V1E1 | 148.823644 | -0.20739841 | 0.26510609 | -0.7823223 | 0.43402517 | 0.74808934 |
| DNAJC21 | 98.9858385 | -0.207146232 | 0.24862599 | -0.833164 | 0.40475222 | 0.73196613 |
| TSPAN7 | 138.61189 | -0.207099654 | 0.89642703 | -0.2310279 | 0.81729313 | 0.93550282 |
| DDB1 | 129.895457 | -0.206993892 | 0.21943495 | -0.9433041 | 0.34552536 | 0.69497757 |
| PIK3C2A | 1434.51472 | -0.206780014 | 0.20016721 | -1.0330364 | 0.30158686 | 0.65926326 |
| ABHD3 | 98.0352216 | -0.206658268 | 0.2872748 | -0.7193749 | 0.47190998 | 0.77855633 |
| 11-Sep | 173.522804 | -0.206658098 | 0.23616406 | -0.8750616 | 0.38154039 | 0.71543195 |
| DUSP3 | 98.4754328 | -0.2066443 | 0.30329579 | -0.6813293 | 0.49566316 | 0.79358475 |
| GALNT10 | 261.968502 | -0.206274541 | 0.26025751 | -0.7925786 | 0.42802336 | 0.74347594 |
| AATF | 158.385137 | -0.206219678 | 0.21569241 | -0.9560822 | 0.33903069 | 0.68955779 |
| POC5 | 57.4530559 | -0.206139163 | 0.25055089 | -0.8227437 | 0.41065377 | 0.73434874 |
| IQCB1 | 173.662864 | -0.206013836 | 0.2345074 | -0.8784961 | 0.37967456 | 0.71418244 |
| ASB8 | 45.6447716 | -0.205707162 | 0.29198291 | -0.7045178 | 0.48111036 | 0.78487362 |
| TMOD2 | 70.675021 | -0.205631809 | 0.46748491 | -0.4398683 | 0.66003247 | 0.87335081 |
| RAB4A | 50.4206239 | -0.20562253 | 0.30788553 | -0.6678538 | 0.5042269 | 0.79776858 |
| ERLEC1 | 83.4516472 | -0.205347267 | 0.2144784 | -0.9574263 | 0.33835211 | 0.68908295 |
| ATMIN | 51.1559196 | -0.205284883 | 0.34525686 | -0.594586 | 0.55212027 | 0.8236419 |
| ADGRA2 | 47.6670963 | -0.204971465 | 0.48258963 | -0.4247324 | 0.67103174 | 0.87730512 |
| RNF41 | 52.5394662 | -0.204752785 | 0.27165114 | -0.7537347 | 0.45100856 | 0.76394184 |
| KIAA0586 | 173.104985 | -0.204516467 | 0.19553686 | -1.0459228 | 0.29559666 | 0.65731032 |
| PAM | 1079.69375 | -0.204425498 | 0.28667046 | -0.7131028 | 0.47578216 | 0.78022377 |
| RRP1B | 75.9468419 | -0.20435702 | 0.22483623 | -0.908915 | 0.36339501 | 0.7056112 |
| SNRNP40 | 95.6687341 | -0.204153544 | 0.17179054 | -1.1883864 | 0.2346812 | 0.59949266 |
| DUSP22 | 122.64481 | -0.204044745 | 0.24101057 | -0.8466215 | 0.3972061 | 0.72680823 |
| COA5 | 47.1701322 | -0.203816921 | 0.34057713 | -0.5984457 | 0.54954258 | 0.82355262 |
| NSUN6 | 159.870979 | -0.20377192 | 0.37191906 | -0.5478932 | 0.58376525 | 0.83850098 |
| CNOT7 | 81.9385562 | -0.203738646 | 0.27433977 | -0.7426508 | 0.45769314 | 0.76905605 |
| UBA2 | 234.502817 | -0.203184413 | 0.24210268 | -0.8392489 | 0.40132964 | 0.72911494 |
| KIF9-AS1 | 118.393777 | -0.20317302 | 0.32761117 | -0.6201651 | 0.53514908 | 0.81602082 |
| SCYL1 | 54.9796843 | -0.203169732 | 0.25044279 | -0.8112421 | 0.41722666 | 0.73821752 |
| E2F6 | 50.812126 | -0.20248102 | 0.29978118 | -0.6754294 | 0.49940299 | 0.79532568 |
| DYNLRB1 | 219.906933 | -0.202476103 | 0.38924221 | -0.5201802 | 0.60293796 | 0.85048808 |
| AC138150.1 | 61.2730509 | -0.20238563 | 0.360632 | -0.5611971 | 0.57466319 | 0.83436282 |
| PDE7A | 156.237992 | -0.202217079 | 0.24055927 | -0.8406123 | 0.40056517 | 0.72864885 |
| SWAP70 | 778.021838 | -0.202178187 | 0.31979793 | -0.6322061 | 0.52725224 | 0.81084479 |
| CEPT1 | 126.880005 | -0.20172561 | 0.26158163 | -0.7711765 | 0.44060231 | 0.75537551 |
| AC009061.2 | 54.9143159 | -0.201641708 | 0.4120321 | -0.4893835 | 0.62457022 | 0.86016045 |
| THOC1 | 165.844767 | -0.201017287 | 0.22612788 | -0.888954 | 0.37402779 | 0.7121148 |
| UBE2D3 | 779.223721 | -0.200796723 | 0.21261026 | -0.9444357 | 0.34494704 | 0.69456241 |
| NFU1 | 79.2024769 | -0.200649793 | 0.22270867 | -0.9009519 | 0.3676139 | 0.70814138 |
| NUBP1 | 45.0418609 | -0.200573921 | 0.25535439 | -0.7854728 | 0.43217644 | 0.74625444 |
| RBL2 | 224.834107 | -0.200424755 | 0.18219868 | -1.1000341 | 0.27131728 | 0.63561102 |
| JAK1 | 646.152956 | -0.20003196 | 0.15639821 | -1.2789914 | 0.20090008 | 0.56551957 |
| CYBRD1 | 92.5795093 | -0.199981341 | 0.43246046 | -0.4624269 | 0.64377523 | 0.86517342 |
| CREB3L2 | 195.928278 | -0.199929496 | 0.27236185 | -0.7340584 | 0.46291317 | 0.77282771 |
| USP31 | 273.706961 | -0.199677908 | 0.36765578 | -0.543111 | 0.58705336 | 0.84027596 |
| SNX17 | 54.4185298 | -0.199408627 | 0.3762873 | -0.5299372 | 0.59615549 | 0.84697301 |
| ATP2A2 | 263.017463 | -0.199100555 | 0.27844657 | -0.7150404 | 0.47458406 | 0.77998019 |
| BAP1 | 39.4183134 | -0.199060245 | 0.34277039 | -0.5807393 | 0.56141615 | 0.82940393 |
| ABRACL | 42.2129939 | -0.198986917 | 0.48607083 | -0.4093784 | 0.68226196 | 0.88454837 |
| PGPEP1 | 56.4284163 | -0.198775336 | 0.44105514 | -0.4506814 | 0.65221921 | 0.8685756 |
| HSF1 | 153.197248 | -0.198624932 | 0.26362827 | -0.7534281 | 0.45119273 | 0.76409563 |
| FGGY | 190.22205 | -0.198599876 | 0.33141785 | -0.5992431 | 0.54901076 | 0.82355262 |
| TRMT1L | 75.730625 | -0.198565805 | 0.25617766 | -0.7751098 | 0.4382748 | 0.75281332 |
| TMEM164 | 128.293006 | -0.198424076 | 0.20054767 | -0.989411 | 0.32246208 | 0.6761919 |
| UBL3 | 275.09688 | -0.198413671 | 0.20944082 | -0.9473496 | 0.34346067 | 0.69361764 |
| TSPAN15 | 140.589294 | -0.198379921 | 0.30862061 | -0.6427954 | 0.52035685 | 0.80691303 |
| SPG21 | 97.8265936 | -0.198350402 | 0.1873499 | -1.0587164 | 0.28972897 | 0.65294745 |
| SDAD1 | 93.0531455 | -0.198350261 | 0.30133356 | -0.6582415 | 0.51038295 | 0.80177277 |
| WBP11 | 78.5981063 | -0.197839521 | 0.18954058 | -1.0437845 | 0.2965851 | 0.65753344 |
| FAM219A | 319.247231 | -0.197524304 | 0.25618262 | -0.7710293 | 0.44068956 | 0.75537551 |
| APTX | 106.702439 | -0.197443613 | 0.26681264 | -0.7400085 | 0.45929485 | 0.76976328 |
| TPR | 291.625629 | -0.197434885 | 0.20019917 | -0.9861923 | 0.32403876 | 0.67764098 |
| ACTR1B | 77.7450085 | -0.197371789 | 0.2380893 | -0.8289822 | 0.40711449 | 0.73411589 |
| DEF8 | 51.3739187 | -0.197080511 | 0.23722078 | -0.8307894 | 0.4060926 | 0.73307328 |
| MAPK1IP1L | 214.157692 | -0.196850555 | 0.24622195 | -0.7994842 | 0.42400972 | 0.74152442 |
| ACBD6 | 388.984732 | -0.196736401 | 0.21508721 | -0.914682 | 0.3603586 | 0.70386371 |

| | | | | | | |
|---|---|---|---|---|---|---|
| PRDX3 | 93.445617 | -0.196664915 | 0.30780755 | -0.6389217 | 0.52287389 | 0.80854347 |
| STMP1 | 115.979454 | -0.196605728 | 0.30522356 | -0.6441368 | 0.51948672 | 0.80617483 |
| ANXA6 | 102.66604 | -0.196526896 | 0.37737745 | -0.5207701 | 0.60252693 | 0.85030923 |
| TAF12 | 60.7401594 | -0.195987839 | 0.22522545 | -0.8701852 | 0.38419923 | 0.71699718 |
| CFAP54 | 106.461373 | -0.1956848 | 0.4537602 | -0.4312516 | 0.66628546 | 0.87587657 |
| ZNF267 | 78.7042931 | -0.19557605 | 0.32938172 | -0.5937672 | 0.55266786 | 0.8236419 |
| CHCHD1 | 56.617565 | -0.195220142 | 0.44244775 | -0.4412276 | 0.65904826 | 0.87335081 |
| ABHD17A | 171.823423 | -0.195173665 | 0.40503287 | -0.4818712 | 0.62989746 | 0.86097142 |
| ECSIT | 56.0090202 | -0.195169609 | 0.29355125 | -0.664857 | 0.50614195 | 0.79890154 |
| ARMCX3 | 97.4938225 | -0.195133999 | 0.301815 | -0.6465351 | 0.51793285 | 0.80546868 |
| CNIH4 | 87.2950821 | -0.195028294 | 0.31818051 | -0.6129486 | 0.53991034 | 0.81900022 |
| HMG20B | 51.1562076 | -0.194968341 | 0.29931522 | -0.6513813 | 0.51480037 | 0.80422048 |
| AKAP1 | 50.2537692 | -0.194891556 | 0.30218412 | -0.6449431 | 0.51896408 | 0.80603851 |
| BCL10 | 51.4567977 | -0.19476843 | 0.27201225 | -0.7160282 | 0.47397396 | 0.77932979 |
| FARP1 | 404.662601 | -0.194474941 | 0.4393165 | -0.4426762 | 0.65799999 | 0.87267648 |
| RNF40 | 53.6447819 | -0.194443204 | 0.28305952 | -0.686934 | 0.49212432 | 0.79094859 |
| ARPC1B | 112.180641 | -0.194133835 | 0.5543414 | -0.3502063 | 0.7261839 | 0.9026831 |
| ZDHHC5 | 106.173202 | -0.194087236 | 0.17504516 | -1.1087838 | 0.26752345 | 0.6327007 |
| CDHR3 | 55.8342526 | -0.19390363 | 0.57496742 | -0.3372428 | 0.73593387 | 0.90649363 |
| ARNTL | 73.5124189 | -0.193893379 | 0.3225749 | -0.6010802 | 0.54778658 | 0.82355262 |
| SHQ1 | 116.76865 | -0.193870803 | 0.25161401 | -0.7705088 | 0.44099815 | 0.75574615 |
| EMP1 | 519.535302 | -0.193416349 | 0.52022434 | -0.3717941 | 0.71004614 | 0.89727558 |
| ZNF75D | 131.855452 | -0.193168625 | 0.28439807 | -0.6792192 | 0.496999 | 0.79463517 |
| FKTN | 99.3158209 | -0.193091297 | 0.26929263 | -0.7170315 | 0.47335467 | 0.77899012 |
| SEL1L3 | 76.7781838 | -0.19307014 | 0.28086875 | -0.6874034 | 0.49182855 | 0.79094859 |
| MAP3K13 | 225.946914 | -0.192772301 | 0.24467739 | -0.7878631 | 0.43077676 | 0.744811 |
| RSAD1 | 46.2147311 | -0.192749274 | 0.35395726 | -0.5445552 | 0.58605946 | 0.83950091 |
| EPM2AIP1 | 81.0882921 | -0.192579222 | 0.230107 | -0.8369116 | 0.40264225 | 0.73037606 |
| GTF2A2 | 102.572274 | -0.192476318 | 0.31293404 | -0.6150699 | 0.53850854 | 0.81807397 |
| DDX27 | 45.4670905 | -0.192351574 | 0.21234551 | -0.9058424 | 0.36501925 | 0.70668224 |
| ARHGAP25 | 106.296594 | -0.192289095 | 0.41146304 | -0.4673302 | 0.64026367 | 0.86442461 |
| ATF4 | 165.091728 | -0.192253135 | 0.26822466 | -0.7167616 | 0.47352121 | 0.77899012 |
| GALM | 61.6642394 | -0.19209853 | 0.34700854 | -0.5535844 | 0.57986327 | 0.83701203 |
| PLOD3 | 94.6638278 | -0.192042873 | 0.30710199 | -0.6253391 | 0.53174854 | 0.81423995 |
| SEC11C | 46.0346536 | -0.191884739 | 0.45904787 | -0.4180059 | 0.67594277 | 0.88033151 |
| ATE1 | 447.383317 | -0.191702011 | 0.20981649 | -0.9136651 | 0.36089282 | 0.70386371 |
| UFL1 | 127.60141 | -0.19158183 | 0.16443685 | -1.1650785 | 0.24398724 | 0.608518 |
| HEBP1 | 140.842886 | -0.191412081 | 0.54340306 | -0.352247 | 0.72465306 | 0.9026193 |
| MRPS27 | 169.886216 | -0.191167726 | 0.22027518 | -0.8678587 | 0.38547171 | 0.71754209 |
| PRKAB1 | 76.7970201 | -0.191127959 | 0.3230878 | -0.5915666 | 0.55414083 | 0.82401134 |
| TOM1L1 | 57.9257972 | -0.190906531 | 0.32409685 | -0.5890416 | 0.55583336 | 0.82487928 |
| NUAK1 | 430.747251 | -0.190762222 | 0.47355281 | -0.402832 | 0.68707182 | 0.88671976 |
| ALDH6A1 | 54.9667329 | -0.190745742 | 0.40071454 | -0.476014 | 0.63406439 | 0.86207417 |
| ANKRD40 | 80.4796711 | -0.190428512 | 0.31639346 | -0.6018725 | 0.54725903 | 0.82355262 |
| SNAP29 | 66.8195061 | -0.190345404 | 0.32092203 | -0.5931204 | 0.55310058 | 0.8236419 |
| SPCS3 | 293.02936 | -0.190222348 | 0.39896597 | -0.4767884 | 0.6335128 | 0.86207417 |
| FBXL18 | 61.8142459 | -0.190009309 | 0.37021293 | -0.5132433 | 0.60778114 | 0.85264411 |
| FAM69A | 51.8741188 | -0.189707909 | 0.28213609 | -0.6723986 | 0.50132997 | 0.79642461 |
| UBAP2 | 253.777373 | -0.189508923 | 0.18924446 | -1.0013975 | 0.31663469 | 0.67161471 |
| CASP4 | 172.836672 | -0.189466939 | 0.29746335 | -0.6369421 | 0.52416254 | 0.80889528 |
| COX5A | 77.804489 | -0.189356482 | 0.29886952 | -0.6335758 | 0.52635772 | 0.81036438 |
| PISD | 86.7149531 | -0.189235289 | 0.30391438 | -0.6226599 | 0.53350805 | 0.81479963 |
| ZNF155 | 46.1558681 | -0.189069027 | 0.40072369 | -0.4718189 | 0.63705603 | 0.86207417 |
| RRAGB | 115.359746 | -0.188433448 | 0.26759038 | -0.7041862 | 0.48131684 | 0.78505396 |
| ACACA | 285.786472 | -0.188274852 | 0.29421367 | -0.6399256 | 0.52222099 | 0.80811921 |
| CA12 | 88.5233502 | -0.188166058 | 0.65474886 | -0.2873866 | 0.77381631 | 0.91726032 |
| DPY30 | 75.7395274 | -0.187920487 | 0.26880526 | -0.6990953 | 0.48449248 | 0.78781193 |
| TRMU | 57.4683946 | -0.187911675 | 0.20976938 | -0.8958012 | 0.37035892 | 0.70984233 |
| WDFY2 | 208.941916 | -0.187891911 | 0.25505941 | -0.7366594 | 0.4613295 | 0.77104969 |
| MTRR | 59.3940084 | -0.187880732 | 0.24666929 | -0.7616705 | 0.44625666 | 0.76093316 |
| CAP1 | 237.335395 | -0.187765107 | 0.31607165 | -0.5940587 | 0.55247287 | 0.8236419 |
| SMARCA4 | 231.006179 | -0.18758632 | 0.32520066 | -0.5768325 | 0.56405259 | 0.83089872 |
| AAMP | 57.8602611 | -0.187557848 | 0.22874985 | -0.8199256 | 0.41225854 | 0.73593186 |
| ZMAT2 | 53.3980083 | -0.187463845 | 0.2833447 | -0.6616106 | 0.50822084 | 0.79974767 |
| ACOT13 | 54.9872215 | -0.187406933 | 0.32852831 | -0.5704438 | 0.56837674 | 0.83354423 |
| SLC27A4 | 79.7579401 | -0.187202727 | 0.34547447 | -0.5418714 | 0.58790711 | 0.8409165 |
| CDC40 | 87.7345289 | -0.187181502 | 0.22692132 | -0.824874 | 0.40944311 | 0.73419295 |
| AP2A1 | 98.1727959 | -0.187113133 | 0.22094232 | -0.8468868 | 0.39705822 | 0.72680823 |
| TTYH3 | 80.4249055 | -0.186898758 | 0.34927575 | -0.5351037 | 0.59257813 | 0.84419687 |
| KYAT1 | 48.9012352 | -0.186809194 | 0.37311205 | -0.5006785 | 0.61659738 | 0.85732696 |
| REST | 136.163201 | -0.186605188 | 0.27639304 | -0.6751443 | 0.49958408 | 0.79534952 |
| C5orf24 | 155.583042 | -0.186518175 | 0.32238121 | -0.578564 | 0.56288337 | 0.83082829 |
| PGAP1 | 100.236617 | -0.186428714 | 0.22644603 | -0.823281 | 0.41034823 | 0.73419295 |
| AKAP17A | 74.3613513 | -0.186184362 | 0.27544685 | -0.6759357 | 0.49908147 | 0.79516816 |
| CEP83 | 147.828997 | -0.186178201 | 0.24457029 | -0.7612462 | 0.44651004 | 0.76094002 |
| AFF4 | 584.072782 | -0.186173566 | 0.23215238 | -0.8019455 | 0.42258446 | 0.74011688 |
| CCDC32 | 96.2144993 | -0.186118984 | 0.41520101 | -0.4482624 | 0.65396386 | 0.86987315 |
| GMDS-DT | 648.686054 | -0.185865851 | 0.39357317 | -0.4722523 | 0.63674668 | 0.86207417 |
| SUPT5H | 110.650045 | -0.185863391 | 0.19379883 | -0.9590532 | 0.33753193 | 0.68826907 |
| ITFG2 | 86.8172763 | -0.185663886 | 0.40916063 | -0.4537677 | 0.64999602 | 0.86766991 |
| HNRNPK | 463.267286 | -0.185662802 | 0.3318388 | -0.559497 | 0.5758226 | 0.83530514 |
| LRRC69 | 86.4744343 | -0.185598378 | 0.22544347 | -0.823259 | 0.4103607 | 0.73419295 |
| ZMYM4 | 702.12204 | -0.184546697 | 0.13529004 | -1.3640819 | 0.17254179 | 0.53461168 |
| LARGE1 | 601.588273 | -0.184343138 | 0.27217207 | -0.6773036 | 0.49821332 | 0.79513036 |
| CCDC57 | 109.224913 | -0.184257462 | 0.29585701 | -0.6227923 | 0.53342103 | 0.81479963 |
| LMCD1 | 391.992905 | -0.184242142 | 0.55081984 | -0.3344871 | 0.73801201 | 0.9078343 |
| PRKCSH | 85.8545238 | -0.184219049 | 0.32213598 | -0.5718674 | 0.56741184 | 0.83329704 |
| C11orf58 | 253.393013 | -0.183933863 | 0.21157728 | -0.869346 | 0.38465791 | 0.71699718 |
| AL359232.1 | 100.068655 | -0.183726059 | 0.45432659 | -0.404392 | 0.68592445 | 0.88671976 |
| TPGS2 | 80.2535799 | -0.183426255 | 0.2657446 | -0.6902351 | 0.49004635 | 0.79046901 |
| ROBO4 | 370.511426 | -0.183132182 | 0.37374173 | -0.4899966 | 0.6241363 | 0.86010565 |
| DNTTIP1 | 42.8036755 | -0.183031272 | 0.23951994 | -0.7641588 | 0.44477261 | 0.75998628 |
| BEX3 | 170.668996 | -0.182710195 | 0.38251347 | -0.4776569 | 0.63289443 | 0.86186971 |
| PDZD8 | 212.809314 | -0.182584946 | 0.16520385 | -1.10521 | 0.26906862 | 0.63347822 |
| RNF167 | 41.3705635 | -0.182350647 | 0.42402124 | -0.4300507 | 0.66715874 | 0.87587657 |
| ADD3 | 479.83761 | -0.182139532 | 0.375784 | -0.4846921 | 0.62789478 | 0.86052754 |
| ADNP2 | 102.944621 | -0.182090632 | 0.20539797 | -0.886526 | 0.37533417 | 0.71257833 |
| FCF1 | 51.1135936 | -0.182090391 | 0.22165329 | -0.82151 | 0.41135584 | 0.73470224 |
| SLC39A9 | 129.574733 | -0.181764051 | 0.17640929 | -1.0303542 | 0.30284378 | 0.65940027 |
| SMARCC1 | 403.41159 | -0.181758058 | 0.25581423 | -0.710508 | 0.47738917 | 0.78098272 |
| ARHGAP4 | 44.4887243 | -0.181619042 | 0.73506689 | -0.2470783 | 0.80484766 | 0.9311439 |
| LGALS3BP | 92.820483 | -0.181095936 | 0.60869844 | -0.2975134 | 0.76607459 | 0.91600937 |
| SASH1 | 2473.19739 | -0.181038354 | 0.28041728 | -0.6456034 | 0.51853622 | 0.8057739 |
| SMARCD1 | 64.2428248 | -0.180846454 | 0.25047401 | -0.7220168 | 0.47028413 | 0.77792258 |
| KPNA2 | 55.6482256 | -0.180829296 | 0.45402754 | -0.3982783 | 0.69042509 | 0.88832346 |
| MMS19 | 128.457778 | -0.180814276 | 0.19216988 | -0.9409085 | 0.34675174 | 0.6956798 |
| HDGFL3 | 340.199041 | -0.180677661 | 0.21164325 | -0.8536897 | 0.39327695 | 0.72360479 |
| CCDC125 | 48.1968939 | -0.180673072 | 0.43769774 | -0.4127805 | 0.67976747 | 0.88266221 |
| MTHFR | 72.7298403 | -0.180611327 | 0.35446413 | -0.5095334 | 0.61037837 | 0.85403069 |
| NUDT19 | 52.7888272 | -0.180487804 | 0.2305203 | -0.7829584 | 0.43365155 | 0.74785472 |
| USP14 | 186.378365 | -0.18030195 | 0.24945639 | -0.7227795 | 0.46981539 | 0.77792258 |
| KMT2E | 1214.4405 | -0.180206852 | 0.37549746 | -0.479915 | 0.63128785 | 0.86097142 |
| TRDMT1 | 50.1825943 | -0.179989132 | 0.39673292 | -0.4536783 | 0.65006036 | 0.86766991 |

| | | | | | | |
|---|---|---|---|---|---|---|
| FGD5-AS1 | 142.604336 | -0.179735974 | 0.28161172 | -0.6382404 | 0.52331721 | 0.80859228 |
| TECPR2 | 160.067995 | -0.179286865 | 0.24277365 | -0.7384939 | 0.46021436 | 0.770424 |
| EIF4A3 | 49.1750394 | -0.17912994 | 0.34022099 | -0.5265105 | 0.59853353 | 0.84839769 |
| TMEM259 | 145.051244 | -0.178945512 | 0.2699022 | -0.6630013 | 0.50732971 | 0.79921563 |
| CASP7 | 106.179119 | -0.178541051 | 0.20751488 | -0.8603771 | 0.3895812 | 0.72202943 |
| GULP1 | 362.620739 | -0.178328416 | 0.47425248 | -0.37602 | 0.70690203 | 0.89558435 |
| DCBLD1 | 116.064599 | -0.177799859 | 0.33381305 | -0.532633 | 0.59428763 | 0.84545474 |
| TNFAIP8L1 | 134.677814 | -0.177765441 | 0.34589641 | -0.5139268 | 0.60730316 | 0.85226578 |
| EIF3L | 142.702274 | -0.17772107 | 0.27560894 | -0.6448306 | 0.51903699 | 0.80603851 |
| SCP2 | 219.949358 | -0.177391872 | 0.2360509 | -0.7514984 | 0.45235277 | 0.76480455 |
| ARPP19 | 189.033215 | -0.177279426 | 0.19189213 | -0.9238494 | 0.35556473 | 0.70018061 |
| ZNF639 | 41.4927647 | -0.177052174 | 0.28910396 | -0.612417 | 0.54026191 | 0.81900022 |
| TMEM267 | 89.4526707 | -0.176954775 | 0.22212604 | -0.7966413 | 0.42565939 | 0.74197066 |
| HLA-DPA1 | 400.076567 | -0.176868637 | 0.4781615 | -0.3698931 | 0.71146215 | 0.89747106 |
| ADSS | 101.285891 | -0.176831705 | 0.2792921 | -0.6331425 | 0.52664057 | 0.81036438 |
| DHX57 | 84.3448924 | -0.176506426 | 0.30663398 | -0.5756258 | 0.56486815 | 0.83113895 |
| GTF2H1 | 118.303706 | -0.17647582 | 0.23243697 | -0.7592416 | 0.44770804 | 0.7614672 |
| FAM89B | 59.7769148 | -0.17579784 | 0.56638601 | -0.3103852 | 0.75626804 | 0.91345267 |
| ZFAT | 89.4147576 | -0.175563765 | 0.26847743 | -0.6539237 | 0.51316095 | 0.80321271 |
| CDK9 | 92.7900017 | -0.174678741 | 0.32742753 | -0.5334883 | 0.59369564 | 0.84502915 |
| POFUT2 | 43.378118 | -0.174564338 | 0.23221597 | -0.7517327 | 0.45221182 | 0.76480455 |
| LETMD1 | 84.567369 | -0.174343463 | 0.23610992 | -0.7383996 | 0.46027167 | 0.770424 |
| KDM3A | 213.488651 | -0.174318713 | 0.37200621 | -0.4685909 | 0.6393621 | 0.86388557 |
| NPHP4 | 43.7030377 | -0.17431577 | 0.26304694 | -0.6626793 | 0.50753595 | 0.79921852 |
| FRY | 361.356572 | -0.174252418 | 0.30956745 | -0.56289 | 0.5735098 | 0.8342245 |
| CBWD5 | 80.1132574 | -0.174107102 | 0.26845045 | -0.6485633 | 0.51662071 | 0.80498569 |
| CAPN15 | 43.9172542 | -0.173866343 | 0.38787812 | -0.4482499 | 0.65397282 | 0.86987315 |
| ZNF131 | 275.073579 | -0.173792671 | 0.15081546 | -1.1523532 | 0.24917597 | 0.61434376 |
| MTCH1 | 180.882578 | -0.173724694 | 0.22141446 | -0.7846131 | 0.43268043 | 0.74680953 |
| SOD2.1 | 63.4301297 | -0.173707989 | 0.49966362 | -0.3476499 | 0.72810315 | 0.9037084 |
| PLBD2 | 41.6775566 | -0.173658425 | 0.22680304 | -0.7656794 | 0.44386707 | 0.75907298 |
| PTEN | 479.298993 | -0.173545251 | 0.1599091 | -1.0852744 | 0.27780014 | 0.64148182 |
| ESD | 100.301415 | -0.17311064 | 0.24755316 | -0.6992867 | 0.48437285 | 0.78781193 |
| AJAP1 | 110.030435 | -0.172973724 | 0.70812189 | -0.2442711 | 0.80702085 | 0.93187644 |
| YY1 | 310.812739 | -0.172730052 | 0.18362001 | -0.940693 | 0.34686223 | 0.6956798 |
| FAM210B | 50.6445659 | -0.172618334 | 0.43421211 | -0.3975438 | 0.69096649 | 0.88832346 |
| CCDC90B | 62.4473838 | -0.17230164 | 0.28976905 | -0.5946171 | 0.55209942 | 0.8236419 |
| LRPPRC | 261.147466 | -0.172301462 | 0.21617907 | -0.7970312 | 0.42543292 | 0.74197066 |
| STIP1 | 84.8636239 | -0.172201559 | 0.32693186 | -0.52672 | 0.59838802 | 0.84839769 |
| XPC | 138.5166 | -0.17217038 | 0.19251034 | -0.8943435 | 0.37113811 | 0.70984233 |
| XPOT | 103.155916 | -0.172031949 | 0.26372874 | -0.6523064 | 0.51420353 | 0.80398873 |
| MAPKAPK2 | 125.339668 | -0.171889358 | 0.22358074 | -0.7688022 | 0.44201076 | 0.75668913 |
| RAMP2 | 457.145431 | -0.171711628 | 0.68328469 | -0.2513032 | 0.8015797 | 0.93040095 |
| TLK1 | 388.055345 | -0.171593174 | 0.15077931 | -1.1380419 | 0.25510299 | 0.6208774 |
| NARS2 | 147.307806 | -0.171367151 | 0.29438925 | -0.5821108 | 0.56049207 | 0.82908652 |
| TRIR | 169.335786 | -0.171197242 | 0.32751606 | -0.522714 | 0.60117326 | 0.8499944 |
| ZNF354B | 44.9890196 | -0.171127511 | 0.28154912 | -0.6078069 | 0.54331552 | 0.82165051 |
| KIF3B | 113.420738 | -0.171118522 | 0.25836477 | -0.6623137 | 0.50777021 | 0.79921852 |
| CHD3 | 112.405144 | -0.171019185 | 0.19901871 | -0.8593121 | 0.39016835 | 0.72234109 |
| ANXA7 | 206.36615 | -0.170904526 | 0.18289799 | -0.9344254 | 0.3500845 | 0.69858573 |
| NECTIN2 | 275.640814 | -0.170645166 | 0.27340619 | -0.6241452 | 0.53253222 | 0.81467773 |
| HMCN2 | 71.2004672 | -0.170534329 | 1.36195499 | -0.1252129 | 0.90035501 | 0.96744565 |
| ATXN2L | 126.83564 | -0.170455805 | 0.2018958 | -0.8442761 | 0.39851512 | 0.72770737 |
| TMCC3 | 253.718556 | -0.170417484 | 0.35508516 | -0.4799341 | 0.63127424 | 0.86097142 |
| YTHDF1 | 92.942231 | -0.170268634 | 0.22838983 | -0.7455176 | 0.45595888 | 0.76740262 |
| TADA3 | 42.0512169 | -0.169667482 | 0.31263071 | -0.5427089 | 0.58733021 | 0.84052521 |
| MYO18A | 120.057708 | -0.169502183 | 0.38288605 | -0.4426962 | 0.65798552 | 0.87267648 |
| NUFIP2 | 344.805756 | -0.1694671 | 0.1726146 | -0.9817658 | 0.32621526 | 0.67997183 |
| LZTS2 | 69.5184841 | -0.169448145 | 0.26088803 | -0.6495052 | 0.51601186 | 0.80483629 |
| CAPN7 | 288.708641 | -0.16934076 | 0.21418978 | -0.7906108 | 0.42917113 | 0.74408005 |
| TSTD2 | 77.8805987 | -0.169313243 | 0.27186767 | -0.6227781 | 0.53343032 | 0.81479963 |
| SLX4IP | 168.526161 | -0.169123102 | 0.30466444 | -0.5551127 | 0.57881757 | 0.83649569 |
| NCOA7 | 1047.89782 | -0.168933459 | 0.3505838 | -0.4818633 | 0.62990308 | 0.86097142 |
| CEBPG | 85.5274561 | -0.168906676 | 0.28398678 | -0.5947695 | 0.55199756 | 0.8236419 |
| KLHDC3 | 48.9669719 | -0.16888427 | 0.27306196 | -0.6184833 | 0.53625679 | 0.81701532 |
| DUSP11 | 157.815756 | -0.168864672 | 0.41459048 | -0.4073048 | 0.68378417 | 0.88565439 |
| FAM3C | 169.935305 | -0.168829764 | 0.34563999 | -0.4884555 | 0.62522722 | 0.86016045 |
| THRA | 68.1410707 | -0.168792158 | 0.46602236 | -0.3621976 | 0.71720441 | 0.9002058 |
| MED27 | 254.546068 | -0.168256876 | 0.27993059 | -0.6010664 | 0.54779576 | 0.82355262 |
| NEMP1 | 42.438199 | -0.168018181 | 0.33153026 | -0.5067959 | 0.61229802 | 0.85573702 |
| DDX46 | 266.89766 | -0.168015981 | 0.18600909 | -0.9032676 | 0.36638389 | 0.70695551 |
| C1orf123 | 40.2942303 | -0.168015928 | 0.35329258 | -0.4755716 | 0.63437959 | 0.86207417 |
| TCF25 | 445.308254 | -0.168000667 | 0.20629594 | -0.8143673 | 0.41543456 | 0.73741887 |
| PTPRF | 44.9429354 | -0.167403455 | 0.39201947 | -0.4270284 | 0.66935863 | 0.8765181 |
| MALAT1 | 73401.3909 | -0.167134913 | 0.28757738 | -0.5811824 | 0.56111753 | 0.82940393 |
| TWSG1 | 52.8772635 | -0.166891754 | 0.27300463 | -0.6113147 | 0.54099123 | 0.81928059 |
| SRRT | 46.6601789 | -0.166826709 | 0.27625916 | -0.6038776 | 0.54592505 | 0.82355262 |
| HLA-DRA | 871.286699 | -0.166606479 | 0.49083676 | -0.3394336 | 0.73428312 | 0.9055183 |
| BAD | 55.7773458 | -0.166093073 | 0.41223851 | -0.4029053 | 0.6870179 | 0.88671976 |
| APOL1 | 184.01114 | -0.165860957 | 0.41697845 | -0.3977687 | 0.69080072 | 0.88832346 |
| TOP1 | 509.27685 | -0.165828174 | 0.26589507 | -0.6236602 | 0.53285076 | 0.81470811 |
| CFAP36 | 91.3755888 | -0.165782427 | 0.23086604 | -0.7180893 | 0.47270224 | 0.77899012 |
| TCTN3 | 78.2692485 | -0.165533592 | 0.29701258 | -0.5573286 | 0.57730297 | 0.83603913 |
| COX19 | 60.2783998 | -0.16547266 | 0.33960437 | -0.4872513 | 0.62608028 | 0.86052754 |
| ATOX1 | 146.834078 | -0.165243212 | 0.33996027 | -0.4860662 | 0.6269202 | 0.86052754 |
| NAA20 | 91.1795668 | -0.165120311 | 0.29681977 | -0.5562982 | 0.57800701 | 0.83613556 |
| FYCO1 | 68.6673849 | -0.165012586 | 0.23841872 | -0.6921125 | 0.48886665 | 0.78955612 |
| APOE | 186.468491 | -0.164726965 | 0.71251252 | -0.2311917 | 0.81716589 | 0.93550282 |
| GLYR1 | 212.533553 | -0.164725437 | 0.20851025 | -0.7900112 | 0.42952122 | 0.74408005 |
| GSPT1 | 221.261053 | -0.164701413 | 0.22239646 | -0.7405757 | 0.45895076 | 0.76976328 |
| MTERF4 | 133.759756 | -0.164678133 | 0.23191054 | -0.7100933 | 0.47764625 | 0.78124711 |
| MFGE8 | 76.6215465 | -0.163995128 | 0.67058464 | -0.2445555 | 0.80680066 | 0.93187644 |
| RNASE1 | 792.200657 | -0.163949825 | 0.67001933 | -0.2446942 | 0.80669324 | 0.93187644 |
| FILIP1L | 191.686384 | -0.163879641 | 0.41074528 | -0.3989812 | 0.68990706 | 0.88817873 |
| PIGB | 51.5208798 | -0.163854266 | 0.35911045 | -0.4562782 | 0.64818995 | 0.86739793 |
| BAALC | 94.4988586 | -0.163703361 | 0.49295627 | -0.332085 | 0.73982511 | 0.90826918 |
| CAVIN2 | 562.234093 | -0.163397797 | 0.51343168 | -0.3182464 | 0.75029802 | 0.91109327 |
| PIH1D1 | 43.0807116 | -0.163237954 | 0.3159911 | -0.5165904 | 0.60544215 | 0.85156118 |
| INTS2 | 66.5236805 | -0.163123363 | 0.31219757 | -0.5225004 | 0.60132196 | 0.8499944 |
| NAA60 | 42.1524927 | -0.163007793 | 0.32799583 | -0.4969813 | 0.61920224 | 0.85855421 |
| NAXD | 42.085863 | -0.162917695 | 0.21700391 | -0.7507593 | 0.45279754 | 0.76507171 |
| FUT11 | 141.70178 | -0.162228858 | 0.28546497 | -0.5682969 | 0.56983338 | 0.83368112 |
| MRPL33 | 234.669448 | -0.162012398 | 0.43009357 | -0.376691 | 0.70640322 | 0.8952296 |
| DDX23 | 63.4336944 | -0.161984257 | 0.21880663 | -0.7403078 | 0.45911324 | 0.76976328 |
| ZNF787 | 53.5886473 | -0.161870319 | 0.28398545 | -0.5699951 | 0.56868101 | 0.83368112 |
| RLIM | 96.8742106 | -0.161865065 | 0.24622201 | -0.6573948 | 0.51092711 | 0.80177277 |
| CCDC149 | 159.359591 | -0.161823507 | 0.21964502 | -0.7367502 | 0.4612743 | 0.77104969 |
| ZSCAN5A | 121.634202 | -0.16181572 | 0.41709446 | -0.3879594 | 0.69804607 | 0.89307552 |
| PRDX5 | 80.3059374 | -0.161553539 | 0.43640244 | -0.3701939 | 0.71123799 | 0.89747106 |
| TSN | 71.3173364 | -0.161540459 | 0.23417636 | -0.6898239 | 0.49030491 | 0.79057439 |
| DAPK1 | 369.129537 | -0.16153133 | 0.37215639 | -0.4340415 | 0.66425828 | 0.8755434 |
| ATP5PD | 53.7365594 | -0.161445201 | 0.35202199 | -0.4586225 | 0.64650529 | 0.86641873 |
| NSFL1C | 97.1842906 | -0.16142711 | 0.2530533 | -0.6379174 | 0.52352743 | 0.80861173 |

| | | | | | | |
|---|---|---|---|---|---|---|
| ERGIC2 | 123.470049 | -0.161393795 | 0.21312791 | -0.7572626 | 0.44889256 | 0.76209291 |
| TMEM39A | 78.3532629 | -0.161246217 | 0.25105113 | -0.6422844 | 0.52068856 | 0.80705591 |
| TIE1 | 439.704313 | -0.161090582 | 0.37186486 | -0.4331966 | 0.66487197 | 0.87568764 |
| NFE2L2 | 747.42371 | -0.161005759 | 0.19657258 | -0.8190652 | 0.41274921 | 0.73622771 |
| MSH3 | 246.986224 | -0.160804964 | 0.17902346 | -0.898234 | 0.36906079 | 0.70942552 |
| ERLIN1 | 60.621418 | -0.160772703 | 0.20171613 | -0.7970245 | 0.42543679 | 0.74197066 |
| ARHGAP1 | 64.7655682 | -0.160598341 | 0.24115193 | -0.6659633 | 0.50543455 | 0.79844998 |
| SLC35G2 | 159.059918 | -0.160560458 | 0.23091881 | -0.6953113 | 0.48686023 | 0.78826504 |
| COL5A2 | 91.126137 | -0.160111617 | 0.51817074 | -0.3089939 | 0.75732614 | 0.913637 |
| AGK | 113.897755 | -0.159872286 | 0.23553093 | -0.6787741 | 0.49728103 | 0.7946635 |
| SNRPC | 52.1017679 | -0.159836559 | 0.34292662 | -0.4660955 | 0.64114714 | 0.86490071 |
| ARFRP1 | 68.9888792 | -0.159739739 | 0.28338778 | -0.563679 | 0.57297262 | 0.8342245 |
| STK38 | 276.235091 | -0.159726339 | 0.25681364 | -0.6219543 | 0.53397194 | 0.81535593 |
| EEF2K | 155.330999 | -0.159694612 | 0.42972729 | -0.3716185 | 0.71017691 | 0.89727558 |
| GIPC2 | 158.491864 | -0.159694367 | 0.33945268 | -0.4704466 | 0.63803596 | 0.862698 |
| PACRGL | 105.399164 | -0.15968759 | 0.25952993 | -0.6152955 | 0.53835962 | 0.81807397 |
| INTS13 | 44.7872917 | -0.159578568 | 0.26942322 | -0.592297 | 0.55365171 | 0.82373309 |
| CACYBP | 53.8256371 | -0.159223644 | 0.32808713 | -0.485309 | 0.62745716 | 0.86052754 |
| LGALS8 | 132.553751 | -0.159195124 | 0.21833609 | -0.7291288 | 0.4659229 | 0.77517793 |
| DNASE1L1 | 49.8649413 | -0.159023579 | 0.33833922 | -0.4700123 | 0.63834627 | 0.86297497 |
| CYYR1 | 826.034787 | -0.159010923 | 0.41107036 | -0.3868217 | 0.69888824 | 0.89364164 |
| SEMA6A-AS1 | 153.536115 | -0.158788969 | 0.23643548 | -0.6715954 | 0.50184133 | 0.79661835 |
| PPP2R3C | 189.202579 | -0.158676192 | 0.17154339 | -0.9249916 | 0.35497028 | 0.70018061 |
| ANXA11 | 200.379417 | -0.158475547 | 0.27237935 | -0.5818192 | 0.56068844 | 0.82917467 |
| ERI3 | 162.330579 | -0.158364453 | 0.26108424 | -0.6065646 | 0.54413991 | 0.82244124 |
| ENSA | 284.572049 | -0.15829893 | 0.19884479 | -0.7960929 | 0.42597802 | 0.74197066 |
| PRDM10 | 63.7194275 | -0.158088002 | 0.36713968 | -0.4305936 | 0.66676389 | 0.87587657 |
| MED15 | 233.840666 | -0.158031353 | 0.2420005 | -0.6530208 | 0.51374289 | 0.80396979 |
| TP53BP1 | 150.69778 | -0.15800862 | 0.20008905 | -0.7896915 | 0.42970797 | 0.74408005 |
| UBE2L3 | 191.495925 | -0.157869312 | 0.2000201 | -0.7892672 | 0.42995583 | 0.74420645 |
| SLC25A36 | 211.582344 | -0.157592115 | 0.23718432 | -0.6644289 | 0.50641584 | 0.7989142 |
| WDR37 | 224.508729 | -0.157297118 | 0.20926296 | -0.7516721 | 0.45224829 | 0.76480455 |
| HADHB | 239.54803 | -0.157208241 | 0.17736809 | -0.8863389 | 0.37543493 | 0.71257833 |
| FBN1 | 475.03788 | -0.15709934 | 0.33148238 | -0.4739297 | 0.63555005 | 0.86207417 |
| AC092069.1 | 61.9033791 | -0.157036323 | 0.7070407 | -0.2221037 | 0.82423319 | 0.93852534 |
| EIF4E2 | 75.6924111 | -0.157029241 | 0.27260493 | -0.5760323 | 0.56459336 | 0.83113895 |
| ABI3 | 59.0892577 | -0.156977897 | 0.41325824 | -0.3798542 | 0.70405361 | 0.89446478 |
| AGO1 | 66.2080328 | -0.156924486 | 0.31785153 | -0.4937037 | 0.62151543 | 0.85896293 |
| MCMBP | 176.894251 | -0.156895875 | 0.24266064 | -0.646565 | 0.51791352 | 0.80546868 |
| TARBP1 | 169.205296 | -0.156869336 | 0.40574135 | -0.386624 | 0.6990346 | 0.89364164 |
| DCAF7 | 198.960257 | -0.156597619 | 0.22608392 | -0.6926526 | 0.48852757 | 0.78941966 |
| MBD2 | 327.576978 | -0.156316331 | 0.1438383 | -1.0867504 | 0.27714715 | 0.64137305 |
| TMEM184C | 69.4453116 | -0.156282923 | 0.23898852 | -0.6539348 | 0.51315379 | 0.80321271 |
| DPYSL2 | 340.292253 | -0.156263244 | 0.40966464 | -0.3814419 | 0.7028754 | 0.89443605 |
| TEX10 | 55.8703382 | -0.156253187 | 0.27787693 | -0.5623108 | 0.57390431 | 0.8342245 |
| ANKS3 | 59.5553673 | -0.155405174 | 0.33799203 | -0.4597895 | 0.64566735 | 0.86572246 |
| TTC9C | 65.5625267 | -0.155209805 | 0.23206402 | -0.6688232 | 0.50360825 | 0.79776858 |
| RNF181 | 40.1048662 | -0.155162857 | 0.38622879 | -0.4017382 | 0.68787671 | 0.88727856 |
| GLUD1 | 227.39366 | -0.15488448 | 0.22086219 | -0.701272 | 0.48313326 | 0.78738886 |
| LAMTOR3 | 45.9046465 | -0.154811753 | 0.24419269 | -0.6339737 | 0.52609796 | 0.81036438 |
| AZI2 | 133.629467 | -0.154796221 | 0.35499881 | -0.4360472 | 0.66280252 | 0.87493354 |
| HSBP1 | 127.446642 | -0.154746579 | 0.40392361 | -0.3831085 | 0.7016393 | 0.89443605 |
| TXN2 | 59.1621142 | -0.154600286 | 0.24368506 | -0.6344266 | 0.52580244 | 0.81036438 |
| TAPT1 | 167.928524 | -0.154545953 | 0.24259441 | -0.6370549 | 0.52408909 | 0.80889528 |
| CREBL2 | 70.9692946 | -0.154526282 | 0.17960583 | -0.8603634 | 0.38958876 | 0.72202943 |
| SORT1 | 111.275914 | -0.154466725 | 0.46635006 | -0.3312248 | 0.74047466 | 0.90826918 |
| CPSF2 | 143.629982 | -0.154392638 | 0.22172587 | -0.6963222 | 0.48622709 | 0.78826504 |
| PLEKHA8 | 81.4403119 | -0.154291193 | 0.22936881 | -0.6726773 | 0.50115261 | 0.79642461 |
| GLG1 | 645.403265 | -0.154169669 | 0.13485168 | -1.1432536 | 0.2529333 | 0.61839055 |
| DENR | 126.289455 | -0.154014002 | 0.30473028 | -0.5054109 | 0.61327027 | 0.85573702 |
| AC025159.1 | 144.380656 | -0.15392444 | 0.3414681 | -0.4507725 | 0.6521535 | 0.8685756 |
| RCC2 | 87.2905033 | -0.153801528 | 0.33436975 | -0.4599744 | 0.64553459 | 0.86572246 |
| NUDT5 | 100.970615 | -0.153742857 | 0.23211338 | -0.662361 | 0.50773987 | 0.79921852 |
| PPP4C | 98.0367198 | -0.153448282 | 0.25427876 | -0.6034648 | 0.54619953 | 0.82355262 |
| ATF7IP2 | 194.592857 | -0.153126713 | 0.35494457 | -0.4314102 | 0.66617013 | 0.87587657 |
| WDR61 | 77.626616 | -0.152696426 | 0.19397402 | -0.7872004 | 0.43116456 | 0.74490865 |
| WDR19 | 94.1259208 | -0.15258068 | 0.3860285 | -0.3952576 | 0.69265282 | 0.88965281 |
| RAE1 | 45.2079944 | -0.15250988 | 0.21948072 | -0.6948669 | 0.48713874 | 0.78826504 |
| AASDHPPT | 79.9320087 | -0.152386577 | 0.23539564 | -0.6473636 | 0.51739663 | 0.80546868 |
| SENP1 | 105.253005 | -0.152318578 | 0.19679591 | -0.7739926 | 0.43893518 | 0.75347316 |
| NAA15 | 261.586866 | -0.152203685 | 0.2190898 | -0.6947091 | 0.48723761 | 0.78826915 |
| EIF2B3 | 115.631245 | -0.152074052 | 0.35540211 | -0.4278929 | 0.66872908 | 0.87625461 |
| BRD4 | 354.009714 | -0.152007881 | 0.20988697 | -0.7242369 | 0.46892034 | 0.77737861 |
| FLVCR1 | 63.4304306 | -0.151748924 | 0.27645174 | -0.5489165 | 0.58306274 | 0.83837681 |
| TACC2 | 133.019375 | -0.151489442 | 0.3110552 | -0.4870179 | 0.62624567 | 0.86052754 |
| TAF1B | 107.73577 | -0.151222071 | 0.33066018 | -0.4573338 | 0.64743115 | 0.86714786 |
| PPP1R8 | 53.499651 | -0.151091129 | 0.34096862 | -0.4431233 | 0.65767657 | 0.87254693 |
| TTF1 | 90.0549598 | -0.150942339 | 0.23504032 | -0.6421976 | 0.52074486 | 0.80705591 |
| CHPT1 | 134.247438 | -0.150857694 | 0.46580132 | -0.323867 | 0.7460387 | 0.91021838 |
| EML2 | 55.8369845 | -0.150821399 | 0.25062266 | -0.6017868 | 0.54731609 | 0.82355262 |
| AP000311.1 | 55.466181 | -0.150709847 | 0.43421487 | -0.3470859 | 0.72852681 | 0.90376371 |
| TMEM214 | 39.7719252 | -0.150622688 | 0.31597807 | -0.4766872 | 0.63358491 | 0.86207417 |
| ITGB1 | 907.344324 | -0.150578179 | 0.29184463 | -0.5159532 | 0.60588708 | 0.85156118 |
| HSDL2 | 167.207141 | -0.150455276 | 0.31039719 | -0.4847186 | 0.627876 | 0.86052754 |
| RAB27A | 186.184284 | -0.150424205 | 0.18290996 | -0.8223948 | 0.41085222 | 0.73454308 |
| MYH10 | 98.6425929 | -0.15024875 | 0.38697429 | -0.3882655 | 0.69781959 | 0.89303379 |
| FYTTD1 | 126.953659 | -0.150225839 | 0.19793475 | -0.7589665 | 0.44787262 | 0.7614672 |
| EML3 | 87.8034089 | -0.150055515 | 0.39072404 | -0.3840447 | 0.70094528 | 0.8944316 |
| TEAD4 | 277.160337 | -0.150045918 | 0.30703912 | -0.4886866 | 0.62506356 | 0.86016045 |
| MTIF3 | 102.868581 | -0.149893979 | 0.23507914 | -0.637632 | 0.52371327 | 0.80874613 |
| FBXW8 | 132.031898 | -0.149889186 | 0.38849141 | -0.3858237 | 0.69962725 | 0.89370119 |
| FAM32A | 49.2967348 | -0.149845341 | 0.39208862 | -0.3821721 | 0.70233369 | 0.89443605 |
| BICD1 | 710.62788 | -0.149644337 | 0.22476163 | -0.6657913 | 0.50554451 | 0.79846955 |
| EHBP1L1 | 178.549753 | -0.149562051 | 0.29495486 | -0.5070676 | 0.6121074 | 0.85563287 |
| TOX4 | 102.988468 | -0.14952245 | 0.21385397 | -0.6991801 | 0.48443946 | 0.78781193 |
| LSG1 | 86.5122906 | -0.149338429 | 0.23943049 | -0.6237235 | 0.53280918 | 0.81470811 |
| KDM5C | 88.6743422 | -0.149283537 | 0.25386757 | -0.5880371 | 0.55650742 | 0.82527172 |
| HDAC4 | 376.565026 | -0.149274489 | 0.35142953 | -0.4247636 | 0.67100898 | 0.87730512 |
| HHEX | 86.0120558 | -0.149026547 | 0.38169016 | -0.3904385 | 0.6962123 | 0.8920567 |
| VPS4A | 61.3005885 | -0.1490054 | 0.27281486 | -0.5461777 | 0.58494378 | 0.83895258 |
| AL645568.1 | 65.6745457 | -0.148638761 | 0.28535552 | -0.5208897 | 0.60244358 | 0.85030923 |
| ZBTB21 | 55.9459814 | -0.14827794 | 0.38911806 | -0.3810616 | 0.70315756 | 0.8944409 |
| CLCN7 | 70.7576068 | -0.148214024 | 0.26899966 | -0.5509822 | 0.58164588 | 0.83789748 |
| UPF3B | 109.07656 | -0.148183292 | 0.30139505 | -0.491658 | 0.62296112 | 0.85940448 |
| SLC35A5 | 53.0489751 | -0.148178242 | 0.23048626 | -0.642894 | 0.52029286 | 0.80691303 |
| NR1D2 | 175.611885 | -0.147930306 | 0.30337675 | -0.4876125 | 0.62582432 | 0.86052754 |
| AC097376.2 | 135.434818 | -0.147713886 | 0.24622729 | -0.5999087 | 0.5485671 | 0.82355262 |
| TPRKB | 89.3054654 | -0.147673353 | 0.43532578 | -0.3392249 | 0.7344403 | 0.9055183 |
| TXNRD2 | 43.8913333 | -0.147613234 | 0.39292256 | -0.3756802 | 0.70715465 | 0.89576571 |
| IMPACT | 43.8670881 | -0.147612918 | 0.32379591 | -0.4558826 | 0.64847441 | 0.86749486 |
| ATP5MG | 179.350918 | -0.147600959 | 0.34563145 | -0.4270472 | 0.66934491 | 0.8765181 |
| RBM17 | 302.783225 | -0.147217326 | 0.24826273 | -0.59299 | 0.55318782 | 0.8236419 |
| ANKRD11 | 1029.47316 | -0.147152039 | 0.27769897 | -0.5298977 | 0.59618288 | 0.84697301 |

| | | | | | | |
|---|---|---|---|---|---|---|
| ESYT1 | 70.9508426 | -0.147136981 | 0.32404668 | -0.4540611 | 0.64978487 | 0.86766991 |
| B4GALT5 | 598.498546 | -0.147063071 | 0.3421653 | -0.4298012 | 0.66734023 | 0.87587657 |
| WDR5 | 42.141735 | -0.146713492 | 0.33456235 | -0.4385236 | 0.66100675 | 0.87369056 |
| TBC1D9B | 111.387514 | -0.146597494 | 0.25507123 | -0.5747316 | 0.56547283 | 0.83179295 |
| CARM1 | 56.3169362 | -0.146461083 | 0.35061083 | -0.4177312 | 0.67614366 | 0.88033151 |
| CYSTM1 | 366.890139 | -0.146398192 | 0.41722947 | -0.3508817 | 0.72567709 | 0.9026193 |
| SMPD1 | 46.8489004 | -0.146102712 | 0.34107005 | -0.4283657 | 0.66838489 | 0.87625461 |
| TUBGCP2 | 109.863258 | -0.145951933 | 0.2007362 | -0.7270833 | 0.46717494 | 0.77590675 |
| RSBN1 | 132.923857 | -0.145662923 | 0.17773108 | -0.8195692 | 0.41246174 | 0.736134 |
| NMT1 | 149.463341 | -0.145479721 | 0.18718121 | -0.7772133 | 0.43703295 | 0.75146894 |
| AK9 | 109.785423 | -0.145371345 | 0.33673148 | -0.431713 | 0.66595004 | 0.87587657 |
| SMAD1 | 429.041194 | -0.145346874 | 0.31441308 | -0.46228 | 0.64388055 | 0.86517342 |
| SRP19 | 94.3544003 | -0.145116853 | 0.25620056 | -0.566419 | 0.57110901 | 0.83368112 |
| ATP9B | 444.242955 | -0.144877617 | 0.25311455 | -0.5723796 | 0.56706481 | 0.83308641 |
| CITED4 | 53.8851667 | -0.144827563 | 0.51531085 | -0.2810489 | 0.77867287 | 0.91979327 |
| EIF4H | 168.428848 | -0.144488692 | 0.29698692 | -0.4865153 | 0.62660183 | 0.86052754 |
| SRPK1 | 154.722729 | -0.144293516 | 0.26426387 | -0.5460206 | 0.58505177 | 0.83895258 |
| PDCL | 76.475167 | -0.144287847 | 0.32687558 | -0.4414152 | 0.65891245 | 0.87335081 |
| FBXO33 | 143.983567 | -0.14422871 | 0.17918185 | -0.8049292 | 0.42086052 | 0.73910721 |
| FBRS | 57.2280423 | -0.144098939 | 0.27186654 | -0.5300356 | 0.59608727 | 0.84697301 |
| ZNF3 | 44.4393 | -0.143891486 | 0.40310884 | -0.3569544 | 0.72112592 | 0.90162538 |
| GALNT2 | 193.415345 | -0.143426098 | 0.21012439 | -0.6825771 | 0.49487411 | 0.79325266 |
| EIF4G2 | 388.901436 | -0.143422594 | 0.30295903 | -0.4734059 | 0.63592362 | 0.86207417 |
| NFYA | 62.9611009 | -0.14334204 | 0.31809065 | -0.4506327 | 0.65225432 | 0.8685756 |
| PRMT9 | 81.2948987 | -0.14331292 | 0.30383223 | -0.4716844 | 0.63715208 | 0.86207417 |
| QRSL1 | 63.743756 | -0.143278988 | 0.25529287 | -0.5612338 | 0.57463816 | 0.83436282 |
| SAMD4B | 195.477145 | -0.143012699 | 0.31037216 | -0.4607781 | 0.64495781 | 0.86558646 |
| LRRC8A | 137.810538 | -0.142955062 | 0.29759343 | -0.4803704 | 0.63096407 | 0.86097142 |
| LMO4 | 86.9247043 | -0.14275591 | 0.28769077 | -0.496213 | 0.61974412 | 0.85855421 |
| ATP8B1 | 806.257254 | -0.142655826 | 0.34231276 | -0.4167412 | 0.67686773 | 0.88085379 |
| STK35 | 47.0964756 | -0.142468255 | 0.24868654 | -0.5728828 | 0.56672402 | 0.83285692 |
| WWOX | 1990.0631 | -0.142465414 | 0.29054292 | -0.4903421 | 0.62389186 | 0.85997304 |
| PSMD4 | 122.564313 | -0.142407365 | 0.23822905 | -0.597775 | 0.54999009 | 0.82355262 |
| ATP5ME | 66.7304566 | -0.142151175 | 0.41720202 | -0.340725 | 0.73331058 | 0.90508002 |
| COMMD4 | 59.6546251 | -0.142119925 | 0.28084031 | -0.5060524 | 0.61281984 | 0.85573702 |
| CNOT6L | 468.033003 | -0.141804105 | 0.29950236 | -0.4734657 | 0.63588094 | 0.86207417 |
| POGK | 61.0749392 | -0.14145947 | 0.25809397 | -0.5480929 | 0.58362813 | 0.83850098 |
| C16orf58 | 42.4089728 | -0.141352496 | 0.23649979 | -0.5976855 | 0.55004981 | 0.82355262 |
| NME3 | 153.849841 | -0.141213299 | 0.43661855 | -0.3234249 | 0.74637349 | 0.91031794 |
| SYTL4 | 115.81495 | -0.141158225 | 0.37204045 | -0.3794163 | 0.70437872 | 0.89446478 |
| SLC20A2 | 274.999522 | -0.140603152 | 0.2299985 | -0.611322 | 0.5409864 | 0.81928059 |
| AKR1E2 | 47.1328208 | -0.140559297 | 0.35614522 | -0.3946685 | 0.69308756 | 0.88975484 |
| PPP2R1A | 169.489172 | -0.140247654 | 0.21693028 | -0.6465103 | 0.51794894 | 0.80546868 |
| GLOD4 | 50.8384966 | -0.140158654 | 0.21597313 | -0.6489634 | 0.51636203 | 0.80483629 |
| KLHL22 | 65.6502918 | -0.139817529 | 0.44644762 | -0.3131779 | 0.75414551 | 0.91262536 |
| KIAA0753 | 73.1467821 | -0.139640626 | 0.30280641 | -0.4611548 | 0.64468755 | 0.86542551 |
| LCA5 | 64.5602151 | -0.139639065 | 0.23952527 | -0.5829826 | 0.55990502 | 0.82851768 |
| ACVR1 | 145.602654 | -0.139340529 | 0.21453778 | -0.6494918 | 0.51602055 | 0.80483629 |
| TBC1D15 | 464.615024 | -0.139260879 | 0.31097558 | -0.4478193 | 0.65428359 | 0.87014506 |
| SRP68 | 60.9618494 | -0.139151208 | 0.22623767 | -0.6150665 | 0.53851081 | 0.81807397 |
| ZNF682 | 47.3214682 | -0.139137187 | 0.23282763 | -0.5975974 | 0.5501086 | 0.82355262 |
| CTR9 | 86.0178714 | -0.139058269 | 0.28427968 | -0.4891601 | 0.62472838 | 0.86016045 |
| MAGT1 | 167.363731 | -0.139049058 | 0.17155008 | -0.810545 | 0.41762702 | 0.73821752 |
| FAM104A | 64.030651 | -0.138876398 | 0.2001406 | -0.6938942 | 0.48774857 | 0.7887839 |
| SLC16A4 | 52.66518 | -0.138836272 | 0.28881674 | -0.4807072 | 0.63072465 | 0.86097142 |
| UCHL3 | 60.2393887 | -0.138815747 | 0.38552097 | -0.3600731 | 0.71879244 | 0.90051656 |
| VMA21 | 62.8623672 | -0.138609024 | 0.41509703 | -0.3339196 | 0.73844025 | 0.90798746 |
| SMYD4 | 46.5799993 | -0.138510334 | 0.23496631 | -0.5894902 | 0.55553249 | 0.82487928 |
| GEMIN5 | 68.710464 | -0.137566859 | 0.23893333 | -0.5757542 | 0.56478136 | 0.83113895 |
| ZBTB8A | 47.3028992 | -0.137459695 | 0.29878703 | -0.4600591 | 0.6454738 | 0.86572246 |
| USP3 | 372.588512 | -0.137374697 | 0.17042394 | -0.8060763 | 0.42019886 | 0.73901389 |
| MTRNR2L12 | 258.707358 | -0.137253885 | 0.42914982 | -0.3198274 | 0.74909915 | 0.91075944 |
| FAM114A1 | 128.424244 | -0.136565605 | 0.283013 | -0.4825418 | 0.6294211 | 0.8608646 |
| STX3 | 91.5316479 | -0.136461202 | 0.36837199 | -0.370444 | 0.71105168 | 0.89747106 |
| H3F3A | 74.8843972 | -0.136340978 | 0.38653283 | -0.3527281 | 0.72429232 | 0.9026193 |
| SNX9 | 586.556492 | -0.136069309 | 0.28098061 | -0.4842658 | 0.62819722 | 0.86052754 |
| CCDC174 | 59.8428568 | -0.136019241 | 0.26602812 | -0.5112965 | 0.60914346 | 0.85323878 |
| AKAP13 | 1400.31788 | -0.136015165 | 0.25274727 | -0.5381469 | 0.59047563 | 0.842228 |
| JADE1 | 95.4987628 | -0.135878263 | 0.24618043 | -0.5519458 | 0.58098547 | 0.83774521 |
| PRCP | 467.078495 | -0.13575479 | 0.43613226 | -0.3112698 | 0.75559554 | 0.91343453 |
| ACLY | 124.685887 | -0.135407984 | 0.27396538 | -0.4942522 | 0.6211281 | 0.85890999 |
| RANBP1 | 59.7161123 | -0.1351803 | 0.36848279 | -0.3668565 | 0.71372607 | 0.89943635 |
| DOT1L | 85.7113661 | -0.134934148 | 0.24891968 | -0.5420791 | 0.58776403 | 0.8409165 |
| FAM172A | 1015.89187 | -0.134863206 | 0.16472605 | -0.8187121 | 0.4129507 | 0.73622771 |
| CSNK1A1 | 1070.97777 | -0.134800302 | 0.2051669 | -0.6570275 | 0.51116321 | 0.80177277 |
| HEXB | 105.210479 | -0.134629607 | 0.2663546 | -0.5054525 | 0.61324103 | 0.85573702 |
| MAPRE1 | 75.5519673 | -0.13438196 | 0.29573092 | -0.4544062 | 0.6495365 | 0.86766991 |
| PTPA | 84.4603833 | -0.134202025 | 0.31715892 | -0.4231381 | 0.6721945 | 0.87803719 |
| RNF126 | 45.949055 | -0.133858234 | 0.30829782 | -0.4341848 | 0.66415423 | 0.8755434 |
| TOR1AIP2 | 187.868076 | -0.133548233 | 0.24245015 | -0.5508276 | 0.58175187 | 0.83789748 |
| MBD1 | 52.3275514 | -0.133402513 | 0.22138725 | -0.6025754 | 0.54679118 | 0.82355262 |
| RHOB | 336.826212 | -0.133271389 | 0.43660924 | -0.3052418 | 0.760182 | 0.91455455 |
| SLC41A3 | 48.9428034 | -0.133251646 | 0.37978288 | -0.3508627 | 0.72569135 | 0.9026193 |
| UBA1 | 130.248016 | -0.133165895 | 0.20267571 | -0.6570393 | 0.51115567 | 0.80177277 |
| TMEM50A | 205.311029 | -0.132603905 | 0.30128272 | -0.4401311 | 0.65984213 | 0.87335081 |
| MAPK3 | 132.017688 | -0.132551066 | 0.39063308 | -0.3393237 | 0.73436588 | 0.9055183 |
| JPT1 | 59.9288862 | -0.132506776 | 0.27992444 | -0.4733662 | 0.63595192 | 0.86207417 |
| ZNF248 | 136.262095 | -0.132290353 | 0.21896194 | -0.6041705 | 0.54573027 | 0.82355262 |
| SEPHS1 | 80.1844192 | -0.132025192 | 0.23725776 | -0.5564631 | 0.57789431 | 0.83613556 |
| ADAM9 | 230.722048 | -0.132010876 | 0.29792165 | -0.443106 | 0.65768905 | 0.87254693 |
| SUCLG2 | 454.620632 | -0.131777989 | 0.11492515 | -1.1466418 | 0.25152968 | 0.61661694 |
| TUBE1 | 80.3589762 | -0.131691004 | 0.34581904 | -0.380809 | 0.70334498 | 0.89446478 |
| INVS | 461.270083 | -0.131687243 | 0.21822191 | -0.6034556 | 0.54620561 | 0.82355262 |
| RARRES3 | 84.7562281 | -0.131679425 | 0.45828415 | -0.2873314 | 0.77385858 | 0.91726032 |
| CCNI | 446.345997 | -0.131643604 | 0.25694282 | -0.5123459 | 0.60840894 | 0.85289019 |
| BOD1L1 | 264.12473 | -0.13161146 | 0.25659824 | -0.5129087 | 0.60801521 | 0.85277972 |
| PLD3 | 177.176062 | -0.131603226 | 0.28834539 | -0.4564083 | 0.64809641 | 0.86739793 |
| RAB8A | 38.5860703 | -0.131464312 | 0.28128451 | -0.4673713 | 0.64023424 | 0.86442461 |
| SRP54 | 196.423377 | -0.131460997 | 0.18710138 | -0.7026191 | 0.48229318 | 0.78648965 |
| SPRYD7 | 47.811867 | -0.131289318 | 0.22870949 | -0.574044 | 0.56593801 | 0.83232759 |
| EIF3M | 107.053378 | -0.131232311 | 0.26510438 | -0.4950213 | 0.62058509 | 0.85890999 |
| PIGG | 95.4119555 | -0.131055069 | 0.23023606 | -0.5692204 | 0.56920657 | 0.83368112 |
| TCP11L2 | 74.6413914 | -0.130922273 | 0.43287869 | -0.3024456 | 0.76231237 | 0.91477008 |
| ZKSCAN1 | 255.64591 | -0.130900325 | 0.18132016 | -0.7219292 | 0.47033799 | 0.77792258 |
| C9orf78 | 64.2500389 | -0.130877634 | 0.34728972 | -0.3768543 | 0.70628186 | 0.89521445 |
| PPP2R5C | 436.938131 | -0.130585819 | 0.27693755 | -0.4715353 | 0.63725854 | 0.86207417 |
| RAP2C-AS1 | 92.4215933 | -0.130521291 | 0.40756344 | -0.3202478 | 0.7487805 | 0.91065347 |
| TNS2 | 341.186065 | -0.130030305 | 0.29888691 | -0.4350485 | 0.66352722 | 0.8755434 |
| UBE3D | 158.944503 | -0.129888115 | 0.27989029 | -0.464068 | 0.64259905 | 0.86490071 |
| C5orf22 | 55.7927966 | -0.129811431 | 0.22951717 | -0.5655848 | 0.57167604 | 0.83368112 |
| POT1 | 135.760229 | -0.129589233 | 0.23296326 | -0.5562647 | 0.57802992 | 0.83613556 |
| TMEM33 | 115.38249 | -0.129465855 | 0.20997742 | -0.6165704 | 0.53751814 | 0.81807397 |
| AC011092.2 | 72.5097095 | -0.129461497 | 0.52077126 | -0.2485957 | 0.80367353 | 0.93081002 |

| | | | | | | |
|---|---|---|---|---|---|---|
| YAP1 | 328.255093 | -0.129259138 | 0.29432972 | -0.4391644 | 0.66054241 | 0.87350009 |
| SETDB1 | 110.886481 | -0.129210865 | 0.28111277 | -0.4596407 | 0.64577415 | 0.86572246 |
| GORASP2 | 106.522347 | -0.128927917 | 0.22235255 | -0.5798356 | 0.5620255 | 0.83015427 |
| KHDC4 | 157.580588 | -0.128913298 | 0.16252815 | -0.7931752 | 0.42767575 | 0.74318766 |
| BAG1 | 69.9181245 | -0.128450744 | 0.33755769 | -0.3805298 | 0.70355221 | 0.89446478 |
| ATF6B | 99.9081818 | -0.128426638 | 0.25650038 | -0.5006879 | 0.61659078 | 0.85732696 |
| HTATIP2 | 41.714333 | -0.128311969 | 0.35529973 | -0.3611373 | 0.71799684 | 0.90051656 |
| SCPEP1 | 57.8144255 | -0.128310097 | 0.34625715 | -0.370563 | 0.71096303 | 0.89747106 |
| CLTC | 454.222579 | -0.128308253 | 0.1760733 | -0.7287207 | 0.46617255 | 0.77518594 |
| SLC30A6 | 124.195397 | -0.128209517 | 0.18975344 | -0.6756637 | 0.49925416 | 0.79528845 |
| TYMP | 57.1523127 | -0.12817925 | 0.99757748 | -0.1284905 | 0.8977608 | 0.96624709 |
| ASCC2 | 89.2236599 | -0.127838486 | 0.31723712 | -0.4029746 | 0.68696694 | 0.88671976 |
| PPP1R16B | 770.255518 | -0.127805705 | 0.30472823 | -0.4194088 | 0.67491739 | 0.87985486 |
| DGCR2 | 137.557277 | -0.12759283 | 0.28051969 | -0.4548445 | 0.64922113 | 0.86766991 |
| POLR3B | 45.298236 | -0.127561427 | 0.36762363 | -0.3469892 | 0.72859943 | 0.90376371 |
| ZNF106 | 204.448262 | -0.127416861 | 0.19300654 | -0.6601686 | 0.50914562 | 0.80091093 |
| SCAP | 95.4970793 | -0.127225263 | 0.25191117 | -0.5050402 | 0.61353061 | 0.85573702 |
| ANKUB1 | 112.941057 | -0.127082177 | 0.42112357 | -0.3017693 | 0.76282791 | 0.91477008 |
| GNG11 | 569.487606 | -0.127072395 | 0.54038049 | -0.2351536 | 0.81408955 | 0.93460727 |
| GID8 | 63.3354472 | -0.126862304 | 0.24185912 | -0.5245297 | 0.59991014 | 0.8497602 |
| CDC42SE2 | 383.118161 | -0.126843979 | 0.28090394 | -0.4515564 | 0.65158856 | 0.8685756 |
| CTDP1 | 54.4948312 | -0.126793038 | 0.27217263 | -0.4658552 | 0.64131914 | 0.86490071 |
| AHCYL1 | 187.358235 | -0.126781601 | 0.25327315 | -0.5005726 | 0.61667194 | 0.85732696 |
| GMFG | 94.3032129 | -0.126628922 | 0.28078551 | -0.450981 | 0.65200326 | 0.8685756 |
| THADA | 478.623064 | -0.126515177 | 0.23019121 | -0.5496091 | 0.58258754 | 0.83813534 |
| NPTN | 364.154955 | -0.126427876 | 0.17932842 | -0.7050075 | 0.48080559 | 0.78473327 |
| ZNF611 | 124.027329 | -0.126427706 | 0.30692954 | -0.4119112 | 0.68040453 | 0.88294217 |
| COPS8 | 61.0927975 | -0.126305903 | 0.2581598 | -0.4892547 | 0.62466136 | 0.86016045 |
| EFCAB11 | 132.689476 | -0.126178963 | 0.2709964 | -0.4656112 | 0.64149382 | 0.86490071 |
| CLEC16A | 335.931413 | -0.12615209 | 0.24070317 | -0.5240982 | 0.60021026 | 0.84976293 |
| EGFL7 | 1657.70085 | -0.126087811 | 0.32036482 | -0.3935757 | 0.69389432 | 0.89026924 |
| RTCB | 107.140881 | -0.125950386 | 0.21122265 | -0.596292 | 0.55098015 | 0.8236419 |
| OSBPL5 | 221.302658 | -0.125938159 | 0.29294409 | -0.4299051 | 0.66726467 | 0.87587657 |
| OGFR | 113.37376 | -0.125794241 | 0.35340482 | -0.3559494 | 0.72187845 | 0.90162538 |
| ESF1 | 128.827496 | -0.125794028 | 0.28774097 | -0.437178 | 0.66198225 | 0.87469736 |
| SELENOF | 240.060575 | -0.125761557 | 0.24123269 | -0.5213288 | 0.60213771 | 0.85030923 |
| OSBPL9 | 336.556234 | -0.125654119 | 0.18701292 | -0.6719007 | 0.50164689 | 0.79661835 |
| IARS2 | 131.461863 | -0.125620977 | 0.19574775 | -0.6417493 | 0.52103598 | 0.80735418 |
| ZFYVE21 | 72.1987507 | -0.125403937 | 0.35740183 | -0.3508766 | 0.72568093 | 0.9026193 |
| ATP6V0A2 | 86.941396 | -0.125366612 | 0.34673814 | -0.3615599 | 0.71768092 | 0.90033983 |
| C1QTNF3-AM | 150.675285 | -0.124920471 | 0.42905911 | -0.2911498 | 0.77093676 | 0.91695334 |
| GABARAPL1 | 211.928182 | -0.124798727 | 0.55303199 | -0.2256628 | 0.82146372 | 0.93726124 |
| TRMT10B | 86.7680988 | -0.124544147 | 0.35219863 | -0.3536191 | 0.72362439 | 0.90258339 |
| DNAJB12 | 62.1558829 | -0.124531599 | 0.27609563 | -0.4510452 | 0.651957 | 0.8685756 |
| KEAP1 | 64.2053815 | -0.124457025 | 0.48609185 | -0.256036 | 0.79792302 | 0.92959949 |
| PKIG | 387.728781 | -0.12414301 | 0.24415848 | -0.5084526 | 0.61113598 | 0.85449687 |
| PIGU | 103.272253 | -0.124058582 | 0.22261063 | -0.5572896 | 0.5773296 | 0.83603913 |
| SF3B2 | 120.58959 | -0.123772829 | 0.1937752 | -0.6387444 | 0.5229892 | 0.80854347 |
| AL445426.1 | 62.5634204 | -0.123704678 | 0.69867821 | -0.1770553 | 0.85946495 | 0.95259403 |
| SLC30A5 | 121.659133 | -0.123559526 | 0.22177424 | -0.557141 | 0.57743108 | 0.83603913 |
| DNAJC10 | 182.412954 | -0.123524164 | 0.24963107 | -0.4948269 | 0.62072233 | 0.85890999 |
| PUDP | 79.3430676 | -0.123272696 | 0.33304397 | -0.3701394 | 0.71127862 | 0.89747106 |
| FNBP4 | 412.570265 | -0.123246015 | 0.21119305 | -0.5835704 | 0.55950937 | 0.82823176 |
| SETD7 | 119.462731 | -0.122971526 | 0.19107846 | -0.6435656 | 0.51985716 | 0.80659673 |
| HAUS1 | 54.032466 | -0.122715854 | 0.31960838 | -0.3839569 | 0.70101037 | 0.8944316 |
| WDFY3 | 940.198343 | -0.12264808 | 0.18117618 | -0.6769548 | 0.49843464 | 0.79513466 |
| UBQLN2 | 38.5725472 | -0.122456494 | 0.34536078 | -0.3545756 | 0.72290758 | 0.90217367 |
| NUPL2 | 86.2123722 | -0.122172966 | 0.28687522 | -0.4258749 | 0.67019897 | 0.87705713 |
| RERE | 1677.75841 | -0.121598533 | 0.15376084 | -0.790829 | 0.42904378 | 0.74408005 |
| PWP1 | 66.1763027 | -0.121354343 | 0.24036466 | -0.504876 | 0.61364595 | 0.85573702 |
| MORF4L2 | 175.579637 | -0.121308236 | 0.30040791 | -0.4038117 | 0.68635118 | 0.88671976 |
| KIAA2013 | 59.8447368 | -0.121179856 | 0.31905142 | -0.3798129 | 0.70408428 | 0.89446478 |
| NDUFV2 | 150.05091 | -0.121135538 | 0.17797285 | -0.6806406 | 0.49609896 | 0.79381653 |
| OSBPL2 | 71.6517573 | -0.121093664 | 0.18756792 | -0.645599 | 0.51853909 | 0.8057739 |
| DSTN | 430.177344 | -0.120875042 | 0.39371815 | -0.3070091 | 0.75883647 | 0.91442519 |
| NUCKS1 | 449.300054 | -0.120802009 | 0.17376624 | -0.6951984 | 0.48693097 | 0.78826504 |
| UBA6 | 283.132143 | -0.120665179 | 0.2599281 | -0.4642252 | 0.6424864 | 0.86490071 |
| MLKL | 201.389509 | -0.12059253 | 0.37369657 | -0.3227017 | 0.74692114 | 0.91031794 |
| CLK2 | 47.8997308 | -0.12052373 | 0.22668975 | -0.5316682 | 0.59495583 | 0.84587674 |
| MAST4 | 954.230215 | -0.120316392 | 0.23206989 | -0.518449 | 0.60414506 | 0.85098921 |
| YIPF6 | 115.055737 | -0.12014201 | 0.23593899 | -0.509208 | 0.61060647 | 0.85411842 |
| CHCHD10 | 70.1074333 | -0.120062137 | 0.28589506 | -0.4199518 | 0.67452069 | 0.87947782 |
| CFL2 | 55.2399067 | -0.119795878 | 0.3012733 | -0.3976319 | 0.69090153 | 0.88832346 |
| PYGB | 52.4232618 | -0.119715821 | 0.35903337 | -0.3334393 | 0.73880273 | 0.90802385 |
| LMNA | 286.802005 | -0.119527328 | 0.41382777 | -0.2888335 | 0.77270879 | 0.91726032 |
| DDAH1 | 224.024494 | -0.119504044 | 0.45827922 | -0.2607669 | 0.79427228 | 0.92810654 |
| RAB34 | 42.0611335 | -0.119324397 | 0.54994824 | -0.2169739 | 0.82822871 | 0.94018985 |
| HMGCL | 64.262043 | -0.119279986 | 0.24277962 | -0.4913097 | 0.6232074 | 0.85940448 |
| ME3 | 175.452767 | -0.119122982 | 0.22980903 | -0.5183564 | 0.60420962 | 0.85098921 |
| KLHDC4 | 82.777149 | -0.118879545 | 0.31774566 | -0.3741343 | 0.70830442 | 0.89680567 |
| ZBTB41 | 59.5209602 | -0.118782865 | 0.21927178 | -0.5417152 | 0.58801469 | 0.8409165 |
| DDX1 | 115.340368 | -0.118673435 | 0.18215642 | -0.6514919 | 0.514729 | 0.80422048 |
| PTPN2 | 162.532584 | -0.118500662 | 0.27303208 | -0.4340174 | 0.66427584 | 0.8755434 |
| WDR36 | 126.158585 | -0.118462065 | 0.25894574 | -0.4574783 | 0.64732728 | 0.86714786 |
| CAPZA1 | 337.562117 | -0.11795082 | 0.19609955 | -0.6014844 | 0.5475174 | 0.82355262 |
| NOL10 | 246.652038 | -0.117754437 | 0.20963878 | -0.5617016 | 0.57431935 | 0.83430769 |
| MID2 | 115.881935 | -0.117481806 | 0.31559113 | -0.3722595 | 0.70969963 | 0.89727558 |
| MYL9 | 85.1890031 | -0.117436788 | 0.48349669 | -0.2428906 | 0.80809017 | 0.93264026 |
| C11orf49 | 248.813477 | -0.117370663 | 0.22541436 | -0.5206885 | 0.6025838 | 0.85030923 |
| MTPAP | 60.055513 | -0.117192192 | 0.19999333 | -0.5859805 | 0.55788862 | 0.82661057 |
| CEBPZOS | 96.5904241 | -0.116755776 | 0.27764754 | -0.420518 | 0.67410712 | 0.87910389 |
| NIPA2 | 96.4030618 | -0.116595841 | 0.22609782 | -0.5156876 | 0.60607263 | 0.85156118 |
| METAP2 | 170.09802 | -0.116580646 | 0.21779663 | -0.535273 | 0.59246112 | 0.84417714 |
| SON | 659.073416 | -0.116385273 | 0.21765152 | -0.5347322 | 0.59283506 | 0.84426894 |
| RNF217 | 312.894773 | -0.116161124 | 0.31982746 | -0.3631993 | 0.71645597 | 0.9001716 |
| DYSF | 971.490795 | -0.116035164 | 0.36452531 | -0.3183185 | 0.75024332 | 0.91109327 |
| PPP1R35 | 53.5993519 | -0.115945499 | 0.38422938 | -0.3017612 | 0.76283415 | 0.91477008 |
| HIPK2 | 2178.16404 | -0.115846019 | 0.40393158 | -0.2867961 | 0.77426841 | 0.91726032 |
| PKN3 | 158.570509 | -0.115684071 | 0.30994662 | -0.3732387 | 0.7089708 | 0.89727558 |
| FAM227B | 62.3169282 | -0.115671642 | 0.44664897 | -0.2589766 | 0.79565328 | 0.92847059 |
| ZNF347 | 116.726091 | -0.115617944 | 0.33272347 | -0.3474896 | 0.72822353 | 0.9037084 |
| PPP2R5E | 519.779829 | -0.115413639 | 0.20882163 | -0.5526901 | 0.58047566 | 0.83760048 |
| ZNF281 | 47.4519389 | -0.115280246 | 0.27399922 | -0.420732 | 0.67395077 | 0.87910389 |
| TRMT2B | 53.618092 | -0.11515079 | 0.36391366 | -0.3164234 | 0.75168118 | 0.9112603 |
| TOMM70 | 79.7054794 | -0.115079143 | 0.23422412 | -0.4913206 | 0.62319969 | 0.85940448 |
| TTTY15 | 44.2431374 | -0.114976329 | 1.85705119 | -0.0619134 | 0.95063181 | 0.98412513 |
| BRMS1 | 38.5605143 | -0.114926578 | 0.4434962 | -0.2591377 | 0.79552901 | 0.92847059 |
| SAMD8 | 177.92242 | -0.114883655 | 0.12939593 | -0.8878459 | 0.37462363 | 0.71257833 |
| MAK | 79.1671982 | -0.114726544 | 0.43918705 | -0.2612248 | 0.79391916 | 0.92790728 |
| ZNF621 | 105.37693 | -0.114693185 | 0.23605594 | -0.4858729 | 0.62705728 | 0.86052754 |
| TIMM23 | 76.6943589 | -0.114595307 | 0.23713609 | -0.483247 | 0.62892035 | 0.86061125 |
| CS | 74.9005004 | -0.114591378 | 0.33568381 | -0.341367 | 0.73282731 | 0.90502956 |
| YKT6 | 37.9596942 | -0.114457409 | 0.40384116 | -0.2834218 | 0.77685348 | 0.91892777 |

| | | | | | | |
|---|---|---|---|---|---|---|
| AP001011.1 | 201.084428 | -0.114352039 | 0.57597143 | -0.1985377 | 0.8426244 | 0.94471448 |
| HARS | 88.5511722 | -0.114114867 | 0.20665731 | -0.5521937 | 0.58081565 | 0.83774521 |
| MANBA | 114.129372 | -0.114076867 | 0.34447277 | -0.3311637 | 0.74052086 | 0.90826918 |
| NDUFA10 | 205.879698 | -0.114066492 | 0.18923386 | -0.6027806 | 0.54665469 | 0.82355262 |
| ARSJ | 66.6930342 | -0.113914184 | 0.55048907 | -0.2069327 | 0.83606243 | 0.9432647 |
| FLNB | 876.683453 | -0.113863409 | 0.32651425 | -0.3487242 | 0.7272964 | 0.90337985 |
| SCAMP2 | 68.0614604 | -0.113846781 | 0.2353946 | -0.4836423 | 0.62863976 | 0.86052754 |
| AP3D1 | 125.109392 | -0.113813279 | 0.22477516 | -0.5063428 | 0.61261603 | 0.85573702 |
| GCLC | 87.4961219 | -0.113812869 | 0.3230218 | -0.352338 | 0.72458476 | 0.9026193 |
| ZBED5 | 97.7785077 | -0.113796509 | 0.20738802 | -0.548713 | 0.58320241 | 0.8384303 |
| KLHL36 | 65.289079 | -0.11370518 | 0.24104734 | -0.4717131 | 0.6371316 | 0.86207417 |
| GGCX | 73.4833637 | -0.113465262 | 0.36913133 | -0.3073845 | 0.7585507 | 0.91442519 |
| CDYL | 348.677849 | -0.11325749 | 0.21426907 | -0.528576 | 0.59709961 | 0.84764654 |
| PSD3 | 185.949355 | -0.113110799 | 0.37592299 | -0.3008882 | 0.76349973 | 0.91490985 |
| KCNMB3 | 46.2930397 | -0.113082603 | 0.39201307 | -0.2884664 | 0.77298974 | 0.91726032 |
| TEP1 | 84.443497 | -0.112992653 | 0.17831163 | -0.6336808 | 0.52628917 | 0.81036438 |
| ERCC4 | 56.6612736 | -0.112867143 | 0.23099763 | -0.4886074 | 0.6251197 | 0.86016045 |
| ZNF12 | 67.4929349 | -0.112827591 | 0.23688583 | -0.4762952 | 0.63386406 | 0.86207417 |
| MRPL19 | 125.929439 | -0.112572429 | 0.26107856 | -0.4311822 | 0.66633589 | 0.87587657 |
| EIF2S3 | 123.633115 | -0.112462638 | 0.17857396 | -0.6297818 | 0.52883733 | 0.81207005 |
| GMEB1 | 72.7563295 | -0.112448593 | 0.34275217 | -0.3280755 | 0.74285457 | 0.90850686 |
| ARL17A | 78.4422585 | -0.112330228 | 0.46181745 | -0.2432351 | 0.80782325 | 0.93248945 |
| POC1B | 133.120965 | -0.112275479 | 0.24047365 | -0.4668931 | 0.64057638 | 0.86456153 |
| ZNF398 | 69.1418122 | -0.112126401 | 0.34682594 | -0.323293 | 0.74647335 | 0.91031794 |
| ALKBH8 | 44.9352349 | -0.112039973 | 0.29255071 | -0.3829762 | 0.70173738 | 0.89443605 |
| RGS3 | 437.819516 | -0.111876834 | 0.36119793 | -0.3097383 | 0.75675997 | 0.91345267 |
| CTSF | 35.9098581 | -0.111702703 | 0.4767004 | -0.2343248 | 0.81473287 | 0.93505738 |
| PTPN11 | 193.083228 | -0.111534173 | 0.19473089 | -0.5727606 | 0.56680682 | 0.83285692 |
| TMA7 | 156.592464 | -0.11142247 | 0.47154937 | -0.2362901 | 0.81320753 | 0.9344584 |
| C2CD3 | 124.183773 | -0.111386238 | 0.22178945 | -0.5022161 | 0.61551551 | 0.85714845 |
| UBAC2 | 380.556924 | -0.111352464 | 0.17375034 | -0.6408762 | 0.5216031 | 0.80772643 |
| BAZ1A | 354.738672 | -0.110938668 | 0.33459108 | -0.3315649 | 0.7402178 | 0.90826918 |
| TMEM222 | 50.2324614 | -0.110599972 | 0.27563428 | -0.4012562 | 0.68823148 | 0.88739126 |
| ASCC1 | 250.502838 | -0.110536281 | 0.15402047 | -0.7176727 | 0.47295913 | 0.77899012 |
| ARL6IP4 | 166.70962 | -0.110474886 | 0.43828377 | -0.2520625 | 0.80099278 | 0.93019678 |
| AL138686.1 | 69.1675697 | -0.110438854 | 0.67627705 | -0.1633042 | 0.87027896 | 0.95689355 |
| PDE2A | 375.678913 | -0.110397689 | 0.31634921 | -0.3489741 | 0.72710874 | 0.90328386 |
| EIF3E | 238.716447 | -0.11021898 | 0.16369297 | -0.6733275 | 0.50073896 | 0.79625863 |
| CAMTA2 | 67.4088537 | -0.110126518 | 0.25523306 | -0.4314744 | 0.6661235 | 0.87587657 |
| PLEC | 228.528403 | -0.109931151 | 0.4021139 | -0.2733831 | 0.78455873 | 0.92302847 |
| UBXN4 | 335.437618 | -0.109915381 | 0.1471548 | -0.7469371 | 0.45510154 | 0.76710126 |
| EIF4G3 | 1603.02666 | -0.109910135 | 0.22776109 | -0.4825676 | 0.62940275 | 0.8608646 |
| MAPK14 | 317.706241 | -0.10977862 | 0.20872524 | -0.525948 | 0.59892435 | 0.84880463 |
| MAOA | 63.0974276 | -0.109671091 | 0.53512016 | -0.2049467 | 0.8376138 | 0.94358503 |
| PCNX4 | 313.539117 | -0.109540658 | 0.25036965 | -0.4375157 | 0.66173738 | 0.87451501 |
| GPR173 | 79.9869285 | -0.109310244 | 0.4380602 | -0.2495325 | 0.80294893 | 0.93081002 |
| RAB7A | 541.042594 | -0.109131859 | 0.21150889 | -0.5159682 | 0.60587663 | 0.85156118 |
| ZNF431 | 157.069559 | -0.109049112 | 0.19574079 | -0.5571098 | 0.57745242 | 0.83603913 |
| DDX6 | 280.143344 | -0.10899024 | 0.14057473 | -0.7753188 | 0.43815128 | 0.75275917 |
| PREPL | 156.789459 | -0.108814932 | 0.33148385 | -0.3282662 | 0.74271043 | 0.90850686 |
| RNF7 | 132.567191 | -0.108666611 | 0.50959 | -0.2132432 | 0.83113726 | 0.94121176 |
| ZBTB38 | 460.990089 | -0.108637223 | 0.22626742 | -0.4801276 | 0.6311367 | 0.86097142 |
| CSNK2A1 | 275.442925 | -0.108630963 | 0.19702403 | -0.551359 | 0.58138763 | 0.83789748 |
| SCAMP4 | 39.723239 | -0.10851527 | 0.25520252 | -0.4252124 | 0.67068186 | 0.87730512 |
| HLCS | 188.569573 | -0.1084538 | 0.31386082 | -0.3455474 | 0.72968286 | 0.90414821 |
| DNAJB6 | 320.055969 | -0.108448723 | 0.22980364 | -0.4719191 | 0.63698454 | 0.86207417 |
| FOXO1 | 1243.86108 | -0.108292457 | 0.30327878 | -0.3570723 | 0.72103767 | 0.90162538 |
| ZNF266 | 78.3158438 | -0.108208386 | 0.30780478 | -0.3515488 | 0.7251767 | 0.9026193 |
| MPHOSPH9 | 104.883368 | -0.107872947 | 0.19859008 | -0.543194 | 0.58699621 | 0.84027596 |
| METTL2B | 79.744842 | -0.107723809 | 0.24558569 | -0.4386404 | 0.66092211 | 0.87369056 |
| CAMK2D | 900.503733 | -0.107663963 | 0.22711304 | -0.4740545 | 0.63546104 | 0.86207417 |
| SLC35E2B | 56.3373981 | -0.10738315 | 0.3123744 | -0.3437642 | 0.73102359 | 0.90430325 |
| CEP104 | 82.771229 | -0.107257332 | 0.25633773 | -0.4184219 | 0.67563864 | 0.88033151 |
| GOLGA5 | 78.7224483 | -0.10680033 | 0.24064795 | -0.4438032 | 0.65718488 | 0.87234121 |
| POLR3C | 52.3575218 | -0.106767806 | 0.26343581 | -0.4052896 | 0.68526462 | 0.88671976 |
| HELQ | 77.6945548 | -0.106672384 | 0.21304559 | -0.5007022 | 0.61658076 | 0.85732696 |
| PRDM5 | 133.331056 | -0.106611521 | 0.28663839 | -0.3719373 | 0.7099395 | 0.89727558 |
| RAD52 | 70.7757061 | -0.106517158 | 0.27922605 | -0.3814728 | 0.70285241 | 0.89443605 |
| NGLY1 | 207.551767 | -0.106511138 | 0.35291768 | -0.3018016 | 0.76280328 | 0.91477008 |
| XPO6 | 167.792077 | -0.106486496 | 0.20223055 | -0.5265599 | 0.59849925 | 0.84839769 |
| SSR3 | 125.600522 | -0.106455654 | 0.39018429 | -0.2728343 | 0.7849806 | 0.92318141 |
| ECI2 | 85.5264818 | -0.10632887 | 0.28742531 | -0.3699357 | 0.71143044 | 0.89747106 |
| AC010618.3 | 56.1783346 | -0.10626916 | 0.30726175 | -0.3458587 | 0.72944888 | 0.90413211 |
| SETD1A | 42.6947231 | -0.106095497 | 0.36120376 | -0.2937276 | 0.7689661 | 0.91633167 |
| ACSL1 | 56.6667547 | -0.106046348 | 0.39246583 | -0.2702053 | 0.78700231 | 0.92401864 |
| AC016831.7 | 2071.59504 | -0.106029192 | 0.33446036 | -0.3170157 | 0.75123168 | 0.9112603 |
| KHSRP | 80.7979782 | -0.106021508 | 0.36979812 | -0.286701 | 0.77434123 | 0.91726032 |
| PRKACA | 82.3115442 | -0.105771668 | 0.32521702 | -0.3252341 | 0.74500389 | 0.90944569 |
| MICU2 | 250.111661 | -0.105578884 | 0.19204094 | -0.5497728 | 0.58247522 | 0.83812111 |
| RRM1 | 50.0569755 | -0.105575262 | 0.32784385 | -0.3220291 | 0.74743064 | 0.91055362 |
| RPE | 77.0879456 | -0.105284776 | 0.21717257 | -0.4847978 | 0.62781982 | 0.86052754 |
| ACADVL | 162.033078 | -0.104915941 | 0.27206571 | -0.3856272 | 0.69977278 | 0.89374756 |
| DNAJC7 | 208.037558 | -0.104294046 | 0.17816106 | -0.5853919 | 0.55828421 | 0.82686692 |
| SUFU | 175.484183 | -0.103869945 | 0.33709278 | -0.3081346 | 0.75797993 | 0.91402148 |
| GOLGA3 | 131.047392 | -0.103861117 | 0.21990676 | -0.4722961 | 0.63671542 | 0.86207417 |
| SERPINB9 | 109.193842 | -0.10366402 | 0.40385742 | -0.2566847 | 0.79742218 | 0.92939834 |
| KSR1 | 263.60841 | -0.103523854 | 0.27264509 | -0.3797019 | 0.70416673 | 0.89446478 |
| NTPCR | 77.4494875 | -0.103419381 | 0.30853633 | -0.3351935 | 0.73747911 | 0.90748745 |
| PDGFC | 113.229863 | -0.103159005 | 0.56682609 | -0.1819941 | 0.85558736 | 0.95134391 |
| SMAP2 | 87.5571552 | -0.103022904 | 0.29874578 | -0.3448514 | 0.73020608 | 0.90426442 |
| WDR1 | 193.820979 | -0.10301585 | 0.26517014 | -0.3884896 | 0.69765373 | 0.89303379 |
| RORA-AS1 | 88.5208405 | -0.10250103 | 0.50724566 | -0.2020737 | 0.83985907 | 0.94377462 |
| GJA4 | 173.127053 | -0.102461696 | 0.64250659 | -0.1594718 | 0.87329716 | 0.9579344 |
| AC098829.1 | 132.557563 | -0.102009219 | 0.23989708 | -0.4252208 | 0.67067575 | 0.87730512 |
| LINC01184 | 229.79671 | -0.101883 | 0.3378642 | -0.3015502 | 0.76299501 | 0.91477008 |
| GRSF1 | 77.0237702 | -0.101787118 | 0.19903328 | -0.5114075 | 0.60906572 | 0.85323878 |
| PAFAH1B2 | 180.866544 | -0.101739187 | 0.15080988 | -0.6746189 | 0.49991795 | 0.79557148 |
| ARL3 | 84.9024381 | -0.10151858 | 0.25998183 | -0.3904834 | 0.69617915 | 0.8920567 |
| SENP5 | 341.968878 | -0.101498963 | 0.27513505 | -0.368906 | 0.71219781 | 0.89798378 |
| TDG | 88.4756185 | -0.101404549 | 0.215031 | -0.4715811 | 0.63722585 | 0.86207417 |
| TMEM165 | 565.166437 | -0.101354741 | 0.24464531 | -0.4142926 | 0.67865984 | 0.88164367 |
| APPL1 | 183.321988 | -0.101201127 | 0.1828045 | -0.553603 | 0.57985054 | 0.83701203 |
| RMND1 | 59.0365067 | -0.101081844 | 0.27790814 | -0.3637239 | 0.71606417 | 0.9001716 |
| SLC41A1 | 56.2298144 | -0.100920582 | 0.35992745 | -0.2803915 | 0.77917719 | 0.91979327 |
| ZFYVE9 | 203.402539 | -0.100837423 | 0.1511499 | -0.6671352 | 0.50468577 | 0.79776858 |
| CWC22 | 109.339154 | -0.100832201 | 0.17464912 | -0.5773416 | 0.56370872 | 0.83089872 |
| MED1 | 106.670848 | -0.100799568 | 0.24651474 | -0.4088987 | 0.68261398 | 0.8848139 |
| SQLE | 108.756228 | -0.100792966 | 0.56243842 | -0.1792071 | 0.85777508 | 0.95213965 |
| ENTPD4 | 119.680729 | -0.100782403 | 0.24239623 | -0.4157755 | 0.67757432 | 0.88143805 |
| SDR39U1 | 43.7949804 | -0.100707542 | 0.26395534 | -0.3815325 | 0.70280815 | 0.89443605 |
| CABIN1 | 198.284972 | -0.100568989 | 0.15374372 | -0.654134 | 0.51302551 | 0.80321271 |
| GGA2 | 117.107128 | -0.100493684 | 0.24397751 | -0.4118973 | 0.68041468 | 0.88294217 |
| GCC2 | 305.144259 | -0.100441227 | 0.2628318 | -0.3821502 | 0.70234996 | 0.89443605 |

| | | | | | | |
|---|---|---|---|---|---|---|
| ZNF213 | 51.2949082 | -0.100304699 | 0.34411816 | -0.2914833 | 0.77068171 | 0.91695334 |
| PON2 | 172.563768 | -0.10001549 | 0.26271274 | -0.3807028 | 0.70342375 | 0.89446478 |
| DHX9 | 212.722485 | -0.099948905 | 0.15196042 | -0.6577299 | 0.51071174 | 0.80177277 |
| DENND6A-A! | 69.2583368 | -0.09992961 | 0.37272445 | -0.2681059 | 0.78861783 | 0.92519853 |
| EBAG9 | 54.5184767 | -0.099794999 | 0.21566775 | -0.4627256 | 0.64356104 | 0.86517342 |
| ZNF770 | 81.3065546 | -0.099597226 | 0.25143421 | -0.3961164 | 0.69201913 | 0.88897842 |
| SKP1 | 248.715235 | -0.099548195 | 0.35098932 | -0.2836217 | 0.77670029 | 0.91892777 |
| ARHGAP23 | 217.246152 | -0.099376383 | 0.38517657 | -0.2580021 | 0.79640525 | 0.92860988 |
| DRAM1 | 197.464141 | -0.099361705 | 0.17065074 | -0.5822518 | 0.56039707 | 0.82908652 |
| PANK3 | 130.211808 | -0.099285725 | 0.17702129 | -0.5608688 | 0.57488695 | 0.834481 |
| USP45 | 139.204999 | -0.099134083 | 0.21693292 | -0.4569804 | 0.64768517 | 0.86714786 |
| RCN2 | 104.321951 | -0.099104325 | 0.33267446 | -0.2979018 | 0.7657781 | 0.91600937 |
| NRAS | 58.1396995 | -0.098904019 | 0.2782563 | -0.3554422 | 0.72225837 | 0.90163848 |
| HMG20A | 157.639559 | -0.098325167 | 0.15624934 | -0.6292837 | 0.52916331 | 0.81225724 |
| NUTM2B-AS | 440.64105 | -0.098225129 | 0.24501964 | -0.4008868 | 0.6885035 | 0.88739126 |
| OTULIN | 109.828897 | -0.098112857 | 0.26022487 | -0.377031 | 0.70615053 | 0.89518665 |
| FLYWCH2 | 45.837287 | -0.097542067 | 0.38735863 | -0.2518133 | 0.80118535 | 0.93020712 |
| PACS2 | 227.108393 | -0.097510169 | 0.25664885 | -0.3799361 | 0.70399283 | 0.89446478 |
| COX10 | 83.3389203 | -0.097398374 | 0.24149547 | -0.4033135 | 0.68671765 | 0.88671976 |
| VAPB | 209.026522 | -0.097130393 | 0.18676797 | -0.5200592 | 0.60302235 | 0.85048808 |
| ELAVL1 | 137.461866 | -0.097048564 | 0.17275396 | -0.5617733 | 0.57427047 | 0.83430769 |
| KIF1C | 73.9152038 | -0.097013529 | 0.2908315 | -0.333573 | 0.73870181 | 0.90802385 |
| RANBP2 | 281.67591 | -0.096944717 | 0.17804599 | -0.5444926 | 0.58610256 | 0.83950091 |
| NBPF1 | 66.3818709 | -0.096701728 | 0.26674264 | -0.3625282 | 0.71695736 | 0.90019386 |
| CD2BP2 | 50.0266194 | -0.096469877 | 0.29374671 | -0.3284118 | 0.74260035 | 0.90850686 |
| GOPC | 103.063353 | -0.096457773 | 0.23204483 | -0.4156859 | 0.67763983 | 0.88143805 |
| SEC11A | 224.259228 | -0.096243457 | 0.1766052 | -0.5449639 | 0.58577836 | 0.83933187 |
| ZBTB1 | 229.500641 | -0.095915577 | 0.30422224 | -0.3152813 | 0.7525481 | 0.91163598 |
| FLI1 | 1151.04673 | -0.095710143 | 0.33593118 | -0.28491 | 0.77571312 | 0.91811196 |
| CCDC122 | 70.7683725 | -0.09514693 | 0.46433569 | -0.2049098 | 0.8376426 | 0.94358503 |
| CASP1 | 41.602884 | -0.094959347 | 0.40787859 | -0.2328128 | 0.81590679 | 0.93550282 |
| MRPS35 | 70.9834505 | -0.094852161 | 0.32497119 | -0.2918787 | 0.77037939 | 0.91695334 |
| PPP4R2 | 273.52514 | -0.094781216 | 0.19598625 | -0.4836116 | 0.62866157 | 0.86052754 |
| TOLLIP | 59.8450496 | -0.094714284 | 0.26048842 | -0.3636027 | 0.71615474 | 0.9001716 |
| ECI1 | 77.1361958 | -0.094660605 | 0.21698694 | -0.4362502 | 0.66265518 | 0.87493354 |
| DDX50 | 136.690677 | -0.094643359 | 0.26682757 | -0.3546986 | 0.72281542 | 0.90217367 |
| AGPS | 196.250751 | -0.094496591 | 0.19501044 | -0.484572 | 0.62798001 | 0.86052754 |
| NSD3 | 429.350277 | -0.094388128 | 0.21616156 | -0.4366555 | 0.66236122 | 0.87491556 |
| SMARCC2 | 152.21048 | -0.09409438 | 0.25696519 | -0.3661756 | 0.71423406 | 0.89971941 |
| DAZAP2 | 233.124638 | -0.093930266 | 0.29243721 | -0.3211981 | 0.74806029 | 0.91065347 |
| NOC3L | 57.2839574 | -0.093925817 | 0.32721901 | -0.2870427 | 0.77407965 | 0.91726032 |
| IFT122 | 101.330311 | -0.09382816 | 0.26782727 | -0.3503309 | 0.7260904 | 0.9026831 |
| HSD17B4 | 194.364548 | -0.093727103 | 0.31606175 | -0.2965468 | 0.76681252 | 0.91603312 |
| ZNF562 | 131.72197 | -0.093716952 | 0.23248187 | -0.4031151 | 0.68686356 | 0.88671976 |
| IL7 | 82.423345 | -0.093663022 | 0.45669306 | -0.2050897 | 0.83750208 | 0.94358503 |
| PDHB | 49.4147041 | -0.093624736 | 0.2899597 | -0.3228888 | 0.74677947 | 0.91031794 |
| INPP5F | 153.497558 | -0.093458359 | 0.31381417 | -0.2978143 | 0.76584487 | 0.91600937 |
| NENF | 77.8444689 | -0.093361735 | 0.36639596 | -0.254811 | 0.79886905 | 0.92962819 |
| CTTN | 290.371679 | -0.093059681 | 0.19860498 | -0.4685667 | 0.63937938 | 0.86388557 |
| RNF187 | 45.6428121 | -0.092880111 | 0.36692691 | -0.2531297 | 0.80016796 | 0.9298524 |
| TBC1D22A | 933.345219 | -0.092803064 | 0.2668453 | -0.3477785 | 0.72800653 | 0.9037084 |
| MCCC2 | 118.622421 | -0.092787879 | 0.24405103 | -0.3801987 | 0.70379794 | 0.89446478 |
| NUP58 | 176.384333 | -0.092786597 | 0.30691414 | -0.302321 | 0.76240736 | 0.91477008 |
| RNF216 | 372.927707 | -0.092472873 | 0.19636412 | -0.4709255 | 0.63769393 | 0.86237803 |
| MIPEP | 100.448641 | -0.092388877 | 0.33350538 | -0.2770236 | 0.78176195 | 0.92124971 |
| HIF1AN | 48.2315301 | -0.092084073 | 0.26683799 | -0.3450936 | 0.73002404 | 0.90426442 |
| ACOT7 | 64.6217966 | -0.092040629 | 0.3016411 | -0.3051329 | 0.76026492 | 0.91455455 |
| ANP32A | 151.855888 | -0.091945015 | 0.25413472 | -0.3617964 | 0.71750422 | 0.90033983 |
| TMOD3 | 533.028115 | -0.091884452 | 0.19876595 | -0.4622746 | 0.64388441 | 0.86517342 |
| RAB10 | 572.663407 | -0.091848175 | 0.23753925 | -0.3866653 | 0.69900404 | 0.89364164 |
| WDR45B | 201.424724 | -0.091709807 | 0.29618672 | -0.3096351 | 0.75683845 | 0.91345267 |
| DBN1 | 56.4640315 | -0.091514664 | 0.35848147 | -0.2552842 | 0.79850358 | 0.92959949 |
| IDH3A | 57.6522784 | -0.091462552 | 0.32826039 | -0.278628 | 0.78053029 | 0.92046106 |
| SUB1 | 288.068997 | -0.090840321 | 0.36051766 | -0.2519719 | 0.80106278 | 0.93019678 |
| IRAK2 | 242.752468 | -0.090766884 | 0.36248347 | -0.2504028 | 0.80227585 | 0.93081002 |
| RHEB | 253.028735 | -0.089998702 | 0.2038553 | -0.4414833 | 0.65886318 | 0.87335081 |
| TAF13 | 59.6976506 | -0.089975285 | 0.30102088 | -0.2989005 | 0.76501598 | 0.9157586 |
| AFAP1L2 | 94.6744339 | -0.089841776 | 0.51209921 | -0.1754382 | 0.86073531 | 0.95284051 |
| SELENOO | 87.453018 | -0.089811252 | 0.23870925 | -0.376237 | 0.70674071 | 0.89551861 |
| DLGAP4-AS1 | 53.2905905 | -0.089794286 | 0.31857937 | -0.2818584 | 0.77805206 | 0.91979327 |
| WDR26 | 315.584259 | -0.089583357 | 0.20870845 | -0.4292273 | 0.66775784 | 0.87587657 |
| SF3A1 | 62.4320708 | -0.089567319 | 0.30175644 | -0.2968199 | 0.766604 | 0.91603312 |
| CALM3 | 89.8665697 | -0.089531638 | 0.41542559 | -0.2155179 | 0.82936358 | 0.94063377 |
| CC2D1B | 47.2737178 | -0.089486872 | 0.40141851 | -0.2229266 | 0.82359262 | 0.93845094 |
| TBCB | 66.0067355 | -0.089248301 | 0.28458939 | -0.3136038 | 0.75382201 | 0.91250377 |
| UBE2I | 160.416478 | -0.089209367 | 0.23334712 | -0.3823033 | 0.70223643 | 0.89443605 |
| CDK11B | 81.9564948 | -0.089149625 | 0.3191752 | -0.2793125 | 0.78000501 | 0.9203826 |
| ZNF516 | 165.768552 | -0.089040024 | 0.22243668 | -0.4002938 | 0.68894013 | 0.88765832 |
| STT3A | 60.7993922 | -0.088979386 | 0.23251305 | -0.3826856 | 0.70195293 | 0.89443605 |
| FOXJ3 | 494.400739 | -0.088953735 | 0.16832395 | -0.5284675 | 0.59717491 | 0.84764654 |
| CRKL | 57.721499 | -0.088928211 | 0.2674761 | -0.3324716 | 0.73953317 | 0.90826918 |
| CTCF | 191.656159 | -0.088834169 | 0.22751149 | -0.3904601 | 0.69619632 | 0.8920567 |
| MMRN2 | 548.468215 | -0.088712713 | 0.49980868 | -0.1774933 | 0.85912089 | 0.95247071 |
| SAT1 | 602.266421 | -0.08866769 | 0.60794357 | -0.1458486 | 0.88404095 | 0.96178798 |
| RAB5A | 254.103993 | -0.088658429 | 0.17178309 | -0.5161068 | 0.60577979 | 0.85156118 |
| CDK16 | 44.6365892 | -0.088507684 | 0.27611682 | -0.3205443 | 0.74855572 | 0.91065347 |
| BABAM2 | 790.913551 | -0.088506995 | 0.24271252 | -0.3646577 | 0.71536692 | 0.9001716 |
| METTL9 | 190.25733 | -0.088467034 | 0.18708571 | -0.472869 | 0.63630664 | 0.86207417 |
| LINC02256 | 188.338914 | -0.088445842 | 0.26503361 | -0.3337156 | 0.7385942 | 0.90802385 |
| ARRDC2 | 92.3984116 | -0.08834722 | 0.31506384 | -0.2804105 | 0.77916256 | 0.91979327 |
| CA8 | 308.113364 | -0.088310152 | 0.65546317 | -0.1347294 | 0.89282584 | 0.96486976 |
| DCAF12 | 71.3250543 | -0.088256318 | 0.2127435 | -0.4148485 | 0.67825282 | 0.88164367 |
| PTPN1 | 336.187722 | -0.087661137 | 0.28872685 | -0.3036127 | 0.76142298 | 0.91477008 |
| CMTM3 | 49.2607278 | -0.087548318 | 0.27397586 | -0.3195476 | 0.74931133 | 0.91075944 |
| ZNF638 | 1022.6906 | -0.087432896 | 0.19846275 | -0.4405507 | 0.65953833 | 0.87335081 |
| BLCAP | 155.49296 | -0.087387149 | 0.24536163 | -0.3561565 | 0.72172334 | 0.90162538 |
| RAP1B | 1235.50726 | -0.087301956 | 0.30377551 | -0.2873897 | 0.77381394 | 0.91726032 |
| RABEP2 | 74.4541975 | -0.087156396 | 0.38329653 | -0.2273863 | 0.82012334 | 0.93710409 |
| PHF21A | 770.782247 | -0.087044396 | 0.29927938 | -0.2908466 | 0.77116863 | 0.91695334 |
| DYNC1H1 | 423.470011 | -0.086946987 | 0.25608781 | -0.3395202 | 0.73421787 | 0.9055183 |
| SLC25A12 | 153.804825 | -0.086872252 | 0.19975595 | -0.4348919 | 0.66364087 | 0.8755434 |
| NDUFAB1 | 75.3267069 | -0.086850234 | 0.34611858 | -0.2509262 | 0.80187114 | 0.93060722 |
| ZNF146 | 120.661114 | -0.086834586 | 0.20928491 | -0.4149109 | 0.67820714 | 0.88164367 |
| PSMD13 | 130.836303 | -0.086686414 | 0.22529651 | -0.3847659 | 0.70041086 | 0.89442292 |
| SRSF3 | 257.444374 | -0.086617952 | 0.25332069 | -0.34193 | 0.73240355 | 0.90471573 |
| TTC27 | 183.424016 | -0.086528392 | 0.26295953 | -0.3290559 | 0.74211341 | 0.90850686 |
| ZNF335 | 50.0765841 | -0.086508644 | 0.37305802 | -0.2318906 | 0.816623 | 0.93550282 |
| MRPS18C | 46.8844484 | -0.086476366 | 0.28052109 | -0.3082705 | 0.75787654 | 0.91402148 |
| ANAPC1 | 191.203034 | -0.0863028 | 0.17924085 | -0.4814907 | 0.6301678 | 0.86097142 |
| BIRC3 | 102.20359 | -0.086096012 | 0.65350782 | -0.1317444 | 0.89518645 | 0.96575575 |
| VPS41 | 228.1419 | -0.086041923 | 0.25599648 | -0.3361059 | 0.73679102 | 0.90708057 |
| MGAT4A | 488.303578 | -0.086024791 | 0.48869166 | -0.1760308 | 0.86026973 | 0.95267061 |
| RARS | 130.127157 | -0.086005664 | 0.21653154 | -0.3971969 | 0.69122224 | 0.88837311 |

| | | | | | | |
|---|---|---|---|---|---|---|
| LINC00843 | 53.686164 | -0.085939786 | 0.26313455 | -0.3266002 | 0.74397032 | 0.90877879 |
| SNRPA1 | 98.1113872 | -0.085881532 | 0.24815334 | -0.3460825 | 0.7292807 | 0.90408479 |
| TRIM47 | 60.2178054 | -0.085802 | 0.42055822 | -0.2040193 | 0.83833841 | 0.94362079 |
| MBIP | 47.0364759 | -0.085741887 | 0.21493421 | -0.3989215 | 0.68995102 | 0.88817873 |
| SYNJ1 | 180.062139 | -0.085694177 | 0.21015225 | -0.4077719 | 0.68344116 | 0.88560554 |
| EXOSC8 | 50.8873545 | -0.08558014 | 0.27109806 | -0.3156796 | 0.75224568 | 0.91163598 |
| ITGB8 | 54.109278 | -0.085313875 | 0.5068896 | -0.1683086 | 0.86634051 | 0.95510194 |
| SLC25A33 | 44.9043751 | -0.084888006 | 0.28142575 | -0.3016355 | 0.76292992 | 0.91477008 |
| TASP1 | 217.52131 | -0.084861995 | 0.29318162 | -0.289452 | 0.77223553 | 0.91702269 |
| HNRNPC | 641.140617 | -0.084621994 | 0.22865615 | -0.3700841 | 0.71131986 | 0.89747106 |
| SREBF2 | 277.917078 | -0.084571444 | 0.29215744 | -0.2894722 | 0.77222008 | 0.91702269 |
| RUFY1 | 142.064555 | -0.084557918 | 0.25676216 | -0.3293239 | 0.74191088 | 0.90850686 |
| ZSWIM6 | 1624.84091 | -0.084465125 | 0.28305496 | -0.2984054 | 0.76539378 | 0.91580893 |
| SZRD1 | 99.180183 | -0.084291571 | 0.19584089 | -0.4304084 | 0.66689855 | 0.87587657 |
| BTBD1 | 168.06332 | -0.083991639 | 0.16635589 | -0.5048913 | 0.6136352 | 0.85573702 |
| ASB13 | 37.9325011 | -0.083927288 | 0.46098335 | -0.1820614 | 0.85553452 | 0.95134391 |
| MLH1 | 67.5980288 | -0.083639565 | 0.24545811 | -0.3407488 | 0.73329267 | 0.90508002 |
| LMTK2 | 169.017566 | -0.083585707 | 0.24991076 | -0.3344622 | 0.7380308 | 0.9078343 |
| SORBS3 | 87.7907876 | -0.083512673 | 0.30332883 | -0.2753206 | 0.78306995 | 0.92204994 |
| USP1 | 57.9982478 | -0.082797218 | 0.30014824 | -0.2758544 | 0.78265989 | 0.92178175 |
| TMEM123 | 300.248381 | -0.082774179 | 0.3200882 | -0.258598 | 0.7959454 | 0.92847059 |
| SGTA | 64.5353092 | -0.082760564 | 0.21713351 | -0.3811506 | 0.70309152 | 0.8944409 |
| USPL1 | 131.493066 | -0.082740706 | 0.24168401 | -0.3423508 | 0.73208693 | 0.90466134 |
| PSMB7 | 182.814239 | -0.082712237 | 0.2278192 | -0.3630609 | 0.71655942 | 0.9001716 |
| HNRNPD | 382.884629 | -0.082654805 | 0.16461991 | -0.5020948 | 0.61560081 | 0.85714845 |
| METTL6 | 47.24576 | -0.082573005 | 0.27902582 | -0.2959332 | 0.76728109 | 0.91611154 |
| RAB3D | 48.0260568 | -0.082548783 | 0.3254266 | -0.2536633 | 0.79975569 | 0.9298524 |
| WARS2 | 63.8547371 | -0.082479932 | 0.34016718 | -0.2424688 | 0.80841692 | 0.93264848 |
| ZMYND8 | 299.821175 | -0.082408329 | 0.27706236 | -0.297436 | 0.76613364 | 0.91600937 |
| PITRM1 | 66.2618288 | -0.082345122 | 0.28311439 | -0.2908546 | 0.77116252 | 0.91695334 |
| MAVS | 139.75548 | -0.082276631 | 0.24481782 | -0.3360729 | 0.7368159 | 0.90708057 |
| GSTCD | 92.9915775 | -0.082093562 | 0.26037105 | -0.3152945 | 0.75253806 | 0.91163598 |
| MFSD10 | 66.6687238 | -0.081928778 | 0.37940636 | -0.2159394 | 0.82903499 | 0.94063377 |
| ZNF254 | 214.317274 | -0.081777023 | 0.14427462 | -0.566815 | 0.57083985 | 0.83368112 |
| CASP8 | 60.3270909 | -0.081697965 | 0.39023191 | -0.2093575 | 0.83416919 | 0.9432647 |
| JKAMP | 72.3696206 | -0.081576581 | 0.28491686 | -0.2863171 | 0.77463523 | 0.91747577 |
| HSPH1 | 172.984849 | -0.081526839 | 0.56929369 | -0.143207 | 0.8861267 | 0.9627377 |
| SREK1IP1 | 117.461437 | -0.081496189 | 0.17947526 | -0.4540803 | 0.649771 | 0.86766991 |
| ZUP1 | 52.8261299 | -0.081445655 | 0.26982987 | -0.3018408 | 0.76277345 | 0.91477008 |
| CEP63 | 108.930744 | -0.081393667 | 0.17990026 | -0.4524377 | 0.65095367 | 0.86839808 |
| SART3 | 78.6223528 | -0.081376586 | 0.17944506 | -0.4534903 | 0.65019576 | 0.86766991 |
| BTBD8 | 55.6299258 | -0.081215226 | 0.4536928 | -0.1790093 | 0.85793041 | 0.9521829 |
| MDH2 | 74.2357661 | -0.081109798 | 0.28987438 | -0.2798102 | 0.77962315 | 0.92018697 |
| ELP3 | 141.447074 | -0.081036849 | 0.17151027 | -0.4724898 | 0.63657725 | 0.86207417 |
| NT5DC1 | 199.276338 | -0.080955027 | 0.18116983 | -0.4468461 | 0.65498621 | 0.87093796 |
| JARID2 | 525.625037 | -0.080953445 | 0.25830187 | -0.3134063 | 0.75397197 | 0.9125503 |
| TRIP11 | 311.247171 | -0.080899392 | 0.17119146 | -0.4725667 | 0.63652232 | 0.86207417 |
| ESM1 | 276.138749 | -0.080690302 | 0.70734757 | -0.1140745 | 0.90917876 | 0.96940376 |
| RAP1GDS1 | 373.547369 | -0.080617909 | 0.19844871 | -0.4062405 | 0.68456588 | 0.88636116 |
| AL139260.1 | 79.7735558 | -0.080321218 | 0.64407005 | -0.1247088 | 0.90075407 | 0.96744565 |
| SPART | 214.973324 | -0.079915788 | 0.2429522 | -0.3289363 | 0.74220387 | 0.90850686 |
| ZNF140 | 49.2773812 | -0.079311088 | 0.30095474 | -0.2635316 | 0.79214086 | 0.92733744 |
| ZNF329 | 41.2656666 | -0.079288654 | 0.31096457 | -0.2549765 | 0.79874126 | 0.92962819 |
| CWC15 | 74.6714714 | -0.07925008 | 0.24104164 | -0.3287817 | 0.7423207 | 0.90850686 |
| CRYBG1 | 101.337101 | -0.079094868 | 0.43105084 | -0.1834931 | 0.85441111 | 0.95098559 |
| RFWD3 | 69.8234085 | -0.079069395 | 0.29884365 | -0.2645845 | 0.79132957 | 0.92665282 |
| CUX1 | 849.704621 | -0.079045715 | 0.30615153 | -0.2581915 | 0.79625914 | 0.92857183 |
| ZNF814 | 108.678833 | -0.078987367 | 0.41175451 | -0.1918312 | 0.84787442 | 0.94731791 |
| DAPK3 | 102.096721 | -0.078923028 | 0.27496129 | -0.2870332 | 0.77408687 | 0.91726032 |
| PPP1R7 | 66.7312209 | -0.078525785 | 0.22219986 | -0.3534016 | 0.72378739 | 0.90258339 |
| LIX1L | 102.391007 | -0.078504828 | 0.25085164 | -0.3129532 | 0.75431621 | 0.91269696 |
| JMJD6 | 46.1234189 | -0.078358436 | 0.26594309 | -0.2946436 | 0.76826613 | 0.91631735 |
| TRAM1 | 246.800081 | -0.07816534 | 0.23783693 | -0.328651 | 0.74241952 | 0.90850686 |
| FBXL15 | 46.3465224 | -0.078074512 | 0.40404749 | -0.193231 | 0.84677804 | 0.94719648 |
| KIAA0100 | 111.152445 | -0.077997127 | 0.24638477 | -0.3165663 | 0.75157268 | 0.9112603 |
| RNF44 | 46.9073183 | -0.077864911 | 0.30078279 | -0.2588742 | 0.79573229 | 0.92847059 |
| RAB35 | 74.7822748 | -0.077681754 | 0.29217355 | -0.2658754 | 0.79033518 | 0.92613348 |
| USP5 | 39.4464204 | -0.077481122 | 0.37675109 | -0.205656 | 0.83705963 | 0.94358503 |
| PGM3 | 55.2834336 | -0.077425301 | 0.31290498 | -0.2474403 | 0.80456749 | 0.93109287 |
| 9-Sep | 427.820085 | -0.07724807 | 0.30509403 | -0.2531943 | 0.80011807 | 0.9298524 |
| BAG4 | 53.3386898 | -0.076891289 | 0.22774075 | -0.3376264 | 0.73564474 | 0.90649363 |
| UHRF1BP1L | 128.465383 | -0.076889404 | 0.22327217 | -0.3443752 | 0.73056412 | 0.90429739 |
| MAF1 | 68.009361 | -0.076709809 | 0.42257242 | -0.1815306 | 0.85595115 | 0.95134391 |
| DNAJC2 | 93.8948813 | -0.076694502 | 0.22536952 | -0.3403056 | 0.73362643 | 0.90519678 |
| PHF20 | 383.686274 | -0.076679548 | 0.18587029 | -0.4125433 | 0.67994123 | 0.88274775 |
| ACSL4 | 335.135759 | -0.076630128 | 0.25305438 | -0.3028208 | 0.76202644 | 0.91477008 |
| ENTPD3-AS1 | 47.0598637 | -0.076523411 | 0.36832307 | -0.2077617 | 0.83541507 | 0.9432647 |
| MAST2 | 155.05868 | -0.076339912 | 0.18090483 | -0.4219894 | 0.67303278 | 0.87837755 |
| BHLHB9 | 44.0491977 | -0.076125416 | 0.41082451 | -0.1852991 | 0.85299444 | 0.95044301 |
| OXSR1 | 223.771027 | -0.076034188 | 0.26169253 | -0.2905478 | 0.77139719 | 0.91695863 |
| WSB1 | 1179.11897 | -0.075891582 | 0.3456989 | -0.2195309 | 0.82623653 | 0.93977669 |
| CPNE3 | 110.689495 | -0.075886304 | 0.28617852 | -0.2651712 | 0.79087757 | 0.92638866 |
| TCN2 | 164.324295 | -0.075852046 | 0.24840353 | -0.3053582 | 0.76009338 | 0.91455455 |
| NDUFB9 | 110.789529 | -0.075792701 | 0.29785628 | -0.2544606 | 0.7991397 | 0.92963087 |
| PPP1R2 | 170.118019 | -0.075786147 | 0.28576007 | -0.265209 | 0.79084845 | 0.92638866 |
| CUTC | 87.3304408 | -0.075605339 | 0.24944288 | -0.3030968 | 0.76181609 | 0.91477008 |
| KAT6A | 1003.27042 | -0.075324255 | 0.21169501 | -0.355815 | 0.72197913 | 0.90162538 |
| ADGRL4 | 1987.63584 | -0.075035035 | 0.25022731 | -0.2998675 | 0.76427823 | 0.9154077 |
| HERC2 | 520.759004 | -0.074915987 | 0.16480209 | -0.4545815 | 0.64941031 | 0.86766991 |
| TRIOBP | 283.237824 | -0.074800314 | 0.24025835 | -0.3113328 | 0.7555476 | 0.91343453 |
| EZR | 92.4461269 | -0.073942518 | 0.33038707 | -0.2238057 | 0.82290848 | 0.93812484 |
| PPIC | 70.8760082 | -0.073771715 | 0.49575916 | -0.1488056 | 0.88170707 | 0.96059232 |
| POLR2A | 187.266644 | -0.07365333 | 0.25163034 | -0.2927045 | 0.76974804 | 0.91695334 |
| ATP5IF1 | 165.511432 | -0.073594478 | 0.4480782 | -0.1642447 | 0.8695385 | 0.95673577 |
| NUS1 | 128.965532 | -0.073474157 | 0.21408023 | -0.3432085 | 0.73144161 | 0.90454688 |
| DMXL2 | 124.924486 | -0.073472626 | 0.29038491 | -0.2530181 | 0.80025426 | 0.9298524 |
| INTS4 | 117.727002 | -0.073384729 | 0.20836707 | -0.3521897 | 0.72469602 | 0.9026193 |
| MAX | 308.416828 | -0.073138762 | 0.27070572 | -0.2701781 | 0.78702323 | 0.92401864 |
| PCNP | 230.735992 | -0.073089088 | 0.19220125 | -0.3802737 | 0.70374223 | 0.89446478 |
| CAV1 | 244.664818 | -0.072977784 | 0.53390789 | -0.1366861 | 0.89127892 | 0.96448584 |
| RAB18 | 197.054046 | -0.07280852 | 0.15179839 | -0.4796396 | 0.63148369 | 0.86109499 |
| CDC42 | 391.527587 | -0.072538246 | 0.21206605 | -0.342055 | 0.73230953 | 0.90471573 |
| NIFK-AS1 | 57.6717058 | -0.072447728 | 0.39901945 | -0.1815644 | 0.85592459 | 0.95134391 |
| UBE2E1 | 680.036747 | -0.072433782 | 0.26588107 | -0.2724293 | 0.78529198 | 0.92328222 |
| SEC16A | 115.744434 | -0.07219123 | 0.20062316 | -0.359835 | 0.71897055 | 0.90051656 |
| UNC50 | 78.3945559 | -0.072181873 | 0.25168883 | -0.2867901 | 0.77427302 | 0.91726032 |
| COQ4 | 65.8450334 | -0.072132479 | 0.30813704 | -0.2340922 | 0.8149134 | 0.93513341 |
| PHTF1 | 118.271992 | -0.07190645 | 0.25632303 | -0.2805306 | 0.77907046 | 0.91979327 |
| UTP18 | 68.6449997 | -0.07167737 | 0.35410946 | -0.2024159 | 0.83959163 | 0.94376667 |
| NFRKB | 55.5044219 | -0.071603423 | 0.24339764 | -0.2941829 | 0.76861815 | 0.91631735 |
| PRKACB | 145.486604 | -0.071526715 | 0.29836999 | -0.2397249 | 0.81054353 | 0.93351089 |
| SHE | 426.476289 | -0.071489166 | 0.35967992 | -0.1987577 | 0.84245226 | 0.94471448 |
| BRIX1 | 46.3985421 | -0.07120117 | 0.28661431 | -0.2484216 | 0.80380826 | 0.93081002 |

| | | | | | | |
|---|---|---|---|---|---|---|
| DCHS1 | 94.5621529 | -0.071183309 | 0.36368096 | -0.1957301 | 0.84482143 | 0.94584399 |
| PIBF1 | 517.273951 | -0.071091101 | 0.19586836 | -0.3629535 | 0.71663964 | 0.9001716 |
| GJA5 | 352.425029 | -0.070920814 | 0.68732905 | -0.1031832 | 0.91781757 | 0.97233474 |
| DEAF1 | 107.956006 | -0.070873304 | 0.20607648 | -0.3439175 | 0.73090835 | 0.9042974 |
| REXO1 | 55.8664747 | -0.070727394 | 0.21619988 | -0.3271389 | 0.74356282 | 0.90856581 |
| ARRB1 | 443.60974 | -0.070687231 | 0.26015827 | -0.2717086 | 0.78584611 | 0.92353565 |
| SUPT7L | 47.8961857 | -0.070655686 | 0.30447902 | -0.2320544 | 0.81649579 | 0.93550282 |
| TUT7 | 196.149429 | -0.070336173 | 0.17577028 | -0.4001596 | 0.68903894 | 0.88765832 |
| QARS | 39.4079787 | -0.070271559 | 0.30626063 | -0.2294502 | 0.81851903 | 0.93612037 |
| R3HCC1L | 205.306617 | -0.070203236 | 0.18584099 | -0.3777597 | 0.70560912 | 0.89491621 |
| NEK11 | 62.845171 | -0.069789579 | 0.43075576 | -0.1620166 | 0.87129279 | 0.95709072 |
| RHBDD1 | 185.192868 | -0.069615757 | 0.29184711 | -0.238535 | 0.81146616 | 0.93351089 |
| MLX | 40.3679914 | -0.069502579 | 0.29993197 | -0.2317278 | 0.81674943 | 0.93550282 |
| ZNF789 | 67.1696774 | -0.069034286 | 0.24901903 | -0.2772249 | 0.78160739 | 0.92120024 |
| FAF2 | 179.302055 | -0.068982152 | 0.17474324 | -0.3947629 | 0.69301787 | 0.88975484 |
| ZFAND5 | 229.696896 | -0.06895005 | 0.29815544 | -0.2312554 | 0.81711641 | 0.93550282 |
| VKORC1L1 | 169.781874 | -0.068889131 | 0.29221579 | -0.2357475 | 0.81362864 | 0.93460727 |
| GLMN | 90.2182153 | -0.068867329 | 0.21479304 | -0.3206218 | 0.74849703 | 0.91065347 |
| CBX5 | 130.922026 | -0.068863753 | 0.24185029 | -0.2847371 | 0.77584556 | 0.91811196 |
| SDCBP | 704.725832 | -0.068854739 | 0.37698243 | -0.1826471 | 0.85507494 | 0.95134391 |
| SLF2 | 184.34409 | -0.06874399 | 0.1933879 | -0.355472 | 0.722236 | 0.90163848 |
| DERL2 | 56.8607755 | -0.068715801 | 0.32110327 | -0.2139991 | 0.83054778 | 0.94075329 |
| TMEM220 | 48.7203453 | -0.068551409 | 0.32758513 | -0.2092629 | 0.83424302 | 0.9432647 |
| RBM12B | 88.449764 | -0.068154068 | 0.19655467 | -0.3467436 | 0.72878398 | 0.9038556 |
| ZNF354C | 44.3613981 | -0.067793497 | 0.28328929 | -0.2393084 | 0.81086648 | 0.93351089 |
| RNF150 | 63.6399047 | -0.067770192 | 0.6454468 | -0.1049973 | 0.91637793 | 0.97233474 |
| KBTBD2 | 233.349611 | -0.067594066 | 0.22770754 | -0.296846 | 0.76658411 | 0.91603312 |
| SHMT1 | 51.7171157 | -0.067585064 | 0.26719404 | -0.2529438 | 0.80031168 | 0.9298524 |
| LSM6 | 69.9733949 | -0.06755164 | 0.30553581 | -0.2210924 | 0.8250205 | 0.93870172 |
| ZNF490 | 58.3143895 | -0.067530929 | 0.28166886 | -0.2397529 | 0.81052181 | 0.93351089 |
| STAU1 | 289.805837 | -0.06752779 | 0.14936515 | -0.4520987 | 0.65119789 | 0.86853188 |
| FAM96A | 44.0958198 | -0.067514714 | 0.23310349 | -0.2896341 | 0.77209619 | 0.91702269 |
| ZNF24 | 115.727728 | -0.067482253 | 0.27172556 | -0.2483471 | 0.80386587 | 0.93081002 |
| PLRG1 | 51.2995263 | -0.067401528 | 0.23612143 | -0.2854528 | 0.77529725 | 0.91786131 |
| ZNF677 | 42.4596604 | -0.067363321 | 0.24818288 | -0.2714261 | 0.7860633 | 0.92365824 |
| PUM1 | 656.244772 | -0.067284473 | 0.16683469 | -0.4033003 | 0.68672736 | 0.88671976 |
| ZNF780B | 114.621391 | -0.067212601 | 0.30784737 | -0.2183309 | 0.82717128 | 0.94018985 |
| DPY19L3 | 114.980166 | -0.067116364 | 0.25600861 | -0.2621645 | 0.79319464 | 0.92777469 |
| UBE4A | 154.207203 | -0.067058164 | 0.15461271 | -0.433717 | 0.66449395 | 0.8755434 |
| NSUN4 | 77.4206928 | -0.066946012 | 0.30278167 | -0.2211033 | 0.82501203 | 0.93870172 |
| RMDN1 | 131.547009 | -0.066869129 | 0.29939847 | -0.2233449 | 0.82326706 | 0.9382722 |
| TBC1D14 | 92.2819173 | -0.066671412 | 0.23160125 | -0.2878716 | 0.77344506 | 0.91726032 |
| HS2ST1 | 191.581491 | -0.066631118 | 0.1647072 | -0.4045428 | 0.68581359 | 0.88671976 |
| NIPSNAP2 | 135.026608 | -0.066418106 | 0.20618694 | -0.3221257 | 0.7473575 | 0.91055362 |
| CDKN1B | 93.0518556 | -0.066333765 | 0.43629444 | -0.152039 | 0.87915618 | 0.95972986 |
| WDR82 | 157.330145 | -0.066147261 | 0.2039465 | -0.3243363 | 0.74568341 | 0.91005223 |
| CNRIP1 | 150.453577 | -0.066133504 | 0.17038089 | -0.388151 | 0.69790432 | 0.89303379 |
| SPEN | 390.699236 | -0.066024029 | 0.19313656 | -0.3418515 | 0.73246263 | 0.90471573 |
| ZNF827 | 127.195926 | -0.065965309 | 0.35521369 | -0.185706 | 0.85267533 | 0.95044301 |
| AC113383.1 | 83.8082118 | -0.065752959 | 1.30349973 | -0.0504434 | 0.95976905 | 0.98739605 |
| GPAA1 | 54.4320038 | -0.065688296 | 0.47302921 | -0.1388673 | 0.889555 | 0.96398417 |
| STAT6 | 183.343341 | -0.065635261 | 0.23887624 | -0.2747668 | 0.78349541 | 0.92222999 |
| HUWE1 | 412.810845 | -0.065378437 | 0.16157451 | -0.4046334 | 0.68574703 | 0.88671976 |
| MTOR | 187.355071 | -0.065228366 | 0.21811688 | -0.2990523 | 0.76490011 | 0.91575385 |
| ANKRD26 | 214.121413 | -0.064860275 | 0.2707321 | -0.2395736 | 0.8106608 | 0.93351089 |
| CD59 | 98.0791046 | -0.064840604 | 0.54036054 | -0.1199951 | 0.90448705 | 0.96889875 |
| SLC25A13 | 449.671914 | -0.064730974 | 0.22669794 | -0.2855384 | 0.77523167 | 0.91786131 |
| MALSU1 | 60.5944723 | -0.064681907 | 0.2368659 | -0.273074 | 0.78479637 | 0.92309741 |
| SUDS3 | 66.5202399 | -0.064608713 | 0.22292215 | -0.2898263 | 0.77194909 | 0.91702269 |
| TMEM126B | 54.7623043 | -0.064581944 | 0.27824162 | -0.2321074 | 0.81645458 | 0.93550282 |
| SERTAD1 | 78.6096734 | -0.064501626 | 0.52339576 | -0.1232368 | 0.90191957 | 0.96801711 |
| PARD3B | 600.046576 | -0.064388073 | 0.21763417 | -0.2958546 | 0.76734111 | 0.91611154 |
| ZNF160 | 233.970597 | -0.064024807 | 0.34060201 | -0.1879754 | 0.85089592 | 0.9493975 |
| AC092683.1 | 161.00787 | -0.063894901 | 0.37108529 | -0.1721839 | 0.86329298 | 0.9538653 |
| FDPS | 268.306868 | -0.063881133 | 0.29852286 | -0.2139908 | 0.83055426 | 0.94075329 |
| MCPH1 | 458.274281 | -0.063859381 | 0.16871396 | -0.3785068 | 0.70505414 | 0.89448961 |
| ANKRD13A | 110.25153 | -0.063419908 | 0.28277523 | -0.2242767 | 0.82254197 | 0.93793807 |
| SCUBE1 | 77.6532964 | -0.063367669 | 0.50298833 | -0.1259824 | 0.89974587 | 0.96723863 |
| ZNF43 | 176.479387 | -0.063282973 | 0.22459968 | -0.281759 | 0.77812835 | 0.91979327 |
| DIS3 | 104.786066 | -0.06303332 | 0.17522372 | -0.3597305 | 0.71904867 | 0.90051656 |
| UBE3C | 477.523602 | -0.062994555 | 0.17686887 | -0.3561653 | 0.72171677 | 0.90162538 |
| CAMTA1 | 571.060946 | -0.062926883 | 0.26369756 | -0.2386328 | 0.81139034 | 0.93351089 |
| ISCA1 | 55.3783497 | -0.062657845 | 0.26530339 | -0.2361743 | 0.81329742 | 0.9344584 |
| KIF13B | 334.267176 | -0.062550956 | 0.20756924 | -0.3013498 | 0.76314774 | 0.91477008 |
| ATG13 | 81.9928612 | -0.062145227 | 0.29014366 | -0.2141878 | 0.83040063 | 0.94075329 |
| ERCC3 | 84.5804753 | -0.062113516 | 0.2209934 | -0.281065 | 0.77866053 | 0.91979327 |
| ARAF | 40.6275187 | -0.062064016 | 0.26620796 | -0.2331411 | 0.81565184 | 0.93550282 |
| BRCA1 | 58.6552024 | -0.062047296 | 0.34924793 | -0.1776597 | 0.8589902 | 0.95247071 |
| HACD2 | 92.7363518 | -0.061912687 | 0.24232712 | -0.2554922 | 0.79834298 | 0.92959949 |
| RREB1 | 597.52388 | -0.061824389 | 0.30571275 | -0.2022303 | 0.83973667 | 0.94376667 |
| NOD1 | 268.26122 | -0.061769632 | 0.27425409 | -0.2252278 | 0.8218021 | 0.9375166 |
| SBF2-AS1 | 47.6931067 | -0.061648281 | 0.3166042 | -0.1947172 | 0.84561435 | 0.94647274 |
| TEX2 | 99.9540028 | -0.061311331 | 0.22226407 | -0.275849 | 0.78266401 | 0.92178175 |
| ST3GAL1 | 415.895818 | -0.061300344 | 0.36288073 | -0.168927 | 0.86585408 | 0.95510194 |
| MBTPS1 | 166.811055 | -0.061226782 | 0.16816149 | -0.3640952 | 0.71578696 | 0.9001716 |
| MTMR1 | 95.6415955 | -0.061131834 | 0.2047755 | -0.298531 | 0.76529792 | 0.91580893 |
| YME1L1 | 257.424936 | -0.060838426 | 0.14384822 | -0.4229349 | 0.67234279 | 0.87803719 |
| SLF1 | 132.533041 | -0.060620118 | 0.369275 | -0.1641598 | 0.86960533 | 0.95673577 |
| PAIP2 | 184.080407 | -0.060593398 | 0.27697253 | -0.2187704 | 0.82682888 | 0.9401057 |
| VRK3 | 71.2663552 | -0.060496034 | 0.31630275 | -0.1912599 | 0.84832197 | 0.9475592 |
| AP3B1 | 562.808009 | -0.060331813 | 0.1557532 | -0.3873552 | 0.69849326 | 0.8933683 |
| ACSL5 | 139.397465 | -0.060262531 | 0.32660688 | -0.1845109 | 0.85361267 | 0.9508242 |
| ATL3 | 215.623801 | -0.060234644 | 0.19444634 | -0.3097752 | 0.75673195 | 0.91345267 |
| SLC35C2 | 60.4270323 | -0.060148886 | 0.32718807 | -0.1838358 | 0.85414226 | 0.95094505 |
| RFTN1 | 338.45093 | -0.060120133 | 0.25512618 | -0.2356486 | 0.81370534 | 0.93460727 |
| UGDH | 46.5775314 | -0.060076328 | 0.33286793 | -0.180481 | 0.85677499 | 0.95134391 |
| UPF1 | 82.8738419 | -0.060034815 | 0.2524289 | -0.2378286 | 0.81201402 | 0.93357139 |
| PEX26 | 80.7788059 | -0.060015509 | 0.27838561 | -0.2155841 | 0.82931196 | 0.94063377 |
| ZNF717 | 66.0505315 | -0.059913145 | 0.40137553 | -0.1492696 | 0.88134094 | 0.96059232 |
| DIP2C | 468.702588 | -0.059890867 | 0.412953 | -0.1450307 | 0.88468662 | 0.96204527 |
| ASXL2 | 292.775341 | -0.059754434 | 0.20818125 | -0.2870308 | 0.77408873 | 0.91726032 |
| RBBP4 | 171.690221 | -0.059630293 | 0.13732477 | -0.4342282 | 0.6641227 | 0.8755434 |
| HPCAL1 | 652.569173 | -0.059580738 | 0.2004021 | -0.297306 | 0.76623294 | 0.91600937 |
| MSL1 | 135.184018 | -0.059487687 | 0.22331767 | -0.2663815 | 0.78994544 | 0.92602336 |
| PLA2G4C | 164.518446 | -0.059353945 | 0.52056488 | -0.1140183 | 0.90922325 | 0.96940376 |
| CMC1 | 177.32008 | -0.05928009 | 0.24038567 | -0.2466041 | 0.80521464 | 0.93131751 |
| ACVR1B | 147.73126 | -0.059127186 | 0.36262422 | -0.1630536 | 0.87047623 | 0.95689355 |
| KAT5 | 52.0958606 | -0.058946664 | 0.34395692 | -0.171378 | 0.86392652 | 0.95405003 |
| PWWP2A | 177.268098 | -0.058762872 | 0.25224875 | -0.232956 | 0.81579554 | 0.93550282 |
| RMDN3 | 59.9691164 | -0.058402021 | 0.28483909 | -0.2050351 | 0.83754467 | 0.94358503 |
| SCMH1 | 418.524391 | -0.058031694 | 0.24459198 | -0.2372592 | 0.81245573 | 0.9338847 |
| WBP4 | 87.9559674 | -0.057923598 | 0.22643632 | -0.2558052 | 0.79810122 | 0.92959949 |
| DHRSX | 234.718279 | -0.057772122 | 0.27865067 | -0.2073281 | 0.83575361 | 0.9432647 |

| | | | | | | |
|---|---|---|---|---|---|---|
| NKAIN2 | 239.492504 | -0.057704969 | 1.12301845 | -0.0513838 | 0.95901968 | 0.98725593 |
| ELF2 | 1096.75824 | -0.057628226 | 0.18493894 | -0.3116068 | 0.75533939 | 0.91339475 |
| UGT2B7 | 91.5914092 | -0.057612711 | 0.61822441 | -0.0931906 | 0.92575213 | 0.97614605 |
| TMX1 | 87.4597774 | -0.057532584 | 0.21871991 | -0.2630423 | 0.79251799 | 0.92735129 |
| SLC7A6OS | 74.0690066 | -0.057339683 | 0.27682793 | -0.2071311 | 0.83590745 | 0.9432647 |
| ST8SIA6 | 283.080077 | -0.057278299 | 1.21911576 | -0.0469835 | 0.9625264 | 0.98828315 |
| SLC17A5 | 116.690277 | -0.057224744 | 0.39997284 | -0.1430716 | 0.88623365 | 0.9627377 |
| NCBP1 | 74.2238161 | -0.057032762 | 0.20602762 | -0.276821 | 0.78191759 | 0.92130045 |
| PMS2 | 51.5708215 | -0.056948058 | 0.23517644 | -0.2421503 | 0.80866367 | 0.93280163 |
| LRSAM1 | 93.6346108 | -0.056927513 | 0.35922753 | -0.158472 | 0.87408486 | 0.95804618 |
| AGO2 | 244.52944 | -0.056775393 | 0.19421045 | -0.2923395 | 0.77002704 | 0.91695334 |
| SNF8 | 79.9328757 | -0.056685484 | 0.26675794 | -0.2124978 | 0.83171867 | 0.94161971 |
| ZNF785 | 59.5594189 | -0.056571341 | 0.2493487 | -0.2268764 | 0.82051984 | 0.93711673 |
| SLC5A4-AS1 | 73.4165853 | -0.056425427 | 0.51961426 | -0.108591 | 0.91352691 | 0.97120835 |
| MGST3 | 332.166991 | -0.056294626 | 0.36422545 | -0.1545598 | 0.87716834 | 0.95947981 |
| SIKE1 | 50.6010673 | -0.056073565 | 0.41556276 | -0.134934 | 0.89266402 | 0.96482231 |
| SLC38A9 | 176.309304 | -0.056059035 | 0.1504476 | -0.372615 | 0.70943498 | 0.89727558 |
| EIF3J-DT | 64.4418307 | -0.056040501 | 0.33310338 | -0.1682376 | 0.86639639 | 0.95510194 |
| EP400 | 226.482148 | -0.05601963 | 0.26913041 | -0.2081505 | 0.83511145 | 0.9432647 |
| CNOT8 | 127.993584 | -0.055822772 | 0.28900916 | -0.1931523 | 0.84683973 | 0.94719648 |
| DHX36 | 291.400808 | -0.055767623 | 0.21226232 | -0.2627297 | 0.79275889 | 0.92739757 |
| HLA-DQB1 | 191.704866 | -0.055632358 | 0.62189988 | -0.0894555 | 0.92871993 | 0.97696444 |
| SKI | 274.635157 | -0.055604212 | 0.34004925 | -0.1635181 | 0.87011051 | 0.956876 |
| TTC23 | 176.477874 | -0.055459122 | 0.23286101 | -0.238164 | 0.81175386 | 0.93357139 |
| DHX15 | 303.389254 | -0.055253044 | 0.16585754 | -0.3331356 | 0.73903196 | 0.908116 |
| MRPL45 | 49.8037873 | -0.054940361 | 0.20977414 | -0.2619024 | 0.79339666 | 0.92787835 |
| PITPNC1 | 2856.64086 | -0.054482192 | 0.38165932 | -0.1427509 | 0.88648695 | 0.9627377 |
| E2F4 | 65.9272313 | -0.054350276 | 0.36378065 | -0.149404 | 0.88123487 | 0.96059232 |
| ATRN | 241.904733 | -0.054267526 | 0.19949007 | -0.2720312 | 0.78559801 | 0.92347689 |
| UBE2R2 | 437.931976 | -0.054228246 | 0.20013597 | -0.270957 | 0.78642409 | 0.92382124 |
| ERG | 1489.57578 | -0.053874592 | 0.18256494 | -0.2950982 | 0.76791883 | 0.91627807 |
| OSBPL1A | 291.758874 | -0.05308171 | 0.4014706 | -0.1322182 | 0.89481173 | 0.96575575 |
| LINC01320 | 56.9947194 | -0.052995561 | 0.39897737 | -0.1328285 | 0.89432902 | 0.965729 |
| TRIM44 | 417.366631 | -0.052980066 | 0.21548073 | -0.2458692 | 0.80578352 | 0.93158047 |
| GPM6A | 2209.73796 | -0.052659601 | 0.6511909 | -0.0808666 | 0.93554803 | 0.98010864 |
| L2HGDH | 43.3510231 | -0.052568537 | 0.33360333 | -0.157578 | 0.87478936 | 0.95841496 |
| VRK1 | 66.9828342 | -0.052441244 | 0.28763258 | -0.1823203 | 0.85533141 | 0.95134391 |
| PUS10 | 87.2363653 | -0.052384844 | 0.24489228 | -0.2139097 | 0.83061744 | 0.94075329 |
| RPAIN | 82.6100714 | -0.052251025 | 0.21353947 | -0.2446902 | 0.80669628 | 0.93187644 |
| AZIN1 | 121.562387 | -0.052246563 | 0.26219805 | -0.1992637 | 0.84205645 | 0.94468713 |
| NSMAF | 118.15449 | -0.052101801 | 0.31841418 | -0.163629 | 0.8700232 | 0.956876 |
| SLC31A1 | 54.4707611 | -0.051996193 | 0.31079383 | -0.1673012 | 0.86713303 | 0.95535286 |
| RPA1 | 111.783375 | -0.051712514 | 0.18877195 | -0.2739417 | 0.7841294 | 0.9227108 |
| MED10 | 94.2119784 | -0.051607113 | 0.2741429 | -0.188249 | 0.8506815 | 0.9492877 |
| PEX2 | 50.9473179 | -0.051562609 | 0.31649025 | -0.1629201 | 0.87058138 | 0.95689355 |
| MESD | 125.654269 | -0.051539393 | 0.19886753 | -0.2591644 | 0.79550836 | 0.92847059 |
| HAUS2 | 48.9302021 | -0.051444144 | 0.22235927 | -0.231356 | 0.81703827 | 0.93550282 |
| PTCD3 | 120.972627 | -0.051181509 | 0.22490057 | -0.2275739 | 0.81997748 | 0.93710409 |
| DCUN1D5 | 75.3975902 | -0.051126481 | 0.19956469 | -0.25619 | 0.79780411 | 0.92959949 |
| BIRC2 | 134.929977 | -0.050942876 | 0.25074112 | -0.2031692 | 0.83900277 | 0.94376667 |
| BST2 | 676.94845 | -0.050884615 | 0.46610353 | -0.1091702 | 0.9130675 | 0.97097223 |
| HNRNPA3 | 499.860813 | -0.050875639 | 0.18003129 | -0.2825933 | 0.7774886 | 0.91952439 |
| MBNL2 | 788.753557 | -0.050645756 | 0.28577634 | -0.1772217 | 0.85933429 | 0.95257823 |
| VDAC3 | 123.055927 | -0.050589377 | 0.22815403 | -0.2217334 | 0.8245214 | 0.93852534 |
| MANEA | 54.0178309 | -0.050536403 | 0.34366146 | -0.1470529 | 0.88309029 | 0.96133136 |
| ORC3 | 128.735983 | -0.05014854 | 0.32172609 | -0.1558734 | 0.87613281 | 0.95883833 |
| BAHCC1 | 44.7359173 | -0.050131146 | 0.33395062 | -0.1501154 | 0.88067353 | 0.96048934 |
| MTFR1 | 160.645976 | -0.049984849 | 0.32547756 | -0.1535739 | 0.87794574 | 0.95969416 |
| UBE2Q1 | 74.6065343 | -0.049918924 | 0.30054609 | -0.1660941 | 0.86808292 | 0.95580259 |
| TBCA | 333.920169 | -0.049720755 | 0.21996725 | -0.2260371 | 0.82117258 | 0.93726124 |
| RABL6 | 124.052328 | -0.04913307 | 0.26343881 | -0.1865066 | 0.85204752 | 0.9501642 |
| PLEKHM2 | 135.9242 | -0.049090377 | 0.18472536 | -0.2657479 | 0.79043337 | 0.92613348 |
| PRRC2A | 103.32707 | -0.048945567 | 0.25995865 | -0.1882821 | 0.8506555 | 0.9492877 |
| ZBTB18 | 57.8012919 | -0.048934292 | 0.36760043 | -0.1331182 | 0.89409993 | 0.96560905 |
| ARPC2 | 431.978881 | -0.048732428 | 0.20440267 | -0.2384139 | 0.81156012 | 0.93351089 |
| DR1 | 85.4906739 | -0.048376335 | 0.21257456 | -0.2275735 | 0.81997783 | 0.93710409 |
| ZBTB7A | 170.999605 | -0.048343031 | 0.21565962 | -0.2241636 | 0.82263003 | 0.93793807 |
| GOLGA1 | 84.6806497 | -0.048037216 | 0.32223814 | -0.1490737 | 0.88149552 | 0.96059232 |
| IRF2 | 294.506156 | -0.047702455 | 0.32331501 | -0.1475417 | 0.88270445 | 0.96103918 |
| DOCK1 | 1455.06427 | -0.047690434 | 0.20878982 | -0.2284136 | 0.81932472 | 0.93691087 |
| GTF2A1 | 130.454566 | -0.047584493 | 0.20588093 | -0.2311263 | 0.8172167 | 0.93550282 |
| COA3 | 63.5921457 | -0.047501749 | 0.4095523 | -0.1159846 | 0.90766477 | 0.96940376 |
| TRPC4AP | 385.661983 | -0.047373021 | 0.16436406 | -0.2882201 | 0.77317829 | 0.91726032 |
| MTDH | 525.653578 | -0.047339376 | 0.16975654 | -0.2788663 | 0.78034745 | 0.92045723 |
| TTI1 | 64.7980722 | -0.047140393 | 0.24144107 | -0.195246 | 0.8452004 | 0.94613884 |
| GPN2 | 39.1792793 | -0.046772074 | 0.25865747 | -0.1808263 | 0.85650391 | 0.95134391 |
| AP5Z1 | 48.6812767 | -0.046663843 | 0.36771581 | -0.1269019 | 0.89901803 | 0.96709143 |
| COPS3 | 48.1947064 | -0.046627259 | 0.32079225 | -0.1453503 | 0.88443428 | 0.96189869 |
| CBR4 | 292.338727 | -0.046439997 | 0.251909 | -0.1843523 | 0.85373711 | 0.9508242 |
| GGA1 | 85.4730076 | -0.046408962 | 0.23899319 | -0.1941853 | 0.8460308 | 0.94667989 |
| SSBP4 | 112.846093 | -0.045963489 | 0.285009 | -0.1612703 | 0.87188051 | 0.95727819 |
| CCNK | 88.8490061 | -0.045943454 | 0.28946564 | -0.1587182 | 0.87389093 | 0.95804618 |
| TOR1AIP1 | 113.390172 | -0.045923425 | 0.20202777 | -0.2273124 | 0.8201808 | 0.93710409 |
| GMPS | 129.660409 | -0.045808765 | 0.17143722 | -0.2672043 | 0.78931186 | 0.92588001 |
| RAB11A | 446.788673 | -0.045790361 | 0.34763815 | -0.1317185 | 0.89520699 | 0.96575575 |
| PHF14 | 619.692992 | -0.045535386 | 0.15135276 | -0.300856 | 0.7635243 | 0.91490985 |
| RNF6 | 63.1815952 | -0.04550328 | 0.32581254 | -0.1396609 | 0.8889279 | 0.96371185 |
| ARHGEF37 | 47.2584743 | -0.045366998 | 0.48795985 | -0.0929728 | 0.92592516 | 0.97614605 |
| KIAA1211L | 146.236649 | -0.045272892 | 0.43315745 | -0.1045183 | 0.91675802 | 0.97233474 |
| FAM196A | 56.6873513 | -0.045037109 | 0.50438195 | -0.0892917 | 0.92885011 | 0.97696444 |
| ACOX1 | 116.033018 | -0.044965749 | 0.2237244 | -0.2009872 | 0.84070856 | 0.94397833 |
| OSBP | 121.222326 | -0.044958107 | 0.16891142 | -0.2661638 | 0.79011305 | 0.92602336 |
| ARID1B | 1745.18553 | -0.044761364 | 0.17581259 | -0.254597 | 0.79903435 | 0.92963087 |
| RBM22 | 67.7168969 | -0.04447117 | 0.21542305 | -0.2064365 | 0.83645 | 0.94344181 |
| SH3BP2 | 148.348187 | -0.044308242 | 0.31181839 | -0.1420963 | 0.88700393 | 0.96290172 |
| YTHDF3 | 206.493758 | -0.044307113 | 0.17902868 | -0.2474861 | 0.80453204 | 0.93109287 |
| LMF2 | 55.8326071 | -0.044221791 | 0.27844261 | -0.1588183 | 0.87381201 | 0.95804618 |
| SEH1L | 76.0494952 | -0.044072737 | 0.25460864 | -0.1730999 | 0.86257288 | 0.95358469 |
| CLCN3 | 244.006691 | -0.044025583 | 0.17773858 | -0.2476985 | 0.80436767 | 0.93109287 |
| DDX42 | 180.170813 | -0.04379045 | 0.15064859 | -0.2906795 | 0.77129649 | 0.91695863 |
| DHX8 | 82.3862079 | -0.043789864 | 0.23614743 | -0.1854344 | 0.85288831 | 0.95044301 |
| KPNA4 | 314.117864 | -0.043428083 | 0.27909898 | -0.155601 | 0.87634753 | 0.95883833 |
| WDPCP | 516.997112 | -0.043412663 | 0.220027 | -0.1973061 | 0.84358803 | 0.94501377 |
| MAPK8 | 472.646691 | -0.043390367 | 0.20271214 | -0.2140492 | 0.83050871 | 0.94075329 |
| BRCC3 | 66.4882516 | -0.043367546 | 0.25005928 | -0.1734291 | 0.86231418 | 0.95356178 |
| NOTCH1 | 378.235626 | -0.043254492 | 0.29213229 | -0.1480647 | 0.88229168 | 0.9608775 |
| TSPAN14 | 613.648609 | -0.043011361 | 0.20894128 | -0.2058538 | 0.8369051 | 0.94356495 |
| SAMD12 | 261.439661 | -0.042667843 | 0.51919497 | -0.0821808 | 0.93450296 | 0.9797614 |
| STUB1 | 86.0614737 | -0.042541513 | 0.48205327 | -0.0882506 | 0.92967746 | 0.97708075 |
| RIOK3 | 189.717342 | -0.042525077 | 0.19778417 | -0.2150075 | 0.82976148 | 0.94063377 |
| ZNF302 | 115.929981 | -0.04250649 | 0.27866038 | -0.1525387 | 0.87876208 | 0.95972986 |
| FEM1C | 69.5787296 | -0.042133901 | 0.34675199 | -0.1215102 | 0.90328694 | 0.96875453 |
| LBR | 145.761318 | -0.042118889 | 0.18984027 | -0.2218649 | 0.82441907 | 0.93852534 |

| | | | | | | | |
|---|---|---|---|---|---|---|---|
| ARHGEF10L | 46.1502646 | | -0.042056873 | 0.46304398 | -0.090827 | 0.9276301 | 0.97688257 |
| NDUFS4 | 228.964086 | | -0.041937912 | 0.15625473 | -0.2683945 | 0.78839566 | 0.92507051 |
| CHIC2 | 371.406772 | | -0.041791769 | 0.31996604 | -0.1306131 | 0.89608135 | 0.96596412 |
| TNKS | 554.944082 | | -0.041775202 | 0.19744878 | -0.2115749 | 0.83243871 | 0.94229436 |
| EIF2AK4 | 169.788398 | | -0.041397065 | 0.24913049 | -0.1661662 | 0.86802616 | 0.95580259 |
| IGBP1 | 79.6265504 | | -0.041277084 | 0.20379773 | -0.2025395 | 0.83949501 | 0.94376667 |
| PRKAR1B | 128.349244 | | -0.041183702 | 0.34629996 | -0.1189249 | 0.90533482 | 0.96904724 |
| TMEM248 | 169.747584 | | -0.041009228 | 0.21638473 | -0.18952 | 0.8496853 | 0.94882298 |
| FAM102B | 61.9324208 | | -0.040880114 | 0.46098826 | -0.0886793 | 0.92933679 | 0.97696444 |
| CEP135 | 148.25602 | | -0.040793012 | 0.18589796 | -0.2194377 | 0.82630914 | 0.93977669 |
| GRPEL2 | 93.347235 | | -0.040756298 | 0.40563014 | -0.1004765 | 0.91996604 | 0.97362676 |
| SMIM12 | 112.775607 | | -0.040746857 | 0.20085058 | -0.2028715 | 0.83923548 | 0.94376667 |
| ZSWIM7 | 83.327683 | | -0.040681611 | 0.21708969 | -0.1873954 | 0.85135062 | 0.94977534 |
| ADRM1 | 48.2161846 | | -0.040666748 | 0.52473702 | -0.0774993 | 0.93822635 | 0.98165276 |
| RNF13 | 368.84275 | | -0.040587739 | 0.18600947 | -0.2182025 | 0.82727131 | 0.94018985 |
| RBM28 | 81.0233142 | | -0.040535301 | 0.29544738 | -0.1371997 | 0.89087292 | 0.96448584 |
| ANKRA2 | 51.0957822 | | -0.040475512 | 0.30814543 | -0.131352 | 0.89549687 | 0.96575575 |
| PSMD1 | 341.06318 | | -0.040376826 | 0.2708379 | -0.1490811 | 0.8814896 | 0.96059232 |
| NRDC | 399.495132 | | -0.040221285 | 0.17361963 | -0.2316632 | 0.81679959 | 0.93550282 |
| CSE1L | 122.642538 | | -0.039548992 | 0.19017234 | -0.207964 | 0.83525711 | 0.9432647 |
| STK24 | 378.436306 | | -0.039467435 | 0.19653339 | -0.200818 | 0.84084092 | 0.94397833 |
| MYADM | 54.3906425 | | -0.039427981 | 0.51857332 | -0.0760316 | 0.93939392 | 0.98171087 |
| IFT52 | 71.9219466 | | -0.039207268 | 0.1931272 | -0.2030127 | 0.83912513 | 0.94376667 |
| DNMBP | 246.522345 | | -0.039088347 | 0.32802871 | -0.1191614 | 0.90514752 | 0.96904724 |
| ME2 | 105.543968 | | -0.038991748 | 0.27619026 | -0.1411771 | 0.88773002 | 0.96330655 |
| PIP5K1A | 273.325831 | | -0.038892536 | 0.17193381 | -0.2262064 | 0.82104086 | 0.93726124 |
| STXBP4 | 119.890546 | | -0.03878915 | 0.24261266 | -0.159881 | 0.87297483 | 0.95783763 |
| MFN2 | 64.1005112 | | -0.038634805 | 0.2303075 | -0.1677531 | 0.86677749 | 0.95526468 |
| HACE1 | 190.503652 | | -0.038565955 | 0.29580546 | -0.1303761 | 0.89626889 | 0.96603904 |
| ARF4 | 162.861005 | | -0.038364273 | 0.31845391 | -0.1204704 | 0.90411052 | 0.9688759 |
| GCN1 | 85.9480137 | | -0.038320503 | 0.22344815 | -0.1714962 | 0.86383364 | 0.95405003 |
| AGPAT1 | 69.3676191 | | -0.038133381 | 0.46701604 | -0.0816533 | 0.93492245 | 0.9799501 |
| CUL1 | 325.276611 | | -0.037962759 | 0.19773777 | -0.1919854 | 0.84775367 | 0.94731791 |
| TPST1 | 333.017204 | | -0.037922354 | 0.23768076 | -0.1595516 | 0.87323428 | 0.9579344 |
| SPICE1 | 91.1803952 | | -0.037775204 | 0.2514767 | -0.1502135 | 0.88059615 | 0.96048934 |
| ETFA | 233.068318 | | -0.03766765 | 0.18240565 | -0.2065048 | 0.83639658 | 0.94344181 |
| ID2 | 274.584914 | | -0.037618232 | 0.84847937 | -0.0443361 | 0.96463653 | 0.9886632 |
| FASTKD2 | 72.14534 | | -0.037587586 | 0.23468591 | -0.1601612 | 0.87275406 | 0.95772381 |
| BCAT2 | 45.5738955 | | -0.037450244 | 0.24434142 | -0.1532701 | 0.87818525 | 0.95969416 |
| GNPDA2 | 58.1121718 | | -0.037442683 | 0.29064196 | -0.1288275 | 0.89749413 | 0.96608714 |
| PDLIM7 | 87.1778446 | | -0.037318186 | 0.36158724 | -0.1032066 | 0.91779901 | 0.97233474 |
| C16orf72 | 160.8639 | | -0.03728266 | 0.17211795 | -0.2166111 | 0.82851143 | 0.94018985 |
| TES | 162.420042 | | -0.037229447 | 0.3396171 | -0.1096218 | 0.9127093 | 0.9709041 |
| OTUD6B-AS1 | 61.1374536 | | -0.037162677 | 0.33113099 | -0.1122295 | 0.91064141 | 0.9700595 |
| NDUFB5 | 70.6657924 | | -0.037014103 | 0.35759656 | -0.103508 | 0.91755981 | 0.97233474 |
| POLR3A | 89.8110259 | | -0.037004357 | 0.19153442 | -0.1931995 | 0.84680272 | 0.94719648 |
| DDX5 | 1142.39375 | | -0.036933363 | 0.24025379 | -0.1537265 | 0.87782543 | 0.95969416 |
| XPO4 | 243.264851 | | -0.036894559 | 0.18321928 | -0.2013683 | 0.84041059 | 0.94397833 |
| ACBD5 | 81.599597 | | -0.036601913 | 0.28149804 | -0.1300255 | 0.89654628 | 0.96608352 |
| SSH2 | 764.87939 | | -0.036585003 | 0.12203827 | -0.299783 | 0.76434265 | 0.9154077 |
| FADS3 | 52.4670324 | | -0.036542751 | 0.56095002 | -0.0651444 | 0.94805903 | 0.98379261 |
| TIMMDC1 | 142.331245 | | -0.036307529 | 0.21459506 | -0.1691909 | 0.8656465 | 0.95510194 |
| PPP5C | 93.8199005 | | -0.036262608 | 0.23410853 | -0.1548966 | 0.87690286 | 0.95931766 |
| BMS1P14 | 245.655847 | | -0.036244139 | 0.6253852 | -0.0579549 | 0.95378455 | 0.98545694 |
| FAM149B1 | 83.2759681 | | -0.036215569 | 0.31193149 | -0.116101 | 0.90757247 | 0.96940376 |
| RAB11FIP2 | 112.783585 | | -0.036197664 | 0.17472102 | -0.2071741 | 0.83587392 | 0.9432647 |
| CARNMT1 | 85.4502742 | | -0.03609828 | 0.23642431 | -0.1526843 | 0.87864725 | 0.95972986 |
| BRD3 | 60.1694096 | | -0.036079561 | 0.29762405 | -0.1212253 | 0.9035126 | 0.96886956 |
| MAGOH | 62.3178249 | | -0.035924783 | 0.30341917 | -0.1183998 | 0.90575085 | 0.96904724 |
| ANKRD36B | 80.538795 | | -0.035788674 | 0.41735163 | -0.0857518 | 0.93166368 | 0.97845553 |
| TMEM127 | 83.9812981 | | -0.035691561 | 0.26060336 | -0.1369574 | 0.89106446 | 0.96448584 |
| TRMT11 | 91.7534824 | | -0.035220588 | 0.27939175 | -0.1260617 | 0.89968311 | 0.96723863 |
| ANAPC7 | 99.3171212 | | -0.035151761 | 0.30720365 | -0.1144249 | 0.90890094 | 0.96940376 |
| FLT4 | 314.091292 | | -0.03508594 | 0.33659815 | -0.1042369 | 0.91698137 | 0.97233474 |
| PDCD2 | 73.1194337 | | -0.035083152 | 0.31346004 | -0.1119222 | 0.91088506 | 0.9700595 |
| GPATCH2 | 269.86839 | | -0.035032207 | 0.29950351 | -0.1169676 | 0.90688573 | 0.96940376 |
| ZNF41 | 50.7654059 | | -0.034962342 | 0.23187106 | -0.1507836 | 0.88014646 | 0.96042318 |
| KRR1 | 81.3088976 | | -0.034885061 | 0.2720918 | -0.1282106 | 0.89798229 | 0.96635837 |
| AKIRIN1 | 94.5089959 | | -0.034884171 | 0.25612373 | -0.1362005 | 0.89166281 | 0.96460264 |
| VTI1A | 982.430186 | | -0.034647059 | 0.22186117 | -0.1561655 | 0.87590257 | 0.95883833 |
| NCOA5 | 94.3048934 | | -0.034633322 | 0.27451053 | -0.1261639 | 0.89960218 | 0.96723863 |
| KPNA6 | 239.814002 | | -0.034531529 | 0.17377573 | -0.1987132 | 0.84248711 | 0.94471448 |
| PSMB2 | 136.636307 | | -0.034443026 | 0.2435894 | -0.1413979 | 0.88755563 | 0.96330655 |
| SCOC | 85.0716602 | | -0.03444228 | 0.34342086 | -0.1002918 | 0.9201127 | 0.97365605 |
| DENND4C | 720.664888 | | -0.0342113 | 0.28459959 | -0.1202085 | 0.90431796 | 0.96889875 |
| ANKS1A | 756.360073 | | -0.033960591 | 0.3605819 | -0.0941827 | 0.924964 | 0.97576555 |
| SFPQ | 423.349244 | | -0.033912039 | 0.21781265 | -0.1556936 | 0.87627453 | 0.95883833 |
| CCNB1IP1 | 46.50234 | | -0.033884531 | 0.27520365 | -0.1231253 | 0.90200788 | 0.96801711 |
| MIR3681HG | 55.3924527 | | -0.03348381 | 0.43527416 | -0.0769258 | 0.93868258 | 0.98171087 |
| DTNB | 153.99701 | | -0.033458894 | 0.35063451 | -0.0954238 | 0.92397817 | 0.97530455 |
| COPS2 | 113.656262 | | -0.033440675 | 0.21429263 | -0.1560514 | 0.87599247 | 0.95883833 |
| NCAPD3 | 47.6469689 | | -0.033419768 | 0.22014885 | -0.1518053 | 0.87934048 | 0.95980301 |
| RECK | 205.939347 | | -0.033156175 | 0.26769747 | -0.1238569 | 0.90142859 | 0.96790318 |
| GNPNAT1 | 60.6211048 | | -0.033088637 | 0.3219889 | -0.1027633 | 0.91815085 | 0.97233474 |
| FAM168B | 161.118013 | | -0.032996824 | 0.20598354 | -0.1601916 | 0.87273018 | 0.95772381 |
| GAB2 | 377.489872 | | -0.032952488 | 0.2502187 | -0.1316947 | 0.89522574 | 0.96575575 |
| SLC25A29 | 108.577132 | | -0.032869078 | 0.3862024 | -0.0851084 | 0.93217519 | 0.97871274 |
| IAH1 | 66.517293 | | -0.032841684 | 0.32296191 | -0.101689 | 0.91900351 | 0.97285973 |
| RER1 | 121.454341 | | -0.032544094 | 0.20885543 | -0.1558212 | 0.87617399 | 0.95883833 |
| MPRIP | 473.347053 | | -0.032164124 | 0.18888696 | -0.1702824 | 0.86478805 | 0.95474375 |
| KLHL42 | 78.1060139 | | -0.032159303 | 0.29176102 | -0.1102248 | 0.91223109 | 0.97083977 |
| SUPV3L1 | 51.209416 | | -0.03190035 | 0.30476558 | -0.1046718 | 0.91663627 | 0.97233474 |
| ATP6V1C1 | 153.225244 | | -0.031514789 | 0.25221186 | -0.1249536 | 0.90056025 | 0.96744565 |
| PMS1 | 155.795731 | | -0.031509993 | 0.16877737 | -0.1866956 | 0.8518993 | 0.9501642 |
| SLC38A10 | 142.858967 | | -0.031280971 | 0.20538657 | -0.1523029 | 0.87894804 | 0.95972986 |
| MPST | 45.5865247 | | -0.031071306 | 0.37109633 | -0.0837284 | 0.93327237 | 0.97952849 |
| PITHD1 | 62.7462875 | | -0.030843211 | 0.27658352 | -0.111515 | 0.91120797 | 0.97018454 |
| USP9X | 454.504629 | | -0.030817198 | 0.24991008 | -0.1233131 | 0.90185913 | 0.96801711 |
| PTBP3 | 471.854453 | | -0.030599206 | 0.2188712 | -0.1398046 | 0.88881436 | 0.96371185 |
| FOXK2 | 313.394375 | | -0.03038773 | 0.18875095 | -0.1609938 | 0.8720983 | 0.95727819 |
| GNA12 | 221.035848 | | -0.030356066 | 0.22631534 | -0.1341317 | 0.89329842 | 0.96512555 |
| ZNF451 | 321.977723 | | -0.030217097 | 0.15387305 | -0.1963768 | 0.84431525 | 0.94540664 |
| ABCA3 | 58.7616878 | | -0.030034152 | 0.28787513 | -0.1043305 | 0.91690709 | 0.97233474 |
| TEAD1 | 962.856767 | | -0.029928633 | 0.27782486 | -0.1077248 | 0.91421398 | 0.97168632 |
| EXOC6 | 1434.6294 | | -0.029896368 | 0.38720956 | -0.0772098 | 0.93845666 | 0.9817035 |
| NOM1 | 50.8622065 | | -0.02975255 | 0.41606407 | -0.0715095 | 0.94299223 | 0.98263114 |
| CSGALNACT2 | 102.054922 | | -0.029748317 | 0.23727367 | -0.1253756 | 0.90022624 | 0.96744565 |
| CDC34 | 41.2932068 | | -0.029552292 | 0.29812585 | -0.0991269 | 0.92103751 | 0.97425672 |
| SAR1A | 118.796968 | | -0.029497696 | 0.25791462 | -0.11437 | 0.90894449 | 0.96940376 |
| METTL21A | 79.6042999 | | -0.029385315 | 0.31942407 | -0.0919947 | 0.92670227 | 0.97645253 |
| INTS8 | 116.35609 | | -0.02924577 | 0.21016118 | -0.1391588 | 0.88932469 | 0.96388662 |
| EXOC1 | 315.752474 | | -0.029104525 | 0.18503299 | -0.1572937 | 0.87501338 | 0.95853206 |
| VGLL4 | 848.00126 | | -0.02902901 | 0.26708752 | -0.1086873 | 0.91345055 | 0.97120835 |

| | | | | | | |
|---|---|---|---|---|---|---|
| REL | 212.081341 | -0.028738663 | 0.37869601 | -0.0758885 | 0.93950782 | 0.98171087 |
| SF1 | 327.141106 | -0.028711662 | 0.26235403 | -0.1094386 | 0.91285461 | 0.9709041 |
| VRK2 | 246.954616 | -0.028514381 | 0.2452337 | -0.1162743 | 0.90743514 | 0.96940376 |
| FSCN1 | 109.891502 | -0.028508207 | 0.45773578 | -0.0622809 | 0.95033912 | 0.98412513 |
| SEC62 | 291.415251 | -0.028445344 | 0.20867102 | -0.1363167 | 0.89157092 | 0.96460264 |
| NOL4L | 155.410371 | -0.028415833 | 0.3522555 | -0.0806682 | 0.93570579 | 0.98010864 |
| ZC3H3 | 66.624691 | -0.028395726 | 0.29494805 | -0.0962737 | 0.92330324 | 0.97530455 |
| AQP1 | 1056.83819 | -0.028394199 | 0.57198095 | -0.0496419 | 0.96040779 | 0.98739605 |
| DDX3X | 365.866047 | -0.027996932 | 0.3047918 | -0.0918559 | 0.92681251 | 0.97645253 |
| ESAM | 398.014711 | -0.027915328 | 0.37584456 | -0.0742736 | 0.94079268 | 0.98256304 |
| ACADSB | 56.5057859 | -0.02768926 | 0.31097546 | -0.08904 | 0.92905011 | 0.97696444 |
| DNPEP | 66.2770494 | -0.027234047 | 0.33899892 | -0.0803367 | 0.93596949 | 0.98010864 |
| USP49 | 65.0362314 | -0.027118851 | 0.33596941 | -0.0807182 | 0.93566605 | 0.98010864 |
| JAM2 | 398.512603 | -0.027081477 | 0.46631332 | -0.0580757 | 0.95368832 | 0.98545694 |
| UBAP2L | 157.574199 | -0.027007423 | 0.21247363 | -0.1271095 | 0.89885371 | 0.9670418 |
| PAK2 | 406.736449 | -0.026794535 | 0.11947336 | -0.224272 | 0.82254563 | 0.93793807 |
| RBPJ | 330.729054 | -0.026783905 | 0.16333929 | -0.1639771 | 0.86974916 | 0.95673577 |
| CCDC148 | 63.2797291 | -0.026778432 | 0.46964166 | -0.0570189 | 0.95453017 | 0.98585389 |
| KDELC2 | 48.3414511 | -0.026570542 | 0.36889748 | -0.0720269 | 0.9425805 | 0.98263114 |
| PINK1 | 160.908226 | -0.026569842 | 0.33889041 | -0.0784025 | 0.93750792 | 0.98137001 |
| MAF | 233.262381 | -0.026507753 | 0.56744257 | -0.0467144 | 0.96274083 | 0.98834629 |
| PLVAP | 1376.68457 | -0.026191285 | 0.4829798 | -0.0542285 | 0.95675309 | 0.98665539 |
| NEPRO | 70.6107504 | -0.025902485 | 0.20754889 | -0.1248018 | 0.90068042 | 0.96744565 |
| VASP | 153.346535 | -0.025900425 | 0.27461892 | -0.0943141 | 0.92485968 | 0.97576555 |
| C1orf21 | 891.599665 | -0.025870814 | 0.48422347 | -0.0534274 | 0.95739135 | 0.98683301 |
| MTA1 | 80.2707091 | -0.025859485 | 0.27792898 | -0.0930435 | 0.92586901 | 0.97614605 |
| CDC16 | 143.153412 | -0.025509165 | 0.1638912 | -0.1556469 | 0.87631132 | 0.95883833 |
| TM9SF3 | 407.010784 | -0.025126556 | 0.21146885 | -0.1188192 | 0.90541861 | 0.96904724 |
| GPD1L | 43.0349745 | -0.025063914 | 0.3884942 | -0.0645155 | 0.94855973 | 0.98379261 |
| RAB11B | 79.647931 | -0.02488039 | 0.32920768 | -0.0755766 | 0.93975597 | 0.98174685 |
| PHF3 | 375.230143 | -0.024757152 | 0.2208623 | -0.1120932 | 0.91074955 | 0.9700595 |
| IPO9-AS1 | 180.481769 | -0.024707325 | 0.35697054 | -0.0692139 | 0.94481936 | 0.98377106 |
| AC006059.1 | 157.810766 | -0.02468192 | 0.51031972 | -0.0483656 | 0.96142487 | 0.98786285 |
| TRABD | 43.3015592 | -0.024493841 | 0.26008121 | -0.0941777 | 0.92496803 | 0.97576555 |
| STK11 | 72.1029723 | -0.0241472 | 0.21323335 | -0.1132431 | 0.90983785 | 0.96980632 |
| ADNP | 380.533309 | -0.024143981 | 0.13322317 | -0.1812296 | 0.85618737 | 0.95134391 |
| DUT | 104.306521 | -0.024062493 | 0.29981351 | -0.0802582 | 0.9360319 | 0.98010864 |
| NEDD4 | 702.949769 | -0.024031719 | 0.27642584 | -0.0869373 | 0.93072134 | 0.97792665 |
| SEC23IP | 125.857459 | -0.024027315 | 0.18637496 | -0.1289192 | 0.89742157 | 0.96608714 |
| XPO1 | 423.945971 | -0.024010362 | 0.38855739 | -0.0617936 | 0.9507272 | 0.98412513 |
| H2AFY | 215.060712 | -0.02399877 | 0.20898975 | -0.1148323 | 0.90857806 | 0.96940376 |
| SH3BP4 | 118.102195 | -0.023968379 | 0.34506348 | -0.0694608 | 0.94462285 | 0.98369162 |
| TFB1M | 84.4585672 | -0.023726753 | 0.25717217 | -0.0922602 | 0.92649132 | 0.97645253 |
| ETFRF1 | 58.1466005 | -0.023692832 | 0.24434179 | -0.0969659 | 0.92275344 | 0.97518761 |
| CDK11A | 42.179953 | -0.023529138 | 0.46567149 | -0.0505273 | 0.95970217 | 0.98739605 |
| CDC5L | 99.1904992 | -0.023524681 | 0.19459384 | -0.1208912 | 0.90377722 | 0.9688759 |
| PAPSS1 | 115.652835 | -0.023459823 | 0.2691626 | -0.0871586 | 0.93054548 | 0.97786743 |
| TAF1C | 75.8707729 | -0.023272571 | 0.32079266 | -0.0725471 | 0.94216654 | 0.98263114 |
| HRAS | 53.8187455 | -0.022934687 | 0.36752025 | -0.0624039 | 0.95024121 | 0.98412513 |
| TTC13 | 109.513015 | -0.022874376 | 0.34306531 | -0.0666764 | 0.94683929 | 0.98379261 |
| MCTS1 | 64.4966697 | -0.022829035 | 0.19049116 | -0.119843 | 0.9046075 | 0.96890094 |
| BRD2 | 188.798211 | -0.022693358 | 0.25039828 | -0.090629 | 0.92778735 | 0.97688257 |
| FAM20B | 110.528108 | -0.022670297 | 0.21883572 | -0.103595 | 0.91749072 | 0.97233474 |
| SF3B3 | 173.619079 | -0.022322897 | 0.20029567 | -0.1114497 | 0.91125973 | 0.97018454 |
| AKAP8L | 154.886669 | -0.022257447 | 0.18987876 | -0.1172193 | 0.90668631 | 0.96940376 |
| AKAP9 | 504.969348 | -0.022219075 | 0.17223679 | -0.1290031 | 0.89735522 | 0.96608714 |
| CUL4A | 249.263906 | -0.022208274 | 0.19426577 | -0.114319 | 0.9089849 | 0.96940376 |
| MORC2 | 58.5258534 | -0.022124489 | 0.22665294 | -0.097614 | 0.92223885 | 0.9749687 |
| C12orf65 | 52.3323411 | -0.022104551 | 0.32550436 | -0.0679086 | 0.94585838 | 0.98379261 |
| HDAC1 | 204.082201 | -0.022102996 | 0.23186251 | -0.095328 | 0.92405428 | 0.97530455 |
| DNAJC9 | 75.719809 | -0.021975474 | 0.28079661 | -0.0782612 | 0.9376203 | 0.98137001 |
| MAPK8IP3 | 142.852828 | -0.021817036 | 0.18727862 | -0.1164951 | 0.90726019 | 0.96940376 |
| KDM1A | 151.283772 | -0.021749736 | 0.1443835 | -0.1506387 | 0.88026077 | 0.96042318 |
| SEC31A | 439.609203 | -0.021397946 | 0.1296057 | -0.1651003 | 0.868865 | 0.95627738 |
| AC027644.4 | 74.4858279 | -0.021141924 | 0.42619689 | -0.049606 | 0.96043636 | 0.98739605 |
| SERGEF | 129.001874 | -0.020794993 | 0.18177756 | -0.114398 | 0.90892228 | 0.96940376 |
| TTN-AS1 | 281.853031 | -0.020717301 | 0.23211676 | -0.0892538 | 0.92888022 | 0.97696444 |
| TGFBR3 | 933.564348 | -0.020572415 | 0.47476306 | -0.043332 | 0.96543691 | 0.98875723 |
| OSGEP | 50.9268784 | -0.020363662 | 0.24339671 | -0.0836645 | 0.93332319 | 0.97952849 |
| DHTKD1 | 64.0529646 | -0.020343546 | 0.31443016 | -0.0646997 | 0.94841308 | 0.98379261 |
| CAPRIN1 | 248.873154 | -0.020206594 | 0.19087999 | -0.1058602 | 0.91569327 | 0.97233474 |
| TTC7B | 271.081945 | -0.020087645 | 0.29838754 | -0.0673207 | 0.94632643 | 0.98379261 |
| CCDC12 | 138.583485 | -0.01994208 | 0.22434734 | -0.0888893 | 0.92916988 | 0.97696444 |
| QSOX1 | 153.768296 | -0.0199233 | 0.46256942 | -0.0430709 | 0.96564498 | 0.98884656 |
| MYNN | 54.620793 | -0.019836993 | 0.27736 | -0.0715207 | 0.94298332 | 0.98263114 |
| PCED1A | 44.6971366 | -0.019797676 | 0.30653486 | -0.0645854 | 0.94850411 | 0.98379261 |
| SAP30BP | 136.699457 | -0.019748255 | 0.19762389 | -0.0999285 | 0.92040111 | 0.97383531 |
| MIA3 | 147.842999 | -0.019711192 | 0.14134644 | -0.139453 | 0.88909216 | 0.96376224 |
| ASXL1 | 655.957115 | -0.019608305 | 0.24324104 | -0.0806126 | 0.93575001 | 0.98010864 |
| NUP153 | 232.785222 | -0.019502944 | 0.16899807 | -0.1154034 | 0.90812542 | 0.96940376 |
| AL117336.3 | 63.8014582 | -0.019443945 | 0.32503146 | -0.0598217 | 0.95229762 | 0.9849155 |
| ZCCHC10 | 92.6410745 | -0.019421809 | 0.23710704 | -0.0819116 | 0.93471704 | 0.9798603 |
| MAPK12 | 56.9249268 | -0.019300054 | 0.36754014 | -0.0525114 | 0.9581212 | 0.98725593 |
| UQCRC2 | 154.330566 | -0.019119678 | 0.18730833 | -0.102076 | 0.91869638 | 0.97266043 |
| KPNB1 | 332.236139 | -0.019080309 | 0.26610196 | -0.071703 | 0.94283827 | 0.98263114 |
| QSER1 | 221.935741 | -0.018706347 | 0.13756244 | -0.1359844 | 0.89183361 | 0.96460264 |
| SECISBP2 | 301.378014 | -0.01832922 | 0.32646709 | -0.0561442 | 0.95522697 | 0.98607572 |
| SIRT2 | 118.090804 | -0.018021001 | 0.25244434 | -0.071386 | 0.94309052 | 0.98263114 |
| NOP53 | 53.0544227 | -0.01794296 | 0.39503735 | -0.0454209 | 0.96377181 | 0.9886632 |
| IWS1 | 197.131441 | -0.017917548 | 0.18273801 | -0.0980505 | 0.92189222 | 0.9749687 |
| CLCN6 | 50.3170992 | -0.017729444 | 0.33703216 | -0.0526046 | 0.95804695 | 0.98725593 |
| TAMM41 | 140.6153 | -0.017666275 | 0.27837885 | -0.0634613 | 0.9493992 | 0.98401602 |
| AP002954.1 | 46.1898534 | -0.017537235 | 0.60317159 | -0.029075 | 0.97680475 | 0.99296979 |
| PRCC | 86.729746 | -0.017510817 | 0.23666143 | -0.073991 | 0.94101754 | 0.98256304 |
| PHKA2 | 65.942127 | -0.01749293 | 0.21245398 | -0.0823375 | 0.93437833 | 0.97975627 |
| NSUN3 | 97.5721217 | -0.017214905 | 0.17979662 | -0.0957465 | 0.92372187 | 0.97530455 |
| CLNS1A | 137.928362 | -0.017122319 | 0.25953142 | -0.065974 | 0.94739855 | 0.98379261 |
| ENOX2 | 185.411663 | -0.017020632 | 0.26264693 | -0.0648042 | 0.94832987 | 0.98379261 |
| VMP1 | 588.955929 | -0.017019851 | 0.36919489 | -0.0460999 | 0.96323062 | 0.98860105 |
| FTO | 803.984334 | -0.016990399 | 0.19779489 | -0.0858991 | 0.93154664 | 0.97845553 |
| AXIN1 | 58.2579645 | -0.016825889 | 0.26416752 | -0.063694 | 0.94921387 | 0.98401602 |
| TXNDC11 | 147.852183 | -0.016560009 | 0.19290146 | -0.085847 | 0.93158806 | 0.97845553 |
| KLHL20 | 177.909005 | -0.016394909 | 0.21371463 | -0.076714 | 0.93885105 | 0.98171087 |
| POM121 | 196.101534 | -0.016171919 | 0.17535265 | -0.0922251 | 0.92651918 | 0.97645253 |
| CNOT9 | 123.199992 | -0.016143174 | 0.20043796 | -0.0805395 | 0.93580818 | 0.98010864 |
| GOT1 | 53.4102282 | -0.016132871 | 0.31156336 | -0.0517804 | 0.95870369 | 0.98725593 |
| CDK7 | 53.7491313 | -0.016020388 | 0.28772186 | -0.0556801 | 0.95559664 | 0.98610144 |
| SLC18B1 | 85.4412578 | -0.016019447 | 0.32534397 | -0.0492385 | 0.96072923 | 0.98739605 |
| YARS2 | 79.4598431 | -0.015925355 | 0.22902155 | -0.0695365 | 0.94456259 | 0.98369162 |
| NBDY | 327.422251 | -0.015856321 | 0.18667378 | -0.0849413 | 0.93230803 | 0.97871412 |
| MCUR1 | 52.4942334 | -0.01580042 | 0.26884877 | -0.0587707 | 0.95313478 | 0.98545694 |
| HAUS6 | 69.6912553 | -0.015761975 | 0.27959517 | -0.0563743 | 0.95504365 | 0.98603425 |
| ENDOV | 44.4348438 | -0.015721939 | 0.28705695 | -0.0547694 | 0.95632217 | 0.98647256 |

| | | | | | | |
|---|---|---|---|---|---|---|
| PTER | 60.984438 | -0.015711152 | 0.43736682 | -0.0359221 | 0.97134444 | 0.99163087 |
| LMBRD2 | 75.1128849 | -0.015481363 | 0.26460235 | -0.058508 | 0.95334396 | 0.98545694 |
| SEC63 | 285.007546 | -0.015426676 | 0.15803755 | -0.097614 | 0.92223881 | 0.9749687 |
| PAXBP1 | 210.855913 | -0.015366697 | 0.1856618 | -0.0827671 | 0.93403669 | 0.97971855 |
| PMPCB | 84.6529048 | -0.015197799 | 0.23481589 | -0.0647222 | 0.94839519 | 0.98379261 |
| PDHA1 | 61.9705459 | -0.015082051 | 0.27629586 | -0.0545866 | 0.95646782 | 0.98648552 |
| ZC3H8 | 73.3993167 | -0.015019108 | 0.2587323 | -0.0580488 | 0.95370973 | 0.98545694 |
| TMEM109 | 123.904448 | -0.014920143 | 0.42823543 | -0.034841 | 0.97220654 | 0.99210665 |
| RBMS3-AS2 | 141.742942 | -0.014890314 | 0.54757462 | -0.0271932 | 0.97830563 | 0.99296979 |
| THOC5 | 50.7857009 | -0.014731157 | 0.24278809 | -0.060675 | 0.95161808 | 0.98446153 |
| GTF2F2 | 180.716299 | -0.014590398 | 0.24786267 | -0.0588648 | 0.95305976 | 0.98545694 |
| EPS15 | 761.271231 | -0.014571026 | 0.1754823 | -0.0830342 | 0.93382437 | 0.97967754 |
| ANGPT2 | 261.644542 | -0.014494174 | 0.87759108 | -0.0165159 | 0.98682285 | 0.99644258 |
| LSM5 | 41.6536896 | -0.014395918 | 0.38151733 | -0.0377333 | 0.96990031 | 0.99121946 |
| SAFB | 171.187507 | -0.014234905 | 0.25998075 | -0.0547537 | 0.9563347 | 0.98647256 |
| AC006978.2 | 108.906686 | -0.014062388 | 0.18463009 | -0.0761652 | 0.93928767 | 0.98171087 |
| SSBP1 | 157.102994 | -0.01380464 | 0.21743485 | -0.0634886 | 0.94937742 | 0.98401602 |
| COPA | 336.376257 | -0.013407536 | 0.21174039 | -0.0633206 | 0.94951118 | 0.98401602 |
| ATF1 | 128.844384 | -0.013394802 | 0.24785843 | -0.0540421 | 0.95690158 | 0.98668418 |
| ZSWIM8 | 100.157431 | -0.013367942 | 0.25030365 | -0.0534069 | 0.95740771 | 0.98683301 |
| DGLUCY | 282.186566 | -0.013330728 | 0.42342143 | -0.0314834 | 0.97488406 | 0.99260131 |
| DCAF1 | 125.603878 | -0.013305706 | 0.25876076 | -0.0514209 | 0.95899015 | 0.98725593 |
| ZCCHC7 | 844.830149 | -0.013202205 | 0.17581893 | -0.0750898 | 0.94014328 | 0.9820261 |
| CLPX | 106.894067 | -0.013041724 | 0.19292731 | -0.0675992 | 0.94610472 | 0.98379261 |
| WDR6 | 83.261437 | -0.012878741 | 0.31377504 | -0.0410445 | 0.96726042 | 0.99000525 |
| ELP2 | 156.07749 | -0.012793242 | 0.19334854 | -0.0661667 | 0.94724508 | 0.98379261 |
| NSRP1 | 413.739033 | -0.01253446 | 0.18587149 | -0.0674362 | 0.94623448 | 0.98379261 |
| EXOC2 | 344.692476 | -0.012478764 | 0.15724484 | -0.0793588 | 0.93674723 | 0.98073216 |
| SNAPC3 | 173.449328 | -0.012406067 | 0.15907297 | -0.0779898 | 0.93783618 | 0.98137001 |
| CCDC88A | 267.598268 | -0.012308423 | 0.24571586 | -0.0500921 | 0.960049 | 0.98739605 |
| PLEKHA6 | 101.377496 | -0.012301466 | 0.45029325 | -0.0273188 | 0.97820547 | 0.99296979 |
| RNF5 | 43.7988172 | -0.012244959 | 0.44040814 | -0.0278037 | 0.97781875 | 0.99296979 |
| ZNF326 | 123.905015 | -0.011894544 | 0.20682114 | -0.0575113 | 0.95413794 | 0.98557318 |
| ACO2 | 111.178312 | -0.011853048 | 0.23643103 | -0.0501332 | 0.96001623 | 0.98739605 |
| SPRED1 | 408.651879 | -0.011834509 | 0.31202621 | -0.0379279 | 0.96974514 | 0.99118467 |
| KANK3 | 487.127252 | -0.011771471 | 0.36720601 | -0.0320569 | 0.97442671 | 0.99244992 |
| NRBP1 | 91.5071058 | -0.011695322 | 0.23114553 | -0.0505972 | 0.95964648 | 0.98739605 |
| SCAPER | 638.937664 | -0.011644389 | 0.20055063 | -0.0580621 | 0.95369917 | 0.98545694 |
| UCHL5 | 89.8443625 | -0.011498352 | 0.20658536 | -0.0556591 | 0.95561339 | 0.98610144 |
| PRPF4B | 307.650639 | -0.011280312 | 0.18139991 | -0.0621848 | 0.95041569 | 0.98412513 |
| NDUFS7 | 51.2284683 | -0.011243736 | 0.25435448 | -0.044205 | 0.96474101 | 0.9886632 |
| TXLNG | 86.039767 | -0.011126904 | 0.33093203 | -0.0336229 | 0.97317784 | 0.99210665 |
| COX20 | 80.5266725 | -0.011013825 | 0.32608354 | -0.0337761 | 0.97305571 | 0.99210665 |
| BAG5 | 72.9920853 | -0.010865056 | 0.26936634 | -0.0403356 | 0.96782556 | 0.99021649 |
| SH3PXD2A | 162.323988 | -0.010808148 | 0.39320398 | -0.0274874 | 0.978071 | 0.99296979 |
| RTCA | 94.7992049 | -0.010578417 | 0.2776015 | -0.0381065 | 0.96960278 | 0.99117915 |
| RAB14 | 229.165487 | -0.010355871 | 0.20015725 | -0.0517387 | 0.95873692 | 0.98725593 |
| GLE1 | 64.2348653 | -0.010275926 | 0.25479447 | -0.0403303 | 0.96782983 | 0.99021649 |
| CDH5 | 511.197786 | -0.010160135 | 0.36842829 | -0.027577 | 0.97799955 | 0.99296979 |
| SFT2D1 | 84.4251848 | -0.009687153 | 0.23189816 | -0.0417733 | 0.96667941 | 0.98953436 |
| UBN1 | 133.768827 | -0.009606823 | 0.32303756 | -0.029739 | 0.97627519 | 0.99296979 |
| CXXC5 | 66.572191 | -0.00955972 | 0.3590133 | -0.0266278 | 0.97875663 | 0.99297384 |
| DYNC1LI1 | 123.483513 | -0.009499602 | 0.21832141 | -0.043512 | 0.9652934 | 0.98875723 |
| ADGRL1 | 64.9902764 | -0.009438051 | 0.28446463 | -0.0331783 | 0.97353241 | 0.99220522 |
| FLOT2 | 155.547395 | -0.009403749 | 0.25717816 | -0.0365651 | 0.97083176 | 0.99142846 |
| AC004889.1 | 84.0633022 | -0.009286666 | 0.41402889 | -0.02243 | 0.98210495 | 0.99474451 |
| SIDT2 | 43.7643705 | -0.009232192 | 0.27367927 | -0.0337336 | 0.97308957 | 0.99210665 |
| POLR1D | 110.226289 | -0.009085877 | 0.28098678 | -0.0323356 | 0.97420442 | 0.99244992 |
| RFX7 | 364.674693 | -0.009034992 | 0.14530587 | -0.0621791 | 0.95042019 | 0.98412513 |
| DIS3L2 | 351.616491 | -0.00891621 | 0.25991791 | -0.0343039 | 0.97263478 | 0.99210665 |
| WDR48 | 133.229678 | -0.008684442 | 0.27168845 | -0.0319647 | 0.9745002 | 0.99244992 |
| RFXAP | 44.3320219 | -0.008621647 | 0.26936056 | -0.0320078 | 0.97446581 | 0.99244992 |
| RABAC1 | 202.44941 | -0.008378413 | 0.53827542 | -0.0155653 | 0.9875812 | 0.99656516 |
| CAB39 | 380.301844 | -0.008294295 | 0.16813363 | -0.0493316 | 0.96065507 | 0.98739605 |
| SH2D3C | 572.916438 | -0.007959507 | 0.29869465 | -0.0266476 | 0.97874078 | 0.99297384 |
| MOB1A | 165.911088 | -0.007724165 | 0.2628601 | -0.0293851 | 0.97655747 | 0.99296979 |
| DCTN5 | 101.163578 | -0.007628902 | 0.21006516 | -0.0363168 | 0.97102973 | 0.9915069 |
| TRIM13 | 134.399339 | -0.007604088 | 0.27095828 | -0.0280637 | 0.97761136 | 0.99296979 |
| PARP6 | 83.8299004 | -0.007530806 | 0.30763039 | -0.02448 | 0.9804697 | 0.99384993 |
| NOL9 | 90.3506282 | -0.00733375 | 0.2841357 | -0.0258107 | 0.9794083 | 0.99326577 |
| PRKN | 868.278646 | -0.007223299 | 0.27746886 | -0.0260328 | 0.97923116 | 0.99326577 |
| TMEM135 | 568.06786 | -0.007122962 | 0.24304199 | -0.0293075 | 0.97661932 | 0.99296979 |
| MMP28 | 46.0591985 | -0.006650869 | 0.71025598 | -0.009364 | 0.99252868 | 0.99826232 |
| GHR | 240.873027 | -0.006604605 | 0.23855334 | -0.0276861 | 0.97791253 | 0.99296979 |
| PQLC3 | 49.7426192 | -0.006333892 | 0.3076089 | -0.0205907 | 0.98357214 | 0.99512497 |
| PCDH12 | 127.709604 | -0.005988835 | 0.38305725 | -0.0156343 | 0.98752614 | 0.99656516 |
| CHTOP | 105.18355 | -0.005956071 | 0.15638222 | -0.0380866 | 0.96961862 | 0.99117915 |
| JDP2 | 69.4614367 | -0.005889588 | 0.4229108 | -0.0139263 | 0.98888877 | 0.99678917 |
| BBIP1 | 77.5027725 | -0.005772588 | 0.23712533 | -0.024344 | 0.98057819 | 0.99384993 |
| GIPC1 | 98.6888803 | -0.005684999 | 0.2604919 | -0.0218241 | 0.98258828 | 0.99474451 |
| DEFB1 | 130.27819 | -0.005684453 | 0.52532953 | -0.0108207 | 0.99136647 | 0.99782925 |
| LSM1 | 45.2106292 | -0.005573646 | 0.33162789 | -0.0168069 | 0.98659064 | 0.99644258 |
| CIAPIN1 | 50.2540822 | -0.005569598 | 0.31216947 | -0.0178416 | 0.98576523 | 0.99635831 |
| LARP7 | 109.596184 | -0.005329517 | 0.21930177 | -0.0243022 | 0.98061155 | 0.99384993 |
| REEP3 | 624.344901 | -0.005300543 | 0.36047257 | -0.0147044 | 0.98826799 | 0.99678917 |
| BTAF1 | 300.00824 | -0.005240216 | 0.31147561 | -0.0168238 | 0.98657715 | 0.99644258 |
| IPO9 | 199.011814 | -0.005046212 | 0.21062962 | -0.0239578 | 0.98088631 | 0.99400528 |
| MGA | 211.934438 | -0.005031625 | 0.22733683 | -0.0221329 | 0.98234194 | 0.99474451 |
| SCARF1 | 180.407297 | -0.004955136 | 0.17610958 | -0.0281367 | 0.97755315 | 0.99296979 |
| TRIQK | 123.149313 | -0.004810162 | 0.4271069 | -0.0112622 | 0.99101426 | 0.99782925 |
| SESTD1 | 302.304867 | -0.004605182 | 0.27915921 | -0.0164966 | 0.9868382 | 0.99644258 |
| C9orf72 | 66.2529902 | -0.004441333 | 0.27318746 | -0.0162575 | 0.987029 | 0.99644258 |
| LMBRD1 | 270.586065 | -0.004410457 | 0.24374588 | -0.0180945 | 0.98556347 | 0.99635831 |
| AKIRIN2 | 71.3168071 | -0.004149213 | 0.25595529 | -0.0162107 | 0.9870663 | 0.99644258 |
| MTCH2 | 97.4032941 | -0.004127234 | 0.30618146 | -0.0134797 | 0.98924508 | 0.99692026 |
| LMBR1L | 108.67281 | -0.004126794 | 0.20029889 | -0.0206032 | 0.9835622 | 0.99512497 |
| IQCG | 59.048037 | -0.004120344 | 0.43588755 | -0.0094528 | 0.99245789 | 0.99826232 |
| USP7 | 178.48552 | -0.003772399 | 0.17514442 | -0.0215388 | 0.98281586 | 0.99485183 |
| MRFAP1L1 | 46.0903747 | -0.003690765 | 0.23891723 | -0.0154479 | 0.98767486 | 0.99656516 |
| UBE2Z | 190.381705 | -0.003625418 | 0.17942773 | -0.0202055 | 0.98387948 | 0.99531287 |
| CYB5R4 | 111.777389 | -0.003505529 | 0.24732392 | -0.0141738 | 0.98869129 | 0.99678917 |
| TMEM209 | 68.1240289 | -0.003295979 | 0.19972867 | -0.0165023 | 0.98683368 | 0.99644258 |
| AC123768.3 | 42.0123068 | -0.003288848 | 0.48505079 | -0.0067804 | 0.99459005 | 0.99898392 |
| RAD51C | 53.0406047 | -0.003229374 | 0.2340902 | -0.0137954 | 0.98899319 | 0.99678917 |
| CAPZA2 | 519.466042 | -0.002992489 | 0.31666754 | -0.0094499 | 0.99246015 | 0.99826232 |
| ATG16L1 | 56.0462142 | -0.002739722 | 0.23368164 | -0.0117242 | 0.99064569 | 0.99771737 |
| YIPF3 | 88.5474003 | -0.0026751 | 0.36802654 | -0.0072688 | 0.99420041 | 0.99896144 |
| ANGPTL2 | 91.4560696 | -0.002598214 | 0.89442549 | -0.0029049 | 0.99768223 | 0.99938628 |
| SDE2 | 57.9727471 | -0.002486061 | 0.41763715 | -0.0059527 | 0.99525048 | 0.99903516 |
| NUDT21 | 111.664769 | -0.002435047 | 0.20280989 | -0.0120066 | 0.99042039 | 0.99770404 |
| PSMG4 | 101.309827 | -0.002105487 | 0.19199447 | -0.0109664 | 0.99125026 | 0.99782925 |
| RBPMS | 1215.52272 | -0.002082421 | 0.25192176 | -0.0082661 | 0.99340465 | 0.99863362 |
| UGGT1 | 173.952816 | -0.001876304 | 0.19403706 | -0.0096698 | 0.99228472 | 0.99826232 |

| | | | | | | |
|---|---|---|---|---|---|---|
| GTF3C5 | 43.2838042 | -0.001821945 | 0.33407876 | -0.0054536 | 0.99564865 | 0.99931226 |
| GUF1 | 50.766997 | -0.001819325 | 0.23233424 | -0.0078306 | 0.99375212 | 0.99863362 |
| DNASE1L3 | 403.741967 | -0.001798694 | 0.44523327 | -0.0040399 | 0.99677664 | 0.99938628 |
| UTP14C | 41.4037743 | -0.00178624 | 0.28022907 | -0.0063742 | 0.99491415 | 0.99898392 |
| TENT4B | 171.843084 | -0.001723689 | 0.26413555 | -0.0065258 | 0.99479322 | 0.99898392 |
| SIN3A | 168.983449 | -0.001566338 | 0.18414985 | -0.0085058 | 0.99321345 | 0.99863362 |
| TCF15 | 42.6046651 | -0.001166711 | 0.71650848 | -0.0016283 | 0.99870078 | 0.99938628 |
| FP700111.1 | 92.2593208 | -0.001063701 | 0.38829049 | -0.0027394 | 0.99781424 | 0.99938628 |
| RNF2 | 132.935524 | -0.000997246 | 0.19773459 | -0.0050434 | 0.995976 | 0.99933015 |
| SLC30A9 | 268.14969 | -0.000969656 | 0.15614384 | -0.00621 | 0.99504516 | 0.99898392 |
| HNRNPL | 121.78486 | -0.000960674 | 0.29037498 | -0.0033084 | 0.99736029 | 0.99938628 |
| HSPA4 | 143.512805 | -0.000860292 | 0.23789461 | -0.0036163 | 0.99711464 | 0.99938628 |
| MKLN1 | 1486.18067 | -0.000809619 | 0.25466145 | -0.0031792 | 0.99746337 | 0.99938628 |
| PEAR1 | 87.1655187 | -0.000731933 | 0.20536486 | -0.0035641 | 0.9971563 | 0.99938628 |
| RHOBTB3 | 69.3499742 | -0.000686766 | 0.36573064 | -0.0018778 | 0.99850174 | 0.99938628 |
| TBC1D9 | 391.965132 | -0.00049742 | 0.26916904 | -0.001848 | 0.99852552 | 0.99938628 |
| CA5B | 125.98088 | -0.000387443 | 0.23651253 | -0.0016381 | 0.99869295 | 0.99938628 |
| FEM1B | 87.1105929 | -0.000320285 | 0.25908411 | -0.0012362 | 0.99901364 | 0.99938628 |
| USP39 | 128.275198 | -0.000313745 | 0.25541343 | -0.0012284 | 0.9990199 | 0.99938628 |
| FAM120A | 396.688824 | -2.74E-05 | 0.1746896 | -0.0001571 | 0.99987465 | 0.99987465 |
| ARHGAP21 | 314.11939 | 8.09E-05 | 0.26054377 | 0.00031063 | 0.99975215 | 0.99987434 |
| FBXO38 | 141.699979 | 0.000132248 | 0.16113486 | 0.00082073 | 0.99934515 | 0.99958946 |
| FBRSL1 | 91.220821 | 0.000240553 | 0.1950346 | 0.00123339 | 0.9990159 | 0.99938628 |
| ABCB10 | 70.3092646 | 0.000431545 | 0.25573301 | 0.00168748 | 0.99865358 | 0.99938628 |
| AC020928.1 | 53.490705 | 0.000468604 | 0.23481553 | 0.00199563 | 0.99840772 | 0.99938628 |
| ORC5 | 124.137038 | 0.000539335 | 0.19107477 | 0.00282264 | 0.99774786 | 0.99938628 |
| DNAJA1 | 249.900598 | 0.000592004 | 0.23153429 | 0.00255687 | 0.99795991 | 0.99938628 |
| OSTM1 | 107.781777 | 0.000951582 | 0.21925974 | 0.00433998 | 0.99653721 | 0.99938628 |
| UBE2J2 | 96.8407577 | 0.001051305 | 0.21144005 | 0.00497212 | 0.99603284 | 0.99933015 |
| MSRA | 361.487451 | 0.001125473 | 0.27003597 | 0.00416786 | 0.99667454 | 0.99938628 |
| FAM149A | 47.4329826 | 0.001236442 | 0.43698173 | 0.00282951 | 0.99774238 | 0.99938628 |
| CCDC85A | 388.770739 | 0.001568114 | 0.61823515 | 0.00253644 | 0.99797622 | 0.99938628 |
| TXNIP | 1544.22815 | 0.00168097 | 0.59648725 | 0.00281812 | 0.99775147 | 0.99938628 |
| TRAPPC9 | 578.924513 | 0.001782928 | 0.26251472 | 0.00679172 | 0.99458103 | 0.99898392 |
| PLGRKT | 60.4914484 | 0.001902153 | 0.36374363 | 0.00522938 | 0.99582758 | 0.99933015 |
| FBXO8 | 53.5108569 | 0.001952917 | 0.31653539 | 0.00616966 | 0.99507735 | 0.99898392 |
| KAT2A | 51.3850634 | 0.002067453 | 0.48983354 | 0.00422073 | 0.99663236 | 0.99938628 |
| ZNF426 | 40.9135789 | 0.002261844 | 0.28624484 | 0.00790178 | 0.99369536 | 0.99863362 |
| POLA1 | 215.580559 | 0.00226362 | 0.23554 | 0.00961034 | 0.99233217 | 0.99826232 |
| RBM47 | 79.8895885 | 0.002765634 | 0.43737144 | 0.00632331 | 0.99495476 | 0.99898392 |
| ARID5B | 399.518937 | 0.003024479 | 0.37306383 | 0.00810713 | 0.99353151 | 0.99863362 |
| POM121C | 156.802194 | 0.00302483 | 0.20801914 | 0.01454112 | 0.98839828 | 0.99678917 |
| AC005332.7 | 58.1628357 | 0.003184393 | 0.3753167 | 0.00848455 | 0.99323039 | 0.99863362 |
| WNK1 | 666.783089 | 0.003630535 | 0.23428709 | 0.0154961 | 0.9876364 | 0.99656516 |
| 6-Mar | 498.358665 | 0.00386613 | 0.20261743 | 0.01908094 | 0.98477654 | 0.99577315 |
| MTMR14 | 105.680415 | 0.003989398 | 0.23967731 | 0.01664487 | 0.98671993 | 0.99644258 |
| FMNL2 | 2176.29721 | 0.004016754 | 0.49970078 | 0.00803832 | 0.99358642 | 0.99863362 |
| ZNF282 | 44.7507236 | 0.004065604 | 0.31676416 | 0.0128348 | 0.98975959 | 0.99731594 |
| TIGAR | 49.0758774 | 0.004128307 | 0.29689891 | 0.01390476 | 0.98890597 | 0.99678917 |
| TOX2 | 232.101042 | 0.004403012 | 0.31068657 | 0.01417188 | 0.98869286 | 0.99678917 |
| RNASEH2C | 68.6731453 | 0.004841413 | 0.44006268 | 0.01100164 | 0.99122214 | 0.99782925 |
| VTA1 | 130.724334 | 0.00487938 | 0.16610132 | 0.02937592 | 0.97656477 | 0.99296979 |
| KLHL8 | 115.592609 | 0.004891981 | 0.34258021 | 0.01427981 | 0.98860675 | 0.99678917 |
| GAPVD1 | 272.418551 | 0.005047811 | 0.13586598 | 0.03715287 | 0.97036312 | 0.99132846 |
| DPP8 | 310.105343 | 0.005068699 | 0.13146375 | 0.03855586 | 0.96924449 | 0.9911089 |
| RBAK-RBAKD | 122.069208 | 0.005191371 | 0.25100439 | 0.02068239 | 0.98349902 | 0.99512497 |
| RNF130 | 286.542155 | 0.005295668 | 0.16174344 | 0.03274116 | 0.973881 | 0.99243689 |
| SORBS1 | 743.88191 | 0.005307476 | 0.54034821 | 0.00982232 | 0.99216304 | 0.99826232 |
| KCNN3 | 304.33765 | 0.005415821 | 0.38970939 | 0.01389708 | 0.98891209 | 0.99678917 |
| CORO1C | 174.956972 | 0.005460389 | 0.3300661 | 0.01654332 | 0.98680094 | 0.99644258 |
| CAST | 564.540747 | 0.005512833 | 0.12414042 | 0.04440804 | 0.96457915 | 0.9886632 |
| GRB2 | 308.056245 | 0.00558842 | 0.18307421 | 0.03052544 | 0.97564801 | 0.99296979 |
| ZNF678 | 112.912753 | 0.005663964 | 0.20302842 | 0.0278974 | 0.97774399 | 0.99296979 |
| CDC25B | 65.7585842 | 0.005669717 | 0.21312302 | 0.02660302 | 0.97877636 | 0.99297384 |
| ARHGAP44 | 46.2554417 | 0.005727513 | 0.45455145 | 0.01260036 | 0.98994663 | 0.99738159 |
| YPEL3 | 101.95971 | 0.005862625 | 0.49292367 | 0.01189358 | 0.99051052 | 0.99770404 |
| CDC42EP4 | 121.132081 | 0.005952534 | 0.20639952 | 0.02883986 | 0.97699231 | 0.99296979 |
| DDX52 | 109.780376 | 0.006127219 | 0.22639369 | 0.02706444 | 0.97840834 | 0.99296979 |
| FOXJ2 | 48.5109073 | 0.006461911 | 0.28792109 | 0.02244334 | 0.98209431 | 0.99474451 |
| GNL1 | 111.496419 | 0.006475931 | 0.21983708 | 0.02945786 | 0.97649942 | 0.99296979 |
| COA1 | 335.105605 | 0.006485633 | 0.18085675 | 0.03586061 | 0.9713935 | 0.99163087 |
| MOB3A | 79.9672157 | 0.0066333 | 0.2957151 | 0.02243139 | 0.98210384 | 0.99474451 |
| LINC00551 | 75.9389994 | 0.006641276 | 0.57435155 | 0.01156309 | 0.9907742 | 0.997724 |
| SMC3 | 147.515814 | 0.006750369 | 0.20310859 | 0.03323527 | 0.97348697 | 0.99220522 |
| FAF1 | 960.503442 | 0.007121848 | 0.19343846 | 0.03681713 | 0.97063082 | 0.99134698 |
| LGMN | 102.693766 | 0.007275478 | 0.3824184 | 0.01902492 | 0.98482123 | 0.99577315 |
| FAM177A1 | 206.290917 | 0.007372091 | 0.16409272 | 0.04492637 | 0.964166 | 0.9886632 |
| GUSB | 78.5953507 | 0.007409453 | 0.22049547 | 0.03360365 | 0.97319321 | 0.99210665 |
| PKD2 | 339.346664 | 0.007493408 | 0.29336092 | 0.02554331 | 0.97962161 | 0.99333874 |
| ZNF521 | 925.687906 | 0.007623496 | 0.33096088 | 0.02303443 | 0.98162281 | 0.99462846 |
| RSPRY1 | 140.915565 | 0.00806793 | 0.17641033 | 0.04573389 | 0.96352235 | 0.9886632 |
| CDK17 | 1644.13792 | 0.008152873 | 0.38470046 | 0.02119278 | 0.98309187 | 0.99500814 |
| MTERF1 | 153.614956 | 0.008226723 | 0.45877887 | 0.01793178 | 0.98569327 | 0.99635831 |
| UBFD1 | 46.3261322 | 0.008269441 | 0.25658139 | 0.03222931 | 0.97428918 | 0.99244992 |
| CPNE8 | 998.63114 | 0.008318582 | 0.1755985 | 0.04737274 | 0.96221616 | 0.98828315 |
| AP002336.2 | 112.97797 | 0.008443721 | 0.34065827 | 0.02478648 | 0.98022527 | 0.9938277 |
| LGR4 | 408.766185 | 0.008513715 | 0.42459041 | 0.02005159 | 0.98400221 | 0.99531398 |
| USF1 | 40.9490123 | 0.008550589 | 0.43695338 | 0.01956865 | 0.98438747 | 0.9955806 |
| PLCG2 | 706.084901 | 0.008639832 | 0.33506288 | 0.0257857 | 0.97942827 | 0.99326577 |
| BPTF | 683.463878 | 0.0086815 | 0.23308417 | 0.0372462 | 0.9702887 | 0.99132846 |
| BECN1 | 74.855608 | 0.008751936 | 0.23562344 | 0.03714374 | 0.9703704 | 0.99132846 |
| AC007216.4 | 80.5865169 | 0.009187731 | 0.3077397 | 0.02985553 | 0.97618228 | 0.99296979 |
| CCNC | 166.731881 | 0.009207154 | 0.22019461 | 0.04181371 | 0.9666472 | 0.98953436 |
| POLDIP3 | 72.1024814 | 0.009331574 | 0.21506312 | 0.04338993 | 0.9653907 | 0.98875723 |
| TRIM4 | 121.060168 | 0.009354587 | 0.29721539 | 0.0314741 | 0.97489145 | 0.99260131 |
| MRPL13 | 55.7263026 | 0.009469048 | 0.27391855 | 0.03456884 | 0.97242355 | 0.99210665 |
| SMG5 | 85.6847905 | 0.009762427 | 0.22397013 | 0.04358808 | 0.96523276 | 0.98875723 |
| RBMS1 | 1944.70934 | 0.009867428 | 0.323794 | 0.0304744 | 0.97568871 | 0.99296979 |
| MPLKIP | 68.5674787 | 0.009898158 | 0.22231685 | 0.04452275 | 0.96448772 | 0.9886632 |
| CLIC2 | 206.180728 | 0.009936909 | 0.22219258 | 0.04472206 | 0.96432885 | 0.9886632 |
| WRN | 265.855282 | 0.010019427 | 0.20592365 | 0.04865603 | 0.96119342 | 0.98774906 |
| ZMPSTE24 | 88.2805045 | 0.010068239 | 0.24713736 | 0.04073944 | 0.96750362 | 0.99013033 |
| GPBP1L1 | 411.614999 | 0.010119212 | 0.1759479 | 0.05751254 | 0.95413691 | 0.98557318 |
| FAM174A | 64.1982183 | 0.010209478 | 0.22936 | 0.0445129 | 0.96449557 | 0.9886632 |
| ETS2 | 963.219016 | 0.010243364 | 0.46684174 | 0.02194183 | 0.98249435 | 0.99474451 |
| GBP2 | 161.898998 | 0.010401721 | 0.33231989 | 0.03130033 | 0.97503003 | 0.9926189 |
| TRIT1 | 71.9858824 | 0.010451734 | 0.2250235 | 0.0464473 | 0.96295374 | 0.98844085 |
| SCGB2B2 | 104.709855 | 0.010530215 | 0.36517222 | 0.0288363 | 0.97699515 | 0.99296979 |
| EXT2 | 231.761719 | 0.010562407 | 0.14694796 | 0.07187855 | 0.94269856 | 0.98263114 |
| RNF219 | 63.4867007 | 0.010657632 | 0.27255937 | 0.03910206 | 0.96880902 | 0.99099174 |
| ZW10 | 43.8186401 | 0.01073617 | 0.24208666 | 0.04434846 | 0.96462665 | 0.9886632 |
| TYW3 | 69.2370829 | 0.010764194 | 0.29130181 | 0.03695203 | 0.97052325 | 0.99134698 |
| SEMA3G | 231.676377 | 0.010920329 | 0.6211978 | 0.01757947 | 0.98597433 | 0.99644258 |

| | | | | | | |
|---|---|---|---|---|---|---|
| IPO11 | 226.869251 | 0.01117042 | 0.2255556 | 0.04952402 | 0.96050169 | 0.98739605 |
| MED14 | 154.684585 | 0.01118964 | 0.18233156 | 0.06136974 | 0.95106475 | 0.98426241 |
| C2CD2 | 192.205807 | 0.011226834 | 0.33000483 | 0.03402021 | 0.97286103 | 0.99210665 |
| IDH3B | 41.3554044 | 0.011289078 | 0.33510199 | 0.03368849 | 0.97312556 | 0.99210665 |
| RBMX | 277.307821 | 0.01141991 | 0.15896706 | 0.07183822 | 0.94273066 | 0.98263114 |
| ITM2B | 1392.07084 | 0.011520244 | 0.38644561 | 0.02981078 | 0.97621796 | 0.99296979 |
| OCIAD1 | 300.384024 | 0.011651634 | 0.18121089 | 0.06429875 | 0.94873235 | 0.98379261 |
| RINT1 | 56.3821841 | 0.011884776 | 0.21664544 | 0.05485819 | 0.95625144 | 0.98647256 |
| PRPF40A | 259.77922 | 0.011962998 | 0.19951746 | 0.05995965 | 0.95218777 | 0.9849155 |
| CCDC126 | 72.9860304 | 0.012137175 | 0.24477262 | 0.04958551 | 0.96045269 | 0.98739605 |
| DENND4A | 570.174284 | 0.01214848 | 0.25879151 | 0.04694312 | 0.96255857 | 0.98828315 |
| PPP4R3B | 325.863265 | 0.012201844 | 0.15958767 | 0.07645856 | 0.93905428 | 0.98171087 |
| KPNA3 | 297.488705 | 0.012332179 | 0.182854 | 0.06744276 | 0.94622923 | 0.98379261 |
| MAP3K14 | 132.207425 | 0.012715018 | 0.32582556 | 0.039024 | 0.96887125 | 0.99099174 |
| NAA50 | 231.850557 | 0.012939935 | 0.16571105 | 0.07808734 | 0.93775858 | 0.98137001 |
| TMEM106B | 140.780175 | 0.012991274 | 0.25041698 | 0.05187857 | 0.95862545 | 0.98725593 |
| GNPTAB | 135.890027 | 0.013070555 | 0.25440089 | 0.05137779 | 0.95902449 | 0.98725593 |
| MROH1 | 115.377917 | 0.01317638 | 0.38517885 | 0.03420847 | 0.97271091 | 0.99210665 |
| AP2A2 | 292.883698 | 0.013226561 | 0.20239746 | 0.06534944 | 0.94789578 | 0.98379261 |
| MAPKAP1 | 433.275476 | 0.013324937 | 0.23574699 | 0.05652219 | 0.95492582 | 0.98603425 |
| BCLAF3 | 89.8554367 | 0.01353147 | 0.24015024 | 0.05634585 | 0.95506629 | 0.98603425 |
| OTUD4 | 222.42534 | 0.013535122 | 0.18941969 | 0.07145573 | 0.94303506 | 0.98263114 |
| PIR | 150.859241 | 0.013797236 | 0.27086314 | 0.05093803 | 0.9593749 | 0.98739605 |
| GORASP1 | 60.2226183 | 0.013958846 | 0.31752716 | 0.04396111 | 0.96493541 | 0.9887386 |
| BOC | 377.646215 | 0.013966648 | 0.29087929 | 0.04801527 | 0.96170407 | 0.98802566 |
| PMPCA | 47.3671689 | 0.014230507 | 0.25551848 | 0.05569267 | 0.95558664 | 0.98610144 |
| DHX16 | 42.0506848 | 0.01498365 | 0.23716277 | 0.06317876 | 0.94962416 | 0.98401602 |
| DUSP16 | 296.080465 | 0.015078239 | 0.42509287 | 0.03547046 | 0.9717046 | 0.99182472 |
| ARL15 | 7198.91035 | 0.015511138 | 0.53538496 | 0.02897193 | 0.97688697 | 0.99296979 |
| TRRAP | 273.544632 | 0.015598063 | 0.18330511 | 0.08509344 | 0.9321871 | 0.97871274 |
| NUDCD3 | 301.76005 | 0.015654289 | 0.168296 | 0.09301641 | 0.92589052 | 0.97614605 |
| TSPAN4 | 115.207414 | 0.015693035 | 0.32960115 | 0.0476122 | 0.96202531 | 0.98823162 |
| SMARCB1 | 80.2486159 | 0.015834542 | 0.23769459 | 0.06661717 | 0.94688647 | 0.98379261 |
| LATS1 | 211.363438 | 0.016035684 | 0.24291661 | 0.06601312 | 0.94736738 | 0.98379261 |
| ZER1 | 149.823774 | 0.01624646 | 0.24718972 | 0.06572466 | 0.94759704 | 0.98379261 |
| ZBED4 | 69.9028956 | 0.016476372 | 0.23094338 | 0.07134377 | 0.94312416 | 0.98263114 |
| SRSF10 | 220.058775 | 0.016549615 | 0.1617913 | 0.10228989 | 0.91852658 | 0.97260649 |
| TMBIM1 | 142.840098 | 0.01681647 | 0.23005228 | 0.07309847 | 0.94172776 | 0.98263114 |
| SLC9A6 | 52.527702 | 0.016879165 | 0.2405476 | 0.07016975 | 0.94405855 | 0.98335427 |
| PLLP | 74.7935901 | 0.016952404 | 0.60378696 | 0.0280768 | 0.9776009 | 0.99296979 |
| EIF4E | 171.528767 | 0.017031246 | 0.15558303 | 0.10946725 | 0.9128319 | 0.9709041 |
| LMF1 | 124.576255 | 0.017193196 | 0.33293138 | 0.05164186 | 0.95881406 | 0.98725593 |
| TCERG1 | 241.682262 | 0.017341658 | 0.24037463 | 0.07214429 | 0.94248708 | 0.98263114 |
| URI1 | 236.1886 | 0.01739186 | 0.2562039 | 0.06788288 | 0.94587886 | 0.98379261 |
| TAF15 | 521.758142 | 0.017553735 | 0.14819678 | 0.11844883 | 0.90571204 | 0.96904724 |
| CYHR1 | 59.382898 | 0.017607013 | 0.39591257 | 0.04447197 | 0.96452819 | 0.9886632 |
| SRRM2 | 638.568779 | 0.017611958 | 0.12997495 | 0.13550271 | 0.89221442 | 0.96471863 |
| RPA2 | 51.5613554 | 0.017713182 | 0.24385023 | 0.0726396 | 0.94209292 | 0.98263114 |
| USP13 | 224.177945 | 0.017872253 | 0.46449612 | 0.03847665 | 0.96930765 | 0.9911089 |
| ZNF720 | 117.643121 | 0.018003979 | 0.24314795 | 0.07404536 | 0.94097429 | 0.98256304 |
| TECR | 53.4426301 | 0.018062723 | 0.38357549 | 0.0470904 | 0.96244118 | 0.98828315 |
| ELMO1 | 2027.6526 | 0.018489463 | 0.14124399 | 0.13090442 | 0.89585092 | 0.96584296 |
| ADH5 | 80.3924818 | 0.018534614 | 0.30407859 | 0.06095337 | 0.95139635 | 0.98446153 |
| MORF4L1 | 402.165171 | 0.019138389 | 0.19649507 | 0.09739883 | 0.92240968 | 0.9749687 |
| MAP2K5 | 433.258049 | 0.019198486 | 0.18438009 | 0.10412451 | 0.91707054 | 0.97233474 |
| SIGIRR | 99.2094257 | 0.019466646 | 0.37354422 | 0.05211336 | 0.95843837 | 0.98725593 |
| TRA2B | 237.633049 | 0.019495144 | 0.21997079 | 0.08862606 | 0.9293791 | 0.97696444 |
| ZNF337 | 82.5667843 | 0.019580843 | 0.22115224 | 0.08854011 | 0.92944741 | 0.97696444 |
| BROX | 101.861136 | 0.019760716 | 0.19877423 | 0.09941287 | 0.92081047 | 0.97414248 |
| BRMS1L | 57.3377248 | 0.01994484 | 0.30273122 | 0.065883 | 0.94747098 | 0.98379261 |
| IMMT | 195.469983 | 0.020031258 | 0.18166819 | 0.11026288 | 0.91220089 | 0.97083977 |
| AL603840.1 | 84.3580426 | 0.020107378 | 0.90889311 | 0.02212293 | 0.9823499 | 0.99474451 |
| SNW1 | 128.329343 | 0.020151044 | 0.20991917 | 0.0959943 | 0.9235251 | 0.97530455 |
| TAGLN | 96.1316319 | 0.020302659 | 0.70802027 | 0.02867525 | 0.9771236 | 0.99296979 |
| NXPE3 | 131.282885 | 0.020651088 | 0.33477043 | 0.06168731 | 0.95081184 | 0.98412513 |
| ABHD15-AS1 | 65.8145584 | 0.021133023 | 0.42299561 | 0.04996038 | 0.96015396 | 0.98739605 |
| GPR75-ASB3 | 81.5318513 | 0.021221369 | 0.34362145 | 0.06175799 | 0.95075556 | 0.98412513 |
| POLR2H | 91.3268683 | 0.021230979 | 0.36209011 | 0.05863452 | 0.95324322 | 0.98545694 |
| MECOM | 3184.68456 | 0.021260296 | 0.39546496 | 0.05376025 | 0.95712618 | 0.98679142 |
| GDI1 | 87.460198 | 0.021290414 | 0.2790439 | 0.07629772 | 0.93918224 | 0.98171087 |
| RIT1 | 87.8924695 | 0.021426958 | 0.32713868 | 0.06549809 | 0.94777743 | 0.98379261 |
| SH3GLB2 | 40.9237827 | 0.021476201 | 0.33172886 | 0.06474023 | 0.94838083 | 0.98379261 |
| SNCAIP | 350.098214 | 0.021555302 | 0.50229674 | 0.04291348 | 0.9657705 | 0.98885135 |
| CTNNA1 | 974.592587 | 0.021777716 | 0.1840495 | 0.11832532 | 0.9058099 | 0.96904724 |
| ANKLE2 | 159.314177 | 0.021824182 | 0.2427164 | 0.08991639 | 0.92835366 | 0.97694418 |
| ITGB1BP1 | 138.259823 | 0.02201804 | 0.178728 | 0.12319301 | 0.90195426 | 0.96801711 |
| UBE2E3 | 294.980682 | 0.022026816 | 0.2086908 | 0.10554762 | 0.91594129 | 0.97233474 |
| ZZEF1 | 246.528029 | 0.022091952 | 0.17130339 | 0.12896389 | 0.89738622 | 0.96608714 |
| CWF19L1 | 85.1857004 | 0.022549967 | 0.17327252 | 0.13014162 | 0.89645438 | 0.96608352 |
| SUGP2 | 160.014097 | 0.022710049 | 0.17289997 | 0.1313479 | 0.8955001 | 0.96575575 |
| UBAC1 | 60.3852555 | 0.022766283 | 0.23281669 | 0.0977863 | 0.92210198 | 0.9749687 |
| NFYB | 52.8766018 | 0.02277131 | 0.28056917 | 0.08116113 | 0.93531381 | 0.98010864 |
| ERCC6L2 | 249.108556 | 0.022819378 | 0.1888679 | 0.1208219 | 0.90383211 | 0.9688759 |
| COL6A1 | 67.4105233 | 0.022887786 | 0.58800984 | 0.03892415 | 0.96895086 | 0.99099174 |
| MGRN1 | 110.596059 | 0.023067032 | 0.25074223 | 0.091995 | 0.92670201 | 0.97645253 |
| NRDE2 | 95.9238714 | 0.023243621 | 0.33770541 | 0.0688281 | 0.94512645 | 0.98379261 |
| MGAT5 | 349.176981 | 0.02365587 | 0.28342702 | 0.08346371 | 0.93348283 | 0.97957041 |
| MXRA7 | 146.832938 | 0.023714103 | 0.33180026 | 0.07147102 | 0.94302289 | 0.98263114 |
| RSRC1 | 409.440981 | 0.02393477 | 0.1702351 | 0.14059833 | 0.88818727 | 0.96341946 |
| TMX3 | 166.457592 | 0.024243809 | 0.2314923 | 0.10472836 | 0.91659136 | 0.97233474 |
| SNX4 | 195.769099 | 0.024283317 | 0.13430082 | 0.18081287 | 0.85651446 | 0.95134391 |
| PTBP1 | 234.485949 | 0.024461258 | 0.34439387 | 0.07102698 | 0.94337628 | 0.98276869 |
| OGFOD1 | 47.4509244 | 0.024468736 | 0.33420765 | 0.07321417 | 0.94163569 | 0.98263114 |
| PARG | 247.447319 | 0.024726166 | 0.14625196 | 0.16906554 | 0.86574509 | 0.95510194 |
| SYT11 | 82.4981839 | 0.024976712 | 0.39953899 | 0.06251383 | 0.95015365 | 0.98412513 |
| PSMD5 | 112.692728 | 0.024987578 | 0.18265728 | 0.13680034 | 0.89118862 | 0.96448584 |
| DCAF8 | 49.9514586 | 0.02566406 | 0.26848302 | 0.09558914 | 0.92384689 | 0.97530455 |
| ZNF766 | 53.3870076 | 0.026310939 | 0.40583511 | 0.0648316 | 0.94830808 | 0.98379261 |
| YWHAZ | 448.351313 | 0.026323951 | 0.22773285 | 0.11559136 | 0.90797641 | 0.96940376 |
| NDUFB1 | 71.3636443 | 0.026388773 | 0.29230783 | 0.09027734 | 0.92806683 | 0.9768936 |
| LINC01197 | 103.580258 | 0.026503113 | 0.34978064 | 0.07577067 | 0.93960155 | 0.98171087 |
| MYO9B | 256.117244 | 0.026725967 | 0.23716618 | 0.11268878 | 0.9102773 | 0.9700595 |
| RBBP5 | 67.5430637 | 0.026764343 | 0.22558584 | 0.11864372 | 0.90555763 | 0.96904724 |
| BORCS5 | 99.1369368 | 0.026790778 | 0.22042537 | 0.12154126 | 0.90326233 | 0.96875453 |
| TRIM56 | 369.408203 | 0.027210701 | 0.35907739 | 0.07577949 | 0.93959453 | 0.98171087 |
| RNF10 | 195.809434 | 0.027429941 | 0.13117233 | 0.20911378 | 0.83435942 | 0.9432647 |
| ELOVL5 | 342.381894 | 0.027459337 | 0.28436487 | 0.09656375 | 0.92307285 | 0.97527503 |
| ZHX3 | 346.611304 | 0.027565045 | 0.24860609 | 0.1108784 | 0.91171278 | 0.97054061 |
| NDUFA5 | 98.4217641 | 0.027741435 | 0.25357667 | 0.10940058 | 0.91288477 | 0.9709041 |
| ATG4A | 56.2484801 | 0.028141952 | 0.22234847 | 0.12656688 | 0.89928322 | 0.96723863 |
| SLC25A25 | 52.5024893 | 0.028191176 | 0.34095135 | 0.08268387 | 0.93410291 | 0.97971855 |
| SLC25A17 | 115.700736 | 0.028309365 | 0.24523541 | 0.11543751 | 0.90809835 | 0.96940376 |
| STX5 | 140.724638 | 0.028502063 | 0.17035809 | 0.16730678 | 0.86712867 | 0.95535286 |

| | | | | | | |
|---|---|---|---|---|---|---|
| GMCL1 | 83.6692072 | 0.028686488 | 0.20089888 | 0.14279068 | 0.8864555 | 0.9627377 |
| SH3BGRL | 324.088311 | 0.028693922 | 0.19241524 | 0.149125 | 0.881455 | 0.96059232 |
| SOAT1 | 130.993272 | 0.028702391 | 0.20175889 | 0.14226085 | 0.88687397 | 0.96288838 |
| CCDC50 | 463.888385 | 0.028977378 | 0.35159409 | 0.08241714 | 0.93431501 | 0.97975627 |
| MFF | 142.669491 | 0.02911615 | 0.2061407 | 0.14124406 | 0.88767714 | 0.96330655 |
| DDHD1 | 195.73947 | 0.029157706 | 0.22606198 | 0.12898103 | 0.89737266 | 0.96608714 |
| PYROXD1 | 56.4733313 | 0.029357352 | 0.1914791 | 0.15331883 | 0.87814685 | 0.95969416 |
| LINC00662 | 90.5485965 | 0.029457283 | 0.30766711 | 0.09574401 | 0.92372388 | 0.97530455 |
| TMEM173 | 127.303591 | 0.029514525 | 0.38262616 | 0.07713671 | 0.93851479 | 0.9817035 |
| NUP155 | 88.2850378 | 0.029581191 | 0.19208327 | 0.15400192 | 0.87760823 | 0.95969416 |
| PPIE | 41.6735876 | 0.02970424 | 0.26462724 | 0.11224937 | 0.91062569 | 0.9700595 |
| TRAPPC12 | 196.600038 | 0.029788755 | 0.23036282 | 0.12931234 | 0.89711051 | 0.96608714 |
| VPS13B | 950.632075 | 0.029824125 | 0.14174064 | 0.21041337 | 0.83334506 | 0.94303136 |
| NUMA1 | 214.492841 | 0.029850594 | 0.22088096 | 0.13514336 | 0.89249853 | 0.96482231 |
| ATM | 261.197198 | 0.030009942 | 0.21850162 | 0.13734425 | 0.89075869 | 0.96448584 |
| DDI2 | 127.37476 | 0.030075302 | 0.22723421 | 0.13235376 | 0.89470449 | 0.96575575 |
| ARFIP1 | 359.995088 | 0.030156997 | 0.24591758 | 0.1226305 | 0.9023997 | 0.96818365 |
| IL15 | 298.058513 | 0.030211217 | 0.352513 | 0.08570242 | 0.93170298 | 0.97845553 |
| ZNF76 | 78.6440819 | 0.030414875 | 0.19895835 | 0.15287057 | 0.87850035 | 0.95972986 |
| CBLL1 | 47.8996799 | 0.03069086 | 0.26380399 | 0.11633964 | 0.90738337 | 0.96940376 |
| SCN8A | 51.6146145 | 0.030871139 | 0.48019406 | 0.06428888 | 0.94874021 | 0.98379261 |
| KDM2B | 87.6819972 | 0.030878664 | 0.29352962 | 0.10519778 | 0.91621888 | 0.97233474 |
| COG7 | 109.168495 | 0.031160912 | 0.2320669 | 0.13427556 | 0.89318468 | 0.96512555 |
| TUBD1 | 51.804914 | 0.031414076 | 0.25601571 | 0.12270371 | 0.90234173 | 0.96818365 |
| ARL13B | 176.465088 | 0.031445432 | 0.18619447 | 0.16888489 | 0.86588719 | 0.95510194 |
| PIK3R4 | 107.712021 | 0.031585337 | 0.18391331 | 0.17174035 | 0.86364166 | 0.95399294 |
| PPARG | 956.135045 | 0.031788848 | 0.47567402 | 0.06682906 | 0.94671779 | 0.98379261 |
| EPS8 | 221.293325 | 0.032394764 | 0.3632124 | 0.08918959 | 0.92893124 | 0.97696444 |
| MRPS5 | 128.267325 | 0.032788914 | 0.19459854 | 0.16849517 | 0.86619375 | 0.95510194 |
| MVB12A | 68.6163949 | 0.032884174 | 0.24991326 | 0.13158235 | 0.89531465 | 0.96575575 |
| ZNF7 | 48.0847486 | 0.032891946 | 0.31031568 | 0.10599511 | 0.91558623 | 0.97233474 |
| PALLD | 72.5202677 | 0.033055034 | 0.32060674 | 0.10310149 | 0.91788242 | 0.97233474 |
| ERMARD | 46.6259813 | 0.033276424 | 0.31548439 | 0.10547724 | 0.91599713 | 0.97233474 |
| DYM | 775.001908 | 0.033329697 | 0.16306711 | 0.20439252 | 0.83804678 | 0.94358503 |
| MAN2C1 | 145.629496 | 0.033365719 | 0.15308626 | 0.21795372 | 0.82746517 | 0.94018985 |
| CLASRP | 102.06866 | 0.033540832 | 0.25128069 | 0.13347954 | 0.89381414 | 0.96542781 |
| DCTN2 | 119.120688 | 0.033571835 | 0.2472486 | 0.1357817 | 0.89199386 | 0.9646076 |
| PPP2R5A | 479.675061 | 0.033590899 | 0.15628792 | 0.21492959 | 0.82982221 | 0.94063377 |
| METTL16 | 162.717528 | 0.033613555 | 0.25632661 | 0.13113564 | 0.89566801 | 0.965773 |
| CIRBP | 213.095243 | 0.033643994 | 0.28830454 | 0.11669602 | 0.90710094 | 0.96940376 |
| MXD4 | 255.472986 | 0.033649429 | 0.35112126 | 0.09583421 | 0.92365225 | 0.97530455 |
| PER3 | 172.072844 | 0.033844403 | 0.55775811 | 0.06067936 | 0.95161457 | 0.98446153 |
| HIRA | 62.3783354 | 0.033921369 | 0.31281375 | 0.10843951 | 0.91364706 | 0.97120992 |
| ING3 | 63.1436837 | 0.033928742 | 0.2922094 | 0.11611106 | 0.90756453 | 0.96940376 |
| GNB4 | 400.976455 | 0.033992718 | 0.29891936 | 0.11371869 | 0.9094608 | 0.96953071 |
| STK25 | 100.932036 | 0.034355914 | 0.22856995 | 0.1503081 | 0.88052154 | 0.96048934 |
| AP4E1 | 107.706479 | 0.034470833 | 0.18268008 | 0.18869509 | 0.8503318 | 0.9492877 |
| NOSIP | 85.9731406 | 0.034485695 | 0.2781357 | 0.12398874 | 0.90132419 | 0.96790318 |
| EPB41L4A | 1721.33537 | 0.034498618 | 0.27498074 | 0.12545831 | 0.90016073 | 0.96744565 |
| TEX9 | 50.173493 | 0.034508777 | 0.53748269 | 0.06420444 | 0.94880744 | 0.98379261 |
| HNRNPDL | 325.045903 | 0.034546462 | 0.18627471 | 0.18545975 | 0.85286844 | 0.95044301 |
| ZNF592 | 126.911064 | 0.035384761 | 0.24790619 | 0.14273448 | 0.88649989 | 0.9627377 |
| ITGAE | 41.8767661 | 0.035536378 | 0.34580925 | 0.10276295 | 0.91815111 | 0.97233474 |
| ACYP2 | 429.516785 | 0.035627977 | 0.21715426 | 0.1640676 | 0.86967793 | 0.95673577 |
| SRSF8 | 73.1782603 | 0.035701648 | 0.30802456 | 0.1159052 | 0.90772768 | 0.96940376 |
| AL078459.1 | 58.5311466 | 0.035715393 | 0.49181726 | 0.07261923 | 0.94210912 | 0.98263114 |
| ANKRD17 | 1233.7247 | 0.035718514 | 0.18609963 | 0.19193221 | 0.8477953 | 0.94731791 |
| ACTL6A | 58.4800109 | 0.035810575 | 0.2674732 | 0.13388472 | 0.89349373 | 0.96520914 |
| AGPAT3 | 210.988504 | 0.035823845 | 0.22662317 | 0.15807671 | 0.87439635 | 0.95824097 |
| LYPLA1 | 210.687398 | 0.035944868 | 0.20615369 | 0.17435957 | 0.86158289 | 0.95324701 |
| EXOC5 | 264.454207 | 0.03598277 | 0.14460656 | 0.24883222 | 0.80349057 | 0.93081002 |
| CTDSPL2 | 299.496105 | 0.036002466 | 0.16820593 | 0.21403804 | 0.8305174 | 0.94075329 |
| SETD5 | 719.456912 | 0.036035722 | 0.17367286 | 0.20749197 | 0.83562567 | 0.9432647 |
| SLC25A46 | 129.567763 | 0.036041675 | 0.21386518 | 0.16852521 | 0.86617011 | 0.95510194 |
| LYL1 | 67.8176223 | 0.036102171 | 0.5531253 | 0.06526943 | 0.94795949 | 0.98379261 |
| ANKRD12 | 931.591466 | 0.036116194 | 0.21744469 | 0.16609371 | 0.8680832 | 0.95580259 |
| FAM174B | 122.654193 | 0.036129698 | 0.39731589 | 0.09093444 | 0.92754468 | 0.97688257 |
| SUCO | 184.445474 | 0.036337154 | 0.20140717 | 0.18041639 | 0.85682569 | 0.95134391 |
| YTHDC1 | 322.266079 | 0.036497153 | 0.22672724 | 0.16097384 | 0.87211401 | 0.95727819 |
| TMED5 | 123.57911 | 0.036889675 | 0.26795139 | 0.13767301 | 0.89049885 | 0.96439281 |
| FERMT2 | 466.08291 | 0.036922762 | 0.19264675 | 0.19166045 | 0.84800819 | 0.94733802 |
| CEP89 | 80.1278742 | 0.036971836 | 0.24976336 | 0.14802746 | 0.8823211 | 0.9608775 |
| CYTH1 | 706.77421 | 0.037041647 | 0.31931227 | 0.11600446 | 0.90764901 | 0.96940376 |
| HAT1 | 121.951407 | 0.03706361 | 0.20151694 | 0.18392305 | 0.85407382 | 0.95094505 |
| TRAP1 | 98.209272 | 0.037267416 | 0.26475876 | 0.1407599 | 0.88805962 | 0.96340871 |
| EIF2AK1 | 98.9506515 | 0.037336237 | 0.16886157 | 0.22110559 | 0.82501022 | 0.93870172 |
| AGFG1 | 391.028122 | 0.037345299 | 0.17958181 | 0.20795703 | 0.83526252 | 0.9432647 |
| PRTG | 139.933327 | 0.037482785 | 0.38491679 | 0.09737893 | 0.92242548 | 0.9749687 |
| DNM2 | 302.417625 | 0.037785289 | 0.2169139 | 0.17419487 | 0.86171232 | 0.95324701 |
| PLAA | 121.099555 | 0.037901332 | 0.29745323 | 0.12741947 | 0.89860841 | 0.96690501 |
| UXS1 | 79.4929367 | 0.037904036 | 0.26338392 | 0.14391173 | 0.88557017 | 0.9626223 |
| B3GALNT2 | 85.7768476 | 0.037972487 | 0.29321777 | 0.12950268 | 0.89695991 | 0.96608714 |
| PLA2G16 | 128.96872 | 0.038083845 | 0.39335505 | 0.09681799 | 0.92287094 | 0.97518761 |
| PRKRIP1 | 358.497702 | 0.03812132 | 0.24998467 | 0.15249463 | 0.87879682 | 0.95972986 |
| ANAPC10 | 202.815061 | 0.038130242 | 0.14917215 | 0.25561235 | 0.79825018 | 0.92959949 |
| TERF1 | 104.312284 | 0.038210117 | 0.21860383 | 0.17479162 | 0.86124338 | 0.95314506 |
| MMADHC | 75.9497523 | 0.038281711 | 0.26195878 | 0.14613639 | 0.88381372 | 0.96173506 |
| PLCB1 | 3111.64614 | 0.038656243 | 0.22309245 | 0.17327455 | 0.86243563 | 0.95356178 |
| LMAN2L | 40.1766175 | 0.038681332 | 0.25422882 | 0.15215164 | 0.87906733 | 0.95972986 |
| PITPNB | 443.446118 | 0.038687493 | 0.26353179 | 0.14680389 | 0.88328681 | 0.96141487 |
| COX7A1 | 57.3593379 | 0.038899898 | 0.59167492 | 0.06574539 | 0.94758053 | 0.98379261 |
| PCYT1A | 86.4620222 | 0.038963576 | 0.24493882 | 0.15907473 | 0.87361 | 0.95804618 |
| CYB5R1 | 39.5017969 | 0.038971609 | 0.37030546 | 0.10524179 | 0.91618395 | 0.97233474 |
| TBK1 | 289.034473 | 0.038985365 | 0.2205816 | 0.17673897 | 0.85971343 | 0.95263718 |
| CRNDE | 139.504849 | 0.039486657 | 0.2771416 | 0.14247827 | 0.88670224 | 0.96282968 |
| SUCLA2 | 162.488196 | 0.039624595 | 0.28322057 | 0.13990719 | 0.88873332 | 0.96371185 |
| QTRT2 | 82.8101572 | 0.039653016 | 0.19225972 | 0.20624714 | 0.83659787 | 0.94347855 |
| TAF2 | 240.586731 | 0.039660618 | 0.16504384 | 0.24030353 | 0.81009496 | 0.93351089 |
| PRRC2B | 453.705761 | 0.039692662 | 0.17900825 | 0.2217365 | 0.82451902 | 0.93852534 |
| IQGAP1 | 625.487768 | 0.039779012 | 0.22573753 | 0.17621798 | 0.86012269 | 0.95267061 |
| DESI2 | 148.079995 | 0.040022449 | 0.13224402 | 0.3026409 | 0.76216354 | 0.91477008 |
| PCSK7 | 182.743153 | 0.040093916 | 0.19941221 | 0.20106049 | 0.84065128 | 0.94397833 |
| TCHP | 108.936988 | 0.040145455 | 0.25660756 | 0.15644689 | 0.87568078 | 0.95883833 |
| HNRNPU | 627.391721 | 0.040496918 | 0.2206885 | 0.18350262 | 0.85440366 | 0.95098559 |
| SH3GLB1 | 348.342115 | 0.040520867 | 0.15484885 | 0.26168013 | 0.79356806 | 0.92790728 |
| ZNRF1 | 329.24681 | 0.040635513 | 0.29287551 | 0.13874671 | 0.88965031 | 0.96398417 |
| TBC1D12 | 132.382619 | 0.040664324 | 0.20102427 | 0.20228564 | 0.83969342 | 0.94376667 |
| CDC123 | 125.875605 | 0.040796784 | 0.22046138 | 0.18505184 | 0.85318837 | 0.95052967 |
| TMEM242 | 77.866129 | 0.040896158 | 0.21227135 | 0.19265981 | 0.8472254 | 0.94731791 |
| DENND3 | 380.04836 | 0.041006698 | 0.39099769 | 0.10487708 | 0.91647334 | 0.97233474 |
| PKNOX1 | 91.5879085 | 0.041116698 | 0.28793538 | 0.14279835 | 0.88644944 | 0.9627377 |
| TIPRL | 106.889206 | 0.041137231 | 0.20541084 | 0.20026807 | 0.84127094 | 0.94419422 |
| JPT2 | 52.175621 | 0.041184578 | 0.23345556 | 0.17641292 | 0.85996955 | 0.95263718 |

| | | | | | | |
|---|---|---|---|---|---|---|
| BCAM | 174.77387 | 0.041278392 | 0.4557187 | 0.09057867 | 0.92782738 | 0.97688257 |
| EPN2 | 555.099307 | 0.041533504 | 0.24061218 | 0.17261596 | 0.8629533 | 0.95381365 |
| IFT74 | 105.525896 | 0.04161647 | 0.22938917 | 0.181423 | 0.85603557 | 0.95134391 |
| ABHD13 | 72.2072947 | 0.041827636 | 0.28909235 | 0.14468607 | 0.88495874 | 0.96208547 |
| CPM | 70.1727759 | 0.042248597 | 0.50746993 | 0.0832534 | 0.93365005 | 0.97962026 |
| CAMKK1 | 69.0810762 | 0.042316175 | 0.39803539 | 0.10631259 | 0.91533434 | 0.97233474 |
| SLC38A2 | 624.631147 | 0.042360371 | 0.5890017 | 0.07191893 | 0.94266643 | 0.98263114 |
| TEFM | 70.8362912 | 0.042410139 | 0.28734774 | 0.14759169 | 0.88266501 | 0.96103918 |
| SGCE | 94.8924611 | 0.04245033 | 0.47165567 | 0.0900028 | 0.92828499 | 0.97694418 |
| MAML1 | 93.9480183 | 0.04245691 | 0.20806285 | 0.2040581 | 0.83830809 | 0.94362079 |
| VPS53 | 419.079299 | 0.042974468 | 0.23938349 | 0.17952143 | 0.85752829 | 0.95199484 |
| AL683813.1 | 59.7991812 | 0.043051398 | 0.42529249 | 0.10122774 | 0.91936967 | 0.97312147 |
| CGRRF1 | 52.943034 | 0.043254877 | 0.25170187 | 0.17184965 | 0.86355573 | 0.95399294 |
| SMARCAD1 | 218.850345 | 0.043411504 | 0.16069614 | 0.27014653 | 0.78704753 | 0.92401864 |
| ATG7 | 732.783279 | 0.043428831 | 0.38811485 | 0.11189686 | 0.91090519 | 0.9700595 |
| DNAJC11 | 117.429669 | 0.043895466 | 0.24246756 | 0.18103645 | 0.85633897 | 0.95134391 |
| BRD8 | 74.7818658 | 0.044141285 | 0.21577291 | 0.20457288 | 0.83790584 | 0.94358503 |
| AC008014.1 | 387.620436 | 0.044176302 | 0.67169858 | 0.06576804 | 0.9475625 | 0.98379261 |
| RAD54L2 | 135.855006 | 0.044256223 | 0.24191678 | 0.18293986 | 0.8548452 | 0.95133936 |
| RSRC2 | 200.025136 | 0.044301993 | 0.22365412 | 0.19808261 | 0.84298043 | 0.94481699 |
| MOSMO | 299.330228 | 0.044429724 | 0.16764572 | 0.26502152 | 0.79099288 | 0.92639112 |
| METTL3 | 47.0003869 | 0.044526903 | 0.25025086 | 0.17792907 | 0.85877868 | 0.95247071 |
| TIMM23B | 202.221677 | 0.044838998 | 0.39261661 | 0.11420556 | 0.90907485 | 0.96940376 |
| SLC38A6 | 83.1987714 | 0.044869192 | 0.31005565 | 0.14471335 | 0.8849372 | 0.96208547 |
| CGGBP1 | 249.50701 | 0.044926065 | 0.276645 | 0.16239608 | 0.87099396 | 0.95709072 |
| MTG2 | 78.8461526 | 0.044989109 | 0.22823177 | 0.19712027 | 0.84373342 | 0.94501377 |
| SP1 | 129.34774 | 0.045151225 | 0.20960689 | 0.21540907 | 0.8294484 | 0.94063377 |
| PRPF6 | 129.393675 | 0.045224147 | 0.19675364 | 0.22985164 | 0.81820706 | 0.93593281 |
| RBM23 | 95.3458981 | 0.045297889 | 0.20871366 | 0.21703366 | 0.82818212 | 0.94018985 |
| VPS29 | 150.863361 | 0.045303079 | 0.19267715 | 0.2351243 | 0.81411226 | 0.93460727 |
| SMARCAL1 | 79.4240976 | 0.045338705 | 0.25770888 | 0.17592993 | 0.86034899 | 0.95267061 |
| AKT2 | 93.6489764 | 0.045368811 | 0.23076244 | 0.19660396 | 0.84413747 | 0.94533692 |
| ASPSCR1 | 44.2474199 | 0.045402712 | 0.43964039 | 0.10327239 | 0.91774679 | 0.97233474 |
| PIGC | 82.4187556 | 0.045459693 | 0.35040119 | 0.12973613 | 0.8967752 | 0.96608714 |
| ZNF790 | 89.2447805 | 0.045498974 | 0.28817441 | 0.15788693 | 0.87454589 | 0.95827652 |
| TAOK1 | 597.253265 | 0.045513089 | 0.15528253 | 0.29309858 | 0.76944681 | 0.916771 |
| CDC26 | 48.1087247 | 0.045683501 | 0.39099365 | 0.1168395 | 0.90698724 | 0.96940376 |
| ANKRD23 | 45.7476668 | 0.045781985 | 0.39858606 | 0.11486098 | 0.90855531 | 0.96940376 |
| ALKBH3 | 64.9113155 | 0.045953137 | 0.27232959 | 0.16874089 | 0.86600046 | 0.95510194 |
| TOPBP1 | 93.7349816 | 0.045985813 | 0.16911501 | 0.27192036 | 0.78568326 | 0.92347689 |
| RPP30 | 41.2633394 | 0.046230861 | 0.33421347 | 0.13832734 | 0.88998172 | 0.96407367 |
| NAE1 | 84.2809808 | 0.046402706 | 0.19883621 | 0.23337151 | 0.81547294 | 0.93550282 |
| CNOT6 | 135.787257 | 0.046420331 | 0.22251649 | 0.20861524 | 0.83474861 | 0.9432647 |
| HIKESHI | 91.2878193 | 0.046625733 | 0.23006772 | 0.20266091 | 0.83940008 | 0.94376667 |
| ZMIZ2 | 101.675698 | 0.046738514 | 0.27850332 | 0.16782032 | 0.86672464 | 0.95526468 |
| YWHAQ | 266.406611 | 0.046818869 | 0.15490523 | 0.30224201 | 0.76246758 | 0.91477008 |
| NMNAT3 | 62.8467172 | 0.046835946 | 0.34269461 | 0.13666963 | 0.89129194 | 0.96448584 |
| RRP7A | 71.2828678 | 0.046869543 | 0.41809709 | 0.11210206 | 0.91074249 | 0.9700595 |
| VPS35 | 111.135018 | 0.046876631 | 0.19054119 | 0.24601836 | 0.80566802 | 0.93158047 |
| API5 | 134.982383 | 0.047094719 | 0.1571688 | 0.2996442 | 0.76444857 | 0.9154077 |
| SUGP1 | 41.9836967 | 0.047171274 | 0.29770283 | 0.15845087 | 0.87410153 | 0.95804618 |
| EXOC7 | 93.654139 | 0.047231526 | 0.19590485 | 0.24109421 | 0.8094821 | 0.93351089 |
| CKAP5 | 116.779595 | 0.047239269 | 0.17707089 | 0.26678167 | 0.78963726 | 0.92602336 |
| SLC35D1 | 72.0490302 | 0.047288228 | 0.25915387 | 0.18247163 | 0.85521262 | 0.95134391 |
| DBT | 79.2686249 | 0.0473973 | 0.20284067 | 0.23366764 | 0.81524301 | 0.93538048 |
| POLR2B | 186.861944 | 0.04747944 | 0.17662758 | 0.26881102 | 0.78807511 | 0.924827 |
| NCOA1 | 1106.48976 | 0.047570785 | 0.16785864 | 0.28339789 | 0.77687185 | 0.91892777 |
| AATK | 42.3847927 | 0.047580481 | 0.29991524 | 0.15864642 | 0.87394745 | 0.95804618 |
| CASC3 | 143.902913 | 0.047865444 | 0.2761708 | 0.17331827 | 0.86240127 | 0.95356178 |
| CCNH | 780.53809 | 0.047879113 | 0.2953053 | 0.16213428 | 0.87120011 | 0.95709072 |
| FIG4 | 87.1342186 | 0.048017947 | 0.24688772 | 0.19449306 | 0.84578983 | 0.94653969 |
| JMY | 126.017051 | 0.048039106 | 0.33018242 | 0.14549262 | 0.88432194 | 0.96189869 |
| SPTLC1 | 192.054258 | 0.048211159 | 0.15654038 | 0.30797906 | 0.75809826 | 0.91402948 |
| MAN1A2 | 353.677524 | 0.04837669 | 0.14996615 | 0.32258407 | 0.74701026 | 0.91031794 |
| PPIG | 294.695417 | 0.048393719 | 0.13502956 | 0.35839352 | 0.72004884 | 0.90114265 |
| RANBP3 | 73.6987382 | 0.048413509 | 0.28511846 | 0.16980138 | 0.86516634 | 0.95503253 |
| PLXNB2 | 102.802215 | 0.048571108 | 0.3995348 | 0.12156915 | 0.90324024 | 0.96875453 |
| RPAP2 | 228.5937 | 0.048693741 | 0.18060296 | 0.26961763 | 0.78745444 | 0.92423116 |
| AC002070.1 | 430.945071 | 0.049122166 | 0.33176749 | 0.14806202 | 0.88229382 | 0.9608775 |
| FAM208B | 550.273549 | 0.049218236 | 0.22714733 | 0.21667979 | 0.8284579 | 0.94018985 |
| LMBR1 | 609.325252 | 0.049310448 | 0.29763328 | 0.16567518 | 0.86841258 | 0.95591135 |
| EHBP1 | 736.355967 | 0.049356754 | 0.21833431 | 0.22606046 | 0.8211544 | 0.93726124 |
| H2AFV | 164.065425 | 0.049416809 | 0.23656561 | 0.20889262 | 0.83453207 | 0.9432647 |
| IL3RA | 160.323689 | 0.049479664 | 0.52465811 | 0.09430839 | 0.92486418 | 0.97576555 |
| LINC00476 | 100.164594 | 0.049557535 | 0.43130489 | 0.1149014 | 0.90852328 | 0.96940376 |
| MACF1 | 3336.8294 | 0.049567778 | 0.30698727 | 0.16146526 | 0.87172697 | 0.95727819 |
| NES | 203.039853 | 0.049919003 | 0.43256781 | 0.11540157 | 0.90812684 | 0.96940376 |
| RNF170 | 103.605802 | 0.050096495 | 0.26860092 | 0.18650902 | 0.8520456 | 0.9501642 |
| AC139887.2 | 220.262135 | 0.050164645 | 0.31963317 | 0.15694443 | 0.87528864 | 0.95870526 |
| PPP6R1 | 71.5502365 | 0.0502626 | 0.31012349 | 0.16207286 | 0.87124848 | 0.95709072 |
| MFSD1 | 64.9327481 | 0.050264202 | 0.2823211 | 0.17803912 | 0.85869225 | 0.95247071 |
| TRERF1 | 85.3856855 | 0.050335338 | 0.42375842 | 0.1187831 | 0.9054472 | 0.96904724 |
| TUT4 | 402.618577 | 0.050414483 | 0.20161325 | 0.25005541 | 0.8025445 | 0.93081002 |
| FAM160B1 | 153.552684 | 0.050678307 | 0.15487531 | 0.32722004 | 0.74350147 | 0.90856581 |
| CIR1 | 131.560936 | 0.050881773 | 0.21235621 | 0.23960577 | 0.81063589 | 0.93351089 |
| PHRF1 | 73.6264854 | 0.050960813 | 0.22960469 | 0.22195022 | 0.82435263 | 0.93852534 |
| ZNF706 | 185.814997 | 0.05101345 | 0.24403109 | 0.20904488 | 0.8344132 | 0.9432647 |
| RBM41 | 99.0714925 | 0.051035472 | 0.24923442 | 0.20476895 | 0.83775264 | 0.94358503 |
| TERF2 | 80.175699 | 0.05104451 | 0.28616021 | 0.17837739 | 0.85842661 | 0.95247071 |
| CTDSP1 | 119.505788 | 0.051053956 | 0.26466529 | 0.19290008 | 0.84703722 | 0.9472879 |
| BCOR | 158.035445 | 0.051373609 | 0.3378236 | 0.15207229 | 0.87912992 | 0.95972986 |
| ERCC6 | 68.1568001 | 0.051441249 | 0.24424061 | 0.2106171 | 0.83318607 | 0.9430099 |
| NCBP3 | 146.217304 | 0.05147683 | 0.18277853 | 0.28163499 | 0.77822341 | 0.91979327 |
| CCPG1 | 241.201636 | 0.051635709 | 0.27724339 | 0.18624686 | 0.85225118 | 0.95026181 |
| ZFAND3 | 2296.80808 | 0.05164234 | 0.26832282 | 0.19246346 | 0.84737919 | 0.94731791 |
| FBXW4 | 220.098054 | 0.051772188 | 0.20377636 | 0.25406376 | 0.7994463 | 0.92963892 |
| NDC1 | 65.2207419 | 0.052066639 | 0.21446703 | 0.24277223 | 0.80818185 | 0.93264026 |
| UQCC1 | 165.943086 | 0.052225685 | 0.14397235 | 0.36274802 | 0.71679312 | 0.9001716 |
| TTLL11 | 71.5944163 | 0.05225261 | 0.34944641 | 0.14952968 | 0.88113569 | 0.96059232 |
| PSMD12 | 135.107944 | 0.052523723 | 0.21269396 | 0.24694506 | 0.80495074 | 0.9311439 |
| FBXO46 | 42.9390234 | 0.052540124 | 0.28284217 | 0.18575775 | 0.85263474 | 0.95044301 |
| PAFAH1B1 | 718.541233 | 0.052680724 | 0.18087946 | 0.29124769 | 0.77086189 | 0.91695334 |
| RAB28 | 246.326634 | 0.052880332 | 0.22131204 | 0.23894015 | 0.81115199 | 0.93351089 |
| VAPA | 425.615294 | 0.053052509 | 0.2308554 | 0.2298084 | 0.81824066 | 0.93593281 |
| DDHD2 | 128.596959 | 0.053065341 | 0.21339692 | 0.24866967 | 0.80361631 | 0.93081002 |
| GOLGA2 | 91.3612906 | 0.053176944 | 0.26539218 | 0.20037117 | 0.8411903 | 0.94419422 |
| CTIF | 303.458314 | 0.053279486 | 0.24350836 | 0.21879941 | 0.82680631 | 0.9401057 |
| LRRFIP2 | 448.480805 | 0.05349063 | 0.22228988 | 0.24063458 | 0.80983835 | 0.93351089 |
| R3HDM1 | 425.756485 | 0.053602517 | 0.148724 | 0.36041606 | 0.71853602 | 0.90051656 |
| TTC28 | 2410.62059 | 0.053676976 | 0.17814997 | 0.30130218 | 0.76318408 | 0.91477008 |
| MED24 | 71.4978789 | 0.054014992 | 0.23315578 | 0.23166911 | 0.81679503 | 0.93550282 |
| AC005261.1 | 113.61839 | 0.054126925 | 0.264867 | 0.20435511 | 0.83807601 | 0.94358503 |
| DCLRE1C | 60.2524282 | 0.054163066 | 0.24874055 | 0.21774925 | 0.82762449 | 0.94018985 |

| | | | | | | |
|---|---|---|---|---|---|---|
| LAMA4 | 174.245084 | 0.054349222 | 0.45867783 | 0.11849106 | 0.90567858 | 0.96904724 |
| C12orf4 | 84.1180575 | 0.054403198 | 0.19514823 | 0.27877885 | 0.78041455 | 0.92045723 |
| YPEL5 | 104.32426 | 0.054514875 | 0.2731274 | 0.19959504 | 0.8417973 | 0.94452589 |
| PSMF1 | 189.397715 | 0.054660643 | 0.30018642 | 0.18208899 | 0.85551289 | 0.95134391 |
| ZNF736 | 99.9793697 | 0.054972201 | 0.21618583 | 0.25428216 | 0.79927758 | 0.92963087 |
| LZIC | 65.8904807 | 0.055181325 | 0.27938845 | 0.19750753 | 0.84343039 | 0.94501377 |
| GOSR1 | 185.849455 | 0.055238132 | 0.23474135 | 0.23531488 | 0.81396435 | 0.93460727 |
| ALKBH1 | 108.730445 | 0.055257958 | 0.39967402 | 0.13825757 | 0.89003686 | 0.96407367 |
| TCF7L2 | 943.734871 | 0.055312592 | 0.27814331 | 0.19886364 | 0.84236941 | 0.94471448 |
| AC001226.2 | 65.7513783 | 0.0555958 | 0.46092208 | 0.12061865 | 0.90399309 | 0.9688759 |
| SUMO1 | 153.307935 | 0.055617878 | 0.28019838 | 0.19849465 | 0.84265807 | 0.94471448 |
| LPP | 2606.57129 | 0.055820153 | 0.19444364 | 0.28707625 | 0.77405393 | 0.91726032 |
| CALM1 | 595.745451 | 0.05592294 | 0.32817086 | 0.170408 | 0.86468928 | 0.95474375 |
| PLK2 | 177.958013 | 0.055953032 | 0.37078213 | 0.15090542 | 0.88005033 | 0.96042318 |
| SCARB1 | 46.7698023 | 0.056345895 | 0.54057872 | 0.10423254 | 0.91698481 | 0.97233474 |
| TJAP1 | 49.0124082 | 0.056384196 | 0.22800275 | 0.24729612 | 0.80467906 | 0.93109287 |
| MRS2 | 48.3907155 | 0.056614878 | 0.26108949 | 0.21684089 | 0.82833234 | 0.94018985 |
| ANKMY1 | 82.4056336 | 0.056655254 | 0.36831621 | 0.15382232 | 0.87774983 | 0.95969416 |
| DYNC1LI2 | 262.892148 | 0.056918496 | 0.17173723 | 0.33142783 | 0.74032135 | 0.90826918 |
| RAB6A | 477.969169 | 0.056948594 | 0.14662524 | 0.38839557 | 0.69772333 | 0.89303379 |
| FAR1 | 419.958858 | 0.056960908 | 0.19269047 | 0.29560834 | 0.76752921 | 0.91611154 |
| FAM50A | 51.540734 | 0.057297417 | 0.3284094 | 0.17446948 | 0.86149652 | 0.95324701 |
| DIS3L | 54.4692899 | 0.057531406 | 0.24743035 | 0.23251556 | 0.8161376 | 0.93550282 |
| FOPNL | 43.065216 | 0.057554907 | 0.22517516 | 0.2556006 | 0.79825925 | 0.92959949 |
| ITGA10 | 56.9782184 | 0.05769221 | 0.55016937 | 0.10486264 | 0.91648481 | 0.97233474 |
| CHM | 223.920988 | 0.057872723 | 0.18851223 | 0.30699717 | 0.75884552 | 0.91442519 |
| REPS1 | 251.461865 | 0.057972696 | 0.20661657 | 0.28058106 | 0.77903174 | 0.91979327 |
| RABL3 | 57.221203 | 0.058187276 | 0.24240471 | 0.24004186 | 0.81029781 | 0.93351089 |
| MRPS14 | 45.1100521 | 0.058258912 | 0.32987939 | 0.1766067 | 0.85981733 | 0.95263718 |
| MYO1D | 547.708708 | 0.058412861 | 0.35857284 | 0.16290375 | 0.87059421 | 0.95689355 |
| MIR29B2CH( | 390.591022 | 0.058464416 | 0.29114429 | 0.20080908 | 0.84084786 | 0.94397833 |
| NHSL1 | 744.325179 | 0.058859423 | 0.43299168 | 0.13593661 | 0.89187139 | 0.96460264 |
| HOXB3 | 292.20229 | 0.058926028 | 0.33196092 | 0.17750947 | 0.85910823 | 0.95247071 |
| SCML1 | 133.678813 | 0.058949026 | 0.34194836 | 0.17239161 | 0.86312967 | 0.95381365 |
| SS18L1 | 89.7892293 | 0.058956829 | 0.23191283 | 0.25421978 | 0.79932577 | 0.92963087 |
| PLXNA2 | 708.651851 | 0.059003093 | 0.35344731 | 0.16693604 | 0.86742038 | 0.95545847 |
| CSK | 40.7346388 | 0.059304908 | 0.34383679 | 0.17247982 | 0.86306033 | 0.95381365 |
| ZNF493 | 39.9873475 | 0.060039666 | 0.25330134 | 0.23702862 | 0.81263459 | 0.93395911 |
| CDC73 | 402.485216 | 0.060051985 | 0.20790395 | 0.28884485 | 0.77270011 | 0.91726032 |
| HSP90AA1 | 842.785408 | 0.060137454 | 0.45523579 | 0.13210177 | 0.8949038 | 0.96575575 |
| DICER1 | 455.251207 | 0.060237465 | 0.19012317 | 0.31683389 | 0.75136965 | 0.9112603 |
| ATP6V1H | 169.853105 | 0.060343793 | 0.18837709 | 0.3203351 | 0.74871432 | 0.91065347 |
| PALM | 227.91651 | 0.060667288 | 0.32159438 | 0.18864536 | 0.85037078 | 0.9492877 |
| WASHC2C | 70.95733 | 0.06068946 | 0.272515 | 0.22270136 | 0.82376795 | 0.93845094 |
| MXD1 | 50.2671714 | 0.061030324 | 0.44162484 | 0.13819495 | 0.89008634 | 0.96407367 |
| CSNK1G3 | 285.314122 | 0.061275762 | 0.20740733 | 0.29543682 | 0.76766021 | 0.91611154 |
| PTPRN2 | 83.7457006 | 0.061312575 | 0.26679669 | 0.2298101 | 0.81823933 | 0.93593281 |
| PPARA | 107.927486 | 0.061343788 | 0.37027403 | 0.16567132 | 0.86841562 | 0.95591135 |
| DAP3 | 186.457901 | 0.061819196 | 0.26369077 | 0.23443822 | 0.81464479 | 0.93505738 |
| TTC17 | 747.44759 | 0.061942243 | 0.19744261 | 0.31372276 | 0.75373161 | 0.91250377 |
| OXCT1 | 114.644077 | 0.061995004 | 0.20215749 | 0.30666687 | 0.75909695 | 0.91442519 |
| ZNF397 | 85.3575691 | 0.062277854 | 0.29970461 | 0.20779745 | 0.83538712 | 0.9432647 |
| NUP133 | 124.169216 | 0.062563308 | 0.17082727 | 0.36623723 | 0.71418806 | 0.89971941 |
| CCDC97 | 41.2543341 | 0.062742137 | 0.31397359 | 0.19983253 | 0.84161156 | 0.94444698 |
| PBX2 | 81.1254374 | 0.062757554 | 0.26644621 | 0.23553554 | 0.81379309 | 0.93460727 |
| BRF1 | 91.9189137 | 0.063050481 | 0.28341135 | 0.22246985 | 0.82394814 | 0.93852534 |
| ERCC8 | 103.838154 | 0.063304036 | 0.22145504 | 0.28585502 | 0.77498917 | 0.91776213 |
| KIF5B | 331.352719 | 0.063348515 | 0.21352463 | 0.29668013 | 0.76671073 | 0.91603312 |
| PPIL2 | 92.3567316 | 0.063578402 | 0.29557647 | 0.21509967 | 0.82968961 | 0.94063377 |
| SPATA5 | 430.00608 | 0.06366525 | 0.19123315 | 0.33291953 | 0.73919502 | 0.908116 |
| ERGIC3 | 70.7793573 | 0.063796666 | 0.26102493 | 0.24440833 | 0.80691459 | 0.93187644 |
| FBXO21 | 154.956327 | 0.063805111 | 0.2122395 | 0.30062788 | 0.76369827 | 0.91498433 |
| WDR13 | 89.2190027 | 0.064240105 | 0.22744751 | 0.28243925 | 0.77760672 | 0.91953119 |
| IQSEC2 | 58.7040636 | 0.064439431 | 0.31358799 | 0.20549075 | 0.83718872 | 0.94358503 |
| ZFPM2 | 514.602625 | 0.064471354 | 0.42037975 | 0.15336456 | 0.87811079 | 0.95969416 |
| CLDND1 | 122.010146 | 0.064500003 | 0.25947524 | 0.24857864 | 0.80368673 | 0.93081002 |
| YIPF4 | 233.970292 | 0.064556569 | 0.11977747 | 0.53897088 | 0.58990695 | 0.8420771 |
| SCAF1 | 45.0254147 | 0.064642482 | 0.32627473 | 0.19812286 | 0.84294894 | 0.94481699 |
| CHMP1A | 74.6335429 | 0.064663015 | 0.26956091 | 0.23988276 | 0.81042114 | 0.93351089 |
| ZNF791 | 132.043938 | 0.064807696 | 0.19197475 | 0.33758448 | 0.73567634 | 0.90649363 |
| ANKRD13C | 289.548309 | 0.064948332 | 0.15867172 | 0.40932519 | 0.68230103 | 0.88454837 |
| NUP35 | 59.1604517 | 0.065054094 | 0.25328815 | 0.25683828 | 0.79730362 | 0.92939834 |
| SCAMP1 | 227.410773 | 0.065059193 | 0.19631026 | 0.33141005 | 0.74033478 | 0.90826918 |
| FAM98B | 58.4104673 | 0.065086614 | 0.22444602 | 0.28998783 | 0.77182555 | 0.91702269 |
| INTS6 | 416.145322 | 0.065253679 | 0.27187079 | 0.24001725 | 0.81031688 | 0.93351089 |
| NELFCD | 67.873302 | 0.065552114 | 0.21372992 | 0.30670538 | 0.75906764 | 0.91442519 |
| NAV1 | 992.37255 | 0.065770997 | 0.25426389 | 0.25867219 | 0.79588818 | 0.92847059 |
| RNF8 | 70.1500016 | 0.065853036 | 0.23590182 | 0.27915442 | 0.78012632 | 0.9203826 |
| PARVA | 218.403429 | 0.065999477 | 0.32056549 | 0.20588453 | 0.83688111 | 0.94356495 |
| ZNF75A | 73.8237913 | 0.066043556 | 0.45304506 | 0.14577701 | 0.88409742 | 0.96178798 |
| SLC48A1 | 72.0695387 | 0.066245951 | 0.2716605 | 0.24385566 | 0.80734261 | 0.93206611 |
| TNPO1 | 423.114076 | 0.066327391 | 0.16628632 | 0.39887461 | 0.6899856 | 0.88817873 |
| TYW1B | 175.319378 | 0.066638915 | 0.16059965 | 0.41493811 | 0.67818721 | 0.88164367 |
| HIBADH | 218.575484 | 0.06670205 | 0.21928393 | 0.3041812 | 0.76098985 | 0.91477008 |
| G2E3 | 96.9693203 | 0.066706787 | 0.20982057 | 0.317923 | 0.75054334 | 0.91109327 |
| WDR70 | 564.20571 | 0.066848758 | 0.15201115 | 0.4397622 | 0.66010935 | 0.87335081 |
| MEX3C | 46.5341417 | 0.066857343 | 0.26844248 | 0.24905649 | 0.80331708 | 0.93081002 |
| AL139383.1 | 61.9706073 | 0.066918697 | 0.65056727 | 0.10286207 | 0.91807244 | 0.97233474 |
| CTTNBP2NL | 554.147194 | 0.067117472 | 0.29488007 | 0.22760939 | 0.81994992 | 0.93710409 |
| THOC3 | 62.8451228 | 0.067141561 | 0.35316235 | 0.19011529 | 0.84921879 | 0.94843147 |
| AUH | 289.749116 | 0.06737057 | 0.22314282 | 0.30191682 | 0.76271548 | 0.91477008 |
| RPRD1A | 312.288215 | 0.067716676 | 0.1654271 | 0.40934451 | 0.68228686 | 0.88454837 |
| TARSL2 | 84.163672 | 0.06773031 | 0.23169209 | 0.29232896 | 0.77003511 | 0.91695334 |
| ITGB5 | 312.593036 | 0.067736776 | 0.3040053 | 0.22281446 | 0.82367991 | 0.93845094 |
| ZNF141 | 135.170793 | 0.068355821 | 0.1835061 | 0.3724989 | 0.70952143 | 0.89727558 |
| CDC42BPA | 1632.15781 | 0.068374803 | 0.18295762 | 0.37371935 | 0.70861313 | 0.89705774 |
| PIK3C3 | 287.087217 | 0.068446037 | 0.2024997 | 0.33800562 | 0.73535895 | 0.90649363 |
| GIMAP8 | 390.167849 | 0.068825524 | 0.3041241 | 0.22630737 | 0.82096237 | 0.93726124 |
| SNX6 | 314.18796 | 0.068990414 | 0.14417669 | 0.47851296 | 0.63228515 | 0.86141143 |
| PEX3 | 119.467521 | 0.069322307 | 0.22786923 | 0.30421969 | 0.76096052 | 0.91477008 |
| 8-Sep | 58.7622155 | 0.069482977 | 0.29025658 | 0.23938467 | 0.81080732 | 0.93351089 |
| DNAJC16 | 90.1750601 | 0.06966251 | 0.21312411 | 0.32686359 | 0.74377106 | 0.90867104 |
| IDS | 75.8853459 | 0.069911431 | 0.34417521 | 0.20312745 | 0.83903542 | 0.94376667 |
| STYXL1 | 59.5764743 | 0.069912597 | 0.40160857 | 0.17408144 | 0.86180146 | 0.95324701 |
| FBH1 | 209.046539 | 0.069919604 | 0.2197854 | 0.3181267 | 0.75038883 | 0.91109327 |
| EIF3K | 118.692085 | 0.069974304 | 0.3465761 | 0.2019017 | 0.83999357 | 0.94379616 |
| DAAM1 | 280.284319 | 0.070109325 | 0.3138042 | 0.22341742 | 0.82321065 | 0.9382722 |
| HSPA1A | 581.701739 | 0.070178487 | 1.02341609 | 0.06857278 | 0.94532969 | 0.98379261 |
| AC009093.2 | 75.7831112 | 0.070280975 | 0.58568569 | 0.11999777 | 0.90448492 | 0.96889875 |
| PDK3 | 65.965079 | 0.070320252 | 0.4991329 | 0.14088483 | 0.88796093 | 0.96340871 |
| RSF1 | 523.386792 | 0.070470588 | 0.17421793 | 0.40449677 | 0.68584746 | 0.88671976 |
| CHP1 | 115.226895 | 0.070506979 | 0.19453968 | 0.36242981 | 0.71703086 | 0.90019386 |
| CDK12 | 219.562454 | 0.070602747 | 0.22280958 | 0.31687482 | 0.75133859 | 0.9112603 |

| | | | | | | |
|---|---|---|---|---|---|---|
| PNKD | 90.3317058 | 0.070618805 | 0.31107299 | 0.22701684 | 0.82041065 | 0.93711673 |
| PPP4R3A | 306.669645 | 0.070706804 | 0.15390039 | 0.45943227 | 0.64592378 | 0.86578121 |
| DPM1 | 102.980183 | 0.070885126 | 0.18363017 | 0.38602113 | 0.69948102 | 0.89370119 |
| TMEM184B | 317.469142 | 0.071506536 | 0.2468255 | 0.28970482 | 0.77204207 | 0.91702269 |
| ZNF708 | 126.458488 | 0.071506768 | 0.30075485 | 0.23775765 | 0.81206906 | 0.93357139 |
| ARL14EP | 60.5327178 | 0.071577899 | 0.26878463 | 0.26630206 | 0.79000659 | 0.92602336 |
| YTHDF2 | 167.643243 | 0.071661843 | 0.16737072 | 0.42816235 | 0.66853293 | 0.87625461 |
| RAB31 | 149.78218 | 0.07176526 | 0.40132791 | 0.17881951 | 0.85807943 | 0.95221914 |
| PAIP1 | 97.1008935 | 0.071879873 | 0.21973263 | 0.32712426 | 0.7435739 | 0.90856581 |
| MTF1 | 61.0967562 | 0.071969513 | 0.38222491 | 0.18829101 | 0.85064853 | 0.9492877 |
| FBXL14 | 49.1961835 | 0.072030424 | 0.3458168 | 0.2082907 | 0.83500199 | 0.9432647 |
| THEM6 | 39.082282 | 0.072034576 | 0.67351391 | 0.10695336 | 0.91482598 | 0.97221052 |
| FDX1 | 81.0245059 | 0.072049737 | 0.20252379 | 0.35575938 | 0.72202077 | 0.90162538 |
| PNPLA6 | 79.9520427 | 0.072100998 | 0.33231922 | 0.21696307 | 0.82823713 | 0.94018985 |
| LSM12 | 97.3302507 | 0.072756703 | 0.23441086 | 0.31038112 | 0.75627115 | 0.91345267 |
| FOXP1 | 1501.92215 | 0.072769706 | 0.24090741 | 0.30206504 | 0.76260248 | 0.91477008 |
| PQLC1 | 134.187435 | 0.073002436 | 0.19122357 | 0.38176483 | 0.70263581 | 0.89443605 |
| UHRF2 | 269.538179 | 0.073149712 | 0.18556702 | 0.39419565 | 0.6934366 | 0.8898215 |
| STARD7 | 135.737282 | 0.073677862 | 0.30104746 | 0.24473836 | 0.80665902 | 0.93187644 |
| FAM91A1 | 203.855672 | 0.074306175 | 0.21589892 | 0.34417113 | 0.73071759 | 0.90429739 |
| GAREM1 | 96.2334769 | 0.074515908 | 0.30079809 | 0.24772733 | 0.80434538 | 0.93109287 |
| C2orf88 | 50.3273599 | 0.074618669 | 0.28033737 | 0.26617453 | 0.7901048 | 0.92602336 |
| CHD8 | 187.787193 | 0.074702397 | 0.23312144 | 0.32044412 | 0.74863168 | 0.91065347 |
| NUP43 | 75.0899041 | 0.074704325 | 0.20537859 | 0.3637396 | 0.71605247 | 0.9001716 |
| COPB1 | 198.280441 | 0.074811139 | 0.20329252 | 0.36799749 | 0.7128751 | 0.89869927 |
| RIPK2 | 71.2762344 | 0.074840987 | 0.40614759 | 0.18427042 | 0.85380132 | 0.9508242 |
| FAM135A | 332.254822 | 0.074863793 | 0.23523116 | 0.31825627 | 0.75029055 | 0.91109327 |
| LRCH2 | 89.588818 | 0.074874543 | 0.42429931 | 0.17646633 | 0.8599276 | 0.95263718 |
| SRP9 | 174.519124 | 0.075020553 | 0.30083763 | 0.24937223 | 0.80307286 | 0.93081002 |
| PLEKHM1 | 128.72195 | 0.075082798 | 0.27383256 | 0.27419237 | 0.78393679 | 0.92261682 |
| SLU7 | 48.939007 | 0.075273118 | 0.24593549 | 0.30606855 | 0.75955245 | 0.91455455 |
| SPG7 | 332.988739 | 0.075331945 | 0.31569134 | 0.23862531 | 0.81139613 | 0.93351089 |
| STXBP5 | 388.254204 | 0.075479983 | 0.19367986 | 0.38971519 | 0.69674716 | 0.89253006 |
| KCTD13 | 64.6524602 | 0.075572094 | 0.3116136 | 0.2425186 | 0.80837834 | 0.93264848 |
| MAP3K20 | 149.537534 | 0.075627735 | 0.30226718 | 0.25020161 | 0.80243144 | 0.93081002 |
| CELF2 | 212.045181 | 0.075749786 | 0.83727619 | 0.09047168 | 0.9279124 | 0.97688257 |
| KMT2D | 101.182021 | 0.075772743 | 0.27143379 | 0.27915737 | 0.78012406 | 0.9203826 |
| SPAG1 | 58.8442186 | 0.075868918 | 0.34970416 | 0.21695171 | 0.82824598 | 0.94018985 |
| ZNF484 | 53.4267326 | 0.07594152 | 0.28495325 | 0.26650519 | 0.78985016 | 0.92602336 |
| SUMO2 | 67.817173 | 0.075958883 | 0.2566473 | 0.29596603 | 0.76725602 | 0.91611154 |
| METTL7A | 72.9588835 | 0.076129446 | 0.57979066 | 0.13130506 | 0.89553399 | 0.96575575 |
| DRAP1 | 57.803211 | 0.076141023 | 0.421765 | 0.1805295 | 0.8567369 | 0.95134391 |
| CARS2 | 104.622779 | 0.076497093 | 0.23027983 | 0.33219189 | 0.73974437 | 0.90826918 |
| PRKD2 | 339.228589 | 0.076510751 | 0.26251835 | 0.29144915 | 0.77070783 | 0.91695334 |
| UBAP1 | 240.54701 | 0.07657091 | 0.29470167 | 0.25982517 | 0.79499864 | 0.92847059 |
| HECTD1 | 523.03613 | 0.077228078 | 0.17917125 | 0.43102942 | 0.66644698 | 0.87587657 |
| PPP2R2D | 206.475449 | 0.077361774 | 0.25359692 | 0.30505802 | 0.76032196 | 0.91455455 |
| AIG1 | 177.118165 | 0.077976307 | 0.30006105 | 0.25986814 | 0.79496549 | 0.92847059 |
| PLA2R1 | 75.4443659 | 0.078158559 | 0.5329308 | 0.14665799 | 0.88340198 | 0.96141487 |
| PPP1R10 | 110.34372 | 0.078224828 | 0.26880847 | 0.29100582 | 0.77104687 | 0.91695334 |
| THRAP3 | 344.422336 | 0.078363151 | 0.16845107 | 0.46519829 | 0.64178947 | 0.86490071 |
| THOC7 | 67.0031036 | 0.078366064 | 0.25898906 | 0.30258446 | 0.76220656 | 0.91477008 |
| ICE2 | 167.239893 | 0.078426367 | 0.18773768 | 0.41774442 | 0.67613399 | 0.88033151 |
| ACTR6 | 53.3866408 | 0.07849174 | 0.22349008 | 0.35120905 | 0.72543152 | 0.9026193 |
| SP110 | 208.69762 | 0.078758597 | 0.52877178 | 0.14894629 | 0.88159601 | 0.96059232 |
| CPSF3 | 50.6038548 | 0.078977828 | 0.23045157 | 0.342709 | 0.73181739 | 0.90460147 |
| EXOG | 77.8092686 | 0.079073291 | 0.24501597 | 0.32272709 | 0.74690193 | 0.91031794 |
| KRBOX4 | 129.145218 | 0.079136264 | 0.26580693 | 0.29772085 | 0.76591623 | 0.91600937 |
| POC1B-AS1 | 48.7008564 | 0.079382953 | 0.30191888 | 0.26292809 | 0.79260601 | 0.92735129 |
| SMU1 | 102.283881 | 0.079449885 | 0.16783122 | 0.47339156 | 0.63593385 | 0.86207417 |
| FUBP3 | 199.770441 | 0.079566483 | 0.19641066 | 0.40510267 | 0.68540204 | 0.88671976 |
| NDUFAF6 | 156.570942 | 0.079571603 | 0.2380732 | 0.33423167 | 0.73820475 | 0.9078343 |
| MTIF2 | 43.6630161 | 0.079986874 | 0.30371439 | 0.26336215 | 0.79227146 | 0.92735129 |
| SHOC2 | 294.159793 | 0.080201762 | 0.14842158 | 0.54036454 | 0.58894565 | 0.84140756 |
| FAM241A | 445.396929 | 0.080240197 | 0.31686641 | 0.25323037 | 0.8000902 | 0.9298524 |
| FBXW5 | 104.0247 | 0.080343825 | 0.32239681 | 0.24920787 | 0.80319999 | 0.93081002 |
| ZFP90 | 65.2532009 | 0.080366955 | 0.28924863 | 0.27784732 | 0.78112957 | 0.92103505 |
| PCBP2 | 331.322043 | 0.080382737 | 0.18654215 | 0.43090924 | 0.66653437 | 0.87587657 |
| TSPYL1 | 54.6735146 | 0.080517061 | 0.35492528 | 0.22685637 | 0.82053543 | 0.93711673 |
| FRG1 | 50.7696009 | 0.080619551 | 0.24021846 | 0.33560932 | 0.7371655 | 0.90723797 |
| SPPL2A | 315.438879 | 0.080696356 | 0.18326201 | 0.44033325 | 0.65969576 | 0.87335081 |
| NDUFV3 | 86.3409064 | 0.080980972 | 0.19144954 | 0.42298859 | 0.67230359 | 0.87803719 |
| MAU2 | 133.686298 | 0.081120911 | 0.21022815 | 0.38587082 | 0.69959234 | 0.89370119 |
| NFATC2 | 140.699112 | 0.081228848 | 0.8981443 | 0.09044075 | 0.92793697 | 0.97688257 |
| UFSP2 | 88.003707 | 0.081381794 | 0.32247732 | 0.25236439 | 0.80075942 | 0.93010849 |
| PDXDC1 | 398.493687 | 0.08156649 | 0.20431166 | 0.39922582 | 0.68972682 | 0.88817873 |
| TYW5 | 51.0471531 | 0.082080548 | 0.2078067 | 0.39498508 | 0.6928539 | 0.88975484 |
| S100A4 | 440.027894 | 0.082318145 | 1.25584904 | 0.0655478 | 0.94773785 | 0.98379261 |
| ARFGEF2 | 355.199423 | 0.082459702 | 0.16450098 | 0.5012718 | 0.61617985 | 0.85732696 |
| SAFB2 | 126.25143 | 0.0825056 | 0.25278597 | 0.3263852 | 0.74413293 | 0.90884176 |
| ACACB | 484.752334 | 0.082558299 | 0.486779 | 0.16960119 | 0.86532378 | 0.95507749 |
| OAZ2 | 127.659831 | 0.082751759 | 0.38101617 | 0.217187 | 0.82806262 | 0.94018985 |
| BTBD10 | 221.511935 | 0.082790665 | 0.18540784 | 0.44653272 | 0.65521248 | 0.87094432 |
| SLC25A26 | 167.270923 | 0.083065592 | 0.33771307 | 0.24596499 | 0.80570933 | 0.93158047 |
| FBXO22 | 125.408344 | 0.083079526 | 0.21446665 | 0.38737737 | 0.69847686 | 0.8933683 |
| ASH2L | 91.9847534 | 0.083086107 | 0.28117046 | 0.29550084 | 0.76761131 | 0.91611154 |
| DDAH2 | 77.4900918 | 0.083176364 | 0.30814875 | 0.26992277 | 0.78721967 | 0.92408816 |
| MOB1B | 167.303613 | 0.083200242 | 0.17184091 | 0.48417017 | 0.6282651 | 0.86052754 |
| FAM160B2 | 73.7961033 | 0.083200365 | 0.25254084 | 0.32945311 | 0.74181323 | 0.90850686 |
| DCP2 | 208.723315 | 0.083210172 | 0.15449392 | 0.53859836 | 0.59016402 | 0.8420771 |
| MIB1 | 410.334679 | 0.083312399 | 0.13003043 | 0.64071463 | 0.52170811 | 0.80772643 |
| GTPBP10 | 70.9550051 | 0.083376956 | 0.2529518 | 0.32961598 | 0.74169015 | 0.90850686 |
| AC087500.1 | 43.298803 | 0.0833988 | 0.42250229 | 0.19739254 | 0.84352037 | 0.94501377 |
| SIPA1 | 97.5425386 | 0.083667667 | 0.27069126 | 0.30908892 | 0.75725389 | 0.913637 |
| USF2 | 72.8840364 | 0.08380853 | 0.34888714 | 0.24021674 | 0.81016224 | 0.93351089 |
| SZT2 | 91.2473195 | 0.083906628 | 0.25805977 | 0.32514417 | 0.74507196 | 0.90944569 |
| ARMC1 | 82.5499475 | 0.083911951 | 0.17626084 | 0.4760669 | 0.63402672 | 0.86207417 |
| FSD1L | 87.4680133 | 0.083918255 | 0.22159631 | 0.37869879 | 0.70491155 | 0.89448961 |
| TSG101 | 180.418599 | 0.083928067 | 0.2384672 | 0.35194805 | 0.72487722 | 0.9026193 |
| NT5C2 | 494.957687 | 0.084022904 | 0.14155113 | 0.59358696 | 0.55278841 | 0.8236419 |
| CRLS1 | 107.213862 | 0.084079905 | 0.27489372 | 0.30586332 | 0.75970871 | 0.91455455 |
| TMUB2 | 48.8109972 | 0.084112435 | 0.36303068 | 0.23169511 | 0.81677483 | 0.93550282 |
| TAP1 | 145.677901 | 0.084125151 | 0.42667273 | 0.19716552 | 0.84369801 | 0.94501377 |
| LNPEP | 258.919918 | 0.084162645 | 0.15671279 | 0.53705025 | 0.59123291 | 0.8428674 |
| MPHOSPH8 | 289.957209 | 0.084262093 | 0.24246262 | 0.34752612 | 0.7281961 | 0.9037084 |
| ZNF236 | 141.920717 | 0.084471476 | 0.16378609 | 0.51574269 | 0.60603414 | 0.85156118 |
| AC074032.1 | 155.191623 | 0.084633069 | 0.17664903 | 0.47910294 | 0.6318654 | 0.86118497 |
| SDHA | 153.218346 | 0.084874265 | 0.149025 | 0.56953039 | 0.56899626 | 0.83368112 |
| ZFP64 | 74.4549233 | 0.084938671 | 0.39441151 | 0.21535546 | 0.82949019 | 0.94063377 |
| LARP4 | 275.511626 | 0.085247432 | 0.1625752 | 0.52435694 | 0.6000303 | 0.84976293 |
| UBR4 | 324.99303 | 0.08532103 | 0.19865416 | 0.4294953 | 0.66756281 | 0.87587657 |
| L3MBTL3 | 78.1706436 | 0.08560088 | 0.28059914 | 0.30506466 | 0.76031691 | 0.91455455 |
| STX7 | 263.198936 | 0.085629621 | 0.21292334 | 0.40216174 | 0.687565 | 0.88715616 |

| | | | | | | | |
|---|---|---|---|---|---|---|---|
| AC005480.1 | 50.1854722 | | 0.085784227 | 0.39092913 | 0.21943678 | 0.82630982 | 0.93977669 |
| SENP6 | 680.975 | | 0.086129662 | 0.16591112 | 0.51913134 | 0.60366916 | 0.85081376 |
| ZNF586 | 182.376279 | | 0.086203486 | 0.374288 | 0.23031325 | 0.81784836 | 0.93587654 |
| APBB1 | 42.7103889 | | 0.086284947 | 0.33253847 | 0.25947358 | 0.79526987 | 0.92847059 |
| SAR1B | 123.070805 | | 0.086329805 | 0.18712881 | 0.46133892 | 0.64455546 | 0.86542551 |
| HGSNAT | 263.073286 | | 0.086450874 | 0.17843384 | 0.4844982 | 0.62803234 | 0.86052754 |
| HSPBAP1 | 77.1176714 | | 0.086475229 | 0.18747029 | 0.4612743 | 0.64460182 | 0.86542551 |
| RBM10 | 88.454117 | | 0.086545319 | 0.2410582 | 0.35902251 | 0.71957825 | 0.90076622 |
| STIM1 | 383.91639 | | 0.086862174 | 0.34560641 | 0.25133265 | 0.80155694 | 0.93040095 |
| RNF135 | 45.7038893 | | 0.08690353 | 0.24383399 | 0.35640449 | 0.72153767 | 0.90162538 |
| ORC2 | 228.733603 | | 0.086955314 | 0.2105613 | 0.41296912 | 0.67962924 | 0.88262277 |
| 8-Mar | 122.895214 | | 0.087390088 | 0.24561758 | 0.35579737 | 0.72199232 | 0.90162538 |
| ICA1L | 52.9008986 | | 0.087621361 | 0.35878805 | 0.24421483 | 0.80706445 | 0.93187644 |
| MIATNB | 111.159858 | | 0.087981876 | 0.38161855 | 0.23054926 | 0.81766499 | 0.93579756 |
| PATL1 | 113.494195 | | 0.088067401 | 0.41446149 | 0.21248633 | 0.83172764 | 0.94161971 |
| UBE2D2 | 400.546896 | | 0.08821487 | 0.20004556 | 0.44097389 | 0.6592319 | 0.87335081 |
| AC007038.1 | 72.1516381 | | 0.088384817 | 0.26051601 | 0.33926828 | 0.73440764 | 0.9055183 |
| ZNF260 | 48.7108757 | | 0.088410917 | 0.21206888 | 0.41689719 | 0.67675361 | 0.88084536 |
| BANP | 148.413414 | | 0.08841795 | 0.29776437 | 0.29693932 | 0.76651284 | 0.91603312 |
| TRUB1 | 42.0418129 | | 0.08848935 | 0.23843734 | 0.37112203 | 0.71054664 | 0.89742293 |
| RBBP6 | 188.11853 | | 0.088599623 | 0.18779296 | 0.47179416 | 0.63707372 | 0.86207417 |
| NUP214 | 259.516957 | | 0.088711634 | 0.16110871 | 0.55063215 | 0.58188587 | 0.83789748 |
| COP1 | 774.531355 | | 0.088813503 | 0.12122626 | 0.73262593 | 0.46378661 | 0.77373412 |
| AL445472.1 | 114.134718 | | 0.088818829 | 0.34851394 | 0.25485015 | 0.79883884 | 0.92962819 |
| ZNF432 | 77.3050942 | | 0.088989332 | 0.27059328 | 0.32886749 | 0.74225585 | 0.90850686 |
| PCNT | 86.6055892 | | 0.089555969 | 0.26083448 | 0.34334405 | 0.73133964 | 0.90454688 |
| IFIT5 | 53.0566477 | | 0.089640162 | 0.34547806 | 0.25946702 | 0.79527492 | 0.92847059 |
| UIMC1 | 283.611784 | | 0.08978681 | 0.22488957 | 0.39924845 | 0.68971015 | 0.88817873 |
| TAX1BP1 | 419.424072 | | 0.08985543 | 0.13866773 | 0.64799091 | 0.51699083 | 0.80535617 |
| PCBD2 | 87.6584172 | | 0.090098929 | 0.51523672 | 0.174869 | 0.86118257 | 0.95314506 |
| RANGAP1 | 51.1948648 | | 0.090157772 | 0.28717457 | 0.31394762 | 0.75356082 | 0.91250377 |
| TSC1 | 101.409844 | | 0.090265661 | 0.19451168 | 0.46406292 | 0.64260266 | 0.86490071 |
| C1orf159 | 54.8618688 | | 0.090290306 | 0.36145223 | 0.24979872 | 0.80274301 | 0.93081002 |
| ZNF783 | 42.2114538 | | 0.090423022 | 0.24335186 | 0.37157317 | 0.71021067 | 0.89727558 |
| SMS | 77.7427035 | | 0.090774512 | 0.26268822 | 0.34555989 | 0.72967349 | 0.90414821 |
| KDM2A | 724.22366 | | 0.090859792 | 0.20101555 | 0.45200381 | 0.65126624 | 0.86853188 |
| CPNE1 | 88.0482871 | | 0.090945275 | 0.23407296 | 0.38853387 | 0.69762099 | 0.89303379 |
| TDRD3 | 328.024731 | | 0.091329997 | 0.16472549 | 0.55443755 | 0.57927943 | 0.83649569 |
| AP3M1 | 64.5201161 | | 0.091348085 | 0.23151851 | 0.39456061 | 0.69316719 | 0.88975484 |
| SEC24B | 489.438352 | | 0.091518876 | 0.18370838 | 0.49817475 | 0.61836088 | 0.85855421 |
| PRPF38A | 74.3529408 | | 0.091611623 | 0.20516127 | 0.44653468 | 0.65521107 | 0.87094432 |
| ERGIC1 | 209.580534 | | 0.091832129 | 0.23452259 | 0.39157051 | 0.69537558 | 0.89183015 |
| CXorf38 | 70.0638136 | | 0.091862888 | 0.20170353 | 0.45543519 | 0.64879618 | 0.86764163 |
| AAGAB | 153.244915 | | 0.0918907 | 0.26481089 | 0.34700499 | 0.72858757 | 0.90376371 |
| AC099669.1 | 92.9476584 | | 0.091892814 | 0.28010669 | 0.32806362 | 0.74286356 | 0.90850686 |
| NADK | 72.2880691 | | 0.091958352 | 0.28814457 | 0.31913964 | 0.74962063 | 0.91078628 |
| BIRC6 | 1244.77436 | | 0.092018431 | 0.15818298 | 0.58172145 | 0.56075432 | 0.82917467 |
| MTA3 | 217.083728 | | 0.092040647 | 0.20183789 | 0.45601273 | 0.64838082 | 0.86749486 |
| NBEA | 609.348315 | | 0.092080119 | 0.284028 | 0.32419381 | 0.74579131 | 0.91005223 |
| MTMR6 | 134.934309 | | 0.092104371 | 0.23430255 | 0.39310016 | 0.6942455 | 0.89058018 |
| TRIM35 | 45.7014257 | | 0.092490002 | 0.3620962 | 0.25542936 | 0.79839149 | 0.92959949 |
| ODR4 | 160.120188 | | 0.092851018 | 0.16804386 | 0.55254039 | 0.58057817 | 0.8376007 |
| MACROD1 | 56.6823099 | | 0.092922095 | 0.29570707 | 0.31423698 | 0.75334106 | 0.9124615 |
| NBPF11 | 64.7846905 | | 0.093011667 | 0.27116448 | 0.34300829 | 0.73159222 | 0.90459642 |
| CCDC82 | 154.033925 | | 0.093017755 | 0.163553 | 0.56873159 | 0.56953831 | 0.83368112 |
| CARD8 | 448.090885 | | 0.093066638 | 0.13576265 | 0.6855099 | 0.49302221 | 0.79136932 |
| ASH1L | 931.037478 | | 0.093412799 | 0.1594211 | 0.58595002 | 0.5579091 | 0.82661057 |
| MTFMT | 69.8319566 | | 0.093463114 | 0.25253012 | 0.3701068 | 0.71130292 | 0.89747106 |
| DNM1L | 235.551031 | | 0.093464793 | 0.17992785 | 0.51945708 | 0.60344204 | 0.85064017 |
| AHCTF1 | 273.306218 | | 0.09355858 | 0.22376259 | 0.41811538 | 0.67586276 | 0.88033151 |
| RASIP1 | 447.168199 | | 0.093561619 | 0.26435579 | 0.3539231 | 0.72339652 | 0.90250857 |
| RAD21 | 210.385399 | | 0.093675615 | 0.25768596 | 0.36352627 | 0.71621179 | 0.9001716 |
| SH3YL1 | 91.7493249 | | 0.093736949 | 0.21008793 | 0.4461796 | 0.65546751 | 0.8710118 |
| KAZN | 357.529463 | | 0.093762979 | 0.57902236 | 0.16193326 | 0.87135841 | 0.95709072 |
| PALD1 | 275.897411 | | 0.093780531 | 0.30624496 | 0.30622718 | 0.75943168 | 0.91455455 |
| TMEM168 | 178.860901 | | 0.093793139 | 0.20194985 | 0.46443778 | 0.64233413 | 0.86490071 |
| MLLT10 | 702.392281 | | 0.093793814 | 0.17159912 | 0.54658679 | 0.58466265 | 0.83895258 |
| SMNDC1 | 78.7008415 | | 0.093918554 | 0.31957697 | 0.29388398 | 0.76884656 | 0.91632266 |
| LHFPL6 | 982.81425 | | 0.093992596 | 0.18329034 | 0.51280714 | 0.60808624 | 0.85277972 |
| SDHB | 94.799216 | | 0.094113589 | 0.19574808 | 0.48078934 | 0.63066623 | 0.86097142 |
| PHIP | 661.550424 | | 0.094156192 | 0.2055036 | 0.45817297 | 0.64682818 | 0.86670951 |
| FAM120B | 470.651966 | | 0.094186281 | 0.24815525 | 0.37954579 | 0.70428261 | 0.89446478 |
| PDSS2 | 433.383834 | | 0.094186485 | 0.24844249 | 0.3791078 | 0.70460782 | 0.89448961 |
| CERS5 | 131.897885 | | 0.094200691 | 0.1901041 | 0.49552161 | 0.62023197 | 0.85877466 |
| SFT2D2 | 209.292021 | | 0.094260312 | 0.19130577 | 0.49272069 | 0.62220996 | 0.85940448 |
| HSPG2 | 1577.52118 | | 0.094394434 | 0.53154593 | 0.17758472 | 0.85904913 | 0.95247071 |
| CEP85L | 258.489875 | | 0.094435461 | 0.26760471 | 0.35289163 | 0.72416969 | 0.9026193 |
| PI4KB | 108.636095 | | 0.094457469 | 0.18358457 | 0.51451747 | 0.60689025 | 0.85197854 |
| DLG5 | 93.5611831 | | 0.094498722 | 0.32401761 | 0.29164687 | 0.77055664 | 0.91695334 |
| RNF115 | 437.482971 | | 0.094520427 | 0.15744168 | 0.60035198 | 0.54827168 | 0.82355262 |
| PCGF2 | 145.72719 | | 0.094541917 | 0.31075002 | 0.30423785 | 0.76094669 | 0.91477008 |
| RBM43 | 41.452781 | | 0.094680743 | 0.39597817 | 0.23910597 | 0.81102341 | 0.93351089 |
| RAB40C | 76.5907746 | | 0.094943537 | 0.26566044 | 0.35738681 | 0.72080225 | 0.90162538 |
| DAG1 | 93.8838773 | | 0.095104007 | 0.28663581 | 0.33179388 | 0.74004491 | 0.90826918 |
| ZNF721 | 329.891618 | | 0.095475425 | 0.28992021 | 0.32931621 | 0.74191669 | 0.90850686 |
| RC3H2 | 258.859437 | | 0.095613187 | 0.17058086 | 0.56051534 | 0.57512798 | 0.834481 |
| GCFC2 | 154.367464 | | 0.095782005 | 0.22536224 | 0.42501355 | 0.6708268 | 0.87730512 |
| LDB2 | 11411.3621 | | 0.095794672 | 0.27064403 | 0.35395081 | 0.72337575 | 0.90250857 |
| C2orf42 | 48.1175826 | | 0.095911945 | 0.27383846 | 0.35025009 | 0.72615102 | 0.9026831 |
| USP38 | 83.7407988 | | 0.095955465 | 0.22136637 | 0.43346902 | 0.66467407 | 0.87556792 |
| KLHL24 | 222.867738 | | 0.096423048 | 0.35389319 | 0.2724637 | 0.7852655 | 0.92328222 |
| MAPKBP1 | 132.689513 | | 0.096499156 | 0.19639259 | 0.49135844 | 0.62317295 | 0.85940448 |
| AMMECR1L | 52.7558496 | | 0.096540349 | 0.24300076 | 0.39728415 | 0.69115794 | 0.88837311 |
| GIGYF2 | 444.658013 | | 0.096646092 | 0.19412417 | 0.49785709 | 0.61858477 | 0.85855421 |
| MAP4 | 1089.74354 | | 0.097077973 | 0.20296378 | 0.47830196 | 0.6324353 | 0.86141143 |
| GLCE | 77.7976906 | | 0.09710274 | 0.28216805 | 0.34413088 | 0.73074786 | 0.90429739 |
| CARD10 | 94.3865911 | | 0.097242316 | 0.3011767 | 0.32287463 | 0.74679019 | 0.91031794 |
| SEC22B | 135.8289 | | 0.097302105 | 0.2677171 | 0.36345121 | 0.71626786 | 0.9001716 |
| SERINC3 | 254.45145 | | 0.097338206 | 0.31388678 | 0.3101061 | 0.75648027 | 0.91345267 |
| MED6 | 77.1571753 | | 0.097365794 | 0.17725635 | 0.54929369 | 0.58280391 | 0.83815192 |
| EXOC4 | 1511.85881 | | 0.097377206 | 0.13813072 | 0.70496418 | 0.48083253 | 0.78473327 |
| GTF2B | 103.372456 | | 0.097459161 | 0.23698656 | 0.41124341 | 0.68089406 | 0.88326957 |
| GABPB1 | 171.278242 | | 0.097790957 | 0.15923584 | 0.61412654 | 0.53913171 | 0.81864721 |
| RYBP | 346.17432 | | 0.097885088 | 0.26770566 | 0.36564445 | 0.7146304 | 0.89980314 |
| WASL | 209.059157 | | 0.097892042 | 0.1989088 | 0.49214536 | 0.62261659 | 0.85940448 |
| RPIA | 70.9242828 | | 0.098060085 | 0.27843607 | 0.35218169 | 0.72470201 | 0.9026193 |
| HMGCR | 54.1247823 | | 0.098151788 | 0.3251648 | 0.30185244 | 0.76276455 | 0.91477008 |
| HEXDC | 74.0110484 | | 0.09821607 | 0.33367945 | 0.29434258 | 0.76849614 | 0.91631735 |
| PDS5A | 589.322852 | | 0.098255393 | 0.18572267 | 0.52904361 | 0.5967752 | 0.84737315 |
| ACIN1 | 242.299985 | | 0.098676283 | 0.231275 | 0.42666212 | 0.66962544 | 0.8767272 |
| ETS1 | 1392.4882 | | 0.098914722 | 0.29013901 | 0.34092183 | 0.73316243 | 0.90508002 |
| FBXL17 | 778.452679 | | 0.099025914 | 0.1815286 | 0.54551137 | 0.58540186 | 0.83908626 |
| STRN4 | 61.0890084 | | 0.099039166 | 0.27839511 | 0.35575038 | 0.72202751 | 0.90162538 |

| | | | | | | |
|---|---|---|---|---|---|---|
| EMSY | 266.389117 | 0.099062667 | 0.24095594 | 0.41112357 | 0.68098193 | 0.88326957 |
| SUGT1 | 124.269247 | 0.099139111 | 0.21363602 | 0.46405615 | 0.64260752 | 0.86490071 |
| N4BP2L1 | 154.352852 | 0.099284345 | 0.36662615 | 0.27080541 | 0.7865407 | 0.92382124 |
| RIPK1 | 185.825264 | 0.099388397 | 0.30201218 | 0.32908738 | 0.74208965 | 0.90850686 |
| GALNT7 | 98.5831562 | 0.099539582 | 0.23641871 | 0.4210309 | 0.67373252 | 0.87910389 |
| ZNF136 | 47.8785104 | 0.09968889 | 0.23861549 | 0.41778047 | 0.67610763 | 0.88033151 |
| SCAF11 | 696.301516 | 0.099819242 | 0.10632782 | 0.93878764 | 0.34783979 | 0.696616 |
| CAMKK2 | 100.748833 | 0.100180468 | 0.36993394 | 0.27080637 | 0.78653996 | 0.92382124 |
| CLDN15 | 140.951373 | 0.100325158 | 0.97566657 | 0.1028273 | 0.91810004 | 0.97233474 |
| TRIO | 1540.17038 | 0.10037495 | 0.20624627 | 0.48667523 | 0.6264885 | 0.86052754 |
| SPOP | 257.813818 | 0.100724755 | 0.17898908 | 0.56274247 | 0.57361026 | 0.8342245 |
| MAP4K5 | 508.055294 | 0.100819484 | 0.18902582 | 0.53336356 | 0.59378194 | 0.84502915 |
| PNRC1 | 213.168383 | 0.100972579 | 0.44697449 | 0.22590233 | 0.82127738 | 0.93726124 |
| RHOT2 | 51.6792747 | 0.101452505 | 0.27774174 | 0.36527641 | 0.71490509 | 0.90001051 |
| TULP3 | 123.922707 | 0.102040718 | 0.25743054 | 0.39638156 | 0.69182357 | 0.88886674 |
| TBCK | 456.854295 | 0.102110444 | 0.23600284 | 0.43266616 | 0.66525732 | 0.87587657 |
| TM7SF3 | 159.550719 | 0.102186992 | 0.3188061 | 0.32053022 | 0.74856642 | 0.91065347 |
| RDH13 | 48.0250329 | 0.102443759 | 0.26799405 | 0.38226131 | 0.70226755 | 0.89443605 |
| MKNK2 | 82.7729284 | 0.102734832 | 0.27941999 | 0.36767174 | 0.71311802 | 0.89886703 |
| TGS1 | 89.0459715 | 0.102783906 | 0.28501814 | 0.36062233 | 0.7183818 | 0.90051656 |
| C2orf49 | 41.0576696 | 0.102946385 | 0.2383395 | 0.4319317 | 0.66579105 | 0.87587657 |
| LTBP4 | 89.0535623 | 0.103103385 | 0.72221769 | 0.14275943 | 0.88648018 | 0.9627377 |
| DNAJC24 | 79.4895655 | 0.103269709 | 0.26902136 | 0.38387178 | 0.70107348 | 0.8944316 |
| TPRA1 | 43.4278416 | 0.10328674 | 0.30622737 | 0.33728775 | 0.7359 | 0.90649363 |
| TET3 | 117.715816 | 0.103344592 | 0.35036954 | 0.29495883 | 0.76802533 | 0.91627807 |
| OGG1 | 65.4826535 | 0.103618433 | 0.2432937 | 0.42589854 | 0.67018178 | 0.87705713 |
| JMJD1C | 3157.81751 | 0.103714312 | 0.36911372 | 0.280982 | 0.7787242 | 0.91979327 |
| RHOA | 620.505149 | 0.103838828 | 0.24188198 | 0.42929542 | 0.66770825 | 0.87587657 |
| RBM39 | 683.401126 | 0.103917623 | 0.22438957 | 0.46311253 | 0.64328371 | 0.86517342 |
| ZCCHC14 | 233.713408 | 0.104082154 | 0.19632551 | 0.53015094 | 0.59600728 | 0.84697301 |
| AK6 | 51.2827551 | 0.104137371 | 0.27611693 | 0.37714953 | 0.70606248 | 0.89518665 |
| ATAD3B | 51.6977469 | 0.104161046 | 0.25719629 | 0.40498658 | 0.68548738 | 0.88671976 |
| C6orf106 | 260.747416 | 0.104166378 | 0.19545325 | 0.53294778 | 0.59406973 | 0.8452917 |
| EHMT1 | 550.062965 | 0.104169396 | 0.21892355 | 0.47582546 | 0.63419873 | 0.86207417 |
| SEC22C | 85.3054542 | 0.104224401 | 0.23745372 | 0.43892511 | 0.6607158 | 0.8735882 |
| ISG15 | 960.572051 | 0.104226658 | 0.9525491 | 0.10941867 | 0.91287042 | 0.9709041 |
| KDM5D | 71.4796708 | 0.104288934 | 2.32578349 | 0.04484034 | 0.96423457 | 0.9886632 |
| MAPK9 | 105.053883 | 0.104290028 | 0.21486 | 0.48538597 | 0.62740258 | 0.86052754 |
| SCRN2 | 39.54423 | 0.104327475 | 0.2920302 | 0.35724892 | 0.72090547 | 0.90162538 |
| ARIH2 | 278.617253 | 0.104344046 | 0.17036885 | 0.61245966 | 0.54023368 | 0.81900022 |
| SETD1B | 51.8115286 | 0.1043519 | 0.30329173 | 0.34406444 | 0.73079783 | 0.90429739 |
| SMAD2 | 580.004513 | 0.104522572 | 0.26698915 | 0.39148621 | 0.69543788 | 0.89183015 |
| UBQLN1 | 237.409736 | 0.104582416 | 0.15157645 | 0.68996482 | 0.49021631 | 0.79057439 |
| PCMT1 | 152.843437 | 0.105176125 | 0.20222356 | 0.52009828 | 0.60299508 | 0.85048808 |
| SNAP23 | 268.363109 | 0.105190251 | 0.30904706 | 0.34036969 | 0.73357814 | 0.90519678 |
| USP25 | 607.075053 | 0.105277822 | 0.20966836 | 0.50211592 | 0.61558598 | 0.85714845 |
| DEPTOR | 128.744289 | 0.10574461 | 0.50713158 | 0.20851514 | 0.83482677 | 0.9432647 |
| FKBP3 | 68.0162931 | 0.10580181 | 0.27912665 | 0.37904589 | 0.70465379 | 0.89448961 |
| DHX30 | 96.8697609 | 0.105900695 | 0.18464436 | 0.57353875 | 0.56627997 | 0.83238172 |
| CUL3 | 502.314125 | 0.106046777 | 0.13688918 | 0.77469071 | 0.43852244 | 0.75300027 |
| CD99L2 | 117.807955 | 0.106139255 | 0.21895813 | 0.48474681 | 0.62785596 | 0.86052754 |
| LMO2 | 537.494804 | 0.10627577 | 0.50535924 | 0.21029747 | 0.83343551 | 0.94303136 |
| STRAP | 134.447391 | 0.106426098 | 0.24231567 | 0.43920435 | 0.66051347 | 0.87350009 |
| TRMT61B | 62.5662915 | 0.106439965 | 0.23282605 | 0.4571652 | 0.64755231 | 0.86714786 |
| STX8 | 526.732476 | 0.10646775 | 0.21039427 | 0.5060392 | 0.61282913 | 0.85573702 |
| CAPN1 | 89.4945008 | 0.106534263 | 0.18572602 | 0.57360979 | 0.56623189 | 0.83238172 |
| MIER1 | 318.963365 | 0.106561844 | 0.1938264 | 0.54977982 | 0.5824704 | 0.83812111 |
| XPO7 | 193.877049 | 0.107063254 | 0.15278796 | 0.70073097 | 0.48347093 | 0.78748452 |
| AL157392.5 | 78.8216 | 0.107259816 | 0.2314703 | 0.46338478 | 0.64308858 | 0.86517342 |
| TJP1 | 1134.7461 | 0.107383936 | 0.287847 | 0.37305908 | 0.70910447 | 0.89727558 |
| CSRP1 | 125.453949 | 0.107857521 | 0.47416617 | 0.22746777 | 0.82006003 | 0.93710409 |
| TBL1X | 340.933035 | 0.107986606 | 0.28209195 | 0.38280641 | 0.70186331 | 0.89443605 |
| KRIT1 | 248.971912 | 0.108315935 | 0.23120232 | 0.46848984 | 0.63943434 | 0.86388557 |
| USP16 | 122.001204 | 0.108393324 | 0.20735986 | 0.5227305 | 0.60116181 | 0.8499944 |
| THAP12 | 61.4442223 | 0.108485642 | 0.2168133 | 0.50036433 | 0.61681856 | 0.85738514 |
| CNOT10 | 171.671414 | 0.108559227 | 0.21834262 | 0.49719669 | 0.61905035 | 0.85855421 |
| DOCK4 | 5089.02163 | 0.108651877 | 0.23585809 | 0.46066631 | 0.64503803 | 0.86558646 |
| TMEM140 | 126.089314 | 0.108659546 | 0.35538442 | 0.30575213 | 0.75979338 | 0.91455455 |
| SGPL1 | 70.6342587 | 0.108689724 | 0.21815098 | 0.49823165 | 0.61832078 | 0.85855421 |
| STARD3 | 77.0689574 | 0.108756445 | 0.21997603 | 0.49440134 | 0.62102277 | 0.85890999 |
| NAPG | 128.257104 | 0.10880281 | 0.18327345 | 0.59366377 | 0.55273702 | 0.8236419 |
| ACSS1 | 64.0123159 | 0.108883328 | 0.30116699 | 0.36153805 | 0.71769726 | 0.90033983 |
| MYO1E | 1156.93752 | 0.108905371 | 0.31612049 | 0.34450589 | 0.73046587 | 0.90429739 |
| NCK2 | 159.789042 | 0.109262447 | 0.33538929 | 0.32577799 | 0.74459234 | 0.90926714 |
| RNPEPL1 | 60.323703 | 0.109285705 | 0.38369299 | 0.28482591 | 0.77577752 | 0.91811196 |
| VPS51 | 138.637338 | 0.109414146 | 0.17740332 | 0.61675364 | 0.53739723 | 0.81807397 |
| SPATA6L | 105.950357 | 0.109427825 | 0.3403118 | 0.32155166 | 0.74779237 | 0.91065347 |
| SPATA7 | 75.6653386 | 0.10947924 | 0.25576292 | 0.42804969 | 0.66861494 | 0.87625461 |
| SBF2 | 1869.23905 | 0.109513387 | 0.2166786 | 0.50541857 | 0.61326488 | 0.85573702 |
| ATP8B2 | 66.9665953 | 0.109763516 | 0.30678731 | 0.35778375 | 0.72050515 | 0.90151279 |
| USP20 | 47.0675628 | 0.109794969 | 0.31684192 | 0.34652918 | 0.72894506 | 0.90391839 |
| CIZ1 | 56.1808012 | 0.10989186 | 0.28816203 | 0.38136135 | 0.70293514 | 0.89443605 |
| SOCS2 | 166.137012 | 0.110018494 | 0.44258438 | 0.24858197 | 0.80368416 | 0.93081002 |
| C9orf85 | 149.188373 | 0.110050674 | 0.19836578 | 0.55478658 | 0.57904065 | 0.83649569 |
| GNE | 99.3477998 | 0.110057934 | 0.26034096 | 0.42274537 | 0.67248105 | 0.87806854 |
| ATG5 | 287.919094 | 0.11008144 | 0.14829103 | 0.74233379 | 0.45788513 | 0.76922069 |
| RBBP7 | 92.2113113 | 0.110160337 | 0.2175179 | 0.50644264 | 0.61254595 | 0.85573702 |
| RARA | 65.0201322 | 0.110221315 | 0.32692453 | 0.33714605 | 0.7360068 | 0.90649363 |
| ZNF761 | 71.6398835 | 0.110987947 | 0.25592063 | 0.43368112 | 0.66452002 | 0.8755434 |
| CDC42EP1 | 74.8402053 | 0.110998122 | 0.36173676 | 0.30684779 | 0.75895923 | 0.91442519 |
| CD47 | 219.941122 | 0.111036076 | 0.2398163 | 0.4630047 | 0.643361 | 0.86517342 |
| CCAR2 | 65.9740953 | 0.111117326 | 0.22820138 | 0.48692662 | 0.62631033 | 0.86052754 |
| PDE5A | 91.7717497 | 0.11124879 | 0.43336089 | 0.25671165 | 0.79740138 | 0.92939834 |
| STX18-AS1 | 83.4592182 | 0.111310564 | 0.49601543 | 0.22440948 | 0.82243869 | 0.93793807 |
| LIN7C | 77.1663208 | 0.11131749 | 0.3184475 | 0.34956309 | 0.72666662 | 0.90290665 |
| PPP1R9B | 48.789618 | 0.111444156 | 0.3203576 | 0.34787424 | 0.72793463 | 0.9037084 |
| SLC13A3 | 73.7242385 | 0.111469688 | 0.53201587 | 0.20952324 | 0.8340398 | 0.9432647 |
| PRR12 | 71.9611214 | 0.111538906 | 0.33657776 | 0.3313912 | 0.74034901 | 0.90826918 |
| MAPKAPK5 | 92.5152796 | 0.111569488 | 0.18227626 | 0.61209006 | 0.54047817 | 0.81902461 |
| TBCD | 668.946918 | 0.111636643 | 0.24209455 | 0.46112828 | 0.64470658 | 0.86542551 |
| ACCS | 50.8994887 | 0.111711776 | 0.55232509 | 0.20225729 | 0.83971559 | 0.94376667 |
| RABGAP1 | 447.441637 | 0.111730071 | 0.18777879 | 0.59500902 | 0.55183744 | 0.8236419 |
| ZFP36L1 | 585.322092 | 0.11180798 | 0.34341554 | 0.32557636 | 0.7447449 | 0.90931774 |
| PDCD6IP | 403.377702 | 0.111855536 | 0.17521632 | 0.63838537 | 0.52322285 | 0.80859228 |
| TMEM128 | 42.6410163 | 0.112032261 | 0.26286778 | 0.42619244 | 0.66996763 | 0.87703489 |
| GLB1 | 130.607357 | 0.112065423 | 0.17606308 | 0.63650723 | 0.52444587 | 0.80889528 |
| ZNF609 | 810.356927 | 0.112088007 | 0.16309747 | 0.68724551 | 0.49192803 | 0.79094859 |
| PAAF1 | 59.3729267 | 0.112103104 | 0.26196326 | 0.42793445 | 0.66869885 | 0.87625461 |
| ATXN7L1 | 299.540216 | 0.112180903 | 0.26019703 | 0.43113829 | 0.66636782 | 0.87587657 |
| NOS3 | 288.208319 | 0.112184118 | 0.37020369 | 0.30303349 | 0.76186434 | 0.91477008 |
| IRAK1BP1 | 69.9169083 | 0.112214686 | 0.35898128 | 0.31259203 | 0.75459063 | 0.91289071 |
| TYW1 | 257.451388 | 0.112264895 | 0.15420675 | 0.72801542 | 0.46660415 | 0.77550392 |
| STRBP | 131.187333 | 0.112483706 | 0.24609365 | 0.45707684 | 0.64761582 | 0.86714786 |

| | | | | | | |
|---|---|---|---|---|---|---|
| SPPL2B | 73.7467397 | 0.112700411 | 0.23815295 | 0.47322703 | 0.63605122 | 0.86207417 |
| AES | 322.750837 | 0.112849887 | 0.29810233 | 0.3785609 | 0.70501396 | 0.89448961 |
| TRIM33 | 344.126812 | 0.11289702 | 0.19146747 | 0.58964072 | 0.55543154 | 0.82487928 |
| ZFYVE26 | 67.0837778 | 0.112912898 | 0.34420695 | 0.32803782 | 0.74288307 | 0.90850686 |
| ARHGEF11 | 185.508512 | 0.113108841 | 0.25999792 | 0.43503749 | 0.66353522 | 0.8755434 |
| PIGN | 208.522555 | 0.113340292 | 0.19250748 | 0.58875785 | 0.55602372 | 0.82500619 |
| ACAD9 | 78.3818797 | 0.11360408 | 0.21035528 | 0.54005813 | 0.58915694 | 0.84152055 |
| LAP3 | 253.093239 | 0.113692166 | 0.40994801 | 0.27733314 | 0.78152432 | 0.92120024 |
| RAB9A | 58.3393271 | 0.113707305 | 0.33169013 | 0.34281184 | 0.73174002 | 0.90460147 |
| PURA | 214.355957 | 0.113926969 | 0.16683242 | 0.68288266 | 0.49468099 | 0.79309846 |
| COL19A1 | 69.354787 | 0.113952448 | 0.31581567 | 0.36081948 | 0.7182344 | 0.90051656 |
| TMEM131 | 420.863176 | 0.11396766 | 0.15883265 | 0.71753295 | 0.47304531 | 0.77899012 |
| CDK8 | 319.848805 | 0.114085908 | 0.21739847 | 0.52477788 | 0.59973761 | 0.8496629 |
| CTDSP2 | 163.339967 | 0.114172897 | 0.22070341 | 0.51731371 | 0.60493719 | 0.85144569 |
| RAB22A | 135.29652 | 0.114178775 | 0.22788559 | 0.50103553 | 0.61634612 | 0.85732696 |
| VEZF1 | 123.722254 | 0.114263709 | 0.21934837 | 0.52092345 | 0.6024201 | 0.85030923 |
| DNAJB14 | 331.773559 | 0.114529333 | 0.18108108 | 0.63247543 | 0.52707626 | 0.81084479 |
| AC108866.1 | 84.8820067 | 0.114700894 | 0.35546944 | 0.32267442 | 0.74694183 | 0.91031794 |
| NPAS2 | 551.41786 | 0.115051895 | 0.95393503 | 0.12060768 | 0.90400178 | 0.9688759 |
| ZNF654 | 310.053998 | 0.115165246 | 0.2536988 | 0.45394478 | 0.64986857 | 0.86766991 |
| CCNT1 | 118.643674 | 0.115235908 | 0.31513861 | 0.36566737 | 0.7146133 | 0.89980314 |
| ADAMTS9-AS | 563.903189 | 0.115320605 | 0.56946574 | 0.20250666 | 0.83952065 | 0.94376667 |
| MINDY2 | 242.892317 | 0.115417341 | 0.35124177 | 0.32859799 | 0.74245957 | 0.90850686 |
| NEIL1 | 46.4023406 | 0.115480553 | 0.32098019 | 0.3597747 | 0.71901562 | 0.90051656 |
| METTL5 | 43.4650576 | 0.115556239 | 0.3207879 | 0.3602263 | 0.71867791 | 0.90051656 |
| ESCO1 | 133.212644 | 0.115637413 | 0.23311897 | 0.49604464 | 0.61986292 | 0.85855421 |
| EPC2 | 285.015376 | 0.115684414 | 0.19688153 | 0.58758389 | 0.55681163 | 0.82543289 |
| B3GLCT | 110.592999 | 0.115834214 | 0.26556058 | 0.43618752 | 0.66270068 | 0.87493354 |
| AC007388.1 | 195.565425 | 0.115843318 | 0.44460532 | 0.26055315 | 0.79443712 | 0.92816661 |
| LSM8 | 110.492946 | 0.11616617 | 0.21941987 | 0.52942412 | 0.59651127 | 0.84714539 |
| PCAT1 | 962.44474 | 0.116315915 | 0.33232088 | 0.35001085 | 0.72633056 | 0.90272827 |
| KDM4C | 630.734799 | 0.116326156 | 0.16295934 | 0.71383545 | 0.47532893 | 0.78022377 |
| BARD1 | 78.6025998 | 0.116399228 | 0.28698043 | 0.40559988 | 0.68503661 | 0.88671976 |
| FBXL13 | 49.650395 | 0.116654328 | 0.26895066 | 0.43373877 | 0.66447815 | 0.8755434 |
| PPM1H | 75.4605106 | 0.116672188 | 0.64616032 | 0.18056229 | 0.85671116 | 0.95134391 |
| ZFP30 | 58.4389598 | 0.116929491 | 0.27642608 | 0.42300456 | 0.67229194 | 0.87803719 |
| ARAP1 | 54.1178416 | 0.117033829 | 0.23554047 | 0.49687355 | 0.61927823 | 0.85855421 |
| MAT2B | 92.5043655 | 0.117404352 | 0.31761278 | 0.36964618 | 0.71164614 | 0.89756479 |
| NECAP2 | 107.360425 | 0.117467948 | 0.22603596 | 0.51968699 | 0.60328176 | 0.85056075 |
| PNN | 209.765021 | 0.117606632 | 0.21461722 | 0.5479832 | 0.58370344 | 0.83850098 |
| MALT1 | 262.204765 | 0.117770418 | 0.23822756 | 0.49436101 | 0.62105125 | 0.85890999 |
| GOLPH3L | 111.487486 | 0.117776615 | 0.18679136 | 0.63052497 | 0.52835117 | 0.81207005 |
| VPS26A | 164.322918 | 0.117784073 | 0.2148994 | 0.54808935 | 0.58363055 | 0.83850098 |
| TXNL1 | 141.818328 | 0.118164301 | 0.25084094 | 0.47107262 | 0.63758887 | 0.86237803 |
| ZBTB43 | 81.8893712 | 0.118271838 | 0.33545853 | 0.35256768 | 0.72441257 | 0.9026193 |
| SLCO2A1 | 2271.29416 | 0.118448928 | 0.49539533 | 0.23909981 | 0.81102819 | 0.93351089 |
| PRPSAP2 | 115.344944 | 0.118572484 | 0.21084027 | 0.56238063 | 0.57385671 | 0.8342245 |
| KMT5A | 65.4064609 | 0.118668119 | 0.23915948 | 0.49618823 | 0.61976161 | 0.85855421 |
| TAZ | 80.8456656 | 0.118933474 | 0.27059096 | 0.43953232 | 0.66027587 | 0.87342991 |
| BCLAF1 | 346.698645 | 0.118951832 | 0.19304644 | 0.61618247 | 0.5377741 | 0.81807397 |
| NDUFV1 | 90.3408424 | 0.119046119 | 0.24163799 | 0.49266309 | 0.62225066 | 0.85940448 |
| MNAT1 | 387.692188 | 0.119099761 | 0.21476912 | 0.55454787 | 0.57920395 | 0.83649569 |
| TROVE2 | 225.530225 | 0.119177999 | 0.17940858 | 0.6642826 | 0.50650945 | 0.7989142 |
| INIP | 54.9375648 | 0.119317404 | 0.28828897 | 0.41388127 | 0.67896106 | 0.88189498 |
| SELENOI | 114.140539 | 0.119495525 | 0.25261863 | 0.47302737 | 0.63619365 | 0.86207417 |
| DCTN4 | 208.693437 | 0.119509401 | 0.19974945 | 0.59829653 | 0.54964209 | 0.82355262 |
| TMTC3 | 104.604653 | 0.119604974 | 0.19069537 | 0.62720439 | 0.53052527 | 0.81320326 |
| TRIP12 | 712.970249 | 0.119751131 | 0.19740945 | 0.60661296 | 0.5441078 | 0.82244124 |
| TTC21B | 234.778476 | 0.119811338 | 0.20758903 | 0.57715639 | 0.56383381 | 0.83089872 |
| GYS1 | 61.9141881 | 0.119848814 | 0.30625911 | 0.39133142 | 0.69555228 | 0.89183709 |
| OPA1 | 283.636624 | 0.120112806 | 0.16665206 | 0.72074001 | 0.47106949 | 0.77826805 |
| FAM102A | 84.6381491 | 0.120235063 | 0.43699859 | 0.27513834 | 0.78320996 | 0.92204994 |
| COQ9 | 43.8967463 | 0.120341219 | 0.2481556 | 0.48494258 | 0.62771708 | 0.86052754 |
| PRR14L | 218.019879 | 0.120404803 | 0.22351401 | 0.53869019 | 0.59010064 | 0.8420771 |
| KIF2A | 186.107357 | 0.120461894 | 0.17812562 | 0.67627493 | 0.49886611 | 0.79513466 |
| SPTY2D1 | 47.0552841 | 0.120609042 | 0.27178726 | 0.44376268 | 0.65721417 | 0.87234121 |
| EIF4A2 | 255.685067 | 0.120809915 | 0.33530559 | 0.36029794 | 0.71862434 | 0.90051656 |
| DRG2 | 37.8446341 | 0.120821919 | 0.28733502 | 0.42049146 | 0.67412647 | 0.87910389 |
| ITFG1 | 669.899555 | 0.120892952 | 0.2312547 | 0.52276972 | 0.60113452 | 0.8499944 |
| CLYBL | 130.911224 | 0.121064369 | 0.3810966 | 0.3176737 | 0.75073246 | 0.91109327 |
| UEVLD | 134.738941 | 0.121107203 | 0.23655604 | 0.51195988 | 0.60867909 | 0.85302637 |
| SMAD3 | 440.477132 | 0.121380138 | 0.5100674 | 0.23796882 | 0.81190527 | 0.93357139 |
| HOXB-AS3 | 53.2847976 | 0.121675083 | 0.4178662 | 0.29118192 | 0.77091219 | 0.91695334 |
| TANGO6 | 234.577962 | 0.122148094 | 0.25003496 | 0.48852407 | 0.62517868 | 0.86016045 |
| FAM13B | 353.504158 | 0.122265699 | 0.12838349 | 0.95234752 | 0.34092076 | 0.69116603 |
| TMEM87B | 328.829091 | 0.122386222 | 0.25273168 | 0.48425358 | 0.62820591 | 0.86052754 |
| RNF149 | 388.648174 | 0.12270792 | 0.40110756 | 0.30592273 | 0.75966348 | 0.91455455 |
| CLUAP1 | 52.5242399 | 0.123049654 | 0.30212666 | 0.40727837 | 0.68380355 | 0.88565439 |
| ZDHHC6 | 79.1561108 | 0.123184506 | 0.202472 | 0.60840268 | 0.54292044 | 0.82150849 |
| CTNNBL1 | 170.060081 | 0.123333034 | 0.22667883 | 0.54408712 | 0.58638152 | 0.83960805 |
| RPP38 | 83.2417675 | 0.123468498 | 0.22149518 | 0.55743199 | 0.57723232 | 0.83603913 |
| CCDC14 | 162.197808 | 0.123486064 | 0.28611098 | 0.43160198 | 0.66603072 | 0.87587657 |
| BBX | 613.231437 | 0.123875262 | 0.22690051 | 0.54594528 | 0.58510355 | 0.83895258 |
| LRP12 | 74.8080802 | 0.124192296 | 0.19490734 | 0.63718633 | 0.52400348 | 0.80889528 |
| HCFC1 | 51.7674669 | 0.124528012 | 0.24922935 | 0.49965227 | 0.61731995 | 0.85790478 |
| KAT7 | 83.8086257 | 0.124531697 | 0.22015294 | 0.56565994 | 0.57162498 | 0.83368112 |
| TAPBP | 355.774515 | 0.124617445 | 0.27689346 | 0.45005558 | 0.65267036 | 0.86875728 |
| LTBP3 | 164.947702 | 0.124940777 | 0.44497953 | 0.2807787 | 0.77888014 | 0.91979327 |
| CEP295 | 97.367481 | 0.124973802 | 0.25490233 | 0.49028113 | 0.62393498 | 0.85997304 |
| FN1 | 1116.45958 | 0.124974944 | 0.49512642 | 0.25241017 | 0.80072404 | 0.93010849 |
| TUBA4A | 57.8945982 | 0.125139514 | 0.57706368 | 0.21685564 | 0.82832085 | 0.94018985 |
| FANCL | 241.51499 | 0.125177673 | 0.27841688 | 0.44960518 | 0.65299515 | 0.86899648 |
| MAEA | 80.4226361 | 0.125443666 | 0.29038655 | 0.43198856 | 0.66574973 | 0.87587657 |
| SLTM | 390.107962 | 0.125742121 | 0.19299681 | 0.65152434 | 0.51470807 | 0.80422048 |
| ZNF274 | 77.1166826 | 0.125831652 | 0.4762579 | 0.26420906 | 0.79161883 | 0.92685891 |
| AL035413.1 | 57.7325043 | 0.126302332 | 0.31342443 | 0.40297539 | 0.68696633 | 0.88671976 |
| CRY2 | 73.8855553 | 0.126519647 | 0.28032239 | 0.45133621 | 0.65174725 | 0.8685756 |
| TADA2A | 110.756213 | 0.126586814 | 0.24630642 | 0.51394037 | 0.60729368 | 0.85226578 |
| KCTD3 | 221.173725 | 0.126985583 | 0.16162843 | 0.78566367 | 0.43206455 | 0.74621871 |
| SPG11 | 367.119898 | 0.12708882 | 0.25821076 | 0.49219026 | 0.62258485 | 0.85940448 |
| HERPUD1 | 296.303392 | 0.127272051 | 0.46018831 | 0.27656516 | 0.78211402 | 0.92139923 |
| LRIG2 | 127.634955 | 0.127287079 | 0.16099836 | 0.79061105 | 0.429171 | 0.74408005 |
| MT1G | 421.873243 | 0.127344897 | 1.02126153 | 0.12469372 | 0.90076603 | 0.96744565 |
| GON4L | 258.720456 | 0.127590283 | 0.22511708 | 0.566773 | 0.57086842 | 0.83368112 |
| AC015923.1 | 60.4662523 | 0.127690361 | 0.52165442 | 0.2447796 | 0.80662709 | 0.93187644 |
| LANCL2 | 59.0602709 | 0.127784528 | 0.46026535 | 0.2776323 | 0.78129463 | 0.92109696 |
| CHCHD6 | 69.200603 | 0.127786353 | 0.4117888 | 0.31032012 | 0.75631753 | 0.91345267 |
| RPRD1B | 135.809942 | 0.127790173 | 0.17069482 | 0.74864704 | 0.45406997 | 0.76611434 |
| CALD1 | 851.900355 | 0.12782897 | 0.30848172 | 0.41438102 | 0.67859509 | 0.88164367 |
| SELENOT | 120.451128 | 0.128032034 | 0.29109288 | 0.43983224 | 0.66005862 | 0.87335081 |
| MED21 | 59.6601033 | 0.128142019 | 0.28123094 | 0.45564695 | 0.64864387 | 0.86757974 |
| NCOR1 | 692.05851 | 0.12816238 | 0.14486595 | 0.88469639 | 0.37632041 | 0.71257833 |
| ZNF420 | 110.072273 | 0.128173869 | 0.40121746 | 0.31946234 | 0.74937594 | 0.91075944 |

| | | | | | | |
|---|---|---|---|---|---|---|
| PPFIA1 | 293.901982 | 0.128267905 | 0.25958376 | 0.49412916 | 0.62121497 | 0.85890999 |
| KIAA0556 | 143.070153 | 0.128280938 | 0.29866801 | 0.42951013 | 0.66755202 | 0.87587657 |
| FAM208A | 266.220472 | 0.128476648 | 0.2022074 | 0.63537064 | 0.52518671 | 0.80979892 |
| CCAR1 | 326.277732 | 0.128645241 | 0.21458884 | 0.59949641 | 0.5488419 | 0.82355262 |
| IBTK | 237.299812 | 0.128674954 | 0.18236935 | 0.70557336 | 0.48045349 | 0.7845841 |
| NPNT | 169.566775 | 0.128789567 | 0.63906852 | 0.20152701 | 0.84028651 | 0.94397833 |
| MIGA1 | 224.310378 | 0.128791869 | 0.18505667 | 0.69595906 | 0.48645447 | 0.78826504 |
| GTF2H2C | 57.5908703 | 0.128957816 | 0.20055521 | 0.64300406 | 0.52022147 | 0.80691303 |
| YLPM1 | 299.954942 | 0.12896934 | 0.14335993 | 0.89961914 | 0.36832297 | 0.70909779 |
| CAT | 105.693689 | 0.129055035 | 0.43683102 | 0.29543469 | 0.76766184 | 0.91611154 |
| SCYL3 | 63.8193327 | 0.129087552 | 0.22700647 | 0.56865141 | 0.56959273 | 0.83368112 |
| ARL2BP | 112.311147 | 0.129156817 | 0.27279324 | 0.47346048 | 0.63588469 | 0.86207417 |
| PHF12 | 139.551414 | 0.129381871 | 0.19882701 | 0.65072583 | 0.51522349 | 0.80444072 |
| AIFM1 | 51.5660217 | 0.12941213 | 0.32265164 | 0.40108933 | 0.68835436 | 0.88739126 |
| CDC42SE1 | 129.100414 | 0.129420343 | 0.26764622 | 0.48355006 | 0.62870522 | 0.86052754 |
| DLL1 | 89.8038507 | 0.129552188 | 0.44064379 | 0.29400661 | 0.76875285 | 0.91632266 |
| MGST2 | 272.33229 | 0.129677992 | 0.34646584 | 0.37428796 | 0.70819009 | 0.89680567 |
| GBF1 | 382.163862 | 0.12978235 | 0.21166293 | 0.61315578 | 0.53977334 | 0.81900022 |
| POR | 99.3617284 | 0.129808653 | 0.20496966 | 0.63330666 | 0.5265334 | 0.81036438 |
| TNFRSF14 | 133.466951 | 0.130072445 | 0.22971156 | 0.56624247 | 0.57122896 | 0.83368112 |
| UBE3B | 87.372168 | 0.130250225 | 0.23039747 | 0.56532837 | 0.57185043 | 0.83368112 |
| GNAQ | 2409.68103 | 0.130275729 | 0.22111311 | 0.58918138 | 0.5557396 | 0.82487928 |
| INTS14 | 58.4868982 | 0.130428374 | 0.23363304 | 0.55826169 | 0.5766657 | 0.83585958 |
| CTC1 | 42.0012859 | 0.130531219 | 0.2822874 | 0.46240541 | 0.64379062 | 0.86517342 |
| CAAP1 | 102.313263 | 0.130601426 | 0.19719272 | 0.66230348 | 0.50777674 | 0.79921852 |
| CD46 | 499.88695 | 0.13072097 | 0.35165788 | 0.37172769 | 0.71009561 | 0.89727558 |
| EHD4 | 653.373675 | 0.130835058 | 0.30997009 | 0.4220893 | 0.67295984 | 0.87837755 |
| HHAT | 64.4778712 | 0.130842581 | 0.52565249 | 0.2489146 | 0.80342684 | 0.93081002 |
| APLF | 100.602037 | 0.130857808 | 0.31579083 | 0.41438128 | 0.67859489 | 0.88164367 |
| CACUL1 | 234.032813 | 0.130968272 | 0.14222942 | 0.92082405 | 0.3571423 | 0.70058911 |
| CYTOR | 110.314145 | 0.131089622 | 0.50156804 | 0.2613596 | 0.79381521 | 0.92790728 |
| MUS81 | 43.6998393 | 0.131262104 | 0.26568854 | 0.49404503 | 0.62127438 | 0.85890999 |
| USP4 | 175.369712 | 0.131287211 | 0.21084793 | 0.62266303 | 0.53350598 | 0.81479963 |
| LYSMD2 | 50.3924468 | 0.13137058 | 0.36576382 | 0.35916778 | 0.71946958 | 0.90076622 |
| MRPS31 | 114.92346 | 0.131407672 | 0.17645631 | 0.74470373 | 0.45645084 | 0.76775688 |
| GTF3C2 | 106.767436 | 0.131479909 | 0.22920672 | 0.57363026 | 0.56621803 | 0.83238172 |
| NUP93 | 125.238952 | 0.131553556 | 0.28979809 | 0.45394901 | 0.64986553 | 0.86766991 |
| HSPB11 | 58.4211274 | 0.131691522 | 0.2237317 | 0.5886136 | 0.55612051 | 0.82500619 |
| PPM1D | 245.267311 | 0.131962203 | 0.15155355 | 0.8707299 | 0.38390165 | 0.71699718 |
| PPA2 | 313.804882 | 0.132096363 | 0.20488692 | 0.64472814 | 0.51910338 | 0.80603851 |
| EDC3 | 67.5567795 | 0.132305425 | 0.30299421 | 0.43665992 | 0.662358 | 0.87491556 |
| NAP1L4 | 244.454196 | 0.132498642 | 0.1677952 | 0.78964503 | 0.4297351 | 0.74408005 |
| ZNF407 | 721.639279 | 0.132599794 | 0.25682539 | 0.51630329 | 0.6056426 | 0.85156118 |
| TRIP10 | 101.998417 | 0.132617853 | 0.23548849 | 0.56316066 | 0.57332549 | 0.8342245 |
| DLD | 102.207307 | 0.132627679 | 0.19856486 | 0.66793127 | 0.50417746 | 0.79776858 |
| FGD5 | 961.150971 | 0.132703349 | 0.28862997 | 0.45976982 | 0.64568145 | 0.86572246 |
| CNOT11 | 63.4996633 | 0.132734292 | 0.24658352 | 0.53829344 | 0.59037448 | 0.842228 |
| NAF1 | 86.5145113 | 0.132753034 | 0.19326011 | 0.68691381 | 0.49213705 | 0.79094859 |
| TRAF6 | 80.710691 | 0.132755293 | 0.22662951 | 0.58578113 | 0.55802261 | 0.82662907 |
| NUP50 | 85.9978529 | 0.132889359 | 0.21464536 | 0.61911125 | 0.53584307 | 0.81653703 |
| NUP188 | 68.838043 | 0.13302942 | 0.25675058 | 0.51812706 | 0.60436962 | 0.85105198 |
| SNX29 | 468.520898 | 0.133130783 | 0.3210049 | 0.41473132 | 0.67833861 | 0.88164367 |
| AGO3 | 495.43704 | 0.133230782 | 0.16777142 | 0.79412084 | 0.42712508 | 0.74286176 |
| SLFN11 | 117.320877 | 0.133242661 | 0.33049837 | 0.40315679 | 0.68683288 | 0.88671976 |
| TMEM88 | 143.284709 | 0.133416031 | 0.98816086 | 0.13501449 | 0.89260042 | 0.96482231 |
| AMFR | 113.05041 | 0.133476469 | 0.17964102 | 0.74301776 | 0.45747093 | 0.76899848 |
| ZNF529 | 69.3574455 | 0.133506815 | 0.2199559 | 0.60697084 | 0.54387026 | 0.82233746 |
| TGFB1 | 232.578224 | 0.133562336 | 0.27613903 | 0.48367787 | 0.6286145 | 0.86052754 |
| ICK | 54.426921 | 0.133567738 | 0.22332683 | 0.59808191 | 0.54978528 | 0.82355262 |
| ZNF710 | 238.28516 | 0.13363428 | 0.48574401 | 0.27511256 | 0.78322976 | 0.92204994 |
| LYPD6B | 378.14079 | 0.133787754 | 0.64620286 | 0.20703677 | 0.83598114 | 0.9432647 |
| USP34 | 1263.52106 | 0.133889966 | 0.19505322 | 0.68642786 | 0.49244334 | 0.79099316 |
| ARHGEF9 | 73.9761462 | 0.133911119 | 0.31141068 | 0.43001454 | 0.66718507 | 0.87587657 |
| NECAP1 | 48.866091 | 0.134030562 | 0.25595213 | 0.52365481 | 0.60051865 | 0.8498698 |
| TTC37 | 321.35491 | 0.134246368 | 0.16168708 | 0.83028507 | 0.40637762 | 0.73327191 |
| ICA1 | 454.660486 | 0.134269087 | 0.302256 | 0.44422306 | 0.65688132 | 0.87218235 |
| TRAPPC13 | 102.101909 | 0.134272336 | 0.19850475 | 0.67641875 | 0.49877481 | 0.79513466 |
| BAZ2A | 228.179591 | 0.134649989 | 0.17801375 | 0.75640219 | 0.44940809 | 0.76249356 |
| EHHADH | 60.5414637 | 0.134666515 | 0.39879378 | 0.33768459 | 0.73560089 | 0.90649363 |
| DMAC2 | 41.5408379 | 0.134785867 | 0.27101668 | 0.49733422 | 0.61895339 | 0.85855421 |
| RASSF3 | 418.414813 | 0.134792385 | 0.23916415 | 0.56359779 | 0.57302789 | 0.8342245 |
| DROSHA | 158.146539 | 0.134855181 | 0.20200393 | 0.66758692 | 0.50439731 | 0.79776858 |
| PPP1R21 | 244.938777 | 0.135281285 | 0.22043991 | 0.6136878 | 0.53942165 | 0.81878823 |
| CARD6 | 76.9236448 | 0.135386848 | 0.27675503 | 0.48919381 | 0.62470449 | 0.86016045 |
| LINC00630 | 130.615093 | 0.135472291 | 0.25483509 | 0.53160769 | 0.59499774 | 0.84587674 |
| TMEM260 | 158.841893 | 0.135625097 | 0.19330436 | 0.70161428 | 0.48291975 | 0.78719766 |
| AC108062.1 | 76.7786348 | 0.135644704 | 0.42220276 | 0.32127858 | 0.74799929 | 0.91065347 |
| FAM122B | 54.8275513 | 0.135655947 | 0.25512855 | 0.53171607 | 0.59492266 | 0.84587674 |
| MRPL16 | 53.1755287 | 0.135816966 | 0.33787568 | 0.40197319 | 0.68770376 | 0.88719531 |
| ZNF26 | 79.4059124 | 0.135855764 | 0.26395916 | 0.51468479 | 0.60677331 | 0.85196053 |
| FOXO3 | 514.133691 | 0.13588734 | 0.28697555 | 0.47351539 | 0.63584552 | 0.86207417 |
| CFDP1 | 389.929909 | 0.135968207 | 0.20643618 | 0.65864523 | 0.51012361 | 0.80177277 |
| TLK2 | 412.479505 | 0.136018711 | 0.17519858 | 0.77636879 | 0.43753127 | 0.75200973 |
| IFRD1 | 167.530426 | 0.136209154 | 0.43862327 | 0.31053791 | 0.75615193 | 0.91345267 |
| RAF1 | 295.930546 | 0.13628332 | 0.16068661 | 0.84813116 | 0.39636493 | 0.7264986 |
| USP24 | 412.159164 | 0.13629198 | 0.17944625 | 0.75951424 | 0.447545 | 0.7614672 |
| OTUD7B | 186.302997 | 0.136508771 | 0.26388937 | 0.51729546 | 0.60494992 | 0.85144569 |
| KLHDC2 | 86.7826724 | 0.136574387 | 0.26456738 | 0.51621778 | 0.60570231 | 0.85156118 |
| GNA11 | 147.537692 | 0.136617797 | 0.3606157 | 0.37884595 | 0.70480226 | 0.89448961 |
| STAG2 | 439.709484 | 0.136886658 | 0.19696033 | 0.69499608 | 0.48705776 | 0.78826504 |
| SMARCD2 | 71.2783091 | 0.136932977 | 0.30786887 | 0.44477695 | 0.65648095 | 0.87193372 |
| GIGYF1 | 83.1082968 | 0.136978775 | 0.24676019 | 0.5551089 | 0.57882018 | 0.83649569 |
| CUL2 | 162.989063 | 0.137026037 | 0.16035651 | 0.85450873 | 0.39282317 | 0.72332854 |
| RBX1 | 139.652447 | 0.137104085 | 0.2436764 | 0.56264819 | 0.57367447 | 0.8342245 |
| SYNRG | 225.230209 | 0.137179474 | 0.15885672 | 0.86354213 | 0.38783946 | 0.72096555 |
| FNIP2 | 472.880038 | 0.137330538 | 0.26003254 | 0.52812828 | 0.5974103 | 0.84775936 |
| ATAD2B | 451.917369 | 0.137462495 | 0.1740704 | 0.78969485 | 0.429706 | 0.74408005 |
| 7-Mar | 504.236326 | 0.137554211 | 0.14330925 | 0.95984183 | 0.33713483 | 0.68780211 |
| VTI1B | 53.2748092 | 0.137693604 | 0.36018408 | 0.38228676 | 0.70224868 | 0.89443605 |
| ACVR2B | 47.9414102 | 0.137917854 | 0.33765082 | 0.40846296 | 0.68293382 | 0.88508828 |
| CD164 | 213.790338 | 0.137944212 | 0.24239378 | 0.56909139 | 0.56929412 | 0.83368112 |
| RWDD4 | 49.1101957 | 0.138363342 | 0.23982075 | 0.57694483 | 0.56397672 | 0.83089872 |
| MRE11 | 122.123838 | 0.138523799 | 0.1939235 | 0.71432187 | 0.47502817 | 0.78022377 |
| PARN | 218.537519 | 0.138775636 | 0.25721486 | 0.53953195 | 0.58951986 | 0.84189197 |
| GPHN | 1145.45058 | 0.138825187 | 0.15526519 | 0.89411662 | 0.3712595 | 0.70984233 |
| ZNF81 | 93.9193463 | 0.138833877 | 0.24544771 | 0.56563525 | 0.57164176 | 0.83368112 |
| RSBN1L | 241.828856 | 0.138955667 | 0.18195367 | 0.76368708 | 0.44505374 | 0.76014919 |
| VPS39 | 144.823029 | 0.139061311 | 0.18760864 | 0.74123086 | 0.45855348 | 0.76968366 |
| RIF1 | 310.31806 | 0.139350375 | 0.2026057 | 0.68779101 | 0.4915844 | 0.79094859 |
| ZNF598 | 38.0623199 | 0.139390413 | 0.39630534 | 0.35172479 | 0.72504467 | 0.9026193 |
| HBS1L | 166.099201 | 0.139478646 | 0.18907962 | 0.73767151 | 0.46071409 | 0.77047321 |
| DERA | 206.556438 | 0.139503001 | 0.19913603 | 0.70054124 | 0.48358936 | 0.78748452 |
| ATP5MD | 58.3038523 | 0.139652777 | 0.38565052 | 0.36212262 | 0.7172604 | 0.9002058 |

| | | | | | | |
|---|---|---|---|---|---|---|
| NUP160 | 225.218018 | 0.139781207 | 0.18749018 | 0.7455388 | 0.45594606 | 0.76740262 |
| GABPA | 146.454673 | 0.139832088 | 0.23519846 | 0.59452807 | 0.55215897 | 0.8236419 |
| ATP11C | 650.580376 | 0.139855479 | 0.35150888 | 0.39787182 | 0.69072468 | 0.88832346 |
| EIF4EBP2 | 109.291219 | 0.139926814 | 0.33754811 | 0.41453888 | 0.6784795 | 0.88164367 |
| CC2D1A | 67.3983535 | 0.139954647 | 0.28352577 | 0.49362232 | 0.62157293 | 0.85896293 |
| SLC26A11 | 53.7076915 | 0.140118889 | 0.26902006 | 0.52084922 | 0.60247181 | 0.85030923 |
| APPBP2 | 363.359439 | 0.140154899 | 0.14190473 | 0.98766896 | 0.3233148 | 0.67664579 |
| IFI6 | 259.036567 | 0.140165628 | 0.86907666 | 0.16128109 | 0.87187202 | 0.95727819 |
| TANC2 | 1019.62892 | 0.140275657 | 0.37609057 | 0.37298371 | 0.70916057 | 0.89727558 |
| MED13L | 1713.05664 | 0.140439492 | 0.25125642 | 0.55894886 | 0.57619662 | 0.83550638 |
| PRKAR2A | 326.177913 | 0.140802482 | 0.17017056 | 0.82741976 | 0.40799919 | 0.73419295 |
| PIGF | 109.599287 | 0.140929208 | 0.24345376 | 0.57887465 | 0.56267376 | 0.83082829 |
| RAB5B | 79.1734093 | 0.141051216 | 0.21826896 | 0.64622663 | 0.5181326 | 0.80557522 |
| BAIAP2 | 88.1274932 | 0.141062167 | 0.43053688 | 0.32764247 | 0.74318201 | 0.90850686 |
| MCM3AP | 156.94381 | 0.141492051 | 0.15770568 | 0.89719061 | 0.36961722 | 0.70984233 |
| CLK3 | 53.2617766 | 0.141494536 | 0.21535178 | 0.65703909 | 0.51115578 | 0.80177277 |
| DCP1A | 237.691348 | 0.141530082 | 0.28620363 | 0.49450834 | 0.62094721 | 0.85890999 |
| GIMAP6 | 114.361817 | 0.141888709 | 0.40767254 | 0.34804579 | 0.7278058 | 0.9037084 |
| SCAF4 | 162.890803 | 0.141976215 | 0.17492737 | 0.8116295 | 0.41700426 | 0.73821752 |
| NHLRC3 | 78.6188096 | 0.141986375 | 0.39126456 | 0.36289097 | 0.71668634 | 0.9001716 |
| LCORL | 393.149775 | 0.142090066 | 0.19905861 | 0.71381019 | 0.47534456 | 0.78022377 |
| SPATS2 | 359.040216 | 0.142253025 | 0.17653191 | 0.80582047 | 0.42034637 | 0.73901389 |
| YWHAH | 126.032213 | 0.142341853 | 0.47514131 | 0.29957794 | 0.76449911 | 0.9154077 |
| PAX8 | 108.376037 | 0.142389331 | 0.3924618 | 0.36281068 | 0.71674632 | 0.9001716 |
| NUP54 | 139.741638 | 0.142488674 | 0.19719676 | 0.72257108 | 0.46994344 | 0.77792258 |
| PRKRA | 107.174586 | 0.142675013 | 0.19743723 | 0.7226348 | 0.46990428 | 0.77792258 |
| CYB5A | 126.696912 | 0.142773477 | 0.38337548 | 0.3724116 | 0.70958642 | 0.89727558 |
| ZKSCAN5 | 41.2219395 | 0.142937518 | 0.28725257 | 0.49760222 | 0.61876444 | 0.85855421 |
| EPHB4 | 295.256337 | 0.143020742 | 0.28799765 | 0.49660386 | 0.61946843 | 0.85855421 |
| MAP1LC3A | 51.7994877 | 0.143046636 | 0.38751285 | 0.36914036 | 0.7120231 | 0.89790184 |
| ATF7IP | 469.749177 | 0.143057987 | 0.27955854 | 0.5117282 | 0.60884125 | 0.85310752 |
| PHTF2 | 174.343436 | 0.143219091 | 0.23771435 | 0.60248398 | 0.54685202 | 0.82355262 |
| TRAM2 | 167.995836 | 0.143445032 | 0.29805501 | 0.48127032 | 0.63032438 | 0.86097142 |
| ZNF169 | 50.5812221 | 0.143688404 | 0.27530366 | 0.52192696 | 0.60172119 | 0.85011818 |
| THSD4 | 1338.47669 | 0.143692457 | 0.44640502 | 0.32188808 | 0.74753748 | 0.91055362 |
| RNF111 | 372.643389 | 0.143694583 | 0.17624034 | 0.81533309 | 0.41488167 | 0.7367571 |
| AGRN | 65.6699905 | 0.143708853 | 0.40911702 | 0.35126589 | 0.72538888 | 0.9026193 |
| CCDC138 | 58.6042194 | 0.14372376 | 0.3380997 | 0.42509284 | 0.670769 | 0.87730512 |
| AC007376.2 | 74.6596532 | 0.143770775 | 0.36126359 | 0.39796641 | 0.69065495 | 0.88832346 |
| FARP2 | 154.506939 | 0.143843136 | 0.23912314 | 0.6015442 | 0.54747758 | 0.82355262 |
| PTBP2 | 660.336701 | 0.143916654 | 0.18283016 | 0.78716034 | 0.43118801 | 0.74490865 |
| LYPLAL1 | 222.210092 | 0.143926438 | 0.31951337 | 0.45045514 | 0.65238229 | 0.86860467 |
| LAMB2 | 216.213275 | 0.14407252 | 0.46110759 | 0.31244881 | 0.75469946 | 0.91289071 |
| FRMD6 | 130.163367 | 0.144188985 | 0.22895377 | 0.62977335 | 0.52884288 | 0.81207005 |
| HLA-DPB1 | 296.162511 | 0.144403304 | 0.5035731 | 0.28675738 | 0.7742981 | 0.91726032 |
| ERO1B | 135.094921 | 0.144464123 | 0.30923912 | 0.46715993 | 0.64038546 | 0.86444642 |
| USP11 | 39.6020757 | 0.144593243 | 0.30409417 | 0.47548837 | 0.63443892 | 0.86207417 |
| ELP1 | 101.267789 | 0.14464187 | 0.19611492 | 0.7375363 | 0.46079627 | 0.77047321 |
| PPP1R12A | 800.740995 | 0.144699581 | 0.20333055 | 0.71164701 | 0.47668339 | 0.78092096 |
| SMAD7 | 182.11167 | 0.144721839 | 0.56012067 | 0.25837618 | 0.7961166 | 0.92853793 |
| WASHC3 | 116.628186 | 0.144821542 | 0.23535524 | 0.61533171 | 0.53833568 | 0.81807397 |
| PPP4R1 | 277.874472 | 0.144892987 | 0.16572126 | 0.87431746 | 0.38194539 | 0.71553552 |
| SAMM50 | 45.6428591 | 0.145129418 | 0.23709805 | 0.61210718 | 0.54046684 | 0.81902461 |
| ZNF91 | 290.02003 | 0.145182154 | 0.20773659 | 0.69887617 | 0.48462942 | 0.78781193 |
| ZNF518A | 238.607481 | 0.145403071 | 0.25649252 | 0.56689009 | 0.57078886 | 0.83368112 |
| SGMS1 | 408.899164 | 0.145425587 | 0.22345906 | 0.65079298 | 0.51518013 | 0.80444072 |
| GALK2 | 274.818138 | 0.145694489 | 0.29062384 | 0.50131637 | 0.61614848 | 0.85732696 |
| ATP11B | 284.637864 | 0.145825675 | 0.23032168 | 0.63313916 | 0.52664277 | 0.81036438 |
| GK5 | 92.6830448 | 0.14583707 | 0.1805894 | 0.80756164 | 0.41934298 | 0.7385888 |
| CNOT4 | 449.120026 | 0.14619674 | 0.16804816 | 0.86996927 | 0.3843172 | 0.71699718 |
| VPS13D | 596.603455 | 0.146300153 | 0.16581954 | 0.88228536 | 0.37762251 | 0.71305115 |
| BRD7 | 127.172982 | 0.146343556 | 0.22525486 | 0.64967989 | 0.51589902 | 0.80483629 |
| CAMK1D | 605.416285 | 0.146701757 | 0.26354015 | 0.55665808 | 0.57776106 | 0.83613556 |
| MAZ | 53.8998813 | 0.146793556 | 0.42518389 | 0.34524722 | 0.72990852 | 0.90426442 |
| TRMT13 | 89.7238336 | 0.147133273 | 0.16904313 | 0.87038894 | 0.38408789 | 0.71699718 |
| EXD2 | 42.3655055 | 0.147204282 | 0.25342438 | 0.58086077 | 0.5613343 | 0.82940393 |
| ABRAXAS2 | 59.3835341 | 0.147242492 | 0.26438655 | 0.55692127 | 0.57758122 | 0.83607769 |
| MDC1 | 47.2267225 | 0.147246882 | 0.28401996 | 0.5184385 | 0.60415236 | 0.85098921 |
| CAPRIN2 | 89.4084728 | 0.147308708 | 0.32494693 | 0.45333159 | 0.65030999 | 0.86768084 |
| ATG4B | 145.009917 | 0.147373777 | 0.18122431 | 0.81321197 | 0.41609654 | 0.73795362 |
| NRF1 | 315.464792 | 0.14745057 | 0.19175002 | 0.7689729 | 0.4419094 | 0.75667391 |
| LINC01004 | 260.436521 | 0.147801321 | 0.25640654 | 0.57643351 | 0.5643222 | 0.83099668 |
| USP32 | 390.785754 | 0.147822449 | 0.22698305 | 0.65124883 | 0.51488587 | 0.80422048 |
| NVL | 149.099893 | 0.147894577 | 0.18318257 | 0.80736161 | 0.41945818 | 0.73863273 |
| DCUN1D4 | 220.352472 | 0.148131517 | 0.15855358 | 0.93426787 | 0.35016573 | 0.69858573 |
| TUBGCP5 | 73.5996349 | 0.148233988 | 0.34539562 | 0.42917159 | 0.66779835 | 0.87587657 |
| ZCCHC8 | 133.725312 | 0.148239548 | 0.22712051 | 0.65269117 | 0.5139554 | 0.80398873 |
| RHOT1 | 214.271086 | 0.148367039 | 0.25182966 | 0.58915633 | 0.55575641 | 0.82487928 |
| SSFA2 | 329.216641 | 0.14851281 | 0.27222955 | 0.54554257 | 0.58538041 | 0.83908626 |
| EDRF1 | 110.408397 | 0.148997152 | 0.16872969 | 0.88305238 | 0.37720797 | 0.71261673 |
| HEPH | 51.2263272 | 0.149312657 | 0.5676762 | 0.26302434 | 0.79253182 | 0.92735129 |
| PHKA1 | 62.1669104 | 0.149380165 | 0.50752719 | 0.29432939 | 0.76850622 | 0.91631735 |
| ZNF430 | 97.9229073 | 0.149452435 | 0.20427744 | 0.73161497 | 0.46440361 | 0.77428988 |
| ANKRD27 | 114.876144 | 0.149777152 | 0.27962667 | 0.53563256 | 0.59221252 | 0.84399017 |
| LPGAT1 | 291.668287 | 0.149826627 | 0.12858798 | 1.16516819 | 0.24395092 | 0.608518 |
| DYNC2LI1 | 76.6662067 | 0.149835143 | 0.2646303 | 0.56620555 | 0.57125406 | 0.83368112 |
| CBWD2 | 48.5542504 | 0.149888063 | 0.39250308 | 0.38187742 | 0.70255229 | 0.89443605 |
| SAMHD1 | 213.167742 | 0.150138113 | 0.47511429 | 0.3160042 | 0.75199932 | 0.91151095 |
| ARID4B | 961.978253 | 0.150411926 | 0.25111881 | 0.59896719 | 0.54919476 | 0.82355262 |
| ERP44 | 213.644692 | 0.150461339 | 0.14199735 | 1.0596067 | 0.28932356 | 0.65239315 |
| HLA-F | 160.822395 | 0.150546771 | 0.3497658 | 0.43042164 | 0.66688895 | 0.87587657 |
| PHC3 | 511.925518 | 0.150636355 | 0.18474982 | 0.8153532 | 0.41487016 | 0.7367571 |
| ZNF529-AS1 | 44.5242499 | 0.150704811 | 0.33760244 | 0.44639728 | 0.6553103 | 0.87094432 |
| AC108134.3 | 70.2325662 | 0.150740742 | 0.3177835 | 0.47435044 | 0.63525003 | 0.86207417 |
| PSMG2 | 115.195645 | 0.151076616 | 0.22501654 | 0.67140226 | 0.50196431 | 0.79665902 |
| AL359220.1 | 106.254601 | 0.151080514 | 0.25432504 | 0.59404499 | 0.55248202 | 0.8236419 |
| SBNO1 | 194.08528 | 0.151111261 | 0.19095553 | 0.79134268 | 0.42874405 | 0.74408005 |
| BDP1 | 356.280039 | 0.15115905 | 0.21836995 | 0.69221544 | 0.48880205 | 0.78955612 |
| DYNC1I2 | 308.081771 | 0.15118932 | 0.14117426 | 1.0709411 | 0.28419591 | 0.64726501 |
| CCDC102B | 62.6168833 | 0.151456999 | 0.31490778 | 0.48095668 | 0.63054729 | 0.86097142 |
| PPP2R2A | 585.11975 | 0.151484407 | 0.21628052 | 0.70040706 | 0.48367313 | 0.78748452 |
| GNAI3 | 185.591667 | 0.151532483 | 0.17624619 | 0.85977737 | 0.38991177 | 0.72202943 |
| GGNBP2 | 226.55222 | 0.151573866 | 0.16896042 | 0.8970969 | 0.36966722 | 0.70984233 |
| FAM192A | 114.008616 | 0.151842911 | 0.22570221 | 0.67275777 | 0.50110141 | 0.79642461 |
| MARK3 | 718.652721 | 0.151921434 | 0.23292253 | 0.65224018 | 0.51424624 | 0.80398873 |
| CREBBP | 460.460684 | 0.152088063 | 0.21296317 | 0.71415194 | 0.47513323 | 0.78022377 |
| PSMD11 | 154.24538 | 0.152615714 | 0.24706907 | 0.61770466 | 0.53677004 | 0.81764505 |
| ARL8B | 260.62919 | 0.152677518 | 0.21328583 | 0.71583525 | 0.47409308 | 0.77932979 |
| GGA3 | 83.6274932 | 0.152713686 | 0.23233349 | 0.6573038 | 0.5109856 | 0.80177277 |
| CRBN | 258.591365 | 0.152833521 | 0.26462445 | 0.57754876 | 0.56356881 | 0.83089872 |
| MICAL3 | 320.868058 | 0.153041681 | 0.23670475 | 0.64655095 | 0.51792261 | 0.80546868 |
| APOL2 | 99.242933 | 0.153387676 | 0.493025 | 0.31111541 | 0.75571289 | 0.91344144 |
| ZNF608 | 263.002301 | 0.153542942 | 0.29786304 | 0.51548169 | 0.60621646 | 0.85161705 |

| | | | | | | |
|---|---|---|---|---|---|---|
| SIK3 | 1219.74067 | 0.153606546 | 0.32087481 | 0.47871176 | 0.6321437 | 0.86141143 |
| TCF3 | 77.1560883 | 0.153837587 | 0.32030246 | 0.4802885 | 0.63102227 | 0.86097142 |
| TCF12 | 2026.66556 | 0.153842697 | 0.18820487 | 0.81742146 | 0.41368761 | 0.73655477 |
| CCNG2 | 98.7036493 | 0.153924934 | 0.46037618 | 0.334346 | 0.73811848 | 0.9078343 |
| ARMC8 | 295.366878 | 0.154087278 | 0.17241537 | 0.89369805 | 0.37148347 | 0.70991341 |
| RNF152 | 197.55271 | 0.154136547 | 0.35018998 | 0.44015122 | 0.65982759 | 0.87335081 |
| SKAP2 | 564.869303 | 0.154359586 | 0.30676241 | 0.50318938 | 0.61483113 | 0.85680098 |
| AC053527.1 | 82.7208482 | 0.154385079 | 0.52959606 | 0.29151478 | 0.77065764 | 0.91695334 |
| BICD2 | 90.2200894 | 0.154522809 | 0.21788164 | 0.70920528 | 0.4781971 | 0.78162783 |
| AFF1 | 1740.94851 | 0.154543737 | 0.21507178 | 0.71856818 | 0.47240703 | 0.77874833 |
| CD55 | 293.696858 | 0.154625983 | 0.44000678 | 0.35141727 | 0.72527532 | 0.9026193 |
| NEDD9 | 1055.70098 | 0.154817289 | 0.51860637 | 0.29852562 | 0.76530202 | 0.91580893 |
| NFATC2IP | 231.355364 | 0.154994502 | 0.24826418 | 0.62431278 | 0.53242219 | 0.8146617 |
| VPS4B | 152.990858 | 0.15514794 | 0.16413956 | 0.94521964 | 0.34454676 | 0.69409801 |
| LANCL1 | 103.710795 | 0.155428378 | 0.17790539 | 0.87365751 | 0.3823048 | 0.71588105 |
| SLC35E2A | 48.2931542 | 0.155443991 | 0.34274547 | 0.45352602 | 0.65017001 | 0.86766991 |
| SHANK3 | 1157.62723 | 0.155937953 | 0.21256846 | 0.73358931 | 0.46319908 | 0.77306915 |
| BAZ2B | 1483.72157 | 0.155944927 | 0.24342822 | 0.64061977 | 0.52176975 | 0.80772643 |
| ZSCAN30 | 92.0117984 | 0.156006402 | 0.28126665 | 0.5546566 | 0.57912957 | 0.83649569 |
| NIPAL3 | 69.5680622 | 0.156154506 | 0.24868506 | 0.62792074 | 0.53005586 | 0.8131655 |
| RNASEH2B | 88.461311 | 0.156328829 | 0.26273179 | 0.59501298 | 0.55183479 | 0.8236419 |
| SMPD4 | 87.2719587 | 0.156367209 | 0.21957948 | 0.71212123 | 0.47638971 | 0.78059611 |
| MAP7D1 | 98.2710251 | 0.156478556 | 0.28419818 | 0.55059662 | 0.58191023 | 0.83789748 |
| ZDHHC8 | 56.4848887 | 0.15673322 | 0.31103606 | 0.5039069 | 0.6143268 | 0.85624871 |
| ZFR | 491.725285 | 0.156740413 | 0.13930234 | 1.12518148 | 0.26051214 | 0.62580682 |
| AC084337.2 | 71.0276395 | 0.156769681 | 0.29390561 | 0.53340145 | 0.59375571 | 0.84502915 |
| DET1 | 45.954803 | 0.156887186 | 0.57405393 | 0.27329695 | 0.78462496 | 0.92302847 |
| FBXO11 | 780.238624 | 0.157000603 | 0.19191863 | 0.81805817 | 0.41332397 | 0.73622771 |
| SUPT3H | 285.783061 | 0.157007625 | 0.3776926 | 0.41570215 | 0.67762797 | 0.88143805 |
| MRPL39 | 52.5501056 | 0.157191874 | 0.36707404 | 0.42822934 | 0.66848416 | 0.87625461 |
| ANKH | 351.375562 | 0.157244774 | 0.170781 | 0.92073925 | 0.35718658 | 0.70058911 |
| TRIM22 | 225.086017 | 0.157417209 | 0.49137276 | 0.3203621 | 0.74869385 | 0.91065347 |
| WSB2 | 66.3990431 | 0.157657883 | 0.276975 | 0.5692134 | 0.56921133 | 0.83368112 |
| KDM5A | 351.57819 | 0.157718173 | 0.19448095 | 0.8109698 | 0.41738302 | 0.73821752 |
| BACE1 | 64.4621222 | 0.157795782 | 0.24064939 | 0.65570821 | 0.51201188 | 0.80233497 |
| ACTR3 | 514.994027 | 0.157855367 | 0.23333543 | 0.67651692 | 0.49871251 | 0.79513466 |
| EXOSC6 | 76.5275061 | 0.157931941 | 0.33190963 | 0.47582814 | 0.63419682 | 0.86207417 |
| UBA5 | 93.1414788 | 0.158022481 | 0.29223434 | 0.54073891 | 0.58868756 | 0.84140756 |
| DDX56 | 41.0378844 | 0.158080457 | 0.30322699 | 0.52132714 | 0.6021389 | 0.85030923 |
| FAM171B | 65.2427875 | 0.158098915 | 0.48255547 | 0.32762848 | 0.74319258 | 0.90850686 |
| JAG2 | 142.95988 | 0.158133657 | 0.64574579 | 0.24488531 | 0.80654524 | 0.93187644 |
| CCDC93 | 186.241284 | 0.158293435 | 0.17881514 | 0.88523508 | 0.37602986 | 0.71257833 |
| INO80 | 289.514038 | 0.1583402 | 0.26415248 | 0.59942728 | 0.54888799 | 0.82355262 |
| LRRC57 | 48.1688583 | 0.158375542 | 0.24126159 | 0.65644738 | 0.51153631 | 0.80205051 |
| NOL11 | 61.4704656 | 0.158429975 | 0.28002999 | 0.56576074 | 0.57155644 | 0.83368112 |
| DVL3 | 75.3690731 | 0.158434683 | 0.22009278 | 0.71985408 | 0.47161485 | 0.77855633 |
| EXOSC10 | 81.7154945 | 0.158682992 | 0.2362081 | 0.6717932 | 0.50171536 | 0.79661835 |
| KCTD2 | 45.3249422 | 0.15886137 | 0.2278518 | 0.69721359 | 0.48566913 | 0.78826504 |
| DCAF5 | 379.562295 | 0.159050691 | 0.17635289 | 0.90188875 | 0.36711597 | 0.7075153 |
| BRAP | 85.3335656 | 0.15906401 | 0.21415471 | 0.74275281 | 0.45763135 | 0.76905605 |
| UBLCP1 | 90.1366941 | 0.159500257 | 0.26368543 | 0.6048884 | 0.54525315 | 0.82355262 |
| AL359915.2 | 149.067363 | 0.159643376 | 0.39505826 | 0.40410084 | 0.68613856 | 0.88671976 |
| PIAS2 | 276.271412 | 0.159731661 | 0.17949743 | 0.88988273 | 0.37352886 | 0.71166161 |
| FHOD1 | 65.7201006 | 0.159797706 | 0.3774787 | 0.42332907 | 0.67205519 | 0.87803719 |
| DELE1 | 181.80844 | 0.160129712 | 0.18264428 | 0.87672998 | 0.38063332 | 0.71529048 |
| ZNHIT6 | 170.049107 | 0.160231557 | 0.25416352 | 0.63042704 | 0.52841522 | 0.81207005 |
| THSD1 | 93.9979308 | 0.160258704 | 0.2809502 | 0.57041677 | 0.56839506 | 0.83354423 |
| SEMA6A | 877.676689 | 0.160321361 | 0.37836656 | 0.42371969 | 0.67177025 | 0.87799009 |
| XIAP | 309.515785 | 0.160428712 | 0.16791237 | 0.95543114 | 0.33935971 | 0.68959039 |
| GARNL3 | 43.5034361 | 0.160445338 | 0.39621567 | 0.40494445 | 0.68551834 | 0.88671976 |
| TP53I11 | 97.5299543 | 0.160838289 | 0.4778955 | 0.33655535 | 0.73645212 | 0.90690559 |
| RCHY1 | 96.2237114 | 0.160956368 | 0.29301091 | 0.54931869 | 0.58278677 | 0.83815192 |
| ZFC3H1 | 348.533255 | 0.161168456 | 0.21583821 | 0.74670956 | 0.4552389 | 0.76710126 |
| APIP | 70.9093037 | 0.161261058 | 0.34742844 | 0.46415619 | 0.64253585 | 0.86490071 |
| HLTF | 88.9028848 | 0.161394884 | 0.35759775 | 0.45133082 | 0.65175114 | 0.8685756 |
| RIPOR1 | 192.509788 | 0.161457555 | 0.25886241 | 0.62371959 | 0.53281175 | 0.81470811 |
| CSNK1G1 | 46.5694188 | 0.161472637 | 0.24305456 | 0.66434728 | 0.50646807 | 0.7989142 |
| TRIP6 | 49.7338501 | 0.1615425 | 0.3483514 | 0.46373433 | 0.6428381 | 0.86504591 |
| C1orf56 | 46.8329823 | 0.161595993 | 0.61882153 | 0.26113505 | 0.79398836 | 0.92790728 |
| TRAPPC4 | 72.4109307 | 0.161654788 | 0.32226731 | 0.50161709 | 0.61593689 | 0.85732465 |
| RABEP1 | 338.688905 | 0.1616773 | 0.2375199 | 0.68068949 | 0.49606799 | 0.79381653 |
| ATF2 | 325.618766 | 0.161700435 | 0.21322658 | 0.75835029 | 0.44824131 | 0.76164496 |
| ARFGEF1 | 672.956184 | 0.161858942 | 0.13474757 | 1.2012012 | 0.22967317 | 0.59648134 |
| SPOPL | 150.027615 | 0.161874878 | 0.14943137 | 1.08327238 | 0.27868754 | 0.64221351 |
| DIAPH1 | 259.632653 | 0.161961296 | 0.34252452 | 0.47284584 | 0.63632317 | 0.86207417 |
| SHROOM4 | 545.776688 | 0.162520327 | 0.21357394 | 0.7609558 | 0.44668347 | 0.76094002 |
| DENND1A | 577.602567 | 0.162796954 | 0.31910167 | 0.51017268 | 0.60993049 | 0.85375662 |
| PRKAA1 | 263.062855 | 0.162816076 | 0.26461811 | 0.61528697 | 0.53836522 | 0.81807397 |
| ARHGAP35 | 207.323199 | 0.163219738 | 0.27300732 | 0.59785847 | 0.54993437 | 0.82355262 |
| SH2B1 | 66.206367 | 0.16334911 | 0.21743019 | 0.75127152 | 0.45248926 | 0.76486669 |
| DCAF16 | 70.497042 | 0.163369965 | 0.22588598 | 0.72324085 | 0.46953193 | 0.77792258 |
| ELMOD2 | 62.1754756 | 0.163419051 | 0.22625646 | 0.72227354 | 0.47012632 | 0.77792258 |
| TRAPPC6B | 87.3263696 | 0.163546471 | 0.19482919 | 0.83943517 | 0.40122515 | 0.72911494 |
| FYN | 1166.70846 | 0.163966338 | 0.3592113 | 0.45646208 | 0.64805773 | 0.86739793 |
| BRAT1 | 43.8093784 | 0.163983461 | 0.28381426 | 0.57778443 | 0.56340967 | 0.83089872 |
| SEC14L1 | 2227.1736 | 0.163989338 | 0.3473674 | 0.4720919 | 0.63686119 | 0.86207417 |
| POU2F1 | 510.165475 | 0.164044633 | 0.18066872 | 0.90798583 | 0.36388571 | 0.70570548 |
| CLOCK | 339.44132 | 0.164278218 | 0.17802563 | 0.92277848 | 0.35612266 | 0.70018061 |
| ELK4 | 167.17872 | 0.16435712 | 0.18102884 | 0.90790572 | 0.36392804 | 0.70570548 |
| CLSTN1 | 173.210177 | 0.164376192 | 0.27583692 | 0.59591803 | 0.55122999 | 0.8236419 |
| XPNPEP3 | 126.603652 | 0.164408536 | 0.2648508 | 0.62075907 | 0.53475816 | 0.81575648 |
| GNB1 | 1049.60744 | 0.164426055 | 0.13661129 | 1.20360513 | 0.22874223 | 0.59497702 |
| TBL1XR1 | 897.609404 | 0.164509243 | 0.24313641 | 0.67661296 | 0.49865156 | 0.79513466 |
| AKAP10 | 275.747452 | 0.164628687 | 0.16645887 | 0.98900518 | 0.32266061 | 0.6761919 |
| BLMH | 106.661414 | 0.164682907 | 0.26485747 | 0.62177936 | 0.53408696 | 0.81537941 |
| IREB2 | 199.32878 | 0.164947459 | 0.20569166 | 0.80191612 | 0.42260148 | 0.74011688 |
| RAP2C | 47.7571995 | 0.165023487 | 0.28411843 | 0.58082641 | 0.56135746 | 0.82940393 |
| ECPAS | 456.462208 | 0.165074019 | 0.27853883 | 0.59264276 | 0.55342026 | 0.82372318 |
| GTF2H2 | 56.3400103 | 0.165478273 | 0.47201057 | 0.35058171 | 0.72590218 | 0.9026831 |
| IMMP2L | 1570.03626 | 0.165483173 | 0.23806624 | 0.69511398 | 0.48698388 | 0.78826504 |
| CDKAL1 | 1036.36706 | 0.16568582 | 0.17123489 | 0.9675938 | 0.33324727 | 0.6852158 |
| RBM4B | 46.6743818 | 0.165820854 | 0.33349966 | 0.49721446 | 0.61903783 | 0.85855421 |
| TCP11L1 | 72.5407945 | 0.165834879 | 0.2671971 | 0.62064626 | 0.5348324 | 0.81575648 |
| TMEM87A | 268.074547 | 0.165959523 | 0.25160636 | 0.65959987 | 0.50951064 | 0.80133107 |
| OTUD6B | 59.1766555 | 0.165993178 | 0.24197185 | 0.68600202 | 0.49271183 | 0.79102628 |
| FRMD4B | 1208.60449 | 0.166004178 | 0.1935996 | 0.85746416 | 0.39118841 | 0.72247779 |
| FBXL7 | 3031.5822 | 0.166011977 | 0.22767035 | 0.72917698 | 0.46589341 | 0.77517793 |
| RNLS | 88.4889199 | 0.166047957 | 0.46222309 | 0.3592377 | 0.71941728 | 0.90076622 |
| LPIN1 | 77.0705235 | 0.166065328 | 0.29072584 | 0.57120939 | 0.56785771 | 0.83343068 |
| PPID | 62.9578385 | 0.166075213 | 0.27070041 | 0.61350188 | 0.53954454 | 0.81882288 |
| GPR107 | 206.026467 | 0.166088653 | 0.17864465 | 0.92971525 | 0.35251853 | 0.69949552 |
| EFHC1 | 82.0144118 | 0.166239545 | 0.30056367 | 0.5530926 | 0.5802 | 0.83735037 |
| VEGFA | 85.5074517 | 0.166312012 | 0.53269925 | 0.31220621 | 0.75488381 | 0.91297875 |

| | | | | | | |
|---|---|---|---|---|---|---|
| KCNQ1 | 161.40556 | 0.1664018 | 0.25098522 | 0.66299442 | 0.50733413 | 0.79921563 |
| DCBLD2 | 58.5252181 | 0.167094987 | 0.40027573 | 0.41744971 | 0.6763495 | 0.88045943 |
| FAM53C | 41.8632469 | 0.167098478 | 0.31549237 | 0.52964348 | 0.59635914 | 0.84707635 |
| LRIG1 | 68.3251925 | 0.167350589 | 0.43778341 | 0.382268 | 0.70226259 | 0.89443605 |
| WIPF2 | 262.162209 | 0.167379048 | 0.23475098 | 0.71300681 | 0.47584154 | 0.78022377 |
| ETHE1 | 45.3975324 | 0.167493189 | 0.2670267 | 0.62725258 | 0.53049368 | 0.81320326 |
| ARHGDIB | 107.542712 | 0.167586185 | 0.42930929 | 0.39036235 | 0.69626862 | 0.8920567 |
| MED4 | 181.709626 | 0.167596762 | 0.13895831 | 1.20609382 | 0.22778132 | 0.59455647 |
| WDR59 | 184.024291 | 0.167603052 | 0.16225942 | 1.03293265 | 0.30163542 | 0.65926326 |
| SECISBP2L | 233.950995 | 0.167852817 | 0.18118599 | 0.92641168 | 0.35423208 | 0.69999543 |
| LYPLA2 | 46.848377 | 0.167867706 | 0.35022514 | 0.47931369 | 0.63171548 | 0.86118497 |
| BCL7B | 60.4899231 | 0.167955573 | 0.22063457 | 0.76123868 | 0.44651452 | 0.76094002 |
| UPF2 | 237.932059 | 0.168116914 | 0.15580606 | 1.07901398 | 0.2805815 | 0.64385822 |
| TSNARE1 | 50.4461745 | 0.168132329 | 0.28300989 | 0.59408641 | 0.55245432 | 0.8236419 |
| TBXAS1 | 110.395914 | 0.168269963 | 0.54521493 | 0.30863051 | 0.75760261 | 0.91383581 |
| APOOL | 102.895223 | 0.168923456 | 0.2056334 | 0.82147871 | 0.41137365 | 0.73470224 |
| ZNF445 | 72.4011707 | 0.168930388 | 0.23264607 | 0.72612613 | 0.46776145 | 0.77656563 |
| TUBGCP3 | 196.632374 | 0.169070981 | 0.24865872 | 0.67993184 | 0.49654762 | 0.79420452 |
| TAF4 | 94.6350242 | 0.169072025 | 0.2867391 | 0.58963714 | 0.55543394 | 0.82487928 |
| CHTF8 | 78.677854 | 0.169168127 | 0.26418362 | 0.64034299 | 0.52194964 | 0.80785207 |
| SFSWAP | 249.168741 | 0.169188341 | 0.22827762 | 0.74115167 | 0.45860148 | 0.76968366 |
| CAMSAP2 | 512.350725 | 0.169209235 | 0.1420944 | 1.19082267 | 0.23372321 | 0.59935976 |
| NNT | 180.799847 | 0.169290856 | 0.22428456 | 0.75480389 | 0.45036666 | 0.76375998 |
| BMPR2 | 2496.13784 | 0.169550115 | 0.32386402 | 0.52352255 | 0.60061067 | 0.8498698 |
| YIF1B | 42.5618462 | 0.169728294 | 0.37836321 | 0.44858562 | 0.65373061 | 0.86983375 |
| MSH2 | 101.264379 | 0.169877523 | 0.23828876 | 0.71290616 | 0.47590383 | 0.78022377 |
| PTPRA | 464.935411 | 0.169974561 | 0.22940173 | 0.74094717 | 0.45872547 | 0.76968435 |
| DCP1B | 68.5165733 | 0.17002973 | 0.27608277 | 0.61586506 | 0.53798358 | 0.81807397 |
| ARFGAP2 | 67.1834829 | 0.170195372 | 0.27893482 | 0.61016181 | 0.54175462 | 0.82004774 |
| PDPK1 | 169.726134 | 0.170402382 | 0.24782838 | 0.6875822 | 0.49171592 | 0.79094859 |
| GATA2-AS1 | 136.434266 | 0.170406688 | 0.34661593 | 0.49162971 | 0.62298114 | 0.85940448 |
| AC118549.1 | 459.542341 | 0.170722917 | 0.16668032 | 1.02425362 | 0.3057155 | 0.66234311 |
| CHFR | 185.331799 | 0.170747524 | 0.2780996 | 0.61397976 | 0.5392287 | 0.81864721 |
| RB1CC1 | 392.984859 | 0.170970853 | 0.15906358 | 1.07485861 | 0.28243804 | 0.64558393 |
| OCRL | 61.7016865 | 0.17139617 | 0.21525992 | 0.79622889 | 0.42589901 | 0.74197066 |
| FBXO42 | 263.03331 | 0.171509574 | 0.19561151 | 0.87678671 | 0.3806025 | 0.71529048 |
| ZNF429 | 72.1878729 | 0.171557226 | 0.29993786 | 0.5719759 | 0.5673383 | 0.83329704 |
| GDAP2 | 117.169827 | 0.171605289 | 0.19388061 | 0.88510805 | 0.37609836 | 0.71257833 |
| NUP88 | 106.772705 | 0.171607584 | 0.21034286 | 0.81584695 | 0.41458767 | 0.73671465 |
| MERTK | 126.467169 | 0.171766365 | 0.56479778 | 0.30412011 | 0.76103638 | 0.91477008 |
| TBC1D10A | 47.6478073 | 0.171780295 | 0.2811339 | 0.61102662 | 0.54118196 | 0.81933246 |
| EFR3A | 307.399497 | 0.172035224 | 0.18847328 | 0.91278311 | 0.36135662 | 0.70386371 |
| ZEB1 | 3385.21289 | 0.172147461 | 0.19805727 | 0.86918022 | 0.38474857 | 0.71699718 |
| MSH6 | 51.4547482 | 0.172155879 | 0.27544739 | 0.62500458 | 0.53196805 | 0.81427134 |
| FAM171A1 | 384.946202 | 0.172586048 | 0.33691915 | 0.51224767 | 0.60847769 | 0.85289019 |
| LPXN | 62.1508071 | 0.172658244 | 0.30400943 | 0.56793714 | 0.57007766 | 0.83368112 |
| CRK | 325.285742 | 0.172686986 | 0.3089468 | 0.5589538 | 0.57619325 | 0.83550638 |
| SPIN1 | 211.113033 | 0.173058787 | 0.15119046 | 1.14464095 | 0.25235792 | 0.61777104 |
| BLZF1 | 97.3755309 | 0.173092991 | 0.34163652 | 0.50665833 | 0.61239457 | 0.85573702 |
| AVL9 | 362.873884 | 0.173195067 | 0.19547031 | 0.88604282 | 0.37559445 | 0.71257833 |
| SCLT1 | 395.438833 | 0.17328151 | 0.35656791 | 0.48597057 | 0.62698803 | 0.86052754 |
| ZKSCAN8 | 55.6276255 | 0.173545256 | 0.25366339 | 0.68415572 | 0.49387684 | 0.79243023 |
| RAP1A | 602.61659 | 0.173706444 | 0.22874853 | 0.75937731 | 0.44762688 | 0.7614672 |
| NAPRT | 37.703263 | 0.17392439 | 0.40058276 | 0.43417842 | 0.66415888 | 0.8755434 |
| ZNF69 | 49.4349161 | 0.173966895 | 0.24511544 | 0.70973455 | 0.47786877 | 0.78129872 |
| ERC1 | 1274.26005 | 0.174010482 | 0.11720951 | 1.48461058 | 0.13764701 | 0.48962215 |
| RBM26 | 508.177845 | 0.174061404 | 0.10050001 | 1.73195408 | 0.08328174 | 0.39543345 |
| DGKH | 969.50615 | 0.174068147 | 0.18670232 | 0.93232986 | 0.35116608 | 0.69858573 |
| SHARPIN | 46.1729474 | 0.174075258 | 0.22817678 | 0.76289647 | 0.44552514 | 0.76047815 |
| DCAF10 | 291.519018 | 0.174306189 | 0.19540116 | 0.89204276 | 0.37237001 | 0.71084822 |
| RNF121 | 80.873112 | 0.174364698 | 0.28126583 | 0.61992848 | 0.53530487 | 0.81602082 |
| NEMF | 402.83197 | 0.174516304 | 0.14990115 | 1.16420926 | 0.24433922 | 0.60883916 |
| PCID2 | 99.7756341 | 0.174835987 | 0.18744615 | 0.93272647 | 0.35096121 | 0.69858573 |
| IMMP1L | 156.410583 | 0.17498226 | 0.21378333 | 0.81850281 | 0.41307014 | 0.73622771 |
| PSMD6 | 47.8786136 | 0.175192899 | 0.31857381 | 0.54992875 | 0.58236825 | 0.83812111 |
| ZFP14 | 93.1857832 | 0.175401104 | 0.26049863 | 0.67332832 | 0.50073844 | 0.79625863 |
| SRPK2 | 910.888433 | 0.175462001 | 0.17925631 | 0.97883306 | 0.32766247 | 0.68017808 |
| MPP5 | 150.635493 | 0.175483837 | 0.17000012 | 1.03225715 | 0.30195167 | 0.65926326 |
| MKNK1 | 73.0713959 | 0.175801103 | 0.18293366 | 0.96101015 | 0.33654706 | 0.68745997 |
| FARS2 | 542.267933 | 0.175878018 | 0.26457653 | 0.66475292 | 0.50620854 | 0.79890154 |
| MBD6 | 49.7304211 | 0.175885962 | 0.31380283 | 0.56049834 | 0.57513957 | 0.834481 |
| VPS35L | 261.116979 | 0.175981765 | 0.21576619 | 0.81561325 | 0.41472137 | 0.7367571 |
| OCLN | 67.3471222 | 0.176221381 | 0.41214874 | 0.42756744 | 0.66896607 | 0.87628452 |
| SMC4 | 145.76439 | 0.176281847 | 0.26865469 | 0.65616517 | 0.51171786 | 0.80218146 |
| NADK2 | 115.151924 | 0.176476952 | 0.49927594 | 0.35346576 | 0.72373929 | 0.90258339 |
| NBAS | 503.675492 | 0.176610274 | 0.18228428 | 0.96887279 | 0.33260866 | 0.6852158 |
| TTC38 | 60.2014229 | 0.176636016 | 0.29522174 | 0.59831642 | 0.54962882 | 0.82355262 |
| FGF2 | 220.911302 | 0.1766727 | 0.44529301 | 0.39675606 | 0.69154735 | 0.88865138 |
| MINDY3 | 247.032634 | 0.176776764 | 0.19227739 | 0.91938404 | 0.35789473 | 0.70147368 |
| ITM2A | 105.425708 | 0.176785245 | 0.91914231 | 0.19233718 | 0.84747809 | 0.94731791 |
| GLT8D1 | 47.7942396 | 0.176849458 | 0.25665411 | 0.68905757 | 0.49078704 | 0.79073252 |
| USP8 | 231.499806 | 0.177193531 | 0.16028672 | 1.10547857 | 0.26895228 | 0.63347822 |
| RALY | 284.511127 | 0.177352142 | 0.20052416 | 0.88444275 | 0.37645725 | 0.71257833 |
| MAP3K11 | 200.028519 | 0.177616298 | 0.34814742 | 0.51017553 | 0.60992849 | 0.85375662 |
| EPS15L1 | 197.086727 | 0.177750654 | 0.24434609 | 0.72745448 | 0.46694759 | 0.77568659 |
| HPF1 | 57.149785 | 0.177875691 | 0.32474124 | 0.54774592 | 0.58386638 | 0.83850098 |
| TRIB2 | 109.211905 | 0.177898036 | 0.37511101 | 0.47425437 | 0.63531852 | 0.86207417 |
| EPHX1 | 112.833764 | 0.177932473 | 0.28356465 | 0.62748468 | 0.53034157 | 0.81320326 |
| HKR1 | 125.962022 | 0.177981208 | 0.22826979 | 0.77969672 | 0.43556941 | 0.74971913 |
| ZCCHC4 | 49.9493104 | 0.17800852 | 0.3043742 | 0.58483446 | 0.55865903 | 0.82727232 |
| IFT81 | 59.9686292 | 0.178022619 | 0.32931602 | 0.54058293 | 0.58879509 | 0.84140756 |
| RNF168 | 215.344661 | 0.178038364 | 0.15769138 | 1.1290304 | 0.25888501 | 0.62362556 |
| AC007319.1 | 1150.39847 | 0.178171698 | 0.5166873 | 0.34483468 | 0.73021866 | 0.90426442 |
| PPP3CC | 442.165273 | 0.178238247 | 0.19935117 | 0.89409179 | 0.37127278 | 0.70984233 |
| PPP6R2 | 383.147183 | 0.178362174 | 0.2360676 | 0.75555549 | 0.44991575 | 0.76319664 |
| HPS3 | 183.599427 | 0.178411908 | 0.37484575 | 0.47596087 | 0.63410226 | 0.86207417 |
| AC092422.1 | 106.04396 | 0.178432022 | 0.36648992 | 0.48686748 | 0.62635225 | 0.86052754 |
| ENTPD5 | 54.9584389 | 0.178508785 | 0.31692424 | 0.5632538 | 0.57326207 | 0.8342245 |
| PDHX | 120.210705 | 0.178517483 | 0.17557332 | 1.01676883 | 0.30926341 | 0.66457524 |
| FAM204A | 132.357333 | 0.178808414 | 0.18065116 | 0.9897994 | 0.32227218 | 0.6761919 |
| TMEM161B | 107.488275 | 0.178838024 | 0.20696711 | 0.8640891 | 0.38753895 | 0.7205706 |
| GAN | 125.304256 | 0.179239038 | 0.23595771 | 0.75962357 | 0.44747963 | 0.7614672 |
| IPPK | 79.5966794 | 0.179247689 | 0.35518781 | 0.50465609 | 0.61380041 | 0.85580656 |
| TMTC1 | 2853.81838 | 0.179407732 | 0.34709473 | 0.51688406 | 0.6052371 | 0.85156118 |
| TTC3 | 327.664467 | 0.179441428 | 0.11695789 | 1.53423955 | 0.12497073 | 0.47038289 |
| HECA | 111.312823 | 0.17946233 | 0.21090006 | 0.8509354 | 0.39480524 | 0.72461789 |
| CTPS2 | 147.746157 | 0.179463639 | 0.30012568 | 0.59796162 | 0.54986554 | 0.82355262 |
| MTF2 | 266.528466 | 0.179588084 | 0.20253943 | 0.88668208 | 0.37525009 | 0.71257833 |
| CKMT2-AS1 | 140.845292 | 0.179682838 | 0.26504249 | 0.67793974 | 0.4978099 | 0.79513036 |
| RBM33 | 428.282302 | 0.179868648 | 0.20359336 | 0.8834701 | 0.37698233 | 0.71257833 |
| CYP20A1 | 207.958933 | 0.180001229 | 0.32015328 | 0.56223452 | 0.57395624 | 0.8342245 |
| OGDH | 216.717595 | 0.180177742 | 0.19418871 | 0.92784869 | 0.35348606 | 0.69977303 |
| ATXN3 | 356.204384 | 0.180214727 | 0.20441147 | 0.88162728 | 0.3779784 | 0.71305115 |

| | | | | | | |
|---|---|---|---|---|---|---|
| ILK | 162.587629 | 0.180667518 | 0.22773337 | 0.79332913 | 0.42758609 | 0.74318766 |
| AL592183.1 | 123.271275 | 0.180770838 | 0.55087807 | 0.32815036 | 0.74279798 | 0.90850686 |
| ORMDL1 | 128.578792 | 0.180801257 | 0.22623116 | 0.79918814 | 0.42418133 | 0.74152442 |
| MCM9 | 166.938419 | 0.180871048 | 0.23762756 | 0.76115348 | 0.4465654 | 0.76094002 |
| SQOR | 68.1794498 | 0.180875153 | 0.29111701 | 0.62131427 | 0.53439287 | 0.81562583 |
| CHSY3 | 219.148067 | 0.181049977 | 0.46885151 | 0.38615632 | 0.6993809 | 0.89370119 |
| AC142391.1 | 45.6061154 | 0.181159505 | 0.49764635 | 0.36403262 | 0.71583365 | 0.9001716 |
| TCAF1 | 148.378434 | 0.181191137 | 0.28832932 | 0.6284173 | 0.5297306 | 0.81297553 |
| TMLHE | 147.019709 | 0.181298127 | 0.36458148 | 0.49727739 | 0.61899345 | 0.85855421 |
| RNF214 | 92.4303094 | 0.181664668 | 0.20195507 | 0.8995301 | 0.36837037 | 0.70909779 |
| TRPM7 | 415.449774 | 0.181687039 | 0.18606203 | 0.9764864 | 0.32882348 | 0.68172348 |
| OGT | 368.687621 | 0.181696393 | 0.24012787 | 0.75666515 | 0.4492505 | 0.76241624 |
| KLHDC10 | 287.84354 | 0.181736712 | 0.20117234 | 0.90338816 | 0.36631992 | 0.70695551 |
| TAF8 | 78.7852043 | 0.181744027 | 0.30312257 | 0.59957274 | 0.54879102 | 0.82355262 |
| MON2 | 612.515856 | 0.18176537 | 0.15105139 | 1.20333467 | 0.22884684 | 0.59505995 |
| NAA30 | 60.1186923 | 0.18184731 | 0.22465288 | 0.80945906 | 0.41825114 | 0.73821752 |
| ZNF33A | 294.989129 | 0.182266471 | 0.27301058 | 0.66761687 | 0.50437819 | 0.79776858 |
| PRDM2 | 267.045491 | 0.182393491 | 0.2120257 | 0.86024238 | 0.38965545 | 0.72202943 |
| PCSK1N | 51.372933 | 0.182395576 | 0.58153 | 0.31364775 | 0.75378859 | 0.91250377 |
| ULK2 | 57.0168226 | 0.182495559 | 0.41404007 | 0.44076786 | 0.65938106 | 0.87335081 |
| SLC35F5 | 241.757299 | 0.182733205 | 0.18462913 | 0.98973115 | 0.32230555 | 0.6761919 |
| ANAPC5 | 250.131022 | 0.182837068 | 0.16705327 | 1.09448363 | 0.27374292 | 0.63946284 |
| STRN3 | 614.907903 | 0.182923368 | 0.21255927 | 0.86057582 | 0.38947171 | 0.72202943 |
| MAMLD1 | 69.8208809 | 0.183040179 | 0.38746037 | 0.47241007 | 0.63663412 | 0.86207417 |
| AGAP1 | 652.011257 | 0.18341591 | 0.27964521 | 0.6558879 | 0.51189626 | 0.80230743 |
| UBR1 | 565.828635 | 0.183685472 | 0.22702654 | 0.8090925 | 0.41846194 | 0.73821752 |
| CDC27 | 563.57707 | 0.183764544 | 0.16834437 | 1.09159896 | 0.27500941 | 0.64017771 |
| PDS5B | 733.280441 | 0.184150222 | 0.2315869 | 0.79516685 | 0.42651645 | 0.7424344 |
| SAP30L-AS1 | 67.3727633 | 0.184283212 | 0.32499568 | 0.56703281 | 0.57069189 | 0.83368112 |
| HELZ | 480.544405 | 0.184335038 | 0.20513412 | 0.89860738 | 0.36886183 | 0.70937634 |
| SNX1 | 211.570126 | 0.184580963 | 0.17016843 | 1.08469568 | 0.27805646 | 0.64148182 |
| MYSM1 | 213.420138 | 0.184623497 | 0.22821181 | 0.80900062 | 0.41851479 | 0.73821752 |
| SNCG | 90.8100838 | 0.184645 | 0.45325512 | 0.40737543 | 0.68373227 | 0.88565439 |
| TET2 | 255.614909 | 0.184901917 | 0.22637356 | 0.81679996 | 0.41404276 | 0.73671465 |
| NSD1 | 536.518429 | 0.1851336 | 0.18849457 | 0.98216941 | 0.32601639 | 0.67986548 |
| WDR7 | 385.605583 | 0.185244007 | 0.15545383 | 1.19163363 | 0.23340493 | 0.59910683 |
| PRMT2 | 247.812119 | 0.185411036 | 0.18525269 | 1.00085476 | 0.31689703 | 0.67180528 |
| DGCR8 | 71.9892605 | 0.185790384 | 0.38660418 | 0.48057003 | 0.63082212 | 0.86097142 |
| CASTOR3 | 62.3594847 | 0.185862979 | 0.37457861 | 0.49619218 | 0.61975883 | 0.85855421 |
| SLC10A7 | 273.95567 | 0.186074153 | 0.17152765 | 1.08480559 | 0.27800777 | 0.64148182 |
| EPB41L2 | 554.934854 | 0.186124493 | 0.18057802 | 1.03071513 | 0.30267443 | 0.65940027 |
| GDAP1 | 103.19477 | 0.18642431 | 0.32391242 | 0.57553924 | 0.56492666 | 0.83113895 |
| PTK2B | 90.5211377 | 0.186532428 | 0.55560818 | 0.33572657 | 0.73707707 | 0.90723797 |
| DLEU2 | 898.289854 | 0.186558034 | 0.31439919 | 0.5933795 | 0.55292721 | 0.8236419 |
| ZNF644 | 537.836842 | 0.186859977 | 0.21009807 | 0.88939408 | 0.37379132 | 0.7119959 |
| ATP5F1C | 149.138192 | 0.186903951 | 0.17146772 | 1.09002414 | 0.27570251 | 0.64048591 |
| ARFGAP1 | 56.3968938 | 0.187049199 | 0.23138258 | 0.80839795 | 0.41886153 | 0.73821752 |
| ZMYM5 | 119.301727 | 0.18742734 | 0.25330102 | 0.73993913 | 0.45933693 | 0.76976328 |
| AGL | 73.530609 | 0.187447978 | 0.30460313 | 0.61538427 | 0.53830097 | 0.81807397 |
| MED16 | 41.290721 | 0.187478429 | 0.32438724 | 0.57794637 | 0.56330033 | 0.83089872 |
| NBN | 112.0046 | 0.187538743 | 0.29960831 | 0.62594641 | 0.53135009 | 0.81393445 |
| SF3B1 | 325.682137 | 0.187806525 | 0.18453195 | 1.01774531 | 0.30879901 | 0.66424338 |
| GSR | 73.2719876 | 0.187907175 | 0.31668895 | 0.59334932 | 0.5529474 | 0.8236419 |
| OSBPL8 | 427.791278 | 0.187913538 | 0.14999788 | 1.25277464 | 0.21028773 | 0.57593532 |
| CMTR2 | 43.7129315 | 0.187976933 | 0.22273366 | 0.84395385 | 0.39869519 | 0.72770737 |
| RUFY3 | 364.688543 | 0.188232889 | 0.21418436 | 0.87883583 | 0.3794903 | 0.71418244 |
| CROT | 64.0572224 | 0.188251698 | 0.28289002 | 0.66545896 | 0.50575698 | 0.79865098 |
| SREK1 | 282.268499 | 0.188406374 | 0.18567088 | 1.01473303 | 0.3102331 | 0.6648266 |
| SNX13 | 414.000147 | 0.188496982 | 0.25543948 | 0.73793205 | 0.46055574 | 0.770424 |
| PSMC6 | 175.35225 | 0.188580085 | 0.22007601 | 0.85688614 | 0.39150782 | 0.72247779 |
| PUM2 | 650.908007 | 0.1887006 | 0.12401636 | 1.52157833 | 0.12811477 | 0.47566387 |
| TMEM9B | 81.2541482 | 0.188960367 | 0.30988544 | 0.60977491 | 0.54201092 | 0.82028396 |
| BBS4 | 68.3136194 | 0.189232238 | 0.26291582 | 0.71974457 | 0.47168228 | 0.77855633 |
| NDUFAF5 | 52.5582596 | 0.18947794 | 0.31728991 | 0.59717607 | 0.55038983 | 0.8236419 |
| LIPA | 173.877955 | 0.189558638 | 0.2455087 | 0.77210559 | 0.44005189 | 0.75475678 |
| ZYG11B | 170.929507 | 0.189639535 | 0.16992331 | 1.11603011 | 0.26440924 | 0.63018314 |
| HMBOX1 | 959.631146 | 0.189875027 | 0.23574863 | 0.80541307 | 0.42058135 | 0.73902022 |
| CNTLN | 361.770466 | 0.189968266 | 0.15238372 | 1.24664408 | 0.21252803 | 0.57720441 |
| DIDO1 | 211.786508 | 0.190134517 | 0.19834766 | 0.95859218 | 0.33776423 | 0.68839968 |
| GATAD2A | 401.064217 | 0.190158057 | 0.26209205 | 0.72553921 | 0.4681213 | 0.77664704 |
| AC079447.1 | 122.1045 | 0.190244904 | 0.32222955 | 0.59040178 | 0.55492131 | 0.82472232 |
| ELK3 | 633.19233 | 0.190339664 | 0.25915916 | 0.73445085 | 0.46267401 | 0.77266559 |
| INTS1 | 73.6313096 | 0.190352298 | 0.26789003 | 0.71056134 | 0.47735611 | 0.78098272 |
| PTGES3 | 391.376967 | 0.190371787 | 0.23020423 | 0.8269691 | 0.40825458 | 0.73419295 |
| SUPT20H | 174.889537 | 0.190400883 | 0.20514674 | 0.92812046 | 0.35334509 | 0.69977303 |
| CLHC1 | 56.2795497 | 0.190491891 | 0.6002128 | 0.31737393 | 0.75095989 | 0.91119584 |
| WIPF1 | 310.521962 | 0.190595638 | 0.29948026 | 0.63642137 | 0.52450181 | 0.80889528 |
| ATG10 | 254.181041 | 0.190686428 | 0.22321739 | 0.85426333 | 0.3929591 | 0.72341604 |
| AKAP11 | 189.873627 | 0.191019716 | 0.1638249 | 1.1659993 | 0.24361473 | 0.608518 |
| RYK | 379.971482 | 0.191133777 | 0.19216344 | 0.99464175 | 0.31991053 | 0.67434487 |
| LTN1 | 167.024448 | 0.191331774 | 0.20786409 | 0.92046576 | 0.35732942 | 0.70070132 |
| 7-Sep | 527.468844 | 0.19133264 | 0.20375629 | 0.93902691 | 0.34771693 | 0.696616 |
| ADAR | 364.87772 | 0.191373535 | 0.3278265 | 0.58376469 | 0.55937864 | 0.82818805 |
| ZFYVE1 | 91.6577262 | 0.191374206 | 0.29982393 | 0.63828863 | 0.52328581 | 0.80859228 |
| ZNF507 | 91.1357687 | 0.191521278 | 0.19783544 | 0.96808377 | 0.33300254 | 0.6852158 |
| UBE2K | 680.846386 | 0.191610328 | 0.20981895 | 0.91321744 | 0.36112819 | 0.70386371 |
| ARL8A | 71.0107798 | 0.1916734 | 0.22796109 | 0.8408163 | 0.40045086 | 0.72864885 |
| STK19 | 46.4500925 | 0.191952918 | 0.22973653 | 0.83553503 | 0.40341653 | 0.73065697 |
| VPS8 | 307.951731 | 0.19200804 | 0.16531294 | 1.16148224 | 0.24544584 | 0.6104646 |
| RILPL2 | 186.691695 | 0.192161134 | 0.32613247 | 0.58921191 | 0.55571913 | 0.82487928 |
| UBE3A | 378.270876 | 0.192213201 | 0.1667552 | 1.15266691 | 0.24904713 | 0.61434376 |
| DLGAP4 | 224.595541 | 0.192384443 | 0.27749409 | 0.69329203 | 0.4881263 | 0.78923879 |
| VPS13A | 455.064428 | 0.192454955 | 0.24562341 | 0.7835367 | 0.43331201 | 0.74751487 |
| NAA16 | 184.963241 | 0.192634397 | 0.18441855 | 1.04455002 | 0.29623099 | 0.65753344 |
| DENND4B | 108.827907 | 0.19263967 | 0.22314128 | 0.86330809 | 0.3879681 | 0.72104087 |
| TRMT9B | 88.5186865 | 0.192723849 | 0.52546036 | 0.36677143 | 0.71378952 | 0.89943635 |
| CUL4B | 91.4644076 | 0.192765538 | 0.21328138 | 0.90380856 | 0.36609692 | 0.70695551 |
| SEMA4C | 128.474618 | 0.192863374 | 0.34060123 | 0.56624391 | 0.57122798 | 0.83368112 |
| ZNF207 | 496.549427 | 0.193077585 | 0.17938025 | 1.07635924 | 0.28176662 | 0.64471757 |
| PPFIBP1 | 1464.72539 | 0.193135942 | 0.28093926 | 0.68746511 | 0.49178968 | 0.79094859 |
| ZFYVE16 | 280.580807 | 0.19326142 | 0.18684625 | 1.03433397 | 0.30098006 | 0.65926326 |
| UBA6-AS1 | 123.448348 | 0.193495709 | 0.50368048 | 0.3841636 | 0.70085719 | 0.8944316 |
| MAPK1 | 447.021315 | 0.193556526 | 0.16604683 | 1.16567432 | 0.24374615 | 0.608518 |
| PTPRK | 932.746094 | 0.193602943 | 0.27221428 | 0.71121522 | 0.47695088 | 0.78098272 |
| KIAA1217 | 1818.99632 | 0.193634357 | 0.48075825 | 0.40276866 | 0.68711842 | 0.88671976 |
| KLHL12 | 51.5918975 | 0.19378993 | 0.27716543 | 0.69918506 | 0.48443639 | 0.78781193 |
| PIP5K1C | 163.388039 | 0.193790214 | 0.26012661 | 0.74498421 | 0.45628127 | 0.76762944 |
| AC004594.1 | 75.4850261 | 0.193819929 | 0.53751034 | 0.36058828 | 0.71840725 | 0.90051656 |
| ZDHHC2 | 124.672918 | 0.194220921 | 0.4851611 | 0.40032253 | 0.68891897 | 0.88765832 |
| MTO1 | 67.9618377 | 0.194545697 | 0.22435141 | 0.8671472 | 0.38586136 | 0.71810405 |
| KCNQ1OT1 | 216.420238 | 0.194606917 | 0.40524833 | 0.48021646 | 0.63107349 | 0.86097142 |
| MTREX | 257.237186 | 0.194688074 | 0.20156995 | 0.96585864 | 0.33411492 | 0.6852158 |
| KATNBL1 | 276.030899 | 0.19471412 | 0.17481076 | 1.11385662 | 0.2653407 | 0.63060082 |

| | | | | | | |
|---|---|---|---|---|---|---|
| STRADB | 53.871626 | 0.194794408 | 0.43750136 | 0.44524297 | 0.65614417 | 0.87162789 |
| TMEM161B-, | 184.724997 | 0.194797269 | 0.32300487 | 0.60307842 | 0.54645652 | 0.82355262 |
| DYNC2H1 | 238.880615 | 0.19544446 | 0.26816631 | 0.72881809 | 0.46611294 | 0.77518594 |
| TOGARAM1 | 191.339867 | 0.195666813 | 0.25822594 | 0.75773491 | 0.44860969 | 0.76192883 |
| RC3H1 | 358.76675 | 0.195780411 | 0.18027795 | 1.085992 | 0.27748253 | 0.64148182 |
| ST3GAL2 | 194.753684 | 0.195816557 | 0.31957214 | 0.61274602 | 0.54004429 | 0.81900022 |
| KIF1BP | 64.486807 | 0.195930834 | 0.24775241 | 0.7908332 | 0.42904133 | 0.74408005 |
| ANKZF1 | 45.9201137 | 0.196047025 | 0.29637904 | 0.66147398 | 0.5083084 | 0.79974767 |
| KLF10 | 159.295396 | 0.19605151 | 0.45321686 | 0.43257771 | 0.66532159 | 0.87587657 |
| EEPD1 | 260.679785 | 0.19621023 | 0.50379557 | 0.38946398 | 0.69693295 | 0.89262832 |
| ATPAF1 | 67.7214922 | 0.196283696 | 0.28272743 | 0.69425064 | 0.48752504 | 0.78857826 |
| SGK3 | 226.858049 | 0.196375974 | 0.21058144 | 0.93254168 | 0.35105666 | 0.69858573 |
| CPSF7 | 116.442872 | 0.196504644 | 0.21644026 | 0.90789323 | 0.36393464 | 0.70570548 |
| YY1AP1 | 179.324468 | 0.196540987 | 0.18108388 | 1.08535883 | 0.27776276 | 0.64148182 |
| RANBP17 | 409.239103 | 0.196676496 | 0.30498418 | 0.64487442 | 0.51900857 | 0.80603851 |
| CDK13 | 648.564341 | 0.196850328 | 0.13027829 | 1.51099867 | 0.13078879 | 0.47936448 |
| AP003086.2 | 106.887442 | 0.197525848 | 0.4256725 | 0.46403244 | 0.64262451 | 0.86490071 |
| ACAD10 | 52.6763356 | 0.197582021 | 0.36514105 | 0.5411115 | 0.58843074 | 0.84136444 |
| SNHG7 | 151.035424 | 0.19800929 | 0.40119808 | 0.49354496 | 0.62162758 | 0.85896293 |
| PCMTD1 | 506.993003 | 0.198125864 | 0.21244838 | 0.93258354 | 0.35103503 | 0.69858573 |
| IP6K1 | 195.930524 | 0.198171437 | 0.23698565 | 0.83621702 | 0.40303283 | 0.73065099 |
| PSME4 | 335.353536 | 0.198220905 | 0.24072382 | 0.82343702 | 0.41025952 | 0.73419295 |
| MEGF9 | 106.201706 | 0.198455376 | 0.24538313 | 0.8087572 | 0.41865482 | 0.73821752 |
| IRF3 | 92.07753 | 0.198463016 | 0.2403511 | 0.82572128 | 0.40896222 | 0.73419295 |
| NDFIP2 | 107.243131 | 0.198640393 | 0.27844852 | 0.71338282 | 0.4756089 | 0.78022377 |
| METTL4 | 100.173655 | 0.198898723 | 0.32924783 | 0.60410032 | 0.54577694 | 0.82355262 |
| NHSL2 | 251.740624 | 0.198982501 | 0.33027116 | 0.60248222 | 0.5468532 | 0.82355262 |
| WDR25 | 90.3582833 | 0.199088794 | 0.33356598 | 0.59684982 | 0.55060765 | 0.8236419 |
| UBXN7 | 166.820016 | 0.199097102 | 0.1915221 | 1.03955158 | 0.29854828 | 0.65856711 |
| NREP | 201.308465 | 0.199123525 | 0.41236558 | 0.48288105 | 0.6291802 | 0.86082287 |
| ARID1A | 387.278664 | 0.199227339 | 0.19474184 | 1.02303305 | 0.30629222 | 0.66324139 |
| ZNF496 | 72.9404322 | 0.199246727 | 0.29234096 | 0.68155596 | 0.49551977 | 0.79351043 |
| UBE4B | 368.575974 | 0.199603657 | 0.20416253 | 0.97767038 | 0.32823738 | 0.6806808 |
| SMARCA5 | 234.607986 | 0.199641305 | 0.21220215 | 0.94080718 | 0.34680368 | 0.6956798 |
| CDK5RAP2 | 157.738396 | 0.199677992 | 0.21856765 | 0.91357521 | 0.3609401 | 0.70386371 |
| TTC33 | 132.8515 | 0.199916518 | 0.26123257 | 0.76528174 | 0.44410379 | 0.75931913 |
| MED13 | 628.230343 | 0.20007965 | 0.15820611 | 1.26467713 | 0.20598709 | 0.57080676 |
| BRWD3 | 274.989657 | 0.200252904 | 0.21443096 | 0.93388056 | 0.35036551 | 0.69858573 |
| SLCO3A1 | 729.218357 | 0.200305813 | 0.33219035 | 0.60298505 | 0.54651864 | 0.82355262 |
| SHPRH | 332.965081 | 0.200448744 | 0.140455 | 1.42713851 | 0.15353997 | 0.5097092 |
| FKBP9 | 203.655309 | 0.200665955 | 0.40901232 | 0.49061102 | 0.62370159 | 0.85994104 |
| SLC5A4 | 132.445778 | 0.200724176 | 0.56017163 | 0.35832621 | 0.72009921 | 0.90114265 |
| TBRG1 | 128.516887 | 0.200948376 | 0.23392432 | 0.85903156 | 0.39032311 | 0.72246414 |
| CRADD | 586.942565 | 0.201456463 | 0.26885149 | 0.74932246 | 0.45366287 | 0.76578691 |
| SEC23A | 291.263243 | 0.201461465 | 0.17080498 | 1.1794824 | 0.23820614 | 0.60143192 |
| TM9SF2 | 252.523807 | 0.20146581 | 0.17279244 | 1.16594111 | 0.24363825 | 0.608518 |
| MTX2 | 243.63877 | 0.201484126 | 0.24252228 | 0.83078604 | 0.40609451 | 0.73307328 |
| NAGK | 51.1723155 | 0.201684531 | 0.26096482 | 0.77284183 | 0.43961599 | 0.75432537 |
| SUSD1 | 83.6915365 | 0.20176971 | 0.2815465 | 0.7166479 | 0.47359138 | 0.77899012 |
| PLEKHA5 | 382.362753 | 0.201788741 | 0.3401419 | 0.5932487 | 0.55301473 | 0.8236419 |
| ANKRD10 | 608.344396 | 0.201799293 | 0.20185678 | 0.99971519 | 0.31744836 | 0.67227741 |
| UNK | 188.300298 | 0.20188623 | 0.25768885 | 0.7834496 | 0.43336314 | 0.74751487 |
| KPNA1 | 314.398842 | 0.201906438 | 0.21638953 | 0.93306934 | 0.35078417 | 0.69858573 |
| TCF20 | 180.867485 | 0.2020159 | 0.23903422 | 0.84513381 | 0.39803613 | 0.72752505 |
| PRKD3 | 328.154469 | 0.202265683 | 0.18539591 | 1.09099321 | 0.27527587 | 0.64017771 |
| PROSER1 | 67.019788 | 0.20239628 | 0.20774996 | 0.97423015 | 0.32994228 | 0.68216508 |
| LRRC6 | 44.5508472 | 0.202433189 | 0.344335 | 0.58789606 | 0.55660206 | 0.82527172 |
| RFX3 | 1176.68128 | 0.202584971 | 0.29349128 | 0.6902589 | 0.49003139 | 0.79046901 |
| EYA3 | 242.78506 | 0.202603782 | 0.23001055 | 0.88084562 | 0.37840139 | 0.71314107 |
| MAP3K2 | 393.275347 | 0.202642885 | 0.14660566 | 1.382231 | 0.16690079 | 0.52764813 |
| ZNF652 | 253.080363 | 0.202659043 | 0.24472955 | 0.82809387 | 0.40761734 | 0.73419295 |
| NBEAL1 | 858.05394 | 0.202805028 | 0.17463393 | 1.16131518 | 0.24551375 | 0.6104646 |
| HDAC7 | 408.655713 | 0.202831682 | 0.21779573 | 0.93129319 | 0.35170193 | 0.69889281 |
| FRYL | 1303.26837 | 0.20286796 | 0.20171312 | 1.00572514 | 0.31454781 | 0.66883106 |
| LNPK | 86.4027477 | 0.202914472 | 0.18591048 | 1.09146333 | 0.27506906 | 0.64017771 |
| UBXN2A | 145.687895 | 0.2030198 | 0.30031874 | 0.67601444 | 0.49903148 | 0.79516816 |
| MOB4 | 57.7118382 | 0.203024927 | 0.42744521 | 0.47497298 | 0.63480623 | 0.86207417 |
| CLIC4 | 1442.43816 | 0.203048277 | 0.34174367 | 0.59415373 | 0.5524093 | 0.8236419 |
| OGA | 528.199939 | 0.203061144 | 0.33221745 | 0.61122962 | 0.54104758 | 0.81928059 |
| WDFY1 | 296.880685 | 0.203424456 | 0.20497195 | 0.9924502 | 0.32097796 | 0.67473727 |
| PPHLN1 | 269.763512 | 0.203478894 | 0.15511881 | 1.31176155 | 0.1896006 | 0.55529769 |
| KLF13 | 126.356342 | 0.203623683 | 0.30716354 | 0.66291618 | 0.50738424 | 0.79921563 |
| CD109 | 402.566464 | 0.203633885 | 0.35697838 | 0.57043759 | 0.56838094 | 0.83354423 |
| RSU1 | 462.981072 | 0.203854719 | 0.14614719 | 1.39485902 | 0.16305836 | 0.52335809 |
| TRAPPC8 | 275.427523 | 0.204185947 | 0.13332405 | 1.53150123 | 0.12564556 | 0.47103973 |
| COMMD10 | 757.456616 | 0.204190635 | 0.18867409 | 1.08223994 | 0.27914593 | 0.64254602 |
| KAT8 | 65.3553961 | 0.204245928 | 0.27198298 | 0.75095113 | 0.45268207 | 0.76503457 |
| NELFA | 84.8369585 | 0.204307349 | 0.18261812 | 1.11876824 | 0.26323902 | 0.62948128 |
| DYNLL2 | 58.1869423 | 0.204320085 | 0.23052612 | 0.88632076 | 0.37544471 | 0.71257833 |
| LARP1B | 142.235415 | 0.204361549 | 0.24648148 | 0.82911523 | 0.40703921 | 0.73411589 |
| TRNAU1AP | 108.435357 | 0.204530693 | 0.20496756 | 0.9978686 | 0.31834308 | 0.67312698 |
| ARMCX4 | 248.088244 | 0.204613429 | 0.38201803 | 0.53561197 | 0.59222676 | 0.84399017 |
| JAG1 | 288.978344 | 0.204853086 | 0.42114411 | 0.48642039 | 0.62666913 | 0.86052754 |
| UBE2V2 | 126.046989 | 0.204901618 | 0.19194969 | 1.06747563 | 0.28575711 | 0.64828124 |
| FBXW11 | 453.865488 | 0.204926848 | 0.19282981 | 1.06273427 | 0.28790247 | 0.65121935 |
| RRN3 | 94.0271855 | 0.204947749 | 0.25734352 | 0.79639755 | 0.425801 | 0.74197066 |
| PEX1 | 76.5397308 | 0.205045844 | 0.20594198 | 0.99564861 | 0.31942091 | 0.67395826 |
| POLR2F | 65.6159621 | 0.20509942 | 0.30170657 | 0.67979765 | 0.4966326 | 0.79420452 |
| KMT2A | 527.423165 | 0.205153435 | 0.19977974 | 1.0268981 | 0.30446845 | 0.66086613 |
| DPF2 | 67.6052751 | 0.205251197 | 0.2043515 | 1.00440269 | 0.31518455 | 0.66938884 |
| GLI3 | 134.764174 | 0.205361076 | 0.33412077 | 0.61463128 | 0.53879825 | 0.81829735 |
| GIT2 | 232.939479 | 0.205379415 | 0.12984553 | 1.58172111 | 0.11371325 | 0.45258538 |
| DTWD1 | 109.806236 | 0.205420798 | 0.25697352 | 0.79938507 | 0.42406716 | 0.74152442 |
| UBE2A | 70.2793229 | 0.205518369 | 0.25009361 | 0.82176578 | 0.41121021 | 0.73470224 |
| HSPA1B | 205.92033 | 0.205594308 | 1.22958497 | 0.16720626 | 0.86720776 | 0.95535286 |
| NCK1 | 266.510922 | 0.206109014 | 0.29598954 | 0.69633884 | 0.48621665 | 0.78826504 |
| TIAL1 | 210.602053 | 0.206255661 | 0.19202759 | 1.07409387 | 0.28278061 | 0.64582577 |
| MPC1 | 86.7054751 | 0.206436636 | 0.23318687 | 0.88528412 | 0.37600342 | 0.71257833 |
| METTL14 | 95.1657552 | 0.206684011 | 0.22329062 | 0.92562784 | 0.35463942 | 0.70011254 |
| CASC4 | 292.87081 | 0.206736525 | 0.16290642 | 1.26905078 | 0.20442295 | 0.56929203 |
| ARHGEF12 | 1163.6534 | 0.206751211 | 0.21327398 | 0.96941602 | 0.33233766 | 0.6852158 |
| PCGF3 | 188.367889 | 0.206789137 | 0.19336666 | 1.06941464 | 0.28488287 | 0.64773452 |
| SP3 | 262.424297 | 0.206793435 | 0.12707269 | 1.62736332 | 0.10365996 | 0.43503085 |
| MARK2 | 183.81425 | 0.20687886 | 0.29023557 | 0.71279636 | 0.47597178 | 0.78022377 |
| LOXL2 | 91.106408 | 0.207037885 | 0.25360191 | 0.81638929 | 0.41427752 | 0.73671465 |
| MCPH1-AS1 | 48.9455467 | 0.207083095 | 0.30734742 | 0.67377529 | 0.50045419 | 0.7961152 |
| NFATC1 | 87.4029259 | 0.207211357 | 0.2798875 | 0.74033801 | 0.45909492 | 0.76976328 |
| NDUFAF7 | 75.8357826 | 0.207634719 | 0.2201308 | 0.9432334 | 0.34556153 | 0.69497757 |
| PHC1 | 61.102123 | 0.207672589 | 0.24798595 | 0.83743693 | 0.40234702 | 0.73035873 |
| CCZ1 | 55.5511114 | 0.207736732 | 0.24710633 | 0.8406775 | 0.40052863 | 0.72864885 |
| STAM | 119.447559 | 0.207767251 | 0.23206114 | 0.89531253 | 0.37062004 | 0.70984233 |
| CCNT2 | 234.639632 | 0.207835683 | 0.24423226 | 0.85097555 | 0.39478293 | 0.72461789 |
| RAB2B | 84.5917685 | 0.207890486 | 0.25255512 | 0.82314898 | 0.41042328 | 0.73419295 |

| | | | | | | |
|---|---|---|---|---|---|---|
| GTF2IRD2B | 44.849088 | 0.208047525 | 0.40610433 | 0.51230068 | 0.60844059 | 0.85289019 |
| TMCO3 | 187.634442 | 0.208135205 | 0.16535762 | 1.25869741 | 0.20813965 | 0.57338509 |
| SIMC1 | 144.997251 | 0.208193023 | 0.26342467 | 0.79033229 | 0.42933373 | 0.74408005 |
| ADK | 537.261358 | 0.208590378 | 0.22952246 | 0.90880158 | 0.36345487 | 0.7056112 |
| SLC4A1AP | 54.7941276 | 0.208609342 | 0.20624764 | 1.01145081 | 0.31180071 | 0.66561244 |
| TBC1D19 | 186.547695 | 0.208675685 | 0.25897044 | 0.80578958 | 0.42036419 | 0.73901389 |
| RBM14-RBM | 65.8876856 | 0.209148845 | 0.30821628 | 0.67857819 | 0.49740517 | 0.7946635 |
| PDPR | 152.533882 | 0.209185193 | 0.34394923 | 0.60818625 | 0.54306395 | 0.82151188 |
| GALC | 57.0933516 | 0.20924633 | 0.26108239 | 0.80145709 | 0.42286707 | 0.74026603 |
| COX11 | 61.7230328 | 0.209987717 | 0.25125774 | 0.83574626 | 0.40329766 | 0.73065697 |
| PNPLA2 | 60.6435585 | 0.210052755 | 0.23769853 | 0.88369394 | 0.37686145 | 0.71257833 |
| AC010997.3 | 81.69052 | 0.210078756 | 0.47734989 | 0.44009386 | 0.65986913 | 0.87335081 |
| CDK10 | 82.3094561 | 0.210260325 | 0.18463201 | 1.13880757 | 0.25478342 | 0.6205038 |
| MAP3K4 | 235.028541 | 0.21029569 | 0.16812999 | 1.25079228 | 0.21101027 | 0.5768065 |
| SETD4 | 45.1622539 | 0.210463679 | 0.2189582 | 0.96120485 | 0.33644918 | 0.68745652 |
| DENND5B | 420.398821 | 0.21072112 | 0.24058072 | 0.8758853 | 0.38109239 | 0.71543195 |
| ERAP1 | 152.07462 | 0.210916923 | 0.19263563 | 1.09490088 | 0.27356006 | 0.63926188 |
| NEK7 | 578.218057 | 0.211075074 | 0.1652097 | 1.27761911 | 0.20138376 | 0.56649134 |
| RAB3GAP2 | 362.712325 | 0.211272325 | 0.18943667 | 1.11526627 | 0.26473634 | 0.63018797 |
| CCDC171 | 80.1086831 | 0.211512452 | 0.4244071 | 0.49837161 | 0.61822214 | 0.85855421 |
| CEP95 | 138.627644 | 0.211626735 | 0.24988669 | 0.84689078 | 0.397056 | 0.72680823 |
| SS18 | 235.240289 | 0.211645281 | 0.2116045 | 1.00019273 | 0.31721724 | 0.67213587 |
| ZBTB8OS | 148.371872 | 0.211799273 | 0.19569561 | 1.08228936 | 0.27912397 | 0.64254602 |
| ACTN4 | 748.735859 | 0.211887651 | 0.30250721 | 0.70043836 | 0.48365359 | 0.78748452 |
| NF1 | 1016.80594 | 0.21197055 | 0.23928937 | 0.88583355 | 0.37570723 | 0.71257833 |
| RASA2 | 505.405678 | 0.211990432 | 0.18662888 | 1.13589298 | 0.25600136 | 0.62095131 |
| MCEE | 47.0175657 | 0.212052791 | 0.2859088 | 0.74167982 | 0.45828135 | 0.76968366 |
| CEP97 | 55.9744051 | 0.212074028 | 0.24050012 | 0.88180426 | 0.37788267 | 0.71305115 |
| PAK4 | 56.0549545 | 0.21210625 | 0.30903901 | 0.68634135 | 0.49249788 | 0.79099316 |
| UBE2D1 | 67.1851223 | 0.212110871 | 0.23537589 | 0.90115802 | 0.36750431 | 0.70809695 |
| AASDH | 66.5014325 | 0.212289814 | 0.19293549 | 1.10031498 | 0.27119491 | 0.63561102 |
| SNRNP35 | 68.0404561 | 0.212434587 | 0.23019463 | 0.92284772 | 0.35608657 | 0.70018061 |
| ZNF195 | 73.7048556 | 0.212447709 | 0.34936511 | 0.60809652 | 0.54312346 | 0.82151188 |
| LRP5 | 250.400807 | 0.212513278 | 0.44344344 | 0.47923423 | 0.631772 | 0.86118497 |
| RNGTT | 373.993339 | 0.212687941 | 0.23655578 | 0.89910269 | 0.36859796 | 0.70919884 |
| YTHDC2 | 293.068594 | 0.212896363 | 0.20425653 | 1.04229894 | 0.2972731 | 0.65795352 |
| FGF12 | 179.334954 | 0.213097033 | 0.67089261 | 0.31763211 | 0.75076402 | 0.91109327 |
| BBS7 | 78.5999744 | 0.213130995 | 0.23397755 | 0.9109036 | 0.36234617 | 0.70497203 |
| STK4 | 154.403134 | 0.213236127 | 0.18743156 | 1.13767464 | 0.25525637 | 0.6208774 |
| GOLPH3 | 624.530285 | 0.213308284 | 0.31412906 | 0.67904665 | 0.4971083 | 0.79465467 |
| CHD6 | 377.06313 | 0.213319171 | 0.16027676 | 1.33094264 | 0.18320789 | 0.54813352 |
| SLMAP | 454.236261 | 0.213732482 | 0.14644027 | 1.45951988 | 0.14442207 | 0.49679586 |
| DENND6A | 299.616059 | 0.21375082 | 0.16359359 | 1.30659657 | 0.19134976 | 0.55590015 |
| ZFAND6 | 382.768002 | 0.213885846 | 0.1560046 | 1.37102267 | 0.17036789 | 0.53193492 |
| EML6 | 50.3236883 | 0.21397443 | 0.25822873 | 0.82862365 | 0.4073174 | 0.73419295 |
| KLHL7 | 99.8930183 | 0.213997251 | 0.19337683 | 1.10663334 | 0.26845249 | 0.63347822 |
| PI4KA | 312.986253 | 0.214417149 | 0.27000322 | 0.79412811 | 0.42712086 | 0.74286176 |
| SYTL2 | 71.518848 | 0.214525528 | 0.55777436 | 0.38460988 | 0.70052647 | 0.89443098 |
| MED28 | 99.7413838 | 0.214590355 | 0.20804237 | 1.03147429 | 0.30231846 | 0.65926326 |
| TNRC6B | 1341.25308 | 0.214761552 | 0.20069801 | 1.07007317 | 0.28458638 | 0.6476002 |
| TRAF5 | 63.9672448 | 0.214872769 | 0.29302212 | 0.7332988 | 0.46337621 | 0.77320707 |
| TYK2 | 74.2761461 | 0.214941847 | 0.24330674 | 0.88341919 | 0.37700983 | 0.71257833 |
| KCTD1 | 79.4845831 | 0.214987596 | 0.37222841 | 0.57756902 | 0.56355513 | 0.83089872 |
| CARMIL1 | 609.688834 | 0.214995256 | 0.33094006 | 0.64965014 | 0.51591824 | 0.80483629 |
| GOLGA8A | 109.769238 | 0.215007353 | 0.25634253 | 0.83875021 | 0.4016095 | 0.72933211 |
| RBMS2 | 893.598475 | 0.215080667 | 0.33485073 | 0.64231805 | 0.5206667 | 0.80705591 |
| LRP6 | 413.742371 | 0.215110976 | 0.20955749 | 1.02650099 | 0.3046555 | 0.66092151 |
| ASTN2 | 49.8713018 | 0.215183792 | 0.44664329 | 0.48177998 | 0.62996226 | 0.86097142 |
| RAMP3 | 1024.14282 | 0.215306763 | 0.29866515 | 0.72089685 | 0.47097298 | 0.77826574 |
| PSKH1 | 42.7318934 | 0.215568663 | 0.25743859 | 0.83735957 | 0.40239049 | 0.73035873 |
| NBPF12 | 57.0365272 | 0.215621304 | 0.30992751 | 0.69571529 | 0.48660715 | 0.78826504 |
| HOOK3 | 352.373489 | 0.216051511 | 0.21284898 | 1.01504603 | 0.31008389 | 0.6648266 |
| PAXIP1 | 61.2576782 | 0.21610822 | 0.27873429 | 0.77531982 | 0.4381507 | 0.75275917 |
| MSN | 1054.37579 | 0.216264808 | 0.25345101 | 0.85328051 | 0.39350377 | 0.72360479 |
| MBTD1 | 256.992929 | 0.216331094 | 0.20652184 | 1.04749741 | 0.29487023 | 0.65671864 |
| CNOT3 | 40.5076621 | 0.216343482 | 0.27093419 | 0.79850934 | 0.42457497 | 0.74157887 |
| AREL1 | 97.3900857 | 0.216540197 | 0.22116507 | 0.97908857 | 0.32753622 | 0.68017808 |
| SHKBP1 | 42.1900028 | 0.216559497 | 0.31019196 | 0.69814671 | 0.48508545 | 0.78805921 |
| FANCM | 65.2645633 | 0.216766574 | 0.27649367 | 0.78398387 | 0.43304958 | 0.74728905 |
| CSPP1 | 255.931331 | 0.216787219 | 0.17178424 | 1.26197385 | 0.2069582 | 0.57162978 |
| NINL | 48.8267906 | 0.216925595 | 0.40654396 | 0.5335846 | 0.59362896 | 0.84502915 |
| NDUFS1 | 227.905525 | 0.217016719 | 0.14510637 | 1.49556989 | 0.13476577 | 0.48474744 |
| COG6 | 167.626308 | 0.217218543 | 0.30184192 | 0.7196434 | 0.47174459 | 0.77855633 |
| PBRM1 | 434.539099 | 0.217245423 | 0.2343186 | 0.92713691 | 0.35385546 | 0.69977303 |
| PELI1 | 947.109268 | 0.217282986 | 0.56286305 | 0.38603171 | 0.69947318 | 0.89370119 |
| RBP1 | 51.3171888 | 0.217638883 | 0.73936375 | 0.29435969 | 0.76848307 | 0.91631735 |
| RILPL1 | 200.601587 | 0.217751466 | 0.33680018 | 0.64653013 | 0.51793609 | 0.80546868 |
| MORC3 | 216.971352 | 0.218025383 | 0.23418747 | 0.93098653 | 0.35186053 | 0.69889281 |
| RSRP1 | 531.697455 | 0.218153368 | 0.28962101 | 0.75323738 | 0.45130728 | 0.76413149 |
| RNF146 | 82.8271175 | 0.218187589 | 0.25856804 | 0.84383048 | 0.39876414 | 0.72770737 |
| UHMK1 | 172.951727 | 0.218308259 | 0.20443575 | 1.06785754 | 0.28558478 | 0.64813609 |
| PLEKHA3 | 99.2246276 | 0.218515757 | 0.22859832 | 0.95589398 | 0.3391258 | 0.68955779 |
| MAP3K7CL | 50.6245147 | 0.218618549 | 0.38651881 | 0.56560908 | 0.57165955 | 0.83368112 |
| PSIP1 | 197.005644 | 0.218706793 | 0.26324489 | 0.83081117 | 0.40608031 | 0.73307328 |
| RNPEP | 88.500461 | 0.218714832 | 0.29082274 | 0.75205547 | 0.4520177 | 0.76480455 |
| SFMBT1 | 48.0222824 | 0.218874157 | 0.36605716 | 0.59792345 | 0.54989101 | 0.82355262 |
| CSTF3 | 133.229109 | 0.218943667 | 0.21625845 | 1.01241669 | 0.31133886 | 0.66561244 |
| TBC1D17 | 46.93567 | 0.218961131 | 0.27792835 | 0.78783301 | 0.43079439 | 0.744811 |
| GIMAP2 | 62.8748613 | 0.219128818 | 0.29252714 | 0.74908885 | 0.45380365 | 0.76582291 |
| ZMYM2 | 662.562819 | 0.219363344 | 0.23747862 | 0.92371829 | 0.35563301 | 0.70018061 |
| SMCHD1 | 1260.80356 | 0.219417167 | 0.46360486 | 0.47328487 | 0.63600995 | 0.86207417 |
| METAP1 | 82.4539835 | 0.219475546 | 0.22369943 | 0.98111805 | 0.32653453 | 0.67997183 |
| AC093110.1 | 84.918413 | 0.219541442 | 0.66471851 | 0.33027731 | 0.74119043 | 0.90850686 |
| MAST3 | 51.155341 | 0.21972879 | 0.38001546 | 0.57821014 | 0.56312225 | 0.83089872 |
| KITLG | 112.752316 | 0.219782317 | 0.38904341 | 0.56493007 | 0.57212133 | 0.83392748 |
| HMGXB3 | 113.392811 | 0.220007346 | 0.29012274 | 0.75832507 | 0.4482564 | 0.76164496 |
| CCP110 | 48.5252882 | 0.220018307 | 0.2453232 | 0.89685078 | 0.36979855 | 0.70984233 |
| WAC | 961.76554 | 0.220065282 | 0.20831654 | 1.0563985 | 0.29078619 | 0.65335074 |
| METTL15 | 430.344739 | 0.220091385 | 0.25753233 | 0.85461652 | 0.39276348 | 0.72332854 |
| PDZD2 | 857.710914 | 0.22016553 | 0.34736923 | 0.63380839 | 0.52620588 | 0.81036438 |
| GLRX3 | 150.348165 | 0.22017116 | 0.27875642 | 0.78983351 | 0.42962501 | 0.74408005 |
| HEATR5B | 241.558572 | 0.220311152 | 0.16896869 | 1.30385788 | 0.19228204 | 0.55659725 |
| EXOC6B | 1191.56673 | 0.220318066 | 0.23515488 | 0.93690622 | 0.3488068 | 0.69804011 |
| ETNK1 | 384.640183 | 0.220362518 | 0.21644407 | 1.01810373 | 0.30862866 | 0.66414532 |
| CTNNBIP1 | 73.0512778 | 0.220701842 | 0.38987427 | 0.56608465 | 0.57133624 | 0.83368112 |
| GABPB2 | 88.9935671 | 0.220796649 | 0.41813241 | 0.52805438 | 0.59746159 | 0.84775936 |
| KDM1B | 66.6534884 | 0.221095338 | 0.23643117 | 0.93513616 | 0.34971813 | 0.69858573 |
| IQCJ-SCHIP1 | 399.060531 | 0.221402002 | 0.48666081 | 0.45494109 | 0.64915162 | 0.86766991 |
| PPP1R12C | 91.7356398 | 0.221450498 | 0.24478959 | 0.9046565 | 0.36564739 | 0.70668224 |
| TBPL1 | 68.6394713 | 0.221458321 | 0.20083016 | 1.10271447 | 0.27015118 | 0.63524343 |
| CEP152 | 67.4905981 | 0.2216223 | 0.27811729 | 0.79686631 | 0.42552868 | 0.74197066 |
| TBC1D23 | 164.392275 | 0.222012177 | 0.23952225 | 0.92689583 | 0.35398062 | 0.69977303 |
| TRIM38 | 178.562294 | 0.222224601 | 0.25137569 | 0.88403377 | 0.37667798 | 0.71257833 |

| | | | | | | |
|---|---|---|---|---|---|---|
| IKBKB | 439.715124 | 0.222572209 | 0.21360369 | 1.04198675 | 0.29741782 | 0.65795352 |
| IQSEC1 | 249.296047 | 0.222818905 | 0.4079532 | 0.54618742 | 0.58493712 | 0.83895258 |
| PITPNA | 257.216398 | 0.222851838 | 0.27719578 | 0.80395104 | 0.42142525 | 0.73939169 |
| KIAA0319L | 327.204393 | 0.223497907 | 0.25408177 | 0.87962985 | 0.37905986 | 0.7138888 |
| TULP4 | 443.755574 | 0.2237897 | 0.20110233 | 1.11281502 | 0.26578788 | 0.63060082 |
| SENP2 | 128.017661 | 0.223792851 | 0.24736816 | 0.90469544 | 0.36562676 | 0.70668224 |
| MEF2C | 2130.29459 | 0.22415414 | 0.26169035 | 0.85656251 | 0.39168671 | 0.72247779 |
| GATAD2B | 538.608514 | 0.224340849 | 0.21936243 | 1.02269493 | 0.30645211 | 0.66324542 |
| CCDC6 | 179.408575 | 0.224452575 | 0.1825832 | 1.22931668 | 0.2189531 | 0.58619521 |
| TMEM243 | 85.2662385 | 0.224538018 | 0.29066552 | 0.77249623 | 0.43982058 | 0.75451819 |
| AP2B1 | 308.268038 | 0.224724447 | 0.23021595 | 0.9761463 | 0.32899197 | 0.68172735 |
| IDH1 | 44.8012064 | 0.224819961 | 0.41729487 | 0.53875564 | 0.59005548 | 0.8420771 |
| THUMPD1 | 102.273343 | 0.224820519 | 0.17499302 | 1.2847399 | 0.19888319 | 0.56402001 |
| MAP2K4 | 349.43765 | 0.225026732 | 0.16213764 | 1.38787475 | 0.16517519 | 0.52531231 |
| ARID2 | 640.500387 | 0.225053851 | 0.18370413 | 1.22508868 | 0.22054181 | 0.58706366 |
| KHNYN | 45.8755882 | 0.225201987 | 0.23611473 | 0.95378202 | 0.34019399 | 0.69059971 |
| CELSR1 | 112.250402 | 0.225296028 | 0.32202609 | 0.69962041 | 0.48416439 | 0.78781193 |
| SEM1 | 296.693662 | 0.22535533 | 0.2778542 | 0.81105606 | 0.41733348 | 0.73821752 |
| AUP1 | 58.1257117 | 0.225474079 | 0.30652148 | 0.73558982 | 0.46198036 | 0.77197984 |
| PEAK1 | 1296.50928 | 0.225530946 | 0.37909496 | 0.5949194 | 0.55189735 | 0.8236419 |
| RPS6KA3 | 531.618525 | 0.225579367 | 0.19157711 | 1.17748601 | 0.23900158 | 0.60290148 |
| PPM1A | 123.397536 | 0.22565686 | 0.22372958 | 1.00861433 | 0.31315963 | 0.66751375 |
| LRRFIP1 | 1062.80966 | 0.225745216 | 0.23727412 | 0.95141103 | 0.34139577 | 0.69116603 |
| BEND7 | 214.455156 | 0.225748634 | 0.21446574 | 1.05260929 | 0.2925201 | 0.65502684 |
| AC005237.1 | 78.2034641 | 0.225803676 | 0.48199322 | 0.46847894 | 0.63944212 | 0.86388557 |
| YEATS2 | 323.586428 | 0.225813531 | 0.26681997 | 0.84631419 | 0.39737749 | 0.72680823 |
| PDCD10 | 183.176797 | 0.225837638 | 0.21413991 | 1.05462658 | 0.29159614 | 0.65416882 |
| NFIA | 1640.33663 | 0.22587542 | 0.24711396 | 0.91405365 | 0.36068865 | 0.70386371 |
| TSC22D1 | 741.641946 | 0.225881774 | 0.25026172 | 0.90258219 | 0.36674768 | 0.70730527 |
| FBXO9 | 125.075018 | 0.226003608 | 0.17764662 | 1.2722089 | 0.20329891 | 0.567308 |
| MRPL1 | 146.114665 | 0.226006256 | 0.24344532 | 0.9283656 | 0.35321796 | 0.69977303 |
| CWC27 | 283.255106 | 0.226017797 | 0.16551282 | 1.36556067 | 0.1720769 | 0.53377758 |
| SCYL2 | 218.652614 | 0.226099023 | 0.23342388 | 0.96861995 | 0.33273485 | 0.6852158 |
| LRIG3 | 73.1564139 | 0.226185228 | 0.43158614 | 0.52407899 | 0.6002236 | 0.84976293 |
| PAXIP1-AS2 | 59.6002695 | 0.226315914 | 0.36797749 | 0.61502651 | 0.53853721 | 0.81807397 |
| PCF11 | 176.259478 | 0.226587198 | 0.18812756 | 1.20443383 | 0.22842194 | 0.59497702 |
| PANK2 | 206.008623 | 0.22675662 | 0.17989617 | 1.26048606 | 0.20749407 | 0.57246257 |
| ZFAND1 | 71.5620198 | 0.226787685 | 0.30479659 | 0.74406241 | 0.45683872 | 0.76825139 |
| ADAM10 | 795.622115 | 0.226847861 | 0.19891389 | 1.14043249 | 0.25410616 | 0.61966901 |
| TRA2A | 441.10149 | 0.226862518 | 0.27539693 | 0.8237656 | 0.41007276 | 0.73419295 |
| SGSM1 | 64.2484759 | 0.227014682 | 0.56583099 | 0.40120581 | 0.6882686 | 0.88739126 |
| ZNF273 | 51.0184429 | 0.227046881 | 0.44468232 | 0.51058221 | 0.60964364 | 0.85375662 |
| AMMECR1 | 133.748398 | 0.227063793 | 0.2062515 | 1.10090734 | 0.27093699 | 0.63561102 |
| PREX2 | 1073.09498 | 0.227264854 | 0.3262512 | 0.6965947 | 0.48605647 | 0.78826504 |
| STX18 | 216.994352 | 0.227295574 | 0.24763766 | 0.91785544 | 0.35869455 | 0.70242891 |
| PUS7L | 75.5380584 | 0.227315774 | 0.20748208 | 1.09559235 | 0.27325721 | 0.63887536 |
| ABCC10 | 39.6716717 | 0.227755318 | 0.24223792 | 0.94021333 | 0.34710814 | 0.69583193 |
| USP40 | 205.542149 | 0.227768436 | 0.29982647 | 0.75966752 | 0.44745335 | 0.7614672 |
| CBFA2T2 | 315.247376 | 0.227889803 | 0.20183477 | 1.12909092 | 0.25885948 | 0.62362556 |
| GTF3C3 | 86.8941041 | 0.227932459 | 0.187822 | 1.21355571 | 0.22491743 | 0.59122266 |
| C6orf62 | 196.736969 | 0.227941614 | 0.2181981 | 1.04465446 | 0.2961827 | 0.65753344 |
| FAM200B | 46.7699114 | 0.228037824 | 0.305718 | 0.74590904 | 0.45572236 | 0.76740262 |
| MYO15B | 233.32393 | 0.228470713 | 0.31496514 | 0.72538412 | 0.46821641 | 0.77664704 |
| CAND1 | 193.450363 | 0.228625585 | 0.15230924 | 1.50106187 | 0.13333956 | 0.48236854 |
| ROCK1 | 657.981699 | 0.228629251 | 0.14684354 | 1.55695817 | 0.11948042 | 0.46293005 |
| CDK19 | 288.476471 | 0.228739251 | 0.32897043 | 0.69531858 | 0.48685568 | 0.78826504 |
| TPST2 | 265.572567 | 0.228770059 | 0.18384856 | 1.24433966 | 0.21337458 | 0.5777777 |
| PRKAA2 | 49.9252799 | 0.229022574 | 0.28702483 | 0.79791903 | 0.42491748 | 0.74186041 |
| ZNRF2 | 118.019898 | 0.229033452 | 0.20994199 | 1.09093686 | 0.27530067 | 0.64017771 |
| ATAD1 | 113.806884 | 0.229214346 | 0.19037281 | 1.2040288 | 0.22857845 | 0.59497702 |
| NAA35 | 195.206344 | 0.229612894 | 0.17627536 | 1.30258081 | 0.19271791 | 0.55703491 |
| MAN2A1 | 480.589288 | 0.229952322 | 0.2314292 | 0.99361847 | 0.32040865 | 0.67435802 |
| GNPAT | 127.055553 | 0.230436855 | 0.18522636 | 1.24408243 | 0.21346923 | 0.57779965 |
| FAM49B | 240.399378 | 0.230471527 | 0.26889402 | 0.85710915 | 0.39138456 | 0.72247779 |
| NPAT | 145.94225 | 0.230852592 | 0.23176664 | 0.99605617 | 0.31922285 | 0.67394237 |
| ANKIB1 | 509.503924 | 0.230958043 | 0.17945024 | 1.28703109 | 0.19808345 | 0.56327864 |
| STAMBP | 85.4435207 | 0.230976203 | 0.25903616 | 0.89167551 | 0.37256688 | 0.71084822 |
| FMR1 | 121.504236 | 0.231155627 | 0.25258419 | 0.91516269 | 0.36010623 | 0.70386371 |
| CNOT1 | 285.748739 | 0.231204214 | 0.16805143 | 1.37579441 | 0.16888529 | 0.53071749 |
| PML | 221.792103 | 0.23125055 | 0.286724 | 0.80652667 | 0.41993924 | 0.73901389 |
| TOP2B | 300.356498 | 0.231272958 | 0.19113753 | 1.20998192 | 0.22628583 | 0.59267552 |
| ING4 | 50.0474145 | 0.231667424 | 0.25511589 | 0.90808699 | 0.36383226 | 0.70570548 |
| AP003108.2 | 105.13406 | 0.231765535 | 0.34725176 | 0.66742797 | 0.5044988 | 0.79776858 |
| TRPM4 | 104.625906 | 0.23183873 | 0.24960782 | 0.92881196 | 0.35298655 | 0.69977303 |
| CSNK2A2 | 113.898759 | 0.231950068 | 0.25649528 | 0.90430539 | 0.36583349 | 0.70687495 |
| PACS1 | 550.233093 | 0.232036866 | 0.21415913 | 1.08347876 | 0.27859597 | 0.64218333 |
| UBE2O | 105.206602 | 0.232500973 | 0.37500514 | 0.6199941 | 0.53526167 | 0.81602082 |
| NSD2 | 163.982744 | 0.232854035 | 0.27141932 | 0.85791252 | 0.39094077 | 0.72247779 |
| NSMCE2 | 763.967534 | 0.233006691 | 0.2454494 | 0.94930641 | 0.34246479 | 0.6921189 |
| ZNF263 | 53.9020168 | 0.233104388 | 0.26816004 | 0.86927338 | 0.38469762 | 0.71699718 |
| SEC22A | 243.265316 | 0.233261932 | 0.23427049 | 0.99569491 | 0.3193984 | 0.67395826 |
| MINK1 | 157.824848 | 0.23328643 | 0.22324112 | 1.04499758 | 0.29602409 | 0.65753344 |
| MPDZ | 656.210823 | 0.233397323 | 0.31921884 | 0.73115146 | 0.46468665 | 0.77460396 |
| CDK5RAP3 | 55.7686969 | 0.233458596 | 0.2317078 | 1.00755604 | 0.31366764 | 0.66807452 |
| LTA4H | 101.073098 | 0.233593805 | 0.2899712 | 0.80557589 | 0.42048743 | 0.73901389 |
| DNAJB2 | 48.9714259 | 0.233717131 | 0.31529382 | 0.74126772 | 0.45853113 | 0.76968366 |
| PSEN1 | 229.649867 | 0.23384106 | 0.32528155 | 0.71888817 | 0.47220982 | 0.77858009 |
| SPC25 | 200.838898 | 0.233925208 | 0.5883585 | 0.39758958 | 0.69093275 | 0.88832346 |
| MTMR9 | 141.911211 | 0.234189224 | 0.26745184 | 0.87563139 | 0.38123045 | 0.71543195 |
| TAF11 | 95.7134411 | 0.234271633 | 0.23699208 | 0.98852094 | 0.32289759 | 0.67633553 |
| TRIM65 | 43.6784544 | 0.234326293 | 0.26288301 | 0.89137101 | 0.37273016 | 0.71084822 |
| ILKAP | 153.652551 | 0.234399983 | 0.26924891 | 0.87056984 | 0.38398907 | 0.71699718 |
| VPS26C | 136.947826 | 0.234523821 | 0.19378413 | 1.21023234 | 0.22618975 | 0.59267075 |
| CARF | 88.4535082 | 0.234620047 | 0.44936522 | 0.52211439 | 0.60159069 | 0.85011818 |
| HDAC3 | 69.9409603 | 0.234620927 | 0.26220494 | 0.8947998 | 0.37089412 | 0.70984233 |
| KDM3B | 213.938181 | 0.234784715 | 0.22516362 | 1.04272934 | 0.29707367 | 0.65795352 |
| CTBP2 | 971.402198 | 0.235323373 | 0.26908571 | 0.87452942 | 0.38183001 | 0.71553552 |
| HIVEP3 | 1119.35153 | 0.235401258 | 0.41929899 | 0.56141623 | 0.57451382 | 0.83436282 |
| KLHL28 | 117.981942 | 0.235429926 | 0.19820035 | 1.18783808 | 0.23489721 | 0.59949266 |
| TUBA1A | 110.188208 | 0.235509046 | 0.55727561 | 0.42260785 | 0.6725814 | 0.87806854 |
| DNMT3A | 167.037862 | 0.235601615 | 0.36866647 | 0.63906439 | 0.52278104 | 0.80854347 |
| APBA1 | 79.6681181 | 0.235655784 | 0.33272397 | 0.70826212 | 0.47878249 | 0.78232371 |
| NDFIP1 | 303.197759 | 0.235740909 | 0.1590988 | 1.48172653 | 0.13841307 | 0.48991657 |
| SLC37A3 | 93.6900928 | 0.235825324 | 0.1919968 | 1.22827738 | 0.21934286 | 0.58626926 |
| CEP128 | 114.395106 | 0.235842366 | 0.36589594 | 0.6445613 | 0.51921152 | 0.80605347 |
| SAP30L | 51.2654799 | 0.236150701 | 0.2903638 | 0.81329251 | 0.41605037 | 0.73795362 |
| WRNIP1 | 104.807995 | 0.236189328 | 0.1747977 | 1.35121531 | 0.17662647 | 0.54037749 |
| ZNF577 | 58.6558269 | 0.236220616 | 0.37247183 | 0.63419727 | 0.52595209 | 0.81036438 |
| AQR | 225.704912 | 0.236429558 | 0.14739393 | 1.60406572 | 0.10869957 | 0.44474428 |
| USP48 | 317.630797 | 0.236607207 | 0.17168158 | 1.3781747 | 0.16814936 | 0.52962517 |
| RBM6 | 890.691448 | 0.236633502 | 0.18971239 | 1.24732763 | 0.21227739 | 0.57690664 |
| TMEM219 | 131.956964 | 0.236860336 | 0.26043646 | 0.90947458 | 0.36309967 | 0.70525625 |
| STK40 | 113.954065 | 0.236992498 | 0.27655204 | 0.85695443 | 0.39147007 | 0.72247779 |

| | | | | | | |
|---|---|---|---|---|---|---|
| BCKDHB | 181.535667 | 0.237053337 | 0.22464636 | 1.05522891 | 0.29132065 | 0.65401285 |
| PRICKLE2 | 111.619625 | 0.237170926 | 0.42092376 | 0.5634534 | 0.57312618 | 0.8342245 |
| ZBTB44 | 182.541374 | 0.237287211 | 0.22721287 | 1.04433879 | 0.29632867 | 0.65753344 |
| PAPOLA | 423.910855 | 0.237364218 | 0.19856609 | 1.19539148 | 0.23193411 | 0.59852313 |
| ATP2C1 | 301.777303 | 0.237504834 | 0.16017902 | 1.48274623 | 0.13814185 | 0.48991657 |
| CASP8AP2 | 134.018842 | 0.237529398 | 0.18046135 | 1.31623417 | 0.18809546 | 0.55388137 |
| CHUK | 67.6861912 | 0.237936002 | 0.32948907 | 0.72213625 | 0.47021071 | 0.77792258 |
| GOLGA8B | 49.4807152 | 0.238179474 | 0.34367671 | 0.6930335 | 0.48828853 | 0.78931968 |
| FANCC | 261.193992 | 0.238291699 | 0.26163431 | 0.91078152 | 0.3624105 | 0.70497203 |
| ZNF251 | 133.379344 | 0.238295782 | 0.27572176 | 0.8642618 | 0.38744409 | 0.72055795 |
| FAM133B | 139.895865 | 0.238331894 | 0.25912825 | 0.91974492 | 0.35770607 | 0.70127187 |
| RPH3AL | 365.933922 | 0.238387661 | 0.42313682 | 0.56338198 | 0.57317481 | 0.8342245 |
| VPS50 | 219.455625 | 0.238760597 | 0.21686819 | 1.10094798 | 0.2709193 | 0.63561102 |
| KIAA1958 | 68.8897463 | 0.238763847 | 0.36009969 | 0.66304929 | 0.50729899 | 0.79921563 |
| GRK2 | 82.086579 | 0.238784004 | 0.28242502 | 0.84547752 | 0.39784428 | 0.72733685 |
| WNT2B | 197.900444 | 0.238894978 | 0.49777734 | 0.47992337 | 0.63128188 | 0.86097142 |
| SNX27 | 133.44988 | 0.238934715 | 0.17441802 | 1.36989694 | 0.17071908 | 0.53193492 |
| CTBP1 | 85.7453221 | 0.239325177 | 0.25845661 | 0.92597815 | 0.35445734 | 0.70004491 |
| CNIH3 | 100.717702 | 0.23933695 | 0.37716038 | 0.63457606 | 0.52570495 | 0.81036438 |
| EIF4ENIF1 | 103.048458 | 0.239531326 | 0.23296583 | 1.02818222 | 0.30386412 | 0.66060576 |
| R3HDM2 | 426.62668 | 0.2396938 | 0.24339523 | 0.98479253 | 0.32472599 | 0.67855791 |
| TRIM52 | 52.6129634 | 0.23979275 | 0.2310212 | 1.03796861 | 0.29928467 | 0.65856711 |
| DPYD | 1482.20486 | 0.239896831 | 0.28013477 | 0.85636222 | 0.39179745 | 0.72247779 |
| PIKFYVE | 296.670542 | 0.240010304 | 0.1726289 | 1.39032516 | 0.16443016 | 0.52418378 |
| SH3BGRL2 | 228.58605 | 0.240062051 | 0.36089575 | 0.66518393 | 0.50593286 | 0.79877457 |
| DNMT1 | 120.837256 | 0.240362853 | 0.21953789 | 1.09485818 | 0.27357877 | 0.63926188 |
| NMNAT1 | 67.765334 | 0.240600246 | 0.29901808 | 0.80463444 | 0.42103066 | 0.73910721 |
| ANKRD13D | 86.904259 | 0.240619073 | 0.29435744 | 0.81743838 | 0.41367795 | 0.73655477 |
| MFSD6 | 89.8102882 | 0.24071405 | 0.39140157 | 0.61500533 | 0.5385512 | 0.81807397 |
| LYST | 402.982848 | 0.240766945 | 0.31719226 | 0.7590568 | 0.44781858 | 0.7614672 |
| ZBTB11 | 96.4327812 | 0.240951677 | 0.16672771 | 1.44518076 | 0.14840712 | 0.50188244 |
| HDX | 116.136136 | 0.241264181 | 0.32039278 | 0.75302628 | 0.45143412 | 0.76418813 |
| PDK4 | 405.522222 | 0.241341475 | 0.92269341 | 0.26156194 | 0.79365919 | 0.92790728 |
| SNX30 | 54.6966262 | 0.241355914 | 0.38485843 | 0.62712907 | 0.53057463 | 0.81320326 |
| PHF6 | 80.4545274 | 0.241513796 | 0.19604343 | 1.23194028 | 0.21797139 | 0.58419738 |
| NSL1 | 140.607175 | 0.241675227 | 0.2417049 | 0.99987724 | 0.31736992 | 0.67227741 |
| TRAF3IP1 | 67.3072097 | 0.241708585 | 0.32745577 | 0.73814117 | 0.46042867 | 0.770424 |
| ITGA2 | 143.332296 | 0.241781886 | 0.51955986 | 0.46535905 | 0.64167435 | 0.86490071 |
| AEBP2 | 332.336449 | 0.241800344 | 0.26579198 | 0.90973528 | 0.36296214 | 0.70525625 |
| SOX17 | 109.893424 | 0.241808799 | 0.61332683 | 0.39425766 | 0.69339082 | 0.8898215 |
| WASHC5 | 112.44774 | 0.241813383 | 0.28170595 | 0.85838935 | 0.39067751 | 0.72247779 |
| ERMP1 | 120.095006 | 0.241865577 | 0.32700306 | 0.73964317 | 0.45951654 | 0.76990661 |
| ENG | 2129.27885 | 0.242158415 | 0.57821224 | 0.41880541 | 0.67535835 | 0.88028948 |
| RFX1 | 53.6140873 | 0.242177309 | 0.28287077 | 0.85614117 | 0.3919197 | 0.72247779 |
| ETFDH | 135.088896 | 0.242403173 | 0.37058942 | 0.65410171 | 0.51304628 | 0.80321271 |
| OSMR-AS1 | 241.891131 | 0.242502156 | 0.35465152 | 0.68377588 | 0.4941167 | 0.7926408 |
| FBXW2 | 139.492478 | 0.242789711 | 0.1906568 | 1.27343853 | 0.20286247 | 0.56730017 |
| FER | 892.024475 | 0.242980257 | 0.20083502 | 1.20985007 | 0.22633643 | 0.59267552 |
| KLF6 | 402.535068 | 0.243232823 | 0.44632314 | 0.54497023 | 0.58577399 | 0.83933187 |
| NXF1 | 47.4542674 | 0.243448169 | 0.3171109 | 0.76770671 | 0.44266145 | 0.75748612 |
| NFIX | 101.303985 | 0.243498854 | 0.39504491 | 0.61638271 | 0.53764196 | 0.81807397 |
| MARK1 | 56.6807065 | 0.243702087 | 0.52353459 | 0.46549377 | 0.6415779 | 0.86490071 |
| WTIP | 39.7460856 | 0.243796956 | 0.44475138 | 0.54816459 | 0.58357889 | 0.83850098 |
| KIAA1468 | 302.897651 | 0.2439054 | 0.18414791 | 1.32450808 | 0.18533436 | 0.55068666 |
| SNX18 | 77.0628217 | 0.244051779 | 0.27887 | 0.87514532 | 0.38149484 | 0.71543195 |
| SLC25A24 | 141.93815 | 0.244270027 | 0.30024863 | 0.81355916 | 0.41589754 | 0.73795362 |
| SMC5 | 428.137278 | 0.244273065 | 0.15329227 | 1.59351196 | 0.11104539 | 0.44762779 |
| EP300 | 314.37141 | 0.244395496 | 0.19059662 | 1.28226561 | 0.19974949 | 0.56449294 |
| REX1BD | 73.2042922 | 0.244436692 | 0.47475112 | 0.51487333 | 0.60664154 | 0.85196053 |
| ODF2 | 95.4164635 | 0.244605552 | 0.28063784 | 0.87160575 | 0.38342349 | 0.71699718 |
| MED25 | 46.4256913 | 0.244653444 | 0.2637153 | 0.92771805 | 0.35355385 | 0.69977303 |
| NPR1 | 242.72297 | 0.244893583 | 0.34281388 | 0.7143631 | 0.47500268 | 0.78022377 |
| CBX7 | 73.206843 | 0.244952923 | 0.36626162 | 0.66879223 | 0.50362803 | 0.79776858 |
| RCSD1 | 136.457903 | 0.244997023 | 0.64952885 | 0.3771919 | 0.706031 | 0.89518665 |
| UQCC2 | 42.768047 | 0.245475793 | 0.3702765 | 0.66295267 | 0.50736086 | 0.79921563 |
| TRPS1 | 509.812047 | 0.245797447 | 0.36053419 | 0.68175906 | 0.49539132 | 0.79346 |
| CAMSAP1 | 100.968601 | 0.24584971 | 0.31939148 | 0.7697441 | 0.44145171 | 0.75620668 |
| PDCD1LG2 | 49.1877116 | 0.24615771 | 0.59876147 | 0.41111147 | 0.6809908 | 0.88326957 |
| RICTOR | 640.96636 | 0.246311873 | 0.19007358 | 1.29587642 | 0.19501806 | 0.56007526 |
| ATXN7L3 | 41.6572172 | 0.246540097 | 0.25184599 | 0.97893198 | 0.32761359 | 0.68017808 |
| PLAGL1 | 62.0245517 | 0.246557723 | 0.28804352 | 0.85597384 | 0.39201225 | 0.72248563 |
| ZFAND4 | 79.2234889 | 0.24662036 | 0.19149573 | 1.28786347 | 0.19779349 | 0.56316776 |
| ALG13 | 358.907763 | 0.247122466 | 0.25189479 | 0.9810543 | 0.32656596 | 0.67997183 |
| WASF2 | 555.29823 | 0.247304783 | 0.1619853 | 1.5267113 | 0.12683282 | 0.47369828 |
| ZNF675 | 45.9118817 | 0.247387212 | 0.29437768 | 0.84037352 | 0.40069899 | 0.72864885 |
| ITCH | 546.615118 | 0.247430893 | 0.17347823 | 1.42629364 | 0.1537836 | 0.50978388 |
| SYT1 | 93.9707949 | 0.247464597 | 1.28947764 | 0.19191073 | 0.84781213 | 0.94731791 |
| EFNB1 | 225.446188 | 0.247761995 | 0.30743344 | 0.8059045 | 0.42029792 | 0.73901389 |
| RNF34 | 86.3670083 | 0.248152337 | 0.3668852 | 0.67637598 | 0.49880196 | 0.79513466 |
| VAMP4 | 95.1548253 | 0.248374654 | 0.23099735 | 1.0752273 | 0.28227297 | 0.64538691 |
| ZNF276 | 70.395713 | 0.248916529 | 0.18931309 | 1.31484053 | 0.1885635 | 0.55388137 |
| BTN2A1 | 76.6257598 | 0.248941136 | 0.32611521 | 0.76335334 | 0.4452527 | 0.76017167 |
| GPR155 | 60.3660439 | 0.249172558 | 0.25792104 | 0.96608078 | 0.33400376 | 0.6852158 |
| STK39 | 144.851268 | 0.249327825 | 0.28603578 | 0.87166656 | 0.38339031 | 0.71699718 |
| SUN1 | 440.633273 | 0.249396327 | 0.25583156 | 0.97484582 | 0.32963674 | 0.68216508 |
| CDKN1C | 98.8521906 | 0.249433343 | 0.48565452 | 0.51360244 | 0.60752998 | 0.85243789 |
| TGFBR1 | 126.392281 | 0.249621723 | 0.25641118 | 0.97352123 | 0.33029432 | 0.68269724 |
| YAF2 | 296.361456 | 0.249763952 | 0.14887812 | 1.67764046 | 0.09341731 | 0.41231599 |
| RP2 | 49.8147278 | 0.249941671 | 0.23323709 | 1.07162061 | 0.28389047 | 0.6470963 |
| TOM1L2 | 229.031641 | 0.250209568 | 0.21061734 | 1.18798179 | 0.23484058 | 0.59949266 |
| INTS3 | 175.024589 | 0.250295266 | 0.26525415 | 0.94360547 | 0.3453713 | 0.69497757 |
| LPAR6 | 243.367486 | 0.250387776 | 0.38753695 | 0.64610039 | 0.51821434 | 0.80557522 |
| RRAS2 | 132.854902 | 0.250735256 | 0.25574236 | 0.9804213 | 0.3268782 | 0.68007671 |
| KANSL3 | 146.10949 | 0.250747209 | 0.19282396 | 1.30039446 | 0.19346581 | 0.55822662 |
| NR2C2 | 315.932268 | 0.250813738 | 0.15100179 | 1.66099844 | 0.09671376 | 0.41851332 |
| KTN1 | 618.258898 | 0.251143032 | 0.22005591 | 1.1412692 | 0.25375791 | 0.61961636 |
| TMEM41B | 128.830982 | 0.251492715 | 0.28815284 | 0.87277543 | 0.3827855 | 0.71637321 |
| PPP1R13L | 71.12242 | 0.251530809 | 0.22021636 | 1.14219854 | 0.2533715 | 0.61909197 |
| TBC1D2B | 167.715424 | 0.251625991 | 0.2809505 | 0.89562392 | 0.37045365 | 0.70984233 |
| NAPA | 93.8480464 | 0.251630992 | 0.31264676 | 0.8048412 | 0.42091133 | 0.73910721 |
| NMD3 | 133.744818 | 0.252280944 | 0.18628942 | 1.35424193 | 0.1756592 | 0.53876285 |
| ZBTB10 | 105.192103 | 0.252293654 | 0.36464181 | 0.69189448 | 0.4890036 | 0.78955612 |
| NTAN1 | 74.2330558 | 0.252540762 | 0.26962873 | 0.93662407 | 0.34895197 | 0.69809042 |
| RETREG3 | 114.329773 | 0.252685505 | 0.24503716 | 1.03121299 | 0.30244095 | 0.65926326 |
| SENP7 | 576.572206 | 0.252724341 | 0.25465906 | 0.9924027 | 0.32100112 | 0.67473727 |
| SART1 | 42.1203423 | 0.252773993 | 0.32745509 | 0.77193483 | 0.44015303 | 0.75477205 |
| AP3S1 | 158.786325 | 0.252965835 | 0.1714565 | 1.47539372 | 0.14010672 | 0.49118733 |
| RING1 | 53.6590691 | 0.253035427 | 0.25829568 | 0.97963476 | 0.32726644 | 0.68017808 |
| HERPUD2 | 145.952787 | 0.253334977 | 0.1903051 | 1.33120432 | 0.18312179 | 0.54813352 |
| PRIMPOL | 101.91879 | 0.253449851 | 0.30138424 | 0.84095257 | 0.40037451 | 0.72864885 |
| BTBD9 | 665.867841 | 0.253770746 | 0.30709839 | 0.82634996 | 0.4086056 | 0.73419295 |
| CHMP5 | 134.977481 | 0.253822464 | 0.31894507 | 0.79581874 | 0.4261374 | 0.74203245 |
| NLRP1 | 405.28305 | 0.25391716 | 0.31434097 | 0.80777621 | 0.41921942 | 0.73853015 |

| | | | | | | |
|---|---|---|---|---|---|---|
| IL6ST | 1188.80111 | 0.254036715 | 0.36558847 | 0.69487069 | 0.48713634 | 0.78826504 |
| LRP11 | 64.965101 | 0.254099164 | 0.44457211 | 0.57155894 | 0.56762082 | 0.83332923 |
| LRRC49 | 113.102283 | 0.25451277 | 0.25895707 | 0.98283769 | 0.32568733 | 0.67964945 |
| CCDC66 | 189.300702 | 0.254569904 | 0.20499287 | 1.24184757 | 0.21429281 | 0.57873203 |
| ADAM15 | 352.890148 | 0.254616188 | 0.41574128 | 0.61243904 | 0.54024732 | 0.81900022 |
| TTC19 | 329.752254 | 0.254692701 | 0.16068472 | 1.58504617 | 0.11295584 | 0.45066683 |
| DIP2B | 525.631899 | 0.254779871 | 0.24686568 | 1.03205868 | 0.30204463 | 0.65926326 |
| SNX24 | 144.124269 | 0.255213386 | 0.26176359 | 0.97497665 | 0.32957184 | 0.68216508 |
| ABHD12 | 102.572296 | 0.255265438 | 0.29836677 | 0.85554245 | 0.39225092 | 0.72276272 |
| FGFR1OP2 | 214.201941 | 0.255290949 | 0.31804372 | 0.80269136 | 0.42215315 | 0.73995396 |
| ATP6V1D | 110.98138 | 0.255354094 | 0.38771014 | 0.65862115 | 0.51013908 | 0.80177277 |
| RPRD2 | 349.034455 | 0.255685592 | 0.19167901 | 1.33392586 | 0.18222815 | 0.54721945 |
| IFT172 | 65.8563875 | 0.255746078 | 0.31133504 | 0.82144971 | 0.41139016 | 0.73470224 |
| ERBIN | 812.380315 | 0.255780474 | 0.1734656 | 1.47453138 | 0.14033858 | 0.49118733 |
| PIGX | 87.0567391 | 0.256059675 | 0.38341775 | 0.66783469 | 0.50423912 | 0.79776858 |
| ATXN7 | 47.1328593 | 0.256110097 | 0.34309159 | 0.74647733 | 0.45537912 | 0.76710126 |
| CCDC92 | 126.101132 | 0.256112369 | 0.21183021 | 1.20904553 | 0.22664536 | 0.59310487 |
| WIPI2 | 116.817844 | 0.256191387 | 0.21909196 | 1.16933266 | 0.24226963 | 0.6070269 |
| UBR5 | 871.822559 | 0.256259864 | 0.2388381 | 1.07294384 | 0.28329631 | 0.6466426 |
| PRKCE | 1608.41537 | 0.256536738 | 0.38385708 | 0.66831316 | 0.50393371 | 0.79776858 |
| STAM2 | 174.265526 | 0.256621829 | 0.19470525 | 1.31800161 | 0.18750311 | 0.55388137 |
| ARIH1 | 758.7744 | 0.256758811 | 0.16483107 | 1.55770883 | 0.11930229 | 0.46290743 |
| SMTN | 142.211521 | 0.256845955 | 0.22315367 | 1.15098242 | 0.24973947 | 0.6149919 |
| ZNF568 | 59.8703872 | 0.25693401 | 0.25067135 | 1.02498357 | 0.30537094 | 0.66194713 |
| PIGQ | 38.9289905 | 0.257277447 | 0.28774308 | 0.8941221 | 0.37125656 | 0.70984233 |
| GSK3B | 1045.51952 | 0.257613068 | 0.20864839 | 1.23467556 | 0.21695127 | 0.58245153 |
| NELFB | 87.8944693 | 0.257658635 | 0.31786159 | 0.8106001 | 0.41759536 | 0.73821752 |
| APOL6 | 178.923032 | 0.258049106 | 0.51095265 | 0.50503527 | 0.61353407 | 0.85573702 |
| GAB3 | 67.7184432 | 0.258808689 | 0.2449194 | 1.05670962 | 0.29064413 | 0.65333738 |
| AC009554.1 | 102.381637 | 0.258816741 | 0.53661188 | 0.48231645 | 0.62958116 | 0.86093961 |
| COG5 | 541.809116 | 0.259031853 | 0.16924359 | 1.53052682 | 0.12588638 | 0.47145458 |
| ANXA1 | 224.252154 | 0.259101393 | 0.47386807 | 0.5467796 | 0.58453016 | 0.83895258 |
| STAG1 | 1598.30444 | 0.259210962 | 0.17526744 | 1.47894536 | 0.13915492 | 0.49082099 |
| HTT | 286.766593 | 0.259469138 | 0.17992978 | 1.44205777 | 0.14928608 | 0.50283523 |
| TFCP2 | 338.67077 | 0.259648384 | 0.22888602 | 1.13440036 | 0.25662665 | 0.62095131 |
| CDC42BPB | 553.945059 | 0.259730842 | 0.22349309 | 1.1621426 | 0.24517754 | 0.6103705 |
| CTNND1 | 432.348917 | 0.25979919 | 0.15255818 | 1.7029516 | 0.08857713 | 0.40312938 |
| IDE | 205.471964 | 0.259805749 | 0.24898353 | 1.04346559 | 0.29673271 | 0.6576825 |
| STXBP5-AS1 | 103.333794 | 0.260126634 | 0.28064336 | 0.92689395 | 0.3539816 | 0.69977303 |
| NR2C1 | 177.581595 | 0.260148706 | 0.16494511 | 1.57718349 | 0.11475331 | 0.45569454 |
| TXNDC16 | 68.0211249 | 0.260226411 | 0.47314075 | 0.54999788 | 0.58232082 | 0.83812111 |
| NIPBL | 1091.53239 | 0.260504721 | 0.17354299 | 1.50109621 | 0.13333068 | 0.48236854 |
| CYP27A1 | 61.3660482 | 0.260545076 | 0.35738191 | 0.72903823 | 0.46597828 | 0.77517793 |
| PTPN4 | 297.112669 | 0.260603566 | 0.1785277 | 1.45973747 | 0.14436224 | 0.49679586 |
| ZBTB40 | 80.5214603 | 0.260671445 | 0.31552812 | 0.82614331 | 0.4087228 | 0.73419295 |
| FAM222B | 400.751036 | 0.260703374 | 0.27515414 | 0.9474812 | 0.34339363 | 0.69361764 |
| SNRNP70 | 289.179143 | 0.2607846 | 0.25052318 | 1.04095997 | 0.29789413 | 0.6579563 |
| SPSB3 | 81.810722 | 0.260797718 | 0.29304619 | 0.88995429 | 0.37349043 | 0.71166161 |
| STT3B | 426.705455 | 0.261213702 | 0.25451367 | 1.02632484 | 0.30473849 | 0.66092634 |
| ANGEL2 | 112.445502 | 0.261215376 | 0.21915355 | 1.19192859 | 0.23328925 | 0.59910683 |
| MAPK6 | 174.889615 | 0.261409115 | 0.22676848 | 1.15275772 | 0.24900984 | 0.61434376 |
| EIF2B1 | 63.5278936 | 0.261444732 | 0.2272963 | 1.15023752 | 0.25004606 | 0.61500658 |
| AHDC1 | 111.373915 | 0.261713409 | 0.24270594 | 1.07831481 | 0.28089329 | 0.64403189 |
| ZBTB4 | 126.97473 | 0.261841449 | 0.29218802 | 0.89614025 | 0.37017786 | 0.70984233 |
| ARHGAP17 | 347.516111 | 0.261881092 | 0.13945861 | 1.87784096 | 0.06040292 | 0.34485038 |
| PPIP5K1 | 62.2095541 | 0.26226123 | 0.46228019 | 0.56732094 | 0.57049615 | 0.83368112 |
| JUND | 344.335174 | 0.262439268 | 0.56415183 | 0.46519262 | 0.64179352 | 0.86490071 |
| ICE1 | 209.617183 | 0.262654854 | 0.21754904 | 1.20733633 | 0.22730265 | 0.59406501 |
| PPP2CB | 250.31966 | 0.262934307 | 0.18545748 | 1.41776062 | 0.15626066 | 0.5124974 |
| VHL | 65.2959043 | 0.263059854 | 0.27437764 | 0.95875108 | 0.33768416 | 0.68839968 |
| CCDC7 | 208.360268 | 0.263312687 | 0.38232029 | 0.68872276 | 0.49099775 | 0.79075666 |
| IFT140 | 106.084143 | 0.264111715 | 0.33037025 | 0.79944157 | 0.42403441 | 0.74152442 |
| ZADH2 | 135.672405 | 0.264488439 | 0.309888 | 0.85349686 | 0.39338383 | 0.72360479 |
| CEP70 | 105.601504 | 0.264698093 | 0.2271774 | 1.1651603 | 0.24395411 | 0.608518 |
| TSPAN12 | 130.070727 | 0.264991107 | 0.55405292 | 0.47827761 | 0.63245263 | 0.86141143 |
| TLR3 | 42.9521595 | 0.265009267 | 0.49153865 | 0.53914227 | 0.58978869 | 0.8420771 |
| PDE8A | 886.86226 | 0.265011509 | 0.30436306 | 0.87070852 | 0.38391332 | 0.71699718 |
| CERS4 | 67.3197346 | 0.265353624 | 0.45864712 | 0.57855727 | 0.56288794 | 0.83082829 |
| UBE2G1 | 254.539781 | 0.265369386 | 0.25906644 | 1.02432945 | 0.30567969 | 0.66234311 |
| HOTAIRM1 | 88.6515222 | 0.266036792 | 0.29378592 | 0.90554644 | 0.36517597 | 0.70668224 |
| TAB3 | 85.8287071 | 0.266096703 | 0.21377979 | 1.24472341 | 0.21323344 | 0.57758664 |
| MACO1 | 225.352588 | 0.266372896 | 0.15530392 | 1.71517174 | 0.08631374 | 0.39864674 |
| CEP78 | 54.2884564 | 0.266578497 | 0.26473885 | 1.00694891 | 0.31395933 | 0.66834786 |
| FAM124A | 62.4167412 | 0.266598823 | 0.30088481 | 0.88604947 | 0.37559087 | 0.71257833 |
| GALNT11 | 173.449519 | 0.266623949 | 0.23297929 | 1.14441053 | 0.25245342 | 0.61777104 |
| ELP4 | 352.178093 | 0.266651693 | 0.20021874 | 1.33180185 | 0.18292531 | 0.5479055 |
| C3orf38 | 58.6146505 | 0.266836671 | 0.23927581 | 1.11518446 | 0.26477138 | 0.63018797 |
| DNAJC17 | 77.0009932 | 0.266941131 | 0.18177 | 1.46856541 | 0.1419507 | 0.49378081 |
| PAN2 | 55.6089773 | 0.266946831 | 0.21183593 | 1.2601584 | 0.20761223 | 0.57254369 |
| ARMH3 | 335.848406 | 0.266954718 | 0.18389358 | 1.45168049 | 0.14659047 | 0.50043796 |
| MFSD12 | 78.5580509 | 0.26743286 | 0.24458478 | 1.09341581 | 0.27421128 | 0.63975272 |
| JAM3 | 296.234524 | 0.267501355 | 0.19825269 | 1.34929499 | 0.17724223 | 0.54037749 |
| PTOV1 | 65.5112592 | 0.267602645 | 0.1931158 | 1.38571078 | 0.16583523 | 0.5261375 |
| ARL16 | 43.2277523 | 0.267829426 | 0.39734335 | 0.67405036 | 0.5002793 | 0.79599174 |
| ZNF862 | 79.835764 | 0.268262919 | 0.22381591 | 1.19858737 | 0.23068843 | 0.59719185 |
| PRELID2 | 308.286974 | 0.268583239 | 0.40853336 | 0.65743282 | 0.51090266 | 0.80177277 |
| SPPL3 | 465.339574 | 0.268759117 | 0.15110863 | 1.77858222 | 0.07530828 | 0.38047182 |
| ARHGEF1 | 109.627722 | 0.268808316 | 0.1902099 | 1.41321935 | 0.15759124 | 0.51438736 |
| TPRG1L | 58.0571772 | 0.2688275 | 0.33661159 | 0.79862818 | 0.42450604 | 0.74157887 |
| DYRK1A | 899.729193 | 0.268892664 | 0.20710878 | 1.29831612 | 0.19417873 | 0.55913011 |
| MLLT6 | 205.149205 | 0.269034112 | 0.24259415 | 1.10898847 | 0.26743516 | 0.6327007 |
| ALG6 | 56.7926494 | 0.269082641 | 0.23778728 | 1.13161075 | 0.25779812 | 0.62258498 |
| AC058822.1 | 174.55827 | 0.269089861 | 0.48266257 | 0.55751135 | 0.5771781 | 0.83603913 |
| ZNF800 | 259.669815 | 0.269237168 | 0.2535824 | 1.06173446 | 0.28835625 | 0.65138125 |
| TM2D3 | 114.266562 | 0.26943342 | 0.22235957 | 1.21170149 | 0.22562667 | 0.59165529 |
| SP140L | 158.835071 | 0.26963681 | 0.24388482 | 1.10559077 | 0.26890369 | 0.63347822 |
| LINC02241 | 39.6641037 | 0.27001942 | 0.35786114 | 0.75453685 | 0.45052693 | 0.76375998 |
| ADGRA3 | 46.6564154 | 0.270539469 | 0.41460258 | 0.65252722 | 0.51406112 | 0.80398873 |
| ZCCHC24 | 144.340096 | 0.270620974 | 0.37506601 | 0.72152893 | 0.47058415 | 0.77797169 |
| HERC4 | 921.745834 | 0.270697883 | 0.19122639 | 1.4155885 | 0.15689602 | 0.51396323 |
| HLA-DQA1 | 162.750447 | 0.270769237 | 0.47629856 | 0.56848637 | 0.56970476 | 0.83368112 |
| TMCC1 | 617.498178 | 0.270841299 | 0.213945 | 1.26593893 | 0.20553495 | 0.57051983 |
| SMAD5 | 220.471096 | 0.271348351 | 0.16039131 | 1.69178962 | 0.09068609 | 0.40729106 |
| MTSS1L | 39.7911271 | 0.271450999 | 0.43296963 | 0.6269516 | 0.53069096 | 0.81322924 |
| KLHL5 | 754.457775 | 0.271476384 | 0.30676799 | 0.88495669 | 0.37617999 | 0.71257833 |
| ALMS1 | 194.998084 | 0.271717809 | 0.22159018 | 1.22621775 | 0.22011674 | 0.58688069 |
| RAB30-AS1 | 99.7408069 | 0.271896389 | 0.31218709 | 0.87094053 | 0.38378662 | 0.71699718 |
| SYNE3 | 47.6050112 | 0.27266747 | 0.34909904 | 0.78106049 | 0.43476692 | 0.74898899 |
| ATP6V1A | 234.590576 | 0.272854552 | 0.17670957 | 1.54408472 | 0.12256781 | 0.46802258 |
| HCG17 | 123.265514 | 0.273045882 | 0.27096131 | 1.00769324 | 0.31360175 | 0.66807452 |
| PDE6D | 113.783219 | 0.273143017 | 0.26308685 | 1.03822376 | 0.29916589 | 0.65856711 |
| RAD17 | 88.0361014 | 0.273169043 | 0.21427991 | 1.2748234 | 0.20237175 | 0.56693091 |
| LONP2 | 425.601076 | 0.273222064 | 0.15935199 | 1.71458202 | 0.08642189 | 0.39864674 |

| | | | | | | |
|---|---|---|---|---|---|---|
| RBM15 | 51.1009325 | 0.273248106 | 0.29569571 | 0.92408544 | 0.35544183 | 0.70018061 |
| EXD3 | 174.046856 | 0.273684162 | 0.24508842 | 1.11667523 | 0.26413321 | 0.63018314 |
| DCAF17 | 73.7629414 | 0.273722047 | 0.20335318 | 1.3460426 | 0.17828878 | 0.54195285 |
| GPATCH8 | 596.736485 | 0.273780914 | 0.23375422 | 1.17123409 | 0.2415047 | 0.60620642 |
| UBR3 | 690.020824 | 0.273867726 | 0.15347205 | 1.78447945 | 0.07434579 | 0.37740174 |
| SNX19 | 133.37765 | 0.274167745 | 0.31532606 | 0.8694738 | 0.38458803 | 0.71699718 |
| NAB1 | 147.277704 | 0.275297059 | 0.16752094 | 1.64335907 | 0.10030866 | 0.42658751 |
| PIEZO1 | 251.544162 | 0.275442656 | 0.15557031 | 1.77053488 | 0.07663808 | 0.38356538 |
| NPLOC4 | 264.656473 | 0.275736636 | 0.22753815 | 1.21182596 | 0.22557901 | 0.59165529 |
| SPATA1 | 59.363325 | 0.276048869 | 0.54186052 | 0.50944636 | 0.61043939 | 0.85403069 |
| BICRAL | 263.418461 | 0.27606239 | 0.28600365 | 0.96524078 | 0.33442422 | 0.68544254 |
| ZNF384 | 53.6135615 | 0.276066658 | 0.31186103 | 0.88522333 | 0.37603619 | 0.71257833 |
| ITPR3 | 193.363829 | 0.276290535 | 0.30071291 | 0.91878508 | 0.358208 | 0.70175151 |
| RAB1A | 597.026626 | 0.276613493 | 0.20974911 | 1.31878265 | 0.18724179 | 0.55388137 |
| NUP205 | 110.500657 | 0.277166065 | 0.23086905 | 1.20053364 | 0.22993215 | 0.59648134 |
| SCARB2 | 341.29398 | 0.277379185 | 0.21760758 | 1.27467614 | 0.2024239 | 0.56693091 |
| LRRC37B | 97.6335543 | 0.277524098 | 0.39051959 | 0.71065347 | 0.477299 | 0.78098272 |
| MRPS9 | 95.1592123 | 0.277789781 | 0.24710487 | 1.12417767 | 0.26093766 | 0.62580682 |
| VPS54 | 459.392244 | 0.277833633 | 0.15939019 | 1.74310376 | 0.08131549 | 0.39187554 |
| DOCK7 | 245.103941 | 0.277923049 | 0.18931927 | 1.46801243 | 0.14210085 | 0.49392944 |
| ZNF33B | 119.124954 | 0.278147245 | 0.23970037 | 1.16039556 | 0.24588779 | 0.61094785 |
| SOX6 | 211.221994 | 0.278194445 | 0.55849682 | 0.49811285 | 0.6184045 | 0.85855421 |
| PHF8 | 126.390428 | 0.278230336 | 0.1435372 | 1.93838482 | 0.05257629 | 0.32082907 |
| SCFD1 | 336.932759 | 0.278273821 | 0.2130008 | 1.30644498 | 0.19140128 | 0.55590015 |
| TM6SF1 | 100.912621 | 0.278407304 | 0.62444418 | 0.44584819 | 0.65570691 | 0.87118845 |
| KCMF1 | 328.147294 | 0.278479172 | 0.19801804 | 1.40633234 | 0.15962546 | 0.51768507 |
| ERICH1 | 324.178715 | 0.278593587 | 0.40773763 | 0.68326681 | 0.49443826 | 0.79286465 |
| RBSN | 77.5261359 | 0.278730205 | 0.21548726 | 1.29348812 | 0.19584229 | 0.56062979 |
| N4BP2 | 58.6827719 | 0.278820224 | 0.23732989 | 1.17482138 | 0.24006618 | 0.60463576 |
| RAB2A | 564.780103 | 0.278867818 | 0.18402687 | 1.5153647 | 0.12968007 | 0.4777902 |
| PRKX | 257.420177 | 0.278928298 | 0.23825081 | 1.17073391 | 0.24170575 | 0.60636513 |
| FLRT2 | 465.151183 | 0.27900615 | 0.41176984 | 0.67757791 | 0.49803936 | 0.79513036 |
| RTN3 | 199.361131 | 0.279715629 | 0.17590483 | 1.5901532 | 0.11180028 | 0.44839019 |
| NDEL1 | 170.891315 | 0.279725277 | 0.23519394 | 1.1893388 | 0.23430637 | 0.59949266 |
| TBC1D8B | 56.9739201 | 0.279776705 | 0.34912565 | 0.80136395 | 0.42292098 | 0.74026603 |
| PLXDC2 | 927.069486 | 0.280072036 | 0.29076585 | 0.9632219 | 0.33543618 | 0.68639016 |
| SLC4A2 | 45.5592694 | 0.280884702 | 0.32654916 | 0.86016055 | 0.38970055 | 0.72202943 |
| QKI | 1766.68303 | 0.280893658 | 0.19400225 | 1.44788869 | 0.14764818 | 0.50136364 |
| MARF1 | 153.215842 | 0.281068165 | 0.23296367 | 1.20648924 | 0.22762891 | 0.59434823 |
| SMG7 | 290.591309 | 0.281121623 | 0.2248384 | 1.25032747 | 0.21117995 | 0.5768065 |
| SMARCA1 | 100.165658 | 0.28118426 | 0.37498733 | 0.74985004 | 0.45334503 | 0.76568057 |
| MBD5 | 1421.20884 | 0.281245391 | 0.21732 | 1.29415325 | 0.19561249 | 0.56046814 |
| MTMR2 | 240.039321 | 0.281437931 | 0.23586625 | 1.19320985 | 0.2327872 | 0.59910683 |
| PPM1L | 83.1597237 | 0.281457017 | 0.44401944 | 0.63388445 | 0.52615623 | 0.81036438 |
| NAPEPLD | 68.2902368 | 0.281473116 | 0.2394985 | 1.17526048 | 0.23989052 | 0.60456313 |
| FBXL20 | 331.949679 | 0.281482738 | 0.2377213 | 1.18408717 | 0.23637856 | 0.60018541 |
| CUL5 | 344.115757 | 0.28179969 | 0.17651446 | 1.59646801 | 0.11038434 | 0.44631216 |
| GNPTG | 50.786666 | 0.28180916 | 0.21220523 | 1.32800289 | 0.18417715 | 0.54912489 |
| AC025682.1 | 59.9103156 | 0.282059588 | 0.39208774 | 0.71937875 | 0.47190758 | 0.77855633 |
| FASTKD1 | 104.57177 | 0.282492206 | 0.24017756 | 1.17618067 | 0.23952269 | 0.60382444 |
| ANKRD52 | 50.7140112 | 0.282499112 | 0.23774984 | 1.18822 | 0.23474675 | 0.59949266 |
| ZNF224 | 67.4203231 | 0.282569468 | 0.26447892 | 1.06840073 | 0.28533979 | 0.64806045 |
| ZNF467 | 57.8821637 | 0.282796278 | 0.40584942 | 0.69680099 | 0.48592735 | 0.78826504 |
| TCEA1 | 139.637508 | 0.282911322 | 0.19382739 | 1.45960447 | 0.14439881 | 0.49679586 |
| RALGAPA1 | 754.680222 | 0.283023659 | 0.22243777 | 1.2723723 | 0.20324087 | 0.567308 |
| VOPP1 | 498.515981 | 0.283077983 | 0.23000701 | 1.23073634 | 0.21842149 | 0.58505502 |
| PHF2 | 115.064184 | 0.28349281 | 0.21886287 | 1.29529881 | 0.19521717 | 0.56007526 |
| DCAF6 | 379.501202 | 0.283783489 | 0.20184767 | 1.40592897 | 0.15974522 | 0.51770104 |
| BRD1 | 122.457055 | 0.283876735 | 0.31376223 | 0.90475113 | 0.36559725 | 0.70668224 |
| COMMD1 | 494.016504 | 0.283911737 | 0.23961963 | 1.18484339 | 0.23607937 | 0.60018541 |
| AC017101.1 | 51.6461231 | 0.284202609 | 0.63155903 | 0.45000166 | 0.65270925 | 0.86875728 |
| CIC | 56.4881056 | 0.284260809 | 0.3801555 | 0.74774878 | 0.4546117 | 0.76671219 |
| SOCS6 | 66.9272635 | 0.284444157 | 0.31179937 | 0.91226662 | 0.36162838 | 0.70406973 |
| MIOS | 57.1399211 | 0.284506293 | 0.23927468 | 1.18903635 | 0.23442537 | 0.59949266 |
| DMPK | 50.4420646 | 0.284549554 | 0.31537074 | 0.90227 | 0.36691346 | 0.70745825 |
| MED12 | 42.4094566 | 0.284556933 | 0.27935686 | 1.01861445 | 0.30838604 | 0.66414532 |
| UFM1 | 145.336954 | 0.284705877 | 0.29009517 | 0.98142232 | 0.32638452 | 0.67997183 |
| RAB11FIP3 | 214.818926 | 0.284944946 | 0.2761598 | 1.03181182 | 0.30216028 | 0.65926326 |
| ZRANB2 | 369.57171 | 0.285123075 | 0.16619288 | 1.71561542 | 0.08623245 | 0.39864674 |
| ZNF512 | 148.428458 | 0.285136486 | 0.28494278 | 1.00067982 | 0.31698163 | 0.67181058 |
| GTF2E2 | 134.372505 | 0.285220089 | 0.25808067 | 1.10515869 | 0.26909084 | 0.63347822 |
| PBX3 | 430.807713 | 0.285515017 | 0.23414296 | 1.21940466 | 0.22269064 | 0.58858855 |
| PPP6C | 181.877714 | 0.285529039 | 0.13634233 | 2.09420689 | 0.03624155 | 0.2678994 |
| PRPSAP1 | 103.298417 | 0.285718551 | 0.23014776 | 1.24145701 | 0.21443697 | 0.5789303 |
| ZFY | 67.4852667 | 0.285744069 | 2.0430137 | 0.139864 | 0.88876745 | 0.96371185 |
| SH3D19 | 735.649765 | 0.285772092 | 0.26216111 | 1.09006289 | 0.27568544 | 0.64048591 |
| LASP1 | 158.53868 | 0.286283678 | 0.21252031 | 1.34708855 | 0.17795172 | 0.54173322 |
| PARD3 | 835.913775 | 0.286287619 | 0.36217897 | 0.79045897 | 0.42925977 | 0.74408005 |
| SLC25A40 | 54.4886839 | 0.286381222 | 0.28307578 | 1.01167689 | 0.31169257 | 0.66561244 |
| EWSR1 | 282.241076 | 0.286409716 | 0.1911473 | 1.49837178 | 0.13403668 | 0.48404962 |
| TRPM3 | 120.623315 | 0.286665667 | 0.38478135 | 0.74500926 | 0.45626612 | 0.76762944 |
| ZNF439 | 45.2087359 | 0.286915691 | 0.24224455 | 1.18440516 | 0.23625272 | 0.60018541 |
| SIAH1 | 118.494049 | 0.287158576 | 0.20204248 | 1.42127819 | 0.1552359 | 0.51129169 |
| MIS18BP1 | 278.325331 | 0.287344911 | 0.30844269 | 0.93159903 | 0.35154379 | 0.69889281 |
| FAM185A | 102.976207 | 0.287473451 | 0.23026235 | 1.24846049 | 0.21186247 | 0.57690664 |
| CMPK1 | 189.80345 | 0.287509532 | 0.22540197 | 1.27554135 | 0.2021177 | 0.56693091 |
| KAT6B | 349.158569 | 0.287676886 | 0.27183535 | 1.05827621 | 0.28992953 | 0.65303974 |
| SETD2 | 655.711764 | 0.287717188 | 0.19038052 | 1.51127431 | 0.13071858 | 0.47936448 |
| PTPN14 | 772.063702 | 0.287948246 | 0.32335514 | 0.89050151 | 0.37319666 | 0.71135994 |
| TRIM37 | 117.861442 | 0.288109307 | 0.20883193 | 1.37962287 | 0.16770279 | 0.52903314 |
| CEP44 | 76.0723372 | 0.288281526 | 0.25410651 | 1.13449091 | 0.25658868 | 0.62095131 |
| CDKN2D | 45.7543022 | 0.288558132 | 0.64953458 | 0.44425368 | 0.65685918 | 0.87218235 |
| TMEM50B | 182.823087 | 0.288615267 | 0.24830455 | 1.16234386 | 0.24509582 | 0.61035273 |
| MYO9A | 651.593403 | 0.288768852 | 0.1881159 | 1.53505814 | 0.12476955 | 0.47038289 |
| AVEN | 166.914736 | 0.288770399 | 0.37050051 | 0.77940622 | 0.43574046 | 0.74971913 |
| KMT5B | 283.828149 | 0.288861675 | 0.29245754 | 0.98770465 | 0.32329732 | 0.67664579 |
| ITSN1 | 192.220159 | 0.289311536 | 0.31023574 | 0.93255384 | 0.35105037 | 0.69858573 |
| OSBPL11 | 72.811726 | 0.28969298 | 0.23848274 | 1.21473354 | 0.22446774 | 0.59083342 |
| ZC3H6 | 174.863531 | 0.289860771 | 0.41078302 | 0.70562987 | 0.48041834 | 0.7845841 |
| MINPP1 | 47.9214508 | 0.289941656 | 0.24568659 | 1.18012815 | 0.23794925 | 0.60096874 |
| RASA1 | 553.143272 | 0.290041511 | 0.16731771 | 1.73347764 | 0.08301081 | 0.39515849 |
| CUL9 | 53.6591258 | 0.290428278 | 0.22817504 | 1.27283105 | 0.203078 | 0.567308 |
| PLCG1 | 155.250564 | 0.290704525 | 0.20942178 | 1.3881294 | 0.16509764 | 0.52526984 |
| OTUD5 | 92.2648426 | 0.290913211 | 0.19436137 | 1.49676458 | 0.13445453 | 0.48474744 |
| USO1 | 241.822268 | 0.290992301 | 0.17074216 | 1.70427909 | 0.08832896 | 0.40244759 |
| SLFN5 | 298.053974 | 0.291034692 | 0.27822536 | 1.04603941 | 0.29554284 | 0.65731032 |
| EZH2 | 49.1088005 | 0.291106973 | 0.30745615 | 0.94682437 | 0.34372828 | 0.69381562 |
| CLIC5 | 52.0741758 | 0.291137235 | 0.66057917 | 0.44073027 | 0.65940828 | 0.87335081 |
| ZNF280C | 47.1308762 | 0.29121341 | 0.24080127 | 1.20935163 | 0.22652778 | 0.59298684 |
| KCTD16 | 173.605669 | 0.291414671 | 1.0392969 | 0.28039598 | 0.77917372 | 0.91979327 |
| STAT1 | 254.955013 | 0.291437736 | 0.57797102 | 0.50424282 | 0.61409075 | 0.85606552 |
| JUP | 234.001948 | 0.291453915 | 0.38036876 | 0.76624042 | 0.44353326 | 0.75866068 |
| HMCES | 83.4702096 | 0.291553149 | 0.22129311 | 1.31749766 | 0.18767187 | 0.55388137 |

| | | | | | | |
|---|---|---|---|---|---|---|
| CSRNP3 | 56.0328654 | 0.291616779 | 0.60095964 | 0.48525185 | 0.6274977 | 0.86052754 |
| C15orf40 | 52.816251 | 0.291957259 | 0.26445438 | 1.10399858 | 0.26959375 | 0.63429721 |
| ARID4A | 248.111245 | 0.292063732 | 0.2937856 | 0.99413905 | 0.32015518 | 0.67434487 |
| SHLD2 | 209.883634 | 0.292115002 | 0.29526784 | 0.98932211 | 0.32250557 | 0.6761919 |
| TBP | 56.0366521 | 0.292166505 | 0.22002337 | 1.32788851 | 0.18421494 | 0.54912489 |
| USP9Y | 192.760579 | 0.292259785 | 2.31384316 | 0.12630925 | 0.89948714 | 0.96723863 |
| RTTN | 154.854349 | 0.292726267 | 0.30432214 | 0.96189605 | 0.33610182 | 0.68706501 |
| GSTA4 | 48.5140807 | 0.293154855 | 0.39727698 | 0.7379105 | 0.46056884 | 0.770424 |
| METTL25 | 118.777255 | 0.293407391 | 0.29970826 | 0.97897664 | 0.32759152 | 0.68017808 |
| ATRX | 947.43061 | 0.29356828 | 0.16914797 | 1.73557078 | 0.08263976 | 0.39477007 |
| MZT2A | 40.9507622 | 0.293624942 | 0.36430316 | 0.80599066 | 0.42024824 | 0.73901389 |
| CEP290 | 186.273877 | 0.293782092 | 0.17629603 | 1.66641358 | 0.09563107 | 0.41580714 |
| TNRC6C | 229.391404 | 0.294218977 | 0.27046861 | 1.08781192 | 0.27667815 | 0.64098811 |
| CAMK2G | 123.698846 | 0.294927695 | 0.38603989 | 0.76398243 | 0.44487772 | 0.76000717 |
| AC007364.1 | 93.1378505 | 0.295241004 | 0.22585498 | 1.30721494 | 0.19113972 | 0.55590015 |
| CDADC1 | 56.7424858 | 0.295288736 | 0.32446022 | 0.91009226 | 0.36277386 | 0.70519842 |
| LONRF1 | 146.308865 | 0.29538466 | 0.41218427 | 0.71663254 | 0.47360086 | 0.77899012 |
| EFCAB7 | 45.7301143 | 0.295451336 | 0.21211236 | 1.39290015 | 0.16364999 | 0.52357299 |
| KIAA0232 | 256.086801 | 0.295529144 | 0.2151613 | 1.3735237 | 0.1695896 | 0.5312985 |
| ORAOV1 | 50.0156032 | 0.295690548 | 0.30994702 | 0.95400352 | 0.34008186 | 0.69059971 |
| CD2AP | 408.756593 | 0.295806376 | 0.16917557 | 1.74851713 | 0.08037452 | 0.38871437 |
| INTS6L | 76.0738404 | 0.295908287 | 0.29986573 | 0.98680263 | 0.3237394 | 0.67718802 |
| RASSF1 | 134.536632 | 0.296159337 | 0.35072487 | 0.84442069 | 0.39843436 | 0.72770737 |
| SGSM2 | 67.4507072 | 0.296210079 | 0.22036818 | 1.34415996 | 0.17889667 | 0.54282239 |
| CEP112 | 935.612158 | 0.296213587 | 0.23562372 | 1.25714671 | 0.20870052 | 0.57372449 |
| ZNF346 | 133.81714 | 0.296242884 | 0.25074309 | 1.18145982 | 0.23742011 | 0.60074482 |
| APPL2 | 208.562922 | 0.296443669 | 0.23131106 | 1.28158017 | 0.19998996 | 0.56449294 |
| MFSD11 | 167.585671 | 0.29655473 | 0.25723477 | 1.15285631 | 0.24896937 | 0.61434376 |
| NCOA6 | 285.356063 | 0.296774825 | 0.17722567 | 1.67455888 | 0.09402081 | 0.41280519 |
| CWF19L2 | 255.49615 | 0.296779438 | 0.18568541 | 1.59829162 | 0.10997809 | 0.44594475 |
| SWT1 | 166.523015 | 0.296905545 | 0.29261952 | 1.0146471 | 0.31027408 | 0.6648266 |
| POLG2 | 50.4036057 | 0.296960307 | 0.24021311 | 1.23623688 | 0.21637052 | 0.58218265 |
| DECR1 | 201.20896 | 0.296993065 | 0.28568004 | 1.03960034 | 0.29852562 | 0.65856711 |
| CDKL3 | 66.6666302 | 0.297022551 | 0.36582099 | 0.81193414 | 0.41682942 | 0.73821752 |
| AFTPH | 251.798519 | 0.297329165 | 0.16893962 | 1.75997294 | 0.07841239 | 0.38566641 |
| ADD1 | 607.725973 | 0.297473948 | 0.25113568 | 1.18451485 | 0.23620932 | 0.60018541 |
| PACSIN2 | 288.773126 | 0.297588869 | 0.24253947 | 1.2269709 | 0.21983353 | 0.58687522 |
| ZNF567 | 44.1317969 | 0.297896703 | 0.30417449 | 0.97936123 | 0.32740153 | 0.68017808 |
| FBXL5 | 290.600613 | 0.297903341 | 0.16012049 | 1.86049477 | 0.06281556 | 0.35158667 |
| C11orf74 | 42.1445723 | 0.297956033 | 0.3054411 | 0.97549423 | 0.32931516 | 0.68205162 |
| TMPO | 166.437852 | 0.297991206 | 0.25213999 | 1.18184824 | 0.23726592 | 0.60074482 |
| EIF2B5 | 47.9851417 | 0.297992036 | 0.26940352 | 1.10611781 | 0.26867553 | 0.63347822 |
| ATL2 | 173.210851 | 0.298105481 | 0.14689277 | 2.02940874 | 0.04241668 | 0.291241 |
| INTS7 | 74.280414 | 0.298310339 | 0.33327876 | 0.89507756 | 0.37074563 | 0.70984233 |
| PGGT1B | 124.530053 | 0.298329218 | 0.19718984 | 1.51290359 | 0.13030415 | 0.47901117 |
| MIA2 | 235.103079 | 0.298480538 | 0.23718179 | 1.25844624 | 0.20823042 | 0.57338509 |
| TRIM23 | 80.1841656 | 0.298649714 | 0.23303302 | 1.2815768 | 0.19999114 | 0.56449294 |
| PRDM11 | 65.3865093 | 0.298829189 | 0.23114958 | 1.29279573 | 0.19608172 | 0.56102682 |
| THOC2 | 409.938171 | 0.298859431 | 0.19433778 | 1.53783497 | 0.12408897 | 0.46936415 |
| MFSD14B | 145.480226 | 0.298936886 | 0.2267813 | 1.31817256 | 0.18744589 | 0.55388137 |
| COPG2 | 75.3618851 | 0.299019846 | 0.39891531 | 0.74958227 | 0.45350633 | 0.76578691 |
| SLC27A3 | 61.5571514 | 0.299043234 | 0.30364773 | 0.98483607 | 0.3247046 | 0.67855791 |
| RNF114 | 144.17113 | 0.299085589 | 0.34651032 | 0.86313616 | 0.38806261 | 0.72105276 |
| FBXL3 | 145.025923 | 0.29925225 | 0.23717956 | 1.26171181 | 0.20705251 | 0.57162978 |
| RCBTB1 | 72.9036068 | 0.299650165 | 0.23998007 | 1.24864606 | 0.21179456 | 0.57690664 |
| KIZ | 100.283095 | 0.299746007 | 0.28105222 | 1.06651359 | 0.28619154 | 0.64883774 |
| VPS13C | 749.910061 | 0.29983525 | 0.22928188 | 1.30771453 | 0.19097015 | 0.55586607 |
| DOCK6 | 724.476768 | 0.300491919 | 0.2169494 | 1.38507838 | 0.1660285 | 0.52618561 |
| ZC3HC1 | 65.3088411 | 0.300981798 | 0.25481255 | 1.18118905 | 0.23752763 | 0.60083109 |
| BRWD1 | 590.065605 | 0.30113564 | 0.18662789 | 1.6135618 | 0.10662251 | 0.43932066 |
| NPEPPS | 431.819842 | 0.301143191 | 0.15584038 | 1.93238227 | 0.05331234 | 0.32315176 |
| MPP3 | 51.1124551 | 0.301348326 | 0.41260382 | 0.73035757 | 0.46517165 | 0.77477727 |
| RANBP9 | 318.661826 | 0.301402733 | 0.20196555 | 1.49234722 | 0.13560814 | 0.48508929 |
| SLC12A6 | 196.511666 | 0.301604826 | 0.23616676 | 1.27708416 | 0.20157254 | 0.56682752 |
| SNRNP48 | 68.2047789 | 0.301897029 | 0.2170105 | 1.39116325 | 0.16417593 | 0.52376282 |
| HEATR5A | 73.1528606 | 0.30273508 | 0.23892319 | 1.26708121 | 0.20512625 | 0.56996541 |
| ENTPD1-AS1 | 390.07211 | 0.302753701 | 0.50468144 | 0.5998907 | 0.54857908 | 0.82355262 |
| DPY19L1 | 149.664547 | 0.302907815 | 0.37699283 | 0.80348429 | 0.42169487 | 0.73954761 |
| INTS11 | 77.5876756 | 0.302971336 | 0.18865321 | 1.60596969 | 0.10828057 | 0.44369549 |
| OPHN1 | 389.849434 | 0.30325444 | 0.16897281 | 1.79469373 | 0.07270251 | 0.37296554 |
| NCKAP1 | 840.028195 | 0.303450645 | 0.21902705 | 1.38544827 | 0.16591544 | 0.5261375 |
| VEZT | 355.794923 | 0.304166031 | 0.18060652 | 1.6841365 | 0.09215529 | 0.41050994 |
| TSEN15 | 92.0083577 | 0.304414521 | 0.19840518 | 1.53430734 | 0.12495406 | 0.47038289 |
| DUSP7 | 42.5661835 | 0.30485208 | 0.51578509 | 0.59104477 | 0.55449042 | 0.82438138 |
| RB1 | 846.951771 | 0.305260223 | 0.19219164 | 1.58831166 | 0.11221588 | 0.44902815 |
| A2M | 947.953196 | 0.305411324 | 0.60804528 | 0.50228385 | 0.61546787 | 0.85714845 |
| CEP350 | 529.373088 | 0.30553958 | 0.1543445 | 1.97959481 | 0.04774908 | 0.30661402 |
| ZNF641 | 38.8426731 | 0.305596494 | 0.35123733 | 0.87005698 | 0.38426927 | 0.71699718 |
| CHD1 | 417.455376 | 0.305642056 | 0.25165704 | 1.21451818 | 0.22454991 | 0.59083342 |
| MYCT1 | 219.22543 | 0.306064762 | 0.27659833 | 1.1065315 | 0.26849654 | 0.63347822 |
| ATG12 | 102.578716 | 0.306077063 | 0.2641706 | 1.15863411 | 0.24660536 | 0.61160108 |
| FKBP15 | 124.826068 | 0.306101532 | 0.27814365 | 1.10051598 | 0.27110737 | 0.63561102 |
| ZNF32 | 44.0499513 | 0.306216509 | 0.27631668 | 1.10820856 | 0.26777176 | 0.63276201 |
| MAFK | 55.5603107 | 0.306231328 | 0.36575115 | 0.83726689 | 0.40244257 | 0.73035873 |
| GRASP | 356.624151 | 0.306271389 | 0.29042777 | 1.0545527 | 0.29162995 | 0.65416882 |
| RMC1 | 56.2609469 | 0.306699048 | 0.26692557 | 1.14900587 | 0.25055356 | 0.61583146 |
| CNTRL | 96.7963086 | 0.307083423 | 0.33208951 | 0.92470075 | 0.35512159 | 0.70018061 |
| NUBPL | 287.887153 | 0.307707109 | 0.3411638 | 0.90193365 | 0.36709212 | 0.7075153 |
| UTP6 | 80.6707271 | 0.307861035 | 0.29558436 | 1.04153357 | 0.29762798 | 0.6579563 |
| IPO8 | 160.273856 | 0.308067066 | 0.17374825 | 1.77306571 | 0.07621782 | 0.38263215 |
| SAE1 | 126.601123 | 0.308134695 | 0.25336679 | 1.21616058 | 0.22392376 | 0.59051502 |
| SETD3 | 248.622258 | 0.308271628 | 0.21525121 | 1.43214819 | 0.15210142 | 0.50814312 |
| PCGF6 | 70.0514742 | 0.308356414 | 0.21534221 | 1.43193671 | 0.15216194 | 0.50814312 |
| DPH6 | 349.743987 | 0.308366331 | 0.29488946 | 1.04570145 | 0.29569889 | 0.65731032 |
| SLC39A10 | 187.255786 | 0.308473661 | 0.58703815 | 0.52547464 | 0.59925328 | 0.84912374 |
| IQCK | 321.746785 | 0.308519093 | 0.30023997 | 1.02757503 | 0.30414978 | 0.66086613 |
| ECD | 97.5974767 | 0.308982077 | 0.27697496 | 1.11555961 | 0.26461069 | 0.63018314 |
| COMMD2 | 86.4095987 | 0.309097108 | 0.28011912 | 1.1034488 | 0.26983231 | 0.634676 |
| GPD2 | 230.286308 | 0.30912652 | 0.28433465 | 1.08719257 | 0.27695171 | 0.64110208 |
| ZNF782 | 122.570601 | 0.309186603 | 0.19363535 | 1.59674665 | 0.11032219 | 0.44631216 |
| ALDH3A2 | 65.6857477 | 0.309202992 | 0.34679857 | 0.89159247 | 0.37261141 | 0.71084822 |
| SLC27A1 | 81.5206138 | 0.309211396 | 0.27501973 | 1.12432439 | 0.26087543 | 0.62580682 |
| AL158152.2 | 124.899398 | 0.309255757 | 0.30350272 | 1.01895547 | 0.30822411 | 0.66414532 |
| TRPC1 | 215.08291 | 0.309448063 | 0.30787345 | 1.00511449 | 0.31484172 | 0.66894474 |
| TMOD1 | 110.706635 | 0.309516085 | 0.49823036 | 0.62123088 | 0.53444772 | 0.81562583 |
| FRS2 | 248.899498 | 0.309678982 | 0.2361983 | 1.31109744 | 0.18982485 | 0.55535814 |
| FEZ2 | 523.368079 | 0.310033388 | 0.41574595 | 0.74572797 | 0.45583176 | 0.76740262 |
| C2orf68 | 46.0095091 | 0.310466769 | 0.33940184 | 0.91474687 | 0.36032453 | 0.70386371 |
| CPQ | 521.157604 | 0.310601732 | 0.23183888 | 1.33973098 | 0.18033282 | 0.5444489 |
| GMPR | 68.9949244 | 0.310790904 | 0.52609898 | 0.59074607 | 0.55469057 | 0.82452915 |
| FUS | 398.153249 | 0.310932504 | 0.29425401 | 1.05668059 | 0.29065739 | 0.65333738 |
| TDP2 | 73.7047698 | 0.311009477 | 0.29470574 | 1.0553221 | 0.29127804 | 0.65401285 |
| CXCL12 | 105.5558 | 0.311174631 | 1.00476435 | 0.30969912 | 0.75678978 | 0.91345267 |

| | | | | | | |
|---|---|---|---|---|---|---|
| ANAPC4 | 105.323321 | 0.311434014 | 0.27646001 | 1.12650656 | 0.25995116 | 0.6252109 |
| RGS12 | 313.264118 | 0.31151618 | 0.28393877 | 1.0971245 | 0.27258697 | 0.63767272 |
| PEX14 | 165.624798 | 0.311654188 | 0.29480907 | 1.05713908 | 0.29044812 | 0.65333738 |
| NISCH | 134.292539 | 0.311669907 | 0.21516291 | 1.44852991 | 0.14746891 | 0.50134527 |
| RYR3 | 236.05233 | 0.311780922 | 1.08599649 | 0.28709202 | 0.77404186 | 0.91726032 |
| CAB39L | 63.8885174 | 0.311968146 | 0.23460816 | 1.32974122 | 0.18360355 | 0.54833133 |
| PKN1 | 231.404526 | 0.312616056 | 0.30019852 | 1.04136442 | 0.29770645 | 0.6579563 |
| SKA2 | 72.325301 | 0.312855232 | 0.18880255 | 1.65704983 | 0.0975094 | 0.42092574 |
| INTS12 | 101.853566 | 0.312907125 | 0.33810452 | 0.92547453 | 0.35471912 | 0.70011254 |
| KATNA1 | 55.2012664 | 0.313011838 | 0.21188436 | 1.47727677 | 0.13960146 | 0.49091481 |
| ARL5A | 103.419739 | 0.313037812 | 0.34561347 | 0.90574541 | 0.36507062 | 0.70668224 |
| FXR1 | 296.415 | 0.313131124 | 0.16901429 | 1.85269019 | 0.06392678 | 0.3533385 |
| SACM1L | 131.378872 | 0.313225879 | 0.18792838 | 1.66673006 | 0.09556809 | 0.41575423 |
| DYNC1I1 | 94.9822906 | 0.313257435 | 0.61619348 | 0.50837512 | 0.6111903 | 0.85449687 |
| RCOR3 | 255.361333 | 0.313305391 | 0.20127898 | 1.55657284 | 0.11957194 | 0.46306538 |
| SUCLG1 | 115.793363 | 0.313401566 | 0.20724254 | 1.51224531 | 0.13047147 | 0.47929621 |
| HIC2 | 46.7097096 | 0.313493152 | 0.28157953 | 1.11333787 | 0.26556335 | 0.63060082 |
| DHX29 | 135.738618 | 0.313597409 | 0.19018036 | 1.64894743 | 0.09915841 | 0.42376187 |
| NSUN2 | 95.3576423 | 0.313696403 | 0.25991634 | 1.20691297 | 0.22746566 | 0.59430125 |
| HIPK1 | 128.162795 | 0.313946441 | 0.16405336 | 1.91368497 | 0.05566042 | 0.32988018 |
| ZDHHC20 | 316.514652 | 0.314051672 | 0.22851184 | 1.37433437 | 0.1693379 | 0.5312985 |
| AGPAT4 | 248.195412 | 0.314293558 | 0.17247488 | 1.82225703 | 0.06841599 | 0.36142549 |
| JPX | 472.231469 | 0.314311522 | 0.22325084 | 1.40788504 | 0.15916511 | 0.51663947 |
| THUMPD3-A! | 168.373651 | 0.31456761 | 0.2528813 | 1.24393385 | 0.21352391 | 0.57779965 |
| CDK5RAP1 | 78.018198 | 0.314761397 | 0.27775204 | 1.13324605 | 0.25711094 | 0.62154766 |
| SAV1 | 150.208403 | 0.314855325 | 0.25547284 | 1.23244146 | 0.21778422 | 0.58393378 |
| CREB1 | 326.444872 | 0.315364466 | 0.16795895 | 1.87762818 | 0.06043205 | 0.34485038 |
| STON2 | 212.565178 | 0.315743017 | 0.3582394 | 0.88137435 | 0.37811523 | 0.71305115 |
| CALCOCO2 | 301.013977 | 0.315873432 | 0.30306407 | 1.04226617 | 0.29728829 | 0.65795352 |
| DLST | 103.86005 | 0.316188625 | 0.26849139 | 1.17764905 | 0.23893655 | 0.60290148 |
| INPP4A | 194.414205 | 0.316385251 | 0.25921374 | 1.2205574 | 0.22225364 | 0.58800568 |
| ZNF569 | 45.2699491 | 0.316912321 | 0.23763047 | 1.33363506 | 0.18232349 | 0.54730487 |
| STK17A | 82.6964026 | 0.317179376 | 0.30795621 | 1.02994959 | 0.30303367 | 0.6595012 |
| ACSS2 | 109.725043 | 0.317192839 | 0.3212751 | 0.98729355 | 0.32349875 | 0.67685765 |
| PARP16 | 43.6031787 | 0.317218638 | 0.30552965 | 1.0382581 | 0.29914991 | 0.65856711 |
| SMC6 | 172.875156 | 0.317373253 | 0.17271555 | 1.83754886 | 0.06612891 | 0.35600847 |
| YPEL2 | 176.303941 | 0.317514229 | 0.20968866 | 1.51421746 | 0.12997069 | 0.47843013 |
| PRKCI | 174.745935 | 0.317528202 | 0.18034671 | 1.76065426 | 0.07829694 | 0.38566641 |
| CSRNP2 | 54.4386205 | 0.31780503 | 0.20448903 | 1.55414219 | 0.1201505 | 0.46382142 |
| HOXB2 | 61.7130853 | 0.317815469 | 0.50807626 | 0.6255271 | 0.53162517 | 0.8142034 |
| COG4 | 87.0935675 | 0.318073941 | 0.25429334 | 1.25081507 | 0.21100196 | 0.5768065 |
| ALG10B | 41.7630164 | 0.318284369 | 0.42640617 | 0.74643471 | 0.45540486 | 0.76710126 |
| NFYC | 179.036492 | 0.318324529 | 0.16871934 | 1.88671038 | 0.05919929 | 0.34138675 |
| DTX2 | 74.5873983 | 0.318410181 | 0.34870408 | 0.91312432 | 0.36117716 | 0.70386371 |
| ZFYVE27 | 65.9472691 | 0.3184688 | 0.24126209 | 1.32001177 | 0.18683109 | 0.55312547 |
| HSPA2 | 62.7131071 | 0.318668646 | 1.02600933 | 0.3105904 | 0.75611203 | 0.91345267 |
| ZNF737 | 50.7934033 | 0.319160577 | 0.28286155 | 1.12832788 | 0.25918147 | 0.62403413 |
| KCTD15 | 87.2362716 | 0.319409146 | 0.45688224 | 0.69910608 | 0.48448574 | 0.78781193 |
| PTPN12 | 1223.81379 | 0.319426371 | 0.13450491 | 2.3748305 | 0.01755701 | 0.18854202 |
| CCNL1 | 351.547848 | 0.319661581 | 0.4237409 | 0.75437982 | 0.45062119 | 0.76375998 |
| ARSB | 199.31061 | 0.320070088 | 0.36326409 | 0.88109476 | 0.37826654 | 0.71305115 |
| EPHX2 | 39.4945335 | 0.320176618 | 0.45049181 | 0.71072684 | 0.47725352 | 0.78098272 |
| POMT1 | 43.9023584 | 0.32022681 | 0.28686894 | 1.11628263 | 0.26430117 | 0.63018314 |
| GNG2 | 148.45175 | 0.320384356 | 0.23383303 | 1.37014155 | 0.17064272 | 0.53193492 |
| LZTR1 | 44.0423214 | 0.320921117 | 0.23274368 | 1.3788607 | 0.16793771 | 0.52916221 |
| SEC24C | 79.5960847 | 0.320945969 | 0.19456091 | 1.64959119 | 0.09902658 | 0.42376187 |
| N4BP2L2 | 1266.16522 | 0.321127737 | 0.21512999 | 1.49271487 | 0.13551184 | 0.48508022 |
| FZR1 | 37.8884743 | 0.321282266 | 0.28923206 | 1.11081139 | 0.26664954 | 0.63206134 |
| RNF20 | 49.3540687 | 0.321975036 | 0.34596454 | 0.93065908 | 0.35202994 | 0.69889281 |
| MAP3K7 | 229.427217 | 0.322024122 | 0.21403193 | 1.50456112 | 0.13243695 | 0.48165847 |
| MVB12B | 135.682213 | 0.32205459 | 0.33338644 | 0.96600987 | 0.33403924 | 0.6852158 |
| SVIP | 92.4573053 | 0.322500573 | 0.25879786 | 1.24614855 | 0.21270986 | 0.57730926 |
| PTPRJ | 172.385652 | 0.322552606 | 0.46616754 | 0.69192421 | 0.48898492 | 0.78955612 |
| GDPD5 | 89.2699727 | 0.322644787 | 0.40651741 | 0.79368013 | 0.42738167 | 0.74309294 |
| PIK3CB | 253.33075 | 0.322692372 | 0.23754314 | 1.35845797 | 0.17431841 | 0.53666198 |
| TRAK2 | 173.912281 | 0.322716127 | 0.23993913 | 1.34499164 | 0.17862793 | 0.54258069 |
| LRCH3 | 453.976368 | 0.323091431 | 0.20518083 | 1.57466676 | 0.11533338 | 0.4568354 |
| DNAJB1 | 78.628017 | 0.323638362 | 0.70362006 | 0.45996182 | 0.64554363 | 0.86572246 |
| SNIP1 | 56.564562 | 0.323847684 | 0.33521059 | 0.96610218 | 0.33399305 | 0.6852158 |
| RBM25 | 346.054529 | 0.323853451 | 0.23031075 | 1.40615863 | 0.15967703 | 0.51768507 |
| IDH3G | 39.8778289 | 0.323978264 | 0.25442544 | 1.27337213 | 0.20288602 | 0.56730017 |
| CYTH2 | 61.5118098 | 0.324599412 | 0.3138907 | 1.03411607 | 0.30108191 | 0.65926326 |
| HDAC6 | 47.566527 | 0.324697872 | 0.20717662 | 1.56725151 | 0.11705591 | 0.45985048 |
| ZNF280D | 229.044824 | 0.325024867 | 0.28291919 | 1.14882582 | 0.25062781 | 0.61583146 |
| CHD9 | 1080.6236 | 0.325365014 | 0.18940043 | 1.71786837 | 0.08582063 | 0.39833817 |
| SMIM8 | 93.6962922 | 0.325700017 | 0.29920707 | 1.08854386 | 0.27635509 | 0.64068436 |
| AC027808.1 | 49.6760044 | 0.325715058 | 0.47239701 | 0.68949432 | 0.49051225 | 0.79073252 |
| ATXN2 | 697.076721 | 0.325845002 | 0.18771441 | 1.73585505 | 0.08258947 | 0.39477007 |
| USP47 | 433.799718 | 0.326315746 | 0.15834784 | 2.06075272 | 0.03932664 | 0.28057468 |
| CREBRF | 273.917279 | 0.326699891 | 0.34933689 | 0.93520009 | 0.34968519 | 0.69858573 |
| RAD9A | 122.274636 | 0.326948341 | 0.248521 | 1.31557629 | 0.1883163 | 0.55388137 |
| SUZ12 | 290.666563 | 0.327089583 | 0.21922672 | 1.4920151 | 0.13569519 | 0.48509992 |
| FUBP1 | 306.472114 | 0.327629614 | 0.19434481 | 1.68581615 | 0.09183122 | 0.40973546 |
| ST7L | 191.587959 | 0.327657369 | 0.19695724 | 1.66359645 | 0.0961931 | 0.41727618 |
| BCL2L13 | 168.275717 | 0.327845324 | 0.34916909 | 0.93892996 | 0.3477667 | 0.696616 |
| UPF3A | 332.483185 | 0.328476075 | 0.19673262 | 1.66965742 | 0.09498716 | 0.41466145 |
| TMLHE-AS1 | 70.7222913 | 0.32854484 | 0.40923736 | 0.80282221 | 0.4220775 | 0.73995396 |
| UBA3 | 143.399769 | 0.328549892 | 0.14955247 | 2.19688707 | 0.02802851 | 0.23214303 |
| PRKCH | 1779.3818 | 0.328584356 | 0.186979 | 1.75733293 | 0.07886108 | 0.38686531 |
| CYFIP1 | 417.305429 | 0.328650206 | 0.24709031 | 1.33008133 | 0.18349147 | 0.54819669 |
| SRBD1 | 365.95101 | 0.32867502 | 0.2232019 | 1.47254577 | 0.14087356 | 0.49249381 |
| EMC2 | 145.743404 | 0.328683646 | 0.23925521 | 1.37377844 | 0.16951048 | 0.5312985 |
| RNF185 | 110.798085 | 0.32871933 | 0.25114891 | 1.30886225 | 0.19058101 | 0.55558405 |
| CATSPER2 | 73.0038661 | 0.328937575 | 0.54511481 | 0.60342806 | 0.54622396 | 0.82355262 |
| KPNA5 | 139.534897 | 0.328993278 | 0.21291257 | 1.54520365 | 0.12229701 | 0.46748939 |
| PPP2R5B | 59.5707053 | 0.329288196 | 0.24403928 | 1.34932459 | 0.17723273 | 0.54037749 |
| CNST | 388.542075 | 0.32934137 | 0.35312833 | 0.93263933 | 0.35100622 | 0.69858573 |
| ZNF532 | 341.186658 | 0.329362792 | 0.30652622 | 1.07450119 | 0.28259811 | 0.64576944 |
| GTPBP2 | 71.528117 | 0.32967703 | 0.34951188 | 0.94324986 | 0.34555312 | 0.69497757 |
| ZXDC | 176.261123 | 0.329688337 | 0.28097935 | 1.17335435 | 0.24065374 | 0.6051624 |
| IL27RA | 40.548032 | 0.329806705 | 0.40756605 | 0.80921044 | 0.41839411 | 0.73821752 |
| ALAS1 | 42.852444 | 0.329985529 | 0.3490155 | 0.94547529 | 0.34441628 | 0.69400602 |
| USP54 | 96.9916478 | 0.330053854 | 0.33102058 | 0.99707957 | 0.31872589 | 0.67376232 |
| WDTC1 | 96.1845699 | 0.330138438 | 0.18871973 | 1.74935838 | 0.08022909 | 0.38871437 |
| CALCRL | 1685.33027 | 0.330639728 | 0.47546077 | 0.69540906 | 0.48679898 | 0.78826504 |
| VWA8 | 322.545279 | 0.330695838 | 0.2656734 | 1.24474576 | 0.21322522 | 0.57758664 |
| ACTR10 | 160.456208 | 0.330783322 | 0.14160515 | 2.3359555 | 0.01949356 | 0.19632601 |
| SFXN3 | 41.9598001 | 0.330923717 | 0.31580214 | 1.04788307 | 0.29469248 | 0.65671864 |
| ATP13A2 | 67.2358521 | 0.331146424 | 0.35618593 | 0.92970101 | 0.35252591 | 0.69949552 |
| SLC37A1 | 77.2278731 | 0.33122748 | 0.35142106 | 0.94253737 | 0.34591759 | 0.69497757 |
| TMEM68 | 75.1134018 | 0.331336701 | 0.27117715 | 1.22184594 | 0.22176589 | 0.58750144 |
| UAP1 | 58.6507985 | 0.331436951 | 0.31239623 | 1.06095054 | 0.28871238 | 0.65160592 |
| ARHGAP31 | 1256.13055 | 0.331533675 | 0.22392782 | 1.4805381 | 0.1387297 | 0.49037803 |

| | | | | | | |
|---|---|---|---|---|---|---|
| LUC7L | 261.470318 | 0.331827596 | 0.19227823 | 1.72576793 | 0.08438917 | 0.39616527 |
| AC245297.3 | 89.6081264 | 0.331999848 | 0.37600341 | 0.88297031 | 0.37725231 | 0.71261673 |
| AC079380.1 | 87.3187676 | 0.332079608 | 0.58197111 | 0.57061184 | 0.56826279 | 0.83354423 |
| USF3 | 111.599509 | 0.33210658 | 0.19117418 | 1.73719369 | 0.08235299 | 0.39457252 |
| CBFB | 362.156634 | 0.332241617 | 0.27182918 | 1.22224411 | 0.22161533 | 0.58750144 |
| PSPC1 | 443.322504 | 0.332276584 | 0.20487452 | 1.62185412 | 0.10483459 | 0.43679299 |
| CHIC1 | 170.228472 | 0.333565601 | 0.33586182 | 0.99316321 | 0.32063042 | 0.67465126 |
| TNPO3 | 241.239517 | 0.333568861 | 0.15754393 | 2.11730697 | 0.0342338 | 0.25808697 |
| C11orf80 | 62.3552169 | 0.333710025 | 0.38011162 | 0.87792639 | 0.37998367 | 0.71447756 |
| KCTD9 | 137.463928 | 0.334063622 | 0.28093181 | 1.18912706 | 0.23438967 | 0.59949266 |
| PTPRM | 6370.50185 | 0.33428615 | 0.23327942 | 1.43298603 | 0.15186183 | 0.50784035 |
| TBC1D5 | 1635.21311 | 0.334513816 | 0.23324991 | 1.43414339 | 0.15153135 | 0.50771689 |
| MICA | 68.3465709 | 0.334588515 | 0.29755155 | 1.12447245 | 0.26081265 | 0.62580682 |
| MICU1 | 376.598011 | 0.334622863 | 0.25496727 | 1.31241498 | 0.18938016 | 0.55524823 |
| CCS | 62.6518853 | 0.33468053 | 0.29179471 | 1.14697259 | 0.25139295 | 0.61661694 |
| CRIM1 | 1530.91326 | 0.335011941 | 0.48451503 | 0.69143767 | 0.48929054 | 0.78971686 |
| PATJ | 230.678572 | 0.335457914 | 0.32922134 | 1.0189434 | 0.30822984 | 0.66414532 |
| VSIR | 77.9046486 | 0.335694731 | 0.50293389 | 0.66747288 | 0.50447013 | 0.79776858 |
| POLG | 60.9814633 | 0.335767943 | 0.24441312 | 1.37377217 | 0.16951242 | 0.5312985 |
| EIF4E3 | 58.5959002 | 0.335778674 | 0.417901 | 0.80348857 | 0.4216924 | 0.73954761 |
| WDR33 | 334.061521 | 0.336238919 | 0.18051721 | 1.86264186 | 0.06251267 | 0.35061082 |
| CDK14 | 299.493257 | 0.336537904 | 0.34691032 | 0.97010058 | 0.33199636 | 0.68483141 |
| TPD52L1 | 249.050773 | 0.337393947 | 0.92958779 | 0.36295006 | 0.71664219 | 0.9001716 |
| GPBP1 | 561.340634 | 0.337421104 | 0.26453141 | 1.27554267 | 0.20211723 | 0.56693091 |
| LARP4B | 463.625957 | 0.337444238 | 0.26845114 | 1.25700429 | 0.20875208 | 0.57372449 |
| ZNF148 | 674.022846 | 0.3376086 | 0.14149045 | 2.38608762 | 0.01702869 | 0.18593697 |
| NAPB | 109.128278 | 0.338030198 | 0.2548923 | 1.32616872 | 0.18478382 | 0.5500631 |
| SLC9A9 | 233.627249 | 0.338185237 | 0.3945552 | 0.85713034 | 0.39137285 | 0.72247779 |
| FAM182B | 65.558284 | 0.338279304 | 0.41795312 | 0.8093714 | 0.41830155 | 0.73821752 |
| ACAT1 | 133.37603 | 0.338405325 | 0.23238068 | 1.45625416 | 0.14532237 | 0.49826376 |
| ZNF25 | 62.8009513 | 0.338827699 | 0.24471494 | 1.38458118 | 0.16618057 | 0.52646365 |
| SYBU | 71.2518797 | 0.339008547 | 0.37395631 | 0.90654588 | 0.36464699 | 0.70668224 |
| THAP4 | 61.2412233 | 0.339132968 | 0.2353144 | 1.44119086 | 0.14953077 | 0.50333619 |
| CHORDC1 | 95.8475413 | 0.339256723 | 0.39394006 | 0.86118869 | 0.38913413 | 0.72202943 |
| RCOR1 | 244.778133 | 0.339341726 | 0.22147395 | 1.53219705 | 0.12547382 | 0.47103973 |
| MRPL48 | 54.9187019 | 0.339743323 | 0.24050454 | 1.41262746 | 0.15776529 | 0.51454498 |
| ETAA1 | 43.2453069 | 0.33975982 | 0.31554311 | 1.07674612 | 0.2815937 | 0.64471757 |
| C16orf70 | 42.5539902 | 0.339826945 | 0.25388618 | 1.33850115 | 0.18073312 | 0.54513054 |
| PTPN9 | 226.261984 | 0.340023546 | 0.19672698 | 1.72840325 | 0.08391595 | 0.39616527 |
| MFSD14C | 302.169312 | 0.340110478 | 0.26802962 | 1.26892872 | 0.20446649 | 0.56929203 |
| RBCK1 | 139.88601 | 0.340232677 | 0.27269049 | 1.24768808 | 0.21214531 | 0.57690664 |
| RPS6KB1 | 110.124877 | 0.340283318 | 0.28800854 | 1.18150425 | 0.23740247 | 0.60074482 |
| RIC8B | 163.246351 | 0.340338731 | 0.17693647 | 1.92350804 | 0.05441628 | 0.32727552 |
| NME7 | 207.988489 | 0.340412446 | 0.22450165 | 1.51630262 | 0.12944285 | 0.47777665 |
| S100PBP | 184.028354 | 0.340457131 | 0.23812446 | 1.42974449 | 0.15279037 | 0.50900346 |
| ING5 | 66.7171288 | 0.340944473 | 0.35632651 | 0.95683161 | 0.33865225 | 0.68935109 |
| ZNF587 | 53.7235419 | 0.341156869 | 0.30865218 | 1.1053117 | 0.26902455 | 0.63347822 |
| SP4 | 89.2865109 | 0.342304375 | 0.2365742 | 1.44692182 | 0.14791882 | 0.50143858 |
| ZMAT1 | 91.7121132 | 0.342359402 | 0.34599008 | 0.98950641 | 0.32241544 | 0.6761919 |
| CKB | 47.3504445 | 0.342948192 | 0.47330707 | 0.72457864 | 0.46871057 | 0.77719525 |
| BMP6 | 283.722844 | 0.342948361 | 0.51247309 | 0.66920267 | 0.5033662 | 0.79776858 |
| ZC3H13 | 246.376376 | 0.34295339 | 0.177399 | 1.93323179 | 0.05320765 | 0.32275626 |
| AP002812.2 | 57.0756119 | 0.343047242 | 0.30160044 | 1.13742289 | 0.25536154 | 0.6208774 |
| USP12 | 253.943717 | 0.343179604 | 0.25451943 | 1.34834342 | 0.17754795 | 0.54070521 |
| GLS | 835.606866 | 0.343195482 | 0.28774743 | 1.19269695 | 0.23298808 | 0.59910683 |
| RNF14 | 86.9657734 | 0.343422788 | 0.19049898 | 1.80275394 | 0.07142687 | 0.36926912 |
| LIN52 | 138.241443 | 0.343633879 | 0.28276482 | 1.21526392 | 0.22426545 | 0.59083342 |
| APBB2 | 2013.92466 | 0.343950322 | 0.22245656 | 1.54614603 | 0.1220693 | 0.46733953 |
| DPH7 | 51.0213333 | 0.344229291 | 0.24094279 | 1.42867647 | 0.15309725 | 0.50905923 |
| TFEB | 66.3088482 | 0.344333057 | 0.37774205 | 0.91155606 | 0.36200246 | 0.70463038 |
| WDR11 | 206.108056 | 0.344627758 | 0.16243548 | 2.12162865 | 0.03386893 | 0.25638251 |
| FBXO7 | 170.854796 | 0.344647424 | 0.21637273 | 1.59284133 | 0.11119579 | 0.44768463 |
| CDC23 | 45.0706874 | 0.344952626 | 0.22961969 | 1.50227806 | 0.13302531 | 0.48236854 |
| FAM117B | 88.3683229 | 0.345012469 | 0.41054226 | 0.84038235 | 0.40069404 | 0.72864885 |
| ELF1 | 639.127419 | 0.345193028 | 0.27838114 | 1.24000148 | 0.21497485 | 0.57982196 |
| SH2B3 | 349.273651 | 0.345415002 | 0.25378126 | 1.36107369 | 0.1734904 | 0.53536344 |
| FAM76A | 58.3662621 | 0.34551705 | 0.32968207 | 1.04803106 | 0.2946243 | 0.65671864 |
| MSL2 | 127.150611 | 0.345521493 | 0.27161525 | 1.27209902 | 0.20333794 | 0.567308 |
| CD40 | 78.1351261 | 0.345592367 | 0.31817588 | 1.0861677 | 0.27740481 | 0.64148182 |
| GTPBP1 | 81.1116085 | 0.34563122 | 0.45424883 | 0.76088521 | 0.44672563 | 0.76094002 |
| WRB | 87.6481284 | 0.345779943 | 0.24354703 | 1.41976664 | 0.15567563 | 0.51180943 |
| HECTD2 | 154.814402 | 0.345932895 | 0.23908081 | 1.44692872 | 0.14791689 | 0.50143858 |
| EML4 | 341.820224 | 0.346043045 | 0.25308676 | 1.36729017 | 0.17153437 | 0.53351036 |
| TTC39B | 149.352226 | 0.346240147 | 0.2177991 | 1.58972257 | 0.11189735 | 0.44839019 |
| DGAT1 | 44.5283059 | 0.346359132 | 0.30771558 | 1.12558206 | 0.26034247 | 0.62580682 |
| ITSN2 | 777.820278 | 0.346442382 | 0.14848216 | 2.3332257 | 0.0196363 | 0.1969165 |
| REV1 | 272.000108 | 0.346565873 | 0.18928846 | 1.83088752 | 0.06711733 | 0.35896806 |
| HS6ST1 | 141.850284 | 0.347022223 | 0.35247807 | 0.98452145 | 0.32485919 | 0.67866294 |
| GPALPP1 | 111.327428 | 0.34711687 | 0.18059207 | 1.92210474 | 0.05459258 | 0.32727552 |
| ABI2 | 308.964176 | 0.347229628 | 0.19150325 | 1.81317882 | 0.06980425 | 0.3652226 |
| MBNL1 | 2190.24357 | 0.347235177 | 0.28982679 | 1.19807828 | 0.23088654 | 0.59732678 |
| HLA-DMB | 64.5646833 | 0.347407915 | 0.45573217 | 0.7623072 | 0.44587668 | 0.76074282 |
| PHKB | 587.680224 | 0.347691779 | 0.19688697 | 1.76594609 | 0.07740489 | 0.38411416 |
| DNAH1 | 50.4448673 | 0.34776447 | 0.34970321 | 0.99445604 | 0.3200009 | 0.67434487 |
| L3MBTL4 | 391.264476 | 0.347987453 | 0.39615126 | 0.87842066 | 0.37971547 | 0.71418244 |
| HCFC2 | 119.440876 | 0.34799384 | 0.20546988 | 1.69364894 | 0.09033201 | 0.4068172 |
| WWP2 | 256.35105 | 0.348134162 | 0.26093515 | 1.33417887 | 0.18214524 | 0.54717126 |
| ZNF701 | 52.0322803 | 0.34826255 | 0.25589852 | 1.36094007 | 0.17353263 | 0.53536344 |
| BRAF | 871.989744 | 0.348287961 | 0.20531381 | 1.69636891 | 0.08981604 | 0.40583359 |
| PNISR | 577.832908 | 0.348377921 | 0.20536179 | 1.69641059 | 0.08980815 | 0.40583359 |
| SRSF2 | 198.774722 | 0.348438402 | 0.27007835 | 1.29013821 | 0.19700268 | 0.56189365 |
| ARMH4 | 118.330172 | 0.348529673 | 0.2544828 | 1.36956082 | 0.17082404 | 0.53193492 |
| U2SURP | 307.059508 | 0.348733237 | 0.18546582 | 1.88031003 | 0.06006584 | 0.34442742 |
| TTBK2 | 294.123017 | 0.348838203 | 0.18269353 | 1.90941739 | 0.05620827 | 0.33113914 |
| ATP2B1 | 231.523061 | 0.349241734 | 0.30635427 | 1.13999304 | 0.2542892 | 0.61966901 |
| FXR2 | 38.8953403 | 0.349870026 | 0.30367153 | 1.1521331 | 0.24926638 | 0.61438157 |
| ZNF680 | 130.838835 | 0.350106875 | 0.30424104 | 1.15075493 | 0.24983307 | 0.61500658 |
| SRSF11 | 496.0315 | 0.35019889 | 0.1475869 | 2.37283175 | 0.0176523 | 0.1890691 |
| ZNF438 | 559.026093 | 0.350227968 | 0.29621079 | 1.18236061 | 0.23706264 | 0.60074482 |
| RALGAPB | 315.172065 | 0.350330402 | 0.16286815 | 2.15100622 | 0.03147571 | 0.24759838 |
| ZFAND2B | 39.6621557 | 0.35061664 | 0.32137973 | 1.09097308 | 0.27528473 | 0.64017771 |
| SNTB2 | 196.123766 | 0.35067109 | 0.29404884 | 1.19256068 | 0.23304147 | 0.59910683 |
| NEDD4L | 443.269228 | 0.350841773 | 0.39051015 | 0.8984191 | 0.36896216 | 0.70940257 |
| BTRC | 321.771211 | 0.35097712 | 0.17359655 | 2.02179783 | 0.04319724 | 0.2945692 |
| COX17 | 110.410045 | 0.351222693 | 0.34492395 | 1.01826126 | 0.30855381 | 0.66414532 |
| CPLANE1 | 275.548586 | 0.351280171 | 0.15327621 | 2.29181148 | 0.02191652 | 0.20902438 |
| FZD4 | 322.896196 | 0.351357826 | 0.31654822 | 1.10996621 | 0.26701359 | 0.63222575 |
| ABHD18 | 240.119092 | 0.351379764 | 0.17789891 | 1.97516536 | 0.04824939 | 0.30798222 |
| DPY19L4 | 154.589033 | 0.351458817 | 0.13733934 | 2.55905423 | 0.01049574 | 0.14485284 |
| KIAA2026 | 460.411677 | 0.351562056 | 0.15329275 | 2.29340309 | 0.02182481 | 0.20902438 |
| MFSD8 | 90.4866975 | 0.351674862 | 0.22661254 | 1.55187732 | 0.12069158 | 0.46476198 |
| TAF3 | 373.209549 | 0.352446297 | 0.2262608 | 1.55769933 | 0.11930454 | 0.46290743 |
| AP1M1 | 65.1907183 | 0.352727206 | 0.21318105 | 1.65458984 | 0.09800773 | 0.42165994 |

| | | | | | | |
|---|---|---|---|---|---|---|
| PIP4K2B | 94.7076015 | 0.352795669 | 0.25040388 | 1.40890653 | 0.15886281 | 0.51606763 |
| STRIP1 | 68.5051265 | 0.352923592 | 0.20136261 | 1.75267689 | 0.07965748 | 0.38738328 |
| GRAMD1C | 115.257887 | 0.352951669 | 0.28833493 | 1.22410306 | 0.22091335 | 0.58706366 |
| KLHL18 | 77.1239143 | 0.353342964 | 0.261536 | 1.35102992 | 0.17668585 | 0.54037749 |
| SOS2 | 491.98762 | 0.353545076 | 0.17648676 | 2.00323854 | 0.04515169 | 0.29989958 |
| WDR20 | 247.433724 | 0.353586234 | 0.26413317 | 1.33866652 | 0.18067926 | 0.54513054 |
| INO80E | 52.4944442 | 0.354019034 | 0.24042473 | 1.47247345 | 0.14089307 | 0.49249381 |
| PDE4A | 41.7120577 | 0.354183168 | 0.36164128 | 0.97937705 | 0.32739371 | 0.68017808 |
| STXBP3 | 180.036322 | 0.354224621 | 0.20869873 | 1.69730125 | 0.08963972 | 0.40583359 |
| ZMYND11 | 408.758345 | 0.354356642 | 0.1960842 | 1.80716567 | 0.07073645 | 0.36751516 |
| PLD2 | 45.4994761 | 0.354552767 | 0.30876323 | 1.14829984 | 0.25084481 | 0.6158605 |
| LSM14A | 506.862013 | 0.355010011 | 0.14718127 | 2.41205974 | 0.01586268 | 0.18072864 |
| NCKAP5L | 56.659824 | 0.355049688 | 0.22691327 | 1.56469336 | 0.11765483 | 0.46042456 |
| ALPL | 87.069048 | 0.35524188 | 0.63567177 | 0.55884483 | 0.57626763 | 0.83550638 |
| PRKAB2 | 40.4735685 | 0.35527984 | 0.24518261 | 1.44904178 | 0.14732592 | 0.50127566 |
| ATG2B | 204.249223 | 0.355354963 | 0.2146482 | 1.65552266 | 0.09781852 | 0.42128893 |
| LRMP | 83.9681101 | 0.355823434 | 0.53039057 | 0.67087059 | 0.50230297 | 0.79691777 |
| CCDC186 | 113.264543 | 0.355843157 | 0.25570313 | 1.39162614 | 0.16403564 | 0.52357299 |
| BMI1 | 45.9391853 | 0.356007938 | 0.4480066 | 0.79464887 | 0.42681778 | 0.74280091 |
| FNTA | 89.6322821 | 0.356127502 | 0.19846141 | 1.79444207 | 0.07274264 | 0.37296554 |
| PLEKHM3 | 174.339301 | 0.356392348 | 0.26652882 | 1.33716252 | 0.18116959 | 0.54524118 |
| HMGN3 | 217.672115 | 0.356765578 | 0.35313532 | 1.01028008 | 0.31236112 | 0.66633239 |
| AC074386.1 | 49.3084707 | 0.356815054 | 0.33174691 | 1.07556406 | 0.28212227 | 0.64522262 |
| RDX | 827.976512 | 0.356961463 | 0.30420207 | 1.17343534 | 0.24062127 | 0.6051624 |
| CENPC | 198.977114 | 0.356977886 | 0.24092832 | 1.48167673 | 0.13842633 | 0.48991657 |
| UQCRC1 | 47.3141505 | 0.357159952 | 0.28280334 | 1.26292693 | 0.20661544 | 0.57138701 |
| BAZ1B | 238.540765 | 0.357181088 | 0.1991985 | 1.79309126 | 0.07295833 | 0.37361976 |
| MTMR3 | 354.351178 | 0.357306974 | 0.1972312 | 1.81161483 | 0.07004573 | 0.36578445 |
| MUC20-OT1 | 431.602816 | 0.35750211 | 0.35105165 | 1.01837469 | 0.30849993 | 0.66414532 |
| FBXL4 | 228.843442 | 0.357957844 | 0.25713082 | 1.39212345 | 0.16388502 | 0.52357299 |
| WDR41 | 131.355205 | 0.358207883 | 0.23696055 | 1.51167729 | 0.13061598 | 0.47929621 |
| SLC2A9 | 166.356266 | 0.358386892 | 0.49974666 | 0.71713714 | 0.47328948 | 0.77899012 |
| ZNF44 | 219.563734 | 0.358446914 | 0.27852191 | 1.2869613 | 0.19810778 | 0.56327864 |
| ULK1 | 89.8217615 | 0.358813973 | 0.25828441 | 1.38922043 | 0.16476573 | 0.52480753 |
| FBXL2 | 66.4425813 | 0.358864017 | 0.3523793 | 1.01840267 | 0.30848663 | 0.66414532 |
| LCLAT1 | 208.708767 | 0.359377207 | 0.27429044 | 1.31020684 | 0.19012587 | 0.55552647 |
| PSMB9 | 166.125745 | 0.35991313 | 0.42705007 | 0.84278906 | 0.39934643 | 0.72820151 |
| MRNIP | 61.8414363 | 0.359936574 | 0.29401808 | 1.22419877 | 0.22087725 | 0.58706366 |
| AP002495.2 | 88.8276549 | 0.359993721 | 0.27160955 | 1.32540894 | 0.18503555 | 0.55006698 |
| VIRMA | 138.287244 | 0.360245126 | 0.18757791 | 1.9205093 | 0.0547936 | 0.32800002 |
| CRTC2 | 41.7636596 | 0.360260666 | 0.27231881 | 1.32293713 | 0.18585629 | 0.55183672 |
| CCNYL1 | 145.472033 | 0.360681104 | 0.22878146 | 1.57653118 | 0.11490344 | 0.45577064 |
| USP15 | 586.973137 | 0.360791424 | 0.26615095 | 1.35558949 | 0.17522982 | 0.53845498 |
| PPP1R13B | 884.623413 | 0.360915683 | 0.27581334 | 1.30855051 | 0.19068666 | 0.55566466 |
| SMAP1 | 352.326923 | 0.361009988 | 0.24179633 | 1.49303337 | 0.13542845 | 0.48508022 |
| DMXL1 | 361.973704 | 0.361301004 | 0.16014165 | 2.25613395 | 0.02406224 | 0.21636008 |
| RXRB | 39.5827242 | 0.36149892 | 0.26370037 | 1.37086999 | 0.17041549 | 0.53193492 |
| FAM45A | 49.621871 | 0.361635329 | 0.2157946 | 1.67583126 | 0.09377125 | 0.41280519 |
| ITGA6 | 455.272714 | 0.362281782 | 0.37770118 | 0.95917568 | 0.33747025 | 0.68826907 |
| ULK4 | 375.892127 | 0.362631336 | 0.32006314 | 1.13299936 | 0.25721452 | 0.62161443 |
| TFDP2 | 325.621147 | 0.362782508 | 0.25484811 | 1.42352445 | 0.15458418 | 0.51037454 |
| GBA2 | 89.1981472 | 0.363444249 | 0.29904007 | 1.21536973 | 0.22422511 | 0.59083342 |
| AC068724.3 | 63.8347429 | 0.36355063 | 0.44755305 | 0.81230735 | 0.4166153 | 0.73807382 |
| KIF27 | 86.8606502 | 0.363916166 | 0.29275016 | 1.24309467 | 0.21383295 | 0.57825348 |
| LRRC28 | 176.189532 | 0.363917822 | 0.1978984 | 1.83891242 | 0.06592807 | 0.35579065 |
| SLC35G1 | 45.1739447 | 0.364117999 | 0.46144572 | 0.78908089 | 0.43006473 | 0.74420645 |
| EFNB2 | 1105.56024 | 0.364387415 | 0.40014788 | 0.91063187 | 0.36248937 | 0.70497203 |
| ARHGEF3 | 1255.79887 | 0.364545391 | 0.32407038 | 1.12489574 | 0.26063322 | 0.62580682 |
| LEMD3 | 212.164771 | 0.364578584 | 0.15906171 | 2.29205752 | 0.02190232 | 0.20902438 |
| CRIP2 | 446.765554 | 0.3646425 | 0.52924981 | 0.68897993 | 0.4908359 | 0.79073252 |
| SPAST | 159.163419 | 0.364648706 | 0.21069376 | 1.73070484 | 0.08350442 | 0.3954379 |
| AP001528.2 | 210.774012 | 0.364662363 | 0.55306074 | 0.65935319 | 0.509669 | 0.80142609 |
| NRP1 | 1012.65222 | 0.364794431 | 0.4854295 | 0.75148797 | 0.45235904 | 0.76480455 |
| FAM76B | 90.6812767 | 0.364959823 | 0.24618999 | 1.4824316 | 0.13822549 | 0.48991657 |
| PPIP5K2 | 253.916808 | 0.365167976 | 0.17058996 | 2.14061821 | 0.03230484 | 0.25128374 |
| UBE2H | 680.198307 | 0.365390389 | 0.25571889 | 1.42887525 | 0.15304009 | 0.50905923 |
| AASS | 115.506272 | 0.365479002 | 0.29598812 | 1.23477593 | 0.21691391 | 0.58245153 |
| ANKRD46 | 62.2158278 | 0.365796137 | 0.20494503 | 1.78485001 | 0.07428565 | 0.37733052 |
| ACSL3 | 299.612158 | 0.366148442 | 0.19399818 | 1.88738078 | 0.05910913 | 0.34110719 |
| KLF3 | 364.845071 | 0.366427682 | 0.24774902 | 1.47902778 | 0.13913289 | 0.49082099 |
| NFATC3 | 340.306901 | 0.366530499 | 0.27976913 | 1.31011775 | 0.19015601 | 0.55552647 |
| NUP85 | 81.466805 | 0.366669856 | 0.23018108 | 1.59296265 | 0.11116857 | 0.44768463 |
| LATS2 | 222.059357 | 0.366759208 | 0.23004953 | 1.59426193 | 0.11087738 | 0.44739133 |
| PKN2 | 579.463196 | 0.366968552 | 0.12465599 | 2.94385023 | 0.00324157 | 0.07557195 |
| AC005154.6 | 91.4621511 | 0.36701085 | 0.30962641 | 1.18533446 | 0.23588523 | 0.60018541 |
| HDGFL2 | 46.6992206 | 0.367054245 | 0.24049991 | 1.52621363 | 0.12695667 | 0.47383808 |
| HIP1 | 1009.78377 | 0.367353875 | 0.31688447 | 1.15926753 | 0.24634715 | 0.61160108 |
| MAD1L1 | 166.803663 | 0.367476592 | 0.31853955 | 1.15362942 | 0.24865213 | 0.61416251 |
| AC015813.2 | 50.5061175 | 0.36748082 | 0.24178604 | 1.51985954 | 0.12854628 | 0.47653642 |
| CLCN5 | 140.31215 | 0.367520601 | 0.30309292 | 1.21256744 | 0.22529524 | 0.59146326 |
| LEO1 | 41.857276 | 0.367579222 | 0.22882127 | 1.60640322 | 0.10818534 | 0.44369549 |
| UGDH-AS1 | 74.5110308 | 0.367833555 | 0.29192757 | 1.26001651 | 0.20766341 | 0.57254369 |
| C1D | 92.2502853 | 0.367848033 | 0.22542827 | 1.63177422 | 0.10272706 | 0.43263794 |
| PNPT1 | 173.149778 | 0.368182447 | 0.45176742 | 0.81498229 | 0.41508245 | 0.73695372 |
| TANGO2 | 52.024928 | 0.368382716 | 0.28875829 | 1.27574769 | 0.20204472 | 0.56693091 |
| POLK | 370.386403 | 0.36880576 | 0.22875822 | 1.61220767 | 0.10691676 | 0.43964815 |
| TBC1D32 | 155.947317 | 0.369126924 | 0.33815681 | 1.09158505 | 0.27501553 | 0.64017771 |
| AC025569.1 | 122.335109 | 0.369181815 | 0.59610238 | 0.61932619 | 0.53570149 | 0.81647333 |
| DACH1 | 394.841205 | 0.369185203 | 0.55207503 | 0.66872288 | 0.50367227 | 0.79776858 |
| SIRT5 | 75.8320748 | 0.369263063 | 0.31935221 | 1.15628781 | 0.24756345 | 0.61258293 |
| KANK2 | 131.463544 | 0.369324514 | 0.34411223 | 1.07326762 | 0.28315106 | 0.64649138 |
| UBTF | 92.4775722 | 0.369547715 | 0.27555202 | 1.34111777 | 0.17988222 | 0.54382715 |
| DNAL1 | 50.956484 | 0.369567038 | 0.27691685 | 1.33457766 | 0.18201461 | 0.54697965 |
| LIN54 | 88.8292328 | 0.370150774 | 0.28275421 | 1.30909023 | 0.19050379 | 0.55558405 |
| SMCO4 | 55.7185999 | 0.370339543 | 0.40896446 | 0.90555434 | 0.36517179 | 0.70668224 |
| SGPP1 | 88.5462616 | 0.370802125 | 0.25130992 | 1.47547748 | 0.14008422 | 0.49118733 |
| NBPF15 | 187.93481 | 0.371166616 | 0.39468362 | 0.94041557 | 0.34700444 | 0.6957945 |
| SLAIN2 | 222.482786 | 0.371240808 | 0.19946468 | 1.86118569 | 0.06271796 | 0.35128067 |
| RNF38 | 570.392814 | 0.372108551 | 0.16404263 | 2.26836494 | 0.02330697 | 0.2129727 |
| CCDC88C | 172.388361 | 0.372254009 | 0.46380202 | 0.80261403 | 0.42219785 | 0.73995396 |
| SMARCA2 | 638.721786 | 0.372275922 | 0.28453197 | 1.30837993 | 0.19074448 | 0.55566466 |
| ZC3H14 | 181.42702 | 0.373289129 | 0.1460724 | 2.55550755 | 0.01060331 | 0.14485284 |
| ABCB7 | 152.190161 | 0.373503291 | 0.17361529 | 2.15132712 | 0.03145039 | 0.24759838 |
| PKD1 | 82.5958104 | 0.373554461 | 0.23118189 | 1.61584656 | 0.10612749 | 0.43901385 |
| ARGLU1 | 1053.60212 | 0.373628888 | 0.25390119 | 1.47155235 | 0.1411418 | 0.49265411 |
| CELF1 | 578.60055 | 0.3736535 | 0.21166535 | 1.7653031 | 0.07751284 | 0.3844167 |
| KIFAP3 | 195.357856 | 0.374359378 | 0.22888934 | 1.63554746 | 0.10193434 | 0.43085158 |
| AFG1L | 108.864502 | 0.375017064 | 0.2899478 | 1.2933951 | 0.19587444 | 0.56062979 |
| NHLRC2 | 233.584954 | 0.375648578 | 0.12569106 | 2.98866581 | 0.00280198 | 0.07081266 |
| KIF16B | 204.806205 | 0.375668646 | 0.20306566 | 1.84998613 | 0.06431555 | 0.35347506 |
| ATG14 | 66.1941956 | 0.37585657 | 0.25534297 | 1.47196758 | 0.14102963 | 0.49255036 |
| RFTN2 | 106.69319 | 0.375863939 | 0.45128595 | 0.83287313 | 0.40491629 | 0.73208793 |
| VAV2 | 114.620251 | 0.376107911 | 0.4809633 | 0.78198879 | 0.43422116 | 0.7482063 |

| | | | | | | |
|---|---|---|---|---|---|---|
| AGBL3 | 92.0178345 | 0.376704077 | 0.38032007 | 0.99049223 | 0.32193359 | 0.6761919 |
| CNP | 62.0613753 | 0.377330682 | 0.283647 | 1.33028265 | 0.18342516 | 0.54819669 |
| AKT3 | 2496.79977 | 0.377371707 | 0.23600431 | 1.59900343 | 0.10981984 | 0.44576178 |
| QRICH1 | 158.381936 | 0.377954841 | 0.18659761 | 2.0255074 | 0.04281529 | 0.29235086 |
| INF2 | 131.547053 | 0.378395899 | 0.19390164 | 1.95148373 | 0.05099953 | 0.31710683 |
| KMT2C | 1120.12748 | 0.378546411 | 0.19266995 | 1.96474025 | 0.04944431 | 0.31166266 |
| IST1 | 190.552159 | 0.378976249 | 0.15064467 | 2.5156964 | 0.01187975 | 0.15189817 |
| WASHC1 | 148.029164 | 0.379025606 | 0.30386711 | 1.24734 | 0.21227286 | 0.57690664 |
| PICALM | 2017.2628 | 0.379048913 | 0.37475628 | 1.01145446 | 0.31179897 | 0.66561244 |
| IP6K2 | 156.522363 | 0.379075371 | 0.28266929 | 1.34105607 | 0.17990225 | 0.54382715 |
| AL031428.1 | 75.2797537 | 0.379387453 | 0.24338262 | 1.55881076 | 0.11904118 | 0.46290743 |
| RAB3GAP1 | 502.362704 | 0.379873311 | 0.17065128 | 2.22602084 | 0.02601279 | 0.22518506 |
| SPRY4-AS1 | 57.9973427 | 0.379895513 | 0.31172617 | 1.21868342 | 0.22296437 | 0.58858855 |
| CDC37L1 | 72.404625 | 0.379913537 | 0.26943608 | 1.41003217 | 0.15853018 | 0.51560115 |
| ARAP2 | 206.03463 | 0.380699036 | 0.36813976 | 1.03411551 | 0.30108217 | 0.65926326 |
| ABI1 | 418.83976 | 0.380983888 | 0.20937192 | 1.81965134 | 0.06881212 | 0.36281544 |
| PIAS4 | 46.5410432 | 0.38106612 | 0.28982715 | 1.31480478 | 0.18857552 | 0.55388137 |
| C18orf25 | 144.608951 | 0.381131002 | 0.20728186 | 1.83870891 | 0.06595801 | 0.35579065 |
| CGAS | 57.5475957 | 0.381508755 | 0.39106022 | 0.97557547 | 0.32927489 | 0.68205162 |
| TAB2 | 305.057508 | 0.381600723 | 0.22270126 | 1.71350953 | 0.08661884 | 0.39932506 |
| AFDN | 839.677309 | 0.382346671 | 0.19619009 | 1.94885827 | 0.05131235 | 0.31806717 |
| UMAD1 | 376.284021 | 0.382642351 | 0.20749215 | 1.84412932 | 0.0651643 | 0.35535588 |
| ADCY4 | 215.813716 | 0.382719452 | 0.24839662 | 1.5407595 | 0.12337533 | 0.46870024 |
| ZRANB1 | 160.692807 | 0.382903682 | 0.15985272 | 2.39535299 | 0.01660437 | 0.18464043 |
| MAP4K4 | 1167.39824 | 0.383179922 | 0.40412146 | 0.94818009 | 0.34303778 | 0.69310572 |
| ARHGAP5 | 497.451798 | 0.383329531 | 0.30226404 | 1.26819431 | 0.20472857 | 0.56960077 |
| GPR137C | 55.9554219 | 0.383644724 | 0.73270975 | 0.52359714 | 0.60055877 | 0.8498698 |
| RAB12 | 192.325407 | 0.383645042 | 0.15969458 | 2.4023673 | 0.01628934 | 0.18309849 |
| AC013394.1 | 243.279377 | 0.383666079 | 0.2514388 | 1.52588253 | 0.12703912 | 0.47383808 |
| PLXND1 | 488.049686 | 0.383895502 | 0.28656654 | 1.33963828 | 0.18036297 | 0.5444489 |
| CLASP1 | 835.43189 | 0.384145487 | 0.23985327 | 1.60158534 | 0.10924734 | 0.44576178 |
| HUS1 | 99.37126 | 0.384167256 | 0.29902103 | 1.28474995 | 0.19887968 | 0.56402001 |
| OFD1 | 102.309974 | 0.3842633 | 0.31005335 | 1.23934573 | 0.21521749 | 0.58008061 |
| TLE4 | 189.000339 | 0.384270655 | 0.28443542 | 1.35099439 | 0.17669723 | 0.54037749 |
| QDPR | 46.5924996 | 0.384285638 | 0.23972908 | 1.60299967 | 0.10893473 | 0.44503888 |
| SUMF1 | 532.053804 | 0.384463121 | 0.34233782 | 1.12305184 | 0.26141548 | 0.62654428 |
| CFAP44 | 128.093888 | 0.384478007 | 0.2748422 | 1.39890456 | 0.1618416 | 0.52098735 |
| MDM4 | 293.509522 | 0.384848708 | 0.19714724 | 1.95208775 | 0.05092779 | 0.31710683 |
| CASP2 | 46.1706056 | 0.38500572 | 0.3118142 | 1.23472798 | 0.21693176 | 0.58245153 |
| PDGFB | 185.105672 | 0.385307273 | 0.25358756 | 1.51942499 | 0.12865556 | 0.4765905 |
| NCEH1 | 99.3021189 | 0.385490034 | 0.35447968 | 1.08748133 | 0.27682414 | 0.64098811 |
| FBXW7 | 662.059269 | 0.385593163 | 0.21646742 | 1.78129883 | 0.07486365 | 0.37933183 |
| ARHGAP6 | 200.574746 | 0.385628384 | 0.38192638 | 1.00969298 | 0.31264241 | 0.66675862 |
| UBQLN4 | 42.6957704 | 0.386095348 | 0.30178067 | 1.27939057 | 0.20075955 | 0.56536969 |
| PXDC1 | 100.963831 | 0.386117793 | 0.19396034 | 1.99070488 | 0.04651334 | 0.30359341 |
| AC242426.2 | 102.691837 | 0.386315602 | 0.29612545 | 1.30456736 | 0.19204021 | 0.55646778 |
| BTBD7 | 542.551446 | 0.386356283 | 0.1965143 | 1.96604671 | 0.04929322 | 0.31155957 |
| CEP192 | 281.672801 | 0.386402064 | 0.229954 | 1.68034504 | 0.0928902 | 0.41198944 |
| ERBB4 | 199.576786 | 0.386509587 | 0.38721276 | 0.99818402 | 0.31819014 | 0.6730872 |
| AL357153.2 | 104.854658 | 0.387336458 | 0.37693965 | 1.02758214 | 0.30414643 | 0.66086613 |
| PIK3CA | 334.380233 | 0.38740293 | 0.2244358 | 1.72611911 | 0.08432599 | 0.39616527 |
| IFI35 | 81.2870009 | 0.388086887 | 0.45638155 | 0.85035621 | 0.39512707 | 0.72496073 |
| CYLD | 167.262739 | 0.388304547 | 0.24722407 | 1.57065832 | 0.11626204 | 0.4592091 |
| DAGLB | 53.8426885 | 0.388654456 | 0.25798618 | 1.5064933 | 0.13194059 | 0.48092078 |
| GABBR1 | 119.37029 | 0.388839818 | 0.34602721 | 1.12372611 | 0.26112923 | 0.62608278 |
| TNKS2 | 295.724957 | 0.389169815 | 0.18478946 | 2.10601733 | 0.03520285 | 0.26331343 |
| KANK1 | 140.40738 | 0.389979462 | 0.40157369 | 0.97112802 | 0.33148453 | 0.68446579 |
| PAPOLG | 94.8377942 | 0.390461013 | 0.28155225 | 1.38681544 | 0.16549805 | 0.52581169 |
| SHLD1 | 124.287136 | 0.390630477 | 0.26177235 | 1.49225262 | 0.13563293 | 0.48508929 |
| DDX17 | 1403.88874 | 0.390754839 | 0.24026316 | 1.62636187 | 0.1038727 | 0.43532543 |
| PARP11 | 112.223365 | 0.390834622 | 0.18693448 | 2.09075722 | 0.03654983 | 0.26964679 |
| FOXN2 | 206.637803 | 0.390953838 | 0.2151877 | 1.81680383 | 0.06924716 | 0.36386894 |
| PPP6R3 | 957.630158 | 0.391878383 | 0.22335566 | 1.75450391 | 0.0793442 | 0.38738328 |
| EBF4 | 105.110206 | 0.392342746 | 0.25589999 | 1.53318781 | 0.12522959 | 0.47093183 |
| BCL9 | 62.8538705 | 0.392409274 | 0.38634865 | 1.01568808 | 0.30977794 | 0.6648266 |
| SCAF8 | 525.092746 | 0.393676814 | 0.1770123 | 2.22400824 | 0.02614789 | 0.22519168 |
| C1GALT1 | 99.158889 | 0.393945632 | 0.29957828 | 1.31500066 | 0.18850968 | 0.55388137 |
| PRKAG2 | 175.735971 | 0.393955489 | 0.43063815 | 0.91481791 | 0.36028723 | 0.70386371 |
| PPWD1 | 132.296617 | 0.394543547 | 0.18893462 | 2.08825439 | 0.03677489 | 0.27037642 |
| AC009690.1 | 139.501898 | 0.395069788 | 0.45817291 | 0.86227225 | 0.38853772 | 0.72177165 |
| PHACTR2 | 466.728456 | 0.395615454 | 0.15356236 | 2.57625271 | 0.00998776 | 0.14189211 |
| ANKRD6 | 154.778147 | 0.395881804 | 0.23855948 | 1.65946794 | 0.09702154 | 0.4194016 |
| WWC2 | 606.807334 | 0.396573578 | 0.1810334 | 2.19061001 | 0.02848003 | 0.23351909 |
| DTX3L | 131.426075 | 0.396714778 | 0.37367072 | 1.06166943 | 0.28838578 | 0.65138125 |
| USP6NL | 252.170191 | 0.396938879 | 0.17761778 | 2.23479252 | 0.02543097 | 0.22138474 |
| RBFOX2 | 1302.05997 | 0.396944302 | 0.1723897 | 2.30259866 | 0.02130143 | 0.20628732 |
| LNX2 | 119.209722 | 0.397357003 | 0.31346125 | 1.26764316 | 0.20492541 | 0.56960077 |
| PRAG1 | 114.615898 | 0.397959131 | 0.58499418 | 0.68027879 | 0.49632795 | 0.79402769 |
| ZNF566 | 73.1771798 | 0.398053699 | 0.25226312 | 1.57793064 | 0.11458154 | 0.45559805 |
| PGM5 | 611.55479 | 0.398516534 | 0.39319396 | 1.01353676 | 0.31080385 | 0.66561244 |
| UVSSA | 96.7248066 | 0.398609027 | 0.30932699 | 1.28863319 | 0.19752564 | 0.56299279 |
| RNF138 | 144.393935 | 0.398926305 | 0.17274745 | 2.3093036 | 0.02092674 | 0.20434787 |
| RGL2 | 132.780448 | 0.39895335 | 0.25722288 | 1.55100255 | 0.12090107 | 0.46513092 |
| SNX2 | 140.423706 | 0.39940644 | 0.19950829 | 2.00195409 | 0.04528967 | 0.30043379 |
| ITPKB | 447.452084 | 0.399592028 | 0.42297476 | 0.94471838 | 0.34480267 | 0.69444259 |
| SLC35B3 | 84.8361599 | 0.399657307 | 0.2925429 | 1.36614938 | 0.17189208 | 0.53377758 |
| MED23 | 150.379591 | 0.399676753 | 0.16385097 | 2.43926999 | 0.01471697 | 0.17529687 |
| HSPB1 | 659.238772 | 0.399920257 | 0.47382772 | 0.84402039 | 0.39865801 | 0.72770737 |
| PCM1 | 739.920246 | 0.399963783 | 0.15675775 | 2.55147687 | 0.01072674 | 0.14556708 |
| LRRC8B | 131.478281 | 0.400373371 | 0.27533509 | 1.45413131 | 0.1459099 | 0.4993646 |
| CRLF3 | 168.624885 | 0.400455852 | 0.21770125 | 1.83947428 | 0.06584546 | 0.35579065 |
| AL365273.2 | 228.91926 | 0.400485532 | 0.45045946 | 0.88906011 | 0.37397078 | 0.7121148 |
| ERBB2 | 46.4092373 | 0.400678613 | 0.36075549 | 1.11066533 | 0.26671243 | 0.63206134 |
| MITD1 | 80.0939403 | 0.401075614 | 0.19345168 | 2.07325988 | 0.0381481 | 0.2764977 |
| MN1 | 123.736553 | 0.401423059 | 0.45850819 | 0.87549812 | 0.38130293 | 0.71543195 |
| STXBP1 | 222.447312 | 0.4014498 | 0.25623988 | 1.56669524 | 0.11718594 | 0.45991971 |
| PSMA3 | 143.551601 | 0.401589363 | 0.21757268 | 1.84577107 | 0.06492545 | 0.35498206 |
| FAM168A | 381.311666 | 0.401639625 | 0.1511518 | 2.65719375 | 0.00787941 | 0.12559463 |
| MEF2A | 2128.9908 | 0.401839285 | 0.34987851 | 1.14851089 | 0.25075772 | 0.61583146 |
| TNRC6A | 724.479065 | 0.402096802 | 0.17862526 | 2.25106356 | 0.02438151 | 0.21755892 |
| NIPAL2 | 99.8177704 | 0.402156434 | 0.30718787 | 1.30915468 | 0.19048196 | 0.55558405 |
| SPTLC3 | 67.6166646 | 0.402513801 | 0.37680868 | 1.06821797 | 0.2854222 | 0.64806045 |
| ZNF138 | 72.6640781 | 0.402521868 | 0.30485784 | 1.32035923 | 0.18671511 | 0.55302731 |
| ZDHHC3 | 106.91669 | 0.403450216 | 0.25119054 | 1.60615212 | 0.10824049 | 0.44369549 |
| AGAP4 | 71.3686171 | 0.403596966 | 0.22129712 | 1.82377863 | 0.06818555 | 0.36142549 |
| C15orf41 | 370.244431 | 0.403650519 | 0.20547204 | 1.96450343 | 0.04947174 | 0.31166266 |
| NSF | 318.528672 | 0.403977832 | 0.31720047 | 1.2735726 | 0.20281493 | 0.56730017 |
| MTM1 | 178.613157 | 0.404313145 | 0.2045146 | 1.97694028 | 0.04804838 | 0.30741196 |
| AP1G1 | 266.647837 | 0.404354405 | 0.16450202 | 2.45805128 | 0.01396932 | 0.16985285 |
| HSF2 | 51.0211105 | 0.404478637 | 0.32729969 | 1.23580512 | 0.21653101 | 0.58218265 |
| ARID3B | 49.7108665 | 0.40464685 | 0.22005508 | 1.83884346 | 0.06593821 | 0.35579065 |
| SOS1 | 828.045344 | 0.405049816 | 0.15761952 | 2.56979483 | 0.01017588 | 0.14236887 |
| FRA10AC1 | 139.566736 | 0.40518828 | 0.22055863 | 1.83710006 | 0.06619513 | 0.35613065 |

| | | | | | | |
|---|---|---|---|---|---|---|
| VCL | 364.731316 | 0.405387824 | 0.31674424 | 1.27985857 | 0.20059488 | 0.56536969 |
| DNAJC13 | 254.049339 | 0.4054339 | 0.1960245 | 2.06828175 | 0.03861354 | 0.2776578 |
| SUGCT | 88.0918504 | 0.405681813 | 0.39326792 | 1.03156599 | 0.30227548 | 0.65926326 |
| ARNT | 181.174013 | 0.406316445 | 0.18281344 | 2.22257429 | 0.02624452 | 0.22535037 |
| PCDH9 | 264.979688 | 0.407168451 | 0.94632263 | 0.43026389 | 0.66700369 | 0.87587657 |
| SFXN5 | 96.7236827 | 0.407307307 | 0.31273653 | 1.3023976 | 0.1927805 | 0.55703491 |
| LIMA1 | 720.548558 | 0.407728711 | 0.19156708 | 2.1283861 | 0.03330508 | 0.25490607 |
| DDX59 | 52.3939097 | 0.40807828 | 0.33596561 | 1.214643 | 0.22450228 | 0.59083342 |
| FAM193B | 183.778535 | 0.408804012 | 0.18826278 | 2.17145421 | 0.02989686 | 0.2403202 |
| DUSP1 | 734.2757 | 0.409206608 | 0.72682881 | 0.56300273 | 0.57343302 | 0.8342245 |
| SLC44A1 | 398.070043 | 0.409352278 | 0.31450059 | 1.30159461 | 0.193055 | 0.55752247 |
| AL139807.1 | 303.798019 | 0.40940033 | 0.68636206 | 0.59647867 | 0.5508555 | 0.8236419 |
| EPC1 | 483.697022 | 0.409605903 | 0.20723878 | 1.97649253 | 0.04809902 | 0.30749554 |
| TSPAN9 | 855.461164 | 0.40997787 | 0.33004928 | 1.24217169 | 0.21417322 | 0.57870917 |
| MADD | 111.569988 | 0.410337371 | 0.21274849 | 1.92874398 | 0.05376265 | 0.32491861 |
| TNRC18 | 382.949742 | 0.410578071 | 0.29270107 | 1.40272143 | 0.16069992 | 0.51944206 |
| SAMD4A | 550.440301 | 0.410643289 | 0.60592768 | 0.67771007 | 0.49795554 | 0.79513036 |
| CCM2 | 258.067276 | 0.41078407 | 0.26939244 | 1.52485372 | 0.12729559 | 0.47449441 |
| DLC1 | 1223.5663 | 0.410935187 | 0.32329283 | 1.2710928 | 0.20369564 | 0.56791871 |
| ERAP2 | 165.778159 | 0.411003218 | 0.62508328 | 0.65751754 | 0.5108482 | 0.80177277 |
| ITGAV | 516.299137 | 0.411010593 | 0.35071694 | 1.17191545 | 0.24123101 | 0.60589113 |
| HLA-DOA | 54.1721672 | 0.411381313 | 0.43639439 | 0.94268241 | 0.34584337 | 0.69497757 |
| SMG1 | 491.116252 | 0.411451762 | 0.21696072 | 1.89643439 | 0.05790263 | 0.33795806 |
| PTK2 | 2178.33604 | 0.412854721 | 0.18168288 | 2.27239201 | 0.02306284 | 0.21247788 |
| BCL2L1 | 371.001458 | 0.413245483 | 0.39699106 | 1.04094405 | 0.29790151 | 0.6579563 |
| SIPA1L1 | 484.950995 | 0.413258841 | 0.30219914 | 1.367505 | 0.17146707 | 0.53350382 |
| COPS7B | 60.6293025 | 0.413437368 | 0.27673374 | 1.49398972 | 0.13517831 | 0.48508022 |
| GNAI1 | 196.76143 | 0.413592619 | 0.39008904 | 1.06025182 | 0.28903005 | 0.65209069 |
| RUFY2 | 212.288938 | 0.413698355 | 0.21642219 | 1.91153389 | 0.05593601 | 0.33024844 |
| GPR180 | 60.237894 | 0.41437989 | 0.26648893 | 1.55496101 | 0.11995536 | 0.46382142 |
| TUSC3 | 65.4991027 | 0.414827723 | 0.75274062 | 0.55108986 | 0.58157208 | 0.83789748 |
| ZNF84 | 83.2046438 | 0.415093939 | 0.28745509 | 1.4440306 | 0.14873037 | 0.50222622 |
| TPCN1 | 167.144105 | 0.415323595 | 0.44452385 | 0.93431117 | 0.3501434 | 0.69858573 |
| KANTR | 179.729185 | 0.415801285 | 0.28365809 | 1.46585381 | 0.14268812 | 0.49496264 |
| ZDHHC17 | 302.286705 | 0.415853743 | 0.20983331 | 1.98182903 | 0.04749838 | 0.30604664 |
| UBR2 | 534.069688 | 0.416345644 | 0.18824111 | 2.21176789 | 0.02698271 | 0.22894365 |
| SLC25A53 | 83.0344356 | 0.416754313 | 0.2152812 | 1.93586021 | 0.05288482 | 0.3215522 |
| BCAP29 | 220.597175 | 0.417621261 | 0.34863422 | 1.19787798 | 0.23096452 | 0.59733966 |
| MPPE1 | 73.5234392 | 0.417707051 | 0.24176465 | 1.72774245 | 0.08403441 | 0.39616527 |
| EVI5 | 405.919873 | 0.417719259 | 0.22335115 | 1.87023552 | 0.06145112 | 0.34778328 |
| POLN | 51.1372668 | 0.417785664 | 0.38357195 | 1.08919764 | 0.27606674 | 0.64068436 |
| ARHGAP32 | 164.002725 | 0.418521573 | 0.27749735 | 1.50820026 | 0.13150328 | 0.48069696 |
| XPR1 | 690.896643 | 0.420449217 | 0.25957487 | 1.61976085 | 0.10528366 | 0.43797647 |
| SERTAD2 | 115.589641 | 0.420576067 | 0.32106926 | 1.30992319 | 0.19022182 | 0.55552647 |
| ZFHX3 | 820.810115 | 0.421144864 | 0.29197227 | 1.44241393 | 0.14918564 | 0.50283523 |
| VPS26B | 57.0172996 | 0.421195962 | 0.31957636 | 1.31798223 | 0.1875096 | 0.55388137 |
| ASAP1 | 2494.60559 | 0.421490232 | 0.24147219 | 1.7455022 | 0.08089749 | 0.39055112 |
| GFM2 | 62.5697125 | 0.421537198 | 0.22615198 | 1.86395538 | 0.06232797 | 0.35005476 |
| DHX35 | 75.0820058 | 0.421721945 | 0.23252767 | 1.81364197 | 0.06973286 | 0.3652226 |
| EPB41L5 | 153.141831 | 0.421832185 | 0.33536932 | 1.25781387 | 0.20845908 | 0.57338509 |
| ELMO2 | 54.8508512 | 0.42186804 | 0.21503544 | 1.96185356 | 0.04977954 | 0.3126216 |
| NEK1 | 257.828633 | 0.422610631 | 0.18909016 | 2.23496886 | 0.02541939 | 0.22138474 |
| KANSL1 | 1310.54369 | 0.423089174 | 0.28753958 | 1.47141193 | 0.14117975 | 0.49265411 |
| NUB1 | 137.547325 | 0.423225693 | 0.29083808 | 1.45519356 | 0.14561568 | 0.49856617 |
| LIPE | 75.3046525 | 0.423519704 | 0.30693417 | 1.37983891 | 0.16763625 | 0.52902716 |
| MAP3K6 | 100.895583 | 0.42375339 | 0.4618703 | 0.91747269 | 0.358895 | 0.70242891 |
| SLC8A1 | 483.024864 | 0.423860866 | 0.39800353 | 1.0649676 | 0.28689059 | 0.64995175 |
| PPP3CB | 285.770298 | 0.424146746 | 0.18008294 | 2.3552855 | 0.01850848 | 0.19394842 |
| OTUD7A | 116.895772 | 0.424199486 | 0.34726515 | 1.22154351 | 0.2218803 | 0.58750144 |
| AC078785.1 | 76.9990589 | 0.424333473 | 0.35475145 | 1.19614303 | 0.23164075 | 0.59795465 |
| SLC2A13 | 212.028858 | 0.424736816 | 0.37746903 | 1.12522294 | 0.26049458 | 0.62580682 |
| RRAS | 136.304162 | 0.425034281 | 0.42978086 | 0.98895582 | 0.32268476 | 0.6761919 |
| RXRA | 141.056807 | 0.425065418 | 0.39473024 | 1.0768504 | 0.2815471 | 0.64471757 |
| RFC1 | 230.728478 | 0.425331391 | 0.3078834 | 1.38146906 | 0.16713479 | 0.52764813 |
| SP2 | 39.2848371 | 0.425752238 | 0.39332659 | 1.0824395 | 0.27905729 | 0.64254602 |
| PRPF39 | 90.1490408 | 0.425917493 | 0.25433874 | 1.67460723 | 0.09401132 | 0.41280519 |
| MMP24OS | 103.140408 | 0.425922111 | 0.26072895 | 1.63358198 | 0.10234665 | 0.43132417 |
| SNX14 | 262.273768 | 0.426452718 | 0.20300096 | 2.10074237 | 0.03566359 | 0.26554608 |
| PAXX | 61.1376287 | 0.426810098 | 0.49636023 | 0.85987973 | 0.38985534 | 0.72202943 |
| LRRC40 | 110.993381 | 0.427273333 | 0.23760719 | 1.798234 | 0.07213993 | 0.37150477 |
| MT1H | 139.518253 | 0.427811722 | 1.29202667 | 0.33111679 | 0.74055627 | 0.90826918 |
| PHF20L1 | 295.994405 | 0.427828532 | 0.18795219 | 2.27626261 | 0.0228303 | 0.21136949 |
| NPIPB5 | 195.430685 | 0.427939853 | 0.25903566 | 1.65204999 | 0.09852436 | 0.42254971 |
| TENT2 | 558.178966 | 0.428062452 | 0.18987799 | 2.25440791 | 0.02417052 | 0.21639754 |
| SLC4A4 | 95.0028059 | 0.428221459 | 0.75413566 | 0.56783081 | 0.57014986 | 0.83368112 |
| LSS | 67.3513384 | 0.428529595 | 0.36526823 | 1.17319154 | 0.24071901 | 0.6051624 |
| SPRED2 | 382.952136 | 0.428535169 | 0.25248096 | 1.69729699 | 0.08964053 | 0.40583359 |
| BCL7C | 94.0122317 | 0.428641928 | 0.24361764 | 1.75948641 | 0.07849493 | 0.38578017 |
| SRRM1 | 231.94426 | 0.428894301 | 0.17844 | 2.4035771 | 0.01623554 | 0.18299648 |
| RBMS3 | 1256.06314 | 0.429872593 | 0.27570281 | 1.55918828 | 0.11895182 | 0.46285439 |
| SOCS7 | 47.9379599 | 0.42989423 | 0.28198991 | 1.52450215 | 0.12738332 | 0.47449441 |
| HID1 | 73.8466463 | 0.430120906 | 0.44150848 | 0.97420757 | 0.32995349 | 0.68216508 |
| CDS2 | 237.429341 | 0.430158761 | 0.23204556 | 1.85376858 | 0.06377228 | 0.3533385 |
| LUC7L3 | 513.574737 | 0.430318548 | 0.16328953 | 2.63531009 | 0.00840605 | 0.13026671 |
| MAPK11 | 73.1934321 | 0.430550583 | 0.29366055 | 1.46615057 | 0.14260727 | 0.49489199 |
| POGZ | 509.644939 | 0.430674997 | 0.18333208 | 2.34915237 | 0.0188162 | 0.19394842 |
| APOL3 | 255.097372 | 0.430684404 | 0.39913942 | 1.07903249 | 0.28057324 | 0.64385822 |
| CLASP2 | 541.213947 | 0.431199435 | 0.22199218 | 1.94240825 | 0.0520877 | 0.32016393 |
| BET1 | 87.6003846 | 0.43139118 | 0.34142726 | 1.26349367 | 0.20641182 | 0.57126492 |
| ZNF573 | 60.1646166 | 0.431681102 | 0.33311472 | 1.29589322 | 0.19501228 | 0.56007526 |
| UNKL | 58.5670122 | 0.431954284 | 0.33877542 | 1.27504611 | 0.20229292 | 0.56693091 |
| IPP | 56.9915012 | 0.432305421 | 0.36348507 | 1.18933474 | 0.23430797 | 0.59949266 |
| WDSUB1 | 110.776565 | 0.433038948 | 0.26354207 | 1.6431492 | 0.10035207 | 0.42658751 |
| STK10 | 339.684635 | 0.433549307 | 0.20073267 | 2.15983426 | 0.0307855 | 0.24387005 |
| DCUN1D1 | 132.233621 | 0.433829397 | 0.21057902 | 2.06017389 | 0.03938192 | 0.28071625 |
| NUCB2 | 205.507193 | 0.434077212 | 0.24779505 | 1.75175903 | 0.07981525 | 0.38738328 |
| ATP9A | 53.5396031 | 0.434291426 | 0.37161757 | 1.1686515 | 0.24254407 | 0.60732501 |
| KRAS | 208.349876 | 0.434579253 | 0.26053375 | 1.66803441 | 0.0953089 | 0.41531149 |
| MAPRE2 | 221.077885 | 0.435286012 | 0.34331714 | 1.26788314 | 0.20483968 | 0.56960077 |
| MAML3 | 1208.76068 | 0.435479952 | 0.31215707 | 1.39506676 | 0.16299571 | 0.52335809 |
| SHC2 | 146.077042 | 0.435528158 | 0.26662154 | 1.63350703 | 0.1023624 | 0.43132417 |
| ZNF366 | 357.561438 | 0.43584408 | 0.30728958 | 1.41834969 | 0.1560887 | 0.51234409 |
| CEP162 | 115.985426 | 0.436051774 | 0.25371723 | 1.71865256 | 0.08567765 | 0.39790026 |
| STARD13 | 849.964504 | 0.436295305 | 0.40029855 | 1.08992477 | 0.27574629 | 0.64048591 |
| FILIP1 | 117.007989 | 0.436390018 | 0.60654889 | 0.71946388 | 0.47185515 | 0.77855633 |
| DLG2 | 676.813043 | 0.437512936 | 0.39985813 | 1.09417042 | 0.27388024 | 0.63960103 |
| TMEM245 | 323.047782 | 0.437588219 | 0.19875576 | 2.20163796 | 0.02769089 | 0.23121895 |
| IGF2BP2 | 273.732751 | 0.437654798 | 0.51945069 | 0.84253387 | 0.39948919 | 0.72822901 |
| DAPK2 | 97.2523881 | 0.438942716 | 0.44738385 | 0.98113224 | 0.32652753 | 0.67997183 |
| DTWD2 | 63.1147836 | 0.438958402 | 0.27062704 | 1.62200496 | 0.10480229 | 0.43679299 |
| TSPAN18 | 199.674416 | 0.439070511 | 0.60155165 | 0.72989661 | 0.46545339 | 0.77477727 |
| VPS52 | 57.494857 | 0.439207512 | 0.19314178 | 2.27401613 | 0.02296502 | 0.21234207 |
| STIM2 | 502.366445 | 0.439381856 | 0.16858511 | 2.60629108 | 0.00915286 | 0.13593083 |

| | | | | | | |
|---|---|---|---|---|---|---|
| RNPC3 | 147.560857 | 0.439743825 | 0.25616719 | 1.7166282 | 0.08604713 | 0.39864674 |
| NT5C3A | 231.165572 | 0.439897897 | 0.30634987 | 1.43593301 | 0.1510214 | 0.50677567 |
| AK3 | 215.298366 | 0.440093002 | 0.26456761 | 1.66344248 | 0.0962239 | 0.41727618 |
| TMEM44 | 240.926283 | 0.440222141 | 0.29061043 | 1.51481877 | 0.12981831 | 0.47808425 |
| NTN4 | 798.164602 | 0.440572866 | 0.43225551 | 1.01924175 | 0.30808821 | 0.66414532 |
| SAMD9 | 99.4080994 | 0.440721543 | 0.39027205 | 1.12926751 | 0.258785 | 0.62362556 |
| TRANK1 | 110.192161 | 0.441026904 | 0.54022057 | 0.81638303 | 0.4142811 | 0.73671465 |
| ZNF318 | 53.7932158 | 0.441045803 | 0.25441617 | 1.73356044 | 0.08299611 | 0.39515849 |
| IQCE | 38.800425 | 0.441241553 | 0.35076298 | 1.25794789 | 0.2084106 | 0.57338509 |
| MIR99AHG | 2887.36984 | 0.441350825 | 0.49096625 | 0.8989433 | 0.36868286 | 0.70919884 |
| MEF2D | 131.593723 | 0.441454323 | 0.25016385 | 1.76466076 | 0.0776208 | 0.38471895 |
| COG2 | 55.5842004 | 0.441534344 | 0.2694797 | 1.63846975 | 0.10132374 | 0.42937968 |
| IKZF5 | 52.8904801 | 0.441568696 | 0.28341173 | 1.55804667 | 0.11922219 | 0.46290743 |
| DIP2A | 338.030896 | 0.441818958 | 0.14482664 | 3.05067464 | 0.00228328 | 0.06286489 |
| NDRG3 | 87.2903048 | 0.442714687 | 0.28512711 | 1.55269237 | 0.12049665 | 0.46432492 |
| FRG1-DT | 77.2946163 | 0.44279621 | 0.45449685 | 0.97425584 | 0.32992953 | 0.68216508 |
| ADGRB3 | 1621.41912 | 0.442836045 | 0.54495276 | 0.81261363 | 0.41643961 | 0.73802299 |
| SFI1 | 90.7608134 | 0.442956091 | 0.37236874 | 1.18956305 | 0.23421818 | 0.59949266 |
| EMP3 | 164.105098 | 0.443338941 | 0.37058965 | 1.19630686 | 0.23157683 | 0.59795465 |
| GTF2IRD1 | 126.561904 | 0.443824187 | 0.33310316 | 1.33239259 | 0.18273122 | 0.54790102 |
| ABCC5 | 75.1823166 | 0.443878399 | 0.26926436 | 1.64848549 | 0.09925308 | 0.42376187 |
| KIDINS220 | 369.803163 | 0.444107366 | 0.16072023 | 2.76323256 | 0.0057232 | 0.10415242 |
| USP28 | 64.4776793 | 0.444142645 | 0.23585349 | 1.8831294 | 0.05968283 | 0.34296331 |
| RAD18 | 69.1523632 | 0.444637423 | 0.20732384 | 2.14465168 | 0.03198071 | 0.24982585 |
| TGFBR2 | 2538.35157 | 0.444653349 | 0.33774579 | 1.31653264 | 0.18799533 | 0.55388137 |
| CRELD1 | 54.9616244 | 0.444917591 | 0.38051386 | 1.16925463 | 0.24230106 | 0.6070269 |
| RMND5A | 139.732716 | 0.445089199 | 0.21735598 | 2.04774303 | 0.04058518 | 0.28580837 |
| PDLIM5 | 1537.56472 | 0.445229042 | 0.29788176 | 1.49465022 | 0.13500575 | 0.484753 |
| PAN3 | 934.112805 | 0.446142572 | 0.166681 | 2.67662518 | 0.00743678 | 0.12171031 |
| CLDN10 | 48.0930117 | 0.446296654 | 0.44966204 | 0.99251575 | 0.320946 | 0.67473727 |
| MRAS | 51.4477258 | 0.447029606 | 0.37504569 | 1.19193373 | 0.23328723 | 0.59910683 |
| PTPN18 | 74.3806936 | 0.448016998 | 0.28059123 | 1.59668925 | 0.11033499 | 0.44631216 |
| RASSF8 | 258.622626 | 0.448080588 | 0.33184538 | 1.35026917 | 0.17692966 | 0.54037749 |
| MIRLET7BHG | 281.355478 | 0.448257806 | 0.34704933 | 1.29162562 | 0.19648683 | 0.56164812 |
| AC083798.2 | 49.7971051 | 0.448592164 | 0.37859029 | 1.1849014 | 0.23605644 | 0.60018541 |
| TMEM131L | 311.351082 | 0.449153978 | 0.26470211 | 1.69682811 | 0.08972916 | 0.40583359 |
| SRGAP2B | 205.230137 | 0.449262054 | 0.22446656 | 2.00146537 | 0.04534226 | 0.30043379 |
| HECTD4 | 388.833222 | 0.449546435 | 0.16606828 | 2.70699754 | 0.00678948 | 0.11693671 |
| ELMOD3 | 93.7222073 | 0.449781081 | 0.24681807 | 1.82231826 | 0.06840671 | 0.36142549 |
| LRCH1 | 1219.99739 | 0.449845202 | 0.23861412 | 1.8852413 | 0.05939727 | 0.34186127 |
| CTNNB1 | 1002.91291 | 0.450696831 | 0.27417838 | 1.64380877 | 0.10021571 | 0.42645093 |
| LCMT1 | 101.883886 | 0.45071476 | 0.22304739 | 2.02071301 | 0.04330948 | 0.29508867 |
| OAS2 | 125.693327 | 0.450856247 | 0.67128105 | 0.6716356 | 0.50181571 | 0.79661835 |
| ZDHHC14 | 697.822609 | 0.451519271 | 0.40209674 | 1.12291204 | 0.26147485 | 0.62654428 |
| ACAP2 | 744.207988 | 0.451666369 | 0.15746028 | 2.86844632 | 0.00412493 | 0.08875226 |
| RNF19A | 505.788631 | 0.451902382 | 0.24482621 | 1.84580887 | 0.06491996 | 0.35498206 |
| KIF1B | 486.324071 | 0.452468057 | 0.2523425 | 1.79307113 | 0.07296155 | 0.37361976 |
| ZNF595 | 83.6982854 | 0.452503755 | 0.21267323 | 2.12769492 | 0.03336239 | 0.25490607 |
| AC104806.2 | 73.0221609 | 0.452615444 | 0.29822357 | 1.51770516 | 0.12908874 | 0.47711526 |
| JADE2 | 114.128579 | 0.452693807 | 0.22058439 | 2.05224773 | 0.04014559 | 0.28442544 |
| ANKAR | 188.526158 | 0.452777073 | 0.30603039 | 1.4795167 | 0.13900227 | 0.49082099 |
| ATP10D | 202.131653 | 0.453114639 | 0.22699241 | 1.99616643 | 0.04591581 | 0.302276 |
| BTN3A2 | 116.727861 | 0.453260095 | 0.56862455 | 0.79711665 | 0.42538329 | 0.74197066 |
| UBTD2 | 122.577326 | 0.453558803 | 0.26161445 | 1.73369173 | 0.08297279 | 0.39515849 |
| SPATS2L | 397.435382 | 0.453643907 | 0.35555428 | 1.27587807 | 0.20199863 | 0.56693091 |
| AL024474.2 | 56.7901913 | 0.45404498 | 0.5119698 | 0.88685891 | 0.37515487 | 0.71257833 |
| NEK4 | 79.5389254 | 0.454364206 | 0.21036077 | 2.15992846 | 0.03077821 | 0.24387005 |
| TBKBP1 | 44.3037016 | 0.45462442 | 0.24699572 | 1.84061657 | 0.06567777 | 0.35579065 |
| SPAG16 | 162.426255 | 0.455080171 | 0.23914088 | 1.90297944 | 0.05704323 | 0.33402904 |
| XPA | 76.162648 | 0.456249392 | 0.23466203 | 1.94428298 | 0.05186134 | 0.31908369 |
| RARS2 | 257.555414 | 0.4563979 | 0.19210417 | 2.37578345 | 0.01751174 | 0.18854202 |
| AL157938.2 | 62.1174102 | 0.456600692 | 0.49577956 | 0.92097522 | 0.35706336 | 0.70058911 |
| EPHA2 | 82.9234653 | 0.456634939 | 0.43530094 | 1.04900976 | 0.29417363 | 0.65671864 |
| SLC25A16 | 168.945083 | 0.456887466 | 0.19853874 | 2.30125101 | 0.02137745 | 0.20628732 |
| SCAI | 152.974859 | 0.456961472 | 0.2494921 | 1.83156692 | 0.06701597 | 0.35889508 |
| PTAR1 | 262.186514 | 0.457085885 | 0.18490622 | 2.47198765 | 0.01343642 | 0.1660879 |
| STRN | 314.874856 | 0.457300585 | 0.15034105 | 3.04175461 | 0.00235204 | 0.06331154 |
| BCORL1 | 43.6633215 | 0.45760071 | 0.39998539 | 1.14404356 | 0.25260557 | 0.61795857 |
| POLI | 65.0816623 | 0.457897683 | 0.29201821 | 1.56804499 | 0.11687063 | 0.4596919 |
| CBL | 277.344849 | 0.458048654 | 0.16387573 | 2.79509763 | 0.0051884 | 0.09919793 |
| MEIS1 | 206.885367 | 0.45808172 | 0.27333587 | 1.67589315 | 0.09375912 | 0.41280519 |
| IGF1R | 986.241489 | 0.458451619 | 0.30636503 | 1.49642282 | 0.13454351 | 0.48474744 |
| C1orf52 | 83.2068732 | 0.458476579 | 0.27151129 | 1.68860963 | 0.09129426 | 0.40862523 |
| ZNF264 | 86.6862242 | 0.458650057 | 0.26636822 | 1.72186479 | 0.08509401 | 0.39686932 |
| MMS22L | 96.4828582 | 0.458735172 | 0.27772422 | 1.65176511 | 0.09858244 | 0.42257734 |
| TAOK2 | 157.217564 | 0.459107131 | 0.24201993 | 1.89698065 | 0.05783049 | 0.33777795 |
| SMYD3 | 1686.13397 | 0.459196434 | 0.16985701 | 2.70342944 | 0.0068628 | 0.11719379 |
| KIAA1328 | 362.086564 | 0.460024138 | 0.26519267 | 1.73467893 | 0.08279769 | 0.39515849 |
| BTD | 40.7943309 | 0.461078303 | 0.27270725 | 1.69074455 | 0.0908856 | 0.40744637 |
| UBP1 | 175.994876 | 0.462186635 | 0.22702904 | 2.035804 | 0.04177004 | 0.28963374 |
| PODXL | 1334.45606 | 0.463400836 | 0.3330056 | 1.39157068 | 0.16405244 | 0.52357299 |
| HSPA12B | 140.834361 | 0.463798325 | 0.40817367 | 1.13627693 | 0.25584068 | 0.62095131 |
| KLF7 | 861.954828 | 0.464128968 | 0.25679452 | 1.80739437 | 0.07070081 | 0.36751516 |
| AL390728.6 | 100.975135 | 0.464179875 | 0.32149059 | 1.44383659 | 0.14878495 | 0.50222622 |
| WASHC2A | 61.6069484 | 0.464336329 | 0.33293099 | 1.3946924 | 0.16310862 | 0.52335809 |
| TM2D1 | 215.716125 | 0.464401024 | 0.17267322 | 2.68947914 | 0.00715636 | 0.12024745 |
| DUSP5 | 192.757846 | 0.464449335 | 0.44379586 | 1.04653824 | 0.2953126 | 0.65720506 |
| TMCO4 | 88.4329438 | 0.464839444 | 0.4374502 | 1.0626111 | 0.28795835 | 0.65121935 |
| HIBCH | 391.331957 | 0.465450198 | 0.23540019 | 1.97727197 | 0.0480109 | 0.30741196 |
| PPM1B | 196.432946 | 0.465619965 | 0.17543843 | 2.65403627 | 0.00795353 | 0.12613123 |
| ZMYM1 | 89.9541781 | 0.465661955 | 0.21270255 | 2.18926363 | 0.02857768 | 0.23392531 |
| HP1BP3 | 427.084027 | 0.465755417 | 0.20393566 | 2.28383506 | 0.02238123 | 0.2108594 |
| PHF1 | 57.0250833 | 0.465854556 | 0.26995331 | 1.72568565 | 0.08440398 | 0.39616527 |
| RBL1 | 51.8939178 | 0.466253946 | 0.25503438 | 1.82820037 | 0.06751949 | 0.3605285 |
| CAPN2 | 265.756008 | 0.466398217 | 0.26403047 | 1.76645606 | 0.07731937 | 0.38392256 |
| KIAA1109 | 812.116064 | 0.467098451 | 0.26986346 | 1.73086956 | 0.08347503 | 0.3954379 |
| NOTCH4 | 625.095328 | 0.467210506 | 0.24370563 | 1.91710999 | 0.05522396 | 0.32937147 |
| FASTK | 36.8511564 | 0.467287829 | 0.32971072 | 1.41726611 | 0.15640514 | 0.5126563 |
| CEP164 | 82.9886717 | 0.467356068 | 0.34125222 | 1.3695327 | 0.17083282 | 0.53193492 |
| TBC1D13 | 52.0617732 | 0.468160864 | 0.23520254 | 1.99045836 | 0.04654047 | 0.30359341 |
| NR6A1 | 141.70028 | 0.468651661 | 0.43039312 | 1.08889209 | 0.27620147 | 0.64068436 |
| ZNF83 | 383.528375 | 0.469075831 | 0.26451111 | 1.7733691 | 0.07616757 | 0.38261461 |
| ARAP3 | 360.811701 | 0.469201808 | 0.20850358 | 2.25032979 | 0.02442802 | 0.21755892 |
| RBM5 | 332.595616 | 0.46928481 | 0.13923144 | 3.37053759 | 0.00075022 | 0.0305424 |
| TMC6 | 85.4638282 | 0.469599014 | 0.30654648 | 1.5319015 | 0.12554674 | 0.47103973 |
| SLC7A6 | 156.031547 | 0.471189139 | 0.29123481 | 1.61790118 | 0.1056839 | 0.43832302 |
| SLC40A1 | 128.687403 | 0.471390273 | 0.52197682 | 0.90308661 | 0.36647993 | 0.70695551 |
| AAMDC | 161.968011 | 0.471519097 | 0.26156244 | 1.80270186 | 0.07143505 | 0.36926912 |
| ZNF235 | 107.090821 | 0.471827898 | 0.36297672 | 1.29988473 | 0.19364048 | 0.55851167 |
| N4BP1 | 263.167249 | 0.471947561 | 0.39321168 | 1.2002379 | 0.23004696 | 0.59648134 |
| NOVA2 | 268.470316 | 0.472003227 | 0.27350962 | 1.72572804 | 0.08439635 | 0.39616527 |
| HNRNPH3 | 226.065285 | 0.472389569 | 0.25145329 | 1.87863747 | 0.06029401 | 0.34478399 |
| TTC14 | 269.289229 | 0.473027804 | 0.19904077 | 2.37653722 | 0.017476 | 0.18854202 |

| | | | | | | |
|---|---|---|---|---|---|---|
| CADPS2 | 1522.44795 | 0.473304911 | 0.20948947 | 2.25932557 | 0.02386314 | 0.2155321 |
| STYX | 76.1281048 | 0.473582875 | 0.18738836 | 2.52728008 | 0.01149498 | 0.15147088 |
| KIAA1147 | 188.215676 | 0.473690445 | 0.32329874 | 1.46517875 | 0.14287216 | 0.49500775 |
| DOPEY2 | 93.7357746 | 0.473988663 | 0.39878506 | 1.1885818 | 0.23460427 | 0.59949266 |
| B3GNT2 | 86.6372707 | 0.474343903 | 0.2745468 | 1.72773419 | 0.08403589 | 0.39616527 |
| UBE2Q2 | 209.210526 | 0.474351258 | 0.2392772 | 1.98243402 | 0.04743068 | 0.30604664 |
| CC2D2B | 175.659956 | 0.474419892 | 0.64918145 | 0.73079705 | 0.46490313 | 0.77464921 |
| CREBZF | 144.710133 | 0.474956184 | 0.22584706 | 2.10299916 | 0.03546584 | 0.26499217 |
| AC091948.1 | 65.9574906 | 0.475193828 | 0.42352177 | 1.12200566 | 0.26186003 | 0.6270738 |
| AHCYL2 | 342.025614 | 0.475708884 | 0.2844276 | 1.67251308 | 0.09442319 | 0.41318982 |
| MOB2 | 112.417685 | 0.475810439 | 0.17909262 | 2.65678411 | 0.00788899 | 0.12559463 |
| EHMT2 | 46.97197 | 0.476304124 | 0.24253928 | 1.9638226 | 0.04955067 | 0.31166266 |
| TLNRD1 | 103.549018 | 0.476387153 | 0.36472532 | 1.30615321 | 0.19150046 | 0.55590015 |
| GATA2 | 140.15823 | 0.476458887 | 0.45347125 | 1.05069259 | 0.2933998 | 0.6556551 |
| SMIM14 | 264.117003 | 0.476659549 | 0.21317714 | 2.23597877 | 0.02535316 | 0.22138474 |
| FCHO2 | 639.513886 | 0.477094743 | 0.21576947 | 2.21113182 | 0.02702671 | 0.22894365 |
| MICU3 | 163.362562 | 0.477295748 | 0.18178677 | 2.62558024 | 0.00865014 | 0.13154285 |
| MSRB3 | 267.552429 | 0.477589792 | 0.39668368 | 1.20395623 | 0.2286065 | 0.59497702 |
| OSTF1 | 246.143822 | 0.477618175 | 0.21561043 | 2.21519048 | 0.02674699 | 0.22775301 |
| NUMB | 948.688635 | 0.477786917 | 0.16919621 | 2.82386307 | 0.00474487 | 0.09401267 |
| SLC44A2 | 349.851173 | 0.47779107 | 0.28872566 | 1.65482716 | 0.09795956 | 0.42165994 |
| CCDC30 | 75.0328296 | 0.477886206 | 0.33568187 | 1.42362827 | 0.15455411 | 0.51037454 |
| CAPS2 | 92.4469742 | 0.477960042 | 0.5726786 | 0.83460434 | 0.40394052 | 0.73096878 |
| RAVER2 | 95.7801903 | 0.478067746 | 0.50245364 | 0.95146637 | 0.34136768 | 0.69116603 |
| AP5M1 | 97.9206363 | 0.4782886 | 0.22700123 | 2.1069868 | 0.03511872 | 0.26292453 |
| MOSPD2 | 105.375817 | 0.47838395 | 0.28776554 | 1.66240874 | 0.09643085 | 0.41773088 |
| ITGA1 | 1000.45828 | 0.478487295 | 0.29862187 | 1.60231831 | 0.10908524 | 0.44543139 |
| EFCAB2 | 191.07933 | 0.478636023 | 0.28958706 | 1.65282256 | 0.09836698 | 0.42231742 |
| MAML2 | 1381.9023 | 0.478845627 | 0.41965213 | 1.14105372 | 0.25384757 | 0.61961636 |
| TMEM150C | 110.230946 | 0.479747913 | 0.30862 | 1.5544939 | 0.12006665 | 0.46382142 |
| ACAP3 | 38.898441 | 0.480234219 | 0.27603084 | 1.73978464 | 0.08189684 | 0.39328747 |
| ABCC1 | 445.888355 | 0.481049577 | 0.30091387 | 1.59862877 | 0.10990311 | 0.44587862 |
| SRGAP2C | 156.475457 | 0.481155597 | 0.21847434 | 2.20234372 | 0.02764104 | 0.23121895 |
| GXYLT1 | 119.582708 | 0.481258791 | 0.22658633 | 2.12395336 | 0.03367404 | 0.25585395 |
| ST7 | 335.755584 | 0.481850808 | 0.28154064 | 1.7114787 | 0.08699278 | 0.39969788 |
| MGP | 975.042843 | 0.481898513 | 1.2752213 | 0.37789403 | 0.70550932 | 0.89491621 |
| MRI1 | 46.4860717 | 0.481961575 | 0.31586161 | 1.5258631 | 0.12704396 | 0.47383808 |
| BCAS3 | 1033.24004 | 0.482232123 | 0.16863808 | 2.85956838 | 0.00424218 | 0.08946844 |
| NMRK1 | 117.004109 | 0.482263277 | 0.40835204 | 1.18099883 | 0.23760319 | 0.60083649 |
| C20orf194 | 523.977842 | 0.482318306 | 0.27819284 | 1.73375526 | 0.08296152 | 0.39515849 |
| IER5 | 78.0965961 | 0.482516866 | 0.37383409 | 1.29072464 | 0.19679918 | 0.56172669 |
| BHMT2 | 56.4382248 | 0.483164315 | 0.5757974 | 0.83912208 | 0.4014008 | 0.72911494 |
| PLEKHF2 | 58.3774121 | 0.483669052 | 0.2342537 | 2.06472323 | 0.0389492 | 0.27982552 |
| AC019197.1 | 171.24113 | 0.483687373 | 0.43387819 | 1.11479992 | 0.26493617 | 0.63020787 |
| SLC12A9 | 94.2555202 | 0.483979649 | 0.31159487 | 1.55323369 | 0.12036732 | 0.46416859 |
| VPS37B | 222.089859 | 0.484133584 | 0.2475625 | 1.95560142 | 0.05051213 | 0.31516299 |
| HERC3 | 105.198854 | 0.484280438 | 0.2890918 | 1.67517874 | 0.09389917 | 0.41280519 |
| PHACTR4 | 462.520617 | 0.484538603 | 0.28220852 | 1.71695242 | 0.08598787 | 0.39864674 |
| YPEL1 | 45.5071934 | 0.484678022 | 0.44661649 | 1.08522197 | 0.27782336 | 0.64148182 |
| ATXN1 | 770.771419 | 0.484697533 | 0.37625311 | 1.28822199 | 0.1976687 | 0.56300834 |
| AVIL | 58.4325552 | 0.484803783 | 0.44690302 | 1.08480758 | 0.27800689 | 0.64148182 |
| DCLK2 | 433.787816 | 0.485386582 | 0.34313854 | 1.41454988 | 0.15720051 | 0.51434298 |
| SETX | 694.586934 | 0.485442272 | 0.28374645 | 1.71083115 | 0.08711229 | 0.39981956 |
| NBPF9 | 50.6317407 | 0.48566235 | 0.46111706 | 1.05323006 | 0.29223557 | 0.65502684 |
| ZC4H2 | 53.9030066 | 0.486519428 | 0.32470223 | 1.49835566 | 0.13404087 | 0.48404962 |
| AC096711.2 | 100.83612 | 0.486554015 | 0.3920106 | 1.24117565 | 0.21454087 | 0.57901976 |
| HIP1R | 94.2227733 | 0.486694766 | 0.28991221 | 1.67876602 | 0.09319765 | 0.41205855 |
| ZNF444 | 50.9937021 | 0.486842903 | 0.22148957 | 2.19803987 | 0.02794626 | 0.23193132 |
| ZC3H7A | 246.392549 | 0.487268698 | 0.22211088 | 2.19380832 | 0.02824919 | 0.23279269 |
| NOSTRIN | 1119.65668 | 0.487273201 | 0.28458731 | 1.71220989 | 0.086858 | 0.39969788 |
| SPTBN1 | 3064.20807 | 0.487877785 | 0.16247725 | 3.00274517 | 0.00267556 | 0.06906668 |
| KATNAL2 | 83.847604 | 0.487880205 | 0.42185327 | 1.15651636 | 0.24747001 | 0.61253693 |
| PHLDB2 | 413.146313 | 0.48812482 | 0.27619772 | 1.76730213 | 0.07717764 | 0.38356763 |
| GAS2L1 | 58.6369017 | 0.488127518 | 0.47136844 | 1.03555409 | 0.30041022 | 0.65926326 |
| ABCD3 | 177.430144 | 0.4881847 | 0.1847584 | 2.64228689 | 0.00823483 | 0.12859845 |
| APC | 242.911719 | 0.488354679 | 0.17192176 | 2.84056348 | 0.00450339 | 0.09141064 |
| SSBP2 | 930.315733 | 0.488666354 | 0.39007877 | 1.25273764 | 0.2103012 | 0.57593532 |
| GTDC1 | 284.885776 | 0.488827504 | 0.13043446 | 3.74768677 | 0.00017847 | 0.01298167 |
| LUC7L2 | 77.2822979 | 0.489407314 | 0.2684309 | 1.82321529 | 0.06827079 | 0.36142549 |
| INPP1 | 147.449835 | 0.489911236 | 0.41754643 | 1.1733096 | 0.24067167 | 0.6051624 |
| CAMKMT | 579.525201 | 0.490057551 | 0.21634313 | 2.26518655 | 0.02350123 | 0.21378373 |
| ATG16L2 | 111.255255 | 0.490274433 | 0.28634725 | 1.71216743 | 0.08686582 | 0.39969788 |
| GANC | 117.426149 | 0.490357072 | 0.35986325 | 1.36262059 | 0.17300212 | 0.53502507 |
| EVA1B | 155.099202 | 0.490839383 | 0.41503202 | 1.18265425 | 0.2369462 | 0.60074482 |
| PPIL4 | 127.155242 | 0.491087941 | 0.23792503 | 2.06404489 | 0.03901346 | 0.28002839 |
| SPACA6 | 66.2596036 | 0.491270082 | 0.40958994 | 1.1994193 | 0.23036495 | 0.5967991 |
| ABI3BP | 322.65871 | 0.491777852 | 0.55765663 | 0.88186497 | 0.37784983 | 0.71305115 |
| VPS36 | 246.997535 | 0.491970765 | 0.17747571 | 2.77204563 | 0.00557052 | 0.10266574 |
| CCNL2 | 290.869238 | 0.492031307 | 0.15917852 | 3.09106596 | 0.00199439 | 0.05746521 |
| FLYWCH1 | 80.34078 | 0.492070994 | 0.25974507 | 1.89443828 | 0.05816686 | 0.33831183 |
| TCF7L1 | 1069.44558 | 0.492132464 | 0.25843199 | 1.90430163 | 0.05687091 | 0.33347614 |
| CRTC1 | 73.2544088 | 0.492371661 | 0.30392749 | 1.62003002 | 0.10522583 | 0.43797647 |
| GPRC5B | 307.291378 | 0.492995489 | 0.28040789 | 1.75813704 | 0.07872419 | 0.38667472 |
| WDR47 | 80.2746491 | 0.493172246 | 0.22678263 | 2.17464738 | 0.02965655 | 0.23885783 |
| SYN3 | 517.8246 | 0.493208556 | 0.60934825 | 0.80940341 | 0.41828314 | 0.73821752 |
| TESK2 | 99.0435628 | 0.493367092 | 0.26815654 | 1.83984731 | 0.06579066 | 0.35579065 |
| MIPOL1 | 264.288992 | 0.493471515 | 0.20614546 | 2.39380249 | 0.01667473 | 0.18464043 |
| POLB | 109.205686 | 0.494052316 | 0.37657381 | 1.31196673 | 0.18953136 | 0.55529769 |
| ANKFY1 | 235.182393 | 0.494452089 | 0.33532885 | 1.47452891 | 0.14033924 | 0.49118733 |
| AL356056.2 | 51.4334619 | 0.494703267 | 0.40836241 | 1.21143194 | 0.22572991 | 0.59165529 |
| CGNL1 | 341.459524 | 0.494711765 | 0.4500248 | 1.09929888 | 0.27163772 | 0.63617958 |
| ST5 | 139.440103 | 0.495314585 | 0.38163099 | 1.29788878 | 0.19432556 | 0.55913011 |
| CHN1 | 677.515946 | 0.495503546 | 0.24812167 | 1.9970184 | 0.04582318 | 0.30190911 |
| FAM199X | 100.00381 | 0.496445392 | 0.21951675 | 2.2615376 | 0.02372599 | 0.21500526 |
| CALCOCO1 | 87.3079068 | 0.498026933 | 0.24044497 | 2.07127197 | 0.03833339 | 0.27652057 |
| ZNF345 | 99.5249333 | 0.498631638 | 0.32744283 | 1.52280519 | 0.12780745 | 0.47495385 |
| CDYL2 | 211.231137 | 0.498854862 | 0.28680618 | 1.73934491 | 0.08197411 | 0.39342766 |
| STARD4 | 158.85902 | 0.498861394 | 0.51769596 | 0.96361847 | 0.33523724 | 0.68631363 |
| ADAM22 | 46.9289998 | 0.499080874 | 0.35337493 | 1.41232677 | 0.15785377 | 0.51462844 |
| PTPDC1 | 89.3957337 | 0.499179971 | 0.22966593 | 2.17350463 | 0.02974236 | 0.23931339 |
| TCIRG1 | 51.3032172 | 0.500034387 | 0.30260315 | 1.65244279 | 0.09844432 | 0.42242782 |
| OVCH1 | 83.4711303 | 0.500316005 | 0.71713275 | 0.69766163 | 0.48538883 | 0.78823909 |
| RIC1 | 348.898305 | 0.500810139 | 0.2238332 | 2.23742566 | 0.02525853 | 0.22129612 |
| EFR3B | 47.8583748 | 0.500940158 | 0.61793671 | 0.8106658 | 0.41755762 | 0.73821752 |
| OXR1 | 423.244293 | 0.501084677 | 0.21484126 | 2.33234848 | 0.01968237 | 0.19704872 |
| ODF2L | 520.099047 | 0.501318166 | 0.30131179 | 1.66378543 | 0.09615532 | 0.41727618 |
| INPP4B | 353.007714 | 0.501347128 | 0.88663634 | 0.56544843 | 0.57176879 | 0.83368112 |
| BTNL9 | 753.119362 | 0.501695824 | 0.5893144 | 0.85132117 | 0.39459097 | 0.72461789 |
| TCEA2 | 99.0212211 | 0.502079506 | 0.26523875 | 1.89293422 | 0.05836662 | 0.33873336 |
| FAM160A2 | 60.3422207 | 0.502305416 | 0.26122922 | 1.92285307 | 0.0544985 | 0.32727552 |
| MAGI3 | 551.887863 | 0.50272391 | 0.25318889 | 1.98556859 | 0.04708125 | 0.30480001 |
| GRK3 | 67.7142685 | 0.503482168 | 0.31183791 | 1.61456368 | 0.10640522 | 0.43901385 |
| IGHMBP2 | 39.3850142 | 0.503516434 | 0.35343761 | 1.42462606 | 0.15426532 | 0.51037454 |

| | | | | | | | |
|---|---|---|---|---|---|---|---|
| CHST11 | 681.450298 | | 0.504060702 | 0.41065249 | 1.22746293 | 0.21964865 | 0.58680538 |
| FHIT | 707.459226 | | 0.504309589 | 0.31700992 | 1.59083222 | 0.11164734 | 0.44839019 |
| ZNF891 | 50.1782729 | | 0.504890022 | 0.24051691 | 2.09918724 | 0.0358004 | 0.26597323 |
| AC124319.2 | 132.043513 | | 0.504908178 | 0.39159683 | 1.28935715 | 0.19727395 | 0.56247134 |
| CCM2L | 204.312952 | | 0.505153725 | 0.33832562 | 1.49309925 | 0.13541121 | 0.48508022 |
| PFDN1 | 125.048356 | | 0.505358007 | 0.17166197 | 2.94391358 | 0.0032409 | 0.07557195 |
| HAGH | 93.0983963 | | 0.505558955 | 0.25046733 | 2.01846267 | 0.0435431 | 0.29561649 |
| FAM214A | 335.569222 | | 0.505724028 | 0.23127953 | 2.18663547 | 0.02876915 | 0.23448 |
| PTK7 | 44.0251475 | | 0.505938001 | 0.41023497 | 1.23328833 | 0.21746821 | 0.58345652 |
| SOX5 | 1208.11224 | | 0.506168993 | 0.54141485 | 0.93490046 | 0.3498396 | 0.69858573 |
| SNX7 | 83.4707764 | | 0.506569099 | 0.23691886 | 2.13815442 | 0.03250421 | 0.25201798 |
| LMCD1-AS1 | 226.152412 | | 0.506822275 | 0.39596319 | 1.27997322 | 0.20055455 | 0.56536969 |
| PTGIS | 138.189328 | | 0.507467979 | 0.58950576 | 0.86083634 | 0.38932819 | 0.72202943 |
| LIPE-AS1 | 410.290013 | | 0.508329047 | 0.34051369 | 1.49282997 | 0.1354817 | 0.48508022 |
| TMEM67 | 55.275178 | | 0.508982861 | 0.37796952 | 1.34662409 | 0.17810133 | 0.54178558 |
| MSANTD2 | 146.145736 | | 0.509363007 | 0.22083196 | 2.30656379 | 0.02107915 | 0.20534606 |
| LRBA | 936.785494 | | 0.509697168 | 0.28505589 | 1.78806048 | 0.07376625 | 0.37609297 |
| KIAA0355 | 805.30374 | | 0.509858142 | 0.27000557 | 1.88832452 | 0.0589824 | 0.34075389 |
| CNNM2 | 140.807083 | | 0.510268301 | 0.31102478 | 1.64060336 | 0.10087978 | 0.42816349 |
| PXK | 123.924153 | | 0.51051949 | 0.28417797 | 1.79647806 | 0.07241852 | 0.37247061 |
| TMEM63A | 91.0443688 | | 0.510791598 | 0.31941448 | 1.59914978 | 0.10978733 | 0.44576178 |
| HNRNPLL | 132.168776 | | 0.511307805 | 0.25714454 | 1.98840625 | 0.04676678 | 0.30359341 |
| KDM7A | 176.803118 | | 0.511575583 | 0.26128568 | 1.95791665 | 0.05023979 | 0.31422181 |
| ACYP1 | 63.0050182 | | 0.512360507 | 0.47776819 | 1.07240398 | 0.28353862 | 0.6468618 |
| PRMT7 | 62.0687516 | | 0.512585051 | 0.41727232 | 1.22841853 | 0.2192899 | 0.58626926 |
| SELENOP | 179.544046 | | 0.513355154 | 0.44475787 | 1.15423513 | 0.24840378 | 0.61373435 |
| ABO | 126.932427 | | 0.513382627 | 0.40989748 | 1.25246593 | 0.21040013 | 0.57601348 |
| PER2 | 49.8908281 | | 0.513410557 | 0.39110357 | 1.31272277 | 0.18927638 | 0.55514288 |
| CPSF6 | 355.272587 | | 0.513575055 | 0.22406107 | 2.29212084 | 0.02189867 | 0.20902438 |
| SLK | 449.395518 | | 0.514300343 | 0.24274524 | 2.11868358 | 0.03411722 | 0.25778503 |
| MEF2C-AS1 | 363.156248 | | 0.515133062 | 0.37353646 | 1.37907038 | 0.16787305 | 0.52916221 |
| ARHGEF2 | 217.005756 | | 0.515256658 | 0.1631155 | 3.15884554 | 0.00158395 | 0.05081457 |
| RPAP3 | 45.3568163 | | 0.515361195 | 0.28599149 | 1.80201582 | 0.07154292 | 0.36936005 |
| TAOK3 | 405.500475 | | 0.515553004 | 0.24269587 | 2.12427598 | 0.03364707 | 0.25585395 |
| WDR44 | 182.766808 | | 0.51630853 | 0.14727602 | 3.50572023 | 0.00045537 | 0.02314486 |
| ACADM | 123.050632 | | 0.516515969 | 0.19350465 | 2.66926907 | 0.00760165 | 0.12366665 |
| POLA2 | 54.7204652 | | 0.517222826 | 0.24650254 | 2.09824542 | 0.03588347 | 0.26597323 |
| ADCY6 | 55.5769732 | | 0.517245229 | 0.2436578 | 2.12283467 | 0.03376771 | 0.2563276 |
| SMAD4 | 121.615644 | | 0.517547429 | 0.18630753 | 2.77792004 | 0.00547081 | 0.10194119 |
| COG3 | 167.757307 | | 0.518325731 | 0.18691376 | 2.77307416 | 0.00555295 | 0.10266574 |
| TIMP3 | 6055.86241 | | 0.518390106 | 0.36993013 | 1.40131895 | 0.16111872 | 0.51944206 |
| FUT10 | 97.8599611 | | 0.518904488 | 0.44791382 | 1.15849181 | 0.24666339 | 0.61160108 |
| RIN2 | 1023.3354 | | 0.519415067 | 0.2109134 | 2.46269354 | 0.01378977 | 0.16892472 |
| PRR14 | 43.4269192 | | 0.519992813 | 0.22317996 | 2.32992605 | 0.01981006 | 0.19704872 |
| MRPS18A | 46.8211899 | | 0.520061386 | 0.28703602 | 1.81183316 | 0.07001198 | 0.36578445 |
| STARD8 | 130.746667 | | 0.520378604 | 0.21184274 | 2.45643826 | 0.01403219 | 0.17036413 |
| RARB | 911.333422 | | 0.520475331 | 0.31165668 | 1.67002786 | 0.09491385 | 0.41466145 |
| NUTM2A-AS | 579.5234 | | 0.521151138 | 0.18795631 | 2.77272489 | 0.00555891 | 0.10266574 |
| FBXO3 | 130.319646 | | 0.521854747 | 0.23301272 | 2.23959767 | 0.02511705 | 0.22029245 |
| OPTN | 223.043506 | | 0.522353323 | 0.34689985 | 1.50577558 | 0.13212479 | 0.48092078 |
| RAD51B | 824.05254 | | 0.522915639 | 0.34337085 | 1.52288887 | 0.12778651 | 0.47495385 |
| TRAPPC10 | 213.144165 | | 0.523540221 | 0.2460456 | 2.12781789 | 0.03335219 | 0.25490607 |
| BBS2 | 69.3544194 | | 0.524433052 | 0.22973538 | 2.28277012 | 0.02244391 | 0.2108594 |
| CCDC191 | 80.5766398 | | 0.524528393 | 0.43035554 | 1.21882571 | 0.22291035 | 0.58858855 |
| ARFGEF3 | 103.23318 | | 0.524891874 | 0.42798628 | 1.22642219 | 0.22003984 | 0.58688069 |
| ALS2 | 139.800373 | | 0.525357284 | 0.20883261 | 2.51568603 | 0.0118801 | 0.15189817 |
| SNX16 | 67.0819248 | | 0.52631431 | 0.29611794 | 1.77738072 | 0.07550562 | 0.38092632 |
| LTBP1 | 384.12475 | | 0.526861034 | 0.56809823 | 0.92741186 | 0.35371274 | 0.69977303 |
| GAS6 | 357.662361 | | 0.52723912 | 0.51005989 | 1.03368081 | 0.30128541 | 0.65926326 |
| AL445250.1 | 82.0203182 | | 0.527261302 | 0.54597645 | 0.9657217 | 0.33418346 | 0.6852158 |
| ZNF667-AS1 | 42.8933265 | | 0.527323725 | 0.25129069 | 2.09846108 | 0.03586443 | 0.26597323 |
| ZNF423 | 317.294896 | | 0.527558345 | 0.35129609 | 1.50174843 | 0.13316209 | 0.48236854 |
| NEK3 | 121.58023 | | 0.527880386 | 0.35055533 | 1.50584041 | 0.13210815 | 0.48092078 |
| CCSER1 | 315.571711 | | 0.52808726 | 0.67695685 | 0.78008999 | 0.43533791 | 0.74965701 |
| USP46 | 59.5454454 | | 0.528309242 | 0.23649868 | 2.23387821 | 0.02549109 | 0.22167223 |
| OGFRL1 | 51.1360272 | | 0.530024167 | 0.27490069 | 1.92805691 | 0.05384805 | 0.32519452 |
| PROSER2 | 47.4545205 | | 0.530475589 | 0.37940469 | 1.39817878 | 0.16205939 | 0.52148327 |
| ARHGAP29 | 1680.68869 | | 0.530697418 | 0.22515483 | 2.35703329 | 0.0184216 | 0.19350952 |
| SAP130 | 107.753893 | | 0.532305693 | 0.34656344 | 1.53595456 | 0.12454953 | 0.47032246 |
| TP53INP1 | 74.9214721 | | 0.532407113 | 0.52177872 | 1.02036954 | 0.30755323 | 0.66400081 |
| AC124312.1 | 221.779922 | | 0.532449572 | 0.20267616 | 2.62709516 | 0.00861172 | 0.1314734 |
| PIP4P2 | 62.2339315 | | 0.532598753 | 0.36284627 | 1.46783582 | 0.14214882 | 0.49392944 |
| SLC14A1 | 492.121333 | | 0.533055184 | 1.15401783 | 0.46191243 | 0.64414412 | 0.86523823 |
| TDRD10 | 140.72775 | | 0.533700225 | 0.27202759 | 1.96193417 | 0.04977015 | 0.3126216 |
| MEAF6 | 81.3328934 | | 0.534533868 | 0.19542687 | 2.73521172 | 0.00623402 | 0.11041766 |
| CSNK1E | 76.5595677 | | 0.534562212 | 0.46133226 | 1.15873582 | 0.24656389 | 0.61160108 |
| GTF2I | 731.441899 | | 0.534880864 | 0.14365556 | 3.72335644 | 0.00019659 | 0.01398878 |
| CASTOR2 | 46.6361593 | | 0.535229181 | 0.40259227 | 1.32945716 | 0.18369719 | 0.54841084 |
| ARL6IP6 | 123.089121 | | 0.535384531 | 0.2597042 | 2.06151664 | 0.03925378 | 0.28057468 |
| DENND5A | 608.072677 | | 0.535469103 | 0.32473573 | 1.64893803 | 0.09916033 | 0.42376187 |
| ZBTB20 | 4960.95318 | | 0.53551928 | 0.29904045 | 1.79079214 | 0.07332666 | 0.37502002 |
| SETDB2 | 80.3128312 | | 0.535642544 | 0.21599738 | 2.47985664 | 0.01314352 | 0.16347517 |
| MICALL1 | 78.1324589 | | 0.535885659 | 0.31130766 | 1.72140208 | 0.08517789 | 0.39686932 |
| ASB7 | 64.1588739 | | 0.535937159 | 0.19240967 | 2.7853962 | 0.00534624 | 0.10080241 |
| TMEM117 | 259.087458 | | 0.536409383 | 0.23122623 | 2.31984662 | 0.02034918 | 0.20014099 |
| NDE1 | 128.718597 | | 0.536658887 | 0.24215618 | 2.21616845 | 0.02667997 | 0.22752964 |
| XRN1 | 610.201408 | | 0.537342654 | 0.36632035 | 1.46686543 | 0.14241267 | 0.49463618 |
| FAM157C | 60.7035852 | | 0.537596725 | 0.31143342 | 1.72620112 | 0.08431124 | 0.39616527 |
| SRSF5 | 382.856574 | | 0.53761665 | 0.26279924 | 2.04573134 | 0.04078281 | 0.28670444 |
| GPIHBP1 | 79.0049406 | | 0.537928662 | 0.73861992 | 0.72828887 | 0.46643678 | 0.77546772 |
| TRIM26 | 93.9572956 | | 0.538092506 | 0.36304406 | 1.48216862 | 0.13829543 | 0.48991657 |
| ATG4C | 133.374609 | | 0.538363632 | 0.29023123 | 1.85494727 | 0.06360376 | 0.3533385 |
| LCOR | 482.195067 | | 0.538575262 | 0.18795819 | 2.86539923 | 0.00416484 | 0.08875226 |
| FTX | 3212.3435 | | 0.538950126 | 0.26180568 | 2.05858838 | 0.03953369 | 0.28130797 |
| MCL1 | 257.797323 | | 0.541364036 | 0.43258017 | 1.25147679 | 0.21076058 | 0.57661444 |
| ZFX | 190.113103 | | 0.541516951 | 0.22940615 | 2.3605163 | 0.01824952 | 0.19288766 |
| MSI2 | 1110.8156 | | 0.541747323 | 0.40347724 | 1.34269612 | 0.17937039 | 0.54322277 |
| WARS | 673.598339 | | 0.541765008 | 0.41859059 | 1.29425987 | 0.19557567 | 0.56046814 |
| DSTYK | 165.271126 | | 0.542107262 | 0.32251978 | 1.68084963 | 0.09279213 | 0.41187311 |
| MTMR4 | 47.3338437 | | 0.542261463 | 0.22917458 | 2.36615017 | 0.01797415 | 0.19138504 |
| ELN | 141.312659 | | 0.542437438 | 0.50607974 | 1.07184183 | 0.28379108 | 0.64704999 |
| ECT2 | 75.7883691 | | 0.542854605 | 0.45026546 | 1.20563235 | 0.22795928 | 0.59483125 |
| RUBCN | 94.6160413 | | 0.542897108 | 0.26621535 | 2.03931554 | 0.04141855 | 0.28844933 |
| EPG5 | 176.207015 | | 0.543216297 | 0.29298998 | 1.85404396 | 0.06373287 | 0.3533385 |
| CSAD | 118.84912 | | 0.543341443 | 0.27846887 | 1.95117478 | 0.05103626 | 0.31710683 |
| NAIP | 332.464137 | | 0.544959806 | 0.46119108 | 1.18163561 | 0.23735032 | 0.60074482 |
| AC021231.1 | 95.9264746 | | 0.544974965 | 0.36887247 | 1.47740754 | 0.13956643 | 0.49091481 |
| NBEAL2 | 86.1616363 | | 0.545481303 | 0.31243412 | 1.74590824 | 0.0808269 | 0.39055112 |
| AQP2 | 72.1553787 | | 0.545984321 | 0.63486948 | 0.85999459 | 0.38979202 | 0.72202943 |
| PLEKHA2 | 158.851024 | | 0.546474297 | 0.29338447 | 1.86265585 | 0.0625107 | 0.35061082 |
| SMURF1 | 448.065168 | | 0.546741279 | 0.25656099 | 2.13103823 | 0.03308599 | 0.25421848 |
| AGO4 | 249.788254 | | 0.54707952 | 0.33478639 | 1.63411516 | 0.10223467 | 0.43132417 |
| ARID3A | 54.4097235 | | 0.547172816 | 0.32039231 | 1.70782132 | 0.0876695 | 0.40026508 |

| | | | | | | |
|---|---|---|---|---|---|---|
| CROCC | 88.1832714 | 0.547242986 | 0.27506894 | 1.98947576 | 0.04664872 | 0.30359341 |
| PPP1R12B | 338.359111 | 0.547600644 | 0.16833187 | 3.25310142 | 0.00114153 | 0.04043775 |
| NKTR | 958.421539 | 0.54799983 | 0.25589613 | 2.14149321 | 0.03223429 | 0.25097351 |
| PTPN21 | 101.070108 | 0.548466669 | 0.24416272 | 2.24631617 | 0.02468376 | 0.21860091 |
| SP100 | 913.297889 | 0.549067333 | 0.30933935 | 1.77496762 | 0.07590323 | 0.38199026 |
| RFX8 | 50.6174472 | 0.549351842 | 0.96612717 | 0.56861235 | 0.56961924 | 0.83368112 |
| AC011447.3 | 52.9762001 | 0.549521567 | 0.24109319 | 2.27929108 | 0.02264977 | 0.21131824 |
| SLC15A3 | 57.476126 | 0.549881545 | 0.41549605 | 1.32343386 | 0.18569114 | 0.55154649 |
| PABPN1 | 97.1058413 | 0.55049715 | 0.27646956 | 1.99116733 | 0.04646249 | 0.30359341 |
| UNC5B | 147.059191 | 0.55116082 | 0.46249466 | 1.19171284 | 0.23337386 | 0.59910683 |
| KIN | 64.0810436 | 0.55134241 | 0.24176579 | 2.28048152 | 0.02257915 | 0.21131824 |
| CLK1 | 276.475034 | 0.551535217 | 0.35286661 | 1.56301332 | 0.11804946 | 0.46065749 |
| ZBTB37 | 88.315447 | 0.552045792 | 0.27745377 | 1.98968565 | 0.04662557 | 0.30359341 |
| HNRNPA2B1 | 899.582211 | 0.552101086 | 0.23496445 | 2.34972177 | 0.01878745 | 0.19394842 |
| ZNF217 | 93.5873189 | 0.552307004 | 0.25255263 | 2.18689864 | 0.02874992 | 0.23448 |
| TMEM233 | 171.581265 | 0.553639437 | 0.50663611 | 1.09277532 | 0.27449246 | 0.63975272 |
| AC005920.2 | 59.7384232 | 0.553664439 | 0.64056491 | 0.86433776 | 0.38740237 | 0.72055795 |
| CUEDC1 | 170.65928 | 0.55399597 | 0.33907829 | 1.63382909 | 0.10229474 | 0.43132417 |
| PGGHG | 52.2074564 | 0.554923139 | 0.39334653 | 1.41077421 | 0.1583112 | 0.51533541 |
| NOS1AP | 190.107604 | 0.557121532 | 0.30863363 | 1.8051226 | 0.0710555 | 0.3687046 |
| FAM66D | 54.9570487 | 0.557176292 | 0.64040729 | 0.87003427 | 0.38428167 | 0.71699718 |
| CCND3 | 325.014001 | 0.557819091 | 0.47246974 | 1.18064511 | 0.23774374 | 0.60083682 |
| VPS45 | 253.816167 | 0.558362671 | 0.17309399 | 3.22577729 | 0.00125631 | 0.04283496 |
| MIER3 | 51.2140606 | 0.558440574 | 0.28175267 | 1.9820241 | 0.04747654 | 0.30604664 |
| TAPT1-AS1 | 100.155685 | 0.559103394 | 0.47237258 | 1.18360679 | 0.23656875 | 0.60027509 |
| FOXN3 | 896.020746 | 0.559248389 | 0.27304267 | 2.0482088 | 0.04053954 | 0.28580837 |
| CCDC84 | 161.452073 | 0.559581212 | 0.27404319 | 2.04194535 | 0.04115695 | 0.28785243 |
| COBLL1 | 500.538384 | 0.560175395 | 0.61817042 | 0.90618278 | 0.36483911 | 0.70668224 |
| PRRG4 | 56.6544436 | 0.560221936 | 0.54941984 | 1.01966091 | 0.30788931 | 0.66414532 |
| RBM45 | 60.3327958 | 0.560419571 | 0.22196271 | 2.52483663 | 0.01157521 | 0.15149764 |
| DNM3OS | 65.744699 | 0.560880037 | 0.3201101 | 1.75214727 | 0.07974849 | 0.38738328 |
| FAM85B | 134.282249 | 0.560916601 | 0.47358931 | 1.18439457 | 0.23625691 | 0.60018541 |
| NIN | 317.443983 | 0.561804196 | 0.27242142 | 2.06226148 | 0.03918285 | 0.28057468 |
| SPAG9 | 970.657809 | 0.561879738 | 0.27352304 | 2.05423186 | 0.03995326 | 0.28379991 |
| DOPEY1 | 270.359565 | 0.562072075 | 0.20204898 | 2.78186044 | 0.00540483 | 0.10143969 |
| PCDH11Y | 285.58056 | 0.562193499 | 1.60844729 | 0.3495256 | 0.72669476 | 0.90290665 |
| COL4A3 | 44.0452953 | 0.562640131 | 0.41502729 | 1.35567021 | 0.17520412 | 0.53845498 |
| NIPA1 | 59.8487734 | 0.563154086 | 0.2970266 | 1.8959719 | 0.05796376 | 0.33805823 |
| NLK | 929.516676 | 0.564299646 | 0.27034462 | 2.08733447 | 0.03685791 | 0.27062343 |
| RUSC2 | 237.250961 | 0.564484107 | 0.25762596 | 2.19109949 | 0.02844459 | 0.2334625 |
| NSMCE4A | 62.6635462 | 0.565320716 | 0.30961789 | 1.82586577 | 0.06787048 | 0.36115704 |
| KLF8 | 47.9164278 | 0.565558849 | 0.42970605 | 1.31615288 | 0.18812274 | 0.55388137 |
| ACVR2A | 152.741491 | 0.565865475 | 0.2090683 | 2.70660577 | 0.00679749 | 0.11693671 |
| DCUN1D2 | 136.803488 | 0.566309193 | 0.24065498 | 2.35319951 | 0.01861264 | 0.19394842 |
| AP1S2 | 266.448425 | 0.566316319 | 0.23518459 | 2.40796528 | 0.01604171 | 0.1817684 |
| CCSER2 | 458.959387 | 0.56649475 | 0.20528785 | 2.75951432 | 0.00578874 | 0.10503153 |
| CD38 | 108.248951 | 0.567021515 | 0.90011896 | 0.62994064 | 0.52873342 | 0.81207005 |
| SRGAP2 | 377.45377 | 0.567150084 | 0.21286928 | 2.66431155 | 0.00771461 | 0.12405 |
| CDC14A | 465.250489 | 0.567244897 | 0.19960752 | 2.84180125 | 0.00448594 | 0.09141064 |
| SMAGP | 73.1393788 | 0.567713822 | 0.32258288 | 1.75990067 | 0.07842465 | 0.38566641 |
| ZCCHC2 | 248.095322 | 0.567920419 | 0.41567613 | 1.366257 | 0.17185831 | 0.53377758 |
| 5-Mar | 193.817468 | 0.568737597 | 0.20690276 | 2.74881592 | 0.0059811 | 0.10663031 |
| FUT8 | 500.411105 | 0.568870718 | 0.33921115 | 1.67704017 | 0.09353463 | 0.41261124 |
| KATNAL1 | 130.388582 | 0.569141499 | 0.21906683 | 2.59802677 | 0.00937612 | 0.13676431 |
| EED | 77.1820366 | 0.569434699 | 0.2492169 | 2.28489599 | 0.02231893 | 0.2108594 |
| WASHC4 | 410.87844 | 0.570110968 | 0.1748765 | 3.26007772 | 0.00111382 | 0.03980071 |
| AC090825.1 | 137.219033 | 0.570538452 | 0.188772 | 3.02236794 | 0.00250805 | 0.06620456 |
| ZNF718 | 196.357759 | 0.57083185 | 0.1844528 | 3.09473134 | 0.00196991 | 0.05722046 |
| USHBP1 | 63.3446427 | 0.57097801 | 0.29453618 | 1.93856664 | 0.05255413 | 0.32082907 |
| POU6F1 | 74.6735602 | 0.571151765 | 0.354488 | 1.61120196 | 0.10713571 | 0.44010619 |
| AR | 218.680276 | 0.572447257 | 0.28205389 | 2.02956693 | 0.04240058 | 0.291241 |
| WDR78 | 56.5037431 | 0.572533069 | 0.47925075 | 1.19464199 | 0.23222694 | 0.59908986 |
| ARL6 | 48.7897966 | 0.573214249 | 0.30937203 | 1.85283153 | 0.06390651 | 0.3533385 |
| FOCAD | 340.324812 | 0.573892752 | 0.29258269 | 1.96147196 | 0.04982399 | 0.31266084 |
| GLUL | 215.853662 | 0.574419221 | 0.374136 | 1.53532196 | 0.12470476 | 0.47038289 |
| FAM126B | 254.238572 | 0.574599244 | 0.21549147 | 2.6664594 | 0.00766549 | 0.12405 |
| MIER2 | 122.891 | 0.574884914 | 0.33201112 | 1.73152309 | 0.08335851 | 0.39543345 |
| KDM6A | 686.703128 | 0.575016739 | 0.34240507 | 1.67934645 | 0.09308454 | 0.41205855 |
| PCMTD2 | 164.609393 | 0.575720894 | 0.33667642 | 1.7100125 | 0.08726356 | 0.39981956 |
| REPS2 | 53.2234262 | 0.575913188 | 0.40972654 | 1.4056038 | 0.15984181 | 0.51780899 |
| KLC1 | 293.795599 | 0.575979354 | 0.19127619 | 3.01124437 | 0.00260179 | 0.0680207 |
| ENAH | 192.342224 | 0.575995229 | 0.32189036 | 1.78941436 | 0.07354811 | 0.375683 |
| MYCBP2 | 1044.93959 | 0.576289487 | 0.22203283 | 2.59551472 | 0.00944494 | 0.13752297 |
| RALGPS1 | 107.126541 | 0.57642598 | 0.45177141 | 1.27592398 | 0.20198239 | 0.56693091 |
| GRB10 | 1825.99345 | 0.577604259 | 0.37721841 | 1.53121968 | 0.12571511 | 0.47103973 |
| RNF213 | 922.060738 | 0.578251916 | 0.4247323 | 1.3614503 | 0.17337143 | 0.53536344 |
| MLLT3 | 699.363055 | 0.578985473 | 0.32684979 | 1.77141148 | 0.0764923 | 0.38307007 |
| CIITA | 274.262638 | 0.580677167 | 0.44162807 | 1.31485566 | 0.18855842 | 0.55388137 |
| KLHDC1 | 127.59068 | 0.581061 | 0.33533956 | 1.73275411 | 0.08313938 | 0.39543345 |
| SESN1 | 325.157409 | 0.581243029 | 0.24088466 | 2.41295165 | 0.01582392 | 0.18072864 |
| COPS4 | 67.8837648 | 0.58218386 | 0.29261473 | 1.98959175 | 0.04663593 | 0.30359341 |
| HIF3A | 415.636877 | 0.582757632 | 0.59326845 | 0.9822832 | 0.32596035 | 0.67986548 |
| DNASE1 | 114.648301 | 0.583747511 | 0.37194307 | 1.56945391 | 0.11654221 | 0.4592091 |
| ITPK1 | 119.909817 | 0.584134202 | 0.28305147 | 2.06370311 | 0.03904587 | 0.28002839 |
| USP42 | 143.208948 | 0.585088962 | 0.27232293 | 2.14851157 | 0.03167314 | 0.24849596 |
| ACOT9 | 120.844324 | 0.585265143 | 0.22808354 | 2.56601223 | 0.01028752 | 0.14331772 |
| NPIPB13 | 54.7728705 | 0.585452972 | 0.35507723 | 1.6488046 | 0.09918767 | 0.42376187 |
| TSPAN13 | 110.095937 | 0.58628406 | 0.52872788 | 1.10885784 | 0.26749151 | 0.6327007 |
| TINAGL1 | 336.968368 | 0.58671646 | 0.49424632 | 1.18709322 | 0.23519085 | 0.59949266 |
| AIF1L | 176.735851 | 0.587346154 | 0.36753573 | 1.59806548 | 0.11002841 | 0.44594475 |
| UVRAG | 839.360393 | 0.588016255 | 0.36223573 | 1.62329721 | 0.10452589 | 0.43661833 |
| SLC35D2 | 47.6793142 | 0.588321393 | 0.28886476 | 2.03666723 | 0.0416834 | 0.28961808 |
| SLC12A2 | 205.362781 | 0.588431563 | 0.30526494 | 1.92760936 | 0.05390373 | 0.32529075 |
| ST6GAL1 | 2076.47745 | 0.58877788 | 0.45758712 | 1.28670115 | 0.19819847 | 0.56334077 |
| GKAP1 | 71.8270658 | 0.5896099 | 0.3206571 | 1.8387552 | 0.0659512 | 0.35579065 |
| ADGRF5 | 1329.26281 | 0.589928284 | 0.30017203 | 1.96530062 | 0.04937945 | 0.31166266 |
| SRSF6 | 100.175237 | 0.590124506 | 0.35153381 | 1.67871339 | 0.09320791 | 0.41205855 |
| SESN3 | 540.211695 | 0.590280275 | 1.23757037 | 0.47696704 | 0.63338559 | 0.86207417 |
| ST3GAL6 | 126.708972 | 0.590408593 | 0.58026428 | 1.01748223 | 0.30892408 | 0.66424338 |
| CSMD1 | 154.407632 | 0.590807795 | 1.27692883 | 0.46267872 | 0.64359468 | 0.86517342 |
| FAXDC2 | 116.458077 | 0.591315392 | 0.54675279 | 1.08150411 | 0.27947293 | 0.64286511 |
| VCPKMT | 51.1652677 | 0.591520433 | 0.27153796 | 2.17840788 | 0.02937568 | 0.23789138 |
| PCED1B | 779.206788 | 0.59153713 | 0.30704206 | 1.92656707 | 0.05403361 | 0.32559425 |
| PRIM2 | 253.893756 | 0.592194001 | 0.22508665 | 2.63096014 | 0.0085144 | 0.13071924 |
| RBM4 | 57.5773633 | 0.592215721 | 0.33107555 | 1.78876311 | 0.07365297 | 0.37598396 |
| RNF19B | 96.6806313 | 0.592932068 | 0.50889224 | 1.16514269 | 0.24396124 | 0.608518 |
| WAPL | 328.274912 | 0.59337606 | 0.13484828 | 4.40032338 | 1.08E-05 | 0.001769 |
| TAB1 | 60.7664693 | 0.59431882 | 0.26498149 | 2.24286917 | 0.02490525 | 0.21937533 |
| AMOTL1 | 403.678279 | 0.595473924 | 0.27138804 | 2.19417897 | 0.02822255 | 0.23279269 |
| TENT4A | 200.98029 | 0.595943755 | 0.16976751 | 3.51035227 | 0.00044751 | 0.02292344 |
| MFN1 | 155.196664 | 0.596386198 | 0.20644938 | 2.88877688 | 0.00386743 | 0.0851496 |
| STN1 | 115.535436 | 0.597271558 | 0.24281137 | 2.45981711 | 0.01390078 | 0.16969375 |
| AL021707.1 | 54.4255409 | 0.597838452 | 0.49489595 | 1.20800838 | 0.22704404 | 0.59376842 |

| | | | | | | |
|---|---|---|---|---|---|---|
| SMIM13 | 69.3256321 | 0.598589362 | 0.19651226 | 3.04606621 | 0.00231857 | 0.06319319 |
| CC2D2A | 85.9574631 | 0.59879972 | 0.52784971 | 1.13441328 | 0.25662123 | 0.62095131 |
| WHAMM | 92.7490967 | 0.600616637 | 0.23647246 | 2.53990105 | 0.01108838 | 0.14826183 |
| SPEF2 | 51.1197213 | 0.601761621 | 0.56809832 | 1.05925612 | 0.28948315 | 0.65257318 |
| JADE3 | 179.32816 | 0.603135353 | 0.18489884 | 3.26197478 | 0.00110639 | 0.03970872 |
| GCH1 | 209.344456 | 0.603751054 | 0.64018646 | 0.94308626 | 0.34563678 | 0.69497757 |
| RGL1 | 403.012238 | 0.60417206 | 0.41819282 | 1.44472126 | 0.1485362 | 0.50205357 |
| HLX | 66.5977791 | 0.604240056 | 0.28537105 | 2.11738389 | 0.03422728 | 0.25808697 |
| PPP1R9A | 147.514402 | 0.605285699 | 0.41925898 | 1.4437036 | 0.14882237 | 0.50222622 |
| GBP1 | 119.157273 | 0.605743852 | 0.64646941 | 0.93700311 | 0.34875696 | 0.69804011 |
| ROCK2 | 835.22668 | 0.606033886 | 0.25233495 | 2.40170409 | 0.0163189 | 0.18317913 |
| TNS3 | 303.459473 | 0.606697116 | 0.37611379 | 1.61306797 | 0.10672974 | 0.43932066 |
| GIT1 | 153.045598 | 0.606907441 | 0.27439684 | 2.21178726 | 0.02698137 | 0.22894365 |
| JHY | 50.6474158 | 0.607536913 | 0.3356997 | 1.80976306 | 0.07033254 | 0.36634702 |
| PLEKHH2 | 69.2787018 | 0.607960626 | 0.51251516 | 1.18622955 | 0.23553166 | 0.60004843 |
| CDC14B | 213.244371 | 0.608447282 | 0.2751298 | 2.21149173 | 0.0270018 | 0.22894365 |
| HDAC9 | 396.445761 | 0.608598066 | 0.58901535 | 1.03324654 | 0.30148854 | 0.65926326 |
| ANTXR2 | 176.773094 | 0.609349304 | 0.38661367 | 1.57611939 | 0.11499829 | 0.45592588 |
| NECTIN3 | 193.245279 | 0.610328411 | 0.339825 | 1.79600797 | 0.07249325 | 0.37262076 |
| PHF11 | 149.52019 | 0.610501972 | 0.33966414 | 1.79736954 | 0.07227697 | 0.37197639 |
| USP33 | 366.081676 | 0.611087419 | 0.20168158 | 3.02996149 | 0.00244585 | 0.06478413 |
| DMTF1 | 294.431018 | 0.611235219 | 0.32946457 | 1.85523808 | 0.06356224 | 0.3533385 |
| ZNF107 | 73.7344385 | 0.612230322 | 0.32812121 | 1.86586633 | 0.06206007 | 0.34926929 |
| CDH13 | 1416.59136 | 0.613863537 | 0.42331136 | 1.45014662 | 0.14701764 | 0.50126889 |
| ENTPD1 | 445.089293 | 0.614815922 | 0.38987842 | 1.5769427 | 0.11480871 | 0.45569454 |
| TSC22D2 | 330.917027 | 0.614977782 | 0.30695936 | 2.0034502 | 0.04512899 | 0.29989958 |
| PHLPP2 | 79.2693758 | 0.61512383 | 0.30203823 | 2.03657605 | 0.04169254 | 0.28961808 |
| TTL | 140.925486 | 0.615394119 | 0.23947851 | 2.5697258 | 0.0101779 | 0.14236887 |
| SMAD9 | 397.993618 | 0.616537 | 0.48425726 | 1.27316006 | 0.20296125 | 0.56730017 |
| TRIP4 | 126.315491 | 0.617554752 | 0.17231125 | 3.58394911 | 0.00033844 | 0.01923221 |
| STX17 | 131.200202 | 0.618669374 | 0.29028714 | 2.13123244 | 0.03307 | 0.25421848 |
| SOX18 | 600.199318 | 0.618912633 | 0.72490963 | 0.85377902 | 0.39322744 | 0.72360479 |
| SLC30A4 | 48.0215553 | 0.619455871 | 0.2764763 | 2.24053876 | 0.02505597 | 0.22005414 |
| TPP2 | 234.039992 | 0.619734936 | 0.18706772 | 3.3128908 | 0.00092337 | 0.0349812 |
| AKR1B1 | 248.063564 | 0.620053941 | 1.1108321 | 0.55818871 | 0.57671553 | 0.83585958 |
| CMTM4 | 91.8415858 | 0.62007729 | 0.19888214 | 3.11781279 | 0.00182198 | 0.05451467 |
| EGFR | 139.171437 | 0.620228098 | 0.41312661 | 1.50130269 | 0.13327729 | 0.48236854 |
| PLPP1 | 5242.1184 | 0.620323245 | 0.42203094 | 1.46985254 | 0.1416017 | 0.49349518 |
| ANK2 | 63.3683167 | 0.621179262 | 0.51908817 | 1.1966739 | 0.23143368 | 0.59795465 |
| ZC3H12C | 174.625204 | 0.62141347 | 0.31382654 | 1.98011768 | 0.04769031 | 0.30655914 |
| PIEZO2 | 194.629204 | 0.621937117 | 0.52631808 | 1.18167538 | 0.23733453 | 0.60074482 |
| C2CD5 | 252.165199 | 0.622823994 | 0.24490693 | 2.54310481 | 0.01098723 | 0.14749418 |
| FHL1 | 109.42409 | 0.623601302 | 0.32556619 | 1.91543628 | 0.05543689 | 0.32988018 |
| DPF3 | 51.1336789 | 0.623732415 | 0.40103919 | 1.55529044 | 0.11987691 | 0.46380746 |
| PDGFD | 545.264998 | 0.623870581 | 0.39082013 | 1.59631128 | 0.11041931 | 0.44631216 |
| CRY1 | 178.182076 | 0.624499056 | 0.36276316 | 1.72150629 | 0.08515899 | 0.39686932 |
| SLC25A27 | 93.71106 | 0.625199599 | 0.51193403 | 1.22125032 | 0.22199126 | 0.58750144 |
| GSE1 | 929.957834 | 0.62557182 | 0.53780434 | 1.16319592 | 0.24475002 | 0.60967714 |
| AP3M2 | 134.105504 | 0.62645989 | 0.31860273 | 1.9662728 | 0.04926711 | 0.31155957 |
| SLC47A1 | 49.4247825 | 0.626563267 | 0.36311425 | 1.7255265 | 0.08443263 | 0.39616527 |
| RAPH1 | 327.843295 | 0.627735928 | 0.339147 | 1.85092577 | 0.06418023 | 0.35347506 |
| MITF | 78.0870776 | 0.628779077 | 0.33191201 | 1.89441494 | 0.05816996 | 0.33831183 |
| MYLK | 152.104276 | 0.629488645 | 0.4528386 | 1.39009493 | 0.16450005 | 0.52418378 |
| IDH2 | 86.462748 | 0.6308076 | 0.30331891 | 2.07968438 | 0.03755449 | 0.27364951 |
| ATP2B4 | 403.657436 | 0.630948728 | 0.22875312 | 2.75820816 | 0.00581192 | 0.10504875 |
| 2-Mar | 109.72462 | 0.631451269 | 0.33818536 | 1.86717505 | 0.06187715 | 0.3489463 |
| ZBTB46 | 663.107946 | 0.632328081 | 0.33762653 | 1.87286256 | 0.06108737 | 0.34737868 |
| AC090517.4 | 210.378085 | 0.633027864 | 0.31017142 | 2.04089685 | 0.04126108 | 0.28808825 |
| AKAP7 | 105.740751 | 0.633216023 | 0.40499737 | 1.56350651 | 0.11793351 | 0.46042456 |
| XAF1 | 823.508598 | 0.633232117 | 0.69130584 | 0.91599417 | 0.35966995 | 0.70360488 |
| HEG1 | 1105.7464 | 0.633713865 | 0.44520624 | 1.4234164 | 0.15461548 | 0.51037454 |
| LRP2 | 100.484321 | 0.63469035 | 1.07187517 | 0.59213084 | 0.55376297 | 0.82374885 |
| TOB2 | 79.0776801 | 0.63709786 | 0.18808799 | 3.38723314 | 0.00070601 | 0.02932644 |
| AC106869.1 | 104.811114 | 0.637504461 | 0.40100496 | 1.58976704 | 0.11188733 | 0.44839019 |
| CEP57L1 | 101.260898 | 0.637559804 | 0.26444441 | 2.41094073 | 0.01591143 | 0.18083787 |
| EZH1 | 158.518021 | 0.638192621 | 0.19699071 | 3.2397093 | 0.00119652 | 0.04184227 |
| RASGRF2 | 1109.52061 | 0.63842958 | 0.30022802 | 2.12648235 | 0.03346312 | 0.25519915 |
| SOGA1 | 144.002239 | 0.639828732 | 0.34512611 | 1.85389836 | 0.0637537 | 0.3533385 |
| KLHL14 | 99.0680487 | 0.640247442 | 0.48694603 | 1.31482217 | 0.18856967 | 0.55388137 |
| ORC4 | 181.581109 | 0.64073376 | 0.22649276 | 2.82893704 | 0.00467029 | 0.09321212 |
| IRF1 | 140.609053 | 0.641246076 | 0.34108475 | 1.88001976 | 0.06010538 | 0.34442742 |
| MACROD2 | 692.831319 | 0.641729794 | 0.45944471 | 1.39675085 | 0.16248851 | 0.52265859 |
| FBXO25 | 135.591346 | 0.642023697 | 0.28092571 | 2.28538603 | 0.0222902 | 0.2108594 |
| RAPGEF1 | 1026.04298 | 0.642423874 | 0.31631116 | 2.03098706 | 0.04225631 | 0.291241 |
| TBX19 | 63.3666631 | 0.642634038 | 0.40823995 | 1.57415764 | 0.11545101 | 0.4568354 |
| KCNK6 | 89.9969205 | 0.642763 | 0.36188451 | 1.77615507 | 0.07570736 | 0.38160963 |
| MTMR8 | 59.66408 | 0.642824136 | 0.26693356 | 2.40818033 | 0.01603226 | 0.1817684 |
| GLIDR | 51.5780073 | 0.642999973 | 0.37597275 | 1.71023026 | 0.0872233 | 0.39981956 |
| PIP4K2A | 1210.85235 | 0.643427249 | 0.3483627 | 1.84700386 | 0.06474658 | 0.35487024 |
| CYP4V2 | 50.9627255 | 0.643754636 | 0.46596505 | 1.38155133 | 0.16710951 | 0.52764813 |
| MDM1 | 46.3068506 | 0.645611937 | 0.33351038 | 1.9358076 | 0.05289127 | 0.3215522 |
| CMC2 | 95.1222532 | 0.645906427 | 0.2449212 | 2.63720097 | 0.00835933 | 0.1297996 |
| PRPF3 | 170.559658 | 0.646391498 | 0.27778071 | 2.32698485 | 0.01996607 | 0.19732169 |
| TPTEP2-CSNI | 423.851546 | 0.64653003 | 0.26528918 | 2.4370765 | 0.01480655 | 0.17549246 |
| SLC25A28 | 101.010005 | 0.646634541 | 0.42088596 | 1.53636519 | 0.12444884 | 0.47015921 |
| HABP4 | 114.665377 | 0.647073382 | 0.34498407 | 1.8756616 | 0.06070176 | 0.34614809 |
| DMGDH | 79.0546642 | 0.648384132 | 0.355977 | 1.82142141 | 0.06854282 | 0.36169608 |
| WWC3 | 313.592764 | 0.648499337 | 0.18973418 | 3.41793632 | 0.00063098 | 0.02761122 |
| FBXO34 | 235.432179 | 0.64935422 | 0.22764542 | 2.85248095 | 0.00433794 | 0.0903241 |
| GCNT2 | 263.258875 | 0.650089099 | 0.35668281 | 1.82259723 | 0.06836441 | 0.36142549 |
| ZNF846 | 91.714831 | 0.650558878 | 0.31590144 | 2.05937293 | 0.03945853 | 0.28101751 |
| CASK | 391.015068 | 0.65116226 | 0.18551974 | 3.5099352 | 0.00044822 | 0.02292344 |
| B4GALT1 | 397.941456 | 0.651592791 | 0.3476025 | 1.87453427 | 0.06085682 | 0.34654933 |
| DSE | 111.798596 | 0.651977104 | 0.35427849 | 1.8402955 | 0.06572487 | 0.35579065 |
| PLS3 | 416.722956 | 0.652961361 | 0.3255737 | 2.00557155 | 0.04490198 | 0.29946125 |
| TTC5 | 49.6690182 | 0.652983415 | 0.24303601 | 2.68677643 | 0.00721452 | 0.12072891 |
| ANKRD36 | 268.792125 | 0.653870362 | 0.268532 | 2.43498113 | 0.01489257 | 0.17555058 |
| CLK4 | 100.274072 | 0.65445309 | 0.24551261 | 2.66565974 | 0.00768374 | 0.12405 |
| C5orf15 | 87.298807 | 0.656557054 | 0.25547379 | 2.56995859 | 0.01017107 | 0.14236887 |
| TNFSF10 | 573.295944 | 0.657518583 | 0.48324371 | 1.36063558 | 0.17362888 | 0.53536344 |
| SPAAR | 243.362553 | 0.657697201 | 0.63489201 | 1.0359198 | 0.30023956 | 0.65926326 |
| AC012358.3 | 64.2523359 | 0.658550951 | 0.32088374 | 2.05230393 | 0.04014013 | 0.28442544 |
| LINC00486 | 207.829946 | 0.659681651 | 0.2058353 | 3.20490053 | 0.00135109 | 0.04507706 |
| RAB30 | 100.217112 | 0.659761816 | 0.32344178 | 2.0398163 | 0.04136863 | 0.28834707 |
| UBN2 | 209.82747 | 0.662717856 | 0.21538993 | 3.07682845 | 0.00209216 | 0.05903489 |
| C9orf3 | 1081.62143 | 0.663101865 | 0.19823025 | 3.34510938 | 0.0008225 | 0.03283184 |
| MAP4K3 | 610.980682 | 0.663115892 | 0.20455625 | 3.2417289 | 0.00118807 | 0.04172522 |
| EPB41 | 210.78375 | 0.664944701 | 0.41670111 | 1.59573536 | 0.11054789 | 0.44631216 |
| AIDA | 86.546337 | 0.665741461 | 0.22688369 | 2.93428523 | 0.00334317 | 0.07718524 |
| ZC2HC1A | 96.5662361 | 0.667141072 | 0.33732002 | 1.977769 | 0.04795477 | 0.30730127 |
| AL021937.3 | 60.6032057 | 0.667699144 | 0.64933312 | 1.02828444 | 0.30381605 | 0.66060576 |
| RABGAP1L | 1723.03644 | 0.667888714 | 0.25859148 | 2.58279476 | 0.00980036 | 0.1397149 |
| COL4A3BP | 389.116527 | 0.668469321 | 0.22393817 | 2.98506198 | 0.00283521 | 0.07116722 |

| | | | | | | |
|---|---|---|---|---|---|---|
| NOX4 | 86.7440439 | 0.668562283 | 0.70664993 | 0.94610112 | 0.34409702 | 0.6938754 |
| LINC02100 | 58.7946713 | 0.668773716 | 0.56645547 | 1.18062893 | 0.23775017 | 0.60083682 |
| NMI | 85.9946635 | 0.670036416 | 0.40936045 | 1.63678834 | 0.1016747 | 0.43042116 |
| DUSP18 | 69.1769848 | 0.67078192 | 0.20820224 | 3.22178052 | 0.00127397 | 0.04315685 |
| NFKBIZ | 208.119588 | 0.671586539 | 0.48800218 | 1.37619579 | 0.16876102 | 0.53071749 |
| STARD9 | 311.382218 | 0.671667931 | 0.34519423 | 1.94576815 | 0.0516826 | 0.31875077 |
| NEDD1 | 100.921197 | 0.67412937 | 0.23439559 | 2.87603262 | 0.00402708 | 0.087386 |
| INO80D | 514.722094 | 0.674421385 | 0.55848235 | 1.20759659 | 0.22720247 | 0.59399292 |
| CPT1A | 247.608147 | 0.675167704 | 0.32585022 | 2.07201859 | 0.03826371 | 0.27652057 |
| KIRREL1 | 145.005467 | 0.675356608 | 0.39871366 | 1.69383863 | 0.09029595 | 0.4068172 |
| OVCH1-AS1 | 144.756857 | 0.67660064 | 0.72685928 | 0.93085506 | 0.35192854 | 0.69889281 |
| MARK4 | 512.968828 | 0.67768754 | 0.33790362 | 2.00556463 | 0.04490272 | 0.29946125 |
| SIRT1 | 100.475791 | 0.677722776 | 0.19068385 | 3.55416971 | 0.00037917 | 0.02110738 |
| TRIM25 | 102.758 | 0.678086406 | 0.40032704 | 1.69383114 | 0.09029738 | 0.4068172 |
| CASKIN2 | 237.196235 | 0.678215964 | 0.38134256 | 1.77849531 | 0.07532254 | 0.38047182 |
| STAT2 | 177.282655 | 0.678409144 | 0.32593742 | 2.08140918 | 0.03739647 | 0.27298421 |
| FGFR1OP | 77.0015442 | 0.678639801 | 0.28248756 | 2.4023706 | 0.0162892 | 0.18309849 |
| AL669831.5 | 68.3277555 | 0.679007791 | 0.34486846 | 1.96888924 | 0.04896581 | 0.31085123 |
| KL | 70.9147365 | 0.67931643 | 0.25397256 | 2.67476304 | 0.00747821 | 0.12214407 |
| CLTCL1 | 52.3270649 | 0.67932461 | 0.31152259 | 2.18065924 | 0.02920863 | 0.23735275 |
| CMTR1 | 77.9588528 | 0.679385823 | 0.37279067 | 1.82243246 | 0.06838939 | 0.36142549 |
| BBS9 | 507.132853 | 0.679689533 | 0.25825714 | 2.63183248 | 0.00849257 | 0.13062917 |
| DPYSL3 | 165.03932 | 0.680346558 | 0.63798098 | 1.06640571 | 0.28624028 | 0.64883774 |
| TLN2 | 95.4736699 | 0.681105326 | 0.45297012 | 1.50364295 | 0.13267333 | 0.48208962 |
| CPEB4 | 282.337452 | 0.681654248 | 0.22579181 | 3.01895028 | 0.00253652 | 0.06674069 |
| ARHGEF40 | 45.330712 | 0.681787245 | 0.31978158 | 2.13204036 | 0.03300353 | 0.25406197 |
| CREB5 | 227.335585 | 0.682271212 | 0.59022215 | 1.15595663 | 0.2476989 | 0.61273279 |
| SUCLG2-AS1 | 119.168927 | 0.682454186 | 0.48687617 | 1.40169972 | 0.16100493 | 0.51944206 |
| PDE3B | 68.5972073 | 0.685009951 | 0.67276058 | 1.01820762 | 0.3085793 | 0.66414532 |
| CDR2 | 477.31327 | 0.685127709 | 0.25354652 | 2.7021775 | 0.0068887 | 0.11719379 |
| KIAA1324L | 141.937445 | 0.685291302 | 0.30899913 | 2.21777743 | 0.02657001 | 0.22752964 |
| OAS3 | 96.6986103 | 0.685511178 | 1.03266731 | 0.66382578 | 0.50680183 | 0.79909604 |
| SEMA6D | 83.2242671 | 0.686619213 | 0.34145968 | 2.01083542 | 0.04434284 | 0.29889414 |
| FAT4 | 541.099523 | 0.687293288 | 0.37152422 | 1.8499286 | 0.06432384 | 0.35347506 |
| COQ8A | 77.4132279 | 0.687315366 | 0.36737494 | 1.87088254 | 0.06136136 | 0.34778328 |
| LINC00278 | 108.169962 | 0.68739414 | 2.09774843 | 0.32768188 | 0.74315221 | 0.90850686 |
| ZRSR2 | 57.5332335 | 0.68864054 | 0.31317663 | 2.19888863 | 0.02788584 | 0.23190345 |
| BTN3A1 | 78.3530889 | 0.688990158 | 0.64410777 | 1.06968148 | 0.2847627 | 0.64764125 |
| DLGAP2 | 109.797816 | 0.689225389 | 0.67915073 | 1.0148342 | 0.31018487 | 0.6648266 |
| DSCAML1 | 181.803875 | 0.689472323 | 0.71235488 | 0.96787758 | 0.33310551 | 0.6852158 |
| FOS | 76.0469642 | 0.690520563 | 1.0036161 | 0.68803257 | 0.49143227 | 0.79094859 |
| MYH11 | 61.9235687 | 0.690889697 | 0.3517351 | 1.96423304 | 0.04950307 | 0.31166266 |
| ID3 | 711.368895 | 0.691011717 | 0.77575534 | 0.89075986 | 0.37305802 | 0.71126136 |
| AC092839.1 | 120.254601 | 0.691415976 | 0.49859555 | 1.38672714 | 0.16552498 | 0.52581169 |
| MAP3K1 | 318.885441 | 0.692439441 | 0.1498559 | 4.62070185 | 3.82E-06 | 0.00074513 |
| SMURF2 | 663.249446 | 0.69268174 | 0.2851388 | 2.42927912 | 0.01512888 | 0.17660432 |
| AGTPBP1 | 537.302031 | 0.693170014 | 0.46934781 | 1.4768792 | 0.13970802 | 0.49107849 |
| KANSL2 | 44.394597 | 0.696308157 | 0.25735032 | 2.70568216 | 0.00681643 | 0.11693671 |
| HNRNPH1 | 548.007646 | 0.696328331 | 0.26755985 | 2.60251432 | 0.0092543 | 0.13620125 |
| SPRY4 | 85.1663658 | 0.696758034 | 0.47363728 | 1.47107939 | 0.14126965 | 0.49275769 |
| ZDHHC7 | 81.7855489 | 0.697989062 | 0.20400547 | 3.42142325 | 0.00062294 | 0.02740614 |
| MDFIC | 389.508805 | 0.698857725 | 0.26525168 | 2.63469667 | 0.00842125 | 0.13026671 |
| ARRDC1 | 58.9385397 | 0.700023996 | 0.28073614 | 2.49353006 | 0.01264799 | 0.15947378 |
| UNC13B | 355.314267 | 0.700730918 | 0.25100162 | 2.79173862 | 0.00524257 | 0.09999986 |
| UHRF1BP1 | 102.016882 | 0.700743813 | 0.277205 | 2.52789025 | 0.01147502 | 0.15145178 |
| FAM189A1 | 75.4222786 | 0.701370911 | 0.43527557 | 1.61132616 | 0.10710865 | 0.44010619 |
| XRRA1 | 210.374531 | 0.702727289 | 0.56817234 | 1.23682065 | 0.21615367 | 0.58202879 |
| LUZP1 | 478.923421 | 0.703314684 | 0.22343103 | 3.14779324 | 0.00164508 | 0.05170301 |
| ANKRD36C | 194.816318 | 0.704433661 | 0.31129719 | 2.26289759 | 0.023642 | 0.21448172 |
| PREX1 | 411.302105 | 0.704962994 | 0.32612982 | 2.16160235 | 0.03064884 | 0.24325845 |
| NDRG2 | 53.6729271 | 0.706755333 | 0.35256718 | 2.00459761 | 0.04500608 | 0.29976402 |
| NMT2 | 112.833083 | 0.707400841 | 0.31508377 | 2.24511993 | 0.02476043 | 0.21904285 |
| KCNC4 | 59.7445835 | 0.708057553 | 0.35497174 | 1.99468713 | 0.04607701 | 0.30284996 |
| CEP120 | 109.788644 | 0.708693704 | 0.30024653 | 2.36037266 | 0.01825659 | 0.19288766 |
| ITIH5 | 184.280933 | 0.70871187 | 1.21484966 | 0.58337414 | 0.55964147 | 0.82827747 |
| ZNF362 | 90.1380803 | 0.708929981 | 0.3467479 | 2.04451125 | 0.04090307 | 0.28670444 |
| FAR2 | 602.11578 | 0.710276087 | 0.20662745 | 3.43747198 | 0.00058717 | 0.02686611 |
| IMPA1 | 69.342194 | 0.710420808 | 0.28221473 | 2.51730594 | 0.01182561 | 0.15189817 |
| ZDHHC21 | 266.732533 | 0.711215177 | 0.37578439 | 1.892615 | 0.05840909 | 0.33873959 |
| BCL9L | 59.6602211 | 0.711527398 | 0.31134781 | 2.28531361 | 0.02229445 | 0.2108594 |
| TMC7 | 83.6173885 | 0.712195232 | 0.39254476 | 1.81430324 | 0.06963105 | 0.36501659 |
| ADAM17 | 344.562812 | 0.712591656 | 0.19344172 | 3.68375365 | 0.00022982 | 0.01505003 |
| CMTM8 | 597.658535 | 0.712968052 | 0.42457703 | 1.6792431 | 0.09310468 | 0.41205855 |
| TBC1D4 | 343.384461 | 0.713218547 | 0.56442651 | 1.26361631 | 0.20636778 | 0.57126492 |
| ADCY9 | 240.905961 | 0.715369595 | 0.38867116 | 1.84055228 | 0.0656872 | 0.35579065 |
| KANSL1L | 377.006956 | 0.715419899 | 0.27207666 | 2.62947916 | 0.00855158 | 0.13104411 |
| ANKMY2 | 45.3370204 | 0.716677758 | 0.29253168 | 2.44991504 | 0.01428899 | 0.17286813 |
| PARP9 | 339.251536 | 0.716838708 | 0.46620581 | 1.5376014 | 0.12414611 | 0.46936415 |
| FXYD2 | 46.8527699 | 0.717600909 | 0.37686089 | 1.90415329 | 0.05689022 | 0.33347614 |
| ZNF133 | 83.7427898 | 0.718120125 | 0.22293169 | 3.22125632 | 0.0012763 | 0.04315685 |
| EXOC3L1 | 68.9310602 | 0.718735777 | 0.35032992 | 2.05159687 | 0.04020886 | 0.28462723 |
| GBP3 | 83.242309 | 0.720105144 | 0.28521464 | 2.52478326 | 0.01157697 | 0.15149764 |
| GPR160 | 50.3411189 | 0.720800622 | 0.44284464 | 1.62766026 | 0.10359694 | 0.43503085 |
| LIMCH1 | 2438.96826 | 0.7221188 | 0.2136584 | 3.379782 | 0.00072543 | 0.02983026 |
| GCA | 132.497485 | 0.724897122 | 0.41152845 | 1.76147512 | 0.07815802 | 0.38566641 |
| FNDC3A | 727.081622 | 0.725223564 | 0.34515772 | 2.10113671 | 0.03562897 | 0.26552991 |
| STOX2 | 1338.91277 | 0.726394779 | 0.26103565 | 2.78274162 | 0.00539017 | 0.10139718 |
| ZEB2 | 209.874932 | 0.726970453 | 0.31800584 | 2.28602861 | 0.02225259 | 0.2108594 |
| UBASH3B | 127.225488 | 0.72770022 | 0.52463032 | 1.38707238 | 0.16541969 | 0.52581169 |
| PDIA5 | 523.421756 | 0.727892027 | 0.3340878 | 2.17874468 | 0.02935064 | 0.23789138 |
| CD58 | 193.108065 | 0.72812685 | 0.29321642 | 2.4832404 | 0.01301932 | 0.16340044 |
| IER2 | 355.188378 | 0.72813018 | 0.43685918 | 1.66673888 | 0.09556634 | 0.41575423 |
| AC034195.1 | 96.3242156 | 0.72820099 | 0.42043839 | 1.73200403 | 0.08327284 | 0.39543345 |
| ITGA3 | 201.781966 | 0.728602349 | 0.32167905 | 2.26499782 | 0.02351281 | 0.21378373 |
| FAM122C | 93.5520218 | 0.728816558 | 0.33869278 | 2.1518515 | 0.03140905 | 0.24759838 |
| KIAA0825 | 222.596702 | 0.729211836 | 0.32623011 | 2.23526831 | 0.02539974 | 0.22138474 |
| GLCCI1 | 106.198209 | 0.729507229 | 0.34457937 | 2.11709488 | 0.0342518 | 0.25808697 |
| UTRN | 4112.57986 | 0.731571481 | 0.30524908 | 2.39663777 | 0.01654627 | 0.18464043 |
| RORA | 1355.25856 | 0.734445514 | 0.40917092 | 1.79496019 | 0.07266004 | 0.37296554 |
| RAI2 | 73.552167 | 0.734896069 | 0.4385807 | 1.67562336 | 0.09381199 | 0.41280519 |
| FAM13A-AS1 | 45.7256267 | 0.737436547 | 0.28257967 | 2.60965888 | 0.00906325 | 0.13583263 |
| TSC2 | 134.234216 | 0.737484801 | 0.28591827 | 2.57935525 | 0.00989849 | 0.14086847 |
| NCOA2 | 900.202911 | 0.73751018 | 0.25398934 | 2.90370529 | 0.00368775 | 0.08250421 |
| TBC1D1 | 786.186102 | 0.737652902 | 0.17390314 | 4.24174564 | 2.22E-05 | 0.00324088 |
| NLRC5 | 235.596495 | 0.73810948 | 0.51878127 | 1.42277589 | 0.15480113 | 0.51078132 |
| RAPGEF2 | 1186.46228 | 0.741618446 | 0.24681234 | 3.00478678 | 0.00265767 | 0.0688219 |
| LEF1 | 109.277917 | 0.741623908 | 0.75654425 | 0.98027829 | 0.32694877 | 0.68007671 |
| BTN3A3 | 62.4229301 | 0.74440106 | 0.5426915 | 1.37168366 | 0.17016194 | 0.53193492 |
| CCDC146 | 141.832542 | 0.745026834 | 0.28459422 | 2.61785649 | 0.0088484 | 0.13359126 |
| KIF3A | 69.2973826 | 0.745929426 | 0.23621884 | 3.15778971 | 0.0015897 | 0.05081457 |
| BTBD3 | 260.090691 | 0.746058361 | 0.17710623 | 4.21249084 | 2.53E-05 | 0.00345277 |
| KHDRBS3 | 210.204217 | 0.746397053 | 0.39738627 | 1.87826582 | 0.06034481 | 0.3448335 |
| LINC00894 | 146.989799 | 0.747359833 | 0.36568532 | 2.0437239 | 0.04098083 | 0.28686581 |

| | | | | | | |
|---|---|---|---|---|---|---|
| HES4 | 120.661348 | 0.748012214 | 0.59064463 | 1.26643361 | 0.20535789 | 0.57041534 |
| ASS1 | 86.3399518 | 0.748431966 | 0.40549565 | 1.84572132 | 0.06493268 | 0.35498206 |
| PTPN13 | 63.0318757 | 0.749082924 | 0.44371091 | 1.6882229 | 0.09136845 | 0.40862523 |
| AP000787.1 | 252.708949 | 0.749348258 | 0.70003407 | 1.07044541 | 0.28441887 | 0.64739905 |
| PTPRB | 3001.72607 | 0.749551148 | 0.32903572 | 2.27802366 | 0.02272517 | 0.21131824 |
| PCNX1 | 745.638163 | 0.749698677 | 0.27740581 | 2.70253416 | 0.00688131 | 0.11719379 |
| PDE4DIP | 52.1244373 | 0.750540696 | 0.29664811 | 2.53007075 | 0.01140395 | 0.15075694 |
| RCBTB2 | 96.280811 | 0.75105291 | 0.21561757 | 3.48326393 | 0.00049534 | 0.02427165 |
| MAP2 | 362.675282 | 0.751132914 | 0.46303632 | 1.62219006 | 0.10476266 | 0.43679299 |
| SIDT1 | 81.4361054 | 0.75160643 | 0.46252537 | 1.62500584 | 0.10416131 | 0.4357628 |
| ZNFX1 | 226.078445 | 0.752709514 | 0.41432651 | 1.81670613 | 0.06926212 | 0.36386894 |
| CD34 | 644.59922 | 0.753708554 | 0.46101452 | 1.63489113 | 0.10207187 | 0.43121018 |
| CLIP2 | 161.363668 | 0.754283238 | 0.30102432 | 2.50572194 | 0.01222017 | 0.15551732 |
| RASSF9 | 95.7457512 | 0.755471391 | 0.39386198 | 1.91811202 | 0.05509681 | 0.32909284 |
| COL15A1 | 98.3708374 | 0.755859834 | 0.88835659 | 0.85085183 | 0.39485166 | 0.72461789 |
| TGFB2 | 65.4640216 | 0.75641908 | 0.50774947 | 1.48974862 | 0.13629034 | 0.48637762 |
| LINC00854 | 179.193655 | 0.756583265 | 0.69167105 | 1.0938484 | 0.27402147 | 0.63974828 |
| STK38L | 341.002232 | 0.758739978 | 0.52670997 | 1.44052707 | 0.14971834 | 0.50376034 |
| EBF1 | 2112.78274 | 0.758956689 | 0.22166123 | 3.42394875 | 0.00061718 | 0.02729949 |
| ATP2A3 | 124.938254 | 0.759945149 | 0.67005716 | 1.13414974 | 0.25673174 | 0.62099788 |
| ANO6 | 689.482527 | 0.761500488 | 0.19687684 | 3.86790284 | 0.00010978 | 0.00935721 |
| ABR | 550.840741 | 0.762489987 | 0.27044512 | 2.81938897 | 0.00481152 | 0.09480113 |
| AC104389.4 | 128.995528 | 0.766613869 | 0.43825919 | 1.74922486 | 0.08025216 | 0.38871437 |
| TMEM47 | 196.92524 | 0.767060884 | 0.50688395 | 1.51328698 | 0.13020678 | 0.47888574 |
| TLE2 | 147.169497 | 0.76720887 | 0.34213319 | 2.24242748 | 0.02493376 | 0.21939025 |
| UBE2J1 | 318.891132 | 0.768712346 | 0.3886786 | 1.97775837 | 0.04795597 | 0.30730127 |
| PPM1K | 97.8321566 | 0.769901794 | 0.48100442 | 1.60061273 | 0.10946272 | 0.44576178 |
| ZNF519 | 47.6596456 | 0.770888626 | 0.32427534 | 2.37726566 | 0.01744152 | 0.18853893 |
| LDLRAD4 | 993.705193 | 0.770966428 | 0.45309037 | 1.70157318 | 0.08883541 | 0.4036314 |
| MTSS1 | 1026.9273 | 0.774477645 | 0.33972451 | 2.2797226 | 0.02262415 | 0.21131824 |
| TIAM2 | 414.034114 | 0.77545029 | 0.23662766 | 3.27709064 | 0.00104883 | 0.03823551 |
| LLGL2 | 70.0790418 | 0.776877708 | 0.26438281 | 2.93845776 | 0.0032985 | 0.07646344 |
| MUM1 | 87.7035648 | 0.777606934 | 0.25593692 | 3.03827573 | 0.00237936 | 0.06383709 |
| ARHGAP10 | 174.731301 | 0.77897392 | 0.2782167 | 2.79988192 | 0.00511213 | 0.0986617 |
| CDK6 | 433.428267 | 0.779421334 | 0.30040869 | 2.59453662 | 0.00947185 | 0.13766993 |
| DANT2 | 90.0864259 | 0.780358194 | 1.02319755 | 0.7626662 | 0.44566249 | 0.76055395 |
| PRKG1 | 783.578889 | 0.780523528 | 0.65250161 | 1.1962017 | 0.23161786 | 0.59795465 |
| GBP4 | 358.522258 | 0.780842414 | 0.52920391 | 1.47550386 | 0.14007713 | 0.49118733 |
| GSN | 769.386208 | 0.781382412 | 0.61595325 | 1.26857422 | 0.20459296 | 0.56945042 |
| CTNNAL1 | 247.347193 | 0.781745221 | 0.44227709 | 1.76754627 | 0.07713679 | 0.38356763 |
| SOX13 | 237.357404 | 0.784123667 | 0.3508077 | 2.23519511 | 0.02540454 | 0.22138474 |
| NIPSNAP3A | 44.6339706 | 0.78580011 | 0.25338751 | 3.10117934 | 0.00192752 | 0.05673689 |
| NR3C2 | 901.057742 | 0.788628028 | 0.32943793 | 2.39385922 | 0.01667215 | 0.18464043 |
| MIGA2 | 41.6906099 | 0.788838078 | 0.2295226 | 3.43686452 | 0.00058849 | 0.02686611 |
| KAT2B | 316.392121 | 0.78908868 | 0.39264649 | 2.00966698 | 0.04446645 | 0.29898844 |
| TAL1 | 103.710615 | 0.789330341 | 0.36268076 | 2.17637774 | 0.02952703 | 0.23828369 |
| GSAP | 98.9509422 | 0.78947655 | 0.43802587 | 1.80235141 | 0.07149014 | 0.36932056 |
| CACNA2D3 | 53.5717157 | 0.791622281 | 0.65242726 | 1.21334948 | 0.22499623 | 0.59122266 |
| LINC02062 | 66.3245222 | 0.792035895 | 0.4302867 | 1.84071666 | 0.06566309 | 0.35579065 |
| STX2 | 99.2932339 | 0.792695762 | 0.32799627 | 2.41678288 | 0.01565835 | 0.17945702 |
| NCOA3 | 843.209656 | 0.794975236 | 0.17834333 | 4.45755513 | 8.29E-06 | 0.00141327 |
| C19orf66 | 142.429688 | 0.796259879 | 0.38849533 | 2.0495996 | 0.04040352 | 0.285758 |
| CTGF | 244.156067 | 0.799331478 | 0.98876539 | 0.80841369 | 0.41885247 | 0.73821752 |
| PLCL2 | 504.683619 | 0.799971745 | 0.2636304 | 3.03444427 | 0.00240979 | 0.06444227 |
| CDKL5 | 150.040466 | 0.800436379 | 0.28279413 | 2.83045616 | 0.00464817 | 0.09299746 |
| INPP5A | 210.857697 | 0.802218965 | 0.2898067 | 2.76811734 | 0.00563812 | 0.10300181 |
| KIAA1549 | 118.482256 | 0.802705015 | 0.5039846 | 1.59271734 | 0.11122362 | 0.44768463 |
| ARMC2 | 67.9644804 | 0.803424663 | 0.45725732 | 1.75705152 | 0.07890903 | 0.38686531 |
| PCGF5 | 410.410079 | 0.804458167 | 0.36445931 | 2.20726472 | 0.02729557 | 0.23022064 |
| MRC2 | 44.846211 | 0.805102895 | 0.49081419 | 1.64034152 | 0.10093418 | 0.42817231 |
| AC119674.2 | 144.804489 | 0.807401702 | 0.44758447 | 1.80390913 | 0.07124555 | 0.36912889 |
| PTPRG | 5333.57268 | 0.807808696 | 0.21632301 | 3.7342707 | 0.00018826 | 0.01351343 |
| AL807752.6 | 40.3232357 | 0.808909995 | 0.62987249 | 1.28424405 | 0.19905658 | 0.56402001 |
| DNAJC1 | 969.581261 | 0.809314682 | 0.42532751 | 1.90280354 | 0.05706619 | 0.33402904 |
| D2HGDH | 87.6332108 | 0.810862406 | 0.34321616 | 2.36254147 | 0.01815011 | 0.192636 |
| PDE9A | 76.4655284 | 0.811324724 | 0.5303754 | 1.52971786 | 0.12608658 | 0.47177251 |
| TBC1D8 | 724.548159 | 0.816398657 | 0.47298096 | 1.72607086 | 0.08433467 | 0.39616527 |
| CFLAR-AS1 | 105.4567 | 0.818399815 | 0.43834362 | 1.86702803 | 0.06189767 | 0.3489463 |
| EDA | 82.6011732 | 0.819082114 | 0.52525185 | 1.55940834 | 0.11889976 | 0.46285439 |
| STPG2 | 114.264379 | 0.821606043 | 0.43094951 | 1.90650187 | 0.05658512 | 0.33287999 |
| AC105052.3 | 73.0894038 | 0.822587841 | 0.45333717 | 1.81451665 | 0.06959822 | 0.36501659 |
| TDRP | 225.528783 | 0.822652311 | 0.19979842 | 4.11741152 | 3.83E-05 | 0.00461078 |
| LENG8 | 196.49548 | 0.822944451 | 0.26729125 | 3.07883044 | 0.00207815 | 0.05884254 |
| JAK2 | 363.176184 | 0.823402228 | 0.38513993 | 2.13793002 | 0.03252242 | 0.25201798 |
| ASAP2 | 717.633764 | 0.824917712 | 0.29917658 | 2.75729377 | 0.0058282 | 0.10504875 |
| AC239800.3 | 37.6721239 | 0.825004298 | 0.44515649 | 1.85329052 | 0.06384073 | 0.3533385 |
| RNF125 | 89.6909755 | 0.827155586 | 0.40991306 | 2.01788055 | 0.0436037 | 0.29561649 |
| AL117381.1 | 64.3381725 | 0.82734859 | 0.586466 | 1.41073582 | 0.15832253 | 0.51533541 |
| DNM3 | 1578.53473 | 0.828072605 | 0.35233001 | 2.35027554 | 0.01875952 | 0.19394842 |
| FMNL3 | 441.186854 | 0.829121003 | 0.22360084 | 3.70804072 | 0.00020887 | 0.01434275 |
| NPIPB4 | 68.0629491 | 0.829375165 | 0.2388043 | 3.47303281 | 0.00051461 | 0.02477102 |
| EXT1 | 1215.80812 | 0.829843595 | 0.27890211 | 2.97539375 | 0.00292613 | 0.07188756 |
| CACNA2D1 | 847.072667 | 0.83031528 | 0.4616678 | 1.79851244 | 0.07209584 | 0.37150477 |
| SLC12A4 | 53.3290964 | 0.831916922 | 0.24472606 | 3.39938023 | 0.00067539 | 0.02848812 |
| HERC1 | 1066.58738 | 0.832255292 | 0.31727396 | 2.62314404 | 0.00871224 | 0.13202275 |
| SEMA3D | 436.802513 | 0.833066402 | 0.63276774 | 1.31654372 | 0.18799161 | 0.55388137 |
| DGKZ | 164.896952 | 0.83310065 | 0.26790214 | 3.10972005 | 0.00187265 | 0.05552128 |
| RPS6KC1 | 440.715674 | 0.834174759 | 0.3492998 | 2.3881341 | 0.01693416 | 0.18591296 |
| TET1 | 58.4852618 | 0.837574853 | 0.4615484 | 1.81470644 | 0.06956903 | 0.36501659 |
| FAM111A | 130.017847 | 0.83934336 | 0.27100643 | 3.09713445 | 0.00195401 | 0.05710601 |
| AC105001.1 | 71.0729182 | 0.839918019 | 0.55701899 | 1.5078804 | 0.13158514 | 0.48069696 |
| AF117829.1 | 242.235059 | 0.840684045 | 0.42920582 | 1.95869675 | 0.05014831 | 0.31421411 |
| MTURN | 90.1537986 | 0.841219317 | 0.33523253 | 2.50936065 | 0.01209499 | 0.15440456 |
| SYNE2 | 2220.61987 | 0.841451513 | 0.17035121 | 4.93951 | 7.83E-07 | 0.00023102 |
| PARP12 | 211.210222 | 0.841554176 | 0.49064365 | 1.71520447 | 0.08630775 | 0.39864674 |
| CHST3 | 67.4845821 | 0.843157183 | 0.31373398 | 2.68749083 | 0.00719911 | 0.12071783 |
| CSF1 | 72.038776 | 0.844129363 | 0.45639926 | 1.84954149 | 0.06437966 | 0.35347506 |
| ARHGEF15 | 251.429016 | 0.844931598 | 0.29982402 | 2.81809178 | 0.004831 | 0.09480113 |
| CFLAR | 1236.04714 | 0.847659512 | 0.21027247 | 4.03124336 | 5.55E-05 | 0.00589628 |
| FGD6 | 283.708111 | 0.849026711 | 0.16555522 | 5.12835962 | 2.92E-07 | 0.00011959 |
| MOV10 | 117.632053 | 0.850353217 | 0.29756346 | 2.85772056 | 0.00426696 | 0.08952957 |
| INPP5K | 73.7771282 | 0.851291183 | 0.32389135 | 2.62832333 | 0.00858069 | 0.13124448 |
| AC093152.1 | 59.3907052 | 0.852175051 | 0.30289199 | 2.81346186 | 0.00490112 | 0.09526334 |
| PAG1 | 66.5374795 | 0.85229239 | 0.45723109 | 1.86402985 | 0.06231751 | 0.35005476 |
| TRIM2 | 169.375962 | 0.852556159 | 0.31827552 | 2.67867333 | 0.00739145 | 0.12155049 |
| DDX60L | 442.831125 | 0.855038843 | 0.65578628 | 1.30383765 | 0.19228894 | 0.55659725 |
| INPPL1 | 51.2058694 | 0.855795055 | 0.27132646 | 3.15411569 | 0.00160985 | 0.05125846 |
| CASP10 | 208.065764 | 0.856877117 | 0.23721353 | 3.61226068 | 0.00030354 | 0.01826369 |
| JAZF1 | 372.004062 | 0.856944666 | 0.36443096 | 2.35145954 | 0.01869992 | 0.19394842 |
| TPCN2 | 51.4044344 | 0.859176776 | 0.43574324 | 1.97175011 | 0.04863814 | 0.30949137 |
| LINC00639 | 138.500597 | 0.85918026 | 1.42975476 | 0.60092842 | 0.54788767 | 0.82355262 |
| SEC16B | 65.1000826 | 0.86024167 | 0.70393245 | 1.22205145 | 0.22168817 | 0.58750144 |
| ANKRD33B | 128.669146 | 0.86086692 | 0.47117912 | 1.8270481 | 0.06769254 | 0.36086517 |

| | | | | | | |
|---|---|---|---|---|---|---|
| MEIS2 | 6554.28123 | 0.86346078 | 0.19178964 | 4.50212426 | 6.73E-06 | 0.00124771 |
| AC009948.1 | 95.499836 | 0.864575077 | 0.31760007 | 2.72221313 | 0.00648463 | 0.11362684 |
| EXOC3L2 | 458.709972 | 0.866890328 | 0.41791644 | 2.07431495 | 0.03805007 | 0.2762766 |
| EIF2AK2 | 334.136178 | 0.867977829 | 0.50376178 | 1.72299262 | 0.08488986 | 0.39686932 |
| MX1 | 387.794449 | 0.870062448 | 0.99635305 | 0.87324713 | 0.38252839 | 0.71613585 |
| TEK | 1585.43649 | 0.87220066 | 0.3004846 | 2.90264675 | 0.00370024 | 0.08250421 |
| TMEM62 | 93.3348031 | 0.872664694 | 0.46805397 | 1.86445315 | 0.06225809 | 0.35005476 |
| SORBS2 | 938.166774 | 0.874059153 | 0.35209577 | 2.4824472 | 0.01304834 | 0.16344193 |
| ARHGEF6 | 181.95316 | 0.874074101 | 0.27458733 | 3.18322812 | 0.00145643 | 0.04767181 |
| VSIG10 | 74.7309158 | 0.874605285 | 0.4174095 | 2.09531714 | 0.03614281 | 0.26741104 |
| ARHGAP27 | 119.29763 | 0.875549983 | 0.37125367 | 2.35836048 | 0.01835586 | 0.19310354 |
| TRAFD1 | 59.8512084 | 0.875849614 | 0.33590148 | 2.60745983 | 0.00912168 | 0.13593083 |
| TMEM204 | 262.383283 | 0.876927654 | 0.43678515 | 2.00768653 | 0.04467661 | 0.29946125 |
| DIRC3 | 139.33102 | 0.878593167 | 0.46986971 | 1.86986551 | 0.0615025 | 0.34780576 |
| ARHGAP12 | 261.545337 | 0.879931403 | 0.23000009 | 3.82578718 | 0.00013035 | 0.01048909 |
| RAP2A | 119.962508 | 0.880112077 | 0.25581314 | 3.44044912 | 0.00058075 | 0.02686611 |
| MYO6 | 646.930535 | 0.880386051 | 0.22481412 | 3.91606205 | 9.00E-05 | 0.0082756 |
| KIAA1841 | 160.956428 | 0.88582111 | 0.30752068 | 2.88052535 | 0.00397013 | 0.08640313 |
| PLCL1 | 534.714327 | 0.887757134 | 0.26866731 | 3.30429907 | 0.00095214 | 0.03585533 |
| DDX60 | 294.33605 | 0.887994935 | 0.57980155 | 1.53154979 | 0.12563357 | 0.47103973 |
| PHLDB1 | 186.235509 | 0.888357186 | 0.23525883 | 3.7760843 | 0.00015931 | 0.01229867 |
| KLF11 | 73.2085368 | 0.888747662 | 0.36045093 | 2.46565507 | 0.0136763 | 0.16829042 |
| SLC26A3 | 767.431721 | 0.888943937 | 0.68478861 | 1.29812898 | 0.19424302 | 0.55913011 |
| LINC00623 | 165.082992 | 0.891838817 | 0.3325885 | 2.68150831 | 0.00732911 | 0.1211598 |
| DENND2C | 76.7401478 | 0.89500639 | 0.38872819 | 2.30239642 | 0.02131283 | 0.20628732 |
| AL022323.4 | 58.5739421 | 0.896764558 | 0.35079954 | 2.55634472 | 0.01057783 | 0.14485284 |
| TSPAN2 | 55.462722 | 0.897398382 | 0.57476849 | 1.56132146 | 0.11844792 | 0.46185998 |
| KIF13A | 782.482785 | 0.89827774 | 0.21923334 | 4.09735923 | 4.18E-05 | 0.00488514 |
| LINGO1 | 1321.5447 | 0.898278715 | 0.6198985 | 1.44907386 | 0.14731696 | 0.50127566 |
| RBMS3-AS3 | 540.490308 | 0.899116985 | 0.44732426 | 2.00998934 | 0.04443232 | 0.29898844 |
| ALS2CL | 67.8099421 | 0.900078658 | 0.40836766 | 2.204089 | 0.02751809 | 0.23083108 |
| LRRK2 | 113.695999 | 0.900959042 | 0.32672803 | 2.75751992 | 0.00582417 | 0.10504875 |
| MYLIP | 133.139932 | 0.903026742 | 0.30412929 | 2.96921992 | 0.00298557 | 0.07188756 |
| CUBN | 154.444748 | 0.908694321 | 0.49233913 | 1.84566746 | 0.0649405 | 0.35498206 |
| RAPGEF3 | 186.97409 | 0.911741505 | 0.37633531 | 2.42268389 | 0.01540633 | 0.17781377 |
| TCIM | 693.446665 | 0.914170432 | 0.65988963 | 1.38533837 | 0.16594903 | 0.5261375 |
| AP4S1 | 69.2546044 | 0.918516808 | 0.3153743 | 2.91246559 | 0.00358588 | 0.08141758 |
| AP002989.1 | 123.063258 | 0.920379834 | 0.57400643 | 1.60343124 | 0.10883948 | 0.44503888 |
| ADARB1 | 366.493526 | 0.920753882 | 0.23211711 | 3.96676437 | 7.29E-05 | 0.00727039 |
| KCNJ15 | 45.2033719 | 0.920857431 | 0.52102075 | 1.76741027 | 0.07715954 | 0.38356763 |
| ID1 | 1606.52163 | 0.924332021 | 0.78095436 | 1.18359288 | 0.23657426 | 0.60027509 |
| LRRC37A2 | 61.114021 | 0.92788925 | 0.30002109 | 3.0927467 | 0.00198313 | 0.05734268 |
| KIFC3 | 301.534357 | 0.928422976 | 0.26997556 | 3.43891488 | 0.00058405 | 0.02686611 |
| MAP2K6 | 91.9987767 | 0.930112239 | 0.64044656 | 1.45228704 | 0.14642181 | 0.50027961 |
| SLC9A3R2 | 787.70678 | 0.93042929 | 0.58501197 | 1.59044486 | 0.11173456 | 0.44839019 |
| RIMKLB | 464.113241 | 0.931028166 | 0.38523056 | 2.41680764 | 0.01565729 | 0.17945702 |
| TDRD7 | 114.105673 | 0.931611682 | 0.3501896 | 2.66030653 | 0.00780696 | 0.12501039 |
| CPNE2 | 145.616142 | 0.932515314 | 0.27911817 | 3.34093377 | 0.00083497 | 0.03297047 |
| TMEM94 | 98.693859 | 0.932694946 | 0.26958677 | 3.45972071 | 0.00054074 | 0.02587627 |
| EMB | 59.8253068 | 0.934728044 | 1.047744 | 0.892134 | 0.37232111 | 0.71084822 |
| MATN2 | 139.191707 | 0.936866699 | 0.64019573 | 1.46340667 | 0.14335613 | 0.49538989 |
| MTMR10 | 213.374456 | 0.939266137 | 0.27280381 | 3.4430096 | 0.00057528 | 0.02686611 |
| EPHA4 | 395.721643 | 0.939746584 | 0.30812602 | 3.04987735 | 0.00228935 | 0.06286489 |
| CPEB3 | 493.32588 | 0.941399922 | 0.24611435 | 3.82505085 | 0.00013075 | 0.01048909 |
| LIMS2 | 740.808199 | 0.944439694 | 0.38771295 | 2.43592505 | 0.01485376 | 0.17549246 |
| ZNF618 | 152.68803 | 0.946443167 | 0.28909632 | 3.27379878 | 0.00106112 | 0.03842105 |
| KIAA1671 | 732.93016 | 0.94644457 | 0.36556254 | 2.58900862 | 0.00962527 | 0.13807701 |
| CDR2L | 48.5630252 | 0.949202375 | 0.26755313 | 3.54771542 | 0.00038859 | 0.02112867 |
| RASAL2 | 2517.28146 | 0.951152204 | 0.23193147 | 4.10100537 | 4.11E-05 | 0.00487848 |
| LNX1 | 239.825304 | 0.953640198 | 0.69353275 | 1.37504711 | 0.16911684 | 0.53103726 |
| MYO1C | 361.481426 | 0.95881526 | 0.29721672 | 3.22598019 | 0.00125542 | 0.04283496 |
| LRRC37A3 | 247.953031 | 0.960241087 | 0.33498708 | 2.86650182 | 0.00415036 | 0.08875226 |
| PXDNL | 60.3748932 | 0.961478742 | 0.33278831 | 2.88916016 | 0.00386272 | 0.0851496 |
| HDAC5 | 93.4734136 | 0.963228318 | 0.20199855 | 4.76849123 | 1.86E-06 | 0.00043396 |
| PRX | 44.9511814 | 0.963608398 | 0.5719847 | 1.68467513 | 0.09205127 | 0.41026989 |
| ABHD6 | 111.720873 | 0.964229453 | 0.34718323 | 2.77729273 | 0.00548138 | 0.10194119 |
| KLHL4 | 94.4088766 | 0.966280549 | 0.52254667 | 1.8491756 | 0.06443246 | 0.35347506 |
| CDH2 | 99.9576244 | 0.967228966 | 1.09064586 | 0.88684054 | 0.37516476 | 0.71257833 |
| NCKIPSD | 53.7227738 | 0.968862637 | 0.32614951 | 2.97060887 | 0.0029721 | 0.07188756 |
| VANGL1 | 63.0695275 | 0.969587594 | 0.31849238 | 3.04430392 | 0.00233219 | 0.06319319 |
| RAMP2-AS1 | 41.5505791 | 0.971725215 | 0.38052695 | 2.55363048 | 0.01066063 | 0.14515134 |
| NAV3 | 412.37026 | 0.979000417 | 0.67002399 | 1.46114234 | 0.14397638 | 0.49679586 |
| TNK2 | 74.7317438 | 0.979090367 | 0.34494389 | 2.83840469 | 0.00453397 | 0.09149853 |
| ZBTB20-AS5 | 104.794809 | 0.980180427 | 0.9390447 | 1.04380593 | 0.29657519 | 0.65753344 |
| UACA | 1078.58525 | 0.980223026 | 0.34805692 | 2.81627219 | 0.00485845 | 0.09504268 |
| CCND1 | 100.859795 | 0.980855359 | 0.57013017 | 1.72040598 | 0.08535867 | 0.39686932 |
| DOK6 | 268.056812 | 0.9839665 | 0.49587931 | 1.98428625 | 0.04722394 | 0.30523972 |
| KCNJ16 | 56.8370893 | 0.985489739 | 0.39467359 | 2.49697415 | 0.01252581 | 0.15866671 |
| PDE1A | 88.3392831 | 0.988921659 | 0.48600618 | 2.03479233 | 0.04187177 | 0.28963374 |
| LRRC8C-DT | 46.9427583 | 0.989207319 | 0.29989159 | 3.29854969 | 0.00097186 | 0.03631372 |
| NRG3 | 1928.38055 | 0.9917955 | 0.78955802 | 1.25614011 | 0.20906518 | 0.57428007 |
| AC116366.3 | 552.864321 | 0.992098911 | 0.44942361 | 2.20749177 | 0.02727972 | 0.23022064 |
| GAB1 | 636.870432 | 0.992526933 | 0.27892114 | 3.55845002 | 0.00037305 | 0.02090867 |
| RIPOR2 | 136.0193 | 0.994362538 | 0.57656139 | 1.72464296 | 0.08459184 | 0.3964812 |
| BACH2 | 165.811835 | 0.994641409 | 0.39514975 | 2.51712523 | 0.01183168 | 0.15189817 |
| LINC02328 | 165.883565 | 0.997326458 | 0.48860859 | 2.04115621 | 0.0412353 | 0.28808825 |
| ISPD | 174.041316 | 1.002851315 | 0.43663502 | 2.2967725 | 0.02163175 | 0.20825014 |
| NFASC | 86.2217294 | 1.0079197 | 0.48611915 | 2.07340054 | 0.03813502 | 0.2764977 |
| HELZ2 | 77.8303892 | 1.008344598 | 0.71314746 | 1.41393563 | 0.1573808 | 0.51438736 |
| KLF12 | 517.576516 | 1.01076375 | 0.4669698 | 2.16451629 | 0.03042475 | 0.24194918 |
| NEBL | 124.857008 | 1.018330256 | 0.4104564 | 2.48097059 | 0.01310252 | 0.16344193 |
| THRB | 337.175687 | 1.020091719 | 0.62827328 | 1.62364333 | 0.10445195 | 0.43653234 |
| CEACAM1 | 1436.24294 | 1.025554368 | 0.34227514 | 2.99628645 | 0.0027329 | 0.06991543 |
| PARP14 | 1217.17298 | 1.025686884 | 0.54732649 | 1.8739946 | 0.06093117 | 0.3467314 |
| BICC1 | 105.941078 | 1.02804629 | 0.46736398 | 2.19966952 | 0.02783035 | 0.23190345 |
| PLA1A | 104.535638 | 1.028597689 | 0.66625876 | 1.54384115 | 0.12262682 | 0.46802951 |
| MEG3 | 51.385686 | 1.029466824 | 0.68636661 | 1.49987895 | 0.13364576 | 0.48326261 |
| RAPGEF4 | 1583.82166 | 1.02993679 | 0.29282908 | 3.51719432 | 0.00043613 | 0.0225879 |
| GPC5 | 1379.73663 | 1.036054849 | 0.74231269 | 1.39571216 | 0.16280119 | 0.52335809 |
| C17orf67 | 79.6977613 | 1.03672956 | 0.40774462 | 2.54259532 | 0.01100326 | 0.14749418 |
| HERC5 | 160.546743 | 1.036796821 | 0.66886731 | 1.55007848 | 0.12112268 | 0.46542849 |
| HLA-DRB1 | 957.234881 | 1.037656069 | 0.6938049 | 1.49560211 | 0.13475737 | 0.48474744 |
| TM4SF18 | 348.234581 | 1.038320482 | 0.67669109 | 1.53440838 | 0.12492922 | 0.47038289 |
| IFIT3 | 207.259333 | 1.039484009 | 0.85695584 | 1.212996 | 0.22513135 | 0.59122266 |
| NPY1R | 92.5541595 | 1.039805343 | 0.67385334 | 1.54307368 | 0.1228129 | 0.46808475 |
| P3H2 | 147.277164 | 1.042340769 | 0.44307396 | 2.35252093 | 0.01864664 | 0.19394842 |
| PLPP3 | 1755.22323 | 1.043300039 | 0.42041915 | 2.48157116 | 0.01308046 | 0.16344193 |
| CADM1 | 148.514582 | 1.043771434 | 0.41208921 | 2.53287736 | 0.01131305 | 0.15028362 |
| SKIL | 441.2756 | 1.044740308 | 0.28454108 | 3.67166773 | 0.00024097 | 0.01540532 |
| PPP1R14A | 66.8632264 | 1.044901505 | 0.77230525 | 1.3529644 | 0.17606701 | 0.5396091 |
| AC098850.3 | 143.925245 | 1.056326324 | 0.55917104 | 1.88909342 | 0.05887931 | 0.34050135 |
| HLA-DRB5 | 356.684833 | 1.05700915 | 0.68884068 | 1.5344755 | 0.12491272 | 0.47038289 |
| PRKD1 | 1104.65689 | 1.057093933 | 0.45736961 | 2.31124654 | 0.02081924 | 0.20354104 |

| | | | | | | |
|---|---|---|---|---|---|---|
| HEY1 | 50.4826666 | 1.05750114 | 0.75474144 | 1.40114361 | 0.16117113 | 0.51944206 |
| ST8SIA4 | 67.3528352 | 1.058268455 | 0.28966917 | 3.65336934 | 0.00025882 | 0.01627997 |
| TBC1D20 | 92.0880607 | 1.059229561 | 0.28594751 | 3.70427967 | 0.00021199 | 0.01434275 |
| A2ML1-AS1 | 61.8202601 | 1.061379186 | 0.52589932 | 2.01821746 | 0.04356862 | 0.29561649 |
| ANK3 | 337.50651 | 1.061979424 | 0.37262284 | 2.85001165 | 0.00437176 | 0.09052099 |
| FRMD4A | 477.677658 | 1.064088016 | 0.52431906 | 2.0294666 | 0.04241079 | 0.291241 |
| SPTAN1 | 436.095633 | 1.064237044 | 0.2619436 | 4.06284806 | 4.85E-05 | 0.00542299 |
| MAPK10 | 62.1430269 | 1.064321466 | 0.41335984 | 2.57480616 | 0.01002963 | 0.14223994 |
| CYTH3 | 385.238616 | 1.06500951 | 0.28836126 | 3.69331692 | 0.00022135 | 0.01472593 |
| GATM | 49.3721595 | 1.0704377 | 0.7158517 | 1.49533444 | 0.13482718 | 0.48474992 |
| RASGEF1B | 1007.20406 | 1.070565353 | 0.61223849 | 1.74860837 | 0.08035874 | 0.38871437 |
| MRPS6 | 121.556842 | 1.077030121 | 0.71689125 | 1.50236193 | 0.13300366 | 0.48236854 |
| DAB1 | 62.3772183 | 1.078107184 | 0.61591344 | 1.75041996 | 0.08004587 | 0.38827231 |
| SMAD6 | 585.816401 | 1.078497583 | 0.68370008 | 1.57744252 | 0.11469374 | 0.45569454 |
| LMO7 | 137.702938 | 1.078917574 | 0.51871543 | 2.07997971 | 0.03752739 | 0.27364951 |
| ATP13A3 | 761.624626 | 1.079194075 | 0.32124125 | 3.35945048 | 0.00078098 | 0.03148143 |
| LPIN2 | 308.896293 | 1.09025135 | 0.38064114 | 2.8642499 | 0.00417998 | 0.08880727 |
| MBOAT2 | 160.521756 | 1.090513862 | 0.56010177 | 1.94699236 | 0.05153565 | 0.31859402 |
| PRKAR2B | 51.85413 | 1.09970135 | 0.74573349 | 1.47465732 | 0.1403047 | 0.49118733 |
| NET1 | 100.601218 | 1.104854943 | 0.29798957 | 3.70769663 | 0.00020915 | 0.01434275 |
| MAP4K2 | 69.5733899 | 1.106595893 | 0.36338796 | 3.04521893 | 0.00232511 | 0.06319319 |
| RUNX1T1 | 690.840909 | 1.1080037 | 0.43503296 | 2.54694199 | 0.01086715 | 0.14698493 |
| IFIH1 | 238.351563 | 1.10855093 | 0.56899856 | 1.94824909 | 0.05138516 | 0.31806717 |
| SH3KBP1 | 252.825781 | 1.11448834 | 0.48962657 | 2.27620068 | 0.022834 | 0.21136949 |
| FNBP1 | 592.896309 | 1.118335944 | 0.23737277 | 4.71130679 | 2.46E-06 | 0.00054188 |
| AC008537.1 | 195.498793 | 1.125477723 | 0.43761494 | 2.57184485 | 0.01011582 | 0.14236887 |
| AC004918.1 | 71.453831 | 1.131954042 | 0.4109641 | 2.75438667 | 0.00588023 | 0.10529077 |
| PBX1 | 4179.25664 | 1.132881746 | 0.31516756 | 3.59453791 | 0.00032497 | 0.01885966 |
| ARHGAP24 | 118.714081 | 1.138742274 | 0.70632195 | 1.6122142 | 0.10691534 | 0.43964815 |
| PARP10 | 56.334731 | 1.138907398 | 0.55233825 | 2.06197453 | 0.03921016 | 0.28057468 |
| ADAMTS10 | 68.9865891 | 1.139782455 | 0.35198193 | 3.23818457 | 0.00120293 | 0.04185098 |
| ZNF503-AS1 | 185.860867 | 1.142370639 | 0.37877495 | 3.01596144 | 0.00256166 | 0.06718606 |
| SEMA5B | 166.378354 | 1.142739554 | 0.5506284 | 2.07533712 | 0.0379553 | 0.27583324 |
| ECHDC2 | 206.299288 | 1.145758248 | 0.38240955 | 2.99615486 | 0.00273408 | 0.06991543 |
| RASGRP3 | 494.365833 | 1.146569478 | 0.27070736 | 4.23545735 | 2.28E-05 | 0.00327445 |
| DHRS3 | 460.892127 | 1.150725675 | 0.44984381 | 2.55805605 | 0.01052591 | 0.14485284 |
| SNED1 | 764.532895 | 1.15118493 | 0.36893544 | 3.12028826 | 0.00180674 | 0.05451467 |
| ANKRD44 | 594.684799 | 1.155071703 | 0.31066047 | 3.71811608 | 0.00020071 | 0.01415899 |
| KDR | 644.564389 | 1.157205885 | 0.32802438 | 3.52780452 | 0.00041902 | 0.02197983 |
| SAMD9L | 204.312101 | 1.160903761 | 0.63050061 | 1.8412413 | 0.06558621 | 0.35579065 |
| ATP1B1 | 354.377483 | 1.166649735 | 0.50075456 | 2.32978354 | 0.01981759 | 0.19704872 |
| VAV3 | 478.204337 | 1.169104514 | 0.51807221 | 2.25664394 | 0.02403033 | 0.21632587 |
| ADAMTS6 | 993.893412 | 1.170512863 | 0.52991901 | 2.20885236 | 0.02718491 | 0.22980799 |
| KCNE4 | 86.4073886 | 1.17138982 | 1.11591977 | 1.04970792 | 0.29385242 | 0.65627576 |
| EMID1 | 186.495564 | 1.172035503 | 0.49899789 | 2.34877846 | 0.01883511 | 0.19394842 |
| DENND1B | 165.198386 | 1.176974083 | 0.23000153 | 5.11724457 | 3.10E-07 | 0.00012081 |
| ACSF2 | 44.5711334 | 1.179360557 | 0.42319082 | 2.78682924 | 0.00532265 | 0.10080241 |
| IFI44L | 616.117113 | 1.181044407 | 0.9608274 | 1.22919518 | 0.21899864 | 0.58619521 |
| MCU | 442.043366 | 1.182047939 | 0.28296821 | 4.17731714 | 2.95E-05 | 0.0038313 |
| CAMK1 | 58.9057696 | 1.184169354 | 0.37559262 | 3.15280247 | 0.00161711 | 0.05129003 |
| NR3C1 | 594.286321 | 1.187153965 | 0.2229078 | 5.32576231 | 1.01E-07 | 4.57E-05 |
| DGKE | 221.785979 | 1.191750638 | 0.34649941 | 3.43940167 | 0.000583 | 0.02686611 |
| LTC4S | 58.0850815 | 1.196603537 | 0.74100029 | 1.61484896 | 0.10634341 | 0.43901385 |
| ZEB1-AS1 | 112.027931 | 1.205477032 | 0.33987378 | 3.54683743 | 0.00038989 | 0.02112867 |
| ABCB1 | 136.84454 | 1.21557737 | 1.11233152 | 1.09281931 | 0.27447314 | 0.63975272 |
| AC010976.2 | 77.894767 | 1.218039176 | 0.94048935 | 1.29511214 | 0.19528155 | 0.56007526 |
| MPPED2 | 174.180805 | 1.219794969 | 0.43145469 | 2.82716821 | 0.00469617 | 0.09350054 |
| NRG2 | 184.42925 | 1.223958023 | 0.63216455 | 1.93613834 | 0.05285076 | 0.3215522 |
| DDX58 | 514.271008 | 1.226838274 | 0.66745004 | 1.83809752 | 0.06604804 | 0.35600847 |
| PARP8 | 137.828596 | 1.228362376 | 0.3941871 | 3.11619125 | 0.00183203 | 0.05451467 |
| EPSTI1 | 366.295239 | 1.235240823 | 0.87116738 | 1.41791445 | 0.15621574 | 0.5124974 |
| TRAK1 | 466.189819 | 1.237446398 | 0.19246189 | 6.42956574 | 1.28E-10 | 2.44E-07 |
| PRICKLE1 | 97.2449005 | 1.240978355 | 0.54263199 | 2.28696128 | 0.02219808 | 0.2108594 |
| UMOD | 248.554682 | 1.241994466 | 0.70752063 | 1.75541803 | 0.07918783 | 0.3870932 |
| CYR61 | 175.479768 | 1.247186047 | 0.87803106 | 1.42043499 | 0.15548108 | 0.51178668 |
| SLFN12 | 57.6803743 | 1.254332198 | 0.33779284 | 3.71331794 | 0.00020456 | 0.01430694 |
| SLC6A6 | 263.352927 | 1.256145992 | 0.3932168 | 3.19453799 | 0.00140055 | 0.04621245 |
| MX2 | 268.419675 | 1.262246486 | 1.16414613 | 1.08426807 | 0.27824596 | 0.64157752 |
| RANBP3L | 378.72121 | 1.262289716 | 1.52599405 | 0.82719177 | 0.40812838 | 0.73419295 |
| FCN3 | 243.393506 | 1.263121563 | 1.17364964 | 1.07623392 | 0.28182265 | 0.64471757 |
| OLFML2A | 58.1543438 | 1.264913528 | 0.48660219 | 2.59948179 | 0.00933646 | 0.13676431 |
| ZC3HAV1 | 552.213886 | 1.271143512 | 0.57881758 | 2.19610384 | 0.02808451 | 0.23222363 |
| RARG | 46.5682674 | 1.287225787 | 0.48050222 | 2.67891747 | 0.00738606 | 0.12155049 |
| PHLPP1 | 526.682952 | 1.300973061 | 0.32042475 | 4.06015156 | 4.90E-05 | 0.00542299 |
| FAM118A | 94.8601791 | 1.302974205 | 0.47863221 | 2.72228693 | 0.00648318 | 0.11362684 |
| EML1 | 619.965077 | 1.306344855 | 0.51077492 | 2.55757439 | 0.0105405 | 0.14485284 |
| NIPSNAP3B | 50.8719741 | 1.311384182 | 0.37007868 | 3.54352803 | 0.00039481 | 0.02125314 |
| PTH2R | 94.5957084 | 1.313700245 | 0.42596436 | 3.08406144 | 0.00204195 | 0.05828136 |
| EGR1 | 91.789625 | 1.324615528 | 1.27153966 | 1.04174142 | 0.29753158 | 0.6579563 |
| GJC1 | 64.6554974 | 1.325625358 | 0.39350665 | 3.3687496 | 0.0007551 | 0.03058902 |
| CFAP161 | 158.033125 | 1.332112188 | 0.45419354 | 2.93291753 | 0.00335793 | 0.07718524 |
| AC020916.1 | 75.7568758 | 1.336286847 | 0.4674799 | 2.85849048 | 0.00425662 | 0.08952957 |
| CCDC68 | 64.9120131 | 1.34301667 | 0.51580045 | 2.60375243 | 0.00922093 | 0.13620125 |
| HECW2 | 545.485185 | 1.343659087 | 0.37085878 | 3.6231017 | 0.00029109 | 0.01764445 |
| NRIR | 86.397777 | 1.344563099 | 0.91843597 | 1.46397043 | 0.14320203 | 0.49538989 |
| CPED1 | 130.979166 | 1.351589515 | 0.55396922 | 2.43982782 | 0.01469426 | 0.17528157 |
| IFIT1 | 262.668838 | 1.351711203 | 0.89002831 | 1.51872832 | 0.1288309 | 0.47679271 |
| HERC6 | 250.220842 | 1.356152462 | 0.9967556 | 1.36056668 | 0.17365067 | 0.53536344 |
| CMKLR1 | 96.9166242 | 1.371998589 | 0.70077655 | 1.95782607 | 0.05025043 | 0.31422181 |
| AC090630.1 | 87.1771942 | 1.37370778 | 1.31598932 | 1.04385937 | 0.29655046 | 0.65753344 |
| LIFR | 950.982591 | 1.390170168 | 0.39886714 | 3.48529632 | 0.00049159 | 0.02423313 |
| SNCA | 92.4696867 | 1.391678721 | 1.49540096 | 0.93063918 | 0.35204024 | 0.69889281 |
| AL050403.2 | 117.736721 | 1.401954676 | 0.53960755 | 2.59810055 | 0.0093741 | 0.13676431 |
| UBA7 | 50.9906829 | 1.404771346 | 0.52761488 | 2.66249379 | 0.0077564 | 0.1244522 |
| FRMD3 | 527.891439 | 1.407194559 | 0.42297036 | 3.32693423 | 0.00087807 | 0.03373359 |
| HIST1H2AC | 290.776374 | 1.42868782 | 0.89123652 | 1.60304004 | 0.10892581 | 0.44503888 |
| SYNPO2 | 70.1083832 | 1.437295077 | 0.43140629 | 3.33165071 | 0.00086333 | 0.03348148 |
| SGK1 | 512.78734 | 1.441608932 | 0.60823835 | 2.37013818 | 0.01778144 | 0.18995498 |
| IFI44 | 462.860121 | 1.44254162 | 0.47273522 | 3.05147907 | 0.00227717 | 0.06286489 |
| CHRM3 | 1359.03014 | 1.450139052 | 0.33850374 | 4.28396756 | 1.84E-05 | 0.00278206 |
| RHOBTB1 | 129.413708 | 1.451695876 | 0.23449504 | 6.19073167 | 5.99E-10 | 7.00E-07 |
| PTGER3 | 49.9582433 | 1.452485312 | 0.5448956 | 2.66562129 | 0.00768462 | 0.12405 |
| SYCP2L | 80.7769041 | 1.45459183 | 0.41184713 | 3.53187318 | 0.00041263 | 0.02178405 |
| DPH6-DT | 366.731806 | 1.47737167 | 0.8688269 | 1.70042118 | 0.08905173 | 0.4043684 |
| MEG8 | 79.1822934 | 1.48051294 | 0.67933671 | 2.17935071 | 0.02930562 | 0.23789138 |
| OAS1 | 98.552085 | 1.489326146 | 0.74873235 | 1.98913023 | 0.04668683 | 0.30359341 |
| AL136164.2 | 120.516911 | 1.498743848 | 0.42315639 | 3.54182019 | 0.00039738 | 0.02125314 |
| AC005162.3 | 86.6622214 | 1.50059766 | 0.71024694 | 2.11278299 | 0.03461935 | 0.25989918 |
| PLAG1 | 68.7652788 | 1.514307481 | 0.49106021 | 3.08375111 | 0.00204409 | 0.05828136 |
| IFIT2 | 220.614786 | 1.517780612 | 1.03691313 | 1.46374905 | 0.14326253 | 0.49538989 |
| MIR222HG | 185.363205 | 1.541561791 | 0.65157049 | 2.36591714 | 0.01798547 | 0.19138504 |
| WDR72 | 63.2656665 | 1.551930548 | 0.59861831 | 2.59252101 | 0.00952754 | 0.13774528 |
| ARHGAP18 | 320.207291 | 1.553528199 | 0.31506291 | 4.93085083 | 8.19E-07 | 0.00023102 |

| | | | | | | | |
|---|---|---|---|---|---|---|---|
| KLF4 | 135.834894 | | 1.560068803 | 0.51435017 | 3.03308699 | 0.00242066 | 0.06452197 |
| GNG7 | 205.930376 | | 1.561102312 | 0.41408448 | 3.77000921 | 0.00016324 | 0.01230754 |
| AL157871.4 | 115.345118 | | 1.562884575 | 0.60299263 | 2.59188007 | 0.0095453 | 0.13775877 |
| CA4 | 74.701573 | | 1.562995003 | 0.98661204 | 1.58420426 | 0.11314724 | 0.45099069 |
| CX3CL1 | 174.897266 | | 1.580323978 | 0.61814739 | 2.55654882 | 0.01057162 | 0.14485284 |
| AMOTL2 | 58.9508649 | | 1.583430915 | 0.28883088 | 5.48220792 | 4.20E-08 | 2.29E-05 |
| CNTN4 | 146.592634 | | 1.590552197 | 0.73502401 | 2.16394589 | 0.0304685 | 0.2420619 |
| XIST | 404.008731 | | 1.615598036 | 3.97593282 | 0.4063444 | 0.68448957 | 0.88636116 |
| ADGRE5 | 67.9903422 | | 1.633889194 | 0.42522623 | 3.84239981 | 0.00012184 | 0.0101734 |
| ITGA8 | 568.744048 | | 1.650970571 | 0.63140693 | 2.61474888 | 0.00892931 | 0.13439403 |
| ZNF704 | 356.167623 | | 1.660701057 | 0.40819675 | 4.0683838 | 4.73E-05 | 0.00538036 |
| ANXA3 | 125.962683 | | 1.666315679 | 1.04182118 | 1.5994258 | 0.10972602 | 0.44576178 |
| ABCG2 | 99.9350478 | | 1.68636217 | 0.60297739 | 2.7967254 | 0.00516234 | 0.09916296 |
| LCAT | 41.831058 | | 1.686909306 | 0.38883208 | 4.3384005 | 1.44E-05 | 0.00221595 |
| SYNE1 | 3348.84709 | | 1.711852191 | 0.37276054 | 4.59236426 | 4.38E-06 | 0.000834 |
| NCKAP5 | 141.370476 | | 1.715221004 | 0.88028526 | 1.94848317 | 0.05135718 | 0.31806717 |
| EHD3 | 164.475001 | | 1.72000141 | 0.64252223 | 2.67695238 | 0.00742952 | 0.12171031 |
| CMPK2 | 88.9105365 | | 1.729307923 | 0.6624566 | 2.61044712 | 0.0090424 | 0.13576866 |
| CHRM3-AS2 | 55.0967873 | | 1.744892242 | 0.55444768 | 3.14708186 | 0.00164909 | 0.05170301 |
| PKHD1 | 124.508754 | | 1.748133058 | 0.56073874 | 3.11755357 | 0.00182359 | 0.05451467 |
| LINC02384 | 186.265114 | | 1.779735722 | 0.64233216 | 2.7707405 | 0.0055929 | 0.10277085 |
| IQGAP2 | 149.452118 | | 1.804797141 | 0.63183534 | 2.8564359 | 0.00428427 | 0.08955895 |
| LINC00882 | 89.0089064 | | 1.831581573 | 0.93156648 | 1.96613082 | 0.0492835 | 0.31155957 |
| NEO1 | 91.0856955 | | 1.88504417 | 0.49202886 | 3.83116585 | 0.00012754 | 0.01043639 |
| ARL17B | 281.518741 | | 1.995380931 | 1.14770822 | 1.73857858 | 0.08210892 | 0.39384365 |
| AL078590.3 | 355.304864 | | 2.013655958 | 1.38086824 | 1.45825351 | 0.14477068 | 0.49729058 |
| KLF2 | 661.087889 | | 2.028285831 | 0.50732978 | 3.99796325 | 6.39E-05 | 0.0067027 |
| SNTB1 | 72.6034174 | | 2.081649441 | 0.52797129 | 3.94273229 | 8.06E-05 | 0.00775542 |
| RSAD2 | 313.84666 | | 2.083459339 | 1.18801087 | 1.7537376 | 0.07947548 | 0.38738328 |
| NCALD | 340.724983 | | 2.112702401 | 0.43568519 | 4.84914896 | 1.24E-06 | 0.0003273 |
| TBX3 | 138.221931 | | 2.114919417 | 0.58692088 | 3.60341483 | 0.00031406 | 0.01861588 |
| RGS9 | 46.3300523 | | 2.123607294 | 0.58879594 | 3.60669485 | 0.00031012 | 0.01852356 |
| SLC12A1 | 137.049839 | | 2.137382388 | 0.5808266 | 3.67989758 | 0.00023333 | 0.01515333 |
| ESRRG | 152.40916 | | 2.157501205 | 0.60804577 | 3.54825461 | 0.00038779 | 0.02112867 |
| ASIC2 | 93.4233053 | | 2.197355447 | 0.9175697 | 2.3947559 | 0.01663143 | 0.18464043 |
| EMCN | 3739.13588 | | 2.200195562 | 0.49727793 | 4.42447862 | 9.67E-06 | 0.00161448 |
| AP001347.1 | 82.4811549 | | 2.256556554 | 0.44629931 | 5.05615072 | 4.28E-07 | 0.00015912 |
| RCAN2 | 327.873528 | | 2.373838617 | 0.42148693 | 5.63205752 | 1.78E-08 | 1.26E-05 |
| RGCC | 500.155789 | | 2.496583004 | 1.0847593 | 2.30150873 | 0.02136289 | 0.20628732 |
| CRHBP | 390.780424 | | 2.510557842 | 1.15683391 | 2.17019732 | 0.0299919 | 0.24082201 |
| AL078590.2 | 146.832232 | | 2.526076509 | 1.32067425 | 1.91271731 | 0.05578425 | 0.32988018 |
| ADCY1 | 53.3114398 | | 2.539783232 | 0.4514263 | 5.6261304 | 1.84E-08 | 1.26E-05 |
| NR4A1 | 192.426338 | | 2.584755469 | 0.66091862 | 3.91085288 | 9.20E-05 | 0.00836219 |
| APLNR | 38.963228 | | 2.625729682 | 1.15235118 | 2.27858463 | 0.02269177 | 0.21131824 |
| FABP5 | 143.780762 | | 2.817052417 | 1.68887185 | 1.66800839 | 0.09531406 | 0.41531149 |
| RXFP1 | 162.658697 | | 3.303970639 | 0.59867874 | 5.51877058 | 3.41E-08 | 2.00E-05 |
| GRIP2 | 50.1489841 | | 3.309375843 | 0.56039408 | 5.90544398 | 3.52E-09 | 3.04E-06 |
| SOST | 67.0762511 | | 3.485854687 | 1.45705296 | 2.39240082 | 0.01673855 | 0.18509671 |
| EGF | 55.4061104 | | 3.737214892 | 0.75711187 | 4.93614624 | 7.97E-07 | 0.00023102 |

**Interstitial cells**

*This means that positive fold changes imply higher mean value in controls than in AKI and vice versa.

| | baseMean | log2 fold change control/AKI* | lfcSE | stat | pvalue | padj |
|---|---|---|---|---|---|---|
| AL669831.5 | 32.6000421 | 0.312520164 | 0.27962588 | 1.11763677 | 0.26372216 | 0.6579701 |
| SAMD11 | 31.8417224 | 0.508679374 | 0.32553225 | 1.56260824 | 0.11814477 | 0.49192402 |
| NOC2L | 22.7672471 | -0.279029661 | 0.32766268 | -0.851576 | 0.39444949 | 0.75988253 |
| HES4 | 66.582799 | -0.226713675 | 0.51369557 | -0.4413386 | 0.6589679 | 0.90831499 |
| ISG15 | 264.626103 | 1.185498214 | 1.21625333 | 0.97471323 | 0.32970253 | 0.71362138 |
| AGRN | 19.6123781 | 0.756050045 | 0.40145172 | 1.88329011 | 0.05966106 | 0.36410704 |
| C1orf159 | 25.08242 | 0.067359556 | 0.29586203 | 0.22767219 | 0.8199011 | 0.96284007 |
| SDF4 | 42.2143163 | -0.083714805 | 0.28307082 | -0.295738 | 0.76743015 | 0.94564792 |
| UBE2J2 | 30.6710557 | 0.22880289 | 0.2652795 | 0.86249743 | 0.38841385 | 0.75549104 |
| ACAP3 | 21.6881338 | 0.840534353 | 0.34510138 | 2.43561574 | 0.01486647 | 0.17735986 |
| INTS11 | 33.02019 | 0.419023919 | 0.23089568 | 1.81477594 | 0.06955835 | 0.39291646 |
| MXRA8 | 39.7021008 | 0.491259728 | 0.37908818 | 1.2958983 | 0.19501053 | 0.59308963 |
| AURKAIP1 | 30.4957617 | -0.01855173 | 0.4207689 | -0.0440901 | 0.96483261 | 0.99275266 |
| CCNL2 | 88.0269672 | -0.066827903 | 0.19371421 | -0.3449819 | 0.73010795 | 0.93455731 |
| MRPL20 | 44.0789191 | -0.225408559 | 0.33025423 | -0.6825304 | 0.49490362 | 0.82044816 |
| ATAD3B | 17.1170556 | 0.298203853 | 0.3305074 | 0.90226074 | 0.36691837 | 0.73896496 |
| SSU72 | 44.2019559 | -0.065113754 | 0.25315126 | -0.2572128 | 0.79701448 | 0.95463848 |
| CDK11B | 18.66467 | -0.035069136 | 0.41955724 | -0.0835861 | 0.93338555 | 0.98725539 |
| SLC35E2B | 23.7810081 | -0.01875987 | 0.32558342 | -0.0576192 | 0.95405192 | 0.99100036 |
| SLC35E2A | 18.68708 | 0.507316627 | 0.30340554 | 1.67207435 | 0.09450966 | 0.44706027 |
| NADK | 28.388152 | -0.378672812 | 0.29551828 | -1.2813854 | 0.20005832 | 0.59922861 |
| GNB1 | 238.014468 | 0.228116092 | 0.13228422 | 1.72443919 | 0.08462859 | 0.42695086 |
| FAAP20 | 26.778249 | -0.095175168 | 0.25426416 | -0.3743161 | 0.70816916 | 0.93009618 |
| SKI | 110.373669 | -0.025589485 | 0.27142092 | -0.0942797 | 0.92488697 | 0.98561736 |
| RER1 | 39.9552718 | -0.306675295 | 0.25425098 | -1.2061912 | 0.22774377 | 0.62183669 |
| TNFRSF14 | 34.0568336 | -0.467086071 | 0.47479513 | -0.9837634 | 0.32523186 | 0.71149668 |
| PRDM16 | 45.0406513 | 0.420234522 | 0.88062729 | 0.47719907 | 0.63322037 | 0.89542746 |
| CEP104 | 27.9483235 | 0.041958283 | 0.26872824 | 0.15613649 | 0.87592544 | 0.9768643 |
| RPL22 | 26.8630813 | -0.035835407 | 0.34787037 | -0.1030137 | 0.91795211 | 0.98477389 |
| ICMT | 18.3536637 | 0.044256947 | 0.30341822 | 0.1458612 | 0.88403096 | 0.97783617 |
| NOL9 | 24.3118447 | 0.012213909 | 0.48303621 | 0.0252857 | 0.97982708 | 0.99503182 |
| DNAJC11 | 39.985956 | 0.038787304 | 0.29852726 | 0.12992885 | 0.89662272 | 0.98040427 |
| CAMTA1 | 166.744809 | -0.151153259 | 0.1781746 | -0.8483435 | 0.39624672 | 0.76020146 |
| VAMP3 | 19.770221 | 0.004194122 | 0.31401622 | 0.01335639 | 0.98934346 | 0.99739981 |
| PER3 | 68.042059 | 0.695631624 | 0.52951912 | 1.31370445 | 0.18894568 | 0.58671246 |
| PARK7 | 62.2451629 | -0.16992855 | 0.22108507 | -0.7686116 | 0.44212392 | 0.79074438 |
| ERRFI1 | 119.071231 | 0.283347153 | 0.85110469 | 0.33291692 | 0.739197 | 0.93594619 |
| RERE | 547.845735 | -0.021946637 | 0.15788053 | -0.1390079 | 0.88944393 | 0.97998852 |
| ENO1 | 149.14189 | -1.500076131 | 0.34920712 | -4.295663 | 1.74E-05 | 0.00271172 |
| MIR34AHG | 36.4421647 | -0.991099365 | 0.40202834 | -2.4652475 | 0.01369186 | 0.17073456 |
| H6PD | 44.4016171 | -0.871357255 | 0.3458534 | -2.5194411 | 0.01175413 | 0.15480487 |
| SPSB1 | 70.7875287 | 0.345883255 | 0.42992977 | 0.80451107 | 0.42110188 | 0.77771095 |
| SLC25A33 | 23.0465022 | 0.218844163 | 0.2919232 | 0.74966349 | 0.4534574 | 0.79534858 |
| CLSTN1 | 55.8615058 | 0.287601578 | 0.20705314 | 1.38902305 | 0.16482574 | 0.55965519 |
| LZIC | 19.0966657 | 0.178180344 | 0.32543362 | 0.54751671 | 0.5840238 | 0.86828629 |
| NMNAT1 | 19.5359129 | -0.216710898 | 0.28142672 | -0.7700438 | 0.44127392 | 0.79053729 |
| UBE4B | 147.717568 | -0.140360994 | 0.21426716 | -0.6550747 | 0.51241966 | 0.82891748 |
| KIF1B | 163.403151 | 0.295991788 | 0.24276624 | 1.2192461 | 0.2227508 | 0.61772146 |
| DFFA | 19.8069651 | -0.155684808 | 0.34425341 | -0.452239 | 0.65109683 | 0.9050963 |
| PEX14 | 53.9474258 | -0.255024921 | 0.19948666 | -1.2784059 | 0.20110634 | 0.60005299 |
| SRM | 28.3859772 | -0.672866624 | 0.5368126 | -1.2534479 | 0.21004275 | 0.60731347 |
| EXOSC10 | 26.3302225 | 0.214770261 | 0.26051042 | 0.82442099 | 0.40970039 | 0.7710341 |
| MTOR | 63.9463686 | 0.227154239 | 0.26069315 | 0.87134716 | 0.38356463 | 0.75256313 |
| MTHFR | 46.8408965 | -0.427773792 | 0.38805475 | -1.1023542 | 0.27030771 | 0.6651094 |
| CLCN6 | 25.6862523 | -0.582797567 | 0.34377655 | -1.6952802 | 0.09002228 | 0.43812831 |
| PLOD1 | 34.8281116 | -0.157785864 | 0.21666232 | -0.7282571 | 0.46645624 | 0.80355999 |
| MFN2 | 28.1685491 | 0.379269894 | 0.28731933 | 1.32002917 | 0.18682528 | 0.58501647 |
| TNFRSF1B | 26.5236968 | -0.455095028 | 0.48681034 | -0.9348508 | 0.3498652 | 0.72803645 |
| VPS13D | 338.231813 | -0.54380686 | 0.25109725 | -2.1657221 | 0.03033243 | 0.26568894 |
| DHRS3 | 195.65804 | -0.321943201 | 0.50364429 | -0.6392273 | 0.52267505 | 0.8350836 |
| PRDM2 | 110.481676 | -0.290025693 | 0.19611544 | -1.4788519 | 0.1391799 | 0.52506833 |
| KAZN | 1241.12909 | -0.8556595 | 0.45922734 | -1.8632591 | 0.06242583 | 0.37164525 |
| TMEM51 | 64.6096464 | -0.092814495 | 0.36740497 | -0.2526218 | 0.8005605 | 0.95648173 |
| DNAJC16 | 34.4718815 | -0.098078384 | 0.27517788 | -0.3564181 | 0.72152746 | 0.93444541 |
| DDI2 | 70.2727118 | -0.266070195 | 0.22136657 | -1.2019439 | 0.22938527 | 0.62342914 |
| PLEKHM2 | 61.711746 | -0.481510164 | 0.25870157 | -1.8612572 | 0.06270787 | 0.37164525 |
| FBLIM1 | 123.394396 | -0.523175381 | 0.30338814 | -1.7244424 | 0.084628 | 0.42695086 |
| SPEN | 144.831286 | -0.161500503 | 0.18428344 | -0.8763701 | 0.38082885 | 0.74991898 |
| EPHA2 | 25.3594236 | 0.421191138 | 0.40961144 | 1.02826994 | 0.30382286 | 0.69579129 |
| FBXO42 | 152.070985 | -0.259879232 | 0.2090952 | -1.2428752 | 0.21391384 | 0.61001472 |
| SZRD1 | 38.1730454 | 0.105099354 | 0.25645135 | 0.40982181 | 0.68193667 | 0.91921361 |
| NECAP2 | 18.1716372 | 0.356948589 | 0.27753266 | 1.28614986 | 0.19839076 | 0.59582089 |
| CROCC | 34.3924373 | 0.441592912 | 0.28815309 | 1.53249412 | 0.12540055 | 0.50440978 |
| SDHB | 36.6131617 | -0.220808862 | 0.22757582 | -0.9702651 | 0.33191439 | 0.71490002 |
| RCC2 | 20.1523215 | -0.587101035 | 0.37067398 | -1.5838744 | 0.1132223 | 0.48363945 |
| ARHGEF10L | 82.5587764 | 1.01334608 | 0.31446283 | 3.22246692 | 0.00127092 | 0.04375316 |
| UBR4 | 135.170468 | -0.159607571 | 0.1981217 | -0.8056037 | 0.42047141 | 0.77771095 |
| AL035413.1 | 22.0230377 | -0.070660369 | 0.25490368 | -0.2772042 | 0.78162332 | 0.94939885 |
| EMC1 | 20.922311 | 0.037000399 | 0.30233694 | 0.12238134 | 0.90259702 | 0.98164203 |
| CAPZB | 296.48487 | -0.571779128 | 0.27790124 | -2.0574903 | 0.03963908 | 0.30093762 |
| NBL1 | 31.6518983 | -0.344287447 | 0.33995731 | -1.0127373 | 0.31118566 | 0.70126123 |
| TMCO4 | 29.0865984 | 0.44403884 | 0.30945837 | 1.43489036 | 0.15131834 | 0.53898138 |
| CAMK2N1 | 31.6096945 | -0.006027229 | 0.25380272 | -0.0237477 | 0.98105386 | 0.99533442 |
| PINK1 | 22.3895564 | 0.305430022 | 0.36059461 | 0.84701772 | 0.39698524 | 0.76043276 |
| DDOST | 19.730064 | -0.175945417 | 0.40859337 | -0.4306125 | 0.66675014 | 0.9133172 |
| HP1BP3 | 152.369288 | 0.359846439 | 0.15678481 | 2.29516136 | 0.02172388 | 0.21750516 |
| EIF4G3 | 387.67391 | -0.298668931 | 0.22330445 | -1.3374966 | 0.1810606 | 0.57953588 |
| ECE1 | 147.54535 | -0.516481872 | 0.35727565 | -1.4456118 | 0.14828613 | 0.53680621 |
| USP48 | 104.586303 | 0.010978259 | 0.23762598 | 0.04619974 | 0.96315105 | 0.99231823 |
| HSPG2 | 115.821127 | 0.964689106 | 0.41177657 | 2.34274891 | 0.01914226 | 0.20295514 |
| CDC42 | 84.5156528 | 0.010129948 | 0.20420462 | 0.04960685 | 0.96043569 | 0.99181157 |
| ZBTB40 | 32.39219 | 0.062124007 | 0.23100683 | 0.26892714 | 0.78798575 | 0.95129454 |
| KDM1A | 57.7492203 | -0.084755247 | 0.2048593 | -0.4137242 | 0.67907611 | 0.91831925 |
| LUZP1 | 73.1888086 | 0.552577928 | 0.16540416 | 3.34077398 | 0.00083545 | 0.03373482 |
| HNRNPR | 124.851477 | 0.024283052 | 0.17878033 | 0.1358262 | 0.89195868 | 0.98023547 |
| ASAP3 | 25.9649012 | 0.464044465 | 0.31316415 | 1.48179308 | 0.13839536 | 0.52264518 |
| ID3 | 153.825374 | 0.550057103 | 0.73108928 | 0.7523802 | 0.45182245 | 0.79455448 |
| RPL11 | 148.173363 | -0.15324141 | 0.27285067 | -0.5616311 | 0.57436742 | 0.86294056 |
| ELOA | 20.1570403 | -0.412140104 | 0.33168467 | -1.242566 | 0.21402779 | 0.61001472 |
| HMGCL | 23.9143118 | -0.250139511 | 0.2896321 | -0.8636457 | 0.38778256 | 0.75457985 |
| PNRC2 | 36.1733058 | 0.265899279 | 0.32642195 | 0.81458762 | 0.4153084 | 0.77523531 |
| SRSF10 | 97.7338424 | 0.2092508 | 0.17625523 | 1.18720332 | 0.23514743 | 0.63100657 |
| SRRM1 | 78.4159457 | 0.193193785 | 0.21656357 | 0.8920881 | 0.37234571 | 0.74463512 |
| CLIC4 | 361.020701 | 0.284332934 | 0.33044553 | 0.86045326 | 0.38953924 | 0.75628551 |
| SYF2 | 21.3270064 | -0.084221982 | 0.31508688 | -0.2672977 | 0.78924 | 0.95162014 |
| RSRP1 | 156.721126 | -0.106690037 | 0.34151853 | -0.312399 | 0.75473734 | 0.94170839 |
| TMEM50A | 50.6782146 | -0.243601698 | 0.22692222 | -1.073503 | 0.28304547 | 0.67727609 |
| MACO1 | 59.9884446 | -0.056166117 | 0.17143875 | -0.3276162 | 0.74320185 | 0.93791562 |
| LDLRAP1 | 22.905408 | 1.218197802 | 0.4898203 | 2.48703006 | 0.01288145 | 0.16578947 |
| MAN1C1 | 95.6716979 | 0.631109551 | 0.44421457 | 1.42073132 | 0.15539488 | 0.54412824 |
| SH3BGRL3 | 40.2576353 | -0.745966383 | 0.40467438 | -1.8433744 | 0.06527437 | 0.38095778 |

| | | | | | | |
|---|---|---|---|---|---|---|
| DHDDS | 25.3608345 | -0.364539138 | 0.31859489 | -1.144209 | 0.25253697 | 0.64724776 |
| HMGN2 | 49.3524602 | 0.151055439 | 0.2979793 | 0.50693267 | 0.61220206 | 0.88282721 |
| ARID1A | 145.632986 | -0.104513319 | 0.17262383 | -0.6054397 | 0.54488689 | 0.84730644 |
| NUDC | 36.4626797 | -0.114758037 | 0.25706008 | -0.446425 | 0.6552903 | 0.90691684 |
| SLC9A1 | 20.8444031 | 0.316711203 | 0.26783719 | 1.18247658 | 0.23701665 | 0.63270446 |
| WDTC1 | 36.4009807 | -0.073220615 | 0.2955142 | -0.2477736 | 0.80430959 | 0.95836286 |
| WASF2 | 139.9855 | 0.153597657 | 0.14838915 | 1.03510029 | 0.30062208 | 0.69191459 |
| AHDC1 | 51.2022639 | 0.552467174 | 0.32212855 | 1.71505187 | 0.08633572 | 0.4307408 |
| IFI6 | 91.3345261 | 0.608862901 | 1.15850811 | 0.52555774 | 0.59919553 | 0.87666493 |
| FAM76A | 23.1815665 | 0.491747883 | 0.24855422 | 1.97843308 | 0.04787987 | 0.33132691 |
| STX12 | 126.751825 | 0.098280678 | 0.32633143 | 0.30116829 | 0.76328617 | 0.94462105 |
| EYA3 | 70.5066503 | -0.007692989 | 0.15758046 | -0.0488194 | 0.96106319 | 0.99181157 |
| DNAJC8 | 28.2487419 | 0.184747172 | 0.28860916 | 0.64012928 | 0.52208856 | 0.8350836 |
| ATP5IF1 | 46.7770456 | -0.186159881 | 0.27037026 | -0.6885368 | 0.49111479 | 0.81720854 |
| PHACTR4 | 124.084357 | 0.171817903 | 0.20764537 | 0.82745838 | 0.4079773 | 0.76956499 |
| RCC1 | 42.0722144 | -0.968110939 | 0.44819072 | -2.1600423 | 0.03076939 | 0.26662968 |
| TRNAU1AP | 43.2067143 | -0.253984056 | 0.27452071 | -0.9251909 | 0.35486663 | 0.73100657 |
| TAF12 | 20.4872007 | 0.29349391 | 0.29902182 | 0.98151335 | 0.32633965 | 0.71149668 |
| GMEB1 | 24.893633 | 0.324027982 | 0.26656571 | 1.21556512 | 0.22415063 | 0.61858591 |
| YTHDF2 | 46.5851848 | -0.128437793 | 0.1952781 | -0.6577174 | 0.51071977 | 0.82780758 |
| EPB41 | 54.6790032 | -0.142506844 | 0.28084689 | -0.5074183 | 0.61186136 | 0.88282721 |
| SRSF4 | 183.31689 | -0.156315155 | 0.20682811 | -0.7557733 | 0.44978516 | 0.79399826 |
| MECR | 17.4119907 | -0.058018928 | 0.36893059 | -0.1572624 | 0.87503801 | 0.9768643 |
| PUM1 | 270.352801 | -0.212012439 | 0.19523635 | -1.0859271 | 0.27751126 | 0.67269898 |
| SNRNP40 | 32.6137768 | -0.048399607 | 0.27269305 | -0.1774875 | 0.85912548 | 0.97397549 |
| ZCCHC17 | 33.2869644 | 0.198249574 | 0.23847988 | 0.83130523 | 0.40580122 | 0.76703528 |
| TINAGL1 | 205.856498 | 0.152772196 | 0.32775129 | 0.46612233 | 0.64112794 | 0.89934133 |
| PEF1 | 19.5491921 | -0.203157745 | 0.26938221 | -0.7541617 | 0.45075214 | 0.79401729 |
| COL16A1 | 69.3432524 | 0.365907634 | 0.34244502 | 1.06851498 | 0.28528828 | 0.6783522 |
| PTP4A2 | 38.0495619 | -0.479845018 | 0.3338233 | -1.4374222 | 0.15059807 | 0.53880252 |
| KHDRBS1 | 117.532864 | 0.085261873 | 0.18035469 | 0.47274553 | 0.63639474 | 0.89741446 |
| KPNA6 | 96.9628467 | -0.013794956 | 0.20827116 | -0.0662356 | 0.94719029 | 0.99060401 |
| TXLNA | 34.9844518 | -0.220856182 | 0.26082806 | -0.8467501 | 0.39713443 | 0.76043276 |
| EIF3I | 27.2392025 | -0.379992215 | 0.40500063 | -0.9382509 | 0.34811548 | 0.72612016 |
| HDAC1 | 31.2652758 | 0.007362161 | 0.24115046 | 0.03052933 | 0.9756449 | 0.99335195 |
| BSDC1 | 44.023378 | 0.14146737 | 0.34864341 | 0.40576522 | 0.68491511 | 0.91977784 |
| ZBTB8OS | 65.0214792 | 0.062722132 | 0.17579055 | 0.35680037 | 0.72124126 | 0.93444541 |
| RBBP4 | 59.4478798 | 0.193903378 | 0.17146002 | 1.1308956 | 0.25809903 | 0.65130862 |
| YARS | 51.0206075 | -1.030605315 | 0.35836652 | -2.8758415 | 0.00402952 | 0.08225816 |
| S100PBP | 50.4433691 | 0.319568749 | 0.23641158 | 1.35174742 | 0.17645613 | 0.5732874 |
| RNF19B | 33.4789965 | -0.081958857 | 0.46138478 | -0.1776367 | 0.85900832 | 0.97397549 |
| AK2 | 45.0850024 | -0.631462974 | 0.29791912 | -2.1195785 | 0.03404161 | 0.2823609 |
| ZNF362 | 38.0406905 | 0.748723661 | 0.32788326 | 2.28350683 | 0.02240053 | 0.22134579 |
| AL513327.1 | 39.4975729 | -1.966409247 | 0.42764877 | -4.5981875 | 4.26E-06 | 0.00087 |
| PHC2 | 238.736882 | -0.985422293 | 0.25709088 | -3.8329726 | 0.0001266 | 0.00930413 |
| SMIM12 | 35.9937653 | -0.040212278 | 0.24426008 | -0.1646289 | 0.86923605 | 0.9768643 |
| GJA4 | 51.819821 | -0.915008297 | 0.57361648 | -1.5951569 | 0.11067715 | 0.47776431 |
| ZMYM1 | 27.525694 | 0.084431136 | 0.27703924 | 0.30476237 | 0.76054714 | 0.94425408 |
| SFPQ | 165.821248 | 0.02994718 | 0.19292112 | 0.15523018 | 0.87663986 | 0.9768643 |
| ZMYM4 | 245.684926 | -0.003729914 | 0.14118764 | -0.0264181 | 0.97892383 | 0.99465087 |
| KIAA0319L | 113.016759 | 0.02325768 | 0.21250544 | 0.10944511 | 0.91284946 | 0.98395113 |
| PSMB2 | 35.8803106 | -0.225969607 | 0.29712978 | -0.7605081 | 0.44695092 | 0.79296348 |
| AGO4 | 67.3255856 | 0.369682986 | 0.32343962 | 1.14297372 | 0.2530495 | 0.64796543 |
| AGO1 | 23.8647889 | 0.522174442 | 0.25670773 | 2.03412046 | 0.04193945 | 0.30821303 |
| AGO3 | 212.194937 | 0.022416589 | 0.12359904 | 0.1813654 | 0.85608077 | 0.97379567 |
| MAP7D1 | 25.1649145 | 0.195049017 | 0.37666521 | 0.51783126 | 0.60457601 | 0.87903643 |
| THRAP3 | 140.287982 | -0.094054536 | 0.14075263 | -0.6682258 | 0.50398948 | 0.82438238 |
| EVA1B | 31.1346893 | 0.106512976 | 0.33693575 | 0.31612252 | 0.75190952 | 0.94055881 |
| STK40 | 41.9420224 | 0.119556967 | 0.3344687 | 0.35745338 | 0.72075242 | 0.93444541 |
| MEAF6 | 21.4216602 | 0.314245743 | 0.26556889 | 1.18329276 | 0.23669314 | 0.63270446 |
| C1orf122 | 22.9887052 | -0.166643457 | 0.34142651 | -0.48808 | 0.62549136 | 0.89069797 |
| MTF1 | 20.3391702 | 0.02821835 | 0.32202325 | 0.0876283 | 0.93017211 | 0.98681039 |
| INPP5B | 88.8273434 | -0.143581175 | 0.17824758 | -0.8055154 | 0.4205223 | 0.77771095 |
| SF3A3 | 32.6798968 | -0.045843998 | 0.34317239 | -0.1335888 | 0.89372772 | 0.98023547 |
| RRAGC | 24.4114062 | -0.109643277 | 0.33566958 | -0.3266405 | 0.74393982 | 0.93793339 |
| AL139260.1 | 35.5870887 | 0.163545253 | 0.4390615 | 0.37248825 | 0.70952936 | 0.93009618 |
| AKIRIN1 | 35.2008459 | -0.298103255 | 0.23809857 | -1.2520161 | 0.21056398 | 0.60731347 |
| NDUFS5 | 59.3777688 | -0.68780001 | 0.3909525 | -1.7592931 | 0.07852775 | 0.41542971 |
| MACF1 | 788.882492 | 0.096596674 | 0.18057606 | 0.53493622 | 0.59269396 | 0.87313824 |
| PABPC4 | 33.9585265 | -0.397690932 | 0.36806551 | -1.0804895 | 0.27992425 | 0.6742587 |
| HEYL | 56.280813 | 0.366177032 | 0.35984476 | 1.01759723 | 0.3088694 | 0.69885506 |
| TRIT1 | 31.2642882 | -0.720228306 | 0.27693314 | -2.6007299 | 0.00930257 | 0.13510782 |
| CAP1 | 77.8477672 | 0.199290903 | 0.26083858 | 0.76403922 | 0.44484387 | 0.79252305 |
| PPT1 | 20.1778875 | 0.226043225 | 0.3116073 | 0.72541055 | 0.46820021 | 0.80374006 |
| RLF | 144.492902 | -0.514421264 | 0.29617344 | -1.7368919 | 0.08240625 | 0.42256146 |
| ZMPSTE24 | 28.0217018 | 0.013256347 | 0.27766588 | 0.04774208 | 0.9619218 | 0.99202404 |
| SMAP2 | 36.140541 | -0.472940161 | 0.30171038 | -1.5675303 | 0.11699079 | 0.48989475 |
| NFYC | 46.2986407 | 0.463122842 | 0.27604485 | 1.67770868 | 0.09340398 | 0.44400122 |
| CTPS1 | 25.0227708 | 0.264880591 | 0.42439203 | 0.62414129 | 0.5325348 | 0.84290291 |
| SCMH1 | 299.741715 | -0.203822464 | 0.1984983 | -1.0268222 | 0.3045042 | 0.69648345 |
| HIVEP3 | 148.354437 | 0.440868569 | 0.24096801 | 1.82957303 | 0.06731381 | 0.38712255 |
| FOXJ3 | 141.621518 | -0.143352504 | 0.18957608 | -0.756174 | 0.44954487 | 0.7937783 |
| CCDC30 | 24.5815705 | 0.457094295 | 0.38702992 | 1.1810309 | 0.23759045 | 0.63302595 |
| YBX1 | 77.8358103 | -0.36250348 | 0.33543554 | -1.0806949 | 0.27983284 | 0.6742587 |
| P3H1 | 19.8364384 | -0.460852549 | 0.34581625 | -1.3326515 | 0.1826462 | 0.58106792 |
| SLC2A1 | 29.9548629 | -1.937105496 | 0.62704297 | -3.0892707 | 0.00200648 | 0.05600918 |
| SZT2 | 35.7151012 | 0.077714799 | 0.2747444 | 0.28286217 | 0.77728249 | 0.94808797 |
| PTPRF | 29.6574001 | -0.193275847 | 0.29822866 | -0.6480794 | 0.51693361 | 0.83252743 |
| ATP6V0B | 44.2370286 | -0.505241734 | 0.33675563 | -1.5003215 | 0.13353113 | 0.5153993 |
| ERI3 | 76.2904291 | -0.375077588 | 0.21001819 | -1.7859291 | 0.07411074 | 0.40373597 |
| RPS8 | 148.539981 | -0.480960378 | 0.33938383 | -1.4171576 | 0.15643685 | 0.54607164 |
| EIF2B3 | 61.5005504 | 0.017968859 | 0.27685115 | 0.06490441 | 0.94825011 | 0.99060401 |
| ZSWIM5 | 63.1197777 | 0.0882124 | 0.25912699 | 0.3404215 | 0.73353913 | 0.93589914 |
| TESK2 | 35.583243 | 0.336852181 | 0.29685616 | 1.13473198 | 0.25648763 | 0.65084527 |
| PRDX1 | 105.391768 | -0.564938948 | 0.36547318 | -1.545774 | 0.12215916 | 0.50069584 |
| AKR1A1 | 20.3501741 | 0.04306392 | 0.32454397 | 0.13269056 | 0.89443811 | 0.98023547 |
| NASP | 45.0896396 | -0.014205381 | 0.24078643 | -0.0589958 | 0.95295548 | 0.99060401 |
| GPBP1L1 | 115.364902 | -0.049310237 | 0.1527015 | -0.3229191 | 0.74675648 | 0.93900586 |
| IPP | 19.896797 | 0.473110803 | 0.37919129 | 1.24768374 | 0.2121469 | 0.60910715 |
| MAST2 | 71.7216081 | -0.071533236 | 0.22541476 | -0.3173405 | 0.75098524 | 0.94052327 |
| TSPAN1 | 22.0394791 | -1.562638723 | 0.36791007 | -4.2473388 | 2.16E-05 | 0.00305725 |
| LRRC41 | 38.0491916 | 0.057190609 | 0.20131672 | 0.28408275 | 0.77634696 | 0.94805375 |
| UQCRH | 47.6161909 | -0.448180375 | 0.24370502 | -1.8390281 | 0.06591105 | 0.38351571 |
| NSUN4 | 26.7290634 | -0.154144172 | 0.29271339 | -0.5266044 | 0.5984683 | 0.87618867 |
| MKNK1 | 29.3293824 | 0.176173032 | 0.25503196 | 0.69078807 | 0.48969874 | 0.81720854 |
| ATPAF1 | 32.5551877 | 0.116456186 | 0.24700856 | 0.4714662 | 0.63730785 | 0.89741446 |
| EFCAB14 | 55.6398663 | -0.056173258 | 0.22370607 | -0.251103 | 0.8017345 | 0.95758343 |
| CYP4X1 | 22.4474949 | 1.34356018 | 0.46330319 | 2.89995882 | 0.00373212 | 0.0794995 |
| PDZK1IP1 | 53.6532816 | -1.169717026 | 0.53088148 | -2.2033487 | 0.02757018 | 0.24860521 |
| CMPK1 | 58.3203187 | -0.232703818 | 0.2930212 | -0.7941535 | 0.42710606 | 0.77987274 |
| TRABD2B | 57.8873153 | -1.035063473 | 0.58948909 | -1.7558654 | 0.0791114 | 0.41722081 |
| SPATA6 | 95.4475466 | -0.057966882 | 0.30392402 | -0.1907282 | 0.84873854 | 0.97092453 |
| FAF1 | 344.751015 | -0.146616814 | 0.13839051 | -1.0594427 | 0.28939822 | 0.67992862 |

| | | | | | | |
|---|---|---|---|---|---|---|
| RNF11 | 50.0425844 | 0.377201547 | 0.18817466 | 2.00452893 | 0.04501343 | 0.31898889 |
| EPS15 | 230.704871 | -0.009581074 | 0.19743798 | -0.048527 | 0.96129624 | 0.99181157 |
| OSBPL9 | 225.156993 | 0.311563713 | 0.25404025 | 1.22643443 | 0.22003524 | 0.61590675 |
| NRDC | 167.235292 | -0.356582527 | 0.13351847 | -2.6706606 | 0.00757021 | 0.12147053 |
| TXNDC12 | 23.8167528 | -0.12076897 | 0.35088376 | -0.3441851 | 0.73070707 | 0.93455731 |
| BTF3L4 | 40.1581619 | -0.36812456 | 0.23900361 | -1.5402468 | 0.1235002 | 0.50097964 |
| ZFYVE9 | 102.027238 | 0.021772739 | 0.17408722 | 0.12506799 | 0.90046972 | 0.98110482 |
| PRPF38A | 19.6137118 | 0.576823375 | 0.28145472 | 2.04943581 | 0.04041952 | 0.30187301 |
| TUT4 | 154.423809 | -0.029540965 | 0.21042683 | -0.1403859 | 0.88835508 | 0.97942933 |
| ZYG11B | 73.1384436 | 0.165989276 | 0.21319965 | 0.77856263 | 0.4362374 | 0.78730567 |
| ECHDC2 | 113.649552 | 0.800285789 | 0.429806 | 1.86196981 | 0.06260735 | 0.37164525 |
| SCP2 | 78.7045295 | -0.050896427 | 0.22891921 | -0.2223336 | 0.82405421 | 0.96413949 |
| PODN | 38.49505 | 1.619156676 | 0.51383048 | 3.15114956 | 0.00162629 | 0.05064245 |
| MAGOH | 18.2038538 | 0.458196547 | 0.32093676 | 1.42768483 | 0.15338259 | 0.54114675 |
| NDC1 | 18.2756406 | 0.248513991 | 0.27279369 | 0.91099611 | 0.36229742 | 0.73489477 |
| YIPF1 | 17.7438897 | -0.195763537 | 0.27159719 | -0.7207863 | 0.47104099 | 0.80492221 |
| HSPB11 | 28.0747156 | 0.221392992 | 0.3177583 | 0.69673395 | 0.48596931 | 0.81594436 |
| TMEM59 | 101.97026 | -0.157479087 | 0.20260177 | -0.7772839 | 0.43699132 | 0.78768252 |
| SSBP3 | 463.129323 | -1.090003522 | 0.35428985 | -3.0765869 | 0.00209385 | 0.05624279 |
| USP24 | 144.959186 | 0.103264524 | 0.17826604 | 0.57927198 | 0.56240567 | 0.85545224 |
| PLPP3 | 178.925179 | -0.397042495 | 0.49993799 | -0.7941835 | 0.42708862 | 0.77987274 |
| PRKAA2 | 43.3256303 | 0.144576011 | 0.4722762 | 0.30612597 | 0.75950874 | 0.94385579 |
| DAB1 | 64.0262935 | 1.492450659 | 0.5180028 | 2.8811633 | 0.0039621 | 0.08179072 |
| MYSM1 | 58.972532 | -0.1991189 | 0.21256335 | -0.9367509 | 0.34888673 | 0.72734588 |
| JUN | 154.517971 | 0.112725426 | 0.47896895 | 0.23535017 | 0.81393695 | 0.96038841 |
| FGGY | 62.6997716 | 0.068881292 | 0.33185523 | 0.20756428 | 0.8355692 | 0.9668633 |
| NFIA | 776.987714 | 0.115480187 | 0.26925736 | 0.42888405 | 0.66800761 | 0.9143573 |
| TM2D1 | 79.1757979 | 0.117223283 | 0.16108396 | 0.72771545 | 0.46678779 | 0.80374006 |
| PATJ | 138.622627 | 0.352951848 | 0.30351043 | 1.16289857 | 0.24487066 | 0.6405276 |
| DOCK7 | 124.021747 | 0.036292516 | 0.23952541 | 0.15151844 | 0.87956677 | 0.9776068 |
| ATG4C | 37.4434088 | -0.353024905 | 0.26287701 | -1.342928 | 0.17929528 | 0.57744792 |
| ITGB3BP | 30.8693919 | -0.964745212 | 0.40247166 | -2.3970513 | 0.01652761 | 0.18980307 |
| EFCAB7 | 20.3925158 | 0.255263494 | 0.25433417 | 1.00365397 | 0.31554543 | 0.70306809 |
| PGM1 | 25.2573524 | -0.936930323 | 0.39038475 | -2.4000178 | 0.01639428 | 0.18884254 |
| ROR1 | 228.855181 | 1.211042726 | 0.36285934 | 3.33749908 | 0.00084536 | 0.03394837 |
| CACHD1 | 109.65643 | -0.3664418 | 0.29424097 | -1.2453799 | 0.21299213 | 0.60976984 |
| JAK1 | 205.264745 | -0.393637274 | 0.18931537 | -2.0792674 | 0.03759278 | 0.29453019 |
| AK4 | 46.9310833 | -1.63453992 | 0.55369107 | -2.9520793 | 0.00315642 | 0.07226331 |
| LEPR | 54.7195525 | -0.312381087 | 0.31446157 | -0.993384 | 0.32052287 | 0.70762743 |
| PDE4B | 100.24819 | -1.061765107 | 0.75862504 | -1.3995914 | 0.1616357 | 0.55507512 |
| SGIP1 | 688.673977 | -1.326368204 | 0.37729539 | -3.5154636 | 0.00043899 | 0.02145813 |
| WDR78 | 35.7059221 | 0.665652932 | 0.41430983 | 1.60665492 | 0.10813009 | 0.47385094 |
| MIER1 | 119.430353 | -0.219160548 | 0.2025316 | -1.0821055 | 0.27920567 | 0.67353297 |
| SLC35D1 | 40.223295 | 0.459365301 | 0.22697328 | 2.02387389 | 0.04298313 | 0.31204692 |
| SERBP1 | 78.9745405 | -0.330505128 | 0.31595873 | -1.0460389 | 0.29554307 | 0.68645575 |
| GADD45A | 29.1187005 | -0.683945877 | 0.45969468 | -1.4878264 | 0.13679668 | 0.52089048 |
| GNG12 | 152.509993 | 0.260210005 | 0.18638883 | 1.39606007 | 0.16269641 | 0.55715559 |
| GNG12-AS1 | 28.086249 | -0.108562235 | 0.35609643 | -0.3048675 | 0.76046706 | 0.94425408 |
| WLS | 45.8196038 | 0.351955278 | 0.2342477 | 1.5024919 | 0.13297012 | 0.51431442 |
| LRRC40 | 32.4492765 | 0.23615574 | 0.25592204 | 0.92276437 | 0.35613001 | 0.73129235 |
| SRSF11 | 217.497324 | -0.057558593 | 0.14170672 | -0.4061811 | 0.68460952 | 0.91977784 |
| ANKRD13C | 113.679903 | -0.002526459 | 0.20006055 | -0.0126285 | 0.9899242 | 0.99739981 |
| PTGER3 | 37.2962687 | 0.582112478 | 0.43402634 | 1.34119161 | 0.17985825 | 0.57744792 |
| ZRANB2 | 178.576773 | -0.08558701 | 0.19462239 | -0.4397593 | 0.66011143 | 0.90913772 |
| NEGR1 | 1257.19346 | 0.180637586 | 0.48608678 | 0.37161592 | 0.71017883 | 0.93009618 |
| CRYZ | 24.9303192 | 0.774421269 | 0.48791196 | 1.58721518 | 0.11246392 | 0.48220381 |
| TYW3 | 23.9433655 | 0.271794143 | 0.28341145 | 0.95900904 | 0.33755419 | 0.71925477 |
| ACADM | 37.3529401 | 0.198837176 | 0.29994093 | 0.66292111 | 0.50738108 | 0.82494327 |
| RABGGTB | 23.0625704 | -0.675686686 | 0.35246848 | -1.9170131 | 0.05523627 | 0.35288127 |
| ST6GALNAC3 | 28.9217063 | -0.073620296 | 0.46033529 | -0.1599276 | 0.87293814 | 0.9768643 |
| PIGK | 37.1217808 | 0.185722913 | 0.27168219 | 0.68360357 | 0.49422553 | 0.81957698 |
| AC118549.1 | 167.843375 | -0.150400629 | 0.15442936 | -0.9739121 | 0.33010016 | 0.71362138 |
| USP33 | 124.40064 | -0.344175204 | 0.22715875 | -1.5151307 | 0.12973931 | 0.51204187 |
| MIGA1 | 75.6082639 | 0.020236403 | 0.16747228 | 0.12083434 | 0.90382225 | 0.98225619 |
| NEXN | 116.431109 | 1.124405398 | 0.54325445 | 2.06975827 | 0.03847499 | 0.29669746 |
| FUBP1 | 119.327626 | -0.017310343 | 0.23277429 | -0.0743654 | 0.94071967 | 0.98895798 |
| IFI44L | 134.219307 | 0.381720264 | 1.34409662 | 0.28399764 | 0.77641219 | 0.94805375 |
| IFI44 | 54.3645479 | 0.880294706 | 1.12895055 | 0.77974603 | 0.43554038 | 0.78682061 |
| ADGRL2 | 229.393435 | 0.269986268 | 0.286997 | 0.94072852 | 0.346844 | 0.7249592 |
| TTLL7 | 64.3576563 | 1.250236059 | 0.43203995 | 2.89379731 | 0.00380614 | 0.07977552 |
| PRKACB | 82.1385562 | 0.348228611 | 0.2631258 | 1.32343015 | 0.18569237 | 0.58501647 |
| GNG5 | 43.1539955 | -0.120035805 | 0.2842485 | -0.4222918 | 0.67281206 | 0.91482259 |
| CTBS | 35.9362944 | -0.244663573 | 0.29220701 | -0.8372953 | 0.40242657 | 0.76472383 |
| SYDE2 | 22.6109793 | 0.831135132 | 0.28893272 | 2.87656973 | 0.00402023 | 0.08225816 |
| C1orf52 | 34.5665815 | -0.040934251 | 0.25822702 | -0.1585204 | 0.87404674 | 0.9768643 |
| AL078459.1 | 25.3471627 | 0.090032988 | 0.46603345 | 0.19318997 | 0.8468102 | 0.97014274 |
| DDAH1 | 210.907859 | 0.399692585 | 0.44476194 | 0.89866634 | 0.36883041 | 0.74123132 |
| CYR61 | 524.671522 | 1.016938597 | 0.73551729 | 1.38261684 | 0.16678238 | 0.56275653 |
| ZNHIT6 | 64.471913 | -0.236941121 | 0.26717039 | -0.886854 | 0.37515752 | 0.74723387 |
| ODF2L | 76.4804992 | -0.400801654 | 0.25477212 | -1.5731771 | 0.11567783 | 0.48836096 |
| SH3GLB1 | 88.0893731 | 0.044042736 | 0.21390162 | 0.20590183 | 0.83686759 | 0.9668633 |
| SELENOF | 84.3146028 | 0.017916341 | 0.17119874 | 0.1046523 | 0.91665172 | 0.98477389 |
| HS2ST1 | 59.6894497 | -0.456128671 | 0.29825492 | -1.5293249 | 0.12618393 | 0.50535459 |
| LMO4 | 101.52057 | 0.244682012 | 0.28161381 | 0.8688566 | 0.38492557 | 0.75305975 |
| PKN2 | 191.559343 | 0.104514206 | 0.16804652 | 0.62193613 | 0.53398386 | 0.84333372 |
| GTF2B | 37.3806072 | 0.027614324 | 0.25832622 | 0.1068971 | 0.91487061 | 0.98439006 |
| KYAT3 | 38.378239 | -0.529095456 | 0.33030021 | -1.6018623 | 0.10918606 | 0.47648954 |
| GBP3 | 20.1379819 | -0.541919277 | 0.42972122 | -1.261095 | 0.20727462 | 0.6059796 |
| GBP1 | 39.490997 | -0.063582439 | 0.89358261 | -0.0711545 | 0.94327478 | 0.99001017 |
| GBP2 | 85.4073811 | -0.694776149 | 0.43364684 | -1.6021704 | 0.10911793 | 0.47647513 |
| GBP4 | 24.612446 | 0.11964524 | 0.65996046 | 0.18129153 | 0.85613876 | 0.97379567 |
| LRRC8B | 33.9464188 | -0.740103181 | 0.43067257 | -1.7184823 | 0.08570868 | 0.42858628 |
| ZNF326 | 46.7282604 | -0.28246399 | 0.21047187 | -1.342051 | 0.17957945 | 0.57744792 |
| ZNF644 | 180.124466 | -0.039266907 | 0.18280713 | -0.2147996 | 0.82992353 | 0.96580172 |
| CDC7 | 20.8804783 | -1.186731829 | 0.51434812 | -2.3072541 | 0.02104066 | 0.21390029 |
| TGFBR3 | 282.899061 | -0.085305987 | 0.38500283 | -0.2215724 | 0.8246468 | 0.96413949 |
| BTBD8 | 28.6368671 | -0.150617906 | 0.55163615 | -0.2730385 | 0.78482362 | 0.95065355 |
| GLMN | 39.6670401 | 0.048765429 | 0.19622411 | 0.24851904 | 0.80373284 | 0.95836189 |
| RPAP2 | 87.6755431 | -0.184120054 | 0.21927691 | -0.8396691 | 0.40109392 | 0.76363711 |
| EVI5 | 216.928586 | 0.158975629 | 0.21771295 | 0.73020749 | 0.46526336 | 0.80342007 |
| RPL5 | 98.4696983 | -0.504701461 | 0.28436406 | -1.7748427 | 0.07592387 | 0.40904738 |
| FAM69A | 77.4843334 | 0.154961798 | 0.39566557 | 0.39164843 | 0.695318 | 0.92604059 |
| MTF2 | 60.9129375 | -0.087870119 | 0.19195022 | -0.4577756 | 0.6471137 | 0.90243971 |
| TMED5 | 51.3433054 | -0.444169151 | 0.26463306 | -1.6784341 | 0.09326239 | 0.44361507 |
| CCDC18 | 21.3603672 | -0.390538685 | 0.32908785 | -1.1867308 | 0.23533383 | 0.63119281 |
| DR1 | 27.1780104 | 0.307096004 | 0.26900267 | 1.14160949 | 0.25361638 | 0.64816854 |
| FNBP1L | 104.36012 | -0.021397613 | 0.34560867 | -0.0619128 | 0.95063224 | 0.99060401 |
| BCAR3 | 83.5681373 | -0.326282285 | 0.42632038 | -0.7653453 | 0.44406597 | 0.79248199 |
| GCLM | 32.4585801 | -0.613812597 | 0.44471485 | -1.3802386 | 0.16751319 | 0.56315392 |
| ARHGAP29 | 176.167666 | 0.050431553 | 0.3986809 | 0.12649604 | 0.89933929 | 0.98058672 |
| ABCD3 | 72.6807753 | 0.141109537 | 0.2220255 | 0.63555553 | 0.52506616 | 0.83726969 |
| F3 | 38.182227 | 0.749940516 | 0.39664315 | 1.89071843 | 0.05866194 | 0.36041506 |
| CNN3 | 254.300775 | -0.404097611 | 0.34850578 | -1.1595148 | 0.24624641 | 0.64218217 |

| | | | | | | |
|---|---|---|---|---|---|---|
| ALG14 | 44.7542986 | -0.103749591 | 0.25599347 | -0.4052822 | 0.6852701 | 0.91977784 |
| PTBP2 | 226.016209 | 0.083046652 | 0.2266132 | 0.36646873 | 0.71401534 | 0.93136293 |
| DPYD | 508.224284 | -0.382479309 | 0.20948586 | -1.8258001 | 0.06788037 | 0.38827623 |
| DPYD-AS1 | 21.3401806 | -0.707303447 | 0.34605727 | -2.0438913 | 0.04096429 | 0.30341448 |
| SNX7 | 24.2791473 | 0.014707205 | 0.44091445 | 0.03335614 | 0.97339058 | 0.99300191 |
| AGL | 29.7648872 | 0.10003778 | 0.29048826 | 0.34437804 | 0.730562 | 0.93455731 |
| TRMT13 | 33.986606 | -0.291125437 | 0.2197297 | -1.3249253 | 0.18519593 | 0.58487533 |
| DBT | 27.9233829 | 0.124750079 | 0.33736362 | 0.36977929 | 0.71154695 | 0.93045525 |
| RTCA | 23.9726779 | -0.081680655 | 0.32928174 | -0.248057 | 0.80409027 | 0.95836189 |
| CDC14A | 47.4445666 | 0.414834431 | 0.25685825 | 1.61503253 | 0.10630365 | 0.46993733 |
| VCAM1 | 53.6529407 | -0.07630667 | 0.79499472 | -0.0959839 | 0.92353338 | 0.98561736 |
| EXTL2 | 23.7054995 | 0.050998515 | 0.47054686 | 0.10838137 | 0.91369318 | 0.98395113 |
| SLC30A7 | 98.2009065 | -0.353076477 | 0.22395662 | -1.5765396 | 0.11490151 | 0.48724723 |
| RNPC3 | 56.2183728 | -0.1394697 | 0.20182069 | -0.6910575 | 0.48952942 | 0.81720854 |
| VAV3 | 37.3980927 | 0.89734956 | 0.2516061 | 3.5664857 | 0.0003618 | 0.01915017 |
| SLC25A24 | 39.7987226 | -0.14994204 | 0.22062323 | -0.6796294 | 0.49673913 | 0.8216376 |
| FAM102B | 73.6326258 | 0.218487654 | 0.27689403 | 0.78906596 | 0.43007345 | 0.78298297 |
| PRPF38B | 55.3120336 | -0.184695115 | 0.28061703 | -0.658175 | 0.51042568 | 0.82751342 |
| STXBP3 | 65.9243377 | 0.091839728 | 0.24368954 | 0.37687185 | 0.70626885 | 0.93006079 |
| SARS | 46.845973 | -0.284532508 | 0.20985366 | -1.3558615 | 0.17514323 | 0.57299474 |
| SORT1 | 104.505962 | -0.235359315 | 0.36501646 | -0.644791 | 0.51906265 | 0.83297983 |
| PSMA5 | 38.6361235 | -0.725982935 | 0.33812823 | -2.1470639 | 0.03178819 | 0.2713257 |
| GNAI3 | 49.1347928 | 0.304835527 | 0.26939954 | 1.13153694 | 0.25782917 | 0.65130862 |
| GSTM3 | 19.0587045 | -0.411096845 | 0.33766892 | -1.2174554 | 0.22343101 | 0.61775564 |
| CSF1 | 45.0400437 | 0.368843555 | 0.41747601 | 0.88350838 | 0.37696165 | 0.74832285 |
| AHCYL1 | 100.546744 | -0.104658853 | 0.23365543 | -0.4479196 | 0.65421121 | 0.90646849 |
| STRIP1 | 21.5995997 | 0.522786228 | 0.29823759 | 1.75291866 | 0.07961597 | 0.41779962 |
| RBM15 | 18.5029464 | -0.205998229 | 0.33921513 | -0.607279 | 0.54366577 | 0.84730644 |
| SLC16A4 | 53.5397636 | -0.328765837 | 0.33102484 | -0.9931757 | 0.32062432 | 0.70762743 |
| LAMTOR5 | 48.340709 | -0.632894463 | 0.33810164 | -1.8719059 | 0.06121962 | 0.36816937 |
| DRAM2 | 27.5864785 | -0.522681594 | 0.2746244 | -1.9032598 | 0.05700665 | 0.35717443 |
| CEPT1 | 37.1685841 | -0.351131403 | 0.22414546 | -1.5665337 | 0.11722373 | 0.49031145 |
| ATP5PB | 40.4605903 | -0.199024909 | 0.24777686 | -0.8032425 | 0.42183458 | 0.77771095 |
| RAP1A | 216.310934 | -0.185738274 | 0.24009908 | -0.7735901 | 0.43917322 | 0.78887428 |
| KCND3 | 28.3048329 | -0.859200688 | 0.54166268 | -1.5862283 | 0.11268753 | 0.48288083 |
| AL445426.1 | 57.889849 | 0.11062364 | 0.52759007 | 0.20967726 | 0.83391957 | 0.9666365 |
| CTTNBP2NL | 41.7648779 | 0.250044627 | 0.21819321 | 1.14597804 | 0.25180424 | 0.64619053 |
| ST7L | 72.3751489 | 0.049327335 | 0.20149553 | 0.2448061 | 0.80660657 | 0.95838958 |
| CAPZA1 | 98.8782789 | -0.300132265 | 0.17926553 | -1.674233 | 0.09408482 | 0.44579582 |
| MOV10 | 24.6014718 | 0.033598674 | 0.43220216 | 0.07773833 | 0.9380362 | 0.98827804 |
| RHOC | 50.4685958 | -0.490360267 | 0.38961287 | -1.2585833 | 0.20818087 | 0.6059796 |
| SLC16A1-AS1 | 90.3950583 | -0.424332887 | 0.31026665 | -1.3676394 | 0.17142499 | 0.56870244 |
| LRIG2 | 41.9602394 | -0.117898436 | 0.26475183 | -0.4453168 | 0.65609083 | 0.90751204 |
| MAGI3 | 133.274924 | 0.096455581 | 0.23620539 | 0.4083547 | 0.68301329 | 0.91933701 |
| PHTF1 | 27.1006739 | 0.019780026 | 0.23019318 | 0.08592794 | 0.9315237 | 0.98681039 |
| RSBN1 | 49.9805277 | -0.281143713 | 0.21697305 | -1.2957541 | 0.19506023 | 0.59308963 |
| HIPK1 | 53.5220756 | 0.076074988 | 0.1965707 | 0.38701082 | 0.6987482 | 0.92769314 |
| TRIM33 | 133.561527 | -0.041960943 | 0.15441965 | -0.2717332 | 0.78582718 | 0.95065355 |
| BCAS2 | 22.7014968 | 0.73909496 | 0.35948245 | 2.05599735 | 0.03978277 | 0.30124947 |
| CSDE1 | 132.581906 | 0.101965868 | 0.17283472 | 0.58996172 | 0.55521631 | 0.85207344 |
| NGF | 125.193383 | -0.976785287 | 0.4520713 | -2.1606886 | 0.03071941 | 0.26662968 |
| SLC22A15 | 22.8714071 | -0.540403366 | 0.37577855 | -1.43809 | 0.15040853 | 0.53880252 |
| ATP1A1 | 163.441945 | -0.255484832 | 0.23904749 | -1.0687618 | 0.28517701 | 0.6783522 |
| CD58 | 21.6064515 | 0.185696325 | 0.32132303 | 0.57791166 | 0.56332376 | 0.85605176 |
| MAN1A2 | 126.666355 | -0.014614466 | 0.19491111 | -0.0749802 | 0.9402305 | 0.9889521 |
| GDAP2 | 41.2654388 | 0.082628051 | 0.24726898 | 0.33416262 | 0.73825685 | 0.93594619 |
| WARS2 | 43.3863117 | 0.04157726 | 0.24501553 | 0.16969234 | 0.8652521 | 0.9762143 |
| AL359915.2 | 87.1027323 | 0.307936966 | 0.32252269 | 0.95477612 | 0.33969093 | 0.71925477 |
| ZNF697 | 20.118781 | -0.294509148 | 0.36662 | -0.803309 | 0.42179616 | 0.77771095 |
| NOTCH2 | 148.876803 | 0.289774986 | 0.2814407 | 1.02961296 | 0.30319173 | 0.69521249 |
| SEC22B | 79.6979368 | -0.402492984 | 0.29990376 | -1.3420738 | 0.17957206 | 0.57744792 |
| NBPF26 | 24.3457872 | 0.051684992 | 0.44386536 | 0.11644295 | 0.9073015 | 0.98306694 |
| LINC00623 | 63.0376444 | 0.101784064 | 0.28691321 | 0.35475558 | 0.72277271 | 0.93455731 |
| SRGAP2C | 34.6979088 | 0.54892314 | 0.26563229 | 2.06647745 | 0.03878342 | 0.29782586 |
| NBPF15 | 24.8093407 | 0.235964804 | 0.27829712 | 0.84788806 | 0.39650031 | 0.76020893 |
| FP700111.1 | 24.9660625 | 0.617093021 | 0.36352978 | 1.69750333 | 0.08960154 | 0.43694872 |
| SRGAP2B | 38.9631284 | 0.182562217 | 0.23389484 | 0.78053119 | 0.43507827 | 0.78656586 |
| RNF115 | 139.558104 | 0.252386895 | 0.1545466 | 1.63307962 | 0.10245225 | 0.4627468 |
| RBM8A | 24.5058091 | 0.074181354 | 0.32808681 | 0.22610282 | 0.82112145 | 0.96294563 |
| LIX1L | 33.8790141 | 0.164868045 | 0.24518005 | 0.67243663 | 0.50130577 | 0.82406312 |
| TXNIP | 354.753646 | 0.155220442 | 0.41710645 | 0.37213628 | 0.70979139 | 0.93009618 |
| NBPF12 | 23.9420246 | 0.134023152 | 0.27739649 | 0.48314652 | 0.6289917 | 0.89288391 |
| PRKAB2 | 26.4320573 | -0.355262976 | 0.35866191 | -0.9905233 | 0.32191841 | 0.70810487 |
| AC242426.2 | 27.5682247 | 0.28576521 | 0.31417352 | 0.90957765 | 0.36304529 | 0.73604674 |
| CHD1L | 57.238305 | -0.692439416 | 0.2510162 | -2.7585447 | 0.00580594 | 0.10256688 |
| BCL9 | 23.1716123 | 0.260089996 | 0.37462888 | 0.6942604 | 0.48751892 | 0.81595238 |
| GJA5 | 41.1170111 | 1.711797736 | 0.55558674 | 3.08106297 | 0.00206263 | 0.05618053 |
| GPR89B | 18.9090478 | -0.00994881 | 0.28328609 | -0.0351193 | 0.97198461 | 0.99300191 |
| NBPF11 | 22.1278325 | 0.258513467 | 0.34688415 | 0.74524438 | 0.456124 | 0.79659108 |
| NBPF14 | 28.2552577 | 0.318304847 | 0.24154147 | 1.31780617 | 0.18756854 | 0.58607194 |
| PDE4DIP | 126.602725 | 0.082996277 | 0.25686028 | 0.32311838 | 0.74660559 | 0.93900586 |
| NBPF9 | 20.6301187 | -0.287011563 | 0.2725588 | -1.0530262 | 0.29232898 | 0.6840363 |
| AC245297.3 | 160.605767 | -0.225773425 | 0.24254645 | -0.9308461 | 0.35193317 | 0.73019673 |
| NBPF19 | 42.2204391 | 0.063647349 | 0.60519467 | 0.10516839 | 0.91624219 | 0.98477389 |
| OTUD7B | 81.7033961 | -0.175995661 | 0.25342108 | -0.6944792 | 0.48738176 | 0.81595238 |
| VPS45 | 108.870309 | 0.388388774 | 0.18761 | 2.07019231 | 0.03843434 | 0.29669534 |
| PLEKHO1 | 34.8871389 | -0.277010541 | 0.27550904 | -1.0054499 | 0.31468026 | 0.70282967 |
| ANP32E | 24.0247273 | -0.504605214 | 0.26798097 | -1.882989 | 0.05970186 | 0.36410704 |
| C1orf54 | 21.4462199 | -0.160588096 | 0.29937945 | -0.5364032 | 0.59167992 | 0.87229579 |
| MRPS21 | 38.6730047 | -0.274925866 | 0.21539637 | -1.2763718 | 0.20182411 | 0.60121822 |
| PRPF3 | 49.9332683 | 0.061852097 | 0.22702233 | 0.2724494 | 0.78527649 | 0.95065355 |
| RPRD2 | 143.545197 | 0.360272159 | 0.13214796 | 2.72627868 | 0.00640529 | 0.10918078 |
| MCL1 | 75.9922865 | -0.225672303 | 0.3344676 | -0.674721 | 0.49985306 | 0.82271448 |
| ENSA | 89.1663307 | 0.343943212 | 0.22704049 | 1.51489812 | 0.12979821 | 0.51204187 |
| GOLPH3L | 27.2616467 | 0.137561039 | 0.26816395 | 0.51297364 | 0.60796976 | 0.88042838 |
| ARNT | 109.757441 | -0.05319877 | 0.1981484 | -0.2684794 | 0.7883303 | 0.95129454 |
| SETDB1 | 37.3317207 | 0.213320161 | 0.23918649 | 0.89185707 | 0.37246955 | 0.74463512 |
| CERS2 | 27.2957275 | -0.252764367 | 0.29518744 | -0.8562843 | 0.39184054 | 0.75901355 |
| CDC42SE1 | 43.2624096 | -0.199660277 | 0.30071119 | -0.6639602 | 0.50671576 | 0.82481485 |
| GABPB2 | 67.6834807 | 0.63221693 | 0.29070964 | 2.1747367 | 0.02964985 | 0.26108772 |
| PIP5K1A | 149.821808 | -0.135699378 | 0.23983968 | -0.565792 | 0.57153516 | 0.86228945 |
| PSMD4 | 36.4377443 | -0.2499184 | 0.2455681 | -1.0177152 | 0.3088133 | 0.69885506 |
| PI4KB | 46.1010669 | -0.016063145 | 0.20600516 | -0.0779745 | 0.93784835 | 0.98827804 |
| PSMB4 | 26.509431 | -0.500699058 | 0.36097019 | -1.3870926 | 0.16541354 | 0.56122997 |
| POGZ | 195.677364 | 0.481291266 | 0.246097 | 1.95569742 | 0.05050081 | 0.34017456 |
| SNX27 | 62.4112716 | -0.026071722 | 0.26177319 | -0.0995966 | 0.92066459 | 0.98528826 |
| THEM4 | 24.3103924 | 0.048950317 | 0.38795775 | 0.12617435 | 0.89959392 | 0.98058672 |
| S100A10 | 85.9386723 | 0.066695721 | 0.44572307 | 0.14963488 | 0.88105268 | 0.97782528 |
| S100A11 | 164.759033 | -0.996602088 | 0.30706463 | -3.2455776 | 0.00117213 | 0.04285549 |
| FLG-AS1 | 44.9050369 | 0.908093568 | 0.35193127 | 2.58031512 | 0.00987102 | 0.14050816 |
| S100A6 | 272.807837 | -1.115440249 | 0.49564947 | -2.2504619 | 0.02441964 | 0.23156121 |
| S100A4 | 63.0682373 | 0.608344874 | 0.79993171 | 0.76049601 | 0.44695815 | 0.79296348 |
| S100A16 | 21.8742518 | -0.296388323 | 0.41063283 | -0.7217843 | 0.47042711 | 0.80457054 |

| | | | | | | |
|---|---|---|---|---|---|---|
| S100A13 | 39.984713 | -0.104133848 | 0.25451754 | -0.4091421 | 0.68243536 | 0.91933701 |
| CHTOP | 29.218578 | 0.207029778 | 0.27707995 | 0.74718426 | 0.45495234 | 0.79609632 |
| ILF2 | 25.3256024 | -0.046625226 | 0.32730562 | -0.1424517 | 0.88672327 | 0.97919298 |
| NPR1 | 31.4296147 | 1.004376469 | 0.3240736 | 3.09922336 | 0.00194029 | 0.05557623 |
| INTS3 | 44.8347924 | 0.342155066 | 0.20466294 | 1.67179785 | 0.09456419 | 0.44706027 |
| GATAD2B | 168.710392 | -0.053252213 | 0.1683351 | -0.3163465 | 0.75173956 | 0.94055881 |
| DENND4B | 22.386551 | 0.00280658 | 0.29677793 | 0.00945683 | 0.99245465 | 0.99759781 |
| SLC39A1 | 24.6314905 | -0.591130167 | 0.3875537 | -1.5252858 | 0.12718782 | 0.50683947 |
| JTB | 28.972905 | -0.427286462 | 0.4641413 | -0.9205956 | 0.35726158 | 0.73129235 |
| RAB13 | 33.1584102 | -0.594953379 | 0.30648171 | -1.9412362 | 0.05222964 | 0.34579783 |
| RPS27 | 41.323994 | -0.693479676 | 0.34327488 | -2.0201876 | 0.04336393 | 0.31335448 |
| TPM3 | 49.0970788 | -0.396041192 | 0.30642332 | -1.2924643 | 0.19619641 | 0.59411229 |
| C1orf43 | 44.479968 | -0.324738307 | 0.2030027 | -1.5996748 | 0.10967075 | 0.47680173 |
| UBAP2L | 67.7794572 | -0.242559322 | 0.24460992 | -0.9916169 | 0.32138445 | 0.70780851 |
| ATP8B2 | 43.81097 | 0.124918276 | 0.33986935 | 0.36754793 | 0.71321034 | 0.93115158 |
| IL6R | 38.0427331 | -0.467992569 | 0.47354091 | -0.9882833 | 0.32301392 | 0.70962823 |
| ADAR | 91.1707383 | -0.121230379 | 0.25649126 | -0.4726492 | 0.6364635 | 0.89741446 |
| PBXIP1 | 18.6054984 | 0.502026556 | 0.39042672 | 1.28584066 | 0.19849868 | 0.59590146 |
| SCAMP3 | 17.6257289 | 0.187980321 | 0.32711301 | 0.57466476 | 0.56551804 | 0.85765509 |
| CLK2 | 19.7085045 | -0.359481402 | 0.26816843 | -1.3405061 | 0.18008087 | 0.57783734 |
| FDPS | 43.9566295 | -0.14792696 | 0.42098932 | -0.3513794 | 0.72530376 | 0.93455731 |
| ASH1L | 415.028322 | -0.093048437 | 0.12046901 | -0.7723848 | 0.43988654 | 0.78933067 |
| YY1AP1 | 76.3933211 | 0.031842023 | 0.19916821 | 0.15987503 | 0.87297952 | 0.9768643 |
| DAP3 | 64.4911743 | -0.279703867 | 0.28061165 | -0.996765 | 0.3188786 | 0.70683013 |
| GON4L | 93.1605053 | -0.244082543 | 0.15136654 | -1.6125264 | 0.10684743 | 0.47094784 |
| SYT11 | 19.6605404 | 0.252730103 | 0.28807509 | 0.87730634 | 0.38032027 | 0.74991898 |
| KHDC4 | 58.0613308 | -0.146854119 | 0.23527772 | -0.6241735 | 0.53251365 | 0.84290291 |
| ARHGEF2 | 56.291566 | 0.104175124 | 0.31068851 | 0.33530408 | 0.73739572 | 0.93594619 |
| SSR2 | 39.5353276 | -0.548082962 | 0.32610011 | -1.6807199 | 0.09281733 | 0.44320781 |
| LMNA | 145.612504 | 1.317075243 | 0.39345159 | 3.34748991 | 0.00081547 | 0.03346039 |
| SMG5 | 29.9751158 | -0.057726928 | 0.29502203 | -0.1956699 | 0.84486855 | 0.96938938 |
| CCT3 | 45.8043901 | -0.310778995 | 0.33699988 | -0.9221932 | 0.35642782 | 0.73129235 |
| MEF2D | 76.94973 | 0.376590714 | 0.19758954 | 1.90592435 | 0.05666002 | 0.35680762 |
| NES | 33.806822 | 1.268585092 | 0.41919809 | 3.02621871 | 0.00247633 | 0.062538 |
| HDGF | 40.0664499 | -0.474281912 | 0.36312432 | -1.3061144 | 0.19151365 | 0.59004802 |
| PRCC | 28.4823107 | 0.181302128 | 0.23366671 | 0.77590056 | 0.4378077 | 0.78836643 |
| PEAR1 | 36.7442826 | 0.755711398 | 0.36795063 | 2.05383911 | 0.03999127 | 0.30130305 |
| ARHGEF11 | 75.35921 | 0.111117304 | 0.22758423 | 0.48824693 | 0.62537494 | 0.89069797 |
| KIRREL1 | 169.126335 | 0.354102877 | 0.2251581 | 1.57268549 | 0.11579166 | 0.48843995 |
| IFI16 | 161.797229 | -1.042042993 | 0.38807541 | -2.685156 | 0.0072496 | 0.11709294 |
| DUSP23 | 29.4886113 | -1.47033319 | 0.4146385 | -3.5460605 | 0.00039104 | 0.02041031 |
| TAGLN2 | 71.584298 | -0.015677714 | 0.39470848 | -0.0397197 | 0.96831657 | 0.99300191 |
| PEA15 | 25.4970137 | 0.099957785 | 0.32778585 | 0.30494845 | 0.76040542 | 0.94425408 |
| DCAF8 | 19.668492 | 0.322202922 | 0.32982578 | 0.97688822 | 0.3286245 | 0.71294249 |
| COPA | 157.036304 | -0.357953197 | 0.19015088 | -1.8824693 | 0.05977232 | 0.3642345 |
| NCSTN | 18.9636176 | 0.167911301 | 0.28822317 | 0.58257392 | 0.56018016 | 0.85423726 |
| PFDN2 | 32.6915578 | -0.701297794 | 0.32438281 | -2.161945 | 0.03062241 | 0.26662968 |
| NIT1 | 25.4107044 | -0.300072073 | 0.33731989 | -0.8895772 | 0.37369296 | 0.74545863 |
| UFC1 | 21.8335823 | -0.417929885 | 0.35390392 | -1.1809134 | 0.23763712 | 0.63302595 |
| ADAMTS4 | 139.298902 | -3.717788199 | 0.86622479 | -4.2919439 | 1.77E-05 | 0.00271172 |
| NDUFS2 | 54.882127 | -1.002484316 | 0.3624476 | -2.7658738 | 0.00567705 | 0.10101851 |
| SDHC | 44.0910014 | 0.149665544 | 0.20247999 | 0.73916214 | 0.45980855 | 0.80029179 |
| ATF6 | 148.1238 | -0.733514092 | 0.26541521 | -2.7636475 | 0.00571593 | 0.10146457 |
| NOS1AP | 35.9370446 | -0.171374312 | 0.41115803 | -0.4168089 | 0.67681822 | 0.91753129 |
| UHMK1 | 47.6286852 | -0.1155301 | 0.21300031 | -0.542394 | 0.58754708 | 0.870249 |
| UAP1 | 72.4419782 | 1.774718592 | 0.49752412 | 3.56710063 | 0.00036095 | 0.01915017 |
| DDR2 | 205.222271 | -0.074404243 | 0.36979724 | -0.2012028 | 0.84054 | 0.96803523 |
| RGS5 | 278.98234 | 0.609146522 | 0.38064544 | 1.60029903 | 0.10953226 | 0.47680173 |
| PBX1 | 946.409462 | 0.068679971 | 0.32724704 | 0.20987194 | 0.83376763 | 0.9666365 |
| MGST3 | 106.224318 | -0.48255252 | 0.25409025 | -1.8991383 | 0.05754629 | 0.35839637 |
| ALDH9A1 | 24.4709884 | -0.052879944 | 0.38422274 | -0.1376284 | 0.89053415 | 0.98023547 |
| TMCO1 | 40.6169738 | -0.115824733 | 0.20622983 | -0.5616294 | 0.57436855 | 0.86294056 |
| UCK2 | 25.029153 | -1.459797383 | 0.45165481 | -3.2321086 | 0.0012288 | 0.0436255 |
| POGK | 18.4211831 | 0.053806712 | 0.31818014 | 0.1691077 | 0.86571193 | 0.9762143 |
| POU2F1 | 211.808495 | -0.239160051 | 0.16074542 | -1.4878188 | 0.1367987 | 0.52089048 |
| CREG1 | 19.5928679 | 0.035433339 | 0.29922887 | 0.11841551 | 0.90573844 | 0.98303609 |
| RCSD1 | 52.5635281 | 1.143969918 | 0.36075069 | 3.17108169 | 0.00151872 | 0.04900622 |
| MPZL1 | 137.979508 | -0.362952974 | 0.29480265 | -1.2311727 | 0.21825828 | 0.61478732 |
| MPC2 | 25.7684142 | -0.490017914 | 0.37895498 | -1.2930768 | 0.19598448 | 0.59411229 |
| DCAF6 | 196.815142 | -0.343640582 | 0.18455854 | -1.8619598 | 0.06260876 | 0.37164525 |
| TIPRL | 34.6602517 | 0.231522879 | 0.2198037 | 1.05331657 | 0.29219593 | 0.6840363 |
| SFT2D2 | 57.2619051 | -0.267458724 | 0.24247612 | -1.1030312 | 0.27001362 | 0.6651094 |
| TBX19 | 18.0935064 | -0.086282941 | 0.38079844 | -0.2265843 | 0.82074702 | 0.96291025 |
| ATP1B1 | 136.198689 | 1.149341394 | 0.32632745 | 3.52204936 | 0.00042822 | 0.02141653 |
| NME7 | 81.6329533 | -0.114522994 | 0.20425545 | -0.5606851 | 0.5750122 | 0.86294056 |
| BLZF1 | 37.4288481 | 0.077380941 | 0.26495655 | 0.29205144 | 0.7702473 | 0.94593455 |
| SLC19A2 | 42.9883919 | -1.23961684 | 0.41897291 | -2.958704 | 0.00308936 | 0.07118649 |
| SCYL3 | 25.3017065 | 0.166119861 | 0.28360634 | 0.58574099 | 0.55804958 | 0.85279817 |
| KIFAP3 | 92.6277969 | 0.113467178 | 0.23647531 | 0.47982674 | 0.6313506 | 0.89436221 |
| PRRX1 | 124.930532 | -0.605199231 | 0.54860797 | -1.1031543 | 0.26996017 | 0.6651094 |
| FMO3 | 33.6119326 | -0.519952591 | 0.97264976 | -0.5345733 | 0.59294495 | 0.87313824 |
| FMO2 | 55.9903287 | -1.566558743 | 0.92031794 | -1.702193 | 0.0887192 | 0.43544541 |
| PRRC2C | 259.377601 | -0.349482898 | 0.20057537 | -1.7424019 | 0.08143815 | 0.42087829 |
| VAMP4 | 28.9840696 | 0.262719916 | 0.32521473 | 0.80783523 | 0.41918544 | 0.77771095 |
| DNM3 | 66.1792902 | 0.10022312 | 0.30952367 | 0.32379791 | 0.74609104 | 0.93894983 |
| DNM3OS | 42.4604492 | 0.657165101 | 0.31033526 | 2.11759725 | 0.03420919 | 0.2833153 |
| SUCO | 65.2294966 | 0.158203045 | 0.22568951 | 0.70097652 | 0.48331767 | 0.81409616 |
| AL645568.1 | 34.2733386 | -0.031442887 | 0.27135378 | -0.1158741 | 0.90775229 | 0.98306694 |
| PRDX6 | 82.861276 | -0.641184515 | 0.31840922 | -2.0137122 | 0.04403976 | 0.3152359 |
| KLHL20 | 49.1004578 | -0.05900375 | 0.26723702 | -0.2207918 | 0.82525453 | 0.96450311 |
| ZBTB37 | 29.9393701 | 0.333207609 | 0.24089015 | 1.3832347 | 0.16659291 | 0.56240432 |
| RC3H1 | 120.127329 | -0.012717162 | 0.14335522 | -0.0887108 | 0.92931172 | 0.98681039 |
| RABGAP1L | 355.836108 | 0.147426595 | 0.16177036 | 0.91133254 | 0.36212019 | 0.73473806 |
| KIAA0040 | 27.1023514 | -0.293175872 | 0.29220556 | -1.0033207 | 0.31570618 | 0.70306809 |
| COP1 | 231.908837 | -0.021102103 | 0.15863609 | -0.1330221 | 0.89417591 | 0.98023547 |
| RASAL2 | 436.053704 | 0.869139971 | 0.33309024 | 2.60932285 | 0.00907216 | 0.13367648 |
| RALGPS2 | 47.0800335 | 0.520832464 | 0.39013923 | 1.33499127 | 0.1818792 | 0.58089103 |
| FAM20B | 35.6906293 | 0.264619543 | 0.247778 | 1.06797029 | 0.28553391 | 0.6783522 |
| ABL2 | 123.351992 | -0.16615686 | 0.22054782 | -0.7533825 | 0.45122012 | 0.79401729 |
| SOAT1 | 50.9126015 | -0.452274207 | 0.20981203 | -2.1556162 | 0.03111365 | 0.26774498 |
| TOR1AIP2 | 52.2173938 | -0.363462614 | 0.18054191 | -2.0131758 | 0.04409614 | 0.3152359 |
| TOR1AIP1 | 51.0988791 | 0.157324251 | 0.25113582 | 0.62645086 | 0.53101926 | 0.84177319 |
| CEP350 | 230.347699 | -0.180991621 | 0.19681339 | -0.9196103 | 0.35777645 | 0.73137668 |
| QSOX1 | 38.4448128 | -0.036372213 | 0.23439209 | -0.1551768 | 0.87668194 | 0.9768643 |
| ACBD6 | 128.650908 | -0.057731079 | 0.16762301 | -0.3444102 | 0.73053779 | 0.93455731 |
| XPR1 | 111.780144 | -0.249362609 | 0.29136965 | -0.855829 | 0.39209235 | 0.75901355 |
| STX6 | 28.8808594 | -0.183724359 | 0.2929527 | -0.6271468 | 0.53056298 | 0.84152078 |
| MR1 | 20.5487137 | -0.267529156 | 0.27261401 | -0.9813478 | 0.32642126 | 0.71149668 |
| GLUL | 103.484097 | -0.780442214 | 0.4246595 | -1.837807 | 0.06609084 | 0.3842576 |
| RGS16 | 62.6315709 | 0.143378824 | 0.81097831 | 0.17679736 | 0.85966756 | 0.97397549 |
| DHX9 | 90.7659946 | -0.123212244 | 0.21152661 | -0.5824905 | 0.56023633 | 0.85423726 |
| LAMC1 | 259.421115 | 0.082525326 | 0.23804714 | 0.3466764 | 0.72883444 | 0.93455731 |
| SMG7 | 96.8673237 | 0.160875909 | 0.21191438 | 0.75915522 | 0.44775971 | 0.79296348 |

| | | | | | | |
|---|---|---|---|---|---|---|
| ARPC5 | 68.3385505 | 0.301677666 | 0.20672657 | 1.45930767 | 0.14448045 | 0.53398616 |
| RGL1 | 115.541762 | 0.214618251 | 0.2517848 | 0.85238763 | 0.39399898 | 0.75988253 |
| TSEN15 | 30.6520105 | -0.070314301 | 0.31091819 | -0.2261505 | 0.82108438 | 0.96294563 |
| C1orf21 | 288.780821 | -0.510251737 | 0.49944946 | -1.0216284 | 0.30695682 | 0.69801431 |
| EDEM3 | 89.4892009 | -0.38278833 | 0.26478017 | -1.4456835 | 0.14826599 | 0.53680621 |
| FAM129A | 115.787429 | 0.780256897 | 0.33504766 | 2.32879372 | 0.01987 | 0.20828536 |
| TRMT1L | 23.7468918 | 0.09410806 | 0.32453996 | 0.28997372 | 0.77183634 | 0.94671883 |
| SWT1 | 36.7943448 | 0.249804084 | 0.32073627 | 0.77884576 | 0.43607058 | 0.78719791 |
| IVNS1ABP | 41.0708732 | -0.258778638 | 0.34671326 | -0.7463765 | 0.45543999 | 0.79609632 |
| HMCN1 | 78.9049077 | 0.490254711 | 0.48620447 | 1.00833032 | 0.31329591 | 0.70282967 |
| TPR | 101.054122 | -0.073502776 | 0.21746913 | -0.3379918 | 0.73536939 | 0.93594619 |
| ODR4 | 56.5575218 | -0.155334812 | 0.19249985 | -0.8069347 | 0.4197041 | 0.77771095 |
| PTGS2 | 72.5119386 | -0.380204004 | 0.82081506 | -0.463203 | 0.64321886 | 0.90063554 |
| PLA2G4A | 225.288005 | -0.140206705 | 0.40706103 | -0.3444366 | 0.73051799 | 0.93455731 |
| AL390957.1 | 59.8898126 | 0.765971986 | 0.47218496 | 1.62218632 | 0.10476346 | 0.46548166 |
| AL136454.1 | 35.0345406 | -0.181302261 | 0.33583609 | -0.5398534 | 0.58929813 | 0.8702694 |
| RGS2 | 65.6466538 | 1.174499565 | 0.4365385 | 2.69048332 | 0.00713486 | 0.11602488 |
| UCHL5 | 35.1214567 | -0.138199862 | 0.26729601 | -0.5170293 | 0.60513573 | 0.87903643 |
| TROVE2 | 70.9871919 | 0.083614738 | 0.22888684 | 0.36531037 | 0.71487974 | 0.93182887 |
| CDC73 | 162.103081 | -0.171894291 | 0.26022232 | -0.660567 | 0.50889001 | 0.82663984 |
| KCNT2 | 243.059656 | -0.279100665 | 0.33023434 | -0.8451594 | 0.39802184 | 0.7611823 |
| CFH | 209.938145 | -0.864524354 | 0.4582644 | -1.8865187 | 0.0592251 | 0.36270436 |
| ZBTB41 | 30.4500179 | 0.133157678 | 0.30481526 | 0.43684715 | 0.6622222 | 0.91084989 |
| DENND1B | 63.4896326 | 0.712357537 | 0.30875412 | 2.30720011 | 0.02104367 | 0.21390029 |
| NEK7 | 235.929256 | 0.227694269 | 0.18326405 | 1.24243827 | 0.2140749 | 0.61001472 |
| MIR181A1HC | 35.7295493 | 0.423581682 | 0.38191527 | 1.10909858 | 0.26738766 | 0.66207274 |
| DDX59 | 18.0007021 | 0.395091234 | 0.37534084 | 1.0526199 | 0.29251523 | 0.6840363 |
| CAMSAP2 | 187.672103 | -0.096586973 | 0.15321767 | -0.6303906 | 0.52843908 | 0.83995114 |
| PHLDA3 | 21.8597629 | 0.17284222 | 0.40909289 | 0.42250115 | 0.67265926 | 0.91482259 |
| CSRP1 | 207.15568 | 1.361627164 | 0.26352353 | 5.16700418 | 2.38E-07 | 0.00010283 |
| NAV1 | 155.469138 | -0.284036144 | 0.39911115 | -0.7116718 | 0.47666805 | 0.81064148 |
| IPO9-AS1 | 42.3942778 | -0.587857979 | 0.4056278 | -1.4492546 | 0.14726648 | 0.53679369 |
| IPO9 | 75.1693086 | -0.464929292 | 0.21842546 | -2.1285489 | 0.0332916 | 0.27874784 |
| LMOD1 | 172.40129 | 0.924478958 | 0.45130624 | 2.04845151 | 0.04051578 | 0.30197815 |
| TIMM17A | 19.2925427 | -0.356610679 | 0.35467994 | -1.0054436 | 0.31468329 | 0.70282967 |
| RNPEP | 28.7130618 | 0.17700438 | 0.29395535 | 0.60214716 | 0.54707618 | 0.84766242 |
| ARL8A | 24.438711 | 0.017200928 | 0.26996453 | 0.06371551 | 0.94919675 | 0.99060401 |
| PPP1R12B | 444.124969 | 1.148795716 | 0.33339183 | 3.44578248 | 0.00056941 | 0.02567229 |
| KDM5B | 115.869913 | -0.542095137 | 0.26726327 | -2.0283189 | 0.04252771 | 0.31098123 |
| BTG2 | 39.4347518 | 0.39482876 | 0.41405025 | 0.95357692 | 0.34029784 | 0.71925477 |
| FMOD | 29.7161115 | -0.522540125 | 0.43282224 | -1.2072858 | 0.22732211 | 0.62173063 |
| PRELP | 67.7236297 | 0.626096313 | 0.30190556 | 2.07381513 | 0.03809648 | 0.29501902 |
| ATP2B4 | 143.155824 | 0.661133077 | 0.30649708 | 2.15706161 | 0.03100086 | 0.26740065 |
| SOX13 | 85.632665 | 0.749135789 | 0.23263786 | 3.22018008 | 0.0012811 | 0.04375316 |
| REN | 643.974268 | -1.832008015 | 1.44111052 | -1.2712474 | 0.20364064 | 0.60367992 |
| PLEKHA6 | 56.9249995 | 0.737340734 | 0.23356994 | 3.15683058 | 0.00159494 | 0.05045611 |
| PPP1R15B | 30.9480427 | -0.182057412 | 0.38224665 | -0.4762826 | 0.63387309 | 0.89608287 |
| PIK3C2B | 48.5619853 | 0.429275491 | 0.42676745 | 1.00587685 | 0.31447482 | 0.70282967 |
| MDM4 | 111.729673 | 0.160326436 | 0.15122721 | 1.06016927 | 0.2890676 | 0.67992862 |
| NFASC | 414.323804 | 0.010814509 | 0.50776258 | 0.02129836 | 0.98300765 | 0.99601865 |
| RBBP5 | 24.8926256 | -0.06001679 | 0.24820042 | -0.2418078 | 0.80892912 | 0.95939969 |
| DSTYK | 60.4537912 | 0.615355426 | 0.21592109 | 2.84990888 | 0.00437318 | 0.08733279 |
| CDK18 | 44.6688315 | 0.150300383 | 0.33903691 | 0.4433157 | 0.6575374 | 0.90772143 |
| ELK4 | 40.3945459 | -0.118980397 | 0.25539032 | -0.4658767 | 0.64130377 | 0.89941629 |
| NUCKS1 | 174.532959 | 0.361756616 | 0.20481958 | 1.76622086 | 0.0773588 | 0.4114983 |
| SRGAP2 | 69.1167494 | 0.006383892 | 0.2067692 | 0.03087448 | 0.97536964 | 0.9933005 |
| RASSF5 | 37.2583792 | 0.155563254 | 0.46879083 | 0.33183937 | 0.74001055 | 0.93603056 |
| MAPKAPK2 | 122.987886 | -0.900938399 | 0.30377824 | -2.9657766 | 0.0030192 | 0.07066267 |
| PIGR | 25.8589822 | -3.369192132 | 0.69173672 | -4.8706279 | 1.11E-06 | 0.00028191 |
| CD55 | 54.1098278 | 0.4242672 | 0.31053511 | 1.36624552 | 0.17186191 | 0.56870244 |
| CD46 | 238.474385 | -0.154301073 | 0.22175856 | -0.6958066 | 0.48654995 | 0.81595238 |
| MIR29B2CHC | 95.3839272 | -0.139820026 | 0.27915448 | -0.5008697 | 0.61646282 | 0.88466808 |
| CD34 | 27.7844397 | 1.557634298 | 0.46914377 | 3.32016408 | 0.00089965 | 0.03554567 |
| PLXNA2 | 49.9618306 | 0.532722776 | 0.31153079 | 1.71001646 | 0.08726283 | 0.43301454 |
| G0S2 | 148.17116 | -2.249771936 | 0.95272201 | -2.3614149 | 0.01820535 | 0.19733204 |
| HHAT | 31.8364735 | 0.247528087 | 0.51668739 | 0.47906741 | 0.63189067 | 0.89436221 |
| RCOR3 | 105.341706 | 0.103845931 | 0.24332554 | 0.42677777 | 0.6695412 | 0.91445841 |
| TRAF5 | 45.8890009 | 0.101796889 | 0.32814691 | 0.31021742 | 0.75639562 | 0.94216126 |
| LPGAT1 | 104.917048 | -0.152699465 | 0.16805574 | -0.9086239 | 0.36354868 | 0.73604674 |
| INTS7 | 29.0195935 | -0.271605025 | 0.35028393 | -0.7753853 | 0.438112 | 0.78836643 |
| PPP2R5A | 57.9831634 | -0.047456448 | 0.23372336 | -0.2030454 | 0.83909956 | 0.96770443 |
| NENF | 29.1065907 | -0.126769131 | 0.24509751 | -0.5172192 | 0.60500317 | 0.87903643 |
| ATF3 | 62.8524529 | 0.931806174 | 0.54468284 | 1.71073165 | 0.08713066 | 0.43290196 |
| NSL1 | 34.5975676 | 0.208229096 | 0.27871522 | 0.74710344 | 0.45500112 | 0.79609632 |
| ANGEL2 | 35.3230077 | -0.049652744 | 0.21185016 | -0.2343767 | 0.81469253 | 0.96040671 |
| RPS6KC1 | 133.3339 | 0.158568828 | 0.24994488 | 0.63441519 | 0.5258099 | 0.83803447 |
| PTPN14 | 145.965807 | -0.043908108 | 0.23931596 | -0.1834734 | 0.8544266 | 0.97379567 |
| KCTD3 | 72.5697388 | -0.01766629 | 0.24703275 | -0.071514 | 0.94298872 | 0.98986203 |
| ESRRG | 75.2202625 | 1.325640328 | 0.38238865 | 3.46673556 | 0.00052682 | 0.0246904 |
| GPATCH2 | 95.2175074 | -0.370644332 | 0.28076347 | -1.3201302 | 0.18679155 | 0.58501647 |
| RRP15 | 23.0511268 | 0.020476009 | 0.29802437 | 0.06870582 | 0.94522378 | 0.99060401 |
| TGFB2 | 29.02693 | 0.848566658 | 0.4908928 | 1.72861907 | 0.0838773 | 0.42540665 |
| LYPLAL1 | 80.1065559 | 0.121881167 | 0.34401411 | 0.35429119 | 0.72312067 | 0.93455731 |
| EPRS | 49.3228349 | -0.39147505 | 0.23452451 | -1.6692287 | 0.09507206 | 0.44764276 |
| IARS2 | 37.5802947 | 0.073495662 | 0.21084437 | 0.34857777 | 0.72740632 | 0.93455731 |
| RAB3GAP2 | 153.722614 | -0.199878175 | 0.22247268 | -0.8984392 | 0.36895145 | 0.74123132 |
| MARK1 | 81.7116897 | 0.126739052 | 0.20832194 | 0.60838072 | 0.542935 | 0.84678041 |
| 2-Mar | 51.5680931 | -0.360291631 | 0.33148949 | -1.086887 | 0.27708673 | 0.67269898 |
| MIA3 | 59.3890253 | -0.279947511 | 0.20646356 | -1.3559173 | 0.17512548 | 0.57299474 |
| AIDA | 23.7505 | -0.099510237 | 0.2799069 | -0.3555119 | 0.72220613 | 0.93444541 |
| BROX | 23.7532586 | -0.282218874 | 0.25705583 | -1.0978894 | 0.27225278 | 0.66723562 |
| AC093152.1 | 22.63705 | 0.64944418 | 0.41846342 | 1.55197359 | 0.12066854 | 0.49791866 |
| CAPN2 | 61.0833534 | 0.457703731 | 0.28184222 | 1.6239715 | 0.10438189 | 0.46422585 |
| TP53BP2 | 42.6118391 | -0.145651898 | 0.29641586 | -0.4913769 | 0.62315992 | 0.88872545 |
| FBXO28 | 68.6827041 | -0.138712101 | 0.29597558 | -0.4686606 | 0.63931223 | 0.89868126 |
| DEGS1 | 34.4531655 | -0.42923854 | 0.29102796 | -1.4749048 | 0.14023814 | 0.52680084 |
| NVL | 48.5116032 | 0.125423269 | 0.22859551 | 0.54866898 | 0.58323264 | 0.86764709 |
| CNIH4 | 27.2510417 | -0.047802108 | 0.27628718 | -0.173016 | 0.86263884 | 0.97561293 |
| WDR26 | 138.421822 | -0.24059279 | 0.19519596 | -1.2325705 | 0.21773603 | 0.61468881 |
| CNIH3 | 42.9266426 | 0.561031002 | 0.37404266 | 1.4999118 | 0.13363725 | 0.51553814 |
| LBR | 40.2358376 | 0.073082966 | 0.40434212 | 0.18074537 | 0.85656744 | 0.97381501 |
| ENAH | 234.660698 | 0.988337834 | 0.38324857 | 2.57884287 | 0.00991319 | 0.14050816 |
| SRP9 | 47.4672641 | -0.067468073 | 0.21457222 | -0.3144306 | 0.75319402 | 0.94096423 |
| EPHX1 | 35.0650699 | 0.319467241 | 0.23061238 | 1.38529964 | 0.16596086 | 0.56208655 |
| AL591895.1 | 17.6310029 | 0.429834484 | 0.40274702 | 1.06725678 | 0.2858559 | 0.6783522 |
| TMEM63A | 48.7890799 | 0.388270551 | 0.31969084 | 1.21451886 | 0.22454965 | 0.61883685 |
| ACBD3 | 82.7317232 | -0.35036839 | 0.26288161 | -1.3327991 | 0.18259773 | 0.58106792 |
| PARP1 | 20.1116806 | 0.059755224 | 0.2843122 | 0.21017467 | 0.83353135 | 0.9666365 |
| ITPKB | 54.9136808 | 0.315411291 | 0.39624403 | 0.79600263 | 0.42603051 | 0.77902418 |
| COQ8A | 22.7641738 | 0.264874168 | 0.2777284 | 0.95371655 | 0.34022714 | 0.71925477 |
| CDC42BPA | 539.905342 | 0.422007549 | 0.18394022 | 2.29426463 | 0.0217753 | 0.2177234 |
| ZNF678 | 35.8608398 | 0.111491017 | 0.21549622 | 0.51736878 | 0.60489875 | 0.87903643 |
| ARF1 | 91.8043285 | -0.727804062 | 0.32140025 | -2.2644788 | 0.02354468 | 0.22729214 |

| | | | | | | |
|---|---|---|---|---|---|---|
| GUK1 | 58.4010303 | -0.405985734 | 0.22502331 | -1.8041941 | 0.07120088 | 0.39729887 |
| OBSCN | 26.8817967 | 0.599553151 | 0.45529861 | 1.31683502 | 0.18789393 | 0.58645909 |
| NUP133 | 43.9873052 | 0.184362101 | 0.23836192 | 0.7734545 | 0.43925344 | 0.78887428 |
| ABCB10 | 25.0781278 | 0.321985035 | 0.32040484 | 1.00493186 | 0.31492966 | 0.70282967 |
| GALNT2 | 114.925618 | -0.600427955 | 0.4065283 | -1.4769647 | 0.1396851 | 0.52592211 |
| COG2 | 25.1020353 | -0.21953837 | 0.28046538 | -0.7827646 | 0.43376536 | 0.78484089 |
| C1orf198 | 43.0825218 | 0.874690085 | 0.38620445 | 2.26483687 | 0.02352269 | 0.22729214 |
| TTC13 | 24.3040232 | 0.054931782 | 0.35031503 | 0.15680681 | 0.87539711 | 0.9768643 |
| FAM89A | 26.3830291 | -0.501594201 | 0.53858422 | -0.9313199 | 0.35168811 | 0.72989437 |
| GNPAT | 32.52922 | 0.084782787 | 0.22319029 | 0.37986772 | 0.70404361 | 0.92936308 |
| EGLN1 | 101.228634 | -0.773659097 | 0.31542356 | -2.4527626 | 0.01417639 | 0.17480247 |
| DISC1 | 54.4581612 | 0.581143571 | 0.30253899 | 1.9208882 | 0.0547458 | 0.3515795 |
| SIPA1L2 | 68.5182818 | 0.400212593 | 0.42487476 | 0.94195426 | 0.34621606 | 0.72436557 |
| PCNX2 | 68.8793094 | -0.164263043 | 0.2085238 | -0.7877424 | 0.43084738 | 0.78340375 |
| TARBP1 | 37.3200822 | 0.323172206 | 0.37160445 | 0.8696672 | 0.3844823 | 0.75288048 |
| IRF2BP2 | 80.6566956 | 0.135611296 | 0.31062492 | 0.43657571 | 0.66241908 | 0.91095019 |
| TOMM20 | 51.8798029 | -0.497615949 | 0.23908565 | -2.0813292 | 0.03740378 | 0.2933622 |
| ARID4B | 324.128672 | -0.070416869 | 0.23974939 | -0.2937103 | 0.76897928 | 0.94579017 |
| B3GALNT2 | 67.6612407 | -0.056565258 | 0.3274785 | -0.1727297 | 0.86286391 | 0.97571732 |
| LYST | 179.03788 | -0.305655465 | 0.21866373 | -1.3978334 | 0.1621631 | 0.5558473 |
| NID1 | 436.871949 | -0.370949585 | 0.49637681 | -0.7473145 | 0.45487374 | 0.79609632 |
| GPR137B | 69.2344069 | 0.059418391 | 0.39939304 | 0.14877173 | 0.88173376 | 0.97782528 |
| ERO1B | 48.8423101 | -0.690100959 | 0.2891376 | -2.3867562 | 0.01699776 | 0.19063814 |
| LGALS8 | 37.5454067 | -0.640239658 | 0.26222747 | -2.4415431 | 0.01462464 | 0.17653018 |
| HEATR1 | 43.0991896 | -0.493648416 | 0.28709714 | -1.7194474 | 0.08553294 | 0.42848827 |
| MTR | 118.709829 | -0.414882222 | 0.19900373 | -2.0847962 | 0.03708781 | 0.29287595 |
| RYR2 | 43.9317272 | 0.614211186 | 0.53102089 | 1.15666105 | 0.24741087 | 0.64317338 |
| PLD5 | 57.9419811 | 2.209725619 | 0.99125699 | 2.22921567 | 0.02579956 | 0.23969779 |
| CEP170 | 105.372317 | -0.18535794 | 0.15498417 | -1.1959798 | 0.23170445 | 0.62741193 |
| SDCCAG8 | 207.730959 | -0.240757197 | 0.17515855 | -1.3745101 | 0.16928338 | 0.56570963 |
| AKT3 | 732.633933 | -0.093465736 | 0.27110277 | -0.3447613 | 0.73027385 | 0.93455731 |
| ADSS | 33.6851169 | 0.042398467 | 0.24831347 | 0.17074574 | 0.8644237 | 0.9762143 |
| DESI2 | 38.6668807 | 0.241058821 | 0.21146292 | 1.13995785 | 0.25430386 | 0.64869111 |
| COX20 | 33.8196413 | 0.04801196 | 0.23961174 | 0.20037399 | 0.8411881 | 0.96803523 |
| HNRNPU | 254.799385 | -0.089765655 | 0.22295771 | -0.4026129 | 0.68723301 | 0.92100905 |
| EFCAB2 | 122.574876 | -0.027320407 | 0.34008123 | -0.0803349 | 0.93597087 | 0.98827804 |
| KIF26B | 187.467562 | -0.39774338 | 0.59388401 | -0.6697324 | 0.50302837 | 0.82430366 |
| SMYD3 | 409.93312 | -0.004420729 | 0.28936662 | -0.0152773 | 0.98781098 | 0.99719329 |
| CNST | 75.9816692 | -0.055113161 | 0.23429078 | -0.235234 | 0.81402711 | 0.96038841 |
| SCCPDH | 20.6091082 | -0.243350263 | 0.37488257 | -0.6491373 | 0.51624963 | 0.83181726 |
| AHCTF1 | 93.7326004 | -0.11913378 | 0.20797813 | -0.5728188 | 0.5667674 | 0.85826779 |
| ZNF496 | 25.1365183 | 0.187209923 | 0.31767011 | 0.58932181 | 0.55564542 | 0.85207344 |
| SH3YL1 | 48.0447431 | 0.173671094 | 0.25114642 | 0.69151331 | 0.48924302 | 0.81720854 |
| ACP1 | 24.2302124 | -0.299573817 | 0.28204154 | -1.062162 | 0.28816213 | 0.67992862 |
| PXDN | 119.415837 | -0.284681239 | 0.31551628 | -0.9022712 | 0.36691284 | 0.73896496 |
| EIPR1 | 26.9339809 | -0.480530709 | 0.30491091 | -1.5759709 | 0.11503251 | 0.48724723 |
| TRAPPC12 | 74.3526324 | 0.083503983 | 0.22087655 | 0.37805726 | 0.70538806 | 0.92946306 |
| ADI1 | 35.9287341 | -0.033704078 | 0.30208944 | -0.1115699 | 0.91116447 | 0.98343817 |
| RPS7 | 113.5642 | -0.395465777 | 0.4042283 | -0.9783228 | 0.32791468 | 0.71294249 |
| RNF144A | 217.531259 | 0.338548893 | 0.46294804 | 0.73128919 | 0.46460253 | 0.80319078 |
| ID2 | 82.3387517 | -1.113045275 | 0.35696729 | -3.1180596 | 0.00182046 | 0.05398305 |
| KIDINS220 | 154.694688 | 0.024908271 | 0.17662111 | 0.14102658 | 0.88784894 | 0.97920559 |
| MBOAT2 | 57.3697391 | 0.584906924 | 0.31968534 | 1.82963326 | 0.0673048 | 0.38712255 |
| ASAP2 | 244.614983 | 0.203550399 | 0.31581687 | 0.64452034 | 0.51923807 | 0.83297983 |
| ITGB1BP1 | 51.989534 | 0.196135299 | 0.22374273 | 0.87661082 | 0.38069806 | 0.74991898 |
| IAH1 | 21.1822132 | 0.060293191 | 0.29156949 | 0.20678841 | 0.83617511 | 0.9668633 |
| ADAM17 | 109.292516 | -0.070326779 | 0.25641301 | -0.2742715 | 0.78387599 | 0.95064024 |
| YWHAQ | 97.8040849 | 0.174031392 | 0.18779946 | 0.92668737 | 0.35408888 | 0.73059415 |
| TAF1B | 49.1867842 | -0.018257391 | 0.27307008 | -0.0668597 | 0.94669337 | 0.99060401 |
| HPCAL1 | 23.2106163 | 0.250658719 | 0.27819172 | 0.90102867 | 0.36757308 | 0.73987799 |
| NOL10 | 83.5425235 | -0.370363728 | 0.23633544 | -1.5671104 | 0.11708888 | 0.4900263 |
| PDIA6 | 59.2548081 | -0.478113941 | 0.33266018 | -1.4372443 | 0.15064859 | 0.53880252 |
| ROCK2 | 485.254925 | 0.264108372 | 0.37722416 | 0.70013642 | 0.48384211 | 0.8144078 |
| LPIN1 | 54.0895403 | -0.24914784 | 0.21725882 | -1.1467789 | 0.25147299 | 0.6460368 |
| TRIB2 | 82.530608 | -0.893752184 | 0.46091295 | -1.9390911 | 0.05249025 | 0.34627545 |
| NBAS | 200.88543 | -0.354339337 | 0.23898998 | -1.4826535 | 0.13816649 | 0.52264518 |
| DDX1 | 47.4913476 | -0.092488091 | 0.19613371 | -0.4715563 | 0.6372435 | 0.89741446 |
| FAM49A | 41.5945774 | -0.104079563 | 0.30155735 | -0.3451402 | 0.72998897 | 0.93455731 |
| SMC6 | 36.6801451 | 0.457037566 | 0.26373681 | 1.73293051 | 0.08310802 | 0.42333667 |
| LINC01376 | 52.3229318 | 0.354250255 | 0.40797668 | 0.86831005 | 0.38522462 | 0.75333042 |
| WDR35 | 40.223963 | -0.343319821 | 0.27261041 | -1.259379 | 0.20789348 | 0.6059796 |
| LAPTM4A | 122.898609 | 0.073279166 | 0.21005602 | 0.34885535 | 0.72719791 | 0.93455731 |
| PUM2 | 219.386781 | 0.181167892 | 0.12322621 | 1.47020585 | 0.14150601 | 0.52895609 |
| RHOB | 111.224117 | 0.030397377 | 0.50182923 | 0.06057315 | 0.95169916 | 0.99060401 |
| HS1BP3 | 19.5808291 | -0.216050764 | 0.33692744 | -0.6412383 | 0.52136789 | 0.83493847 |
| LDAH | 70.9831372 | -0.219680681 | 0.18862008 | -1.1646728 | 0.24415144 | 0.63966808 |
| KLHL29 | 201.864851 | -0.346509988 | 0.22428192 | -1.5449751 | 0.12235227 | 0.50092862 |
| ATAD2B | 168.880204 | -0.02038825 | 0.20833418 | -0.0978632 | 0.92204092 | 0.98561736 |
| UBXN2A | 52.3130682 | 0.271151285 | 0.21965136 | 1.23446215 | 0.21703074 | 0.61410359 |
| SF3B6 | 25.0178679 | -0.314699163 | 0.32828942 | -0.9586028 | 0.33775887 | 0.71925477 |
| FAM228B | 36.7177454 | 0.464699974 | 0.34702915 | 1.33908053 | 0.18054446 | 0.57863987 |
| ITSN2 | 167.727523 | -0.024082478 | 0.2293648 | -0.1049964 | 0.91637867 | 0.98477389 |
| NCOA1 | 449.844235 | 0.254091956 | 0.23390703 | 1.08629466 | 0.27734865 | 0.67269898 |
| ADCY3 | 230.050975 | -0.70936626 | 0.33772073 | -2.1004522 | 0.03568908 | 0.28821871 |
| DNAJC27 | 26.3687404 | 0.295526056 | 0.38972494 | 0.75829393 | 0.44827504 | 0.79296348 |
| DNMT3A | 58.8307149 | 0.542776894 | 0.26061419 | 2.0826836 | 0.03728007 | 0.29301102 |
| DTNB | 46.2801778 | 0.16433 | 0.31707404 | 0.51827012 | 0.60426982 | 0.87903643 |
| ASXL2 | 75.6491342 | -0.219437185 | 0.26222935 | -0.836814 | 0.40269714 | 0.76472383 |
| RAB10 | 178.929032 | -0.277821859 | 0.2216485 | -1.2534344 | 0.21004765 | 0.60731347 |
| HADHA | 63.5911905 | -0.24997854 | 0.20058328 | -1.2462581 | 0.21266964 | 0.60955897 |
| HADHB | 70.9051032 | -0.163220578 | 0.24200206 | -0.6744595 | 0.50001925 | 0.82280373 |
| SELENOI | 35.4060882 | -0.249260513 | 0.29353175 | -0.8491774 | 0.39578261 | 0.76020146 |
| KCNK3 | 20.220417 | 1.175743981 | 0.59720095 | 1.96875771 | 0.04898092 | 0.33550435 |
| MAPRE3 | 22.2954258 | -0.285925382 | 0.26317706 | -1.0864373 | 0.27728555 | 0.67269898 |
| TMEM214 | 24.6479044 | -0.093034921 | 0.4429752 | -0.2100229 | 0.83364983 | 0.9666365 |
| OST4 | 36.2206847 | -0.684367283 | 0.50260918 | -1.3616291 | 0.17331497 | 0.57065039 |
| EMILIN1 | 113.39151 | -0.298679892 | 0.3747529 | -0.7970049 | 0.42544819 | 0.77861803 |
| ATRAID | 24.2520832 | -0.471517175 | 0.3209819 | -1.4689837 | 0.14183722 | 0.52938635 |
| GTF3C2 | 30.9844639 | 0.132076596 | 0.34888455 | 0.3785682 | 0.70500854 | 0.92946306 |
| PPM1G | 51.0672103 | -0.748855814 | 0.42802777 | -1.7495496 | 0.08019606 | 0.41887764 |
| NRBP1 | 29.8192532 | 0.42447382 | 0.2414839 | 1.75777278 | 0.07878618 | 0.41594801 |
| IFT172 | 28.9610554 | -0.011517536 | 0.29393318 | -0.0391842 | 0.96874353 | 0.99300191 |
| ZNF512 | 48.9564643 | -0.215045664 | 0.22875341 | -0.9400763 | 0.34717842 | 0.72524566 |
| MRPL33 | 51.1885993 | -0.060421122 | 0.2526277 | -0.2391706 | 0.81097329 | 0.95967025 |
| BABAM2 | 253.065125 | -0.425357418 | 0.21983723 | -1.9348744 | 0.05300571 | 0.34749239 |
| FOSL2 | 128.841184 | -0.77501452 | 0.39052252 | -1.9845578 | 0.04719369 | 0.32874541 |
| PPP1CB | 111.808459 | -0.177077059 | 0.45670261 | -0.3877295 | 0.69821625 | 0.92737958 |
| TRMT61B | 22.1669079 | 0.074576474 | 0.2647748 | 0.28166002 | 0.77820421 | 0.94820337 |
| WDR43 | 77.1041694 | -0.501907218 | 0.33098999 | -1.5163819 | 0.12942282 | 0.51204187 |
| CLIP4 | 101.959286 | -0.52097465 | 0.33975332 | -1.5333909 | 0.12517957 | 0.50407927 |
| YPEL5 | 38.3531569 | -0.229236397 | 0.28562574 | -0.8025761 | 0.42221977 | 0.77771095 |
| LBH | 81.6420524 | 0.907477313 | 0.35890538 | 2.52845836 | 0.01145647 | 0.1525246 |
| LCLAT1 | 105.225845 | -0.017847072 | 0.17903355 | -0.0996856 | 0.92059391 | 0.98528826 |

| | | | | | | |
|---|---|---|---|---|---|---|
| EHD3 | 23.6863983 | 1.562104981 | 0.37264585 | 4.19192914 | 2.77E-05 | 0.00369578 |
| DPY30 | 23.1297271 | -0.274573893 | 0.30269252 | -0.907105 | 0.36435128 | 0.73648283 |
| SPAST | 37.9564769 | 0.738014777 | 0.28490235 | 2.59041309 | 0.00958608 | 0.1379822 |
| SLC30A6 | 43.0775109 | -0.198288821 | 0.2815254 | -0.7043372 | 0.48122278 | 0.8133639 |
| YIPF4 | 76.4779152 | 0.143573795 | 0.17934644 | 0.80053885 | 0.42339867 | 0.77827834 |
| BIRC6 | 450.220203 | -0.125696149 | 0.12911738 | -0.9735029 | 0.33030341 | 0.71362138 |
| TTC27 | 66.4569563 | -0.220279724 | 0.19287146 | -1.1421064 | 0.2534098 | 0.64812282 |
| LINC00486 | 72.5289974 | 0.73024331 | 0.18183344 | 4.01600128 | 5.92E-05 | 0.00557713 |
| LTBP1 | 585.59405 | 0.998126744 | 0.52121844 | 1.91498741 | 0.05549411 | 0.35370877 |
| FAM98A | 18.2486682 | -0.181475949 | 0.29456742 | -0.6160761 | 0.5378443 | 0.84439601 |
| LINC01320 | 32.6322505 | -0.314928431 | 0.31334117 | -1.0050656 | 0.31486526 | 0.70282967 |
| CRIM1 | 678.198702 | 1.139362139 | 0.43030474 | 2.64780288 | 0.00810168 | 0.12587571 |
| FEZ2 | 86.6076488 | 0.378292901 | 0.26163763 | 1.44586582 | 0.14821484 | 0.53680621 |
| STRN | 70.5034666 | 0.356849222 | 0.2264014 | 1.57617941 | 0.11498446 | 0.48724723 |
| HEATR5B | 69.6387386 | -0.02387834 | 0.23006193 | -0.1037909 | 0.91733527 | 0.98477389 |
| EIF2AK2 | 92.9484311 | 0.285199784 | 0.49218905 | 0.57945171 | 0.56228442 | 0.85545224 |
| CEBPZOS | 35.3736082 | -0.225693199 | 0.23636998 | -0.9548302 | 0.33966356 | 0.71925477 |
| CEBPZ | 42.2850025 | -0.445992582 | 0.24193407 | -1.8434468 | 0.0652638 | 0.38095778 |
| NDUFAF7 | 27.0687839 | -0.234646521 | 0.24695439 | -0.9501614 | 0.34203027 | 0.7206366 |
| PRKD3 | 96.7092142 | -0.213748373 | 0.29509428 | -0.7243393 | 0.46885748 | 0.80374006 |
| CDC42EP3 | 56.6193879 | -0.137377009 | 0.46605288 | -0.294767 | 0.76817188 | 0.94576899 |
| RMDN2 | 37.0974838 | 0.393881308 | 0.39448907 | 0.99845937 | 0.31805666 | 0.7065896 |
| CYP1B1 | 120.193468 | -0.172366479 | 0.42849295 | -0.4022621 | 0.68749113 | 0.92100905 |
| CYP1B1-AS1 | 72.3100721 | 0.648556743 | 0.34543316 | 1.87751735 | 0.06044722 | 0.36535567 |
| ATL2 | 75.6797012 | -0.111056415 | 0.17053164 | -0.6512364 | 0.51489388 | 0.83109052 |
| HNRNPLL | 88.8242673 | 0.418188477 | 0.38798499 | 1.07784706 | 0.28110202 | 0.6759878 |
| GALM | 27.6235256 | 0.654485926 | 0.31017307 | 2.1100669 | 0.03485259 | 0.28490735 |
| SRSF7 | 66.2597529 | -0.120593258 | 0.40516127 | -0.2976426 | 0.76597595 | 0.9447922 |
| DHX57 | 34.6148419 | -0.247444682 | 0.28715321 | -0.8617166 | 0.3888435 | 0.75549104 |
| SOS1 | 212.651922 | 0.055711506 | 0.22589335 | 0.24662747 | 0.80519654 | 0.95836441 |
| MAP4K3 | 331.886255 | 0.1292263 | 0.27797453 | 0.4648854 | 0.64201353 | 0.90015315 |
| AC007388.1 | 99.5640304 | 0.130536059 | 0.32584091 | 0.40061286 | 0.68870517 | 0.92207949 |
| THUMPD2 | 45.9603969 | -0.413183419 | 0.19936577 | -2.0724893 | 0.03821983 | 0.29566057 |
| SLC8A1 | 414.553064 | 1.600117751 | 0.39719009 | 4.0285943 | 5.61E-05 | 0.00542582 |
| PKDCC | 67.3916755 | 0.022697856 | 0.29528814 | 0.07686681 | 0.9387295 | 0.98827804 |
| EML4 | 275.348628 | -0.133287794 | 0.30084608 | -0.4430431 | 0.65773453 | 0.90772143 |
| COX7A2L | 44.3861392 | -0.052433764 | 0.2635849 | -0.1989255 | 0.84232101 | 0.96828048 |
| MTA3 | 106.851276 | 0.248675824 | 0.18080004 | 1.37541906 | 0.16900156 | 0.56570963 |
| HAAO | 17.7419156 | 0.880613903 | 0.27027001 | 3.25827454 | 0.00112092 | 0.04160421 |
| ZFP36L2 | 129.29479 | 0.312645844 | 0.31042054 | 1.00716867 | 0.31385373 | 0.70282967 |
| THADA | 169.855422 | -0.259843037 | 0.22053889 | -1.1782187 | 0.23870944 | 0.63422836 |
| PLEKHH2 | 121.641977 | -0.011247452 | 0.35550281 | -0.0316382 | 0.97476062 | 0.99300191 |
| DYNC2LI1 | 64.7756702 | 0.115253141 | 0.31471175 | 0.36621811 | 0.71420233 | 0.93144151 |
| LRPPRC | 114.176834 | -0.390684465 | 0.22951385 | -1.7022261 | 0.08871299 | 0.43544541 |
| PPM1B | 105.362766 | -0.009847693 | 0.21458639 | -0.0458915 | 0.96339672 | 0.99231823 |
| SLC3A1 | 29.4579683 | -0.163865991 | 0.63594408 | -0.2576736 | 0.79665883 | 0.95463848 |
| PREPL | 82.4194102 | 0.078363601 | 0.27558118 | 0.28435759 | 0.77613636 | 0.94805375 |
| CAMKMT | 216.327379 | -0.053087462 | 0.23531376 | -0.2256029 | 0.8215103 | 0.96303704 |
| SRBD1 | 105.653219 | -0.326482385 | 0.20740739 | -1.5741117 | 0.11546164 | 0.48836096 |
| PRKCE | 294.29896 | -0.099185309 | 0.14413077 | -0.6881619 | 0.49135081 | 0.81732392 |
| EPAS1 | 635.057486 | -0.130546783 | 0.38762255 | -0.3367884 | 0.73627641 | 0.93594619 |
| RHOQ | 188.54925 | -0.919141139 | 0.32327541 | -2.8432139 | 0.00446611 | 0.08834811 |
| AC018682.1 | 34.5431817 | -1.943537744 | 0.49918375 | -3.8934315 | 9.88E-05 | 0.00789507 |
| PIGF | 32.8946467 | -0.177292839 | 0.27417486 | -0.6466415 | 0.517864 | 0.83295744 |
| SOCS5 | 141.934374 | -0.364530683 | 0.37240642 | -0.9788518 | 0.32765323 | 0.71282522 |
| MCFD2 | 43.8893564 | -0.19062928 | 0.38241616 | -0.4984865 | 0.61814121 | 0.88534783 |
| TTC7A | 46.6484175 | 0.631464432 | 0.316802 | 1.99324633 | 0.04623448 | 0.32328943 |
| EPCAM | 25.5064504 | -0.466784867 | 0.38352802 | -1.2170815 | 0.2235732 | 0.61791629 |
| FBXO11 | 250.392794 | -0.045883002 | 0.15404351 | -0.2978574 | 0.765812 | 0.9447922 |
| FOXN2 | 66.4404376 | 0.440520896 | 0.2673671 | 1.64762566 | 0.09942951 | 0.45779653 |
| PPP1R21 | 99.8697532 | 0.106922801 | 0.32043564 | 0.33367949 | 0.73862143 | 0.93594619 |
| STON1-GTF2 | 31.3665601 | -1.389751617 | 0.50408054 | -2.7570031 | 0.00583338 | 0.10280457 |
| AC009975.1 | 46.3693717 | -2.704241388 | 1.13851058 | -2.3752448 | 0.01753731 | 0.19333326 |
| GPR75-ASB3 | 29.976582 | -0.532015597 | 0.32509315 | -1.636502 | 0.10173456 | 0.46169744 |
| ERLEC1 | 33.675846 | -0.230447062 | 0.25435205 | -0.9060161 | 0.36492732 | 0.73726023 |
| PSME4 | 159.513291 | -0.039052228 | 0.26277637 | -0.1486139 | 0.88185828 | 0.97782528 |
| ACYP2 | 226.657448 | 0.24492013 | 0.19490366 | 1.2566215 | 0.20889073 | 0.6059796 |
| SPTBN1 | 544.946578 | 0.002967557 | 0.26930916 | 0.01101915 | 0.99120817 | 0.99759781 |
| RTN4 | 342.266722 | 0.101427599 | 0.2855958 | 0.35514388 | 0.72248181 | 0.93444541 |
| CLHC1 | 27.5669432 | -0.15230769 | 0.46895279 | -0.3247826 | 0.74534564 | 0.93891908 |
| RPS27A | 163.029993 | -0.151952057 | 0.39406792 | -0.3855986 | 0.69979394 | 0.92769314 |
| CCDC88A | 90.2562044 | 0.311390874 | 0.17787521 | 1.75061427 | 0.08001237 | 0.41881121 |
| CFAP36 | 32.6201759 | -0.655345902 | 0.24693689 | -2.6539003 | 0.00795673 | 0.1246781 |
| PPP4R3B | 109.82569 | -0.033939636 | 0.17002408 | -0.1996166 | 0.84178041 | 0.96803523 |
| PNPT1 | 59.7795441 | 0.013108127 | 0.53448919 | 0.02452459 | 0.98043417 | 0.99519485 |
| EFEMP1 | 120.829967 | -0.104897008 | 0.50808503 | -0.2064556 | 0.83643503 | 0.9668633 |
| VRK2 | 62.9969975 | -0.031838175 | 0.23788422 | -0.133839 | 0.89352992 | 0.98023547 |
| FANCL | 37.9094023 | -0.2028347 | 0.40345092 | -0.5027494 | 0.61514048 | 0.88360697 |
| PAPOLG | 33.9899521 | -0.043413243 | 0.22579403 | -0.1922692 | 0.84753132 | 0.97044434 |
| REL | 53.8590476 | -0.579622459 | 0.46946054 | -1.2346564 | 0.21695841 | 0.61410359 |
| PEX13 | 19.7904336 | -0.368147722 | 0.28938736 | -1.2721624 | 0.20331543 | 0.60367992 |
| KIAA1841 | 30.1831044 | 0.658991503 | 0.36672358 | 1.79697062 | 0.07234028 | 0.40009608 |
| USP34 | 459.668615 | -0.248996216 | 0.18317371 | -1.3593447 | 0.17403738 | 0.57154863 |
| XPO1 | 179.265246 | -0.15121968 | 0.36608573 | -0.4130718 | 0.67955403 | 0.91852907 |
| CCT4 | 40.9872968 | 0.053198265 | 0.25917485 | 0.20526013 | 0.83736889 | 0.96697423 |
| COMMD1 | 161.933407 | -0.005885454 | 0.17969283 | -0.0327529 | 0.97387167 | 0.99300191 |
| B3GNT2 | 23.271296 | 0.15024944 | 0.33845842 | 0.44392289 | 0.65709834 | 0.90772143 |
| EHBP1 | 355.36564 | 0.333784226 | 0.22952388 | 1.4542462 | 0.14587806 | 0.53576106 |
| WDPCP | 168.138059 | -0.059392797 | 0.19840217 | -0.2993556 | 0.76466875 | 0.94462105 |
| MDH1 | 36.9196727 | -0.10259896 | 0.32552346 | -0.3151815 | 0.75262388 | 0.94058012 |
| UGP2 | 114.211677 | -0.840566404 | 0.35664409 | -2.3568774 | 0.01842933 | 0.19887981 |
| VPS54 | 142.39841 | 0.162152861 | 0.12338379 | 1.31421531 | 0.18877376 | 0.58671246 |
| PELI1 | 85.1463632 | -0.003068681 | 0.33590394 | -0.0091356 | 0.99271095 | 0.99759781 |
| AFTPH | 76.4379728 | 0.244901817 | 0.19483378 | 1.25697822 | 0.20876152 | 0.6059796 |
| SERTAD2 | 53.7880529 | 0.066539239 | 0.23869846 | 0.27875856 | 0.78043012 | 0.94916975 |
| CEP68 | 26.7981227 | 0.420352126 | 0.34203752 | 1.22896497 | 0.21908494 | 0.61522936 |
| RAB1A | 276.107646 | -0.499436653 | 0.28985818 | -1.723038 | 0.08488166 | 0.42746464 |
| ACTR2 | 103.046567 | -0.099610464 | 0.21034833 | -0.47355 | 0.6358208 | 0.89741446 |
| SPRED2 | 194.255477 | 0.153418194 | 0.23457734 | 0.65401966 | 0.51309914 | 0.82983397 |
| MEIS1 | 745.299484 | -0.35931769 | 0.23813126 | -1.508906 | 0.13132281 | 0.51290135 |
| LINC01798 | 64.5115407 | 0.304097042 | 0.38851663 | 0.78271307 | 0.43379563 | 0.78484089 |
| LINC01828 | 29.0470714 | 0.125300757 | 0.34866843 | 0.35936938 | 0.71931878 | 0.93444541 |
| C1D | 31.8653524 | -0.026696657 | 0.28413011 | -0.0939593 | 0.92514151 | 0.98561736 |
| CNRIP1 | 22.0366896 | -0.537104611 | 0.2663506 | -2.0165324 | 0.04374433 | 0.31491241 |
| APLF | 37.2236332 | -0.130848559 | 0.32372214 | -0.4042002 | 0.6860655 | 0.92055785 |
| ARHGAP25 | 31.615214 | -0.447560304 | 0.8065379 | -0.5549154 | 0.57895253 | 0.86486944 |
| ANTXR1 | 231.076496 | 0.422373813 | 0.21555744 | 1.95944905 | 0.05006022 | 0.33782605 |
| GFPT1 | 131.205433 | -0.060784413 | 0.15501033 | -0.3921314 | 0.69496115 | 0.92589762 |
| NFU1 | 34.2009299 | 0.447712676 | 0.31878998 | 1.40441265 | 0.16019601 | 0.55310356 |
| AAK1 | 216.47765 | -0.172490316 | 0.18671334 | -0.9238243 | 0.3555778 | 0.73125149 |
| ANXA4 | 73.159653 | -0.010751303 | 0.28623633 | -0.0375609 | 0.97003776 | 0.99300191 |
| GMCL1 | 49.425463 | -0.365363157 | 0.19884923 | -1.8373878 | 0.06615266 | 0.38431299 |
| MXD1 | 23.2699427 | -0.689858882 | 0.32872326 | -2.0986007 | 0.03585211 | 0.28882238 |

| | | | | | | |
|---|---|---|---|---|---|---|
| PCBP1 | 55.558212 | -0.310173889 | 0.29190955 | -1.0625685 | 0.28797769 | 0.67992862 |
| C2orf42 | 18.4400393 | 0.397320565 | 0.31324924 | 1.26838476 | 0.20466058 | 0.60370888 |
| PCYOX1 | 45.4104673 | 0.543588413 | 0.4134491 | 1.31476501 | 0.18858889 | 0.58671246 |
| SNRPG | 22.0970431 | -0.690154542 | 0.38576888 | -1.7890363 | 0.07360897 | 0.40189622 |
| MCEE | 33.9774749 | -0.379311253 | 0.27755091 | -1.3666367 | 0.17173921 | 0.56870244 |
| MPHOSPH10 | 22.0544046 | -0.033200902 | 0.33291756 | -0.0997271 | 0.92056099 | 0.98528826 |
| ZNF638 | 440.892587 | -0.505146729 | 0.18181184 | -2.7784039 | 0.00546267 | 0.09815437 |
| EXOC6B | 361.720947 | -0.006496623 | 0.26095371 | -0.0248957 | 0.98013816 | 0.99503182 |
| SFXN5 | 49.1467708 | 0.572572326 | 0.25694862 | 2.22835338 | 0.02585696 | 0.23992777 |
| CCT7 | 38.8453927 | -0.515775797 | 0.28664778 | -1.7993365 | 0.07196548 | 0.39925993 |
| ALMS1 | 84.9544108 | 0.096394173 | 0.1643117 | 0.58665435 | 0.55743587 | 0.85274692 |
| DUSP11 | 23.8167786 | -0.301511428 | 0.31257287 | -0.9646117 | 0.33473936 | 0.716992 |
| STAMBP | 32.3369857 | 0.004185789 | 0.25269458 | 0.01656462 | 0.98678395 | 0.99719329 |
| DGUOK | 42.0885881 | 0.039484935 | 0.3059667 | 0.12904978 | 0.89731826 | 0.98040427 |
| TET3 | 34.5935616 | -0.210327139 | 0.28788455 | -0.7305954 | 0.4650263 | 0.80342007 |
| MOB1A | 45.933736 | -0.089911802 | 0.23128948 | -0.3887414 | 0.69746743 | 0.92702563 |
| MTHFD2 | 45.2829929 | -1.106283473 | 0.35229477 | -3.1402211 | 0.0016882 | 0.05147971 |
| AC006030.1 | 21.5129654 | -0.800797311 | 0.34384632 | -2.3289396 | 0.01986227 | 0.20828536 |
| LOXL3 | 21.5263931 | -1.283969165 | 0.49968536 | -2.5695553 | 0.01018291 | 0.14209086 |
| MRPL19 | 44.5321533 | -0.07149891 | 0.33914066 | -0.2108238 | 0.8330248 | 0.9666365 |
| GCFC2 | 59.2811614 | -0.20654921 | 0.24763827 | -0.8340763 | 0.40423799 | 0.7660508 |
| SUCLG1 | 41.0412019 | -0.0540797 | 0.23055309 | -0.2345651 | 0.81454633 | 0.96038841 |
| TMSB10 | 211.105956 | -0.669867158 | 0.37156488 | -1.8028269 | 0.0714154 | 0.39729887 |
| KCMF1 | 105.124114 | -0.075104565 | 0.17621413 | -0.4262119 | 0.66995343 | 0.91445841 |
| TCF7L1 | 232.170481 | 0.462099381 | 0.18636434 | 2.47954831 | 0.01315489 | 0.16754817 |
| TGOLN2 | 64.1027436 | 0.321096201 | 0.2457318 | 1.30669373 | 0.19131675 | 0.58975955 |
| ELMOD3 | 34.4406524 | 0.119101007 | 0.30200327 | 0.39436992 | 0.69330794 | 0.9248682 |
| MAT2A | 78.8849775 | -0.244237742 | 0.28891205 | -0.8453706 | 0.39790398 | 0.7611823 |
| GGCX | 22.1817772 | -0.410970057 | 0.26960788 | -1.5243251 | 0.12742753 | 0.50683947 |
| VAMP5 | 24.6455013 | -0.342217708 | 0.25294888 | -1.3529125 | 0.17608357 | 0.57311399 |
| USP39 | 33.9939483 | 0.025658178 | 0.24742476 | 0.10370093 | 0.91740669 | 0.98477389 |
| ATOH8 | 29.3027126 | 0.298533031 | 0.40477367 | 0.73753076 | 0.46079965 | 0.80039155 |
| POLR1A | 28.9216605 | -0.388006546 | 0.28608151 | -1.3562797 | 0.17501018 | 0.57299474 |
| PTCD3 | 51.3071445 | -0.002345728 | 0.1982011 | -0.0118351 | 0.99055718 | 0.99759781 |
| IMMT | 66.7616396 | -0.150230869 | 0.20821064 | -0.7215331 | 0.47058158 | 0.80457054 |
| KDM3A | 70.5135515 | -0.642476622 | 0.41312799 | -1.5551515 | 0.11990999 | 0.49618158 |
| CHMP3 | 19.4833722 | -0.064844159 | 0.35985025 | -0.1801976 | 0.85699743 | 0.97381501 |
| RMND5A | 33.4489385 | -0.019083691 | 0.31302071 | -0.0609662 | 0.95138611 | 0.99060401 |
| CYTOR | 98.2731695 | 1.161456804 | 0.5087388 | 2.28301205 | 0.02242966 | 0.22134579 |
| KRCC1 | 46.5986929 | 0.233142594 | 0.26717569 | 0.87261904 | 0.38287076 | 0.75179833 |
| EIF2AK3 | 97.2879383 | -0.953379365 | 0.31516234 | -3.0250421 | 0.00248598 | 0.06256679 |
| MRPS5 | 43.5220734 | -0.354300624 | 0.3014236 | -1.1754243 | 0.23982501 | 0.63451757 |
| ANKRD36C | 67.1841462 | 0.637460265 | 0.29685022 | 2.14741385 | 0.03176035 | 0.2713257 |
| STARD7 | 31.4567489 | 0.089464861 | 0.25335967 | 0.35311406 | 0.72400293 | 0.93455731 |
| TMEM127 | 22.7291992 | 0.295904426 | 0.29764231 | 0.99416118 | 0.3201444 | 0.70728138 |
| CIAO1 | 28.5935725 | -0.303980644 | 0.36352271 | -0.8362081 | 0.40303783 | 0.76511661 |
| SNRNP200 | 45.5145575 | -0.008842588 | 0.33731568 | -0.0262146 | 0.97908619 | 0.99465087 |
| ARID5A | 27.8251575 | 0.062487308 | 0.40457928 | 0.15445009 | 0.87725486 | 0.9768643 |
| KANSL3 | 65.1551046 | 0.230923409 | 0.20240379 | 1.14090458 | 0.25390963 | 0.64832175 |
| LMAN2L | 19.7309153 | -0.012230295 | 0.35190127 | -0.0347549 | 0.97227518 | 0.99300191 |
| ANKRD23 | 18.4838349 | 0.216526067 | 0.29362936 | 0.73741287 | 0.46087131 | 0.80039155 |
| SEMA4C | 21.9613699 | -0.116513163 | 0.32666282 | -0.3566772 | 0.72133351 | 0.93444541 |
| ANKRD36 | 78.1287155 | 0.848223128 | 0.33519979 | 2.53050014 | 0.01139 | 0.15215018 |
| AC092683.1 | 41.7676151 | -0.043299427 | 0.30856803 | -0.1403238 | 0.88840419 | 0.97942933 |
| ANKRD36B | 20.937241 | 0.546356689 | 0.36023248 | 1.51667802 | 0.129348 | 0.51204187 |
| COX5B | 47.6063001 | -0.515933033 | 0.40090431 | -1.2869231 | 0.19812108 | 0.59549768 |
| ACTR1B | 21.2911265 | -0.012593032 | 0.28271443 | -0.0445433 | 0.96447134 | 0.99275266 |
| TMEM131 | 186.372708 | -0.100931008 | 0.18953446 | -0.5325206 | 0.59436546 | 0.87394794 |
| INPP4A | 89.2712542 | -0.111217291 | 0.31162816 | -0.356891 | 0.72117339 | 0.93444541 |
| UNC50 | 23.6161245 | -0.100239589 | 0.24130278 | -0.41541 | 0.67784178 | 0.91799007 |
| MGAT4A | 21.5383866 | 0.458917905 | 0.29824003 | 1.53875356 | 0.12386447 | 0.50097964 |
| AC079447.1 | 48.0617018 | 0.495506902 | 0.4143972 | 1.19572936 | 0.23180219 | 0.62744541 |
| MITD1 | 23.5724493 | -0.618322025 | 0.31774435 | -1.9459733 | 0.05165795 | 0.34512206 |
| EIF5B | 87.2015099 | -0.123846413 | 0.32420577 | -0.3819994 | 0.7024618 | 0.92865476 |
| REV1 | 106.686145 | 0.406793809 | 0.19796185 | 2.05491014 | 0.03988769 | 0.30126886 |
| AFF3 | 395.421203 | 0.603897179 | 0.48635729 | 1.24167396 | 0.21435688 | 0.61009832 |
| NPAS2 | 284.069244 | -1.177234149 | 0.48043568 | -2.450347 | 0.01427186 | 0.17509832 |
| RPL31 | 50.471434 | -0.620627531 | 0.2669896 | -2.3245382 | 0.02009666 | 0.20948991 |
| TBC1D8 | 64.9695973 | -0.664555403 | 0.28546591 | -2.3279677 | 0.01991382 | 0.20828536 |
| CNOT11 | 19.0928866 | 0.113329882 | 0.28058675 | 0.40390319 | 0.68628391 | 0.9206249 |
| RNF149 | 114.108421 | -0.33559368 | 0.23783219 | -1.4110524 | 0.15822917 | 0.55005969 |
| MAP4K4 | 261.444639 | -0.36980463 | 0.30421574 | -1.2155999 | 0.22413736 | 0.61858591 |
| IL1R1 | 226.047078 | -1.105902007 | 0.39471878 | -2.8017466 | 0.00508268 | 0.09338149 |
| IL18R1 | 115.600793 | -1.088695754 | 0.77542121 | -1.4040056 | 0.16031717 | 0.55313187 |
| MRPS9 | 31.0533017 | 0.087659634 | 0.27901279 | 0.31417783 | 0.75338598 | 0.94096423 |
| FHL2 | 141.815786 | 0.369129638 | 0.44639978 | 0.82690371 | 0.40829164 | 0.76963974 |
| NCK2 | 104.962322 | 0.070650352 | 0.23584196 | 0.2995665 | 0.76450784 | 0.94462105 |
| UXS1 | 39.2003654 | 0.298562414 | 0.27956995 | 1.06793458 | 0.28555002 | 0.6783522 |
| GCC2 | 141.487998 | -0.08610717 | 0.22662559 | -0.3799534 | 0.70397998 | 0.92936308 |
| LIMS1 | 247.092241 | -0.343548276 | 0.17976917 | -1.9110522 | 0.05599787 | 0.35446025 |
| RANBP2 | 116.150736 | 0.06426609 | 0.20694669 | 0.31054418 | 0.75614717 | 0.94216126 |
| SH3RF3 | 412.333294 | -1.292108536 | 0.30460236 | -4.2419519 | 2.22E-05 | 0.00307249 |
| 10-Sep | 80.5645382 | -0.027070553 | 0.23377291 | -0.1157985 | 0.90781223 | 0.98306694 |
| MIR4435-2HG | 383.084085 | -0.009423724 | 0.3954486 | -0.0238305 | 0.98098784 | 0.99533442 |
| BCL2L11 | 22.515085 | -0.579010477 | 0.4719303 | -1.2268983 | 0.21986082 | 0.61590675 |
| AC017002.5 | 35.630185 | 0.17299822 | 0.80066231 | 0.21606889 | 0.82893405 | 0.96580172 |
| ANAPC1 | 65.4693884 | -0.230430579 | 0.17448177 | -1.320657 | 0.18661575 | 0.58501647 |
| MERTK | 41.3086283 | -0.35931805 | 0.37208004 | -0.965701 | 0.33419384 | 0.71686823 |
| TMEM87B | 81.0399888 | -0.379683798 | 0.24822608 | -1.5295887 | 0.12611857 | 0.50535459 |
| ZC3H8 | 29.0921181 | -0.081270637 | 0.23148133 | -0.3510894 | 0.7255213 | 0.93455731 |
| ZC3H6 | 63.3390666 | 0.498821333 | 0.34323282 | 1.45330313 | 0.14613961 | 0.53584686 |
| TTL | 28.1337314 | -0.022486079 | 0.32198476 | -0.0698358 | 0.94432431 | 0.99055658 |
| SLC20A1 | 20.5462213 | -0.335935058 | 0.35143276 | -0.9559014 | 0.33912206 | 0.71925477 |
| PAX8 | 52.2528454 | -0.613174475 | 0.31675704 | -1.935788 | 0.05289367 | 0.34706747 |
| CBWD2 | 19.5241645 | 0.419143595 | 0.28785016 | 1.45611727 | 0.1453602 | 0.53538077 |
| SLC35F5 | 109.56279 | -0.022582381 | 0.29148479 | -0.0774736 | 0.93824678 | 0.98827804 |
| ACTR3 | 180.228323 | 0.054102668 | 0.23742376 | 0.22787386 | 0.8197443 | 0.96284007 |
| DDX18 | 41.0207986 | -0.741143628 | 0.27514807 | -2.6936174 | 0.00706813 | 0.11576428 |
| CCDC93 | 53.8385956 | -0.050251507 | 0.22354808 | -0.2247906 | 0.82214218 | 0.96331679 |
| INSIG2 | 37.7646735 | -0.87014443 | 0.34742786 | -2.5045327 | 0.01226133 | 0.15976684 |
| DBI | 53.6737317 | -0.808821653 | 0.28745363 | -2.8137465 | 0.00489678 | 0.09203696 |
| PTPN4 | 73.7355755 | 0.176383043 | 0.18004143 | 0.97968029 | 0.32724396 | 0.71256766 |
| EPB41L5 | 68.9256691 | 0.151457133 | 0.32734601 | 0.46268209 | 0.64359226 | 0.90063554 |
| RALB | 20.5228647 | 0.228170115 | 0.3871093 | 0.58942039 | 0.5555793 | 0.85207344 |
| GLI2 | 128.981197 | 0.276714438 | 0.287842 | 0.96134141 | 0.33638053 | 0.71883121 |
| CLASP1 | 308.568461 | -0.098207586 | 0.2677412 | -0.3668004 | 0.71376789 | 0.93136293 |
| GYPC | 51.7370376 | -0.631542345 | 0.38000092 | -1.6619495 | 0.0965229 | 0.45210121 |
| BIN1 | 23.2089841 | -0.005214075 | 0.26686324 | -0.0195384 | 0.98441162 | 0.9964038 |
| ERCC3 | 30.2752437 | 0.085069179 | 0.25044108 | 0.33967741 | 0.73409948 | 0.93594619 |
| MAP3K2 | 144.541619 | 0.014510613 | 0.16028185 | 0.09053186 | 0.92786458 | 0.98681039 |
| IWS1 | 75.4490004 | 0.144934043 | 0.16954054 | 0.85486362 | 0.39262665 | 0.75931928 |
| LIMS2 | 30.8274955 | 1.466703956 | 0.46410482 | 3.16028601 | 0.00157614 | 0.05036121 |
| WDR33 | 109.942108 | 0.027771996 | 0.19131579 | 0.14516311 | 0.88458209 | 0.97815133 |

| | | | | | | |
|---|---|---|---|---|---|---|
| AMMECR1L | 20.1979563 | -0.521396692 | 0.27867866 | -1.8709603 | 0.06135059 | 0.36839337 |
| SAP130 | 36.3866018 | -0.015458351 | 0.22670603 | -0.0681868 | 0.94563696 | 0.99060401 |
| UGGT1 | 57.762467 | -0.124176139 | 0.25035622 | -0.4959978 | 0.61989595 | 0.88658432 |
| HS6ST1 | 28.0300322 | -0.521586355 | 0.39997406 | -1.3040504 | 0.19221638 | 0.59129268 |
| SMPD4 | 30.6180179 | -0.216314927 | 0.22904568 | -0.9444183 | 0.34495594 | 0.72325086 |
| MZT2B | 37.9385065 | -0.437730343 | 0.3504753 | -1.248962 | 0.21167896 | 0.60910715 |
| FAM168B | 60.6893219 | 0.077258668 | 0.192215 | 0.40193881 | 0.68772906 | 0.92101828 |
| PLEKHB2 | 42.4475396 | -0.420058369 | 0.2652367 | -1.5837113 | 0.11325942 | 0.48363945 |
| NCKAP5 | 135.82669 | 0.131413346 | 0.89623525 | 0.14662818 | 0.88342551 | 0.97782528 |
| MGAT5 | 158.500036 | -0.460685588 | 0.24676188 | -1.8669236 | 0.06191226 | 0.37021413 |
| CCNT2 | 83.09207 | -0.146200437 | 0.22765148 | -0.6422117 | 0.52073576 | 0.83447168 |
| RAB3GAP1 | 160.285361 | 0.205912555 | 0.21447261 | 0.96008787 | 0.33701099 | 0.71925477 |
| ZRANB3 | 72.8112048 | -0.041289892 | 0.20259938 | -0.2038007 | 0.83850926 | 0.96753094 |
| R3HDM1 | 140.758014 | -0.196048409 | 0.24102593 | -0.8133913 | 0.41599372 | 0.77573656 |
| UBXN4 | 110.036318 | -0.393040325 | 0.1773405 | -2.2163032 | 0.02667075 | 0.24439315 |
| DARS | 99.911166 | -0.650506948 | 0.26772662 | -2.4297432 | 0.01510952 | 0.17917181 |
| HNMT | 44.5523517 | 0.487597301 | 0.4710644 | 1.0350969 | 0.30062366 | 0.69191459 |
| AC114763.1 | 27.1604612 | -0.103218116 | 0.27793863 | -0.3713702 | 0.71036184 | 0.93009618 |
| SPOPL | 56.0549881 | 0.148431141 | 0.22144239 | 0.67029235 | 0.50267144 | 0.82430366 |
| LRP1B | 39.7662427 | -0.176155494 | 0.62041013 | -0.2839339 | 0.77646101 | 0.94805375 |
| ARHGAP15 | 145.485999 | 0.255279961 | 0.42171832 | 0.60533287 | 0.54495784 | 0.84730644 |
| GTDC1 | 229.811062 | 0.185444293 | 0.23915185 | 0.77542487 | 0.43808865 | 0.78836643 |
| ZEB2 | 700.675032 | 0.196781789 | 0.2165407 | 0.90875197 | 0.36348107 | 0.73604674 |
| TEX41 | 100.895678 | 1.406717231 | 0.50047411 | 2.81076923 | 0.00494232 | 0.09242016 |
| ACVR2A | 88.5088188 | 0.064115288 | 0.26014807 | 0.2464569 | 0.80532856 | 0.95836441 |
| ORC4 | 65.2515376 | 0.17353551 | 0.29574777 | 0.58676861 | 0.55735912 | 0.85274692 |
| MBD5 | 502.308227 | -0.091841892 | 0.23721798 | -0.3871624 | 0.69863595 | 0.92769314 |
| EPC2 | 130.121408 | 0.043264667 | 0.15194867 | 0.28473212 | 0.77584938 | 0.94805375 |
| MMADHC | 23.5855842 | -0.391836205 | 0.29236233 | -1.3402418 | 0.18016676 | 0.57783734 |
| AC007364.1 | 38.8542692 | 0.424316491 | 0.37923437 | 1.11887668 | 0.26319275 | 0.65753753 |
| RND3 | 99.8693433 | -1.490793246 | 0.48928994 | -3.0468504 | 0.00231253 | 0.05984073 |
| RIF1 | 128.211458 | -0.126905614 | 0.18039425 | -0.7034904 | 0.48175021 | 0.81369393 |
| ARL5A | 35.1884394 | 0.055655046 | 0.35323443 | 0.15755838 | 0.8748048 | 0.9768643 |
| STAM2 | 70.5877979 | 0.132483515 | 0.20800673 | 0.63691937 | 0.52417736 | 0.83697897 |
| FMNL2 | 291.983122 | -0.669476378 | 0.23386855 | -2.8626182 | 0.00420156 | 0.08482775 |
| PRPF40A | 84.0978227 | -0.080405554 | 0.17745473 | -0.4531046 | 0.65047342 | 0.9050963 |
| ARL6IP6 | 27.0391206 | 0.422969725 | 0.37129089 | 1.13918691 | 0.2546252 | 0.64906023 |
| NR4A2 | 57.0623848 | 1.110274977 | 0.43407455 | 2.55779791 | 0.01053373 | 0.14415712 |
| GPD2 | 91.7405266 | -0.266973964 | 0.24495161 | -1.0899049 | 0.27575503 | 0.67125661 |
| ACVR1 | 65.452067 | 0.035229721 | 0.28825523 | 0.12221711 | 0.90272708 | 0.98164203 |
| PKP4 | 127.088366 | 0.247709509 | 0.13959197 | 1.77452543 | 0.07597628 | 0.40904738 |
| TANC1 | 405.815603 | -0.110547743 | 0.36878025 | -0.2997659 | 0.76435573 | 0.94462105 |
| WDSUB1 | 39.5684882 | 0.02600808 | 0.35288989 | 0.07370027 | 0.94124888 | 0.98895798 |
| BAZ2B | 373.961996 | 0.061368465 | 0.28592011 | 0.21463501 | 0.8300519 | 0.96580172 |
| 7-Mar | 179.526206 | -0.026945334 | 0.19110744 | -0.1409957 | 0.88787331 | 0.97920559 |
| CD302 | 22.3416862 | -0.088641139 | 0.27565742 | -0.3215627 | 0.74778402 | 0.93903859 |
| PLA2R1 | 159.484179 | 0.485360458 | 0.40661699 | 1.19365513 | 0.2326129 | 0.62871357 |
| ITGB6 | 31.4803223 | -2.431117511 | 0.62064186 | -3.917102 | 8.96E-05 | 0.00748427 |
| RBMS1 | 819.048664 | -0.148226123 | 0.26044883 | -0.569118 | 0.56927605 | 0.85958696 |
| TANK | 208.561798 | -0.71954161 | 0.2424203 | -2.9681574 | 0.00299591 | 0.0705671 |
| PSMD14 | 63.9875219 | -0.449062597 | 0.30302614 | -1.4819269 | 0.13835973 | 0.52264518 |
| IFIH1 | 33.9126122 | 0.239563055 | 1.02969118 | 0.23265525 | 0.81602912 | 0.96132648 |
| FIGN | 283.213402 | 0.898380278 | 0.36585285 | 2.45557816 | 0.01406582 | 0.17402303 |
| COBLL1 | 177.704626 | 0.577698761 | 0.35415545 | 1.63120108 | 0.1028479 | 0.46284703 |
| AC019197.1 | 57.1763676 | 0.255599871 | 0.35217407 | 0.72577709 | 0.46797543 | 0.80374006 |
| SLC38A11 | 88.7326077 | -0.541025622 | 0.92810973 | -0.5829328 | 0.55993853 | 0.85423726 |
| CSRNP3 | 47.6370931 | 0.917769678 | 0.35786589 | 2.56456317 | 0.01033058 | 0.14280022 |
| TTC21B | 61.9429521 | 0.025074891 | 0.16635266 | 0.15073334 | 0.88018608 | 0.97781634 |
| STK39 | 176.688666 | -0.474156131 | 0.26770898 | -1.7711626 | 0.07653366 | 0.41054445 |
| CERS6 | 296.584292 | -0.053616677 | 0.47893945 | -0.1119488 | 0.91086404 | 0.98343817 |
| LRP2 | 45.5569191 | 0.853973164 | 0.81133649 | 1.05255116 | 0.29254675 | 0.6840363 |
| PPIG | 110.261259 | 0.114850991 | 0.17440545 | 0.65852867 | 0.51019848 | 0.82732759 |
| KLHL23 | 54.2572752 | 0.147531442 | 0.36664854 | 0.40237837 | 0.68740558 | 0.92100905 |
| SSB | 46.0124024 | 0.043857891 | 0.21952005 | 0.19978991 | 0.84164489 | 0.96803523 |
| UBR3 | 329.679362 | -0.06074711 | 0.21409297 | -0.2837417 | 0.77660831 | 0.94805375 |
| GORASP2 | 42.0340759 | -0.187872242 | 0.25622449 | -0.733233 | 0.46341635 | 0.80225845 |
| TLK1 | 109.579606 | -0.082701633 | 0.17813879 | -0.4642539 | 0.64246583 | 0.90036905 |
| METTL8 | 41.1776934 | -0.331143158 | 0.26688141 | -1.2407876 | 0.21468421 | 0.61009832 |
| DCAF17 | 26.0774914 | 0.024096918 | 0.26619824 | 0.09052245 | 0.92787205 | 0.98681039 |
| CYBRD1 | 95.851131 | -0.026567613 | 0.3160419 | -0.0840636 | 0.93300588 | 0.98725539 |
| DYNC1I2 | 93.1021474 | 0.224041771 | 0.17194563 | 1.30298034 | 0.19258147 | 0.59142551 |
| SLC25A12 | 74.1015646 | -0.43177574 | 0.19362843 | -2.2299191 | 0.02575281 | 0.23956637 |
| HAT1 | 45.3685067 | -0.265155579 | 0.21362568 | -1.2412158 | 0.21452603 | 0.61009832 |
| METAP1D | 20.7889151 | -0.141400886 | 0.30572785 | -0.4625057 | 0.64371869 | 0.90063554 |
| ITGA6 | 24.8414339 | 1.000135552 | 0.47126422 | 2.12223953 | 0.03381763 | 0.28113773 |
| PDK1 | 80.8494645 | -2.380352323 | 0.74837617 | -3.1806896 | 0.00146925 | 0.04777661 |
| RAPGEF4 | 80.227213 | 0.360670928 | 0.38039268 | 0.94815423 | 0.34305095 | 0.72177972 |
| MAP3K20 | 172.678697 | -0.221404967 | 0.16291636 | -1.3590101 | 0.1741434 | 0.57154863 |
| SP3 | 111.715369 | -0.219310033 | 0.19112373 | -1.1474767 | 0.25118465 | 0.6460368 |
| OLA1 | 106.006285 | -0.63901509 | 0.2274873 | -2.8090143 | 0.00496934 | 0.09251529 |
| CIR1 | 47.3820628 | -0.394711696 | 0.27135444 | -1.4545983 | 0.14578051 | 0.53567049 |
| GPR155 | 27.7933524 | 0.360415031 | 0.29300606 | 1.23005999 | 0.21867464 | 0.61522936 |
| AC010894.3 | 55.3239014 | -0.599780926 | 0.35249856 | -1.7015131 | 0.08884668 | 0.43544541 |
| WIPF1 | 79.2974348 | -0.045730752 | 0.16145656 | -0.2832387 | 0.77699384 | 0.94805375 |
| CHN1 | 38.1964811 | -0.307485821 | 0.29014004 | -1.0597842 | 0.2892428 | 0.67992862 |
| ATF2 | 108.000488 | -0.192442024 | 0.22667095 | -0.8489929 | 0.39588526 | 0.76020146 |
| ATP5MC3 | 55.1842968 | -0.450149711 | 0.36040255 | -1.2490192 | 0.21165805 | 0.60910715 |
| LNPK | 49.8303892 | -0.264977493 | 0.23859968 | -1.1105526 | 0.26676099 | 0.66188434 |
| HOXD10 | 73.0168046 | -0.093727651 | 0.19267054 | -0.4864659 | 0.62663687 | 0.89160148 |
| HOXD9 | 16.6774526 | 0.132886975 | 0.37660693 | 0.35285324 | 0.72419847 | 0.93455731 |
| HOXD3 | 30.8099391 | 0.314737906 | 0.24676849 | 1.27543801 | 0.20215425 | 0.60171389 |
| MTX2 | 79.6313413 | 0.196761018 | 0.19749255 | 0.99629588 | 0.3191064 | 0.70683013 |
| HNRNPA3 | 151.964103 | 0.078737114 | 0.1638414 | 0.48056911 | 0.63082278 | 0.89436221 |
| NFE2L2 | 212.575888 | -0.004846373 | 0.17445972 | -0.0277793 | 0.97783816 | 0.99407008 |
| AGPS | 80.9441478 | 0.324694273 | 0.27776279 | 1.16896246 | 0.24241876 | 0.63672023 |
| PRKRA | 44.8378561 | 0.138790154 | 0.19001599 | 0.73041301 | 0.46513777 | 0.80342007 |
| PLEKHA3 | 36.0680153 | -0.212712642 | 0.20536449 | -1.035781 | 0.3003043 | 0.69191459 |
| TTN-AS1 | 122.804886 | -0.138160956 | 0.24098386 | -0.5733204 | 0.5664278 | 0.85825441 |
| TTN | 54.0086895 | -1.275544798 | 0.93908723 | -1.3582815 | 0.17437438 | 0.57183281 |
| SESTD1 | 124.499765 | -0.292647517 | 0.27924737 | -1.0479866 | 0.29464476 | 0.68566952 |
| CWC22 | 40.4278192 | 0.002000122 | 0.20899086 | 0.00957038 | 0.99236406 | 0.99759781 |
| UBE2E3 | 96.5694173 | 0.071540831 | 0.18622051 | 0.38417267 | 0.70085047 | 0.92852895 |
| SSFA2 | 93.4376056 | -0.12874184 | 0.31595437 | -0.4074697 | 0.68366303 | 0.91950744 |
| PDE1A | 481.590514 | 0.052153681 | 0.25763939 | 0.20242899 | 0.83958137 | 0.96770443 |
| DNAJC10 | 66.6337878 | -0.165427595 | 0.27908563 | -0.5927485 | 0.55334946 | 0.85188037 |
| FRZB | 48.527021 | 1.466413025 | 0.26084958 | 5.62168066 | 1.89E-08 | 1.37E-05 |
| NCKAP1 | 246.166563 | -0.191486513 | 0.25759279 | -0.7433691 | 0.45725827 | 0.79762426 |
| NUP35 | 19.8833011 | -0.084680766 | 0.27146803 | -0.3119364 | 0.75508884 | 0.94189722 |
| ZC3H15 | 54.9136727 | 0.070799881 | 0.22960419 | 0.30835622 | 0.75781129 | 0.94301125 |
| ITGAV | 165.822647 | -0.320839007 | 0.29138185 | -1.1010947 | 0.27085545 | 0.66572465 |
| AC007319.1 | 123.683361 | -0.032125067 | 0.35632742 | -0.090156 | 0.92816322 | 0.98681039 |
| CALCRL | 66.9816643 | 0.282118695 | 0.72557804 | 0.38881923 | 0.69740987 | 0.92702563 |
| TFPI | 324.532528 | -0.494259621 | 0.22811676 | -2.1666958 | 0.03025806 | 0.26535376 |

| | | | | | | |
|---|---|---|---|---|---|---|
| GULP1 | 281.737067 | 0.244701444 | 0.24840448 | 0.98509272 | 0.32457853 | 0.71118891 |
| COL3A1 | 407.890953 | -0.005046237 | 0.4577145 | -0.0110249 | 0.99120361 | 0.99759781 |
| COL5A2 | 218.191906 | -0.037863821 | 0.35478674 | -0.1067228 | 0.91500892 | 0.98439476 |
| WDR75 | 47.009406 | -0.501281236 | 0.21257513 | -2.3581367 | 0.01836693 | 0.19849785 |
| SLC40A1 | 81.0671507 | 0.201712473 | 0.40005666 | 0.50420976 | 0.61411399 | 0.88360697 |
| ANKAR | 82.4351048 | 0.54446167 | 0.30689921 | 1.77407321 | 0.07605104 | 0.40915015 |
| ORMDL1 | 38.3903357 | -0.3340051 | 0.25852343 | -1.2919723 | 0.19636675 | 0.59411229 |
| PMS1 | 45.5416027 | -0.13295272 | 0.19167347 | -0.6936418 | 0.4879069 | 0.81595238 |
| HIBCH | 98.0558397 | 0.460779344 | 0.19306594 | 2.38664231 | 0.01700303 | 0.19063814 |
| INPP1 | 27.6383135 | 0.333310455 | 0.29047238 | 1.14747727 | 0.25118442 | 0.6460368 |
| MFSD6 | 26.2594906 | -0.038904072 | 0.35876764 | -0.1084381 | 0.9136482 | 0.98395113 |
| NAB1 | 52.1176353 | 0.44223182 | 0.18933665 | 2.33569055 | 0.01950738 | 0.20568106 |
| GLS | 364.797816 | 0.487773008 | 0.30695615 | 1.58906415 | 0.11204591 | 0.4809788 |
| STAT1 | 93.9601904 | -0.52284162 | 0.59310886 | -0.8815272 | 0.37803252 | 0.74933113 |
| STAT4 | 54.221357 | -0.306746105 | 0.36005177 | -0.85195 | 0.39424184 | 0.75988253 |
| MYO1B | 357.690648 | -0.502428617 | 0.32698063 | -1.53657 | 0.12439865 | 0.50231081 |
| NABP1 | 98.9160198 | -2.690365255 | 0.73630932 | -3.653852 | 0.00025834 | 0.01573508 |
| SLC39A10 | 32.1342492 | 0.004937848 | 0.33494172 | 0.01474241 | 0.98823768 | 0.99719329 |
| STK17B | 100.912387 | 0.20312362 | 0.40084462 | 0.50673904 | 0.61233793 | 0.88282721 |
| HECW2 | 49.5959611 | 0.371664465 | 0.4850001 | 0.76631833 | 0.44348691 | 0.7920275 |
| GTF3C3 | 28.9297767 | 0.25250956 | 0.22004972 | 1.14751141 | 0.25117032 | 0.6460368 |
| PGAP1 | 79.9332709 | 0.219442693 | 0.45268948 | 0.48475324 | 0.62785139 | 0.89212682 |
| ANKRD44 | 189.537988 | 0.408773927 | 0.21799189 | 1.87517953 | 0.06076803 | 0.36605266 |
| SF3B1 | 122.331702 | 0.05677419 | 0.17186496 | 0.33034187 | 0.74114166 | 0.93711378 |
| COQ10B | 24.715806 | -0.438691537 | 0.29382265 | -1.4930487 | 0.13542443 | 0.51914127 |
| HSPD1 | 76.9492298 | -0.125981292 | 0.30306285 | -0.4156936 | 0.67763422 | 0.91799007 |
| RFTN2 | 48.2796601 | 0.518222797 | 0.36944324 | 1.40271288 | 0.16070248 | 0.55420108 |
| PLCL1 | 725.936511 | 0.736084707 | 0.42176079 | 1.74526586 | 0.0809386 | 0.42036593 |
| SPATS2L | 145.610979 | 0.302009741 | 0.42540672 | 0.70993176 | 0.47774645 | 0.81138532 |
| AOX1 | 52.0672541 | -0.276253959 | 0.41708804 | -0.6623397 | 0.50775356 | 0.82518374 |
| BZW1 | 24.3161848 | 0.016630649 | 0.3329299 | 0.0499524 | 0.96016032 | 0.99181157 |
| CLK1 | 97.5103832 | 0.154520914 | 0.24040885 | 0.6427422 | 0.52039139 | 0.83428367 |
| ORC2 | 62.7077968 | -0.266316657 | 0.22722284 | -1.1720506 | 0.24117675 | 0.63527165 |
| FAM126B | 77.3563561 | 0.110708358 | 0.16768359 | 0.66022178 | 0.50911152 | 0.82663984 |
| NDUFB3 | 23.5054362 | -0.390305745 | 0.45141551 | -0.8646264 | 0.3872439 | 0.75443982 |
| CFLAR | 235.296956 | -0.50992254 | 0.26931357 | -1.8934157 | 0.05830261 | 0.36041506 |
| TRAK2 | 68.827658 | -0.088305931 | 0.30061102 | -0.2937548 | 0.76894527 | 0.94579017 |
| STRADB | 30.1521615 | -0.051328955 | 0.48752902 | -0.1052839 | 0.91615054 | 0.98477389 |
| ALS2 | 56.9401964 | -0.120882846 | 0.26954607 | -0.4484682 | 0.65381532 | 0.90646849 |
| SUMO1 | 36.6804353 | -0.095058553 | 0.22499044 | -0.4225004 | 0.6726598 | 0.91482259 |
| NOP58 | 80.3477944 | -0.479141038 | 0.21262853 | -2.2534184 | 0.02423278 | 0.23086389 |
| BMPR2 | 477.512319 | 0.143941116 | 0.30048522 | 0.47902894 | 0.63191804 | 0.89436221 |
| FAM117B | 37.723965 | -0.0761856 | 0.32414275 | -0.2350372 | 0.81417987 | 0.96038841 |
| ICA1L | 27.448439 | 0.998516782 | 0.4194937 | 2.3802903 | 0.017299 | 0.19236057 |
| WDR12 | 25.9055797 | -0.397982849 | 0.30263116 | -1.3150756 | 0.1884845 | 0.58668894 |
| CARF | 29.0017538 | 0.422837687 | 0.36552065 | 1.15680929 | 0.24735028 | 0.64317338 |
| NBEAL1 | 752.720386 | -0.072635082 | 0.22772104 | -0.3189652 | 0.74975291 | 0.94004394 |
| CYP20A1 | 93.4855299 | 0.441157447 | 0.22841001 | 1.93142781 | 0.05343017 | 0.34871963 |
| ABI2 | 141.933579 | 0.301789491 | 0.23258548 | 1.29754227 | 0.19444467 | 0.59272783 |
| RAPH1 | 258.445582 | 0.591670742 | 0.23092278 | 2.56220174 | 0.01040109 | 0.14309893 |
| PARD3B | 460.340479 | 0.637053303 | 0.27989657 | 2.27603108 | 0.02284415 | 0.22440695 |
| NRP2 | 58.705038 | -0.453412123 | 0.75148578 | -0.6033542 | 0.54627306 | 0.84758329 |
| INO80D | 181.991863 | -0.230519431 | 0.53420605 | -0.4315178 | 0.6660919 | 0.91292603 |
| NDUFS1 | 89.2134446 | -0.055413495 | 0.20389448 | -0.2717754 | 0.78579475 | 0.95065355 |
| EEF1B2 | 42.7556771 | -0.389729255 | 0.35887269 | -1.0859819 | 0.27748699 | 0.67269898 |
| ZDBF2 | 46.3613945 | -0.283179239 | 0.44553616 | -0.635592 | 0.52504236 | 0.83726969 |
| FASTKD2 | 25.08684 | 0.009908701 | 0.24124538 | 0.04107312 | 0.9672376 | 0.99300191 |
| KLF7 | 133.279108 | 0.290811051 | 0.29483243 | 0.98636046 | 0.32395626 | 0.71051224 |
| CREB1 | 86.9615394 | 0.36945469 | 0.18362592 | 2.01199637 | 0.04422032 | 0.31581172 |
| CCNYL1 | 18.9890096 | 0.013285991 | 0.38862458 | 0.03418721 | 0.97272786 | 0.99300191 |
| PLEKHM3 | 57.0504244 | 0.201880718 | 0.1951464 | 1.03450904 | 0.30089825 | 0.69211307 |
| PIKFYVE | 109.028264 | -0.375425228 | 0.21137934 | -1.7760734 | 0.07572082 | 0.40887016 |
| PTH2R | 46.8256124 | 0.917897161 | 0.32288645 | 2.84278629 | 0.0044721 | 0.08834811 |
| MAP2 | 118.875053 | 1.177514828 | 0.41674213 | 2.82552385 | 0.00472034 | 0.09152971 |
| RPE | 28.1261878 | -0.068949924 | 0.26632622 | -0.2588927 | 0.795718 | 0.95463848 |
| KANSL1L | 148.6562 | 0.269273581 | 0.23679689 | 1.13715 | 0.25547559 | 0.65030654 |
| AC007038.1 | 22.6913979 | 0.03123808 | 0.32538137 | 0.09600451 | 0.92351699 | 0.98561736 |
| LANCL1 | 36.928318 | -0.003029603 | 0.20371122 | -0.014872 | 0.98813426 | 0.99719329 |
| ERBB4 | 159.426579 | 0.35226852 | 0.34248055 | 1.02857964 | 0.30367725 | 0.69579129 |
| IKZF2 | 33.1527519 | -0.724175713 | 0.32570141 | -2.2234344 | 0.02618653 | 0.24176479 |
| SPAG16 | 83.187237 | -0.175551267 | 0.21676277 | -0.8098774 | 0.41801065 | 0.77722447 |
| BARD1 | 24.2745888 | 0.255742983 | 0.42788023 | 0.5976976 | 0.55004173 | 0.84939203 |
| ATIC | 21.6955951 | -0.173764981 | 0.39210132 | -0.4431635 | 0.6576475 | 0.90772143 |
| FN1 | 366.301619 | -0.067554844 | 0.61792651 | -0.109325 | 0.91294468 | 0.98395113 |
| XRCC5 | 165.555102 | -0.262723547 | 0.15398748 | -1.7061358 | 0.0879828 | 0.43541117 |
| SMARCAL1 | 20.5385207 | 0.144180795 | 0.26713048 | 0.53973921 | 0.5893769 | 0.8702694 |
| RPL37A | 113.14986 | -0.491437503 | 0.3083724 | -1.5936494 | 0.11101458 | 0.47822164 |
| IGFBP2 | 109.185534 | -0.295014589 | 0.7655946 | -0.3853405 | 0.69998518 | 0.92771705 |
| IGFBP5 | 522.195913 | 0.845136742 | 0.48464522 | 1.7438256 | 0.08118951 | 0.42077692 |
| AC007563.2 | 27.6781918 | 0.370522673 | 0.485852 | 0.76262457 | 0.44568733 | 0.79296348 |
| DIRC3 | 115.062643 | 0.266619918 | 0.43747215 | 0.60945575 | 0.54222239 | 0.84643373 |
| TNS1 | 537.885748 | -0.14327947 | 0.26799084 | -0.5346432 | 0.59289664 | 0.87313824 |
| ARPC2 | 106.698403 | -0.030550216 | 0.21643735 | -0.1411504 | 0.88775114 | 0.97920559 |
| AAMP | 17.9762566 | 0.27243874 | 0.33174236 | 0.82123591 | 0.41151191 | 0.77167671 |
| PNKD | 39.8962042 | -0.348646512 | 0.28314957 | -1.2313157 | 0.21820481 | 0.61478732 |
| TMBIM1 | 41.3170213 | -0.567338719 | 0.24461488 | -2.319314 | 0.02037802 | 0.21063016 |
| CTDSP1 | 37.5612481 | 0.364293998 | 0.27602871 | 1.31976849 | 0.18691232 | 0.58501647 |
| USP37 | 92.0235201 | -0.790298452 | 0.33358669 | -2.3690946 | 0.01783169 | 0.19558968 |
| CNOT9 | 36.4688838 | -0.44776805 | 0.2947582 | -1.5191029 | 0.12873659 | 0.51084514 |
| CYP27A1 | 20.7976 | -0.092238486 | 0.334245 | -0.2759607 | 0.78257825 | 0.95014325 |
| DNAJB2 | 19.734913 | 0.288565542 | 0.29733193 | 0.97051648 | 0.33178911 | 0.71487809 |
| DNPEP | 20.452259 | -0.306058758 | 0.39050162 | -0.783758 | 0.43318214 | 0.78484089 |
| DES | 48.0079167 | 1.32042165 | 1.41647725 | 0.93218698 | 0.3512399 | 0.72936942 |
| SPEG | 26.0820488 | 0.767037838 | 0.62213058 | 1.23292098 | 0.21760525 | 0.61468881 |
| OBSL1 | 19.8539484 | 0.334182803 | 0.262429 | 1.27342177 | 0.20286842 | 0.60335087 |
| FARSB | 43.2882101 | -0.105783118 | 0.30688748 | -0.3446968 | 0.73002236 | 0.93455731 |
| ACSL3 | 131.228247 | 0.267688362 | 0.17037267 | 1.57119306 | 0.11613781 | 0.48882756 |
| KCNE4 | 106.937198 | -0.72204809 | 0.39345493 | -1.8351481 | 0.06648374 | 0.38523695 |
| WDFY1 | 79.7442972 | -0.22016481 | 0.25163604 | -0.8749335 | 0.38161007 | 0.75065642 |
| SERPINE2 | 72.9602554 | -1.465779532 | 0.72137972 | -2.0319112 | 0.04216265 | 0.3087494 |
| CUL3 | 180.146935 | -0.050260471 | 0.18614144 | -0.2700123 | 0.78715082 | 0.95129454 |
| DOCK10 | 125.32322 | 0.946487333 | 0.20907833 | 4.52695092 | 5.98E-06 | 0.00109943 |
| IRS1 | 76.0430253 | -0.143577903 | 0.36391927 | -0.3945323 | 0.69318809 | 0.9248682 |
| RHBDD1 | 77.0410176 | -0.388666905 | 0.17905754 | -2.1706257 | 0.02995948 | 0.26304922 |
| COL4A4 | 356.37759 | -0.08177195 | 0.3482677 | -0.2347962 | 0.81436688 | 0.96038841 |
| COL4A3 | 216.752636 | 0.125760329 | 0.41929503 | 0.29993279 | 0.76422842 | 0.94462105 |
| AC097662.1 | 35.0221287 | 0.076321657 | 0.60187708 | 0.12680605 | 0.8990939 | 0.98058672 |
| MFF | 48.1726038 | -0.123384205 | 0.22644937 | -0.5448644 | 0.58584677 | 0.86977533 |
| AGFG1 | 113.777713 | -0.055690272 | 0.25825903 | -0.2156373 | 0.8292705 | 0.96580172 |
| PID1 | 386.984853 | -0.657068358 | 0.53067555 | -1.2381734 | 0.21565177 | 0.61213784 |
| TRIP12 | 236.018896 | -0.198382519 | 0.2140452 | -0.9268253 | 0.35401722 | 0.73059415 |
| SP110 | 37.9170958 | -0.603964816 | 0.56716484 | -1.0648841 | 0.28692838 | 0.67921285 |
| SP140L | 29.0589661 | -0.2578828 | 0.3356415 | -0.7683281 | 0.44229228 | 0.79085304 |

| | | | | | | |
|---|---|---|---|---|---|---|
| SP100 | 187.536218 | -0.590107437 | 0.36707405 | -1.6075978 | 0.10792329 | 0.47350942 |
| CAB39 | 105.640176 | -0.000240012 | 0.21238275 | -0.0011301 | 0.99909832 | 0.99969762 |
| ITM2C | 35.2605438 | 0.144525748 | 0.40526737 | 0.35661827 | 0.7213776 | 0.93444541 |
| PSMD1 | 115.531142 | -0.118865469 | 0.20924224 | -0.5680759 | 0.56998344 | 0.86047829 |
| ARMC9 | 118.775367 | -0.534465142 | 0.29920112 | -1.7863073 | 0.07404952 | 0.40373597 |
| NCL | 71.0029433 | -0.443591212 | 0.41971736 | -1.0568808 | 0.290566 | 0.68179104 |
| PTMA | 402.6873 | -0.717536097 | 0.39978456 | -1.7948069 | 0.07268447 | 0.40072954 |
| PDE6D | 21.9702323 | -0.130618098 | 0.3452113 | -0.3783714 | 0.70515468 | 0.92946306 |
| COPS7B | 19.5857242 | -0.017822597 | 0.26070884 | -0.0683621 | 0.94549741 | 0.99060401 |
| DIS3L2 | 105.094766 | 0.032571249 | 0.22987252 | 0.14169266 | 0.88732277 | 0.97920559 |
| EIF4E2 | 26.4481715 | 0.163774533 | 0.24583699 | 0.66619157 | 0.50528866 | 0.82481485 |
| EFHD1 | 58.3593072 | 0.958598206 | 0.35188017 | 2.72421778 | 0.0064454 | 0.10918078 |
| GIGYF2 | 194.082028 | -0.308538879 | 0.15139627 | -2.0379556 | 0.04155437 | 0.30568877 |
| DGKD | 97.4125272 | -0.46299308 | 0.36041988 | -1.2845936 | 0.19893433 | 0.59696545 |
| USP40 | 82.541617 | 0.156036042 | 0.30613226 | 0.50970141 | 0.61026067 | 0.88163721 |
| SH3BP4 | 22.7353111 | 0.039478638 | 0.3836556 | 0.10290124 | 0.91804135 | 0.98477389 |
| AGAP1 | 532.213146 | -0.121866842 | 0.23475463 | -0.5191243 | 0.60367404 | 0.87903643 |
| COPS8 | 19.5838215 | 0.246396748 | 0.26709315 | 0.92251243 | 0.35626135 | 0.73129235 |
| COL6A3 | 288.861425 | -0.324701305 | 0.47642474 | -0.6815374 | 0.49553148 | 0.82093346 |
| LRRFIP1 | 264.681453 | -0.225319695 | 0.27419014 | -0.8217644 | 0.41121101 | 0.77166383 |
| RAMP1 | 94.6776831 | -0.407689523 | 0.45705317 | -0.8919958 | 0.37239517 | 0.74463512 |
| ILKAP | 61.1823002 | 0.279426604 | 0.18910387 | 1.47763559 | 0.13950534 | 0.52576347 |
| PER2 | 28.9735618 | 0.278390294 | 0.37663299 | 0.73915537 | 0.45981266 | 0.80029179 |
| TRAF3IP1 | 31.6979049 | 0.067284635 | 0.26122321 | 0.25757526 | 0.79673473 | 0.95463848 |
| TWIST2 | 52.5556971 | 0.223284033 | 0.42576953 | 0.52442465 | 0.59998322 | 0.87764265 |
| HDAC4 | 143.241604 | -0.126062562 | 0.30563383 | -0.4124627 | 0.6800003 | 0.91860979 |
| NDUFA10 | 69.1014797 | 0.137084602 | 0.19896429 | 0.68899098 | 0.49082894 | 0.81720854 |
| COPS9 | 21.7090816 | -0.565853209 | 0.33293551 | -1.699588 | 0.08920845 | 0.43560988 |
| GPC1 | 36.1172286 | -1.273509991 | 0.43283806 | -2.942232 | 0.00325856 | 0.07300955 |
| ANKMY1 | 24.3123391 | 0.091419574 | 0.27749051 | 0.32945117 | 0.74181469 | 0.9373446 |
| SNED1 | 217.430956 | -0.349774563 | 0.39282052 | -0.8904182 | 0.37324135 | 0.74512029 |
| AC005237.1 | 27.0567925 | -0.822802539 | 0.39640365 | -2.0756684 | 0.03792463 | 0.29501902 |
| MTERF4 | 36.7213754 | -0.064114723 | 0.26814765 | -0.2391023 | 0.81102625 | 0.95967025 |
| PPP1R7 | 22.8092299 | 0.052814573 | 0.28953866 | 0.1824094 | 0.85526145 | 0.97379567 |
| HDLBP | 149.136439 | -0.264370529 | 0.19296006 | -1.370079 | 0.17066224 | 0.56802392 |
| 2-Sep | 200.326788 | -0.046186297 | 0.17671952 | -0.2613537 | 0.79381978 | 0.95416774 |
| FARP2 | 91.6017609 | 0.21150761 | 0.15645355 | 1.35188753 | 0.1764113 | 0.5732874 |
| STK25 | 25.2455432 | -0.010180795 | 0.26841263 | -0.0379296 | 0.96974378 | 0.99300191 |
| THAP4 | 23.9708391 | -0.242395499 | 0.28115032 | -0.8621562 | 0.38860156 | 0.75549104 |
| ATG4B | 47.2386885 | -0.291249546 | 0.24070748 | -1.209973 | 0.22628926 | 0.62072404 |
| ING5 | 37.6204791 | 0.130213079 | 0.22602984 | 0.57608801 | 0.56455569 | 0.85721483 |
| D2HGDH | 47.1292886 | 0.126582596 | 0.3115039 | 0.40635958 | 0.68447841 | 0.91977784 |
| LINC01266 | 75.0498984 | 0.563524338 | 0.40571517 | 1.38896542 | 0.16484327 | 0.55965519 |
| CNTN4 | 307.44155 | 0.813177099 | 0.45611859 | 1.78281947 | 0.0746157 | 0.4046869 |
| TRNT1 | 32.678445 | -0.33598846 | 0.29619853 | -1.1343353 | 0.25665391 | 0.65084527 |
| CRBN | 45.1574896 | -0.120657256 | 0.29568395 | -0.4080616 | 0.68322848 | 0.91933701 |
| AC034195.1 | 34.7709645 | 0.386162878 | 0.23763801 | 1.62500465 | 0.10416157 | 0.46422585 |
| SUMF1 | 183.549337 | 0.011642291 | 0.23428462 | 0.04969294 | 0.96036708 | 0.99181157 |
| ITPR1 | 313.327295 | 0.05891704 | 0.28523975 | 0.2065527 | 0.8363592 | 0.9668633 |
| AC018816.1 | 25.8357124 | -0.352551578 | 0.39071561 | -0.9023227 | 0.36688546 | 0.73896496 |
| BHLHE40 | 99.5856254 | -1.107404828 | 0.48990238 | -2.2604602 | 0.02379271 | 0.22886465 |
| ARL8B | 101.882221 | -0.306643159 | 0.20612625 | -1.4876473 | 0.13684392 | 0.52089048 |
| EDEM1 | 28.8852572 | -0.603327132 | 0.37644524 | -1.6026956 | 0.10900187 | 0.47636267 |
| LMCD1-AS1 | 123.988601 | -0.240528664 | 0.38097798 | -0.6313453 | 0.52781477 | 0.83947444 |
| LMCD1 | 244.597057 | -0.868917495 | 0.51054527 | -1.7019402 | 0.08876659 | 0.43544541 |
| RAD18 | 24.0377588 | 0.102338632 | 0.26789431 | 0.38201122 | 0.70245304 | 0.92865476 |
| THUMPD3-A | 67.7286399 | 0.153991318 | 0.21296466 | 0.72308391 | 0.46962833 | 0.80412363 |
| SETD5 | 306.451143 | -0.387486782 | 0.17746512 | -2.1834531 | 0.02900246 | 0.25741431 |
| MTMR14 | 29.3001114 | -0.188531001 | 0.33364894 | -0.565058 | 0.57203432 | 0.86236719 |
| OGG1 | 19.6121498 | 0.17906044 | 0.37449604 | 0.47813707 | 0.63255265 | 0.8948276 |
| TADA3 | 18.7814344 | 0.48416374 | 0.29747943 | 1.62755366 | 0.10361956 | 0.4637638 |
| EMC3 | 34.2861064 | -0.238595763 | 0.24412132 | -0.9773655 | 0.32838822 | 0.71294249 |
| BRK1 | 35.7342726 | -0.362075632 | 0.3698675 | -0.9789333 | 0.32761292 | 0.71282522 |
| VHL | 18.0992148 | -0.246504355 | 0.28145859 | -0.8758104 | 0.38113313 | 0.74991898 |
| IRAK2 | 25.2018737 | -0.766399067 | 0.60780152 | -1.2609364 | 0.20733176 | 0.6059796 |
| SEC13 | 44.4412599 | -0.267656872 | 0.28679686 | -0.9332629 | 0.35068424 | 0.72904624 |
| ATG7 | 197.90911 | -0.383691331 | 0.25630633 | -1.4970029 | 0.13439249 | 0.51662128 |
| VGLL4 | 182.053498 | 0.120237044 | 0.19956675 | 0.60249038 | 0.54684777 | 0.84758329 |
| TAMM41 | 41.219748 | 0.075243522 | 0.24154622 | 0.31150776 | 0.75541464 | 0.94189722 |
| PPARG | 229.40748 | 0.05057783 | 0.24519441 | 0.20627644 | 0.83657498 | 0.9668633 |
| TSEN2 | 39.2152292 | -0.375293109 | 0.32088499 | -1.1695564 | 0.24217952 | 0.63672023 |
| RAF1 | 130.128882 | 0.060774812 | 0.16477631 | 0.36883222 | 0.71225279 | 0.93104691 |
| RPL32 | 158.009742 | -0.33999822 | 0.33388205 | -1.0183184 | 0.30852668 | 0.69885506 |
| IQSEC1 | 40.4878658 | 0.400359815 | 0.29130418 | 1.3743703 | 0.16932675 | 0.56570963 |
| XPC | 51.1121155 | 0.119033051 | 0.19811493 | 0.60082829 | 0.54795436 | 0.84785354 |
| SLC6A6 | 25.4717339 | -0.96525302 | 0.43661375 | -2.210771 | 0.0270517 | 0.24634813 |
| GRIP2 | 22.6314501 | 1.299993837 | 0.43474618 | 2.99023638 | 0.00278762 | 0.0676112 |
| CCDC174 | 23.3756011 | 0.2263247 | 0.31026784 | 0.72944943 | 0.46572679 | 0.80355051 |
| FGD5 | 43.681471 | -0.152693925 | 0.56023997 | -0.2725509 | 0.78519844 | 0.95065355 |
| FGD5-AS1 | 43.5976904 | 0.44717149 | 0.30871331 | 1.44850086 | 0.14747703 | 0.53679369 |
| NR2C2 | 83.8772651 | 0.176299197 | 0.19137876 | 0.92120568 | 0.35694305 | 0.73129235 |
| MRPS25 | 40.3965261 | -0.294358637 | 0.30702984 | -0.9587297 | 0.33769492 | 0.71925477 |
| RBSN | 32.4027888 | -0.001648809 | 0.2045429 | -0.0080609 | 0.99356836 | 0.99805002 |
| CAPN7 | 120.57078 | -0.236324403 | 0.16261728 | -1.4532552 | 0.14615291 | 0.53584686 |
| SH3BP5 | 205.882824 | -1.546220235 | 0.4604024 | -3.3584105 | 0.00078392 | 0.03268107 |
| METTL6 | 21.4611982 | -0.319817603 | 0.37609935 | -0.8503541 | 0.39512827 | 0.76015646 |
| COLQ | 23.0372593 | -0.659047563 | 0.5385949 | -1.2236424 | 0.22108715 | 0.61653656 |
| HACL1 | 30.3551948 | -0.149957992 | 0.25314387 | -0.5923825 | 0.5535945 | 0.85188037 |
| ANKRD28 | 105.110509 | 0.215819199 | 0.388883 | 0.55497205 | 0.57891377 | 0.86486944 |
| OXNAD1 | 40.9394403 | -0.393753356 | 0.24020917 | -1.6392104 | 0.10116946 | 0.46151108 |
| RFTN1 | 277.413664 | 0.048643079 | 0.30866624 | 0.15759119 | 0.87477895 | 0.9768643 |
| PLCL2 | 143.906885 | 0.195946858 | 0.2632623 | 0.74430276 | 0.45669333 | 0.79732984 |
| TBC1D5 | 371.804628 | 0.188820248 | 0.25108997 | 0.75200235 | 0.45204965 | 0.79476384 |
| RAB5A | 71.8586019 | -0.127888288 | 0.19979719 | -0.6400905 | 0.52211375 | 0.8350836 |
| KAT2B | 41.6562098 | 0.702254368 | 0.37068164 | 1.89449462 | 0.05815939 | 0.36041506 |
| ZNF385D | 41.6043777 | 0.522980245 | 0.48601565 | 1.0760564 | 0.28190203 | 0.676814 |
| UBE2E2 | 583.657888 | -0.11949102 | 0.22032034 | -0.5423513 | 0.58757654 | 0.870249 |
| UBE2E1 | 189.240978 | -0.098446872 | 0.17830367 | -0.5521304 | 0.58085903 | 0.86639598 |
| RPL15 | 190.78595 | -0.660516239 | 0.34869274 | -1.8942644 | 0.05818993 | 0.36041506 |
| NR1D2 | 52.1098554 | 0.428959232 | 0.42829687 | 1.0015465 | 0.31656267 | 0.70433517 |
| THRB | 228.285467 | 0.614205909 | 0.35422446 | 1.73394548 | 0.08292776 | 0.42321951 |
| RARB | 253.217727 | 0.229493144 | 0.46466532 | 0.49388912 | 0.62138449 | 0.88729641 |
| TOP2B | 112.845103 | -0.022814639 | 0.15356395 | -0.1485677 | 0.88189478 | 0.97782528 |
| NGLY1 | 63.0082121 | -1.043437184 | 0.35689616 | -2.9236436 | 0.00345961 | 0.07625361 |
| SLC4A7 | 122.413459 | -0.400594357 | 0.39535337 | -1.0132565 | 0.31093767 | 0.70126123 |
| CMC1 | 76.4457561 | -0.243449524 | 0.34062421 | -0.7147159 | 0.47478463 | 0.80851131 |
| AZI2 | 108.74391 | -1.140146029 | 0.46455069 | -2.4542984 | 0.01411598 | 0.17435012 |
| RBMS3 | 825.268541 | -0.371523488 | 0.24546908 | -1.5135246 | 0.13014646 | 0.51204187 |
| RBMS3-AS3 | 310.651479 | -0.408558911 | 0.47781213 | -0.8550618 | 0.39251693 | 0.75930689 |
| RBMS3-AS2 | 89.89444 | -0.574386578 | 0.47877212 | -1.1997077 | 0.2302529 | 0.62447176 |
| TGFBR2 | 288.570395 | -0.098807859 | 0.29223094 | -0.3381157 | 0.73527603 | 0.93594619 |
| STT3B | 162.846124 | 0.217473393 | 0.23829118 | 0.91263718 | 0.36143339 | 0.73354709 |
| OSBPL10 | 219.278038 | 0.561924619 | 0.39358042 | 1.42772503 | 0.15337102 | 0.54114675 |

| | | | | | | |
|---|---|---|---|---|---|---|
| CMTM8 | 48.238028 | -1.57770463 | 0.5418085 | -2.911923 | 0.00359211 | 0.07758886 |
| CMTM7 | 30.6493812 | -0.222934258 | 0.3397953 | -0.656084 | 0.51177008 | 0.82882457 |
| CMTM6 | 39.9859304 | 0.011081962 | 0.23934462 | 0.04630128 | 0.96307012 | 0.99231823 |
| DYNC1LI1 | 37.8680542 | 0.262789395 | 0.22724048 | 1.15643741 | 0.24750229 | 0.64317338 |
| CNOT10 | 57.6427918 | 0.067284085 | 0.221768 | 0.30339853 | 0.76158616 | 0.94425408 |
| GLB1 | 43.6223403 | 0.061718798 | 0.27630477 | 0.22337218 | 0.82324586 | 0.96413949 |
| CRTAP | 42.9092639 | -0.195618065 | 0.28178124 | -0.6942196 | 0.48754449 | 0.81595238 |
| FBXL2 | 37.4639699 | 0.849457855 | 0.42847393 | 1.98251934 | 0.04742114 | 0.3297075 |
| UBP1 | 51.7032288 | 0.062495382 | 0.2990291 | 0.20899431 | 0.83445268 | 0.9666365 |
| CLASP2 | 230.415796 | 0.306854377 | 0.29411818 | 1.04330301 | 0.29680798 | 0.68830604 |
| PDCD6IP | 105.609228 | -0.031371417 | 0.20226665 | -0.1550993 | 0.87674303 | 0.9768643 |
| STAC | 54.5655034 | 2.03357989 | 0.40952521 | 4.96570134 | 6.85E-07 | 0.00018632 |
| TRANK1 | 70.4973724 | -0.216191273 | 0.7518981 | -0.2875274 | 0.77370856 | 0.94750611 |
| EPM2AIP1 | 37.0036524 | -0.189878061 | 0.19658956 | -0.9658604 | 0.33411406 | 0.71686823 |
| MLH1 | 29.6664825 | -0.053388446 | 0.23112655 | -0.2309923 | 0.81732081 | 0.96134613 |
| LRRFIP2 | 175.520089 | 0.216691257 | 0.28184151 | 0.76884082 | 0.44198781 | 0.79069339 |
| GOLGA4 | 366.431203 | -0.533745774 | 0.21602455 | -2.4707645 | 0.01348246 | 0.16974928 |
| ITGA9 | 105.362263 | 0.798875882 | 0.43501836 | 1.83641879 | 0.06629574 | 0.38483998 |
| CTDSPL | 136.821109 | 0.285891319 | 0.30829256 | 0.92733773 | 0.35375121 | 0.73059415 |
| ACAA1 | 20.7260597 | -0.218873913 | 0.33798198 | -0.6475905 | 0.51724985 | 0.83269861 |
| OXSR1 | 68.8536437 | -0.195988011 | 0.2735931 | -0.7163485 | 0.47377618 | 0.80789916 |
| EXOG | 28.8988346 | 0.0235974 | 0.22888046 | 0.10309923 | 0.91788422 | 0.98477389 |
| WDR48 | 36.3002466 | 0.154716095 | 0.31314109 | 0.49407792 | 0.62125115 | 0.88729641 |
| RPSA | 90.5994957 | -0.360227381 | 0.3212855 | -1.1212065 | 0.26219999 | 0.65630111 |
| EIF1B | 28.4581746 | -0.397186415 | 0.31592174 | -1.2572304 | 0.20867021 | 0.6059796 |
| RPL14 | 157.074004 | -0.405257891 | 0.39197774 | -1.0338798 | 0.30119234 | 0.69248569 |
| ZNF621 | 36.9257151 | -0.236164802 | 0.27451072 | -0.8603118 | 0.38961722 | 0.75628551 |
| CTNNB1 | 166.931152 | -0.082093887 | 0.24064892 | -0.3411355 | 0.73300158 | 0.93574285 |
| ULK4 | 109.57767 | -0.200484497 | 0.33888219 | -0.5916053 | 0.55411493 | 0.85191704 |
| TRAK1 | 134.857193 | 0.592431377 | 0.29221854 | 2.02735723 | 0.04262589 | 0.31108012 |
| SEC22C | 24.4749889 | 0.203834022 | 0.32255758 | 0.63193064 | 0.5274322 | 0.83938843 |
| NKTR | 202.169218 | 0.112124778 | 0.18317317 | 0.61212446 | 0.54045542 | 0.84578923 |
| AC006059.1 | 37.1143707 | -0.57203257 | 0.3957805 | -1.4453278 | 0.14836582 | 0.53680621 |
| SNRK | 131.691433 | 0.983290524 | 0.2351023 | 4.18239433 | 2.88E-05 | 0.00375164 |
| ANO10 | 110.13766 | 0.397723918 | 0.26593239 | 1.49558283 | 0.1347624 | 0.51743411 |
| ABHD5 | 27.3411875 | -0.043705998 | 0.38817918 | -0.1125923 | 0.91035377 | 0.98343817 |
| TCAIM | 56.6542428 | -0.379306703 | 0.30007994 | -1.2640189 | 0.20622325 | 0.6050285 |
| ZNF445 | 28.4327939 | -0.55075883 | 0.31875492 | -1.7278442 | 0.08401617 | 0.4256693 |
| AC099669.1 | 36.6162314 | 0.081644638 | 0.26481512 | 0.30830807 | 0.75784793 | 0.94301125 |
| ZNF197 | 34.592158 | -0.077175141 | 0.36603511 | -0.2108408 | 0.83301149 | 0.9666365 |
| ZDHHC3 | 33.3964887 | 0.164299685 | 0.22412634 | 0.73306727 | 0.4635174 | 0.80225845 |
| LARS2 | 42.3883385 | -0.49329386 | 0.27547231 | -1.7907203 | 0.07333819 | 0.40189622 |
| LIMD1 | 51.6824679 | -0.164894467 | 0.25944924 | -0.6355558 | 0.52506599 | 0.83726969 |
| SACM1L | 44.820671 | 0.10462886 | 0.26903765 | 0.38890044 | 0.6973498 | 0.92702563 |
| FYCO1 | 29.0108657 | -0.466407841 | 0.25734626 | -1.8123747 | 0.06992833 | 0.39397538 |
| LTF | 29.6227315 | -4.276859097 | 1.21603402 | -3.5170555 | 0.00043636 | 0.02145813 |
| PTH1R | 101.95796 | 0.416678744 | 0.32572826 | 1.27922197 | 0.2008189 | 0.59975737 |
| CCDC12 | 34.3791158 | -0.039915077 | 0.24003269 | -0.1662902 | 0.8679286 | 0.97648611 |
| SETD2 | 238.193715 | -0.047247797 | 0.14389484 | -0.3283495 | 0.74264744 | 0.93759079 |
| KIF9-AS1 | 46.4231824 | -0.171595944 | 0.25005749 | -0.686226 | 0.49257063 | 0.818797 |
| KLHL18 | 23.7447594 | 0.504024312 | 0.27950681 | 1.8032631 | 0.07134691 | 0.39729887 |
| PTPN23 | 19.8374331 | -0.18345059 | 0.30555049 | -0.6003937 | 0.54824388 | 0.84786286 |
| SCAP | 32.8131283 | 0.101139258 | 0.2462533 | 0.4107123 | 0.68128351 | 0.91886128 |
| SMARCC1 | 131.096829 | 0.019955169 | 0.20062113 | 0.09946693 | 0.92076754 | 0.98528826 |
| DHX30 | 34.5880694 | 0.206875222 | 0.20449667 | 1.01163123 | 0.31171441 | 0.70140514 |
| MAP4 | 373.349329 | 0.006735541 | 0.2238366 | 0.03009133 | 0.97599422 | 0.99343234 |
| PLXNB1 | 25.2405977 | 0.595026213 | 0.28678104 | 2.07484503 | 0.0380009 | 0.29501902 |
| TMA7 | 42.4870045 | 0.018607799 | 0.37080155 | 0.05018264 | 0.95997685 | 0.99181157 |
| SHISA5 | 54.8668707 | -0.137620524 | 0.25166081 | -0.5468493 | 0.5844823 | 0.86845136 |
| UQCRC1 | 19.062994 | 0.140553628 | 0.34889776 | 0.40285048 | 0.68705823 | 0.92100905 |
| NCKIPSD | 19.6879982 | -0.197832101 | 0.30028068 | -0.6588239 | 0.51000883 | 0.82720258 |
| IP6K2 | 52.4687692 | 0.146442638 | 0.2609808 | 0.56112419 | 0.57471288 | 0.86294056 |
| PRKAR2A | 113.265236 | 0.113278548 | 0.22784024 | 0.49718411 | 0.61905923 | 0.8859718 |
| ARIH2 | 95.8184646 | -0.221147917 | 0.19093031 | -1.1582651 | 0.24675587 | 0.64247811 |
| P4HTM | 18.629156 | 0.236531367 | 0.25754633 | 0.91840319 | 0.35840782 | 0.73144657 |
| WDR6 | 31.5342132 | 0.337462846 | 0.26637976 | 1.26684867 | 0.20520941 | 0.60370888 |
| QRICH1 | 51.4255816 | 0.009328958 | 0.22707315 | 0.04108349 | 0.96722933 | 0.99300191 |
| LAMB2 | 63.6222168 | 0.376131987 | 0.28189765 | 1.33428566 | 0.18211025 | 0.58094117 |
| USP4 | 54.14094 | 0.449525212 | 0.2490928 | 1.80464957 | 0.07112954 | 0.39721198 |
| RHOA | 138.424027 | 0.188839693 | 0.26917353 | 0.70155374 | 0.48295751 | 0.81409616 |
| DAG1 | 36.3672421 | 0.330301661 | 0.30938946 | 1.06759182 | 0.28570468 | 0.6783522 |
| IP6K1 | 68.9981601 | -0.030652479 | 0.23143162 | -0.1324472 | 0.89463055 | 0.98023547 |
| RBM6 | 306.095854 | 0.090339408 | 0.13430751 | 0.67263107 | 0.50118203 | 0.82406312 |
| RBM5 | 110.87633 | 0.261105678 | 0.21096781 | 1.23765647 | 0.21584347 | 0.61220904 |
| GNAI2 | 64.6716324 | 0.226807876 | 0.33252603 | 0.68207555 | 0.49519118 | 0.82073974 |
| RASSF1 | 32.4479833 | 0.033457119 | 0.26340277 | 0.12701886 | 0.89892547 | 0.98058672 |
| DOCK3 | 28.1324805 | -0.348687628 | 0.34462759 | -1.0117809 | 0.31164281 | 0.70140514 |
| MANF | 21.6869736 | -0.337203885 | 0.45186381 | -0.7462511 | 0.45551572 | 0.79609632 |
| DCAF1 | 40.6531751 | -0.009415397 | 0.35759038 | -0.0263301 | 0.97899403 | 0.99465087 |
| RAD54L2 | 49.3869978 | -0.019409587 | 0.24418676 | -0.0794866 | 0.93664555 | 0.98827804 |
| RPL29 | 200.757957 | -0.803976969 | 0.35141037 | -2.2878578 | 0.0221458 | 0.21963494 |
| WDR82 | 52.2171572 | -0.020260993 | 0.23531405 | -0.0861019 | 0.93138539 | 0.98681039 |
| NISCH | 57.4217652 | 0.183490096 | 0.18988267 | 0.96633409 | 0.33387703 | 0.71681633 |
| NT5DC2 | 29.5752206 | 0.244322788 | 0.51922339 | 0.47055428 | 0.63795907 | 0.89779507 |
| SMIM4 | 24.8709781 | -0.339882725 | 0.24856242 | -1.3673938 | 0.1715019 | 0.56870244 |
| PBRM1 | 149.083909 | -0.026923662 | 0.19611054 | -0.1372882 | 0.890803 | 0.98023547 |
| GNL3 | 31.3857947 | -0.986526258 | 0.33501925 | -2.9446853 | 0.00323283 | 0.07300955 |
| SPCS1 | 40.1640617 | -0.16195531 | 0.28702292 | -0.5642592 | 0.57257777 | 0.86236719 |
| NEK4 | 30.1551435 | 0.074879791 | 0.26869603 | 0.27867844 | 0.78049161 | 0.94916975 |
| TKT | 40.6378797 | -0.717079949 | 0.45924925 | -1.5614178 | 0.11842521 | 0.49281248 |
| DCP1A | 86.1601582 | -0.016155004 | 0.18521735 | -0.0872219 | 0.93049515 | 0.98681039 |
| CACNA1D | 49.086269 | 1.0871167 | 0.56372212 | 1.92846202 | 0.05379768 | 0.34959616 |
| SELENOK | 26.1200888 | -0.503390503 | 0.31814895 | -1.5822479 | 0.11359299 | 0.48395978 |
| CACNA2D3 | 70.0457348 | 1.013993133 | 0.37019354 | 2.7390892 | 0.00616097 | 0.10632774 |
| CCDC66 | 68.6814894 | 0.253632557 | 0.18665922 | 1.35880007 | 0.17420995 | 0.57154863 |
| FAM208A | 92.1050076 | 0.31894999 | 0.23451098 | 1.36006421 | 0.17380961 | 0.57125528 |
| ARHGEF3 | 70.2104573 | 0.129859389 | 0.29952026 | 0.43355794 | 0.66460948 | 0.91201464 |
| APPL1 | 54.7742846 | 0.27291097 | 0.21021528 | 1.29824519 | 0.19420309 | 0.59269042 |
| ARF4 | 76.6355566 | -0.27983346 | 0.27336386 | -1.0236666 | 0.30599276 | 0.69728397 |
| DENND6A | 98.8500601 | 0.240634389 | 0.19051921 | 1.26304526 | 0.20657292 | 0.60530477 |
| DENND6A-AS | 20.9852435 | -0.964765325 | 0.39080137 | -2.4686846 | 0.01356107 | 0.17006877 |
| SLMAP | 177.347417 | 0.16239744 | 0.16277559 | 0.99767689 | 0.31843606 | 0.70683013 |
| FLNB | 147.115963 | 1.018190222 | 0.46181584 | 2.20475378 | 0.02747138 | 0.24847431 |
| PXK | 73.4491776 | -0.497708453 | 0.34709901 | -1.4339092 | 0.15159819 | 0.53898138 |
| FHIT | 344.323492 | -0.479180692 | 0.40731831 | -1.1764281 | 0.23942386 | 0.63451757 |
| PTPRG | 1491.83183 | -0.40708701 | 0.27197112 | -1.4968023 | 0.1344447 | 0.51662128 |
| THOC7 | 22.9214642 | -0.198183817 | 0.27425473 | -0.7226268 | 0.46990921 | 0.80441714 |
| PRICKLE2 | 258.782296 | -0.428811483 | 0.34526035 | -1.2419946 | 0.21423856 | 0.61001472 |
| ADAMTS9 | 1316.19094 | -3.438362637 | 0.5731897 | -5.998647 | 1.99E-09 | 2.44E-06 |
| ADAMTS9-AS | 79.7142397 | -1.423060362 | 0.39531313 | -3.5998308 | 0.00031842 | 0.01772803 |
| ADAMTS9-AS | 756.577248 | 0.240174843 | 0.48701611 | 0.49315585 | 0.62190247 | 0.88762114 |
| MAGI1 | 513.733744 | 0.34079671 | 0.29713131 | 1.14695658 | 0.25139956 | 0.6460368 |
| SLC25A26 | 57.9526315 | -0.224977671 | 0.25604538 | -0.8786633 | 0.37958387 | 0.74991898 |

| | | | | | | |
|---|---|---|---|---|---|---|
| LRIG1 | 41.5959721 | 0.658663911 | 0.42113211 | 1.56403158 | 0.11781015 | 0.49136596 |
| SUCLG2 | 174.262848 | -0.183886394 | 0.17361142 | -1.0591837 | 0.28951611 | 0.67997888 |
| SUCLG2-AS1 | 49.4902192 | 0.770425272 | 0.38441709 | 2.00413899 | 0.04505517 | 0.31898889 |
| EOGT | 34.3213009 | -0.261799039 | 0.38698152 | -0.6765156 | 0.49871334 | 0.82271448 |
| TMF1 | 82.8841953 | -0.306498396 | 0.17841768 | -1.7178701 | 0.08582031 | 0.4287515 |
| UBA3 | 44.0484164 | 0.057844411 | 0.24065054 | 0.24036685 | 0.81004588 | 0.95954661 |
| ARL6IP5 | 55.0150912 | -0.054677024 | 0.32073215 | -0.1704757 | 0.86463608 | 0.9762143 |
| FRMD4B | 41.5574447 | 0.531948802 | 0.3941042 | 1.34976691 | 0.17709076 | 0.57388115 |
| MITF | 196.081991 | -0.511768645 | 0.4795126 | -1.0672684 | 0.28585066 | 0.6783522 |
| FOXP1 | 397.791151 | -0.143922163 | 0.20521836 | -0.7013123 | 0.48310813 | 0.81409616 |
| RYBP | 93.4800751 | -0.001996776 | 0.31462675 | -0.0063465 | 0.99493627 | 0.99854015 |
| SHQ1 | 50.3524302 | -0.557445247 | 0.24690976 | -2.2576881 | 0.02396511 | 0.23001094 |
| GXYLT2 | 68.1875166 | -1.079731796 | 0.3564911 | -3.0287763 | 0.00245546 | 0.062538 |
| PPP4R2 | 104.207341 | -0.309466851 | 0.17400317 | -1.7785127 | 0.07531969 | 0.40760265 |
| PDZRN3 | 370.349675 | 0.053976497 | 0.30344043 | 0.17788169 | 0.85881589 | 0.97397549 |
| CNTN3 | 59.9761564 | 1.916758989 | 0.31727559 | 6.04130622 | 1.53E-09 | 2.25E-06 |
| ZNF717 | 31.0029861 | -0.074407767 | 0.29920703 | -0.2486832 | 0.80360583 | 0.95836189 |
| ROBO2 | 312.871939 | 0.939020118 | 0.47659694 | 1.9702605 | 0.04880852 | 0.33522789 |
| ROBO1 | 686.543007 | 0.28326642 | 0.55614087 | 0.50934293 | 0.61051187 | 0.88163721 |
| GBE1 | 408.134616 | -1.900900308 | 0.41538582 | -4.5762282 | 4.73E-06 | 0.00094034 |
| VGLL3 | 66.6346405 | 1.125988859 | 0.85736275 | 1.31331675 | 0.18907624 | 0.58671246 |
| CHMP2B | 27.9409648 | -0.287947356 | 0.27455288 | -1.0487865 | 0.29427639 | 0.68566952 |
| CGGBP1 | 73.3820228 | -0.070907437 | 0.19670586 | -0.3604745 | 0.71849236 | 0.93432517 |
| ZNF654 | 111.506685 | -0.565402992 | 0.29105905 | -1.9425714 | 0.05206797 | 0.34566169 |
| C3orf38 | 20.6522797 | -0.044442002 | 0.25233072 | -0.176126 | 0.86019495 | 0.97397549 |
| EPHA3 | 68.224183 | 0.628959564 | 0.43519265 | 1.44524401 | 0.14838936 | 0.53680621 |
| PROS1 | 88.4284069 | -0.970409661 | 0.37634616 | -2.5785029 | 0.00992295 | 0.14050816 |
| ARL13B | 63.3436273 | 0.072279165 | 0.24645527 | 0.29327499 | 0.76931197 | 0.94579017 |
| NSUN3 | 34.1541194 | -0.351015014 | 0.26848889 | -1.3073726 | 0.19108621 | 0.58975955 |
| MTRNR2L12 | 110.390965 | -0.762117173 | 0.35629523 | -2.1390047 | 0.03243529 | 0.27525049 |
| ARL6 | 24.7798865 | 0.705497683 | 0.31827338 | 2.21664053 | 0.02664767 | 0.24439315 |
| CRYBG3 | 110.951366 | -0.568478919 | 0.26986174 | -2.1065562 | 0.03515607 | 0.28643233 |
| CLDND1 | 36.7691346 | 0.165873983 | 0.28061901 | 0.59110032 | 0.55445321 | 0.85191704 |
| DCBLD2 | 102.716321 | 0.470423701 | 0.26939577 | 1.74621785 | 0.0807731 | 0.42027326 |
| COL8A1 | 131.552142 | -1.159878715 | 0.40229197 | -2.8831764 | 0.00393687 | 0.08179072 |
| CMSS1 | 194.651627 | -0.77913758 | 0.3153662 | -2.4705805 | 0.0134894 | 0.16974928 |
| FILIP1L | 204.640719 | 0.578118178 | 0.42396904 | 1.36358582 | 0.17269796 | 0.56912883 |
| TBC1D23 | 73.8916348 | -0.199900342 | 0.2608303 | -0.7664 | 0.44343833 | 0.7920275 |
| TOMM70 | 36.4238489 | 0.035630046 | 0.24766988 | 0.14386104 | 0.8856102 | 0.97867793 |
| TMEM45A | 50.6221228 | -1.836532883 | 0.52316398 | -3.5104345 | 0.00044738 | 0.02157489 |
| TFG | 106.458059 | -0.360212694 | 0.22179748 | -1.6240613 | 0.10436273 | 0.46422585 |
| ABI3BP | 116.018436 | -0.173864024 | 0.88295358 | -0.1969119 | 0.84389652 | 0.9691438 |
| SENP7 | 120.711069 | 0.322809344 | 0.22708966 | 1.42150615 | 0.15516967 | 0.54412824 |
| PCNP | 77.1348311 | -0.242995898 | 0.19843951 | -1.2245339 | 0.2207509 | 0.61590675 |
| ZBTB11 | 29.2102302 | -0.080124233 | 0.24783429 | -0.3232976 | 0.74646987 | 0.93900586 |
| RPL24 | 72.0436646 | -0.277573578 | 0.32598577 | -0.8514899 | 0.3944973 | 0.75988253 |
| CEP97 | 18.5158428 | -0.03160133 | 0.28250932 | -0.1118594 | 0.91093487 | 0.98343817 |
| NFKBIZ | 64.8238659 | -0.394707706 | 0.41317969 | -0.9552931 | 0.3394295 | 0.71925477 |
| ALCAM | 68.8607792 | -0.314232211 | 0.3432433 | -0.9154795 | 0.35993995 | 0.73273991 |
| CBLB | 806.634651 | -0.582970563 | 0.34691913 | -1.6804221 | 0.09287522 | 0.44320781 |
| LINC00882 | 152.385195 | 1.202167067 | 0.46204533 | 2.60183795 | 0.00927257 | 0.13493878 |
| DUBR | 35.770605 | 0.586411285 | 0.42014195 | 1.39574562 | 0.16279111 | 0.55718935 |
| BBX | 282.250827 | -0.439277314 | 0.22404541 | -1.960662 | 0.04991847 | 0.33748407 |
| CD47 | 68.4070305 | -0.099104445 | 0.32457343 | -0.3053375 | 0.76010911 | 0.94422614 |
| IFT57 | 36.5148899 | -0.025203493 | 0.32578773 | -0.0773617 | 0.93833581 | 0.98827804 |
| DZIP3 | 42.586098 | 0.136009202 | 0.34803374 | 0.39079315 | 0.69595014 | 0.92648248 |
| NECTIN3 | 49.8863241 | 0.258167622 | 0.27954093 | 0.92354139 | 0.35572514 | 0.73125149 |
| PHLDB2 | 322.957093 | 0.43918074 | 0.35084812 | 1.25176882 | 0.21065411 | 0.60733506 |
| ATG3 | 19.8104439 | -0.388380883 | 0.2773245 | -1.4004564 | 0.16137668 | 0.55470404 |
| CCDC80 | 116.402825 | 0.8318167 | 0.41526426 | 2.00310208 | 0.04516633 | 0.31946813 |
| NEPRO | 31.5433754 | -0.299939664 | 0.27428171 | -1.093546 | 0.27415414 | 0.66930898 |
| BOC | 52.2937027 | -0.855169877 | 0.59420431 | -1.4391849 | 0.15009814 | 0.53880252 |
| CFAP44 | 35.5900521 | 0.241447004 | 0.24265656 | 0.99501536 | 0.31972879 | 0.70688535 |
| SPICE1 | 27.680948 | -0.625628034 | 0.2933271 | -2.1328681 | 0.03293554 | 0.27693743 |
| USF3 | 35.9469164 | 0.281312237 | 0.2179226 | 1.29088141 | 0.1967448 | 0.59452202 |
| NAA50 | 77.4257713 | -0.144231599 | 0.21058625 | -0.6849051 | 0.49340381 | 0.81908904 |
| ATP6V1A | 67.9679582 | -0.115818454 | 0.20682407 | -0.5599854 | 0.57548941 | 0.86294056 |
| ZBTB20 | 2127.7386 | 0.221975157 | 0.33144942 | 0.6697105 | 0.50304236 | 0.82430366 |
| ZBTB20-AS5 | 35.2611028 | -0.370422853 | 0.59189787 | -0.6258222 | 0.53143154 | 0.84190241 |
| LSAMP | 319.511104 | 1.20879115 | 0.63363112 | 1.9077206 | 0.05642734 | 0.35564709 |
| AC092691.1 | 109.8068 | 2.403139036 | 0.88093891 | 2.72792927 | 0.00637333 | 0.10918078 |
| ARHGAP31 | 51.6843608 | 0.102875928 | 0.3041501 | 0.33824066 | 0.73518184 | 0.93594619 |
| TMEM39A | 38.9526364 | -0.737544889 | 0.25388293 | -2.905059 | 0.00367184 | 0.0789016 |
| POGLUT1 | 25.2632748 | -0.175068039 | 0.3123579 | -0.5604726 | 0.57515713 | 0.86294056 |
| TIMMDC1 | 34.1352029 | -0.040659974 | 0.24340952 | -0.1670435 | 0.86733584 | 0.97634923 |
| COX17 | 19.0081812 | 0.099773742 | 0.27824492 | 0.35858243 | 0.71990749 | 0.93444541 |
| GSK3B | 241.803181 | -0.174305482 | 0.14260614 | -1.2222859 | 0.22159952 | 0.61733695 |
| LRRC58 | 45.1844261 | -0.000999145 | 0.21147492 | -0.0047247 | 0.99623029 | 0.99854015 |
| FSTL1 | 270.467506 | 0.002931891 | 0.30560023 | 0.00959388 | 0.99234531 | 0.99759781 |
| NDUFB4 | 36.3787228 | -0.478748094 | 0.36481141 | -1.3123167 | 0.18941331 | 0.58678196 |
| RABL3 | 20.8477679 | -0.490857944 | 0.3252385 | -1.5092246 | 0.1312414 | 0.51290135 |
| HCLS1 | 25.5704498 | -1.02281227 | 0.6001454 | -1.7042741 | 0.08832989 | 0.43544541 |
| GOLGB1 | 151.277046 | -0.351760891 | 0.19295073 | -1.8230607 | 0.0682942 | 0.38936702 |
| IQCB1 | 60.7162769 | -0.290132968 | 0.23667146 | -1.2258891 | 0.22024041 | 0.61590675 |
| CASR | 41.0997923 | 1.623926515 | 0.93188766 | 1.74262047 | 0.08139993 | 0.42087829 |
| FAM162A | 62.8510844 | -0.842355261 | 0.51516356 | -1.635122 | 0.10202348 | 0.46169744 |
| AC083798.2 | 21.5212947 | 0.011239481 | 0.34218312 | 0.03284639 | 0.97379708 | 0.99300191 |
| KPNA1 | 103.242722 | 0.093296744 | 0.17993994 | 0.51848826 | 0.60411765 | 0.87903643 |
| PARP9 | 71.5027086 | -0.23774834 | 0.73892732 | -0.3217479 | 0.74764366 | 0.93903859 |
| DTX3L | 26.5719767 | -0.420718444 | 0.49313763 | -0.8531461 | 0.3935783 | 0.75988253 |
| PARP14 | 211.886132 | -0.13568791 | 0.83017535 | -0.1634449 | 0.87016817 | 0.9768643 |
| HSPBAP1 | 29.0765411 | -0.795127467 | 0.26874774 | -2.9586387 | 0.00309001 | 0.07118649 |
| DIRC2 | 42.557835 | -0.591725166 | 0.34414904 | -1.7193864 | 0.08554404 | 0.42848827 |
| PDIA5 | 43.81716 | -0.184105662 | 0.33589634 | -0.5481026 | 0.58362144 | 0.86804978 |
| SEC22A | 86.2294124 | -0.036556616 | 0.21382128 | -0.1709681 | 0.86424885 | 0.9762143 |
| ADCY5 | 88.9281506 | 0.469519253 | 0.27233165 | 1.72407157 | 0.08469492 | 0.42695086 |
| HACD2 | 30.6936405 | -0.42597268 | 0.29672066 | -1.4356017 | 0.15111571 | 0.53884006 |
| MYLK | 329.259989 | 0.787832122 | 0.38481886 | 2.04728044 | 0.04063056 | 0.30252681 |
| CCDC14 | 90.6089524 | 0.038328607 | 0.29169886 | 0.13139787 | 0.89546058 | 0.98023547 |
| KALRN | 532.822928 | -0.089950827 | 0.34018356 | -0.2644185 | 0.79145746 | 0.95257889 |
| ITGB5 | 95.7785251 | 0.551048145 | 0.22692235 | 2.42835556 | 0.01516747 | 0.17920531 |
| HEG1 | 41.2359645 | 1.055784531 | 0.32998588 | 3.19948398 | 0.00137674 | 0.045575 |
| ZNF148 | 213.867894 | 0.049554963 | 0.12595932 | 0.39342038 | 0.69400902 | 0.92496777 |
| SNX4 | 75.3950356 | 0.26805954 | 0.20944916 | 1.27983106 | 0.20060456 | 0.59975737 |
| OSBPL11 | 23.6205461 | 0.193611246 | 0.27277855 | 0.70977445 | 0.47784401 | 0.81138532 |
| ZXDC | 60.9158965 | -0.080340486 | 0.2562703 | -0.313499 | 0.75390157 | 0.94128825 |
| CHCHD6 | 19.1677202 | 0.164641796 | 0.31722832 | 0.51900093 | 0.60376009 | 0.87903643 |
| ABTB1 | 22.0763026 | 0.568309127 | 0.30803019 | 1.84497868 | 0.06504064 | 0.38086347 |
| MGLL | 105.444182 | 0.33923357 | 0.32584124 | 1.0411008 | 0.29782877 | 0.68850067 |
| SEC61A1 | 82.6705005 | -0.584154411 | 0.35747575 | -1.6341092 | 0.10223592 | 0.46235803 |
| RUVBL1 | 30.8799848 | -0.437390839 | 0.30138416 | -1.4512735 | 0.14670372 | 0.53611418 |
| EEFSEC | 52.789724 | 0.13501734 | 0.24248415 | 0.55680892 | 0.57765799 | 0.86425255 |
| RPN1 | 42.1298167 | -0.238200065 | 0.25748351 | -0.925108 | 0.35490971 | 0.73100657 |

| | | | | | | |
|---|---|---|---|---|---|---|
| RAB7A | 170.422187 | -0.54464701 | 0.22628292 | -2.4069294 | 0.01608728 | 0.18618176 |
| ACAD9 | 26.3285024 | 0.128807102 | 0.23288693 | 0.55308859 | 0.58020275 | 0.86612025 |
| CNBP | 109.89874 | -0.746598214 | 0.42268804 | -1.7663103 | 0.07734381 | 0.4114983 |
| COPG1 | 34.9055499 | -0.219773831 | 0.32123363 | -0.6841557 | 0.49387683 | 0.81930042 |
| HMCES | 19.9414319 | 0.064942298 | 0.34881576 | 0.18617937 | 0.8523041 | 0.97289794 |
| H1FX | 44.2939159 | 0.335621283 | 0.33348901 | 1.00639384 | 0.31422615 | 0.70282967 |
| MBD4 | 32.6157489 | -0.274108767 | 0.22382759 | -1.2246425 | 0.22070996 | 0.61590675 |
| IFT122 | 36.6214163 | -0.356934771 | 0.23639546 | -1.5099054 | 0.13106757 | 0.51290135 |
| PLXND1 | 84.7857748 | 0.127704584 | 0.23808786 | 0.53637587 | 0.5916988 | 0.87229579 |
| TMCC1 | 222.503038 | 0.238946612 | 0.21309841 | 1.12129701 | 0.26216146 | 0.65630111 |
| PIK3R4 | 28.2585305 | -0.24305086 | 0.23655412 | -1.0274641 | 0.30420199 | 0.69600884 |
| ATP2C1 | 94.1292434 | -0.124809285 | 0.18432688 | -0.6771084 | 0.49833716 | 0.82271448 |
| NEK11 | 26.0631006 | 0.519237677 | 0.36612769 | 1.4181874 | 0.15613606 | 0.54562239 |
| MRPL3 | 41.8721209 | -0.302507023 | 0.27038768 | -1.11879 | 0.26322975 | 0.65753753 |
| DNAJC13 | 79.3133251 | -0.317409723 | 0.22535422 | -1.4084925 | 0.15898529 | 0.55091387 |
| UBA5 | 37.3612069 | -0.60612778 | 0.27242484 | -2.2249358 | 0.02608555 | 0.24143916 |
| CDV3 | 45.5863016 | -0.611088959 | 0.36882705 | -1.6568442 | 0.09755098 | 0.45507247 |
| TOPBP1 | 25.9542248 | 0.035565986 | 0.23929889 | 0.14862579 | 0.88184892 | 0.97782528 |
| SLCO2A1 | 69.3233463 | 0.797855515 | 0.76121779 | 1.04813041 | 0.29457853 | 0.68566952 |
| RYK | 122.095053 | -0.126271833 | 0.17860294 | -0.7069975 | 0.47956803 | 0.81206116 |
| AMOTL2 | 41.8550409 | 1.461130623 | 0.40563542 | 3.60207847 | 0.00031568 | 0.01770958 |
| CEP63 | 46.061762 | 0.02727344 | 0.23516806 | 0.11597425 | 0.90767295 | 0.98306694 |
| EPHB1 | 37.8499518 | -0.820329703 | 0.43212038 | -1.8983824 | 0.05764573 | 0.35871167 |
| PPP2R3A | 85.2492397 | -0.182418197 | 0.27260423 | -0.6691686 | 0.50338791 | 0.82430366 |
| MSL2 | 45.1284549 | -0.040239192 | 0.20163274 | -0.1995668 | 0.84181943 | 0.96803523 |
| PCCB | 23.2952432 | 0.040431878 | 0.27657324 | 0.14618868 | 0.88377244 | 0.97782528 |
| STAG1 | 570.566365 | -0.037925722 | 0.17549889 | -0.2161024 | 0.82890797 | 0.96580172 |
| NCK1 | 50.4587143 | -0.676369065 | 0.21752363 | -3.109405 | 0.00187465 | 0.05488753 |
| DZIP1L | 18.5338501 | 0.004129894 | 0.32738953 | 0.01261462 | 0.98993526 | 0.99739981 |
| ARMC8 | 105.990169 | -0.206536445 | 0.23579268 | -0.8759239 | 0.3810714 | 0.74991898 |
| MRAS | 24.1207713 | 0.406165676 | 0.34059279 | 1.19252578 | 0.23305515 | 0.62939306 |
| CEP70 | 40.9334787 | 0.147258323 | 0.23912989 | 0.61580894 | 0.53802062 | 0.84449243 |
| PIK3CB | 59.63002 | 0.039966424 | 0.19186597 | 0.20830387 | 0.83499171 | 0.96665943 |
| COPB2 | 71.085866 | -0.46823369 | 0.19132344 | -2.4473409 | 0.01439146 | 0.17568582 |
| AC097103.2 | 43.7705935 | 0.22457016 | 0.4296861 | 0.52263771 | 0.60122639 | 0.87876148 |
| CLSTN2 | 118.49006 | 0.264991691 | 0.56226967 | 0.47128933 | 0.63743414 | 0.89741446 |
| SLC25A36 | 114.115749 | -0.26883467 | 0.30801551 | -0.8727959 | 0.38277435 | 0.75179833 |
| ZBTB38 | 207.67144 | -0.556103389 | 0.2353341 | -2.3630378 | 0.01812582 | 0.19729344 |
| RASA2 | 141.773754 | 0.254488918 | 0.15933993 | 1.59714464 | 0.11023347 | 0.47706445 |
| RNF7 | 26.073893 | -0.296455814 | 0.28435799 | -1.0425443 | 0.29715938 | 0.68849737 |
| ATP1B3 | 214.521659 | -0.662877017 | 0.61054042 | -1.0857218 | 0.27760211 | 0.67269898 |
| TFDP2 | 136.792736 | 0.072502772 | 0.27437687 | 0.26424521 | 0.79159098 | 0.95257935 |
| GK5 | 51.9295408 | -0.915359449 | 0.31599136 | -2.8967863 | 0.00377006 | 0.07961552 |
| XRN1 | 203.101176 | -0.154404243 | 0.28015395 | -0.5511407 | 0.58153724 | 0.86687975 |
| ATR | 105.003357 | -0.457184866 | 0.26078829 | -1.7530881 | 0.07958688 | 0.41779962 |
| TRPC1 | 107.644582 | 0.050089251 | 0.34893709 | 0.14354809 | 0.88585734 | 0.97867793 |
| PCOLCE2 | 31.5552329 | 0.001644748 | 0.34308021 | 0.00479406 | 0.99617491 | 0.99854015 |
| U2SURP | 107.361652 | -0.034811775 | 0.18260161 | -0.1906433 | 0.84880506 | 0.97092453 |
| SLC9A9 | 122.694792 | -0.753905144 | 0.38982734 | -1.9339463 | 0.05311973 | 0.34792949 |
| C3orf58 | 25.6937028 | 0.106872579 | 0.28846468 | 0.37048758 | 0.71101923 | 0.93009618 |
| PLOD2 | 279.448347 | -1.938354987 | 0.5281924 | -3.6697896 | 0.00024275 | 0.01499135 |
| PLSCR4 | 245.228992 | 0.052310695 | 0.38381252 | 0.13629231 | 0.8915902 | 0.98023547 |
| PLSCR1 | 205.819477 | -0.972823336 | 0.52439358 | -1.8551397 | 0.06357628 | 0.37532828 |
| AGTR1 | 120.447836 | 0.619328732 | 0.4656506 | 1.33002886 | 0.18350876 | 0.58298863 |
| GYG1 | 24.7655048 | 0.209230763 | 0.23443346 | 0.89249529 | 0.37212752 | 0.74463512 |
| HLTF | 43.6326546 | 0.295797662 | 0.35084021 | 0.84311221 | 0.39916569 | 0.76213787 |
| HPS3 | 30.6665858 | 0.174145459 | 0.38791238 | 0.44892988 | 0.65348224 | 0.90646849 |
| CP | 73.0169974 | -2.796498537 | 0.97445731 | -2.869801 | 0.0041073 | 0.08338278 |
| TM4SF1 | 68.3841092 | 1.014457519 | 0.53772719 | 1.8865654 | 0.0592188 | 0.36270436 |
| WWTR1 | 232.681601 | 0.367272775 | 0.20932489 | 1.75455856 | 0.07933485 | 0.41737582 |
| COMMD2 | 20.0151911 | 0.263992263 | 0.26224611 | 1.00665845 | 0.31409893 | 0.70282967 |
| ANKUB1 | 49.8824803 | -0.045252228 | 0.41575451 | -0.1088436 | 0.91332652 | 0.98395113 |
| RNF13 | 96.1663109 | -0.294866839 | 0.18210225 | -1.6192378 | 0.10539611 | 0.46772706 |
| TSC22D2 | 158.079306 | 0.339144241 | 0.24556855 | 1.38105733 | 0.16726133 | 0.56282213 |
| SERP1 | 66.7765305 | -0.38250183 | 0.22118435 | -1.7293349 | 0.08374917 | 0.42505016 |
| EIF2A | 42.2151575 | -0.50810094 | 0.25757429 | -1.9726385 | 0.04853677 | 0.33413582 |
| SELENOT | 40.5746884 | -0.136877886 | 0.26352792 | -0.5194056 | 0.60347791 | 0.87903643 |
| MBNL1 | 796.902545 | 0.046016837 | 0.34643183 | 0.13283086 | 0.89432715 | 0.98023547 |
| ARHGEF26 | 45.2697408 | -0.069559279 | 0.5334934 | -0.1303845 | 0.89626222 | 0.98040427 |
| DHX36 | 110.869202 | -0.042407177 | 0.18547763 | -0.2286377 | 0.81915053 | 0.96272785 |
| GMPS | 48.8280832 | -0.168788411 | 0.18022592 | -0.9365379 | 0.34899629 | 0.72734588 |
| KCNAB1 | 152.826832 | 2.096710511 | 0.50595454 | 4.14406897 | 3.41E-05 | 0.00424993 |
| SSR3 | 57.7230776 | -0.438419734 | 0.30856717 | -1.4208243 | 0.15536785 | 0.54412824 |
| TIPARP | 80.5972137 | -1.349276993 | 0.84648136 | -1.5939831 | 0.11093982 | 0.4781799 |
| CCNL1 | 152.613014 | -0.681754124 | 0.32709803 | -2.0842502 | 0.03713742 | 0.29287595 |
| RSRC1 | 131.42908 | -0.115642752 | 0.18028513 | -0.6414437 | 0.52123448 | 0.83490675 |
| GFM1 | 40.6141522 | -0.359653399 | 0.24981173 | -1.4396978 | 0.14995291 | 0.53880252 |
| LXN | 39.7228794 | -0.905789986 | 0.51122291 | -1.7718102 | 0.07642606 | 0.41026672 |
| RARRES1 | 338.404614 | -1.787950505 | 0.57812093 | -3.0926929 | 0.00198349 | 0.05584938 |
| MFSD1 | 32.1032835 | -0.197368639 | 0.31912072 | -0.6184764 | 0.53626134 | 0.84423863 |
| IQCJ-SCHIP1 | 85.824679 | 0.805603768 | 0.4395654 | 1.83272789 | 0.06684305 | 0.3864906 |
| KPNA4 | 90.5356132 | -0.017756046 | 0.24029188 | -0.0738937 | 0.941095 | 0.98895798 |
| PPM1L | 130.00728 | 0.107189974 | 0.44541488 | 0.24065198 | 0.80982486 | 0.95953411 |
| NMD3 | 104.336954 | 0.347449184 | 0.3204768 | 1.08416328 | 0.27829241 | 0.67319648 |
| PDCD10 | 45.1665711 | -0.21791567 | 0.21361265 | -1.020144 | 0.30766016 | 0.69801431 |
| GOLIM4 | 65.4988767 | 0.032419175 | 0.20298489 | 0.15971226 | 0.87310774 | 0.9768643 |
| MECOM | 1139.01894 | 0.450916526 | 0.42227979 | 1.06781461 | 0.28560415 | 0.6783522 |
| SEC62 | 80.4411604 | -0.159447417 | 0.19177217 | -0.8314419 | 0.40572403 | 0.76703528 |
| PHC3 | 229.263262 | -0.15771849 | 0.19790968 | -0.7969215 | 0.4254966 | 0.77861803 |
| PRKCI | 78.2111292 | 0.295523757 | 0.24439196 | 1.20922046 | 0.22657816 | 0.62085119 |
| SKIL | 98.7488882 | 0.423292211 | 0.28158571 | 1.50324465 | 0.13277597 | 0.51393747 |
| TNIK | 35.4074747 | 0.44046399 | 0.33727579 | 1.30594605 | 0.19157091 | 0.59004802 |
| PLD1 | 152.618004 | 0.099829723 | 0.37690973 | 0.26486374 | 0.79111443 | 0.95257889 |
| FNDC3B | 1224.92905 | -0.741253326 | 0.28294028 | -2.6198226 | 0.00879755 | 0.1324861 |
| TNFSF10 | 48.5626123 | -1.300667809 | 0.54514978 | -2.3858907 | 0.01703781 | 0.19063814 |
| NCEH1 | 27.5918745 | 0.596497939 | 0.46164726 | 1.2921076 | 0.19631988 | 0.59411229 |
| NLGN1 | 177.365652 | -0.61849715 | 0.30953235 | -1.9981664 | 0.04569862 | 0.32106995 |
| NAALADL2 | 416.893086 | 0.34799486 | 0.29633307 | 1.17433689 | 0.24026011 | 0.63451757 |
| TBL1XR1 | 335.291439 | -0.021427594 | 0.23265533 | -0.0921002 | 0.92661846 | 0.98677279 |
| KCNMB2-AS1 | 33.2930441 | -0.663553567 | 0.43927089 | -1.5105794 | 0.13089564 | 0.51290135 |
| ZMAT3 | 50.9844177 | -0.641080743 | 0.3701723 | -1.7318442 | 0.08330131 | 0.42348124 |
| PIK3CA | 138.080077 | -0.268514101 | 0.30116472 | -0.8915855 | 0.37261514 | 0.7447236 |
| MFN1 | 56.990237 | 0.089082517 | 0.25001332 | 0.35631108 | 0.72160761 | 0.93444541 |
| GNB4 | 59.0724959 | 0.33771833 | 0.23040394 | 1.4657663 | 0.14271196 | 0.53022827 |
| NDUFB5 | 20.1139191 | -0.020425064 | 0.27521659 | -0.0742145 | 0.9408397 | 0.98895798 |
| USP13 | 26.8593736 | 0.223118839 | 0.26764582 | 0.83363469 | 0.40448687 | 0.76622334 |
| TTC14 | 91.4385397 | 0.115963211 | 0.15881513 | 0.73017736 | 0.46528178 | 0.80342007 |
| FXR1 | 105.234102 | -0.10596558 | 0.20778249 | -0.5099832 | 0.61006323 | 0.88163721 |
| ATP11B | 130.777933 | 0.114820325 | 0.22641403 | 0.50712549 | 0.61206677 | 0.88282721 |
| DCUN1D1 | 42.1140086 | 0.458001644 | 0.21870044 | 2.09419622 | 0.0362425 | 0.28969304 |
| MCCC1 | 53.7162875 | -0.169136113 | 0.27723996 | -0.6100712 | 0.54181464 | 0.84641877 |
| MCF2L2 | 27.8994428 | 0.451273698 | 0.41394854 | 1.0901686 | 0.27563888 | 0.6711962 |
| KLHL24 | 123.386834 | -0.515856777 | 0.38990499 | -1.323032 | 0.18582474 | 0.58501647 |

| | | | | | | |
|---|---|---|---|---|---|---|
| YEATS2 | 111.764001 | -0.841240263 | 0.36524141 | -2.3032445 | 0.02126509 | 0.21525781 |
| ABCC5 | 40.2137489 | 0.017152202 | 0.28460322 | 0.06026707 | 0.95194293 | 0.99060401 |
| DVL3 | 34.341355 | 0.070465557 | 0.30889281 | 0.22812301 | 0.81955061 | 0.96284007 |
| AP2M1 | 60.9084465 | -0.472008384 | 0.31328868 | -1.5066244 | 0.13190695 | 0.51290135 |
| PSMD2 | 28.379542 | -0.398262113 | 0.36221422 | -1.0995209 | 0.27154092 | 0.66674046 |
| EIF4G1 | 57.986276 | -0.157957351 | 0.33788143 | -0.4674934 | 0.64014689 | 0.89914713 |
| POLR2H | 27.3434932 | -0.338340424 | 0.31929082 | -1.0596622 | 0.28929829 | 0.67992862 |
| VPS8 | 98.9945056 | 0.016378619 | 0.27216727 | 0.0601785 | 0.95201347 | 0.99060401 |
| C3orf70 | 20.4149062 | 0.532122089 | 0.45292983 | 1.17484443 | 0.24005696 | 0.63451757 |
| EHHADH | 19.1405554 | 0.322337559 | 0.35290058 | 0.91339481 | 0.36103493 | 0.73314333 |
| MAP3K13 | 102.478382 | -0.000999702 | 0.25828356 | -0.0038706 | 0.99691175 | 0.9989507 |
| SENP2 | 50.514242 | 0.279719316 | 0.23234709 | 1.20388558 | 0.22863381 | 0.62213472 |
| IGF2BP2 | 48.1486067 | -0.196517418 | 0.31965099 | -0.6147875 | 0.53869509 | 0.8449569 |
| TRA2B | 93.3674822 | -0.179775617 | 0.2240046 | -0.8025532 | 0.422233 | 0.77771095 |
| ETV5 | 22.0077675 | -0.062700899 | 0.32016724 | -0.195838 | 0.84473699 | 0.96938938 |
| DGKG | 110.227516 | 0.983591121 | 0.4854656 | 2.0260779 | 0.0427568 | 0.31172594 |
| EIF4A2 | 88.753796 | -0.0514575 | 0.32299213 | -0.159315 | 0.87342068 | 0.9768643 |
| ST6GAL1 | 46.9617305 | -0.057643846 | 0.42172145 | -0.136687 | 0.89127819 | 0.98023547 |
| BCL6 | 223.12291 | -1.487621168 | 0.35999599 | -4.1323271 | 3.59E-05 | 0.0042566 |
| LPP | 1731.69614 | -0.000575858 | 0.3881981 | -0.0014834 | 0.99881641 | 0.99969762 |
| TPRG1 | 75.5158059 | -0.303686736 | 0.28904017 | -1.0506731 | 0.29340874 | 0.68480256 |
| P3H2 | 148.110038 | 0.0013935 | 0.55807727 | 0.00249697 | 0.99800771 | 0.99950377 |
| CCDC50 | 103.69033 | 0.048127091 | 0.19656903 | 0.24483557 | 0.80658375 | 0.95838958 |
| MB21D2 | 60.0470073 | -0.520890233 | 0.53310466 | -0.9770881 | 0.32852552 | 0.71294249 |
| OPA1 | 90.0458834 | -0.059006323 | 0.18509764 | -0.3187848 | 0.74988967 | 0.94004394 |
| HES1 | 39.2049071 | -0.058010632 | 0.44238106 | -0.1311327 | 0.89567032 | 0.98023547 |
| ATP13A3 | 272.287472 | -0.833367357 | 0.42959141 | -1.939907 | 0.052391 | 0.34611061 |
| LSG1 | 22.6916306 | -0.206879407 | 0.33942902 | -0.6094924 | 0.54219812 | 0.84643373 |
| XXYLT1 | 37.2893401 | 0.444566234 | 0.28253329 | 1.57350038 | 0.115603 | 0.48836096 |
| ACAP2 | 186.712173 | -0.079187082 | 0.14026292 | -0.5645618 | 0.57237187 | 0.86236719 |
| PPP1R2 | 40.8409017 | 0.101493279 | 0.20013226 | 0.50713104 | 0.61206288 | 0.88282721 |
| MUC20-OT1 | 175.698902 | 0.174917942 | 0.37086733 | 0.4716456 | 0.63717978 | 0.89741446 |
| TNK2 | 17.4211182 | 0.546206804 | 0.3667817 | 1.48918773 | 0.13643794 | 0.52087399 |
| TFRC | 29.9869327 | -0.796207943 | 0.35151455 | -2.2650782 | 0.02350788 | 0.22729214 |
| PCYT1A | 24.9692577 | -0.191263881 | 0.23597934 | -0.8105111 | 0.41764648 | 0.77722447 |
| UBXN7 | 63.346678 | 0.035663822 | 0.21544803 | 0.16553329 | 0.86852425 | 0.97670768 |
| RNF168 | 57.5181792 | -0.217574208 | 0.2567232 | -0.847505 | 0.39671368 | 0.76041962 |
| PAK2 | 92.1882795 | 0.09428411 | 0.18947638 | 0.49760349 | 0.61876354 | 0.88593086 |
| SENP5 | 104.190754 | -0.011367472 | 0.26360854 | -0.0431225 | 0.96560385 | 0.99293098 |
| DLG1 | 270.948793 | -0.30318832 | 0.20570005 | -1.4739341 | 0.14049934 | 0.52680084 |
| RUBCN | 32.8166225 | 0.049626317 | 0.24057266 | 0.20628411 | 0.83656899 | 0.9668633 |
| FYTTD1 | 40.1276681 | -0.13270734 | 0.21401082 | -0.6200964 | 0.53519433 | 0.84377224 |
| LRCH3 | 191.8798 | -0.151819883 | 0.16369552 | -0.9274529 | 0.35369144 | 0.73059415 |
| IQCG | 20.4093909 | -0.196296787 | 0.28512913 | -0.6884487 | 0.49117024 | 0.81720854 |
| RPL35A | 89.3407791 | -0.52537581 | 0.31256986 | -1.6808268 | 0.09279656 | 0.44320781 |
| ZNF595 | 27.0579881 | 0.189306273 | 0.26070954 | 0.72611948 | 0.46776553 | 0.80374006 |
| ZNF718 | 54.8664166 | 0.330610233 | 0.25252403 | 1.30922287 | 0.19045887 | 0.58909185 |
| ZNF141 | 57.2864108 | -0.643544318 | 0.20007931 | -3.2164461 | 0.00129789 | 0.04375316 |
| ZNF721 | 115.030432 | 0.354338712 | 0.22950635 | 1.54391678 | 0.12260849 | 0.50097964 |
| PIGG | 32.2780757 | 0.175064791 | 0.29061281 | 0.60239875 | 0.54690874 | 0.84758329 |
| ATP5ME | 22.9808834 | -0.01017457 | 0.31495277 | -0.0323051 | 0.97422877 | 0.99300191 |
| PCGF3 | 46.4064891 | -0.070320615 | 0.19770055 | -0.3556926 | 0.72207081 | 0.93444541 |
| AC139887.2 | 84.4979917 | 0.039869545 | 0.35431114 | 0.11252693 | 0.91040562 | 0.98343817 |
| GAK | 57.7562184 | -0.052681087 | 0.22082758 | -0.2385621 | 0.81144517 | 0.9598303 |
| AC092535.3 | 20.7267101 | -0.204979037 | 0.33199257 | -0.6174206 | 0.53695737 | 0.84423863 |
| SPON2 | 42.9732104 | -0.431912858 | 0.52290577 | -0.825986 | 0.40881202 | 0.77015112 |
| CTBP1 | 35.420034 | 0.106694865 | 0.29029831 | 0.36753527 | 0.71321979 | 0.93115158 |
| MAEA | 28.8492788 | 0.059774264 | 0.27496812 | 0.21738616 | 0.82790742 | 0.96580172 |
| UVSSA | 37.8652433 | 0.123148149 | 0.25685965 | 0.47943751 | 0.63162742 | 0.89436221 |
| SLBP | 33.03908 | -0.016659689 | 0.27460217 | -0.0606685 | 0.95162326 | 0.99060401 |
| LETM1 | 18.0684816 | 0.046652022 | 0.30555611 | 0.15267907 | 0.87865137 | 0.97718052 |
| NSD2 | 40.0369533 | 0.230650283 | 0.30416141 | 0.75831541 | 0.44826219 | 0.79296348 |
| NELFA | 22.9131096 | 0.081102891 | 0.26859998 | 0.30194675 | 0.76269266 | 0.94462105 |
| MXD4 | 52.4892763 | 0.268912972 | 0.31682369 | 0.84877798 | 0.39600484 | 0.76020146 |
| ZFYVE28 | 26.4689549 | -0.568394712 | 0.42164398 | -1.3480442 | 0.17764418 | 0.57486 |
| RNF4 | 39.1386864 | 0.005243088 | 0.28146749 | 0.01862769 | 0.98513812 | 0.99680298 |
| FAM193A | 128.617021 | 0.011899746 | 0.20188075 | 0.05894443 | 0.95299637 | 0.99060401 |
| TNIP2 | 18.3934092 | -0.934458043 | 0.390633 | -2.3921636 | 0.01674938 | 0.19005424 |
| SH3BP2 | 21.0770187 | -0.340420357 | 0.28968611 | -1.1751352 | 0.23994062 | 0.63451757 |
| ADD1 | 186.873363 | 0.312387712 | 0.14764031 | 2.11587013 | 0.03435585 | 0.2833153 |
| MFSD10 | 21.7041059 | 0.159202263 | 0.33046859 | 0.48174704 | 0.62998566 | 0.89365764 |
| GRK4 | 20.4396996 | 0.433304783 | 0.42257977 | 1.02537986 | 0.30518399 | 0.69673722 |
| HTT | 101.908144 | 0.313745609 | 0.15846621 | 1.9798897 | 0.04771592 | 0.3305036 |
| RGS12 | 38.8369788 | 0.324336542 | 0.2343777 | 1.38381999 | 0.16641358 | 0.56228662 |
| LRPAP1 | 44.2482753 | -0.393977368 | 0.20572581 | -1.9150605 | 0.05548479 | 0.35370877 |
| ADRA2C | 32.04177 | -0.976569387 | 0.33374561 | -2.9260891 | 0.00343253 | 0.07621037 |
| STX18 | 125.925283 | 0.120717179 | 0.30466108 | 0.39623433 | 0.69193217 | 0.92471531 |
| STX18-AS1 | 44.0123122 | 0.695844496 | 0.48362708 | 1.43880383 | 0.15020611 | 0.53880252 |
| EVC2 | 34.9621342 | -0.422338626 | 0.29346876 | -1.4391264 | 0.15011472 | 0.53880252 |
| EVC | 24.7055028 | 0.297892961 | 0.27743063 | 1.07375655 | 0.28293181 | 0.67727609 |
| MAN2B2 | 18.4801521 | 0.263688892 | 0.30503086 | 0.86446627 | 0.3873318 | 0.75443982 |
| MRFAP1 | 49.283759 | 0.134361343 | 0.38434943 | 0.34958122 | 0.72665301 | 0.93455731 |
| KIAA0232 | 99.8734904 | 0.143870993 | 0.19414935 | 0.74103256 | 0.4586737 | 0.79933436 |
| TBC1D14 | 38.7808103 | -0.037826392 | 0.2929234 | -0.1291341 | 0.89725156 | 0.98040427 |
| SORCS2 | 27.1637764 | 0.427009931 | 0.34052605 | 1.2539714 | 0.2098524 | 0.60731347 |
| AFAP1 | 78.804706 | -0.070833824 | 0.31958065 | -0.2216462 | 0.82458935 | 0.96413949 |
| ABLIM2 | 61.6307868 | -0.496153679 | 0.43726656 | -1.134671 | 0.2565132 | 0.65084527 |
| ACOX3 | 25.6581294 | -0.003827612 | 0.35915451 | -0.0106573 | 0.99149688 | 0.99759781 |
| SLC2A9 | 24.6165106 | 0.042819067 | 0.3712753 | 0.11532969 | 0.9081838 | 0.98306694 |
| WDR1 | 69.0395453 | 0.374605244 | 0.29078568 | 1.28825205 | 0.19765824 | 0.59530234 |
| RAB28 | 104.338756 | 0.093506227 | 0.16482193 | 0.56731666 | 0.57049906 | 0.86107981 |
| BOD1L1 | 83.0544731 | -0.03933079 | 0.23697189 | -0.1659724 | 0.86817868 | 0.97656528 |
| CPEB2 | 60.8939569 | 0.168455783 | 0.30149405 | 0.55873667 | 0.57634145 | 0.86325068 |
| AC098829.1 | 137.704391 | 0.623959871 | 0.37801772 | 1.6506101 | 0.09881822 | 0.45673904 |
| C1QTNF7 | 26.2736403 | 1.008873196 | 0.44749455 | 2.25449271 | 0.02416519 | 0.23086389 |
| CC2D2A | 48.1359715 | 0.148651302 | 0.27889166 | 0.53300733 | 0.5940285 | 0.87394794 |
| FBXL5 | 101.980674 | -0.01953477 | 0.2028639 | -0.096295 | 0.92328632 | 0.98561736 |
| TAPT1 | 64.0312078 | 0.001675816 | 0.18147021 | 0.00923466 | 0.99263191 | 0.99759781 |
| TAPT1-AS1 | 56.2180917 | 0.217712393 | 0.39408588 | 0.55244911 | 0.5806407 | 0.86624614 |
| LDB2 | 544.219951 | 0.996103629 | 0.42992166 | 2.3169422 | 0.02050688 | 0.21107152 |
| LAP3 | 35.070534 | -0.889918813 | 0.55738613 | -1.596593 | 0.11035645 | 0.47706445 |
| MED28 | 28.8397004 | -0.119056319 | 0.2250969 | -0.5289114 | 0.5968669 | 0.8753492 |
| DCAF16 | 28.2322303 | -0.115738006 | 0.29958962 | -0.3863218 | 0.69925834 | 0.92769314 |
| LCORL | 81.8109811 | 0.038123382 | 0.20996653 | 0.18156885 | 0.8559211 | 0.97379567 |
| AC024230.1 | 18.8563091 | -0.041193438 | 0.36730705 | -0.1121499 | 0.91070458 | 0.98343817 |
| SLIT2 | 161.844208 | 0.576218433 | 0.31510123 | 1.82867721 | 0.06744798 | 0.38712255 |
| PACRGL | 45.0777085 | -0.0106321 | 0.2756659 | -0.0385688 | 0.96923419 | 0.99300191 |
| KCNIP4 | 172.683599 | 0.079652437 | 0.34118738 | 0.23345658 | 0.81540688 | 0.96094054 |
| ADGRA3 | 47.4414129 | 0.56698202 | 0.31679789 | 1.7897279 | 0.07349766 | 0.40189622 |
| PPARGC1A | 64.2212764 | -0.565114307 | 0.38554206 | -1.4657657 | 0.14271214 | 0.53022827 |
| DHX15 | 115.955117 | -0.160828374 | 0.20470254 | -0.7856687 | 0.43206161 | 0.78417671 |
| SOD3 | 104.338094 | -0.344149086 | 0.38865901 | -0.8854782 | 0.37589876 | 0.74787433 |
| CCDC149 | 33.2826357 | 0.110803828 | 0.43138761 | 0.25685445 | 0.79729113 | 0.95463848 |

| | | | | | | |
|---|---|---|---|---|---|---|
| ANAPC4 | 37.8043278 | 0.141240138 | 0.27529965 | 0.51304148 | 0.60792231 | 0.88042838 |
| RBPJ | 160.335632 | -0.32866603 | 0.26760367 | -1.2281821 | 0.21937862 | 0.61567703 |
| TBC1D19 | 62.6920189 | 0.016290127 | 0.2360911 | 0.06899933 | 0.94499016 | 0.99060401 |
| STIM2 | 52.2730298 | 0.435129427 | 0.22117338 | 1.96736797 | 0.0491408 | 0.33550435 |
| PCDH7 | 144.48129 | 1.490364864 | 0.80067498 | 1.86138558 | 0.06268974 | 0.37164525 |
| ARAP2 | 23.2484016 | -0.248003494 | 0.34003899 | -0.7293384 | 0.46579469 | 0.80355051 |
| C4orf19 | 27.2729965 | -0.362447745 | 0.28127993 | -1.288566 | 0.19754903 | 0.59530234 |
| RELL1 | 59.7549911 | 0.328235403 | 0.25179993 | 1.30355636 | 0.19238489 | 0.59142551 |
| PGM2 | 32.6361422 | -0.902652848 | 0.22234018 | -4.0597828 | 4.91E-05 | 0.00494481 |
| TBC1D1 | 429.811431 | -0.232116616 | 0.268661 | -0.8639758 | 0.38760116 | 0.75457985 |
| KLF3 | 41.913896 | 0.409390007 | 0.32277909 | 1.26832878 | 0.20468056 | 0.60370888 |
| FAM114A1 | 70.4508436 | -0.311715471 | 0.29307045 | -1.0636196 | 0.28750106 | 0.67963037 |
| KLHL5 | 68.9701944 | -0.320323554 | 0.40012193 | -0.8005648 | 0.42338361 | 0.77827834 |
| WDR19 | 46.2808002 | -0.555150293 | 0.23524564 | -2.359875 | 0.01828109 | 0.19786121 |
| RFC1 | 61.1177021 | -0.071516543 | 0.24467336 | -0.2922939 | 0.77006189 | 0.94593455 |
| UGDH | 52.7967233 | -0.173444865 | 0.56798668 | -0.3053678 | 0.76008601 | 0.94422614 |
| UGDH-AS1 | 34.3031521 | 0.001828688 | 0.37452494 | 0.00488269 | 0.99610419 | 0.99854015 |
| SMIM14 | 107.839835 | 0.041958373 | 0.23495464 | 0.17858074 | 0.85826692 | 0.97397549 |
| UBE2K | 218.595799 | -0.039242227 | 0.19580617 | -0.2004136 | 0.84115709 | 0.96803523 |
| PDS5A | 198.953825 | -0.189576311 | 0.1228489 | -1.5431665 | 0.12279039 | 0.50097964 |
| RBM47 | 44.6487042 | -0.343570616 | 0.32109679 | -1.0699908 | 0.28462345 | 0.6783522 |
| APBB2 | 637.98181 | 0.553119264 | 0.24735306 | 2.23615292 | 0.02534176 | 0.23785002 |
| LIMCH1 | 220.768193 | 0.592435868 | 0.49847262 | 1.18850231 | 0.23463557 | 0.63080829 |
| TMEM33 | 31.7537274 | -0.060007605 | 0.24946115 | -0.2405489 | 0.80990476 | 0.95953411 |
| SLC30A9 | 96.8995101 | 0.03845302 | 0.18357049 | 0.20947278 | 0.83407918 | 0.9666365 |
| ATP8A1 | 28.9895993 | -0.205904381 | 0.46680139 | -0.4410963 | 0.65914326 | 0.90831499 |
| GUF1 | 20.1122834 | -0.275588523 | 0.30752416 | -0.8961524 | 0.37017135 | 0.74264779 |
| GNPDA2 | 20.4179042 | -0.469191352 | 0.30632742 | -1.5316662 | 0.12560483 | 0.50440978 |
| ATP10D | 221.37537 | 0.898378031 | 0.52223105 | 1.72026928 | 0.08538351 | 0.4283163 |
| NFXL1 | 42.306242 | -0.323305991 | 0.28292876 | -1.1427117 | 0.25315833 | 0.6480183 |
| SLAIN2 | 87.0264252 | 0.23555161 | 0.20210817 | 1.165473 | 0.24382758 | 0.63964329 |
| FRYL | 253.566736 | 0.075339479 | 0.20187672 | 0.37319547 | 0.70900296 | 0.93009618 |
| OCIAD1 | 91.8141286 | 0.018429579 | 0.14579346 | 0.12640882 | 0.89940832 | 0.98058672 |
| DCUN1D4 | 89.7988576 | -0.114263901 | 0.27096493 | -0.4216926 | 0.67324941 | 0.91505229 |
| SGCB | 22.9637407 | 0.147305495 | 0.26161466 | 0.56306285 | 0.57339209 | 0.86261176 |
| USP46 | 34.0775796 | 0.607518669 | 0.24838783 | 2.44584713 | 0.01445123 | 0.17612282 |
| SCFD2 | 242.132908 | -0.548679946 | 0.34758171 | -1.5785639 | 0.11443612 | 0.48612197 |
| AC058822.1 | 70.0058609 | -0.485387756 | 0.44628265 | -1.0876241 | 0.27676109 | 0.67259169 |
| CHIC2 | 198.309748 | -0.680185374 | 0.28920725 | -2.351896 | 0.018678 | 0.20009416 |
| PDGFRA | 115.735557 | -0.189507559 | 0.4088848 | -0.4634742 | 0.6430245 | 0.90063554 |
| TMEM165 | 228.290094 | -0.655853942 | 0.25526216 | -2.5693347 | 0.0101894 | 0.14209086 |
| CLOCK | 104.828176 | 0.005326224 | 0.21381495 | 0.02491044 | 0.9801264 | 0.99503182 |
| EXOC1 | 85.1098453 | 0.085659545 | 0.24908516 | 0.34389662 | 0.73092404 | 0.93467213 |
| CEP135 | 24.15904 | 0.296803247 | 0.35372781 | 0.83907242 | 0.40142867 | 0.76402268 |
| KIAA1211 | 60.2532575 | -0.069123954 | 0.40614542 | -0.1701951 | 0.86485672 | 0.9762143 |
| AASDH | 24.2052622 | 0.102566211 | 0.30264382 | 0.33890072 | 0.73468452 | 0.93594619 |
| PAICS | 20.9888511 | -1.019982512 | 0.36034003 | -2.8306112 | 0.00464592 | 0.0906884 |
| SRP72 | 52.3154852 | -0.186107018 | 0.21803687 | -0.8535576 | 0.39335018 | 0.75988253 |
| REST | 44.3016171 | 0.056320393 | 0.24719269 | 0.22784004 | 0.81977059 | 0.96284007 |
| POLR2B | 63.0960418 | -0.142421178 | 0.17817973 | -0.7993119 | 0.42410958 | 0.77857941 |
| IGFBP7 | 1910.15949 | 0.143605804 | 0.28042885 | 0.51209354 | 0.60858555 | 0.8807592 |
| ADGRL3 | 239.982964 | 2.073020746 | 0.41717483 | 4.96918941 | 6.72E-07 | 0.00018632 |
| AC104806.2 | 18.6212461 | 0.437617347 | 0.33435641 | 1.30883493 | 0.19059027 | 0.58925027 |
| CENPC | 82.2272387 | 0.453780312 | 0.34393258 | 1.31938741 | 0.18703963 | 0.5851657 |
| UBA6 | 66.8477131 | 0.078086008 | 0.19808585 | 0.39420285 | 0.69343128 | 0.9248682 |
| UBA6-AS1 | 64.4347932 | 0.593489797 | 0.4176057 | 1.42117264 | 0.15526658 | 0.54412824 |
| YTHDC1 | 109.586401 | -0.091596392 | 0.26074737 | -0.351284 | 0.72537526 | 0.93455731 |
| UGT2B7 | 40.7284427 | -0.383895568 | 0.54199828 | -0.7082967 | 0.47876106 | 0.81162976 |
| RUFY3 | 190.81554 | 0.299612392 | 0.25500883 | 1.17490987 | 0.24003078 | 0.63451757 |
| GRSF1 | 32.4994095 | -0.149402014 | 0.23344529 | -0.6399873 | 0.52218088 | 0.8350836 |
| MOB1B | 63.6436572 | -0.165050027 | 0.21864247 | -0.7548855 | 0.4503177 | 0.79399826 |
| SLC4A4 | 40.4025934 | 0.706185983 | 0.40021116 | 1.76453344 | 0.07764221 | 0.412576 |
| ADAMTS3 | 86.1280166 | -0.004413415 | 0.72665845 | -0.0060736 | 0.99515402 | 0.99854015 |
| ANKRD17 | 442.108863 | -0.362276812 | 0.21910211 | -1.6534611 | 0.09823706 | 0.45606074 |
| AC053527.1 | 44.2361411 | 0.522628328 | 0.42517876 | 1.22919671 | 0.21899806 | 0.61522936 |
| MTHFD2L | 30.7985141 | -0.662866464 | 0.33455909 | -1.9813135 | 0.04755612 | 0.3297075 |
| PARM1 | 152.008614 | 0.222403676 | 0.32068033 | 0.69353701 | 0.48797261 | 0.81595238 |
| RCHY1 | 35.019607 | -0.074573768 | 0.28378167 | -0.2627857 | 0.79271575 | 0.95346449 |
| THAP6 | 22.1804245 | 0.222127788 | 0.30280347 | 0.73357081 | 0.46321035 | 0.80225845 |
| G3BP2 | 61.4676322 | -0.048724116 | 0.31810938 | -0.1531678 | 0.87826594 | 0.9770476 |
| USO1 | 146.196261 | -0.228060164 | 0.21409489 | -1.0652294 | 0.28677215 | 0.67921285 |
| SDAD1 | 36.653819 | -0.297311626 | 0.2857739 | -1.0403736 | 0.29816636 | 0.68868829 |
| NUP54 | 48.2818514 | -0.166576704 | 0.18875078 | -0.882522 | 0.37749461 | 0.749056 |
| SCARB2 | 106.951043 | -0.069651701 | 0.24135022 | -0.2885918 | 0.77289375 | 0.9472495 |
| SHROOM3 | 43.212386 | 0.992425203 | 0.34198922 | 2.90191954 | 0.00370884 | 0.07946428 |
| 11-Sep | 237.575939 | 0.618082855 | 0.42577475 | 1.45166631 | 0.14659441 | 0.53611418 |
| CCNI | 164.680126 | 0.254734563 | 0.17943617 | 1.41963888 | 0.15571284 | 0.54466143 |
| CCNG2 | 34.1558955 | -0.220916904 | 0.38484536 | -0.5740407 | 0.56594026 | 0.85807612 |
| CNOT6L | 91.2786195 | -0.069310348 | 0.23615126 | -0.2934998 | 0.76914015 | 0.94579017 |
| MRPL1 | 46.5731475 | -0.185740772 | 0.21757536 | -0.8536848 | 0.39327966 | 0.75988253 |
| FRAS1 | 103.928583 | 1.091273899 | 0.4766373 | 2.28952687 | 0.02204876 | 0.21901484 |
| BMP2K | 77.2267029 | 0.392953398 | 0.2718415 | 1.44552395 | 0.14831077 | 0.53680621 |
| LINC01088 | 72.8583436 | 0.664741497 | 0.88205761 | 0.75362595 | 0.45107386 | 0.79401729 |
| LINC00989 | 48.5119955 | 0.095729968 | 0.52614205 | 0.181947 | 0.85562432 | 0.97379567 |
| ANTXR2 | 210.575861 | -0.082774342 | 0.33754348 | -0.2452257 | 0.80628166 | 0.95838958 |
| RASGEF1B | 422.560747 | -0.021039779 | 0.55581686 | -0.0378538 | 0.96980425 | 0.99300191 |
| HNRNPD | 120.899062 | -0.178486094 | 0.18639725 | -0.9575576 | 0.33828591 | 0.71925477 |
| HNRNPDL | 124.65825 | -0.211682232 | 0.21519085 | -0.9836953 | 0.32526534 | 0.71149668 |
| ENOPH1 | 18.32872 | 0.004171934 | 0.2978905 | 0.01400492 | 0.98882605 | 0.99723928 |
| SEC31A | 249.840564 | -0.315899552 | 0.18701043 | -1.6892082 | 0.09117954 | 0.44115857 |
| LIN54 | 26.6978516 | 0.029262947 | 0.28190098 | 0.10380577 | 0.91732349 | 0.98477389 |
| COPS4 | 30.331578 | 0.406830836 | 0.26115874 | 1.55779139 | 0.11928271 | 0.4941424 |
| HELQ | 30.1001768 | 0.112603503 | 0.2947589 | 0.382019 | 0.70244727 | 0.92865476 |
| WDFY3 | 370.382976 | 0.127875712 | 0.21584366 | 0.59244599 | 0.55355198 | 0.85188037 |
| ARHGAP24 | 418.117375 | -0.410365945 | 0.49019006 | -0.8371568 | 0.40250444 | 0.76472383 |
| MAPK10 | 487.012754 | 0.380547475 | 0.31429305 | 1.21080463 | 0.22597029 | 0.62072404 |
| PTPN13 | 133.644719 | 0.057677583 | 0.20592161 | 0.28009486 | 0.77940473 | 0.94894721 |
| AFF1 | 455.888629 | -0.283622441 | 0.25542244 | -1.1104054 | 0.26682439 | 0.66188434 |
| KLHL8 | 34.0368171 | -0.338086542 | 0.23919422 | -1.4134394 | 0.15752656 | 0.54865532 |
| HSD17B11 | 34.8691408 | -0.21598411 | 0.270326 | -0.7989765 | 0.42430406 | 0.77857941 |
| NUDT9 | 32.4622591 | -0.288603645 | 0.23742422 | -1.2155611 | 0.22415217 | 0.61858591 |
| SPARCL1 | 317.002507 | -0.331354498 | 0.42350619 | -0.7824077 | 0.43397502 | 0.78490415 |
| SPP1 | 1102.4676 | -3.12439034 | 0.50565683 | -6.1788749 | 6.46E-10 | 2.25E-06 |
| PKD2 | 229.514804 | 0.419202204 | 0.23939227 | 1.75111003 | 0.07992696 | 0.41881121 |
| PPM1K | 53.2177319 | 0.533655045 | 0.43172391 | 1.23610258 | 0.21642044 | 0.61337207 |
| HERC3 | 33.770327 | 0.465554877 | 0.30247233 | 1.53916517 | 0.12376398 | 0.50097964 |
| FAM13A | 393.092802 | -1.858726235 | 0.55179868 | -3.3684862 | 0.00075582 | 0.03182184 |
| GPRIN3 | 107.161157 | 0.117368047 | 0.4230672 | 0.27742176 | 0.78145627 | 0.94939885 |
| CCSER1 | 79.7050674 | 0.507719755 | 0.30620855 | 1.65808486 | 0.09730034 | 0.45487289 |
| GRID2 | 142.894688 | 1.405814119 | 0.49948994 | 2.81449938 | 0.00488533 | 0.09203696 |
| SMARCAD1 | 76.8174723 | -0.166874629 | 0.22089204 | -0.7554579 | 0.4499743 | 0.79399826 |
| PDLIM5 | 1083.22498 | 0.61563258 | 0.48600963 | 1.2667086 | 0.2052595 | 0.60370888 |
| BMPR1B | 234.591233 | 0.313456026 | 0.90965972 | 0.34458602 | 0.73040562 | 0.93455731 |

| | | | | | | |
|---|---|---|---|---|---|---|
| UNC5C | 363.087398 | 0.751134344 | 0.19310187 | 3.88983465 | 0.00010031 | 0.00791782 |
| STPG2 | 25.533253 | 0.872185192 | 0.53126107 | 1.64172616 | 0.10064677 | 0.4602695 |
| RAP1GDS1 | 90.238988 | 0.369984287 | 0.23282588 | 1.58910292 | 0.11203716 | 0.4809788 |
| TSPAN5 | 33.5701642 | 1.030374867 | 0.33133929 | 3.10972738 | 0.0018726 | 0.05488753 |
| EIF4E | 54.618297 | 0.030601664 | 0.29595764 | 0.1033988 | 0.91764647 | 0.98477389 |
| METAP1 | 33.4406981 | 0.105516286 | 0.25652986 | 0.41132166 | 0.68083669 | 0.91860979 |
| ADH5 | 53.4681603 | -0.014832966 | 0.23119899 | -0.0641567 | 0.94884545 | 0.99060401 |
| AP002026.1 | 39.8856383 | 0.654532554 | 0.45679132 | 1.43289183 | 0.15188875 | 0.53898138 |
| ADH1B | 223.696121 | -0.191808684 | 0.59752815 | -0.3210036 | 0.74820767 | 0.93912522 |
| DNAJB14 | 94.4744474 | 0.099962322 | 0.218626 | 0.4572298 | 0.64750588 | 0.90256375 |
| H2AFZ | 26.3820904 | -0.635005377 | 0.39322714 | -1.6148565 | 0.10634178 | 0.46993733 |
| EMCN | 246.548398 | 0.973205242 | 0.43347611 | 2.24511851 | 0.02476053 | 0.23328859 |
| PPP3CA | 238.317636 | -0.485784997 | 0.2524368 | -1.9243826 | 0.05430664 | 0.35039464 |
| NFKB1 | 199.786231 | -0.908107172 | 0.43433246 | -2.0908112 | 0.03654499 | 0.28984581 |
| MANBA | 41.7913206 | -0.50437315 | 0.28046084 | -1.7983728 | 0.07211795 | 0.39969442 |
| UBE2D3 | 254.014758 | -0.328521736 | 0.18469483 | -1.7787273 | 0.07528448 | 0.40760265 |
| SLC9B2 | 20.0773288 | -0.049706323 | 0.38274641 | -0.1298675 | 0.89667125 | 0.98040427 |
| TET2 | 121.046445 | -0.201111175 | 0.27753093 | -0.7246442 | 0.46867034 | 0.80374006 |
| PPA2 | 109.648075 | 0.025633542 | 0.18490458 | 0.13863119 | 0.88974159 | 0.98016953 |
| INTS12 | 24.6312189 | -0.105198119 | 0.30161444 | -0.3487834 | 0.7272519 | 0.93455731 |
| GSTCD | 30.7233556 | 0.050043388 | 0.36842375 | 0.13583106 | 0.89195484 | 0.98023547 |
| NPNT | 46.4843662 | 0.495116371 | 0.37011289 | 1.3377442 | 0.18097984 | 0.57952978 |
| TBCK | 193.257244 | -0.1217578 | 0.29448918 | -0.4134542 | 0.67927384 | 0.91831925 |
| AIMP1 | 27.0203592 | -0.269134728 | 0.24406874 | -1.1027005 | 0.27015725 | 0.6651094 |
| PAPSS1 | 41.17637 | 0.368372122 | 0.22333079 | 1.64944619 | 0.09905626 | 0.45678074 |
| SGMS2 | 29.7355727 | 1.138043107 | 0.43674471 | 2.60573986 | 0.00916761 | 0.13415695 |
| RPL34 | 71.1541163 | -0.429648159 | 0.3452139 | -1.2445853 | 0.21328422 | 0.61001472 |
| OSTC | 35.3564487 | -0.612524203 | 0.3192811 | -1.9184481 | 0.05505422 | 0.35260065 |
| COL25A1 | 405.458717 | 1.822069606 | 0.58325559 | 3.12396426 | 0.00178432 | 0.05374171 |
| SEC24B | 147.676733 | -0.054848185 | 0.20372972 | -0.2692203 | 0.78776013 | 0.95129454 |
| EGF | 22.6648697 | 2.24943073 | 0.44851717 | 5.01526117 | 5.30E-07 | 0.00016922 |
| ELOVL6 | 30.0483557 | -0.676400196 | 0.32666792 | -2.0706049 | 0.03839574 | 0.29669534 |
| ENPEP | 58.7111477 | 0.382948959 | 0.45963942 | 0.83315082 | 0.40475967 | 0.76624906 |
| FAM241A | 58.2010666 | -0.125077297 | 0.27280161 | -0.4584918 | 0.64659916 | 0.90219427 |
| AP1AR | 24.4391352 | -0.03076811 | 0.34470383 | -0.0892596 | 0.92887564 | 0.98681039 |
| ALPK1 | 110.410296 | -1.021550539 | 0.35004397 | -2.9183492 | 0.0035189 | 0.07673709 |
| LARP7 | 37.294761 | 0.322100584 | 0.30543057 | 1.05457873 | 0.29161803 | 0.68334882 |
| ANK2 | 143.195184 | 0.889082736 | 0.28811256 | 3.08588679 | 0.00202946 | 0.05607827 |
| CAMK2D | 460.480854 | -0.521228616 | 0.22707233 | -2.2954299 | 0.0217085 | 0.21750516 |
| ARSJ | 38.6331253 | -0.218013159 | 0.33231682 | -0.6560401 | 0.51179833 | 0.82882457 |
| METTL14 | 36.5994459 | 0.318262477 | 0.25120155 | 1.26696065 | 0.20516936 | 0.60370888 |
| SEC24D | 233.097771 | -0.715705549 | 0.18897746 | -3.7872536 | 0.00015232 | 0.01046179 |
| SYNPO2 | 519.259071 | 0.411753348 | 0.49941028 | 0.82447913 | 0.40966737 | 0.7710341 |
| USP53 | 809.638375 | -0.368540481 | 0.47111304 | -0.7822761 | 0.43405232 | 0.78490415 |
| C4orf3 | 59.8393454 | -0.2712609 | 0.29469717 | -0.9204734 | 0.35732544 | 0.73129235 |
| PDE5A | 211.042852 | 0.654102295 | 0.38750241 | 1.68799544 | 0.0914121 | 0.44124451 |
| AC108866.1 | 45.5451736 | 0.23868954 | 0.34769465 | 0.68649185 | 0.492403 | 0.81870355 |
| PRDM5 | 53.4974371 | 0.094101466 | 0.25743899 | 0.3655292 | 0.71471642 | 0.93179044 |
| ANXA5 | 131.057731 | -0.153183039 | 0.31521594 | -0.4859622 | 0.62699396 | 0.89177058 |
| BBS7 | 27.9739699 | -0.029042676 | 0.30747941 | -0.0944541 | 0.92474848 | 0.98561736 |
| KIAA1109 | 223.661663 | 0.357938927 | 0.21133242 | 1.69372467 | 0.09031761 | 0.43898422 |
| FGF2 | 41.2346712 | -0.342171367 | 0.28386847 | -1.205387 | 0.22805393 | 0.62188065 |
| SPATA5 | 120.548999 | -0.115785952 | 0.20275319 | -0.5710684 | 0.56795325 | 0.85953221 |
| SPRY1 | 56.9612654 | 0.206379264 | 0.35436979 | 0.58238392 | 0.56030811 | 0.85423726 |
| ANKRD50 | 31.5540194 | 0.835262557 | 0.25808208 | 3.23642214 | 0.00121038 | 0.04338282 |
| FAT4 | 101.205724 | 0.428128221 | 0.56758675 | 0.75429566 | 0.45067171 | 0.79401729 |
| INTU | 40.4784075 | 0.249618985 | 0.29439082 | 0.847917 | 0.3964842 | 0.76020893 |
| HSPA4L | 21.7559057 | -0.663912316 | 0.34287074 | -1.9363341 | 0.05282679 | 0.34706747 |
| MFSD8 | 28.8269794 | 0.077893297 | 0.25907039 | 0.30066461 | 0.76367026 | 0.94462105 |
| ABHD18 | 62.1316605 | 0.113877929 | 0.24253683 | 0.4695284 | 0.63869199 | 0.89815297 |
| LARP1B | 88.9166991 | -0.066361745 | 0.23141894 | -0.2867602 | 0.77429593 | 0.9475938 |
| PGRMC2 | 42.2165553 | -0.227520125 | 0.23457577 | -0.9699217 | 0.33208554 | 0.71505908 |
| JADE1 | 65.5040745 | -0.575279347 | 0.41485198 | -1.3867099 | 0.16553024 | 0.56136675 |
| SCLT1 | 117.035393 | -0.167876308 | 0.2568624 | -0.6535651 | 0.51339202 | 0.82994235 |
| AC096711.2 | 35.0525699 | 0.278931598 | 0.24265261 | 1.14950998 | 0.25034575 | 0.6460368 |
| PCDH10 | 24.2304989 | 0.027692138 | 0.52291987 | 0.05295675 | 0.95776637 | 0.99181157 |
| PCDH18 | 41.0646489 | 0.378622706 | 0.47345723 | 0.79969781 | 0.4238859 | 0.77857941 |
| AC109927.1 | 33.1206058 | -0.745180526 | 0.33691445 | -2.2117797 | 0.02698189 | 0.24601728 |
| ELF2 | 299.627939 | -0.192078523 | 0.19385535 | -0.9908342 | 0.32176653 | 0.7079827 |
| NDUFC1 | 20.6173504 | -0.306000871 | 0.29830524 | -1.0257978 | 0.30498689 | 0.69650363 |
| NAA15 | 106.027231 | 0.083422207 | 0.24056906 | 0.3467703 | 0.72876389 | 0.93455731 |
| AC097376.2 | 51.9638552 | 0.161289483 | 0.32126334 | 0.50204758 | 0.61563405 | 0.8836513 |
| SETD7 | 44.0523882 | -0.243177771 | 0.22475019 | -1.0819914 | 0.27925636 | 0.67353297 |
| MGST2 | 22.506842 | 0.145823336 | 0.36510283 | 0.39940346 | 0.68959594 | 0.92259978 |
| MAML3 | 407.456605 | 0.810532763 | 0.30730175 | 2.63757935 | 0.00835001 | 0.12814321 |
| SCOC | 26.5871006 | -0.41852164 | 0.30600449 | -1.3676977 | 0.17140671 | 0.56870244 |
| ELMOD2 | 20.2677222 | 0.190197773 | 0.28805478 | 0.66028335 | 0.50907201 | 0.82663984 |
| TBC1D9 | 33.1963129 | 0.154865086 | 0.23245225 | 0.66622321 | 0.50526844 | 0.82481485 |
| RNF150 | 179.480995 | 0.66521625 | 0.37914322 | 1.75452499 | 0.07934059 | 0.41737582 |
| ZNF330 | 21.1524762 | -0.72651374 | 0.38625727 | -1.8809063 | 0.05998466 | 0.36492325 |
| IL15 | 24.6853322 | -0.267016489 | 0.40034505 | -0.6669659 | 0.50479393 | 0.82481485 |
| INPP4B | 996.410237 | 0.012053723 | 0.40367261 | 0.02986014 | 0.97617859 | 0.99348241 |
| AC139720.1 | 37.613256 | -1.075787353 | 0.52505563 | -2.0489017 | 0.04047173 | 0.3019561 |
| AC104596.1 | 29.6407729 | 0.819180707 | 0.58448125 | 1.40155173 | 0.16104915 | 0.55470404 |
| USP38 | 27.9081145 | -0.286477328 | 0.30124815 | -0.9509679 | 0.34162067 | 0.72039319 |
| GAB1 | 214.148124 | 0.387674688 | 0.3394769 | 1.14197662 | 0.25346374 | 0.64812282 |
| SMARCA5 | 77.5582624 | -0.008932017 | 0.18001893 | -0.0496171 | 0.96042752 | 0.99181157 |
| ANAPC10 | 63.0269305 | -0.055769903 | 0.20428735 | -0.2729973 | 0.78485526 | 0.95065355 |
| ABCE1 | 43.9595174 | -0.539988186 | 0.35067326 | -1.5398613 | 0.12359416 | 0.50097964 |
| OTUD4 | 62.2447407 | 0.06541904 | 0.22388509 | 0.29219919 | 0.77013433 | 0.94593455 |
| SMAD1 | 39.5182871 | 0.123481147 | 0.31613497 | 0.39059629 | 0.69609566 | 0.92648248 |
| ZNF827 | 41.4280617 | 1.000015752 | 0.31692988 | 3.15532181 | 0.00160321 | 0.05045611 |
| SLC10A7 | 100.075611 | -0.474624589 | 0.24796511 | -1.9140781 | 0.05561017 | 0.35414139 |
| EDNRA | 178.228225 | -0.062443896 | 0.27531409 | -0.2268097 | 0.82057175 | 0.96291025 |
| TMEM184C | 26.1816857 | 0.067719644 | 0.31575862 | 0.21446649 | 0.83018329 | 0.96580172 |
| PRMT9 | 32.0977589 | 0.30971456 | 0.21288924 | 1.45481548 | 0.14572035 | 0.53567049 |
| ARHGAP10 | 412.0294 | 0.067231253 | 0.41888175 | 0.16050175 | 0.87248584 | 0.9768643 |
| NR3C2 | 189.967598 | 0.694091453 | 0.37902065 | 1.83127611 | 0.06705934 | 0.38712255 |
| DCLK2 | 66.0586197 | 0.464897488 | 0.33427542 | 1.39076183 | 0.16429766 | 0.55949572 |
| LRBA | 246.834237 | 0.016343292 | 0.27485117 | 0.05946233 | 0.95258387 | 0.99060401 |
| RPS3A | 58.6750758 | 0.071260602 | 0.37187869 | 0.19162325 | 0.84803733 | 0.97075177 |
| SH3D19 | 324.040125 | 1.12351748 | 0.30870851 | 3.6394121 | 0.00027326 | 0.0161874 |
| FBXW7 | 193.807535 | 0.09957631 | 0.17708138 | 0.5623195 | 0.57389836 | 0.86294056 |
| ARFIP1 | 99.0360953 | 0.133981759 | 0.15256325 | 0.87820469 | 0.37983265 | 0.74991898 |
| TRIM2 | 76.1992566 | 0.178109195 | 0.28770609 | 0.61906648 | 0.53587257 | 0.84412047 |
| TMEM131L | 244.533803 | 0.281537173 | 0.21447491 | 1.31268115 | 0.18929041 | 0.58671246 |
| MAP9 | 19.9655022 | -0.024703727 | 0.33251647 | -0.0742932 | 0.94077705 | 0.98895798 |
| GUCY1A1 | 180.78896 | 0.389559966 | 0.27118329 | 1.43651905 | 0.1508547 | 0.53880252 |
| GUCY1B1 | 76.7048906 | 0.277438802 | 0.17539042 | 1.58183556 | 0.11368711 | 0.48395978 |
| CTSO | 19.9662865 | 0.27195004 | 0.35745821 | 0.76078835 | 0.44678349 | 0.79296348 |
| PDGFC | 150.877233 | 0.307348338 | 0.23785547 | 1.29216427 | 0.19630026 | 0.59411229 |
| FAM198B | 39.1040887 | -0.357454325 | 0.47254546 | -0.7564443 | 0.44938285 | 0.7936829 |
| RXFP1 | 89.1633287 | -0.04151247 | 0.8272147 | -0.0501834 | 0.95997622 | 0.99181157 |

| | | | | | | |
|---|---|---|---|---|---|---|
| ETFDH | 59.8232865 | -0.157173031 | 0.23969846 | -0.6557115 | 0.51200979 | 0.82885603 |
| PPID | 21.0894811 | -0.147520029 | 0.32891121 | -0.4485102 | 0.65378504 | 0.90646849 |
| FNIP2 | 170.319061 | 0.159157181 | 0.27240764 | 0.58426108 | 0.55904467 | 0.85343152 |
| RAPGEF2 | 328.850853 | 0.811386495 | 0.22101873 | 3.67112091 | 0.00024149 | 0.01499135 |
| NAF1 | 64.4424582 | 0.158249444 | 0.2172905 | 0.72828516 | 0.46643904 | 0.80355999 |
| NPY1R | 183.411781 | 0.364293618 | 0.39555605 | 0.92096586 | 0.35706825 | 0.73129235 |
| TMA16 | 47.7885449 | -0.275635125 | 0.27948894 | -0.9862112 | 0.32402947 | 0.71051224 |
| TMEM192 | 28.4239099 | -0.095272889 | 0.38800373 | -0.2455463 | 0.80603345 | 0.95838958 |
| KLHL2 | 56.5580764 | -0.55666394 | 0.31855354 | -1.7474737 | 0.0805552 | 0.42015625 |
| MSMO1 | 21.4230523 | -0.070242265 | 0.41095184 | -0.1709258 | 0.86428212 | 0.9762143 |
| CPE | 177.920055 | 0.014652235 | 0.46563171 | 0.03146743 | 0.97489676 | 0.99300296 |
| DDX60 | 62.8195146 | 0.132460728 | 0.90185939 | 0.14687514 | 0.88323057 | 0.97782528 |
| DDX60L | 49.9782251 | 1.04995238 | 1.07762737 | 0.97431859 | 0.32989838 | 0.71362138 |
| PALLD | 770.71159 | 0.916262214 | 0.55656459 | 1.64628192 | 0.09970573 | 0.45779653 |
| CBR4 | 138.652211 | -0.026837408 | 0.34774839 | -0.0771748 | 0.9384845 | 0.98827804 |
| SH3RF1 | 178.439034 | -0.005717868 | 0.15191361 | -0.0376389 | 0.96997556 | 0.99300191 |
| NEK1 | 118.538298 | 0.308937104 | 0.27388263 | 1.12799087 | 0.25932378 | 0.65213055 |
| CLCN3 | 67.0782847 | -0.195942464 | 0.20446604 | -0.958313 | 0.33790494 | 0.71925477 |
| GALNT7 | 39.9332896 | 0.355996604 | 0.23085387 | 1.54208633 | 0.12305262 | 0.50097964 |
| CEP44 | 31.7697381 | -0.057636916 | 0.28471168 | -0.2024396 | 0.83957308 | 0.96770443 |
| SPCS3 | 62.170157 | -0.317135498 | 0.27675217 | -1.1459187 | 0.25182878 | 0.64619053 |
| TENM3 | 338.450474 | -0.991803204 | 0.49602849 | -1.9994884 | 0.04555554 | 0.32037095 |
| DCTD | 22.6252681 | -0.770117364 | 0.33450591 | -2.3022534 | 0.02132089 | 0.21552573 |
| WWC2 | 187.295672 | 0.142838965 | 0.17763554 | 0.80411255 | 0.42133198 | 0.77771095 |
| TRAPPC11 | 47.4112564 | -0.228808348 | 0.2048954 | -1.1167081 | 0.26411917 | 0.6579701 |
| STOX2 | 26.7692407 | 0.501854772 | 0.47432226 | 1.05804599 | 0.29003447 | 0.68076121 |
| IRF2 | 81.5886875 | -0.471942139 | 0.28833175 | -1.6368025 | 0.10167174 | 0.46169744 |
| PRIMPOL | 28.1345289 | 0.101855025 | 0.27685628 | 0.36789855 | 0.71294888 | 0.93115158 |
| ACSL1 | 60.7584557 | -0.84684762 | 0.41649226 | -2.0332854 | 0.0420237 | 0.30845286 |
| SLC25A4 | 30.632015 | -0.620545889 | 0.44447478 | -1.3961329 | 0.16267447 | 0.55715559 |
| CFAP97 | 42.4405856 | -0.240752997 | 0.24650493 | -0.976666 | 0.32873451 | 0.71294249 |
| SNX25 | 134.165087 | -0.103746299 | 0.31204829 | -0.3324687 | 0.73953534 | 0.93594619 |
| ANKRD37 | 65.1129243 | -3.274909053 | 1.36886177 | -2.3924323 | 0.01673712 | 0.19005424 |
| UFSP2 | 34.506376 | -0.141501538 | 0.34210461 | -0.4136207 | 0.67915194 | 0.91831925 |
| PDLIM3 | 158.444204 | -0.034642793 | 0.55033314 | -0.0629488 | 0.9498073 | 0.99060401 |
| SORBS2 | 310.775924 | 0.878751883 | 0.33305894 | 2.63842753 | 0.00832915 | 0.12814321 |
| CYP4V2 | 21.6202113 | 0.816076134 | 0.40580356 | 2.01101274 | 0.04432411 | 0.31594364 |
| FAT1 | 76.5103005 | 0.62931692 | 0.34917587 | 1.8022921 | 0.07149946 | 0.39746561 |
| SDHA | 58.6444658 | -0.291921714 | 0.25082101 | -1.1638647 | 0.24447886 | 0.64029762 |
| PDCD6 | 28.2800477 | -0.324310868 | 0.33750931 | -0.9608946 | 0.33660518 | 0.71910216 |
| EXOC3 | 27.190887 | -0.106256837 | 0.32386952 | -0.3280853 | 0.74284715 | 0.93768184 |
| BRD9 | 19.5285195 | 0.267988737 | 0.38092698 | 0.70351734 | 0.48173341 | 0.81369393 |
| SLC12A7 | 30.2834026 | -0.565603049 | 0.33513643 | -1.68768 | 0.09147268 | 0.44124451 |
| CLPTM1L | 26.9432596 | -0.344791323 | 0.36607489 | -0.9418601 | 0.34626429 | 0.72436557 |
| LPCAT1 | 32.2177733 | -0.317058042 | 0.25501024 | -1.2433149 | 0.21375181 | 0.61001472 |
| NDUFS6 | 30.8767898 | -0.307020561 | 0.30597607 | -1.0034136 | 0.31566132 | 0.70306809 |
| ICE1 | 67.9144661 | 0.234613377 | 0.17570936 | 1.33523552 | 0.18179927 | 0.5808882 |
| MED10 | 20.3091265 | -0.41814364 | 0.2551756 | -1.6386506 | 0.10128606 | 0.46169744 |
| NSUN2 | 32.5874183 | 0.259155045 | 0.30109853 | 0.86069847 | 0.38940414 | 0.75627141 |
| TENT4A | 62.4195111 | -0.115887481 | 0.20651484 | -0.5611581 | 0.57468974 | 0.86294056 |
| ADCY2 | 34.8886628 | -1.122144847 | 0.42313153 | -2.6520001 | 0.00800165 | 0.12511519 |
| MTRR | 25.2407753 | -0.450147124 | 0.31342279 | -1.4362297 | 0.15093698 | 0.53880252 |
| SEMA5A | 199.169606 | 0.646719722 | 0.25281773 | 2.55804735 | 0.01052618 | 0.14415712 |
| CCT5 | 35.1888124 | -0.226904757 | 0.36720174 | -0.6179294 | 0.53662186 | 0.84423863 |
| CMBL | 28.5756847 | -0.968958718 | 1.02696743 | -0.9435146 | 0.34541778 | 0.72341842 |
| 6-Mar | 164.963235 | -0.015895453 | 0.21766505 | -0.0730271 | 0.94178453 | 0.98916315 |
| DAP | 48.5026282 | -0.252414815 | 0.26493103 | -0.9527567 | 0.34071335 | 0.71952201 |
| TRIO | 432.826315 | 0.202535246 | 0.23506472 | 0.8616148 | 0.38889952 | 0.75549104 |
| OTULIN | 27.4825798 | -0.175020579 | 0.31882035 | -0.548963 | 0.58303083 | 0.86764709 |
| ANKH | 122.880523 | -0.051124925 | 0.33861783 | -0.1509812 | 0.87999055 | 0.97778206 |
| FBXL7 | 649.338445 | 0.760292987 | 0.28553672 | 2.66268028 | 0.0077521 | 0.1230458 |
| RETREG1 | 28.5628923 | -1.966316281 | 0.41606224 | -4.7260148 | 2.29E-06 | 0.00052584 |
| MYO10 | 217.943973 | -0.039585646 | 0.3151712 | -0.1256005 | 0.9000482 | 0.98079096 |
| BASP1 | 79.5269008 | -0.735931942 | 0.34496848 | -2.133331 | 0.03289759 | 0.27693513 |
| CDH6 | 45.1466468 | 0.859410584 | 0.32342075 | 2.65725248 | 0.00787804 | 0.12441768 |
| DROSHA | 49.7677034 | -0.090996826 | 0.18294649 | -0.4973959 | 0.61890992 | 0.88593086 |
| PDZD2 | 168.857476 | -0.025496647 | 0.51048527 | -0.0499459 | 0.9601655 | 0.99181157 |
| GOLPH3 | 137.740805 | -0.176562245 | 0.25453465 | -0.6936668 | 0.48789117 | 0.81595238 |
| MTMR12 | 44.1742263 | -0.17849748 | 0.23583993 | -0.7568586 | 0.44913459 | 0.79362589 |
| ZFR | 181.507839 | -0.025429385 | 0.2193016 | -0.1159562 | 0.90768724 | 0.98306694 |
| SUB1 | 75.3661983 | 0.010294788 | 0.2703193 | 0.03808381 | 0.96962086 | 0.99300191 |
| NPR3 | 35.5745951 | 1.386268562 | 0.78343667 | 1.76947113 | 0.07681528 | 0.41115479 |
| TARS | 30.8908305 | -0.506871833 | 0.3250345 | -1.5594401 | 0.11889225 | 0.493917 |
| ADAMTS12 | 172.445968 | -0.035468829 | 0.48980866 | -0.0724136 | 0.94227273 | 0.9893931 |
| C1QTNF3-AN | 45.675961 | 0.482275334 | 0.39594341 | 1.21804108 | 0.22320836 | 0.61772146 |
| RAI14 | 172.313259 | 0.326084028 | 0.31776073 | 1.0261936 | 0.30480034 | 0.69650244 |
| DNAJC21 | 31.2315542 | 0.325254811 | 0.27374132 | 1.1881831 | 0.23476128 | 0.63080829 |
| SPEF2 | 29.1958621 | 0.942605119 | 0.34996901 | 2.693396 | 0.00707282 | 0.11576428 |
| LMBRD2 | 27.2007209 | 0.158975049 | 0.29380304 | 0.54109395 | 0.58844283 | 0.8702694 |
| NADK2 | 42.9604945 | 0.04712545 | 0.34874331 | 0.13512933 | 0.89250962 | 0.98023547 |
| RANBP3L | 95.3120839 | 1.488160302 | 0.97148408 | 1.53184219 | 0.12556138 | 0.50440978 |
| NIPBL | 402.202906 | -0.104831775 | 0.18667573 | -0.5615715 | 0.57440799 | 0.86294056 |
| CPLANE1 | 117.34332 | 0.321615496 | 0.21602575 | 1.48878315 | 0.13654448 | 0.52089048 |
| NUP155 | 25.496579 | -0.154075042 | 0.29726692 | -0.5183054 | 0.60424522 | 0.87903643 |
| WDR70 | 201.623114 | -0.14741912 | 0.13754497 | -1.0717885 | 0.28381502 | 0.67741668 |
| LIFR | 210.760003 | 0.168650906 | 0.41344293 | 0.40791823 | 0.68333371 | 0.91933701 |
| OSMR-AS1 | 157.118352 | -0.711941247 | 0.2763401 | -2.5763226 | 0.00998574 | 0.14112543 |
| OSMR | 374.725256 | -1.030480951 | 0.33798675 | -3.0488797 | 0.00229696 | 0.05984073 |
| RICTOR | 186.040004 | -0.248367103 | 0.25419259 | -0.9770824 | 0.32852836 | 0.71294249 |
| TTC33 | 48.5836013 | -0.150855161 | 0.23374472 | -0.6453842 | 0.51867821 | 0.83297983 |
| PTGER4 | 34.8551115 | 0.935953882 | 0.30163115 | 3.10297483 | 0.00191586 | 0.05543169 |
| PRKAA1 | 79.514238 | -0.265569447 | 0.2470501 | -1.0749619 | 0.28239178 | 0.6768745 |
| RPL37 | 74.9359412 | -0.550976026 | 0.26325835 | -2.09291 | 0.03635719 | 0.28969304 |
| C7 | 1142.63976 | 0.033839515 | 0.27569184 | 0.12274398 | 0.90230983 | 0.98164203 |
| PLCXD3 | 215.145273 | -0.781580605 | 0.38991519 | -2.0044887 | 0.04501773 | 0.31898889 |
| OXCT1 | 50.9939856 | 0.374735593 | 0.25204731 | 1.48676686 | 0.1370764 | 0.52089048 |
| GHR | 370.461266 | 0.393994664 | 0.4469913 | 0.88143698 | 0.37808135 | 0.74933113 |
| SELENOP | 120.071175 | 0.942055627 | 0.37233349 | 2.53013936 | 0.01140172 | 0.15215018 |
| ZNF131 | 95.7654353 | 0.031538922 | 0.23903527 | 0.13194255 | 0.89502973 | 0.98023547 |
| HMGCS1 | 42.510418 | 0.230618292 | 0.38888879 | 0.59301861 | 0.5531687 | 0.85188037 |
| TMEM267 | 31.8491969 | -0.508640193 | 0.29535223 | -1.7221478 | 0.08504275 | 0.42746464 |
| PAIP1 | 29.4293843 | -0.008679624 | 0.21287451 | -0.0407734 | 0.96747652 | 0.99300191 |
| NNT-AS1 | 22.840583 | 0.217309099 | 0.31791783 | 0.68353856 | 0.49426659 | 0.81957698 |
| NNT | 72.4000268 | -0.160051976 | 0.21395085 | -0.7480782 | 0.45441296 | 0.79609632 |
| PARP8 | 97.8999299 | -0.53489834 | 0.30562498 | -1.7501787 | 0.08008748 | 0.41886353 |
| ITGA1 | 807.964304 | 0.529916202 | 0.37703292 | 1.40549052 | 0.15987547 | 0.5530386 |
| ITGA2 | 75.4378721 | -0.280076183 | 0.5617401 | -0.4985868 | 0.61807053 | 0.88534783 |
| NDUFS4 | 86.0015165 | -0.052979023 | 0.22316614 | -0.2373972 | 0.81234864 | 0.96026221 |
| ARL15 | 275.347312 | -0.051406536 | 0.24996248 | -0.205657 | 0.83705884 | 0.96692006 |
| SNX18 | 35.5622765 | -0.000664027 | 0.34720988 | -0.0019125 | 0.99847408 | 0.99969762 |
| DHX29 | 57.3028351 | -0.089845793 | 0.20879628 | -0.4303036 | 0.6669748 | 0.91345468 |
| MTREX | 126.424838 | -0.318855802 | 0.1957992 | -1.6284837 | 0.10342237 | 0.46372851 |
| PLPP1 | 611.632503 | -0.081397476 | 0.42300282 | -0.1924277 | 0.84740717 | 0.97044434 |

| | | | | | | |
|---|---|---|---|---|---|---|
| SLC38A9 | 62.8942697 | -0.212554756 | 0.21700774 | -0.9794801 | 0.32734283 | 0.71257181 |
| IL6ST | 333.240787 | -0.263809753 | 0.26852537 | -0.9824388 | 0.32588371 | 0.71149668 |
| MAP3K1 | 63.8547877 | 0.107898098 | 0.24624026 | 0.43818219 | 0.66125422 | 0.91019989 |
| MIER3 | 19.6648097 | 0.305472541 | 0.27138712 | 1.12559706 | 0.26033611 | 0.65328956 |
| GPBP1 | 191.838934 | -0.121532337 | 0.26033193 | -0.4668361 | 0.64061716 | 0.89931147 |
| PDE4D | 1525.99087 | -0.81997642 | 0.36420426 | -2.2514191 | 0.02435901 | 0.23128467 |
| ERCC8 | 28.6474519 | 0.034936196 | 0.3006238 | 0.11621234 | 0.90748426 | 0.98306694 |
| NDUFAF2 | 112.339046 | -0.685280828 | 0.238469 | -2.8736684 | 0.00405735 | 0.08259679 |
| ZSWIM6 | 411.421608 | 0.451804594 | 0.19327401 | 2.33763764 | 0.01940605 | 0.20490674 |
| KIF2A | 64.3088337 | 0.180997482 | 0.18416321 | 0.98281019 | 0.32570087 | 0.71149668 |
| DIMT1 | 21.1526992 | -0.533007568 | 0.28393455 | -1.8772198 | 0.06048797 | 0.36535567 |
| IPO11 | 38.0972314 | -0.205048211 | 0.2172261 | -0.9439391 | 0.34520077 | 0.72325086 |
| RNF180 | 132.737443 | 0.562676521 | 0.21585109 | 2.60678096 | 0.00913978 | 0.13406836 |
| RGS7BP | 50.0343413 | 0.455570236 | 0.50818724 | 0.89646138 | 0.3700064 | 0.74253878 |
| SREK1IP1 | 38.786184 | -0.200706538 | 0.25200381 | -0.7964425 | 0.42577491 | 0.77875057 |
| CWC27 | 97.2242176 | -0.354612935 | 0.17096295 | -2.0742093 | 0.03805988 | 0.29501902 |
| ADAMTS6 | 47.9546295 | 1.579260288 | 0.54875364 | 2.87790401 | 0.00400327 | 0.08225816 |
| PPWD1 | 41.8408747 | 0.090386451 | 0.25956409 | 0.34822403 | 0.72767195 | 0.93455731 |
| TRIM23 | 32.8241524 | 0.026154266 | 0.22717114 | 0.11513023 | 0.9083419 | 0.98306694 |
| TRAPPC13 | 32.7097692 | -0.051441691 | 0.24260477 | -0.2120391 | 0.83207655 | 0.96643177 |
| ERBIN | 268.337605 | -0.033050435 | 0.26544968 | -0.1245073 | 0.90091359 | 0.98118567 |
| SREK1 | 99.1300512 | 0.002739638 | 0.16719631 | 0.01638576 | 0.98692664 | 0.99719329 |
| MAST4 | 165.641716 | 0.071359521 | 0.2423178 | 0.29448733 | 0.76838555 | 0.94579017 |
| PIK3R1 | 193.872133 | -0.107597997 | 0.40465391 | -0.2659013 | 0.79031523 | 0.95194192 |
| SLC30A5 | 47.2818381 | -0.033502545 | 0.21499344 | -0.1558305 | 0.87616659 | 0.9768643 |
| CDK7 | 20.3438971 | -0.186137256 | 0.27544255 | -0.6757752 | 0.49918336 | 0.82271448 |
| RAD17 | 24.3881008 | 0.376138412 | 0.34905488 | 1.07759103 | 0.28121631 | 0.67604143 |
| GTF2H2C | 17.4712883 | 0.594789569 | 0.3227755 | 1.84273457 | 0.06536777 | 0.38095778 |
| NAIP | 60.5791217 | 0.77563396 | 0.28461296 | 2.7252236 | 0.0064258 | 0.10918078 |
| GTF2H2 | 19.0074615 | 0.177668423 | 0.36623091 | 0.48512678 | 0.62758642 | 0.89192276 |
| BDP1 | 121.962379 | 0.117834678 | 0.15376763 | 0.76631654 | 0.44348798 | 0.7920275 |
| MCCC2 | 36.0273738 | -0.108996348 | 0.21110879 | -0.5163042 | 0.60564198 | 0.87944337 |
| MAP1B | 210.818218 | 0.270987456 | 0.27165428 | 0.99754534 | 0.31849988 | 0.70683013 |
| MRPS27 | 68.1041265 | -0.051169855 | 0.18385572 | -0.2783153 | 0.78077036 | 0.9491946 |
| TNPO1 | 152.573269 | 0.227438243 | 0.17053036 | 1.33371112 | 0.18229855 | 0.58106792 |
| FCHO2 | 112.547244 | -0.018479306 | 0.21502649 | -0.0859397 | 0.93151438 | 0.98681039 |
| BTF3 | 54.2534783 | -0.425408202 | 0.31487629 | -1.3510328 | 0.17668492 | 0.57377707 |
| ANKRA2 | 22.3928964 | 0.046255324 | 0.29800949 | 0.15521426 | 0.8766524 | 0.9768643 |
| ARHGEF28 | 68.9192565 | 0.101570555 | 0.33810198 | 0.30041396 | 0.76386141 | 0.94462105 |
| HEXB | 27.3042725 | -0.233563077 | 0.29017037 | -0.804917 | 0.42086756 | 0.77771095 |
| GFM2 | 22.2027625 | 0.065544345 | 0.33827088 | 0.19376289 | 0.84636154 | 0.97014274 |
| NSA2 | 21.7750352 | -0.454925832 | 0.37915083 | -1.1998545 | 0.23019584 | 0.62447176 |
| HMGCR | 19.2002321 | -0.082462538 | 0.43221808 | -0.1907892 | 0.84869076 | 0.97092453 |
| COL4A3BP | 148.490124 | -0.575804079 | 0.27747614 | -2.0751481 | 0.03797281 | 0.29501902 |
| POLK | 117.888633 | -0.139258319 | 0.25790458 | -0.5399606 | 0.58922418 | 0.8702694 |
| POC5 | 19.2575493 | -0.29733772 | 0.29857705 | -0.9958492 | 0.31932341 | 0.70683013 |
| IQGAP2 | 25.4177294 | -0.335399381 | 0.53074571 | -0.6319399 | 0.52742616 | 0.83938843 |
| F2R | 42.8213142 | 0.78435675 | 0.29145547 | 2.69117184 | 0.00712015 | 0.11602488 |
| AC022414.1 | 28.2908173 | -0.606572589 | 0.49523445 | -1.2248191 | 0.2206434 | 0.61590675 |
| PDE8B | 23.5989342 | -0.069991473 | 0.59806614 | -0.1170297 | 0.90683656 | 0.98306694 |
| WDR41 | 111.306945 | 0.025175478 | 0.41647151 | 0.06044946 | 0.95179767 | 0.99060401 |
| TBCA | 124.750301 | -0.352193349 | 0.16115993 | -2.1853654 | 0.02886206 | 0.25647799 |
| AP3B1 | 176.736849 | -0.406308179 | 0.18093462 | -2.2456077 | 0.02472915 | 0.23328859 |
| SCAMP1 | 55.9504783 | 0.114439982 | 0.27159137 | 0.42136826 | 0.67348619 | 0.91505229 |
| LHFPL2 | 80.1561965 | -0.884704458 | 0.46253421 | -1.912733 | 0.05578224 | 0.35425851 |
| ARSB | 34.9218905 | -0.263962856 | 0.38859849 | -0.6792689 | 0.49696751 | 0.82183039 |
| DMGDH | 18.499284 | 0.571381417 | 0.2931919 | 1.94883083 | 0.05131563 | 0.34408629 |
| BHMT2 | 27.1620164 | 0.063251151 | 0.52980267 | 0.11938625 | 0.90496936 | 0.98265659 |
| JMY | 52.2633132 | -0.023607191 | 0.27512858 | -0.0858042 | 0.93162206 | 0.98681039 |
| HOMER1 | 36.7943172 | 0.239627272 | 0.43194855 | 0.55475882 | 0.57905963 | 0.86486944 |
| TENT2 | 196.402934 | 0.185181598 | 0.21821807 | 0.84860801 | 0.39609945 | 0.76020146 |
| SERINC5 | 82.9309608 | 0.625130076 | 0.24777371 | 2.52298795 | 0.01163624 | 0.15416382 |
| ZFYVE16 | 110.678935 | -0.071059457 | 0.25151276 | -0.2825282 | 0.77753849 | 0.94808867 |
| DHFR | 158.434461 | -0.822230866 | 0.24757434 | -3.3211473 | 0.00089648 | 0.03554567 |
| MSH3 | 75.6684709 | 0.046024027 | 0.23193984 | 0.19843088 | 0.84270796 | 0.96844084 |
| RASGRF2 | 28.2279742 | 0.474333119 | 0.48924138 | 0.96952779 | 0.33228192 | 0.71527236 |
| CKMT2-AS1 | 45.6635857 | 0.100188122 | 0.2978086 | 0.33641782 | 0.73655581 | 0.93594619 |
| SSBP2 | 584.531835 | 0.181791352 | 0.28606023 | 0.63550027 | 0.52510219 | 0.83726969 |
| ATG10 | 70.9130764 | 0.392767381 | 0.29911477 | 1.31309924 | 0.18914951 | 0.58671246 |
| RPS23 | 143.637837 | -0.348051086 | 0.35279252 | -0.9865603 | 0.32385826 | 0.71051224 |
| ATP6AP1L | 22.818009 | 0.68788171 | 0.44701406 | 1.53883684 | 0.12384413 | 0.50097964 |
| TMEM167A | 31.7096138 | -0.530845916 | 0.25291877 | -2.0988791 | 0.03582756 | 0.28882238 |
| XRCC4 | 40.3602263 | -0.859741328 | 0.25727478 | -3.3417241 | 0.0008326 | 0.03373482 |
| VCAN | 213.350928 | -0.633092883 | 0.48213611 | -1.3130999 | 0.18914928 | 0.58671246 |
| EDIL3 | 182.379889 | 0.402886877 | 0.43489214 | 0.92640644 | 0.3542348 | 0.73059415 |
| AC113383.1 | 30.5880961 | 0.667912843 | 0.39962873 | 1.6713334 | 0.09465584 | 0.44706027 |
| COX7C | 54.8064927 | -0.711657436 | 0.4028815 | -1.7664188 | 0.07732562 | 0.4114983 |
| RASA1 | 145.501655 | 0.00905203 | 0.2449259 | 0.03695824 | 0.9705183 | 0.99300191 |
| CCNH | 346.834973 | -0.372016693 | 0.29294166 | -1.2699344 | 0.20410799 | 0.60367992 |
| TMEM161B | 39.1013307 | -0.449943951 | 0.21285531 | -2.1138488 | 0.03452818 | 0.2833153 |
| TMEM161B-. | 78.1799691 | -0.056736979 | 0.24524058 | -0.2313523 | 0.8170411 | 0.96134613 |
| MEF2C | 262.214664 | 0.538938168 | 0.29695056 | 1.81490872 | 0.06953794 | 0.39291646 |
| MEF2C-AS1 | 81.3347102 | 0.835732458 | 0.44812279 | 1.86496308 | 0.06218658 | 0.37064816 |
| LYSMD3 | 21.5024637 | -0.440566534 | 0.38434867 | -1.1462679 | 0.25168433 | 0.64619053 |
| ARRDC3 | 121.818795 | -1.446497525 | 0.44542096 | -3.2474842 | 0.0011643 | 0.04285549 |
| AC124854.1 | 63.0512394 | -0.808318259 | 0.45945444 | -1.7593001 | 0.07852655 | 0.41542971 |
| NR2F1-AS1 | 368.352041 | 0.012855569 | 0.2369124 | 0.05426297 | 0.95672565 | 0.99181157 |
| NR2F1 | 277.236476 | -0.307658485 | 0.36398339 | -0.8452542 | 0.39796894 | 0.7611823 |
| FAM172A | 300.760703 | -0.126603601 | 0.19965121 | -0.6341239 | 0.52599997 | 0.83815563 |
| KIAA0825 | 79.9242113 | 0.240136785 | 0.20291497 | 1.18343552 | 0.23663659 | 0.63270446 |
| TTC37 | 119.272472 | -0.489343141 | 0.23742154 | -2.0610731 | 0.03929607 | 0.30005099 |
| RHOBTB3 | 174.291027 | -0.978707046 | 0.39675527 | -2.4667777 | 0.0136335 | 0.17065692 |
| GLRX | 62.5518686 | -1.581152554 | 0.49456519 | -3.1970559 | 0.00138838 | 0.04575429 |
| ELL2 | 546.038592 | -1.869564575 | 0.63959031 | -2.9230658 | 0.00346603 | 0.07625361 |
| AC104123.1 | 81.5942484 | -0.191795384 | 0.34278393 | -0.5595227 | 0.57580505 | 0.86295198 |
| CAST | 188.864276 | -0.262804801 | 0.27962988 | -0.9398309 | 0.34730431 | 0.72530246 |
| ERAP1 | 43.6572104 | -0.392421657 | 0.32312984 | -1.2144395 | 0.22457992 | 0.61883685 |
| ERAP2 | 18.5757801 | 0.740168298 | 0.50121361 | 1.4767522 | 0.13974207 | 0.52592211 |
| LNPEP | 97.1816681 | -0.097726772 | 0.22466475 | -0.4349894 | 0.66357016 | 0.91141325 |
| LIX1-AS1 | 26.8902174 | -0.32582998 | 0.41540263 | -0.7843715 | 0.43282216 | 0.78461027 |
| CHD1 | 129.190364 | -0.093015346 | 0.26948066 | -0.3451652 | 0.72997018 | 0.93455731 |
| LINC02062 | 29.9987604 | -0.249365446 | 0.26999714 | -0.9235855 | 0.35570216 | 0.73125149 |
| FAM174A | 20.2866212 | -0.07226986 | 0.30126432 | -0.2398885 | 0.81041666 | 0.95967025 |
| PAM | 320.579679 | -0.504789236 | 0.27701698 | -1.8222321 | 0.06841977 | 0.38947862 |
| PPIP5K2 | 86.0005331 | 0.242499603 | 0.1924309 | 1.26019055 | 0.20760063 | 0.6059796 |
| EFNA5 | 140.440806 | 0.017000351 | 0.45484048 | 0.03737651 | 0.9701848 | 0.99300191 |
| FBXL17 | 301.164034 | -0.010773257 | 0.17142373 | -0.0628458 | 0.94988932 | 0.99060401 |
| FER | 382.632174 | 0.244691747 | 0.30695472 | 0.79715908 | 0.42535865 | 0.77861803 |
| PJA2 | 157.552715 | -0.100687793 | 0.20022983 | -0.5028611 | 0.61506193 | 0.88360697 |
| MAN2A1 | 124.502693 | 0.309652919 | 0.29217786 | 1.05980965 | 0.2892312 | 0.67992862 |
| TMEM232 | 28.278575 | 0.596353486 | 0.44434854 | 1.34208493 | 0.17956846 | 0.57744792 |
| SLC25A46 | 46.8617001 | -0.143341552 | 0.22066662 | -0.6495842 | 0.51596084 | 0.83174649 |
| WDR36 | 35.442015 | -0.208976789 | 0.24830823 | -0.8416023 | 0.40001058 | 0.7630739 |

| | | | | | | |
|---|---|---|---|---|---|---|
| CAMK4 | 43.2616398 | 1.260856831 | 0.90646471 | 1.39096075 | 0.16423733 | 0.55949572 |
| STARD4 | 24.8684081 | -0.014954308 | 0.42238355 | -0.0354046 | 0.97175714 | 0.99300191 |
| NREP | 23.6397967 | 1.54399223 | 0.52037809 | 2.96705852 | 0.00300664 | 0.07059354 |
| EPB41L4A | 82.8927191 | 0.510930322 | 0.27967913 | 1.82684464 | 0.06772313 | 0.38791683 |
| APC | 69.1836861 | 0.477231379 | 0.25112971 | 1.90033821 | 0.05738875 | 0.358022 |
| SRP19 | 29.6510142 | 0.001201057 | 0.33294796 | 0.00360734 | 0.99712177 | 0.99902493 |
| REEP5 | 77.8898954 | -0.015547908 | 0.16966976 | -0.0916363 | 0.92698702 | 0.98678435 |
| DCP2 | 57.2143448 | 0.207456531 | 0.20557793 | 1.00913813 | 0.31290839 | 0.70282967 |
| MCC | 189.54446 | -0.628206213 | 0.4160712 | -1.5098527 | 0.13108102 | 0.51290135 |
| YTHDC2 | 106.050182 | 0.186457254 | 0.2271471 | 0.82086564 | 0.4117228 | 0.77172894 |
| PGGT1B | 32.1366379 | -0.223798577 | 0.22578694 | -0.9911936 | 0.32159105 | 0.70780851 |
| ATG12 | 41.2278589 | -0.377976976 | 0.22519133 | -1.6784704 | 0.09325531 | 0.44361507 |
| AP3S1 | 122.586933 | -0.487982587 | 0.26110508 | -1.8689127 | 0.06163497 | 0.36915678 |
| COMMD10 | 183.263684 | -0.31636139 | 0.18590561 | -1.7017313 | 0.08880575 | 0.43544541 |
| SEMA6A | 24.7978111 | 0.180930509 | 0.26756771 | 0.67620458 | 0.49891077 | 0.82271448 |
| SEMA6A-AS1 | 24.2413511 | 0.000937364 | 0.32536331 | 0.00288098 | 0.99770132 | 0.99945221 |
| DTWD2 | 24.0311876 | -0.374972851 | 0.3043791 | -1.2319271 | 0.21797633 | 0.61478732 |
| DMXL1 | 161.442933 | 0.132825727 | 0.15950515 | 0.83273629 | 0.40499347 | 0.76629687 |
| TNFAIP8 | 78.1351058 | -0.670527983 | 0.41446062 | -1.6178328 | 0.10569864 | 0.4682214 |
| HSD17B4 | 99.6026758 | -0.083815358 | 0.24585431 | -0.3409147 | 0.73316777 | 0.93574285 |
| PRR16 | 338.148116 | -0.460934324 | 0.32802425 | -1.4051837 | 0.15996666 | 0.5530386 |
| SRFBP1 | 46.3572905 | -0.580808404 | 0.29816614 | -1.9479355 | 0.05142268 | 0.34417603 |
| LOX | 35.594011 | -0.9462858 | 0.64689896 | -1.4628031 | 0.14352127 | 0.53188996 |
| SNCAIP | 49.2393521 | -1.091077537 | 0.41888499 | -2.6047186 | 0.00919497 | 0.13415695 |
| SNX2 | 47.7544098 | 0.178563738 | 0.2354225 | 0.75848203 | 0.44816247 | 0.79296348 |
| SNX24 | 49.3021606 | 0.298067869 | 0.298281 | 0.99928547 | 0.31765642 | 0.70591383 |
| CEP120 | 36.4053641 | -0.079201252 | 0.32783445 | -0.2415892 | 0.80909852 | 0.95939969 |
| CSNK1G3 | 71.6156506 | -0.253731456 | 0.22636195 | -1.1209104 | 0.26232602 | 0.65630111 |
| ZNF608 | 97.9452817 | -0.346092822 | 0.71441123 | -0.4844448 | 0.62807022 | 0.89226523 |
| GRAMD2B | 304.639513 | -0.214929652 | 0.40104762 | -0.5359205 | 0.59201348 | 0.87258028 |
| AC093535.1 | 32.057697 | -0.326237133 | 0.63412093 | -0.5144715 | 0.6069224 | 0.88000768 |
| ALDH7A1 | 91.9263841 | 0.305173944 | 0.49753918 | 0.61336665 | 0.53963393 | 0.84521947 |
| PHAX | 19.6978016 | -0.015875685 | 0.31694727 | -0.0500894 | 0.96005118 | 0.99181157 |
| 3-Mar | 138.812313 | -1.932998678 | 0.77237497 | -2.5026687 | 0.01232609 | 0.16004312 |
| PRRC1 | 41.5349482 | -0.182180287 | 0.29056173 | -0.6269934 | 0.53066357 | 0.84152078 |
| LINC01184 | 75.0890709 | 0.192698243 | 0.23531277 | 0.81890262 | 0.41284198 | 0.77260518 |
| SLC12A2 | 60.4256622 | 0.214095929 | 0.33205711 | 0.64475635 | 0.5190851 | 0.83297983 |
| CHSY3 | 97.9409997 | -0.336536366 | 0.29306126 | -1.1483482 | 0.25082486 | 0.6460368 |
| HINT1 | 88.6232062 | -1.14970718 | 0.30799311 | -3.7328991 | 0.00018929 | 0.01276221 |
| CDC42SE2 | 72.79987 | -0.682296111 | 0.28416582 | -2.4010491 | 0.01634814 | 0.18860673 |
| P4HA2 | 23.6714224 | -0.110588229 | 0.33600012 | -0.3291315 | 0.74205628 | 0.93748868 |
| AC116366.3 | 119.179412 | 0.299656758 | 0.43088375 | 0.69544688 | 0.48677529 | 0.81595238 |
| IRF1 | 29.3446078 | 0.365301936 | 0.52425699 | 0.69679937 | 0.48592836 | 0.81594436 |
| KIF3A | 29.4138521 | 0.475788518 | 0.35164339 | 1.35304269 | 0.17604199 | 0.57311399 |
| 8-Sep | 22.3763902 | 0.200249785 | 0.35144393 | 0.56979157 | 0.56881908 | 0.85958696 |
| UQCRQ | 31.9275542 | -0.523171898 | 0.34584161 | -1.51275 | 0.13034317 | 0.51204187 |
| AFF4 | 261.203993 | -0.206346181 | 0.18439781 | -1.1190273 | 0.2631285 | 0.65753753 |
| ZCCHC10 | 20.6639887 | -0.194773666 | 0.27208618 | -0.7158528 | 0.47408222 | 0.80792204 |
| HSPA4 | 50.4986289 | 0.161185254 | 0.24089425 | 0.66911209 | 0.50342398 | 0.82430366 |
| C5orf15 | 32.6341503 | -0.15761 | 0.2577207 | -0.6115535 | 0.5408332 | 0.84601601 |
| VDAC1 | 50.4820289 | -0.422343306 | 0.24848992 | -1.6996396 | 0.08919873 | 0.43560988 |
| SKP1 | 88.947697 | -0.071232718 | 0.23273618 | -0.3060664 | 0.75955411 | 0.94385579 |
| CDKL3 | 29.1767887 | 0.17889833 | 0.31349292 | 0.57066147 | 0.56822914 | 0.8595777 |
| UBE2B | 115.341775 | -0.526006413 | 0.17482826 | -3.0087035 | 0.00262365 | 0.06512862 |
| JADE2 | 24.8184154 | 0.093681795 | 0.33800251 | 0.27716301 | 0.78165494 | 0.94939885 |
| SAR1B | 39.7532437 | -0.336219238 | 0.22162919 | -1.517035 | 0.12925786 | 0.51204187 |
| SEC24A | 77.5956685 | -0.453119801 | 0.26490386 | -1.7105066 | 0.08717223 | 0.43290196 |
| CAMLG | 23.9910809 | 0.180029832 | 0.26280346 | 0.685036 | 0.49332121 | 0.81908904 |
| DDX46 | 92.2239788 | -0.024495294 | 0.26942104 | -0.0909183 | 0.92755753 | 0.98678435 |
| C5orf24 | 38.2836569 | -0.067426003 | 0.27471478 | -0.24544 | 0.80611574 | 0.95838958 |
| TXNDC15 | 36.977064 | -0.204051425 | 0.25651723 | -0.7954687 | 0.42634091 | 0.77939785 |
| PCBD2 | 27.2877755 | 0.639219372 | 0.42257427 | 1.51267934 | 0.13036113 | 0.51204187 |
| H2AFY | 60.4958346 | -0.238298241 | 0.31188067 | -0.7640687 | 0.44482633 | 0.79252305 |
| CXCL14 | 40.5553856 | -0.831494356 | 0.77505705 | -1.072817 | 0.28335325 | 0.67727609 |
| TGFBI | 94.6639737 | -1.011560496 | 0.3510596 | -2.8814495 | 0.00395851 | 0.08179072 |
| SMAD5 | 99.880822 | -0.264912326 | 0.19877022 | -1.3327566 | 0.18261168 | 0.58106792 |
| SPOCK1 | 277.745249 | -1.35429033 | 0.4388724 | -3.0858407 | 0.00202978 | 0.05607827 |
| HNRNPA0 | 35.12181 | -0.340999181 | 0.37205882 | -0.9165195 | 0.35939447 | 0.73264631 |
| FAM13B | 142.518988 | 0.225232873 | 0.16002113 | 1.40751955 | 0.15927338 | 0.55160231 |
| BRD8 | 29.6090935 | 0.041039834 | 0.24970374 | 0.1643541 | 0.8694524 | 0.9768643 |
| KDM3B | 75.1742908 | 0.038586846 | 0.23807705 | 0.16207713 | 0.87124511 | 0.9768643 |
| EGR1 | 123.736033 | 0.745234748 | 0.86957812 | 0.85700724 | 0.39144088 | 0.75882328 |
| ETF1 | 63.3921223 | -0.44163921 | 0.28478217 | -1.5507965 | 0.12095045 | 0.49849255 |
| HSPA9 | 92.0551735 | -0.564974465 | 0.2349477 | -2.4046818 | 0.01618655 | 0.18703614 |
| CTNNA1 | 305.245665 | -0.062092194 | 0.20256205 | -0.3065342 | 0.75919795 | 0.94385579 |
| SIL1 | 122.61461 | -0.224855288 | 0.22131576 | -1.0159931 | 0.30963267 | 0.69972033 |
| PAIP2 | 63.3694825 | -0.113635355 | 0.21466449 | -0.5293626 | 0.59655392 | 0.87523952 |
| TMEM173 | 35.6551942 | 0.057979572 | 0.28602473 | 0.20270825 | 0.83936308 | 0.96770443 |
| UBE2D2 | 129.948105 | 0.037465588 | 0.18771752 | 0.19958493 | 0.84180522 | 0.96803523 |
| CXXC5 | 35.1143044 | 0.221309635 | 0.36395866 | 0.60806256 | 0.54314598 | 0.84692974 |
| PURA | 73.156909 | 0.01598565 | 0.18745 | 0.08527954 | 0.93203916 | 0.98710992 |
| CYSTM1 | 116.449038 | 0.053302239 | 0.28864873 | 0.18466126 | 0.85349473 | 0.97320912 |
| PFDN1 | 29.6165985 | 0.108487454 | 0.30012487 | 0.36147438 | 0.71774485 | 0.93372582 |
| SRA1 | 24.6179551 | -0.721062813 | 0.29679274 | -2.4295163 | 0.01511898 | 0.17917181 |
| NDUFA2 | 20.5060789 | -0.428098666 | 0.30600328 | -1.3990003 | 0.1618129 | 0.55516481 |
| HARS | 30.8024785 | -0.290142157 | 0.22966265 | -1.2633406 | 0.20646679 | 0.60523513 |
| TAF7 | 33.1177602 | 0.164108837 | 0.29272125 | 0.56063178 | 0.57504858 | 0.86294056 |
| DIAPH1 | 80.0326311 | 0.456047771 | 0.23196001 | 1.96606205 | 0.04929144 | 0.33589213 |
| HDAC3 | 27.9583152 | -0.115673888 | 0.3656026 | -0.3163924 | 0.75170467 | 0.94055881 |
| ARAP3 | 20.3828236 | 0.562383959 | 0.36627023 | 1.53543452 | 0.12467713 | 0.50315883 |
| DELE1 | 32.6972932 | 0.038908696 | 0.2920523 | 0.1332251 | 0.89401536 | 0.98023547 |
| RNF14 | 26.2362446 | 0.151980686 | 0.22660369 | 0.67068935 | 0.50241845 | 0.82430366 |
| NDFIP1 | 125.045346 | 0.049091113 | 0.17009202 | 0.28861503 | 0.77287599 | 0.9472495 |
| SPRY4 | 54.454943 | -0.498857447 | 0.40689736 | -1.2260031 | 0.22019749 | 0.61590675 |
| SPRY4-AS1 | 117.376754 | 0.374512819 | 0.34077386 | 1.09900689 | 0.27176506 | 0.66686313 |
| FGF1 | 42.7483249 | 0.238790003 | 0.3819579 | 0.62517362 | 0.53185711 | 0.84237455 |
| ARHGAP26 | 148.925483 | 0.161068534 | 0.27067248 | 0.595068 | 0.55179802 | 0.85138854 |
| NR3C1 | 288.960526 | -0.043579789 | 0.32312105 | -0.1348714 | 0.89271355 | 0.98023547 |
| YIPF5 | 24.6501715 | -0.765084221 | 0.40446923 | -1.8915758 | 0.05854752 | 0.36041506 |
| PRELID2 | 41.1547904 | 0.848476462 | 0.40471327 | 2.09648788 | 0.03603893 | 0.28882238 |
| LARS | 79.3680272 | -0.285597821 | 0.20385553 | -1.4009814 | 0.16121962 | 0.55470404 |
| TCERG1 | 90.5769046 | -0.088583004 | 0.28109621 | -0.3151341 | 0.75265983 | 0.94058012 |
| DPYSL3 | 55.2781461 | 0.94052748 | 0.55055836 | 1.70831568 | 0.08757778 | 0.43393454 |
| FBXO38 | 44.648251 | 0.081924058 | 0.21206574 | 0.38631444 | 0.69926381 | 0.92769314 |
| ABLIM3 | 23.4200859 | -1.461537654 | 0.45623451 | -3.2034789 | 0.00135778 | 0.04535601 |
| CARMN | 625.763473 | 0.753299159 | 0.40944725 | 1.83979536 | 0.06579829 | 0.38316292 |
| AC131025.2 | 53.6497949 | 0.171435231 | 0.34582909 | 0.49572241 | 0.62009027 | 0.88658432 |
| CSNK1A1 | 410.483261 | -0.309929853 | 0.23362211 | -1.3266289 | 0.18463146 | 0.58363936 |
| PPARGC1B | 24.1404545 | 0.13077396 | 0.53968597 | 0.24231491 | 0.80853616 | 0.95936772 |
| SLC26A2 | 18.959539 | -0.089583877 | 0.42985603 | -0.2084044 | 0.83491324 | 0.96665943 |
| HMGXB3 | 38.1440545 | -0.593474477 | 0.27290421 | -2.1746623 | 0.02965543 | 0.26108772 |
| CSF1R | 19.3312701 | 0.330205711 | 0.54073478 | 0.61066113 | 0.54142395 | 0.84641877 |
| PDGFRB | 476.661082 | 0.332623713 | 0.24647331 | 1.3495324 | 0.17716602 | 0.57388115 |

| | | | | | | |
|---|---|---|---|---|---|---|
| CD74 | 81.8253919 | -0.564695522 | 0.38624432 | -1.4620164 | 0.14373672 | 0.532104 |
| RPS14 | 136.150278 | -0.227679502 | 0.28865813 | -0.7887514 | 0.43025733 | 0.78298297 |
| NDST1 | 37.5841566 | 0.535407771 | 0.28137718 | 1.90281166 | 0.05706512 | 0.35717443 |
| SYNPO | 48.7746195 | -0.122924782 | 0.38551018 | -0.3188626 | 0.7498307 | 0.94004394 |
| DCTN4 | 81.0609133 | 0.003749007 | 0.23682175 | 0.0158305 | 0.98736961 | 0.99719329 |
| SMIM3 | 59.5776651 | -1.334806736 | 0.27643597 | -4.828629 | 1.37E-06 | 0.00032974 |
| GPX3 | 638.555217 | -1.001810044 | 0.44060066 | -2.2737371 | 0.0229818 | 0.22440695 |
| TNIP1 | 62.440561 | -1.354177795 | 0.58368107 | -2.3200646 | 0.02033738 | 0.21050622 |
| ANXA6 | 91.9415693 | 0.393772222 | 0.28728299 | 1.3706771 | 0.17047564 | 0.56801217 |
| SLC36A1 | 18.3784233 | 0.003685843 | 0.36168647 | 0.01019071 | 0.99186913 | 0.99759781 |
| SPARC | 300.551982 | 0.564384875 | 0.35798684 | 1.57655202 | 0.11489864 | 0.48724723 |
| ATOX1 | 32.1452716 | -0.229640881 | 0.29530336 | -0.777644 | 0.43677893 | 0.78768252 |
| G3BP1 | 92.7534916 | -0.240497758 | 0.21173037 | -1.135868 | 0.2560118 | 0.65078891 |
| FAM114A2 | 20.1324343 | -0.537919767 | 0.27826694 | -1.933107 | 0.05322302 | 0.34798571 |
| MFAP3 | 24.2066113 | -0.179599314 | 0.39120853 | -0.4590884 | 0.64617065 | 0.90193887 |
| GALNT10 | 54.4513226 | -0.163119799 | 0.46558153 | -0.3503571 | 0.72607071 | 0.93455731 |
| LARP1 | 109.862999 | 0.027559848 | 0.23302529 | 0.11826977 | 0.90585391 | 0.98303609 |
| FAXDC2 | 35.8700905 | 0.355150506 | 0.49229873 | 0.7214126 | 0.4706557 | 0.80457054 |
| CNOT8 | 30.543138 | -0.576256689 | 0.2513921 | -2.2922625 | 0.0218905 | 0.21819108 |
| GEMIN5 | 20.0916951 | -0.277810486 | 0.28301559 | -0.9816084 | 0.3262928 | 0.71149668 |
| SGCD | 60.8841686 | 1.56070581 | 0.52978566 | 2.94591932 | 0.00321996 | 0.07300955 |
| CYFIP2 | 23.4224785 | 0.751965029 | 0.33715235 | 2.23034194 | 0.02572475 | 0.23956637 |
| ADAM19 | 56.7957665 | -0.054762517 | 0.36927173 | -0.1482987 | 0.88210704 | 0.97782528 |
| CLINT1 | 105.053373 | -0.171436007 | 0.18274693 | -0.9381061 | 0.34818988 | 0.72612016 |
| EBF1 | 1348.24024 | 0.312562494 | 0.19503703 | 1.60258027 | 0.10902735 | 0.47636267 |
| RNF145 | 108.609362 | -0.29910513 | 0.25111697 | -1.1910988 | 0.2336148 | 0.62956917 |
| UBLCP1 | 25.8179523 | 0.030599813 | 0.27568319 | 0.1109963 | 0.91161928 | 0.98343817 |
| ADRA1B | 27.7713191 | 0.992909379 | 0.56964043 | 1.74304583 | 0.08132561 | 0.42087829 |
| TTC1 | 46.0245054 | -0.090693237 | 0.26109782 | -0.3473535 | 0.72832578 | 0.93455731 |
| PWWP2A | 83.3962682 | 0.054584243 | 0.17918443 | 0.30462604 | 0.76065098 | 0.94425408 |
| MAT2B | 22.1886742 | 0.262386863 | 0.2448128 | 1.0717857 | 0.2838163 | 0.67741668 |
| WWC1 | 22.4943272 | -0.567293236 | 0.49961536 | -1.13546 | 0.25618265 | 0.65084527 |
| RARS | 47.4959595 | -0.106459343 | 0.23846902 | -0.4464284 | 0.65528783 | 0.90691684 |
| PANK3 | 48.9106256 | -0.066665963 | 0.21218402 | -0.3141894 | 0.75337722 | 0.94096423 |
| SLIT3 | 1449.2016 | 0.472605346 | 0.30927617 | 1.5281014 | 0.12648736 | 0.50599102 |
| KCNIP1 | 71.2576059 | 0.689177634 | 1.1554655 | 0.59645021 | 0.55087451 | 0.85014212 |
| NPM1 | 51.3898115 | -0.372167517 | 0.27261917 | -1.3651553 | 0.17220423 | 0.56882549 |
| FBXW11 | 169.407166 | -0.000112544 | 0.19599415 | -0.0005742 | 0.99954184 | 0.99969762 |
| STK10 | 61.6391815 | 0.548391446 | 0.20343976 | 2.69559623 | 0.00702628 | 0.11576428 |
| UBTD2 | 52.8396853 | 0.346413337 | 0.21473901 | 1.61318307 | 0.10670474 | 0.47073496 |
| SH3PXD2B | 78.7462299 | -0.269101384 | 0.3364909 | -0.7997286 | 0.42386807 | 0.77857941 |
| NEURL1B | 114.895203 | 0.443712194 | 0.4561279 | 0.97278022 | 0.33066255 | 0.71362138 |
| DUSP1 | 215.326483 | 1.644545423 | 0.59070721 | 2.78402802 | 0.00536884 | 0.09670491 |
| ERGIC1 | 81.0415035 | 0.297782679 | 0.22400283 | 1.32937015 | 0.18372588 | 0.58324038 |
| ATP6V0E1 | 107.777853 | -0.687731086 | 0.17381734 | -3.9566311 | 7.60E-05 | 0.00674861 |
| CREBRF | 99.5073928 | -0.050572376 | 0.33695055 | -0.1500884 | 0.88069486 | 0.97782528 |
| CPEB4 | 142.481213 | -0.224690832 | 0.25429324 | -0.8835895 | 0.37691785 | 0.74832285 |
| SFXN1 | 53.5550237 | -0.364240249 | 0.28207893 | -1.2912707 | 0.19660984 | 0.59435859 |
| HRH2 | 43.0422432 | -1.308705794 | 0.63037256 | -2.0760831 | 0.03788627 | 0.29501902 |
| SIMC1 | 49.2997085 | -0.062629083 | 0.28986459 | -0.2160632 | 0.82893845 | 0.96580172 |
| KIAA1191 | 24.1009798 | -0.385069897 | 0.25821988 | -1.4912481 | 0.13589638 | 0.51988679 |
| CLTB | 36.8989622 | 0.05869244 | 0.36303088 | 0.16167341 | 0.87156305 | 0.9768643 |
| FAF2 | 64.5125235 | -0.115922246 | 0.21907645 | -0.5291406 | 0.59670791 | 0.87529071 |
| UIMC1 | 88.8894512 | 0.104675821 | 0.16867215 | 0.62058745 | 0.5348711 | 0.84361209 |
| ZNF346 | 40.0603703 | 0.231707898 | 0.21006983 | 1.10300415 | 0.27002536 | 0.6651094 |
| NSD1 | 198.493794 | 0.027618602 | 0.15125003 | 0.18260229 | 0.85511009 | 0.97379567 |
| PRELID1 | 27.4421442 | -0.975169834 | 0.31997698 | -3.047625 | 0.00230658 | 0.05984073 |
| LMAN2 | 32.319779 | -0.067799207 | 0.27520012 | -0.2463633 | 0.80540102 | 0.95836441 |
| PDLIM7 | 79.3149192 | 0.869155748 | 0.36768855 | 2.36383683 | 0.01808678 | 0.19729344 |
| FAM193B | 73.8838281 | 0.046812671 | 0.2062531 | 0.22696712 | 0.82044931 | 0.96291025 |
| TMED9 | 45.8281454 | -0.316943453 | 0.42589283 | -0.7441859 | 0.456764 | 0.79732984 |
| RMND5B | 23.9526675 | 0.200806131 | 0.32645657 | 0.61510826 | 0.53848323 | 0.8449569 |
| NHP2 | 21.388639 | -0.922600417 | 0.39200864 | -2.3535206 | 0.01859658 | 0.20009416 |
| HNRNPAB | 37.898849 | -0.665332151 | 0.40966184 | -1.6241009 | 0.10435429 | 0.46422585 |
| PHYKPL | 91.0777018 | -0.121502592 | 0.34728239 | -0.3498668 | 0.72643864 | 0.93455731 |
| COL23A1 | 101.62614 | 0.153048292 | 0.56201978 | 0.27231834 | 0.78537726 | 0.95065355 |
| CLK4 | 41.1723152 | -0.022146228 | 0.29515457 | -0.0750326 | 0.94018874 | 0.9889521 |
| ADAMTS2 | 182.101135 | -0.375334151 | 0.3322198 | -1.1297766 | 0.25857038 | 0.6514155 |
| RUFY1 | 55.8777187 | 0.05762794 | 0.2561377 | 0.22498813 | 0.82198851 | 0.96331679 |
| HNRNPH1 | 175.040995 | 0.104723077 | 0.22927564 | 0.45675622 | 0.64784628 | 0.90256375 |
| CANX | 126.188991 | -0.307261168 | 0.24229849 | -1.2681101 | 0.20475864 | 0.60370888 |
| MAML1 | 41.8483945 | -0.184322744 | 0.22321886 | -0.825749 | 0.4089465 | 0.77020702 |
| SQSTM1 | 197.028794 | -0.781904769 | 0.32721653 | -2.3895638 | 0.01686839 | 0.19013163 |
| MRNIP | 24.4228587 | 0.260324778 | 0.32308623 | 0.80574396 | 0.4203905 | 0.77771095 |
| TBC1D9B | 32.6436646 | 0.036890272 | 0.25065494 | 0.14717552 | 0.88299348 | 0.97782528 |
| RNF130 | 72.2233375 | 0.021416581 | 0.20793808 | 0.10299499 | 0.91796695 | 0.98477389 |
| MAPK9 | 26.5992783 | 0.292326415 | 0.28533662 | 1.02449667 | 0.30560074 | 0.69722386 |
| GFPT2 | 43.54058 | 0.183455367 | 0.62885735 | 0.29172811 | 0.77049452 | 0.94593455 |
| CNOT6 | 44.9134198 | 0.011085503 | 0.2130174 | 0.05204036 | 0.95849653 | 0.99181157 |
| MGAT1 | 38.0147909 | -0.617808729 | 0.29270117 | -2.1107149 | 0.03479682 | 0.28488012 |
| RACK1 | 99.4697531 | -0.643955887 | 0.35084268 | -1.8354548 | 0.06643833 | 0.38523695 |
| DUSP22 | 25.5235876 | -0.416708718 | 0.29865761 | -1.3952724 | 0.16293372 | 0.55718935 |
| EXOC2 | 89.7446886 | -0.030387381 | 0.18406307 | -0.1650922 | 0.8688714 | 0.9768643 |
| GMDS | 172.398498 | -0.515853208 | 0.28658148 | -1.8000228 | 0.07185704 | 0.39915144 |
| GMDS-DT | 169.150879 | 0.037282345 | 0.34291739 | 0.10872107 | 0.91342373 | 0.98395113 |
| WRNIP1 | 32.1302546 | 0.120132776 | 0.20555432 | 0.58443323 | 0.55892888 | 0.85343152 |
| SERPINB1 | 18.6749711 | -0.197603212 | 0.33716851 | -0.5860666 | 0.55783074 | 0.85279817 |
| SERPINB9 | 22.3731314 | 0.225399795 | 0.31548165 | 0.71446246 | 0.47494126 | 0.80851131 |
| SERPINB6 | 85.0827006 | -0.392466017 | 0.27542947 | -1.4249238 | 0.15417922 | 0.54343931 |
| NQO2 | 31.2403006 | 0.07566093 | 0.24894502 | 0.30392626 | 0.76118407 | 0.94425408 |
| RIPK1 | 49.275446 | -0.080745724 | 0.23021928 | -0.350734 | 0.72578793 | 0.93455731 |
| PSMG4 | 19.0721533 | -0.023062272 | 0.41018614 | -0.0562239 | 0.95516343 | 0.99181157 |
| PXDC1 | 31.6004888 | 0.401760438 | 0.30378645 | 1.32250939 | 0.18599858 | 0.58501647 |
| PRPF4B | 97.2751199 | -0.328774665 | 0.19911088 | -1.6512139 | 0.0986949 | 0.45657782 |
| ECI2 | 32.943574 | 0.103738962 | 0.42356963 | 0.24491596 | 0.8065215 | 0.95838958 |
| CDYL | 90.9770013 | -0.111039013 | 0.1882351 | -0.5898954 | 0.55526081 | 0.85207344 |
| LYRM4 | 66.1513776 | -0.748864215 | 0.39132588 | -1.9136588 | 0.05566377 | 0.35417578 |
| FARS2 | 169.871465 | 0.038746358 | 0.19404651 | 0.19967562 | 0.84173428 | 0.96803523 |
| RREB1 | 139.626568 | -0.464063986 | 0.23187921 | -2.0013178 | 0.04535816 | 0.31959454 |
| SSR1 | 60.6630718 | -0.149236494 | 0.32008527 | -0.4662398 | 0.64104386 | 0.89934133 |
| SNRNP48 | 17.8120955 | -0.027240426 | 0.31269277 | -0.0871156 | 0.93057961 | 0.98681039 |
| SLC35B3 | 31.1605664 | -0.133113414 | 0.26843999 | -0.4958777 | 0.61998067 | 0.88658432 |
| TMEM14C | 25.9347686 | 0.196274958 | 0.32177459 | 0.60997656 | 0.54187733 | 0.84641877 |
| MAK | 32.5516565 | 0.087697826 | 0.29903149 | 0.29327288 | 0.76931358 | 0.94579017 |
| AL139807.1 | 34.4231147 | -0.11489023 | 0.5387769 | -0.2132427 | 0.83113768 | 0.96621638 |
| NEDD9 | 74.8642342 | -0.234540303 | 0.43612191 | -0.5377861 | 0.59072472 | 0.87190922 |
| HIVEP1 | 105.22611 | 0.130102191 | 0.36887224 | 0.35270259 | 0.72431142 | 0.93455731 |
| PHACTR1 | 65.9340842 | 0.50487655 | 0.56257507 | 0.89743854 | 0.36948496 | 0.74169488 |
| GFOD1 | 24.2367974 | 1.68243135 | 0.41357384 | 4.06803135 | 4.74E-05 | 0.00488287 |
| NOL7 | 25.3944514 | -0.441657804 | 0.29602957 | -1.4919381 | 0.13571537 | 0.51955288 |
| RANBP9 | 126.102612 | -0.14529021 | 0.23523391 | -0.6176414 | 0.53681173 | 0.84423863 |
| JARID2 | 174.55573 | -0.314465567 | 0.2441121 | -1.2882015 | 0.19767584 | 0.59530234 |
| DTNBP1 | 55.1158298 | -0.21762674 | 0.27679629 | -0.7862343 | 0.43173023 | 0.78398454 |

| | | | | | | |
|---|---|---|---|---|---|---|
| MYLIP | 33.769105 | 0.811680889 | 0.42071011 | 1.92931159 | 0.05369219 | 0.34959616 |
| ATXN1 | 387.678019 | 0.44567695 | 0.27232125 | 1.63658526 | 0.10171716 | 0.46169744 |
| CAP2 | 44.8209399 | 0.285002138 | 0.35698888 | 0.79835017 | 0.42466731 | 0.77861803 |
| NUP153 | 112.346563 | -0.030630428 | 0.31726982 | -0.0965438 | 0.9230887 | 0.98561736 |
| KIF13A | 258.639164 | 0.889643903 | 0.23370364 | 3.8067182 | 0.00014082 | 0.00985627 |
| DEK | 62.6578028 | -0.023653565 | 0.22379116 | -0.1056948 | 0.91582449 | 0.98477389 |
| AL022068.1 | 23.378313 | 0.516428159 | 0.52469503 | 0.98424443 | 0.32499534 | 0.71149668 |
| ID4 | 55.4018387 | 0.625878693 | 0.46849706 | 1.33592875 | 0.18157256 | 0.580649 |
| MBOAT1 | 31.1852534 | 0.155119467 | 0.36648019 | 0.42326835 | 0.67209949 | 0.91482259 |
| E2F3 | 45.1120063 | -0.606114268 | 0.2955735 | -2.0506381 | 0.0403022 | 0.30130305 |
| CDKAL1 | 318.91281 | -0.056539865 | 0.13259244 | -0.4264185 | 0.66980295 | 0.91445841 |
| SOX4 | 40.59966 | 0.299073064 | 0.4530962 | 0.66006527 | 0.50921194 | 0.82663984 |
| CASC15 | 205.795787 | -0.009140011 | 0.57355078 | -0.0159358 | 0.98728558 | 0.99719329 |
| DCDC2 | 23.1257163 | -0.650023976 | 0.36203956 | -1.7954501 | 0.07258201 | 0.40072954 |
| MRS2 | 18.9699192 | 0.02893644 | 0.29108051 | 0.09941043 | 0.9208124 | 0.98528826 |
| TDP2 | 22.4641066 | 0.145375841 | 0.32891678 | 0.44198366 | 0.65850103 | 0.9081111 |
| C6orf62 | 68.6538445 | 0.039037818 | 0.22058215 | 0.17697633 | 0.85952699 | 0.97397549 |
| RIPOR2 | 29.7133015 | 0.760141337 | 0.40510975 | 1.87638373 | 0.06060261 | 0.36535567 |
| CARMIL1 | 201.206266 | -0.193520619 | 0.35816465 | -0.5403119 | 0.58898196 | 0.8702694 |
| TRIM38 | 75.9348702 | 0.047798664 | 0.35563227 | 0.13440474 | 0.89308253 | 0.98023547 |
| HIST1H4C | 19.1391896 | -1.563667081 | 0.90830333 | -1.7215252 | 0.08515556 | 0.42746464 |
| HIST1H2AC | 88.1887982 | -0.149646953 | 0.77738269 | -0.192501 | 0.84734978 | 0.97044434 |
| BTN2A1 | 20.2824203 | 0.722203366 | 0.34049263 | 2.12105432 | 0.03391723 | 0.28164716 |
| TRIM27 | 40.3420194 | -0.179257419 | 0.27454189 | -0.6529328 | 0.51379957 | 0.83023594 |
| GABBR1 | 66.5973335 | 0.347990821 | 0.21734828 | 1.60107465 | 0.10936039 | 0.47668415 |
| HLA-A | 189.504929 | -0.457953022 | 0.2985316 | -1.5340186 | 0.12502508 | 0.50400948 |
| TRIM26 | 25.7793962 | 0.071397918 | 0.29211881 | 0.24441397 | 0.80691022 | 0.95845858 |
| HCG17 | 76.1323642 | 0.373605149 | 0.32130102 | 1.16278855 | 0.2449153 | 0.6405276 |
| HLA-E | 112.917264 | -0.386717988 | 0.32030733 | -1.2073342 | 0.22730348 | 0.62173063 |
| GNL1 | 34.3020135 | 0.494092756 | 0.23943255 | 2.06359891 | 0.03905576 | 0.29897999 |
| ABCF1 | 31.2062387 | -0.277719981 | 0.35944841 | -0.7726282 | 0.43974245 | 0.78933067 |
| PPP1R10 | 45.7776104 | 0.230811965 | 0.27350592 | 0.84390117 | 0.39872463 | 0.76149359 |
| MRPS18B | 20.9299323 | -0.013162339 | 0.31713261 | -0.0415042 | 0.96689394 | 0.99300191 |
| PPP1R18 | 20.9004329 | -0.124374511 | 0.46731442 | -0.2661474 | 0.7901257 | 0.95194192 |
| MDC1 | 17.3878336 | -0.207299668 | 0.31830924 | -0.6512524 | 0.51488355 | 0.83109052 |
| TUBB | 81.7035834 | -0.403596353 | 0.49133243 | -0.8214324 | 0.41140004 | 0.77166383 |
| FLOT1 | 81.8143295 | -0.280854971 | 0.22849212 | -1.229167 | 0.2190092 | 0.61522936 |
| IER3 | 123.184096 | -2.107827172 | 0.84844608 | -2.4843384 | 0.01297924 | 0.1665263 |
| HLA-C | 112.501365 | -0.234797975 | 0.44465418 | -0.5280463 | 0.59746723 | 0.87535619 |
| HLA-B | 162.864091 | -0.769402289 | 0.45449429 | -1.6928756 | 0.09047915 | 0.43947871 |
| MICA | 22.28303 | 0.644079116 | 0.47181528 | 1.36510866 | 0.1722189 | 0.56882549 |
| PRRC2A | 38.358893 | -0.051210538 | 0.30691178 | -0.1668575 | 0.86748216 | 0.97634923 |
| BAG6 | 21.5202666 | -0.155934804 | 0.32829537 | -0.4749833 | 0.6347989 | 0.89672816 |
| DDAH2 | 19.0043853 | 1.015352851 | 0.36096486 | 2.81288558 | 0.00490991 | 0.09204832 |
| CLIC1 | 75.5566361 | -0.274569935 | 0.373617 | -0.7348968 | 0.46240235 | 0.80221787 |
| HSPA1A | 165.196969 | 0.491965659 | 0.75268189 | 0.65361697 | 0.51335861 | 0.82994235 |
| HSPA1B | 76.2231952 | 0.36929155 | 0.88381125 | 0.41783984 | 0.67606422 | 0.91718589 |
| C6orf48 | 25.5620314 | -0.002999449 | 0.31235173 | -0.0096028 | 0.9923382 | 0.99759781 |
| TNXB | 111.010449 | 0.581904914 | 0.48316054 | 1.20437176 | 0.22844592 | 0.62213472 |
| ATF6B | 26.0195114 | 0.191749183 | 0.27602393 | 0.69468318 | 0.48725387 | 0.81595238 |
| PBX2 | 22.5027992 | 0.297555435 | 0.25762259 | 1.15500521 | 0.24808829 | 0.64333128 |
| AL662796.1 | 62.389912 | 1.311677222 | 0.30845349 | 4.25243113 | 2.11E-05 | 0.00305725 |
| HLA-DRA | 31.9761164 | -0.316760981 | 0.47271458 | -0.6700893 | 0.50280087 | 0.82430366 |
| HLA-DRB1 | 26.1174855 | 0.9785211 | 0.64868502 | 1.50846879 | 0.13143459 | 0.51290135 |
| BRD2 | 60.5239771 | -0.106147698 | 0.21213664 | -0.5003742 | 0.61681163 | 0.88482308 |
| SLC39A7 | 26.7514446 | -0.154806897 | 0.41767096 | -0.3706432 | 0.71090331 | 0.93009618 |
| RING1 | 21.9319679 | 0.098598515 | 0.25737187 | 0.38309748 | 0.70164749 | 0.92865476 |
| VPS52 | 17.0914018 | 0.139003156 | 0.27869728 | 0.49876036 | 0.61794821 | 0.88534783 |
| RPS18 | 94.368971 | -0.519950865 | 0.33800958 | -1.5382726 | 0.12398198 | 0.5011791 |
| RGL2 | 17.4690587 | 0.812817834 | 0.36442069 | 2.23043823 | 0.02571836 | 0.23956637 |
| TAPBP | 74.7438559 | -0.071122569 | 0.30275643 | -0.2349168 | 0.81427332 | 0.96038841 |
| PHF1 | 25.0629431 | 0.507725199 | 0.25105741 | 2.02234698 | 0.04314052 | 0.31204692 |
| CUTA | 18.7793244 | 0.594199517 | 0.36919756 | 1.60943511 | 0.10752123 | 0.47265636 |
| SYNGAP1 | 34.6850902 | -0.25726859 | 0.31631846 | -0.8133215 | 0.41603378 | 0.77573656 |
| LEMD2 | 20.6036258 | -0.189339005 | 0.31517153 | -0.6007491 | 0.54800712 | 0.84785354 |
| NUDT3 | 19.703557 | 0.534072252 | 0.49785906 | 1.07273784 | 0.28338876 | 0.67727609 |
| C6orf106 | 90.1275409 | -0.144010019 | 0.17932369 | -0.8030731 | 0.42193251 | 0.77771095 |
| UHRF1BP1 | 33.1298046 | -0.391651985 | 0.33173368 | -1.1806217 | 0.23775304 | 0.63306054 |
| TAF11 | 27.6608548 | 0.103016662 | 0.22819025 | 0.45145076 | 0.6516647 | 0.90548004 |
| ANKS1A | 129.755682 | 0.1105216 | 0.21644653 | 0.5106185 | 0.60961822 | 0.88163721 |
| ZNF76 | 22.6664596 | 0.362708727 | 0.29112989 | 1.24586566 | 0.21281372 | 0.60956449 |
| PPARD | 59.0506956 | -0.495474513 | 0.25834714 | -1.9178634 | 0.05512834 | 0.35260065 |
| RPL10A | 102.267011 | -0.334187564 | 0.35521279 | -0.9408095 | 0.34680249 | 0.7249592 |
| TEAD3 | 64.3515458 | 0.034689369 | 0.27246104 | 0.12731864 | 0.89868821 | 0.98058672 |
| FKBP5 | 467.03695 | -1.511579394 | 0.62527923 | -2.417447 | 0.01562981 | 0.18290362 |
| SRPK1 | 31.9173848 | -0.33856707 | 0.24023546 | -1.4093134 | 0.1587425 | 0.55091387 |
| MAPK14 | 100.223617 | -0.55078586 | 0.25486735 | -2.1610687 | 0.03069003 | 0.26662968 |
| KCTD20 | 60.8941821 | -0.317874483 | 0.29045199 | -1.0944132 | 0.27377382 | 0.66906109 |
| STK38 | 72.6795719 | 0.035476882 | 0.21055955 | 0.1684886 | 0.86619891 | 0.9762143 |
| SRSF3 | 85.3467772 | -0.04123823 | 0.23616156 | -0.1746187 | 0.86137923 | 0.97470567 |
| CDKN1A | 48.3420511 | -2.485969443 | 0.49435485 | -5.0287146 | 4.94E-07 | 0.00016494 |
| C6orf89 | 46.8148448 | -0.017352031 | 0.2319587 | -0.0748066 | 0.94036863 | 0.9889521 |
| MTCH1 | 67.0971276 | -0.239950981 | 0.23425277 | -1.0243251 | 0.30568176 | 0.69722386 |
| PIM1 | 24.9801898 | -0.214761419 | 0.59889831 | -0.3585941 | 0.71989874 | 0.93444541 |
| TBC1D22B | 36.9560957 | -0.195097134 | 0.20903257 | -0.9333337 | 0.35064771 | 0.72904624 |
| RNF8 | 20.5924836 | 0.161823407 | 0.28030908 | 0.57730348 | 0.56373447 | 0.85632174 |
| CMTR1 | 21.3111349 | 0.040506646 | 0.31414338 | 0.12894318 | 0.89740261 | 0.98040427 |
| ZFAND3 | 716.338192 | -0.262553029 | 0.22871566 | -1.1479452 | 0.2509912 | 0.6460368 |
| BTBD9 | 377.152914 | -0.583665743 | 0.26770725 | -2.1802389 | 0.02923976 | 0.25858362 |
| GLO1 | 37.4105003 | -0.095142026 | 0.29796294 | -0.3193082 | 0.74949278 | 0.94004394 |
| DAAM2 | 191.765915 | 0.335959584 | 0.31221665 | 1.07604635 | 0.28190653 | 0.676814 |
| MOCS1 | 30.3913941 | -0.628564568 | 0.36191966 | -1.7367517 | 0.08243102 | 0.42256146 |
| NFYA | 23.6182611 | -0.522582415 | 0.26793478 | -1.950409 | 0.05112738 | 0.34374561 |
| TFEB | 19.5269445 | 0.331391609 | 0.37606881 | 0.88119941 | 0.3782099 | 0.74938381 |
| USP49 | 28.8116571 | -0.007894898 | 0.2240284 | -0.0352406 | 0.97188788 | 0.99300191 |
| CCND3 | 142.181049 | -0.244644797 | 0.43342277 | -0.5644484 | 0.57244899 | 0.86236719 |
| TAF8 | 22.6844123 | -0.237639312 | 0.35620852 | -0.6671354 | 0.50468565 | 0.82481485 |
| TRERF1 | 302.342862 | 0.768972479 | 0.31657898 | 2.42900676 | 0.01514025 | 0.17917181 |
| UBR2 | 220.994387 | -0.205839036 | 0.1897063 | -1.0850406 | 0.27790367 | 0.67269898 |
| BICRAL | 82.6133148 | 0.762938854 | 0.32510414 | 2.3467522 | 0.01893785 | 0.20213493 |
| RPL7L1 | 39.4683311 | -0.13462649 | 0.38039247 | -0.3539147 | 0.7234028 | 0.93455731 |
| PEX6 | 20.3652265 | -0.199305585 | 0.3406721 | -0.5850364 | 0.55852323 | 0.85316716 |
| KLHDC3 | 18.9450413 | 0.03920074 | 0.26370665 | 0.14865283 | 0.88182758 | 0.97782528 |
| RRP36 | 26.8306335 | -0.264568196 | 0.30498035 | -0.8674926 | 0.38567217 | 0.75340372 |
| PTK7 | 50.3812651 | 0.400073082 | 0.54592261 | 0.73283846 | 0.46365696 | 0.80227475 |
| ZNF318 | 18.0211777 | 0.452084921 | 0.26577161 | 1.70102786 | 0.08893776 | 0.43544541 |
| TJAP1 | 18.2581049 | -0.036919469 | 0.26369316 | -0.1400092 | 0.88865272 | 0.97942933 |
| POLR1C | 22.6559743 | -0.721908409 | 0.35607285 | -2.0274177 | 0.04261971 | 0.31108012 |
| YIPF3 | 21.9482728 | 0.124633717 | 0.28854238 | 0.4319425 | 0.66578321 | 0.91284091 |
| XPO5 | 75.9392643 | -0.220377919 | 0.31659061 | -0.6960975 | 0.4863678 | 0.81595238 |
| GTPBP2 | 18.200862 | -0.082127534 | 0.33357961 | -0.2462007 | 0.80552687 | 0.95836441 |
| VEGFA | 129.857716 | -1.906058245 | 0.75407361 | -2.5276819 | 0.01148183 | 0.15258586 |
| MRPL14 | 19.5619763 | -1.386464654 | 0.34390581 | -4.0315244 | 5.54E-05 | 0.00542582 |

| | | | | | | |
|---|---|---|---|---|---|---|
| TMEM63B | 21.4058786 | -0.651378235 | 0.36805426 | -1.7697886 | 0.07676236 | 0.41115479 |
| HSP90AB1 | 156.884634 | -0.131104079 | 0.28946568 | -0.4529175 | 0.65060813 | 0.9050963 |
| CDC5L | 32.5023413 | 0.048486595 | 0.20885612 | 0.23215309 | 0.81641911 | 0.96132648 |
| SUPT3H | 175.46817 | 0.08419613 | 0.22508648 | 0.37406125 | 0.70835875 | 0.93009618 |
| RUNX2 | 91.5552742 | -0.148349511 | 0.23665177 | -0.6268684 | 0.53074552 | 0.84152078 |
| CLIC5 | 28.7309377 | 0.91131107 | 0.89363524 | 1.01977969 | 0.30783296 | 0.69801431 |
| RCAN2 | 350.611497 | 0.636601628 | 0.26913251 | 2.36538363 | 0.0180114 | 0.19726646 |
| SLC25A27 | 61.2356356 | -0.332877431 | 0.37405645 | -0.8899123 | 0.37351298 | 0.7453019 |
| ADGRF5 | 191.056098 | 0.40277957 | 0.24804379 | 1.62382446 | 0.10441328 | 0.46422585 |
| TNFRSF21 | 32.2964496 | -0.495541759 | 0.44927806 | -1.1029734 | 0.2700387 | 0.6651094 |
| CD2AP | 117.352656 | 0.040268079 | 0.21173745 | 0.19017929 | 0.84916864 | 0.97113917 |
| MUT | 27.7854408 | 0.047568136 | 0.30617568 | 0.15536223 | 0.87653576 | 0.9768643 |
| PKHD1 | 58.2315038 | 1.125255994 | 0.42367975 | 2.65591168 | 0.00790943 | 0.12446771 |
| EFHC1 | 46.2254278 | -0.102492635 | 0.34230284 | -0.2994209 | 0.76461889 | 0.94462105 |
| TRAM2 | 121.352775 | -0.311815879 | 0.43199449 | -0.7218052 | 0.47041423 | 0.80457054 |
| GSTA4 | 31.1075955 | 0.140951846 | 0.34473491 | 0.40887024 | 0.68263489 | 0.91933701 |
| ICK | 21.6354649 | 0.145417714 | 0.39122713 | 0.3716964 | 0.71011891 | 0.93009618 |
| FBXO9 | 59.2034173 | -0.015570515 | 0.17867437 | -0.0871446 | 0.93055654 | 0.98681039 |
| ELOVL5 | 165.216217 | -0.416210196 | 0.26313529 | -1.5817346 | 0.11371017 | 0.48395978 |
| GCLC | 24.3541948 | -0.470866781 | 0.41787513 | -1.1268122 | 0.2598219 | 0.65257388 |
| COL21A1 | 57.4535954 | 1.39388641 | 0.29693473 | 4.69425184 | 2.68E-06 | 0.0005959 |
| DST | 604.393977 | 0.491506219 | 0.29405486 | 1.67147794 | 0.09462731 | 0.44706027 |
| ZNF451 | 126.946856 | -0.448809213 | 0.27927664 | -1.6070417 | 0.10804522 | 0.47376152 |
| BAG2 | 30.8996336 | -0.176080464 | 0.50478119 | -0.3488253 | 0.72722045 | 0.93455731 |
| RAB23 | 37.2561051 | 0.124437768 | 0.34195555 | 0.36390042 | 0.71593237 | 0.93237409 |
| PRIM2 | 102.834979 | -0.105920711 | 0.29602499 | -0.35781 | 0.72048548 | 0.93444541 |
| AL445250.1 | 39.4888833 | 0.053535438 | 0.57052644 | 0.09383516 | 0.9252401 | 0.98561736 |
| PTP4A1 | 50.6079626 | -1.349414319 | 0.46915368 | -2.8762736 | 0.00402401 | 0.08225816 |
| PHF3 | 139.818726 | -0.128084745 | 0.21042393 | -0.6086986 | 0.54272426 | 0.84678041 |
| ADGRB3 | 529.575509 | -0.866451927 | 0.51374381 | -1.6865447 | 0.09169092 | 0.44124451 |
| LMBRD1 | 92.347838 | 0.178536862 | 0.2375983 | 0.75142314 | 0.45239804 | 0.79504127 |
| COL19A1 | 31.2076139 | 0.190962389 | 0.27062727 | 0.70562878 | 0.48041902 | 0.81237906 |
| FAM135A | 121.441145 | 0.077716814 | 0.17193795 | 0.45200501 | 0.65126538 | 0.9050963 |
| SMAP1 | 108.269059 | -0.009263349 | 0.19761213 | -0.0468764 | 0.96261172 | 0.99231823 |
| OGFRL1 | 38.8095854 | -0.196962096 | 0.24167917 | -0.8149734 | 0.41508754 | 0.7750199 |
| AL136164.2 | 31.8077916 | 0.076886781 | 0.38054222 | 0.20204533 | 0.83988128 | 0.96789831 |
| KCNQ5 | 96.3734611 | 1.414989634 | 0.49522467 | 2.85726804 | 0.00427305 | 0.08579954 |
| OOEP | 86.2643101 | -0.451648143 | 0.30369963 | -1.4871541 | 0.13697412 | 0.52089048 |
| MTO1 | 23.5069324 | 0.292644507 | 0.25422302 | 1.15113299 | 0.24967753 | 0.64565668 |
| EEF1A1 | 407.76805 | -0.166685452 | 0.33099213 | -0.5035934 | 0.61454713 | 0.88360697 |
| SLC17A5 | 41.1501072 | -0.33384547 | 0.36519679 | -0.9141522 | 0.36063685 | 0.73273991 |
| COL12A1 | 470.956735 | 0.633534042 | 0.42266463 | 1.4989048 | 0.13389833 | 0.51600358 |
| COX7A2 | 40.6870085 | -0.489104522 | 0.37638132 | -1.299492 | 0.19377513 | 0.59261481 |
| TMEM30A | 79.5746174 | -0.361416143 | 0.29381581 | -1.2300772 | 0.21866818 | 0.61522936 |
| FILIP1 | 203.875154 | -0.270267283 | 0.39231195 | -0.6889091 | 0.49088045 | 0.81720854 |
| SENP6 | 320.782221 | 0.210473841 | 0.15035724 | 1.39982509 | 0.1615657 | 0.55507512 |
| MYO6 | 118.463271 | 0.0879961 | 0.22346628 | 0.39377798 | 0.69374496 | 0.9248682 |
| PHIP | 241.578494 | -0.059552572 | 0.18536999 | -0.3212633 | 0.74801088 | 0.93903859 |
| HMGN3 | 64.8910693 | 0.415061886 | 0.2146971 | 1.93324402 | 0.05320614 | 0.34798571 |
| LCA5 | 18.2297321 | 0.152852969 | 0.29501681 | 0.51811613 | 0.60437724 | 0.87903643 |
| BCKDHB | 147.42832 | 0.014607933 | 0.37414498 | 0.03904351 | 0.9688557 | 0.99300191 |
| TENT5A | 177.133774 | 0.280899246 | 0.45507306 | 0.61726186 | 0.53706202 | 0.84423863 |
| IBTK | 96.3335728 | 0.037696904 | 0.18464379 | 0.20416015 | 0.83822835 | 0.96735869 |
| UBE3D | 53.6739962 | -0.032925596 | 0.19628133 | -0.167747 | 0.86678236 | 0.97624269 |
| DOPEY1 | 93.4022223 | 0.125585457 | 0.24978226 | 0.50277974 | 0.61511914 | 0.88360697 |
| PGM3 | 38.7348742 | -0.262368855 | 0.25051975 | -1.0472981 | 0.29496213 | 0.68597363 |
| ME1 | 31.1042038 | -0.695004736 | 0.55202465 | -1.2590103 | 0.2080266 | 0.6059796 |
| CYB5R4 | 34.9413827 | 0.249693379 | 0.25876903 | 0.96492761 | 0.33458107 | 0.71690491 |
| CEP162 | 36.5668552 | -0.091342 | 0.26284053 | -0.3475187 | 0.72820167 | 0.93455731 |
| SNX14 | 73.5969246 | 0.270154316 | 0.21661533 | 1.24716156 | 0.21233827 | 0.60910715 |
| SYNCRIP | 38.3413235 | -0.013598657 | 0.22634339 | -0.0600798 | 0.95209211 | 0.99060401 |
| ZNF292 | 266.824513 | -0.859842861 | 0.3052615 | -2.8167419 | 0.00485135 | 0.09203696 |
| SMIM8 | 29.4423747 | 0.197138508 | 0.29503018 | 0.66819778 | 0.50400735 | 0.82438238 |
| RARS2 | 96.9827052 | -0.083192382 | 0.23085636 | -0.3603643 | 0.71857475 | 0.93432517 |
| ORC3 | 56.8136054 | 0.062044594 | 0.25325384 | 0.24498974 | 0.80646437 | 0.95838958 |
| AKIRIN2 | 22.9918258 | 0.056501763 | 0.31922572 | 0.17699627 | 0.85951132 | 0.97397549 |
| RNGTT | 139.161706 | -0.279444702 | 0.28165395 | -0.9921562 | 0.32112135 | 0.70780851 |
| PNRC1 | 81.8008805 | -0.387347501 | 0.29914188 | -1.2948621 | 0.19536779 | 0.59328838 |
| PM20D2 | 19.5952863 | 0.051761409 | 0.37158075 | 0.13930057 | 0.88921264 | 0.9798806 |
| UBE2J1 | 50.2865662 | -0.229957979 | 0.22413059 | -1.026 | 0.3048916 | 0.69650244 |
| ANKRD6 | 53.6884307 | 0.853949932 | 0.403969 | 2.11389964 | 0.03452384 | 0.2833153 |
| LYRM2 | 23.6575876 | -0.113851191 | 0.29779415 | -0.3823151 | 0.70222768 | 0.92865476 |
| MDN1 | 90.3892162 | -0.453612678 | 0.28244967 | -1.6059947 | 0.10827507 | 0.47420349 |
| CASP8AP2 | 38.1097758 | 0.131802423 | 0.20114928 | 0.6552468 | 0.51230887 | 0.82891748 |
| BACH2 | 87.9437961 | 0.015102069 | 0.37881275 | 0.03986684 | 0.96819929 | 0.99300191 |
| MAP3K7 | 76.4844128 | -0.012920941 | 0.25755143 | -0.0501684 | 0.9599882 | 0.99181157 |
| UFL1 | 54.8962043 | -0.155820821 | 0.17522828 | -0.8892447 | 0.37387157 | 0.74561254 |
| FHL5 | 117.411079 | 0.789238974 | 0.47681549 | 1.6552293 | 0.09787799 | 0.45529992 |
| MMS22L | 30.274453 | 0.204392145 | 0.37650894 | 0.54286133 | 0.58722528 | 0.870249 |
| FBXL4 | 74.7954917 | -0.33415578 | 0.27757707 | -1.2038307 | 0.22865504 | 0.62213472 |
| PNISR | 287.120284 | 0.017375044 | 0.18401078 | 0.09442405 | 0.92477231 | 0.98561736 |
| USP45 | 46.7168339 | -0.459315085 | 0.27415005 | -1.6754149 | 0.09385285 | 0.44536803 |
| CCNC | 49.6355756 | -0.408663645 | 0.19967845 | -2.0466087 | 0.04069652 | 0.30271128 |
| ASCC3 | 228.375348 | -0.508380351 | 0.22453875 | -2.2641097 | 0.02356737 | 0.22729214 |
| GRIK2 | 90.2055528 | 2.545664108 | 0.44641943 | 5.70240436 | 1.18E-08 | 9.65E-06 |
| HACE1 | 54.9107731 | -0.226005065 | 0.30436263 | -0.742552 | 0.45775298 | 0.79810834 |
| PREP | 43.2715538 | -0.836184424 | 0.3089414 | -2.7066118 | 0.00679737 | 0.11301781 |
| ATG5 | 71.2096471 | -0.024950815 | 0.18184658 | -0.1372081 | 0.89086634 | 0.98023547 |
| QRSL1 | 23.1078071 | -0.212875904 | 0.28180155 | -0.7554107 | 0.45000259 | 0.79399826 |
| LINC02532 | 20.5739295 | -0.110031096 | 0.43475598 | -0.253087 | 0.80020097 | 0.95636314 |
| CD24 | 66.3487146 | -0.892650802 | 0.26135224 | -3.4155086 | 0.00063663 | 0.02784879 |
| PDSS2 | 132.752983 | -0.006947492 | 0.17515325 | -0.0396652 | 0.96836003 | 0.99300191 |
| SOBP | 165.721525 | 0.075610608 | 0.29295118 | 0.25809969 | 0.79632997 | 0.95463848 |
| SEC63 | 136.813187 | -0.206521748 | 0.17478539 | -1.1815733 | 0.23737505 | 0.63302595 |
| OSTM1 | 44.6722984 | -0.242573266 | 0.31214387 | -0.7771201 | 0.43708794 | 0.78768252 |
| SNX3 | 66.9233888 | -0.514036529 | 0.20987022 | -2.4493067 | 0.01431315 | 0.17531225 |
| AFG1L | 34.2248073 | 0.14061691 | 0.24687417 | 0.5695894 | 0.56895622 | 0.85958696 |
| FOXO3 | 444.963348 | -0.521261598 | 0.33537122 | -1.5542824 | 0.12011706 | 0.49647936 |
| ARMC2 | 51.7607175 | 0.227042849 | 0.37394649 | 0.60715332 | 0.54374917 | 0.84730644 |
| SESN1 | 141.644855 | -0.263589435 | 0.22805603 | -1.15581 | 0.24775889 | 0.64333128 |
| CEP57L1 | 36.5381302 | 0.172556772 | 0.25456051 | 0.6778615 | 0.49785951 | 0.82238021 |
| CD164 | 70.3535483 | -0.207584495 | 0.22598621 | -0.9185715 | 0.35831974 | 0.73144657 |
| FIG4 | 32.2307417 | -0.11101925 | 0.22076477 | -0.5028848 | 0.61504526 | 0.88360697 |
| WASF1 | 34.9639334 | -0.460166014 | 0.36570937 | -1.2582834 | 0.20828929 | 0.6059796 |
| CDC40 | 33.5379992 | -0.276568542 | 0.27290162 | -1.0134368 | 0.31085157 | 0.70126123 |
| METTL24 | 32.6431879 | 0.367217938 | 0.43797176 | 0.83845118 | 0.40177736 | 0.7641692 |
| CDK19 | 97.2420713 | 0.043007291 | 0.30722234 | 0.13998751 | 0.88866986 | 0.97942933 |
| AMD1 | 44.7320403 | -0.247992828 | 0.33182062 | -0.7473701 | 0.45484016 | 0.79609632 |
| GTF3C6 | 24.3875753 | -0.71772107 | 0.33926195 | -2.1155366 | 0.03438424 | 0.2833153 |
| RPF2 | 28.5649642 | -0.594951254 | 0.24545039 | -2.4239165 | 0.01535413 | 0.18089458 |
| REV3L | 1025.62617 | -1.554103912 | 0.44752216 | -3.4726859 | 0.00051528 | 0.02458947 |
| TRAF3IP2-AS | 231.174001 | -0.6799897 | 0.32755044 | -2.0759847 | 0.03789537 | 0.29501902 |
| TRAF3IP2 | 33.8513423 | -1.070540217 | 0.25173518 | -4.2526445 | 2.11E-05 | 0.00305725 |

| | | | | | | |
|---|---|---|---|---|---|---|
| FYN | 193.950824 | -0.565528744 | 0.3398351 | -1.6641269 | 0.09608707 | 0.4510884 |
| TUBE1 | 29.6920989 | -0.384766944 | 0.36251903 | -1.0613703 | 0.28852163 | 0.67992862 |
| LAMA4 | 160.400474 | 0.561073411 | 0.46009526 | 1.21947226 | 0.222665 | 0.61772146 |
| MARCKS | 85.3391794 | 0.190299217 | 0.39961544 | 0.47620587 | 0.63392772 | 0.89608287 |
| HDAC2 | 37.3421543 | 0.026227101 | 0.21943017 | 0.11952368 | 0.90486048 | 0.98265659 |
| NT5DC1 | 126.032436 | -0.097563433 | 0.35545716 | -0.2744731 | 0.78372106 | 0.95064024 |
| DSE | 54.8389847 | 0.39243032 | 0.30125008 | 1.30267291 | 0.19268646 | 0.59150074 |
| TSPYL1 | 28.6022321 | 0.442650687 | 0.26302116 | 1.68294704 | 0.09238534 | 0.44238068 |
| RWDD1 | 22.4422402 | 0.269847386 | 0.29084297 | 0.92781129 | 0.35350547 | 0.73059415 |
| KPNA5 | 52.3185713 | 0.426974336 | 0.21724909 | 1.96536768 | 0.0493717 | 0.33595611 |
| FAM162B | 26.2325484 | 0.607827226 | 0.57249393 | 1.06171821 | 0.28836363 | 0.67992862 |
| DCBLD1 | 40.9242711 | -0.606901813 | 0.30852162 | -1.9671289 | 0.04916835 | 0.33550435 |
| GOPC | 37.7759477 | 0.238989146 | 0.25318695 | 0.94392365 | 0.34520867 | 0.72325086 |
| NUS1 | 45.4832912 | -0.005415956 | 0.26788008 | -0.0202178 | 0.9838696 | 0.99620525 |
| SLC35F1 | 157.488362 | 0.538200524 | 0.45966176 | 1.17086208 | 0.24165422 | 0.63584563 |
| CEP85L | 213.869729 | -0.113303874 | 0.22125148 | -0.5121045 | 0.60857789 | 0.8807592 |
| PLN | 27.6812305 | 0.353769424 | 0.38920303 | 0.90895855 | 0.363372 | 0.73604674 |
| MCM9 | 47.53764 | -0.178729481 | 0.25211158 | -0.7089301 | 0.47836787 | 0.81152481 |
| MAN1A1 | 126.685222 | -0.619327394 | 0.30610702 | -2.0232381 | 0.0430486 | 0.31204692 |
| TBC1D32 | 63.2419514 | -0.205350022 | 0.339668 | -0.604561 | 0.54547073 | 0.84758329 |
| GJA1 | 25.551719 | -1.136396704 | 0.3333347 | -3.4091762 | 0.00065159 | 0.02800329 |
| SERINC1 | 79.8245907 | 0.25153992 | 0.18952017 | 1.32724617 | 0.18442726 | 0.58363936 |
| RNF217 | 117.80118 | -0.384413392 | 0.39106141 | -0.9830001 | 0.3256074 | 0.71149668 |
| HDDC2 | 21.6177784 | -0.228806902 | 0.31611591 | -0.723807 | 0.46918424 | 0.80388079 |
| NCOA7 | 148.342065 | -1.086988772 | 0.34955386 | -3.109646 | 0.00187312 | 0.05488753 |
| TRMT11 | 58.1003139 | -0.462399206 | 0.23995997 | -1.9269848 | 0.05398153 | 0.34959616 |
| RSPO3 | 60.0454945 | -0.910289937 | 0.85168364 | -1.0688123 | 0.28515426 | 0.6783522 |
| RNF146 | 35.4414863 | -0.173161211 | 0.29481972 | -0.5873461 | 0.55697128 | 0.85256862 |
| ECHDC1 | 62.9365582 | -0.260881177 | 0.18105435 | -1.4408998 | 0.14961299 | 0.53844556 |
| PTPRK | 381.159716 | 0.192970971 | 0.23444821 | 0.82308572 | 0.41045925 | 0.77166383 |
| LAMA2 | 819.048325 | -1.158641073 | 0.31870964 | -3.6354127 | 0.00027754 | 0.0161874 |
| ARHGAP18 | 64.0075619 | 0.971439365 | 0.32298112 | 3.00772801 | 0.00263209 | 0.06512862 |
| L3MBTL3 | 33.679803 | -0.011576474 | 0.26341051 | -0.0439484 | 0.96494552 | 0.99275266 |
| SAMD3 | 105.471838 | 0.656450753 | 0.40292681 | 1.62920596 | 0.10326943 | 0.46365765 |
| TMEM200A | 77.3094445 | 0.11485492 | 0.4869208 | 0.23588009 | 0.81352572 | 0.96038841 |
| EPB41L2 | 448.364172 | 0.343553417 | 0.20433827 | 1.68129749 | 0.09270515 | 0.44320781 |
| AKAP7 | 33.5394088 | -0.242219826 | 0.25091255 | -0.9653556 | 0.33436674 | 0.71690491 |
| MED23 | 49.3681049 | 0.299638142 | 0.21866003 | 1.37033798 | 0.17058142 | 0.56801217 |
| ENPP1 | 54.1210628 | 0.564035422 | 0.53630281 | 1.05171073 | 0.29293228 | 0.6842846 |
| CTGF | 229.463067 | 2.719110157 | 0.84487652 | 3.21835216 | 0.00128929 | 0.04375316 |
| AL133346.1 | 50.4762772 | 0.436936614 | 0.84874023 | 0.51480606 | 0.60668855 | 0.88000768 |
| MOXD1 | 359.225319 | 0.013593578 | 0.42837418 | 0.03173295 | 0.97468501 | 0.99300191 |
| STX7 | 72.3874792 | 0.031519254 | 0.30538674 | 0.10321095 | 0.91779555 | 0.98477389 |
| RPS12 | 90.1487346 | -0.280321437 | 0.37310005 | -0.7513305 | 0.45245379 | 0.79504127 |
| EYA4 | 238.9966 | -0.184629718 | 0.27423116 | -0.6732631 | 0.50077993 | 0.8236866 |
| TARID | 76.5586671 | -0.076449045 | 0.55962279 | -0.1366082 | 0.89134052 | 0.98023547 |
| TCF21 | 43.9961853 | 0.814473202 | 0.58610548 | 1.38963587 | 0.16463948 | 0.55965519 |
| TBPL1 | 26.3532436 | 0.197103857 | 0.26854892 | 0.73395885 | 0.46297382 | 0.80225845 |
| SGK1 | 44.4288822 | 0.869120676 | 0.39976829 | 2.17406107 | 0.02970055 | 0.26108772 |
| HBS1L | 55.4297205 | -0.085980553 | 0.19178985 | -0.4483061 | 0.65393232 | 0.90646849 |
| AHI1 | 295.188952 | -0.819720835 | 0.17995132 | -4.5552365 | 5.23E-06 | 0.00101197 |
| PDE7B | 540.410609 | 0.567665208 | 0.36187762 | 1.56866625 | 0.11672573 | 0.48989475 |
| AL138828.1 | 92.9706623 | -0.098176203 | 0.51368967 | -0.1911197 | 0.84843183 | 0.97092453 |
| BCLAF1 | 135.207305 | -0.074331181 | 0.1802403 | -0.4124004 | 0.68004594 | 0.91860979 |
| MAP7 | 31.9704802 | -0.126312815 | 0.29912854 | -0.4222693 | 0.67282843 | 0.91482259 |
| MAP3K5 | 746.555468 | -0.833227227 | 0.27316835 | -3.0502334 | 0.00228664 | 0.05980244 |
| PEX7 | 23.4986348 | -0.265054767 | 0.27225088 | -0.9735681 | 0.33027105 | 0.71362138 |
| IFNGR1 | 55.5754551 | -0.514276024 | 0.23781674 | -2.1624887 | 0.03058053 | 0.26662968 |
| TNFAIP3 | 66.8628567 | -2.256635324 | 1.01884009 | -2.2149063 | 0.0267665 | 0.24496511 |
| HEBP2 | 24.6785178 | -0.72597475 | 0.26734692 | -2.7154783 | 0.00661801 | 0.11104057 |
| NHSL1 | 146.52433 | -0.185464334 | 0.3930763 | -0.4718278 | 0.63704969 | 0.89741446 |
| REPS1 | 120.594328 | 0.135242319 | 0.2939994 | 0.46000883 | 0.64550988 | 0.90150535 |
| HECA | 25.9281948 | 0.246741349 | 0.25423463 | 0.97052612 | 0.33178431 | 0.71487809 |
| CITED2 | 24.0139161 | 0.553317356 | 0.45936651 | 1.20452263 | 0.22838764 | 0.62213472 |
| VTA1 | 34.1105433 | 0.096671551 | 0.21054908 | 0.45914022 | 0.64613347 | 0.90193887 |
| HIVEP2 | 373.794511 | -0.60263276 | 0.28882556 | -2.0864939 | 0.03693391 | 0.29217146 |
| AIG1 | 70.97379 | 0.156935199 | 0.22109893 | 0.70979628 | 0.47783048 | 0.81138532 |
| PEX3 | 35.6304944 | -0.068705705 | 0.25763571 | -0.2666777 | 0.78971731 | 0.9517272 |
| PHACTR2 | 127.217973 | 0.695074075 | 0.28781147 | 2.41503254 | 0.0157338 | 0.18382786 |
| PLAGL1 | 114.698624 | -0.035862817 | 0.31187461 | -0.1149911 | 0.90845214 | 0.98306694 |
| SF3B5 | 27.2830205 | -0.696221111 | 0.36591066 | -1.9027079 | 0.05707867 | 0.35717443 |
| UTRN | 987.517783 | 0.41937712 | 0.34428868 | 1.21809733 | 0.22318699 | 0.61772146 |
| EPM2A | 26.9931233 | -0.083866211 | 0.34007973 | -0.2466075 | 0.805212 | 0.95836441 |
| SHPRH | 125.315071 | -0.116513349 | 0.14496318 | -0.8037444 | 0.42154458 | 0.77771095 |
| STXBP5 | 134.17422 | -0.125900667 | 0.18463762 | -0.6818798 | 0.49531494 | 0.82075974 |
| SAMD5 | 37.29667 | 1.416242522 | 0.39413006 | 3.59333804 | 0.00032647 | 0.01803923 |
| SASH1 | 129.605713 | 0.128685327 | 0.35339701 | 0.36413813 | 0.71575487 | 0.93237409 |
| UST | 152.601706 | 0.254463664 | 0.43173073 | 0.58940364 | 0.55559054 | 0.85207344 |
| TAB2 | 101.671967 | -0.023624347 | 0.22831902 | -0.1034708 | 0.91758934 | 0.98477389 |
| PPIL4 | 46.0667198 | 0.102920226 | 0.25286952 | 0.40700922 | 0.68400122 | 0.91963501 |
| GINM1 | 49.2255094 | 0.133585304 | 0.20574799 | 0.64926664 | 0.51616605 | 0.83181726 |
| KATNA1 | 22.4704013 | 0.233972416 | 0.28178968 | 0.83030867 | 0.40636429 | 0.76770466 |
| LATS1 | 75.4640243 | 0.158039641 | 0.1720276 | 0.91868769 | 0.35825895 | 0.73144657 |
| NUP43 | 23.0231405 | -0.050719526 | 0.26397903 | -0.1921347 | 0.84763671 | 0.97044434 |
| PCMT1 | 50.3181362 | -0.079448486 | 0.19738176 | -0.4025118 | 0.68730741 | 0.92100905 |
| PLEKHG1 | 119.188205 | -0.857108378 | 0.48726608 | -1.7590151 | 0.07857495 | 0.41542971 |
| MTHFD1L | 367.034001 | -1.177508504 | 0.46694834 | -2.5217104 | 0.01167858 | 0.15416382 |
| AKAP12 | 1148.88038 | -0.541165312 | 0.58757721 | -0.9210114 | 0.35704448 | 0.73129235 |
| RMND1 | 24.2229344 | -0.28471075 | 0.24156549 | -1.1786069 | 0.23855475 | 0.63422836 |
| CCDC170 | 21.3065438 | 0.130327489 | 0.40930809 | 0.31840927 | 0.75017451 | 0.94004394 |
| ESR1 | 161.603763 | 0.600029204 | 0.40091079 | 1.49666516 | 0.13448041 | 0.51662128 |
| SYNE1 | 827.131443 | 0.53460344 | 0.37859355 | 1.41207752 | 0.15792714 | 0.54952961 |
| CNKSR3 | 111.827431 | 0.641145533 | 0.31261975 | 2.05087979 | 0.04027865 | 0.30130305 |
| SCAF8 | 203.176506 | 0.094090937 | 0.17291473 | 0.54414646 | 0.58634069 | 0.87014172 |
| TIAM2 | 124.106035 | -0.001985436 | 0.15682419 | -0.0126603 | 0.98989884 | 0.99739981 |
| TFB1M | 29.8343069 | -0.241201175 | 0.27076853 | -0.8908021 | 0.37303535 | 0.74512029 |
| ARID1B | 706.432638 | -0.125408391 | 0.14808687 | -0.8468569 | 0.39707488 | 0.76043276 |
| TMEM242 | 27.2303967 | 0.238980876 | 0.29847415 | 0.80067529 | 0.42331965 | 0.77827834 |
| ZDHHC14 | 226.066169 | 0.30617954 | 0.23186849 | 1.32048792 | 0.18667217 | 0.58501647 |
| SNX9 | 201.391456 | -0.242314124 | 0.2614749 | -0.9267204 | 0.35407172 | 0.73059415 |
| SYNJ2 | 111.72544 | 0.713979669 | 0.34655864 | 2.0601987 | 0.03937955 | 0.30020778 |
| TULP4 | 156.516462 | 0.007305415 | 0.25984889 | 0.02811409 | 0.97757116 | 0.99407008 |
| TMEM181 | 59.9162524 | -0.453227274 | 0.28464103 | -1.5922767 | 0.11132255 | 0.47870652 |
| DYNLT1 | 47.1739919 | -0.493634049 | 0.25496941 | -1.9360521 | 0.05286132 | 0.34706747 |
| EZR | 40.7811933 | 0.0869315 | 0.2501202 | 0.34755889 | 0.72817148 | 0.93455731 |
| AL035530.2 | 21.1511768 | 0.639039048 | 0.37806527 | 1.69028764 | 0.09097294 | 0.44071202 |
| WTAP | 136.029356 | -0.952955984 | 0.3919478 | -2.4313339 | 0.01504334 | 0.17917181 |
| TCP1 | 29.0460046 | -0.513445118 | 0.28311779 | -1.8135389 | 0.06974875 | 0.39338722 |
| IGF2R | 192.450631 | -0.216407658 | 0.15896547 | -1.3613501 | 0.17340309 | 0.57068486 |
| SLC22A3 | 70.0297558 | -0.524890929 | 0.73673467 | -0.7124559 | 0.47618254 | 0.81006145 |
| MAP3K4 | 100.005297 | -0.100221626 | 0.25648847 | -0.3907452 | 0.69598562 | 0.92648248 |
| AGPAT4 | 40.711212 | 0.300519743 | 0.29860368 | 1.00641673 | 0.31421515 | 0.70282967 |
| PRKN | 311.280239 | 0.328170171 | 0.21055443 | 1.55860017 | 0.11909104 | 0.49395083 |

| | | | | | | |
|---|---|---|---|---|---|---|
| QKI | 411.80493 | -0.398495588 | 0.21195399 | -1.8801042 | 0.06009388 | 0.36498338 |
| PDE10A | 369.22426 | -0.848414354 | 0.49107038 | -1.7276838 | 0.08404492 | 0.4256693 |
| SFT2D1 | 33.1288915 | -0.076562146 | 0.22750921 | -0.3365233 | 0.7364763 | 0.93594619 |
| MPC1 | 37.2737195 | 0.086303231 | 0.2581566 | 0.33430573 | 0.73814887 | 0.93594619 |
| RPS6KA2 | 76.4943703 | -0.274285063 | 0.39503104 | -0.694338 | 0.48747026 | 0.81595238 |
| FGFR1OP | 37.2750429 | 0.297758901 | 0.25102582 | 1.18616841 | 0.2355558 | 0.63132734 |
| AFDN | 122.534095 | 0.103891466 | 0.30427546 | 0.34143886 | 0.73277322 | 0.93574285 |
| SMOC2 | 87.0188173 | 0.070582249 | 0.42250862 | 0.16705517 | 0.86732664 | 0.97634923 |
| BX322234.1 | 26.5848615 | -1.865828239 | 0.48820292 | -3.8218294 | 0.00013247 | 0.00953067 |
| THBS2 | 198.420781 | -1.537697611 | 0.43456442 | -3.5384803 | 0.00040244 | 0.0206819 |
| WDR27 | 59.665715 | 0.529617844 | 0.23613094 | 2.24289895 | 0.02490333 | 0.234034 |
| PHF10 | 35.7219881 | -0.355388102 | 0.32844601 | -1.082029 | 0.27923965 | 0.67353297 |
| ERMARD | 17.1355512 | 0.298859018 | 0.31663369 | 0.9438636 | 0.34523935 | 0.72325086 |
| FAM120B | 134.956964 | 0.052944193 | 0.20515236 | 0.25807256 | 0.79635091 | 0.95463848 |
| PSMB1 | 63.7691204 | -0.564867074 | 0.28838399 | -1.9587324 | 0.05014413 | 0.33808186 |
| PDCD2 | 17.9147865 | 0.063247391 | 0.30394687 | 0.208087 | 0.83516104 | 0.96670318 |
| FAM20C | 61.2670621 | -0.410788975 | 0.36193966 | -1.1349654 | 0.25638983 | 0.65084527 |
| PDGFA | 44.1886082 | -0.096780486 | 0.2757445 | -0.3509788 | 0.72560422 | 0.93455731 |
| PRKAR1B | 19.4863415 | 0.546796086 | 0.34493327 | 1.58522282 | 0.11291572 | 0.48301375 |
| DNAAF5 | 17.7292661 | -0.172652559 | 0.30872123 | -0.5592507 | 0.57599065 | 0.86298783 |
| SUN1 | 110.326816 | 0.217916501 | 0.3510974 | 0.6206725 | 0.53481513 | 0.84361209 |
| C7orf50 | 38.2968845 | 0.554693633 | 0.21508148 | 2.57899299 | 0.00990888 | 0.14050816 |
| MICALL2 | 35.3563276 | -0.181229397 | 0.39095742 | -0.4635528 | 0.6429682 | 0.90063554 |
| INTS1 | 22.8690345 | 0.55837005 | 0.30918784 | 1.80592502 | 0.07093005 | 0.39670088 |
| MAFK | 23.3886845 | 0.573795097 | 0.34023386 | 1.68647264 | 0.0917048 | 0.44124451 |
| MAD1L1 | 48.1689179 | -0.066787136 | 0.30158326 | -0.221455 | 0.82473813 | 0.96413949 |
| SNX8 | 19.6947595 | -0.665523519 | 0.491574 | -1.3538623 | 0.17578031 | 0.57311399 |
| EIF3B | 60.9295351 | -0.324071949 | 0.29254247 | -1.1077775 | 0.26795794 | 0.66303801 |
| CHST12 | 19.8223148 | 0.394008165 | 0.31523494 | 1.24988734 | 0.2113407 | 0.60907562 |
| TTYH3 | 18.566433 | -0.109043541 | 0.28749415 | -0.3792896 | 0.70447282 | 0.92936308 |
| GNA12 | 111.128378 | -0.224594523 | 0.25744513 | -0.8723976 | 0.38299149 | 0.75179833 |
| SDK1 | 592.86647 | -0.780426435 | 0.48806695 | -1.5990151 | 0.10981724 | 0.47680173 |
| FOXK1 | 90.9530498 | -0.104918334 | 0.33807917 | -0.3103366 | 0.75630501 | 0.94216126 |
| RBAK-RBAKI | 48.5479682 | 0.175186682 | 0.2816559 | 0.62198832 | 0.53394955 | 0.84333372 |
| WIPI2 | 40.3900307 | -0.074770936 | 0.25621942 | -0.2918238 | 0.77042131 | 0.94593455 |
| TNRC18 | 153.645012 | 0.667130699 | 0.27886142 | 2.39233775 | 0.01674143 | 0.19005424 |
| FBXL18 | 18.1351456 | -0.022853893 | 0.39427558 | -0.0579643 | 0.9537771 | 0.99100036 |
| ACTB | 921.92965 | 0.674282793 | 0.31588557 | 2.13457927 | 0.03279539 | 0.27660385 |
| RNF216 | 129.027503 | 0.053584533 | 0.23138053 | 0.23158619 | 0.81685944 | 0.96134613 |
| PMS2 | 17.9823525 | -0.221705572 | 0.31077837 | -0.713388 | 0.47560567 | 0.80926744 |
| EIF2AK1 | 36.4971657 | -0.565268104 | 0.24027725 | -2.3525661 | 0.01864438 | 0.20009416 |
| USP42 | 52.7625326 | 0.319900676 | 0.21788386 | 1.46821648 | 0.14204543 | 0.52954815 |
| CYTH3 | 158.628109 | 0.567752486 | 0.34014064 | 1.66916981 | 0.09508373 | 0.44764276 |
| RAC1 | 117.359016 | 0.196436809 | 0.25379 | 0.77401318 | 0.438923 | 0.78887428 |
| ZNF12 | 26.7635611 | -0.47312247 | 0.24907405 | -1.8995253 | 0.05749544 | 0.35838335 |
| CCZ1B | 23.2218294 | 0.387969705 | 0.29730642 | 1.30494896 | 0.19191023 | 0.59084553 |
| C1GALT1 | 50.6889194 | -1.100797605 | 0.33938298 | -3.2435263 | 0.0011806 | 0.0429516 |
| MIOS | 20.4417171 | -0.194392221 | 0.27399536 | -0.7094727 | 0.4780312 | 0.81139113 |
| UMAD1 | 154.916502 | 0.229839227 | 0.17682354 | 1.29982257 | 0.19366179 | 0.59251477 |
| GLCCI1 | 79.1690645 | 0.893219016 | 0.28897324 | 3.09100942 | 0.00199477 | 0.05595262 |
| NDUFA4 | 84.453849 | -0.533229604 | 0.32295708 | -1.6510851 | 0.09872121 | 0.45657782 |
| PHF14 | 204.312129 | -0.074727161 | 0.16178473 | -0.4618926 | 0.64415837 | 0.90084107 |
| THSD7A | 132.397392 | -0.074682436 | 0.46434219 | -0.1608349 | 0.87222343 | 0.9768643 |
| TMEM106B | 41.3292798 | -0.074478267 | 0.23007135 | -0.3237181 | 0.74615145 | 0.93894983 |
| ETV1 | 25.2063249 | -0.048625309 | 0.37917434 | -0.12824 | 0.89795906 | 0.98040427 |
| ISPD | 46.4202254 | 0.385397648 | 0.29200664 | 1.31982495 | 0.18689347 | 0.58501647 |
| BZW2 | 23.0090684 | -0.342816478 | 0.41729218 | -0.8215263 | 0.41134658 | 0.77166383 |
| SNX13 | 143.544191 | -0.294539049 | 0.18052102 | -1.6316053 | 0.10276266 | 0.4627468 |
| HDAC9 | 163.94054 | 0.414905982 | 0.51521739 | 0.80530275 | 0.42064499 | 0.77771095 |
| ITGB8 | 87.8667271 | -0.762733884 | 0.5156125 | -1.4792773 | 0.13906621 | 0.52490887 |
| SP4 | 32.9460874 | 0.336937943 | 0.29216847 | 1.1532317 | 0.2488153 | 0.64453423 |
| RAPGEF5 | 520.704392 | -0.981331963 | 0.56240908 | -1.7448722 | 0.08100712 | 0.42042465 |
| STEAP1B | 72.8704074 | -0.548467273 | 0.28979685 | -1.8925922 | 0.05841212 | 0.36041506 |
| TOMM7 | 40.0628632 | -0.10097858 | 0.32911976 | -0.3068141 | 0.75898488 | 0.94385579 |
| FAM126A | 88.4071176 | -0.178983632 | 0.21622403 | -0.8277694 | 0.40780113 | 0.76944531 |
| KLHL7 | 36.1759997 | 0.101906665 | 0.25768671 | 0.39546729 | 0.69249806 | 0.9248682 |
| NUPL2 | 28.6153612 | -0.310782479 | 0.34018438 | -0.9135707 | 0.36094247 | 0.73314333 |
| MALSU1 | 22.0667212 | 0.014876172 | 0.25033642 | 0.05942472 | 0.95261382 | 0.99060401 |
| TRA2A | 161.866096 | -0.05365613 | 0.26792453 | -0.2002658 | 0.84127268 | 0.96803523 |
| OSBPL3 | 33.3799867 | 0.555619091 | 0.58694347 | 0.94663136 | 0.34382666 | 0.72261198 |
| CYCS | 31.9473845 | -0.275913707 | 0.31478625 | -0.8765113 | 0.38075214 | 0.74991898 |
| HNRNPA2B1 | 283.669385 | 0.214755741 | 0.18298155 | 1.173647 | 0.24053645 | 0.63466484 |
| CBX3 | 38.9157198 | -0.539605564 | 0.25039901 | -2.1549828 | 0.03116318 | 0.26776774 |
| SKAP2 | 206.612637 | -0.399090779 | 0.34729735 | -1.1491328 | 0.25050122 | 0.6460368 |
| HOTAIRM1 | 38.1441129 | 0.441518974 | 0.35792463 | 1.23355293 | 0.21736954 | 0.61440337 |
| HOXA3 | 52.3272954 | 0.445688747 | 0.23744503 | 1.87701866 | 0.06051554 | 0.36535567 |
| HIBADH | 113.468506 | -0.148471878 | 0.22850865 | -0.6497429 | 0.51585829 | 0.83174649 |
| TAX1BP1 | 173.63389 | 0.054733432 | 0.16661991 | 0.32849274 | 0.74253913 | 0.93759079 |
| JAZF1 | 125.704889 | 0.499699071 | 0.3033038 | 1.64751999 | 0.09945121 | 0.45779653 |
| CREB5 | 81.1357711 | 1.211032057 | 0.42582433 | 2.84397103 | 0.00445551 | 0.08834811 |
| SCRN1 | 24.2151859 | 0.360494371 | 0.26733554 | 1.34847153 | 0.17750677 | 0.5746684 |
| FKBP14 | 19.8284409 | -0.964459154 | 0.38836815 | -2.4833632 | 0.01301483 | 0.16653913 |
| PLEKHA8 | 36.8633169 | -0.418843611 | 0.29521429 | -1.4187782 | 0.15596368 | 0.54527928 |
| MTURN | 31.5342935 | 0.047378067 | 0.21861297 | 0.2167212 | 0.82842562 | 0.96580172 |
| ZNRF2 | 59.0867764 | -0.540491522 | 0.31111143 | -1.7372924 | 0.08233557 | 0.42256146 |
| AC006978.2 | 40.0875101 | -0.671372897 | 0.35179562 | -1.9084174 | 0.05633728 | 0.35564709 |
| NOD1 | 62.9147884 | -0.941490371 | 0.37567307 | -2.5061428 | 0.01220563 | 0.15932359 |
| AC005154.6 | 29.985193 | 0.109485054 | 0.27585947 | 0.39688706 | 0.69145075 | 0.92440814 |
| GARS | 41.3624585 | -0.999405369 | 0.34671739 | -2.8824783 | 0.0039456 | 0.08179072 |
| AQP1 | 77.1352833 | 0.464547433 | 0.75327404 | 0.61670442 | 0.5374297 | 0.84423863 |
| PDE1C | 187.79549 | -0.91659734 | 0.52471631 | -1.7468436 | 0.08066447 | 0.42022908 |
| AVL9 | 131.002095 | -0.252667276 | 0.17863226 | -1.4144549 | 0.15722839 | 0.54813635 |
| KBTBD2 | 32.4634386 | -0.186386954 | 0.27809538 | -0.6702267 | 0.50271327 | 0.82430366 |
| FKBP9 | 36.4639562 | 0.641202593 | 0.23366281 | 2.74413631 | 0.00606703 | 0.10540575 |
| NT5C3A | 58.1819304 | -0.316067054 | 0.2793662 | -1.1313719 | 0.25789861 | 0.65130862 |
| BBS9 | 141.233536 | 0.242379488 | 0.26765715 | 0.90555953 | 0.36516904 | 0.73726023 |
| DPY19L1 | 34.465459 | -0.022291478 | 0.28905538 | -0.0771184 | 0.93852939 | 0.98827804 |
| HERPUD2 | 53.1241714 | 0.265354091 | 0.20953473 | 1.26639668 | 0.2053711 | 0.60370888 |
| 7-Sep | 258.294499 | 0.31634167 | 0.20128316 | 1.5716251 | 0.11603752 | 0.48868753 |
| EEPD1 | 146.480692 | -0.531497401 | 0.49441955 | -1.0749927 | 0.282378 | 0.6768745 |
| ELMO1 | 85.0284792 | 0.552991957 | 0.37663175 | 1.46825633 | 0.14203461 | 0.52954815 |
| STARD3NL | 26.4290209 | -0.429307635 | 0.28039725 | -1.5310694 | 0.12575225 | 0.504726 |
| AMPH | 41.4143166 | -0.3959656 | 0.41417103 | -0.9560437 | 0.33905016 | 0.71925477 |
| VPS41 | 94.1989279 | -0.227994412 | 0.30014852 | -0.7596053 | 0.44749053 | 0.79296348 |
| RALA | 59.507477 | 0.067556081 | 0.24178501 | 0.27940558 | 0.77993359 | 0.94911939 |
| CDK13 | 249.555755 | 0.134803543 | 0.14242543 | 0.9464851 | 0.34390122 | 0.72261198 |
| MPLKIP | 18.3890802 | -0.136939698 | 0.40892812 | -0.3348747 | 0.73771958 | 0.93594619 |
| SUGCT | 141.204224 | -0.129182139 | 0.42152794 | -0.3064616 | 0.75925319 | 0.94385579 |
| INHBA | 79.3019805 | -1.187957945 | 0.82017812 | -1.4484146 | 0.14750114 | 0.53679369 |
| GLI3 | 104.051561 | 0.438149181 | 0.20892102 | 2.09720009 | 0.03597587 | 0.28882238 |
| MRPL32 | 25.6075823 | -0.080691566 | 0.25960944 | -0.3108191 | 0.75593816 | 0.94216126 |
| STK17A | 24.1679222 | 0.42573023 | 0.41723336 | 1.0203648 | 0.30755548 | 0.69801431 |
| COA1 | 107.792397 | -0.164331992 | 0.22622802 | -0.7263998 | 0.4675937 | 0.80374006 |

| | | | | | | |
|---|---|---|---|---|---|---|
| DBNL | 21.6956928 | 0.202413091 | 0.25209941 | 0.80290983 | 0.42202686 | 0.77771095 |
| AEBP1 | 221.220647 | 0.201307378 | 0.39762393 | 0.50627581 | 0.61266304 | 0.88300857 |
| NUDCD3 | 85.8211189 | -0.160483479 | 0.16938782 | -0.9474322 | 0.34341857 | 0.72231915 |
| TMED4 | 25.3410245 | 0.28629531 | 0.32873463 | 0.87090098 | 0.38380822 | 0.75256313 |
| OGDH | 69.1252024 | 0.334895177 | 0.21945634 | 1.52602185 | 0.12700442 | 0.50643981 |
| ZMIZ2 | 30.5496693 | -0.00520032 | 0.28442093 | -0.0182839 | 0.98541238 | 0.996806 |
| PPIA | 124.870727 | -0.368405282 | 0.36972532 | -0.9964297 | 0.31904142 | 0.70683013 |
| H2AFV | 55.7790944 | 0.33378933 | 0.27414126 | 1.21758151 | 0.22338305 | 0.61775564 |
| PURB | 36.4311188 | -0.048467377 | 0.26078316 | -0.1858532 | 0.85255991 | 0.97289794 |
| CCM2 | 66.4111365 | 0.323066239 | 0.20780966 | 1.55462571 | 0.12003524 | 0.49642035 |
| ADCY1 | 33.4672027 | 0.353512037 | 0.64962725 | 0.54417674 | 0.58631986 | 0.87014172 |
| IGFBP3 | 73.1679937 | 0.496409918 | 0.71874046 | 0.69066644 | 0.48977518 | 0.81720854 |
| TNS3 | 382.846394 | 0.311520725 | 0.30985959 | 1.00536094 | 0.31472308 | 0.70282967 |
| HUS1 | 26.7568497 | -0.148049482 | 0.25061459 | -0.5907457 | 0.55469085 | 0.85191704 |
| UPP1 | 25.8475455 | -1.636638069 | 0.65552088 | -2.4966986 | 0.01253554 | 0.1619046 |
| GRB10 | 90.19987 | -0.732180451 | 0.2844088 | -2.5743945 | 0.01004157 | 0.14150196 |
| SEC61G | 29.1739551 | -0.872208076 | 0.48225951 | -1.8085866 | 0.07051525 | 0.3958874 |
| EGFR | 304.632182 | -0.394331936 | 0.39067034 | -1.0093726 | 0.31279598 | 0.70282967 |
| LANCL2 | 26.9525006 | 0.180393196 | 0.28976616 | 0.62254749 | 0.53358193 | 0.84333372 |
| VOPP1 | 103.598785 | -0.077587412 | 0.2841216 | -0.2730782 | 0.78479311 | 0.95065355 |
| NIPSNAP2 | 53.1039978 | 0.366331852 | 0.26464032 | 1.38426318 | 0.16627788 | 0.56208655 |
| CCT6A | 40.633905 | -0.196206179 | 0.34243249 | -0.5729777 | 0.5666598 | 0.85826779 |
| SUMF2 | 39.2601361 | -0.431119909 | 0.26684749 | -1.6156042 | 0.10617992 | 0.46978703 |
| CHCHD2 | 75.1336834 | -0.318505595 | 0.3306657 | -0.9632254 | 0.33543443 | 0.71785313 |
| ZNF736 | 33.9083999 | 0.102465922 | 0.22901241 | 0.44742519 | 0.65456809 | 0.90646849 |
| ZNF680 | 45.0118349 | -0.297169978 | 0.32465651 | -0.9153366 | 0.36001496 | 0.73273991 |
| ZNF138 | 19.4492601 | -0.056885139 | 0.31677274 | -0.1795771 | 0.85748456 | 0.97383002 |
| ZNF117 | 18.3288192 | 0.281249029 | 0.35299553 | 0.79674956 | 0.4255965 | 0.77861803 |
| VKORC1L1 | 73.1411872 | 0.043279774 | 0.26505826 | 0.163284 | 0.87029483 | 0.9768643 |
| TPST1 | 212.189703 | 0.00154825 | 0.30051851 | 0.00515193 | 0.99588937 | 0.99854015 |
| AC027644.4 | 50.1364629 | 0.211430965 | 0.25814742 | 0.81903187 | 0.41276823 | 0.77260518 |
| TMEM248 | 52.1446543 | -0.014719081 | 0.23565812 | -0.0624595 | 0.95019694 | 0.99060401 |
| SBDS | 52.4009779 | 0.121876806 | 0.3482964 | 0.34992267 | 0.72639673 | 0.93455731 |
| TYW1 | 101.932221 | 0.071303089 | 0.16747683 | 0.42574898 | 0.67029077 | 0.91445841 |
| AUTS2 | 685.930026 | 0.319446217 | 0.33974445 | 0.94025442 | 0.34708707 | 0.72524566 |
| GALNT17 | 48.5260061 | 0.878250825 | 0.46429852 | 1.89156498 | 0.05854897 | 0.36041506 |
| TYW1B | 60.426583 | 0.270410245 | 0.20570117 | 1.31457807 | 0.18865175 | 0.58671246 |
| POM121 | 81.7096788 | -0.491068423 | 0.24896405 | -1.9724471 | 0.04855859 | 0.33413582 |
| BAZ1B | 97.3750023 | 0.22823404 | 0.16842529 | 1.35510554 | 0.17538393 | 0.57311399 |
| BUD23 | 21.1306643 | -0.088232287 | 0.33704308 | -0.2617834 | 0.79348843 | 0.95416774 |
| ELN | 72.7881302 | 2.274290062 | 0.45995478 | 4.94459486 | 7.63E-07 | 0.00020027 |
| EIF4H | 49.4369093 | -0.525404393 | 0.30562421 | -1.719119 | 0.0855927 | 0.42848827 |
| CLIP2 | 104.601583 | 0.266585171 | 0.28179977 | 0.9460092 | 0.34414389 | 0.72261198 |
| GTF2IRD1 | 56.3298501 | 0.0421704 | 0.34158759 | 0.12345413 | 0.90174749 | 0.98164203 |
| GTF2I | 231.995421 | -0.011755826 | 0.13998869 | -0.083977 | 0.93307474 | 0.98725539 |
| GTF2IRD2 | 25.7869882 | 0.326230166 | 0.47260741 | 0.6902773 | 0.49001982 | 0.81720854 |
| CASTOR2 | 23.6618936 | -0.064929113 | 0.35799366 | -0.1813694 | 0.8560776 | 0.97379567 |
| RCC1L | 27.3743719 | -0.212067867 | 0.27439513 | -0.7728558 | 0.43960773 | 0.78931767 |
| GTF2IRD2B | 41.2733249 | 0.729136599 | 0.44021374 | 1.65632405 | 0.09765621 | 0.45507247 |
| POM121C | 66.5724845 | -0.382209633 | 0.24795334 | -1.5414579 | 0.12320539 | 0.50097964 |
| HIP1 | 404.881107 | 0.64444148 | 0.27885676 | 2.31101257 | 0.02083216 | 0.21233779 |
| RHBDD2 | 23.3394498 | -0.28407947 | 0.29059328 | -0.9775844 | 0.3282799 | 0.71294249 |
| POR | 59.1752781 | -0.507045191 | 0.22866541 | -2.217411 | 0.02659502 | 0.24430847 |
| STYXL1 | 22.3446713 | -0.042342775 | 0.37999983 | -0.1114284 | 0.91127664 | 0.98343817 |
| MDH2 | 24.6749692 | -0.37048978 | 0.32257459 | -1.1485399 | 0.25074577 | 0.6460368 |
| HSPB1 | 216.26129 | 0.887750199 | 0.42940734 | 2.06738477 | 0.03869791 | 0.29748008 |
| YWHAG | 63.4374791 | -0.191349506 | 0.24554205 | -0.7792942 | 0.43580641 | 0.78710773 |
| DTX2 | 28.9417018 | 0.406840216 | 0.3686496 | 1.10359599 | 0.26976843 | 0.6651094 |
| POMZP3 | 18.2961279 | -0.372984836 | 0.33990205 | -1.0973304 | 0.27249701 | 0.6673044 |
| CCDC146 | 60.6381354 | 0.052452178 | 0.27781438 | 0.18880296 | 0.85024725 | 0.97191897 |
| GSAP | 28.9451596 | 0.458733677 | 0.31947283 | 1.43590829 | 0.15102844 | 0.53880252 |
| PTPN12 | 283.729921 | -0.171162289 | 0.19992742 | -0.8561221 | 0.39193024 | 0.75901355 |
| RSBN1L | 110.066693 | 0.231565506 | 0.20677412 | 1.11989597 | 0.2627581 | 0.65702935 |
| PHTF2 | 119.206301 | 0.742377166 | 0.33624705 | 2.20783249 | 0.02725595 | 0.24775789 |
| MAGI2 | 572.448411 | -0.101657589 | 0.22094709 | -0.4600992 | 0.64544499 | 0.90150535 |
| GNAI1 | 211.932952 | 0.162902123 | 0.2922284 | 0.55744795 | 0.57722141 | 0.86377523 |
| SEMA3C | 153.637054 | 1.493191885 | 0.47448306 | 3.14698671 | 0.00164962 | 0.05093736 |
| HGF | 158.240475 | -0.47444419 | 0.31615813 | -1.5006547 | 0.1334449 | 0.51533712 |
| CACNA2D1 | 364.982107 | 0.112442917 | 0.23260638 | 0.48340426 | 0.62880872 | 0.89279662 |
| KIAA1324L | 27.035525 | 0.643428553 | 0.31985986 | 2.01159516 | 0.04426263 | 0.31581172 |
| DMTF1 | 125.919258 | 0.105684911 | 0.30948729 | 0.34148385 | 0.73273936 | 0.93574285 |
| TMEM243 | 25.5484963 | 0.045117708 | 0.28934817 | 0.15592878 | 0.87608916 | 0.9768643 |
| ADAM22 | 57.9101865 | 0.644918111 | 0.54921233 | 1.17426007 | 0.24029087 | 0.63451757 |
| SRI | 25.5527099 | -0.121084426 | 0.35897798 | -0.3373032 | 0.73588834 | 0.93594619 |
| STEAP4 | 117.29805 | -2.288267306 | 0.40801761 | -5.6082562 | 2.04E-08 | 1.37E-05 |
| STEAP2 | 39.0984566 | -1.431360294 | 0.40641159 | -3.5219475 | 0.00042839 | 0.02141653 |
| CFAP69 | 42.2616081 | -1.159407262 | 0.43837512 | -2.6447834 | 0.00817432 | 0.12673647 |
| GTPBP10 | 29.0645214 | -0.477485246 | 0.3011106 | -1.5857471 | 0.1127967 | 0.48301375 |
| CLDN12 | 22.9302803 | -0.192343308 | 0.33253407 | -0.5784169 | 0.56298272 | 0.85571045 |
| CDK14 | 271.16698 | -0.424196973 | 0.37495621 | -1.131324 | 0.25791873 | 0.65130862 |
| FZD1 | 28.2793741 | 0.796847334 | 0.38232877 | 2.08419404 | 0.03714252 | 0.29287595 |
| MTERF1 | 54.3920094 | 0.151175378 | 0.28720395 | 0.52636943 | 0.59863155 | 0.87618867 |
| AKAP9 | 248.409121 | 0.225170307 | 0.19930343 | 1.12978643 | 0.25856623 | 0.6514155 |
| KRIT1 | 89.9752964 | -0.205952928 | 0.2051128 | -1.0040959 | 0.31533238 | 0.70306809 |
| ANKIB1 | 173.918091 | 0.025979506 | 0.18978598 | 0.13688844 | 0.89111898 | 0.98023547 |
| GATAD1 | 31.75209 | 0.111383856 | 0.36106009 | 0.30849119 | 0.7577086 | 0.94301125 |
| AC007566.1 | 18.9080463 | 0.034896778 | 0.40281855 | 0.08663151 | 0.93096442 | 0.98681039 |
| PEX1 | 28.5780713 | -0.036944809 | 0.26125021 | -0.1414154 | 0.88754177 | 0.97920559 |
| FAM133B | 51.17839 | -0.112936889 | 0.20216335 | -0.5586418 | 0.57640624 | 0.86325068 |
| CDK6 | 110.782311 | 0.682601268 | 0.16630076 | 4.10461893 | 4.05E-05 | 0.00444211 |
| SAMD9L | 29.1055958 | 0.429243608 | 0.96836332 | 0.44326711 | 0.65757254 | 0.90772143 |
| VPS50 | 60.5032602 | -0.008470105 | 0.24030336 | -0.0352476 | 0.97188235 | 0.99300191 |
| GNG11 | 19.8749298 | 0.677197997 | 0.33630148 | 2.01366348 | 0.04404488 | 0.3152359 |
| BET1 | 23.7372711 | -0.289161328 | 0.33985739 | -0.8508314 | 0.39486303 | 0.75988253 |
| AC002074.1 | 30.6275981 | -0.118075604 | 0.40783649 | -0.289517 | 0.77218576 | 0.94690358 |
| COL1A2 | 448.936815 | 0.25020442 | 0.36041866 | 0.69420495 | 0.48755369 | 0.81595238 |
| CASD1 | 50.9854509 | 0.276495712 | 0.2217154 | 1.24707493 | 0.21237002 | 0.60910715 |
| SGCE | 50.2873569 | 0.118403982 | 0.31756062 | 0.37285474 | 0.70925656 | 0.93009618 |
| PPP1R9A | 31.3438307 | 0.340728945 | 0.36883984 | 0.92378563 | 0.35559794 | 0.73125149 |
| PON2 | 34.4692961 | -0.087384589 | 0.24968522 | -0.349979 | 0.72635444 | 0.93455731 |
| PDK4 | 281.214078 | -1.857876302 | 0.59871396 | -3.1031117 | 0.00191497 | 0.05543169 |
| DYNC1I1 | 87.3921502 | -0.297193421 | 0.34609789 | -0.8586976 | 0.39050739 | 0.75729345 |
| SLC25A13 | 185.686358 | -0.573879882 | 0.21992424 | -2.6094435 | 0.00906896 | 0.13367648 |
| SEM1 | 109.361159 | 0.409555553 | 0.29575262 | 1.38479098 | 0.16611639 | 0.56208655 |
| LMTK2 | 49.5922571 | -0.239242419 | 0.21166281 | -1.1302997 | 0.25834994 | 0.6514155 |
| TECPR1 | 17.4045191 | 0.718030282 | 0.47798943 | 1.50218861 | 0.1330484 | 0.51434652 |
| BRI3 | 84.2537201 | -0.466552595 | 0.27979275 | -1.6674935 | 0.09541631 | 0.44884198 |
| BAIAP2L1 | 113.570164 | -0.014464586 | 0.35571519 | -0.0406634 | 0.96756425 | 0.99300191 |
| TRRAP | 102.99757 | 0.085535345 | 0.15965443 | 0.53575302 | 0.59212925 | 0.87258028 |
| SMURF1 | 291.198424 | 0.081633621 | 0.28974771 | 0.28174035 | 0.77814261 | 0.94820337 |
| PDAP1 | 26.1844617 | -0.035148008 | 0.24708061 | -0.1422532 | 0.88688001 | 0.97920559 |
| ZNF789 | 23.6280789 | -0.149271634 | 0.33453106 | -0.4462116 | 0.65544442 | 0.90695933 |
| TRIM4 | 40.920448 | 0.004625613 | 0.28525126 | 0.01621592 | 0.98706213 | 0.99719329 |

| | | | | | | |
|---|---|---|---|---|---|---|
| ZKSCAN1 | 96.1401609 | -0.243196667 | 0.2359834 | -1.0305668 | 0.30274399 | 0.6944025 |
| COPS6 | 27.8857153 | -0.245065897 | 0.29918589 | -0.8191091 | 0.41272415 | 0.77260518 |
| LAMTOR4 | 44.5742822 | -0.010571968 | 0.2413418 | -0.043805 | 0.96505987 | 0.99275266 |
| CASTOR3 | 47.7096205 | -0.122543223 | 0.37776814 | -0.3243874 | 0.74564476 | 0.93891908 |
| SPDYE3 | 21.9488959 | -0.065753386 | 0.35201123 | -0.1867934 | 0.8518226 | 0.97285733 |
| PILRB | 48.1015981 | -0.273958225 | 0.36676547 | -0.7469575 | 0.45508921 | 0.79609632 |
| TSC22D4 | 26.2248041 | -0.241800813 | 0.29640451 | -0.8157798 | 0.41462609 | 0.77476594 |
| PCOLCE | 40.7270197 | 0.058186879 | 0.37894126 | 0.15355119 | 0.87796363 | 0.9768643 |
| GNB2 | 52.4084418 | -0.284543998 | 0.28697177 | -0.99154 | 0.32142195 | 0.70780851 |
| GIGYF1 | 33.7719383 | 0.037171471 | 0.24122476 | 0.15409475 | 0.87753502 | 0.9768643 |
| TRIP6 | 24.7359672 | 0.536962967 | 0.24930078 | 2.15387598 | 0.0312499 | 0.26776774 |
| SRRT | 19.921979 | -0.030811133 | 0.33699559 | -0.0914289 | 0.92715181 | 0.98678435 |
| TRIM56 | 86.012971 | -0.165871187 | 0.31843077 | -0.5209019 | 0.60243512 | 0.87903643 |
| SERPINE1 | 850.987379 | -2.231429528 | 0.99347128 | -2.2460936 | 0.02469801 | 0.23328859 |
| PLOD3 | 23.5015779 | -0.375260986 | 0.30185874 | -1.2431676 | 0.2138061 | 0.61001472 |
| ZNHIT1 | 22.3925308 | -0.010414635 | 0.35331598 | -0.0294768 | 0.9764843 | 0.99365593 |
| FIS1 | 41.4807119 | -0.455499676 | 0.31503949 | -1.4458495 | 0.14821943 | 0.53680621 |
| CUX1 | 488.683048 | -0.422645346 | 0.25413544 | -1.6630712 | 0.09629818 | 0.45133629 |
| PRKRIP1 | 110.178893 | -0.37417091 | 0.23206892 | -1.6123267 | 0.10689087 | 0.47094784 |
| AC105052.3 | 27.1813912 | 0.779240293 | 0.28165375 | 2.76666047 | 0.00566337 | 0.10101851 |
| FAM185A | 51.7743195 | 0.033759754 | 0.26090894 | 0.12939286 | 0.8970468 | 0.98040427 |
| FBXL13 | 22.9076808 | 0.223436966 | 0.34019849 | 0.65678412 | 0.51131974 | 0.8285973 |
| ARMC10 | 30.4537684 | -0.327267411 | 0.23535527 | -1.3905251 | 0.16436947 | 0.55949572 |
| NAPEPLD | 41.017654 | 0.185991504 | 0.27542584 | 0.67528705 | 0.49949341 | 0.82271448 |
| PMPCB | 27.837913 | -0.144398254 | 0.30418261 | -0.4747091 | 0.63499433 | 0.89672816 |
| DNAJC2 | 24.9670496 | -0.10459818 | 0.28868659 | -0.3623243 | 0.71710967 | 0.93308056 |
| PSMC2 | 28.8511847 | -0.217097832 | 0.22565512 | -0.962078 | 0.33601041 | 0.71883121 |
| ORC5 | 44.3470972 | -0.302084438 | 0.25337485 | -1.1922432 | 0.2331659 | 0.62939306 |
| LINC01004 | 112.82728 | 0.430554345 | 0.27497682 | 1.56578415 | 0.11739916 | 0.49048688 |
| KMT2E | 406.317797 | -0.116665776 | 0.32762941 | -0.3560907 | 0.72177266 | 0.93444541 |
| SRPK2 | 263.11988 | 0.013660378 | 0.1801072 | 0.07584581 | 0.93954177 | 0.98864439 |
| PUS7 | 41.0581248 | -0.760989983 | 0.45050885 | -1.6891788 | 0.09118518 | 0.44115857 |
| RINT1 | 23.9704899 | -0.366661546 | 0.30486502 | -1.2027013 | 0.22909194 | 0.62286225 |
| ATXN7L1 | 131.291373 | 0.132673947 | 0.44597771 | 0.29749009 | 0.76609237 | 0.9447922 |
| CDHR3 | 27.734044 | 0.34090745 | 0.42480988 | 0.80249416 | 0.42226717 | 0.77771095 |
| SYPL1 | 51.4997765 | 0.05000297 | 0.20768722 | 0.24076094 | 0.8097404 | 0.95953411 |
| NAMPT | 883.324841 | -2.855260168 | 0.54912839 | -5.1996222 | 2.00E-07 | 9.17E-05 |
| CCDC71L | 46.1253188 | -1.280573651 | 0.46629185 | -2.7462922 | 0.00602731 | 0.10521299 |
| PRKAR2B | 62.2455831 | -0.288252008 | 0.7757818 | -0.3715633 | 0.71021805 | 0.93009618 |
| HBP1 | 82.344437 | -0.416976875 | 0.34855128 | -1.1963143 | 0.23157393 | 0.62728965 |
| COG5 | 221.118865 | 0.033493924 | 0.23411612 | 0.14306543 | 0.8862385 | 0.97895187 |
| BCAP29 | 61.5189962 | -0.12352965 | 0.2311047 | -0.5345181 | 0.59298312 | 0.87313824 |
| SLC26A3 | 262.906993 | 0.102820751 | 0.61277222 | 0.16779604 | 0.86674374 | 0.97624269 |
| DLD | 39.0709034 | -0.167563837 | 0.24293604 | -0.6897447 | 0.49035477 | 0.81720854 |
| LAMB1 | 172.763789 | -1.012477699 | 0.4044921 | -2.503084 | 0.01231163 | 0.16004312 |
| NRCAM | 33.7604452 | -0.426624819 | 0.78998029 | -0.5400449 | 0.58916608 | 0.8702694 |
| PNPLA8 | 74.6822036 | -0.441879169 | 0.22683611 | -1.9480107 | 0.05141368 | 0.34417603 |
| IMMP2L | 989.62786 | -0.128416402 | 0.2473833 | -0.5190989 | 0.60369176 | 0.87903643 |
| DOCK4 | 586.61038 | -0.035072392 | 0.29931157 | -0.1171769 | 0.9067199 | 0.98306694 |
| ZNF277 | 66.4475185 | -0.246226972 | 0.22891448 | -1.0756287 | 0.28209337 | 0.676814 |
| IFRD1 | 80.8946488 | -0.385641127 | 0.37810816 | -1.0199228 | 0.30776508 | 0.69801431 |
| TMEM168 | 65.9679179 | -0.025236746 | 0.18926719 | -0.1333393 | 0.89392508 | 0.98023547 |
| BMT2 | 54.9613184 | -0.24466061 | 0.20078868 | -1.218498 | 0.22303479 | 0.61772146 |
| FOXP2 | 75.4167264 | 0.531722595 | 0.54183337 | 0.9813397 | 0.32642525 | 0.71149668 |
| MDFIC | 66.7711259 | -0.545932438 | 0.38011769 | -1.4362195 | 0.1509399 | 0.53880252 |
| TES | 35.2017642 | 0.989087463 | 0.46510004 | 2.1266123 | 0.03345231 | 0.2790477 |
| CAV2 | 34.5065096 | 0.765629634 | 0.33722445 | 2.27038588 | 0.02318418 | 0.22566959 |
| CAV1 | 44.4354666 | 1.284198125 | 0.38784144 | 3.31114212 | 0.00092916 | 0.03612909 |
| CAPZA2 | 86.3962803 | -0.178729288 | 0.17305205 | -1.0328066 | 0.30169444 | 0.69264368 |
| ST7 | 121.475806 | 0.093407316 | 0.17600379 | 0.53071195 | 0.5956184 | 0.87456537 |
| CTTNBP2 | 36.322919 | -0.013152106 | 0.46583617 | -0.0282333 | 0.97747606 | 0.99407008 |
| LSM8 | 26.1035485 | 0.260559012 | 0.28957334 | 0.89980319 | 0.368225 | 0.74098727 |
| ING3 | 18.6893125 | 0.466032993 | 0.26091093 | 1.78617661 | 0.07407067 | 0.40373597 |
| CPED1 | 406.209996 | 0.035373203 | 0.37355923 | 0.09469235 | 0.92455919 | 0.98561736 |
| FAM3C | 76.1123571 | 0.294100439 | 0.16751243 | 1.7556932 | 0.07914081 | 0.41722081 |
| AASS | 120.890074 | 0.139476173 | 0.42607486 | 0.32735133 | 0.74340217 | 0.93791562 |
| CADPS2 | 177.590365 | -0.418629179 | 0.19980027 | -2.0952383 | 0.03614982 | 0.28939542 |
| NDUFA5 | 33.5986744 | 0.125042289 | 0.2561735 | 0.48811564 | 0.62546794 | 0.89069797 |
| WASL | 88.6809102 | 0.055453831 | 0.1899364 | 0.29196 | 0.7703172 | 0.94593455 |
| POT1 | 48.5596388 | -0.195288382 | 0.28938376 | -0.6748422 | 0.49977601 | 0.82271448 |
| ZNF800 | 58.038414 | -0.290426321 | 0.22329058 | -1.3006653 | 0.19337303 | 0.59237115 |
| SND1 | 277.989039 | -0.982214723 | 0.33420695 | -2.9389416 | 0.00329335 | 0.07334193 |
| RBM28 | 31.6598708 | -0.157792225 | 0.28170082 | -0.5601412 | 0.57538315 | 0.86294056 |
| HILPDA | 92.554275 | -4.276774066 | 0.85972049 | -4.9746099 | 6.54E-07 | 0.00018632 |
| METTL2B | 25.3068771 | -0.145266305 | 0.42932667 | -0.3383584 | 0.73509309 | 0.93594619 |
| CALU | 127.122618 | -0.204044149 | 0.30263327 | -0.6742291 | 0.50016568 | 0.82286044 |
| ATP6V1F | 29.4748163 | -0.588367948 | 0.35289289 | -1.6672706 | 0.09546061 | 0.44884198 |
| TNPO3 | 72.0977096 | 0.246901663 | 0.16908212 | 1.460247 | 0.1442222 | 0.53341166 |
| AHCYL2 | 132.217635 | 0.381773369 | 0.20415379 | 1.87002832 | 0.06147989 | 0.36882912 |
| NRF1 | 107.872179 | -0.005439124 | 0.16471733 | -0.033021 | 0.97365787 | 0.99300191 |
| UBE2H | 200.254261 | 0.095950403 | 0.27155933 | 0.35333127 | 0.7238401 | 0.93455731 |
| KLHDC10 | 82.5272235 | 0.159306212 | 0.19366883 | 0.8225702 | 0.41075245 | 0.77166383 |
| COPG2 | 27.2970213 | 0.479683733 | 0.30560282 | 1.56963122 | 0.11650093 | 0.48979711 |
| AC016831.7 | 1071.00628 | -0.118421086 | 0.27281993 | -0.4340632 | 0.66424256 | 0.9117517 |
| LINC00513 | 80.1266899 | -0.661649842 | 0.40274538 | -1.642849 | 0.10041418 | 0.45949178 |
| MKLN1 | 712.023888 | -0.127592505 | 0.29983877 | -0.425537 | 0.67044522 | 0.91445841 |
| PODXL | 24.8712065 | 0.197405269 | 0.57352783 | 0.34419475 | 0.73069983 | 0.93455731 |
| PLXNA4 | 73.3943233 | -0.600476068 | 0.44573343 | -1.3471641 | 0.17792739 | 0.57526987 |
| CHCHD3 | 223.127719 | -0.154181474 | 0.24818453 | -0.6212373 | 0.53444353 | 0.84361209 |
| EXOC4 | 483.184292 | -0.240776303 | 0.19708416 | -1.2216928 | 0.22182381 | 0.61772146 |
| SLC35B4 | 19.9023541 | 0.019957294 | 0.39500746 | 0.05052384 | 0.95970495 | 0.99181157 |
| AKR1B1 | 26.3077125 | 0.229238434 | 0.50045521 | 0.45805984 | 0.64690945 | 0.90243971 |
| CALD1 | 2372.97465 | 0.322672846 | 0.24635103 | 1.30980918 | 0.1902604 | 0.58890833 |
| AGBL3 | 29.2267055 | 0.466974428 | 0.23493974 | 1.98763487 | 0.04685209 | 0.32667551 |
| CYREN | 21.6114894 | 0.497515411 | 0.3230461 | 1.54007557 | 0.12354193 | 0.50097964 |
| WDR91 | 18.708931 | 0.032099932 | 0.3154002 | 0.10177524 | 0.91893508 | 0.98528826 |
| CNOT4 | 162.991043 | 0.047297199 | 0.18420311 | 0.25676656 | 0.79735899 | 0.95463848 |
| NUP205 | 35.3203392 | 0.072199856 | 0.25361949 | 0.28467786 | 0.77589095 | 0.94805375 |
| STMP1 | 47.1850639 | 0.010592181 | 0.22010121 | 0.04812414 | 0.96161731 | 0.99198843 |
| MTPN | 48.2058149 | -0.170487492 | 0.21040386 | -0.8102869 | 0.4177753 | 0.77722447 |
| PTN | 56.6351878 | 0.657741299 | 0.45152124 | 1.45672282 | 0.14519291 | 0.53538077 |
| CREB3L2 | 273.004328 | 0.362561215 | 0.36572535 | 0.99134833 | 0.32151552 | 0.70780851 |
| TRIM24 | 43.9626864 | 0.175191692 | 0.23262643 | 0.75310313 | 0.45138794 | 0.79401729 |
| KIAA1549 | 32.9578986 | 0.39540292 | 0.51951698 | 0.76109721 | 0.44659901 | 0.79296348 |
| ZC3HAV1 | 67.7494383 | 0.228851612 | 0.50517914 | 0.45301081 | 0.65054096 | 0.9050963 |
| UBN2 | 89.5224737 | 0.050642733 | 0.18909076 | 0.26782235 | 0.78883606 | 0.95129454 |
| LUC7L2 | 25.5980652 | 0.302049497 | 0.37325089 | 0.80923985 | 0.4183772 | 0.77722447 |
| HIPK2 | 452.212321 | -1.431182982 | 0.30858847 | -4.6378368 | 3.52E-06 | 0.000761 |
| PARP12 | 33.6671842 | 0.223485427 | 0.79420266 | 0.28139597 | 0.77840671 | 0.94820337 |
| KDM7A | 86.4537401 | -1.098548461 | 0.28529944 | -3.8505104 | 0.00011787 | 0.00879608 |
| SLC37A3 | 53.7698718 | 0.094400603 | 0.21076382 | 0.44789756 | 0.65422713 | 0.90646849 |
| MKRN1 | 32.540517 | 0.116722054 | 0.24240033 | 0.48152598 | 0.63014272 | 0.89365764 |
| DENND2A | 119.593341 | -0.233177552 | 0.2958966 | -0.7880373 | 0.4306749 | 0.78340375 |

| | | | | | | |
|---|---|---|---|---|---|---|
| NDUFB2 | 39.5323738 | -0.066787788 | 0.25590081 | -0.2609909 | 0.79409951 | 0.95434788 |
| BRAF | 309.101843 | 0.118068899 | 0.15442495 | 0.76457138 | 0.44452682 | 0.79252305 |
| AGK | 33.723707 | 0.012681601 | 0.24457279 | 0.05185205 | 0.95864658 | 0.99181157 |
| KIAA1147 | 26.2190909 | 0.136569997 | 0.31629962 | 0.43177414 | 0.66590558 | 0.91284091 |
| SSBP1 | 39.7114714 | -0.307729383 | 0.22915374 | -1.3428949 | 0.17930602 | 0.57744792 |
| GSTK1 | 25.4480403 | -0.330525939 | 0.27065494 | -1.2212078 | 0.22200733 | 0.61772146 |
| ZYX | 27.1755507 | 0.07095397 | 0.30513031 | 0.23253662 | 0.81612125 | 0.96132648 |
| TCAF2 | 38.0023518 | -1.422940613 | 0.54402941 | -2.6155583 | 0.00890817 | 0.1336044 |
| TCAF1 | 49.6850386 | -0.195965278 | 0.2467734 | -0.7941102 | 0.42713128 | 0.77987274 |
| AC004889.1 | 26.4479432 | 1.090436024 | 0.26765872 | 4.0739791 | 4.62E-05 | 0.00485209 |
| TPK1 | 41.8129221 | -0.016360176 | 0.23043119 | -0.0709981 | 0.94339927 | 0.99001017 |
| CUL1 | 144.28309 | -0.187076592 | 0.23535591 | -0.7948668 | 0.42669101 | 0.77966505 |
| PDIA4 | 28.9407717 | -0.394455858 | 0.30788203 | -1.2811915 | 0.2001264 | 0.59922861 |
| ZNF398 | 28.8847676 | 0.060400776 | 0.30646977 | 0.19708559 | 0.84376056 | 0.9691438 |
| ZNF783 | 22.5677507 | 0.374202841 | 0.23855367 | 1.5686317 | 0.11673378 | 0.48989475 |
| ZNF862 | 25.0128914 | -0.024606781 | 0.26960058 | -0.0912712 | 0.92727706 | 0.98678435 |
| RARRES2 | 52.3240696 | 0.440247933 | 0.43120644 | 1.0209679 | 0.30726965 | 0.69801431 |
| TMEM176B | 31.4907128 | -0.495614835 | 0.45036944 | -1.1004628 | 0.27113051 | 0.6661779 |
| AGAP3 | 30.2519769 | 0.082432594 | 0.22643401 | 0.36404688 | 0.715823 | 0.93237409 |
| CHPF2 | 17.7899255 | -0.412865693 | 0.44622973 | -0.9252313 | 0.35484563 | 0.73100657 |
| SMARCD3 | 42.0603267 | -0.394777649 | 0.27591057 | -1.4308174 | 0.15248256 | 0.54004545 |
| NUB1 | 37.2256572 | -0.161354895 | 0.2802389 | -0.5757762 | 0.56476645 | 0.8572637 |
| WDR86 | 31.1818347 | 0.46581349 | 0.37738579 | 1.23431644 | 0.21708501 | 0.61410359 |
| RHEB | 116.547685 | 0.010586514 | 0.21800922 | 0.04855994 | 0.96127 | 0.99181157 |
| PRKAG2 | 89.024174 | 0.077587588 | 0.33469578 | 0.23181525 | 0.81668151 | 0.96134613 |
| GALNT11 | 70.6953063 | 0.231558192 | 0.27974236 | 0.82775519 | 0.40780916 | 0.76944531 |
| KMT2C | 407.989221 | 0.037783101 | 0.20191093 | 0.18712757 | 0.85156062 | 0.97281501 |
| ACTR3B | 35.6778593 | -0.121144534 | 0.28648612 | -0.4228635 | 0.67239483 | 0.91482259 |
| DPP6 | 68.7204495 | 0.747038058 | 0.56601634 | 1.31981713 | 0.18689608 | 0.58501647 |
| PAXIP1 | 19.7172814 | 0.254668964 | 0.26978313 | 0.94397659 | 0.34518161 | 0.72325086 |
| INSIG1 | 23.0404198 | -0.956868654 | 0.63489027 | -1.5071402 | 0.13177472 | 0.51290135 |
| RBM33 | 149.440924 | -0.210099085 | 0.15497529 | -1.3556941 | 0.17519652 | 0.57299474 |
| LMBR1 | 85.1626604 | -0.043831179 | 0.24481369 | -0.1790389 | 0.85790714 | 0.97397549 |
| UBE3C | 157.551773 | -0.14259164 | 0.18623489 | -0.7656548 | 0.44388172 | 0.79248199 |
| DNAJB6 | 88.9181105 | 0.064433464 | 0.20013008 | 0.32195792 | 0.74748457 | 0.93903859 |
| ESYT2 | 350.784303 | -0.107528826 | 0.45624153 | -0.235684 | 0.81367789 | 0.96038841 |
| WDR60 | 106.970763 | -0.347656946 | 0.20618215 | -1.6861642 | 0.09176418 | 0.44124451 |
| VIPR2 | 49.7385308 | -0.057828841 | 0.49989334 | -0.1156824 | 0.90790429 | 0.98306694 |
| GTPBP6 | 26.3217349 | 0.057492897 | 0.26827447 | 0.21430626 | 0.83030824 | 0.96580172 |
| SLC25A6 | 78.6980928 | -0.618141578 | 0.38062584 | -1.6240137 | 0.10437289 | 0.46422585 |
| ASMTL | 23.5950616 | -0.117024126 | 0.27225187 | -0.4298377 | 0.66731369 | 0.91374851 |
| AKAP17A | 20.4493856 | -0.22596901 | 0.26907757 | -0.8397913 | 0.4010254 | 0.76363711 |
| DHRSX | 107.301666 | -0.025475595 | 0.15839244 | -0.1608384 | 0.87222064 | 0.9768643 |
| CD99 | 87.2314183 | 0.511036543 | 0.33980199 | 1.50392449 | 0.13260081 | 0.51369707 |
| PRKX | 51.2274732 | -0.224877296 | 0.26626446 | -0.8445637 | 0.39835449 | 0.7611823 |
| TBL1X | 283.273283 | 0.247051768 | 0.17507568 | 1.41111415 | 0.15821096 | 0.55005969 |
| SHROOM2 | 25.7351616 | 0.084777667 | 0.42178535 | 0.20099718 | 0.84070078 | 0.96803523 |
| WWC3 | 56.7677752 | 0.02459402 | 0.20015878 | 0.12287255 | 0.90220802 | 0.98164203 |
| MID1 | 159.480343 | -0.200031142 | 0.39563926 | -0.5055897 | 0.6131447 | 0.88331141 |
| ARHGAP6 | 348.017354 | 1.03441814 | 0.36655709 | 2.82198375 | 0.00477276 | 0.09157966 |
| TMSB4X | 661.913199 | -0.018375495 | 0.4069692 | -0.0451521 | 0.96398611 | 0.99274284 |
| RAB9A | 28.0103667 | -0.340864294 | 0.36880383 | -0.9242428 | 0.35535991 | 0.73125149 |
| OFD1 | 40.5798779 | 0.118356569 | 0.25301126 | 0.46779171 | 0.63993355 | 0.89903874 |
| GPM6B | 86.1467244 | 0.406885691 | 0.50556454 | 0.80481454 | 0.42092671 | 0.77771095 |
| MOSPD2 | 24.9171518 | 0.274342341 | 0.36716212 | 0.74719674 | 0.45494481 | 0.79609632 |
| PIR | 36.7513864 | -0.029538505 | 0.27607695 | -0.1069937 | 0.91479395 | 0.98439006 |
| CLTRN | 22.6428248 | -1.585280009 | 0.83571399 | -1.8969169 | 0.0578389 | 0.35960922 |
| CA5B | 46.075494 | 0.016238156 | 0.29598484 | 0.05486144 | 0.95624885 | 0.99181157 |
| ZRSR2 | 19.2033554 | 0.10394675 | 0.39369777 | 0.26402677 | 0.79175929 | 0.95262591 |
| CTPS2 | 36.1211561 | 0.183029533 | 0.30916445 | 0.59201352 | 0.55384153 | 0.85191704 |
| SYAP1 | 38.933283 | -0.13283988 | 0.24250751 | -0.5477763 | 0.58384548 | 0.86820729 |
| TXLNG | 44.318255 | -0.460907414 | 0.34136414 | -1.3501928 | 0.17695414 | 0.57388115 |
| RBBP7 | 43.962873 | 0.127908964 | 0.2282705 | 0.56033943 | 0.57524793 | 0.86294056 |
| NHS | 520.271286 | 0.208567284 | 0.26898561 | 0.77538454 | 0.43811247 | 0.78836643 |
| SCML1 | 66.1298522 | -0.769866339 | 0.42584854 | -1.8078408 | 0.07063129 | 0.3959339 |
| CDKL5 | 40.4428496 | 0.488863904 | 0.34782737 | 1.40547853 | 0.15987903 | 0.5530386 |
| PHKA2 | 47.3190258 | 0.258845797 | 0.38980263 | 0.66404323 | 0.50666264 | 0.82481485 |
| ADGRG2 | 24.3010886 | 0.047099075 | 0.47121002 | 0.09995347 | 0.92038127 | 0.98528826 |
| PDHA1 | 24.1400763 | 0.167962817 | 0.27820306 | 0.60374181 | 0.54601531 | 0.84758329 |
| SH3KBP1 | 94.7557291 | 0.31213501 | 0.30939971 | 1.00884068 | 0.31305105 | 0.70282967 |
| BCLAF3 | 31.1498631 | 0.4644902 | 0.23982786 | 1.93676496 | 0.05277408 | 0.34706747 |
| EIF1AX | 64.6064882 | 0.156043906 | 0.26402622 | 0.5910167 | 0.55450923 | 0.85191704 |
| RPS6KA3 | 158.018581 | -0.209410371 | 0.26726013 | -0.7835451 | 0.43330707 | 0.78484089 |
| MBTPS2 | 29.5826522 | -0.361852994 | 0.27953278 | -1.2944922 | 0.19549547 | 0.5934309 |
| SMS | 32.4883903 | -0.231382348 | 0.42020431 | -0.5506425 | 0.58187878 | 0.86703713 |
| ACOT9 | 33.5069074 | 0.513841962 | 0.28559286 | 1.79921154 | 0.07198522 | 0.39925993 |
| SAT1 | 143.254969 | -1.309948455 | 0.40572057 | -3.2286962 | 0.00124356 | 0.04375316 |
| APOO | 222.571818 | -0.116031672 | 0.30934581 | -0.3750873 | 0.70759557 | 0.93009618 |
| EIF2S3 | 52.3163895 | -0.555891189 | 0.3186967 | -1.744264 | 0.08111306 | 0.42067742 |
| ZFX | 90.8048111 | -0.088776623 | 0.29892711 | -0.2969842 | 0.76647858 | 0.9447922 |
| PDK3 | 53.4140681 | -0.873645591 | 0.48619984 | -1.7968858 | 0.07235375 | 0.40009608 |
| POLA1 | 61.8641579 | -0.019104051 | 0.21825772 | -0.0875298 | 0.93025041 | 0.98681039 |
| IL1RAPL1 | 96.1891964 | 1.757281905 | 0.35170166 | 4.99651297 | 5.84E-07 | 0.00017875 |
| TAB3 | 27.2669128 | -0.07974779 | 0.24537658 | -0.3250016 | 0.74517983 | 0.93891908 |
| DMD | 719.001968 | 0.307522074 | 0.43924636 | 0.70011296 | 0.48385676 | 0.8144078 |
| TMEM47 | 55.3468478 | 0.903031762 | 0.2785806 | 3.24154572 | 0.00118883 | 0.04303812 |
| DYNLT3 | 27.2170931 | 0.213014303 | 0.27687559 | 0.76935025 | 0.44168541 | 0.79053729 |
| RPGR | 24.6545172 | -0.182685641 | 0.33660794 | -0.5427253 | 0.58731896 | 0.870249 |
| BCOR | 48.5773857 | 0.261123691 | 0.3063269 | 0.85243474 | 0.39397284 | 0.75988253 |
| ATP6AP2 | 63.2203637 | -0.313157758 | 0.28276732 | -1.1074751 | 0.26808859 | 0.66313802 |
| CXorf38 | 19.105537 | -0.530731489 | 0.28048363 | -1.8922013 | 0.05846416 | 0.36041506 |
| MED14 | 66.7896774 | 0.034176856 | 0.19451644 | 0.17570163 | 0.86052835 | 0.97397549 |
| USP9X | 198.747099 | -0.031770936 | 0.23188415 | -0.1370121 | 0.89102122 | 0.98023547 |
| DDX3X | 108.440001 | -0.190209448 | 0.26391508 | -0.7207222 | 0.47108047 | 0.80492221 |
| CASK | 128.235904 | 0.017949884 | 0.22216621 | 0.08079484 | 0.93560511 | 0.98827804 |
| MAOA | 52.6484196 | -1.012339792 | 0.46377435 | -2.1828283 | 0.02904846 | 0.25751158 |
| MAOB | 34.651914 | 0.750138607 | 0.58143284 | 1.29015521 | 0.19699677 | 0.59479429 |
| KDM6A | 230.501507 | -0.120994354 | 0.33902245 | -0.3568919 | 0.72117276 | 0.93444541 |
| MIR222HG | 44.7754974 | -0.078273736 | 0.59522632 | -0.1315025 | 0.89537783 | 0.98023547 |
| KRBOX4 | 29.8062009 | -0.119215153 | 0.23398626 | -0.5094964 | 0.61040433 | 0.88163721 |
| JADE3 | 24.6094578 | 0.402693208 | 0.25500161 | 1.57917909 | 0.11429499 | 0.48580326 |
| NDUFB11 | 31.1789465 | -0.35274588 | 0.33551033 | -1.0513711 | 0.29308816 | 0.68443118 |
| RBM10 | 31.6490224 | 0.079204846 | 0.23071159 | 0.34330675 | 0.7313677 | 0.93507677 |
| UBA1 | 44.3722203 | 0.163347731 | 0.2230394 | 0.73237163 | 0.46394177 | 0.80227475 |
| USP11 | 33.3088502 | 0.155228822 | 0.26405663 | 0.58786187 | 0.55662501 | 0.85243179 |
| ARAF | 17.3413495 | 0.104648585 | 0.32005287 | 0.32697281 | 0.74368845 | 0.93791562 |
| TIMP1 | 588.196824 | -1.829714382 | 0.67933256 | -2.6934001 | 0.00707273 | 0.11576428 |
| ZNF81 | 37.0608121 | -0.072884705 | 0.24528323 | -0.2971451 | 0.76635575 | 0.9447922 |
| RBM3 | 52.8860367 | -0.288826153 | 0.35819085 | -0.8063471 | 0.42004274 | 0.77771095 |
| WDR13 | 26.6675572 | 0.168154505 | 0.3928695 | 0.42801619 | 0.66863933 | 0.91445841 |
| HDAC6 | 19.4782137 | 0.176159006 | 0.26098261 | 0.6749837 | 0.49968612 | 0.82271448 |
| PCSK1N | 32.2281008 | -0.568096297 | 0.40538918 | -1.4013603 | 0.16110636 | 0.55470404 |
| OTUD5 | 40.8559361 | 0.056285369 | 0.23222018 | 0.24237932 | 0.80848626 | 0.95936772 |

| | | | | | | |
|---|---|---|---|---|---|---|
| GRIPAP1 | 31.2714134 | -0.076940526 | 0.23124967 | -0.3327163 | 0.73934848 | 0.93594619 |
| TFE3 | 28.2359439 | -0.288767396 | 0.24855251 | -1.1617963 | 0.2453182 | 0.64112498 |
| WDR45 | 22.1125752 | -0.404070003 | 0.2498065 | -1.617532 | 0.1057635 | 0.46822648 |
| CLCN5 | 43.9777027 | -0.161000502 | 0.28878108 | -0.5575175 | 0.57717392 | 0.86377523 |
| SHROOM4 | 35.1135477 | 0.073808026 | 0.32818068 | 0.22490059 | 0.82205661 | 0.96331679 |
| MAGED1 | 96.8962176 | -0.016550212 | 0.17374334 | -0.0952567 | 0.92411096 | 0.98561736 |
| GPR173 | 50.9321979 | 0.198826233 | 0.25757714 | 0.77190947 | 0.44016805 | 0.78935944 |
| TSPYL2 | 39.4810221 | 0.042839913 | 0.25500014 | 0.16799956 | 0.86658362 | 0.97624269 |
| KANTR | 80.3745422 | 0.127050562 | 0.26272873 | 0.48358078 | 0.62868342 | 0.8927912 |
| KDM5C | 37.233118 | -0.503288036 | 0.25673316 | -1.9603546 | 0.04995436 | 0.33748407 |
| SMC1A | 37.4749976 | 0.031837938 | 0.24742299 | 0.12867817 | 0.89761231 | 0.98040427 |
| HUWE1 | 161.702007 | -0.031564541 | 0.18152694 | -0.1738835 | 0.86195701 | 0.97514133 |
| PHF8 | 49.1898005 | 0.126698915 | 0.24429534 | 0.51863009 | 0.60401872 | 0.87903643 |
| FAM120C | 27.7896927 | 0.254098101 | 0.26226281 | 0.96886821 | 0.33261095 | 0.71542878 |
| MAGED2 | 29.1674708 | 0.682717503 | 0.29392311 | 2.32277588 | 0.0201912 | 0.20958347 |
| RRAGB | 45.6864855 | -0.321585703 | 0.24768336 | -1.2983743 | 0.19415875 | 0.59269042 |
| AL445472.1 | 39.2300243 | 0.135593624 | 0.31470467 | 0.43085991 | 0.66657024 | 0.9133172 |
| UBQLN2 | 17.5245592 | 0.025168308 | 0.30286982 | 0.08309943 | 0.93377248 | 0.98752252 |
| NBDY | 71.2660345 | -0.267935821 | 0.22652931 | -1.1827866 | 0.23689375 | 0.63270446 |
| ARHGEF9 | 38.9889228 | 0.280545962 | 0.29816256 | 0.94091612 | 0.34674785 | 0.7249592 |
| ZC4H2 | 37.0824113 | 0.190675049 | 0.31678437 | 0.601908 | 0.54723538 | 0.84773035 |
| ZC3H12B | 26.8193786 | 0.181715086 | 0.3204331 | 0.56709212 | 0.5706516 | 0.86113318 |
| MSN | 174.352349 | 0.049137669 | 0.22391904 | 0.2194439 | 0.82630427 | 0.96557642 |
| HEPH | 41.3514975 | 0.233966111 | 0.36321461 | 0.64415391 | 0.51947564 | 0.83317906 |
| AR | 70.5504825 | 0.74210441 | 0.31315425 | 2.36977278 | 0.01779902 | 0.19558968 |
| OPHN1 | 86.693549 | 0.629801277 | 0.20443631 | 3.08067228 | 0.00206534 | 0.05618053 |
| YIPF6 | 37.1016202 | 0.25981565 | 0.23289417 | 1.11559533 | 0.26459539 | 0.6584868 |
| STARD8 | 23.8015599 | 0.297540411 | 0.43108477 | 0.69021323 | 0.4900601 | 0.81720854 |
| EFNB1 | 24.2686647 | 0.454921431 | 0.33649906 | 1.35192483 | 0.17639936 | 0.5732874 |
| EDA | 71.476365 | 0.813250707 | 0.39180489 | 2.07565227 | 0.03792612 | 0.29501902 |
| IGBP1 | 25.4418817 | -0.128004672 | 0.2831294 | -0.4521066 | 0.65119219 | 0.9050963 |
| NONO | 47.7503259 | -0.182040996 | 0.27088075 | -0.6720337 | 0.50156224 | 0.82406312 |
| TAF1 | 93.1672762 | -0.030093975 | 0.25502584 | -0.1180036 | 0.90606478 | 0.98306694 |
| OGT | 144.826167 | -0.073509137 | 0.21509027 | -0.3417595 | 0.73253191 | 0.93574285 |
| NHSL2 | 49.6505572 | 0.061705967 | 0.28759014 | 0.21456218 | 0.83010868 | 0.96580172 |
| PIN4 | 28.6021175 | 0.089509458 | 0.23085448 | 0.38773109 | 0.69821504 | 0.92737958 |
| RPS4X | 129.699652 | -0.172506569 | 0.2396158 | -0.7199299 | 0.47156818 | 0.80556824 |
| HDAC8 | 180.988368 | -0.144238069 | 0.18919628 | -0.7623727 | 0.44583762 | 0.79296348 |
| CHIC1 | 63.2046342 | -0.031115897 | 0.25619666 | -0.1214532 | 0.9033321 | 0.98189434 |
| XIST | 300.946339 | -0.904146556 | 3.14718342 | -0.2872875 | 0.77389217 | 0.94757307 |
| JPX | 189.195704 | 0.043199488 | 0.29170732 | 0.14809189 | 0.88227025 | 0.97782528 |
| FTX | 1476.39826 | 0.09551895 | 0.22614337 | 0.42238226 | 0.67274602 | 0.91482259 |
| SLC16A2 | 25.3282072 | 0.427667529 | 0.30454447 | 1.404286 | 0.1602337 | 0.55310356 |
| RLIM | 26.7972979 | -0.31621663 | 0.33268784 | -0.9504905 | 0.34186308 | 0.7204909 |
| NEXMIF | 28.1116529 | 0.200728626 | 0.30154967 | 0.66565692 | 0.50563042 | 0.82481485 |
| ABCB7 | 51.5263246 | -0.051993673 | 0.19952827 | -0.260583 | 0.79441411 | 0.95454757 |
| ATRX | 314.539848 | 0.139435994 | 0.15250065 | 0.9143305 | 0.36054321 | 0.73273991 |
| MAGT1 | 51.7189158 | -0.015204616 | 0.25478502 | -0.0596763 | 0.95241348 | 0.99060401 |
| COX7B | 47.6012716 | -0.774914114 | 0.25301936 | -3.0626673 | 0.00219374 | 0.05799201 |
| PGK1 | 115.327632 | -1.577841036 | 0.44533047 | -3.543079 | 0.00039548 | 0.0204677 |
| BRWD3 | 111.065277 | -0.33313461 | 0.29557644 | -1.1270675 | 0.25971392 | 0.65252568 |
| SH3BGRL | 137.200347 | 0.006916526 | 0.24332331 | 0.02842525 | 0.97732298 | 0.99407008 |
| HDX | 41.2204436 | -0.195624834 | 0.28961414 | -0.6754671 | 0.49937903 | 0.82271448 |
| APOOL | 33.0356943 | 0.152343966 | 0.22956213 | 0.66362848 | 0.50692812 | 0.82481485 |
| CHM | 104.546941 | -0.166134288 | 0.25963618 | -0.6398734 | 0.5222549 | 0.8350836 |
| DACH2 | 143.023526 | 0.529790285 | 1.31696242 | 0.40228201 | 0.6874765 | 0.92100905 |
| DIAPH2 | 449.310607 | -0.79958044 | 0.30464845 | -2.6246004 | 0.00867507 | 0.13117922 |
| SRPX2 | 115.927409 | -0.45009733 | 0.45821247 | -0.9822896 | 0.32595722 | 0.71149668 |
| SYTL4 | 78.4934703 | 0.23803831 | 0.50162155 | 0.47453765 | 0.63511656 | 0.89672845 |
| ARMCX4 | 103.489369 | 0.334788009 | 0.27553939 | 1.21502779 | 0.22435549 | 0.61883685 |
| ARMCX3 | 34.6838038 | 0.26267022 | 0.30087245 | 0.87302849 | 0.38264755 | 0.75179833 |
| ZMAT1 | 20.2286502 | 0.927973193 | 0.33880574 | 2.73895359 | 0.00616351 | 0.10632774 |
| LINC00630 | 31.9172589 | 0.030185006 | 0.3197454 | 0.09440325 | 0.92478883 | 0.98561736 |
| BEX4 | 18.6918401 | -0.046986558 | 0.2864709 | -0.1640186 | 0.8697165 | 0.9768643 |
| TCEAL9 | 49.1009857 | -0.324595975 | 0.28925311 | -1.1221866 | 0.2617831 | 0.65593044 |
| BEX3 | 58.9076039 | 0.164511636 | 0.24931992 | 0.65984154 | 0.50935552 | 0.82669032 |
| TCEAL4 | 75.4810413 | -0.237359171 | 0.2887762 | -0.8219485 | 0.41110619 | 0.77166383 |
| TCEAL3 | 27.5922795 | 0.102692753 | 0.26074838 | 0.39383851 | 0.69370027 | 0.9248682 |
| MORF4L2 | 149.556251 | -0.49577028 | 0.36110593 | -1.372922 | 0.16977659 | 0.56661587 |
| SLC25A53 | 23.7647238 | -0.010077549 | 0.2871884 | -0.0350904 | 0.97200767 | 0.99300191 |
| FAM199X | 31.8138168 | 0.385625946 | 0.26380227 | 1.46179922 | 0.14379624 | 0.532104 |
| TBC1D8B | 27.1978198 | -0.40116535 | 0.35541685 | -1.1287179 | 0.25901687 | 0.65211201 |
| MORC4 | 27.8491314 | -0.131434243 | 0.323965 | -0.4057051 | 0.68495931 | 0.91977784 |
| RBM41 | 34.0046892 | -0.07606172 | 0.34181527 | -0.2225229 | 0.82390686 | 0.96413949 |
| TSC22D3 | 80.450469 | 0.037895005 | 0.58901693 | 0.06433602 | 0.94870267 | 0.99060401 |
| MID2 | 29.7548078 | 0.446044784 | 0.26931683 | 1.65620835 | 0.09767963 | 0.45507247 |
| COL4A5 | 97.1008131 | 0.461054561 | 0.39751639 | 1.15983788 | 0.24611482 | 0.64218217 |
| ACSL4 | 252.662116 | -1.378704608 | 0.48844056 | -2.8226661 | 0.00476261 | 0.09157966 |
| TMEM164 | 43.9698123 | 0.288635986 | 0.27189755 | 1.06156155 | 0.28843478 | 0.67992862 |
| AMMECR1 | 42.4760714 | -0.007543223 | 0.34443642 | -0.0219002 | 0.98252757 | 0.99599681 |
| CHRDL1 | 26.2893976 | 1.173473886 | 0.36145921 | 3.24649051 | 0.00116837 | 0.04285549 |
| PAK3 | 58.9683383 | 0.812022417 | 0.55910275 | 1.45236705 | 0.14639957 | 0.53585673 |
| ALG13 | 115.454404 | 0.044301907 | 0.25963453 | 0.1706318 | 0.86451329 | 0.9762143 |
| LRCH2 | 55.9955123 | -0.438956458 | 0.45071059 | -0.9739209 | 0.33009582 | 0.71362138 |
| PLS3 | 197.87298 | 0.567107452 | 0.28348634 | 2.00047538 | 0.04544896 | 0.31992756 |
| DANT2 | 65.6908016 | 0.646510336 | 0.35070906 | 1.8434378 | 0.06526512 | 0.38095778 |
| KLHL13 | 143.06901 | -0.103204769 | 0.38698309 | -0.2666906 | 0.78970736 | 0.9517272 |
| WDR44 | 75.6864911 | 0.181712265 | 0.20210518 | 0.89909751 | 0.36860072 | 0.74123132 |
| DOCK11 | 50.9733291 | 0.335840095 | 0.2962814 | 1.13351729 | 0.25699708 | 0.65130862 |
| IL13RA1 | 53.7040416 | 0.323011266 | 0.26911287 | 1.20028174 | 0.23002994 | 0.62447176 |
| PGRMC1 | 39.264894 | 0.162660861 | 0.28835346 | 0.56410233 | 0.57268449 | 0.86236719 |
| SLC25A43 | 51.6452465 | 0.022363953 | 0.25433063 | 0.0879326 | 0.92993025 | 0.98681039 |
| SLC25A5 | 38.0119242 | -0.74579566 | 0.44296503 | -1.6836445 | 0.09225038 | 0.44238068 |
| 6-Sep | 21.8651753 | -0.368995501 | 0.29467535 | -1.2522103 | 0.21049325 | 0.60731347 |
| UPF3B | 25.3806526 | -0.058152927 | 0.32290847 | -0.1800911 | 0.85708109 | 0.97381501 |
| NDUFA1 | 32.7618076 | -0.415503311 | 0.33876758 | -1.2265144 | 0.22000517 | 0.61590675 |
| LAMP2 | 71.1857268 | -0.645598821 | 0.23492535 | -2.7481019 | 0.00599414 | 0.10497927 |
| CUL4B | 39.3145214 | -0.30334403 | 0.32395196 | -0.9363859 | 0.34907456 | 0.72734588 |
| MCTS1 | 21.9506108 | 0.022316913 | 0.26985867 | 0.08269852 | 0.93409126 | 0.98757541 |
| THOC2 | 169.477695 | -0.155157082 | 0.24147837 | -0.6425299 | 0.52052917 | 0.83432254 |
| XIAP | 88.3923898 | 8.14E-05 | 0.14153058 | 0.00057539 | 0.99954091 | 0.99969762 |
| STAG2 | 170.332243 | -0.231010958 | 0.23755979 | -0.9724329 | 0.33083522 | 0.71362138 |
| SMARCA1 | 90.3693812 | 0.427848167 | 0.31137883 | 1.37404385 | 0.16942807 | 0.56570963 |
| OCRL | 24.4087613 | 0.347609421 | 0.30193419 | 1.15127544 | 0.24961894 | 0.64565668 |
| ZDHHC9 | 26.388269 | -1.908100817 | 0.48325318 | -3.9484496 | 7.87E-05 | 0.0068121 |
| BCORL1 | 22.0480295 | 0.541306597 | 0.24227583 | 2.23425756 | 0.02546613 | 0.2387125 |
| AIFM1 | 20.2089982 | -0.084168304 | 0.37132678 | -0.2266691 | 0.82068106 | 0.96291025 |
| ENOX2 | 48.3306703 | -0.460812902 | 0.3475515 | -1.3258838 | 0.18487819 | 0.58412288 |
| RAP2C-AS1 | 43.391215 | 0.479394279 | 0.38431623 | 1.24739536 | 0.21225257 | 0.60910715 |
| GPC4 | 47.148748 | -0.288861488 | 0.36628109 | -0.7886334 | 0.43032633 | 0.78298297 |
| GPC3 | 110.090859 | 1.148831014 | 0.72460899 | 1.58544957 | 0.11286423 | 0.48301375 |
| PHF6 | 28.674924 | -0.246236433 | 0.32238739 | -0.7637905 | 0.44499209 | 0.79259498 |
| HPRT1 | 42.2008629 | -0.252982818 | 0.25724423 | -0.9834344 | 0.32539371 | 0.71149668 |

| | | | | | | |
|---|---|---|---|---|---|---|
| FAM122C | 20.2648802 | -0.121453773 | 0.53731083 | -0.2260401 | 0.82117026 | 0.96294563 |
| RTL8C | 35.2033723 | -0.551775889 | 0.28886169 | -1.9101733 | 0.0561109 | 0.35487005 |
| ZNF75D | 44.0726476 | 0.085949573 | 0.32505689 | 0.26441394 | 0.79146097 | 0.95257889 |
| INTS6L | 50.1276885 | 0.250660968 | 0.28953569 | 0.86573427 | 0.3866359 | 0.75402666 |
| FHL1 | 115.805997 | 1.61241106 | 0.53138418 | 3.03436027 | 0.00241046 | 0.06193881 |
| MAP7D3 | 26.6487257 | 0.246178712 | 0.30668122 | 0.80271857 | 0.42213742 | 0.77771095 |
| ARHGEF6 | 58.4407106 | 0.996422338 | 0.28836165 | 3.45546066 | 0.00054935 | 0.02523249 |
| AL683813.1 | 28.3851211 | 0.358270435 | 0.27224025 | 1.31600832 | 0.18817125 | 0.58645909 |
| RBMX | 118.012608 | -0.002245902 | 0.1676266 | -0.0133982 | 0.98931007 | 0.99739981 |
| FGF13 | 52.8810614 | -0.041402209 | 0.4554521 | -0.0909035 | 0.92756923 | 0.98678435 |
| ATP11C | 271.557453 | -0.444258203 | 0.3206638 | -1.385433 | 0.16592011 | 0.56208655 |
| FMR1 | 49.845121 | -0.010225045 | 0.3323902 | -0.0307622 | 0.97545921 | 0.9933005 |
| IDS | 42.2285831 | 0.273896973 | 0.28038707 | 0.97685308 | 0.3286419 | 0.71294249 |
| LINC00894 | 96.7130469 | 0.531848704 | 0.30663922 | 1.73444449 | 0.08283925 | 0.42306159 |
| MAMLD1 | 150.191174 | -0.559336965 | 0.42518439 | -1.3155162 | 0.18833647 | 0.58668894 |
| MTM1 | 44.7732969 | 0.063362913 | 0.21134943 | 0.29980168 | 0.76432843 | 0.94462105 |
| MTMR1 | 40.5913841 | -0.04953066 | 0.23045329 | -0.2149271 | 0.82982415 | 0.96580172 |
| CD99L2 | 37.2116947 | 0.095979241 | 0.26863109 | 0.35729015 | 0.72087461 | 0.93444541 |
| GABRE | 25.1614004 | -1.071681308 | 0.55812402 | -1.9201491 | 0.05483907 | 0.35166873 |
| BGN | 331.143221 | -0.017258703 | 0.50191116 | -0.034386 | 0.97256937 | 0.99300191 |
| BCAP31 | 31.0586073 | 0.172032338 | 0.32495918 | 0.52939676 | 0.59653024 | 0.87523952 |
| SSR4 | 26.6403411 | -0.606682802 | 0.38558215 | -1.5734204 | 0.11562152 | 0.48836096 |
| MECP2 | 107.369691 | -0.096404229 | 0.26019436 | -0.3705085 | 0.71100363 | 0.93009618 |
| FLNA | 474.827414 | 0.77167188 | 0.26054237 | 2.96179033 | 0.00305856 | 0.07113088 |
| EMD | 27.4931705 | 0.019953671 | 0.33425042 | 0.05969677 | 0.95239714 | 0.99060401 |
| RPL10 | 238.013183 | -0.227546108 | 0.35240356 | -0.6456975 | 0.51847526 | 0.83297983 |
| TAZ | 20.2781457 | 0.006512379 | 0.29186883 | 0.02231269 | 0.98219853 | 0.99588534 |
| ATP6AP1 | 26.3186984 | -0.112545136 | 0.32875227 | -0.3423403 | 0.73209484 | 0.93568087 |
| GDI1 | 34.609411 | 0.015179902 | 0.30931356 | 0.0490761 | 0.96085865 | 0.99181157 |
| FAM50A | 20.5571249 | 0.121589587 | 0.35665623 | 0.34091536 | 0.7331673 | 0.93574285 |
| FUNDC2 | 26.982786 | -0.655509279 | 0.28598456 | -2.2921142 | 0.02189905 | 0.21819108 |
| BRCC3 | 23.9719457 | -0.657824322 | 0.40116243 | -1.6397954 | 0.10104771 | 0.46131565 |
| TMLHE | 46.5925567 | -0.029391658 | 0.35066673 | -0.0838165 | 0.93320233 | 0.98725539 |
| VAMP7 | 23.606895 | -0.355509022 | 0.29109151 | -1.2212964 | 0.2219738 | 0.61772146 |
| FBXO25 | 34.7605266 | -0.136252023 | 0.26197441 | -0.5200967 | 0.60299619 | 0.87903643 |
| TDRP | 24.6396077 | -0.55208794 | 0.34433362 | -1.6033518 | 0.108857 | 0.47618458 |
| ERICH1 | 61.0908974 | -0.174029756 | 0.29441322 | -0.5911071 | 0.55444864 | 0.85191704 |
| MYOM2 | 45.4211598 | -1.674218912 | 0.51147681 | -3.2733036 | 0.00106298 | 0.0400608 |
| CSMD1 | 98.110486 | -0.088658512 | 0.5240016 | -0.1691951 | 0.86564317 | 0.9762143 |
| MCPH1 | 136.061965 | -0.22190507 | 0.22819065 | -0.9724547 | 0.33082438 | 0.71362138 |
| ANGPT2 | 72.1263568 | 0.166383562 | 0.29792358 | 0.55847733 | 0.57651848 | 0.86325068 |
| AGPAT5 | 40.956739 | -1.328496212 | 0.31779085 | -4.1804105 | 2.91E-05 | 0.00375164 |
| DEFB1 | 64.1927138 | -0.107409814 | 0.43815628 | -0.2451404 | 0.80634771 | 0.95838958 |
| FAM85B | 106.789926 | 0.475670801 | 0.43819338 | 1.08552713 | 0.27768826 | 0.67269898 |
| PRAG1 | 24.3138696 | 0.326193509 | 0.36299985 | 0.89860509 | 0.36886305 | 0.74123132 |
| MFHAS1 | 95.5242573 | -0.145077159 | 0.38968324 | -0.3722951 | 0.70967315 | 0.93009618 |
| ERI1 | 29.9699768 | -0.194524237 | 0.29122023 | -0.6679627 | 0.50415743 | 0.82444437 |
| PPP1R3B | 30.196538 | -0.608698157 | 0.49759981 | -1.2232685 | 0.22122832 | 0.61653656 |
| TNKS | 223.77139 | -0.158248292 | 0.15126045 | -1.0461974 | 0.2954699 | 0.68645575 |
| MSRA | 76.6899388 | 0.254600442 | 0.2941749 | 0.86547302 | 0.38677922 | 0.75402666 |
| XKR6 | 56.1180661 | -0.332230167 | 0.36976323 | -0.8984943 | 0.36892207 | 0.74123132 |
| MTMR9 | 35.4172017 | 0.349803952 | 0.23519924 | 1.48726648 | 0.13694444 | 0.52089048 |
| FDFT1 | 48.890471 | -0.111744909 | 0.34190755 | -0.3268278 | 0.7437981 | 0.93791562 |
| CTSB | 89.8207116 | -0.492244969 | 0.36376074 | -1.3532108 | 0.17598829 | 0.57311399 |
| FAM66D | 31.0299519 | 0.998762704 | 0.57580215 | 1.73455883 | 0.08281898 | 0.42306159 |
| LONRF1 | 110.200189 | 0.716365844 | 0.54422004 | 1.31631655 | 0.18806782 | 0.58645909 |
| DLC1 | 1023.22359 | 0.092869245 | 0.2555719 | 0.36337815 | 0.71632242 | 0.93247094 |
| TUSC3 | 73.3733058 | -0.075218631 | 0.35273045 | -0.2132468 | 0.83113449 | 0.96621638 |
| MICU3 | 118.23666 | 0.209752738 | 0.23355781 | 0.89807632 | 0.36914486 | 0.74141722 |
| VPS37A | 49.0085519 | -0.393412824 | 0.24846607 | -1.5833664 | 0.11333798 | 0.48369387 |
| SLC7A2 | 388.294279 | -1.607035709 | 0.42093513 | -3.8177752 | 0.00013466 | 0.00953067 |
| MTUS1 | 142.433438 | -0.876476285 | 0.42366021 | -2.068819 | 0.03856307 | 0.29706503 |
| AC027117.2 | 25.421728 | 0.373524287 | 0.54111389 | 0.69028775 | 0.49001325 | 0.81720854 |
| PCM1 | 235.379327 | 0.211009419 | 0.17133926 | 1.23152989 | 0.21812474 | 0.61478732 |
| ASAH1 | 53.2556075 | -0.236295813 | 0.33704003 | -0.7010912 | 0.48324607 | 0.81409616 |
| PSD3 | 300.16761 | 0.460623115 | 0.31346198 | 1.46947046 | 0.14170523 | 0.52921602 |
| CSGALNACT1 | 102.88835 | 0.397697615 | 0.44806459 | 0.88758991 | 0.37476138 | 0.74698166 |
| INTS10 | 60.3867888 | -0.3807558 | 0.24734 | -1.5394024 | 0.12370608 | 0.50097964 |
| ATP6V1B2 | 41.7430518 | -0.495191475 | 0.250839 | -1.9741407 | 0.04836574 | 0.33343326 |
| LZTS1 | 54.5099364 | -0.269190781 | 0.30849375 | -0.8725972 | 0.38288267 | 0.75179833 |
| XPO7 | 66.7498427 | -0.331761036 | 0.20257529 | -1.6377172 | 0.10148071 | 0.46169744 |
| FAM160B2 | 24.2032328 | -0.027017645 | 0.27225818 | -0.0992354 | 0.92095138 | 0.98528826 |
| BMP1 | 40.2050635 | -0.30999345 | 0.21295422 | -1.4556812 | 0.14548078 | 0.53538077 |
| SLC39A14 | 100.202667 | -1.949148162 | 0.52302024 | -3.7267165 | 0.00019399 | 0.01296033 |
| PPP3CC | 146.94804 | -0.00873956 | 0.1696586 | -0.0515126 | 0.95891704 | 0.99181157 |
| SORBS3 | 61.5438939 | 0.568734553 | 0.27710345 | 2.05242681 | 0.0401282 | 0.30130305 |
| BIN3 | 31.7562159 | -0.252720511 | 0.33316282 | -0.7585496 | 0.44812205 | 0.79296348 |
| RHOBTB2 | 18.8990552 | -0.491769523 | 0.4619237 | -1.064612 | 0.28705154 | 0.67921285 |
| TNFRSF10B | 32.1174011 | -0.927511248 | 0.29536281 | -3.1402438 | 0.00168807 | 0.05147971 |
| TNFRSF10D | 23.0598575 | -1.399686207 | 0.33912824 | -4.1273065 | 3.67E-05 | 0.00428152 |
| LOXL2 | 58.4244107 | -1.733188922 | 0.50077376 | -3.4610219 | 0.00053813 | 0.02502981 |
| ENTPD4 | 42.2241618 | -0.503964408 | 0.21269994 | -2.3693679 | 0.01781852 | 0.19558968 |
| SLC25A37 | 59.5509319 | -0.363937495 | 0.2539511 | -1.4331007 | 0.15182906 | 0.53898138 |
| DOCK5 | 78.3108282 | -0.042955689 | 0.37977595 | -0.113108 | 0.90994495 | 0.98343817 |
| KCTD9 | 46.8632468 | 0.163518706 | 0.35824861 | 0.45643919 | 0.64807419 | 0.90270986 |
| PPP2R2A | 148.18971 | -0.050439312 | 0.26038925 | -0.1937073 | 0.84640504 | 0.97014274 |
| BNIP3L | 204.894418 | -1.144396613 | 0.44458585 | -2.5740734 | 0.01005089 | 0.14150196 |
| DPYSL2 | 90.0813495 | -0.102601637 | 0.2765606 | -0.3709915 | 0.71064385 | 0.93009618 |
| ADRA1A | 38.5151298 | 1.110149726 | 0.58589579 | 1.8947904 | 0.05812018 | 0.36041506 |
| CLU | 107.079571 | -1.206002188 | 0.43933521 | -2.7450615 | 0.00604996 | 0.1053581 |
| CCDC25 | 20.8312613 | 0.176191217 | 0.29714638 | 0.59294417 | 0.55321851 | 0.85188037 |
| ELP3 | 52.9034228 | -0.492700169 | 0.19539709 | -2.5215327 | 0.01168448 | 0.15416382 |
| ZNF395 | 20.3449994 | -1.169879805 | 0.47462528 | -2.4648493 | 0.01370709 | 0.17073456 |
| FZD3 | 23.9992512 | -0.602750896 | 0.31275847 | -1.9272089 | 0.05395361 | 0.34959616 |
| EXTL3 | 28.9336508 | 0.146675869 | 0.26033184 | 0.56341886 | 0.5731497 | 0.86260027 |
| INTS9 | 46.856473 | -0.456697598 | 0.24222809 | -1.8854031 | 0.05937544 | 0.36322722 |
| HMBOX1 | 175.956754 | -0.168949227 | 0.19892219 | -0.8493232 | 0.39570149 | 0.76020146 |
| KIF13B | 76.9371888 | 0.161521215 | 0.24817141 | 0.65084538 | 0.5151463 | 0.83131536 |
| SARAF | 71.166481 | -0.194287023 | 0.19970978 | -0.9728468 | 0.33062944 | 0.71362138 |
| LEPROTL1 | 30.440947 | -0.293541428 | 0.39609362 | -0.741091 | 0.45863825 | 0.79933436 |
| RBPMS | 1819.04818 | -0.383679013 | 0.23505275 | -1.6323102 | 0.10261415 | 0.4627468 |
| GTF2E2 | 52.8460618 | -0.20987723 | 0.20799243 | -1.0090619 | 0.31294495 | 0.70282967 |
| GSR | 26.7527883 | 0.328523183 | 0.26921326 | 1.22030834 | 0.22234801 | 0.61772146 |
| PPP2CB | 129.227313 | 0.534300615 | 0.14454827 | 3.69634727 | 0.00021872 | 0.0139774 |
| WRN | 106.787531 | -0.245944902 | 0.19996816 | -1.2299203 | 0.21872695 | 0.61522936 |
| FUT10 | 29.5970352 | 0.155326085 | 0.23907079 | 0.6497075 | 0.51588118 | 0.83174649 |
| RNF122 | 19.567358 | -1.984845086 | 0.58411776 | -3.3980222 | 0.00067875 | 0.02893045 |
| ADGRA2 | 66.3800672 | 0.686565318 | 0.28326157 | 2.4237856 | 0.01535967 | 0.18089458 |
| RAB11FIP1 | 22.9736926 | -0.343385933 | 0.57159221 | -0.6007534 | 0.54800428 | 0.84785354 |
| ASH2L | 30.343557 | 0.419647407 | 0.25744983 | 1.63001624 | 0.10309806 | 0.46365765 |
| BAG4 | 23.6230095 | 0.163093908 | 0.299186 | 0.54512547 | 0.58566722 | 0.86968447 |
| DDHD2 | 40.818763 | 0.370676205 | 0.23419675 | 1.58275553 | 0.1134772 | 0.48395978 |
| PLPP5 | 20.5916461 | -0.288480645 | 0.43436477 | -0.6641437 | 0.50659832 | 0.82481485 |

| | | | | | | |
|---|---|---|---|---|---|---|
| NSD3 | 165.970281 | -0.202131923 | 0.17038478 | -1.1863261 | 0.23549353 | 0.63132734 |
| FGFR1 | 134.097546 | 0.056140657 | 0.43885439 | 0.12792547 | 0.89820795 | 0.98053034 |
| TACC1 | 503.589838 | -0.388663527 | 0.46648015 | -0.8331834 | 0.40474129 | 0.76624906 |
| PLEKHA2 | 150.003639 | 0.169936562 | 0.31416339 | 0.54091777 | 0.58856426 | 0.8702694 |
| ADAM9 | 122.361337 | -0.017110484 | 0.30664554 | -0.0557989 | 0.95550201 | 0.99181157 |
| TCIM | 60.113573 | 1.684587229 | 0.83180617 | 2.02521608 | 0.04284518 | 0.31175172 |
| SFRP1 | 113.381868 | -0.629052113 | 0.41006381 | -1.5340347 | 0.12502111 | 0.50400948 |
| GOLGA7 | 20.1309792 | -0.154141535 | 0.27523125 | -0.5600437 | 0.57544962 | 0.86294056 |
| GPAT4 | 24.4655269 | -0.271409529 | 0.25454871 | -1.0662381 | 0.28631603 | 0.67897273 |
| KAT6A | 215.488981 | -0.030720823 | 0.14582031 | -0.2106759 | 0.83314019 | 0.9666365 |
| AP3M2 | 29.2941577 | 0.470716686 | 0.32771565 | 1.43635704 | 0.15090077 | 0.53880252 |
| PLAT | 41.6666115 | 1.113531874 | 0.30254656 | 3.68053056 | 0.00023275 | 0.01461943 |
| IKBKB | 57.7236148 | -0.052701475 | 0.22041522 | -0.2391009 | 0.81102735 | 0.95967025 |
| VDAC3 | 45.2907199 | -0.021365255 | 0.18601461 | -0.1148579 | 0.90855772 | 0.98306694 |
| SLC20A2 | 165.467329 | 0.350797107 | 0.28933111 | 1.21244172 | 0.22534334 | 0.62047516 |
| SMIM19 | 20.5645389 | -0.064866015 | 0.26874207 | -0.241369 | 0.80926911 | 0.95939969 |
| RNF170 | 30.2819163 | 0.482145726 | 0.3010953 | 1.60130605 | 0.10930915 | 0.47668415 |
| HOOK3 | 115.841436 | 0.038238148 | 0.18034899 | 0.21202307 | 0.83208903 | 0.96643177 |
| FNTA | 25.8633513 | 0.117239806 | 0.23760043 | 0.49343264 | 0.62170692 | 0.88751441 |
| HGSNAT | 96.6295854 | 0.285835771 | 0.22610753 | 1.26415856 | 0.20617312 | 0.6050285 |
| SPIDR | 470.528582 | -0.744953464 | 0.31753428 | -2.3460568 | 0.01897322 | 0.20213493 |
| CEBPD | 323.396338 | -2.197627277 | 0.53146855 | -4.135009 | 3.55E-05 | 0.0042566 |
| PRKDC | 130.526872 | -0.211992513 | 0.27084798 | -0.7826993 | 0.43380374 | 0.78484089 |
| UBE2V2 | 47.7442667 | -0.23763462 | 0.25618238 | -0.9275994 | 0.35361541 | 0.73059415 |
| PCMTD1 | 130.930939 | -0.175224109 | 0.15575443 | -1.1250024 | 0.26058802 | 0.65360455 |
| RB1CC1 | 148.110493 | -0.168360381 | 0.15401249 | -1.0931606 | 0.27432331 | 0.66930898 |
| ATP6V1H | 70.5079257 | -0.133658075 | 0.25834291 | -0.5173669 | 0.60490005 | 0.87903643 |
| TCEA1 | 55.5082946 | 0.065154547 | 0.20639864 | 0.31567334 | 0.75225046 | 0.94058012 |
| LYPLA1 | 41.9753701 | -0.113853208 | 0.24620139 | -0.4624393 | 0.6437663 | 0.90063554 |
| TMEM68 | 19.4496662 | 0.522300456 | 0.29398476 | 1.77662425 | 0.07563008 | 0.4086805 |
| TGS1 | 34.0761602 | -0.203664959 | 0.26407902 | -0.7712273 | 0.4405722 | 0.78962177 |
| LYN | 45.98036 | -0.443978431 | 0.26349739 | -1.6849443 | 0.09199932 | 0.44160875 |
| RPS20 | 69.4244136 | -0.382353843 | 0.30104608 | -1.2700841 | 0.20405467 | 0.60367992 |
| IMPAD1 | 44.5170561 | -0.194495171 | 0.2616339 | -0.7433867 | 0.45724757 | 0.79762426 |
| FAM110B | 108.157688 | -0.020067052 | 0.41069569 | -0.0488611 | 0.96102997 | 0.99181157 |
| SDCBP | 71.6468117 | -0.459638466 | 0.30921177 | -1.4864844 | 0.13715105 | 0.52089048 |
| NSMAF | 54.0038498 | -0.28638258 | 0.22312738 | -1.2834937 | 0.19931917 | 0.59763223 |
| TOX | 49.4332746 | 1.186854309 | 0.35241466 | 3.36777796 | 0.00075777 | 0.03182184 |
| CA8 | 82.1863226 | -0.725064176 | 1.39308966 | -0.520472 | 0.60273463 | 0.87903643 |
| RAB2A | 211.971665 | -0.214087075 | 0.14834124 | -1.4432067 | 0.14896225 | 0.53689238 |
| CHD7 | 33.8355669 | -0.398857092 | 0.32442612 | -1.2294235 | 0.21891307 | 0.61522936 |
| ASPH | 145.451442 | -0.449657182 | 0.17060051 | -2.6357317 | 0.00839561 | 0.12814321 |
| YTHDF3 | 76.2817952 | -0.053804899 | 0.19631656 | -0.2740721 | 0.78402918 | 0.95064024 |
| CYP7B1 | 322.080069 | -1.580841008 | 0.47854908 | -3.3034042 | 0.00095519 | 0.0366558 |
| ARMC1 | 28.5845023 | -0.042402094 | 0.27375533 | -0.1548905 | 0.87690766 | 0.9768643 |
| MTFR1 | 40.7182546 | -0.299552103 | 0.31317117 | -0.9565124 | 0.33881343 | 0.71925477 |
| PDE7A | 54.6453192 | 0.110060552 | 0.24757017 | 0.44456305 | 0.65663556 | 0.90772143 |
| VCPIP1 | 20.7125998 | -0.394410914 | 0.30404555 | -1.29721 | 0.19455895 | 0.59272783 |
| SGK3 | 44.1059828 | 0.407876418 | 0.29735693 | 1.37167281 | 0.17016532 | 0.56765543 |
| CSPP1 | 90.2547182 | 0.293900539 | 0.28386361 | 1.03535828 | 0.30050162 | 0.69191459 |
| ARFGEF1 | 186.932385 | -0.027440509 | 0.14524259 | -0.1889288 | 0.85014862 | 0.97191897 |
| PREX2 | 77.1669115 | 0.194486259 | 0.50043854 | 0.38863166 | 0.69754864 | 0.92702563 |
| SULF1 | 212.05484 | -0.288170577 | 0.49705182 | -0.5797596 | 0.56207673 | 0.85545224 |
| NCOA2 | 306.691567 | 0.192209296 | 0.27176866 | 0.70725335 | 0.47940905 | 0.81197905 |
| TRAM1 | 74.1691374 | -0.396015839 | 0.21767431 | -1.8193045 | 0.06886499 | 0.39050064 |
| MSC-AS1 | 194.827161 | -0.74893979 | 0.55191931 | -1.3569733 | 0.17478968 | 0.57293905 |
| TERF1 | 47.3300984 | 0.069334932 | 0.23645343 | 0.29322871 | 0.76934734 | 0.94579017 |
| SBSPON | 33.032123 | 1.103878301 | 0.3519013 | 3.13689749 | 0.00170746 | 0.05185169 |
| RPL7 | 111.074092 | -0.073419434 | 0.39105188 | -0.1877486 | 0.85107375 | 0.97256119 |
| RDH10 | 23.6135631 | -0.880324413 | 0.42744115 | -2.0595219 | 0.03944427 | 0.30038956 |
| STAU2 | 72.8682415 | -0.303280336 | 0.26734329 | -1.1344229 | 0.25661722 | 0.65084527 |
| ELOC | 35.2581831 | -0.575366038 | 0.24851273 | -2.3152377 | 0.02059993 | 0.21114239 |
| GDAP1 | 22.2907749 | -0.225750959 | 0.3682965 | -0.6129598 | 0.5399029 | 0.84546056 |
| PI15 | 114.33977 | -4.660543399 | 0.88108071 | -5.289576 | 1.23E-07 | 6.44E-05 |
| ZFHX4 | 55.2793368 | 0.21095392 | 0.52494411 | 0.40185977 | 0.68778723 | 0.92101828 |
| PEX2 | 21.6599821 | 0.045571385 | 0.29638814 | 0.15375577 | 0.87780231 | 0.9768643 |
| ZC2HC1A | 36.8548595 | 0.551117319 | 0.29145249 | 1.89093366 | 0.0586332 | 0.36041506 |
| ZBTB10 | 42.660846 | 0.161608694 | 0.21406105 | 0.75496542 | 0.45026973 | 0.79399826 |
| ZNF704 | 150.655782 | 0.959418858 | 0.37413969 | 2.5643333 | 0.01033742 | 0.14280022 |
| PAG1 | 264.613835 | -0.342618194 | 0.57193257 | -0.5990535 | 0.54913723 | 0.84855413 |
| ZFAND1 | 25.6850784 | 0.07035746 | 0.27528469 | 0.25558072 | 0.79827461 | 0.95499269 |
| C8orf59 | 18.8412457 | -0.591044226 | 0.2736373 | -2.1599549 | 0.03077616 | 0.26662968 |
| WWP1 | 96.2976958 | -0.341363898 | 0.20695189 | -1.6494843 | 0.09904846 | 0.45678074 |
| RMDN1 | 42.741376 | -0.095390744 | 0.22307906 | -0.4276096 | 0.66893539 | 0.91445841 |
| CPNE3 | 36.4557524 | 0.10888838 | 0.25691549 | 0.42382956 | 0.67169012 | 0.91482259 |
| AF117829.1 | 103.091393 | 0.034122572 | 0.38007887 | 0.08977761 | 0.92846394 | 0.98681039 |
| RIPK2 | 23.3658991 | -0.737871557 | 0.34257092 | -2.1539235 | 0.03124617 | 0.26776774 |
| NBN | 36.9436176 | 0.324405485 | 0.33980418 | 0.95468362 | 0.33973772 | 0.71925477 |
| DECR1 | 69.0601245 | -0.423481451 | 0.33086963 | -1.2799042 | 0.20057881 | 0.59975737 |
| PIP4P2 | 45.3430561 | 0.188270519 | 0.29173906 | 0.64533875 | 0.51870768 | 0.83297983 |
| OTUD6B-AS1 | 19.6877456 | 0.33606603 | 0.38829165 | 0.865499 | 0.38676497 | 0.75402666 |
| OTUD6B | 18.3278095 | 0.030149498 | 0.34662803 | 0.0869794 | 0.93068789 | 0.98681039 |
| LRRC69 | 23.4954124 | -0.271690939 | 0.31857211 | -0.8528397 | 0.39374821 | 0.75988253 |
| RUNX1T1 | 280.833369 | 0.178130401 | 0.29763058 | 0.59849495 | 0.54950973 | 0.84874884 |
| TRIQK | 46.7562833 | -0.001314904 | 0.26082006 | -0.0050414 | 0.99597754 | 0.99854015 |
| FAM92A | 23.768385 | -0.115967329 | 0.27577837 | -0.4205091 | 0.67411355 | 0.91555359 |
| RBM12B | 33.4636476 | -0.385187684 | 0.23319778 | -1.6517639 | 0.09858269 | 0.45657782 |
| TMEM67 | 32.9735906 | -0.03908554 | 0.26759533 | -0.1460621 | 0.88387235 | 0.97782528 |
| GEM | 362.829244 | -0.226943928 | 0.4625705 | -0.4906148 | 0.62369893 | 0.88932158 |
| VIRMA | 49.1803877 | 0.135480121 | 0.21975906 | 0.61649389 | 0.5375686 | 0.84423863 |
| DPY19L4 | 49.9044469 | 0.07201405 | 0.29216774 | 0.24648187 | 0.80530924 | 0.95836441 |
| INTS8 | 39.4577972 | -0.153103189 | 0.27139872 | -0.5641264 | 0.5726681 | 0.86236719 |
| NDUFAF6 | 33.7334039 | 0.264918096 | 0.36137771 | 0.73307814 | 0.46351077 | 0.80225845 |
| TP53INP1 | 25.1785563 | -0.156991595 | 0.49417507 | -0.3176842 | 0.75072452 | 0.94050927 |
| UQCRB | 51.3835769 | -0.484109874 | 0.36475388 | -1.3272234 | 0.18443481 | 0.58363936 |
| SDC2 | 124.241961 | 0.688456088 | 0.40274181 | 1.70942294 | 0.08737264 | 0.43326687 |
| CPQ | 175.999149 | 0.384550573 | 0.32180699 | 1.19497271 | 0.23209769 | 0.62798586 |
| MTDH | 207.729158 | -0.069301034 | 0.18638492 | -0.3718167 | 0.71002929 | 0.93009618 |
| LAPTM4B | 51.06983 | -0.593809802 | 0.27508038 | -2.1586774 | 0.0308752 | 0.26662968 |
| MATN2 | 90.431467 | 0.705149722 | 0.45696211 | 1.54312515 | 0.12280041 | 0.50097964 |
| RPL30 | 127.122988 | -0.191443384 | 0.31040936 | -0.6167449 | 0.53740301 | 0.84423863 |
| NIPAL2 | 64.8644651 | -0.013896542 | 0.42356219 | -0.0328087 | 0.97382711 | 0.99300191 |
| STK3 | 474.95153 | -0.234216061 | 0.2335329 | -1.0029253 | 0.31589689 | 0.7032797 |
| AC104986.2 | 20.1938349 | 0.067919791 | 0.3186108 | 0.21317479 | 0.83119063 | 0.96621638 |
| VPS13B | 421.579546 | -0.274563422 | 0.19333232 | -1.4201631 | 0.15556021 | 0.54438666 |
| COX6C | 59.3300611 | -0.439399106 | 0.22855742 | -1.9224889 | 0.05454427 | 0.35100338 |
| RNF19A | 110.794065 | 0.290217462 | 0.24696781 | 1.17512264 | 0.23994566 | 0.63451757 |
| PABPC1 | 143.029458 | -0.354597939 | 0.27909454 | -1.2705298 | 0.20389597 | 0.60367992 |
| YWHAZ | 146.465537 | 0.40760395 | 0.16733797 | 2.43581272 | 0.01485838 | 0.17735986 |
| ZNF706 | 57.0921716 | 0.15287533 | 0.29296851 | 0.52181489 | 0.60179922 | 0.87903643 |
| NCALD | 230.3602 | 1.416403101 | 0.5001722 | 2.83183091 | 0.00462823 | 0.0906884 |
| RRM2B | 27.6727231 | -0.039864558 | 0.29576871 | -0.1347829 | 0.89278354 | 0.98023547 |
| UBR5 | 346.711567 | -0.158993516 | 0.26268178 | -0.6052704 | 0.54499932 | 0.84730644 |

| | | | | | | |
|---|---|---|---|---|---|---|
| KLF10 | 30.931747 | 0.769187671 | 0.44824253 | 1.7160078 | 0.08616061 | 0.43015921 |
| AZIN1 | 42.2310424 | -0.237027676 | 0.22088113 | -1.0731006 | 0.28322597 | 0.67727609 |
| AZIN1-AS1 | 40.7759831 | 0.368925431 | 0.37604348 | 0.98107122 | 0.32655762 | 0.71149668 |
| ATP6V1C1 | 60.4676932 | -0.533356816 | 0.26471557 | -2.0148298 | 0.04392248 | 0.3152359 |
| SLC25A32 | 37.1177261 | -0.458522048 | 0.22476151 | -2.0400381 | 0.04134653 | 0.30452956 |
| LRP12 | 29.6876252 | -0.185408705 | 0.26437222 | -0.7013169 | 0.48310527 | 0.81409616 |
| ZFPM2 | 535.220392 | 0.42253377 | 0.30985971 | 1.36362927 | 0.17268428 | 0.56912883 |
| OXR1 | 296.924032 | 0.207695614 | 0.26330058 | 0.78881563 | 0.43021978 | 0.78298297 |
| ANGPT1 | 184.449242 | 1.048988646 | 0.51422285 | 2.03994951 | 0.04135535 | 0.30452956 |
| EIF3E | 98.521886 | -0.246629526 | 0.22876891 | -1.0780727 | 0.2810013 | 0.67596679 |
| EMC2 | 51.8497592 | -0.254483127 | 0.24317405 | -1.0465061 | 0.29532743 | 0.68638877 |
| NUDCD1 | 43.6456928 | -0.67061752 | 0.29291699 | -2.2894456 | 0.02205347 | 0.21901484 |
| ENY2 | 34.8709339 | -0.341300188 | 0.29085218 | -1.173449 | 0.24061581 | 0.63466484 |
| TRPS1 | 160.108191 | -0.087481631 | 0.23359415 | -0.3745027 | 0.70803038 | 0.93009618 |
| EIF3H | 101.351597 | -0.102156312 | 0.2502298 | -0.40825 | 0.68309016 | 0.91933701 |
| UTP23 | 19.2999867 | -0.231088378 | 0.26356117 | -0.8767922 | 0.38059951 | 0.74991898 |
| RAD21 | 63.3723716 | -0.054318935 | 0.19044859 | -0.2852157 | 0.77547887 | 0.94805375 |
| EXT1 | 1057.82444 | -0.138319678 | 0.3956635 | -0.3495892 | 0.72664703 | 0.93455731 |
| SAMD12 | 143.030216 | 0.855515726 | 0.36938624 | 2.31604657 | 0.02055573 | 0.21114239 |
| ENPP2 | 24.5365493 | -0.729254908 | 0.47341613 | -1.5404099 | 0.12346046 | 0.50097964 |
| TAF2 | 85.1744946 | -0.15253902 | 0.18669065 | -0.8170684 | 0.41388937 | 0.77396258 |
| DEPTOR | 54.5582648 | 0.112391142 | 0.26461908 | 0.42472803 | 0.67103495 | 0.91475345 |
| COL14A1 | 173.405591 | 0.625126556 | 0.50077634 | 1.24831488 | 0.21191577 | 0.60910715 |
| MRPL13 | 22.1513304 | -0.446655733 | 0.26757136 | -1.6692958 | 0.09505878 | 0.44764276 |
| SNTB1 | 89.6798035 | 0.516195956 | 0.22730405 | 2.27094924 | 0.02315005 | 0.22563624 |
| ZHX2 | 272.971284 | -0.608761005 | 0.35746427 | -1.7029982 | 0.08856841 | 0.43544541 |
| DERL1 | 30.8149733 | -0.728838924 | 0.22614189 | -3.2229276 | 0.00126888 | 0.04375316 |
| ATAD2 | 24.4449123 | 0.398406091 | 0.33813267 | 1.17825375 | 0.23869546 | 0.63422836 |
| FBXO32 | 71.3698911 | -0.971807683 | 0.45976941 | -2.113685 | 0.03454218 | 0.2833153 |
| FAM91A1 | 70.833024 | -0.081276296 | 0.27635544 | -0.2941006 | 0.76868105 | 0.94579017 |
| TMEM65 | 45.0388157 | 0.553680842 | 0.31892946 | 1.73606054 | 0.08255313 | 0.42256146 |
| TATDN1 | 40.3204492 | -0.115505926 | 0.27071743 | -0.426666 | 0.66962261 | 0.91445841 |
| NDUFB9 | 34.8482497 | -0.36375333 | 0.33663372 | -1.0805612 | 0.27989237 | 0.6742587 |
| MTSS1 | 311.429832 | 0.205108412 | 0.39876337 | 0.51436122 | 0.60699947 | 0.88000768 |
| SQLE | 21.5109529 | -0.472162123 | 0.45917078 | -1.0282931 | 0.303812 | 0.69579129 |
| WASHC5 | 32.8138921 | 0.148881857 | 0.28772655 | 0.51744219 | 0.60484751 | 0.87903643 |
| NSMCE2 | 268.299086 | -0.333042041 | 0.19858538 | -1.6770723 | 0.09352834 | 0.44430497 |
| PVT1 | 101.050548 | -1.548865519 | 0.40471153 | -3.8270853 | 0.00012967 | 0.00943507 |
| FAM49B | 38.0736507 | -0.048522145 | 0.30586889 | -0.1586371 | 0.87395482 | 0.9768643 |
| ASAP1 | 601.6166 | -0.337926216 | 0.33710353 | -1.0024405 | 0.31613091 | 0.70358754 |
| EFR3A | 112.784508 | -0.086823108 | 0.22868301 | -0.3796658 | 0.70419354 | 0.92936308 |
| KCNQ3 | 49.5223534 | 1.253050635 | 1.0894781 | 1.15013843 | 0.25008686 | 0.6460368 |
| PHF20L1 | 116.878809 | 0.011475052 | 0.19845442 | 0.05782211 | 0.95389033 | 0.99100036 |
| TG | 28.8077111 | 0.122185579 | 0.43886149 | 0.2784149 | 0.78069388 | 0.9491946 |
| SLA | 55.1893663 | 1.041168349 | 0.5283492 | 1.97060648 | 0.0487689 | 0.33522789 |
| WISP1 | 279.279597 | -0.816746199 | 0.57318736 | -1.4249201 | 0.15418029 | 0.54343931 |
| NDRG1 | 161.488789 | -0.797651846 | 0.44493869 | -1.7927231 | 0.07301722 | 0.40164935 |
| ST3GAL1 | 163.875054 | -0.560753762 | 0.3660451 | -1.5319253 | 0.12554086 | 0.50440978 |
| ZFAT | 34.3865048 | -0.04331872 | 0.20949075 | -0.2067811 | 0.83618085 | 0.9668633 |
| KHDRBS3 | 63.7705796 | 0.75094484 | 0.33092103 | 2.26925692 | 0.02325271 | 0.22603722 |
| TRAPPC9 | 168.169634 | -0.113478013 | 0.21551774 | -0.5265368 | 0.59851532 | 0.87618867 |
| AGO2 | 148.203275 | -0.307723678 | 0.20081787 | -1.5323521 | 0.12543558 | 0.50440978 |
| PTK2 | 500.898106 | 0.11370154 | 0.12895688 | 0.88170201 | 0.37793797 | 0.74933113 |
| DENND3 | 52.9163219 | -1.218555562 | 0.49788834 | -2.4474475 | 0.01438721 | 0.17568582 |
| PTP4A3 | 100.183836 | -1.470171155 | 0.44321509 | -3.3170602 | 0.0009097 | 0.03556056 |
| LY6E | 84.2709318 | -0.487194077 | 0.5449406 | -0.8940315 | 0.37130502 | 0.74392601 |
| ZC3H3 | 26.0118633 | -0.021249523 | 0.32601495 | -0.0651796 | 0.94803101 | 0.99060401 |
| GSDMD | 21.6967663 | -0.078974761 | 0.36660124 | -0.2154242 | 0.82943664 | 0.96580172 |
| EEF1D | 93.9877665 | -0.053284679 | 0.28826368 | -0.184847 | 0.85334904 | 0.97319399 |
| PUF60 | 18.6143901 | -0.129834703 | 0.43947487 | -0.2954315 | 0.7676643 | 0.94576899 |
| NRBP2 | 19.7096556 | 0.305257311 | 0.37957352 | 0.8042113 | 0.42127495 | 0.77771095 |
| PLEC | 91.6837211 | 0.048964464 | 0.31399712 | 0.15593921 | 0.87608094 | 0.9768643 |
| GRINA | 32.3504997 | -0.368524485 | 0.24787702 | -1.4867231 | 0.13708796 | 0.52089048 |
| GPAA1 | 18.8337998 | 0.196639437 | 0.29604528 | 0.66422082 | 0.50654899 | 0.82481485 |
| CYC1 | 18.3494127 | -0.34088226 | 0.41648024 | -0.8184836 | 0.41308108 | 0.77264771 |
| MAF1 | 20.0071307 | -0.075226211 | 0.32421713 | -0.2320242 | 0.81651924 | 0.96132648 |
| MROH1 | 32.506932 | 0.339534414 | 0.340624 | 0.9968012 | 0.31886101 | 0.70683013 |
| HSF1 | 35.0374365 | 0.014579864 | 0.24611347 | 0.05924041 | 0.95276062 | 0.99060401 |
| VPS28 | 41.2141423 | -0.09329988 | 0.26946877 | -0.3462363 | 0.7291651 | 0.93455731 |
| CYHR1 | 21.4032141 | 0.331443045 | 0.24857736 | 1.33335974 | 0.18241378 | 0.58106792 |
| ZNF251 | 37.0477443 | 0.124385625 | 0.27991231 | 0.44437355 | 0.65677254 | 0.90772143 |
| RPL8 | 154.84274 | -0.277389759 | 0.37455869 | -0.7405775 | 0.45894962 | 0.79943607 |
| WASHC1 | 51.4178888 | 0.421596502 | 0.26531223 | 1.58905793 | 0.11204731 | 0.4809788 |
| DOCK8 | 172.733027 | 0.459764102 | 0.31832851 | 1.44430704 | 0.14865263 | 0.53682956 |
| KANK1 | 127.318096 | 1.268964025 | 0.32899502 | 3.85709189 | 0.00011474 | 0.00878389 |
| SMARCA2 | 142.700757 | 0.459337964 | 0.27015225 | 1.70029296 | 0.08907583 | 0.43554112 |
| VLDLR-AS1 | 41.7519952 | 2.199694624 | 0.51763496 | 4.24950941 | 2.14E-05 | 0.00305725 |
| PUM3 | 124.07803 | -0.531825191 | 0.26040448 | -2.0423043 | 0.04112135 | 0.30341448 |
| RFX3 | 214.141462 | 0.142081769 | 0.33571926 | 0.42321602 | 0.67213766 | 0.91482259 |
| RFX3-AS1 | 62.6373042 | 0.046761845 | 0.2971184 | 0.15738455 | 0.87494179 | 0.9768643 |
| GLIS3 | 639.81472 | -1.337412938 | 0.33367524 | -4.0081276 | 6.12E-05 | 0.00569334 |
| SLC1A1 | 49.8022736 | -0.600786623 | 0.39416674 | -1.5241941 | 0.12746023 | 0.50683947 |
| CDC37L1 | 49.334149 | 0.156119654 | 0.32565949 | 0.47939537 | 0.63165738 | 0.89436221 |
| AK3 | 76.7102906 | 0.388726357 | 0.21252331 | 1.82909986 | 0.06738465 | 0.38712255 |
| RCL1 | 56.7136035 | -0.364129685 | 0.3466368 | -1.0504646 | 0.29350456 | 0.68480256 |
| JAK2 | 190.212695 | -0.007619676 | 0.36889832 | -0.0206552 | 0.98352069 | 0.99620525 |
| PLGRKT | 25.7163924 | -0.384097736 | 0.29008522 | -1.3240859 | 0.18547453 | 0.585001 |
| RIC1 | 165.238556 | -0.243766533 | 0.22815849 | -1.0684088 | 0.28533615 | 0.6783522 |
| ERMP1 | 34.0630821 | 0.508798086 | 0.32969382 | 1.54324422 | 0.12277153 | 0.50097964 |
| KIAA2026 | 136.065888 | 0.523748405 | 0.22587636 | 2.3187394 | 0.02040917 | 0.21065589 |
| IL33 | 43.2777804 | -1.436532216 | 0.89021279 | -1.6136953 | 0.10659353 | 0.47073496 |
| UHRF2 | 72.2546553 | -0.304044975 | 0.2041335 | -1.4894418 | 0.13637106 | 0.52087399 |
| KDM4C | 230.32411 | -0.677692012 | 0.22004728 | -3.0797563 | 0.0020717 | 0.05618053 |
| PTPRD | 176.808918 | 1.174120642 | 0.41579886 | 2.82377069 | 0.00474623 | 0.09157966 |
| LURAP1L | 51.0814357 | -0.143073011 | 0.39763601 | -0.359809 | 0.71898998 | 0.93444541 |
| MPDZ | 183.268457 | 0.050296341 | 0.27959507 | 0.17988994 | 0.85723897 | 0.97381501 |
| NFIB | 319.915908 | 0.258013753 | 0.27683845 | 0.93200116 | 0.35133593 | 0.72936942 |
| ZDHHC21 | 110.7639 | 0.51533164 | 0.28581344 | 1.80303502 | 0.07138271 | 0.39729887 |
| FREM1 | 91.9312964 | 1.409497433 | 0.56043041 | 2.5150267 | 0.01190234 | 0.15619692 |
| TTC39B | 124.225167 | 0.062779322 | 0.24308211 | 0.25826385 | 0.79620328 | 0.95463848 |
| SNAPC3 | 66.0606463 | -0.133633358 | 0.21234812 | -0.6293127 | 0.52914438 | 0.8404327 |
| PSIP1 | 48.6087395 | 0.235088962 | 0.23384917 | 1.00530165 | 0.31475162 | 0.70282967 |
| CCDC171 | 53.2368871 | 0.769118831 | 0.36154327 | 2.12732165 | 0.03339337 | 0.2790477 |
| BNC2 | 277.411398 | -0.195524922 | 0.42208305 | -0.463238 | 0.64319375 | 0.90063554 |
| CNTLN | 95.8431103 | -0.112081242 | 0.19890114 | -0.5635023 | 0.57309293 | 0.86260027 |
| ADAMTSL1 | 524.820242 | 0.390375253 | 0.39332324 | 0.99250492 | 0.32095128 | 0.70780851 |
| PLIN2 | 162.088209 | -3.52697053 | 0.5905762 | -5.9720837 | 2.34E-09 | 2.46E-06 |
| DENND4C | 191.795717 | -0.229702586 | 0.25324283 | -0.9070448 | 0.36438312 | 0.73648283 |
| RPS6 | 169.303829 | -0.599010162 | 0.33781093 | -1.7732113 | 0.0761937 | 0.40931834 |
| ACER2 | 77.8715749 | -0.661960573 | 0.71893266 | -0.9207546 | 0.35717855 | 0.73129235 |
| MLLT3 | 189.298786 | 0.075568534 | 0.2866979 | 0.26358245 | 0.79210169 | 0.95288186 |
| FOCAD | 131.04176 | 0.001446268 | 0.23519368 | 0.00614927 | 0.99509363 | 0.99854015 |
| CAAP1 | 31.6962405 | 0.037230752 | 0.23926382 | 0.15560544 | 0.87634403 | 0.9768643 |

| | | | | | | |
|---|---|---|---|---|---|---|
| PLAA | 37.4075109 | 0.098774691 | 0.20211845 | 0.48869705 | 0.6250562 | 0.89069797 |
| IFT74 | 42.0351052 | -0.240691303 | 0.25806333 | -0.9326831 | 0.35098362 | 0.72929137 |
| MOB3B | 57.8717122 | 0.410444078 | 0.2300485 | 1.7841632 | 0.07439715 | 0.40409804 |
| ACO1 | 94.7097135 | -0.537322854 | 0.27940996 | -1.9230627 | 0.05447218 | 0.35100338 |
| DDX58 | 132.31238 | 0.790122722 | 1.02464495 | 0.77111855 | 0.44063667 | 0.78962177 |
| NDUFB6 | 28.2569197 | 0.084934588 | 0.28598123 | 0.29699357 | 0.76647141 | 0.9447922 |
| APTX | 43.1955904 | -0.334407379 | 0.27631822 | -1.2102256 | 0.22619234 | 0.62072404 |
| DNAJA1 | 52.3072336 | -0.103036231 | 0.33973516 | -0.303284 | 0.76167345 | 0.94425408 |
| SMU1 | 33.5718852 | -0.150751188 | 0.23857199 | -0.6318897 | 0.52745894 | 0.83938843 |
| B4GALT1 | 155.4248 | -0.317155325 | 0.41122717 | -0.7712412 | 0.44056399 | 0.78962177 |
| BAG1 | 23.5999473 | 0.185903182 | 0.35828422 | 0.51887069 | 0.60385092 | 0.87903643 |
| CHMP5 | 29.0180116 | 0.311732453 | 0.37798866 | 0.82471376 | 0.40953412 | 0.7710341 |
| NFX1 | 105.986602 | -0.098578823 | 0.22456784 | -0.4389712 | 0.66068238 | 0.90958315 |
| UBE2R2 | 163.576088 | -0.07085623 | 0.16999451 | -0.4168148 | 0.67681387 | 0.91753129 |
| UBAP2 | 75.6373833 | -0.157749228 | 0.21747518 | -0.7253666 | 0.46822718 | 0.80374006 |
| DCAF12 | 17.5077167 | 0.094943648 | 0.36978781 | 0.2567517 | 0.79737046 | 0.95463848 |
| UBAP1 | 118.766021 | -0.305091922 | 0.24586812 | -1.2408763 | 0.21465145 | 0.61009832 |
| FAM219A | 65.3163869 | -0.90910783 | 0.22948568 | -3.9615014 | 7.45E-05 | 0.00674861 |
| VCP | 70.5079783 | -0.217593821 | 0.281821 | -0.7720994 | 0.44005556 | 0.78935034 |
| UNC13B | 107.714247 | 0.537440302 | 0.222045 | 2.42041159 | 0.01550295 | 0.18198087 |
| RUSC2 | 105.604003 | 0.304867311 | 0.23601308 | 1.29173906 | 0.19644752 | 0.59411229 |
| CCDC107 | 37.4078327 | -0.488606784 | 0.43211374 | -1.1307365 | 0.25816601 | 0.65130862 |
| TPM2 | 140.455267 | 0.671229724 | 0.33480148 | 2.00485887 | 0.04497814 | 0.31898889 |
| TLN1 | 95.8201645 | 0.302873643 | 0.20580179 | 1.4716764 | 0.14110828 | 0.52800651 |
| GBA2 | 20.5910766 | 0.134179622 | 0.38186565 | 0.35137914 | 0.72530393 | 0.93455731 |
| RECK | 55.0523581 | 0.034253851 | 0.28597234 | 0.11978029 | 0.90465719 | 0.98265659 |
| CLTA | 76.8892054 | -0.07282572 | 0.24446433 | -0.2978992 | 0.76578013 | 0.9447922 |
| GNE | 28.0829633 | 0.071993527 | 0.26799692 | 0.26863565 | 0.78821008 | 0.95129454 |
| RNF38 | 142.157644 | 0.35018338 | 0.18969782 | 1.8460064 | 0.06489128 | 0.38029186 |
| EBLN3P | 26.5676984 | -0.591096468 | 0.31560707 | -1.8728873 | 0.06108395 | 0.36765437 |
| ZCCHC7 | 383.04135 | -0.350708643 | 0.20057767 | -1.7484929 | 0.08037871 | 0.41953348 |
| GRHPR | 24.6492528 | -0.183751662 | 0.29528415 | -0.6222876 | 0.53375277 | 0.84333372 |
| AL138752.2 | 27.8598548 | -0.138810828 | 0.27839642 | -0.4986085 | 0.6180552 | 0.88534783 |
| TRMT10B | 31.387798 | -0.16475632 | 0.28401003 | -0.5801074 | 0.56184219 | 0.85545224 |
| DCAF10 | 102.943964 | -0.0297554 | 0.17540057 | -0.1696425 | 0.86529126 | 0.9762143 |
| GLIDR | 28.2385124 | 0.597226162 | 0.41205715 | 1.44937702 | 0.14723232 | 0.53679369 |
| CBWD5 | 28.8569937 | 0.010195192 | 0.28900946 | 0.03527633 | 0.9718594 | 0.99300191 |
| PGM5 | 63.2721431 | 0.832411246 | 0.47908521 | 1.73750145 | 0.0822987 | 0.42256146 |
| PIP5K1B | 539.679486 | -0.274394404 | 0.44126479 | -0.6218362 | 0.5340496 | 0.84333372 |
| APBA1 | 80.4542805 | 0.275909801 | 0.46254435 | 0.59650453 | 0.55083823 | 0.85014212 |
| PTAR1 | 72.0844883 | -0.12355747 | 0.17457688 | -0.7077539 | 0.4790981 | 0.81163945 |
| MAMDC2 | 112.675535 | 0.055860644 | 0.4129001 | 0.13528852 | 0.89238376 | 0.98023547 |
| SMC5-AS1 | 23.9851294 | -0.099688732 | 0.35333765 | -0.2821345 | 0.7778404 | 0.94820337 |
| SMC5 | 210.228244 | -0.088780859 | 0.1819084 | -0.4880526 | 0.62551261 | 0.89069797 |
| KLF9 | 288.058135 | -1.062006705 | 0.45711175 | -2.3232977 | 0.02016316 | 0.20958347 |
| TRPM3 | 312.589597 | 0.092524293 | 0.49881666 | 0.18548758 | 0.85284662 | 0.97307403 |
| CEMIP2 | 62.0996908 | -0.5956546 | 0.38262211 | -1.5567699 | 0.11952512 | 0.49486766 |
| ABHD17B | 21.3808211 | -0.025116223 | 0.38592902 | -0.0650799 | 0.94811038 | 0.99060401 |
| C9orf85 | 43.5675875 | 0.252985919 | 0.25422475 | 0.99512703 | 0.31967448 | 0.70688535 |
| ZFAND5 | 131.39756 | -0.092531249 | 0.38534999 | -0.2401226 | 0.8102352 | 0.9596162 |
| ALDH1A1 | 65.8839581 | 0.718768564 | 0.3550434 | 2.02445269 | 0.0429236 | 0.31201336 |
| ANXA1 | 48.7653785 | 0.94510849 | 0.44693785 | 2.11463069 | 0.03446144 | 0.2833153 |
| CARNMT1 | 26.4062467 | 0.004376338 | 0.30749175 | 0.01423238 | 0.98864459 | 0.99722663 |
| NMRK1 | 19.2753307 | 0.43145563 | 0.34632789 | 1.24580101 | 0.21283746 | 0.60956449 |
| OSTF1 | 48.7119066 | 0.097036479 | 0.23950092 | 0.4051612 | 0.68535902 | 0.91977784 |
| PCSK5 | 168.994669 | -0.574963605 | 0.36743256 | -1.564814 | 0.11762654 | 0.49087872 |
| PRUNE2 | 62.1740062 | 0.040378858 | 0.83240603 | 0.04850861 | 0.9613109 | 0.99181157 |
| VPS13A | 123.080126 | -0.194033952 | 0.32739258 | -0.5926645 | 0.55340572 | 0.85188037 |
| GNA14 | 358.626402 | 0.175760116 | 0.46876746 | 0.37494095 | 0.70770438 | 0.93009618 |
| GNA14-AS1 | 52.2060954 | -0.175745227 | 0.5889122 | -0.2984235 | 0.76537997 | 0.9447922 |
| GNAQ | 335.024524 | 0.134456312 | 0.18327986 | 0.73361204 | 0.46318522 | 0.80225845 |
| TLE4 | 118.13737 | 0.290127018 | 0.2364475 | 1.2270251 | 0.21981316 | 0.61590675 |
| TLE1 | 113.747377 | 0.321107512 | 0.35331927 | 0.90883101 | 0.36343933 | 0.73604674 |
| FRMD3 | 122.984022 | 0.110329993 | 0.37758006 | 0.29220291 | 0.77013149 | 0.94593455 |
| UBQLN1 | 84.1438985 | 0.09565718 | 0.20437023 | 0.46805829 | 0.63974291 | 0.89903874 |
| KIF27 | 28.8140837 | 0.091001587 | 0.24504816 | 0.37136205 | 0.71036789 | 0.93009618 |
| HNRNPK | 130.696534 | -0.046321279 | 0.22834611 | -0.2028556 | 0.83924792 | 0.96770443 |
| NTRK2 | 78.97381 | 0.711851946 | 0.32297663 | 2.20403543 | 0.02752185 | 0.24847431 |
| AGTPBP1 | 63.8220623 | 0.330669874 | 0.26281909 | 1.25816538 | 0.20833195 | 0.6059796 |
| NAA35 | 87.3765198 | 0.283317797 | 0.17392728 | 1.62894395 | 0.10332488 | 0.46365765 |
| GOLM1 | 21.108006 | -0.144309484 | 0.32191482 | -0.4482847 | 0.65394775 | 0.90646849 |
| ISCA1 | 23.0031694 | -0.342967704 | 0.29867891 | -1.1482823 | 0.25085205 | 0.6460368 |
| TUT7 | 73.9075147 | -0.243376316 | 0.26274293 | -0.9262906 | 0.35429497 | 0.73059415 |
| GAS1 | 31.7068046 | 0.846645844 | 0.40825579 | 2.07381221 | 0.03809675 | 0.29501902 |
| DAPK1 | 51.6282029 | -0.477824243 | 0.39188784 | -1.2192883 | 0.22273481 | 0.61772146 |
| CTSL | 48.8600312 | -0.389469405 | 0.40649061 | -0.9581264 | 0.337999 | 0.71925477 |
| SPIN1 | 123.324584 | 0.411153911 | 0.25074674 | 1.63971785 | 0.10106385 | 0.46131565 |
| S1PR3 | 120.212314 | -0.554401763 | 0.37565547 | -1.4758251 | 0.13999086 | 0.52623673 |
| SHC3 | 44.6101301 | -0.568347147 | 0.81195914 | -0.6999701 | 0.48394596 | 0.8144078 |
| SECISBP2 | 97.852518 | -0.119540152 | 0.20406844 | -0.5857846 | 0.55802027 | 0.85279817 |
| SEMA4D | 20.2355632 | -0.417358254 | 0.38233708 | -1.0915976 | 0.27500999 | 0.67029497 |
| AUH | 130.685262 | -0.537115047 | 0.3176117 | -1.691106 | 0.09081657 | 0.44024469 |
| NFIL3 | 57.0373194 | -1.388928019 | 0.45167523 | -3.0750591 | 0.00210461 | 0.05624279 |
| ROR2 | 126.820496 | -0.760204208 | 0.74360447 | -1.0223233 | 0.30662789 | 0.69801431 |
| SPTLC1 | 63.6657672 | 0.182640598 | 0.23883296 | 0.76472107 | 0.44443766 | 0.79252305 |
| IARS | 77.7190758 | -0.508637372 | 0.36055016 | -1.4107257 | 0.15832552 | 0.55013438 |
| NOL8 | 30.4014356 | -0.486322148 | 0.2480887 | -1.9602753 | 0.04996362 | 0.33748407 |
| CENPP | 129.05069 | 0.02851979 | 0.37179063 | 0.07670928 | 0.93885483 | 0.98827804 |
| ASPN | 24.217348 | 1.271300411 | 0.53194008 | 2.38993161 | 0.01685151 | 0.19013163 |
| ECM2 | 48.1592786 | 1.129514846 | 0.49677267 | 2.27370569 | 0.02298369 | 0.22440695 |
| IPPK | 21.618305 | -0.268674388 | 0.25777841 | -1.0422688 | 0.29728709 | 0.68849737 |
| BICD2 | 33.1699215 | 0.077532311 | 0.28920034 | 0.26809205 | 0.78862847 | 0.95129454 |
| NINJ1 | 18.1182739 | -1.112742278 | 0.5476808 | -2.031735 | 0.04218049 | 0.3087494 |
| FAM120A | 126.508357 | -0.203230929 | 0.2385887 | -0.8518045 | 0.3943226 | 0.75988253 |
| PHF2 | 39.2349147 | 0.225623823 | 0.21278695 | 1.06032737 | 0.28899569 | 0.67992862 |
| PTPDC1 | 40.9132477 | -0.284812061 | 0.39859085 | -0.7145474 | 0.47488874 | 0.80851131 |
| AL158152.2 | 54.2951193 | 0.084246532 | 0.29374854 | 0.28679813 | 0.7742669 | 0.9475938 |
| MFSD14B | 44.1829988 | 0.148318827 | 0.24702507 | 0.60042013 | 0.54822627 | 0.84786286 |
| C9orf3 | 667.041947 | 0.328611354 | 0.30071709 | 1.09275915 | 0.27449956 | 0.66930898 |
| FANCC | 94.0953009 | -0.174575844 | 0.23682106 | -0.7371635 | 0.46102292 | 0.80039155 |
| PTCH1 | 52.0946548 | 0.566519465 | 0.42413486 | 1.33570596 | 0.1816454 | 0.580649 |
| LINC00476 | 49.4837547 | 0.589445278 | 0.42701455 | 1.38038686 | 0.16746756 | 0.56315392 |
| ERCC6L2 | 126.954981 | 0.087417762 | 0.18538926 | 0.47153629 | 0.63725781 | 0.89741446 |
| HABP4 | 32.9886591 | 0.735173319 | 0.2597194 | 2.83064466 | 0.00464543 | 0.0906884 |
| CDC14B | 60.1145669 | 0.640444492 | 0.41936684 | 1.52717009 | 0.12671872 | 0.50611733 |
| ZNF782 | 38.6068029 | -0.032657938 | 0.24777531 | -0.1318047 | 0.89513881 | 0.98023547 |
| MFSD14C | 88.5315519 | 0.312844833 | 0.19161328 | 1.63268864 | 0.1025345 | 0.4627468 |
| TDRD7 | 21.6177628 | 0.25882555 | 0.46924301 | 0.55158105 | 0.58123542 | 0.86660562 |
| TMOD1 | 52.9277442 | -0.811558784 | 0.57998425 | -1.3992773 | 0.16172985 | 0.55513904 |
| TSTD2 | 37.8856158 | -0.264531306 | 0.20864489 | -1.2678542 | 0.20485002 | 0.60370888 |
| NCBP1 | 22.3656947 | -0.04437362 | 0.23792937 | -0.1864991 | 0.85205336 | 0.97285733 |
| XPA | 26.5177436 | 0.298624615 | 0.23349771 | 1.27891879 | 0.20092565 | 0.59975737 |
| ANP32B | 38.4952546 | -0.041085412 | 0.31303391 | -0.1312491 | 0.89557827 | 0.98023547 |

| | | | | | | |
|---|---|---|---|---|---|---|
| COL15A1 | 160.900491 | -0.86525957 | 0.79074337 | -1.0942356 | 0.27385165 | 0.66906109 |
| TGFBR1 | 69.6276882 | 0.374441915 | 0.28780073 | 1.30104575 | 0.19324279 | 0.59237115 |
| SEC61B | 57.5020145 | -0.606368395 | 0.3006163 | -2.0170843 | 0.04368672 | 0.31491241 |
| AL359710.1 | 32.5131852 | 0.718959802 | 0.50217024 | 1.43170533 | 0.15222817 | 0.53966466 |
| NR4A3 | 129.850553 | -0.791496066 | 1.04844433 | -0.7549243 | 0.45029443 | 0.79399826 |
| STX17 | 53.411643 | 0.081709775 | 0.28036246 | 0.29144335 | 0.77071226 | 0.94604383 |
| ERP44 | 77.3162655 | -0.116165055 | 0.2123347 | -0.5470847 | 0.58432057 | 0.86838662 |
| INVS | 180.37488 | -0.453413297 | 0.16481705 | -2.7510097 | 0.00594119 | 0.10445406 |
| TEX10 | 29.8309964 | -0.769561429 | 0.29971341 | -2.5676576 | 0.01023882 | 0.14224028 |
| ALDOB | 74.0105096 | 0.003623349 | 0.86501885 | 0.00418875 | 0.99665787 | 0.99883249 |
| ABCA1 | 242.214814 | -1.593775892 | 0.54141952 | -2.9436986 | 0.00324316 | 0.07300955 |
| SLC44A1 | 91.6546319 | 0.030289652 | 0.29220704 | 0.10365819 | 0.91744061 | 0.98477389 |
| FSD1L | 31.3097945 | 0.193600032 | 0.22914299 | 0.84488744 | 0.39817369 | 0.7611823 |
| FKTN | 37.1782009 | 0.030508804 | 0.24727525 | 0.12337993 | 0.90180625 | 0.98164203 |
| TMEM38B | 31.4526966 | -0.568779276 | 0.32205854 | -1.7660742 | 0.07738341 | 0.4114983 |
| ZNF462 | 78.5979518 | 0.128865333 | 0.22630992 | 0.56941973 | 0.56907133 | 0.85958696 |
| RAD23B | 101.274697 | -0.456940335 | 0.20474653 | -2.2317366 | 0.02563237 | 0.23956637 |
| ELP1 | 33.681054 | 0.315993293 | 0.23339991 | 1.3538707 | 0.17577763 | 0.57311399 |
| CTNNAL1 | 70.5419771 | -0.247199426 | 0.31419054 | -0.7867819 | 0.43140956 | 0.78389374 |
| TMEM245 | 97.095132 | 0.267856395 | 0.18586326 | 1.4411476 | 0.14954299 | 0.53844556 |
| PTPN3 | 25.6315546 | 0.058076072 | 0.32230546 | 0.18018954 | 0.85700377 | 0.97381501 |
| PALM2-AKAF | 80.0549074 | -0.136920668 | 0.34042108 | -0.4022097 | 0.68752969 | 0.92100905 |
| TXN | 47.4679041 | -0.558389185 | 0.44280049 | -1.2610401 | 0.20729439 | 0.6059796 |
| SVEP1 | 512.599941 | -0.562824251 | 0.44440121 | -1.2664778 | 0.20534208 | 0.60370888 |
| LPAR1 | 516.140551 | -1.164615581 | 0.41303368 | -2.8196625 | 0.00480742 | 0.09200446 |
| ECPAS | 182.407378 | -0.33008543 | 0.22953649 | -1.4380521 | 0.15041928 | 0.53880252 |
| ZNF483 | 30.5860445 | 0.057817135 | 0.40211134 | 0.1437839 | 0.88567112 | 0.97867793 |
| PTGR1 | 77.9043366 | -0.836709092 | 0.24978539 | -3.3497119 | 0.00080896 | 0.033399 |
| C9orf84 | 28.8480282 | -0.676645358 | 0.43372505 | -1.560079 | 0.11874121 | 0.49356852 |
| UGCG | 205.30521 | -1.943991585 | 0.42783244 | -4.5438153 | 5.52E-06 | 0.00104102 |
| SUSD1 | 47.2026047 | 0.46923933 | 0.33130827 | 1.4163224 | 0.15668113 | 0.54648773 |
| PTBP3 | 142.417345 | -0.324274568 | 0.2333757 | -1.3894958 | 0.16468203 | 0.55965519 |
| HSDL2 | 70.524526 | 0.074047112 | 0.31463653 | 0.23534175 | 0.81394349 | 0.96038841 |
| KIAA1958 | 77.6527683 | -0.388979851 | 0.23168002 | -1.6789529 | 0.09316122 | 0.44361507 |
| FKBP15 | 36.1624594 | -0.139780864 | 0.27835902 | -0.5021603 | 0.61555473 | 0.8836513 |
| SLC31A1 | 25.8755849 | -0.363100233 | 0.30019169 | -1.2095613 | 0.22644729 | 0.62072404 |
| RGS3 | 30.2690939 | 0.181944184 | 0.28398359 | 0.64068556 | 0.521727 | 0.8350836 |
| ZNF618 | 92.059807 | 0.795682567 | 0.26212668 | 3.03548865 | 0.00240146 | 0.06192403 |
| COL27A1 | 181.097626 | -0.220866866 | 0.4327869 | -0.5103363 | 0.60981588 | 0.88163721 |
| AKNA | 28.3813368 | 0.285445434 | 0.26686884 | 1.06960944 | 0.28479515 | 0.6783522 |
| ATP6V1G1 | 56.8293207 | -0.516484762 | 0.3103969 | -1.6639495 | 0.09612252 | 0.4510884 |
| TNC | 720.737063 | -0.221844021 | 1.13575201 | -0.1953279 | 0.84513628 | 0.96952174 |
| PAPPA | 190.289931 | -2.995540415 | 0.62067008 | -4.8263007 | 1.39E-06 | 0.00032974 |
| ASTN2 | 43.1666167 | 0.288401535 | 0.26653144 | 1.08205446 | 0.27922833 | 0.67353297 |
| CDK5RAP2 | 60.6044993 | 0.272968723 | 0.21958968 | 1.2430854 | 0.21383637 | 0.61001472 |
| MEGF9 | 33.9719663 | 0.230050311 | 0.35398958 | 0.64987877 | 0.51577054 | 0.83174649 |
| FBXW2 | 39.1379452 | 0.092789639 | 0.29222881 | 0.31752393 | 0.75084609 | 0.94050927 |
| PSMD5 | 42.2930886 | -0.488384835 | 0.20671651 | -2.3625826 | 0.0181481 | 0.19729344 |
| CNTRL | 48.1907915 | 0.256658923 | 0.24378073 | 1.05282694 | 0.29242032 | 0.6840363 |
| RAB14 | 70.4733417 | 0.071438815 | 0.21670786 | 0.32965494 | 0.7416607 | 0.93731118 |
| GSN | 206.060234 | 0.96652331 | 0.27502518 | 3.51430849 | 0.0004409 | 0.02145813 |
| STOM | 127.470548 | -0.945456457 | 0.31500742 | -3.0013784 | 0.0026876 | 0.06583733 |
| DAB2IP | 118.84623 | -0.178077598 | 0.32212642 | -0.552819 | 0.58038737 | 0.8662199 |
| TTLL11 | 28.4730076 | 0.156401174 | 0.34334692 | 0.45551937 | 0.64873563 | 0.90328878 |
| MRRF | 30.5843056 | -0.526229605 | 0.24962207 | -2.1081053 | 0.03502188 | 0.28565573 |
| RC3H2 | 79.5051894 | 0.097392451 | 0.18906006 | 0.51514028 | 0.606455 | 0.88000768 |
| RABGAP1 | 224.709424 | 0.169852857 | 0.22090699 | 0.76888856 | 0.44195947 | 0.79069339 |
| STRBP | 23.8310969 | 0.104804373 | 0.43148472 | 0.24289243 | 0.80808873 | 0.95923826 |
| DENND1A | 245.079238 | -0.281456685 | 0.22389615 | -1.2570859 | 0.20872254 | 0.6059796 |
| NEK6 | 50.0853881 | -1.08899021 | 0.29077513 | -3.7451284 | 0.0001803 | 0.01226885 |
| PSMB7 | 57.4342466 | -0.250609732 | 0.18888826 | -1.3267618 | 0.18458749 | 0.58363936 |
| NR6A1 | 35.2558096 | 0.408726444 | 0.44126621 | 0.92625819 | 0.35431182 | 0.73059415 |
| RPL35 | 81.5081022 | -0.398515615 | 0.31649498 | -1.2591531 | 0.20797506 | 0.6059796 |
| GOLGA1 | 28.2427345 | -0.024238205 | 0.28019581 | -0.0865045 | 0.93106536 | 0.98681039 |
| SCAI | 37.7006626 | 0.210336965 | 0.28148068 | 0.74725188 | 0.45491153 | 0.79609632 |
| PPP6C | 62.6798698 | 0.069159568 | 0.18403637 | 0.37579294 | 0.70707084 | 0.93009618 |
| RABEPK | 20.8495826 | 0.058770958 | 0.33121668 | 0.17743961 | 0.8591631 | 0.97397549 |
| HSPA5 | 145.870363 | -1.123720238 | 0.52796071 | -2.1284164 | 0.03330257 | 0.27874784 |
| GAPVD1 | 106.918653 | -0.20802389 | 0.17166617 | -1.2117931 | 0.22559158 | 0.62063573 |
| MAPKAP1 | 133.554862 | -0.143441184 | 0.18237871 | -0.7865018 | 0.43157356 | 0.78389374 |
| PBX3 | 160.267253 | 0.272409303 | 0.23697922 | 1.14950714 | 0.25034693 | 0.6460368 |
| MVB12B | 34.9471352 | 0.105631903 | 0.34501799 | 0.30616347 | 0.75948019 | 0.94385579 |
| ZBTB43 | 26.161996 | -0.537832931 | 0.37956169 | -1.4169843 | 0.15648753 | 0.54607164 |
| RALGPS1 | 21.3481169 | 0.99728813 | 0.53828636 | 1.85270926 | 0.06392404 | 0.37702872 |
| ANGPTL2 | 23.8988012 | 0.022521493 | 0.30747784 | 0.0732459 | 0.94161044 | 0.98912166 |
| GARNL3 | 82.1314215 | 0.676713756 | 0.35595343 | 1.90113005 | 0.05728498 | 0.35767826 |
| RPL12 | 38.4054488 | -0.711062815 | 0.40968177 | -1.7356467 | 0.08262632 | 0.42256146 |
| LRSAM1 | 34.5059303 | 0.39099018 | 0.24528466 | 1.5940262 | 0.11093017 | 0.4781799 |
| FAM129B | 79.1925857 | -0.202319999 | 0.38874225 | -0.5204477 | 0.60275159 | 0.87903643 |
| STXBP1 | 72.1317273 | 0.310612009 | 0.33708661 | 0.92146054 | 0.35681004 | 0.73129235 |
| CDK9 | 20.6724229 | -0.124967655 | 0.26804239 | -0.4662235 | 0.64105554 | 0.89934133 |
| FPGS | 34.8870187 | -0.673527201 | 0.31483164 | -2.139325 | 0.03240935 | 0.27525049 |
| ENG | 368.243558 | 0.285199778 | 0.41421366 | 0.68853301 | 0.49111719 | 0.81720854 |
| SLC25A25 | 42.1846291 | -0.206514544 | 0.27175249 | -0.7599362 | 0.44729274 | 0.79296348 |
| PTGES2 | 19.4385731 | -0.646053305 | 0.39148614 | -1.6502584 | 0.09889009 | 0.45678074 |
| C9orf16 | 23.9862834 | -0.945586991 | 0.39798524 | -2.3759348 | 0.01750455 | 0.19333326 |
| CIZ1 | 22.1734794 | 0.178522086 | 0.29764725 | 0.59977737 | 0.54865462 | 0.84831954 |
| GOLGA2 | 49.4685116 | 0.077794595 | 0.28543965 | 0.27254306 | 0.78520449 | 0.95065355 |
| COQ4 | 24.8785986 | 0.251342902 | 0.30036217 | 0.83679947 | 0.4027053 | 0.76472383 |
| SLC27A4 | 25.7277915 | 0.117259277 | 0.25905479 | 0.45264276 | 0.650806 | 0.9050963 |
| ODF2 | 27.6614776 | -0.27188257 | 0.34080435 | -0.7977673 | 0.42500553 | 0.77861803 |
| GLE1 | 20.5769369 | 0.246002275 | 0.353512 | 0.69588097 | 0.48650337 | 0.81595238 |
| SPTAN1 | 113.17866 | 0.541035699 | 0.26975362 | 2.00566616 | 0.04489188 | 0.31898889 |
| SET | 56.2768214 | -0.292942871 | 0.32978688 | -0.8882793 | 0.37439051 | 0.7464449 |
| ZER1 | 55.7394859 | 0.06488806 | 0.34236651 | 0.18952806 | 0.84967897 | 0.97157161 |
| KYAT1 | 18.9790724 | 0.016056305 | 0.2949012 | 0.05444639 | 0.95657952 | 0.99181157 |
| LRRC8A | 23.830006 | -0.533242358 | 0.43113832 | -1.2368243 | 0.21615231 | 0.61284851 |
| NUP188 | 23.8699942 | -0.293585486 | 0.31632451 | -0.9281149 | 0.35334799 | 0.73059415 |
| SH3GLB2 | 16.6183425 | 0.574587742 | 0.29399386 | 1.95442086 | 0.05065147 | 0.34087694 |
| PTPA | 34.333073 | -0.31906736 | 0.35229304 | -0.9056874 | 0.36510134 | 0.73726023 |
| IER5L | 68.1645582 | -0.409770723 | 0.39692888 | -1.032353 | 0.30190678 | 0.69291471 |
| NTMT1 | 22.8705686 | -1.0762573 | 0.39270718 | -2.7406102 | 0.00613252 | 0.10629223 |
| C9orf78 | 19.9938935 | 0.200927709 | 0.34838865 | 0.57673437 | 0.56411891 | 0.85672864 |
| USP20 | 24.7406975 | 0.299261068 | 0.33798349 | 0.88543102 | 0.37592421 | 0.74787433 |
| FNBP1 | 200.2273 | 0.319961739 | 0.28407674 | 1.1263215 | 0.26002946 | 0.65287206 |
| GPR107 | 81.3413419 | 0.019307221 | 0.18581202 | 0.10390728 | 0.91724293 | 0.98477389 |
| NCS1 | 18.8050543 | 0.948016062 | 0.33682083 | 2.81460042 | 0.00488379 | 0.09203696 |
| ASS1 | 18.5634948 | 0.476577014 | 0.37935463 | 1.25628365 | 0.20901315 | 0.6059796 |
| FUBP3 | 67.8151761 | 0.04632547 | 0.1881389 | 0.24623015 | 0.80550408 | 0.95836441 |
| ABL1 | 244.947718 | -0.33501419 | 0.25629831 | -1.307126 | 0.19116991 | 0.58975955 |
| LAMC3 | 129.728445 | -0.112816102 | 0.52417615 | -0.2152256 | 0.82959147 | 0.96580172 |
| AIF1L | 21.7067779 | 0.687545461 | 0.38926514 | 1.76626516 | 0.07735137 | 0.4114983 |
| NUP214 | 80.2166579 | 0.01358263 | 0.25116755 | 0.05407797 | 0.95687305 | 0.99181157 |

| | | | | | | |
|---|---|---|---|---|---|---|
| PRRC2B | 159.575677 | -0.015813311 | 0.19395986 | -0.0815288 | 0.93502144 | 0.98827449 |
| RAPGEF1 | 82.993956 | 0.174519666 | 0.29392272 | 0.59376038 | 0.55267239 | 0.85188037 |
| MED27 | 114.248394 | -0.381848628 | 0.30129391 | -1.2673626 | 0.20502567 | 0.60370888 |
| SETX | 175.714469 | -0.043823097 | 0.16567339 | -0.264515 | 0.79138311 | 0.95257889 |
| TTF1 | 20.8580899 | -0.004198399 | 0.28324671 | -0.0148224 | 0.98817386 | 0.99719329 |
| DDX31 | 23.188216 | -0.165420924 | 0.32690389 | -0.5060231 | 0.61284043 | 0.88309104 |
| TSC1 | 43.5079227 | -0.36460483 | 0.25081296 | -1.4536921 | 0.14603168 | 0.53584686 |
| RPL7A | 91.0885153 | -0.207730098 | 0.34300675 | -0.6056152 | 0.54477031 | 0.84730644 |
| SURF4 | 63.4986233 | -0.591746067 | 0.39318755 | -1.504997 | 0.13232486 | 0.51310477 |
| ADAMTSL2 | 95.7698873 | 0.700674604 | 0.57163896 | 1.22572926 | 0.22030057 | 0.61590675 |
| VAV2 | 68.7951591 | -0.097013686 | 0.27205354 | -0.3565978 | 0.72139294 | 0.93444541 |
| BRD3 | 22.499394 | 0.29309016 | 0.27955865 | 1.04840313 | 0.29445291 | 0.68566952 |
| WDR5 | 21.9122938 | -0.22285501 | 0.27817085 | -0.8011444 | 0.42304808 | 0.77827834 |
| RXRA | 45.7995549 | 0.2129285 | 0.35299052 | 0.60321308 | 0.54636694 | 0.84758329 |
| COL5A1 | 141.121206 | -0.084750074 | 0.42473775 | -0.1995351 | 0.84184422 | 0.96803523 |
| CAMSAP1 | 25.83964 | 0.31952806 | 0.23590221 | 1.3544937 | 0.17557892 | 0.57311399 |
| NACC2 | 44.4546123 | 0.076495672 | 0.27428212 | 0.27889412 | 0.78032608 | 0.94916975 |
| ENTR1 | 18.1674696 | -0.25839382 | 0.33250279 | -0.7771177 | 0.43708931 | 0.78768252 |
| SEC16A | 45.6888703 | -0.377096409 | 0.25701919 | -1.4671916 | 0.14232393 | 0.52958915 |
| NOTCH1 | 19.7600756 | 0.568404474 | 0.29654419 | 1.91676144 | 0.05526825 | 0.35288127 |
| RABL6 | 45.1203304 | -0.105149952 | 0.2979718 | -0.3528856 | 0.72417422 | 0.93455731 |
| PHPT1 | 38.1187906 | -0.383421575 | 0.35039538 | -1.0942541 | 0.27384356 | 0.66906109 |
| EDF1 | 54.1695715 | -0.074562832 | 0.32861635 | -0.2268993 | 0.82050202 | 0.96291025 |
| FBXW5 | 38.9243823 | -0.128562876 | 0.33520765 | -0.3835321 | 0.70132531 | 0.92865476 |
| MAN1B1 | 22.3849002 | 0.36139096 | 0.24950749 | 1.4484173 | 0.14750038 | 0.53679369 |
| DPP7 | 19.547975 | -0.290343002 | 0.28794914 | -1.0083135 | 0.31330398 | 0.70282967 |
| TUBB4B | 43.2764115 | -0.164102673 | 0.50834791 | -0.3228157 | 0.74683484 | 0.93900586 |
| NELFB | 25.9562174 | 0.659315728 | 0.26540286 | 2.48420728 | 0.01298402 | 0.1665263 |
| EXD3 | 32.3854086 | 0.317841972 | 0.32670827 | 0.97286171 | 0.33062204 | 0.71362138 |
| MRPL41 | 38.4780985 | -0.487782111 | 0.25803083 | -1.8904025 | 0.05870415 | 0.36041506 |
| EHMT1 | 164.147173 | 0.154554366 | 0.18000264 | 0.85862277 | 0.39054867 | 0.75729345 |
| PSMD13 | 30.6375605 | -0.233081475 | 0.343657 | -0.6782387 | 0.49762036 | 0.82235486 |
| PGGHG | 18.6020019 | -0.119634197 | 0.38736535 | -0.3088407 | 0.75744269 | 0.94298599 |
| IFITM2 | 262.540443 | -1.155970188 | 0.32736008 | -3.5311886 | 0.0004137 | 0.02111289 |
| IFITM3 | 443.186394 | -1.166810611 | 0.31983188 | -3.6481998 | 0.00026408 | 0.01590783 |
| PTDSS2 | 23.2227923 | -0.153747574 | 0.27183942 | -0.5655823 | 0.57167773 | 0.86232751 |
| RNH1 | 37.8370052 | -0.320205875 | 0.2314953 | -1.3832068 | 0.16660147 | 0.56240432 |
| PHRF1 | 25.8318049 | -0.362780891 | 0.25042982 | -1.4486329 | 0.14744012 | 0.53679369 |
| DEAF1 | 36.4036689 | 0.075024512 | 0.24438713 | 0.30699044 | 0.75885065 | 0.94385579 |
| TALDO1 | 29.1476806 | -0.564933165 | 0.35911947 | -1.5731065 | 0.11569417 | 0.48836096 |
| RPLP2 | 102.634619 | -0.351531645 | 0.35306387 | -0.9956602 | 0.31941527 | 0.70683013 |
| PNPLA2 | 23.576198 | -0.99268368 | 0.35489774 | -2.7970978 | 0.00515639 | 0.0940306 |
| CD151 | 88.9556374 | -0.058675283 | 0.34815781 | -0.1685307 | 0.86616579 | 0.9762143 |
| POLR2L | 56.1647728 | -0.579235623 | 0.29781095 | -1.9449776 | 0.05177768 | 0.34513804 |
| TSPAN4 | 36.5339965 | 0.57651964 | 0.35328822 | 1.63186772 | 0.10270736 | 0.4627468 |
| AP2A2 | 80.1281907 | -0.07409423 | 0.26906899 | -0.2753726 | 0.78302998 | 0.9503695 |
| TOLLIP | 21.5137927 | -0.234407294 | 0.28465704 | -0.8234727 | 0.41023925 | 0.77165299 |
| MOB2 | 29.2925951 | 0.686540256 | 0.28843412 | 2.38023247 | 0.01730172 | 0.19236057 |
| CD81 | 228.215926 | 0.017342167 | 0.21896123 | 0.079202 | 0.93687195 | 0.98827804 |
| KCNQ1 | 34.1792978 | -0.424673331 | 0.37024927 | -1.146993 | 0.25138451 | 0.6460368 |
| KCNQ1OT1 | 161.758675 | -0.130478557 | 0.33273152 | -0.3921437 | 0.69495206 | 0.92589762 |
| CDKN1C | 48.5544662 | -0.630636806 | 0.78056332 | -0.8079252 | 0.41913362 | 0.77771095 |
| NAP1L4 | 91.1738373 | -0.016566583 | 0.19280738 | -0.085923 | 0.93152765 | 0.98681039 |
| CARS | 31.2106155 | -0.661633849 | 0.30169745 | -2.1930376 | 0.02830467 | 0.25305477 |
| OSBPL5 | 82.2233176 | -0.568312837 | 0.23279516 | -2.4412571 | 0.01463623 | 0.17653018 |
| ZNF195 | 19.3532691 | -0.240481598 | 0.38493512 | -0.6247328 | 0.53214644 | 0.84265119 |
| NUP98 | 132.840464 | -0.031071159 | 0.23338449 | -0.1331329 | 0.89408826 | 0.98023547 |
| STIM1 | 109.051313 | 0.337007744 | 0.22504217 | 1.49753152 | 0.13425501 | 0.51662128 |
| AC104389.4 | 59.9896043 | 0.629462787 | 0.77197259 | 0.81539526 | 0.4148461 | 0.77476594 |
| TRIM22 | 66.6362856 | -0.202660162 | 0.91552341 | -0.2213599 | 0.82481222 | 0.96413949 |
| CAVIN3 | 36.9364383 | 0.233005473 | 0.55953213 | 0.41642912 | 0.67709602 | 0.91767419 |
| APBB1 | 18.4731814 | -0.013617095 | 0.30258032 | -0.0450032 | 0.96410473 | 0.99274284 |
| AC084337.2 | 24.3764229 | -0.54182256 | 0.33620802 | -1.6115694 | 0.10705567 | 0.47139134 |
| DNHD1 | 36.9884364 | -0.260004961 | 0.30560187 | -0.8507964 | 0.39488248 | 0.75988253 |
| ILK | 72.3240265 | 0.545398518 | 0.2994275 | 1.82147105 | 0.06853528 | 0.38953271 |
| TPP1 | 27.7356424 | -0.583249953 | 0.2843444 | -2.0512095 | 0.04024655 | 0.30130305 |
| PPFIBP2 | 25.6310507 | 0.799831226 | 0.41119479 | 1.9451395 | 0.05175819 | 0.34513804 |
| EIF3F | 28.8131487 | 0.129303996 | 0.25380425 | 0.50946347 | 0.6104274 | 0.88163721 |
| STK33 | 28.9794566 | 0.585577405 | 0.36178802 | 1.61856496 | 0.1055409 | 0.46789211 |
| TRIM66 | 36.5607399 | 0.458362588 | 0.28997777 | 1.58068182 | 0.11395081 | 0.48462065 |
| RPL27A | 115.754755 | -0.4556754 | 0.27011141 | -1.6869905 | 0.09160517 | 0.44124451 |
| ST5 | 222.736012 | 0.525699705 | 0.31143125 | 1.68801207 | 0.09140891 | 0.44124451 |
| TMEM9B | 17.925252 | -0.028690813 | 0.27927795 | -0.1027321 | 0.91817559 | 0.98477414 |
| DENND5A | 168.046214 | -0.24991719 | 0.31161081 | -0.8020171 | 0.42254308 | 0.77787302 |
| TMEM41B | 56.3761958 | -0.045387288 | 0.31827143 | -0.1426056 | 0.88660168 | 0.97919298 |
| IPO7 | 97.24457 | -0.252137625 | 0.3348138 | -0.7530682 | 0.45140893 | 0.79401729 |
| ZNF143 | 29.931045 | -0.012286489 | 0.24255514 | -0.0506544 | 0.9596009 | 0.99181157 |
| WEE1 | 21.6355163 | 0.992841877 | 0.53067235 | 1.87091314 | 0.06135712 | 0.36839337 |
| SWAP70 | 71.1395906 | 0.06883749 | 0.25106991 | 0.27417659 | 0.78394892 | 0.95064024 |
| SBF2 | 649.062663 | -0.093022008 | 0.21854168 | -0.4256488 | 0.67036376 | 0.91445841 |
| AC011092.2 | 23.1991831 | -0.586648516 | 0.38959272 | -1.5057995 | 0.13211865 | 0.51310477 |
| ADM | 74.7984808 | -2.282988512 | 0.80888902 | -2.8223754 | 0.00476693 | 0.09157966 |
| MTRNR2L8 | 47.2138654 | -0.416709175 | 0.55687843 | -0.7482947 | 0.45428242 | 0.79609632 |
| MRVI1 | 118.376325 | 0.511426433 | 0.39119655 | 1.30733881 | 0.19109767 | 0.58975955 |
| CTR9 | 29.677346 | 0.342808618 | 0.24606097 | 1.39318569 | 0.16356364 | 0.55830433 |
| EIF4G2 | 108.791519 | 0.165985394 | 0.23155344 | 0.71683408 | 0.47347648 | 0.80770164 |
| ZBED5 | 40.2087112 | 0.068055463 | 0.26697368 | 0.25491451 | 0.79878913 | 0.95529721 |
| GALNT18 | 55.0420228 | 0.487754232 | 0.32237662 | 1.51299505 | 0.13028092 | 0.51204187 |
| USP47 | 231.42809 | -0.097819851 | 0.1536148 | -0.6367866 | 0.52426383 | 0.83697897 |
| DKK3 | 52.5921136 | 0.567422604 | 0.36413625 | 1.55827003 | 0.11916925 | 0.49395083 |
| MICAL2 | 163.3092 | 0.541282083 | 0.29096707 | 1.86028642 | 0.06284501 | 0.37215794 |
| PARVA | 224.099395 | 0.125750287 | 0.21367224 | 0.58851954 | 0.55618362 | 0.85207344 |
| TEAD1 | 319.850806 | 0.035340921 | 0.27874034 | 0.12678797 | 0.89910822 | 0.98058672 |
| ARNTL | 35.7064221 | -1.121586245 | 0.42194168 | -2.6581546 | 0.00785698 | 0.12441768 |
| BTBD10 | 45.2667881 | 0.270469037 | 0.29310271 | 0.92277905 | 0.35612236 | 0.73129235 |
| FAR1 | 100.46439 | 0.353151582 | 0.20226017 | 1.74602629 | 0.08080638 | 0.42027326 |
| SPON1 | 187.536915 | -0.963783621 | 0.56087043 | -1.7183713 | 0.08572892 | 0.42858628 |
| RRAS2 | 68.4647881 | 0.340636956 | 0.38021837 | 0.89589821 | 0.37030713 | 0.74264779 |
| COPB1 | 81.5064855 | -0.030460357 | 0.18785018 | -0.1621524 | 0.87118584 | 0.9768643 |
| SOX6 | 152.184618 | 1.081554758 | 0.33538184 | 3.22484593 | 0.0012604 | 0.04375316 |
| C11orf58 | 89.9412027 | 0.04743042 | 0.17305351 | 0.2740795 | 0.78402353 | 0.95064024 |
| PLEKHA7 | 133.893803 | -0.460698442 | 0.26577697 | -1.7334024 | 0.08302417 | 0.42333667 |
| RPS13 | 90.7646821 | -0.476317551 | 0.35891238 | -1.3271137 | 0.18447106 | 0.58363936 |
| PIK3C2A | 156.635325 | -0.300026398 | 0.15770193 | -1.9024903 | 0.05710708 | 0.35717443 |
| NUCB2 | 28.2081557 | -0.243226959 | 0.288663 | -0.8425983 | 0.39945314 | 0.7624886 |
| SERGEF | 56.7433338 | -0.470472658 | 0.26934422 | -1.7467338 | 0.08068353 | 0.42022908 |
| HPS5 | 27.4594134 | -0.144978131 | 0.24114874 | -0.601198 | 0.54770814 | 0.84785354 |
| GTF2H1 | 40.7779498 | 0.065332069 | 0.21045292 | 0.31043555 | 0.75622976 | 0.94216126 |
| LDHA | 137.369756 | -1.75871088 | 0.3370103 | -5.2185672 | 1.80E-07 | 8.83E-05 |
| TSG101 | 53.4050035 | 0.167023152 | 0.19272783 | 0.86662703 | 0.3861464 | 0.75392929 |
| UEVLD | 34.6585641 | -0.246013484 | 0.2533957 | -0.9708669 | 0.33161459 | 0.71487809 |
| NAV2 | 799.761807 | -0.53306686 | 0.463395 | -1.1503509 | 0.24999937 | 0.6460368 |
| PRMT3 | 52.9950426 | -0.611496509 | 0.21606322 | -2.830174 | 0.00465227 | 0.0906884 |

| | | | | | | |
|---|---|---|---|---|---|---|
| GAS2 | 49.0201183 | 3.061499477 | 0.55892754 | 5.47745329 | 4.31E-08 | 2.64E-05 |
| AC099687.1 | 23.9311805 | 1.607468577 | 0.5637736 | 2.85126613 | 0.00435455 | 0.0871978 |
| ANO3 | 136.450691 | 2.351247462 | 0.38735284 | 6.07004065 | 1.28E-09 | 2.25E-06 |
| LGR4 | 52.3341644 | 0.179257278 | 0.24311929 | 0.73732233 | 0.46092636 | 0.80039155 |
| LIN7C | 23.0974302 | 0.090848876 | 0.25884416 | 0.35097904 | 0.72560407 | 0.93455731 |
| BDNF-AS | 65.5563661 | -0.111053641 | 0.2501993 | -0.4438607 | 0.65714329 | 0.90772143 |
| METTL15 | 182.61973 | 0.272103906 | 0.21615859 | 1.25881605 | 0.20809678 | 0.6059796 |
| ARL14EP | 21.536278 | 0.011737805 | 0.33119931 | 0.03544031 | 0.97172864 | 0.99300191 |
| DNAJC24 | 24.3621944 | 0.040470794 | 0.28573198 | 0.14163901 | 0.88736515 | 0.97920559 |
| IMMP1L | 43.5471901 | -0.043404279 | 0.24420416 | -0.1777377 | 0.858929 | 0.97397549 |
| ELP4 | 100.375936 | 0.137275617 | 0.15518456 | 0.88459586 | 0.37637464 | 0.74787433 |
| RCN1 | 31.054509 | -0.334972152 | 0.41171579 | -0.8136005 | 0.41587388 | 0.77573656 |
| WT1 | 57.822986 | 0.885500229 | 0.47473463 | 1.86525308 | 0.06214594 | 0.37064816 |
| EIF3M | 39.3652693 | -0.140480488 | 0.29312988 | -0.4792432 | 0.63176565 | 0.89436221 |
| QSER1 | 60.3004973 | -0.134944221 | 0.2031654 | -0.6642087 | 0.50655677 | 0.82481485 |
| TCP11L1 | 28.0587866 | 0.126770786 | 0.27633483 | 0.45875789 | 0.64640803 | 0.90209887 |
| CSTF3 | 46.7246791 | -0.044661371 | 0.20772449 | -0.2150029 | 0.82976504 | 0.96580172 |
| HIPK3 | 270.978195 | -0.352469312 | 0.31244186 | -1.1281117 | 0.25927276 | 0.65213055 |
| FBXO3 | 40.5834694 | 0.133697469 | 0.2299626 | 0.58138789 | 0.56097906 | 0.85477139 |
| CAPRIN1 | 83.5411486 | -0.101621422 | 0.17151351 | -0.5924981 | 0.55351712 | 0.85188037 |
| NAT10 | 28.4183056 | -0.520074285 | 0.37305027 | -1.3941132 | 0.16328344 | 0.55760686 |
| ABTB2 | 77.8874928 | -0.372239484 | 0.78960119 | -0.4714272 | 0.63733569 | 0.89741446 |
| CAT | 41.30266 | 0.293571308 | 0.37223736 | 0.78866696 | 0.43030669 | 0.78298297 |
| APIP | 28.1199792 | 0.247605134 | 0.35248326 | 0.70245928 | 0.48239279 | 0.8140309 |
| PDHX | 67.7941972 | -0.237885385 | 0.27777472 | -0.8563968 | 0.39177832 | 0.75901355 |
| CD44 | 226.208383 | -0.075295784 | 0.35194549 | -0.2139416 | 0.83059259 | 0.96597958 |
| PAMR1 | 69.0483828 | -0.18455777 | 0.78019489 | -0.2365534 | 0.81300326 | 0.96038841 |
| TRIM44 | 169.181266 | -0.17217791 | 0.13375131 | -1.2872989 | 0.19799014 | 0.59534761 |
| LDLRAD3 | 345.378348 | -1.192826643 | 0.31404277 | -3.7982936 | 0.0001457 | 0.01010111 |
| PRR5L | 37.7621068 | 0.1780046 | 0.34093957 | 0.52210015 | 0.6016006 | 0.87903643 |
| TRAF6 | 18.5000124 | -0.198822245 | 0.27942173 | -0.711549 | 0.47674411 | 0.81064148 |
| API5 | 46.5842915 | -0.086769016 | 0.21925104 | -0.3957519 | 0.69228806 | 0.9248682 |
| TTC17 | 242.370115 | -0.003571505 | 0.13287789 | -0.0268781 | 0.97855696 | 0.99465087 |
| HSD17B12 | 167.347439 | 0.079251241 | 0.20756647 | 0.38181139 | 0.70260127 | 0.92867207 |
| ALKBH3 | 19.4996566 | 0.263723871 | 0.32092083 | 0.82177238 | 0.41120646 | 0.77166383 |
| C11orf96 | 486.599633 | -0.824638234 | 0.48201427 | -1.7108171 | 0.08711489 | 0.43290196 |
| EXT2 | 74.4590722 | -0.039116661 | 0.18704258 | -0.2091324 | 0.83434489 | 0.9666365 |
| TP53I11 | 28.4201817 | 0.041130531 | 0.40747177 | 0.10094081 | 0.91959745 | 0.98528826 |
| PRDM11 | 31.2378035 | 0.418535879 | 0.22392701 | 1.86907279 | 0.06161269 | 0.36915678 |
| CRY2 | 34.5823998 | 0.110520032 | 0.26922022 | 0.41051906 | 0.68142523 | 0.91886128 |
| PHF21A | 358.42952 | -0.893922969 | 0.22732307 | -3.93239 | 8.41E-05 | 0.0071045 |
| DGKZ | 18.5213623 | 0.705626443 | 0.31630417 | 2.23084775 | 0.02569122 | 0.23956637 |
| AMBRA1 | 287.191062 | -0.31572913 | 0.18042743 | -1.7498954 | 0.08013636 | 0.41886353 |
| ATG13 | 46.7079534 | -0.009992546 | 0.24067909 | -0.0415181 | 0.96688284 | 0.99300191 |
| ARHGAP1 | 62.4533204 | -0.082110385 | 0.37614737 | -0.2182931 | 0.82720073 | 0.96580172 |
| CKAP5 | 49.5812228 | 0.131854876 | 0.1895225 | 0.69572147 | 0.48660327 | 0.81595238 |
| LRP4 | 46.3019659 | -0.421135746 | 0.41261211 | -1.0206577 | 0.30741662 | 0.69801431 |
| C11orf49 | 98.1041519 | -0.156674683 | 0.26541658 | -0.5902973 | 0.55499136 | 0.85207344 |
| ARFGAP2 | 21.5290526 | 0.408687649 | 0.29732399 | 1.37455323 | 0.16927 | 0.56570963 |
| DDB2 | 30.5381103 | -0.200363 | 0.38957581 | -0.5143107 | 0.60703479 | 0.88000768 |
| MADD | 31.304696 | 0.247317053 | 0.23510504 | 1.05194281 | 0.29282579 | 0.6842846 |
| CELF1 | 198.273303 | 0.183842536 | 0.1370169 | 1.34175084 | 0.1796768 | 0.57744792 |
| MTCH2 | 36.3173746 | -0.298868272 | 0.23861469 | -1.2525142 | 0.21038257 | 0.60731347 |
| FNBP4 | 170.917192 | -0.385486487 | 0.20273401 | -1.9014397 | 0.05724444 | 0.35767826 |
| NUP160 | 78.6463542 | 0.115901902 | 0.18462588 | 0.62776627 | 0.53015707 | 0.84131382 |
| PTPRJ | 63.0267405 | 1.065587968 | 0.25914902 | 4.11187347 | 3.92E-05 | 0.00443201 |
| TNKS1BP1 | 45.3714988 | 0.087368012 | 0.20659134 | 0.42290259 | 0.67236633 | 0.91482259 |
| SSRP1 | 20.0731437 | -0.439439654 | 0.45995018 | -0.9554071 | 0.33937188 | 0.71925477 |
| SERPING1 | 227.936212 | -0.737885711 | 0.28717462 | -2.569467 | 0.01018551 | 0.14209086 |
| ZDHHC5 | 32.1828129 | -0.20957933 | 0.23717267 | -0.8836571 | 0.37688132 | 0.74832285 |
| CTNND1 | 94.6178081 | 0.217605023 | 0.27021145 | 0.80531386 | 0.42063859 | 0.77771095 |
| LPXN | 21.4114223 | 0.286341367 | 0.28805418 | 0.99405386 | 0.32019664 | 0.70728138 |
| FAM111A | 27.7465522 | 0.240473343 | 0.4746563 | 0.50662626 | 0.61241708 | 0.88282721 |
| DTX4 | 18.7485424 | -0.610177369 | 0.31425257 | -1.9416782 | 0.05217607 | 0.34575469 |
| OSBP | 51.5242353 | -0.223097419 | 0.21990793 | -1.0145038 | 0.31034243 | 0.70067789 |
| PATL1 | 39.491602 | -0.828232364 | 0.33962486 | -2.4386683 | 0.01474149 | 0.17699624 |
| STX3 | 20.4719629 | -0.067783766 | 0.46525994 | -0.1456901 | 0.88416604 | 0.97783841 |
| PRPF19 | 17.5579246 | -0.525574905 | 0.45120522 | -1.1648245 | 0.24409003 | 0.63966808 |
| TMEM109 | 21.8224943 | 0.447575616 | 0.30549706 | 1.46507339 | 0.1429009 | 0.53039327 |
| VPS37C | 18.7127736 | -0.08092702 | 0.28137363 | -0.2876141 | 0.77364216 | 0.94750611 |
| DDB1 | 56.8994844 | -0.055543094 | 0.18714823 | -0.2967866 | 0.7666294 | 0.94481963 |
| CPSF7 | 49.525414 | 0.006293065 | 0.26359233 | 0.02387424 | 0.98095293 | 0.99533442 |
| AP003108.2 | 38.4791063 | -0.065263965 | 0.30254438 | -0.215717 | 0.82920835 | 0.96580172 |
| FADS2 | 54.0156798 | -0.041381877 | 0.34604011 | -0.1195869 | 0.90481037 | 0.98265659 |
| FADS1 | 44.9225822 | -0.472443525 | 0.48840653 | -0.9673162 | 0.33338601 | 0.71622463 |
| FADS3 | 34.601254 | 0.044104633 | 0.35785904 | 0.12324582 | 0.90191244 | 0.98164203 |
| FTH1 | 701.537203 | -1.332284517 | 0.43024137 | -3.0965979 | 0.00195755 | 0.05557623 |
| AHNAK | 171.783147 | 0.768211184 | 0.23870214 | 3.2182836 | 0.0012896 | 0.04375316 |
| EML3 | 19.8396663 | -0.146791928 | 0.33918153 | -0.4327828 | 0.66517258 | 0.91235816 |
| B3GAT3 | 21.7717786 | -0.309023349 | 0.27717484 | -1.114904 | 0.26489155 | 0.65892534 |
| GANAB | 68.0097467 | -0.052771299 | 0.23055366 | -0.2288894 | 0.81895484 | 0.96265181 |
| UBXN1 | 23.153187 | -0.068779153 | 0.38123651 | -0.1804107 | 0.85683014 | 0.97381501 |
| TTC9C | 20.1411342 | 0.034012831 | 0.29414093 | 0.11563447 | 0.90794224 | 0.98306694 |
| NXF1 | 20.2297512 | 0.089430881 | 0.3495323 | 0.2558587 | 0.79805995 | 0.95489133 |
| STX5 | 53.4988777 | 0.054956257 | 0.18381244 | 0.29898007 | 0.76495525 | 0.9447922 |
| SLC3A2 | 40.8388522 | -0.917706308 | 0.29188972 | -3.1440172 | 0.00166646 | 0.0512418 |
| RARRES3 | 29.5979883 | -0.304526111 | 0.59687543 | -0.5102004 | 0.60991104 | 0.88163721 |
| PLA2G16 | 20.5170034 | -0.458741337 | 0.32060922 | -1.4308426 | 0.15247533 | 0.54004545 |
| ATL3 | 88.0723668 | -0.108942601 | 0.26637549 | -0.4089813 | 0.68255336 | 0.91933701 |
| RTN3 | 79.4078448 | 0.343102687 | 0.19517863 | 1.75789066 | 0.07876612 | 0.41594801 |
| MARK2 | 52.615802 | 0.168126375 | 0.21967176 | 0.7653527 | 0.44406155 | 0.79248199 |
| MACROD1 | 21.8746895 | 0.332836785 | 0.29409455 | 1.13173393 | 0.25774631 | 0.65130862 |
| STIP1 | 26.1530756 | 0.327313773 | 0.25493927 | 1.28388919 | 0.19918073 | 0.59746089 |
| VEGFB | 32.0151297 | -0.0972367 | 0.308147 | -0.315553 | 0.75234185 | 0.94058012 |
| FKBP2 | 28.3538336 | 0.181161825 | 0.33463999 | 0.54136335 | 0.58825717 | 0.8702694 |
| ESRRA | 22.5832126 | -0.111844783 | 0.38804225 | -0.2882284 | 0.77317194 | 0.94730145 |
| TRMT112 | 30.3414529 | -0.325144172 | 0.29748853 | -1.0929637 | 0.27440973 | 0.66930898 |
| PRDX5 | 28.0763995 | -0.187777018 | 0.32460657 | -0.5784757 | 0.56294299 | 0.85571045 |
| NRXN2 | 18.3987275 | -0.731313804 | 0.57318963 | -1.2758671 | 0.20200249 | 0.6015058 |
| RASGRP2 | 29.510636 | 0.938810681 | 0.26477575 | 3.54568227 | 0.0003916 | 0.02041031 |
| SF1 | 134.149543 | 0.039279601 | 0.15878126 | 0.24738185 | 0.80461272 | 0.95836286 |
| EHD1 | 23.3529729 | 0.879359386 | 0.45102289 | 1.94970012 | 0.05121187 | 0.34374561 |
| ATG2A | 18.2745709 | -0.500272721 | 0.36500667 | -1.3705851 | 0.17050432 | 0.56801217 |
| VPS51 | 42.6155241 | 0.049607671 | 0.19783488 | 0.2507529 | 0.80200516 | 0.95758343 |
| FAU | 97.4210229 | -0.227494974 | 0.37277918 | -0.6102674 | 0.5416847 | 0.84641877 |
| SYVN1 | 19.7770914 | -0.209720155 | 0.34591457 | -0.6062773 | 0.54433062 | 0.84730644 |
| CAPN1 | 31.7455249 | 0.393960693 | 0.30411838 | 1.29541888 | 0.19517577 | 0.5931955 |
| DPF2 | 27.1812106 | 0.167605483 | 0.23634046 | 0.70916967 | 0.47821919 | 0.8114599 |
| FRMD8 | 18.3309919 | -0.282213598 | 0.28566808 | -0.9879074 | 0.32319802 | 0.70964513 |
| NEAT1 | 4726.0414 | -1.431000566 | 0.3993596 | -3.5832382 | 0.00033936 | 0.0184738 |
| MALAT1 | 41344.0507 | -0.413350794 | 0.24430122 | -1.6919719 | 0.09065133 | 0.43973376 |
| SCYL1 | 27.8999077 | -0.471208334 | 0.30325852 | -1.5538173 | 0.120228 | 0.49665855 |
| LTBP3 | 84.7873672 | 0.735236786 | 0.23575088 | 3.11870226 | 0.00181649 | 0.05398305 |

| | | | | | | |
|---|---|---|---|---|---|---|
| EHBP1L1 | 61.298144 | 0.640299619 | 0.3463082 | 1.84892998 | 0.06446792 | 0.37871685 |
| RELA | 22.8502932 | -0.230665306 | 0.35183749 | -0.6556018 | 0.51208034 | 0.82885603 |
| KAT5 | 24.3144146 | 0.385937488 | 0.28509192 | 1.35372999 | 0.17582253 | 0.57311399 |
| RNASEH2C | 23.0976848 | -0.197656164 | 0.33569737 | -0.5887927 | 0.55600034 | 0.85207344 |
| CFL1 | 101.623977 | -0.331038447 | 0.38147661 | -0.8677818 | 0.38551382 | 0.75340372 |
| EFEMP2 | 26.8679337 | 0.342767344 | 0.33724625 | 1.01637112 | 0.30945269 | 0.69972033 |
| CCDC85B | 43.2696517 | -0.449810829 | 0.37495449 | -1.1996411 | 0.23027874 | 0.62447176 |
| DRAP1 | 19.6046424 | 0.293364174 | 0.36158617 | 0.81132576 | 0.41717862 | 0.77722447 |
| BANF1 | 21.8175946 | 0.218629104 | 0.40047953 | 0.5459183 | 0.5851221 | 0.86905058 |
| SF3B2 | 43.2541162 | 0.118399705 | 0.25135388 | 0.47104784 | 0.63760657 | 0.89748529 |
| PACS1 | 178.937108 | 0.11539691 | 0.18643822 | 0.61895522 | 0.53594586 | 0.84412047 |
| RAB1B | 22.5169568 | 0.288792025 | 0.2911701 | 0.99183271 | 0.32127914 | 0.70780851 |
| CD248 | 40.1931355 | 0.814135551 | 0.56406969 | 1.44332441 | 0.14892911 | 0.53689238 |
| CCS | 31.4252221 | 0.219375754 | 0.31435783 | 0.69785364 | 0.48526873 | 0.81569988 |
| RBM14-RBM | 27.3598441 | 0.473984903 | 0.24840403 | 1.90812081 | 0.0563756 | 0.35564709 |
| RBM4 | 22.5040051 | 0.433383 | 0.30492172 | 1.42129265 | 0.1552317 | 0.54412824 |
| C11orf80 | 21.678035 | 1.247549942 | 0.33791303 | 3.69192612 | 0.00022256 | 0.01410008 |
| KDM2A | 287.217717 | 0.00701566 | 0.15298191 | 0.04585941 | 0.96342231 | 0.99231823 |
| GRK2 | 22.5463541 | -0.063819811 | 0.26345024 | -0.2422462 | 0.80858942 | 0.95936772 |
| ANKRD13D | 30.0538293 | 0.008493748 | 0.29218501 | 0.02906976 | 0.97680895 | 0.99384868 |
| RAD9A | 49.8082876 | 0.241078971 | 0.23326104 | 1.03351581 | 0.30136258 | 0.69253083 |
| PPP1CA | 25.3268216 | -0.05182938 | 0.38837631 | -0.1334514 | 0.89383636 | 0.98023547 |
| CDK2AP2 | 19.3584383 | -0.501044518 | 0.39462156 | -1.2696836 | 0.20419736 | 0.60367992 |
| GSTP1 | 120.97753 | -0.486808173 | 0.28763908 | -1.6924271 | 0.09056457 | 0.43960307 |
| NDUFV1 | 31.4339257 | 0.389376951 | 0.26351479 | 1.47762848 | 0.13950725 | 0.52576347 |
| NDUFS8 | 27.1344952 | -0.537378206 | 0.3092026 | -1.7379485 | 0.08221988 | 0.42256146 |
| CHKA | 37.4701265 | -0.482729443 | 0.23534817 | -2.0511289 | 0.04025439 | 0.30130305 |
| KMT5B | 97.0305201 | 0.491608741 | 0.28871454 | 1.70275018 | 0.08861483 | 0.43544541 |
| LRP5 | 42.7770459 | 0.262959786 | 0.40640795 | 0.64703406 | 0.5176099 | 0.83277421 |
| PPP6R3 | 339.426685 | 0.083166112 | 0.21513692 | 0.38657294 | 0.6990724 | 0.92769314 |
| CPT1A | 66.9810829 | 1.024992981 | 0.37678849 | 2.72034044 | 0.00652147 | 0.10992273 |
| MRGPRF | 23.0440787 | 0.294366518 | 0.42125411 | 0.6987861 | 0.48468571 | 0.81546596 |
| TPCN2 | 26.5605142 | 0.399837845 | 0.28456676 | 1.40507573 | 0.15999876 | 0.5530386 |
| CCND1 | 20.5015555 | 1.022838996 | 0.41384187 | 2.47156964 | 0.01345213 | 0.16974928 |
| ORAOV1 | 17.2643674 | 0.345400881 | 0.31514099 | 1.09602016 | 0.27306995 | 0.6682621 |
| ANO1 | 70.5136957 | 0.721602579 | 0.38992019 | 1.85064176 | 0.06422111 | 0.37847708 |
| PPFIA1 | 117.236635 | -0.317127941 | 0.2147588 | -1.4766703 | 0.13976403 | 0.52592211 |
| AP002336.2 | 38.8736255 | -0.372472486 | 0.27739094 | -1.3427709 | 0.17934616 | 0.57744792 |
| CTTN | 92.5062281 | -0.198697154 | 0.23522314 | -0.8447177 | 0.39826847 | 0.7611823 |
| AP002495.2 | 42.0982303 | 0.318113311 | 0.24872225 | 1.27899015 | 0.20090053 | 0.59975737 |
| RNF121 | 33.449295 | 0.147387536 | 0.24146568 | 0.6103871 | 0.54160541 | 0.84641877 |
| NUMA1 | 78.4426636 | 0.14047885 | 0.24169238 | 0.58122996 | 0.56108548 | 0.85477139 |
| LAMTOR1 | 33.2761052 | -0.657486841 | 0.27567829 | -2.3849786 | 0.01708011 | 0.19063814 |
| INPPL1 | 24.2660922 | 0.476410382 | 0.25850124 | 1.84297134 | 0.0653332 | 0.38095778 |
| CLPB | 23.9380256 | -0.148498483 | 0.36412564 | -0.4078221 | 0.68340429 | 0.91933701 |
| ARAP1 | 33.4197741 | 0.373460224 | 0.28072429 | 1.33034526 | 0.18340454 | 0.58297577 |
| ATG16L2 | 23.7650057 | 0.411133116 | 0.30466228 | 1.34947167 | 0.17718551 | 0.57388115 |
| FCHSD2 | 465.988037 | 0.273625277 | 0.29822195 | 0.91752226 | 0.35886903 | 0.73218448 |
| ARHGEF17 | 180.532353 | 0.678675235 | 0.27048877 | 2.50906995 | 0.01210495 | 0.15829055 |
| FAM168A | 135.261337 | -0.2743257 | 0.23293706 | -1.1776817 | 0.23892354 | 0.63433855 |
| RAB6A | 145.652924 | 0.041082755 | 0.15926667 | 0.25794948 | 0.79644589 | 0.95463848 |
| COA4 | 18.6943809 | -1.139422117 | 0.47974862 | -2.3750399 | 0.01754705 | 0.19333326 |
| PAAF1 | 25.8633633 | 0.093447723 | 0.34693764 | 0.2693502 | 0.7876602 | 0.95129454 |
| C2CD3 | 45.5115083 | -0.269155411 | 0.22063684 | -1.2199024 | 0.22250188 | 0.61772146 |
| PPME1 | 81.4605976 | -0.776762205 | 0.34967833 | -2.2213621 | 0.02632644 | 0.24214398 |
| PGM2L1 | 25.2758438 | -0.151484908 | 0.3904978 | -0.3879277 | 0.69806955 | 0.92737958 |
| POLD3 | 23.5633976 | 0.133245557 | 0.3161259 | 0.42149522 | 0.6733935 | 0.91505229 |
| RNF169 | 76.2329925 | -0.095002531 | 0.17700938 | -0.536709 | 0.59146862 | 0.87229579 |
| XRRA1 | 85.5210346 | -0.071476498 | 0.48810177 | -0.1464377 | 0.88357587 | 0.97782528 |
| SPCS2 | 62.7229534 | -0.499236988 | 0.27372502 | -1.8238632 | 0.06817276 | 0.38897639 |
| ARRB1 | 58.2381667 | 0.405460315 | 0.29683848 | 1.36592909 | 0.17196122 | 0.56874123 |
| RPS3 | 159.154318 | -0.30429733 | 0.32152256 | -0.946426 | 0.34393133 | 0.72261198 |
| SERPINH1 | 81.7769222 | -0.276377331 | 0.43204152 | -0.6397009 | 0.5223671 | 0.8350836 |
| UVRAG | 224.999229 | 0.009426758 | 0.18825432 | 0.05007459 | 0.96006295 | 0.99181157 |
| THAP12 | 18.5773386 | 0.13079665 | 0.26288362 | 0.49754584 | 0.61880419 | 0.88593086 |
| EMSY | 115.785813 | -0.387074989 | 0.18806069 | -2.0582451 | 0.03956661 | 0.30069808 |
| LRRC32 | 74.2776323 | -0.118002415 | 0.42766336 | -0.2759236 | 0.78260674 | 0.95014325 |
| TSKU | 23.5048876 | -0.414905484 | 0.50398251 | -0.8232537 | 0.41036371 | 0.77166383 |
| ACER3 | 106.717077 | 0.26582338 | 0.26429855 | 1.00576933 | 0.31452654 | 0.70282967 |
| PAK1 | 32.0441026 | 0.385510076 | 0.24582459 | 1.56823237 | 0.11682691 | 0.48989475 |
| CLNS1A | 47.8344069 | 0.029784929 | 0.2622403 | 0.11357876 | 0.90957172 | 0.98343817 |
| RSF1 | 176.85402 | -0.140377555 | 0.15308812 | -0.9169722 | 0.35915719 | 0.73256903 |
| AAMDC | 48.3575073 | 0.340451183 | 0.26961319 | 1.26273935 | 0.20668287 | 0.60538557 |
| INTS4 | 38.6130023 | 0.125743971 | 0.23719922 | 0.53011967 | 0.59602896 | 0.87481862 |
| ALG8 | 23.2749936 | -0.447889511 | 0.33246337 | -1.3471846 | 0.1779208 | 0.57526987 |
| GAB2 | 278.674716 | -0.392377458 | 0.26301397 | -1.4918503 | 0.1357384 | 0.51955288 |
| AP003086.2 | 70.8965166 | -0.650327812 | 0.4806953 | -1.3528899 | 0.17609081 | 0.57311399 |
| NARS2 | 73.0950092 | -0.000374421 | 0.30826623 | -0.0012146 | 0.99903089 | 0.99969762 |
| TENM4 | 34.0042978 | -0.336217916 | 0.49085668 | -0.6849615 | 0.49336823 | 0.81908904 |
| PRCP | 30.8212245 | -0.192839885 | 0.21063354 | -0.9155232 | 0.35991704 | 0.73273991 |
| RAB30 | 40.6068522 | 0.289417488 | 0.26255911 | 1.10229459 | 0.27033362 | 0.6651094 |
| RAB30-AS1 | 47.6394374 | 0.22104085 | 0.30286691 | 0.72982833 | 0.46549513 | 0.80355051 |
| PCF11 | 60.0704299 | -0.006050825 | 0.21682504 | -0.0279065 | 0.97773674 | 0.99407008 |
| ANKRD42 | 23.2979305 | -0.029488706 | 0.27826206 | -0.1059746 | 0.91560252 | 0.98477389 |
| DLG2 | 789.360324 | 0.182636943 | 0.38043622 | 0.48007244 | 0.63117589 | 0.89436221 |
| CREBZF | 64.847651 | 0.024429445 | 0.24682725 | 0.09897386 | 0.92115903 | 0.98528826 |
| SYTL2 | 178.54137 | 0.937787371 | 0.51282285 | 1.82867704 | 0.067448 | 0.38712255 |
| PICALM | 587.083034 | 0.3006044 | 0.30185621 | 0.99585296 | 0.3193216 | 0.70683013 |
| EED | 27.2854387 | 0.123243746 | 0.30190687 | 0.40821776 | 0.68311381 | 0.91933701 |
| HIKESHI | 32.9314507 | 0.035607744 | 0.24228651 | 0.14696544 | 0.8831593 | 0.97782528 |
| ME3 | 58.1689695 | 0.595249613 | 0.28272573 | 2.10539594 | 0.03525686 | 0.28693537 |
| PRSS23 | 177.192132 | 1.362543787 | 0.37607572 | 3.62305709 | 0.00029114 | 0.01671562 |
| AP001528.2 | 39.1590925 | 0.694314672 | 0.55142209 | 1.25913466 | 0.2079817 | 0.6059796 |
| FZD4 | 30.8322357 | -0.050514131 | 0.39211724 | -0.128824 | 0.89749688 | 0.98040427 |
| TMEM135 | 180.278192 | -0.258067012 | 0.29454431 | -0.8761568 | 0.38094478 | 0.74991898 |
| CTSC | 58.2738987 | -0.947288271 | 0.53131255 | -1.7829209 | 0.07459918 | 0.4046869 |
| NOX4 | 88.9287538 | 0.202170699 | 0.44709222 | 0.45219015 | 0.651132 | 0.9050963 |
| NAALAD2 | 31.4369327 | 1.034942349 | 0.62987019 | 1.6431042 | 0.10036138 | 0.45949178 |
| CHORDC1 | 34.7380399 | 0.075680351 | 0.24154011 | 0.31332415 | 0.7540344 | 0.94129418 |
| FAT3 | 44.7629716 | 1.319872674 | 0.50593008 | 2.60880452 | 0.00908591 | 0.13367648 |
| SLC36A4 | 40.9805941 | -0.595287683 | 0.29393759 | -2.025218 | 0.04284499 | 0.31175172 |
| CEP295 | 31.6912006 | -0.150732783 | 0.30454985 | -0.4949363 | 0.62064507 | 0.8870324 |
| TAF1D | 98.3771573 | -0.399475479 | 0.24542567 | -1.6276841 | 0.10359188 | 0.4637638 |
| PANX1 | 21.7082876 | -1.422482302 | 0.44592566 | -3.1899539 | 0.00142295 | 0.04668435 |
| MRE11 | 35.8637994 | -0.279586045 | 0.2381368 | -1.1740564 | 0.24037242 | 0.63451757 |
| AMOTL1 | 99.3416723 | 0.093384373 | 0.21918872 | 0.42604552 | 0.67007468 | 0.91445841 |
| CWC15 | 20.9512492 | 0.070119427 | 0.25070597 | 0.27968791 | 0.77971696 | 0.9490129 |
| SRSF8 | 19.260919 | -0.248762045 | 0.3333173 | -0.746322 | 0.4554729 | 0.79609632 |
| ENDOD1 | 48.904123 | 0.18049854 | 0.29386918 | 0.61421391 | 0.53907399 | 0.8449569 |
| SESN3 | 71.4050086 | 0.614388028 | 0.38409485 | 1.5995737 | 0.10969319 | 0.47680173 |
| FAM76B | 25.4065743 | 0.104231091 | 0.24325929 | 0.42847733 | 0.66830363 | 0.9143573 |
| CEP57 | 46.8705386 | -0.215303317 | 0.22258746 | -0.9672751 | 0.33340651 | 0.71622463 |
| MTMR2 | 84.1086725 | 0.206636482 | 0.16825796 | 1.22809339 | 0.21941191 | 0.61567703 |

| | | | | | | |
|---|---|---|---|---|---|---|
| MAML2 | 1277.90411 | -0.27215055 | 0.38770729 | -0.7019485 | 0.48271129 | 0.81409616 |
| CCDC82 | 51.4109112 | -0.379273409 | 0.22247154 | -1.7048177 | 0.08822843 | 0.43544541 |
| CNTN5 | 142.031976 | 1.378551276 | 0.5798021 | 2.37762381 | 0.01742459 | 0.19325071 |
| ARHGAP42 | 483.909382 | 0.181093256 | 0.30754962 | 0.58882613 | 0.55597791 | 0.85207344 |
| PGR | 51.5028643 | 0.446111015 | 0.27724502 | 1.60908576 | 0.10759759 | 0.47265636 |
| TRPC6 | 43.4211499 | 1.172208355 | 0.39414711 | 2.9740377 | 0.00293909 | 0.06945135 |
| CEP126 | 49.6364524 | 0.046675266 | 0.29604686 | 0.15766175 | 0.87472334 | 0.9768643 |
| YAP1 | 420.44393 | 0.058542565 | 0.27240509 | 0.21490995 | 0.82983753 | 0.96580172 |
| BIRC3 | 27.2794122 | -1.151653099 | 0.81064005 | -1.4206714 | 0.15541232 | 0.54412824 |
| BIRC2 | 50.3794644 | -0.208102906 | 0.24311957 | -0.8559694 | 0.39201472 | 0.75901355 |
| TMEM123 | 73.4507625 | -0.280592609 | 0.23789665 | -1.1794727 | 0.23820999 | 0.63385933 |
| DCUN1D5 | 26.4805202 | -0.209161937 | 0.26005274 | -0.8043058 | 0.42122037 | 0.77771095 |
| DYNC2H1 | 178.532014 | -0.35381282 | 0.31141101 | -1.1361603 | 0.25588949 | 0.65070308 |
| PDGFD | 75.5724256 | -0.457669627 | 0.50052472 | -0.9143797 | 0.36051738 | 0.73273991 |
| CASP4 | 43.5972219 | -0.932503007 | 0.35740824 | -2.6090697 | 0.00907888 | 0.13367648 |
| AASDHPPT | 35.0171923 | -0.01709525 | 0.27591002 | -0.0619595 | 0.95059507 | 0.99060401 |
| GUCY1A2 | 354.452111 | 0.413666922 | 0.45226338 | 0.91465933 | 0.3603705 | 0.73273991 |
| CWF19L2 | 79.9534237 | 0.113215259 | 0.19547304 | 0.57918606 | 0.56246363 | 0.85545224 |
| CUL5 | 120.930673 | 0.302402987 | 0.13779593 | 2.19457125 | 0.02819437 | 0.25237565 |
| ACAT1 | 43.9247205 | 0.080112642 | 0.24862121 | 0.3222277 | 0.7472802 | 0.93903859 |
| NPAT | 45.2772571 | -0.501125674 | 0.2769836 | -1.8092251 | 0.07041604 | 0.39563263 |
| ATM | 131.466482 | -0.176554293 | 0.21856728 | -0.8077801 | 0.41921721 | 0.77771095 |
| KDELC2 | 29.1663752 | 0.33443969 | 0.28435807 | 1.17612166 | 0.23954626 | 0.63451757 |
| DDX10 | 120.887226 | -0.392217909 | 0.31323037 | -1.2521708 | 0.21050765 | 0.60731347 |
| RDX | 86.147956 | 0.00192404 | 0.21687052 | 0.00887184 | 0.99292139 | 0.99767286 |
| ZC3H12C | 36.5836976 | -0.761189905 | 0.35789396 | -2.1268588 | 0.03343182 | 0.2790477 |
| FDX1 | 30.0935612 | -0.025650308 | 0.35577933 | -0.0720961 | 0.94252542 | 0.98951704 |
| ARHGAP20 | 41.2047227 | 1.653641395 | 0.56589431 | 2.92217356 | 0.00347598 | 0.07625361 |
| SIK2 | 326.348444 | -0.062614043 | 0.40181875 | -0.1558266 | 0.87616972 | 0.9768643 |
| PPP2R1B | 75.689688 | 0.293265213 | 0.3188397 | 0.91978888 | 0.35768309 | 0.73137668 |
| CRYAB | 305.426719 | -1.49140569 | 0.60115321 | -2.4809078 | 0.01310483 | 0.1672003 |
| DIXDC1 | 47.5620707 | 0.391114038 | 0.37453642 | 1.04426171 | 0.29636432 | 0.68771122 |
| PTS | 57.0164666 | -0.050435883 | 0.2414032 | -0.208928 | 0.83450446 | 0.9666365 |
| AP002884.1 | 24.9892118 | -0.170731787 | 0.35741496 | -0.4776851 | 0.63287437 | 0.89511043 |
| NCAM1 | 71.3185023 | 0.695745494 | 0.76695673 | 0.90715091 | 0.364327 | 0.73648283 |
| USP28 | 27.6770757 | -0.290455357 | 0.32110546 | -0.9045482 | 0.3657048 | 0.73753145 |
| ZBTB16 | 726.139574 | -0.826802941 | 0.71342009 | -1.1589286 | 0.24648529 | 0.64225758 |
| AP002518.2 | 88.78272 | -1.939742034 | 0.88070438 | -2.2024894 | 0.02763076 | 0.24884614 |
| NNMT | 258.528705 | -1.977262135 | 0.47775607 | -4.1386437 | 3.49E-05 | 0.0042566 |
| REXO2 | 36.6741009 | -0.074445556 | 0.48822589 | -0.1524818 | 0.87880696 | 0.97720568 |
| CADM1 | 29.2923111 | 0.768908339 | 0.36920865 | 2.08258485 | 0.03728908 | 0.29301102 |
| SIK3 | 605.827356 | 0.176748943 | 0.2997969 | 0.58956227 | 0.55548415 | 0.85207344 |
| PAFAH1B2 | 55.042108 | -0.238677115 | 0.21103663 | -1.1309748 | 0.25806568 | 0.65130862 |
| SIDT2 | 41.4128882 | 0.147652976 | 0.34468409 | 0.42837189 | 0.66838038 | 0.9143573 |
| TAGLN | 922.468189 | 0.381080773 | 0.39680696 | 0.96036817 | 0.33686995 | 0.71924965 |
| PCSK7 | 83.294581 | 0.051473053 | 0.22813196 | 0.22562841 | 0.82149044 | 0.96303704 |
| RNF214 | 31.7867671 | 0.113816287 | 0.27092915 | 0.42009613 | 0.67441523 | 0.91562489 |
| BACE1 | 31.9134593 | 0.599542246 | 0.24679044 | 2.42935763 | 0.0151256 | 0.17917181 |
| CEP164 | 36.6850025 | -0.015368144 | 0.20161247 | -0.0762262 | 0.93923917 | 0.98846752 |
| DSCAML1 | 50.1004907 | 0.585276911 | 0.8751143 | 0.66880053 | 0.50362273 | 0.82430366 |
| FXYD2 | 22.1896403 | -0.301111699 | 0.44300968 | -0.6796955 | 0.49669727 | 0.8216376 |
| UBE4A | 50.6648196 | -0.012854175 | 0.20474831 | -0.0627804 | 0.9499414 | 0.99060401 |
| ATP5MG | 56.3544929 | -0.205059693 | 0.28281423 | -0.7250685 | 0.46840999 | 0.80374006 |
| KMT2A | 187.098235 | 0.206453279 | 0.15022653 | 1.37427978 | 0.16935484 | 0.56570963 |
| ARCN1 | 43.619663 | -0.211541023 | 0.20778037 | -1.0180992 | 0.30863082 | 0.69885506 |
| PHLDB1 | 94.7801516 | 0.320881754 | 0.31690042 | 1.01256337 | 0.31126876 | 0.70126123 |
| DDX6 | 110.284056 | -0.075931344 | 0.16624052 | -0.4567559 | 0.64784648 | 0.90256375 |
| BCL9L | 39.3179923 | -0.072204458 | 0.3342888 | -0.2159942 | 0.82899223 | 0.96580172 |
| CCDC84 | 62.1751298 | 0.156414492 | 0.27850487 | 0.56162211 | 0.57437351 | 0.86294056 |
| RPS25 | 102.894484 | -0.569904696 | 0.32646762 | -1.7456699 | 0.08086833 | 0.42029797 |
| TRAPPC4 | 20.7159207 | -0.129186997 | 0.30350512 | -0.4256501 | 0.6703628 | 0.91445841 |
| HYOU1 | 19.2087095 | -0.234420485 | 0.46456394 | -0.5046033 | 0.61383751 | 0.88360697 |
| CBL | 76.2354118 | 0.465230484 | 0.2288433 | 2.03296529 | 0.04205603 | 0.30845286 |
| MCAM | 187.43115 | 0.264206844 | 0.43026526 | 0.61405572 | 0.53917851 | 0.8449569 |
| THY1 | 22.3719653 | 1.02772884 | 0.92974127 | 1.1053923 | 0.26898964 | 0.66469564 |
| OAF | 42.141301 | 0.367778753 | 0.41932453 | 0.87707427 | 0.3804463 | 0.74991898 |
| ARHGEF12 | 353.807194 | -0.040508133 | 0.22250882 | -0.1820518 | 0.85554207 | 0.97379567 |
| SORL1 | 32.3175468 | 0.890746097 | 0.39520875 | 2.2538623 | 0.02420483 | 0.23086389 |
| UBASH3B | 28.0266075 | 0.243484445 | 0.52118945 | 0.46717071 | 0.64037775 | 0.89914713 |
| HSPA8 | 81.2215549 | 0.440358348 | 0.33004949 | 1.33421913 | 0.18213205 | 0.58094117 |
| CLMP | 205.520654 | -0.179920659 | 0.59170841 | -0.3040698 | 0.76107471 | 0.94425408 |
| TBRG1 | 43.3511662 | -0.324058662 | 0.24547673 | -1.3201197 | 0.18679506 | 0.58501647 |
| SIAE | 20.4171373 | 0.495035255 | 0.27260638 | 1.81593419 | 0.06938047 | 0.39251505 |
| NRGN | 39.5830232 | 1.854078172 | 0.4744314 | 3.90800057 | 9.31E-05 | 0.00759912 |
| ESAM | 21.6919576 | 0.819609095 | 0.29106871 | 2.81586119 | 0.00486467 | 0.09203696 |
| MSANTD2 | 43.8398798 | 0.485237063 | 0.22762517 | 2.13173725 | 0.03302845 | 0.27740126 |
| PKNOX2 | 78.2402783 | 0.238185415 | 0.35899574 | 0.663477 | 0.50702511 | 0.82481485 |
| EI24 | 26.6773162 | -0.118438606 | 0.35094895 | -0.337481 | 0.73575433 | 0.93594619 |
| STT3A | 29.3427542 | -0.157413417 | 0.23713531 | -0.6638126 | 0.50681023 | 0.82481485 |
| FAM118B | 19.9289088 | 0.095248384 | 0.27421841 | 0.34734496 | 0.72833218 | 0.93455731 |
| SRPRA | 27.6511075 | -0.35176276 | 0.31708462 | -1.1093656 | 0.26727251 | 0.66201068 |
| ST3GAL4 | 38.4613454 | -0.450192103 | 0.41021108 | -1.0974645 | 0.27243839 | 0.6673044 |
| ETS1 | 207.199097 | 0.234104782 | 0.26023955 | 0.89957418 | 0.3683469 | 0.74102967 |
| FLI1 | 167.09341 | 0.298040518 | 0.31399194 | 0.94919798 | 0.34251993 | 0.72125471 |
| ARHGAP32 | 156.920152 | -0.110211773 | 0.24979591 | -0.4412073 | 0.65906295 | 0.90831499 |
| NFRKB | 20.8260188 | 0.238067757 | 0.25175485 | 0.94563326 | 0.34433568 | 0.72280002 |
| PRDM10 | 28.0277945 | -0.137317737 | 0.27498018 | -0.4993732 | 0.61751648 | 0.88534783 |
| APLP2 | 127.156547 | 0.047948465 | 0.17583205 | 0.27269468 | 0.78508792 | 0.95065355 |
| ZBTB44 | 71.3831959 | 0.041283325 | 0.24360523 | 0.16946814 | 0.86542843 | 0.9762143 |
| SNX19 | 56.4947794 | -0.058021404 | 0.27300393 | -0.2125296 | 0.83169392 | 0.96634287 |
| NTM | 72.2353422 | 0.60006085 | 0.43424065 | 1.38186247 | 0.16701393 | 0.56278128 |
| JAM3 | 68.0997959 | 0.206029761 | 0.22392836 | 0.92006999 | 0.35753618 | 0.73129235 |
| VPS26B | 21.7179588 | 0.436984176 | 0.30467331 | 1.43427129 | 0.15149486 | 0.53898138 |
| ZMYND11 | 134.992885 | 0.069379124 | 0.17117521 | 0.40531059 | 0.68524922 | 0.91977784 |
| DIP2C | 304.218118 | -0.731299937 | 0.27723913 | -2.6377948 | 0.00834471 | 0.12814321 |
| LARP4B | 146.824383 | -0.190029604 | 0.19639374 | -0.967595 | 0.33324666 | 0.71622463 |
| GTPBP4 | 26.9594026 | -0.500729396 | 0.40071881 | -1.249578 | 0.21145375 | 0.60910715 |
| IDI1 | 20.5859142 | 0.012244826 | 0.36583337 | 0.03347105 | 0.97329895 | 0.99300191 |
| WDR37 | 52.2677647 | -0.045614006 | 0.1835683 | -0.2484852 | 0.80375901 | 0.95836189 |
| PITRM1 | 36.9498656 | -0.581077471 | 0.32410908 | -1.7928454 | 0.07299765 | 0.40164935 |
| KLF6 | 108.286999 | 0.522703509 | 0.36965117 | 1.41404534 | 0.15734859 | 0.5482953 |
| NET1 | 36.079053 | 0.516075082 | 0.41201589 | 1.25256111 | 0.21036548 | 0.60731347 |
| FAM208B | 169.686192 | -0.297686078 | 0.25960214 | -1.1467012 | 0.25150514 | 0.6460368 |
| GDI2 | 92.5104489 | -0.252275789 | 0.24300413 | -1.0381543 | 0.29919821 | 0.69014677 |
| ANKRD16 | 25.5276795 | -0.014397275 | 0.36011606 | -0.0399795 | 0.96810944 | 0.99300191 |
| FBH1 | 85.1573137 | -0.540298403 | 0.24601735 | -2.1961801 | 0.02807906 | 0.25226528 |
| IL15RA | 29.5659737 | -1.875640573 | 0.58298636 | -3.2172975 | 0.00129404 | 0.04375316 |
| RBM17 | 63.8397162 | -0.429495838 | 0.31475681 | -1.3645323 | 0.17240009 | 0.56904425 |
| PFKFB3 | 94.0232933 | -0.463733268 | 0.314743 | -1.4733712 | 0.14065097 | 0.52710045 |
| SFMBT2 | 194.625592 | -1.045694225 | 0.50973045 | -2.0514651 | 0.04022168 | 0.30130305 |
| ITIH5 | 64.3000219 | 1.090506968 | 0.4796506 | 2.27354445 | 0.02299339 | 0.22440695 |
| KIN | 17.6222782 | 0.015591953 | 0.3034785 | 0.05137745 | 0.95902475 | 0.99181157 |
| ATP5F1C | 50.2962989 | -0.197509742 | 0.23283924 | -0.8482666 | 0.39628954 | 0.76020146 |

| | | | | | | |
|---|---|---|---|---|---|---|
| TAF3 | 124.835425 | -0.087204546 | 0.17905017 | -0.4870397 | 0.62623017 | 0.8912584 |
| GATA3 | 85.0348857 | 1.214380653 | 0.58718335 | 2.06814559 | 0.03862633 | 0.29724077 |
| CELF2 | 106.657705 | 1.247401536 | 0.39631779 | 3.14747803 | 0.00164685 | 0.05093736 |
| USP6NL | 80.7872175 | 0.062523659 | 0.2490223 | 0.25107654 | 0.80175493 | 0.95758343 |
| PROSER2 | 31.7851345 | -0.721033731 | 0.59201569 | -1.2179301 | 0.22325054 | 0.61772146 |
| UPF2 | 75.3846713 | 0.11424547 | 0.16680915 | 0.68488729 | 0.49341505 | 0.81908904 |
| DHTKD1 | 28.2235571 | 0.503836144 | 0.26981828 | 1.86731659 | 0.06185739 | 0.37018726 |
| SEC61A2 | 17.6009044 | 0.330838749 | 0.31771527 | 1.04130578 | 0.29773366 | 0.68849737 |
| NUDT5 | 31.0662358 | -0.590383446 | 0.25700039 | -2.2972083 | 0.02160689 | 0.21698983 |
| CDC123 | 32.7052637 | -0.40599483 | 0.2627237 | -1.54533 | 0.12226645 | 0.50085628 |
| CAMK1D | 238.388078 | 0.176270535 | 0.35349826 | 0.49864612 | 0.61802871 | 0.88534783 |
| CCDC3 | 72.9625333 | 1.22815987 | 0.4844628 | 2.53509636 | 0.01124164 | 0.15130918 |
| OPTN | 70.7572532 | -0.185405465 | 0.26851771 | -0.6904776 | 0.48989389 | 0.81720854 |
| BEND7 | 35.7525127 | -0.01402868 | 0.32595657 | -0.0430385 | 0.96567085 | 0.99293098 |
| AL157392.5 | 22.3722729 | -0.021738081 | 0.2796095 | -0.0777444 | 0.93803135 | 0.98827804 |
| FRMD4A | 367.718117 | 0.561559123 | 0.37501009 | 1.49745071 | 0.13427602 | 0.51662128 |
| FAM107B | 89.8367456 | 0.626003455 | 0.30619617 | 2.04445227 | 0.04090889 | 0.30336974 |
| DCLRE1C | 22.2154861 | 0.167638468 | 0.26402822 | 0.63492634 | 0.52547646 | 0.83768471 |
| RPP38 | 28.4115295 | 0.124346589 | 0.30528826 | 0.40730878 | 0.68378121 | 0.91950744 |
| NMT2 | 34.5795742 | 0.839038343 | 0.28408487 | 2.95347776 | 0.00314215 | 0.0721615 |
| FAM171A1 | 34.2534267 | 0.416177282 | 0.29717249 | 1.40045694 | 0.16137653 | 0.55470404 |
| ITGA8 | 345.971714 | 1.023727661 | 0.50753965 | 2.01703977 | 0.04369136 | 0.31491241 |
| MINDY3 | 81.6954387 | -0.225435681 | 0.18667475 | -1.2076388 | 0.22718622 | 0.62173063 |
| RSU1 | 167.921863 | 0.248731521 | 0.2981189 | 0.83433663 | 0.40409132 | 0.76597036 |
| CUBN | 47.8603239 | 1.311714507 | 0.51577753 | 2.54317884 | 0.0109849 | 0.14812481 |
| VIM | 332.103477 | 0.35174658 | 0.34055697 | 1.03285681 | 0.30167092 | 0.69264368 |
| STAM | 44.0021957 | -0.244855028 | 0.19988852 | -1.224958 | 0.22059106 | 0.61590675 |
| CACNB2 | 430.344468 | -0.237022076 | 0.33729749 | -0.7027093 | 0.48223694 | 0.81402096 |
| AL450384.2 | 24.6249509 | -0.203734803 | 0.30652185 | -0.6646665 | 0.50626384 | 0.82481485 |
| NSUN6 | 48.8215928 | -0.226038144 | 0.2042783 | -1.1065206 | 0.26850126 | 0.66393532 |
| ARL5B | 28.1686121 | -1.108733291 | 0.45817948 | -2.4198668 | 0.01552619 | 0.18198087 |
| MALRD1 | 46.9735871 | 0.306127004 | 0.82912412 | 0.36921734 | 0.71196573 | 0.93083724 |
| PLXDC2 | 641.887814 | 0.722933487 | 0.38192072 | 1.89288889 | 0.05837265 | 0.36041506 |
| NEBL | 47.8149825 | 0.316897515 | 0.82430191 | 0.3844435 | 0.70064976 | 0.92843042 |
| MLLT10 | 237.765875 | -0.011221492 | 0.21926876 | -0.0511769 | 0.95918457 | 0.99181157 |
| DNAJC1 | 257.837377 | -0.517585667 | 0.25224394 | -2.051925 | 0.04017695 | 0.30130305 |
| PIP4K2A | 259.545264 | 0.280963134 | 0.27710913 | 1.0139079 | 0.3106267 | 0.70110431 |
| MSRB2 | 29.4577208 | -0.01015522 | 0.26969762 | -0.0376541 | 0.96996348 | 0.99300191 |
| KIAA1217 | 997.648772 | 1.107131643 | 0.3955316 | 2.79909784 | 0.00512456 | 0.09368258 |
| ARHGAP21 | 195.359834 | 0.476731068 | 0.17880328 | 2.66623218 | 0.00767067 | 0.12228146 |
| ABI1 | 127.613186 | 0.010584316 | 0.26664124 | 0.03969497 | 0.96833631 | 0.99300191 |
| ANKRD26 | 67.6749833 | -0.306997438 | 0.2232498 | -1.3751297 | 0.16909124 | 0.56570963 |
| YME1L1 | 91.4907974 | -0.157157678 | 0.18090087 | -0.8687503 | 0.38498373 | 0.75305975 |
| ACBD5 | 22.4059401 | -0.079496989 | 0.27595222 | -0.2880824 | 0.77328363 | 0.94730145 |
| RAB18 | 72.3738421 | -0.187567995 | 0.19033472 | -0.9854639 | 0.32439625 | 0.71100151 |
| MPP7 | 35.4892082 | 1.228066174 | 0.48684707 | 2.52248859 | 0.01165277 | 0.15416382 |
| WAC-AS1 | 21.3256372 | 0.123393433 | 0.38054912 | 0.324251 | 0.74574802 | 0.93891908 |
| WAC | 314.626183 | -0.170567127 | 0.18300154 | -0.932053 | 0.35130915 | 0.72936942 |
| SVIL | 244.773624 | 0.0408235 | 0.4307443 | 0.09477432 | 0.92449408 | 0.98561736 |
| JCAD | 19.4751754 | -0.157044757 | 0.3612995 | -0.4346664 | 0.66380458 | 0.91141325 |
| MTPAP | 19.893444 | -0.234070396 | 0.26671917 | -0.8775912 | 0.3801656 | 0.74991898 |
| MAP3K8 | 98.6666914 | -0.851455316 | 0.36694669 | -2.3203788 | 0.02032039 | 0.21050622 |
| ZNF438 | 78.4738645 | 0.049680598 | 0.26841527 | 0.18508857 | 0.85315957 | 0.97312893 |
| ZEB1-AS1 | 26.3774254 | 0.734862554 | 0.28371798 | 2.59011622 | 0.00959435 | 0.1379822 |
| ZEB1 | 400.716102 | -0.036124667 | 0.23498066 | -0.1537346 | 0.87781897 | 0.9768643 |
| ARHGAP12 | 90.6107693 | 0.258657093 | 0.31719424 | 0.81545332 | 0.41481287 | 0.77476594 |
| KIF5B | 114.816492 | 0.220009434 | 0.19273555 | 1.14150938 | 0.25365801 | 0.64816854 |
| EPC1 | 209.242693 | 0.266646811 | 0.189221 | 1.40918193 | 0.15878138 | 0.55091387 |
| CCDC7 | 89.8115108 | 0.380895583 | 0.25211519 | 1.51079983 | 0.13083946 | 0.51290135 |
| ITGB1 | 338.444287 | 0.381627535 | 0.20866723 | 1.82888105 | 0.06741743 | 0.38712255 |
| NRP1 | 979.043201 | -0.301142972 | 0.35012552 | -0.8601 | 0.38973391 | 0.75631224 |
| PARD3 | 871.855193 | 0.015472692 | 0.31432196 | 0.04922562 | 0.9607395 | 0.99181157 |
| CUL2 | 42.1816427 | 0.132609253 | 0.19619165 | 0.6759169 | 0.4990934 | 0.82271448 |
| CREM | 95.8783439 | 0.036385091 | 0.28196831 | 0.12903965 | 0.89732628 | 0.98040427 |
| CCNY | 247.768935 | 0.107615444 | 0.15188986 | 0.70850973 | 0.47862877 | 0.81159272 |
| AL117336.3 | 19.6518317 | -0.039558528 | 0.42897793 | -0.0922158 | 0.92652661 | 0.98677279 |
| ZNF248 | 61.6188829 | 0.129307559 | 0.22904244 | 0.5645572 | 0.57237498 | 0.86236719 |
| ZNF25 | 19.937468 | 0.243040065 | 0.26386231 | 0.9210867 | 0.35700516 | 0.73129235 |
| ZNF33A | 77.7603401 | -0.044557743 | 0.28455733 | -0.1565862 | 0.875571 | 0.9768643 |
| ZNF33B | 29.18472 | 0.811033797 | 0.39394608 | 2.05874316 | 0.03951885 | 0.30064598 |
| BMS1 | 38.8713449 | 0.102220427 | 0.25545106 | 0.40015659 | 0.68904119 | 0.92217934 |
| CSGALNACT2 | 58.195527 | -0.620630911 | 0.19649052 | -3.1585793 | 0.0015854 | 0.05043773 |
| HNRNPF | 66.2934612 | -0.355950983 | 0.2816078 | -1.2639955 | 0.20623165 | 0.6050285 |
| CXCL12 | 89.6951069 | 0.70553154 | 0.28865751 | 2.44418219 | 0.01451809 | 0.17653018 |
| RASSF4 | 23.1087743 | -0.002104526 | 0.27883699 | -0.0075475 | 0.99397801 | 0.99819806 |
| DEPP1 | 103.61596 | -2.160858379 | 0.29199754 | -7.4002622 | 1.36E-13 | 9.99E-10 |
| 8-Mar | 41.1044156 | -0.140504661 | 0.2735791 | -0.5135797 | 0.6075459 | 0.88042838 |
| ZFAND4 | 27.9937868 | 0.103770234 | 0.27349683 | 0.37942025 | 0.70437582 | 0.92936308 |
| WASHC2C | 23.6683547 | -0.076230625 | 0.32611356 | -0.2337548 | 0.81517531 | 0.9608217 |
| AGAP4 | 26.376086 | 0.452612562 | 0.26058073 | 1.73693794 | 0.08239813 | 0.42256146 |
| TIMM23 | 26.7917022 | -0.030557037 | 0.25744577 | -0.1186931 | 0.9055185 | 0.98303609 |
| NCOA4 | 30.9681132 | -0.016347949 | 0.26874071 | -0.0608317 | 0.95149326 | 0.99060401 |
| AGAP9 | 18.5363737 | -0.139609181 | 0.39840544 | -0.3504199 | 0.72602362 | 0.93455731 |
| MAPK8 | 157.44417 | 0.164753027 | 0.1634303 | 1.0080935 | 0.31340958 | 0.70282967 |
| VSTM4 | 50.8702594 | 0.128132278 | 0.40079056 | 0.31969885 | 0.74919663 | 0.94004394 |
| ERCC6 | 41.1821226 | -0.579842935 | 0.33899866 | -1.7104579 | 0.08718123 | 0.43290196 |
| PARG | 87.5632666 | -0.104721679 | 0.16909125 | -0.6193205 | 0.53570524 | 0.84412047 |
| TIMM23B | 81.3266794 | -0.017920779 | 0.29936868 | -0.0598619 | 0.95226562 | 0.99060401 |
| LINC00843 | 20.2008134 | 0.162237713 | 0.36275887 | 0.44723294 | 0.65470688 | 0.90646849 |
| AGAP6 | 28.5867986 | 0.009480401 | 0.29868104 | 0.03174089 | 0.97467869 | 0.99300191 |
| WASHC2A | 20.0400757 | -0.109623781 | 0.40293016 | -0.2720665 | 0.78557092 | 0.95065355 |
| SGMS1 | 165.919283 | 0.203684649 | 0.17317417 | 1.17618377 | 0.23952145 | 0.63451757 |
| PRKG1 | 3030.40969 | 0.610053106 | 0.29728288 | 2.05209632 | 0.0401603 | 0.30130305 |
| IPMK | 34.5868035 | -0.847599837 | 0.32477809 | -2.6097815 | 0.00906001 | 0.13367648 |
| CISD1 | 33.0900144 | -0.790335077 | 0.30246723 | -2.612961 | 0.00897616 | 0.13367648 |
| UBE2D1 | 25.1857936 | -0.364870722 | 0.28131106 | -1.2970365 | 0.19461862 | 0.59272783 |
| BICC1 | 413.89479 | -0.364779643 | 0.2804365 | -1.3007567 | 0.19334176 | 0.59237115 |
| FAM13C | 50.5987518 | 0.349434554 | 0.91366265 | 0.38245468 | 0.70212414 | 0.92865476 |
| CCDC6 | 78.8057108 | 0.157763009 | 0.2902531 | 0.54353601 | 0.58676081 | 0.870249 |
| ANK3 | 283.11314 | 0.421687515 | 0.43289762 | 0.9741045 | 0.33000466 | 0.71362138 |
| RHOBTB1 | 71.5064597 | 0.691614258 | 0.26289318 | 2.63078054 | 0.0085189 | 0.12961784 |
| TMEM26 | 24.7588383 | 0.053707324 | 0.42519949 | 0.12631089 | 0.89948584 | 0.98058672 |
| ARID5B | 485.778531 | -0.365719554 | 0.30687932 | -1.1917374 | 0.23336424 | 0.62956917 |
| JMJD1C | 818.101857 | 0.204694621 | 0.23510664 | 0.87064584 | 0.38394756 | 0.75263553 |
| REEP3 | 124.530132 | 0.153074749 | 0.23220669 | 0.65921766 | 0.50975601 | 0.82715763 |
| CTNNA3 | 83.9379934 | 1.122026027 | 0.30861898 | 3.63563522 | 0.0002773 | 0.0161874 |
| SIRT1 | 39.5610612 | 0.387786751 | 0.26426879 | 1.46739518 | 0.14226859 | 0.52958915 |
| HERC4 | 301.426679 | 0.002295709 | 0.12966965 | 0.01770429 | 0.98587476 | 0.99713647 |
| HNRNPH3 | 86.1079894 | 0.123474469 | 0.24547841 | 0.50299524 | 0.61496762 | 0.88360697 |
| RUFY2 | 76.2147324 | 0.383060051 | 0.22588629 | 1.69580923 | 0.08992201 | 0.43806588 |
| SLC25A16 | 53.8109547 | 0.046270083 | 0.17187612 | 0.269206 | 0.78777116 | 0.95129454 |
| TET1 | 39.6936322 | 0.228819173 | 0.38312732 | 0.59724056 | 0.55034679 | 0.84968456 |
| CCAR1 | 119.749094 | -0.108333778 | 0.15511539 | -0.6984077 | 0.48492227 | 0.81549057 |

| | | | | | | |
|---|---|---|---|---|---|---|
| DDX50 | 56.0528244 | -0.139465401 | 0.24326523 | -0.5733059 | 0.56643756 | 0.85825441 |
| DDX21 | 72.9433048 | -1.368472681 | 0.45379156 | -3.0156416 | 0.00256436 | 0.0643191 |
| SRGN | 152.053365 | -0.241644821 | 0.5368724 | -0.4500973 | 0.65264028 | 0.90646849 |
| VPS26A | 54.2490597 | -0.150094694 | 0.24467968 | -0.6134334 | 0.53958979 | 0.84521947 |
| SUPV3L1 | 19.8391098 | -0.167221797 | 0.27717062 | -0.6033172 | 0.5462977 | 0.84758329 |
| HK1 | 107.830351 | -0.376071855 | 0.31570871 | -1.1911989 | 0.23357553 | 0.62956917 |
| SAR1A | 49.2718103 | -0.3600885 | 0.26306185 | -1.3688359 | 0.17105058 | 0.56870244 |
| PPA1 | 52.8240683 | -1.195804064 | 0.35068901 | -3.4098704 | 0.00064994 | 0.02800329 |
| EIF4EBP2 | 43.2095269 | 0.01266667 | 0.20514374 | 0.06174534 | 0.95076563 | 0.99060401 |
| PALD1 | 22.3473355 | 0.611843039 | 0.39261289 | 1.55838755 | 0.1191414 | 0.49395083 |
| SGPL1 | 51.4711619 | 0.299858909 | 0.2309648 | 1.29828834 | 0.19418827 | 0.59269042 |
| UNC5B | 43.058491 | 0.942967953 | 0.4048657 | 2.32908829 | 0.01985439 | 0.20828536 |
| VSIR | 20.013918 | 0.891793489 | 0.33838057 | 2.63547489 | 0.00840196 | 0.12814321 |
| PSAP | 189.781856 | -0.126245477 | 0.20562218 | -0.6139682 | 0.53923634 | 0.8449569 |
| CHST3 | 27.0116883 | 0.253523301 | 0.41564806 | 0.60994703 | 0.5418969 | 0.84641877 |
| ASCC1 | 74.2529822 | -0.294159279 | 0.23866913 | -1.2324982 | 0.21776302 | 0.61468881 |
| ANAPC16 | 106.449787 | -0.098693131 | 0.24286229 | -0.4063749 | 0.68446719 | 0.91977784 |
| DDIT4 | 131.735072 | -3.024862256 | 0.87761974 | -3.4466661 | 0.00056755 | 0.02567229 |
| DNAJB12 | 22.3371148 | -0.033679312 | 0.31065827 | -0.1084127 | 0.9136683 | 0.98395113 |
| MICU1 | 173.037346 | 0.01675145 | 0.26929035 | 0.06220591 | 0.95039886 | 0.99060401 |
| MCU | 170.685464 | 0.600108436 | 0.40713721 | 1.47397099 | 0.1404894 | 0.52680084 |
| P4HA1 | 228.310755 | -1.147787309 | 0.52279739 | -2.1954725 | 0.02812972 | 0.25237565 |
| ECD | 31.3178043 | -0.319927936 | 0.31864439 | -1.0040281 | 0.31536505 | 0.70306809 |
| FAM149B1 | 29.9601085 | 0.280556693 | 0.34801241 | 0.80616864 | 0.42014562 | 0.77771095 |
| DNAJC9 | 28.4702454 | 0.305512323 | 0.31344977 | 0.97467713 | 0.32972044 | 0.71362138 |
| MRPS16 | 22.7653671 | -0.548418201 | 0.31119626 | -1.7622904 | 0.07802024 | 0.41398608 |
| ANXA7 | 67.0038179 | 0.105493334 | 0.23657036 | 0.44592793 | 0.6556493 | 0.90707205 |
| PPP3CB | 110.515705 | 0.472402854 | 0.22093595 | 2.13818916 | 0.03250139 | 0.27542099 |
| USP54 | 68.4689074 | 0.187997634 | 0.26529982 | 0.7086233 | 0.47855827 | 0.81159272 |
| SEC24C | 34.5334323 | -0.137143978 | 0.2276479 | -0.602439 | 0.54688194 | 0.84758329 |
| FUT11 | 25.1641029 | -0.288931131 | 0.35078021 | -0.8236814 | 0.41012061 | 0.77162734 |
| ZSWIM8 | 44.2803798 | -0.257848664 | 0.22462624 | -1.1479009 | 0.25100948 | 0.6460368 |
| CAMK2G | 49.8063229 | 1.329304674 | 0.44492173 | 2.987727 | 0.0028106 | 0.06794452 |
| VCL | 430.872233 | 0.911315953 | 0.33812764 | 2.69518326 | 0.00703499 | 0.11576428 |
| AP3M1 | 19.0262141 | 0.17212535 | 0.30153664 | 0.57082732 | 0.5681167 | 0.8595777 |
| ADK | 223.693941 | 0.056759774 | 0.22927365 | 0.24756344 | 0.8044722 | 0.95836286 |
| KAT6B | 187.497769 | 0.464199621 | 0.2252476 | 2.06084162 | 0.03931815 | 0.30005099 |
| SAMD8 | 81.675011 | -0.088932543 | 0.22558411 | -0.3942323 | 0.69340954 | 0.9248682 |
| VDAC2 | 31.8125158 | -0.31383947 | 0.25466014 | -1.2323855 | 0.2178051 | 0.61468881 |
| LRMDA | 149.618925 | 0.336360469 | 0.28214878 | 1.19213868 | 0.23320687 | 0.62939306 |
| KCNMA1 | 102.922044 | 2.195404027 | 0.86712896 | 2.53180798 | 0.01134761 | 0.15215018 |
| DLG5 | 88.6975766 | -0.093663342 | 0.24749746 | -0.3784416 | 0.70510255 | 0.92946306 |
| POLR3A | 30.5989436 | -0.291548174 | 0.27236206 | -1.0704434 | 0.28441977 | 0.6783522 |
| RPS24 | 168.1264 | -0.312585781 | 0.29781175 | -1.0496086 | 0.29389809 | 0.68545131 |
| ZMIZ1 | 156.483621 | -0.592639219 | 0.28181036 | -2.1029717 | 0.03546824 | 0.28738272 |
| ZCCHC24 | 79.7014214 | -0.34502524 | 0.20741583 | -1.663447 | 0.09622299 | 0.45127171 |
| NUTM2B-AS | 153.020064 | -0.440200663 | 0.24867559 | -1.7701804 | 0.07669709 | 0.41112103 |
| PLAC9 | 33.4401513 | 0.267992958 | 0.4255556 | 0.6297484 | 0.52885921 | 0.84016133 |
| ANXA11 | 83.1553642 | 0.301970641 | 0.2168323 | 1.392646 | 0.16372687 | 0.55847737 |
| FAM213A | 42.8264915 | -0.321486627 | 0.48361596 | -0.664756 | 0.50620655 | 0.82481485 |
| TSPAN14 | 59.8613348 | 0.255207809 | 0.24349375 | 1.04810825 | 0.29458874 | 0.68566952 |
| NRG3 | 98.8155093 | -0.56283952 | 0.88860845 | -0.6333943 | 0.52647619 | 0.83873261 |
| GHITM | 34.2595541 | -0.405104448 | 0.29143268 | -1.3900447 | 0.16451531 | 0.55965519 |
| CCSER2 | 150.23127 | 0.034551401 | 0.16377843 | 0.2109643 | 0.83291513 | 0.9666365 |
| GRID1 | 68.8267014 | -0.35585244 | 0.31852845 | -1.1171763 | 0.26391895 | 0.6579701 |
| WAPL | 102.88207 | 0.115909145 | 0.17122601 | 0.67693654 | 0.49844622 | 0.82271448 |
| LDB3 | 43.2426795 | 0.847024415 | 0.43957975 | 1.92689591 | 0.0539926 | 0.34959616 |
| BMPR1A | 236.415365 | 0.452127207 | 0.24086525 | 1.87709605 | 0.06050493 | 0.36535567 |
| ADIRF | 344.598615 | -0.258402444 | 0.50801738 | -0.5086488 | 0.61099839 | 0.88216644 |
| GLUD1 | 74.853303 | 0.170012467 | 0.35260548 | 0.48216059 | 0.62969187 | 0.89365764 |
| SHLD2 | 88.3771621 | 0.429312382 | 0.25318271 | 1.69566234 | 0.08994985 | 0.43806588 |
| NUTM2A-AS | 134.671767 | -0.208845075 | 0.1761309 | -1.1857379 | 0.23572583 | 0.63155273 |
| PAPSS2 | 90.6873064 | -0.484602687 | 0.40024871 | -1.2107539 | 0.22598974 | 0.62072404 |
| ATAD1 | 58.6703049 | -0.168851327 | 0.21734835 | -0.7768696 | 0.4372357 | 0.78775317 |
| PTEN | 688.053289 | -0.774395281 | 0.31759618 | -2.4383017 | 0.01475645 | 0.17699624 |
| RNLS | 70.7489436 | 0.287504723 | 0.26431229 | 1.08774634 | 0.2767071 | 0.67259169 |
| STAMBPL1 | 37.9415529 | 0.300892685 | 0.32687463 | 0.92051405 | 0.3573042 | 0.73129235 |
| ACTA2 | 605.625497 | 1.305932105 | 0.42284378 | 3.08845053 | 0.00201203 | 0.05600918 |
| FAS | 50.0611634 | -0.940036594 | 0.40457494 | -2.3235166 | 0.02015142 | 0.20958347 |
| LIPA | 39.7309704 | 0.32112412 | 0.2237821 | 1.43498576 | 0.15129116 | 0.53898138 |
| IFIT3 | 47.7059503 | 1.027255433 | 1.11646512 | 0.9200963 | 0.35752244 | 0.73129235 |
| RPP30 | 17.7330153 | 0.102205092 | 0.35842041 | 0.28515422 | 0.77552599 | 0.94805375 |
| PCGF5 | 269.950027 | 0.179219227 | 0.26546146 | 0.67512333 | 0.49959741 | 0.82271448 |
| HECTD2 | 86.7901352 | 0.146812548 | 0.28054832 | 0.52330575 | 0.6007615 | 0.87825667 |
| TNKS2 | 109.745923 | -0.061455701 | 0.18479368 | -0.3325639 | 0.73946352 | 0.93594619 |
| BTAF1 | 146.332279 | -0.506222684 | 0.26083498 | -1.9407776 | 0.05228526 | 0.34585451 |
| CPEB3 | 184.606839 | 0.21727569 | 0.24743843 | 0.87810001 | 0.37988945 | 0.74991898 |
| 5-Mar | 64.6307438 | -0.278436623 | 0.17013375 | -1.6365749 | 0.10171933 | 0.46169744 |
| IDE | 60.918439 | -0.078243682 | 0.27112048 | -0.2885938 | 0.77289226 | 0.9472495 |
| EXOC6 | 45.2163312 | 0.635331889 | 0.29620367 | 2.14491566 | 0.03195959 | 0.27215647 |
| MYOF | 245.033052 | -0.392718338 | 0.51441404 | -0.7634285 | 0.44520789 | 0.79268341 |
| FRA10AC1 | 43.2105915 | -0.060548961 | 0.2946748 | -0.2054772 | 0.83719929 | 0.96693031 |
| PLCE1 | 147.915278 | -1.167988836 | 0.32416577 | -3.6030604 | 0.00031449 | 0.01770958 |
| NOC3L | 23.3431493 | -0.21562638 | 0.35718673 | -0.6036797 | 0.54605661 | 0.84758329 |
| TBC1D12 | 70.3989279 | 0.203156567 | 0.19508641 | 1.0413671 | 0.2977052 | 0.68849737 |
| PDLIM1 | 115.259391 | 0.483664871 | 0.37726259 | 1.28203774 | 0.19982941 | 0.59891775 |
| SORBS1 | 924.392706 | 0.043438088 | 0.43789746 | 0.09919694 | 0.9209819 | 0.98528826 |
| ALDH18A1 | 29.1443382 | -0.908519203 | 0.25888391 | -3.5093692 | 0.00044917 | 0.02157489 |
| TCTN3 | 32.9904018 | 0.063082407 | 0.32203712 | 0.19588552 | 0.84469977 | 0.96938938 |
| ENTPD1 | 31.8783791 | 1.580867308 | 0.44799696 | 3.52874561 | 0.00041753 | 0.02116179 |
| ENTPD1-AS1 | 110.822269 | 0.447641178 | 0.23790147 | 1.88162426 | 0.05988705 | 0.36463126 |
| AL365273.2 | 21.5107052 | 0.512064003 | 0.32664962 | 1.56762465 | 0.11696876 | 0.48989475 |
| ZNF518A | 80.6301969 | 0.409744507 | 0.24680511 | 1.6601946 | 0.09687531 | 0.45317419 |
| TM9SF3 | 142.445326 | 0.169063966 | 0.18477928 | 0.91495088 | 0.36021742 | 0.73273991 |
| LCOR | 176.116975 | 0.110876062 | 0.17806829 | 0.62266035 | 0.53350775 | 0.84333372 |
| PGAM1 | 51.3209215 | -0.303696208 | 0.28959112 | -1.0487069 | 0.29431304 | 0.68566952 |
| EXOSC1 | 22.5172664 | -0.098733083 | 0.25952237 | -0.3804415 | 0.7036177 | 0.92936308 |
| MMS19 | 40.4744884 | 0.260881636 | 0.19760644 | 1.32020815 | 0.18676553 | 0.58501647 |
| UBTD1 | 20.6640377 | 0.112869019 | 0.26485638 | 0.42615178 | 0.66999725 | 0.91445841 |
| MARVELD1 | 21.9186568 | 0.779886611 | 0.36486264 | 2.13748004 | 0.03255897 | 0.27542099 |
| ZFYVE27 | 22.5390951 | -0.363637565 | 0.23603315 | -1.5406207 | 0.12340912 | 0.50097964 |
| R3HCC1L | 52.1641084 | -0.577329535 | 0.19210834 | -3.0052288 | 0.00265381 | 0.06544583 |
| HPS1 | 25.9610372 | -0.424962192 | 0.23242782 | -1.828362 | 0.06749525 | 0.38712255 |
| HPSE2 | 338.546166 | 0.424787432 | 0.58309787 | 0.72850109 | 0.4663069 | 0.80355999 |
| GOT1 | 21.7879365 | -0.460717918 | 0.30988831 | -1.4867225 | 0.13708813 | 0.52089048 |
| SLC25A28 | 31.3869096 | -0.948473425 | 0.39399215 | -2.4073409 | 0.01606916 | 0.18618176 |
| CUTC | 28.0730527 | -0.373123032 | 0.25605063 | -1.4572236 | 0.14505466 | 0.53538077 |
| COX15 | 22.9842925 | -0.286221442 | 0.25537489 | -1.1207893 | 0.26237755 | 0.65630111 |
| DNMBP | 64.0676234 | 0.142114625 | 0.32009314 | 0.44397898 | 0.65705778 | 0.90772143 |
| ERLIN1 | 20.6637482 | -0.043346324 | 0.24417979 | -0.1775181 | 0.85910148 | 0.97397549 |
| CHUK | 27.1493273 | -0.044087312 | 0.31285554 | -0.1409191 | 0.88793388 | 0.97920559 |
| CWF19L1 | 28.8792813 | -0.108852677 | 0.24340109 | -0.4472152 | 0.65471965 | 0.90646849 |
| HIF1AN | 19.2743581 | 0.018293361 | 0.28670502 | 0.06380551 | 0.94912509 | 0.99060401 |

| | | | | | | |
|---|---|---|---|---|---|---|
| SLF2 | 93.0094384 | -0.08058247 | 0.29662233 | -0.2716669 | 0.78587815 | 0.95065355 |
| LZTS2 | 38.4971853 | -0.246909327 | 0.25907926 | -0.9530262 | 0.3405768 | 0.71952201 |
| BTRC | 131.599279 | 0.136854817 | 0.21521247 | 0.6359056 | 0.52483795 | 0.83726969 |
| FBXW4 | 107.54095 | -0.12153761 | 0.22139438 | -0.5489643 | 0.58302995 | 0.86764709 |
| OGA | 200.698187 | -0.138563452 | 0.26092784 | -0.5310413 | 0.59539019 | 0.87454826 |
| ARMH3 | 109.637453 | 0.068836337 | 0.15045863 | 0.45751007 | 0.64730446 | 0.90249298 |
| LDB1 | 20.0587597 | -0.153989626 | 0.37226796 | -0.4136526 | 0.67912852 | 0.91831925 |
| PPRC1 | 25.2448589 | -0.488922248 | 0.47643344 | -1.0262131 | 0.30479114 | 0.69650244 |
| NOLC1 | 19.7124653 | -0.654606817 | 0.40682215 | -1.6090737 | 0.10760023 | 0.47265636 |
| GBF1 | 180.904564 | -0.128746697 | 0.16829376 | -0.7650117 | 0.44426458 | 0.79252305 |
| NFKB2 | 23.3175615 | -0.85276155 | 0.55334944 | -1.5410905 | 0.12329477 | 0.50097964 |
| ACTR1A | 26.4847552 | 0.205651456 | 0.29768857 | 0.69082751 | 0.48967395 | 0.81720854 |
| SUFU | 42.7365837 | 0.140902886 | 0.24934073 | 0.56510177 | 0.57200454 | 0.86236719 |
| TRIM8 | 35.2301008 | -0.097535084 | 0.28623371 | -0.3407533 | 0.7332893 | 0.93574285 |
| ARL3 | 32.1941141 | 0.096609481 | 0.24817928 | 0.38927296 | 0.69707424 | 0.92702563 |
| WBP1L | 123.49861 | -0.964839793 | 0.23483306 | -4.1086199 | 3.98E-05 | 0.00443201 |
| CNNM2 | 122.783358 | 0.535518113 | 0.27608531 | 1.93968349 | 0.05241817 | 0.34611061 |
| NT5C2 | 188.513942 | -0.173818341 | 0.15776516 | -1.1017536 | 0.27056879 | 0.66546521 |
| PCGF6 | 27.1539157 | 0.199592493 | 0.25793828 | 0.77379943 | 0.43904941 | 0.78887428 |
| ATP5MD | 20.6809828 | -0.07563845 | 0.31849695 | -0.2374856 | 0.81228006 | 0.96026221 |
| PDCD11 | 23.2304409 | -0.432564273 | 0.35844082 | -1.2067941 | 0.22751144 | 0.62183669 |
| SH3PXD2A | 140.103302 | -0.500587259 | 0.30419389 | -1.645619 | 0.09984222 | 0.45801526 |
| SLK | 78.8669148 | -0.092691319 | 0.19314101 | -0.4799153 | 0.63128765 | 0.89436221 |
| GSTO1 | 36.9363534 | -0.794016222 | 0.31563017 | -2.5156538 | 0.01188118 | 0.15619692 |
| ITPRIP | 39.4817962 | -0.554110944 | 0.36496483 | -1.5182585 | 0.12894926 | 0.51141289 |
| XPNPEP1 | 38.8587238 | -0.423827986 | 0.26866623 | -1.5775261 | 0.11467452 | 0.4868533 |
| ADD3 | 172.077384 | 0.38986967 | 0.28268573 | 1.37916289 | 0.16784454 | 0.56323722 |
| MXI1 | 83.9405128 | -1.240748662 | 0.43881647 | -2.8274888 | 0.00469146 | 0.09121052 |
| SMNDC1 | 21.5118322 | 0.413795786 | 0.29454695 | 1.40485508 | 0.16006438 | 0.5530386 |
| SMC3 | 39.1618455 | 0.044035774 | 0.22088284 | 0.19936258 | 0.84197913 | 0.96803889 |
| RBM20 | 39.2372838 | 0.025312081 | 0.48710613 | 0.0519642 | 0.95855722 | 0.99181157 |
| PDCD4 | 53.5499892 | -0.196312445 | 0.30788424 | -0.6376177 | 0.52372256 | 0.83652187 |
| BBIP1 | 33.0962814 | 0.319753558 | 0.33561205 | 0.95274754 | 0.340718 | 0.71952201 |
| SHOC2 | 103.049447 | -0.005119762 | 0.23541889 | -0.0217475 | 0.98264941 | 0.99599681 |
| ZDHHC6 | 21.6159006 | 0.051333509 | 0.26494325 | 0.19375285 | 0.84636941 | 0.97014274 |
| VTI1A | 259.551995 | -0.038086678 | 0.18253187 | -0.2086577 | 0.83471548 | 0.96664419 |
| TCF7L2 | 359.874788 | 0.197548517 | 0.27168012 | 0.72713645 | 0.46714237 | 0.80374006 |
| CASP7 | 22.7665852 | 0.05108085 | 0.34366026 | 0.14863764 | 0.88183957 | 0.97782528 |
| NHLRC2 | 66.5166678 | 0.324984628 | 0.21497149 | 1.51175686 | 0.13059573 | 0.51241218 |
| CCDC186 | 36.9262952 | -0.035222908 | 0.23576931 | -0.1493956 | 0.88124145 | 0.97782528 |
| AFAP1L2 | 45.2747305 | 1.560228239 | 0.25739876 | 6.06152204 | 1.35E-09 | 2.25E-06 |
| ABLIM1 | 54.6352851 | 0.226791446 | 0.35477666 | 0.63925131 | 0.52265946 | 0.8350836 |
| FAM160B1 | 56.1776192 | -0.116148868 | 0.26715478 | -0.4347625 | 0.66373485 | 0.91141325 |
| TRUB1 | 19.4041811 | -0.182291958 | 0.3524688 | -0.5171861 | 0.60502626 | 0.87903643 |
| ATRNL1 | 479.326459 | 0.667646442 | 0.5471429 | 1.22024144 | 0.22237336 | 0.61772146 |
| GFRA1 | 72.4276693 | 0.665589683 | 0.50468511 | 1.31882172 | 0.18722872 | 0.58550803 |
| HSPA12A | 66.1621219 | 0.317233621 | 0.27713388 | 1.14469446 | 0.25233575 | 0.64703956 |
| PDZD8 | 70.5614075 | -0.105862418 | 0.20765182 | -0.5098073 | 0.61018646 | 0.88163721 |
| RAB11FIP2 | 49.6048582 | 0.257155421 | 0.27664247 | 0.92955871 | 0.35259961 | 0.73059415 |
| FAM204A | 45.570127 | 0.04267852 | 0.21510235 | 0.19841029 | 0.84272407 | 0.96844084 |
| CACUL1 | 82.9463748 | -0.191540733 | 0.19687157 | -0.9729223 | 0.33059193 | 0.71362138 |
| EIF3A | 110.532426 | -0.240395498 | 0.33832437 | -0.7105474 | 0.47736475 | 0.81132136 |
| PRDX3 | 34.6667356 | 0.095265354 | 0.22237308 | 0.42840327 | 0.66835754 | 0.9143573 |
| GRK5 | 574.017592 | -0.320182964 | 0.36429209 | -0.8789182 | 0.37944561 | 0.74991898 |
| TIAL1 | 74.7524904 | -0.246058091 | 0.23802262 | -1.0337593 | 0.30124871 | 0.69248569 |
| BAG3 | 26.921218 | -0.467585369 | 0.30605121 | -1.5278011 | 0.12656194 | 0.50599102 |
| INPP5F | 34.1353367 | 0.13431981 | 0.37311147 | 0.35999914 | 0.71884777 | 0.93444541 |
| MCMBP | 42.9127808 | -0.454484531 | 0.23392762 | -1.9428426 | 0.05203518 | 0.34566169 |
| SEC23IP | 53.9020634 | -0.088229686 | 0.24875779 | -0.3546811 | 0.72282851 | 0.93455731 |
| WDR11 | 65.0906798 | 0.052184989 | 0.19410284 | 0.26885227 | 0.78804337 | 0.95129454 |
| ATE1 | 124.871415 | -0.258093761 | 0.18424695 | -1.4008035 | 0.16127285 | 0.55470404 |
| TACC2 | 49.2193313 | -0.098063481 | 0.30041681 | -0.3264247 | 0.74410301 | 0.93797823 |
| PLEKHA1 | 78.3057163 | -0.44647385 | 0.24923132 | -1.7914035 | 0.07322857 | 0.40189622 |
| HTRA1 | 35.5837307 | 0.940949178 | 0.34013816 | 2.76637342 | 0.00566836 | 0.10101851 |
| IKZF5 | 18.6879355 | -0.27766547 | 0.33117317 | -0.8384298 | 0.40178933 | 0.7641692 |
| ACADSB | 27.2771818 | 0.413723948 | 0.35680828 | 1.15951329 | 0.24624702 | 0.64218217 |
| BUB3 | 22.9886273 | -0.225662783 | 0.27164062 | -0.8307402 | 0.40612042 | 0.76744123 |
| CPXM2 | 29.1160315 | 1.062482967 | 0.497098 | 2.13737122 | 0.03256781 | 0.27542099 |
| CHST15 | 118.3348 | -0.473191971 | 0.39812329 | -1.1885564 | 0.23461429 | 0.63080829 |
| OAT | 39.2857067 | -0.322920243 | 0.30117507 | -1.0722011 | 0.28362972 | 0.67741138 |
| ZRANB1 | 71.075946 | 0.167278769 | 0.21192725 | 0.78932165 | 0.42992403 | 0.78298297 |
| CTBP2 | 320.286011 | -0.017171801 | 0.19390039 | -0.0885599 | 0.92943167 | 0.98681039 |
| EDRF1 | 40.3949273 | 0.251362795 | 0.26306251 | 0.95552497 | 0.33931229 | 0.71925477 |
| BCCIP | 29.7622627 | -0.081377068 | 0.22897351 | -0.3553995 | 0.72229033 | 0.93444541 |
| DHX32 | 55.2081212 | -0.685732194 | 0.31062695 | -2.2075747 | 0.02727393 | 0.24775789 |
| ADAM12 | 136.335584 | -0.24726937 | 0.39717468 | -0.6225708 | 0.53356658 | 0.84333372 |
| DOCK1 | 376.085987 | 0.037250537 | 0.17577164 | 0.21192576 | 0.83216496 | 0.96643177 |
| PTPRE | 94.9956612 | -1.496550955 | 0.84861102 | -1.7635299 | 0.07781115 | 0.41317496 |
| MGMT | 76.5763377 | 0.412059222 | 0.20461265 | 2.01385016 | 0.04402527 | 0.3152359 |
| GLRX3 | 52.6574883 | -0.115617409 | 0.23182444 | -0.4987283 | 0.6179708 | 0.88534783 |
| PPP2R2D | 79.1028039 | -0.079021754 | 0.15652141 | -0.5048623 | 0.61365558 | 0.88357266 |
| BNIP3 | 58.7477461 | -2.033479322 | 0.52394402 | -3.8811003 | 0.00010398 | 0.00804406 |
| INPP5A | 118.941417 | 0.619517387 | 0.32148403 | 1.92705492 | 0.05397279 | 0.34959616 |
| TUBGCP2 | 34.4615103 | -0.017400496 | 0.35110709 | -0.0495589 | 0.96047386 | 0.99181157 |
| ECHS1 | 36.2026536 | -0.547044236 | 0.38248946 | -1.4302204 | 0.15265376 | 0.54039138 |
| KDM5A | 120.084287 | 0.042881097 | 0.17206985 | 0.2492075 | 0.80320028 | 0.95836189 |
| WNK1 | 208.928785 | -0.050778 | 0.23401585 | -0.2169853 | 0.8282198 | 0.96580172 |
| RAD52 | 26.8361843 | -0.131987197 | 0.30177045 | -0.4373761 | 0.66183858 | 0.91076998 |
| ERC1 | 330.076468 | 0.003151037 | 0.19670663 | 0.01601897 | 0.98721926 | 0.99719329 |
| WNT5B | 33.6485955 | 0.177800569 | 0.33032457 | 0.53826019 | 0.59039743 | 0.87160119 |
| ADIPOR2 | 76.1908278 | -0.168860266 | 0.1822136 | -0.9267161 | 0.35407398 | 0.73059415 |
| DCP1B | 44.2727934 | 0.074300931 | 0.34441703 | 0.21572955 | 0.82919856 | 0.96580172 |
| CACNA1C | 1269.47262 | 0.138185515 | 0.22791085 | 0.60631389 | 0.54430634 | 0.84730644 |
| FKBP4 | 19.290997 | -0.128054826 | 0.38335108 | -0.3340406 | 0.73834892 | 0.93594619 |
| ITFG2 | 44.9227429 | 0.170600235 | 0.41190249 | 0.41417626 | 0.67874502 | 0.91831925 |
| NRIP2 | 21.0899029 | 0.532952467 | 0.43688176 | 1.21990094 | 0.22250243 | 0.61772146 |
| TULP3 | 79.2212848 | 0.192207727 | 0.18839085 | 1.02026042 | 0.30760497 | 0.69801431 |
| TEAD4 | 32.1096806 | -0.309252977 | 0.43901262 | -0.7044284 | 0.48116599 | 0.8133639 |
| TSPAN9 | 210.95879 | -0.142931407 | 0.22631528 | -0.6315588 | 0.52767521 | 0.83947444 |
| PARP11 | 39.6217179 | 0.126288271 | 0.21374716 | 0.59083016 | 0.55463422 | 0.85191704 |
| CCND2 | 48.7405553 | 0.227128997 | 0.43674039 | 0.52005494 | 0.60302528 | 0.87903643 |
| C12orf4 | 29.8054446 | -0.087741587 | 0.23462804 | -0.3739604 | 0.70843381 | 0.93009618 |
| NTF3 | 56.7482001 | 0.241924745 | 0.47048259 | 0.51420552 | 0.60710831 | 0.88000768 |
| CD9 | 87.8582188 | 0.269080463 | 0.43801729 | 0.61431471 | 0.53900738 | 0.8449569 |
| TNFRSF1A | 117.942724 | -0.09945903 | 0.25263159 | -0.393692 | 0.69380847 | 0.9248682 |
| LTBR | 43.7875352 | -0.322500019 | 0.31492101 | -1.0240664 | 0.30580392 | 0.69728397 |
| MRPL51 | 22.9392139 | -0.575231235 | 0.31008417 | -1.855081 | 0.06358467 | 0.37532828 |
| GAPDH | 265.988751 | -1.232430605 | 0.39825124 | -3.0946058 | 0.00197075 | 0.05570389 |
| CHD4 | 22.2507886 | 0.153689787 | 0.39864005 | 0.38553524 | 0.6998409 | 0.92769314 |
| COPS7A | 18.8744082 | 0.259083478 | 0.2921116 | 0.8869332 | 0.37511486 | 0.74723387 |
| MLF2 | 31.7139963 | -0.345156141 | 0.3619174 | -0.9536876 | 0.34024179 | 0.71925477 |
| PTMS | 87.4355404 | 0.407229906 | 0.27469095 | 1.48250208 | 0.13820675 | 0.52264518 |
| P3H3 | 22.3478614 | -0.05267513 | 0.32509193 | -0.1620315 | 0.87128105 | 0.9768643 |

| | | | | | | |
|---|---|---|---|---|---|---|
| TPI1 | 80.8821609 | -0.945197796 | 0.33384486 | -2.8312486 | 0.00463667 | 0.0906884 |
| LRRC23 | 18.1457293 | 0.402971688 | 0.41898566 | 0.9617792 | 0.33616053 | 0.71883121 |
| ATN1 | 59.9123207 | -0.173289436 | 0.20877977 | -0.8300107 | 0.40653275 | 0.76782554 |
| C12orf57 | 39.0714217 | -0.432937966 | 0.36861665 | -1.1744938 | 0.2401973 | 0.63451757 |
| PHB2 | 28.2063411 | -0.224615381 | 0.32269812 | -0.6960542 | 0.48639489 | 0.81595238 |
| LPCAT3 | 67.7637448 | -0.699199357 | 0.28513703 | -2.4521521 | 0.01420046 | 0.17480605 |
| C1S | 276.285755 | -0.653963767 | 0.32674465 | -2.0014521 | 0.04534369 | 0.31959454 |
| C1R | 319.675383 | -0.579137487 | 0.30833925 | -1.8782477 | 0.06034728 | 0.36535567 |
| C1RL | 92.2905944 | -1.120400133 | 0.32644211 | -3.4321556 | 0.0005988 | 0.0265097 |
| C1RL-AS1 | 18.761896 | -0.659682884 | 0.58400518 | -1.129584 | 0.25865157 | 0.6514155 |
| CLSTN3 | 22.9420792 | -1.026268547 | 0.42914846 | -2.3914068 | 0.01678394 | 0.19005424 |
| SLC2A3 | 478.130565 | -0.35525651 | 0.62270368 | -0.5705065 | 0.5683342 | 0.8595777 |
| RIMKLB | 55.1375202 | 0.762623949 | 0.28599053 | 2.66660559 | 0.00766215 | 0.12228146 |
| PHC1 | 24.0676796 | 0.22174107 | 0.27078185 | 0.81889192 | 0.41284808 | 0.77260518 |
| M6PR | 29.663056 | -0.363365559 | 0.24503431 | -1.4829171 | 0.13809644 | 0.52264518 |
| A2M | 814.343216 | 0.107014873 | 0.43233046 | 0.24753026 | 0.80449787 | 0.95836286 |
| CLEC2D | 32.3414846 | -0.246615148 | 0.26357701 | -0.9356474 | 0.34945475 | 0.7277254 |
| GABARAPL1 | 68.2764915 | -0.503933764 | 0.39348849 | -1.2806824 | 0.20030526 | 0.59936617 |
| YBX3 | 152.899445 | -1.151030513 | 0.28390477 | -4.0542838 | 5.03E-05 | 0.00499416 |
| AC244131.2 | 25.550105 | -0.300580141 | 0.4530389 | -0.6634753 | 0.50702617 | 0.82481485 |
| ETV6 | 511.930993 | 0.055844949 | 0.3668593 | 0.15222443 | 0.87900993 | 0.9772835 |
| BCL2L14 | 29.730868 | -0.341577277 | 0.29557499 | -1.1556366 | 0.24782983 | 0.64333128 |
| LRP6 | 92.8338026 | 0.392513739 | 0.21723552 | 1.80685801 | 0.07078442 | 0.3961879 |
| BORCS5 | 34.4515748 | -0.085807084 | 0.23001256 | -0.3730539 | 0.70910832 | 0.93009618 |
| DUSP16 | 89.3335125 | -0.314802549 | 0.29854353 | -1.0544611 | 0.29167185 | 0.68334882 |
| CREBL2 | 21.4437177 | -0.021630476 | 0.36240985 | -0.0596851 | 0.95240642 | 0.99060401 |
| CDKN1B | 23.4491379 | -0.321456102 | 0.4026106 | -0.7984293 | 0.4246214 | 0.77861803 |
| APOLD1 | 45.4485675 | -2.023449087 | 0.82313144 | -2.4582333 | 0.01396224 | 0.17332523 |
| HEBP1 | 19.1955888 | 0.207747613 | 0.29759274 | 0.6980937 | 0.4851186 | 0.81563409 |
| EMP1 | 52.8705844 | 1.059223818 | 0.81535822 | 1.29909013 | 0.193913 | 0.59269042 |
| ATF7IP | 161.191868 | -0.030365855 | 0.26849016 | -0.1130986 | 0.9099524 | 0.98343817 |
| H2AFJ | 95.8226224 | -0.856664562 | 0.40616794 | -2.1091388 | 0.0349326 | 0.28524412 |
| WBP11 | 30.9713417 | -0.394996785 | 0.25884206 | -1.5260147 | 0.1270062 | 0.50643981 |
| C12orf60 | 33.1137453 | -0.774027419 | 0.46801173 | -1.6538633 | 0.09815529 | 0.45596917 |
| ART4 | 44.337248 | -0.603702802 | 0.59127658 | -1.0210159 | 0.3072469 | 0.69801431 |
| MGP | 2198.25889 | -0.974164126 | 0.49632748 | -1.9627447 | 0.04967585 | 0.33731209 |
| ARHGDIB | 32.5123818 | 0.648441059 | 0.33068856 | 1.96088145 | 0.04989285 | 0.33748407 |
| RERG | 553.290243 | -0.921091232 | 0.34951109 | -2.6353705 | 0.00840455 | 0.12814321 |
| EPS8 | 829.674955 | -0.372096231 | 0.38993749 | -0.9542458 | 0.33995922 | 0.71925477 |
| STRAP | 49.693859 | 0.065573125 | 0.24672554 | 0.26577356 | 0.79041361 | 0.95194192 |
| DERA | 57.0388117 | -0.184898961 | 0.27154103 | -0.6809246 | 0.49591921 | 0.82114698 |
| MGST1 | 84.3516763 | -0.767710123 | 0.21527461 | -3.5661899 | 0.00036221 | 0.01915017 |
| LMO3 | 41.920949 | 0.119194079 | 0.50576055 | 0.23567295 | 0.81368647 | 0.96038841 |
| RERGL | 38.0023247 | 1.005036857 | 0.41964673 | 2.39495936 | 0.01662221 | 0.19005424 |
| PLEKHA5 | 333.544917 | 0.4697831 | 0.29577553 | 1.58830954 | 0.11221636 | 0.48142325 |
| AEBP2 | 286.716993 | 0.155208315 | 0.30626153 | 0.50678357 | 0.61230668 | 0.88282721 |
| AC024901.1 | 31.1331148 | -0.106693073 | 0.32596338 | -0.3273161 | 0.74342879 | 0.93791562 |
| PDE3A | 1118.33669 | -0.071888558 | 0.27726416 | -0.2592782 | 0.79542058 | 0.95463848 |
| PYROXD1 | 19.3368258 | -0.298175856 | 0.28862713 | -1.0330833 | 0.30156494 | 0.69264368 |
| RECQL | 31.5018009 | -0.202179622 | 0.27921278 | -0.7241059 | 0.46900072 | 0.8037981 |
| GOLT1B | 30.3156392 | -0.228010845 | 0.21915858 | -1.0403921 | 0.29815779 | 0.68868829 |
| LDHB | 91.4273053 | -0.515665616 | 0.28794974 | -1.7908181 | 0.07332248 | 0.40189622 |
| KCNJ8 | 28.4727212 | 0.249260524 | 0.42808553 | 0.58226804 | 0.56038615 | 0.85423726 |
| ABCC9 | 243.079282 | -0.134588304 | 0.39680794 | -0.3391774 | 0.73447606 | 0.93594619 |
| CMAS | 20.9871104 | 0.307850478 | 0.27883412 | 1.10406317 | 0.26956574 | 0.6651094 |
| ST8SIA1 | 36.0733049 | 0.971983229 | 0.56425206 | 1.72260465 | 0.08496004 | 0.42746464 |
| C2CD5 | 88.4150345 | 0.342094969 | 0.26312527 | 1.30012206 | 0.19355914 | 0.59251477 |
| ETNK1 | 196.053022 | -0.347629468 | 0.33094447 | -1.0504163 | 0.29352675 | 0.68480256 |
| SOX5 | 1046.0803 | 0.12939711 | 0.45266044 | 0.28585911 | 0.77498603 | 0.94805375 |
| KRAS | 35.6333065 | 0.098203198 | 0.29066587 | 0.33785597 | 0.73547173 | 0.93594619 |
| RASSF8-AS1 | 26.8072157 | -0.516110359 | 0.40643253 | -1.2698549 | 0.20413631 | 0.60367992 |
| RASSF8 | 141.92263 | -0.303746864 | 0.23896706 | -1.2710826 | 0.20369928 | 0.60367992 |
| SSPN | 30.2275455 | 0.581452096 | 0.27840266 | 2.08852927 | 0.03675012 | 0.29103084 |
| ITPR2 | 348.541335 | -0.145372665 | 0.30904211 | -0.4703976 | 0.63807099 | 0.89779507 |
| FGFR1OP2 | 70.3193004 | 0.099574307 | 0.33191093 | 0.3000031 | 0.76417479 | 0.94462105 |
| TM7SF3 | 40.6177151 | 0.223129442 | 0.33279041 | 0.67048038 | 0.50255161 | 0.82430366 |
| STK38L | 88.5561885 | 0.053356814 | 0.45068695 | 0.11838997 | 0.90575867 | 0.98303609 |
| PPFIBP1 | 246.810671 | 0.344961529 | 0.20122327 | 1.71432227 | 0.08646956 | 0.43111585 |
| MRPS35 | 30.2042452 | -0.192076446 | 0.28891508 | -0.6648197 | 0.50616581 | 0.82481485 |
| KLHL42 | 49.5740599 | 0.044767753 | 0.20127275 | 0.22242331 | 0.82398436 | 0.96413949 |
| CCDC91 | 207.585268 | -0.184025136 | 0.16512447 | -1.1144631 | 0.26508056 | 0.65897365 |
| ERGIC2 | 44.6965652 | -0.099066056 | 0.19166278 | -0.5168769 | 0.60524212 | 0.87903643 |
| TMTC1 | 1053.93339 | 0.353376102 | 0.37124568 | 0.9518659 | 0.341165 | 0.71963879 |
| IPO8 | 48.8377398 | 0.010708148 | 0.29296391 | 0.03655108 | 0.97084295 | 0.99300191 |
| CAPRIN2 | 27.5102834 | 0.057555943 | 0.27454279 | 0.20964289 | 0.8339464 | 0.9666365 |
| SINHCAF | 28.7946131 | -0.684815159 | 0.30530492 | -2.2430531 | 0.02489339 | 0.234034 |
| DENND5B | 212.320936 | 0.019370391 | 0.37517968 | 0.05162964 | 0.9588238 | 0.99181157 |
| AMN1 | 38.3024855 | 0.304619008 | 0.32541275 | 0.9361004 | 0.3492215 | 0.7274458 |
| KIAA1551 | 76.2223076 | -0.898655749 | 0.30504588 | -2.9459692 | 0.00321944 | 0.07300955 |
| BICD1 | 151.610058 | -0.578920292 | 0.39053419 | -1.4823805 | 0.13823907 | 0.52264518 |
| FGD4 | 214.345567 | -0.85664989 | 0.32075155 | -2.6707584 | 0.00756801 | 0.12147053 |
| DNM1L | 86.0295456 | 0.088193027 | 0.17859356 | 0.49381976 | 0.62143348 | 0.88729641 |
| YARS2 | 21.7441042 | -0.14923007 | 0.26888384 | -0.5549983 | 0.57889584 | 0.86486944 |
| CPNE8 | 109.000917 | -0.897516742 | 0.24123036 | -3.7205796 | 0.00019877 | 0.01312136 |
| KIF21A | 25.1906555 | 0.443187996 | 0.45112518 | 0.98240581 | 0.32589996 | 0.71149668 |
| SLC2A13 | 108.373621 | -0.550563624 | 0.2800647 | -1.9658444 | 0.04931659 | 0.33589213 |
| LRRK2 | 50.3122167 | -0.141294171 | 0.38761627 | -0.3645207 | 0.71546919 | 0.93237409 |
| CNTN1 | 167.05156 | 1.346612429 | 0.49716803 | 2.70856599 | 0.00675747 | 0.11286506 |
| PDZRN4 | 182.357103 | 1.298476857 | 0.57102237 | 2.27395094 | 0.02296894 | 0.22440695 |
| GXYLT1 | 27.6194983 | 0.151389677 | 0.24732007 | 0.61212047 | 0.54045805 | 0.84578923 |
| YAF2 | 128.046286 | -0.370386152 | 0.17523084 | -2.1137041 | 0.03454055 | 0.2833153 |
| PPHLN1 | 104.290943 | -0.059987322 | 0.16544867 | -0.3625736 | 0.71692342 | 0.9330034 |
| PRICKLE1 | 132.662556 | -0.055987049 | 0.45873877 | -0.1220456 | 0.9028629 | 0.98164203 |
| PUS7L | 19.2233686 | 0.208884576 | 0.24810676 | 0.84191408 | 0.39983605 | 0.76302133 |
| IRAK4 | 25.0332661 | -0.476324618 | 0.25647479 | -1.8571986 | 0.06328291 | 0.37444936 |
| TMEM117 | 99.0371896 | 0.103444385 | 0.36480372 | 0.28356176 | 0.77674625 | 0.94805375 |
| ANO6 | 256.521789 | 0.456148523 | 0.29841229 | 1.52858488 | 0.12636739 | 0.50581368 |
| ARID2 | 211.010463 | -0.2117351 | 0.183823 | -1.1518422 | 0.24938591 | 0.64565668 |
| SCAF11 | 232.800715 | -0.041398285 | 0.18648681 | -0.2219904 | 0.82432134 | 0.96413949 |
| SLC38A1 | 49.9888132 | -0.382671157 | 0.40121799 | -0.9537737 | 0.34019822 | 0.71925477 |
| SLC38A2 | 377.644102 | 0.533957532 | 0.47845021 | 1.11601483 | 0.26441579 | 0.65826275 |
| AC008014.1 | 189.746402 | 0.212361888 | 0.48838544 | 0.43482436 | 0.66368992 | 0.91141325 |
| PCED1B | 49.3220817 | 1.134770354 | 0.37870227 | 2.99647098 | 0.00273124 | 0.06668405 |
| RAPGEF3 | 26.4870152 | 0.833548264 | 0.34774451 | 2.39701341 | 0.01652932 | 0.18980307 |
| HDAC7 | 83.5370461 | 0.00460837 | 0.27709915 | 0.01663076 | 0.98673118 | 0.99719329 |
| SENP1 | 29.2077028 | 0.243259098 | 0.31327938 | 0.77649253 | 0.43745823 | 0.78796092 |
| ZNF641 | 20.5016126 | 0.977364546 | 0.31726442 | 3.08059924 | 0.00206584 | 0.05618053 |
| CCNT1 | 40.6869275 | -0.196620958 | 0.2499455 | -0.7866553 | 0.43148366 | 0.78389374 |
| ADCY6 | 64.1608457 | 0.432823714 | 0.27690163 | 1.56309559 | 0.11803011 | 0.49172522 |
| DDX23 | 18.4240158 | 0.007154842 | 0.27502389 | 0.02601535 | 0.9792451 | 0.99467481 |
| CCDC65 | 20.5798212 | -0.623538986 | 0.36631717 | -1.7021834 | 0.088721 | 0.43544541 |
| PRKAG1 | 26.4426263 | 0.217316039 | 0.32690587 | 0.6647664 | 0.50619992 | 0.82481485 |
| KMT2D | 33.7219337 | -0.008635847 | 0.30978395 | -0.027877 | 0.97776025 | 0.99407008 |

| | | | | | | |
|---|---|---|---|---|---|---|
| LMBR1L | 38.0244563 | 0.132233534 | 0.26846378 | 0.49255632 | 0.62232612 | 0.88788092 |
| TUBA1B | 94.7731134 | -0.203048397 | 0.53531035 | -0.3793097 | 0.70445792 | 0.92936308 |
| TUBA1A | 46.3232838 | 1.274242213 | 0.49616343 | 2.56819053 | 0.01022309 | 0.14224028 |
| TUBA1C | 34.7894225 | -0.158471453 | 0.38530833 | -0.4112848 | 0.68086372 | 0.91860979 |
| SPATS2 | 92.7542197 | -0.314529091 | 0.16518273 | -1.9041281 | 0.0568935 | 0.35705408 |
| PRPF40B | 40.1950843 | 0.089796814 | 0.34936855 | 0.25702604 | 0.79715867 | 0.95463848 |
| FMNL3 | 75.3796728 | 0.495144085 | 0.30280018 | 1.63521725 | 0.10200351 | 0.46169744 |
| TMBIM6 | 122.790876 | -0.156621709 | 0.24978873 | -0.6270167 | 0.53064828 | 0.84152078 |
| NCKAP5L | 30.3356456 | 0.364899186 | 0.24925546 | 1.46395664 | 0.1432058 | 0.5309886 |
| AQP2 | 32.2329427 | 0.206210034 | 0.59053776 | 0.34919026 | 0.72694648 | 0.93455731 |
| AC074032.1 | 59.2293963 | -0.282758655 | 0.25122743 | -1.1255087 | 0.26037354 | 0.65328956 |
| CERS5 | 54.007939 | 0.15524937 | 0.20695354 | 0.75016532 | 0.45315514 | 0.79534858 |
| LIMA1 | 298.034228 | 0.54642612 | 0.30445049 | 1.79479469 | 0.07268642 | 0.40072954 |
| LARP4 | 133.268426 | -0.51205082 | 0.23026071 | -2.2237872 | 0.02616276 | 0.24176479 |
| DIP2B | 160.582441 | -0.377539761 | 0.26011935 | -1.4514097 | 0.1466658 | 0.53611418 |
| ATF1 | 49.7727938 | 0.010434899 | 0.27237796 | 0.03831037 | 0.96944023 | 0.99300191 |
| METTL7A | 45.7469959 | 0.278209804 | 0.32518679 | 0.85553846 | 0.39225312 | 0.75901355 |
| SLC11A2 | 46.3559448 | -0.767754637 | 0.3560479 | -2.156324 | 0.03105837 | 0.26758262 |
| LETMD1 | 35.2029709 | -0.261709148 | 0.2583827 | -1.0128741 | 0.31112029 | 0.70126123 |
| CSRNP2 | 19.0376888 | 0.101771481 | 0.27995665 | 0.36352586 | 0.7162121 | 0.93247094 |
| TFCP2 | 108.994665 | 0.464802454 | 0.17599043 | 2.64106667 | 0.00826455 | 0.12786557 |
| POU6F1 | 27.2160403 | 0.407027206 | 0.27974517 | 1.45499278 | 0.14567126 | 0.53567049 |
| DAZAP2 | 61.2303825 | 0.009912216 | 0.28022784 | 0.03537199 | 0.97178312 | 0.99300191 |
| ACVR1B | 71.9975019 | -0.762917961 | 0.23243597 | -3.2822715 | 0.00102974 | 0.03921031 |
| NR4A1 | 271.276196 | 1.732481138 | 0.44490668 | 3.89403267 | 9.86E-05 | 0.00789507 |
| KRT8 | 19.8671893 | -0.912742271 | 0.31269157 | -2.9189858 | 0.00351172 | 0.07673709 |
| KRT18 | 22.0542195 | -0.752934703 | 0.47054532 | -1.6001322 | 0.10956927 | 0.47680173 |
| EIF4B | 74.0238977 | -0.24285242 | 0.21872295 | -1.1103198 | 0.26686124 | 0.66188434 |
| AC068888.1 | 40.2096389 | -0.172686093 | 0.25829817 | -0.6685533 | 0.50378047 | 0.82437825 |
| TNS2 | 105.948016 | 0.315515826 | 0.23218308 | 1.35890964 | 0.17417522 | 0.57154863 |
| IGFBP6 | 145.565589 | 1.033151616 | 0.87014928 | 1.18732687 | 0.23509871 | 0.63100657 |
| CSAD | 54.6896729 | 0.10427387 | 0.26713137 | 0.39034678 | 0.69628014 | 0.92648248 |
| PFDN5 | 58.6447847 | 0.056463015 | 0.27992994 | 0.20170409 | 0.84014806 | 0.96803523 |
| SP1 | 44.773414 | 0.331232698 | 0.26361078 | 1.25652184 | 0.20892683 | 0.6059796 |
| PRR13 | 31.6939871 | -0.658995185 | 0.30525495 | -2.1588354 | 0.03086294 | 0.26662968 |
| PCBP2 | 121.135319 | -0.014726614 | 0.1991689 | -0.0739403 | 0.94105787 | 0.98895798 |
| MAP3K12 | 17.862668 | -0.166602415 | 0.27028204 | -0.6164021 | 0.53762917 | 0.84423863 |
| ATP5MC2 | 68.890944 | -0.198319605 | 0.20963982 | -0.9460016 | 0.34414777 | 0.72261198 |
| CALCOCO1 | 43.5136663 | -0.205085371 | 0.3240878 | -0.6328081 | 0.526859 | 0.83897872 |
| HOXC4 | 33.6631416 | 0.833788313 | 0.28027556 | 2.97488764 | 0.00293096 | 0.06945135 |
| LINC02381 | 17.8090933 | 0.345435117 | 0.27145497 | 1.27253192 | 0.20318419 | 0.60367992 |
| CBX5 | 60.0722608 | 0.271858177 | 0.20803638 | 1.30678188 | 0.1912868 | 0.58975955 |
| HNRNPA1 | 53.4633262 | -0.114956705 | 0.22829624 | -0.5035418 | 0.61458338 | 0.88360697 |
| COPZ1 | 31.8180066 | -0.345366666 | 0.24275605 | -1.4226902 | 0.15482597 | 0.54412824 |
| ITGA5 | 55.0744732 | -0.388114791 | 0.47897066 | -0.8103102 | 0.41776194 | 0.77722447 |
| ITGA7 | 71.1772401 | 0.56117858 | 0.30829941 | 1.82023892 | 0.06872263 | 0.39029567 |
| CD63 | 126.910875 | -0.248969174 | 0.35161411 | -0.708075 | 0.47889866 | 0.81163945 |
| GDF11 | 19.7801714 | 0.660337392 | 0.30175925 | 2.18829213 | 0.02864833 | 0.25557869 |
| RAB5B | 34.2500078 | 0.182753612 | 0.21162136 | 0.86358773 | 0.38781441 | 0.75457985 |
| RPS26 | 20.7631374 | -0.909146574 | 0.37747145 | -2.4085174 | 0.01601746 | 0.18618176 |
| PA2G4 | 18.7390646 | -0.68398208 | 0.41323645 | -1.6551833 | 0.09788731 | 0.45529992 |
| RPL41 | 246.150428 | -0.498036367 | 0.33436114 | -1.4895163 | 0.13635146 | 0.52087399 |
| ESYT1 | 21.3211367 | 0.326601447 | 0.26987457 | 1.2101972 | 0.22620323 | 0.62072404 |
| MYL6 | 227.669683 | 0.498213616 | 0.36345994 | 1.37075248 | 0.17045213 | 0.56801217 |
| SMARCC2 | 51.4707416 | -0.152700581 | 0.26080841 | -0.5854895 | 0.55821864 | 0.85287916 |
| RNF41 | 18.49108 | -0.46809668 | 0.30225027 | -1.5487056 | 0.12145252 | 0.49924175 |
| ANKRD52 | 18.7283284 | 0.407146576 | 0.34193848 | 1.19070123 | 0.23377089 | 0.62975891 |
| CS | 26.6800093 | -0.074723202 | 0.31257983 | -0.2390532 | 0.81106435 | 0.95967025 |
| STAT2 | 107.584587 | -0.257362773 | 0.4413259 | -0.5831581 | 0.55978688 | 0.85420967 |
| RBMS2 | 111.882781 | 0.639487033 | 0.3462241 | 1.8470321 | 0.06474249 | 0.380026 |
| BAZ2A | 81.3477236 | 0.023662716 | 0.17478985 | 0.13537809 | 0.89231295 | 0.98023547 |
| ATP5F1B | 75.5685849 | -0.342027703 | 0.21118111 | -1.6195942 | 0.10531947 | 0.46766937 |
| PTGES3 | 121.152935 | 0.111250542 | 0.16546376 | 0.67235595 | 0.50135712 | 0.82406312 |
| NACA | 73.3392622 | -0.510720897 | 0.28038833 | -1.821477 | 0.06853438 | 0.38953271 |
| NAB2 | 18.9751213 | 0.016908328 | 0.36781485 | 0.04596967 | 0.96333442 | 0.99231823 |
| STAT6 | 89.17701 | -0.554765308 | 0.22423903 | -2.4739908 | 0.01336131 | 0.16900567 |
| LRP1 | 166.815586 | 0.030721358 | 0.19552407 | 0.15712315 | 0.87514779 | 0.9768643 |
| NDUFA4L2 | 43.9467813 | -1.564399802 | 0.50069489 | -3.1244573 | 0.00178133 | 0.05374171 |
| R3HDM2 | 162.875792 | 0.058267458 | 0.19340523 | 0.30127137 | 0.76320757 | 0.94462105 |
| MARS | 28.1964434 | -0.153954278 | 0.29383536 | -0.5239474 | 0.60031512 | 0.87795339 |
| DDIT3 | 23.2732757 | -1.286049788 | 0.53930113 | -2.3846599 | 0.01709492 | 0.19063814 |
| DCTN2 | 31.7194732 | 0.261736946 | 0.3123985 | 0.83783035 | 0.40212599 | 0.76441384 |
| DTX3 | 24.9329601 | 0.86146562 | 0.33761969 | 2.55158588 | 0.01072339 | 0.14566763 |
| OS9 | 54.0430685 | -0.046679757 | 0.26209537 | -0.1781022 | 0.85864273 | 0.97397549 |
| AVIL | 24.9155712 | -0.054364661 | 0.4153648 | -0.1308841 | 0.89586697 | 0.9803047 |
| CTDSP2 | 72.1632851 | -0.129065762 | 0.19455342 | -0.663395 | 0.50707763 | 0.82481485 |
| LRIG3 | 31.9691916 | 0.475228005 | 0.42321033 | 1.12291212 | 0.26147482 | 0.65560507 |
| SLC16A7 | 88.5629398 | -0.6030003 | 0.36685808 | -1.6436882 | 0.10024062 | 0.45949178 |
| USP15 | 186.121578 | -0.092037639 | 0.21615451 | -0.4257956 | 0.67025679 | 0.91445841 |
| MON2 | 306.287882 | -0.106550224 | 0.19250735 | -0.5534865 | 0.57993031 | 0.86588945 |
| AC048341.1 | 24.9440782 | 0.029683939 | 0.34234326 | 0.08670812 | 0.93090353 | 0.98681039 |
| DPY19L2 | 37.4138716 | 0.685561206 | 0.45669961 | 1.50112063 | 0.13332437 | 0.51514235 |
| SRGAP1 | 249.966575 | -0.362238186 | 0.42759002 | -0.8471624 | 0.39690461 | 0.76043276 |
| XPOT | 36.6598919 | -0.452261779 | 0.32491227 | -1.3919505 | 0.1639374 | 0.55880147 |
| TBK1 | 82.9328665 | -0.592372437 | 0.27986302 | -2.1166514 | 0.03428945 | 0.2833153 |
| RASSF3 | 107.621104 | 0.728485601 | 0.34741654 | 2.09686504 | 0.03600552 | 0.28882238 |
| GNS | 54.1087314 | -0.235638632 | 0.22618702 | -1.0417867 | 0.29751058 | 0.68849737 |
| LEMD3 | 93.0641463 | -0.031878951 | 0.20464675 | -0.1557755 | 0.87620998 | 0.9768643 |
| MSRB3 | 359.420411 | 0.662473988 | 0.3213369 | 2.06161815 | 0.0392441 | 0.30005099 |
| IRAK3 | 196.007152 | -1.720794141 | 0.47104809 | -3.6531178 | 0.00025908 | 0.01573508 |
| HELB | 23.6166248 | -0.770684241 | 0.3564913 | -2.1618599 | 0.03062898 | 0.26662968 |
| GRIP1 | 39.306359 | 0.205753726 | 0.29144989 | 0.70596604 | 0.48020926 | 0.81237906 |
| CAND1 | 80.403646 | -0.107682644 | 0.16202644 | -0.6645992 | 0.50630688 | 0.82481485 |
| MDM1 | 18.6798959 | 0.597988447 | 0.29767166 | 2.00888606 | 0.04454922 | 0.31724051 |
| RAP1B | 336.926082 | -0.180393959 | 0.23814347 | -0.7575012 | 0.44874966 | 0.79313643 |
| NUP107 | 44.825791 | 0.085231593 | 0.32907728 | 0.25900176 | 0.79563389 | 0.95463848 |
| SLC35E3 | 37.0094099 | -0.242908041 | 0.34556598 | -0.7029281 | 0.48210056 | 0.81402096 |
| MDM2 | 96.7019158 | -0.852778655 | 0.28489343 | -2.993325 | 0.00275956 | 0.06715226 |
| CPM | 221.599211 | -0.418420689 | 0.39186549 | -1.0677661 | 0.28562604 | 0.6783522 |
| CPSF6 | 175.4166 | 0.302290559 | 0.26710685 | 1.13172148 | 0.25775155 | 0.65130862 |
| FRS2 | 105.000153 | 0.137124477 | 0.17179031 | 0.79820843 | 0.42474955 | 0.77861803 |
| CCT2 | 28.9983067 | -0.440679771 | 0.32718627 | -1.3468773 | 0.17801974 | 0.57531535 |
| AC025159.1 | 51.0729806 | 0.578935928 | 0.43201207 | 1.34009203 | 0.18021543 | 0.57783734 |
| CNOT2 | 157.937342 | -0.293116813 | 0.25547611 | -1.1473355 | 0.25124298 | 0.6460368 |
| ZFC3H1 | 133.271796 | -0.232344197 | 0.22601148 | -1.0280194 | 0.30394069 | 0.69584049 |
| RAB21 | 96.0267355 | -0.456138569 | 0.21793346 | -2.0930176 | 0.03634758 | 0.28969304 |
| TBC1D15 | 108.644584 | -0.8602809 | 0.26047995 | -3.302676 | 0.00095767 | 0.0366558 |
| ATXN7L3B | 24.9315251 | 0.179585747 | 0.29884493 | 0.60093288 | 0.54788469 | 0.84785354 |
| CAPS2 | 77.5479835 | -0.037854062 | 0.39114488 | -0.0967776 | 0.92290301 | 0.98561736 |
| GLIPR1 | 25.1168138 | 0.395254658 | 0.53500914 | 0.73878113 | 0.46003991 | 0.80038667 |
| KRR1 | 25.8342454 | -0.11886049 | 0.30785411 | -0.3860936 | 0.69942737 | 0.92769314 |
| PHLDA1 | 64.1518452 | -0.548197177 | 0.505462 | -1.0845468 | 0.27812244 | 0.67300686 |
| NAP1L1 | 177.688841 | -0.309773334 | 0.36406233 | -0.8508799 | 0.39483605 | 0.75988253 |

| | | | | | | |
|---|---|---|---|---|---|---|
| OSBPL8 | 262.579129 | -0.785327731 | 0.23592678 | -3.3286926 | 0.00087255 | 0.0348497 |
| ZDHHC17 | 100.572678 | 0.073335411 | 0.22085734 | 0.33204878 | 0.73985243 | 0.93603056 |
| CSRP2 | 57.5070843 | -0.028092367 | 0.33304082 | -0.0843511 | 0.93277727 | 0.98725539 |
| NAV3 | 162.862703 | -0.295251164 | 0.40430019 | -0.7302771 | 0.46522083 | 0.80342007 |
| SYT1 | 237.166497 | 0.958091499 | 0.63598709 | 1.50646376 | 0.13194816 | 0.51290135 |
| AC027288.3 | 36.7777697 | 0.758916455 | 0.44431509 | 1.70805913 | 0.08762537 | 0.43393454 |
| PAWR | 245.061473 | 0.830330046 | 0.20319107 | 4.08644958 | 4.38E-05 | 0.00466528 |
| PPP1R12A | 360.890465 | 0.407190907 | 0.23921889 | 1.70216871 | 0.08872375 | 0.43544541 |
| LIN7A | 71.8374228 | -1.002060641 | 0.46283199 | -2.1650635 | 0.03038282 | 0.26581354 |
| ACSS3 | 166.484104 | -0.533673364 | 0.28309302 | -1.885152 | 0.05940932 | 0.36322722 |
| PPFIA2 | 77.1458176 | -0.017861804 | 0.40178696 | -0.0444559 | 0.964541 | 0.99275266 |
| METTL25 | 35.9454826 | 0.231295568 | 0.30671608 | 0.75410317 | 0.45078727 | 0.79401729 |
| TMTC2 | 113.289834 | 0.283173259 | 0.25988604 | 1.08960552 | 0.27588695 | 0.67135536 |
| ALX1 | 27.2800002 | -0.054420294 | 0.61658712 | -0.0882605 | 0.92966963 | 0.98681039 |
| CEP290 | 62.4517176 | -0.062800365 | 0.19802776 | -0.3171291 | 0.75114564 | 0.94055881 |
| TMTC3 | 43.6069143 | 0.219203911 | 0.22343363 | 0.98106947 | 0.32655848 | 0.71149668 |
| KITLG | 61.4722215 | 0.806912049 | 0.23266945 | 3.46806191 | 0.00052423 | 0.0246904 |
| DUSP6 | 25.6985985 | -0.264376156 | 0.46008577 | -0.5746236 | 0.56554586 | 0.85765509 |
| POC1B | 38.6954368 | -0.316801697 | 0.21802198 | -1.4530723 | 0.14620369 | 0.53584686 |
| ATP2B1 | 112.318709 | 0.248775669 | 0.22657695 | 1.09797429 | 0.27221571 | 0.66723562 |
| LUM | 141.219111 | -0.302382548 | 0.47124818 | -0.6416631 | 0.52109198 | 0.83486047 |
| DCN | 402.118081 | 0.771035945 | 0.35236412 | 2.1881795 | 0.02865653 | 0.25557869 |
| BTG1 | 205.324036 | -0.914971247 | 0.41480635 | -2.2057793 | 0.02739946 | 0.24828436 |
| EEA1 | 170.394544 | 0.085184992 | 0.20135982 | 0.42304861 | 0.67225979 | 0.91482259 |
| AC138123.1 | 40.9000788 | 0.592372188 | 0.45507687 | 1.30169698 | 0.19301999 | 0.59223254 |
| NUDT4 | 80.255837 | -0.489808171 | 0.31803366 | -1.5401143 | 0.1235325 | 0.50097964 |
| UBE2N | 49.0364983 | -0.032215363 | 0.29098383 | -0.1107119 | 0.91184484 | 0.98343817 |
| MRPL42 | 37.5375235 | 0.017758101 | 0.28582183 | 0.06212997 | 0.95045933 | 0.99060401 |
| SOCS2 | 26.8316436 | -0.012214715 | 0.37864523 | -0.032259 | 0.97426551 | 0.99300191 |
| CRADD | 182.300341 | -0.076889 | 0.24726128 | -0.3109626 | 0.75582909 | 0.94216126 |
| CEP83 | 63.0393666 | -0.462761555 | 0.16977917 | -2.7256674 | 0.00641716 | 0.10918078 |
| NDUFA12 | 19.7728064 | -0.310289368 | 0.29990449 | -1.0346273 | 0.300843 | 0.69211307 |
| NR2C1 | 54.6390409 | -0.012067964 | 0.19652632 | -0.0614064 | 0.95103559 | 0.99060401 |
| FGD6 | 34.7433176 | -0.288241472 | 0.4296952 | -0.6708045 | 0.50234508 | 0.82430366 |
| VEZT | 128.088882 | -0.103179637 | 0.28344888 | -0.364015 | 0.71584682 | 0.93237409 |
| METAP2 | 58.4749341 | -0.015851135 | 0.18664885 | -0.0849249 | 0.93232109 | 0.98725539 |
| NTN4 | 95.2908951 | 0.680418475 | 0.41206712 | 1.65123215 | 0.09869118 | 0.45657782 |
| LTA4H | 21.3589234 | -0.156397111 | 0.26923333 | -0.580898 | 0.56130918 | 0.85493496 |
| ELK3 | 37.9123871 | -0.244221622 | 0.40081226 | -0.6093167 | 0.54231451 | 0.84643373 |
| CDK17 | 143.337942 | -0.129904787 | 0.36482308 | -0.3560761 | 0.72178356 | 0.93444541 |
| NEDD1 | 21.4591591 | 0.259842902 | 0.27551705 | 0.94311005 | 0.34562461 | 0.72364538 |
| TMPO | 31.0182613 | 0.245568083 | 0.24565848 | 0.99963202 | 0.31748862 | 0.70575435 |
| SLC25A3 | 76.4721841 | -0.200094006 | 0.3317823 | -0.6030882 | 0.54644999 | 0.84758329 |
| IKBIP | 26.9749932 | -0.641560912 | 0.33002064 | -1.9440024 | 0.05189516 | 0.34513804 |
| APAF1 | 26.5513287 | -0.123686594 | 0.26124687 | -0.4734472 | 0.63589417 | 0.89741446 |
| ANKS1B | 81.6644492 | 1.094062472 | 0.46328454 | 2.36153458 | 0.01819947 | 0.19733204 |
| UHRF1BP1L | 76.9262344 | -0.24384819 | 0.26950787 | -0.9047906 | 0.36557633 | 0.73747473 |
| ACTR6 | 20.0301068 | -0.067356124 | 0.26046447 | -0.2586 | 0.79594385 | 0.95463848 |
| SCYL2 | 71.724482 | -0.143679868 | 0.18895125 | -0.7604071 | 0.44701131 | 0.79296348 |
| UTP20 | 35.1794834 | -0.381588833 | 0.32265532 | -1.1826516 | 0.23694725 | 0.63270446 |
| ARL1 | 33.2328094 | -0.148556159 | 0.30716405 | -0.4836378 | 0.62864292 | 0.8927912 |
| CHPT1 | 103.706903 | -0.334360477 | 0.36340038 | -0.9200884 | 0.35752656 | 0.73129235 |
| GNPTAB | 56.5428806 | -0.005263589 | 0.25978641 | -0.0202612 | 0.98383499 | 0.99620525 |
| DRAM1 | 104.710783 | -0.923112519 | 0.27173545 | -3.3971 | 0.00068104 | 0.02893045 |
| WASHC3 | 38.0524408 | 0.437292263 | 0.25323662 | 1.72681288 | 0.08420127 | 0.42616747 |
| IGF1 | 112.223089 | -0.636823356 | 0.75457234 | -0.8439527 | 0.39869585 | 0.76149359 |
| HSP90B1 | 184.430979 | -0.47905239 | 0.37682583 | -1.2712833 | 0.20362787 | 0.60367992 |
| TDG | 31.5530867 | -0.371355871 | 0.24310166 | -1.5275744 | 0.12661825 | 0.50599102 |
| HCFC2 | 40.3050341 | -0.03251278 | 0.20761633 | -0.1566003 | 0.87555987 | 0.9768643 |
| NFYB | 22.6670355 | -0.278995986 | 0.25651641 | -1.0876341 | 0.27675669 | 0.67259169 |
| TXNRD1 | 300.145884 | -0.577810068 | 0.26512535 | -2.1793844 | 0.02930312 | 0.2588325 |
| CHST11 | 190.341385 | -1.542696317 | 0.30423805 | -5.0706883 | 3.96E-07 | 0.00015332 |
| SLC41A2 | 37.7928027 | -0.093403279 | 0.31398425 | -0.2974776 | 0.76610191 | 0.9447922 |
| WASHC4 | 153.259139 | 0.000432653 | 0.29186827 | 0.00148236 | 0.99881725 | 0.99969762 |
| APPL2 | 58.645468 | -0.135038037 | 0.30485183 | -0.4429628 | 0.65779261 | 0.90772143 |
| C12orf75 | 32.5072201 | 0.761186911 | 0.37258374 | 2.04299553 | 0.04105288 | 0.30341448 |
| NUAK1 | 123.35508 | 0.467921586 | 0.31884526 | 1.46755069 | 0.14222631 | 0.52958915 |
| CKAP4 | 39.6167195 | -0.374171755 | 0.41434503 | -0.9030439 | 0.3665026 | 0.73873494 |
| TCP11L2 | 23.5967505 | 0.351583472 | 0.31551904 | 1.1143019 | 0.26514969 | 0.65897365 |
| POLR3B | 30.772329 | 0.154152884 | 0.26800782 | 0.57518054 | 0.5651692 | 0.8576148 |
| RIC8B | 66.4082541 | 0.139334756 | 0.17209473 | 0.80963988 | 0.41814718 | 0.77722447 |
| TMEM263 | 51.648781 | -0.142761295 | 0.22033445 | -0.64793 | 0.51703023 | 0.83252743 |
| CRY1 | 74.0360508 | 0.156756222 | 0.38234144 | 0.40999014 | 0.68181318 | 0.91921361 |
| PWP1 | 22.5425949 | -0.026953991 | 0.28689161 | -0.0939518 | 0.92514743 | 0.98561736 |
| PRDM4 | 19.1001857 | 0.172132128 | 0.3302902 | 0.52115421 | 0.60225935 | 0.87903643 |
| SART3 | 30.4053545 | -0.290799956 | 0.24496164 | -1.1871245 | 0.23517853 | 0.63100657 |
| ISCU | 41.7687201 | -0.12126657 | 0.31420655 | -0.3859454 | 0.69953711 | 0.92769314 |
| CORO1C | 110.865206 | 0.377131527 | 0.34498409 | 1.09318527 | 0.27431246 | 0.66930898 |
| SSH1 | 121.998068 | -0.789657478 | 0.34598495 | -2.2823463 | 0.0224689 | 0.22134579 |
| ACACB | 144.333071 | 0.139901178 | 0.46697322 | 0.29959144 | 0.76448882 | 0.94462105 |
| KCTD10 | 38.2557881 | -0.069807058 | 0.24322319 | -0.2870082 | 0.77410601 | 0.9475938 |
| UBE3B | 32.0179029 | -0.048795148 | 0.23668982 | -0.2061565 | 0.83666866 | 0.9668633 |
| MMAB | 19.27229 | 0.051803309 | 0.47779626 | 0.10842134 | 0.91366148 | 0.98395113 |
| GLTP | 28.0727855 | -0.158658787 | 0.36672735 | -0.4326342 | 0.66528055 | 0.91235816 |
| TCHP | 37.1860029 | -0.355908968 | 0.24528381 | -1.4510088 | 0.1467774 | 0.53611686 |
| GIT2 | 84.6988812 | -0.701263018 | 0.24217057 | -2.8957401 | 0.00378266 | 0.07965254 |
| ANKRD13A | 55.3632833 | -0.780859136 | 0.49806679 | -1.56778 | 0.11693249 | 0.48989475 |
| IFT81 | 54.7121938 | -0.106203856 | 0.25610785 | -0.4146841 | 0.67837316 | 0.91825081 |
| ATP2A2 | 202.943035 | -0.138142127 | 0.21211597 | -0.6512576 | 0.51488024 | 0.83109052 |
| ANAPC7 | 39.0717363 | -0.013878768 | 0.3664828 | -0.0378702 | 0.96979119 | 0.99300191 |
| ARPC3 | 72.1216778 | -0.333250848 | 0.20854124 | -1.5980093 | 0.11004091 | 0.47680173 |
| VPS29 | 40.1135908 | -0.457683944 | 0.24241336 | -1.888031 | 0.05902178 | 0.36206268 |
| PPTC7 | 37.1645674 | -0.235894755 | 0.26670069 | -0.8844925 | 0.37643042 | 0.74787433 |
| PPP1CC | 36.1449159 | 0.090836445 | 0.2175123 | 0.41761522 | 0.67622847 | 0.91723939 |
| SH2B3 | 50.9841016 | 0.340101254 | 0.36643647 | 0.92813157 | 0.35333932 | 0.73059415 |
| ATXN2 | 295.029493 | 0.061590994 | 0.15074197 | 0.40858558 | 0.68284381 | 0.91933701 |
| BRAP | 26.7536884 | 0.097687962 | 0.28491583 | 0.34286604 | 0.73169924 | 0.93533792 |
| ALDH2 | 41.8293928 | -0.343892349 | 0.32694177 | -1.0518459 | 0.29287027 | 0.6842846 |
| MAPKAPK5 | 28.3845197 | 0.088694996 | 0.28632685 | 0.30976835 | 0.75673712 | 0.94226721 |
| ERP29 | 29.326391 | 0.145246183 | 0.28897124 | 0.50263197 | 0.61522305 | 0.88360697 |
| NAA25 | 78.2805911 | -0.32031872 | 0.26290441 | -1.2183848 | 0.2230778 | 0.61772146 |
| TRAFD1 | 25.9868804 | -0.537053114 | 0.30914472 | -1.7372223 | 0.08234793 | 0.42256146 |
| HECTD4 | 164.378419 | 0.184149546 | 0.15642385 | 1.17724728 | 0.23909682 | 0.63451757 |
| RPL6 | 112.472547 | -0.597524153 | 0.33817374 | -1.7669147 | 0.07724251 | 0.4114983 |
| PTPN11 | 80.7862488 | -0.170854277 | 0.21755861 | -0.7853253 | 0.43226288 | 0.78417671 |
| DDX54 | 18.3637899 | -0.576426331 | 0.36664491 | -1.5721651 | 0.11591227 | 0.48843995 |
| TPCN1 | 114.954679 | 0.206832859 | 0.26724578 | 0.77394247 | 0.43896482 | 0.78887428 |
| SLC8B1 | 32.7817338 | -0.031365452 | 0.33389983 | -0.0939367 | 0.92515943 | 0.98561736 |
| PLBD2 | 22.6558298 | -0.474070959 | 0.35248545 | -1.3449377 | 0.17864537 | 0.57682988 |
| RBM19 | 30.1272592 | -0.487245024 | 0.33896858 | -1.4374342 | 0.15059466 | 0.53880252 |
| TBX3 | 115.697919 | 0.153092786 | 0.43201265 | 0.35437107 | 0.72306081 | 0.93455731 |
| MED13L | 758.824881 | -0.209025585 | 0.2578542 | -0.8106348 | 0.41757543 | 0.77722447 |
| FBXW8 | 55.1632393 | 0.019155221 | 0.27346733 | 0.07004574 | 0.94415726 | 0.99052272 |

| | | | | | | |
|---|---|---|---|---|---|---|
| FBXO21 | 52.2920251 | -0.012426938 | 0.23871322 | -0.052058 | 0.95848246 | 0.99181157 |
| WSB2 | 25.3791534 | 0.236809706 | 0.24248736 | 0.97658577 | 0.32877427 | 0.71294249 |
| VSIG10 | 29.7188143 | 0.659102649 | 0.45583905 | 1.4459109 | 0.1482022 | 0.53680621 |
| PEBP1 | 126.576535 | 0.008788892 | 0.37474634 | 0.02345291 | 0.981289 | 0.99533442 |
| TAOK3 | 151.740312 | 0.08069797 | 0.22017736 | 0.36651347 | 0.71398196 | 0.93136293 |
| SUDS3 | 23.6111079 | 0.130630573 | 0.23680585 | 0.55163576 | 0.58119793 | 0.86660562 |
| HSPB8 | 24.4466441 | -0.13456198 | 0.42488726 | -0.3167004 | 0.75147092 | 0.94055881 |
| RAB35 | 18.4021876 | 0.028131791 | 0.33961093 | 0.08283535 | 0.93398246 | 0.98757541 |
| GCN1 | 36.1018339 | 0.146844489 | 0.27046226 | 0.54293893 | 0.58717185 | 0.870249 |
| RPLP0 | 171.140673 | -0.865836136 | 0.33799867 | -2.5616554 | 0.01041746 | 0.14309893 |
| PXN | 36.6126304 | 0.317841866 | 0.34673809 | 0.91666267 | 0.35931943 | 0.73264631 |
| SRSF9 | 35.4773382 | -0.27550452 | 0.28933324 | -0.9522049 | 0.3409931 | 0.71963879 |
| DYNLL1 | 73.1275426 | -0.005748456 | 0.37117622 | -0.0154871 | 0.98764355 | 0.99719329 |
| RNF10 | 74.2167228 | 0.003020572 | 0.17342394 | 0.01741727 | 0.98610373 | 0.99719329 |
| MLEC | 46.3283071 | 0.108841 | 0.32068036 | 0.33940651 | 0.73430352 | 0.93594619 |
| SPPL3 | 162.006928 | 0.129395671 | 0.17919776 | 0.72208308 | 0.4702434 | 0.80457054 |
| P2RX4 | 20.2877045 | 0.616094683 | 0.45522518 | 1.35338445 | 0.17593284 | 0.57311399 |
| CAMKK2 | 37.5256278 | 0.435318923 | 0.26263457 | 1.65750808 | 0.0974168 | 0.45507247 |
| ANAPC5 | 89.3064497 | 0.142177978 | 0.21255887 | 0.66888752 | 0.50356723 | 0.82430366 |
| RNF34 | 22.566097 | 0.216259021 | 0.32560461 | 0.66417678 | 0.50657718 | 0.82481485 |
| KDM2B | 37.7536325 | 0.310800775 | 0.36988792 | 0.84025662 | 0.40076452 | 0.76340551 |
| TMEM120B | 42.6030208 | 0.258729732 | 0.35077478 | 0.73759502 | 0.46076058 | 0.80039155 |
| SETD1B | 27.051615 | 0.099250241 | 0.39991081 | 0.24818094 | 0.80399441 | 0.95836189 |
| WDR66 | 39.6204897 | 0.356332387 | 0.49785561 | 0.7157344 | 0.47415536 | 0.80792204 |
| MLXIP | 127.777252 | -0.088288797 | 0.28333659 | -0.3116039 | 0.75534154 | 0.94189722 |
| CLIP1 | 154.054869 | 0.039653721 | 0.25274808 | 0.15689029 | 0.87533131 | 0.9768643 |
| ZCCHC8 | 48.787321 | 0.012610055 | 0.20730604 | 0.06082821 | 0.95149602 | 0.99060401 |
| RSRC2 | 67.6913535 | -0.00105627 | 0.19527226 | -0.0054092 | 0.99568409 | 0.99854015 |
| DENR | 41.9933119 | 0.002261149 | 0.2651785 | 0.00852689 | 0.99319661 | 0.99781297 |
| VPS37B | 48.5244429 | -0.013989074 | 0.26256007 | -0.0532795 | 0.9575092 | 0.99181157 |
| ARL6IP4 | 54.8690328 | 0.124758444 | 0.33149135 | 0.37635506 | 0.70665295 | 0.93009618 |
| PITPNM2 | 34.422788 | 0.214165012 | 0.28175183 | 0.76011933 | 0.44718326 | 0.79296348 |
| MPHOSPH9 | 41.4770915 | 0.171269229 | 0.24111073 | 0.71033434 | 0.47749683 | 0.8113582 |
| C12orf65 | 19.5089045 | 0.199675813 | 0.27197184 | 0.73417825 | 0.46284011 | 0.80225845 |
| SBNO1 | 81.1355187 | -0.174628496 | 0.1787023 | -0.9772034 | 0.32846846 | 0.71294249 |
| KMT5A | 19.0982615 | 0.343131858 | 0.33146896 | 1.03518548 | 0.3005823 | 0.69191459 |
| RILPL2 | 29.7382504 | 0.055553852 | 0.3837144 | 0.14477917 | 0.88488523 | 0.97833933 |
| SNRNP35 | 21.5656328 | 0.222098894 | 0.32108285 | 0.69171833 | 0.48911423 | 0.81720854 |
| RILPL1 | 57.9217985 | 0.208953784 | 0.25016947 | 0.83524894 | 0.40357756 | 0.76539855 |
| TMED2 | 30.9881541 | -0.29063662 | 0.28860647 | -1.0070343 | 0.31391828 | 0.70282967 |
| DDX55 | 19.5793914 | -0.11182445 | 0.33575023 | -0.3330584 | 0.73909016 | 0.93594619 |
| EIF2B1 | 19.6018654 | -0.195880699 | 0.27375395 | -0.7155356 | 0.47427815 | 0.80794393 |
| ATP6V0A2 | 30.1181087 | -0.22125387 | 0.39499981 | -0.5601366 | 0.57538623 | 0.86294056 |
| CCDC92 | 59.5813 | 0.391130024 | 0.24685506 | 1.58445215 | 0.11309086 | 0.48348153 |
| NCOR2 | 168.408714 | 0.054335032 | 0.22721226 | 0.23913777 | 0.81099876 | 0.95967025 |
| SCARB1 | 21.5691097 | -0.431121221 | 0.32781657 | -1.3151294 | 0.1884664 | 0.58668894 |
| UBC | 321.282803 | -0.534894023 | 0.31784695 | -1.6828666 | 0.09240092 | 0.44238068 |
| TMEM132C | 56.4337513 | -0.347494162 | 0.56360862 | -0.6165522 | 0.5375301 | 0.84423863 |
| SLC15A4 | 26.9561206 | -0.275500527 | 0.26519962 | -1.0388421 | 0.29887819 | 0.68984164 |
| STX2 | 26.989765 | 0.343623783 | 0.312466 | 1.09971577 | 0.27145598 | 0.66674046 |
| RAN | 40.7107443 | -0.558634409 | 0.41580414 | -1.3435037 | 0.17910892 | 0.57744792 |
| ADGRD1 | 53.7109599 | 0.803063303 | 0.56999146 | 1.4089041 | 0.15886353 | 0.55091387 |
| SFSWAP | 90.5405037 | -0.071080533 | 0.1797167 | -0.3955143 | 0.69246334 | 0.9248682 |
| ULK1 | 38.9767743 | 0.464080707 | 0.3018296 | 1.53755865 | 0.12415656 | 0.5016089 |
| EP400 | 70.6602665 | 0.011237555 | 0.20674715 | 0.0543541 | 0.95665305 | 0.99181157 |
| FBRSL1 | 36.0495512 | -0.188311014 | 0.31773195 | -0.5926726 | 0.5534003 | 0.85188037 |
| ANKLE2 | 54.4006821 | -0.319013157 | 0.27713578 | -1.1511078 | 0.24968789 | 0.64565668 |
| GOLGA3 | 63.3544961 | -0.621912816 | 0.32520737 | -1.9123577 | 0.05583033 | 0.35425851 |
| CHFR | 56.6273185 | -0.144848437 | 0.26074154 | -0.555525 | 0.57853563 | 0.86486944 |
| ZNF26 | 28.8432334 | 0.131983431 | 0.26390113 | 0.50012454 | 0.61698739 | 0.88490248 |
| ZNF84 | 30.9032249 | -0.011066306 | 0.36545191 | -0.0302812 | 0.97584283 | 0.99341584 |
| ZNF140 | 18.6284087 | 0.134172687 | 0.29916681 | 0.44848788 | 0.65380113 | 0.90646849 |
| MPHOSPH8 | 104.646672 | 0.051579897 | 0.28339184 | 0.18200911 | 0.85557557 | 0.97379567 |
| PSPC1 | 160.197585 | 0.119113888 | 0.16374006 | 0.72745723 | 0.46694591 | 0.80374006 |
| ZMYM5 | 43.1722748 | 0.17482335 | 0.22948221 | 0.76181656 | 0.44616949 | 0.79296348 |
| ZMYM2 | 259.90511 | -0.054412418 | 0.27877291 | -0.1951855 | 0.84524776 | 0.96952174 |
| CRYL1 | 26.6104057 | -0.078512889 | 0.33058161 | -0.2374993 | 0.81226949 | 0.96026221 |
| IFT88 | 81.173764 | 0.036008468 | 0.23151103 | 0.15553672 | 0.8763982 | 0.9768643 |
| XPO4 | 88.3776059 | 0.012227484 | 0.19942403 | 0.061314 | 0.95110914 | 0.99060401 |
| LATS2 | 93.8962773 | 0.611180676 | 0.29230244 | 2.09091883 | 0.03653534 | 0.28984581 |
| SAP18 | 39.5002475 | -0.341151406 | 0.30910448 | -1.1036767 | 0.26973341 | 0.6651094 |
| MRPL57 | 24.4674015 | -0.974931149 | 0.3009372 | -3.2396498 | 0.00119677 | 0.0431129 |
| ZDHHC20 | 55.4512092 | -0.195958467 | 0.27161896 | -0.7214462 | 0.47063503 | 0.80457054 |
| MICU2 | 106.661868 | 0.07918813 | 0.20725342 | 0.38208359 | 0.70239936 | 0.92865476 |
| SACS | 23.6615569 | 0.181996254 | 0.31051511 | 0.58611079 | 0.55780107 | 0.85279817 |
| MIPEP | 34.0422246 | -0.51460035 | 0.39794076 | -1.2931582 | 0.19595635 | 0.59411229 |
| PARP4 | 51.5269855 | -0.291736069 | 0.28422807 | -1.0264154 | 0.30469582 | 0.69650244 |
| MTMR6 | 34.3607845 | 0.05414978 | 0.2497969 | 0.21677523 | 0.82838352 | 0.96580172 |
| NUP58 | 85.4194942 | -0.667814387 | 0.29738881 | -2.2455935 | 0.02473005 | 0.23328859 |
| CDK8 | 149.367097 | 0.035780325 | 0.23757765 | 0.15060476 | 0.88028751 | 0.97781634 |
| WASF3 | 80.8580432 | -0.488302195 | 0.47330128 | -1.0316942 | 0.30221539 | 0.69340645 |
| USP12 | 111.476826 | 0.218504129 | 0.20858517 | 1.04755353 | 0.29484436 | 0.6859168 |
| RPL21 | 64.2917779 | 0.118042339 | 0.34627902 | 0.34088793 | 0.73318795 | 0.93574285 |
| GTF3A | 19.5745557 | -0.506584729 | 0.36655933 | -1.3819993 | 0.16697192 | 0.56278128 |
| MTIF3 | 27.6882071 | 0.042025716 | 0.24812989 | 0.16936982 | 0.86550576 | 0.9762143 |
| POLR1D | 26.6278886 | 0.253568688 | 0.23923307 | 1.05992323 | 0.28917953 | 0.67992862 |
| PAN3 | 306.486358 | 0.121892184 | 0.15729401 | 0.77493212 | 0.43837977 | 0.78846132 |
| FLT1 | 29.6681207 | -0.408079895 | 0.5612445 | -0.7270983 | 0.46716577 | 0.80374006 |
| POMP | 50.710421 | -0.596068447 | 0.28860609 | -2.0653356 | 0.03889125 | 0.29803111 |
| UBL3 | 80.6871305 | 0.04011783 | 0.18564455 | 0.21610023 | 0.82890962 | 0.96580172 |
| KATNAL1 | 65.2041138 | 0.495417904 | 0.3398395 | 1.45779967 | 0.14489577 | 0.53509498 |
| HMGB1 | 153.544561 | 0.376971536 | 0.18892492 | 1.99535106 | 0.0460046 | 0.32236114 |
| USPL1 | 36.2481529 | -0.235851647 | 0.19908738 | -1.184664 | 0.23615034 | 0.63222238 |
| HSPH1 | 42.7055603 | 0.462356686 | 0.33995148 | 1.36006669 | 0.17380882 | 0.57125528 |
| B3GLCT | 29.342177 | 0.759944558 | 0.33618095 | 2.26052237 | 0.02378885 | 0.22886465 |
| FRY | 323.388005 | 1.294783014 | 0.34814865 | 3.71905222 | 0.00019997 | 0.01312136 |
| N4BP2L1 | 35.4789109 | -0.440589594 | 0.38701548 | -1.1384289 | 0.25494145 | 0.64964102 |
| N4BP2L2 | 441.715611 | -0.139919297 | 0.13980316 | -1.0008307 | 0.31690864 | 0.70489153 |
| PDS5B | 298.072553 | -0.091256696 | 0.27215404 | -0.3353127 | 0.73738925 | 0.93594619 |
| STARD13 | 513.12434 | -0.416907479 | 0.31472931 | -1.3246541 | 0.1852859 | 0.5849081 |
| AL139383.1 | 90.4286467 | 0.839230667 | 0.54868433 | 1.52953277 | 0.12613242 | 0.50535459 |
| RFC3 | 25.7123377 | 0.244698234 | 0.3009466 | 0.8130952 | 0.41616348 | 0.77573656 |
| NBEA | 324.036046 | 0.591626012 | 0.24739365 | 2.39143572 | 0.01678262 | 0.19005424 |
| DCLK1 | 156.945953 | 1.24721868 | 0.48287737 | 2.58288908 | 0.00979768 | 0.14035704 |
| SPART | 128.072627 | -0.458526442 | 0.30393815 | -1.5086176 | 0.13139653 | 0.51290135 |
| SMAD9 | 28.4486746 | 0.333341545 | 0.34195593 | 0.97480848 | 0.32965527 | 0.71362138 |
| ALG5 | 28.0097732 | 0.139636101 | 0.24959183 | 0.55945783 | 0.5758493 | 0.86295198 |
| SUPT20H | 71.9314914 | 0.069097902 | 0.24217324 | 0.28532427 | 0.77539572 | 0.94805375 |
| POSTN | 178.237435 | 0.052182887 | 0.93978649 | 0.05552632 | 0.95571916 | 0.99181157 |
| TRPC4 | 323.42944 | -1.867839369 | 0.43079614 | -4.3357849 | 1.45E-05 | 0.00232038 |
| UFM1 | 70.6864624 | -0.696276533 | 0.23664227 | -2.9423168 | 0.00325766 | 0.07300955 |
| NHLRC3 | 33.5642818 | 0.148347039 | 0.38922894 | 0.38113055 | 0.70310638 | 0.9291726 |
| LHFPL6 | 913.98732 | 0.152351931 | 0.38186673 | 0.39896623 | 0.68991808 | 0.92269481 |

| | | | | | | |
|---|---|---|---|---|---|---|
| COG6 | 65.6888735 | -0.09154355 | 0.3045725 | -0.3005641 | 0.76374693 | 0.94462105 |
| FOXO1 | 210.024358 | -0.234821025 | 0.33255576 | -0.7061102 | 0.48011958 | 0.81237906 |
| MRPS31 | 33.1842374 | -0.12204692 | 0.22983714 | -0.5310148 | 0.59540853 | 0.87454826 |
| ELF1 | 176.39257 | 0.003097005 | 0.26448445 | 0.01170959 | 0.99065731 | 0.99759781 |
| WBP4 | 37.248869 | 0.190764868 | 0.24076404 | 0.79233122 | 0.42816757 | 0.78079491 |
| NAA16 | 68.0344903 | 0.014461257 | 0.22989821 | 0.06290287 | 0.94984385 | 0.99060401 |
| RGCC | 44.1031945 | -0.347286556 | 0.94466241 | -0.3676303 | 0.7131489 | 0.93115158 |
| VWA8 | 210.52942 | 0.19550809 | 0.2224306 | 0.87896221 | 0.37942176 | 0.74991898 |
| DGKH | 301.146149 | 0.282144914 | 0.24275444 | 1.16226468 | 0.24512797 | 0.64085572 |
| AKAP11 | 78.2845541 | 0.049409328 | 0.19024399 | 0.25971557 | 0.79508318 | 0.95463848 |
| EPSTI1 | 84.7667195 | -0.099056082 | 0.98437371 | -0.1006285 | 0.91984535 | 0.98528826 |
| DNAJC15 | 54.1227077 | -0.473699176 | 0.25995475 | -1.8222371 | 0.06841902 | 0.38947862 |
| ENOX1 | 102.175805 | 0.577123156 | 0.29601398 | 1.94964832 | 0.05121805 | 0.34374561 |
| CCDC122 | 25.6341148 | -0.223032461 | 0.48884323 | -0.4562454 | 0.64821354 | 0.90273286 |
| TSC22D1 | 292.654659 | 0.285068221 | 0.21813429 | 1.30684736 | 0.19126456 | 0.58975955 |
| GPALPP1 | 38.3181223 | 0.124116002 | 0.29649038 | 0.41861729 | 0.67549585 | 0.91675328 |
| GTF2F2 | 64.3040419 | -0.091320471 | 0.26299674 | -0.3472304 | 0.72841821 | 0.93455731 |
| TPT1 | 317.063674 | -0.356124159 | 0.36197394 | -0.9838392 | 0.32519457 | 0.71149668 |
| COG3 | 69.0438445 | -0.211208249 | 0.19681577 | -1.0731266 | 0.2832143 | 0.67727609 |
| ZC3H13 | 95.7824798 | -0.008781985 | 0.18369323 | -0.0478079 | 0.96186935 | 0.99202404 |
| LRCH1 | 226.882675 | 0.226018936 | 0.22424531 | 1.00790931 | 0.313498 | 0.70282967 |
| ESD | 42.2427402 | -0.130346443 | 0.22323374 | -0.5839012 | 0.5592868 | 0.85362382 |
| SUCLA2 | 59.5433758 | -0.108958693 | 0.34580348 | -0.3150885 | 0.75269447 | 0.94058012 |
| MED4 | 50.6741505 | 0.022143274 | 0.17428997 | 0.12704847 | 0.89890203 | 0.98058672 |
| ITM2B | 353.205117 | 0.259592808 | 0.19166437 | 1.35441347 | 0.1756045 | 0.57311399 |
| RB1 | 227.557592 | -0.264333283 | 0.11355654 | -2.327768 | 0.01992443 | 0.20828536 |
| LPAR6 | 38.6640392 | -0.353781866 | 0.3838459 | -0.9216768 | 0.35669717 | 0.73129235 |
| RCBTB2 | 20.0106076 | 0.482512896 | 0.27182682 | 1.77507463 | 0.07588557 | 0.40904738 |
| FNDC3A | 415.328546 | 0.190390334 | 0.2947009 | 0.64604598 | 0.51824958 | 0.83297983 |
| CDADC1 | 20.5192604 | 0.049754606 | 0.29832038 | 0.16678245 | 0.86754122 | 0.97634923 |
| CAB39L | 36.5354807 | 0.572900971 | 0.22644689 | 2.52995731 | 0.01140764 | 0.15215018 |
| SETDB2 | 25.9320086 | -0.173236252 | 0.40848344 | -0.4240961 | 0.6714957 | 0.91482259 |
| PHF11 | 40.3611226 | -0.229943367 | 0.34337928 | -0.6696483 | 0.50308198 | 0.82430366 |
| RCBTB1 | 23.5244154 | 0.444893696 | 0.37568012 | 1.18423541 | 0.23631989 | 0.63222238 |
| KPNA3 | 71.4919712 | 0.153168994 | 0.15147998 | 1.01115005 | 0.31194462 | 0.7017083 |
| DLEU2 | 330.829375 | 0.114755206 | 0.34063106 | 0.33689003 | 0.7361998 | 0.93594619 |
| TRIM13 | 42.9296242 | 0.04162311 | 0.27237823 | 0.15281365 | 0.87854524 | 0.97718052 |
| RNASEH2B | 49.7502616 | -0.186110845 | 0.26350613 | -0.7062866 | 0.48000993 | 0.81237906 |
| INTS6 | 156.70399 | -0.233368168 | 0.26562744 | -0.8785544 | 0.3796429 | 0.74991898 |
| WDFY2 | 130.402445 | 0.07699405 | 0.29563221 | 0.26043864 | 0.79452544 | 0.95454757 |
| VPS36 | 61.6171354 | 0.228912742 | 0.17501515 | 1.30795958 | 0.19088702 | 0.58975955 |
| SUGT1 | 38.816004 | 0.029690076 | 0.22286788 | 0.13321828 | 0.89402075 | 0.98023547 |
| PCDH17 | 46.5186068 | 0.230414682 | 0.60928184 | 0.37817421 | 0.70530118 | 0.92946306 |
| TDRD3 | 120.394365 | 0.186528952 | 0.21719441 | 0.85881103 | 0.39044478 | 0.75729345 |
| PCDH9 | 383.101567 | 1.310819483 | 0.4668177 | 2.80799014 | 0.00498518 | 0.09251529 |
| DACH1 | 612.039034 | 0.258401228 | 0.48873825 | 0.52871087 | 0.59700603 | 0.87535619 |
| DIS3 | 34.8134297 | -0.110023967 | 0.21191615 | -0.5191863 | 0.60363081 | 0.87903643 |
| PIBF1 | 168.592978 | -0.123886024 | 0.1802659 | -0.6872405 | 0.4919312 | 0.81810418 |
| KLF12 | 296.643195 | 0.577694083 | 0.29151539 | 1.98169325 | 0.04751358 | 0.3297075 |
| TBC1D4 | 178.083378 | -0.148940661 | 0.31752938 | -0.469061 | 0.63902601 | 0.89845076 |
| COMMD6 | 29.4547534 | 0.099452928 | 0.35335258 | 0.28145522 | 0.77836127 | 0.94820337 |
| UCHL3 | 25.6828884 | 0.177193047 | 0.28496505 | 0.62180625 | 0.53406928 | 0.84333372 |
| LMO7 | 166.275294 | 1.344432745 | 0.4017902 | 3.34610632 | 0.00081955 | 0.03346039 |
| AC001226.2 | 26.750668 | -0.894906414 | 0.3865538 | -2.3150889 | 0.02060807 | 0.21114239 |
| FBXL3 | 75.3797095 | -0.305953275 | 0.29307602 | -1.0439383 | 0.29651394 | 0.68784121 |
| MYCBP2 | 383.595681 | -0.245540194 | 0.251823 | -0.9750507 | 0.32953511 | 0.71362138 |
| EDNRB | 180.121589 | -0.732561654 | 0.45841183 | -1.5980426 | 0.11003349 | 0.47680173 |
| RBM26 | 186.773684 | 0.225673564 | 0.19261261 | 1.1716448 | 0.2413397 | 0.6354731 |
| NDFIP2 | 43.3251454 | 0.273695135 | 0.29539735 | 0.92653212 | 0.35416951 | 0.73059415 |
| SPRY2 | 34.5463371 | -0.12287077 | 0.59678357 | -0.2058883 | 0.83687814 | 0.9668633 |
| GPC5 | 36.3246148 | 1.241087885 | 0.44557712 | 2.78534921 | 0.00534701 | 0.09654835 |
| GPC6 | 760.92312 | 0.35478671 | 0.32434409 | 1.09385902 | 0.27401682 | 0.66924213 |
| GPR180 | 23.2257064 | -0.231170438 | 0.28324999 | -0.8161357 | 0.4144225 | 0.7747624 |
| ABCC4 | 88.3601462 | -0.147783968 | 0.7222191 | -0.2046248 | 0.83786525 | 0.96735869 |
| DZIP1 | 56.2853081 | 0.023745225 | 0.20209886 | 0.11749312 | 0.9064693 | 0.98306694 |
| DNAJC3 | 109.020245 | -0.174039023 | 0.23890706 | -0.72848 | 0.46631979 | 0.80355999 |
| UGGT2 | 272.376998 | -0.522318334 | 0.2439478 | -2.141107 | 0.03226541 | 0.27444273 |
| HS6ST3 | 252.821319 | 0.222099973 | 0.62469306 | 0.35553456 | 0.72218916 | 0.93444541 |
| MBNL2 | 418.23701 | 0.303951684 | 0.29273978 | 1.03829991 | 0.29913045 | 0.69014677 |
| RAP2A | 32.7821362 | 0.238393591 | 0.34319633 | 0.69462745 | 0.48728881 | 0.81595238 |
| IPO5 | 55.4368133 | 0.207267693 | 0.26929695 | 0.76966223 | 0.44150028 | 0.79053729 |
| FARP1 | 246.783432 | 0.489257297 | 0.16524783 | 2.96074872 | 0.00306892 | 0.07114671 |
| STK24 | 112.30639 | -0.589623992 | 0.21915645 | -2.6904251 | 0.00713611 | 0.11602488 |
| DOCK9 | 93.1376422 | 0.289659692 | 0.23287486 | 1.2438427 | 0.21355746 | 0.61001472 |
| UBAC2 | 114.335688 | -0.366957256 | 0.18334638 | -2.0014426 | 0.04534471 | 0.31959454 |
| TM9SF2 | 79.1502012 | -0.132638085 | 0.17610425 | -0.7531794 | 0.45134214 | 0.79401729 |
| CLYBL | 34.1766069 | -0.023173439 | 0.36212831 | -0.0639923 | 0.94897632 | 0.99060401 |
| PCCA | 253.546987 | -0.006064087 | 0.18201493 | -0.0333164 | 0.97342225 | 0.99300191 |
| GGACT | 21.8734531 | -0.691852259 | 0.2927499 | -2.3632878 | 0.0181136 | 0.19729344 |
| ITGBL1 | 54.2849481 | 1.599483311 | 0.90193034 | 1.77340005 | 0.07616244 | 0.40931834 |
| FGF14 | 160.158174 | 1.180336003 | 0.56187901 | 2.10069423 | 0.03566782 | 0.28821871 |
| TPP2 | 88.1800758 | -0.002174287 | 0.18692747 | -0.0116317 | 0.99071945 | 0.99759781 |
| EFNB2 | 23.5367933 | 0.398791649 | 0.53847482 | 0.74059479 | 0.45893917 | 0.79943607 |
| ARGLU1 | 183.284131 | 0.154736035 | 0.13840656 | 1.11798198 | 0.26357469 | 0.6579701 |
| FAM155A | 55.3563654 | 1.335618408 | 0.53119498 | 2.51436564 | 0.01192467 | 0.15621109 |
| ABHD13 | 19.7037874 | -0.017187199 | 0.37088253 | -0.0463414 | 0.96303818 | 0.99231823 |
| IRS2 | 55.0978245 | -0.605384306 | 0.23899377 | -2.5330547 | 0.01130733 | 0.1519151 |
| COL4A1 | 1460.23409 | -0.788328717 | 0.54135381 | -1.4562172 | 0.14533259 | 0.53538077 |
| COL4A2 | 1161.32251 | -0.700666594 | 0.39020217 | -1.7956502 | 0.07255017 | 0.40072954 |
| RAB20 | 47.4644916 | -0.744961103 | 0.45107737 | -1.6515151 | 0.09863343 | 0.45657782 |
| NAXD | 27.9039305 | -0.640777186 | 0.25807207 | -2.4829389 | 0.01303034 | 0.16653913 |
| CARS2 | 54.4148119 | 0.157124205 | 0.23781802 | 0.66069091 | 0.50881055 | 0.82663984 |
| ANKRD10 | 259.45902 | -0.274874636 | 0.24365031 | -1.1281522 | 0.25925564 | 0.65213055 |
| ARHGEF7 | 244.888979 | -0.196055423 | 0.23575379 | -0.8316109 | 0.4056286 | 0.76703528 |
| TUBGCP3 | 47.2279548 | 0.061583752 | 0.2404534 | 0.25611513 | 0.79786194 | 0.95489133 |
| ATP11A | 112.108125 | -0.716986684 | 0.34863881 | -2.0565315 | 0.03973132 | 0.30124947 |
| PCID2 | 34.4785212 | -0.056845537 | 0.22263489 | -0.2553308 | 0.79846764 | 0.95506814 |
| CUL4A | 80.2610244 | -0.059487959 | 0.20521418 | -0.2898823 | 0.77190628 | 0.94671883 |
| LAMP1 | 71.2295847 | 0.184706192 | 0.18293105 | 1.00970389 | 0.31263718 | 0.70282967 |
| DCUN1D2 | 27.7418241 | -0.018884168 | 0.24062116 | -0.0784809 | 0.93744551 | 0.98827804 |
| TMCO3 | 56.0294604 | -0.019635825 | 0.25390292 | -0.077336 | 0.93835629 | 0.98827804 |
| TFDP1 | 26.1701619 | -0.421819861 | 0.33096268 | -1.2745239 | 0.20247779 | 0.60243292 |
| GAS6 | 69.310391 | -0.049038608 | 0.27807805 | -0.1763484 | 0.86002027 | 0.97397549 |
| RASA3 | 99.9597006 | -0.785219177 | 0.38438465 | -2.0427954 | 0.0410727 | 0.30341448 |
| CDC16 | 49.1093253 | 0.094297291 | 0.17256256 | 0.54645279 | 0.58475473 | 0.86868052 |
| UPF3A | 64.7063517 | 0.475060511 | 0.18418421 | 2.57926842 | 0.00990098 | 0.14050816 |
| CCNB1IP1 | 30.3180297 | 0.055912239 | 0.32033506 | 0.17454299 | 0.86143875 | 0.97470567 |
| TEP1 | 25.0791539 | -0.072590909 | 0.24151386 | -0.3005662 | 0.76374529 | 0.94462105 |
| NDRG2 | 49.2051495 | 0.346929673 | 0.36545773 | 0.94930177 | 0.34246715 | 0.72125471 |
| ARHGEF40 | 26.3613567 | 0.472525858 | 0.33048968 | 1.42977493 | 0.15278163 | 0.54042019 |
| HNRNPC | 245.751711 | -0.425723189 | 0.21225381 | -2.005727 | 0.04488538 | 0.31898889 |
| SUPT16H | 54.161246 | 0.040072051 | 0.21812378 | 0.18371244 | 0.85423905 | 0.97375567 |
| CHD8 | 65.5298268 | -0.079120682 | 0.1891363 | -0.4183263 | 0.67570859 | 0.91687267 |

| | | | | | | |
|---|---|---|---|---|---|---|
| RAB2B | 21.6698726 | 0.58953966 | 0.29451731 | 2.00171483 | 0.04531541 | 0.31959454 |
| TOX4 | 33.9989607 | -0.269816538 | 0.21797563 | -1.2378289 | 0.21577951 | 0.61220904 |
| DAD1 | 55.6405892 | -0.351347363 | 0.36616783 | -0.9595255 | 0.33729409 | 0.71925477 |
| OXA1L | 19.0042946 | -0.561929906 | 0.37304006 | -1.5063527 | 0.13197666 | 0.51290135 |
| MMP14 | 42.4999422 | -0.587383262 | 0.34028298 | -1.7261611 | 0.08431843 | 0.42646669 |
| LRP10 | 37.9757972 | -0.190997417 | 0.29704673 | -0.6429878 | 0.52023203 | 0.83421016 |
| RBM23 | 34.5598358 | 0.517692785 | 0.26927542 | 1.92254007 | 0.05453784 | 0.35100338 |
| PSMB5 | 35.5569327 | -0.830190255 | 0.35634366 | -2.3297461 | 0.01981958 | 0.20828536 |
| ACIN1 | 100.156512 | -0.20334125 | 0.18350147 | -1.1081178 | 0.26781094 | 0.66289747 |
| SLC7A8 | 28.5333478 | 0.097353247 | 0.36342017 | 0.26788069 | 0.78879116 | 0.95129454 |
| PABPN1 | 38.8301851 | 0.428534119 | 0.28577879 | 1.4995309 | 0.13373596 | 0.51564825 |
| PSME1 | 28.0244708 | -0.493840383 | 0.38354312 | -1.2875746 | 0.19789408 | 0.59530234 |
| NFATC4 | 66.0674838 | -0.509347265 | 0.33590632 | -1.5163372 | 0.1294341 | 0.51204187 |
| KHNYN | 17.7499116 | 0.250819479 | 0.29090854 | 0.8621936 | 0.38858099 | 0.75549104 |
| NOVA1 | 88.7903822 | -0.335878435 | 0.26031087 | -1.2902974 | 0.19694743 | 0.59479429 |
| PRKD1 | 308.935288 | -0.020599618 | 0.21525983 | -0.0956965 | 0.9237616 | 0.98561736 |
| G2E3 | 40.868469 | -0.420352251 | 0.21799529 | -1.928263 | 0.05382242 | 0.34959616 |
| SCFD1 | 173.333219 | -0.184376452 | 0.15694471 | -1.174786 | 0.24008032 | 0.63451757 |
| STRN3 | 266.167395 | -0.015218093 | 0.21534424 | -0.0706687 | 0.94366145 | 0.99014392 |
| AP4S1 | 19.3817729 | 0.507039906 | 0.29437511 | 1.72242788 | 0.08499204 | 0.42746464 |
| HECTD1 | 183.134016 | 0.074076948 | 0.13710136 | 0.54030791 | 0.5889847 | 0.8702694 |
| HEATR5A | 41.4120523 | 0.32311026 | 0.2209474 | 1.46238544 | 0.14363562 | 0.53204544 |
| NUBPL | 111.6647 | 0.107872834 | 0.28197002 | 0.38256845 | 0.70203976 | 0.92865476 |
| ARHGAP5 | 272.70483 | -0.050735122 | 0.17759913 | -0.2856721 | 0.77512926 | 0.94805375 |
| AKAP6 | 125.286427 | 0.402035775 | 0.51346105 | 0.78299177 | 0.43363194 | 0.78484089 |
| NPAS3 | 202.332171 | 0.78698373 | 0.40762381 | 1.93066181 | 0.05352489 | 0.34902787 |
| EAPP | 42.483959 | 0.000104559 | 0.28195579 | 0.00037084 | 0.99970412 | 0.99970412 |
| SNX6 | 89.7597421 | 0.009877882 | 0.22406116 | 0.04408565 | 0.96483613 | 0.99275266 |
| CFL2 | 19.746169 | 0.425845424 | 0.31127477 | 1.36806921 | 0.17129041 | 0.56870244 |
| BAZ1A | 81.0440376 | -0.677298862 | 0.32185582 | -2.1043549 | 0.0353475 | 0.28703733 |
| SRP54 | 85.649456 | -0.302594523 | 0.205269 | -1.4741365 | 0.14044484 | 0.52680084 |
| FAM177A1 | 45.413697 | -0.012452749 | 0.26390112 | -0.0471872 | 0.96236405 | 0.99231823 |
| PPP2R3C | 44.503053 | 0.033520008 | 0.23333778 | 0.14365444 | 0.88577335 | 0.97867793 |
| NFKBIA | 126.599633 | -0.774209722 | 0.55923731 | -1.3844028 | 0.16623514 | 0.56208655 |
| RALGAPA1 | 288.359343 | -0.036934598 | 0.18141846 | -0.2035879 | 0.83867557 | 0.96757092 |
| BRMS1L | 26.8079981 | 0.459013602 | 0.27484543 | 1.67007909 | 0.09490372 | 0.44764276 |
| MBIP | 19.7858132 | 0.075869006 | 0.2694412 | 0.28157908 | 0.77826628 | 0.94820337 |
| MIPOL1 | 119.316271 | -0.025309013 | 0.23385505 | -0.1082252 | 0.91381704 | 0.98395113 |
| SEC23A | 118.795818 | -0.227415262 | 0.20956002 | -1.0852035 | 0.27783155 | 0.67269898 |
| TRAPPC6B | 32.6674839 | 0.341121876 | 0.24690519 | 1.38159052 | 0.16709747 | 0.56278128 |
| PNN | 67.089794 | 0.230765581 | 0.25194241 | 0.91594575 | 0.35969535 | 0.73273991 |
| MIA2 | 71.7369746 | -0.21965212 | 0.1745336 | -1.2585091 | 0.20820771 | 0.6059796 |
| FBXO33 | 48.7825802 | 0.092548416 | 0.2232519 | 0.41454705 | 0.67847352 | 0.91825081 |
| LRFN5 | 66.8110234 | -0.320750668 | 0.53544989 | -0.5990302 | 0.54915273 | 0.84855413 |
| KLHL28 | 38.4399304 | 0.16706203 | 0.30521699 | 0.54735494 | 0.5841349 | 0.86828629 |
| TOGARAM1 | 66.6811215 | -0.013820048 | 0.21120439 | -0.0654345 | 0.94782808 | 0.99060401 |
| PRPF39 | 40.5982926 | -0.37858838 | 0.32753119 | -1.155885 | 0.24772821 | 0.64333128 |
| FKBP3 | 20.4519132 | -0.037664731 | 0.28473105 | -0.1322818 | 0.89476142 | 0.98023547 |
| MIS18BP1 | 32.2076583 | -0.227838139 | 0.27833236 | -0.818583 | 0.41302437 | 0.77264771 |
| MDGA2 | 62.3433449 | 0.433305487 | 0.54224131 | 0.79910084 | 0.42423195 | 0.77857941 |
| RPS29 | 61.0803461 | -0.261658135 | 0.23561402 | -1.1105372 | 0.26676761 | 0.66188434 |
| RPL36AL | 71.1810776 | -0.856525402 | 0.29602098 | -2.8934618 | 0.00381021 | 0.07977552 |
| KLHDC1 | 62.3261951 | 0.426555151 | 0.32882958 | 1.29719214 | 0.19456509 | 0.59272783 |
| KLHDC2 | 43.023896 | 0.195845656 | 0.29878017 | 0.65548412 | 0.51215611 | 0.82885603 |
| NEMF | 138.149362 | 0.018930074 | 0.14745559 | 0.12837814 | 0.89784973 | 0.98040427 |
| SOS2 | 202.983132 | 0.449222263 | 0.1719625 | 2.6123269 | 0.00899282 | 0.13367648 |
| L2HGDH | 18.9838957 | 0.192491261 | 0.32382399 | 0.59443176 | 0.55222337 | 0.85186599 |
| ATP5S | 18.5225747 | 0.314226986 | 0.30297739 | 1.03713015 | 0.29967521 | 0.69081339 |
| CDKL1 | 35.9568558 | -0.059432173 | 0.42024169 | -0.1414238 | 0.88753516 | 0.97920559 |
| MAP4K5 | 176.985244 | -0.100325342 | 0.23039743 | -0.4354447 | 0.66323966 | 0.91141325 |
| ATL1 | 21.7406234 | 0.310007245 | 0.39876226 | 0.77742374 | 0.43690881 | 0.78768252 |
| SAV1 | 55.5506349 | -0.382661881 | 0.32120096 | -1.1913473 | 0.23351729 | 0.62956917 |
| NIN | 78.7850329 | 0.134016855 | 0.30351592 | 0.44154802 | 0.65881631 | 0.90831499 |
| PYGL | 45.1466982 | -0.494662374 | 0.37107274 | -1.3330604 | 0.18251199 | 0.58106792 |
| TRIM9 | 52.1144628 | 0.991127358 | 0.46671329 | 2.12363217 | 0.03370091 | 0.28048473 |
| TMX1 | 29.1539305 | -0.526472721 | 0.29397159 | -1.7908966 | 0.07330989 | 0.40189622 |
| FRMD6 | 98.0350331 | 0.459049312 | 0.24093972 | 1.9052455 | 0.05674817 | 0.35705408 |
| GNG2 | 30.8780503 | 0.5571286 | 0.35576388 | 1.56600665 | 0.11734706 | 0.49048688 |
| RTRAF | 35.6803611 | -0.351091845 | 0.25463227 | -1.3788191 | 0.16795053 | 0.56333568 |
| NID2 | 80.5808941 | 0.273730276 | 0.34530972 | 0.79270944 | 0.42794713 | 0.7807804 |
| TXNDC16 | 33.259586 | -0.144590626 | 0.44021617 | -0.3284537 | 0.74256865 | 0.93759079 |
| ERO1A | 119.691666 | -2.307611949 | 0.74010866 | -3.1179367 | 0.00182122 | 0.05398305 |
| PSMC6 | 55.8578862 | -0.223699018 | 0.20994785 | -1.065498 | 0.28665062 | 0.67921285 |
| FERMT2 | 257.593362 | 0.340409369 | 0.29832196 | 1.14108048 | 0.25383643 | 0.64832175 |
| DDHD1 | 63.159521 | -0.2182425 | 0.20257083 | -1.0773639 | 0.28131772 | 0.67606407 |
| CNIH1 | 26.8761713 | -0.435877111 | 0.2453145 | -1.7768094 | 0.0755996 | 0.4086805 |
| GMFB | 20.7357876 | -0.301698381 | 0.26676986 | -1.1309313 | 0.25808401 | 0.65130862 |
| SAMD4A | 444.860911 | -0.290459169 | 0.47563006 | -0.6106829 | 0.5414095 | 0.84641877 |
| MAPK1IP1L | 76.6116447 | -0.276616519 | 0.21253606 | -1.3015039 | 0.19308602 | 0.59223254 |
| LGALS3 | 46.9104934 | -0.67872768 | 0.27571893 | -2.4616651 | 0.01382937 | 0.17196626 |
| FBXO34 | 77.4699312 | 0.040635719 | 0.27459683 | 0.1479832 | 0.88235602 | 0.97782528 |
| ATG14 | 26.7855374 | -0.130475331 | 0.48718169 | -0.2678166 | 0.78884051 | 0.95129454 |
| KTN1 | 179.307399 | 0.279140177 | 0.17247129 | 1.61847329 | 0.10556064 | 0.46789211 |
| PELI2 | 309.620047 | 0.15609617 | 0.46035651 | 0.33907671 | 0.73455194 | 0.93594619 |
| TMEM260 | 57.4070484 | 0.064026802 | 0.20654663 | 0.30998715 | 0.75657073 | 0.94221967 |
| EXOC5 | 63.0301604 | 0.110258687 | 0.18179889 | 0.60648713 | 0.54419132 | 0.84730644 |
| AP5M1 | 28.8218377 | 0.197062135 | 0.23614489 | 0.83449671 | 0.40400114 | 0.76597036 |
| NAA30 | 20.237099 | 0.200545639 | 0.25598043 | 0.78344128 | 0.43336802 | 0.78484089 |
| ARMH4 | 47.8415224 | 0.431178512 | 0.21076826 | 2.04574686 | 0.04078128 | 0.30303503 |
| ACTR10 | 50.563633 | -0.010280126 | 0.17165723 | -0.0598875 | 0.95224522 | 0.99060401 |
| PSMA3 | 53.6807534 | -0.139463841 | 0.27189437 | -0.5129339 | 0.60799756 | 0.88042838 |
| ARID4A | 79.203461 | -0.064711098 | 0.30174945 | -0.2144531 | 0.83019375 | 0.96580172 |
| TIMM9 | 22.1732225 | -0.599854121 | 0.25788295 | -2.3260713 | 0.02001475 | 0.20893236 |
| KIAA0586 | 57.7538803 | -0.425211323 | 0.2453836 | -1.7328433 | 0.08312353 | 0.42333667 |
| DACT1 | 25.7397868 | 0.771783733 | 0.43135722 | 1.7891986 | 0.07358284 | 0.40189622 |
| DAAM1 | 278.628797 | 0.44342794 | 0.46350742 | 0.95667927 | 0.33872916 | 0.71925477 |
| GPR135 | 25.5273917 | -0.433469267 | 0.42712336 | -1.0148573 | 0.31017384 | 0.70051247 |
| L3HYPDH | 21.7547062 | 0.057923127 | 0.34040889 | 0.1701575 | 0.86488627 | 0.9762143 |
| JKAMP | 27.0850538 | 0.021993702 | 0.27840904 | 0.0789978 | 0.93703437 | 0.98827804 |
| PCNX4 | 118.749945 | -0.015175005 | 0.22647298 | -0.0670058 | 0.94657708 | 0.99060401 |
| DHRS7 | 21.6637126 | 0.437900978 | 0.33088699 | 1.32341551 | 0.18569724 | 0.58501647 |
| PPM1A | 52.8355976 | 0.241854075 | 0.21407106 | 1.12978408 | 0.25856722 | 0.6514155 |
| MNAT1 | 157.747977 | -0.184317793 | 0.19025436 | -0.9687967 | 0.33264664 | 0.71542878 |
| SLC38A6 | 42.4909995 | -0.584222248 | 0.25891596 | -2.2564165 | 0.02404456 | 0.23038262 |
| PRKCH | 74.4584745 | -0.440399043 | 0.33461535 | -1.3161352 | 0.18812866 | 0.58645909 |
| HIF1A | 755.382023 | -1.239841219 | 0.2818325 | -4.399213 | 1.09E-05 | 0.00177428 |
| HIF1A-AS2 | 268.023071 | -2.753228341 | 0.61219227 | -4.4973262 | 6.88E-06 | 0.00117607 |
| SNAPC1 | 40.3327377 | -1.322974323 | 0.31434219 | -4.2087074 | 2.57E-05 | 0.00349534 |
| RHOJ | 65.1941559 | 0.8548658 | 0.3693184 | 2.31471217 | 0.02062869 | 0.21114239 |
| PPP2R5E | 171.292193 | -0.076822236 | 0.19920469 | -0.3856447 | 0.69975981 | 0.92769314 |
| SGPP1 | 42.7653716 | 0.336305479 | 0.20302578 | 1.65646684 | 0.09762731 | 0.45507247 |
| SYNE2 | 580.656115 | 0.183974057 | 0.33300553 | 0.55246547 | 0.58062949 | 0.86624614 |
| ESR2 | 93.9217994 | -0.910585909 | 0.47016265 | -1.9367466 | 0.05277633 | 0.34706747 |

| | | | | | | |
|---|---|---|---|---|---|---|
| ZBTB25 | 39.841213 | -0.931204629 | 0.30073501 | -3.0964291 | 0.00195867 | 0.05557623 |
| ZBTB1 | 101.319144 | -0.661233321 | 0.29766835 | -2.221376 | 0.02632551 | 0.24214398 |
| PLEKHG3 | 42.8166798 | -0.15787769 | 0.32419515 | -0.4869835 | 0.62627002 | 0.8912584 |
| MAX | 73.6717721 | -0.709422753 | 0.30287615 | -2.3422866 | 0.01916599 | 0.20295514 |
| FUT8 | 98.8103069 | -0.222540079 | 0.22179516 | -1.0033586 | 0.31568787 | 0.70306809 |
| AL359232.1 | 31.4424041 | -0.379805207 | 0.46715727 | -0.8130136 | 0.41621027 | 0.77573656 |
| GPHN | 502.420801 | -0.368831516 | 0.21066439 | -1.7508014 | 0.07998012 | 0.41881121 |
| MPP5 | 54.7275968 | 0.356029503 | 0.27539019 | 1.29281841 | 0.19607387 | 0.59411229 |
| ATP6V1D | 36.7973685 | 0.058635838 | 0.28691917 | 0.20436361 | 0.83806936 | 0.96735869 |
| EIF2S1 | 32.2124492 | -0.316593015 | 0.26064634 | -1.2146459 | 0.22450117 | 0.61883685 |
| ZFYVE26 | 25.344908 | -0.073180081 | 0.33428112 | -0.2189178 | 0.82671409 | 0.96580172 |
| RAD51B | 205.308615 | 0.609389876 | 0.38985812 | 1.5631068 | 0.11802748 | 0.49172522 |
| ZFP36L1 | 434.67109 | -0.346648066 | 0.40738914 | -0.8509016 | 0.39482402 | 0.75988253 |
| ACTN1 | 403.415247 | 0.020411104 | 0.21083837 | 0.09680925 | 0.92287788 | 0.98561736 |
| DCAF5 | 138.212435 | -0.271448193 | 0.23159616 | -1.1720756 | 0.24116672 | 0.63527165 |
| GALNT16 | 22.3314469 | 0.196537692 | 0.56346991 | 0.34879891 | 0.72724028 | 0.93455731 |
| ERH | 39.7470223 | -0.242195525 | 0.27541532 | -0.8793829 | 0.37919369 | 0.74991898 |
| SLC39A9 | 52.0637175 | -0.200430463 | 0.21358001 | -0.9384327 | 0.34802209 | 0.72612016 |
| SUSD6 | 115.031066 | -0.842473324 | 0.31624602 | -2.6639808 | 0.0077222 | 0.12283642 |
| SRSF5 | 148.437796 | 0.247144233 | 0.21672018 | 1.14038402 | 0.25412635 | 0.64846337 |
| AL357153.2 | 38.255303 | 0.10402095 | 0.34340617 | 0.30290938 | 0.76195892 | 0.94444866 |
| MED6 | 29.5140061 | 0.015768519 | 0.28268069 | 0.05578209 | 0.95551541 | 0.99181157 |
| PCNX1 | 268.251844 | -0.657508431 | 0.25788644 | -2.5496045 | 0.01078452 | 0.14622774 |
| SIPA1L1 | 203.087753 | 0.118100039 | 0.22245232 | 0.53090047 | 0.59548775 | 0.87454826 |
| RGS6 | 134.318042 | 1.041112131 | 0.34877486 | 2.98505495 | 0.00283527 | 0.06800182 |
| ZFYVE1 | 31.7440793 | -0.04287543 | 0.27923631 | -0.1535453 | 0.87796825 | 0.9768643 |
| RBM25 | 129.579352 | 0.135070751 | 0.17429002 | 0.77497697 | 0.43835327 | 0.78846132 |
| PSEN1 | 66.7654181 | -0.041170419 | 0.16899211 | -0.2436233 | 0.80752256 | 0.95887596 |
| PAPLN | 23.3455704 | -0.378404213 | 0.4344181 | -0.87106 | 0.38372141 | 0.75256313 |
| NUMB | 197.074709 | -0.060779404 | 0.12802226 | -0.4747565 | 0.63496052 | 0.89672816 |
| DNAL1 | 18.8631741 | -0.216824064 | 0.2768029 | -0.7833157 | 0.43344174 | 0.78484089 |
| ELMSAN1 | 59.9566279 | -0.207833575 | 0.32219338 | -0.6450585 | 0.51888928 | 0.83297983 |
| AC005480.1 | 22.6526202 | -0.124781672 | 0.41034837 | -0.3040872 | 0.76106148 | 0.94425408 |
| ENTPD5 | 21.7274785 | 0.170973736 | 0.34791596 | 0.49142251 | 0.62312764 | 0.88872545 |
| BBOF1 | 22.4762494 | -0.258063829 | 0.32000433 | -0.8064386 | 0.41999002 | 0.77771095 |
| ALDH6A1 | 25.5970798 | 0.553550659 | 0.38398876 | 1.44158039 | 0.14942078 | 0.53828105 |
| LIN52 | 50.5565969 | -0.143574552 | 0.23566279 | -0.6092373 | 0.54236718 | 0.84643373 |
| ABCD4 | 34.9024468 | 0.278006399 | 0.24492918 | 1.13504809 | 0.25635517 | 0.65084527 |
| NPC2 | 48.7812709 | -0.783444315 | 0.3637631 | -2.1537213 | 0.03126204 | 0.26776774 |
| LTBP2 | 27.1928399 | 1.01879558 | 0.5765913 | 1.76692847 | 0.07724021 | 0.4114983 |
| AREL1 | 34.1197724 | 0.264616507 | 0.24381136 | 1.08533295 | 0.27777422 | 0.67269898 |
| YLPM1 | 100.466149 | 0.185758208 | 0.14617983 | 1.2707513 | 0.20381714 | 0.60367992 |
| DLST | 37.1614661 | -0.229963592 | 0.22105074 | -1.0403204 | 0.29819107 | 0.68868829 |
| PGF | 25.1737775 | 0.098934444 | 0.39832199 | 0.24837806 | 0.80384191 | 0.95836189 |
| MLH3 | 58.2843617 | 0.143744204 | 0.2084782 | 0.68949272 | 0.49051325 | 0.81720854 |
| NEK9 | 53.9257111 | 0.110311928 | 0.21184299 | 0.52072495 | 0.60255839 | 0.87903643 |
| TMED10 | 93.3927383 | -0.187288241 | 0.21574711 | -0.8680916 | 0.38534421 | 0.75336383 |
| FOS | 115.112711 | 2.808811143 | 0.77168361 | 3.63984814 | 0.0002728 | 0.0161874 |
| JDP2 | 20.2933682 | 1.211419988 | 0.26821515 | 4.51659799 | 6.28E-06 | 0.00111215 |
| FLVCR2 | 46.308704 | -2.03198549 | 0.6225132 | -3.2641645 | 0.00109787 | 0.04106353 |
| TTLL5 | 106.02892 | -0.417252872 | 0.31598534 | -1.3204817 | 0.18667423 | 0.58501647 |
| IFT43 | 112.417164 | -0.408578531 | 0.24057492 | -1.6983422 | 0.0894432 | 0.43646619 |
| TGFB3 | 65.7736541 | -0.50208069 | 0.50488688 | -0.994442 | 0.32000775 | 0.70728138 |
| GPATCH2L | 99.6571204 | -0.244738351 | 0.17194118 | -1.4233841 | 0.15462483 | 0.54412824 |
| IRF2BPL | 29.6350599 | -0.383210567 | 0.3728376 | -1.0278217 | 0.30403372 | 0.69584049 |
| TMED8 | 32.493129 | -0.46466937 | 0.27277847 | -1.7034679 | 0.08848053 | 0.43544541 |
| AHSA1 | 24.1619678 | 0.126470099 | 0.37190692 | 0.34005848 | 0.73381249 | 0.93594619 |
| SPTLC2 | 54.0558519 | 0.359741062 | 0.24552818 | 1.4651722 | 0.14287394 | 0.53039327 |
| ALKBH1 | 23.9640874 | 0.192562378 | 0.33698276 | 0.57143094 | 0.56770756 | 0.85933735 |
| SLIRP | 29.9730312 | -0.410281532 | 0.31050529 | -1.3213351 | 0.18638966 | 0.58501647 |
| SNW1 | 40.4418554 | -0.291372506 | 0.20506613 | -1.4208709 | 0.15535429 | 0.54412824 |
| NRXN3 | 237.016404 | 0.448272688 | 0.50206328 | 0.89286093 | 0.37193165 | 0.74463512 |
| CEP128 | 35.5106078 | 0.42231555 | 0.44098373 | 0.95766697 | 0.33823072 | 0.71925477 |
| GTF2A1 | 44.5721471 | 0.038840932 | 0.20971117 | 0.18521156 | 0.8530631 | 0.97312893 |
| SEL1L | 74.6564402 | -0.235941196 | 0.22249697 | -1.0604243 | 0.2889516 | 0.67992862 |
| FLRT2 | 158.478121 | -0.110817381 | 0.48336663 | -0.2292615 | 0.81866564 | 0.9626198 |
| GALC | 27.2592156 | -0.05441184 | 0.2986278 | -0.1822062 | 0.8554209 | 0.97379567 |
| SPATA7 | 31.3071198 | 0.338938762 | 0.34968322 | 0.96927375 | 0.33240862 | 0.71533557 |
| PTPN21 | 55.7638163 | 1.020472354 | 0.3650174 | 2.79568141 | 0.00517904 | 0.09420986 |
| ZC3H14 | 73.9909436 | 0.09140522 | 0.15980522 | 0.57197894 | 0.56733624 | 0.85895221 |
| TTC8 | 24.2230885 | 0.485877634 | 0.26616408 | 1.82548163 | 0.06792838 | 0.38827623 |
| FOXN3 | 399.615014 | -0.263185261 | 0.29501375 | -0.8921118 | 0.37233299 | 0.74463512 |
| PSMC1 | 31.9627891 | -0.108078304 | 0.3013896 | -0.3586 | 0.71989436 | 0.93444541 |
| NRDE2 | 32.524343 | 0.367061843 | 0.37665709 | 0.97452524 | 0.32979581 | 0.71362138 |
| CALM1 | 97.81632 | 0.50775596 | 0.23790603 | 2.1342711 | 0.0328206 | 0.27660385 |
| TTC7B | 90.5796384 | 0.264593774 | 0.19361977 | 1.36656382 | 0.17176206 | 0.56870244 |
| RPS6KA5 | 55.939268 | 0.874201011 | 0.36592982 | 2.38898544 | 0.01689497 | 0.19013958 |
| DGLUCY | 49.8135101 | -0.196984116 | 0.3633291 | -0.5421644 | 0.58770523 | 0.870249 |
| PPP4R3A | 93.5926411 | 0.071465397 | 0.16818007 | 0.42493379 | 0.67088494 | 0.91475345 |
| CATSPERB | 27.2399013 | 0.814180798 | 0.33154231 | 2.45573726 | 0.01405959 | 0.17402303 |
| TC2N | 58.322557 | 0.677081091 | 0.44663927 | 1.51594616 | 0.12953297 | 0.51204187 |
| FBLN5 | 255.863291 | 1.006507592 | 0.40756618 | 2.46955623 | 0.01352808 | 0.169945 |
| TRIP11 | 114.948471 | 0.103652155 | 0.1527758 | 0.67845925 | 0.49748055 | 0.82230872 |
| ATXN3 | 102.270503 | 0.281171127 | 0.23533285 | 1.19478062 | 0.23217276 | 0.62798586 |
| NDUFB1 | 22.0619828 | -0.258805041 | 0.29213105 | -0.885921 | 0.37566008 | 0.74775892 |
| CPSF2 | 40.1588612 | -0.191971548 | 0.21871161 | -0.8777383 | 0.38008576 | 0.74991898 |
| RIN3 | 29.6164838 | -0.316556457 | 0.29496323 | -1.0732065 | 0.28317847 | 0.67727609 |
| LGMN | 36.9297372 | -0.120352905 | 0.32468246 | -0.3706788 | 0.71087678 | 0.93009618 |
| GOLGA5 | 31.554357 | -0.458422705 | 0.25511242 | -1.7969439 | 0.07234453 | 0.40009608 |
| ITPK1 | 30.1215077 | 0.480489502 | 0.31921205 | 1.50523611 | 0.13226338 | 0.51310477 |
| BTBD7 | 115.996961 | 0.032288329 | 0.19466861 | 0.16586304 | 0.86826473 | 0.97656528 |
| DDX24 | 82.8307572 | -0.316856806 | 0.1800021 | -1.7602951 | 0.07835779 | 0.41517764 |
| IFI27 | 74.1577879 | -0.378111093 | 0.95371814 | -0.39646 | 0.69176571 | 0.924661 |
| SERPINA1 | 44.5349063 | -2.50604595 | 0.83436534 | -3.0035355 | 0.00266862 | 0.06559102 |
| DICER1 | 193.364502 | -0.503465943 | 0.183225 | -2.7478015 | 0.00599963 | 0.10497927 |
| CLMN | 327.80131 | -0.480930303 | 0.25149356 | -1.9122967 | 0.05583815 | 0.35425851 |
| GLRX5 | 17.8263626 | -0.211808611 | 0.35142956 | -0.6027057 | 0.54670446 | 0.84758329 |
| C14orf132 | 25.361384 | 0.455002123 | 0.45200965 | 1.00662038 | 0.31411724 | 0.70282967 |
| ATG2B | 78.6765319 | -0.080114382 | 0.19442546 | -0.4120571 | 0.6802976 | 0.91860979 |
| PAPOLA | 132.779255 | 0.001957699 | 0.20464893 | 0.00956613 | 0.99236745 | 0.99759781 |
| SETD3 | 76.1612792 | -0.168409019 | 0.22222709 | -0.757824 | 0.44855636 | 0.7929855 |
| CCNK | 28.4808879 | -0.165852496 | 0.21862397 | -0.7586199 | 0.44807997 | 0.79296348 |
| EML1 | 40.3660934 | 1.811841062 | 0.44301355 | 4.08980958 | 4.32E-05 | 0.00466528 |
| EVL | 131.281093 | -0.676744201 | 0.4161461 | -1.6262178 | 0.10390333 | 0.46422585 |
| YY1 | 92.6073593 | -0.066659958 | 0.21075724 | -0.3162879 | 0.75178402 | 0.94055881 |
| SLC25A29 | 18.2464537 | -0.362180466 | 0.33769765 | -1.0724992 | 0.28349587 | 0.67731182 |
| WARS | 48.4868137 | -0.804959583 | 0.49229839 | -1.635105 | 0.10202703 | 0.46169744 |
| WDR25 | 33.5784357 | -0.207673317 | 0.30756129 | -0.6752258 | 0.49953231 | 0.82271448 |
| MEG3 | 73.4014973 | -0.437583647 | 0.34701349 | -1.260999 | 0.20730922 | 0.6059796 |
| MEG8 | 98.0752905 | 0.414006581 | 0.39184922 | 1.05654563 | 0.290719 | 0.68193232 |
| PPP2R5C | 114.911189 | 0.062261257 | 0.18719324 | 0.33260419 | 0.73943308 | 0.93594619 |
| DYNC1H1 | 127.492838 | -0.032747182 | 0.21142729 | -0.1548863 | 0.87691099 | 0.9768643 |
| HSP90AA1 | 249.543745 | 0.546602329 | 0.34208309 | 1.59786422 | 0.1100732 | 0.47680173 |

| | | | | | | |
|---|---|---|---|---|---|---|
| WDR20 | 81.2327565 | 0.145886935 | 0.17780009 | 0.82051101 | 0.41192485 | 0.77186021 |
| MOK | 31.0561639 | 0.584600103 | 0.46518047 | 1.25671678 | 0.20885621 | 0.6059796 |
| TECPR2 | 49.7985204 | 0.212309851 | 0.24885815 | 0.85313601 | 0.39358388 | 0.75988253 |
| RCOR1 | 86.9973063 | 0.055139334 | 0.19364865 | 0.28473906 | 0.77584406 | 0.94805375 |
| TRAF3 | 95.4875126 | -0.732588825 | 0.43786689 | -1.6730857 | 0.09431043 | 0.44657689 |
| CDC42BPB | 122.001481 | 0.357094332 | 0.16375959 | 2.18060101 | 0.02921294 | 0.25858362 |
| TNFAIP2 | 53.7784776 | -1.104870533 | 0.89017389 | -1.2411851 | 0.21453739 | 0.61009832 |
| EIF5 | 55.545548 | -0.045964502 | 0.25302078 | -0.181663 | 0.85584724 | 0.97379567 |
| MARK3 | 301.584994 | -0.113706573 | 0.21385542 | -0.5316983 | 0.59493494 | 0.87429987 |
| CKB | 40.749023 | 0.612023591 | 0.34918327 | 1.75272886 | 0.07964856 | 0.41779962 |
| KLC1 | 97.1476039 | 0.251435127 | 0.19271646 | 1.30468944 | 0.19199862 | 0.59087012 |
| ZFYVE21 | 25.397651 | 0.398399712 | 0.27112764 | 1.46941755 | 0.14171957 | 0.52921602 |
| PPP1R13B | 46.6512821 | 0.382746983 | 0.29151712 | 1.31294857 | 0.18920028 | 0.58671246 |
| ATP5MPL | 31.6636839 | -0.150222661 | 0.23903088 | -0.6284655 | 0.52969904 | 0.84095015 |
| INF2 | 36.8075324 | 0.653460952 | 0.20713819 | 3.15471011 | 0.00160658 | 0.05045611 |
| SIVA1 | 23.8264454 | -0.337869251 | 0.27551782 | -1.2263063 | 0.22008342 | 0.61590675 |
| AKT1 | 21.9444711 | -0.318002745 | 0.38769515 | -0.8202392 | 0.41207978 | 0.77195369 |
| BRF1 | 29.9731306 | -0.080248647 | 0.23235843 | -0.3453658 | 0.72981942 | 0.93455731 |
| PACS2 | 43.9153443 | 0.421574632 | 0.25609686 | 1.6461531 | 0.09973224 | 0.45779653 |
| MTA1 | 23.9865202 | 0.371291443 | 0.29206469 | 1.27126442 | 0.2036346 | 0.60367992 |
| CRIP2 | 49.6826334 | 1.170498176 | 0.45168285 | 2.59141602 | 0.00955819 | 0.1379822 |
| NIPA2 | 22.6996352 | -0.124159999 | 0.24703327 | -0.5026044 | 0.61524246 | 0.88360697 |
| CYFIP1 | 102.862208 | 0.371751277 | 0.21335071 | 1.74244217 | 0.0814311 | 0.42087829 |
| TUBGCP5 | 27.7064013 | 0.065181777 | 0.26642063 | 0.24465739 | 0.80672172 | 0.95838958 |
| PWRN1 | 29.9935827 | 1.484654659 | 0.6200917 | 2.39425018 | 0.01665438 | 0.19005424 |
| AC124312.1 | 116.564322 | 0.539711279 | 0.19242095 | 2.80484677 | 0.00503405 | 0.09281959 |
| PWAR6 | 16.8399266 | 0.613624666 | 0.31302016 | 1.96033591 | 0.04995654 | 0.33748407 |
| UBE3A | 142.595342 | 0.193691482 | 0.19042249 | 1.01716707 | 0.30907396 | 0.69910265 |
| ATP10A | 144.484495 | 0.320064358 | 0.29574549 | 1.08222904 | 0.27915077 | 0.67353297 |
| HERC2 | 149.59477 | 0.07424197 | 0.19880152 | 0.3734477 | 0.70881527 | 0.93009618 |
| TJP1 | 339.754049 | -0.001904803 | 0.25343248 | -0.007516 | 0.99400314 | 0.99819806 |
| FAN1 | 20.7776038 | -0.235228424 | 0.27231303 | -0.8638163 | 0.38768884 | 0.75457985 |
| MTMR10 | 61.4628039 | 0.0264729 | 0.21268783 | 0.12446833 | 0.90094447 | 0.98118567 |
| KLF13 | 50.6615938 | 0.298997563 | 0.25047129 | 1.19373987 | 0.23257974 | 0.62871357 |
| OTUD7A | 57.4958561 | -0.134971852 | 0.20480926 | -0.6590125 | 0.50988777 | 0.82718879 |
| LINC02256 | 96.964481 | -0.051421878 | 0.25170144 | -0.2042971 | 0.83812133 | 0.96735869 |
| AVEN | 51.7178212 | 0.034018446 | 0.30367286 | 0.11202333 | 0.91080491 | 0.98343817 |
| EMC7 | 27.2046455 | -0.64959123 | 0.25316387 | -2.5658923 | 0.01029108 | 0.14269647 |
| KATNBL1 | 88.3010117 | -0.102424146 | 0.17620007 | -0.5812946 | 0.56104193 | 0.85477139 |
| SLC12A6 | 32.7719705 | 0.028414108 | 0.35437107 | 0.08018179 | 0.93609267 | 0.98827804 |
| NOP10 | 28.7819129 | -0.504845823 | 0.44411543 | -1.1367446 | 0.25564507 | 0.65030654 |
| GOLGA8A | 41.0491136 | 0.117617856 | 0.31957059 | 0.36804969 | 0.71283618 | 0.93115158 |
| GOLGA8B | 20.8322769 | -0.152252667 | 0.36230672 | -0.4202314 | 0.6743164 | 0.91562489 |
| AQR | 63.1354534 | 0.320099102 | 0.19352901 | 1.65401096 | 0.09812529 | 0.45596917 |
| ZNF770 | 25.2597095 | -0.479608468 | 0.27708864 | -1.7308847 | 0.08347234 | 0.42393795 |
| AC021231.1 | 51.9301923 | 0.519378696 | 0.35022804 | 1.48297292 | 0.13808161 | 0.52264518 |
| DPH6 | 155.046571 | 0.419389978 | 0.30929006 | 1.35597628 | 0.17510672 | 0.57299474 |
| DPH6-DT | 109.50324 | 0.42526364 | 0.47968672 | 0.88654453 | 0.37532417 | 0.74729269 |
| C15orf41 | 77.0150173 | 0.084579387 | 0.18218748 | 0.46424369 | 0.64247316 | 0.90036905 |
| MEIS2 | 1085.25955 | 0.097211934 | 0.26256309 | 0.37024219 | 0.71120204 | 0.93016975 |
| SPRED1 | 176.296497 | 0.099413625 | 0.21470129 | 0.46303227 | 0.64334124 | 0.90063554 |
| FAM98B | 18.4663228 | 0.398251677 | 0.27411795 | 1.45284787 | 0.146266 | 0.53584686 |
| THBS1 | 597.908846 | -0.518344209 | 0.5466517 | -0.9482166 | 0.3430192 | 0.72177972 |
| FSIP1 | 47.1893232 | -0.420896905 | 0.63079057 | -0.667253 | 0.50461054 | 0.82481485 |
| GPR176 | 71.0158013 | 0.413030869 | 0.72563729 | 0.56919741 | 0.56922218 | 0.85958696 |
| EIF2AK4 | 50.5368061 | 0.179775294 | 0.18699888 | 0.96137098 | 0.33636567 | 0.71883121 |
| SRP14 | 102.608503 | -0.530428622 | 0.29644179 | -1.789318 | 0.07356362 | 0.40189622 |
| IVD | 28.9944173 | -0.114701345 | 0.38401792 | -0.2986875 | 0.76517851 | 0.9447922 |
| CCDC32 | 28.0561632 | 0.167672315 | 0.33771317 | 0.49649326 | 0.61954644 | 0.88632408 |
| DNAJC17 | 29.0441206 | 0.245566258 | 0.28232794 | 0.86979084 | 0.38441472 | 0.75288048 |
| INO80 | 104.365524 | 0.133435706 | 0.22067372 | 0.6046742 | 0.5453955 | 0.84758329 |
| CHP1 | 33.8191954 | -0.148628218 | 0.2291227 | -0.6486839 | 0.51654268 | 0.83210701 |
| RTF1 | 52.5186983 | 0.13719503 | 0.25749754 | 0.53280133 | 0.59417111 | 0.87394794 |
| MGA | 80.593102 | -0.029620723 | 0.19221826 | -0.1540994 | 0.87753134 | 0.9768643 |
| MAPKBP1 | 38.6133785 | -0.030776806 | 0.32780832 | -0.0938866 | 0.92519925 | 0.98561736 |
| VPS39 | 51.1694394 | -0.218684153 | 0.2093439 | -1.0446168 | 0.29620012 | 0.68754727 |
| TMEM87A | 105.283926 | -0.014992094 | 0.21703259 | -0.0690776 | 0.94492784 | 0.99060401 |
| GANC | 45.3046333 | -0.259985958 | 0.30601497 | -0.8495857 | 0.39555544 | 0.76020146 |
| ZNF106 | 94.2281399 | 0.00961574 | 0.24431913 | 0.0393573 | 0.96860553 | 0.99300191 |
| SNAP23 | 64.1157455 | -0.310165931 | 0.275004 | -1.1278597 | 0.25937918 | 0.65213055 |
| STARD9 | 88.8177286 | 0.836926284 | 0.33483379 | 2.49952757 | 0.0124359 | 0.16118421 |
| TTBK2 | 96.2317708 | 0.018084343 | 0.20437718 | 0.08848514 | 0.9294911 | 0.98681039 |
| UBR1 | 214.093771 | -0.059530855 | 0.21209261 | -0.2806833 | 0.77895331 | 0.94855475 |
| AC068724.3 | 20.5419099 | 0.079952478 | 0.40628778 | 0.1967878 | 0.8439936 | 0.9691438 |
| TMEM62 | 24.8911756 | 0.700411358 | 0.43867447 | 1.59665402 | 0.11034285 | 0.47706445 |
| TP53BP1 | 65.2314923 | 0.428515177 | 0.26262821 | 1.63164187 | 0.10275495 | 0.4627468 |
| CATSPER2 | 18.7227837 | 0.461613093 | 0.42524088 | 1.08553319 | 0.27768557 | 0.67269898 |
| PDIA3 | 110.56459 | -0.045903599 | 0.27361249 | -0.1677687 | 0.86676528 | 0.97624269 |
| SERF2 | 229.654433 | -0.53698433 | 0.31579653 | -1.7004124 | 0.08905338 | 0.43554112 |
| CASC4 | 139.936935 | 0.213349671 | 0.15321169 | 1.39251558 | 0.16376633 | 0.55847737 |
| CTDSPL2 | 106.201204 | -0.149084248 | 0.15607913 | -0.9551837 | 0.33948478 | 0.71925477 |
| EIF3J-DT | 23.8677188 | 0.212665312 | 0.33978647 | 0.62587928 | 0.53139412 | 0.84190241 |
| EIF3J | 52.7719966 | -0.092107013 | 0.28412254 | -0.3241806 | 0.74580132 | 0.93891908 |
| SPG11 | 110.428021 | -0.157642794 | 0.20562827 | -0.7666397 | 0.44329577 | 0.7920275 |
| B2M | 692.142505 | -0.173764878 | 0.38845794 | -0.4473197 | 0.65464426 | 0.90646849 |
| GATM | 23.3933739 | 1.09278657 | 0.54885214 | 1.99104002 | 0.04647649 | 0.32467273 |
| SEMA6D | 22.7811579 | 0.479832213 | 0.51397363 | 0.93357361 | 0.35052388 | 0.72904624 |
| MYEF2 | 22.3108708 | 0.117773306 | 0.35333109 | 0.33332279 | 0.73889064 | 0.93594619 |
| SLC12A1 | 63.3974416 | 1.902918069 | 0.49848745 | 3.81738408 | 0.00013487 | 0.00953067 |
| DUT | 27.5657189 | 0.390133694 | 0.23235085 | 1.67907152 | 0.09313811 | 0.44361507 |
| FBN1 | 305.834172 | 0.48237175 | 0.35007774 | 1.37789896 | 0.16823448 | 0.56403067 |
| EID1 | 71.7017611 | -0.175561797 | 0.37352774 | -0.4700101 | 0.63834783 | 0.89801267 |
| SECISBP2L | 96.3736539 | 0.00103318 | 0.17669866 | 0.00584713 | 0.99533469 | 0.99854015 |
| COPS2 | 43.5533151 | -0.15597408 | 0.26576764 | -0.5868814 | 0.55728337 | 0.85274692 |
| GALK2 | 90.0817577 | -0.605491163 | 0.231842 | -2.6116543 | 0.00901053 | 0.13367648 |
| FAM227B | 45.8396641 | -1.089263014 | 0.32831711 | -3.3177163 | 0.00090757 | 0.03556056 |
| FGF7 | 261.393099 | -1.89757531 | 0.48097731 | -3.9452491 | 7.97E-05 | 0.0068121 |
| DTWD1 | 99.6943983 | -0.506441562 | 0.34446465 | -1.470228 | 0.14150003 | 0.52895609 |
| GABPB1 | 53.1327218 | -0.286431005 | 0.3165136 | -0.9049564 | 0.36548849 | 0.73747473 |
| GABPB1-AS1 | 76.7984651 | 0.069156515 | 0.32538497 | 0.21253752 | 0.83168771 | 0.96634287 |
| USP8 | 82.285386 | 0.016833174 | 0.17226296 | 0.0977179 | 0.92215631 | 0.98561736 |
| TRPM7 | 182.432793 | -0.402817078 | 0.21256202 | -1.8950567 | 0.05808489 | 0.36041506 |
| SPPL2A | 102.279507 | -0.127932844 | 0.24273076 | -0.5270566 | 0.5981543 | 0.87601354 |
| AP4E1 | 38.8920667 | -0.146943834 | 0.195612 | -0.7512005 | 0.45253199 | 0.79504127 |
| DMXL2 | 98.7203292 | -0.920429609 | 0.35124416 | -2.6204837 | 0.00878051 | 0.1324861 |
| TMOD3 | 129.997114 | -0.079856604 | 0.20214606 | -0.3950441 | 0.69281035 | 0.9248682 |
| MAPK6 | 57.4561702 | -0.205806271 | 0.24885676 | -0.827007 | 0.40823312 | 0.76963974 |
| GNB5 | 33.1201863 | 0.035572882 | 0.29168188 | 0.12195781 | 0.90293243 | 0.98164203 |
| MYO5A | 61.555794 | 0.198958506 | 0.28045514 | 0.70941296 | 0.47806825 | 0.81139113 |
| ARPP19 | 50.9270075 | -0.058064229 | 0.22344928 | -0.2598542 | 0.79497626 | 0.95463848 |
| FAM214A | 130.358408 | -0.362240346 | 0.34664856 | -1.0449787 | 0.29603283 | 0.68737608 |
| WDR72 | 27.9573932 | 0.840177845 | 0.5073428 | 1.65603581 | 0.09771457 | 0.45507247 |
| RSL24D1 | 22.4504357 | -0.072589689 | 0.35826245 | -0.202616 | 0.83943522 | 0.96770443 |

| | | | | | | |
|---|---|---|---|---|---|---|
| RAB27A | 60.7618655 | -0.452971655 | 0.29358787 | -1.5428827 | 0.12285923 | 0.50097964 |
| PIGB | 22.1612762 | 0.414485895 | 0.38298535 | 1.08225 | 0.27914146 | 0.67353297 |
| CCPG1 | 75.3623699 | 0.461072144 | 0.27097707 | 1.7015172 | 0.08884591 | 0.43544541 |
| PYGO1 | 20.8304027 | 0.357051701 | 0.30910893 | 1.15509991 | 0.24804952 | 0.64333128 |
| NEDD4 | 99.8491991 | -0.278304552 | 0.1895772 | -1.4680275 | 0.14209674 | 0.52954815 |
| RFX7 | 124.037788 | -0.129850075 | 0.17395949 | -0.7464386 | 0.45540253 | 0.79609632 |
| TEX9 | 22.7985808 | 0.003557001 | 0.36925808 | 0.00963283 | 0.99231423 | 0.99759781 |
| ZNF280D | 73.9816893 | 0.285473469 | 0.2597598 | 1.09899016 | 0.27177236 | 0.66686313 |
| AC090517.4 | 60.6220062 | 0.582189812 | 0.36426505 | 1.59825878 | 0.1099854 | 0.47680173 |
| TCF12 | 666.979451 | -0.053684444 | 0.18306705 | -0.2932502 | 0.76933094 | 0.94579017 |
| CGNL1 | 79.5744812 | 0.864568971 | 0.29802727 | 2.90097269 | 0.00372006 | 0.07947309 |
| POLR2M | 20.5141153 | -0.287317622 | 0.34008044 | -0.8448519 | 0.39819351 | 0.7611823 |
| ALDH1A2 | 123.312537 | 0.113830676 | 0.27618971 | 0.41214669 | 0.68023191 | 0.91860979 |
| ADAM10 | 199.188008 | -0.385180868 | 0.21866047 | -1.7615478 | 0.07814573 | 0.41435277 |
| MINDY2 | 100.170653 | 0.154003675 | 0.33694997 | 0.45705205 | 0.64763363 | 0.90256375 |
| RNF111 | 112.184378 | -0.253055362 | 0.22909231 | -1.1046 | 0.26933296 | 0.6651094 |
| SLTM | 138.181892 | -0.024180049 | 0.15985628 | -0.1512612 | 0.8797697 | 0.97768448 |
| MYO1E | 395.251581 | 0.736367162 | 0.30875162 | 2.38498235 | 0.01707994 | 0.19063814 |
| GTF2A2 | 29.6852179 | -0.400714197 | 0.27368563 | -1.4641405 | 0.14315558 | 0.5309886 |
| BNIP2 | 49.3390162 | 0.079733855 | 0.23677026 | 0.3367562 | 0.73630069 | 0.93594619 |
| ANXA2 | 111.849131 | 0.415818952 | 0.42046489 | 0.98895047 | 0.32268737 | 0.70937168 |
| ICE2 | 56.1241277 | 0.071483514 | 0.21284919 | 0.33584114 | 0.73699067 | 0.93594619 |
| RORA-AS1 | 36.3726261 | -0.710817481 | 0.60979899 | -1.1656587 | 0.24375248 | 0.63964329 |
| RORA | 1352.51862 | 0.19764921 | 0.47474923 | 0.41632339 | 0.67717338 | 0.91767419 |
| AC012404.1 | 41.1624381 | 0.151336812 | 0.56881286 | 0.2660573 | 0.79019508 | 0.95194192 |
| AC009554.1 | 25.4701677 | -0.093613275 | 0.57600919 | -0.1625205 | 0.87089602 | 0.9768643 |
| VPS13C | 224.075547 | -0.271029301 | 0.17135861 | -1.5816497 | 0.11372956 | 0.48395978 |
| TLN2 | 179.980775 | 0.237512061 | 0.35138282 | 0.67593532 | 0.49908171 | 0.82271448 |
| TPM1 | 509.42666 | 0.948426404 | 0.32545108 | 2.91419036 | 0.00356612 | 0.07730809 |
| LACTB | 32.5587751 | -0.675806423 | 0.29155062 | -2.3179729 | 0.0204508 | 0.21078948 |
| RPS27L | 58.9816878 | -0.57161449 | 0.26476415 | -2.1589573 | 0.03085348 | 0.26662968 |
| RAB8B | 144.97734 | -0.733377163 | 0.36876801 | -1.9887223 | 0.04673187 | 0.32614671 |
| APH1B | 32.2947198 | 0.041263752 | 0.43989553 | 0.09380353 | 0.92526523 | 0.98561736 |
| CA12 | 122.392481 | -1.028633094 | 0.78347543 | -1.3129105 | 0.18921312 | 0.58671246 |
| USP3 | 132.194579 | -0.402290625 | 0.19364479 | -2.0774668 | 0.0377585 | 0.29501902 |
| USP3-AS1 | 29.9478336 | -0.243136551 | 0.29168986 | -0.8335448 | 0.40453757 | 0.76622334 |
| HERC1 | 343.0487 | 0.503564159 | 0.18884031 | 2.6666137 | 0.00766197 | 0.12228146 |
| DAPK2 | 122.366557 | 1.185080649 | 0.43452392 | 2.7273082 | 0.00638534 | 0.10918078 |
| FAM96A | 18.1690682 | -0.306558027 | 0.33093821 | -0.9263301 | 0.35427445 | 0.73059415 |
| SNX1 | 89.8462121 | 0.192198782 | 0.17865439 | 1.07581338 | 0.28201073 | 0.676814 |
| PPIB | 107.96863 | -0.449609183 | 0.36303957 | -1.2384578 | 0.21554636 | 0.61207504 |
| TRIP4 | 36.8246063 | 0.323391473 | 0.24057498 | 1.34424402 | 0.17886949 | 0.57704649 |
| ZNF609 | 298.029636 | -0.031388465 | 0.20125198 | -0.155966 | 0.87605983 | 0.9768643 |
| OAZ2 | 42.4595694 | 0.350442708 | 0.28463518 | 1.23119957 | 0.21824823 | 0.61478732 |
| RBPMS2 | 25.3465733 | 1.071925469 | 0.41677052 | 2.57198005 | 0.01011187 | 0.14181707 |
| ANKDD1A | 26.559388 | 0.264566051 | 0.32440647 | 0.81553876 | 0.41476399 | 0.77476594 |
| SPG21 | 32.9210252 | -0.052423287 | 0.24395649 | -0.2148879 | 0.82985475 | 0.96580172 |
| MTFMT | 25.3616783 | -0.243896502 | 0.4043944 | -0.6031154 | 0.5464319 | 0.84758329 |
| RASL12 | 26.5403267 | 0.733462619 | 0.30851242 | 2.37741682 | 0.01743437 | 0.19325071 |
| PDCD7 | 21.851523 | -0.169478432 | 0.38773542 | -0.4370981 | 0.66204018 | 0.91076998 |
| CLPX | 43.4757974 | -0.193802102 | 0.24262983 | -0.7987563 | 0.42443174 | 0.77861803 |
| DPP8 | 69.114827 | 0.234776549 | 0.21954174 | 1.06939368 | 0.28489232 | 0.6783522 |
| HACD3 | 36.4976781 | 0.108447846 | 0.25383937 | 0.4272302 | 0.66921167 | 0.91445841 |
| DENND4A | 186.244754 | -0.335158841 | 0.18498452 | -1.811821 | 0.07001386 | 0.39397538 |
| RAB11A | 86.844813 | -0.519811212 | 0.2356124 | -2.2062134 | 0.02736906 | 0.24828436 |
| DIS3L | 18.3615034 | 0.240758806 | 0.3935622 | 0.6117427 | 0.540708 | 0.84600024 |
| MAP2K1 | 82.2495475 | -0.457032784 | 0.36222338 | -1.2617429 | 0.20704131 | 0.6059796 |
| RPL4 | 105.643281 | -0.298046916 | 0.36828713 | -0.8092787 | 0.41835488 | 0.77722447 |
| SMAD6 | 29.6084131 | 1.294117576 | 0.46303301 | 2.79487109 | 0.00519204 | 0.09421312 |
| SMAD3 | 269.347936 | -0.289951019 | 0.37668219 | -0.7697497 | 0.44144836 | 0.79053729 |
| AAGAB | 45.7542287 | 0.113008206 | 0.30716654 | 0.36790532 | 0.71294383 | 0.93115158 |
| MAP2K5 | 138.818085 | -0.036307753 | 0.20644043 | -0.1758752 | 0.86039198 | 0.97397549 |
| PIAS1 | 220.61237 | -0.078122968 | 0.23531663 | -0.3319909 | 0.73989617 | 0.93603056 |
| FEM1B | 43.0221841 | -0.096871016 | 0.27352721 | -0.354155 | 0.72322276 | 0.93455731 |
| ITGA11 | 199.449719 | 0.751304304 | 0.26789239 | 2.80450036 | 0.00503946 | 0.09281959 |
| CORO2B | 56.4802023 | -0.126326123 | 0.44122908 | -0.2863051 | 0.77464447 | 0.94786251 |
| ANP32A | 41.2361222 | -0.222247997 | 0.26785027 | -0.8297471 | 0.40668176 | 0.76790963 |
| GLCE | 35.4135838 | 0.344865147 | 0.50862883 | 0.67802909 | 0.49775325 | 0.82238021 |
| RPLP1 | 253.010021 | -0.147048331 | 0.3510852 | -0.4188394 | 0.67533348 | 0.9167022 |
| TLE3 | 27.1168789 | -0.390038621 | 0.35354801 | -1.1032126 | 0.26993485 | 0.6651094 |
| UACA | 333.356268 | 0.649513097 | 0.32892463 | 1.9746563 | 0.04830717 | 0.33343326 |
| LRRC49 | 37.1363852 | -0.01221659 | 0.26392837 | -0.0462875 | 0.96308108 | 0.99231823 |
| THSD4 | 776.509008 | 0.760957913 | 0.41211685 | 1.84646155 | 0.06482522 | 0.38020792 |
| THSD4-AS1 | 53.0635322 | 0.278310861 | 0.30020744 | 0.92706184 | 0.35389442 | 0.73059415 |
| MYO9A | 306.616165 | 0.426983573 | 0.19405694 | 2.20030048 | 0.02778558 | 0.24993421 |
| PKM | 129.524227 | -1.210398981 | 0.29769208 | -4.0659428 | 4.78E-05 | 0.00488287 |
| PARP6 | 38.2482888 | 0.168335419 | 0.25049083 | 0.67202228 | 0.50156951 | 0.82406312 |
| AC009690.1 | 41.1578844 | 0.134074215 | 0.40173068 | 0.33374153 | 0.73857461 | 0.93594619 |
| ARIH1 | 292.693616 | 0.033342697 | 0.19335571 | 0.17244227 | 0.86308985 | 0.97582266 |
| BBS4 | 38.9195387 | -0.103746531 | 0.23551397 | -0.4405112 | 0.65956693 | 0.90855808 |
| ADPGK | 36.9608351 | -0.598959791 | 0.3209214 | -1.8663754 | 0.06198887 | 0.37037087 |
| NEO1 | 166.074558 | 0.673232807 | 0.24036935 | 2.80082635 | 0.00509719 | 0.09341466 |
| NPTN | 159.835198 | 0.210927858 | 0.22105472 | 0.95418844 | 0.33998827 | 0.71925477 |
| LOXL1 | 30.1301107 | 0.654351553 | 0.44122 | 1.48305051 | 0.13806099 | 0.52264518 |
| PML | 49.7658515 | -0.020110439 | 0.29351606 | -0.0685156 | 0.94537518 | 0.99060401 |
| ISLR | 42.7036903 | 0.119008871 | 0.35899172 | 0.33150868 | 0.74026028 | 0.93618531 |
| EDC3 | 21.3056738 | 0.372481912 | 0.29988406 | 1.24208638 | 0.21420469 | 0.61001472 |
| SCAMP2 | 19.0707798 | -0.0708456 | 0.29343206 | -0.2414378 | 0.80921579 | 0.95939969 |
| COX5A | 22.0663094 | -0.395720487 | 0.27804066 | -1.4232468 | 0.15466461 | 0.54412824 |
| PPCDC | 57.5645527 | 0.101256023 | 0.42987104 | 0.23554977 | 0.81378206 | 0.96038841 |
| MAN2C1 | 37.6759649 | 0.183942027 | 0.23061681 | 0.79760895 | 0.42509746 | 0.77861803 |
| SIN3A | 63.5168346 | 0.124928855 | 0.18662587 | 0.66940803 | 0.50323523 | 0.82430366 |
| PTPN9 | 103.931247 | 0.326222125 | 0.19384766 | 1.68287884 | 0.09239855 | 0.44238068 |
| SNX33 | 17.3290218 | 0.086620426 | 0.2878514 | 0.30092063 | 0.76347502 | 0.94462105 |
| CSPG4 | 53.7599067 | 0.325569463 | 0.37165096 | 0.87600867 | 0.38102532 | 0.74991898 |
| UBE2Q2 | 50.231392 | 0.214109556 | 0.2320433 | 0.92271379 | 0.35615638 | 0.73129235 |
| FBXO22 | 40.530048 | 0.096388628 | 0.24591181 | 0.39196421 | 0.69508466 | 0.92589762 |
| ETFA | 84.4209528 | -0.091161441 | 0.19613014 | -0.4648008 | 0.64207413 | 0.90015315 |
| SCAPER | 139.282428 | 0.17340437 | 0.29440125 | 0.58900691 | 0.55585664 | 0.85207344 |
| RCN2 | 41.3207231 | 0.341930098 | 0.23614134 | 1.44798913 | 0.14762009 | 0.53679369 |
| TSPAN3 | 52.3031707 | 0.057736967 | 0.191265 | 0.30186897 | 0.76275195 | 0.94462105 |
| PEAK1 | 305.683372 | 0.500667237 | 0.28055918 | 1.78453346 | 0.07433702 | 0.40409804 |
| HMG20A | 58.6064465 | 0.083520075 | 0.26496517 | 0.31521152 | 0.75260106 | 0.94058012 |
| LINGO1 | 478.091138 | 0.215602625 | 0.57945342 | 0.3720793 | 0.70983381 | 0.93009618 |
| TBC1D2B | 140.200665 | -0.199257434 | 0.20869119 | -0.9547956 | 0.33968106 | 0.71925477 |
| IDH3A | 30.9427817 | -0.069698437 | 0.31311197 | -0.2225991 | 0.82384755 | 0.96413949 |
| WDR61 | 29.0403741 | -0.112136038 | 0.25198562 | -0.4450097 | 0.65631276 | 0.90764819 |
| IREB2 | 85.7611645 | 0.006329309 | 0.15950302 | 0.03968144 | 0.9683471 | 0.99300191 |
| PSMA4 | 34.8348771 | -0.300699749 | 0.25298967 | -1.1885851 | 0.23460298 | 0.63080829 |
| MORF4L1 | 132.907919 | 0.144499981 | 0.18280269 | 0.79046968 | 0.42925352 | 0.78220055 |
| CTSH | 27.5545945 | -0.14652233 | 0.26652246 | -0.549756 | 0.58248672 | 0.867524 |
| TMED3 | 23.7184411 | -1.096471687 | 0.41409501 | -2.6478747 | 0.00809995 | 0.12587571 |
| ZFAND6 | 117.0981 | -0.121709112 | 0.19009031 | -0.6402699 | 0.52199712 | 0.8350836 |

| | | | | | | |
|---|---|---|---|---|---|---|
| ARNT2 | 24.0443209 | -0.112863989 | 0.38270814 | -0.2949088 | 0.76806357 | 0.94576899 |
| MESD | 32.7954138 | -0.053413136 | 0.28381065 | -0.1881999 | 0.85071995 | 0.97230808 |
| EFL1 | 27.3007516 | -0.392884324 | 0.3539652 | -1.1099518 | 0.26701978 | 0.66201068 |
| WHAMM | 31.1834627 | -0.28891398 | 0.24926181 | -1.1590784 | 0.24642424 | 0.64225758 |
| BTBD1 | 45.8721 | -0.002851096 | 0.22361949 | -0.0127498 | 0.98982744 | 0.99739981 |
| HDGFL3 | 69.7249706 | 0.249621644 | 0.18097398 | 1.37932341 | 0.16779506 | 0.56323722 |
| ADAMTSL3 | 127.257351 | 0.857720354 | 0.75190531 | 1.14072921 | 0.25398263 | 0.64832175 |
| WDR73 | 36.0105489 | -0.194728676 | 0.28077493 | -0.6935401 | 0.48797067 | 0.81595238 |
| SEC11A | 62.2315404 | 0.103833894 | 0.26317737 | 0.3945396 | 0.6931827 | 0.9248682 |
| ZNF592 | 46.5355571 | 0.173457487 | 0.21479137 | 0.80756265 | 0.4193424 | 0.77771095 |
| ALPK3 | 23.3143874 | -0.476833764 | 0.42774356 | -1.1147655 | 0.26495093 | 0.65892534 |
| PDE8A | 426.654465 | -0.928517559 | 0.24103524 | -3.8522066 | 0.00011706 | 0.00879608 |
| AKAP13 | 618.743334 | -0.147676201 | 0.31062903 | -0.4754102 | 0.63449465 | 0.89653935 |
| AC087286.2 | 43.9313704 | -0.406760121 | 0.56145976 | -0.7244689 | 0.46877793 | 0.80374006 |
| KLHL25 | 23.7048223 | -1.040887205 | 0.43205609 | -2.4091483 | 0.0159898 | 0.18618176 |
| NTRK3 | 402.844626 | -0.330726443 | 0.37294863 | -0.8867882 | 0.37519295 | 0.74723387 |
| DET1 | 21.4148827 | 0.431149167 | 0.567809 | 0.75932077 | 0.44766069 | 0.79296348 |
| MFGE8 | 178.285329 | 0.305880794 | 0.25759005 | 1.18747133 | 0.23504176 | 0.63100657 |
| ABHD2 | 133.538851 | 0.154088126 | 0.3068564 | 0.50215061 | 0.61556158 | 0.8836513 |
| POLG | 17.5392362 | -0.231608416 | 0.35932228 | -0.6445702 | 0.51920578 | 0.83297983 |
| KIF7 | 17.4397748 | 0.070265762 | 0.2965355 | 0.23695565 | 0.81269119 | 0.96038841 |
| ANPEP | 24.0964238 | -0.015424933 | 0.48612963 | -0.0317301 | 0.97468731 | 0.99300191 |
| ZNF710 | 30.1463955 | 0.534779262 | 0.37579758 | 1.42305139 | 0.15472126 | 0.54412824 |
| IDH2 | 25.7063788 | 0.056221781 | 0.28364337 | 0.19821292 | 0.84287848 | 0.96846684 |
| SEMA4B | 63.202739 | -1.470374328 | 0.25574408 | -5.7493973 | 8.96E-09 | 8.23E-06 |
| CIB1 | 31.269333 | -0.50424867 | 0.26819197 | -1.8801781 | 0.06008381 | 0.36498338 |
| IQGAP1 | 194.694543 | 0.276224825 | 0.18350024 | 1.50531047 | 0.13224428 | 0.51310477 |
| CRTC3 | 146.400885 | 0.051658886 | 0.18273571 | 0.28269727 | 0.77740891 | 0.94808797 |
| UNC45A | 25.45902 | 0.334514278 | 0.24521643 | 1.36415931 | 0.17251743 | 0.56904425 |
| SLCO3A1 | 496.022478 | 0.004842 | 0.47601906 | 0.01017186 | 0.99188417 | 0.99759781 |
| FAM174B | 23.7015357 | 0.331030523 | 0.24928854 | 1.32790111 | 0.18421078 | 0.58363936 |
| AC013394.1 | 104.17932 | 0.240189568 | 0.29642322 | 0.8102927 | 0.41777197 | 0.77722447 |
| CHD2 | 166.442682 | -0.309167886 | 0.21526605 | -1.4362129 | 0.15094177 | 0.53880252 |
| MCTP2 | 39.129626 | -1.274647774 | 0.59412306 | -2.1454272 | 0.03191871 | 0.27212364 |
| LINC01197 | 115.091851 | 0.607855885 | 0.42385029 | 1.43412875 | 0.15153553 | 0.53898138 |
| LINC00924 | 131.125643 | 0.1645225 | 0.43650241 | 0.37691086 | 0.70623985 | 0.93006079 |
| AC015574.1 | 43.6738588 | -1.348114504 | 0.81181768 | -1.6606124 | 0.09679132 | 0.45306967 |
| AC012409.2 | 239.278875 | 0.105245222 | 0.44318727 | 0.23747348 | 0.81228949 | 0.96026221 |
| NR2F2-AS1 | 1333.3954 | 0.428228869 | 0.40848074 | 1.04834531 | 0.29447954 | 0.68566952 |
| NR2F2 | 484.062794 | 0.047676246 | 0.22700119 | 0.21002642 | 0.83364705 | 0.9666365 |
| AC087477.2 | 86.9898869 | 0.330297856 | 0.40171727 | 0.82221473 | 0.4109547 | 0.77166383 |
| LINC00923 | 28.9871672 | 0.265555358 | 0.38064038 | 0.69765421 | 0.48539347 | 0.81570449 |
| IGF1R | 431.923116 | -0.140827177 | 0.26209667 | -0.5373101 | 0.59105346 | 0.87221925 |
| SYNM | 30.0246493 | -0.310010288 | 0.60616558 | -0.5114284 | 0.60905112 | 0.88125943 |
| TTC23 | 57.3970136 | -0.262816563 | 0.20586655 | -1.2766356 | 0.20173093 | 0.60118434 |
| LRRC28 | 49.9339525 | 0.075536121 | 0.24085602 | 0.31361525 | 0.75381327 | 0.94128825 |
| MEF2A | 336.58218 | -0.133162618 | 0.35354779 | -0.3766467 | 0.70643616 | 0.93006079 |
| AC090825.1 | 48.3884395 | 0.674468237 | 0.40892961 | 1.64935044 | 0.09907586 | 0.45678074 |
| ASB7 | 19.2653002 | 0.155177403 | 0.28676919 | 0.541123 | 0.58842281 | 0.8702694 |
| LRRK1 | 52.9973145 | -0.263020898 | 0.36599494 | -0.7186463 | 0.4723589 | 0.80671473 |
| CHSY1 | 140.814466 | -0.944363745 | 0.42276894 | -2.2337586 | 0.02549896 | 0.23871578 |
| SELENOS | 48.0761591 | -0.385949131 | 0.3393551 | -1.1373017 | 0.25541219 | 0.65030654 |
| SNRPA1 | 35.2502148 | -0.336901369 | 0.24156081 | -1.3946855 | 0.16311069 | 0.5572759 |
| TM2D3 | 34.4117271 | 0.093040651 | 0.28960512 | 0.32126728 | 0.74800785 | 0.93903859 |
| TARSL2 | 45.8858204 | -0.058637819 | 0.23406696 | -0.2505173 | 0.80218734 | 0.95758343 |
| RHBDF1 | 18.4862102 | 0.455336325 | 0.30097642 | 1.51286376 | 0.13031427 | 0.51204187 |
| MPG | 31.2639613 | -0.562648662 | 0.29213988 | -1.9259564 | 0.05410982 | 0.34985531 |
| NPRL3 | 44.5432536 | -0.21059307 | 0.34098805 | -0.6175966 | 0.53684126 | 0.84423863 |
| LUC7L | 105.932945 | 0.105956634 | 0.17414303 | 0.60844604 | 0.54289169 | 0.84678041 |
| AXIN1 | 22.2685443 | 0.362282108 | 0.4131467 | 0.8768849 | 0.38054916 | 0.74991898 |
| RAB11FIP3 | 75.8045778 | 0.233063489 | 0.34405092 | 0.67740988 | 0.49814593 | 0.82266842 |
| RAB40C | 27.1975619 | -0.115904424 | 0.28165037 | -0.4115188 | 0.68069215 | 0.91860979 |
| RHOT2 | 21.5010654 | -0.149547695 | 0.3252007 | -0.4598628 | 0.64561473 | 0.90150535 |
| STUB1 | 28.3034152 | 0.313180681 | 0.3175902 | 0.9861157 | 0.32407634 | 0.71051224 |
| JMJD8 | 18.1641304 | 0.369484093 | 0.33858163 | 1.09127036 | 0.27515394 | 0.67029497 |
| LMF1 | 35.0091897 | 0.397924177 | 0.38167263 | 1.0425798 | 0.29714295 | 0.68849737 |
| CACNA1H | 27.6109877 | 0.473735479 | 0.39131003 | 1.21063977 | 0.2260335 | 0.62072404 |
| UBE2I | 57.5208078 | -0.00396011 | 0.27204324 | -0.0145569 | 0.98838567 | 0.9972057 |
| GNPTG | 19.2239823 | 0.183947431 | 0.33164193 | 0.55465673 | 0.57912948 | 0.86486944 |
| UNKL | 23.9854117 | 0.507771091 | 0.42118373 | 1.20558098 | 0.2279791 | 0.62188065 |
| CLCN7 | 24.8574376 | -0.231636235 | 0.30877895 | -0.7501685 | 0.45315324 | 0.79534858 |
| IFT140 | 32.7148703 | -0.460181697 | 0.2793941 | -1.6470702 | 0.09954361 | 0.45779653 |
| TMEM204 | 36.4833793 | 0.03306372 | 0.33429694 | 0.09890524 | 0.92121351 | 0.98528826 |
| MAPK8IP3 | 56.4775337 | -0.178196682 | 0.24822815 | -0.7178746 | 0.47283461 | 0.80716876 |
| NME3 | 44.280763 | 0.361586352 | 0.38283451 | 0.94449779 | 0.34491533 | 0.72325086 |
| MRPS34 | 21.7327553 | -0.174142102 | 0.39178133 | -0.444488 | 0.6566898 | 0.90772143 |
| SPSB3 | 23.9301605 | 0.41774634 | 0.27722746 | 1.50687218 | 0.13184342 | 0.51290135 |
| HAGH | 26.7265376 | 0.222222499 | 0.28008066 | 0.79342322 | 0.42753129 | 0.7804092 |
| NDUFB10 | 29.2424976 | 0.000198919 | 0.36201967 | 0.00054947 | 0.99956159 | 0.99969762 |
| RPS2 | 109.98948 | -0.514305072 | 0.33748073 | -1.5239539 | 0.12752023 | 0.50683947 |
| SLC9A3R2 | 25.0507371 | 1.173118633 | 0.39281845 | 2.98641427 | 0.0028227 | 0.06800182 |
| TSC2 | 35.5851517 | 0.320094468 | 0.26533719 | 1.20636867 | 0.22767537 | 0.62183669 |
| PKD1 | 60.4316762 | 0.461341731 | 0.20083704 | 2.2970949 | 0.02161336 | 0.21698983 |
| PGP | 20.4314175 | 0.029768305 | 0.44885366 | 0.06632074 | 0.94712247 | 0.99060401 |
| ECI1 | 25.517463 | -0.1625117 | 0.33749513 | -0.4815231 | 0.63014477 | 0.89365764 |
| RNPS1 | 38.4655141 | 0.032931712 | 0.25078492 | 0.13131456 | 0.89552647 | 0.98023547 |
| PDPK1 | 70.8123687 | -0.008282272 | 0.20373527 | -0.0406521 | 0.96757323 | 0.99300191 |
| SRRM2 | 250.340163 | -0.140449778 | 0.17002419 | -0.8260576 | 0.40877139 | 0.77015112 |
| ELOB | 44.3117236 | -0.404790616 | 0.39713809 | -1.0192692 | 0.30807519 | 0.69834811 |
| FLYWCH1 | 31.2043236 | 0.062711946 | 0.28262742 | 0.22188911 | 0.82440021 | 0.96413949 |
| TNFRSF12A | 31.1403486 | -0.758108046 | 0.59364006 | -1.27705 | 0.20158459 | 0.60099196 |
| HCFC1R1 | 32.5756917 | 0.363790631 | 0.41459757 | 0.87745481 | 0.38023965 | 0.74991898 |
| IL32 | 26.0585247 | -0.731091969 | 0.4709715 | -1.5523062 | 0.12058898 | 0.49786991 |
| ZNF213 | 20.8480051 | -0.147560171 | 0.27305801 | -0.5403986 | 0.58892217 | 0.8702694 |
| ZNF263 | 19.3611243 | 0.135967111 | 0.31430189 | 0.43260036 | 0.66530513 | 0.91235816 |
| ZNF75A | 24.9389796 | 0.415370301 | 0.4361924 | 0.95226395 | 0.34096314 | 0.71963879 |
| CLUAP1 | 22.0388623 | -0.158359675 | 0.28376884 | -0.5580587 | 0.5768043 | 0.8635027 |
| DNASE1 | 39.910084 | 0.328456881 | 0.30842339 | 1.0649545 | 0.28689652 | 0.67921285 |
| TRAP1 | 33.7985895 | -0.095601464 | 0.27552688 | -0.3469769 | 0.72860867 | 0.93455731 |
| CREBBP | 192.767102 | 0.169962254 | 0.20497522 | 0.8291844 | 0.40700007 | 0.76831326 |
| ADCY9 | 121.205892 | 0.375823252 | 0.21698087 | 1.73205706 | 0.0832634 | 0.42348124 |
| VASN | 48.2197785 | 0.487756903 | 0.32217472 | 1.51395152 | 0.13003813 | 0.51204187 |
| DNAJA3 | 24.3539783 | -0.329436601 | 0.2640697 | -1.2475365 | 0.21220083 | 0.60910715 |
| NMRAL1 | 23.1504905 | -0.581506445 | 0.34183929 | -1.7011106 | 0.08892221 | 0.43544541 |
| HMOX2 | 53.6734115 | -1.226875106 | 0.38316065 | -3.2019862 | 0.00136484 | 0.04538541 |
| CDIP1 | 19.2188896 | -0.205309719 | 0.33088246 | -0.6204914 | 0.53493432 | 0.84361209 |
| MGRN1 | 55.705786 | 0.057889365 | 0.25962338 | 0.22297439 | 0.82355544 | 0.96413949 |
| ANKS3 | 30.2812707 | 0.027930629 | 0.24785452 | 0.11268961 | 0.91027664 | 0.98343817 |
| GLYR1 | 71.2015913 | -0.034141376 | 0.21425621 | -0.1593484 | 0.87339442 | 0.9768643 |
| UBN1 | 33.432354 | -0.276654833 | 0.2258863 | -1.2247526 | 0.22066845 | 0.61590675 |
| RBFOX1 | 21.9552775 | 0.541019242 | 0.3229367 | 1.67531049 | 0.09387333 | 0.44536803 |
| PMM2 | 66.6298613 | -1.079170342 | 0.27279417 | -3.9559876 | 7.62E-05 | 0.00674861 |
| CARHSP1 | 55.2805383 | -0.312015246 | 0.24987686 | -1.248676 | 0.21178359 | 0.60910715 |

| | | | | | | |
|---|---|---|---|---|---|---|
| USP7 | 62.6527397 | 0.230179574 | 0.26705434 | 0.86192037 | 0.38873134 | 0.75549104 |
| C16orf72 | 51.7848187 | -0.01094018 | 0.2220436 | -0.0492704 | 0.9607038 | 0.99181157 |
| GRIN2A | 147.328575 | -0.090693064 | 1.1261098 | -0.0805366 | 0.93581048 | 0.98827804 |
| ATF7IP2 | 61.527501 | 0.488435804 | 0.37651732 | 1.29724657 | 0.19454636 | 0.59272783 |
| EMP2 | 47.6469871 | 0.297938158 | 0.41029183 | 0.72616157 | 0.46773973 | 0.80374006 |
| CLEC16A | 122.59454 | -0.306537987 | 0.2360899 | -1.2983952 | 0.19415157 | 0.59269042 |
| AC099489.1 | 24.0402376 | -0.950160199 | 0.65272927 | -1.4556727 | 0.14548311 | 0.53538077 |
| LITAF | 184.281185 | -0.899229263 | 0.34960337 | -2.5721413 | 0.01010716 | 0.14181707 |
| TXNDC11 | 68.3223391 | -0.206553129 | 0.161666 | -1.2776535 | 0.20137163 | 0.60060069 |
| ZC3H7A | 98.8413171 | -0.026781715 | 0.17919871 | -0.1494526 | 0.8811965 | 0.97782528 |
| RSL1D1 | 23.9104122 | -0.326355431 | 0.35241646 | -0.9260505 | 0.35441974 | 0.7306117 |
| AC007216.4 | 34.749939 | 0.143086221 | 0.24854507 | 0.57569526 | 0.56482118 | 0.8572637 |
| GSPT1 | 84.2046867 | -0.083023462 | 0.20829796 | -0.3985803 | 0.69020248 | 0.92290721 |
| SNX29 | 158.021923 | -0.030115478 | 0.22096163 | -0.1362928 | 0.89158982 | 0.98023547 |
| MKL2 | 175.908697 | -0.014970475 | 0.19313433 | -0.0775133 | 0.93821523 | 0.98827804 |
| PARN | 80.3902617 | -0.124740406 | 0.20376711 | -0.6121714 | 0.54040434 | 0.84578923 |
| BFAR | 20.9077251 | -0.056127392 | 0.31916946 | -0.1758545 | 0.86040824 | 0.97397549 |
| PDXDC1 | 197.140132 | -0.007072902 | 0.21640515 | -0.0326836 | 0.97392689 | 0.99300191 |
| RRN3 | 33.5965114 | 0.026134495 | 0.231234 | 0.11302185 | 0.91001323 | 0.98343817 |
| C16orf45 | 95.6836441 | -0.452518604 | 0.29992488 | -1.5087731 | 0.13135677 | 0.51290135 |
| MARF1 | 47.4458743 | 0.065114586 | 0.31143981 | 0.20907598 | 0.83438892 | 0.9666365 |
| NDE1 | 33.2146406 | 0.357171121 | 0.32431586 | 1.10130636 | 0.27076334 | 0.66572091 |
| MYH11 | 436.75616 | 1.695864286 | 0.53787344 | 3.1529058 | 0.00161654 | 0.05055298 |
| ABCC1 | 126.807577 | -0.72194178 | 0.26667226 | -2.7072249 | 0.00678483 | 0.11301781 |
| XYLT1 | 125.257259 | 0.396503563 | 0.34146899 | 1.16117004 | 0.24557275 | 0.64133411 |
| RPS15A | 27.1939933 | -0.148506613 | 0.41502523 | -0.3578255 | 0.7204739 | 0.93444541 |
| SMG1 | 173.318863 | -0.116243659 | 0.15332283 | -0.7581628 | 0.44835355 | 0.79296348 |
| CCP110 | 30.8544937 | -0.218668046 | 0.42609793 | -0.5131873 | 0.60782031 | 0.88042838 |
| VPS35L | 87.4221731 | -0.0711292 | 0.21651588 | -0.3285172 | 0.74252061 | 0.93759079 |
| IQCK | 34.234004 | 0.491619614 | 0.35150378 | 1.39861827 | 0.16192748 | 0.55529869 |
| UMOD | 143.712794 | 0.563117139 | 0.59157269 | 0.95189848 | 0.34114847 | 0.71963879 |
| THUMPD1 | 29.9643644 | 0.251648444 | 0.25266229 | 0.99598737 | 0.31925628 | 0.70683013 |
| METTL9 | 41.9652628 | -0.361115919 | 0.2453095 | -1.4720829 | 0.1409985 | 0.52800651 |
| NPIPB4 | 20.0418059 | 0.702654709 | 0.35713044 | 1.96750158 | 0.04912541 | 0.33550435 |
| UQCRC2 | 57.2156223 | -0.202804714 | 0.1879314 | -1.0791422 | 0.28052433 | 0.67548272 |
| MOSMO | 106.790195 | -0.1531984 | 0.24709886 | -0.6199883 | 0.5352655 | 0.84377224 |
| EEF2K | 42.3090186 | 0.839255616 | 0.32926374 | 2.54888565 | 0.01080677 | 0.14625962 |
| POLR3E | 20.7013558 | -0.670286829 | 0.32266882 | -2.0773213 | 0.03777191 | 0.29501902 |
| CDR2 | 129.929182 | 0.702420568 | 0.24937796 | 2.81669063 | 0.00485212 | 0.09203696 |
| NPIPB5 | 72.6524928 | 0.086040647 | 0.22603682 | 0.38064881 | 0.70346386 | 0.92936308 |
| USP31 | 62.2046652 | -0.037663041 | 0.38568392 | -0.0976526 | 0.92220815 | 0.98561736 |
| COG7 | 46.9250237 | -0.329637998 | 0.2124706 | -1.5514523 | 0.12079334 | 0.49815388 |
| GGA2 | 43.9286268 | -0.338936986 | 0.30638823 | -1.1062337 | 0.26862539 | 0.66401883 |
| UBFD1 | 18.5627718 | -0.361984131 | 0.27951682 | -1.2950352 | 0.19530808 | 0.59328838 |
| NDUFAB1 | 24.5224714 | -0.223050664 | 0.32324036 | -0.6900458 | 0.49016536 | 0.81720854 |
| DCTN5 | 25.6007796 | -0.187974039 | 0.28770639 | -0.6533537 | 0.51352826 | 0.82998003 |
| PRKCB | 65.9659423 | 0.1136994 | 1.05711703 | 0.10755611 | 0.91434781 | 0.98411571 |
| RBBP6 | 73.9701623 | -0.094805227 | 0.21052998 | -0.450317 | 0.65248189 | 0.90644412 |
| TNRC6A | 410.340894 | 0.134903285 | 0.26218848 | 0.5145279 | 0.60688296 | 0.88000768 |
| ARHGAP17 | 145.605795 | 0.198593977 | 0.23883056 | 0.83152666 | 0.40567617 | 0.76703528 |
| LCMT1 | 39.9677966 | 0.312660471 | 0.24299976 | 1.28666987 | 0.19820938 | 0.59551951 |
| NSMCE1 | 25.4553899 | 0.2085567 | 0.32234792 | 0.64699255 | 0.51763677 | 0.83277421 |
| IL4R | 149.735179 | -0.725104801 | 0.30935684 | -2.3439107 | 0.01908274 | 0.20295087 |
| GTF3C1 | 42.3616537 | 0.060490859 | 0.2176891 | 0.2778773 | 0.78110655 | 0.94939885 |
| KIAA0556 | 37.7038744 | 0.026630369 | 0.32469553 | 0.08201643 | 0.93463365 | 0.98800671 |
| XPO6 | 79.7536663 | -0.344646865 | 0.21474272 | -1.6049292 | 0.10850939 | 0.47494671 |
| NUPR1 | 73.7936096 | -0.991169714 | 0.31926746 | -3.1045122 | 0.00190593 | 0.05543169 |
| ATXN2L | 41.5113586 | -0.207225747 | 0.28589226 | -0.7248386 | 0.46855104 | 0.80374006 |
| TUFM | 34.8692635 | -0.152170796 | 0.34012345 | -0.4473987 | 0.6545872 | 0.90646849 |
| SH2B1 | 24.9535749 | 0.443659126 | 0.25766974 | 1.72181306 | 0.08510339 | 0.42746464 |
| NFATC2IP | 56.3677062 | 0.027288007 | 0.22154377 | 0.12317208 | 0.90197083 | 0.98164203 |
| AC009093.2 | 29.2846951 | -0.296386834 | 0.39732272 | -0.7459599 | 0.45569161 | 0.79621437 |
| MAZ | 17.5179986 | -0.093810471 | 0.3311712 | -0.2832688 | 0.77697078 | 0.94805375 |
| MVP | 67.0261395 | -0.643972286 | 0.30511974 | -2.1105559 | 0.0348105 | 0.28488012 |
| CDIPT | 21.6377721 | -0.112613901 | 0.37074308 | -0.3037519 | 0.76131694 | 0.94425408 |
| KCTD13 | 18.666036 | -0.231450358 | 0.33996626 | -0.6808039 | 0.49599557 | 0.82114698 |
| TMEM219 | 28.0735232 | 0.418980375 | 0.27669186 | 1.51424899 | 0.1299627 | 0.51204187 |
| INO80E | 15.8961053 | 0.418025114 | 0.33031703 | 1.26552697 | 0.20568249 | 0.60431091 |
| PPP4C | 24.3792774 | -0.200520413 | 0.44574988 | -0.4498496 | 0.65281888 | 0.90646849 |
| YPEL3 | 36.1747434 | 0.014981432 | 0.25944041 | 0.05774518 | 0.95395161 | 0.99100036 |
| MAPK3 | 20.6624779 | 0.214991744 | 0.25189317 | 0.85350367 | 0.39338005 | 0.75988253 |
| SEPHS2 | 26.1699662 | -0.675808586 | 0.43312966 | -1.5602916 | 0.11869098 | 0.49356852 |
| ZNF785 | 17.6786132 | 0.003248065 | 0.44098315 | 0.00736551 | 0.99412323 | 0.99819806 |
| FBRS | 20.9381456 | -0.036677139 | 0.33956492 | -0.1080122 | 0.91398605 | 0.98395113 |
| RNF40 | 18.7061181 | 0.275010103 | 0.34498298 | 0.79717005 | 0.42535228 | 0.77861803 |
| BCL7C | 37.9063157 | 0.808532121 | 0.26285678 | 3.07594166 | 0.00209839 | 0.05624279 |
| KAT8 | 25.3310239 | -0.036977891 | 0.23724109 | -0.1558663 | 0.87613841 | 0.9768643 |
| FUS | 152.957088 | 0.333883239 | 0.18261819 | 1.82831312 | 0.06750257 | 0.38712255 |
| TGFB1I1 | 37.2596961 | 0.285534888 | 0.3524812 | 0.81007126 | 0.41789922 | 0.77722447 |
| C16orf58 | 20.9054722 | -0.180400569 | 0.29132826 | -0.6192347 | 0.53576175 | 0.84412047 |
| ZNF720 | 35.9211389 | -0.327796075 | 0.25882663 | -1.2664697 | 0.20534498 | 0.60370888 |
| VPS35 | 33.2170163 | 0.092327218 | 0.21858041 | 0.42239476 | 0.6727369 | 0.91482259 |
| DNAJA2 | 21.0789031 | -0.345668688 | 0.25603646 | -1.350076 | 0.17699159 | 0.57388115 |
| ITFG1 | 231.061086 | 0.034280329 | 0.14835032 | 0.23107688 | 0.81725508 | 0.96134613 |
| PHKB | 166.957962 | 0.197000015 | 0.24838948 | 0.79310933 | 0.42771413 | 0.78054907 |
| LONP2 | 148.340507 | 0.030777785 | 0.1593505 | 0.19314521 | 0.84684525 | 0.97014274 |
| SIAH1 | 35.3954547 | -0.122568352 | 0.23556077 | -0.5203258 | 0.6028365 | 0.87903643 |
| N4BP1 | 67.5161617 | -0.127631637 | 0.26245397 | -0.486301 | 0.62675374 | 0.89160148 |
| ZNF423 | 74.0629078 | 0.788875243 | 0.44443133 | 1.77502168 | 0.07589431 | 0.40904738 |
| TENT4B | 65.4579093 | 0.108561295 | 0.28378443 | 0.38254846 | 0.70205459 | 0.92865476 |
| BRD7 | 36.9578437 | 0.124951535 | 0.20055213 | 0.62303768 | 0.53325976 | 0.84333372 |
| NKD1 | 60.0199433 | -0.13355407 | 0.5006692 | -0.2667511 | 0.78966079 | 0.9517272 |
| CYLD | 49.7234367 | -0.163835068 | 0.22961265 | -0.7135281 | 0.47551905 | 0.80926744 |
| CHD9 | 402.866918 | 0.212679208 | 0.24038147 | 0.88475709 | 0.37628766 | 0.74787433 |
| RBL2 | 70.6804557 | -0.045010836 | 0.21553335 | -0.2088347 | 0.8345773 | 0.9666365 |
| FTO | 432.528647 | -0.177138296 | 0.19860865 | -0.8918962 | 0.37244858 | 0.74463512 |
| MMP2 | 80.0285817 | -0.285575205 | 0.32973547 | -0.8660737 | 0.38644977 | 0.75402666 |
| LPCAT2 | 22.0800808 | 0.739227557 | 0.30272199 | 2.44193545 | 0.01460876 | 0.17653018 |
| GNAO1 | 27.1214438 | 0.044680799 | 0.42996256 | 0.10391788 | 0.91723451 | 0.98477389 |
| AMFR | 40.5233504 | 0.325948264 | 0.23596425 | 1.38134596 | 0.16717261 | 0.56278128 |
| NUDT21 | 33.5196499 | -0.03349987 | 0.21809715 | -0.1536007 | 0.8779246 | 0.9768643 |
| OGFOD1 | 20.9107483 | -0.221594714 | 0.35172307 | -0.6300261 | 0.5286775 | 0.84005426 |
| BBS2 | 28.6602426 | 0.418542406 | 0.23850284 | 1.75487385 | 0.07928089 | 0.41737582 |
| MT2A | 638.234056 | -2.975820124 | 0.58457932 | -5.0905327 | 3.57E-07 | 0.00014578 |
| MT1E | 136.781208 | -1.372560446 | 0.53125276 | -2.5836298 | 0.00977667 | 0.14032956 |
| MT1M | 46.5703747 | -2.28016565 | 0.57340141 | -3.9765609 | 6.99E-05 | 0.00642294 |
| MT1G | 181.906412 | -1.148809979 | 0.94302393 | -1.2182193 | 0.22314064 | 0.61772146 |
| MT1H | 54.7122132 | -1.498071299 | 1.2011338 | -1.2472143 | 0.21231892 | 0.60910715 |
| MT1X | 106.817076 | -2.065293553 | 0.59580308 | -3.4664029 | 0.00052747 | 0.0246904 |
| NUP93 | 29.1664876 | 0.057925216 | 0.28837833 | 0.20086536 | 0.84080386 | 0.96803523 |
| HERPUD1 | 82.5070661 | -0.994923524 | 0.49181458 | -2.0229647 | 0.04307679 | 0.31204692 |
| NLRC5 | 29.3481036 | 0.092842632 | 0.39559306 | 0.23469226 | 0.81444759 | 0.96038841 |
| FAM192A | 36.427105 | -0.241758569 | 0.21925903 | -1.1026163 | 0.27019384 | 0.6651094 |

| | | | | | | |
|---|---|---|---|---|---|---|
| RSPRY1 | 53.2203386 | -0.070771121 | 0.25751276 | -0.2748257 | 0.78345016 | 0.95064024 |
| ARL2BP | 23.4341823 | 0.119744169 | 0.32555153 | 0.3678194 | 0.7130079 | 0.93115158 |
| COQ9 | 20.0557678 | -0.258695495 | 0.31766156 | -0.8143746 | 0.41543041 | 0.77526615 |
| ADGRG1 | 23.6895898 | -0.460749683 | 0.52213996 | -0.8824256 | 0.37754668 | 0.749056 |
| KIFC3 | 53.8147988 | 0.698694343 | 0.28330476 | 2.46622873 | 0.01365441 | 0.17065692 |
| CSNK2A2 | 28.9849127 | 0.090369781 | 0.2339517 | 0.38627538 | 0.69929273 | 0.92769314 |
| CNOT1 | 103.984229 | -0.14927402 | 0.17077685 | -0.8740881 | 0.38207026 | 0.75115954 |
| GOT2 | 20.7158965 | -0.006338338 | 0.31210176 | -0.0203086 | 0.98379723 | 0.99620525 |
| CDH11 | 378.028723 | 0.101110059 | 0.3229884 | 0.31304548 | 0.75424611 | 0.94139855 |
| CMTM4 | 39.9397316 | 0.410121679 | 0.28480579 | 1.44000472 | 0.14986606 | 0.53880252 |
| DYNC1LI2 | 73.5355189 | 0.068145756 | 0.20307271 | 0.33557319 | 0.73719274 | 0.93594619 |
| NAE1 | 32.6427771 | 0.015288621 | 0.27444383 | 0.05570765 | 0.9555747 | 0.99181157 |
| CDH16 | 29.6223421 | -0.27891695 | 0.38085925 | -0.732336 | 0.46396349 | 0.80227475 |
| RRAD | 29.5491315 | 0.489026141 | 0.40720347 | 1.20093806 | 0.22977523 | 0.62425809 |
| FAM96B | 27.0927425 | -0.204860325 | 0.26526028 | -0.7722993 | 0.43993719 | 0.78933067 |
| CES2 | 18.3378704 | 0.110832038 | 0.34198142 | 0.32408789 | 0.74587149 | 0.93891908 |
| CBFB | 106.538301 | 0.093784087 | 0.1932549 | 0.48528698 | 0.62747279 | 0.89192276 |
| KIAA0895L | 25.0790562 | -0.59575732 | 0.41233511 | -1.4448377 | 0.14850348 | 0.53680621 |
| FHOD1 | 22.9545488 | 0.820962305 | 0.44398515 | 1.84907606 | 0.06444683 | 0.37871685 |
| ATP6V0D1 | 31.8362073 | 0.229057611 | 0.31531369 | 0.72644361 | 0.46756687 | 0.80374006 |
| RIPOR1 | 32.5628657 | 0.07002497 | 0.2969822 | 0.23578844 | 0.81359684 | 0.96038841 |
| CTCF | 58.5184522 | 0.238689674 | 0.23732439 | 1.00575282 | 0.31453449 | 0.70282967 |
| GFOD2 | 22.7639589 | -0.915576649 | 0.46527083 | -1.9678359 | 0.04908692 | 0.33550435 |
| RANBP10 | 49.6418874 | -0.390436996 | 0.28263058 | -1.3814393 | 0.16714392 | 0.56278128 |
| LCAT | 19.2629018 | 0.872673308 | 0.45416941 | 1.92147089 | 0.05467237 | 0.3515199 |
| SLC12A4 | 22.6121047 | 0.358984417 | 0.32350831 | 1.10966058 | 0.26714531 | 0.66201068 |
| NFATC3 | 119.069052 | 0.218723516 | 0.18162451 | 1.20426214 | 0.22848828 | 0.62213472 |
| SLC7A6 | 92.0713881 | -0.817074288 | 0.44003907 | -1.8568221 | 0.06333647 | 0.37446475 |
| SLC7A6OS | 27.2671776 | -0.723152275 | 0.30739976 | -2.3524816 | 0.01864861 | 0.20009416 |
| PRMT7 | 28.4328252 | 0.418478744 | 0.48099983 | 0.87001848 | 0.38429031 | 0.75288048 |
| ZFP90 | 25.7097333 | -0.019869927 | 0.29306218 | -0.0678011 | 0.945944 | 0.99060401 |
| TANGO6 | 81.6823354 | -0.145709506 | 0.18263297 | -0.797827 | 0.42497091 | 0.77861803 |
| CHTF8 | 21.8025875 | 0.297694013 | 0.39112518 | 0.76112208 | 0.44658415 | 0.79296348 |
| UTP4 | 19.8203697 | -0.202207434 | 0.34358695 | -0.588519 | 0.55618401 | 0.85207344 |
| SNTB2 | 44.8464487 | 0.200308575 | 0.29453991 | 0.68007277 | 0.49645839 | 0.82154305 |
| VPS4A | 19.1977091 | 0.313998103 | 0.2595664 | 1.20970245 | 0.22639309 | 0.62072404 |
| TERF2 | 25.0094378 | -0.230180991 | 0.26321813 | -0.8744876 | 0.38185277 | 0.75093284 |
| CYB5B | 59.7748892 | -0.160004438 | 0.25477726 | -0.6280169 | 0.52999284 | 0.84123485 |
| NFAT5 | 529.827069 | 0.005805916 | 0.18078661 | 0.03211474 | 0.97438054 | 0.99300191 |
| NQO1 | 23.5234102 | -0.810421732 | 0.64925162 | -1.2482398 | 0.21194324 | 0.60910715 |
| WWP2 | 78.015203 | 0.54071023 | 0.25967525 | 2.08225555 | 0.03731913 | 0.29301102 |
| PDPR | 45.1547705 | -0.089561876 | 0.2755162 | -0.3250694 | 0.74512857 | 0.93891908 |
| EXOSC6 | 17.6580083 | -0.187381974 | 0.35456448 | -0.5284849 | 0.59716283 | 0.87535619 |
| AARS | 31.4499545 | 0.087785022 | 0.25675434 | 0.34190278 | 0.73242406 | 0.93574285 |
| ST3GAL2 | 47.924349 | 0.492739423 | 0.31992916 | 1.54015163 | 0.1235234 | 0.50097964 |
| COG4 | 25.990288 | 0.125384445 | 0.24470683 | 0.51238638 | 0.60838062 | 0.8807592 |
| SF3B3 | 71.6366983 | -0.203224516 | 0.21740341 | -0.9347807 | 0.34990131 | 0.72803645 |
| IL34 | 34.6998644 | -0.622419961 | 0.53517046 | -1.1630312 | 0.24481683 | 0.6405276 |
| MTSS1L | 25.2297238 | 0.358930724 | 0.33715586 | 1.06458396 | 0.28706424 | 0.67921285 |
| VAC14 | 27.5658923 | -0.406401759 | 0.42434064 | -0.9577253 | 0.33820131 | 0.71925477 |
| HYDIN | 27.7043307 | 0.512046625 | 0.38896043 | 1.31644913 | 0.18802334 | 0.58645909 |
| PHLPP2 | 39.1091408 | 0.367115828 | 0.2921402 | 1.2566426 | 0.20888308 | 0.6059796 |
| AP1G1 | 94.4626822 | -0.031831224 | 0.19553332 | -0.1627918 | 0.87068235 | 0.9768643 |
| IST1 | 67.5554698 | -0.046344118 | 0.2178378 | -0.212746 | 0.8315251 | 0.96634287 |
| DHX38 | 18.6881351 | 0.020437876 | 0.31861154 | 0.06414669 | 0.94885343 | 0.99060401 |
| AC004943.2 | 21.377185 | -0.202234691 | 0.40826025 | -0.4953573 | 0.62034793 | 0.88678019 |
| ZFHX3 | 475.717866 | 0.058278318 | 0.29615099 | 0.19678583 | 0.84399514 | 0.9691438 |
| PSMD7 | 38.6900317 | -0.482596238 | 0.31685601 | -1.5230775 | 0.12773933 | 0.50743585 |
| GLG1 | 215.348987 | 0.1591557 | 0.17584563 | 0.90508759 | 0.36541899 | 0.73747473 |
| RFWD3 | 21.5130539 | -0.157446397 | 0.38991999 | -0.4037916 | 0.68636601 | 0.9206249 |
| WDR59 | 70.4830041 | -0.202492884 | 0.21010707 | -0.9637604 | 0.33516604 | 0.71748769 |
| ZNRF1 | 97.3531749 | -0.150969921 | 0.16066945 | -0.9396305 | 0.34740711 | 0.72531103 |
| BCAR1 | 58.8724171 | 0.234341603 | 0.1999135 | 1.17221499 | 0.24111075 | 0.63527165 |
| CFDP1 | 137.063438 | 0.00812364 | 0.194877 | 0.04168599 | 0.96674902 | 0.99300191 |
| GABARAPL2 | 40.318899 | 0.121823002 | 0.30939286 | 0.39374859 | 0.69376666 | 0.9248682 |
| KARS | 22.9467035 | -0.342679434 | 0.28482795 | -1.2031103 | 0.22893365 | 0.62266226 |
| TERF2IP | 96.4428455 | -0.172392441 | 0.24426962 | -0.7057465 | 0.48034577 | 0.81237906 |
| WWOX | 674.499796 | -0.057508874 | 0.25186982 | -0.2283278 | 0.81939143 | 0.96284007 |
| MAF | 35.9002558 | 0.771642695 | 0.2912386 | 2.64952067 | 0.0080606 | 0.12576938 |
| CDYL2 | 36.8145458 | -0.105015964 | 0.32687719 | -0.3212704 | 0.74800549 | 0.93903859 |
| CMC2 | 34.2390566 | 0.289737145 | 0.28964977 | 1.00030165 | 0.31716455 | 0.70524728 |
| GAN | 36.6094809 | 0.128894399 | 0.22231998 | 0.57976976 | 0.5620699 | 0.85545224 |
| CMIP | 170.566716 | 0.007862902 | 0.29474236 | 0.0266772 | 0.97871719 | 0.99465087 |
| PLCG2 | 87.4574432 | -0.668978896 | 0.43168775 | -1.5496824 | 0.12121777 | 0.49897659 |
| CDH13 | 177.835706 | -0.189600934 | 0.57969738 | -0.3270688 | 0.74361584 | 0.93791562 |
| HSBP1 | 32.4725444 | -0.276779775 | 0.27102308 | -1.0212406 | 0.30714045 | 0.69801431 |
| MBTPS1 | 62.2823541 | -0.10486104 | 0.20117249 | -0.5212494 | 0.60219304 | 0.87903643 |
| WFDC1 | 86.0541134 | -0.044858473 | 0.5528687 | -0.0811377 | 0.93533248 | 0.98827804 |
| COTL1 | 23.0679406 | -0.027657147 | 0.34431495 | -0.0803251 | 0.93597866 | 0.98827804 |
| KLHL36 | 28.6295331 | -0.366006262 | 0.25610579 | -1.4291214 | 0.15296935 | 0.54048732 |
| USP10 | 71.0070644 | -0.500561147 | 0.25260316 | -1.9816108 | 0.04752282 | 0.3297075 |
| CRISPLD2 | 579.897013 | -0.814974359 | 0.5644968 | -1.4437183 | 0.14881824 | 0.53689238 |
| ZDHHC7 | 32.5190148 | -0.202608646 | 0.21140232 | -0.9584031 | 0.33785952 | 0.71925477 |
| GSE1 | 287.269811 | 0.472784435 | 0.37274633 | 1.2683812 | 0.20466185 | 0.60370888 |
| COX4I1 | 85.1628876 | -0.300335327 | 0.26251397 | -1.1440737 | 0.25259308 | 0.64724776 |
| FOXC2 | 22.252355 | 0.409976576 | 0.27855231 | 1.47181179 | 0.14107171 | 0.52800651 |
| FBXO31 | 25.9662817 | -0.113996425 | 0.23834308 | -0.4782871 | 0.63244586 | 0.8948276 |
| MAP1LC3B | 37.0601208 | -0.399568702 | 0.39031154 | -1.0237174 | 0.30596879 | 0.69728397 |
| ZCCHC14 | 87.0924962 | 0.170529521 | 0.33899477 | 0.5030447 | 0.61493284 | 0.88360697 |
| KLHDC4 | 28.6195706 | 0.119979137 | 0.33655142 | 0.35649571 | 0.72146937 | 0.93444541 |
| BANP | 52.2289281 | -0.305233538 | 0.23856777 | -1.2794416 | 0.20074159 | 0.59975737 |
| ZFPM1 | 17.6103637 | 0.012738185 | 0.3425141 | 0.03719025 | 0.97033331 | 0.99300191 |
| ZC3H18 | 29.2205298 | 0.25634967 | 0.33798876 | 0.75845619 | 0.44817793 | 0.79296348 |
| CYBA | 41.5413258 | -0.921308105 | 0.37744949 | -2.4408778 | 0.01465161 | 0.17653018 |
| PIEZO1 | 36.8904568 | -0.385127384 | 0.24838218 | -1.5505435 | 0.12101112 | 0.49849255 |
| APRT | 26.6740469 | -0.747809199 | 0.33651494 | -2.2222169 | 0.02626865 | 0.24214398 |
| GALNS | 22.8302125 | -0.24844115 | 0.34109345 | -0.7283668 | 0.46638907 | 0.80355999 |
| ACSF3 | 23.4181729 | 0.106985445 | 0.30716005 | 0.34830521 | 0.72761099 | 0.93455731 |
| ANKRD11 | 326.077538 | -0.163752514 | 0.18345406 | -0.8926078 | 0.37206726 | 0.74463512 |
| SPG7 | 72.9019193 | -0.040465691 | 0.34847516 | -0.1161222 | 0.90755572 | 0.98306694 |
| RPL13 | 298.118216 | -0.343431916 | 0.29485479 | -1.1647493 | 0.24412048 | 0.63966808 |
| CHMP1A | 20.87801 | -0.104109299 | 0.25623444 | -0.4063049 | 0.68451861 | 0.91977784 |
| TCF25 | 166.100713 | -0.169367599 | 0.26870949 | -0.6303 | 0.52849831 | 0.83995114 |
| FAM157C | 23.430229 | 0.376631397 | 0.31227549 | 1.20608695 | 0.22778397 | 0.62183669 |
| VPS53 | 115.882013 | -0.268253697 | 0.21737463 | -1.2340617 | 0.21717992 | 0.61410359 |
| NXN | 262.515288 | -1.61061327 | 0.52371041 | -3.0753891 | 0.00210228 | 0.05624279 |
| ABR | 83.8989429 | -0.019160429 | 0.35324581 | -0.0542411 | 0.9567431 | 0.99181157 |
| YWHAE | 139.083829 | -0.032220444 | 0.1407215 | -0.228966 | 0.81889531 | 0.96265181 |
| CRK | 64.3753979 | 0.115249459 | 0.28809165 | 0.40004443 | 0.68912379 | 0.92217934 |
| MYO1C | 75.250093 | 0.772904691 | 0.25533293 | 3.02704666 | 0.00246956 | 0.062538 |
| PITPNA | 63.5352179 | 0.01559343 | 0.23294512 | 0.06694036 | 0.94662918 | 0.99060401 |
| SLC43A2 | 24.8062643 | -0.672135905 | 0.59962707 | -1.1209232 | 0.26232054 | 0.65630111 |
| PRPF8 | 67.8802115 | 0.109427116 | 0.23734004 | 0.46105628 | 0.64475823 | 0.90107359 |

| | | | | | | |
|---|---|---|---|---|---|---|
| MIR22HG | 23.2086684 | -0.201966635 | 0.54234212 | -0.3723971 | 0.70959721 | 0.93009618 |
| SERPINF1 | 54.8858158 | -0.745075484 | 0.4554634 | -1.6358625 | 0.10186838 | 0.46169744 |
| SMYD4 | 22.2048239 | 0.036225365 | 0.29713648 | 0.12191491 | 0.90296641 | 0.98164203 |
| RPA1 | 37.385082 | -0.040117653 | 0.24099109 | -0.1664694 | 0.86778752 | 0.97647688 |
| HIC1 | 40.9920398 | -0.749569762 | 0.46053803 | -1.6275958 | 0.10361062 | 0.4637638 |
| SMG6 | 334.389172 | -0.497779037 | 0.38295563 | -1.2998348 | 0.19365761 | 0.59251477 |
| SGSM2 | 24.7059984 | 0.286882714 | 0.24578049 | 1.16723145 | 0.24311691 | 0.63832305 |
| MNT | 18.3275781 | -0.510679214 | 0.35630986 | -1.4332447 | 0.15178792 | 0.53898138 |
| METTL16 | 70.9627975 | -0.018753391 | 0.1976782 | -0.0948683 | 0.92441945 | 0.98561736 |
| PAFAH1B1 | 263.71106 | 0.033713804 | 0.13343144 | 0.25266761 | 0.80052508 | 0.95648173 |
| ASPA | 37.3440299 | 0.977960087 | 0.54500666 | 1.79440024 | 0.07274931 | 0.40077561 |
| NCBP3 | 43.8496611 | 0.284538764 | 0.18810292 | 1.51267592 | 0.130362 | 0.51204187 |
| ATP2A3 | 20.9030665 | 1.763924603 | 0.55679574 | 3.16799226 | 0.00153496 | 0.04925935 |
| ZZEF1 | 95.32143 | 0.092881951 | 0.18785826 | 0.49442569 | 0.62100557 | 0.88720256 |
| ANKFY1 | 63.9418512 | -0.140978951 | 0.27380436 | -0.5148893 | 0.60663034 | 0.88000768 |
| UBE2G1 | 82.4898418 | 0.18791574 | 0.17348599 | 1.08317534 | 0.27873061 | 0.67353297 |
| PSMB6 | 21.291832 | -0.404605419 | 0.44894354 | -0.901239 | 0.36746127 | 0.73985558 |
| MINK1 | 29.7626331 | 0.271877032 | 0.29942162 | 0.90800736 | 0.36387434 | 0.73626445 |
| PFN1 | 109.103911 | -0.373007917 | 0.41062049 | -0.9084007 | 0.36366658 | 0.73604674 |
| SPAG7 | 18.1163589 | 0.162559644 | 0.37990346 | 0.42789725 | 0.66872593 | 0.91445841 |
| CAMTA2 | 19.7665293 | -0.195728611 | 0.33043352 | -0.5923388 | 0.55362371 | 0.85188037 |
| KIF1C | 27.5315933 | 0.291721318 | 0.40735332 | 0.71613831 | 0.47390595 | 0.80789916 |
| AC087500.1 | 24.782603 | 0.466675685 | 0.28236466 | 1.6527411 | 0.09838357 | 0.45645254 |
| RABEP1 | 121.031543 | 0.154608523 | 0.20678603 | 0.74767393 | 0.45465686 | 0.79609632 |
| NUP88 | 44.0299844 | -0.032996258 | 0.21742193 | -0.1517614 | 0.87937512 | 0.97754164 |
| RPAIN | 26.8778889 | -0.001706186 | 0.31678694 | -0.0053859 | 0.99570269 | 0.99854015 |
| C1QBP | 25.5387741 | -0.672255258 | 0.3206467 | -2.0965607 | 0.03603248 | 0.28882238 |
| DERL2 | 21.2143266 | -0.096160987 | 0.31847666 | -0.3019405 | 0.76269745 | 0.94462105 |
| KIAA0753 | 33.779287 | -0.232425088 | 0.24849635 | -0.935326 | 0.34962034 | 0.72786398 |
| XAF1 | 116.795375 | -0.156766842 | 1.13791466 | -0.1377668 | 0.89042474 | 0.98023547 |
| DLG4 | 18.7955586 | 0.2103942 | 0.3827787 | 0.54964971 | 0.58255966 | 0.867524 |
| ACADVL | 81.7386586 | -0.226593833 | 0.29684315 | -0.7633453 | 0.44525747 | 0.79268341 |
| EIF5A | 32.135019 | -0.518699267 | 0.4664089 | -1.1121127 | 0.2660897 | 0.66086285 |
| GPS2 | 25.8401056 | -0.751650375 | 0.35711853 | -2.1047644 | 0.03531182 | 0.28703733 |
| ZBTB4 | 54.0436978 | 0.359642805 | 0.25155481 | 1.42967968 | 0.15280898 | 0.54042019 |
| POLR2A | 64.2238169 | 0.021626473 | 0.31782512 | 0.0680452 | 0.94574966 | 0.99060401 |
| EIF4A1 | 48.8698287 | -0.972656065 | 0.48748216 | -1.9952649 | 0.04601399 | 0.32236114 |
| FXR2 | 18.3524063 | 0.147192344 | 0.28532617 | 0.51587397 | 0.60594244 | 0.87970584 |
| SAT2 | 35.6687555 | -0.344082155 | 0.30412473 | -1.131385 | 0.25789309 | 0.65130862 |
| TP53 | 18.7389787 | -0.133822299 | 0.36611509 | -0.3655198 | 0.71472346 | 0.93179044 |
| KDM6B | 32.3176896 | -0.412540892 | 0.44484544 | -0.9273803 | 0.35372912 | 0.73059415 |
| NAA38 | 61.5071936 | -0.108694732 | 0.2566291 | -0.423548 | 0.67189551 | 0.91482259 |
| CHD3 | 55.0860505 | 0.31853166 | 0.30695815 | 1.03770388 | 0.29940794 | 0.69041385 |
| PER1 | 73.7396401 | -0.485718886 | 0.4291874 | -1.1317175 | 0.25775322 | 0.65130862 |
| NDEL1 | 63.4672857 | 0.228045253 | 0.27112481 | 0.84110803 | 0.40028742 | 0.76328807 |
| MYH10 | 283.37979 | 1.226963675 | 0.26498044 | 4.63039346 | 3.65E-06 | 0.00076634 |
| STX8 | 143.649225 | -0.188503886 | 0.18558577 | -1.0157238 | 0.30976092 | 0.69979495 |
| GAS7 | 65.8226522 | 1.369268983 | 0.25300002 | 5.41212998 | 6.23E-08 | 3.52E-05 |
| SCO1 | 24.4429226 | -0.018551162 | 0.28760522 | -0.0645022 | 0.94857038 | 0.99060401 |
| TMEM220 | 28.2929321 | 0.567757926 | 0.34063552 | 1.66676081 | 0.09556198 | 0.44903131 |
| MAP2K4 | 142.202095 | 0.005755969 | 0.15004474 | 0.03836168 | 0.96939931 | 0.99300191 |
| MYOCD | 71.1259112 | 0.351691026 | 0.39177684 | 0.89768202 | 0.3693551 | 0.74163678 |
| ARHGAP44 | 105.217751 | 0.565941561 | 0.29292411 | 1.93204159 | 0.05335437 | 0.34853448 |
| HS3ST3A1 | 49.3109327 | 1.689906835 | 0.40600866 | 4.16224336 | 3.15E-05 | 0.00399299 |
| COX10 | 26.3793921 | -0.216220556 | 0.27353676 | -0.7904625 | 0.4292577 | 0.78220055 |
| HS3ST3B1 | 27.0447372 | 0.355583111 | 0.60144387 | 0.59121579 | 0.55437584 | 0.85191704 |
| PMP22 | 54.6342726 | 0.192123412 | 0.26355898 | 0.72895795 | 0.46602739 | 0.80355999 |
| ZSWIM7 | 32.6644658 | -0.110345437 | 0.26804243 | -0.4116715 | 0.6805802 | 0.91860979 |
| TTC19 | 77.7512194 | -0.180031069 | 0.19385846 | -0.9286727 | 0.35305871 | 0.73059415 |
| NCOR1 | 266.116744 | 0.111143673 | 0.1536762 | 0.72323283 | 0.46953685 | 0.80412363 |
| PIGL | 80.9047777 | -0.040933416 | 0.28028482 | -0.1460422 | 0.88388806 | 0.97782528 |
| UBB | 100.154249 | -0.113229974 | 0.34060917 | -0.3324337 | 0.73956178 | 0.93594619 |
| LRRC75A | 30.3274519 | -0.068380541 | 0.28666051 | -0.2385419 | 0.81146083 | 0.9598303 |
| AC098850.3 | 64.4266703 | -0.199274528 | 0.56054686 | -0.3555002 | 0.72221488 | 0.93444541 |
| MPRIP | 280.699791 | -0.264054141 | 0.30081473 | -0.8777966 | 0.38005411 | 0.74991898 |
| RAI1 | 58.5802829 | 0.089272033 | 0.26444034 | 0.33758856 | 0.73567326 | 0.93594619 |
| SREBF1 | 45.1661032 | -0.41070197 | 0.30367831 | -1.3524244 | 0.17623958 | 0.5732874 |
| TOM1L2 | 110.236938 | 0.50267751 | 0.15795036 | 3.18250311 | 0.00146008 | 0.04768944 |
| ALKBH5 | 43.7744919 | -0.192262172 | 0.26531643 | -0.7246523 | 0.46866535 | 0.80374006 |
| FLII | 21.6509765 | 0.210290167 | 0.29971591 | 0.70163165 | 0.48290891 | 0.81409616 |
| TOP3A | 24.8584269 | -0.15205833 | 0.28331112 | -0.5367185 | 0.59146205 | 0.87229579 |
| PRPSAP2 | 38.7671754 | 0.161120123 | 0.23653196 | 0.68117698 | 0.49575951 | 0.82112613 |
| EPN2 | 152.01228 | 0.170656201 | 0.16422164 | 1.03918216 | 0.29872003 | 0.6896932 |
| MFAP4 | 33.174492 | 0.997020879 | 0.33506822 | 2.97557574 | 0.00292439 | 0.06945135 |
| SLC47A1 | 23.313476 | -0.000197772 | 0.32764147 | -0.0006036 | 0.99951838 | 0.99969762 |
| ALDH3A2 | 40.490301 | 0.553248183 | 0.31307343 | 1.76715149 | 0.07720286 | 0.4114983 |
| ULK2 | 67.09787 | 0.272704219 | 0.34064865 | 0.80054396 | 0.42339571 | 0.77827834 |
| AKAP10 | 110.564878 | -0.144284252 | 0.23491569 | -0.6141959 | 0.53908589 | 0.8449569 |
| SPECC1 | 134.833856 | 0.689318537 | 0.83870583 | 0.82188356 | 0.41114317 | 0.77166383 |
| USP22 | 100.724512 | -0.134457855 | 0.15240179 | -0.882259 | 0.37763675 | 0.749056 |
| DHRS7B | 47.5786741 | -0.240458379 | 0.32162208 | -0.7476426 | 0.45467574 | 0.79609632 |
| MAP2K3 | 30.6014602 | 0.129827449 | 0.38434336 | 0.33779027 | 0.73552124 | 0.93594619 |
| WSB1 | 378.259259 | -0.309059349 | 0.41612156 | -0.7427141 | 0.45765479 | 0.79810834 |
| KSR1 | 120.846437 | -1.144810586 | 0.31918042 | -3.5867193 | 0.00033486 | 0.01836507 |
| NLK | 127.149085 | 0.083146437 | 0.14505814 | 0.57319386 | 0.56651344 | 0.85825441 |
| SARM1 | 21.8471883 | 0.420561327 | 0.39541978 | 1.06358191 | 0.28751814 | 0.67963037 |
| KIAA0100 | 35.0783266 | 0.243558205 | 0.28806712 | 0.84549116 | 0.39783666 | 0.7611823 |
| SUPT6H | 54.9492294 | -0.175326721 | 0.23769437 | -0.7376141 | 0.46074898 | 0.80039155 |
| RAB34 | 29.4864889 | 0.219273403 | 0.36986739 | 0.5928433 | 0.55328603 | 0.85188037 |
| RPL23A | 115.191498 | -0.206118989 | 0.27341984 | -0.7538553 | 0.45093613 | 0.79401729 |
| FAM222B | 101.823112 | 0.054761393 | 0.19481337 | 0.28109669 | 0.77863624 | 0.94832577 |
| FLOT2 | 36.0543043 | -0.364826023 | 0.3157618 | -1.1553837 | 0.24793335 | 0.64333128 |
| PHF12 | 47.3734827 | -0.291610122 | 0.23268623 | -1.2532332 | 0.21012084 | 0.60731347 |
| MYO18A | 25.6964632 | 0.155185382 | 0.43903277 | 0.35347107 | 0.72373532 | 0.93455731 |
| NUFIP2 | 112.746692 | -0.164721566 | 0.16519765 | -0.9971181 | 0.31870718 | 0.70683013 |
| TAOK1 | 224.161838 | 0.116758981 | 0.17202671 | 0.67872589 | 0.49731156 | 0.82221432 |
| SSH2 | 134.248576 | -0.128507087 | 0.20356503 | -0.6312827 | 0.52785568 | 0.83947444 |
| NSRP1 | 134.310801 | -0.03787668 | 0.16926401 | -0.2237728 | 0.8229341 | 0.96393731 |
| BLMH | 25.973538 | 0.165171847 | 0.24076006 | 0.6860434 | 0.49268574 | 0.81880315 |
| CPD | 79.9128695 | -0.810783179 | 0.26451699 | -3.0651459 | 0.00217564 | 0.05772117 |
| GOSR1 | 61.68892 | 0.024898153 | 0.18184022 | 0.13692325 | 0.89109146 | 0.98023547 |
| CRLF3 | 49.4899039 | -0.017805081 | 0.25587238 | -0.0695858 | 0.94452335 | 0.99060356 |
| TEFM | 22.8389097 | 0.02034403 | 0.25588399 | 0.0795049 | 0.93663104 | 0.98827804 |
| NF1 | 400.732662 | -0.038376765 | 0.19548206 | -0.1963186 | 0.8443608 | 0.96926078 |
| UTP6 | 29.4550508 | -0.020290278 | 0.25299197 | -0.0802013 | 0.93607718 | 0.98827804 |
| SUZ12 | 83.5201704 | 0.112480553 | 0.19239184 | 0.58464305 | 0.55878775 | 0.85339385 |
| LRRC37B | 33.4582278 | 0.050666183 | 0.32684255 | 0.1550171 | 0.87680784 | 0.9768643 |
| RHOT1 | 74.0952508 | 0.236288778 | 0.22242084 | 1.06235001 | 0.28807682 | 0.67992862 |
| ZNF207 | 171.163064 | -0.12405906 | 0.15405992 | -0.805265 | 0.42066677 | 0.77771095 |
| PSMD11 | 51.8710976 | -0.316078108 | 0.22441652 | -1.408444 | 0.15899963 | 0.55091387 |
| MYO1D | 428.420846 | 0.541131902 | 0.31248371 | 1.73171236 | 0.08332479 | 0.42348124 |
| TMEM98 | 19.3351457 | 0.551545368 | 0.28850273 | 1.91175093 | 0.05590815 | 0.35425851 |
| CCL2 | 111.340584 | -1.270798205 | 1.0231941 | -1.2419913 | 0.21423976 | 0.61001472 |

| | | | | | | |
|---|---|---|---|---|---|---|
| SLFN5 | 46.227259 | -0.135140565 | 0.41821961 | -0.323133 | 0.74659451 | 0.93900586 |
| SLFN11 | 32.2555688 | -0.531236181 | 0.47552821 | -1.1171497 | 0.26393034 | 0.6579701 |
| AP2B1 | 105.092826 | -0.129482025 | 0.21837774 | -0.5929269 | 0.5532301 | 0.85188037 |
| TAF15 | 173.872288 | -0.210996236 | 0.21003001 | -1.0046004 | 0.31508928 | 0.70297242 |
| MYO19 | 30.1394454 | -0.344362852 | 0.33474046 | -1.0287458 | 0.30359912 | 0.69579129 |
| GGNBP2 | 90.4296668 | 0.015824631 | 0.18093056 | 0.08746246 | 0.93030393 | 0.98681039 |
| AATF | 58.5807154 | -0.368719978 | 0.19146207 | -1.9258121 | 0.05412784 | 0.34985531 |
| ACACA | 123.049205 | -0.306644092 | 0.31134766 | -0.9848929 | 0.32467669 | 0.71119194 |
| TADA2A | 32.5980738 | -0.08351917 | 0.23441169 | -0.3562927 | 0.72162139 | 0.93444541 |
| SYNRG | 68.9147147 | 0.146321433 | 0.22435599 | 0.6521842 | 0.51428235 | 0.83083337 |
| DDX52 | 34.1102156 | -0.392438038 | 0.24133632 | -1.6261043 | 0.10392746 | 0.46422585 |
| SOCS7 | 27.9634573 | 0.408814014 | 0.22362557 | 1.82811838 | 0.06753179 | 0.38712255 |
| ARHGAP23 | 33.8159207 | 0.901488566 | 0.32634046 | 2.76241741 | 0.00573751 | 0.10160227 |
| MLLT6 | 82.8634432 | -0.217501419 | 0.29174684 | -0.7455142 | 0.45596092 | 0.79649555 |
| PCGF2 | 19.856201 | 0.334887126 | 0.34916838 | 0.95909923 | 0.33750876 | 0.71925477 |
| PSMB3 | 29.0405899 | -0.46326077 | 0.38431339 | -1.2054245 | 0.22803947 | 0.62188065 |
| PIP4K2B | 29.9188951 | 0.494712223 | 0.24221541 | 2.04244735 | 0.04110718 | 0.30341448 |
| CWC25 | 23.7553598 | 0.085986061 | 0.25718274 | 0.33433839 | 0.73812423 | 0.93594619 |
| RPL23 | 75.2468194 | -0.226143226 | 0.26903655 | -0.8405669 | 0.4005906 | 0.76337055 |
| LASP1 | 70.5184381 | 1.00070568 | 0.30664328 | 3.26341959 | 0.00110076 | 0.04106353 |
| PLXDC1 | 23.7097705 | 0.945221839 | 0.58028859 | 1.62888233 | 0.10333793 | 0.46365765 |
| RPL19 | 132.346692 | -0.498370999 | 0.34315668 | -1.4523133 | 0.14641452 | 0.53585673 |
| FBXL20 | 122.70647 | 0.178001768 | 0.24526555 | 0.7257512 | 0.46799131 | 0.80374006 |
| MED1 | 27.6640286 | -0.028144808 | 0.27866074 | -0.1010003 | 0.91955025 | 0.98528826 |
| CDK12 | 77.627155 | 0.088978559 | 0.19042983 | 0.46725116 | 0.64032019 | 0.89914713 |
| ERBB2 | 24.1849205 | 0.967304505 | 0.28173635 | 3.43336778 | 0.00059613 | 0.0265097 |
| PSMD3 | 24.5635738 | -0.314683245 | 0.36782583 | -0.8555224 | 0.392262 | 0.75901355 |
| MED24 | 22.4313574 | -0.195251889 | 0.25716739 | -0.7592405 | 0.44770872 | 0.79296348 |
| THRA | 27.3308196 | 0.164445636 | 0.38707818 | 0.4248383 | 0.67095456 | 0.91475345 |
| MSL1 | 49.2690135 | -0.162148787 | 0.18611933 | -0.8712087 | 0.38364019 | 0.75256313 |
| CASC3 | 55.7824643 | -0.144714639 | 0.27650481 | -0.5233711 | 0.600716 | 0.87825667 |
| WIPF2 | 66.5751428 | 0.422501598 | 0.19432041 | 2.17425227 | 0.02968619 | 0.26108772 |
| RARA | 21.88809 | 0.306908251 | 0.31059808 | 0.98812025 | 0.32309376 | 0.70962823 |
| IGFBP4 | 484.783236 | -1.255593803 | 0.36513913 | -3.4386723 | 0.00058457 | 0.02619535 |
| KRT10 | 30.6456879 | -0.505557524 | 0.54716261 | -0.9239621 | 0.35550604 | 0.73125149 |
| EIF1 | 156.858453 | -0.359583526 | 0.31556826 | -1.1394794 | 0.25450326 | 0.64897448 |
| JUP | 37.4404371 | 0.650056169 | 0.39557665 | 1.6433128 | 0.10031823 | 0.45949178 |
| FKBP10 | 57.7232208 | -0.869559942 | 0.34798379 | -2.4988519 | 0.01245964 | 0.16120752 |
| ACLY | 37.4750517 | -0.128284599 | 0.27806158 | -0.4613532 | 0.64454522 | 0.90107359 |
| DNAJC7 | 74.41771 | 0.033071677 | 0.20622591 | 0.16036626 | 0.87259257 | 0.9768643 |
| KAT2A | 18.7004186 | -0.288665464 | 0.37001617 | -0.7801428 | 0.43530681 | 0.78678547 |
| STAT5B | 119.59916 | 0.005295392 | 0.25985954 | 0.0203779 | 0.98374191 | 0.99620525 |
| STAT5A | 18.6627306 | -0.366061397 | 0.33840731 | -1.0817184 | 0.27937769 | 0.67360455 |
| STAT3 | 484.51711 | -0.737793148 | 0.29622088 | -2.4906859 | 0.01274968 | 0.16438139 |
| CAVIN1 | 104.753981 | 1.165490067 | 0.37381136 | 3.11785619 | 0.00182172 | 0.05398305 |
| ATP6V0A1 | 99.0192627 | 0.017336251 | 0.22025412 | 0.07871023 | 0.93726311 | 0.98827804 |
| NAGLU | 28.7275149 | 0.185380046 | 0.42188506 | 0.43940889 | 0.66036529 | 0.90931694 |
| RETREG3 | 44.9432935 | 0.262847109 | 0.2475931 | 1.06160917 | 0.28841315 | 0.67992862 |
| TUBG2 | 22.2503524 | 0.428000103 | 0.35293392 | 1.21269188 | 0.22524764 | 0.62044413 |
| PLEKHH3 | 31.5367897 | 0.882473959 | 0.2783161 | 3.17076141 | 0.0015204 | 0.04900622 |
| AC100793.2 | 21.5183124 | 0.846623427 | 0.57748941 | 1.46604148 | 0.14263699 | 0.53022827 |
| EZH1 | 49.1774877 | -0.113102653 | 0.26351407 | -0.4292092 | 0.66777102 | 0.9142044 |
| COA3 | 21.8681331 | -0.150794342 | 0.36307806 | -0.4153221 | 0.67790612 | 0.91799007 |
| BECN1 | 17.2843405 | -0.209454283 | 0.32477563 | -0.6449199 | 0.51897907 | 0.83297983 |
| AOC3 | 26.9715362 | 0.152807076 | 0.31905167 | 0.47894147 | 0.63198027 | 0.89436221 |
| RPL27 | 66.3141705 | -0.836351442 | 0.34735039 | -2.4078034 | 0.01604882 | 0.18618176 |
| NBR1 | 63.4219789 | -0.257493022 | 0.26602502 | -0.9679279 | 0.3330804 | 0.7161521 |
| LINC00854 | 97.8208005 | 1.041561783 | 0.81969453 | 1.27067065 | 0.20384584 | 0.60367992 |
| DHX8 | 23.1682942 | 0.368194648 | 0.25642149 | 1.43589624 | 0.15103187 | 0.53880252 |
| DUSP3 | 27.9256204 | -0.242075062 | 0.29491919 | -0.8208183 | 0.41174978 | 0.77172894 |
| LSM12 | 29.8475899 | -0.139920827 | 0.27609769 | -0.5067801 | 0.61230909 | 0.88282721 |
| HDAC5 | 44.8852637 | 0.551673018 | 0.38051892 | 1.4497913 | 0.14711673 | 0.53679369 |
| UBTF | 22.2785446 | 0.867028085 | 0.30241863 | 2.86697977 | 0.0041441 | 0.08389795 |
| SLC25A39 | 19.6685382 | -0.412072638 | 0.40196528 | -1.0251449 | 0.30529485 | 0.69677386 |
| GRN | 34.3828998 | -0.444700744 | 0.31117505 | -1.4291016 | 0.15297505 | 0.54048732 |
| GPATCH8 | 247.594516 | 0.2436847 | 0.19356806 | 1.25890964 | 0.20806297 | 0.6059796 |
| GJC1 | 38.5733051 | 0.912520681 | 0.59213244 | 1.5410753 | 0.12329846 | 0.50097964 |
| EFTUD2 | 31.1471373 | -0.117000116 | 0.3114735 | -0.3756343 | 0.70718882 | 0.93009618 |
| DCAKD | 25.7383526 | 0.381701201 | 0.28450589 | 1.34162849 | 0.17971648 | 0.57744792 |
| NMT1 | 59.0471075 | -0.227653853 | 0.17932232 | -1.2695232 | 0.20425451 | 0.60367992 |
| AC138150.1 | 30.2290532 | 0.479033787 | 0.38216242 | 1.25348219 | 0.21003028 | 0.60731347 |
| MAP3K14 | 33.7370234 | 0.299728376 | 0.39773651 | 0.75358526 | 0.4510983 | 0.79401729 |
| PLEKHM1 | 36.9019827 | -0.455775082 | 0.21733095 | -2.0971476 | 0.03598051 | 0.28882238 |
| MAPT | 24.5034756 | 1.073791931 | 0.58042304 | 1.85001603 | 0.06431124 | 0.37870457 |
| KANSL1 | 462.661761 | 0.344198615 | 0.23057251 | 1.49279986 | 0.13548958 | 0.51914127 |
| ARL17B | 106.700831 | 1.816278058 | 0.97576458 | 1.86138962 | 0.06268918 | 0.37164525 |
| LRRC37A2 | 23.8362556 | 0.564766086 | 0.30062945 | 1.87861199 | 0.06029749 | 0.36535567 |
| ARL17A | 26.4697708 | -0.245689664 | 0.37055599 | -0.6630298 | 0.50731147 | 0.82494327 |
| NSF | 145.110685 | -0.555468419 | 0.24102001 | -2.3046569 | 0.0211858 | 0.21475093 |
| CDC27 | 168.115172 | -0.208942069 | 0.19032621 | -1.0978103 | 0.27228733 | 0.66723562 |
| AC068234.1 | 26.0047855 | -0.649439771 | 0.595152 | -1.0912166 | 0.27517757 | 0.67029497 |
| EFCAB13 | 42.1325229 | -0.385952313 | 0.32665843 | -1.1815165 | 0.23739761 | 0.63302595 |
| NPEPPS | 185.369574 | -0.150771083 | 0.18739731 | -0.8045531 | 0.42107762 | 0.77771095 |
| AC025682.1 | 18.3607486 | 0.337779292 | 0.3172332 | 1.06476653 | 0.2869816 | 0.67921285 |
| KPNB1 | 98.2739099 | -0.084818971 | 0.20933114 | -0.4051904 | 0.68533755 | 0.91977784 |
| CDK5RAP3 | 19.4026282 | 0.057580942 | 0.29657577 | 0.19415255 | 0.84605644 | 0.97014274 |
| NFE2L1 | 51.7697471 | -0.125387572 | 0.26566021 | -0.4719848 | 0.63693766 | 0.89741446 |
| CBX1 | 18.957123 | 0.212603445 | 0.29590558 | 0.71848408 | 0.47245886 | 0.80671473 |
| HOXB3 | 42.5952433 | 0.371749032 | 0.34827219 | 1.06740945 | 0.28578698 | 0.6783522 |
| CALCOCO2 | 80.5651593 | -0.075074514 | 0.28709672 | -0.2614955 | 0.79371039 | 0.95416774 |
| ATP5MC1 | 28.3918969 | -0.425245203 | 0.32109843 | -1.3243453 | 0.18538838 | 0.58498035 |
| UBE2Z | 49.7193801 | -0.309595122 | 0.24386823 | -1.269518 | 0.20425636 | 0.60367992 |
| SNF8 | 20.9843114 | -0.094321706 | 0.29520735 | -0.31951 | 0.74933979 | 0.94004394 |
| ZNF652 | 83.540644 | 0.022287189 | 0.17861851 | 0.12477536 | 0.90070139 | 0.98118567 |
| PHB | 24.0805956 | -0.240733557 | 0.32788514 | -0.7342009 | 0.46282632 | 0.80225845 |
| SPOP | 92.8091691 | 0.328557632 | 0.18403327 | 1.78531647 | 0.07421 | 0.40397725 |
| KAT7 | 30.6050946 | 0.075200768 | 0.23187938 | 0.32430986 | 0.74570345 | 0.93891908 |
| ITGA3 | 78.7132742 | 0.852481489 | 0.40730407 | 2.09298542 | 0.03635046 | 0.28969304 |
| COL1A1 | 178.161407 | -0.038185124 | 0.48281628 | -0.0790883 | 0.93696238 | 0.98827804 |
| MRPL27 | 19.4373115 | -0.444986328 | 0.29475361 | -1.5096891 | 0.13112276 | 0.51290135 |
| LRRC59 | 20.3499641 | -0.711798673 | 0.41261196 | -1.7251043 | 0.08450868 | 0.42695086 |
| ABCC3 | 26.179924 | -0.698566192 | 0.37499723 | -1.8628569 | 0.0624824 | 0.37164525 |
| ANKRD40 | 19.8748596 | -0.179573208 | 0.32724667 | -0.5487396 | 0.58318416 | 0.86764709 |
| LUC7L3 | 216.673281 | 0.387904597 | 0.14243721 | 2.72333755 | 0.0064626 | 0.10918078 |
| TOB1 | 61.1534596 | 0.524894778 | 0.27564693 | 1.90422861 | 0.05688042 | 0.35705408 |
| SPAG9 | 312.585683 | 0.05572018 | 0.2292305 | 0.24307489 | 0.80794738 | 0.95922541 |
| MBTD1 | 91.4137292 | -0.145670384 | 0.2288063 | -0.6366537 | 0.52435041 | 0.83697897 |
| UTP18 | 24.6700843 | -0.246297284 | 0.29033404 | -0.8483238 | 0.39625766 | 0.76020146 |
| TOM1L1 | 31.4462111 | -0.112426242 | 0.25772551 | -0.4362247 | 0.66267368 | 0.91112982 |
| COX11 | 21.9649771 | -0.110574636 | 0.30166994 | -0.3665418 | 0.71396084 | 0.93136293 |
| STXBP4 | 128.889464 | -0.89271929 | 0.30312505 | -2.9450529 | 0.00322899 | 0.07300955 |
| DGKE | 25.5513072 | 0.435059539 | 0.35387557 | 1.22941387 | 0.21891667 | 0.61522936 |
| TRIM25 | 26.1975573 | -0.082106756 | 0.41009161 | -0.2002156 | 0.84131193 | 0.96803523 |

| | | | | | | |
|---|---|---|---|---|---|---|
| SCPEP1 | 33.9569972 | -0.375344816 | 0.22267078 | -1.6856491 | 0.0918634 | 0.44124451 |
| AKAP1 | 21.1730995 | 0.254051587 | 0.33584462 | 0.75645572 | 0.44937601 | 0.7936829 |
| MSI2 | 353.166199 | -0.544728489 | 0.33257273 | -1.6379229 | 0.10143778 | 0.46169744 |
| CUEDC1 | 70.9433345 | 0.346439817 | 0.30870563 | 1.12223357 | 0.26176314 | 0.65593044 |
| VEZF1 | 39.1562781 | 0.371452251 | 0.20750164 | 1.79011718 | 0.07343508 | 0.40189622 |
| AC015813.2 | 23.0875603 | 0.095600734 | 0.32411708 | 0.29495741 | 0.76802641 | 0.94576899 |
| DYNLL2 | 21.1874735 | 0.3302874 | 0.2768155 | 1.193168 | 0.23280358 | 0.62899763 |
| MTMR4 | 21.3670287 | 0.041761503 | 0.30357011 | 0.1375679 | 0.89058193 | 0.98023547 |
| 4-Sep | 21.7663595 | 0.960213784 | 0.33138761 | 2.89755488 | 0.00376084 | 0.07961552 |
| TRIM37 | 58.9244155 | 0.513791464 | 0.2809162 | 1.82898482 | 0.06740188 | 0.38712255 |
| SKA2 | 29.9352461 | 0.625586177 | 0.23894514 | 2.6181164 | 0.00884166 | 0.13287809 |
| YPEL2 | 124.775338 | 0.060812135 | 0.36055993 | 0.16866027 | 0.86606387 | 0.9762143 |
| DHX40 | 33.9780578 | -0.336080711 | 0.30895548 | -1.0877966 | 0.27668492 | 0.67259169 |
| CLTC | 196.39397 | -0.164368847 | 0.189931 | -0.8654135 | 0.38681188 | 0.75402666 |
| PTRH2 | 33.1164403 | -0.641880733 | 0.29242423 | -2.1950327 | 0.02816125 | 0.25237565 |
| VMP1 | 737.943011 | -1.276422122 | 0.52075166 | -2.4511148 | 0.01424145 | 0.17501743 |
| TUBD1 | 25.0797078 | 0.359573934 | 0.40140339 | 0.89579197 | 0.37036388 | 0.74264779 |
| RPS6KB1 | 50.3442157 | -0.035201787 | 0.26942047 | -0.1306574 | 0.89604631 | 0.98035497 |
| USP32 | 206.128755 | -0.461506752 | 0.30604783 | -1.5079563 | 0.13156571 | 0.51290135 |
| APPBP2 | 114.348556 | 0.275992806 | 0.19403644 | 1.42237616 | 0.15491708 | 0.54412824 |
| PPM1D | 39.7569958 | 0.249952923 | 0.28652314 | 0.87236558 | 0.38300897 | 0.75179833 |
| BCAS3 | 345.152808 | 0.337543409 | 0.20212516 | 1.66997225 | 0.09492485 | 0.44764276 |
| TBX2 | 113.971792 | 0.410380533 | 0.30749766 | 1.334581 | 0.18201352 | 0.58094117 |
| INTS2 | 20.7391031 | -0.005970201 | 0.2755668 | -0.0216652 | 0.98271505 | 0.99599681 |
| MED13 | 235.742183 | -0.092714041 | 0.16709838 | -0.554847 | 0.5789993 | 0.86486944 |
| TLK2 | 140.553178 | -0.123626054 | 0.21342357 | -0.5792521 | 0.56241908 | 0.85545224 |
| MRC2 | 116.283098 | 0.300597693 | 0.34283789 | 0.87679252 | 0.38059934 | 0.74991898 |
| TANC2 | 148.336085 | -0.272803887 | 0.43373366 | -0.6289664 | 0.52937107 | 0.84061106 |
| DCAF7 | 63.1915267 | -0.140686539 | 0.29741593 | -0.4730296 | 0.63619206 | 0.89741446 |
| MAP3K3 | 74.8885905 | 0.410072161 | 0.22369545 | 1.83317165 | 0.06677705 | 0.38641301 |
| CCDC47 | 25.5352051 | -0.973935311 | 0.31444066 | -3.0973581 | 0.00195254 | 0.05557623 |
| DDX42 | 77.9904571 | 0.202065794 | 0.15194615 | 1.32985137 | 0.18356725 | 0.58298863 |
| SMARCD2 | 23.154995 | 0.231159264 | 0.31277881 | 0.73905027 | 0.45987647 | 0.80029179 |
| ERN1 | 55.9222258 | -0.967016021 | 0.42671159 | -2.2662052 | 0.02343882 | 0.22729214 |
| SNHG25 | 23.4120661 | -0.37870371 | 0.4164046 | -0.9094609 | 0.36310688 | 0.73604674 |
| TEX2 | 38.4768716 | -0.011804 | 0.19810029 | -0.059586 | 0.95248538 | 0.99060401 |
| PECAM1 | 41.9119623 | -0.99528267 | 0.53344526 | -1.8657635 | 0.06207447 | 0.37058104 |
| POLG2 | 22.5184802 | -0.290328636 | 0.30089376 | -0.9648875 | 0.33460115 | 0.71690491 |
| DDX5 | 325.981219 | -0.336339887 | 0.24112078 | -1.3949021 | 0.16304535 | 0.5572759 |
| CEP95 | 71.4128283 | 0.11557039 | 0.21411463 | 0.53975942 | 0.58936296 | 0.8702694 |
| SMURF2 | 131.027978 | 0.51452199 | 0.18819059 | 2.73404735 | 0.0062561 | 0.1076724 |
| LRRC37A3 | 85.3806611 | 0.904153524 | 0.3331881 | 2.71364294 | 0.00665479 | 0.11140322 |
| GNA13 | 58.8889465 | -0.458358224 | 0.20701635 | -2.214116 | 0.0268208 | 0.24515678 |
| CEP112 | 176.07957 | 0.250013379 | 0.21214442 | 1.17850553 | 0.23859513 | 0.63422836 |
| PRKCA | 730.667021 | 0.35160669 | 0.31475692 | 1.11707375 | 0.26396279 | 0.6579701 |
| PRKCA-AS1 | 31.247789 | -0.096979588 | 0.59960486 | -0.1617392 | 0.87151126 | 0.9768643 |
| HELZ | 191.851258 | -0.452697146 | 0.23067812 | -1.9624625 | 0.04970867 | 0.33731209 |
| PSMD12 | 48.1702999 | -0.380451134 | 0.24579995 | -1.547808 | 0.12166852 | 0.49924175 |
| PITPNC1 | 552.968031 | -1.335906851 | 0.48103455 | -2.7771537 | 0.00548372 | 0.09829242 |
| NOL11 | 22.1368301 | 0.42134232 | 0.28000076 | 1.50478991 | 0.1323781 | 0.51310477 |
| BPTF | 246.353071 | 0.426828062 | 0.22132591 | 1.92850474 | 0.05379238 | 0.34959616 |
| ARSG | 20.3550342 | -0.330104325 | 0.33661567 | -0.9806564 | 0.32676219 | 0.71172951 |
| WIPI1 | 39.3672186 | -0.889411621 | 0.27561017 | -3.2270639 | 0.00125068 | 0.04375316 |
| PRKAR1A | 133.84813 | -0.307541173 | 0.26896903 | -1.1434074 | 0.25286949 | 0.64773018 |
| FAM20A | 31.4520339 | -0.829170553 | 0.52782853 | -1.5709089 | 0.1162038 | 0.48882756 |
| LINC01482 | 29.4686476 | -0.091051099 | 0.82041344 | -0.110982 | 0.91163064 | 0.98343817 |
| ABCA8 | 206.090172 | 0.219767258 | 0.44915848 | 0.48928667 | 0.62463875 | 0.89031618 |
| ABCA9 | 158.170365 | 0.341651491 | 0.4500453 | 0.75914911 | 0.44776336 | 0.79296348 |
| ABCA6 | 172.809613 | 0.193124261 | 0.51457626 | 0.37530736 | 0.70743189 | 0.93009618 |
| ABCA10 | 126.256106 | 0.400395926 | 0.44772732 | 0.89428523 | 0.3711693 | 0.74385689 |
| ABCA5 | 80.0871991 | -0.755810576 | 0.28663785 | -2.6368136 | 0.00836888 | 0.12814321 |
| KCNJ16 | 31.1999463 | 0.561021806 | 0.35113479 | 1.59773916 | 0.11010104 | 0.47680173 |
| SLC39A11 | 123.679995 | 0.341599887 | 0.21215786 | 1.61012131 | 0.10737138 | 0.47249835 |
| SSTR2 | 41.7891651 | -0.551995195 | 0.60072558 | -0.9188808 | 0.35815793 | 0.73144657 |
| FAM104A | 18.8281094 | -0.162318678 | 0.26135065 | -0.6210762 | 0.53454947 | 0.84361209 |
| CDC42EP4 | 43.8599794 | -0.388765229 | 0.23609719 | -1.6466322 | 0.09963367 | 0.45779653 |
| RPL38 | 46.5360703 | -0.429046178 | 0.37267144 | -1.1512719 | 0.24962038 | 0.64565668 |
| GPRC5C | 32.3609391 | -0.165088368 | 0.34024932 | -0.4851982 | 0.62753573 | 0.89192276 |
| ATP5PD | 20.0885603 | 0.000596237 | 0.35503117 | 0.00167939 | 0.99866004 | 0.99969762 |
| NT5C | 22.8599476 | -0.111915747 | 0.30806075 | -0.3632912 | 0.71638739 | 0.93247094 |
| SUMO2 | 19.1253918 | -0.054276604 | 0.3084955 | -0.1759397 | 0.86034132 | 0.97397549 |
| NUP85 | 22.5311805 | -0.148809507 | 0.27861029 | -0.5341135 | 0.59326304 | 0.87337541 |
| GGA3 | 27.7948574 | 0.231505066 | 0.25352266 | 0.91315336 | 0.36116189 | 0.73319855 |
| GRB2 | 79.698774 | -0.180165687 | 0.22454687 | -0.8023523 | 0.42234922 | 0.77771095 |
| TMEM94 | 28.2867915 | 0.3074755 | 0.29516557 | 1.04170515 | 0.2975484 | 0.68849737 |
| MYO15B | 39.7727959 | 0.394663051 | 0.4313186 | 0.91501514 | 0.36018368 | 0.73273991 |
| SAP30BP | 31.3185216 | -0.031052979 | 0.24565719 | -0.1264078 | 0.89940915 | 0.98058672 |
| H3F3B | 262.922791 | -0.146069335 | 0.27419772 | -0.5327154 | 0.59423063 | 0.87394794 |
| UNK | 70.4360951 | 0.292605527 | 0.24187322 | 1.20974752 | 0.22637579 | 0.62072404 |
| WBP2 | 34.7179923 | -0.325706415 | 0.23826959 | -1.3669659 | 0.17163598 | 0.56870244 |
| ACOX1 | 36.1248162 | 0.162020857 | 0.24323536 | 0.66610734 | 0.50534249 | 0.82481485 |
| SRP68 | 25.1262579 | 0.087006542 | 0.31081926 | 0.27992648 | 0.77953391 | 0.94894728 |
| EXOC7 | 29.6420073 | 0.686291652 | 0.37974057 | 1.80726452 | 0.07072105 | 0.39613488 |
| PRPSAP1 | 40.594761 | 0.235536369 | 0.23178841 | 1.01616973 | 0.30954857 | 0.69972033 |
| UBE2O | 41.7354555 | -0.394401634 | 0.2729369 | -1.4450286 | 0.14844985 | 0.53680621 |
| MXRA7 | 57.0102623 | 0.275452189 | 0.24662929 | 1.11686729 | 0.26405107 | 0.6579701 |
| SRSF2 | 57.7324479 | 0.008908882 | 0.20762021 | 0.04290951 | 0.96577366 | 0.99293098 |
| MFSD11 | 43.4215331 | 0.166100469 | 0.29129343 | 0.57021701 | 0.56853052 | 0.85958696 |
| SEC14L1 | 106.144643 | -0.239310825 | 0.22696163 | -1.0544109 | 0.29169482 | 0.68334882 |
| 9-Sep | 187.92281 | -0.049400765 | 0.3370803 | -0.1465549 | 0.88348336 | 0.97782528 |
| TNRC6C | 145.897183 | 0.053608296 | 0.30067149 | 0.17829524 | 0.85849112 | 0.97397549 |
| SYNGR2 | 21.3652273 | -0.250219168 | 0.3415731 | -0.7325494 | 0.4638333 | 0.80227475 |
| SOCS3 | 52.4466586 | -1.739627545 | 0.50443038 | -3.448697 | 0.0005633 | 0.02567229 |
| PGS1 | 31.4886242 | -0.872608017 | 0.35201643 | -2.4788843 | 0.0131794 | 0.16756996 |
| CYTH1 | 68.8636857 | -0.299439931 | 0.25615116 | -1.168997 | 0.24240486 | 0.63672023 |
| USP36 | 60.3754952 | -0.303572621 | 0.24130619 | -1.2580391 | 0.20837762 | 0.6059796 |
| TIMP2 | 205.278867 | 0.696663244 | 0.30600631 | 2.27663031 | 0.02280831 | 0.22438859 |
| LGALS3BP | 84.9244991 | 0.082464505 | 0.52654497 | 0.15661436 | 0.87554878 | 0.9768643 |
| C1QTNF1 | 64.0197769 | -1.050985787 | 0.54230597 | -1.9379941 | 0.05262394 | 0.34684602 |
| TBC1D16 | 87.3881413 | -0.411196166 | 0.26269247 | -1.5653138 | 0.11750934 | 0.49066825 |
| EIF4A3 | 21.9465398 | 0.253873494 | 0.44560988 | 0.56972143 | 0.56886665 | 0.85958696 |
| RNF213 | 195.177722 | -0.049571692 | 0.45893531 | -0.1080146 | 0.91398415 | 0.98395113 |
| AC124319.2 | 36.5559943 | 0.262390824 | 0.35499401 | 0.73914156 | 0.45982104 | 0.80029179 |
| RPTOR | 105.720081 | -0.344986313 | 0.23277286 | -1.4820728 | 0.13832093 | 0.52264518 |
| SLC38A10 | 47.4406621 | -0.176288484 | 0.25046458 | -0.703846 | 0.48152871 | 0.81369383 |
| BAHCC1 | 24.0352378 | 0.503870351 | 0.41331859 | 1.21908466 | 0.22281206 | 0.61772146 |
| ACTG1 | 356.395968 | 0.521763833 | 0.29152376 | 1.78978151 | 0.07348904 | 0.40189622 |
| NPLOC4 | 85.0819528 | 0.101736544 | 0.19099453 | 0.53266733 | 0.59426389 | 0.87394794 |
| HGS | 29.0436637 | -0.343807171 | 0.26009246 | -1.3218652 | 0.18621304 | 0.58501647 |
| MCRIP1 | 27.7887586 | 0.263576207 | 0.35159821 | 0.74965172 | 0.45346449 | 0.79534858 |
| P4HB | 134.51655 | -0.728470568 | 0.37846452 | -1.9248054 | 0.0542537 | 0.3503607 |
| ARHGDIA | 63.8641798 | -0.436848622 | 0.37334757 | -1.1700856 | 0.24196652 | 0.6364395 |
| ANAPC11 | 32.1251189 | -0.555484871 | 0.36525404 | -1.5208179 | 0.12830554 | 0.50940975 |

| | | | | | | |
|---|---|---|---|---|---|---|
| ASPSCR1 | 16.2934574 | 0.038153686 | 0.45427535 | 0.08398802 | 0.93306596 | 0.98725539 |
| DUS1L | 20.7411889 | -0.418620347 | 0.32504385 | -1.2878889 | 0.19778466 | 0.59530234 |
| CCDC57 | 65.5400018 | -0.14125163 | 0.23321082 | -0.6056821 | 0.54472584 | 0.84730644 |
| SLC16A3 | 23.4641302 | -1.075189306 | 0.43446318 | -2.4747536 | 0.01333281 | 0.16893595 |
| CSNK1D | 64.1023531 | -0.476468021 | 0.18351851 | -2.5962941 | 0.00942354 | 0.13659482 |
| HEXDC | 26.4878527 | 0.196880208 | 0.30458926 | 0.64637936 | 0.51803371 | 0.83297983 |
| NARF | 29.708268 | -0.521107765 | 0.24903963 | -2.0924692 | 0.03639656 | 0.28969304 |
| FOXK2 | 124.380617 | -0.290216624 | 0.24398087 | -1.1895057 | 0.23424074 | 0.63076093 |
| WDR45B | 71.484497 | -0.490452158 | 0.33261804 | -1.4745206 | 0.14034147 | 0.52680084 |
| TBCD | 55.2951167 | 0.01392951 | 0.28168403 | 0.04945083 | 0.96056002 | 0.99181157 |
| USP14 | 70.1148148 | 0.113351043 | 0.19347099 | 0.58588134 | 0.55795526 | 0.85279817 |
| THOC1 | 60.3277748 | -0.032076711 | 0.23616805 | -0.1358216 | 0.89196235 | 0.98023547 |
| COLEC12 | 76.1168307 | 0.047603678 | 0.7618328 | 0.06248573 | 0.95017603 | 0.99060401 |
| ENOSF1 | 104.349974 | -0.713366164 | 0.2763251 | -2.5816192 | 0.0098338 | 0.14050816 |
| YES1 | 60.3877938 | -0.061324255 | 0.2034113 | -0.3014791 | 0.76304918 | 0.94462105 |
| METTL4 | 19.7052074 | 0.357206004 | 0.2611822 | 1.36765062 | 0.17142146 | 0.56870244 |
| SMCHD1 | 288.492539 | 0.057774881 | 0.35228258 | 0.16400153 | 0.86972994 | 0.9768643 |
| AP001011.1 | 40.3223074 | -0.423457289 | 0.4392585 | -0.9640275 | 0.33503212 | 0.71740999 |
| LPIN2 | 175.887609 | 0.669620189 | 0.3758653 | 1.78154299 | 0.07482379 | 0.40551624 |
| MYOM1 | 87.3547534 | 1.459934774 | 0.54997763 | 2.65453484 | 0.00794178 | 0.1246781 |
| MYL12A | 106.989438 | 0.336033829 | 0.33987328 | 0.98870329 | 0.32280833 | 0.70942537 |
| MYL12B | 123.222416 | 0.056202726 | 0.31028118 | 0.18113482 | 0.85626175 | 0.97379567 |
| DLGAP1 | 132.359281 | 0.670013273 | 0.78727991 | 0.85104836 | 0.39474249 | 0.75988253 |
| LINC00667 | 19.6001396 | 0.46629067 | 0.35536625 | 1.31214111 | 0.18947253 | 0.58678196 |
| EPB41L3 | 38.6631533 | 0.616898203 | 0.37928914 | 1.62645894 | 0.10385206 | 0.46422585 |
| L3MBTL4 | 172.852884 | 0.299453045 | 0.43749601 | 0.68447034 | 0.49367821 | 0.81915583 |
| ARHGAP28 | 30.2440379 | 1.852385969 | 0.41034758 | 4.51418759 | 6.36E-06 | 0.00111215 |
| PTPRM | 399.565544 | 0.042542341 | 0.22822533 | 0.18640499 | 0.85212718 | 0.97285733 |
| RAB12 | 46.9746534 | -0.172008963 | 0.22937566 | -0.7499007 | 0.45331453 | 0.79534858 |
| NDUFV2 | 53.0442543 | -0.283878072 | 0.25105278 | -1.1307506 | 0.25816009 | 0.65130862 |
| ANKRD12 | 329.998215 | -0.331639092 | 0.1605718 | -2.0653632 | 0.03888865 | 0.29803111 |
| TWSG1 | 50.3620793 | 0.243361146 | 0.1921038 | 1.26682111 | 0.20521926 | 0.60370888 |
| RALBP1 | 42.8468761 | 0.301520691 | 0.21608316 | 1.39539192 | 0.16289769 | 0.55718935 |
| PPP4R1 | 70.4546894 | 0.080032423 | 0.21461477 | 0.37291201 | 0.70921393 | 0.93009618 |
| RAB31 | 142.584947 | -0.186396039 | 0.35150286 | -0.5302831 | 0.59591564 | 0.87481862 |
| VAPA | 164.311193 | -0.062684823 | 0.25032115 | -0.2504176 | 0.80226442 | 0.95758343 |
| APCDD1 | 28.9598909 | -0.272558189 | 0.36352689 | -0.7497607 | 0.45339882 | 0.79534858 |
| NAPG | 44.9811867 | -0.064699177 | 0.29016874 | -0.2229709 | 0.82355818 | 0.96413949 |
| PIEZO2 | 185.429657 | 1.07879706 | 0.31623631 | 3.41136364 | 0.00064639 | 0.02800329 |
| MPPE1 | 20.6815376 | 0.468753581 | 0.25688013 | 1.82479505 | 0.06803196 | 0.38847467 |
| IMPA2 | 24.5206264 | 0.104994922 | 0.40578681 | 0.25874405 | 0.79583273 | 0.95463848 |
| TUBB6 | 94.0713153 | 0.113734816 | 0.30594459 | 0.37174972 | 0.7100792 | 0.93009618 |
| AFG3L2 | 46.3758847 | 0.032195971 | 0.28434441 | 0.11322878 | 0.90984918 | 0.98343817 |
| SPIRE1 | 139.115416 | 0.250895352 | 0.23149365 | 1.08381096 | 0.27844862 | 0.67335273 |
| PSMG2 | 34.4045012 | -0.021773246 | 0.21895792 | -0.0994403 | 0.92078867 | 0.98528826 |
| PTPN2 | 80.1910762 | -0.644213177 | 0.23979435 | -2.6865236 | 0.00721998 | 0.11687151 |
| SEH1L | 38.8155098 | -0.409099984 | 0.33764502 | -1.2116275 | 0.22565501 | 0.62063573 |
| CEP192 | 79.7681695 | -0.023569476 | 0.2116126 | -0.1113803 | 0.91131478 | 0.98343817 |
| LDLRAD4 | 432.237642 | -0.096274565 | 0.35455835 | -0.2715338 | 0.78598053 | 0.95065355 |
| FAM210A | 33.2039971 | -1.548705735 | 0.48279582 | -3.2077861 | 0.00133761 | 0.04488626 |
| RNMT | 79.9896306 | -1.129806431 | 0.39487239 | -2.8611938 | 0.00422049 | 0.08497639 |
| ROCK1 | 354.931808 | 0.590598379 | 0.23117588 | 2.55475775 | 0.01062617 | 0.14488263 |
| GREB1L | 39.6054292 | 0.718917508 | 0.59542176 | 1.20740887 | 0.22727472 | 0.62173063 |
| ESCO1 | 50.5927732 | -0.086450867 | 0.196248 | -0.4405185 | 0.65956164 | 0.90855808 |
| ABHD3 | 45.6424741 | -0.115864152 | 0.30539996 | -0.379385 | 0.70440201 | 0.92936308 |
| MIB1 | 159.174306 | 0.044267882 | 0.18146664 | 0.24394501 | 0.80727341 | 0.95873502 |
| GATA6 | 62.7064543 | -0.499855893 | 0.44204131 | -1.13079 | 0.25814349 | 0.65130862 |
| RBBP8 | 29.7174979 | -0.279491462 | 0.32169973 | -0.8687961 | 0.38495868 | 0.75305975 |
| CABLES1 | 97.2165059 | 0.925073578 | 0.40006112 | 2.31233063 | 0.02075947 | 0.21189076 |
| TMEM241 | 61.6172728 | -0.05548543 | 0.22202454 | -0.2499067 | 0.80265947 | 0.95789939 |
| RIOK3 | 75.0398319 | -0.292212357 | 0.2309312 | -1.2653654 | 0.20574036 | 0.60431091 |
| NPC1 | 57.7485541 | -0.36692626 | 0.31589963 | -1.161528 | 0.24542725 | 0.64118197 |
| TTC39C | 34.7092848 | -0.738054701 | 0.36601336 | -2.0164693 | 0.04375093 | 0.31491241 |
| OSBPL1A | 186.881792 | -0.446135059 | 0.31244822 | -1.4278688 | 0.15332964 | 0.54114675 |
| ZNF521 | 29.8738123 | 0.661480591 | 0.33178048 | 1.99372969 | 0.0461816 | 0.32322724 |
| SS18 | 109.18652 | -0.207266924 | 0.22223437 | -0.9326502 | 0.35100062 | 0.72929137 |
| TAF4B | 21.6233353 | -0.529660035 | 0.27614115 | -1.9180772 | 0.05510123 | 0.35260065 |
| KCTD1 | 46.6743336 | -0.47317294 | 0.30175598 | -1.5680648 | 0.116866 | 0.48989475 |
| TRAPPC8 | 91.9731447 | 0.007338283 | 0.21151401 | 0.03469407 | 0.97232369 | 0.99300191 |
| RNF138 | 32.8149789 | 0.014893748 | 0.33580279 | 0.04435266 | 0.96462329 | 0.99275266 |
| GAREM1 | 162.999944 | 0.781652273 | 0.25569219 | 3.05700492 | 0.00223561 | 0.05888698 |
| DTNA | 60.7410823 | 0.683117572 | 0.53813606 | 1.26941422 | 0.20429337 | 0.60367992 |
| MAPRE2 | 114.271258 | 0.007429549 | 0.3817438 | 0.01946213 | 0.98447244 | 0.9964038 |
| ZNF397 | 33.3909813 | 0.136071534 | 0.25013636 | 0.54398941 | 0.58644876 | 0.87014172 |
| ZSCAN30 | 23.7339506 | -0.300791187 | 0.28278241 | -1.0636842 | 0.28747177 | 0.67963037 |
| ZNF24 | 37.7896743 | -0.181042412 | 0.25965141 | -0.6972518 | 0.48564524 | 0.81577299 |
| GALNT1 | 73.6264924 | -0.002714872 | 0.26409845 | -0.0102798 | 0.99179807 | 0.99759781 |
| RPRD1A | 95.0177224 | -0.00347804 | 0.16292986 | -0.0213469 | 0.98296897 | 0.99601865 |
| SLC39A6 | 23.6143246 | -0.076019467 | 0.27228139 | -0.2791945 | 0.78009556 | 0.94915932 |
| ELP2 | 75.7924254 | -0.250136958 | 0.25576186 | -0.9780073 | 0.32807073 | 0.71294249 |
| FHOD3 | 59.4185458 | 0.370967622 | 0.56545793 | 0.65604814 | 0.51179315 | 0.82882457 |
| TPGS2 | 29.9166926 | 0.209304384 | 0.22141902 | 0.94528637 | 0.3445127 | 0.72296511 |
| KIAA1328 | 160.597053 | 0.360929016 | 0.28730934 | 1.25623835 | 0.20902957 | 0.6059796 |
| AC016205.1 | 19.2175555 | -0.435024387 | 0.33039678 | -1.3166726 | 0.18794839 | 0.58645909 |
| PIK3C3 | 119.648352 | -0.582110721 | 0.25836463 | -2.2530589 | 0.02425544 | 0.23086389 |
| SETBP1 | 1049.35055 | -0.356924486 | 0.36677627 | -0.9731395 | 0.33048396 | 0.71362138 |
| SLC14A2-AS1 | 30.7865761 | 0.592918708 | 0.83757755 | 0.70789708 | 0.47900917 | 0.81163945 |
| EPG5 | 73.1280512 | -0.054222064 | 0.18652394 | -0.2906976 | 0.7712826 | 0.94654641 |
| PSTPIP2 | 24.9911607 | -0.414360838 | 0.31011349 | -1.3361587 | 0.18149741 | 0.580649 |
| ATP5F1A | 52.9980221 | -0.07194676 | 0.20596425 | -0.3493167 | 0.72685153 | 0.93455731 |
| C18orf25 | 45.4223834 | -0.071574671 | 0.22428626 | -0.319122 | 0.74963403 | 0.94004394 |
| PIAS2 | 81.323852 | -0.190633763 | 0.19643231 | -0.9704807 | 0.33180693 | 0.71487809 |
| KATNAL2 | 21.9430676 | 0.800068915 | 0.37822668 | 2.11531592 | 0.03440303 | 0.2833153 |
| SMAD2 | 127.110777 | -0.183371544 | 0.19775076 | -0.9272862 | 0.35377797 | 0.73059415 |
| ZBTB7C | 81.2529354 | 0.309322415 | 0.2914253 | 1.06141237 | 0.28850254 | 0.67992862 |
| CTIF | 103.318965 | 0.0113348 | 0.28148111 | 0.04026842 | 0.96787913 | 0.99300191 |
| SMAD7 | 44.3693499 | 0.396136491 | 0.24395586 | 1.62380397 | 0.10441765 | 0.46422585 |
| DYM | 268.558943 | -0.442223365 | 0.17322783 | -2.5528425 | 0.01068478 | 0.14541195 |
| SNHG22 | 23.1282243 | -0.072400408 | 0.41155237 | -0.1759203 | 0.86035657 | 0.97397549 |
| ME2 | 28.8227522 | 0.070823019 | 0.2934006 | 0.24138675 | 0.80925538 | 0.95939969 |
| SMAD4 | 56.3336637 | 0.107803014 | 0.22818003 | 0.47244718 | 0.63660763 | 0.89741446 |
| DCC | 45.1068352 | 2.187576908 | 0.80312573 | 2.72382871 | 0.006453 | 0.10918078 |
| MBD2 | 98.7852924 | -0.016912038 | 0.1381479 | -0.1224198 | 0.90256656 | 0.98164203 |
| POLI | 25.6910233 | 0.280242011 | 0.38126438 | 0.73503328 | 0.46231922 | 0.80221787 |
| TCF4 | 406.005163 | 0.114108858 | 0.26791563 | 0.42591341 | 0.67017094 | 0.91445841 |
| TXNL1 | 45.9872722 | 0.030306705 | 0.19563144 | 0.15491735 | 0.87688647 | 0.9768643 |
| WDR7 | 140.168971 | 0.000883489 | 0.16361988 | 0.00539965 | 0.99569173 | 0.99854015 |
| NARS | 37.5268308 | -0.550959874 | 0.30525231 | -1.8049327 | 0.07108522 | 0.39721198 |
| AC027097.2 | 54.9422066 | -0.207267266 | 0.27330493 | -0.7583737 | 0.44822731 | 0.79296348 |
| ATP8B1 | 137.344333 | 0.027853712 | 0.41725565 | 0.06675455 | 0.94677711 | 0.99060401 |
| NEDD4L | 50.9163331 | -0.331217088 | 0.25719246 | -1.287818 | 0.19780931 | 0.59530234 |
| MALT1 | 50.6052479 | -0.398971627 | 0.33574858 | -1.1883047 | 0.23471337 | 0.63080829 |
| ZNF532 | 138.29005 | 0.003398646 | 0.30745201 | 0.01105423 | 0.99118018 | 0.99759781 |

| | | | | | | |
|---|---|---|---|---|---|---|
| LMAN1 | 83.0261525 | -0.73325387 | 0.32250128 | -2.2736464 | 0.02298726 | 0.22440695 |
| CCBE1 | 284.405971 | -0.393260857 | 0.49625293 | -0.7924605 | 0.4280922 | 0.78079491 |
| RNF152 | 281.169263 | -0.793368416 | 0.59704349 | -1.3288285 | 0.18390455 | 0.58355551 |
| PIGN | 100.87372 | 0.389323513 | 0.29494409 | 1.31999088 | 0.18683806 | 0.58501647 |
| KIAA1468 | 96.0822561 | 0.035931769 | 0.20293219 | 0.17706293 | 0.85945896 | 0.97397549 |
| ZCCHC2 | 59.0902472 | -0.334101684 | 0.39703396 | -0.841494 | 0.40007126 | 0.7630739 |
| PHLPP1 | 105.807385 | 0.453783369 | 0.26134887 | 1.73631271 | 0.08250856 | 0.42256146 |
| BCL2 | 142.823466 | 0.454741059 | 0.34722737 | 1.30963483 | 0.1903194 | 0.58890833 |
| KDSR | 53.0976155 | -0.434200606 | 0.19848735 | -2.187548 | 0.02870254 | 0.25567874 |
| VPS4B | 49.3723514 | 0.258975188 | 0.2362613 | 1.09613883 | 0.27301802 | 0.6682621 |
| TMX3 | 63.8617125 | -0.261824937 | 0.22226693 | -1.1779752 | 0.23880648 | 0.63425689 |
| CCDC102B | 303.280862 | -0.323303948 | 0.40461457 | -0.7990418 | 0.42426619 | 0.77857941 |
| DOK6 | 157.685907 | -0.187321603 | 0.28745528 | -0.6516548 | 0.5146239 | 0.83109052 |
| RTTN | 41.9629335 | 0.175767524 | 0.23639442 | 0.74353499 | 0.45715784 | 0.79762426 |
| CYB5A | 87.416954 | -0.257596164 | 0.32043011 | -0.8039075 | 0.4214504 | 0.77771095 |
| CNDP2 | 40.6997328 | -0.787770731 | 0.3296555 | -2.3896787 | 0.01686312 | 0.19013163 |
| ZNF407 | 213.044085 | -0.118874595 | 0.13521228 | -0.8791701 | 0.37930905 | 0.74991898 |
| ZADH2 | 23.1848023 | 0.295186815 | 0.36926201 | 0.79939665 | 0.42406046 | 0.77857941 |
| TSHZ1 | 43.1338018 | 0.360097247 | 0.23776005 | 1.51454058 | 0.12988879 | 0.51204187 |
| ZNF516 | 130.328872 | -0.446291055 | 0.41498856 | -1.0754298 | 0.28218235 | 0.676814 |
| ZNF236 | 45.5930853 | 0.297718583 | 0.2112055 | 1.40961567 | 0.1586532 | 0.55091387 |
| MBP | 82.8188425 | -0.229682886 | 0.44234368 | -0.5192408 | 0.60359283 | 0.87903643 |
| ATP9B | 173.085552 | 0.023340632 | 0.17542193 | 0.13305424 | 0.89415048 | 0.98023547 |
| NFATC1 | 37.1733577 | 0.215058693 | 0.52002332 | 0.41355586 | 0.6791994 | 0.91831925 |
| PQLC1 | 36.8237817 | -0.443669102 | 0.26282083 | -1.6881048 | 0.09139111 | 0.44124451 |
| TXNL4A | 41.8979332 | -0.183547016 | 0.24213847 | -0.758025 | 0.44843601 | 0.79296348 |
| ADNP2 | 36.2959674 | 0.031750744 | 0.23908204 | 0.13280272 | 0.89434941 | 0.98023547 |
| PARD6G | 23.9875894 | 0.566314506 | 0.3758658 | 1.50669334 | 0.13188928 | 0.51290135 |
| NRSN2-AS1 | 19.9352686 | 0.621641849 | 0.41606296 | 1.49410524 | 0.13514811 | 0.51849793 |
| RBCK1 | 35.8801248 | -0.154117848 | 0.34844442 | -0.4423025 | 0.6582703 | 0.9079633 |
| TBC1D20 | 26.4934861 | -0.206198603 | 0.26068003 | -0.7910027 | 0.42894243 | 0.78201388 |
| CSNK2A1 | 101.269651 | -0.208420155 | 0.17241681 | -1.2088158 | 0.22673363 | 0.62104564 |
| PSMF1 | 67.3792567 | -0.441304597 | 0.22353221 | -1.9742327 | 0.04835529 | 0.33343326 |
| AL136531.2 | 38.4380779 | -1.114306258 | 0.30704014 | -3.6291876 | 0.00028431 | 0.01645219 |
| SDCBP2-AS1 | 29.438062 | -0.202874283 | 0.23384924 | -0.867543 | 0.38564454 | 0.75340372 |
| NSFL1C | 29.3542261 | 0.087070168 | 0.23403519 | 0.37203879 | 0.70986397 | 0.93009618 |
| STK35 | 18.8907612 | 0.186111523 | 0.32237612 | 0.57731176 | 0.56372887 | 0.85632174 |
| SNRPB | 27.8395324 | -0.610583521 | 0.40509743 | -1.507251 | 0.13174633 | 0.51290135 |
| NOP56 | 24.9941286 | -0.169796645 | 0.44923209 | -0.3779709 | 0.70545223 | 0.92946306 |
| EBF4 | 41.0842066 | -0.137487015 | 0.2376121 | -0.5786196 | 0.56284589 | 0.85571045 |
| PCED1A | 27.5021856 | -0.91195298 | 0.43588903 | -2.0921678 | 0.03642351 | 0.28969304 |
| PTPRA | 159.212658 | -0.013333215 | 0.19200815 | -0.0694409 | 0.94463869 | 0.99060356 |
| ITPA | 25.8815742 | -0.588302946 | 0.3591414 | -1.6380817 | 0.10140465 | 0.46169744 |
| C20orf194 | 165.58244 | 0.231566002 | 0.26740564 | 0.86597275 | 0.3865051 | 0.75402666 |
| ATRN | 87.7167394 | -0.185430849 | 0.18159452 | -1.0211258 | 0.30719485 | 0.69801431 |
| MAVS | 47.441752 | -0.1548355 | 0.33498631 | -0.4622144 | 0.64392756 | 0.9006897 |
| PANK2 | 66.8647296 | 0.065139783 | 0.17564468 | 0.37086112 | 0.71074097 | 0.93009618 |
| RNF24 | 134.024312 | -0.949902556 | 0.25560005 | -3.7163629 | 0.00020211 | 0.01314438 |
| PRNP | 53.5425602 | -0.275728945 | 0.31154275 | -0.8850437 | 0.37613307 | 0.74787433 |
| SLC23A2 | 78.5030411 | 0.006752195 | 0.26793245 | 0.02520111 | 0.97989455 | 0.99503182 |
| TMEM230 | 43.6398434 | 0.207405869 | 0.23863748 | 0.86912529 | 0.3847786 | 0.75305975 |
| CDS2 | 79.118098 | 0.684168516 | 0.26886495 | 2.54465493 | 0.01093858 | 0.14777134 |
| GPCPD1 | 131.564288 | -0.200760493 | 0.42908375 | -0.4678818 | 0.6398691 | 0.89903874 |
| SHLD1 | 25.534982 | 0.521345609 | 0.36414286 | 1.43170625 | 0.15222791 | 0.53966466 |
| CRLS1 | 44.7110065 | -0.137512164 | 0.29273215 | -0.4697542 | 0.63853062 | 0.8980979 |
| TMX4 | 34.7913117 | -0.099173639 | 0.26727743 | -0.3710513 | 0.71059932 | 0.93009618 |
| PLCB1 | 522.179066 | -0.558203681 | 0.5472136 | -1.0200837 | 0.30768876 | 0.69801431 |
| PLCB4 | 96.6157382 | 0.537262255 | 0.54097383 | 0.99313909 | 0.32064218 | 0.70762743 |
| SLX4IP | 44.8737878 | -0.037966391 | 0.29603541 | -0.1282495 | 0.89795153 | 0.98040427 |
| JAG1 | 251.106845 | 0.318576536 | 0.30057946 | 1.05987461 | 0.28920165 | 0.67992862 |
| AL050403.2 | 40.1415911 | 0.214523455 | 0.43162514 | 0.49701335 | 0.61917964 | 0.8859718 |
| BTBD3 | 29.7103501 | 0.310618353 | 0.24093343 | 1.28922894 | 0.19731851 | 0.59530234 |
| SPTLC3 | 21.3777805 | -0.200476255 | 0.39275774 | -0.5104323 | 0.6097486 | 0.88163721 |
| TASP1 | 115.036777 | -0.613882008 | 0.21572936 | -2.8456118 | 0.00443262 | 0.08828001 |
| ESF1 | 51.8047136 | -0.636887379 | 0.24911106 | -2.5566403 | 0.01056884 | 0.14436882 |
| NDUFAF5 | 23.8461069 | 0.339086034 | 0.28633117 | 1.18424423 | 0.2363164 | 0.63222238 |
| MACROD2 | 327.088928 | 1.444303515 | 0.51780443 | 2.78928381 | 0.00528248 | 0.095618 |
| KIF16B | 70.4239722 | 0.382304314 | 0.23998863 | 1.59301013 | 0.11115792 | 0.47827842 |
| SNRPB2 | 19.0145213 | -0.153270062 | 0.34133808 | -0.4490271 | 0.65341208 | 0.90646849 |
| DSTN | 290.845679 | 0.563933257 | 0.26486804 | 2.12911025 | 0.03324514 | 0.27874784 |
| RRBP1 | 137.91506 | -0.421558663 | 0.37038379 | -1.1381672 | 0.25505069 | 0.64969412 |
| SNX5 | 34.2926703 | -0.372360174 | 0.28069061 | -1.3265858 | 0.18464573 | 0.58363936 |
| ZNF133 | 29.0527257 | -0.211511202 | 0.31587521 | -0.6696037 | 0.50311047 | 0.82430366 |
| SEC23B | 35.1670185 | -0.684141083 | 0.29082596 | -2.3524072 | 0.01865234 | 0.20009416 |
| SLC24A3 | 220.030683 | -0.965609348 | 0.51077323 | -1.8904854 | 0.05869307 | 0.36041506 |
| AL121761.1 | 42.699266 | -0.272617595 | 0.38011407 | -0.7171994 | 0.47325105 | 0.80758763 |
| RIN2 | 107.120885 | 0.295639193 | 0.29662109 | 0.99668974 | 0.31891513 | 0.70683013 |
| NAA20 | 23.8585634 | -0.114625568 | 0.25532662 | -0.448937 | 0.65347711 | 0.90646849 |
| RALGAPA2 | 107.221031 | -0.006258991 | 0.19306973 | -0.0324183 | 0.97413848 | 0.99300191 |
| KIZ | 76.3492995 | -0.168898097 | 0.28002007 | -0.6031642 | 0.54639943 | 0.84758329 |
| XRN2 | 116.565886 | -0.733963951 | 0.20883968 | -3.5144851 | 0.00044061 | 0.02145813 |
| NAPB | 43.41524 | 0.145648789 | 0.27371543 | 0.53211756 | 0.59464458 | 0.87418344 |
| CST3 | 228.517565 | -0.130553989 | 0.31629323 | -0.4127625 | 0.67978062 | 0.91860979 |
| SYNDIG1 | 42.6953697 | 0.648349732 | 0.49758325 | 1.3029975 | 0.19257561 | 0.59142551 |
| APMAP | 21.2676645 | -0.334883459 | 0.29027362 | -1.153682 | 0.24863055 | 0.64428277 |
| ACSS1 | 34.5852936 | -0.133694633 | 0.27032265 | -0.4945743 | 0.62090065 | 0.88720256 |
| PYGB | 20.0482807 | 0.458574545 | 0.41877408 | 1.09504041 | 0.27349893 | 0.66906109 |
| ABHD12 | 34.9016039 | -0.025338876 | 0.26907485 | -0.0941704 | 0.92497383 | 0.98561736 |
| NINL | 33.5621079 | -0.023355276 | 0.26258729 | -0.0889429 | 0.92912728 | 0.98681039 |
| ZNF337 | 20.6545275 | 0.068949318 | 0.25234645 | 0.27323276 | 0.7846743 | 0.95065355 |
| FAM182B | 40.6727983 | -0.035801219 | 0.31908829 | -0.1121985 | 0.91066604 | 0.98343817 |
| HM13 | 102.902779 | -0.740058152 | 0.30941543 | -2.3917946 | 0.01676622 | 0.19005424 |
| ID1 | 94.8203083 | -0.402427453 | 0.5693554 | -0.7068124 | 0.47968307 | 0.81206886 |
| BCL2L1 | 86.0838349 | 0.096519796 | 0.25451343 | 0.37923262 | 0.70451513 | 0.92936308 |
| TM9SF4 | 57.1732002 | -0.283641227 | 0.18985597 | -1.493981 | 0.13518057 | 0.51849793 |
| POFUT1 | 19.6446467 | -0.268731747 | 0.37059386 | -0.7251381 | 0.46836728 | 0.80374006 |
| KIF3B | 29.0799916 | -0.450354481 | 0.29785557 | -1.5119895 | 0.13053655 | 0.51241218 |
| ASXL1 | 196.873798 | -0.109275657 | 0.20102924 | -0.5435809 | 0.58672991 | 0.870249 |
| MAPRE1 | 27.6936539 | 0.29004734 | 0.25329684 | 1.14508867 | 0.25217243 | 0.64684649 |
| CDK5RAP1 | 27.2721296 | -0.139983139 | 0.24820094 | -0.5639912 | 0.57276014 | 0.86236719 |
| CBFA2T2 | 130.695081 | 0.136941251 | 0.22152171 | 0.61818434 | 0.53645383 | 0.84423863 |
| CHMP4B | 37.3603382 | -0.192044641 | 0.28930172 | -0.6638213 | 0.5068047 | 0.82481485 |
| RALY | 80.7290056 | 0.364983329 | 0.19800207 | 1.84333087 | 0.06528072 | 0.38095778 |
| EIF2S2 | 55.2094079 | -0.373514828 | 0.31668998 | -1.1794337 | 0.23822554 | 0.63385933 |
| ITCH | 216.152879 | -0.002697221 | 0.14712722 | -0.0183326 | 0.98537354 | 0.996806 |
| DYNLRB1 | 63.2953095 | -0.151105929 | 0.27158673 | -0.5563819 | 0.57794984 | 0.8645132 |
| MAP1LC3A | 23.1686143 | 0.208754962 | 0.38497659 | 0.54225365 | 0.58764378 | 0.870249 |
| PIGU | 36.7048114 | 0.350140135 | 0.33049737 | 1.05943395 | 0.2894022 | 0.67992862 |
| NCOA6 | 104.976924 | 0.02964122 | 0.17452103 | 0.16984325 | 0.86513341 | 0.9762143 |
| GGT7 | 31.1407954 | -0.227946364 | 0.27121146 | -0.8404747 | 0.40064229 | 0.76337055 |
| ACSS2 | 53.6698366 | -0.63310505 | 0.45903595 | -1.3792058 | 0.16783131 | 0.56323722 |
| TRPC4AP | 145.498498 | -0.146333758 | 0.13806289 | -1.0599065 | 0.28918712 | 0.67992862 |
| EDEM2 | 21.3090868 | -0.661323238 | 0.27139268 | -2.4367762 | 0.01481885 | 0.17735986 |

| | | | | | | |
|---|---|---|---|---|---|---|
| PROCR | 53.9061426 | 0.141357041 | 0.42310794 | 0.33409215 | 0.73831002 | 0.93594619 |
| MMP24OS | 38.4429964 | 0.204794402 | 0.23975107 | 0.85419598 | 0.39299641 | 0.75983441 |
| UQCC1 | 64.7233545 | 0.034467346 | 0.21030158 | 0.16389485 | 0.86981392 | 0.9768643 |
| CEP250 | 17.8429012 | -0.221728496 | 0.26816565 | -0.8268341 | 0.40833111 | 0.76963974 |
| ERGIC3 | 24.7316365 | 0.430406466 | 0.2760746 | 1.55902236 | 0.11899109 | 0.49395083 |
| CPNE1 | 33.1088427 | 0.121464248 | 0.35930383 | 0.33805442 | 0.73532218 | 0.93594619 |
| ROMO1 | 22.2552589 | -0.487875559 | 0.45241404 | -1.0783829 | 0.28086293 | 0.67585515 |
| RBM39 | 258.681224 | -0.001116658 | 0.16864676 | -0.0066213 | 0.99471702 | 0.99854015 |
| PHF20 | 129.48831 | 0.264512797 | 0.17478826 | 1.51333278 | 0.13019515 | 0.51204187 |
| SCAND1 | 36.3519972 | 0.028078576 | 0.38144625 | 0.07361083 | 0.94132005 | 0.98895798 |
| NORAD | 92.8111858 | 0.170871675 | 0.24497929 | 0.69749437 | 0.48549346 | 0.81570449 |
| EPB41L1 | 32.7239274 | 0.622734935 | 0.32030443 | 1.94419706 | 0.05187169 | 0.34513804 |
| DLGAP4 | 93.0669525 | 0.20777373 | 0.26674211 | 0.77893111 | 0.4360203 | 0.78719791 |
| DLGAP4-AS1 | 28.5820291 | 0.070151387 | 0.27176663 | 0.25813098 | 0.79630582 | 0.95463848 |
| MYL9 | 393.974504 | 0.233926101 | 0.39826462 | 0.5873635 | 0.5569596 | 0.85256862 |
| NDRG3 | 25.5148292 | 0.248562415 | 0.3841648 | 0.64702028 | 0.51761882 | 0.83277421 |
| SOGA1 | 51.1364025 | 0.655784292 | 0.27915097 | 2.34921016 | 0.01881328 | 0.20125012 |
| SAMHD1 | 100.268331 | 0.030695786 | 0.59531721 | 0.05156207 | 0.95887765 | 0.99181157 |
| RPN2 | 62.544401 | -0.1441855 | 0.2529924 | -0.5699203 | 0.56873177 | 0.85958696 |
| MANBAL | 16.7579628 | -0.014672055 | 0.33960888 | -0.0432028 | 0.96553987 | 0.99293098 |
| SRC | 18.0287196 | -0.185380555 | 0.33041667 | -0.5610509 | 0.57476286 | 0.86294056 |
| BLCAP | 45.7307814 | 0.194007143 | 0.21220856 | 0.91422862 | 0.36059673 | 0.73273991 |
| CTNNBL1 | 53.4982643 | -0.095410507 | 0.23852601 | -0.4000004 | 0.6891562 | 0.92217934 |
| TTI1 | 20.1614845 | -0.202489686 | 0.28818148 | -0.7026464 | 0.48227613 | 0.81402096 |
| RPRD1B | 55.6427178 | 0.067312381 | 0.19157404 | 0.35136483 | 0.72531466 | 0.93455731 |
| TGM2 | 210.495571 | -1.371219741 | 0.52186843 | -2.62752 | 0.00860098 | 0.13046619 |
| RALGAPB | 81.060991 | 0.122078628 | 0.1456993 | 0.83788069 | 0.40209771 | 0.76441384 |
| DHX35 | 27.247241 | 0.18401276 | 0.26493991 | 0.69454526 | 0.48734033 | 0.81595238 |
| TOP1 | 208.89054 | -0.46658915 | 0.26150219 | -1.7842648 | 0.07438065 | 0.40409804 |
| PLCG1 | 26.3151467 | 0.146471004 | 0.27849242 | 0.52594252 | 0.59892814 | 0.87644821 |
| ZHX3 | 111.565568 | -0.533176006 | 0.28829826 | -1.8493903 | 0.06440148 | 0.37871685 |
| CHD6 | 154.493409 | -0.002262497 | 0.17508431 | -0.0129223 | 0.98968976 | 0.99739981 |
| SRSF6 | 40.7486176 | 0.354948458 | 0.34848582 | 1.01854492 | 0.30841907 | 0.69885506 |
| IFT52 | 27.5816713 | 0.468268905 | 0.2581981 | 1.8136032 | 0.06973884 | 0.39338722 |
| JPH2 | 28.4576295 | 1.056596284 | 0.52243111 | 2.02246049 | 0.0431288 | 0.31204692 |
| SERINC3 | 62.5686516 | 0.200147664 | 0.27949204 | 0.71611221 | 0.47392207 | 0.80789916 |
| PKIG | 98.3145534 | 0.198527719 | 0.22440662 | 0.88467854 | 0.37633004 | 0.74787433 |
| YWHAB | 111.529319 | -0.096241814 | 0.37610255 | -0.2558925 | 0.79803386 | 0.95489133 |
| STK4 | 58.4945096 | -0.307351233 | 0.19869227 | -1.5468706 | 0.12189445 | 0.4998897 |
| SLPI | 93.0040468 | -2.047270468 | 0.87268189 | -2.3459527 | 0.01897851 | 0.20213493 |
| SDC4 | 111.697116 | -0.997423038 | 0.37309879 | -2.6733483 | 0.00750982 | 0.12103002 |
| PIGT | 25.8927103 | 0.514269903 | 0.41951744 | 1.22586059 | 0.22025114 | 0.61590675 |
| WFDC2 | 115.961357 | -2.554536413 | 0.50669591 | -5.0415572 | 4.62E-07 | 0.00016494 |
| SNX21 | 22.3912403 | 0.708463711 | 0.43420938 | 1.6316177 | 0.10276005 | 0.4627468 |
| CTSA | 34.0729097 | 0.13411676 | 0.26533453 | 0.5054629 | 0.61323375 | 0.88331141 |
| PLTP | 29.2130344 | -0.926805717 | 0.42850047 | -2.1629048 | 0.0305485 | 0.26662968 |
| ZNF335 | 19.1018009 | -0.110353336 | 0.33424017 | -0.3301618 | 0.74127771 | 0.93711378 |
| NCOA5 | 35.0708484 | 0.123216459 | 0.26405758 | 0.46662724 | 0.64076659 | 0.89934133 |
| SLC35C2 | 26.5030946 | -0.048099543 | 0.2617191 | -0.1837831 | 0.85418363 | 0.97375567 |
| ELMO2 | 31.312533 | -0.583183224 | 0.34596136 | -1.6856889 | 0.09185573 | 0.44124451 |
| SLC13A3 | 42.5159588 | -0.061352617 | 0.25444887 | -0.2411196 | 0.8094624 | 0.95947406 |
| EYA2 | 60.5143027 | 0.406495826 | 0.43724312 | 0.92967918 | 0.35253722 | 0.73059415 |
| ZMYND8 | 112.920568 | -0.131605357 | 0.24677956 | -0.5332912 | 0.59383205 | 0.87394794 |
| NCOA3 | 176.82996 | 0.276156864 | 0.30490938 | 0.90570145 | 0.3650939 | 0.73726023 |
| SULF2 | 32.4422078 | -0.77121097 | 0.31958234 | -2.413184 | 0.01581384 | 0.18446965 |
| PREX1 | 23.5490826 | -0.163748439 | 0.48978627 | -0.3343263 | 0.73813333 | 0.93594619 |
| ARFGEF2 | 100.484289 | -0.142327385 | 0.1866453 | -0.7625554 | 0.44572859 | 0.79296348 |
| CSE1L | 33.0555411 | 0.048447643 | 0.28760091 | 0.16845441 | 0.8662258 | 0.9762143 |
| STAU1 | 96.9224284 | -0.04245365 | 0.15805497 | -0.2686005 | 0.7882371 | 0.95129454 |
| DDX27 | 19.5768948 | -0.079681689 | 0.29416632 | -0.2708729 | 0.78648877 | 0.95095525 |
| ZNFX1 | 67.9261324 | 0.181874699 | 0.42225868 | 0.43071867 | 0.66667294 | 0.9133172 |
| ZFAS1 | 41.4952474 | -0.816992271 | 0.33509063 | -2.4381233 | 0.01476374 | 0.17699624 |
| PTGIS | 197.980668 | 0.02038142 | 0.61324685 | 0.03323526 | 0.97348698 | 0.99300191 |
| B4GALT5 | 126.231503 | -0.357605592 | 0.21291472 | -1.6795719 | 0.09304063 | 0.44361507 |
| SLC9A8 | 38.5388501 | -0.029049074 | 0.33176788 | -0.0875584 | 0.93022764 | 0.98681039 |
| RNF114 | 48.947119 | -0.319804353 | 0.35756079 | -0.8944055 | 0.37110495 | 0.74385689 |
| CEBPB | 145.59583 | -1.757487317 | 0.42676834 | -4.1181296 | 3.82E-05 | 0.00438597 |
| PTPN1 | 203.284008 | -0.979824699 | 0.33950997 | -2.8859968 | 0.00390176 | 0.08146035 |
| RIPOR3 | 312.610192 | -0.137330857 | 0.44583505 | -0.3080306 | 0.75805902 | 0.94311422 |
| PARD6B | 27.2888107 | 0.023694203 | 0.47717199 | 0.04965548 | 0.96039694 | 0.99181157 |
| ADNP | 147.759134 | 0.01037111 | 0.13535509 | 0.0766215 | 0.93892465 | 0.98827804 |
| DPM1 | 36.172615 | 0.033630547 | 0.2134829 | 0.15753275 | 0.874825 | 0.9768643 |
| NFATC2 | 52.0275891 | 2.30452093 | 0.9512525 | 2.42261748 | 0.01540914 | 0.18118686 |
| ATP9A | 63.3433522 | -0.055813774 | 0.34848117 | -0.160163 | 0.87275271 | 0.9768643 |
| ZFP64 | 25.2804778 | -0.119961356 | 0.31839926 | -0.3767639 | 0.70634905 | 0.93006079 |
| TSHZ2 | 896.691264 | -0.647342233 | 0.33932167 | -1.9077539 | 0.05642303 | 0.35564709 |
| AL109930.1 | 72.8236388 | -1.186578864 | 0.35926033 | -3.3028386 | 0.00095711 | 0.0366558 |
| ZNF217 | 47.6716164 | -0.558733561 | 0.19895701 | -2.808313 | 0.00498018 | 0.09251529 |
| DOK5 | 30.8974292 | -0.499829641 | 0.61728807 | -0.8097186 | 0.41810192 | 0.77722447 |
| FAM210B | 19.2386163 | 0.341227415 | 0.34279927 | 0.99541465 | 0.31953463 | 0.70688141 |
| RTF2 | 26.5389465 | -0.008876867 | 0.27283462 | -0.0325357 | 0.97404484 | 0.99300191 |
| PCK1 | 93.6364928 | -0.016181206 | 1.085237 | -0.0149103 | 0.98810375 | 0.99719329 |
| PMEPA1 | 53.7300759 | 0.413870906 | 0.30388855 | 1.36191673 | 0.17322416 | 0.57060708 |
| RAB22A | 58.4715074 | 0.266783608 | 0.21429882 | 1.24491405 | 0.21316335 | 0.61001472 |
| VAPB | 68.6167888 | -0.119034298 | 0.17279301 | -0.6888838 | 0.49089642 | 0.81720854 |
| GNAS | 279.279726 | 0.373687019 | 0.28164308 | 1.32681059 | 0.18457135 | 0.58363936 |
| NELFCD | 26.2406106 | 0.087645308 | 0.25231099 | 0.34737016 | 0.72831325 | 0.93455731 |
| CTSZ | 31.8389564 | 0.280559589 | 0.30920159 | 0.90736786 | 0.3642123 | 0.73648283 |
| ATP5F1E | 97.5488199 | -0.366513865 | 0.34342663 | -1.0672261 | 0.28586975 | 0.6783522 |
| PHACTR3 | 43.6808534 | -0.581507354 | 0.85486429 | -0.6802335 | 0.4963566 | 0.82154305 |
| MIR646HG | 81.8728676 | -0.476639364 | 0.50373753 | -0.9462058 | 0.34404364 | 0.72261198 |
| TAF4 | 27.4387008 | 0.189173689 | 0.30652192 | 0.61716203 | 0.53712786 | 0.84423863 |
| PSMA7 | 59.1168339 | -0.312530104 | 0.36808947 | -0.8490602 | 0.39584781 | 0.76020146 |
| SS18L1 | 25.6445462 | -0.116159669 | 0.24198038 | -0.4800376 | 0.63120069 | 0.89436221 |
| MTG2 | 26.4104504 | -0.014874474 | 0.29684766 | -0.0501081 | 0.96003624 | 0.99181157 |
| OSBPL2 | 23.4819626 | -0.089544078 | 0.29455067 | -0.3040023 | 0.76112614 | 0.94425408 |
| LAMA5 | 41.4653673 | 0.811164315 | 0.40919452 | 1.98234401 | 0.04744075 | 0.3297075 |
| OGFR | 27.7989599 | -0.623808666 | 0.28003176 | -2.2276354 | 0.02590484 | 0.24006894 |
| DIDO1 | 71.4350836 | 0.06552827 | 0.18447872 | 0.35520773 | 0.72243397 | 0.93444541 |
| GID8 | 21.3662909 | 0.301575034 | 0.25464949 | 1.18427506 | 0.2363042 | 0.63222238 |
| YTHDF1 | 24.2726749 | 0.049033976 | 0.25366469 | 0.19330233 | 0.8467222 | 0.97014274 |
| ARFGAP1 | 27.3675339 | -0.314769506 | 0.40107237 | -0.7848197 | 0.43255928 | 0.7843272 |
| PPDPF | 65.1494186 | -0.461116819 | 0.28910238 | -1.5949949 | 0.11071338 | 0.47776431 |
| ARFRP1 | 17.2580165 | 0.125668346 | 0.31485327 | 0.39913305 | 0.68979517 | 0.92269481 |
| ZBTB46 | 221.591106 | 0.599770706 | 0.30849023 | 1.94421294 | 0.05186978 | 0.34513804 |
| TPD52L2 | 31.75515 | -0.005542831 | 0.28140011 | -0.0196973 | 0.98428482 | 0.9964038 |
| DNAJC5 | 58.4357259 | -0.479445921 | 0.3106935 | -1.5431476 | 0.12279497 | 0.50097964 |
| PRPF6 | 41.8484616 | 0.260346714 | 0.26977106 | 0.96506538 | 0.33451206 | 0.71690491 |
| PCMTD2 | 53.3022061 | 0.03378037 | 0.37098383 | 0.09105618 | 0.92744795 | 0.98678435 |
| SHC2 | 21.5476737 | -0.046690158 | 0.35336673 | -0.1321295 | 0.89488189 | 0.98023547 |
| BSG | 69.4832364 | -0.303414294 | 0.23014256 | -1.3183754 | 0.18737801 | 0.58572565 |
| FSTL3 | 38.5734945 | -0.831710599 | 0.45707047 | -1.8196551 | 0.06881155 | 0.39049889 |
| PALM | 27.3957107 | 0.464531791 | 0.25316546 | 1.83489406 | 0.06652139 | 0.38523695 |

| | | | | | | |
|---|---|---|---|---|---|---|
| PTBP1 | 56.7931028 | 0.06665537 | 0.24055776 | 0.27708675 | 0.78171349 | 0.94939885 |
| ARID3A | 42.0926742 | -0.492829844 | 0.28315379 | -1.7405024 | 0.08177084 | 0.4223007 |
| TMEM259 | 50.2059723 | -0.075976832 | 0.32703358 | -0.2323212 | 0.81628855 | 0.96132648 |
| CNN2 | 40.2249186 | 0.851421412 | 0.41420014 | 2.05557974 | 0.03982304 | 0.30124947 |
| POLR2E | 19.4612343 | 0.088587816 | 0.37430408 | 0.23667339 | 0.81291018 | 0.96038841 |
| GPX4 | 61.9338342 | -0.305384604 | 0.29320879 | -1.0415261 | 0.29763145 | 0.68849737 |
| SBNO2 | 67.3132816 | -1.271418325 | 0.35136004 | -3.6185627 | 0.00029624 | 0.01687671 |
| STK11 | 30.6369536 | 0.067587774 | 0.26033726 | 0.25961622 | 0.79515983 | 0.95463848 |
| ATP5F1D | 46.1926316 | -0.121909049 | 0.26441055 | -0.4610597 | 0.64475577 | 0.90107359 |
| MIDN | 62.1530334 | -0.005946571 | 0.37969805 | -0.0156613 | 0.98750459 | 0.99719329 |
| CIRBP | 92.6371401 | 0.218023166 | 0.29713864 | 0.73374222 | 0.46310586 | 0.80225845 |
| MUM1 | 37.855555 | 0.37441625 | 0.27844727 | 1.34465764 | 0.17873582 | 0.57686849 |
| DAZAP1 | 59.2565912 | -0.561277661 | 0.23952782 | -2.3432671 | 0.01911569 | 0.20295514 |
| RPS15 | 172.015872 | -0.593681565 | 0.30178221 | -1.9672517 | 0.0491542 | 0.33550435 |
| C19orf25 | 22.4946008 | -0.354143876 | 0.32832241 | -1.0786467 | 0.28074526 | 0.6757933 |
| MBD3 | 21.2813819 | -0.133291891 | 0.4266959 | -0.3123815 | 0.75475063 | 0.94170839 |
| TCF3 | 20.7413526 | 0.306800968 | 0.32270297 | 0.95072249 | 0.34174527 | 0.72044921 |
| ABHD17A | 24.5355943 | -0.375479725 | 0.34858654 | -1.0771492 | 0.2814136 | 0.6760734 |
| CSNK1G2 | 29.1013403 | -0.421899487 | 0.43923767 | -0.9605267 | 0.33679022 | 0.71924965 |
| BTBD2 | 28.1595578 | 0.412298612 | 0.33065528 | 1.24691375 | 0.21242912 | 0.60910715 |
| MOB3A | 27.9704163 | 0.198307925 | 0.34497138 | 0.57485327 | 0.56539053 | 0.85765509 |
| AP3D1 | 46.6478125 | -0.086541115 | 0.23951328 | -0.3613207 | 0.71785969 | 0.93372582 |
| DOT1L | 43.5489607 | 0.018692593 | 0.32007822 | 0.05840008 | 0.95342995 | 0.99091454 |
| LSM7 | 24.9142854 | -0.40390576 | 0.3247296 | -1.2438218 | 0.21356515 | 0.61001472 |
| SPPL2B | 26.2312374 | -0.065312874 | 0.30391689 | -0.2149037 | 0.82984238 | 0.96580172 |
| GADD45B | 130.510638 | -1.990487521 | 0.50884219 | -3.9117974 | 9.16E-05 | 0.00756466 |
| GNG7 | 36.141443 | 0.543439344 | 0.26988557 | 2.01359173 | 0.04405242 | 0.3152359 |
| SGTA | 23.9743589 | 0.012022817 | 0.24111926 | 0.04986253 | 0.96023194 | 0.99181157 |
| TLE2 | 117.29342 | -0.511744926 | 0.2948233 | -1.7357682 | 0.08260482 | 0.42256146 |
| AES | 115.588009 | 0.109810237 | 0.17351295 | 0.63286478 | 0.52682195 | 0.83897872 |
| GNA11 | 46.381104 | -0.048342342 | 0.26642164 | -0.1814505 | 0.85601397 | 0.97379567 |
| NCLN | 19.1254547 | 0.025647653 | 0.45480309 | 0.05639287 | 0.95502883 | 0.99181157 |
| NFIC | 187.448696 | 0.005866307 | 0.16912397 | 0.03468643 | 0.97232978 | 0.99300191 |
| MFSD12 | 23.9163524 | 0.032839354 | 0.34587779 | 0.09494496 | 0.92435854 | 0.98561736 |
| HMG20B | 16.9287191 | 0.365362226 | 0.34668977 | 1.05385927 | 0.29194735 | 0.68372247 |
| PIP5K1C | 39.346124 | 0.254795313 | 0.21753438 | 1.17128754 | 0.24148322 | 0.63562327 |
| DAPK3 | 44.3640586 | 0.517275782 | 0.31408719 | 1.64691781 | 0.09957494 | 0.45779653 |
| EEF2 | 155.482515 | -0.442169445 | 0.36361942 | -1.2160226 | 0.2239763 | 0.61858591 |
| ZBTB7A | 50.5363728 | 0.152763826 | 0.34015159 | 0.44910514 | 0.65335581 | 0.90646849 |
| MAP2K2 | 43.6239417 | -0.023839919 | 0.24141565 | -0.0987505 | 0.92133636 | 0.98528826 |
| SH3GL1 | 42.3172026 | -0.216755745 | 0.2726993 | -0.7948526 | 0.42669925 | 0.77966505 |
| UBXN6 | 19.3828691 | -0.648521662 | 0.33041589 | -1.9627436 | 0.04967598 | 0.33731209 |
| MYDGF | 47.0582456 | -0.535765561 | 0.40035597 | -1.338223 | 0.18082376 | 0.57928238 |
| DPP9 | 24.3182776 | 0.007394322 | 0.3234219 | 0.02286277 | 0.98175973 | 0.9955778 |
| PLIN3 | 24.7960492 | -0.061697576 | 0.36129884 | -0.1707661 | 0.86440772 | 0.9762143 |
| KDM4B | 106.797212 | -0.372133092 | 0.30138601 | -1.2347391 | 0.21692762 | 0.61410359 |
| PTPRS | 119.863418 | 0.45505806 | 0.35093204 | 1.29671276 | 0.19473004 | 0.59277982 |
| SAFB2 | 52.2357175 | -0.161079469 | 0.20031905 | -0.8041146 | 0.42133079 | 0.77771095 |
| SAFB | 67.1465724 | -0.192434413 | 0.20011503 | -0.961619 | 0.33624103 | 0.71883121 |
| RPL36 | 96.4223757 | -0.816213951 | 0.27106128 | -3.0111787 | 0.00260236 | 0.06505007 |
| C19orf70 | 22.1537118 | -0.161639012 | 0.3063312 | -0.5276609 | 0.59773468 | 0.87557349 |
| LONP1 | 34.2739093 | -0.635854794 | 0.32746285 | -1.9417616 | 0.05216597 | 0.34575469 |
| RANBP3 | 23.4044593 | -0.092256313 | 0.26606966 | -0.3467374 | 0.72878858 | 0.93455731 |
| RFX2 | 69.7962345 | 1.558757553 | 0.53786795 | 2.8980302 | 0.00375514 | 0.07961552 |
| MLLT1 | 47.0661211 | -0.934448892 | 0.30442757 | -3.0695278 | 0.00214397 | 0.0570872 |
| KHSRP | 29.9005102 | 0.027813698 | 0.33152806 | 0.08389546 | 0.93313955 | 0.98725539 |
| C3 | 103.49143 | 0.260068749 | 0.58762241 | 0.442578 | 0.658071 | 0.90785879 |
| TRIP10 | 36.8020759 | 0.269804599 | 0.28734258 | 0.93896491 | 0.34774876 | 0.72581813 |
| INSR | 445.731882 | -0.940158459 | 0.59798671 | -1.5722063 | 0.11590272 | 0.48843995 |
| ZNF358 | 17.5203745 | 0.31284043 | 0.37218835 | 0.84054331 | 0.40060383 | 0.76337055 |
| PNPLA6 | 19.0685658 | 0.364141877 | 0.30990898 | 1.1749962 | 0.23999624 | 0.63451757 |
| MAP2K7 | 28.8768847 | -0.074908716 | 0.38146655 | -0.1963703 | 0.84432031 | 0.96926078 |
| ELAVL1 | 42.7569364 | 0.193098349 | 0.24964449 | 0.77349335 | 0.43923046 | 0.78887428 |
| CERS4 | 45.252249 | 0.414317597 | 0.3382252 | 1.22497554 | 0.22058444 | 0.61590675 |
| RPS28 | 40.50804 | -0.615707971 | 0.29450802 | -2.0906323 | 0.03656104 | 0.28984581 |
| ANGPTL4 | 65.6036501 | -2.138844256 | 0.79573115 | -2.6878981 | 0.00719033 | 0.11664848 |
| RAB11B | 26.514262 | 0.004517319 | 0.31832873 | 0.01419074 | 0.98867781 | 0.99722663 |
| HNRNPM | 115.102631 | -0.035425783 | 0.2420365 | -0.1463655 | 0.88363289 | 0.97782528 |
| ADAMTS10 | 39.0281754 | 0.365645341 | 0.25384433 | 1.44043137 | 0.14974539 | 0.53865829 |
| ZNF266 | 22.1086197 | 0.148412169 | 0.30797152 | 0.48190225 | 0.62987539 | 0.89365764 |
| ZNF121 | 22.7030733 | -0.653662527 | 0.34702845 | -1.8835992 | 0.05961921 | 0.36410704 |
| ZNF562 | 33.7813344 | 0.105213176 | 0.2421067 | 0.43457358 | 0.66387197 | 0.91141325 |
| ZNF846 | 38.9324852 | 1.146868663 | 0.29051576 | 3.94769864 | 7.89E-05 | 0.0068121 |
| COL5A3 | 36.60783 | 0.295654816 | 0.44134584 | 0.66989374 | 0.50292553 | 0.82430366 |
| C19orf66 | 34.9828421 | -0.230436265 | 0.42516882 | -0.5419877 | 0.58782699 | 0.87025394 |
| EIF3G | 18.9078554 | -0.029215165 | 0.36734365 | -0.0795309 | 0.93661037 | 0.98827804 |
| DNMT1 | 38.1636139 | 0.059538919 | 0.31239687 | 0.19058744 | 0.84884883 | 0.97092453 |
| ICAM1 | 45.7641514 | -1.19959798 | 0.86625646 | -1.384807 | 0.1661115 | 0.56208655 |
| TYK2 | 23.8366957 | 0.217786529 | 0.26924767 | 0.80887063 | 0.41858956 | 0.77741084 |
| CDC37 | 48.5949579 | -0.390032722 | 0.28082802 | -1.3888668 | 0.16487325 | 0.55965519 |
| PDE4A | 19.6156734 | 0.380410024 | 0.35588435 | 1.06891474 | 0.28510809 | 0.6783522 |
| SLC44A2 | 57.2624699 | 0.185664349 | 0.19469424 | 0.95362013 | 0.34027596 | 0.71925477 |
| ILF3 | 66.9598055 | -0.301074687 | 0.2648317 | -1.1368529 | 0.25559979 | 0.65030654 |
| QTRT1 | 16.5694545 | 0.575977708 | 0.37199522 | 1.54834706 | 0.12153876 | 0.49924175 |
| DNM2 | 88.5467367 | -0.088211829 | 0.24692352 | -0.3572435 | 0.72090951 | 0.93444541 |
| CARM1 | 28.0475421 | -0.004383619 | 0.2275055 | -0.0192682 | 0.98462716 | 0.99642316 |
| SMARCA4 | 76.5787239 | 0.093816639 | 0.30087349 | 0.31181424 | 0.7551817 | 0.94189722 |
| LDLR | 69.3907192 | -0.769751457 | 0.49677789 | -1.5494882 | 0.12126442 | 0.49897659 |
| KANK2 | 122.512 | 0.822451893 | 0.2694821 | 3.05197224 | 0.00227343 | 0.05966945 |
| DOCK6 | 92.5537349 | 0.475604008 | 0.22627679 | 2.10186826 | 0.03556482 | 0.28784784 |
| RGL3 | 94.0536863 | -0.531381343 | 0.31724395 | -1.6749928 | 0.09393564 | 0.44537614 |
| PRKCSH | 26.1900079 | 0.329030444 | 0.29069124 | 1.13188977 | 0.25768078 | 0.65130862 |
| ECSIT | 18.5386964 | -0.044825292 | 0.35683802 | -0.125618 | 0.90003428 | 0.98079096 |
| CNN1 | 68.6842252 | 1.679289661 | 0.3745659 | 4.48329557 | 7.35E-06 | 0.0012276 |
| ZNF439 | 28.3348626 | -0.07538066 | 0.26130775 | -0.2884746 | 0.77298344 | 0.9472495 |
| ZNF69 | 20.2332374 | 0.633604846 | 0.33516874 | 1.89040553 | 0.05870374 | 0.36041506 |
| ZNF44 | 68.3064271 | 0.326565974 | 0.21804542 | 1.497697 | 0.13421199 | 0.51662128 |
| ZNF791 | 40.7636621 | -0.031456163 | 0.27361819 | -0.1149637 | 0.90847389 | 0.98306694 |
| TNPO2 | 35.2912876 | -0.076559217 | 0.26363112 | -0.2904028 | 0.77150809 | 0.94654641 |
| TRIR | 40.6208618 | 0.049966148 | 0.29039294 | 0.17206392 | 0.86338728 | 0.97600878 |
| HOOK2 | 31.7171296 | -0.172510882 | 0.26558295 | -0.6495556 | 0.51597935 | 0.83174649 |
| JUNB | 132.188023 | 0.012280984 | 0.52000777 | 0.02361692 | 0.98115817 | 0.99533442 |
| CALR | 102.767915 | -0.080338285 | 0.35774746 | -0.224567 | 0.82231612 | 0.96336699 |
| RAD23A | 32.6922726 | -0.418516672 | 0.34745764 | -1.2045114 | 0.22839199 | 0.62213472 |
| GADD45GIP1 | 42.2470431 | -0.698661094 | 0.35891193 | -1.9466087 | 0.05158166 | 0.34492595 |
| NFIX | 88.4967565 | -0.129846586 | 0.33844704 | -0.3836541 | 0.70123486 | 0.92865476 |
| IER2 | 61.6948945 | 1.005403239 | 0.45428242 | 2.21316784 | 0.02688608 | 0.24544817 |
| CCDC130 | 29.0425185 | 0.108649919 | 0.2615938 | 0.41533828 | 0.67789428 | 0.91799007 |
| MRI1 | 16.9975558 | 0.705930313 | 0.35846999 | 1.96928707 | 0.04892014 | 0.33550435 |
| C19orf53 | 38.9644086 | -1.213647503 | 0.3700374 | -3.2797969 | 0.00103882 | 0.03935194 |
| ZSWIM4 | 23.8658179 | -0.071265742 | 0.27998215 | -0.2545367 | 0.79908093 | 0.95549068 |
| AC020916.1 | 25.3241226 | 1.120447311 | 0.52485069 | 2.13479251 | 0.03277796 | 0.27660385 |
| RFX1 | 21.0723656 | 0.386245759 | 0.26712606 | 1.44593066 | 0.14819665 | 0.53680621 |

| | | | | | | |
|---|---|---|---|---|---|---|
| PRKACA | 30.4351661 | 0.262948773 | 0.25958425 | 1.01296121 | 0.31107869 | 0.70126123 |
| ADGRL1 | 33.3526059 | 0.277964102 | 0.2731393 | 1.01766427 | 0.30883753 | 0.69885506 |
| ADGRE5 | 98.7494151 | 0.023502481 | 0.34919248 | 0.06730523 | 0.94633871 | 0.99060401 |
| PKN1 | 34.4306123 | 0.578382067 | 0.38940958 | 1.48527951 | 0.1374698 | 0.52183137 |
| GIPC1 | 33.2434643 | 0.419317317 | 0.27076351 | 1.5486478 | 0.12146641 | 0.49924175 |
| DNAJB1 | 21.5673242 | 1.014952507 | 0.36170549 | 2.80601908 | 0.00501577 | 0.09281959 |
| TECR | 18.393056 | 0.115629255 | 0.34509548 | 0.33506453 | 0.73757641 | 0.93594619 |
| NDUFB7 | 35.651743 | 0.075533414 | 0.2350785 | 0.32131146 | 0.74797438 | 0.93903859 |
| ZNF333 | 24.6965186 | 0.257204499 | 0.32733895 | 0.78574365 | 0.43201769 | 0.78417671 |
| ADGRE2 | 55.7003149 | -2.536354425 | 0.85566421 | -2.9641937 | 0.00303477 | 0.07080172 |
| SYDE1 | 24.0274971 | 0.266369758 | 0.2644874 | 1.00711702 | 0.31387855 | 0.70282967 |
| NOTCH3 | 149.268236 | 0.431272185 | 0.25571866 | 1.68651044 | 0.09169753 | 0.44124451 |
| BRD4 | 179.671871 | -0.36599354 | 0.15274706 | -2.3960759 | 0.01657166 | 0.18999241 |
| AKAP8 | 37.2489507 | -0.244335368 | 0.26420908 | -0.9247804 | 0.35508017 | 0.73115274 |
| AKAP8L | 78.5218052 | 0.110880868 | 0.22044561 | 0.50298514 | 0.61497472 | 0.88360697 |
| TPM4 | 292.482858 | 0.528275533 | 0.27743333 | 1.90415312 | 0.05689024 | 0.35705408 |
| AP1M1 | 24.0328956 | -0.352889359 | 0.26561784 | -1.3285605 | 0.18399303 | 0.58358427 |
| KLF2 | 52.6258238 | 1.676312601 | 0.48983896 | 3.42217081 | 0.00062123 | 0.02733796 |
| EPS15L1 | 57.6358479 | 0.109391297 | 0.22189888 | 0.49297814 | 0.62202803 | 0.88762797 |
| SIN3B | 33.2315665 | 0.062998026 | 0.22190821 | 0.28389228 | 0.77649294 | 0.94805375 |
| MYO9B | 112.666151 | 0.320094961 | 0.24987013 | 1.28104533 | 0.20017775 | 0.59922861 |
| MRPL34 | 19.5222336 | -0.665669575 | 0.33345964 | -1.9962523 | 0.04590647 | 0.32222219 |
| BST2 | 88.2569021 | -0.183609541 | 0.54637618 | -0.3360497 | 0.7368334 | 0.93594619 |
| SLC27A1 | 33.5545547 | 0.55756644 | 0.26773672 | 2.08251765 | 0.03729521 | 0.29301102 |
| AC010618.3 | 20.0937757 | 0.221950218 | 0.34702414 | 0.63958149 | 0.52244472 | 0.8350836 |
| PGLS | 21.7442053 | -0.213219 | 0.29463527 | -0.723671 | 0.46926773 | 0.80388079 |
| COLGALT1 | 21.3536644 | 0.344691578 | 0.32645187 | 1.05587257 | 0.29102644 | 0.68243563 |
| JAK3 | 28.1586671 | -0.733505366 | 0.51757846 | -1.4171868 | 0.15642832 | 0.54607164 |
| ARRDC2 | 25.3241978 | -0.284768666 | 0.38629231 | -0.7371844 | 0.46101022 | 0.80039155 |
| JUND | 136.066096 | 0.365879459 | 0.47554187 | 0.76939483 | 0.44165896 | 0.79053729 |
| LSM4 | 18.8830606 | -0.781565289 | 0.42816685 | -1.8253755 | 0.06794438 | 0.38827623 |
| PGPEP1 | 28.5446384 | 0.082230656 | 0.32335356 | 0.25430571 | 0.79925939 | 0.9555486 |
| GDF15 | 39.670255 | -3.233068054 | 0.64221474 | -5.0342477 | 4.80E-07 | 0.00016494 |
| SSBP4 | 44.1425953 | -0.025276134 | 0.33790846 | -0.0748017 | 0.94037247 | 0.9889521 |
| ISYNA1 | 43.3073089 | 1.190238318 | 0.35452411 | 3.35728457 | 0.00078712 | 0.03268107 |
| ELL | 89.2071823 | -1.298368341 | 0.33730675 | -3.8492213 | 0.00011849 | 0.00879608 |
| FKBP8 | 36.7023367 | 0.377424748 | 0.20872938 | 1.80820135 | 0.07057517 | 0.39592131 |
| KXD1 | 22.136643 | -0.239972936 | 0.38346429 | -0.6258026 | 0.53144445 | 0.84190241 |
| UBA52 | 68.3134033 | -0.099840156 | 0.38541979 | -0.2590426 | 0.79560235 | 0.95463848 |
| CRLF1 | 26.3766041 | -0.973031849 | 0.55482099 | -1.7537762 | 0.07946886 | 0.41775156 |
| REX1BD | 16.7344057 | 0.575467997 | 0.28048316 | 2.05170248 | 0.04019859 | 0.30130305 |
| CRTC1 | 47.7186534 | -0.018560502 | 0.29765386 | -0.062356 | 0.95027934 | 0.99060401 |
| UPF1 | 30.4935394 | 0.039793597 | 0.23841779 | 0.16690699 | 0.86744323 | 0.97634923 |
| COPE | 23.1738605 | -0.168156492 | 0.31628595 | -0.5316597 | 0.59496171 | 0.87429987 |
| SUGP2 | 74.0016789 | -0.187030159 | 0.15604424 | -1.1985714 | 0.23069465 | 0.62536886 |
| SLC25A42 | 25.5338779 | 0.035157322 | 0.32027034 | 0.10977389 | 0.9125887 | 0.98395113 |
| MAU2 | 54.1854112 | -0.173744204 | 0.26224883 | -0.6625166 | 0.5076402 | 0.82518199 |
| GATAD2A | 128.221892 | 0.120911325 | 0.17262004 | 0.70044778 | 0.48364771 | 0.8144078 |
| ZNF90 | 75.7441527 | -0.253774804 | 0.2980353 | -0.8514924 | 0.39449586 | 0.75988253 |
| ZNF430 | 23.4738079 | -0.332313717 | 0.24088117 | -1.3795753 | 0.16771745 | 0.56323722 |
| ZNF431 | 44.5779967 | 0.048553549 | 0.20915787 | 0.23213829 | 0.81643061 | 0.96132648 |
| ZNF708 | 42.7149756 | 0.41520159 | 0.24246942 | 1.71238743 | 0.08682529 | 0.43259599 |
| AC010615.4 | 21.0802658 | -0.219335679 | 0.59805755 | -0.3667468 | 0.71380791 | 0.93136293 |
| ZNF429 | 26.5007439 | 0.150620445 | 0.26743365 | 0.56320679 | 0.57329408 | 0.86261176 |
| ZNF100 | 21.6719764 | -0.344762195 | 0.29980516 | -1.1499542 | 0.25016274 | 0.6460368 |
| ZNF43 | 38.1121447 | 0.466766931 | 0.29264093 | 1.59501588 | 0.11070868 | 0.47776431 |
| ZNF208 | 23.4288758 | -0.22651499 | 0.47603112 | -0.4758407 | 0.63418786 | 0.89627818 |
| ZNF91 | 104.033 | -0.028143015 | 0.15828726 | -0.1777971 | 0.85888233 | 0.97397549 |
| ZNF254 | 89.572921 | 0.01546749 | 0.19299913 | 0.0801428 | 0.93612368 | 0.98827804 |
| LINC00662 | 29.9179157 | -0.033153117 | 0.25635317 | -0.129326 | 0.89709974 | 0.98040427 |
| UQCRFS1 | 27.5547271 | -0.519494648 | 0.44134408 | -1.177074 | 0.23916597 | 0.63451757 |
| C19orf12 | 21.8324884 | 0.007532037 | 0.32280009 | 0.02333344 | 0.9813843 | 0.99533442 |
| URI1 | 112.321522 | -0.053217998 | 0.15374088 | -0.3461538 | 0.7292271 | 0.93455731 |
| TSHZ3 | 82.7806587 | -0.33528749 | 0.26708694 | -1.2553496 | 0.20935188 | 0.60667466 |
| ZNF507 | 34.8480304 | 0.054232745 | 0.23534525 | 0.23043908 | 0.81775059 | 0.96169773 |
| DPY19L3 | 47.9646826 | -0.00776867 | 0.23240099 | -0.0334279 | 0.97333338 | 0.99300191 |
| ANKRD27 | 29.0604577 | 0.295692432 | 0.25568268 | 1.15648204 | 0.24748404 | 0.64317338 |
| CEP89 | 29.0905755 | 0.338591219 | 0.24783193 | 1.36621304 | 0.17187211 | 0.56870244 |
| PEPD | 38.2262347 | -0.115548181 | 0.31098161 | -0.3715595 | 0.71022082 | 0.93009618 |
| LSM14A | 190.875179 | 0.117484694 | 0.12649012 | 0.92880532 | 0.35298998 | 0.73059415 |
| KIAA0355 | 76.1194488 | 0.446699235 | 0.19787007 | 2.25753813 | 0.02397447 | 0.23001094 |
| GPI | 88.2553311 | -0.978645386 | 0.42316263 | -2.3126933 | 0.02073951 | 0.21189076 |
| UBA2 | 163.947537 | 0.029193563 | 0.21307571 | 0.13701028 | 0.89102267 | 0.98023547 |
| WTIP | 48.1072512 | 1.398310901 | 0.35969272 | 3.88751514 | 0.00010128 | 0.00791782 |
| SCGB2B2 | 29.056896 | 0.074365025 | 0.42808546 | 0.17371537 | 0.86208916 | 0.97514133 |
| ZNF302 | 41.9882781 | -0.183080293 | 0.26713567 | -0.6853457 | 0.49312577 | 0.81908904 |
| GRAMD1A | 36.4357018 | -0.324163164 | 0.27445137 | -1.1811315 | 0.23755047 | 0.63302595 |
| USF2 | 21.515444 | 0.278494802 | 0.33196131 | 0.83893753 | 0.40150436 | 0.76402268 |
| COX6B1 | 48.6292217 | -0.182040888 | 0.30412078 | -0.5985809 | 0.5494524 | 0.84874884 |
| KMT2B | 20.3974832 | -0.019035939 | 0.30117051 | -0.0632065 | 0.94960205 | 0.99060401 |
| HSPB6 | 46.4300131 | -0.328394437 | 0.42234351 | -0.7775529 | 0.43683262 | 0.78768252 |
| CAPNS1 | 60.853476 | -0.173839918 | 0.32174255 | -0.5403075 | 0.58898498 | 0.8702694 |
| ZNF565 | 44.5456623 | -0.203885014 | 0.27944801 | -0.7295991 | 0.46563528 | 0.80355051 |
| ZNF146 | 50.5501461 | -0.088929302 | 0.18317841 | -0.4854792 | 0.62733649 | 0.89192276 |
| ZFP14 | 22.2959807 | 0.288796527 | 0.29878702 | 0.96656317 | 0.33376246 | 0.71677974 |
| ZNF566 | 27.9592797 | -0.202339867 | 0.2272412 | -0.8904189 | 0.37324099 | 0.74512029 |
| ZNF529 | 25.8096246 | 0.204393209 | 0.23574867 | 0.86699622 | 0.38594408 | 0.75373453 |
| AC020928.1 | 19.8826357 | 0.048544803 | 0.26686558 | 0.18190733 | 0.85565546 | 0.97379567 |
| ZNF345 | 28.2702059 | 0.565467098 | 0.41411384 | 1.36548707 | 0.17210001 | 0.56882549 |
| ZNF568 | 20.1915984 | -0.376058534 | 0.31619676 | -1.1893181 | 0.23431451 | 0.63076093 |
| ZNF420 | 20.8786272 | 0.39752231 | 0.28139492 | 1.41268476 | 0.15774844 | 0.54916781 |
| HKR1 | 42.1556329 | 0.078904876 | 0.19214087 | 0.4106616 | 0.68132069 | 0.91886128 |
| ZFP30 | 19.2637652 | 0.392415567 | 0.32774576 | 1.19731698 | 0.23118303 | 0.62646168 |
| ZNF573 | 20.6098112 | 0.04686373 | 0.28569958 | 0.16403149 | 0.86970635 | 0.9768643 |
| SIPA1L3 | 79.6899891 | 0.03132199 | 0.27573563 | 0.11359428 | 0.90955942 | 0.98343817 |
| PPP1R14A | 56.6440476 | 0.680461392 | 0.57996968 | 1.17327064 | 0.24068729 | 0.63466484 |
| KCNK6 | 22.8511388 | 0.054082148 | 0.53431894 | 0.10121698 | 0.91937822 | 0.98528826 |
| PSMD8 | 41.0373633 | -0.738191117 | 0.34747134 | -2.1244662 | 0.03363118 | 0.28022171 |
| EIF3K | 33.9450178 | 0.1626243 | 0.26996192 | 0.60239719 | 0.54690978 | 0.84758329 |
| ACTN4 | 420.25279 | 0.971597807 | 0.31540555 | 3.08047148 | 0.00206673 | 0.05618053 |
| ECH1 | 21.8152136 | 0.406181639 | 0.4071046 | 0.99773286 | 0.31840891 | 0.70683013 |
| HNRNPL | 46.0531545 | -0.000724761 | 0.2652912 | -0.0027319 | 0.99782022 | 0.99945221 |
| SIRT2 | 23.0880999 | 0.307496132 | 0.30122437 | 1.02082089 | 0.30733931 | 0.69801431 |
| FBXO17 | 31.7333957 | 0.406899458 | 0.3284643 | 1.23879354 | 0.21542197 | 0.61195827 |
| SAMD4B | 92.7048374 | -0.030981366 | 0.31662728 | -0.0978481 | 0.92205294 | 0.98561736 |
| ZFP36 | 162.202101 | -0.440505607 | 0.7493825 | -0.5878248 | 0.55664991 | 0.85243179 |
| PLEKHG2 | 51.2894992 | 0.617584869 | 0.22714656 | 2.71888281 | 0.00655028 | 0.11015564 |
| RPS16 | 85.4312839 | -0.352479109 | 0.31646094 | -1.1138155 | 0.26535834 | 0.65926925 |
| SUPT5H | 41.6224208 | -0.444255794 | 0.2065843 | -2.1504818 | 0.03151712 | 0.26963833 |
| ZNF780B | 42.1693932 | -0.039651371 | 0.2116835 | -0.1873144 | 0.85141412 | 0.97279888 |
| AKT2 | 43.45475 | 0.148677886 | 0.23034574 | 0.64545532 | 0.51863216 | 0.83297983 |
| PLD3 | 73.8555969 | 0.076499388 | 0.22518288 | 0.33972116 | 0.73406653 | 0.93594619 |
| BLVRB | 32.3817619 | -0.256396697 | 0.31870601 | -0.8044928 | 0.42111242 | 0.77771095 |

| | | | | | | |
|---|---|---|---|---|---|---|
| LTBP4 | 116.36491 | 1.030496898 | 0.34031395 | 3.02807717 | 0.00246115 | 0.062538 |
| NUMBL | 27.5366147 | 0.020205565 | 0.27902851 | 0.07241398 | 0.94227246 | 0.9893931 |
| ITPKC | 31.7102091 | -0.829476819 | 0.28492401 | -2.9112212 | 0.00360019 | 0.07758886 |
| SNRPA | 20.9484703 | -0.41289267 | 0.30784211 | -1.3412482 | 0.17983988 | 0.57744792 |
| AC008537.1 | 32.0579335 | 0.701460603 | 0.34299955 | 2.04507733 | 0.04084724 | 0.30321852 |
| AXL | 168.398504 | -0.091835466 | 0.26469078 | -0.3469538 | 0.72862605 | 0.93455731 |
| HNRNPUL1 | 53.1473765 | -0.030926345 | 0.26507647 | -0.1166695 | 0.90712194 | 0.98306694 |
| TGFB1 | 50.5582967 | 0.213262449 | 0.27164257 | 0.78508478 | 0.43240386 | 0.78423889 |
| RPS19 | 218.55756 | -0.789273547 | 0.3004288 | -2.6271568 | 0.00861017 | 0.13046619 |
| ARHGEF1 | 21.4105839 | -0.199785136 | 0.29182317 | -0.6846103 | 0.4935899 | 0.81915583 |
| RABAC1 | 55.0760855 | -0.058894359 | 0.32762273 | -0.1797627 | 0.85733884 | 0.97381501 |
| ETHE1 | 23.1822781 | -0.332002149 | 0.36283094 | -0.9150326 | 0.36017451 | 0.73273991 |
| ZNF428 | 21.9060528 | 0.128309742 | 0.30497458 | 0.42072274 | 0.67395755 | 0.91551091 |
| PLAUR | 22.7764137 | -0.797081743 | 0.52272977 | -1.5248448 | 0.12729782 | 0.50683947 |
| ZNF235 | 33.3871721 | 0.597817289 | 0.37992732 | 1.57350435 | 0.11560208 | 0.48836096 |
| BCL3 | 23.4156155 | -0.870734907 | 0.51400051 | -1.6940351 | 0.09025861 | 0.43898422 |
| BCAM | 70.37452 | 0.332331109 | 0.28429643 | 1.16895984 | 0.24241981 | 0.63672023 |
| NECTIN2 | 66.6324355 | 0.357030602 | 0.19730634 | 1.80952424 | 0.0703696 | 0.39563263 |
| APOE | 127.481444 | -1.316005186 | 0.4409516 | -2.9844663 | 0.00284073 | 0.06800182 |
| CLPTM1 | 32.9864335 | 0.109291488 | 0.32288505 | 0.3384842 | 0.73499834 | 0.93594619 |
| RELB | 49.2950617 | -0.529829789 | 0.54841134 | -0.9661175 | 0.33398539 | 0.71683955 |
| CLASRP | 49.3987821 | 0.252804561 | 0.20345614 | 1.24255067 | 0.21403346 | 0.61001472 |
| MARK4 | 56.5672799 | 0.425157298 | 0.311621 | 1.36434098 | 0.17246027 | 0.56904425 |
| ERCC1 | 38.4560966 | 1.230980965 | 0.4138556 | 2.97442141 | 0.00293542 | 0.06945135 |
| VASP | 43.061537 | 0.597692839 | 0.28409653 | 2.10383715 | 0.03539266 | 0.28708678 |
| OPA3 | 32.2207975 | -0.519764291 | 0.35689779 | -1.4563393 | 0.14529884 | 0.53538077 |
| EML2 | 25.216037 | -0.043173424 | 0.25260059 | -0.1709158 | 0.86429 | 0.9762143 |
| SNRPD2 | 32.3822859 | -0.730610768 | 0.31775134 | -2.2993161 | 0.021487 | 0.21631225 |
| DMPK | 36.2507896 | 0.822070943 | 0.4156404 | 1.97784178 | 0.04794656 | 0.33147625 |
| SYMPK | 68.8937225 | -0.203748198 | 0.19976967 | -1.0199156 | 0.30776851 | 0.69801431 |
| HIF3A | 79.8399385 | 0.635269977 | 0.82806164 | 0.76717716 | 0.44297619 | 0.79188325 |
| PPP5C | 33.3858444 | -0.133690461 | 0.24306193 | -0.5500263 | 0.58230132 | 0.86749086 |
| CALM3 | 28.1096513 | 0.142570333 | 0.29134546 | 0.48935148 | 0.62459288 | 0.89031618 |
| PTGIR | 61.3140456 | -1.24599933 | 0.41141948 | -3.0285375 | 0.00245741 | 0.062538 |
| DACT3 | 34.5011311 | 0.336838511 | 0.3835119 | 0.87830003 | 0.37978092 | 0.74991898 |
| PRKD2 | 34.5593417 | 0.414057436 | 0.37095049 | 1.11620673 | 0.26433365 | 0.65826275 |
| SLC1A5 | 19.8196295 | -0.096631263 | 0.35282939 | -0.2738753 | 0.78418044 | 0.95065355 |
| AP2S1 | 40.2225186 | -0.419244474 | 0.36179223 | -1.158799 | 0.24653812 | 0.64225758 |
| ARHGAP35 | 87.47999 | 0.400773651 | 0.19904704 | 2.013462 | 0.04406605 | 0.3152359 |
| ZC3H4 | 23.6951906 | 0.267519404 | 0.32512522 | 0.82281959 | 0.41061059 | 0.77166383 |
| SAE1 | 50.2974106 | 0.288061915 | 0.29124693 | 0.98906422 | 0.32263172 | 0.70937168 |
| INAFM1 | 21.658101 | -0.258627891 | 0.42729565 | -0.6052668 | 0.5450017 | 0.84730644 |
| MEIS3 | 22.7630328 | 0.066922891 | 0.41346168 | 0.16185996 | 0.87141614 | 0.9768643 |
| NAPA | 44.1944155 | -0.203520492 | 0.26625971 | -0.7643683 | 0.44464777 | 0.79252305 |
| EHD2 | 53.2742056 | 0.313264546 | 0.253831 | 1.23414611 | 0.21714846 | 0.61410359 |
| NOP53 | 16.8638073 | 0.213669515 | 0.38786012 | 0.55089323 | 0.58170687 | 0.86695676 |
| SELENOW | 80.6122159 | 0.31953056 | 0.28594682 | 1.11744751 | 0.26380303 | 0.6579701 |
| PLA2G4C | 54.7509655 | -0.241109388 | 0.5780477 | -0.4171099 | 0.67659806 | 0.91753129 |
| CARD8 | 28.5015026 | 0.185932464 | 0.26148414 | 0.71106592 | 0.47704339 | 0.81096272 |
| EMP3 | 30.5003963 | 0.515050407 | 0.48645692 | 1.05877908 | 0.2897004 | 0.68019434 |
| KDELR1 | 28.6875282 | -0.29203088 | 0.30622218 | -0.9536568 | 0.34025737 | 0.71925477 |
| RPL18 | 105.776254 | -0.188824666 | 0.29932955 | -0.6308253 | 0.52815473 | 0.8397683 |
| PLEKHA4 | 51.9817847 | 0.186415727 | 0.4288857 | 0.4346513 | 0.66381555 | 0.91141325 |
| PPP1R15A | 35.1726306 | -0.17059503 | 0.3774078 | -0.4520178 | 0.65125618 | 0.9050963 |
| NUCB1 | 60.2739059 | -0.241349153 | 0.24455587 | -0.9868876 | 0.32369774 | 0.71051224 |
| FTL | 912.380886 | -1.12130527 | 0.51304372 | -2.185594 | 0.02884532 | 0.25647799 |
| GYS1 | 61.0196917 | -0.962140692 | 0.36885291 | -2.6084671 | 0.00909488 | 0.13367648 |
| SNRNP70 | 136.85693 | 0.487767133 | 0.29002294 | 1.68182258 | 0.09260325 | 0.44306073 |
| SLC6A16 | 47.0238529 | 0.129869716 | 0.3154196 | 0.41173635 | 0.68053268 | 0.91860979 |
| TEAD2 | 25.3669706 | 0.155011729 | 0.2790651 | 0.55546799 | 0.5785746 | 0.86486944 |
| RPL13A | 146.960357 | -0.339008838 | 0.33506022 | -1.0117848 | 0.31164096 | 0.70140514 |
| RPS11 | 126.382282 | -0.292463043 | 0.29475439 | -0.9922263 | 0.32108717 | 0.70780851 |
| FCGRT | 45.6078635 | 0.680438344 | 0.27854274 | 2.4428508 | 0.01457176 | 0.17653018 |
| RCN3 | 27.3056265 | -0.388307225 | 0.42738906 | -0.9085568 | 0.36358413 | 0.73604674 |
| NOSIP | 20.3153513 | 0.049240766 | 0.30551779 | 0.16117152 | 0.87195831 | 0.9768643 |
| PRR12 | 20.7339704 | 0.499242504 | 0.3782629 | 1.31982942 | 0.18689198 | 0.58501647 |
| RRAS | 38.9736343 | 0.381554823 | 0.36440195 | 1.04707131 | 0.29506669 | 0.68599972 |
| IRF3 | 30.9408107 | -0.016682309 | 0.26782146 | -0.0622889 | 0.95033275 | 0.99060401 |
| PRMT1 | 28.3055033 | -0.273613616 | 0.38973355 | -0.702053 | 0.4826461 | 0.81409616 |
| AP2A1 | 50.8093116 | -0.110789476 | 0.27063657 | -0.4093662 | 0.68227091 | 0.91933701 |
| MED25 | 19.1078554 | -0.400865346 | 0.33222583 | -1.206605 | 0.2275843 | 0.62183669 |
| PTOV1 | 23.9258592 | 0.042120321 | 0.28331822 | 0.14866788 | 0.8818157 | 0.97782528 |
| PNKP | 17.7632285 | -0.006094703 | 0.58275803 | -0.0104584 | 0.99165557 | 0.99759781 |
| TBC1D17 | 19.0666403 | 0.131228576 | 0.3323034 | 0.39490591 | 0.69291233 | 0.9248682 |
| VRK3 | 20.9993003 | 0.238192043 | 0.26735569 | 0.89091816 | 0.37297307 | 0.74512029 |
| NR1H2 | 20.1463812 | 0.078486226 | 0.31315771 | 0.25062843 | 0.8021014 | 0.95758343 |
| EMC10 | 25.6529917 | -0.134575852 | 0.4181835 | -0.3218105 | 0.74759624 | 0.93903859 |
| CLEC11A | 27.3989986 | 0.473831074 | 0.29145109 | 1.62576531 | 0.10399959 | 0.46422585 |
| ETFB | 30.3015395 | 0.220348804 | 0.35701796 | 0.61719249 | 0.53710777 | 0.84423863 |
| SPACA6 | 27.0048319 | 0.659022251 | 0.34473537 | 1.91167577 | 0.05591779 | 0.35425851 |
| ZNF577 | 22.1111301 | 0.349002634 | 0.32160264 | 1.08519828 | 0.27783384 | 0.67269898 |
| ZNF350-AS1 | 23.027795 | 0.730610105 | 0.38877627 | 1.87925591 | 0.06020956 | 0.36535567 |
| ZNF432 | 43.8156403 | -0.333029617 | 0.21993335 | -1.5142297 | 0.12996758 | 0.51204187 |
| PPP2R1A | 58.0205579 | -0.182592146 | 0.21843543 | -0.835909 | 0.40320609 | 0.76511661 |
| ZNF83 | 92.0591592 | 0.342475454 | 0.22461149 | 1.52474592 | 0.12732248 | 0.50683947 |
| ZNF611 | 36.3755572 | -0.287473999 | 0.26531285 | -1.0835284 | 0.27857397 | 0.67343425 |
| ZNF160 | 42.7817635 | -0.423924696 | 0.29558992 | -1.4341649 | 0.15152521 | 0.53898138 |
| ZNF761 | 24.3369325 | -0.648720356 | 0.31562352 | -2.0553612 | 0.03984413 | 0.30124947 |
| ZNF331 | 110.329653 | -1.007122026 | 0.78203217 | -1.2878268 | 0.19780625 | 0.59530234 |
| MYADM | 47.5218265 | 1.050239632 | 0.40215629 | 2.61152105 | 0.00901404 | 0.13367648 |
| PRPF31 | 20.8639866 | -0.022512679 | 0.26925185 | -0.083612 | 0.93336495 | 0.98725539 |
| MBOAT7 | 20.2814943 | -0.244509987 | 0.33351078 | -0.7331397 | 0.46347324 | 0.80225845 |
| RPS9 | 198.58791 | -0.448604182 | 0.30995105 | -1.4473388 | 0.14780206 | 0.53680621 |
| LENG8 | 90.218287 | 0.435454551 | 0.30143818 | 1.44458988 | 0.14857312 | 0.53680621 |
| CDC42EP5 | 20.0208216 | -0.02758043 | 0.2740585 | -0.100637 | 0.91983861 | 0.98528826 |
| PPP1R12C | 29.5271412 | 0.423691797 | 0.29795312 | 1.42200827 | 0.15502385 | 0.54412824 |
| PPP6R1 | 21.676278 | 0.095963613 | 0.32182155 | 0.29818889 | 0.765559 | 0.9447922 |
| RPL28 | 142.235734 | -0.765221316 | 0.26173573 | -2.9236411 | 0.00345963 | 0.07625361 |
| SSC5D | 22.7003821 | 0.115566402 | 0.45031056 | 0.25663711 | 0.79745892 | 0.95463848 |
| U2AF2 | 21.031852 | -0.136500761 | 0.28926161 | -0.4718938 | 0.63700259 | 0.89741446 |
| EPN1 | 22.8839916 | 0.119541639 | 0.25833416 | 0.46274034 | 0.6435505 | 0.90063554 |
| ZSCAN5A | 51.7396227 | -0.142370819 | 0.37670362 | -0.3779385 | 0.70547625 | 0.92946306 |
| ZNF264 | 26.500807 | 0.162722111 | 0.26773621 | 0.60777027 | 0.54333985 | 0.8470523 |
| AC005261.1 | 42.5961626 | -0.18220838 | 0.3028619 | -0.601622 | 0.54742579 | 0.8478466 |
| ZNF586 | 45.1219996 | 0.273714877 | 0.37066855 | 0.73843567 | 0.46024975 | 0.80039155 |
| ZNF814 | 41.1043903 | -0.145956364 | 0.31886712 | -0.4577341 | 0.64714346 | 0.90243971 |
| ZNF587 | 22.2791194 | 0.255182608 | 0.25993965 | 0.98169945 | 0.32624794 | 0.71149668 |
| ZSCAN18 | 30.2717192 | -0.061273943 | 0.32942743 | -0.1860013 | 0.85244371 | 0.97289794 |
| ZNF274 | 24.17247 | 0.316610374 | 0.25238973 | 1.2544503 | 0.20967838 | 0.60731347 |
| RPS5 | 64.533332 | -0.631178275 | 0.33634605 | -1.8765741 | 0.0605765 | 0.36535567 |
| TRIM28 | 30.5851155 | -0.050962376 | 0.34467648 | -0.1478557 | 0.88245667 | 0.97782528 |
| CHMP2A | 21.5767655 | 0.485451587 | 0.35475134 | 1.36842778 | 0.17117821 | 0.56870244 |
| MZF1 | 30.6533325 | -0.506194568 | 0.26355538 | -1.9206383 | 0.05477732 | 0.3515795 |

| | | | | | | |
|---|---|---|---|---|---|---|
| RPS4Y1 | 48.8144023 | -0.399866803 | 1.84687445 | -0.21651 | 0.82859022 | 0.96580172 |
| ZFY | 23.861884 | 0.695145567 | 1.64978355 | 0.42135562 | 0.67349542 | 0.91505229 |
| LINC00278 | 45.0509816 | 0.506976247 | 1.53615013 | 0.3300304 | 0.74137699 | 0.93711378 |
| TTTY15 | 22.1415396 | 0.346693767 | 1.65526872 | 0.20944863 | 0.83409803 | 0.9666365 |
| USP9Y | 93.2000972 | 0.150605674 | 1.7122071 | 0.08795996 | 0.9299085 | 0.98681039 |
| DDX3Y | 34.0553234 | 0.027926561 | 1.68338611 | 0.01658952 | 0.98676409 | 0.99719329 |
| UTY | 165.6505 | 0.204434216 | 1.41773763 | 0.1441975 | 0.88534452 | 0.97867793 |
| NLGN4Y | 51.7611925 | -0.279132065 | 1.35562693 | -0.2059063 | 0.83686414 | 0.9668633 |
| TTTY14 | 181.744516 | 0.118639306 | 1.32891756 | 0.08927514 | 0.92886325 | 0.98681039 |
| KDM5D | 37.5730949 | 1.014559215 | 1.95980014 | 0.51768504 | 0.60467804 | 0.87903643 |
| TTTY10 | 20.0893119 | 0.85038835 | 1.5479376 | 0.54936862 | 0.5827525 | 0.86763536 |
| IL17RA | 26.1121838 | -0.779560411 | 0.29931179 | -2.6045096 | 0.00920059 | 0.13415695 |
| HDHD5 | 21.7477796 | -0.232844655 | 0.29647106 | -0.7853875 | 0.43222643 | 0.78417671 |
| CECR2 | 32.13235 | -1.226961216 | 0.94635094 | -1.2965182 | 0.19479702 | 0.59277982 |
| ATP6V1E1 | 47.9460584 | -0.710405483 | 0.22008919 | -3.2278073 | 0.00124743 | 0.04375316 |
| BCL2L13 | 80.8016436 | -0.082859991 | 0.19116052 | -0.4334576 | 0.66468233 | 0.91201464 |
| MICAL3 | 101.374252 | 0.378218438 | 0.28423727 | 1.33064338 | 0.18330638 | 0.58291588 |
| PEX26 | 28.1102449 | -0.030087685 | 0.2997058 | -0.1003907 | 0.92003413 | 0.98528826 |
| DGCR2 | 51.896213 | 0.163794065 | 0.2767053 | 0.59194407 | 0.55388804 | 0.85191704 |
| CLTCL1 | 52.3162801 | -0.088546085 | 0.31896712 | -0.2776025 | 0.78131748 | 0.94939885 |
| HIRA | 21.8362626 | -0.602578537 | 0.40459901 | -1.4893228 | 0.13640238 | 0.52087399 |
| UFD1 | 16.9643541 | -0.277807728 | 0.36596859 | -0.7591027 | 0.44779115 | 0.79296348 |
| TXNRD2 | 22.9061587 | 0.544959633 | 0.33889362 | 1.60805517 | 0.1078231 | 0.47335241 |
| COMT | 62.4666426 | 0.093713266 | 0.19224581 | 0.48746584 | 0.62592824 | 0.89111714 |
| ARVCF | 16.4069886 | 1.801061959 | 0.44830878 | 4.01745859 | 5.88E-05 | 0.00557713 |
| TANGO2 | 20.2695616 | -0.208996711 | 0.30202047 | -0.6919952 | 0.48894036 | 0.81720854 |
| DGCR8 | 24.2590661 | 0.345679314 | 0.30257729 | 1.14244964 | 0.25326717 | 0.64807117 |
| KLHL22 | 44.015301 | 0.173152705 | 0.28158276 | 0.61492652 | 0.53860325 | 0.8449569 |
| MED15 | 86.2495142 | -0.183056846 | 0.19177876 | -0.954521 | 0.33982 | 0.71925477 |
| PI4KA | 141.483951 | -0.033341542 | 0.1530738 | -0.2178135 | 0.82757441 | 0.96580172 |
| SNAP29 | 22.1744029 | -0.111146872 | 0.25096023 | -0.4428864 | 0.65784791 | 0.90772143 |
| CRKL | 18.946609 | -0.30545148 | 0.34309957 | -0.8902706 | 0.37332058 | 0.74512029 |
| LZTR1 | 17.3147651 | 0.128183358 | 0.32826898 | 0.3904827 | 0.69617965 | 0.92648248 |
| HIC2 | 23.2321769 | 0.384634359 | 0.26053111 | 1.47634715 | 0.13985072 | 0.525979 |
| UBE2L3 | 56.3265615 | -0.085100225 | 0.24635473 | -0.3454378 | 0.7297653 | 0.93455731 |
| PPIL2 | 32.7296149 | 0.406214379 | 0.24938628 | 1.62885613 | 0.10334348 | 0.46365765 |
| MAPK1 | 155.404951 | 0.121257209 | 0.17586906 | 0.68947437 | 0.4905248 | 0.81720854 |
| PPM1F | 17.9520281 | 0.538690571 | 0.37864157 | 1.42269262 | 0.15482528 | 0.54412824 |
| BCR | 59.0464546 | -0.357271375 | 0.35981225 | -0.9929383 | 0.32074001 | 0.70763084 |
| CHCHD10 | 43.6094408 | -0.5081173 | 0.22427748 | -2.2655743 | 0.02347746 | 0.22729214 |
| SMARCB1 | 31.423221 | 0.373599665 | 0.27453003 | 1.36086994 | 0.17355479 | 0.57092846 |
| DDT | 30.5074803 | -0.453790681 | 0.30189498 | -1.5031408 | 0.13280273 | 0.51393747 |
| CABIN1 | 73.5810166 | 0.071395689 | 0.20321718 | 0.35132703 | 0.72534302 | 0.93455731 |
| GGT5 | 139.07665 | 0.179260837 | 0.23444843 | 0.76460668 | 0.44450579 | 0.79252305 |
| SNRPD3 | 36.545221 | 0.134583824 | 0.25089856 | 0.53640732 | 0.59167708 | 0.87229579 |
| KIAA1671 | 45.9392346 | 1.025119834 | 0.36404226 | 2.81593637 | 0.00486353 | 0.09203696 |
| GRK3 | 95.6412348 | -0.696394868 | 0.40912282 | -1.7021658 | 0.0887243 | 0.43544541 |
| HPS4 | 20.0285347 | 0.068786353 | 0.33404018 | 0.20592239 | 0.83685153 | 0.9668633 |
| MIATNB | 29.5578681 | 0.52484025 | 0.38867704 | 1.35032481 | 0.17691182 | 0.57388115 |
| PITPNB | 194.908059 | -0.218470239 | 0.29124293 | -0.7501306 | 0.45317604 | 0.79534858 |
| TTC28 | 673.291848 | -0.104132657 | 0.17161918 | -0.6067658 | 0.54400632 | 0.84730644 |
| HSCB | 30.7022416 | -0.167062838 | 0.36313457 | -0.4600577 | 0.64547484 | 0.90150535 |
| XBP1 | 40.2073736 | -0.706837181 | 0.38894157 | -1.8173351 | 0.06916581 | 0.39160212 |
| Z93930.2 | 52.4671755 | -0.090728133 | 0.32898306 | -0.2757836 | 0.78271428 | 0.95014325 |
| ZNRF3 | 66.9138135 | 0.265926984 | 0.19324835 | 1.37608928 | 0.16879399 | 0.56539064 |
| KREMEN1 | 28.7477338 | 0.09680339 | 0.38220961 | 0.25327304 | 0.80005723 | 0.95634687 |
| EMID1 | 55.581295 | 1.218715368 | 0.51535789 | 2.36479425 | 0.01804009 | 0.19728664 |
| EWSR1 | 114.666606 | 0.040613783 | 0.17559705 | 0.23128967 | 0.81708977 | 0.96134613 |
| AP1B1 | 22.4695405 | 0.132867048 | 0.30487331 | 0.43581069 | 0.66297409 | 0.91137235 |
| THOC5 | 19.5557208 | 0.26548857 | 0.24898948 | 1.06626419 | 0.28630423 | 0.67897273 |
| NF2 | 34.1612734 | -0.390435104 | 0.27047838 | -1.4434984 | 0.14888013 | 0.53689238 |
| UQCR10 | 44.5459293 | -0.859916206 | 0.37375074 | -2.3007746 | 0.02140438 | 0.21577607 |
| ASCC2 | 28.7932578 | 0.097320456 | 0.33311947 | 0.2921488 | 0.77017285 | 0.94593455 |
| MTMR3 | 142.614055 | 0.257481734 | 0.14934981 | 1.72401778 | 0.08470463 | 0.42695086 |
| RNF215 | 24.0344576 | -0.442164041 | 0.41169395 | -1.0740115 | 0.28281753 | 0.67727609 |
| MORC2 | 26.4597902 | -0.192133921 | 0.26399353 | -0.7277978 | 0.46673734 | 0.80374006 |
| SMTN | 92.9620832 | 0.406438064 | 0.3216979 | 1.26341533 | 0.20643996 | 0.60523513 |
| SELENOM | 113.497827 | -0.274871755 | 0.38329927 | -0.7171205 | 0.47329976 | 0.80758763 |
| RNF185 | 36.9895838 | -0.230878413 | 0.27162187 | -0.8499993 | 0.39532545 | 0.76020146 |
| LIMK2 | 29.1464377 | 0.063436773 | 0.34020549 | 0.18646605 | 0.8520793 | 0.97285733 |
| DRG1 | 36.7689673 | -0.176513654 | 0.2040872 | -0.8648933 | 0.38709735 | 0.75438304 |
| EIF4ENIF1 | 32.6547145 | 0.217650665 | 0.23681977 | 0.91905613 | 0.35806622 | 0.73144657 |
| SFI1 | 25.3207975 | 0.278190008 | 0.35673192 | 0.7798293 | 0.43549136 | 0.78682061 |
| PISD | 33.1022625 | 0.223760055 | 0.23744469 | 0.94236706 | 0.34600475 | 0.72423494 |
| PRR14L | 69.9711154 | -0.021803652 | 0.1815715 | -0.120083 | 0.90441739 | 0.982638 |
| YWHAH | 20.0052779 | 0.117308372 | 0.39280105 | 0.29864577 | 0.76521034 | 0.9447922 |
| RTCB | 33.0391419 | -0.203928215 | 0.23448248 | -0.8696949 | 0.38446717 | 0.75288048 |
| FBXO7 | 43.3854613 | 0.226374906 | 0.21734016 | 1.04156962 | 0.29761126 | 0.68849737 |
| SYN3 | 106.191654 | 0.136206217 | 0.87465106 | 0.15572635 | 0.87624873 | 0.9768643 |
| TIMP3 | 1163.80541 | 0.476494738 | 0.38952437 | 1.22327325 | 0.22122651 | 0.61653656 |
| LARGE1 | 507.346778 | -1.002630637 | 0.37142606 | -2.6994084 | 0.00694629 | 0.11497356 |
| HMGXB4 | 35.246914 | -0.407253772 | 0.31250589 | -1.3031875 | 0.19251075 | 0.59142551 |
| TOM1 | 31.3497824 | -1.005451875 | 0.3447116 | -2.9167915 | 0.00353652 | 0.07689317 |
| HMOX1 | 34.4595589 | -3.490575007 | 0.94381402 | -3.6983716 | 0.00021699 | 0.0139774 |
| APOL6 | 49.7614566 | -0.498168346 | 0.45506449 | -1.0947203 | 0.27363918 | 0.66906109 |
| RBFOX2 | 397.238526 | 0.364488485 | 0.14932859 | 2.44084857 | 0.0146528 | 0.17653018 |
| APOL3 | 20.3443889 | -0.692427382 | 0.4830196 | -1.4335389 | 0.1517039 | 0.53898138 |
| APOL2 | 22.0426207 | -1.136869324 | 0.43866074 | -2.5916824 | 0.00955079 | 0.1379822 |
| APOL1 | 26.7005703 | -0.63589682 | 0.38777042 | -1.6398796 | 0.1010302 | 0.46131565 |
| MYH9 | 397.079777 | 0.304451949 | 0.19890683 | 1.5306259 | 0.12586188 | 0.50489024 |
| TXN2 | 18.0968866 | -0.494233853 | 0.31928833 | -1.5479233 | 0.12164075 | 0.49924175 |
| EIF3D | 32.177933 | -0.336674602 | 0.31356543 | -1.0736981 | 0.28295803 | 0.67727609 |
| CDC42EP1 | 20.2053231 | 0.296665934 | 0.35491605 | 0.83587634 | 0.40322447 | 0.76511661 |
| GGA1 | 26.64178 | 0.13177135 | 0.2858523 | 0.46097704 | 0.64481508 | 0.90107359 |
| LGALS1 | 150.103061 | 0.153690151 | 0.54301247 | 0.28303245 | 0.77715196 | 0.94808797 |
| TRIOBP | 44.7550064 | 0.366008667 | 0.22897474 | 1.59846741 | 0.10993899 | 0.47680173 |
| H1F0 | 30.6093908 | 0.31714598 | 0.60041532 | 0.52821101 | 0.59735289 | 0.87535619 |
| EIF3L | 51.1146304 | 0.0278408 | 0.30418471 | 0.09152597 | 0.92707467 | 0.98678435 |
| POLR2F | 19.7426001 | 0.327537587 | 0.34151465 | 0.95907331 | 0.33752182 | 0.71925477 |
| MAFF | 24.861179 | -0.369778454 | 0.28717439 | -1.2876442 | 0.19786983 | 0.59530234 |
| TMEM184B | 66.387325 | 0.129385989 | 0.28596647 | 0.45245161 | 0.65094367 | 0.9050963 |
| CSNK1E | 30.1910503 | -0.030892667 | 0.30590532 | -0.1009877 | 0.91956024 | 0.98528826 |
| TPTEP2-CSNI | 57.2810166 | 0.251615559 | 0.24610143 | 1.02240591 | 0.30658883 | 0.69801431 |
| DDX17 | 496.035391 | 0.215459242 | 0.23850756 | 0.90336441 | 0.36633252 | 0.7385947 |
| CBY1 | 17.6360344 | 0.137392032 | 0.43112281 | 0.31868421 | 0.74996599 | 0.94004394 |
| JOSD1 | 28.8667985 | -0.2058224 | 0.26129836 | -0.7876911 | 0.43087739 | 0.78340375 |
| GTPBP1 | 25.1514286 | -0.118059828 | 0.35207011 | -0.3353304 | 0.73737583 | 0.93594619 |
| SUN2 | 39.9821995 | 0.204498228 | 0.32783237 | 0.62378901 | 0.53276616 | 0.84290603 |
| CBX6 | 47.2783913 | 0.340210931 | 0.42430346 | 0.80181041 | 0.42266264 | 0.77789825 |
| CBX7 | 34.0566647 | 0.576968987 | 0.42145582 | 1.36899045 | 0.17100226 | 0.56870244 |
| RPL3 | 110.627996 | 0.283124024 | 0.32223863 | 0.87861603 | 0.3796095 | 0.74991898 |
| TAB1 | 18.4600494 | 0.614273926 | 0.26801897 | 2.29190463 | 0.02191115 | 0.21819108 |
| ATF4 | 57.5686396 | 0.031952811 | 0.2647393 | 0.12069538 | 0.90393232 | 0.98225619 |

| | | | | | | |
|---|---|---|---|---|---|---|
| RPS19BP1 | 34.8222378 | -0.344618194 | 0.30572484 | -1.1272169 | 0.2596508 | 0.65252568 |
| TNRC6B | 402.294679 | 0.086510004 | 0.22563647 | 0.38340434 | 0.70141998 | 0.92865476 |
| MKL1 | 350.531635 | -0.010996453 | 0.31404531 | -0.0350155 | 0.97206738 | 0.99300191 |
| SLC25A17 | 41.2279041 | 0.156429557 | 0.19330692 | 0.80922897 | 0.41838346 | 0.77722447 |
| ST13 | 73.097429 | -0.077896928 | 0.19977166 | -0.3899298 | 0.69658843 | 0.92672491 |
| XPNPEP3 | 43.7424311 | 0.20086922 | 0.27637411 | 0.72680188 | 0.46734733 | 0.80374006 |
| RBX1 | 47.2478615 | -0.059368692 | 0.21949009 | -0.2704846 | 0.78678746 | 0.9511599 |
| EP300 | 129.742149 | 0.171042817 | 0.13217224 | 1.29409033 | 0.19563422 | 0.5936069 |
| ZC3H7B | 36.2285414 | -0.184053792 | 0.32111173 | -0.5731768 | 0.56652499 | 0.85825441 |
| TOB2 | 37.8116666 | 0.595446266 | 0.26043833 | 2.28632345 | 0.02223534 | 0.22022579 |
| ACO2 | 42.6886808 | 0.183421197 | 0.30248561 | 0.60637992 | 0.5442625 | 0.84730644 |
| CSDC2 | 20.4091139 | 0.496103957 | 0.42571559 | 1.1653413 | 0.24388087 | 0.63964329 |
| XRCC6 | 72.9064754 | 0.12744273 | 0.2052754 | 0.62083782 | 0.53470634 | 0.84361209 |
| SNU13 | 55.3856585 | -0.503724129 | 0.22854064 | -2.2040899 | 0.02751802 | 0.24847431 |
| SREBF2 | 147.936109 | -0.106338657 | 0.33624424 | -0.3162542 | 0.75180956 | 0.94055881 |
| SMDT1 | 23.6164822 | -0.601649885 | 0.33363911 | -1.8032955 | 0.07134183 | 0.39729887 |
| TCF20 | 74.5800509 | -0.065863099 | 0.2267602 | -0.2904526 | 0.77146998 | 0.94654641 |
| RRP7A | 20.9325492 | 0.06285884 | 0.4241932 | 0.14818446 | 0.8821972 | 0.97782528 |
| POLDIP3 | 22.6136456 | 0.325698415 | 0.27135296 | 1.20027588 | 0.23003221 | 0.62447176 |
| CYB5R3 | 70.2985496 | -0.319418938 | 0.22782273 | -1.4020504 | 0.1609002 | 0.5546227 |
| A4GALT | 30.2419163 | -0.867458909 | 0.37691176 | -2.3014907 | 0.02136391 | 0.21566399 |
| ARFGAP3 | 105.127332 | -0.60051491 | 0.33039272 | -1.8175791 | 0.06912848 | 0.39160212 |
| PACSIN2 | 119.652396 | 0.011064257 | 0.31237419 | 0.03541988 | 0.97174493 | 0.99300191 |
| TSPO | 42.2967321 | -0.405966301 | 0.36590928 | -1.1094725 | 0.26722638 | 0.66201068 |
| EFCAB6 | 21.2087093 | 0.043675085 | 0.379955 | 0.11494805 | 0.9084863 | 0.98306694 |
| NUP50-DT | 35.308108 | -1.538482589 | 0.44509999 | -3.4564876 | 0.00054726 | 0.02523249 |
| NUP50 | 24.1731727 | -0.276163865 | 0.28141242 | -0.9813493 | 0.32642054 | 0.71149668 |
| FAM118A | 49.5568373 | 0.387777609 | 0.46558327 | 0.83288562 | 0.40490924 | 0.76629687 |
| FBLN1 | 209.980147 | 0.394134153 | 0.34131682 | 1.15474577 | 0.24819455 | 0.64337981 |
| ATXN10 | 209.982924 | -0.060673719 | 0.15613372 | -0.388601 | 0.69757133 | 0.92702563 |
| MIRLET7BHC | 97.1989147 | 0.387618162 | 0.33468367 | 1.15816277 | 0.24679762 | 0.64247811 |
| PPARA | 62.1124009 | 0.441531074 | 0.25414039 | 1.73735108 | 0.08232522 | 0.42256146 |
| TTC38 | 24.3706538 | -0.217643951 | 0.43087992 | -0.5051151 | 0.613478 | 0.88349007 |
| TRMU | 22.6132797 | -0.324932265 | 0.2415362 | -1.3452736 | 0.17853691 | 0.57673307 |
| GRAMD4 | 22.9413229 | -0.658274742 | 0.26570397 | -2.4774743 | 0.0132316 | 0.167943 |
| CERK | 37.4952434 | -0.36144106 | 0.26244215 | -1.3772218 | 0.16844368 | 0.5644745 |
| TBC1D22A | 197.13545 | -0.34886912 | 0.28787268 | -1.2118869 | 0.22555569 | 0.62063573 |
| BRD1 | 44.1339009 | 0.059570093 | 0.30775788 | 0.19356155 | 0.8465192 | 0.97014274 |
| ZBED4 | 22.90145 | -0.07537031 | 0.30276589 | -0.2489392 | 0.80340778 | 0.95836189 |
| PIM3 | 39.2738671 | -0.918283088 | 0.39818832 | -2.3061527 | 0.0211021 | 0.21419798 |
| SELENOO | 29.8158591 | -0.101700079 | 0.24511867 | -0.4149014 | 0.67821408 | 0.9182379 |
| PLXNB2 | 65.7001911 | 0.472932116 | 0.27639583 | 1.71106817 | 0.08706853 | 0.43290196 |
| PPP6R2 | 144.636631 | 0.123566 | 0.16796408 | 0.73566922 | 0.46193203 | 0.80178046 |
| FP671120.1 | 548.58525 | -0.479725029 | 0.40565357 | -1.1825978 | 0.23696857 | 0.63270446 |
| FP236383.1 | 317.79829 | -0.429992865 | 0.44900471 | -0.9576578 | 0.33823534 | 0.71925477 |
| HSPA13 | 22.4821157 | -0.601727867 | 0.3678051 | -1.6359965 | 0.10184031 | 0.46169744 |
| NRIP1 | 102.336833 | -0.678886191 | 0.22561329 | -3.00907 | 0.00262049 | 0.06512862 |
| USP25 | 130.209607 | -0.066731793 | 0.24735914 | -0.2697769 | 0.78733186 | 0.95129454 |
| MIR99AHG | 583.656173 | 0.957696617 | 0.41953468 | 2.28275912 | 0.02244456 | 0.22134579 |
| BTG3 | 28.4042948 | -0.646843985 | 0.35699774 | -1.8118994 | 0.07000174 | 0.39397538 |
| ATP5PF | 35.0721023 | -0.322725367 | 0.4252356 | -0.7589331 | 0.44789258 | 0.79296348 |
| GABPA | 32.7347132 | -0.025488679 | 0.24600232 | -0.1036115 | 0.91747763 | 0.98477389 |
| APP | 379.285851 | -0.178309426 | 0.18806456 | -0.9481288 | 0.34306389 | 0.72177972 |
| ADAMTS1 | 327.127658 | -1.636987175 | 0.4584851 | -3.5704261 | 0.0003564 | 0.01915017 |
| ADAMTS5 | 62.7399947 | -0.094556705 | 0.52314594 | -0.1807463 | 0.8565667 | 0.97381501 |
| LTN1 | 53.596071 | -0.060255997 | 0.1984741 | -0.3035963 | 0.76143549 | 0.94425408 |
| USP16 | 47.0954423 | 0.243510679 | 0.19433075 | 1.25307334 | 0.21017902 | 0.60731347 |
| CCT8 | 29.8272329 | -0.28870251 | 0.31426696 | -0.9186537 | 0.35827673 | 0.73144657 |
| MAP3K7CL | 22.3126828 | 0.842374615 | 0.28647214 | 2.94051145 | 0.00327671 | 0.07319311 |
| BACH1 | 263.854884 | -0.695593333 | 0.22425749 | -3.1017619 | 0.00192373 | 0.05544103 |
| SOD1 | 53.9595304 | -0.139241962 | 0.22316529 | -0.623941 | 0.53266636 | 0.84290603 |
| SCAF4 | 67.8345525 | -0.148574871 | 0.17788238 | -0.8352422 | 0.40358136 | 0.76539855 |
| URB1 | 51.6105322 | -0.483072744 | 0.32545621 | -1.4842941 | 0.1377309 | 0.52255261 |
| EVA1C | 159.164525 | -0.403893255 | 0.2767997 | -1.4591535 | 0.14452286 | 0.53398616 |
| SYNJ1 | 42.2956114 | 0.238507166 | 0.23718 | 1.00559561 | 0.31461013 | 0.70282967 |
| PAXBP1 | 91.3934892 | -0.227388272 | 0.22477293 | -1.0116355 | 0.31171236 | 0.70140514 |
| IFNAR1 | 48.614879 | -0.053748696 | 0.23680847 | -0.2269712 | 0.82044616 | 0.96291025 |
| IFNGR2 | 63.8304418 | -0.349271506 | 0.25229213 | -1.3843932 | 0.1662381 | 0.56208655 |
| TMEM50B | 57.3019686 | 0.578594729 | 0.24739375 | 2.33876048 | 0.01934783 | 0.20458593 |
| GART | 33.9892476 | -0.156830802 | 0.24536405 | -0.639176 | 0.52270847 | 0.8350836 |
| SON | 241.103463 | -0.146302962 | 0.20943317 | -0.6985663 | 0.48482309 | 0.81549057 |
| AP000311.1 | 79.2084385 | -0.053001757 | 0.59548084 | -0.0890067 | 0.92907662 | 0.98681039 |
| ITSN1 | 292.453622 | -0.17374566 | 0.31983857 | -0.5432292 | 0.58697198 | 0.870249 |
| MRPS6 | 128.699377 | 0.507327498 | 0.75065803 | 0.6758437 | 0.49913989 | 0.82271448 |
| RCAN1 | 54.8611121 | -0.765918497 | 0.34046912 | -2.2495976 | 0.0244745 | 0.23178238 |
| RUNX1 | 184.228062 | -1.038601517 | 0.5983918 | -1.7356547 | 0.08262491 | 0.42256146 |
| CBR1 | 25.1972431 | -0.608821561 | 0.37741434 | -1.6131384 | 0.10671444 | 0.47073496 |
| DOPEY2 | 20.6206944 | 0.490670931 | 0.33880278 | 1.44824943 | 0.1475473 | 0.53679369 |
| MORC3 | 92.8786881 | 0.115978304 | 0.20718584 | 0.55977911 | 0.57563012 | 0.86295198 |
| HLCS | 100.975646 | 0.049012654 | 0.18198234 | 0.26932643 | 0.7876785 | 0.95129454 |
| TTC3 | 221.969832 | 0.118116613 | 0.15005108 | 0.78717604 | 0.43117882 | 0.78375789 |
| VPS26C | 35.8445206 | 0.049619589 | 0.33776289 | 0.14690657 | 0.88320576 | 0.97782528 |
| DYRK1A | 311.862844 | -0.022860303 | 0.19939672 | -0.1146473 | 0.90872466 | 0.98310283 |
| KCNJ15 | 17.4794062 | 0.808360284 | 0.46490644 | 1.73875907 | 0.08207715 | 0.42256146 |
| ERG | 237.733887 | -0.676035097 | 0.44286468 | -1.5265049 | 0.12688418 | 0.50643981 |
| ETS2 | 107.79257 | -0.813133274 | 0.36062598 | -2.2547828 | 0.02414696 | 0.23086389 |
| BRWD1 | 197.862117 | -0.12547219 | 0.20936324 | -0.5993038 | 0.54897031 | 0.84855413 |
| HMGN1 | 46.5562567 | 0.00451161 | 0.28249639 | 0.01597051 | 0.98725792 | 0.99719329 |
| WRB | 21.6464994 | 0.217850474 | 0.29507271 | 0.73829422 | 0.46033568 | 0.80039155 |
| BACE2 | 86.8813186 | -0.207993961 | 0.31059592 | -0.6696609 | 0.50307395 | 0.82430366 |
| MX2 | 59.7642201 | 0.406831965 | 1.27793558 | 0.31835092 | 0.75021877 | 0.94004394 |
| MX1 | 85.544602 | 0.688944666 | 1.30408772 | 0.52829626 | 0.59729373 | 0.87535619 |
| C2CD2 | 196.528546 | -0.98220353 | 0.43602765 | -2.2526175 | 0.02428327 | 0.23086389 |
| ZBTB21 | 26.6050102 | -0.761517927 | 0.28663299 | -2.65677 | 0.00788932 | 0.12441768 |
| SLC37A1 | 17.4494313 | -0.115589269 | 0.44205597 | -0.2614811 | 0.79372154 | 0.95416774 |
| PDE9A | 32.4230015 | 0.561044888 | 0.36584628 | 1.53355363 | 0.1251395 | 0.50407927 |
| NDUFV3 | 38.8228562 | -0.268549976 | 0.23084173 | -1.1633511 | 0.2446871 | 0.6405276 |
| PKNOX1 | 51.5705853 | 0.215978759 | 0.19671061 | 1.09795174 | 0.27222556 | 0.66723562 |
| RRP1B | 26.5348209 | -0.107987559 | 0.24702572 | -0.4371511 | 0.66200178 | 0.91076998 |
| PDXK | 56.8418277 | -0.255201038 | 0.22052525 | -1.1572418 | 0.24717357 | 0.64317338 |
| CSTB | 47.0329719 | -0.602406887 | 0.25356815 | -2.3757199 | 0.01751475 | 0.19333326 |
| AGPAT3 | 64.2239155 | 0.084739203 | 0.28742667 | 0.29482025 | 0.7681312 | 0.94576899 |
| TRAPPC10 | 60.1676375 | 0.058668016 | 0.1669035 | 0.35150861 | 0.72520681 | 0.93455731 |
| PFKL | 53.7572231 | -0.821829182 | 0.25403634 | -3.2350851 | 0.00121606 | 0.04338282 |
| UBE2G2 | 78.0642992 | -0.16650142 | 0.24161702 | -0.689113 | 0.49075218 | 0.81720854 |
| SUMO3 | 24.0035343 | -0.096919773 | 0.35780446 | -0.2708736 | 0.78648827 | 0.95095525 |
| PTTG1IP | 77.1117162 | -0.240911639 | 0.20401179 | -1.1808712 | 0.23765391 | 0.63302595 |
| ADARB1 | 158.812606 | 0.028852772 | 0.372215 | 0.07751641 | 0.93821273 | 0.98827804 |
| POFUT2 | 18.9692464 | -0.156515136 | 0.31278037 | -0.5003995 | 0.61679381 | 0.88482308 |
| COL18A1 | 483.736321 | -0.829733616 | 0.30697149 | -2.7029664 | 0.00687237 | 0.11400688 |
| PCBP3 | 61.6507199 | -2.108311589 | 0.82292288 | -2.5619795 | 0.01040774 | 0.14309893 |
| COL6A1 | 136.834909 | 0.449512935 | 0.25430906 | 1.76758521 | 0.07713027 | 0.4114983 |
| COL6A2 | 243.052385 | 0.36903289 | 0.30270683 | 1.2191099 | 0.22280248 | 0.61772146 |

| | | | | | | | |
|---|---|---|---|---|---|---|---|
| LSS | 30.0309164 | | -0.468848351 | 0.40431973 | -1.159598 | 0.24621251 | 0.64218217 |
| MCM3AP | 48.7587858 | | -0.020180611 | 0.18700008 | -0.1079177 | 0.91406101 | 0.98395113 |
| PCNT | 33.4832188 | | 0.038360785 | 0.34608142 | 0.11084324 | 0.91174066 | 0.98343817 |
| DIP2A | 71.9836819 | | 0.283639147 | 0.17803597 | 1.59315643 | 0.1111251 | 0.47827842 |
| PRMT2 | 64.3083631 | | 0.640351326 | 0.41539208 | 1.54155882 | 0.12318084 | 0.50097964 |
| AL592183.1 | 34.6911875 | | 0.148897321 | 0.4950688 | 0.30076086 | 0.76359685 | 0.94462105 |

**Leukocytes**

*This means that positive fold changes imply higher mean value in controls than in AKI and vice versa.

| | baseMean | log2 fold change control/AKI* | lfcSE | stat | pvalue | padj |
|---|---|---|---|---|---|---|
| AL669831.5 | 11.4426808 | -0.187821471 | 0.47567375 | -0.3948536 | 0.69295097 | 0.99945981 |
| NOC2L | 9.32746858 | -0.519692113 | 0.55286045 | -0.940006 | 0.34721451 | 0.99945981 |
| ISG15 | 79.0322025 | 0.634167652 | 0.99833072 | 0.63522803 | 0.52527971 | 0.99945981 |
| C1orf159 | 8.14888496 | 0.563024355 | 0.52972508 | 1.06286142 | 0.2878448 | 0.99945981 |
| SDF4 | 16.5523511 | 0.11742375 | 0.43027479 | 0.27290409 | 0.78492694 | 0.99945981 |
| UBE2J2 | 18.5593821 | 0.515481164 | 0.40402479 | 1.2758652 | 0.20200317 | 0.99945981 |
| INTS11 | 8.08083264 | 0.188321631 | 0.50267766 | 0.37463696 | 0.70793048 | 0.99945981 |
| AURKAIP1 | 12.0702703 | -0.281045474 | 0.6086943 | -0.4617186 | 0.64428314 | 0.99945981 |
| CCNL2 | 26.7462897 | -0.080663938 | 0.3237464 | -0.2491578 | 0.80323873 | 0.99945981 |
| MRPL20 | 18.4340156 | -0.274914534 | 0.35800776 | -0.767901 | 0.44254597 | 0.99945981 |
| SSU72 | 19.2629869 | -0.063339861 | 0.40091324 | -0.157989 | 0.87446551 | 0.99945981 |
| CDK11B | 12.4634196 | -0.119010768 | 0.39607565 | -0.3004748 | 0.76381499 | 0.99945981 |
| SLC35E2B | 14.5438172 | -0.121953292 | 0.53170888 | -0.229361 | 0.81858833 | 0.99945981 |
| SLC35E2A | 9.09848858 | -0.582351741 | 0.52497201 | -1.1093006 | 0.26730054 | 0.99945981 |
| NADK | 25.0722065 | 0.281300059 | 0.44642598 | 0.63011579 | 0.52861883 | 0.99945981 |
| GNB1 | 165.529027 | -0.120019417 | 0.24216838 | -0.4956032 | 0.62017442 | 0.99945981 |
| PRKCZ | 8.47675968 | -0.530821219 | 0.58557686 | -0.9064928 | 0.36467506 | 0.99945981 |
| FAAP20 | 8.28089249 | -0.417032982 | 0.50055437 | -0.8331422 | 0.40476452 | 0.99945981 |
| SKI | 41.6208551 | -0.896839671 | 0.37553279 | -2.3881794 | 0.01693207 | 0.74280553 |
| RER1 | 18.6174104 | 0.053551773 | 0.40458502 | 0.13236223 | 0.89469779 | 0.99945981 |
| TNFRSF14 | 19.1694129 | 0.22975323 | 0.33850167 | 0.67873589 | 0.49730522 | 0.99945981 |
| CEP104 | 17.5953909 | -0.680688769 | 0.42457956 | -1.6032066 | 0.10888904 | 0.99945981 |
| KCNAB2 | 59.5555324 | -0.025150444 | 0.38293936 | -0.0656774 | 0.9476347 | 0.99945981 |
| RPL22 | 9.59351648 | 0.401511324 | 0.41887615 | 0.95854425 | 0.33778839 | 0.99945981 |
| NOL9 | 13.0176432 | 0.000881799 | 0.4649188 | 0.00189667 | 0.99848667 | 0.9997934 |
| DNAJC11 | 15.7121 | -0.154632379 | 0.39272268 | -0.3937445 | 0.69376971 | 0.99945981 |
| CAMTA1 | 54.3191471 | -0.764866572 | 0.33631164 | -2.2742792 | 0.0229492 | 0.80320673 |
| VAMP3 | 7.39451092 | -0.03410529 | 0.50106776 | -0.0680652 | 0.94573371 | 0.99945981 |
| PARK7 | 24.5277181 | -0.594386214 | 0.3714121 | -1.6003415 | 0.10952284 | 0.99945981 |
| RERE | 194.975072 | -0.459499604 | 0.33249325 | -1.3819818 | 0.1669773 | 0.99945981 |
| ENO1 | 72.9643657 | -0.538765797 | 0.43153519 | -1.2484864 | 0.211853 | 0.99945981 |
| MIR34AHG | 12.7678359 | -0.554035985 | 0.81957144 | -0.6760069 | 0.49903624 | 0.99945981 |
| H6PD | 12.2948002 | -0.184561796 | 0.4259778 | -0.4332662 | 0.66482137 | 0.99945981 |
| PIK3CD | 80.1269624 | 0.320671695 | 0.36978933 | 0.86717401 | 0.38584668 | 0.99945981 |
| CLSTN1 | 29.2897297 | -0.193825905 | 0.393644 | -0.4923888 | 0.62244451 | 0.99945981 |
| CTNNBIP1 | 11.0994501 | -0.463984935 | 0.47117209 | -0.9847462 | 0.32474874 | 0.99945981 |
| LZIC | 9.19056564 | -0.134047537 | 0.5585542 | -0.2399902 | 0.81033785 | 0.99945981 |
| UBE4B | 54.3482911 | -0.10021149 | 0.27720318 | -0.3615092 | 0.71771885 | 0.99945981 |
| KIF1B | 56.344207 | 0.514132868 | 0.38968946 | 1.31933993 | 0.18705549 | 0.99945981 |
| PGD | 21.3976891 | 0.3020626 | 0.47084798 | 0.64152893 | 0.52117909 | 0.99945981 |
| DFFA | 7.69328501 | 0.10692121 | 0.49228798 | 0.2171924 | 0.8280584 | 0.99945981 |
| PEX14 | 38.1458986 | 0.024945185 | 0.41524679 | 0.06007316 | 0.95209737 | 0.99945981 |
| SRM | 10.3698894 | -0.558226586 | 0.54847389 | -1.0177815 | 0.3087818 | 0.99945981 |
| EXOSC10 | 12.1929609 | -0.268041959 | 0.38994771 | -0.6873792 | 0.49184378 | 0.99945981 |
| MTOR | 29.2796481 | 0.083728415 | 0.30821453 | 0.27165628 | 0.78588632 | 0.99945981 |
| AGTRAP | 12.2219895 | 0.164143268 | 0.58262072 | 0.28173263 | 0.77814853 | 0.99945981 |
| MTHFR | 19.2484408 | -0.908342118 | 0.53223963 | -1.7066413 | 0.08788875 | 0.99640647 |
| CLCN6 | 12.2077155 | -0.354159294 | 0.47578366 | -0.7443704 | 0.4566524 | 0.99945981 |
| KIAA2013 | 9.11884155 | 0.20060353 | 0.53233738 | 0.37683533 | 0.70629599 | 0.99945981 |
| MFN2 | 12.0313549 | -0.134671263 | 0.41308546 | -0.3260131 | 0.74441446 | 0.99945981 |
| TNFRSF1B | 152.007501 | 0.195687299 | 0.57112574 | 0.34263436 | 0.73187355 | 0.99945981 |
| VPS13D | 77.2872679 | -0.085547853 | 0.20890916 | -0.4094978 | 0.68217434 | 0.99945981 |
| DHRS3 | 21.5546907 | -0.161790018 | 0.59616673 | -0.2713838 | 0.78609583 | 0.99945981 |
| PRDM2 | 89.1391901 | -0.025375891 | 0.27095085 | -0.093655 | 0.92538325 | 0.99945981 |
| KAZN | 29.0286656 | -1.357944946 | 0.58497489 | -2.321373 | 0.02026672 | 0.78749881 |
| TMEM51 | 25.3421033 | -0.132684197 | 0.67214983 | -0.1974027 | 0.84351241 | 0.99945981 |
| EFHD2 | 35.3320172 | 0.301087347 | 0.65962576 | 0.45645177 | 0.64806515 | 0.99945981 |
| DNAJC16 | 14.7800157 | -0.146441723 | 0.3459047 | -0.4233586 | 0.67203366 | 0.99945981 |
| DDI2 | 27.16173 | -0.047887324 | 0.35825055 | -0.1336699 | 0.89366359 | 0.99945981 |
| PLEKHM2 | 27.1594918 | -0.703860848 | 0.48320434 | -1.4566526 | 0.14521231 | 0.99945981 |
| SPEN | 54.5565015 | -0.049537682 | 0.32197779 | -0.1538543 | 0.87772458 | 0.99945981 |
| FBXO42 | 56.7566239 | -0.00638915 | 0.30040229 | -0.0212686 | 0.98303136 | 0.99945981 |
| SZRD1 | 15.3024133 | -0.3489734 | 0.42618227 | -0.818836 | 0.41287997 | 0.99945981 |
| NECAP2 | 16.6162389 | 0.651735286 | 0.47096236 | 1.38383731 | 0.16640827 | 0.99945981 |
| NBPF1 | 8.82126879 | -0.402992676 | 0.45587152 | -0.8840049 | 0.37669355 | 0.99945981 |
| CROCC | 10.6168775 | 0.32534824 | 0.54532042 | 0.59661848 | 0.55076213 | 0.99945981 |
| ATP13A2 | 9.15146132 | -0.707485043 | 0.57251207 | -1.2357557 | 0.21654939 | 0.99945981 |
| SDHB | 18.8385209 | 0.11597571 | 0.38250568 | 0.30319997 | 0.76173747 | 0.99945981 |
| RCC2 | 20.3706556 | 0.201942321 | 0.38105322 | 0.52995831 | 0.59614084 | 0.99945981 |
| ARHGEF10L | 20.6704898 | 0.52604209 | 0.52843944 | 0.99546334 | 0.31951096 | 0.99945981 |
| IGSF21 | 17.568416 | -0.700471071 | 0.64162692 | -1.0917108 | 0.27496022 | 0.99945981 |
| UBR4 | 53.4780318 | -0.047843271 | 0.26877928 | -0.1780021 | 0.85872134 | 0.99945981 |
| CAPZB | 146.432898 | -0.019504919 | 0.29352744 | -0.0664501 | 0.94701951 | 0.99945981 |
| TMCO4 | 17.5854487 | -0.304097677 | 0.40910707 | -0.7433205 | 0.45728765 | 0.99945981 |
| DDOST | 11.747545 | 0.012410466 | 0.4699081 | 0.02641041 | 0.97892999 | 0.99945981 |
| HP1BP3 | 67.1086686 | 0.06121226 | 0.28729071 | 0.21306732 | 0.83127446 | 0.99945981 |
| EIF4G3 | 247.165416 | -0.225637982 | 0.27660689 | -0.8157352 | 0.41465159 | 0.99945981 |
| ECE1 | 31.6549194 | 0.501202315 | 0.40347047 | 1.242228 | 0.21415245 | 0.99945981 |
| USP48 | 63.5701566 | -0.363473597 | 0.4113791 | -0.883549 | 0.37693972 | 0.99945981 |
| CDC42 | 69.1786181 | 0.108201148 | 0.263441 | 0.41072251 | 0.68127602 | 0.99945981 |
| ZBTB40 | 15.9394907 | -0.215983952 | 0.42972976 | -0.5026041 | 0.61524263 | 0.99945981 |
| C1QA | 88.0504087 | -1.112851429 | 0.91379524 | -1.2178346 | 0.22328685 | 0.99945981 |
| C1QC | 56.3120088 | -0.781143395 | 0.9582856 | -0.8151468 | 0.41498831 | 0.99945981 |
| C1QB | 104.417503 | -1.410791162 | 0.88602838 | -1.5922641 | 0.11132538 | 0.99945981 |
| KDM1A | 19.5796636 | 0.018476627 | 0.44595476 | 0.04143162 | 0.96695181 | 0.99945981 |
| LUZP1 | 28.4821868 | -0.465013676 | 0.50481508 | -0.9211565 | 0.35696874 | 0.99945981 |
| HNRNPR | 49.3046147 | -0.062911114 | 0.25320707 | -0.2484572 | 0.8037807 | 0.99945981 |
| RPL11 | 126.496084 | -0.012848909 | 0.54836607 | -0.0234313 | 0.98130627 | 0.99945981 |
| PITHD1 | 9.7083548 | 0.262122008 | 0.49055988 | 0.53433234 | 0.59311162 | 0.99945981 |
| HMGCL | 8.49034038 | -0.832666931 | 0.46257179 | -1.8000815 | 0.07184777 | 0.959601 |
| PNRC2 | 14.4876644 | 0.156023372 | 0.36460517 | 0.42792419 | 0.66870632 | 0.99945981 |
| SRSF10 | 31.3416759 | -0.130655925 | 0.34149823 | -0.3825962 | 0.70201919 | 0.99945981 |
| NIPAL3 | 19.1939655 | -0.868172064 | 0.38893011 | -2.2322058 | 0.02560136 | 0.81367289 |
| RCAN3 | 14.6884008 | -0.29900985 | 0.59590355 | -0.5017756 | 0.61582539 | 0.99945981 |
| SRRM1 | 39.3332312 | 0.159739445 | 0.24469402 | 0.65281304 | 0.51387682 | 0.99945981 |
| CLIC4 | 44.097297 | -0.580024341 | 0.63960043 | -0.9068542 | 0.3644839 | 0.99945981 |
| RUNX3 | 55.8993265 | -0.937745008 | 0.31123618 | -3.0129692 | 0.00258705 | 0.45570351 |
| SYF2 | 11.9427526 | -0.123245311 | 0.42501473 | -0.2899789 | 0.77183237 | 0.99945981 |
| RSRP1 | 102.7066 | 0.160318909 | 0.31376272 | 0.51095588 | 0.60938195 | 0.99945981 |
| TMEM50A | 24.0996974 | -0.041499859 | 0.33717281 | -0.1230819 | 0.90204227 | 0.99945981 |
| MACO1 | 30.1121445 | -0.353615797 | 0.3184572 | -1.1104029 | 0.26682544 | 0.99945981 |
| MAN1C1 | 22.2685189 | -0.377425797 | 0.31594656 | -1.1945875 | 0.23224825 | 0.99945981 |
| SH3BGRL3 | 97.068866 | -0.710766311 | 0.74025124 | -0.960169 | 0.33697015 | 0.99945981 |
| UBXN11 | 9.06085099 | 0.673401921 | 0.52334947 | 1.28671559 | 0.19819344 | 0.99945981 |
| CD52 | 42.9218707 | -0.624733254 | 0.70700254 | -0.8836365 | 0.37689247 | 0.99945981 |
| DHDDS | 14.344464 | 0.108275726 | 0.58503247 | 0.18507644 | 0.85316908 | 0.99945981 |
| HMGN2 | 25.0925107 | 0.249068924 | 0.50227384 | 0.49588273 | 0.61997715 | 0.99945981 |
| RPS6KA1 | 31.6052088 | 0.527368899 | 0.35247007 | 1.49620901 | 0.1345992 | 0.99945981 |
| ARID1A | 94.233715 | -0.001554818 | 0.27696587 | -0.0056138 | 0.9955209 | 0.99945981 |
| ZDHHC18 | 9.98572724 | 0.205956051 | 0.4850315 | 0.42462408 | 0.67111074 | 0.99945981 |
| NUDC | 11.7456177 | 0.166843509 | 0.41605828 | 0.40100994 | 0.6884128 | 0.99945981 |

| | | | | | | |
|---|---|---|---|---|---|---|
| SLC9A1 | 9.00055935 | 0.512859503 | 0.5171096 | 0.99178104 | 0.32130435 | 0.99945981 |
| WDTC1 | 21.6027267 | 0.071523325 | 0.35793793 | 0.19982047 | 0.84162099 | 0.99945981 |
| SYTL1 | 11.1472321 | 0.810236587 | 0.41616192 | 1.94692632 | 0.05154357 | 0.9230968 |
| WASF2 | 94.0285596 | 0.350802972 | 0.19988187 | 1.75505152 | 0.0792505 | 0.9846219 |
| AHDC1 | 10.2340879 | -0.193170769 | 0.54691416 | -0.3532013 | 0.72393757 | 0.99945981 |
| FGR | 52.1709259 | 0.452717722 | 0.71966391 | 0.62906826 | 0.52930437 | 0.99945981 |
| IFI6 | 32.9386249 | 0.680744119 | 1.00336159 | 0.6784634 | 0.49747792 | 0.99945981 |
| FAM76A | 9.86488353 | 0.382811179 | 0.44282825 | 0.86446874 | 0.38733044 | 0.99945981 |
| STX12 | 61.0765832 | 0.116724005 | 0.40084085 | 0.29119788 | 0.77089998 | 0.99945981 |
| THEMIS2 | 43.4914606 | 0.217882711 | 0.52487301 | 0.4151151 | 0.67805765 | 0.99945981 |
| RPA2 | 15.975956 | -0.2940705 | 0.411284 | -0.7150059 | 0.47460538 | 0.99945981 |
| EYA3 | 46.4308372 | 0.035666185 | 0.25688929 | 0.13883874 | 0.88957758 | 0.99945981 |
| PTAFR | 17.7361701 | 0.182732536 | 0.69535366 | 0.26279079 | 0.79271184 | 0.99945981 |
| DNAJC8 | 15.0078233 | 0.10293861 | 0.45256427 | 0.22745633 | 0.82006892 | 0.99945981 |
| ATP5IF1 | 24.9676514 | -0.313318735 | 0.47498789 | -0.6596352 | 0.50948796 | 0.99945981 |
| PHACTR4 | 56.6181023 | -0.244269112 | 0.32992639 | -0.7403746 | 0.45907273 | 0.99945981 |
| RCC1 | 15.975387 | -1.048152979 | 0.53071273 | -1.9749912 | 0.04826914 | 0.91035074 |
| TRNAU1AP | 19.743156 | -0.295765663 | 0.36685776 | -0.8062135 | 0.42011978 | 0.99945981 |
| TAF12 | 9.60163256 | 0.140382646 | 0.46675393 | 0.30076372 | 0.76359468 | 0.99945981 |
| GMEB1 | 14.4550946 | 0.319508182 | 0.43545231 | 0.73373863 | 0.46310805 | 0.99945981 |
| YTHDF2 | 28.1648464 | -0.336184079 | 0.36006156 | -0.933685 | 0.3504664 | 0.99945981 |
| EPB41 | 110.016164 | 0.268302179 | 0.43657425 | 0.61456254 | 0.53884366 | 0.99945981 |
| SRSF4 | 66.9291157 | -0.215907155 | 0.32863923 | -0.6569732 | 0.51119817 | 0.99945981 |
| LAPTM5 | 154.067318 | -0.617921488 | 0.61342106 | -1.0073366 | 0.31377305 | 0.99945981 |
| SDC3 | 10.5372664 | -2.171748831 | 0.90778539 | -2.3923593 | 0.01674044 | 0.74280553 |
| PUM1 | 120.781076 | 0.111482137 | 0.18625101 | 0.59855857 | 0.54946729 | 0.99945981 |
| SNRNP40 | 12.334022 | 0.363602101 | 0.38993834 | 0.9324605 | 0.35109859 | 0.99945981 |
| ZCCHC17 | 15.9695042 | -0.59795908 | 0.40674692 | -1.4701011 | 0.14153439 | 0.99945981 |
| PTP4A2 | 36.9837784 | 0.539189095 | 0.474884 | 1.13541222 | 0.25620264 | 0.99945981 |
| KHDRBS1 | 53.7081518 | 0.013315105 | 0.23910805 | 0.05568656 | 0.95559151 | 0.99945981 |
| KPNA6 | 38.1880509 | -0.43803988 | 0.33459167 | -1.3091775 | 0.19047424 | 0.99945981 |
| TXLNA | 11.5454802 | -0.501708541 | 0.454025 | -1.105024 | 0.26914917 | 0.99945981 |
| EIF3I | 8.96206963 | -1.015369507 | 0.67973826 | -1.4937654 | 0.13523694 | 0.99945981 |
| LCK | 35.5289067 | -0.149296802 | 0.35972096 | -0.415035 | 0.67811626 | 0.99945981 |
| HDAC1 | 21.3605272 | 0.040716783 | 0.35224304 | 0.11559287 | 0.90797522 | 0.99945981 |
| BSDC1 | 16.7130589 | -0.28978559 | 0.40470149 | -0.7160478 | 0.47396186 | 0.99945981 |
| ZBTB8OS | 33.0059037 | -0.093631283 | 0.40746914 | -0.2297874 | 0.81825696 | 0.99945981 |
| RBBP4 | 30.7224452 | -0.089337934 | 0.34364354 | -0.2599727 | 0.79488484 | 0.99945981 |
| YARS | 27.020604 | -0.774546363 | 0.44190888 | -1.7527287 | 0.0796486 | 0.9846219 |
| S100PBP | 39.4503877 | -0.266835259 | 0.40546301 | -0.6581001 | 0.51047379 | 0.99945981 |
| RNF19B | 19.0216133 | 0.911896488 | 0.56813711 | 1.60506412 | 0.10847969 | 0.99945981 |
| AK2 | 15.8893459 | -0.368976425 | 0.41813764 | -0.8824281 | 0.37754532 | 0.99945981 |
| ZNF362 | 8.84029961 | -0.802801964 | 0.43635784 | -1.839779 | 0.0658007 | 0.94594189 |
| PHC2 | 55.0683115 | -0.070694432 | 0.46281247 | -0.1527496 | 0.87859573 | 0.99945981 |
| SMIM12 | 10.8978917 | -0.341186546 | 0.50342538 | -0.6777301 | 0.49794282 | 0.99945981 |
| ZMYM1 | 12.9121974 | 0.248322165 | 0.39066063 | 0.63564677 | 0.52500668 | 0.99945981 |
| SFPQ | 62.9158796 | -0.050623491 | 0.35033421 | -0.1445006 | 0.88510521 | 0.99945981 |
| ZMYM4 | 94.4810469 | -0.033083425 | 0.19586236 | -0.1689116 | 0.86586618 | 0.99945981 |
| KIAA0319L | 60.0302011 | -0.340108259 | 0.3412768 | -0.996576 | 0.31897037 | 0.99945981 |
| PSMB2 | 17.6096466 | -0.05809032 | 0.38228728 | -0.1519546 | 0.87922272 | 0.99945981 |
| AGO4 | 53.7112547 | -0.043049455 | 0.35372277 | -0.1217039 | 0.90313349 | 0.99945981 |
| AGO1 | 20.4784145 | 0.283963853 | 0.4261785 | 0.66630263 | 0.50521768 | 0.99945981 |
| AGO3 | 87.8336192 | 0.001092886 | 0.30767582 | 0.00355207 | 0.99716586 | 0.99945981 |
| TRAPPC3 | 8.7109115 | -0.48553029 | 0.47624209 | -1.0195031 | 0.30796418 | 0.99945981 |
| MAP7D1 | 15.200763 | 0.480893551 | 0.52206596 | 0.92113562 | 0.35697962 | 0.99945981 |
| THRAP3 | 61.6064652 | -0.215653521 | 0.220584 | -0.9776481 | 0.32824842 | 0.99945981 |
| STK40 | 29.8645943 | -0.091595354 | 0.39297755 | -0.2330804 | 0.815699 | 0.99945981 |
| CSF3R | 48.6934842 | 0.427339003 | 0.64466184 | 0.66288863 | 0.50740189 | 0.99945981 |
| MEAF6 | 12.2387433 | 0.095580176 | 0.43757674 | 0.21843066 | 0.82709358 | 0.99945981 |
| C1orf122 | 9.61796298 | 0.202405956 | 0.54455067 | 0.37169352 | 0.71012105 | 0.99945981 |
| MTF1 | 16.253026 | 0.503165109 | 0.58443382 | 0.86094455 | 0.38926858 | 0.99945981 |
| INPP5B | 29.053023 | 0.099096205 | 0.30090331 | 0.32932906 | 0.74190698 | 0.99945981 |
| SF3A3 | 14.5216194 | 0.524730662 | 0.38163185 | 1.37496558 | 0.16914212 | 0.99945981 |
| RRAGC | 12.5459324 | -0.621132135 | 0.61428449 | -1.0111473 | 0.31194591 | 0.99945981 |
| AKIRIN1 | 22.9775539 | 0.110689763 | 0.32121324 | 0.344599 | 0.73039586 | 0.99945981 |
| NDUFS5 | 25.4190064 | -0.324249697 | 0.5212159 | -0.6221025 | 0.53387449 | 0.99945981 |
| MACF1 | 270.927133 | -0.095257531 | 0.26116897 | -0.3647353 | 0.71530903 | 0.99945981 |
| PABPC4 | 16.0170343 | 0.169411236 | 0.3740931 | 0.45285849 | 0.65065065 | 0.99945981 |
| TRIT1 | 12.7258617 | 0.070818458 | 0.47092084 | 0.15038293 | 0.88046251 | 0.99945981 |
| CAP1 | 65.0872025 | 0.46694094 | 0.44748386 | 1.04348108 | 0.29672554 | 0.99945981 |
| PPT1 | 21.0471197 | 0.699966161 | 0.54827895 | 1.27666066 | 0.20172208 | 0.99945981 |
| RLF | 71.374952 | -0.285224735 | 0.28433812 | -1.0031182 | 0.31580384 | 0.99945981 |
| ZMPSTE24 | 11.4310536 | -0.022750289 | 0.45403899 | -0.0501065 | 0.96003755 | 0.99945981 |
| SMAP2 | 202.071812 | 0.05219573 | 0.41948444 | 0.12442829 | 0.90097618 | 0.99945981 |
| NFYC | 32.5676681 | 0.200423921 | 0.35043071 | 0.57193595 | 0.56736537 | 0.99945981 |
| SCMH1 | 54.2654392 | -0.179764119 | 0.51456619 | -0.3493508 | 0.72682595 | 0.99945981 |
| HIVEP3 | 76.8733243 | 0.227033203 | 0.44168589 | 0.51401507 | 0.60724145 | 0.99945981 |
| FOXJ3 | 66.2788647 | -0.081327645 | 0.22371552 | -0.3635315 | 0.71620786 | 0.99945981 |
| YBX1 | 46.7405708 | -0.010067185 | 0.46454932 | -0.0216709 | 0.98271051 | 0.99945981 |
| SZT2 | 16.5081555 | -0.226156916 | 0.48863892 | -0.4628303 | 0.64348598 | 0.99945981 |
| ATP6V0B | 39.3740093 | 0.060863328 | 0.41687844 | 0.14599778 | 0.88392314 | 0.99945981 |
| ERI3 | 23.0845996 | -0.227405786 | 0.32912696 | -0.6909364 | 0.48960553 | 0.99945981 |
| RNF220 | 38.3162665 | -3.565851661 | 1.59517304 | -2.2354012 | 0.02539102 | 0.81367289 |
| RPS8 | 108.483554 | -0.461736014 | 0.67119291 | -0.6879334 | 0.49149473 | 0.99945981 |
| EIF2B3 | 23.2740635 | -0.373290966 | 0.39588851 | -0.9429194 | 0.34572211 | 0.99945981 |
| ZSWIM5 | 12.9518311 | -0.94281894 | 0.65461394 | -1.4402671 | 0.14979184 | 0.99945981 |
| TESK2 | 22.2006587 | -0.200033028 | 0.44540127 | -0.4491075 | 0.65335415 | 0.99945981 |
| PRDX1 | 43.3536367 | -0.725800563 | 0.66140572 | -1.0973606 | 0.2724838 | 0.99945981 |
| AKR1A1 | 17.7271582 | -0.510611401 | 0.50155483 | -1.018057 | 0.30865087 | 0.99945981 |
| NASP | 23.3103105 | -0.358791784 | 0.33472433 | -1.0719023 | 0.28376392 | 0.99945981 |
| GPBP1L1 | 75.8613177 | -0.041105708 | 0.31103687 | -0.132157 | 0.89486009 | 0.99945981 |
| MAST2 | 25.5723648 | -0.13877898 | 0.42943769 | -0.3231644 | 0.74657073 | 0.99945981 |
| LRRC41 | 10.3705604 | -0.732149695 | 0.42778824 | -1.7114769 | 0.08699311 | 0.99640647 |
| UQCRH | 26.1384858 | 0.526509395 | 0.42633121 | 1.23497736 | 0.21683893 | 0.99945981 |
| NSUN4 | 9.78609336 | -1.426645517 | 0.5669842 | -2.5161998 | 0.01186279 | 0.67187199 |
| MKNK1 | 40.3666471 | 0.321379513 | 0.40144984 | 0.80054712 | 0.42339387 | 0.99945981 |
| ATPAF1 | 9.89067872 | -0.24197003 | 0.46707657 | -0.5180522 | 0.60442187 | 0.99945981 |
| EFCAB14 | 29.8891152 | -0.30486973 | 0.39335424 | -0.7750513 | 0.43830932 | 0.99945981 |
| PDZK1IP1 | 21.1861592 | -0.496764035 | 0.76410629 | -0.6501243 | 0.51561196 | 0.99945981 |
| CMPK1 | 23.8468564 | 0.207890215 | 0.33667593 | 0.6174787 | 0.53691902 | 0.99945981 |
| SPATA6 | 21.1272686 | 0.522738179 | 0.63550373 | 0.82255722 | 0.41075984 | 0.99945981 |
| FAF1 | 160.369988 | -0.071059635 | 0.21356998 | -0.332723 | 0.73934342 | 0.99945981 |
| RNF11 | 16.7574524 | -0.080095403 | 0.44363404 | -0.1805439 | 0.85672562 | 0.99945981 |
| EPS15 | 161.547108 | -0.131303868 | 0.25729309 | -0.510328 | 0.60982169 | 0.99945981 |
| OSBPL9 | 63.983745 | 0.13237685 | 0.34558982 | 0.38304615 | 0.70168555 | 0.99945981 |
| NRDC | 82.3545639 | -0.126591238 | 0.2800777 | -0.4519861 | 0.65127897 | 0.99945981 |
| TXNDC12 | 17.222034 | -0.36901009 | 0.45311687 | -0.8143817 | 0.41542632 | 0.99945981 |
| BTF3L4 | 20.6247081 | 0.034706002 | 0.32814716 | 0.10576353 | 0.91576997 | 0.99945981 |
| ZFYVE9 | 11.5658061 | -1.527319139 | 0.8633267 | -1.7691091 | 0.07687567 | 0.9821652 |
| PRPF38A | 10.1739405 | -0.1137792 | 0.56781801 | -0.2003797 | 0.84118364 | 0.99945981 |
| TUT4 | 98.2931239 | -0.734240624 | 0.39950761 | -1.8378639 | 0.06608246 | 0.94594189 |
| SHISAL2A | 12.3753773 | -0.078052865 | 0.41594647 | -0.1876512 | 0.85115007 | 0.99945981 |
| ZYG11B | 22.6471897 | 0.202354758 | 0.32835194 | 0.61627399 | 0.5377137 | 0.99945981 |

| | | | | | | |
|---|---|---|---|---|---|---|
| ECHDC2 | 19.3728374 | 0.132782935 | 0.49446215 | 0.26854014 | 0.78828358 | 0.99945981 |
| SCP2 | 30.1049437 | 0.082492299 | 0.31358745 | 0.26305995 | 0.79250437 | 0.99945981 |
| NDC1 | 13.3974733 | 0.503701195 | 0.40770247 | 1.2354627 | 0.21665835 | 0.99945981 |
| HSPB11 | 15.1002323 | -0.371990941 | 0.39075914 | -0.9519699 | 0.34111225 | 0.99945981 |
| TMEM59 | 37.2692598 | -0.113014566 | 0.35293341 | -0.320215 | 0.74880536 | 0.99945981 |
| SSBP3 | 49.2810884 | 0.367263335 | 0.42072212 | 0.87293565 | 0.38269815 | 0.99945981 |
| USP24 | 115.332536 | -0.154904494 | 0.37817222 | -0.4096136 | 0.6820894 | 0.99945981 |
| DAB1 | 14.8214986 | -0.34554677 | 0.73326791 | -0.4712422 | 0.6374678 | 0.99945981 |
| MYSM1 | 33.0288468 | 0.110489704 | 0.28058413 | 0.39378458 | 0.69374009 | 0.99945981 |
| JUN | 23.6337813 | -0.695719201 | 0.53383841 | -1.3032393 | 0.19249307 | 0.99945981 |
| FGGY | 26.3514743 | 0.210356504 | 0.48138281 | 0.43698383 | 0.66212307 | 0.99945981 |
| NFIA | 69.5725418 | 0.844440902 | 0.59346839 | 1.42289112 | 0.15476772 | 0.99945981 |
| TM2D1 | 35.0164132 | 0.359869264 | 0.27036381 | 1.33105558 | 0.18317072 | 0.99945981 |
| PATJ | 48.8296581 | -0.251855491 | 0.45028601 | -0.5593234 | 0.57594105 | 0.99945981 |
| USP1 | 9.11886308 | 0.279977341 | 0.45052801 | 0.62144269 | 0.53430839 | 0.99945981 |
| DOCK7 | 28.7975813 | -0.357931674 | 0.32803642 | -1.091134 | 0.27521395 | 0.99945981 |
| ATG4C | 28.6396536 | 0.417478813 | 0.41929544 | 0.99566742 | 0.31941176 | 0.99945981 |
| ALG6 | 12.7873534 | 0.790559413 | 0.42717094 | 1.85068633 | 0.06421469 | 0.93776345 |
| ITGB3BP | 16.7490141 | 0.498165595 | 0.46841283 | 1.06351826 | 0.28754699 | 0.99945981 |
| PGM1 | 10.2203339 | -1.129681575 | 0.51632664 | -2.1879204 | 0.0286754 | 0.84763724 |
| JAK1 | 140.998808 | 0.054065495 | 0.26024645 | 0.20774729 | 0.83542629 | 0.99945981 |
| AK4 | 11.3179759 | -1.192652806 | 0.85942043 | -1.3877408 | 0.16521598 | 0.99945981 |
| LEPR | 14.8298424 | -0.245889908 | 0.72230963 | -0.3404218 | 0.73353894 | 0.99945981 |
| PDE4B | 218.767956 | 0.028259166 | 0.67566396 | 0.04182429 | 0.96663877 | 0.99945981 |
| MIER1 | 58.1054909 | -0.032468172 | 0.27069815 | -0.1199423 | 0.90452882 | 0.99945981 |
| SLC35D1 | 15.6988624 | -1.310419174 | 0.78504267 | -1.669233 | 0.0950712 | 0.99945981 |
| IL12RB2 | 18.3556597 | -0.543374507 | 0.6565595 | -0.8276089 | 0.40789201 | 0.99945981 |
| SERBP1 | 40.7280313 | -0.072824584 | 0.32288862 | -0.2255409 | 0.82155852 | 0.99945981 |
| WLS | 15.3656549 | 0.967982284 | 0.76449303 | 1.26617542 | 0.20545029 | 0.99945981 |
| LRRC40 | 13.8669278 | -0.237098001 | 0.53979722 | -0.4392353 | 0.66049104 | 0.99945981 |
| SRSF11 | 82.1349094 | 0.26362209 | 0.29648885 | 0.88914672 | 0.37392423 | 0.99945981 |
| ANKRD13C | 42.5993696 | -0.001870499 | 0.3100281 | -0.0060333 | 0.99518614 | 0.99945981 |
| ZRANB2 | 56.5046915 | -0.180799761 | 0.29017835 | -0.6230643 | 0.5332423 | 0.99945981 |
| NEGR1 | 27.5805707 | -0.845843389 | 0.96403961 | -0.8773948 | 0.38027221 | 0.99945981 |
| ACADM | 21.0631293 | 0.399215995 | 0.38150143 | 1.04643382 | 0.29536079 | 0.99945981 |
| RABGGTB | 12.6918137 | 0.494090377 | 0.53956684 | 0.91571673 | 0.35981549 | 0.99945981 |
| ST6GALNAC3 | 58.3419551 | 0.606339619 | 0.5964095 | 1.01664983 | 0.30932004 | 0.99945981 |
| PIGK | 13.8978346 | 0.309716925 | 0.51112637 | 0.60594981 | 0.54454809 | 0.99945981 |
| AC118549.1 | 65.3176953 | -0.035429479 | 0.33295654 | -0.1064087 | 0.91525807 | 0.99945981 |
| USP33 | 66.9158925 | 0.272713827 | 0.22866475 | 1.19263605 | 0.23301194 | 0.99945981 |
| MIGA1 | 33.2430318 | -0.364437141 | 0.33507808 | -1.0876186 | 0.27676353 | 0.99945981 |
| FUBP1 | 62.4329902 | 0.269995068 | 0.25290916 | 1.06755751 | 0.28572016 | 0.99945981 |
| IFI44L | 48.1091392 | 1.707240693 | 1.47751822 | 1.1554786 | 0.24789449 | 0.99945981 |
| IFI44 | 34.5728539 | 1.743182228 | 0.86212116 | 2.02196897 | 0.04317956 | 0.89562588 |
| PRKACB | 57.7552097 | -0.262216486 | 0.35744688 | -0.7335817 | 0.46320372 | 0.99945981 |
| GNG5 | 24.2454869 | -0.108346565 | 0.39783581 | -0.2723399 | 0.78536068 | 0.99945981 |
| CTBS | 15.3223628 | -0.311872681 | 0.38797395 | -0.8038495 | 0.42148387 | 0.99945981 |
| MCOLN2 | 24.4334572 | -0.226228089 | 0.67592154 | -0.3346958 | 0.73785458 | 0.99945981 |
| C1orf52 | 12.2030921 | 0.434748175 | 0.40927698 | 1.06223462 | 0.28812919 | 0.99945981 |
| ZNHIT6 | 19.8364293 | -0.047588428 | 0.43999029 | -0.1081579 | 0.91387043 | 0.99945981 |
| ODF2L | 34.3665906 | -0.312557967 | 0.39047164 | -0.8004627 | 0.42344279 | 0.99945981 |
| SH3GLB1 | 45.5765722 | -0.212277186 | 0.34007386 | -0.6242091 | 0.5324903 | 0.99945981 |
| SELENOF | 44.83374 | 0.067175664 | 0.3132253 | 0.21446436 | 0.83018495 | 0.99945981 |
| HS2ST1 | 34.7151394 | 0.140049543 | 0.33575728 | 0.41711544 | 0.67659397 | 0.99945981 |
| LMO4 | 25.8326515 | -0.301493786 | 0.40427402 | -0.7457659 | 0.45580883 | 0.99945981 |
| PKN2 | 102.425869 | 0.453838837 | 0.23851806 | 1.90274409 | 0.05707395 | 0.92445778 |
| GTF2B | 23.271507 | -0.038965551 | 0.38550416 | -0.1010769 | 0.91948944 | 0.99945981 |
| KYAT3 | 23.1104154 | -0.165195285 | 0.30889574 | -0.534793 | 0.59279299 | 0.99945981 |
| GBP3 | 11.1871085 | -0.565162133 | 0.50087913 | -1.1283403 | 0.25917621 | 0.99945981 |
| GBP1 | 16.5178085 | 1.062637074 | 0.78176572 | 1.35927817 | 0.17405846 | 0.99945981 |
| GBP2 | 41.0588466 | 0.458693286 | 0.42421716 | 1.08127 | 0.27957703 | 0.99945981 |
| GBP4 | 16.4582521 | 0.672981668 | 0.67676626 | 0.99440783 | 0.32002436 | 0.99945981 |
| GBP5 | 21.0561769 | -1.039812058 | 0.43689447 | -2.3800074 | 0.01731229 | 0.74280553 |
| LRRC8B | 21.6785474 | 0.133907556 | 0.42954473 | 0.31174299 | 0.75523586 | 0.99945981 |
| ZNF326 | 16.9876734 | -0.239604758 | 0.50783769 | -0.4718137 | 0.6370598 | 0.99945981 |
| ZNF644 | 112.025102 | 0.242008726 | 0.23506314 | 1.02954775 | 0.30322235 | 0.99945981 |
| TGFBR3 | 51.2016277 | 0.375233263 | 0.56891111 | 0.65956395 | 0.50953369 | 0.99945981 |
| GLMN | 24.3283306 | -0.527205556 | 0.46405653 | -1.1360804 | 0.2559229 | 0.99945981 |
| RPAP2 | 39.4746673 | -0.321806674 | 0.42356729 | -0.7597534 | 0.44740202 | 0.99945981 |
| EVI5 | 79.0148529 | 0.313760167 | 0.20753282 | 1.51185807 | 0.13056997 | 0.99945981 |
| RPL5 | 57.7807791 | -0.385602801 | 0.51820986 | -0.7441055 | 0.45681266 | 0.99945981 |
| FAM69A | 59.7773056 | -0.26482606 | 0.44483716 | -0.5953326 | 0.55162118 | 0.99945981 |
| MTF2 | 47.2964386 | 0.161948835 | 0.25235734 | 0.6417441 | 0.52103935 | 0.99945981 |
| TMED5 | 28.8681498 | 0.211737205 | 0.36555636 | 0.57921904 | 0.56244139 | 0.99945981 |
| CCDC18 | 29.9801799 | 0.079154572 | 0.38854366 | 0.20372118 | 0.83857139 | 0.99945981 |
| DR1 | 15.9355719 | -0.286132601 | 0.34881558 | -0.8202977 | 0.41204644 | 0.99945981 |
| FNBP1L | 15.1181926 | 0.577996589 | 0.47939484 | 1.20567962 | 0.22794104 | 0.99945981 |
| BCAR3 | 21.6812947 | -1.724724116 | 0.94484096 | -1.8254121 | 0.06793887 | 0.94796811 |
| GCLM | 15.8537864 | -0.342993322 | 0.43449821 | -0.789401 | 0.42987767 | 0.99945981 |
| ARHGAP29 | 10.4238406 | 0.491587608 | 0.42924841 | 1.14522871 | 0.25211443 | 0.99945981 |
| ABCD3 | 24.1333437 | 0.590513326 | 0.32272037 | 1.82979875 | 0.06728004 | 0.94796811 |
| CNN3 | 10.0885385 | -1.001861767 | 0.49179565 | -2.0371505 | 0.04163496 | 0.89562588 |
| PTBP2 | 89.8559427 | -0.146675177 | 0.3883061 | -0.3777308 | 0.70563058 | 0.99945981 |
| DPYD | 644.195236 | 0.071816292 | 0.22437177 | 0.32007723 | 0.74890979 | 0.99945981 |
| DPYD-AS1 | 27.5368739 | -0.230430492 | 0.66779713 | -0.3450606 | 0.7300488 | 0.99945981 |
| AGL | 17.3004535 | -0.276683169 | 0.4454317 | -0.6211573 | 0.53449611 | 0.99945981 |
| TRMT13 | 14.9721974 | -0.175187256 | 0.37435682 | -0.4679686 | 0.63980701 | 0.99945981 |
| DBT | 13.6296456 | -0.820900857 | 0.44662798 | -1.837997 | 0.06606285 | 0.94594189 |
| RTCA | 12.9541162 | -0.343189269 | 0.4090183 | -0.839056 | 0.40143787 | 0.99945981 |
| CDC14A | 89.878033 | 0.050712509 | 0.27412717 | 0.18499629 | 0.85323195 | 0.99945981 |
| VCAM1 | 9.54889251 | -1.90862426 | 0.75819073 | -2.5173405 | 0.01182445 | 0.67187199 |
| SLC30A7 | 49.5462595 | 0.088425834 | 0.4091302 | 0.21613128 | 0.82888542 | 0.99945981 |
| AC093157.1 | 13.2878907 | -0.385900804 | 0.61418936 | -0.6283092 | 0.52980143 | 0.99945981 |
| RNPC3 | 31.3223069 | 0.105316959 | 0.3126438 | 0.33685926 | 0.736223 | 0.99945981 |
| VAV3 | 219.629163 | -0.197836994 | 0.39859701 | -0.4963334 | 0.61965923 | 0.99945981 |
| SLC25A24 | 24.9319009 | 0.460787197 | 0.39774687 | 1.15849359 | 0.24666266 | 0.99945981 |
| FAM102B | 52.1193145 | 0.292137273 | 0.4794256 | 0.60934851 | 0.54229346 | 0.99945981 |
| PRPF38B | 42.0573975 | -0.163460735 | 0.31587128 | -0.5174916 | 0.60481302 | 0.99945981 |
| STXBP3 | 37.0313125 | 0.047809768 | 0.34505513 | 0.1385569 | 0.88980031 | 0.99945981 |
| WDR47 | 12.2678618 | 0.990178858 | 0.41117028 | 2.40819655 | 0.01603155 | 0.74280553 |
| SARS | 16.6922299 | 0.158046011 | 0.35468129 | 0.44560008 | 0.65588615 | 0.99945981 |
| SORT1 | 22.1129891 | 0.266235711 | 0.46257354 | 0.57555326 | 0.56491718 | 0.99945981 |
| PSMA5 | 29.1331804 | -0.197169143 | 0.35835286 | -0.5502095 | 0.58217569 | 0.99945981 |
| GNAI3 | 52.3851225 | 0.656904039 | 0.45370927 | 1.44785236 | 0.14765835 | 0.99945981 |
| AMPD2 | 9.46492365 | 0.380146798 | 0.80915526 | 0.46980699 | 0.63849292 | 0.99945981 |
| AHCYL1 | 22.3395774 | -0.443399095 | 0.37674128 | -1.1769326 | 0.23922241 | 0.99945981 |
| STRIP1 | 9.58815944 | -0.182971174 | 0.51852752 | -0.3528669 | 0.72418826 | 0.99945981 |
| RBM15 | 11.6688004 | 0.248414766 | 0.51233428 | 0.48486852 | 0.62776961 | 0.99945981 |
| LAMTOR5 | 22.6076928 | -0.675657606 | 0.389823 | -1.733242 | 0.08305266 | 0.98529784 |
| CD53 | 124.609341 | -0.353239756 | 0.2782187 | -1.2696478 | 0.20421012 | 0.99945981 |
| LRIF1 | 8.85643186 | -0.100205068 | 0.49634233 | -0.201887 | 0.84000506 | 0.99945981 |
| DRAM2 | 33.4489707 | -0.049364177 | 0.31386217 | -0.1572798 | 0.87502434 | 0.99945981 |
| CEPT1 | 41.3114498 | 0.249102693 | 0.33646653 | 0.7403491 | 0.4590882 | 0.99945981 |

| | | | | | | |
|---|---|---|---|---|---|---|
| DENND2D | 19.6020236 | 0.013521401 | 0.40652449 | 0.03326098 | 0.97346647 | 0.99945981 |
| ATP5PB | 18.9895213 | -0.326775521 | 0.38015478 | -0.8595855 | 0.39001759 | 0.99945981 |
| C1orf162 | 27.3834325 | 0.286812684 | 0.71508672 | 0.40108797 | 0.68835536 | 0.99945981 |
| RAP1A | 177.7917 | 0.069166627 | 0.22858337 | 0.30258818 | 0.76220372 | 0.99945981 |
| INKA2 | 14.5838321 | 0.024448557 | 0.67267574 | 0.03634523 | 0.97100708 | 0.99945981 |
| CTTNBP2NL | 27.7339451 | 0.33594694 | 0.48415917 | 0.69387706 | 0.48775931 | 0.99945981 |
| ST7L | 17.325787 | 0.109556416 | 0.46541075 | 0.23539726 | 0.81390041 | 0.99945981 |
| CAPZA1 | 100.002221 | -0.087604785 | 0.25143025 | -0.3484258 | 0.72752043 | 0.99945981 |
| RHOC | 16.6232085 | 0.355156316 | 0.59757107 | 0.59433319 | 0.55228929 | 0.99945981 |
| SLC16A1-AS1 | 26.04322 | -0.739568901 | 0.40107527 | -1.8439654 | 0.06518819 | 0.9441285 |
| LRIG2 | 23.4718945 | 0.249741583 | 0.30934049 | 0.80733558 | 0.41947317 | 0.99945981 |
| MAGI3 | 26.8498069 | 0.054123433 | 0.608837 | 0.08889642 | 0.92916422 | 0.99945981 |
| PHTF1 | 20.1704053 | -0.520177027 | 0.47663652 | -1.0913495 | 0.27511912 | 0.99945981 |
| RSBN1 | 28.1922591 | -0.500641247 | 0.30984798 | -1.6157641 | 0.10614533 | 0.99945981 |
| PTPN22 | 79.5270741 | -0.653122226 | 0.39630829 | -1.6480156 | 0.09934948 | 0.99945981 |
| HIPK1 | 42.8619748 | 0.154406928 | 0.29300063 | 0.52698497 | 0.59820402 | 0.99945981 |
| TRIM33 | 76.7976247 | 0.231237414 | 0.22600551 | 1.02314945 | 0.30623719 | 0.99945981 |
| CSDE1 | 60.2162431 | 0.30526496 | 0.23807509 | 1.28222131 | 0.19976502 | 0.99945981 |
| SLC22A15 | 21.9449292 | -0.09005955 | 0.44287409 | -0.2033525 | 0.83885953 | 0.99945981 |
| ATP1A1 | 77.6144127 | 0.056421981 | 0.38480813 | 0.14662367 | 0.88342907 | 0.99945981 |
| CD58 | 50.2377351 | -0.144698137 | 0.41172874 | -0.3514405 | 0.72525793 | 0.99945981 |
| CD2 | 44.6038332 | -0.565769425 | 0.47234901 | -1.1977784 | 0.23100331 | 0.99945981 |
| TTF2 | 11.6249283 | -0.190501096 | 0.44288203 | -0.4301396 | 0.6670941 | 0.99945981 |
| MAN1A2 | 71.3540585 | 0.076069152 | 0.31741188 | 0.23965439 | 0.81059819 | 0.99945981 |
| TENT5C | 25.309316 | -0.081896528 | 1.02641926 | -0.0797886 | 0.93640541 | 0.99945981 |
| GDAP2 | 26.9045058 | -0.155784199 | 0.31931523 | -0.4878696 | 0.6256422 | 0.99945981 |
| AL359915.2 | 16.5355015 | 0.409281189 | 0.66210786 | 0.61814881 | 0.53647725 | 0.99945981 |
| NOTCH2 | 75.2832814 | 0.498851345 | 0.32783309 | 1.52166256 | 0.12809365 | 0.99945981 |
| SEC22B | 23.712585 | -0.052267964 | 0.39773716 | -0.1314133 | 0.89544835 | 0.99945981 |
| NBPF26 | 13.9685371 | 1.154006715 | 0.66283044 | 1.74102855 | 0.08167857 | 0.9846219 |
| LINC00623 | 69.6192033 | 0.23148397 | 0.39644231 | 0.5839033 | 0.55928537 | 0.99945981 |
| SRGAP2C | 30.3148762 | 0.468679797 | 0.37264408 | 1.25771433 | 0.20849509 | 0.99945981 |
| NBPF15 | 8.41450811 | -0.663907443 | 0.47572477 | -1.3955705 | 0.16284388 | 0.99945981 |
| SRGAP2B | 39.3215351 | 0.67651936 | 0.4986793 | 1.3566221 | 0.17490132 | 0.99945981 |
| RNF115 | 75.7546089 | -0.545313912 | 0.30344502 | -1.7970765 | 0.07232348 | 0.96034943 |
| POLR3C | 11.2410383 | -0.905659737 | 0.46875492 | -1.9320538 | 0.05335286 | 0.92445778 |
| RBM8A | 12.9617225 | 0.030660067 | 0.47958149 | 0.06393088 | 0.94902526 | 0.99945981 |
| LIX1L | 9.91034744 | -0.010920646 | 0.53611244 | -0.0203701 | 0.98374816 | 0.99945981 |
| TXNIP | 199.797611 | 0.435423841 | 0.46845662 | 0.92948594 | 0.35263731 | 0.99945981 |
| NBPF12 | 15.0990964 | -0.104130665 | 0.41009271 | -0.2539198 | 0.7995575 | 0.99945981 |
| AC242426.2 | 13.1591287 | -0.14688702 | 0.51743375 | -0.283876 | 0.77650542 | 0.99945981 |
| CHD1L | 27.4851698 | -0.164196704 | 0.27924006 | -0.5880127 | 0.55652377 | 0.99945981 |
| GPR89B | 10.9994514 | -0.033172002 | 0.42875821 | -0.0773676 | 0.9383311 | 0.99945981 |
| NBPF11 | 11.9038419 | -0.088890468 | 0.53687613 | -0.1655698 | 0.86849552 | 0.99945981 |
| NBPF14 | 17.3913997 | 0.922461092 | 0.44291147 | 2.08272118 | 0.03727665 | 0.87920079 |
| PDE4DIP | 69.3844882 | -1.80176877 | 0.66052636 | -2.7277772 | 0.00637627 | 0.52554078 |
| NBPF9 | 9.46016948 | 0.410814511 | 0.4563115 | 0.90029401 | 0.36796381 | 0.99945981 |
| AC245297.3 | 87.2727329 | 0.296409649 | 0.37637998 | 0.78752767 | 0.43097304 | 0.99945981 |
| NBPF19 | 34.423779 | 0.672044552 | 0.8392102 | 0.80080599 | 0.42324397 | 0.99945981 |
| OTUD7B | 9.91737349 | 0.1520961 | 0.50766583 | 0.29959885 | 0.76448316 | 0.99945981 |
| VPS45 | 39.0830203 | -0.012028376 | 0.2472589 | -0.0486469 | 0.9612007 | 0.99945981 |
| PLEKHO1 | 41.0059221 | -0.068567363 | 0.50459953 | -0.1358847 | 0.89191242 | 0.99945981 |
| ANP32E | 17.0464884 | 0.905351517 | 0.52239068 | 1.73309278 | 0.08307918 | 0.98529784 |
| APH1A | 12.1574911 | -0.112159142 | 0.47448312 | -0.2363817 | 0.81313647 | 0.99945981 |
| MRPS21 | 16.5924927 | -0.746884667 | 0.41428327 | -1.8028357 | 0.07141402 | 0.95583904 |
| PRPF3 | 22.5845281 | 0.374294514 | 0.33589442 | 1.11432191 | 0.26514111 | 0.99945981 |
| RPRD2 | 62.9831565 | -0.038784553 | 0.24940295 | -0.1555096 | 0.87641959 | 0.99945981 |
| MCL1 | 73.4687545 | 0.978014789 | 0.57308528 | 1.70657811 | 0.0879005 | 0.99640647 |
| ENSA | 38.838521 | -0.138078032 | 0.2865986 | -0.4817819 | 0.62996086 | 0.99945981 |
| GOLPH3L | 18.0959273 | 0.504833666 | 0.48685021 | 1.03693838 | 0.29976457 | 0.99945981 |
| CTSS | 149.764976 | 1.073543204 | 0.618689 | 1.73519039 | 0.08270709 | 0.98529784 |
| ARNT | 34.3444922 | 0.091221578 | 0.30792012 | 0.29625079 | 0.76703856 | 0.99945981 |
| SETDB1 | 22.7668626 | 0.015949101 | 0.36855456 | 0.04327473 | 0.96548253 | 0.99945981 |
| CERS2 | 12.092786 | -0.041492567 | 0.43638756 | -0.0950819 | 0.92424976 | 0.99945981 |
| C1orf56 | 10.6391025 | 0.63333405 | 0.767012 | 0.82571596 | 0.40896523 | 0.99945981 |
| CDC42SE1 | 34.7661616 | 0.059884367 | 0.33359203 | 0.17951379 | 0.85753429 | 0.99945981 |
| GABPB2 | 16.6294441 | 0.5007948 | 0.42317065 | 1.18343464 | 0.23663694 | 0.99945981 |
| PIP5K1A | 48.1725397 | 0.03763418 | 0.31311388 | 0.12019326 | 0.90433005 | 0.99945981 |
| PSMD4 | 17.6177014 | -0.046928435 | 0.41046025 | -0.1143313 | 0.90897521 | 0.99945981 |
| PI4KB | 20.9803003 | 0.538803048 | 0.34055619 | 1.58212671 | 0.11362065 | 0.99945981 |
| PSMB4 | 9.36658087 | -0.185949896 | 0.51625172 | -0.3601923 | 0.71870333 | 0.99945981 |
| POGZ | 59.7932947 | -0.082581209 | 0.27917271 | -0.2958069 | 0.76737757 | 0.99945981 |
| SNX27 | 45.3788171 | 0.488258339 | 0.41020694 | 1.19027324 | 0.23393902 | 0.99945981 |
| THEM4 | 11.1804261 | -1.15705635 | 0.52628806 | -2.1985229 | 0.02791187 | 0.84763724 |
| S100A10 | 79.2212199 | -0.334660335 | 0.4913003 | -0.6811727 | 0.49576224 | 0.99945981 |
| S100A11 | 131.448577 | -0.069472101 | 0.55153123 | -0.1259622 | 0.89976183 | 0.99945981 |
| S100A9 | 462.948593 | 0.565577937 | 1.53029194 | 0.36958826 | 0.7116893 | 0.99945981 |
| S100A8 | 325.902216 | 0.120207711 | 1.40242085 | 0.08571444 | 0.93169343 | 0.99945981 |
| S100A6 | 146.641257 | -0.833531551 | 0.59140401 | -1.4094114 | 0.15871355 | 0.99945981 |
| S100A4 | 154.746597 | 0.118537126 | 0.94336718 | 0.12565322 | 0.90000643 | 0.99945981 |
| CHTOP | 13.7091203 | 0.18695781 | 0.39524829 | 0.47301358 | 0.63620349 | 0.99945981 |
| ILF2 | 11.3335401 | 0.196532835 | 0.49633211 | 0.39597042 | 0.69212685 | 0.99945981 |
| INTS3 | 13.3843672 | 0.450285298 | 0.42545203 | 1.05836914 | 0.28988718 | 0.99945981 |
| GATAD2B | 101.722809 | 0.071422533 | 0.21225765 | 0.3364898 | 0.73650154 | 0.99945981 |
| DENND4B | 33.9727385 | 0.204436985 | 0.37780879 | 0.5411123 | 0.58843018 | 0.99945981 |
| CRTC2 | 10.2047546 | 0.473777197 | 0.50022443 | 0.94712926 | 0.34357292 | 0.99945981 |
| JTB | 16.499584 | -0.050912543 | 0.5177472 | -0.0983348 | 0.92166648 | 0.99945981 |
| RPS27 | 32.8387773 | 0.177728937 | 0.51963095 | 0.34202916 | 0.73232895 | 0.99945981 |
| TPM3 | 69.2238753 | -0.26026121 | 0.36055119 | -0.7218426 | 0.47039125 | 0.99945981 |
| C1orf43 | 19.3114407 | -0.613042311 | 0.38767459 | -1.5813322 | 0.1138021 | 0.99945981 |
| UBAP2L | 28.6676992 | -0.021377419 | 0.34544294 | -0.0618841 | 0.95065514 | 0.99945981 |
| ATP8B2 | 12.8276511 | -0.424617572 | 0.44410298 | -0.9561241 | 0.33900953 | 0.99945981 |
| IL6R | 30.1495788 | 0.511020058 | 0.4039164 | 1.26516292 | 0.20581293 | 0.99945981 |
| ADAR | 53.7690686 | 0.282331627 | 0.34189679 | 0.82578028 | 0.40892874 | 0.99945981 |
| PBXIP1 | 18.4619639 | -0.142821468 | 0.47346548 | -0.3016513 | 0.76291791 | 0.99945981 |
| ZBTB7B | 8.58476604 | 0.513020472 | 0.68125741 | 0.75304938 | 0.45142024 | 0.99945981 |
| SCAMP3 | 10.8032038 | -0.812974584 | 0.48976995 | -1.6599111 | 0.09693233 | 0.99945981 |
| FDPS | 21.8819231 | -0.481758969 | 0.37312606 | -1.2911427 | 0.19665421 | 0.99945981 |
| ASH1L | 191.864082 | 0.134136413 | 0.16934181 | 0.79210453 | 0.42829973 | 0.99945981 |
| YY1AP1 | 32.3960039 | -0.054736742 | 0.31242688 | -0.1751986 | 0.86092361 | 0.99945981 |
| DAP3 | 35.2872422 | 0.319875046 | 0.28960724 | 1.1045133 | 0.26937054 | 0.99945981 |
| GON4L | 42.298198 | -0.087175357 | 0.3842362 | -0.2268796 | 0.82051736 | 0.99945981 |
| RIT1 | 10.7190144 | -0.639991312 | 0.54061376 | -1.1838236 | 0.23648291 | 0.99945981 |
| KHDC4 | 23.000616 | -0.340423834 | 0.34733487 | -0.9801027 | 0.32703544 | 0.99945981 |
| ARHGEF2 | 42.304744 | -0.120148437 | 0.45704423 | -0.2628814 | 0.79264198 | 0.99945981 |
| SSR2 | 26.2145649 | -0.206887572 | 0.47505178 | -0.4355053 | 0.66319569 | 0.99945981 |
| LMNA | 26.1217897 | 0.862242225 | 0.94295096 | 0.91440835 | 0.36050232 | 0.99945981 |
| SEMA4A | 12.9832295 | 0.665720974 | 0.50623819 | 1.31503508 | 0.18849811 | 0.99945981 |
| SMG5 | 16.6987225 | -0.194771079 | 0.42055902 | -0.4631242 | 0.64327531 | 0.99945981 |
| CCT3 | 16.0504409 | -0.305816862 | 0.47512053 | -0.6436616 | 0.51979487 | 0.99945981 |
| MEF2D | 29.3775111 | -0.34083107 | 0.3011024 | -1.131944 | 0.25765796 | 0.99945981 |
| HDGF | 12.9938265 | 0.342266202 | 0.57652883 | 0.5936671 | 0.55273479 | 0.99945981 |
| PRCC | 12.7093114 | -0.465381663 | 0.46092227 | -1.0096749 | 0.31265105 | 0.99945981 |

| | | | | | | |
|---|---|---|---|---|---|---|
| SH2D2A | 13.9872094 | -1.38765542 | 0.70504483 | -1.9681804 | 0.04904728 | 0.91888954 |
| ARHGEF11 | 21.6334036 | -0.560500613 | 0.37423762 | -1.4977132 | 0.13420778 | 0.99945981 |
| FCRL5 | 35.358338 | -3.586742784 | 1.25060646 | -2.8680028 | 0.00413072 | 0.507383 |
| MNDA | 51.7258839 | 0.73659419 | 0.41649871 | 1.76853894 | 0.07697085 | 0.9821652 |
| PYHIN1 | 49.7107203 | -0.783843088 | 0.2886408 | -2.7156351 | 0.00661488 | 0.52803642 |
| IFI16 | 126.562046 | 0.028659838 | 0.33597776 | 0.08530278 | 0.93202068 | 0.99945981 |
| DUSP23 | 13.1012391 | -0.882269227 | 0.61348141 | -1.4381352 | 0.15039569 | 0.99945981 |
| TAGLN2 | 34.8991108 | -0.187509079 | 0.61810697 | -0.3033602 | 0.76161534 | 0.99945981 |
| COPA | 70.7097448 | -0.03597908 | 0.26994452 | -0.1332832 | 0.89396938 | 0.99945981 |
| NCSTN | 9.216499 | 0.190899196 | 0.5443168 | 0.3507134 | 0.72580337 | 0.99945981 |
| SLAMF6 | 35.2712495 | -1.249909842 | 0.47211449 | -2.6474719 | 0.00810961 | 0.58811673 |
| CD84 | 35.4957016 | -0.941370077 | 0.44491384 | -2.115848 | 0.03435773 | 0.87717522 |
| SLAMF1 | 14.3538544 | -0.890159509 | 0.56796569 | -1.5672769 | 0.11704998 | 0.99945981 |
| CD48 | 40.6779828 | 0.289953545 | 0.55271081 | 0.52460263 | 0.59985946 | 0.99945981 |
| SLAMF7 | 20.7104603 | 0.269262993 | 0.42725447 | 0.63021692 | 0.52855267 | 0.99945981 |
| LY9 | 15.3439708 | -0.146627918 | 0.69604732 | -0.210658 | 0.83315417 | 0.99945981 |
| CD244 | 13.6369205 | 0.608738098 | 0.77906236 | 0.78137275 | 0.4345833 | 0.99945981 |
| ARHGAP30 | 53.6729412 | 0.236123436 | 0.32209939 | 0.73307632 | 0.46351188 | 0.99945981 |
| PFDN2 | 14.3145589 | -0.467917903 | 0.47485758 | -0.9853858 | 0.32443462 | 0.99945981 |
| NIT1 | 11.7850405 | -0.299634613 | 0.40951881 | -0.7316749 | 0.46436705 | 0.99945981 |
| UFC1 | 11.3066063 | 0.006276046 | 0.48446909 | 0.01295448 | 0.98966411 | 0.99945981 |
| NDUFS2 | 11.2320292 | 0.014217635 | 0.40315386 | 0.03526603 | 0.97186761 | 0.99945981 |
| FCER1G | 54.4077935 | 0.81868648 | 0.8788495 | 0.93154344 | 0.35157253 | 0.99945981 |
| SDHC | 20.3215381 | -0.229278293 | 0.32929089 | -0.6962789 | 0.4862542 | 0.99945981 |
| FCGR2A | 60.3551218 | 0.2858473 | 0.65057461 | 0.43937666 | 0.66038864 | 0.99945981 |
| HSPA6 | 9.83682219 | 0.20251772 | 0.76454342 | 0.26488714 | 0.7910964 | 0.99945981 |
| FCGR3A | 68.5003436 | 0.702402026 | 0.92015754 | 0.76334975 | 0.44525484 | 0.99945981 |
| FCGR2B | 15.717372 | -1.03925848 | 0.51390048 | -2.0222952 | 0.04314586 | 0.89562588 |
| ATF6 | 93.7361119 | 0.017767975 | 0.33741824 | 0.05265861 | 0.95800392 | 0.99945981 |
| OLFML2B | 13.582608 | -2.242572765 | 0.85453053 | -2.6243331 | 0.00868188 | 0.59011016 |
| UHMK1 | 32.1512869 | -0.157932032 | 0.34455926 | -0.4583596 | 0.64669414 | 0.99945981 |
| MGST3 | 26.9367609 | -0.308488362 | 0.39590452 | -0.7791989 | 0.43586257 | 0.99945981 |
| ALDH9A1 | 11.8325544 | -0.068186052 | 0.58915016 | -0.1157363 | 0.90786155 | 0.99945981 |
| TMCO1 | 20.1976856 | -0.033973129 | 0.38042891 | -0.0893022 | 0.92884177 | 0.99945981 |
| UCK2 | 10.318952 | -0.499847979 | 0.63671101 | -0.7850469 | 0.4324261 | 0.99945981 |
| POGK | 8.14302277 | -0.191715637 | 0.47746698 | -0.4015265 | 0.68803255 | 0.99945981 |
| POU2F1 | 82.383964 | 0.073440023 | 0.41019578 | 0.17903651 | 0.85790903 | 0.99945981 |
| CD247 | 150.637198 | -0.037328857 | 0.42820358 | -0.0871755 | 0.93053202 | 0.99945981 |
| CREG1 | 19.46721 | -0.629263343 | 0.44749388 | -1.4061943 | 0.15966644 | 0.99945981 |
| RCSD1 | 121.668491 | -0.707482052 | 0.36681843 | -1.9286982 | 0.05376834 | 0.92445778 |
| MPZL1 | 23.2757647 | -0.387322092 | 0.31956319 | -1.212036 | 0.22549859 | 0.99945981 |
| MPC2 | 9.45497328 | -0.46042772 | 0.60924939 | -0.7557295 | 0.44981141 | 0.99945981 |
| DCAF6 | 70.4345034 | 0.117708896 | 0.26800608 | 0.43920233 | 0.66051493 | 0.99945981 |
| TIPRL | 13.2787907 | -0.337720294 | 0.37630516 | -0.8974639 | 0.36947143 | 0.99945981 |
| SFT2D2 | 39.5446155 | 0.12801403 | 0.33306384 | 0.38435283 | 0.70071695 | 0.99945981 |
| TBX19 | 10.1690919 | -0.202167099 | 0.50670217 | -0.3989861 | 0.68990348 | 0.99945981 |
| ATP1B1 | 39.9468215 | 0.857736124 | 0.4513776 | 1.90026294 | 0.05739862 | 0.92445778 |
| NME7 | 21.3899561 | -0.117383939 | 0.40430775 | -0.2903331 | 0.77156138 | 0.99945981 |
| BLZF1 | 14.5501142 | 0.442663163 | 0.60136682 | 0.73609508 | 0.46167283 | 0.99945981 |
| C1orf112 | 11.6168412 | 0.67892223 | 0.44611477 | 1.52185551 | 0.12804529 | 0.99945981 |
| SELL | 20.0182313 | 1.481531053 | 0.54043679 | 2.74135862 | 0.00611857 | 0.52554078 |
| SCYL3 | 10.548279 | -0.59739873 | 0.54671157 | -1.0927128 | 0.27451992 | 0.99945981 |
| KIFAP3 | 29.610281 | 0.343399049 | 0.34755701 | 0.9880366 | 0.32313472 | 0.99945981 |
| PRRC2C | 129.244309 | -0.000743328 | 0.23171075 | -0.003208 | 0.99744039 | 0.99947866 |
| VAMP4 | 14.4938964 | 0.358558192 | 0.47633512 | 0.75274356 | 0.45160403 | 0.99945981 |
| SUCO | 47.0208633 | 0.338392645 | 0.33326979 | 1.01537149 | 0.30992878 | 0.99945981 |
| AL645568.1 | 28.7032377 | 0.177318695 | 0.37723662 | 0.47004635 | 0.63832191 | 0.99945981 |
| PRDX6 | 22.2907805 | -0.417959004 | 0.39733431 | -1.0519077 | 0.29284192 | 0.99945981 |
| KLHL20 | 23.353551 | -0.156266302 | 0.40105062 | -0.3896423 | 0.69680104 | 0.99945981 |
| ZBTB37 | 15.4879349 | -0.030833644 | 0.38562657 | -0.0799573 | 0.93627125 | 0.99945981 |
| RC3H1 | 80.6892404 | 0.226734881 | 0.25937953 | 0.8741433 | 0.38204022 | 0.99945981 |
| RABGAP1L | 456.730322 | 0.239158229 | 0.22820145 | 1.04801364 | 0.29463232 | 0.99945981 |
| CACYBP | 10.6377599 | -0.348442053 | 0.49823234 | -0.6993566 | 0.48432923 | 0.99945981 |
| KIAA0040 | 11.1451201 | -0.632042871 | 0.58273713 | -1.0846106 | 0.27809416 | 0.99945981 |
| COP1 | 210.885155 | 0.224347171 | 0.22000518 | 1.01973588 | 0.30785374 | 0.99945981 |
| RASAL2 | 50.7754584 | -0.83824428 | 0.59037269 | -1.4198561 | 0.15564957 | 0.99945981 |
| RALGPS2 | 92.1859446 | -1.314149831 | 0.77832253 | -1.6884386 | 0.09132706 | 0.99640647 |
| FAM20B | 13.2680424 | -0.685542883 | 0.53293021 | -1.2863652 | 0.19831563 | 0.99945981 |
| TOR3A | 8.64443457 | 0.248064717 | 0.51112564 | 0.48533021 | 0.62744213 | 0.99945981 |
| ABL2 | 41.8357339 | -0.184805517 | 0.32019308 | -0.577169 | 0.5638253 | 0.99945981 |
| SOAT1 | 47.5916331 | 0.036432172 | 0.39078865 | 0.0932273 | 0.92572299 | 0.99945981 |
| TOR1AIP2 | 24.0167721 | -0.285445878 | 0.35280819 | -0.8090682 | 0.41847594 | 0.99945981 |
| TOR1AIP1 | 20.8912269 | -0.072911724 | 0.33607089 | -0.2169534 | 0.82824466 | 0.99945981 |
| CEP350 | 107.097683 | 0.262607961 | 0.22663752 | 1.15871351 | 0.24657298 | 0.99945981 |
| QSOX1 | 15.7462381 | 0.216224207 | 0.4259643 | 0.5076111 | 0.61172611 | 0.99945981 |
| ACBD6 | 57.5197268 | -0.085631095 | 0.26660389 | -0.3211922 | 0.74806473 | 0.99945981 |
| XPR1 | 67.678346 | 0.215380784 | 0.31896894 | 0.67524063 | 0.4995229 | 0.99945981 |
| STX6 | 20.7638784 | -0.274100981 | 0.43023947 | -0.6370893 | 0.52406665 | 0.99945981 |
| MR1 | 16.7694558 | -0.228989796 | 0.48534969 | -0.4718037 | 0.63706688 | 0.99945981 |
| IER5 | 13.2484561 | 0.881621943 | 0.72095388 | 1.22285485 | 0.22138453 | 0.99945981 |
| GLUL | 73.2220782 | -0.18831377 | 0.58850373 | -0.3199874 | 0.74897789 | 0.99945981 |
| NPL | 51.1514535 | -0.26029781 | 0.48736089 | -0.5340966 | 0.59327468 | 0.99945981 |
| DHX9 | 46.3631762 | 0.434675508 | 0.28875125 | 1.50536319 | 0.13223073 | 0.99945981 |
| LAMC1 | 13.1046455 | -0.793618149 | 0.75267397 | -1.0543983 | 0.2917006 | 0.99945981 |
| SMG7 | 64.6215456 | 0.530426249 | 0.23507342 | 2.25642798 | 0.02404384 | 0.80320673 |
| NCF2 | 35.8603452 | 0.835262862 | 0.61281248 | 1.3629991 | 0.17288279 | 0.99945981 |
| ARPC5 | 62.3212024 | 0.128652876 | 0.46486493 | 0.27675324 | 0.78196959 | 0.99945981 |
| RGL1 | 93.0529153 | 0.733583988 | 0.70522719 | 1.04020945 | 0.2982426 | 0.99945981 |
| TSEN15 | 13.3904689 | 0.001446856 | 0.41741449 | 0.00346623 | 0.99723435 | 0.99945981 |
| C1orf21 | 41.2763618 | -0.314485626 | 0.59627593 | -0.5274163 | 0.59790454 | 0.99945981 |
| EDEM3 | 44.5751925 | -0.002327762 | 0.32779394 | -0.0071013 | 0.99433403 | 0.99945981 |
| FAM129A | 105.450823 | -0.520988275 | 0.4554331 | -1.1439403 | 0.2526484 | 0.99945981 |
| RNF2 | 18.0914179 | 0.532526889 | 0.33284858 | 1.59990736 | 0.10961914 | 0.99945981 |
| TRMT1L | 14.4230697 | -0.08898179 | 0.47442303 | -0.1875579 | 0.85122323 | 0.99945981 |
| SWT1 | 13.1165504 | -0.748501303 | 0.47347413 | -1.5808705 | 0.11390764 | 0.99945981 |
| IVNS1ABP | 59.9052599 | 0.445492914 | 0.41296402 | 1.07876931 | 0.28069058 | 0.99945981 |
| TPR | 45.6607506 | -0.178501433 | 0.25210156 | -0.7080537 | 0.47891193 | 0.99945981 |
| ODR4 | 21.5661089 | 0.069839204 | 0.32932455 | 0.21206801 | 0.83205397 | 0.99945981 |
| PLA2G4A | 25.3067183 | 1.123246988 | 0.46446779 | 2.41835282 | 0.01559095 | 0.7371189 |
| RGS18 | 16.0449719 | 1.653279008 | 0.69715417 | 2.37146829 | 0.01771757 | 0.74759796 |
| AL390957.1 | 20.9867331 | -0.935926687 | 0.74877752 | -1.2499396 | 0.2113216 | 0.99945981 |
| RGS1 | 67.0903263 | 0.734961783 | 0.53995419 | 1.36115581 | 0.17346446 | 0.99945981 |
| AL136454.1 | 31.1904517 | -0.031199352 | 0.54899815 | -0.0568296 | 0.95468093 | 0.99945981 |
| RGS2 | 27.284564 | 1.266392701 | 0.83698698 | 1.51303752 | 0.13027013 | 0.99945981 |
| UCHL5 | 17.397712 | 0.217495407 | 0.44316715 | 0.49077511 | 0.62358551 | 0.99945981 |
| TROVE2 | 30.0201701 | -0.229862458 | 0.31969966 | -0.718995 | 0.472144 | 0.99945981 |
| CDC73 | 74.3544382 | 0.434799614 | 0.23815546 | 1.82569659 | 0.06789598 | 0.94796811 |
| ZBTB41 | 13.9401568 | 0.097860926 | 0.55766455 | 0.1754835 | 0.86069974 | 0.99945981 |
| DENND1B | 108.270864 | 0.226063861 | 0.40267175 | 0.56140978 | 0.57451822 | 0.99945981 |
| NEK7 | 125.321133 | 0.12449301 | 0.29257486 | 0.42550823 | 0.67046622 | 0.99945981 |
| PTPRC | 704.641993 | 0.387119458 | 0.33680979 | 1.14937115 | 0.25040297 | 0.99945981 |
| AL157402.2 | 12.259415 | -0.273868166 | 0.45881988 | -0.5968969 | 0.55057621 | 0.99945981 |
| MIR181A1HG | 90.6101158 | -1.159767014 | 0.82275241 | -1.4096185 | 0.15865237 | 0.99945981 |

| | | | | | | |
|---|---|---|---|---|---|---|
| CAMSAP2 | 23.9261732 | -0.073281486 | 0.48769549 | -0.1502607 | 0.8805589 | 0.99945981 |
| KIF21B | 11.6336881 | -0.813110293 | 0.58101872 | -1.3994563 | 0.1616762 | 0.99945981 |
| NAV1 | 29.0145351 | 0.417717455 | 0.67147131 | 0.62209278 | 0.53388086 | 0.99945981 |
| IPO9 | 28.691699 | -0.649429963 | 0.3711803 | -1.7496348 | 0.08018136 | 0.9846219 |
| RNPEP | 18.8981826 | 0.615747635 | 0.55443081 | 1.1105942 | 0.26674306 | 0.99945981 |
| ARL8A | 14.246451 | 0.292274455 | 0.43286993 | 0.67520156 | 0.49954772 | 0.99945981 |
| PTPN7 | 21.705755 | -0.582385122 | 0.40790536 | -1.4277457 | 0.15336507 | 0.99945981 |
| PPP1R12B | 58.1510543 | -0.009068407 | 0.26368516 | -0.034391 | 0.97256533 | 0.99945981 |
| KDM5B | 39.6755908 | -1.010413668 | 0.4389331 | -2.3019765 | 0.0213365 | 0.79606018 |
| KLHL12 | 9.2706419 | -0.448014982 | 0.5212238 | -0.8595444 | 0.39004025 | 0.99945981 |
| ADIPOR1 | 11.5994636 | 0.421766101 | 0.52287183 | 0.80663382 | 0.41987748 | 0.99945981 |
| BTG2 | 37.6635481 | 0.234661492 | 0.62416755 | 0.37595913 | 0.70694728 | 0.99945981 |
| ATP2B4 | 54.3876905 | -0.310227738 | 0.36667035 | -0.8460672 | 0.39751528 | 0.99945981 |
| PLEKHA6 | 10.5839847 | -0.009492775 | 0.60994654 | -0.0155633 | 0.98758279 | 0.99945981 |
| PPP1R15B | 11.0166404 | 0.206299072 | 0.52142663 | 0.39564353 | 0.69236802 | 0.99945981 |
| PIK3C2B | 12.4247239 | -0.422079014 | 0.47823888 | -0.8825694 | 0.37746895 | 0.99945981 |
| MDM4 | 63.9194448 | 0.21204921 | 0.25588179 | 0.82869989 | 0.40727425 | 0.99945981 |
| RBBP5 | 12.0194348 | 0.175832361 | 0.42333437 | 0.41535102 | 0.67788496 | 0.99945981 |
| DSTYK | 18.08037 | -0.432498533 | 0.42622781 | -1.0147121 | 0.31024307 | 0.99945981 |
| ELK4 | 28.2159733 | -0.05228064 | 0.32021806 | -0.1632657 | 0.87030921 | 0.99945981 |
| NUCKS1 | 54.9860585 | -0.256415227 | 0.34414107 | -0.7450875 | 0.4562188 | 0.99945981 |
| RAB29 | 23.0775934 | -0.446798149 | 0.55207199 | -0.8093114 | 0.41833606 | 0.99945981 |
| RHEX | 38.9167613 | -2.015724182 | 1.1309366 | -1.7823494 | 0.07469227 | 0.97114097 |
| SRGAP2 | 71.9567099 | 0.780937817 | 0.33277982 | 2.34671025 | 0.01893998 | 0.76950676 |
| IKBKE | 12.4560469 | 0.01398681 | 0.49166893 | 0.02844762 | 0.97730515 | 0.99945981 |
| RASSF5 | 35.2497902 | 0.128090794 | 0.33491421 | 0.38245852 | 0.70212129 | 0.99945981 |
| MAPKAPK2 | 38.0372607 | -0.507424701 | 0.26961272 | -1.8820503 | 0.05982918 | 0.92445778 |
| FCMR | 22.1290258 | -1.827749135 | 0.97806732 | -1.8687355 | 0.06165962 | 0.92949575 |
| PIGR | 12.7329455 | -2.526132804 | 0.73073405 | -3.4569797 | 0.00054627 | 0.23820088 |
| CD55 | 82.0651248 | 1.150301586 | 0.45710344 | 2.51650168 | 0.01185263 | 0.67187199 |
| CR1 | 54.1133399 | -0.425100991 | 0.61368984 | -0.6926968 | 0.48849985 | 0.99945981 |
| CD46 | 49.8354741 | 0.212077404 | 0.2647555 | 0.80103116 | 0.42311361 | 0.99945981 |
| MIR29B2CHC | 45.4254945 | -0.11331665 | 0.37948314 | -0.2986079 | 0.76523926 | 0.99945981 |
| TRAF3IP3 | 59.692954 | 0.387981614 | 0.38038755 | 1.01996403 | 0.30774552 | 0.99945981 |
| HHAT | 11.4749832 | 0.111989044 | 0.75901807 | 0.14754463 | 0.88270215 | 0.99945981 |
| RCOR3 | 49.0217658 | 0.087236825 | 0.27085889 | 0.32207481 | 0.74739602 | 0.99945981 |
| TRAF5 | 25.5375126 | 0.260384587 | 0.5306975 | 0.49064596 | 0.62367687 | 0.99945981 |
| LPGAT1 | 61.6441412 | 0.222990717 | 0.31088233 | 0.71728334 | 0.47319929 | 0.99945981 |
| INTS7 | 19.0580137 | 0.062263839 | 0.45113352 | 0.13801643 | 0.89022743 | 0.99945981 |
| PPP2R5A | 68.0988617 | -0.035966143 | 0.27136501 | -0.1325379 | 0.89455886 | 0.99945981 |
| AL360091.3 | 11.4981419 | -0.687618307 | 0.77696853 | -0.8850015 | 0.37615583 | 0.99945981 |
| TMEM206 | 13.8970572 | 0.244169546 | 0.46691571 | 0.52294138 | 0.60101505 | 0.99945981 |
| NSL1 | 25.4188818 | 0.097706467 | 0.33058142 | 0.29555947 | 0.76756653 | 0.99945981 |
| TATDN3 | 16.121956 | 0.19270679 | 0.42213659 | 0.4565034 | 0.64802802 | 0.99945981 |
| FLVCR1 | 10.8211534 | 0.23787429 | 0.44255421 | 0.53750316 | 0.5909201 | 0.99945981 |
| ANGEL2 | 17.6425832 | 0.403948075 | 0.35494058 | 1.13807239 | 0.25509026 | 0.99945981 |
| RPS6KC1 | 85.1358507 | -0.020777643 | 0.21253377 | -0.0977616 | 0.9221216 | 0.99945981 |
| SMYD2 | 11.0905172 | 0.033955917 | 0.44370902 | 0.07652744 | 0.93899948 | 0.99945981 |
| ESRRG | 25.2876016 | 1.658775622 | 0.78685283 | 2.10811419 | 0.03502111 | 0.87920079 |
| GPATCH2 | 32.7404196 | -0.201981439 | 0.28419967 | -0.7107026 | 0.47726857 | 0.99945981 |
| LYPLAL1 | 27.3733465 | 0.275389032 | 0.42509436 | 0.64783036 | 0.51709467 | 0.99945981 |
| EPRS | 23.3123637 | -0.254917045 | 0.32016672 | -0.7962009 | 0.42591525 | 0.99945981 |
| IARS2 | 15.7064958 | -0.227167857 | 0.39362803 | -0.577113 | 0.56386311 | 0.99945981 |
| RAB3GAP2 | 58.6877926 | -0.077778114 | 0.27751335 | -0.280268 | 0.77927189 | 0.99945981 |
| DUSP10 | 11.9707222 | -0.952110319 | 0.66305886 | -1.4359364 | 0.15102045 | 0.99945981 |
| MIA3 | 23.0475799 | -0.571106935 | 0.33244931 | -1.7178767 | 0.0858191 | 0.99193047 |
| AIDA | 15.2208956 | 0.378656359 | 0.50720914 | 0.74654877 | 0.45533598 | 0.99945981 |
| BROX | 19.9231246 | -0.277739405 | 0.39628846 | -0.7008516 | 0.48339563 | 0.99945981 |
| AC093152.1 | 15.7882669 | -0.240184899 | 0.4046561 | -0.5935531 | 0.55281103 | 0.99945981 |
| TLR5 | 9.91835778 | 0.754445174 | 0.80479479 | 0.93743794 | 0.34853333 | 0.99945981 |
| CAPN2 | 32.6396792 | 0.31480438 | 0.28835653 | 1.09171925 | 0.27495652 | 0.99945981 |
| TP53BP2 | 21.3779995 | 0.407537152 | 0.35791366 | 1.13864654 | 0.25485061 | 0.99945981 |
| FBXO28 | 34.2572699 | -0.486442976 | 0.35003446 | -1.3897002 | 0.16461992 | 0.99945981 |
| DEGS1 | 14.4610183 | -0.282017887 | 0.37973161 | -0.7426769 | 0.45767733 | 0.99945981 |
| NVL | 26.2042065 | -0.025820885 | 0.40003525 | -0.0645465 | 0.94853506 | 0.99945981 |
| CNIH4 | 19.5062192 | -0.436589373 | 0.4448197 | -0.9814974 | 0.32634752 | 0.99945981 |
| WDR26 | 74.0547456 | -0.083102347 | 0.3506991 | -0.236962 | 0.8126863 | 0.99945981 |
| LBR | 38.6565489 | 0.299635834 | 0.3906185 | 0.7670805 | 0.44303366 | 0.99945981 |
| SRP9 | 18.9522016 | -0.216325453 | 0.3749154 | -0.576998 | 0.56394078 | 0.99945981 |
| TMEM63A | 25.965832 | 0.36668396 | 0.43886187 | 0.83553387 | 0.40341718 | 0.99945981 |
| H3F3A | 24.3493317 | 0.531659107 | 0.32215143 | 1.65033912 | 0.09887359 | 0.99945981 |
| ACBD3 | 30.6536401 | -0.030533162 | 0.33224815 | -0.0918987 | 0.92677855 | 0.99945981 |
| PARP1 | 27.9723527 | -1.310070807 | 0.68545044 | -1.9112553 | 0.05597178 | 0.92445778 |
| ITPKB | 50.5907384 | -0.392182585 | 0.32083249 | -1.2223905 | 0.22155999 | 0.99945981 |
| COQ8A | 10.4894399 | -0.403928601 | 0.47122074 | -0.8571961 | 0.39133649 | 0.99945981 |
| CDC42BPA | 20.71633 | -0.020510856 | 0.44463621 | -0.0461295 | 0.96320701 | 0.99945981 |
| ZNF678 | 20.4096515 | -0.687534488 | 0.47774464 | -1.4391255 | 0.15011497 | 0.99945981 |
| ARF1 | 42.0287256 | -0.26024836 | 0.32568684 | -0.7990755 | 0.42424666 | 0.99945981 |
| GUK1 | 36.4460455 | -0.411945269 | 0.34273462 | -1.2019366 | 0.22938811 | 0.99945981 |
| RAB4A | 13.4228376 | -0.468108999 | 0.40693575 | -1.1503266 | 0.2500094 | 0.99945981 |
| NUP133 | 20.1931654 | -0.021096787 | 0.3254583 | -0.0648218 | 0.9483159 | 0.99945981 |
| ABCB10 | 16.0745391 | 0.308586278 | 0.3559646 | 0.86690158 | 0.38599594 | 0.99945981 |
| GALNT2 | 50.6513628 | 0.118343127 | 0.3232117 | 0.36614741 | 0.71425509 | 0.99945981 |
| COG2 | 14.1344357 | -0.11364726 | 0.38348391 | -0.2963547 | 0.76695922 | 0.99945981 |
| TTC13 | 43.9430184 | -0.028797817 | 0.26802186 | -0.1074458 | 0.91443534 | 0.99945981 |
| GNPAT | 21.833306 | 0.21237298 | 0.33128374 | 0.64106068 | 0.52148326 | 0.99945981 |
| EGLN1 | 32.6293951 | -0.811342267 | 0.40488767 | -2.00387 | 0.04508398 | 0.90096666 |
| DISC1 | 100.9011 | 0.404120011 | 0.46708643 | 0.86519322 | 0.38693274 | 0.99945981 |
| SIPA1L2 | 26.6595349 | -0.656221231 | 0.52613642 | -1.2472454 | 0.21230753 | 0.99945981 |
| PCNX2 | 69.7478942 | -0.026645314 | 0.32764505 | -0.0813237 | 0.93518451 | 0.99945981 |
| TARBP1 | 17.7802788 | -0.353326411 | 0.49284644 | -0.7169097 | 0.4734298 | 0.99945981 |
| IRF2BP2 | 27.4395412 | -0.500785147 | 0.33548175 | -1.4927344 | 0.13550673 | 0.99945981 |
| TOMM20 | 17.4479246 | -0.218111607 | 0.52356985 | -0.4165855 | 0.67698162 | 0.99945981 |
| ARID4B | 188.119833 | 0.191160741 | 0.24697118 | 0.77402043 | 0.43891871 | 0.99945981 |
| B3GALNT2 | 13.5008116 | 0.441907042 | 0.53689189 | 0.82308385 | 0.41046031 | 0.99945981 |
| LYST | 245.689259 | 0.957869077 | 0.38241886 | 2.50476419 | 0.01225331 | 0.67187199 |
| GPR137B | 35.7435053 | -0.173533467 | 0.40699542 | -0.426377 | 0.66983319 | 0.99945981 |
| ERO1B | 28.7183054 | -0.198054603 | 0.42379735 | -0.4673333 | 0.64026146 | 0.99945981 |
| LGALS8 | 22.056132 | -0.446834809 | 0.37724629 | -1.1844644 | 0.23622927 | 0.99945981 |
| HEATR1 | 16.6569832 | -0.119694576 | 0.46544631 | -0.2571609 | 0.7970546 | 0.99945981 |
| MTR | 42.1674077 | -0.12036446 | 0.34361436 | -0.3502894 | 0.72612153 | 0.99945981 |
| KMO | 22.7551496 | -0.544081523 | 0.58615588 | -0.9282199 | 0.35329354 | 0.99945981 |
| CEP170 | 112.607101 | -0.090176514 | 0.3757201 | -0.2400098 | 0.81032264 | 0.99945981 |
| SDCCAG8 | 111.546104 | -0.133612413 | 0.41968996 | -0.3183598 | 0.75021203 | 0.99945981 |
| AKT3 | 180.890269 | -0.198156865 | 0.50476674 | -0.3925712 | 0.69463624 | 0.99945981 |
| ZBTB18 | 11.8980102 | 0.595620602 | 0.46209386 | 1.28896023 | 0.19741191 | 0.99945981 |
| ADSS | 26.184058 | -0.071152987 | 0.40727643 | -0.1747044 | 0.8613119 | 0.99945981 |
| DESI2 | 20.233335 | -0.227171012 | 0.41495794 | -0.5474555 | 0.58406582 | 0.99945981 |
| COX20 | 15.840534 | -1.011591267 | 0.44748274 | -2.2606263 | 0.02378241 | 0.80320673 |
| HNRNPU | 111.162154 | 0.295341962 | 0.22476248 | 1.31401811 | 0.18884011 | 0.99945981 |
| EFCAB2 | 15.8255486 | -0.289092438 | 0.93811345 | -0.3081636 | 0.75795783 | 0.99945981 |
| SMYD3 | 207.020099 | 0.207667419 | 0.36441825 | 0.5698601 | 0.56877259 | 0.99945981 |
| CNST | 47.667736 | 0.256260317 | 0.46063021 | 0.55632547 | 0.57798838 | 0.99945981 |

| | | | | | | |
|---|---|---|---|---|---|---|
| AHCTF1 | 48.8092753 | -1.004050687 | 0.55934564 | -1.7950452 | 0.07264651 | 0.96092152 |
| ZNF124 | 22.9034358 | -0.308462151 | 0.3585303 | -0.8603517 | 0.3895952 | 0.99945981 |
| AL390728.6 | 14.4796506 | 0.657735896 | 0.4505512 | 1.45984717 | 0.14433208 | 0.99945981 |
| NLRP3 | 12.0358788 | 0.925894618 | 0.93293309 | 0.99245555 | 0.32097535 | 0.99945981 |
| SH3YL1 | 11.7755066 | 0.211636778 | 0.42343464 | 0.49980978 | 0.61720902 | 0.99945981 |
| ACP1 | 9.48838957 | -0.040405852 | 0.45566141 | -0.0886752 | 0.92934007 | 0.99945981 |
| EIPR1 | 22.2980055 | -0.49984373 | 0.39582545 | -1.2627882 | 0.20666529 | 0.99945981 |
| TRAPPC12 | 40.4864839 | 0.421903363 | 0.32482091 | 1.29887993 | 0.19398514 | 0.99945981 |
| ADI1 | 10.6977195 | -1.069394856 | 0.70635239 | -1.5139679 | 0.13003398 | 0.99945981 |
| RPS7 | 75.011404 | -0.272761224 | 0.56372473 | -0.4838553 | 0.62848853 | 0.99945981 |
| RNF144A | 18.3890443 | -0.551108088 | 0.68124513 | -0.8089718 | 0.41853138 | 0.99945981 |
| LINC00299 | 13.2599224 | 0.323635162 | 0.99671007 | 0.32470342 | 0.74540555 | 0.99945981 |
| ID2 | 40.356566 | -0.148460644 | 0.64468264 | -0.2302848 | 0.81787043 | 0.99945981 |
| KIDINS220 | 55.4340182 | 0.369996252 | 0.26607087 | 1.39059286 | 0.16434892 | 0.99945981 |
| ITGB1BP1 | 16.288185 | 0.106421555 | 0.39219865 | 0.27134605 | 0.78612489 | 0.99945981 |
| CPSF3 | 8.54398977 | 0.140678477 | 0.43761915 | 0.32146326 | 0.74785935 | 0.99945981 |
| IAH1 | 11.2584455 | -0.10571702 | 0.4208142 | -0.2512202 | 0.80164388 | 0.99945981 |
| ADAM17 | 65.3944995 | 0.283086215 | 0.34634188 | 0.81736064 | 0.41372236 | 0.99945981 |
| YWHAQ | 35.0506888 | 0.130701082 | 0.35809058 | 0.36499447 | 0.71511553 | 0.99945981 |
| TAF1B | 26.1125193 | -0.141429983 | 0.44854252 | -0.3153101 | 0.75252622 | 0.99945981 |
| HPCAL1 | 56.0370067 | 0.128710163 | 0.30778218 | 0.41818589 | 0.67581121 | 0.99945981 |
| ODC1 | 10.3060976 | -0.903451439 | 0.55326362 | -1.6329493 | 0.10247966 | 0.99945981 |
| NOL10 | 28.0390817 | -0.177087631 | 0.31336047 | -0.5651243 | 0.57198919 | 0.99945981 |
| PDIA6 | 32.677512 | -0.39820589 | 0.42109951 | -0.9456337 | 0.34433544 | 0.99945981 |
| PQLC3 | 8.66648534 | 0.066033681 | 0.49895481 | 0.13234401 | 0.8947122 | 0.99945981 |
| ROCK2 | 61.1000213 | 0.236103927 | 0.28109338 | 0.83994837 | 0.40093733 | 0.99945981 |
| LPIN1 | 32.6393329 | -0.649459523 | 0.27958004 | -2.3229824 | 0.0201801 | 0.78749881 |
| TRIB2 | 7.70341432 | -0.226696561 | 0.62012099 | -0.3655683 | 0.71468725 | 0.99945981 |
| NBAS | 75.7751279 | -0.004678976 | 0.24322463 | -0.0192373 | 0.98465183 | 0.99945981 |
| DDX1 | 18.305314 | -0.514330194 | 0.3679888 | -1.3976789 | 0.1622095 | 0.99945981 |
| FAM49A | 125.585437 | 0.21707474 | 0.59067296 | 0.36750411 | 0.71324303 | 0.99945981 |
| SMC6 | 33.3136004 | -0.699610815 | 0.42599354 | -1.6423038 | 0.10052705 | 0.99945981 |
| GEN1 | 14.1551861 | -1.555005095 | 0.78946494 | -1.9696949 | 0.04887334 | 0.91888954 |
| LAPTM4A | 33.4814654 | -0.266024889 | 0.43660738 | -0.6093 | 0.54232559 | 0.99945981 |
| PUM2 | 147.508267 | 0.318992097 | 0.20241524 | 1.57592927 | 0.1150421 | 0.99945981 |
| RHOB | 18.0527503 | 1.131946602 | 0.42965777 | 2.63453072 | 0.00842537 | 0.58811673 |
| HS1BP3 | 9.08928691 | -0.690986981 | 0.4828977 | -1.4309179 | 0.15245373 | 0.99945981 |
| LDAH | 28.8321353 | -0.465319504 | 0.40848941 | -1.1391226 | 0.25465204 | 0.99945981 |
| ATAD2B | 96.2328509 | 0.081985677 | 0.23065185 | 0.35545207 | 0.72225095 | 0.99945981 |
| UBXN2A | 19.7277429 | -0.486658525 | 0.47562605 | -1.0231957 | 0.30621533 | 0.99945981 |
| SF3B6 | 12.0165246 | -0.121078388 | 0.54266916 | -0.2231164 | 0.82344492 | 0.99945981 |
| FAM228B | 14.597938 | -1.195748112 | 0.54648541 | -2.1880696 | 0.02866453 | 0.84763724 |
| ITSN2 | 143.905064 | -0.112917729 | 0.30499613 | -0.3702268 | 0.71121354 | 0.99945981 |
| NCOA1 | 179.688199 | 0.337812952 | 0.26838789 | 1.25867434 | 0.20814798 | 0.99945981 |
| ADCY3 | 12.6984351 | -1.100467208 | 0.44883075 | -2.4518534 | 0.01421226 | 0.7227195 |
| DNMT3A | 35.2123816 | -0.074222888 | 0.45716612 | -0.1623543 | 0.87102685 | 0.99945981 |
| DTNB | 57.6049565 | -0.191741193 | 0.4031266 | -0.4756352 | 0.63433431 | 0.99945981 |
| ASXL2 | 69.8327263 | -0.129129169 | 0.22207065 | -0.5814779 | 0.56091843 | 0.99945981 |
| RAB10 | 163.257672 | 0.507160806 | 0.26572296 | 1.9086074 | 0.05631276 | 0.92445778 |
| HADHA | 33.1183611 | -0.256419685 | 0.35353589 | -0.7253003 | 0.46826783 | 0.99945981 |
| HADHB | 43.4489666 | -0.058801694 | 0.32778122 | -0.1793931 | 0.85762903 | 0.99945981 |
| SELENOI | 22.2198545 | 0.099912658 | 0.34794076 | 0.28715422 | 0.77399424 | 0.99945981 |
| OST4 | 18.4227099 | -0.528998285 | 0.57742417 | -0.9161346 | 0.35959628 | 0.99945981 |
| ATRAID | 8.8757926 | -0.631057521 | 0.55458154 | -1.1378985 | 0.25516286 | 0.99945981 |
| GTF3C2 | 19.9132244 | -0.186577248 | 0.38602942 | -0.4833239 | 0.62886576 | 0.99945981 |
| PPM1G | 32.3057937 | -0.229753283 | 0.4491856 | -0.5114885 | 0.60900902 | 0.99945981 |
| NRBP1 | 15.2907519 | -0.076215006 | 0.50425087 | -0.151145 | 0.87986132 | 0.99945981 |
| ZNF512 | 20.4988459 | 0.409274643 | 0.32507679 | 1.25900913 | 0.20802703 | 0.99945981 |
| SLC4A1AP | 7.95614281 | -0.032072372 | 0.45008803 | -0.071258 | 0.94319242 | 0.99945981 |
| MRPL33 | 15.9903556 | -0.184847084 | 0.49174588 | -0.3758996 | 0.70699153 | 0.99945981 |
| RBKS | 10.6405778 | -0.313277399 | 0.67783639 | -0.4621726 | 0.64395756 | 0.99945981 |
| BABAM2 | 126.99163 | -0.278075619 | 0.21510639 | -1.2927353 | 0.19610262 | 0.99945981 |
| FOSL2 | 53.5517967 | 0.731859 | 0.66836129 | 1.09500506 | 0.27351442 | 0.99945981 |
| PLB1 | 10.730892 | -0.155694468 | 0.71844708 | -0.2167097 | 0.82843457 | 0.99945981 |
| PPP1CB | 55.4158902 | 0.659001883 | 0.51342435 | 1.28354233 | 0.19930214 | 0.99945981 |
| TRMT61B | 8.96940521 | -1.116971882 | 0.63593371 | -1.7564282 | 0.07901533 | 0.98446503 |
| WDR43 | 25.0631843 | -0.544590768 | 0.36123353 | -1.5075864 | 0.1316604 | 0.99945981 |
| CLIP4 | 48.4360394 | 0.385573042 | 0.57107339 | 0.67517249 | 0.49956619 | 0.99945981 |
| YPEL5 | 21.0473566 | -0.002327168 | 0.51160684 | -0.0045487 | 0.99637064 | 0.99945981 |
| LBH | 24.5765958 | -1.223285359 | 0.68612115 | -1.7828999 | 0.07460259 | 0.97114097 |
| LCLAT1 | 31.8035446 | -0.684436358 | 0.59337783 | -1.1534579 | 0.24872248 | 0.99945981 |
| DPY30 | 8.9021375 | -0.118632329 | 0.54689539 | -0.2169196 | 0.828271 | 0.99945981 |
| SPAST | 30.4570665 | 0.242044377 | 0.32642971 | 0.74149004 | 0.45839637 | 0.99945981 |
| SLC30A6 | 17.7025544 | 0.125750244 | 0.3901219 | 0.32233578 | 0.74719833 | 0.99945981 |
| NLRC4 | 16.2840611 | 0.819929214 | 0.52817597 | 1.55237887 | 0.1205716 | 0.99945981 |
| YIPF4 | 34.9439313 | 0.003430794 | 0.30657635 | 0.01119067 | 0.99107133 | 0.99945981 |
| BIRC6 | 243.553963 | -0.074897772 | 0.18434673 | -0.4062875 | 0.68453137 | 0.99945981 |
| TTC27 | 40.4101679 | -1.336265409 | 1.02188989 | -1.3076413 | 0.19099501 | 0.99945981 |
| LINC00486 | 29.3342793 | 0.663856624 | 0.37081777 | 1.79025031 | 0.07341368 | 0.96364229 |
| RASGRP3 | 24.649573 | -0.531237836 | 1.06871067 | -0.4970829 | 0.61913057 | 0.99945981 |
| LINC01320 | 9.68296443 | -0.229498854 | 0.55563075 | -0.413042 | 0.67957582 | 0.99945981 |
| CRIM1 | 25.9808108 | -0.106579335 | 0.46830522 | -0.2275852 | 0.81996873 | 0.99945981 |
| FEZ2 | 42.7977759 | 0.038965811 | 0.36700058 | 0.1061737 | 0.91544453 | 0.99945981 |
| STRN | 47.4846661 | 0.52566069 | 0.25560742 | 2.0565158 | 0.03973283 | 0.89132455 |
| HEATR5B | 59.7691708 | 0.092737811 | 0.41594068 | 0.22295922 | 0.82356724 | 0.99945981 |
| EIF2AK2 | 37.8424917 | 0.718051092 | 0.65079104 | 1.10335123 | 0.26987467 | 0.99945981 |
| CEBPZOS | 11.0106891 | -0.878859345 | 0.46556071 | -1.8877438 | 0.05906035 | 0.92445778 |
| CEBPZ | 14.6367625 | 0.103181732 | 0.38811834 | 0.26585121 | 0.7903538 | 0.99945981 |
| NDUFAF7 | 11.4738944 | -0.301305139 | 0.44046648 | -0.6840592 | 0.4939378 | 0.99945981 |
| PRKD3 | 44.1512213 | -1.059907018 | 0.67793378 | -1.5634374 | 0.11794975 | 0.99945981 |
| AC006369.1 | 18.2339704 | 1.401039719 | 0.60742649 | 2.30651731 | 0.02108174 | 0.79476944 |
| CDC42EP3 | 24.6766734 | 1.301767768 | 0.62255685 | 2.09100224 | 0.03652786 | 0.87920079 |
| CYP1B1 | 17.3699679 | -0.825091531 | 1.10729111 | -0.7451442 | 0.45618456 | 0.99945981 |
| ATL2 | 28.8066596 | -0.31049916 | 0.38654293 | -0.8032721 | 0.42181747 | 0.99945981 |
| HNRNPLL | 37.6624757 | 0.109003269 | 0.74146487 | 0.1470107 | 0.88312357 | 0.99945981 |
| GALM | 36.3673384 | -0.501816831 | 0.54198655 | -0.9258843 | 0.35450611 | 0.99945981 |
| SRSF7 | 47.4176779 | 0.559828529 | 0.25633387 | 2.1839819 | 0.02896358 | 0.84763724 |
| DHX57 | 16.6558464 | 0.492754447 | 0.39262812 | 1.25501567 | 0.20947307 | 0.99945981 |
| SOS1 | 162.101147 | -0.060468844 | 0.19044454 | -0.3175142 | 0.75085348 | 0.99945981 |
| MAP4K3 | 83.2396692 | -0.545586607 | 0.5365721 | -1.0168002 | 0.3092485 | 0.99945981 |
| AC007388.1 | 22.3211832 | -1.218684495 | 0.72323674 | -1.6850423 | 0.09198041 | 0.99640647 |
| THUMPD2 | 14.7877301 | -0.002176536 | 0.46375432 | -0.0046933 | 0.99625531 | 0.99945981 |
| SLC8A1-AS1 | 15.0149565 | 0.728054038 | 0.53355217 | 1.36454143 | 0.17239722 | 0.99945981 |
| SLC8A1 | 400.040273 | 1.21252714 | 0.66187764 | 1.83195059 | 0.06695878 | 0.94594189 |
| EML4 | 201.787796 | -0.352774309 | 0.3948788 | -0.8933736 | 0.37165712 | 0.99945981 |
| COX7A2L | 22.5148993 | 0.089351763 | 0.40472459 | 0.22077177 | 0.82527015 | 0.99945981 |
| MTA3 | 23.0682427 | -0.292054739 | 0.40745945 | -0.7167701 | 0.47351598 | 0.99945981 |
| ZFP36L2 | 119.551777 | -0.124375485 | 0.39687613 | -0.3133862 | 0.7539873 | 0.99945981 |
| THADA | 73.2344808 | -0.387017255 | 0.31537586 | -1.227162 | 0.21976172 | 0.99945981 |
| LRPPRC | 39.919065 | 0.177541072 | 0.36690093 | 0.48389377 | 0.62846126 | 0.99945981 |
| PPM1B | 70.6600406 | -0.170233385 | 0.26084643 | -0.6526192 | 0.51400182 | 0.99945981 |
| SLC3A1 | 11.9328999 | -0.437439384 | 0.69218855 | -0.6319656 | 0.52740933 | 0.99945981 |
| PREPL | 20.6483868 | 0.216014538 | 0.3848618 | 0.5612782 | 0.5746079 | 0.99945981 |

| | | | | | | |
|---|---|---|---|---|---|---|
| CAMKMT | 112.490248 | 0.278154728 | 0.29645454 | 0.93827111 | 0.3481051 | 0.99945981 |
| SRBD1 | 77.7385937 | 0.012947023 | 0.32005656 | 0.0404523 | 0.96773254 | 0.99945981 |
| PRKCE | 133.77145 | -0.788666149 | 0.53714082 | -1.468267 | 0.14203171 | 0.99945981 |
| EPAS1 | 20.1222434 | -0.681823184 | 0.45126514 | -1.5109148 | 0.13081016 | 0.99945981 |
| RHOQ | 56.5981797 | 0.208170389 | 0.32269308 | 0.64510336 | 0.51886021 | 0.99945981 |
| AC018682.1 | 11.8213327 | 0.332143251 | 0.45721076 | 0.72645546 | 0.4675596 | 0.99945981 |
| PIGF | 19.278127 | -0.509602989 | 0.3922475 | -1.2991873 | 0.19387964 | 0.99945981 |
| SOCS5 | 14.2745699 | -0.149365268 | 0.44552215 | -0.3352589 | 0.73742979 | 0.99945981 |
| MCFD2 | 12.8493974 | -0.687474639 | 0.46243169 | -1.4866512 | 0.13710695 | 0.99945981 |
| TTC7A | 45.7318095 | 0.232845621 | 0.38834248 | 0.59958834 | 0.54878062 | 0.99945981 |
| EPCAM | 9.24726217 | -0.445481688 | 0.64669356 | -0.6888606 | 0.49091102 | 0.99945981 |
| MSH2 | 10.2701355 | 0.584524248 | 0.6717445 | 0.87015859 | 0.38421374 | 0.99945981 |
| FBXO11 | 142.638455 | 0.035075961 | 0.20380372 | 0.17210658 | 0.86335374 | 0.99945981 |
| FOXN2 | 59.2177453 | 0.08205676 | 0.27914939 | 0.29395285 | 0.76879393 | 0.99945981 |
| PPP1R21 | 57.9419532 | -0.015786878 | 0.29588848 | -0.0533541 | 0.95744974 | 0.99945981 |
| GPR75-ASB3 | 12.9477232 | -1.045620452 | 0.60419583 | -1.7305986 | 0.08352338 | 0.9868985 |
| ERLEC1 | 12.1242895 | -1.213845553 | 0.52474029 | -2.3132311 | 0.02070994 | 0.78985396 |
| PSME4 | 84.4375578 | 0.182475437 | 0.22536648 | 0.80968311 | 0.41812233 | 0.99945981 |
| ACYP2 | 83.0853832 | 0.045996199 | 0.38603551 | 0.11915018 | 0.90515638 | 0.99945981 |
| SPTBN1 | 35.5046876 | -0.028720127 | 0.40915712 | -0.0701934 | 0.94403973 | 0.99945981 |
| RTN4 | 118.819507 | 0.214557526 | 0.23872441 | 0.89876659 | 0.368777 | 0.99945981 |
| RPS27A | 133.397218 | 0.139742058 | 0.5944028 | 0.23509657 | 0.81413378 | 0.99945981 |
| AC012358.3 | 10.9569722 | 0.694604436 | 0.6692894 | 1.03782376 | 0.29935211 | 0.99945981 |
| CCDC88A | 139.48766 | -0.258058853 | 0.40926622 | -0.6305403 | 0.52834114 | 0.99945981 |
| PPP4R3B | 54.504997 | 0.052755423 | 0.27281676 | 0.1933731 | 0.84666678 | 0.99945981 |
| PNPT1 | 20.2674813 | 0.78056268 | 0.68874587 | 1.13331014 | 0.25708404 | 0.99945981 |
| VRK2 | 54.8396234 | 0.309270441 | 0.36155884 | 0.85538067 | 0.39234044 | 0.99945981 |
| FANCL | 15.9183748 | -0.038972779 | 0.56714637 | -0.0687173 | 0.94521463 | 0.99945981 |
| BCL11A | 32.9577677 | -0.28373264 | 0.75000185 | -0.3783093 | 0.70520087 | 0.99945981 |
| PAPOLG | 22.5620057 | 0.185003482 | 0.30678046 | 0.60304845 | 0.54647646 | 0.99945981 |
| REL | 90.9557035 | 0.358367306 | 0.35197188 | 1.01817029 | 0.30859704 | 0.99945981 |
| PUS10 | 13.0695219 | 0.467303981 | 0.4084996 | 1.1439521 | 0.2526435 | 0.99945981 |
| PEX13 | 12.133221 | -0.293662378 | 0.46129451 | -0.636605 | 0.52438219 | 0.99945981 |
| KIAA1841 | 12.5359617 | -0.235014926 | 0.50893765 | -0.4617755 | 0.64424234 | 0.99945981 |
| USP34 | 205.795181 | 0.093553933 | 0.21111304 | 0.44314616 | 0.65766002 | 0.99945981 |
| XPO1 | 97.6796802 | -0.045218546 | 0.43580123 | -0.1037596 | 0.91736016 | 0.99945981 |
| CCT4 | 20.3899727 | -0.030053035 | 0.33133857 | -0.0907019 | 0.92772947 | 0.99945981 |
| COMMD1 | 57.3867612 | 0.001146141 | 0.26456391 | 0.00433219 | 0.99654342 | 0.99945981 |
| B3GNT2 | 9.84672967 | 1.000665839 | 0.44123811 | 2.26785906 | 0.0233378 | 0.80320673 |
| EHBP1 | 48.9628522 | 0.343650758 | 0.46601216 | 0.73742873 | 0.46086167 | 0.99945981 |
| WDPCP | 55.6375833 | 0.135037373 | 0.36195847 | 0.37307422 | 0.70909321 | 0.99945981 |
| MDH1 | 17.4338002 | 0.030364546 | 0.47157959 | 0.06438902 | 0.94866048 | 0.99945981 |
| UGP2 | 50.9621378 | -1.014027023 | 0.58931724 | -1.7206811 | 0.08530871 | 0.99193047 |
| VPS54 | 55.511662 | 0.313623179 | 0.30638896 | 1.02361123 | 0.30601894 | 0.99945981 |
| PELI1 | 120.336463 | 1.43436008 | 0.629339 | 2.27915332 | 0.02265795 | 0.80320673 |
| AFTPH | 70.5101743 | 0.108712868 | 0.26351863 | 0.41254339 | 0.67994119 | 0.99945981 |
| SERTAD2 | 25.961536 | -0.080468937 | 0.38124666 | -0.2110679 | 0.83283428 | 0.99945981 |
| CEP68 | 13.6191672 | 0.858401862 | 0.76354834 | 1.12422726 | 0.26091663 | 0.99945981 |
| RAB1A | 118.011862 | 0.510828638 | 0.24914031 | 2.05036526 | 0.0403288 | 0.89562588 |
| ACTR2 | 143.881971 | 0.053653392 | 0.31627733 | 0.16964033 | 0.865293 | 0.99945981 |
| SPRED2 | 27.7767351 | -0.219166805 | 0.53373205 | -0.4106308 | 0.6813433 | 0.99945981 |
| C1D | 10.7938114 | 0.891962535 | 0.39519578 | 2.25701432 | 0.02400718 | 0.80320673 |
| PLEK | 47.2204075 | 0.672769157 | 0.54024212 | 1.24531046 | 0.21301765 | 0.99945981 |
| APLF | 14.8945854 | -0.623731032 | 0.45462901 | -1.3719561 | 0.17007711 | 0.99945981 |
| ARHGAP25 | 102.444597 | -0.360766086 | 0.40731106 | -0.8857262 | 0.37576507 | 0.99945981 |
| GFPT1 | 37.3490906 | -0.533646371 | 0.40653114 | -1.3126826 | 0.18928991 | 0.99945981 |
| NFU1 | 8.97113363 | -0.229629379 | 0.47022545 | -0.4883389 | 0.62530982 | 0.99945981 |
| AAK1 | 170.875464 | -0.194895129 | 0.52634572 | -0.3702797 | 0.71117411 | 0.99945981 |
| ANXA4 | 40.2028161 | 0.180739101 | 0.40040593 | 0.45138967 | 0.65170873 | 0.99945981 |
| GMCL1 | 25.4887959 | -0.113386175 | 0.34036354 | -0.3331325 | 0.73903423 | 0.99945981 |
| MXD1 | 52.5431422 | -0.04411141 | 0.59911387 | -0.0736278 | 0.94130658 | 0.99945981 |
| PCBP1 | 36.5120492 | -0.208001245 | 0.50362086 | -0.4130116 | 0.67959813 | 0.99945981 |
| SNRPG | 14.2139377 | 0.509617832 | 0.44298909 | 1.15040718 | 0.24997621 | 0.99945981 |
| NAGK | 27.7623164 | 0.550564092 | 0.38174406 | 1.44223355 | 0.1492365 | 0.99945981 |
| MCEE | 11.079867 | -0.412299664 | 0.47484378 | -0.8682849 | 0.38523841 | 0.99945981 |
| MPHOSPH10 | 11.018364 | -0.119838179 | 0.40307415 | -0.2973105 | 0.76622946 | 0.99945981 |
| ZNF638 | 148.226919 | -0.059957815 | 0.22256733 | -0.2693918 | 0.78762819 | 0.99945981 |
| DYSF | 32.7799639 | -0.795108658 | 0.58468411 | -1.3598944 | 0.17386334 | 0.99945981 |
| EXOC6B | 94.1211266 | 0.611188531 | 0.4167911 | 1.46641455 | 0.14253538 | 0.99945981 |
| SFXN5 | 16.4599916 | -0.09906267 | 0.45916506 | -0.2157452 | 0.82918635 | 0.99945981 |
| CCT7 | 13.7129469 | -0.074293466 | 0.40161228 | -0.184988 | 0.85323842 | 0.99945981 |
| ALMS1 | 22.5970558 | -0.030426513 | 0.44627636 | -0.0681786 | 0.94564344 | 0.99945981 |
| DUSP11 | 8.35068979 | 0.123434384 | 0.47741265 | 0.25854863 | 0.79598353 | 0.99945981 |
| STAMBP | 17.0499946 | -0.308965196 | 0.39763154 | -0.7770138 | 0.43715062 | 0.99945981 |
| DGUOK | 19.5320453 | -0.524114743 | 0.37869946 | -1.383986 | 0.16636275 | 0.99945981 |
| TET3 | 77.9238442 | 0.139180404 | 0.2873 | 0.48444276 | 0.62807168 | 0.99945981 |
| MOB1A | 48.8485158 | 0.260959197 | 0.33957567 | 0.76848613 | 0.44219843 | 0.99945981 |
| MTHFD2 | 11.613223 | -0.497030241 | 0.55321792 | -0.8984348 | 0.36895379 | 0.99945981 |
| MOGS | 8.56873691 | -0.472415473 | 0.55644961 | -0.8489816 | 0.39589154 | 0.99945981 |
| AUP1 | 9.5655338 | 0.052967766 | 0.60148522 | 0.08806163 | 0.9298277 | 0.99945981 |
| HK2 | 19.1340205 | -1.4024657 | 1.09226987 | -1.2839919 | 0.19914478 | 0.99945981 |
| POLE4 | 10.1039452 | -0.399816887 | 0.51740353 | -0.7727371 | 0.43967801 | 0.99945981 |
| MRPL19 | 9.98315181 | -0.118235966 | 0.46912381 | -0.2520357 | 0.80101344 | 0.99945981 |
| GCFC2 | 11.6811873 | -0.151512684 | 0.4245023 | -0.3569184 | 0.72115289 | 0.99945981 |
| SUCLG1 | 19.2373438 | 0.181519425 | 0.36076139 | 0.50315646 | 0.61485428 | 0.99945981 |
| TMSB10 | 266.670412 | 0.375111188 | 0.57135397 | 0.65653029 | 0.51148299 | 0.99945981 |
| KCMF1 | 56.5043073 | -0.18416278 | 0.27663742 | -0.665719 | 0.50559073 | 0.99945981 |
| TGOLN2 | 50.2763743 | -0.085754493 | 0.35383451 | -0.2423576 | 0.80850306 | 0.99945981 |
| ELMOD3 | 16.9988017 | 0.180843333 | 0.35184839 | 0.51398085 | 0.60726538 | 0.99945981 |
| CAPG | 11.2277707 | -0.507740244 | 0.56057325 | -0.9057518 | 0.36506723 | 0.99945981 |
| MAT2A | 36.028713 | 1.164251096 | 0.61946583 | 1.87944361 | 0.06018395 | 0.92445778 |
| VAMP8 | 32.5938516 | -0.267824821 | 0.37974347 | -0.7052783 | 0.48063707 | 0.99945981 |
| VAMP5 | 17.4834989 | 0.171111781 | 0.43410079 | 0.39417524 | 0.69345166 | 0.99945981 |
| USP39 | 24.2771606 | 0.382471919 | 0.32818 | 1.16543337 | 0.24384362 | 0.99945981 |
| C2orf68 | 8.74059422 | 0.198598905 | 0.49492851 | 0.40126786 | 0.68822292 | 0.99945981 |
| GNLY | 65.0321973 | -0.408297426 | 1.17906536 | -0.3462891 | 0.7291255 | 0.99945981 |
| ST3GAL5 | 33.487516 | -0.233326328 | 0.33221781 | -0.7023294 | 0.48247377 | 0.99945981 |
| POLR1A | 9.82163306 | -0.053913837 | 0.44510367 | -0.1211265 | 0.90359086 | 0.99945981 |
| PTCD3 | 22.944222 | -0.214841636 | 0.44813532 | -0.4794124 | 0.63164526 | 0.99945981 |
| IMMT | 34.0764012 | 0.025906504 | 0.36168943 | 0.07162638 | 0.94289925 | 0.99945981 |
| KDM3A | 37.0168729 | -0.076911895 | 0.44971094 | -0.1710252 | 0.86420397 | 0.99945981 |
| RMND5A | 29.0421804 | 0.574153428 | 0.30624073 | 1.87484345 | 0.06081426 | 0.92445778 |
| CD8A | 16.0164298 | -1.421527311 | 0.5882927 | -2.4163606 | 0.01567653 | 0.7371189 |
| CYTOR | 33.2367961 | 0.046118963 | 0.40510234 | 0.11384522 | 0.90936049 | 0.99945981 |
| KRCC1 | 21.9868238 | -0.190536281 | 0.45514701 | -0.4186258 | 0.67548963 | 0.99945981 |
| EIF2AK3 | 32.4775961 | 0.224308885 | 0.43280343 | 0.51826966 | 0.60427014 | 0.99945981 |
| IGKC | 439.7482 | -1.472397381 | 1.75579268 | -0.8385941 | 0.40169712 | 0.99945981 |
| MRPS5 | 19.3477967 | 0.254439799 | 0.32871181 | 0.77405128 | 0.43890047 | 0.99945981 |
| ANKRD36C | 34.7501906 | -0.119168499 | 0.42120743 | -0.2829212 | 0.77723726 | 0.99945981 |
| DUSP2 | 16.2174816 | -0.689336627 | 0.79235931 | -0.8699798 | 0.38431142 | 0.99945981 |
| STARD7 | 17.4926176 | 0.273115345 | 0.3634435 | 0.75146576 | 0.4523724 | 0.99945981 |
| TMEM127 | 15.0919075 | 0.267773562 | 0.53803151 | 0.49769122 | 0.61870169 | 0.99945981 |
| CIAO1 | 13.3939925 | -0.405940068 | 0.4532256 | -0.8956689 | 0.37042963 | 0.99945981 |

| | | | | | | |
|---|---|---|---|---|---|---|
| SNRNP200 | 30.9649671 | -0.425129683 | 0.28673089 | -1.4826783 | 0.1381599 | 0.99945981 |
| ARID5A | 17.1134836 | -0.824693482 | 0.41948129 | -1.9659839 | 0.04930047 | 0.91969484 |
| KANSL3 | 24.5101051 | -0.22159242 | 0.42545706 | -0.5208338 | 0.60248255 | 0.99945981 |
| ANKRD23 | 9.49980809 | -0.569290906 | 0.64155925 | -0.8873552 | 0.37488772 | 0.99945981 |
| ANKRD36 | 48.7582231 | 0.047215175 | 0.42338766 | 0.1115176 | 0.91120591 | 0.99945981 |
| AC092683.1 | 25.4244674 | -0.847305594 | 0.39773553 | -2.1303241 | 0.03314486 | 0.8663391 |
| ANKRD36B | 14.1240734 | -0.063691737 | 0.49162301 | -0.129554 | 0.89691928 | 0.99945981 |
| COX5B | 21.7021284 | 0.087536248 | 0.38907323 | 0.22498656 | 0.82198973 | 0.99945981 |
| ZAP70 | 21.8458785 | 0.620829764 | 0.41972646 | 1.47912944 | 0.13910572 | 0.99945981 |
| TMEM131 | 104.946175 | 0.140866157 | 0.27887475 | 0.50512337 | 0.61347219 | 0.99945981 |
| INPP4A | 134.270313 | 0.263650213 | 0.35839634 | 0.7356387 | 0.46195061 | 0.99945981 |
| UNC50 | 9.42682177 | 0.163440196 | 0.51173843 | 0.3193823 | 0.74943663 | 0.99945981 |
| MGAT4A | 99.956817 | 0.139391675 | 0.42163705 | 0.33059636 | 0.74094939 | 0.99945981 |
| TSGA10 | 10.4870479 | -0.255023659 | 0.74677813 | -0.3414986 | 0.73272829 | 0.99945981 |
| AC079447.1 | 14.471729 | -0.059508586 | 0.6407144 | -0.0928785 | 0.92600009 | 0.99945981 |
| MITD1 | 16.14673 | 0.237214547 | 0.35512934 | 0.66796663 | 0.50415489 | 0.99945981 |
| EIF5B | 39.2816737 | -0.486513135 | 0.39638677 | -1.2273698 | 0.21968364 | 0.99945981 |
| REV1 | 39.7577079 | 0.060275754 | 0.30026132 | 0.20074432 | 0.8408985 | 0.99945981 |
| AFF3 | 153.70899 | -0.520024488 | 0.86472458 | -0.6013759 | 0.54758967 | 0.99945981 |
| RPL31 | 27.3900527 | -0.275512955 | 0.45507152 | -0.6054278 | 0.54489477 | 0.99945981 |
| TBC1D8 | 71.9519445 | 0.977561707 | 0.61839621 | 1.58080157 | 0.11392341 | 0.99945981 |
| CNOT11 | 14.2223547 | -0.154821818 | 0.40702743 | -0.380372 | 0.70366933 | 0.99945981 |
| RNF149 | 263.477877 | 0.422682698 | 0.44811041 | 0.9432557 | 0.34555013 | 0.99945981 |
| MAP4K4 | 113.957867 | 0.068900116 | 0.32707938 | 0.21065258 | 0.83315838 | 0.99945981 |
| IL1R2 | 86.2571899 | -2.609387302 | 1.56286594 | -1.6696168 | 0.0949952 | 0.99945981 |
| IL1R1 | 19.6465468 | -1.830553216 | 0.71227168 | -2.5700211 | 0.01016923 | 0.6404751 |
| IL18R1 | 38.4518155 | -1.259436557 | 0.41184385 | -3.0580439 | 0.00222787 | 0.45570351 |
| IL18RAP | 16.40735 | -1.207055288 | 0.66956186 | -1.802754 | 0.07142686 | 0.95583904 |
| MRPS9 | 12.95744 | -0.260838223 | 0.46481576 | -0.5611648 | 0.57468523 | 0.99945981 |
| TGFBRAP1 | 9.85729945 | -0.580371593 | 0.56309637 | -1.030679 | 0.30269138 | 0.99945981 |
| NCK2 | 64.4025792 | -0.371971634 | 0.32484744 | -1.1450656 | 0.25218198 | 0.99945981 |
| UXS1 | 25.5106022 | -0.336585377 | 0.43236141 | -0.7784815 | 0.43628518 | 0.99945981 |
| GCC2 | 70.0790467 | -0.439489323 | 0.42158548 | -1.0424679 | 0.29719481 | 0.99945981 |
| LIMS1 | 98.8427005 | 0.068750439 | 0.28822066 | 0.23853404 | 0.81146691 | 0.99945981 |
| RANBP2 | 60.4502204 | 0.669778269 | 0.42728017 | 1.56753885 | 0.11698879 | 0.99945981 |
| CCDC138 | 11.7361656 | -0.817882638 | 0.532199 | -1.5367985 | 0.12434265 | 0.99945981 |
| SH3RF3 | 35.2351645 | -0.111219958 | 0.56190795 | -0.1979327 | 0.84309772 | 0.99945981 |
| 10-Sep | 11.025887 | -0.876235691 | 0.44747845 | -1.9581629 | 0.0502109 | 0.92165202 |
| MIR4435-2H | 89.0346721 | 0.127238243 | 0.42822859 | 0.29712692 | 0.76636961 | 0.99945981 |
| BCL2L11 | 31.6963154 | -0.023316901 | 0.68987873 | -0.0337985 | 0.97303779 | 0.99945981 |
| ANAPC1 | 31.3683208 | 0.077563837 | 0.42728912 | 0.18152542 | 0.85595518 | 0.99945981 |
| MERTK | 86.7731638 | -0.593759512 | 0.59162222 | -1.0036126 | 0.31556538 | 0.99945981 |
| TMEM87B | 33.7498177 | -0.041536139 | 0.27436235 | -0.1513915 | 0.87966686 | 0.99945981 |
| ZC3H8 | 16.0840911 | -1.058905923 | 0.42173532 | -2.5108305 | 0.01204475 | 0.67187199 |
| ZC3H6 | 24.9752548 | -0.190760228 | 0.43295114 | -0.4406045 | 0.65949933 | 0.99945981 |
| TTL | 19.5669727 | 0.099404927 | 0.52397473 | 0.18971321 | 0.84953387 | 0.99945981 |
| SLC20A1 | 36.7704628 | 0.407192613 | 0.41214475 | 0.98798448 | 0.32316025 | 0.99945981 |
| PSD4 | 15.2154949 | 0.053315153 | 0.36700733 | 0.14526999 | 0.8844977 | 0.99945981 |
| PAX8 | 25.6039762 | -0.41139457 | 0.44530803 | -0.9238427 | 0.35556822 | 0.99945981 |
| CBWD2 | 14.5202128 | 1.094148884 | 0.45065591 | 2.42790313 | 0.0151864 | 0.73270878 |
| SLC35F5 | 40.452372 | -0.06885569 | 0.30091472 | -0.2288213 | 0.81900783 | 0.99945981 |
| ACTR3 | 139.280416 | 0.397741298 | 0.2830696 | 1.4051007 | 0.15999134 | 0.99945981 |
| DDX18 | 19.1193571 | -0.368137433 | 0.44713597 | -0.8233232 | 0.4103242 | 0.99945981 |
| CCDC93 | 38.2139757 | 0.080209791 | 0.34808512 | 0.23043154 | 0.81775645 | 0.99945981 |
| INSIG2 | 19.3356786 | -0.064426456 | 0.40359962 | -0.1596296 | 0.87317284 | 0.99945981 |
| MARCO | 28.4351789 | -0.091576619 | 1.23423741 | -0.0741969 | 0.94085369 | 0.99945981 |
| DBI | 29.5011633 | -0.660025115 | 0.5362986 | -1.2307045 | 0.21843339 | 0.99945981 |
| PTPN4 | 67.9919926 | 0.281572826 | 0.29141384 | 0.96623012 | 0.33392904 | 0.99945981 |
| EPB41L5 | 13.5787367 | -0.26207142 | 0.50339869 | -0.5206041 | 0.6026426 | 0.99945981 |
| RALB | 17.1707254 | 0.103205527 | 0.46243217 | 0.22317982 | 0.82339556 | 0.99945981 |
| CLASP1 | 146.620562 | -0.11106894 | 0.29085512 | -0.3818703 | 0.70255755 | 0.99945981 |
| NIFK-AS1 | 8.73548662 | -1.434251008 | 0.63490159 | -2.2590131 | 0.02388257 | 0.80320673 |
| TSN | 9.87888925 | 0.332222813 | 0.44553089 | 0.74567851 | 0.45586164 | 0.99945981 |
| GYPC | 20.7938408 | -0.690850436 | 0.55699054 | -1.2403271 | 0.21485445 | 0.99945981 |
| BIN1 | 11.0666977 | 0.085140574 | 0.43755044 | 0.19458459 | 0.84571817 | 0.99945981 |
| ERCC3 | 14.4914477 | -0.098452817 | 0.42766176 | -0.2302119 | 0.81792713 | 0.99945981 |
| MAP3K2 | 91.2750189 | 0.23117799 | 0.2730891 | 0.84652953 | 0.39725741 | 0.99945981 |
| IWS1 | 45.5223428 | -0.019725953 | 0.33660944 | -0.0586019 | 0.9532692 | 0.99945981 |
| WDR33 | 58.5835812 | 0.143761689 | 0.24928213 | 0.57670276 | 0.56414027 | 0.99945981 |
| SAP130 | 18.1936178 | -0.040569864 | 0.34095933 | -0.1189874 | 0.90528534 | 0.99945981 |
| UGGT1 | 35.206043 | -0.004404255 | 0.39120253 | -0.0112582 | 0.99101741 | 0.99945981 |
| HS6ST1 | 10.2480848 | -0.910595808 | 0.64946902 | -1.402062 | 0.16089675 | 0.99945981 |
| SMPD4 | 10.3520593 | 0.099404513 | 0.46336083 | 0.21452938 | 0.83013426 | 0.99945981 |
| MZT2B | 20.3410701 | 0.10972058 | 0.51811261 | 0.21176975 | 0.83228667 | 0.99945981 |
| PTPN18 | 28.4300046 | 0.734114414 | 0.51387438 | 1.4285873 | 0.15312289 | 0.99945981 |
| FAM168B | 30.321372 | -0.28729078 | 0.39276131 | -0.7314641 | 0.46449575 | 0.99945981 |
| PLEKHB2 | 29.3584731 | 0.055784527 | 0.4172442 | 0.13369755 | 0.89364174 | 0.99945981 |
| MZT2A | 10.1466507 | -0.137967934 | 0.53463503 | -0.25806 | 0.79636058 | 0.99945981 |
| NCKAP5 | 54.8835342 | -0.307922969 | 0.62976988 | -0.4889452 | 0.62488049 | 0.99945981 |
| MGAT5 | 175.407897 | -0.040527575 | 0.28210576 | -0.1436609 | 0.88576823 | 0.99945981 |
| TMEM163 | 25.6230399 | -1.862241822 | 0.90364887 | -2.0608025 | 0.03932189 | 0.89132455 |
| CCNT2 | 48.5436427 | -0.034222657 | 0.2826218 | -0.1210899 | 0.9036198 | 0.99945981 |
| RAB3GAP1 | 69.2974375 | 0.190786452 | 0.25320803 | 0.75347711 | 0.45116326 | 0.99945981 |
| ZRANB3 | 22.0681286 | 0.131439807 | 0.41621968 | 0.31579431 | 0.75215864 | 0.99945981 |
| R3HDM1 | 67.6200284 | -0.136571546 | 0.29190533 | -0.4678625 | 0.63988295 | 0.99945981 |
| UBXN4 | 55.4149952 | -0.06935377 | 0.23745311 | -0.2920735 | 0.77023039 | 0.99945981 |
| DARS | 47.9915486 | -0.554860966 | 0.29529129 | -1.8790292 | 0.0602405 | 0.92445778 |
| CXCR4 | 72.4402183 | -0.131469472 | 0.45698377 | -0.2876896 | 0.77358436 | 0.99945981 |
| HNMT | 27.903789 | 0.388769043 | 0.49437247 | 0.78638894 | 0.43163966 | 0.99945981 |
| SPOPL | 54.7322535 | 0.364455497 | 0.28704034 | 1.26970131 | 0.20419105 | 0.99945981 |
| KYNU | 172.199647 | 0.644562899 | 0.54678472 | 1.17882391 | 0.2384683 | 0.99945981 |
| ARHGAP15 | 592.629446 | 0.256353104 | 0.24213709 | 1.05871062 | 0.28973159 | 0.99945981 |
| GTDC1 | 93.1720335 | -0.285923696 | 0.30967988 | -0.9232879 | 0.35585717 | 0.99945981 |
| ZEB2 | 534.127513 | 0.511305707 | 0.223053 | 2.29230589 | 0.021888 | 0.79988354 |
| ACVR2A | 29.2629688 | 0.583771682 | 0.45181738 | 1.29205229 | 0.19633904 | 0.99945981 |
| ORC4 | 31.6509827 | 0.566364383 | 0.27434786 | 2.06440237 | 0.03897958 | 0.8905562 |
| MBD5 | 176.207435 | -0.059277898 | 0.29615914 | -0.2001556 | 0.84135892 | 0.99945981 |
| EPC2 | 60.1715261 | 0.030962003 | 0.28834614 | 0.1073779 | 0.91448919 | 0.99945981 |
| MMADHC | 13.2368767 | -0.055816552 | 0.40238641 | -0.1387138 | 0.8896763 | 0.99945981 |
| NMI | 14.0748506 | 0.427543658 | 0.62677479 | 0.68213283 | 0.49515497 | 0.99945981 |
| RIF1 | 48.2447818 | 0.144185718 | 0.29544859 | 0.48802303 | 0.62553353 | 0.99945981 |
| ARL5A | 17.8699748 | 0.401077278 | 0.54028709 | 0.74234103 | 0.45788074 | 0.99945981 |
| STAM2 | 32.4430112 | 0.061476955 | 0.3727073 | 0.16494701 | 0.86898569 | 0.99945981 |
| FMNL2 | 223.635221 | 0.191908249 | 0.49404818 | 0.38844035 | 0.69769019 | 0.99945981 |
| PRPF40A | 54.9925972 | 0.18725829 | 0.24355291 | 0.76886081 | 0.44197595 | 0.99945981 |
| ARL6IP6 | 23.5870512 | 0.600173469 | 0.37939852 | 1.58190776 | 0.11367063 | 0.99945981 |
| NR4A2 | 20.5016705 | 2.076794824 | 0.90760557 | 2.28821295 | 0.02212512 | 0.79988354 |
| GPD2 | 65.0624287 | 0.215775265 | 0.35263182 | 0.61189958 | 0.5406042 | 0.99945981 |
| CYTIP | 120.776662 | 0.726797613 | 0.43025503 | 1.68922515 | 0.09117629 | 0.99640647 |
| ACVR1 | 28.8260791 | -0.045323037 | 0.31016389 | -0.1461261 | 0.88382185 | 0.99945981 |
| PKP4 | 40.8319066 | 0.420521566 | 0.47584812 | 0.88373064 | 0.37684163 | 0.99945981 |
| WDSUB1 | 18.2550924 | -0.368342799 | 0.39318325 | -0.9368222 | 0.34885002 | 0.99945981 |
| BAZ2B | 156.968476 | 0.191367453 | 0.2392551 | 0.7998469 | 0.4237995 | 0.99945981 |

| | | | | | | |
|---|---|---|---|---|---|---|
| 7-Mar | 89.9794622 | 0.278110794 | 0.22259304 | 1.24941372 | 0.21151379 | 0.99945981 |
| ITGB6 | 10.1887631 | -2.686786117 | 0.9388539 | -2.8617723 | 0.00421279 | 0.507383 |
| RBMS1 | 183.4935 | 0.40146078 | 0.37451983 | 1.07193464 | 0.28374939 | 0.99945981 |
| TANK | 141.393594 | 0.378525175 | 0.26733855 | 1.41590197 | 0.15680421 | 0.99945981 |
| PSMD14 | 28.4715323 | -0.456891555 | 0.36424684 | -1.254346 | 0.20971628 | 0.99945981 |
| DPP4 | 14.4786234 | 0.291973423 | 0.47664104 | 0.61256459 | 0.54016428 | 0.99945981 |
| IFIH1 | 18.5020225 | 1.113674526 | 0.81093941 | 1.37331411 | 0.16965472 | 0.99945981 |
| GCA | 30.5610438 | 0.086168302 | 0.48535682 | 0.17753599 | 0.8590874 | 0.99945981 |
| COBLL1 | 35.1281004 | 0.706222313 | 0.60425507 | 1.16874868 | 0.2425049 | 0.99945981 |
| AC019197.1 | 18.8067927 | 0.559545239 | 0.77442959 | 0.72252564 | 0.46997137 | 0.99945981 |
| GALNT3 | 14.9260757 | 0.332064611 | 0.42362044 | 0.78387297 | 0.43311465 | 0.99945981 |
| TTC21B | 23.6999933 | 0.0906215 | 0.35442351 | 0.25568705 | 0.7981925 | 0.99945981 |
| STK39 | 82.1018752 | 0.30352741 | 0.35009633 | 0.86698255 | 0.38595157 | 0.99945981 |
| CERS6 | 103.30343 | 0.535489375 | 0.40769845 | 1.31344471 | 0.18903314 | 0.99945981 |
| LRP2 | 19.1609398 | 0.46009236 | 0.94947232 | 0.48457691 | 0.6279765 | 0.99945981 |
| FASTKD1 | 18.1962268 | 0.079566395 | 0.5272744 | 0.15090131 | 0.88005357 | 0.99945981 |
| PPIG | 55.0997053 | 0.44679902 | 0.23164329 | 1.9288235 | 0.05375278 | 0.92445778 |
| SSB | 15.3829053 | 0.16469087 | 0.41425606 | 0.39755814 | 0.69095592 | 0.99945981 |
| UBR3 | 100.433183 | 0.283909625 | 0.29556796 | 0.96055616 | 0.33677538 | 0.99945981 |
| GORASP2 | 18.5061054 | -0.418329943 | 0.40825535 | -1.0246772 | 0.30551553 | 0.99945981 |
| TLK1 | 99.1477028 | -0.210146459 | 0.22654928 | -0.9275971 | 0.3536166 | 0.99945981 |
| METTL8 | 18.767157 | -0.404141424 | 0.38406342 | -1.0522778 | 0.2926721 | 0.99945981 |
| DCAF17 | 12.0501781 | 0.25183494 | 0.41484236 | 0.60706178 | 0.54380991 | 0.99945981 |
| CYBRD1 | 12.0415952 | -0.334649397 | 0.66091953 | -0.5063391 | 0.61261862 | 0.99945981 |
| DYNC1I2 | 31.793974 | 0.284259693 | 0.28261017 | 1.00583674 | 0.31449411 | 0.99945981 |
| SLC25A12 | 26.081528 | 0.140700138 | 0.44884383 | 0.31347237 | 0.75392181 | 0.99945981 |
| HAT1 | 20.5706646 | 0.085790846 | 0.43567276 | 0.19691579 | 0.84389344 | 0.99945981 |
| ITGA6 | 27.5762262 | -0.504629561 | 0.3960418 | -1.2741826 | 0.20259871 | 0.99945981 |
| AC078883.1 | 11.3319867 | -0.761902698 | 0.69438967 | -1.0972264 | 0.27254242 | 0.99945981 |
| PDK1 | 26.7901608 | -0.351682358 | 0.43834882 | -0.8022888 | 0.4223859 | 0.99945981 |
| MAP3K20 | 28.3922248 | -0.343911337 | 0.49315231 | -0.6973735 | 0.4855691 | 0.99945981 |
| SP3 | 68.4369794 | 0.087744255 | 0.24498727 | 0.35815843 | 0.72022476 | 0.99945981 |
| OLA1 | 53.0025148 | 0.007571172 | 0.29134372 | 0.02598708 | 0.97926765 | 0.99945981 |
| CIR1 | 27.8753912 | 0.279284619 | 0.35409072 | 0.78873747 | 0.43026546 | 0.99945981 |
| GPR155 | 52.5821234 | -0.311872437 | 0.55609814 | -0.5608227 | 0.57491844 | 0.99945981 |
| AC010894.3 | 61.2837676 | -0.568462056 | 0.43965747 | -1.2929658 | 0.1960229 | 0.99945981 |
| WIPF1 | 173.809365 | -0.25920243 | 0.22268235 | -1.1640008 | 0.2444237 | 0.99945981 |
| ATF2 | 38.5176432 | -0.307253029 | 0.30338538 | -1.0127483 | 0.3111804 | 0.99945981 |
| ATP5MC3 | 26.1447898 | -0.475171975 | 0.52836021 | -0.8993334 | 0.36847511 | 0.99945981 |
| LNPK | 21.510378 | -0.259614553 | 0.49882668 | -0.5204504 | 0.60274968 | 0.99945981 |
| MTX2 | 26.9875683 | 0.75034959 | 0.35734838 | 2.09977048 | 0.03574904 | 0.87920079 |
| HNRNPA3 | 73.3696643 | 0.314389554 | 0.28634285 | 1.09794796 | 0.27222721 | 0.99945981 |
| NFE2L2 | 112.221037 | 0.256036991 | 0.26034069 | 0.98346897 | 0.32537668 | 0.99945981 |
| AGPS | 42.9200366 | 0.245340632 | 0.33715149 | 0.72768664 | 0.46680543 | 0.99945981 |
| RBM45 | 9.71768661 | -0.12555034 | 0.45577328 | -0.2754667 | 0.78295774 | 0.99945981 |
| AC009948.1 | 12.7855636 | -0.068893789 | 0.6062038 | -0.1136479 | 0.90951691 | 0.99945981 |
| PRKRA | 13.1405855 | -1.26276843 | 0.44860783 | -2.8148604 | 0.00487984 | 0.50757754 |
| PLEKHA3 | 12.5264058 | -0.113602609 | 0.43963296 | -0.2584033 | 0.79609567 | 0.99945981 |
| TTN-AS1 | 35.9281853 | -0.474044914 | 0.33284135 | -1.4242368 | 0.15437792 | 0.99945981 |
| TTN | 64.0620665 | 0.105392794 | 0.80288455 | 0.13126768 | 0.89556356 | 0.99945981 |
| CCDC141 | 30.4885426 | 0.860743844 | 0.94755864 | 0.90838056 | 0.36367719 | 0.99945981 |
| SESTD1 | 59.9875951 | 0.406369352 | 0.38408158 | 1.05802874 | 0.29004234 | 0.99945981 |
| CWC22 | 18.1181073 | 0.079280971 | 0.4388752 | 0.18064582 | 0.85664559 | 0.99945981 |
| UBE2E3 | 43.1273428 | 0.152324898 | 0.44434628 | 0.34280673 | 0.73174386 | 0.99945981 |
| LINC01934 | 73.9256619 | -0.124257659 | 0.57273587 | -0.2169546 | 0.82824376 | 0.99945981 |
| ITGA4 | 165.09909 | -0.131983922 | 0.43170251 | -0.3057289 | 0.75981109 | 0.99945981 |
| SSFA2 | 24.0252111 | 0.594368702 | 0.35238443 | 1.68670537 | 0.09166002 | 0.99640647 |
| PDE1A | 15.4688363 | 1.499835031 | 0.71597557 | 2.09481313 | 0.03618761 | 0.87920079 |
| DNAJC10 | 22.8143499 | 0.186009224 | 0.36307099 | 0.51232191 | 0.60842573 | 0.99945981 |
| NCKAP1 | 10.4909244 | -0.441801382 | 0.58813443 | -0.7511912 | 0.45253762 | 0.99945981 |
| ZNF804A | 34.3040647 | 0.152720556 | 0.68253941 | 0.22375346 | 0.82294914 | 0.99945981 |
| ZC3H15 | 20.4446199 | 0.432865493 | 0.30997355 | 1.39645947 | 0.16257618 | 0.99945981 |
| ITGAV | 37.751946 | 0.193629471 | 0.32345853 | 0.59862224 | 0.54942482 | 0.99945981 |
| AC007319.1 | 18.0504371 | 0.702245118 | 1.07691909 | 0.65208717 | 0.51434494 | 0.99945981 |
| TFPI | 21.3411632 | -2.11087037 | 0.8415442 | -2.5083298 | 0.01213034 | 0.67187199 |
| WDR75 | 17.1150954 | 0.284752266 | 0.3602067 | 0.79052462 | 0.42922145 | 0.99945981 |
| SLC40A1 | 12.7152492 | 0.022071821 | 0.85378001 | 0.02585188 | 0.97937548 | 0.99945981 |
| ANKAR | 22.8665251 | 0.342133273 | 0.59525041 | 0.57477201 | 0.5654455 | 0.99945981 |
| ORMDL1 | 33.0595205 | -0.066409387 | 0.4007557 | -0.1657104 | 0.86838486 | 0.99945981 |
| PMS1 | 26.3410543 | 0.075763875 | 0.34495718 | 0.21963269 | 0.82615723 | 0.99945981 |
| HIBCH | 39.9091346 | -0.029654427 | 0.43598716 | -0.0680167 | 0.94577231 | 0.99945981 |
| MFSD6 | 23.4062566 | -0.486701081 | 0.39975388 | -1.2175018 | 0.22341334 | 0.99945981 |
| NAB1 | 17.4558524 | 0.187726639 | 0.37024536 | 0.50703306 | 0.61213162 | 0.99945981 |
| GLS | 137.213683 | 0.161468703 | 0.32889107 | 0.49094889 | 0.6234626 | 0.99945981 |
| STAT1 | 65.1010597 | 0.370478037 | 0.64686171 | 0.57273144 | 0.56682655 | 0.99945981 |
| STAT4 | 143.329864 | -0.387465323 | 0.64397094 | -0.6016814 | 0.54738625 | 0.99945981 |
| NABP1 | 59.3583852 | -0.064824215 | 0.43038341 | -0.1506197 | 0.88027573 | 0.99945981 |
| SLC39A10 | 33.0860098 | 0.288796242 | 0.4789356 | 0.60299598 | 0.54651136 | 0.99945981 |
| STK17B | 169.56801 | -0.014021944 | 0.32699595 | -0.0428811 | 0.96579632 | 0.99945981 |
| AC114760.2 | 13.4199949 | -0.767957219 | 0.53657288 | -1.4312263 | 0.15236536 | 0.99945981 |
| GTF3C3 | 13.197995 | 0.094934953 | 0.47192139 | 0.20116688 | 0.84056809 | 0.99945981 |
| ANKRD44 | 530.007821 | 0.067179433 | 0.30748683 | 0.21847906 | 0.82705588 | 0.99945981 |
| SF3B1 | 63.8511135 | 0.009539267 | 0.30436059 | 0.03134199 | 0.9749968 | 0.99945981 |
| COQ10B | 11.60419 | -0.24300523 | 0.41931726 | -0.5795259 | 0.56223435 | 0.99945981 |
| HSPD1 | 30.9893752 | 0.116183259 | 0.38319252 | 0.30319814 | 0.76173887 | 0.99945981 |
| PLCL1 | 61.3266496 | 0.686093614 | 0.43023402 | 1.59469866 | 0.11077964 | 0.99945981 |
| SPATS2L | 39.9094471 | 0.787340831 | 0.61122543 | 1.28813494 | 0.197699 | 0.99945981 |
| BZW1 | 13.779663 | 0.402476939 | 0.52023478 | 0.77364482 | 0.43914086 | 0.99945981 |
| CLK1 | 52.2972005 | 0.709200749 | 0.33767933 | 2.10021963 | 0.03570953 | 0.87920079 |
| ORC2 | 28.7418527 | -0.213253893 | 0.30186393 | -0.706457 | 0.47990397 | 0.99945981 |
| FAM126B | 49.5081221 | -0.015730679 | 0.31340984 | -0.050192 | 0.95996936 | 0.99945981 |
| NDUFB3 | 12.8627244 | -0.275537504 | 0.56204676 | -0.4902395 | 0.62396446 | 0.99945981 |
| CFLAR | 147.41786 | 0.436264887 | 0.27626915 | 1.57912992 | 0.11430626 | 0.99945981 |
| CFLAR-AS1 | 14.2878425 | 0.015287515 | 0.54779649 | 0.02790729 | 0.97773609 | 0.99945981 |
| CASP10 | 24.3515024 | 0.147977278 | 0.55806067 | 0.26516343 | 0.79088357 | 0.99945981 |
| CASP8 | 47.6292298 | 0.278317518 | 0.35487444 | 0.7842704 | 0.43288146 | 0.99945981 |
| TRAK2 | 21.8540563 | 0.07351961 | 0.36635698 | 0.20067752 | 0.84095074 | 0.99945981 |
| ALS2 | 11.9384318 | -0.878800717 | 0.54197143 | -1.621489 | 0.10491281 | 0.99945981 |
| SUMO1 | 18.3633264 | 0.021231829 | 0.38458866 | 0.05520659 | 0.95597388 | 0.99945981 |
| NOP58 | 29.8049276 | -0.562624537 | 0.3988721 | -1.4105387 | 0.15838068 | 0.99945981 |
| BMPR2 | 73.8854255 | 0.055574746 | 0.3393663 | 0.16376035 | 0.8699198 | 0.99945981 |
| FAM117B | 30.5461167 | 0.845059049 | 0.32308214 | 2.61561669 | 0.00890665 | 0.59184657 |
| WDR12 | 9.7995878 | 0.265089039 | 0.43398953 | 0.61081897 | 0.54131943 | 0.99945981 |
| CARF | 13.7373601 | -0.202998545 | 0.44371029 | -0.4575025 | 0.64730994 | 0.99945981 |
| NBEAL1 | 95.8974222 | -0.038224568 | 0.31748534 | -0.1203979 | 0.90416796 | 0.99945981 |
| CYP20A1 | 33.5627955 | 0.121624783 | 0.32698739 | 0.37195557 | 0.70992593 | 0.99945981 |
| ABI2 | 26.0653266 | 0.50150655 | 0.41119829 | 1.21962217 | 0.22260814 | 0.99945981 |
| RAPH1 | 30.6190217 | 1.288699008 | 0.57507161 | 2.24093658 | 0.02503018 | 0.81052928 |
| CD28 | 12.4313278 | 0.672116694 | 0.67004286 | 1.00309508 | 0.31581499 | 0.99945981 |
| PARD3B | 12.0586278 | 0.690437377 | 0.50121724 | 1.37752119 | 0.16835117 | 0.99945981 |
| NRP2 | 25.3253385 | 0.124541365 | 0.62095133 | 0.20056542 | 0.8410384 | 0.99945981 |
| INO80D | 84.652268 | 0.747719808 | 0.65667767 | 1.13864053 | 0.25485312 | 0.99945981 |
| NDUFS1 | 32.3889581 | 0.007676981 | 0.33673843 | 0.02279805 | 0.98181136 | 0.99945981 |

| | | | | | | |
|---|---|---|---|---|---|---|
| EEF1B2 | 32.761796 | 0.003354506 | 0.53619925 | 0.00625608 | 0.9950084 | 0.99945981 |
| FASTKD2 | 11.0115692 | -0.393361259 | 0.53949219 | -0.7291324 | 0.46592065 | 0.99945981 |
| KLF7 | 47.2503733 | 0.500303704 | 0.30165581 | 1.65852503 | 0.09721154 | 0.99945981 |
| CREB1 | 57.9889058 | 0.168105395 | 0.26141341 | 0.6430634 | 0.52018296 | 0.99945981 |
| METTL21A | 14.4168056 | 0.24527466 | 0.45839161 | 0.53507668 | 0.59259683 | 0.99945981 |
| CCNYL1 | 14.851578 | -0.358270368 | 0.38471486 | -0.9312621 | 0.351718 | 0.99945981 |
| PLEKHM3 | 37.1012242 | 0.040834077 | 0.29558447 | 0.1381469 | 0.89012432 | 0.99945981 |
| IDH1 | 18.8323957 | -0.791705048 | 0.49862057 | -1.5877906 | 0.1123337 | 0.99945981 |
| PIKFYVE | 71.3283739 | -0.619606359 | 0.38852732 | -1.5947562 | 0.11076677 | 0.99945981 |
| PTH2R | 15.8132278 | 1.000480459 | 0.48106404 | 2.07972407 | 0.03755085 | 0.87920079 |
| RPE | 17.3557945 | 0.41604133 | 0.37510184 | 1.10914233 | 0.26736878 | 0.99945981 |
| KANSL1L | 38.4482439 | 0.506864739 | 0.46771192 | 1.08371139 | 0.27849278 | 0.99945981 |
| AC007038.1 | 11.7690764 | 0.532584655 | 0.48803421 | 1.0912855 | 0.27514728 | 0.99945981 |
| LANCL1 | 13.264449 | -0.451369907 | 0.39044906 | -1.1560276 | 0.24766985 | 0.99945981 |
| ERBB4 | 29.6248466 | 0.670711224 | 0.66750477 | 1.00480364 | 0.31499141 | 0.99945981 |
| IKZF2 | 46.6069589 | 0.445545955 | 0.62926689 | 0.70803973 | 0.47892059 | 0.99945981 |
| SPAG16 | 10.4294886 | -0.490820928 | 0.56092339 | -0.8750231 | 0.38156133 | 0.99945981 |
| BARD1 | 18.7749906 | 0.310196764 | 0.39365297 | 0.7879955 | 0.43069934 | 0.99945981 |
| ATIC | 9.20994954 | -0.047623797 | 0.53700459 | -0.0886842 | 0.92933293 | 0.99945981 |
| XRCC5 | 92.3231895 | 0.081533622 | 0.23496823 | 0.34699849 | 0.72859245 | 0.99945981 |
| SMARCAL1 | 15.6558614 | -0.000867011 | 0.37965464 | -0.0022837 | 0.99817789 | 0.9997934 |
| RPL37A | 58.7188098 | -0.111716605 | 0.42695454 | -0.2616592 | 0.79358416 | 0.99945981 |
| IGFBP5 | 52.3943393 | 0.950563616 | 1.12824529 | 0.84251503 | 0.39949973 | 0.99945981 |
| DIRC3 | 65.0129771 | -0.147339398 | 0.72944989 | -0.201987 | 0.83992688 | 0.99945981 |
| TNS1 | 21.9034568 | -0.897645659 | 0.62543741 | -1.4352286 | 0.15122196 | 0.99945981 |
| ARPC2 | 120.21059 | -0.172538388 | 0.33220897 | -0.519367 | 0.60350482 | 0.99945981 |
| PNKD | 16.4563367 | 0.068277775 | 0.49155882 | 0.13890052 | 0.88952876 | 0.99945981 |
| TMBIM1 | 19.1944861 | -0.009263456 | 0.47577157 | -0.0194704 | 0.98446586 | 0.99945981 |
| SLC11A1 | 113.75396 | -0.179384105 | 0.5394677 | -0.3325206 | 0.73949621 | 0.99945981 |
| CTDSP1 | 15.7418461 | -0.158165525 | 0.43026642 | -0.367599 | 0.71317223 | 0.99945981 |
| USP37 | 31.3169949 | -0.02183933 | 0.37649036 | -0.0580077 | 0.95374252 | 0.99945981 |
| CNOT9 | 15.6238454 | 0.151767966 | 0.48027832 | 0.31600004 | 0.75200248 | 0.99945981 |
| RETREG2 | 11.4577338 | 0.231277916 | 0.4204787 | 0.5500348 | 0.5822955 | 0.99945981 |
| ANKZF1 | 11.0766243 | -0.176424555 | 0.42302897 | -0.4170508 | 0.67664128 | 0.99945981 |
| TUBA4A | 14.0980478 | -0.708100926 | 0.56809293 | -1.2464526 | 0.21259827 | 0.99945981 |
| FARSB | 14.0744642 | 0.182995802 | 0.4726013 | 0.38720968 | 0.69860098 | 0.99945981 |
| ACSL3 | 51.458336 | 0.246869898 | 0.51069958 | 0.48339554 | 0.62881491 | 0.99945981 |
| AP1S3 | 10.0485863 | -0.136877927 | 0.52386532 | -0.2612846 | 0.79387306 | 0.99945981 |
| WDFY1 | 36.881423 | 0.156917698 | 0.32903182 | 0.47690736 | 0.63342809 | 0.99945981 |
| CUL3 | 92.5046585 | 0.130327844 | 0.25992688 | 0.50140196 | 0.61608826 | 0.99945981 |
| DOCK10 | 275.049861 | 0.00291279 | 0.31341153 | 0.00929382 | 0.99258471 | 0.99945981 |
| RHBDD1 | 44.5161833 | 0.508155961 | 0.45197066 | 1.12431184 | 0.26088076 | 0.99945981 |
| COL4A3 | 9.87196292 | 0.636982175 | 0.55256262 | 1.15277826 | 0.24900141 | 0.99945981 |
| MFF | 24.0259575 | -0.301300272 | 0.37375368 | -0.8061466 | 0.42015831 | 0.99945981 |
| AGFG1 | 131.915383 | 0.197863874 | 0.42166414 | 0.4692452 | 0.63889438 | 0.99945981 |
| PID1 | 125.742242 | 1.234818401 | 0.81758497 | 1.51032425 | 0.13096071 | 0.99945981 |
| TRIP12 | 119.432795 | 0.301611495 | 0.19610115 | 1.5380404 | 0.12403874 | 0.99945981 |
| SP110 | 63.0313322 | 0.245677593 | 0.54279144 | 0.45261877 | 0.65082328 | 0.99945981 |
| SP140 | 42.8125619 | -0.407432808 | 0.38811737 | -1.049767 | 0.29382525 | 0.99945981 |
| SP140L | 36.585538 | -0.022279941 | 0.35130791 | -0.06342 | 0.94943208 | 0.99945981 |
| SP100 | 165.859056 | 0.277907967 | 0.3083941 | 0.90114555 | 0.36751094 | 0.99945981 |
| CAB39 | 139.152233 | -0.124464707 | 0.21910262 | -0.5680658 | 0.56999027 | 0.99945981 |
| ITM2C | 9.79699798 | -1.544734502 | 0.76940907 | -2.0076895 | 0.0446763 | 0.90039076 |
| PSMD1 | 48.2831385 | -0.180771822 | 0.29784877 | -0.6069249 | 0.54390078 | 0.99945981 |
| ARMC9 | 31.1888338 | -0.586455104 | 0.52662365 | -1.1136133 | 0.26544511 | 0.99945981 |
| NCL | 31.0859226 | -0.851644386 | 0.39764606 | -2.1417146 | 0.03221645 | 0.8663391 |
| PTMA | 165.252537 | -0.347710713 | 0.49030389 | -0.7091739 | 0.47821658 | 0.99945981 |
| PDE6D | 16.8573477 | -0.462984587 | 0.44355548 | -1.0438031 | 0.2965765 | 0.99945981 |
| DIS3L2 | 47.2856189 | -0.062670144 | 0.32887699 | -0.190558 | 0.84887189 | 0.99945981 |
| EIF4E2 | 13.5790253 | -0.39040074 | 0.42632567 | -0.9157336 | 0.35980664 | 0.99945981 |
| GIGYF2 | 73.2667806 | 0.027137196 | 0.25639627 | 0.10584084 | 0.91570863 | 0.99945981 |
| INPP5D | 182.034177 | 0.042824056 | 0.29279032 | 0.14626186 | 0.88371468 | 0.99945981 |
| ATG16L1 | 21.153729 | -0.691611307 | 0.34827897 | -1.9857969 | 0.04705588 | 0.90884462 |
| DGKD | 92.0946676 | -0.464548354 | 0.60394383 | -0.7691913 | 0.44177974 | 0.99945981 |
| USP40 | 19.7504031 | 0.071056282 | 0.43772923 | 0.16232931 | 0.87104654 | 0.99945981 |
| ARL4C | 50.4768456 | -0.041342127 | 0.3995286 | -0.1034773 | 0.91758419 | 0.99945981 |
| AGAP1 | 31.6897367 | -0.22878484 | 0.38166147 | -0.5994444 | 0.54887656 | 0.99945981 |
| LRRFIP1 | 171.873164 | -0.013933018 | 0.30036358 | -0.0463872 | 0.96300166 | 0.99945981 |
| ILKAP | 21.5095221 | -0.688443876 | 0.36263694 | -1.8984384 | 0.05763836 | 0.92445778 |
| HDAC4 | 80.0829226 | -0.695406111 | 0.30727428 | -2.2631445 | 0.02362678 | 0.80320673 |
| NDUFA10 | 32.9570493 | -0.319025535 | 0.37466353 | -0.8514988 | 0.39449235 | 0.99945981 |
| COPS9 | 10.6882181 | -0.392395341 | 0.58905688 | -0.6661417 | 0.50532055 | 0.99945981 |
| ANKMY1 | 23.5934852 | -0.383893832 | 0.58359099 | -0.6578132 | 0.5106582 | 0.99945981 |
| RNPEPL1 | 16.0249206 | 0.145048989 | 0.54212698 | 0.26755538 | 0.78904158 | 0.99945981 |
| SNED1 | 19.8874866 | 0.141962909 | 0.68403497 | 0.2075375 | 0.83559011 | 0.99945981 |
| MTERF4 | 19.8776318 | -0.233084195 | 0.40116005 | -0.5810254 | 0.56122331 | 0.99945981 |
| PPP1R7 | 11.3342936 | -0.26253806 | 0.45620026 | -0.5754886 | 0.56496088 | 0.99945981 |
| HDLBP | 50.3395873 | -0.187664081 | 0.29072186 | -0.6455107 | 0.51859626 | 0.99945981 |
| 2-Sep | 68.9878748 | -0.410096968 | 0.29179885 | -1.4054098 | 0.15989945 | 0.99945981 |
| FARP2 | 31.1891158 | -0.829563551 | 0.53642952 | -1.546454 | 0.12199497 | 0.99945981 |
| STK25 | 10.3631655 | 0.157713775 | 0.42231583 | 0.37344983 | 0.70881368 | 0.99945981 |
| ATG4B | 23.2408121 | -0.189446991 | 0.31167561 | -0.6078339 | 0.54329767 | 0.99945981 |
| ING5 | 14.9348252 | 0.083874216 | 0.40453356 | 0.20733562 | 0.83574776 | 0.99945981 |
| D2HGDH | 19.3479093 | -0.018101202 | 0.45008396 | -0.0402174 | 0.96791981 | 0.99945981 |
| TRNT1 | 16.9936868 | -0.227453286 | 0.60204915 | -0.3777985 | 0.70558026 | 0.99945981 |
| CRBN | 29.1198863 | 0.165180756 | 0.48713915 | 0.33908331 | 0.73454697 | 0.99945981 |
| SUMF1 | 127.134303 | 0.440667223 | 0.28353399 | 1.55419542 | 0.12013781 | 0.99945981 |
| ITPR1 | 94.8022677 | 0.123994615 | 0.41248181 | 0.30060626 | 0.76371476 | 0.99945981 |
| AC018816.1 | 12.1314286 | 0.384160691 | 0.60766412 | 0.63219249 | 0.52726111 | 0.99945981 |
| BHLHE40 | 21.4777617 | -0.595295536 | 0.65624412 | -0.9071251 | 0.36434063 | 0.99945981 |
| ARL8B | 65.148572 | 0.630075549 | 0.2996016 | 2.10304465 | 0.03546187 | 0.87920079 |
| AC026202.2 | 13.562478 | 0.101372467 | 0.75232301 | 0.13474593 | 0.89281275 | 0.99945981 |
| EDEM1 | 34.428918 | -0.009586709 | 0.53124854 | -0.0180456 | 0.98560246 | 0.99945981 |
| RAD18 | 11.1495839 | 0.588240646 | 0.54383512 | 1.08165254 | 0.27940695 | 0.99945981 |
| SRGAP3 | 20.8138354 | -0.47146194 | 0.60442206 | -0.7800211 | 0.43537847 | 0.99945981 |
| THUMPD3-A | 30.9411438 | -0.208633674 | 0.38665321 | -0.5395886 | 0.58948077 | 0.99945981 |
| SETD5 | 96.3102835 | -0.310951198 | 0.20964225 | -1.4832468 | 0.13800886 | 0.99945981 |
| MTMR14 | 31.9608521 | -0.171322647 | 0.34054169 | -0.5030886 | 0.61490197 | 0.99945981 |
| OGG1 | 10.0681822 | 0.383635036 | 0.47114577 | 0.81425975 | 0.41549616 | 0.99945981 |
| TADA3 | 8.38250103 | -0.736668146 | 0.52639063 | -1.3994705 | 0.16167195 | 0.99945981 |
| EMC3 | 14.0712942 | -0.296989989 | 0.47562163 | -0.6244249 | 0.53234858 | 0.99945981 |
| BRK1 | 19.0937709 | -0.020610787 | 0.46310219 | -0.0445059 | 0.96450114 | 0.99945981 |
| VHL | 12.647085 | -0.227029808 | 0.43149308 | -0.5261494 | 0.59878444 | 0.99945981 |
| IRAK2 | 19.3508037 | 1.18839667 | 0.39308994 | 3.0232182 | 0.00250102 | 0.45570351 |
| SEC13 | 18.6648651 | -0.063839514 | 0.40098404 | -0.1592071 | 0.87350569 | 0.99945981 |
| ATG7 | 201.844858 | 0.302008841 | 0.40308062 | 0.7492517 | 0.45370551 | 0.99945981 |
| VGLL4 | 40.8530438 | -0.458484329 | 0.28600464 | -1.6030661 | 0.10892007 | 0.99945981 |
| TAMM41 | 19.2358871 | -0.387296898 | 0.35958204 | -1.0770752 | 0.28144667 | 0.99945981 |
| PPARG | 37.9913141 | 0.257393378 | 0.79761623 | 0.32270329 | 0.74691996 | 0.99945981 |
| RAF1 | 64.67508 | 0.146626124 | 0.20562945 | 0.71305994 | 0.47580867 | 0.99945981 |
| RPL32 | 111.327657 | -0.325959759 | 0.51678493 | -0.6307455 | 0.52820696 | 0.99945981 |
| IQSEC1 | 75.0131465 | -0.585337061 | 0.41406068 | -1.4136504 | 0.15746456 | 0.99945981 |
| NUP210 | 49.0220879 | -0.047699511 | 0.30893782 | -0.1543984 | 0.8772956 | 0.99945981 |

| | | | | | | |
|---|---|---|---|---|---|---|
| XPC | 20.4180871 | -0.399422124 | 0.4002994 | -0.9978085 | 0.31837225 | 0.99945981 |
| SLC6A6 | 42.6584919 | 0.648746757 | 0.29967002 | 2.16487044 | 0.03039761 | 0.85704136 |
| CCDC174 | 12.0250514 | 0.039557687 | 0.44467144 | 0.08895936 | 0.92911421 | 0.99945981 |
| FGD5-AS1 | 17.696495 | 0.225153834 | 0.41866745 | 0.53778682 | 0.59072423 | 0.99945981 |
| NR2C2 | 52.7159852 | 0.384858309 | 0.25320669 | 1.51993736 | 0.12852672 | 0.99945981 |
| MRPS25 | 12.2399955 | 0.224385976 | 0.42975815 | 0.52212152 | 0.60158573 | 0.99945981 |
| RBSN | 13.278352 | 0.219122902 | 0.39184141 | 0.55921323 | 0.5760162 | 0.99945981 |
| CAPN7 | 47.448539 | -0.170910072 | 0.38602967 | -0.4427382 | 0.65795513 | 0.99945981 |
| SH3BP5 | 53.0261614 | -0.167807456 | 0.37070878 | -0.4526665 | 0.65078889 | 0.99945981 |
| METTL6 | 9.48080495 | -0.324287499 | 0.51225197 | -0.6330625 | 0.52669285 | 0.99945981 |
| HACL1 | 13.5229217 | 0.395839669 | 0.43590896 | 0.90807875 | 0.36383661 | 0.99945981 |
| ANKRD28 | 88.4331585 | -0.811835784 | 0.74793947 | -1.0854298 | 0.27773135 | 0.99945981 |
| OXNAD1 | 57.2138719 | -0.478563055 | 0.36956781 | -1.2949262 | 0.19534568 | 0.99945981 |
| RFTN1 | 103.276603 | -1.154290407 | 0.59690597 | -1.9337894 | 0.05313903 | 0.92445778 |
| PLCL2 | 152.536609 | 0.162316771 | 0.21969929 | 0.73881335 | 0.46002034 | 0.99945981 |
| TBC1D5 | 284.420781 | 0.198623083 | 0.23490981 | 0.84552911 | 0.39781549 | 0.99945981 |
| RAB5A | 33.3087871 | 0.266421681 | 0.31503731 | 0.84568294 | 0.39772964 | 0.99945981 |
| KAT2B | 98.8159104 | 0.047707498 | 0.2786893 | 0.17118525 | 0.86407811 | 0.99945981 |
| UBE2E2 | 183.950551 | 0.33692098 | 0.38834812 | 0.86757463 | 0.38562724 | 0.99945981 |
| UBE2E1 | 81.9581087 | 0.238344053 | 0.19977631 | 1.19305461 | 0.23284798 | 0.99945981 |
| RPL15 | 128.826296 | -0.652835081 | 0.51663577 | -1.2636273 | 0.20636382 | 0.99945981 |
| NR1D2 | 19.0050278 | -0.445577104 | 0.34799132 | -1.2804259 | 0.20039539 | 0.99945981 |
| THRB | 35.0194911 | 1.28056487 | 0.7324734 | 1.74827492 | 0.08041643 | 0.9846219 |
| TOP2B | 65.5479878 | 0.129077785 | 0.34387298 | 0.37536472 | 0.70738924 | 0.99945981 |
| NGLY1 | 49.4520107 | -0.377577363 | 0.32086639 | -1.1767433 | 0.239298 | 0.99945981 |
| NEK10 | 20.5671139 | -2.283387849 | 1.20582469 | -1.8936317 | 0.05827391 | 0.92445778 |
| SLC4A7 | 95.4553391 | 0.202817559 | 0.4337788 | 0.46755987 | 0.64009937 | 0.99945981 |
| CMC1 | 43.2900166 | 0.034305091 | 0.47047467 | 0.07291591 | 0.94187303 | 0.99945981 |
| AZI2 | 25.2042879 | 0.327783382 | 0.39406488 | 0.83180054 | 0.40552154 | 0.99945981 |
| TGFBR2 | 126.184444 | -0.071519318 | 0.31336291 | -0.2282316 | 0.8194662 | 0.99945981 |
| STT3B | 83.7456304 | 0.349121448 | 0.27212252 | 1.28295682 | 0.1995072 | 0.99945981 |
| OSBPL10 | 38.6423816 | -1.209316992 | 0.95703818 | -1.2636037 | 0.2063723 | 0.99945981 |
| GPD1L | 10.5251846 | 0.032278698 | 0.41024144 | 0.0786822 | 0.93728541 | 0.99945981 |
| CMTM7 | 51.3683686 | -0.352374228 | 0.38171623 | -0.9231314 | 0.35593874 | 0.99945981 |
| CMTM6 | 57.3554057 | 0.488772737 | 0.46445577 | 1.05235583 | 0.29263632 | 0.99945981 |
| DYNC1LI1 | 24.2305076 | 0.215309949 | 0.37042133 | 0.58125688 | 0.56106733 | 0.99945981 |
| CNOT10 | 29.1058922 | -0.182011869 | 0.34503687 | -0.5275143 | 0.59783651 | 0.99945981 |
| GLB1 | 36.2244787 | 0.182839794 | 0.35747993 | 0.5114687 | 0.6090229 | 0.99945981 |
| CRTAP | 12.8261597 | -0.134989468 | 0.37070275 | -0.3641448 | 0.7157499 | 0.99945981 |
| UBP1 | 29.8101788 | 0.545091015 | 0.31317429 | 1.74053565 | 0.08176501 | 0.9846219 |
| CLASP2 | 100.037614 | -0.0984983 | 0.27359422 | -0.360016 | 0.71883515 | 0.99945981 |
| PDCD6IP | 73.0195429 | 0.242727642 | 0.25212653 | 0.96272153 | 0.33568729 | 0.99945981 |
| TRANK1 | 40.0435908 | 0.330803909 | 0.4235978 | 0.78093869 | 0.43483855 | 0.99945981 |
| EPM2AIP1 | 15.869975 | -0.309231782 | 0.37780632 | -0.8184929 | 0.41307581 | 0.99945981 |
| MLH1 | 12.7072156 | -0.204922706 | 0.40579242 | -0.5049939 | 0.61356311 | 0.99945981 |
| LRRFIP2 | 61.815233 | -0.244791454 | 0.28067801 | -0.8721433 | 0.3831302 | 0.99945981 |
| GOLGA4 | 122.46856 | -0.34571492 | 0.18291186 | -1.8900629 | 0.05874955 | 0.92445778 |
| ITGA9 | 25.0889717 | -0.569440126 | 0.50953098 | -1.1175771 | 0.26374767 | 0.99945981 |
| ACAA1 | 17.2194281 | 0.175556034 | 0.60850461 | 0.28850403 | 0.77296094 | 0.99945981 |
| MYD88 | 11.2425007 | 0.943230381 | 0.63839936 | 1.47749268 | 0.13954362 | 0.99945981 |
| OXSR1 | 43.9650404 | 0.099485827 | 0.25762843 | 0.38616012 | 0.69937809 | 0.99945981 |
| EXOG | 14.9038194 | -0.333080008 | 0.38953017 | -0.8550814 | 0.39250608 | 0.99945981 |
| WDR48 | 20.4611415 | 0.017236484 | 0.44550313 | 0.03868993 | 0.9691376 | 0.99945981 |
| CX3CR1 | 18.4400913 | 2.29534093 | 0.9232893 | 2.48604735 | 0.01291708 | 0.69277921 |
| RPSA | 62.9938338 | -0.446704823 | 0.53856 | -0.829443 | 0.40685378 | 0.99945981 |
| EIF1B | 14.0724724 | -0.507405336 | 0.44880227 | -1.1305766 | 0.25823335 | 0.99945981 |
| RPL14 | 115.521505 | -0.484451103 | 0.62705309 | -0.7725839 | 0.43976869 | 0.99945981 |
| ZNF621 | 9.87278524 | 0.010467835 | 0.43485315 | 0.02407211 | 0.98079509 | 0.99945981 |
| CTNNB1 | 99.4840368 | 0.521583508 | 0.38299365 | 1.36185941 | 0.17324226 | 0.99945981 |
| ULK4 | 44.375055 | -0.309598639 | 0.27576661 | -1.1226836 | 0.26157191 | 0.99945981 |
| TRAK1 | 82.1308469 | 0.125432047 | 0.33794417 | 0.37116204 | 0.71051685 | 0.99945981 |
| NKTR | 132.045676 | 0.22947742 | 0.23675367 | 0.96926656 | 0.33241221 | 0.99945981 |
| AC006059.1 | 20.4248264 | 0.142079043 | 0.44682009 | 0.31797819 | 0.75050148 | 0.99945981 |
| SNRK | 54.3198116 | 0.112088555 | 0.3299842 | 0.33967855 | 0.73409861 | 0.99945981 |
| ANO10 | 35.1636187 | -0.06613667 | 0.35729966 | -0.1851014 | 0.8531495 | 0.99945981 |
| ABHD5 | 14.7133768 | 0.668578668 | 0.57475206 | 1.16324709 | 0.24472926 | 0.99945981 |
| TCAIM | 25.7657256 | -0.474072872 | 0.30303217 | -1.5644309 | 0.11771642 | 0.99945981 |
| ZNF445 | 13.3107656 | -1.500467289 | 0.47552908 | -3.1553639 | 0.00160298 | 0.42324844 |
| ZDHHC3 | 16.7195698 | -0.132214347 | 0.41277073 | -0.3203094 | 0.7487338 | 0.99945981 |
| CDCP1 | 12.3076589 | -0.270075375 | 0.95162375 | -0.2838048 | 0.77655999 | 0.99945981 |
| LARS2 | 19.9286844 | 0.054532184 | 0.37906962 | 0.14385796 | 0.88561263 | 0.99945981 |
| LIMD1 | 60.8971575 | -0.156510371 | 0.36757206 | -0.4257951 | 0.67025718 | 0.99945981 |
| SACM1L | 48.4575438 | -0.233881172 | 0.43921629 | -0.5324966 | 0.59438211 | 0.99945981 |
| FYCO1 | 17.4640088 | -0.087715279 | 0.39885221 | -0.2199192 | 0.82593405 | 0.99945981 |
| CCR1 | 14.5326682 | 0.559438773 | 0.74982492 | 0.74609253 | 0.45561152 | 0.99945981 |
| LTF | 22.128374 | -3.5299588 | 1.19645771 | -2.9503415 | 0.00317423 | 0.46723702 |
| CCDC12 | 31.1424089 | -0.344317011 | 0.48450377 | -0.7106591 | 0.47729552 | 0.99945981 |
| NBEAL2 | 27.4479929 | 0.179169288 | 0.72114314 | 0.24845177 | 0.80378489 | 0.99945981 |
| SETD2 | 123.598832 | -0.249715559 | 0.24700874 | -1.0109584 | 0.31203633 | 0.99945981 |
| KIF9-AS1 | 13.686665 | 0.413014369 | 0.47812447 | 0.86382185 | 0.38768576 | 0.99945981 |
| KLHL18 | 20.3723099 | -0.083646865 | 0.41649994 | -0.2008328 | 0.84082929 | 0.99945981 |
| SCAP | 15.5518834 | 0.30244712 | 0.40360984 | 0.74935517 | 0.45364316 | 0.99945981 |
| SMARCC1 | 73.5918947 | -0.06251594 | 0.2771369 | -0.2255778 | 0.82152979 | 0.99945981 |
| DHX30 | 10.5605287 | -0.121953964 | 0.50740688 | -0.2403475 | 0.81006089 | 0.99945981 |
| MAP4 | 92.5315952 | -0.203122792 | 0.28842347 | -0.7042519 | 0.48127589 | 0.99945981 |
| ZNF589 | 10.0854869 | -0.061588496 | 0.42202422 | -0.1459359 | 0.88397197 | 0.99945981 |
| TMA7 | 24.0350828 | 0.345932776 | 0.65244243 | 0.53021195 | 0.59596498 | 0.99945981 |
| SHISA5 | 20.9224049 | 0.156779208 | 0.39313527 | 0.39879202 | 0.69004646 | 0.99945981 |
| PFKFB4 | 16.6277018 | -0.132792786 | 0.7372 | -0.1801313 | 0.8570495 | 0.99945981 |
| UQCRC1 | 8.69190593 | 0.437314216 | 0.59999183 | 0.72886695 | 0.46608305 | 0.99945981 |
| IP6K2 | 16.896699 | -0.584169303 | 0.4663132 | -1.2527402 | 0.21030026 | 0.99945981 |
| PRKAR2A | 44.7880203 | 0.05397923 | 0.34203817 | 0.15781639 | 0.87460148 | 0.99945981 |
| SLC25A20 | 8.9437772 | 1.098762394 | 0.61499222 | 1.78662812 | 0.07399761 | 0.96760689 |
| ARIH2 | 43.7660065 | -0.002423716 | 0.29381545 | -0.0082491 | 0.99341824 | 0.99945981 |
| WDR6 | 13.3311884 | 0.205850214 | 0.40942049 | 0.50278435 | 0.6151159 | 0.99945981 |
| QRICH1 | 29.125554 | -0.544058755 | 0.31500415 | -1.7271479 | 0.0841411 | 0.99078675 |
| USP4 | 43.240622 | -0.181342784 | 0.39989044 | -0.4534812 | 0.6502023 | 0.99945981 |
| RHOA | 92.9577533 | -0.173906423 | 0.35480113 | -0.4901518 | 0.62402647 | 0.99945981 |
| IP6K1 | 32.4386422 | -0.184661917 | 0.32190898 | -0.5736464 | 0.56620713 | 0.99945981 |
| RBM6 | 158.904411 | -0.134895758 | 0.28215569 | -0.4780898 | 0.63258629 | 0.99945981 |
| RBM5 | 52.5725553 | 0.370438286 | 0.23537258 | 1.57383792 | 0.11552492 | 0.99945981 |
| GNAI2 | 76.3058003 | -0.016717761 | 0.54295885 | -0.0307901 | 0.97543693 | 0.99945981 |
| RASSF1 | 31.4685717 | -0.127959946 | 0.3918491 | -0.3265541 | 0.74400513 | 0.99945981 |
| MAPKAPK3 | 18.3170622 | 0.558816819 | 0.53199633 | 1.05041479 | 0.29352745 | 0.99945981 |
| MANF | 11.3721988 | -0.67740205 | 0.66116471 | -1.0245587 | 0.30557147 | 0.99945981 |
| DCAF1 | 21.7849065 | 0.293564115 | 0.35083264 | 0.83676397 | 0.40272525 | 0.99945981 |
| RAD54L2 | 19.9888259 | -0.030064515 | 0.4242474 | -0.0708655 | 0.94350478 | 0.99945981 |
| TEX264 | 8.83889156 | -0.720613757 | 0.48486784 | -1.4862065 | 0.1372245 | 0.99945981 |
| RPL29 | 140.850963 | -1.009119485 | 0.54157161 | -1.8633168 | 0.0624177 | 0.92949575 |
| WDR82 | 30.6753604 | 0.359285019 | 0.38782061 | 0.92642063 | 0.35422742 | 0.99945981 |
| DNAH1 | 10.8337232 | -0.374854726 | 0.43980987 | -0.8523109 | 0.39404158 | 0.99945981 |
| NISCH | 18.0439047 | 0.108097296 | 0.36367693 | 0.29723441 | 0.76628755 | 0.99945981 |
| STAB1 | 41.2572311 | 0.016259057 | 0.51778687 | 0.03140106 | 0.9749497 | 0.99945981 |

| | | | | | | |
|---|---|---|---|---|---|---|
| SMIM4 | 11.4846614 | -0.268130724 | 0.65390473 | -0.4100455 | 0.68177254 | 0.99945981 |
| PBRM1 | 91.9905441 | 0.00116715 | 0.22265259 | 0.00524202 | 0.99581749 | 0.99945981 |
| GNL3 | 11.8270508 | -0.326977297 | 0.42916162 | -0.7618978 | 0.446121 | 0.99945981 |
| SPCS1 | 20.0203597 | -0.315143107 | 0.46886572 | -0.6721394 | 0.50149498 | 0.99945981 |
| NEK4 | 12.9688567 | -0.505406701 | 0.60085172 | -0.8411505 | 0.40026365 | 0.99945981 |
| SFMBT1 | 15.8773464 | -0.472379839 | 0.5816843 | -0.8120897 | 0.41674015 | 0.99945981 |
| PRKCD | 25.8573601 | 0.775964318 | 0.40430976 | 1.9192322 | 0.05495495 | 0.92445778 |
| TKT | 29.3821254 | 0.044868997 | 0.46272814 | 0.09696622 | 0.92275322 | 0.99945981 |
| DCP1A | 36.2624615 | -0.314950583 | 0.37394956 | -0.8422275 | 0.39966065 | 0.99945981 |
| SELENOK | 14.2013908 | -0.458191397 | 0.52193007 | -0.8778789 | 0.38000944 | 0.99945981 |
| CACNA2D3 | 12.325075 | 0.711370121 | 0.50192673 | 1.41727883 | 0.15640142 | 0.99945981 |
| CCDC66 | 31.9360507 | -0.418860329 | 0.42991965 | -0.9742758 | 0.3299196 | 0.99945981 |
| FAM208A | 48.058658 | 0.055453411 | 0.36200318 | 0.15318487 | 0.87825249 | 0.99945981 |
| ARHGEF3 | 102.935753 | 0.052121136 | 0.27262115 | 0.19118523 | 0.84838048 | 0.99945981 |
| APPL1 | 31.0671268 | 0.177718794 | 0.33377049 | 0.53245807 | 0.59440877 | 0.99945981 |
| ARF4 | 21.6172813 | 0.049625608 | 0.36807232 | 0.1348257 | 0.89274969 | 0.99945981 |
| DENND6A | 50.6857957 | -0.266879687 | 0.24068223 | -1.1088467 | 0.26749633 | 0.99945981 |
| DENND6A-A! | 11.4813672 | -0.087928553 | 0.57350042 | -0.1533191 | 0.87814666 | 0.99945981 |
| SLMAP | 76.3906951 | 0.108993896 | 0.22678716 | 0.48059994 | 0.63080086 | 0.99945981 |
| FLNB | 18.8492612 | 0.040365686 | 0.41503101 | 0.09725944 | 0.92252036 | 0.99945981 |
| ABHD6 | 17.5361685 | -0.009857076 | 0.6185672 | -0.0159353 | 0.98728598 | 0.99945981 |
| PXK | 38.8905785 | -0.880209099 | 0.43728421 | -2.0128993 | 0.04412522 | 0.89803991 |
| FHIT | 171.227666 | 0.082358759 | 0.28641802 | 0.28754741 | 0.77369321 | 0.99945981 |
| PTPRG | 31.1146133 | -0.441807954 | 0.78038943 | -0.5661378 | 0.57130011 | 0.99945981 |
| THOC7 | 10.2600015 | -0.030800203 | 0.53502112 | -0.0575682 | 0.95409258 | 0.99945981 |
| ATXN7 | 10.968241 | 0.237455712 | 0.42453426 | 0.55933227 | 0.57593497 | 0.99945981 |
| ADAMTS9-A! | 10.2453985 | 0.739272796 | 0.73375674 | 1.00751755 | 0.31368613 | 0.99945981 |
| MAGI1 | 41.0002612 | 0.537062245 | 0.70219779 | 0.76483044 | 0.44437252 | 0.99945981 |
| SLC25A26 | 31.529475 | -0.005022721 | 0.33270572 | -0.0150966 | 0.98795512 | 0.99945981 |
| LRIG1 | 18.8613392 | -0.643589038 | 0.66964422 | -0.961091 | 0.33650641 | 0.99945981 |
| SUCLG2 | 118.052291 | -0.082161855 | 0.22312231 | -0.3682368 | 0.71269664 | 0.99945981 |
| FAM19A1 | 20.1349692 | -0.09182299 | 0.82315672 | -0.1115498 | 0.91118036 | 0.99945981 |
| TMF1 | 35.3349361 | -0.446039153 | 0.29713272 | -1.5011445 | 0.13331818 | 0.99945981 |
| UBA3 | 35.3111772 | 0.103595963 | 0.2663635 | 0.38892702 | 0.69733013 | 0.99945981 |
| ARL6IP5 | 42.1601429 | -0.149271527 | 0.4585516 | -0.3255283 | 0.74478126 | 0.99945981 |
| FRMD4B | 134.967574 | 0.968314751 | 0.66080783 | 1.46535 | 0.14282545 | 0.99945981 |
| MITF | 59.0344998 | 0.835560266 | 0.59204581 | 1.41131015 | 0.15815318 | 0.99945981 |
| FOXP1 | 461.609556 | 0.266502101 | 0.2526298 | 1.05491156 | 0.29146577 | 0.99945981 |
| EIF4E3 | 25.4160346 | 0.01015768 | 0.47544565 | 0.02136454 | 0.98295486 | 0.99945981 |
| RYBP | 57.4031204 | 0.324147803 | 0.33299967 | 0.9734178 | 0.3303457 | 0.99945981 |
| SHQ1 | 14.2815971 | -0.15251003 | 0.40747184 | -0.3742836 | 0.70819333 | 0.99945981 |
| GXYLT2 | 10.6495715 | -0.266990295 | 0.5196888 | -0.5137503 | 0.60742656 | 0.99945981 |
| PPP4R2 | 44.9611783 | -0.087940274 | 0.29065409 | -0.3025599 | 0.76222528 | 0.99945981 |
| GBE1 | 57.9630622 | -0.059613816 | 0.43797336 | -0.1361129 | 0.89173204 | 0.99945981 |
| CHMP2B | 10.8897807 | 0.102828675 | 0.4108942 | 0.25025585 | 0.8023895 | 0.99945981 |
| CGGBP1 | 39.6440111 | 0.232072041 | 0.27535076 | 0.84282332 | 0.39932726 | 0.99945981 |
| ZNF654 | 45.0131986 | -0.446977377 | 0.3336078 | -1.3398289 | 0.18030097 | 0.99945981 |
| C3orf38 | 11.987341 | 0.177733959 | 0.43297336 | 0.41049629 | 0.68144192 | 0.99945981 |
| ARL13B | 12.9751474 | -0.297003438 | 0.451139 | -0.6583413 | 0.51031884 | 0.99945981 |
| NSUN3 | 24.8225905 | 0.046398941 | 0.33242938 | 0.13957533 | 0.88899553 | 0.99945981 |
| MTRNR2L12 | 58.282646 | 0.13113592 | 0.52335433 | 0.25056814 | 0.80214802 | 0.99945981 |
| CRYBG3 | 19.1956939 | 0.25617624 | 0.44110441 | 0.58076101 | 0.56140154 | 0.99945981 |
| CLDND1 | 20.425447 | 0.556261553 | 0.5095963 | 1.09157298 | 0.27502084 | 0.99945981 |
| ST3GAL6 | 27.2621711 | 0.01020853 | 0.71825091 | 0.01421304 | 0.98866002 | 0.99945981 |
| CMSS1 | 52.3281336 | 0.406436475 | 0.38672977 | 1.05095729 | 0.29327821 | 0.99945981 |
| FILIP1L | 24.7094307 | 0.288697634 | 0.49495978 | 0.58327494 | 0.55970823 | 0.99945981 |
| TBC1D23 | 27.3123104 | 0.356317377 | 0.31187822 | 1.14248881 | 0.2532509 | 0.99945981 |
| TOMM70 | 12.0764984 | -0.038985997 | 0.40168496 | -0.0970562 | 0.9226818 | 0.99945981 |
| TFG | 31.5919516 | 0.050814807 | 0.27000539 | 0.18819923 | 0.85072047 | 0.99945981 |
| SENP7 | 78.6931068 | 0.164674155 | 0.27921835 | 0.58976839 | 0.55534594 | 0.99945981 |
| PCNP | 31.8649755 | 0.043991349 | 0.27928369 | 0.15751492 | 0.87483905 | 0.99945981 |
| ZBTB11 | 21.9599017 | 0.177941552 | 0.38592824 | 0.46107419 | 0.64474538 | 0.99945981 |
| RPL24 | 43.4108708 | 0.096499126 | 0.52626924 | 0.18336456 | 0.85451198 | 0.99945981 |
| CEP97 | 9.066315 | -0.702838822 | 0.52152547 | -1.3476596 | 0.17776788 | 0.99945981 |
| NXPE3 | 15.6859653 | 0.179037073 | 0.60558632 | 0.29564253 | 0.76750309 | 0.99945981 |
| NFKBIZ | 38.5460791 | 1.340780737 | 0.44920771 | 2.98476783 | 0.00283794 | 0.45570351 |
| ALCAM | 94.0772415 | 0.095990448 | 0.48758034 | 0.19687104 | 0.84392846 | 0.99945981 |
| CBLB | 284.90565 | -0.599079991 | 0.35594667 | -1.6830611 | 0.09236326 | 0.99640647 |
| BBX | 133.973626 | 0.001051292 | 0.23417183 | 0.00448941 | 0.99641798 | 0.99945981 |
| CD47 | 87.8663955 | 0.12326315 | 0.26862625 | 0.45886487 | 0.64633121 | 0.99945981 |
| IFT57 | 12.2268583 | -0.539168643 | 0.67196356 | -0.8023778 | 0.42233445 | 0.99945981 |
| DZIP3 | 13.0009878 | -0.264749596 | 0.42468697 | -0.6233994 | 0.5330221 | 0.99945981 |
| TRAT1 | 12.9788603 | -0.669862141 | 0.6462666 | -1.0365105 | 0.29996402 | 0.99945981 |
| CD96 | 92.6210055 | -0.277947652 | 0.49970668 | -0.5562216 | 0.57805937 | 0.99945981 |
| PHLDB2 | 12.2359345 | 0.33121316 | 0.46040157 | 0.71940058 | 0.47189414 | 0.99945981 |
| ATG3 | 26.0491952 | 0.425734566 | 0.37980739 | 1.12092227 | 0.26232095 | 0.99945981 |
| SLC35A5 | 7.55846031 | -1.0457168 | 0.63030732 | -1.6590586 | 0.09710398 | 0.99945981 |
| CD200R1 | 16.4180441 | 0.22936907 | 0.56529659 | 0.40574996 | 0.68492633 | 0.99945981 |
| NEPRO | 12.4919944 | -0.696501462 | 0.48702015 | -1.4301286 | 0.1526801 | 0.99945981 |
| CFAP44 | 9.24965728 | -0.502295478 | 0.47340932 | -1.0610173 | 0.28868204 | 0.99945981 |
| SPICE1 | 13.6619456 | 0.043374148 | 0.50645463 | 0.08564271 | 0.93175044 | 0.99945981 |
| SIDT1 | 36.3740091 | -0.017665442 | 0.61403027 | -0.0287697 | 0.9770483 | 0.99945981 |
| USF3 | 26.0320512 | -0.054758485 | 0.47718308 | -0.1147536 | 0.90864041 | 0.99945981 |
| NAA50 | 51.6970926 | -0.217121784 | 0.43712194 | -0.4967076 | 0.61939527 | 0.99945981 |
| ATP6V1A | 44.4341952 | 0.005730968 | 0.32781205 | 0.01748248 | 0.98605171 | 0.99945981 |
| QTRT2 | 9.7777861 | 0.067416989 | 0.45266688 | 0.1489329 | 0.88160658 | 0.99945981 |
| TIGIT | 8.55308601 | -0.647821063 | 0.7517639 | -0.8617347 | 0.38883351 | 0.99945981 |
| ZBTB20 | 401.231086 | 0.184471428 | 0.24283399 | 0.75966066 | 0.44745745 | 0.99945981 |
| ARHGAP31 | 45.1210657 | 0.596194314 | 0.48656485 | 1.22531316 | 0.22045725 | 0.99945981 |
| TMEM39A | 19.5428349 | -0.325956132 | 0.50371421 | -0.6471053 | 0.5175638 | 0.99945981 |
| POGLUT1 | 14.0045747 | 0.447428002 | 0.6001495 | 0.74552758 | 0.45595284 | 0.99945981 |
| TIMMDC1 | 14.273679 | 0.093472387 | 0.40698914 | 0.22966801 | 0.81834975 | 0.99945981 |
| COX17 | 10.375074 | 0.508627843 | 0.44837411 | 1.13438273 | 0.25663404 | 0.99945981 |
| GSK3B | 133.38086 | -0.003391891 | 0.22362972 | -0.0151674 | 0.9878986 | 0.99945981 |
| LRRC58 | 19.2818748 | -0.83936957 | 0.45028424 | -1.8640883 | 0.06230931 | 0.92949575 |
| NDUFB4 | 10.3351967 | -0.678680256 | 0.63669933 | -1.0659352 | 0.28645292 | 0.99945981 |
| HCLS1 | 78.3071558 | 0.231734121 | 0.29088222 | 0.79665962 | 0.42564874 | 0.99945981 |
| GOLGB1 | 56.0304407 | -0.570176008 | 0.29762109 | -1.9157783 | 0.05539332 | 0.92445778 |
| IQCB1 | 44.7081811 | 0.012323774 | 0.38401696 | 0.03209174 | 0.97439889 | 0.99945981 |
| EAF2 | 35.9534925 | 0.697652174 | 0.46011289 | 1.51626304 | 0.12945285 | 0.99945981 |
| SLC15A2 | 17.1318876 | 0.10956455 | 0.38574317 | 0.28403497 | 0.77638358 | 0.99945981 |
| CD86 | 61.4468971 | 1.089065865 | 0.69931067 | 1.55734198 | 0.11938931 | 0.99945981 |
| FAM162A | 15.8629091 | -0.127541708 | 0.39616462 | -0.3219412 | 0.74749725 | 0.99945981 |
| KPNA1 | 46.605819 | -0.015804669 | 0.25758835 | -0.0613563 | 0.95107545 | 0.99945981 |
| PARP9 | 51.7778262 | 0.332027377 | 0.54068804 | 0.61408308 | 0.53916043 | 0.99945981 |
| DTX3L | 19.1364358 | 0.26845309 | 0.62161897 | 0.43186116 | 0.66584233 | 0.99945981 |
| PARP15 | 37.436694 | -0.278679786 | 0.62096423 | -0.4487856 | 0.65358635 | 0.99945981 |
| PARP14 | 148.967803 | 0.350901112 | 0.68047365 | 0.51567186 | 0.60608361 | 0.99945981 |
| HSPBAP1 | 27.5820312 | 0.485869233 | 0.45685274 | 1.06351388 | 0.28754897 | 0.99945981 |
| DIRC2 | 41.98656 | 0.493694541 | 0.53861181 | 0.91660549 | 0.35934941 | 0.99945981 |
| SEC22A | 32.9150243 | -0.364741052 | 0.31799699 | -1.1469953 | 0.25138356 | 0.99945981 |
| HACD2 | 18.9431372 | 0.118560827 | 0.36319616 | 0.32643744 | 0.74409341 | 0.99945981 |
| CCDC14 | 28.802435 | -0.079664688 | 0.49376084 | -0.1613427 | 0.87182352 | 0.99945981 |

| | | | | | | |
|---|---|---|---|---|---|---|
| ITGB5 | 16.9989377 | 0.518920772 | 0.55942216 | 0.92760139 | 0.35361438 | 0.99945981 |
| HEG1 | 26.4280983 | 0.215114965 | 0.40260241 | 0.53431117 | 0.59312627 | 0.99945981 |
| ZNF148 | 116.604701 | 0.083716862 | 0.23906775 | 0.3501805 | 0.72620324 | 0.99945981 |
| SNX4 | 27.4040702 | 0.156482843 | 0.29582069 | 0.52897869 | 0.59682023 | 0.99945981 |
| OSBPL11 | 31.498072 | 0.493901121 | 0.36869256 | 1.33960154 | 0.18037492 | 0.99945981 |
| ZXDC | 38.641072 | -0.10172776 | 0.41518344 | -0.2450188 | 0.80644185 | 0.99945981 |
| MGLL | 17.2595328 | -0.030299089 | 0.51638723 | -0.0586751 | 0.95321087 | 0.99945981 |
| SEC61A1 | 26.6505277 | -0.17218368 | 0.53311074 | -0.3229792 | 0.74671099 | 0.99945981 |
| RUVBL1 | 12.6953202 | -0.388551036 | 0.4137847 | -0.9390174 | 0.34772181 | 0.99945981 |
| EEFSEC | 37.7212483 | 0.135066824 | 0.39954331 | 0.33805302 | 0.73532323 | 0.99945981 |
| RPN1 | 23.7423561 | -0.238606996 | 0.46860341 | -0.5091875 | 0.61062081 | 0.99945981 |
| RAB7A | 95.904748 | 0.302489974 | 0.25144718 | 1.2029961 | 0.22897783 | 0.99945981 |
| ACAD9 | 8.20355605 | -0.204311896 | 0.4707266 | -0.4340352 | 0.66426291 | 0.99945981 |
| CNBP | 46.3770429 | -0.68831428 | 0.43062311 | -1.5984146 | 0.10995073 | 0.99945981 |
| COPG1 | 13.933036 | -0.227395739 | 0.45155665 | -0.5035819 | 0.61455524 | 0.99945981 |
| HMCES | 10.2462202 | -0.496693729 | 0.520008 | -0.9551656 | 0.33949398 | 0.99945981 |
| H1FX | 22.6278213 | 0.267104132 | 0.5246392 | 0.50911966 | 0.61066835 | 0.99945981 |
| MBD4 | 20.0665367 | 0.167482668 | 0.39746366 | 0.42137857 | 0.67347866 | 0.99945981 |
| PLXND1 | 28.3679432 | -0.189101454 | 0.43970742 | -0.430062 | 0.66715054 | 0.99945981 |
| TMCC1 | 103.390116 | 0.250657153 | 0.21157455 | 1.18472262 | 0.23612714 | 0.99945981 |
| PIK3R4 | 18.6470748 | 0.312951895 | 0.56896195 | 0.55004011 | 0.58229186 | 0.99945981 |
| ATP2C1 | 54.0688382 | 0.150082177 | 0.30552399 | 0.49122879 | 0.62326464 | 0.99945981 |
| MRPL3 | 15.8639302 | -0.059494783 | 0.36821291 | -0.1615771 | 0.87163887 | 0.99945981 |
| ACPP | 11.6149494 | 1.225521933 | 0.69513991 | 1.762986 | 0.07790285 | 0.98289661 |
| DNAJC13 | 56.8052603 | 0.231308606 | 0.34313067 | 0.6741123 | 0.50023992 | 0.99945981 |
| UBA5 | 12.7738874 | 0.455819667 | 0.57340516 | 0.79493472 | 0.42665148 | 0.99945981 |
| CDV3 | 37.2668669 | -0.265717152 | 0.39653581 | -0.6700962 | 0.50279645 | 0.99945981 |
| TOPBP1 | 24.4507432 | 0.375549864 | 0.31009491 | 1.2110804 | 0.22586459 | 0.99945981 |
| RYK | 36.0132088 | -0.029504151 | 0.3121599 | -0.0945161 | 0.92469916 | 0.99945981 |
| CEP63 | 22.7677676 | -0.111130497 | 0.40638466 | -0.2734614 | 0.7844986 | 0.99945981 |
| PPP2R3A | 15.5077986 | -0.580150625 | 0.52043904 | -1.1147331 | 0.26496481 | 0.99945981 |
| MSL2 | 35.9559164 | -0.06469889 | 0.31024203 | -0.2085433 | 0.83480479 | 0.99945981 |
| PCCB | 11.5249208 | -0.633491424 | 0.41425186 | -1.5292422 | 0.12620441 | 0.99945981 |
| STAG1 | 226.71539 | 0.166974265 | 0.23136705 | 0.72168559 | 0.47048781 | 0.99945981 |
| NCK1 | 39.6513744 | 0.024767712 | 0.35909986 | 0.06897166 | 0.94501218 | 0.99945981 |
| ARMC8 | 53.6762787 | 0.054268178 | 0.2574063 | 0.21082692 | 0.83302233 | 0.99945981 |
| PIK3CB | 70.8438456 | -0.003085437 | 0.42008882 | -0.0073447 | 0.99413981 | 0.99945981 |
| COPB2 | 25.4010486 | 0.251093766 | 0.34264678 | 0.73280643 | 0.4636765 | 0.99945981 |
| AC097103.2 | 13.2516977 | -0.204320051 | 0.6083712 | -0.3358477 | 0.73698574 | 0.99945981 |
| SLC25A36 | 32.3765445 | -0.043423423 | 0.40991635 | -0.1059324 | 0.91563599 | 0.99945981 |
| ZBTB38 | 37.1054613 | -0.587111873 | 0.29307464 | -2.0032845 | 0.04514676 | 0.90096666 |
| RASA2 | 186.305347 | -0.026742489 | 0.35005446 | -0.0763952 | 0.93910467 | 0.99945981 |
| RNF7 | 13.7769426 | -0.551473309 | 0.5097091 | -1.0819374 | 0.27928037 | 0.99945981 |
| ATP1B3 | 94.2403625 | 0.763630624 | 0.56141486 | 1.36018955 | 0.17376995 | 0.99945981 |
| TFDP2 | 62.0364493 | 0.035668432 | 0.4408849 | 0.08090191 | 0.93551996 | 0.99945981 |
| GK5 | 28.743946 | 0.384119956 | 0.52130574 | 0.73684199 | 0.46121844 | 0.99945981 |
| XRN1 | 115.887243 | 0.266922422 | 0.40663485 | 0.65641797 | 0.51155523 | 0.99945981 |
| ATR | 61.9355471 | 0.148350752 | 0.22809526 | 0.65038946 | 0.51544068 | 0.99945981 |
| U2SURP | 43.9251732 | 0.148936243 | 0.30432407 | 0.48940013 | 0.62455844 | 0.99945981 |
| SLC9A9 | 213.703036 | 0.552261207 | 0.44665236 | 1.23644528 | 0.2162931 | 0.99945981 |
| C3orf58 | 11.2382454 | -0.158048184 | 0.48627128 | -0.3250206 | 0.74516548 | 0.99945981 |
| PLSCR1 | 81.2810067 | 0.282936174 | 0.6341429 | 0.44617101 | 0.65547373 | 0.99945981 |
| GYG1 | 13.837395 | -1.357853286 | 0.48235506 | -2.8150493 | 0.00487697 | 0.50757754 |
| HLTF | 11.6135486 | 0.751640701 | 0.46454753 | 1.61800604 | 0.1056613 | 0.99945981 |
| HPS3 | 26.0094444 | -0.342389735 | 0.44555275 | -0.7684606 | 0.44221359 | 0.99945981 |
| WWTR1 | 12.7473098 | -0.511217541 | 0.60162566 | -0.849727 | 0.39547691 | 0.99945981 |
| ANKUB1 | 48.7822151 | 1.001826879 | 0.74596803 | 1.34298903 | 0.17927553 | 0.99945981 |
| RNF13 | 131.417733 | 0.483930701 | 0.4591453 | 1.0539816 | 0.29189134 | 0.99945981 |
| TSC22D2 | 52.0947393 | 0.386656526 | 0.2538084 | 1.52341892 | 0.12765393 | 0.99945981 |
| SERP1 | 46.4106345 | 0.464149921 | 0.50601194 | 0.91727068 | 0.35900082 | 0.99945981 |
| EIF2A | 19.2088773 | -0.082724374 | 0.42368552 | -0.1952495 | 0.84519765 | 0.99945981 |
| SELENOT | 30.3482919 | -0.279857381 | 0.34937913 | -0.8010135 | 0.42312386 | 0.99945981 |
| SIAH2 | 12.9592969 | 0.171050277 | 0.58429356 | 0.29274716 | 0.76971542 | 0.99945981 |
| GPR171 | 15.9142055 | -1.35607446 | 0.55570638 | -2.4402715 | 0.01467623 | 0.73008906 |
| MBNL1 | 850.449995 | 0.386478143 | 0.22441864 | 1.72213034 | 0.08504591 | 0.99193047 |
| RAP2B | 18.7925465 | -0.238973981 | 0.39059017 | -0.6118279 | 0.5406516 | 0.99945981 |
| DHX36 | 56.8032809 | 0.034771913 | 0.31530779 | 0.11027927 | 0.9121879 | 0.99945981 |
| GMPS | 19.7377121 | 0.294474336 | 0.3269099 | 0.90078132 | 0.3677046 | 0.99945981 |
| SSR3 | 21.4376458 | -0.401362963 | 0.47462837 | -0.8456363 | 0.39775569 | 0.99945981 |
| TIPARP | 14.777378 | 0.392395175 | 0.48720304 | 0.80540379 | 0.42058671 | 0.99945981 |
| LINC00886 | 10.7280851 | 0.088755371 | 0.54719522 | 0.16220056 | 0.87114792 | 0.99945981 |
| CCNL1 | 64.3694993 | 0.986668325 | 0.36840246 | 2.67823492 | 0.00740113 | 0.57509411 |
| RSRC1 | 99.2296454 | 0.196069284 | 0.22810218 | 0.85956777 | 0.39002735 | 0.99945981 |
| GFM1 | 17.0231693 | 0.238379905 | 0.42896938 | 0.55570378 | 0.57841337 | 0.99945981 |
| MFSD1 | 38.0573351 | 0.351424279 | 0.47837552 | 0.73462012 | 0.46257088 | 0.99945981 |
| SMC4 | 26.6272824 | -0.130109816 | 0.54945643 | -0.2367973 | 0.81281402 | 0.99945981 |
| KPNA4 | 54.6128673 | 0.146107371 | 0.32020954 | 0.45628676 | 0.64818378 | 0.99945981 |
| PPM1L | 85.2453512 | 1.723172652 | 0.5157805 | 3.34090304 | 0.00083506 | 0.27121928 |
| NMD3 | 17.7489075 | 0.520510664 | 0.53817119 | 0.96718419 | 0.33345196 | 0.99945981 |
| PDCD10 | 34.6992732 | 0.185770204 | 0.33861814 | 0.54861268 | 0.58327128 | 0.99945981 |
| GOLIM4 | 22.2875412 | 0.362089801 | 0.43711781 | 0.82835747 | 0.40746809 | 0.99945981 |
| MECOM | 67.261176 | 0.550296389 | 0.5372109 | 1.0243582 | 0.30566612 | 0.99945981 |
| SEC62 | 42.144197 | -0.098576493 | 0.31986513 | -0.3081814 | 0.75794428 | 0.99945981 |
| GPR160 | 8.75251039 | -0.859725297 | 0.61224449 | -1.4042189 | 0.16025367 | 0.99945981 |
| PHC3 | 124.036744 | -0.014069232 | 0.25865829 | -0.0543931 | 0.95662196 | 0.99945981 |
| PRKCI | 19.3184909 | 0.21461328 | 0.3640775 | 0.58947142 | 0.55554508 | 0.99945981 |
| SKIL | 42.8902529 | 0.454501418 | 0.37383526 | 1.21577996 | 0.22406876 | 0.99945981 |
| RPL22L1 | 9.48846823 | -0.613881027 | 0.60317123 | -1.0177558 | 0.30879401 | 0.99945981 |
| TNIK | 107.726812 | 0.352507022 | 0.50696651 | 0.69532606 | 0.48685099 | 0.99945981 |
| PLD1 | 38.0234859 | 0.142016814 | 0.73850209 | 0.19230388 | 0.84750418 | 0.99945981 |
| FNDC3B | 344.399906 | 0.067374315 | 0.49498861 | 0.13611286 | 0.89173206 | 0.99945981 |
| TNFSF10 | 36.9319645 | 0.833096784 | 0.45556216 | 1.82872252 | 0.06744119 | 0.94796811 |
| NCEH1 | 22.7087086 | -0.412982075 | 0.42934074 | -0.9618982 | 0.33610075 | 0.99945981 |
| NLGN1 | 10.3215 | 0.33339288 | 0.53210637 | 0.62655307 | 0.53095224 | 0.99945981 |
| NAALADL2 | 30.4839278 | 0.097506467 | 0.6325706 | 0.15414322 | 0.87749681 | 0.99945981 |
| TBL1XR1 | 189.6259 | 0.127617509 | 0.30244792 | 0.4219487 | 0.67306246 | 0.99945981 |
| ZMAT3 | 19.0110934 | -0.040610596 | 0.38922973 | -0.1043358 | 0.91690287 | 0.99945981 |
| PIK3CA | 54.8263167 | 0.045292972 | 0.31859412 | 0.14216512 | 0.88694958 | 0.99945981 |
| MFN1 | 26.2294247 | -0.045340715 | 0.37902768 | -0.1196238 | 0.9047812 | 0.99945981 |
| GNB4 | 35.7951351 | 0.469054546 | 0.49851005 | 0.94091292 | 0.34674949 | 0.99945981 |
| NDUFB5 | 10.4222541 | 0.105185807 | 0.45380925 | 0.23178418 | 0.81670564 | 0.99945981 |
| USP13 | 8.22233243 | 0.511606156 | 0.43678157 | 1.17130893 | 0.24147462 | 0.99945981 |
| TTC14 | 48.1552556 | 0.435615077 | 0.24968666 | 1.74464698 | 0.08104634 | 0.9846219 |
| FXR1 | 42.6240551 | -0.002002321 | 0.32894067 | -0.0060872 | 0.99514316 | 0.99945981 |
| ATP11B | 99.1118398 | 0.040852113 | 0.23090331 | 0.17692303 | 0.85956885 | 0.99945981 |
| DCUN1D1 | 27.6073024 | -0.255696222 | 0.31373686 | -0.8150022 | 0.41507107 | 0.99945981 |
| MCCC1 | 21.844993 | -0.425553263 | 0.38006333 | -1.1196904 | 0.26284572 | 0.99945981 |
| MCF2L2 | 15.647443 | 0.087504568 | 0.40407827 | 0.21655351 | 0.82855632 | 0.99945981 |
| B3GNT5 | 14.0282817 | 0.891997989 | 0.51472189 | 1.73297076 | 0.08310087 | 0.98529784 |
| KLHL6 | 34.0141433 | -0.263721586 | 0.43896743 | -0.6007771 | 0.54798845 | 0.99945981 |
| KLHL24 | 48.2497176 | -0.380078115 | 0.37633918 | -1.009935 | 0.31252643 | 0.99945981 |
| YEATS2 | 48.476692 | -0.206196789 | 0.41012033 | -0.5027714 | 0.61512498 | 0.99945981 |
| ABCC5 | 28.2161702 | -0.156566312 | 0.41390336 | -0.3782678 | 0.70523167 | 0.99945981 |

| | | | | | | |
|---|---|---|---|---|---|---|
| DVL3 | 9.5272426 | -0.133771467 | 0.50525221 | -0.2647618 | 0.79119299 | 0.99945981 |
| AP2M1 | 17.2583235 | 0.115748085 | 0.42693 | 0.27111724 | 0.78630086 | 0.99945981 |
| PSMD2 | 11.4514248 | 0.245253491 | 0.40831215 | 0.60065195 | 0.54807183 | 0.99945981 |
| EIF4G1 | 24.8320435 | -0.483724773 | 0.50451164 | -0.9587981 | 0.33766049 | 0.99945981 |
| POLR2H | 9.92160449 | -0.161210833 | 0.53104819 | -0.303571 | 0.76145475 | 0.99945981 |
| VPS8 | 91.7564653 | 0.064262422 | 0.27764501 | 0.23145535 | 0.81696107 | 0.99945981 |
| MAP3K13 | 16.4636563 | -0.214721107 | 0.60386944 | -0.3555754 | 0.72215858 | 0.99945981 |
| SENP2 | 19.4745396 | -0.269205124 | 0.38842497 | -0.6930685 | 0.48826655 | 0.99945981 |
| IGF2BP2 | 16.4939513 | -0.784086722 | 1.09883356 | -0.7135628 | 0.47549758 | 0.99945981 |
| TRA2B | 47.436118 | 0.224631522 | 0.27244849 | 0.82449171 | 0.40966022 | 0.99945981 |
| ETV5 | 16.101558 | 0.02622456 | 0.6806117 | 0.03853087 | 0.96926442 | 0.99945981 |
| DGKG | 24.1633722 | -1.022133532 | 0.9593384 | -1.0654567 | 0.28666931 | 0.99945981 |
| EIF4A2 | 41.7127123 | 0.083169401 | 0.3108454 | 0.26755873 | 0.789039 | 0.99945981 |
| ST6GAL1 | 181.055742 | -0.19793734 | 0.3978257 | -0.4975479 | 0.61880274 | 0.99945981 |
| BCL6 | 56.8834395 | 0.039297911 | 0.62913033 | 0.06246386 | 0.95019344 | 0.99945981 |
| LPP | 319.171814 | 0.253653117 | 0.27690878 | 0.91601689 | 0.35965804 | 0.99945981 |
| TPRG1 | 40.572276 | -0.202524324 | 0.56181828 | -0.3604801 | 0.71848812 | 0.99945981 |
| P3H2 | 15.0309802 | 1.708621467 | 0.47064358 | 3.63039363 | 0.00028299 | 0.17661106 |
| IL1RAP | 33.1251506 | -0.419687944 | 0.70304767 | -0.5969552 | 0.55053731 | 0.99945981 |
| CCDC50 | 30.5844203 | 0.346417829 | 0.40249971 | 0.86066603 | 0.38942201 | 0.99945981 |
| MB21D2 | 35.9940128 | -0.486569634 | 0.85668121 | -0.5679705 | 0.57005502 | 0.99945981 |
| OPA1 | 47.7677187 | -0.026474242 | 0.28063203 | -0.0943379 | 0.92484073 | 0.99945981 |
| ATP13A3 | 71.6779194 | 0.120185718 | 0.36672601 | 0.32772619 | 0.7431187 | 0.99945981 |
| LSG1 | 9.65200368 | -0.269157263 | 0.43451685 | -0.6194403 | 0.53562632 | 0.99945981 |
| XXYLT1 | 11.7860814 | -0.297957355 | 0.42710879 | -0.6976147 | 0.48541821 | 0.99945981 |
| ACAP2 | 191.568577 | 0.409043296 | 0.19318221 | 2.11739634 | 0.03422622 | 0.87717522 |
| PPP1R2 | 38.714643 | 0.119919808 | 0.40514867 | 0.29598964 | 0.76723799 | 0.99945981 |
| MUC20-OT1 | 72.0774889 | 0.227149918 | 0.38756689 | 0.58609217 | 0.55781358 | 0.99945981 |
| TNK2 | 25.4675387 | 0.53478108 | 0.34356048 | 1.55658498 | 0.11956905 | 0.99945981 |
| TFRC | 69.8291986 | 1.976982133 | 1.01692603 | 1.94407663 | 0.05188621 | 0.9230968 |
| PCYT1A | 17.1012208 | 0.05264933 | 0.33727649 | 0.15610139 | 0.8759531 | 0.99945981 |
| UBXN7 | 33.8997788 | -0.221070466 | 0.3576781 | -0.618071 | 0.53652853 | 0.99945981 |
| RNF168 | 32.8298369 | -0.241166796 | 0.29202564 | -0.8258412 | 0.40889419 | 0.99945981 |
| PIGX | 19.8090109 | 0.059513775 | 0.36063236 | 0.16502616 | 0.86892339 | 0.99945981 |
| PAK2 | 93.443467 | 0.07930782 | 0.25505696 | 0.3109416 | 0.75584502 | 0.99945981 |
| SENP5 | 51.8413835 | -0.346595541 | 0.44407943 | -0.780481 | 0.43510782 | 0.99945981 |
| NCBP2 | 12.9536164 | 0.219819292 | 0.40854514 | 0.53805387 | 0.59053986 | 0.99945981 |
| DLG1 | 105.180261 | 0.017965185 | 0.23179246 | 0.07750547 | 0.93822144 | 0.99945981 |
| RUBCN | 36.7132228 | -0.00418888 | 0.34008435 | -0.0123172 | 0.99017256 | 0.99945981 |
| FYTTD1 | 16.9739784 | -0.333602663 | 0.39944509 | -0.8351653 | 0.40362467 | 0.99945981 |
| LRCH3 | 76.6515511 | 0.037112945 | 0.20881665 | 0.17772982 | 0.85893517 | 0.99945981 |
| IQCG | 12.268366 | -0.446155172 | 0.47840378 | -0.9325912 | 0.35103106 | 0.99945981 |
| RPL35A | 67.455313 | -0.005829823 | 0.46967276 | -0.0124125 | 0.9900965 | 0.99945981 |
| ZNF595 | 10.459181 | 1.175498239 | 0.67565646 | 1.73978688 | 0.08189645 | 0.9846219 |
| ZNF718 | 28.5549827 | -0.333009476 | 0.3805367 | -0.8751048 | 0.3815169 | 0.99945981 |
| ZNF141 | 25.0698415 | -0.040891616 | 0.3042899 | -0.1343837 | 0.89309914 | 0.99945981 |
| ZNF721 | 42.0432541 | -0.204485569 | 0.30370025 | -0.6733138 | 0.50074767 | 0.99945981 |
| PIGG | 16.2177586 | -0.742330011 | 0.42095061 | -1.7634611 | 0.07782276 | 0.98289661 |
| ATP5ME | 8.53008741 | -0.062274769 | 0.52679089 | -0.1182154 | 0.90589703 | 0.99945981 |
| PCGF3 | 21.6266329 | 0.226210028 | 0.35876615 | 0.63052222 | 0.52835297 | 0.99945981 |
| AC139887.2 | 32.6130762 | -0.471240714 | 0.37582207 | -1.2538931 | 0.20988085 | 0.99945981 |
| GAK | 42.3592141 | -0.247595928 | 0.30326665 | -0.8164298 | 0.41425436 | 0.99945981 |
| SPON2 | 19.208608 | 0.079144596 | 0.61306842 | 0.12909586 | 0.8972818 | 0.99945981 |
| CTBP1 | 19.254417 | -0.090480509 | 0.37829779 | -0.239178 | 0.81096756 | 0.99945981 |
| MAEA | 16.7771179 | -0.402285958 | 0.36297198 | -1.1083113 | 0.26772738 | 0.99945981 |
| UVSSA | 16.6630102 | 0.282731034 | 0.36298081 | 0.77891454 | 0.43603006 | 0.99945981 |
| SLBP | 18.9937241 | -0.085615405 | 0.36844266 | -0.232371 | 0.81624984 | 0.99945981 |
| TACC3 | 14.8597132 | 0.211114481 | 0.37401715 | 0.56445133 | 0.57244701 | 0.99945981 |
| NSD2 | 40.8874604 | -0.206767158 | 0.31700696 | -0.652248 | 0.51424119 | 0.99945981 |
| NELFA | 11.4857928 | 0.149308541 | 0.42418484 | 0.35198933 | 0.72484626 | 0.99945981 |
| MXD4 | 16.8168771 | 0.054227705 | 0.44586931 | 0.12162242 | 0.90319806 | 0.99945981 |
| ZFYVE28 | 38.7225302 | -0.725806243 | 0.43289735 | -1.6766244 | 0.09361596 | 0.99945981 |
| RNF4 | 28.8827539 | -0.200680075 | 0.31254902 | -0.6420755 | 0.52082415 | 0.99945981 |
| FAM193A | 52.5370183 | 0.146496209 | 0.28216333 | 0.5191894 | 0.60362867 | 0.99945981 |
| TNIP2 | 11.8963935 | -0.157986843 | 0.52220443 | -0.3025383 | 0.76224174 | 0.99945981 |
| SH3BP2 | 39.5274082 | 0.282786895 | 0.44794681 | 0.6312957 | 0.5278472 | 0.99945981 |
| ADD1 | 84.4105816 | -0.042864493 | 0.3148907 | -0.136125 | 0.89172247 | 0.99945981 |
| MFSD10 | 10.5801702 | 0.012389799 | 0.5736865 | 0.02159681 | 0.98276958 | 0.99945981 |
| NOP14 | 7.03661123 | -0.781856884 | 0.59736546 | -1.3088418 | 0.19058795 | 0.99945981 |
| HTT | 68.6977751 | -0.253393792 | 0.23992511 | -1.056137 | 0.29090561 | 0.99945981 |
| RGS12 | 12.9257258 | 0.101441819 | 0.5792321 | 0.17513156 | 0.86097626 | 0.99945981 |
| LRPAP1 | 17.238197 | -0.734404234 | 0.45892773 | -1.6002612 | 0.10954065 | 0.99945981 |
| LYAR | 8.59580848 | -0.236348085 | 0.57378411 | -0.4119112 | 0.68040453 | 0.99945981 |
| STX18 | 40.0064186 | -0.119345655 | 0.32708055 | -0.3648815 | 0.71519983 | 0.99945981 |
| STX18-AS1 | 12.9023232 | -0.771007004 | 0.54124021 | -1.4245191 | 0.15429627 | 0.99945981 |
| JAKMIP1 | 20.1730955 | -1.202802994 | 0.76240976 | -1.5776333 | 0.11464989 | 0.99945981 |
| MAN2B2 | 10.7865282 | 0.233806227 | 0.50173938 | 0.46599138 | 0.64122167 | 0.99945981 |
| MRFAP1 | 14.120458 | -0.389926692 | 0.45827243 | -0.8508622 | 0.3948459 | 0.99945981 |
| KIAA0232 | 54.9383096 | 0.061302175 | 0.26513715 | 0.2312093 | 0.8171522 | 0.99945981 |
| TBC1D14 | 80.5713212 | -0.321077169 | 0.45568207 | -0.7046079 | 0.4810543 | 0.99945981 |
| AFAP1 | 9.25189665 | -0.79658125 | 0.57378404 | -1.3882945 | 0.16504738 | 0.99945981 |
| SH3TC1 | 30.041759 | -0.539665994 | 0.44313768 | -1.2178292 | 0.2232889 | 0.99945981 |
| ACOX3 | 9.10490148 | -0.826270282 | 0.58811385 | -1.4049495 | 0.16003629 | 0.99945981 |
| SLC2A9 | 23.0781418 | -0.410449123 | 0.55914889 | -0.7340605 | 0.46291186 | 0.99945981 |
| WDR1 | 34.7674175 | 0.2244047 | 0.32919395 | 0.68167929 | 0.49544177 | 0.99945981 |
| RAB28 | 34.4984019 | -0.039393599 | 0.28464033 | -0.1383978 | 0.88992603 | 0.99945981 |
| BOD1L1 | 48.0251827 | -0.024755945 | 0.34057765 | -0.0726881 | 0.94205431 | 0.99945981 |
| AC092546.1 | 19.8668411 | 1.681800504 | 0.67084873 | 2.50697426 | 0.01217696 | 0.67187199 |
| CPEB2 | 31.4606114 | 0.708198425 | 0.32805514 | 2.1587786 | 0.03086735 | 0.85791231 |
| AC098829.1 | 13.428729 | 0.633253801 | 0.71565642 | 0.88485729 | 0.37623361 | 0.99945981 |
| FBXL5 | 71.7750342 | 0.754354067 | 0.35551916 | 2.12183804 | 0.03385134 | 0.87694135 |
| FAM200B | 10.0231831 | 0.606449142 | 0.47640323 | 1.27297444 | 0.20302711 | 0.99945981 |
| CD38 | 43.0218922 | 0.046430985 | 0.76260553 | 0.06088467 | 0.95145106 | 0.99945981 |
| TAPT1 | 32.7851252 | -0.1158885 | 0.34948757 | -0.3315955 | 0.74019473 | 0.99945981 |
| TAPT1-AS1 | 11.8612681 | 0.933924127 | 0.47467104 | 1.96751866 | 0.04912345 | 0.91888954 |
| LDB2 | 11.9366744 | 0.417084361 | 0.71242717 | 0.5854414 | 0.55825096 | 0.99945981 |
| LAP3 | 26.6073154 | 0.237287079 | 0.60967061 | 0.38920538 | 0.69712423 | 0.99945981 |
| MED28 | 17.6862541 | 0.233932293 | 0.33191228 | 0.70480155 | 0.48093374 | 0.99945981 |
| DCAF16 | 11.5503844 | 0.132100401 | 0.55286474 | 0.23893801 | 0.81115365 | 0.99945981 |
| LCORL | 75.9735716 | 0.159190472 | 0.301214 | 0.52849625 | 0.59715495 | 0.99945981 |
| AC024230.1 | 11.4734672 | 0.38687126 | 0.80057124 | 0.48324401 | 0.62892249 | 0.99945981 |
| PACRGL | 11.2306169 | -0.219113974 | 0.46258402 | -0.4736739 | 0.63573247 | 0.99945981 |
| KCNIP4 | 41.2594958 | -0.676883806 | 0.94550206 | -0.7158988 | 0.47405383 | 0.99945981 |
| DHX15 | 45.6775763 | 0.035046731 | 0.2599489 | 0.13482162 | 0.89275291 | 0.99945981 |
| CCDC149 | 10.4319216 | 0.531979392 | 0.44090453 | 1.20656369 | 0.22760022 | 0.99945981 |
| SEPSECS | 11.0636482 | 0.598588119 | 0.53831054 | 1.11197548 | 0.26614869 | 0.99945981 |
| ANAPC4 | 20.4221138 | 0.18309254 | 0.38544789 | 0.47501243 | 0.63477811 | 0.99945981 |
| SEL1L3 | 87.5809626 | -1.127855048 | 0.40035273 | -2.8171534 | 0.00484514 | 0.50757754 |
| RBPJ | 315.102998 | 0.18677541 | 0.6319536 | 0.29555241 | 0.76757192 | 0.99945981 |
| TBC1D19 | 18.3610359 | -0.608367806 | 0.46757746 | -1.3011059 | 0.1932222 | 0.99945981 |
| STIM2 | 81.851012 | -0.250410125 | 0.32839796 | -0.7625203 | 0.44574951 | 0.99945981 |
| ARAP2 | 163.553677 | -0.314536709 | 0.3657811 | -0.8599042 | 0.38984184 | 0.99945981 |
| DTHD1 | 38.3316177 | -1.346202077 | 0.52120005 | -2.5828894 | 0.00979767 | 0.62860768 |

| | | | | | | |
|---|---|---|---|---|---|---|
| C4orf19 | 12.3877189 | -0.055294638 | 0.68312703 | -0.0809434 | 0.93548695 | 0.99945981 |
| RELL1 | 29.6181134 | -0.192076226 | 0.54314647 | -0.3536362 | 0.72361158 | 0.99945981 |
| TBC1D1 | 102.310407 | 0.18474071 | 0.25131679 | 0.73509101 | 0.46228407 | 0.99945981 |
| KLF3 | 38.1192221 | 0.053538191 | 0.36813468 | 0.14543099 | 0.8843706 | 0.99945981 |
| TLR1 | 25.8368493 | 0.41101103 | 0.43808119 | 0.93820744 | 0.34813782 | 0.99945981 |
| TMEM156 | 31.0767771 | -0.511081162 | 0.45644119 | -1.1197087 | 0.26283792 | 0.99945981 |
| KLHL5 | 43.7464225 | -0.552724071 | 0.41781847 | -1.3228809 | 0.18587499 | 0.99945981 |
| RFC1 | 28.3636192 | -0.151207396 | 0.34599134 | -0.4370265 | 0.66209211 | 0.99945981 |
| SMIM14 | 33.4151836 | 0.00788459 | 0.31916363 | 0.02470391 | 0.98029114 | 0.99945981 |
| UBE2K | 113.662036 | 0.200987468 | 0.18714651 | 1.07395786 | 0.28284157 | 0.99945981 |
| PDS5A | 117.550254 | 0.176478901 | 0.28179864 | 0.62625889 | 0.53114515 | 0.99945981 |
| N4BP2 | 29.9357259 | -0.263755549 | 0.41950681 | -0.6287277 | 0.52952734 | 0.99945981 |
| RHOH | 96.7607428 | -1.76528147 | 0.54272928 | -3.2526004 | 0.00114354 | 0.34132242 |
| RBM47 | 249.898191 | 0.343799743 | 0.45098508 | 0.76233063 | 0.4458627 | 0.99945981 |
| APBB2 | 14.0431271 | 0.588294001 | 0.76545212 | 0.76855754 | 0.44215602 | 0.99945981 |
| TMEM33 | 22.1361897 | 0.196738426 | 0.36687215 | 0.53625882 | 0.59177969 | 0.99945981 |
| SLC30A9 | 36.8870908 | -0.039916728 | 0.33514943 | -0.1191013 | 0.90519511 | 0.99945981 |
| ATP8A1 | 102.853545 | -0.782592962 | 0.49005905 | -1.5969361 | 0.11027996 | 0.99945981 |
| GUF1 | 10.4044022 | 0.282577949 | 0.44946992 | 0.62869157 | 0.529551 | 0.99945981 |
| ATP10D | 28.644789 | 0.148380884 | 0.35510237 | 0.41785383 | 0.67605399 | 0.99945981 |
| NFXL1 | 18.3839188 | -1.963384533 | 0.68875653 | -2.850622 | 0.00436338 | 0.50757754 |
| TXK | 58.9253032 | 0.765474638 | 0.39881572 | 1.91936929 | 0.05493761 | 0.92445778 |
| TEC | 25.5201655 | -0.806909122 | 0.50854646 | -1.586697 | 0.11258129 | 0.99945981 |
| SLAIN2 | 39.8068108 | 0.26320517 | 0.27143515 | 0.96967976 | 0.33220614 | 0.99945981 |
| FRYL | 153.496859 | -0.179199158 | 0.26262901 | -0.6823281 | 0.49503149 | 0.99945981 |
| OCIAD1 | 38.1202566 | -0.026094671 | 0.31217251 | -0.0835905 | 0.93338198 | 0.99945981 |
| DCUN1D4 | 20.6340968 | 0.344869536 | 0.37388545 | 0.92239358 | 0.35632332 | 0.99945981 |
| SCFD2 | 70.6560323 | -0.257347629 | 0.4804456 | -0.5356436 | 0.59220487 | 0.99945981 |
| AC058822.1 | 19.6433902 | -0.024925127 | 0.66667689 | -0.0373871 | 0.97017634 | 0.99945981 |
| CHIC2 | 57.0901527 | 0.044991783 | 0.30533676 | 0.14735135 | 0.8828547 | 0.99945981 |
| TMEM165 | 75.8665467 | -0.067166978 | 0.21431724 | -0.3133998 | 0.75397694 | 0.99945981 |
| CLOCK | 37.7558153 | 0.240025141 | 0.24529921 | 0.97849945 | 0.32782736 | 0.99945981 |
| EXOC1 | 43.2618609 | -0.088405168 | 0.31419673 | -0.2813688 | 0.77842752 | 0.99945981 |
| CEP135 | 26.1757685 | 0.281865099 | 0.43105604 | 0.65389432 | 0.51317989 | 0.99945981 |
| AASDH | 11.4670977 | -0.201294683 | 0.48584938 | -0.414315 | 0.67864343 | 0.99945981 |
| SRP72 | 20.6853156 | -0.249156676 | 0.39979617 | -0.6232093 | 0.53314701 | 0.99945981 |
| REST | 21.3686522 | -0.573186808 | 0.42965844 | -1.3340522 | 0.18218674 | 0.99945981 |
| POLR2B | 37.2577837 | 0.121603487 | 0.30341838 | 0.40077825 | 0.68858339 | 0.99945981 |
| IGFBP7 | 49.7887295 | -0.494931256 | 0.48582127 | -1.0187517 | 0.30832084 | 0.99945981 |
| CENPC | 34.3826422 | -0.106184905 | 0.33847869 | -0.3137122 | 0.75373961 | 0.99945981 |
| UBA6 | 52.9250572 | 0.039128115 | 0.27396315 | 0.14282255 | 0.88643033 | 0.99945981 |
| UBA6-AS1 | 24.7587418 | 0.177465079 | 0.80363447 | 0.22082811 | 0.82522628 | 0.99945981 |
| YTHDC1 | 54.1246645 | 0.339281153 | 0.28163769 | 1.20467239 | 0.2283298 | 0.99945981 |
| UGT2B7 | 20.0060297 | -1.1289917 | 0.68901932 | -1.6385487 | 0.10130729 | 0.99945981 |
| RUFY3 | 61.049524 | 0.623384036 | 0.25964516 | 2.40090755 | 0.01635447 | 0.74280553 |
| GRSF1 | 14.4300258 | 0.053264013 | 0.37465706 | 0.14216738 | 0.8869478 | 0.99945981 |
| MOB1B | 50.1205778 | 0.580609541 | 0.29342022 | 1.97876459 | 0.04784252 | 0.90884462 |
| DCK | 9.17417533 | 0.709058282 | 0.46197691 | 1.53483489 | 0.12482439 | 0.99945981 |
| SLC4A4 | 17.1488656 | 0.943401095 | 0.41986243 | 2.24692905 | 0.02464456 | 0.80325576 |
| ANKRD17 | 173.231241 | 0.048577618 | 0.20641767 | 0.23533653 | 0.81394754 | 0.99945981 |
| MTHFD2L | 12.7507849 | -0.281250642 | 0.72788977 | -0.3863918 | 0.69920652 | 0.99945981 |
| RCHY1 | 21.3067824 | -0.373615975 | 0.38998069 | -0.9580371 | 0.33804404 | 0.99945981 |
| G3BP2 | 33.0344915 | -0.298534882 | 0.27864381 | -1.0713853 | 0.28399623 | 0.99945981 |
| USO1 | 47.1345772 | -0.208030915 | 0.27786187 | -0.7486846 | 0.4540473 | 0.99945981 |
| NAAA | 38.5082427 | 0.782082173 | 0.64869568 | 1.20562259 | 0.22796304 | 0.99945981 |
| SDAD1 | 13.1648314 | -0.085745338 | 0.46090876 | -0.1860354 | 0.85241701 | 0.99945981 |
| NUP54 | 20.8856908 | -0.250269624 | 0.40554568 | -0.6171182 | 0.53715676 | 0.99945981 |
| SCARB2 | 41.3983868 | 0.311145268 | 0.32355452 | 0.96164712 | 0.33622689 | 0.99945981 |
| 11-Sep | 37.5462053 | 0.470255081 | 0.52774252 | 0.89106916 | 0.37289207 | 0.99945981 |
| CCNI | 77.0774023 | 0.054604305 | 0.29666519 | 0.18406037 | 0.85396609 | 0.99945981 |
| CCNG2 | 17.2861637 | 0.346050407 | 0.42951545 | 0.80567627 | 0.42042954 | 0.99945981 |
| CNOT6L | 167.952736 | -0.387136405 | 0.47931488 | -0.807687 | 0.41927078 | 0.99945981 |
| MRPL1 | 22.5922411 | -0.571213959 | 0.4890392 | -1.1680331 | 0.24279343 | 0.99945981 |
| BMP2K | 146.940194 | -0.368171403 | 0.38415153 | -0.9584015 | 0.33786034 | 0.99945981 |
| PAQR3 | 8.66341666 | -0.533726566 | 0.45660699 | -1.168897 | 0.24244511 | 0.99945981 |
| ANTXR2 | 74.7377082 | 0.176198707 | 0.4805811 | 0.36663678 | 0.71388997 | 0.99945981 |
| RASGEF1B | 183.87793 | 1.68099098 | 0.57843148 | 2.90611945 | 0.00365942 | 0.48727327 |
| HNRNPD | 62.1477082 | 0.01735363 | 0.22812764 | 0.07606982 | 0.93936355 | 0.99945981 |
| HNRNPDL | 55.1966881 | 0.197911349 | 0.37513408 | 0.52757497 | 0.59779437 | 0.99945981 |
| ENOPH1 | 11.4527542 | -0.123947095 | 0.52962943 | -0.2340261 | 0.81496474 | 0.99945981 |
| SEC31A | 59.0858865 | -0.186647542 | 0.32498673 | -0.5743236 | 0.56574885 | 0.99945981 |
| LIN54 | 19.8509803 | -0.131563952 | 0.44534084 | -0.2954231 | 0.76767072 | 0.99945981 |
| PLAC8 | 37.5797729 | 0.548458443 | 0.5553158 | 0.98765143 | 0.32332339 | 0.99945981 |
| HELQ | 14.8549741 | 0.358924264 | 0.3714741 | 0.96621611 | 0.33393605 | 0.99945981 |
| GPAT3 | 24.691434 | 1.742996396 | 0.63605418 | 2.74032694 | 0.00613781 | 0.52554078 |
| WDFY3 | 132.775556 | 0.520839408 | 0.57707092 | 0.90255702 | 0.36676105 | 0.99945981 |
| ARHGAP24 | 203.177144 | -0.656224274 | 0.45681929 | -1.4365074 | 0.15085802 | 0.99945981 |
| MAPK10 | 34.0832308 | -3.425956369 | 1.17996629 | -2.9034358 | 0.00369093 | 0.48727327 |
| PTPN13 | 20.5939586 | 0.628361446 | 0.86426485 | 0.72704732 | 0.46719697 | 0.99945981 |
| AFF1 | 223.682189 | -0.268685443 | 0.41319576 | -0.6502619 | 0.51552309 | 0.99945981 |
| KLHL8 | 20.1462279 | -0.334039464 | 0.4256446 | -0.7847849 | 0.43257968 | 0.99945981 |
| HSD17B11 | 25.501707 | 0.246494895 | 0.3091766 | 0.79726246 | 0.42529862 | 0.99945981 |
| SPP1 | 477.097064 | -2.200500661 | 0.73063092 | -3.0117815 | 0.0025972 | 0.45570351 |
| PKD2 | 39.8622555 | -0.633063804 | 0.62814352 | -1.0078331 | 0.31353461 | 0.99945981 |
| PPM1K | 30.7813289 | -0.10407592 | 0.40567838 | -0.2565479 | 0.79752783 | 0.99945981 |
| HERC6 | 13.6822171 | 0.551436061 | 0.88552637 | 0.62272122 | 0.53346774 | 0.99945981 |
| HERC5 | 25.3808106 | 1.020246984 | 0.83775355 | 1.21783665 | 0.22328605 | 0.99945981 |
| HERC3 | 29.3575131 | 0.03871411 | 0.29669236 | 0.1304857 | 0.89618217 | 0.99945981 |
| FAM13A-AS1 | 11.0580712 | 0.550455479 | 0.44310096 | 1.24228005 | 0.21413325 | 0.99945981 |
| FAM13A | 62.8574049 | -0.012750459 | 0.70854823 | -0.0179952 | 0.98564269 | 0.99945981 |
| GPRIN3 | 131.510595 | -0.446631963 | 0.38631687 | -1.1561286 | 0.24762858 | 0.99945981 |
| SNCA | 21.6511035 | 0.71739604 | 0.7108787 | 1.00916801 | 0.31289407 | 0.99945981 |
| CCSER1 | 96.1129834 | 0.067158917 | 0.86720158 | 0.07744326 | 0.93827093 | 0.99945981 |
| SMARCAD1 | 26.4760773 | 0.046469454 | 0.33973094 | 0.13678311 | 0.89120224 | 0.99945981 |
| PDLIM5 | 69.7599629 | 0.301122225 | 0.41497163 | 0.72564532 | 0.46805623 | 0.99945981 |
| RAP1GDS1 | 97.2568893 | -0.211821207 | 0.25642704 | -0.8260487 | 0.40887649 | 0.99945981 |
| TSPAN5 | 52.4446675 | -0.839559808 | 0.34700378 | -2.4194543 | 0.01554381 | 0.7371189 |
| EIF4E | 35.9976908 | 0.201237741 | 0.42851789 | 0.4696134 | 0.63863125 | 0.99945981 |
| METAP1 | 18.3493436 | 0.356091194 | 0.35610535 | 0.99996025 | 0.31732974 | 0.99945981 |
| ADH5 | 12.9109749 | -0.063012851 | 0.40941499 | -0.1539095 | 0.8776811 | 0.99945981 |
| DAPP1 | 32.6402549 | -0.461607469 | 0.93377799 | -0.4943439 | 0.62106331 | 0.99945981 |
| DNAJB14 | 62.9224178 | -0.191058179 | 0.22844249 | -0.8363513 | 0.40295728 | 0.99945981 |
| H2AFZ | 25.2467328 | -0.12649434 | 0.62234325 | -0.2032549 | 0.83893577 | 0.99945981 |
| PPP3CA | 212.486858 | -0.486312065 | 0.37861155 | -1.2844618 | 0.19898043 | 0.99945981 |
| BANK1 | 134.434464 | -2.633784841 | 1.18125336 | -2.2296528 | 0.0257705 | 0.81444982 |
| SLC39A8 | 41.7811245 | -0.027249489 | 0.83691642 | -0.0325594 | 0.97402595 | 0.99945981 |
| NFKB1 | 82.5116718 | 0.143900027 | 0.37236158 | 0.3864524 | 0.69916165 | 0.99945981 |
| MANBA | 97.3927467 | -0.001480578 | 0.37696622 | -0.0039276 | 0.99686622 | 0.99945981 |
| UBE2D3 | 118.401818 | 0.199330737 | 0.1927633 | 1.03406994 | 0.30110347 | 0.99945981 |
| CISD2 | 9.7290814 | -0.415293684 | 0.5006915 | -0.8294403 | 0.40685533 | 0.99945981 |
| TET2 | 127.148996 | 0.430926562 | 0.2850887 | 1.51155259 | 0.13064772 | 0.99945981 |
| PPA2 | 48.1152901 | -0.015638825 | 0.3680543 | -0.0424905 | 0.96610766 | 0.99945981 |
| GSTCD | 13.7454745 | -0.551688131 | 0.44835147 | -1.2304814 | 0.2185169 | 0.99945981 |

| | | | | | | |
|---|---|---|---|---|---|---|
| TBCK | 70.2847289 | -0.078853188 | 0.3534908 | -0.22307 | 0.82348104 | 0.99945981 |
| AIMP1 | 13.0561719 | -0.842410689 | 0.43890647 | -1.9193399 | 0.05494134 | 0.92445778 |
| PAPSS1 | 32.7954957 | -0.178865927 | 0.42600575 | -0.4198674 | 0.67458232 | 0.99945981 |
| SGMS2 | 14.0971071 | 0.24976324 | 0.65636034 | 0.38052762 | 0.7035538 | 0.99945981 |
| LEF1 | 27.1759892 | 1.242388016 | 0.63786684 | 1.94772315 | 0.0514481 | 0.9230968 |
| RPL34 | 65.5988504 | 0.196909604 | 0.51050049 | 0.38571874 | 0.69970498 | 0.99945981 |
| OSTC | 14.7696451 | -0.126333823 | 0.59925937 | -0.2108166 | 0.83303038 | 0.99945981 |
| SEC24B | 68.6702353 | 0.233694421 | 0.23390998 | 0.99907844 | 0.3177567 | 0.99945981 |
| MCUB | 55.4292641 | 0.049013532 | 0.33468802 | 0.14644543 | 0.88356976 | 0.99945981 |
| ELOVL6 | 13.0234684 | -0.123160759 | 0.48458291 | -0.2541583 | 0.79937328 | 0.99945981 |
| FAM241A | 12.9624565 | 0.322464586 | 0.55489695 | 0.58112517 | 0.5611561 | 0.99945981 |
| AP1AR | 13.0413359 | 0.088909747 | 0.43234833 | 0.20564378 | 0.83706917 | 0.99945981 |
| ALPK1 | 43.757152 | 0.083550602 | 0.46023539 | 0.18153885 | 0.85594465 | 0.99945981 |
| LARP7 | 19.8348121 | -0.239548763 | 0.46775713 | -0.5121221 | 0.60856556 | 0.99945981 |
| ANK2 | 28.7083149 | -2.300411655 | 0.97547439 | -2.3582491 | 0.01836137 | 0.759342 |
| CAMK2D | 167.370713 | -0.509347477 | 0.57638981 | -0.8836858 | 0.37686586 | 0.99945981 |
| SNHG8 | 10.2175161 | -0.420296014 | 0.4718601 | -0.8907217 | 0.37307851 | 0.99945981 |
| METTL14 | 11.530071 | -0.336410047 | 0.50500479 | -0.6661522 | 0.50531383 | 0.99945981 |
| SEC24D | 47.4895332 | -0.36977844 | 0.39571089 | -0.9344662 | 0.35006348 | 0.99945981 |
| USP53 | 54.1833366 | -0.768754727 | 0.44776654 | -1.7168651 | 0.08600383 | 0.99204414 |
| C4orf3 | 23.5047795 | -0.31446047 | 0.42284268 | -0.7436819 | 0.45706892 | 0.99945981 |
| PDE5A | 9.85430783 | 0.335035843 | 0.62197398 | 0.53866536 | 0.59011778 | 0.99945981 |
| ANXA5 | 44.6993236 | 0.25234587 | 0.52750612 | 0.47837525 | 0.63238315 | 0.99945981 |
| KIAA1109 | 115.312655 | 0.158252109 | 0.28667688 | 0.55202258 | 0.58093289 | 0.99945981 |
| SPATA5 | 63.6598926 | -0.076831859 | 0.33110966 | -0.2320435 | 0.81650419 | 0.99945981 |
| MFSD8 | 17.670169 | -0.354193386 | 0.41652881 | -0.8503455 | 0.39513305 | 0.99945981 |
| ABHD18 | 44.4908428 | -0.225731341 | 0.28273998 | -0.7983708 | 0.42465535 | 0.99945981 |
| LARP1B | 22.0558274 | -0.677565576 | 0.48573584 | -1.394926 | 0.16303814 | 0.99945981 |
| PGRMC2 | 10.8358037 | -0.48453919 | 0.44750044 | -1.0827681 | 0.27891138 | 0.99945981 |
| JADE1 | 20.9785965 | -0.229939602 | 0.37904307 | -0.6066319 | 0.54409525 | 0.99945981 |
| SCLT1 | 112.952029 | 0.332815981 | 0.38600786 | 0.8622 | 0.38857747 | 0.99945981 |
| AC096711.2 | 13.3808823 | -0.411682451 | 0.45423691 | -0.9063166 | 0.36476831 | 0.99945981 |
| ELF2 | 152.728751 | 0.199429111 | 0.41517036 | 0.48035489 | 0.63097507 | 0.99945981 |
| NDUFC1 | 9.70631113 | -0.340890558 | 0.65943265 | -0.5169452 | 0.6051944 | 0.99945981 |
| NAA15 | 42.8533684 | 0.243714051 | 0.25434722 | 0.95819427 | 0.3379648 | 0.99945981 |
| AC097376.2 | 26.9944125 | -0.501363347 | 0.48263522 | -1.0388039 | 0.29889595 | 0.99945981 |
| SETD7 | 10.0955996 | 0.680049831 | 0.55162642 | 1.23280866 | 0.21764716 | 0.99945981 |
| MGST2 | 13.2847873 | 0.365376759 | 0.74325451 | 0.49159037 | 0.62300895 | 0.99945981 |
| MAML3 | 162.005381 | 0.266597448 | 0.36911992 | 0.72225159 | 0.47013981 | 0.99945981 |
| ELMOD2 | 9.69981101 | -0.349689436 | 0.47036159 | -0.7434481 | 0.45721043 | 0.99945981 |
| TBC1D9 | 46.4950815 | -0.031581513 | 0.47546923 | -0.0664218 | 0.94704203 | 0.99945981 |
| RNF150 | 22.492939 | 0.254648689 | 0.64826585 | 0.39281522 | 0.69445596 | 0.99945981 |
| ZNF330 | 11.7629747 | 0.248127335 | 0.53622505 | 0.46272984 | 0.64355802 | 0.99945981 |
| IL15 | 32.5895463 | 0.631117975 | 0.4353167 | 1.44979041 | 0.14711698 | 0.99945981 |
| INPP4B | 121.126013 | 0.026520003 | 0.49871091 | 0.05317711 | 0.9575908 | 0.99945981 |
| USP38 | 12.4053395 | 0.403606686 | 0.39478252 | 1.02235197 | 0.30661434 | 0.99945981 |
| GAB1 | 53.223786 | 0.381658041 | 0.35221055 | 1.08360764 | 0.2785388 | 0.99945981 |
| SMARCA5 | 32.9264672 | 0.067248091 | 0.35950428 | 0.18705783 | 0.85161529 | 0.99945981 |
| ANAPC10 | 23.4366766 | -0.109345778 | 0.32218031 | -0.3393931 | 0.73431361 | 0.99945981 |
| ABCE1 | 12.0819835 | -0.216593237 | 0.44617492 | -0.4854447 | 0.62736095 | 0.99945981 |
| OTUD4 | 28.5068839 | 0.142780926 | 0.30823613 | 0.46321931 | 0.64320718 | 0.99945981 |
| ZNF827 | 20.2646494 | 0.052505804 | 0.53801502 | 0.09759171 | 0.92225651 | 0.99945981 |
| LSM6 | 10.9131678 | -0.059259656 | 0.47037956 | -0.1259826 | 0.89974567 | 0.99945981 |
| SLC10A7 | 43.4573991 | -0.249070055 | 0.36086517 | -0.6902025 | 0.49006683 | 0.99945981 |
| TMEM184C | 12.8463507 | -0.170471748 | 0.43871394 | -0.3885715 | 0.69759313 | 0.99945981 |
| PRMT9 | 13.7466508 | -0.009514721 | 0.46606863 | -0.0204148 | 0.98371244 | 0.99945981 |
| ARHGAP10 | 36.9240497 | 0.567359255 | 0.52678436 | 1.07702374 | 0.28146966 | 0.99945981 |
| NR3C2 | 53.2741013 | 1.007676917 | 0.62720125 | 1.60662454 | 0.10813676 | 0.99945981 |
| LRBA | 238.243002 | -0.099446206 | 0.26629443 | -0.3734446 | 0.7088176 | 0.99945981 |
| RPS3A | 48.0673243 | -0.390188655 | 0.56993383 | -0.684621 | 0.49358313 | 0.99945981 |
| FBXW7 | 218.90369 | 0.205350791 | 0.37494495 | 0.5476825 | 0.58390993 | 0.99945981 |
| TMEM154 | 26.3642636 | 0.306289607 | 0.58598342 | 0.5226933 | 0.6011877 | 0.99945981 |
| ARFIP1 | 34.6007946 | 0.081419744 | 0.27655131 | 0.29441098 | 0.76844387 | 0.99945981 |
| TMEM131L | 139.055851 | 0.511029954 | 0.30716295 | 1.6637096 | 0.09617048 | 0.99945981 |
| TLR2 | 115.624626 | 0.434193626 | 0.52060164 | 0.83402278 | 0.40426815 | 0.99945981 |
| FGB | 8.9282793 | -4.974742521 | 2.04913357 | -2.4277297 | 0.01519366 | 0.73270878 |
| PDGFC | 59.8680669 | 0.488594618 | 0.86346931 | 0.56585059 | 0.57149535 | 0.99945981 |
| FAM198B | 18.9780732 | 0.370249772 | 0.87610811 | 0.4226074 | 0.67258173 | 0.99945981 |
| FAM198B-AS | 14.0091379 | 0.827227412 | 0.68543986 | 1.20685629 | 0.22748749 | 0.99945981 |
| TMEM144 | 12.9444671 | 0.869888456 | 0.71355132 | 1.21909726 | 0.22280728 | 0.99945981 |
| ETFDH | 16.7036923 | -0.016632762 | 0.59426261 | -0.0279889 | 0.977671 | 0.99945981 |
| PPID | 10.4102955 | 0.450353394 | 0.40234727 | 1.11931515 | 0.26300571 | 0.99945981 |
| FNIP2 | 67.6180363 | -0.005716672 | 0.39265703 | -0.0145589 | 0.98838405 | 0.99945981 |
| RAPGEF2 | 104.058848 | 0.207602731 | 0.41814091 | 0.49648988 | 0.61954883 | 0.99945981 |
| NAF1 | 10.3104156 | 0.619497301 | 0.44830331 | 1.3818709 | 0.16701135 | 0.99945981 |
| TMA16 | 9.81588589 | 0.64846904 | 0.46876198 | 1.38336525 | 0.1665529 | 0.99945981 |
| 1-Mar | 172.594947 | -0.501633768 | 0.53570694 | -0.9363959 | 0.3490694 | 0.99945981 |
| TMEM192 | 12.1618823 | -0.368781345 | 0.47421434 | -0.7776681 | 0.43676473 | 0.99945981 |
| KLHL2 | 63.7960179 | -0.538936791 | 0.49962827 | -1.0786755 | 0.28073239 | 0.99945981 |
| DDX60 | 29.8106526 | 0.624155071 | 0.9117314 | 0.68458218 | 0.49360761 | 0.99945981 |
| DDX60L | 82.1221148 | 0.961425232 | 0.69298961 | 1.38735879 | 0.16533238 | 0.99945981 |
| PALLD | 18.5831666 | -0.07171067 | 0.35260407 | -0.2033745 | 0.83884234 | 0.99945981 |
| CBR4 | 37.0907508 | 0.334605637 | 0.33630887 | 0.9949355 | 0.31976763 | 0.99945981 |
| SH3RF1 | 26.2511742 | -0.643381555 | 0.68193564 | -0.9434637 | 0.34544375 | 0.99945981 |
| NEK1 | 33.4816607 | 0.120739899 | 0.3261922 | 0.37014956 | 0.71127106 | 0.99945981 |
| CLCN3 | 31.9422962 | -0.212200561 | 0.31269589 | -0.6786164 | 0.49738094 | 0.99945981 |
| GALNT7 | 40.5280088 | 0.3721703 | 0.29262521 | 1.27183265 | 0.20343259 | 0.99945981 |
| HMGB2 | 32.4871498 | -0.098145107 | 0.52978525 | -0.1852545 | 0.85302942 | 0.99945981 |
| SAP30 | 14.1007903 | -2.07698677 | 0.76245565 | -2.7240755 | 0.00644818 | 0.52554078 |
| CEP44 | 13.0491896 | 0.396478265 | 0.51999126 | 0.76247101 | 0.44577894 | 0.99945981 |
| SPCS3 | 35.8943388 | 0.15420802 | 0.38734117 | 0.39811936 | 0.6905422 | 0.99945981 |
| DCTD | 10.2380605 | -0.079293511 | 0.44150875 | -0.1795967 | 0.8574692 | 0.99945981 |
| WWC2 | 25.8682216 | 0.365352972 | 0.33360958 | 1.09515133 | 0.27345034 | 0.99945981 |
| TRAPPC11 | 28.8404705 | -0.190356521 | 0.33728297 | -0.5643823 | 0.57249401 | 0.99945981 |
| IRF2 | 53.9479984 | 0.151267056 | 0.26321592 | 0.57468809 | 0.56550226 | 0.99945981 |
| CASP3 | 21.3253835 | -0.823196408 | 0.62258541 | -1.3222224 | 0.18609409 | 0.99945981 |
| PRIMPOL | 14.6556169 | -0.368646358 | 0.6573852 | -0.5607768 | 0.57494971 | 0.99945981 |
| ACSL1 | 173.812103 | 0.122860279 | 0.50779218 | 0.24194993 | 0.80881896 | 0.99945981 |
| CFAP97 | 12.9757136 | -0.685423476 | 0.4076567 | -1.6813742 | 0.09269025 | 0.99640647 |
| SNX25 | 36.0667301 | -0.473077693 | 0.42722879 | -1.107317 | 0.26815691 | 0.99945981 |
| UFSP2 | 10.6800476 | 0.0095243 | 0.44153661 | 0.0215708 | 0.98279032 | 0.99945981 |
| SDHA | 25.1731987 | -0.008691181 | 0.32792215 | -0.0265038 | 0.97885551 | 0.99945981 |
| PDCD6 | 14.6488442 | -0.661415569 | 0.4188648 | -1.579067 | 0.1143207 | 0.99945981 |
| EXOC3 | 15.3992827 | -0.187030606 | 0.34883423 | -0.536159 | 0.59184868 | 0.99945981 |
| BRD9 | 11.6557168 | -0.085518893 | 0.42592742 | -0.2007828 | 0.84086842 | 0.99945981 |
| SLC12A7 | 14.1326976 | 0.213586497 | 0.61073853 | 0.34971839 | 0.72655005 | 0.99945981 |
| CLPTM1L | 18.0102898 | -0.242590421 | 0.64911024 | -0.3737276 | 0.70860698 | 0.99945981 |
| LPCAT1 | 35.8596979 | -0.086552352 | 0.39318412 | -0.2201319 | 0.82576846 | 0.99945981 |
| NDUFS6 | 17.9457835 | 0.510504977 | 0.41160368 | 1.24028283 | 0.2148708 | 0.99945981 |
| ICE1 | 28.3383503 | -0.216581632 | 0.27978852 | -0.7740905 | 0.43887728 | 0.99945981 |
| MED10 | 11.2298152 | 0.328819353 | 0.40547999 | 0.81093854 | 0.41740097 | 0.99945981 |
| NSUN2 | 19.6691479 | 0.47446244 | 0.39747355 | 1.19369564 | 0.23259705 | 0.99945981 |
| SRD5A1 | 13.369455 | 1.18262654 | 0.58777348 | 2.01204474 | 0.04421522 | 0.89803991 |

| | | | | | | |
|---|---|---|---|---|---|---|
| TENT4A | 41.4329575 | 0.366444393 | 0.45309997 | 0.80874955 | 0.41865922 | 0.99945981 |
| MTRR | 12.8239586 | 0.248744917 | 0.39711841 | 0.62637468 | 0.53106921 | 0.99945981 |
| CCT5 | 15.8524488 | 0.202882382 | 0.44212257 | 0.45888266 | 0.64631843 | 0.99945981 |
| 6-Mar | 77.0870039 | 0.087132108 | 0.30539898 | 0.28530583 | 0.77540985 | 0.99945981 |
| DAP | 26.2509788 | -0.293033974 | 0.45165236 | -0.6488043 | 0.5164649 | 0.99945981 |
| TRIO | 140.51239 | -0.372576018 | 0.30412025 | -1.2250944 | 0.22053965 | 0.99945981 |
| OTULINL | 25.4148354 | 0.44905252 | 0.53900949 | 0.83310689 | 0.40478445 | 0.99945981 |
| OTULIN | 26.1556729 | -0.44580483 | 0.34149813 | -1.3054386 | 0.19174355 | 0.99945981 |
| ANKH | 74.5167015 | 0.619832011 | 0.60095914 | 1.03140459 | 0.30235113 | 0.99945981 |
| RETREG1 | 27.3496617 | -1.234674834 | 0.5175515 | -2.3856077 | 0.01705093 | 0.74280553 |
| MYO10 | 11.9989306 | -0.609392803 | 0.44505323 | -1.3692583 | 0.17091856 | 0.99945981 |
| BASP1 | 49.6877688 | 0.894675199 | 0.61496036 | 1.4548502 | 0.14571074 | 0.99945981 |
| LINC02100 | 13.695485 | 0.093201787 | 0.43996234 | 0.21184038 | 0.83223157 | 0.99945981 |
| LINC02241 | 8.96699672 | 0.104415516 | 0.49664318 | 0.21024252 | 0.83347839 | 0.99945981 |
| CDH6 | 8.09958154 | -1.347477208 | 0.98864712 | -1.3629506 | 0.17289807 | 0.99945981 |
| DROSHA | 23.3967393 | 0.258732056 | 0.37416401 | 0.69149371 | 0.48925533 | 0.99945981 |
| PDZD2 | 12.8938307 | -0.85804897 | 0.89662606 | -0.9569753 | 0.33857974 | 0.99945981 |
| GOLPH3 | 59.5245613 | 0.063919717 | 0.26745343 | 0.23899382 | 0.81111038 | 0.99945981 |
| MTMR12 | 20.4782549 | -0.834610064 | 0.40442577 | -2.0636916 | 0.03904697 | 0.8905562 |
| ZFR | 81.4520893 | -0.021576277 | 0.24505992 | -0.0880449 | 0.92984099 | 0.99945981 |
| SUB1 | 64.044472 | 0.352127231 | 0.48621438 | 0.72422217 | 0.46892935 | 0.99945981 |
| TARS | 12.1050565 | -0.188380573 | 0.42266463 | -0.4456975 | 0.65581577 | 0.99945981 |
| C1QTNF3-AM | 23.7521962 | -0.378938095 | 0.5004533 | -0.7571897 | 0.4489362 | 0.99945981 |
| DNAJC21 | 13.5268716 | 0.551153493 | 0.45545531 | 1.21011543 | 0.2262346 | 0.99945981 |
| IL7R | 97.7139425 | -0.309676384 | 0.60947392 | -0.5081044 | 0.61138013 | 0.99945981 |
| LMBRD2 | 17.2874566 | 0.351438823 | 0.45054711 | 0.7800268 | 0.4353751 | 0.99945981 |
| NADK2 | 9.68688424 | -0.215081084 | 0.62261093 | -0.3454502 | 0.72975592 | 0.99945981 |
| SLC1A3 | 79.3473452 | -0.859677826 | 0.89289961 | -0.9627934 | 0.33565123 | 0.99945981 |
| NIPBL | 214.413354 | 0.271289446 | 0.17302381 | 1.56793126 | 0.11689718 | 0.99945981 |
| CPLANE1 | 23.5616542 | -0.406130922 | 0.31175378 | -1.3027297 | 0.19266704 | 0.99945981 |
| NUP155 | 11.717961 | 0.271674201 | 0.44175619 | 0.61498675 | 0.53856347 | 0.99945981 |
| WDR70 | 117.571363 | 0.448060236 | 0.29437463 | 1.5220749 | 0.12799031 | 0.99945981 |
| RICTOR | 124.677839 | 0.205696193 | 0.25284453 | 0.81352835 | 0.4159152 | 0.99945981 |
| FYB1 | 222.440889 | 0.416460026 | 0.48566126 | 0.85751132 | 0.39116236 | 0.99945981 |
| TTC33 | 18.0986104 | 0.134340083 | 0.5523596 | 0.24321128 | 0.80784173 | 0.99945981 |
| PTGER4 | 17.2308119 | 0.254199166 | 0.50211582 | 0.50625604 | 0.61267692 | 0.99945981 |
| PRKAA1 | 25.8356403 | 0.482398417 | 0.35168903 | 1.37166182 | 0.17016874 | 0.99945981 |
| RPL37 | 42.8223759 | -0.040829127 | 0.40187212 | -0.1015973 | 0.91907631 | 0.99945981 |
| C7 | 11.3256973 | -0.910922306 | 0.65530671 | -1.3900702 | 0.16450757 | 0.99945981 |
| OXCT1 | 25.056435 | -0.623743168 | 0.38534202 | -1.6186741 | 0.10551739 | 0.99945981 |
| SELENOP | 54.3893194 | 0.060067681 | 0.73033337 | 0.08224693 | 0.93445035 | 0.99945981 |
| ZNF131 | 46.5613076 | -0.164393058 | 0.30448375 | -0.5399075 | 0.58926083 | 0.99945981 |
| HMGCS1 | 13.067852 | 0.010269505 | 0.3855161 | 0.02663833 | 0.9787482 | 0.99945981 |
| TMEM267 | 13.9526723 | -0.246624294 | 0.35642506 | -0.6919387 | 0.48897583 | 0.99945981 |
| PAIP1 | 13.0436424 | -0.266834829 | 0.45871652 | -0.5816988 | 0.56076961 | 0.99945981 |
| NNT-AS1 | 9.98432891 | -0.429040348 | 0.49534576 | -0.8661432 | 0.38641165 | 0.99945981 |
| NNT | 44.4237909 | 0.306639947 | 0.2569284 | 1.19348404 | 0.23267986 | 0.99945981 |
| EMB | 107.497467 | -0.093649706 | 0.58115696 | -0.1611436 | 0.87198033 | 0.99945981 |
| PARP8 | 353.715122 | -0.073414146 | 0.51699173 | -0.1420026 | 0.88707799 | 0.99945981 |
| ITGA1 | 16.5998786 | -0.766944567 | 0.55643482 | -1.3783188 | 0.16810489 | 0.99945981 |
| NDUFS4 | 37.6624003 | 0.283679144 | 0.25424564 | 1.11576796 | 0.26452147 | 0.99945981 |
| ARL15 | 227.375134 | 0.448460906 | 0.26420289 | 1.69741104 | 0.08961898 | 0.99640647 |
| SNX18 | 26.9075817 | -0.185993665 | 0.50118489 | -0.3711079 | 0.71055718 | 0.99945981 |
| GZMK | 10.0742143 | -2.007376505 | 0.9139182 | -2.196451 | 0.02805968 | 0.84763724 |
| GZMA | 23.9616988 | -0.293722845 | 0.56794146 | -0.517171 | 0.6050368 | 0.99945981 |
| DHX29 | 20.7692663 | -0.578048428 | 0.40258453 | -1.4358436 | 0.15104684 | 0.99945981 |
| MTREX | 47.8521118 | 0.048306467 | 0.26760897 | 0.18051138 | 0.85675112 | 0.99945981 |
| PLPP1 | 20.9353535 | -1.569674086 | 0.49292568 | -3.1844032 | 0.00145053 | 0.3983151 |
| SLC38A9 | 38.510185 | -0.227852772 | 0.29350251 | -0.7763231 | 0.43755825 | 0.99945981 |
| IL6ST | 47.9263826 | 0.512888102 | 0.39069444 | 1.31276018 | 0.18926377 | 0.99945981 |
| MAP3K1 | 143.835456 | 0.230562538 | 0.25554853 | 0.90222603 | 0.36693681 | 0.99945981 |
| GPBP1 | 104.890498 | 0.293608943 | 0.28336171 | 1.03616309 | 0.30012607 | 0.99945981 |
| PDE4D | 212.72001 | -0.493728992 | 0.51657469 | -0.9557746 | 0.3391861 | 0.99945981 |
| ERCC8 | 19.7590386 | 0.57792282 | 0.53910988 | 1.07199448 | 0.28372251 | 0.99945981 |
| NDUFAF2 | 34.9641032 | 0.78580798 | 0.61982362 | 1.2677929 | 0.20487192 | 0.99945981 |
| ZSWIM6 | 281.590894 | 0.827183261 | 0.33673802 | 2.45645936 | 0.01403137 | 0.7227195 |
| KIF2A | 81.6393812 | 0.116806328 | 0.23703403 | 0.49278295 | 0.62216596 | 0.99945981 |
| IPO11 | 14.802917 | 0.50342709 | 0.348825 | 1.44320815 | 0.14896184 | 0.99945981 |
| SREK1IP1 | 15.2856904 | -0.140931897 | 0.34793872 | -0.405048 | 0.6854422 | 0.99945981 |
| CWC27 | 37.635473 | -0.185742943 | 0.2879252 | -0.6451083 | 0.518857 | 0.99945981 |
| CENPK | 11.3005887 | 1.86432745 | 0.66797753 | 2.79100325 | 0.00525449 | 0.5234882 |
| PPWD1 | 21.7333138 | 0.093164976 | 0.35544118 | 0.26211081 | 0.79323601 | 0.99945981 |
| TRIM23 | 13.2263431 | 0.126527802 | 0.48350566 | 0.26168836 | 0.79356172 | 0.99945981 |
| TRAPPC13 | 13.5574184 | 0.047849233 | 0.37288654 | 0.12832116 | 0.89789482 | 0.99945981 |
| SGTB | 17.8070269 | 0.592453821 | 0.37055166 | 1.5988427 | 0.10985556 | 0.99945981 |
| NLN | 14.9919563 | -0.177947443 | 0.57643056 | -0.3087058 | 0.75754535 | 0.99945981 |
| ERBIN | 173.097944 | 0.669452741 | 0.24510641 | 2.73127394 | 0.006309 | 0.52554078 |
| SREK1 | 33.4776885 | 0.265124808 | 0.35777584 | 0.74103609 | 0.45867156 | 0.99945981 |
| MAST4 | 41.8356715 | -0.324109588 | 0.45757773 | -0.7083159 | 0.4787491 | 0.99945981 |
| PIK3R1 | 128.495747 | -0.171833347 | 0.46456677 | -0.3698787 | 0.71147289 | 0.99945981 |
| SLC30A5 | 17.2507745 | -0.040952511 | 0.51670151 | -0.0792576 | 0.93682775 | 0.99945981 |
| CDK7 | 11.5555595 | 0.240267175 | 0.47823526 | 0.50240373 | 0.61538356 | 0.99945981 |
| CCDC125 | 10.3911672 | -0.415326587 | 0.45271613 | -0.9174106 | 0.35892751 | 0.99945981 |
| RAD17 | 11.2151303 | 0.017677968 | 0.40166776 | 0.04401142 | 0.9648953 | 0.99945981 |
| GTF2H2C | 9.1771133 | 0.560434403 | 0.48419369 | 1.15745911 | 0.24708483 | 0.99945981 |
| NAIP | 134.106822 | 0.496481197 | 0.56075863 | 0.88537416 | 0.37595486 | 0.99945981 |
| GTF2H2 | 8.93745032 | 0.480023505 | 0.60632321 | 0.79169574 | 0.4285381 | 0.99945981 |
| BDP1 | 57.1691033 | -0.048576388 | 0.41432905 | -0.1172411 | 0.90666901 | 0.99945981 |
| MCCC2 | 18.4972398 | 0.020597481 | 0.37954492 | 0.05426889 | 0.95672094 | 0.99945981 |
| MRPS27 | 26.305937 | -0.180503243 | 0.49687965 | -0.3632736 | 0.71640053 | 0.99945981 |
| TNPO1 | 56.9999982 | -0.091911071 | 0.34366386 | -0.2674447 | 0.7891268 | 0.99945981 |
| FCHO2 | 82.0972025 | -0.528346537 | 0.4873786 | -1.0840577 | 0.2783392 | 0.99945981 |
| BTF3 | 30.5759329 | -0.484291295 | 0.53531307 | -0.904688 | 0.36563072 | 0.99945981 |
| ANKRA2 | 9.64025999 | -0.108194081 | 0.52535267 | -0.2059456 | 0.83683339 | 0.99945981 |
| HEXB | 24.2745889 | -0.130842945 | 0.37554786 | -0.3484055 | 0.72753566 | 0.99945981 |
| GFM2 | 12.4057947 | 0.097266967 | 0.41978007 | 0.23170935 | 0.81676377 | 0.99945981 |
| NSA2 | 17.710729 | -0.367643787 | 0.49088396 | -0.7489423 | 0.45389195 | 0.99945981 |
| HMGCR | 9.53434243 | 0.702629538 | 0.46575004 | 1.50859793 | 0.13140156 | 0.99945981 |
| COL4A3BP | 95.1700735 | 0.193757596 | 0.23456171 | 0.82604102 | 0.40878082 | 0.99945981 |
| POLK | 52.0343887 | 0.271600396 | 0.27172919 | 0.999526 | 0.31753995 | 0.99945981 |
| POC5 | 16.5635358 | 0.5322862 | 0.34460351 | 1.5446337 | 0.12243489 | 0.99945981 |
| IQGAP2 | 265.158475 | -0.236532675 | 0.24992523 | -0.9464137 | 0.3439376 | 0.99945981 |
| F2R | 10.7900196 | -0.280464874 | 0.56527299 | -0.4961583 | 0.61978274 | 0.99945981 |
| S100Z | 30.4771951 | 1.730201966 | 0.89515301 | 1.93285611 | 0.05325393 | 0.92445778 |
| WDR41 | 26.1929761 | 0.1924084 | 0.4069965 | 0.47275198 | 0.63639014 | 0.99945981 |
| TBCA | 57.9821805 | -0.154819446 | 0.29771975 | -0.5200174 | 0.60305145 | 0.99945981 |
| AP3B1 | 128.097341 | 0.282893452 | 0.23753488 | 1.19095542 | 0.23367109 | 0.99945981 |
| SCAMP1 | 25.0299861 | 0.008672556 | 0.33424346 | 0.02594682 | 0.97929975 | 0.99945981 |
| LHFPL2 | 31.665166 | -0.037883044 | 0.62231053 | -0.0608748 | 0.9514589 | 0.99945981 |
| ARSB | 37.0787937 | -0.227413709 | 0.36691996 | -0.619791 | 0.53539536 | 0.99945981 |
| JMY | 20.7253206 | -0.070598338 | 0.41341575 | -0.1707684 | 0.86440589 | 0.99945981 |
| TENT2 | 139.330233 | 0.296789048 | 0.20922796 | 1.41849609 | 0.15604598 | 0.99945981 |
| SERINC5 | 85.4416139 | 0.119416782 | 0.53001108 | 0.22530997 | 0.82173814 | 0.99945981 |

| | | | | | | |
|---|---|---|---|---|---|---|
| ZFYVE16 | 45.0395058 | -0.030284798 | 0.31095198 | -0.0973938 | 0.92241366 | 0.99945981 |
| DHFR | 86.0616458 | -0.152333677 | 0.31516744 | -0.4833421 | 0.62885288 | 0.99945981 |
| MSH3 | 48.2744334 | -0.082792211 | 0.27395251 | -0.3022137 | 0.76248914 | 0.99945981 |
| RASGRF2 | 21.2111207 | 0.68839866 | 0.560971 | 1.22715552 | 0.21976414 | 0.99945981 |
| CKMT2-AS1 | 12.4441233 | 0.069023641 | 0.38223613 | 0.18057853 | 0.85669841 | 0.99945981 |
| SSBP2 | 100.074527 | -0.243634969 | 0.462093 | -0.5272423 | 0.59802535 | 0.99945981 |
| ATG10 | 28.3062114 | -0.237905593 | 0.38014547 | -0.6258278 | 0.53142792 | 0.99945981 |
| RPS23 | 107.379124 | -0.047818966 | 0.57233994 | -0.0835499 | 0.93341427 | 0.99945981 |
| TMEM167A | 20.1597463 | 0.152179102 | 0.48212582 | 0.31564189 | 0.75227434 | 0.99945981 |
| XRCC4 | 42.5848378 | 0.353526938 | 0.33183817 | 1.06535947 | 0.28671329 | 0.99945981 |
| VCAN | 311.556393 | 0.022420542 | 1.63567548 | 0.01370721 | 0.98906357 | 0.99945981 |
| COX7C | 28.512443 | -0.72043288 | 0.55413497 | -1.3001036 | 0.19356546 | 0.99945981 |
| RASA1 | 124.498258 | 0.378314045 | 0.28731711 | 1.31671255 | 0.187935 | 0.99945981 |
| CCNH | 109.191676 | -0.135534665 | 0.35795617 | -0.3786348 | 0.70495907 | 0.99945981 |
| TMEM161B | 21.0880214 | 0.145726526 | 0.45856297 | 0.31778956 | 0.75064457 | 0.99945981 |
| TMEM161B-, | 24.9341171 | 0.051522886 | 0.34267134 | 0.15035657 | 0.88048331 | 0.99945981 |
| MEF2C | 220.52322 | -0.127358978 | 0.56921016 | -0.2237468 | 0.82295429 | 0.99945981 |
| MEF2C-AS1 | 49.4440015 | -2.117912551 | 1.04617409 | -2.0244361 | 0.0429253 | 0.89562588 |
| LYSMD3 | 9.57864186 | -0.725306433 | 0.54869166 | -1.3218835 | 0.18620696 | 0.99945981 |
| ARRDC3 | 38.6836455 | 0.442918877 | 0.70249924 | 0.63049019 | 0.52837392 | 0.99945981 |
| FAM172A | 165.288884 | 0.09115446 | 0.20289703 | 0.44926464 | 0.65324076 | 0.99945981 |
| KIAA0825 | 33.3311052 | 0.248527002 | 0.42100571 | 0.59031742 | 0.55497786 | 0.99945981 |
| SLF1 | 26.2894358 | -0.465153226 | 0.37315167 | -1.2465527 | 0.21256155 | 0.99945981 |
| MCTP1 | 145.883678 | 0.802895284 | 0.31425321 | 2.55493104 | 0.01062088 | 0.6628397 |
| TTC37 | 42.3743475 | 0.244658791 | 0.31522903 | 0.77613028 | 0.43767207 | 0.99945981 |
| RHOBTB3 | 26.8318111 | -0.733140809 | 0.51393064 | -1.4265365 | 0.15371354 | 0.99945981 |
| GLRX | 29.7840415 | 0.416301788 | 0.57601118 | 0.72273213 | 0.46984447 | 0.99945981 |
| ELL2 | 119.741924 | -0.805563242 | 0.70104143 | -1.1490951 | 0.25051679 | 0.99945981 |
| AC104123.1 | 17.4941051 | -0.730606112 | 0.60481321 | -1.2079864 | 0.22705251 | 0.99945981 |
| CAST | 93.6453365 | 0.19968371 | 0.29830217 | 0.6694008 | 0.50323984 | 0.99945981 |
| ERAP1 | 29.0656343 | 0.315155436 | 0.3835369 | 0.82170824 | 0.41124297 | 0.99945981 |
| ERAP2 | 36.4585201 | 0.236382127 | 0.41526237 | 0.5692356 | 0.56919626 | 0.99945981 |
| LNPEP | 106.660972 | 0.33611158 | 0.21822558 | 1.54020248 | 0.12351101 | 0.99945981 |
| CHD1 | 101.621613 | 0.577911263 | 0.35813403 | 1.61367313 | 0.10659835 | 0.99945981 |
| ST8SIA4 | 122.488355 | 0.264294978 | 0.3948784 | 0.66930725 | 0.5032995 | 0.99945981 |
| PAM | 82.0928225 | -0.1306882 | 0.42796935 | -0.3053681 | 0.76008579 | 0.99945981 |
| PPIP5K2 | 52.7311376 | 0.46700907 | 0.3835989 | 1.21744111 | 0.22343643 | 0.99945981 |
| EFNA5 | 18.3872477 | -0.237848795 | 0.76166513 | -0.3122748 | 0.75483172 | 0.99945981 |
| FBXL17 | 161.24509 | 0.053194185 | 0.16978563 | 0.31330205 | 0.75405119 | 0.99945981 |
| FER | 85.3065421 | -0.156355587 | 0.25810918 | -0.605773 | 0.54466547 | 0.99945981 |
| PJA2 | 62.6212185 | -0.408091772 | 0.23750765 | -1.7182258 | 0.08575543 | 0.99193047 |
| MAN2A1 | 101.341147 | 0.467260983 | 0.29789228 | 1.56855689 | 0.11675122 | 0.99945981 |
| SLC25A46 | 21.8643522 | -0.737461494 | 0.39743687 | -1.8555437 | 0.06351862 | 0.93775336 |
| WDR36 | 17.2487947 | 0.28488125 | 0.35390207 | 0.80497198 | 0.42083585 | 0.99945981 |
| CAMK4 | 83.2805783 | -0.340943627 | 0.55998345 | -0.6088459 | 0.54262659 | 0.99945981 |
| STARD4 | 10.5151052 | 0.160432517 | 0.45059303 | 0.35604749 | 0.721805 | 0.99945981 |
| EPB41L4A | 15.0812336 | -0.195983555 | 0.55841714 | -0.3509626 | 0.72561638 | 0.99945981 |
| APC | 46.9967978 | 0.397754021 | 0.2992546 | 1.32914925 | 0.18379874 | 0.99945981 |
| SRP19 | 13.329531 | -0.057058457 | 0.44075845 | -0.1294552 | 0.8969975 | 0.99945981 |
| REEP5 | 37.6322162 | 0.19188286 | 0.36621556 | 0.52396151 | 0.60030531 | 0.99945981 |
| DCP2 | 47.9220062 | 0.479633395 | 0.30708445 | 1.56189412 | 0.11831293 | 0.99945981 |
| MCC | 16.5492769 | 0.183402447 | 0.6337965 | 0.28937119 | 0.77229734 | 0.99945981 |
| YTHDC2 | 49.7137475 | 0.00414091 | 0.30457768 | 0.01359558 | 0.98915263 | 0.99945981 |
| PGGT1B | 26.4455019 | 0.357510673 | 0.41803253 | 0.85522214 | 0.39242818 | 0.99945981 |
| CCDC112 | 12.402253 | -0.34110102 | 0.62842683 | -0.5427856 | 0.58727744 | 0.99945981 |
| FEM1C | 10.6473719 | -0.505955509 | 0.50779904 | -0.9963696 | 0.31907062 | 0.99945981 |
| ATG12 | 20.3252735 | -0.300972901 | 0.3396944 | -0.8860108 | 0.37561171 | 0.99945981 |
| AP3S1 | 27.0709494 | 0.440563196 | 0.44212369 | 0.99647045 | 0.31902162 | 0.99945981 |
| COMMD10 | 107.946729 | -0.036239758 | 0.32033262 | -0.1131316 | 0.90992618 | 0.99945981 |
| SEMA6A-AS1 | 12.2810626 | -0.119052984 | 0.50567214 | -0.2354351 | 0.81387102 | 0.99945981 |
| DTWD2 | 14.123663 | 0.332571994 | 0.56836661 | 0.5851364 | 0.558456 | 0.99945981 |
| DMXL1 | 88.53972 | 0.036878134 | 0.25655113 | 0.14374575 | 0.88570124 | 0.99945981 |
| TNFAIP8 | 165.510855 | 0.015856128 | 0.28925897 | 0.05481637 | 0.95628475 | 0.99945981 |
| HSD17B4 | 38.8608087 | -0.75194217 | 0.73071549 | -1.0290492 | 0.30345657 | 0.99945981 |
| SRFBP1 | 16.236356 | 0.238424824 | 0.43608372 | 0.54674094 | 0.58455672 | 0.99945981 |
| SNX2 | 62.4266931 | -0.465014514 | 0.48905654 | -0.95084 | 0.34168561 | 0.99945981 |
| SNX24 | 34.0291628 | 0.29767221 | 0.39457828 | 0.75440597 | 0.45060549 | 0.99945981 |
| CEP120 | 23.5189634 | -0.057388195 | 0.3134739 | -0.1830717 | 0.85474177 | 0.99945981 |
| CSNK1G3 | 44.9006932 | -0.220011015 | 0.36579294 | -0.6014633 | 0.54753147 | 0.99945981 |
| GRAMD2B | 17.3935011 | 0.103372319 | 0.50979868 | 0.20277087 | 0.83931413 | 0.99945981 |
| LMNB1 | 13.4857542 | -0.090640087 | 0.59584659 | -0.1521198 | 0.87909242 | 0.99945981 |
| 3-Mar | 36.3654243 | -1.156399703 | 0.54924576 | -2.1054322 | 0.03525371 | 0.87920079 |
| PRRC1 | 11.44125 | 0.137548855 | 0.48485637 | 0.2836899 | 0.77664804 | 0.99945981 |
| LINC01184 | 30.3412614 | -0.278431841 | 0.46248244 | -0.6020376 | 0.54714908 | 0.99945981 |
| SLC12A2 | 16.1767163 | 0.253852384 | 0.36850553 | 0.68886994 | 0.49090512 | 0.99945981 |
| HINT1 | 50.1620642 | -0.600030165 | 0.41740069 | -1.43754 | 0.15056463 | 0.99945981 |
| CDC42SE2 | 263.003013 | -0.249942588 | 0.22008637 | -1.1356568 | 0.25610022 | 0.99945981 |
| AC116366.3 | 65.6290148 | 0.197182852 | 0.43544082 | 0.45283502 | 0.65066754 | 0.99945981 |
| IRF1 | 35.249491 | -0.006976028 | 0.42451895 | -0.0164328 | 0.98688913 | 0.99945981 |
| KIF3A | 10.9462042 | -0.055597036 | 0.48458104 | -0.1147322 | 0.90865741 | 0.99945981 |
| UQCRQ | 13.0547446 | -0.518402958 | 0.6785636 | -0.7639711 | 0.44488448 | 0.99945981 |
| AFF4 | 72.805894 | -0.101681803 | 0.26478577 | -0.3840154 | 0.70096706 | 0.99945981 |
| ZCCHC10 | 17.9913614 | -0.033973834 | 0.44023812 | -0.0771715 | 0.93848712 | 0.99945981 |
| HSPA4 | 26.2238947 | 0.129219576 | 0.29278744 | 0.44134263 | 0.65896497 | 0.99945981 |
| C5orf15 | 13.7219398 | 0.10082741 | 0.49026131 | 0.20566055 | 0.83705608 | 0.99945981 |
| VDAC1 | 18.9007447 | -0.204217689 | 0.39077718 | -0.5225937 | 0.60125703 | 0.99945981 |
| TCF7 | 18.7841717 | 0.291256301 | 0.35351533 | 0.82388592 | 0.41000438 | 0.99945981 |
| SKP1 | 27.165429 | -0.260215327 | 0.38075228 | -0.6834242 | 0.49433882 | 0.99945981 |
| UBE2B | 48.8146655 | -0.003594074 | 0.23447395 | -0.0153282 | 0.98777031 | 0.99945981 |
| JADE2 | 18.6165731 | -0.310218798 | 0.36245419 | -0.8558842 | 0.39206186 | 0.99945981 |
| SAR1B | 17.976626 | 0.114661747 | 0.43887431 | 0.26126329 | 0.79388947 | 0.99945981 |
| SEC24A | 24.7724466 | -0.338983396 | 0.59771227 | -0.5671347 | 0.57062264 | 0.99945981 |
| CAMLG | 9.99677074 | -0.346714827 | 0.46300179 | -0.7488412 | 0.45395291 | 0.99945981 |
| DDX46 | 40.6591139 | -0.09401753 | 0.30138397 | -0.3119527 | 0.7550765 | 0.99945981 |
| C5orf24 | 10.2334736 | -0.317331762 | 0.45289901 | -0.7006678 | 0.48351034 | 0.99945981 |
| TXNDC15 | 14.8711369 | -0.382422662 | 0.45040748 | -0.8490593 | 0.39584829 | 0.99945981 |
| PCBD2 | 21.847962 | -0.056411364 | 0.47709457 | -0.1182394 | 0.90587799 | 0.99945981 |
| H2AFY | 68.921302 | 0.117692927 | 0.33497063 | 0.35135297 | 0.72532356 | 0.99945981 |
| CXCL14 | 12.9787264 | -1.335519569 | 1.06239963 | -1.2570783 | 0.20872527 | 0.99945981 |
| TGFBI | 63.60344 | 0.573982642 | 0.60397448 | 0.95034254 | 0.34193823 | 0.99945981 |
| SMAD5 | 21.9558648 | 0.144259412 | 0.34637241 | 0.41648644 | 0.67705409 | 0.99945981 |
| HNRNPA0 | 14.0339002 | -0.65007383 | 0.47631642 | -1.3647941 | 0.17231778 | 0.99945981 |
| FAM13B | 95.7543884 | 0.124767652 | 0.23004589 | 0.54235983 | 0.58757065 | 0.99945981 |
| BRD8 | 10.21043 | -0.368245402 | 0.63998939 | -0.575393 | 0.56502555 | 0.99945981 |
| FAM53C | 11.6571776 | 0.609261148 | 0.47599053 | 1.27998586 | 0.20055011 | 0.99945981 |
| KDM3B | 49.0944672 | -0.081976524 | 0.25270422 | -0.3243971 | 0.74563739 | 0.99945981 |
| ETF1 | 24.0871665 | 0.231688026 | 0.37912768 | 0.61110817 | 0.54112797 | 0.99945981 |
| HSPA9 | 33.7528502 | -0.130798842 | 0.29331088 | -0.4459393 | 0.65564111 | 0.99945981 |
| CTNNA1 | 67.7011554 | 0.025937643 | 0.2846427 | 0.09112351 | 0.92739445 | 0.99945981 |
| SIL1 | 48.3383728 | -0.044876857 | 0.3503016 | -0.1281092 | 0.89806256 | 0.99945981 |
| PAIP2 | 30.8417923 | 0.090359392 | 0.44916948 | 0.20116993 | 0.84056571 | 0.99945981 |
| TMEM173 | 10.2034109 | -0.084036069 | 0.58877255 | -0.142731 | 0.88650267 | 0.99945981 |
| UBE2D2 | 86.1359194 | -0.358066618 | 0.36441107 | -0.9825899 | 0.32580934 | 0.99945981 |

| | | | | | | |
|---|---|---|---|---|---|---|
| CXXC5 | 8.34050135 | 0.130449278 | 0.50443855 | 0.25860291 | 0.79594164 | 0.99945981 |
| PURA | 22.944631 | -0.155251274 | 0.35708569 | -0.4347732 | 0.66372709 | 0.99945981 |
| CYSTM1 | 41.1144688 | -0.364139276 | 0.35160451 | -1.0356502 | 0.30036537 | 0.99945981 |
| PFDN1 | 15.9238163 | 0.11741286 | 0.48334138 | 0.24291911 | 0.80806806 | 0.99945981 |
| SRA1 | 10.716302 | -0.121093193 | 0.48396023 | -0.2502131 | 0.80242255 | 0.99945981 |
| CD14 | 70.4148548 | 0.283746571 | 0.79766826 | 0.35572002 | 0.72205025 | 0.99945981 |
| NDUFA2 | 12.2469557 | 0.08539905 | 0.48232383 | 0.1770575 | 0.85946323 | 0.99945981 |
| IK | 7.30561481 | -0.737575356 | 0.58383306 | -1.2633326 | 0.20646966 | 0.99945981 |
| HARS | 14.2440285 | -0.217031405 | 0.38759726 | -0.5599405 | 0.57552002 | 0.99945981 |
| TAF7 | 15.2841977 | -0.426242726 | 0.44732484 | -0.9528707 | 0.3406556 | 0.99945981 |
| DIAPH1 | 111.302607 | 0.27058419 | 0.25473279 | 1.06222757 | 0.28813239 | 0.99945981 |
| HDAC3 | 12.6605335 | 0.016890024 | 0.41866625 | 0.04034246 | 0.96782011 | 0.99945981 |
| DELE1 | 13.8762717 | 0.190797544 | 0.48629035 | 0.39235313 | 0.69479731 | 0.99945981 |
| RNF14 | 8.43376327 | -0.629534447 | 0.47908836 | -1.3140258 | 0.18883753 | 0.99945981 |
| GNPDA1 | 10.9912594 | -0.074122106 | 0.47515843 | -0.1559945 | 0.87603735 | 0.99945981 |
| NDFIP1 | 52.9475344 | -0.012200814 | 0.27055452 | -0.0450956 | 0.96403112 | 0.99945981 |
| ARHGAP26 | 342.549961 | 0.310064139 | 0.43464252 | 0.71337737 | 0.47561227 | 0.99945981 |
| NR3C1 | 173.975557 | -0.194935701 | 0.26098914 | -0.7469112 | 0.45511718 | 0.99945981 |
| LARS | 26.0701869 | 0.321840985 | 0.36520033 | 0.88127243 | 0.37817039 | 0.99945981 |
| TCERG1 | 41.9229327 | -0.383968233 | 0.31935642 | -1.2023188 | 0.22924003 | 0.99945981 |
| PPP2R2B | 91.1469028 | -1.846584768 | 0.70867311 | -2.6056933 | 0.00916885 | 0.59381294 |
| JAKMIP2 | 21.5604767 | -0.296034366 | 0.57471662 | -0.5150962 | 0.60648577 | 0.99945981 |
| FBXO38 | 32.1807841 | -0.007290917 | 0.38336433 | -0.0190182 | 0.98482655 | 0.99945981 |
| CSNK1A1 | 135.79668 | 0.084365009 | 0.21953304 | 0.38429299 | 0.7007613 | 0.99945981 |
| PPARGC1B | 13.3565696 | 1.17150027 | 0.49145432 | 2.38374192 | 0.01713762 | 0.74280553 |
| SLC26A2 | 9.70930105 | -0.699765531 | 0.49243942 | -1.4210185 | 0.15531138 | 0.99945981 |
| HMGXB3 | 17.6737581 | -0.260659787 | 0.39358161 | -0.6622763 | 0.50779414 | 0.99945981 |
| CSF1R | 55.0565548 | 0.245987684 | 0.59174812 | 0.4156966 | 0.67763203 | 0.99945981 |
| TCOF1 | 8.85705298 | -0.2760141 | 0.49987123 | -0.5521704 | 0.58083161 | 0.99945981 |
| CD74 | 276.786657 | -0.544150238 | 0.68610157 | -0.7931045 | 0.42771695 | 0.99945981 |
| RPS14 | 107.639091 | -0.031222702 | 0.42823362 | -0.0729104 | 0.94187739 | 0.99945981 |
| DCTN4 | 44.6458627 | 0.115829238 | 0.34088674 | 0.33978804 | 0.73401615 | 0.99945981 |
| GPX3 | 150.810931 | -0.766559858 | 0.7636857 | -1.0037635 | 0.31549261 | 0.99945981 |
| TNIP1 | 34.4871374 | 0.327472787 | 0.36597606 | 0.89479291 | 0.3708978 | 0.99945981 |
| ANXA6 | 33.4528983 | -0.391733976 | 0.41860593 | -0.9358061 | 0.34937304 | 0.99945981 |
| CCDC69 | 29.4699639 | 0.322916625 | 0.48325309 | 0.6682143 | 0.50399681 | 0.99945981 |
| GM2A | 11.1742595 | -0.615327547 | 0.64342726 | -0.9563281 | 0.33890651 | 0.99945981 |
| SLC36A1 | 22.9923802 | -0.648532643 | 0.56019372 | -1.1576935 | 0.24698913 | 0.99945981 |
| SPARC | 10.2928456 | 0.594173173 | 0.81214071 | 0.73161358 | 0.46440446 | 0.99945981 |
| ATOX1 | 17.5831061 | 0.256336514 | 0.41629671 | 0.61575436 | 0.53805665 | 0.99945981 |
| G3BP1 | 39.9868831 | -0.162928582 | 0.27088717 | -0.6014629 | 0.54753172 | 0.99945981 |
| MFAP3 | 9.94038387 | -0.317180089 | 0.46084723 | -0.6882543 | 0.49129265 | 0.99945981 |
| GALNT10 | 55.723987 | -0.091065449 | 0.36663846 | -0.2483794 | 0.80384085 | 0.99945981 |
| SAP30L-AS1 | 12.8625334 | -0.485186601 | 0.41460042 | -1.1702511 | 0.24189994 | 0.99945981 |
| LARP1 | 46.4264126 | -0.078997435 | 0.29384644 | -0.2688392 | 0.78805344 | 0.99945981 |
| CNOT8 | 17.3969723 | -0.053498417 | 0.41365301 | -0.1293316 | 0.89709524 | 0.99945981 |
| HAVCR2 | 38.6184479 | 0.159951862 | 0.53980003 | 0.29631688 | 0.7669881 | 0.99945981 |
| ITK | 67.4755133 | -0.266542732 | 0.33021648 | -0.8071757 | 0.41956524 | 0.99945981 |
| AC010609.1 | 9.8014504 | -0.79535169 | 0.73650041 | -1.0799067 | 0.28018375 | 0.99945981 |
| CYFIP2 | 69.9246345 | 0.197337115 | 0.3114524 | 0.63360281 | 0.52634007 | 0.99945981 |
| ADAM19 | 54.8652208 | -1.098908397 | 0.60165614 | -1.8264725 | 0.06777912 | 0.94796811 |
| CLINT1 | 55.4567833 | -0.161831776 | 0.28118573 | -0.5755334 | 0.56493062 | 0.99945981 |
| EBF1 | 39.3294236 | -0.380144118 | 0.94829472 | -0.4008713 | 0.6885149 | 0.99945981 |
| RNF145 | 67.2014085 | -0.102806336 | 0.27262551 | -0.3770973 | 0.70610132 | 0.99945981 |
| UBLCP1 | 14.3949132 | 0.253125989 | 0.36012095 | 0.7028916 | 0.48212331 | 0.99945981 |
| TTC1 | 20.3760227 | -0.036169737 | 0.35206354 | -0.1027364 | 0.91817219 | 0.99945981 |
| PWWP2A | 25.7774942 | 0.045559905 | 0.32785946 | 0.13896169 | 0.88948042 | 0.99945981 |
| SLU7 | 9.48763136 | -0.360764203 | 0.49060171 | -0.7353505 | 0.46212608 | 0.99945981 |
| CCNG1 | 10.8744684 | -0.124947651 | 0.52091817 | -0.2398604 | 0.81043846 | 0.99945981 |
| MAT2B | 17.0518928 | 0.465180267 | 0.35922842 | 1.29494281 | 0.19533996 | 0.99945981 |
| RARS | 21.1238506 | -0.154591824 | 0.35268049 | -0.4383339 | 0.66114424 | 0.99945981 |
| PANK3 | 18.5232168 | -0.173989724 | 0.38082129 | -0.4568802 | 0.64775712 | 0.99945981 |
| DOCK2 | 321.693287 | 0.101415857 | 0.26328434 | 0.38519518 | 0.70009281 | 0.99945981 |
| FAM196B | 15.5415854 | 0.058054553 | 0.66879534 | 0.08680466 | 0.93082678 | 0.99945981 |
| LCP2 | 105.775632 | 0.218848942 | 0.36210233 | 0.60437595 | 0.54559372 | 0.99945981 |
| NPM1 | 22.5091521 | -0.373217932 | 0.3626357 | -1.0291814 | 0.30339443 | 0.99945981 |
| FBXW11 | 79.0378033 | -0.232287964 | 0.30846358 | -0.7530483 | 0.45142091 | 0.99945981 |
| STK10 | 157.168087 | 0.029044967 | 0.31506865 | 0.09218615 | 0.92655014 | 0.99945981 |
| UBTD2 | 14.4560322 | 0.049301921 | 0.39753902 | 0.12401782 | 0.90130117 | 0.99945981 |
| SH3PXD2B | 38.8660009 | -2.051100575 | 0.71284619 | -2.8773396 | 0.00401044 | 0.507383 |
| DUSP1 | 52.1321504 | 2.033076191 | 0.60677104 | 3.35064804 | 0.00080623 | 0.27121928 |
| ERGIC1 | 44.3329882 | -0.316677018 | 0.29092643 | -1.0885124 | 0.27636898 | 0.99945981 |
| ATP6V0E1 | 65.1629107 | -0.130304415 | 0.24474096 | -0.5324177 | 0.59443674 | 0.99945981 |
| CREBRF | 51.6403788 | 0.13345142 | 0.32568371 | 0.40975773 | 0.68198368 | 0.99945981 |
| CPEB4 | 99.8632804 | -0.457909854 | 0.51972154 | -0.8810677 | 0.37828119 | 0.99945981 |
| SFXN1 | 23.8602297 | -0.556841432 | 0.41428593 | -1.3440993 | 0.17891627 | 0.99945981 |
| HRH2 | 24.7677766 | 0.91945554 | 0.66025444 | 1.3925776 | 0.16374756 | 0.99945981 |
| THOC3 | 24.2712553 | 0.514821814 | 0.41596271 | 1.23766337 | 0.21584091 | 0.99945981 |
| SIMC1 | 27.262123 | -0.165613924 | 0.28358827 | -0.5839943 | 0.55922417 | 0.99945981 |
| KIAA1191 | 9.64592881 | -0.240641834 | 0.49825938 | -0.482965 | 0.6291206 | 0.99945981 |
| CLTB | 9.71509268 | 0.271532936 | 0.48180323 | 0.56357641 | 0.57304244 | 0.99945981 |
| FAF2 | 24.2266895 | -0.47687613 | 0.31186613 | -1.5291052 | 0.12623836 | 0.99945981 |
| RNF44 | 10.3333801 | 0.913543663 | 0.68991394 | 1.32414148 | 0.18545606 | 0.99945981 |
| HK3 | 11.4040488 | 0.089083893 | 0.9375531 | 0.09501744 | 0.92430098 | 0.99945981 |
| UIMC1 | 47.450386 | 0.022447327 | 0.31868643 | 0.07043703 | 0.94384582 | 0.99945981 |
| ZNF346 | 14.5462922 | 0.3967574 | 0.39211501 | 1.01183936 | 0.31161486 | 0.99945981 |
| NSD1 | 102.236038 | 0.003115685 | 0.20357126 | 0.01530513 | 0.98778875 | 0.99945981 |
| PRELID1 | 21.5103297 | 0.139556087 | 0.67684381 | 0.20618655 | 0.8366452 | 0.99945981 |
| LMAN2 | 18.0773882 | -0.212691833 | 0.57039328 | -0.3728863 | 0.70923308 | 0.99945981 |
| RGS14 | 9.90270921 | 0.335736241 | 0.55695688 | 0.60280473 | 0.5466386 | 0.99945981 |
| GRK6 | 12.9681026 | 0.327882223 | 0.44693302 | 0.73362721 | 0.46317597 | 0.99945981 |
| DOK3 | 17.7762454 | 0.405207715 | 0.38959175 | 1.0400829 | 0.29830139 | 0.99945981 |
| FAM193B | 25.8575736 | 0.011517078 | 0.28391268 | 0.04056556 | 0.96764224 | 0.99945981 |
| TMED9 | 17.7547943 | -0.074825126 | 0.59157716 | -0.1264841 | 0.89934871 | 0.99945981 |
| RMND5B | 9.74304522 | -0.156055504 | 0.47764347 | -0.3267196 | 0.74387994 | 0.99945981 |
| HNRNPAB | 16.2299193 | 0.118986274 | 0.47158167 | 0.25231319 | 0.800799 | 0.99945981 |
| PHYKPL | 52.9999603 | -0.211804756 | 0.48399158 | -0.4376207 | 0.66166123 | 0.99945981 |
| CLK4 | 16.1604974 | 0.35992287 | 0.38141967 | 0.94364002 | 0.34535363 | 0.99945981 |
| ADAMTS2 | 20.4375237 | -0.809067341 | 1.07962984 | -0.7493933 | 0.4536202 | 0.99945981 |
| RUFY1 | 32.3852824 | 0.202058783 | 0.43614722 | 0.46328114 | 0.64316286 | 0.99945981 |
| HNRNPH1 | 102.007701 | 0.825143728 | 0.27409212 | 3.01046132 | 0.00260851 | 0.45570351 |
| CANX | 50.6069074 | -0.464125052 | 0.39002327 | -1.1899932 | 0.23404905 | 0.99945981 |
| MAML1 | 16.2763517 | -0.055973145 | 0.34366177 | -0.1628728 | 0.87061861 | 0.99945981 |
| SQSTM1 | 53.6204665 | -0.841193496 | 0.382443 | -2.1995265 | 0.02784051 | 0.84763724 |
| MRNIP | 9.57602712 | 0.353752764 | 0.5349016 | 0.66134175 | 0.50839317 | 0.99945981 |
| TBC1D9B | 19.9262457 | -0.96666244 | 0.38625163 | -2.5026754 | 0.01232585 | 0.67187199 |
| RNF130 | 116.376545 | -0.074243245 | 0.41057137 | -0.1808291 | 0.85650173 | 0.99945981 |
| MAPK9 | 17.6024099 | -0.463515935 | 0.37632832 | -1.2316797 | 0.21806877 | 0.99945981 |
| CNOT6 | 26.8519239 | -0.193188031 | 0.33690784 | -0.5734151 | 0.5663637 | 0.99945981 |
| MGAT1 | 59.6037243 | 0.050139192 | 0.35908712 | 0.1396296 | 0.88895265 | 0.99945981 |
| RACK1 | 57.2372627 | -0.414999352 | 0.49373659 | -0.8405278 | 0.4006125 | 0.99945981 |
| TRIM52 | 10.0683811 | 0.093400156 | 0.41379536 | 0.22571581 | 0.82142246 | 0.99945981 |
| DUSP22 | 20.3519505 | -0.366598509 | 0.38953399 | -0.9411207 | 0.346643 | 0.99945981 |

| | | | | | | |
|---|---|---|---|---|---|---|
| EXOC2 | 58.8653258 | 0.147243812 | 0.24354369 | 0.6045889 | 0.54545218 | 0.99945981 |
| GMDS | 113.678727 | -0.082373767 | 0.39263559 | -0.209797 | 0.83382612 | 0.99945981 |
| GMDS-DT | 84.9383141 | -0.211835101 | 0.51475941 | -0.4115225 | 0.68068942 | 0.99945981 |
| WRNIP1 | 14.3705477 | 0.248006841 | 0.39234677 | 0.63211133 | 0.52731413 | 0.99945981 |
| SERPINB1 | 33.6796482 | -0.558295848 | 0.42060121 | -1.3273758 | 0.18438442 | 0.99945981 |
| SERPINB9 | 56.0630352 | 0.889739873 | 0.59592359 | 1.49304356 | 0.13542578 | 0.99945981 |
| SERPINB6 | 31.8127046 | -0.241787972 | 0.42340797 | -0.571052 | 0.56796439 | 0.99945981 |
| NQO2 | 14.1271867 | -0.367251867 | 0.50416595 | -0.7284345 | 0.46634766 | 0.99945981 |
| RIPK1 | 25.0942529 | 0.259011472 | 0.31835607 | 0.8135905 | 0.41587958 | 0.99945981 |
| PSMG4 | 9.65375637 | 0.04762805 | 0.44350903 | 0.10738913 | 0.91448028 | 0.99945981 |
| SLC22A23 | 16.8641315 | 0.76831577 | 0.61737223 | 1.24449357 | 0.21331796 | 0.99945981 |
| PRPF4B | 60.0628411 | 0.267380258 | 0.28415229 | 0.94097519 | 0.34671757 | 0.99945981 |
| CDYL | 69.0327727 | 0.17356581 | 0.23943012 | 0.72491219 | 0.4685059 | 0.99945981 |
| LYRM4 | 24.4709348 | -0.475824943 | 0.48015784 | -0.9909761 | 0.32169726 | 0.99945981 |
| FARS2 | 89.6367305 | 0.175107452 | 0.28711176 | 0.60989301 | 0.54193269 | 0.99945981 |
| F13A1 | 193.342038 | -0.940215638 | 0.93146916 | -1.00939 | 0.31278764 | 0.99945981 |
| LY86 | 26.8860999 | 0.744272921 | 0.80271021 | 0.92720001 | 0.3538227 | 0.99945981 |
| RREB1 | 115.461927 | 0.390916457 | 0.29803151 | 1.3116615 | 0.18963437 | 0.99945981 |
| SSR1 | 27.6068213 | -0.205660779 | 0.4959693 | -0.4146643 | 0.67838765 | 0.99945981 |
| SNRNP48 | 10.0491848 | 0.248411657 | 0.44007867 | 0.56447103 | 0.57243361 | 0.99945981 |
| SLC35B3 | 11.3044193 | 0.601034843 | 0.42668725 | 1.40860745 | 0.15895128 | 0.99945981 |
| GCNT2 | 22.2221592 | 0.553672807 | 0.34208423 | 1.61852772 | 0.10554892 | 0.99945981 |
| MAK | 9.04459796 | -0.97031567 | 0.58065605 | -1.6710679 | 0.09470827 | 0.99945981 |
| AL139807.1 | 11.7691419 | 0.538806219 | 0.43570074 | 1.23664287 | 0.2162197 | 0.99945981 |
| NEDD9 | 150.100015 | 0.000371033 | 0.46477273 | 0.00079831 | 0.99936304 | 0.9997934 |
| TMEM170B | 29.1821359 | 1.040896031 | 0.69129586 | 1.50571715 | 0.1321398 | 0.99945981 |
| HIVEP1 | 56.031392 | 0.047367783 | 0.39580121 | 0.11967569 | 0.90474006 | 0.99945981 |
| PHACTR1 | 44.9512261 | -0.431305531 | 0.52737792 | -0.8178301 | 0.41345422 | 0.99945981 |
| GFOD1 | 36.5097265 | 0.600452822 | 0.43637785 | 1.3759929 | 0.16882383 | 0.99945981 |
| SIRT5 | 12.8028912 | 0.338158352 | 0.52845897 | 0.63989519 | 0.52224074 | 0.99945981 |
| NOL7 | 15.5218572 | -0.125099943 | 0.46129627 | -0.2711922 | 0.78624322 | 0.99945981 |
| RANBP9 | 68.7024527 | 0.515016215 | 0.28401095 | 1.81336746 | 0.06977516 | 0.95505283 |
| CD83 | 52.7932256 | 1.969079999 | 0.61643728 | 3.19429092 | 0.00140175 | 0.3983151 |
| JARID2 | 184.817755 | -0.005920997 | 0.23171669 | -0.0255527 | 0.97961408 | 0.99945981 |
| DTNBP1 | 56.4304262 | -0.014938611 | 0.35734314 | -0.0418047 | 0.96665441 | 0.99945981 |
| MYLIP | 13.5608595 | -0.422771388 | 0.50390515 | -0.83899 | 0.40147491 | 0.99945981 |
| ATXN1 | 343.279912 | -0.391087402 | 0.270549 | -1.4455326 | 0.14830834 | 0.99945981 |
| NUP153 | 50.8943357 | 0.548841596 | 0.34030007 | 1.61281658 | 0.10678436 | 0.99945981 |
| KIF13A | 53.9942778 | 0.760343047 | 0.40078914 | 1.89711488 | 0.05781278 | 0.92445778 |
| TPMT | 9.92525782 | 0.474626286 | 0.5411099 | 0.87713473 | 0.38041346 | 0.99945981 |
| KDM1B | 26.2010485 | 1.204116183 | 0.46075663 | 2.61334531 | 0.00896607 | 0.59184657 |
| DEK | 46.1853345 | 0.103829139 | 0.25732341 | 0.40349666 | 0.68658289 | 0.99945981 |
| RNF144B | 123.274427 | 0.8751238 | 0.78196293 | 1.1191372 | 0.26308161 | 0.99945981 |
| MBOAT1 | 18.8568605 | 0.286215486 | 0.43382296 | 0.65975182 | 0.50941311 | 0.99945981 |
| E2F3 | 54.1602167 | 0.008688702 | 0.26146794 | 0.03323047 | 0.9734908 | 0.99945981 |
| CDKAL1 | 162.665284 | 0.074722418 | 0.2821268 | 0.26485402 | 0.79112192 | 0.99945981 |
| SOX4 | 17.5898257 | -2.686150029 | 1.06160502 | -2.5302725 | 0.0113974 | 0.67187199 |
| DCDC2 | 9.38478051 | -0.391656505 | 0.63311473 | -0.6186185 | 0.53616769 | 0.99945981 |
| TDP2 | 16.6894155 | -0.387577559 | 0.45946818 | -0.8435351 | 0.39892922 | 0.99945981 |
| C6orf62 | 53.8844743 | 0.426607668 | 0.29848603 | 1.42923833 | 0.15293575 | 0.99945981 |
| RIPOR2 | 325.675058 | 0.805169113 | 0.32210459 | 2.49971324 | 0.01242939 | 0.67187199 |
| CARMIL1 | 65.3431689 | -1.047415927 | 0.54580946 | -1.9190139 | 0.05498258 | 0.92445778 |
| TRIM38 | 56.8299347 | -0.062722239 | 0.35108039 | -0.1786549 | 0.85820867 | 0.99945981 |
| HIST1H1C | 10.1446942 | -1.223579338 | 0.81781934 | -1.4961487 | 0.13461491 | 0.99945981 |
| HIST1H4C | 10.0342285 | -2.844576139 | 0.7420659 | -3.8333201 | 0.00012643 | 0.12398707 |
| HIST1H2AC | 83.9940107 | 0.523165559 | 0.65072189 | 0.80397719 | 0.42141015 | 0.99945981 |
| HIST1H1E | 10.6633259 | -0.980028955 | 0.86940303 | -1.1272435 | 0.25963952 | 0.99945981 |
| HIST1H1D | 10.5060449 | -0.79286054 | 0.65656848 | -1.2075824 | 0.22720793 | 0.99945981 |
| BTN3A2 | 30.7711559 | -0.23289465 | 0.41523556 | -0.5608736 | 0.57488373 | 0.99945981 |
| BTN3A1 | 30.9167704 | -0.041346522 | 0.55592407 | -0.0743744 | 0.94071247 | 0.99945981 |
| BTN3A3 | 12.5050576 | -0.06120615 | 0.41521243 | -0.1474092 | 0.88280901 | 0.99945981 |
| BTN2A1 | 16.690995 | 0.319294456 | 0.3450546 | 0.92534473 | 0.35478662 | 0.99945981 |
| HMGN4 | 8.50976436 | -0.135668374 | 0.56735242 | -0.2391254 | 0.81100835 | 0.99945981 |
| TRIM27 | 24.8658022 | 0.165076834 | 0.31443769 | 0.52499061 | 0.59958972 | 0.99945981 |
| GABBR1 | 13.5769344 | 0.970555962 | 0.7522467 | 1.29020966 | 0.19697787 | 0.99945981 |
| HLA-F | 30.5772487 | -0.24644904 | 0.43850185 | -0.5620251 | 0.57409893 | 0.99945981 |
| HLA-A | 231.457775 | -0.272082186 | 0.45817805 | -0.593835 | 0.55262244 | 0.99945981 |
| TRIM26 | 17.9694165 | 0.01317786 | 0.36966285 | 0.03564832 | 0.97156278 | 0.99945981 |
| HCG17 | 14.4619163 | -0.535371783 | 0.545601 | -0.9812515 | 0.32646875 | 0.99945981 |
| HLA-E | 155.794746 | -0.149969322 | 0.37001364 | -0.4053075 | 0.68525146 | 0.99945981 |
| GNL1 | 9.9458403 | -1.171282234 | 0.48969362 | -2.3918675 | 0.01676289 | 0.74280553 |
| ABCF1 | 12.7305663 | -0.251141312 | 0.44517299 | -0.5641432 | 0.57265668 | 0.99945981 |
| PPP1R10 | 22.7625118 | 0.395407089 | 0.34602067 | 1.14272679 | 0.25315204 | 0.99945981 |
| PPP1R18 | 21.7249989 | -0.250308671 | 0.53677635 | -0.4663184 | 0.64098763 | 0.99945981 |
| TUBB | 25.948939 | -0.316974508 | 0.57347368 | -0.5527272 | 0.58045022 | 0.99945981 |
| FLOT1 | 31.099321 | -0.446861873 | 0.4460144 | -1.0019001 | 0.31639184 | 0.99945981 |
| IER3 | 23.3354584 | -0.480818768 | 0.6002668 | -0.8010084 | 0.42312676 | 0.99945981 |
| HLA-C | 139.906178 | 0.058613409 | 0.55147425 | 0.10628494 | 0.91535628 | 0.99945981 |
| HLA-B | 274.103497 | 0.080161203 | 0.49889305 | 0.16067813 | 0.87234691 | 0.99945981 |
| LTB | 31.6892296 | 0.44488646 | 0.68926213 | 0.64545322 | 0.51863353 | 0.99945981 |
| LST1 | 27.9245944 | 1.321714237 | 0.80387535 | 1.64417808 | 0.10013942 | 0.99945981 |
| AIF1 | 31.0703139 | 0.365470215 | 0.96779399 | 0.37763224 | 0.70570381 | 0.99945981 |
| PRRC2A | 18.3931077 | -0.015142591 | 0.41080821 | -0.0368605 | 0.97059624 | 0.99945981 |
| DDAH2 | 11.403356 | -0.101563006 | 0.5255959 | -0.193234 | 0.8467757 | 0.99945981 |
| CLIC1 | 71.5999122 | -0.576811439 | 0.58583451 | -0.9845979 | 0.32482161 | 0.99945981 |
| HSPA1A | 66.2806653 | 1.206431836 | 0.70824651 | 1.70340669 | 0.08849199 | 0.99640647 |
| HSPA1B | 31.4375959 | 1.436220051 | 0.73303007 | 1.95929214 | 0.05007858 | 0.92165202 |
| C6orf48 | 9.04378791 | 0.228005518 | 0.54813922 | 0.41596279 | 0.67743723 | 0.99945981 |
| ATF6B | 15.3722359 | -0.181146235 | 0.40968191 | -0.4421631 | 0.65837116 | 0.99945981 |
| PBX2 | 10.0918715 | 0.699140725 | 0.42373963 | 1.64992998 | 0.09895726 | 0.99945981 |
| GPSM3 | 23.5426477 | 0.352094588 | 0.68510147 | 0.51393057 | 0.60730053 | 0.99945981 |
| HLA-DRA | 173.660105 | -0.18501584 | 0.67694343 | -0.2733106 | 0.78461444 | 0.99945981 |
| HLA-DRB5 | 46.5360586 | 0.502184625 | 0.73194127 | 0.68609962 | 0.49265029 | 0.99945981 |
| HLA-DRB1 | 125.576694 | 0.269473256 | 0.75304426 | 0.35784518 | 0.72045918 | 0.99945981 |
| HLA-DQA1 | 41.5763087 | -0.713875006 | 0.75626661 | -0.9439462 | 0.34519713 | 0.99945981 |
| HLA-DQB1 | 55.694168 | -0.015312268 | 0.84346883 | -0.0181539 | 0.98551606 | 0.99945981 |
| PSMB8 | 20.2129663 | 0.111014892 | 0.43156615 | 0.25723725 | 0.79699563 | 0.99945981 |
| PSMB9 | 26.2450417 | 0.008638833 | 0.47328553 | 0.0182529 | 0.9854371 | 0.99945981 |
| TAP1 | 21.6100413 | 0.734391635 | 0.5041119 | 1.4568028 | 0.14517082 | 0.99945981 |
| HLA-DMB | 16.1601636 | 0.175555003 | 0.58047195 | 0.30243494 | 0.76232052 | 0.99945981 |
| BRD2 | 30.9205622 | 0.165749689 | 0.36397434 | 0.45538839 | 0.64882984 | 0.99945981 |
| HLA-DOA | 10.6173011 | -0.743648578 | 0.79474739 | -0.9357043 | 0.34942544 | 0.99945981 |
| HLA-DPA1 | 70.8631977 | -0.306086936 | 0.67339899 | -0.4545402 | 0.64944004 | 0.99945981 |
| HLA-DPB1 | 82.2602683 | 0.236279846 | 0.64917386 | 0.36397005 | 0.71588037 | 0.99945981 |
| RPS18 | 63.357152 | -0.30564869 | 0.64751053 | -0.4720366 | 0.63690064 | 0.99945981 |
| RGL2 | 9.15591392 | 0.291351825 | 0.43435415 | 0.67077022 | 0.50236692 | 0.99945981 |
| TAPBP | 55.9863611 | 0.063667525 | 0.35567214 | 0.17900622 | 0.85793282 | 0.99945981 |
| CUTA | 9.14972782 | -0.288855305 | 0.53739431 | -0.5375109 | 0.59091475 | 0.99945981 |
| ITPR3 | 10.8451592 | 0.046644386 | 0.55097763 | 0.0846575 | 0.93253369 | 0.99945981 |
| LEMD2 | 12.434982 | -0.151454484 | 0.41305263 | -0.3666712 | 0.71386433 | 0.99945981 |
| NUDT3 | 13.8388643 | -0.539701487 | 0.61141778 | -0.8827049 | 0.37739572 | 0.99945981 |
| C6orf106 | 48.798264 | -0.43647583 | 0.36730653 | -1.1883149 | 0.23470936 | 0.99945981 |
| SNRPC | 8.40009153 | -1.481307424 | 0.58953716 | -2.5126617 | 0.01198242 | 0.67187199 |

| | | | | | | |
|---|---|---|---|---|---|---|
| UHRF1BP1 | 26.7314659 | 0.414973574 | 0.41872984 | 0.99102938 | 0.32167123 | 0.99945981 |
| TAF11 | 13.2976316 | 0.027049737 | 0.4022844 | 0.06724033 | 0.94639038 | 0.99945981 |
| ANKS1A | 93.0658466 | 0.002488604 | 0.29972374 | 0.00830299 | 0.99337525 | 0.99945981 |
| ZNF76 | 13.7706737 | -0.175165629 | 0.38704589 | -0.4525707 | 0.6508579 | 0.99945981 |
| DEF6 | 26.3369467 | -0.134482769 | 0.34068789 | -0.3947389 | 0.69303558 | 0.99945981 |
| PPARD | 54.6021352 | -0.322434522 | 0.54195054 | -0.5949519 | 0.5518756 | 0.99945981 |
| RPL10A | 57.6201127 | -0.666511089 | 0.63158539 | -1.0552985 | 0.29128885 | 0.99945981 |
| FKBP5 | 354.201408 | -0.602101539 | 0.65563624 | -0.9183469 | 0.35843726 | 0.99945981 |
| SRPK1 | 31.8925238 | -0.041985714 | 0.31595606 | -0.1328847 | 0.8942846 | 0.99945981 |
| MAPK14 | 88.4743746 | -0.170799965 | 0.33807856 | -0.5052079 | 0.61341281 | 0.99945981 |
| KCTD20 | 33.565393 | -0.396209642 | 0.40716277 | -0.9730989 | 0.33050415 | 0.99945981 |
| STK38 | 109.813994 | 0.690128942 | 0.39412614 | 1.75103569 | 0.07993976 | 0.9846219 |
| SRSF3 | 35.9649468 | 0.034364378 | 0.31636381 | 0.10862297 | 0.91350154 | 0.99945981 |
| CDKN1A | 13.2501926 | 0.52626299 | 0.73035188 | 0.72056087 | 0.47117973 | 0.99945981 |
| C6orf89 | 21.6367049 | 0.06098801 | 0.3173026 | 0.19220772 | 0.8475795 | 0.99945981 |
| MTCH1 | 27.7645884 | 0.351355364 | 0.37401708 | 0.93940995 | 0.34752031 | 0.99945981 |
| FGD2 | 52.1403389 | 0.849114854 | 0.44541216 | 1.9063576 | 0.05660383 | 0.92445778 |
| PIM1 | 20.2000084 | 0.001811082 | 0.53048045 | 0.00341404 | 0.997276 | 0.99945981 |
| TBC1D22B | 15.7099945 | -0.26707018 | 0.44320081 | -0.6025941 | 0.54677877 | 0.99945981 |
| CMTR1 | 11.73743 | 0.214825482 | 0.46326163 | 0.46372388 | 0.64284559 | 0.99945981 |
| ZFAND3 | 381.809562 | 0.422070875 | 0.17004133 | 2.48216636 | 0.01305863 | 0.69426883 |
| BTBD9 | 163.854821 | 0.012624389 | 0.26676172 | 0.04732459 | 0.96225453 | 0.99945981 |
| GLO1 | 15.4986924 | -0.581452906 | 0.69644595 | -0.8348859 | 0.40378195 | 0.99945981 |
| OARD1 | 11.9513725 | 0.234110604 | 0.48415987 | 0.48353988 | 0.62871245 | 0.99945981 |
| NFYA | 10.2448875 | -0.278154555 | 0.45306869 | -0.6139346 | 0.53925853 | 0.99945981 |
| TFEB | 30.2566198 | -0.818281711 | 0.78929377 | -1.0367264 | 0.29986337 | 0.99945981 |
| CCND3 | 255.810263 | -0.960649925 | 0.56661804 | -1.69541 | 0.08999766 | 0.99640647 |
| TAF8 | 11.2816815 | -0.635023206 | 0.50806362 | -1.2498892 | 0.21134004 | 0.99945981 |
| TRERF1 | 89.5238297 | -0.190797624 | 0.28457719 | -0.67046 | 0.50256459 | 0.99945981 |
| UBR2 | 111.898197 | 0.158394234 | 0.23963689 | 0.66097602 | 0.5086277 | 0.99945981 |
| BICRAL | 51.989455 | -0.278758346 | 0.36480643 | -0.7641267 | 0.44479172 | 0.99945981 |
| RPL7L1 | 14.4684242 | -0.776326596 | 0.41038614 | -1.8916979 | 0.05853124 | 0.92445778 |
| CNPY3 | 22.2493107 | 0.395011471 | 0.5267097 | 0.74996051 | 0.45327849 | 0.99945981 |
| PEX6 | 8.45340706 | -0.985946946 | 0.48915014 | -2.0156326 | 0.04383841 | 0.89568661 |
| KLHDC3 | 8.13581382 | -0.33602338 | 0.51794154 | -0.648767 | 0.51648899 | 0.99945981 |
| RRP36 | 11.3507988 | -0.398989824 | 0.42465369 | -0.9395652 | 0.34744064 | 0.99945981 |
| CUL9 | 8.71602597 | -0.258477529 | 0.50924637 | -0.5075687 | 0.61175583 | 0.99945981 |
| ZNF318 | 12.9515105 | -1.204033622 | 0.93551782 | -1.2870237 | 0.19808602 | 0.99945981 |
| ABCC10 | 8.96824014 | -0.47852941 | 0.46527939 | -1.0284776 | 0.30372524 | 0.99945981 |
| POLR1C | 10.1209358 | -0.430491192 | 0.57301016 | -0.7512802 | 0.45248404 | 0.99945981 |
| XPO5 | 51.5639993 | -0.110934915 | 0.49223048 | -0.2253719 | 0.82168997 | 0.99945981 |
| POLH | 10.4997037 | -0.951110047 | 0.68565664 | -1.3871521 | 0.16539539 | 0.99945981 |
| GTPBP2 | 10.2222222 | -0.003557763 | 0.48443334 | -0.0073442 | 0.99414025 | 0.99945981 |
| VEGFA | 19.517127 | 0.178113681 | 0.84829105 | 0.20996765 | 0.83369292 | 0.99945981 |
| MRPL14 | 8.32777617 | -0.402785286 | 0.73506281 | -0.5479604 | 0.58371912 | 0.99945981 |
| HSP90AB1 | 61.2725793 | -0.166646298 | 0.43253671 | -0.3852767 | 0.70003245 | 0.99945981 |
| CDC5L | 16.5053064 | 0.026199324 | 0.51142781 | 0.05122781 | 0.959144 | 0.99945981 |
| SUPT3H | 78.0337595 | -0.06264913 | 0.42287939 | -0.1481489 | 0.88222523 | 0.99945981 |
| RUNX2 | 63.694542 | 0.248519503 | 0.42460992 | 0.58528897 | 0.55835343 | 0.99945981 |
| ENPP4 | 11.1393299 | -0.210695318 | 0.56945497 | -0.3699947 | 0.71138644 | 0.99945981 |
| CD2AP | 55.0171207 | 0.248285008 | 0.35562151 | 0.69817208 | 0.48506958 | 0.99945981 |
| PKHD1 | 20.5590595 | 1.54925382 | 0.72318716 | 2.14225848 | 0.03217269 | 0.8663391 |
| TRAM2 | 15.8191475 | -0.001028675 | 0.70851368 | -0.0014519 | 0.99884157 | 0.9997934 |
| FBXO9 | 21.1353681 | -0.06841535 | 0.33946122 | -0.201541 | 0.84027559 | 0.99945981 |
| ELOVL5 | 71.6379408 | 0.406279911 | 0.39911795 | 1.01794447 | 0.30870435 | 0.99945981 |
| GCLC | 18.6936941 | -0.573519926 | 0.4199704 | -1.3656199 | 0.1720583 | 0.99945981 |
| DST | 53.8809453 | -0.480933777 | 0.49279707 | -0.9759266 | 0.32910084 | 0.99945981 |
| ZNF451 | 52.9658139 | 0.145056265 | 0.29755567 | 0.48749286 | 0.6259091 | 0.99945981 |
| PRIM2 | 70.7219652 | 0.221885668 | 0.25810049 | 0.85968713 | 0.38996153 | 0.99945981 |
| KHDRBS2 | 36.6730213 | 1.036124591 | 0.65327293 | 1.58605161 | 0.1127276 | 0.99945981 |
| PTP4A1 | 18.5993254 | 0.347989842 | 0.42829962 | 0.81249159 | 0.41650961 | 0.99945981 |
| PHF3 | 73.0144529 | 0.089222685 | 0.25885933 | 0.34467634 | 0.73033771 | 0.99945981 |
| LMBRD1 | 67.8578206 | 0.258786652 | 0.22076763 | 1.17221284 | 0.24111161 | 0.99945981 |
| COL19A1 | 14.2177862 | 0.546413219 | 0.56936022 | 0.95969686 | 0.3372078 | 0.99945981 |
| FAM135A | 45.2120537 | -0.004152729 | 0.41572179 | -0.0099892 | 0.9920299 | 0.99945981 |
| SMAP1 | 64.2436271 | 0.14427955 | 0.32104433 | 0.44940694 | 0.65313813 | 0.99945981 |
| OGFRL1 | 41.896639 | 0.486972497 | 0.42101389 | 1.15666611 | 0.2474088 | 0.99945981 |
| KCNQ5 | 79.6902776 | 0.089100126 | 0.5822131 | 0.15303696 | 0.87836913 | 0.99945981 |
| OOEP | 81.5071698 | -0.599139215 | 0.61808497 | -0.9693477 | 0.33237176 | 0.99945981 |
| CGAS | 17.130529 | 0.065774231 | 0.38856276 | 0.1692757 | 0.86557979 | 0.99945981 |
| MTO1 | 12.5083637 | 0.07906366 | 0.42147881 | 0.18758632 | 0.85120095 | 0.99945981 |
| EEF1A1 | 261.597146 | -0.198216722 | 0.4908686 | -0.4038081 | 0.68635383 | 0.99945981 |
| SLC17A5 | 24.9540198 | 0.11268228 | 0.40294388 | 0.27964757 | 0.7797479 | 0.99945981 |
| COX7A2 | 15.3125793 | 0.401731178 | 0.4309786 | 0.93213717 | 0.35126564 | 0.99945981 |
| TMEM30A | 29.8226187 | -0.123270186 | 0.47657139 | -0.2586605 | 0.79589721 | 0.99945981 |
| SENP6 | 118.178278 | 0.092288216 | 0.23566124 | 0.39161389 | 0.69534353 | 0.99945981 |
| MYO6 | 21.6115182 | -0.289471196 | 0.40186112 | -0.7203265 | 0.47132401 | 0.99945981 |
| PHIP | 144.233459 | 0.342876991 | 0.25434714 | 1.34806702 | 0.17763683 | 0.99945981 |
| HMGN3 | 29.4037872 | 0.113010556 | 0.37209056 | 0.30371788 | 0.76134283 | 0.99945981 |
| BCKDHB | 27.2579302 | -0.175162449 | 0.41310403 | -0.4240154 | 0.67155462 | 0.99945981 |
| TENT5A | 28.5660385 | 0.308133198 | 0.7372279 | 0.41796193 | 0.67597495 | 0.99945981 |
| IBTK | 47.0249074 | 0.083003021 | 0.27240711 | 0.30470211 | 0.76059304 | 0.99945981 |
| UBE3D | 19.9787546 | -0.39939818 | 0.39818367 | -1.0030501 | 0.31583667 | 0.99945981 |
| DOPEY1 | 46.5752019 | 0.30130818 | 0.29783357 | 1.01166628 | 0.31169764 | 0.99945981 |
| PGM3 | 10.0773219 | -0.316858138 | 0.66382405 | -0.4773225 | 0.63313251 | 0.99945981 |
| ME1 | 37.2145287 | 0.603412357 | 0.69116442 | 0.87303736 | 0.38264271 | 0.99945981 |
| CYB5R4 | 54.3982789 | 0.219557276 | 0.33656233 | 0.65235249 | 0.51417381 | 0.99945981 |
| CEP162 | 17.3351438 | 0.398324603 | 0.36728731 | 1.08450412 | 0.27814134 | 0.99945981 |
| SNX14 | 47.4250846 | 0.486561048 | 0.27607981 | 1.76239272 | 0.07800297 | 0.98289661 |
| SYNCRIP | 17.878435 | -0.374899832 | 0.46559833 | -0.8052001 | 0.4207042 | 0.99945981 |
| ZNF292 | 133.471666 | -0.323683216 | 0.37043263 | -0.8737978 | 0.38222838 | 0.99945981 |
| SMIM8 | 10.747061 | 0.619608443 | 0.5041658 | 1.22897753 | 0.21908023 | 0.99945981 |
| RARS2 | 38.3062242 | 0.062075994 | 0.37315902 | 0.16635266 | 0.86787943 | 0.99945981 |
| ORC3 | 20.4827317 | 0.158686173 | 0.36057276 | 0.44009473 | 0.6598685 | 0.99945981 |
| AKIRIN2 | 26.7832739 | 0.366100987 | 0.32237215 | 1.13564706 | 0.2561043 | 0.99945981 |
| RNGTT | 75.7801108 | -0.242602305 | 0.31648026 | -0.7665638 | 0.44334091 | 0.99945981 |
| PNRC1 | 39.9688555 | 0.202091899 | 0.27754306 | 0.72814612 | 0.46652415 | 0.99945981 |
| UBE2J1 | 25.7751662 | 0.386876799 | 0.45155609 | 0.85676356 | 0.39157557 | 0.99945981 |
| ANKRD6 | 15.5554637 | 0.521082449 | 0.37847816 | 1.37678341 | 0.16857922 | 0.99945981 |
| LYRM2 | 10.4519041 | -0.011335154 | 0.48649301 | -0.0232997 | 0.98141119 | 0.99945981 |
| MDN1 | 28.34198 | -0.247220033 | 0.29621534 | -0.8345957 | 0.40394541 | 0.99945981 |
| CASP8AP2 | 21.3963986 | 0.036706745 | 0.32687292 | 0.11229668 | 0.91058817 | 0.99945981 |
| BACH2 | 173.540538 | -1.961218492 | 0.98790873 | -1.9852223 | 0.04711974 | 0.90884462 |
| MAP3K7 | 34.128355 | -0.107094235 | 0.32936325 | -0.3251554 | 0.74506347 | 0.99945981 |
| UFL1 | 24.8566682 | -0.063242003 | 0.45864727 | -0.1378881 | 0.89032885 | 0.99945981 |
| MMS22L | 23.9914699 | 0.704170455 | 0.45733254 | 1.53973399 | 0.12362521 | 0.99945981 |
| FBXL4 | 31.5092488 | -0.003076473 | 0.37878294 | -0.008122 | 0.99351966 | 0.99945981 |
| PNISR | 102.091478 | 0.218054568 | 0.34797707 | 0.62663487 | 0.53089861 | 0.99945981 |
| USP45 | 15.2787601 | -0.199719355 | 0.44519527 | -0.4486107 | 0.65371253 | 0.99945981 |
| CCNC | 21.5512718 | 0.45539286 | 0.50476137 | 0.90219435 | 0.36695364 | 0.99945981 |
| ASCC3 | 97.4502034 | 0.092222925 | 0.25328758 | 0.36410362 | 0.71578063 | 0.99945981 |
| HACE1 | 21.264732 | -0.204516752 | 0.55925318 | -0.3656962 | 0.7145918 | 0.99945981 |
| PREP | 31.0862316 | -0.292493553 | 0.34772523 | -0.8411629 | 0.4002567 | 0.99945981 |

| | | | | | | |
|---|---|---|---|---|---|---|
| ATG5 | 61.5641523 | 0.109199026 | 0.29421461 | 0.37115432 | 0.71052259 | 0.99945981 |
| PRDM1 | 31.7360859 | -0.348911587 | 0.64570508 | -0.5403575 | 0.58895051 | 0.99945981 |
| CRYBG1 | 61.5443679 | -0.180973963 | 0.49253903 | -0.3674307 | 0.71329777 | 0.99945981 |
| QRSL1 | 10.0695277 | 0.295013216 | 0.48445287 | 0.60896164 | 0.54254987 | 0.99945981 |
| CD24 | 31.2074438 | -0.639163071 | 0.35361999 | -1.8074857 | 0.07068659 | 0.95505283 |
| PDSS2 | 76.739964 | 0.024944804 | 0.22653176 | 0.11011615 | 0.91231726 | 0.99945981 |
| SCML4 | 37.0388675 | -0.182287416 | 0.57005625 | -0.3197709 | 0.74914199 | 0.99945981 |
| SEC63 | 52.1770991 | 0.05477509 | 0.25303127 | 0.21647557 | 0.82861707 | 0.99945981 |
| OSTM1 | 20.18335 | 0.026375703 | 0.3998235 | 0.06596837 | 0.94740301 | 0.99945981 |
| SNX3 | 43.2505473 | -0.583141026 | 0.40948097 | -1.424098 | 0.15441811 | 0.99945981 |
| AFG1L | 15.3702207 | -0.039849743 | 0.40860043 | -0.0975274 | 0.92230756 | 0.99945981 |
| FOXO3 | 127.333784 | 0.40763094 | 0.4860475 | 0.83866482 | 0.40165742 | 0.99945981 |
| ARMC2 | 17.8471475 | 0.258286298 | 0.56242459 | 0.45923721 | 0.64606383 | 0.99945981 |
| SESN1 | 62.0757455 | 0.447325462 | 0.37302053 | 1.19919796 | 0.23045098 | 0.99945981 |
| CD164 | 49.4389027 | 0.115980625 | 0.26218778 | 0.4423571 | 0.65823082 | 0.99945981 |
| MICAL1 | 14.4294749 | -0.182101116 | 0.55351682 | -0.3289893 | 0.74216377 | 0.99945981 |
| AK9 | 13.8374609 | -0.0452613 | 0.63581771 | -0.071186 | 0.94324975 | 0.99945981 |
| FIG4 | 32.0265029 | 0.040030042 | 0.2884911 | 0.13875659 | 0.8896425 | 0.99945981 |
| CDC40 | 21.3423606 | 0.267702562 | 0.38021697 | 0.70407842 | 0.48138394 | 0.99945981 |
| CDK19 | 82.0119567 | 0.485759894 | 0.39828672 | 1.21962363 | 0.22260758 | 0.99945981 |
| AMD1 | 27.7393822 | 0.158470145 | 0.43275224 | 0.36619139 | 0.71422226 | 0.99945981 |
| GTF3C6 | 11.5606112 | 0.170070368 | 0.52879591 | 0.32161816 | 0.74774198 | 0.99945981 |
| SLC16A10 | 152.796587 | -0.300544378 | 0.98097073 | -0.3063745 | 0.75931955 | 0.99945981 |
| REV3L | 91.0439167 | 0.131873439 | 0.35430423 | 0.37220396 | 0.709741 | 0.99945981 |
| TRAF3IP2-AS | 41.9148729 | -0.109892953 | 0.34019702 | -0.3230274 | 0.7466745 | 0.99945981 |
| FYN | 317.404468 | -0.35412378 | 0.5137298 | -0.6893191 | 0.49062246 | 0.99945981 |
| MARCKS | 38.9891842 | 0.315787868 | 0.60312841 | 0.52358314 | 0.60056851 | 0.99945981 |
| HDAC2 | 17.983914 | -0.091328082 | 0.35019898 | -0.2607891 | 0.79425514 | 0.99945981 |
| NT5DC1 | 48.7824889 | 0.259011996 | 0.33797426 | 0.76636605 | 0.44345853 | 0.99945981 |
| DSE | 19.6693959 | 0.110950044 | 0.5468835 | 0.20287693 | 0.83923123 | 0.99945981 |
| TSPYL1 | 10.2562087 | 0.192359236 | 0.4701149 | 0.40917494 | 0.68241128 | 0.99945981 |
| RWDD1 | 8.82463584 | -1.009121148 | 0.57763973 | -1.7469732 | 0.08064198 | 0.9846219 |
| ZUP1 | 11.1787499 | 0.230735967 | 0.44595171 | 0.51740123 | 0.6048761 | 0.99945981 |
| KPNA5 | 30.8452414 | -0.01163849 | 0.39487347 | -0.029474 | 0.97648658 | 0.99945981 |
| GOPC | 19.7148506 | -0.648998555 | 0.33417867 | -1.9420706 | 0.05212856 | 0.9230968 |
| NUS1 | 20.2254045 | 0.061584015 | 0.33260669 | 0.18515567 | 0.85310694 | 0.99945981 |
| CEP85L | 117.84348 | 0.330020105 | 0.23260208 | 1.41881839 | 0.15595197 | 0.99945981 |
| MCM9 | 30.8973975 | -0.439326671 | 0.28162867 | -1.5599501 | 0.11877168 | 0.99945981 |
| MAN1A1 | 156.483051 | -0.074676985 | 0.42633083 | -0.1751621 | 0.8609523 | 0.99945981 |
| TBC1D32 | 22.6283023 | -0.227362937 | 0.47489942 | -0.4787602 | 0.63210924 | 0.99945981 |
| SERINC1 | 38.997569 | -0.103740188 | 0.304581 | -0.3405997 | 0.73340498 | 0.99945981 |
| RNF217 | 18.9510752 | -0.022625907 | 0.46641969 | -0.0485098 | 0.96130999 | 0.99945981 |
| NCOA7 | 51.4944326 | -0.342054551 | 0.36474605 | -0.9377882 | 0.34835325 | 0.99945981 |
| TRMT11 | 13.1045175 | -0.033412822 | 0.6346437 | -0.0526482 | 0.95801225 | 0.99945981 |
| RNF146 | 14.8929803 | -0.099794425 | 0.40455271 | -0.2466784 | 0.80515711 | 0.99945981 |
| ECHDC1 | 35.1780956 | 0.096660343 | 0.29843503 | 0.32389074 | 0.74602076 | 0.99945981 |
| THEMIS | 161.679702 | -0.373376114 | 0.48005254 | -0.7777818 | 0.43669768 | 0.99945981 |
| PTPRK | 13.9397183 | 0.808763429 | 0.54046615 | 1.4964183 | 0.13454468 | 0.99945981 |
| ARHGAP18 | 105.544359 | -0.22133262 | 0.35747696 | -0.6191521 | 0.53581614 | 0.99945981 |
| L3MBTL3 | 31.9551724 | 0.396335258 | 0.44218843 | 0.89630399 | 0.37009043 | 0.99945981 |
| SAMD3 | 74.8149188 | 0.022795933 | 0.48031893 | 0.04745999 | 0.96214662 | 0.99945981 |
| EPB41L2 | 56.2732656 | 0.069449074 | 0.51179688 | 0.13569656 | 0.89206117 | 0.99945981 |
| AKAP7 | 26.0155559 | 0.465367175 | 0.51943332 | 0.89591321 | 0.37029912 | 0.99945981 |
| MED23 | 26.6105653 | 0.391023401 | 0.40776709 | 0.95893811 | 0.33758992 | 0.99945981 |
| STX7 | 101.403181 | -0.189438481 | 0.40083125 | -0.472614 | 0.63648856 | 0.99945981 |
| SLC18B1 | 12.5098618 | 0.209521034 | 0.50662228 | 0.41356458 | 0.67919302 | 0.99945981 |
| RPS12 | 66.6271368 | 0.286532865 | 0.52111556 | 0.54984515 | 0.58242559 | 0.99945981 |
| TBPL1 | 12.7329345 | -0.229976951 | 0.46160911 | -0.4982071 | 0.61833805 | 0.99945981 |
| SGK1 | 42.8804506 | -0.629916781 | 0.96449436 | -0.6531057 | 0.51368813 | 0.99945981 |
| HBS1L | 28.1856069 | 0.249367021 | 0.28904474 | 0.86272812 | 0.38828697 | 0.99945981 |
| AHI1 | 40.3934999 | -0.746232317 | 0.36918382 | -2.0213029 | 0.04324842 | 0.89562588 |
| PDE7B | 26.5804866 | -0.177551823 | 0.69933446 | -0.2538869 | 0.79958297 | 0.99945981 |
| BCLAF1 | 66.887017 | -0.000433881 | 0.26221467 | -0.0016547 | 0.99867976 | 0.9997934 |
| MAP7 | 12.7429578 | -1.267664188 | 0.85116664 | -1.4893255 | 0.13640168 | 0.99945981 |
| MAP3K5 | 158.757317 | -0.118962913 | 0.38722006 | -0.307223 | 0.75867363 | 0.99945981 |
| IFNGR1 | 67.2867174 | 0.535329885 | 0.38213281 | 1.40090007 | 0.16124396 | 0.99945981 |
| AL357060.1 | 18.5458353 | -1.044229293 | 0.67340876 | -1.5506619 | 0.12098273 | 0.99945981 |
| TNFAIP3 | 43.2769913 | -0.155122353 | 0.60969243 | -0.2544272 | 0.79916552 | 0.99945981 |
| HEBP2 | 11.7031681 | -0.538937192 | 0.43409973 | -1.2415055 | 0.21441907 | 0.99945981 |
| NHSL1 | 54.3326888 | 0.374738948 | 0.72807144 | 0.5147008 | 0.60676212 | 0.99945981 |
| REPS1 | 50.2966575 | -0.115260852 | 0.393781 | -0.2927029 | 0.76974923 | 0.99945981 |
| ABRACL | 15.476891 | 0.61869338 | 0.47152934 | 1.31209944 | 0.18948658 | 0.99945981 |
| HECA | 37.9505633 | 0.281374652 | 0.31057652 | 0.90597528 | 0.36494894 | 0.99945981 |
| AL031772.1 | 12.6735398 | -1.162535718 | 0.97449844 | -1.192958 | 0.23288582 | 0.99945981 |
| VTA1 | 24.6159534 | 0.045335434 | 0.33026923 | 0.13726811 | 0.89081887 | 0.99945981 |
| HIVEP2 | 103.645534 | -0.01363563 | 0.36237049 | -0.037629 | 0.96998351 | 0.99945981 |
| AIG1 | 25.4808491 | 0.545458371 | 0.44616225 | 1.22255608 | 0.22149741 | 0.99945981 |
| PEX3 | 12.4768302 | -0.67495067 | 0.4089883 | -1.6502933 | 0.09888296 | 0.99945981 |
| PHACTR2 | 116.695797 | -0.038592141 | 0.33635649 | -0.1147358 | 0.90865451 | 0.99945981 |
| PLAGL1 | 30.2484518 | -0.040030272 | 0.57148227 | -0.0700464 | 0.94415673 | 0.99945981 |
| SF3B5 | 15.5457554 | -0.174712377 | 0.56481042 | -0.3093292 | 0.75707109 | 0.99945981 |
| STX11 | 22.521771 | 1.100258726 | 0.62185372 | 1.76932081 | 0.07684035 | 0.9821652 |
| UTRN | 550.428259 | 0.544278781 | 0.39859701 | 1.36548637 | 0.17210023 | 0.99945981 |
| SHPRH | 50.2722312 | 0.262367402 | 0.32692952 | 0.80251977 | 0.42225236 | 0.99945981 |
| STXBP5 | 75.7486006 | -0.555186482 | 0.41372917 | -1.341908 | 0.17962583 | 0.99945981 |
| SASH1 | 67.4809935 | 1.208436354 | 0.58975019 | 2.0490648 | 0.04045578 | 0.89562588 |
| UST | 61.4658283 | -0.294163855 | 0.52710927 | -0.55807 | 0.57679661 | 0.99945981 |
| TAB2 | 99.5315679 | 0.002540238 | 0.24769697 | 0.01025542 | 0.9918175 | 0.99945981 |
| ZC3H12D | 13.0233029 | -1.109323919 | 0.9590669 | -1.15667 | 0.24740721 | 0.99945981 |
| PPIL4 | 20.7721774 | -0.144437395 | 0.38552523 | -0.374651 | 0.70792007 | 0.99945981 |
| GINM1 | 13.6687906 | -0.659732338 | 0.37903387 | -1.740563 | 0.0817602 | 0.9846219 |
| KATNA1 | 12.3440098 | -0.454828154 | 0.40654833 | -1.1187554 | 0.26324449 | 0.99945981 |
| LATS1 | 31.3391957 | -0.317238484 | 0.33519899 | -0.9464184 | 0.34393524 | 0.99945981 |
| NUP43 | 12.9567655 | 0.283362702 | 0.36513451 | 0.77605017 | 0.43771936 | 0.99945981 |
| PCMT1 | 36.5378254 | 0.293943626 | 0.26243792 | 1.12005014 | 0.2626924 | 0.99945981 |
| PLEKHG1 | 35.9424111 | -1.306139729 | 0.67932992 | -1.9226884 | 0.0545192 | 0.92445778 |
| MTHFD1L | 38.9858573 | -0.588772493 | 0.59577484 | -0.9882466 | 0.32303187 | 0.99945981 |
| AKAP12 | 11.6703669 | -1.838442146 | 0.66107973 | -2.7809689 | 0.00541969 | 0.5234882 |
| RMND1 | 13.317476 | -0.520655809 | 0.51301339 | -1.0148971 | 0.31015488 | 0.99945981 |
| ESR1 | 31.1089517 | -0.985641714 | 0.54079736 | -1.8225712 | 0.06836836 | 0.9520259 |
| SYNE1 | 130.216463 | 0.441945034 | 0.32195117 | 1.37270826 | 0.16984306 | 0.99945981 |
| OPRM1 | 10.9892542 | 0.231099585 | 0.58750505 | 0.39335762 | 0.69405537 | 0.99945981 |
| IPCEF1 | 49.1181695 | 1.019702031 | 0.33261178 | 3.06574238 | 0.0021713 | 0.45570351 |
| SCAF8 | 91.5407352 | 0.185377099 | 0.24631506 | 0.75260157 | 0.45168938 | 0.99945981 |
| TIAM2 | 41.53556 | 0.152423558 | 0.34345198 | 0.44379875 | 0.65718809 | 0.99945981 |
| TFB1M | 22.0613219 | -0.17657012 | 0.3283808 | -0.5376993 | 0.59078468 | 0.99945981 |
| ARID1B | 300.417401 | -0.184259576 | 0.22237788 | -0.8285877 | 0.40733776 | 0.99945981 |
| ZDHHC14 | 46.063387 | 0.224609833 | 0.3540663 | 0.63437225 | 0.52583792 | 0.99945981 |
| SNX9 | 88.2524926 | -0.450320858 | 0.57543594 | -0.7825734 | 0.43387767 | 0.99945981 |
| SYNJ2 | 16.968342 | -1.964104285 | 0.80021878 | -2.4544591 | 0.01410967 | 0.7227195 |
| TULP4 | 67.7319532 | 0.255170774 | 0.2516433 | 1.01401776 | 0.31057428 | 0.99945981 |
| TMEM181 | 59.9854386 | -0.27706856 | 0.27989874 | -0.9898885 | 0.3222286 | 0.99945981 |
| DYNLT1 | 30.4310859 | -0.489841657 | 0.47356249 | -1.034376 | 0.30096044 | 0.99945981 |

| | | | | | | |
|---|---|---|---|---|---|---|
| SYTL3 | 148.875997 | -1.370278462 | 0.64366293 | -2.1288758 | 0.03326453 | 0.8663391 |
| EZR | 82.7816352 | -0.28539822 | 0.56944751 | -0.5011844 | 0.61624134 | 0.99945981 |
| AL035530.2 | 11.8248447 | 0.964412336 | 0.5816046 | 1.65819241 | 0.09727864 | 0.99945981 |
| TAGAP | 17.1577001 | 0.940363089 | 0.43549604 | 2.15929195 | 0.03082752 | 0.85791231 |
| WTAP | 48.754519 | 0.094530769 | 0.26173677 | 0.36116733 | 0.71797437 | 0.99945981 |
| TCP1 | 12.9482562 | 0.341214775 | 0.39269245 | 0.86891095 | 0.38489584 | 0.99945981 |
| IGF2R | 104.694134 | 0.0437048 | 0.20232729 | 0.21601041 | 0.82897963 | 0.99945981 |
| MAP3K4 | 33.604401 | 0.011144062 | 0.32508373 | 0.03428059 | 0.9726534 | 0.99945981 |
| AGPAT4 | 40.7809614 | -0.404057304 | 0.65584126 | -0.6160901 | 0.53783506 | 0.99945981 |
| PRKN | 72.7912435 | 0.242344284 | 0.46748223 | 0.5184032 | 0.60417698 | 0.99945981 |
| QKI | 282.656087 | 0.519203607 | 0.42771405 | 1.21390356 | 0.22478455 | 0.99945981 |
| SFT2D1 | 22.3447302 | 0.093155213 | 0.33036152 | 0.28197961 | 0.77795914 | 0.99945981 |
| MPC1 | 21.1717573 | 0.063996473 | 0.34900206 | 0.1833699 | 0.85450779 | 0.99945981 |
| RPS6KA2 | 53.6176053 | -0.609068004 | 0.49005256 | -1.2428626 | 0.21391847 | 0.99945981 |
| AL159163.1 | 10.4242872 | 0.205668296 | 0.56701327 | 0.36272219 | 0.71681242 | 0.99945981 |
| Z94721.1 | 17.9126541 | -0.494770836 | 0.45567485 | -1.085798 | 0.27756838 | 0.99945981 |
| FGFR1OP | 18.6954311 | -0.245147501 | 0.43361092 | -0.5653628 | 0.571827 | 0.99945981 |
| AFDN | 21.1280077 | -0.09998895 | 0.33588229 | -0.2976905 | 0.76593943 | 0.99945981 |
| WDR27 | 15.4230481 | -0.119016745 | 0.42756195 | -0.2783614 | 0.78073495 | 0.99945981 |
| PHF10 | 15.6852318 | -0.183327132 | 0.43850102 | -0.4180769 | 0.67589092 | 0.99945981 |
| FAM120B | 63.4791095 | 0.01449552 | 0.42196942 | 0.03435206 | 0.97259641 | 0.99945981 |
| PSMB1 | 25.4624506 | -0.401104505 | 0.46743155 | -0.8581032 | 0.39083548 | 0.99945981 |
| PDCD2 | 8.73151104 | -0.039058111 | 0.4880616 | -0.080027 | 0.93621577 | 0.99945981 |
| FAM20C | 16.7797279 | -0.319122625 | 0.62945928 | -0.506979 | 0.61216957 | 0.99945981 |
| SUN1 | 20.4721164 | -1.533151935 | 0.76912888 | -1.9933615 | 0.04622187 | 0.90884462 |
| COX19 | 14.3108656 | -0.148151645 | 0.39668809 | -0.3734714 | 0.70879765 | 0.99945981 |
| C7orf50 | 13.730529 | -0.405026173 | 0.38582321 | -1.0497714 | 0.29382322 | 0.99945981 |
| INTS1 | 10.5198896 | 0.012778076 | 0.51275671 | 0.02492035 | 0.9801185 | 0.99945981 |
| MAD1L1 | 49.3865923 | -0.059258195 | 0.34751511 | -0.1705198 | 0.86460139 | 0.99945981 |
| SNX8 | 36.6109995 | -1.380379673 | 0.83520014 | -1.6527532 | 0.09838111 | 0.99945981 |
| EIF3B | 21.9599128 | 0.090570322 | 0.42530085 | 0.21295589 | 0.83136136 | 0.99945981 |
| CHST12 | 16.6516004 | -0.431202387 | 0.45345278 | -0.9509312 | 0.34163931 | 0.99945981 |
| LFNG | 16.6003358 | 0.860231928 | 0.54148984 | 1.5886391 | 0.1121419 | 0.99945981 |
| TTYH3 | 34.3947665 | 0.18513867 | 0.35450542 | 0.52224496 | 0.60149979 | 0.99945981 |
| GNA12 | 45.1369463 | 0.166302496 | 0.60300269 | 0.27579064 | 0.78270887 | 0.99945981 |
| CARD11 | 100.534427 | -0.579501665 | 0.36390017 | -1.5924743 | 0.11127817 | 0.99945981 |
| SDK1 | 37.4795697 | -1.068218966 | 0.57303271 | -1.8641501 | 0.06230063 | 0.92949575 |
| FOXK1 | 38.6541482 | -0.448719902 | 0.34611155 | -1.2964604 | 0.19481693 | 0.99945981 |
| AP5Z1 | 9.55238086 | -0.070235172 | 0.42557251 | -0.1650369 | 0.86891493 | 0.99945981 |
| RBAK-RBAKD | 18.7250909 | -1.019727351 | 0.41923109 | -2.4323753 | 0.01500016 | 0.73270878 |
| WIPI2 | 26.3929304 | 0.242596499 | 0.34351136 | 0.70622555 | 0.48004788 | 0.99945981 |
| TNRC18 | 66.6666362 | 0.213739105 | 0.38922095 | 0.54914595 | 0.5829053 | 0.99945981 |
| FBXL18 | 7.04032171 | -0.090007954 | 0.54565874 | -0.1649528 | 0.86898111 | 0.99945981 |
| ACTB | 468.761614 | 0.152818523 | 0.68108818 | 0.22437406 | 0.82246625 | 0.99945981 |
| RNF216 | 64.2009039 | -0.17245033 | 0.23252526 | -0.7416413 | 0.4583047 | 0.99945981 |
| EIF2AK1 | 20.9445843 | 0.133548555 | 0.33853485 | 0.39448983 | 0.69321943 | 0.99945981 |
| USP42 | 19.6105695 | -0.25237609 | 0.34531084 | -0.7308664 | 0.46486077 | 0.99945981 |
| CYTH3 | 21.2420618 | 0.834879747 | 0.39129999 | 2.13360535 | 0.0328751 | 0.8663391 |
| RAC1 | 62.9226408 | -0.020013111 | 0.33556033 | -0.0596409 | 0.95244167 | 0.99945981 |
| DAGLB | 16.1258793 | 0.032790913 | 0.40605429 | 0.080755 | 0.9356368 | 0.99945981 |
| ZNF12 | 10.266526 | -0.346291558 | 0.4324219 | -0.8008187 | 0.4232366 | 0.99945981 |
| CCZ1B | 12.5864569 | 0.140047489 | 0.47071833 | 0.29751866 | 0.76607056 | 0.99945981 |
| C1GALT1 | 46.926596 | 0.154523318 | 0.35600984 | 0.43404226 | 0.66425775 | 0.99945981 |
| MIOS | 24.684081 | -0.067932028 | 0.40213861 | -0.1689269 | 0.86585414 | 0.99945981 |
| RPA3 | 9.9487264 | 0.028612787 | 0.54301714 | 0.05269224 | 0.95797713 | 0.99945981 |
| UMAD1 | 75.4648583 | 0.070750981 | 0.24623618 | 0.28732975 | 0.77385985 | 0.99945981 |
| GLCCI1 | 103.674995 | -0.57606765 | 0.47306877 | -1.217725 | 0.22332851 | 0.99945981 |
| ICA1 | 13.5141585 | -0.297268134 | 0.47927118 | -0.6202504 | 0.53509295 | 0.99945981 |
| NDUFA4 | 27.5014176 | -0.264208634 | 0.42586991 | -0.6203975 | 0.5349961 | 0.99945981 |
| PHF14 | 71.5841917 | 0.256786882 | 0.32762405 | 0.7837852 | 0.43316616 | 0.99945981 |
| THSD7A | 14.6640495 | -0.428026387 | 0.72867463 | -0.587404 | 0.55693241 | 0.99945981 |
| TMEM106B | 17.3224727 | 0.098381894 | 0.38874826 | 0.25307353 | 0.80021139 | 0.99945981 |
| ANKMY2 | 10.8747675 | 0.329088066 | 0.46090674 | 0.71400141 | 0.47522631 | 0.99945981 |
| BZW2 | 11.0276007 | -0.037007007 | 0.59898388 | -0.061783 | 0.95073566 | 0.99945981 |
| SNX13 | 83.2189445 | 0.274645517 | 0.31725503 | 0.86569319 | 0.38665843 | 0.99945981 |
| HDAC9 | 206.97598 | 0.091964454 | 0.85066731 | 0.1081086 | 0.91390954 | 0.99945981 |
| MACC1 | 13.9458433 | -0.196483565 | 0.46575203 | -0.421863 | 0.67312499 | 0.99945981 |
| ITGB8 | 13.1237508 | -1.637047878 | 0.85897016 | -1.9058262 | 0.05667275 | 0.92445778 |
| SP4 | 27.6572779 | 0.126741466 | 0.38615002 | 0.3282182 | 0.74274668 | 0.99945981 |
| STEAP1B | 49.7288494 | -0.339660498 | 0.38608882 | -0.8797471 | 0.37899635 | 0.99945981 |
| TOMM7 | 24.2335045 | 0.082090342 | 0.48892654 | 0.16789913 | 0.86666263 | 0.99945981 |
| FAM126A | 54.3936081 | -1.0914956 | 0.83080623 | -1.3137788 | 0.18892066 | 0.99945981 |
| KLHL7 | 11.3576884 | -0.284636483 | 0.47306144 | -0.6016903 | 0.5473803 | 0.99945981 |
| NUPL2 | 17.4256935 | -0.499484695 | 0.36151959 | -1.3816255 | 0.16708671 | 0.99945981 |
| TRA2A | 74.6660383 | 0.04718637 | 0.26793431 | 0.17611171 | 0.86020618 | 0.99945981 |
| CCDC126 | 13.89594 | 0.27184761 | 0.49769025 | 0.54621847 | 0.58491577 | 0.99945981 |
| MPP6 | 24.3956436 | -2.865511328 | 0.82995218 | -3.4526222 | 0.00055517 | 0.23820088 |
| OSBPL3 | 83.4405151 | 0.149033971 | 0.4187374 | 0.35591273 | 0.72190592 | 0.99945981 |
| CYCS | 15.1147806 | 0.982865359 | 0.46969135 | 2.09257708 | 0.03638692 | 0.87920079 |
| HNRNPA2B1 | 155.534989 | 0.528547932 | 0.19354186 | 2.73092311 | 0.00631572 | 0.52554078 |
| CBX3 | 27.2433447 | -0.30434604 | 0.40590376 | -0.7497985 | 0.45337606 | 0.99945981 |
| SNX10 | 43.8924036 | 0.702255676 | 0.40113532 | 1.75067025 | 0.08000273 | 0.9846219 |
| SKAP2 | 111.8013 | 0.41575617 | 0.32313231 | 1.28664376 | 0.19821848 | 0.99945981 |
| HOTAIRM1 | 8.37172766 | -0.383815348 | 0.62038758 | -0.6186703 | 0.5361336 | 0.99945981 |
| HIBADH | 31.7140122 | -0.165050095 | 0.50601875 | -0.3261739 | 0.74429281 | 0.99945981 |
| TAX1BP1 | 92.2074507 | -0.06599973 | 0.23716245 | -0.2782891 | 0.78079043 | 0.99945981 |
| JAZF1 | 187.068294 | -0.005180702 | 0.47838809 | -0.0108295 | 0.99135948 | 0.99945981 |
| CREB5 | 41.2028354 | 0.141998843 | 0.5455523 | 0.26028456 | 0.79464428 | 0.99945981 |
| CPVL | 82.5661244 | -0.027643296 | 0.64036363 | -0.0431681 | 0.96556752 | 0.99945981 |
| SCRN1 | 10.5859471 | -0.031699918 | 0.49298054 | -0.0643026 | 0.9487293 | 0.99945981 |
| PLEKHA8 | 12.5887614 | -0.025794222 | 0.40670981 | -0.0634217 | 0.94943072 | 0.99945981 |
| MTURN | 10.1545712 | 0.564894584 | 0.53102344 | 1.06378465 | 0.28742626 | 0.99945981 |
| ZNRF2 | 45.6502595 | -0.162673733 | 0.29834746 | -0.5452493 | 0.58558209 | 0.99945981 |
| AC006978.2 | 38.412694 | 0.503288488 | 0.39736976 | 1.26654954 | 0.20531641 | 0.99945981 |
| NOD1 | 13.8562107 | 0.075030685 | 0.53572846 | 0.14005357 | 0.88861766 | 0.99945981 |
| AC005154.6 | 17.2199544 | 0.753283717 | 0.41695697 | 1.80662218 | 0.07082121 | 0.95505283 |
| GARS | 13.3412568 | -0.178865255 | 0.4094107 | -0.4368847 | 0.662195 | 0.99945981 |
| AQP1 | 10.2007199 | -1.216489502 | 0.90726715 | -1.3408283 | 0.1799762 | 0.99945981 |
| AVL9 | 72.7698172 | 0.153342923 | 0.22512355 | 0.68115007 | 0.49577654 | 0.99945981 |
| KBTBD2 | 23.4049688 | -0.154542015 | 0.44257381 | -0.3491892 | 0.72694724 | 0.99945981 |
| NT5C3A | 43.419011 | 0.086314336 | 0.40648857 | 0.21234136 | 0.83184073 | 0.99945981 |
| BBS9 | 44.8895496 | 0.032081504 | 0.4521174 | 0.07095835 | 0.94343091 | 0.99945981 |
| DPY19L1 | 18.4102106 | 0.86412765 | 0.47169931 | 1.83194596 | 0.06695947 | 0.94594189 |
| HERPUD2 | 38.6563104 | -0.111577156 | 0.27591631 | -0.4043877 | 0.68592766 | 0.99945981 |
| 7-Sep | 103.645611 | -0.111681634 | 0.23549879 | -0.4742344 | 0.63533274 | 0.99945981 |
| EEPD1 | 28.9668866 | 1.069737542 | 0.60672789 | 1.76312572 | 0.07787929 | 0.98289661 |
| AOAH | 390.017759 | -0.02588865 | 0.66545396 | -0.0389037 | 0.96896713 | 0.99945981 |
| ELMO1 | 325.231941 | -0.070239467 | 0.23405438 | -0.3000989 | 0.7641017 | 0.99945981 |
| ELMO1-AS1 | 11.1314287 | -0.809208281 | 0.52765841 | -1.5335836 | 0.12513213 | 0.99945981 |
| STARD3NL | 15.587312 | -0.024755463 | 0.44548238 | -0.05557 | 0.95568435 | 0.99945981 |
| TRGC2 | 22.2830989 | 0.461436573 | 0.47653362 | 0.96831903 | 0.33288506 | 0.99945981 |
| TRGC1 | 8.74110884 | -0.110755235 | 0.91637253 | -0.1208627 | 0.90379981 | 0.99945981 |
| TRG-AS1 | 50.7712876 | -0.26508665 | 0.5941153 | -0.4461872 | 0.65546202 | 0.99945981 |

| | | | | | | |
|---|---|---|---|---|---|---|
| VPS41 | 62.2885213 | 0.159127935 | 0.26142881 | 0.60868553 | 0.5427329 | 0.99945981 |
| RALA | 40.4925217 | -0.766909903 | 0.34444645 | -2.2264996 | 0.02598073 | 0.81444982 |
| CDK13 | 125.093416 | 0.115895856 | 0.21645231 | 0.53543368 | 0.59235001 | 0.99945981 |
| STK17A | 66.5703035 | -0.356639405 | 0.3658178 | -0.9749099 | 0.32960495 | 0.99945981 |
| COA1 | 61.2244715 | -0.044854039 | 0.27027772 | -0.1659554 | 0.86819207 | 0.99945981 |
| BLVRA | 15.2893775 | 0.072193469 | 0.44821716 | 0.16106806 | 0.87203979 | 0.99945981 |
| DBNL | 23.2279221 | 0.170027409 | 0.42104447 | 0.40382292 | 0.68634294 | 0.99945981 |
| NUDCD3 | 37.0053743 | -0.547784808 | 0.27413474 | -1.998232 | 0.04569151 | 0.90395454 |
| TMED4 | 11.4541659 | -0.648990017 | 0.42657391 | -1.5214011 | 0.12815921 | 0.99945981 |
| OGDH | 70.5562408 | -0.306381038 | 0.44191888 | -0.693297 | 0.48812318 | 0.99945981 |
| ZMIZ2 | 18.5995743 | 0.72078303 | 0.4284657 | 1.68224208 | 0.09252191 | 0.99640647 |
| PPIA | 64.4570445 | -0.613739081 | 0.48101081 | -1.2759362 | 0.20197809 | 0.99945981 |
| H2AFV | 31.6282265 | -0.062034525 | 0.35520541 | -0.1746441 | 0.8613593 | 0.99945981 |
| PURB | 13.563221 | -0.180984554 | 0.39509646 | -0.4580769 | 0.64689721 | 0.99945981 |
| MYO1G | 31.8751891 | 0.320125523 | 0.5471473 | 0.58508106 | 0.55849321 | 0.99945981 |
| CCM2 | 87.4695615 | -0.498898395 | 0.32261366 | -1.5464267 | 0.12200154 | 0.99945981 |
| TNS3 | 99.8683414 | -0.217962031 | 0.43797346 | -0.4976604 | 0.61872345 | 0.99945981 |
| HUS1 | 16.64676 | 0.349624736 | 0.41310413 | 0.84633561 | 0.39736555 | 0.99945981 |
| UPP1 | 25.4974702 | 0.452241695 | 0.62066809 | 0.72863694 | 0.46622378 | 0.99945981 |
| IKZF1 | 254.883712 | -0.151744797 | 0.35569498 | -0.426615 | 0.66965981 | 0.99945981 |
| GRB10 | 19.2896115 | -1.348401015 | 0.90670582 | -1.4871428 | 0.1369771 | 0.99945981 |
| VOPP1 | 128.697309 | -0.693279041 | 0.46048705 | -1.5055343 | 0.13218677 | 0.99945981 |
| NIPSNAP2 | 21.5732838 | 0.095719802 | 0.3706203 | 0.25826918 | 0.79619917 | 0.99945981 |
| CCT6A | 14.0676028 | -0.304636655 | 0.38566787 | -0.7898938 | 0.42958979 | 0.99945981 |
| SUMF2 | 16.725033 | -0.728159783 | 0.41157516 | -1.7692025 | 0.07686009 | 0.9821652 |
| CHCHD2 | 37.2500718 | -0.180023501 | 0.49996029 | -0.3600756 | 0.7187906 | 0.99945981 |
| ZNF736 | 18.3155209 | -0.094287453 | 0.45121794 | -0.2089621 | 0.83447782 | 0.99945981 |
| ZNF680 | 16.2373929 | -0.455243866 | 0.48622491 | -0.9362825 | 0.34912776 | 0.99945981 |
| ZNF107 | 23.9369803 | 0.2712543 | 0.58648326 | 0.46250988 | 0.64371572 | 0.99945981 |
| ZNF138 | 11.2574203 | -0.298707229 | 0.47516118 | -0.628644 | 0.52958214 | 0.99945981 |
| ZNF117 | 9.08003272 | 0.808716577 | 0.55975748 | 1.44476243 | 0.14852463 | 0.99945981 |
| VKORC1L1 | 27.5791756 | 0.058984067 | 0.31436052 | 0.18763192 | 0.8511652 | 0.99945981 |
| GUSB | 13.5228519 | -0.080574539 | 0.42891906 | -0.1878549 | 0.85099041 | 0.99945981 |
| TPST1 | 38.5390534 | -0.096231304 | 0.63146261 | -0.1523943 | 0.87887595 | 0.99945981 |
| AC027644.4 | 17.2291918 | -0.380823666 | 0.55424469 | -0.6871039 | 0.49201728 | 0.99945981 |
| TMEM248 | 27.1238973 | -0.019407817 | 0.30800715 | -0.0630109 | 0.9497578 | 0.99945981 |
| SBDS | 12.2194539 | -0.340111057 | 0.47702253 | -0.7129874 | 0.47585355 | 0.99945981 |
| TYW1 | 44.7899173 | 0.173411817 | 0.31710246 | 0.54686368 | 0.58447239 | 0.99945981 |
| AUTS2 | 164.023202 | 0.037137341 | 0.38701759 | 0.09595776 | 0.92355412 | 0.99945981 |
| TYW1B | 26.2861653 | 0.343585233 | 0.35685797 | 0.96280666 | 0.33564456 | 0.99945981 |
| POM121 | 32.1112361 | 0.000395303 | 0.32213468 | 0.00122714 | 0.99902089 | 0.9997934 |
| BAZ1B | 41.8143033 | 0.041984937 | 0.28300784 | 0.14835256 | 0.88206454 | 0.99945981 |
| BUD23 | 10.5962812 | 0.150777454 | 0.47034318 | 0.32056903 | 0.74853701 | 0.99945981 |
| LIMK1 | 12.0561862 | -1.153571972 | 0.6512802 | -1.7712376 | 0.0765212 | 0.9821652 |
| EIF4H | 22.4226656 | -0.015997503 | 0.36997516 | -0.0432394 | 0.9655107 | 0.99945981 |
| LAT2 | 27.8926134 | -0.066268468 | 0.50789341 | -0.1304771 | 0.89618896 | 0.99945981 |
| CLIP2 | 18.0573283 | -0.393738889 | 0.63936289 | -0.6158301 | 0.53800668 | 0.99945981 |
| GTF2I | 96.5389669 | -0.121310545 | 0.26857937 | -0.4516748 | 0.65150325 | 0.99945981 |
| NCF1 | 22.3838485 | 0.650512209 | 0.61446151 | 1.05867039 | 0.28974992 | 0.99945981 |
| RCC1L | 9.47760866 | -0.369617148 | 0.48143435 | -0.7677415 | 0.44264076 | 0.99945981 |
| GTF2IRD2B | 8.17786232 | -0.567558503 | 0.58169645 | -0.9756953 | 0.32921548 | 0.99945981 |
| POM121C | 29.0723974 | -0.785345648 | 0.3379866 | -2.3236 | 0.02014694 | 0.78749881 |
| HIP1 | 67.8373982 | 0.291017208 | 0.47035146 | 0.61872287 | 0.53609894 | 0.99945981 |
| POR | 19.3792649 | -0.168715847 | 0.42536988 | -0.3966333 | 0.69163791 | 0.99945981 |
| STYXL1 | 14.387177 | -0.005447762 | 0.51615691 | -0.0105545 | 0.99157891 | 0.99945981 |
| MDH2 | 11.0536869 | -0.818394243 | 0.45107459 | -1.8143213 | 0.06962827 | 0.95505283 |
| HSPB1 | 21.4510932 | -0.041926348 | 0.47074418 | -0.089064 | 0.92903107 | 0.99945981 |
| YWHAG | 26.8258194 | 0.118367104 | 0.33613033 | 0.35214646 | 0.72472843 | 0.99945981 |
| DTX2 | 23.2044814 | 0.107733532 | 0.38780634 | 0.2778024 | 0.78116405 | 0.99945981 |
| CCDC146 | 29.7940289 | -0.139324716 | 0.46234383 | -0.3013444 | 0.7631519 | 0.99945981 |
| FGL2 | 46.5031029 | 1.33082276 | 0.82571155 | 1.6117284 | 0.10702106 | 0.99945981 |
| GSAP | 79.7942678 | 0.339520076 | 0.36504641 | 0.93007373 | 0.35233291 | 0.99945981 |
| PTPN12 | 101.622347 | 0.430762153 | 0.32931472 | 1.30805616 | 0.19085426 | 0.99945981 |
| RSBN1L | 56.0480463 | -0.097759393 | 0.25430091 | -0.3844241 | 0.70066416 | 0.99945981 |
| PHTF2 | 91.8920903 | -0.024830054 | 0.38951171 | -0.0637466 | 0.94917198 | 0.99945981 |
| MAGI2 | 15.2878277 | -0.51565432 | 0.55210842 | -0.9339729 | 0.35031786 | 0.99945981 |
| CD36 | 47.7170259 | 2.981509607 | 1.01776387 | 2.92947088 | 0.0033954 | 0.4856124 |
| DMTF1 | 54.1479049 | 0.028443096 | 0.29712858 | 0.09572656 | 0.92373775 | 0.99945981 |
| TMEM243 | 21.3820686 | -0.196655182 | 0.43274775 | -0.4544337 | 0.64951667 | 0.99945981 |
| ABCB1 | 14.8994895 | -0.410735781 | 0.56900341 | -0.7218512 | 0.47038598 | 0.99945981 |
| SLC25A40 | 18.4441669 | -0.166428941 | 0.38220843 | -0.4354403 | 0.66324289 | 0.99945981 |
| DBF4 | 12.8335899 | -0.101963164 | 0.59446102 | -0.171522 | 0.86381331 | 0.99945981 |
| SRI | 11.9303938 | 0.062448709 | 0.44885302 | 0.13912953 | 0.88934779 | 0.99945981 |
| GTPBP10 | 12.494159 | -0.26493337 | 0.48925659 | -0.5415019 | 0.58816169 | 0.99945981 |
| CDK14 | 175.087786 | -2.397189549 | 0.9880882 | -2.4260886 | 0.01526254 | 0.73270878 |
| MTERF1 | 18.6970339 | 0.170709947 | 0.48067582 | 0.35514569 | 0.72248045 | 0.99945981 |
| AKAP9 | 107.173479 | -0.020694434 | 0.33017135 | -0.0626779 | 0.95002303 | 0.99945981 |
| KRIT1 | 36.778843 | -0.384885268 | 0.32884038 | -1.1704319 | 0.24182721 | 0.99945981 |
| ANKIB1 | 75.9026118 | 0.099323334 | 0.26567812 | 0.37384838 | 0.70851712 | 0.99945981 |
| GATAD1 | 11.1682456 | 0.022299041 | 0.51167496 | 0.04358048 | 0.96523881 | 0.99945981 |
| PEX1 | 9.56433456 | -0.15556262 | 0.45806658 | -0.339607 | 0.7341525 | 0.99945981 |
| FAM133B | 25.4138185 | 0.246783393 | 0.34812968 | 0.70888353 | 0.47839676 | 0.99945981 |
| CDK6 | 84.5828291 | 0.050106955 | 0.52398698 | 0.09562634 | 0.92381735 | 0.99945981 |
| SAMD9 | 24.001693 | 1.206599304 | 0.68543378 | 1.7603441 | 0.07834948 | 0.98326702 |
| SAMD9L | 44.2568486 | 0.539815226 | 0.62117141 | 0.86902781 | 0.38483192 | 0.99945981 |
| VPS50 | 43.8577956 | 0.288476356 | 0.28749274 | 1.00342136 | 0.3156576 | 0.99945981 |
| BET1 | 8.89820149 | 0.776509757 | 0.55534599 | 1.398245 | 0.16203951 | 0.99945981 |
| CASD1 | 30.1108147 | -0.011291738 | 0.39720931 | -0.0284277 | 0.97732105 | 0.99945981 |
| PPP1R9A | 10.5517141 | -0.193734309 | 0.62686843 | -0.309051 | 0.75728274 | 0.99945981 |
| PON2 | 10.0681934 | -0.130392897 | 0.46210214 | -0.2821733 | 0.77781061 | 0.99945981 |
| PDK4 | 65.618899 | 1.992582264 | 1.07523306 | 1.85316313 | 0.06385898 | 0.93776345 |
| SLC25A13 | 63.0902397 | 0.125803144 | 0.34509796 | 0.36454328 | 0.71545236 | 0.99945981 |
| SEM1 | 24.8976679 | -0.438021659 | 0.34206855 | -1.2805084 | 0.20036639 | 0.99945981 |
| LMTK2 | 23.7181219 | -0.023359292 | 0.3479399 | -0.067136 | 0.94647344 | 0.99945981 |
| TECPR1 | 17.3391644 | 0.093141829 | 0.37937877 | 0.24551144 | 0.80606045 | 0.99945981 |
| BRI3 | 55.0629227 | 0.174761347 | 0.40368782 | 0.43291212 | 0.66507862 | 0.99945981 |
| BAIAP2L1 | 90.0513472 | -0.071123106 | 0.64621627 | -0.1100608 | 0.91236112 | 0.99945981 |
| TRRAP | 48.3214633 | 0.060591574 | 0.28211025 | 0.21477977 | 0.82993902 | 0.99945981 |
| SMURF1 | 55.951556 | 0.406691597 | 0.24078394 | 1.68903127 | 0.09121344 | 0.99640647 |
| ARPC1B | 25.817225 | 0.50112989 | 0.60685926 | 0.82577612 | 0.4089311 | 0.99945981 |
| PDAP1 | 10.0116901 | -0.442335357 | 0.50068363 | -0.8834628 | 0.37698628 | 0.99945981 |
| ZNF789 | 13.8548601 | -0.818129388 | 0.50749056 | -1.6121076 | 0.10693853 | 0.99945981 |
| ZNF394 | 14.7450576 | 0.010136594 | 0.39020575 | 0.02597756 | 0.97927523 | 0.99945981 |
| TRIM4 | 16.4630761 | -0.39208912 | 0.40061391 | -0.9787207 | 0.32771801 | 0.99945981 |
| ZKSCAN1 | 33.401985 | -0.676359949 | 0.33534475 | -2.0169093 | 0.04370498 | 0.89562588 |
| COPS6 | 8.88668987 | -0.482888816 | 0.6789526 | -0.7112261 | 0.47694413 | 0.99945981 |
| LAMTOR4 | 30.4052967 | -0.249821194 | 0.40902621 | -0.6107706 | 0.54135144 | 0.99945981 |
| CASTOR3 | 19.9713684 | -1.041954228 | 0.44923312 | -2.3194065 | 0.020373 | 0.78749881 |
| PILRB | 21.2511596 | -0.515889921 | 0.35075547 | -1.4707965 | 0.14134615 | 0.99945981 |
| PILRA | 39.3841503 | 0.692780203 | 0.61397152 | 1.12835886 | 0.25916839 | 0.99945981 |
| TSC22D4 | 19.1204581 | -0.124601999 | 0.36684618 | -0.3396573 | 0.73411459 | 0.99945981 |
| LRCH4 | 9.94320415 | 0.390123825 | 0.43632496 | 0.894113 | 0.37126143 | 0.99945981 |
| GNB2 | 24.1858141 | -0.119656496 | 0.56296128 | -0.2125484 | 0.83167925 | 0.99945981 |

| | | | | | | |
|---|---|---|---|---|---|---|
| GIGYF1 | 12.8790112 | -0.479179175 | 0.4063972 | -1.1790907 | 0.23836205 | 0.99945981 |
| SLC12A9 | 16.7461313 | -0.326149451 | 0.47872803 | -0.6812834 | 0.49569219 | 0.99945981 |
| SRRT | 9.53967272 | -0.388251042 | 0.61108559 | -0.6353464 | 0.5252025 | 0.99945981 |
| TRIM56 | 42.5172885 | 0.058565117 | 0.32514044 | 0.18012252 | 0.85705638 | 0.99945981 |
| PLOD3 | 7.79025075 | -0.191361503 | 0.52193444 | -0.366639 | 0.71388834 | 0.99945981 |
| ZNHIT1 | 10.7334799 | -0.279236469 | 0.50756718 | -0.5501468 | 0.58221867 | 0.99945981 |
| FIS1 | 17.0840023 | -0.575785959 | 0.3956948 | -1.4551264 | 0.14563426 | 0.99945981 |
| CUX1 | 199.712738 | -0.186305545 | 0.35451367 | -0.5255243 | 0.59921877 | 0.99945981 |
| SH2B2 | 11.3955427 | -0.893915484 | 0.77447064 | -1.1542277 | 0.24840682 | 0.99945981 |
| PRKRIP1 | 41.035335 | -0.288388728 | 0.30086256 | -0.9585398 | 0.33779065 | 0.99945981 |
| ORAI2 | 18.0800181 | -1.523252681 | 0.79709641 | -1.9110018 | 0.05600435 | 0.92445778 |
| AC105052.3 | 9.479229 | 0.468277659 | 0.70638933 | 0.66291723 | 0.50738356 | 0.99945981 |
| FAM185A | 15.7392134 | 0.700006949 | 0.50977037 | 1.37318093 | 0.16969611 | 0.99945981 |
| ARMC10 | 10.0378295 | -0.574776911 | 0.500487 | -1.1484352 | 0.25078893 | 0.99945981 |
| NAPEPLD | 11.3517196 | -0.172943218 | 0.41924604 | -0.4125101 | 0.6799656 | 0.99945981 |
| PMPCB | 11.5440275 | -0.455478734 | 0.46165219 | -0.9866275 | 0.32382529 | 0.99945981 |
| DNAJC2 | 15.3535599 | 0.05956913 | 0.36811842 | 0.16182056 | 0.87144716 | 0.99945981 |
| PSMC2 | 10.9186003 | -0.219642708 | 0.45325216 | -0.4845927 | 0.62796526 | 0.99945981 |
| ORC5 | 20.5304108 | 0.372352969 | 0.34807255 | 1.06975676 | 0.28472881 | 0.99945981 |
| AC007384.1 | 18.9161813 | -0.064526776 | 0.70964643 | -0.0909281 | 0.92754975 | 0.99945981 |
| LINC01004 | 38.8324997 | 0.303427457 | 0.45549149 | 0.66615395 | 0.5053127 | 0.99945981 |
| KMT2E | 185.601379 | 0.10651241 | 0.22637018 | 0.47052316 | 0.63798129 | 0.99945981 |
| SRPK2 | 183.60364 | 0.309523639 | 0.18916531 | 1.63626007 | 0.10178517 | 0.99945981 |
| RINT1 | 10.1829985 | -0.346815681 | 0.49007493 | -0.7076789 | 0.47914469 | 0.99945981 |
| ATXN7L1 | 63.2247607 | 0.347921762 | 0.42718423 | 0.81445366 | 0.41538511 | 0.99945981 |
| SYPL1 | 16.6794908 | -0.469722194 | 0.54116179 | -0.8679885 | 0.38540064 | 0.99945981 |
| NAMPT | 282.047856 | 1.139514184 | 0.70403942 | 1.61853748 | 0.10554682 | 0.99945981 |
| PIK3CG | 21.6478616 | 0.280371373 | 0.44412451 | 0.63129003 | 0.52785091 | 0.99945981 |
| HBP1 | 33.6615824 | -0.026941969 | 0.32168742 | -0.083752 | 0.9332536 | 0.99945981 |
| COG5 | 145.037774 | -0.073920538 | 0.28612818 | -0.2583476 | 0.79613863 | 0.99945981 |
| BCAP29 | 16.6364856 | -0.219715122 | 0.38051116 | -0.577421 | 0.56365511 | 0.99945981 |
| SLC26A3 | 104.871665 | 0.916909601 | 0.72255869 | 1.26897596 | 0.20444964 | 0.99945981 |
| DLD | 19.0898078 | 0.143293771 | 0.38484763 | 0.37233897 | 0.70964049 | 0.99945981 |
| PNPLA8 | 33.6005917 | 0.182537886 | 0.34814149 | 0.52432098 | 0.6000553 | 0.99945981 |
| IMMP2L | 178.40668 | 0.318212501 | 0.38300413 | 0.83083309 | 0.40606793 | 0.99945981 |
| DOCK4 | 326.417723 | -0.258426168 | 0.63190036 | -0.4089666 | 0.68256414 | 0.99945981 |
| ZNF277 | 33.0534613 | 0.198501069 | 0.32245784 | 0.61558766 | 0.53816669 | 0.99945981 |
| IFRD1 | 17.6935099 | 0.249125073 | 0.36000487 | 0.69200474 | 0.48893435 | 0.99945981 |
| TMEM168 | 27.5729326 | -0.08946104 | 0.34209239 | -0.2615113 | 0.79369821 | 0.99945981 |
| BMT2 | 35.0927966 | -0.112580864 | 0.35758526 | -0.3148364 | 0.75288585 | 0.99945981 |
| MDFIC | 96.3503119 | -0.227007363 | 0.32151481 | -0.7060557 | 0.48015351 | 0.99945981 |
| TFEC | 69.4689336 | -0.877120561 | 0.67837599 | -1.2929711 | 0.19602105 | 0.99945981 |
| TES | 42.0291838 | -0.00317774 | 0.43893109 | -0.0072397 | 0.99422359 | 0.99945981 |
| CAPZA2 | 62.6094713 | 0.388045521 | 0.317175 | 1.22344297 | 0.22116243 | 0.99945981 |
| ST7 | 32.8365477 | 0.385597467 | 0.48718643 | 0.79147826 | 0.42866496 | 0.99945981 |
| CTTNBP2 | 13.4186515 | -0.009254312 | 0.69017758 | -0.0134086 | 0.98930181 | 0.99945981 |
| LSM8 | 20.0399995 | 0.12575904 | 0.33943986 | 0.37048991 | 0.7110175 | 0.99945981 |
| CPED1 | 27.802782 | 1.412813518 | 0.53011565 | 2.66510433 | 0.00769644 | 0.57727153 |
| FAM3C | 53.5794336 | -2.719654887 | 0.99854066 | -2.7236296 | 0.00645689 | 0.52554078 |
| CADPS2 | 16.5906902 | -0.621635551 | 0.62131711 | -1.0005125 | 0.31706254 | 0.99945981 |
| NDUFA5 | 12.4700024 | 0.203880047 | 0.41743072 | 0.48841649 | 0.62525486 | 0.99945981 |
| WASL | 25.1123195 | 0.514397929 | 0.31223546 | 1.64746798 | 0.09946189 | 0.99945981 |
| POT1 | 16.4887795 | 0.03994477 | 0.4044268 | 0.09876885 | 0.9213218 | 0.99945981 |
| ZNF800 | 45.8361078 | 0.309097342 | 0.35047724 | 0.88193271 | 0.3778132 | 0.99945981 |
| SND1 | 147.249327 | -0.123094727 | 0.22617431 | -0.5442472 | 0.5862714 | 0.99945981 |
| RBM28 | 12.0387622 | 0.044277384 | 0.44266904 | 0.10002367 | 0.92032553 | 0.99945981 |
| IMPDH1 | 10.1335132 | 0.626981648 | 0.51373894 | 1.22042851 | 0.22230248 | 0.99945981 |
| METTL2B | 9.39369729 | 0.187996893 | 0.50559743 | 0.37183119 | 0.71001854 | 0.99945981 |
| CALU | 19.3154507 | -0.485530782 | 0.4247785 | -1.1430211 | 0.25302984 | 0.99945981 |
| ATP6V1F | 18.8861899 | -0.789362398 | 0.40597211 | -1.944376 | 0.05185013 | 0.9230968 |
| TNPO3 | 52.1533239 | 0.168393383 | 0.27504066 | 0.61224905 | 0.54037299 | 0.99945981 |
| TSPAN33 | 10.8047395 | -0.802987811 | 0.85846654 | -0.9353746 | 0.34959527 | 0.99945981 |
| AHCYL2 | 67.9091901 | 0.849017168 | 0.30774679 | 2.7588173 | 0.0058011 | 0.52554078 |
| NRF1 | 54.8054197 | -0.131491264 | 0.24608019 | -0.5343431 | 0.59310415 | 0.99945981 |
| UBE2H | 91.7704253 | 0.095917936 | 0.25842738 | 0.37116011 | 0.71051828 | 0.99945981 |
| KLHDC10 | 35.8746985 | -0.039301875 | 0.34065025 | -0.1153731 | 0.9081494 | 0.99945981 |
| TMEM209 | 9.17253471 | 0.47934188 | 0.43668196 | 1.09769105 | 0.27233942 | 0.99945981 |
| COPG2 | 14.859375 | 0.037680964 | 0.42007414 | 0.08970075 | 0.92852502 | 0.99945981 |
| AC016831.7 | 378.079526 | -0.255012615 | 0.38702668 | -0.6589019 | 0.50995876 | 0.99945981 |
| LINC00513 | 32.5970449 | -1.18309715 | 0.50074767 | -2.3626613 | 0.01814424 | 0.75650017 |
| MKLN1 | 163.491379 | 0.236722055 | 0.21781492 | 1.08680369 | 0.27712358 | 0.99945981 |
| CHCHD3 | 125.308294 | -0.124009634 | 0.32115332 | -0.3861384 | 0.69939416 | 0.99945981 |
| EXOC4 | 227.045469 | 0.068489077 | 0.21259662 | 0.32215507 | 0.74733522 | 0.99945981 |
| SLC35B4 | 10.5714983 | 0.222029864 | 0.50881676 | 0.43636508 | 0.66257187 | 0.99945981 |
| AKR1B1 | 11.0547723 | -0.79241016 | 0.47761264 | -1.6591064 | 0.09709436 | 0.99945981 |
| CALD1 | 10.9628622 | -0.489176026 | 0.54320416 | -0.9005381 | 0.36783397 | 0.99945981 |
| CYREN | 16.1647156 | -0.048578038 | 0.61823442 | -0.0785754 | 0.93737033 | 0.99945981 |
| TMEM140 | 15.5714397 | -0.889999743 | 0.45786634 | -1.9437981 | 0.0519198 | 0.9230968 |
| WDR91 | 11.4322788 | -0.511164153 | 0.48601307 | -1.0517498 | 0.29291435 | 0.99945981 |
| CNOT4 | 83.6433718 | 0.119596403 | 0.22455 | 0.53260479 | 0.59430719 | 0.99945981 |
| NUP205 | 22.5480414 | 0.556993622 | 0.38122112 | 1.46107755 | 0.14399416 | 0.99945981 |
| STMP1 | 17.2714212 | -0.013382744 | 0.48292477 | -0.0277119 | 0.97789196 | 0.99945981 |
| MTPN | 52.3710769 | 0.020928158 | 0.5125799 | 0.04082906 | 0.96743217 | 0.99945981 |
| AC024084.1 | 12.3447615 | 0.974275229 | 0.51162506 | 1.90427582 | 0.05687427 | 0.92445778 |
| AC078845.1 | 12.9757772 | 1.237260157 | 0.82591993 | 1.49803887 | 0.13412315 | 0.99945981 |
| CREB3L2 | 45.2681295 | -0.434149777 | 0.55372926 | -0.784047 | 0.43301252 | 0.99945981 |
| TRIM24 | 27.8626866 | -0.015765947 | 0.49215784 | -0.0320343 | 0.97444467 | 0.99945981 |
| ZC3HAV1 | 114.587447 | 0.316166039 | 0.4397952 | 0.7188938 | 0.47220636 | 0.99945981 |
| UBN2 | 27.9906231 | 0.186962999 | 0.29436979 | 0.63512972 | 0.52534382 | 0.99945981 |
| HIPK2 | 110.337407 | 0.125245629 | 0.35569076 | 0.35211944 | 0.72474869 | 0.99945981 |
| TBXAS1 | 204.6286 | 0.523095382 | 0.59878012 | 0.87360179 | 0.38233515 | 0.99945981 |
| PARP12 | 29.8113161 | 0.674798707 | 0.70051024 | 0.96329599 | 0.335399 | 0.99945981 |
| KDM7A | 55.6191358 | 0.229404268 | 0.3322055 | 0.69054927 | 0.48984884 | 0.99945981 |
| SLC37A3 | 12.0664959 | 0.020326499 | 0.391057 | 0.05197835 | 0.95854594 | 0.99945981 |
| MKRN1 | 21.653761 | 0.32214698 | 0.38195408 | 0.84341808 | 0.39899466 | 0.99945981 |
| NDUFB2 | 19.5771151 | -0.323172928 | 0.45545202 | -0.7095653 | 0.47797377 | 0.99945981 |
| BRAF | 125.024952 | 0.194538642 | 0.22063156 | 0.88173534 | 0.37791995 | 0.99945981 |
| AGK | 19.0784428 | -0.035928882 | 0.41375518 | -0.0868361 | 0.9308018 | 0.99945981 |
| KIAA1147 | 20.3739457 | -0.095190792 | 0.3487082 | -0.2729812 | 0.78486765 | 0.99945981 |
| SSBP1 | 19.7116955 | -0.091820576 | 0.4571742 | -0.2008437 | 0.84082076 | 0.99945981 |
| TRBC1 | 11.568159 | -0.344658157 | 0.82650479 | -0.4170068 | 0.6766734 | 0.99945981 |
| TRBC2 | 16.9543578 | -0.105626875 | 0.58949054 | -0.1791833 | 0.85779376 | 0.99945981 |
| GSTK1 | 24.7627258 | -0.263373045 | 0.41787786 | -0.6302632 | 0.52852239 | 0.99945981 |
| CASP2 | 12.5468939 | -0.440648903 | 0.43387255 | -1.0156183 | 0.30981119 | 0.99945981 |
| ZYX | 20.1409761 | 0.297138427 | 0.4188086 | 0.70948502 | 0.47802354 | 0.99945981 |
| EPHA1-AS1 | 14.7764131 | -0.966860102 | 0.57529818 | -1.6806243 | 0.09283591 | 0.99640647 |
| TCAF2 | 21.9003012 | -1.084009808 | 0.48179599 | -2.2499353 | 0.02445305 | 0.80320673 |
| TCAF1 | 12.0954597 | -0.446897156 | 0.60911258 | -0.7336856 | 0.46314035 | 0.99945981 |
| TPK1 | 67.4788766 | -0.089673313 | 0.25244872 | -0.355214 | 0.72242929 | 0.99945981 |
| CUL1 | 50.2218008 | -0.295787247 | 0.32866228 | -0.8999732 | 0.36813451 | 0.99945981 |
| EZH2 | 23.3812378 | 0.782010633 | 0.62730422 | 1.24662103 | 0.21253649 | 0.99945981 |
| PDIA4 | 13.3137024 | -0.476639051 | 0.62047317 | -0.7681864 | 0.44237645 | 0.99945981 |
| ZNF398 | 12.7216623 | -0.482046849 | 0.41835719 | -1.1522375 | 0.24922348 | 0.99945981 |

| | | | | | | |
|---|---|---|---|---|---|---|
| ZNF862 | 11.9745205 | -0.094387149 | 0.5682746 | -0.1660943 | 0.86808277 | 0.99945981 |
| GIMAP8 | 20.2314485 | 0.6913696 | 0.6682292 | 1.03462943 | 0.30084201 | 0.99945981 |
| GIMAP7 | 34.9977759 | 0.033837809 | 0.51478773 | 0.06573158 | 0.94759153 | 0.99945981 |
| GIMAP4 | 26.4020116 | -0.009004681 | 0.5129985 | -0.017553 | 0.98599542 | 0.99945981 |
| GIMAP2 | 11.2534165 | -0.115238426 | 0.53000394 | -0.2174294 | 0.82787374 | 0.99945981 |
| TMEM176B | 21.5333707 | -0.330879193 | 0.75228115 | -0.4398345 | 0.66005699 | 0.99945981 |
| AGAP3 | 15.474571 | 0.182183103 | 0.43717153 | 0.4167314 | 0.67687489 | 0.99945981 |
| NUB1 | 22.2889058 | 0.675367023 | 0.4826716 | 1.39922678 | 0.16174499 | 0.99945981 |
| RHEB | 36.4949506 | 0.361127987 | 0.27736788 | 1.30198202 | 0.19292253 | 0.99945981 |
| PRKAG2 | 128.534028 | -0.018165506 | 0.49498659 | -0.036699 | 0.97072502 | 0.99945981 |
| GALNT11 | 43.1585204 | 0.250766472 | 0.38170457 | 0.65696482 | 0.51120354 | 0.99945981 |
| KMT2C | 256.761643 | 0.239175368 | 0.1476376 | 1.62001663 | 0.10522871 | 0.99945981 |
| INSIG1 | 18.041322 | 0.755903761 | 0.62658092 | 1.20639449 | 0.22766542 | 0.99945981 |
| RBM33 | 83.8262247 | -0.024243159 | 0.19697268 | -0.1230788 | 0.90204471 | 0.99945981 |
| LMBR1 | 58.0058722 | 0.222984427 | 0.34541586 | 0.64555352 | 0.51856855 | 0.99945981 |
| UBE3C | 81.7386033 | 0.021563852 | 0.22789007 | 0.09462392 | 0.92461355 | 0.99945981 |
| DNAJB6 | 62.3859903 | -0.184870907 | 0.26279046 | -0.7034917 | 0.48174938 | 0.99945981 |
| ESYT2 | 92.4011889 | -0.045911496 | 0.43907007 | -0.1045653 | 0.91672075 | 0.99945981 |
| WDR60 | 17.7606518 | -0.397378398 | 0.51393294 | -0.7732106 | 0.43939775 | 0.99945981 |
| GTPBP6 | 12.8346344 | 0.01471977 | 0.43417375 | 0.03390295 | 0.97295454 | 0.99945981 |
| CSF2RA | 43.4489752 | 0.319145325 | 0.65627416 | 0.48629878 | 0.62675534 | 0.99945981 |
| IL3RA | 14.0697458 | -0.679738358 | 0.84653356 | -0.8029668 | 0.42199392 | 0.99945981 |
| SLC25A6 | 39.8226773 | -0.321689027 | 0.54726818 | -0.5878088 | 0.55666067 | 0.99945981 |
| LINC00106 | 8.26337099 | -0.786160761 | 0.65503549 | -1.2001804 | 0.23006928 | 0.99945981 |
| ASMTL | 10.6400726 | -0.795907249 | 0.42315464 | -1.8808898 | 0.05998691 | 0.92445778 |
| P2RY8 | 63.4247368 | -0.543298071 | 0.30831571 | -1.7621485 | 0.0780442 | 0.98289661 |
| AKAP17A | 16.8044039 | 0.130318427 | 0.38058163 | 0.34241912 | 0.7320355 | 0.99945981 |
| DHRSX | 68.342027 | 0.466523627 | 0.54635093 | 0.85389006 | 0.39316591 | 0.99945981 |
| CD99 | 58.8349555 | -1.038410715 | 0.54662547 | -1.8996749 | 0.05747579 | 0.92445778 |
| PRKX | 56.7821221 | -0.42036123 | 0.35186948 | -1.194651 | 0.23222342 | 0.99945981 |
| PUDP | 10.9482619 | -0.00605619 | 0.41244616 | -0.0146836 | 0.98828461 | 0.99945981 |
| TBL1X | 125.42744 | -0.56508431 | 0.65199774 | -0.8666967 | 0.3861082 | 0.99945981 |
| WWC3 | 22.1288505 | -0.188440728 | 0.45432993 | -0.4147663 | 0.67831302 | 0.99945981 |
| ARHGAP6 | 52.2752315 | 0.553909832 | 0.38523414 | 1.43785239 | 0.15047594 | 0.99945981 |
| MSL3 | 13.3459328 | 0.257032165 | 0.38248776 | 0.67200102 | 0.50158304 | 0.99945981 |
| TMSB4X | 636.948678 | 0.088900777 | 0.6321465 | 0.14063319 | 0.88815973 | 0.99945981 |
| RAB9A | 17.6354864 | 0.033651886 | 0.3963708 | 0.08490001 | 0.93234088 | 0.99945981 |
| OFD1 | 30.9847898 | 0.509231371 | 0.35679708 | 1.42722967 | 0.15351371 | 0.99945981 |
| MOSPD2 | 14.1730312 | 0.455287658 | 0.43772109 | 1.04013189 | 0.29827863 | 0.99945981 |
| CA5B | 30.4898532 | -0.365671011 | 0.51292288 | -0.7129162 | 0.47589764 | 0.99945981 |
| ZRSR2 | 24.326071 | 0.929986059 | 0.42386644 | 2.19405449 | 0.02823149 | 0.84763724 |
| AP1S2 | 32.3622375 | 0.545599374 | 0.42223395 | 1.29217315 | 0.19629719 | 0.99945981 |
| CTPS2 | 13.970722 | -0.611896999 | 0.46837027 | -1.3064386 | 0.19140344 | 0.99945981 |
| SYAP1 | 24.8696257 | -0.480319529 | 0.47102827 | -1.0197255 | 0.30785867 | 0.99945981 |
| TXLNG | 11.7622394 | -0.26926599 | 0.49491944 | -0.5440602 | 0.58640002 | 0.99945981 |
| RBBP7 | 17.7568519 | -0.452809355 | 0.56821817 | -0.7968935 | 0.4255129 | 0.99945981 |
| REPS2 | 16.5926263 | 0.253459417 | 0.55544802 | 0.45631528 | 0.64816328 | 0.99945981 |
| NHS | 18.2418404 | 0.06498337 | 0.35541287 | 0.1828391 | 0.85492427 | 0.99945981 |
| SCML1 | 11.514231 | 0.052801941 | 0.67418655 | 0.07831948 | 0.93757392 | 0.99945981 |
| CDKL5 | 17.8032905 | 0.434647429 | 0.43946637 | 0.98903455 | 0.32264624 | 0.99945981 |
| PHKA2 | 16.5591977 | -0.167469009 | 0.46307167 | -0.3616481 | 0.71761499 | 0.99945981 |
| PDHA1 | 9.82809085 | 0.118149906 | 0.44926145 | 0.26298696 | 0.79256063 | 0.99945981 |
| SH3KBP1 | 172.162502 | 0.431224085 | 0.35484544 | 1.21524483 | 0.22427273 | 0.99945981 |
| BCLAF3 | 18.5674803 | 0.742359787 | 0.45611622 | 1.62756719 | 0.10361669 | 0.99945981 |
| EIF1AX | 26.1093557 | 0.363642437 | 0.37014452 | 0.98243365 | 0.32588625 | 0.99945981 |
| RPS6KA3 | 150.629634 | -0.267521027 | 0.24371807 | -1.0976659 | 0.27235039 | 0.99945981 |
| SMS | 26.1173819 | -0.231961435 | 0.34438623 | -0.6735503 | 0.50059728 | 0.99945981 |
| ACOT9 | 25.2717276 | 0.714191492 | 0.40051501 | 1.78318282 | 0.07455655 | 0.97114097 |
| SAT1 | 300.203218 | 0.992443767 | 0.46493601 | 2.1345814 | 0.03279522 | 0.8663391 |
| APOO | 133.694376 | -0.131881647 | 0.53408759 | -0.2469289 | 0.80496327 | 0.99945981 |
| EIF2S3 | 23.8181843 | 0.049574455 | 0.39773568 | 0.12464171 | 0.90080721 | 0.99945981 |
| ZFX | 39.1912055 | 0.57358787 | 0.40308163 | 1.42300674 | 0.1547342 | 0.99945981 |
| PDK3 | 34.5937793 | -0.118680012 | 0.44755734 | -0.2651727 | 0.79087639 | 0.99945981 |
| POLA1 | 26.1696558 | -0.618222495 | 0.46801158 | -1.3209555 | 0.18651619 | 0.99945981 |
| GK | 67.8341178 | -0.286724636 | 0.62818277 | -0.4564351 | 0.64807717 | 0.99945981 |
| TAB3 | 19.0837087 | 0.326886388 | 0.39538951 | 0.82674522 | 0.40838149 | 0.99945981 |
| DMD | 11.5189014 | -0.089521758 | 0.45394186 | -0.1972097 | 0.8436634 | 0.99945981 |
| RPGR | 9.88201077 | -0.425425062 | 0.55567208 | -0.7656045 | 0.44391164 | 0.99945981 |
| BCOR | 19.2085222 | 0.156056703 | 0.35802723 | 0.43587943 | 0.66292421 | 0.99945981 |
| ATP6AP2 | 27.541671 | -0.279773001 | 0.34018137 | -0.8224231 | 0.41083616 | 0.99945981 |
| CXorf38 | 19.1837701 | 0.350248401 | 0.43607894 | 0.80317661 | 0.42187266 | 0.99945981 |
| MED14 | 30.2793823 | -0.04657555 | 0.29696098 | -0.1568406 | 0.87537044 | 0.99945981 |
| USP9X | 89.9091464 | 0.079068649 | 0.36188188 | 0.21849298 | 0.82704503 | 0.99945981 |
| DDX3X | 55.6639919 | 0.63475313 | 0.3609269 | 1.75867506 | 0.07863272 | 0.98326702 |
| CASK | 100.374276 | 0.085297109 | 0.35377718 | 0.24110405 | 0.80947448 | 0.99945981 |
| KDM6A | 132.236593 | 0.768957609 | 0.37091842 | 2.07311788 | 0.03816131 | 0.87920079 |
| KRBOX4 | 9.32228663 | -0.460269599 | 0.48499858 | -0.9490123 | 0.34261438 | 0.99945981 |
| SLC9A7 | 19.9206833 | -0.116146978 | 0.63820755 | -0.1819894 | 0.85559108 | 0.99945981 |
| RP2 | 23.3937554 | 0.348484793 | 0.47574191 | 0.73250808 | 0.46385851 | 0.99945981 |
| NDUFB11 | 13.4486601 | -0.776024921 | 0.64292215 | -1.2070278 | 0.22742143 | 0.99945981 |
| RBM10 | 14.6188953 | 0.068919811 | 0.382965 | 0.17996373 | 0.85718104 | 0.99945981 |
| UBA1 | 21.0837959 | 0.078531438 | 0.40087794 | 0.19589863 | 0.84468951 | 0.99945981 |
| ZNF41 | 8.18836138 | 0.15206415 | 0.44463581 | 0.34199708 | 0.73235309 | 0.99945981 |
| TIMP1 | 59.665006 | -0.210237959 | 0.74665265 | -0.281574 | 0.7782702 | 0.99945981 |
| ZNF81 | 16.7496965 | -0.317307843 | 0.39319473 | -0.8069992 | 0.41966693 | 0.99945981 |
| RBM3 | 22.5825537 | -0.262127276 | 0.46781441 | -0.5603232 | 0.57525899 | 0.99945981 |
| WAS | 15.1588606 | 0.234413919 | 0.4172531 | 0.5618027 | 0.57425045 | 0.99945981 |
| OTUD5 | 20.070814 | -0.388476476 | 0.36446974 | -1.0658676 | 0.28648352 | 0.99945981 |
| GRIPAP1 | 20.0253209 | -0.031674213 | 0.39712374 | -0.0797591 | 0.93642889 | 0.99945981 |
| TFE3 | 11.6313765 | 0.707862545 | 0.45655079 | 1.5504574 | 0.12103177 | 0.99945981 |
| PLP2 | 11.4684337 | -0.470464768 | 0.64447058 | -0.7300019 | 0.46538901 | 0.99945981 |
| CLCN5 | 25.3327926 | 0.850150628 | 0.45001842 | 1.88914628 | 0.05887223 | 0.92445778 |
| MAGED1 | 33.7334219 | 0.077472628 | 0.30931476 | 0.25046535 | 0.8022275 | 0.99945981 |
| KANTR | 21.3479435 | -0.634969285 | 0.33425808 | -1.8996378 | 0.05748067 | 0.92445778 |
| KDM5C | 16.7274587 | 0.289523964 | 0.53814754 | 0.53800109 | 0.5905763 | 0.99945981 |
| IQSEC2 | 11.0178747 | 1.334345816 | 0.45844681 | 2.91057936 | 0.00360759 | 0.48727327 |
| SMC1A | 21.8387514 | -0.064181035 | 0.34370246 | -0.1867343 | 0.85186897 | 0.99945981 |
| HUWE1 | 72.5978359 | -0.10754958 | 0.27069416 | -0.3973103 | 0.69113865 | 0.99945981 |
| PHF8 | 19.3225459 | 0.367140408 | 0.31894488 | 1.15110929 | 0.24968728 | 0.99945981 |
| FAM120C | 10.7342769 | -0.315022027 | 0.46882863 | -0.6719343 | 0.50162554 | 0.99945981 |
| RRAGB | 12.1141931 | -0.144667016 | 0.39691311 | -0.3644803 | 0.71549937 | 0.99945981 |
| AL445472.1 | 13.6068548 | -0.932771376 | 0.62960564 | -1.4815169 | 0.13846889 | 0.99945981 |
| NBDY | 39.3817352 | 0.183961592 | 0.3067661 | 0.59968032 | 0.54871931 | 0.99945981 |
| FAAH2 | 14.444326 | -0.316934362 | 0.6173344 | -0.5133917 | 0.60767735 | 0.99945981 |
| ARHGEF9 | 17.001384 | 0.072356643 | 0.45904732 | 0.15762349 | 0.87475349 | 0.99945981 |
| MTMR8 | 11.5687981 | -0.321050405 | 0.47305956 | -0.678668 | 0.49734822 | 0.99945981 |
| MSN | 130.908948 | 0.061238781 | 0.30197023 | 0.20279741 | 0.83929338 | 0.99945981 |
| AL034397.3 | 35.3661141 | 0.933225946 | 0.65128434 | 1.43290093 | 0.15188615 | 0.99945981 |
| VSIG4 | 28.0281792 | -0.401645833 | 0.79213168 | -0.5070443 | 0.61212375 | 0.99945981 |
| OPHN1 | 17.4929606 | 0.855011422 | 0.45526533 | 1.87805081 | 0.06037421 | 0.92445778 |
| YIPF6 | 15.6252925 | 0.225525437 | 0.41356738 | 0.54531728 | 0.58553532 | 0.99945981 |
| EDA | 41.4069162 | 1.209774923 | 0.65449 | 1.84842383 | 0.06454105 | 0.93871681 |
| IGBP1 | 15.9909424 | -0.25698545 | 0.3767147 | -0.6821753 | 0.49512813 | 0.99945981 |

| | | | | | | |
|---|---|---|---|---|---|---|
| NONO | 21.5776155 | 0.192550717 | 0.34750045 | 0.55410206 | 0.579509 | 0.99945981 |
| TAF1 | 45.874878 | -0.015823398 | 0.34991313 | -0.0452209 | 0.96393121 | 0.99945981 |
| OGT | 96.842484 | 0.518661484 | 0.31889538 | 1.62643149 | 0.1038579 | 0.99945981 |
| NHSL2 | 16.6160376 | 0.211867981 | 0.45510005 | 0.46554155 | 0.64154369 | 0.99945981 |
| PIN4 | 14.2208781 | -0.502495112 | 0.57102855 | -0.8799825 | 0.3788688 | 0.99945981 |
| RPS4X | 97.6230462 | -0.320300704 | 0.48539781 | -0.6598726 | 0.5093356 | 0.99945981 |
| HDAC8 | 70.5810469 | -0.168292565 | 0.28270174 | -0.5953008 | 0.55164244 | 0.99945981 |
| CHIC1 | 21.3828549 | -0.108891624 | 0.60875945 | -0.1788746 | 0.85803614 | 0.99945981 |
| XIST | 90.1917128 | 4.89585773 | 4.12699295 | 1.18630145 | 0.23550327 | 0.99945981 |
| JPX | 63.0652257 | 0.065973103 | 0.50576502 | 0.1304422 | 0.89621658 | 0.99945981 |
| FTX | 332.137486 | -0.191087767 | 0.26470861 | -0.7218797 | 0.47036845 | 0.99945981 |
| RLIM | 14.8426259 | 0.269618258 | 0.41693766 | 0.64666323 | 0.51784992 | 0.99945981 |
| ABCB7 | 29.1888477 | 0.1001571 | 0.31198041 | 0.3210365 | 0.74818273 | 0.99945981 |
| ATRX | 139.832794 | 0.044702402 | 0.20919605 | 0.21368664 | 0.83079143 | 0.99945981 |
| MAGT1 | 24.1672687 | -0.306015865 | 0.38565986 | -0.7934864 | 0.42749447 | 0.99945981 |
| COX7B | 18.9568894 | -0.411433222 | 0.58992082 | -0.697438 | 0.4855287 | 0.99945981 |
| PGK1 | 57.7090015 | -0.237204578 | 0.37358368 | -0.6349436 | 0.52546518 | 0.99945981 |
| CYSLTR1 | 11.221438 | 0.472878714 | 0.5538729 | 0.85376756 | 0.39323379 | 0.99945981 |
| GPR174 | 32.0595454 | -0.188799531 | 0.41984472 | -0.449689 | 0.65293474 | 0.99945981 |
| BRWD3 | 52.2383951 | 0.101078965 | 0.28265429 | 0.35760633 | 0.72063794 | 0.99945981 |
| SH3BGRL | 92.6087937 | 0.179072344 | 0.2230547 | 0.80281806 | 0.4220799 | 0.99945981 |
| APOOL | 12.4156653 | 0.172242779 | 0.43586151 | 0.39517777 | 0.6927117 | 0.99945981 |
| CHM | 32.5177019 | 0.211330586 | 0.33841053 | 0.62447993 | 0.53231245 | 0.99945981 |
| DIAPH2 | 186.466993 | 0.574691347 | 0.27735809 | 2.07201943 | 0.03826363 | 0.87920079 |
| TRMT2B | 13.4223625 | 0.75664716 | 0.38968066 | 1.94171086 | 0.05217211 | 0.9230968 |
| BTK | 14.9749131 | -0.131500983 | 0.56987164 | -0.2307554 | 0.8175048 | 0.99945981 |
| ARMCX4 | 12.8918635 | 0.115087101 | 0.5230959 | 0.22001148 | 0.82586222 | 0.99945981 |
| ARMCX3 | 12.1275305 | -0.491075909 | 0.42227841 | -1.1629198 | 0.24486206 | 0.99945981 |
| ZMAT1 | 13.2956285 | 0.011906332 | 0.52245464 | 0.02278922 | 0.98181841 | 0.99945981 |
| MORF4L2 | 23.397522 | 0.242211453 | 0.46208478 | 0.52417102 | 0.60015959 | 0.99945981 |
| SLC25A53 | 14.5823135 | -0.185039059 | 0.40263519 | -0.45957 | 0.64582489 | 0.99945981 |
| FAM199X | 13.0632451 | 0.12815977 | 0.3859239 | 0.33208561 | 0.73982462 | 0.99945981 |
| RBM41 | 16.1347864 | -0.144836189 | 0.39325418 | -0.3683017 | 0.71264828 | 0.99945981 |
| TSC22D3 | 54.4923482 | -0.360870821 | 0.57352719 | -0.6292131 | 0.52920956 | 0.99945981 |
| ACSL4 | 83.6072734 | -0.049849978 | 0.21202938 | -0.2351088 | 0.81412427 | 0.99945981 |
| TMEM164 | 42.1400235 | -0.259286122 | 0.67640171 | -0.3833316 | 0.70147394 | 0.99945981 |
| AMMECR1 | 21.7919822 | 0.173328619 | 0.42429933 | 0.40850552 | 0.68290258 | 0.99945981 |
| ALG13 | 52.7355093 | 0.438081391 | 0.35708621 | 1.2268225 | 0.21988931 | 0.99945981 |
| WDR44 | 40.0474504 | 0.084520548 | 0.36445263 | 0.23191093 | 0.8166072 | 0.99945981 |
| DOCK11 | 148.726415 | 0.50711705 | 0.18761831 | 2.70291875 | 0.00687335 | 0.54236298 |
| IL13RA1 | 29.0180797 | -0.159992594 | 0.38366222 | -0.4170142 | 0.67666802 | 0.99945981 |
| SLC25A43 | 21.2470257 | 0.261856839 | 0.56199898 | 0.46593828 | 0.64125968 | 0.99945981 |
| SLC25A5 | 17.9198026 | -0.827271292 | 0.59008876 | -1.4019438 | 0.16093202 | 0.99945981 |
| UBE2A | 12.193701 | 0.183349188 | 0.52485578 | 0.34933252 | 0.72683969 | 0.99945981 |
| 6-Sep | 90.3782773 | -0.167341569 | 0.31549861 | -0.5304035 | 0.5958322 | 0.99945981 |
| UPF3B | 14.4533631 | 0.224546683 | 0.4349437 | 0.51626609 | 0.60566858 | 0.99945981 |
| NDUFA1 | 16.101788 | 0.300142543 | 0.60860065 | 0.49316829 | 0.62189368 | 0.99945981 |
| LAMP2 | 33.5677603 | -0.088360124 | 0.36563334 | -0.2416632 | 0.80904114 | 0.99945981 |
| CUL4B | 13.6502582 | 0.435750224 | 0.43004874 | 1.01325777 | 0.31093705 | 0.99945981 |
| MCTS1 | 7.84133564 | 0.02248097 | 0.47843189 | 0.04698886 | 0.96252211 | 0.99945981 |
| THOC2 | 58.1678716 | 0.358000676 | 0.2586308 | 1.38421517 | 0.16629258 | 0.99945981 |
| XIAP | 40.5661781 | 0.17984764 | 0.28827457 | 0.62387618 | 0.5327089 | 0.99945981 |
| STAG2 | 100.398618 | 0.092090357 | 0.25659648 | 0.35889174 | 0.71967608 | 0.99945981 |
| SASH3 | 30.6266036 | 0.666624067 | 0.43215834 | 1.54254589 | 0.122941 | 0.99945981 |
| ELF4 | 14.5642754 | 0.425605836 | 0.46392878 | 0.91739477 | 0.35893581 | 0.99945981 |
| AIFM1 | 8.05418777 | -0.335297858 | 0.59592804 | -0.5626482 | 0.57367444 | 0.99945981 |
| ENOX2 | 33.8529256 | -0.040757169 | 0.37937718 | -0.1074318 | 0.91444644 | 0.99945981 |
| STK26 | 17.2385933 | -0.426448982 | 0.52693205 | -0.8093055 | 0.41833947 | 0.99945981 |
| RAP2C | 13.5807828 | 0.863263569 | 0.60090635 | 1.4366025 | 0.15083097 | 0.99945981 |
| RAP2C-AS1 | 20.9677352 | 0.040935558 | 0.49009896 | 0.08352509 | 0.93343403 | 0.99945981 |
| MBNL3 | 23.2505687 | 0.169123601 | 0.50959667 | 0.33187737 | 0.73998186 | 0.99945981 |
| PHF6 | 14.615915 | -0.816098074 | 0.58808892 | -1.387712 | 0.16522475 | 0.99945981 |
| HPRT1 | 12.5931275 | -0.504777106 | 0.4888233 | -1.0326372 | 0.30177373 | 0.99945981 |
| FAM122B | 12.3065991 | 0.691444345 | 0.44212497 | 1.56391155 | 0.11783834 | 0.99945981 |
| ZNF75D | 23.6959113 | -0.170357622 | 0.4276489 | -0.3983586 | 0.69036586 | 0.99945981 |
| INTS6L | 29.6554297 | 0.080022689 | 0.41242932 | 0.19402764 | 0.84615424 | 0.99945981 |
| SLC9A6 | 10.5521851 | 0.091417976 | 0.41582588 | 0.21984677 | 0.8259905 | 0.99945981 |
| HTATSF1 | 9.16673061 | 0.268231112 | 0.58512066 | 0.45842017 | 0.6466506 | 0.99945981 |
| ARHGEF6 | 50.0888732 | -0.11715718 | 0.31146822 | -0.3761449 | 0.70680918 | 0.99945981 |
| RBMX | 36.0945077 | -0.409118159 | 0.29608308 | -1.3817681 | 0.16704291 | 0.99945981 |
| ATP11C | 43.8296911 | 0.362962546 | 0.39430068 | 0.92052224 | 0.35729991 | 0.99945981 |
| FMR1 | 20.6788513 | 0.261363219 | 0.39772209 | 0.65715038 | 0.51108423 | 0.99945981 |
| IDS | 18.7092802 | 0.071851506 | 0.41179833 | 0.17448227 | 0.86148647 | 0.99945981 |
| LINC00894 | 21.6839349 | 0.396602634 | 0.39420113 | 1.00609208 | 0.31437128 | 0.99945981 |
| MTM1 | 43.7508865 | 0.399667619 | 0.34841421 | 1.14710482 | 0.2513383 | 0.99945981 |
| MTMR1 | 30.4436179 | -0.302565704 | 0.42747936 | -0.7077902 | 0.47907555 | 0.99945981 |
| CD99L2 | 15.5623704 | -0.733257263 | 0.63863861 | -1.1481568 | 0.25090384 | 0.99945981 |
| BCAP31 | 16.7016286 | 0.24333035 | 0.57821351 | 0.42083132 | 0.67387826 | 0.99945981 |
| SSR4 | 14.8403147 | 0.034425128 | 0.62822684 | 0.05479729 | 0.95629996 | 0.99945981 |
| ARHGAP4 | 20.7441962 | -0.354241845 | 0.4084359 | -0.8673132 | 0.38577043 | 0.99945981 |
| IRAK1 | 10.4252233 | -0.435452142 | 0.57435934 | -0.7581528 | 0.44835951 | 0.99945981 |
| MECP2 | 71.5250922 | 0.263402024 | 0.24831903 | 1.06074039 | 0.2888079 | 0.99945981 |
| FLNA | 71.8600148 | 0.459341587 | 0.50366236 | 0.912003 | 0.36176714 | 0.99945981 |
| EMD | 8.6548973 | 0.558903819 | 0.49177985 | 1.1364919 | 0.25575076 | 0.99945981 |
| RPL10 | 176.600465 | -0.267810769 | 0.56737013 | -0.4720213 | 0.63691161 | 0.99945981 |
| TAZ | 15.6146043 | -0.108785418 | 0.35297126 | -0.3081991 | 0.75793085 | 0.99945981 |
| ATP6AP1 | 16.8877165 | 0.289505945 | 0.41038317 | 0.70545277 | 0.48052851 | 0.99945981 |
| GDI1 | 11.5922076 | 0.317068533 | 0.4328673 | 0.73248438 | 0.46387297 | 0.99945981 |
| GAB3 | 40.7785459 | 0.478486095 | 0.47792313 | 1.00117795 | 0.31674079 | 0.99945981 |
| MPP1 | 16.4992546 | 0.562315937 | 0.48702817 | 1.15458607 | 0.24825997 | 0.99945981 |
| BRCC3 | 12.8922369 | -0.443223084 | 0.44751323 | -0.9904134 | 0.32197212 | 0.99945981 |
| TMLHE-AS1 | 12.3473684 | -0.62941085 | 0.58739446 | -1.0715301 | 0.28393114 | 0.99945981 |
| TMLHE | 33.6111424 | 0.152096505 | 0.29045695 | 0.5236456 | 0.60052506 | 0.99945981 |
| VAMP7 | 12.1397713 | -0.057916569 | 0.41350451 | -0.1400627 | 0.88861043 | 0.99945981 |
| FBXO25 | 10.6824595 | -0.542765778 | 0.50189491 | -1.0814331 | 0.2795045 | 0.99945981 |
| ERICH1 | 58.1415942 | 0.034413249 | 0.32780724 | 0.10498014 | 0.91639157 | 0.99945981 |
| CSMD1 | 17.7926918 | 2.008824649 | 1.11020738 | 1.80941389 | 0.07038673 | 0.95505283 |
| MCPH1 | 61.3804221 | -0.176459614 | 0.26044029 | -0.6775435 | 0.4980612 | 0.99945981 |
| ANGPT2 | 10.4859843 | -0.272615346 | 0.66505123 | -0.4099163 | 0.68186734 | 0.99945981 |
| AGPAT5 | 18.2990311 | -0.14015947 | 0.39275673 | -0.3568608 | 0.72119604 | 0.99945981 |
| DEFB1 | 24.0952247 | -0.668307546 | 0.62031551 | -1.0773671 | 0.28131628 | 0.99945981 |
| FAM85B | 18.6142956 | -2.031201798 | 0.80309283 | -2.5292242 | 0.0114315 | 0.67187199 |
| PRAG1 | 28.5074417 | -1.426462005 | 0.68852179 | -2.0717747 | 0.03828646 | 0.87920079 |
| MFHAS1 | 36.3995638 | 0.040418084 | 0.38475347 | 0.1050493 | 0.91633669 | 0.99945981 |
| ERI1 | 18.1079715 | -0.120700352 | 0.48117193 | -0.2508466 | 0.8019327 | 0.99945981 |
| TNKS | 80.7404117 | 0.038766231 | 0.22821184 | 0.1698695 | 0.86511277 | 0.99945981 |
| MSRA | 55.4201732 | -0.149036346 | 0.43908272 | -0.3394266 | 0.7342884 | 0.99945981 |
| XKR6 | 86.785894 | -0.361307296 | 0.31289392 | -1.1547278 | 0.24820193 | 0.99945981 |
| MTMR9 | 23.0339944 | 0.164144161 | 0.36026644 | 0.4556188 | 0.64866412 | 0.99945981 |
| BLK | 32.700154 | -1.821456275 | 1.11120419 | -1.6391733 | 0.10117717 | 0.99945981 |
| FDFT1 | 22.9284843 | -0.253783405 | 0.34703478 | -0.731291 | 0.46460144 | 0.99945981 |
| CTSB | 195.121873 | -0.335084075 | 0.46989285 | -0.7131074 | 0.47577929 | 0.99945981 |
| LONRF1 | 14.0068463 | -0.360799787 | 0.6397395 | -0.5639792 | 0.57276827 | 0.99945981 |

| | | | | | | |
|---|---|---|---|---|---|---|
| MSR1 | 124.434204 | 0.621962822 | 0.55866502 | 1.11330189 | 0.2655788 | 0.99945981 |
| ZDHHC2 | 26.5197784 | -1.54839024 | 0.58584054 | -2.6430234 | 0.00821694 | 0.58811673 |
| CNOT7 | 11.8408788 | -0.022312219 | 0.40867124 | -0.054597 | 0.95645954 | 0.99945981 |
| VPS37A | 22.7216987 | -0.227562828 | 0.39759922 | -0.5723422 | 0.56709014 | 0.99945981 |
| MTUS1 | 11.0188376 | 0.17327618 | 0.57051122 | 0.3037209 | 0.76134054 | 0.99945981 |
| PCM1 | 121.166311 | -0.046497446 | 0.23753382 | -0.1957508 | 0.84480519 | 0.99945981 |
| ASAH1 | 65.8874034 | 0.440499492 | 0.5037138 | 0.87450353 | 0.3818441 | 0.99945981 |
| PSD3 | 150.788093 | -2.558060941 | 1.10995594 | -2.3046509 | 0.02118613 | 0.79476944 |
| AC100849.1 | 39.6201773 | 0.307790688 | 1.15884359 | 0.26560158 | 0.79054607 | 0.99945981 |
| CSGALNACT1 | 82.6051347 | -0.427417028 | 0.435812 | -0.9807372 | 0.32672237 | 0.99945981 |
| INTS10 | 32.2940995 | -0.041885205 | 0.28981202 | -0.1445254 | 0.88508558 | 0.99945981 |
| ATP6V1B2 | 36.5291077 | 0.53366832 | 0.41312803 | 1.29177466 | 0.19643519 | 0.99945981 |
| DOK2 | 14.5413341 | 0.439581865 | 0.82589655 | 0.53224809 | 0.59455418 | 0.99945981 |
| XPO7 | 32.3486276 | -0.214891459 | 0.29869286 | -0.7194395 | 0.47187014 | 0.99945981 |
| SLC39A14 | 10.9095588 | -0.717959112 | 0.56555255 | -1.2694826 | 0.204269 | 0.99945981 |
| PPP3CC | 86.8634175 | -0.492862114 | 0.51783811 | -0.9517687 | 0.34121429 | 0.99945981 |
| BIN3 | 12.8481535 | 0.08624097 | 0.69509579 | 0.12407062 | 0.90125936 | 0.99945981 |
| TNFRSF10B | 25.7335385 | 1.102732574 | 0.51291234 | 2.14994353 | 0.03155968 | 0.86065765 |
| TNFRSF10D | 8.99597939 | -0.17117647 | 0.52309759 | -0.3272362 | 0.74348924 | 0.99945981 |
| TNFRSF10A | 11.5225679 | 0.482701624 | 0.57713233 | 0.83637945 | 0.40294147 | 0.99945981 |
| CHMP7 | 10.9484456 | 0.334948589 | 0.42750432 | 0.78349755 | 0.43333499 | 0.99945981 |
| ENTPD4 | 27.9972093 | 0.075390556 | 0.33815825 | 0.2229446 | 0.82357862 | 0.99945981 |
| SLC25A37 | 34.9021642 | 0.216669919 | 0.51094831 | 0.42405448 | 0.67152608 | 0.99945981 |
| ADAM28 | 69.0305779 | -0.196653989 | 0.61192559 | -0.3213691 | 0.74793068 | 0.99945981 |
| AC120193.1 | 24.4325809 | -0.692330353 | 0.58588077 | -1.1816915 | 0.23732811 | 0.99945981 |
| DOCK5 | 101.475295 | 0.23475436 | 0.48027194 | 0.48879466 | 0.62498708 | 0.99945981 |
| KCTD9 | 15.1572976 | -0.509777594 | 0.38351016 | -1.3292414 | 0.18376833 | 0.99945981 |
| PPP2R2A | 83.9297025 | 0.275812478 | 0.29998696 | 0.91941488 | 0.35787861 | 0.99945981 |
| BNIP3L | 45.6834957 | -0.751039932 | 0.42823647 | -1.7537972 | 0.07946526 | 0.9846219 |
| DPYSL2 | 25.3970533 | 0.538069591 | 0.44277052 | 1.21523356 | 0.22427702 | 0.99945981 |
| PTK2B | 113.113142 | 0.164214371 | 0.31507178 | 0.5211967 | 0.60222976 | 0.99945981 |
| CLU | 21.6022357 | -1.627194883 | 0.4463679 | -3.645412 | 0.00026696 | 0.17661106 |
| ELP3 | 30.525126 | 0.334144406 | 0.29386536 | 1.13706634 | 0.25551055 | 0.99945981 |
| INTS9 | 23.3436304 | 0.160512778 | 0.30380653 | 0.5283388 | 0.59726421 | 0.99945981 |
| HMBOX1 | 82.9616876 | -0.136718442 | 0.28191006 | -0.4849718 | 0.62769632 | 0.99945981 |
| KIF13B | 76.9429764 | 0.004762452 | 0.27493348 | 0.0173222 | 0.98617958 | 0.99945981 |
| SARAF | 93.1606815 | 0.013874146 | 0.48015532 | 0.02889512 | 0.97694824 | 0.99945981 |
| LEPROTL1 | 48.2195919 | -0.308965239 | 0.48886086 | -0.6320106 | 0.52737996 | 0.99945981 |
| RBPMS | 21.0154879 | -0.155536764 | 0.49701429 | -0.3129422 | 0.75432455 | 0.99945981 |
| GTF2E2 | 24.0086654 | -0.490053462 | 0.38271 | -1.2804825 | 0.2003755 | 0.99945981 |
| GSR | 19.4258717 | 0.179530599 | 0.35796921 | 0.50152525 | 0.61600151 | 0.99945981 |
| PPP2CB | 22.1803089 | -0.21238975 | 0.44757566 | -0.4745337 | 0.63511935 | 0.99945981 |
| WRN | 30.5783326 | -0.110156453 | 0.28489503 | -0.3866563 | 0.69901069 | 0.99945981 |
| NRG1 | 28.2011549 | -1.545763627 | 0.66388177 | -2.3283719 | 0.01989237 | 0.78749881 |
| FUT10 | 9.54177108 | -0.048263554 | 0.5266075 | -0.09165 | 0.92697616 | 0.99945981 |
| RAB11FIP1 | 39.0895138 | -0.512676239 | 0.47609829 | -1.0768286 | 0.28155686 | 0.99945981 |
| ASH2L | 15.5053551 | 0.263950618 | 0.39001368 | 0.67677271 | 0.49855018 | 0.99945981 |
| BAG4 | 11.1895829 | 0.034390273 | 0.44944638 | 0.07651697 | 0.93900782 | 0.99945981 |
| DDHD2 | 11.9405345 | -0.465862313 | 0.44993839 | -1.0353913 | 0.30048619 | 0.99945981 |
| NSD3 | 82.7055927 | 0.006828398 | 0.22204075 | 0.0307529 | 0.9754666 | 0.99945981 |
| TACC1 | 86.4883776 | -0.056373934 | 0.37538526 | -0.1501762 | 0.8806256 | 0.99945981 |
| PLEKHA2 | 108.725474 | -0.750817463 | 0.37610013 | -1.9963233 | 0.04589874 | 0.90544495 |
| ADAM9 | 47.3245467 | 0.632474969 | 0.55022032 | 1.14949402 | 0.25035233 | 0.99945981 |
| GOLGA7 | 10.443836 | -0.101194079 | 0.62821846 | -0.161081 | 0.87202958 | 0.99945981 |
| GPAT4 | 21.9885347 | -0.192255302 | 0.4461626 | -0.4309086 | 0.66653483 | 0.99945981 |
| KAT6A | 143.967165 | -0.006128943 | 0.23660149 | -0.0259041 | 0.97933385 | 0.99945981 |
| AP3M2 | 9.5855388 | -0.522089942 | 0.48685176 | -1.0723797 | 0.28354953 | 0.99945981 |
| IKBKB | 52.7754957 | -0.239779025 | 0.25755776 | -0.9309719 | 0.35186812 | 0.99945981 |
| POLB | 13.0569286 | -0.043419021 | 0.39310799 | -0.1104506 | 0.91205201 | 0.99945981 |
| VDAC3 | 20.8735319 | -0.219409777 | 0.34118162 | -0.6430879 | 0.52016705 | 0.99945981 |
| SLC20A2 | 36.7707356 | -0.193381243 | 0.43652392 | -0.4430026 | 0.65776384 | 0.99945981 |
| RNF170 | 14.2722395 | -0.047347505 | 0.41063377 | -0.1153035 | 0.90820458 | 0.99945981 |
| HOOK3 | 49.8538386 | -0.146939226 | 0.28027641 | -0.5242654 | 0.60009395 | 0.99945981 |
| FNTA | 10.7359945 | -0.273723908 | 0.45351079 | -0.6035665 | 0.54613191 | 0.99945981 |
| HGSNAT | 32.8137123 | -0.386478763 | 0.42752102 | -0.9039994 | 0.3659957 | 0.99945981 |
| SPIDR | 235.703833 | -0.169522518 | 0.3146566 | -0.538754 | 0.59005658 | 0.99945981 |
| CEBPD | 55.0980663 | -0.208161611 | 0.53437197 | -0.3895444 | 0.69687347 | 0.99945981 |
| PRKDC | 63.8063299 | 0.138883157 | 0.25970513 | 0.53477248 | 0.5928072 | 0.99945981 |
| UBE2V2 | 19.4597049 | 0.09473478 | 0.35288262 | 0.26845975 | 0.78834545 | 0.99945981 |
| PCMTD1 | 68.0650913 | -0.034786651 | 0.26927772 | -0.129185 | 0.89721124 | 0.99945981 |
| ST18 | 22.8418209 | 0.297422844 | 0.823736 | 0.36106574 | 0.71805031 | 0.99945981 |
| RB1CC1 | 66.0150994 | 0.253761406 | 0.26324298 | 0.96398165 | 0.33505512 | 0.99945981 |
| ATP6V1H | 50.1146827 | 0.075879832 | 0.32786486 | 0.2314363 | 0.81697586 | 0.99945981 |
| TCEA1 | 21.8932967 | -0.201317512 | 0.33454497 | -0.6017652 | 0.54733046 | 0.99945981 |
| LYPLA1 | 38.9100304 | 0.037844024 | 0.29084971 | 0.13011539 | 0.89647513 | 0.99945981 |
| TMEM68 | 8.81738787 | -0.556563271 | 0.50753131 | -1.0966088 | 0.27281246 | 0.99945981 |
| TGS1 | 16.2994681 | -0.083894039 | 0.39279479 | -0.2135824 | 0.83087276 | 0.99945981 |
| LYN | 278.687201 | 0.950928523 | 0.40928521 | 2.32338846 | 0.02015829 | 0.78749881 |
| RPS20 | 45.5694883 | 0.390946377 | 0.41531253 | 0.94133057 | 0.34653548 | 0.99945981 |
| IMPAD1 | 14.8906578 | 0.307929795 | 0.38139037 | 0.80738743 | 0.41944331 | 0.99945981 |
| FAM110B | 22.0900044 | -0.836690653 | 0.94183395 | -0.8883632 | 0.37434539 | 0.99945981 |
| UBXN2B | 28.3942903 | 0.117299578 | 0.4410121 | 0.26597814 | 0.79025604 | 0.99945981 |
| SDCBP | 74.0355038 | 0.503107218 | 0.31234948 | 1.61071895 | 0.10724099 | 0.99945981 |
| NSMAF | 40.7190644 | -0.229489516 | 0.55731003 | -0.4117807 | 0.68050017 | 0.99945981 |
| TOX | 183.944743 | -1.659092484 | 0.55879403 | -2.9690591 | 0.00298713 | 0.45570351 |
| RAB2A | 105.975689 | 0.185370404 | 0.21116243 | 0.87785693 | 0.38002137 | 0.99945981 |
| CHD7 | 41.8551585 | -0.223329577 | 0.29058249 | -0.7685583 | 0.44215558 | 0.99945981 |
| ASPH | 63.6107251 | -0.501755633 | 0.38878265 | -1.2905813 | 0.19684889 | 0.99945981 |
| YTHDF3 | 44.3899909 | 0.255216844 | 0.3118689 | 0.81834657 | 0.41315932 | 0.99945981 |
| ARMC1 | 12.9647793 | 0.137246386 | 0.44473131 | 0.30860518 | 0.75762188 | 0.99945981 |
| MTFR1 | 27.3718378 | -0.621486972 | 0.44403166 | -1.3996456 | 0.16161946 | 0.99945981 |
| PDE7A | 179.228621 | -0.321107262 | 0.32965781 | -0.9740623 | 0.33002559 | 0.99945981 |
| MYBL1 | 45.1912715 | -0.116409909 | 0.66772034 | -0.1743393 | 0.8615988 | 0.99945981 |
| VCPIP1 | 20.4263162 | 0.197196774 | 0.33842558 | 0.58268873 | 0.56010286 | 0.99945981 |
| SGK3 | 56.024821 | 0.379699302 | 0.47214008 | 0.80420901 | 0.42127628 | 0.99945981 |
| CSPP1 | 31.9246956 | 0.68513122 | 0.39752496 | 1.72349234 | 0.08479953 | 0.99193047 |
| ARFGEF1 | 118.025396 | 0.102889502 | 0.23426696 | 0.43919767 | 0.66051832 | 0.99945981 |
| NCOA2 | 220.308738 | 0.484038876 | 0.22207868 | 2.17958286 | 0.0292884 | 0.84837487 |
| TRAM1 | 35.8659646 | -0.147809784 | 0.42263509 | -0.3497338 | 0.72653847 | 0.99945981 |
| LACTB2 | 8.82996205 | -0.333392949 | 0.58306653 | -0.5717923 | 0.56746269 | 0.99945981 |
| MSC-AS1 | 14.9013682 | -1.76726106 | 0.58776286 | -3.0067587 | 0.00264049 | 0.45570351 |
| TERF1 | 18.381824 | -0.029160612 | 0.38957825 | -0.0748517 | 0.94033267 | 0.99945981 |
| RPL7 | 65.9913937 | -0.020119158 | 0.60560923 | -0.0332214 | 0.97349807 | 0.99945981 |
| STAU2 | 54.8158962 | -0.918584603 | 0.52907632 | -1.7362043 | 0.08252771 | 0.98529784 |
| ELOC | 15.4381727 | -0.08963813 | 0.3823765 | -0.2344237 | 0.81465603 | 0.99945981 |
| LY96 | 21.5913039 | 0.466123919 | 0.49453207 | 0.9425555 | 0.34590831 | 0.99945981 |
| GDAP1 | 12.9739059 | 0.06098874 | 0.55330034 | 0.11022719 | 0.9122292 | 0.99945981 |
| PEX2 | 11.8663259 | -0.188407685 | 0.43455804 | -0.4335616 | 0.66460681 | 0.99945981 |
| ZC2HC1A | 9.57874015 | 0.671074998 | 0.48556666 | 1.38204506 | 0.16695787 | 0.99945981 |
| IL7 | 14.2546305 | -1.14094355 | 0.54616663 | -2.0890027 | 0.03670748 | 0.87920079 |
| TPD52 | 44.1120645 | -0.405269171 | 0.4077005 | -0.9940365 | 0.32020511 | 0.99945981 |
| ZNF704 | 9.12170005 | -0.47730184 | 0.6398613 | -0.7459458 | 0.45570017 | 0.99945981 |
| PAG1 | 183.745015 | 0.806550422 | 0.58088566 | 1.38848395 | 0.16498973 | 0.99945981 |

| | | | | | | |
|---|---|---|---|---|---|---|
| IMPA1 | 9.82358786 | 0.163956914 | 0.4690426 | 0.34955655 | 0.72667152 | 0.99945981 |
| ZFAND1 | 9.67475706 | -0.189534159 | 0.54081112 | -0.3504628 | 0.72599143 | 0.99945981 |
| C8orf59 | 10.9391744 | -0.137423443 | 0.49803362 | -0.2759321 | 0.78260025 | 0.99945981 |
| WWP1 | 89.7844187 | 0.328677459 | 0.33869819 | 0.97041399 | 0.33184018 | 0.99945981 |
| RMDN1 | 20.3882044 | -0.26073818 | 0.37344608 | -0.698195 | 0.48505527 | 0.99945981 |
| CPNE3 | 17.1441488 | -0.131168189 | 0.37866405 | -0.3463973 | 0.72904418 | 0.99945981 |
| AF117829.1 | 18.9997648 | -0.660401302 | 0.57356867 | -1.1513901 | 0.24957177 | 0.99945981 |
| RIPK2 | 22.3011829 | 1.067749841 | 0.63048958 | 1.69352496 | 0.09035559 | 0.99640647 |
| NBN | 20.8316204 | -0.021560237 | 0.32367008 | -0.0666118 | 0.94689078 | 0.99945981 |
| DECR1 | 46.73362 | 0.425779658 | 0.29499179 | 1.44336104 | 0.14891879 | 0.99945981 |
| PIP4P2 | 14.2273292 | 0.97964288 | 0.4932492 | 1.9861013 | 0.04702208 | 0.90884462 |
| LRRC69 | 13.1376627 | 0.242881513 | 0.5011799 | 0.48461942 | 0.62794634 | 0.99945981 |
| TRIQK | 12.7435403 | -0.348205299 | 0.59376629 | -0.5864349 | 0.55758327 | 0.99945981 |
| RBM12B | 15.0913361 | 0.345191113 | 0.46924185 | 0.73563582 | 0.46195235 | 0.99945981 |
| VIRMA | 22.5519287 | -0.05781272 | 0.41189365 | -0.1403584 | 0.88837685 | 0.99945981 |
| DPY19L4 | 15.0654697 | -0.164839001 | 0.49502703 | -0.3329899 | 0.73914191 | 0.99945981 |
| INTS8 | 23.3539841 | -0.11324728 | 0.37977858 | -0.2981929 | 0.76555593 | 0.99945981 |
| NDUFAF6 | 29.2863626 | -0.203982165 | 0.53451061 | -0.3816242 | 0.70274016 | 0.99945981 |
| TP53INP1 | 28.7326467 | -0.521049188 | 0.65992269 | -0.789561 | 0.42978422 | 0.99945981 |
| PLEKHF2 | 11.2350102 | 0.107933759 | 0.44461888 | 0.24275568 | 0.80819467 | 0.99945981 |
| UQCRB | 33.3647757 | 0.259739486 | 0.51607982 | 0.50329324 | 0.61475812 | 0.99945981 |
| PTDSS1 | 17.9358866 | 0.018331209 | 0.38068479 | 0.04815325 | 0.96159411 | 0.99945981 |
| SDC2 | 25.597924 | -0.505417532 | 0.68422903 | -0.7386672 | 0.46010912 | 0.99945981 |
| CPQ | 103.222608 | 0.251092589 | 0.38392759 | 0.65401027 | 0.51310519 | 0.99945981 |
| MTDH | 102.383769 | -0.047138865 | 0.20910324 | -0.2254335 | 0.82164209 | 0.99945981 |
| RPL30 | 99.4904182 | 0.070418544 | 0.60304976 | 0.1167707 | 0.90704176 | 0.99945981 |
| NIPAL2 | 14.5660541 | -0.052209774 | 0.67748689 | -0.0770639 | 0.93857272 | 0.99945981 |
| STK3 | 82.3878421 | 0.16667958 | 0.30418627 | 0.54795235 | 0.58372462 | 0.99945981 |
| AC104986.2 | 9.75967033 | 0.069642996 | 0.48778465 | 0.14277406 | 0.88646862 | 0.99945981 |
| VPS13B | 224.877123 | 0.129275594 | 0.17885346 | 0.72280176 | 0.46980169 | 0.99945981 |
| COX6C | 27.0051264 | 0.297669092 | 0.40594484 | 0.73327473 | 0.46339089 | 0.99945981 |
| RNF19A | 89.5454717 | -0.213367403 | 0.30926886 | -0.6899091 | 0.49025134 | 0.99945981 |
| ANKRD46 | 9.62428186 | 0.239251926 | 0.51271649 | 0.46663591 | 0.64076039 | 0.99945981 |
| PABPC1 | 144.219305 | 0.056050503 | 0.40436416 | 0.13861392 | 0.88975524 | 0.99945981 |
| YWHAZ | 110.68789 | 0.063963476 | 0.30315716 | 0.21099115 | 0.83289418 | 0.99945981 |
| ZNF706 | 36.2105234 | 0.291045467 | 0.42283764 | 0.68831495 | 0.49125447 | 0.99945981 |
| NCALD | 75.0804825 | 0.338832914 | 0.40830427 | 0.82985395 | 0.40662136 | 0.99945981 |
| RRM2B | 11.6399964 | -0.052005441 | 0.51369277 | -0.1012384 | 0.9193612 | 0.99945981 |
| UBR5 | 134.538895 | 0.294943285 | 0.25771871 | 1.14443876 | 0.25244172 | 0.99945981 |
| KLF10 | 11.9175254 | 1.059419893 | 0.57068924 | 1.85638665 | 0.06339847 | 0.93775336 |
| AZIN1 | 28.4071238 | 0.350232911 | 0.35241559 | 0.99380653 | 0.32031707 | 0.99945981 |
| ATP6V1C1 | 33.7965507 | 0.201118818 | 0.41920309 | 0.47976463 | 0.63139476 | 0.99945981 |
| SLC25A32 | 12.7094006 | 0.020563185 | 0.41184925 | 0.04992891 | 0.96017904 | 0.99945981 |
| OXR1 | 69.2427127 | 0.759687372 | 0.25492005 | 2.9801005 | 0.00288154 | 0.45570351 |
| EIF3E | 38.3968556 | 0.142803218 | 0.3965605 | 0.3601045 | 0.71876899 | 0.99945981 |
| EMC2 | 14.7441528 | -0.257737158 | 0.43411194 | -0.5937113 | 0.55270523 | 0.99945981 |
| NUDCD1 | 12.6962655 | -0.470508272 | 0.41545084 | -1.1325245 | 0.25741397 | 0.99945981 |
| ENY2 | 19.6694131 | 0.43274922 | 0.37624361 | 1.15018356 | 0.25006827 | 0.99945981 |
| TRPS1 | 297.281327 | 0.605997369 | 0.47011412 | 1.28904312 | 0.1973831 | 0.99945981 |
| EIF3H | 58.5318123 | 0.092332679 | 0.34289508 | 0.26927386 | 0.78771895 | 0.99945981 |
| RAD21 | 39.7884391 | -0.018739709 | 0.35537694 | -0.0527319 | 0.9579455 | 0.99945981 |
| EXT1 | 174.996923 | 0.723153186 | 0.72633117 | 0.99562461 | 0.31943257 | 0.99945981 |
| SAMD12 | 29.9650903 | 0.726670696 | 0.62001191 | 1.172027 | 0.24118622 | 0.99945981 |
| ENPP2 | 12.0586306 | -0.060190961 | 0.66012234 | -0.0911815 | 0.92734836 | 0.99945981 |
| TAF2 | 39.1104146 | 0.125932092 | 0.33558637 | 0.37525985 | 0.70746722 | 0.99945981 |
| DEPTOR | 11.8694732 | 1.0066393 | 0.58483153 | 1.72124664 | 0.08520608 | 0.99193047 |
| SNTB1 | 87.6502987 | -0.197330252 | 0.5746258 | -0.3434065 | 0.73129265 | 0.99945981 |
| ZHX2 | 116.546017 | -1.02334845 | 0.47057599 | -2.174672 | 0.0296547 | 0.85472619 |
| DERL1 | 19.5542923 | -0.084811233 | 0.45473885 | -0.1865054 | 0.85204848 | 0.99945981 |
| TBC1D31 | 16.7975161 | 0.253769431 | 0.50037148 | 0.50716206 | 0.61204111 | 0.99945981 |
| ATAD2 | 20.7791225 | 0.277484584 | 0.54081794 | 0.51308318 | 0.60789314 | 0.99945981 |
| FBXO32 | 10.090127 | -1.774592 | 0.76558923 | -2.3179427 | 0.02045243 | 0.78749881 |
| FAM91A1 | 42.1095697 | 0.584748133 | 0.27638693 | 2.11568662 | 0.03437147 | 0.87717522 |
| TMEM65 | 55.3802207 | -0.330256089 | 0.34495084 | -0.9574005 | 0.33836517 | 0.99945981 |
| TATDN1 | 14.8156929 | -0.171391875 | 0.46244092 | -0.3706244 | 0.71091733 | 0.99945981 |
| NDUFB9 | 18.791396 | 0.548251078 | 0.5803809 | 0.94464011 | 0.34484264 | 0.99945981 |
| MTSS1 | 313.609248 | -0.536524115 | 0.85464991 | -0.6277706 | 0.53015421 | 0.99945981 |
| WASHC5 | 19.3003722 | -0.168235782 | 0.39108489 | -0.4301772 | 0.66706678 | 0.99945981 |
| NSMCE2 | 131.869931 | 0.060214037 | 0.19565265 | 0.30775988 | 0.75826505 | 0.99945981 |
| LINC00861 | 29.3083804 | 0.3042138 | 0.66130965 | 0.46001718 | 0.64550389 | 0.99945981 |
| PCAT1 | 44.2638582 | 0.656932763 | 0.53388725 | 1.23047097 | 0.21852079 | 0.99945981 |
| PVT1 | 65.1951124 | -0.276233195 | 0.50599634 | -0.5459194 | 0.58512137 | 0.99945981 |
| CCDC26 | 30.9560183 | 0.81588498 | 0.69438191 | 1.17498018 | 0.24000265 | 0.99945981 |
| FAM49B | 350.328357 | 0.388192395 | 0.2971783 | 1.30626089 | 0.19146385 | 0.99945981 |
| ASAP1 | 205.764546 | 0.327853773 | 0.32514852 | 1.00832004 | 0.31330084 | 0.99945981 |
| EFR3A | 77.058748 | 0.082978982 | 0.31956463 | 0.2596626 | 0.79512404 | 0.99945981 |
| KCNQ3 | 28.9637163 | 0.364884861 | 1.08850449 | 0.33521668 | 0.73746164 | 0.99945981 |
| TMEM71 | 35.7531695 | -0.317233935 | 0.4433612 | -0.7155203 | 0.47428761 | 0.99945981 |
| PHF20L1 | 59.2389612 | 0.080326635 | 0.41836318 | 0.19200216 | 0.84774051 | 0.99945981 |
| TG | 31.1826151 | -0.452862209 | 0.5379894 | -0.8417679 | 0.39991788 | 0.99945981 |
| SLA | 104.510473 | -0.436909019 | 0.35277748 | -1.2384833 | 0.21553691 | 0.99945981 |
| NDRG1 | 28.1627715 | -0.453108494 | 0.55826236 | -0.8116408 | 0.41699779 | 0.99945981 |
| ST3GAL1 | 106.225114 | -0.394065919 | 0.35189816 | -1.1198294 | 0.26278646 | 0.99945981 |
| ZFAT | 22.8646037 | 0.024251064 | 0.40151836 | 0.06039839 | 0.95183834 | 0.99945981 |
| TRAPPC9 | 140.20633 | -0.535928602 | 0.21103834 | -2.5394845 | 0.0111016 | 0.67187199 |
| AGO2 | 54.2005202 | -0.431070107 | 0.31689456 | -1.3602951 | 0.17373657 | 0.99945981 |
| PTK2 | 62.6868592 | -0.7183216 | 0.69806828 | -1.0290134 | 0.30347339 | 0.99945981 |
| DENND3 | 51.825641 | -0.518515397 | 0.52546542 | -0.9867736 | 0.32375365 | 0.99945981 |
| SLC45A4 | 18.0709422 | -0.395222411 | 0.44884882 | -0.8805246 | 0.37857521 | 0.99945981 |
| LY6E | 33.9432352 | 0.047725623 | 0.6628961 | 0.07199563 | 0.94260538 | 0.99945981 |
| ZC3H3 | 16.7940292 | -0.503936023 | 0.44875678 | -1.1229602 | 0.26145438 | 0.99945981 |
| GSDMD | 13.6933994 | -0.221095199 | 0.44569628 | -0.496067 | 0.61984717 | 0.99945981 |
| EEF1D | 64.8522624 | -0.269434526 | 0.3630754 | -0.7420897 | 0.45803297 | 0.99945981 |
| PUF60 | 9.07045072 | -0.425814579 | 0.52323386 | -0.8138131 | 0.41575202 | 0.99945981 |
| PLEC | 36.7578561 | -0.067969059 | 0.54797728 | -0.1240363 | 0.90128656 | 0.99945981 |
| PARP10 | 12.4905332 | 0.725753986 | 0.51606697 | 1.40631745 | 0.15962988 | 0.99945981 |
| GRINA | 20.5492088 | 0.53055804 | 0.68540169 | 0.77408335 | 0.4388815 | 0.99945981 |
| CYC1 | 8.32429709 | -0.249991211 | 0.49164093 | -0.5084833 | 0.61111444 | 0.99945981 |
| MAF1 | 10.3041611 | -1.096244111 | 0.65083026 | -1.6843779 | 0.09210866 | 0.99640647 |
| MROH1 | 23.9118136 | 0.063590359 | 0.34660957 | 0.18346394 | 0.85443401 | 0.99945981 |
| HSF1 | 16.8570925 | -0.0868457 | 0.38860882 | -0.2234785 | 0.82316315 | 0.99945981 |
| VPS28 | 18.4966376 | 0.062954424 | 0.36675334 | 0.17165331 | 0.8637101 | 0.99945981 |
| CYHR1 | 8.76099707 | -0.197594764 | 0.45630164 | -0.4330354 | 0.66498905 | 0.99945981 |
| ZNF251 | 13.154837 | -0.273507187 | 0.44051926 | -0.6208745 | 0.53468218 | 0.99945981 |
| RPL8 | 96.1107168 | -0.318484262 | 0.5492392 | -0.5798644 | 0.56200606 | 0.99945981 |
| WASHC1 | 29.7031283 | 0.077894587 | 0.37807268 | 0.20603072 | 0.83676691 | 0.99945981 |
| DOCK8 | 345.114105 | 0.404526296 | 0.24891687 | 1.62514616 | 0.10413142 | 0.99945981 |
| KANK1 | 21.8373522 | -0.369911146 | 0.58425976 | -0.6331279 | 0.52665013 | 0.99945981 |
| SMARCA2 | 104.529069 | -0.306298808 | 0.2996598 | -1.0221551 | 0.30670748 | 0.99945981 |
| PUM3 | 28.5409314 | -0.229470502 | 0.33924317 | -0.6764189 | 0.4987747 | 0.99945981 |
| RFX3 | 109.949954 | 0.019717879 | 0.34428156 | 0.05727254 | 0.95432809 | 0.99945981 |
| RFX3-AS1 | 32.8041129 | -0.066004746 | 0.51991785 | -0.1269523 | 0.89897818 | 0.99945981 |
| GLIS3 | 28.1660618 | 0.209368665 | 0.46536321 | 0.44990378 | 0.65277982 | 0.99945981 |

| | | | | | | |
|---|---|---|---|---|---|---|
| CDC37L1 | 11.5406941 | 0.083000935 | 0.4177416 | 0.19868965 | 0.84250552 | 0.99945981 |
| AK3 | 22.5284536 | -0.108015789 | 0.41746962 | -0.2587393 | 0.79583641 | 0.99945981 |
| RCL1 | 13.3034253 | -0.154485524 | 0.41444789 | -0.3727502 | 0.70933438 | 0.99945981 |
| JAK2 | 98.8163932 | 0.730831251 | 0.24610149 | 2.96963363 | 0.00298155 | 0.45570351 |
| PLGRKT | 10.2984163 | -1.015787048 | 0.4416942 | -2.2997519 | 0.02146228 | 0.7964247 |
| RIC1 | 88.715504 | 0.448007722 | 0.32668444 | 1.3713776 | 0.17025728 | 0.99945981 |
| ERMP1 | 14.7596705 | 0.159080738 | 0.37202521 | 0.42760741 | 0.66893697 | 0.99945981 |
| KIAA2026 | 75.3970507 | 0.28315977 | 0.25992039 | 1.08940962 | 0.27597329 | 0.99945981 |
| UHRF2 | 53.2939448 | -0.206128208 | 0.36998296 | -0.5571289 | 0.57743935 | 0.99945981 |
| KDM4C | 179.036192 | 0.002683883 | 0.2305316 | 0.01164215 | 0.99071112 | 0.99945981 |
| NFIB | 13.4988378 | 0.678967453 | 0.60761436 | 1.11743154 | 0.26380986 | 0.99945981 |
| ZDHHC21 | 45.7921993 | -0.407614668 | 0.564184 | -0.7224853 | 0.46999613 | 0.99945981 |
| TTC39B | 30.1108222 | 0.619959276 | 0.41680742 | 1.48739981 | 0.13690925 | 0.99945981 |
| SNAPC3 | 27.9474088 | 0.164188222 | 0.34795551 | 0.47186557 | 0.63702275 | 0.99945981 |
| PSIP1 | 25.7540592 | 0.067177274 | 0.42004338 | 0.15992937 | 0.87293671 | 0.99945981 |
| CCDC171 | 11.1099275 | -0.567345358 | 0.62140643 | -0.9130021 | 0.36124145 | 0.99945981 |
| BNC2 | 66.1861389 | -0.023298474 | 0.44997846 | -0.0517769 | 0.95870649 | 0.99945981 |
| CNTLN | 35.7339863 | 0.417855076 | 0.3751103 | 1.11395254 | 0.26529955 | 0.99945981 |
| HAUS6 | 14.8955723 | -0.071476772 | 0.51561785 | -0.1386235 | 0.88974764 | 0.99945981 |
| PLIN2 | 50.3064808 | -1.534255565 | 0.54280751 | -2.8265187 | 0.0047057 | 0.50757754 |
| DENND4C | 83.5310567 | -0.315735651 | 0.36972271 | -0.8539796 | 0.3931163 | 0.99945981 |
| RPS6 | 124.133819 | -0.41805862 | 0.57172631 | -0.7312216 | 0.46464382 | 0.99945981 |
| MLLT3 | 82.9695523 | 0.124616513 | 0.31677895 | 0.39338635 | 0.69403415 | 0.99945981 |
| FOCAD | 39.6551188 | 0.146202794 | 0.43565864 | 0.33559026 | 0.73717987 | 0.99945981 |
| HACD4 | 14.0632405 | -0.111109936 | 0.48135923 | -0.2308254 | 0.81745045 | 0.99945981 |
| CAAP1 | 16.614219 | 9.51E-05 | 0.3409946 | 0.00027892 | 0.99977745 | 0.9997934 |
| PLAA | 16.901148 | 0.505154286 | 0.35106486 | 1.43892012 | 0.15017316 | 0.99945981 |
| IFT74 | 8.47800397 | -0.421394604 | 0.52643304 | -0.8004714 | 0.42343772 | 0.99945981 |
| MOB3B | 53.0742456 | -0.702454728 | 0.42985181 | -1.6341788 | 0.10222131 | 0.99945981 |
| C9orf72 | 57.7784394 | 1.21476651 | 0.5067424 | 2.39720717 | 0.01652058 | 0.74280553 |
| ACO1 | 11.8159481 | 0.394125269 | 0.55299135 | 0.71271507 | 0.47602209 | 0.99945981 |
| DDX58 | 42.7970903 | 0.779967876 | 0.70915149 | 1.09986074 | 0.2713928 | 0.99945981 |
| APTX | 15.5787989 | -0.431534608 | 0.45810938 | -0.9419903 | 0.34619759 | 0.99945981 |
| DNAJA1 | 30.9978333 | 0.606415103 | 0.42622992 | 1.42274174 | 0.15481104 | 0.99945981 |
| SMU1 | 15.8915042 | 0.655515867 | 0.33424847 | 1.96116338 | 0.04985997 | 0.92165202 |
| B4GALT1 | 70.8984798 | 0.110629222 | 0.36973742 | 0.29921024 | 0.76477964 | 0.99945981 |
| BAG1 | 9.88429627 | 0.147832888 | 0.43758163 | 0.33784071 | 0.73548323 | 0.99945981 |
| CHMP5 | 13.0019671 | 0.450008935 | 0.56378914 | 0.7981866 | 0.42476221 | 0.99945981 |
| NFX1 | 49.1363824 | -0.067430257 | 0.33886749 | -0.1989871 | 0.84227284 | 0.99945981 |
| UBE2R2 | 106.636696 | 0.595543922 | 0.28553565 | 2.08570779 | 0.0370051 | 0.87920079 |
| UBAP2 | 38.9959484 | 0.136261565 | 0.30688121 | 0.44402056 | 0.65702772 | 0.99945981 |
| DCAF12 | 13.8642054 | 0.136575046 | 0.46245515 | 0.29532604 | 0.76774483 | 0.99945981 |
| UBAP1 | 53.274089 | 0.468328761 | 0.29857034 | 1.56857095 | 0.11674795 | 0.99945981 |
| FAM219A | 12.775082 | -0.186925965 | 0.62113535 | -0.3009424 | 0.76345842 | 0.99945981 |
| VCP | 26.8667963 | -0.277565079 | 0.36377294 | -0.7630174 | 0.44545303 | 0.99945981 |
| UNC13B | 14.9669879 | -0.071633164 | 0.65383234 | -0.1095589 | 0.91275919 | 0.99945981 |
| RUSC2 | 12.0638979 | -0.21022063 | 0.65351513 | -0.3216768 | 0.74769758 | 0.99945981 |
| CD72 | 14.2909528 | -1.744108001 | 1.02207317 | -1.7064414 | 0.08792592 | 0.99640647 |
| CCDC107 | 11.8324002 | -0.511199154 | 0.4933957 | -1.0360835 | 0.30016318 | 0.99945981 |
| TLN1 | 43.2364748 | -0.099228959 | 0.44088252 | -0.2250689 | 0.82192565 | 0.99945981 |
| GBA2 | 9.03587989 | 0.390206316 | 0.58565169 | 0.66627711 | 0.505234 | 0.99945981 |
| RECK | 14.1193224 | -0.468556392 | 0.59606393 | -0.7860841 | 0.43181821 | 0.99945981 |
| GLIPR2 | 24.3873637 | 0.502879793 | 0.42019291 | 1.19678315 | 0.23139109 | 0.99945981 |
| CLTA | 41.1227898 | -0.434986587 | 0.31640362 | -1.3747838 | 0.16919847 | 0.99945981 |
| GNE | 11.7020002 | -0.312004545 | 0.46199013 | -0.6753489 | 0.4994541 | 0.99945981 |
| RNF38 | 100.886205 | -0.091934196 | 0.24912102 | -0.3690343 | 0.71210217 | 0.99945981 |
| EBLN3P | 15.9706499 | -0.164901701 | 0.40595464 | -0.4062072 | 0.68459036 | 0.99945981 |
| ZCCHC7 | 209.530987 | -0.581775874 | 0.47443955 | -1.2262382 | 0.22010907 | 0.99945981 |
| GRHPR | 7.98812698 | -0.21739534 | 0.46148264 | -0.4710802 | 0.63758345 | 0.99945981 |
| AL138752.2 | 17.0904284 | -0.525306055 | 0.54166424 | -0.9698001 | 0.33214612 | 0.99945981 |
| TRMT10B | 16.4973801 | -0.26552822 | 0.3785381 | -0.701457 | 0.48301783 | 0.99945981 |
| DCAF10 | 38.7035702 | 0.162150283 | 0.35862423 | 0.45214536 | 0.65116427 | 0.99945981 |
| CBWD5 | 16.2605418 | 0.645213537 | 0.45347943 | 1.42280662 | 0.15479222 | 0.99945981 |
| PIP5K1B | 24.5332057 | -1.299836322 | 1.06015817 | -1.2260777 | 0.22016942 | 0.99945981 |
| APBA1 | 13.9591774 | 0.084538993 | 0.53542952 | 0.15789005 | 0.87454344 | 0.99945981 |
| PTAR1 | 35.7580547 | 0.011549699 | 0.30722306 | 0.03759386 | 0.97001151 | 0.99945981 |
| MAMDC2 | 83.7225794 | 0.53625367 | 0.94379677 | 0.56818765 | 0.56990756 | 0.99945981 |
| SMC5 | 64.9897155 | -0.454101433 | 0.27330243 | -1.6615346 | 0.09660613 | 0.99945981 |
| KLF9 | 21.2070681 | 0.042161875 | 0.61478977 | 0.06857934 | 0.94532447 | 0.99945981 |
| CEMIP2 | 149.173308 | -1.318108058 | 0.44712652 | -2.9479532 | 0.00319886 | 0.46723702 |
| ABHD17B | 21.3874248 | -0.598573674 | 0.42733714 | -1.400706 | 0.16130202 | 0.99945981 |
| C9orf85 | 19.9334157 | -0.003447748 | 0.33729582 | -0.0102217 | 0.99184438 | 0.99945981 |
| ZFAND5 | 46.8198784 | 0.659445645 | 0.30646942 | 2.15175021 | 0.03141703 | 0.86065765 |
| ALDH1A1 | 11.206652 | -0.607279672 | 0.84011223 | -0.7228554 | 0.46976872 | 0.99945981 |
| ANXA1 | 55.4394285 | 0.382238146 | 0.53653582 | 0.71241869 | 0.47620554 | 0.99945981 |
| CARNMT1 | 19.9493509 | -0.467682895 | 0.347603 | -1.3454513 | 0.17847955 | 0.99945981 |
| NMRK1 | 16.501314 | 0.213294644 | 0.5349938 | 0.3986862 | 0.69012444 | 0.99945981 |
| OSTF1 | 50.7445504 | 0.117320274 | 0.33984523 | 0.34521677 | 0.72993141 | 0.99945981 |
| PCSK5 | 25.6778945 | 0.253463344 | 0.41470641 | 0.61118743 | 0.54107551 | 0.99945981 |
| GCNT1 | 20.0379608 | -0.618007176 | 0.53316706 | -1.1591248 | 0.24640531 | 0.99945981 |
| PRUNE2 | 12.4371873 | -0.288852706 | 0.55583415 | -0.5196743 | 0.60329062 | 0.99945981 |
| VPS13A | 41.6138207 | -0.143181734 | 0.26401081 | -0.5423328 | 0.58758923 | 0.99945981 |
| GNAQ | 292.661169 | 0.331770361 | 0.38903885 | 0.85279493 | 0.39377304 | 0.99945981 |
| CEP78 | 14.9869989 | 0.296451385 | 0.52868875 | 0.56072951 | 0.57498194 | 0.99945981 |
| TLE4 | 84.6514601 | 0.602750709 | 0.33378736 | 1.8057925 | 0.07095076 | 0.95505283 |
| TLE1 | 11.8607831 | 0.010778681 | 0.64182697 | 0.01679375 | 0.98660116 | 0.99945981 |
| FRMD3 | 19.9153087 | 0.430879311 | 0.53363437 | 0.80744295 | 0.41941133 | 0.99945981 |
| UBQLN1 | 38.6764436 | 0.052010498 | 0.26758142 | 0.19437261 | 0.84588414 | 0.99945981 |
| GKAP1 | 12.0333835 | -0.594269271 | 0.47131823 | -1.2608663 | 0.20735702 | 0.99945981 |
| HNRNPK | 59.050504 | 0.435477671 | 0.31074726 | 1.40138861 | 0.1610979 | 0.99945981 |
| AGTPBP1 | 164.360939 | 0.344658355 | 0.30458914 | 1.13155168 | 0.25782297 | 0.99945981 |
| NAA35 | 25.743836 | -0.326538091 | 0.36387317 | -0.8973954 | 0.36950799 | 0.99945981 |
| TUT7 | 84.69431 | 0.47169418 | 0.30791934 | 1.5318758 | 0.12555309 | 0.99945981 |
| DAPK1 | 114.529955 | -0.075730848 | 0.28028836 | -0.2701891 | 0.78701481 | 0.99945981 |
| CTSL | 58.2092674 | -0.612905804 | 0.73052357 | -0.8389953 | 0.40147197 | 0.99945981 |
| SPIN1 | 41.9342735 | -0.307584778 | 0.31739973 | -0.969077 | 0.33250678 | 0.99945981 |
| SECISBP2 | 44.9900475 | -0.521139662 | 0.34632681 | -1.5047627 | 0.13238509 | 0.99945981 |
| SEMA4D | 93.9806884 | -0.066098602 | 0.39579514 | -0.1670021 | 0.86736843 | 0.99945981 |
| SYK | 113.704075 | 0.119724905 | 0.31782981 | 0.37669501 | 0.70640027 | 0.99945981 |
| AL158071.4 | 15.8863765 | -0.454909756 | 0.7228185 | -0.6293554 | 0.52911638 | 0.99945981 |
| AUH | 53.2371807 | -0.104922044 | 0.42468869 | -0.2470564 | 0.8048646 | 0.99945981 |
| NFIL3 | 15.7596441 | 0.168878272 | 0.53283527 | 0.31694274 | 0.75128705 | 0.99945981 |
| SPTLC1 | 29.805997 | 0.038005022 | 0.30259807 | 0.12559572 | 0.90005195 | 0.99945981 |
| IARS | 26.7021705 | -0.618081055 | 0.49520683 | -1.2481271 | 0.21198453 | 0.99945981 |
| NOL8 | 15.9371589 | 0.148354139 | 0.40371729 | 0.36747037 | 0.71326819 | 0.99945981 |
| CENPP | 51.5313699 | 0.584453118 | 0.49862905 | 1.17212008 | 0.24114885 | 0.99945981 |
| IPPK | 8.8450192 | 0.143972863 | 0.62268329 | 0.23121363 | 0.81714884 | 0.99945981 |
| BICD2 | 17.974613 | 0.130508906 | 0.38938706 | 0.33516498 | 0.73750064 | 0.99945981 |
| FGD3 | 62.6870577 | 0.300272461 | 0.32953373 | 0.91120402 | 0.36218789 | 0.99945981 |
| CARD19 | 9.70862007 | 0.612552929 | 0.46098529 | 1.32879062 | 0.18391706 | 0.99945981 |
| NINJ1 | 16.4195973 | 0.149438262 | 0.53666204 | 0.27845879 | 0.7806602 | 0.99945981 |
| FAM120A | 73.2111854 | 0.329485227 | 0.22872678 | 1.44051881 | 0.14972067 | 0.99945981 |
| PHF2 | 16.0858402 | 0.184449848 | 0.4269105 | 0.43205741 | 0.66569969 | 0.99945981 |

| | | | | | | |
|---|---|---|---|---|---|---|
| AL158152.2 | 12.699406 | 0.176272253 | 0.44147371 | 0.39928143 | 0.68968585 | 0.99945981 |
| ZNF169 | 11.7493212 | -0.449579819 | 0.50179769 | -0.8959384 | 0.37028566 | 0.99945981 |
| MFSD14B | 30.7734053 | 0.61154048 | 0.35787552 | 1.70880782 | 0.08748655 | 0.99640647 |
| FBP1 | 11.9550649 | 0.205697396 | 0.70840078 | 0.29036868 | 0.7715342 | 0.99945981 |
| C9orf3 | 58.1021198 | 0.058274789 | 0.3802087 | 0.15327053 | 0.87818494 | 0.99945981 |
| FANCC | 17.409787 | 0.141840733 | 0.4711495 | 0.3010525 | 0.76337446 | 0.99945981 |
| LINC00476 | 14.5193225 | 0.159190816 | 0.58831231 | 0.27058896 | 0.78670719 | 0.99945981 |
| ERCC6L2 | 35.4404619 | 0.236812208 | 0.31469992 | 0.75250164 | 0.45174944 | 0.99945981 |
| HABP4 | 11.9244025 | -1.041644141 | 0.45187645 | -2.3051525 | 0.02115803 | 0.79476944 |
| ZNF782 | 17.1793889 | 0.072058873 | 0.46513681 | 0.15491974 | 0.87688459 | 0.99945981 |
| MFSD14C | 59.1178521 | 0.152267754 | 0.32670691 | 0.46606836 | 0.64116657 | 0.99945981 |
| TDRD7 | 18.0924705 | 0.255400974 | 0.44281416 | 0.57676786 | 0.56409628 | 0.99945981 |
| TSTD2 | 13.2317378 | -0.280872476 | 0.68399977 | -0.4106324 | 0.68134209 | 0.99945981 |
| NCBP1 | 12.6139409 | 0.129514528 | 0.43845352 | 0.29538942 | 0.76769642 | 0.99945981 |
| XPA | 13.2604704 | -0.27220837 | 0.55635591 | -0.4892702 | 0.62465041 | 0.99945981 |
| ANP32B | 27.3145645 | 0.105067693 | 0.32890537 | 0.31944657 | 0.7493879 | 0.99945981 |
| NANS | 9.49193179 | 0.197760187 | 0.49277277 | 0.40132125 | 0.68818362 | 0.99945981 |
| TRIM14 | 47.5018104 | 0.76224072 | 0.38555769 | 1.97698228 | 0.04804363 | 0.90884462 |
| CORO2A | 8.31421849 | -0.783697495 | 0.47304837 | -1.6566963 | 0.09758088 | 0.99945981 |
| TBC1D2 | 21.1986322 | -0.083483695 | 0.46239509 | -0.1805462 | 0.85672375 | 0.99945981 |
| TGFBR1 | 58.9247157 | -0.195759735 | 0.40743897 | -0.480464 | 0.63089753 | 0.99945981 |
| SEC61B | 31.6611543 | -0.276509746 | 0.47491003 | -0.5822361 | 0.56040768 | 0.99945981 |
| STX17 | 17.1189335 | -0.146652617 | 0.3912219 | -0.3748579 | 0.70776616 | 0.99945981 |
| ERP44 | 46.4417477 | -0.124773547 | 0.27714228 | -0.4502148 | 0.65255559 | 0.99945981 |
| INVS | 60.2937049 | -0.287679108 | 0.30223756 | -0.9518311 | 0.34118265 | 0.99945981 |
| TEX10 | 10.9455476 | -0.239048507 | 0.58110303 | -0.4113703 | 0.68080105 | 0.99945981 |
| ALDOB | 29.7691237 | -0.331514427 | 1.06293063 | -0.3118872 | 0.75512627 | 0.99945981 |
| RNF20 | 10.4867362 | 0.415422236 | 0.4147211 | 1.00169062 | 0.31649304 | 0.99945981 |
| ABCA1 | 93.6942928 | 0.064664079 | 0.6225373 | 0.10387181 | 0.91727108 | 0.99945981 |
| SLC44A1 | 48.6281755 | -0.656088593 | 0.42424789 | -1.5464746 | 0.12198999 | 0.99945981 |
| FSD1L | 15.6568246 | -0.641995509 | 0.48503142 | -1.3236163 | 0.1856305 | 0.99945981 |
| FKTN | 11.6853483 | -0.294739788 | 0.42409359 | -0.6949876 | 0.48706307 | 0.99945981 |
| RAD23B | 44.5714801 | -0.214371363 | 0.27791351 | -0.77136 | 0.44049359 | 0.99945981 |
| ELP1 | 13.4303343 | 0.10518619 | 0.38978969 | 0.2698537 | 0.78727281 | 0.99945981 |
| TMEM245 | 41.8941835 | 0.296934485 | 0.27897333 | 1.06438306 | 0.28715521 | 0.99945981 |
| TXN | 21.4615143 | -0.972502953 | 0.5024137 | -1.9356617 | 0.05290915 | 0.92445778 |
| LPAR1 | 14.5072781 | 1.231522336 | 0.89313027 | 1.37888322 | 0.16793076 | 0.99945981 |
| ECPAS | 50.5610947 | -0.190103946 | 0.32998856 | -0.5760925 | 0.56455264 | 0.99945981 |
| PTGR1 | 12.2163683 | -1.540989403 | 0.67439732 | -2.2849874 | 0.02231357 | 0.80200327 |
| C9orf84 | 16.3478758 | -0.45154632 | 0.95151119 | -0.474557 | 0.63510274 | 0.99945981 |
| UGCG | 62.4906285 | 0.173594394 | 0.58758614 | 0.2954365 | 0.76766045 | 0.99945981 |
| SUSD1 | 43.223059 | 0.622079459 | 0.40045827 | 1.55341894 | 0.12032308 | 0.99945981 |
| PTBP3 | 112.268931 | -0.01246056 | 0.21366483 | -0.0583183 | 0.95349513 | 0.99945981 |
| HSDL2 | 26.7684743 | 0.139722593 | 0.38457629 | 0.36331567 | 0.71636909 | 0.99945981 |
| KIAA1958 | 29.55564 | 0.108691409 | 0.48596369 | 0.22366159 | 0.82302064 | 0.99945981 |
| INIP | 13.6525755 | 0.244359991 | 0.44192677 | 0.55294227 | 0.58030294 | 0.99945981 |
| SNX30 | 33.5300538 | 0.387353913 | 0.30873726 | 1.25463934 | 0.20960972 | 0.99945981 |
| FKBP15 | 44.2829069 | 0.396407917 | 0.37841617 | 1.04754486 | 0.29484836 | 0.99945981 |
| SLC31A1 | 16.7792883 | -0.034980762 | 0.54497958 | -0.0641873 | 0.9488211 | 0.99945981 |
| CDC26 | 8.59231158 | -0.094961642 | 0.7259853 | -0.1308038 | 0.89593051 | 0.99945981 |
| RGS3 | 10.7098347 | 0.140479931 | 0.5809048 | 0.24182952 | 0.80891226 | 0.99945981 |
| ZNF618 | 15.0614546 | 0.99800924 | 0.63613417 | 1.56886595 | 0.11667918 | 0.99945981 |
| AKNA | 93.7709809 | -0.692191663 | 0.43022697 | -1.6088988 | 0.10763847 | 0.99945981 |
| ATP6V1G1 | 28.6925473 | -0.544268628 | 0.47448136 | -1.1470812 | 0.25134808 | 0.99945981 |
| TMEM268 | 12.9196881 | -0.434197874 | 0.76001082 | -0.5713049 | 0.56779301 | 0.99945981 |
| TNFSF8 | 19.0949341 | 0.410074316 | 0.45819356 | 0.89498053 | 0.3707975 | 0.99945981 |
| ASTN2 | 11.3075622 | 0.331302946 | 0.56552491 | 0.58583263 | 0.55798799 | 0.99945981 |
| CDK5RAP2 | 43.5804668 | -0.012719538 | 0.2459823 | -0.0517092 | 0.95876044 | 0.99945981 |
| MEGF9 | 78.4103772 | 0.677351928 | 0.38888959 | 1.74175895 | 0.08155063 | 0.9846219 |
| FBXW2 | 23.7196581 | -0.09723136 | 0.41471806 | -0.2344517 | 0.81463431 | 0.99945981 |
| PSMD5 | 17.7465756 | -0.347314347 | 0.3261584 | -1.064864 | 0.28693747 | 0.99945981 |
| TRAF1 | 17.513269 | -0.931995589 | 0.39987452 | -2.3307201 | 0.01976812 | 0.78749881 |
| CNTRL | 74.3840827 | -0.051593149 | 0.25361012 | -0.2034349 | 0.83879512 | 0.99945981 |
| RAB14 | 36.1930928 | 0.276643419 | 0.28618728 | 0.96665168 | 0.33371819 | 0.99945981 |
| GSN | 39.3651204 | 0.428484401 | 0.49385917 | 0.86762467 | 0.38559984 | 0.99945981 |
| STOM | 38.9386042 | 0.028693086 | 0.28640719 | 0.10018284 | 0.92019917 | 0.99945981 |
| MRRF | 12.1865809 | 0.050882832 | 0.39118154 | 0.13007473 | 0.8965073 | 0.99945981 |
| RC3H2 | 46.6295014 | -0.296619319 | 0.24628982 | -1.2043507 | 0.22845406 | 0.99945981 |
| RABGAP1 | 63.0633255 | 0.115789636 | 0.25982118 | 0.44565127 | 0.65584917 | 0.99945981 |
| STRBP | 47.0375394 | -1.006556117 | 0.7209348 | -1.3961819 | 0.16265973 | 0.99945981 |
| DENND1A | 186.976339 | 0.528781661 | 0.25871811 | 2.04385253 | 0.04096812 | 0.89562588 |
| NEK6 | 26.5577246 | -0.383754778 | 0.43567817 | -0.8808217 | 0.37841435 | 0.99945981 |
| PSMB7 | 25.7808864 | 0.039785259 | 0.35792241 | 0.1111561 | 0.91149256 | 0.99945981 |
| NR6A1 | 28.6066666 | 0.853372547 | 0.47059342 | 1.81339669 | 0.06977066 | 0.95505283 |
| RPL35 | 45.3148299 | 0.092466836 | 0.55574459 | 0.16638369 | 0.86785501 | 0.99945981 |
| ARPC5L | 16.5098644 | -0.649089511 | 0.53541392 | -1.2123135 | 0.22539241 | 0.99945981 |
| GOLGA1 | 15.2386347 | 0.07961149 | 0.37954273 | 0.20975633 | 0.83385786 | 0.99945981 |
| SCAI | 27.0115478 | 0.29736463 | 0.37549525 | 0.79192647 | 0.42840355 | 0.99945981 |
| PPP6C | 25.361445 | 0.237552384 | 0.34900415 | 0.68065775 | 0.49608807 | 0.99945981 |
| HSPA5 | 39.3932389 | -0.89532694 | 0.60692989 | -1.4751736 | 0.14016588 | 0.99945981 |
| GAPVD1 | 48.4553481 | 0.146896705 | 0.2401958 | 0.61157066 | 0.54082185 | 0.99945981 |
| MAPKAP1 | 83.7746551 | 0.019603062 | 0.19709911 | 0.09945789 | 0.92077472 | 0.99945981 |
| PBX3 | 108.072153 | -0.230746036 | 0.63180406 | -0.3652177 | 0.71494889 | 0.99945981 |
| MVB12B | 61.8080545 | -0.339387124 | 0.48810803 | -0.6953115 | 0.48686011 | 0.99945981 |
| ZBTB43 | 19.2424824 | -0.413998086 | 0.43476216 | -0.9522404 | 0.34097509 | 0.99945981 |
| RALGPS1 | 16.8632922 | 0.412837241 | 0.43124862 | 0.95730682 | 0.33841242 | 0.99945981 |
| RPL12 | 20.9261003 | 0.032112582 | 0.43472528 | 0.07386868 | 0.94111488 | 0.99945981 |
| LRSAM1 | 19.7313358 | -0.615975251 | 0.34461299 | -1.7874406 | 0.0738663 | 0.96760689 |
| FAM129B | 14.8913625 | -0.279931413 | 0.56261016 | -0.4975584 | 0.61879533 | 0.99945981 |
| SH2D3C | 35.6090262 | 1.030070962 | 0.44885386 | 2.29489162 | 0.02173934 | 0.79988354 |
| CDK9 | 9.90111094 | 0.350446746 | 0.52955805 | 0.6617721 | 0.50811729 | 0.99945981 |
| FPGS | 11.5052376 | -0.148823915 | 0.59467886 | -0.2502593 | 0.80238683 | 0.99945981 |
| ENG | 24.250327 | 0.08269672 | 0.36642029 | 0.22568816 | 0.82144397 | 0.99945981 |
| FAM102A | 15.2461841 | -0.173318223 | 0.51271136 | -0.3380425 | 0.73533117 | 0.99945981 |
| SLC25A25 | 17.8859977 | 0.438091289 | 0.42614859 | 1.02802473 | 0.3039382 | 0.99945981 |
| C9orf16 | 13.3062774 | -0.695419719 | 0.49530075 | -1.4040353 | 0.16030835 | 0.99945981 |
| GOLGA2 | 12.2140481 | -0.714816639 | 0.40254963 | -1.775723 | 0.07577858 | 0.97969864 |
| SLC27A4 | 12.4903926 | 0.160949231 | 0.47385416 | 0.33965985 | 0.7341127 | 0.99945981 |
| URM1 | 9.10114036 | -0.304864002 | 0.48060257 | -0.634337 | 0.5258609 | 0.99945981 |
| ODF2 | 8.09880437 | -0.309041103 | 0.47584142 | -0.6494624 | 0.51603955 | 0.99945981 |
| GLE1 | 13.4551417 | -0.551561545 | 0.39380317 | -1.4006021 | 0.1613331 | 0.99945981 |
| SPTAN1 | 34.9932286 | -0.050940011 | 0.33476058 | -0.1521685 | 0.87905405 | 0.99945981 |
| SET | 34.9839309 | -0.186907776 | 0.42793291 | -0.4367689 | 0.66227897 | 0.99945981 |
| ZER1 | 19.268749 | -0.18356073 | 0.42576123 | -0.4311354 | 0.66636994 | 0.99945981 |
| TBC1D13 | 8.18380401 | -0.408112371 | 0.4812111 | -0.8480943 | 0.39638548 | 0.99945981 |
| NUP188 | 11.7343635 | -0.219064462 | 0.40843681 | -0.5363485 | 0.59171773 | 0.99945981 |
| PTPA | 11.0414015 | 0.278091956 | 0.46306566 | 0.6005454 | 0.54814281 | 0.99945981 |
| C9orf78 | 16.1696661 | -0.231703738 | 0.37541058 | -0.6172009 | 0.53710224 | 0.99945981 |
| USP20 | 14.7009951 | -0.612507044 | 0.39138965 | -1.5649546 | 0.11759355 | 0.99945981 |
| FNBP1 | 250.413522 | -0.544746308 | 0.24076242 | -2.2625886 | 0.02366106 | 0.80320673 |
| GPR107 | 37.2600933 | -0.131834344 | 0.26305519 | -0.5011661 | 0.61625423 | 0.99945981 |
| FUBP3 | 26.1319697 | 0.104951837 | 0.28144498 | 0.37290356 | 0.70922022 | 0.99945981 |
| ABL1 | 62.9014684 | -0.510874193 | 0.23567084 | -2.1677446 | 0.03017813 | 0.8560861 |

| | | | | | |
|---|---|---|---|---|---|
| NUP214 | 42.8747024 | 0.144383882 | 0.35086335 | 0.4115103 | 0.68069839 | 0.99945981 |
| AL157938.2 | 9.90859799 | -0.135351723 | 0.51732664 | -0.2616369 | 0.79360142 | 0.99945981 |
| FAM78A | 14.7366654 | 0.233615248 | 0.40456362 | 0.57744996 | 0.56363553 | 0.99945981 |
| PRRC2B | 72.5999112 | -0.286172052 | 0.25607767 | -1.1175205 | 0.26377183 | 0.99945981 |
| RAPGEF1 | 232.111138 | -0.636580323 | 0.30597976 | -2.0804654 | 0.03748286 | 0.87920079 |
| MED27 | 42.2260339 | -0.023926821 | 0.32030041 | -0.0747012 | 0.94045246 | 0.99945981 |
| SETX | 125.007781 | 0.109668065 | 0.22649477 | 0.4841969 | 0.62824614 | 0.99945981 |
| TTF1 | 16.3843526 | -0.201445237 | 0.35326421 | -0.5702396 | 0.56851521 | 0.99945981 |
| DDX31 | 8.72822157 | -0.690299913 | 0.54343291 | -1.2702578 | 0.2039928 | 0.99945981 |
| TSC1 | 15.8203003 | 0.371434137 | 0.40512737 | 0.91683299 | 0.35923016 | 0.99945981 |
| RALGDS | 10.6027876 | -0.38131068 | 0.47975751 | -0.7947988 | 0.42673057 | 0.99945981 |
| RPL7A | 58.5298024 | -0.260006785 | 0.53906139 | -0.4823324 | 0.62956982 | 0.99945981 |
| SURF4 | 21.9050917 | -0.529792907 | 0.39023971 | -1.3576089 | 0.17458782 | 0.99945981 |
| VAV2 | 21.1845055 | 0.094158839 | 0.48143318 | 0.19558029 | 0.84493869 | 0.99945981 |
| BRD3 | 10.6524949 | 0.262376483 | 0.45954307 | 0.5709508 | 0.568033 | 0.99945981 |
| RXRA | 27.1017968 | 0.477905179 | 0.52350161 | 0.91290106 | 0.36129458 | 0.99945981 |
| FCN1 | 71.8768484 | 1.27486352 | 1.18706591 | 1.07396187 | 0.28283977 | 0.99945981 |
| CAMSAP1 | 15.8315534 | 0.340371456 | 0.46889025 | 0.72590859 | 0.46789481 | 0.99945981 |
| UBAC1 | 11.3981183 | 0.398006457 | 0.55417153 | 0.71820084 | 0.47263346 | 0.99945981 |
| NACC2 | 21.3621911 | 0.401617866 | 0.42524507 | 0.94443862 | 0.34494555 | 0.99945981 |
| SEC16A | 20.2835349 | 0.09891627 | 0.36727123 | 0.26932758 | 0.78767761 | 0.99945981 |
| NOTCH1 | 12.9861156 | 0.452345757 | 0.72812282 | 0.62124925 | 0.53443564 | 0.99945981 |
| AGPAT2 | 10.4572115 | -0.107506744 | 0.54263726 | -0.198119 | 0.84295197 | 0.99945981 |
| SNHG7 | 11.0589069 | -0.223574323 | 0.47819684 | -0.4675362 | 0.64011632 | 0.99945981 |
| RABL6 | 22.4855706 | -0.227808613 | 0.37423087 | -0.6087382 | 0.542698 | 0.99945981 |
| PHPT1 | 13.6646892 | 0.17044058 | 0.48940995 | 0.34825729 | 0.72764697 | 0.99945981 |
| EDF1 | 22.4098363 | -0.201841555 | 0.56176648 | -0.359298 | 0.71937219 | 0.99945981 |
| FBXW5 | 17.2089119 | 0.045548586 | 0.44447804 | 0.10247657 | 0.91837841 | 0.99945981 |
| PAXX | 9.65036867 | -0.048938272 | 0.60152498 | -0.081357 | 0.93515804 | 0.99945981 |
| MAN1B1 | 11.380595 | -0.379217242 | 0.38643626 | -0.981319 | 0.32643546 | 0.99945981 |
| DPP7 | 13.5100586 | -0.341410586 | 0.50659661 | -0.6739299 | 0.5003559 | 0.99945981 |
| SSNA1 | 8.58888747 | -0.456113847 | 0.69067618 | -0.6603874 | 0.50900525 | 0.99945981 |
| TUBB4B | 11.5639386 | -0.813593548 | 0.60287201 | -1.3495295 | 0.17716696 | 0.99945981 |
| NELFB | 14.3543758 | 0.591778051 | 0.42803658 | 1.38254083 | 0.1668057 | 0.99945981 |
| EXD3 | 10.2281196 | -0.549921011 | 0.50036768 | -1.0990338 | 0.27175331 | 0.99945981 |
| MRPL41 | 18.2319104 | -0.210311744 | 0.4196801 | -0.5011239 | 0.6162839 | 0.99945981 |
| ARRDC1 | 12.0095773 | 0.320813552 | 0.40464795 | 0.79282139 | 0.42788189 | 0.99945981 |
| EHMT1 | 116.185995 | -0.487404037 | 0.35515819 | -1.3723576 | 0.16995215 | 0.99945981 |
| PSMD13 | 16.2151344 | 0.334743204 | 0.40554205 | 0.82542168 | 0.40913223 | 0.99945981 |
| PGGHG | 14.1591887 | 0.701489596 | 0.44943888 | 1.56081198 | 0.11856812 | 0.99945981 |
| IFITM2 | 88.1629998 | 0.02096829 | 0.57059192 | 0.03674831 | 0.97068569 | 0.99945981 |
| IFITM3 | 71.4424707 | -0.557862219 | 0.53189107 | -1.0488279 | 0.29425732 | 0.99945981 |
| SIGIRR | 13.9954731 | 0.484647334 | 0.37173944 | 1.30372857 | 0.19232614 | 0.99945981 |
| RNH1 | 16.132513 | -0.007667624 | 0.48219923 | -0.0159014 | 0.98731308 | 0.99945981 |
| PHRF1 | 15.6596236 | -0.471221771 | 0.57009796 | -0.8265628 | 0.40848491 | 0.99945981 |
| DEAF1 | 16.5308674 | 0.10902306 | 0.41879085 | 0.26032818 | 0.79461064 | 0.99945981 |
| TALDO1 | 27.4542924 | -0.410495631 | 0.54614091 | -0.7516295 | 0.45227389 | 0.99945981 |
| RPLP2 | 64.9045331 | 0.339580671 | 0.49934914 | 0.68004657 | 0.49647498 | 0.99945981 |
| PNPLA2 | 9.4579923 | 0.015890296 | 0.50550547 | 0.03143447 | 0.97492305 | 0.99945981 |
| CD151 | 13.695014 | -0.482788917 | 0.55200769 | -0.8746054 | 0.38178864 | 0.99945981 |
| POLR2L | 18.9480775 | -0.746090893 | 0.55708777 | -1.3392699 | 0.18048282 | 0.99945981 |
| AP2A2 | 63.5249469 | 0.457994707 | 0.39510574 | 1.15916997 | 0.24638691 | 0.99945981 |
| TOLLIP | 12.3994076 | -0.168449866 | 0.57266525 | -0.2941507 | 0.76864278 | 0.99945981 |
| MOB2 | 9.33779823 | -0.371681806 | 0.40948777 | -0.907675 | 0.36404997 | 0.99945981 |
| LSP1 | 27.1955662 | 0.148243448 | 0.50729477 | 0.29222349 | 0.77011575 | 0.99945981 |
| TSPAN32 | 10.9265434 | 0.764304143 | 0.61306745 | 1.2466885 | 0.21251174 | 0.99945981 |
| CD81 | 48.7072052 | -0.293240443 | 0.3845217 | -0.7626109 | 0.44569548 | 0.99945981 |
| KCNQ1 | 53.1912948 | 0.649093757 | 0.35353421 | 1.83601401 | 0.06635558 | 0.94594189 |
| KCNQ1OT1 | 25.6234715 | 0.348952476 | 0.74291109 | 0.46970961 | 0.6385625 | 0.99945981 |
| CDKN1C | 13.9929982 | 1.331999354 | 0.83194523 | 1.60106615 | 0.10936227 | 0.99945981 |
| NAP1L4 | 73.8464478 | -0.106674643 | 0.24720224 | -0.4315278 | 0.66608463 | 0.99945981 |
| CARS | 18.8733728 | -0.287010074 | 0.37713461 | -0.7610282 | 0.44664022 | 0.99945981 |
| OSBPL5 | 22.4306874 | 0.215304199 | 1.01385815 | 0.21236126 | 0.8318252 | 0.99945981 |
| ZNF195 | 12.6989053 | 0.090534262 | 0.59475009 | 0.15222236 | 0.87901156 | 0.99945981 |
| NUP98 | 60.1304534 | -0.076125427 | 0.2137927 | -0.3560712 | 0.72178723 | 0.99945981 |
| RHOG | 22.8120314 | 0.351126922 | 0.62818114 | 0.55895808 | 0.57619033 | 0.99945981 |
| STIM1 | 119.54472 | -0.023507063 | 0.23314239 | -0.1008271 | 0.91968774 | 0.99945981 |
| HBB | 29.4108007 | -0.496279203 | 1.74066106 | -0.2851096 | 0.77556016 | 0.99945981 |
| TRIM5 | 13.0413301 | -0.346593936 | 0.58502395 | -0.592444 | 0.5535533 | 0.99945981 |
| TRIM22 | 58.5262252 | 0.567475937 | 0.45767839 | 1.2399011 | 0.21501198 | 0.99945981 |
| AC084337.2 | 9.80916429 | -0.380260124 | 0.4467495 | -0.8511708 | 0.39467448 | 0.99945981 |
| DNHD1 | 16.3338852 | -0.314540607 | 0.46559728 | -0.6755637 | 0.49931771 | 0.99945981 |
| ILK | 13.3973283 | -0.668728678 | 0.50141568 | -1.3336812 | 0.18230835 | 0.99945981 |
| TAF10 | 9.50702856 | -0.157288984 | 0.48620202 | -0.3235054 | 0.74631251 | 0.99945981 |
| TPP1 | 25.7382299 | -0.155963829 | 0.44656227 | -0.3492544 | 0.72689834 | 0.99945981 |
| PPFIBP2 | 21.1213621 | 0.430851343 | 0.56194989 | 0.76670777 | 0.44325529 | 0.99945981 |
| EIF3F | 17.6571297 | 0.237762111 | 0.47210214 | 0.50362431 | 0.61452541 | 0.99945981 |
| TRIM66 | 17.6521152 | 0.308964567 | 0.4048419 | 0.7631734 | 0.44535999 | 0.99945981 |
| RPL27A | 72.5501931 | -0.153064796 | 0.57028375 | -0.2684011 | 0.78839058 | 0.99945981 |
| TMEM9B | 11.6664384 | -0.115038515 | 0.47396612 | -0.2427146 | 0.80822647 | 0.99945981 |
| DENND5A | 74.0800182 | 0.747822284 | 0.39405557 | 1.89775843 | 0.05772791 | 0.92445778 |
| TMEM41B | 22.9090824 | -0.99640798 | 0.35780294 | -2.7847954 | 0.00535615 | 0.5234882 |
| IPO7 | 36.3158839 | -0.162955548 | 0.39844778 | -0.4089759 | 0.68255733 | 0.99945981 |
| ZNF143 | 18.9980404 | 0.016828008 | 0.40325886 | 0.04173004 | 0.96671391 | 0.99945981 |
| SWAP70 | 54.4727897 | -0.594474483 | 0.53123936 | -1.1190332 | 0.26312597 | 0.99945981 |
| SBF2 | 205.755622 | 0.351256576 | 0.40148047 | 0.87490326 | 0.38162654 | 0.99945981 |
| AMPD3 | 22.9668917 | -0.107036115 | 0.45904997 | -0.2331688 | 0.81563036 | 0.99945981 |
| MTRNR2L8 | 19.7981106 | -0.421374573 | 0.77055058 | -0.5468487 | 0.58448269 | 0.99945981 |
| RNF141 | 10.7796801 | -1.182645281 | 0.72505224 | -1.6311173 | 0.10286556 | 0.99945981 |
| LYVE1 | 28.9710757 | 2.185477303 | 1.03422051 | 2.11316375 | 0.03458675 | 0.87920079 |
| CTR9 | 18.688953 | 0.11566788 | 0.44213907 | 0.26160972 | 0.79362235 | 0.99945981 |
| EIF4G2 | 48.3921259 | 0.070495017 | 0.35647755 | 0.19775444 | 0.8432372 | 0.99945981 |
| ZBED5 | 16.3540819 | 0.172941661 | 0.41723168 | 0.41449792 | 0.67850949 | 0.99945981 |
| USP47 | 91.9293535 | 0.053500187 | 0.22168184 | 0.24133771 | 0.80929339 | 0.99945981 |
| MICAL2 | 23.9390476 | -0.144713539 | 0.62875476 | -0.230159 | 0.81796825 | 0.99945981 |
| ARNTL | 34.9611898 | 0.152295908 | 0.3589087 | 0.4243305 | 0.6713248 | 0.99945981 |
| BTBD10 | 30.5654658 | 0.279138971 | 0.33001497 | 0.84583731 | 0.39764351 | 0.99945981 |
| FAR1 | 77.9521152 | 0.324379963 | 0.25607418 | 1.26674218 | 0.20524749 | 0.99945981 |
| RRAS2 | 30.3406086 | 0.010912595 | 0.45300781 | 0.0240892 | 0.98078146 | 0.99945981 |
| COPB1 | 35.6025756 | 0.105241598 | 0.29530587 | 0.35638167 | 0.72155475 | 0.99945981 |
| PSMA1 | 41.4630038 | -0.889908861 | 0.52685954 | -1.6890818 | 0.09120375 | 0.99640647 |
| PDE3B | 240.194385 | -1.050427547 | 0.51652731 | -2.0336341 | 0.0419885 | 0.89562588 |
| SOX6 | 10.5094022 | 0.491889306 | 0.49054236 | 1.00274584 | 0.31598351 | 0.99945981 |
| C11orf58 | 35.7659616 | 0.147993039 | 0.25178353 | 0.58777887 | 0.55668073 | 0.99945981 |
| PLEKHA7 | 91.3959531 | -0.912094074 | 0.60559899 | -1.5061024 | 0.1320409 | 0.99945981 |
| RPS13 | 58.6653076 | -0.223507008 | 0.60581805 | -0.3689342 | 0.71217676 | 0.99945981 |
| PIK3C2A | 51.147669 | 0.126126858 | 0.26285863 | 0.47982772 | 0.6313499 | 0.99945981 |
| NUCB2 | 15.5016216 | -0.824681875 | 0.65687134 | -1.2554694 | 0.2093084 | 0.99945981 |
| SERGEF | 40.1437086 | 0.206038738 | 0.37758168 | 0.54567991 | 0.58528598 | 0.99945981 |
| HPS5 | 22.1433742 | 0.025580211 | 0.51485471 | 0.04968433 | 0.96037395 | 0.99945981 |
| GTF2H1 | 20.0634029 | 0.273305196 | 0.44424746 | 0.61520936 | 0.53841647 | 0.99945981 |
| LDHA | 41.6651762 | -1.279537455 | 0.38029431 | -3.364598 | 0.00076655 | 0.27121928 |
| TSG101 | 26.4178048 | -0.708063806 | 0.33226536 | -2.131019 | 0.03308758 | 0.8663391 |

| | | | | | | |
|---|---|---|---|---|---|---|
| UEVLD | 18.7189932 | -0.294166473 | 0.39717054 | -0.7406553 | 0.45890246 | 0.99945981 |
| SPTY2D1 | 11.9098668 | 0.087295836 | 0.43095106 | 0.20256554 | 0.83947462 | 0.99945981 |
| ZDHHC13 | 16.3909532 | 0.355486835 | 0.35785532 | 0.99338145 | 0.3205241 | 0.99945981 |
| NAV2 | 28.2803388 | 0.595042567 | 0.63226714 | 0.94112525 | 0.34664068 | 0.99945981 |
| PRMT3 | 25.2612897 | 0.015542846 | 0.37948011 | 0.04095826 | 0.96732917 | 0.99945981 |
| LGR4 | 11.0171174 | 0.783105587 | 0.62245397 | 1.25809397 | 0.20835777 | 0.99945981 |
| BDNF-AS | 30.8365353 | -0.313351081 | 0.56338002 | -0.5561984 | 0.57807522 | 0.99945981 |
| METTL15 | 52.3152205 | -0.113442797 | 0.34165007 | -0.3320438 | 0.73985616 | 0.99945981 |
| ARL14EP | 12.0440059 | -0.270225917 | 0.48356543 | -0.5588198 | 0.57628474 | 0.99945981 |
| DNAJC24 | 16.3429971 | -0.173738015 | 0.38132014 | -0.4556224 | 0.6486615 | 0.99945981 |
| IMMP1L | 19.6238725 | -0.103982523 | 0.37838697 | -0.2748047 | 0.78346628 | 0.99945981 |
| ELP4 | 55.125005 | 0.267010947 | 0.24311278 | 1.09830073 | 0.27207319 | 0.99945981 |
| EIF3M | 20.8307825 | 0.4309649 | 0.445827 | 0.96666398 | 0.33371204 | 0.99945981 |
| QSER1 | 30.4534441 | 0.288743415 | 0.3416144 | 0.84523199 | 0.39798133 | 0.99945981 |
| TCP11L1 | 16.2814903 | 0.237024111 | 0.37089134 | 0.63906619 | 0.52277988 | 0.99945981 |
| CSTF3 | 25.5795898 | 0.290488922 | 0.35370586 | 0.82127257 | 0.41149103 | 0.99945981 |
| HIPK3 | 86.635501 | 0.09647292 | 0.35029475 | 0.27540498 | 0.78300512 | 0.99945981 |
| FBXO3 | 16.3898811 | -0.220682839 | 0.35123172 | -0.6283112 | 0.52980007 | 0.99945981 |
| LMO2 | 13.5551984 | 1.112591108 | 0.72196311 | 1.54106364 | 0.1233013 | 0.99945981 |
| CAPRIN1 | 42.7429485 | 0.189599109 | 0.30917464 | 0.61324275 | 0.53971584 | 0.99945981 |
| NAT10 | 10.1367977 | -0.469065329 | 0.51344953 | -0.9135568 | 0.36094975 | 0.99945981 |
| ABTB2 | 21.2544662 | -1.886082214 | 0.93256974 | -2.022457 | 0.04312916 | 0.89562588 |
| CAT | 34.6969021 | -0.578265463 | 0.68836718 | -0.8400538 | 0.40087823 | 0.99945981 |
| APIP | 12.325192 | 0.697817517 | 0.48410913 | 1.44144672 | 0.14945852 | 0.99945981 |
| PDHX | 18.6434621 | 0.26920459 | 0.34770798 | 0.77422609 | 0.4387971 | 0.99945981 |
| CD44 | 207.2445 | 0.099300846 | 0.32935087 | 0.30150473 | 0.76302964 | 0.99945981 |
| TRIM44 | 60.4650922 | 0.211647413 | 0.38059026 | 0.55610307 | 0.5781404 | 0.99945981 |
| LDLRAD3 | 35.0017938 | -0.180565552 | 0.69618005 | -0.2593662 | 0.79535273 | 0.99945981 |
| PRR5L | 31.7421878 | 0.207126055 | 0.5184899 | 0.39947944 | 0.68953997 | 0.99945981 |
| TRAF6 | 10.0176128 | 0.49531827 | 0.41055066 | 1.20647295 | 0.22763518 | 0.99945981 |
| API5 | 22.1935161 | -0.090452465 | 0.37364968 | -0.2420783 | 0.80871952 | 0.99945981 |
| TTC17 | 136.553433 | -0.017570001 | 0.23078062 | -0.0761329 | 0.93931335 | 0.99945981 |
| HSD17B12 | 54.8854018 | -0.53088142 | 0.38990849 | -1.3615539 | 0.17333872 | 0.99945981 |
| EXT2 | 22.5376479 | -0.120627824 | 0.31592396 | -0.3818255 | 0.7025908 | 0.99945981 |
| CD82 | 25.124635 | -1.788125482 | 0.73011768 | -2.4490921 | 0.01432168 | 0.72292893 |
| PHF21A | 181.669308 | -0.174405471 | 0.2526983 | -0.6901727 | 0.49008558 | 0.99945981 |
| DGKZ | 27.4847585 | 0.030151094 | 0.40524972 | 0.07440127 | 0.9406911 | 0.99945981 |
| AMBRA1 | 153.296796 | -0.149461183 | 0.33035458 | -0.4524266 | 0.65096172 | 0.99945981 |
| ATG13 | 18.6123364 | -0.134364578 | 0.46020846 | -0.2919646 | 0.77031369 | 0.99945981 |
| ARHGAP1 | 15.0052745 | 0.071880105 | 0.4657874 | 0.15431956 | 0.87735778 | 0.99945981 |
| CKAP5 | 25.6167062 | -0.482345278 | 0.34082278 | -1.4152378 | 0.15699878 | 0.99945981 |
| LRP4 | 17.2704205 | -0.627400527 | 0.60734422 | -1.033023 | 0.30159315 | 0.99945981 |
| C11orf49 | 37.3469915 | -0.54638852 | 0.41136207 | -1.3282423 | 0.18409806 | 0.99945981 |
| ARFGAP2 | 11.4744179 | -0.082645488 | 0.52406037 | -0.1577022 | 0.87469144 | 0.99945981 |
| DDB2 | 24.7312936 | -0.084630276 | 0.52339817 | -0.1616939 | 0.87154693 | 0.99945981 |
| MADD | 24.1366359 | -0.310852408 | 0.3404064 | -0.9131803 | 0.36114774 | 0.99945981 |
| SPI1 | 50.9852978 | 1.038220688 | 0.53641705 | 1.93547294 | 0.05293229 | 0.92445778 |
| CELF1 | 122.173295 | 0.223855629 | 0.20875746 | 1.07232396 | 0.28357455 | 0.99945981 |
| MTCH2 | 19.714644 | -0.134332698 | 0.41565411 | -0.3231839 | 0.74655601 | 0.99945981 |
| FNBP4 | 84.6356647 | -0.111795648 | 0.24141683 | -0.4630814 | 0.64330601 | 0.99945981 |
| NUP160 | 39.7182801 | -0.080965868 | 0.40239671 | -0.2012091 | 0.8405351 | 0.99945981 |
| PTPRJ | 294.113539 | 0.681512305 | 0.54559712 | 1.24911272 | 0.21162385 | 0.99945981 |
| SERPING1 | 14.6354721 | -2.169600483 | 0.72478721 | -2.993431 | 0.0027586 | 0.45570351 |
| ZDHHC5 | 18.3641163 | -0.388907541 | 0.43654514 | -0.8908759 | 0.37299576 | 0.99945981 |
| CTNND1 | 23.4366228 | 0.457795409 | 0.48608189 | 0.94180716 | 0.34629138 | 0.99945981 |
| LPXN | 40.6123798 | 0.099098856 | 0.33656017 | 0.29444618 | 0.76841699 | 0.99945981 |
| AP001636.3 | 13.1103054 | 1.292764548 | 1.04277766 | 1.23973173 | 0.21507464 | 0.99945981 |
| FAM111A | 21.6758835 | 0.055255856 | 0.32408231 | 0.17049945 | 0.86461737 | 0.99945981 |
| MPEG1 | 38.5443434 | 0.427331485 | 0.90453805 | 0.47243064 | 0.63661944 | 0.99945981 |
| OSBP | 23.172587 | -0.371349816 | 0.44182165 | -0.8404971 | 0.40062972 | 0.99945981 |
| PATL1 | 22.2276768 | 0.343518741 | 0.3338183 | 1.02905905 | 0.30345193 | 0.99945981 |
| STX3 | 15.6422278 | -0.094533303 | 0.46175155 | -0.2047276 | 0.83778493 | 0.99945981 |
| MS4A6A | 66.8576735 | -0.111248602 | 0.73125897 | -0.152133 | 0.87908206 | 0.99945981 |
| MS4A4E | 52.7640839 | -0.024740146 | 1.14588683 | -0.0215904 | 0.9827747 | 0.99945981 |
| MS4A4A | 37.458857 | 0.293215497 | 0.69156717 | 0.42398701 | 0.67157529 | 0.99945981 |
| MS4A14 | 10.9046405 | 0.893420115 | 0.67002239 | 1.33341831 | 0.18239457 | 0.99945981 |
| MS4A1 | 33.978681 | -3.211892172 | 1.54588983 | -2.077698 | 0.03773719 | 0.87920079 |
| PRPF19 | 7.87166271 | 0.032293406 | 0.50869431 | 0.06348293 | 0.94938195 | 0.99945981 |
| SLC15A3 | 11.853577 | 0.313166059 | 0.70315474 | 0.44537289 | 0.6560503 | 0.99945981 |
| CD6 | 26.4890846 | -0.546635083 | 0.44347989 | -1.232604 | 0.21772355 | 0.99945981 |
| CD5 | 6.86562219 | -0.891080363 | 0.70878497 | -1.2571942 | 0.20868332 | 0.99945981 |
| VPS37C | 9.76064455 | 0.47441864 | 0.63314876 | 0.74930043 | 0.45367615 | 0.99945981 |
| DDB1 | 17.6880753 | -0.402587716 | 0.34949697 | -1.1519062 | 0.24935963 | 0.99945981 |
| CPSF7 | 19.0315012 | -0.243282121 | 0.38243098 | -0.6361465 | 0.52468096 | 0.99945981 |
| AP003108.2 | 11.5819701 | -0.52760408 | 0.54299649 | -0.9716528 | 0.33122328 | 0.99945981 |
| TMEM258 | 9.79425577 | -0.263776351 | 0.79250848 | -0.3328373 | 0.73925713 | 0.99945981 |
| FADS1 | 10.2732568 | -1.386389405 | 0.50950119 | -2.7210719 | 0.00650706 | 0.52554078 |
| FTH1 | 499.540758 | -0.376711259 | 0.52849653 | -0.712798 | 0.47597077 | 0.99945981 |
| AHNAK | 98.7596863 | -0.184785432 | 0.36262888 | -0.5095717 | 0.61035152 | 0.99945981 |
| B3GAT3 | 10.2673002 | -0.492562227 | 0.49306433 | -0.9989817 | 0.31780357 | 0.99945981 |
| GANAB | 19.4530807 | -0.089797543 | 0.34089783 | -0.2634148 | 0.79223086 | 0.99945981 |
| UBXN1 | 16.5387861 | 0.173301936 | 0.45285628 | 0.3826864 | 0.7019523 | 0.99945981 |
| TTC9C | 10.6205146 | -0.396801456 | 0.40446702 | -0.9810477 | 0.3265692 | 0.99945981 |
| TMEM179B | 11.6243087 | -0.08947724 | 0.41358652 | -0.2163447 | 0.82871909 | 0.99945981 |
| NXF1 | 10.4085945 | 0.621018413 | 0.49757172 | 1.2480983 | 0.21199506 | 0.99945981 |
| STX5 | 20.4735842 | -0.153132065 | 0.36959497 | -0.414324 | 0.67863684 | 0.99945981 |
| SLC3A2 | 20.1730804 | -0.667847621 | 0.44601386 | -1.4973696 | 0.1342971 | 0.99945981 |
| RARRES3 | 22.3924387 | 0.38999274 | 0.75290048 | 0.51798711 | 0.60446727 | 0.99945981 |
| PLA2G16 | 10.0576013 | -1.097466189 | 0.61618832 | -1.7810565 | 0.07490323 | 0.97204281 |
| ATL3 | 22.4975343 | -0.370152646 | 0.47555911 | -0.7783526 | 0.4363612 | 0.99945981 |
| RTN3 | 63.6541245 | 0.122262085 | 0.36047372 | 0.33917059 | 0.73448122 | 0.99945981 |
| MARK2 | 56.3584417 | -0.045399751 | 0.25196154 | -0.1801852 | 0.85700715 | 0.99945981 |
| OTUB1 | 11.2100285 | -0.173593656 | 0.47155783 | -0.368128 | 0.71277777 | 0.99945981 |
| STIP1 | 16.6005842 | 0.84742252 | 0.36060531 | 2.35000009 | 0.01877341 | 0.76950676 |
| FERMT3 | 26.9116261 | 0.542076856 | 0.48943074 | 1.10756601 | 0.26804929 | 0.99945981 |
| VEGFB | 10.0400676 | -0.953712476 | 0.51556928 | -1.8498241 | 0.06433891 | 0.93776345 |
| FKBP2 | 9.4955837 | -0.394044605 | 0.58741483 | -0.6708115 | 0.50234064 | 0.99945981 |
| ESRRA | 16.3049069 | -0.296696568 | 0.54268158 | -0.5467231 | 0.58456897 | 0.99945981 |
| TRMT112 | 14.5915862 | -0.000158581 | 0.61244694 | -0.0002589 | 0.9997934 | 0.9997934 |
| PRDX5 | 11.6050058 | -0.219354399 | 0.50228173 | -0.4367159 | 0.66231742 | 0.99945981 |
| CCDC88B | 23.9033333 | -0.054111412 | 0.60321801 | -0.0897046 | 0.92852198 | 0.99945981 |
| RASGRP2 | 19.6070719 | 0.494654053 | 0.56888599 | 0.8695135 | 0.38456633 | 0.99945981 |
| SF1 | 54.9089694 | -0.174374589 | 0.31840156 | -0.5476562 | 0.58392799 | 0.99945981 |
| MAP4K2 | 10.4322647 | 0.053546099 | 0.41976074 | 0.12756338 | 0.89849451 | 0.99945981 |
| EHD1 | 16.6121386 | 0.612067778 | 0.40267174 | 1.52001673 | 0.12850677 | 0.99945981 |
| ATG2A | 9.72775461 | -1.369812491 | 0.60871164 | -2.2503471 | 0.02442692 | 0.80320673 |
| VPS51 | 19.0242325 | -0.602752441 | 0.36897461 | -1.6335879 | 0.10234542 | 0.99945981 |
| FAU | 67.4821751 | -0.070124067 | 0.61212121 | -0.1145591 | 0.90879458 | 0.99945981 |
| CAPN1 | 15.1309031 | -0.752392371 | 0.53652316 | -1.4023484 | 0.16081125 | 0.99945981 |
| POLA2 | 11.528863 | 0.053326923 | 0.43042217 | 0.12389446 | 0.90139884 | 0.99945981 |
| DPF2 | 11.9578444 | -0.384597218 | 0.40843126 | -0.9416449 | 0.34637447 | 0.99945981 |
| FRMD8 | 10.4129658 | -0.023823262 | 0.44913747 | -0.0530422 | 0.95769825 | 0.99945981 |
| NEAT1 | 1691.76238 | -0.621319798 | 0.47032286 | -1.3210495 | 0.18648485 | 0.99945981 |

| | | | | | | |
|---|---|---|---|---|---|---|
| MALAT1 | 12108.6361 | -0.378016645 | 0.28104734 | -1.3450284 | 0.17861608 | 0.99945981 |
| SCYL1 | 10.9542533 | -0.638386405 | 0.45809843 | -1.3935573 | 0.16345133 | 0.99945981 |
| LTBP3 | 9.30468755 | -0.093125671 | 0.52160805 | -0.1785357 | 0.85830227 | 0.99945981 |
| EHBP1L1 | 42.7486213 | 0.230362961 | 0.57146769 | 0.40310759 | 0.68686908 | 0.99945981 |
| MAP3K11 | 14.0575521 | 0.697022029 | 0.58414391 | 1.19323683 | 0.23277663 | 0.99945981 |
| SIPA1 | 17.924987 | 0.364922451 | 0.5215254 | 0.69972134 | 0.48410135 | 0.99945981 |
| RELA | 7.97516388 | 0.008938737 | 0.45462346 | 0.01966185 | 0.98431313 | 0.99945981 |
| CFL1 | 83.4393768 | -0.03914277 | 0.61775084 | -0.0633634 | 0.94947716 | 0.99945981 |
| CTSW | 9.29236555 | -0.051714227 | 0.65364977 | -0.0791161 | 0.93694027 | 0.99945981 |
| CCDC85B | 16.7700557 | -0.382600674 | 0.44987242 | -0.8504648 | 0.3950667 | 0.99945981 |
| DRAP1 | 12.7723837 | 0.497308575 | 0.57786079 | 0.86060273 | 0.38945688 | 0.99945981 |
| SART1 | 7.80204666 | -0.842086674 | 0.48497673 | -1.7363445 | 0.08250294 | 0.98529784 |
| BANF1 | 9.37711287 | -0.437667466 | 0.51615176 | -0.8479434 | 0.3964695 | 0.99945981 |
| SF3B2 | 21.6298707 | -0.06162219 | 0.39791088 | -0.1548643 | 0.8769283 | 0.99945981 |
| PACS1 | 160.583737 | -0.071169222 | 0.28641336 | -0.2484843 | 0.80375972 | 0.99945981 |
| RAB1B | 10.7567609 | -0.331567887 | 0.64670475 | -0.5127037 | 0.60815863 | 0.99945981 |
| ZDHHC24 | 7.92327442 | -0.85875535 | 0.50614922 | -1.6966446 | 0.08976387 | 0.99640647 |
| CCS | 11.2854579 | 0.115153873 | 0.41517431 | 0.27736271 | 0.78150161 | 0.99945981 |
| RBM14-RBM | 16.1253492 | 0.235401493 | 0.41371928 | 0.56898845 | 0.56936398 | 0.99945981 |
| RBM4 | 10.7382345 | -0.586590452 | 0.54648449 | -1.073389 | 0.28309661 | 0.99945981 |
| RBM4B | 7.70718361 | 0.126187478 | 0.59387795 | 0.21248049 | 0.8317322 | 0.99945981 |
| C11orf80 | 21.9683792 | -0.622634784 | 0.52185349 | -1.1931218 | 0.23282167 | 0.99945981 |
| KDM2A | 174.270427 | 0.025863551 | 0.18429549 | 0.14033741 | 0.88839341 | 0.99945981 |
| GRK2 | 36.5723665 | 0.385297683 | 0.31739511 | 1.21393706 | 0.22477176 | 0.99945981 |
| ANKRD13D | 43.570106 | 0.015322017 | 0.37087181 | 0.04131351 | 0.96704596 | 0.99945981 |
| RAD9A | 14.8322939 | 0.19326995 | 0.39849846 | 0.48499548 | 0.62767955 | 0.99945981 |
| PPP1CA | 19.5439482 | 0.134013276 | 0.58051324 | 0.23085309 | 0.81742893 | 0.99945981 |
| TBC1D10C | 16.6759739 | -0.047232259 | 0.48344342 | -0.0976997 | 0.92217078 | 0.99945981 |
| CORO1B | 9.48313667 | -1.090499739 | 0.52337405 | -2.0835953 | 0.03719699 | 0.87920079 |
| AIP | 8.61274135 | 0.319650542 | 0.53945756 | 0.59254066 | 0.5534886 | 0.99945981 |
| CDK2AP2 | 10.4012968 | -0.125008955 | 0.66386239 | -0.1883055 | 0.85063716 | 0.99945981 |
| GSTP1 | 42.8756514 | -0.304863477 | 0.60648719 | -0.5026709 | 0.61519565 | 0.99945981 |
| NDUFV1 | 12.4156296 | -0.105579112 | 0.39819433 | -0.2651447 | 0.790898 | 0.99945981 |
| UNC93B1 | 21.5889842 | 0.238381625 | 0.46916454 | 0.50809813 | 0.61138453 | 0.99945981 |
| NDUFS8 | 12.4057138 | -0.48422576 | 0.5910047 | -0.8193264 | 0.41260021 | 0.99945981 |
| TCIRG1 | 18.3910208 | 0.607823712 | 0.55874199 | 1.08784326 | 0.27666431 | 0.99945981 |
| CHKA | 31.9200147 | -0.096463299 | 0.42997584 | -0.2243459 | 0.8224882 | 0.99945981 |
| KMT5B | 38.9823755 | 0.227884196 | 0.28328347 | 0.80443872 | 0.42114365 | 0.99945981 |
| PPP6R3 | 146.542505 | 0.178277537 | 0.22070863 | 0.80775064 | 0.41923414 | 0.99945981 |
| CPT1A | 32.6326335 | 1.192552847 | 0.58737699 | 2.03030228 | 0.04232582 | 0.89562588 |
| TPCN2 | 12.6671235 | -0.428736429 | 0.62620867 | -0.6846543 | 0.49356212 | 0.99945981 |
| ORAOV1 | 8.55316742 | -0.625263672 | 0.49843315 | -1.2544584 | 0.20967543 | 0.99945981 |
| PPFIA1 | 53.3927642 | -0.060367482 | 0.25145127 | -0.2400763 | 0.81027113 | 0.99945981 |
| AP002336.2 | 20.1580518 | -0.292720628 | 0.49314832 | -0.5935752 | 0.55279625 | 0.99945981 |
| NADSYN1 | 15.4286949 | -0.084372138 | 0.37322113 | -0.2260647 | 0.82115107 | 0.99945981 |
| AP002495.2 | 11.9930823 | 0.138803428 | 0.42986086 | 0.32290316 | 0.74676859 | 0.99945981 |
| RNF121 | 12.7655376 | -0.355255447 | 0.48661479 | -0.7300548 | 0.46535671 | 0.99945981 |
| NUMA1 | 29.9715424 | 0.046472056 | 0.30303508 | 0.15335537 | 0.87811804 | 0.99945981 |
| LAMTOR1 | 19.5877714 | -0.690488366 | 0.38271856 | -1.8041675 | 0.07120506 | 0.95583904 |
| INPPL1 | 14.4128962 | -0.079688823 | 0.4984774 | -0.1598645 | 0.87298784 | 0.99945981 |
| CLPB | 12.2996516 | -0.403348084 | 0.54116969 | -0.7453265 | 0.45607439 | 0.99945981 |
| ARAP1 | 40.7165692 | 0.534014699 | 0.48679335 | 1.09700491 | 0.27263925 | 0.99945981 |
| ATG16L2 | 34.2024782 | 0.743052505 | 0.39661613 | 1.87348028 | 0.06100209 | 0.92445778 |
| FCHSD2 | 324.913026 | -0.283152578 | 0.35599156 | -0.7953913 | 0.42638594 | 0.99945981 |
| RELT | 14.1472544 | 0.378862666 | 0.44684866 | 0.84785454 | 0.39651898 | 0.99945981 |
| FAM168A | 69.9449134 | 0.30903852 | 0.34850252 | 0.88676121 | 0.37520748 | 0.99945981 |
| AP000763.3 | 9.4041461 | 0.611675641 | 0.72932926 | 0.83868243 | 0.40164754 | 0.99945981 |
| RAB6A | 56.4418228 | 0.07783791 | 0.2244847 | 0.34674039 | 0.72878637 | 0.99945981 |
| MRPL48 | 8.68202183 | 0.218090038 | 0.60607012 | 0.35984291 | 0.71896461 | 0.99945981 |
| COA4 | 8.82071172 | -0.754409031 | 0.66668997 | -1.131574 | 0.25781358 | 0.99945981 |
| PAAF1 | 10.5950507 | -0.220066349 | 0.48989144 | -0.4492145 | 0.65327691 | 0.99945981 |
| UCP2 | 22.6185426 | 0.051633839 | 0.67629985 | 0.07634755 | 0.93914259 | 0.99945981 |
| C2CD3 | 23.9751028 | -0.376873182 | 0.32274789 | -1.1677015 | 0.2429272 | 0.99945981 |
| PPME1 | 14.6544365 | -0.679519349 | 0.43441471 | -1.5642181 | 0.11776636 | 0.99945981 |
| PGM2L1 | 10.4933189 | 0.260297303 | 0.50759789 | 0.51280218 | 0.60808971 | 0.99945981 |
| POLD3 | 13.6440804 | 0.407036434 | 0.41325566 | 0.98495067 | 0.3246483 | 0.99945981 |
| RNF169 | 37.7808124 | 0.090820811 | 0.26014241 | 0.34911959 | 0.72699953 | 0.99945981 |
| XRRA1 | 29.846073 | 0.574113922 | 0.68102465 | 0.84301489 | 0.39922011 | 0.99945981 |
| SPCS2 | 33.153687 | -0.361044526 | 0.48601369 | -0.742869 | 0.45756097 | 0.99945981 |
| NEU3 | 9.78717557 | -0.290552321 | 0.53829048 | -0.5397686 | 0.5893566 | 0.99945981 |
| SLCO2B1 | 71.7884739 | -0.795489546 | 0.5827302 | -1.3651078 | 0.17221917 | 0.99945981 |
| ARRB1 | 59.167821 | 1.076334401 | 0.59937545 | 1.7957599 | 0.07253271 | 0.96092152 |
| RPS3 | 139.428798 | -0.378893508 | 0.60787836 | -0.6233048 | 0.53308424 | 0.99945981 |
| UVRAG | 133.345136 | 0.47656928 | 0.29505556 | 1.61518486 | 0.10627067 | 0.99945981 |
| THAP12 | 10.8008962 | 0.298104574 | 0.48944971 | 0.60906068 | 0.54248422 | 0.99945981 |
| EMSY | 40.6276194 | -0.215102325 | 0.34081323 | -0.6311443 | 0.52794618 | 0.99945981 |
| ACER3 | 81.6699412 | 0.245728606 | 0.40716138 | 0.60351649 | 0.54616515 | 0.99945981 |
| PAK1 | 39.1553995 | 0.775003939 | 0.55058797 | 1.4075933 | 0.15925153 | 0.99945981 |
| CLNS1A | 22.780045 | -0.510525245 | 0.40920182 | -1.2476124 | 0.21217305 | 0.99945981 |
| RSF1 | 90.1830556 | 0.08377212 | 0.25026111 | 0.33473886 | 0.73782208 | 0.99945981 |
| AAMDC | 9.89970576 | -0.07216789 | 0.49028465 | -0.1471959 | 0.8829774 | 0.99945981 |
| INTS4 | 27.2630868 | 0.398851135 | 0.28390051 | 1.40489756 | 0.16005174 | 0.99945981 |
| GAB2 | 155.891101 | 0.046502629 | 0.49214442 | 0.0944898 | 0.92472008 | 0.99945981 |
| AP003086.2 | 43.3188119 | 0.248675109 | 0.66619829 | 0.37327492 | 0.70894384 | 0.99945981 |
| NARS2 | 21.9911029 | 0.138244679 | 0.3737376 | 0.3698977 | 0.71145871 | 0.99945981 |
| TENM4 | 17.5842809 | -0.397519988 | 1.05186454 | -0.3779194 | 0.70549049 | 0.99945981 |
| PRCP | 20.3217855 | 0.572044467 | 0.50099725 | 1.1418116 | 0.25353234 | 0.99945981 |
| RAB30 | 29.9612445 | -0.274436261 | 0.68253154 | -0.4020858 | 0.68762086 | 0.99945981 |
| RAB30-AS1 | 19.5403165 | 0.025996522 | 0.45351752 | 0.05732198 | 0.95428871 | 0.99945981 |
| PCF11 | 43.4969468 | 0.407523373 | 0.44556946 | 0.91461245 | 0.36039512 | 0.99945981 |
| DLG2 | 35.7681781 | -0.571968224 | 0.8949041 | -0.6391391 | 0.52273243 | 0.99945981 |
| CREBZF | 23.1887814 | 0.302633066 | 0.4760438 | 0.63572525 | 0.52495551 | 0.99945981 |
| SYTL2 | 45.5755952 | -0.789977002 | 0.66653929 | -1.185192 | 0.23594156 | 0.99945981 |
| PICALM | 350.92711 | 0.445181845 | 0.24639663 | 1.80676922 | 0.07079827 | 0.95505283 |
| EED | 15.4128711 | 0.469553031 | 0.43367974 | 1.0827184 | 0.27893343 | 0.99945981 |
| HIKESHI | 13.3880376 | -0.399636977 | 0.44337572 | -0.9013506 | 0.36740193 | 0.99945981 |
| TMEM135 | 49.1992175 | -0.108373811 | 0.3529761 | -0.3070288 | 0.75882149 | 0.99945981 |
| CTSC | 59.9893883 | 0.029821104 | 0.36640198 | 0.08138904 | 0.93513257 | 0.99945981 |
| CHORDC1 | 20.075711 | 0.115947286 | 0.56460692 | 0.20535931 | 0.83729141 | 0.99945981 |
| SLC36A4 | 38.3464049 | 0.215475532 | 0.43251228 | 0.49819518 | 0.61834648 | 0.99945981 |
| SMCO4 | 15.9645985 | 1.049129078 | 0.6735425 | 1.55762863 | 0.11932131 | 0.99945981 |
| CEP295 | 19.2971279 | -0.000362447 | 0.48737343 | -0.0007437 | 0.99940663 | 0.9997934 |
| TAF1D | 51.2289843 | -0.51813119 | 0.34149747 | -1.5172329 | 0.1292079 | 0.99945981 |
| MRE11 | 23.3556316 | 0.29016397 | 0.37442524 | 0.77495836 | 0.43836426 | 0.99945981 |
| CWC15 | 9.47985271 | -0.018471229 | 0.437788 | -0.0421922 | 0.9663455 | 0.99945981 |
| AP000787.1 | 11.0892177 | -0.272075846 | 0.76639684 | -0.3550065 | 0.72258473 | 0.99945981 |
| SESN3 | 50.2623685 | 0.235210883 | 0.41329987 | 0.56910467 | 0.56928511 | 0.99945981 |
| FAM76B | 12.1179881 | 0.19714657 | 0.44975526 | 0.43834189 | 0.66113846 | 0.99945981 |
| CEP57 | 27.8350108 | 0.044439107 | 0.3879791 | 0.11453995 | 0.90880978 | 0.99945981 |
| MTMR2 | 14.1701674 | -0.605304978 | 0.51469823 | -1.1760386 | 0.23957946 | 0.99945981 |
| MAML2 | 350.659079 | 0.551017838 | 0.38777937 | 1.42095707 | 0.15532924 | 0.99945981 |
| CCDC82 | 21.2164481 | -0.35713931 | 0.35385259 | -1.0092884 | 0.31283635 | 0.99945981 |
| BIRC3 | 44.6760079 | -0.127459326 | 0.57876891 | -0.2202249 | 0.825696 | 0.99945981 |

| | | | | | | |
|---|---|---|---|---|---|---|
| BIRC2 | 19.7178892 | -0.177292446 | 0.36681459 | -0.4833299 | 0.62886154 | 0.99945981 |
| TMEM123 | 50.8638144 | 0.263792666 | 0.34378661 | 0.76731513 | 0.44289417 | 0.99945981 |
| DCUN1D5 | 10.1929041 | -0.636176858 | 0.53342203 | -1.1926333 | 0.23301303 | 0.99945981 |
| DYNC2H1 | 12.4412912 | 0.316160279 | 0.59206751 | 0.53399363 | 0.59334594 | 0.99945981 |
| PDGFD | 33.5319131 | -0.483882549 | 0.60921561 | -0.7942714 | 0.42703743 | 0.99945981 |
| CASP4 | 31.0199322 | 0.234913012 | 0.30959356 | 0.75877874 | 0.44798492 | 0.99945981 |
| CASP1 | 17.3579447 | 1.051597584 | 0.45919759 | 2.29007643 | 0.02201689 | 0.79988354 |
| AASDHPPT | 11.985012 | 0.529020401 | 0.47912109 | 1.10414759 | 0.26952912 | 0.99945981 |
| CWF19L2 | 36.5465527 | 0.201548351 | 0.32716743 | 0.61604038 | 0.53786787 | 0.99945981 |
| CUL5 | 58.0773256 | -0.120764527 | 0.24470941 | -0.4935018 | 0.6216581 | 0.99945981 |
| ACAT1 | 17.2346972 | -0.406314259 | 0.46325944 | -0.8770771 | 0.38044475 | 0.99945981 |
| NPAT | 29.2063056 | 0.162193596 | 0.31002141 | 0.52316902 | 0.60085664 | 0.99945981 |
| ATM | 186.706374 | 0.363160401 | 0.31839144 | 1.14060981 | 0.25403233 | 0.99945981 |
| C11orf65 | 13.1129499 | 0.189524258 | 0.88137845 | 0.21503164 | 0.82974265 | 0.99945981 |
| DDX10 | 44.7014399 | 0.407272056 | 0.33063423 | 1.23179035 | 0.21802741 | 0.99945981 |
| RDX | 34.8593137 | 0.175943361 | 0.39386896 | 0.44670532 | 0.65508784 | 0.99945981 |
| FDX1 | 15.643731 | -0.619472508 | 0.51840263 | -1.194964 | 0.23210108 | 0.99945981 |
| SIK2 | 53.7658701 | 0.215817969 | 0.34999651 | 0.61662891 | 0.53747952 | 0.99945981 |
| PPP2R1B | 11.1300689 | 0.339774298 | 0.42782552 | 0.79418895 | 0.42708544 | 0.99945981 |
| CRYAB | 93.9784124 | -2.524224199 | 1.12153166 | -2.2506937 | 0.02440495 | 0.80320673 |
| IL18 | 28.1502284 | 0.007670068 | 0.46086394 | 0.0166428 | 0.98672158 | 0.99945981 |
| PTS | 15.1179416 | -0.981588263 | 0.57529185 | -1.706244 | 0.08796266 | 0.99640647 |
| NCAM1 | 25.5859209 | -0.059829243 | 0.76898071 | -0.0778033 | 0.93798452 | 0.99945981 |
| USP28 | 22.7241015 | 0.04847228 | 0.44300999 | 0.10941577 | 0.91287272 | 0.99945981 |
| ZBTB16 | 60.0290831 | -1.019960761 | 0.58920749 | -1.7310723 | 0.08343887 | 0.9868985 |
| NNMT | 16.1230424 | -4.503227139 | 1.0683186 | -4.2152473 | 2.50E-05 | 0.03425698 |
| CADM1 | 44.4330697 | -0.410137402 | 0.44584577 | -0.9199087 | 0.35762047 | 0.99945981 |
| SIK3 | 308.97692 | 0.36051133 | 0.26895263 | 1.34042684 | 0.18010661 | 0.99945981 |
| PAFAH1B2 | 27.2003625 | -0.043737479 | 0.37326861 | -0.1171743 | 0.90672195 | 0.99945981 |
| SIDT2 | 21.5709404 | 0.193294515 | 0.63598658 | 0.30392861 | 0.76118228 | 0.99945981 |
| TAGLN | 14.2946133 | 0.511115532 | 0.63075529 | 0.81032301 | 0.41775455 | 0.99945981 |
| PCSK7 | 56.3707804 | -0.073237725 | 0.21124897 | -0.3466891 | 0.72882487 | 0.99945981 |
| RNF214 | 13.2025623 | -0.360521561 | 0.4061884 | -0.8875723 | 0.37477088 | 0.99945981 |
| CEP164 | 9.23208195 | 0.366688592 | 0.44853859 | 0.81751849 | 0.41363219 | 0.99945981 |
| IL10RA | 61.3747827 | 0.007748585 | 0.32965071 | 0.02350544 | 0.9812471 | 0.99945981 |
| JAML | 43.8286055 | 0.238796416 | 0.73181862 | 0.32630546 | 0.74419325 | 0.99945981 |
| MPZL3 | 10.9773518 | -0.101298529 | 0.51229434 | -0.197735 | 0.84325239 | 0.99945981 |
| CD3E | 17.6212111 | -0.2417104 | 0.49928093 | -0.484117 | 0.62830281 | 0.99945981 |
| CD3D | 14.7856099 | -0.998196393 | 0.76263886 | -1.3088717 | 0.19057783 | 0.99945981 |
| CD3G | 16.3484636 | -0.325254617 | 0.55761477 | -0.5832963 | 0.55969387 | 0.99945981 |
| UBE4A | 35.7510062 | 0.193888101 | 0.29153875 | 0.66505087 | 0.50601796 | 0.99945981 |
| ATP5MG | 34.0542136 | 0.053801468 | 0.42825464 | 0.12562962 | 0.90002512 | 0.99945981 |
| KMT2A | 84.3572191 | -0.155046727 | 0.22319808 | -0.6946598 | 0.48726856 | 0.99945981 |
| ARCN1 | 16.548551 | -0.055366465 | 0.46568373 | -0.1188928 | 0.90536025 | 0.99945981 |
| AP002954.1 | 8.80365072 | -0.494193494 | 0.75264064 | -0.6566128 | 0.5114299 | 0.99945981 |
| DDX6 | 65.6513929 | -0.238373288 | 0.2270506 | -1.0498686 | 0.29377854 | 0.99945981 |
| BCL9L | 11.5104924 | 0.698797639 | 0.42728975 | 1.63541867 | 0.10196131 | 0.99945981 |
| CCDC84 | 25.9808698 | 0.096068111 | 0.33879154 | 0.28356113 | 0.77674673 | 0.99945981 |
| RPS25 | 76.8717382 | -0.278301202 | 0.52639701 | -0.5286907 | 0.59702003 | 0.99945981 |
| CBL | 89.1790442 | 0.780411579 | 0.23217364 | 3.3613272 | 0.00077569 | 0.27121928 |
| ARHGEF12 | 40.05255 | -0.313263144 | 0.29947513 | -1.0460406 | 0.29554229 | 0.99945981 |
| SORL1 | 118.954679 | 0.278125833 | 0.31515271 | 0.88251131 | 0.37750037 | 0.99945981 |
| UBASH3B | 116.756925 | -0.69592927 | 0.78639477 | -0.8849617 | 0.37617727 | 0.99945981 |
| HSPA8 | 41.130633 | 0.062233188 | 0.3736934 | 0.16653542 | 0.86773561 | 0.99945981 |
| GRAMD1B | 42.3179731 | -0.163933137 | 0.60989924 | -0.2687872 | 0.78809341 | 0.99945981 |
| TBRG1 | 24.7498811 | -0.009629122 | 0.38286914 | -0.0251499 | 0.9799354 | 0.99945981 |
| MSANTD2 | 12.7598694 | 0.039565621 | 0.41391 | 0.09558991 | 0.92384628 | 0.99945981 |
| EI24 | 9.21788688 | 0.506531606 | 0.55885475 | 0.90637435 | 0.36473775 | 0.99945981 |
| STT3A | 11.7583988 | -0.014450019 | 0.49118764 | -0.0294185 | 0.97653079 | 0.99945981 |
| FAM118B | 14.3298403 | -0.234414803 | 0.44500539 | -0.5267685 | 0.59835439 | 0.99945981 |
| SRPRA | 11.3959305 | -0.008087333 | 0.50633385 | -0.0159723 | 0.98725646 | 0.99945981 |
| ETS1 | 132.915534 | -0.625382411 | 0.35157697 | -1.7787923 | 0.07527382 | 0.97500897 |
| FLI1 | 176.741366 | 0.348169622 | 0.31265618 | 1.11358626 | 0.26545672 | 0.99945981 |
| ARHGAP32 | 24.8586122 | 0.49127382 | 0.47800091 | 1.02776753 | 0.30405919 | 0.99945981 |
| NFRKB | 10.0371195 | -0.194913591 | 0.49570088 | -0.3932081 | 0.6941658 | 0.99945981 |
| PRDM10 | 15.1580069 | 0.376094803 | 0.48345372 | 0.77793341 | 0.43660827 | 0.99945981 |
| APLP2 | 75.3007451 | 0.294769915 | 0.29501804 | 0.99915896 | 0.31771769 | 0.99945981 |
| ST14 | 12.4959964 | 0.155137033 | 0.6026395 | 0.25742925 | 0.79684743 | 0.99945981 |
| ZBTB44 | 40.6513137 | 0.484037649 | 0.33690064 | 1.43673709 | 0.15079271 | 0.99945981 |
| SNX19 | 24.0268598 | 0.356611289 | 0.50584854 | 0.70497642 | 0.48082491 | 0.99945981 |
| NCAPD3 | 13.6382067 | 0.053639819 | 0.43929712 | 0.12210373 | 0.90281686 | 0.99945981 |
| VPS26B | 9.07995806 | -0.242997661 | 0.43632561 | -0.5569182 | 0.57758334 | 0.99945981 |
| ZMYND11 | 45.5367965 | -0.286286795 | 0.2834184 | -1.0101207 | 0.31243745 | 0.99945981 |
| DIP2C | 14.7731663 | -0.289752491 | 0.37348549 | -0.7758066 | 0.43786321 | 0.99945981 |
| LARP4B | 102.61368 | 0.214930232 | 0.22869897 | 0.93979536 | 0.34732254 | 0.99945981 |
| GTPBP4 | 11.2061372 | -0.724904585 | 0.48728733 | -1.4876327 | 0.13684777 | 0.99945981 |
| IDI1 | 13.7241967 | -0.431235004 | 0.61290324 | -0.7035939 | 0.48168569 | 0.99945981 |
| WDR37 | 55.7102626 | 0.088558203 | 0.28759945 | 0.30792202 | 0.75814167 | 0.99945981 |
| PFKP | 13.2334267 | -0.939013844 | 0.4287306 | -2.1902189 | 0.02850837 | 0.84763724 |
| PITRM1 | 10.2206173 | 0.191372548 | 0.48143557 | 0.39750396 | 0.69099586 | 0.99945981 |
| KLF6 | 74.2693303 | 0.136420619 | 0.43942645 | 0.31045154 | 0.75621761 | 0.99945981 |
| FAM208B | 77.4620944 | 0.209891687 | 0.24246617 | 0.86565348 | 0.38668022 | 0.99945981 |
| GDI2 | 71.9797383 | -0.592567043 | 0.52220341 | -1.1347437 | 0.25648271 | 0.99945981 |
| FBH1 | 23.2572161 | 0.33240576 | 0.41699508 | 0.79714552 | 0.42536652 | 0.99945981 |
| IL15RA | 11.0545694 | -0.254275039 | 0.56859038 | -0.4472025 | 0.65472886 | 0.99945981 |
| IL2RA | 28.9366188 | -2.632801588 | 0.94849331 | -2.7757724 | 0.00550707 | 0.5234882 |
| RBM17 | 25.1593702 | -0.39566316 | 0.37826291 | -1.0460004 | 0.29556084 | 0.99945981 |
| PFKFB3 | 89.7685199 | 0.080921765 | 0.65807804 | 0.12296682 | 0.90213337 | 0.99945981 |
| AL137145.2 | 9.39325928 | -1.543400623 | 0.64956705 | -2.3760451 | 0.01749932 | 0.74616677 |
| PRKCQ | 70.0020341 | -0.452464847 | 0.34925997 | -1.2954958 | 0.19514923 | 0.99945981 |
| SFMBT2 | 188.661269 | 0.352761072 | 0.52144795 | 0.67650294 | 0.49872138 | 0.99945981 |
| KIN | 13.4377174 | 0.746910717 | 0.39199477 | 1.90540992 | 0.05672681 | 0.92445778 |
| ATP5F1C | 22.7048773 | -0.135360203 | 0.3427406 | -0.3949348 | 0.69289098 | 0.99945981 |
| TAF3 | 58.0617016 | -0.185791301 | 0.31206465 | -0.5953616 | 0.55160181 | 0.99945981 |
| GATA3 | 10.5882197 | 0.442017534 | 0.54764323 | 0.80712681 | 0.41959343 | 0.99945981 |
| CELF2 | 435.950668 | 0.008569504 | 0.23705283 | 0.03615019 | 0.97116261 | 0.99945981 |
| USP6NL | 43.3649396 | -0.841451982 | 0.7234204 | -1.1631577 | 0.24476554 | 0.99945981 |
| UPF2 | 49.1434301 | -0.121312758 | 0.28653361 | -0.4233806 | 0.67201763 | 0.99945981 |
| DHTKD1 | 17.2483888 | -0.114420363 | 0.47037461 | -0.2432537 | 0.80780887 | 0.99945981 |
| NUDT5 | 13.8840535 | -0.172904953 | 0.39255161 | -0.4404643 | 0.6596009 | 0.99945981 |
| CDC123 | 28.4608776 | -0.707106623 | 0.4177006 | -1.6928552 | 0.09048304 | 0.99640647 |
| CAMK1D | 209.443803 | -0.302931723 | 0.54857667 | -0.552214 | 0.58080174 | 0.99945981 |
| OPTN | 21.3218233 | -0.207991187 | 0.33270085 | -0.6251598 | 0.5318662 | 0.99945981 |
| AL157392.5 | 10.6404755 | -0.166990582 | 0.41998088 | -0.3976147 | 0.69091421 | 0.99945981 |
| FRMD4A | 118.553706 | -0.190566294 | 0.61322672 | -0.3107599 | 0.75598313 | 0.99945981 |
| FAM107B | 100.158049 | -0.139381352 | 0.51320387 | -0.2715906 | 0.78593682 | 0.99945981 |
| DCLRE1C | 21.8903987 | -0.185855721 | 0.49562297 | -0.3749942 | 0.70766481 | 0.99945981 |
| RPP38 | 12.1725558 | 0.083760081 | 0.43344855 | 0.19324111 | 0.84677014 | 0.99945981 |
| NMT2 | 12.5843786 | 0.319819714 | 0.48925187 | 0.65369134 | 0.51331068 | 0.99945981 |
| MINDY3 | 38.329643 | 0.034283896 | 0.29856601 | 0.11482853 | 0.90858104 | 0.99945981 |
| PTER | 15.2681099 | 0.175571734 | 0.36698839 | 0.47841223 | 0.63235683 | 0.99945981 |
| RSU1 | 57.5058743 | 0.301625971 | 0.2315064 | 1.30288394 | 0.19261439 | 0.99945981 |
| CUBN | 23.6484157 | 1.19496961 | 0.57576695 | 2.07543976 | 0.0379458 | 0.87920079 |

| | | | | | | |
|---|---|---|---|---|---|---|
| VIM-AS1 | 10.326345 | 0.565254194 | 0.53113584 | 1.06423659 | 0.28722154 | 0.99945981 |
| VIM | 141.514884 | -0.589002144 | 0.56855115 | -1.0359704 | 0.30021597 | 0.99945981 |
| ST8SIA6 | 33.1495335 | -1.583180931 | 1.01050476 | -1.5667229 | 0.11717948 | 0.99945981 |
| STAM | 31.2304531 | -0.102297807 | 0.4067246 | -0.2515162 | 0.80141507 | 0.99945981 |
| MRC1 | 131.571996 | 0.349314199 | 1.08744541 | 0.32122458 | 0.7480402 | 0.99945981 |
| NSUN6 | 22.1045114 | -0.398238473 | 0.37934656 | -1.0498012 | 0.29380953 | 0.99945981 |
| ARL5B | 8.27088033 | -0.17358627 | 0.51817641 | -0.3349945 | 0.7376292 | 0.99945981 |
| PLXDC2 | 420.925667 | 0.555486072 | 0.5873666 | 0.94572295 | 0.34428991 | 0.99945981 |
| NEBL | 10.7422982 | -1.5624682 | 0.93844993 | -1.6649457 | 0.09592359 | 0.99945981 |
| MLLT10 | 104.231134 | 0.076322773 | 0.22005406 | 0.34683646 | 0.72871418 | 0.99945981 |
| DNAJC1 | 101.42457 | -0.052857715 | 0.40288578 | -0.1311978 | 0.89561886 | 0.99945981 |
| PIP4K2A | 308.018225 | -0.03297287 | 0.24174076 | -0.1363976 | 0.89150694 | 0.99945981 |
| ARHGAP21 | 52.7660543 | 0.287920077 | 0.35763531 | 0.80506613 | 0.42078152 | 0.99945981 |
| APBB1IP | 207.400496 | -0.072774934 | 0.28830715 | -0.2524215 | 0.80071526 | 0.99945981 |
| ABI1 | 109.60893 | 0.376163864 | 0.1875188 | 2.00600615 | 0.04485559 | 0.90039076 |
| ANKRD26 | 17.9424312 | 0.215518627 | 0.45626318 | 0.47235595 | 0.63667274 | 0.99945981 |
| YME1L1 | 45.4112294 | -0.083367122 | 0.27243953 | -0.3060023 | 0.7596029 | 0.99945981 |
| ACBD5 | 18.6874283 | 0.175680368 | 0.42686613 | 0.41155846 | 0.68066308 | 0.99945981 |
| RAB18 | 35.3442509 | 0.072797049 | 0.36762764 | 0.19801843 | 0.84303064 | 0.99945981 |
| MPP7 | 44.2739824 | 0.191033356 | 0.4906197 | 0.38937155 | 0.69700131 | 0.99945981 |
| WAC-AS1 | 12.722039 | -0.200174014 | 0.53837979 | -0.3718082 | 0.71003567 | 0.99945981 |
| WAC | 179.596476 | 0.521191337 | 0.18358644 | 2.83894248 | 0.00452633 | 0.50757754 |
| SVIL | 64.164446 | 1.194288991 | 0.54954313 | 2.17323979 | 0.02976227 | 0.85472619 |
| MAP3K8 | 73.0512653 | -0.454556127 | 0.3568431 | -1.2738263 | 0.20272498 | 0.99945981 |
| ZNF438 | 68.6054661 | 0.058180607 | 0.44687414 | 0.13019462 | 0.89641245 | 0.99945981 |
| ZEB1 | 92.0983966 | -0.481135352 | 0.3838663 | -1.2533931 | 0.21006269 | 0.99945981 |
| ARHGAP12 | 50.4914727 | 0.151660324 | 0.37608651 | 0.40325914 | 0.6867576 | 0.99945981 |
| KIF5B | 55.4798384 | 0.194039075 | 0.28212418 | 0.68777896 | 0.49159198 | 0.99945981 |
| EPC1 | 102.840349 | -0.100402519 | 0.20226774 | -0.4963842 | 0.61962334 | 0.99945981 |
| CCDC7 | 31.6799451 | -0.079450203 | 0.53378392 | -0.1488434 | 0.88167722 | 0.99945981 |
| ITGB1 | 74.5054367 | -0.011802306 | 0.27584334 | -0.0427863 | 0.96587191 | 0.99945981 |
| NRP1 | 79.6093975 | -0.016149172 | 0.58957459 | -0.0273912 | 0.9781477 | 0.99945981 |
| PARD3 | 28.6976012 | 0.114870905 | 0.69373444 | 0.1655834 | 0.86848481 | 0.99945981 |
| CUL2 | 25.1339977 | -0.053016837 | 0.30986853 | -0.1710946 | 0.86414937 | 0.99945981 |
| CREM | 82.3466407 | -0.141627396 | 0.76550864 | -0.1850108 | 0.85322053 | 0.99945981 |
| CCNY | 145.887266 | 0.245125838 | 0.25770947 | 0.95117123 | 0.34151746 | 0.99945981 |
| ZNF248 | 13.6881126 | -0.248250519 | 0.44279578 | -0.5606434 | 0.57504068 | 0.99945981 |
| ZNF33A | 58.7853583 | -0.054825647 | 0.29012963 | -0.1889695 | 0.85011673 | 0.99945981 |
| ZNF33B | 21.6676286 | -0.121530819 | 0.47048474 | -0.2583098 | 0.79616782 | 0.99945981 |
| BMS1 | 13.3197247 | -0.260871923 | 0.5286728 | -0.4934468 | 0.62169689 | 0.99945981 |
| CSGALNACT2 | 68.6381157 | 0.665031764 | 0.29495243 | 2.25470854 | 0.02415163 | 0.80320673 |
| HNRNPF | 35.0198857 | -0.369466805 | 0.30669289 | -1.20468 | 0.22832685 | 0.99945981 |
| RASSF4 | 33.6142596 | 0.086017804 | 0.74845321 | 0.11492743 | 0.90850264 | 0.99945981 |
| ALOX5 | 60.3776068 | 0.30196514 | 0.61091223 | 0.49428563 | 0.62110447 | 0.99945981 |
| 8-Mar | 32.0063234 | -0.076984414 | 0.3618587 | -0.2127472 | 0.83152417 | 0.99945981 |
| ZFAND4 | 10.7714275 | -0.148280674 | 0.44241689 | -0.3351605 | 0.737504 | 0.99945981 |
| WASHC2C | 19.1293139 | -0.053618905 | 0.44866412 | -0.1195079 | 0.90487298 | 0.99945981 |
| AGAP4 | 10.0323273 | 0.193832061 | 0.49548218 | 0.39119885 | 0.69565026 | 0.99945981 |
| TIMM23 | 12.0832711 | 0.817958826 | 0.48491621 | 1.68680445 | 0.09164096 | 0.99640647 |
| NCOA4 | 28.4138755 | 0.148371452 | 0.40459351 | 0.36671733 | 0.71382988 | 0.99945981 |
| MAPK8 | 58.8193422 | -0.032580009 | 0.34926228 | -0.0932824 | 0.92567925 | 0.99945981 |
| ARHGAP22 | 27.7392377 | 1.073011576 | 0.49843202 | 2.15277417 | 0.03133643 | 0.86065765 |
| WDFY4 | 81.0409948 | -1.089135344 | 0.7347106 | -1.4824005 | 0.13823377 | 0.99945981 |
| TMEM273 | 9.85939077 | -1.151157806 | 0.63619397 | -1.8094447 | 0.07038195 | 0.95505283 |
| ERCC6 | 15.4937147 | -0.806258502 | 0.48612616 | -1.6585376 | 0.09720901 | 0.99945981 |
| PARG | 40.3420588 | -0.132250893 | 0.33614485 | -0.3934342 | 0.69399878 | 0.99945981 |
| TIMM23B | 40.753842 | 0.033383407 | 0.4695298 | 0.07109966 | 0.94331844 | 0.99945981 |
| AGAP6 | 12.3334174 | -0.464771987 | 0.42627416 | -1.0903124 | 0.27557557 | 0.99945981 |
| WASHC2A | 9.56558904 | -0.114587432 | 0.53474047 | -0.2142861 | 0.83032398 | 0.99945981 |
| SGMS1 | 121.523842 | 0.175255548 | 0.26372333 | 0.66454322 | 0.5063427 | 0.99945981 |
| PRKG1 | 11.934853 | 0.60267964 | 0.45858785 | 1.31420761 | 0.18877635 | 0.99945981 |
| AC074327.1 | 44.6230582 | -2.719021764 | 1.03235684 | -2.6338003 | 0.00844351 | 0.58811673 |
| IPMK | 34.1374706 | 0.304171766 | 0.31536623 | 0.96450327 | 0.33479367 | 0.99945981 |
| CISD1 | 10.0235869 | -0.005648684 | 0.59780546 | -0.009449 | 0.99246087 | 0.99945981 |
| UBE2D1 | 18.4198254 | 0.990546723 | 0.46496371 | 2.13037426 | 0.03314073 | 0.8663391 |
| BICC1 | 20.6266598 | 1.001284877 | 0.51845344 | 1.93129181 | 0.05344698 | 0.92445778 |
| CCDC6 | 41.2822883 | -0.973757113 | 0.39268213 | -2.4797592 | 0.01314712 | 0.69426883 |
| ANK3 | 179.607043 | -0.032123157 | 0.58072853 | -0.0553153 | 0.95588729 | 0.99945981 |
| ARID5B | 198.788429 | -0.498610621 | 0.29056095 | -1.7160276 | 0.08615698 | 0.99204414 |
| NRBF2 | 10.0983257 | -0.905320959 | 0.51523284 | -1.7571104 | 0.078899 | 0.98446503 |
| JMJD1C | 269.693332 | 0.212868839 | 0.26770437 | 0.79516386 | 0.42651819 | 0.99945981 |
| REEP3 | 28.1862459 | 0.03838731 | 0.32230495 | 0.11910245 | 0.90519419 | 0.99945981 |
| SIRT1 | 19.4440563 | -0.12457392 | 0.39105462 | -0.3185589 | 0.75006104 | 0.99945981 |
| HERC4 | 158.567893 | 0.132929608 | 0.24753321 | 0.53701726 | 0.59125569 | 0.99945981 |
| HNRNPH3 | 41.6189852 | 0.554071234 | 0.35403914 | 1.56499993 | 0.11758293 | 0.99945981 |
| RUFY2 | 19.0604874 | 0.528402836 | 0.3236236 | 1.6327698 | 0.10251742 | 0.99945981 |
| SLC25A16 | 19.911504 | -0.183222111 | 0.40678368 | -0.4504166 | 0.6524101 | 0.99945981 |
| CCAR1 | 52.7462732 | -0.151274755 | 0.36694433 | -0.4122553 | 0.68015229 | 0.99945981 |
| DDX50 | 19.3240424 | -1.057276825 | 0.36874411 | -2.8672372 | 0.00414073 | 0.507383 |
| DDX21 | 29.5069101 | -0.205091808 | 0.33167486 | -0.618352 | 0.53634334 | 0.99945981 |
| SRGN | 340.429049 | 1.096882769 | 0.60662279 | 1.8081793 | 0.0705786 | 0.95505283 |
| VPS26A | 42.0185327 | 0.329793636 | 0.29800154 | 1.10668433 | 0.26843043 | 0.99945981 |
| SUPV3L1 | 9.93943824 | -0.081594788 | 0.64167242 | -0.1271596 | 0.89881411 | 0.99945981 |
| HK1 | 48.7012756 | 0.539767455 | 0.41535936 | 1.29951918 | 0.19376582 | 0.99945981 |
| SAR1A | 20.5338593 | 0.659942433 | 0.3746071 | 1.76169228 | 0.0781213 | 0.98289661 |
| PPA1 | 19.9177488 | -0.842679976 | 0.56397568 | -1.4941779 | 0.13512914 | 0.99945981 |
| EIF4EBP2 | 20.5168478 | -0.105344946 | 0.37561159 | -0.2804625 | 0.77912273 | 0.99945981 |
| PALD1 | 13.3765952 | 0.230795686 | 0.9695263 | 0.23804995 | 0.81184235 | 0.99945981 |
| PRF1 | 34.5329804 | -0.331445925 | 0.66541756 | -0.4981022 | 0.61841204 | 0.99945981 |
| SGPL1 | 31.3606757 | -0.408972986 | 0.33626947 | -1.2162061 | 0.22390641 | 0.99945981 |
| CDH23 | 16.0329648 | 1.432601503 | 0.47707074 | 3.00291209 | 0.0026741 | 0.45570351 |
| VSIR | 56.6648039 | -0.073953703 | 0.33874436 | -0.2183171 | 0.82718202 | 0.99945981 |
| PSAP | 205.929678 | 0.666986244 | 0.61557183 | 1.08352302 | 0.27857634 | 0.99945981 |
| SPOCK2 | 30.1855225 | -0.92541982 | 0.3994984 | -2.3164544 | 0.02053347 | 0.78749881 |
| ASCC1 | 29.1075946 | -0.188931497 | 0.37755351 | -0.5004099 | 0.61678651 | 0.99945981 |
| ANAPC16 | 53.4543308 | -0.487227107 | 0.250572 | -1.9444595 | 0.05184007 | 0.9230968 |
| DDIT4 | 35.4602481 | -2.304712423 | 0.48508523 | -4.7511494 | 2.02E-06 | 0.00462847 |
| DNAJB12 | 9.41682669 | -0.292362336 | 0.48232336 | -0.6061542 | 0.54441236 | 0.99945981 |
| MICU1 | 78.0625073 | 0.084876571 | 0.44590009 | 0.19034885 | 0.84903577 | 0.99945981 |
| MCU | 79.8227244 | -0.952982739 | 0.79475727 | -1.1990865 | 0.2304943 | 0.99945981 |
| P4HA1 | 64.6946525 | -0.590608115 | 0.54510122 | -1.0834834 | 0.27859392 | 0.99945981 |
| ECD | 13.9272792 | 0.054618706 | 0.40108403 | 0.13617771 | 0.89168079 | 0.99945981 |
| FAM149B1 | 14.2310509 | -0.082168234 | 0.55168896 | -0.1489394 | 0.88160143 | 0.99945981 |
| DNAJC9 | 14.0736058 | -0.137451275 | 0.51476695 | -0.2670165 | 0.78945645 | 0.99945981 |
| MRPS16 | 8.92092467 | -0.706403589 | 0.51016203 | -1.3846652 | 0.16615488 | 0.99945981 |
| ANXA7 | 25.9810419 | -0.054423724 | 0.39334447 | -0.1383615 | 0.88995474 | 0.99945981 |
| PPP3CB | 52.3579637 | -0.365137949 | 0.37223002 | -0.9809471 | 0.32661884 | 0.99945981 |
| USP54 | 11.1824747 | -1.091432628 | 0.81747173 | -1.3351319 | 0.18183316 | 0.99945981 |
| SEC24C | 13.5917962 | 0.046052064 | 0.3656999 | 0.12592857 | 0.89978847 | 0.99945981 |
| FUT11 | 10.2115006 | -0.778171592 | 0.47743052 | -1.6299159 | 0.10311928 | 0.99945981 |
| ZSWIM8 | 15.3506911 | -0.284847133 | 0.35030918 | -0.8131307 | 0.41614315 | 0.99945981 |
| CAMK2G | 27.3486453 | -0.082778495 | 0.34306381 | -0.2412918 | 0.80932894 | 0.99945981 |
| VCL | 50.8659324 | 1.15636592 | 0.42320604 | 2.73239465 | 0.00628758 | 0.52554078 |

| | | | | | | |
|---|---|---|---|---|---|---|
| AP3M1 | 14.4051193 | -0.38282288 | 0.56539896 | -0.6770845 | 0.49835234 | 0.99945981 |
| ADK | 172.818377 | -0.142483725 | 0.41954229 | -0.3396171 | 0.73414492 | 0.99945981 |
| KAT6B | 78.0277712 | 0.053707763 | 0.28639208 | 0.18753229 | 0.85124331 | 0.99945981 |
| SAMD8 | 31.0347788 | -0.141560653 | 0.31563357 | -0.4484968 | 0.65379468 | 0.99945981 |
| VDAC2 | 12.2076801 | -0.559192648 | 0.52906511 | -1.0569449 | 0.29053676 | 0.99945981 |
| LRMDA | 544.179214 | 0.120852997 | 0.5669049 | 0.21318037 | 0.83118628 | 0.99945981 |
| KCNMA1 | 154.987474 | -2.62921778 | 0.98260099 | -2.6757736 | 0.0074557 | 0.57509411 |
| POLR3A | 11.7615671 | 0.146348623 | 0.39775064 | 0.36794063 | 0.7129175 | 0.99945981 |
| RPS24 | 119.154228 | 0.111036273 | 0.55293364 | 0.20081302 | 0.84084478 | 0.99945981 |
| ZMIZ1 | 52.7452795 | 0.060164446 | 0.4328224 | 0.13900493 | 0.88944626 | 0.99945981 |
| NUTM2B-AS | 72.541594 | -0.308070953 | 0.28913501 | -1.0654917 | 0.28665348 | 0.99945981 |
| ANXA11 | 42.4581029 | -0.088400632 | 0.25554412 | -0.345931 | 0.72939457 | 0.99945981 |
| TSPAN14 | 63.6716935 | 0.485332566 | 0.43764852 | 1.10895511 | 0.26744955 | 0.99945981 |
| GHITM | 17.3339635 | -0.253640691 | 0.41845133 | -0.6061414 | 0.54442083 | 0.99945981 |
| CCSER2 | 100.931127 | -0.413887428 | 0.24635838 | -1.6800217 | 0.09295309 | 0.99640647 |
| WAPL | 71.2147149 | 0.254995063 | 0.22895186 | 1.11374968 | 0.26538659 | 0.99945981 |
| BMPR1A | 25.4064371 | -0.173625103 | 0.42309626 | -0.4103678 | 0.68153613 | 0.99945981 |
| GLUD1 | 41.741087 | 0.240347599 | 0.31127691 | 0.77213437 | 0.44003485 | 0.99945981 |
| SHLD2 | 48.4411563 | -0.124698143 | 0.388446 | -0.321018 | 0.74819678 | 0.99945981 |
| NUTM2A-AS | 62.1684793 | -0.099232781 | 0.29365156 | -0.337927 | 0.73541822 | 0.99945981 |
| PAPSS2 | 39.7654871 | 1.691800942 | 0.91370592 | 1.85158146 | 0.06408595 | 0.93776345 |
| ATAD1 | 29.807244 | 0.062424511 | 0.27807976 | 0.2244842 | 0.82238056 | 0.99945981 |
| PTEN | 151.617228 | -0.219535557 | 0.29861143 | -0.735188 | 0.46222498 | 0.99945981 |
| RNLS | 12.8891173 | 0.025116553 | 0.56756362 | 0.04425328 | 0.96470251 | 0.99945981 |
| STAMBPL1 | 24.016776 | -0.888922194 | 0.60585696 | -1.4672146 | 0.14231768 | 0.99945981 |
| FAS | 9.72522271 | 0.390024252 | 0.54880748 | 0.71067591 | 0.47728509 | 0.99945981 |
| LIPA | 27.009692 | 0.283604246 | 0.67815446 | 0.41820007 | 0.67580084 | 0.99945981 |
| KIF20B | 17.9312569 | 0.11976314 | 0.58287792 | 0.20546865 | 0.83720599 | 0.99945981 |
| PCGF5 | 78.2174428 | 0.190997703 | 0.34191979 | 0.55860383 | 0.57643213 | 0.99945981 |
| HECTD2 | 11.7286127 | 0.348719838 | 0.54957477 | 0.63452665 | 0.52573718 | 0.99945981 |
| TNKS2 | 57.3894761 | 0.058275098 | 0.26402087 | 0.22072156 | 0.82530924 | 0.99945981 |
| BTAF1 | 65.2075622 | 0.00237347 | 0.26652495 | 0.00890524 | 0.99289474 | 0.99945981 |
| CPEB3 | 64.1784379 | 0.48719439 | 0.28951651 | 1.6827862 | 0.09241649 | 0.99640647 |
| 5-Mar | 26.4434515 | -0.118190628 | 0.39865119 | -0.2964763 | 0.76686637 | 0.99945981 |
| IDE | 24.3446459 | 0.07059176 | 0.33250196 | 0.21230479 | 0.83186926 | 0.99945981 |
| EXOC6 | 62.2062658 | -0.305653462 | 0.27061581 | -1.1294738 | 0.25869802 | 0.99945981 |
| MYOF | 44.0005655 | -0.575280746 | 1.0381837 | -0.5541223 | 0.57949515 | 0.99945981 |
| FRA10AC1 | 16.4510529 | 0.210478019 | 0.36830635 | 0.57147541 | 0.56767743 | 0.99945981 |
| TBC1D12 | 26.4340134 | -0.34786783 | 0.4640559 | -0.7496248 | 0.45348069 | 0.99945981 |
| TCTN3 | 12.5395978 | 0.051350071 | 0.4492949 | 0.11429035 | 0.90900763 | 0.99945981 |
| ENTPD1 | 74.1185713 | 0.545737206 | 0.56549964 | 0.96505315 | 0.33451819 | 0.99945981 |
| ENTPD1-AS1 | 53.0689925 | 0.359505328 | 0.31790122 | 1.13087118 | 0.25810932 | 0.99945981 |
| AL365273.2 | 15.6997981 | 0.076371772 | 0.4869423 | 0.15683947 | 0.87537136 | 0.99945981 |
| ZNF518A | 43.9965345 | 0.064368702 | 0.38737609 | 0.16616591 | 0.86802638 | 0.99945981 |
| BLNK | 24.5504564 | -0.027481999 | 0.4149281 | -0.0662332 | 0.9471922 | 0.99945981 |
| TM9SF3 | 60.9174604 | -0.249789787 | 0.26808024 | -0.9317725 | 0.35145413 | 0.99945981 |
| PIK3AP1 | 113.125889 | 0.481432391 | 0.55013452 | 0.87511758 | 0.38150993 | 0.99945981 |
| LCOR | 115.897157 | 0.148054694 | 0.23332142 | 0.63455253 | 0.5257203 | 0.99945981 |
| PGAM1 | 29.2838431 | -0.168243506 | 0.44701887 | -0.3763678 | 0.70664349 | 0.99945981 |
| EXOSC1 | 12.791866 | -0.110512577 | 0.43814685 | -0.2522273 | 0.80086541 | 0.99945981 |
| MMS19 | 21.6933193 | -0.001897944 | 0.32937011 | -0.0057623 | 0.99540234 | 0.99945981 |
| ZFYVE27 | 11.6350306 | 0.071179122 | 0.39761677 | 0.17901439 | 0.8579264 | 0.99945981 |
| R3HCC1L | 54.1889076 | 0.098711204 | 0.24126965 | 0.4091323 | 0.68244258 | 0.99945981 |
| HPS1 | 16.7683968 | 0.226777392 | 0.33208166 | 0.68289647 | 0.49467227 | 0.99945981 |
| SLC25A28 | 11.1502681 | 0.372375642 | 0.57070868 | 0.65247937 | 0.51409198 | 0.99945981 |
| CUTC | 15.2514171 | -0.460289697 | 0.41026557 | -1.1219311 | 0.26189174 | 0.99945981 |
| COX15 | 11.8780571 | -0.269084217 | 0.42501305 | -0.6331199 | 0.52665535 | 0.99945981 |
| DNMBP | 21.8902101 | 0.276714462 | 0.61254903 | 0.45174255 | 0.65145446 | 0.99945981 |
| ERLIN1 | 12.1658026 | -0.024023159 | 0.79233941 | -0.0303193 | 0.97581242 | 0.99945981 |
| CHUK | 11.6827267 | 0.619081056 | 0.41346673 | 1.49729352 | 0.1343169 | 0.99945981 |
| CWF19L1 | 19.2795474 | -0.405705825 | 0.32489701 | -1.2487213 | 0.21176701 | 0.99945981 |
| BLOC1S2 | 10.2149129 | -0.893769873 | 0.55181321 | -1.6196964 | 0.10529751 | 0.99945981 |
| HIF1AN | 8.99497025 | -0.196742605 | 0.44866604 | -0.4385057 | 0.66101976 | 0.99945981 |
| SLF2 | 28.1104398 | -0.683465525 | 0.31208652 | -2.1899874 | 0.02852515 | 0.84763724 |
| BTRC | 45.1363302 | -0.025199437 | 0.30515326 | -0.0825796 | 0.93418581 | 0.99945981 |
| FBXW4 | 47.3578121 | -0.216915849 | 0.30178148 | -0.7187845 | 0.47227371 | 0.99945981 |
| OGA | 105.929508 | 0.050705217 | 0.27156738 | 0.18671321 | 0.8518855 | 0.99945981 |
| ARMH3 | 79.6243858 | 0.366366233 | 0.27410019 | 1.33661432 | 0.18134856 | 0.99945981 |
| GBF1 | 73.6989879 | 0.123377478 | 0.25839209 | 0.47748164 | 0.63301919 | 0.99945981 |
| NFKB2 | 9.23789251 | -0.264192241 | 0.70365567 | -0.3754567 | 0.70732084 | 0.99945981 |
| ACTR1A | 12.2709578 | 0.044444165 | 0.42377386 | 0.10487708 | 0.91647335 | 0.99945981 |
| SUFU | 30.5088134 | -0.23721657 | 0.32071752 | -0.7396433 | 0.45951646 | 0.99945981 |
| TRIM8 | 15.9736668 | 0.065102721 | 0.48609771 | 0.13392929 | 0.89345849 | 0.99945981 |
| ARL3 | 10.5878255 | -0.26829668 | 0.42572343 | -0.6302136 | 0.52855486 | 0.99945981 |
| WBP1L | 44.0593175 | -0.297141234 | 0.29745601 | -0.9989418 | 0.31782289 | 0.99945981 |
| CNNM2 | 23.6310546 | 0.321457908 | 0.55870371 | 0.57536383 | 0.56504526 | 0.99945981 |
| NT5C2 | 115.172169 | 0.224827258 | 0.22495244 | 0.99944353 | 0.31757988 | 0.99945981 |
| ATP5MD | 8.55805682 | 0.112565215 | 0.5765705 | 0.19523235 | 0.84521105 | 0.99945981 |
| PDCD11 | 9.37455642 | -0.186198909 | 0.4782281 | -0.3893517 | 0.69701602 | 0.99945981 |
| SH3PXD2A | 16.3603993 | -0.072384683 | 0.46661458 | -0.1551274 | 0.87672092 | 0.99945981 |
| STN1 | 10.2471343 | -0.127878801 | 0.41279875 | -0.3097849 | 0.75672457 | 0.99945981 |
| SLK | 31.2530914 | -0.078972114 | 0.34492099 | -0.2289571 | 0.81890225 | 0.99945981 |
| GSTO1 | 26.0167695 | -0.428719566 | 0.47977481 | -0.893585 | 0.37154398 | 0.99945981 |
| ITPRIP | 11.5231135 | -0.617456704 | 0.43819051 | -1.4091056 | 0.15880395 | 0.99945981 |
| XPNPEP1 | 18.324709 | 0.062990761 | 0.37430051 | 0.16828928 | 0.86635571 | 0.99945981 |
| ADD3 | 102.56975 | 0.113895337 | 0.27355818 | 0.41634776 | 0.67715555 | 0.99945981 |
| MXI1 | 24.3939158 | -1.43558444 | 0.38844698 | -3.6957024 | 0.00021928 | 0.17661106 |
| SMNDC1 | 11.7024522 | 0.400475515 | 0.43771398 | 0.91492512 | 0.36023094 | 0.99945981 |
| SMC3 | 23.1714069 | 0.080557024 | 0.32316957 | 0.24927169 | 0.80315063 | 0.99945981 |
| PDCD4 | 73.5946304 | 0.248539692 | 0.45663929 | 0.54428013 | 0.58624872 | 0.99945981 |
| BBIP1 | 12.8367259 | -0.094592304 | 0.48509189 | -0.1949987 | 0.84539394 | 0.99945981 |
| SHOC2 | 60.5441151 | 0.310916294 | 0.27746049 | 1.12057863 | 0.26246727 | 0.99945981 |
| ACSL5 | 16.4119306 | 0.100800119 | 0.39330523 | 0.2562898 | 0.79772707 | 0.99945981 |
| ZDHHC6 | 11.2449571 | -0.488279678 | 0.43794813 | -1.1149258 | 0.26488221 | 0.99945981 |
| VTI1A | 187.273962 | -0.053823251 | 0.18443638 | -0.2918256 | 0.77042 | 0.99945981 |
| TCF7L2 | 161.670607 | 1.774978018 | 0.6275983 | 2.82820719 | 0.00468095 | 0.50757754 |
| CASP7 | 11.6846821 | 0.430713051 | 0.44399411 | 0.9700873 | 0.33200298 | 0.99945981 |
| NHLRC2 | 30.0087873 | 0.119610126 | 0.37900667 | 0.31558844 | 0.75231491 | 0.99945981 |
| CCDC186 | 30.365426 | 0.501157131 | 0.51011117 | 0.98244689 | 0.32587974 | 0.99945981 |
| ABLIM1 | 30.6558322 | -0.087267192 | 0.45264165 | -0.1927953 | 0.84711927 | 0.99945981 |
| FAM160B1 | 20.757955 | -0.422589672 | 0.3955054 | -1.0684802 | 0.28530397 | 0.99945981 |
| SHTN1 | 37.5783194 | 0.596554355 | 0.51963464 | 1.14802654 | 0.25095761 | 0.99945981 |
| PDZD8 | 39.7538333 | -0.810863264 | 0.59390404 | -1.3653102 | 0.17215556 | 0.99945981 |
| RAB11FIP2 | 29.4848336 | -0.029865625 | 0.3244637 | -0.0920461 | 0.9266614 | 0.99945981 |
| FAM204A | 20.4838029 | -0.053738589 | 0.33288695 | -0.1614319 | 0.87175321 | 0.99945981 |
| CACUL1 | 56.7661257 | 0.255593079 | 0.23863674 | 1.07105501 | 0.2841447 | 0.99945981 |
| EIF3A | 67.4845181 | -0.110549057 | 0.3045246 | -0.3630218 | 0.71658862 | 0.99945981 |
| FAM45A | 14.9011521 | 0.560718364 | 0.53195744 | 1.05406621 | 0.2918526 | 0.99945981 |
| PRDX3 | 13.5780794 | -0.122029229 | 0.4256855 | -0.2866652 | 0.77436867 | 0.99945981 |
| GRK5 | 75.1479382 | -0.74755166 | 0.50805727 | -1.4713925 | 0.141185 | 0.99945981 |
| RGS10 | 24.8576583 | 0.734304834 | 0.3936396 | 1.86542418 | 0.06212197 | 0.92949575 |
| TIAL1 | 36.3318679 | -0.050874249 | 0.35296682 | -0.1441332 | 0.88539526 | 0.99945981 |
| INPP5F | 30.095531 | 0.253615365 | 0.5861571 | 0.43267473 | 0.66525109 | 0.99945981 |

| | | | | | | |
|---|---|---|---|---|---|---|
| MCMBP | 24.4767268 | 0.036156631 | 0.36799578 | 0.09825284 | 0.92173152 | 0.99945981 |
| SEC23IP | 21.3215555 | -0.196532936 | 0.34694759 | -0.5664629 | 0.57107915 | 0.99945981 |
| WDR11 | 40.2503515 | 0.17310608 | 0.2949707 | 0.58685856 | 0.55729871 | 0.99945981 |
| ATE1 | 50.3596546 | -0.249302137 | 0.26745013 | -0.9321444 | 0.35126191 | 0.99945981 |
| PLEKHA1 | 39.823377 | -0.133893197 | 0.32422874 | -0.4129591 | 0.6796366 | 0.99945981 |
| IKZF5 | 9.71050465 | -0.521997595 | 0.44803059 | -1.1650937 | 0.24398109 | 0.99945981 |
| ACADSB | 9.7337183 | -0.52039219 | 0.54878171 | -0.9482681 | 0.34299298 | 0.99945981 |
| BUB3 | 17.8401171 | -0.133340958 | 0.37355745 | -0.356949 | 0.72112999 | 0.99945981 |
| CHST15 | 92.1387092 | 0.151011679 | 0.57560417 | 0.26235335 | 0.79304904 | 0.99945981 |
| OAT | 8.70175066 | -0.402138515 | 0.57904959 | -0.6944803 | 0.48738107 | 0.99945981 |
| LHPP | 8.60384247 | -0.516219651 | 0.56554441 | -0.9127836 | 0.36135637 | 0.99945981 |
| ABRAXAS2 | 9.6916314 | -0.435635149 | 0.5386899 | -0.8086937 | 0.41869133 | 0.99945981 |
| ZRANB1 | 22.2572125 | -0.647302143 | 0.33561464 | -1.9287065 | 0.05376731 | 0.92445778 |
| CTBP2 | 89.9641338 | 0.075169036 | 0.39241389 | 0.19155549 | 0.84809041 | 0.99945981 |
| EDRF1 | 19.313306 | -0.02933216 | 0.33189588 | -0.0883776 | 0.92957657 | 0.99945981 |
| BCCIP | 10.7346434 | 0.04020206 | 0.4980286 | 0.08072239 | 0.93566273 | 0.99945981 |
| DHX32 | 16.5666061 | -0.287507646 | 0.50457756 | -0.5697987 | 0.56881423 | 0.99945981 |
| DOCK1 | 24.2571656 | -0.114548411 | 0.56656355 | -0.202181 | 0.83977519 | 0.99945981 |
| PTPRE | 106.204828 | 0.119104334 | 0.48940581 | 0.24336518 | 0.80772251 | 0.99945981 |
| MGMT | 33.7072586 | -0.022552328 | 0.37991626 | -0.0593613 | 0.95266433 | 0.99945981 |
| GLRX3 | 17.1599221 | -0.901680174 | 0.44131893 | -2.0431486 | 0.04103773 | 0.89562588 |
| PPP2R2D | 40.4816617 | 0.115961415 | 0.23992802 | 0.48331751 | 0.62887031 | 0.99945981 |
| BNIP3 | 12.002386 | -1.46430362 | 0.8685302 | -1.6859559 | 0.09180429 | 0.99640647 |
| INPP5A | 28.9152577 | -0.323511977 | 0.45329148 | -0.7136953 | 0.47541564 | 0.99945981 |
| ADAM8 | 11.9502168 | 0.519079205 | 0.58103524 | 0.89336957 | 0.37165929 | 0.99945981 |
| TUBGCP2 | 22.8009677 | -0.869638257 | 0.32634717 | -2.6647642 | 0.00770423 | 0.57727153 |
| ECHS1 | 10.784611 | -0.214745093 | 0.51528405 | -0.4167509 | 0.67686062 | 0.99945981 |
| KDM5A | 78.0086247 | -0.048937153 | 0.243752 | -0.2007662 | 0.84088143 | 0.99945981 |
| NINJ2 | 8.0807273 | -0.521721494 | 0.55729945 | -0.9361601 | 0.34919078 | 0.99945981 |
| WNK1 | 171.780233 | -0.210537543 | 0.21668101 | -0.9716474 | 0.33122599 | 0.99945981 |
| RAD52 | 9.44349386 | 0.224466837 | 0.480596 | 0.46705931 | 0.64045744 | 0.99945981 |
| ERC1 | 132.290433 | 0.219373302 | 0.22400203 | 0.97933623 | 0.32741387 | 0.99945981 |
| ADIPOR2 | 38.6948558 | 0.043000335 | 0.31281939 | 0.13746058 | 0.89066675 | 0.99945981 |
| DCP1B | 11.0563397 | -0.620627171 | 0.60659196 | -1.0231378 | 0.30624269 | 0.99945981 |
| ITFG2 | 23.4412275 | -0.252241658 | 0.44478622 | -0.5671076 | 0.57064106 | 0.99945981 |
| TULP3 | 8.69112521 | -0.513284248 | 0.46334861 | -1.1077712 | 0.26796064 | 0.99945981 |
| CRACR2A | 23.5227557 | -0.274074785 | 0.37669002 | -0.7275871 | 0.46686641 | 0.99945981 |
| PARP11 | 25.1203648 | 0.663149261 | 0.43172901 | 1.53603127 | 0.12453071 | 0.99945981 |
| CCND2 | 31.1182271 | -0.463015056 | 0.39104226 | -1.1840538 | 0.23639179 | 0.99945981 |
| TIGAR | 15.0248864 | -0.157878831 | 0.50517471 | -0.3125232 | 0.75464291 | 0.99945981 |
| C12orf4 | 14.5820111 | 0.369901379 | 0.39046701 | 0.94733068 | 0.3434703 | 0.99945981 |
| CD9 | 19.0523639 | -0.230744082 | 0.47257342 | -0.4882714 | 0.62535762 | 0.99945981 |
| TNFRSF1A | 41.0070689 | 0.263883349 | 0.36421099 | 0.72453428 | 0.46873779 | 0.99945981 |
| LTBR | 13.0526459 | 0.125244517 | 0.48046431 | 0.26067392 | 0.79434398 | 0.99945981 |
| MRPL51 | 10.8510646 | -0.827728498 | 0.60087475 | -1.3775392 | 0.16834562 | 0.99945981 |
| NCAPD2 | 12.630865 | 0.175343454 | 0.64937469 | 0.27001892 | 0.7871457 | 0.99945981 |
| GAPDH | 119.11859 | -1.000869062 | 0.52072236 | -1.9220781 | 0.05459593 | 0.92445778 |
| IFFO1 | 8.98895195 | -0.328863776 | 0.51464355 | -0.6390127 | 0.52281466 | 0.99945981 |
| CHD4 | 9.95126687 | -0.675887771 | 0.45349984 | -1.4903815 | 0.13612395 | 0.99945981 |
| ING4 | 9.96143569 | 0.197493624 | 0.54135701 | 0.36481216 | 0.71525162 | 0.99945981 |
| MLF2 | 11.0818419 | -1.029253848 | 0.54506637 | -1.8883092 | 0.05898445 | 0.92445778 |
| PTMS | 15.4440215 | -0.31275179 | 0.4125524 | -0.7580899 | 0.44839719 | 0.99945981 |
| CD4 | 34.8343123 | 0.709527015 | 0.55934288 | 1.26850102 | 0.20461909 | 0.99945981 |
| TPI1 | 29.6395869 | -0.093808909 | 0.55255606 | -0.1697727 | 0.86518893 | 0.99945981 |
| PTPN6 | 21.5272419 | 0.070165731 | 0.70923412 | 0.09893169 | 0.92119251 | 0.99945981 |
| PHB2 | 13.3537097 | 0.221988379 | 0.58433789 | 0.37989729 | 0.70402166 | 0.99945981 |
| LPCAT3 | 22.293551 | 0.049474347 | 0.48683213 | 0.10162507 | 0.91905428 | 0.99945981 |
| C1S | 12.7929686 | -0.658785569 | 0.58610602 | -1.1240041 | 0.26101129 | 0.99945981 |
| C1RL | 15.6589839 | -0.342875235 | 0.54970719 | -0.6237416 | 0.5327973 | 0.99945981 |
| CLSTN3 | 9.80404072 | -0.744201264 | 0.50220418 | -1.4818699 | 0.13837491 | 0.99945981 |
| CD163L1 | 30.5737189 | 0.151729307 | 0.62941254 | 0.24106495 | 0.80950477 | 0.99945981 |
| CD163 | 128.074147 | -0.581502963 | 0.73353829 | -0.792737 | 0.42793105 | 0.99945981 |
| SLC2A3 | 165.384405 | -0.138013052 | 0.71531282 | -0.1929408 | 0.84700531 | 0.99945981 |
| C3AR1 | 18.5962644 | 0.754062688 | 0.48319846 | 1.56056517 | 0.11862638 | 0.99945981 |
| NECAP1 | 12.0953275 | 0.248234757 | 0.4000375 | 0.62052871 | 0.53490976 | 0.99945981 |
| LINC00937 | 14.1882076 | 0.015357543 | 0.71678765 | 0.02142551 | 0.98290622 | 0.99945981 |
| RIMKLB | 12.148865 | -0.580335493 | 0.49387876 | -1.1750566 | 0.23997207 | 0.99945981 |
| M6PR | 34.9043121 | 0.654821134 | 0.34001375 | 1.92586664 | 0.05412103 | 0.92445778 |
| KLRG1 | 17.2825829 | 0.643173545 | 0.79766237 | 0.80632304 | 0.42005661 | 0.99945981 |
| A2M | 30.6051363 | -0.127976109 | 0.60615414 | -0.211128 | 0.83278739 | 0.99945981 |
| PZP | 15.6039415 | 0.249584621 | 1.27954767 | 0.19505692 | 0.84534839 | 0.99945981 |
| KLRB1 | 11.5096093 | 0.820222308 | 0.94766663 | 0.86551777 | 0.38675467 | 0.99945981 |
| CLEC2D | 123.060291 | -0.526215486 | 0.31746561 | -1.6575512 | 0.0974081 | 0.99945981 |
| CD69 | 45.2674441 | 0.416730162 | 0.60908037 | 0.68419569 | 0.4938516 | 0.99945981 |
| KLRF1 | 19.4238809 | 0.396857645 | 0.98795771 | 0.40169497 | 0.68790853 | 0.99945981 |
| CLEC2B | 39.9734058 | 0.799250423 | 0.59943759 | 1.33333384 | 0.18242227 | 0.99945981 |
| CLEC12A | 29.5892859 | 2.418590655 | 0.78358158 | 3.08658435 | 0.0020247 | 0.45570351 |
| CLEC7A | 48.6739755 | 1.384665058 | 0.61987322 | 2.2337875 | 0.02549706 | 0.81367289 |
| GABARAPL1 | 18.5506554 | -0.230366682 | 0.43809224 | -0.5258406 | 0.59899897 | 0.99945981 |
| KLRD1 | 45.1752489 | 0.706188653 | 0.81026686 | 0.87155071 | 0.38345353 | 0.99945981 |
| AC022075.1 | 20.3731863 | -0.834151394 | 0.54880683 | -1.5199362 | 0.128527 | 0.99945981 |
| KLRC4-KLRK1 | 13.1028794 | 0.297013923 | 0.70170416 | 0.42327513 | 0.67209454 | 0.99945981 |
| YBX3 | 34.7195947 | -1.258525783 | 0.60673457 | -2.0742609 | 0.03805509 | 0.87920079 |
| AC244131.2 | 16.3966231 | -1.029611813 | 0.87955325 | -1.1706077 | 0.2417565 | 0.99945981 |
| ETV6 | 354.033378 | -0.087394146 | 0.45093922 | -0.1938047 | 0.8463288 | 0.99945981 |
| BORCS5 | 34.7271446 | -0.224203879 | 0.36915823 | -0.6073382 | 0.54362652 | 0.99945981 |
| DUSP16 | 46.9014728 | -0.979264321 | 0.46594258 | -2.1016845 | 0.03558092 | 0.87920079 |
| CREBL2 | 21.2725265 | -0.463555354 | 0.41793284 | -1.1091623 | 0.26736017 | 0.99945981 |
| CDKN1B | 24.3605193 | 0.085963635 | 0.54339376 | 0.15819769 | 0.87430103 | 0.99945981 |
| APOLD1 | 20.4150583 | -0.296899353 | 0.72434673 | -0.4098857 | 0.68188981 | 0.99945981 |
| ATF7IP | 169.03557 | 0.056947018 | 0.3624661 | 0.15710991 | 0.87515822 | 0.99945981 |
| PLBD1 | 16.9030988 | 0.548236397 | 0.81630849 | 0.67160443 | 0.50183556 | 0.99945981 |
| H2AFJ | 18.7469486 | -0.001225937 | 0.61789342 | -0.0019841 | 0.99841695 | 0.9997934 |
| WBP11 | 12.4400828 | 0.84044909 | 0.46385757 | 1.81186888 | 0.07000646 | 0.95505283 |
| MGP | 22.3096795 | 0.019810098 | 0.78366224 | 0.02527887 | 0.97983253 | 0.99945981 |
| ARHGDIB | 61.3711271 | -0.607468883 | 0.51248581 | -1.185338 | 0.23588385 | 0.99945981 |
| PTPRO | 14.1183658 | -0.78137851 | 0.74816132 | -1.0443984 | 0.29630109 | 0.99945981 |
| EPS8 | 50.310638 | 0.787562689 | 0.36408221 | 2.16314519 | 0.03053001 | 0.85704136 |
| STRAP | 16.8718042 | -0.183292127 | 0.3691307 | -0.4965508 | 0.61950589 | 0.99945981 |
| DERA | 31.8737855 | 0.419698063 | 0.29663946 | 1.41484231 | 0.15711473 | 0.99945981 |
| MGST1 | 23.4379326 | -1.452702149 | 0.8456612 | -1.71783 | 0.08582763 | 0.99193047 |
| PLEKHA5 | 47.195009 | 0.030042599 | 0.41792867 | 0.07188451 | 0.94269382 | 0.99945981 |
| AEBP2 | 62.4231149 | 0.00368446 | 0.35120243 | 0.01049099 | 0.99162956 | 0.99945981 |
| PYROXD1 | 14.6786561 | 0.15092362 | 0.35953555 | 0.41977384 | 0.67465068 | 0.99945981 |
| RECQL | 20.7452105 | 0.052380233 | 0.35336191 | 0.14823395 | 0.88215814 | 0.99945981 |
| GOLT1B | 11.7110793 | 0.308930526 | 0.45512796 | 0.6787773 | 0.49727898 | 0.99945981 |
| LDHB | 32.4488356 | -0.580279151 | 0.67698184 | -0.8571562 | 0.39135859 | 0.99945981 |
| ST8SIA1 | 25.5662597 | -0.236123179 | 0.68809658 | -0.3431541 | 0.73148252 | 0.99945981 |
| C2CD5 | 63.8446585 | 0.211240768 | 0.42131802 | 0.50138081 | 0.61610314 | 0.99945981 |
| ETNK1 | 85.4822305 | 0.265715743 | 0.26272399 | 1.01138744 | 0.31183103 | 0.99945981 |
| SOX5 | 230.82018 | -5.129661011 | 1.70470099 | -3.0091265 | 0.00262 | 0.45570351 |
| BCAT1 | 35.7466402 | -0.011974719 | 0.61662915 | -0.0194196 | 0.98450634 | 0.99945981 |
| LRMP | 63.309734 | -1.235514576 | 0.74934316 | -1.6487968 | 0.09918928 | 0.99945981 |

| | | | | | | |
|---|---|---|---|---|---|---|
| KRAS | 39.3781 | 0.176806946 | 0.33557574 | 0.52687642 | 0.59827941 | 0.99945981 |
| RASSF8 | 15.8589951 | -0.360195106 | 0.51811336 | -0.6952052 | 0.48692671 | 0.99945981 |
| ITPR2 | 266.154236 | 0.212224511 | 0.31458114 | 0.67462567 | 0.49991362 | 0.99945981 |
| INTS13 | 7.49706823 | -0.063638147 | 0.53146014 | -0.1197421 | 0.90468746 | 0.99945981 |
| FGFR1OP2 | 43.6616823 | -0.040586378 | 0.3293088 | -0.1232472 | 0.90191137 | 0.99945981 |
| TM7SF3 | 26.2116374 | -0.528793718 | 0.54084139 | -0.9777242 | 0.32821074 | 0.99945981 |
| MED21 | 9.73666193 | 0.102291334 | 0.5177392 | 0.19757309 | 0.84337909 | 0.99945981 |
| STK38L | 32.0463215 | 0.765319676 | 0.35916083 | 2.13085506 | 0.03310108 | 0.8663391 |
| PPFIBP1 | 11.4247558 | -0.861735666 | 0.66693891 | -1.2920759 | 0.19633088 | 0.99945981 |
| MRPS35 | 11.8870963 | 0.580813261 | 0.44863662 | 1.29461847 | 0.19545188 | 0.99945981 |
| KLHL42 | 10.829643 | -0.017947239 | 0.52676736 | -0.0340705 | 0.97282092 | 0.99945981 |
| CCDC91 | 176.971996 | -0.071327453 | 0.24655565 | -0.2892956 | 0.77235521 | 0.99945981 |
| FAR2 | 53.0613825 | 0.840662772 | 0.48285486 | 1.7410258 | 0.08167906 | 0.9846219 |
| ERGIC2 | 16.2990271 | -0.27999533 | 0.40421398 | -0.6926909 | 0.48850357 | 0.99945981 |
| TMTC1 | 24.138884 | 0.618255747 | 1.02226805 | 0.60478829 | 0.54531967 | 0.99945981 |
| IPO8 | 24.5208461 | 0.095510563 | 0.35433177 | 0.26955123 | 0.78750553 | 0.99945981 |
| CAPRIN2 | 11.5200856 | -0.367014549 | 0.48978841 | -0.7493329 | 0.45365661 | 0.99945981 |
| SINHCAF | 25.4332151 | 0.16440823 | 0.39042085 | 0.42110515 | 0.6736783 | 0.99945981 |
| DENND5B | 36.7242187 | 0.186845449 | 0.80698932 | 0.23153398 | 0.81689999 | 0.99945981 |
| AMN1 | 13.0789916 | -0.342879813 | 0.56261361 | -0.609441 | 0.54223216 | 0.99945981 |
| KIAA1551 | 108.533985 | -0.100826134 | 0.2897143 | -0.3480192 | 0.72782578 | 0.99945981 |
| BICD1 | 43.8839926 | -0.546256957 | 0.60586384 | -0.9016167 | 0.36726051 | 0.99945981 |
| FGD4 | 190.389197 | 0.985136976 | 0.51661566 | 1.90690498 | 0.05653289 | 0.92445778 |
| DNM1L | 41.7608632 | -0.073149651 | 0.34102655 | -0.2144984 | 0.83015841 | 0.99945981 |
| YARS2 | 10.8388538 | 0.228578267 | 0.5001265 | 0.4570409 | 0.64764165 | 0.99945981 |
| CPNE8 | 59.136108 | 0.229520757 | 0.40310909 | 0.56937629 | 0.5691008 | 0.99945981 |
| KIF21A | 17.656156 | -0.471233696 | 0.47838792 | -0.9850451 | 0.3246019 | 0.99945981 |
| SLC2A13 | 38.6420489 | -0.187762875 | 0.39683997 | -0.4731451 | 0.63610968 | 0.99945981 |
| LRRK2 | 94.7023692 | 0.875226299 | 0.58725196 | 1.49037611 | 0.13612537 | 0.99945981 |
| GXYLT1 | 17.1664065 | -0.276904432 | 0.45983007 | -0.6021886 | 0.5470486 | 0.99945981 |
| YAF2 | 50.1776112 | -0.144557702 | 0.27650274 | -0.5228075 | 0.60110823 | 0.99945981 |
| PPHLN1 | 70.6623148 | -0.068611458 | 0.25692608 | -0.2670475 | 0.78943262 | 0.99945981 |
| PUS7L | 13.4752229 | -0.020687401 | 0.48153968 | -0.0429609 | 0.96573266 | 0.99945981 |
| IRAK4 | 24.2891001 | 0.177543385 | 0.36977795 | 0.48013514 | 0.63113131 | 0.99945981 |
| TMEM117 | 58.3702531 | -0.451790825 | 0.39302448 | -1.1495234 | 0.25034024 | 0.99945981 |
| NELL2 | 26.2497151 | 0.500570849 | 0.99095125 | 0.50514175 | 0.61345928 | 0.99945981 |
| ANO6 | 67.4805513 | 0.401256451 | 0.29141474 | 1.37692571 | 0.16853522 | 0.99945981 |
| ARID2 | 114.093641 | 0.110069966 | 0.21187204 | 0.51951152 | 0.60340408 | 0.99945981 |
| SCAF11 | 139.563999 | -0.014687362 | 0.16785078 | -0.0875025 | 0.9302721 | 0.99945981 |
| SLC38A1 | 115.235202 | -1.360616834 | 0.47739229 | -2.8501022 | 0.00437052 | 0.50757754 |
| SLC38A2 | 59.9660047 | 0.488182702 | 0.54205539 | 0.90061406 | 0.36779356 | 0.99945981 |
| AC008014.1 | 66.2987795 | 0.240043662 | 0.55281741 | 0.43421871 | 0.66412963 | 0.99945981 |
| PCED1B | 94.0198239 | 0.025492263 | 0.39179113 | 0.06506595 | 0.94812149 | 0.99945981 |
| RPAP3 | 13.9849733 | 0.52341712 | 0.40499156 | 1.29241489 | 0.1962135 | 0.99945981 |
| HDAC7 | 19.2033218 | 0.004205037 | 0.45829028 | 0.00917549 | 0.99267912 | 0.99945981 |
| VDR | 10.7379188 | -0.059730074 | 0.7001044 | -0.085316 | 0.93201021 | 0.99945981 |
| SENP1 | 16.6435781 | 0.202169232 | 0.43123149 | 0.46881835 | 0.63919949 | 0.99945981 |
| ZNF641 | 10.4138175 | 0.57391014 | 0.44526256 | 1.28892523 | 0.19742408 | 0.99945981 |
| CCNT1 | 18.6903719 | 0.534024277 | 0.41318763 | 1.29244981 | 0.19620142 | 0.99945981 |
| DDX23 | 9.97504432 | 0.332414468 | 0.56937068 | 0.58382787 | 0.55933613 | 0.99945981 |
| CCDC65 | 10.1616795 | 0.058253956 | 0.59954555 | 0.09716352 | 0.92259654 | 0.99945981 |
| PRKAG1 | 13.6083492 | -0.793503847 | 0.4941124 | -1.6059177 | 0.10829199 | 0.99945981 |
| KMT2D | 23.3204157 | 0.151256941 | 0.42611641 | 0.35496624 | 0.72261488 | 0.99945981 |
| LMBR1L | 13.5034391 | -0.271093013 | 0.49326084 | -0.5495936 | 0.58259813 | 0.99945981 |
| TUBA1B | 40.2051413 | -0.023876106 | 0.53897835 | -0.0442988 | 0.96466621 | 0.99945981 |
| TUBA1A | 20.497914 | 0.487588256 | 0.8412861 | 0.57957484 | 0.56220136 | 0.99945981 |
| TUBA1C | 13.8662418 | 0.165145353 | 0.5309743 | 0.31102325 | 0.75578295 | 0.99945981 |
| SPATS2 | 38.6462933 | -0.416950649 | 0.57927468 | -0.7197806 | 0.47166012 | 0.99945981 |
| PRPF40B | 11.041084 | -0.314295218 | 0.52388004 | -0.5999374 | 0.54854797 | 0.99945981 |
| FMNL3 | 36.1843291 | 0.347243063 | 0.37142862 | 0.93488506 | 0.34984754 | 0.99945981 |
| TMBIM6 | 71.3793207 | 0.101718745 | 0.36907045 | 0.27560794 | 0.78284921 | 0.99945981 |
| NCKAP5L | 11.5343204 | -0.113146497 | 0.42471398 | -0.2664063 | 0.78992628 | 0.99945981 |
| AQP2 | 14.7191696 | 0.352795468 | 0.89915985 | 0.39236124 | 0.69479132 | 0.99945981 |
| AC074032.1 | 25.2920407 | -0.643093954 | 0.42524515 | -1.5122899 | 0.13046013 | 0.99945981 |
| CERS5 | 21.2833597 | -0.408190337 | 0.36737893 | -1.1110881 | 0.26653045 | 0.99945981 |
| LIMA1 | 13.5835396 | -0.33829051 | 0.48721748 | -0.6943316 | 0.48747425 | 0.99945981 |
| LARP4 | 40.0398471 | 0.10799195 | 0.29761604 | 0.36285661 | 0.716712 | 0.99945981 |
| DIP2B | 133.149432 | -0.016005835 | 0.42787174 | -0.037408 | 0.97015968 | 0.99945981 |
| ATF1 | 19.5856082 | 0.550134002 | 0.41061071 | 1.33979459 | 0.18031214 | 0.99945981 |
| METTL7A | 17.9604216 | 0.265466649 | 0.41267742 | 0.64327883 | 0.52004319 | 0.99945981 |
| SLC11A2 | 33.3305218 | 0.001821175 | 0.37784487 | 0.0048199 | 0.99615429 | 0.99945981 |
| LETMD1 | 15.9816572 | -0.11939434 | 0.3834624 | -0.3113587 | 0.75552797 | 0.99945981 |
| TFCP2 | 64.6974 | 0.216659921 | 0.25889657 | 0.83685897 | 0.40267184 | 0.99945981 |
| DAZAP2 | 36.5560483 | 0.066613096 | 0.41686682 | 0.15979467 | 0.87304282 | 0.99945981 |
| BIN2 | 60.6014369 | 0.328640286 | 0.4294911 | 0.76518533 | 0.44416119 | 0.99945981 |
| GALNT6 | 9.79269939 | 0.282117831 | 0.43448494 | 0.64931556 | 0.51613443 | 0.99945981 |
| SLC4A8 | 18.2062629 | -0.285514611 | 0.40006867 | -0.713664 | 0.47543496 | 0.99945981 |
| ACVR1B | 18.7128525 | -0.214932314 | 0.49633984 | -0.4330346 | 0.66498964 | 0.99945981 |
| NR4A1 | 24.8151388 | 1.608751997 | 0.58728638 | 2.73929731 | 0.00615707 | 0.52554078 |
| EIF4B | 46.7401926 | -0.24713455 | 0.27732567 | -0.8911348 | 0.37285686 | 0.99945981 |
| AC068888.1 | 14.2103063 | -0.20589342 | 0.49975725 | -0.4119869 | 0.68034905 | 0.99945981 |
| CSAD | 11.082133 | 0.3768773 | 0.47908442 | 0.78666157 | 0.43148001 | 0.99945981 |
| PFDN5 | 46.0794665 | 0.702683238 | 0.57366431 | 1.22490317 | 0.2206117 | 0.99945981 |
| SP1 | 23.9173869 | 0.174764625 | 0.3283811 | 0.53220063 | 0.59458705 | 0.99945981 |
| PRR13 | 23.8162963 | 0.011315712 | 0.4222017 | 0.02680167 | 0.97861792 | 0.99945981 |
| PCBP2 | 57.7409567 | 0.132952753 | 0.25396771 | 0.52350259 | 0.60062455 | 0.99945981 |
| ATP5MC2 | 37.2144865 | -0.118569309 | 0.3957293 | -0.2996223 | 0.7644653 | 0.99945981 |
| CALCOCO1 | 11.0526425 | 0.107981319 | 0.41403729 | 0.26080095 | 0.79424601 | 0.99945981 |
| CBX5 | 17.7614896 | -0.289371494 | 0.38110017 | -0.7593056 | 0.44766977 | 0.99945981 |
| HNRNPA1 | 24.562582 | -0.146481513 | 0.48944797 | -0.299279 | 0.76472715 | 0.99945981 |
| COPZ1 | 12.7122105 | -0.210602597 | 0.37376534 | -0.5634621 | 0.57312028 | 0.99945981 |
| ITGA5 | 14.9989772 | -0.075732444 | 0.54768407 | -0.1382776 | 0.89002102 | 0.99945981 |
| NCKAP1L | 83.3902078 | -0.161878295 | 0.31280393 | -0.5175072 | 0.60480212 | 0.99945981 |
| TESPA1 | 15.5354211 | 0.243825635 | 0.65588229 | 0.37175212 | 0.71007741 | 0.99945981 |
| CD63 | 40.9538052 | -0.922977682 | 0.5321717 | -1.7343607 | 0.0828541 | 0.98529784 |
| DGKA | 19.9790476 | 0.279601602 | 0.43808433 | 0.63823693 | 0.52331946 | 0.99945981 |
| RAB5B | 12.3594018 | 0.445927573 | 0.39753824 | 1.12172247 | 0.26198046 | 0.99945981 |
| RPS26 | 10.9377068 | -0.917584021 | 0.49243476 | -1.8633616 | 0.06241141 | 0.92949575 |
| PA2G4 | 8.73068306 | -0.228651402 | 0.61968527 | -0.3689799 | 0.71214272 | 0.99945981 |
| RPL41 | 172.679934 | -0.421747895 | 0.50568686 | -0.83401 | 0.40427535 | 0.99945981 |
| ESYT1 | 15.0297878 | -0.27916202 | 0.37566722 | -0.7431099 | 0.45741516 | 0.99945981 |
| MYL6 | 68.528614 | 0.294797655 | 0.57197692 | 0.51540131 | 0.60627261 | 0.99945981 |
| SMARCC2 | 30.8558552 | -0.429248653 | 0.41429704 | -1.0360891 | 0.30016058 | 0.99945981 |
| RNF41 | 8.6263469 | -0.306429154 | 0.54221654 | -0.5651417 | 0.57197741 | 0.99945981 |
| CS | 12.726024 | 0.161919505 | 0.48355543 | 0.334852 | 0.73773673 | 0.99945981 |
| STAT2 | 37.1341733 | 1.025026449 | 0.47235908 | 2.17001535 | 0.03000568 | 0.85472619 |
| RBMS2 | 13.7488589 | -0.79244703 | 0.52183725 | -1.5185712 | 0.12887046 | 0.99945981 |
| BAZ2A | 31.9398086 | 0.168131955 | 0.30173299 | 0.55722099 | 0.57737645 | 0.99945981 |
| ATP5F1B | 31.0468928 | 0.172863625 | 0.44737371 | 0.38639648 | 0.69920306 | 0.99945981 |
| PTGES3 | 66.5083289 | 0.454154578 | 0.29601488 | 1.53422888 | 0.12497335 | 0.99945981 |
| NACA | 49.1332802 | -0.120244517 | 0.52126672 | -0.2306775 | 0.81756533 | 0.99945981 |
| NEMP1 | 10.0785448 | 0.585725974 | 0.5810259 | 1.00808927 | 0.31341161 | 0.99945981 |
| STAT6 | 46.740252 | 0.010884868 | 0.28853661 | 0.03772439 | 0.96990743 | 0.99945981 |

| | | | | | | |
|---|---|---|---|---|---|---|
| LRP1 | 46.8415416 | 0.245429393 | 0.58980499 | 0.41611956 | 0.67732252 | 0.99945981 |
| R3HDM2 | 97.6232469 | 0.359322205 | 0.31514415 | 1.14018364 | 0.2542098 | 0.99945981 |
| ARHGAP9 | 22.3387597 | 0.380360871 | 0.6075407 | 0.62606649 | 0.53127133 | 0.99945981 |
| MARS | 22.9824912 | 0.008039906 | 0.35444004 | 0.0226834 | 0.98190282 | 0.99945981 |
| DCTN2 | 10.9298123 | 0.418835277 | 0.4640668 | 0.9025323 | 0.36677417 | 0.99945981 |
| OS9 | 30.7319174 | 0.11460241 | 0.42483744 | 0.26975591 | 0.78734805 | 0.99945981 |
| CTDSP2 | 16.1068804 | 0.131808614 | 0.46194885 | 0.28533162 | 0.77539009 | 0.99945981 |
| SLC16A7 | 63.0593778 | 0.66391415 | 0.50550648 | 1.31336428 | 0.18906023 | 0.99945981 |
| FAM19A2 | 40.1042713 | -0.973645714 | 0.96186887 | -1.0122437 | 0.31142154 | 0.99945981 |
| USP15 | 251.578195 | 0.471147829 | 0.27052756 | 1.74158904 | 0.08158038 | 0.9846219 |
| MON2 | 107.366945 | 0.134942132 | 0.23798571 | 0.56701778 | 0.5707021 | 0.99945981 |
| PPM1H | 12.6274328 | -0.690507158 | 0.86156432 | -0.8014575 | 0.42286686 | 0.99945981 |
| SRGAP1 | 59.4127409 | -1.603298228 | 0.44137485 | -3.6325093 | 0.00028068 | 0.17661106 |
| XPOT | 10.1818868 | 0.125074789 | 0.46111395 | 0.27124486 | 0.78620271 | 0.99945981 |
| TBK1 | 39.7704566 | -0.272166817 | 0.26990653 | -1.0083744 | 0.31327478 | 0.99945981 |
| RASSF3 | 107.447937 | 0.801956548 | 0.38702871 | 2.07208544 | 0.03825747 | 0.87920079 |
| GNS | 46.5758734 | 0.700528144 | 0.45943151 | 1.52477167 | 0.12731606 | 0.99945981 |
| LEMD3 | 44.4984096 | 0.662824448 | 0.32630928 | 2.03127672 | 0.04222693 | 0.89562588 |
| IRAK3 | 146.113412 | 0.531065948 | 0.57706351 | 0.9202903 | 0.35742108 | 0.99945981 |
| HELB | 33.1219986 | -0.161824286 | 0.40101241 | -0.4035393 | 0.6865515 | 0.99945981 |
| GRIP1 | 14.114319 | 0.097922299 | 0.61190392 | 0.16002888 | 0.87285833 | 0.99945981 |
| CAND1 | 29.7069131 | 0.16068933 | 0.35267253 | 0.45563325 | 0.64865373 | 0.99945981 |
| DYRK2 | 13.8223144 | 0.394544427 | 0.59380639 | 0.66443277 | 0.50641336 | 0.99945981 |
| IFNG-AS1 | 47.6580617 | 0.275434524 | 0.81048787 | 0.33983794 | 0.73397858 | 0.99945981 |
| LINC02384 | 25.0566593 | 0.843058961 | 0.85350154 | 0.98776501 | 0.32326775 | 0.99945981 |
| RAP1B | 198.694431 | 0.523827054 | 0.22322407 | 2.34664234 | 0.01894343 | 0.76950676 |
| NUP107 | 28.7101316 | -0.061088521 | 0.31582197 | -0.1934271 | 0.84662451 | 0.99945981 |
| SLC35E3 | 23.7251523 | -0.385402493 | 0.44694969 | -0.862295 | 0.38852519 | 0.99945981 |
| MDM2 | 55.5448123 | 0.676105929 | 0.34669717 | 1.95013398 | 0.05116015 | 0.9230968 |
| CPM | 34.4421139 | -0.507854805 | 0.60654355 | -0.8372932 | 0.40242776 | 0.99945981 |
| CPSF6 | 44.8970853 | 0.006690833 | 0.39649145 | 0.0168751 | 0.98653626 | 0.99945981 |
| LYZ | 161.380421 | 0.994118371 | 1.44639533 | 0.68730751 | 0.49188897 | 0.99945981 |
| AC020656.1 | 49.2695519 | 0.802545879 | 1.99303572 | 0.40267511 | 0.68718725 | 0.99945981 |
| FRS2 | 45.8248079 | 0.728582128 | 0.38836461 | 1.87602606 | 0.0606517 | 0.92445778 |
| CCT2 | 13.8350526 | -0.595039444 | 0.41787232 | -1.4239743 | 0.15445391 | 0.99945981 |
| CNOT2 | 85.7268003 | 0.628599925 | 0.3358826 | 1.87148704 | 0.0612776 | 0.92658757 |
| ZFC3H1 | 82.1714179 | 0.167571173 | 0.22241222 | 0.75342612 | 0.45119389 | 0.99945981 |
| RAB21 | 37.2975143 | 0.284182683 | 0.31746319 | 0.89516735 | 0.37069764 | 0.99945981 |
| TBC1D15 | 41.0580627 | 0.001530034 | 0.33388838 | 0.00458247 | 0.99634373 | 0.99945981 |
| ATXN7L3B | 10.1396828 | -0.178507066 | 0.43171084 | -0.4134876 | 0.67924943 | 0.99945981 |
| CAPS2 | 12.8491298 | -0.235777625 | 0.59817105 | -0.3941642 | 0.6934598 | 0.99945981 |
| GLIPR1 | 28.8803274 | 0.210753838 | 0.46239066 | 0.45579173 | 0.64853975 | 0.99945981 |
| KRR1 | 11.7543477 | -0.006634971 | 0.46120162 | -0.0143863 | 0.98852181 | 0.99945981 |
| NAP1L1 | 51.0191716 | 0.548787398 | 0.34387986 | 1.59586954 | 0.11051792 | 0.99945981 |
| OSBPL8 | 151.891095 | 0.175936857 | 0.30303253 | 0.58058737 | 0.56151859 | 0.99945981 |
| ZDHHC17 | 40.9221191 | 0.089461773 | 0.26286065 | 0.34033917 | 0.73360113 | 0.99945981 |
| SYT1 | 105.625889 | -7.191398272 | 1.68459192 | -4.268926 | 1.96E-05 | 0.03370996 |
| PAWR | 7.78018253 | 0.204964223 | 0.59691266 | 0.34337389 | 0.7313172 | 0.99945981 |
| PPP1R12A | 155.61179 | 0.438187107 | 0.22165758 | 1.97686496 | 0.0480569 | 0.90884462 |
| METTL25 | 20.8240562 | 0.110550494 | 0.36275408 | 0.30475328 | 0.76055407 | 0.99945981 |
| TMTC2 | 66.8789465 | 0.750085565 | 0.42477464 | 1.76584355 | 0.0774221 | 0.98289661 |
| CEP290 | 20.8028595 | -0.521856195 | 0.40868799 | -1.2769061 | 0.20163539 | 0.99945981 |
| TMTC3 | 12.6335707 | -0.317919661 | 0.61935778 | -0.5133053 | 0.60773774 | 0.99945981 |
| DUSP6 | 36.5431244 | 1.124534537 | 0.67201683 | 1.67337258 | 0.09425397 | 0.99945981 |
| POC1B | 28.2889945 | -0.295320611 | 0.28205805 | -1.0470207 | 0.29509004 | 0.99945981 |
| ATP2B1 | 165.947216 | 0.082884308 | 0.437131 | 0.18960977 | 0.84961493 | 0.99945981 |
| BTG1 | 123.763998 | -0.754496454 | 0.28663153 | -2.632287 | 0.00848122 | 0.58811673 |
| LINC02391 | 16.8389316 | 0.567066992 | 0.58941631 | 0.96208229 | 0.33600827 | 0.99945981 |
| EEA1 | 59.3902327 | -0.280789803 | 0.43406582 | -0.646883 | 0.51770767 | 0.99945981 |
| NUDT4 | 19.1509355 | -0.158708429 | 0.46394514 | -0.3420845 | 0.73228732 | 0.99945981 |
| UBE2N | 22.1042169 | -0.289963698 | 0.3540561 | -0.8189767 | 0.4127997 | 0.99945981 |
| MRPL42 | 14.0375001 | 0.445555119 | 0.42638241 | 1.044966 | 0.29603869 | 0.99945981 |
| CRADD | 59.3471254 | -0.523146316 | 0.27569081 | -1.8975835 | 0.05775097 | 0.92445778 |
| PLXNC1 | 93.1132088 | -0.153477916 | 0.5509556 | -0.2785668 | 0.78057733 | 0.99945981 |
| CEP83 | 18.6291791 | -0.178586986 | 0.44871331 | -0.397998 | 0.69063168 | 0.99945981 |
| TMCC3 | 27.161074 | -0.098360754 | 0.64173279 | -0.1532737 | 0.87818245 | 0.99945981 |
| NDUFA12 | 14.2678449 | 0.085998496 | 0.55108962 | 0.15605174 | 0.87599224 | 0.99945981 |
| NR2C1 | 18.8657801 | -0.03390472 | 0.39762182 | -0.0852688 | 0.93204773 | 0.99945981 |
| FGD6 | 14.6924513 | 0.198577409 | 0.59542947 | 0.33350282 | 0.73875476 | 0.99945981 |
| VEZT | 32.1430977 | -0.297990183 | 0.2936221 | -1.0148765 | 0.31016468 | 0.99945981 |
| METAP2 | 23.3162523 | 0.070793758 | 0.4270499 | 0.16577397 | 0.86833483 | 0.99945981 |
| LTA4H | 36.4372739 | 1.767377004 | 0.74829198 | 2.36188154 | 0.01818245 | 0.75650017 |
| ELK3 | 27.3529889 | -0.464821026 | 0.50781429 | -0.9153366 | 0.36001493 | 0.99945981 |
| CDK17 | 93.921937 | 0.414120434 | 0.27766323 | 1.49144861 | 0.13584375 | 0.99945981 |
| NEDD1 | 17.9480256 | 0.386190605 | 0.40607917 | 0.95102294 | 0.34159273 | 0.99945981 |
| TMPO | 46.8168803 | 0.646845418 | 0.43541481 | 1.48558433 | 0.1373891 | 0.99945981 |
| SLC25A3 | 36.2059479 | -0.125752968 | 0.37208808 | -0.3379656 | 0.73538912 | 0.99945981 |
| IKBIP | 17.5584825 | -0.499783598 | 0.5411394 | -0.9235764 | 0.35570688 | 0.99945981 |
| APAF1 | 42.0345645 | 0.187277105 | 0.37667531 | 0.49718444 | 0.61905899 | 0.99945981 |
| UHRF1BP1L | 36.657604 | 0.215413987 | 0.30921369 | 0.69665088 | 0.48602131 | 0.99945981 |
| ACTR6 | 9.03443644 | 0.224005596 | 0.50381925 | 0.444615 | 0.65659801 | 0.99945981 |
| SCYL2 | 36.4475371 | 0.090736785 | 0.26068358 | 0.3480725 | 0.72778574 | 0.99945981 |
| UTP20 | 11.7665867 | -0.375947014 | 0.60449182 | -0.6219224 | 0.53399288 | 0.99945981 |
| ARL1 | 10.6456525 | -0.719519477 | 0.467195 | -1.5400838 | 0.12353992 | 0.99945981 |
| CHPT1 | 29.8909316 | -0.318296041 | 0.39185772 | -0.8122745 | 0.41663413 | 0.99945981 |
| GNPTAB | 55.1390097 | 0.14653129 | 0.26796342 | 0.54683318 | 0.58449334 | 0.99945981 |
| DRAM1 | 38.3124547 | 0.05490393 | 0.28048373 | 0.19574729 | 0.84480797 | 0.99945981 |
| WASHC3 | 22.8391619 | 0.081775728 | 0.31112705 | 0.26283709 | 0.79267614 | 0.99945981 |
| C12orf42 | 17.2080108 | 0.844294133 | 0.65661847 | 1.28582148 | 0.19850537 | 0.99945981 |
| HSP90B1 | 59.0772011 | 0.029859859 | 0.58694927 | 0.05087298 | 0.95942674 | 0.99945981 |
| TDG | 16.7334449 | -0.029042627 | 0.36155462 | -0.0803271 | 0.93597712 | 0.99945981 |
| HCFC2 | 12.9862866 | -0.13062566 | 0.49213068 | -0.2654288 | 0.79067914 | 0.99945981 |
| TXNRD1 | 150.975559 | -0.490773645 | 0.48613706 | -1.0095376 | 0.31271687 | 0.99945981 |
| CHST11 | 421.83079 | -0.151172276 | 0.27425011 | -0.5512205 | 0.58148255 | 0.99945981 |
| SLC41A2 | 19.4599582 | 0.328099627 | 0.56276151 | 0.58301718 | 0.55988174 | 0.99945981 |
| WASHC4 | 90.3028352 | 0.505743582 | 0.30903911 | 1.63650348 | 0.10173426 | 0.99945981 |
| APPL2 | 33.5308723 | 0.043925362 | 0.39008215 | 0.11260541 | 0.91034339 | 0.99945981 |
| C12orf75 | 21.9636411 | 0.279876723 | 0.53306134 | 0.52503661 | 0.59955774 | 0.99945981 |
| TCP11L2 | 11.0605051 | 0.533989418 | 0.52592084 | 1.01534181 | 0.30994292 | 0.99945981 |
| RIC8B | 20.8340453 | -0.191535859 | 0.39078524 | -0.4901307 | 0.62404139 | 0.99945981 |
| TMEM263 | 9.66259747 | -0.705723676 | 0.63787933 | -1.1063592 | 0.26857107 | 0.99945981 |
| CRY1 | 17.3334664 | 0.479347519 | 0.4732738 | 1.01283342 | 0.31113974 | 0.99945981 |
| BTBD11 | 31.1538837 | -1.141104436 | 0.60787187 | -1.8772121 | 0.06048903 | 0.92445778 |
| PWP1 | 10.3887011 | 0.318902949 | 0.4488861 | 0.71043178 | 0.47743642 | 0.99945981 |
| PRDM4 | 11.9248309 | -0.557830809 | 0.44866503 | -1.2433124 | 0.21375273 | 0.99945981 |
| CMKLR1 | 27.9122208 | 0.278825395 | 0.52084591 | 0.53533184 | 0.59242042 | 0.99945981 |
| SART3 | 15.8837869 | -0.11189352 | 0.39113459 | -0.2860742 | 0.77482128 | 0.99945981 |
| ISCU | 16.4222034 | -0.61129025 | 0.41842927 | -1.4609165 | 0.14403834 | 0.99945981 |
| SELPLG | 18.4540902 | 0.477971606 | 0.65398845 | 0.73085634 | 0.46486691 | 0.99945981 |
| CORO1C | 45.3028191 | 0.375401825 | 0.34748677 | 1.08033415 | 0.27999341 | 0.99945981 |
| SSH1 | 37.6472248 | -0.169386662 | 0.47352295 | -0.3577158 | 0.72055598 | 0.99945981 |
| ACACB | 10.0507644 | -0.211841486 | 0.47811981 | -0.443072 | 0.65771368 | 0.99945981 |
| UBE3B | 13.4096307 | 0.139974966 | 0.41896918 | 0.3340937 | 0.73830885 | 0.99945981 |

| | | | | | | |
|---|---|---|---|---|---|---|
| GLTP | 11.3132455 | 0.343162241 | 0.46213041 | 0.7425658 | 0.4577446 | 0.99945981 |
| TCHP | 21.0771218 | -1.007605971 | 0.54044168 | -1.864412 | 0.06226386 | 0.92949575 |
| GIT2 | 64.2014545 | 0.090648109 | 0.2888186 | 0.31385828 | 0.75362868 | 0.99945981 |
| ANKRD13A | 32.205054 | 0.256855779 | 0.33586796 | 0.76475225 | 0.44441909 | 0.99945981 |
| ATP2A2 | 58.248362 | 0.486586361 | 0.28795952 | 1.68977346 | 0.0910713 | 0.99640647 |
| ANAPC7 | 16.750205 | -0.261762793 | 0.3932446 | -0.6656488 | 0.50563561 | 0.99945981 |
| ARPC3 | 83.0484967 | 0.175328307 | 0.39743864 | 0.44114559 | 0.6591076 | 0.99945981 |
| VPS29 | 30.1887328 | 0.093639552 | 0.31889149 | 0.2936408 | 0.76903239 | 0.99945981 |
| PPTC7 | 29.5139794 | -0.124391978 | 0.35646279 | -0.348962 | 0.72711781 | 0.99945981 |
| HVCN1 | 16.3465852 | 0.135331398 | 0.77014125 | 0.17572283 | 0.8605117 | 0.99945981 |
| PPP1CC | 21.4024572 | -0.029330897 | 0.34597264 | -0.0847781 | 0.93243782 | 0.99945981 |
| SH2B3 | 64.9418839 | 0.570090219 | 0.43053211 | 1.32415261 | 0.18545237 | 0.99945981 |
| ATXN2 | 82.2388423 | 0.175865564 | 0.28841303 | 0.60976983 | 0.54201429 | 0.99945981 |
| BRAP | 13.8159243 | -0.192828048 | 0.43265216 | -0.4456884 | 0.65582235 | 0.99945981 |
| ACAD10 | 9.13476425 | -0.383965302 | 0.49895912 | -0.7695326 | 0.44157721 | 0.99945981 |
| ALDH2 | 19.3697127 | -0.369304896 | 0.6751677 | -0.5469825 | 0.58439077 | 0.99945981 |
| MAPKAPK5 | 16.6471153 | 0.006919486 | 0.57009203 | 0.01213749 | 0.99031592 | 0.99945981 |
| TMEM116 | 14.0788587 | 0.66120335 | 0.55615757 | 1.18887773 | 0.23448779 | 0.99945981 |
| ERP29 | 27.8700248 | -0.610509859 | 0.4609206 | -1.3245445 | 0.18532226 | 0.99945981 |
| NAA25 | 30.1153423 | 0.119809939 | 0.47217716 | 0.25373938 | 0.79969691 | 0.99945981 |
| TRAFD1 | 10.5643066 | 0.19216273 | 0.50995459 | 0.37682322 | 0.70630499 | 0.99945981 |
| HECTD4 | 60.0780963 | -0.5890334 | 0.34804045 | -1.692428 | 0.0905644 | 0.99640647 |
| RPL6 | 79.243972 | -0.218867537 | 0.58459378 | -0.3743925 | 0.70811231 | 0.99945981 |
| PTPN11 | 31.2913537 | 0.056069962 | 0.30180667 | 0.18578106 | 0.85261646 | 0.99945981 |
| OAS1 | 19.2220461 | 1.460064761 | 0.81490382 | 1.79170195 | 0.07318072 | 0.96364229 |
| OAS2 | 14.4901927 | 0.59895571 | 0.66310893 | 0.90325387 | 0.36639117 | 0.99945981 |
| DDX54 | 8.80499286 | -0.133159668 | 0.53072496 | -0.2509015 | 0.80189029 | 0.99945981 |
| TPCN1 | 33.1591716 | -0.148912091 | 0.39798113 | -0.3741687 | 0.70827879 | 0.99945981 |
| SLC8B1 | 17.9928691 | 0.017954282 | 0.47799851 | 0.03756138 | 0.9700374 | 0.99945981 |
| PLBD2 | 9.83374614 | -0.036073795 | 0.46288965 | -0.0779317 | 0.93788235 | 0.99945981 |
| RBM19 | 13.5757455 | -0.504390811 | 0.41158612 | -1.2254806 | 0.22039419 | 0.99945981 |
| MED13L | 401.428493 | -0.004693373 | 0.34685056 | -0.0135314 | 0.98920384 | 0.99945981 |
| FBXW8 | 20.4015238 | -0.942759183 | 0.43811262 | -2.1518649 | 0.03140799 | 0.86065765 |
| TESC | 10.4042534 | 0.182443424 | 0.47822009 | 0.38150514 | 0.70282845 | 0.99945981 |
| FBXO21 | 15.2232287 | -0.231331687 | 0.39657251 | -0.5833276 | 0.55967279 | 0.99945981 |
| WSB2 | 10.9820898 | 0.384163263 | 0.53503558 | 0.71801443 | 0.47274839 | 0.99945981 |
| PEBP1 | 36.9702718 | -0.851259071 | 0.47527963 | -1.7910699 | 0.07328207 | 0.96364229 |
| TAOK3 | 237.451345 | 0.281172193 | 0.20987721 | 1.33969857 | 0.18034336 | 0.99945981 |
| SUDS3 | 10.211453 | -0.126083939 | 0.43435366 | -0.2902794 | 0.77160246 | 0.99945981 |
| BICDL1 | 25.8142284 | -0.768420264 | 0.57055259 | -1.3468001 | 0.17804464 | 0.99945981 |
| RAB35 | 11.5942589 | -0.35945361 | 0.50499786 | -0.7117923 | 0.47659337 | 0.99945981 |
| GCN1 | 13.866392 | 0.137400643 | 0.44387265 | 0.30954969 | 0.75690342 | 0.99945981 |
| RPLP0 | 88.5927128 | -0.497718519 | 0.55747307 | -0.8928118 | 0.37195798 | 0.99945981 |
| PXN | 32.3667406 | -0.044358515 | 0.41910177 | -0.1058419 | 0.91570782 | 0.99945981 |
| SRSF9 | 23.9649836 | 0.23064657 | 0.5232502 | 0.44079595 | 0.65936072 | 0.99945981 |
| DYNLL1 | 27.343396 | -0.051391072 | 0.50292304 | -0.1021848 | 0.91861002 | 0.99945981 |
| RNF10 | 34.6383557 | -0.029725907 | 0.31564137 | -0.0941762 | 0.92496919 | 0.99945981 |
| MLEC | 20.1998949 | -0.400566417 | 0.55362402 | -0.7235351 | 0.46935119 | 0.99945981 |
| SPPL3 | 123.282934 | 0.365275182 | 0.24594975 | 1.48516183 | 0.13750096 | 0.99945981 |
| P2RX7 | 21.6692115 | 0.888041488 | 0.53396324 | 1.66311353 | 0.09628971 | 0.99945981 |
| P2RX4 | 18.190941 | 0.477355898 | 0.32774745 | 1.45647481 | 0.14526141 | 0.99945981 |
| CAMKK2 | 19.0704689 | 0.945723594 | 0.66552666 | 1.42101534 | 0.1553123 | 0.99945981 |
| ANAPC5 | 39.2520157 | 0.150511448 | 0.35214525 | 0.42741298 | 0.66907856 | 0.99945981 |
| RNF34 | 19.9779037 | -0.359965684 | 0.46187361 | -0.7793597 | 0.43576785 | 0.99945981 |
| KDM2B | 45.5300572 | -0.035126819 | 0.37998217 | -0.0924433 | 0.92634581 | 0.99945981 |
| ORAI1 | 10.1199005 | -0.226567686 | 0.63934945 | -0.3543722 | 0.72305995 | 0.99945981 |
| TMEM120B | 14.5175261 | -0.457506234 | 0.42787143 | -1.069261 | 0.28495208 | 0.99945981 |
| RHOF | 10.1666506 | -0.212674596 | 0.49768285 | -0.4273296 | 0.6691393 | 0.99945981 |
| SETD1B | 11.4278595 | -0.575816858 | 0.40270353 | -1.4298778 | 0.15275208 | 0.99945981 |
| MLXIP | 41.1463769 | -0.285587516 | 0.30548115 | -0.9348777 | 0.34985133 | 0.99945981 |
| CLIP1 | 62.0695158 | -0.766286765 | 0.39461794 | -1.9418447 | 0.0521559 | 0.9230968 |
| ZCCHC8 | 21.2930214 | -0.367615761 | 0.29475111 | -1.2472074 | 0.21232147 | 0.99945981 |
| RSRC2 | 36.2668022 | -0.076245039 | 0.26683722 | -0.2857362 | 0.77508021 | 0.99945981 |
| KNTC1 | 11.7312453 | 0.216783639 | 0.64912632 | 0.33396218 | 0.7384081 | 0.99945981 |
| DENR | 24.1745602 | -0.146272633 | 0.38067075 | -0.3842497 | 0.70079336 | 0.99945981 |
| VPS37B | 50.9183219 | -0.074013605 | 0.59082109 | -0.1252725 | 0.90030786 | 0.99945981 |
| ARL6IP4 | 25.1331628 | 0.138513054 | 0.46372863 | 0.29869421 | 0.76517338 | 0.99945981 |
| MPHOSPH9 | 30.3195404 | -0.219988353 | 0.43424562 | -0.5065989 | 0.61243628 | 0.99945981 |
| C12orf65 | 11.0547689 | -0.364591856 | 0.56428288 | -0.6461154 | 0.51820463 | 0.99945981 |
| SBNO1 | 35.0000338 | -0.110498727 | 0.29237049 | -0.3779408 | 0.7054746 | 0.99945981 |
| RILPL2 | 33.1091721 | 0.327595884 | 0.40672516 | 0.80544779 | 0.42056132 | 0.99945981 |
| SNRNP35 | 8.07450508 | -0.006443555 | 0.46913357 | -0.013735 | 0.98904139 | 0.99945981 |
| TMED2 | 13.4229809 | -0.027019758 | 0.53840895 | -0.0501845 | 0.9599754 | 0.99945981 |
| EIF2B1 | 8.22167902 | 0.478753907 | 0.49616175 | 0.96491499 | 0.33458739 | 0.99945981 |
| ATP6V0A2 | 18.8061946 | -0.529281566 | 0.52585984 | -1.0065069 | 0.31417179 | 0.99945981 |
| CCDC92 | 15.2376084 | -0.387135613 | 0.43649475 | -0.8869193 | 0.37512235 | 0.99945981 |
| NCOR2 | 67.8455062 | -0.322643498 | 0.31763605 | -1.0157647 | 0.30974144 | 0.99945981 |
| SCARB1 | 20.3339132 | 0.72534795 | 0.803593 | 0.90263099 | 0.36672177 | 0.99945981 |
| UBC | 105.349317 | -0.31601423 | 0.2781555 | -1.1361063 | 0.25591206 | 0.99945981 |
| BRI3BP | 12.2596478 | 0.15950712 | 0.54676727 | 0.29172763 | 0.77049488 | 0.99945981 |
| LINC00944 | 11.8197797 | -0.898527375 | 0.88810502 | -1.0117355 | 0.31166454 | 0.99945981 |
| SLC15A4 | 22.4782684 | -0.912985165 | 0.65819602 | -1.3871022 | 0.1654106 | 0.99945981 |
| GLT1D1 | 27.3208562 | 0.467375119 | 0.85429435 | 0.54708909 | 0.58431753 | 0.99945981 |
| STX2 | 15.6528689 | 0.819850729 | 0.60565963 | 1.35364929 | 0.17584829 | 0.99945981 |
| RAN | 20.4417099 | -0.62972644 | 0.49493753 | -1.2723352 | 0.20325405 | 0.99945981 |
| SFSWAP | 41.9033491 | 0.068327178 | 0.28344581 | 0.24105905 | 0.80950935 | 0.99945981 |
| ULK1 | 14.5046413 | 1.148702558 | 0.52572194 | 2.18500023 | 0.02888883 | 0.84763724 |
| EP400 | 38.2449425 | -0.118981918 | 0.27056228 | -0.439758 | 0.66011241 | 0.99945981 |
| FBRSL1 | 19.6174429 | -0.115807255 | 0.32898631 | -0.3520124 | 0.72482897 | 0.99945981 |
| ANKLE2 | 28.9349162 | -0.33106852 | 0.41124828 | -0.8050332 | 0.42080052 | 0.99945981 |
| GOLGA3 | 25.224231 | -0.298285306 | 0.49017154 | -0.6085325 | 0.54283437 | 0.99945981 |
| CHFR | 38.1977985 | -0.453400903 | 0.32475733 | -1.3961222 | 0.16267771 | 0.99945981 |
| ZNF26 | 12.4691642 | 0.12572248 | 0.43211494 | 0.29094685 | 0.77109197 | 0.99945981 |
| ZNF84 | 11.5485057 | -0.270784288 | 0.53728926 | -0.5039823 | 0.6142738 | 0.99945981 |
| ZNF140 | 8.01332601 | -0.254394267 | 0.51062908 | -0.4981978 | 0.61834466 | 0.99945981 |
| MPHOSPH8 | 47.0161383 | -0.067436042 | 0.2825838 | -0.2386409 | 0.81138407 | 0.99945981 |
| PSPC1 | 86.4685917 | 0.028506702 | 0.24281274 | 0.11740201 | 0.9065415 | 0.99945981 |
| ZMYM5 | 25.8390205 | -0.052536066 | 0.36212376 | -0.1450777 | 0.88464956 | 0.99945981 |
| ZMYM2 | 149.378066 | -0.088836857 | 0.26273473 | -0.3381238 | 0.73526992 | 0.99945981 |
| CRYL1 | 18.6403404 | -0.905746053 | 0.38048312 | -2.3805158 | 0.01728842 | 0.74280553 |
| IFT88 | 16.7700127 | -0.747986927 | 0.48578173 | -1.5397593 | 0.12361904 | 0.99945981 |
| XPO4 | 45.8640772 | -0.22967784 | 0.3188953 | -0.7202296 | 0.47138364 | 0.99945981 |
| LATS2 | 23.1717043 | 0.4634557 | 0.50905088 | 0.910431 | 0.36259525 | 0.99945981 |
| SAP18 | 15.1173819 | -0.595825989 | 0.46498292 | -1.2813933 | 0.20005557 | 0.99945981 |
| MRPL57 | 11.5859034 | -0.517044364 | 0.46249137 | -1.1179546 | 0.26358638 | 0.99945981 |
| ZDHHC20 | 89.9536062 | -0.032647128 | 0.29430375 | -0.11093 | 0.91167182 | 0.99945981 |
| MICU2 | 75.5428431 | -0.159534885 | 0.23678896 | -0.6737429 | 0.50047477 | 0.99945981 |
| SACS | 12.3049795 | -0.34925463 | 0.55034418 | -0.6346113 | 0.52568196 | 0.99945981 |
| MIPEP | 14.4393455 | -1.192802273 | 0.74688144 | -1.5970437 | 0.11025598 | 0.99945981 |
| PARP4 | 85.7834609 | -0.147761164 | 0.34499225 | -0.4283029 | 0.66843064 | 0.99945981 |
| MTMR6 | 23.4245323 | -0.136577167 | 0.39428113 | -0.3463954 | 0.72904558 | 0.99945981 |
| NUP58 | 32.3387428 | -0.227348461 | 0.38321028 | -0.5932734 | 0.5529982 | 0.99945981 |
| CDK8 | 53.4654405 | 0.035804867 | 0.27670513 | 0.12939719 | 0.89704337 | 0.99945981 |

| | | | | | | |
|---|---|---|---|---|---|---|
| USP12 | 36.4282524 | -0.310850323 | 0.28475176 | -1.0916537 | 0.27498533 | 0.99945981 |
| RPL21 | 40.3152037 | -0.022671531 | 0.59576229 | -0.0380547 | 0.9696441 | 0.99945981 |
| GTF3A | 20.5541654 | 0.071474803 | 0.44214211 | 0.16165572 | 0.87157697 | 0.99945981 |
| MTIF3 | 18.8144899 | -0.141747723 | 0.36883778 | -0.3843091 | 0.70074935 | 0.99945981 |
| LNX2 | 11.7695641 | 0.14740234 | 0.50731579 | 0.29055343 | 0.77139288 | 0.99945981 |
| POLR1D | 19.7881336 | 0.080784627 | 0.36874475 | 0.21908007 | 0.82658768 | 0.99945981 |
| FLT3 | 40.6103678 | -2.323155804 | 1.55776595 | -1.4913382 | 0.13587274 | 0.99945981 |
| PAN3 | 233.49035 | 0.079722368 | 0.26803419 | 0.29743357 | 0.76613552 | 0.99945981 |
| FLT1 | 18.970013 | 0.353822929 | 0.59326862 | 0.59639583 | 0.55091083 | 0.99945981 |
| POMP | 19.06428 | 0.276944459 | 0.42305067 | 0.65463662 | 0.51270174 | 0.99945981 |
| SLC46A3 | 11.7554405 | -0.363140471 | 0.5236142 | -0.6935268 | 0.48797902 | 0.99945981 |
| SLC7A1 | 10.2046408 | -0.49775113 | 0.49179102 | -1.0121192 | 0.31148107 | 0.99945981 |
| UBL3 | 57.9056762 | -0.060104034 | 0.30043914 | -0.2000539 | 0.84143839 | 0.99945981 |
| KATNAL1 | 22.0706726 | -0.796203763 | 0.55018117 | -1.4471665 | 0.1478503 | 0.99945981 |
| LINC00426 | 16.4469957 | -0.742207451 | 0.72124025 | -1.029071 | 0.3034463 | 0.99945981 |
| HMGB1 | 62.0447311 | 0.124911743 | 0.26099628 | 0.47859588 | 0.63222615 | 0.99945981 |
| USPL1 | 25.1185292 | -0.489104514 | 0.32938515 | -1.4849015 | 0.13756991 | 0.99945981 |
| ALOX5AP | 27.7496675 | -0.659230714 | 0.45957317 | -1.4344413 | 0.15144638 | 0.99945981 |
| HSPH1 | 37.7892182 | 0.295683771 | 0.66523888 | 0.44447758 | 0.65669733 | 0.99945981 |
| FRY | 60.4307538 | 0.394664352 | 0.79720298 | 0.49506131 | 0.62055684 | 0.99945981 |
| N4BP2L1 | 31.512158 | -0.459331134 | 0.44662541 | -1.0284483 | 0.30373901 | 0.99945981 |
| N4BP2L2 | 208.014628 | 0.012514638 | 0.22613751 | 0.05534083 | 0.95586693 | 0.99945981 |
| PDS5B | 94.8940734 | 0.299816921 | 0.27488618 | 1.09069477 | 0.27540721 | 0.99945981 |
| STARD13 | 111.128321 | 0.171976351 | 0.65652959 | 0.2619476 | 0.79336184 | 0.99945981 |
| NBEA | 26.3462643 | -0.058078774 | 0.38661655 | -0.1502232 | 0.88058853 | 0.99945981 |
| SPART | 24.3464042 | -0.455783217 | 0.47543247 | -0.9586708 | 0.33772462 | 0.99945981 |
| ALG5 | 12.5488156 | -0.44842208 | 0.38643203 | -1.1604164 | 0.2458793 | 0.99945981 |
| EXOSC8 | 8.13011476 | -0.418090574 | 0.67871397 | -0.6160041 | 0.53789183 | 0.99945981 |
| SUPT20H | 33.2519742 | -0.041314134 | 0.39686681 | -0.1041008 | 0.91708939 | 0.99945981 |
| UFM1 | 23.1075837 | 0.36410971 | 0.44750172 | 0.81364985 | 0.41584557 | 0.99945981 |
| PROSER1 | 12.3368957 | 0.014476648 | 0.48783297 | 0.02967542 | 0.97632592 | 0.99945981 |
| NHLRC3 | 25.8635659 | -0.070397378 | 0.50275348 | -0.1400237 | 0.8886413 | 0.99945981 |
| COG6 | 24.4610269 | -0.458640312 | 0.416642 | -1.1008019 | 0.27098287 | 0.99945981 |
| FOXO1 | 130.34143 | -0.693578557 | 0.36725786 | -1.8885329 | 0.05895445 | 0.92445778 |
| MRPS31 | 22.2243791 | 0.359164306 | 0.36259597 | 0.99053584 | 0.32191228 | 0.99945981 |
| ELF1 | 190.24259 | 0.107712174 | 0.24884203 | 0.43285363 | 0.66512111 | 0.99945981 |
| WBP4 | 12.1832247 | -0.053064345 | 0.47110809 | -0.1126373 | 0.91031811 | 0.99945981 |
| NAA16 | 26.6785373 | 0.206778963 | 0.44303167 | 0.46673631 | 0.64068855 | 0.99945981 |
| VWA8 | 66.4315842 | 0.338363707 | 0.32451985 | 1.0426595 | 0.29710602 | 0.99945981 |
| DGKH | 43.1283501 | -0.094693779 | 0.35256626 | -0.2685844 | 0.78824956 | 0.99945981 |
| AKAP11 | 33.5656677 | 0.280610851 | 0.31712181 | 0.8848677 | 0.37622799 | 0.99945981 |
| EPSTI1 | 88.3244323 | 0.775724514 | 0.79898835 | 0.97088339 | 0.33160635 | 0.99945981 |
| DNAJC15 | 23.3272199 | -0.087861095 | 0.41587897 | -0.211266 | 0.8326797 | 0.99945981 |
| TSC22D1 | 53.2199633 | -0.103518415 | 0.35597672 | -0.2908011 | 0.77120346 | 0.99945981 |
| GPALPP1 | 14.4275735 | -0.271927816 | 0.36646598 | -0.7420275 | 0.4580707 | 0.99945981 |
| GTF2F2 | 42.6918209 | 0.285764388 | 0.27085577 | 1.05504263 | 0.29140583 | 0.99945981 |
| TPT1 | 222.846728 | -0.229000877 | 0.45870229 | -0.4992364 | 0.61761286 | 0.99945981 |
| COG3 | 24.6055871 | 0.065479568 | 0.31483235 | 0.20798234 | 0.83524276 | 0.99945981 |
| ZC3H13 | 41.5474412 | -0.012136422 | 0.24608509 | -0.049318 | 0.96066588 | 0.99945981 |
| LCP1 | 183.479778 | 0.215099614 | 0.40974824 | 0.52495555 | 0.59961409 | 0.99945981 |
| RUBCNL | 27.7041667 | 0.105286765 | 0.63040418 | 0.1670147 | 0.86735848 | 0.99945981 |
| LRCH1 | 75.1239981 | 0.372202253 | 0.44634675 | 0.83388587 | 0.4043453 | 0.99945981 |
| ESD | 9.64375382 | 0.478646617 | 0.41293494 | 1.15913323 | 0.24640188 | 0.99945981 |
| SUCLA2 | 22.3803641 | 0.329198829 | 0.49219247 | 0.66884167 | 0.50359649 | 0.99945981 |
| MED4 | 20.8862785 | 0.166938142 | 0.40893355 | 0.40822804 | 0.68310626 | 0.99945981 |
| ITM2B | 132.155795 | 0.063334879 | 0.31057933 | 0.20392497 | 0.83841214 | 0.99945981 |
| RB1 | 161.825124 | -0.069356646 | 0.37576684 | -0.1845736 | 0.85356348 | 0.99945981 |
| LPAR6 | 40.4158545 | 0.945906858 | 0.51402692 | 1.84018932 | 0.06574045 | 0.94594189 |
| RCBTB2 | 34.3666128 | 0.663266634 | 0.39364571 | 1.68493295 | 0.09200151 | 0.99640647 |
| FNDC3A | 190.648453 | 0.40043302 | 0.40118255 | 0.99813171 | 0.3182155 | 0.99945981 |
| CAB39L | 11.29334 | -0.118969088 | 0.53172325 | -0.2237425 | 0.82295768 | 0.99945981 |
| SETDB2 | 28.9559963 | -0.069571476 | 0.45991453 | -0.1512704 | 0.87976238 | 0.99945981 |
| PHF11 | 23.6108701 | 0.201644821 | 0.40971941 | 0.49215344 | 0.62261087 | 0.99945981 |
| RCBTB1 | 11.6559128 | 0.425504371 | 0.41604903 | 1.0227265 | 0.30643718 | 0.99945981 |
| KPNA3 | 42.6736236 | 0.141985689 | 0.26190259 | 0.54213167 | 0.5877278 | 0.99945981 |
| DLEU2 | 288.420682 | 0.677529709 | 0.46717531 | 1.45026867 | 0.14698361 | 0.99945981 |
| TRIM13 | 24.9752233 | 0.060657093 | 0.35167159 | 0.17248221 | 0.86305845 | 0.99945981 |
| RNASEH2B | 37.9479653 | 0.167351708 | 0.32842997 | 0.50955067 | 0.61036629 | 0.99945981 |
| INTS6 | 73.4584026 | 0.500891275 | 0.33724384 | 1.48524958 | 0.13747772 | 0.99945981 |
| WDFY2 | 92.1292358 | 0.373880348 | 0.31282585 | 1.1951709 | 0.23202026 | 0.99945981 |
| VPS36 | 35.4837296 | 0.124704612 | 0.27969106 | 0.44586556 | 0.65569436 | 0.99945981 |
| SUGT1 | 18.9338757 | 0.310751491 | 0.36201413 | 0.85839604 | 0.39067381 | 0.99945981 |
| TDRD3 | 41.0593419 | -0.263333567 | 0.29843688 | -0.8823761 | 0.37757346 | 0.99945981 |
| PCDH9 | 31.1460419 | -2.994692094 | 1.25481589 | -2.386559 | 0.01700688 | 0.74280553 |
| DACH1 | 34.9328219 | -0.971322364 | 0.84133827 | -1.1544968 | 0.24829654 | 0.99945981 |
| DIS3 | 17.1142416 | 0.230190385 | 0.39226302 | 0.58682662 | 0.55732016 | 0.99945981 |
| PIBF1 | 69.7675727 | -0.262553318 | 0.28495354 | -0.9213899 | 0.3568469 | 0.99945981 |
| KLF12 | 136.594313 | 0.066655657 | 0.27205838 | 0.24500497 | 0.80645258 | 0.99945981 |
| TBC1D4 | 34.9742267 | 1.111639675 | 0.55063047 | 2.01884883 | 0.04350293 | 0.89562588 |
| COMMD6 | 18.7697408 | -0.27741647 | 0.4999421 | -0.5548972 | 0.57896498 | 0.99945981 |
| UCHL3 | 15.7375721 | 0.535929376 | 0.57152437 | 0.9377192 | 0.34838873 | 0.99945981 |
| LMO7 | 17.3008032 | 0.284855298 | 0.55757089 | 0.51088625 | 0.60943071 | 0.99945981 |
| KCTD12 | 36.926383 | 0.183598376 | 0.54971842 | 0.33398622 | 0.73838996 | 0.99945981 |
| AC001226.2 | 14.5179254 | -0.446182194 | 0.79084244 | -0.5641859 | 0.57262759 | 0.99945981 |
| FBXL3 | 23.6243118 | 0.155605497 | 0.30617088 | 0.50823088 | 0.61129143 | 0.99945981 |
| MYCBP2 | 222.511534 | 0.246914284 | 0.19166014 | 1.2882923 | 0.19764423 | 0.99945981 |
| RNF219 | 12.9996876 | 0.20924384 | 0.4336929 | 0.48247006 | 0.62947206 | 0.99945981 |
| RBM26 | 92.4309579 | 0.23518893 | 0.28932261 | 0.81289509 | 0.41627821 | 0.99945981 |
| NDFIP2 | 9.90916523 | 0.028022627 | 0.49050147 | 0.05713057 | 0.95444119 | 0.99945981 |
| GPC5 | 19.4979937 | 0.767907036 | 0.94560133 | 0.81208329 | 0.41674384 | 0.99945981 |
| GPC6 | 23.4454653 | -2.494517551 | 1.73262658 | -1.4397318 | 0.1499433 | 0.99945981 |
| ABCC4 | 32.5777368 | 0.587174434 | 0.57313206 | 1.02450111 | 0.30559864 | 0.99945981 |
| DNAJC3 | 51.0082544 | 0.055971331 | 0.36266638 | 0.15433284 | 0.87734731 | 0.99945981 |
| UGGT2 | 36.3840625 | -0.199101032 | 0.33149225 | -0.6006205 | 0.5480928 | 0.99945981 |
| MBNL2 | 76.1069497 | -0.356495756 | 0.36612889 | -0.9736892 | 0.33021087 | 0.99945981 |
| RAP2A | 16.3696301 | 0.015837447 | 0.39806648 | 0.03978593 | 0.96826379 | 0.99945981 |
| IPO5 | 14.3216807 | -0.008726467 | 0.53382963 | -0.0163469 | 0.98695763 | 0.99945981 |
| FARP1 | 31.2108899 | -0.889080589 | 0.49032227 | -1.8132576 | 0.0697921 | 0.95505283 |
| STK24 | 65.6866809 | -0.257193479 | 0.26643259 | -0.9653229 | 0.33438312 | 0.99945981 |
| DOCK9 | 44.7804756 | -0.330760454 | 0.3071518 | -1.0768632 | 0.28154141 | 0.99945981 |
| UBAC2 | 132.239646 | 0.098995849 | 0.19402758 | 0.51021536 | 0.60990059 | 0.99945981 |
| GPR183 | 28.0944864 | 1.094852284 | 0.72930264 | 1.50123176 | 0.13329563 | 0.99945981 |
| TM9SF2 | 59.3442317 | 0.06607332 | 0.25171466 | 0.26249293 | 0.79294143 | 0.99945981 |
| CLYBL | 32.1167663 | -0.232429618 | 0.49427575 | -0.4702428 | 0.63818155 | 0.99945981 |
| PCCA | 94.71634 | 0.323169096 | 0.24282421 | 1.33087674 | 0.18322957 | 0.99945981 |
| GGACT | 18.7639383 | -0.785223912 | 0.47965822 | -1.6370488 | 0.10162027 | 0.99945981 |
| TMTC4 | 13.0900598 | -0.22003078 | 0.43748028 | -0.5029502 | 0.61499931 | 0.99945981 |
| TPP2 | 54.5822482 | 0.139546496 | 0.31175468 | 0.44761637 | 0.65443009 | 0.99945981 |
| ARGLU1 | 73.3251309 | -0.265896833 | 0.30993678 | -0.8579067 | 0.390944 | 0.99945981 |
| ABHD13 | 14.0177039 | 1.062884447 | 0.37722329 | 2.81765333 | 0.0048376 | 0.50757754 |
| TNFSF13B | 46.2158207 | 0.846276058 | 0.86747532 | 0.97556212 | 0.32928151 | 0.99945981 |
| IRS2 | 27.8045117 | 0.084139682 | 0.91214118 | 0.09224414 | 0.92650407 | 0.99945981 |
| RAB20 | 36.03018 | 0.570973262 | 0.76331454 | 0.74801832 | 0.45444911 | 0.99945981 |

| | | | | | | |
|---|---|---|---|---|---|---|
| CARS2 | 24.9508503 | 0.292891962 | 0.36510469 | 0.80221363 | 0.42242939 | 0.99945981 |
| ANKRD10 | 73.6786192 | -0.082479533 | 0.20987253 | -0.3929982 | 0.69432079 | 0.99945981 |
| ARHGEF7 | 60.2667182 | -0.050669425 | 0.26637738 | -0.1902167 | 0.84913933 | 0.99945981 |
| TUBGCP3 | 39.4326388 | -0.528804735 | 0.37152776 | -1.4233249 | 0.15464198 | 0.99945981 |
| ATP11A | 84.4043414 | 0.350004136 | 0.28856816 | 1.21289938 | 0.22516829 | 0.99945981 |
| PCID2 | 17.2968289 | -0.173260198 | 0.38157374 | -0.4540674 | 0.64978031 | 0.99945981 |
| CUL4A | 52.9090955 | -0.045501202 | 0.35841765 | -0.1269502 | 0.89897979 | 0.99945981 |
| LAMP1 | 34.0401053 | -0.140281843 | 0.3929587 | -0.3569888 | 0.72110021 | 0.99945981 |
| DCUN1D2 | 13.2208241 | 0.125730493 | 0.41855397 | 0.30039254 | 0.76387775 | 0.99945981 |
| TMCO3 | 21.3304673 | -0.317221679 | 0.45844544 | -0.6919508 | 0.48896824 | 0.99945981 |
| TFDP1 | 12.2603426 | -0.517233864 | 0.54836283 | -0.9432329 | 0.34556178 | 0.99945981 |
| GAS6 | 13.2284852 | -0.367530942 | 0.54624637 | -0.67283 | 0.50105545 | 0.99945981 |
| RASA3 | 73.4089287 | 0.199652725 | 0.34056738 | 0.5862356 | 0.5577172 | 0.99945981 |
| CDC16 | 23.9411511 | -0.222471174 | 0.3120188 | -0.7130057 | 0.47584224 | 0.99945981 |
| UPF3A | 22.47426 | -0.623849013 | 0.4140909 | -1.5065509 | 0.13192581 | 0.99945981 |
| TEP1 | 18.7448289 | 0.137459905 | 0.36434938 | 0.37727498 | 0.70596926 | 0.99945981 |
| OSGEP | 10.0411817 | -0.07810159 | 0.43340717 | -0.1802037 | 0.85699263 | 0.99945981 |
| APEX1 | 9.61985683 | -0.406722161 | 0.47105226 | -0.8634332 | 0.38789933 | 0.99945981 |
| PNP | 9.52460911 | -1.052174855 | 0.76516054 | -1.3751034 | 0.16909939 | 0.99945981 |
| RNASE6 | 10.3129454 | -0.405515087 | 0.75635746 | -0.5361421 | 0.59186034 | 0.99945981 |
| RNASE1 | 50.5296098 | 0.895862869 | 0.83895568 | 1.06783098 | 0.28559676 | 0.99945981 |
| HNRNPC | 150.417454 | 0.275952068 | 0.20557755 | 1.34232592 | 0.17949034 | 0.99945981 |
| SUPT16H | 19.2831236 | -0.110103532 | 0.42941415 | -0.2564041 | 0.79763885 | 0.99945981 |
| CHD8 | 36.1147594 | -0.150868293 | 0.27976116 | -0.5392753 | 0.58969689 | 0.99945981 |
| RAB2B | 11.1338112 | -0.485010006 | 0.55969445 | -0.8665621 | 0.38618198 | 0.99945981 |
| TOX4 | 16.8361081 | -0.505642447 | 0.38257723 | -1.3216742 | 0.18627667 | 0.99945981 |
| TRAC | 18.4401283 | -0.833967718 | 0.61673291 | -1.3522348 | 0.17630021 | 0.99945981 |
| DAD1 | 20.952942 | -0.709642906 | 0.40611082 | -1.7474119 | 0.08056591 | 0.9846219 |
| OXA1L | 11.1696093 | 0.02060931 | 0.51594675 | 0.03994465 | 0.96813726 | 0.99945981 |
| SLC7A7 | 31.3290644 | 0.528532913 | 0.64765805 | 0.81606785 | 0.41446133 | 0.99945981 |
| LRP10 | 15.0343268 | -0.501983549 | 0.45178616 | -1.1111087 | 0.26652155 | 0.99945981 |
| RBM23 | 17.0149716 | 0.133441041 | 0.42753031 | 0.31212065 | 0.75494883 | 0.99945981 |
| PSMB5 | 9.52249699 | -0.905146305 | 0.49768143 | -1.8187263 | 0.0689532 | 0.95436228 |
| ACIN1 | 42.3364885 | 0.12329032 | 0.25973317 | 0.47468069 | 0.63501458 | 0.99945981 |
| SLC7A8 | 15.6312412 | 0.363519639 | 0.73278312 | 0.4960808 | 0.6198374 | 0.99945981 |
| PABPN1 | 12.9072079 | 0.487521096 | 0.50006152 | 0.97492224 | 0.32959883 | 0.99945981 |
| AP1G2 | 7.46734879 | -0.072564303 | 0.60339741 | -0.1202596 | 0.90427754 | 0.99945981 |
| DHRS4L2 | 8.52689017 | 0.151217905 | 0.47297477 | 0.31971664 | 0.74918314 | 0.99945981 |
| PSME1 | 21.0760716 | -0.41282063 | 0.55208035 | -0.7477546 | 0.45460818 | 0.99945981 |
| LTB4R | 10.8384815 | 0.416576786 | 0.5449313 | 0.76445744 | 0.44459469 | 0.99945981 |
| GZMH | 12.4095114 | -0.85271745 | 0.95244926 | -0.8952891 | 0.37063256 | 0.99945981 |
| GZMB | 19.7700534 | -1.113208042 | 0.86882189 | -1.2812845 | 0.20009375 | 0.99945981 |
| PRKD1 | 15.4769534 | 0.332605662 | 0.65263624 | 0.50963407 | 0.61030785 | 0.99945981 |
| G2E3 | 24.5902633 | 0.140878231 | 0.39676597 | 0.35506631 | 0.72253991 | 0.99945981 |
| SCFD1 | 84.1587903 | 0.177131564 | 0.29281339 | 0.60492986 | 0.5452256 | 0.99945981 |
| STRN3 | 97.8243322 | -0.047434786 | 0.26243549 | -0.1807484 | 0.85656509 | 0.99945981 |
| AP4S1 | 9.68305108 | 0.393328573 | 0.43921807 | 0.89552001 | 0.37050917 | 0.99945981 |
| HECTD1 | 79.2935222 | -0.049517508 | 0.29829109 | -0.166004 | 0.86815382 | 0.99945981 |
| HEATR5A | 13.8123522 | -0.206694209 | 0.40595284 | -0.5091582 | 0.61064135 | 0.99945981 |
| NUBPL | 39.4600788 | -0.306668217 | 0.39003794 | -0.7862523 | 0.43171971 | 0.99945981 |
| ARHGAP5 | 61.4382281 | -0.074489919 | 0.47840277 | -0.1557055 | 0.8762652 | 0.99945981 |
| NPAS3 | 20.5078118 | 0.261688133 | 0.87760278 | 0.29818517 | 0.76556184 | 0.99945981 |
| EAPP | 14.7847366 | -0.479677456 | 0.37846887 | -1.2674159 | 0.20500663 | 0.99945981 |
| SNX6 | 81.7317246 | 0.063449656 | 0.32087151 | 0.19774163 | 0.84324721 | 0.99945981 |
| BAZ1A | 114.551639 | 0.339721914 | 0.2517586 | 1.34939547 | 0.17720997 | 0.99945981 |
| SRP54 | 35.8899019 | -0.139105981 | 0.41079339 | -0.3386276 | 0.73489029 | 0.99945981 |
| FAM177A1 | 23.1949706 | -0.278139145 | 0.358752 | -0.7752964 | 0.43816452 | 0.99945981 |
| PPP2R3C | 39.5994942 | 0.292820838 | 0.37832281 | 0.77399731 | 0.43893238 | 0.99945981 |
| NFKBIA | 38.5624533 | 0.417997321 | 0.59978157 | 0.69691591 | 0.48585542 | 0.99945981 |
| RALGAPA1 | 86.6152786 | -0.244967351 | 0.30962592 | -0.791172 | 0.42884364 | 0.99945981 |
| SEC23A | 34.8341194 | 0.032640214 | 0.34816528 | 0.09374919 | 0.92530839 | 0.99945981 |
| TRAPPC6B | 15.8455529 | -0.223151504 | 0.37547655 | -0.5943154 | 0.55230117 | 0.99945981 |
| PNN | 32.2990768 | -0.070469551 | 0.30739929 | -0.2292443 | 0.818679 | 0.99945981 |
| MIA2 | 39.4508687 | -0.160068043 | 0.27086878 | -0.5909431 | 0.55455853 | 0.99945981 |
| FBXO33 | 35.4412063 | -0.121598209 | 0.28946806 | -0.4200747 | 0.67443088 | 0.99945981 |
| KLHL28 | 23.7737703 | -0.07216502 | 0.35860477 | -0.2012383 | 0.84051223 | 0.99945981 |
| TOGARAM1 | 22.9084634 | -0.142087255 | 0.42699388 | -0.3327618 | 0.73931409 | 0.99945981 |
| PRPF39 | 13.6319524 | 0.17077816 | 0.41190573 | 0.41460497 | 0.67843111 | 0.99945981 |
| FKBP3 | 11.0605186 | -0.617774969 | 0.43792834 | -1.4106759 | 0.15834019 | 0.99945981 |
| FANCM | 11.9923684 | 0.07053265 | 0.56981666 | 0.12378131 | 0.90148843 | 0.99945981 |
| MIS18BP1 | 59.485716 | 0.248279629 | 0.34317137 | 0.72348586 | 0.46938144 | 0.99945981 |
| RPS29 | 48.3643652 | 0.084953445 | 0.46128939 | 0.18416518 | 0.85388387 | 0.99945981 |
| RPL36AL | 60.5038687 | -0.60901179 | 0.45602919 | -1.3354667 | 0.18172365 | 0.99945981 |
| KLHDC1 | 10.4752662 | -0.232253908 | 0.76414067 | -0.3039413 | 0.76117261 | 0.99945981 |
| KLHDC2 | 12.914517 | -0.611891157 | 0.51611076 | -1.1855811 | 0.23578778 | 0.99945981 |
| NEMF | 67.856932 | -0.170160716 | 0.25290312 | -0.6728296 | 0.50105568 | 0.99945981 |
| ARF6 | 22.5742078 | 0.354277904 | 0.57729439 | 0.61368672 | 0.53942236 | 0.99945981 |
| LINC01588 | 18.2116067 | -0.443633455 | 0.51527919 | -0.8609574 | 0.38926148 | 0.99945981 |
| VCPKMT | 10.6721711 | 0.145195806 | 0.4608452 | 0.31506416 | 0.75271294 | 0.99945981 |
| SOS2 | 79.5266727 | -0.025416065 | 0.24465857 | -0.1038838 | 0.91726155 | 0.99945981 |
| ATP5S | 10.7763305 | 0.248146789 | 0.43399196 | 0.57177738 | 0.5674728 | 0.99945981 |
| MAP4K5 | 49.9601532 | -0.374822927 | 0.32710157 | -1.1458916 | 0.25184003 | 0.99945981 |
| NIN | 89.1876268 | -0.219640606 | 0.28783648 | -0.7630742 | 0.44541912 | 0.99945981 |
| PYGL | 23.0942417 | -0.222716568 | 0.59850114 | -0.3721239 | 0.70980062 | 0.99945981 |
| TMX1 | 15.1155251 | 0.007864485 | 0.39354061 | 0.01998392 | 0.9840562 | 0.99945981 |
| GNG2 | 148.473155 | 0.441085441 | 0.50188553 | 0.87885667 | 0.379479 | 0.99945981 |
| RTRAF | 13.90957 | 0.349450041 | 0.44758611 | 0.78074371 | 0.43495325 | 0.99945981 |
| PTGDR | 19.6545684 | 1.106147647 | 0.56305081 | 1.96456098 | 0.04946507 | 0.92026482 |
| TXNDC16 | 15.8494935 | 0.433909698 | 0.56248496 | 0.77141565 | 0.4404606 | 0.99945981 |
| ERO1A | 39.0163244 | -0.475094169 | 0.47507868 | -1.0000326 | 0.31729473 | 0.99945981 |
| PSMC6 | 25.5236625 | 0.285339567 | 0.34189077 | 0.83459277 | 0.40394704 | 0.99945981 |
| STYX | 13.6913694 | 0.209919818 | 0.50559969 | 0.41518977 | 0.67800299 | 0.99945981 |
| DDHD1 | 50.9646325 | 0.037664747 | 0.3102708 | 0.12139314 | 0.90337965 | 0.99945981 |
| CNIH1 | 10.8822654 | -0.84979592 | 0.57608388 | -1.4751253 | 0.14017885 | 0.99945981 |
| GMFB | 11.6130296 | -0.302056816 | 0.45507104 | -0.6637575 | 0.50684553 | 0.99945981 |
| SAMD4A | 47.4443866 | 0.354054784 | 0.59898551 | 0.59109073 | 0.55445963 | 0.99945981 |
| GCH1 | 38.6982752 | 2.090016563 | 0.5978022 | 3.49616742 | 0.00047199 | 0.23820088 |
| SOCS4 | 10.9266164 | -0.204732641 | 0.47732813 | -0.4289138 | 0.66798593 | 0.99945981 |
| MAPK1IP1L | 34.1764221 | -0.090966852 | 0.38188401 | -0.2382055 | 0.81172175 | 0.99945981 |
| LGALS3 | 19.3358167 | 0.064448049 | 0.54228888 | 0.1188445 | 0.90539856 | 0.99945981 |
| FBXO34 | 62.3379346 | 0.237270927 | 0.50848049 | 0.46662739 | 0.64076649 | 0.99945981 |
| ATG14 | 15.0855251 | 0.079449949 | 0.55267567 | 0.1437551 | 0.88569386 | 0.99945981 |
| KTN1 | 74.8822068 | 0.068943928 | 0.23197196 | 0.29720803 | 0.76630769 | 0.99945981 |
| PELI2 | 58.3275315 | 0.616770174 | 0.44769192 | 1.37766652 | 0.16830627 | 0.99945981 |
| TMEM260 | 28.2298548 | 0.042257054 | 0.30979444 | 0.13640352 | 0.89150229 | 0.99945981 |
| EXOC5 | 33.268343 | 0.189901526 | 0.3015265 | 0.62980045 | 0.52882515 | 0.99945981 |
| AP5M1 | 15.0965797 | -0.594279704 | 0.36662817 | -1.620933 | 0.105032 | 0.99945981 |
| ARMH4 | 15.971099 | -0.977558351 | 0.4351974 | -2.2462412 | 0.02468856 | 0.80325576 |
| ACTR10 | 23.6176292 | 0.104740897 | 0.34717154 | 0.30169782 | 0.76288243 | 0.99945981 |
| PSMA3 | 21.7870964 | 0.223565647 | 0.4150202 | 0.53868618 | 0.59010341 | 0.99945981 |
| ARID4A | 54.2159095 | 0.032675162 | 0.3473511 | 0.09406955 | 0.92505391 | 0.99945981 |
| KIAA0586 | 20.5555435 | -0.43666761 | 0.34923145 | -1.2503674 | 0.21116538 | 0.99945981 |
| DAAM1 | 28.7458096 | -0.027751794 | 0.42563109 | -0.0652015 | 0.94801355 | 0.99945981 |

| | | | | | | |
|---|---|---|---|---|---|---|
| JKAMP | 14.6324744 | 0.819368741 | 0.37436784 | 2.18867291 | 0.02862062 | 0.84763724 |
| RTN1 | 61.5004982 | 0.642649341 | 0.81282601 | 0.7906358 | 0.42915655 | 0.99945981 |
| PCNX4 | 43.2302406 | -0.50735936 | 0.34929518 | -1.4525232 | 0.14635618 | 0.99945981 |
| DHRS7 | 11.2526346 | -0.178489351 | 0.42166782 | -0.4232937 | 0.67208096 | 0.99945981 |
| PPM1A | 27.4547975 | -0.045172619 | 0.39780068 | -0.1135559 | 0.90958984 | 0.99945981 |
| MNAT1 | 55.0306799 | -0.60326527 | 0.33994504 | -1.7745965 | 0.07596454 | 0.98025673 |
| SLC38A6 | 23.6984303 | -0.069763989 | 0.46348194 | -0.1505215 | 0.8803532 | 0.99945981 |
| PRKCH | 225.217575 | -0.046273694 | 0.22506645 | -0.2056001 | 0.83710327 | 0.99945981 |
| HIF1A | 255.329779 | -0.086374722 | 0.55881566 | -0.1545675 | 0.87716232 | 0.99945981 |
| HIF1A-AS2 | 41.8765512 | -0.579875558 | 0.79211644 | -0.7320585 | 0.46413287 | 0.99945981 |
| SNAPC1 | 11.0412677 | 0.478515287 | 0.45719294 | 1.04663753 | 0.29526679 | 0.99945981 |
| PPP2R5E | 89.2662683 | 0.231390989 | 0.20248165 | 1.1427751 | 0.25313198 | 0.99945981 |
| SGPP1 | 39.9498788 | 0.036446416 | 0.48053361 | 0.07584572 | 0.93954184 | 0.99945981 |
| SYNE2 | 165.996714 | -0.178316611 | 0.24402145 | -0.7307415 | 0.46493704 | 0.99945981 |
| ESR2 | 14.7319714 | -0.096452644 | 0.69649837 | -0.1384822 | 0.88985932 | 0.99945981 |
| ZBTB25 | 21.4236781 | -0.427468208 | 0.45400695 | -0.9415455 | 0.34642537 | 0.99945981 |
| ZBTB1 | 43.1798354 | -0.525074366 | 0.23259539 | -2.2574582 | 0.02397946 | 0.80320673 |
| MAX | 55.2105169 | -0.04082166 | 0.32028203 | -0.1274554 | 0.89858001 | 0.99945981 |
| FUT8 | 92.7566688 | -0.002563534 | 0.36435411 | -0.0070358 | 0.99438626 | 0.99945981 |
| GPHN | 83.4434306 | 0.207191843 | 0.38918345 | 0.53237577 | 0.59446576 | 0.99945981 |
| MPP5 | 18.1415174 | -0.070631944 | 0.36440991 | -0.1938255 | 0.84631249 | 0.99945981 |
| ATP6V1D | 14.3471839 | 0.020479407 | 0.4693993 | 0.04362897 | 0.96520016 | 0.99945981 |
| EIF2S1 | 15.8844658 | -0.016944133 | 0.37460499 | -0.045232 | 0.96392239 | 0.99945981 |
| VTI1B | 9.48432348 | 0.253304022 | 0.53296384 | 0.47527431 | 0.63459146 | 0.99945981 |
| ZFYVE26 | 16.6264764 | -0.172945043 | 0.51059958 | -0.3387097 | 0.73482841 | 0.99945981 |
| RAD51B | 172.070097 | 0.496148644 | 0.48128681 | 1.03087938 | 0.30259739 | 0.99945981 |
| ZFP36L1 | 107.397662 | 0.027700542 | 0.44500059 | 0.06224833 | 0.95036508 | 0.99945981 |
| ACTN1 | 22.8622353 | -0.270126686 | 0.44868414 | -0.602042 | 0.54714619 | 0.99945981 |
| DCAF5 | 56.2781272 | -0.314403784 | 0.25567315 | -1.2297098 | 0.2188058 | 0.99945981 |
| ERH | 16.8319506 | 0.137206001 | 0.44389512 | 0.30909554 | 0.75724885 | 0.99945981 |
| SLC39A9 | 20.1785659 | 0.122843963 | 0.3515545 | 0.34943078 | 0.72676593 | 0.99945981 |
| SUSD6 | 91.5158373 | -0.110319997 | 0.42609177 | -0.2589114 | 0.79570364 | 0.99945981 |
| SRSF5 | 56.1188605 | 0.820707879 | 0.3461766 | 2.37077804 | 0.01775069 | 0.74759796 |
| MED6 | 10.0806711 | 0.056123463 | 0.41280038 | 0.13595788 | 0.89185458 | 0.99945981 |
| PCNX1 | 227.196337 | -0.021179093 | 0.29191122 | -0.0725532 | 0.94216167 | 0.99945981 |
| SIPA1L1 | 453.184659 | -0.230333841 | 0.53918006 | -0.4271928 | 0.66923891 | 0.99945981 |
| ZFYVE1 | 14.8788841 | -0.403745836 | 0.42758465 | -0.9442477 | 0.34504307 | 0.99945981 |
| RBM25 | 58.0263428 | 0.030343791 | 0.28044175 | 0.10819998 | 0.91383706 | 0.99945981 |
| PSEN1 | 61.905939 | 0.291132511 | 0.25662112 | 1.13448383 | 0.25659165 | 0.99945981 |
| NUMB | 154.914125 | 0.475742279 | 0.36454645 | 1.30502514 | 0.19188429 | 0.99945981 |
| ELMSAN1 | 24.5764115 | -0.071702172 | 0.44136765 | -0.1624545 | 0.87094794 | 0.99945981 |
| LIN52 | 25.7992363 | -0.166821143 | 0.4466422 | -0.3735006 | 0.70877588 | 0.99945981 |
| ABCD4 | 14.8310434 | -0.786998259 | 0.38822012 | -2.027196 | 0.04264238 | 0.89562588 |
| NPC2 | 41.4937057 | -0.356506046 | 0.47713393 | -0.7471823 | 0.45495351 | 0.99945981 |
| AREL1 | 19.6107725 | 0.045312326 | 0.34458593 | 0.1314979 | 0.89538145 | 0.99945981 |
| YLPM1 | 39.8977185 | -0.422288851 | 0.30466783 | -1.3860631 | 0.16572763 | 0.99945981 |
| DLST | 13.9993463 | -0.00430109 | 0.38422405 | -0.0111942 | 0.99106849 | 0.99945981 |
| MLH3 | 17.0643062 | 0.159891856 | 0.37770372 | 0.42332614 | 0.67205733 | 0.99945981 |
| NEK9 | 24.2119581 | 0.111608699 | 0.33567905 | 0.33248634 | 0.73952205 | 0.99945981 |
| TMED10 | 34.7968931 | -0.357700663 | 0.32822739 | -1.0897953 | 0.27580333 | 0.99945981 |
| FOS | 43.1826593 | 2.979125935 | 0.58252175 | 5.11418832 | 3.15E-07 | 0.00108156 |
| JDP2 | 24.2180172 | 1.347388619 | 0.6708347 | 2.00852554 | 0.04458748 | 0.90039076 |
| BATF | 9.7225494 | -0.14255979 | 0.49186248 | -0.2898367 | 0.77194119 | 0.99945981 |
| FLVCR2 | 18.0315753 | -0.329652915 | 0.70594335 | -0.4669679 | 0.64052281 | 0.99945981 |
| TTLL5 | 32.3736336 | -0.702068762 | 0.37336171 | -1.8803984 | 0.0600538 | 0.92445778 |
| IFT43 | 12.4449525 | 0.507604353 | 0.51003197 | 0.99524026 | 0.31961943 | 0.99945981 |
| GPATCH2L | 54.3276046 | -0.122320459 | 0.26443416 | -0.4625744 | 0.6436695 | 0.99945981 |
| VASH1 | 13.0390917 | 0.701171146 | 0.82765472 | 0.84717833 | 0.39689573 | 0.99945981 |
| ANGEL1 | 9.25883693 | 0.126694661 | 0.47307122 | 0.26781308 | 0.7888432 | 0.99945981 |
| TMED8 | 19.9816857 | -0.563565871 | 0.35667162 | -1.5800693 | 0.114091 | 0.99945981 |
| AHSA1 | 10.4062046 | -0.162604426 | 0.55188753 | -0.2946333 | 0.76827405 | 0.99945981 |
| SPTLC2 | 79.1766939 | 0.633297979 | 0.38324704 | 1.65245367 | 0.0984421 | 0.99945981 |
| ALKBH1 | 10.2761882 | -0.575922085 | 0.46521788 | -1.2379621 | 0.21573013 | 0.99945981 |
| SLIRP | 14.9439939 | 0.075012839 | 0.44840053 | 0.16728981 | 0.86714202 | 0.99945981 |
| SNW1 | 22.7226655 | 0.131181676 | 0.37935535 | 0.34580157 | 0.72949184 | 0.99945981 |
| CEP128 | 69.6699633 | -0.296027032 | 0.75067535 | -0.3943476 | 0.69332442 | 0.99945981 |
| GTF2A1 | 19.8530482 | 0.173044915 | 0.47857273 | 0.3615854 | 0.71766187 | 0.99945981 |
| STON2 | 20.7836658 | 1.235044889 | 0.65133235 | 1.89618232 | 0.05793594 | 0.92445778 |
| SEL1L | 44.58525 | 0.007180298 | 0.28735804 | 0.02498729 | 0.9800651 | 0.99945981 |
| LINC02328 | 46.5788726 | -0.256744914 | 0.40568671 | -0.632865 | 0.52682181 | 0.99945981 |
| GALC | 23.0190084 | 0.067110666 | 0.58329032 | 0.11505534 | 0.90840126 | 0.99945981 |
| GPR65 | 19.0148735 | 0.018914975 | 0.88152187 | 0.02145718 | 0.98288096 | 0.99945981 |
| LINC01146 | 12.5398219 | 0.240037241 | 0.50771618 | 0.47277839 | 0.63637129 | 0.99945981 |
| ZC3H14 | 21.2853235 | 0.265080672 | 0.4617025 | 0.5741374 | 0.56587483 | 0.99945981 |
| FOXN3 | 376.765009 | 0.350715612 | 0.25127521 | 1.39574298 | 0.16279191 | 0.99945981 |
| TDP1 | 14.9844603 | -0.202377572 | 0.44774254 | -0.4519954 | 0.65127229 | 0.99945981 |
| KCNK13 | 11.2656386 | 0.558658486 | 0.6949683 | 0.80386183 | 0.42147678 | 0.99945981 |
| PSMC1 | 13.528524 | 0.053307568 | 0.49043323 | 0.10869486 | 0.91344452 | 0.99945981 |
| NRDE2 | 11.5804283 | 0.066099043 | 0.40322488 | 0.163926 | 0.8697894 | 0.99945981 |
| CALM1 | 70.4737095 | -0.06093962 | 0.34874778 | -0.1747384 | 0.86128521 | 0.99945981 |
| TTC7B | 20.4300697 | -0.062848374 | 0.50196962 | -0.1252035 | 0.90036241 | 0.99945981 |
| RPS6KA5 | 62.5313806 | 0.219818269 | 0.28724142 | 0.76527358 | 0.44410865 | 0.99945981 |
| DGLUCY | 56.1804509 | 0.594254281 | 0.42151491 | 1.40980608 | 0.15859695 | 0.99945981 |
| CCDC88C | 84.8653157 | 0.059478561 | 0.22989364 | 0.25872209 | 0.79584968 | 0.99945981 |
| PPP4R3A | 61.4054302 | -0.040532405 | 0.25122876 | -0.1613366 | 0.87182827 | 0.99945981 |
| CATSPERB | 16.5852666 | -1.0509542 | 0.74394283 | -1.4126814 | 0.15774942 | 0.99945981 |
| TC2N | 65.2094166 | -0.400392932 | 0.53021111 | -0.7551576 | 0.45015445 | 0.99945981 |
| TRIP11 | 39.6653237 | -0.00447706 | 0.30773239 | -0.0145485 | 0.98839235 | 0.99945981 |
| ATXN3 | 32.4556963 | 0.432953162 | 0.30755012 | 1.40774833 | 0.1592056 | 0.99945981 |
| NDUFB1 | 11.3532345 | 0.096623865 | 0.41755758 | 0.23140249 | 0.81700213 | 0.99945981 |
| CPSF2 | 21.5714041 | 0.19700017 | 0.35969808 | 0.54768202 | 0.58391026 | 0.99945981 |
| SLC24A4 | 16.0689121 | 0.393307836 | 0.9052771 | 0.43446127 | 0.66395352 | 0.99945981 |
| RIN3 | 132.057212 | -0.285076918 | 0.3611701 | -0.7893148 | 0.42992802 | 0.99945981 |
| LGMN | 76.560986 | 1.833342392 | 0.87593323 | 2.09301614 | 0.03634771 | 0.87920079 |
| GOLGA5 | 11.6882495 | -0.064362279 | 0.56859758 | -0.1131948 | 0.90987613 | 0.99945981 |
| ITPK1 | 35.8529205 | 0.472151698 | 0.39520322 | 1.19470609 | 0.23220188 | 0.99945981 |
| BTBD7 | 66.0675661 | -0.242028585 | 0.30539066 | -0.7925212 | 0.42805682 | 0.99945981 |
| DDX24 | 36.402098 | -0.933589674 | 0.2986628 | -3.1258987 | 0.00177263 | 0.45070656 |
| SERPINA1 | 37.2192432 | 0.107063303 | 0.75606979 | 0.14160505 | 0.88739198 | 0.99945981 |
| DICER1 | 75.1707012 | 0.09636438 | 0.29690584 | 0.32456209 | 0.74551252 | 0.99945981 |
| CLMN | 19.8957132 | 0.31107947 | 0.37646441 | 0.8263184 | 0.4086235 | 0.99945981 |
| SYNE3 | 23.1292421 | -0.523444575 | 0.58855751 | -0.8893686 | 0.37380501 | 0.99945981 |
| ATG2B | 28.4626047 | 0.132210531 | 0.40606534 | 0.32558931 | 0.7447351 | 0.99945981 |
| PAPOLA | 71.6202855 | 0.275216608 | 0.32058498 | 0.85848255 | 0.39062606 | 0.99945981 |
| VRK1 | 21.1533751 | -0.413807621 | 0.42290083 | -0.978498 | 0.32782807 | 0.99945981 |
| BCL11B | 75.6486739 | -0.293587593 | 0.28943163 | -1.0143591 | 0.31041145 | 0.99945981 |
| SETD3 | 40.9342499 | 0.179804316 | 0.27324408 | 0.65803554 | 0.5105153 | 0.99945981 |
| CCNK | 17.9719827 | 0.111385705 | 0.35931953 | 0.30999068 | 0.75656804 | 0.99945981 |
| EVL | 91.4901308 | 0.045753626 | 0.42033575 | 0.10885019 | 0.91332132 | 0.99945981 |
| YY1 | 44.4220415 | 0.01908293 | 0.27732594 | 0.06881048 | 0.94514048 | 0.99945981 |
| SLC25A29 | 11.4037468 | -1.280131661 | 0.56240256 | -2.2761839 | 0.022835 | 0.80320673 |
| WARS | 48.2030743 | 0.725784803 | 0.68150328 | 1.06497624 | 0.28688668 | 0.99945981 |
| WDR25 | 13.6231918 | 0.17266852 | 0.63451544 | 0.27212658 | 0.78552469 | 0.99945981 |

| | | | | | | |
|---|---|---|---|---|---|---|
| PPP2R5C | 191.167542 | -0.342154495 | 0.37090571 | -0.9224838 | 0.3562763 | 0.99945981 |
| DYNC1H1 | 71.2417624 | -0.127540258 | 0.26263705 | -0.4856141 | 0.62724079 | 0.99945981 |
| HSP90AA1 | 150.967638 | 0.560574793 | 0.47244557 | 1.18653835 | 0.23540976 | 0.99945981 |
| WDR20 | 48.9081795 | 0.021688241 | 0.28139323 | 0.0770745 | 0.93856428 | 0.99945981 |
| TECPR2 | 16.7750077 | 0.559122842 | 0.4818056 | 1.16047394 | 0.2458559 | 0.99945981 |
| RCOR1 | 75.5515074 | 0.469893829 | 0.27620387 | 1.70125725 | 0.08889469 | 0.99640647 |
| TRAF3 | 80.6793876 | -0.062108671 | 0.36668396 | -0.1693793 | 0.86549831 | 0.99945981 |
| CDC42BPB | 20.8159432 | -0.040543087 | 0.36561932 | -0.1108888 | 0.91170453 | 0.99945981 |
| TNFAIP2 | 28.3748207 | 0.7568893 | 0.77216806 | 0.98021317 | 0.32698091 | 0.99945981 |
| EIF5 | 29.3693085 | -0.514234541 | 0.29868655 | -1.7216528 | 0.08513243 | 0.99193047 |
| MARK3 | 113.808426 | 0.044594756 | 0.17958272 | 0.24832431 | 0.80388349 | 0.99945981 |
| CKB | 9.40986197 | 0.263482393 | 0.58576429 | 0.44980959 | 0.65284775 | 0.99945981 |
| BAG5 | 9.14575633 | 0.185808821 | 0.47471505 | 0.39141127 | 0.69549326 | 0.99945981 |
| KLC1 | 32.6547019 | -0.002401112 | 0.36630043 | -0.006555 | 0.99476988 | 0.99945981 |
| PPP1R13B | 15.7031532 | 0.029280661 | 0.43421301 | 0.06743386 | 0.94623631 | 0.99945981 |
| ATP5MPL | 15.4828723 | 0.344988191 | 0.52821025 | 0.65312665 | 0.51367463 | 0.99945981 |
| SIVA1 | 12.7145754 | 0.254535879 | 0.47266542 | 0.53851175 | 0.5902238 | 0.99945981 |
| AKT1 | 12.5184122 | -0.052847497 | 0.41744545 | -0.1265974 | 0.89925908 | 0.99945981 |
| GPR132 | 15.3892404 | -0.711026742 | 0.47273138 | -1.504082 | 0.13256026 | 0.99945981 |
| BRF1 | 14.0575067 | -0.146526639 | 0.50872924 | -0.2880248 | 0.77332775 | 0.99945981 |
| PACS2 | 18.1122629 | -0.411903025 | 0.42124612 | -0.9778203 | 0.32816318 | 0.99945981 |
| IGHG3 | 46.3137805 | -3.005442792 | 1.79388338 | -1.6753836 | 0.09385899 | 0.99945981 |
| IGHM | 188.599027 | -1.097726284 | 2.07278157 | -0.5295909 | 0.5963956 | 0.99945981 |
| NIPA1 | 8.5465588 | -1.586902747 | 0.60589821 | -2.6190913 | 0.00881643 | 0.59184657 |
| NIPA2 | 19.5648117 | -0.526348207 | 0.45816595 | -1.1488156 | 0.25063201 | 0.99945981 |
| CYFIP1 | 35.5344103 | 0.238240442 | 0.46232115 | 0.51531375 | 0.60633379 | 0.99945981 |
| AC124312.1 | 24.944095 | 0.099414476 | 0.42122182 | 0.23601455 | 0.81342138 | 0.99945981 |
| UBE3A | 64.945464 | 0.240463268 | 0.2997993 | 0.8020808 | 0.42250622 | 0.99945981 |
| ATP10A | 48.9587531 | -0.291305081 | 0.82527822 | -0.352978 | 0.72410491 | 0.99945981 |
| HERC2 | 59.7825195 | -0.401580806 | 0.3568599 | -1.1253178 | 0.26045438 | 0.99945981 |
| APBA2 | 31.8323235 | -0.262659535 | 0.68387178 | -0.3840772 | 0.70092125 | 0.99945981 |
| AC091057.6 | 9.5864029 | -0.521157446 | 0.51717497 | -1.0077004 | 0.3135983 | 0.99945981 |
| FAN1 | 8.32843233 | -0.18700585 | 0.45376786 | -0.4121179 | 0.68025302 | 0.99945981 |
| MTMR10 | 36.4481944 | -0.096900981 | 0.35927778 | -0.2697105 | 0.787383 | 0.99945981 |
| KLF13 | 46.0309561 | -0.060503649 | 0.32536267 | -0.1859576 | 0.85247804 | 0.99945981 |
| LINC02256 | 30.7787544 | -0.724293795 | 0.5228036 | -1.3854032 | 0.16592921 | 0.99945981 |
| FMN1 | 189.97215 | -1.171875496 | 0.69275502 | -1.691616 | 0.09071921 | 0.99640647 |
| AVEN | 18.8038945 | -0.362759798 | 0.54444907 | -0.6662878 | 0.50522713 | 0.99945981 |
| KATNBL1 | 44.0820067 | -0.284164266 | 0.31605127 | -0.8991081 | 0.36859507 | 0.99945981 |
| SLC12A6 | 69.8481032 | -0.368054825 | 0.54674894 | -0.6731697 | 0.50083933 | 0.99945981 |
| NOP10 | 21.6583805 | -0.424175207 | 0.60466918 | -0.7014996 | 0.48299127 | 0.99945981 |
| GOLGA8A | 18.0116573 | 0.014215799 | 0.39424497 | 0.03605829 | 0.97123588 | 0.99945981 |
| GOLGA8B | 19.0126407 | -0.680255837 | 0.40583043 | -1.6762071 | 0.09369764 | 0.99945981 |
| AQR | 34.4163192 | 0.084927756 | 0.31531001 | 0.26934684 | 0.78766279 | 0.99945981 |
| ZNF770 | 11.1203899 | 0.569155701 | 0.40704313 | 1.39826876 | 0.16203237 | 0.99945981 |
| DPH6 | 33.4966241 | -0.476005572 | 0.46007367 | -1.034629 | 0.3008422 | 0.99945981 |
| C15orf41 | 15.3154627 | 0.278202086 | 0.36410695 | 0.76406696 | 0.44482734 | 0.99945981 |
| SPRED1 | 103.221487 | 0.540684646 | 0.65031124 | 0.83142442 | 0.40573391 | 0.99945981 |
| FAM98B | 8.91194349 | -0.121380743 | 0.431624 | -0.2812187 | 0.77854266 | 0.99945981 |
| RASGRP1 | 59.0871592 | -0.53536589 | 0.39963511 | -1.3396368 | 0.18036346 | 0.99945981 |
| C15orf53 | 28.0171567 | -0.895897468 | 0.92280199 | -0.9708447 | 0.3316256 | 0.99945981 |
| THBS1 | 79.8729285 | -1.124975368 | 1.1306312 | -0.9949976 | 0.31973742 | 0.99945981 |
| EIF2AK4 | 28.6521987 | 0.01861914 | 0.33371014 | 0.05579435 | 0.95550564 | 0.99945981 |
| SRP14 | 46.2426128 | -0.841662047 | 0.46403537 | -1.8137886 | 0.06971028 | 0.95505283 |
| PLCB2 | 20.3506008 | 0.635883537 | 0.48437106 | 1.3128025 | 0.18924951 | 0.99945981 |
| IVD | 15.2548253 | -0.183911163 | 0.3656599 | -0.5029569 | 0.61499459 | 0.99945981 |
| CCDC32 | 17.4418769 | -0.279925228 | 0.44294703 | -0.631961 | 0.52741239 | 0.99945981 |
| DNAJC17 | 10.3364226 | -0.249765118 | 0.53060136 | -0.4707208 | 0.6378401 | 0.99945981 |
| INO80 | 44.3057317 | 0.081283188 | 0.28370109 | 0.28650996 | 0.77448757 | 0.99945981 |
| CHP1 | 15.5097667 | -0.002387122 | 0.42506027 | -0.005616 | 0.99551913 | 0.99945981 |
| RTF1 | 24.0059889 | -0.495181697 | 0.35168937 | -1.4080087 | 0.15912848 | 0.99945981 |
| MGA | 47.832938 | -0.478707632 | 0.38645741 | -1.2387073 | 0.2154539 | 0.99945981 |
| MAPKBP1 | 14.0960009 | 0.280403472 | 0.37239426 | 0.75297475 | 0.45146509 | 0.99945981 |
| EHD4 | 19.082159 | 0.062000498 | 0.38160534 | 0.16247283 | 0.87093352 | 0.99945981 |
| VPS39 | 28.1113669 | 0.153996733 | 0.28246362 | 0.54519138 | 0.5856219 | 0.99945981 |
| TMEM87A | 47.2822159 | -0.04899895 | 0.30181004 | -0.1623503 | 0.87103001 | 0.99945981 |
| GANC | 23.0502122 | -0.009454347 | 0.44542462 | -0.0212255 | 0.9830658 | 0.99945981 |
| ZNF106 | 52.5478895 | 0.243985074 | 0.32956796 | 0.74031794 | 0.4591071 | 0.99945981 |
| SNAP23 | 38.6284053 | 0.258594261 | 0.29316267 | 0.88208455 | 0.37773108 | 0.99945981 |
| STARD9 | 12.6982825 | -0.303256244 | 0.66102849 | -0.4587643 | 0.64640347 | 0.99945981 |
| TTBK2 | 52.0960884 | 0.421696872 | 0.42209553 | 0.99905552 | 0.3177678 | 0.99945981 |
| UBR1 | 71.733329 | -0.117710381 | 0.22476057 | -0.5237146 | 0.60047709 | 0.99945981 |
| AC068724.3 | 14.3117372 | -0.210786524 | 0.54369632 | -0.3876917 | 0.69824423 | 0.99945981 |
| TMEM62 | 20.564664 | -0.085440624 | 0.37892926 | -0.2254791 | 0.82160659 | 0.99945981 |
| CCNDBP1 | 11.8483758 | -0.526799765 | 0.52147962 | -1.010202 | 0.31239852 | 0.99945981 |
| TUBGCP4 | 9.84430329 | 0.232609396 | 0.44770101 | 0.51956415 | 0.60336739 | 0.99945981 |
| TP53BP1 | 17.0768367 | -0.614492332 | 0.4297701 | -1.4298164 | 0.15276973 | 0.99945981 |
| CATSPER2 | 9.19748651 | 0.008373899 | 0.61922287 | 0.01352324 | 0.98921035 | 0.99945981 |
| PDIA3 | 42.9732309 | -0.073217575 | 0.33651647 | -0.217575 | 0.82776026 | 0.99945981 |
| SERF2 | 121.833928 | -0.095650836 | 0.46847701 | -0.204174 | 0.83821751 | 0.99945981 |
| MFAP1 | 8.4184274 | -0.539528248 | 0.7558249 | -0.713827 | 0.47533414 | 0.99945981 |
| CASC4 | 58.9003625 | -0.277866992 | 0.28447605 | -0.9767676 | 0.32868421 | 0.99945981 |
| CTDSPL2 | 53.2626612 | -0.078717541 | 0.29060622 | -0.2708736 | 0.78648828 | 0.99945981 |
| EIF3J | 20.7849102 | -0.058584184 | 0.35805268 | -0.1636189 | 0.87003117 | 0.99945981 |
| SPG11 | 100.462931 | 0.024513781 | 0.35922207 | 0.0682413 | 0.94559355 | 0.99945981 |
| PATL2 | 16.3850216 | -0.175421142 | 0.71770354 | -0.2444201 | 0.80690551 | 0.99945981 |
| B2M | 817.639008 | 0.185331483 | 0.47417308 | 0.39085197 | 0.69590666 | 0.99945981 |
| GATM | 9.99052841 | 0.740525026 | 0.6621954 | 1.11828779 | 0.2634441 | 0.99945981 |
| SQOR | 14.1004792 | -0.045354311 | 0.62015765 | -0.0731335 | 0.94169987 | 0.99945981 |
| MYEF2 | 12.9913415 | -1.549064247 | 0.68313763 | -2.2675727 | 0.02335526 | 0.80320673 |
| SLC12A1 | 24.8538436 | 2.120256923 | 0.73004683 | 2.90427522 | 0.00368104 | 0.48727327 |
| DUT | 11.9153429 | 0.266935188 | 0.49958341 | 0.53431555 | 0.59312323 | 0.99945981 |
| CEP152 | 14.8094168 | 0.854731598 | 0.51512911 | 1.65925704 | 0.09706401 | 0.99945981 |
| EID1 | 18.4530578 | -1.032016308 | 0.41051507 | -2.5139547 | 0.01193858 | 0.67187199 |
| SECISBP2L | 34.4405879 | -0.014658462 | 0.31259361 | -0.046893 | 0.96259848 | 0.99945981 |
| COPS2 | 15.0486556 | 0.096684784 | 0.48338782 | 0.20001494 | 0.8414689 | 0.99945981 |
| GALK2 | 39.9995371 | 0.089495736 | 0.43028159 | 0.20799341 | 0.83523411 | 0.99945981 |
| DTWD1 | 17.6619504 | -0.136834052 | 0.51193544 | -0.2672877 | 0.78924765 | 0.99945981 |
| ATP8B4 | 198.669451 | -0.263993248 | 0.9333488 | -0.2828452 | 0.77729549 | 0.99945981 |
| GABPB1 | 37.9964883 | -0.14005971 | 0.32750013 | -0.4276631 | 0.66889644 | 0.99945981 |
| GABPB1-AS1 | 33.1091711 | 0.022033915 | 0.5779072 | 0.03812708 | 0.96958636 | 0.99945981 |
| USP8 | 52.0636926 | -0.1373348 | 0.2682145 | -0.5120335 | 0.60862759 | 0.99945981 |
| TRPM7 | 93.3395335 | 0.147161483 | 0.29328425 | 0.50177083 | 0.61582873 | 0.99945981 |
| SPPL2A | 56.2095194 | -0.016601104 | 0.32805416 | -0.0506048 | 0.95964047 | 0.99945981 |
| AP4E1 | 18.3951718 | -0.079809394 | 0.41521233 | -0.1922134 | 0.84757501 | 0.99945981 |
| DMXL2 | 166.214139 | 1.22080212 | 0.61322719 | 1.99078276 | 0.04650478 | 0.90884462 |
| TMOD2 | 13.2637687 | -0.87581903 | 0.45041048 | -1.9444908 | 0.0518363 | 0.9230968 |
| TMOD3 | 69.6311404 | -0.190678637 | 0.30327327 | -0.6287354 | 0.52952231 | 0.99945981 |
| MAPK6 | 26.1893482 | 0.036653495 | 0.33372664 | 0.10983089 | 0.91254349 | 0.99945981 |
| GNB5 | 15.9162466 | -0.572993164 | 0.41825012 | -1.3699773 | 0.170694 | 0.99945981 |
| MYO5A | 146.371838 | 0.281487817 | 0.29783084 | 0.94512649 | 0.3445943 | 0.99945981 |
| ARPP19 | 27.6782365 | -0.115359567 | 0.31038119 | -0.3716706 | 0.71013811 | 0.99945981 |
| FAM214A | 73.9162291 | -0.032141642 | 0.45710457 | -0.0703157 | 0.94394236 | 0.99945981 |

| | | | | | | |
|---|---|---|---|---|---|---|
| RSL24D1 | 12.4107168 | -0.179644572 | 0.50192461 | -0.3579115 | 0.72040957 | 0.99945981 |
| RAB27A | 71.1809395 | -0.375422136 | 0.36003216 | -1.0427461 | 0.29706589 | 0.99945981 |
| PIGB | 17.6694991 | -0.102982842 | 0.42309232 | -0.2434051 | 0.80769157 | 0.99945981 |
| CCPG1 | 38.4508583 | -0.206417301 | 0.32206218 | -0.6409238 | 0.52157222 | 0.99945981 |
| NEDD4 | 10.7576258 | 0.280357413 | 0.58099854 | 0.48254409 | 0.62941949 | 0.99945981 |
| RFX7 | 80.2548982 | -0.17323164 | 0.29783823 | -0.58163 | 0.56081595 | 0.99945981 |
| ZNF280D | 18.0017505 | 0.091079195 | 0.3872595 | 0.23518905 | 0.814062 | 0.99945981 |
| AC090517.4 | 14.2219364 | 0.575884681 | 0.61750619 | 0.93259742 | 0.35102787 | 0.99945981 |
| TCF12 | 268.250966 | 0.060042807 | 0.22556528 | 0.26618816 | 0.7900943 | 0.99945981 |
| CGNL1 | 11.5393344 | 0.788936795 | 0.46891485 | 1.68247349 | 0.09247706 | 0.99640647 |
| ALDH1A2 | 18.5461291 | 0.553682712 | 0.66878997 | 0.82788729 | 0.40773434 | 0.99945981 |
| AQP9 | 27.9679035 | 1.066998762 | 0.84263365 | 1.26626649 | 0.20541769 | 0.99945981 |
| ADAM10 | 143.593326 | 0.537678431 | 0.28588735 | 1.88073531 | 0.06000793 | 0.92445778 |
| MINDY2 | 29.4161156 | -0.116667035 | 0.35614203 | -0.3275857 | 0.74322494 | 0.99945981 |
| RNF111 | 78.3373147 | 0.03530432 | 0.28060687 | 0.12581417 | 0.89987903 | 0.99945981 |
| SLTM | 87.6301405 | 0.056546586 | 0.24783796 | 0.2281595 | 0.81952224 | 0.99945981 |
| MYO1E | 38.5086741 | 0.085473474 | 0.68752213 | 0.12432105 | 0.90106108 | 0.99945981 |
| GTF2A2 | 11.5252935 | 0.243314124 | 0.42302261 | 0.57517995 | 0.5651696 | 0.99945981 |
| BNIP2 | 33.6124833 | -0.011156443 | 0.31656437 | -0.0352423 | 0.97188657 | 0.99945981 |
| ANXA2 | 66.7390824 | 0.010602049 | 0.46606895 | 0.02274781 | 0.98185144 | 0.99945981 |
| ICE2 | 23.8754654 | 0.191754466 | 0.3661855 | 0.5236539 | 0.60051928 | 0.99945981 |
| RORA-AS1 | 17.799255 | -0.734989169 | 0.45291408 | -1.6228004 | 0.10463207 | 0.99945981 |
| RORA | 109.339664 | 0.107168137 | 0.37988286 | 0.28210837 | 0.77786041 | 0.99945981 |
| AC009554.1 | 21.9072855 | -0.263444716 | 0.59715819 | -0.441164 | 0.65909425 | 0.99945981 |
| VPS13C | 172.557502 | 0.23215112 | 0.2307288 | 1.00616448 | 0.31433646 | 0.99945981 |
| TLN2 | 16.0602523 | -0.763148455 | 0.80642482 | -0.9463355 | 0.34397748 | 0.99945981 |
| TPM1 | 13.4736105 | -0.590725151 | 0.39328033 | -1.502046 | 0.13308522 | 0.99945981 |
| LACTB | 16.8774936 | -0.228874242 | 0.53146852 | -0.430645 | 0.66672654 | 0.99945981 |
| RPS27L | 23.3727607 | -0.814696816 | 0.37901285 | -2.149523 | 0.03159297 | 0.86065765 |
| RAB8B | 113.036043 | 0.246850118 | 0.32666615 | 0.75566482 | 0.44985018 | 0.99945981 |
| APH1B | 17.464737 | -0.123628156 | 0.45473917 | -0.2718661 | 0.78572499 | 0.99945981 |
| CA12 | 16.0984677 | -0.093466626 | 0.67868055 | -0.1377181 | 0.89046318 | 0.99945981 |
| USP3 | 134.063533 | 0.449102844 | 0.22676096 | 1.98051218 | 0.04764601 | 0.90884462 |
| USP3-AS1 | 26.2393022 | 0.364809223 | 0.46583423 | 0.783131 | 0.43355019 | 0.99945981 |
| HERC1 | 152.452888 | 0.247423369 | 0.22710417 | 1.08947081 | 0.27594632 | 0.99945981 |
| DAPK2 | 10.3222929 | -0.98928567 | 0.56426135 | -1.7532402 | 0.07956079 | 0.9846219 |
| FAM96A | 10.0760521 | -0.198283189 | 0.46627296 | -0.4252513 | 0.67065348 | 0.99945981 |
| SNX1 | 50.5941352 | -0.133138537 | 0.30727923 | -0.4332819 | 0.66480996 | 0.99945981 |
| PPIB | 38.3060728 | -0.427145263 | 0.61500781 | -0.6945363 | 0.48734593 | 0.99945981 |
| TRIP4 | 25.3157772 | 0.105787087 | 0.3157538 | 0.33503029 | 0.73760224 | 0.99945981 |
| ZNF609 | 153.790484 | 0.015133668 | 0.30699217 | 0.04929659 | 0.96068294 | 0.99945981 |
| OAZ2 | 15.2625457 | 0.392167062 | 0.4793488 | 0.81812463 | 0.41328602 | 0.99945981 |
| AC069368.1 | 14.0242111 | -0.464360277 | 0.65157789 | -0.7126704 | 0.47604973 | 0.99945981 |
| ANKDD1A | 11.3366411 | -0.976173178 | 0.85392669 | -1.1431581 | 0.25297297 | 0.99945981 |
| SPG21 | 20.9572362 | 0.68229908 | 0.35094851 | 1.94415722 | 0.0518765 | 0.9230968 |
| MTFMT | 9.62917741 | -0.86493603 | 0.4687564 | -1.8451717 | 0.06501257 | 0.94357565 |
| PDCD7 | 9.12709917 | -0.342141048 | 0.45567293 | -0.7508479 | 0.45274422 | 0.99945981 |
| CLPX | 16.2970147 | -0.146962623 | 0.38852646 | -0.3782564 | 0.70524013 | 0.99945981 |
| DPP8 | 44.3579202 | -0.252628913 | 0.25188157 | -1.002967 | 0.31587676 | 0.99945981 |
| HACD3 | 11.9997388 | -0.268060687 | 0.48636588 | -0.5511503 | 0.58153067 | 0.99945981 |
| DENND4A | 126.486788 | 0.251466737 | 0.2725797 | 0.92254389 | 0.35624495 | 0.99945981 |
| RAB11A | 58.9560797 | 0.001330313 | 0.48214329 | 0.00275917 | 0.99779851 | 0.99969159 |
| DIS3L | 12.1249002 | -0.165566997 | 0.41399221 | -0.3999278 | 0.68920969 | 0.99945981 |
| MAP2K1 | 88.9123511 | 0.195513044 | 0.39801613 | 0.4912189 | 0.62327163 | 0.99945981 |
| RPL4 | 61.5090147 | -0.156310271 | 0.44323625 | -0.3526568 | 0.72434576 | 0.99945981 |
| SMAD3 | 59.2976226 | 0.636179571 | 0.44945595 | 1.41544365 | 0.15693846 | 0.99945981 |
| AAGAB | 27.4844323 | 0.339091695 | 0.36555311 | 0.92761267 | 0.35360852 | 0.99945981 |
| IQCH-AS1 | 9.3862015 | -0.255574988 | 0.4945502 | -0.5167827 | 0.60530786 | 0.99945981 |
| MAP2K5 | 69.6716021 | 0.098275034 | 0.29099785 | 0.33771739 | 0.73557617 | 0.99945981 |
| PIAS1 | 188.954785 | 0.661267827 | 0.25178742 | 2.62629414 | 0.00863202 | 0.59011016 |
| FEM1B | 13.3654598 | 0.372982705 | 0.45841537 | 0.81363482 | 0.41585418 | 0.99945981 |
| ANP32A | 41.8369819 | 0.131684238 | 0.32470627 | 0.4055488 | 0.68507416 | 0.99945981 |
| GLCE | 11.6225574 | 0.077503154 | 0.47950772 | 0.16163067 | 0.8715967 | 0.99945981 |
| RPLP1 | 156.10966 | 0.07985741 | 0.60649524 | 0.1316703 | 0.89524508 | 0.99945981 |
| TLE3 | 27.2672176 | -0.135716444 | 0.55595742 | -0.244113 | 0.80714329 | 0.99945981 |
| UACA | 14.9372535 | -0.170835882 | 0.50478783 | -0.3384311 | 0.73503837 | 0.99945981 |
| THSD4-AS1 | 22.5297667 | -0.269560762 | 0.45778662 | -0.588835 | 0.555972 | 0.99945981 |
| MYO9A | 76.710956 | -0.056936229 | 0.34389999 | -0.1655604 | 0.86850289 | 0.99945981 |
| PKM | 79.8617556 | -0.611675506 | 0.40337886 | -1.5163797 | 0.12942338 | 0.99945981 |
| PARP6 | 14.0391401 | -0.64652834 | 0.39842364 | -1.6227158 | 0.10465017 | 0.99945981 |
| AC009690.1 | 24.6034308 | -0.282194934 | 0.58050574 | -0.4861191 | 0.62688272 | 0.99945981 |
| ARIH1 | 128.677959 | 0.171844657 | 0.18904296 | 0.90902439 | 0.36333725 | 0.99945981 |
| BBS4 | 7.95066931 | -0.964044975 | 0.54809068 | -1.7589151 | 0.07859193 | 0.98326702 |
| ADPGK | 42.0577224 | -0.060372253 | 0.37817099 | -0.1596427 | 0.87316251 | 0.99945981 |
| NEO1 | 12.1709379 | 0.148631975 | 0.59284107 | 0.25071133 | 0.8020373 | 0.99945981 |
| NPTN | 46.8703779 | 0.175726208 | 0.29896671 | 0.58777851 | 0.55668097 | 0.99945981 |
| PML | 14.9601984 | 0.142015835 | 0.58298603 | 0.24360075 | 0.80754005 | 0.99945981 |
| ARID3B | 18.4732695 | 0.300060771 | 0.40501477 | 0.74086378 | 0.45877604 | 0.99945981 |
| CLK3 | 9.87909739 | -0.089828363 | 0.48522018 | -0.1851291 | 0.8531278 | 0.99945981 |
| EDC3 | 8.1736213 | -0.501959224 | 0.48115254 | -1.0432434 | 0.29683557 | 0.99945981 |
| CSK | 36.8278765 | -0.4573616 | 0.65184728 | -0.7016392 | 0.4829042 | 0.99945981 |
| SCAMP2 | 19.6894728 | -0.005957084 | 0.42057266 | -0.0141642 | 0.98869897 | 0.99945981 |
| COX5A | 13.2564689 | -0.182492991 | 0.47524171 | -0.3840004 | 0.70097818 | 0.99945981 |
| PPCDC | 22.8179382 | -0.263591158 | 0.33404919 | -0.7890789 | 0.43006591 | 0.99945981 |
| MAN2C1 | 15.0012874 | 0.542506426 | 0.417547 | 1.29927034 | 0.19385117 | 0.99945981 |
| SIN3A | 31.0434761 | -0.136072529 | 0.33167751 | -0.4102555 | 0.68161851 | 0.99945981 |
| PTPN9 | 39.1309208 | -0.187031641 | 0.35495123 | -0.5269221 | 0.5982477 | 0.99945981 |
| UBE2Q2 | 28.8759151 | -0.424313237 | 0.51077411 | -0.8307258 | 0.40612854 | 0.99945981 |
| FBXO22 | 19.1226017 | -0.88476074 | 0.46124511 | -1.9182008 | 0.05508556 | 0.92445778 |
| ETFA | 42.2333873 | 0.388387895 | 0.27487443 | 1.41296478 | 0.15766608 | 0.99945981 |
| SCAPER | 84.9991885 | -0.054891185 | 0.31777721 | -0.1727348 | 0.86285988 | 0.99945981 |
| RCN2 | 14.06159 | -0.193882138 | 0.48298148 | -0.4014277 | 0.68810527 | 0.99945981 |
| PSTPIP1 | 22.7414224 | -0.204912239 | 0.33367339 | -0.6141102 | 0.53914249 | 0.99945981 |
| TSPAN3 | 16.9604988 | -0.993970633 | 0.61868923 | -1.606575 | 0.10814763 | 0.99945981 |
| PEAK1 | 107.237853 | 0.900394675 | 0.40440188 | 2.22648488 | 0.02598172 | 0.81444982 |
| HMG20A | 21.475871 | -0.577209848 | 0.35799329 | -1.6123482 | 0.10688618 | 0.99945981 |
| LINGO1 | 197.23615 | 1.108062099 | 0.54246279 | 2.04265087 | 0.04108701 | 0.89562588 |
| TBC1D2B | 53.5609551 | -0.439665926 | 0.39860748 | -1.1030047 | 0.27002512 | 0.99945981 |
| IDH3A | 11.300316 | 0.101545478 | 0.57872828 | 0.17546313 | 0.86071574 | 0.99945981 |
| IREB2 | 39.3326725 | -0.201259434 | 0.32254696 | -0.6239694 | 0.53264768 | 0.99945981 |
| PSMA4 | 22.3264392 | 0.559625368 | 0.41558548 | 1.3465951 | 0.17811067 | 0.99945981 |
| MORF4L1 | 53.6616353 | 0.0238566 | 0.22736584 | 0.10492605 | 0.91643449 | 0.99945981 |
| CTSH | 28.0978557 | 0.278560097 | 0.46441466 | 0.59980901 | 0.54863353 | 0.99945981 |
| TMED3 | 12.8298809 | -0.121569921 | 0.65224487 | -0.1863869 | 0.85214134 | 0.99945981 |
| BCL2A1 | 14.5357791 | 1.381456439 | 0.82663161 | 1.67118752 | 0.09468464 | 0.99945981 |
| ZFAND6 | 97.8902713 | 0.025668515 | 0.27395517 | 0.09369604 | 0.92535061 | 0.99945981 |
| MESD | 16.8338446 | -0.753736728 | 0.35415702 | -2.1282558 | 0.03331588 | 0.8663391 |
| IL16 | 16.8500512 | 0.238976608 | 0.41015357 | 0.58265153 | 0.56012791 | 0.99945981 |
| EFL1 | 21.6789818 | -0.247138802 | 0.33136228 | -0.7458266 | 0.45577217 | 0.99945981 |
| WHAMM | 17.465854 | 0.104897501 | 0.3735397 | 0.28082022 | 0.7788483 | 0.99945981 |
| BTBD1 | 30.631134 | -0.35209575 | 0.30191158 | -1.1662214 | 0.24352493 | 0.99945981 |
| TM6SF1 | 12.0251187 | 0.080059285 | 0.47448743 | 0.16872794 | 0.86601064 | 0.99945981 |
| WDR73 | 12.265804 | -0.193546074 | 0.46598964 | -0.4153442 | 0.67788998 | 0.99945981 |

| | | | | | | | |
|---|---|---|---|---|---|---|---|
| SEC11A | 47.8624714 | | 0.018899125 | 0.2519118 | 0.07502279 | 0.94019658 | 0.99945981 |
| ZNF592 | 27.8906528 | | 0.021581631 | 0.30333607 | 0.07114759 | 0.94328029 | 0.99945981 |
| PDE8A | 84.7218239 | | -0.486574127 | 0.36297699 | -1.3405096 | 0.18007973 | 0.99945981 |
| AKAP13 | 398.73628 | | -0.152092619 | 0.32716303 | -0.4648833 | 0.64201505 | 0.99945981 |
| AC087286.2 | 21.4572746 | | 0.695098234 | 0.51788094 | 1.34219698 | 0.17953214 | 0.99945981 |
| ISG20 | 42.8057982 | | -1.099264134 | 0.52294089 | -2.102081 | 0.03554618 | 0.87920079 |
| ABHD2 | 53.339091 | | -0.141787347 | 0.2985171 | -0.4749723 | 0.63480673 | 0.99945981 |
| POLG | 12.6374572 | | 0.216532946 | 0.41450929 | 0.52238382 | 0.60140312 | 0.99945981 |
| ANPEP | 24.5856241 | | 0.662613986 | 0.59068721 | 1.12176796 | 0.26196111 | 0.99945981 |
| ZNF710 | 87.5855456 | | 0.421823555 | 0.40836766 | 1.03295045 | 0.30162709 | 0.99945981 |
| IDH2 | 21.4114116 | | -0.627807595 | 0.39352068 | -1.5953611 | 0.1106315 | 0.99945981 |
| SEMA4B | 18.2297936 | | -0.363960493 | 0.6307897 | -0.5769918 | 0.56394499 | 0.99945981 |
| CIB1 | 25.0638411 | | -0.813105288 | 0.5625705 | -1.4453394 | 0.14836259 | 0.99945981 |
| IQGAP1 | 164.937948 | | 0.159741925 | 0.26799159 | 0.59607066 | 0.55112802 | 0.99945981 |
| CRTC3 | 59.6539227 | | -0.363483735 | 0.25475772 | -1.426782 | 0.15364273 | 0.99945981 |
| FES | 15.129853 | | -0.470556835 | 0.49189702 | -0.9566166 | 0.33876082 | 0.99945981 |
| MAN2A2 | 12.0564353 | | -0.122843316 | 0.41611635 | -0.2952139 | 0.76783051 | 0.99945981 |
| UNC45A | 8.77325256 | | 0.062673158 | 0.46453688 | 0.13491536 | 0.8926788 | 0.99945981 |
| SLCO3A1 | 128.139175 | | 0.280091762 | 0.32056692 | 0.87373881 | 0.38226051 | 0.99945981 |
| AC013394.1 | 54.7760221 | | 0.426721968 | 0.28712674 | 1.4861798 | 0.13723157 | 0.99945981 |
| CHD2 | 87.9259931 | | -0.034195351 | 0.2245202 | -0.1523041 | 0.87894708 | 0.99945981 |
| MCTP2 | 133.784781 | | -0.198235739 | 0.37734536 | -0.525343 | 0.59934478 | 0.99945981 |
| IGF1R | 60.5689685 | | 0.174121795 | 0.42634954 | 0.40840151 | 0.68297893 | 0.99945981 |
| LRRC28 | 22.1929634 | | -0.118691112 | 0.34033974 | -0.348743 | 0.72728223 | 0.99945981 |
| MEF2A | 253.652034 | | -0.127709558 | 0.44306943 | -0.2882383 | 0.77316437 | 0.99945981 |
| AC090825.1 | 17.3838983 | | -0.16646277 | 0.44007843 | -0.3782571 | 0.70523965 | 0.99945981 |
| ADAMTS17 | 15.6252069 | | -0.218181854 | 0.4129719 | -0.5283213 | 0.59727634 | 0.99945981 |
| LINS1 | 10.9040293 | | 0.289833902 | 0.53311934 | 0.5436567 | 0.58667773 | 0.99945981 |
| ASB7 | 13.6104934 | | 0.033158353 | 0.4130476 | 0.08027732 | 0.9360167 | 0.99945981 |
| LRRK1 | 46.2787564 | | 0.423373461 | 0.4841407 | 0.87448435 | 0.38185454 | 0.99945981 |
| CHSY1 | 50.6305476 | | 0.209222198 | 0.39804543 | 0.52562392 | 0.59914954 | 0.99945981 |
| SELENOS | 19.4970619 | | -0.426611947 | 0.47568091 | -0.8968448 | 0.36980175 | 0.99945981 |
| SNRPA1 | 12.8055624 | | 0.197375678 | 0.43713828 | 0.45151772 | 0.65161646 | 0.99945981 |
| TM2D3 | 13.8982002 | | 0.25060057 | 0.42198686 | 0.59385871 | 0.55260662 | 0.99945981 |
| TARSL2 | 24.3667458 | | -0.372018819 | 0.59196525 | -0.6284471 | 0.52971112 | 0.99945981 |
| MPG | 11.3135766 | | -1.530541672 | 0.49664319 | -3.0817732 | 0.00205772 | 0.45570351 |
| NPRL3 | 15.2897071 | | -0.036672899 | 0.41035503 | -0.0893687 | 0.92878889 | 0.99945981 |
| LUC7L | 38.3450321 | | 0.080286816 | 0.31517887 | 0.25473413 | 0.79892845 | 0.99945981 |
| AXIN1 | 18.8270044 | | 0.009697686 | 0.39754404 | 0.02439399 | 0.98053834 | 0.99945981 |
| TMEM8A | 9.45245564 | | -0.746522197 | 0.47498708 | -1.5716684 | 0.11602747 | 0.99945981 |
| RAB11FIP3 | 24.1174185 | | 0.463472236 | 0.33668661 | 1.37656865 | 0.16864565 | 0.99945981 |
| CAPN15 | 10.8984287 | | 0.179772642 | 0.50912378 | 0.35310203 | 0.72401195 | 0.99945981 |
| RAB40C | 10.6753573 | | -0.408928487 | 0.62570555 | -0.6535478 | 0.51340318 | 0.99945981 |
| RHOT2 | 11.1746443 | | 0.713851505 | 0.45924483 | 1.55440293 | 0.12008834 | 0.99945981 |
| STUB1 | 11.2295653 | | -0.18904807 | 0.51498938 | -0.3670912 | 0.71355099 | 0.99945981 |
| LMF1 | 17.4553797 | | -0.295002156 | 0.54156642 | -0.5447202 | 0.58594599 | 0.99945981 |
| UBE2I | 31.9753248 | | -0.460173777 | 0.44038368 | -1.0449383 | 0.29605148 | 0.99945981 |
| GNPTG | 9.09043461 | | -0.46179165 | 0.50541114 | -0.913695 | 0.36087711 | 0.99945981 |
| UNKL | 11.8002188 | | -1.166575289 | 0.43823426 | -2.6619902 | 0.00776801 | 0.57727153 |
| CLCN7 | 19.4310289 | | 0.251428184 | 0.39779589 | 0.63205325 | 0.52735208 | 0.99945981 |
| MAPK8IP3 | 30.5880976 | | -0.745796224 | 0.45915457 | -1.6242814 | 0.10431576 | 0.99945981 |
| NME3 | 15.0153298 | | 0.137957586 | 0.38454472 | 0.35875563 | 0.71977791 | 0.99945981 |
| MRPS34 | 11.5883989 | | -0.139629045 | 0.71888649 | -0.1942296 | 0.8459961 | 0.99945981 |
| SPSB3 | 13.9834168 | | 0.247974346 | 0.40743292 | 0.6086262 | 0.54277224 | 0.99945981 |
| HAGH | 16.1458215 | | -0.297029822 | 0.44283506 | -0.6707459 | 0.50238239 | 0.99945981 |
| NDUFB10 | 12.530701 | | -0.519010814 | 0.50163974 | -1.0346286 | 0.3008424 | 0.99945981 |
| RPS2 | 89.8855591 | | 0.103315388 | 0.49947295 | 0.20684882 | 0.83612793 | 0.99945981 |
| TSC2 | 15.1449597 | | 0.029520586 | 0.35378295 | 0.08344265 | 0.93349958 | 0.99945981 |
| PKD1 | 10.7684132 | | -0.471016206 | 0.48980815 | -0.9616341 | 0.33623345 | 0.99945981 |
| PGP | 13.217688 | | -0.259494125 | 0.40787672 | -0.6362072 | 0.52464135 | 0.99945981 |
| ECI1 | 11.926597 | | -0.022288591 | 0.4906656 | -0.0454252 | 0.96376838 | 0.99945981 |
| RNPS1 | 19.9473542 | | -0.057076102 | 0.38787551 | -0.1471506 | 0.88301317 | 0.99945981 |
| PDPK1 | 50.1431392 | | -0.359589832 | 0.28526683 | -1.2605386 | 0.20747515 | 0.99945981 |
| SRRM2 | 109.41223 | | 0.111402314 | 0.24774698 | 0.44966164 | 0.65295444 | 0.99945981 |
| ELOB | 18.4460224 | | 0.039315804 | 0.60909961 | 0.06454741 | 0.94853436 | 0.99945981 |
| MMP25-AS1 | 7.38207502 | | -0.612090653 | 0.81972608 | -0.7467015 | 0.45524379 | 0.99945981 |
| IL32 | 34.7149218 | | -0.802319691 | 0.57104182 | -1.4050104 | 0.16001819 | 0.99945981 |
| ZNF263 | 9.08176495 | | -0.010898184 | 0.54070465 | -0.0201555 | 0.98391931 | 0.99945981 |
| ZNF75A | 12.0173644 | | -0.97052867 | 0.71944852 | -1.3489897 | 0.17734026 | 0.99945981 |
| NLRC3 | 17.1231418 | | -0.093753033 | 0.40777521 | -0.2299135 | 0.81815897 | 0.99945981 |
| DNASE1 | 19.459753 | | -0.136063914 | 0.4164732 | -0.3267051 | 0.74389095 | 0.99945981 |
| TRAP1 | 9.03743192 | | -1.069930242 | 0.45710925 | -2.3406445 | 0.01925049 | 0.77737999 |
| CREBBP | 91.1425699 | | 0.099687213 | 0.25341097 | 0.39338161 | 0.69403765 | 0.99945981 |
| ADCY9 | 26.1566517 | | -0.036531133 | 0.4083715 | -0.0894556 | 0.92871981 | 0.99945981 |
| DNAJA3 | 10.6969128 | | -0.507375024 | 0.45637819 | -1.1117425 | 0.26624889 | 0.99945981 |
| NMRAL1 | 9.27658866 | | -0.593077132 | 0.46433377 | -1.2772647 | 0.20150881 | 0.99945981 |
| HMOX2 | 15.3183426 | | -0.488717365 | 0.4513476 | -1.082796 | 0.27889899 | 0.99945981 |
| MGRN1 | 25.7992515 | | -0.340242383 | 0.33945525 | -1.0023188 | 0.31618964 | 0.99945981 |
| ANKS3 | 9.24886415 | | -0.190466057 | 0.43122657 | -0.4416844 | 0.65871758 | 0.99945981 |
| GLYR1 | 34.7416904 | | -0.276033808 | 0.39650672 | -0.6961643 | 0.48632596 | 0.99945981 |
| UBN1 | 18.4986177 | | -0.347907775 | 0.36075969 | -0.9643754 | 0.33485775 | 0.99945981 |
| METTL22 | 8.52760462 | | -0.259403006 | 0.50797576 | -0.5106602 | 0.60958901 | 0.99945981 |
| ABAT | 16.3189161 | | 0.690345163 | 0.46585467 | 1.48188953 | 0.13836969 | 0.99945981 |
| PMM2 | 24.135294 | | -0.651616783 | 0.30133775 | -2.1624134 | 0.03058633 | 0.85704136 |
| CARHSP1 | 18.498971 | | -0.830615736 | 0.45313909 | -1.833026 | 0.0667987 | 0.94594189 |
| USP7 | 33.85513 | | 0.265736452 | 0.30558438 | 0.8696009 | 0.38451854 | 0.99945981 |
| C16orf72 | 29.1819604 | | 0.10803613 | 0.29072328 | 0.37161155 | 0.71018209 | 0.99945981 |
| ATF7IP2 | 51.4023391 | | -0.332573079 | 0.33506834 | -0.992553 | 0.32092785 | 0.99945981 |
| NUBP1 | 8.60013813 | | -0.824110336 | 0.58990253 | -1.397028 | 0.16240516 | 0.99945981 |
| CIITA | 71.0047228 | | -0.402114259 | 0.56849917 | -0.707326 | 0.47936389 | 0.99945981 |
| DEXI | 7.83709498 | | -0.728929004 | 0.4558318 | -1.5991184 | 0.10979431 | 0.99945981 |
| CLEC16A | 59.1790151 | | -0.216153577 | 0.35707893 | -0.6053384 | 0.54495419 | 0.99945981 |
| LITAF | 85.8936517 | | -0.509604371 | 0.42009413 | -1.2130719 | 0.22510234 | 0.99945981 |
| TXNDC11 | 43.7980565 | | -0.537700662 | 0.46289731 | -1.1615982 | 0.24539873 | 0.99945981 |
| ZC3H7A | 41.7418562 | | 0.214943964 | 0.25682209 | 0.83693722 | 0.40262786 | 0.99945981 |
| RSL1D1 | 11.8604417 | | 0.067767462 | 0.46778732 | 0.1448681 | 0.88481501 | 0.99945981 |
| AC007216.4 | 12.6996026 | | -0.035529693 | 0.41496414 | -0.0856211 | 0.93176761 | 0.99945981 |
| GSPT1 | 32.5340724 | | -0.361925628 | 0.27883947 | -1.2979713 | 0.1942972 | 0.99945981 |
| SNX29 | 134.473497 | | 0.320150102 | 0.2928105 | 1.09336962 | 0.27423155 | 0.99945981 |
| CPPED1 | 54.449002 | | 1.018123801 | 0.46887254 | 2.17142978 | 0.0298987 | 0.85472619 |
| MKL2 | 27.5264883 | | 0.020650467 | 0.35359245 | 0.05840189 | 0.95342851 | 0.99945981 |
| PARN | 44.9986262 | | -0.072527814 | 0.24042192 | -0.3016689 | 0.76290449 | 0.99945981 |
| BFAR | 13.6201292 | | -0.209882428 | 0.47073392 | -0.4458621 | 0.65569684 | 0.99945981 |
| PDXDC1 | 66.698101 | | 0.008710162 | 0.30850676 | 0.0282333 | 0.97747608 | 0.99945981 |
| RRN3 | 14.8504352 | | -0.17591668 | 0.37053672 | -0.4747618 | 0.63495677 | 0.99945981 |
| C16orf45 | 13.6920464 | | -1.239478412 | 0.75235104 | -1.6474735 | 0.09946076 | 0.99945981 |
| MARF1 | 45.1028889 | | -0.734678659 | 0.70598444 | -1.0406443 | 0.29804068 | 0.99945981 |
| NDE1 | 25.867259 | | 0.131297418 | 0.45419899 | 0.28907466 | 0.77252424 | 0.99945981 |
| ABCC1 | 60.1860691 | | -0.030326044 | 0.47457282 | -0.0639018 | 0.94904844 | 0.99945981 |
| XYLT1 | 133.299798 | | -0.05274642 | 0.49805964 | -0.1059038 | 0.91565866 | 0.99945981 |
| RPS15A | 23.8567166 | | -0.459509521 | 0.84144906 | -0.5460931 | 0.58500195 | 0.99945981 |
| SMG1 | 99.7943194 | | 0.086137256 | 0.21726188 | 0.39646741 | 0.69176025 | 0.99945981 |
| VPS35L | 36.9345478 | | 0.030881826 | 0.32365986 | 0.09541444 | 0.92398564 | 0.99945981 |

| | | | | | | |
|---|---|---|---|---|---|---|
| UMOD | 48.0203775 | 1.441798894 | 0.77828248 | 1.85253931 | 0.06394842 | 0.93776345 |
| ACSM3 | 39.7773711 | -3.25831628 | 1.67721584 | -1.9426935 | 0.0520532 | 0.9230968 |
| THUMPD1 | 13.8639461 | -0.588327565 | 0.45988808 | -1.2792842 | 0.20079698 | 0.99945981 |
| METTL9 | 37.0334553 | 0.131640657 | 0.34139192 | 0.3855998 | 0.69979308 | 0.99945981 |
| IGSF6 | 15.1923673 | 0.565239013 | 0.62081183 | 0.91048364 | 0.3625675 | 0.99945981 |
| NPIPB4 | 14.6369978 | 0.519378333 | 0.50496744 | 1.02853827 | 0.3036967 | 0.99945981 |
| UQCRC2 | 28.9795412 | 0.247462644 | 0.30228658 | 0.81863591 | 0.41299418 | 0.99945981 |
| MOSMO | 35.5515454 | -0.296446138 | 0.40379877 | -0.7341432 | 0.46286144 | 0.99945981 |
| EEF2K | 17.6467605 | -0.030932863 | 0.42655035 | -0.0725187 | 0.94218915 | 0.99945981 |
| POLR3E | 9.65598965 | -0.641419917 | 0.42848082 | -1.496963 | 0.1344029 | 0.99945981 |
| CDR2 | 53.0237233 | -0.270810227 | 0.47181764 | -0.5739722 | 0.56598659 | 0.99945981 |
| NPIPB5 | 42.149498 | 0.001848487 | 0.2794261 | 0.0066153 | 0.9947218 | 0.99945981 |
| USP31 | 8.74324519 | -0.483343662 | 0.54753669 | -0.8827603 | 0.37736578 | 0.99945981 |
| COG7 | 15.2466061 | 0.123221217 | 0.38281102 | 0.32188524 | 0.74753964 | 0.99945981 |
| GGA2 | 44.7883963 | -0.776676102 | 0.46241939 | -1.6795924 | 0.09303664 | 0.99640647 |
| NDUFAB1 | 8.47469825 | -0.166536102 | 0.51220231 | -0.3251374 | 0.74507712 | 0.99945981 |
| DCTN5 | 14.8222671 | -0.263587047 | 0.4362453 | -0.6042175 | 0.54569905 | 0.99945981 |
| PRKCB | 314.215269 | 0.760710832 | 0.31534959 | 2.41227786 | 0.01585319 | 0.74035497 |
| RBBP6 | 30.645798 | 0.019094144 | 0.36405713 | 0.05244821 | 0.95817156 | 0.99945981 |
| TNRC6A | 98.1227314 | 0.154881276 | 0.19706111 | 0.78595559 | 0.43189351 | 0.99945981 |
| ARHGAP17 | 46.4446814 | 0.74471546 | 0.33841139 | 2.20062176 | 0.02776281 | 0.84763724 |
| LCMT1 | 15.4488473 | -0.111885479 | 0.42016474 | -0.2662896 | 0.79001622 | 0.99945981 |
| NSMCE1 | 7.97318889 | -0.270698989 | 0.59614117 | -0.4540854 | 0.64976738 | 0.99945981 |
| IL4R | 40.5766923 | -0.412851475 | 0.43229864 | -0.9550145 | 0.33957035 | 0.99945981 |
| IL21R | 30.2149698 | -1.321971864 | 0.84130885 | -1.5713277 | 0.11610656 | 0.99945981 |
| GTF3C1 | 17.5255838 | 0.262213844 | 0.38856071 | 0.67483365 | 0.49978146 | 0.99945981 |
| KIAA0556 | 17.3358809 | -0.173449194 | 0.39219296 | -0.4422547 | 0.65830488 | 0.99945981 |
| XPO6 | 78.7443323 | 0.042579403 | 0.35324196 | 0.12053891 | 0.90405626 | 0.99945981 |
| NUPR1 | 11.8397982 | -2.333957615 | 1.10183394 | -2.1182481 | 0.03415406 | 0.87717522 |
| ATXN2L | 23.3634746 | 0.422901142 | 0.34637576 | 1.22093168 | 0.22211189 | 0.99945981 |
| TUFM | 17.1873377 | 0.333687342 | 0.47293312 | 0.70556983 | 0.48045569 | 0.99945981 |
| RABEP2 | 19.172835 | -1.909197819 | 0.94646163 | -2.0171952 | 0.04367515 | 0.89562588 |
| NFATC2IP | 45.8787783 | 0.232669934 | 0.28876655 | 0.80573715 | 0.42039442 | 0.99945981 |
| SPNS1 | 9.38522982 | -0.106800899 | 0.48123304 | -0.2219318 | 0.824367 | 0.99945981 |
| LAT | 8.10566508 | 0.932565435 | 0.57890499 | 1.61091276 | 0.10719874 | 0.99945981 |
| AC009093.2 | 66.7651017 | 0.150846687 | 0.42413358 | 0.35565844 | 0.72209637 | 0.99945981 |
| MAZ | 10.3841631 | 0.094344795 | 0.48335381 | 0.19518786 | 0.84524588 | 0.99945981 |
| MVP | 30.052073 | 0.19509934 | 0.38676368 | 0.50444069 | 0.61395173 | 0.99945981 |
| KCTD13 | 15.2005551 | 0.81134638 | 0.38513407 | 2.1066596 | 0.0351471 | 0.87920079 |
| TMEM219 | 12.7766188 | -0.123529049 | 0.59114174 | -0.2089669 | 0.83447409 | 0.99945981 |
| PPP4C | 14.0580613 | 0.29579877 | 0.38995515 | 0.75854561 | 0.44812442 | 0.99945981 |
| YPEL3 | 24.5128438 | -0.34188863 | 0.45116549 | -0.7577898 | 0.4485768 | 0.99945981 |
| CORO1A | 42.1636793 | -0.043607944 | 0.56250464 | -0.0775246 | 0.93820623 | 0.99945981 |
| NPIPB13 | 12.8880345 | -0.124181234 | 0.46904308 | -0.2647544 | 0.79119864 | 0.99945981 |
| CD2BP2 | 9.05268062 | 0.915924816 | 0.55916136 | 1.6380331 | 0.10141479 | 0.99945981 |
| 1-Sep | 10.0468282 | -0.837858795 | 0.47344946 | -1.76969 | 0.07677879 | 0.9821652 |
| SEPHS2 | 11.9216895 | 0.027416279 | 0.64295498 | 0.04264106 | 0.96598767 | 0.99945981 |
| ITGAL | 92.5147781 | 1.008346839 | 0.5139737 | 1.96186467 | 0.04977824 | 0.92165202 |
| FBRS | 11.1257218 | 0.346990006 | 0.48439101 | 0.71634279 | 0.47377971 | 0.99945981 |
| RNF40 | 7.56012179 | -0.243122975 | 0.4582641 | -0.5305303 | 0.59574433 | 0.99945981 |
| BCL7C | 9.37471381 | -0.162276726 | 0.46276877 | -0.3506648 | 0.72583982 | 0.99945981 |
| STX4 | 10.4764571 | 0.662180838 | 0.53775206 | 1.23138689 | 0.2181782 | 0.99945981 |
| KAT8 | 13.1418389 | -0.043559158 | 0.46224754 | -0.0942334 | 0.92492375 | 0.99945981 |
| FUS | 74.6533159 | 0.191479256 | 0.29760509 | 0.64340047 | 0.51996428 | 0.99945981 |
| PYCARD | 18.7611667 | 0.083368675 | 0.68229123 | 0.12218928 | 0.90274912 | 0.99945981 |
| ITGAM | 40.9343572 | 1.308452237 | 0.63876351 | 2.0484142 | 0.04051943 | 0.89562588 |
| ITGAX | 62.3770137 | -0.080340051 | 0.60083872 | -0.1337132 | 0.89362939 | 0.99945981 |
| ZNF720 | 14.2722911 | -0.076964032 | 0.41355025 | -0.1861056 | 0.85236192 | 0.99945981 |
| ZNF267 | 26.0192135 | 0.040876285 | 0.49317066 | 0.08288467 | 0.93394325 | 0.99945981 |
| VPS35 | 30.2954939 | -0.097237227 | 0.35373879 | -0.2748843 | 0.78340515 | 0.99945981 |
| C16orf87 | 9.6431534 | -0.148220242 | 0.46278791 | -0.3202768 | 0.74875849 | 0.99945981 |
| DNAJA2 | 9.65144725 | -0.037247007 | 0.52043638 | -0.0715688 | 0.94294507 | 0.99945981 |
| NETO2 | 12.6777809 | 0.0993987 | 0.54926512 | 0.18096671 | 0.85639371 | 0.99945981 |
| ITFG1 | 126.747485 | 0.19399127 | 0.2452664 | 0.79094109 | 0.42897837 | 0.99945981 |
| PHKB | 105.127012 | 0.182277429 | 0.19550709 | 0.93233153 | 0.35116522 | 0.99945981 |
| LONP2 | 55.0803608 | -0.076553328 | 0.23097571 | -0.3314345 | 0.74031628 | 0.99945981 |
| SIAH1 | 13.9786884 | -0.017685103 | 0.46036995 | -0.038415 | 0.96935682 | 0.99945981 |
| N4BP1 | 61.6077138 | 0.227432535 | 0.45088691 | 0.50441148 | 0.61397225 | 0.99945981 |
| CNEP1R1 | 9.84259165 | -0.010028429 | 0.4607944 | -0.0217633 | 0.98263673 | 0.99945981 |
| HEATR3 | 26.6465182 | 0.006085649 | 0.4860377 | 0.01252094 | 0.99001 | 0.99945981 |
| TENT4B | 45.1262192 | -0.377394378 | 0.27636054 | -1.365587 | 0.17206861 | 0.99945981 |
| ADCY7 | 52.3675878 | 0.302291845 | 0.40301507 | 0.75007579 | 0.45320906 | 0.99945981 |
| BRD7 | 20.1184531 | -0.026840363 | 0.30679688 | -0.0874858 | 0.93028539 | 0.99945981 |
| SNX20 | 9.34094176 | 0.305677875 | 0.45493447 | 0.67191628 | 0.501637 | 0.99945981 |
| NOD2 | 12.7239879 | 0.558402391 | 0.60545683 | 0.92228275 | 0.35638111 | 0.99945981 |
| CYLD | 57.6046719 | 0.514439242 | 0.25279163 | 2.03503275 | 0.04184758 | 0.89562588 |
| CHD9 | 133.331932 | 0.181304868 | 0.23274512 | 0.77898462 | 0.43598878 | 0.99945981 |
| RBL2 | 73.9415399 | -0.209313875 | 0.26528159 | -0.7890253 | 0.43009725 | 0.99945981 |
| FTO | 98.9790458 | -0.215756636 | 0.38307923 | -0.5632167 | 0.5732873 | 0.99945981 |
| LPCAT2 | 36.4623398 | 0.678496479 | 0.35960817 | 1.88676605 | 0.0591918 | 0.92445778 |
| GNAO1 | 10.5525533 | -2.18404127 | 0.62907402 | -3.4718351 | 0.00051691 | 0.23820088 |
| AMFR | 21.6390722 | 0.075419457 | 0.4291478 | 0.17574238 | 0.86049634 | 0.99945981 |
| NUDT21 | 12.0567891 | -0.168486405 | 0.41655781 | -0.404473 | 0.6858649 | 0.99945981 |
| BBS2 | 11.8131841 | 0.725797296 | 0.54599783 | 1.32930435 | 0.18374758 | 0.99945981 |
| MT2A | 141.052476 | -0.427023456 | 0.68737632 | -0.6212368 | 0.53444384 | 0.99945981 |
| MT1E | 30.662057 | -0.678505543 | 0.89809846 | -0.7554913 | 0.44995427 | 0.99945981 |
| MT1G | 100.674739 | 0.341251431 | 1.01159851 | 0.3373388 | 0.73586151 | 0.99945981 |
| MT1H | 33.4269541 | 1.497542308 | 0.95583724 | 1.56673359 | 0.11717698 | 0.99945981 |
| MT1X | 36.5590114 | -0.087967131 | 0.72491598 | -0.121348 | 0.90341537 | 0.99945981 |
| NUP93 | 21.496669 | 0.502978144 | 0.36538217 | 1.37658098 | 0.16864184 | 0.99945981 |
| HERPUD1 | 80.3640584 | 0.548534589 | 0.40085229 | 1.36842074 | 0.17118041 | 0.99945981 |
| NLRC5 | 72.4327906 | 0.3777212 | 0.42470596 | 0.88937108 | 0.37380368 | 0.99945981 |
| FAM192A | 25.2268584 | 0.068673101 | 0.34990468 | 0.19626231 | 0.84440486 | 0.99945981 |
| RSPRY1 | 17.8161599 | -0.315833152 | 0.4358957 | -0.7245613 | 0.46872121 | 0.99945981 |
| ARL2BP | 10.1771515 | 0.193211081 | 0.43793513 | 0.44118653 | 0.65907796 | 0.99945981 |
| ADGRG5 | 14.6781854 | 0.471276707 | 0.52264124 | 0.90172124 | 0.36720497 | 0.99945981 |
| ADGRG1 | 13.0626531 | -0.012370561 | 0.59678851 | -0.0207286 | 0.98346219 | 0.99945981 |
| KIFC3 | 14.919397 | 0.562102867 | 0.76005965 | 0.73955099 | 0.45957249 | 0.99945981 |
| USB1 | 8.84937916 | -0.846526527 | 0.54249496 | -1.5604321 | 0.11865781 | 0.99945981 |
| CSNK2A2 | 14.8686559 | 0.152536987 | 0.40591567 | 0.37578492 | 0.7070768 | 0.99945981 |
| CNOT1 | 66.1607863 | 0.425354033 | 0.24748872 | 1.71868052 | 0.08567256 | 0.99193047 |
| GOT2 | 9.29716035 | -0.17881245 | 0.52115743 | -0.3431064 | 0.73151841 | 0.99945981 |
| CMTM3 | 11.2729574 | -0.587721798 | 0.53991704 | -1.0885409 | 0.27635638 | 0.99945981 |
| CMTM4 | 10.2972922 | -0.226585533 | 0.61224076 | -0.3700922 | 0.71131379 | 0.99945981 |
| DYNC1LI2 | 30.0605427 | -0.212308069 | 0.35546282 | -0.5972722 | 0.55032565 | 0.99945981 |
| NAE1 | 10.609639 | -0.115573672 | 0.42281963 | -0.2733404 | 0.78459159 | 0.99945981 |
| CDH16 | 12.5805883 | -0.411761752 | 0.54924225 | -0.7496906 | 0.45344107 | 0.99945981 |
| FAM96B | 10.6139717 | -0.107321982 | 0.46359625 | -0.2314988 | 0.81692731 | 0.99945981 |
| CBFB | 67.2132337 | 0.084141313 | 0.22162358 | 0.37965866 | 0.70419881 | 0.99945981 |
| C16orf70 | 13.3601929 | 0.438835797 | 0.4517348 | 0.97144562 | 0.33132642 | 0.99945981 |
| TRADD | 8.41138078 | -0.276975848 | 0.54096777 | -0.5120006 | 0.60865056 | 0.99945981 |
| E2F4 | 8.1009764 | -0.470113728 | 0.51285602 | -0.9166583 | 0.35932172 | 0.99945981 |
| FHOD1 | 8.43928363 | 0.651775098 | 0.5338783 | 1.22083085 | 0.22215007 | 0.99945981 |

| | | | | | | |
|---|---|---|---|---|---|---|
| ATP6V0D1 | 28.5557498 | -0.063639431 | 0.44173506 | -0.144067 | 0.88544759 | 0.99945981 |
| AC009061.2 | 10.0719641 | -0.207643251 | 0.64751313 | -0.3206781 | 0.74845438 | 0.99945981 |
| RIPOR1 | 14.1167326 | -0.421652703 | 0.60970416 | -0.6915693 | 0.48920782 | 0.99945981 |
| CTCF | 36.6063931 | -0.149697431 | 0.29180098 | -0.5130121 | 0.60794286 | 0.99945981 |
| RANBP10 | 20.8715584 | -0.071961697 | 0.37844393 | -0.1901515 | 0.84919038 | 0.99945981 |
| DPEP2 | 16.7298256 | 0.43420747 | 0.47501883 | 0.91408475 | 0.36067232 | 0.99945981 |
| DUS2 | 10.3598793 | 0.457053512 | 0.51611681 | 0.88556215 | 0.37585352 | 0.99945981 |
| NFATC3 | 117.94565 | -0.040659191 | 0.24486095 | -0.1660501 | 0.8681175 | 0.99945981 |
| SLC7A6 | 27.1063751 | 0.353529859 | 0.32212312 | 1.09749919 | 0.27242324 | 0.99945981 |
| SLC7A6OS | 14.9577631 | -0.745719953 | 0.45339305 | -1.6447538 | 0.10002059 | 0.99945981 |
| PRMT7 | 11.7304684 | 0.469749394 | 0.56646396 | 0.82926616 | 0.40695382 | 0.99945981 |
| ZFP90 | 11.9301533 | -0.475441189 | 0.40401969 | -1.1767773 | 0.23928442 | 0.99945981 |
| TANGO6 | 39.8595563 | 0.083569614 | 0.27385493 | 0.30516016 | 0.76024417 | 0.99945981 |
| CHTF8 | 8.62046196 | 0.091601 | 0.56651782 | 0.16169129 | 0.87154896 | 0.99945981 |
| SNTB2 | 20.2016467 | 0.414834254 | 0.44996137 | 0.92193304 | 0.3565635 | 0.99945981 |
| TERF2 | 16.0975895 | -0.383278361 | 0.45992435 | -0.8333509 | 0.40464687 | 0.99945981 |
| CYB5B | 24.1470399 | 0.163065967 | 0.32999542 | 0.49414615 | 0.62120297 | 0.99945981 |
| NFAT5 | 159.248643 | 0.14657289 | 0.22830405 | 0.6420074 | 0.52086838 | 0.99945981 |
| WWP2 | 63.0167097 | 0.22071293 | 0.37122181 | 0.59455808 | 0.55213891 | 0.99945981 |
| PDPR | 20.4278875 | 0.295247637 | 0.39023597 | 0.75658744 | 0.44929707 | 0.99945981 |
| AARS | 8.6280916 | -0.401493902 | 0.50776976 | -0.7907007 | 0.42911867 | 0.99945981 |
| ST3GAL2 | 25.7845599 | 0.13785649 | 0.48179135 | 0.28613317 | 0.77477612 | 0.99945981 |
| COG4 | 17.7301073 | -0.290389941 | 0.35778372 | -0.8116354 | 0.41700087 | 0.99945981 |
| SF3B3 | 27.862 | -0.163812902 | 0.3322756 | -0.4930031 | 0.6220104 | 0.99945981 |
| VAC14 | 27.9056782 | -0.066943849 | 0.46727861 | -0.1432632 | 0.88608228 | 0.99945981 |
| CMTR2 | 8.41195138 | 0.450761256 | 0.45167765 | 0.99797113 | 0.31829336 | 0.99945981 |
| PHLPP2 | 11.3671269 | -0.068634314 | 0.40389715 | -0.1699302 | 0.86506505 | 0.99945981 |
| AP1G1 | 46.4806273 | 0.467417594 | 0.26837644 | 1.74164916 | 0.08156985 | 0.9846219 |
| IST1 | 38.6398887 | 0.432594644 | 0.26342349 | 1.6422022 | 0.10054811 | 0.99945981 |
| DHX38 | 9.51124068 | -0.395530326 | 0.44223936 | -0.8943807 | 0.37111826 | 0.99945981 |
| LINC01572 | 16.8056021 | 0.641819621 | 0.65411308 | 0.98120591 | 0.32649121 | 0.99945981 |
| ZFHX3 | 84.0416346 | 0.867031715 | 0.51506739 | 1.68333645 | 0.09230998 | 0.99640647 |
| PSMD7 | 14.8331381 | -0.24671478 | 0.49569477 | -0.4977151 | 0.61868485 | 0.99945981 |
| GLG1 | 130.169714 | 0.411995478 | 0.28413938 | 1.44997669 | 0.14706502 | 0.99945981 |
| RFWD3 | 15.6512927 | -0.155437428 | 0.3717063 | -0.4181727 | 0.67582085 | 0.99945981 |
| MLKL | 19.5025852 | 0.231978535 | 0.48839278 | 0.47498355 | 0.6347987 | 0.99945981 |
| WDR59 | 26.131096 | -0.198694895 | 0.43576597 | -0.455967 | 0.64841372 | 0.99945981 |
| ZNRF1 | 19.6402787 | -0.520836452 | 0.7746481 | -0.6723523 | 0.50135942 | 0.99945981 |
| CFDP1 | 42.190445 | 0.236141809 | 0.29419163 | 0.80268023 | 0.42215958 | 0.99945981 |
| GABARAPL2 | 16.7706306 | -1.103355327 | 0.45908282 | -2.4033906 | 0.01624383 | 0.74280553 |
| ADAT1 | 10.0392374 | -0.26821909 | 0.54183138 | -0.4950232 | 0.62058376 | 0.99945981 |
| KARS | 11.8214013 | -0.301441032 | 0.52536691 | -0.5737724 | 0.56612183 | 0.99945981 |
| TERF2IP | 36.2136877 | -0.365345171 | 0.35490513 | -1.0294164 | 0.30328404 | 0.99945981 |
| WWOX | 310.529732 | -1.185679831 | 0.56913141 | -2.0833147 | 0.03722255 | 0.87920079 |
| MAF | 37.4454637 | 0.56647629 | 0.63102943 | 0.89770185 | 0.36934452 | 0.99945981 |
| CDYL2 | 33.7640508 | 0.463883777 | 0.32177528 | 1.44163896 | 0.14940425 | 0.99945981 |
| CMC2 | 19.7602169 | 0.022602601 | 0.41187654 | 0.05487713 | 0.95623636 | 0.99945981 |
| GAN | 12.8692279 | 0.303223435 | 0.40396503 | 0.75061802 | 0.45288257 | 0.99945981 |
| CMIP | 236.53457 | -0.572614916 | 0.47002394 | -1.2182676 | 0.22312232 | 0.99945981 |
| PLCG2 | 120.500549 | -0.548633729 | 0.36591208 | -1.4993594 | 0.13378041 | 0.99945981 |
| MPHOSPH6 | 13.8719544 | -0.620754129 | 0.52985541 | -1.1715538 | 0.24137624 | 0.99945981 |
| HSBP1 | 15.4079177 | 0.265236445 | 0.46684773 | 0.56814337 | 0.56993762 | 0.99945981 |
| MBTPS1 | 30.1626865 | -0.452507834 | 0.31629505 | -1.430651 | 0.15253026 | 0.99945981 |
| TLDC1 | 9.81749577 | -1.268193159 | 0.56154404 | -2.2584037 | 0.0239205 | 0.80320673 |
| COTL1 | 104.587118 | 0.913479135 | 0.71381685 | 1.2797108 | 0.20064687 | 0.99945981 |
| KLHL36 | 11.7915861 | -0.533685282 | 0.42987583 | -1.2414871 | 0.21442587 | 0.99945981 |
| USP10 | 43.0234963 | -0.468618069 | 0.38411928 | -1.2199806 | 0.22247223 | 0.99945981 |
| ZDHHC7 | 25.0597105 | 0.800824249 | 0.39161559 | 2.04492433 | 0.04086232 | 0.89562588 |
| KIAA0513 | 20.8820089 | 0.941328832 | 0.63939662 | 1.47221427 | 0.14096303 | 0.99945981 |
| GSE1 | 45.8526666 | 0.156958235 | 0.4236695 | 0.37047329 | 0.71102987 | 0.99945981 |
| COX4I1 | 63.3693065 | -0.020006535 | 0.50504446 | -0.0396134 | 0.96840133 | 0.99945981 |
| IRF8 | 29.5786236 | 0.540081114 | 0.59310039 | 0.91060657 | 0.36250271 | 0.99945981 |
| MAP1LC3B | 21.2767893 | -0.096926812 | 0.38937546 | -0.2489289 | 0.80341576 | 0.99945981 |
| ZCCHC14 | 18.6976223 | -0.098693736 | 0.33945922 | -0.2907381 | 0.77125161 | 0.99945981 |
| KLHDC4 | 29.7824555 | -0.201839317 | 0.37674129 | -0.5357505 | 0.59213103 | 0.99945981 |
| SLC7A5 | 14.9947806 | -1.38685749 | 0.69758725 | -1.9880775 | 0.04680312 | 0.90884462 |
| BANP | 44.757167 | 0.09392026 | 0.25213588 | 0.37249859 | 0.70952166 | 0.99945981 |
| ZFPM1 | 19.4191622 | -1.767351497 | 0.94980706 | -1.8607479 | 0.06277978 | 0.93084929 |
| ZC3H18 | 21.3326919 | -0.780790352 | 0.39367784 | -1.9833231 | 0.04733135 | 0.90884462 |
| CYBA | 134.045435 | -0.122853834 | 0.60692409 | -0.2024204 | 0.83958806 | 0.99945981 |
| MVD | 8.47598226 | -0.992822996 | 0.84703543 | -1.1721151 | 0.24115086 | 0.99945981 |
| RNF166 | 24.8197349 | 0.223086262 | 0.37540363 | 0.59425707 | 0.55234018 | 0.99945981 |
| PIEZO1 | 23.4721368 | 0.355884011 | 0.42615864 | 0.83509748 | 0.40366282 | 0.99945981 |
| APRT | 17.4532038 | -0.717585057 | 0.58711651 | -1.2222192 | 0.22162476 | 0.99945981 |
| GALNS | 9.81070825 | -0.454242954 | 0.46698126 | -0.972722 | 0.33069148 | 0.99945981 |
| CBFA2T3 | 17.0090526 | 0.930459777 | 0.90458986 | 1.0285985 | 0.30366838 | 0.99945981 |
| ACSF3 | 16.8234615 | -0.355177033 | 0.37785339 | -0.9399864 | 0.34722456 | 0.99945981 |
| ANKRD11 | 221.079001 | -0.039527824 | 0.2340442 | -0.1688904 | 0.86588283 | 0.99945981 |
| SPG7 | 40.7084301 | -0.576825829 | 0.3959 | -1.4569988 | 0.14511671 | 0.99945981 |
| RPL13 | 202.973126 | -0.030720727 | 0.50742302 | -0.0605426 | 0.95172346 | 0.99945981 |
| CHMP1A | 8.64387847 | -0.204884569 | 0.60388943 | -0.339275 | 0.7344026 | 0.99945981 |
| CDK10 | 8.39051703 | 0.113055475 | 0.53158397 | 0.21267661 | 0.83157921 | 0.99945981 |
| VPS9D1 | 14.3235417 | -0.068320259 | 0.4236498 | -0.1612659 | 0.87188399 | 0.99945981 |
| ZNF276 | 16.2727427 | -0.171644062 | 0.39260076 | -0.4371975 | 0.66196814 | 0.99945981 |
| TCF25 | 80.848036 | -0.54876002 | 0.26458956 | -2.0740048 | 0.03807887 | 0.87920079 |
| DEF8 | 8.94421157 | 0.119684359 | 0.45233072 | 0.26459481 | 0.79132162 | 0.99945981 |
| FAM157C | 26.7426551 | 0.743169587 | 0.6674273 | 1.11348395 | 0.26550064 | 0.99945981 |
| RPH3AL | 13.2430837 | 0.288580283 | 0.63810472 | 0.45224597 | 0.6510918 | 0.99945981 |
| RFLNB | 10.3843102 | 0.06431109 | 0.76009078 | 0.08460975 | 0.93257165 | 0.99945981 |
| VPS53 | 68.2348959 | 0.328116692 | 0.30891595 | 1.06215523 | 0.28816522 | 0.99945981 |
| GLOD4 | 8.93339215 | 0.015063565 | 0.45447034 | 0.03314532 | 0.9735587 | 0.99945981 |
| ABR | 117.443902 | 0.001787869 | 0.35234378 | 0.00507422 | 0.99595138 | 0.99945981 |
| YWHAE | 56.3189311 | -0.074859254 | 0.28203311 | -0.2654272 | 0.7906804 | 0.99945981 |
| CRK | 35.0742738 | -0.298945234 | 0.41663353 | -0.7175256 | 0.47304983 | 0.99945981 |
| MYO1C | 11.5329839 | -0.95693922 | 0.75947723 | -1.2599972 | 0.20767036 | 0.99945981 |
| PITPNA | 39.2248296 | -0.237505573 | 0.41061994 | -0.5784073 | 0.56298916 | 0.99945981 |
| SLC43A2 | 42.8864452 | 0.783614645 | 0.46193942 | 1.69635804 | 0.0898181 | 0.99640647 |
| PRPF8 | 32.9372943 | -0.218370539 | 0.33153437 | -0.6586664 | 0.51011003 | 0.99945981 |
| WDR81 | 11.5520919 | -0.180206445 | 0.55947541 | -0.322099 | 0.74737773 | 0.99945981 |
| SMYD4 | 11.1146031 | -0.600790877 | 0.43014446 | -1.3967188 | 0.16249814 | 0.99945981 |
| RPA1 | 23.2346754 | 0.625713322 | 0.34094091 | 1.83525444 | 0.066468 | 0.94594189 |
| SMG6 | 71.7245823 | -0.105647647 | 0.25298911 | -0.4175976 | 0.67624134 | 0.99945981 |
| SGSM2 | 10.3781743 | -0.844361513 | 0.4638785 | -1.8202213 | 0.06872532 | 0.95312989 |
| METTL16 | 26.5149216 | -0.348139315 | 0.32402735 | -1.0744134 | 0.28263746 | 0.99945981 |
| PAFAH1B1 | 120.019358 | 0.126660608 | 0.22700007 | 0.55797608 | 0.57686072 | 0.99945981 |
| RAP1GAP2 | 101.371149 | 1.299102984 | 0.46860529 | 2.77227553 | 0.00556659 | 0.5234882 |
| ITGAE | 28.5663568 | -2.196561262 | 0.71927481 | -3.0538554 | 0.00225921 | 0.45570351 |
| NCBP3 | 21.2440338 | 0.357336778 | 0.43572956 | 0.82008845 | 0.41216569 | 0.99945981 |
| P2RX1 | 23.7079952 | -0.389657175 | 0.63069678 | -0.6178201 | 0.5366939 | 0.99945981 |
| ATP2A3 | 31.1170282 | -0.419247391 | 0.4564986 | -0.918398 | 0.35841055 | 0.99945981 |
| ZZEF1 | 55.4452077 | 0.11188949 | 0.22901529 | 0.48856777 | 0.62514773 | 0.99945981 |
| ANKFY1 | 51.0829451 | 0.403724689 | 0.39375261 | 1.02532575 | 0.30520951 | 0.99945981 |
| UBE2G1 | 57.9897554 | -0.001995641 | 0.40636538 | -0.004911 | 0.99608164 | 0.99945981 |

| | | | | | | |
|---|---|---|---|---|---|---|
| ARRB2 | 40.5870375 | -0.102695879 | 0.40067271 | -0.2563086 | 0.79771253 | 0.99945981 |
| CXCL16 | 14.9521865 | 0.290767694 | 0.6389707 | 0.45505638 | 0.64906868 | 0.99945981 |
| MINK1 | 25.3721995 | -0.307424917 | 0.34544529 | -0.8899381 | 0.37349913 | 0.99945981 |
| RNF167 | 9.04709035 | -0.363199061 | 0.43755173 | -0.8300711 | 0.40649856 | 0.99945981 |
| PFN1 | 93.3244932 | -0.106992498 | 0.6036406 | -0.1772454 | 0.85931567 | 0.99945981 |
| SPAG7 | 11.479003 | 0.380890487 | 0.49657699 | 0.76703209 | 0.44306243 | 0.99945981 |
| CAMTA2 | 11.1363772 | 0.354607465 | 0.40703935 | 0.87118719 | 0.38365195 | 0.99945981 |
| AC087500.1 | 18.625725 | -0.618719739 | 0.76969664 | -0.8038488 | 0.42148429 | 0.99945981 |
| SCIMP | 29.7809919 | -0.881627276 | 0.83725777 | -1.0529938 | 0.29234381 | 0.99945981 |
| RABEP1 | 77.5039097 | -0.117686709 | 0.44435714 | -0.2648471 | 0.79112724 | 0.99945981 |
| NUP88 | 24.1551317 | -0.054121703 | 0.39499895 | -0.1370173 | 0.89101709 | 0.99945981 |
| RPAIN | 13.9619495 | -0.17430781 | 0.42965293 | -0.4056944 | 0.68496712 | 0.99945981 |
| DERL2 | 9.68073694 | -0.18457154 | 0.56844409 | -0.324696 | 0.74541114 | 0.99945981 |
| NLRP1 | 59.2747437 | 0.254156704 | 0.36451923 | 0.69723813 | 0.48565378 | 0.99945981 |
| KIAA0753 | 10.7386979 | -0.086392013 | 0.48557971 | -0.1779152 | 0.85878957 | 0.99945981 |
| XAF1 | 53.7915549 | 0.686403227 | 0.92316725 | 0.74353074 | 0.45716042 | 0.99945981 |
| ACADVL | 29.1654817 | 0.433426665 | 0.48026051 | 0.90248243 | 0.36680065 | 0.99945981 |
| EIF5A | 13.2903405 | 0.01458014 | 0.53203795 | 0.02740432 | 0.97813725 | 0.99945981 |
| GPS2 | 9.35655503 | -0.211117005 | 0.69178673 | -0.3051764 | 0.76023178 | 0.99945981 |
| ACAP1 | 42.7354216 | -0.439365609 | 0.30522058 | -1.4395019 | 0.15000836 | 0.99945981 |
| ZBTB4 | 17.9664987 | 0.020233175 | 0.34782859 | 0.05816996 | 0.95361325 | 0.99945981 |
| POLR2A | 27.6708989 | -0.41856786 | 0.43243402 | -0.9679346 | 0.33307702 | 0.99945981 |
| EIF4A1 | 13.7739963 | -0.567955611 | 0.58657922 | -0.9682505 | 0.33291929 | 0.99945981 |
| CD68 | 29.062268 | 0.310403389 | 0.77270566 | 0.40170974 | 0.68789766 | 0.99945981 |
| SAT2 | 15.16974 | -0.242709197 | 0.52194748 | -0.4650069 | 0.64192649 | 0.99945981 |
| TP53 | 10.580805 | -0.456464163 | 0.51195888 | -0.8916032 | 0.37260567 | 0.99945981 |
| KDM6B | 11.6815827 | 0.243270724 | 0.44110432 | 0.55150384 | 0.58128834 | 0.99945981 |
| NAA38 | 19.5974305 | 0.001410535 | 0.36602326 | 0.00385368 | 0.99692522 | 0.99945981 |
| CHD3 | 24.0168637 | 0.016986925 | 0.32291761 | 0.05260452 | 0.95804702 | 0.99945981 |
| PER1 | 17.0938602 | -0.693441568 | 0.51000463 | -1.359677 | 0.17393216 | 0.99945981 |
| CTC1 | 11.4784116 | 0.118124623 | 0.41452005 | 0.28496721 | 0.77566926 | 0.99945981 |
| NDEL1 | 39.0665509 | 0.223168969 | 0.42070418 | 0.53046529 | 0.59578937 | 0.99945981 |
| PIK3R5 | 135.10719 | -0.017276979 | 0.23273619 | -0.0742342 | 0.94082405 | 0.99945981 |
| STX8 | 90.6625245 | 0.216767635 | 0.21816439 | 0.99359769 | 0.32041877 | 0.99945981 |
| GAS7 | 134.723574 | -0.265570547 | 0.60312691 | -0.4403228 | 0.65970331 | 0.99945981 |
| SCO1 | 9.44714246 | 0.151430715 | 0.49754645 | 0.30435493 | 0.7608575 | 0.99945981 |
| MAP2K4 | 56.1022272 | -0.031495123 | 0.25998289 | -0.1211431 | 0.90357772 | 0.99945981 |
| COX10 | 18.7153159 | -0.093532109 | 0.39027568 | -0.2396565 | 0.81059655 | 0.99945981 |
| HS3ST3B1 | 11.8761195 | -0.044972093 | 0.65452292 | -0.0687097 | 0.94522067 | 0.99945981 |
| PMP22 | 16.5039603 | -1.647024851 | 0.73709115 | -2.2344928 | 0.02545067 | 0.81367289 |
| ZSWIM7 | 15.8205856 | -0.418031424 | 0.37619576 | -1.1112072 | 0.26647919 | 0.99945981 |
| TTC19 | 32.6453337 | -0.215474552 | 0.37626002 | -0.5726746 | 0.56686504 | 0.99945981 |
| NCOR1 | 123.862902 | 0.083455074 | 0.18661587 | 0.44720245 | 0.65472889 | 0.99945981 |
| PIGL | 26.4382309 | -0.085589306 | 0.28560516 | -0.299677 | 0.76442352 | 0.99945981 |
| UBB | 47.5151179 | -0.748202206 | 0.37278202 | -2.007077 | 0.04474147 | 0.90039076 |
| TRPV2 | 11.4869229 | -0.044857193 | 0.45013772 | -0.0996522 | 0.92062049 | 0.99945981 |
| AC098850.3 | 18.5274718 | 0.769413897 | 0.73354795 | 1.0488938 | 0.294227 | 0.99945981 |
| MPRIP | 38.5369294 | -0.433423263 | 0.44207756 | -0.9804236 | 0.32687708 | 0.99945981 |
| COPS3 | 10.8415012 | -0.687328372 | 0.47569959 | -1.4448791 | 0.14849186 | 0.99945981 |
| RAI1 | 21.0202742 | -0.250839134 | 0.43076372 | -0.5823126 | 0.56035615 | 0.99945981 |
| SREBF1 | 16.2301142 | -0.852583009 | 0.58053816 | -1.468608 | 0.14193914 | 0.99945981 |
| TOM1L2 | 31.7017459 | 0.079304501 | 0.31984989 | 0.24794287 | 0.80417861 | 0.99945981 |
| ALKBH5 | 15.8925619 | -0.441173824 | 0.36512907 | -1.2082682 | 0.22694413 | 0.99945981 |
| FLII | 15.6573851 | 0.285344281 | 0.42594689 | 0.66990577 | 0.50291786 | 0.99945981 |
| TOP3A | 16.2516971 | 0.193428017 | 0.3625662 | 0.5334971 | 0.59368952 | 0.99945981 |
| SHMT1 | 9.70790288 | -0.65580571 | 0.51204081 | -1.2807685 | 0.20027501 | 0.99945981 |
| PRPSAP2 | 23.0450446 | -0.338605537 | 0.48279652 | -0.7013421 | 0.48308953 | 0.99945981 |
| GRAPL | 12.6725812 | -1.701959382 | 0.86837267 | -1.9599412 | 0.05000266 | 0.92165202 |
| EPN2 | 15.5656873 | -0.31871643 | 0.50805415 | -0.6273277 | 0.53044447 | 0.99945981 |
| AC004448.2 | 8.61045723 | -0.604067518 | 0.59130942 | -1.021576 | 0.30698162 | 0.99945981 |
| SLC47A1 | 9.73380042 | 0.21414009 | 0.63223868 | 0.33870134 | 0.73483473 | 0.99945981 |
| ALDH3A2 | 9.289399 | 0.031891996 | 0.56111232 | 0.0568371 | 0.95467496 | 0.99945981 |
| ULK2 | 19.472319 | 1.124714088 | 0.51548739 | 2.18184598 | 0.0291209 | 0.84763724 |
| AKAP10 | 78.9867038 | 0.174735523 | 0.30099851 | 0.58051956 | 0.5615643 | 0.99945981 |
| SPECC1 | 21.8881213 | 0.676257195 | 0.79338826 | 0.85236603 | 0.39401097 | 0.99945981 |
| USP22 | 29.7125492 | 0.046464568 | 0.27773847 | 0.16729612 | 0.86713706 | 0.99945981 |
| DHRS7B | 18.6554137 | -0.340124184 | 0.60319275 | -0.5638731 | 0.57284048 | 0.99945981 |
| MAP2K3 | 22.3484426 | 0.2690816 | 0.41451165 | 0.64915329 | 0.5162393 | 0.99945981 |
| WSB1 | 169.536846 | 0.532329746 | 0.43298055 | 1.22945417 | 0.21890157 | 0.99945981 |
| KSR1 | 39.854937 | 0.172897564 | 0.48935612 | 0.35331644 | 0.72385122 | 0.99945981 |
| LGALS9 | 15.7642172 | 0.255113839 | 0.7492537 | 0.3404906 | 0.7334871 | 0.99945981 |
| NLK | 83.9333685 | -0.506729697 | 0.49958425 | -1.0143028 | 0.31043829 | 0.99945981 |
| SARM1 | 10.4201711 | -0.464002012 | 0.57520905 | -0.8066667 | 0.41985851 | 0.99945981 |
| KIAA0100 | 22.7202977 | -0.221539122 | 0.37856755 | -0.5852037 | 0.55841077 | 0.99945981 |
| SDF2 | 9.85679013 | -0.129621417 | 0.46558297 | -0.2784067 | 0.78070018 | 0.99945981 |
| SUPT6H | 27.6959631 | -0.491859843 | 0.32781519 | -1.5004181 | 0.13350613 | 0.99945981 |
| RPL23A | 78.6029374 | -0.084668661 | 0.56924099 | -0.1487396 | 0.88175913 | 0.99945981 |
| FAM222B | 43.1406826 | -0.244135637 | 0.28476836 | -0.8573131 | 0.39127189 | 0.99945981 |
| FLOT2 | 29.5190452 | -0.035841671 | 0.38187368 | -0.0938574 | 0.92522243 | 0.99945981 |
| PHF12 | 33.54183 | -0.118225517 | 0.35041729 | -0.3373849 | 0.73582673 | 0.99945981 |
| MYO18A | 13.9622739 | -0.044939022 | 0.51788917 | -0.0867734 | 0.9308516 | 0.99945981 |
| NUFIP2 | 71.700363 | -0.06033412 | 0.25595914 | -0.2357178 | 0.81365168 | 0.99945981 |
| TAOK1 | 126.400324 | 0.200979972 | 0.2211129 | 0.90894729 | 0.36337795 | 0.99945981 |
| ABHD15-AS1 | 44.6346318 | 0.053959906 | 0.6020025 | 0.08963402 | 0.92857805 | 0.99945981 |
| SSH2 | 430.968303 | -0.085544489 | 0.28802732 | -0.2970013 | 0.76646551 | 0.99945981 |
| NSRP1 | 43.7780911 | -0.20310409 | 0.24453278 | -0.8305802 | 0.40621081 | 0.99945981 |
| BLMH | 10.8822485 | -0.549750284 | 0.62664454 | -0.8772921 | 0.38032802 | 0.99945981 |
| CPD | 42.4107604 | -0.861087244 | 0.56099109 | -1.5349392 | 0.12479875 | 0.99945981 |
| GOSR1 | 26.6102597 | -0.297782121 | 0.28936963 | -1.0290718 | 0.30344595 | 0.99945981 |
| CRLF3 | 57.130684 | 0.256208884 | 0.31287329 | 0.81889023 | 0.41284904 | 0.99945981 |
| TEFM | 8.94572285 | -0.600966922 | 0.47498655 | -1.2652294 | 0.20578912 | 0.99945981 |
| ADAP2 | 41.6053871 | 0.617210265 | 0.46772027 | 1.3196141 | 0.18696389 | 0.99945981 |
| RNF135 | 15.016846 | 0.452873427 | 0.38365562 | 1.1804165 | 0.2378346 | 0.99945981 |
| NF1 | 127.799594 | -0.446745367 | 0.30194195 | -1.4795737 | 0.13898704 | 0.99945981 |
| EVI2B | 19.8616569 | -0.134811506 | 0.53054766 | -0.2540988 | 0.79941926 | 0.99945981 |
| RAB11FIP4 | 9.39044598 | 0.056864524 | 0.55773818 | 0.10195559 | 0.91879193 | 0.99945981 |
| UTP6 | 18.4133314 | 0.262413245 | 0.35229214 | 0.74487397 | 0.45634791 | 0.99945981 |
| SUZ12 | 51.8620864 | 0.238372382 | 0.29609383 | 0.80505689 | 0.42078685 | 0.99945981 |
| LRRC37B | 21.3739384 | -0.320351423 | 0.3566373 | -0.8982555 | 0.36904934 | 0.99945981 |
| RHOT1 | 51.6855644 | 0.28726596 | 0.36572331 | 0.78547347 | 0.43217602 | 0.99945981 |
| ZNF207 | 93.0610763 | 0.299925575 | 0.24096399 | 1.24469046 | 0.21324555 | 0.99945981 |
| PSMD11 | 20.7495278 | -0.194952087 | 0.5414479 | -0.360057 | 0.71880453 | 0.99945981 |
| MYO1D | 54.3469529 | -0.727672721 | 0.6147863 | -1.183619 | 0.23656393 | 0.99945981 |
| SLFN5 | 37.1419911 | 0.12108448 | 0.46456413 | 0.26064105 | 0.79436933 | 0.99945981 |
| SLFN11 | 19.4296036 | 0.457641094 | 0.35686125 | 1.28240626 | 0.19970017 | 0.99945981 |
| SLFN12L | 110.040358 | -0.015010638 | 0.43869902 | -0.0342163 | 0.9727047 | 0.99945981 |
| AP2B1 | 49.1536021 | -0.121882773 | 0.28592438 | -0.4262763 | 0.66990655 | 0.99945981 |
| TAF15 | 86.5603894 | -0.048348768 | 0.20626075 | -0.2344061 | 0.81466976 | 0.99945981 |
| CCL5 | 68.0787518 | -1.11285463 | 0.54735515 | -2.0331491 | 0.04203746 | 0.89562588 |
| CCL4 | 26.261983 | -0.754425449 | 0.87382536 | -0.8633595 | 0.38793983 | 0.99945981 |
| AC243829.2 | 16.3598429 | -0.192816638 | 0.51964215 | -0.3710566 | 0.71059539 | 0.99945981 |
| MYO19 | 12.870713 | -0.280961756 | 0.42380542 | -0.6629499 | 0.50736265 | 0.99945981 |
| GGNBP2 | 47.977263 | -0.455352242 | 0.31474961 | -1.4467126 | 0.14797743 | 0.99945981 |

| | | | | | | |
|---|---|---|---|---|---|---|
| AATF | 38.7046668 | 0.210629002 | 0.35685891 | 0.59023047 | 0.55503615 | 0.99945981 |
| ACACA | 35.2264541 | -0.184020383 | 0.42905226 | -0.4288997 | 0.66799622 | 0.99945981 |
| TADA2A | 17.011827 | 0.101074098 | 0.35870388 | 0.28177587 | 0.77811537 | 0.99945981 |
| SYNRG | 80.5709332 | -0.062385353 | 0.19979167 | -0.312252 | 0.754849 | 0.99945981 |
| DDX52 | 14.2251639 | -0.197561253 | 0.41846729 | -0.4721068 | 0.63685056 | 0.99945981 |
| SOCS7 | 9.11435934 | -0.339664383 | 0.71361773 | -0.4759753 | 0.634092 | 0.99945981 |
| MLLT6 | 37.4919506 | -0.157333087 | 0.29175973 | -0.5392557 | 0.58971045 | 0.99945981 |
| PSMB3 | 14.3190727 | 0.659736978 | 0.4444912 | 1.48425206 | 0.13774206 | 0.99945981 |
| PIP4K2B | 10.5525956 | -0.424330837 | 0.53428945 | -0.7941965 | 0.42708102 | 0.99945981 |
| CWC25 | 8.97316463 | -0.761663177 | 0.44528786 | -1.7104962 | 0.08717416 | 0.99640647 |
| RPL23 | 40.3967547 | -0.287228914 | 0.5344852 | -0.5373936 | 0.59099578 | 0.99945981 |
| LASP1 | 31.7836884 | -0.166392161 | 0.41380632 | -0.4021015 | 0.68760929 | 0.99945981 |
| RPL19 | 117.977894 | -0.434795049 | 0.56577029 | -0.768501 | 0.44218958 | 0.99945981 |
| FBXL20 | 76.5773502 | 0.078667582 | 0.35271095 | 0.22303697 | 0.82350673 | 0.99945981 |
| MED1 | 14.6504099 | 0.031200733 | 0.52509966 | 0.05941869 | 0.95261863 | 0.99945981 |
| CDK12 | 42.7407522 | -0.144301365 | 0.3181489 | -0.4535655 | 0.65014159 | 0.99945981 |
| STARD3 | 14.5606936 | -0.547751881 | 0.43468506 | -1.2601121 | 0.20762894 | 0.99945981 |
| IKZF3 | 94.9123518 | -1.006924352 | 0.52344513 | -1.9236483 | 0.05439868 | 0.92445778 |
| PSMD3 | 8.75341539 | -0.753907309 | 0.52799357 | -1.4278721 | 0.15332866 | 0.99945981 |
| MED24 | 9.18382699 | 0.711153465 | 0.57133618 | 1.24471981 | 0.21323476 | 0.99945981 |
| MSL1 | 18.4367203 | -0.211996579 | 0.35941252 | -0.5898419 | 0.55529662 | 0.99945981 |
| CASC3 | 22.7135702 | -0.152389442 | 0.32414825 | -0.4701227 | 0.63826738 | 0.99945981 |
| WIPF2 | 36.0251982 | 0.103327827 | 0.27360728 | 0.37765013 | 0.70569052 | 0.99945981 |
| RARA | 16.1120058 | 0.790144134 | 0.61502441 | 1.28473623 | 0.19888447 | 0.99945981 |
| IGFBP4 | 14.042449 | -0.266329007 | 0.49041684 | -0.5430666 | 0.58708394 | 0.99945981 |
| KRT10 | 10.9211212 | -0.373459583 | 0.52463387 | -0.711848 | 0.4765589 | 0.99945981 |
| EIF1 | 85.3215234 | -0.13300743 | 0.39596287 | -0.3359088 | 0.73693961 | 0.99945981 |
| ACLY | 16.1267511 | -0.521135162 | 0.42872974 | -1.215533 | 0.22416286 | 0.99945981 |
| DNAJC7 | 40.4150288 | 0.466518442 | 0.36159853 | 1.2901558 | 0.19699657 | 0.99945981 |
| STAT5B | 74.7100866 | -0.346382821 | 0.26916675 | -1.2868708 | 0.19813934 | 0.99945981 |
| STAT5A | 19.5295363 | -0.080094529 | 0.42776769 | -0.1872384 | 0.85147374 | 0.99945981 |
| STAT3 | 102.411156 | -0.534149161 | 0.27002231 | -1.9781667 | 0.0479099 | 0.90884462 |
| ATP6V0A1 | 44.2361217 | -0.98275871 | 0.52247507 | -1.8809676 | 0.05997632 | 0.92445778 |
| MLX | 10.5328011 | -0.516299659 | 0.47763756 | -1.0809444 | 0.27972183 | 0.99945981 |
| RETREG3 | 26.8852425 | 0.002072909 | 0.32694699 | 0.0063402 | 0.99494129 | 0.99945981 |
| EZH1 | 20.803381 | -0.323751983 | 0.41791918 | -0.7746761 | 0.4385311 | 0.99945981 |
| BECN1 | 11.1974075 | 0.067144321 | 0.48219323 | 0.13924775 | 0.88925438 | 0.99945981 |
| RPL27 | 32.8687191 | -0.12771098 | 0.59724647 | -0.213833 | 0.83067732 | 0.99945981 |
| IFI35 | 11.9952059 | 0.749912299 | 0.7156741 | 1.04784048 | 0.29471211 | 0.99945981 |
| BRCA1 | 10.1352067 | -0.362502575 | 0.59054242 | -0.6138468 | 0.53931657 | 0.99945981 |
| NBR1 | 24.2662901 | -0.108348666 | 0.30660703 | -0.3533796 | 0.72380389 | 0.99945981 |
| TMEM106A | 18.9514239 | -0.249675899 | 0.41826577 | -0.5969312 | 0.5505533 | 0.99945981 |
| LINC00854 | 24.7102579 | 0.287773692 | 0.48833109 | 0.58930036 | 0.5556598 | 0.99945981 |
| DHX8 | 14.7249545 | -0.017207933 | 0.40487969 | -0.0425014 | 0.96609904 | 0.99945981 |
| DUSP3 | 8.82332471 | -0.620982052 | 0.45518987 | -1.3642264 | 0.17249632 | 0.99945981 |
| LSM12 | 12.8615983 | -0.038484 | 0.39189637 | -0.0981994 | 0.92177394 | 0.99945981 |
| HDAC5 | 10.5397633 | -0.121728223 | 0.63256346 | -0.1924364 | 0.8474004 | 0.99945981 |
| TMUB2 | 9.02961031 | 0.123095523 | 0.48998368 | 0.25122372 | 0.80164115 | 0.99945981 |
| UBTF | 13.0877866 | -0.220458731 | 0.55513859 | -0.3971238 | 0.69127619 | 0.99945981 |
| SLC25A39 | 9.70331983 | -0.395094494 | 0.46741684 | -0.8452723 | 0.39795884 | 0.99945981 |
| GRN | 54.942761 | -0.069046584 | 0.56778358 | -0.1216072 | 0.90321009 | 0.99945981 |
| GPATCH8 | 118.736898 | -0.076977312 | 0.23536918 | -0.3270492 | 0.74363064 | 0.99945981 |
| EFTUD2 | 20.1061666 | 0.345733606 | 0.33742936 | 1.02461034 | 0.30554709 | 0.99945981 |
| DCAKD | 8.28702176 | -0.209907861 | 0.57940262 | -0.3622832 | 0.71714038 | 0.99945981 |
| NMT1 | 22.8259314 | -0.443372106 | 0.32479957 | -1.3650637 | 0.17223303 | 0.99945981 |
| AC138150.1 | 12.1126889 | -0.343032228 | 0.56234984 | -0.6099979 | 0.54186319 | 0.99945981 |
| AC008105.3 | 31.0341549 | 0.18383019 | 0.36997809 | 0.49686777 | 0.6192823 | 0.99945981 |
| FMNL1 | 66.3584677 | 0.16818662 | 0.37332444 | 0.45051061 | 0.65234231 | 0.99945981 |
| MAP3K14 | 40.0666072 | 0.107821934 | 0.38945955 | 0.27685015 | 0.78189517 | 0.99945981 |
| ARHGAP27 | 22.3562205 | 0.438432805 | 0.46322692 | 0.94647522 | 0.34390625 | 0.99945981 |
| PLEKHM1 | 29.9575969 | 0.01560057 | 0.32651533 | 0.04777898 | 0.96189239 | 0.99945981 |
| KANSL1 | 256.990144 | 0.200867639 | 0.19380976 | 1.03641654 | 0.30000786 | 0.99945981 |
| ARL17B | 41.3336638 | 2.263893931 | 0.91952308 | 2.46203057 | 0.01381529 | 0.72008995 |
| LRRC37A2 | 12.3115391 | 0.521691497 | 0.50818201 | 1.02658396 | 0.30461641 | 0.99945981 |
| ARL17A | 15.3971789 | -0.323014151 | 0.48406333 | -0.6672973 | 0.50458225 | 0.99945981 |
| NSF | 105.974276 | 0.395832368 | 0.33980737 | 1.16487281 | 0.24407048 | 0.99945981 |
| AC005670.2 | 8.10838976 | -0.568388965 | 0.47641945 | -1.1930432 | 0.23285247 | 0.99945981 |
| CDC27 | 67.7492625 | -0.004578055 | 0.21315049 | -0.021478 | 0.98286432 | 0.99945981 |
| EFCAB13 | 14.5742035 | -0.177952925 | 0.47691273 | -0.3731352 | 0.70904783 | 0.99945981 |
| NPEPPS | 77.999869 | 0.173470739 | 0.20962012 | 0.82754815 | 0.40792644 | 0.99945981 |
| AC025682.1 | 9.67364198 | -0.162421771 | 0.44564736 | -0.3644625 | 0.71551264 | 0.99945981 |
| KPNB1 | 66.1667193 | -0.238688513 | 0.28831163 | -0.8278837 | 0.40773634 | 0.99945981 |
| TBX21 | 9.92000195 | 0.316064807 | 0.49636792 | 0.63675511 | 0.52428437 | 0.99945981 |
| SP2 | 9.78688848 | 0.078844205 | 0.56295767 | 0.14005352 | 0.8886177 | 0.99945981 |
| CDK5RAP3 | 12.1379747 | -0.16209294 | 0.53447265 | -0.3032764 | 0.76167923 | 0.99945981 |
| NFE2L1 | 12.8632936 | 0.07870103 | 0.52767959 | 0.14914549 | 0.88143883 | 0.99945981 |
| SKAP1 | 241.149171 | 0.125233587 | 0.41563546 | 0.30130631 | 0.76318093 | 0.99945981 |
| CALCOCO2 | 32.7836203 | -0.287650268 | 0.4265386 | -0.6743827 | 0.50006802 | 0.99945981 |
| ATP5MC1 | 9.31761092 | -0.453554895 | 0.5020622 | -0.9033839 | 0.3663222 | 0.99945981 |
| UBE2Z | 26.8188727 | -0.122595916 | 0.42004508 | -0.2918637 | 0.77039083 | 0.99945981 |
| SNF8 | 11.7443972 | -0.193937158 | 0.53157328 | -0.3648362 | 0.7152337 | 0.99945981 |
| ABI3 | 14.763985 | 0.191208736 | 0.78333292 | 0.24409639 | 0.80715617 | 0.99945981 |
| ZNF652 | 65.4879309 | 0.348852487 | 0.32673012 | 1.06770839 | 0.28565207 | 0.99945981 |
| PHB | 13.2544397 | 0.087418251 | 0.46886084 | 0.18644818 | 0.85209331 | 0.99945981 |
| SPOP | 40.7257733 | -0.098651294 | 0.26396276 | -0.3737319 | 0.70860381 | 0.99945981 |
| FAM117A | 22.8185843 | -0.828168103 | 0.718476 | -1.1526733 | 0.2490445 | 0.99945981 |
| KAT7 | 14.8924878 | -0.254125517 | 0.41366719 | -0.6143236 | 0.53900151 | 0.99945981 |
| PPP1R9B | 10.4381975 | 0.606674255 | 0.45505169 | 1.33319855 | 0.18246666 | 0.99945981 |
| ACSF2 | 10.4602883 | 0.867780016 | 0.43404911 | 1.99926689 | 0.04557948 | 0.90395454 |
| ABCC3 | 15.6747071 | 1.467201411 | 0.72003529 | 2.03767986 | 0.04158196 | 0.89562588 |
| LUC7L3 | 84.5512863 | 0.252955844 | 0.26009706 | 0.97254404 | 0.33077997 | 0.99945981 |
| TOB1 | 9.66005601 | 0.133316221 | 0.50324234 | 0.26491455 | 0.79107529 | 0.99945981 |
| SPAG9 | 90.9564046 | 0.153814493 | 0.28569893 | 0.53837966 | 0.59031497 | 0.99945981 |
| MBTD1 | 47.1918582 | -0.250722749 | 0.3414204 | -0.734352 | 0.46273424 | 0.99945981 |
| UTP18 | 14.0631192 | -0.379014428 | 0.37875548 | -1.0006837 | 0.31697976 | 0.99945981 |
| STXBP4 | 17.9662892 | -0.584013352 | 0.49938139 | -1.1694736 | 0.24221287 | 0.99945981 |
| DGKE | 15.5093701 | -0.379697832 | 0.41928333 | -0.9055877 | 0.36515411 | 0.99945981 |
| TRIM25 | 20.0257453 | -0.01968724 | 0.46747155 | -0.0421143 | 0.96640758 | 0.99945981 |
| SCPEP1 | 22.1312055 | -0.170278089 | 0.54581449 | -0.3119706 | 0.75506285 | 0.99945981 |
| MSI2 | 136.071206 | -0.75026818 | 0.63309401 | -1.1850818 | 0.23598511 | 0.99945981 |
| CUEDC1 | 17.3458236 | 0.635751033 | 0.45426767 | 1.39950754 | 0.16166084 | 0.99945981 |
| VEZF1 | 14.5289683 | 0.482510749 | 0.39619995 | 1.21784658 | 0.22328228 | 0.99945981 |
| DYNLL2 | 9.43389555 | -0.168453473 | 0.58351073 | -0.2886896 | 0.77281893 | 0.99945981 |
| TSPOAP1-AS | 24.7669941 | -0.411278109 | 0.55345809 | -0.7431061 | 0.45741742 | 0.99945981 |
| AC004687.1 | 16.46824 | 0.125364787 | 0.69433793 | 0.18055299 | 0.85671846 | 0.99945981 |
| MTMR4 | 9.10589748 | -0.422873727 | 0.51048597 | -0.8283748 | 0.40745826 | 0.99945981 |
| TRIM37 | 20.0670801 | 0.064291918 | 0.34094421 | 0.1885702 | 0.85042969 | 0.99945981 |
| SKA2 | 13.1487533 | 0.063834201 | 0.39914044 | 0.15992917 | 0.87293687 | 0.99945981 |
| YPEL2 | 36.4195486 | 0.647024206 | 0.35319918 | 1.83189612 | 0.0669669 | 0.94594189 |
| DHX40 | 11.7803877 | -0.163268681 | 0.39719801 | -0.4110511 | 0.68103506 | 0.99945981 |
| CLTC | 95.4112282 | -0.021647198 | 0.25185415 | -0.0859513 | 0.93150511 | 0.99945981 |
| PTRH2 | 14.9536426 | -0.604434652 | 0.48648297 | -1.242458 | 0.21406763 | 0.99945981 |
| VMP1 | 264.050575 | 0.863324717 | 0.32459086 | 2.65973203 | 0.00782028 | 0.57727153 |

| | | | | | | |
|---|---|---|---|---|---|---|
| RPS6KB1 | 18.2006375 | 0.054006817 | 0.34820726 | 0.15509963 | 0.87674277 | 0.99945981 |
| RNFT1 | 9.32449183 | -0.173020573 | 0.62898659 | -0.2750783 | 0.78325607 | 0.99945981 |
| USP32 | 113.284512 | 0.203036943 | 0.19286496 | 1.05274148 | 0.29245949 | 0.99945981 |
| APPBP2 | 38.5947512 | -0.081187186 | 0.28227605 | -0.2876163 | 0.77364049 | 0.99945981 |
| PPM1D | 26.4010914 | 0.312944354 | 0.35183282 | 0.88946891 | 0.37375112 | 0.99945981 |
| BCAS3 | 153.006727 | 0.330771142 | 0.22414798 | 1.47568201 | 0.14002928 | 0.99945981 |
| INTS2 | 11.9258788 | -0.103887567 | 0.41816797 | -0.248435 | 0.80379784 | 0.99945981 |
| MED13 | 116.917942 | 0.103188483 | 0.18753485 | 0.55023631 | 0.5821573 | 0.99945981 |
| TLK2 | 73.3501069 | 0.222713611 | 0.25654219 | 0.86813638 | 0.38531968 | 0.99945981 |
| TANC2 | 81.0006235 | -0.203037315 | 0.47252582 | -0.4296851 | 0.66742471 | 0.99945981 |
| DCAF7 | 48.0249517 | -0.384974556 | 0.34612618 | -1.1122376 | 0.266036 | 0.99945981 |
| MAP3K3 | 63.1989415 | -0.047948004 | 0.32202415 | -0.1488957 | 0.88163595 | 0.99945981 |
| CCDC47 | 9.54999014 | -0.369704247 | 0.53846458 | -0.6865897 | 0.4923413 | 0.99945981 |
| DDX42 | 28.1176043 | -0.122794287 | 0.32462764 | -0.378262 | 0.705236 | 0.99945981 |
| SMARCD2 | 11.3234284 | 0.290547046 | 0.47229644 | 0.61517942 | 0.53843624 | 0.99945981 |
| ICAM2 | 12.9198834 | 0.027519661 | 0.52989411 | 0.05193426 | 0.95858107 | 0.99945981 |
| ERN1 | 61.3416927 | 0.153992402 | 0.78265863 | 0.19675551 | 0.84401886 | 0.99945981 |
| TEX2 | 16.6995996 | 0.730771954 | 0.61299469 | 1.19213423 | 0.23320862 | 0.99945981 |
| PECAM1 | 80.2495792 | 1.088827709 | 0.53366285 | 2.04029137 | 0.04132131 | 0.89562588 |
| MILR1 | 8.70799623 | 0.172592583 | 0.59580327 | 0.28968049 | 0.77206068 | 0.99945981 |
| POLG2 | 10.5392754 | -0.123944886 | 0.47778259 | -0.2594169 | 0.79531358 | 0.99945981 |
| DDX5 | 147.553338 | -0.078158082 | 0.27154832 | -0.2878238 | 0.77348158 | 0.99945981 |
| CEP95 | 28.3787227 | -0.183146824 | 0.36388095 | -0.5033152 | 0.61474266 | 0.99945981 |
| SMURF2 | 83.0735131 | -0.080464421 | 0.23601489 | -0.3409294 | 0.73315671 | 0.99945981 |
| LRRC37A3 | 40.6557492 | 0.496256975 | 0.36431931 | 1.36214841 | 0.17315105 | 0.99945981 |
| GNA13 | 98.9300452 | 0.604051842 | 0.51299556 | 1.17749916 | 0.23899633 | 0.99945981 |
| PRKCA | 145.343853 | 0.137040313 | 0.51619955 | 0.26547934 | 0.79064023 | 0.99945981 |
| HELZ | 139.281526 | 0.267033299 | 0.27414521 | 0.97405787 | 0.33002781 | 0.99945981 |
| PSMD12 | 19.8228417 | 0.076714412 | 0.32317712 | 0.23737575 | 0.8123653 | 0.99945981 |
| PITPNC1 | 227.599765 | -0.178051546 | 0.44225 | -0.4026038 | 0.68723969 | 0.99945981 |
| NOL11 | 11.1187957 | 0.485987327 | 0.44364822 | 1.09543396 | 0.27332657 | 0.99945981 |
| BPTF | 141.816943 | -0.457900073 | 0.31616689 | -1.448286 | 0.14753709 | 0.99945981 |
| ARSG | 24.816907 | 0.088292534 | 0.56073182 | 0.15745947 | 0.87488275 | 0.99945981 |
| SLC16A6 | 12.3543472 | 0.405075952 | 0.86404439 | 0.46881382 | 0.63920272 | 0.99945981 |
| WIPI1 | 26.0325121 | -0.591606641 | 0.51012045 | -1.1597391 | 0.24615504 | 0.99945981 |
| PRKAR1A | 52.123791 | -0.199645212 | 0.33981451 | -0.5875123 | 0.55685968 | 0.99945981 |
| FAM20A | 33.1995737 | -0.244998826 | 0.83487551 | -0.2934555 | 0.769174 | 0.99945981 |
| ABCA5 | 17.7054639 | 0.104089121 | 0.52255504 | 0.19919264 | 0.84211205 | 0.99945981 |
| MAP2K6 | 24.5776312 | 0.503746322 | 0.46268296 | 1.08875053 | 0.27626391 | 0.99945981 |
| KCNJ16 | 10.4370259 | 0.521735753 | 0.65922041 | 0.79144357 | 0.4286852 | 0.99945981 |
| SLC39A11 | 114.146686 | -0.28816612 | 0.33876426 | -0.8506391 | 0.39496987 | 0.99945981 |
| FAM104A | 11.4240257 | 0.201722498 | 0.43095541 | 0.46808206 | 0.63972591 | 0.99945981 |
| RPL38 | 29.3925877 | 0.113164507 | 0.54738255 | 0.20673752 | 0.83621486 | 0.99945981 |
| TTYH2 | 7.87144195 | 0.721871727 | 0.67604141 | 1.06779217 | 0.28561427 | 0.99945981 |
| CD300A | 23.6296652 | 0.993182151 | 0.45207403 | 2.19694584 | 0.02802431 | 0.84763724 |
| CD300E | 28.7010027 | 1.110512765 | 0.85392511 | 1.30048028 | 0.19343641 | 0.99945981 |
| RAB37 | 8.94928931 | -0.159736561 | 0.64343391 | -0.2482564 | 0.80393606 | 0.99945981 |
| CD300LF | 11.951535 | 0.313446601 | 0.7330605 | 0.42758626 | 0.66895237 | 0.99945981 |
| SLC9A3R1 | 21.8835236 | 0.230923812 | 0.33606237 | 0.68714569 | 0.49199092 | 0.99945981 |
| TMEM104 | 11.8189888 | -0.050109075 | 0.47606703 | -0.1052563 | 0.9161724 | 0.99945981 |
| NT5C | 10.0071002 | 0.029559567 | 0.51782448 | 0.05708414 | 0.95447817 | 0.99945981 |
| JPT1 | 16.9752051 | -0.129067069 | 0.43860342 | -0.2942683 | 0.76855292 | 0.99945981 |
| SUMO2 | 10.0822319 | -0.144914735 | 0.4331239 | -0.3345803 | 0.73794169 | 0.99945981 |
| NUP85 | 10.2919256 | 0.99912525 | 0.43668243 | 2.28799052 | 0.02213807 | 0.79988354 |
| GGA3 | 19.3104418 | -0.277552292 | 0.41895407 | -0.6624886 | 0.50765814 | 0.99945981 |
| GRB2 | 163.114934 | 0.103542769 | 0.30524826 | 0.33920839 | 0.73445275 | 0.99945981 |
| AC011933.4 | 30.4236874 | -0.504481685 | 0.52391537 | -0.9629068 | 0.33559429 | 0.99945981 |
| TMEM94 | 9.41577971 | -0.114769963 | 0.44341213 | -0.2588336 | 0.79576362 | 0.99945981 |
| TSEN54 | 13.1785077 | -0.640294109 | 0.60486583 | -1.0585721 | 0.28979469 | 0.99945981 |
| LLGL2 | 17.2239916 | -0.863664722 | 0.42023386 | -2.0552002 | 0.03985967 | 0.89132455 |
| MYO15B | 18.9599375 | -0.554213185 | 0.41610744 | -1.3318993 | 0.1828933 | 0.99945981 |
| SAP30BP | 17.25499 | -0.145046703 | 0.35813137 | -0.4050098 | 0.68547033 | 0.99945981 |
| H3F3B | 110.712598 | 0.053351994 | 0.36903374 | 0.14457213 | 0.8850487 | 0.99945981 |
| UNK | 27.4643333 | 0.231633164 | 0.35538984 | 0.65177205 | 0.51454823 | 0.99945981 |
| UNC13D | 23.4196935 | 0.044838368 | 0.41908513 | 0.10699107 | 0.91479606 | 0.99945981 |
| WBP2 | 14.8129724 | -0.236186511 | 0.46293862 | -0.5101897 | 0.60991857 | 0.99945981 |
| ACOX1 | 15.9079667 | 0.109368092 | 0.35547709 | 0.30766565 | 0.75833676 | 0.99945981 |
| SRP68 | 9.09170198 | -0.245458825 | 0.44997178 | -0.5454983 | 0.58541087 | 0.99945981 |
| EXOC7 | 15.9091517 | -0.57279522 | 0.40452024 | -1.4159866 | 0.15677944 | 0.99945981 |
| RNF157 | 14.4498812 | -0.705854334 | 0.56907278 | -1.2403586 | 0.21484277 | 0.99945981 |
| UBALD2 | 20.5293629 | -0.741212623 | 0.49308689 | -1.503209 | 0.13278517 | 0.99945981 |
| PRPSAP1 | 15.7110223 | 0.379144894 | 0.40120766 | 0.94500912 | 0.34465422 | 0.99945981 |
| UBE2O | 21.0426409 | -0.119924097 | 0.46943319 | -0.2554657 | 0.7983634 | 0.99945981 |
| RHBDF2 | 46.6733923 | -0.120769441 | 0.41680701 | -0.2897491 | 0.77200822 | 0.99945981 |
| SRSF2 | 31.5396745 | 0.055434685 | 0.36597485 | 0.1514713 | 0.87960395 | 0.99945981 |
| MFSD11 | 25.6421342 | 0.167407845 | 0.34087413 | 0.49111338 | 0.62334626 | 0.99945981 |
| SEC14L1 | 75.1444351 | -0.066894075 | 0.33409798 | -0.2002229 | 0.84130624 | 0.99945981 |
| 9-Sep | 85.6867454 | 0.089336932 | 0.36791128 | 0.24282195 | 0.80814333 | 0.99945981 |
| TNRC6C | 44.9385639 | -0.283651619 | 0.38430541 | -0.7380891 | 0.46046033 | 0.99945981 |
| TMC6 | 35.7881154 | 0.334451489 | 0.4405586 | 0.75915324 | 0.4477609 | 0.99945981 |
| TMC8 | 60.2164347 | -0.03892315 | 0.32104871 | -0.1212375 | 0.90350291 | 0.99945981 |
| SYNGR2 | 21.7940896 | -0.080522578 | 0.62031625 | -0.1298089 | 0.89671761 | 0.99945981 |
| AFMID | 9.2301642 | -0.626553611 | 0.46088685 | -1.3594521 | 0.17400336 | 0.99945981 |
| SOCS3 | 11.2297876 | -0.281490992 | 0.58187766 | -0.4837632 | 0.62855394 | 0.99945981 |
| PGS1 | 28.6643174 | -0.144819206 | 0.31569517 | -0.4587311 | 0.64642724 | 0.99945981 |
| CYTH1 | 212.520371 | 0.367730487 | 0.396708 | 0.92695507 | 0.35394986 | 0.99945981 |
| USP36 | 40.4099263 | -0.059716074 | 0.61207507 | -0.0975633 | 0.92227905 | 0.99945981 |
| TIMP2 | 45.5843076 | 0.619090408 | 0.40022931 | 1.54683925 | 0.12190201 | 0.99945981 |
| TBC1D16 | 17.4949717 | -2.037902256 | 0.78177867 | -2.6067509 | 0.00914058 | 0.59381294 |
| GAA | 15.1713065 | 0.477799152 | 0.54999212 | 0.86873818 | 0.38499035 | 0.99945981 |
| RNF213 | 232.04533 | 0.517347932 | 0.47748011 | 1.0834963 | 0.27858819 | 0.99945981 |
| AC124319.2 | 18.1452752 | 0.51916154 | 0.51690333 | 1.00436873 | 0.31520092 | 0.99945981 |
| RPTOR | 51.5271549 | 0.056892372 | 0.27717587 | 0.2052573 | 0.8373711 | 0.99945981 |
| SLC38A10 | 28.6420049 | 0.41752337 | 0.34132344 | 1.22324844 | 0.22123587 | 0.99945981 |
| ACTG1 | 143.473672 | -0.259489291 | 0.50243769 | -0.5164606 | 0.60553273 | 0.99945981 |
| NPLOC4 | 34.514488 | 0.226570175 | 0.34782635 | 0.65138876 | 0.51479557 | 0.99945981 |
| CCDC137 | 10.0262683 | -0.10379558 | 0.44139427 | -0.2351539 | 0.81408929 | 0.99945981 |
| HGS | 15.6705882 | 0.314172258 | 0.41281364 | 0.76105105 | 0.44662657 | 0.99945981 |
| MCRIP1 | 10.8065849 | -0.366556689 | 0.63721396 | -0.575249 | 0.56512291 | 0.99945981 |
| P4HB | 43.1336096 | -0.418143626 | 0.43881524 | -0.9528922 | 0.34064468 | 0.99945981 |
| ARHGDIA | 31.3156882 | -0.543241558 | 0.55990283 | -0.9702426 | 0.3319256 | 0.99945981 |
| ANAPC11 | 13.0113627 | -0.02179183 | 0.49581894 | -0.0439512 | 0.96494332 | 0.99945981 |
| SIRT7 | 11.8571929 | 0.139981541 | 0.52293835 | 0.26768268 | 0.78894358 | 0.99945981 |
| CCDC57 | 19.2345709 | -0.356524312 | 0.33859456 | -1.0529535 | 0.29236232 | 0.99945981 |
| SLC16A3 | 27.6219942 | 0.141788414 | 0.54161479 | 0.2617883 | 0.79348466 | 0.99945981 |
| CSNK1D | 34.9477445 | -0.450854499 | 0.37621903 | -1.198383 | 0.23076793 | 0.99945981 |
| CD7 | 19.5652926 | -1.111277575 | 0.78288218 | -1.4194698 | 0.15576211 | 0.99945981 |
| HEXDC | 14.372365 | -0.324655324 | 0.56359643 | -0.5760422 | 0.56458666 | 0.99945981 |
| CYBC1 | 22.384381 | 0.125709562 | 0.48720354 | 0.25802268 | 0.7963894 | 0.99945981 |
| NARF | 22.3925971 | -0.6702589 | 0.33221577 | -2.0175409 | 0.0436391 | 0.89562588 |
| FOXK2 | 47.2237687 | -0.176951561 | 0.26751147 | -0.6614728 | 0.50830915 | 0.99945981 |
| WDR45B | 23.6354726 | 0.211385052 | 0.41351521 | 0.51119051 | 0.60921766 | 0.99945981 |
| TBCD | 33.2926908 | -0.510542649 | 0.30251458 | -1.6876629 | 0.09147595 | 0.99640647 |

| | | | | | | |
|---|---|---|---|---|---|---|
| B3GNTL1 | 14.7773075 | -0.304007804 | 0.3866071 | -0.7863482 | 0.43166351 | 0.99945981 |
| METRNL | 32.2839524 | -0.557698149 | 0.72430935 | -0.7699723 | 0.44131631 | 0.99945981 |
| USP14 | 28.2546803 | -0.291668755 | 0.37120489 | -0.7857352 | 0.43202263 | 0.99945981 |
| THOC1 | 32.1970336 | -0.111697091 | 0.37368733 | -0.2989052 | 0.76501237 | 0.99945981 |
| COLEC12 | 77.1246666 | 1.242742321 | 0.79358074 | 1.56599354 | 0.11735013 | 0.99945981 |
| ENOSF1 | 28.4663279 | -0.643797718 | 0.46295269 | -1.3906339 | 0.16433647 | 0.99945981 |
| YES1 | 19.6195485 | -0.405713457 | 0.3585058 | -1.1316789 | 0.25776945 | 0.99945981 |
| METTL4 | 10.5863678 | -0.578023082 | 0.54836593 | -1.0540828 | 0.29184502 | 0.99945981 |
| SMCHD1 | 301.416501 | 0.218674128 | 0.34658992 | 0.63093043 | 0.52808602 | 0.99945981 |
| AP001011.1 | 55.1143235 | 0.243391221 | 0.61344023 | 0.39676436 | 0.69154123 | 0.99945981 |
| EMILIN2 | 47.146922 | 0.298325538 | 0.49370986 | 0.60425275 | 0.54567562 | 0.99945981 |
| LPIN2 | 59.9184395 | 0.339299006 | 0.2891912 | 1.17326879 | 0.24068804 | 0.99945981 |
| MYL12A | 61.8359591 | -0.561376777 | 0.48488734 | -1.1577468 | 0.24696737 | 0.99945981 |
| MYL12B | 54.2811961 | -0.468972912 | 0.4004541 | -1.1711028 | 0.24155746 | 0.99945981 |
| TGIF1 | 11.3621045 | -0.483748617 | 0.63731035 | -0.7590472 | 0.44782434 | 0.99945981 |
| EPB41L3 | 142.471887 | -0.343041015 | 0.73559958 | -0.466342 | 0.64097069 | 0.99945981 |
| L3MBTL4 | 52.389077 | 0.210243722 | 0.52386385 | 0.40133275 | 0.68817515 | 0.99945981 |
| PTPRM | 99.8994735 | -1.028758939 | 0.71545218 | -1.4379143 | 0.15045837 | 0.99945981 |
| RAB12 | 36.5558679 | -0.095231829 | 0.48329526 | -0.1970469 | 0.84379084 | 0.99945981 |
| NDUFV2 | 33.483899 | -0.453839504 | 0.57536592 | -0.7887841 | 0.43023819 | 0.99945981 |
| ANKRD12 | 233.521574 | -0.31360449 | 0.27799764 | -1.1280833 | 0.25928475 | 0.99945981 |
| TWSG1 | 15.1945026 | -2.288067498 | 0.88878282 | -2.5743831 | 0.0100419 | 0.63831153 |
| RALBP1 | 23.4566893 | -0.165233482 | 0.40710816 | -0.4058712 | 0.68483723 | 0.99945981 |
| PPP4R1 | 35.8249339 | 0.207069424 | 0.41297687 | 0.50140684 | 0.61608483 | 0.99945981 |
| RAB31 | 154.918695 | 0.257842425 | 0.56881576 | 0.45329691 | 0.65033496 | 0.99945981 |
| VAPA | 63.1142712 | 0.200540272 | 0.24769122 | 0.80963819 | 0.41814815 | 0.99945981 |
| NAPG | 22.6124621 | 0.035157798 | 0.41147067 | 0.08544424 | 0.93190823 | 0.99945981 |
| MPPE1 | 17.4819467 | -0.395765014 | 0.37620545 | -1.0519917 | 0.29280336 | 0.99945981 |
| IMPA2 | 13.8378304 | 0.281885046 | 0.6419205 | 0.43912766 | 0.66056904 | 0.99945981 |
| AFG3L2 | 16.324792 | -0.981176465 | 0.49573198 | -1.9792479 | 0.04778811 | 0.90884462 |
| SPIRE1 | 25.5252303 | -0.53042085 | 0.44948096 | -1.1800741 | 0.23797073 | 0.99945981 |
| PSMG2 | 13.0714956 | -0.046568464 | 0.4377579 | -0.1063795 | 0.91528126 | 0.99945981 |
| PTPN2 | 86.5160241 | -0.439651745 | 0.3325997 | -1.3218645 | 0.18621326 | 0.99945981 |
| SEH1L | 11.586263 | -0.148544299 | 0.41725496 | -0.3560037 | 0.72183782 | 0.99945981 |
| CEP192 | 66.5192621 | 0.280955949 | 0.25465115 | 1.10329738 | 0.26989804 | 0.99945981 |
| LDLRAD4 | 235.613634 | 0.354448153 | 0.37048184 | 0.95672206 | 0.33870756 | 0.99945981 |
| FAM210A | 14.3135886 | -0.723603022 | 0.66960261 | -1.0806455 | 0.27985485 | 0.99945981 |
| RNMT | 34.6917985 | 0.111804071 | 0.33906359 | 0.32974366 | 0.74159366 | 0.99945981 |
| ROCK1 | 149.357769 | 0.249823163 | 0.19278223 | 1.29588271 | 0.1950159 | 0.99945981 |
| ESCO1 | 28.5236212 | -0.023443119 | 0.36146529 | -0.0648558 | 0.94828882 | 0.99945981 |
| SNRPD1 | 8.60266976 | -0.608686432 | 0.67824164 | -0.8974477 | 0.36948004 | 0.99945981 |
| ABHD3 | 68.3424245 | 0.127012284 | 0.29651921 | 0.42834421 | 0.66840054 | 0.99945981 |
| MIB1 | 60.0933812 | 0.13455628 | 0.32550323 | 0.41337924 | 0.67932878 | 0.99945981 |
| RBBP8 | 13.1576042 | 0.340538599 | 0.54330785 | 0.62678755 | 0.5307985 | 0.99945981 |
| CABLES1 | 17.8569881 | -0.817694538 | 0.62680878 | -1.3045359 | 0.19205094 | 0.99945981 |
| TMEM241 | 24.2469685 | -1.299563 | 0.68600781 | -1.8943851 | 0.05817391 | 0.92445778 |
| RIOK3 | 40.362621 | -0.182103641 | 0.3266646 | -0.5574637 | 0.57721068 | 0.99945981 |
| RMC1 | 21.4652657 | 0.163667545 | 0.5410028 | 0.30252625 | 0.76225093 | 0.99945981 |
| NPC1 | 46.3364174 | -0.568295267 | 0.38754032 | -1.4664159 | 0.14253503 | 0.99945981 |
| TTC39C | 84.7035212 | -0.073693302 | 0.37809381 | -0.1949075 | 0.8454654 | 0.99945981 |
| OSBPL1A | 24.4707335 | -0.121241207 | 0.47407272 | -0.2557439 | 0.79814859 | 0.99945981 |
| SS18 | 44.5071086 | 0.301991432 | 0.30182884 | 1.00053868 | 0.31704989 | 0.99945981 |
| TAF4B | 11.2448919 | 0.240562742 | 0.47658465 | 0.50476393 | 0.61372465 | 0.99945981 |
| TRAPPC8 | 60.34847 | 0.351234965 | 0.26210028 | 1.34007857 | 0.1802198 | 0.99945981 |
| RNF125 | 37.7885245 | -0.385127041 | 0.56396795 | -0.6828882 | 0.49467751 | 0.99945981 |
| RNF138 | 32.8427512 | -0.004270424 | 0.27923307 | -0.0152934 | 0.98779811 | 0.99945981 |
| GAREM1 | 15.3583282 | 0.118218155 | 0.39125416 | 0.30215181 | 0.76253634 | 0.99945981 |
| DTNA | 41.9284971 | 0.356867315 | 0.4794527 | 0.74432226 | 0.45668154 | 0.99945981 |
| MAPRE2 | 50.6096792 | 0.055256989 | 0.24905168 | 0.22186957 | 0.82441542 | 0.99945981 |
| ZNF397 | 10.1390659 | -0.086855771 | 0.53681943 | -0.161797 | 0.87146572 | 0.99945981 |
| ZNF24 | 15.650759 | 0.148555218 | 0.45272571 | 0.32813515 | 0.74280948 | 0.99945981 |
| GALNT1 | 49.5527161 | -0.302682686 | 0.45432448 | -0.6662258 | 0.50526679 | 0.99945981 |
| RPRD1A | 33.9678751 | -0.035534676 | 0.29924547 | -0.1187476 | 0.90547534 | 0.99945981 |
| ELP2 | 24.6921994 | 0.141143596 | 0.38574533 | 0.3658984 | 0.71444089 | 0.99945981 |
| TPGS2 | 11.9264977 | -0.427400798 | 0.41798875 | -1.0225175 | 0.30653605 | 0.99945981 |
| KIAA1328 | 45.3678985 | 0.216456532 | 0.46258631 | 0.4679268 | 0.63983694 | 0.99945981 |
| AC016205.1 | 26.8240103 | -0.107073611 | 0.49244718 | -0.2174317 | 0.82787195 | 0.99945981 |
| PIK3C3 | 49.8175129 | -0.094216926 | 0.29749206 | -0.316704 | 0.75146821 | 0.99945981 |
| SETBP1 | 141.727904 | -1.729892303 | 0.94162006 | -1.8371447 | 0.06618854 | 0.94594189 |
| SLC14A1 | 10.8082204 | -0.540691635 | 0.76129488 | -0.7102263 | 0.47756382 | 0.99945981 |
| EPG5 | 38.3162087 | -0.031877552 | 0.29940936 | -0.1064681 | 0.91521095 | 0.99945981 |
| PSTPIP2 | 40.7832503 | 0.420065516 | 0.4470439 | 0.9396516 | 0.3473963 | 0.99945981 |
| ATP5F1A | 26.6201054 | -0.121494684 | 0.32828356 | -0.3700907 | 0.71131493 | 0.99945981 |
| C18orf25 | 34.3590972 | 0.44609285 | 0.32634336 | 1.36694323 | 0.1716431 | 0.99945981 |
| PIAS2 | 41.9725272 | 0.190826154 | 0.32054972 | 0.59530906 | 0.5516369 | 0.99945981 |
| SMAD2 | 84.7928714 | 0.30846617 | 0.25072469 | 1.23029833 | 0.21858541 | 0.99945981 |
| CTIF | 18.9947246 | 0.335592842 | 0.42715819 | 0.78564065 | 0.43207804 | 0.99945981 |
| SMAD7 | 10.1051382 | 0.065261058 | 0.49139878 | 0.13280671 | 0.89434624 | 0.99945981 |
| DYM | 120.233592 | 0.247842863 | 0.18222956 | 1.36005853 | 0.1738114 | 0.99945981 |
| ACAA2 | 10.3534201 | 0.728033897 | 0.53090636 | 1.37130379 | 0.17028028 | 0.99945981 |
| MBD1 | 11.5736082 | 0.229598006 | 0.49785913 | 0.46117063 | 0.64467619 | 0.99945981 |
| ME2 | 47.9377105 | -0.201940182 | 0.31984222 | -0.6313744 | 0.52779577 | 0.99945981 |
| SMAD4 | 19.6379292 | 0.622438108 | 0.32433246 | 1.91913602 | 0.05496712 | 0.92445778 |
| MBD2 | 83.3368273 | 0.719874044 | 0.25801997 | 2.78999352 | 0.00527091 | 0.5234882 |
| POLI | 9.72601908 | -0.08849008 | 0.44818808 | -0.1974396 | 0.84348353 | 0.99945981 |
| TCF4 | 148.528742 | 0.750382836 | 0.73950519 | 1.01470936 | 0.31024439 | 0.99945981 |
| TXNL1 | 15.6021738 | -0.389321021 | 0.36025869 | -1.0806707 | 0.27984361 | 0.99945981 |
| WDR7 | 75.4478286 | -0.084788394 | 0.21309583 | -0.3978886 | 0.69071233 | 0.99945981 |
| NARS | 15.4459631 | -0.70235388 | 0.50366327 | -1.394491 | 0.1631694 | 0.99945981 |
| AC027097.2 | 33.9654167 | -0.025097101 | 0.49243977 | -0.0509648 | 0.95935356 | 0.99945981 |
| ATP8B1 | 26.1847946 | 0.397754528 | 0.35588045 | 1.11766332 | 0.26371081 | 0.99945981 |
| NEDD4L | 41.3751974 | 0.279783995 | 0.44413893 | 0.62994701 | 0.52872925 | 0.99945981 |
| MALT1 | 86.2763312 | 0.57834103 | 0.44255529 | 1.3068221 | 0.19127314 | 0.99945981 |
| AC104365.1 | 13.91426 | 0.694401999 | 0.64272452 | 1.08040378 | 0.27996241 | 0.99945981 |
| ZNF532 | 20.463452 | 0.451755143 | 0.57247257 | 0.78912977 | 0.43003617 | 0.99945981 |
| LMAN1 | 21.4955379 | -0.058200519 | 0.53183203 | -0.109434 | 0.91285825 | 0.99945981 |
| PIGN | 34.3021405 | -0.245380322 | 0.29963938 | -0.8189188 | 0.41283274 | 0.99945981 |
| KIAA1468 | 76.5649677 | 0.311290265 | 0.27999859 | 1.11175656 | 0.26624283 | 0.99945981 |
| ZCCHC2 | 44.6423967 | 0.141508249 | 0.39539589 | 0.35789004 | 0.72042561 | 0.99945981 |
| PHLPP1 | 53.086232 | -0.568802436 | 0.64159433 | -0.8865453 | 0.37532376 | 0.99945981 |
| BCL2 | 244.09271 | -0.052798922 | 0.45687004 | -0.1155666 | 0.90799603 | 0.99945981 |
| KDSR | 11.6516698 | -0.764144995 | 0.43739199 | -1.7470484 | 0.08062894 | 0.9846219 |
| VPS4B | 32.9238139 | 0.11589863 | 0.35778759 | 0.32393139 | 0.74598998 | 0.99945981 |
| TMX3 | 28.071275 | 0.171958162 | 0.40779948 | 0.42167332 | 0.67326348 | 0.99945981 |
| CD226 | 42.1028225 | 0.057939752 | 0.5842352 | 0.09917196 | 0.92100174 | 0.99945981 |
| RTTN | 27.4667756 | -0.246847098 | 0.41005658 | -0.601983 | 0.54718544 | 0.99945981 |
| CYB5A | 18.8406028 | -0.158070913 | 0.69364146 | -0.2278856 | 0.81973516 | 0.99945981 |
| CNDP2 | 30.675116 | -0.009464978 | 0.42418 | -0.0223136 | 0.98219781 | 0.99945981 |
| ZNF407 | 116.287557 | 0.117558478 | 0.27403115 | 0.42899677 | 0.66792558 | 0.99945981 |
| ZADH2 | 10.7446725 | -0.182470118 | 0.48810429 | -0.3738343 | 0.70852761 | 0.99945981 |
| TSHZ1 | 19.7963652 | -0.117626369 | 0.39881342 | -0.2949409 | 0.76803906 | 0.99945981 |
| ZNF516 | 46.7941635 | 0.339017934 | 0.38117072 | 0.88941231 | 0.37378153 | 0.99945981 |
| ZNF236 | 29.4046367 | -0.164641897 | 0.28763749 | -0.5723937 | 0.56705526 | 0.99945981 |

| | | | | | | |
|---|---|---|---|---|---|---|
| MBP | 144.57577 | -0.208989015 | 0.2638429 | -0.7920964 | 0.42830445 | 0.99945981 |
| ATP9B | 97.0188574 | -0.384044593 | 0.25713359 | -1.4935606 | 0.13529051 | 0.99945981 |
| NFATC1 | 27.8585817 | -0.38831414 | 0.54340922 | -0.7145888 | 0.47486317 | 0.99945981 |
| CTDP1 | 15.2368143 | -0.34589453 | 0.38471402 | -0.8990952 | 0.36860195 | 0.99945981 |
| PQLC1 | 18.4640846 | -0.33907559 | 0.41734921 | -0.8124505 | 0.41653316 | 0.99945981 |
| TXNL4A | 23.2422703 | -0.033798196 | 0.34710741 | -0.097371 | 0.92243177 | 0.99945981 |
| ADNP2 | 14.7446603 | -0.415246174 | 0.43313206 | -0.9587057 | 0.33770703 | 0.99945981 |
| RBCK1 | 18.6697739 | 0.27386003 | 0.51535972 | 0.53139588 | 0.59514448 | 0.99945981 |
| TBC1D20 | 12.2095052 | -0.312161677 | 0.39842522 | -0.7834888 | 0.43334016 | 0.99945981 |
| CSNK2A1 | 43.7106596 | 0.20447241 | 0.27194256 | 0.75189557 | 0.45211386 | 0.99945981 |
| PSMF1 | 27.3855459 | -0.115786736 | 0.37834982 | -0.3060309 | 0.75958112 | 0.99945981 |
| AL136531.2 | 9.45306385 | -0.619146669 | 0.4797162 | -1.290652 | 0.19682438 | 0.99945981 |
| SDCBP2-AS1 | 10.7842159 | 0.409654472 | 0.43847416 | 0.93427277 | 0.3501632 | 0.99945981 |
| NSFL1C | 18.8471078 | 0.685962427 | 0.36048377 | 1.90289409 | 0.05705437 | 0.92445778 |
| SIRPB2 | 23.7578799 | 0.32356053 | 0.56942504 | 0.56822322 | 0.5698834 | 0.99945981 |
| SIRPA | 25.4242313 | 0.589047363 | 0.61271906 | 0.96136615 | 0.3363681 | 0.99945981 |
| STK35 | 9.21204959 | 0.203436762 | 0.5180062 | 0.39273036 | 0.69451864 | 0.99945981 |
| SNRPB | 17.8853419 | -0.899439943 | 0.44063892 | -2.0412176 | 0.0412292 | 0.89562588 |
| NOP56 | 10.2616088 | -0.316689573 | 0.50983893 | -0.6211561 | 0.5344969 | 0.99945981 |
| PTPRA | 83.782321 | -0.35212112 | 0.24011275 | -1.4664824 | 0.14251691 | 0.99945981 |
| ITPA | 11.1728629 | 0.273123429 | 0.41033204 | 0.66561566 | 0.5056568 | 0.99945981 |
| C20orf194 | 89.291205 | 0.05056696 | 0.29097345 | 0.17378548 | 0.86203406 | 0.99945981 |
| ATRN | 54.8985084 | 0.165069438 | 0.30467846 | 0.54178244 | 0.58796839 | 0.99945981 |
| SIGLEC1 | 26.8619719 | 0.787290493 | 0.88614961 | 0.8884397 | 0.37430428 | 0.99945981 |
| CDC25B | 9.40760301 | 0.301352961 | 0.56696083 | 0.53152342 | 0.59505611 | 0.99945981 |
| MAVS | 18.3981674 | -0.435577598 | 0.41798273 | -1.0420947 | 0.29736776 | 0.99945981 |
| PANK2 | 35.2143129 | 0.326644232 | 0.28292717 | 1.15451703 | 0.24828826 | 0.99945981 |
| RNF24 | 50.0833115 | -0.18482342 | 0.68557705 | -0.2695881 | 0.78747715 | 0.99945981 |
| PRNP | 21.50391 | -0.216774243 | 0.41048664 | -0.5280909 | 0.59743627 | 0.99945981 |
| RASSF2 | 52.614881 | 0.18232079 | 0.44859893 | 0.40642271 | 0.68443204 | 0.99945981 |
| SLC23A2 | 41.6922916 | 0.313520513 | 0.32005263 | 0.9795905 | 0.3272883 | 0.99945981 |
| TMEM230 | 17.2502582 | 0.048707145 | 0.35262185 | 0.13812855 | 0.89013882 | 0.99945981 |
| CDS2 | 34.1726705 | -0.238330446 | 0.27362097 | -0.871024 | 0.38374103 | 0.99945981 |
| GPCPD1 | 110.170103 | 0.428034637 | 0.30704463 | 1.39404696 | 0.16330342 | 0.99945981 |
| SHLD1 | 29.1193315 | -0.47028396 | 0.4841931 | -0.9712736 | 0.33141207 | 0.99945981 |
| CRLS1 | 16.5351202 | 0.208652765 | 0.46691418 | 0.44687605 | 0.65496456 | 0.99945981 |
| TMX4 | 20.1129811 | -0.671765292 | 0.46053863 | -1.4586514 | 0.14466109 | 0.99945981 |
| PLCB1 | 126.221646 | 0.087996834 | 0.53489718 | 0.16451168 | 0.86932835 | 0.99945981 |
| SLX4IP | 35.4397586 | -0.073516085 | 0.47107215 | -0.1560612 | 0.87598478 | 0.99945981 |
| SPTLC3 | 13.5204761 | -1.204553986 | 0.64269498 | -1.8742234 | 0.06089964 | 0.92445778 |
| TASP1 | 34.6102662 | 0.249449446 | 0.34905074 | 0.71465097 | 0.47482474 | 0.99945981 |
| ESF1 | 17.4033012 | -0.718875138 | 0.46604236 | -1.5425103 | 0.12294964 | 0.99945981 |
| NDUFAF5 | 9.87405246 | -0.084495359 | 0.50754729 | -0.1664778 | 0.86778095 | 0.99945981 |
| MACROD2 | 28.4335008 | -0.248947244 | 0.70606326 | -0.3525849 | 0.72439966 | 0.99945981 |
| KIF16B | 44.0435601 | 0.321203362 | 0.41246011 | 0.77875013 | 0.43612692 | 0.99945981 |
| SNRPB2 | 9.54169468 | -0.413735786 | 0.46518599 | -0.8893986 | 0.37378888 | 0.99945981 |
| DSTN | 29.100281 | -0.172343318 | 0.46546389 | -0.3702614 | 0.71118772 | 0.99945981 |
| RRBP1 | 37.0042485 | -0.142173989 | 0.49995032 | -0.2843762 | 0.77612207 | 0.99945981 |
| SNX5 | 26.3098794 | 0.129797892 | 0.36516994 | 0.35544517 | 0.72225611 | 0.99945981 |
| MGME1 | 13.435447 | -0.553103736 | 0.54009355 | -1.0240888 | 0.30579335 | 0.99945981 |
| SEC23B | 24.8820604 | -0.285433137 | 0.31164773 | -0.9158839 | 0.35972779 | 0.99945981 |
| SLC24A3 | 67.3683537 | -5.51080457 | 2.23903808 | -2.4612375 | 0.01384587 | 0.72008995 |
| RIN2 | 79.9469718 | -0.82052978 | 0.76905985 | -1.0669258 | 0.28600535 | 0.99945981 |
| NAA20 | 10.8352797 | -0.547672362 | 0.43298463 | -1.2648771 | 0.20591537 | 0.99945981 |
| RALGAPA2 | 91.8744552 | 0.145928363 | 0.38648462 | 0.3775787 | 0.70574359 | 0.99945981 |
| KIZ | 21.3604643 | -0.051044793 | 0.3794933 | -0.1345078 | 0.89300108 | 0.99945981 |
| XRN2 | 57.7256216 | -0.04894046 | 0.29765216 | -0.1644217 | 0.86939922 | 0.99945981 |
| CD93 | 20.7446307 | -0.113051345 | 0.70086045 | -0.1613036 | 0.87185425 | 0.99945981 |
| NAPB | 14.2618271 | -0.634351681 | 0.44519572 | -1.4248827 | 0.15419111 | 0.99945981 |
| CST3 | 124.974434 | 0.27407835 | 0.66573531 | 0.41169268 | 0.68056469 | 0.99945981 |
| CST7 | 23.0549858 | -1.129187268 | 0.78689395 | -1.4349929 | 0.15128911 | 0.99945981 |
| APMAP | 25.2495291 | 0.422065425 | 0.60204725 | 0.70105034 | 0.4832716 | 0.99945981 |
| ACSS1 | 18.5738991 | 0.207618315 | 0.39053978 | 0.53161887 | 0.59498999 | 0.99945981 |
| ENTPD6 | 9.44103437 | -0.275303666 | 0.45105249 | -0.6103584 | 0.54162444 | 0.99945981 |
| PYGB | 12.3589656 | 0.288247625 | 0.40676268 | 0.70863833 | 0.47854894 | 0.99945981 |
| ABHD12 | 30.7659662 | 0.002895872 | 0.48819418 | 0.0059318 | 0.99526713 | 0.99945981 |
| ZNF337 | 10.0701084 | -0.556218293 | 0.44061835 | -1.2623584 | 0.20681984 | 0.99945981 |
| HM13 | 49.546703 | 0.183711853 | 0.46044998 | 0.3989833 | 0.68990551 | 0.99945981 |
| BCL2L1 | 37.2276256 | 0.037208933 | 0.45128338 | 0.08245137 | 0.93428779 | 0.99945981 |
| AL117381.1 | 13.2783983 | -0.652085814 | 0.58369956 | -1.11716 | 0.26392591 | 0.99945981 |
| HCK | 82.491455 | 0.864228456 | 0.6293372 | 1.37323594 | 0.16967901 | 0.99945981 |
| TM9SF4 | 34.5172373 | 0.189151416 | 0.2734195 | 0.69179929 | 0.48906338 | 0.99945981 |
| KIF3B | 15.3859208 | -0.585163541 | 0.52258036 | -1.119758 | 0.2628169 | 0.99945981 |
| ASXL1 | 69.1629833 | -0.125684369 | 0.24984535 | -0.5030487 | 0.61493006 | 0.99945981 |
| NOL4L | 33.0353435 | 1.189589814 | 0.65643862 | 1.81218743 | 0.06995724 | 0.95505283 |
| MAPRE1 | 16.6839533 | 0.349279315 | 0.42259228 | 0.82651609 | 0.40851139 | 0.99945981 |
| CDK5RAP1 | 13.9844285 | -0.371811587 | 0.43420855 | -0.8562972 | 0.39183339 | 0.99945981 |
| CBFA2T2 | 65.3350114 | 0.361759508 | 0.26699588 | 1.35492544 | 0.17544131 | 0.99945981 |
| CHMP4B | 19.8975981 | 0.418238038 | 0.35118372 | 1.19093801 | 0.23367792 | 0.99945981 |
| RALY | 57.4088705 | 0.016608978 | 0.36994178 | 0.0448962 | 0.96419005 | 0.99945981 |
| EIF2S2 | 24.6840554 | -0.471117579 | 0.36172889 | -1.3024052 | 0.19277791 | 0.99945981 |
| ITCH | 120.044523 | 0.069020373 | 0.29029633 | 0.23775834 | 0.81206853 | 0.99945981 |
| DYNLRB1 | 23.1039558 | -0.232665163 | 0.39121644 | -0.5947224 | 0.55202908 | 0.99945981 |
| PIGU | 16.6913828 | -0.22207375 | 0.39267166 | -0.5655457 | 0.57170268 | 0.99945981 |
| NCOA6 | 46.9089913 | -0.188975903 | 0.25267644 | -0.7478968 | 0.45452241 | 0.99945981 |
| ACSS2 | 17.3681635 | 0.470218624 | 0.47014867 | 1.0001488 | 0.3172385 | 0.99945981 |
| TRPC4AP | 69.9745888 | -0.147890009 | 0.2322847 | -0.6366756 | 0.52433614 | 0.99945981 |
| EDEM2 | 13.4122017 | -0.748982065 | 0.40992277 | -1.8271297 | 0.06768027 | 0.94796811 |
| MMP24OS | 17.4018249 | -0.058304747 | 0.41154093 | -0.1416742 | 0.88733733 | 0.99945981 |
| MMP24 | 9.55949494 | -0.537650683 | 0.6184281 | -0.8693827 | 0.38463785 | 0.99945981 |
| UQCC1 | 24.4937974 | 0.191802786 | 0.3613571 | 0.53078461 | 0.59556804 | 0.99945981 |
| CEP250 | 12.6952621 | -0.686084291 | 0.43301525 | -1.5844345 | 0.11309488 | 0.99945981 |
| FO393401.1 | 16.5712132 | -0.678193081 | 0.39553623 | -1.7146168 | 0.0864155 | 0.99204414 |
| ERGIC3 | 11.5137117 | 0.269821548 | 0.45313803 | 0.59545112 | 0.55154196 | 0.99945981 |
| CPNE1 | 17.9893385 | 0.406724533 | 0.45352428 | 0.89680872 | 0.36982099 | 0.99945981 |
| RBM39 | 127.607542 | 0.189508634 | 0.19041164 | 0.99525763 | 0.31961098 | 0.99945981 |
| PHF20 | 91.43482 | -0.066980268 | 0.24590753 | -0.2723799 | 0.78532993 | 0.99945981 |
| SCAND1 | 16.7136842 | -0.040218591 | 0.48932808 | -0.0821915 | 0.93449446 | 0.99945981 |
| NORAD | 28.8052198 | -0.465869081 | 0.54553366 | -0.8539694 | 0.39312192 | 0.99945981 |
| DLGAP4 | 22.9293525 | -0.608454396 | 0.41480825 | -1.4668329 | 0.14242151 | 0.99945981 |
| RAB5IF | 11.2960786 | 0.108490336 | 0.62171725 | 0.17450109 | 0.86147168 | 0.99945981 |
| SLA2 | 17.3309687 | -1.49196081 | 0.5964018 | -2.5016034 | 0.01236323 | 0.67187199 |
| NDRG3 | 12.4839842 | -0.036122518 | 0.4351694 | -0.0830079 | 0.93384522 | 0.99945981 |
| SOGA1 | 15.6765102 | -0.610073504 | 0.44826036 | -1.3609803 | 0.17351993 | 0.99945981 |
| SAMHD1 | 118.848282 | 1.04613862 | 0.41621935 | 2.51343103 | 0.01195632 | 0.67187199 |
| RBL1 | 24.9882086 | -0.254271867 | 0.34368819 | -0.739833 | 0.45940133 | 0.99945981 |
| RPN2 | 32.832715 | -0.228792986 | 0.45003057 | -0.5083943 | 0.61117683 | 0.99945981 |
| BLCAP | 21.9799552 | -0.109868893 | 0.32207084 | -0.3411327 | 0.73300369 | 0.99945981 |
| CTNNBL1 | 31.5730076 | 0.390205942 | 0.35500052 | 1.09917007 | 0.27169389 | 0.99945981 |
| TTI1 | 9.88174628 | -1.08782093 | 0.44565797 | -2.4409323 | 0.0146494 | 0.73008906 |
| RPRD1B | 22.1238362 | 0.206011574 | 0.39610329 | 0.52009559 | 0.60299695 | 0.99945981 |
| TGM2 | 10.425232 | -2.117224323 | 0.63584631 | -3.3297737 | 0.00086917 | 0.27121928 |
| RALGAPB | 48.8483162 | 0.053522277 | 0.35229928 | 0.15192275 | 0.87924786 | 0.99945981 |

| | | | | | | |
|---|---|---|---|---|---|---|
| PPP1R16B | 86.609716 | -1.129910501 | 0.37208704 | -3.0366833 | 0.00239197 | 0.45570351 |
| DHX35 | 10.9159186 | -0.376075536 | 0.42495173 | -0.8849841 | 0.37616519 | 0.99945981 |
| MAFB | 34.8431881 | 0.449810635 | 0.49759841 | 0.90396316 | 0.36601493 | 0.99945981 |
| TOP1 | 83.5956303 | 0.166719313 | 0.25832458 | 0.64538694 | 0.51867646 | 0.99945981 |
| PLCG1 | 12.2049804 | -0.534330796 | 0.44728927 | -1.1945978 | 0.23224419 | 0.99945981 |
| ZHX3 | 15.3039794 | -0.072387025 | 0.42367644 | -0.1708545 | 0.86433818 | 0.99945981 |
| CHD6 | 66.0151918 | -0.335255192 | 0.30055242 | -1.1154633 | 0.26465194 | 0.99945981 |
| SRSF6 | 14.3282836 | 0.883304303 | 0.42636582 | 2.07170522 | 0.03829294 | 0.87920079 |
| SERINC3 | 25.8218953 | 0.258203417 | 0.35945348 | 0.7183222 | 0.47255864 | 0.99945981 |
| YWHAB | 59.6012623 | -0.265258875 | 0.42526326 | -0.6237522 | 0.53279037 | 0.99945981 |
| STK4 | 163.504695 | -0.019586744 | 0.26851426 | -0.0729449 | 0.94184997 | 0.99945981 |
| SLPI | 29.7278489 | -4.255776099 | 1.07723479 | -3.9506486 | 7.79E-05 | 0.089176 |
| SDC4 | 10.5968926 | -1.302610956 | 0.59709118 | -2.1815947 | 0.02913946 | 0.84763724 |
| PIGT | 9.55173773 | 0.270512618 | 0.47624902 | 0.56800667 | 0.57003044 | 0.99945981 |
| WFDC2 | 44.742615 | -3.417344767 | 0.65866189 | -5.1883141 | 2.12E-07 | 0.00108156 |
| DNTTIP1 | 9.59155637 | 0.068078839 | 0.45757129 | 0.14878302 | 0.88172485 | 0.99945981 |
| CTSA | 21.1500823 | 0.026142992 | 0.5077543 | 0.05148749 | 0.95893707 | 0.99945981 |
| PLTP | 31.8274813 | -1.02345801 | 0.93243288 | -1.0976211 | 0.27236997 | 0.99945981 |
| ZNF335 | 13.9314566 | -0.328047162 | 0.36178877 | -0.9067367 | 0.36454604 | 0.99945981 |
| NCOA5 | 15.3789223 | -0.409978134 | 0.46150839 | -0.8883438 | 0.37435582 | 0.99945981 |
| SLC35C2 | 14.4398156 | 0.27143038 | 0.46671086 | 0.58158145 | 0.56084864 | 0.99945981 |
| ELMO2 | 16.7171825 | 0.34965141 | 0.38152244 | 0.91646355 | 0.35942381 | 0.99945981 |
| SLC13A3 | 10.4683764 | 0.528050177 | 0.59112433 | 0.89329799 | 0.37169762 | 0.99945981 |
| ZMYND8 | 78.1151147 | -0.265701895 | 0.29536882 | -0.8995597 | 0.3683546 | 0.99945981 |
| NCOA3 | 185.877345 | 0.002243252 | 0.39370221 | 0.00569784 | 0.99545381 | 0.99945981 |
| SULF2 | 47.9018318 | 0.707350371 | 0.95876452 | 0.73777278 | 0.46065253 | 0.99945981 |
| PREX1 | 144.567199 | -0.050838819 | 0.32877056 | -0.1546331 | 0.87711055 | 0.99945981 |
| ARFGEF2 | 55.9539009 | -0.047294339 | 0.35174748 | -0.1344554 | 0.8930425 | 0.99945981 |
| CSE1L | 22.0401226 | 0.093924971 | 0.35627618 | 0.26362967 | 0.79206529 | 0.99945981 |
| STAU1 | 50.8572022 | 0.276308889 | 0.32029438 | 0.86267167 | 0.38831801 | 0.99945981 |
| DDX27 | 13.834231 | -0.592392836 | 0.38815865 | -1.5261616 | 0.12696961 | 0.99945981 |
| ZNFX1 | 27.548261 | 0.176321782 | 0.51171923 | 0.34456743 | 0.73041959 | 0.99945981 |
| ZFAS1 | 29.0048419 | -0.07858423 | 0.30523273 | -0.2574568 | 0.79682619 | 0.99945981 |
| B4GALT5 | 73.438484 | 0.231960777 | 0.39576287 | 0.5861105 | 0.55780126 | 0.99945981 |
| SLC9A8 | 27.3603968 | 0.083552138 | 0.47446232 | 0.17609857 | 0.8602165 | 0.99945981 |
| RNF114 | 25.8553931 | 0.670127977 | 0.45662694 | 1.4675612 | 0.14222345 | 0.99945981 |
| CEBPB | 81.0561923 | 1.026399909 | 0.69641605 | 1.47383149 | 0.14052697 | 0.99945981 |
| PTPN1 | 94.152081 | -0.599994672 | 0.29581702 | -2.0282628 | 0.04253343 | 0.89562588 |
| ADNP | 69.1576848 | 0.030256465 | 0.3153527 | 0.09594484 | 0.92356438 | 0.99945981 |
| DPM1 | 19.2194007 | -0.313840779 | 0.41332301 | -0.7593112 | 0.44766643 | 0.99945981 |
| NFATC2 | 85.0128848 | -0.749519389 | 0.44067795 | -1.7008325 | 0.08897444 | 0.99640647 |
| ZFP64 | 9.5641102 | -0.339481686 | 0.44627283 | -0.7607044 | 0.44683367 | 0.99945981 |
| TSHZ2 | 26.0432399 | 0.139923569 | 0.82954524 | 0.16867503 | 0.86605226 | 0.99945981 |
| ZNF217 | 38.1761567 | 0.250801853 | 0.37723266 | 0.66484661 | 0.50614861 | 0.99945981 |
| CASS4 | 13.8310665 | 0.603731417 | 0.69557863 | 0.86795567 | 0.3854186 | 0.99945981 |
| RTF2 | 18.6251502 | 0.067021754 | 0.34659553 | 0.19337167 | 0.8466679 | 0.99945981 |
| RBM38 | 12.7567846 | -0.575438558 | 0.68253964 | -0.8430844 | 0.39918122 | 0.99945981 |
| PCK1 | 21.4991055 | 0.128367555 | 1.19816852 | 0.10713648 | 0.91468071 | 0.99945981 |
| ZBP1 | 12.0877386 | -0.602299764 | 0.86935464 | -0.6928125 | 0.48842722 | 0.99945981 |
| RAB22A | 27.8329707 | 0.177815713 | 0.36644096 | 0.48525065 | 0.62749856 | 0.99945981 |
| VAPB | 28.0076965 | -0.354319407 | 0.32711042 | -1.0831798 | 0.27872861 | 0.99945981 |
| GNAS | 93.2498604 | -0.543975973 | 0.30243149 | -1.798675 | 0.0720701 | 0.96000932 |
| NELFCD | 11.6713924 | -0.1363331 | 0.42353547 | -0.321893 | 0.74753376 | 0.99945981 |
| CTSZ | 45.5670933 | 0.341876353 | 0.43842754 | 0.77977845 | 0.43552129 | 0.99945981 |
| ATP5F1E | 63.4763222 | -0.334186971 | 0.49434726 | -0.6760166 | 0.49903009 | 0.99945981 |
| ZNF831 | 38.1870236 | -0.547703667 | 0.40299557 | -1.3590811 | 0.17412089 | 0.99945981 |
| MIR646HG | 37.2678105 | 1.034000127 | 0.7532964 | 1.37263384 | 0.16986621 | 0.99945981 |
| TAF4 | 16.5584751 | 0.575234144 | 0.35615696 | 1.61511416 | 0.10628598 | 0.99945981 |
| PSMA7 | 30.1506063 | -0.413248653 | 0.42160456 | -0.9801807 | 0.32699692 | 0.99945981 |
| SS18L1 | 12.2524211 | 0.196543401 | 0.4257743 | 0.46161406 | 0.64435811 | 0.99945981 |
| MTG2 | 10.6257674 | -0.070601552 | 0.47392573 | -0.1489718 | 0.88157591 | 0.99945981 |
| OSBPL2 | 15.4842528 | -0.145576333 | 0.35451319 | -0.4106373 | 0.68133852 | 0.99945981 |
| OGFR | 21.873141 | 0.173966927 | 0.50465876 | 0.3447219 | 0.73030345 | 0.99945981 |
| DIDO1 | 41.1651271 | 0.049339468 | 0.25090265 | 0.19664786 | 0.84410312 | 0.99945981 |
| GID8 | 11.7969772 | 0.151961605 | 0.42672544 | 0.35611096 | 0.72175747 | 0.99945981 |
| YTHDF1 | 9.64516422 | -0.306500158 | 0.51267977 | -0.5978394 | 0.54994711 | 0.99945981 |
| ARFGAP1 | 11.9070313 | -0.641806256 | 0.66612053 | -0.9634987 | 0.33529732 | 0.99945981 |
| PPDPF | 23.6273511 | -0.622291688 | 0.52068108 | -1.1951494 | 0.23202866 | 0.99945981 |
| ZBTB46 | 15.926921 | -0.903144955 | 0.79743913 | -1.1325566 | 0.2574005 | 0.99945981 |
| TPD52L2 | 18.221343 | -0.181064674 | 0.44727447 | -0.4048178 | 0.68561143 | 0.99945981 |
| DNAJC5 | 66.0712173 | 0.140165182 | 0.34448559 | 0.40688257 | 0.68409424 | 0.99945981 |
| PRPF6 | 17.461214 | -0.527274824 | 0.38142766 | -1.3823718 | 0.16685759 | 0.99945981 |
| RGS19 | 17.2570534 | 0.343993119 | 0.48703821 | 0.70629596 | 0.4800041 | 0.99945981 |
| PCMTD2 | 17.271212 | 0.083742161 | 0.38668194 | 0.216566 | 0.82854658 | 0.99945981 |
| BSG | 21.6307872 | -0.208802066 | 0.46858359 | -0.4456026 | 0.65588434 | 0.99945981 |
| RNF126 | 13.7750413 | -0.383288137 | 0.51454065 | -0.7449132 | 0.45632418 | 0.99945981 |
| PTBP1 | 24.2256211 | -0.014607697 | 0.4097685 | -0.0356487 | 0.97156251 | 0.99945981 |
| CFD | 19.8540108 | 0.327851739 | 0.87011185 | 0.37679264 | 0.70632771 | 0.99945981 |
| R3HDM4 | 9.23204061 | 0.231602138 | 0.6295335 | 0.36789486 | 0.71295163 | 0.99945981 |
| ARID3A | 31.1685431 | 0.318931155 | 0.41610562 | 0.76646682 | 0.44339858 | 0.99945981 |
| TMEM259 | 28.5694906 | 0.068995859 | 0.32692977 | 0.21104183 | 0.83285463 | 0.99945981 |
| CNN2 | 18.0913224 | -0.038438795 | 0.40958153 | -0.0938489 | 0.92522915 | 0.99945981 |
| ABCA7 | 15.6911648 | 0.592173299 | 0.48838149 | 1.212522 | 0.22531263 | 0.99945981 |
| ARHGAP45 | 36.9819793 | 0.405436069 | 0.34556384 | 1.17325954 | 0.24069174 | 0.99945981 |
| POLR2E | 12.9969254 | -0.277721373 | 0.46273613 | -0.6001722 | 0.54839146 | 0.99945981 |
| GPX4 | 29.4224227 | -0.348615629 | 0.34439725 | -1.0122486 | 0.31141921 | 0.99945981 |
| SBNO2 | 21.3752469 | -0.184999653 | 0.51146881 | -0.3617027 | 0.7175742 | 0.99945981 |
| STK11 | 12.9356564 | -0.031734297 | 0.48191637 | -0.0658502 | 0.94749708 | 0.99945981 |
| ATP5F1D | 20.5729057 | 0.305790023 | 0.41026646 | 0.74534492 | 0.45606323 | 0.99945981 |
| MIDN | 34.5318569 | 0.264963529 | 0.46863314 | 0.56539648 | 0.57180412 | 0.99945981 |
| CIRBP | 34.2750371 | -0.270578736 | 0.50150285 | -0.5395358 | 0.58951721 | 0.99945981 |
| MUM1 | 10.3758653 | -0.455134498 | 0.48024951 | -0.9477042 | 0.34328004 | 0.99945981 |
| DAZAP1 | 23.2560928 | 0.177479341 | 0.33870497 | 0.5239939 | 0.60028278 | 0.99945981 |
| RPS15 | 137.962826 | -0.26937992 | 0.47486544 | -0.5672763 | 0.57052646 | 0.99945981 |
| C19orf25 | 14.441836 | -0.795069599 | 0.40134773 | -1.9809994 | 0.04759134 | 0.90884462 |
| TCF3 | 16.5579962 | -0.982850652 | 0.74058952 | -1.3271193 | 0.18446922 | 0.99945981 |
| REXO1 | 9.64883216 | -0.195954758 | 0.53947076 | -0.3632352 | 0.71642921 | 0.99945981 |
| ABHD17A | 22.460893 | -0.017127001 | 0.60440019 | -0.0283372 | 0.97739322 | 0.99945981 |
| CSNK1G2 | 23.6326588 | -0.217755232 | 0.39087548 | -0.5570962 | 0.57746173 | 0.99945981 |
| BTBD2 | 10.1284332 | -1.117339727 | 0.45892904 | -2.4346677 | 0.01490547 | 0.73270878 |
| MKNK2 | 12.3715167 | -0.122636871 | 0.47122516 | -0.2602511 | 0.79467008 | 0.99945981 |
| MOB3A | 39.2035349 | -0.855492901 | 0.57399517 | -1.4904183 | 0.13611428 | 0.99945981 |
| AP3D1 | 24.8181787 | 0.062971355 | 0.42804329 | 0.14711445 | 0.88304168 | 0.99945981 |
| DOT1L | 18.2826363 | 0.104577408 | 0.57333351 | 0.1824024 | 0.85526694 | 0.99945981 |
| PLEKHJ1 | 10.6204347 | 0.447489952 | 0.48885618 | 0.91538159 | 0.35999134 | 0.99945981 |
| AC005258.1 | 11.0547456 | -1.254034561 | 0.60945635 | -2.0576282 | 0.03962584 | 0.89132455 |
| LSM7 | 17.7384027 | -0.564493314 | 0.60966437 | -0.9259083 | 0.35449362 | 0.99945981 |
| SPPL2B | 9.66666268 | -0.308393252 | 0.48168939 | -0.6402326 | 0.52202139 | 0.99945981 |
| GADD45B | 18.9755892 | 0.14892503 | 0.60174309 | 0.24748939 | 0.8045295 | 0.99945981 |
| GNG7 | 31.8455337 | 0.29170126 | 0.54452415 | 0.5356994 | 0.59216632 | 0.99945981 |
| SGTA | 8.24784764 | -0.276850427 | 0.50071204 | -0.5529135 | 0.58032267 | 0.99945981 |
| AES | 45.7026308 | -0.373336354 | 0.46429536 | -0.8040924 | 0.42134363 | 0.99945981 |
| GNA15 | 13.1181186 | -0.299174287 | 0.77795006 | -0.3845675 | 0.7005579 | 0.99945981 |

| | | | | | | |
|---|---|---|---|---|---|---|
| S1PR4 | 12.2824111 | 0.077624536 | 0.5646602 | 0.13747124 | 0.89065833 | 0.99945981 |
| NCLN | 9.96929175 | 0.203069906 | 0.41984566 | 0.48367751 | 0.62861475 | 0.99945981 |
| NFIC | 33.0343862 | -0.292878439 | 0.30956506 | -0.9460966 | 0.34409933 | 0.99945981 |
| MFSD12 | 19.0583935 | 0.000234109 | 0.42807235 | 0.00054689 | 0.99956364 | 0.9997934 |
| PIP5K1C | 13.2744286 | 0.5055206 | 0.50789206 | 0.99533077 | 0.31957541 | 0.99945981 |
| EEF2 | 69.6887249 | -0.317606628 | 0.52836199 | -0.6011156 | 0.547763 | 0.99945981 |
| ZBTB7A | 28.6751624 | 0.071427936 | 0.48364636 | 0.14768629 | 0.88259035 | 0.99945981 |
| MAP2K2 | 23.2743261 | -0.478206368 | 0.37854276 | -1.2632823 | 0.20648775 | 0.99945981 |
| SH3GL1 | 20.8999202 | -0.276938741 | 0.38665475 | -0.716243 | 0.47384135 | 0.99945981 |
| MYDGF | 24.2115712 | -0.35604793 | 0.53311227 | -0.6678667 | 0.50421869 | 0.99945981 |
| DPP9 | 14.8627589 | -0.514683338 | 0.43759593 | -1.1761612 | 0.23953049 | 0.99945981 |
| PLIN3 | 10.3494226 | -0.42355502 | 0.58719135 | -0.7213237 | 0.4707104 | 0.99945981 |
| ARRDC5 | 10.8501382 | -0.293551441 | 0.65184184 | -0.4503415 | 0.65246421 | 0.99945981 |
| KDM4B | 48.7852457 | -0.789725869 | 0.35917121 | -2.1987449 | 0.02789607 | 0.84763724 |
| SAFB2 | 25.7629781 | -0.384947746 | 0.34288181 | -1.1226835 | 0.26157195 | 0.99945981 |
| SAFB | 36.6176501 | -0.187675256 | 0.28655743 | -0.6549307 | 0.51251238 | 0.99945981 |
| RPL36 | 60.028814 | -0.366978528 | 0.52486801 | -0.6991825 | 0.48443799 | 0.99945981 |
| LONP1 | 14.3466487 | -0.097069649 | 0.3907645 | -0.2484096 | 0.80381752 | 0.99945981 |
| RANBP3 | 12.6333909 | -0.118080549 | 0.39627263 | -0.2979781 | 0.76571991 | 0.99945981 |
| RFX2 | 15.7535206 | -0.195893688 | 0.82296404 | -0.2380343 | 0.81185447 | 0.99945981 |
| MLLT1 | 9.19995768 | -0.16576726 | 0.43459551 | -0.3814288 | 0.70288507 | 0.99945981 |
| KHSRP | 11.6077957 | 0.006118742 | 0.51326732 | 0.01192116 | 0.99048852 | 0.99945981 |
| DENND1C | 14.3680275 | -0.490579088 | 0.4400518 | -1.1148212 | 0.26492704 | 0.99945981 |
| C3 | 15.8298738 | 0.337485906 | 1.16345113 | 0.29007313 | 0.7717603 | 0.99945981 |
| VAV1 | 70.9774705 | 0.19702488 | 0.34722193 | 0.56743213 | 0.57042063 | 0.99945981 |
| INSR | 37.8841666 | -0.068997551 | 0.41609661 | -0.165821 | 0.86829783 | 0.99945981 |
| AC119396.1 | 22.8367392 | 0.494847097 | 0.47076749 | 1.05114967 | 0.29318985 | 0.99945981 |
| AC008878.3 | 11.9292478 | 1.575163275 | 0.46572696 | 3.38216036 | 0.00071918 | 0.27121928 |
| MCOLN1 | 12.8522411 | 0.216312958 | 0.63414439 | 0.34110995 | 0.73302081 | 0.99945981 |
| PNPLA6 | 18.0165357 | 0.125915438 | 0.40218131 | 0.31308127 | 0.75421892 | 0.99945981 |
| STXBP2 | 13.300885 | 1.476698923 | 0.74366551 | 1.98570312 | 0.0470663 | 0.90884462 |
| MAP2K7 | 10.6780099 | -0.697226425 | 0.54421603 | -1.2811574 | 0.20013837 | 0.99945981 |
| ELAVL1 | 19.1306291 | -0.293676452 | 0.39429063 | -0.7448223 | 0.45637915 | 0.99945981 |
| CERS4 | 17.7105441 | -0.255877331 | 0.40802338 | -0.6271144 | 0.53058426 | 0.99945981 |
| RPS28 | 24.5294714 | 0.128563437 | 0.72778406 | 0.17665053 | 0.8597829 | 0.99945981 |
| RAB11B | 10.6496307 | -0.317679845 | 0.59523749 | -0.5337027 | 0.59354725 | 0.99945981 |
| 2-Mar | 12.3925211 | -0.625879373 | 0.5938499 | -1.0539353 | 0.29191254 | 0.99945981 |
| HNRNPM | 59.712767 | 0.115744457 | 0.23290988 | 0.49694953 | 0.61922464 | 0.99945981 |
| PRAM1 | 22.9383041 | 0.5675037 | 0.7334383 | 0.77375793 | 0.43907395 | 0.99945981 |
| MYO1F | 140.991163 | 0.245741813 | 0.38678043 | 0.63535225 | 0.5251987 | 0.99945981 |
| ZNF317 | 8.66407895 | -0.335628271 | 0.51689031 | -0.649322 | 0.51613024 | 0.99945981 |
| ZNF266 | 14.3552891 | -0.258079481 | 0.37408834 | -0.6898891 | 0.4902639 | 0.99945981 |
| ZNF121 | 11.6680355 | -0.429259465 | 0.51442465 | -0.8344458 | 0.40402985 | 0.99945981 |
| ZNF562 | 18.3839186 | -0.169698981 | 0.49134107 | -0.3453792 | 0.72980932 | 0.99945981 |
| ZNF846 | 15.1706536 | 0.074332817 | 0.3962679 | 0.18758223 | 0.85120415 | 0.99945981 |
| UBL5 | 11.1394567 | 0.008412211 | 0.66103453 | 0.01272583 | 0.98984653 | 0.99945981 |
| PIN1 | 9.26741069 | 0.116327673 | 0.46670042 | 0.24925556 | 0.80316311 | 0.99945981 |
| C19orf66 | 17.5088716 | 0.335249326 | 0.56326396 | 0.59519044 | 0.55171618 | 0.99945981 |
| EIF3G | 9.75093681 | 0.342442446 | 0.50276762 | 0.68111476 | 0.49579888 | 0.99945981 |
| DNMT1 | 25.611321 | 0.40432519 | 0.30353328 | 1.33206212 | 0.18283978 | 0.99945981 |
| ICAM1 | 10.1917751 | -0.476499809 | 0.76820382 | -0.6202778 | 0.53507488 | 0.99945981 |
| ICAM3 | 10.2870462 | 0.06875255 | 0.799813 | 0.08596078 | 0.93149759 | 0.99945981 |
| TYK2 | 17.6937253 | 0.241068354 | 0.34079734 | 0.70736572 | 0.47933924 | 0.99945981 |
| CDC37 | 22.2775737 | -0.595289807 | 0.34301749 | -1.7354503 | 0.08266107 | 0.98529784 |
| PDE4A | 23.6392764 | -1.05653388 | 0.74373675 | -1.4205751 | 0.15544033 | 0.99945981 |
| KRI1 | 9.23077303 | -0.90115439 | 0.58643053 | -1.5366772 | 0.12437239 | 0.99945981 |
| CDKN2D | 11.7461181 | -0.299187981 | 0.80253372 | -0.3728042 | 0.70929414 | 0.99945981 |
| SLC44A2 | 27.7287732 | -0.370871356 | 0.54018836 | -0.6865593 | 0.49236047 | 0.99945981 |
| ILF3 | 35.3446452 | 0.418741242 | 0.3870025 | 1.08201173 | 0.27924732 | 0.99945981 |
| DNM2 | 82.2706672 | -0.010649497 | 0.2754883 | -0.0386568 | 0.96916401 | 0.99945981 |
| CARM1 | 9.81195688 | 0.040320281 | 0.60419329 | 0.06673408 | 0.94679341 | 0.99945981 |
| SMARCA4 | 33.5774903 | -0.007158139 | 0.37707274 | -0.0189834 | 0.98485431 | 0.99945981 |
| LDLR | 9.2352389 | -0.612143184 | 0.65771195 | -0.9307162 | 0.35200037 | 0.99945981 |
| PRKCSH | 10.9500322 | -0.140192069 | 0.48485972 | -0.2891394 | 0.77247467 | 0.99945981 |
| ZNF439 | 10.9077386 | -1.132733982 | 0.55019186 | -2.0587981 | 0.03951358 | 0.89132455 |
| ZNF44 | 29.8491221 | 0.206179259 | 0.46973612 | 0.4389257 | 0.66071537 | 0.99945981 |
| ZNF490 | 12.0547146 | -0.164002599 | 0.42563935 | -0.3853088 | 0.70000862 | 0.99945981 |
| ZNF791 | 17.6530283 | -0.462959053 | 0.43056274 | -1.0752418 | 0.28226649 | 0.99945981 |
| MAN2B1 | 17.7794464 | 0.152340925 | 0.56713816 | 0.26861343 | 0.78822718 | 0.99945981 |
| TNPO2 | 11.4745299 | -0.377357286 | 0.53875441 | -0.7004254 | 0.48366166 | 0.99945981 |
| TRIR | 32.2942173 | -0.213742726 | 0.50334678 | -0.4246431 | 0.67109689 | 0.99945981 |
| HOOK2 | 9.56368295 | -0.458036349 | 0.48967816 | -0.9353824 | 0.34959125 | 0.99945981 |
| JUNB | 37.1840094 | 1.253056018 | 0.56499343 | 2.21782405 | 0.02656683 | 0.8290059 |
| CALR | 40.3513339 | -0.304118376 | 0.53088101 | -0.572856 | 0.5667422 | 0.99945981 |
| RAD23A | 14.5056953 | -0.60914879 | 0.4700037 | -1.2960511 | 0.19495789 | 0.99945981 |
| GADD45GIP1 | 15.5148395 | -0.599346855 | 0.52931312 | -1.1323106 | 0.25750388 | 0.99945981 |
| LYL1 | 11.6223968 | 0.172068602 | 0.61001299 | 0.28207367 | 0.77788702 | 0.99945981 |
| IER2 | 25.0830881 | 0.354737769 | 0.32908792 | 1.07794225 | 0.28105953 | 0.99945981 |
| CCDC130 | 12.6873443 | -0.47163811 | 0.46733058 | -1.0092173 | 0.31287043 | 0.99945981 |
| C19orf53 | 17.2510834 | -0.204486865 | 0.58970636 | -0.3467605 | 0.72877126 | 0.99945981 |
| CC2D1A | 8.58761829 | 0.481034279 | 0.45315024 | 1.06153376 | 0.2884474 | 0.99945981 |
| RFX1 | 12.4671913 | 0.915008116 | 0.42790219 | 2.1383581 | 0.03248769 | 0.8663391 |
| IL27RA | 12.9564015 | 0.489184765 | 0.59616004 | 0.82055946 | 0.41189724 | 0.99945981 |
| PRKACA | 13.4561184 | 0.008661054 | 0.58631019 | 0.01477214 | 0.98821397 | 0.99945981 |
| ADGRE5 | 98.9967138 | -0.292817904 | 0.56355682 | -0.519589 | 0.60335009 | 0.99945981 |
| DDX39A | 9.00485703 | 0.462050892 | 0.44765466 | 1.03215923 | 0.30199753 | 0.99945981 |
| PKN1 | 32.4431989 | 0.005158939 | 0.47554342 | 0.01084851 | 0.99134431 | 0.99945981 |
| DNAJB1 | 15.0160949 | 0.536753673 | 0.76267467 | 0.70377802 | 0.48157102 | 0.99945981 |
| TECR | 12.490858 | 0.552899418 | 0.48271379 | 1.14539803 | 0.25204431 | 0.99945981 |
| NDUFB7 | 13.8418882 | -0.050608571 | 0.48094923 | -0.1052264 | 0.91619614 | 0.99945981 |
| ZNF333 | 10.4614181 | 0.259760247 | 0.45958554 | 0.56520544 | 0.57193404 | 0.99945981 |
| ADGRE2 | 53.7870068 | 0.106000426 | 0.52152751 | 0.20324992 | 0.8389397 | 0.99945981 |
| BRD4 | 63.9911404 | 0.005156469 | 0.27030757 | 0.0190763 | 0.98478023 | 0.99945981 |
| AKAP8 | 14.5410308 | -0.056996919 | 0.38220104 | -0.1491281 | 0.88145254 | 0.99945981 |
| AKAP8L | 28.2730766 | -0.394125967 | 0.40189745 | -0.980663 | 0.32675894 | 0.99945981 |
| RASAL3 | 27.6943892 | 0.088304258 | 0.32105025 | 0.27504809 | 0.78327929 | 0.99945981 |
| TPM4 | 58.9889862 | -0.284646013 | 0.45802175 | -0.6214683 | 0.53429152 | 0.99945981 |
| RAB8A | 11.9284289 | -0.316704898 | 0.48495191 | -0.6530645 | 0.51371467 | 0.99945981 |
| HSH2D | 29.4065699 | -0.190523535 | 0.4728609 | -0.4029167 | 0.68700953 | 0.99945981 |
| AP1M1 | 10.1604646 | 0.247804843 | 0.51546492 | 0.48074046 | 0.63070097 | 0.99945981 |
| KLF2 | 45.9772739 | 0.601236733 | 0.66926445 | 0.8983545 | 0.36899659 | 0.99945981 |
| EPS15L1 | 26.5192983 | -0.565175661 | 0.40535939 | -1.3942582 | 0.16323964 | 0.99945981 |
| SIN3B | 14.1496063 | 0.216889652 | 0.3556446 | 0.60984941 | 0.54196157 | 0.99945981 |
| MYO9B | 138.791321 | -0.12404425 | 0.47517744 | -0.2610483 | 0.79405528 | 0.99945981 |
| BST2 | 26.3102375 | 0.227997823 | 0.53425277 | 0.42676021 | 0.66955399 | 0.99945981 |
| SLC27A1 | 9.40381292 | -0.279571994 | 0.44146291 | -0.6332854 | 0.52654732 | 0.99945981 |
| PGLS | 14.0160471 | -0.133883324 | 0.54387715 | -0.2461646 | 0.80555478 | 0.99945981 |
| COLGALT1 | 21.4801696 | 0.117970094 | 0.54088708 | 0.21810485 | 0.82734742 | 0.99945981 |
| FCHO1 | 12.1965013 | 0.522767165 | 0.51891889 | 1.00741594 | 0.31373494 | 0.99945981 |
| JAK3 | 38.9378133 | -0.249127293 | 0.47913893 | -0.5199479 | 0.60309987 | 0.99945981 |
| RPL18A | 10.4889781 | -0.483952918 | 0.57694008 | -0.838827 | 0.4015664 | 0.99945981 |
| ARRDC2 | 15.2614583 | -0.038067792 | 0.35657774 | -0.1067587 | 0.91498038 | 0.99945981 |
| IL12RB1 | 10.2677035 | -0.011172949 | 0.47146072 | -0.0236986 | 0.98109304 | 0.99945981 |

| | | | | | | |
|---|---|---|---|---|---|---|
| MAST3 | 19.8963036 | -0.289339494 | 0.6085154 | -0.4754843 | 0.63444185 | 0.99945981 |
| JUND | 55.9291037 | 0.317731159 | 0.54306078 | 0.58507476 | 0.55849745 | 0.99945981 |
| LSM4 | 14.78379 | -0.188090244 | 0.41773394 | -0.4502633 | 0.65252063 | 0.99945981 |
| PGPEP1 | 15.3570428 | -0.528238449 | 0.51433645 | -1.027029 | 0.30440681 | 0.99945981 |
| LRRC25 | 18.0873861 | 0.981877492 | 0.73630803 | 1.33351458 | 0.18236299 | 0.99945981 |
| SSBP4 | 22.5953045 | -0.439651346 | 0.43291196 | -1.0155676 | 0.30983535 | 0.99945981 |
| ELL | 30.2447053 | -0.196543871 | 0.31808484 | -0.6178976 | 0.53664282 | 0.99945981 |
| FKBP8 | 14.5356069 | 0.491437483 | 0.50527248 | 0.97261873 | 0.33074283 | 0.99945981 |
| KXD1 | 8.10223498 | -0.34288846 | 0.49993805 | -0.6858619 | 0.4928002 | 0.99945981 |
| UBA52 | 47.0233384 | 0.386174572 | 0.56866901 | 0.67908496 | 0.49708403 | 0.99945981 |
| REX1BD | 10.4923287 | 0.264051418 | 0.45095377 | 0.58553989 | 0.55818475 | 0.99945981 |
| CRTC1 | 21.213854 | 0.410152468 | 0.42989469 | 0.95407661 | 0.34004486 | 0.99945981 |
| UPF1 | 14.2129474 | 0.093840943 | 0.38732988 | 0.24227654 | 0.80856589 | 0.99945981 |
| COPE | 14.5997853 | 0.094074561 | 0.3762339 | 0.25004276 | 0.80255428 | 0.99945981 |
| SUGP2 | 36.5059422 | -0.185536927 | 0.28288333 | -0.6558779 | 0.51190269 | 0.99945981 |
| SLC25A42 | 12.6464453 | -0.594760023 | 0.43524415 | -1.3664975 | 0.17178288 | 0.99945981 |
| SUGP1 | 7.98064724 | -0.188485198 | 0.45269588 | -0.4163616 | 0.6771454 | 0.99945981 |
| MAU2 | 23.4849867 | -0.269929355 | 0.29480619 | -0.9156163 | 0.35986817 | 0.99945981 |
| GATAD2A | 71.1396703 | 0.00031852 | 0.32997447 | 0.00096529 | 0.99922981 | 0.9997934 |
| PBX4 | 18.7709706 | -1.456868753 | 0.50616681 | -2.8782384 | 0.00399903 | 0.507383 |
| GMIP | 16.0978447 | 0.171591026 | 0.40560039 | 0.42305439 | 0.67225558 | 0.99945981 |
| ZNF253 | 10.2921354 | -0.702838679 | 0.54819194 | -1.2821033 | 0.19980642 | 0.99945981 |
| ZNF90 | 47.5923892 | -0.189208711 | 0.3944176 | -0.4797167 | 0.63142885 | 0.99945981 |
| ZNF430 | 26.7094122 | -0.304081092 | 0.37501442 | -0.8108517 | 0.41745082 | 0.99945981 |
| ZNF431 | 32.5218194 | -0.289483585 | 0.41263196 | -0.701554 | 0.48295738 | 0.99945981 |
| ZNF708 | 23.0441422 | 0.157874378 | 0.36885808 | 0.42800846 | 0.66864496 | 0.99945981 |
| ZNF429 | 9.93503241 | -0.240366785 | 0.74267474 | -0.3236501 | 0.74620292 | 0.99945981 |
| ZNF100 | 20.6295289 | -0.919595861 | 0.48644107 | -1.8904569 | 0.05869688 | 0.92445778 |
| ZNF43 | 17.1334377 | -0.143466062 | 0.41312807 | -0.3472678 | 0.72839016 | 0.99945981 |
| ZNF91 | 37.1863256 | -0.345138176 | 0.36652006 | -0.9416624 | 0.34636549 | 0.99945981 |
| ZNF675 | 10.7008405 | -0.317805272 | 0.53101997 | -0.5984808 | 0.54951915 | 0.99945981 |
| ZNF254 | 34.1965525 | -0.195106768 | 0.40376565 | -0.4832178 | 0.62894107 | 0.99945981 |
| LINC00662 | 10.5357707 | -0.511814133 | 0.5057046 | -1.0120812 | 0.31149921 | 0.99945981 |
| UQCRFS1 | 12.0415641 | -0.209968349 | 0.58441833 | -0.3592775 | 0.71938751 | 0.99945981 |
| C19orf12 | 10.8549905 | -0.227483083 | 0.58928837 | -0.3860302 | 0.69947433 | 0.99945981 |
| URI1 | 64.5757537 | -0.19336107 | 0.33828708 | -0.5715887 | 0.56760065 | 0.99945981 |
| TSHZ3 | 21.6061046 | -0.334872568 | 0.51879392 | -0.6454828 | 0.51861434 | 0.99945981 |
| ZNF507 | 11.7583981 | -0.375347615 | 0.6101949 | -0.6151274 | 0.53847058 | 0.99945981 |
| DPY19L3 | 12.2482257 | -0.459811242 | 0.43809969 | -1.0495585 | 0.29392116 | 0.99945981 |
| ANKRD27 | 21.0404994 | -0.460576648 | 0.50799389 | -0.9066579 | 0.36458776 | 0.99945981 |
| CEP89 | 10.2905248 | 0.055489684 | 0.51580358 | 0.1075791 | 0.91432958 | 0.99945981 |
| PEPD | 16.3513741 | 0.046374985 | 0.59916016 | 0.07739998 | 0.93830536 | 0.99945981 |
| LSM14A | 89.8406269 | -0.11597248 | 0.28239102 | -0.4106805 | 0.68130684 | 0.99945981 |
| KIAA0355 | 25.0349774 | 0.189372932 | 0.36211838 | 0.52295862 | 0.60100305 | 0.99945981 |
| GPI | 22.5418618 | -1.187501233 | 0.44969444 | -2.6406847 | 0.00827387 | 0.58811673 |
| UBA2 | 34.1096823 | 0.117124061 | 0.27798487 | 0.4213325 | 0.6735123 | 0.99945981 |
| ZNF302 | 15.5306897 | -0.026438874 | 0.48312559 | -0.0547246 | 0.95635784 | 0.99945981 |
| GRAMD1A | 21.1010735 | -0.28031004 | 0.52481657 | -0.5341105 | 0.59326509 | 0.99945981 |
| FXYD5 | 27.8369786 | 0.167919282 | 0.6156105 | 0.27276871 | 0.78503101 | 0.99945981 |
| USF2 | 19.4658883 | -0.201934974 | 0.47961096 | -0.4210391 | 0.67372652 | 0.99945981 |
| COX6B1 | 25.427546 | 0.419037528 | 0.46890186 | 0.89365721 | 0.37150533 | 0.99945981 |
| KMT2B | 14.1350097 | -0.508266766 | 0.39789733 | -1.2773817 | 0.20146752 | 0.99945981 |
| NFKBID | 25.6829705 | 0.921293858 | 0.57451703 | 1.60359713 | 0.10880288 | 0.99945981 |
| HCST | 24.6350977 | -0.231094071 | 0.61509292 | -0.375706 | 0.70713551 | 0.99945981 |
| TYROBP | 54.8152855 | 0.880949783 | 0.74395639 | 1.18414169 | 0.23635698 | 0.99945981 |
| TBCB | 8.6479501 | 0.006922156 | 0.4933155 | 0.0140319 | 0.98880453 | 0.99945981 |
| CAPNS1 | 26.9450867 | -0.298125141 | 0.42425054 | -0.7027101 | 0.48223643 | 0.99945981 |
| ZNF565 | 15.1592101 | -0.075905919 | 0.4997423 | -0.1518901 | 0.8792736 | 0.99945981 |
| ZNF146 | 14.9588354 | 0.380095667 | 0.42910962 | 0.88577755 | 0.37573741 | 0.99945981 |
| ZFP14 | 17.1775262 | -0.072188079 | 0.41855281 | -0.1724707 | 0.86306753 | 0.99945981 |
| ZNF566 | 8.65375013 | -0.854695967 | 0.60869268 | -1.4041502 | 0.16027412 | 0.99945981 |
| ZNF529 | 12.5646466 | -0.140488913 | 0.56664941 | -0.2479292 | 0.80418922 | 0.99945981 |
| ZNF420 | 9.72113664 | -0.029707639 | 0.46104933 | -0.0644348 | 0.94862399 | 0.99945981 |
| HKR1 | 13.1832461 | -0.291123819 | 0.41292483 | -0.7050286 | 0.48079243 | 0.99945981 |
| SIPA1L3 | 98.4161992 | -0.422980493 | 0.50116516 | -0.8439942 | 0.39867264 | 0.99945981 |
| KCNK6 | 12.8123271 | 0.071688129 | 0.63561672 | 0.11278515 | 0.91020089 | 0.99945981 |
| PSMD8 | 15.5043584 | -0.430902832 | 0.46199581 | -0.9326986 | 0.35097562 | 0.99945981 |
| RYR1 | 11.7015147 | 0.440680445 | 0.60345555 | 0.73026165 | 0.46523026 | 0.99945981 |
| MAP4K1 | 10.8372612 | -0.357374534 | 0.56038235 | -0.6377334 | 0.52364725 | 0.99945981 |
| EIF3K | 18.0702218 | 0.055011684 | 0.47410097 | 0.11603369 | 0.90762584 | 0.99945981 |
| ACTN4 | 65.3956298 | -0.262753086 | 0.27056224 | -0.9711373 | 0.3314799 | 0.99945981 |
| ECH1 | 11.0399771 | -0.681141943 | 0.48298472 | -1.4102764 | 0.15845809 | 0.99945981 |
| HNRNPL | 23.1278837 | 0.078374061 | 0.32023796 | 0.24473695 | 0.80666012 | 0.99945981 |
| SIRT2 | 12.4531332 | -0.30685003 | 0.43775321 | -0.7009658 | 0.48332436 | 0.99945981 |
| GMFG | 23.5044624 | 0.137817635 | 0.33385392 | 0.41280819 | 0.67974715 | 0.99945981 |
| SAMD4B | 26.0778295 | -0.58680654 | 0.36202895 | -1.6208829 | 0.10504275 | 0.99945981 |
| MED29 | 9.50175131 | -0.535484475 | 0.66657736 | -0.8033343 | 0.42178152 | 0.99945981 |
| ZFP36 | 45.509434 | 0.709698907 | 0.59275756 | 1.19728361 | 0.23119603 | 0.99945981 |
| RPS16 | 60.6286036 | 0.141689825 | 0.55352984 | 0.25597504 | 0.79797011 | 0.99945981 |
| SUPT5H | 21.1235737 | 0.134760854 | 0.35606723 | 0.37847025 | 0.70508129 | 0.99945981 |
| ZNF780B | 23.3245608 | 0.05733443 | 0.31390983 | 0.18264617 | 0.85507565 | 0.99945981 |
| AKT2 | 17.5847926 | -0.169003526 | 0.44857702 | -0.3767548 | 0.70635586 | 0.99945981 |
| PLD3 | 31.7635516 | -0.591527936 | 0.41088051 | -1.4396593 | 0.14996382 | 0.99945981 |
| BLVRB | 22.2862529 | -0.450310772 | 0.38184263 | -1.1793099 | 0.23827482 | 0.99945981 |
| SHKBP1 | 12.1251082 | 0.47700944 | 0.51867859 | 0.91966287 | 0.35774896 | 0.99945981 |
| SNRPA | 11.484031 | -0.190519505 | 0.39691858 | -0.4799964 | 0.63122993 | 0.99945981 |
| AXL | 12.0979598 | 0.377786692 | 0.67783872 | 0.55734009 | 0.57729509 | 0.99945981 |
| HNRNPUL1 | 26.0748069 | -0.229291294 | 0.35158457 | -0.6521654 | 0.51429447 | 0.99945981 |
| TGFB1 | 58.5256443 | 0.166828528 | 0.34438869 | 0.4844193 | 0.62808833 | 0.99945981 |
| RPS19 | 254.054996 | -0.094015058 | 0.60857693 | -0.1544834 | 0.87722857 | 0.99945981 |
| ARHGEF1 | 43.2434992 | 0.126457742 | 0.27870337 | 0.45373597 | 0.65001888 | 0.99945981 |
| RABAC1 | 22.2550244 | 0.200117306 | 0.64531264 | 0.31010908 | 0.75647801 | 0.99945981 |
| POU2F2 | 126.936333 | -0.676766794 | 0.95662458 | -0.7074529 | 0.4792851 | 0.99945981 |
| LIPE-AS1 | 14.2559972 | -0.24868422 | 0.48917756 | -0.5083721 | 0.61119242 | 0.99945981 |
| PLAUR | 65.4725239 | 1.233587942 | 0.83542157 | 1.47660533 | 0.13978146 | 0.99945981 |
| SMG9 | 10.8574882 | -0.171592164 | 0.66048075 | -0.2597989 | 0.79501891 | 0.99945981 |
| ZNF224 | 11.7675121 | -0.164184907 | 0.41407998 | -0.3965053 | 0.6917323 | 0.99945981 |
| ZNF235 | 17.6768954 | -0.052276203 | 0.4384222 | -0.1192371 | 0.90508749 | 0.99945981 |
| BCL3 | 8.91640087 | -0.690885624 | 0.53667417 | -1.2873465 | 0.19797354 | 0.99945981 |
| NECTIN2 | 12.4010287 | -0.291403029 | 0.67339931 | -0.4327344 | 0.66520775 | 0.99945981 |
| APOE | 97.5159736 | -1.862220881 | 1.01499757 | -1.8347048 | 0.06654945 | 0.94594189 |
| CLPTM1 | 12.4210027 | -0.470129851 | 0.40578258 | -1.1585757 | 0.24662917 | 0.99945981 |
| RELB | 19.9762701 | 0.027277192 | 0.71073915 | 0.03837863 | 0.9693858 | 0.99945981 |
| CLASRP | 25.2740196 | 0.138444822 | 0.37886006 | 0.3654247 | 0.71479441 | 0.99945981 |
| MARK4 | 20.7439693 | -0.422041143 | 0.39109731 | -1.0791205 | 0.280534 | 0.99945981 |
| ERCC1 | 7.75538134 | 0.208089019 | 0.52065126 | 0.39967063 | 0.68939912 | 0.99945981 |
| VASP | 35.6180083 | 0.489559712 | 0.50128747 | 0.97660472 | 0.32876488 | 0.99945981 |
| OPA3 | 9.10109221 | -0.393551069 | 0.56015826 | -0.7025712 | 0.48232301 | 0.99945981 |
| EML2 | 11.799935 | -0.410795872 | 0.42618314 | -0.9638952 | 0.33509848 | 0.99945981 |
| SNRPD2 | 17.1561323 | -0.466516322 | 0.64472862 | -0.7235856 | 0.46932021 | 0.99945981 |
| SYMPK | 30.3610104 | 0.162536771 | 0.37638149 | 0.43184051 | 0.66585734 | 0.99945981 |
| PPP5C | 12.4825686 | 0.29442176 | 0.42197123 | 0.69772946 | 0.4853464 | 0.99945981 |
| CALM3 | 16.5843949 | 0.280750734 | 0.63550519 | 0.44177568 | 0.65865154 | 0.99945981 |

| | | | | | | |
|---|---|---|---|---|---|---|
| PRKD2 | 28.0369487 | -0.76424448 | 0.52272713 | -1.4620333 | 0.14373208 | 0.99945981 |
| AP2S1 | 21.3821152 | 0.360620919 | 0.53840669 | 0.66979278 | 0.50298989 | 0.99945981 |
| ARHGAP35 | 39.7063777 | -0.365326941 | 0.31334376 | -1.1658983 | 0.24365559 | 0.99945981 |
| TMEM160 | 12.8383144 | -0.259389059 | 0.54584122 | -0.4752097 | 0.63463749 | 0.99945981 |
| ZC3H4 | 13.1686784 | 0.288646862 | 0.40625148 | 0.71051276 | 0.47738622 | 0.99945981 |
| SAE1 | 17.6449511 | -0.126869656 | 0.5379998 | -0.2358173 | 0.81357445 | 0.99945981 |
| C5AR1 | 44.8228985 | 1.043196838 | 0.65634672 | 1.58939904 | 0.11197033 | 0.99945981 |
| DHX34 | 13.7128996 | 0.58966554 | 0.43560266 | 1.35367756 | 0.17583927 | 0.99945981 |
| NAPA | 18.4803918 | -0.078216333 | 0.54588718 | -0.143283 | 0.88606671 | 0.99945981 |
| NOP53 | 11.82462 | 0.146572 | 0.56920114 | 0.25750476 | 0.79678914 | 0.99945981 |
| SELENOW | 19.2646005 | -0.245852589 | 0.38518362 | -0.6382737 | 0.52329551 | 0.99945981 |
| CARD8 | 74.511901 | 0.037592687 | 0.25736537 | 0.14606739 | 0.88386819 | 0.99945981 |
| EMP3 | 59.6781395 | -0.375184808 | 0.81956888 | -0.4577831 | 0.64710825 | 0.99945981 |
| KDELR1 | 9.08827787 | -0.437044872 | 0.50731243 | -0.8614906 | 0.38896791 | 0.99945981 |
| RPL18 | 76.5064335 | 0.213056946 | 0.52275048 | 0.40756911 | 0.68359005 | 0.99945981 |
| NUCB1 | 20.2408472 | -0.577765646 | 0.4368767 | -1.3224913 | 0.18600459 | 0.99945981 |
| BAX | 17.8098587 | -0.016676806 | 0.50077624 | -0.0333019 | 0.97343383 | 0.99945981 |
| FTL | 806.861412 | -0.413982995 | 0.66261645 | -0.6247702 | 0.53212191 | 0.99945981 |
| GYS1 | 14.4831094 | -0.501807421 | 0.50529155 | -0.9931047 | 0.32065893 | 0.99945981 |
| SNRNP70 | 46.6439867 | 0.467687248 | 0.28662561 | 1.63170085 | 0.10274252 | 0.99945981 |
| CD37 | 40.3453711 | 0.117899839 | 0.62689655 | 0.18806905 | 0.85082252 | 0.99945981 |
| PIH1D1 | 8.9167849 | 0.381121337 | 0.52500768 | 0.72593479 | 0.46787875 | 0.99945981 |
| RPL13A | 106.402612 | -0.202090355 | 0.47377893 | -0.4265499 | 0.6697072 | 0.99945981 |
| RPS11 | 78.9815612 | -0.112616242 | 0.5611553 | -0.2006864 | 0.84094379 | 0.99945981 |
| FCGRT | 39.3779201 | 0.503316863 | 0.48351449 | 1.04095508 | 0.2978964 | 0.99945981 |
| NOSIP | 11.777514 | 0.466142903 | 0.46005529 | 1.01323235 | 0.31094919 | 0.99945981 |
| PRR12 | 13.1745559 | -0.192744903 | 0.46978896 | -0.4102798 | 0.68160074 | 0.99945981 |
| IRF3 | 21.5331142 | 0.212935727 | 0.33374876 | 0.63801205 | 0.52346584 | 0.99945981 |
| PRMT1 | 9.60679497 | -0.225158244 | 0.45123935 | -0.4989774 | 0.6177953 | 0.99945981 |
| AP2A1 | 18.8874921 | -0.192409745 | 0.52031377 | -0.3697956 | 0.71153479 | 0.99945981 |
| MED25 | 11.222086 | 0.217407158 | 0.49459079 | 0.43956976 | 0.66024874 | 0.99945981 |
| PNKP | 11.3555728 | -0.959338005 | 0.59924741 | -1.6009047 | 0.10939803 | 0.99945981 |
| VRK3 | 11.7785315 | 0.024764637 | 0.47102028 | 0.05257658 | 0.95806928 | 0.99945981 |
| NR1H2 | 9.88071188 | -0.372109579 | 0.47321926 | -0.7863365 | 0.43167037 | 0.99945981 |
| EMC10 | 18.9033772 | 0.2630852 | 0.40674064 | 0.64681316 | 0.51775288 | 0.99945981 |
| SIGLEC9 | 12.3413673 | 0.231446032 | 0.68822265 | 0.33629528 | 0.73664821 | 0.99945981 |
| ETFB | 14.5043032 | -0.507831085 | 0.50204153 | -1.011532 | 0.31176187 | 0.99945981 |
| NKG7 | 50.7548383 | -0.143374885 | 0.9014206 | -0.1590544 | 0.87362604 | 0.99945981 |
| SIGLEC10 | 18.8942396 | -0.130464137 | 0.65266968 | -0.1998931 | 0.84156422 | 0.99945981 |
| FPR1 | 21.4613297 | 0.348321522 | 0.6421466 | 0.54243302 | 0.58752024 | 0.99945981 |
| FPR3 | 30.5545493 | 0.50433236 | 0.8142167 | 0.61940803 | 0.5356476 | 0.99945981 |
| ZNF577 | 8.70389996 | -1.239965862 | 0.79994588 | -1.5500622 | 0.12112659 | 0.99945981 |
| PPP2R1A | 22.6770142 | -0.538495819 | 0.38954596 | -1.3823679 | 0.16685877 | 0.99945981 |
| ZNF766 | 9.18845973 | 0.303670454 | 0.48041294 | 0.63210298 | 0.52731959 | 0.99945981 |
| ZNF83 | 32.1274541 | 0.286147488 | 0.33227074 | 0.86118773 | 0.38913466 | 0.99945981 |
| ZNF611 | 22.1328148 | -0.437027664 | 0.30308593 | -1.4419266 | 0.14932308 | 0.99945981 |
| ZNF160 | 17.832572 | 0.000567651 | 0.5341259 | 0.00106277 | 0.99915203 | 0.9997934 |
| ZNF761 | 11.3721051 | -0.256500182 | 0.39911685 | -0.6426694 | 0.52043865 | 0.99945981 |
| ZNF331 | 42.748328 | -0.301993701 | 0.88836736 | -0.3399424 | 0.73389993 | 0.99945981 |
| MYADM | 14.9793476 | 0.308826942 | 0.78256409 | 0.3946347 | 0.6931125 | 0.99945981 |
| OSCAR | 11.049784 | 0.473072612 | 0.72517417 | 0.65235723 | 0.51417075 | 0.99945981 |
| PRPF31 | 9.7875479 | -0.050599335 | 0.54179414 | -0.0933922 | 0.925592 | 0.99945981 |
| MBOAT7 | 19.0538917 | -0.46099442 | 0.40350806 | -1.1424664 | 0.25326018 | 0.99945981 |
| RPS9 | 163.188673 | -0.018584867 | 0.47886987 | -0.0388098 | 0.96904199 | 0.99945981 |
| LILRB3 | 16.7607133 | -0.043355761 | 0.59228145 | -0.0732013 | 0.94164595 | 0.99945981 |
| LILRA6 | 12.6979771 | 1.451986121 | 0.89901421 | 1.61508696 | 0.10629186 | 0.99945981 |
| LILRB2 | 30.8553083 | 0.037285532 | 0.91840918 | 0.04059795 | 0.96761642 | 0.99945981 |
| LAIR1 | 22.5925992 | 0.277096607 | 0.5044161 | 0.54934133 | 0.58277123 | 0.99945981 |
| LENG8 | 34.2868218 | 0.479890182 | 0.4043726 | 1.18675247 | 0.23532527 | 0.99945981 |
| LILRB1 | 42.0346912 | -0.057910199 | 0.69317256 | -0.0835437 | 0.93341923 | 0.99945981 |
| LILRB4 | 10.8934043 | -0.452733997 | 0.72371094 | -0.625573 | 0.53159508 | 0.99945981 |
| PPP1R12C | 15.2015816 | 0.073020635 | 0.38834806 | 0.18802884 | 0.85085404 | 0.99945981 |
| PPP6R1 | 30.3109691 | 0.020957462 | 0.32488694 | 0.06450694 | 0.94856658 | 0.99945981 |
| RPL28 | 121.905804 | -0.288437539 | 0.44974288 | -0.6413388 | 0.52130262 | 0.99945981 |
| U2AF2 | 9.81678034 | 0.485777414 | 0.4775339 | 1.01726267 | 0.30902849 | 0.99945981 |
| EPN1 | 13.6342253 | 0.28285675 | 0.42730338 | 0.66195767 | 0.50799835 | 0.99945981 |
| ZNF787 | 10.3792675 | 0.053439066 | 0.42627018 | 0.12536431 | 0.90023514 | 0.99945981 |
| ZSCAN5A | 16.9417725 | -0.520213473 | 0.63831771 | -0.8149758 | 0.41508617 | 0.99945981 |
| ZNF264 | 9.9597382 | 0.135420833 | 0.41352306 | 0.32748073 | 0.74330432 | 0.99945981 |
| AC005261.1 | 11.3465823 | 0.259485224 | 0.42463391 | 0.61107985 | 0.54114672 | 0.99945981 |
| ZNF586 | 24.2947387 | -0.158042022 | 0.47830665 | -0.3304199 | 0.74108273 | 0.99945981 |
| ZNF814 | 16.786129 | 0.415448893 | 0.43796149 | 0.94859686 | 0.34282569 | 0.99945981 |
| ZNF274 | 12.7632363 | -0.780666721 | 0.45814758 | -1.7039634 | 0.08838792 | 0.99640647 |
| RPS5 | 40.477814 | -0.680478605 | 0.67788387 | -1.0038277 | 0.31546167 | 0.99945981 |
| TRIM28 | 11.7224761 | -0.339213094 | 0.57378913 | -0.5911808 | 0.55439932 | 0.99945981 |
| CHMP2A | 7.59080368 | -0.214769268 | 0.59369191 | -0.3617521 | 0.71753733 | 0.99945981 |
| MZF1 | 9.66460134 | -0.535457012 | 0.50985906 | -1.0502059 | 0.29362344 | 0.99945981 |
| RPS4Y1 | 25.2262138 | -0.809453056 | 1.86006621 | -0.4351743 | 0.6634359 | 0.99945981 |
| ZFY | 12.2147676 | -0.697466355 | 1.20105834 | -0.5807098 | 0.56143605 | 0.99945981 |
| LINC00278 | 90.8510803 | 0.16661683 | 2.09720083 | 0.07944725 | 0.93667689 | 0.99945981 |
| TTTY15 | 9.34704079 | -0.575804313 | 1.22884358 | -0.4685741 | 0.63937407 | 0.99945981 |
| USP9Y | 46.2959158 | -0.727879444 | 1.90528348 | -0.3820321 | 0.70243756 | 0.99945981 |
| DDX3Y | 17.0475245 | -0.098207593 | 1.69893771 | -0.0578053 | 0.95390372 | 0.99945981 |
| UTY | 101.332309 | -0.891235548 | 2.09086837 | -0.4262514 | 0.66992467 | 0.99945981 |
| TTTY14 | 29.9801024 | 0.048150497 | 0.94973703 | 0.05069877 | 0.95956556 | 0.99945981 |
| KDM5D | 14.4716788 | -0.377816311 | 1.7521284 | -0.2156328 | 0.829274 | 0.99945981 |
| IL17RA | 55.6728486 | 0.164864962 | 0.55982975 | 0.29449125 | 0.76838255 | 0.99945981 |
| HDHD5 | 10.120865 | 0.200384211 | 0.56278945 | 0.35605538 | 0.72179909 | 0.99945981 |
| ADA2 | 33.2988863 | 0.677593979 | 0.52884164 | 1.28127955 | 0.20009549 | 0.99945981 |
| ATP6V1E1 | 19.3102737 | -0.605230453 | 0.39750155 | -1.5225864 | 0.12786222 | 0.99945981 |
| BCL2L13 | 24.5159885 | 0.48100494 | 0.40222822 | 1.19585081 | 0.23175478 | 0.99945981 |
| BID | 30.2622214 | 0.908112766 | 0.48373333 | 1.8773004 | 0.06047693 | 0.92445778 |
| MICAL3 | 29.479556 | -0.737803227 | 0.49846242 | -1.4801582 | 0.13883104 | 0.99945981 |
| PEX26 | 11.5415801 | -0.401592607 | 0.55041605 | -0.7296165 | 0.46562466 | 0.99945981 |
| DGCR2 | 28.5332041 | -0.415758921 | 0.31030574 | -1.3398364 | 0.18029854 | 0.99945981 |
| HIRA | 12.0540559 | -0.084319339 | 0.59721127 | -0.1411885 | 0.88772106 | 0.99945981 |
| UFD1 | 11.6241958 | -0.26343233 | 0.44916383 | -0.586495 | 0.55754294 | 0.99945981 |
| TXNRD2 | 10.0669921 | -0.198887848 | 0.40148773 | -0.4953772 | 0.62033392 | 0.99945981 |
| COMT | 25.3338259 | -0.031580721 | 0.43395916 | -0.0727735 | 0.94198637 | 0.99945981 |
| TANGO2 | 20.8763168 | 0.391844229 | 0.36259637 | 1.08066232 | 0.27984735 | 0.99945981 |
| DGCR8 | 8.85229157 | -0.261102893 | 0.4666127 | -0.5595709 | 0.57577215 | 0.99945981 |
| KLHL22 | 19.9508234 | -0.546814925 | 0.33367723 | -1.6387541 | 0.10126448 | 0.99945981 |
| MED15 | 50.1318495 | 0.116773978 | 0.27928612 | 0.41811594 | 0.67586235 | 0.99945981 |
| PI4KA | 61.2246906 | 0.029146784 | 0.24354926 | 0.11967511 | 0.90474052 | 0.99945981 |
| SNAP29 | 11.7947179 | -0.748438281 | 0.39065088 | -1.9158751 | 0.055381 | 0.92445778 |
| CRKL | 11.1781156 | -0.284649769 | 0.40480872 | -0.703171 | 0.48194917 | 0.99945981 |
| UBE2L3 | 37.8627683 | -0.115309079 | 0.30896355 | -0.3732126 | 0.70899024 | 0.99945981 |
| SDF2L1 | 12.2439717 | -0.736126258 | 0.82795329 | -0.8890915 | 0.37395389 | 0.99945981 |
| PPIL2 | 22.2583461 | -0.296676071 | 0.34779935 | -0.8530093 | 0.39365416 | 0.99945981 |
| YPEL1 | 15.179688 | -0.823237916 | 0.45919669 | -1.7927784 | 0.07300836 | 0.96364229 |
| MAPK1 | 103.777859 | 0.303653073 | 0.29701318 | 1.02235556 | 0.30661264 | 0.99945981 |
| PPM1F | 18.840886 | 0.728483577 | 0.67653271 | 1.07678988 | 0.28157415 | 0.99945981 |
| IGLC2 | 164.887789 | -1.32243561 | 2.4174232 | -0.5470435 | 0.58434886 | 0.99945981 |

| | | | | | | |
|---|---|---|---|---|---|---|
| BCR | 37.1289888 | -0.723255303 | 0.51775747 | -1.3968998 | 0.16244371 | 0.99945981 |
| CHCHD10 | 16.7366423 | -0.568811901 | 0.46427565 | -1.2251599 | 0.22051498 | 0.99945981 |
| SMARCB1 | 13.3169225 | -0.534271494 | 0.54120671 | -0.9871856 | 0.32355164 | 0.99945981 |
| DDT | 14.4418572 | -0.441052299 | 0.61770035 | -0.7140231 | 0.47521291 | 0.99945981 |
| CABIN1 | 31.5187987 | -0.287764361 | 0.28242325 | -1.0189117 | 0.30824487 | 0.99945981 |
| SNRPD3 | 15.9696478 | 0.001672926 | 0.39453563 | 0.00424024 | 0.99661679 | 0.99945981 |
| KIAA1671 | 10.7483885 | -0.505980917 | 0.58097687 | -0.870914 | 0.38380109 | 0.99945981 |
| LRP5L | 9.64159825 | -0.922895348 | 0.47379294 | -1.9478875 | 0.05142843 | 0.9230968 |
| GRK3 | 154.723383 | 0.786898579 | 0.37338495 | 2.10747267 | 0.03507663 | 0.87920079 |
| HPS4 | 11.1075952 | -0.686244228 | 0.44730605 | -1.5341716 | 0.12498745 | 0.99945981 |
| TPST2 | 36.2747429 | 0.775446914 | 0.38442003 | 2.01718656 | 0.04367605 | 0.89562588 |
| MIAT | 12.5331059 | -1.232443679 | 0.57688546 | -2.136375 | 0.03264886 | 0.8663391 |
| MIATNB | 27.452459 | -0.131452767 | 0.36677875 | -0.358398 | 0.7200455 | 0.99945981 |
| PITPNB | 42.7770066 | -0.221235726 | 0.26202126 | -0.8443426 | 0.39847796 | 0.99945981 |
| TTC28 | 40.1001122 | -0.908935581 | 0.76242547 | -1.1921632 | 0.23319726 | 0.99945981 |
| HSCB | 9.92626822 | -0.736662541 | 0.54593653 | -1.3493557 | 0.17722276 | 0.99945981 |
| XBP1 | 40.400191 | -0.949857039 | 0.93516678 | -1.0157087 | 0.30976812 | 0.99945981 |
| Z93930.2 | 18.4205371 | -0.627035054 | 0.35599014 | -1.7613832 | 0.07817357 | 0.98289661 |
| EWSR1 | 46.0951872 | 0.165773131 | 0.28937659 | 0.57286297 | 0.56673748 | 0.99945981 |
| AP1B1 | 37.5778288 | 0.023526428 | 0.31011384 | 0.07586384 | 0.93952742 | 0.99945981 |
| NF2 | 15.7354208 | -0.58160285 | 0.38991687 | -1.4916073 | 0.13580213 | 0.99945981 |
| UQCR10 | 26.5655589 | -0.342629762 | 0.54730359 | -0.6260324 | 0.53129371 | 0.99945981 |
| ASCC2 | 17.9298842 | -0.062893152 | 0.43206084 | -0.1455655 | 0.88426441 | 0.99945981 |
| MTMR3 | 99.3031316 | 0.456794251 | 0.22835459 | 2.00037252 | 0.04546005 | 0.90395454 |
| TBC1D10A | 9.02879587 | -0.188065459 | 0.51362558 | -0.3661528 | 0.71425104 | 0.99945981 |
| SF3A1 | 11.0952523 | 0.584343588 | 0.41406689 | 1.41122993 | 0.15817683 | 0.99945981 |
| DUSP18 | 11.6405569 | 0.126559705 | 0.56011894 | 0.22595148 | 0.82123916 | 0.99945981 |
| RNF185 | 14.1448689 | -0.225080289 | 0.42103395 | -0.5345894 | 0.59293381 | 0.99945981 |
| LIMK2 | 38.4347045 | -0.349434448 | 0.50584917 | -0.6907878 | 0.48969889 | 0.99945981 |
| PIK3IP1 | 22.2850756 | -1.871643758 | 0.5355494 | -3.4948107 | 0.0004744 | 0.23820088 |
| PIK3IP1-AS1 | 11.4869346 | -0.46457668 | 0.51968282 | -0.893962 | 0.37134223 | 0.99945981 |
| DRG1 | 14.7985445 | -0.110151457 | 0.39616523 | -0.2780442 | 0.78097841 | 0.99945981 |
| EIF4ENIF1 | 15.8800399 | -0.651581907 | 0.41550257 | -1.5681778 | 0.11683965 | 0.99945981 |
| SFI1 | 28.3203343 | -0.150730446 | 0.31792358 | -0.4741091 | 0.63542214 | 0.99945981 |
| PISD | 20.4998855 | 0.054241309 | 0.40384807 | 0.13431118 | 0.89315651 | 0.99945981 |
| PRR14L | 47.5897245 | -0.179203879 | 0.27824845 | -0.6440427 | 0.51954776 | 0.99945981 |
| YWHAH | 11.3270308 | -1.046330463 | 1.0378494 | -1.0081718 | 0.31337201 | 0.99945981 |
| RTCB | 21.3426364 | 0.505227797 | 0.40028137 | 1.26218165 | 0.20688343 | 0.99945981 |
| FBXO7 | 25.1468136 | 0.326587966 | 0.31942663 | 1.02241936 | 0.30658246 | 0.99945981 |
| TIMP3 | 15.1075271 | 0.248717525 | 0.54276263 | 0.45824364 | 0.64677741 | 0.99945981 |
| LARGE1 | 42.1304307 | -2.160669893 | 1.11749999 | -1.9334854 | 0.05317643 | 0.92445778 |
| HMGXB4 | 13.4962199 | -0.130885549 | 0.36830514 | -0.3553726 | 0.72231049 | 0.99945981 |
| TOM1 | 22.6379274 | 0.078327135 | 0.41617973 | 0.18820507 | 0.8507159 | 0.99945981 |
| HMOX1 | 52.1977821 | -0.665599354 | 0.64613236 | -1.0301285 | 0.30294969 | 0.99945981 |
| APOL6 | 32.2931373 | 0.008879046 | 0.47730485 | 0.01860246 | 0.98515824 | 0.99945981 |
| RBFOX2 | 17.182884 | -0.131678282 | 0.44373102 | -0.2967525 | 0.76665548 | 0.99945981 |
| APOL3 | 14.1487484 | 0.189104302 | 0.4772884 | 0.39620553 | 0.69195341 | 0.99945981 |
| APOL2 | 13.0479213 | 0.239120591 | 0.56479477 | 0.42337607 | 0.6720209 | 0.99945981 |
| MYH9 | 116.192728 | -0.236848436 | 0.32449023 | -0.7299093 | 0.46544563 | 0.99945981 |
| TXN2 | 9.53007393 | -0.354881719 | 0.50625005 | -0.7010009 | 0.48330248 | 0.99945981 |
| EIF3D | 20.5712259 | -0.053148633 | 0.35582646 | -0.1493667 | 0.88126427 | 0.99945981 |
| NCF4 | 16.293769 | -0.074834347 | 0.50528778 | -0.1481024 | 0.88226193 | 0.99945981 |
| IL2RB | 13.7901517 | -0.03122692 | 0.65654374 | -0.0475626 | 0.96206485 | 0.99945981 |
| RAC2 | 36.0404739 | -0.075030482 | 0.58843725 | -0.127508 | 0.89853831 | 0.99945981 |
| CYTH4 | 49.9911606 | 0.047377466 | 0.46939664 | 0.10093269 | 0.91960389 | 0.99945981 |
| MFNG | 8.79980223 | 0.739962196 | 0.4779534 | 1.548189 | 0.1215768 | 0.99945981 |
| GGA1 | 24.7206425 | -0.225242027 | 0.36048734 | -0.6248265 | 0.53208497 | 0.99945981 |
| LGALS1 | 49.8045515 | -0.398394088 | 0.78213517 | -0.5093673 | 0.61049479 | 0.99945981 |
| TRIOBP | 15.6881113 | -0.252663199 | 0.39142467 | -0.6454964 | 0.51860558 | 0.99945981 |
| EIF3L | 28.9634477 | 0.524586066 | 0.35454679 | 1.47959615 | 0.13898105 | 0.99945981 |
| POLR2F | 8.43170989 | 0.077707539 | 0.4794107 | 0.1620897 | 0.87123521 | 0.99945981 |
| TMEM184B | 13.4740375 | -0.880749395 | 0.54493304 | -1.6162525 | 0.10603973 | 0.99945981 |
| TPTEP2-CSNI | 36.3585052 | -0.21577447 | 0.45625004 | -0.4729303 | 0.63626291 | 0.99945981 |
| DDX17 | 181.412545 | 0.259968987 | 0.22994285 | 1.13058088 | 0.25823154 | 0.99945981 |
| JOSD1 | 19.894369 | -0.258418645 | 0.45076652 | -0.5732871 | 0.5664503 | 0.99945981 |
| GTPBP1 | 12.9785444 | -0.004189062 | 0.47107264 | -0.0088926 | 0.99290482 | 0.99945981 |
| SUN2 | 45.5913453 | 0.109522931 | 0.38811561 | 0.28219151 | 0.77779667 | 0.99945981 |
| AL021707.1 | 11.1156748 | -1.181692237 | 0.49413764 | -2.3914232 | 0.01678319 | 0.74280553 |
| CBX6 | 12.9811777 | -0.36646288 | 0.48546342 | -0.7548723 | 0.45032561 | 0.99945981 |
| APOBEC3G | 12.6110093 | -0.660237507 | 0.4474589 | -1.4755266 | 0.14007102 | 0.99945981 |
| CBX7 | 12.093087 | -0.488128698 | 0.47455566 | -1.0286016 | 0.30366694 | 0.99945981 |
| RPL3 | 63.9369394 | -0.122927544 | 0.57795461 | -0.2126941 | 0.83156556 | 0.99945981 |
| TAB1 | 7.3832188 | 0.108831476 | 0.452286 | 0.24062535 | 0.8098455 | 0.99945981 |
| ATF4 | 19.4794558 | -0.25491969 | 0.45046736 | -0.5659005 | 0.57146144 | 0.99945981 |
| RPS19BP1 | 13.2054004 | -0.018742323 | 0.51691926 | -0.0362577 | 0.97107685 | 0.99945981 |
| GRAP2 | 77.0581675 | -0.742104768 | 0.55012407 | -1.3489771 | 0.17734432 | 0.99945981 |
| TNRC6B | 282.503119 | -0.105914014 | 0.18483756 | -0.5730113 | 0.56663702 | 0.99945981 |
| MKL1 | 119.767749 | 0.2987249 | 0.20648191 | 1.44673641 | 0.14797076 | 0.99945981 |
| SLC25A17 | 14.6597018 | -0.24662113 | 0.47385371 | -0.5204584 | 0.60274414 | 0.99945981 |
| ST13 | 23.0536786 | -0.233575402 | 0.40615439 | -0.5750902 | 0.56523031 | 0.99945981 |
| XPNPEP3 | 12.8723861 | -0.160062717 | 0.48302287 | -0.3313771 | 0.74035966 | 0.99945981 |
| RBX1 | 23.0437637 | 0.497862411 | 0.32167996 | 1.54769481 | 0.12169579 | 0.99945981 |
| EP300 | 79.8768768 | 0.249212497 | 0.21893645 | 1.13828694 | 0.25500069 | 0.99945981 |
| L3MBTL2 | 8.08996115 | -0.007938712 | 0.51194196 | -0.0155071 | 0.98762766 | 0.99945981 |
| RANGAP1 | 10.7365817 | -0.106454037 | 0.47656148 | -0.2233794 | 0.8232402 | 0.99945981 |
| ZC3H7B | 14.3424155 | -0.392996734 | 0.38869026 | -1.0110795 | 0.3119784 | 0.99945981 |
| TOB2 | 12.186261 | -0.091861019 | 0.47084346 | -0.1950989 | 0.84531556 | 0.99945981 |
| ACO2 | 23.7151266 | -0.036787849 | 0.38455331 | -0.0956638 | 0.92378756 | 0.99945981 |
| DESI1 | 11.2567833 | 0.149008302 | 0.52182471 | 0.28555241 | 0.77522096 | 0.99945981 |
| XRCC6 | 35.0483531 | -0.033605231 | 0.32730715 | -0.1026719 | 0.91822341 | 0.99945981 |
| SNU13 | 27.6025867 | -0.220836352 | 0.34555942 | -0.6390691 | 0.52277797 | 0.99945981 |
| SREBF2 | 80.927733 | -0.913997948 | 0.54114036 | -1.6890219 | 0.09121523 | 0.99640647 |
| SMDT1 | 11.765362 | -0.351277655 | 0.5452679 | -0.6442295 | 0.51942663 | 0.99945981 |
| TCF20 | 39.4170135 | -0.056351613 | 0.28706858 | -0.1963002 | 0.84437522 | 0.99945981 |
| NFAM1 | 22.1887027 | 1.107765256 | 0.74372535 | 1.48948164 | 0.13636058 | 0.99945981 |
| RRP7A | 7.72146859 | 0.049482249 | 0.50544831 | 0.09789774 | 0.92201349 | 0.99945981 |
| POLDIP3 | 13.8278402 | 0.286346247 | 0.42870136 | 0.66793874 | 0.5041727 | 0.99945981 |
| CYB5R3 | 15.9621079 | -0.445851697 | 0.45178787 | -0.9868607 | 0.32371093 | 0.99945981 |
| ARFGAP3 | 32.63561 | -0.344379841 | 0.27753824 | -1.2408375 | 0.2146658 | 0.99945981 |
| PACSIN2 | 67.2413057 | 0.056500608 | 0.31385406 | 0.18002191 | 0.85713537 | 0.99945981 |
| TSPO | 48.3662884 | 0.329341085 | 0.78862075 | 0.41761656 | 0.67622749 | 0.99945981 |
| PARVB | 31.1615208 | 0.037618822 | 0.29244378 | 0.12863608 | 0.89764561 | 0.99945981 |
| PARVG | 30.5391932 | 0.116193234 | 0.3119423 | 0.37248309 | 0.7095332 | 0.99945981 |
| PRR5 | 10.3603096 | 0.38832416 | 0.51845554 | 0.74900185 | 0.45385609 | 0.99945981 |
| NUP50-DT | 26.7345013 | -0.001881359 | 0.34221365 | -0.0054976 | 0.99561356 | 0.99945981 |
| NUP50 | 19.5847585 | 0.243425196 | 0.32948358 | 0.73880828 | 0.46002343 | 0.99945981 |
| KIAA0930 | 23.1844766 | 0.357668784 | 0.44666457 | 0.80075477 | 0.42327363 | 0.99945981 |
| FAM118A | 21.4834869 | 0.602643557 | 0.68446772 | 0.88045578 | 0.37861245 | 0.99945981 |
| ATXN10 | 62.3586745 | -0.22131433 | 0.22742211 | -0.9731434 | 0.33048203 | 0.99945981 |
| MIRLET7BHG | 20.9623476 | -1.087346807 | 0.63281018 | -1.7182827 | 0.08574506 | 0.99193047 |
| PPARA | 23.7588524 | -0.403108002 | 0.39285486 | -1.0260991 | 0.30484489 | 0.99945981 |
| TTC38 | 13.0470139 | 0.470647379 | 0.47512278 | 0.99058054 | 0.32189044 | 0.99945981 |
| TRMU | 9.13147688 | -0.696813611 | 0.48142655 | -1.4473934 | 0.14778678 | 0.99945981 |

| | | | | | | |
|---|---|---|---|---|---|---|
| GRAMD4 | 27.0226648 | -0.846045703 | 0.47051686 | -1.7981198 | 0.07215802 | 0.96000932 |
| CERK | 26.1880796 | -0.252765536 | 0.36954846 | -0.6839848 | 0.49398475 | 0.99945981 |
| TBC1D22A | 238.383719 | -0.504437156 | 0.41624362 | -1.2118796 | 0.22555847 | 0.99945981 |
| C22orf34 | 51.3900508 | -0.483992325 | 0.47562274 | -1.0175971 | 0.30886946 | 0.99945981 |
| BRD1 | 26.263947 | -0.46290958 | 0.3394192 | -1.3638285 | 0.17262154 | 0.99945981 |
| ZBED4 | 19.4697638 | -0.580003191 | 0.38989771 | -1.4875778 | 0.13686227 | 0.99945981 |
| PIM3 | 15.8954749 | 0.395912398 | 0.56630752 | 0.69911203 | 0.48448202 | 0.99945981 |
| TRABD | 19.5503332 | -0.212834673 | 0.52763099 | -0.4033779 | 0.68667025 | 0.99945981 |
| SELENOO | 19.7505816 | -0.361552804 | 0.41918818 | -0.8625072 | 0.3884085 | 0.99945981 |
| PLXNB2 | 23.7142512 | 0.756733436 | 0.53152256 | 1.42370896 | 0.15453074 | 0.99945981 |
| PPP6R2 | 72.6568083 | 0.208202948 | 0.29620111 | 0.70291077 | 0.48211136 | 0.99945981 |
| LMF2 | 7.15909203 | 0.171035798 | 0.67021654 | 0.25519483 | 0.79857262 | 0.99945981 |
| TYMP | 33.0426412 | -0.070189165 | 0.84288264 | -0.0832728 | 0.93363466 | 0.99945981 |
| FP671120.1 | 220.216089 | -0.671583833 | 0.4784074 | -1.4037907 | 0.1603812 | 0.99945981 |
| FP236383.1 | 117.551931 | -0.996552177 | 0.58632663 | -1.6996536 | 0.0891961 | 0.99640647 |
| SAMSN1 | 198.139512 | 0.527883423 | 0.69811083 | 0.75615991 | 0.44955333 | 0.99945981 |
| NRIP1 | 28.7707934 | 0.125754336 | 0.38747582 | 0.32454757 | 0.74552351 | 0.99945981 |
| USP25 | 114.128004 | 0.649629727 | 0.34428319 | 1.88690515 | 0.05917308 | 0.92445778 |
| MIR99AHG | 46.7845402 | 1.006702348 | 0.54072366 | 1.86176862 | 0.06263571 | 0.93072329 |
| LINC01684 | 21.7836713 | -0.072185738 | 0.5311845 | -0.1358958 | 0.89190366 | 0.99945981 |
| ATP5PF | 15.9177267 | 0.109484322 | 0.5394614 | 0.20295117 | 0.8391732 | 0.99945981 |
| GABPA | 12.5035188 | 0.021658634 | 0.41539991 | 0.05213923 | 0.95841775 | 0.99945981 |
| APP | 73.6584919 | 0.395815103 | 0.40201071 | 0.98458844 | 0.32482627 | 0.99945981 |
| ADAMTS1 | 12.1504476 | -1.916290759 | 0.68566515 | -2.794791 | 0.00519333 | 0.5234882 |
| AF165147.1 | 30.559713 | -0.029856407 | 0.51344005 | -0.0581497 | 0.95362935 | 0.99945981 |
| LTN1 | 25.3412526 | -0.020453466 | 0.30074228 | -0.0680099 | 0.94577772 | 0.99945981 |
| USP16 | 23.78734 | 0.103940367 | 0.32288643 | 0.32190999 | 0.74752088 | 0.99945981 |
| CCT8 | 16.6704596 | 0.742834653 | 0.50655444 | 1.46644585 | 0.14252686 | 0.99945981 |
| BACH1 | 215.012951 | 0.505106795 | 0.32380843 | 1.55989391 | 0.11878495 | 0.99945981 |
| TIAM1 | 43.9474999 | 0.830818484 | 0.45614054 | 1.821409 | 0.06854471 | 0.95254943 |
| SOD1 | 19.2857676 | -0.28133052 | 0.50826125 | -0.5535156 | 0.57991042 | 0.99945981 |
| SCAF4 | 30.7669169 | 0.509205234 | 0.31084824 | 1.63811522 | 0.10139766 | 0.99945981 |
| URB1 | 9.35275023 | 0.324205976 | 0.43430786 | 0.74648885 | 0.45537216 | 0.99945981 |
| EVA1C | 18.1147671 | -0.510499424 | 0.48546413 | -1.0515698 | 0.29299696 | 0.99945981 |
| SYNJ1 | 33.4600073 | 0.072778297 | 0.37992917 | 0.19155754 | 0.8480888 | 0.99945981 |
| PAXBP1 | 38.5522751 | 0.269247321 | 0.30732528 | 0.87609885 | 0.3809763 | 0.99945981 |
| IFNAR1 | 31.6315568 | -0.258491927 | 0.3905701 | -0.6618324 | 0.50807867 | 0.99945981 |
| IFNGR2 | 69.8748609 | 0.018233225 | 0.40103418 | 0.04546551 | 0.96373626 | 0.99945981 |
| TMEM50B | 19.543704 | -0.213529957 | 0.43627351 | -0.4894406 | 0.62452982 | 0.99945981 |
| GART | 13.7100618 | 0.533984766 | 0.43217328 | 1.23558023 | 0.21661464 | 0.99945981 |
| SON | 92.4970806 | 0.111666144 | 0.21893614 | 0.5100398 | 0.61002358 | 0.99945981 |
| AP000311.1 | 20.7286102 | 1.064312745 | 0.79786147 | 1.33395681 | 0.18221801 | 0.99945981 |
| ITSN1 | 92.2000646 | 1.039004444 | 0.60595168 | 1.71466551 | 0.08640657 | 0.99204414 |
| LINC00649 | 15.9126566 | 0.0266315 | 0.55798259 | 0.04772819 | 0.96193287 | 0.99945981 |
| MRPS6 | 45.1401267 | 1.141884138 | 0.616085 | 1.85345226 | 0.06381756 | 0.93776345 |
| KCNE1 | 36.431599 | 0.435220861 | 0.8580474 | 0.5072224 | 0.61199878 | 0.99945981 |
| RCAN1 | 15.1025831 | 0.743578509 | 0.59634915 | 1.24688449 | 0.21243985 | 0.99945981 |
| RUNX1 | 387.742991 | -0.284543671 | 0.49316679 | -0.5769725 | 0.56395804 | 0.99945981 |
| CBR1 | 12.1061152 | -0.044253668 | 0.59060327 | -0.0749296 | 0.94027072 | 0.99945981 |
| DOPEY2 | 33.6016136 | -0.099873258 | 0.66006965 | -0.1513072 | 0.87973343 | 0.99945981 |
| MORC3 | 55.031495 | 0.476057988 | 0.23153553 | 2.05609041 | 0.0397738 | 0.89132455 |
| HLCS | 38.3189814 | -0.387239417 | 0.3972747 | -0.9747397 | 0.3296894 | 0.99945981 |
| TTC3 | 85.074778 | -0.08548913 | 0.29402865 | -0.290751 | 0.77124175 | 0.99945981 |
| VPS26C | 23.3885633 | -0.577845293 | 0.37138143 | -1.5559348 | 0.1197236 | 0.99945981 |
| DYRK1A | 163.023119 | 0.12005433 | 0.24713735 | 0.48577979 | 0.62712331 | 0.99945981 |
| KCNJ15 | 14.5082082 | 1.21382948 | 0.48123365 | 2.52232878 | 0.01165807 | 0.67187199 |
| ETS2 | 41.7777239 | 0.037187494 | 0.60525134 | 0.06144141 | 0.95100768 | 0.99945981 |
| BRWD1 | 97.1391925 | 0.408135201 | 0.24063243 | 1.69609392 | 0.0898681 | 0.99640647 |
| HMGN1 | 30.5006215 | 0.011871939 | 0.45343732 | 0.02618209 | 0.9791121 | 0.99945981 |
| MX2 | 52.6007528 | 0.676395227 | 0.95085018 | 0.71135836 | 0.47686219 | 0.99945981 |
| MX1 | 52.0697485 | 1.061019464 | 1.07238396 | 0.98940259 | 0.32246621 | 0.99945981 |
| C2CD2 | 9.05950074 | 0.153263681 | 0.516306 | 0.2968466 | 0.76658363 | 0.99945981 |
| ABCG1 | 22.8497428 | -0.885924259 | 0.60137596 | -1.4731621 | 0.14070734 | 0.99945981 |
| UBASH3A | 7.83150704 | -1.144305309 | 0.66477398 | -1.7213449 | 0.08518825 | 0.99193047 |
| SLC37A1 | 15.2921976 | 0.025007531 | 0.45290087 | 0.05521635 | 0.95596611 | 0.99945981 |
| NDUFV3 | 15.0356787 | 0.661359231 | 0.38262144 | 1.72849495 | 0.08389953 | 0.98963959 |
| PKNOX1 | 16.7968748 | 0.396261938 | 0.46555594 | 0.85115859 | 0.39468126 | 0.99945981 |
| RRP1B | 11.4605346 | -0.701587201 | 0.53289214 | -1.3165651 | 0.18798444 | 0.99945981 |
| PDXK | 26.3338453 | -0.480405513 | 0.40329402 | -1.1912042 | 0.23357344 | 0.99945981 |
| CSTB | 39.0548795 | -0.445531601 | 0.59634806 | -0.7470999 | 0.45500323 | 0.99945981 |
| AGPAT3 | 56.4626119 | 0.511922358 | 0.34663322 | 1.47684159 | 0.13971811 | 0.99945981 |
| TRAPPC10 | 61.2842463 | -0.204572206 | 0.27361763 | -0.7476573 | 0.45466692 | 0.99945981 |
| PFKL | 24.2652945 | -0.563720744 | 0.4563398 | -1.2353092 | 0.21671545 | 0.99945981 |
| TRPM2 | 29.1691869 | -0.289999134 | 0.63088922 | -0.4596673 | 0.64575506 | 0.99945981 |
| UBE2G2 | 31.3763919 | 0.283893165 | 0.31916244 | 0.88949428 | 0.37373749 | 0.99945981 |
| SUMO3 | 14.5162766 | 0.404878699 | 0.47726608 | 0.84832909 | 0.39625472 | 0.99945981 |
| PTTG1IP | 34.9795905 | -0.376951308 | 0.36675148 | -1.0278113 | 0.30403861 | 0.99945981 |
| ITGB2 | 72.202681 | 0.287894441 | 0.58072813 | 0.49574737 | 0.62007266 | 0.99945981 |
| ITGB2-AS1 | 11.344053 | 1.222043176 | 0.54048662 | 2.26100541 | 0.02375892 | 0.80320673 |
| ADARB1 | 33.2350593 | -1.453099337 | 0.52553991 | -2.7649648 | 0.00569289 | 0.52554078 |
| COL18A1 | 9.43687807 | -0.565176405 | 0.43933543 | -1.2864348 | 0.19829134 | 0.99945981 |
| LSS | 9.55779445 | -0.503061632 | 0.52924661 | -0.950524 | 0.34184604 | 0.99945981 |
| MCM3AP | 30.6808037 | 0.280424314 | 0.31729469 | 0.88379769 | 0.37680543 | 0.99945981 |
| PCNT | 19.3937687 | -0.396149176 | 0.35085781 | -1.1290875 | 0.25886092 | 0.99945981 |
| DIP2A | 67.2506565 | 0.428341856 | 0.38734196 | 1.10584936 | 0.26879173 | 0.99945981 |
| PRMT2 | 64.4217366 | -0.310122405 | 0.25893037 | -1.1977058 | 0.23103157 | 0.99945981 |
| AL592183.1 | 14.5049358 | 0.061735969 | 0.53911741 | 0.11451303 | 0.90883112 | 0.99945981 |

**Bulk RNA sequencing**

*This means that positive fold changes imply higher mean value in controls than in AKI and vice versa.

| | baseMean | log2 fold change control/AKI* | lfcSE | stat | pvalue | padj |
|---|---|---|---|---|---|---|
| MIR1302-2H( | 1.82201478 | 1.302330707 | 0.94476245 | 1.37847425 | 0.16805691 | 0.33938281 |
| FAM138A | 0 | NA | NA | NA | NA | NA |
| OR4F5 | 0 | NA | NA | NA | NA | NA |
| AL627309.1 | 6.71541223 | 0.14517263 | 0.5295211 | 0.27415835 | 0.78396294 | 0.8819583 |
| AL627309.3 | 0.66757808 | -1.343133912 | 1.63867467 | -0.8196465 | 0.41241767 | NA |
| AL627309.2 | 1.12832931 | 0.415459761 | 1.39851893 | 0.29707125 | 0.76641212 | 0.8711864 |
| AL627309.4 | 0 | NA | NA | NA | NA | NA |
| AL732372.1 | 0.34346317 | -1.70467338 | 2.84472907 | -0.5992393 | 0.54901334 | NA |
| OR4F29 | 0 | NA | NA | NA | NA | NA |
| AC114498.1 | 2.31952407 | -3.581899256 | 1.35084467 | -2.6515997 | 0.00801115 | 0.03816003 |
| OR4F16 | 0 | NA | NA | NA | NA | NA |
| AL669831.2 | 0 | NA | NA | NA | NA | NA |
| AL669831.5 | 95.6793077 | 0.358878943 | 0.26053471 | 1.37747072 | 0.16836676 | 0.33977435 |
| FAM87B | 7.93755327 | 0.673695109 | 0.47674791 | 1.41310553 | 0.1576247 | 0.32475926 |
| LINC00115 | 207.785736 | -0.463293988 | 0.24726676 | -1.8736606 | 0.06097722 | 0.16736033 |
| FAM41C | 1.79012884 | 0.624263384 | 1.0793319 | 0.57837944 | 0.56300797 | 0.73032032 |
| AL645608.7 | 21.8246601 | 0.384700949 | 0.55291122 | 0.69577346 | 0.48657071 | 0.67061463 |
| AL645608.3 | 48.3714125 | 0.828827741 | 0.42296803 | 1.95955173 | 0.05004821 | 0.14571984 |
| AL645608.5 | 11.8170414 | 0.032815951 | 0.46500608 | 0.07057101 | 0.94373918 | 0.97315146 |
| AL645608.1 | 806.913711 | 0.190838233 | 0.41763567 | 0.45694907 | 0.64770765 | 0.79272233 |
| SAMD11 | 1837.94619 | -0.131087525 | 0.25611268 | -0.5118353 | 0.60876625 | 0.76476423 |
| NOC2L | 4798.10953 | -0.189058523 | 0.1190958 | -1.5874492 | 0.11241095 | 0.25794264 |
| KLHL17 | 1327.65945 | -0.773022354 | 0.27335402 | -2.8279165 | 0.0046852 | 0.02559536 |
| PLEKHN1 | 381.51643 | -2.555382798 | 0.29853162 | -8.5598396 | 1.13E-17 | 1.23E-14 |
| PERM1 | 198.587514 | -1.163884529 | 0.40233403 | -2.8928314 | 0.00381786 | 0.02201071 |
| AL645608.8 | 212.571458 | 0.552014038 | 0.60558837 | 0.91153342 | NA | NA |
| HES4 | 545.637929 | -1.112258235 | 0.37238736 | -2.9868314 | 0.00281885 | 0.01749426 |
| ISG15 | 805.25462 | -0.808695295 | 0.37801448 | -2.1393236 | 0.03240947 | 0.10649692 |
| AL645608.2 | 21.0729292 | -0.629021281 | 0.78998853 | -0.796241 | 0.42589195 | 0.61826429 |
| AGRN | 25714.1041 | -0.60782672 | 0.29211022 | -2.0808129 | 0.03745104 | 0.11792046 |
| AL645608.9 | 105.51929 | -2.698625009 | 0.47931549 | -5.6301644 | 1.80E-08 | 1.14E-06 |
| RNF223 | 65.2594112 | -2.099891529 | 0.73656553 | -2.8509229 | 0.00435925 | 0.02432581 |
| C1orf159 | 1280.11683 | -0.565750853 | 0.27216941 | -2.0786717 | 0.03764753 | 0.11839778 |
| LINC01342 | 59.2371236 | -0.400145638 | 0.55997751 | -0.7145745 | 0.47487202 | 0.6602887 |
| AL390719.2 | 843.291855 | -0.605521935 | 0.28825017 | -2.100682 | 0.03566889 | 0.1141117 |
| TTLL10-AS1 | 10.3873564 | -0.157276287 | 0.4225788 | -0.3721822 | 0.70975724 | 0.8350644 |
| TTLL10 | 49.2714732 | -0.412034396 | 0.27559444 | -1.4950751 | 0.13489483 | 0.29233919 |
| TNFRSF18 | 53.073584 | -1.317911068 | 0.65288889 | -2.018584 | 0.04353048 | 0.1318363 |
| TNFRSF4 | 161.328477 | -1.258076915 | 0.52766178 | -2.3842487 | 0.01711403 | 0.06697539 |
| SDF4 | 10801.9906 | -0.119814064 | 0.10017377 | -1.1960622 | 0.23167227 | 0.41970307 |
| B3GALT6 | 902.70958 | -0.354087763 | 0.20485719 | -1.7284615 | 0.08390552 | 0.20953107 |
| C1QTNF12 | 277.200523 | 0.808337556 | 0.22869725 | 3.53453121 | 0.0004085 | 0.00388325 |
| AL162741.1 | 42.7628953 | 0.704152507 | 0.30903063 | 2.27858484 | 0.02269176 | 0.08232997 |
| UBE2J2 | 2824.53912 | -0.71448504 | 0.16789149 | -4.2556357 | 2.08E-05 | 0.00036647 |
| LINC01786 | 73.3669028 | 0.949439888 | 0.36545336 | 2.59797829 | 0.00937744 | 0.0429311 |
| SCNN1D | 693.81577 | 0.637244609 | 0.33399858 | 1.90792609 | 0.05640077 | 0.15840719 |
| ACAP3 | 5195.39751 | -0.191224632 | 0.14959732 | -1.2782624 | 0.20115692 | 0.38294166 |
| PUSL1 | 846.744755 | -0.722281889 | 0.17924475 | -4.0295847 | 5.59E-05 | 0.00080825 |
| INTS11 | 6963.89168 | -0.216460445 | 0.16487397 | -1.3128843 | 0.18922194 | 0.36749025 |
| CPTP | 2518.83099 | -0.270007362 | 0.18418722 | -1.4659397 | 0.14266471 | 0.30393184 |
| TAS1R3 | 81.372033 | -0.368897246 | 0.25554005 | -1.4435986 | 0.14885193 | 0.31289161 |
| DVL1 | 4702.75056 | -0.477459344 | 0.16236468 | -2.9406601 | 0.00327514 | 0.01963295 |
| MXRA8 | 2768.5168 | 0.368393021 | 0.22572059 | 1.63207537 | 0.10266361 | 0.24176097 |
| AURKAIP1 | 4485.34385 | -0.117175216 | 0.12378902 | -0.946572 | 0.34385694 | 0.54151789 |
| CCNL2 | 13383.612 | -0.484868004 | 0.18987193 | -2.5536582 | 0.01065978 | 0.04720378 |
| MRPL20 | 3284.00876 | -0.079337452 | 0.10621965 | -0.7469188 | 0.4551126 | 0.64453917 |
| AL391244.3 | 250.349791 | -0.204755146 | 0.22490058 | -0.9104252 | 0.36259833 | 0.55991227 |
| ANKRD65 | 218.038812 | -0.071360819 | 0.41794128 | -0.1707436 | 0.86442534 | 0.92964904 |
| AL391244.2 | 9.78710634 | 0.150234668 | 0.74606679 | 0.20136893 | 0.84041011 | 0.91505077 |
| TMEM88B | 12.84155 | -1.008856141 | 0.67165672 | -1.5020413 | 0.13308644 | 0.28956183 |
| LINC01770 | 76.041177 | -1.372150991 | 0.43265968 | -3.1714325 | 0.00151689 | 0.01091714 |
| VWA1 | 4397.11815 | -0.83592351 | 0.38924173 | -2.1475691 | 0.031748 | 0.10505031 |
| ATAD3C | 499.500055 | -0.360292505 | 0.32484436 | -1.1091235 | 0.26737693 | 0.46149558 |
| ATAD3B | 3117.94675 | -0.636344236 | 0.19285103 | -3.2996673 | 0.000968 | 0.00764253 |
| ATAD3A | 1591.93778 | -0.50402562 | 0.16459554 | -3.062207 | 0.00219711 | 0.01450987 |
| TMEM240 | 77.5663507 | -0.620288352 | 0.4082265 | -1.5194711 | 0.12864396 | 0.28286011 |
| SSU72 | 5462.67487 | -0.351216945 | 0.20495872 | -1.7135984 | 0.0866025 | 0.21440156 |
| AL645728.1 | 37.7434305 | 0.22974815 | 0.44339983 | 0.51815119 | 0.60435279 | 0.76143254 |
| FNDC10 | 242.793572 | 0.770730328 | 0.46177438 | 1.66906255 | 0.09510498 | 0.22953674 |
| AL691432.2 | 576.636156 | 0.026338227 | 0.22675799 | 0.11615126 | 0.90753266 | 0.95385641 |
| MIB2 | 4457.15522 | -0.081308024 | 0.1727368 | -0.4707047 | 0.63785163 | 0.78562859 |
| MMP23B | 170.557836 | -0.087183165 | 0.37055877 | -0.2352749 | 0.8139954 | 0.89962526 |
| CDK11B | 1814.74917 | -0.419060271 | 0.20341999 | -2.0600742 | 0.03939145 | 0.12245109 |
| FO704657.1 | 653.922596 | -0.173055866 | 0.46807588 | -0.3697175 | 0.71159296 | 0.83606132 |
| SLC35E2B | 3019.77297 | 0.179282019 | 0.27671921 | 0.64788425 | 0.51705982 | 0.69438111 |
| CDK11A | 1731.29867 | -0.428792454 | 0.31284599 | -1.3706184 | 0.17049396 | 0.34259892 |
| SLC35E2A | 1619.33605 | 0.603442989 | 0.42280659 | 1.42723176 | 0.1535131 | 0.3191533 |
| NADK | 3859.65708 | -0.556092635 | 0.16689124 | -3.332066 | 0.00086204 | 0.00699444 |
| GNB1 | 16546.6671 | -0.28082307 | 0.17359985 | -1.6176458 | 0.10573894 | 0.24677792 |
| AL109917.1 | 22.3440978 | -0.032701875 | 0.32967761 | -0.0991935 | 0.92098464 | 0.96149352 |
| CALML6 | 41.030256 | 0.747525218 | 0.25563462 | 2.92419403 | 0.00345349 | 0.02038554 |
| TMEM52 | 83.3221168 | 0.553343963 | 0.50302926 | 1.10002341 | 0.27132192 | 0.46598666 |
| CFAP74 | 11.7925799 | 0.596116118 | 0.92190876 | 0.64661075 | 0.5178839 | 0.69488619 |
| AL391845.2 | 2.12669278 | -1.357311187 | 1.19723822 | -1.1337019 | 0.25691963 | 0.44958173 |
| GABRD | 21.5602146 | -0.158745515 | 0.63252525 | -0.250971 | 0.80183651 | 0.89258426 |
| AL391845.1 | 3.14209311 | 1.725432974 | 0.72686025 | 2.37381668 | 0.01760529 | 0.06831246 |
| PRKCZ | 2496.70653 | 0.021493931 | 0.27147985 | 0.07917321 | 0.93689486 | 0.96969938 |
| AL590822.2 | 261.496924 | 0.558483243 | 0.34117912 | 1.63692096 | 0.10164699 | 0.24024657 |
| PRKCZ-AS1 | 351.726471 | -0.17597955 | 0.28447402 | -0.6186138 | 0.53617081 | 0.70988757 |
| FAAP20 | 3896.72319 | -0.323161478 | 0.12994668 | -2.4868776 | 0.01288697 | 0.05438142 |
| AL590822.1 | 26.9365375 | 1.713672322 | 0.45846259 | 3.73786731 | 0.00018559 | 0.00205888 |
| SKI | 4239.33383 | 0.021255768 | 0.24275874 | 0.08755923 | 0.93022701 | 0.96634079 |
| MORN1 | 664.890606 | 0.602847256 | 0.20961266 | 2.876006 | 0.00402742 | 0.02286888 |
| RER1 | 9098.18275 | -0.211009749 | 0.10747749 | -1.9632924 | 0.04961221 | 0.14489688 |
| PEX10 | 4037.92318 | -0.025223607 | 0.10407498 | -0.24236 | 0.80850126 | 0.89648382 |
| PLCH2 | 3251.54012 | 2.806682009 | 0.56401493 | 4.97625481 | 6.48E-07 | 2.26E-05 |
| AL139246.1 | 21.0830983 | 1.44236716 | 0.682939 | 2.11199998 | 0.03468645 | 0.11192875 |
| AL139246.4 | 22.8023415 | 1.268021112 | 0.70566281 | 1.79692211 | 0.07234799 | 0.18893012 |
| PANK4 | 1709.00386 | -0.069030905 | 0.11989356 | -0.5757683 | 0.56477182 | 0.73164184 |
| HES5 | 3.1428926 | 0.762771288 | 1.13688369 | 0.67093169 | 0.50226405 | 0.68263189 |
| AL139246.5 | 241.280973 | -1.105695517 | 0.33294338 | -3.3209716 | 0.00089705 | 0.00720475 |
| TNFRSF14-A | 4015.38772 | -0.509536576 | 0.26315798 | -1.9362384 | 0.05283851 | 0.15143479 |
| TNFRSF14 | 16057.0369 | -0.564364489 | 0.26010295 | -2.1697735 | 0.03002401 | 0.10090874 |
| AL139246.3 | 36.9421176 | -1.535122112 | 0.51915448 | -2.9569659 | 0.00310682 | 0.0188598 |
| FAM213B | 2808.86001 | -0.379995822 | 0.21774595 | -1.7451338 | 0.08096157 | 0.2041458 |
| MMEL1 | 830.123314 | -0.69097618 | 0.26778945 | -2.5802965 | 0.00987155 | 0.04466216 |
| AL831784.1 | 8.63813768 | 0.486220312 | 0.60535179 | 0.8032029 | 0.42185747 | 0.61446342 |
| TTC34 | 214.203149 | 0.720560239 | 0.42694907 | 1.68769599 | 0.0914696 | 0.22313168 |
| AC242022.1 | 5.99991809 | 1.126460735 | 0.70413909 | 1.59977021 | 0.10964957 | 0.25335933 |
| AL592464.2 | 0 | NA | NA | NA | NA | NA |
| AL592464.1 | 0 | NA | NA | NA | NA | NA |

| | | | | | | |
|---|---|---|---|---|---|---|
| AL589702.1 | 2.7272554 | -3.79967769 | 1.86462103 | -2.0377748 | 0.04157246 | 0.12742832 |
| ACTRT2 | 0 | NA | NA | NA | NA | NA |
| LINC00982 | 6398.40518 | 2.038127394 | 0.44635293 | 4.56617907 | 4.97E-06 | 0.00011667 |
| PRDM16 | 1208.28914 | 0.737287827 | 0.31970703 | 2.30613582 | 0.02110304 | 0.07792848 |
| AL008733.1 | 0.80050411 | 2.000787548 | 1.5966287 | 1.25313265 | 0.21015744 | NA |
| AL590438.1 | 4.91168474 | -1.103060855 | 0.86985244 | -1.2681011 | 0.20476184 | 0.38711193 |
| ARHGEF16 | 4370.13224 | -0.320890015 | 0.14265788 | -2.2493675 | 0.02448912 | 0.08691496 |
| AL512413.1 | 22.6206388 | -0.427693926 | 0.48310875 | -0.8852953 | 0.37599736 | 0.57184583 |
| MEGF6 | 1250.69624 | 0.437484061 | 0.29449851 | 1.48552214 | 0.13740556 | 0.29604946 |
| AL513320.1 | 142.327126 | 0.525997433 | 0.38062459 | 1.38193234 | 0.16699248 | 0.33798385 |
| TPRG1L | 5629.42138 | 0.526660906 | 0.27651081 | 1.90466659 | 0.05682342 | 0.15927037 |
| WRAP73 | 1484.97212 | -0.210098084 | 0.12691283 | -1.6554519 | 0.09783287 | 0.23406082 |
| TP73 | 75.0331013 | -0.450981057 | 0.21689044 | -2.0793035 | 0.03758946 | 0.1182857 |
| AL136528.1 | 0.8786774 | -0.283613187 | 1.41508877 | -0.2004208 | 0.84115151 | NA |
| AL136528.2 | 0 | NA | NA | NA | NA | NA |
| CCDC27 | 70.5756492 | 0.960364064 | 0.2509349 | 3.82714432 | 0.00012964 | 0.00155248 |
| SMIM1 | 189.86685 | 0.679516246 | 0.34045003 | 1.99593532 | 0.04594096 | 0.13690397 |
| LRRC47 | 3081.02247 | 0.088119843 | 0.12932689 | 0.68137292 | 0.49563555 | 0.67769982 |
| AL365330.1 | 465.26162 | 0.338991012 | 0.15771682 | 2.14936497 | 0.03160548 | 0.10474256 |
| CEP104 | 1859.34888 | 0.078045998 | 0.21227851 | 0.36765849 | 0.7131279 | 0.83695061 |
| DFFB | 467.14535 | 0.716609147 | 0.15201707 | 4.71400441 | 2.43E-06 | 6.65E-05 |
| C1orf174 | 1546.35789 | -0.054147123 | 0.21773243 | -0.2486865 | 0.80360327 | 0.89345721 |
| LINC01134 | 88.2588615 | 0.051705695 | 0.53057691 | 0.09745184 | 0.92236758 | 0.96203347 |
| LINC01346 | 1.5765006 | 2.7051429 | 1.24775897 | 2.16800116 | 0.0301586 | 0.1012067 |
| LINC01345 | 1.14212318 | 2.117989678 | 1.30015651 | 1.6290267 | 0.10330737 | 0.24295151 |
| AL805961.1 | 0 | NA | NA | NA | NA | NA |
| AL805961.2 | 0.28299204 | 1.12993107 | 2.97686151 | 0.37957126 | 0.7042637 | NA |
| LINC01777 | 0.10178723 | 1.087385526 | 3.28420134 | 0.33109588 | 0.74057207 | NA |
| AL355602.1 | 0 | NA | NA | NA | NA | NA |
| Z98747.1 | 0 | NA | NA | NA | NA | NA |
| LINC01646 | 0.1357163 | 1.313406857 | 3.28103699 | 0.40030236 | 0.68893383 | NA |
| AJAP1 | 655.702361 | 0.647664877 | 0.22280547 | 2.90686255 | 0.00365074 | 0.02127982 |
| BX005132.2 | 0 | NA | NA | NA | NA | NA |
| BX005132.1 | 0.03392908 | 0.380740593 | 3.29738709 | 0.11546736 | 0.90807469 | NA |
| AL691497.1 | 0 | NA | NA | NA | NA | NA |
| AL139823.1 | 2.59354494 | 2.400468051 | 0.86725632 | 2.76788766 | 0.00564209 | 0.02951015 |
| Z98259.3 | 8.25645562 | 3.065799594 | 1.26521688 | 2.42314156 | 0.01538693 | 0.06200293 |
| Z98259.2 | 1.90694741 | 3.991753719 | 1.67566005 | 2.38219783 | 0.01720965 | 0.06723236 |
| Z98259.1 | 0.30684769 | 1.415461554 | 2.2752879 | 0.62210218 | 0.53387468 | NA |
| AL365255.1 | 0.41623257 | -0.602926659 | 1.95031701 | -0.3091429 | 0.75721283 | NA |
| NPHP4 | 1052.99415 | 0.016386788 | 0.13175577 | 0.12437246 | 0.90102038 | 0.9498264 |
| KCNAB2 | 3703.18714 | 1.626297915 | 0.41490434 | 3.91969366 | 8.87E-05 | 0.00115229 |
| CHD5 | 146.24609 | 0.244193767 | 0.51951968 | 0.47003757 | 0.63832818 | 0.78602447 |
| RPL22 | 13173.5718 | -0.005815536 | 0.21172538 | -0.0274674 | 0.97808698 | 0.99002586 |
| AL031847.1 | 68.837031 | 0.323322812 | 0.40433651 | 0.79963794 | 0.4239206 | 0.61642052 |
| RNF207 | 1922.88218 | 0.125682876 | 0.22783383 | 0.55164273 | 0.58119316 | 0.74384312 |
| ICMT | 5353.92324 | 0.245533333 | 0.20533688 | 1.19575855 | 0.2317908 | 0.4197736 |
| LINC00337 | 14.6440328 | 1.161065253 | 0.65010479 | 1.78596631 | 0.07410471 | 0.19212873 |
| HES3 | 0.08393208 | -0.642905668 | 3.30558948 | -0.1944905 | 0.84579185 | NA |
| GPR153 | 905.44634 | -0.816974054 | 0.35371862 | -2.3096722 | 0.02090631 | 0.07739709 |
| ACOT7 | 3103.09082 | 1.348589797 | 0.46266538 | 2.91482754 | 0.00355885 | 0.02082737 |
| AL031848.2 | 14.74857 | 0.626391701 | 0.34214501 | 1.83077841 | 0.06713362 | 0.1788506 |
| HES2 | 222.644314 | 0.148316209 | 0.40693382 | 0.36447255 | 0.71550517 | 0.83812505 |
| ESPN | 3649.78638 | 1.364540053 | 0.40389744 | 3.37843207 | 0.000729 | 0.00610544 |
| AL031848.1 | 160.725059 | 1.514768251 | 0.44947739 | 3.37006554 | 0.0007515 | 0.00624949 |
| TNFRSF25 | 1167.63756 | 0.202015555 | 0.30727229 | 0.657448 | 0.5108929 | 0.68920615 |
| PLEKHG5 | 1418.9095 | 0.299846265 | 0.30807103 | 0.97330237 | 0.33040305 | 0.52823623 |
| NOL9 | 1102.54021 | 0.129682792 | 0.14319271 | 0.90565222 | 0.36511996 | 0.56210014 |
| TAS1R1 | 10.3552436 | 1.420628837 | 0.75960085 | 1.87023071 | 0.06145179 | 0.1681676 |
| ZBTB48 | 2078.6986 | 0.157249881 | 0.12432619 | 1.26481705 | 0.20593692 | 0.388563 |
| KLHL21 | 5457.08515 | -0.293632319 | 0.49744963 | -0.5902755 | 0.55500598 | 0.72445835 |
| PHF13 | 1201.18515 | -0.534478756 | 0.17302222 | -3.0890759 | 0.0020078 | 0.01354457 |
| THAP3 | 4425.67856 | 0.41294104 | 0.14287573 | 2.89021119 | 0.00384983 | 0.02213396 |
| DNAJC11 | 6104.75889 | 0.889048787 | 0.15474109 | 5.74539568 | 9.17E-09 | 6.43E-07 |
| LINC01672 | 0.20176029 | 1.652197309 | 3.26794479 | 0.50557687 | 0.61315372 | NA |
| AL591163.1 | 0 | NA | NA | NA | NA | NA |
| CAMTA1-DT | 22.9793308 | 0.347322871 | 0.4725045 | 0.73506786 | 0.46229816 | 0.65052308 |
| CAMTA1 | 1878.22905 | -0.199672626 | 0.14327365 | -1.3936452 | 0.16342478 | 0.33267806 |
| AL512330.1 | 0.78113846 | 3.006059616 | 1.59654477 | 1.88285332 | 0.05972025 | NA |
| Z97987.1 | 0.13390214 | 1.04981649 | 3.28477129 | 0.31960109 | 0.74927075 | NA |
| AL365194.1 | 0.68223863 | 0.252450329 | 1.64104836 | 0.15383479 | 0.87774 | NA |
| AL359881.3 | 14.7244534 | -0.150456201 | 0.63695071 | -0.2362133 | 0.81326719 | 0.89913137 |
| AL359881.2 | 11.3512632 | -0.15144215 | 0.47379712 | -0.319635 | 0.74924503 | 0.85996045 |
| AL359881.1 | 14.5918812 | -0.450077696 | 0.46372873 | -0.9705625 | 0.33176621 | 0.5296793 |
| VAMP3 | 5178.03379 | -0.127803593 | 0.16672274 | -0.7665637 | 0.44334099 | 0.63444412 |
| PER3 | 1117.55845 | 0.922023429 | 0.39266449 | 2.34812023 | 0.01886843 | 0.0719122 |
| Z98884.1 | 56.0815185 | 0.685499895 | 0.44772669 | 1.53106776 | 0.12575265 | 0.27855712 |
| UTS2 | 284.66133 | 1.10230028 | 0.45552665 | 2.41983707 | 0.01552746 | 0.06239275 |
| TNFRSF9 | 19.0268226 | -0.649651825 | 0.63483005 | -1.0233476 | 0.30614351 | 0.50305458 |
| PARK7 | 7615.23686 | -0.051981185 | 0.14452799 | -0.3596617 | 0.71910012 | 0.84021138 |
| AL034417.4 | 187.851998 | -0.106312694 | 0.57638602 | -0.184447 | 0.85366277 | 0.92345602 |
| AL034417.3 | 5.92396478 | -1.58379006 | 0.61548598 | -2.573235 | 0.01007528 | 0.04530782 |
| ERRFI1 | 9907.793 | -1.295802148 | 0.5717591 | -2.2663429 | 0.0234304 | 0.08417706 |
| AL034417.2 | 16.9675912 | 0.036313693 | 0.50355206 | 0.07211507 | 0.94251033 | 0.97275716 |
| LINC01714 | 15.6930691 | 0.770195828 | 0.85465858 | 0.90117369 | 0.36749598 | 0.56438321 |
| AL358876.2 | 0.37327355 | 0.785291144 | 3.27645337 | 0.23967719 | 0.81058052 | NA |
| SLC45A1 | 376.901785 | -0.115655142 | 0.3070305 | -0.3766894 | 0.70640443 | 0.83294065 |
| RERE | 11913.2677 | -0.107499951 | 0.18255367 | -0.5888677 | 0.55595006 | 0.72518812 |
| AL096855.1 | 32.2398476 | -0.337300823 | 0.40477154 | -0.8333116 | 0.40466901 | 0.59945083 |
| AL357552.2 | 0.32825158 | -0.279886877 | 2.93064906 | -0.0955034 | 0.92391501 | NA |
| ENO1 | 94369.4398 | -0.845340383 | 0.22489842 | -3.7587654 | 0.00017075 | 0.00193501 |
| ENO1-AS1 | 33.5671492 | 0.660397753 | 0.45039993 | 1.46624745 | 0.14258089 | 0.30378955 |
| CA6 | 0.3320302 | -0.541119081 | 2.88239456 | -0.1877325 | 0.85108637 | NA |
| SLC2A7 | 2.17209838 | 1.374930273 | 1.02179423 | 1.34560387 | 0.1784303 | 0.35385479 |
| SLC2A5 | 2400.23625 | 1.908974706 | 0.61674677 | 3.09523261 | 0.00196659 | 0.01331764 |
| GPR157 | 661.065797 | -0.226969746 | 0.20843784 | -1.0889085 | 0.27619422 | 0.47149878 |
| MIR34AHG | 515.414242 | -1.223093908 | 0.44135245 | -2.7712408 | 0.00558431 | 0.02927982 |
| LINC01759 | 45.8266624 | -1.201596088 | 0.61085833 | -1.9670618 | 0.04917609 | 0.14405392 |
| H6PD | 4415.11358 | -0.678655458 | 0.19040525 | -3.5642686 | 0.00036487 | 0.00354782 |
| SPSB1 | 3992.69968 | -2.684479798 | 0.47146605 | -5.6938984 | 1.24E-08 | 8.40E-07 |
| BX323043.1 | 72.7821129 | -0.060906694 | 0.68810223 | -0.088514 | 0.92946814 | 0.96587622 |
| AL928921.2 | 105.641768 | 0.932675972 | 0.47755716 | 1.95301431 | 0.0508179 | 0.14730879 |
| AL928921.1 | 13.4578393 | 0.749020103 | 0.60994118 | 1.22802023 | 0.21943937 | 0.40497 |
| SLC25A33 | 1020.5575 | 0.003153434 | 0.37770639 | 0.0083489 | 0.99333862 | 0.99693014 |
| TMEM201 | 1154.39447 | 0.252647551 | 0.18034638 | 1.40090172 | 0.16124346 | 0.32994158 |
| PIK3CD | 2255.49221 | -0.436685081 | 0.28294818 | -1.5433394 | 0.12274844 | 0.27398222 |
| PIK3CD-AS1 | 2.87586878 | -0.835191597 | 1.0507904 | -0.7948223 | 0.4267169 | 0.61900696 |
| PIK3CD-AS2 | 121.750654 | 0.501044342 | 0.26975769 | 1.85738666 | 0.06325616 | 0.17159398 |
| CLSTN1 | 10842.9401 | 0.318055598 | 0.11756514 | 2.70535644 | 0.00682311 | 0.03392261 |
| CTNNBIP1 | 1489.29039 | 0.464725774 | 0.17814872 | 2.60863944 | 0.0090903 | 0.04195889 |
| AL357140.1 | 521.291797 | 0.561251842 | 0.20693121 | 2.7122629 | 0.00668256 | 0.03340012 |
| LZIC | 668.879381 | 0.463009016 | 0.19385222 | 2.38846388 | 0.01691897 | 0.06645181 |
| NMNAT1 | 950.569319 | 0.42396389 | 0.22274697 | 1.90334306 | 0.0569958 | 0.15956886 |

| | | | | | | |
|---|---|---|---|---|---|---|
| RBP7 | 528.693407 | -0.582536633 | 0.53613151 | -1.0865555 | 0.27723331 | 0.47266066 |
| UBE4B | 2284.53276 | 0.196994114 | 0.14947854 | 1.31787556 | 0.18754531 | 0.36558026 |
| KIF1B | 2916.31406 | 0.279496398 | 0.1703875 | 1.64035741 | 0.10093088 | 0.23902534 |
| AL139424.3 | 5.49953949 | 0.720812906 | 0.56121799 | 1.28437241 | 0.19901168 | 0.3800809 |
| AL139424.2 | 145.369182 | 0.378605168 | 0.28612425 | 1.32321945 | 0.18576241 | 0.36314347 |
| PGD | 5192.44174 | -0.455894287 | 0.21176954 | -2.1527849 | 0.03133558 | 0.10413602 |
| AL139424.1 | 9.03143214 | 1.422592052 | 0.42207775 | 3.37045021 | 0.00075045 | 0.00624668 |
| CENPS-CORT | 874.838804 | 0.942914743 | 0.20299282 | 4.6450644 | 3.40E-06 | 8.65E-05 |
| CENPS | 871.97163 | 0.952817439 | 0.21916807 | 4.34742807 | 1.38E-05 | 0.00026436 |
| CORT | 141.048594 | 0.367295117 | 0.13581433 | 2.70439147 | 0.00684296 | 0.03398284 |
| DFFA | 2494.81793 | 0.162611842 | 0.14278911 | 1.13882524 | 0.25477605 | 0.44753011 |
| AL354956.1 | 185.038438 | 0.235383797 | 0.22247828 | 1.05800798 | 0.29005181 | 0.48615433 |
| PEX14 | 1842.57117 | -0.056730159 | 0.13353318 | -0.4248394 | 0.67095375 | 0.80780558 |
| CASZ1 | 2098.84024 | 0.102021686 | 0.19390685 | 0.5261376 | 0.59879261 | 0.75677743 |
| AL139423.1 | 919.427599 | 0.052873726 | 0.20699516 | 0.2554346 | 0.79838745 | 0.8908508 |
| C1orf127 | 10.3813675 | 0.256384869 | 0.7030122 | 0.36469476 | 0.71533927 | 0.8380436 |
| TARDBP | 8301.10686 | -0.335577771 | 0.14416658 | -2.3277085 | 0.01992758 | 0.07471782 |
| AL109811.4 | 4302.94514 | -0.107455049 | 0.14065606 | -0.763956 | 0.44489344 | 0.63551603 |
| MASP2 | 306.251611 | 0.506324455 | 0.22515984 | 2.24873346 | 0.02452946 | 0.08701128 |
| AL109811.3 | 438.148993 | 0.039673135 | 0.22379549 | 0.17727406 | 0.85929313 | 0.9265427 |
| SRM | 1934.88355 | -0.995324376 | 0.27864454 | -3.5720218 | 0.00035424 | 0.00346741 |
| EXOSC10 | 3162.1456 | -0.259905733 | 0.16575354 | -1.5680252 | 0.11687525 | 0.26520955 |
| AL109811.1 | 1330.1432 | -0.576682161 | 0.17790718 | -3.2414778 | 0.00118912 | 0.00898482 |
| AL109811.2 | 103.262425 | 0.614559632 | 0.3066543 | 2.00407962 | 0.04506153 | 0.13492365 |
| MTOR | 3048.0153 | 0.246755359 | 0.11194737 | 2.20420863 | 0.02750968 | 0.09483987 |
| MTOR-AS1 | 27.6270666 | 1.747189386 | 0.35578982 | 4.91073461 | 9.07E-07 | 2.99E-05 |
| ANGPTL7 | 71.9728191 | -0.694357792 | 1.0493461 | -0.6617052 | 0.50816016 | 0.68701274 |
| UBIAD1 | 895.771587 | 0.091093936 | 0.27961987 | 0.32577776 | 0.7445925 | 0.85675592 |
| AL031291.1 | 0 | NA | NA | NA | NA | NA |
| DISP3 | 14.8477833 | -0.108307045 | 0.44603107 | -0.242824 | 0.80814175 | 0.89624087 |
| LINC01647 | 0.15318056 | -1.003574173 | 3.30558948 | -0.3035992 | 0.7614333 | NA |
| AL031731.1 | 17.0445742 | -1.048221904 | 0.5700008 | -1.8389832 | 0.06591765 | 0.17661537 |
| FBXO2 | 1504.6415 | -1.041224781 | 0.32088753 | -3.2448278 | 0.00117522 | 0.00890732 |
| FBXO44 | 3838.44705 | -0.141944881 | 0.25891136 | -0.5482374 | 0.58352894 | 0.74575043 |
| FBXO6 | 778.047317 | -0.797890594 | 0.35320871 | -2.2589777 | 0.02388477 | 0.08535593 |
| MAD2L2 | 1236.25366 | -0.911102483 | 0.15302851 | -5.9538086 | 2.62E-09 | 2.25E-07 |
| DRAXIN | 28.8088429 | -0.191495015 | 0.40941739 | -0.4677256 | 0.6399808 | 0.78705286 |
| AGTRAP | 3597.36523 | -0.657623206 | 0.2346029 | -2.8031333 | 0.00506087 | 0.02711174 |
| C1orf167 | 972.857207 | 0.359492979 | 0.28399846 | 1.26582723 | 0.20557494 | 0.38815776 |
| AL953897.1 | 0.57261988 | 2.514002035 | 1.57598581 | 1.59519332 | 0.11066901 | NA |
| MTHFR | 3159.89472 | 0.156564234 | 0.23623222 | 0.66275563 | 0.50748707 | 0.68645469 |
| CLCN6 | 2348.59884 | -0.134261434 | 0.28677883 | -0.4681707 | 0.63966256 | 0.78704498 |
| NPPA-AS1 | 506.063113 | -0.865228307 | 0.57980976 | -1.4922624 | 0.13563037 | 0.29333001 |
| NPPA | 221.501785 | -1.448617249 | 0.63292534 | -2.2887648 | 0.02209302 | 0.08072583 |
| NPPB | 8.9759613 | -5.58626466 | 1.69909743 | -3.2877836 | 0.00100979 | 0.00790157 |
| KIAA2013 | 4162.11518 | 0.130238903 | 0.14322616 | 0.90932342 | 0.36317943 | 0.56029027 |
| PLOD1 | 7671.82877 | 0.0679347 | 0.26246688 | 0.25883152 | 0.79576524 | 0.88920418 |
| MFN2 | 6276.266 | 0.266324623 | 0.19389763 | 1.37353214 | 0.16958698 | 0.3413747 |
| MIIP | 1292.437 | -0.54571909 | 0.14039821 | -3.8869378 | 0.00010152 | 0.00128086 |
| TNFRSF8 | 40.5442129 | 0.45845888 | 0.45398381 | 1.00985732 | 0.31256365 | 0.50941908 |
| TNFRSF1B | 2269.33014 | -0.936361788 | 0.47510812 | -1.9708394 | 0.04874225 | 0.1431144 |
| VPS13D | 3321.19142 | -0.017495456 | 0.10822615 | -0.1616565 | 0.8715764 | 0.93317628 |
| AC243836.1 | 4.65159862 | 1.691102368 | 0.62223291 | 2.7177964 | 0.00657183 | 0.03295299 |
| DHRS3 | 5595.30419 | -0.126394001 | 0.25775928 | -0.4903567 | 0.62388148 | 0.77584318 |
| AC254633.1 | 53.4436579 | -0.223798632 | 0.38851797 | -0.5760316 | 0.56459382 | 0.73156144 |
| AADACL4 | 26.5581583 | -0.311094354 | 0.60952767 | -0.5103859 | 0.6097811 | 0.76577463 |
| AADACL3 | 0.16223974 | -1.003574173 | 3.30558948 | -0.3035992 | 0.7614333 | NA |
| C1orf158 | 3.66175679 | -5.090778665 | 2.19716252 | -2.3169787 | 0.02050489 | 0.07631815 |
| PRAMEF12 | 0.16354576 | -1.003574173 | 3.30558948 | -0.3035992 | 0.7614333 | NA |
| PRAMEF1 | 0.22977084 | -1.364242679 | 3.30558948 | -0.4127078 | 0.67982068 | NA |
| LINC01784 | 0 | NA | NA | NA | NA | NA |
| PRAMEF11 | 0 | NA | NA | NA | NA | NA |
| HNRNPCL1 | 4.00602165 | -0.569578072 | 0.70120552 | -0.8122841 | 0.41662865 | 0.60981891 |
| PRAMEF2 | 0.30636111 | -1.6406616 | 3.30406282 | -0.4965588 | 0.61950019 | NA |
| PRAMEF4 | 0 | NA | NA | NA | NA | NA |
| PRAMEF10 | 0 | NA | NA | NA | NA | NA |
| PRAMEF7 | 0 | NA | NA | NA | NA | NA |
| PRAMEF6 | 0 | NA | NA | NA | NA | NA |
| PRAMEF27 | 0 | NA | NA | NA | NA | NA |
| HNRNPCL3 | 0.21786011 | 1.725828273 | 3.04405575 | 0.56695029 | 0.57074796 | NA |
| PRAMEF25 | 0 | NA | NA | NA | NA | NA |
| HNRNPCL2 | 2.65864974 | -1.402919399 | 0.92430422 | -1.5178113 | 0.12906197 | 0.28361369 |
| AC245056.1 | 0 | NA | NA | NA | NA | NA |
| PRAMEF26 | 0 | NA | NA | NA | NA | NA |
| HNRNPCL4 | 2.38994011 | -0.178848555 | 0.74633439 | -0.239636 | 0.81061249 | 0.89757931 |
| PRAMEF9 | 0.08695548 | -0.642905668 | 3.30558948 | -0.1944905 | 0.84579185 | NA |
| PRAMEF13 | 0.15318056 | -1.003574173 | 3.30558948 | -0.3035992 | 0.7614333 | NA |
| PRAMEF18 | 0.08393208 | -0.642905668 | 3.30558948 | -0.1944905 | 0.84579185 | NA |
| PRAMEF5 | 0 | NA | NA | NA | NA | NA |
| PRAMEF8 | 0.07659028 | -0.642905668 | 3.30558948 | -0.1944905 | 0.84579185 | NA |
| PRAMEF33 | 0.10721287 | -0.642905668 | 3.30558948 | -0.1944905 | 0.84579185 | NA |
| PRAMEF15 | 0 | NA | NA | NA | NA | NA |
| AC243961.1 | 0 | NA | NA | NA | NA | NA |
| PRAMEF14 | 0 | NA | NA | NA | NA | NA |
| PRAMEF19 | 0 | NA | NA | NA | NA | NA |
| PRAMEF17 | 0 | NA | NA | NA | NA | NA |
| PRAMEF20 | 0 | NA | NA | NA | NA | NA |
| LRRC38 | 2.89512443 | -0.254318365 | 0.82472016 | -0.3083693 | 0.75780135 | 0.86608684 |
| AL354712.1 | 0.44330188 | -0.923985434 | 2.82903947 | -0.3266075 | 0.74396479 | NA |
| PDPN | 251.266764 | -0.28433534 | 0.22040724 | -1.2900454 | 0.1970349 | 0.37771229 |
| AL359771.1 | 0.19978473 | -1.003574173 | 3.30558948 | -0.3035992 | 0.7614333 | NA |
| PRDM2 | 1708.84648 | -0.360068763 | 0.14090406 | -2.555418 | 0.01060604 | 0.0470606 |
| KAZN | 3524.03782 | -0.132632499 | 0.1650546 | -0.8035674 | 0.42164685 | 0.61421392 |
| AL357873.1 | 0.92658576 | -1.111249454 | 1.44273378 | -0.7702387 | 0.44115829 | NA |
| KAZN-AS1 | 0.50120839 | -0.00774142 | 1.85362503 | -0.0041764 | 0.99666775 | NA |
| TMEM51-AS | 843.119557 | -0.15283551 | 0.2133353 | -0.7164098 | 0.47373832 | 0.65959453 |
| TMEM51 | 3334.29595 | -0.846513625 | 0.25932141 | -3.2643415 | 0.00109719 | 0.00842541 |
| C1orf195 | 5.36028298 | -0.710690479 | 0.54189124 | -1.3115002 | 0.18968882 | 0.36820716 |
| FHAD1 | 97.0704606 | -0.157031353 | 0.34672141 | -0.4529035 | 0.6506182 | 0.7947011 |
| AL031283.2 | 32.5942943 | 0.205675565 | 0.38152504 | 0.53908799 | 0.58982614 | 0.74983272 |
| AL031283.3 | 3.53662912 | -0.923212229 | 0.74742348 | -1.235193 | 0.2167587 | 0.40159677 |
| AL031283.1 | 6.3907668 | 0.770431635 | 0.66006006 | 1.16721445 | 0.24312378 | 0.43368841 |
| EFHD2 | 2299.59707 | -0.568841102 | 0.22456477 | -2.5330825 | 0.01130643 | 0.04940359 |
| CTRC | 7.9223693 | -0.903377286 | 0.58274955 | -1.5501982 | 0.12109396 | 0.27158086 |
| CELA2A | 2.20703192 | -0.165539837 | 0.8201238 | -0.2018474 | 0.84003604 | 0.91492195 |
| CELA2B | 29.8706678 | 0.261700661 | 0.23517354 | 1.11279808 | 0.26579516 | 0.45996556 |
| CASP9 | 1581.24843 | 0.07553141 | 0.26151991 | 0.28881705 | 0.77272138 | 0.8747977 |
| DNAJC16 | 4119.12717 | 1.667161574 | 0.34822446 | 4.78760622 | 1.69E-06 | 4.97E-05 |
| AL121992.3 | 133.669675 | 1.866574077 | 0.4925098 | 3.78992268 | 0.00015069 | 0.00176066 |
| AGMAT | 7044.60928 | 2.807206855 | 0.65221795 | 4.30409322 | 1.68E-05 | 0.00030833 |
| AL121992.1 | 15.6768378 | 1.016069612 | 0.42969162 | 2.36464845 | 0.01804719 | 0.06955634 |
| DDI2 | 2334.7978 | 0.031984465 | 0.2213178 | 0.14451827 | 0.88509123 | 0.94051643 |
| RSC1A1 | 343.728024 | 0.125257486 | 0.22480366 | 0.55718614 | 0.57740026 | 0.7411933 |

| | | | | | | |
|---|---|---|---|---|---|---|
| AL121992.2 | 1.71171109 | -2.079710638 | 1.2490836 | -1.6649892 | 0.09591492 | 0.2309179 |
| PLEKHM2 | 6051.64059 | -0.845693074 | 0.29921275 | -2.8263939 | 0.00470753 | 0.02569606 |
| AL450998.3 | 890.661054 | -0.703961361 | 0.28692372 | -2.4534791 | 0.01414818 | 0.05837565 |
| SLC25A34 | 838.424868 | 1.412365543 | 0.47157893 | 2.99497167 | 0.0027447 | 0.01717556 |
| SLC25A34-AS | 91.2643724 | 1.235743862 | 0.44807743 | 2.75788017 | 0.00581775 | 0.03011072 |
| TMEM82 | 885.289504 | 1.345722611 | 0.28293898 | 4.75622914 | 1.97E-06 | 5.60E-05 |
| FBLIM1 | 3528.78023 | -0.990169101 | 0.35786505 | -2.7668785 | 0.00565958 | 0.02956192 |
| UQCRHL | 314.934418 | -0.902862261 | 0.32364724 | -2.7896492 | 0.00527652 | 0.02799634 |
| AL450998.2 | 284.204578 | 0.365155474 | 0.24691275 | 1.47888466 | 0.13917114 | 0.29873366 |
| SPEN | 3641.98716 | -0.221489107 | 0.26651671 | -0.8310515 | 0.40594454 | 0.60081706 |
| ZBTB17 | 2027.31209 | -0.391792915 | 0.19746708 | -1.9840923 | 0.04724555 | 0.13974931 |
| SRARP | 99.2747616 | 0.182207873 | 0.40652552 | 0.44820771 | 0.6540033 | 0.79691637 |
| AL355994.3 | 68.8796962 | 0.119562839 | 0.43688668 | 0.27367014 | 0.78433813 | 0.88217189 |
| HSPB7 | 2285.40538 | 0.348190592 | 0.74504812 | 0.46733974 | 0.64025683 | 0.78714964 |
| CLCNKA | 10130.8215 | 2.278789456 | 0.59322858 | 3.84133459 | 0.00012237 | 0.0014836 |
| CLCNKB | 14696.6381 | 1.428623691 | 0.55493783 | 2.57438513 | 0.01004184 | 0.045196 |
| FAM131C | 732.864335 | -0.084315316 | 0.52271328 | -0.1613032 | 0.87185462 | 0.93318205 |
| EPHA2 | 2775.51221 | -0.591246191 | 0.28920185 | -2.0444067 | 0.04091339 | 0.12606506 |
| AL451042.2 | 35.9820238 | -1.200922023 | 0.39228126 | -3.06138 | 0.00220319 | 0.01454637 |
| AL451042.1 | 19.8339127 | -0.802163651 | 0.42595513 | -1.8832116 | 0.05967169 | 0.16495886 |
| ARHGEF19-A | 97.7525809 | -0.652039954 | 0.36955681 | -1.7643835 | 0.07766743 | 0.19836392 |
| ARHGEF19 | 566.628415 | -0.219874289 | 0.33269999 | -0.6608786 | 0.50869019 | 0.68744736 |
| CPLANE2 | 270.722787 | 0.307664306 | 0.27773138 | 1.10777655 | 0.26795833 | 0.46209809 |
| FBXO42 | 1773.92251 | -0.441353914 | 0.16994602 | -2.5970241 | 0.00940353 | 0.04301322 |
| SZRD1 | 7562.77159 | -0.40012271 | 0.1666518 | -2.4009504 | 0.01635255 | 0.06482513 |
| SPATA21 | 2487.97415 | -0.22937957 | 0.17875888 | -1.2831786 | 0.19942952 | 0.38058738 |
| NECAP2 | 3002.41577 | -0.559162724 | 0.12847826 | -4.3521972 | 1.35E-05 | 0.00025981 |
| LINC01772 | 118.046283 | 0.2460526 | 0.24438005 | 1.00684406 | 0.31400972 | 0.51094247 |
| AL137802.2 | 16.026493 | 0.900944948 | 0.63635396 | 1.41579216 | 0.15683637 | 0.32372702 |
| AL137802.1 | 0.48237114 | -0.150743152 | 3.27672046 | -0.0460043 | 0.96330684 | NA |
| LINC01783 | 15.1217449 | 1.472748219 | 0.34190589 | 4.3074666 | 1.65E-05 | 0.00030452 |
| NBPF1 | 487.096878 | 0.238011943 | 0.19477449 | 1.22198727 | 0.22171244 | 0.40769633 |
| AL137798.1 | 2.49846776 | 0.236428936 | 0.82838704 | 0.28540878 | 0.77533098 | 0.87673576 |
| AL137798.2 | 0.41004266 | 1.035826527 | 2.21180262 | 0.4683178 | 0.63955734 | NA |
| AL021920.1 | 4.98938083 | 2.147768925 | 0.99814254 | 2.15176575 | 0.03141581 | 0.10433689 |
| AL021920.3 | 0.73435737 | -0.655127466 | 1.65401957 | -0.3960821 | 0.6920445 | NA |
| CROCC | 2476.89863 | 0.163850391 | 0.21694081 | 0.75527694 | 0.45008283 | 0.63999865 |
| BX284668.1 | 0.33829879 | -1.723487886 | 3.30203076 | -0.5219479 | 0.60170663 | NA |
| BX284668.2 | 30.1013675 | 0.842922462 | 0.39093443 | 2.15617352 | 0.03107012 | 0.10348857 |
| BX284668.5 | 297.691796 | -0.563019124 | 0.4870099 | -1.1560733 | 0.24765119 | 0.43919542 |
| BX284668.6 | 5.04777746 | -0.688145199 | 0.55670162 | -1.2361114 | 0.21641717 | 0.40127742 |
| MFAP2 | 214.986999 | -0.006474171 | 0.40113686 | -0.0161396 | 0.98712306 | 0.99410333 |
| AL049569.1 | 200.188632 | 0.074437009 | 0.18482268 | 0.40274823 | 0.68713345 | 0.81934022 |
| ATP13A2 | 2968.46329 | -0.296945323 | 0.15107915 | -1.965495 | 0.04935697 | 0.14434327 |
| SDHB | 6813.3236 | 0.352058411 | 0.19978572 | 1.76218004 | 0.07803888 | 0.1988504 |
| PADI2 | 1177.25099 | 1.968429339 | 0.52982555 | 3.7152405 | 0.00020301 | 0.00220758 |
| AL590644.1 | 23.2707149 | -3.23763022 | 0.75866479 | -4.2675372 | 1.98E-05 | 0.00035049 |
| AL590644.3 | 6.95927004 | -4.388730513 | 1.18866524 | -3.6921501 | 0.00022237 | 0.00237786 |
| PADI1 | 19.629828 | -2.407960236 | 0.67574339 | -3.563424 | 0.00036605 | 0.00355663 |
| PADI3 | 0.79320204 | -2.882788706 | 3.26206038 | -0.8837325 | 0.37684064 | NA |
| PADI4 | 40.169526 | -1.690534716 | 0.5915892 | -2.8576159 | 0.00426837 | 0.02390249 |
| PADI6 | 0.40883395 | -1.945586661 | 3.29439637 | -0.5905746 | 0.55480552 | NA |
| AC004824.1 | 1272.40904 | -0.997263732 | 0.21489976 | -4.6405995 | 3.47E-06 | 8.80E-05 |
| RCC2 | 3738.04083 | -0.482027182 | 0.16978173 | -2.8390992 | 0.00452411 | 0.02496103 |
| ARHGEF10L | 10030.2334 | 0.124409617 | 0.19400735 | 0.64126239 | 0.52135222 | 0.69769389 |
| AL136529.1 | 0.43423061 | -0.916903581 | 3.28947206 | -0.2787388 | 0.78044526 | NA |
| ACTL8 | 0.41067337 | -0.54357759 | 2.75989125 | -0.1969562 | 0.84386184 | NA |
| AL357509.1 | 0 | NA | NA | NA | NA | NA |
| LINC01654 | 0 | NA | NA | NA | NA | NA |
| IGSF21 | 117.177795 | -0.844988948 | 0.5796785 | -1.4576855 | 0.14492725 | 0.30708972 |
| IGSF21-AS1 | 0.08666371 | -0.642905668 | 3.30558948 | -0.1944905 | 0.84579185 | NA |
| AL591896.1 | 0.15543392 | 1.368031615 | 3.28033652 | 0.41704002 | 0.67664913 | NA |
| KLHDC7A | 3581.70375 | 1.628313265 | 0.44216065 | 3.68262815 | 0.00023084 | 0.00244962 |
| PAX7 | 0.03392908 | 0.380740593 | 3.29738709 | 0.11546736 | 0.90807469 | NA |
| TAS1R2 | 1.18700673 | -0.194314072 | 1.89358492 | -0.102617 | 0.91826692 | 0.96022749 |
| AL080251.1 | 10977.8831 | 1.923147391 | 0.39232304 | 4.90194862 | 9.49E-07 | 3.11E-05 |
| ALDH4A1 | 25196.4037 | 1.626573863 | 0.36016941 | 4.51613546 | 6.30E-06 | 0.00014145 |
| IFFO2 | 1479.33225 | -0.269439085 | 0.2082101 | -1.2940731 | 0.19564017 | 0.37588571 |
| UBR4 | 10116.5143 | -0.117697295 | 0.16425104 | -0.7165696 | 0.47363973 | 0.65956659 |
| AL035413.1 | 102.181096 | 0.704722627 | 0.1996598 | 3.5296171 | 0.00041616 | 0.0039334 |
| EMC1 | 2580.52883 | 0.013562545 | 0.14691295 | 0.09231688 | 0.92644628 | 0.96437374 |
| MRTO4 | 1447.81274 | -0.501894412 | 0.219111 | -2.2905943 | 0.02198689 | 0.08045993 |
| AKR7A3 | 3768.66051 | 2.243735334 | 0.2915703 | 7.69534942 | 1.41E-14 | 6.53E-12 |
| AKR7A2 | 4269.23658 | 1.04481758 | 0.30347123 | 3.44288846 | 0.00057554 | 0.00506234 |
| PQLC2 | 1493.22116 | -0.632456102 | 0.33617834 | -1.8813113 | 0.05992959 | 0.16539438 |
| CAPZB | 9522.77532 | -0.297160979 | 0.14077623 | -2.1108747 | 0.03478308 | 0.11218566 |
| AL031727.2 | 3.27063816 | 0.82768428 | 1.14796624 | 0.72100054 | 0.47090918 | 0.65748881 |
| MINOS1 | 3123.95843 | -0.109689789 | 0.13346372 | -0.8218697 | 0.41115106 | 0.60543073 |
| NBL1 | 5725.97336 | -0.766258036 | 0.14445536 | -5.3044626 | 1.13E-07 | 5.32E-06 |
| MINOS1-NBI | 3657.64655 | -0.385501763 | 0.13470247 | -2.8618759 | 0.00421142 | 0.02366398 |
| HTR6 | 1.52200943 | -0.361695011 | 1.22648036 | -0.2949049 | 0.76806656 | 0.87195714 |
| TMCO4 | 2261.47928 | -0.047448357 | 0.20894297 | -0.2270876 | 0.82035563 | 0.90356178 |
| RNF186 | 1305.61303 | 0.468347873 | 0.72302554 | 0.64776118 | 0.51713943 | 0.69438193 |
| AL391883.1 | 572.885925 | 0.43758898 | 0.7196015 | 0.60809904 | 0.54312179 | 0.71536126 |
| OTUD3 | 981.057227 | -0.273427226 | 0.239439 | -1.1419494 | 0.25347504 | 0.44608495 |
| PLA2G2E | 0 | NA | NA | NA | NA | NA |
| PLA2G2A | 345.965089 | -1.143826859 | 1.60006603 | -0.7148623 | 0.47469414 | 0.66018061 |
| PLA2G5 | 7.32387807 | -0.308071464 | 1.00059912 | -0.307887 | 0.75816832 | 0.86624344 |
| PLA2G2D | 13.0503182 | 2.933556539 | 1.01552102 | 2.88872065 | 0.00386813 | 0.02221308 |
| PLA2G2F | 0.9667549 | -3.16668931 | 2.1397906 | -1.4799062 | 0.13889829 | NA |
| Z98257.1 | 6.51872969 | -1.423490902 | 0.91355197 | -1.5581937 | 0.11918734 | 0.26862994 |
| PLA2G2C | 2.69039513 | 1.128779404 | 0.75189936 | 1.50123734 | 0.13329418 | 0.28991815 |
| UBXN10-AS1 | 21.5860171 | -0.624586656 | 0.28486946 | -2.1925364 | 0.0283408 | 0.09698188 |
| UBXN10 | 456.419856 | -0.375162463 | 0.26572144 | -1.4118637 | 0.1579901 | 0.32512868 |
| LINC01757 | 0 | NA | NA | NA | NA | NA |
| Z98257.2 | 0 | NA | NA | NA | NA | NA |
| VWA5B1 | 70.7072755 | 0.713481733 | 0.53807063 | 1.32600015 | 0.18483965 | 0.36207983 |
| AL020998.1 | 0.51989216 | -1.188532423 | 2.24582597 | -0.5292184 | 0.59665396 | NA |
| LINC01141 | 5.51074924 | 0.101541411 | 0.73431942 | 0.13827962 | 0.89001943 | 0.94301721 |
| AL139254.1 | 0 | NA | NA | NA | NA | NA |
| AL139254.2 | 2.03514243 | 3.624286566 | 1.26543027 | 2.86407451 | 0.0041823 | 0.02356563 |
| AL139254.3 | 4.65834518 | 2.628625023 | 1.29625126 | 2.02786689 | 0.04257383 | 0.12963714 |
| CAMK2N1 | 5520.48878 | 0.260504858 | 0.24869813 | 1.04747412 | 0.29488096 | 0.4916937 |
| MUL1 | 2946.02929 | -0.342186616 | 0.26030524 | -1.3145591 | 0.18865813 | 0.36687944 |
| FAM43B | 111.549087 | 0.810063799 | 0.62025155 | 1.30602463 | 0.19154418 | 0.37072406 |
| CDA | 366.350862 | -0.559348504 | 0.3042996 | -1.8381506 | 0.06604021 | 0.17681799 |
| PINK1 | 10594.1631 | 0.342479442 | 0.20960722 | 1.63391054 | 0.10227764 | 0.24117511 |
| PINK1-AS | 9796.52323 | 0.209444589 | 0.17873118 | 1.17184138 | 0.24126075 | 0.43140822 |
| DDOST | 9830.76095 | -0.102787727 | 0.13448465 | -0.7643082 | 0.44468359 | 0.63537947 |
| KIF17 | 278.321333 | -0.727224548 | 0.23505389 | -3.0938631 | 0.00197569 | 0.0133621 |
| SH2D5 | 506.477176 | -4.403694798 | 0.77024334 | -5.7172774 | 1.08E-08 | 7.41E-07 |
| AL663074.1 | 2.54506568 | -3.128299863 | 1.30719287 | -2.3931433 | 0.01670472 | 0.06584504 |
| HP1BP3 | 7474.74088 | 0.217115398 | 0.09371896 | 2.31666462 | 0.02052201 | 0.07636029 |

| | | | | | | | |
|---|---|---|---|---|---|---|---|
| EIF4G3 | 3007.01766 | | -0.512678525 | 0.21892288 | -2.3418225 | 0.01918984 | 0.07268545 |
| ECE1 | 7057.02797 | | -0.362537451 | 0.19377555 | -1.8709143 | 0.06135696 | 0.16799005 |
| AL031005.1 | 1.4048761 | | -0.183539314 | 1.37712765 | -0.1332769 | 0.89397439 | 0.9455183 |
| AL031728.1 | 17.6831167 | | -0.035072227 | 0.42447376 | -0.0826252 | 0.93414957 | 0.96837347 |
| AL592309.2 | 0 | NA | | NA | NA | NA | NA |
| NBPF3 | 309.699787 | | 0.845855371 | 0.33163348 | 2.55057293 | 0.0107546 | 0.04755976 |
| ALPL | 5350.7797 | | 1.037995605 | 0.40350682 | 2.57243635 | 0.01009855 | 0.04538927 |
| AL359815.1 | 9.64921231 | | 0.992643741 | 0.70316489 | 1.4116799 | 0.15804423 | 0.32516038 |
| RAP1GAP | 15675.0054 | | 1.115839273 | 0.40636956 | 2.74587318 | 0.00603501 | 0.03092561 |
| USP48 | 2308.1067 | | 0.042836002 | 0.09834942 | 0.43554909 | 0.66316392 | 0.80203781 |
| LDLRAD2 | 6951.32793 | | -0.512087551 | 0.48484299 | -1.0561925 | 0.29088026 | 0.48690769 |
| HSPG2 | 18901.5392 | | 0.115627822 | 0.43903553 | 0.2633678 | 0.79226711 | 0.8873257 |
| CELA3B | 0.18692855 | | 0.18657164 | 3.29377195 | 0.05664376 | 0.95482897 | NA |
| CELA3A | 0.23387521 | | 1.816617118 | 3.25499482 | 0.55810139 | 0.57677515 | NA |
| LINC01635 | 20.3491065 | | 1.503746899 | 0.43088024 | 3.48994169 | 0.00048313 | 0.00440832 |
| CDC42 | 12682.9922 | | -0.397405902 | 0.18407917 | -2.1588858 | 0.03085903 | 0.10294151 |
| AL031281.2 | 26.105159 | | -1.051595533 | 0.36955896 | -2.845542 | 0.00443359 | 0.02462097 |
| WNT4 | 85.917627 | | -0.690396627 | 0.36800441 | -1.8760553 | 0.06064768 | 0.16676532 |
| ZBTB40 | 1918.11044 | | -0.01777928 | 0.24936029 | -0.0712996 | 0.94315934 | 0.97315146 |
| EPHA8 | 6.71046884 | | -1.174676167 | 0.8972647 | -1.3091746 | 0.19047521 | 0.3691901 |
| C1QA | 4819.40334 | | -1.668230231 | 0.57056957 | -2.9237981 | 0.00345789 | 0.02039923 |
| C1QC | 4433.91432 | | -1.685601397 | 0.54684633 | -3.0824041 | 0.00205336 | 0.01376037 |
| C1QB | 5127.72019 | | -1.939544966 | 0.56016975 | -3.4624236 | 0.00053533 | 0.00478513 |
| EPHB2 | 595.799384 | | -0.920943327 | 0.39920108 | -2.306966 | 0.02105672 | 0.07782279 |
| AL512444.1 | 0.09644756 | | -0.642905668 | 3.30558948 | -0.1944905 | 0.84579185 | NA |
| LACTBL1 | 0 | NA | | NA | NA | NA | NA |
| TEX46 | 1.30970347 | | 1.48476655 | 1.11785027 | 1.32823384 | 0.18410087 | 0.3611083 |
| KDM1A | 3298.6217 | | -0.169165147 | 0.1087255 | -1.5558921 | 0.11973376 | 0.26953896 |
| AL031428.1 | 57.281714 | | 1.167135005 | 0.3386486 | 3.44644863 | 0.00056801 | 0.00501451 |
| LUZP1 | 1611.50645 | | 0.054529591 | 0.16254289 | 0.33547818 | 0.7372644 | 0.85244277 |
| HTR1D | 1.8755364 | | 1.008938242 | 1.04396587 | 0.96644754 | 0.33382029 | 0.53186158 |
| LINC01355 | 680.757839 | | -0.094814199 | 0.23890906 | -0.3968631 | 0.69146839 | 0.82241065 |
| AL109936.6 | 11.7811559 | | 0.155308034 | 0.66291756 | 0.23427956 | 0.81476795 | 0.90017726 |
| HNRNPR | 5563.85576 | | -0.30590251 | 0.23704481 | -1.2904839 | 0.1968827 | 0.37750286 |
| ZNF436 | 767.872282 | | -0.081169788 | 0.18156217 | -0.4470633 | 0.65482934 | 0.79747572 |
| ZNF436-AS1 | 326.941692 | | 0.094915213 | 0.28194977 | 0.33663873 | 0.73638925 | 0.85207763 |
| AL109936.2 | 31.1116689 | | 1.340271352 | 0.48009016 | 2.7917076 | 0.00524307 | 0.02786372 |
| TCEA3 | 2696.56586 | | 1.3437132 | 0.33839252 | 3.97087152 | 7.16E-05 | 0.00097238 |
| ASAP3 | 2768.95114 | | -0.29616834 | 0.30396545 | -0.9743487 | 0.32988344 | 0.52775746 |
| E2F2 | 34.663652 | | -1.422921558 | 0.58007726 | -2.4529863 | 0.01416757 | 0.0584287 |
| AL021154.1 | 9.96511473 | | 0.536040929 | 0.47827148 | 1.12078798 | 0.26237812 | 0.45603853 |
| ID3 | 3716.40038 | | 0.310043562 | 0.41009709 | 0.75602477 | 0.44963434 | 0.63970098 |
| MDS2 | 5.15825959 | | 0.739381926 | 0.80988266 | 0.91294944 | 0.36126913 | 0.55878832 |
| RPL11 | 26951.4274 | | -0.316212991 | 0.16136375 | -1.9596284 | 0.05003924 | 0.14571984 |
| ELOA | 2542.78417 | | -0.362952218 | 0.18459541 | -1.966204 | 0.04927505 | 0.14425776 |
| ELOA-AS1 | 762.007327 | | -0.054187029 | 0.10783893 | -0.5024811 | 0.61532912 | 0.77003744 |
| PITHD1 | 2212.04011 | | -0.403198427 | 0.21469294 | -1.8780237 | 0.06037792 | 0.16628364 |
| LYPLA2 | 6277.3223 | | -0.611814842 | 0.10943032 | -5.5909082 | 2.26E-08 | 1.39E-06 |
| GALE | 3633.70752 | | -0.435678918 | 0.23303828 | -1.8695594 | 0.06154502 | 0.16833353 |
| HMGCL | 5628.05246 | | 0.696434265 | 0.28110008 | 2.47753139 | 0.01322948 | 0.05543641 |
| FUCA1 | 5271.93508 | | -0.045694706 | 0.18594527 | -0.2457428 | 0.80588135 | 0.89485815 |
| CNR2 | 19.6940642 | | 0.100466614 | 0.7590622 | 0.13235623 | 0.89470253 | 0.94588886 |
| AL590609.3 | 0.07830766 | | -0.642905668 | 3.30558948 | -0.1944905 | 0.84579185 | NA |
| PNRC2 | 6294.38638 | | -0.278386782 | 0.16392605 | -1.6982461 | 0.08946132 | 0.21963457 |
| SRSF10 | 4547.09082 | | -0.101935897 | 0.14925641 | -0.6829583 | 0.49463322 | 0.67694119 |
| AL591178.1 | 327.034493 | | 2.313291575 | 0.49083514 | 4.71297057 | 2.44E-06 | 6.67E-05 |
| MYOM3 | 3903.14763 | | 1.705875946 | 0.41323704 | 4.12808098 | 3.66E-05 | 0.00057624 |
| AL591178.2 | 10.2577113 | | 0.411972451 | 0.50031799 | 0.82342122 | 0.4102685 | 0.60463759 |
| IL22RA1 | 528.440975 | | -1.060856696 | 0.6061831 | -1.7500598 | 0.08010799 | 0.20261727 |
| IFNLR1 | 543.52836 | | 0.489377028 | 0.19229093 | 2.54498243 | 0.01092832 | 0.04812623 |
| AL590683.1 | 0.58162973 | | -2.421131678 | 3.27556873 | -0.7391485 | 0.45981684 | NA |
| AL138902.1 | 1.07265237 | | 0.755568296 | 1.27879736 | 0.59084287 | 0.55462571 | 0.72417116 |
| GRHL3 | 42.3969467 | | -1.311377324 | 0.50713321 | -2.5858636 | 0.00971354 | 0.04410163 |
| STPG1 | 734.73397 | | -0.203025834 | 0.13457334 | -1.5086631 | 0.1313849 | 0.28713905 |
| NIPAL3 | 1626.5668 | | 0.097353852 | 0.18937189 | 0.51408818 | 0.60719034 | 0.76354837 |
| RCAN3 | 818.354852 | | -0.554461475 | 0.25002055 | -2.2176636 | 0.02657778 | 0.09242438 |
| AL445686.2 | 184.45524 | | -0.204812145 | 0.26800521 | -0.7642096 | 0.44474237 | 0.63538066 |
| NCMAP | 390.623556 | | -2.263571027 | 0.41472449 | -5.4580115 | 4.81E-08 | 2.58E-06 |
| SRRM1 | 3315.75489 | | -0.01453269 | 0.12033332 | -0.1207703 | 0.90387298 | 0.95167363 |
| AL445648.1 | 0.26775652 | | -1.364242679 | 3.30558948 | -0.4127078 | 0.67982068 | NA |
| CLIC4 | 17182.4122 | | 0.956308815 | 0.2379975 | 4.01814654 | 5.87E-05 | 0.00083623 |
| RUNX3 | 264.917778 | | -0.584730862 | 0.56339675 | -1.0378669 | 0.299332 | 0.49609609 |
| AL445471.1 | 1.9173351 | | 0.992290504 | 1.05838902 | 0.93754799 | 0.34847675 | 0.54600902 |
| AL445471.2 | 0 | NA | | NA | NA | NA | NA |
| AL050344.1 | 0.16052235 | | -1.003574173 | 3.30558948 | -0.3035992 | 0.7614333 | NA |
| AL031432.1 | 0.64098057 | | -1.787886417 | 2.29935127 | -0.7775612 | 0.43682772 | NA |
| SYF2 | 2848.87431 | | -0.367672086 | 0.14185125 | -2.5919551 | 0.00954322 | 0.04350878 |
| AL031432.5 | 10.4994796 | | 0.538529537 | 0.47884997 | 1.12463104 | 0.26074542 | 0.45391184 |
| RSRP1 | 8275.88134 | | -0.111669771 | 0.19653081 | -0.5682049 | 0.56989583 | 0.73521253 |
| AL031432.3 | 11.989325 | | 0.550591021 | 0.41275881 | 1.33392919 | 0.18222707 | 0.35888956 |
| RHD | 79.3640776 | | -0.370358851 | 1.08368068 | -0.3417601 | 0.73253142 | 0.8492009 |
| TMEM50A | 6317.73224 | | -0.600990382 | 0.12066908 | -4.9804835 | 6.34E-07 | 2.22E-05 |
| RHCE | 257.63445 | | -0.712379323 | 0.17710872 | -4.0222713 | 5.76E-05 | 0.00082758 |
| MACO1 | 1838.69611 | | -0.197293238 | 0.2548414 | -0.7741805 | 0.43882407 | 0.63037483 |
| LDLRAP1 | 1743.04989 | | 0.308392379 | 0.11105022 | 2.77705319 | 0.00548542 | 0.02888389 |
| AL606491.1 | 15.3440739 | | -0.57959555 | 0.54713432 | -1.0593296 | 0.28944972 | 0.48570087 |
| MAN1C1 | 3624.91881 | | 1.164305907 | 0.3779999 | 3.08017514 | 0.00206879 | 0.01381813 |
| AL031280.1 | 19.9521586 | | 0.383898263 | 0.44012186 | 0.87225448 | 0.38306957 | 0.57895142 |
| SELENON | 7760.40258 | | -0.014311274 | 0.20373703 | -0.0702439 | 0.94399957 | 0.97322897 |
| AL020996.2 | 2211.88895 | | 0.745283875 | 0.10288221 | 7.24405017 | 4.35E-13 | 1.30E-10 |
| AL020996.1 | 2012.2302 | | -0.022948092 | 0.20474143 | -0.1120833 | 0.91075737 | 0.9558494 |
| MTFR1L | 5764.41358 | | 0.762999657 | 0.15501176 | 4.92220509 | 8.56E-07 | 2.84E-05 |
| AL020996.3 | 1447.57934 | | 0.754695717 | 0.15693706 | 4.80890681 | 1.52E-06 | 4.57E-05 |
| AUNIP | 1539.11072 | | 0.750510943 | 0.15519017 | 4.8360728 | 1.32E-06 | 4.09E-05 |
| AL033528.2 | 1164.1145 | | 1.344515268 | 0.32737201 | 4.10699521 | 4.01E-05 | 0.00062353 |
| PAQR7 | 4414.70732 | | 1.078180905 | 0.31375245 | 3.43640628 | 0.00058949 | 0.00515239 |
| STMN1 | 2281.1208 | | -0.10787092 | 0.28835466 | -0.3740911 | 0.70833651 | 0.83417046 |
| PAFAH2 | 1923.03167 | | 0.495151587 | 0.22722138 | 2.17915932 | 0.02931983 | 0.0992381 |
| EXTL1 | 48.8546882 | | 0.189121701 | 0.36703739 | 0.5152655 | 0.6063675 | 0.76308872 |
| SLC30A2 | 3444.42687 | | -0.450838178 | 0.48526166 | -0.929062 | 0.35285695 | 0.5505121 |
| TRIM63 | 68.1488809 | | 1.583801406 | 0.56391976 | 2.8085581 | 0.00497639 | 0.02677067 |
| PDIK1L | 463.270493 | | 0.294174925 | 0.21057116 | 1.39703332 | 0.16240355 | 0.33138509 |
| FAM110D | 278.282063 | | -0.553652951 | 0.4428369 | -1.2502412 | 0.21121144 | 0.39461916 |
| C1orf232 | 0.08666371 | | -0.642905668 | 3.30558948 | -0.1944905 | 0.84579185 | NA |
| AL391650.1 | 816.424435 | | -0.774955167 | 0.1554922 | -4.9838846 | 6.23E-07 | 2.20E-05 |
| ZNF593 | 811.200013 | | -0.794632299 | 0.15524665 | -5.1185149 | 3.08E-07 | 1.22E-05 |
| CNKSR1 | 1182.59187 | | -0.124084167 | 0.23171229 | -0.5355096 | 0.59229749 | 0.75170299 |
| CATSPER4 | 2.87058177 | | 0.569735816 | 0.71299454 | 0.79907458 | 0.42424717 | 0.61669135 |
| CEP85 | 806.69476 | | -0.641970129 | 0.32759972 | -1.9596175 | 0.05004052 | 0.14571984 |
| SH3BGRL3 | 5645.44804 | | -1.123444111 | 0.25691779 | -4.3727766 | 1.23E-05 | 0.00024232 |
| UBXN11 | 5656.29483 | | -0.485716264 | 0.15379651 | -3.1581748 | 0.0015876 | 0.01128743 |
| CD52 | 288.007197 | | 0.551603632 | 0.69752071 | 0.7908061 | 0.42905715 | 0.62112297 |
| CRYBG2 | 138.074418 | | -1.336691507 | 0.55896467 | -2.3913703 | 0.01678561 | 0.06610464 |

| | | | | | | |
|---|---|---|---|---|---|---|
| ZNF683 | 31.4859578 | 0.16590216 | 0.84938461 | 0.19532042 | 0.84514211 | 0.91828433 |
| LIN28A | 0.37042038 | -0.03213209 | 2.27356282 | -0.0141329 | 0.98872393 | NA |
| DHDDS | 2311.11608 | 0.117055371 | 0.20193039 | 0.57968179 | 0.56212923 | 0.72980721 |
| AL513365.2 | 15.2772725 | 0.597194055 | 0.31575176 | 1.89134041 | 0.05857892 | 0.16274742 |
| HMGN2 | 6177.90176 | -0.080540654 | 0.25205712 | -0.3195333 | 0.74932211 | 0.85998627 |
| RPS6KA1 | 4732.23174 | -0.50813981 | 0.23129617 | -2.1969227 | 0.02802597 | 0.09616003 |
| AL512408.1 | 41.3568208 | 1.009705776 | 0.50734463 | 1.99017733 | 0.0465714 | 0.13828262 |
| ARID1A | 3399.90883 | 0.201160629 | 0.09626993 | 2.08954779 | 0.03665844 | 0.11628571 |
| PIGV | 960.515077 | 0.607179162 | 0.21447762 | 2.83096739 | 0.00464075 | 0.02545024 |
| ZDHHC18 | 2251.12627 | -1.201316448 | 0.35028202 | -3.4295692 | 0.00060454 | 0.00524917 |
| SFN | 619.44783 | -4.478204426 | 0.46427624 | -9.6455602 | 5.13E-22 | 1.35E-18 |
| AL034380.1 | 169.6825 | -0.550518658 | 0.19130303 | -2.8777311 | 0.00400546 | 0.02277576 |
| GPN2 | 2096.33552 | -0.304742176 | 0.14389523 | -2.1178059 | 0.03419151 | 0.11074278 |
| GPATCH3 | 919.450542 | -0.389538997 | 0.25789831 | -1.5104364 | 0.1309321 | 0.28650565 |
| NUDC | 5226.82542 | -0.486509332 | 0.13014779 | -3.7381298 | 0.00018539 | 0.00205846 |
| NR0B2 | 582.229463 | -0.092325251 | 0.42851606 | -0.2154534 | 0.82941383 | 0.90934819 |
| KDF1 | 832.885073 | -0.352698021 | 0.15789846 | -2.2337014 | 0.02550272 | 0.08958397 |
| TRNP1 | 1853.76587 | -1.974142861 | 0.32694137 | -6.0382168 | 1.56E-09 | 1.47E-07 |
| TENT5B | 402.411121 | 0.009362858 | 0.54756546 | 0.01709907 | 0.98635758 | 0.99391317 |
| SLC9A1 | 2226.92252 | -0.18381943 | 0.17082507 | -1.0760683 | 0.28189673 | 0.47776958 |
| AL590640.1 | 0 | NA | NA | NA | NA | NA |
| WDTC1 | 5408.27094 | 0.163429483 | 0.13307838 | 1.22806941 | 0.21942091 | 0.40496427 |
| TMEM222 | 2854.6874 | -0.193354814 | 0.18733379 | -1.0321406 | 0.30200625 | 0.49883526 |
| SYTL1 | 430.418499 | -0.244373889 | 0.4772009 | -0.5120986 | 0.60858204 | 0.76467853 |
| MAP3K6 | 3371.10946 | -0.677527903 | 0.24207882 | -2.7987905 | 0.00512944 | 0.02740343 |
| FCN3 | 1400.24129 | -0.331976636 | 0.64983381 | -0.5108639 | 0.60944636 | 0.76547276 |
| CD164L2 | 38.1324003 | -1.868982499 | 0.53324809 | -3.5049024 | 0.00045677 | 0.00422785 |
| GPR3 | 93.8930118 | -1.352099192 | 0.51492601 | -2.6258126 | 0.00864424 | 0.04033775 |
| WASF2 | 7124.01971 | -0.120489251 | 0.13888469 | -0.8675489 | 0.38564136 | 0.58143868 |
| FO393419.2 | 0.33992608 | -0.54125709 | 2.87508463 | -0.1882578 | 0.85067457 | NA |
| BX293535.1 | 4.12409456 | 0.368708423 | 0.77979789 | 0.47282563 | 0.63633759 | 0.78445244 |
| AHDC1 | 3459.98646 | 0.065211746 | 0.30493873 | 0.21385197 | 0.8306625 | 0.91033489 |
| FGR | 798.479526 | -0.506465551 | 0.39888371 | -1.2697073 | 0.20418892 | 0.38650935 |
| LINC02574 | 2.51430719 | -2.169156381 | 1.7777599 | -1.2201627 | 0.22240319 | 0.40839745 |
| IFI6 | 2840.84487 | -1.482939947 | 0.68786753 | -2.1558511 | 0.0310953 | 0.10355936 |
| AL445490.1 | 8.4503598 | 1.152740231 | 0.77199679 | 1.49319303 | 0.13538667 | 0.2929471 |
| FAM76A | 811.306123 | 0.117392764 | 0.12835581 | 0.9145886 | 0.36040764 | 0.5580355 |
| STX12 | 3557.09536 | -0.189536226 | 0.13568114 | -1.3969239 | 0.16243647 | 0.33138509 |
| PPP1R8 | 1776.3278 | -0.163293565 | 0.17328714 | -0.9423294 | 0.34602404 | 0.54355477 |
| THEMIS2 | 1322.7532 | -1.630171996 | 0.43527932 | -3.745117 | 0.00018031 | 0.00201217 |
| RPA2 | 1733.77245 | -0.134013341 | 0.1869359 | -0.7168946 | 0.47343913 | 0.65944379 |
| SMPDL3B | 647.024558 | -0.660054417 | 0.28155973 | -2.3442785 | 0.01906393 | 0.07244789 |
| AL512288.1 | 289.746257 | -0.682752099 | 0.29276461 | -2.3320855 | 0.01969619 | 0.074038 |
| XKR8 | 812.359071 | -0.354999633 | 0.1716773 | -2.0678309 | 0.03865592 | 0.12076819 |
| EYA3 | 2052.85735 | 0.002471243 | 0.19702608 | 0.01254272 | 0.98999262 | 0.99527661 |
| PTAFR | 401.712508 | -1.025542642 | 0.35705742 | -2.8722065 | 0.00407617 | 0.0230761 |
| DNAJC8 | 4225.45042 | -0.323248395 | 0.16765773 | -1.9280256 | 0.05385194 | 0.153506 |
| ATP5IF1 | 7068.05164 | 0.544307305 | 0.1503826 | 3.61948333 | 0.00029519 | 0.00299278 |
| AL353622.1 | 418.163581 | 1.245493441 | 0.32531625 | 3.82856199 | 0.00012889 | 0.00154429 |
| AL353622.2 | 5.91178904 | 1.400163092 | 0.75987565 | 1.84262134 | 0.06538432 | 0.17552524 |
| SESN2 | 2758.63407 | 0.45851372 | 0.2415204 | 1.89844717 | 0.0576372 | 0.16090334 |
| MED18 | 569.288355 | 0.006236094 | 0.17842331 | 0.03495111 | 0.97211872 | 0.98686367 |
| PHACTR4 | 2620.59369 | 0.081805109 | 0.17422729 | 0.46953097 | 0.63869016 | 0.78618782 |
| RCC1 | 1770.13297 | -1.551858307 | 0.27343546 | -5.6754099 | 1.38E-08 | 9.17E-07 |
| TRNAU1AP | 1073.81323 | -0.758085501 | 0.15042491 | -5.0396275 | 4.66E-07 | 1.71E-05 |
| SNHG12 | 1880.62303 | -1.720869398 | 0.24755551 | -6.9514484 | 3.62E-12 | 8.59E-10 |
| TAF12 | 758.489615 | -0.245404 | 0.09899462 | -2.4789631 | 0.01317649 | 0.05529066 |
| RAB42 | 159.026615 | -2.289720657 | 0.68162674 | -3.3592002 | 0.00078168 | 0.00645779 |
| LINC01715 | 16.9983151 | 0.735242446 | 0.40862154 | 1.79932374 | 0.07196748 | 0.18808549 |
| AL360012.1 | 1.91407092 | -1.688885474 | 1.06890148 | -1.5800198 | 0.11410234 | 0.2606218 |
| GMEB1 | 651.594988 | -0.237412698 | 0.23702224 | -1.0016474 | 0.31651394 | 0.51369231 |
| YTHDF2 | 3550.21774 | -0.298214905 | 0.2019897 | -1.4763867 | 0.13984011 | 0.29975481 |
| OPRD1 | 24.9731376 | -0.479446606 | 0.37934804 | -1.26387 | 0.2062767 | 0.38889801 |
| AL009181.1 | 9.58450254 | 0.286134321 | 0.48284448 | 0.59260141 | 0.55344793 | 0.72299723 |
| EPB41 | 2750.85836 | -0.424237966 | 0.15913397 | -2.6659171 | 0.00767786 | 0.03704762 |
| TMEM200B | 1520.88273 | -0.497513953 | 0.1720868 | -2.891064 | 0.0038394 | 0.02210918 |
| AL357500.1 | 0.39457337 | -0.543269465 | 2.77439999 | -0.1958151 | 0.84475488 | NA |
| SRSF4 | 6054.58623 | -0.529610988 | 0.14145175 | -3.7441105 | 0.00018103 | 0.00201684 |
| MECR | 1188.88928 | 0.111001116 | 0.16598253 | 0.66875178 | 0.50365383 | 0.68364658 |
| AL590729.1 | 6.12337451 | 0.610043356 | 0.75798477 | 0.80482271 | 0.42092199 | 0.61354889 |
| PTPRU | 1325.43146 | -1.109131188 | 0.25966723 | -4.271356 | 1.94E-05 | 0.00034641 |
| LINC01756 | 0 | NA | NA | NA | NA | NA |
| AC092265.1 | 0 | NA | NA | NA | NA | NA |
| AL645944.2 | 0.6432772 | -2.554755786 | 3.27121203 | -0.7809814 | 0.43481343 | NA |
| LINC01648 | 1.8353303 | -2.842936546 | 1.42721468 | -1.9919474 | 0.04637683 | 0.13787565 |
| AL137076.1 | 0 | NA | NA | NA | NA | NA |
| AL161638.2 | 0 | NA | NA | NA | NA | NA |
| AL161638.1 | 0 | NA | NA | NA | NA | NA |
| MATN1 | 41.897777 | -0.048622227 | 0.32102565 | -0.151459 | 0.87961364 | 0.93749138 |
| MATN1-AS1 | 98.8296562 | -0.379388101 | 0.28690409 | -1.3223517 | 0.18605107 | 0.36355441 |
| LAPTM5 | 4213.88145 | -1.099520188 | 0.41757224 | -2.6331257 | 0.0084603 | 0.03969013 |
| AL137027.1 | 0 | NA | NA | NA | NA | NA |
| LINC01778 | 1.41332528 | 0.182157298 | 0.97671682 | 0.1864996 | 0.85205299 | 0.92250934 |
| AL445235.1 | 0 | NA | NA | NA | NA | NA |
| SDC3 | 3020.87864 | -0.700496669 | 0.22215921 | -3.1531291 | 0.0016153 | 0.01144734 |
| PUM1 | 4021.36352 | -0.072347133 | 0.12159643 | -0.5949774 | 0.55185854 | 0.72178454 |
| NKAIN1 | 30.9924572 | -2.006673565 | 0.72194767 | -2.7795277 | 0.0054438 | 0.02871615 |
| SNRNP40 | 1389.13108 | -0.287760317 | 0.13421867 | -2.1439664 | 0.03203558 | 0.10572943 |
| ZCCHC17 | 1492.98702 | -0.612905664 | 0.12633053 | -4.8516037 | 1.22E-06 | 3.84E-05 |
| AL451070.1 | 501.043283 | -0.394153703 | 0.15991939 | -2.4647024 | 0.01371271 | 0.05691865 |
| FABP3 | 4516.7989 | 0.3802587 | 0.39725355 | 0.95721913 | 0.33845666 | 0.53651565 |
| SERINC2 | 14027.6625 | -0.788434292 | 0.28810111 | -2.7366583 | 0.00620667 | 0.03165016 |
| AC114488.1 | 19.6640346 | 0.319292267 | 0.35084477 | 0.91006704 | 0.36278715 | 0.5600455 |
| TINAGL1 | 17382.5269 | -0.622475889 | 0.24911918 | -2.4987072 | 0.01246473 | 0.0530507 |
| HCRTR1 | 559.658266 | -0.21201802 | 0.15018769 | -1.4116871 | 0.15804213 | 0.32516038 |
| PEF1 | 4568.31132 | -0.430422071 | 0.11648121 | -3.6952059 | 0.00021971 | 0.00235421 |
| AC114488.2 | 550.434143 | -0.332351043 | 0.24742082 | -1.3432622 | 0.17918707 | 0.3548671 |
| AC114488.3 | 158.452854 | -0.149203801 | 0.22806036 | -0.6542294 | 0.51296401 | 0.69102737 |
| COL16A1 | 3548.25513 | -0.091060606 | 0.31730399 | -0.2869822 | 0.77412593 | 0.87574864 |
| ADGRB2 | 220.72702 | 0.531253978 | 0.42426043 | 1.25218838 | 0.21050123 | 0.39390558 |
| AL354919.2 | 13.2615262 | -3.274124019 | 0.86969967 | -3.7646605 | 0.00016678 | 0.00189809 |
| SPOCD1 | 73.944235 | -2.899271025 | 0.58651737 | -4.9431972 | 7.69E-07 | 2.60E-05 |
| AL354919.1 | 3.5853483 | 3.810591599 | 1.01572783 | 3.75158727 | 0.00017572 | 0.00197937 |
| AL136115.2 | 9.01164712 | -0.851980101 | 0.51748358 | -1.6463906 | 0.09968337 | 0.23696479 |
| PTP4A2 | 9216.46787 | -0.927539153 | 0.23530036 | -3.9419369 | 8.08E-05 | 0.00107105 |
| AL136115.1 | 1.40820214 | -0.909553081 | 1.07800384 | -0.8437383 | 0.39881569 | 0.59366467 |
| AL136115.3 | 0 | NA | NA | NA | NA | NA |
| KHDRBS1 | 5610.63994 | -0.109034704 | 0.16333819 | -0.6675396 | 0.50442753 | 0.68392997 |
| AL445248.1 | 1.97234093 | 0.243730365 | 0.83295708 | 0.29260855 | 0.76982138 | 0.87275106 |
| TMEM39B | 712.036806 | -0.330167592 | 0.17766586 | -1.8583626 | 0.06311754 | 0.17130603 |
| KPNA6 | 3753.48697 | -0.149431503 | 0.15846239 | -0.9430093 | 0.34567616 | 0.54333196 |
| TXLNA | 4449.24477 | -0.449321562 | 0.18779506 | -2.3926165 | 0.01672872 | 0.06590025 |
| CCDC28B | 430.711248 | 0.064360169 | 0.21877971 | 0.29417796 | 0.76862192 | 0.87217134 |

| | | | | | | |
|---|---|---|---|---|---|---|
| AL049795.1 | 295.28595 | 0.272849262 | 0.1834647 | 1.48720306 | 0.13696119 | 0.29536036 |
| IQCC | 331.06215 | 0.345160782 | 0.17928188 | 1.92524074 | 0.05419925 | 0.15419508 |
| DCDC2B | 456.528158 | -0.347717549 | 0.29459207 | -1.1803358 | 0.2378667 | 0.42725597 |
| TMEM234 | 1409.17019 | -0.474185365 | 0.22000413 | -2.1553475 | 0.03113465 | 0.10365114 |
| EIF3I | 7233.89263 | -0.289441991 | 0.15548334 | -1.8615627 | 0.06266476 | 0.17048072 |
| FAM167B | 832.560159 | -0.409116075 | 0.46874224 | -0.8727954 | 0.3827746 | 0.5788687 |
| LCK | 299.109329 | -0.083372114 | 0.67024147 | -0.1243912 | 0.90100557 | 0.9498264 |
| HDAC1 | 4361.2974 | -0.139063363 | 0.12679969 | -1.0967169 | 0.27276517 | 0.46762862 |
| MARCKSL1 | 2231.39923 | -1.007311458 | 0.3378671 | -2.9813837 | 0.00286949 | 0.01773341 |
| AL109945.1 | 6.64028087 | 0.885661755 | 0.76684954 | 1.15493551 | 0.24811684 | 0.43979781 |
| TSSK3 | 579.357747 | -0.237251454 | 0.25433653 | -0.9328249 | 0.35091037 | 0.54857388 |
| FAM229A | 957.891954 | -0.340061483 | 0.22818682 | -1.4902766 | 0.13615152 | 0.29423993 |
| BSDC1 | 4826.90582 | -0.001378777 | 0.11308725 | -0.0121922 | 0.99027231 | 0.99533983 |
| AL033529.1 | 94.0410167 | 0.191130733 | 0.25728863 | 0.74286507 | 0.45756338 | 0.64679422 |
| ZBTB8B | 14.7821667 | 0.301031118 | 0.40975149 | 0.73466755 | 0.46254199 | 0.65076201 |
| ZBTB8A | 361.022712 | -0.082064348 | 0.21987193 | -0.373237 | 0.70897203 | 0.83447565 |
| ZBTB8OS | 1124.62462 | -0.409754579 | 0.14124949 | -2.9009278 | 0.0037206 | 0.02158208 |
| RBBP4 | 6119.10178 | -0.060862489 | 0.10896982 | -0.5585261 | 0.57648518 | 0.74050923 |
| SYNC | 945.736177 | -0.223645253 | 0.11914208 | -1.8771307 | 0.06050018 | 0.16648477 |
| AC114489.1 | 0.3503936 | -1.726457206 | 2.8412215 | -0.6076461 | 0.54342221 | NA |
| KIAA1522 | 11670.5472 | -0.373647222 | 0.19272183 | -1.9387904 | 0.05252687 | 0.15073822 |
| YARS | 3503.87473 | -0.865074479 | 0.1890115 | -4.5768351 | 4.72E-06 | 0.00011255 |
| S100PBP | 1446.54475 | -0.212606321 | 0.14908975 | -1.4260291 | 0.15385994 | 0.3195972 |
| FNDC5 | 71.2000443 | 1.623781597 | 0.32858787 | 4.94169664 | 7.74E-07 | 2.62E-05 |
| HPCA | 1017.64141 | -1.629579748 | 0.38403127 | -4.2433518 | 2.20E-05 | 0.0003838 |
| TMEM54 | 2572.71574 | -1.730392117 | 0.39137989 | -4.4212596 | 9.81E-06 | 0.00020043 |
| RNF19B | 2799.4655 | -0.522759145 | 0.29748965 | -1.7572347 | 0.07887781 | 0.20046337 |
| AL020995.1 | 96.3702512 | 0.638714704 | 0.17903813 | 3.56747868 | 0.00036043 | 0.00350983 |
| AK2 | 10181.7314 | 0.158988404 | 0.22230089 | 0.71519465 | 0.47448878 | 0.66007262 |
| AZIN2 | 655.646234 | -0.287865343 | 0.20653406 | -1.3937911 | 0.16338068 | 0.33263964 |
| TRIM62 | 537.381326 | -0.278137244 | 0.21097834 | -1.3183213 | 0.1873961 | 0.36534343 |
| AL662907.3 | 0.35848737 | -0.557500269 | 2.27381927 | -0.2451823 | 0.80631527 | NA |
| ZNF362 | 1748.39225 | 0.662417333 | 0.19692838 | 3.3637474 | 0.00076892 | 0.00636621 |
| AL513327.3 | 3.99200237 | -0.429957336 | 0.63134411 | -0.681019 | 0.49585946 | 0.67772047 |
| A3GALT2 | 13.590637 | -0.211853644 | 0.58735982 | -0.360688 | 0.71833269 | 0.839869 |
| AL513327.1 | 561.567297 | -1.136367848 | 0.30305111 | -3.7497565 | 0.00017701 | 0.00198878 |
| PHC2 | 16873.1038 | -1.511668722 | 0.26875435 | -5.6247228 | 1.86E-08 | 1.17E-06 |
| AL513327.2 | 1.79563226 | -1.758969202 | 1.96043697 | -0.8972332 | 0.36959448 | 0.56638748 |
| ZSCAN20 | 134.73362 | -0.413168273 | 0.27311419 | -1.5128041 | 0.13032942 | 0.28551856 |
| CSMD2 | 39.2762904 | 0.108209893 | 0.42462937 | 0.25483375 | 0.79885151 | 0.89107318 |
| HMGB4 | 0 | NA | NA | NA | NA | NA |
| CSMD2-AS1 | 0 | NA | NA | NA | NA | NA |
| C1orf94 | 0.39177108 | 0.994474711 | 2.22381451 | 0.44719319 | 0.65473558 | NA |
| SMIM12 | 2579.7348 | -0.716406607 | 0.24142602 | -2.967396 | 0.00300334 | 0.01837541 |
| GJB5 | 4.88398582 | 0.122498556 | 0.74613123 | 0.1641783 | 0.86959078 | 0.93213734 |
| GJB4 | 7.68485645 | -2.206529634 | 1.0967818 | -2.0118219 | 0.04423872 | 0.13324671 |
| AL121988.1 | 2.52298728 | -1.518561022 | 1.73336946 | -0.8760746 | 0.38098946 | 0.57703131 |
| GJB3 | 45.9816805 | -2.017053805 | 0.71771408 | -2.8103863 | 0.00494821 | 0.02667298 |
| GJA4 | 1625.51351 | -0.600358256 | 0.44228132 | -1.3574126 | 0.17465013 | 0.34859424 |
| AL122010.1 | 157.624709 | -0.379443848 | 0.38693873 | -0.9806303 | 0.32677508 | 0.52469456 |
| DLGAP3 | 92.9709766 | -0.19755082 | 0.28756221 | -0.6869846 | 0.49209241 | 0.6750144 |
| AC114490.3 | 699.268423 | -0.270790252 | 0.20444499 | -1.324514 | 0.18533239 | 0.36268759 |
| AC114490.2 | 1746.60598 | -0.16073903 | 0.10547673 | -1.5239289 | 0.12752648 | 0.28120021 |
| TMEM35B | 1365.53572 | -0.32033132 | 0.198616 | -1.6128173 | 0.10678421 | 0.24857872 |
| ZMYM6 | 1328.78491 | 0.094041118 | 0.12870026 | 0.73069874 | 0.46496318 | 0.65209958 |
| ZMYM1 | 473.890591 | 0.243058801 | 0.1663246 | 1.46135205 | 0.14391885 | 0.30561747 |
| AL590434.1 | 36.3912128 | 1.708995972 | 0.51306808 | 3.33093411 | 0.00086555 | 0.00702079 |
| SFPQ | 12777.4686 | -0.087776281 | 0.17113869 | -0.5128956 | 0.60802436 | 0.76426914 |
| ZMYM4 | 2234.16151 | -0.213346806 | 0.13668642 | -1.5608486 | 0.11855947 | 0.26765102 |
| ZMYM4-AS1 | 9.4023593 | 0.247924953 | 0.38345759 | 0.64655117 | 0.51792247 | 0.69490259 |
| KIAA0319L | 5035.16637 | 0.18072489 | 0.16753416 | 1.07873456 | 0.28070608 | 0.4763946 |
| NCDN | 2080.65339 | -0.041856597 | 0.15259619 | -0.2742965 | 0.7838568 | 0.88187652 |
| AC004865.2 | 18.9118395 | -0.514188263 | 0.43050226 | -1.1943916 | 0.23232484 | 0.42042314 |
| TFAP2E | 77.0009089 | -0.561192138 | 0.27487056 | -2.0416597 | 0.0411853 | 0.1266095 |
| PSMB2 | 4154.88032 | -0.631917726 | 0.16905687 | -3.7379002 | 0.00018556 | 0.00205888 |
| C1orf216 | 593.292095 | -1.06931281 | 0.24020351 | -4.4516951 | 8.52E-06 | 0.00017958 |
| CLSPN | 9.4192455 | -0.303958024 | 0.57970975 | -0.5243279 | 0.60005046 | 0.75778606 |
| AL354864.1 | 2.05777455 | -1.651607827 | 0.94262217 | -1.7521419 | 0.07974941 | 0.20202197 |
| AGO4 | 1445.83582 | -0.524038189 | 0.12154143 | -4.3116012 | 1.62E-05 | 0.00030035 |
| AGO1 | 1708.62681 | 0.331450381 | 0.13794623 | 2.40275053 | 0.01627228 | 0.06458461 |
| AL139286.2 | 27.5159364 | 0.11310012 | 0.26801073 | 0.42199848 | 0.67302613 | 0.809264 |
| AGO3 | 1413.06379 | -0.138307248 | 0.1133133 | -1.2205738 | 0.22224743 | 0.40826643 |
| AL138787.2 | 34.44662 | -0.176287499 | 0.44393843 | -0.397099 | 0.69129447 | 0.82241065 |
| TEKT2 | 277.506147 | -0.776354168 | 0.50902269 | -1.5251858 | 0.12721277 | 0.28079682 |
| ADPRHL2 | 2936.72076 | -1.15289624 | 0.24473338 | -4.7108255 | 2.47E-06 | 6.73E-05 |
| COL8A2 | 211.627114 | -0.88565418 | 0.42308766 | -2.0933113 | 0.03632137 | 0.11558876 |
| TRAPPC3 | 3467.77132 | -0.387521185 | 0.21026525 | -1.8430111 | 0.06532739 | 0.17540813 |
| MAP7D1 | 4712.2924 | -0.812539833 | 0.30036311 | -2.7051919 | 0.00682649 | 0.03393302 |
| THRAP3 | 5470.04391 | -0.057876765 | 0.1390321 | -0.4162835 | 0.67720259 | 0.81234738 |
| SH3D21 | 2552.24575 | -0.09948607 | 0.18059332 | -0.5508845 | 0.58171283 | 0.74428759 |
| EVA1B | 1429.5768 | -0.557780962 | 0.14334397 | -3.8912064 | 9.97E-05 | 0.00126376 |
| AL591845.1 | 842.753436 | 0.071237956 | 0.27969524 | 0.25469849 | 0.79895598 | 0.89108294 |
| STK40 | 5929.62857 | -0.800672538 | 0.31545678 | -2.538137 | 0.01114443 | 0.04882517 |
| LSM10 | 1131.16263 | -0.267610144 | 0.16040068 | -1.6683854 | 0.09523925 | 0.22975593 |
| OSCP1 | 712.20649 | 0.210718914 | 0.1622177 | 1.29898846 | 0.19394789 | 0.37377811 |
| MRPS15 | 3205.39283 | -0.172883875 | 0.15226129 | -1.1354421 | 0.25619014 | 0.44900532 |
| CSF3R | 902.988753 | -0.818705697 | 0.41236308 | -1.9854001 | 0.04709998 | 0.13939038 |
| AL596257.1 | 0.69809255 | 3.383681486 | 1.79476911 | 1.88530183 | 0.0593891 | NA |
| AC117945.2 | 0 | NA | NA | NA | NA | NA |
| AC117945.1 | 0 | NA | NA | NA | NA | NA |
| GRIK3 | 18.6047035 | 0.773219869 | 0.84264578 | 0.91760962 | 0.35882328 | 0.5564673 |
| AL031430.1 | 0 | NA | NA | NA | NA | NA |
| AL353604.1 | 0 | NA | NA | NA | NA | NA |
| LINC01137 | 331.629806 | -0.740503151 | 0.2291655 | -3.2313029 | 0.00123227 | 0.00924731 |
| ZC3H12A | 2313.09184 | -0.9011784 | 0.46335915 | -1.944881 | 0.0517893 | 0.14945107 |
| MEAF6 | 2547.39725 | -0.46745332 | 0.0939011 | -4.9781452 | 6.42E-07 | 2.24E-05 |
| SNIP1 | 793.532304 | -0.254001707 | 0.19136346 | -1.3273261 | 0.18440085 | 0.36153535 |
| DNALI1 | 1477.35526 | 0.155835619 | 0.25184329 | 0.6187801 | 0.53606123 | 0.70984909 |
| GNL2 | 1847.75018 | -0.605101603 | 0.24207038 | -2.4996929 | 0.0124301 | 0.05296199 |
| RSPO1 | 24.7937504 | -0.004070451 | 0.79996467 | -0.0050883 | 0.99594015 | 0.99802173 |
| C1orf109 | 783.680909 | 0.177366929 | 0.19707842 | 0.89998149 | 0.3681301 | 0.56492032 |
| CDCA8 | 70.1109222 | -0.97068651 | 0.3775635 | -2.5709226 | 0.0101428 | 0.04554933 |
| EPHA10 | 53.7318082 | -1.110185585 | 0.6386592 | -1.7383067 | 0.08215678 | 0.20627196 |
| MANEAL | 273.441264 | 0.388894664 | 0.43809523 | 0.88769435 | 0.37470518 | 0.5707258 |
| AL929472.3 | 174.738041 | 0.311399025 | 0.45827891 | 0.67949674 | 0.49682317 | 0.67865838 |
| YRDC | 1008.10486 | -0.492631551 | 0.26695733 | -1.8453569 | 0.06498564 | 0.17478766 |
| C1orf122 | 2240.20937 | -0.445464072 | 0.16091677 | -2.7682886 | 0.00563515 | 0.02948392 |
| MTF1 | 987.952184 | -0.482517197 | 0.21072773 | -2.289766 | 0.02203489 | 0.08057966 |
| AL929472.2 | 545.978016 | -0.675563739 | 0.20202606 | -3.3439436 | 0.00082597 | 0.00674743 |
| INPP5B | 1977.48801 | -0.232620991 | 0.1780494 | -1.3064969 | 0.19138363 | 0.37051511 |
| SF3A3 | 2912.62304 | -0.212805905 | 0.15960582 | -1.3333217 | 0.18242625 | 0.35910573 |
| FHL3 | 684.118698 | -1.079314956 | 0.3697585 | -2.9189727 | 0.00351187 | 0.02062539 |
| UTP11 | 1015.71094 | -0.509580872 | 0.22893323 | -2.2258929 | 0.02602135 | 0.09100949 |

| | | | | | | |
|---|---|---|---|---|---|---|
| POU3F1 | 6.16271332 | 0.882239652 | 0.83433698 | 1.05741406 | 0.29032266 | 0.48645371 |
| AL139158.2 | 2.63129526 | 0.0001087 | 1.00602355 | 0.00010805 | 0.99991379 | 0.99993924 |
| AL139158.3 | 1.60763466 | 1.244506279 | 1.26393581 | 0.98462776 | 0.32480695 | 0.52236136 |
| LINC01343 | 0.38747328 | 0.706820657 | 2.28672457 | 0.30909742 | 0.75724742 | NA |
| LINC01685 | 0.90075495 | 3.282232326 | 1.38381406 | 2.37187382 | 0.01769813 | NA |
| AL354702.2 | 7.41982709 | 2.875348195 | 1.22200312 | 2.35297943 | 0.01862366 | 0.0712574 |
| RRAGC | 2884.95078 | -0.866966091 | 0.43639674 | -1.9866466 | 0.04696158 | 0.13914345 |
| AL139260.1 | 97.2201262 | -0.829348743 | 0.44995147 | -1.843196 | 0.0653004 | 0.17536884 |
| MYCBP | 1387.44454 | -0.294952824 | 0.17090244 | -1.7258549 | 0.08437352 | 0.21041513 |
| AL139260.3 | 570.583958 | -0.277473996 | 0.14614408 | -1.8986332 | 0.05761272 | 0.16088429 |
| GJA9 | 57.559749 | -0.444994322 | 0.46569318 | -0.9555526 | 0.33929833 | 0.53743974 |
| RHBDL2 | 111.862034 | -1.173477432 | 0.2944419 | -3.9854294 | 6.74E-05 | 0.00093079 |
| AKIRIN1 | 4203.18564 | -0.340431688 | 0.1537321 | -2.2144477 | 0.026798 | 0.09292086 |
| NDUFS5 | 5183.30501 | -0.630061371 | 0.21174545 | -2.9755604 | 0.00292454 | 0.01798509 |
| MACF1 | 9760.75518 | -0.224891637 | 0.20059436 | -1.1211264 | 0.26223404 | 0.45590135 |
| AL442071.1 | 4.00204393 | 0.053889567 | 0.74428227 | 0.07240474 | 0.94227981 | 0.97267355 |
| BMP8A | 238.952132 | -0.144148196 | 0.21985082 | -0.6556637 | 0.51204055 | 0.69020648 |
| PABPC4 | 5113.77328 | -0.407638713 | 0.16032846 | -2.5425225 | 0.01100555 | 0.04838553 |
| AL365277.1 | 855.583532 | -0.073280766 | 0.20922306 | -0.3502519 | 0.72614969 | 0.84507557 |
| HEYL | 1080.39421 | 0.064461382 | 0.50643879 | 0.12728366 | 0.89871589 | 0.94852624 |
| AL035404.2 | 1.20893176 | 0.827468401 | 1.55081984 | 0.53356836 | 0.5936402 | 0.7528212 |
| NT5C1A | 8.45035317 | -0.225558878 | 0.72106742 | -0.3128125 | 0.75442314 | 0.86329626 |
| HPCAL4 | 616.20038 | -0.045559184 | 0.61869372 | -0.0736377 | 0.94129867 | 0.97207934 |
| PPIE | 2005.42769 | -0.092260277 | 0.18905394 | -0.4880103 | 0.62554252 | 0.77702936 |
| BMP8B | 309.914959 | -0.210895034 | 0.24670804 | -0.8548365 | 0.39264168 | 0.58819298 |
| OXCT2 | 8.97030556 | -1.488819889 | 0.56878327 | -2.6175522 | 0.00885629 | 0.04113077 |
| AL033527.3 | 10.7006461 | -1.081492958 | 0.45139572 | -2.3958866 | 0.01658022 | 0.06547175 |
| AL033527.5 | 43.851051 | -2.1492284 | 0.49764287 | -4.3188169 | 1.57E-05 | 0.00029379 |
| AL033527.4 | 13.4486353 | -2.710926877 | 0.61260344 | -4.4252557 | 9.63E-06 | 0.00019768 |
| TRIT1 | 1261.9106 | -0.307990785 | 0.12621686 | -2.4401716 | 0.01468029 | 0.05994413 |
| MYCL | 125.628127 | -0.662387367 | 0.31518199 | -2.1016029 | 0.03558808 | 0.11395964 |
| AL033527.2 | 10.919329 | -0.161265897 | 0.4261093 | -0.3784613 | 0.70508791 | 0.83164862 |
| MFSD2A | 1803.35611 | -3.618631052 | 0.53485596 | -6.7656179 | 1.33E-11 | 2.65E-09 |
| CAP1 | 12059.4312 | -0.359271063 | 0.18761273 | -1.914961 | 0.05549747 | 0.15665472 |
| PPT1 | 11927.198 | -0.352694587 | 0.15331015 | -2.3005299 | 0.02141822 | 0.07886044 |
| RLF | 1111.17215 | -0.678707873 | 0.19575115 | -3.4671974 | 0.00052592 | 0.00471393 |
| TMCO2 | 1.70978178 | -0.446010922 | 1.10033224 | -0.405342 | 0.6852261 | 0.81810054 |
| AL050341.2 | 472.284942 | 0.363040672 | 0.29448436 | 1.23280118 | 0.21764995 | 0.40265393 |
| ZMPSTE24 | 3632.11057 | 0.136127304 | 0.19159339 | 0.71050103 | 0.47739349 | 0.66250218 |
| COL9A2 | 1545.97592 | 0.87417167 | 0.41485944 | 2.10715143 | 0.03510446 | 0.11290456 |
| SMAP2 | 2093.25621 | -0.500663179 | 0.25904 | -1.932764 | 0.05326528 | 0.15229377 |
| AL031985.4 | 0.38168934 | -1.028785817 | 2.27758123 | -0.451701 | 0.65148437 | NA |
| ZFP69B | 27.4169122 | -0.236642085 | 0.43853033 | -0.5396254 | 0.58945543 | 0.74961886 |
| ZFP69 | 185.508376 | 0.07970582 | 0.20255642 | 0.39349934 | 0.69395071 | 0.82378141 |
| AL603839.3 | 26.8192141 | 0.420485394 | 0.47133206 | 0.89212134 | 0.3723279 | 0.56878538 |
| EXO5 | 319.524287 | 0.172238208 | 0.24448612 | 0.70449076 | 0.4811272 | 0.66575665 |
| AL603839.1 | 124.702927 | 0.260978003 | 0.28544877 | 0.91427266 | 0.36057359 | 0.55819692 |
| AL603839.2 | 71.0333307 | -0.123372945 | 0.27719861 | -0.4450706 | 0.65626875 | 0.79827515 |
| ZNF684 | 515.973924 | 0.631615242 | 0.25743247 | 2.45351813 | 0.01414664 | 0.05837565 |
| RIMS3 | 354.887966 | -0.780951768 | 0.36684152 | -2.1288532 | 0.0332664 | 0.1085687 |
| AL031289.1 | 18.5657108 | 0.542254503 | 0.42005514 | 1.29091267 | 0.19673396 | 0.37732741 |
| NFYC-AS1 | 360.567489 | 0.62433535 | 0.17326389 | 3.60337843 | 0.00031411 | 0.00314346 |
| NFYC | 2939.72066 | 0.436676553 | 0.124464 | 3.50845663 | 0.00045071 | 0.0041834 |
| KCNQ4 | 111.328754 | -0.39800288 | 0.3736257 | -1.0652449 | 0.28676511 | 0.48300946 |
| CITED4 | 869.381551 | -1.673238367 | 0.23438178 | -7.1389437 | 9.41E-13 | 2.51E-10 |
| AC119677.1 | 0.5606987 | -1.283648767 | 2.21587629 | -0.5792962 | 0.56238931 | NA |
| CTPS1 | 1731.88156 | -0.445833507 | 0.29248403 | -1.5243003 | 0.1274337 | 0.28104794 |
| SLFNL1-AS1 | 697.890614 | -0.465861328 | 0.16589447 | -2.8081788 | 0.00498226 | 0.02679525 |
| SLFNL1 | 133.239158 | -0.298507338 | 0.27116444 | -1.1008351 | 0.27096842 | 0.46554185 |
| SCMH1 | 3185.63643 | -0.38035536 | 0.14384851 | -2.6441384 | 0.00818992 | 0.03878041 |
| AC093151.3 | 15.1560149 | -1.117563159 | 0.62348499 | -1.792446 | 0.07306156 | 0.19013454 |
| FOXO6 | 75.1050764 | -0.983731752 | 0.52751523 | -1.8648405 | 0.06220377 | 0.16952851 |
| AC093151.2 | 9.53399543 | -0.70477481 | 0.68136711 | -1.034354 | 0.30097069 | 0.49768474 |
| EDN2 | 363.689464 | -1.133772715 | 0.96542573 | -1.1743759 | 0.24024449 | 0.43032517 |
| HIVEP3 | 396.733047 | -0.527346552 | 0.22364932 | -2.3579171 | 0.0183778 | 0.0705624 |
| AL445933.2 | 0 | NA | NA | NA | NA | NA |
| AC119676.1 | 1.55188899 | -3.138133257 | 1.67622312 | -1.8721453 | 0.0611865 | 0.16768739 |
| GUCA2B | 3.34845703 | 0.1727851 | 1.27893915 | 0.13510033 | 0.89253255 | 0.94442773 |
| GUCA2A | 0.73228724 | -1.978516175 | 1.90251609 | -1.0399471 | 0.29836446 | NA |
| FOXJ3 | 2535.69039 | -0.163088449 | 0.16943254 | -0.9625569 | 0.33576996 | 0.53393559 |
| RIMKLA | 373.509238 | -1.137303999 | 0.41241537 | -2.7576664 | 0.00582156 | 0.0301245 |
| ZMYND12 | 328.775413 | 1.267467172 | 0.37928166 | 3.34175706 | 0.0008325 | 0.00678819 |
| PPCS | 3263.02727 | 0.262874351 | 0.24217057 | 1.08549255 | 0.27770356 | 0.4732072 |
| CCDC30 | 561.945412 | 0.338079655 | 0.24905225 | 1.35746477 | 0.17463358 | 0.34859424 |
| PPIH | 634.224471 | -0.34406647 | 0.13620929 | -2.5260133 | 0.01153651 | 0.05012043 |
| AC098484.2 | 14.9928999 | 0.451384021 | 0.3182273 | 1.41843276 | 0.15606446 | 0.32257562 |
| YBX1 | 16227.1275 | -0.405912198 | 0.24424414 | -1.6619117 | 0.09653047 | 0.23186833 |
| CLDN19 | 1451.21925 | 2.428725815 | 0.65621086 | 3.70113629 | 0.00021464 | 0.00231302 |
| P3H1 | 2035.21127 | -0.89838995 | 0.26593497 | -3.3782317 | 0.00072954 | 0.00610544 |
| C1orf50 | 570.662456 | 0.048803788 | 0.1763989 | 0.27666718 | 0.78203567 | 0.88076709 |
| AC098484.3 | 457.856586 | -0.115799744 | 0.2129023 | -0.5439102 | 0.58650324 | 0.74768058 |
| AC098484.1 | 30.2919146 | 0.547802136 | 0.22314834 | 2.45487879 | 0.01409321 | 0.05820485 |
| TMEM269 | 278.208723 | -0.340235318 | 0.18243406 | -1.8649769 | 0.06218464 | 0.16952851 |
| SVBP | 574.080383 | -0.32535067 | 0.15325773 | -2.122899 | 0.03376232 | 0.10976604 |
| ERMAP | 1133.48807 | 0.386577182 | 0.17060292 | 2.26594705 | 0.02345463 | 0.08424114 |
| AL512353.1 | 433.287513 | 0.484085216 | 0.20843575 | 2.32246733 | 0.02020779 | 0.07542882 |
| ZNF691 | 762.644282 | -0.080497221 | 0.13183455 | -0.6105928 | 0.54146921 | 0.7142843 |
| SLC2A1 | 7171.73848 | -2.173835015 | 0.40176529 | -5.4107088 | 6.28E-08 | 3.21E-06 |
| SLC2A1-AS1 | 55.0448033 | -1.021408767 | 0.57010735 | -1.7916078 | 0.07319582 | 0.19036309 |
| AC099795.1 | 21.6935546 | -0.7602904 | 0.45515639 | -1.6703938 | 0.09484148 | 0.22913618 |
| FAM183A | 53.9819356 | -0.441795677 | 0.41538151 | -1.0635901 | 0.28751441 | 0.48378822 |
| EBNA1BP2 | 3390.52315 | 0.375843016 | 0.18586709 | 2.0221063 | 0.04316537 | 0.13100136 |
| CFAP57 | 823.594771 | 0.992023712 | 0.23854755 | 4.15859937 | 3.20E-05 | 0.00051707 |
| TMEM125 | 962.072161 | 0.149257594 | 0.28214676 | 0.52900694 | 0.59680064 | 0.7553822 |
| C1orf210 | 1450.281 | 0.178538462 | 0.36344397 | 0.49124067 | 0.62325624 | 0.77535816 |
| TIE1 | 2528.01525 | -0.106456077 | 0.35111513 | -0.3031942 | 0.76174186 | 0.86791967 |
| MPL | 19.8859525 | 1.155115571 | 0.30519871 | 3.78479838 | 0.00015383 | 0.00178704 |
| AL139289.2 | 32.3016388 | 0.523274414 | 0.40797316 | 1.28261969 | 0.19962535 | 0.3808784 |
| CDC20 | 158.149392 | -1.12519615 | 0.38567315 | -2.9174863 | 0.00352865 | 0.0207009 |
| ELOVL1 | 5014.84226 | -0.518387946 | 0.15587411 | -3.3256833 | 0.00088202 | 0.00711479 |
| MED8 | 1959.65211 | -0.285026693 | 0.15901548 | -1.7924461 | 0.07306153 | 0.19013454 |
| AL139289.1 | 486.27024 | -0.167315128 | 0.16976828 | -0.98555 | 0.32435399 | 0.52189213 |
| SZT2 | 5993.10125 | 0.156132253 | 0.09038429 | 1.72742686 | 0.08409103 | 0.2098893 |
| SZT2-AS1 | 508.674445 | 0.129803263 | 0.15263011 | 0.85044336 | 0.39507864 | 0.59041733 |
| HYI | 3394.28968 | 0.017306693 | 0.13814646 | 0.12527786 | 0.90030358 | 0.94951124 |
| PTPRF | 7503.94688 | -0.099532338 | 0.1284167 | -0.7750732 | 0.43829642 | 0.63008807 |
| AL451062.3 | 1593.40925 | 0.515175871 | 0.11288628 | 4.56367132 | 5.03E-06 | 0.00011751 |
| KDM4A | 1984.85124 | 0.383936548 | 0.118662 | 3.23554761 | 0.0012141 | 0.00913169 |
| KDM4A-AS1 | 785.795637 | 0.140086419 | 0.13867866 | 1.01015121 | 0.31242285 | 0.50924654 |
| ST3GAL3 | 812.956505 | 0.275327252 | 0.1756196 | 1.56774786 | 0.11693999 | 0.26528798 |
| AL451062.1 | 0.69536255 | 0.929273804 | 1.52605346 | 0.60893922 | 0.54256473 | NA |
| ARTN | 338.314655 | -1.897554746 | 0.25726105 | -7.3759892 | 1.63E-13 | 5.87E-11 |
| AL357079.1 | 24.5975809 | 0.728580345 | 0.41086345 | 1.77329074 | 0.07618054 | 0.19589282 |

| | | | | | | |
|---|---|---|---|---|---|---|
| IPO13 | 4161.58115 | 0.29687253 | 0.23691237 | 1.25309001 | 0.21017295 | 0.3934588 |
| DPH2 | 1751.58048 | -0.248576647 | 0.22953599 | -1.0829528 | 0.27882937 | 0.47446154 |
| ATP6V0B | 10415.9243 | -0.59587247 | 0.16700128 | -3.5680713 | 0.00035962 | 0.00350319 |
| B4GALT2 | 2144.98888 | -0.534110376 | 0.18947785 | -2.8188539 | 0.00481954 | 0.02613182 |
| CCDC24 | 1842.54306 | 0.024512475 | 0.29685786 | 0.0825731 | 0.93419099 | 0.96837347 |
| SLC6A9 | 818.241869 | -0.15053518 | 0.23562116 | -0.6388865 | 0.52289677 | 0.69862624 |
| AL139220.2 | 23.6135231 | -0.111575331 | 0.24626432 | -0.4530714 | 0.6504973 | 0.79460075 |
| KLF17 | 0.24156165 | -1.364244367 | 3.02237876 | -0.451381 | 0.65171497 | NA |
| KLF18 | 0 | NA | NA | NA | NA | NA |
| DMAP1 | 2493.1494 | 0.194865227 | 0.11790612 | 1.65271513 | 0.09838885 | 0.23497691 |
| ERI3 | 3691.74463 | 0.218055077 | 0.18676393 | 1.16754385 | 0.24299081 | 0.43356571 |
| RNF220 | 3370.96013 | -0.236867425 | 0.10901537 | -2.1727894 | 0.02979617 | 0.10042499 |
| TMEM53 | 1884.53222 | -0.041597596 | 0.12535813 | -0.3318301 | 0.74001758 | 0.85412805 |
| ARMH1 | 176.892861 | -0.179867215 | 0.43351796 | -0.4149014 | 0.67821407 | 0.81313471 |
| KIF2C | 80.6406685 | -0.687130059 | 0.46418864 | -1.4802819 | 0.13879802 | 0.29815117 |
| AL592166.1 | 3147.06776 | -0.394095609 | 0.21225725 | -1.8566886 | 0.06335547 | 0.17177501 |
| RPS8 | 26313.0449 | -0.682029336 | 0.19017746 | -3.5862784 | 0.00033543 | 0.00331533 |
| BEST4 | 51.5205001 | -1.540992619 | 0.38656046 | -3.9864207 | 6.71E-05 | 0.00092987 |
| PLK3 | 1098.23833 | -0.609200387 | 0.42887175 | -1.4204722 | 0.15547026 | 0.32175096 |
| TCTEX1D4 | 55.4660664 | -0.838959964 | 0.32904862 | -2.5496535 | 0.010783 | 0.04764828 |
| BTBD19 | 675.797055 | 0.077777392 | 0.25023755 | 0.31081423 | 0.75594185 | 0.86456115 |
| PTCH2 | 209.259644 | 0.165303805 | 0.35602907 | 0.46429861 | 0.64243382 | 0.78833864 |
| EIF2B3 | 867.8032 | -0.133224966 | 0.15721344 | -0.8474146 | 0.39676408 | 0.5917799 |
| HECTD3 | 4052.41834 | -0.153153275 | 0.16908523 | -0.9057756 | 0.36505464 | 0.56205495 |
| UROD | 4426.44696 | 0.081003448 | 0.14659203 | 0.55257742 | 0.58055281 | 0.74350374 |
| ZSWIM5 | 496.29635 | 0.421441889 | 0.19132602 | 2.20274212 | 0.02761293 | 0.09510734 |
| LINC01144 | 100.140781 | 0.85090968 | 0.32612974 | 2.609114 | 0.0090777 | 0.0419301 |
| HPDL | 30.5001571 | 0.143350604 | 0.31794491 | 0.45086617 | 0.65208601 | 0.79564631 |
| MUTYH | 1346.52299 | 0.26029905 | 0.17641873 | 1.47546157 | 0.14008849 | 0.30016524 |
| TOE1 | 734.728862 | 0.269695934 | 0.18804695 | 1.43419464 | 0.15151673 | 0.31657328 |
| TESK2 | 390.347531 | 0.151635907 | 0.2249228 | 0.67416868 | 0.50020408 | 0.68135656 |
| CCDC163 | 183.793493 | 0.328934446 | 0.66390528 | 0.49545388 | 0.62027977 | 0.77307768 |
| MMACHC | 13183.327 | -0.64520733 | 0.28465448 | -2.2666333 | 0.02341263 | 0.08413616 |
| PRDX1 | 29046.0314 | -0.63570321 | 0.29650218 | -2.1440086 | 0.0320322 | 0.10572943 |
| AKR1A1 | 13281.9425 | 0.958801137 | 0.29774879 | 3.220168 | 0.00128115 | 0.00953544 |
| NASP | 3999.84999 | -0.568475494 | 0.20346416 | -2.7939835 | 0.00520631 | 0.02770125 |
| CCDC17 | 499.309808 | -1.011548432 | 0.39159414 | -2.5831552 | 0.00979013 | 0.04436737 |
| GPBP1L1 | 5895.7974 | -0.05576847 | 0.1195565 | -0.4664612 | 0.6408854 | 0.78744515 |
| TMEM69 | 1238.53187 | -0.236334412 | 0.13174592 | -1.7938651 | 0.0728347 | 0.18975113 |
| IPP | 538.134021 | 0.561903775 | 0.1843062 | 3.04875135 | 0.00229795 | 0.01503644 |
| MAST2 | 2724.03817 | -0.347795773 | 0.23170725 | -1.5010138 | 0.133352 | 0.29001998 |
| PIK3R3 | 1775.73439 | -0.236286352 | 0.31887057 | -0.7410102 | 0.45868724 | 0.64757615 |
| AL358075.4 | 119.550914 | 0.220815521 | 0.41740335 | 0.52902192 | 0.59679024 | 0.7553822 |
| AL358075.1 | 2.40154942 | 0.278914445 | 0.74075188 | 0.37652884 | 0.70652378 | 0.83304414 |
| AL358075.2 | 3.09359367 | 0.614005182 | 0.83113952 | 0.73875103 | 0.46005819 | 0.6486998 |
| TSPAN1 | 24634.4079 | -0.538777078 | 0.23090297 | -2.3333484 | 0.01962987 | 0.07388518 |
| AL672043.1 | 2.21196622 | -0.017810287 | 1.28884916 | -0.0138188 | 0.98897458 | 0.99503126 |
| POMGNT1 | 5736.17611 | 0.113423762 | 0.12866876 | 0.88151751 | 0.37803778 | 0.57397837 |
| LURAP1 | 314.756077 | -0.123624762 | 0.34991079 | -0.3533037 | 0.72386081 | 0.84360319 |
| RAD54L | 375.561312 | -1.003386449 | 0.23228817 | -4.3195762 | 1.56E-05 | 0.0002932 |
| LRRC41 | 5737.94157 | -0.46536668 | 0.18410615 | -2.5277084 | 0.01148097 | 0.04997424 |
| UQCRH | 6346.37457 | -0.11095619 | 0.17183936 | -0.6456972 | 0.51847549 | 0.69536155 |
| NSUN4 | 1408.34125 | -0.024812174 | 0.13674368 | -0.1814502 | 0.85601419 | 0.92460093 |
| FAAH | 3458.7395 | 1.248247248 | 0.31159975 | 4.00593148 | 6.18E-05 | 0.00086836 |
| DMBX1 | 1.64045938 | -3.209540747 | 1.38701386 | -2.3139933 | 0.02066809 | 0.07674142 |
| AL136373.1 | 0 | NA | NA | NA | NA | NA |
| MKNK1-AS1 | 447.491507 | -0.170219151 | 0.15339988 | -1.1096433 | 0.26715275 | 0.46124006 |
| KNCN | 0.25087713 | 1.011917048 | 2.94486708 | 0.34362062 | 0.73113162 | NA |
| MKNK1 | 2099.8289 | -0.252033512 | 0.13441225 | -1.8750784 | 0.06078194 | 0.16699518 |
| MOB3C | 1478.25161 | -0.047481686 | 0.3034539 | -0.1564708 | 0.8756619 | 0.93543465 |
| ATPAF1 | 3767.38634 | 0.724939155 | 0.21123638 | 3.43188589 | 0.0005994 | 0.00521033 |
| TEX38 | 5.59823646 | 0.788190316 | 0.58425903 | 1.34904258 | 0.17732329 | 0.35240147 |
| EFCAB14-AS | 88.9951777 | 0.948013021 | 0.35706162 | 2.65504039 | 0.00792989 | 0.03791681 |
| EFCAB14 | 6265.37257 | 0.184874919 | 0.10933587 | 1.69088993 | 0.09085783 | 0.22213248 |
| CYP4B1 | 136.706885 | -0.406263928 | 0.7084679 | -0.5734401 | 0.56634672 | 0.73284279 |
| CYP4A11 | 27374.7257 | 1.592509972 | 0.68886698 | 2.31178153 | 0.02078973 | 0.07707368 |
| CYP4X1 | 323.276027 | 1.734161504 | 0.35440704 | 4.89313506 | 9.92E-07 | 3.23E-05 |
| CYP4Z1 | 20.8075066 | 1.350148516 | 0.51334606 | 2.63009425 | 0.00853612 | 0.03996759 |
| CYP4A22-AS | 100.72002 | 1.000743862 | 0.80619018 | 1.2413248 | 0.21448579 | 0.39864605 |
| CYP4A22 | 4391.76047 | 1.258184042 | 0.82039744 | 1.53362745 | 0.12512133 | 0.27747452 |
| LINC00853 | 112.007497 | 0.320114365 | 0.6349012 | 0.50419556 | 0.61412396 | 0.76907674 |
| PDZK1IP1 | 48920.6067 | 0.421467852 | 0.31060257 | 1.35693614 | 0.1748015 | 0.34865396 |
| TAL1 | 440.329072 | -0.250771662 | 0.35523165 | -0.7059384 | 0.48022645 | 0.66492893 |
| AL135960.1 | 79.9645364 | 0.267889608 | 0.35423503 | 0.75624822 | 0.44950039 | 0.63959456 |
| STIL | 141.31174 | -0.820388179 | 0.25749074 | -3.1860881 | 0.00144211 | 0.01049347 |
| CMPK1 | 7022.31152 | -0.251181389 | 0.20005924 | -1.2555351 | 0.20928458 | 0.3924644 |
| LINC01389 | 30.0901495 | -0.707541937 | 0.36255939 | -1.9515201 | 0.05099521 | 0.14772532 |
| FOXE3 | 4.7147269 | -0.147774557 | 0.64106381 | -0.2305146 | 0.81769193 | 0.90201305 |
| FOXD2-AS1 | 135.516805 | -0.607690517 | 0.26554042 | -2.2885048 | 0.02210814 | 0.08075796 |
| FOXD2 | 138.613552 | 0.276716132 | 0.28350527 | 0.97605286 | 0.32903827 | 0.52688482 |
| LINC01738 | 4.66451834 | 1.723132175 | 0.97271106 | 1.77147381 | 0.07648194 | 0.19632326 |
| TRABD2B | 3967.01052 | -0.436995381 | 0.25457289 | -1.7165825 | 0.08605548 | 0.21339881 |
| AL691459.1 | 642.599899 | -0.314722178 | 0.30653119 | -1.0267215 | 0.3045516 | 0.5015003 |
| AC096541.1 | 22.750565 | -0.717857214 | 0.44715046 | -1.6054042 | 0.10840488 | 0.25139605 |
| AL356289.1 | 13.8943292 | -0.093192842 | 0.51438581 | -0.181173 | 0.85623175 | 0.92471767 |
| AL356289.2 | 0.91011852 | -0.761463804 | 1.60451866 | -0.4745746 | 0.63509021 | NA |
| AL109659.1 | 6.81358388 | 0.472643948 | 0.46273198 | 1.02142053 | 0.30705524 | 0.50401785 |
| SLC5A9 | 1228.82047 | 1.57086836 | 0.52848129 | 2.97242002 | 0.00295462 | 0.01813126 |
| AL109659.2 | 200.578821 | 0.955892519 | 0.33772057 | 2.83042432 | 0.00464863 | 0.02548002 |
| SPATA6 | 770.152637 | 0.076169769 | 0.18921322 | 0.40256051 | 0.68727157 | 0.81935672 |
| AGBL4 | 108.030161 | 1.68317495 | 0.4324338 | 3.8923298 | 9.93E-05 | 0.00125874 |
| BEND5 | 529.165759 | 0.522881246 | 0.26057467 | 2.0066465 | 0.04478732 | 0.13440782 |
| AC099788.1 | 0.06785815 | 0.795868262 | 3.28898349 | 0.24198001 | 0.80879565 | NA |
| AL590432.1 | 0.03392908 | 0.380740593 | 3.29738709 | 0.11546736 | 0.90807469 | NA |
| ELAVL4 | 3.49247927 | 1.44382187 | 0.72519302 | 1.99094838 | 0.04648656 | 0.13813958 |
| AL592182.2 | 0 | NA | NA | NA | NA | NA |
| AL592182.1 | 0.09989236 | -0.642905668 | 3.30558948 | -0.1944905 | 0.84579185 | NA |
| AL596275.2 | 0.71040915 | 2.811161836 | 1.60694984 | 1.74937747 | 0.08022579 | NA |
| AL592182.3 | 0.10721287 | -0.642905668 | 3.30558948 | -0.1944905 | 0.84579185 | NA |
| DMRTA2 | 0.99851597 | -1.173681305 | 1.3441603 | -0.8731706 | 0.38257008 | NA |
| FAF1 | 2053.58808 | -0.001235891 | 0.1133972 | -0.0108988 | 0.99130421 | 0.99583588 |
| AL049637.1 | 0.51676656 | -0.048478335 | 1.93980127 | -0.0249914 | 0.98006183 | NA |
| CDKN2C | 492.131798 | 0.013254381 | 0.29187968 | 0.04541043 | 0.96378017 | 0.98305633 |
| C1orf185 | 0.49431582 | 1.489051476 | 1.57421375 | 0.94590171 | 0.34419872 | NA |
| LINC01562 | 7.96049602 | 1.061470997 | 0.45180372 | 2.34940736 | 0.01880332 | 0.0717574 |
| RNF11 | 6778.16436 | -0.141371235 | 0.18894138 | -0.748228 | 0.45432262 | 0.64389264 |
| AL162430.2 | 6.15250807 | -0.276068421 | 0.76526041 | -0.3607509 | 0.71828563 | 0.83985952 |
| TTC39A | 5456.77012 | -2.54727467 | 0.44631592 | -5.7073354 | 1.15E-08 | 7.80E-07 |
| TTC39A-AS1 | 99.5064451 | -2.715390006 | 0.48762374 | -5.5686173 | 2.57E-08 | 1.54E-06 |
| EPS15 | 3059.08866 | 0.001787921 | 0.08245992 | 0.02168231 | 0.98270138 | 0.99232988 |
| AC104170.2 | 2.6144602 | -0.17721134 | 0.87595332 | -0.2023068 | 0.83967686 | 0.91484357 |
| AC104170.1 | 14.851189 | 0.091905962 | 0.50161949 | 0.18321848 | 0.85462659 | 0.92399813 |
| OSBPL9 | 7110.17346 | 0.135697731 | 0.13987352 | 0.97014595 | 0.33197375 | 0.52994417 |

| | | | | | | |
|---|---|---|---|---|---|---|
| NRDC | 5809.19305 | -0.425439943 | 0.15630674 | -2.7218272 | 0.00649221 | 0.03266543 |
| AL050343.1 | 980.342391 | -0.525829821 | 0.18421566 | -2.8544252 | 0.00431148 | 0.02409275 |
| AL050343.2 | 9.2237559 | 0.014724306 | 0.52753621 | 0.02791146 | 0.97773277 | 0.9897433 |
| RAB3B | 124.807307 | -0.260550673 | 0.54985674 | -0.4738519 | 0.6356055 | 0.78387977 |
| TXNDC12 | 2958.51699 | -0.558974105 | 0.12029153 | -4.6468283 | 3.37E-06 | 8.59E-05 |
| KTI12 | 457.595909 | -0.420864404 | 0.29392587 | -1.4318726 | 0.15218029 | 0.31725782 |
| AL445685.1 | 86.7017993 | -0.271165755 | 0.30093346 | -0.9010821 | 0.36754467 | 0.56438321 |
| TXNDC12-AS | 1.05900102 | -0.368384498 | 1.28285871 | -0.2871591 | 0.77399053 | 0.8756706 |
| BTF3L4 | 2043.12045 | -0.414220701 | 0.10795518 | -3.8369693 | 0.00012456 | 0.00150398 |
| ZFYVE9 | 1296.29001 | 0.461709665 | 0.12193911 | 3.78639528 | 0.00015285 | 0.00177952 |
| CC2D1B | 4734.35282 | -0.541663967 | 0.22874078 | -2.3680254 | 0.01788331 | 0.0690256 |
| AL513218.1 | 627.60827 | -0.769475665 | 0.23484266 | -3.2765584 | 0.00105081 | 0.00815168 |
| ORC1 | 69.2517328 | 0.588906717 | 0.23879692 | 2.46614029 | 0.01365778 | 0.05675868 |
| PRPF38A | 1886.67966 | -0.009644345 | 0.11190672 | -0.086182 | 0.93132174 | 0.96690482 |
| TUT4 | 1780.19842 | -0.058778918 | 0.18271653 | -0.3216946 | 0.74768409 | 0.85884766 |
| AL591167.1 | 0.65381436 | 0.993961142 | 1.74773641 | 0.56871341 | 0.56955064 | NA |
| GPX7 | 465.076701 | -0.214951723 | 0.22565469 | -0.9525693 | 0.34080835 | 0.53886055 |
| SHISAL2A | 24.0926745 | 0.092536161 | 0.50500131 | 0.18323945 | 0.85461014 | 0.92399813 |
| COA7 | 931.699058 | 0.36189026 | 0.32313716 | 1.11992771 | 0.26274457 | 0.45630758 |
| ZYG11B | 2115.7864 | 0.361569602 | 0.17422113 | 2.0753488 | 0.03795422 | 0.1191066 |
| ZYG11A | 219.553823 | 1.999564425 | 0.47489219 | 4.21056499 | 2.55E-05 | 0.00043103 |
| AC099677.4 | 21.0075516 | 2.577014609 | 0.5768436 | 4.46744073 | 7.92E-06 | 0.00016958 |
| ECHDC2 | 10759.5255 | 0.525825411 | 0.22614014 | 2.32521928 | 0.02006024 | 0.07507642 |
| SCP2 | 9108.25303 | 0.721169484 | 0.23600891 | 3.05568754 | 0.00224545 | 0.01477728 |
| PODN | 575.467366 | -0.230550709 | 0.53303214 | -0.4325268 | 0.66535855 | 0.80373362 |
| AL445183.1 | 163.366738 | -0.508451383 | 0.55672723 | -0.9132864 | 0.36109196 | 0.55878717 |
| SLC1A7 | 87.7899374 | 0.833325904 | 0.79758662 | 1.04480928 | 0.29611112 | 0.49280217 |
| AL445183.2 | 7.61180164 | 0.608905485 | 1.0022436 | 0.6075424 | 0.54349101 | 0.71573402 |
| CPT2 | 3227.55492 | 0.738538117 | 0.30616187 | 2.41224718 | 0.01585453 | 0.06338446 |
| AL606760.2 | 2006.68502 | 0.795234872 | 0.31045059 | 2.56155052 | 0.01042061 | 0.04650388 |
| C1orf123 | 2011.62371 | 0.11536554 | 0.11871888 | 0.97175397 | 0.33117296 | 0.52905016 |
| AL606760.3 | 1.61610362 | 0.663563345 | 1.36271156 | 0.48694336 | 0.62629847 | 0.77760208 |
| MAGOH | 1005.6086 | -0.328823911 | 0.18423924 | -1.7847659 | 0.07429929 | 0.1924754 |
| AL606760.1 | 104.44581 | 0.027275115 | 0.35469231 | 0.07689796 | 0.93870472 | 0.97076919 |
| LRP8 | 464.509497 | -1.486617848 | 0.42209304 | -3.5220146 | 0.00042828 | 0.00402486 |
| AL355483.4 | 0.90673911 | -0.135843207 | 1.57174424 | -0.0864283 | 0.93112594 | NA |
| AL355483.2 | 4.3465957 | -1.396732435 | 0.8100766 | -1.724198 | 0.08467211 | 0.21076114 |
| AL355483.3 | 0.34471008 | 0.328499558 | 2.81543776 | 0.11667797 | 0.90711525 | NA |
| LINC01771 | 0.22673191 | 0.689158461 | 3.03348976 | 0.22718338 | 0.82028115 | NA |
| AC119428.1 | 0 | NA | NA | NA | NA | NA |
| AC119428.2 | 1.88512584 | 2.687176521 | 1.30814171 | 2.05419375 | 0.03995694 | 0.12377124 |
| DMRTB1 | 0.1357163 | 1.313406857 | 3.28103699 | 0.40030236 | 0.68893383 | NA |
| GLIS1 | 463.226802 | 1.199139291 | 0.43423394 | 2.7615052 | 0.00575356 | 0.02990173 |
| NDC1 | 908.513739 | -0.065807177 | 0.18698601 | -0.3519364 | 0.72488599 | 0.84398179 |
| YIPF1 | 1980.41069 | -0.710962377 | 0.14650691 | -4.8527567 | 1.22E-06 | 3.82E-05 |
| DIO1 | 3736.54724 | 2.343576631 | 0.65910027 | 3.55572096 | 0.00037694 | 0.00364236 |
| HSPB11 | 808.982388 | -0.230059193 | 0.10821415 | -2.1259622 | 0.03350641 | 0.10920331 |
| LRRC42 | 1437.76137 | -0.798714966 | 0.17767544 | -4.4953595 | 6.95E-06 | 0.00015313 |
| LDLRAD1 | 1.95790656 | -0.286187991 | 1.74443944 | -0.1640573 | 0.86968605 | 0.93214735 |
| TMEM59 | 18423.9509 | 0.00023233 | 0.11777583 | 0.00197265 | 0.99842606 | 0.99922179 |
| TCEANC2 | 452.581677 | 0.143456457 | 0.18053023 | 0.79463954 | 0.42682321 | 0.61907648 |
| AL357673.1 | 77.0605129 | 0.208269748 | 0.32422985 | 0.64235218 | 0.52064454 | 0.69724222 |
| CDCP2 | 17.6281813 | -0.721482595 | 0.50014701 | -1.4425411 | 0.1491498 | 0.31330365 |
| AL357673.2 | 11.0646006 | -0.247376665 | 0.45359749 | -0.545366 | 0.5855018 | 0.74706935 |
| CYB5RL | 306.098855 | 0.817535943 | 0.22112282 | 3.69720295 | 0.00021799 | 0.00234122 |
| MRPL37 | 3328.16372 | 0.13096747 | 0.1751275 | 0.74784069 | 0.45455626 | 0.64404672 |
| SSBP3 | 3658.00538 | -0.51192001 | 0.20553894 | -2.4906229 | 0.01275194 | 0.05395837 |
| SSBP3-AS1 | 48.6758463 | -0.320418571 | 0.32539849 | -0.9846959 | 0.32477345 | 0.52234961 |
| AL161644.1 | 0.23320561 | -1.364242679 | 3.30558948 | -0.4127078 | 0.67982068 | NA |
| AL035415.1 | 0.09997307 | 0.547239394 | 3.29377202 | 0.16614368 | 0.86804388 | NA |
| AC099796.1 | 0.08666371 | -0.642905668 | 3.30558948 | -0.1944905 | 0.84579185 | NA |
| ACOT11 | 6038.57121 | 2.353946326 | 0.46580223 | 5.05353171 | 4.34E-07 | 1.61E-05 |
| FAM151A | 9459.54431 | 2.950637101 | 0.70379576 | 4.19246216 | 2.76E-05 | 0.00045967 |
| MROH7 | 1020.12794 | 1.154223592 | 0.30637644 | 3.76733792 | 0.000165 | 0.00187948 |
| MROH7-TTC4 | 2512.36318 | 0.220954561 | 0.18801121 | 1.17522012 | 0.23990666 | 0.42986605 |
| TTC4 | 1663.09008 | -0.362264995 | 0.21461572 | -1.6879705 | 0.0914169 | 0.22302373 |
| PARS2 | 433.040742 | 0.106774859 | 0.21728083 | 0.49141408 | 0.6231336 | 0.77531533 |
| TTC22 | 2386.26075 | -1.095595376 | 0.42781265 | -2.5609233 | 0.01043944 | 0.04656428 |
| AC096536.2 | 0.29032414 | -1.364245079 | 2.89454325 | -0.4713162 | 0.63741496 | NA |
| LEXM | 12.8107062 | -0.229804715 | 0.4250651 | -0.5406342 | 0.58875977 | 0.74926862 |
| DHCR24 | 7712.98097 | -0.769725324 | 0.27034499 | -2.8471966 | 0.00441061 | 0.024522 |
| AC096536.3 | 117.027077 | -0.539282495 | 0.37106524 | -1.4533361 | 0.14613047 | 0.30886987 |
| AC096536.1 | 25.3492998 | -1.334053321 | 0.35338072 | -3.7751163 | 0.00015993 | 0.00184167 |
| TMEM61 | 730.158788 | -0.554115692 | 0.4341986 | -1.2761803 | 0.2018918 | 0.38378654 |
| AL590440.1 | 519.641091 | -0.567689967 | 0.42508585 | -1.3354713 | 0.18172213 | 0.35814742 |
| BSND | 1232.70593 | 0.565508071 | 0.42757556 | 1.32259214 | 0.18597105 | 0.36347885 |
| PCSK9 | 38.8135161 | 0.045597665 | 0.63365254 | 0.07196005 | 0.9426337 | 0.97275716 |
| USP24 | 3808.32498 | 0.139537036 | 0.19849103 | 0.70298912 | 0.48206253 | 0.66659617 |
| MIR4422HG | 3.00126155 | 0.26646201 | 1.08662113 | 0.24522071 | 0.80628554 | 0.8951185 |
| AL603840.1 | 2.87107283 | -0.068818639 | 0.97160516 | -0.0708298 | 0.94353318 | 0.97315146 |
| LINC01753 | 0.09635527 | 0.547239394 | 3.29377202 | 0.16614368 | 0.86804388 | NA |
| LINC01755 | 0.17828073 | 0.18657164 | 3.29377195 | 0.05664376 | 0.95482897 | NA |
| AL353681.1 | 5.53812795 | -0.016332859 | 0.68083703 | -0.0239894 | 0.98086108 | 0.99132745 |
| AC119674.1 | 63.206281 | -0.973051462 | 0.42609439 | -2.2836524 | 0.02239197 | 0.08152371 |
| AC119674.2 | 2747.77304 | 0.130284743 | 0.45764891 | 0.28468273 | 0.77588722 | 0.87702628 |
| LINC01767 | 4.86114647 | -0.83992511 | 1.03793647 | -0.8092259 | 0.4183852 | 0.61113582 |
| PLPP3 | 13098.6155 | -0.225471218 | 0.40747853 | -0.5533328 | 0.58003558 | 0.7431151 |
| PRKAA2 | 2455.55908 | 0.49874779 | 0.15949507 | 3.12704201 | 0.00176575 | 0.0122695 |
| FYB2 | 677.749362 | -0.240196564 | 0.35175841 | -0.6828453 | 0.49470463 | 0.67694261 |
| AL360295.1 | 2.58786185 | -2.120587415 | 1.22191702 | -1.7354594 | 0.08265946 | 0.20727748 |
| C8A | 19.72561 | -3.706495637 | 0.652139 | -5.6835976 | 1.32E-08 | 8.85E-07 |
| C8B | 6.88484443 | -1.446222759 | 1.04069638 | -1.3896683 | 0.16462963 | 0.33458831 |
| AL161740.1 | 5.55493502 | -0.914637003 | 1.02983387 | -0.8881403 | 0.37446527 | 0.5705785 |
| DAB1 | 499.023222 | 0.603655592 | 0.32387903 | 1.86383044 | 0.06234552 | 0.16982247 |
| AL390243.1 | 0 | NA | NA | NA | NA | NA |
| DAB1-AS1 | 0.2441911 | -0.174096114 | 3.29377188 | -0.0528562 | 0.95784652 | NA |
| AL445193.2 | 0.16421342 | 1.36803136 | 3.28033652 | 0.41703994 | 0.67664919 | NA |
| OMA1 | 1182.23781 | 0.719854637 | 0.35097965 | 2.05098682 | 0.04026823 | 0.12441397 |
| TACSTD2 | 10861.0094 | -0.565865572 | 0.48226601 | -1.1733474 | 0.24065652 | 0.43079984 |
| MYSM1 | 1764.53409 | 0.056974979 | 0.1337026 | 0.42613217 | 0.67001154 | 0.80733565 |
| AL136985.3 | 77.5399062 | -0.490120305 | 0.37776114 | -1.2974344 | 0.19448176 | 0.37447857 |
| JUN | 12326.0104 | -0.498351167 | 0.3383884 | -1.4727194 | 0.14082671 | 0.30109945 |
| LINC01135 | 28.0084216 | 1.190508128 | 0.36274925 | 3.28190378 | 0.00103109 | 0.00804194 |
| AL136985.2 | 8.43561297 | 0.167754848 | 0.511287 | 0.3281031 | 0.74283371 | 0.85562255 |
| AL136985.1 | 0.42149465 | -0.032077937 | 1.87730314 | -0.0170872 | 0.98636702 | NA |
| AC105277.1 | 96.4616253 | 0.235755961 | 0.34614038 | 0.68109927 | 0.49580868 | 0.67772047 |
| LINC01358 | 34.3467364 | -1.140352839 | 0.43860554 | -2.5999508 | 0.00932371 | 0.04272221 |
| AL592431.2 | 0.41459949 | -1.973379568 | 2.21435362 | -0.8911763 | 0.37283457 | NA |
| AL592431.1 | 0.85543166 | -1.473612215 | 1.67620149 | -0.8791379 | 0.37932652 | NA |
| AC093424.1 | 2.194346 | -2.906210271 | 1.26531785 | -2.2968223 | 0.02162891 | 0.07948955 |
| FGGY | 1073.53834 | 0.909573966 | 0.34648161 | 2.62517242 | 0.00866051 | 0.04039223 |
| AL035416.1 | 4.01583036 | 0.630966936 | 0.79610887 | 0.79256363 | 0.42803211 | 0.62042202 |
| HOOK1 | 2520.41576 | -0.313093483 | 0.18472614 | -1.6949062 | 0.09009321 | 0.22081588 |

| | | | | | | |
|---|---|---|---|---|---|---|
| CYP2J2 | 288.486522 | 0.146233049 | 0.60684379 | 0.24097313 | 0.80957594 | 0.89703256 |
| C1orf87 | 5.67222049 | -0.775755274 | 1.04831579 | -0.7400015 | 0.45929908 | 0.64801707 |
| AC097066.1 | 5.95827365 | -0.029896459 | 1.05486521 | -0.0283415 | 0.97738978 | 0.98958605 |
| LINC01748 | 6.56387214 | -1.562488988 | 0.87875369 | -1.7780739 | 0.07539172 | 0.19443322 |
| AC099792.1 | 8.41934976 | -0.791607912 | 0.76871826 | -1.0297764 | 0.30311499 | 0.49991508 |
| NFIA | 1503.5256 | 0.458424853 | 0.23439423 | 1.95578559 | 0.05049042 | 0.14666582 |
| NFIA-AS2 | 6.23848426 | -1.902133581 | 0.99934404 | -1.9033821 | 0.0569907 | 0.15956886 |
| AC099791.3 | 0 | NA | NA | NA | NA | NA |
| TM2D1 | 1516.07117 | -0.054212474 | 0.09129974 | -0.5937856 | 0.5526555 | 0.72260638 |
| AC099791.2 | 14.5913871 | -0.188802076 | 0.47310982 | -0.3990661 | 0.68984452 | 0.82134731 |
| PATJ | 5790.76526 | 0.362477244 | 0.20329273 | 1.78303104 | 0.07458125 | 0.19298475 |
| L1TD1 | 14.4697056 | 0.254666333 | 0.643758 | 0.39559327 | 0.6924051 | 0.82302108 |
| KANK4 | 64.5998689 | 3.972708333 | 0.61845131 | 6.42363961 | 1.33E-10 | 1.83E-08 |
| USP1 | 1315.13936 | -0.089981708 | 0.19257955 | -0.4672444 | 0.64032506 | 0.78716012 |
| DOCK7 | 1370.60052 | 0.037885986 | 0.1455047 | 0.26037637 | 0.79457347 | 0.88857752 |
| ANGPTL3 | 1587.22063 | 3.507721031 | 0.56986881 | 6.15531319 | 7.49E-10 | 8.13E-08 |
| AC103923.1 | 7.78611419 | 0.311878442 | 0.60284244 | 0.51734653 | 0.60491428 | 0.76184877 |
| ATG4C | 626.010466 | 0.382422952 | 0.16222883 | 2.35730567 | 0.01840809 | 0.07064783 |
| AC099794.1 | 0.87378556 | -0.037258267 | 1.51696229 | -0.0245611 | 0.98040505 | NA |
| LINC01739 | 15.2203475 | 1.250616439 | 0.85496839 | 1.46276337 | 0.14353214 | 0.30511427 |
| AL162400.1 | 0 | NA | NA | NA | NA | NA |
| AL162400.2 | 2.5826867 | 2.964199605 | 1.24973379 | 2.37186482 | 0.01769856 | 0.06855334 |
| AC096543.1 | 0.44586572 | -0.886821371 | 2.76734578 | -0.3204592 | 0.74862027 | NA |
| AC096543.2 | 0 | NA | NA | NA | NA | NA |
| FOXD3-AS1 | 5.29964054 | -0.017602165 | 1.130632 | -0.0155684 | 0.98757869 | 0.99440414 |
| FOXD3 | 6.80911955 | 0.703890833 | 1.03030821 | 0.68318473 | 0.49449012 | 0.67689496 |
| ALG6 | 613.982498 | -0.008582231 | 0.16048227 | -0.0534778 | 0.95735126 | 0.97951358 |
| ITGB3BP | 1060.81032 | 0.026667193 | 0.22821427 | 0.11685156 | 0.90697769 | 0.95356011 |
| EFCAB7 | 434.421957 | 0.354575565 | 0.22563253 | 1.57147358 | 0.11607269 | 0.26397902 |
| PGM1 | 5057.3151 | -0.005729822 | 0.19145571 | -0.0299277 | 0.97612474 | 0.98894663 |
| ROR1 | 164.301906 | 0.925200412 | 0.24846906 | 3.72360409 | 0.0001964 | 0.0021501 |
| ROR1-AS1 | 4.36342442 | -0.619470146 | 0.89697556 | -0.690621 | 0.48980376 | 0.67311398 |
| AL445205.1 | 55.8329469 | 1.568595338 | 0.68705883 | 2.28305825 | 0.02242694 | 0.08161721 |
| UBE2U | 12.4662971 | 1.420705812 | 1.13147253 | 1.25562554 | 0.20925177 | 0.39243077 |
| CACHD1 | 700.004295 | 0.906012579 | 0.22163939 | 4.08777789 | 4.36E-05 | 0.00066486 |
| RAVER2 | 1000.61516 | 0.164021256 | 0.16517409 | 0.99302052 | 0.32069996 | 0.51819065 |
| JAK1 | 8199.03769 | -0.324963792 | 0.13120803 | -2.4767066 | 0.01326009 | 0.05552055 |
| AL357078.3 | 15.7395453 | -0.390078871 | 0.53469968 | -0.7295289 | 0.46567819 | 0.65288037 |
| AL357078.1 | 2.98170294 | 0.194387785 | 0.66470885 | 0.29244049 | 0.76994985 | 0.87279244 |
| AK4 | 15691.7351 | 1.129753885 | 0.47921697 | 2.35749977 | 0.01839847 | 0.07062119 |
| DNAJC6 | 829.776719 | 0.778835156 | 0.35187461 | 2.2133883 | 0.02687089 | 0.09311239 |
| AL139294.1 | 0.22838051 | 0.077317824 | 3.11260685 | 0.02484022 | 0.98018241 | NA |
| LEPROT | 11790.7047 | -0.302900138 | 0.17704354 | -1.7108793 | 0.08710339 | 0.21514231 |
| LEPR | 1460.1954 | -0.03000943 | 0.22940074 | -0.1308166 | 0.89592037 | 0.94664504 |
| AL513493.1 | 0.03392908 | 0.380740593 | 3.29738709 | 0.11546736 | 0.90807469 | NA |
| PDE4B | 741.449236 | -0.162044012 | 0.49304873 | -0.3286572 | 0.74241481 | 0.8554347 |
| AL590783.1 | 0.17466293 | 0.18657164 | 3.29377195 | 0.05664376 | 0.95482897 | NA |
| SGIP1 | 1421.30822 | 0.02587107 | 0.39063575 | 0.06622812 | 0.94719621 | 0.97455008 |
| AL139147.1 | 33.317935 | 0.019932718 | 0.48920577 | 0.04074506 | 0.96749914 | 0.98467589 |
| TCTEX1D1 | 88.1015309 | -1.994756536 | 0.65566748 | -3.0423295 | 0.00234755 | 0.01529334 |
| INSL5 | 6.82631795 | -0.512582066 | 0.60577153 | -0.846164 | 0.39746126 | 0.5923507 |
| WDR78 | 495.436624 | 0.562278105 | 0.31103843 | 1.80774483 | 0.07064622 | 0.18549646 |
| AL592161.1 | 53.0368782 | 0.653688656 | 0.36886037 | 1.77218458 | 0.07636392 | 0.1961526 |
| MIER1 | 2150.45975 | -0.412299563 | 0.12492696 | -3.3003249 | 0.00096573 | 0.00764032 |
| SLC35D1 | 1293.82835 | 0.583654631 | 0.22031428 | 2.64919108 | 0.00806847 | 0.03834304 |
| C1orf141 | 1.1799357 | 1.288027263 | 1.27482755 | 1.01035412 | 0.31232566 | 0.50918254 |
| IL23R | 3.00576511 | 2.003688117 | 1.10056043 | 1.82060709 | 0.0686666 | 0.18160759 |
| IL12RB2 | 57.7662977 | 0.642496896 | 0.4781578 | 1.34369219 | 0.17904794 | 0.35475951 |
| SERBP1 | 9402.45127 | -0.070651567 | 0.12860323 | -0.5493763 | 0.58274723 | 0.74511123 |
| LINC01702 | 127.842108 | 1.222008155 | 0.58937797 | 2.07338622 | 0.03813635 | 0.11957854 |
| GADD45A | 8118.79423 | -0.833486288 | 0.27016441 | -3.0851076 | 0.00203479 | 0.0136775 |
| GNG12 | 7481.24202 | -0.07180283 | 0.21036488 | -0.3413252 | 0.7328588 | 0.84935993 |
| GNG12-AS1 | 313.328591 | -0.054608499 | 0.23555494 | -0.2318291 | 0.81667073 | 0.90138837 |
| DIRAS3 | 451.209867 | 0.220595914 | 0.26292034 | 0.83902187 | 0.40145704 | 0.59635095 |
| WLS | 6512.89102 | 0.421805645 | 0.17356542 | 2.43024013 | 0.01508882 | 0.06113894 |
| RPE65 | 0.15543392 | 1.368031615 | 3.28033652 | 0.41704002 | 0.67664913 | NA |
| DEPDC1 | 13.2432885 | -1.38078782 | 0.85052741 | -1.6234489 | 0.10449347 | 0.24491092 |
| AL138789.1 | 0.76617325 | 0.009430136 | 1.61915298 | 0.00582412 | 0.99535305 | NA |
| DEPDC1-AS1 | 0.08695548 | -0.642905668 | 3.30558948 | -0.1944905 | 0.84579185 | NA |
| AL035412.1 | 0 | NA | NA | NA | NA | NA |
| AL033530.1 | 0.2657758 | -0.179000381 | 2.96268942 | -0.0604182 | 0.95182256 | NA |
| AL691520.1 | 0.24327904 | -1.364242679 | 3.30558948 | -0.4127078 | 0.67982068 | NA |
| LINC01707 | 0.17828073 | 0.18657164 | 3.29377195 | 0.05664376 | 0.95482897 | NA |
| AL512271.1 | 0 | NA | NA | NA | NA | NA |
| LINC01758 | 17.4694451 | 5.267406764 | 1.0094033 | 5.21833717 | 1.81E-07 | 7.87E-06 |
| LRRC7 | 99.7229006 | 1.283933085 | 0.25819915 | 4.9726464 | 6.60E-07 | 2.28E-05 |
| AL158840.1 | 0 | NA | NA | NA | NA | NA |
| LRRC40 | 865.035421 | 0.380549231 | 0.14114906 | 2.6960806 | 0.00701607 | 0.03468574 |
| SRSF11 | 8405.32664 | -0.323536778 | 0.13641968 | -2.3716284 | 0.01770989 | 0.06857359 |
| AL353771.1 | 0.70509342 | -0.713442481 | 1.52668871 | -0.4673137 | 0.64027549 | NA |
| ANKRD13C | 1203.85699 | -0.038724183 | 0.19320922 | -0.2004262 | 0.8411473 | 0.91521102 |
| HHLA3 | 453.580591 | -0.202961463 | 0.31695653 | -0.6403448 | 0.52194847 | 0.69790813 |
| AL158839.1 | 3.122489 | 2.303489329 | 0.9481742 | 2.42939465 | 0.01512406 | 0.06124561 |
| CTH | 670.37542 | 0.301129767 | 0.38802969 | 0.77604827 | 0.43772049 | 0.62968137 |
| AL354872.2 | 39.3262752 | 0.363823215 | 0.46951267 | 0.77489541 | 0.43840146 | 0.6301797 |
| LINC01788 | 174.755263 | 1.444115863 | 0.53793757 | 2.68454175 | 0.00726293 | 0.03558593 |
| AL513285.1 | 0.03392908 | 0.380740593 | 3.29738709 | 0.11546736 | 0.90807469 | NA |
| PTGER3 | 7046.16527 | 0.622610246 | 0.30125962 | 2.06668997 | 0.03876338 | 0.1210263 |
| AL031429.1 | 35.3286235 | 1.22854968 | 0.44756749 | 2.74494843 | 0.00605204 | 0.03098278 |
| ZRANB2-AS1 | 382.576239 | 0.488224565 | 0.18723968 | 2.60748445 | 0.00912102 | 0.04207179 |
| ZRANB2 | 4350.19425 | -0.034980586 | 0.16077367 | -0.2175766 | 0.82775904 | 0.90828919 |
| NEGR1 | 647.715682 | 1.303178966 | 0.36540436 | 3.56640238 | 0.00036192 | 0.00352297 |
| AL354949.1 | 1.13893478 | 1.067041891 | 1.35344697 | 0.7883884 | 0.43046955 | 0.62251837 |
| AL583808.1 | 0.12088456 | 0.02007176 | 3.29738704 | 0.00608717 | 0.99514317 | NA |
| AL583808.2 | 0.09635527 | 0.547239394 | 3.29377202 | 0.16614368 | 0.86804388 | NA |
| LINC01360 | 0 | NA | NA | NA | NA | NA |
| LINC02238 | 0 | NA | NA | NA | NA | NA |
| LRRIQ3 | 81.8243807 | 1.597532147 | 0.43696696 | 3.65595642 | 0.00025622 | 0.0026685 |
| FPGT | 763.597016 | 0.54938567 | 0.1594993 | 3.44443944 | 0.00057224 | 0.0050418 |
| FPGT-TNNI3I | 409.434039 | 1.85423131 | 0.31985396 | 5.79711843 | 6.75E-09 | 4.92E-07 |
| TNNI3K | 262.27803 | 2.26175518 | 0.37797595 | 5.98386011 | 2.18E-09 | 1.92E-07 |
| AC093158.1 | 34.7403835 | 2.228155023 | 0.40094837 | 5.55721187 | 2.74E-08 | 1.62E-06 |
| AC105271.1 | 2.98314053 | 1.577523855 | 0.86997103 | 1.8133062 | 0.06978461 | 0.18378309 |
| LRRC53 | 0.41418845 | 2.629243739 | 2.05846617 | 1.27728295 | 0.20150237 | NA |
| ERICH3 | 16.0033458 | -0.863790501 | 0.78461403 | -1.1009114 | 0.27093524 | 0.46553813 |
| ERICH3-AS1 | 0.30370406 | -0.699693739 | 2.92440877 | -0.2392599 | 0.81090406 | NA |
| AC135803.1 | 81.2730627 | 0.416421965 | 0.39000735 | 1.06772851 | 0.28564299 | 0.48167322 |
| CRYZ | 12009.5533 | 0.507082206 | 0.32738722 | 1.54887601 | 0.12141153 | 0.2719958 |
| TYW3 | 1338.35366 | 0.275509284 | 0.15977658 | 1.72434084 | 0.08464633 | 0.21073676 |
| AC133865.1 | 0 | NA | NA | NA | NA | NA |
| AC099786.3 | 0 | NA | NA | NA | NA | NA |
| AC099786.1 | 0 | NA | NA | NA | NA | NA |
| LHX8 | 0.17332742 | -1.003574173 | 3.30558948 | -0.3035992 | 0.7614333 | NA |

| | | | | | | |
|---|---|---|---|---|---|---|
| AC099786.2 | 0 | NA | NA | NA | NA | NA |
| SLC44A5 | 133.408506 | -0.423801467 | 0.83906879 | -0.5050855 | 0.6134988 | 0.76843981 |
| AL357314.1 | 1.47939889 | 1.317995214 | 1.28199773 | 1.02807921 | 0.30391257 | 0.50076071 |
| ACADM | 9956.29986 | 1.734005106 | 0.33632306 | 5.15577224 | 2.53E-07 | 1.03E-05 |
| RABGGTB | 4606.67935 | -0.357097998 | 0.25194471 | -1.4173665 | 0.1563758 | 0.32306721 |
| MSH4 | 15.1570265 | -0.237241264 | 0.61044017 | -0.3886397 | 0.69754272 | 0.82618584 |
| ASB17 | 9.28139386 | 1.314698643 | 0.57887743 | 2.27111747 | 0.02313987 | 0.08358881 |
| AC092813.1 | 1.12991508 | -0.476304281 | 1.22803062 | -0.3878603 | 0.69811944 | 0.82660903 |
| AC092813.2 | 55.8537606 | -0.277718131 | 0.88675185 | -0.3131858 | 0.75413947 | 0.86318765 |
| ST6GALNAC3 | 786.381289 | 0.049449291 | 0.29673037 | 0.16664722 | 0.86764764 | 0.93109788 |
| AC104458.1 | 4.48718899 | 0.411837266 | 0.71376704 | 0.57699115 | 0.56394543 | 0.73102497 |
| LINC02567 | 0 | NA | NA | NA | NA | NA |
| ST6GALNAC5 | 19.3130027 | 1.951966128 | 0.53637502 | 3.63918164 | 0.00027351 | 0.00282616 |
| AL035409.1 | 0.53375756 | 2.463056813 | 1.94219803 | 1.26818006 | 0.20473366 | NA |
| PIGK | 1957.61606 | 0.143390221 | 0.20184181 | 0.71040892 | 0.47745059 | 0.66254655 |
| AK5 | 180.099083 | -1.281295868 | 0.56987736 | -2.2483712 | 0.02455253 | 0.08706968 |
| AC095030.1 | 0.78007395 | 0.952094902 | 1.88012893 | 0.50639873 | 0.61257677 | NA |
| AC118549.1 | 1656.28778 | 0.048728145 | 0.14307726 | 0.34057225 | 0.73342563 | 0.84975544 |
| USP33 | 3152.57317 | 0.021675142 | 0.1239817 | 0.17482533 | 0.86121688 | 0.92774579 |
| MIGA1 | 1119.10031 | 0.640321937 | 0.16712915 | 3.83130025 | 0.00012747 | 0.00153136 |
| NEXN-AS1 | 40.6025041 | 0.705378449 | 0.43145062 | 1.63489961 | 0.10207009 | 0.24087957 |
| NEXN | 503.144308 | -0.018559306 | 0.53783347 | -0.0345075 | 0.97247243 | 0.98710875 |
| FUBP1 | 4043.23478 | -0.176305182 | 0.14890477 | -1.184013 | 0.23640791 | 0.42570859 |
| DNAJB4 | 1578.27057 | 0.152387586 | 0.2124309 | 0.71735133 | 0.47315734 | 0.65922361 |
| GIPC2 | 2229.9243 | 1.500789672 | 0.42775182 | 3.50855239 | 0.00045055 | 0.0041834 |
| AC103591.3 | 7.645201 | -0.390657442 | 0.54924606 | -0.7112613 | 0.47692235 | 0.66209226 |
| PTGFR | 322.511647 | 0.675648116 | 0.26782412 | 2.52273066 | 0.01164476 | 0.05047514 |
| IFI44L | 764.545852 | -0.374556547 | 0.57473474 | -0.6517033 | 0.51459257 | 0.6924077 |
| IFI44 | 1134.80405 | -0.551281993 | 0.42174402 | -1.3071483 | 0.19116234 | 0.37023627 |
| ADGRL4 | 1769.31391 | -0.715815202 | 0.3200375 | -2.2366604 | 0.02530854 | 0.08917704 |
| AL353651.2 | 0 | NA | NA | NA | NA | NA |
| AL353651.1 | 0 | NA | NA | NA | NA | NA |
| AL353651.3 | 0 | NA | NA | NA | NA | NA |
| AC099671.1 | 0 | NA | NA | NA | NA | NA |
| AC098657.2 | 0.19994613 | 1.506623855 | 3.27645372 | 0.45983371 | 0.64563559 | NA |
| AL606519.1 | 1.27907876 | 1.576148499 | 1.26142917 | 1.24949425 | 0.21148435 | 0.39498849 |
| AL136234.1 | 1.77631412 | 0.507183576 | 1.07074156 | 0.47367506 | 0.63573164 | 0.78399866 |
| LINC01781 | 136.715168 | -0.294187198 | 0.587487 | -0.5007553 | 0.61654338 | 0.7706427 |
| AC093154.1 | 0 | NA | NA | NA | NA | NA |
| ADGRL2 | 2605.00843 | -0.184306186 | 0.1772555 | -1.039777 | 0.29844353 | 0.49530098 |
| AL138799.4 | 0.28934269 | -1.364242679 | 3.30558948 | -0.4127078 | 0.67982068 | NA |
| AC117944.1 | 0.61821446 | 0.041200007 | 2.7688506 | 0.01487982 | 0.98812806 | NA |
| AC093580.1 | 0 | NA | NA | NA | NA | NA |
| LINC01362 | 0.78286505 | -0.037145912 | 1.30238933 | -0.0285214 | 0.97724633 | NA |
| LINC01361 | 6.25936777 | 0.443240786 | 0.4927089 | 0.89959971 | 0.36833331 | 0.56515711 |
| LINC01712 | 0 | NA | NA | NA | NA | NA |
| LINC01725 | 23.6954101 | 0.229230882 | 0.33813626 | 0.67792458 | 0.49781951 | 0.6795476 |
| AL035706.1 | 3.2446957 | -0.157590971 | 0.73028371 | -0.2157942 | 0.82914819 | 0.90920816 |
| TTLL7 | 471.908922 | -0.863576203 | 0.26521324 | -3.2561579 | 0.00112931 | 0.00862935 |
| AC104454.2 | 0 | NA | NA | NA | NA | NA |
| AL359504.2 | 105.82835 | -0.128005708 | 0.29073021 | -0.4402904 | 0.65972682 | 0.79999826 |
| PRKACB | 1743.4743 | 0.563993202 | 0.16738411 | 3.36945494 | 0.00075317 | 0.00625743 |
| SAMD13 | 67.1157683 | 0.50127958 | 0.42476035 | 1.18014682 | 0.23794183 | 0.42730359 |
| DNASE2B | 1.33195318 | 0.117175565 | 1.04889594 | 0.11171324 | 0.91105078 | 0.95600514 |
| AL844170.1 | 0.35176226 | 2.413587081 | 2.07427612 | 1.16358042 | 0.24459409 | NA |
| RPF1 | 1307.95715 | -0.417295759 | 0.13439381 | -3.1050221 | 0.00190265 | 0.01296456 |
| GNG5 | 4793.58548 | -0.859794151 | 0.16708075 | -5.1459797 | 2.66E-07 | 1.07E-05 |
| SPATA1 | 1297.70189 | -0.140299705 | 0.13772087 | -1.0187251 | 0.30833349 | 0.50529825 |
| CTBS | 2053.58388 | -0.016584578 | 0.17877406 | -0.0927684 | 0.92608758 | 0.9641707 |
| AL359762.2 | 2.48094034 | 0.621428237 | 0.71252818 | 0.87214548 | 0.38312902 | 0.57900809 |
| LINC01555 | 14.9854312 | 1.209007288 | 0.95846822 | 1.26139528 | 0.20716647 | 0.38993432 |
| SSX2IP | 1270.48061 | 0.434620738 | 0.26739147 | 1.62540987 | 0.10407526 | 0.24425636 |
| AC104169.1 | 0 | NA | NA | NA | NA | NA |
| LPAR3 | 5.04721746 | 1.491854748 | 0.99142115 | 1.50476389 | 0.13238479 | 0.28867832 |
| MCOLN2 | 191.807464 | 0.144759756 | 0.3851243 | 0.37587801 | 0.70700759 | 0.83325953 |
| WDR63 | 53.5654734 | 0.137875301 | 0.56335375 | 0.24474019 | 0.80665761 | 0.89545616 |
| MCOLN3 | 630.989811 | -0.148000105 | 0.23582796 | -0.6275766 | 0.53028135 | 0.70472412 |
| SYDE2 | 170.696752 | 1.263602458 | 0.20691102 | 6.10698467 | 1.02E-09 | 1.04E-07 |
| C1orf52 | 897.779356 | -0.497031347 | 0.1238448 | -4.0133403 | 5.99E-05 | 0.00084828 |
| BCL10 | 953.950914 | -0.64455469 | 0.17285044 | -3.7289733 | 0.00019226 | 0.00211424 |
| AL078459.1 | 31.7765518 | -0.300871859 | 0.30988006 | -0.97093 | 0.33158313 | 0.52953852 |
| DDAH1 | 9036.39921 | 0.656063753 | 0.20866605 | 3.14408473 | 0.00166607 | 0.01171116 |
| AL360219.1 | 8.2008824 | 0.877087662 | 0.51514408 | 1.7026065 | 0.08864173 | 0.21813013 |
| AC092807.3 | 20.3037897 | 0.524115944 | 0.55857433 | 0.93831011 | 0.34808506 | 0.54549255 |
| AC092807.2 | 4.50987096 | -0.259218838 | 0.56371333 | -0.4598416 | 0.64562992 | 0.79091523 |
| CYR61 | 15367.1571 | 0.500288193 | 0.62884646 | 0.79556494 | 0.42628495 | 0.61866451 |
| AC092807.1 | 1.08904749 | 0.167405251 | 1.42999727 | 0.11706683 | 0.90680709 | 0.95344908 |
| ZNHIT6 | 913.996928 | -0.204935262 | 0.20170462 | -1.0160167 | 0.30962145 | 0.50688268 |
| COL24A1 | 104.813054 | 1.001805952 | 0.25191214 | 3.97680699 | 6.98E-05 | 0.00095433 |
| AL136382.1 | 67.1321489 | -0.633531407 | 0.44364512 | -1.4280139 | 0.15328786 | 0.31886099 |
| ODF2L | 1638.80367 | -0.447611848 | 0.2267534 | -1.9740028 | 0.04838143 | 0.14229514 |
| CLCA2 | 2.7371488 | -0.295582812 | 1.05407475 | -0.2804192 | 0.7791559 | 0.87898785 |
| CLCA1 | 0.10721287 | -0.642905668 | 3.30558948 | -0.1944905 | 0.84579185 | NA |
| CLCA4 | 1.01816867 | -0.238755329 | 1.32406843 | -0.1803195 | 0.85690176 | 0.92513816 |
| AL356270.1 | 4.79317307 | -2.119548273 | 0.90503318 | -2.3419564 | 0.01918295 | 0.0726698 |
| AL049597.2 | 23.3938225 | -0.477018203 | 0.39979667 | -1.193152 | 0.23280984 | 0.42092531 |
| SH3GLB1 | 4799.09493 | -0.462746831 | 0.16979711 | -2.7252927 | 0.00642445 | 0.03241109 |
| AL355981.1 | 0 | NA | NA | NA | NA | NA |
| SELENOF | 7643.25102 | -0.294092433 | 0.13617954 | -2.1595934 | 0.03080416 | 0.10282094 |
| HS2ST1 | 1644.53635 | 0.071531014 | 0.14316561 | 0.49963823 | 0.61732983 | 0.77126014 |
| AL121989.1 | 1.56946849 | -2.668382766 | 1.75604652 | -1.5195399 | 0.12862665 | 0.28284563 |
| AC093155.3 | 362.248873 | 0.163481102 | 0.21380055 | 0.76464305 | 0.44448413 | 0.63537357 |
| LINC01140 | 150.937211 | -0.515742691 | 0.42839953 | -1.2038825 | 0.228635 | 0.41588639 |
| AC099062.1 | 0.40510741 | -1.952265886 | 2.21477099 | -0.8814753 | 0.37806063 | NA |
| LMO4 | 1878.64643 | -0.206167993 | 0.2009978 | -1.0257227 | 0.30502233 | 0.50208732 |
| LINC01364 | 0.4938063 | 1.565735756 | 2.09367217 | 0.74784189 | 0.45455553 | NA |
| PKN2-AS1 | 12.5426635 | 0.122471803 | 0.54606189 | 0.22428191 | 0.82253795 | 0.9051968 |
| PKN2 | 2311.71831 | 0.080332321 | 0.11939089 | 0.67285133 | 0.50104188 | 0.6819342 |
| GTF2B | 1355.58995 | -0.342115078 | 0.12682728 | -2.6974881 | 0.00698648 | 0.03457834 |
| KYAT3 | 1854.11183 | -0.044180844 | 0.21139752 | -0.2089941 | 0.83445281 | 0.9121801 |
| RBMXL1 | 951.089623 | -0.222277959 | 0.18299485 | -1.2146678 | 0.22449281 | 0.41060288 |
| GBP3 | 2512.43809 | -0.85523024 | 0.38310367 | -2.2323728 | 0.02559033 | 0.08979601 |
| GBP1 | 2272.77784 | -0.952701402 | 0.62057472 | -1.5351921 | 0.12473665 | 0.27692419 |
| GBP2 | 5313.11949 | -2.332843413 | 0.33751652 | -6.9117903 | 4.79E-12 | 1.08E-09 |
| GBP7 | 53.3030292 | 0.231254354 | 0.58312013 | 0.39658099 | 0.69167648 | 0.82245158 |
| GBP4 | 1699.42089 | -0.053838177 | 0.52787812 | -0.1019898 | 0.91876478 | 0.96046034 |
| AC099063.4 | 27.8490017 | 0.977881815 | 0.5436673 | 1.79867691 | 0.0720698 | 0.1882596 |
| GBP5 | 359.395862 | -0.322442462 | 0.73592704 | -0.4381446 | 0.66128146 | 0.80071801 |
| AC099063.1 | 48.1635675 | -0.756438097 | 0.71279036 | -1.061235 | 0.28858311 | 0.48477109 |
| GBP6 | 2.23349541 | 1.464659904 | 0.90497742 | 1.61844911 | 0.10556585 | 0.24656965 |
| LRRC8B | 647.004662 | -0.206789919 | 0.10980186 | -1.8833007 | 0.05965963 | 0.16495701 |
| LRRC8C-DT | 95.4978679 | 0.337338837 | 0.21954416 | 1.53654209 | 0.12440548 | 0.2763984 |
| LRRC8C | 484.653238 | 0.142902326 | 0.25053984 | 0.57037766 | 0.56842158 | 0.73404883 |

| | | | | | | |
|---|---|---|---|---|---|---|
| AC093423.3 | 127.823653 | 1.493196427 | 0.38926201 | 3.83596755 | 0.00012507 | 0.00150806 |
| AC099568.2 | 65.8034818 | 0.176710809 | 0.44811849 | 0.39433948 | 0.69333042 | 0.82352701 |
| LRRC8D | 2031.41575 | 0.652616803 | 0.28092285 | 2.32311755 | 0.02017284 | 0.07537091 |
| AL391497.1 | 1.85280107 | 0.706118369 | 0.93592656 | 0.75445916 | 0.45057356 | 0.64041528 |
| ZNF326 | 1884.41796 | 0.061588301 | 0.13885069 | 0.44355774 | 0.65736236 | 0.79897887 |
| AC092783.1 | 0.09997307 | 0.547239394 | 3.29377202 | 0.16614368 | 0.86804388 | NA |
| BARHL2 | 0 | NA | NA | NA | NA | NA |
| LINC01763 | 0.38295139 | -1.873172457 | 3.29726093 | -0.5680996 | 0.56996737 | NA |
| AC091614.1 | 0.91399279 | 0.549085053 | 1.42601832 | 0.38504768 | 0.70020209 | NA |
| ZNF644 | 1155.38856 | 0.027328093 | 0.13186676 | 0.2072402 | 0.83582227 | 0.91280681 |
| HFM1 | 100.56925 | 0.239208103 | 0.34607662 | 0.69119984 | 0.48943996 | 0.67284599 |
| CDC7 | 406.026782 | -1.368125322 | 0.44106766 | -3.1018491 | 0.00192316 | 0.01307713 |
| TGFBR3 | 3135.8509 | 0.354930437 | 0.17336938 | 2.04724983 | 0.04063356 | 0.12526385 |
| BRDT | 0.9635366 | -2.246806177 | 1.72532522 | -1.3022508 | 0.19283066 | NA |
| EPHX4 | 61.5975462 | -1.999383243 | 0.41410563 | -4.8281963 | 1.38E-06 | 4.22E-05 |
| SETSIP | 2.54310023 | 0.272208768 | 0.78195981 | 0.34811095 | 0.72775686 | 0.84611931 |
| BTBD8 | 188.422924 | 0.000341847 | 0.38143859 | 0.0008962 | 0.99928493 | 0.99973988 |
| AC104836.1 | 22.3061654 | 0.02603778 | 0.47690093 | 0.05459788 | 0.95645883 | 0.97925995 |
| C1orf146 | 4.58060684 | 0.305308 | 0.55682527 | 0.54830127 | 0.58348506 | 0.74575043 |
| GLMN | 423.018681 | -0.409873505 | 0.13428599 | -3.0522432 | 0.00227138 | 0.01489587 |
| RPAP2 | 936.75703 | 0.227574101 | 0.11522143 | 1.97510227 | 0.04825654 | 0.14197535 |
| GFI1 | 63.9724527 | -0.619915169 | 0.58920489 | -1.0521216 | 0.29274379 | 0.48914292 |
| EVI5 | 932.624957 | -0.055855779 | 0.16546058 | -0.3375776 | 0.73568156 | 0.85154608 |
| RPL5 | 26226.0443 | -0.217888039 | 0.23071145 | -0.944418 | 0.3449561 | 0.54262064 |
| FAM69A | 612.170209 | -0.134937472 | 0.18734323 | -0.7202687 | 0.47135955 | 0.65783283 |
| MTF2 | 1333.23682 | -0.315939187 | 0.1805794 | -1.749586 | 0.08018978 | 0.20277947 |
| TMED5 | 6727.65331 | -0.182325922 | 0.20529529 | -0.8881154 | 0.37447866 | 0.5705785 |
| CCDC18 | 912.464611 | -0.003159683 | 0.21407735 | -0.0147595 | 0.98822402 | 0.99471209 |
| DR1 | 2547.62034 | 0.022729157 | 0.11795733 | 0.19268966 | 0.84720202 | 0.91978745 |
| FNBP1L | 4241.55884 | 0.071206678 | 0.12327716 | 0.57761452 | 0.5635244 | 0.73062286 |
| BCAR3 | 2056.10539 | -0.012761307 | 0.33441607 | -0.03816 | 0.96956014 | 0.98592327 |
| AL109613.1 | 273.028755 | 0.735417059 | 0.27853884 | 2.64026754 | 0.00828406 | 0.03907883 |
| DNTTIP2 | 2292.7582 | -0.791886249 | 0.37369042 | -2.119097 | 0.03408227 | 0.11049292 |
| GCLM | 3625.29342 | -1.992218509 | 0.51343904 | -3.8801461 | 0.00010439 | 0.00131026 |
| ABCA4 | 357.765294 | 0.125990551 | 0.74880071 | 0.16825645 | 0.86638153 | 0.93064813 |
| AC093579.1 | 0.18037964 | -1.003574173 | 3.30558948 | -0.3035992 | 0.7614333 | NA |
| ARHGAP29 | 5267.19296 | -0.226857478 | 0.25788746 | -0.8796763 | 0.37903471 | 0.57504694 |
| AC097059.1 | 16.7835802 | -0.033889268 | 0.57046015 | -0.0594069 | 0.95262802 | 0.97717271 |
| AC097059.2 | 28.762496 | -1.65786749 | 0.76742292 | -2.1603049 | 0.03074907 | 0.10269163 |
| ABCD3 | 4703.99497 | 0.782977082 | 0.25530286 | 3.06685594 | 0.00216323 | 0.01432556 |
| F3 | 1430.73887 | -0.994511343 | 0.43055916 | -2.3098135 | 0.02089848 | 0.07737897 |
| AC093117.1 | 0.63342468 | -1.527610692 | 2.21631976 | -0.6892555 | 0.49066247 | NA |
| SLC44A3 | 2611.25537 | 0.14193685 | 0.34643446 | 0.40970765 | 0.68202042 | 0.81588709 |
| CNN3 | 20054.06 | -0.775849291 | 0.17780337 | -4.3635241 | 1.28E-05 | 0.00024926 |
| AC105942.1 | 755.582891 | -0.169081542 | 0.23871089 | -0.708311 | 0.47875216 | 0.66367039 |
| ALG14 | 586.27618 | 0.099819837 | 0.24211057 | 0.41229029 | 0.68012666 | 0.8145211 |
| AC095033.1 | 2.8624268 | -0.664154706 | 0.7362453 | -0.9020835 | 0.36701254 | 0.56395482 |
| TMEM56 | 706.274891 | 0.442336525 | 0.23513414 | 1.88120924 | 0.05994346 | 0.16540189 |
| TMEM56-RW | 780.577324 | 0.043019592 | 0.14539427 | 0.29588231 | 0.76731996 | 0.87152635 |
| AC092802.1 | 20.1532372 | 0.878863179 | 0.41859638 | 2.09954795 | 0.03576863 | 0.11428797 |
| RWDD3 | 908.455072 | -0.04365169 | 0.15446521 | -0.2825988 | 0.77748437 | 0.87819167 |
| AC092802.2 | 15.0101632 | 0.198196257 | 0.53275132 | 0.37202396 | 0.70987501 | 0.83512844 |
| LINC01760 | 0 | NA | NA | NA | NA | NA |
| LINC01650 | 0 | NA | NA | NA | NA | NA |
| AL445218.1 | 0.09997307 | 0.547239394 | 3.29377202 | 0.16614368 | 0.86804388 | NA |
| LINC01761 | 0.10178723 | 1.087385526 | 3.28420134 | 0.33109588 | 0.74057207 | NA |
| AL356479.1 | 0.50026919 | 0.838063145 | 2.12587333 | 0.39422064 | 0.69341815 | NA |
| AL683887.1 | 0 | NA | NA | NA | NA | NA |
| AC099566.1 | 0 | NA | NA | NA | NA | NA |
| LINC01787 | 0 | NA | NA | NA | NA | NA |
| PTBP2 | 1536.47836 | -0.315501291 | 0.16559478 | -1.9052611 | 0.05674613 | 0.15912331 |
| DPYD | 1164.62508 | 0.098928091 | 0.25277784 | 0.39136378 | 0.69552836 | 0.8247678 |
| DPYD-AS1 | 67.8097669 | 0.513321246 | 0.2737201 | 1.87535094 | 0.06074446 | 0.16691996 |
| DPYD-AS2 | 0.38194116 | -0.558425783 | 2.24981117 | -0.2482101 | 0.80397188 | NA |
| MIR137HG | 0.55060704 | -2.343997332 | 2.21442606 | -1.0585124 | 0.28982192 | NA |
| BX005019.1 | 0.19632833 | 1.506623996 | 3.27645372 | 0.45983375 | 0.64563556 | NA |
| LINC01776 | 0 | NA | NA | NA | NA | NA |
| SNX7 | 1486.22655 | -0.215034037 | 0.15975788 | -1.3459996 | 0.17830266 | 0.35377423 |
| PLPPR5 | 8.20829062 | -1.097967529 | 0.89808522 | -1.222565 | 0.22149405 | 0.40759273 |
| AL445433.2 | 1.01837714 | -0.849834911 | 1.39630109 | -0.608633 | 0.54276773 | 0.71520975 |
| PLPPR4 | 24.1033029 | 3.714918222 | 0.84316471 | 4.40592231 | 1.05E-05 | 0.00021318 |
| LINC01708 | 0.17820003 | -1.003574173 | 3.30558948 | -0.3035992 | 0.7614333 | NA |
| PALMD | 640.354156 | 0.233561138 | 0.42602864 | 0.54822872 | 0.58353486 | 0.74575043 |
| FRRS1 | 123.795283 | 0.334965181 | 0.28437664 | 1.17789275 | 0.23883937 | 0.42841943 |
| AGL | 1533.35086 | 0.44532885 | 0.13588513 | 3.27724485 | 0.00104825 | 0.00813732 |
| AC118553.1 | 61.5504628 | 0.959513499 | 0.43231847 | 2.21945986 | 0.02645545 | 0.09207211 |
| SLC35A3 | 1785.66295 | 0.400329957 | 0.14203286 | 2.81857275 | 0.00482377 | 0.02614651 |
| AC118553.2 | 3545.9899 | -0.290044949 | 0.15236575 | -1.9036098 | 0.05696102 | 0.15950536 |
| MFSD14A | 3181.6755 | -0.39929311 | 0.16530118 | -2.4155491 | 0.0157115 | 0.06295584 |
| AC093019.2 | 55.0042396 | -0.384872916 | 0.33370484 | -1.1533333 | 0.24877359 | 0.44051569 |
| SASS6 | 214.503572 | -0.056182584 | 0.24252822 | -0.2316538 | 0.81680692 | 0.90145455 |
| TRMT13 | 943.086432 | 0.126520488 | 0.13672733 | 0.92534896 | 0.35478442 | 0.55230354 |
| LRRC39 | 399.616211 | 0.139231163 | 0.13485014 | 1.03248808 | 0.30184353 | 0.49870254 |
| DBT | 1885.80947 | 1.215198912 | 0.20294268 | 5.98789223 | 2.13E-09 | 1.91E-07 |
| RTCA-AS1 | 75.183044 | 1.696170768 | 0.30992 | 5.472931 | 4.43E-08 | 2.40E-06 |
| RTCA | 2507.07925 | -0.005401955 | 0.25219944 | -0.0214194 | 0.98291111 | 0.99239475 |
| AC104457.2 | 0.71418241 | 2.306372618 | 1.98226202 | 1.16350543 | 0.2446245 | NA |
| CDC14A | 579.091146 | 0.521331662 | 0.25388603 | 2.05340819 | 0.04003301 | 0.12387589 |
| AL589990.1 | 26.604403 | 1.717046373 | 0.49444184 | 3.47269632 | 0.00051526 | 0.00464049 |
| GPR88 | 21.8038235 | 1.359331277 | 0.69191318 | 1.96459805 | 0.04946078 | 0.14455067 |
| LINC01349 | 7.86203104 | 0.517168494 | 0.60355012 | 0.85687746 | 0.39151261 | 0.5872047 |
| VCAM1 | 11611.9361 | -1.195950227 | 0.48266805 | -2.4777903 | 0.01321988 | 0.0554138 |
| EXTL2 | 934.78168 | 0.552549213 | 0.12275339 | 4.50129488 | 6.75E-06 | 0.0001503 |
| AC104506.1 | 27.6510073 | -0.2088146 | 0.5335635 | -0.3913585 | 0.69553228 | 0.8247678 |
| SLC30A7 | 1774.31142 | -0.432449886 | 0.19750131 | -2.1896051 | 0.02855289 | 0.09745529 |
| DPH5 | 889.269122 | -0.004619229 | 0.17585888 | -0.0262667 | 0.97904463 | 0.99050094 |
| AC093157.2 | 10.2313219 | -0.018972066 | 0.57407571 | -0.033048 | 0.97363629 | 0.98779582 |
| AC093157.1 | 259.115179 | 0.012224792 | 0.20833548 | 0.0586784 | 0.95320826 | 0.9774119 |
| AL109741.1 | 19.6061473 | -0.964674957 | 0.53249093 | -1.811627 | 0.07004385 | 0.18431856 |
| S1PR1 | 2069.70989 | -0.572233374 | 0.3352982 | -1.7066401 | 0.08788896 | 0.21666185 |
| LINC01307 | 0.20754827 | -0.339942428 | 3.12492333 | -0.1087842 | 0.91337362 | NA |
| LINC01709 | 8.7229532 | -0.404995628 | 0.54811773 | -0.7388844 | 0.45997721 | 0.64867635 |
| OLFM3 | 11.2704578 | 1.354269961 | 0.84639469 | 1.60004543 | 0.10958851 | 0.2532779 |
| AC114485.1 | 0 | NA | NA | NA | NA | NA |
| AC099567.1 | 0 | NA | NA | NA | NA | NA |
| COL11A1 | 18.7911593 | 0.680870496 | 0.5617119 | 1.21213472 | 0.22546081 | 0.41190257 |
| AC095032.2 | 0 | NA | NA | NA | NA | NA |
| RNPC3 | 1818.57148 | 0.668420578 | 0.17629269 | 3.79153875 | 0.00014972 | 0.00175079 |
| AMY2B | 2952.80223 | 0.306018999 | 0.24813829 | 1.23325988 | 0.21747882 | 0.40250677 |
| AMY2A | 8.1569485 | 0.712758913 | 0.66782691 | 1.06728091 | 0.285845 | 0.48183954 |
| AMY1A | 0.36539036 | -0.032132692 | 2.27971556 | -0.014095 | 0.98875415 | NA |
| AMY1B | 0.6409388 | 2.20050573 | 1.61561171 | 1.36202635 | 0.17318957 | NA |
| AMY1C | 0.03392908 | 0.380740593 | 3.29738709 | 0.11546736 | 0.90807469 | NA |

| | | | | | | |
|---|---|---|---|---|---|---|
| LINC01676 | 0.37038051 | -1.761428036 | 2.82286449 | -0.623986 | 0.53263675 | NA |
| LINC01677 | 0 | NA | NA | NA | NA | NA |
| AL355306.2 | 0.19478864 | 0.18657164 | 3.29377195 | 0.05664376 | 0.95482897 | NA |
| LINC01661 | 0 | NA | NA | NA | NA | NA |
| PRMT6 | 1174.65413 | 0.190459339 | 0.22457114 | 0.84810249 | 0.3963809 | 0.59135803 |
| NTNG1 | 500.52435 | 0.148686522 | 0.63011714 | 0.23596648 | 0.81345868 | 0.89922052 |
| VAV3 | 6589.79304 | 0.907490717 | 0.21067863 | 4.30746452 | 1.65E-05 | 0.00030452 |
| VAV3-AS1 | 1.63222066 | -0.384604845 | 1.32849393 | -0.2895044 | 0.77219541 | 0.87442749 |
| AL390036.1 | 5.35233325 | -0.252037021 | 0.52236932 | -0.4824882 | 0.6294592 | 0.77986909 |
| SLC25A24 | 1549.46011 | -0.018579726 | 0.19241591 | -0.0965602 | 0.92307564 | 0.96245718 |
| AL359258.2 | 105.618369 | 0.551617431 | 0.2747314 | 2.00784269 | 0.04466001 | 0.13406179 |
| NBPF4 | 0.26148673 | -0.174096114 | 3.29377188 | -0.0528562 | 0.95784652 | NA |
| AL359258.1 | 0 | NA | NA | NA | NA | NA |
| AL390038.1 | 0 | NA | NA | NA | NA | NA |
| NBPF6 | 0.07659028 | -0.642905668 | 3.30558948 | -0.1944905 | 0.84579185 | NA |
| FAM102B | 459.317155 | 0.16187589 | 0.29333043 | 0.55185508 | 0.58104764 | 0.7438335 |
| HENMT1 | 137.364112 | 0.119206059 | 0.2649206 | 0.449969 | 0.65273279 | 0.79594194 |
| PRPF38B | 2619.19728 | -0.179072851 | 0.13770272 | -1.3004307 | 0.19345339 | 0.37306875 |
| FNDC7 | 2.01664053 | -1.487335119 | 1.12344476 | -1.3239059 | 0.18553431 | 0.36286362 |
| AL591719.2 | 0 | NA | NA | NA | NA | NA |
| STXBP3 | 2977.82442 | 0.074786345 | 0.10606764 | 0.70508163 | 0.48075944 | 0.66542234 |
| AKNAD1 | 3.68256838 | 0.747604135 | 0.73201075 | 1.02130213 | 0.30711132 | 0.50405799 |
| AL449266.1 | 38.160784 | 0.236438391 | 0.2459417 | 0.96135951 | 0.33637144 | 0.53442067 |
| SPATA42 | 0.71696073 | -0.387942656 | 1.85328512 | -0.209327 | 0.83419296 | NA |
| GPSM2 | 1184.22142 | -0.237635397 | 0.12559687 | -1.8920487 | 0.05848449 | 0.16254596 |
| CLCC1 | 1504.89555 | -0.073879902 | 0.1037968 | -0.7117743 | 0.47660453 | 0.66179458 |
| WDR47 | 737.398126 | 0.012047041 | 0.15760678 | 0.07643732 | 0.93907118 | 0.97099584 |
| TAF13 | 1446.14921 | -0.69419662 | 0.32381514 | -2.1438053 | 0.03204849 | 0.10575881 |
| AL356488.3 | 218.177163 | 0.543108208 | 0.2237662 | 2.42712354 | 0.01521907 | 0.0614872 |
| TMEM167B | 2860.03934 | -0.158528608 | 0.12732538 | -1.2450668 | 0.21310719 | 0.3970244 |
| AL356488.2 | 8.16737611 | -0.525995337 | 0.49292486 | -1.0670903 | 0.28593106 | 0.48194325 |
| C1orf194 | 15.406143 | -2.641420423 | 0.91375116 | -2.8907437 | 0.00384331 | 0.02211877 |
| KIAA1324 | 39.6402638 | 0.807254514 | 0.31661603 | 2.54963251 | 0.01078365 | 0.04764828 |
| SARS | 6252.32363 | -0.360125027 | 0.13511737 | -2.6652755 | 0.00769253 | 0.03709123 |
| CELSR2 | 3094.89394 | 0.377651305 | 0.30959303 | 1.21983141 | 0.22252879 | 0.40849873 |
| PSRC1 | 268.35625 | -0.525660991 | 0.31600743 | -1.663445 | 0.09622338 | 0.23138597 |
| MYBPHL | 3.94536356 | 0.457579705 | 0.63200012 | 0.72401838 | 0.46905445 | 0.65569182 |
| SORT1 | 7196.17724 | -0.084357296 | 0.18962967 | -0.4448528 | 0.65642611 | 0.79835608 |
| PSMA5 | 4275.52865 | -0.523402366 | 0.1459901 | -3.5851909 | 0.00033683 | 0.00332544 |
| SYPL2 | 1542.86366 | 0.494544703 | 0.27180739 | 1.81946747 | 0.06884014 | 0.1819593 |
| ATXN7L2 | 1525.94009 | -0.315914755 | 0.22861435 | -1.3818676 | 0.16701236 | 0.33799816 |
| CYB561D1 | 1118.00572 | 0.145868727 | 0.14954396 | 0.97542371 | 0.32935013 | 0.52712812 |
| AMIGO1 | 771.696921 | 0.084765943 | 0.2237036 | 0.37892077 | 0.7047467 | 0.83139491 |
| GPR61 | 22.994837 | 0.506394208 | 0.4577452 | 1.10627967 | 0.26860549 | 0.46279822 |
| AL355310.3 | 1.34759915 | 0.229969201 | 1.15666479 | 0.19882096 | 0.8424028 | 0.91630899 |
| GNAI3 | 4096.53904 | -0.44399015 | 0.1803982 | -2.4611673 | 0.01384858 | 0.05740128 |
| GNAT2 | 66.6473956 | 0.245325317 | 0.31457822 | 0.77985473 | 0.43547639 | 0.62754863 |
| AMPD2 | 4385.3489 | -0.399569397 | 0.22550908 | -1.771855 | 0.07641863 | 0.1962219 |
| AL355310.2 | 952.59903 | -0.299045659 | 0.22499353 | -1.32913 | 0.18380508 | 0.36082334 |
| GSTM4 | 4040.16789 | 0.766140372 | 0.21974164 | 3.48655076 | 0.00048929 | 0.00445533 |
| GSTM2 | 2990.1215 | 0.908971436 | 0.33741002 | 2.69396689 | 0.00706072 | 0.0348542 |
| GSTM1 | 409.190282 | 0.455031007 | 1.2922682 | 0.35211809 | 0.7247497 | 0.84389349 |
| AC000032.1 | 145.541111 | 0.35904778 | 1.24728894 | 0.28786255 | 0.77345195 | 0.87536172 |
| GSTM5 | 185.352855 | 0.737411324 | 0.46885832 | 1.57278071 | 0.1157696 | 0.26344878 |
| AL158847.1 | 1106.85967 | 0.220303043 | 0.34542492 | 0.63777403 | 0.52362079 | 0.69892989 |
| GSTM3 | 5661.8223 | 0.36430338 | 0.36290433 | 1.00385513 | 0.31544844 | 0.51256204 |
| EPS8L3 | 22.8113433 | -1.561312925 | 0.8787158 | -1.7768122 | 0.07559915 | 0.19483479 |
| LINC01768 | 2.84331153 | 0.766851394 | 0.71514842 | 1.07229684 | 0.28358672 | 0.47952433 |
| AL450468.1 | 0.38636858 | -0.032082483 | 1.90201856 | -0.0168676 | 0.98654224 | NA |
| AL450468.2 | 1.02861197 | 0.194653184 | 1.31917787 | 0.14755644 | 0.88269284 | 0.93902736 |
| CSF1 | 2843.44564 | -0.524602092 | 0.28307741 | -1.8532107 | 0.06385216 | 0.17263173 |
| AHCYL1 | 12008.4757 | -0.056526859 | 0.21820244 | -0.2590569 | 0.7955913 | 0.88912284 |
| STRIP1 | 1662.24674 | 0.006765341 | 0.16199742 | 0.04176203 | 0.9666884 | 0.98419262 |
| AL160006.1 | 951.614281 | 0.010003384 | 0.15886085 | 0.06296947 | 0.94979081 | 0.97588092 |
| ALX3 | 380.150325 | 0.171589967 | 0.1653836 | 1.03752709 | 0.29949028 | 0.49614701 |
| LINC01397 | 1.01063646 | -0.833288736 | 1.53185702 | -0.5439729 | 0.58646011 | 0.74768058 |
| UBL4B | 2.14766899 | -1.381907248 | 1.33705111 | -1.0335486 | 0.30134726 | 0.49812118 |
| SLC6A17 | 412.817723 | 0.062144629 | 0.42142049 | 0.14746466 | 0.88276527 | 0.93903649 |
| AL355990.1 | 18.158895 | 0.751632735 | 0.62960321 | 1.19381973 | 0.23254849 | 0.42059697 |
| AL355990.2 | 2.18555134 | 0.36693056 | 0.9829312 | 0.37330238 | 0.7089234 | 0.83447565 |
| LINC02586 | 41.251285 | -0.028915739 | 0.42989165 | -0.0672629 | 0.94637245 | 0.97427563 |
| KCNC4 | 1860.58414 | -0.138531496 | 0.26910088 | -0.5147939 | 0.60669703 | 0.76318279 |
| RBM15-AS1 | 28.8543 | -0.496638533 | 0.47912886 | -1.0365448 | 0.29994804 | 0.49671125 |
| RBM15 | 502.626994 | -1.003914367 | 0.26960299 | -3.7236767 | 0.00019634 | 0.0021501 |
| SLC16A4 | 6768.88242 | 0.79010536 | 0.38210698 | 2.06775957 | 0.03866264 | 0.12076922 |
| AL355488.1 | 712.301866 | 0.089489161 | 0.21982224 | 0.40709784 | 0.68393613 | 0.81748139 |
| LAMTOR5 | 7595.79696 | 0.630353034 | 0.22781507 | 2.76695052 | 0.00565833 | 0.02956192 |
| PROK1 | 4.67305602 | 0.348487146 | 0.75584105 | 0.46105877 | 0.64475644 | 0.79009304 |
| AL358215.1 | 3.26027024 | 0.1462881 | 0.81700294 | 0.17905456 | 0.85789486 | 0.92575552 |
| AL358215.2 | 0 | NA | NA | NA | NA | NA |
| CYMP-AS1 | 0 | NA | NA | NA | NA | NA |
| KCNA10 | 0.13382144 | 0.02007176 | 3.29738704 | 0.00608717 | 0.99514317 | NA |
| KCNA2 | 11.9577213 | 1.810451911 | 0.56629204 | 3.19702873 | 0.00138851 | 0.01018983 |
| AL365361.1 | 71.3744952 | 0.311181101 | 0.60737548 | 0.51233728 | 0.60841498 | 0.76457791 |
| KCNA3 | 11.5235478 | 0.132210175 | 0.65346087 | 0.20232302 | 0.83966421 | 0.91484357 |
| CD53 | 1309.57159 | -0.771480553 | 0.47551423 | -1.6224132 | 0.10471491 | 0.24520711 |
| AL360270.1 | 70.2385128 | 0.623553614 | 0.34089085 | 1.82918847 | 0.06737138 | 0.1792356 |
| LRIF1 | 570.896476 | -0.010431581 | 0.24825845 | -0.042019 | 0.96648352 | 0.98410057 |
| AL360270.3 | 85.0398698 | 0.291422534 | 0.33450685 | 0.87120049 | 0.38364469 | 0.57938895 |
| DRAM2 | 2831.9476 | -0.5709081 | 0.22008141 | -2.5940768 | 0.00948453 | 0.04332358 |
| CEPT1 | 2689.25279 | -0.138595712 | 0.1487968 | -0.9314429 | 0.35162453 | 0.54930479 |
| AL355816.1 | 40.4247145 | 0.08705032 | 0.25934417 | 0.33565559 | 0.7371306 | 0.8524157 |
| AL355816.2 | 838.53361 | -0.502989696 | 0.19893199 | -2.5284506 | 0.01145672 | 0.04989492 |
| DENND2D | 2699.94282 | -0.790006736 | 0.23250794 | -3.3977625 | 0.00067939 | 0.0057551 |
| CHI3L2 | 903.59808 | -4.780528344 | 0.74482501 | -6.4183241 | 1.38E-10 | 1.87E-08 |
| CHIA | 7.48546594 | 1.359747614 | 0.91023656 | 1.49383982 | 0.13521749 | 0.29265309 |
| AL356387.1 | 1.89643766 | 2.099577173 | 1.27859441 | 1.64209789 | 0.10056972 | 0.23848911 |
| PIFO | 478.199515 | -0.212191342 | 0.34733236 | -0.6109173 | 0.54125434 | 0.71410802 |
| OVGP1 | 673.189986 | -1.142314451 | 0.34362992 | -3.3242578 | 0.00088654 | 0.00714272 |
| WDR77 | 2279.7091 | -0.403731817 | 0.15898803 | -2.539385 | 0.01110476 | 0.04868476 |
| ATP5PB | 9474.26758 | 0.206847252 | 0.17120086 | 1.20821387 | 0.22696501 | 0.41379753 |
| C1orf162 | 1048.43621 | -1.296744691 | 0.48674599 | -2.6641097 | 0.00771924 | 0.03720642 |
| TMIGD3 | 138.794508 | -2.353670772 | 0.53580984 | -4.3927352 | 1.12E-05 | 0.00022327 |
| ADORA3 | 205.704648 | -2.291659838 | 0.57028524 | -4.0184449 | 5.86E-05 | 0.00083596 |
| RAP1A | 3980.86647 | -0.445245248 | 0.1586983 | -2.8056082 | 0.00502217 | 0.02697244 |
| LINC01160 | 56.5182476 | -0.449148304 | 0.43240593 | -1.0387191 | 0.29893539 | 0.49578153 |
| INKA2 | 641.590829 | 0.216821716 | 0.35461892 | 0.61142174 | 0.54092041 | 0.71384602 |
| AL049557.1 | 631.738602 | 0.227896117 | 0.35460275 | 0.64268007 | 0.52043172 | 0.69699386 |
| DDX20 | 779.216099 | 0.004230007 | 0.22140176 | 0.01910557 | 0.98475689 | 0.99345955 |
| KCND3 | 363.764357 | -0.416919653 | 0.29026304 | -1.4363512 | 0.15090244 | 0.31566678 |
| KCND3-AS1 | 0 | NA | NA | NA | NA | NA |
| LINC01750 | 39.3331688 | -1.291878578 | 0.37810635 | -3.4167069 | 0.00063383 | 0.00545093 |
| AL445426.1 | 5.2720014 | -0.317406774 | 0.94459515 | -0.3360241 | 0.73685266 | 0.85226566 |

| | | | | | | |
|---|---|---|---|---|---|---|
| CTTNBP2NL | 1477.73451 | -0.276783214 | 0.16868243 | -1.6408538 | 0.10082777 | 0.23888843 |
| WNT2B | 586.399581 | -0.132599453 | 0.14528766 | -0.9126684 | 0.36141699 | 0.55891192 |
| AL354760.1 | 27.4652446 | -0.574446239 | 0.34731403 | -1.6539678 | 0.09813405 | 0.2346355 |
| ST7L | 878.108237 | -0.159340136 | 0.1389819 | -1.1464812 | 0.25159608 | 0.44387427 |
| CAPZA1 | 4844.01974 | -0.719044669 | 0.11068885 | -6.4960895 | 8.24E-11 | 1.22E-08 |
| MOV10 | 5803.75462 | -0.352753545 | 0.38507667 | -0.9160605 | 0.35963514 | 0.55726752 |
| AL603832.1 | 326.417469 | 0.241206594 | 0.35725735 | 0.67516201 | 0.49957284 | 0.68095398 |
| RHOC | 16366.6574 | -0.557951671 | 0.19765839 | -2.822808 | 0.00476051 | 0.02592618 |
| AL603832.3 | 8498.26589 | -0.463944007 | 0.20131746 | -2.3045394 | 0.02119238 | 0.07820483 |
| PPM1J | 280.758555 | -0.675875288 | 0.47300492 | -1.428897 | 0.15303385 | 0.31857242 |
| AL603832.2 | 0.89464232 | 0.814038171 | 2.19994364 | 0.37002683 | 0.7113625 | NA |
| FAM19A3 | 1.65345865 | -0.351828593 | 1.07546425 | -0.3271411 | 0.74356114 | 0.85615699 |
| LINC01356 | 4.58472109 | 0.701888754 | 0.67737263 | 1.03619296 | 0.30011213 | 0.49682698 |
| AL390729.1 | 2.88152443 | -0.570799685 | 0.79595232 | -0.717128 | 0.47329513 | 0.65931286 |
| SLC16A1 | 1403.45393 | -0.271342538 | 0.33155704 | -0.8183887 | 0.41313526 | 0.607085 |
| SLC16A1-AS1 | 96.8624051 | -0.898724547 | 0.26944674 | -3.3354442 | 0.00085163 | 0.00692493 |
| AL357055.3 | 21.6725473 | 1.031686639 | 0.46211142 | 2.23254953 | 0.02557867 | 0.08977897 |
| LRIG2 | 1277.36931 | 0.181609409 | 0.13737481 | 1.32199932 | 0.18616838 | 0.36370278 |
| MAGI3 | 792.718165 | 0.370333962 | 0.16039534 | 2.30888226 | 0.02095012 | 0.07750485 |
| PHTF1 | 1062.58183 | -0.638985495 | 0.21587908 | -2.9599231 | 0.00307716 | 0.01872281 |
| RSBN1 | 994.417033 | -0.134117633 | 0.12620244 | -1.0627182 | 0.28790974 | 0.4841334 |
| AL137856.1 | 19.8146658 | -0.918528861 | 0.60423467 | -1.5201525 | 0.12847265 | 0.28267198 |
| PTPN22 | 121.604115 | -0.496908508 | 0.60183348 | -0.8256578 | 0.40899824 | 0.60343935 |
| AP4B1-AS1 | 328.407185 | 0.152765675 | 0.18266251 | 0.83632749 | 0.40297069 | 0.59801276 |
| BCL2L15 | 137.467948 | -1.354304311 | 0.51026158 | -2.6541373 | 0.00795115 | 0.03799778 |
| AP4B1 | 1032.09698 | -0.060204382 | 0.14322902 | -0.4203365 | 0.67423967 | 0.81027171 |
| DCLRE1B | 254.654137 | -0.252209336 | 0.27595151 | -0.9139625 | 0.36073655 | 0.55838359 |
| HIPK1-AS1 | 15.9869406 | 0.786860857 | 0.42848026 | 1.83639931 | 0.06629862 | 0.17726497 |
| HIPK1 | 2798.2139 | -0.023182851 | 0.11207629 | -0.2068489 | 0.8361279 | 0.91295155 |
| OLFML3 | 1345.2615 | 1.425286046 | 0.41273771 | 3.45324893 | 0.00055388 | 0.00491942 |
| SYT6 | 228.818118 | 1.711712315 | 0.68277795 | 2.50698241 | 0.01217668 | 0.05210954 |
| AL121999.1 | 6.32525591 | 3.582812266 | 0.69447032 | 5.15905743 | 2.48E-07 | 1.01E-05 |
| TRIM33 | 2172.53527 | -0.063068531 | 0.23666921 | -0.2664839 | 0.78986657 | 0.88584109 |
| BCAS2 | 1536.2477 | -0.189045636 | 0.18713983 | -1.0101839 | 0.31240721 | 0.50924654 |
| DENND2C | 106.483679 | -0.049226175 | 0.28678046 | -0.1716511 | 0.86371185 | 0.92906716 |
| AMPD1 | 7.91543996 | 1.45439528 | 0.97285553 | 1.4949756 | 0.13492081 | 0.2923469 |
| NRAS | 2749.31844 | -0.671988881 | 0.31705143 | -2.1194949 | 0.03404867 | 0.11042857 |
| CSDE1 | 18140.4849 | 0.323367553 | 0.1740944 | 1.85742647 | 0.0632505 | 0.17159398 |
| SIKE1 | 1625.54517 | -0.024647562 | 0.14097074 | -0.1748417 | 0.86120403 | 0.92774579 |
| SYCP1 | 0.61429748 | 0.263488397 | 1.65875598 | 0.15884699 | 0.87378942 | NA |
| TSHB | 2.05517189 | 1.866788103 | 0.89263543 | 2.09132199 | 0.03649921 | 0.11594798 |
| TSPAN2 | 345.972843 | 1.180853062 | 0.27977807 | 4.2206777 | 2.44E-05 | 0.00041626 |
| LINC01765 | 1.21664883 | -2.592557368 | 1.91782056 | -1.3518248 | 0.17643137 | 0.35118473 |
| AL512638.2 | 1.38233486 | 0.063197951 | 1.3053456 | 0.04841473 | 0.96138572 | 0.98156291 |
| NGF | 201.635972 | -0.978275489 | 0.44265344 | -2.2100257 | 0.02710338 | 0.09374555 |
| VANGL1 | 957.554795 | 0.174548319 | 0.18943046 | 0.92143745 | 0.35682208 | 0.55430918 |
| CASQ2 | 178.510396 | 0.188170734 | 0.66643109 | 0.28235588 | 0.77767064 | 0.87832683 |
| NHLH2 | 0.61834888 | -1.499993182 | 1.8555679 | -0.8083742 | 0.41887521 | NA |
| LINC01649 | 2.43277034 | 0.64526777 | 0.83740705 | 0.7705545 | 0.44097104 | 0.63221913 |
| SLC22A15 | 525.293761 | -0.021719072 | 0.24253566 | -0.08955 | 0.92864481 | 0.96531469 |
| AL365318.1 | 0.91114845 | -0.682971701 | 1.45328874 | -0.4699491 | 0.6383914 | NA |
| MAB21L3 | 79.2832389 | -1.66054544 | 0.61626517 | -2.6945308 | 0.00704878 | 0.03481483 |
| LINC01779 | 1.40173774 | 1.681582654 | 1.27583894 | 1.31802111 | 0.18749658 | 0.3655123 |
| ATP1A1 | 153137.211 | 0.041818001 | 0.21841599 | 0.19146034 | 0.84816495 | 0.92016931 |
| LINC01762 | 534.264826 | 0.62018467 | 0.45457253 | 1.36432501 | 0.1724653 | 0.34553334 |
| AL136376.1 | 6.15262137 | 1.388991122 | 0.5313684 | 2.61398895 | 0.00894919 | 0.04147456 |
| AL390066.1 | 228.766689 | 1.013501496 | 0.36558465 | 2.77227588 | 0.00556658 | 0.02924702 |
| CD58 | 1035.29969 | -0.694429651 | 0.19150045 | -3.6262559 | 0.00028756 | 0.00293231 |
| IGSF3 | 3931.59378 | -0.593064348 | 0.20712585 | -2.8633044 | 0.00419248 | 0.023604 |
| C1orf137 | 0.08757577 | 0.547239394 | 3.29377202 | 0.16614368 | 0.86804388 | NA |
| CD2 | 204.064479 | -0.433895409 | 0.75387031 | -0.5755571 | 0.56491458 | 0.73174269 |
| AL157904.1 | 6.67874339 | -0.305047818 | 0.58734884 | -0.519364 | 0.60350696 | 0.76073856 |
| PTGFRN | 1305.60894 | -0.884456002 | 0.26336423 | -3.3582997 | 0.00078424 | 0.00646865 |
| CD101 | 54.3631366 | -0.149464433 | 0.21396704 | -0.6985395 | 0.48483985 | 0.6691144 |
| AL445231.1 | 26.2438521 | -0.020950553 | 0.27038068 | -0.0774854 | 0.93823741 | 0.97060692 |
| TTF2 | 552.9742 | -0.495455347 | 0.14599379 | -3.3936742 | 0.00068962 | 0.00582672 |
| TRIM45 | 408.189636 | 0.67747002 | 0.35348362 | 1.91655281 | 0.05529477 | 0.15633355 |
| VTCN1 | 1051.73987 | 0.299487465 | 0.64275736 | 0.46594172 | 0.64125722 | 0.78758115 |
| LINC01525 | 0 | NA | NA | NA | NA | NA |
| AL358072.1 | 18.6339314 | 0.956342949 | 0.32036443 | 2.98517208 | 0.00283419 | 0.01756395 |
| MAN1A2 | 2114.91291 | 0.032138333 | 0.16003558 | 0.20081993 | 0.84083938 | 0.91512686 |
| AL157902.1 | 0.55394496 | -0.078588551 | 1.94448 | -0.0404162 | 0.96776129 | NA |
| TENT5C | 887.185425 | 0.966346444 | 0.39273473 | 2.46055766 | 0.01387213 | 0.05748083 |
| GDAP2 | 833.293251 | -0.103592716 | 0.0904652 | -1.1451112 | 0.2521631 | 0.44443341 |
| WDR3 | 1137.78738 | 0.009638858 | 0.16913098 | 0.05699049 | 0.95455277 | 0.97829602 |
| SPAG17 | 430.293438 | -0.153706862 | 0.31825917 | -0.4829613 | 0.62912322 | 0.77967757 |
| TBX15 | 56.2561894 | -1.695634055 | 0.90110901 | -1.8817191 | 0.05987416 | 0.16529331 |
| AL139420.1 | 0.67947767 | 0.302946011 | 1.89191729 | 0.16012646 | 0.87278146 | NA |
| AL139420.2 | 0.75606795 | 0.061781202 | 1.91467869 | 0.03226714 | 0.97425902 | NA |
| WARS2 | 877.348664 | 0.408367938 | 0.20237926 | 2.01783496 | 0.04360845 | 0.13201181 |
| WARS2-IT1 | 8.10590936 | 0.940125543 | 0.74871895 | 1.25564545 | 0.20924454 | 0.39243077 |
| AL359915.2 | 459.500486 | -0.043711326 | 0.25364116 | -0.1723353 | 0.86317393 | 0.92871584 |
| AL359915.1 | 10.2203222 | 2.439682824 | 0.55890747 | 4.36509255 | 1.27E-05 | 0.00024784 |
| LINC01780 | 29.8674101 | 3.191701331 | 0.77522685 | 4.11711918 | 3.84E-05 | 0.0006011 |
| HAO2 | 5652.20572 | 3.924190616 | 0.83062794 | 4.72436627 | 2.31E-06 | 6.38E-05 |
| HSD3B2 | 59.6561993 | 1.739495641 | 0.81457686 | 2.13545918 | 0.03272352 | 0.10735497 |
| HSD3B1 | 2.38475507 | 2.804940291 | 1.14040477 | 2.45960064 | 0.01390917 | 0.05759812 |
| ZNF697 | 1331.09966 | 0.158684172 | 0.40905775 | 0.38792608 | 0.69807074 | 0.82658847 |
| PHGDH | 2979.35947 | 1.162342518 | 0.29212963 | 3.97885868 | 6.92E-05 | 0.00094852 |
| HMGCS2 | 3778.19756 | 1.695775485 | 0.56144925 | 3.02035397 | 0.00252479 | 0.01611298 |
| REG4 | 2.59994374 | 1.760455293 | 0.87679699 | 2.00782544 | 0.04466184 | 0.13406179 |
| ADAM30 | 0.08666371 | -0.642905668 | 3.30558948 | -0.1944905 | 0.84579185 | NA |
| NOTCH2 | 3004.31301 | -0.127182562 | 0.11214821 | -1.1340579 | 0.25677025 | 0.44945587 |
| AC245008.1 | 0.07659028 | -0.642905668 | 3.30558948 | -0.1944905 | 0.84579185 | NA |
| SEC22B | 3688.66944 | -0.216385826 | 0.1405997 | -1.5390205 | 0.12379929 | 0.27542002 |
| NBPF26 | 182.103314 | -0.242497575 | 0.33917049 | -0.7149725 | 0.47462604 | 0.66013204 |
| PPIAL4A | 0.30744063 | -0.695202747 | 2.31888909 | -0.2997999 | 0.76432978 | NA |
| LINC00623 | 285.430331 | -0.805369177 | 0.31018831 | -2.5963879 | 0.00942096 | 0.04306307 |
| FCGR1B | 42.2031977 | -0.58919364 | 0.64200054 | -0.9177463 | 0.35875169 | 0.55642171 |
| AC244453.3 | 7.08911761 | -0.32133241 | 0.49879835 | -0.6442131 | 0.51943729 | 0.69628475 |
| AC244453.2 | 5.1274661 | 0.10553491 | 0.63204722 | 0.16697314 | 0.86739118 | 0.93105 |
| AC244453.1 | 3.35741477 | -1.194418873 | 0.96693936 | -1.2352573 | 0.21673477 | 0.40158066 |
| FAM72B | 18.0870833 | -0.089536818 | 0.54410562 | -0.1645578 | 0.86929206 | 0.932003 |
| SRGAP2C | 537.444592 | 0.041442066 | 0.24464045 | 0.16939989 | 0.86548211 | 0.93006085 |
| SRGAP2-AS1 | 1.27441756 | 1.621111632 | 1.86263649 | 0.87033172 | 0.38411915 | 0.57967391 |
| AC244021.1 | 33.216457 | 0.352276397 | 0.51037024 | 0.69023695 | 0.49004519 | 0.67336208 |
| AC244021.2 | 3.30431768 | 0.24562769 | 0.78632105 | 0.31237583 | 0.75475492 | 0.86361569 |
| AL592494.1 | 0.70860307 | 0.98670004 | 1.85289065 | 0.5325193 | 0.59436637 | NA |
| AL592494.3 | 7.8918469 | 0.746468887 | 0.78379152 | 0.95238194 | 0.34090331 | 0.53891376 |
| LINC01691 | 0.7854621 | 0.805619186 | 1.75376172 | 0.45936639 | 0.64597108 | NA |
| AC239859.3 | 0 | NA | NA | NA | NA | NA |
| AC239800.2 | 13.5915008 | 0.17777912 | 0.72979199 | 0.24360245 | 0.80753873 | 0.89601937 |
| AC239800.3 | 1.32759856 | -0.794550126 | 1.44895485 | -0.5483609 | 0.58344415 | 0.74574292 |

| | | | | | | |
|---|---|---|---|---|---|---|
| AC239798.2 | 3.70901761 | -0.512049582 | 0.91978563 | -0.5567053 | 0.57772876 | 0.74134829 |
| AC246680.1 | 3.35479709 | 0.561247721 | 0.79034694 | 0.71012829 | 0.47762458 | 0.66275312 |
| HIST2H3PS2 | 4.00959034 | 0.622559102 | 0.7143269 | 0.8715325 | 0.38346347 | 0.57928115 |
| FAM72C | 3.00186315 | -1.204699879 | 0.88712017 | -1.3579895 | 0.17446702 | 0.3484054 |
| IGKV1OR1-1 | 0 | NA | NA | NA | NA | NA |
| AC245595.1 | 344.287196 | -0.855448227 | 0.52274938 | -1.6364405 | 0.10174744 | 0.24039429 |
| PPIAL4E | 0 | NA | NA | NA | NA | NA |
| AC246785.3 | 1.88033757 | -0.81235393 | 0.92786386 | -0.8755098 | 0.38129656 | 0.57729749 |
| NBPF15 | 844.448493 | -0.265025654 | 0.21758178 | -1.2180507 | 0.22320469 | 0.40924076 |
| PPIAL4F | 0.35126719 | -1.740327409 | 2.84550584 | -0.6116056 | 0.54079871 | NA |
| LINC01632 | 0 | NA | NA | NA | NA | NA |
| FP700111.1 | 0 | NA | NA | NA | NA | NA |
| SRGAP2B | 235.858294 | 0.225149704 | 0.28891551 | 0.77929256 | 0.4358074 | 0.62792269 |
| FAM72D | 8.6457578 | -1.043148486 | 0.48021914 | -2.1722343 | 0.02983799 | 0.10050171 |
| PPIAL4D | 0 | NA | NA | NA | NA | NA |
| AC245014.3 | 5.54425774 | -0.076878077 | 0.78800159 | -0.0975608 | 0.92228104 | 0.96203347 |
| AC245014.1 | 2.00352405 | 0.796571716 | 0.8808671 | 0.90430408 | 0.36583418 | 0.56283824 |
| NBPF20 | 228.594807 | -0.676978873 | 0.31816672 | -2.1277489 | 0.03335791 | 0.10880001 |
| GPR89A | 500.979154 | 0.432793155 | 0.17285065 | 2.50385615 | 0.0122848 | 0.05247855 |
| PDZK1 | 18362.6047 | 1.487975151 | 0.4454888 | 3.34009553 | 0.0008375 | 0.0068205 |
| CD160 | 65.4731511 | -0.364290879 | 0.18345215 | -1.9857542 | 0.04706063 | 0.13931171 |
| RNF115 | 2651.43496 | -0.263537777 | 0.15041 | -1.7521294 | 0.07975156 | 0.20202197 |
| POLR3C | 1074.78106 | -0.451185374 | 0.18654188 | -2.4186814 | 0.01557688 | 0.06252089 |
| NUDT17 | 721.633491 | -0.533075472 | 0.19577709 | -2.7228695 | 0.00647176 | 0.03259365 |
| PIAS3 | 2258.06948 | -0.355486545 | 0.16686429 | -2.1303931 | 0.03313917 | 0.10834118 |
| ANKRD35 | 57.9049473 | 0.05061519 | 0.40422581 | 0.12521514 | 0.90035323 | 0.94951124 |
| ITGA10 | 354.204287 | -0.283096951 | 0.40530075 | -0.6984861 | 0.48487323 | 0.6691144 |
| AC243547.1 | 29.4351198 | -0.501942327 | 0.43621087 | -1.1506873 | 0.24986089 | 0.44194046 |
| PEX11B | 1718.60966 | 0.145853793 | 0.15409596 | 0.94651274 | 0.34388713 | 0.54151789 |
| RBM8A | 4024.88176 | -0.214724908 | 0.1386333 | -1.5488696 | 0.12141306 | 0.2719958 |
| LIX1L-AS1 | 1478.56949 | -0.660061732 | 0.1541656 | -4.281511 | 1.86E-05 | 0.00033459 |
| AC243547.3 | 1031.16124 | -0.320895018 | 0.14132714 | -2.2705831 | 0.02317223 | 0.08364841 |
| LIX1L | 1303.03416 | -0.224584575 | 0.16663677 | -1.3477492 | 0.17773906 | 0.35293516 |
| ANKRD34A | 77.5326068 | -0.204667847 | 0.35502431 | -0.5764897 | 0.56428425 | 0.73132527 |
| AC243547.2 | 10.7392447 | 0.327761792 | 0.38554277 | 0.85013082 | 0.39525236 | 0.59054817 |
| POLR3GL | 1008.11816 | -0.088179741 | 0.1932597 | -0.4562759 | 0.6481916 | 0.79283884 |
| TXNIP | 54115.2464 | -0.27603745 | 0.42020395 | -0.656913 | 0.51123684 | 0.68936978 |
| HJV | 10.9895466 | 2.061338641 | 0.52896997 | 3.8968916 | 9.74E-05 | 0.00123825 |
| LINC01719 | 27.8813997 | 0.94534525 | 0.3564755 | 2.65192205 | 0.0080035 | 0.03813739 |
| NBPF10 | 192.762938 | -0.21261598 | 0.15405449 | -1.3801349 | 0.16754511 | 0.33876434 |
| AC239799.1 | 103.184276 | -0.612123166 | 0.32241909 | -1.8985327 | 0.05762595 | 0.16088897 |
| NOTCH2NL | 108.84964 | -0.444674023 | 0.32567474 | -1.365393 | 0.17212956 | 0.34506776 |
| AC239799.2 | 1.06685495 | 1.076392485 | 1.30240878 | 0.82646286 | 0.40854158 | 0.60300243 |
| AC243756.1 | 1.64347887 | -0.329721675 | 1.13964227 | -0.2893203 | 0.77233626 | 0.87454943 |
| AC245407.2 | 14.4643573 | -0.374354766 | 0.54334388 | -0.6889831 | 0.49083389 | 0.67391907 |
| NBPF12 | 785.553575 | -0.064553579 | 0.14441 | -0.447016 | 0.65486351 | 0.79747572 |
| AC244394.2 | 0.28652914 | -0.179717927 | 2.92211968 | -0.0615026 | 0.95095895 | NA |
| PRKAB2 | 1359.52348 | 0.016697605 | 0.15254235 | 0.10946209 | 0.91283599 | 0.95719548 |
| AC242426.2 | 198.720398 | 0.331691247 | 0.20109596 | 1.64941772 | 0.09906209 | 0.23602605 |
| FMO5 | 1322.24713 | 1.915616687 | 0.45170956 | 4.24081506 | 2.23E-05 | 0.00038737 |
| CHD1L | 1771.41166 | -0.222455533 | 0.15570703 | -1.4286801 | 0.15309621 | 0.318588 |
| LINC00624 | 70.4576926 | 0.796575238 | 0.23591412 | 3.37654753 | 0.00073402 | 0.00613698 |
| BCL9 | 1789.25297 | 0.060949906 | 0.19959085 | 0.30537426 | 0.76008113 | 0.86700095 |
| ACP6 | 1934.48177 | 0.073450933 | 0.20565775 | 0.3571513 | 0.72097854 | 0.84122492 |
| AC241644.1 | 0.18311127 | -1.003574173 | 3.30558948 | -0.3035992 | 0.7614333 | NA |
| GJA5 | 1669.31741 | 0.397817178 | 0.51569308 | 0.77142238 | 0.44045662 | 0.63178083 |
| AC241644.3 | 686.803208 | 0.441028157 | 0.5103297 | 0.86420241 | 0.38747671 | 0.5830724 |
| AC241644.2 | 11.5213835 | -0.477367118 | 0.54144096 | -0.8816605 | 0.37796042 | 0.57390102 |
| GJA8 | 0.47835553 | 1.489856711 | 1.58105509 | 0.94231803 | 0.34602984 | NA |
| GPR89B | 635.42828 | 1.226397197 | 0.20032086 | 6.12216434 | 9.23E-10 | 9.67E-08 |
| AC239803.2 | 20.6940094 | 0.428546642 | 0.44789863 | 0.95679382 | 0.33867133 | 0.53670451 |
| AC239803.3 | 31.5399787 | 0.278995077 | 0.36167668 | 0.7713936 | 0.44047367 | 0.63178083 |
| NBPF11 | 629.757411 | 0.176208139 | 0.12621835 | 1.39605796 | 0.16269704 | 0.33170875 |
| AC239809.3 | 17.5264341 | -0.095315856 | 0.39720927 | -0.2399638 | 0.8103583 | 0.89747017 |
| AC239803.1 | 3.14533154 | -4.019187947 | 1.72131648 | -2.33495 | 0.01954603 | 0.0736748 |
| LINC01731 | 1.7999555 | -0.481378899 | 1.0755296 | -0.4475738 | 0.65446079 | 0.79728574 |
| AC245100.6 | 3.39055312 | -2.18208008 | 0.92084039 | -2.3696616 | 0.01780437 | 0.06880149 |
| AC245100.1 | 12.7924461 | -0.652413152 | 0.84707749 | -0.770193 | 0.44118542 | 0.63242332 |
| AC245100.7 | 11.4487039 | -0.594375046 | 0.61813063 | -0.9615687 | 0.33626632 | 0.53433132 |
| PPIAL4G | 0.86449779 | -0.221578395 | 1.37488881 | -0.161161 | 0.87196663 | NA |
| NBPF14 | 602.062752 | -0.375074027 | 0.18235962 | -2.0567822 | 0.03970718 | 0.12311327 |
| NUDT4B | 0 | NA | NA | NA | NA | NA |
| PDE4DIP | 3809.24895 | 0.306684947 | 0.22650488 | 1.35398827 | 0.17574012 | 0.35002016 |
| AC239802.1 | 7.3903839 | 1.755502686 | 0.43972192 | 3.99230199 | 6.54E-05 | 0.00091065 |
| AC239802.2 | 0.53371791 | 1.53570896 | 1.77425743 | 0.86555025 | 0.38673685 | NA |
| AC239804.1 | 1.10568154 | 0.461878333 | 1.20234945 | 0.3841465 | 0.70086987 | 0.82856354 |
| NBPF9 | 770.477207 | -0.451070878 | 0.24970995 | -1.8063793 | 0.07085912 | 0.18594437 |
| AC245297.3 | 252.723413 | -0.853985975 | 0.34581981 | -2.4694536 | 0.01353196 | 0.05641099 |
| AC245297.2 | 7.62228511 | -1.26266527 | 0.72359303 | -1.7449937 | 0.08098597 | 0.20418778 |
| NBPF19 | 672.68289 | -0.805067727 | 0.51485216 | -1.5636872 | 0.11789105 | 0.26659695 |
| PPIAL4C | 0.48321246 | 1.448695286 | 2.16510349 | 0.66911134 | 0.50342446 | NA |
| FCGR1A | 213.200735 | -1.315864059 | 0.59996103 | -2.1932492 | 0.02828943 | 0.09685695 |
| HIST2H2BF | 173.175368 | 0.711212853 | 0.56256394 | 1.2642347 | 0.2061458 | 0.38879021 |
| AC243772.2 | 12.5964237 | -0.68828195 | 0.64078883 | -1.0741166 | 0.28277041 | 0.47875729 |
| HIST2H3D | 3.1714301 | 0.065471392 | 0.9192691 | 0.07122114 | 0.94322176 | 0.97315146 |
| HIST2H4A | 11.5569516 | -0.683713135 | 0.63910529 | -1.0697973 | 0.28471054 | 0.4808388 |
| HIST2H3C | 0 | NA | NA | NA | NA | NA |
| HIST2H2AA3 | 1.76351434 | -1.077733807 | 1.20914625 | -0.891318 | 0.3727586 | 0.56894393 |
| AC239868.3 | 169.550769 | -0.365246654 | 0.34659366 | -1.0538181 | 0.29196622 | 0.48818396 |
| HIST2H2AA4 | 4.45709411 | -2.383941209 | 1.01950488 | -2.3383323 | 0.01937002 | 0.07321016 |
| HIST2H3A | 0 | NA | NA | NA | NA | NA |
| HIST2H4B | 0.71423544 | -1.504930455 | 2.19236969 | -0.6864401 | 0.49243563 | NA |
| AC239868.2 | 0.07659028 | -0.642905668 | 3.30558948 | -0.1944905 | 0.84579185 | NA |
| HIST2H2BE | 1356.60871 | 0.421879287 | 0.3709895 | 1.13717313 | 0.25546592 | 0.44814219 |
| HIST2H2AC | 21.1311743 | -1.336612059 | 0.45183095 | -2.9582127 | 0.00309429 | 0.01880099 |
| HIST2H2AB | 1.22696043 | -1.683671782 | 1.40328977 | -1.1998034 | 0.23021572 | 0.41778188 |
| BOLA1 | 706.016802 | 0.272992841 | 0.14870297 | 1.83582637 | 0.06638334 | 0.17739448 |
| SV2A | 285.738447 | -0.210889974 | 0.43609612 | -0.483586 | 0.62867972 | 0.77927443 |
| SF3B4 | 3293.83662 | -0.473739328 | 0.23174555 | -2.0442219 | 0.04093163 | 0.12606506 |
| MTMR11 | 2389.1092 | 0.449439748 | 0.31864032 | 1.41049238 | 0.15839434 | 0.32565561 |
| OTUD7B | 1622.89713 | -0.532316532 | 0.27219677 | -1.9556313 | 0.0505086 | 0.14670248 |
| VPS45 | 2017.95895 | -0.008226166 | 0.18106939 | -0.045431 | 0.96376376 | 0.98305633 |
| PLEKHO1 | 1755.72319 | -0.865817513 | 0.35758604 | -2.4212844 | 0.01546577 | 0.06223442 |
| ANP32E | 2635.50678 | -0.047859245 | 0.19780571 | -0.2419508 | 0.8088183 | 0.89660916 |
| AC242988.1 | 0.22147791 | 1.745651581 | 3.03531355 | 0.57511409 | 0.56521414 | NA |
| CA14 | 49.3475295 | 0.05824344 | 0.43173733 | 0.1349048 | 0.89268714 | 0.94447761 |
| APH1A | 9043.93954 | -0.134672326 | 0.08588571 | -1.568041 | 0.11687155 | 0.26520955 |
| C1orf54 | 1575.86948 | 0.026922538 | 0.09556501 | 0.28171961 | 0.77815852 | 0.87857682 |
| CIART | 502.055689 | -1.344746339 | 0.41805345 | -3.2166852 | 0.00129681 | 0.00964922 |
| MRPS21 | 3264.66882 | -0.423008527 | 0.10134915 | -4.1737749 | 3.00E-05 | 0.00048933 |
| PRPF3 | 2722.39829 | -0.24297699 | 0.18193997 | -1.3354789 | 0.18171965 | 0.35814742 |
| RPRD2 | 2636.19891 | 0.392300185 | 0.12648645 | 3.10151956 | 0.0019253 | 0.0130851 |
| TARS2 | 1701.02584 | 0.582786626 | 0.19025722 | 3.06315111 | 0.00219019 | 0.01446871 |

| | | | | | | |
|---|---|---|---|---|---|---|
| ECM1 | 1180.97421 | 0.01485724 | 0.30007858 | 0.04951117 | 0.96051194 | 0.98098634 |
| FALEC | 5.03626377 | 0.84703135 | 0.86788827 | 0.9759682 | 0.32908022 | 0.52689812 |
| AL356356.1 | 241.377728 | -1.206608739 | 0.42939083 | -2.8100477 | 0.00495342 | 0.02669014 |
| ADAMTSL4 | 1996.47629 | -1.577474477 | 0.44288139 | -3.5618441 | 0.00036826 | 0.00357548 |
| MCL1 | 17066.3432 | -0.636751239 | 0.24698117 | -2.5781367 | 0.00993347 | 0.0448436 |
| ENSA | 7174.94969 | -0.154454835 | 0.12291878 | -1.2565601 | 0.20891299 | 0.39215799 |
| GOLPH3L | 1599.07125 | -0.03407496 | 0.14792489 | -0.2303531 | 0.81781738 | 0.90211011 |
| HORMAD1 | 2.29615251 | -0.049524979 | 1.32539691 | -0.0373661 | 0.97019306 | 0.98622863 |
| CTSS | 1803.34303 | -0.439523837 | 0.4030457 | -1.0905062 | 0.27549022 | 0.4708151 |
| CTSK | 812.994618 | -0.543748003 | 0.28760636 | -1.890598 | 0.05867803 | 0.16287786 |
| ARNT | 2463.33714 | 0.058084362 | 0.13211071 | 0.43966429 | 0.66018027 | 0.8000806 |
| CTXND2 | 0.2926029 | 1.145957498 | 3.27645355 | 0.34975545 | 0.72652223 | NA |
| SETDB1 | 2184.86983 | 0.073434168 | 0.1058546 | 0.69372676 | 0.48785358 | 0.67152554 |
| CERS2 | 21436.7689 | -0.030724481 | 0.20532261 | -0.14964 | 0.88104863 | 0.93836253 |
| AL590133.2 | 12585.829 | -0.194868595 | 0.21374962 | -0.9116676 | 0.36194375 | 0.55929686 |
| AL590133.1 | 10.3980839 | 1.298478255 | 0.4982909 | 2.60586387 | 0.00916429 | 0.04220553 |
| ANXA9 | 1623.2706 | 0.811829246 | 0.18688465 | 4.34401248 | 1.40E-05 | 0.00026792 |
| MINDY1 | 2738.31713 | 0.186468838 | 0.25616007 | 0.72793875 | 0.46665108 | 0.65385702 |
| PRUNE1 | 1826.4936 | 0.042203874 | 0.20350612 | 0.20738381 | 0.83571013 | 0.91280681 |
| BNIPL | 258.709293 | 0.083182371 | 0.51898458 | 0.16027908 | 0.87266124 | 0.93366644 |
| C1orf56 | 2415.36396 | -0.683591998 | 0.152966 | -4.4689146 | 7.86E-06 | 0.0001688 |
| CDC42SE1 | 6802.78909 | -0.581075863 | 0.2039668 | -2.8488748 | 0.00438741 | 0.02444456 |
| MLLT11 | 622.441597 | -0.240230288 | 0.25323475 | -0.9486466 | 0.34280037 | 0.54048703 |
| GABPB2 | 1188.31664 | 0.422768561 | 0.2699488 | 1.56610644 | 0.1173237 | 0.26579261 |
| SEMA6C | 1697.96847 | 1.343671166 | 0.28195024 | 4.7656322 | 1.88E-06 | 5.42E-05 |
| TNFAIP8L2 | 105.900079 | -0.35153022 | 0.44035374 | -0.7982905 | 0.42470192 | 0.61708025 |
| SCNM1 | 1080.59452 | -0.543628139 | 0.15774599 | -3.4462248 | 0.00056848 | 0.00501698 |
| LYSMD1 | 597.379912 | -0.062728706 | 0.19344452 | -0.3242723 | 0.74573187 | 0.85745833 |
| TMOD4 | 59.9542016 | 0.415845369 | 0.2812113 | 1.4787648 | 0.13920319 | 0.29877813 |
| VPS72 | 1630.85706 | -0.253151954 | 0.10492071 | -2.412793 | 0.01583081 | 0.06330883 |
| PIP5K1A | 3413.73615 | -0.269264557 | 0.12481455 | -2.157317 | 0.03098097 | 0.1032438 |
| PSMD4 | 5276.5637 | -0.273067603 | 0.11656313 | -2.3426585 | 0.0191469 | 0.07257773 |
| AL391069.2 | 44.6057473 | 0.030413302 | 0.21056639 | 0.14443569 | 0.88515644 | 0.94054782 |
| ZNF687 | 2306.66568 | 0.063900528 | 0.1529517 | 0.4177824 | 0.67610622 | 0.81149917 |
| PI4KB | 5153.89862 | 0.046126048 | 0.11840277 | 0.389569 | 0.69685528 | 0.82577961 |
| AL391069.4 | 3.61410727 | 2.027711919 | 1.04691305 | 1.93684846 | 0.05276387 | 0.15128664 |
| RFX5 | 2272.26508 | 0.096903323 | 0.18780271 | 0.5159847 | 0.6058651 | 0.76260919 |
| AL391069.1 | 416.592111 | -0.287695052 | 0.16631997 | -1.7297686 | 0.08367163 | 0.20907997 |
| AL391069.3 | 20.2812921 | -0.211519232 | 0.3730163 | -0.5670509 | 0.5706796 | 0.73576038 |
| SELENBP1 | 6461.96639 | 1.55320205 | 0.35767884 | 4.34244883 | 1.41E-05 | 0.00026925 |
| PSMB4 | 10137.9681 | -0.409797726 | 0.16760633 | -2.4450015 | 0.01448515 | 0.0593403 |
| POGZ | 4501.19223 | 0.244679972 | 0.19777306 | 1.23717546 | 0.21602195 | 0.4006822 |
| CGN | 4227.77124 | 0.158091316 | 0.28779329 | 0.54932244 | 0.58278419 | 0.74511607 |
| TUFT1 | 1632.5722 | -0.245495949 | 0.21691533 | -1.1317594 | 0.25773562 | 0.45029234 |
| SNX27 | 3520.17431 | 0.281022448 | 0.18471713 | 1.52136651 | 0.12816789 | 0.28228407 |
| AL391335.1 | 9.68890991 | 0.328507004 | 0.48564817 | 0.67643002 | 0.49876766 | 0.68013772 |
| AL589765.1 | 5.7257839 | -0.549488818 | 0.6727529 | -0.8167766 | 0.41405611 | 0.60790934 |
| CELF3 | 28.5845555 | 2.086219774 | 0.51744992 | 4.03173276 | 5.54E-05 | 0.00080401 |
| RIIAD1 | 10.8689431 | -0.798232957 | 0.50855459 | -1.5696112 | 0.1165056 | 0.26469648 |
| AL589765.7 | 577.815796 | -0.684284061 | 0.16721534 | -4.0922325 | 4.27E-05 | 0.00065426 |
| MRPL9 | 2100.21616 | -0.429404105 | 0.13132663 | -3.2697413 | 0.00107646 | 0.00830735 |
| OAZ3 | 682.038334 | -0.346316952 | 0.15747209 | -2.1992275 | 0.02786175 | 0.09577818 |
| AL589765.4 | 275.085396 | -0.432763027 | 0.17388223 | -2.4888284 | 0.01281648 | 0.05416578 |
| AL589765.5 | 17.0527847 | -0.576835825 | 0.40121148 | -1.4377351 | 0.15050923 | 0.31514439 |
| TDRKH | 479.06374 | -0.581358809 | 0.17894144 | -3.2488775 | 0.00115861 | 0.00880987 |
| TDRKH-AS1 | 60.9161615 | -1.073197469 | 0.22010171 | -4.8759161 | 1.08E-06 | 3.48E-05 |
| AL589765.6 | 19.9688339 | -0.558924222 | 0.42888871 | -1.3031917 | 0.19250931 | 0.37187535 |
| LINGO4 | 75.3909837 | -0.019730492 | 0.62278454 | -0.0316811 | 0.97472638 | 0.98843899 |
| RORC | 3625.64853 | 0.758296856 | 0.39545537 | 1.91752827 | 0.05517085 | 0.15605013 |
| C2CD4D | 36.2368962 | -0.902081367 | 0.25275291 | -3.5690246 | 0.00035831 | 0.00349693 |
| AL450992.2 | 93.4866777 | -0.29010335 | 0.27428749 | -1.0576616 | 0.29020973 | 0.4862954 |
| THEM5 | 130.704598 | -0.258318352 | 0.23332214 | -1.1071318 | 0.26823696 | 0.46243518 |
| THEM4 | 1919.72699 | -0.01231555 | 0.24778408 | -0.0497027 | 0.96035927 | 0.98093621 |
| S100A10 | 11222.3942 | -1.554409441 | 0.34380352 | -4.5212145 | 6.15E-06 | 0.00013929 |
| AL450992.1 | 3.00471046 | -0.110927908 | 0.70309607 | -0.1577706 | 0.87463754 | 0.93482598 |
| S100A11 | 15201.7842 | -2.079722218 | 0.32482259 | -6.4026404 | 1.53E-10 | 2.05E-08 |
| TCHHL1 | 0 | NA | NA | NA | NA | NA |
| TCHH | 8.43698007 | -0.73747918 | 0.63824346 | -1.1554826 | 0.24789287 | 0.43948736 |
| AL589986.1 | 0 | NA | NA | NA | NA | NA |
| RPTN | 0.08695548 | -0.642905668 | 3.30558948 | -0.1944905 | 0.84579185 | NA |
| FLG-AS1 | 327.863545 | 0.802336813 | 0.46421852 | 1.72836019 | 0.08392367 | 0.20953107 |
| AL589986.2 | 2.19269758 | 0.777226455 | 0.84133296 | 0.92380364 | 0.35558856 | 0.55305686 |
| HRNR | 41.3175807 | -0.71970584 | 0.52234657 | -1.377832 | 0.16825515 | 0.33960109 |
| FLG | 48.4535467 | 1.850813 | 0.46305663 | 3.9969474 | 6.42E-05 | 0.00089682 |
| FLG2 | 2.25260249 | 1.625729344 | 0.76176008 | 2.13417503 | 0.03282846 | 0.10756541 |
| CRNN | 0.61232958 | 1.954218185 | 2.0496316 | 0.9534485 | 0.34036288 | NA |
| LCE5A | 3.40299956 | 0.29344205 | 1.02750532 | 0.28558689 | 0.77519454 | 0.87661907 |
| CRCT1 | 0 | NA | NA | NA | NA | NA |
| LCE3E | 0 | NA | NA | NA | NA | NA |
| LCE3D | 0.27046685 | -1.364244813 | 2.94293477 | -0.4635661 | 0.64295864 | NA |
| LCE3C | 0 | NA | NA | NA | NA | NA |
| LCE3B | 0 | NA | NA | NA | NA | NA |
| LCE3A | 0 | NA | NA | NA | NA | NA |
| LINC00302 | 0.17361919 | -1.003574173 | 3.30558948 | -0.3035992 | 0.7614333 | NA |
| LCE2D | 2.80053208 | 3.721147537 | 1.42852298 | 2.60489162 | 0.00919033 | 0.04229478 |
| LCE2C | 0.36028579 | -1.693887584 | 2.81438567 | -0.6018676 | 0.54726226 | NA |
| LCE2B | 0 | NA | NA | NA | NA | NA |
| LCE2A | 0 | NA | NA | NA | NA | NA |
| LCE4A | 0 | NA | NA | NA | NA | NA |
| C1orf68 | 0.07830766 | -0.642905668 | 3.30558948 | -0.1944905 | 0.84579185 | NA |
| KPRP | 0.40557068 | -1.788706826 | 3.29999086 | -0.5420339 | 0.58779517 | NA |
| LCE1F | 0 | NA | NA | NA | NA | NA |
| LCE1E | 0.08666371 | -0.642905668 | 3.30558948 | -0.1944905 | 0.84579185 | NA |
| LCE1D | 0 | NA | NA | NA | NA | NA |
| LCE1C | 15.2022464 | -1.050058127 | 1.31727998 | -0.7971412 | 0.42536904 | 0.61770921 |
| LCE1B | 0.09989236 | -0.642905668 | 3.30558948 | -0.1944905 | 0.84579185 | NA |
| LCE1A | 0 | NA | NA | NA | NA | NA |
| LCE6A | 0 | NA | NA | NA | NA | NA |
| SMCP | 0 | NA | NA | NA | NA | NA |
| IVL | 4.08221365 | -2.440183024 | 2.00594556 | -1.2164752 | 0.22380395 | 0.40990863 |
| LINC01527 | 0 | NA | NA | NA | NA | NA |
| SPRR5 | 0 | NA | NA | NA | NA | NA |
| SPRR4 | 0 | NA | NA | NA | NA | NA |
| SPRR1A | 0.65569856 | -0.681194998 | 3.25600078 | -0.2092122 | 0.83428261 | NA |
| SPRR3 | 0.08695548 | -0.642905668 | 3.30558948 | -0.1944905 | 0.84579185 | NA |
| SPRR1B | 0.8231141 | -2.931945933 | 3.26085739 | -0.8991334 | 0.36858159 | NA |
| SPRR2D | 2.27724128 | -2.000140498 | 1.75198379 | -1.1416433 | 0.25360233 | 0.4462494 |
| SPRR2A | 6.87476765 | -3.070218233 | 1.02764277 | -2.9876318 | 0.00281148 | 0.01746084 |
| SPRR2B | 0.12088456 | 0.02007176 | 3.29738704 | 0.00608717 | 0.99514317 | NA |
| SPRR2E | 0 | NA | NA | NA | NA | NA |
| SPRR2F | 0.24118557 | 0.074529887 | 3.28898342 | 0.02266046 | 0.98192111 | NA |
| SPRR2G | 0.10721287 | -0.642905668 | 3.30558948 | -0.1944905 | 0.84579185 | NA |
| AL161636.1 | 0 | NA | NA | NA | NA | NA |

| | | | | | | |
|---|---|---|---|---|---|---|
| LELP1 | 0 | NA | NA | NA | NA | NA |
| PRR9 | 0 | NA | NA | NA | NA | NA |
| LOR | 0.43543444 | 0.363068068 | 2.82007149 | 0.12874428 | 0.89756 | NA |
| PGLYRP3 | 0.25722115 | 2.020122844 | 3.24078384 | 0.6233439 | 0.53305855 | NA |
| PGLYRP4 | 0.87127105 | 0.447489426 | 1.79707813 | 0.24900944 | 0.80335348 | NA |
| S100A9 | 3121.26889 | -2.840665628 | 0.59491719 | -4.7748925 | 1.80E-06 | 5.22E-05 |
| S100A12 | 286.460124 | -3.866594 | 0.85022993 | -4.5477039 | 5.42E-06 | 0.00012501 |
| S100A8 | 1445.59448 | -3.185304705 | 0.61073776 | -5.2155031 | 1.83E-07 | 7.95E-06 |
| S100A7A | 0.09989236 | -0.642905668 | 3.30558948 | -0.1944905 | 0.84579185 | NA |
| S100A7L2 | 0 | NA | NA | NA | NA | NA |
| S100A7 | 0.66950706 | -2.623931078 | 2.68976122 | -0.9755257 | 0.32929958 | NA |
| BX470102.1 | 345.541359 | 0.548100397 | 0.38827099 | 1.41164395 | 0.15805482 | 0.32516038 |
| S100A6 | 28819.5249 | -1.107013789 | 0.39997638 | -2.7676979 | 0.00564538 | 0.02951091 |
| S100A5 | 27.5467263 | 0.597061198 | 0.63058533 | 0.94683649 | 0.3437221 | 0.54151789 |
| S100A4 | 2168.25617 | -0.081939216 | 0.51543545 | -0.1589709 | 0.87369183 | 0.93417196 |
| S100A3 | 225.100065 | -2.153931938 | 0.56198821 | -3.8326995 | 0.00012674 | 0.00152475 |
| S100A2 | 3925.65965 | -1.375314782 | 0.67793542 | -2.0286811 | 0.04249078 | 0.12946065 |
| S100A16 | 6529.51 | -1.703734747 | 0.35082497 | -4.8563668 | 1.20E-06 | 3.76E-05 |
| S100A14 | 1314.19474 | -1.839571277 | 0.35554321 | -5.1739739 | 2.29E-07 | 9.55E-06 |
| S100A13 | 8139.20172 | -1.275623462 | 0.26950485 | -4.7332116 | 2.21E-06 | 6.15E-05 |
| S100A1 | 5072.14404 | -1.29767225 | 0.35988779 | -3.6057691 | 0.00031123 | 0.00312053 |
| AL162258.2 | 4278.01696 | -1.087116825 | 0.3229278 | -3.3664392 | 0.00076145 | 0.00631233 |
| CHTOP | 4663.40406 | -0.128025578 | 0.15573365 | -0.8220804 | 0.41103115 | 0.60533693 |
| SNAPIN | 1505.53694 | -0.193238518 | 0.1502456 | -1.2861509 | 0.1983904 | 0.37934558 |
| ILF2 | 8131.8652 | -0.364756946 | 0.2052387 | -1.7772328 | 0.07552995 | 0.19467546 |
| NPR1 | 5760.16855 | -0.927349598 | 0.30602144 | -3.0303419 | 0.00244277 | 0.01570735 |
| INTS3 | 7583.18578 | -0.301498789 | 0.15473483 | -1.9484869 | 0.05135673 | 0.14852786 |
| AL513523.4 | 164.598127 | 1.090458615 | 0.30635764 | 3.55943018 | 0.00037166 | 0.00360187 |
| AL513523.3 | 115.167371 | 0.892589452 | 0.34757996 | 2.56801186 | 0.01022837 | 0.0458399 |
| SLC27A3 | 2328.03752 | -0.874798116 | 0.17135106 | -5.1052972 | 3.30E-07 | 1.29E-05 |
| GATAD2B | 2289.05005 | -0.269235363 | 0.12792812 | -2.1045831 | 0.03532762 | 0.11344185 |
| AL358472.5 | 173.981722 | 0.437473033 | 0.19367946 | 2.25874765 | 0.02389909 | 0.08539552 |
| DENND4B | 3519.83735 | -0.133349122 | 0.13185745 | -1.0113128 | 0.31186676 | 0.50890634 |
| CRTC2 | 4414.96474 | -0.725795859 | 0.1820819 | -3.9860955 | 6.72E-05 | 0.00092987 |
| SLC39A1 | 11230.8099 | -1.130227229 | 0.27681923 | -4.0829072 | 4.45E-05 | 0.00067559 |
| AL358472.4 | 37.6299801 | -0.866315122 | 0.28361203 | -3.0545782 | 0.00225377 | 0.01480249 |
| AL358472.3 | 34.5857386 | -1.301109406 | 0.38213098 | -3.4048781 | 0.00066194 | 0.00563618 |
| CREB3L4 | 728.634228 | -0.061892958 | 0.2247657 | -0.2753666 | 0.78303463 | 0.88125233 |
| JTB | 5412.42966 | -0.359080646 | 0.15681872 | -2.2897817 | 0.02203397 | 0.08057966 |
| AL358472.2 | 120.435742 | -0.139792498 | 0.26498125 | -0.5275562 | 0.5978074 | 0.75633738 |
| RAB13 | 6356.75416 | -0.859882154 | 0.17836572 | -4.8208936 | 1.43E-06 | 4.35E-05 |
| RPS27 | 25610.137 | -0.442236792 | 0.14545772 | -3.0403115 | 0.00236334 | 0.015359 |
| NUP210L | 23.4306994 | -1.111581863 | 0.75386106 | -1.4745182 | 0.14034212 | 0.30048893 |
| TPM3 | 11175.6623 | -0.680109693 | 0.21228965 | -3.2036875 | 0.0013568 | 0.00999965 |
| C1orf189 | 5.04477895 | -0.642278873 | 1.28152578 | -0.501183 | 0.61624237 | 0.77049015 |
| C1orf43 | 10328.1814 | 0.110127245 | 0.17886139 | 0.61571279 | 0.53808409 | 0.71154757 |
| UBAP2L | 6281.41914 | -0.205007939 | 0.16484252 | -1.2436594 | 0.21362495 | 0.39757852 |
| HAX1 | 3141.97076 | 0.12624656 | 0.15132603 | 0.83426863 | 0.40412962 | 0.59897135 |
| AQP10 | 2.48645965 | -0.387154159 | 0.85750798 | -0.4514875 | 0.65163821 | 0.79524711 |
| ATP8B2 | 1227.68497 | -0.103539102 | 0.32697704 | -0.3166556 | 0.75150496 | 0.86135902 |
| IL6R-AS1 | 14.6043497 | 0.70385115 | 0.58367408 | 1.20589755 | 0.22785699 | 0.41487185 |
| IL6R | 2961.41263 | -1.106316115 | 0.49971944 | -2.2138745 | 0.02683742 | 0.09302085 |
| SHE | 905.019456 | -0.041148549 | 0.33889733 | -0.1214189 | 0.90335923 | 0.95141786 |
| AL162591.2 | 223.776452 | 0.089447095 | 0.36685415 | 0.24382195 | 0.80736872 | 0.89590613 |
| TDRD10 | 197.650247 | -0.443840106 | 0.29645916 | -1.4971375 | 0.1343575 | 0.29158153 |
| UBE2Q1 | 4408.13058 | -0.299514303 | 0.19295838 | -1.5522223 | 0.12060903 | 0.27086188 |
| UBE2Q1-AS1 | 579.389714 | -0.066305795 | 0.21346667 | -0.3106143 | 0.75609387 | 0.86466002 |
| CHRNB2 | 19.688744 | 0.164528684 | 0.47364071 | 0.34737023 | 0.72831319 | 0.84636078 |
| AL592078.1 | 6.22338486 | -0.371009366 | 0.91444739 | -0.4057198 | 0.68494853 | 0.81791843 |
| ADAR | 10366.1682 | -0.233965245 | 0.29413429 | -0.7954368 | 0.42635944 | 0.61867047 |
| KCNN3 | 325.98634 | 1.324512446 | 0.30886603 | 4.28830728 | 1.80E-05 | 0.0003272 |
| PMVK | 2726.48154 | 0.505957484 | 0.28529903 | 1.77342868 | 0.0761577 | 0.19585319 |
| AL451085.1 | 6.12793165 | 1.733180552 | 0.60780391 | 2.85154557 | 0.00435072 | 0.02428633 |
| PBXIP1 | 9166.57923 | 0.152120183 | 0.23531535 | 0.64645244 | 0.51798639 | 0.69495299 |
| PYGO2 | 4204.45605 | -0.424380275 | 0.13306008 | -3.1893884 | 0.00142574 | 0.01040598 |
| AL451085.2 | 1683.42541 | -1.506982881 | 0.2488118 | -6.0567179 | 1.39E-09 | 1.35E-07 |
| SHC1 | 7649.37822 | -1.036523576 | 0.21943337 | -4.7236369 | 2.32E-06 | 6.39E-05 |
| CKS1B | 512.241932 | -0.37116275 | 0.16319189 | -2.2743945 | 0.02294228 | 0.08300011 |
| FLAD1 | 1778.638 | -0.226596073 | 0.12020199 | -1.8851275 | 0.05941263 | 0.1644323 |
| LENEP | 4.80297656 | 0.817343906 | 0.58517449 | 1.39675246 | 0.16248802 | 0.33138509 |
| ZBTB7B | 4258.87706 | -0.309132343 | 0.17655439 | -1.7509185 | 0.07995995 | 0.20235547 |
| DCST2 | 100.411446 | -0.069921964 | 0.3232623 | -0.216301 | 0.82875312 | 0.90888832 |
| DCST1 | 51.7560593 | -0.035751232 | 0.23351895 | -0.1530978 | 0.87832117 | 0.93677394 |
| DCST1-AS1 | 29.2061412 | -0.66885097 | 0.47005687 | -1.422915 | 0.15476079 | 0.32088146 |
| ADAM15 | 4672.72543 | -0.581334311 | 0.29586795 | -1.9648438 | 0.04943231 | 0.1444835 |
| EFNA4 | 393.498533 | -0.941929933 | 0.19787395 | -4.7602524 | 1.93E-06 | 5.52E-05 |
| AL691442.1 | 192.670513 | -1.49177597 | 0.31776427 | -4.6945995 | 2.67E-06 | 7.16E-05 |
| EFNA3 | 222.826851 | -2.070486804 | 0.41130793 | -5.0339092 | 4.81E-07 | 1.75E-05 |
| EFNA1 | 11379.4514 | -2.003654109 | 0.37044272 | -5.4088096 | 6.34E-08 | 3.23E-06 |
| SLC50A1 | 2742.48763 | -1.008596579 | 0.13676416 | -7.3747143 | 1.65E-13 | 5.87E-11 |
| DPM3 | 974.354179 | -0.434685149 | 0.15370438 | -2.8280597 | 0.00468311 | 0.02558921 |
| KRTCAP2 | 3073.1486 | -0.592393109 | 0.13562911 | -4.3677432 | 1.26E-05 | 0.00024647 |
| AL713999.1 | 8034.83165 | -1.232511608 | 0.30258312 | -4.0732993 | 4.64E-05 | 0.00069846 |
| TRIM46 | 133.647779 | -0.829358069 | 0.36023851 | -2.3022471 | 0.02132124 | 0.07861268 |
| MUC1 | 25315.636 | -1.567826703 | 0.42364292 | -3.7008212 | 0.0002149 | 0.00231495 |
| AC234582.1 | 1191.52266 | 0.091398759 | 0.26700602 | 0.34230973 | 0.73211782 | 0.84898243 |
| THBS3 | 2601.08811 | -0.084373937 | 0.27785005 | -0.3036672 | 0.76138147 | 0.8677512 |
| MTX1 | 1564.89562 | -0.039800171 | 0.08779503 | -0.4533306 | 0.65031072 | 0.79447272 |
| AC234582.2 | 184.627144 | -0.085973768 | 0.1359706 | -0.6322967 | 0.527193 | 0.70191253 |
| GBA | 3607.99317 | 0.112017921 | 0.18376099 | 0.60958489 | 0.54213682 | 0.71466072 |
| FAM189B | 1895.9458 | -0.56382888 | 0.21406681 | -2.6338921 | 0.00844123 | 0.0396359 |
| SCAMP3 | 3755.89496 | -0.315637323 | 0.15295834 | -2.0635509 | 0.03906032 | 0.12167992 |
| CLK2 | 4316.53678 | -0.20114262 | 0.15760315 | -1.2762602 | 0.20186356 | 0.38376051 |
| HCN3 | 1540.11409 | 0.527202667 | 0.30362249 | 1.73637552 | 0.08249746 | 0.20696965 |
| PKLR | 2797.79996 | 2.484926496 | 0.45432297 | 5.46951538 | 4.51E-08 | 2.44E-06 |
| FDPS | 5968.23618 | -0.238636497 | 0.20406463 | -1.1694163 | 0.24223595 | 0.43262866 |
| RUSC1-AS1 | 4758.64478 | -0.23285696 | 0.17182995 | -1.3551594 | 0.17536679 | 0.34951419 |
| RUSC1 | 3042.13553 | -0.411320279 | 0.13974817 | -2.9432962 | 0.00324738 | 0.01951533 |
| ASH1L | 2123.66524 | 0.319188159 | 0.10545408 | 3.02679759 | 0.00247159 | 0.01585728 |
| ASH1L-AS1 | 179.784615 | -0.250038674 | 0.30899539 | -0.8091987 | 0.41840084 | 0.61113582 |
| AL353807.2 | 72.0922651 | 0.477285749 | 0.34434467 | 1.38606979 | 0.1657256 | 0.33621972 |
| MSTO1 | 1434.28038 | 0.090325212 | 0.20260354 | 0.44582247 | 0.65572549 | 0.79807876 |
| YY1AP1 | 4181.86562 | -0.33777377 | 0.12551008 | -2.6912082 | 0.00711937 | 0.03506498 |
| DAP3 | 4355.35198 | -0.313840403 | 0.13088871 | -2.3977653 | 0.01649543 | 0.06522485 |
| AL162734.1 | 0.46234773 | -2.085786846 | 2.21991478 | -0.9395797 | 0.3474332 | NA |
| GON4L | 1792.65469 | 0.146452095 | 0.12940009 | 1.13177734 | 0.25772806 | 0.45029234 |
| SYT11 | 971.744977 | -0.12642597 | 0.27171543 | -0.4652882 | 0.64172512 | 0.78770616 |
| RIT1 | 2935.47501 | -0.715183779 | 0.22986327 | -3.1113443 | 0.00186238 | 0.01274614 |
| KHDC4 | 4082.12172 | -0.126291481 | 0.14432178 | -0.8750688 | 0.3815365 | 0.57756127 |
| RXFP4 | 4.26098861 | 1.592622997 | 0.69167641 | 2.30255503 | 0.02130389 | 0.07857114 |
| ARHGEF2 | 3098.10601 | -0.221116353 | 0.2385222 | -0.9270263 | 0.35391287 | 0.55148822 |
| AL355388.2 | 70.778766 | -0.571225402 | 0.50005028 | -1.1423359 | 0.25331442 | 0.44588738 |
| AL355388.1 | 0.63006139 | 2.022563065 | 2.1415485 | 0.94443953 | 0.34494509 | NA |

| | | | | | | |
|---|---|---|---|---|---|---|
| SSR2 | 8128.12379 | -0.24856873 | 0.15878171 | -1.5654745 | 0.11747168 | 0.26601409 |
| UBQLN4 | 3052.28276 | -0.375766032 | 0.12712446 | -2.9558908 | 0.00311767 | 0.01890825 |
| LAMTOR2 | 2186.5066 | -0.168528915 | 0.1227066 | -1.37343 | 0.16961872 | 0.34140975 |
| RAB25 | 1870.82344 | -0.636825249 | 0.27790185 | -2.2915473 | 0.02193178 | 0.08031398 |
| MEX3A | 223.922195 | -0.929702826 | 0.34785611 | -2.672665 | 0.00752514 | 0.03654473 |
| LMNA | 16780.3931 | -0.28600434 | 0.2998007 | -0.9539815 | 0.34009298 | 0.53820287 |
| SEMA4A | 2415.43762 | -0.448672665 | 0.43772196 | -1.0250175 | 0.30535494 | 0.50240696 |
| SLC25A44 | 3673.13839 | 0.653712402 | 0.23080304 | 2.83233884 | 0.00462089 | 0.02539846 |
| PMF1-BGLAF | 2167.10938 | -0.032182921 | 0.14246755 | -0.2258965 | 0.82128192 | 0.9041919 |
| PMF1 | 2950.46461 | -0.040800815 | 0.13999369 | -0.2914475 | 0.77070907 | 0.87338196 |
| BGLAP | 319.349255 | -0.46330529 | 0.20561458 | -2.2532706 | 0.02424209 | 0.0862588 |
| PAQR6 | 603.230873 | -0.905359768 | 0.37316677 | -2.4261533 | 0.01525982 | 0.06160465 |
| SMG5 | 9373.53591 | -0.466272732 | 0.22646435 | -2.0589233 | 0.03950158 | 0.1226633 |
| TMEM79 | 2494.87893 | -0.343493669 | 0.2552705 | -1.3456066 | 0.17842942 | 0.35385479 |
| GLMP | 5528.02424 | -0.216364673 | 0.29158603 | -0.7420269 | 0.45807104 | 0.64706542 |
| VHLL | 1.07658806 | -0.931243117 | 1.42056401 | -0.6555446 | 0.51211716 | 0.69027446 |
| CCT3 | 10017.7239 | -0.178961657 | 0.1522461 | -1.1754761 | 0.23980428 | 0.42971181 |
| TSACC | 25.8364559 | 0.344570979 | 0.34150116 | 1.00898919 | 0.31297982 | 0.50986215 |
| RHBG | 1725.99626 | 1.290535655 | 0.5031106 | 2.5651132 | 0.01031421 | 0.04610711 |
| AL139130.1 | 4.34190139 | 0.562713736 | 0.9137066 | 0.61585824 | 0.53798809 | 0.71150739 |
| C1orf61 | 4.37217937 | -0.904089224 | 1.05099323 | -0.8602236 | 0.38966577 | 0.5852977 |
| AL139412.1 | 3.77131094 | -0.04297861 | 0.83117001 | -0.0517086 | 0.95876091 | 0.98017606 |
| MEF2D | 4070.34303 | -0.088355455 | 0.18255234 | -0.4840007 | 0.6283854 | 0.77902222 |
| AL365181.1 | 0.11223674 | 0.02007176 | 3.29738704 | 0.00608717 | 0.99514317 | NA |
| IQGAP3 | 130.154419 | -0.989304145 | 0.43872014 | -2.2549777 | 0.02413473 | 0.08599281 |
| TTC24 | 13.8185394 | 0.3246564 | 0.7155394 | 0.4537226 | 0.65002849 | 0.79420152 |
| NAXE | 4231.00646 | -0.095615454 | 0.18318873 | -0.5219505 | 0.60170478 | 0.75903919 |
| GPATCH4 | 1411.78227 | -0.441008217 | 0.23685776 | -1.8619116 | 0.06261555 | 0.17038201 |
| AL365181.4 | 6.15705369 | -0.076355247 | 0.76015969 | -0.1004463 | 0.91998999 | 0.96091164 |
| HAPLN2 | 40.5847772 | -0.272282444 | 0.57538384 | -0.4732188 | 0.6360571 | 0.7841886 |
| AL365181.2 | 31.4235375 | -1.895504094 | 0.57570364 | -3.2924997 | 0.00099301 | 0.00779335 |
| BCAN | 59.2697119 | -0.635264513 | 0.48443653 | -1.3113472 | 0.18974047 | 0.36824723 |
| AL365181.3 | 68.4285036 | -1.675233605 | 0.52676652 | -3.1802204 | 0.00147163 | 0.01064661 |
| AL590666.2 | 16.2643321 | -0.864957011 | 0.69712085 | -1.2407562 | 0.21469583 | 0.39886776 |
| NES | 5603.00296 | 0.702246577 | 0.35108167 | 2.0002371 | 0.04547467 | 0.13588294 |
| AL590666.3 | 1.85218533 | -2.682551859 | 1.50243395 | -1.7854707 | 0.07418499 | 0.19228015 |
| CRABP2 | 338.268612 | -1.010141911 | 0.46142306 | -2.1891882 | 0.02858316 | 0.09749554 |
| AL590666.1 | 16.7277954 | -0.54030287 | 0.53160951 | -1.0163529 | 0.30946136 | 0.50667429 |
| ISG20L2 | 1492.81668 | -0.032173272 | 0.17850169 | -0.1802407 | 0.8569636 | 0.92514304 |
| RRNAD1 | 1966.26256 | 0.007942079 | 0.13441489 | 0.0590863 | 0.95288337 | 0.97723075 |
| MRPL24 | 2652.02256 | 0.160803137 | 0.14659213 | 1.09694247 | 0.27266654 | 0.46759554 |
| HDGF | 17660.2554 | -0.595586844 | 0.18963317 | -3.1407314 | 0.00168527 | 0.01181612 |
| PRCC | 2834.59459 | -0.354170122 | 0.2072666 | -1.708766 | 0.0874943 | 0.21593119 |
| SH2D2A | 182.287192 | -1.645156065 | 0.67053987 | -2.4534799 | 0.01414815 | 0.05837565 |
| NTRK1 | 236.385072 | 0.828963606 | 0.5664834 | 1.46335021 | 0.14337157 | 0.30496235 |
| INSRR | 461.287679 | 1.39272841 | 0.54312302 | 2.5642964 | 0.01033852 | 0.0462001 |
| PEAR1 | 1780.97979 | 0.458481641 | 0.30636516 | 1.49652016 | 0.13451816 | 0.29185814 |
| LRRC71 | 32.0525205 | -0.46545574 | 0.49369357 | -0.9428029 | 0.34578172 | 0.54340071 |
| ARHGEF11 | 2947.82843 | 0.023839075 | 0.14674736 | 0.16244977 | 0.87095168 | 0.93297298 |
| ETV3L | 0.27059475 | 1.100737003 | 2.99309129 | 0.36775925 | 0.71305276 | NA |
| ETV3 | 979.03799 | -0.22552506 | 0.18399939 | -1.2256838 | 0.22031771 | 0.40616456 |
| AL138900.1 | 0 | NA | NA | NA | NA | NA |
| AL138900.3 | 8.27557928 | 1.10850071 | 1.05101354 | 1.05469689 | 0.29156398 | 0.48775881 |
| AL138900.2 | 12.4781367 | -2.355902002 | 1.32944218 | -1.7720981 | 0.07637827 | 0.1961526 |
| FCRL5 | 154.137544 | -0.666705577 | 0.88157706 | -0.7562647 | NA | NA |
| FCRL4 | 0.68324079 | 0.265795203 | 2.25189138 | 0.11803198 | 0.90604232 | NA |
| FCRL3 | 58.4955888 | 0.07723514 | 0.84462126 | 0.09144352 | 0.92714018 | 0.96467356 |
| AL356276.1 | 3.53300485 | 0.369176651 | 0.87451997 | 0.42214776 | 0.67291717 | 0.80923138 |
| FCRL2 | 43.4301522 | 1.410827206 | 0.85892457 | 1.64255077 | NA | NA |
| FCRL1 | 43.8775557 | 2.191143983 | 1.02206806 | 2.14383374 | NA | NA |
| CD5L | 7.85646621 | -3.032440819 | 1.26296517 | -2.4010487 | 0.01634816 | 0.06482513 |
| KIRREL1 | 2474.83425 | -0.185946526 | 0.23051331 | -0.8066629 | 0.41986075 | 0.61261525 |
| LINC01704 | 3.24916978 | -3.431403133 | 1.83961844 | -1.8652798 | 0.06214219 | 0.16947871 |
| CD1D | 184.565788 | -0.779984861 | 0.41632173 | -1.8735146 | 0.06099735 | 0.16736033 |
| AL138899.1 | 0.09635527 | 0.547239394 | 3.29377202 | 0.16614368 | 0.86804388 | NA |
| CD1A | 2.26584307 | 1.596158146 | 1.32993837 | 1.20017452 | 0.23007157 | 0.41761051 |
| CD1C | 58.0434675 | 2.944487555 | 0.8412316 | 3.50021037 | NA | NA |
| CD1B | 2.65319031 | 2.737519765 | 1.51822495 | 1.8031055 | 0.07137165 | 0.18691731 |
| CD1E | 24.0691005 | 2.259573548 | 2.02812998 | 1.11411673 | 0.26522911 | 0.45930973 |
| OR10T2 | 0 | NA | NA | NA | NA | NA |
| OR10K2 | 0.09635527 | 0.547239394 | 3.29377202 | 0.16614368 | 0.86804388 | NA |
| OR10K1 | 0 | NA | NA | NA | NA | NA |
| OR10R2 | 0 | NA | NA | NA | NA | NA |
| AL365440.2 | 0 | NA | NA | NA | NA | NA |
| OR6Y1 | 0 | NA | NA | NA | NA | NA |
| OR6P1 | 0 | NA | NA | NA | NA | NA |
| OR10X1 | 0 | NA | NA | NA | NA | NA |
| OR10Z1 | 11.0050426 | -3.223014177 | 1.03297321 | -3.1201334 | 0.00180769 | 0.0124657 |
| SPTA1 | 24.1488774 | -3.197561307 | 0.88064831 | -3.6309174 | 0.00028242 | 0.0028944 |
| OR6K2 | 0 | NA | NA | NA | NA | NA |
| OR6K3 | 0.18009489 | 0.726716446 | 3.28420131 | 0.22127646 | 0.82487718 | NA |
| OR6N1 | 0.51220377 | 0.909221628 | 2.13029126 | 0.42680625 | 0.66952045 | NA |
| OR6K6 | 0 | NA | NA | NA | NA | NA |
| OR6N2 | 0 | NA | NA | NA | NA | NA |
| MNDA | 298.236742 | -0.155097388 | 0.40560111 | -0.382389 | 0.70217288 | 0.82947179 |
| PYHIN1 | 67.511464 | -0.301078132 | 0.70538382 | -0.4268288 | 0.66950402 | 0.80704064 |
| IFI16 | 2795.20036 | -1.328175982 | 0.35797784 | -3.7102185 | 0.00020708 | 0.00224259 |
| AIM2 | 43.4898735 | -0.85613314 | 0.75370897 | -1.1358935 | 0.25600113 | 0.4487337 |
| CADM3 | 392.101582 | -1.965147012 | 0.72564461 | -2.7081397 | 0.00676615 | 0.03368387 |
| CADM3-AS1 | 303.110146 | -1.730573301 | 0.76228482 | -2.270245 | 0.02319272 | 0.08367794 |
| ACKR1 | 2804.99706 | -2.495592814 | 0.86648573 | -2.8801315 | 0.00397509 | 0.02264614 |
| FCER1A | 115.792313 | 2.555474834 | 1.31645602 | 1.94117752 | 0.05223675 | 0.15020333 |
| OR10J3 | 0 | NA | NA | NA | NA | NA |
| AL513323.1 | 0.08666371 | -0.642905668 | 3.30558948 | -0.1944905 | 0.84579185 | NA |
| OR10J1 | 0.09989236 | -0.642905668 | 3.30558948 | -0.1944905 | 0.84579185 | NA |
| AL663023.1 | 0.08757577 | 0.547239394 | 3.29377202 | 0.16614368 | 0.86804388 | NA |
| OR10J5 | 0.06785815 | 0.795868262 | 3.28898349 | 0.24198001 | 0.80879565 | NA |
| APCS | 62.8326734 | -4.906416729 | 1.48793968 | -3.2974567 | NA | NA |
| CRP | 31.7519919 | -2.486748569 | 0.74917675 | -3.3193082 | 0.00090241 | 0.00724178 |
| DUSP23 | 7467.63985 | -2.057390051 | 0.42870383 | -4.7990941 | 1.59E-06 | 4.76E-05 |
| FCRL6 | 91.4212751 | -0.786948904 | 0.53386239 | -1.4740669 | 0.14046357 | 0.300627 |
| SLAMF8 | 657.141206 | -0.564270874 | 0.28715045 | -1.9650705 | 0.04940608 | 0.14444674 |
| AL590560.1 | 874.323355 | 0.022365916 | 0.23256851 | 0.09616915 | 0.92338623 | 0.96266683 |
| SNHG28 | 425.545424 | 0.198859071 | 0.22193217 | 0.89603535 | 0.37023388 | 0.56696095 |
| VSIG8 | 149.677447 | 1.047736185 | 0.53698302 | 1.9511533 | 0.05103881 | 0.14780293 |
| AL590560.2 | 381.311024 | 0.292606993 | 0.459765 | 0.63642729 | 0.52449796 | 0.69951978 |
| CFAP45 | 123.946731 | -0.685292688 | 0.35053502 | -1.9549906 | 0.05058418 | 0.14682495 |
| TAGLN2 | 17489.9834 | -1.138231064 | 0.33885286 | -3.3590717 | 0.00078205 | 0.00645869 |
| IGSF9 | 116.19943 | 1.298469088 | 0.53587289 | 2.42309156 | 0.01538905 | 0.06200293 |
| SLAMF9 | 2.82867149 | -2.105709584 | 1.17027491 | -1.799329 | 0.07196665 | 0.18808549 |
| LINC01133 | 3.17159584 | -0.125365191 | 1.46495924 | -0.0855759 | 0.93180357 | 0.96704554 |
| KCNJ10 | 2073.86561 | 2.401617845 | 0.44177792 | 5.43625596 | 5.44E-08 | 2.83E-06 |
| PIGM | 835.143585 | 0.098318263 | 0.11416378 | 0.8612036 | 0.38912592 | 0.58468663 |

| | | | | | | |
|---|---|---|---|---|---|---|
| AL121987.1 | 4.58928193 | 1.057038363 | 0.71982431 | 1.46846716 | 0.14197737 | 0.30285902 |
| KCNJ9 | 9.55551997 | 1.860909863 | 0.55735594 | 3.33881767 | 0.00084136 | 0.00684772 |
| IGSF8 | 4329.88444 | -0.194220802 | 0.16734666 | -1.1605897 | 0.24580881 | 0.43705606 |
| ATP1A2 | 292.321839 | 0.976175424 | 0.58080027 | 1.68074202 | 0.09281304 | 0.22549282 |
| ATP1A4 | 2.97027685 | 0.368876904 | 0.70897455 | 0.52029639 | 0.60285701 | 0.76012907 |
| CASQ1 | 24.4803439 | -0.056355048 | 0.74629419 | -0.0755132 | 0.93980641 | 0.97115066 |
| AL121987.2 | 532.779827 | -0.583111711 | 0.38864942 | -1.500354 | 0.13352273 | 0.29029553 |
| PEA15 | 9416.23427 | -0.87586306 | 0.37288492 | -2.3488831 | 0.01882982 | 0.07181694 |
| DCAF8 | 8159.43775 | 0.267392607 | 0.10666433 | 2.50686063 | 0.01218087 | 0.05211904 |
| AL139011.2 | 4969.77734 | 0.527913142 | 0.13166983 | 4.00937058 | 6.09E-05 | 0.00086082 |
| AL139011.1 | 300.257595 | 1.052247254 | 0.23768403 | 4.42708436 | 9.55E-06 | 0.00019624 |
| PEX19 | 3946.48999 | 0.31302887 | 0.14397396 | 2.17420472 | 0.02968976 | 0.10015595 |
| COPA | 7403.75774 | 0.127067108 | 0.15099522 | 0.84153067 | 0.40005071 | 0.59481541 |
| NCSTN | 6793.69029 | 0.132872809 | 0.16764698 | 0.79257505 | 0.42802545 | 0.62042202 |
| NHLH1 | 23.637762 | -0.67818785 | 0.48550179 | -1.3968802 | 0.16244959 | 0.33138509 |
| VANGL2 | 722.61942 | 0.006852695 | 0.39802601 | 0.0172167 | 0.98626374 | 0.99391317 |
| SLAMF6 | 131.536399 | -0.671852762 | 0.75422009 | -0.8907914 | 0.3730411 | 0.56921038 |
| AL138930.1 | 0.52394735 | -0.059717422 | 1.93215475 | -0.0309072 | 0.97534358 | NA |
| CD84 | 221.681959 | -0.209247354 | 0.47356662 | -0.4418541 | 0.65859478 | 0.79946078 |
| SLAMF1 | 166.81062 | -1.301058452 | 0.71451139 | -1.8209065 | 0.06862107 | 0.18156552 |
| AL121985.1 | 216.635902 | -0.050569336 | 0.6107396 | -0.0828002 | 0.93401044 | 0.96826888 |
| CD48 | 300.582425 | 0.045048131 | 0.6252715 | 0.07204571 | 0.94256553 | 0.97275716 |
| SLAMF7 | 181.098662 | 0.205461891 | 0.69017716 | 0.29769442 | 0.7659364 | 0.87093023 |
| LY9 | 58.9236527 | 0.374212895 | 0.85216328 | 0.43913286 | 0.66056527 | 0.80025296 |
| CD244 | 35.0312615 | 0.232877817 | 0.61031156 | 0.38157202 | 0.70277884 | 0.8298902 |
| ITLN1 | 127.988123 | 3.17241576 | 0.72688712 | 4.36438574 | 1.27E-05 | 0.00024846 |
| AL354714.3 | 0.91954489 | -0.60315123 | 1.66089879 | -0.3631475 | 0.71649471 | NA |
| AL354714.1 | 2.18324105 | 1.387033736 | 0.94439891 | 1.46869477 | 0.1419156 | 0.30275174 |
| ITLN2 | 5.71936503 | 1.990895195 | 0.85112517 | 2.33913327 | 0.01932854 | 0.0730953 |
| F11R | 10019.8356 | -0.500363039 | 0.16683536 | -2.9991426 | 0.00270741 | 0.01698247 |
| AL591806.3 | 5054.10706 | -0.120843198 | 0.1742511 | -0.6935004 | 0.4879956 | 0.67152554 |
| TSTD1 | 2276.72826 | 0.137167885 | 0.20673274 | 0.66350346 | 0.50700817 | 0.68603166 |
| USF1 | 3590.80281 | -0.490231872 | 0.16777211 | -2.9220105 | 0.0034778 | 0.02048391 |
| ARHGAP30 | 775.516806 | -0.026047925 | 0.45435112 | -0.0573299 | 0.95428237 | 0.97809492 |
| NECTIN4 | 604.1146 | -1.889397641 | 0.38744203 | -4.8765944 | 1.08E-06 | 3.47E-05 |
| AL591806.1 | 0.28660984 | 1.145957499 | 3.27645355 | 0.34975545 | 0.72652223 | NA |
| KLHDC9 | 660.898059 | 0.449502486 | 0.3685287 | 1.21972178 | 0.22257036 | 0.4085466 |
| PFDN2 | 1664.90864 | -0.814740645 | 0.1494416 | -5.4518998 | 4.98E-08 | 2.65E-06 |
| NIT1 | 4701.21744 | 0.163524819 | 0.15337744 | 1.06615953 | 0.28635153 | 0.48249776 |
| DEDD | 2570.01118 | -0.259220988 | 0.1280993 | -2.0235941 | 0.04301193 | 0.13064093 |
| UFC1 | 4189.56233 | -0.232512246 | 0.14214277 | -1.6357655 | 0.10188867 | 0.24057953 |
| AL590714.1 | 945.299389 | -0.107507022 | 0.18522526 | -0.5804123 | 0.56163661 | 0.72942161 |
| USP21 | 2101.63864 | 0.035105216 | 0.16488395 | 0.21290863 | 0.83139823 | 0.91040231 |
| PPOX | 1904.36171 | 0.074435338 | 0.20243065 | 0.36770785 | 0.71309108 | 0.83695061 |
| B4GALT3 | 2695.80217 | -0.43848214 | 0.12845105 | -3.4136128 | 0.00064108 | 0.00549583 |
| ADAMTS4 | 4869.65853 | -2.785714484 | 0.89277348 | -3.1202926 | 0.00180671 | 0.01246479 |
| NDUFS2 | 10885.6324 | 0.070323714 | 0.1297802 | 0.54186783 | 0.58790956 | 0.74877861 |
| FCER1G | 1210.1872 | -1.517669507 | 0.50375689 | -3.0127022 | 0.00258933 | 0.0164294 |
| APOA2 | 12.8800281 | 1.396131705 | 0.98608825 | 1.41582835 | 0.15682577 | 0.32372702 |
| TOMM40L | 1516.71275 | 0.255066328 | 0.12908577 | 1.97594464 | 0.04816105 | 0.14175765 |
| NR1I3 | 1094.36489 | 1.094933872 | 0.301527 | 3.63129625 | 0.000282 | 0.00289128 |
| PCP4L1 | 26.5132706 | -0.023190881 | 0.36470064 | -0.0635888 | 0.94929764 | 0.97552721 |
| MPZ | 302.23351 | -0.561415444 | 0.24277374 | -2.3125048 | 0.02074988 | 0.07696925 |
| SDHC | 8448.9444 | 0.592191718 | 0.24142953 | 2.45285541 | 0.01417273 | 0.0584326 |
| CFAP126 | 342.11888 | 0.806527573 | 0.37479941 | 2.15189126 | 0.03140592 | 0.10431718 |
| AL592295.4 | 33.3753652 | 0.220670609 | 0.29165962 | 0.75660324 | 0.4492876 | 0.63948368 |
| AL592295.3 | 2.05154287 | -0.260301959 | 0.8559089 | -0.3041234 | 0.76103385 | 0.8676003 |
| AL831711.1 | 38.3436374 | 0.023723952 | 0.27930629 | 0.08493884 | 0.93231001 | 0.96734562 |
| FCGR2A | 1363.66098 | -0.818747045 | 0.44969333 | -1.8206786 | 0.06865572 | 0.18160759 |
| HSPA6 | 412.622088 | -1.647700002 | 0.3696938 | -4.4569316 | 8.31E-06 | 0.00017655 |
| FCGR3A | 1490.04015 | -1.566776849 | 0.55565836 | -2.8196766 | 0.00480721 | 0.02609431 |
| AL590385.1 | 0 | NA | NA | NA | NA | NA |
| FCGR3B | 133.538352 | 0.332494292 | 0.68227153 | 0.48733426 | 0.62602147 | 0.77729476 |
| FCGR2B | 294.077849 | -0.196826774 | 0.59784907 | -0.3292249 | 0.74198574 | 0.85508976 |
| AL451067.1 | 55.7603387 | -0.077372852 | 0.67070369 | -0.1153607 | 0.90815922 | 0.95414792 |
| FCRLA | 42.6733552 | 0.283411284 | 1.15276487 | 0.24585351 | 0.80579564 | 0.89480475 |
| FCRLB | 227.269498 | 0.237275725 | 0.45908405 | 0.51684593 | 0.60526371 | 0.76217966 |
| DUSP12 | 750.714565 | -0.435345979 | 0.18229533 | -2.3881357 | 0.01693409 | 0.06649075 |
| AL359541.1 | 5.3771546 | 0.554102252 | 0.78725412 | 0.70384167 | 0.48153138 | 0.66600139 |
| ATF6 | 3373.06867 | -0.238709988 | 0.13470256 | -1.7721266 | 0.07637354 | 0.1961526 |
| OLFML2B | 506.280834 | -0.533484923 | 0.39469353 | -1.3516434 | 0.17648941 | 0.35127374 |
| AL590408.1 | 0.21442573 | -1.003574173 | 3.30558948 | -0.3035992 | 0.7614333 | NA |
| NOS1AP | 106.87577 | 0.459995728 | 0.40837227 | 1.12641273 | 0.25999086 | 0.45292965 |
| AL512785.1 | 6.54346573 | -1.990640021 | 0.77600511 | -2.5652409 | 0.01031042 | 0.04609796 |
| AL512785.2 | 119.04255 | 0.902728178 | 0.39223461 | 2.30150054 | 0.02136335 | 0.07872438 |
| C1orf226 | 1231.54816 | -0.269807588 | 0.23719741 | -1.1374812 | 0.25533718 | 0.44795699 |
| SPATA46 | 0.83652695 | -0.91538766 | 1.89327217 | -0.483495 | 0.6287443 | NA |
| SH2D1B | 29.487994 | -0.267641733 | 0.33792626 | -0.7920122 | 0.42835354 | 0.62074424 |
| UHMK1 | 4107.96336 | 0.082240604 | 0.12875592 | 0.63873261 | 0.5229969 | 0.69862624 |
| AL596325.1 | 0.81050239 | 0.629624174 | 1.85649185 | 0.33914729 | 0.73449878 | NA |
| UAP1 | 1824.13704 | -0.100358118 | 0.22296599 | -0.4501051 | 0.6526347 | 0.79590266 |
| DDR2 | 1111.06928 | 0.245422165 | 0.32638094 | 0.75195006 | 0.45208109 | 0.6417624 |
| HSD17B7 | 798.170247 | -0.630232698 | 0.18912839 | -3.3323008 | 0.00086131 | 0.00699221 |
| CCDC190 | 1.16753727 | 0.932155458 | 1.15477096 | 0.80722107 | 0.41953913 | 0.61243738 |
| RGS4 | 120.441211 | 0.52547456 | 0.43830414 | 1.19888113 | 0.23057417 | 0.4182309 |
| RGS5 | 9236.74697 | 0.264882682 | 0.58853359 | 0.45007232 | 0.65265829 | 0.79590266 |
| AL499616.1 | 1.44775732 | 0.239477711 | 1.0675573 | 0.22432305 | 0.82250594 | 0.9051968 |
| AL592435.1 | 1.61758304 | 1.350016253 | 1.14841141 | 1.17555107 | 0.23977432 | 0.42969352 |
| NUF2 | 32.0856279 | -0.956622299 | 0.50368344 | -1.899253 | 0.05753122 | 0.16074361 |
| PBX1 | 2704.64374 | 1.101289839 | 0.18112171 | 6.08038556 | 1.20E-09 | 1.20E-07 |
| AL357568.1 | 0.36402593 | 1.795827672 | 2.11634291 | 0.84855231 | 0.39613045 | NA |
| LMX1A | 0.49567509 | 1.690623022 | 2.1009352 | 0.80470022 | 0.42099269 | NA |
| AL390730.1 | 0 | NA | NA | NA | NA | NA |
| AL390730.2 | 0 | NA | NA | NA | NA | NA |
| RXRG | 17.3811407 | 0.552087913 | 0.71638212 | 0.77066121 | 0.44090777 | 0.63219716 |
| AL157714.2 | 4.60210702 | 0.417497494 | 0.88679963 | 0.47079124 | 0.63778982 | 0.78561835 |
| LRRC52 | 7.98572202 | 4.022616299 | 1.16042096 | 3.46651467 | 0.00052725 | 0.00472111 |
| AL356441.1 | 1.26828208 | 0.262741261 | 1.1839177 | 0.22192528 | 0.82437205 | 0.90604306 |
| MGST3 | 10147.2136 | -0.514184471 | 0.1649888 | -3.116481 | 0.00183023 | 0.01258167 |
| ALDH9A1 | 7688.14515 | 0.85758694 | 0.25841398 | 3.31865532 | 0.00090452 | 0.00725432 |
| AL451074.6 | 3.81504483 | -0.333092075 | 0.782249 | -0.4258134 | 0.67024385 | 0.80742522 |
| TMCO1 | 4781.28188 | -0.218417616 | 0.12044016 | -1.8134949 | 0.06975553 | 0.1837482 |
| AL451074.2 | 22.1731674 | 0.586018787 | 0.43106536 | 1.3594662 | 0.17399891 | 0.34788772 |
| UCK2 | 963.442269 | -0.73350657 | 0.25792357 | -2.8438912 | 0.00445663 | 0.0247101 |
| AL358115.1 | 123.253075 | -0.92041365 | 0.36018053 | -2.5554231 | 0.01060588 | 0.0470606 |
| FAM78B | 90.2724534 | 1.1266964 | 0.29198456 | 3.85875339 | 0.00011397 | 0.00140304 |
| AL626787.1 | 4.69191353 | 4.988977774 | 1.25394352 | 3.97863037 | 6.93E-05 | 0.00094852 |
| AL390115.1 | 0.19718306 | -0.340099831 | 3.16382928 | -0.1074963 | 0.91439529 | NA |
| AL583804.1 | 0 | NA | NA | NA | NA | NA |
| LINC01675 | 0.72746216 | -1.763275821 | 1.63254888 | -1.0800754 | 0.28010862 | NA |
| POGK | 2593.95281 | 0.096823925 | 0.19284668 | 0.50207721 | 0.61561321 | 0.77018414 |
| TADA1 | 865.595722 | 0.158834713 | 0.14434264 | 1.1004005 | 0.27115766 | 0.46582589 |
| ILDR2 | 568.155212 | 0.746190701 | 0.39017814 | 1.91243595 | 0.0558203 | 0.15727977 |

| | | | | | | |
|---|---|---|---|---|---|---|
| MAEL | 84.8495046 | 0.45270548 | 0.60424153 | 0.74921279 | 0.45372896 | 0.64340895 |
| AL158837.1 | 0 | NA | NA | NA | NA | NA |
| GPA33 | 5.04690167 | 1.337026641 | 0.95442996 | 1.40086408 | 0.16125472 | 0.32994158 |
| DUSP27 | 6.28924038 | 0.269220678 | 0.94388186 | 0.28522709 | 0.77547016 | 0.87674575 |
| LINC01363 | 1.04666622 | -1.491348745 | 1.36231811 | -1.094714 | 0.27364194 | 0.46868914 |
| AL451050.2 | 30.6102011 | -0.936221282 | 0.38176464 | -2.452352 | 0.01419257 | 0.05848698 |
| POU2F1 | 1127.44285 | -0.37318419 | 0.15924778 | -2.3434185 | 0.01910794 | 0.07253232 |
| CD247 | 209.899146 | -0.464744241 | 0.63450871 | -0.7324474 | 0.46389555 | 0.65155396 |
| AL359962.2 | 6.86496121 | 1.389529249 | 0.50290989 | 2.76297858 | 0.00572765 | 0.02981409 |
| CREG1 | 7595.10719 | -0.33286861 | 0.26885751 | -1.2380856 | 0.21568432 | 0.4002812 |
| AL031733.2 | 32.6771012 | -0.07066943 | 0.45428593 | -0.1555616 | 0.87637862 | 0.93555133 |
| RCSD1 | 544.886385 | 0.178645989 | 0.28946888 | 0.61715093 | 0.53713518 | 0.71080718 |
| MPZL1 | 4766.6955 | -0.943200315 | 0.17969324 | -5.2489473 | 1.53E-07 | 6.81E-06 |
| ADCY10 | 36.1906481 | 1.182170003 | 0.36512898 | 3.2376778 | 0.00120507 | 0.00907673 |
| Z99943.1 | 4.68438893 | -0.403882148 | 0.64104884 | -0.6300333 | 0.52867277 | 0.70327906 |
| MPC2 | 6436.54378 | 0.443250187 | 0.1886041 | 2.35016201 | 0.01876524 | 0.07166052 |
| DCAF6 | 6014.64679 | 0.032225333 | 0.14841645 | 0.21712777 | 0.82810877 | 0.90848393 |
| GPR161 | 241.539709 | 0.053315821 | 0.25915924 | 0.2057261 | 0.83700487 | 0.91368209 |
| TIPRL | 1912.75118 | -0.267824974 | 0.17908203 | -1.4955436 | 0.13477264 | 0.29218265 |
| SFT2D2 | 5014.75796 | -0.494414842 | 0.30413462 | -1.6256447 | 0.10402526 | 0.24416075 |
| TBX19 | 351.321567 | -0.73153629 | 0.25639699 | -2.8531392 | 0.00432897 | 0.02418022 |
| AL023755.1 | 7.79872997 | -0.084602299 | 0.72224622 | -0.1171378 | 0.90675089 | 0.95343797 |
| AL022100.2 | 1.45215363 | -0.238047369 | 1.67601514 | -0.1420318 | 0.88705492 | 0.94150285 |
| XCL2 | 14.3386557 | 0.820893236 | 0.84118702 | 0.97587482 | 0.3291265 | 0.52689812 |
| XCL1 | 24.748943 | -0.26310049 | 0.76579558 | -0.3435649 | 0.73117352 | 0.84815227 |
| DPT | 498.018947 | 1.908140645 | 0.73717668 | 2.58844411 | 0.00964106 | 0.04384869 |
| AL031275.1 | 16.230019 | 2.435137725 | 0.68961717 | 3.53114426 | 0.00041377 | 0.003922 |
| LINC00626 | 7.68772139 | 0.613494209 | 0.63110981 | 0.9720879 | 0.33100682 | 0.52888087 |
| LINC00970 | 10.4403002 | 2.248713935 | 0.51562894 | 4.36110886 | 1.29E-05 | 0.00025147 |
| AL031726.1 | 50.8843457 | 3.738429824 | 0.84441917 | 4.42722047 | 9.55E-06 | 0.00019624 |
| ATP1B1 | 53581.9215 | 0.995523994 | 0.23381241 | 4.25778944 | 2.06E-05 | 0.00036368 |
| NME7 | 6467.36486 | 0.433121871 | 0.25523185 | 1.69697423 | 0.08970153 | 0.22003995 |
| Z99758.1 | 0.39103762 | 0.924887234 | 2.72438688 | 0.33948454 | 0.73424474 | NA |
| BLZF1 | 1110.19022 | -0.171162506 | 0.21639637 | -0.7909675 | 0.42896293 | 0.62112297 |
| CCDC181 | 265.658982 | -0.088891082 | 0.27850174 | -0.319176 | 0.74959303 | 0.86013525 |
| SLC19A2 | 1586.64567 | -0.413749792 | 0.18905699 | -2.1884925 | 0.02863375 | 0.09760496 |
| AL021068.1 | 14.6880892 | 1.751922463 | 0.52675214 | 3.3258953 | 0.00088135 | 0.00711368 |
| F5 | 279.460603 | -0.020105858 | 0.38910533 | -0.051672 | 0.95879003 | 0.98017606 |
| SELP | 844.841501 | -0.247109854 | 0.62134919 | -0.3976988 | 0.69085219 | 0.82202745 |
| C1orf112 | 580.482222 | -0.068913668 | 0.13761749 | -0.5007624 | 0.61653834 | 0.7706427 |
| SELL | 387.593464 | -0.015348724 | 0.64033251 | -0.0239699 | 0.9808766 | 0.99132745 |
| SELE | 521.225927 | 0.586897578 | 0.79195718 | 0.74107236 | 0.45864957 | 0.64757615 |
| AL021940.1 | 0.09635527 | 0.547239394 | 3.29377202 | 0.16614368 | 0.86804388 | NA |
| METTL18 | 273.438311 | 0.003153758 | 0.13994944 | 0.02253498 | 0.98202121 | 0.99187603 |
| SCYL3 | 1063.10368 | -0.015632877 | 0.11579516 | -0.1350046 | 0.89260825 | 0.94443203 |
| KIFAP3 | 1908.58952 | 0.212719634 | 0.16237414 | 1.3100586 | 0.19017601 | 0.36885418 |
| AL356475.1 | 102.547054 | 0.717385555 | 0.22583896 | 3.17653583 | 0.00149045 | 0.01075623 |
| METTL11B | 0.35085019 | 1.644831918 | 2.15817781 | 0.7621392 | 0.44597693 | NA |
| LINC01681 | 0.26075328 | 0.074529887 | 3.28898342 | 0.02266046 | 0.98192111 | NA |
| LINC01142 | 0.25337868 | 0.078649477 | 3.03843147 | 0.02588489 | 0.97934915 | NA |
| AL354732.1 | 9.85072925 | -0.518472673 | 0.44358817 | -1.1688154 | 0.24247803 | 0.43288509 |
| GORAB | 517.443779 | -0.219237981 | 0.22860849 | -0.9590107 | 0.33755336 | 0.53552018 |
| AL162399.1 | 0 | NA | NA | NA | NA | NA |
| AL023495.1 | 0.65089782 | 0.761282017 | 2.0773734 | 0.36646374 | 0.71401907 | NA |
| PRRX1 | 375.15218 | -1.117920875 | 0.6407357 | -1.7447457 | 0.08102914 | 0.2042526 |
| Z97200.1 | 9.97607226 | -1.759761581 | 0.93126334 | -1.8896498 | 0.05880481 | 0.16314767 |
| MROH9 | 0.99895425 | -0.789709444 | 1.67880378 | -0.4704001 | 0.6380692 | NA |
| FMO3 | 698.467431 | 1.128563775 | 0.57327974 | 1.9686092 | 0.04899799 | 0.14372379 |
| FMO2 | 1025.1915 | 0.990940422 | 0.5524462 | 1.79373201 | 0.07285595 | 0.18978677 |
| AL021026.1 | 75.3494185 | 2.187193534 | 0.62029848 | 3.52603399 | 0.00042183 | 0.00397367 |
| AL445673.1 | 3.17166918 | -0.363586216 | 0.85268165 | -0.4264032 | 0.66981404 | 0.80715607 |
| FMO1 | 4787.79095 | 3.364719994 | 0.50994371 | 6.59821844 | 4.16E-11 | 6.90E-09 |
| FMO4 | 1494.19875 | 2.438214552 | 0.51058615 | 4.77532449 | 1.79E-06 | 5.22E-05 |
| PRRC2C | 5175.90915 | -0.343890021 | 0.15156384 | -2.268945 | 0.02327167 | 0.08382379 |
| MYOCOS | 90.8647401 | -2.157102799 | 0.8560491 | -2.5198354 | 0.01174097 | 0.05076397 |
| MYOC | 62.3347034 | 1.078930766 | 0.98348872 | 1.09704437 | 0.27262199 | 0.46759554 |
| VAMP4 | 1208.0457 | -0.080011718 | 0.1514897 | -0.5281661 | 0.59738408 | 0.75595249 |
| METTL13 | 1574.42005 | -0.049496337 | 0.13363467 | -0.3703855 | 0.71109531 | 0.83587657 |
| DNM3 | 397.291926 | 0.117712389 | 0.35277603 | 0.33367457 | 0.73862514 | 0.85330522 |
| DNM3OS | 303.642533 | -0.251774899 | 0.30363601 | -0.8291997 | 0.4069914 | 0.60168362 |
| PIGC | 1254.67673 | -0.320641889 | 0.13005702 | -2.4653946 | 0.01368624 | 0.05683562 |
| C1orf105 | 7.62113106 | 1.734660267 | 1.13509974 | 1.52820075 | 0.1264627 | 0.27971997 |
| SUCO | 2286.03464 | -0.795160185 | 0.25803499 | -3.0815982 | 0.00205893 | 0.01377665 |
| FASLG | 20.8770024 | -0.650973196 | 0.61403496 | -1.0601566 | 0.28907338 | 0.4852447 |
| AL031599.1 | 0.27059475 | 1.100737003 | 2.99309129 | 0.36775925 | 0.71305276 | NA |
| Z97198.1 | 0.3108733 | -1.364242679 | 3.30558948 | -0.4127078 | 0.67982068 | NA |
| TNFSF18 | 10.2931866 | 1.756383022 | 0.78910239 | 2.22579864 | 0.02602767 | 0.09100949 |
| AL022310.1 | 1.49393658 | 0.743296931 | 1.10596727 | 0.67207859 | 0.50153366 | 0.68221617 |
| TNFSF4 | 81.1776758 | 0.676633476 | 0.28949285 | 2.33730631 | 0.01942326 | 0.07332733 |
| AL645568.1 | 75.3734066 | 0.166440849 | 0.25297741 | 0.65792772 | 0.51058458 | 0.68898349 |
| PRDX6 | 15198.8176 | -0.299293128 | 0.18802404 | -1.5917812 | 0.11143388 | 0.25626482 |
| SLC9C2 | 15.2137202 | 0.020217541 | 0.64570948 | 0.03131058 | 0.97502185 | 0.98855497 |
| AL139142.1 | 2.26258568 | 0.726801394 | 1.13799765 | 0.63866686 | 0.52303967 | 0.69862624 |
| ANKRD45 | 58.2491335 | 0.167058757 | 0.46483488 | 0.35939376 | 0.71930054 | 0.84021138 |
| TEX50 | 0 | NA | NA | NA | NA | NA |
| KLHL20 | 788.364202 | 0.495017984 | 0.1393433 | 3.55250665 | 0.00038158 | 0.00367638 |
| CENPL | 162.608367 | 0.094113505 | 0.29006231 | 0.32445962 | 0.74559009 | 0.85740777 |
| DARS2 | 1427.10862 | 0.691077588 | 0.20196479 | 3.42177263 | 0.00062214 | 0.00536492 |
| GAS5-AS1 | 341.313868 | -0.740954686 | 0.12462525 | -5.9454618 | 2.76E-09 | 2.35E-07 |
| ZBTB37 | 973.83802 | 0.362110993 | 0.20970358 | 1.72677543 | 0.084208 | 0.2101216 |
| SERPINC1 | 145.102299 | 1.069143186 | 0.65442123 | 1.63372325 | 0.10231697 | 0.24120317 |
| RC3H1 | 1696.15862 | -0.263578666 | 0.14156053 | -1.8619503 | 0.0626101 | 0.17038201 |
| AL121983.1 | 3.50262149 | 0.660983945 | 0.68282973 | 0.96800698 | 0.33304088 | 0.53100478 |
| AL121983.2 | 20.2742504 | 0.77249736 | 0.56926599 | 1.357006 | 0.1747793 | 0.34865396 |
| RABGAP1L | 1611.94845 | 0.078007387 | 0.20708527 | 0.37669211 | 0.70640242 | 0.83294065 |
| GPR52 | 1.18854917 | 0.799320311 | 1.10625942 | 0.72254328 | 0.46996053 | 0.65659606 |
| Z99127.1 | 1.58937669 | 0.042084952 | 1.07343303 | 0.03920594 | 0.9687262 | 0.98550634 |
| CACYBP | 2704.87264 | 0.009480991 | 0.18734085 | 0.05060824 | 0.95963769 | 0.98050555 |
| MRPS14 | 1296.43841 | 0.293268078 | 0.09930595 | 2.95317731 | 0.00314521 | 0.01904898 |
| TNN | 32.0530071 | 3.227095083 | 0.58706046 | 5.4970404 | 3.86E-08 | 2.14E-06 |
| KIAA0040 | 2554.50249 | -0.428315278 | 0.22115898 | -1.936685 | 0.05278387 | 0.15131107 |
| AL008626.1 | 0 | NA | NA | NA | NA | NA |
| Z94057.1 | 0.90723234 | 0.558257297 | 1.42407036 | 0.39201525 | 0.69504695 | NA |
| TNR | 4.03495032 | 0.69274696 | 0.79345059 | 0.87308141 | 0.3826187 | 0.57875886 |
| LINC01657 | 0 | NA | NA | NA | NA | NA |
| AL513329.1 | 23.8882679 | -0.905787759 | 0.49063682 | -1.8461471 | 0.06487084 | 0.1745718 |
| COP1 | 1618.73783 | -0.263125552 | 0.12074403 | -2.1792013 | 0.02931672 | 0.0992381 |
| AL590723.1 | 7.78981676 | -0.016233851 | 0.57394124 | -0.0282849 | 0.97743495 | 0.98959378 |
| AL359265.3 | 12.9690631 | 1.004661762 | 0.42040912 | 2.38972402 | 0.01686104 | 0.06631254 |
| PAPPA2 | 1806.22583 | -0.487787175 | 0.7206514 | -0.6768698 | 0.49848856 | 0.6799953 |
| ASTN1 | 13.9534831 | 1.100206703 | 0.5791779 | 1.89960064 | 0.05748555 | 0.16065005 |
| BRINP2 | 4.63222998 | -1.400035165 | 0.75543612 | -1.8532807 | 0.06384214 | 0.17263173 |
| LINC01645 | 1.97864262 | 1.528290069 | 1.39176114 | 1.09809796 | 0.27216172 | 0.46706394 |

| | | | | | | |
|---|---|---|---|---|---|---|
| AL122019.1 | 8.22513998 | 1.324995156 | 0.78702898 | 1.68354049 | 0.09227051 | 0.22456607 |
| LINC01741 | 0 | NA | NA | NA | NA | NA |
| SEC16B | 1407.67449 | 0.877271822 | 0.47082845 | 1.8632515 | 0.06242689 | 0.16999144 |
| RASAL2-AS1 | 55.2740845 | 0.152977605 | 0.40901941 | 0.37401063 | 0.70839641 | 0.83417046 |
| RASAL2 | 1487.09151 | 0.181806307 | 0.22131048 | 0.82149886 | 0.41136218 | 0.60560633 |
| CLEC20A | 35.3716511 | 0.564719462 | 0.32825305 | 1.72037842 | 0.08536368 | 0.2119424 |
| TEX35 | 16.7701485 | 1.767035268 | 0.45074861 | 3.92022345 | 8.85E-05 | 0.00115033 |
| C1orf220 | 41.4822481 | 1.567067179 | 0.40859954 | 3.83521522 | 0.00012545 | 0.00151199 |
| AL137796.1 | 2.33458731 | 0.97174626 | 1.34390718 | 0.72307543 | 0.46963354 | 0.65627365 |
| AL449106.1 | 28.2738968 | 0.726509049 | 0.27374026 | 2.6540088 | 0.00795418 | 0.03800536 |
| RALGPS2 | 1073.80882 | -0.323791448 | 0.11003642 | -2.9425843 | 0.00325485 | 0.01955082 |
| ANGPTL1 | 605.905679 | 1.136757432 | 0.49788455 | 2.28317475 | 0.02242008 | 0.08160351 |
| FAM20B | 2334.41327 | 0.272200034 | 0.14486894 | 1.87893987 | 0.0602527 | 0.16604282 |
| TOR3A | 2006.73217 | -0.478849004 | 0.20547526 | -2.330446 | 0.01978259 | 0.07430094 |
| ABL2 | 1298.81317 | -0.851524694 | 0.21090558 | -4.0374687 | 5.40E-05 | 0.00079199 |
| SOAT1 | 2894.89959 | -0.893816664 | 0.1727963 | -5.172661 | 2.31E-07 | 9.60E-06 |
| AXDND1 | 2836.40262 | 0.267931853 | 0.81963574 | 0.32689138 | 0.74375004 | 0.85622495 |
| AL160286.3 | 28.7270615 | 1.214691589 | 0.7625002 | 1.59303773 | 0.11115173 | 0.25576536 |
| NPHS2 | 5132.14524 | 0.210070468 | 1.09896377 | 0.19115322 | 0.84840555 | 0.92025018 |
| TDRD5 | 105.95637 | 0.540194479 | 0.38692065 | 1.39613764 | 0.16267305 | 0.33168548 |
| AL359853.3 | 0 | NA | NA | NA | NA | NA |
| FAM163A | 9.31600756 | -1.575857066 | 0.83397839 | -1.8895658 | 0.05881605 | 0.16314767 |
| AL359853.1 | 2.72158644 | -3.830857009 | 1.48607154 | -2.5778416 | 0.00994196 | 0.04486129 |
| AL359853.2 | 11.3098643 | -1.008934288 | 0.51171467 | -1.9716736 | 0.04864688 | 0.14285281 |
| TOR1AIP2 | 3525.67578 | -0.556683511 | 0.17343724 | -3.2097116 | 0.00132868 | 0.00984472 |
| AL353708.3 | 166.46883 | 1.154008031 | 0.23996496 | 4.80906885 | 1.52E-06 | 4.57E-05 |
| TOR1AIP1 | 2916.97838 | 0.461733661 | 0.14404873 | 3.20539905 | 0.00134875 | 0.00994965 |
| AL353708.2 | 0.03392908 | 0.380740593 | 3.29738709 | 0.11546736 | 0.90807469 | NA |
| AL353708.1 | 8.18818373 | 0.258563961 | 0.47116777 | 0.5487726 | 0.58316152 | 0.7454178 |
| CEP350 | 1375.64675 | -0.023369265 | 0.0849675 | -0.2750377 | 0.78328728 | 0.88142381 |
| AL390718.1 | 0 | NA | NA | NA | NA | NA |
| QSOX1 | 6022.64324 | -0.600324003 | 0.23306563 | -2.5757723 | 0.01000165 | 0.04507665 |
| LHX4 | 231.794437 | 0.967168025 | 0.35493706 | 2.72490012 | 0.00643209 | 0.03244344 |
| ACBD6 | 1840.40341 | -0.103338275 | 0.09403269 | -1.0989612 | 0.271785 | 0.46656926 |
| OVAAL | 0.45473089 | 1.479757406 | 2.10007464 | 0.70462134 | 0.48104591 | NA |
| XPR1 | 2679.44755 | 0.054994106 | 0.19272353 | 0.28535232 | 0.77537423 | 0.87674575 |
| AL162431.1 | 8.85527397 | -0.542853465 | 0.45263647 | -1.1993145 | 0.23040569 | 0.41798281 |
| KIAA1614 | 831.137546 | 0.532336228 | 0.26099048 | 2.03967684 | 0.04138252 | 0.12704736 |
| AL162431.2 | 412.64143 | 0.102056433 | 0.11358235 | 0.89852372 | 0.36890641 | 0.56571656 |
| KIAA1614-AS | 263.12836 | 0.323973251 | 0.30198557 | 1.07281036 | 0.28335621 | 0.47925747 |
| STX6 | 1783.87606 | -0.355862677 | 0.13367384 | -2.6621715 | 0.00776383 | 0.0373704 |
| MR1 | 1302.20801 | -0.196447985 | 0.2088353 | -0.9406838 | 0.34686691 | 0.54448958 |
| IER5 | 749.667593 | -0.453181395 | 0.22499568 | -2.0141782 | 0.04399082 | 0.13284906 |
| LINC01732 | 0.18393104 | 1.469068379 | 3.27783996 | 0.44818185 | 0.65402196 | NA |
| AL358393.1 | 0.82136938 | 1.301244427 | 1.51414277 | 0.85939348 | 0.39012346 | NA |
| LINC01699 | 0.09644756 | -0.642905668 | 3.30558948 | -0.1944905 | 0.84579185 | NA |
| CACNA1E | 420.766549 | 0.683587906 | 0.6033806 | 1.13292988 | 0.2572437 | 0.44992482 |
| AL161734.1 | 1.58727856 | 3.686954964 | 1.47431932 | 2.50078453 | 0.01239185 | 0.05284172 |
| ZNF648 | 2.05402288 | 0.742548468 | 0.97047557 | 0.76513876 | 0.44418892 | 0.63510611 |
| AL355482.1 | 4.45207484 | 1.183782372 | 0.99085107 | 1.19471271 | 0.23219929 | 0.42025363 |
| AL355482.2 | 1.46153943 | -1.004665669 | 1.17526755 | -0.85484 | 0.39263975 | 0.58819298 |
| AL390856.1 | 0 | NA | NA | NA | NA | NA |
| LINC01344 | 0.26494444 | -0.178773086 | 2.97589501 | -0.0600737 | 0.95209692 | NA |
| GLUL | 16746.2675 | -0.89408532 | 0.17958666 | -4.9785731 | 6.41E-07 | 2.24E-05 |
| TEDDM1 | 2.32694448 | -0.110711178 | 0.9381067 | -0.1180155 | 0.90605534 | 0.95300065 |
| LINC00272 | 0 | NA | NA | NA | NA | NA |
| RGSL1 | 0.65348178 | -0.05041375 | 1.67100857 | -0.0301697 | 0.97593175 | NA |
| RNASEL | 786.073972 | 0.018748585 | 0.31340502 | 0.05982222 | 0.95229723 | 0.97717271 |
| RGS16 | 804.425258 | -0.204979814 | 0.55727931 | -0.3678224 | 0.71300566 | 0.83694455 |
| LINC01686 | 10.657645 | -0.531823676 | 0.65503104 | -0.8119061 | 0.41684553 | 0.60989939 |
| RGS8 | 3.04637742 | 2.259771476 | 0.77247413 | 2.92536847 | 0.00344049 | 0.02032657 |
| LINC01688 | 0 | NA | NA | NA | NA | NA |
| NPL | 2868.74386 | 0.858713019 | 0.50072775 | 1.71492996 | 0.08635807 | 0.21398819 |
| DHX9 | 7047.88594 | 0.124285903 | 0.15841401 | 0.78456386 | 0.43270933 | 0.62472187 |
| SHCBP1L | 0.27330048 | -0.174096114 | 3.29377188 | -0.0528562 | 0.95784652 | NA |
| LAMC1 | 6390.13857 | -0.810237168 | 0.17590301 | -4.6061587 | 4.10E-06 | 0.00010062 |
| LAMC1-AS1 | 12.4177178 | -0.780304094 | 0.66287359 | -1.1771537 | 0.23913417 | 0.42888988 |
| LAMC2 | 2187.16577 | -1.823056442 | 0.40916272 | -4.4555781 | 8.37E-06 | 0.00017721 |
| NMNAT2 | 43.4514677 | 0.314164034 | 0.36169235 | 0.86859463 | 0.38506889 | 0.58085962 |
| SMG7 | 3506.05243 | -0.160335506 | 0.14620994 | -1.0966115 | 0.27281124 | 0.4676612 |
| NCF2 | 388.569105 | -0.354003229 | 0.36031551 | -0.9824813 | 0.32586282 | 0.52354856 |
| ARPC5 | 5143.53708 | -0.735419482 | 0.1912108 | -3.8461189 | 0.00012 | 0.00146572 |
| RGL1 | 2342.51959 | 1.001817152 | 0.2459559 | 4.07315768 | 4.64E-05 | 0.00069848 |
| APOBEC4 | 0.76989996 | -2.842884152 | 3.26306715 | -0.8712307 | 0.38362822 | NA |
| AL590422.1 | 0 | NA | NA | NA | NA | NA |
| COLGALT2 | 95.5407825 | 0.629346224 | 0.29783019 | 2.11310417 | 0.03459185 | 0.11173287 |
| TSEN15 | 1107.8499 | 0.133306105 | 0.15591551 | 0.85498938 | 0.39255702 | 0.58819298 |
| AL445228.1 | 0 | NA | NA | NA | NA | NA |
| AL445228.2 | 8.13158345 | -0.425528771 | 0.48041958 | -0.885744 | 0.37575548 | 0.57159919 |
| C1orf21 | 917.101045 | -0.613795698 | 0.2688324 | -2.2831909 | 0.02241913 | 0.08160351 |
| EDEM3 | 1817.33808 | -0.361795355 | 0.16750292 | -2.1599347 | 0.03077773 | 0.10274828 |
| FAM129A | 739.076122 | -0.244745175 | 0.44520754 | -0.5497328 | 0.58250268 | 0.74490659 |
| LINC01633 | 0 | NA | NA | NA | NA | NA |
| RNF2 | 769.366254 | -0.270065665 | 0.13062351 | -2.0675119 | 0.03868595 | 0.12081798 |
| TRMT1L | 1240.0501 | 0.406850467 | 0.18665322 | 2.1797131 | 0.02927873 | 0.09918811 |
| SWT1 | 450.515074 | 0.229269981 | 0.17749559 | 1.29169399 | 0.19646314 | 0.37702736 |
| IVNS1ABP | 24399.0085 | -0.359340818 | 0.36303649 | -0.9898201 | 0.32226206 | 0.5200447 |
| AL078644.2 | 23.3496865 | -0.795232715 | 0.43756672 | -1.8173976 | 0.06915625 | 0.18254758 |
| LINC01350 | 2.37531858 | 1.144590154 | 1.11668658 | 1.02498783 | 0.30536893 | 0.50240696 |
| AL133383.1 | 19.2756999 | 0.99941751 | 0.58702337 | 1.70251742 | 0.08865842 | 0.21815083 |
| HMCN1 | 235.974495 | 1.019723997 | 0.47404554 | 2.15110977 | 0.03146754 | 0.10445606 |
| AL133553.1 | 0.06785815 | 0.795868262 | 3.28898349 | 0.24198001 | 0.80879565 | NA |
| PRG4 | 576.351 | -0.260629709 | 0.20588393 | -1.265906 | 0.20554673 | 0.38814058 |
| TPR | 3719.64841 | -0.056451303 | 0.11055726 | -0.5106069 | 0.60962633 | 0.76562587 |
| ODR4 | 1718.08969 | -0.015887674 | 0.13088984 | -0.121382 | 0.90338845 | 0.95141786 |
| OCLM | 3.48682722 | -0.135474374 | 0.66925017 | -0.2024271 | 0.83958284 | 0.91484357 |
| AL596220.1 | 20.5054025 | 0.235764285 | 0.37270391 | 0.63257798 | 0.52700927 | 0.7017742 |
| PDC | 2.58310019 | 0.181238856 | 0.98351004 | 0.18427759 | 0.8537957 | 0.92346684 |
| AL096803.3 | 4.11968163 | 0.620494254 | 0.7260808 | 0.85458017 | 0.39278361 | 0.58833876 |
| PTGS2 | 734.085357 | 0.003659919 | 0.7011998 | 0.00521951 | 0.99583545 | 0.99795473 |
| PACERR | 22.3728272 | 0.246363383 | 0.42448528 | 0.58038146 | 0.56165741 | 0.72942161 |
| PLA2G4A | 561.543446 | 0.340223074 | 0.3400471 | 1.00051749 | 0.31706014 | 0.51438882 |
| ERVMER61-1 | 21.848465 | 0.458685036 | 0.33250146 | 1.3794978 | 0.16774133 | 0.33890122 |
| AL596211.1 | 0 | NA | NA | NA | NA | NA |
| AL929288.1 | 0 | NA | NA | NA | NA | NA |
| AL691515.1 | 0 | NA | NA | NA | NA | NA |
| AL691515.2 | 0 | NA | NA | NA | NA | NA |
| LINC01035 | 0 | NA | NA | NA | NA | NA |
| LINC01701 | 0.27330048 | -0.174096114 | 3.29377188 | -0.0528562 | 0.95784652 | NA |
| AL591504.1 | 0 | NA | NA | NA | NA | NA |
| BRINP3 | 73.7311296 | 0.699993379 | 0.43353023 | 1.61463567 | 0.10638962 | 0.24783518 |
| AL354771.1 | 6.87025545 | 1.102935952 | 0.6459976 | 1.70733755 | 0.08775933 | 0.21648228 |
| LINC01351 | 2.05082668 | 1.143978653 | 0.76557796 | 1.49426801 | 0.13510558 | 0.29250691 |

| | | | | | | | |
|---|---|---|---|---|---|---|---|
| AL391645.1 | 0.96403153 | | 1.760122551 | 1.1554725 | 1.52329246 | 0.12768555 | NA |
| LINC01720 | 0 | NA | | NA | NA | NA | NA |
| AL713866.1 | 0 | NA | | NA | NA | NA | NA |
| LINC01680 | 0 | NA | | NA | NA | NA | NA |
| AL359081.1 | 0 | NA | | NA | NA | NA | NA |
| AL954650.1 | 0.16783122 | | 1.368031398 | 3.28033652 | 0.41703996 | 0.67664918 | NA |
| RGS18 | 73.076651 | | 0.098098768 | 0.58185284 | 0.16859722 | 0.86611347 | 0.9305118 |
| AL390957.1 | 36.8895782 | | -0.274547968 | 0.33742006 | -0.8136682 | 0.41583506 | 0.60913368 |
| RGS21 | 0.16223974 | | -1.003574173 | 3.30558948 | -0.3035992 | 0.7614333 | NA |
| AL136987.1 | 1.81442808 | | 0.400946441 | 1.01207245 | 0.39616377 | 0.69198422 | 0.82270621 |
| RGS1 | 1584.54743 | | -0.364421742 | 0.76688524 | -0.4751972 | 0.6346464 | 0.78339304 |
| RGS13 | 14.8879317 | | -1.061604386 | 1.00365285 | -1.0577406 | 0.29017371 | 0.48626594 |
| AL136454.1 | 50.7822635 | | -0.66264015 | 0.30350584 | -2.1832864 | 0.02901473 | 0.09855963 |
| RGS2 | 2165.26816 | | -0.124607526 | 0.44119879 | -0.2824294 | 0.77761424 | 0.87830075 |
| UCHL5 | 1506.85306 | | 0.026742888 | 0.15649451 | 0.17088707 | 0.86431257 | 0.92963749 |
| TROVE2 | 2070.21114 | | -0.06155473 | 0.09401554 | -0.6547293 | 0.51264206 | 0.69080541 |
| GLRX2 | 1615.01743 | | -0.282095608 | 0.13893844 | -2.030364 | 0.04231956 | 0.12916281 |
| CDC73 | 1512.12341 | | -0.065604252 | 0.13816133 | -0.474838 | 0.63490244 | 0.78341572 |
| B3GALT2 | 59.0134932 | | 1.120941827 | 0.61586306 | 1.82011537 | 0.06874144 | 0.18177175 |
| LINC01031 | 6.43576989 | | -0.158887503 | 0.60095926 | -0.2643898 | 0.79147957 | 0.88679396 |
| AL357793.1 | 0 | NA | | NA | NA | NA | NA |
| AL357793.2 | 0 | NA | | NA | NA | NA | NA |
| AL356272.1 | 0.67963309 | | 1.47134626 | 1.5580485 | 0.94435202 | 0.34498979 | NA |
| AL513348.1 | 0 | NA | | NA | NA | NA | NA |
| LINC01724 | 0.29665639 | | 2.176798933 | 3.23107108 | 0.67370815 | 0.50049688 | NA |
| KCNT2 | 175.517431 | | -0.519516011 | 0.37144348 | -1.3986408 | 0.16192072 | 0.33068846 |
| CFH | 3025.79364 | | -0.277511682 | 0.35254664 | -0.787163 | 0.43118643 | 0.62323976 |
| CFHR3 | 26.172125 | | -1.212083597 | 0.90467233 | -1.339804 | 0.18030908 | 0.35621444 |
| BX248415.1 | 0.09644756 | | -0.642905668 | 3.30558948 | -0.1944905 | 0.84579185 | NA |
| CFHR1 | 97.5106296 | | -0.879680023 | 0.70532895 | -1.2471911 | 0.21232742 | 0.39611419 |
| CFHR4 | 11.7657589 | | -0.528825094 | 1.04193105 | -0.5075433 | 0.61177368 | 0.76749276 |
| CFHR2 | 0.09635527 | | 0.547239394 | 3.29377202 | 0.16614368 | 0.86804388 | NA |
| CFHR5 | 0.07659028 | | -0.642905668 | 3.30558948 | -0.1944905 | 0.84579185 | NA |
| F13B | 0 | NA | | NA | NA | NA | NA |
| ASPM | 29.5016096 | | -0.900315175 | 0.63612358 | -1.4153149 | 0.15697619 | 0.32390157 |
| ZBTB41 | 853.094659 | | 0.080434971 | 0.0986699 | 0.81519255 | 0.4149621 | 0.60849924 |
| CRB1 | 21.1776507 | | 0.476959085 | 0.70840093 | 0.67328975 | 0.50076297 | 0.68176568 |
| AL136322.1 | 1.33267071 | | 0.738623991 | 1.14184729 | 0.64686758 | 0.51771765 | 0.69480454 |
| DENND1B | 761.252863 | | 0.578170408 | 0.14521655 | 3.98143596 | 6.85E-05 | 0.00094278 |
| AL365258.1 | 0.26421942 | | -0.174096114 | 3.29377188 | -0.0528562 | 0.95784652 | NA |
| C1orf53 | 92.0894293 | | -0.35293542 | 0.32862931 | -1.0739621 | 0.28283968 | 0.47881378 |
| LHX9 | 2.31096164 | | -0.246877425 | 0.83108861 | -0.2970531 | 0.76642599 | 0.8711864 |
| NEK7 | 2267.29881 | | 0.297349616 | 0.09300722 | 3.19705942 | 0.00138836 | 0.01018983 |
| ATP6V1G3 | 261.469784 | | 0.950357441 | 0.62413661 | 1.52267536 | 0.12783994 | 0.2817265 |
| AL157402.1 | 0.60628042 | | -0.320395556 | 1.49328144 | -0.2145581 | 0.8301119 | NA |
| PTPRC | 1123.4515 | | 0.203088791 | 0.54981569 | 0.36937613 | 0.71184739 | 0.83618048 |
| AL157402.2 | 0 | NA | | NA | NA | NA | NA |
| MIR181A1H( | 2.03629077 | | 0.157780081 | 0.92670865 | 0.17025856 | 0.8648068 | 0.92971402 |
| AC096631.1 | 0 | NA | | NA | NA | NA | NA |
| LINC01222 | 0 | NA | | NA | NA | NA | NA |
| LINC01221 | 0 | NA | | NA | NA | NA | NA |
| AC099673.1 | 0 | NA | | NA | NA | NA | NA |
| NR5A2 | 92.7208912 | | -0.657464981 | 0.46049833 | -1.427725 | 0.15337103 | 0.31898367 |
| LINC00862 | 14.2460271 | | -0.254368064 | 0.51674276 | -0.4922528 | 0.62254065 | 0.77481557 |
| AC097065.2 | 0.40713206 | | 1.80386971 | 3.23179521 | 0.55816337 | 0.57673283 | NA |
| AC104461.1 | 20.2904814 | | 0.567605607 | 0.49922424 | 1.13697526 | 0.25554863 | 0.44821175 |
| ZNF281 | 684.847395 | | 0.126049993 | 0.27712065 | 0.45485601 | 0.64921283 | 0.79357263 |
| AL359834.1 | 2.23777136 | | 1.906571797 | 1.00420314 | 1.89859175 | 0.05761817 | 0.16088429 |
| KIF14 | 19.3588554 | | -1.81102316 | 0.62431506 | -2.9008161 | 0.00372192 | 0.02158502 |
| DDX59 | 892.08492 | | -0.095622391 | 0.16983305 | -0.5630376 | 0.5734093 | 0.73796013 |
| AL445483.1 | 9.25451154 | | -0.376816027 | 0.63975526 | -0.5890003 | 0.55586109 | 0.7251438 |
| CAMSAP2 | 1526.28154 | | -0.22591763 | 0.16649154 | -1.3569316 | 0.17480296 | 0.34865396 |
| GPR25 | 1.24809965 | | 1.706478338 | 1.79894725 | 0.94859832 | 0.34282495 | 0.54049181 |
| INAVA | 2061.13236 | | -0.115973154 | 0.36612686 | -0.3167568 | 0.75142818 | 0.86135902 |
| KIF21B | 254.239241 | | -0.601557083 | 0.3546066 | -1.6964069 | 0.08980885 | 0.22026221 |
| AL358473.1 | 2.28555999 | | 0.021900208 | 0.76792631 | 0.02851863 | 0.97724851 | 0.98951899 |
| AL358473.2 | 0.71351497 | | 0.65655731 | 1.54651219 | 0.42454066 | 0.67117156 | NA |
| CACNA1S | 19.8389258 | | 1.536166147 | 0.77357285 | 1.98580671 | 0.04705479 | 0.13931171 |
| ASCL5 | 5.09306168 | | -0.556814185 | 0.67699819 | -0.8224751 | 0.41080652 | 0.60515994 |
| TMEM9 | 5122.61323 | | -0.008439505 | 0.19081019 | -0.0442298 | 0.96472119 | 0.98354657 |
| IGFN1 | 228.125285 | | -1.456615751 | 0.6126863 | -2.3774251 | 0.01743399 | 0.06782683 |
| AC103925.1 | 18.1362093 | | -1.597797726 | 0.71960068 | -2.220395 | 0.02639197 | 0.09197254 |
| PKP1 | 57.2987319 | | 1.039177928 | 0.53046913 | 1.95897907 | 0.05011524 | 0.14586521 |
| TNNT2 | 884.128261 | | 0.440475475 | 0.56648556 | 0.77755817 | 0.43682953 | 0.62891432 |
| LAD1 | 4377.67019 | | 0.073839847 | 0.20074165 | 0.36783521 | 0.71299611 | 0.83694455 |
| AC119427.1 | 95.5425427 | | 1.335825444 | 0.34493746 | 3.87265987 | 0.00010765 | 0.00134351 |
| TNNI1 | 546.81359 | | 0.871712855 | 0.44595794 | 1.95469747 | 0.05061879 | 0.14686415 |
| PHLDA3 | 991.339928 | | -0.943057012 | 0.3408889 | -2.7664644 | 0.00566678 | 0.02957896 |
| CSRP1 | 11256.5989 | | 0.177829056 | 0.3538957 | 0.50249001 | 0.61532288 | 0.77003744 |
| AC096677.1 | 201.75452 | | 0.142075548 | 0.33304008 | 0.42660195 | 0.66966928 | 0.80710223 |
| NAV1 | 1110.95384 | | -0.65858925 | 0.21941921 | -3.0015114 | 0.00268643 | 0.01688704 |
| AC092800.1 | 0.2768272 | | -1.364242679 | 3.30558948 | -0.4127078 | 0.67982068 | NA |
| IPO9-AS1 | 220.467616 | | -0.417601106 | 0.20962838 | -1.9921019 | 0.04635987 | 0.13787173 |
| AL645504.1 | 1.8818613 | | 1.850523885 | 0.97686255 | 1.89435442 | 0.05817798 | 0.16203563 |
| IPO9 | 2706.32156 | | 0.065492966 | 0.11214976 | 0.58397776 | 0.55923528 | 0.72772184 |
| SHISA4 | 679.204955 | | -0.281646315 | 0.26231997 | -1.0736747 | 0.28296851 | 0.4789089 |
| AL513217.1 | 434.92088 | | 0.730800244 | 0.48299225 | 1.51306826 | 0.13026232 | 0.2854427 |
| LMOD1 | 1591.83259 | | 0.384311862 | 0.50340236 | 0.76342881 | 0.4452077 | 0.63573887 |
| TIMM17A | 3025.49124 | | -0.423617774 | 0.20280307 | -2.0888135 | 0.03672452 | 0.11642751 |
| RNPEP | 4187.40088 | | -0.27117619 | 0.13362466 | -2.0293873 | 0.04241885 | 0.12937046 |
| AC099676.1 | 657.131458 | | -0.021539339 | 0.17596954 | -0.1224038 | 0.90257924 | 0.9508995 |
| ELF3-AS1 | 133.741196 | | -0.731630138 | 0.38393102 | -1.9056291 | 0.05669834 | 0.15905049 |
| ELF3 | 16237.1372 | | -1.435845311 | 0.34549768 | -4.1558754 | 3.24E-05 | 0.00052199 |
| GPR37L1 | 214.471617 | | -0.986925039 | 0.17987558 | -5.4867093 | 4.09E-08 | 2.25E-06 |
| ARL8A | 3307.07954 | | -0.612302074 | 0.15255818 | -4.0135645 | 5.98E-05 | 0.00084828 |
| PTPN7 | 339.470895 | | -0.10606921 | 0.43922964 | -0.2414892 | 0.80917599 | 0.89689256 |
| LGR6 | 122.93375 | | 0.686013469 | 0.44162768 | 1.55337516 | 0.12033353 | 0.2704634 |
| UBE2T | 129.872062 | | -0.960213144 | 0.23919555 | -4.0143436 | 5.96E-05 | 0.0008465 |
| PPP1R12B | 2494.03016 | | 0.274791747 | 0.38452042 | 0.71463499 | 0.47483462 | 0.66027151 |
| SYT2 | 102.755957 | | 0.09514977 | 0.29685008 | 0.32053139 | 0.74856554 | 0.85955468 |
| AC104463.2 | 27.1226654 | | -0.157936339 | 0.36863271 | -0.4284382 | 0.66833211 | 0.80610702 |
| AC104463.1 | 0.20816856 | | 0.689155606 | 3.09692809 | 0.22252877 | 0.82390228 | NA |
| KDM5B | 2181.87968 | | -0.781240259 | 0.18937634 | -4.1253319 | 3.70E-05 | 0.00058108 |
| AC098934.4 | 1.06962523 | | -1.362004331 | 2.01507457 | -0.6759077 | 0.49909928 | 0.680414 |
| PCAT6 | 164.378831 | | -1.093070274 | 0.43438166 | -2.5163822 | 0.01185665 | 0.05112164 |
| RABIF | 707.050418 | | -0.456334133 | 0.30342451 | -1.5039462 | 0.13259523 | 0.28894606 |
| KLHL12 | 2152.72533 | | 0.141610841 | 0.19496315 | 0.72634671 | 0.46762625 | 0.65456549 |
| ADIPOR1 | 5518.34326 | | 0.028700741 | 0.12051547 | 0.23814984 | 0.81176488 | 0.89840265 |
| CYB5R1 | 3886.68007 | | 0.297838595 | 0.16066252 | 1.85381503 | 0.06376563 | 0.17247919 |
| TMEM183A | 1984.05376 | | -0.210382773 | 0.15570031 | -1.3512033 | 0.17663032 | 0.35147462 |
| PPFIA4 | 672.755013 | | -0.756247904 | 0.39168803 | -1.9307404 | 0.05351516 | 0.15284251 |
| MYOG | 1.65686105 | | 1.681792853 | 1.17765045 | 1.42809172 | 0.15326546 | 0.31883956 |
| ADORA1 | 506.602266 | | 0.43522565 | 0.34677793 | 1.25505578 | 0.20945851 | 0.39262304 |

| | | | | | | |
|---|---|---|---|---|---|---|
| AC105940.2 | 18.1066736 | 0.996580665 | 0.40856948 | 2.43919505 | 0.01472002 | 0.06002272 |
| AC105940.1 | 0.38163648 | -0.55883963 | 2.23931134 | -0.2495587 | 0.80292864 | NA |
| MYBPH | 64.0473398 | 0.736679762 | 0.67976574 | 1.08372593 | 0.27848633 | 0.47412258 |
| CHI3L1 | 3840.40435 | -1.314739003 | 0.62906246 | -2.0899975 | 0.03661802 | 0.11621341 |
| CHIT1 | 41.2351858 | -1.186818296 | 0.86762339 | -1.3678957 | 0.17134473 | 0.34386116 |
| LINC01353 | 13.533415 | -0.363285893 | 0.39995532 | -0.9083162 | 0.36371119 | 0.56082876 |
| LINC01136 | 5.25699203 | -0.399548 | 0.6401863 | -0.6241121 | 0.53255399 | 0.70676641 |
| BTG2 | 8127.1015 | 0.675569143 | 0.42702635 | 1.58203152 | 0.11364237 | 0.25984127 |
| FMOD | 1620.28555 | 0.035067555 | 0.42746976 | 0.08203517 | 0.93461874 | 0.968671 |
| PRELP | 2496.73295 | 0.750989633 | 0.52609265 | 1.42748552 | 0.15344 | 0.319048 |
| OPTC | 1.21111654 | 0.90554552 | 1.25238523 | 0.72305669 | 0.46964506 | 0.65627365 |
| ATP2B4 | 2606.53675 | -0.281083789 | 0.24782777 | -1.13419 | 0.25671485 | 0.44944698 |
| LAX1 | 36.4565295 | 0.731648934 | 0.74405846 | 0.98332184 | 0.32544906 | 0.52301126 |
| ZC3H11A | 7533.22306 | -0.108886281 | 0.18545493 | -0.5871307 | 0.55711593 | 0.72602654 |
| ZBED6 | 7529.9552 | -0.109065056 | 0.18537145 | -0.5883595 | 0.556291 | 0.7253727 |
| SNRPE | 1740.4719 | -0.551928637 | 0.13988813 | -3.9455002 | 7.96E-05 | 0.00105684 |
| LINC00303 | 0.43283449 | 0.298097038 | 1.94030823 | 0.15363386 | 0.87789844 | NA |
| SOX13 | 4092.93304 | 0.318042023 | 0.26539445 | 1.19837479 | 0.23077114 | 0.41844427 |
| AL592146.1 | 2568.04762 | 0.689335296 | 0.54018153 | 1.27611785 | 0.20191386 | 0.3838008 |
| ETNK2 | 6228.52549 | 0.780883387 | 0.5507178 | 1.41793745 | 0.15620903 | 0.32279853 |
| ERLNC1 | 586.64264 | 1.124268655 | 0.58217345 | 1.93115756 | 0.05346357 | 0.15274478 |
| REN | 4411.04354 | -1.28828696 | 0.82886162 | -1.5542847 | 0.12011651 | 0.2700778 |
| KISS1 | 40.9281978 | -1.916348364 | 0.52883619 | -3.6237088 | 0.00029041 | 0.00295907 |
| GOLT1A | 604.164173 | -1.152592921 | 0.29854019 | -3.8607629 | 0.00011303 | 0.00139508 |
| PLEKHA6 | 5903.05708 | 0.501950128 | 0.20325356 | 2.46957604 | 0.01352733 | 0.05640664 |
| AL592114.3 | 9.64059803 | 3.419141195 | 0.62115246 | 5.504512 | 3.70E-08 | 2.08E-06 |
| LINC00628 | 6.92259401 | -0.863491456 | 0.71433178 | -1.2088101 | 0.22673582 | 0.41360152 |
| AL606489.1 | 31.4262565 | 0.753242364 | 0.72603228 | 1.03747778 | 0.29951324 | 0.49614701 |
| PPP1R15B | 3109.25007 | -0.359550892 | 0.20319669 | -1.7694722 | 0.07681511 | 0.19685902 |
| PIK3C2B | 1464.63991 | -0.204416912 | 0.25281361 | -0.8085677 | 0.41876386 | 0.61158634 |
| MDM4 | 4073.12213 | -0.158014833 | 0.15742021 | -1.0037773 | 0.31548597 | 0.51256204 |
| AL512306.3 | 7.56392953 | 1.00342404 | 0.52396171 | 1.91507133 | 0.05548341 | 0.15663178 |
| LRRN2 | 1762.30318 | 0.355156462 | 0.50734279 | 0.70003254 | 0.48390698 | 0.66841062 |
| AL512306.2 | 166.775249 | 0.291795006 | 0.50080933 | 0.5826469 | 0.56013102 | 0.72838393 |
| AL161793.1 | 0.53806857 | 1.106640639 | 2.09519962 | 0.5281791 | 0.59737504 | NA |
| NFASC | 5048.19888 | 0.032276688 | 0.33875543 | 0.09528021 | 0.92409226 | 0.96312021 |
| CNTN2 | 13.5574772 | -1.090571224 | 0.7472472 | -1.4594517 | 0.14444082 | 0.3063544 |
| TMEM81 | 286.90071 | -0.512363315 | 0.33081275 | -1.5488016 | 0.12142943 | 0.27200934 |
| RBBP5 | 1023.90765 | 0.504173824 | 0.15899952 | 3.17091421 | 0.0015196 | 0.01093366 |
| DSTYK | 1200.70885 | 0.284947916 | 0.12283131 | 2.31983124 | 0.02035001 | 0.07584872 |
| TMCC2 | 252.145571 | 0.05038918 | 0.34985424 | 0.14402907 | 0.88547751 | 0.94073175 |
| AC093422.2 | 0.43913079 | 0.455879164 | 2.75396612 | 0.1655355 | 0.86852251 | NA |
| NUAK2 | 3124.6437 | 0.456253933 | 0.33432799 | 1.36468959 | 0.17235063 | 0.34540859 |
| KLHDC8A | 16.0800588 | 0.763375913 | 0.41079776 | 1.85827672 | 0.06312973 | 0.17132149 |
| LEMD1-AS1 | 2.34773666 | -0.722030293 | 1.07798393 | -0.6697969 | 0.50298727 | 0.68324186 |
| LEMD1 | 100.440389 | -3.528366987 | 0.477094 | -7.3955383 | 1.41E-13 | 5.23E-11 |
| BLACAT1 | 207.766914 | -3.756510738 | 0.38426023 | -9.7759551 | 1.43E-22 | 5.38E-19 |
| AC098936.1 | 0.47788515 | -2.122351396 | 2.22314387 | -0.9546622 | 0.33974856 | NA |
| CDK18 | 6219.98871 | 0.011859091 | 0.32521612 | 0.03646526 | 0.97091138 | 0.98654951 |
| MFSD4A | 4616.51086 | 2.626648079 | 0.5561387 | 4.72300903 | 2.32E-06 | 6.40E-05 |
| ELK4 | 3835.01798 | -0.081379968 | 0.20995407 | -0.3876084 | 0.69830583 | 0.8267926 |
| SLC45A3 | 917.789276 | 0.041608217 | 0.27366799 | 0.15203903 | 0.87915615 | 0.93719318 |
| NUCKS1 | 8609.99588 | 0.313993931 | 0.17429029 | 1.80155718 | 0.0716151 | 0.1874339 |
| RAB29 | 5068.59295 | 0.890891785 | 0.35511123 | 2.50876829 | 0.01211529 | 0.05191425 |
| SLC41A1 | 3291.40958 | 0.782488399 | 0.20681882 | 3.78344871 | 0.00015467 | 0.00179518 |
| PM20D1 | 332.410114 | 3.034391152 | 0.87737671 | 3.45848152 | 0.00054323 | 0.00483951 |
| SLC26A9 | 814.720859 | 3.870816781 | 0.36650279 | 10.5614932 | 4.49E-26 | 3.95E-22 |
| AL713965.1 | 0 | NA | NA | NA | NA | NA |
| RAB7B | 81.9383934 | 0.674870551 | 0.57375804 | 1.17622848 | 0.23950358 | 0.42934803 |
| CTSE | 24.1212568 | -0.601339202 | 0.77253357 | -0.7783988 | 0.43633395 | 0.62847532 |
| RHEX | 1351.17167 | -0.122093189 | 0.3994271 | -0.3056708 | 0.75985533 | 0.86700095 |
| AVPR1B | 18.0365711 | 1.220796427 | 0.62641911 | 1.94884928 | 0.05131343 | 0.14843517 |
| AC244035.1 | 11.2137091 | 0.498984232 | 0.66753159 | 0.74750655 | 0.45475785 | 0.64420975 |
| AC244035.2 | 0 | NA | NA | NA | NA | NA |
| FAM72A | 20.5351257 | 0.035285986 | 0.38047428 | 0.09274211 | 0.92610845 | 0.9641707 |
| SRGAP2 | 1706.81229 | -0.240877106 | 0.1793359 | -1.3431617 | 0.17921962 | 0.3548671 |
| IKBKE | 456.775359 | -0.614309452 | 0.38128728 | -1.6111459 | 0.10714792 | 0.24913947 |
| C1orf147 | 58.4597007 | -0.574640545 | 0.39203081 | -1.4658045 | 0.14270154 | 0.30398576 |
| AC244034.2 | 4.85518993 | -0.912372894 | 1.04197102 | -0.8756221 | 0.38123549 | 0.57725981 |
| RASSF5 | 606.709096 | -0.010717209 | 0.416703 | -0.0257191 | 0.97948142 | 0.99069759 |
| EIF2D | 2541.85461 | 0.343812953 | 0.17005734 | 2.02174718 | 0.04320248 | 0.13106002 |
| AL591846.2 | 14.3124852 | -0.277763656 | 0.33321184 | -0.8335948 | 0.40450936 | 0.59933206 |
| DYRK3 | 302.571245 | -0.500577243 | 0.25286779 | -1.9796006 | 0.04774842 | 0.14087334 |
| MAPKAPK2 | 10693.9789 | -0.746071683 | 0.26366027 | -2.8296705 | 0.0046596 | 0.02551214 |
| IL10 | 40.6447698 | -1.614087057 | 0.69464517 | -2.3236137 | 0.02014621 | 0.07532041 |
| IL19 | 22.0777364 | -2.67528149 | 0.88066547 | -3.0377954 | 0.00238316 | 0.01543359 |
| IL20 | 3.49134755 | -1.464376712 | 0.83756752 | -1.7483686 | 0.08040023 | 0.20307577 |
| IL24 | 85.1298756 | -1.010131956 | 0.63373548 | -1.5939331 | 0.11095102 | 0.25539462 |
| FCMR | 276.796264 | 0.166836397 | 0.63189434 | 0.26402578 | 0.79176005 | 0.88698384 |
| PIGR | 23758.4048 | -0.94185186 | 0.44521417 | -2.1155029 | 0.03438711 | 0.11122139 |
| FCAMR | 1845.50848 | 2.989801431 | 0.47753485 | 6.26090729 | 3.83E-10 | 4.49E-08 |
| C1orf116 | 924.92427 | -0.144502391 | 0.2805446 | -0.5150781 | 0.60649842 | 0.76313157 |
| PFKFB2 | 4944.80575 | -0.611838416 | 0.49512397 | -1.2357277 | 0.21655978 | 0.40142567 |
| YOD1 | 1247.35424 | -1.155165067 | 0.41944255 | -2.7540483 | 0.00588631 | 0.03037251 |
| C4BPB | 28.6861008 | -1.176271049 | 0.44218747 | -2.6601184 | 0.00781132 | 0.03755414 |
| C4BPA | 113.755381 | -3.913000272 | 0.8894156 | -4.3995184 | 1.08E-05 | 0.00021806 |
| AL445493.3 | 2.48775015 | 0.534843107 | 0.86967755 | 0.61499012 | 0.53856124 | 0.71192638 |
| AL445493.2 | 0.20366043 | -1.003574173 | 3.30558948 | -0.3035992 | 0.7614333 | NA |
| AL596218.1 | 0.24111501 | 0.689160557 | 2.98884029 | 0.23057791 | 0.81764273 | NA |
| CD55 | 2143.41336 | -0.848559145 | 0.20814957 | -4.0766798 | 4.57E-05 | 0.00069075 |
| AL391597.1 | 0 | NA | NA | NA | NA | NA |
| CR2 | 279.330117 | 1.683317457 | 0.94072415 | 1.78938475 | 0.07355287 | 0.1911302 |
| CR1 | 355.379131 | 0.144344598 | 0.42987105 | 0.33578581 | 0.73703239 | 0.85239876 |
| AL137789.1 | 1.45397531 | -0.487601493 | 1.07998497 | -0.4514891 | 0.65163704 | 0.79524711 |
| CR1L | 56.0626402 | -1.406170052 | 0.68211677 | -2.06148 | 0.03925727 | 0.12216359 |
| AL137789.2 | 0.93319786 | -2.221291546 | 1.52894636 | -1.452825 | 0.14627234 | NA |
| CD46 | 14619.9554 | 0.253881425 | 0.13307918 | 1.90774708 | 0.05642391 | 0.15843878 |
| MIR29B2CHC | 991.473863 | -0.438147847 | 0.31403102 | -1.3952375 | 0.16294424 | 0.33198174 |
| CD34 | 4321.23444 | 0.450833894 | 0.37517377 | 1.20166688 | 0.22949262 | 0.41695744 |
| AL356275.1 | 99.9870845 | 1.018509723 | 0.38991588 | 2.61212678 | 0.00899809 | 0.04166454 |
| PLXNA2 | 1240.68405 | 0.1686776 | 0.24217107 | 0.69652251 | 0.48610166 | 0.67017833 |
| LINC01735 | 0.08757577 | 0.547239394 | 3.29377202 | 0.16614368 | 0.86804388 | NA |
| AC099793.1 | 0.44979913 | 0.340928972 | 2.27090816 | 0.15012891 | 0.88066291 | NA |
| LINC01717 | 0 | NA | NA | NA | NA | NA |
| LINC01774 | 0 | NA | NA | NA | NA | NA |
| AC092810.3 | 0.08393208 | -0.642905668 | 3.30558948 | -0.1944905 | 0.84579185 | NA |
| LINC01696 | 0 | NA | NA | NA | NA | NA |
| LINC01698 | 0.03392908 | 0.380740593 | 3.29738709 | 0.11546736 | 0.90807469 | NA |
| AL023754.1 | 1.10498915 | -0.238677719 | 1.87298534 | -0.1274317 | 0.89859872 | 0.9484869 |
| CAMK1G | 288.940053 | 0.406162712 | 0.54816665 | 0.74094751 | 0.45872527 | 0.64757615 |
| LAMB3 | 3263.48841 | -2.405316489 | 0.66426257 | -3.6210327 | 0.00029343 | 0.00298533 |
| AL031316.1 | 63.0853673 | 0.880043516 | 0.61642064 | 1.42766718 | 0.15338768 | 0.31899314 |
| G0S2 | 6435.75657 | -0.771027324 | 0.48096859 | -1.6030721 | 0.10891874 | 0.25230439 |

| | | | | | | |
|---|---|---|---|---|---|---|
| HSD11B1 | 242.736357 | -0.265013658 | 0.58928646 | -0.4497196 | 0.65291265 | 0.79602192 |
| TRAF3IP3 | 472.934544 | 0.733439338 | 0.46214777 | 1.58702344 | 0.11250734 | 0.25810331 |
| C1orf74 | 405.086687 | 0.728175234 | 0.33058348 | 2.20269697 | 0.02761611 | 0.09510734 |
| IRF6 | 2327.91765 | 0.010800983 | 0.1562807 | 0.06911271 | 0.9448999 | 0.9737607 |
| UTP25 | 1013.05429 | -0.385441729 | 0.29348381 | -1.3133322 | 0.18907104 | 0.36733245 |
| SYT14 | 38.2667022 | -0.240349815 | 0.55977918 | -0.4293654 | 0.66765732 | 0.80551419 |
| SERTAD4-AS | 132.028487 | 0.455995175 | 0.48076437 | 0.94847955 | 0.34288538 | 0.54049181 |
| SERTAD4 | 177.082363 | 1.263695184 | 0.53793246 | 2.34917071 | 0.01881528 | 0.07177186 |
| HHAT | 477.016922 | 0.293179366 | 0.35632544 | 0.82278539 | 0.41063005 | 0.60498364 |
| KCNH1 | 37.1331018 | 1.067898801 | 0.45036411 | 2.37118983 | 0.01773092 | 0.0686274 |
| AL590132.1 | 4.55786326 | 0.666882483 | 0.56109827 | 1.18853063 | 0.23462442 | 0.42334841 |
| AC092017.3 | 0.18746814 | 0.18657164 | 3.29377195 | 0.05664376 | 0.95482897 | NA |
| RCOR3 | 2646.07646 | 0.250983557 | 0.13387357 | 1.87478048 | 0.06082293 | 0.16705558 |
| TRAF5 | 1085.75715 | -0.400475777 | 0.24835907 | -1.612487 | 0.106856 | 0.24863608 |
| LINC00467 | 454.927999 | 0.337457979 | 0.30398453 | 1.11011564 | 0.2669492 | 0.4611392 |
| RD3 | 3.04647876 | -1.446332061 | 1.22497167 | -1.1807065 | 0.23771932 | 0.42708476 |
| AC105275.1 | 0.09644756 | -0.642905668 | 3.30558948 | -0.1944905 | 0.84579185 | NA |
| SLC30A1 | 3499.6235 | -0.399473245 | 0.39642712 | -1.0076839 | 0.31360622 | 0.51067166 |
| AL356310.1 | 2.08705769 | 1.803958047 | 1.30768758 | 1.37950232 | 0.16773993 | 0.33890122 |
| LINC01693 | 13.5980374 | -2.507777757 | 0.81984601 | -3.05884 | 0.00222196 | 0.01464094 |
| NEK2 | 22.3083905 | -1.152290226 | 0.70944933 | -1.6242037 | 0.10433234 | 0.24460631 |
| AC096637.2 | 7.1308299 | -0.937549623 | 0.67560912 | -1.3877101 | 0.16522533 | 0.33554038 |
| AC096637.1 | 0 | NA | NA | NA | NA | NA |
| LPGAT1 | 4466.50778 | -1.418784307 | 0.33915236 | -4.1833243 | 2.87E-05 | 0.00047486 |
| AL445488.1 | 1.40401884 | -0.632963646 | 1.10383264 | -0.5734236 | 0.56635794 | 0.73284279 |
| INTS7 | 594.694043 | -0.277735421 | 0.20443179 | -1.3585726 | 0.17428207 | 0.34824908 |
| DTL | 34.6905235 | -1.047036351 | 0.49078037 | -2.1334112 | 0.03289101 | 0.10771682 |
| AL606468.1 | 1.03162757 | -1.967414976 | 1.33492245 | -1.4738047 | 0.14053418 | 0.30069693 |
| AL451060.1 | 16.8206337 | -1.425672036 | 0.49798245 | -2.8628962 | 0.00419788 | 0.02361308 |
| PPP2R5A | 3914.93106 | -0.653572893 | 0.11417906 | -5.7241045 | 1.04E-08 | 7.16E-07 |
| AL360091.3 | 0.67660057 | -0.134741129 | 1.64502836 | -0.0819081 | 0.9347198 | NA |
| TMEM206 | 900.849452 | 0.532047326 | 0.34843776 | 1.52695084 | 0.12677324 | 0.28014835 |
| NENF | 2385.50243 | -0.107992086 | 0.08178767 | -1.3203956 | 0.18670296 | 0.36453111 |
| AC092803.1 | 0.73701653 | -0.644777325 | 1.98612883 | -0.3246402 | 0.74545337 | NA |
| LINC01740 | 0.96728364 | 0.684662019 | 1.30199275 | 0.52585701 | 0.59898756 | NA |
| AC092803.2 | 129.059039 | 0.366046087 | 0.29953938 | 1.22202994 | 0.2216963 | 0.40769633 |
| ATF3 | 3783.26104 | 0.030948383 | 0.5438009 | 0.05691124 | 0.9546159 | 0.9783227 |
| AL590648.2 | 2.45319364 | -3.126151844 | 1.48834409 | -2.1004228 | 0.03569167 | 0.11413888 |
| FAM71A | 2.45319364 | -3.126151844 | 1.48834409 | -2.1004228 | 0.03569167 | 0.11413888 |
| AL590648.3 | 5.36257959 | -0.391030641 | 0.76874622 | -0.5086602 | 0.61099039 | 0.7667912 |
| BATF3 | 208.908363 | -1.066853956 | 0.45904841 | -2.3240554 | 0.02012252 | 0.0752668 |
| NSL1 | 1022.11721 | 0.071418577 | 0.11708025 | 0.6099968 | 0.54186393 | 0.71448332 |
| TATDN3 | 1004.04319 | 0.251057999 | 0.17130582 | 1.46555439 | 0.14276972 | 0.30408602 |
| SPATA45 | 7.23297729 | 0.847975401 | 0.58965183 | 1.4380951 | 0.15040707 | 0.31500319 |
| FLVCR1-DT | 181.976437 | 0.299294695 | 0.52429427 | 0.5708525 | 0.56809963 | 0.73384869 |
| FLVCR1 | 664.805499 | -0.455628664 | 0.14912909 | -3.0552635 | 0.00224863 | 0.0147945 |
| AC104333.4 | 0.06785815 | 0.795868262 | 3.28898349 | 0.24198001 | 0.80879565 | NA |
| VASH2 | 60.3168549 | 0.58041131 | 0.2447348 | 2.37159292 | 0.01771159 | 0.06857359 |
| ANGEL2 | 1287.84929 | 0.205923067 | 0.1175877 | 1.7512297 | 0.07990635 | 0.20227805 |
| RPS6KC1 | 892.237509 | -0.167321221 | 0.20589382 | -0.8126578 | 0.41641428 | 0.60966391 |
| AL592402.1 | 2.62665821 | -0.550359995 | 1.20128063 | -0.4581444 | 0.6468487 | 0.7919677 |
| AC096639.1 | 2.5149476 | 0.449875145 | 1.72151434 | 0.26132524 | 0.7938417 | 0.88844572 |
| LINC00538 | 0.23111214 | 0.077467214 | 3.10399257 | 0.02495728 | 0.98008904 | NA |
| PROX1 | 495.047578 | 1.42119202 | 0.59788833 | 2.37701916 | 0.01745318 | 0.06787205 |
| AC011700.1 | 15.1982192 | 2.076596353 | 0.69410166 | 2.99177553 | 0.0027736 | 0.01729893 |
| AL606537.1 | 0.16783122 | 1.368031398 | 3.28033652 | 0.41703996 | 0.67664918 | NA |
| AL592287.1 | 0.18663678 | 0.18657164 | 3.29377195 | 0.05664376 | 0.95482897 | NA |
| SMYD2 | 1767.11169 | -0.349103232 | 0.0928971 | -3.7579561 | 0.00017131 | 0.00193958 |
| PTPN14 | 2853.59948 | -0.339736968 | 0.2015153 | -1.6859115 | 0.09181285 | 0.22380336 |
| CENPF | 74.8403361 | -0.97664509 | 0.67575232 | -1.4452708 | 0.14838184 | 0.31224398 |
| KCNK2 | 50.4222988 | -0.652335919 | 1.17411283 | -0.555599 | 0.57848501 | 0.74199376 |
| AL450990.1 | 0 | NA | NA | NA | NA | NA |
| KCTD3 | 3127.90261 | -0.101858677 | 0.13977257 | -0.7287458 | 0.46615715 | 0.65340797 |
| USH2A | 29.2367813 | 1.697798282 | 0.47031473 | 3.60991945 | 0.00030629 | 0.00308397 |
| AL358452.1 | 0.18331075 | 0.18657164 | 3.29377195 | 0.05664376 | 0.95482897 | NA |
| AC093581.1 | 0.03392908 | 0.380740593 | 3.29738709 | 0.11546736 | 0.90807469 | NA |
| AC138024.1 | 0 | NA | NA | NA | NA | NA |
| ESRRG | 1517.15895 | 1.525900399 | 0.17834689 | 8.55580042 | 1.17E-17 | 1.23E-14 |
| GPATCH2 | 760.781829 | -0.12132177 | 0.17042556 | -0.7118755 | 0.4765419 | 0.66179458 |
| SPATA17 | 243.937412 | 0.961706072 | 0.36292648 | 2.64986467 | 0.0080524 | 0.03828738 |
| SPATA17-AS | 55.882777 | 1.297451688 | 0.40476537 | 3.20544142 | 0.00134855 | 0.00994965 |
| LINC00210 | 0 | NA | NA | NA | NA | NA |
| AL355526.1 | 0 | NA | NA | NA | NA | NA |
| LINC01653 | 0 | NA | NA | NA | NA | NA |
| RRP15 | 716.138439 | -0.221803117 | 0.22578964 | -0.9823441 | 0.32593036 | 0.52359327 |
| TGFB2-AS1 | 4.95486964 | 0.76752603 | 0.67431921 | 1.13822358 | 0.25502713 | 0.44771091 |
| TGFB2 | 243.729306 | 0.229778113 | 0.39790946 | 0.57746331 | 0.56362652 | 0.73071933 |
| C1orf143 | 1.67794507 | 3.310191071 | 1.26979104 | 2.60687859 | 0.00913718 | 0.04211629 |
| LINC01710 | 0.17391096 | -1.003574173 | 3.30558948 | -0.3035992 | 0.7614333 | NA |
| LYPLAL1-DT | 192.67944 | 0.866706355 | 0.30818736 | 2.81227092 | 0.0049193 | 0.02654975 |
| LYPLAL1 | 2394.87719 | 0.76423443 | 0.23151187 | 3.30105939 | 0.0009632 | 0.00762754 |
| AL360093.1 | 2.93787141 | -0.21164253 | 0.67326984 | -0.3143502 | 0.75325505 | 0.8625291 |
| AC096642.1 | 81.6421131 | 0.535328538 | 0.28897871 | 1.85248436 | 0.0639563 | 0.17287081 |
| AL513283.1 | 64.4896888 | 2.059188515 | 0.6453086 | 3.19101358 | 0.00141775 | 0.01035622 |
| AL356364.1 | 26.4024398 | 2.07038401 | 0.81618232 | 2.53666853 | 0.01119129 | 0.04898974 |
| ZC3H11B | 9.34764443 | -0.319799625 | 0.45074965 | -0.7094839 | 0.47802424 | 0.66299369 |
| SLC30A10 | 1.67745107 | -1.008281423 | 1.33711378 | -0.754073 | 0.45080538 | 0.64064962 |
| EPRS | 3477.79546 | 0.0642326 | 0.13861511 | 0.46338814 | 0.64308618 | 0.78889944 |
| BPNT1 | 2156.58206 | -0.286750724 | 0.22795095 | -1.2579492 | 0.20841011 | 0.39152061 |
| IARS2 | 5109.64919 | 0.494167441 | 0.18645073 | 2.65039151 | 0.00803985 | 0.03824841 |
| RAB3GAP2 | 1835.28491 | -0.118520361 | 0.11222536 | -1.0560925 | 0.29092596 | 0.48690769 |
| AL451081.1 | 0 | NA | NA | NA | NA | NA |
| AC096644.4 | 0 | NA | NA | NA | NA | NA |
| AC096644.1 | 0.08393208 | -0.642905668 | 3.30558948 | -0.1944905 | 0.84579185 | NA |
| MARK1 | 398.681862 | -0.375157866 | 0.2695208 | -1.391944 | 0.16393935 | 0.33341303 |
| C1orf115 | 4573.27222 | 0.74387844 | 0.22193728 | 3.35175075 | 0.00080302 | 0.00659885 |
| 2-Mar | 4836.25555 | 1.631467948 | 0.27028123 | 6.03618665 | 1.58E-09 | 1.48E-07 |
| 1-Mar | 614.368757 | 0.35113538 | 0.24430835 | 1.4372631 | 0.15064325 | 0.31532485 |
| RNU6ATAC3 | 5.88571192 | 0.965803647 | 0.70156968 | 1.37663253 | 0.16862589 | 0.34024521 |
| AL445423.1 | 5.62617378 | 1.163944862 | 0.72676747 | 1.60153682 | 0.10925807 | 0.25273117 |
| LINC01352 | 8.75271784 | 1.696808213 | 0.59286016 | 2.86207158 | 0.00420882 | 0.02365442 |
| HLX-AS1 | 38.2538385 | 0.03234467 | 0.45935635 | 0.07041303 | 0.94386492 | 0.97318192 |
| HLX | 433.253998 | 0.042942568 | 0.28085753 | 0.15289805 | 0.87847867 | 0.93677394 |
| AL445470.1 | 2.59091273 | 0.410893805 | 0.92522727 | 0.44410041 | 0.65696999 | 0.7986941 |
| AL360013.2 | 0 | NA | NA | NA | NA | NA |
| DUSP10 | 131.986422 | -0.230824299 | 0.29676227 | -0.7778088 | 0.43668175 | 0.62877147 |
| LINC01655 | 0.46797215 | -2.1071007 | 2.74195262 | -0.7684672 | 0.44220966 | NA |
| LINC02257 | 2.02233863 | -4.222325648 | 2.5712659 | -1.6421194 | 0.10056526 | 0.23848911 |
| LINC02474 | 0 | NA | NA | NA | NA | NA |
| LINC01705 | 8.60992406 | -4.583209391 | 1.58145025 | -2.8981053 | 0.00375424 | 0.02175334 |
| AL356108.1 | 0.51897318 | 2.36637478 | 1.95926159 | 1.20778909 | 0.2271284 | NA |
| AL513314.1 | 0.07830766 | -0.642905668 | 3.30558948 | -0.1944905 | 0.84579185 | NA |
| AL513314.2 | 29.2669963 | 1.217994051 | 0.3441736 | 3.53889451 | 0.00040181 | 0.00383209 |

| | | | | | | |
|---|---|---|---|---|---|---|
| HHIPL2 | 9.16236189 | -1.083437427 | 0.48316759 | -2.2423636 | 0.02493788 | 0.08817537 |
| TAF1A | 138.30115 | -0.495767033 | 0.19969109 | -2.4826697 | 0.01304019 | 0.05481745 |
| TAF1A-AS1 | 85.0602186 | 0.892588371 | 0.44926757 | 1.98676342 | 0.04694862 | 0.13913633 |
| MIA3 | 2864.4924 | 0.058884199 | 0.13915872 | 0.42314415 | 0.67219009 | 0.80872583 |
| AL592148.3 | 588.347944 | 0.006991568 | 0.15292488 | 0.04571897 | 0.96353424 | 0.98295478 |
| AIDA | 1628.99046 | -0.349364886 | 0.1433784 | -2.4366633 | 0.01482348 | 0.06032331 |
| BROX | 2463.57568 | -0.256424256 | 0.12684806 | -2.0215071 | 0.0432273 | 0.13108371 |
| FAM177B | 103.484454 | -2.707320316 | 0.48587046 | -5.5721031 | 2.52E-08 | 1.53E-06 |
| AL392172.1 | 533.587337 | -1.034407041 | 0.60740935 | -1.7029818 | 0.08857148 | 0.21803865 |
| AC093152.1 | 178.474765 | -0.170754738 | 0.44584097 | -0.3829947 | 0.70172368 | 0.82913351 |
| DISP1 | 1147.54296 | -0.048292884 | 0.26580738 | -0.1816838 | 0.85583091 | 0.92454691 |
| TLR5 | 584.904569 | -0.234164696 | 0.33312978 | -0.7029233 | 0.48210352 | 0.6666179 |
| AL359979.2 | 1.35145757 | 2.100117561 | 1.18233099 | 1.77625181 | 0.07569142 | 0.19501542 |
| AL359979.1 | 0.47165954 | -0.831446369 | 1.91147281 | -0.4349768 | 0.66357925 | NA |
| SUSD4 | 1105.01329 | 0.983281151 | 0.35777119 | 2.74835196 | 0.00598957 | 0.03073456 |
| CCDC185 | 5.52409147 | 1.396569007 | 0.71916564 | 1.94192955 | 0.05214563 | 0.15008516 |
| CAPN8 | 227.722798 | -1.992517091 | 0.75003704 | -2.6565583 | 0.00789428 | 0.03781511 |
| CAPN2 | 15473.6706 | -0.066511374 | 0.22530215 | -0.2952097 | 0.76783371 | 0.87183611 |
| TP53BP2 | 1985.91687 | -0.653562763 | 0.277351 | -2.3564464 | 0.01845073 | 0.07077315 |
| AC138393.3 | 14.584072 | 0.877535081 | 0.34352137 | 2.55452838 | 0.01063318 | 0.04712552 |
| FBXO28 | 1886.16817 | 0.169619934 | 0.20701108 | 0.81937611 | 0.41257186 | 0.60640544 |
| DEGS1 | 4332.45289 | -0.943192265 | 0.24556581 | -3.8408941 | 0.00012259 | 0.00148462 |
| AC092809.4 | 175.871528 | -0.528882617 | 0.20329994 | -2.6014892 | 0.009282 | 0.04256805 |
| NVL | 806.388828 | -0.130907146 | 0.11183098 | -1.1705804 | 0.2417675 | 0.43204402 |
| CNIH4 | 2151.84342 | -0.797772147 | 0.19328813 | -4.1273728 | 3.67E-05 | 0.00057767 |
| WDR26 | 6080.56539 | -0.425339091 | 0.16818141 | -2.5290494 | 0.01143719 | 0.0498181 |
| CNIH3 | 192.480522 | 0.892104357 | 0.32017808 | 2.78627556 | 0.00533175 | 0.02823826 |
| AL596330.1 | 7.22868643 | 1.380565941 | 0.59787861 | 2.30910744 | 0.02093762 | 0.07749123 |
| AL391811.1 | 0.2839041 | 2.085287735 | 2.91926552 | 0.71431931 | 0.47502975 | NA |
| DNAH14 | 354.883923 | -0.386280175 | 0.18633174 | -2.0730777 | 0.03816505 | 0.11961198 |
| LBR | 3140.93623 | -0.583564321 | 0.24030299 | -2.4284521 | 0.01516343 | 0.06132578 |
| AC092811.2 | 0.51269238 | 1.652014849 | 2.04378599 | 0.80831107 | 0.41891153 | NA |
| ENAH | 1761.94215 | -0.307787865 | 0.30082775 | -1.0231366 | 0.30624328 | 0.50312351 |
| AC099066.2 | 1.12676378 | -0.047708998 | 1.33414528 | -0.03576 | 0.97147375 | 0.98670705 |
| SRP9 | 7994.1483 | 0.037414592 | 0.12941037 | 0.28911588 | 0.7724927 | 0.87466993 |
| EPHX1 | 13865.5176 | -0.079580936 | 0.36147618 | -0.2201554 | 0.82575012 | 0.90695123 |
| AL591895.1 | 1243.46244 | -0.305398853 | 0.32248419 | -0.9470196 | 0.34362878 | 0.54143708 |
| TMEM63A | 7276.18958 | -0.186085064 | 0.33375879 | -0.5575436 | 0.57715611 | 0.74104619 |
| LEFTY1 | 81.9336541 | 0.386586685 | 0.51837616 | 0.74576479 | 0.45580951 | 0.64521428 |
| AL117348.2 | 2466.37861 | -0.335847843 | 0.14229376 | -2.3602429 | 0.01826297 | 0.07021352 |
| PYCR2 | 4463.18835 | -0.483242496 | 0.13862485 | -3.4859731 | 0.00049035 | 0.00446034 |
| LEFTY2 | 19.9604541 | 1.298023018 | 0.76926435 | 1.68735625 | 0.09153487 | 0.22324959 |
| SDE2 | 922.056527 | 0.114457452 | 0.19803513 | 0.5779654 | 0.56328748 | 0.7305312 |
| AL512343.2 | 11.1402219 | 1.073605852 | 0.43222361 | 2.48391299 | 0.01299476 | 0.05471367 |
| H3F3A | 4224.72488 | -0.122840959 | 0.19684926 | -0.6240357 | 0.53260416 | 0.70679743 |
| LINC01703 | 15.9895957 | 0.348406316 | 0.51124127 | 0.68149098 | 0.49556087 | 0.67769982 |
| ACBD3 | 2198.60067 | -0.384486509 | 0.19188384 | -2.0037461 | 0.04509726 | 0.13499261 |
| ACBD3-AS1 | 173.519839 | 0.446673747 | 0.32535634 | 1.3728755 | 0.16979106 | 0.34157691 |
| MIXL1 | 2.34600298 | -0.277720266 | 0.98224182 | -0.2827412 | 0.7773752 | 0.87810598 |
| LIN9 | 131.705568 | 0.382237962 | 0.16603558 | 2.30214493 | 0.021327 | 0.07861268 |
| PARP1 | 4051.92284 | -0.056651692 | 0.12834268 | -0.4414096 | 0.65891648 | 0.79962689 |
| STUM | 1095.72727 | 0.682390312 | 0.44701707 | 1.52654194 | 0.12687495 | 0.28032614 |
| ITPKB | 2560.46055 | 0.476396578 | 0.14231908 | 3.3473838 | 0.00081578 | 0.00668287 |
| ITPKB-AS1 | 0.19478864 | 0.18657164 | 3.29377195 | 0.05664376 | 0.95482897 | NA |
| PSEN2 | 1234.80623 | -0.097170402 | 0.20798872 | -0.4671907 | 0.64036342 | 0.78717058 |
| COQ8A | 4234.13319 | 0.482382236 | 0.17428397 | 2.76779456 | 0.0056437 | 0.02951091 |
| CDC42BPA | 3532.66225 | 0.314821614 | 0.16112127 | 1.95394197 | 0.05070808 | 0.14703894 |
| AL353689.2 | 399.785111 | 0.758241133 | 0.21705651 | 3.49328914 | 0.00047711 | 0.00437309 |
| AL451047.1 | 1.48704161 | -0.396017683 | 1.13443107 | -0.3490892 | 0.72702231 | 0.84575518 |
| LINC01641 | 2.39374021 | -0.573952002 | 0.85683448 | -0.6698517 | 0.50295236 | 0.68322963 |
| ZNF678 | 259.107896 | -0.135333466 | 0.17920254 | -0.7551984 | 0.45012997 | 0.64002628 |
| SNAP47 | 2364.13034 | 0.059514038 | 0.10773002 | 0.55243688 | 0.58064907 | 0.74350374 |
| JMJD4 | 1168.71314 | 0.010273622 | 0.17054525 | 0.06023986 | 0.9519646 | 0.97699287 |
| SNAP47-AS1 | 2.37808917 | 0.924960755 | 0.77561874 | 1.19254566 | 0.23304736 | 0.42124864 |
| PRSS38 | 0.17648264 | -1.003574173 | 3.30558948 | -0.3035992 | 0.7614333 | NA |
| WNT9A | 92.1462515 | -0.638973462 | 0.42604753 | -1.4997704 | 0.1336739 | 0.29048535 |
| WNT3A | 7.25153448 | 0.765665577 | 0.48558983 | 1.57677434 | 0.11484746 | 0.26205348 |
| AL136379.1 | 3.55941935 | -0.237126442 | 0.64083305 | -0.3700284 | 0.71136131 | 0.8359819 |
| ARF1 | 21739.748 | -0.35788833 | 0.14273346 | -2.5073891 | 0.01216268 | 0.05205807 |
| C1orf35 | 988.081677 | -0.516091989 | 0.12990329 | -3.9728939 | 7.10E-05 | 0.00096565 |
| MRPL55 | 1755.21245 | -0.16878316 | 0.1181336 | -1.4287481 | 0.15307664 | 0.31857242 |
| GUK1 | 10631.9888 | -0.378545968 | 0.11752592 | -3.2209573 | 0.00127763 | 0.00951996 |
| GJC2 | 70.5205165 | -0.389634574 | 0.35048193 | -1.1117109 | 0.26626249 | 0.46044605 |
| IBA57-DT | 19.865925 | 0.104235577 | 0.40152363 | 0.25960011 | 0.79517225 | 0.88891824 |
| IBA57 | 693.089851 | 0.515038325 | 0.24806391 | 2.0762324 | 0.03787246 | 0.11894538 |
| OBSCN-AS1 | 223.211117 | 0.429856941 | 0.25108322 | 1.71200985 | 0.08689485 | 0.21485064 |
| OBSCN | 3282.96115 | 0.277511972 | 0.18032668 | 1.5389402 | 0.1238189 | 0.27542002 |
| AL353593.1 | 40.7689429 | 0.645698985 | 0.22483918 | 2.87182596 | 0.00408108 | 0.02308408 |
| AL353593.2 | 19.4451795 | 0.758896317 | 0.35888881 | 2.11457227 | 0.03446642 | 0.11142325 |
| AL353593.3 | 5.82204288 | 0.960581847 | 0.52891781 | 1.81612687 | 0.06935091 | 0.18293324 |
| AL670729.3 | 1.92315942 | -0.478702879 | 0.9325365 | -0.5133342 | 0.60771757 | 0.76405917 |
| TRIM11 | 1515.8169 | -0.878080797 | 0.14690413 | -5.9772369 | 2.27E-09 | 1.98E-07 |
| AL670729.2 | 452.785714 | -0.752765577 | 0.14957425 | -5.0327217 | 4.84E-07 | 1.75E-05 |
| AL670729.1 | 72.376553 | -1.294392662 | 0.29856622 | -4.3353621 | 1.46E-05 | 0.00027667 |
| TRIM17 | 252.83333 | -1.642091943 | 0.30757488 | -5.3388362 | 9.35E-08 | 4.52E-06 |
| HIST3H3 | 0.57646586 | 1.117063453 | 1.73640399 | 0.64332002 | 0.52001647 | NA |
| HIST3H2A | 69.6549684 | -0.928703544 | 0.3332501 | -2.7868065 | 0.00532303 | 0.02819771 |
| HIST3H2BB | 16.735129 | 0.120186272 | 0.78501869 | 0.15309989 | 0.8783195 | 0.93677394 |
| RNF187 | 6274.33352 | -0.29505839 | 0.11360173 | -2.5973054 | 0.00939583 | 0.04298545 |
| RHOU | 1124.76066 | -0.992778394 | 0.30133753 | -3.2945727 | 0.00098571 | 0.0077576 |
| AL137793.2 | 0.18655607 | -1.003574173 | 3.30558948 | -0.3035992 | 0.7614333 | NA |
| AL137793.1 | 1.58319204 | 1.375939324 | 1.12077585 | 1.22766682 | 0.21957207 | 0.40512982 |
| AL162595.2 | 0.07659028 | -0.642905668 | 3.30558948 | -0.1944905 | 0.84579185 | NA |
| TMEM78 | 0.08695548 | -0.642905668 | 3.30558948 | -0.1944905 | 0.84579185 | NA |
| RAB4A | 1717.11697 | 0.075278586 | 0.16165922 | 0.46566218 | 0.64145733 | 0.78758115 |
| SPHAR | 148.409128 | 0.228940832 | 0.22328633 | 1.025324 | 0.30521034 | 0.50224003 |
| AL117350.1 | 23.3114433 | 0.468458566 | 0.28845165 | 1.62404535 | 0.10436613 | 0.24465595 |
| CCSAP | 231.216627 | 0.024500737 | 0.26469641 | 0.09256165 | 0.92625181 | 0.96428189 |
| ACTA1 | 67.9227874 | 2.900567319 | 0.88585226 | 3.274324 | NA | NA |
| NUP133 | 1743.35011 | 0.148584271 | 0.11355197 | 1.30851335 | 0.19069925 | 0.36951511 |
| AL121990.1 | 1.58924417 | 1.163508199 | 0.99668368 | 1.1673796 | 0.24305711 | 0.43362532 |
| ABCB10 | 1057.71914 | 0.298432709 | 0.13507086 | 2.20945288 | 0.02714316 | 0.09385852 |
| TAF5L | 929.129967 | -0.21574534 | 0.25247532 | -0.8545205 | 0.39281664 | 0.58835482 |
| URB2 | 302.396235 | -0.343183255 | 0.27251887 | -1.2593009 | 0.20792168 | 0.39107665 |
| LINC01682 | 0.03392908 | 0.380740593 | 3.29738709 | 0.11546736 | 0.90807469 | NA |
| LINC01736 | 2.32141154 | -0.126634867 | 1.28097965 | -0.0988578 | 0.92125115 | 0.9615743 |
| GALNT2 | 6516.99418 | -0.930602967 | 0.29029758 | -3.2056862 | 0.00134741 | 0.00994965 |
| AL136988.2 | 1.07984636 | -0.052261273 | 1.24288015 | -0.0420485 | 0.96646002 | 0.98410057 |
| AL136988.1 | 2581.95912 | -1.111871177 | 0.28647208 | -3.8812549 | 0.00010392 | 0.0013061 |
| PGBD5 | 327.292218 | -0.11641787 | 0.24480677 | -0.47555 | 0.63439498 | 0.78326602 |
| AL133516.1 | 0.13028434 | 1.049816488 | 3.28477129 | 0.31960109 | 0.74927075 | NA |
| LINC01737 | 0 | NA | NA | NA | NA | NA |
| COG2 | 1452.18496 | 0.446672125 | 0.11788249 | 3.78913045 | 0.00015118 | 0.00176394 |

| | | | | | | | |
|---|---|---|---|---|---|---|---|
| AGT | 11564.7488 | | -1.356399309 | 0.49522809 | -2.7389386 | 0.00616379 | 0.03146324 |
| AL512328.1 | 1532.59337 | | -1.160578418 | 0.52422038 | -2.2139132 | 0.02683475 | 0.09302085 |
| CAPN9 | 5.41886701 | | 0.520576588 | 0.70679496 | 0.73653127 | 0.46140745 | 0.64985923 |
| AL118511.2 | 0 | NA | | NA | NA | NA | NA |
| C1orf198 | 2460.08785 | | -0.329074217 | 0.19793539 | -1.6625335 | 0.09640585 | 0.23176426 |
| AL118511.1 | 7.4699298 | | -0.68943916 | 0.53152788 | -1.2970894 | 0.19460044 | 0.37459768 |
| TTC13 | 678.788477 | | -0.302588872 | 0.17665495 | -1.7128808 | 0.08673447 | 0.2145781 |
| ARV1 | 1045.6131 | | 0.135246339 | 0.10909257 | 1.23973923 | 0.21507186 | 0.39939753 |
| FAM89A | 692.442157 | | -0.908784978 | 0.20598841 | -4.4118258 | 1.03E-05 | 0.00020808 |
| TRIM67 | 6.89549956 | | 1.014520369 | 0.57164186 | 1.77474821 | 0.07593947 | 0.19550169 |
| AL109810.2 | 0 | NA | | NA | NA | NA | NA |
| C1orf131 | 522.709567 | | -0.156866698 | 0.12865489 | -1.2192828 | 0.22273686 | 0.40873842 |
| GNPAT | 2445.02945 | | -0.133029447 | 0.13414539 | -0.9916811 | 0.32135312 | 0.51899079 |
| EXOC8 | 694.150618 | | -0.133001455 | 0.23591043 | -0.5637795 | 0.57290423 | 0.73753036 |
| SPRTN | 457.629831 | | 0.038288975 | 0.30668727 | 0.12484697 | 0.9006447 | 0.94969868 |
| EGLN1 | 4850.61906 | | -0.573777488 | 0.19625401 | -2.9236472 | 0.00345957 | 0.02040455 |
| AL445524.1 | 188.469815 | | -1.370118421 | 0.39782588 | -3.4440153 | 0.00057314 | 0.00504767 |
| TSNAX | 2057.29646 | | 0.250027094 | 0.09912525 | 2.52233512 | 0.01165786 | 0.05051228 |
| TSNAX-DISC1 | 536.604994 | | 0.640880065 | 0.20237051 | 3.16686497 | 0.00154092 | 0.01103885 |
| LINC00582 | 2.33439571 | | 1.024557518 | 0.90288601 | 1.13475844 | 0.25647654 | 0.44921834 |
| DISC1 | 235.575175 | | -0.389996897 | 0.27029295 | -1.4428674 | 0.14905782 | 0.31317463 |
| AL353052.1 | 0 | NA | | NA | NA | NA | NA |
| AL353052.2 | 0 | NA | | NA | NA | NA | NA |
| SIPA1L2 | 932.722047 | | -0.847360617 | 0.31773319 | -2.6668936 | 0.00765559 | 0.03697398 |
| LINC01745 | 0 | NA | | NA | NA | NA | NA |
| LINC01744 | 0.10721287 | | -0.642905668 | 3.30558948 | -0.1944905 | 0.84579185 | NA |
| MAP10 | 85.6371145 | | 0.497441903 | 0.31284677 | 1.59004965 | 0.11182361 | 0.25701659 |
| NTPCR | 1873.51383 | | 0.147453858 | 0.16610499 | 0.88771483 | 0.37469417 | 0.5707258 |
| PCNX2 | 811.162827 | | -0.328502574 | 0.19896011 | -1.6510977 | 0.09871862 | 0.23558262 |
| MAP3K21 | 1803.69835 | | 0.104215168 | 0.23634668 | 0.44094196 | 0.65925502 | 0.7999151 |
| KCNK1 | 1079.09149 | | -1.056419165 | 0.18726272 | -5.6413746 | 1.69E-08 | 1.08E-06 |
| SLC35F3 | 52.9950726 | | 0.204301952 | 0.48971429 | 0.41718601 | 0.67654235 | 0.81185083 |
| AL122008.3 | 3.62830493 | | 1.250532103 | 0.7579105 | 1.64997333 | 0.09894839 | 0.23585547 |
| AL122008.4 | 0 | NA | | NA | NA | NA | NA |
| AL355472.3 | 7.63495771 | | -0.949153041 | 0.61684296 | -1.5387272 | 0.12387091 | 0.27548621 |
| AL355472.4 | 23.6026191 | | -0.035704819 | 0.37170816 | -0.0960561 | 0.92347605 | 0.96272242 |
| AL355472.1 | 478.571803 | | -0.333277769 | 0.1903336 | -1.7510191 | 0.07994261 | 0.202331 |
| COA6 | 1058.69147 | | -0.540777428 | 0.20764915 | -2.6042843 | 0.00920663 | 0.0423395 |
| TARBP1 | 1639.65021 | | 0.312946722 | 0.2560188 | 1.22235836 | 0.22157214 | 0.40759381 |
| LINC01354 | 15.3799219 | | 1.301229916 | 0.45955954 | 2.83147186 | 0.00463343 | 0.02544217 |
| AL161640.1 | 0.03392908 | | 0.380740593 | 3.29738709 | 0.11546736 | 0.90807469 | NA |
| IRF2BP2 | 7546.27729 | | -0.203504947 | 0.20558511 | -0.9898818 | 0.32223192 | 0.52002789 |
| AL160408.2 | 2864.37292 | | -0.21798866 | 0.19813476 | -1.100204 | 0.27124325 | 0.46594257 |
| LINC00184 | 3.10637647 | | 1.087717794 | 0.74913301 | 1.45196885 | 0.14651027 | 0.30947419 |
| AL160408.5 | 0.92168238 | | 2.235860361 | 1.56824451 | 1.42570903 | 0.15395235 | NA |
| AL160408.1 | 3.67884864 | | 1.878854426 | 0.88213684 | 2.12988998 | 0.0331807 | 0.10838288 |
| AL160408.4 | 2.35499412 | | 1.231698065 | 1.21968893 | 1.00984606 | 0.31256905 | 0.50941908 |
| AL160408.3 | 10.0612712 | | 0.776947961 | 1.23717943 | 0.62799942 | 0.53000432 | 0.70458506 |
| AL160408.6 | 6.44685453 | | 1.913131262 | 1.33544138 | 1.43258349 | 0.1519769 | 0.31705941 |
| LINC01132 | 50.249641 | | 0.236703263 | 0.40415676 | 0.58567191 | 0.55809601 | 0.72665737 |
| AL360294.1 | 1.15699 | | 0.014590887 | 1.7626047 | 0.00827803 | 0.99339517 | 0.99694729 |
| AL391832.1 | 9.93710233 | | -0.172780793 | 0.60420141 | -0.2859656 | 0.7749045 | 0.87636625 |
| AL391832.2 | 15.7980455 | | 0.115795916 | 0.55389283 | 0.20905834 | 0.83440269 | 0.9121801 |
| AL391832.3 | 5.40012685 | | -0.332848405 | 0.53747976 | -0.6192762 | 0.53573444 | 0.70948761 |
| LINC01348 | 66.5665727 | | -0.848205959 | 0.3717298 | -2.2817809 | 0.02250228 | 0.08182365 |
| AL732292.2 | 7.03370843 | | -0.445099107 | 0.66079911 | -0.673577 | 0.50058025 | 0.6815873 |
| TOMM20 | 8146.36329 | | 0.308138539 | 0.21506753 | 1.43275246 | 0.15192859 | 0.31700878 |
| RBM34 | 1643.89059 | | -0.427406989 | 0.16139862 | -2.6481452 | 0.00809347 | 0.03842725 |
| ARID4B | 3312.00735 | | -0.353881173 | 0.10257672 | -3.4499171 | 0.00056076 | 0.00496548 |
| GGPS1 | 1529.92699 | | -0.254385361 | 0.11454218 | -2.2208881 | 0.02635854 | 0.09192422 |
| TBCE | 2720.41442 | | -0.081312285 | 0.15602824 | -0.5211383 | 0.60227045 | 0.75957108 |
| AL357556.4 | 2.03691663 | | 0.471352245 | 0.90132301 | 0.52295597 | 0.6010049 | 0.75862808 |
| B3GALNT2 | 212.339544 | | 0.616350419 | 0.18218953 | 3.38301776 | 0.00071694 | 0.00602476 |
| GNG4 | 128.941158 | | -1.173671686 | 0.46653836 | -2.5157024 | 0.01187954 | 0.05119522 |
| LYST | 967.146243 | | -0.0183424 | 0.12284357 | -0.1493151 | 0.881305 | 0.93845958 |
| LYST-AS1 | 0.25481963 | | -1.364242679 | 3.30558948 | -0.4127078 | 0.67982068 | NA |
| AL139161.2 | 0.10721287 | | -0.642905668 | 3.30558948 | -0.1944905 | 0.84579185 | NA |
| AL139161.1 | 2.0671712 | | 0.819278547 | 0.86109067 | 0.95144284 | 0.34137962 | 0.53937972 |
| NID1 | 5359.90143 | | -0.601338687 | 0.2725014 | -2.2067361 | 0.0273325 | 0.09437711 |
| GPR137B | 3486.74337 | | 0.339062543 | 0.47024305 | 0.7210368 | 0.47088687 | 0.65748881 |
| ERO1B | 1277.47849 | | 0.024445543 | 0.16524118 | 0.14793856 | 0.88239125 | 0.93897169 |
| EDARADD | 96.0515779 | | 0.1315238 | 0.20456758 | 0.64293569 | 0.52026583 | 0.6969232 |
| LGALS8 | 4886.52145 | | -0.738214473 | 0.19304115 | -3.8241301 | 0.00013123 | 0.00156733 |
| LGALS8-AS1 | 105.795411 | | -0.64669409 | 0.30966423 | -2.0883719 | 0.0367643 | 0.11652338 |
| AL359921.2 | 82.6687399 | | -0.409478254 | 0.33830111 | -1.2103958 | 0.22612704 | 0.41281666 |
| AL359921.1 | 273.474102 | | -0.187277396 | 0.18096235 | -1.034897 | 0.30071702 | 0.4974536 |
| HEATR1 | 2734.79831 | | -0.492569705 | 0.16552423 | -2.9758163 | 0.0029221 | 0.01797847 |
| ACTN2 | 106.206984 | | -0.293766376 | 0.44883152 | -0.6545137 | 0.5127809 | 0.69088659 |
| MTR | 4253.85152 | | 0.072756033 | 0.22443866 | 0.32416889 | 0.74581017 | 0.85747359 |
| MT1HL1 | 4.97702212 | | 0.071299815 | 0.68974133 | 0.10337182 | 0.91766788 | 0.96005997 |
| RYR2 | 32.2724046 | | 0.263519358 | 0.74739766 | 0.35258253 | 0.72440144 | 0.84387647 |
| AL359924.1 | 0.76633462 | | -0.722499033 | 2.25281725 | -0.3207091 | 0.74843084 | NA |
| ZP4 | 0 | NA | | NA | NA | NA | NA |
| MTRNR2L11 | 0 | NA | | NA | NA | NA | NA |
| AL356010.2 | 0 | NA | | NA | NA | NA | NA |
| LINC01139 | 654.739259 | | 0.448700517 | 0.48451722 | 0.92607754 | 0.35440568 | 0.55199607 |
| AL359551.1 | 3.03972019 | | 2.289906867 | 1.0923308 | 2.09634926 | 0.03605122 | 0.11496803 |
| AC093426.1 | 0.03392908 | | 0.380740593 | 3.29738709 | 0.11546736 | 0.90807469 | NA |
| CHRM3 | 163.946606 | | 1.030127987 | 0.35215201 | 2.92523673 | 0.00344194 | 0.02032789 |
| CHRM3-AS2 | 128.640176 | | 2.036215276 | 0.58174047 | 3.50021251 | 0.00046489 | 0.0042924 |
| CHRM3-AS1 | 21.7508047 | | 2.699708211 | 0.46782178 | 5.77080492 | 7.89E-09 | 5.62E-07 |
| AL356361.2 | 15.7633297 | | 1.475375064 | 0.55511382 | 2.65778838 | 0.00786553 | 0.03771162 |
| FMN2 | 192.338785 | | 0.224730234 | 0.61064887 | 0.36801875 | 0.71285925 | 0.83684719 |
| AL359918.2 | 7.14483989 | | 1.607541563 | 0.7320948 | 2.19581067 | 0.0281055 | 0.0963899 |
| AL590490.1 | 0 | NA | | NA | NA | NA | NA |
| GREM2 | 59.83057 | | 0.794382468 | 0.73009105 | 1.08805945 | 0.27656886 | 0.47192487 |
| AL358176.1 | 0 | NA | | NA | NA | NA | NA |
| AL365184.2 | 0.08666371 | | -0.642905668 | 3.30558948 | -0.1944905 | 0.84579185 | NA |
| AL365184.1 | 17.051067 | | 3.137676624 | 0.71874575 | 4.36548896 | 1.27E-05 | 0.00024757 |
| RGS7 | 45.7904177 | | 3.20583161 | 0.64433599 | 4.9754036 | 6.51E-07 | 2.26E-05 |
| AL359764.1 | 2.54980905 | | 0.808647531 | 0.76053546 | 1.06326078 | 0.28766371 | 0.4838718 |
| FH | 4549.68671 | | 0.609929001 | 0.22006498 | 2.77158597 | 0.00557839 | 0.02926964 |
| KMO | 2281.16813 | | 0.837127929 | 0.3189785 | 2.62440235 | 0.00868012 | 0.04045937 |
| OPN3 | 1775.95668 | | 0.146417902 | 0.22848765 | 0.64081319 | 0.52164406 | 0.69783654 |
| CHML | 622.849433 | | -0.179298091 | 0.27238638 | -0.6582491 | 0.51037807 | 0.68891634 |
| WDR64 | 5.84958945 | | 0.824781251 | 0.6524614 | 1.26410734 | 0.2061915 | 0.38882079 |
| EXO1 | 17.5292515 | | -0.180974473 | 0.56924605 | -0.3179196 | 0.75054593 | 0.8607792 |
| BECN2 | 0.15318056 | | -1.003574173 | 3.30558948 | -0.3035992 | 0.7614333 | NA |
| MAP1LC3C | 19.0951356 | | -0.811713144 | 0.55006067 | -1.4756793 | 0.14003 | 0.3000643 |
| PLD5 | 40.0760962 | | 2.646980387 | 0.63244609 | 4.18530593 | 2.85E-05 | 0.00047142 |
| AL591686.2 | 0.92087757 | | 1.236731663 | 1.54860985 | 0.79860765 | 0.42451795 | NA |
| AL591686.1 | 1.0565098 | | -1.231303601 | 2.82060087 | -0.4365395 | 0.66244537 | 0.80142609 |
| AL606534.3 | 0 | NA | | NA | NA | NA | NA |

| | | | | | | |
|---|---|---|---|---|---|---|
| CEP170 | 804.700093 | -0.334286606 | 0.19384855 | -1.7244731 | 0.08462247 | 0.2107331 |
| AL606534.1 | 44.3039416 | 0.449667216 | 0.33531201 | 1.3410412 | 0.17990708 | 0.35574007 |
| AL606534.2 | 7.73601688 | 0.994823743 | 0.54615079 | 1.82151847 | 0.06852808 | 0.18136145 |
| SDCCAG8 | 1388.05022 | -0.072651656 | 0.12190922 | -0.5959488 | 0.55120942 | 0.72143103 |
| AKT3 | 2366.07254 | -0.159574109 | 0.21834418 | -0.7308375 | 0.46487843 | 0.65205955 |
| AL591721.1 | 28.1662732 | 0.526860253 | 0.46659221 | 1.1291664 | 0.25882764 | 0.4516467 |
| AL591885.1 | 5.3886841 | 1.74352257 | 0.60325637 | 2.8901851 | 0.00385015 | 0.02213396 |
| ZBTB18 | 1648.42249 | 0.418699579 | 0.14045247 | 2.98107656 | 0.00287237 | 0.01773543 |
| AL590483.1 | 0.76314378 | 2.360258026 | 1.71086561 | 1.3795695 | 0.16771923 | NA |
| AL590483.2 | 0.16964538 | 1.509808164 | 3.2763376 | 0.46082191 | 0.64492638 | NA |
| C1orf100 | 12.9938123 | -0.408813282 | 0.42509153 | -0.9617065 | 0.33619706 | 0.53426037 |
| AL645465.1 | 724.862673 | -0.403089697 | 0.15641244 | -2.5770949 | 0.00996346 | 0.04495063 |
| ADSS | 2890.4781 | -0.199560689 | 0.14953148 | -1.3345731 | 0.18201611 | 0.35851245 |
| CATSPERE | 151.786363 | 0.171671486 | 0.25071662 | 0.68472321 | 0.4935186 | 0.67619153 |
| DESI2 | 1612.17979 | -0.420792548 | 0.1750921 | -2.403264 | 0.01624945 | 0.06451341 |
| AL451007.3 | 1.85561138 | 1.451107517 | 0.98772117 | 1.46914693 | 0.14179294 | 0.30251455 |
| AL451007.2 | 2.04639385 | -1.004619269 | 1.37128484 | -0.7326117 | 0.46379532 | 0.65155199 |
| COX20 | 6331.22031 | 0.055648861 | 0.10752051 | 0.51756509 | 0.60476174 | 0.76172942 |
| HNRNPU | 22604.4057 | -0.057601553 | 0.11707823 | -0.491992 | 0.622725 | 0.77488006 |
| BX323046.1 | 1.17304414 | 0.941386253 | 1.18468065 | 0.79463293 | 0.42682706 | 0.61907648 |
| AL356512.1 | 139.138555 | 0.343251106 | 0.22793324 | 1.50592824 | 0.1320856 | 0.2882334 |
| EFCAB2 | 731.780937 | 0.08687329 | 0.21827545 | 0.39799844 | 0.69063133 | 0.82183882 |
| KIF26B | 351.476936 | -0.416235805 | 0.39462963 | -1.0547505 | 0.29153945 | 0.48774871 |
| KIF26B-AS1 | 0.88968815 | -1.431928059 | 1.90329952 | -0.7523398 | 0.45184672 | NA |
| AC104462.2 | 0 | NA | NA | NA | NA | NA |
| AC104462.1 | 0.33510098 | 0.669740542 | 2.23157452 | 0.30012018 | 0.76408549 | NA |
| SMYD3 | 526.395956 | 0.099100383 | 0.21774784 | 0.45511534 | 0.64902627 | 0.79349874 |
| AC118555.1 | 0.23845394 | 0.077863861 | 3.08148914 | 0.02526826 | 0.97984099 | NA |
| AC092801.1 | 0.36945157 | 0.704334393 | 2.29204019 | 0.30729583 | 0.75861821 | NA |
| LINC01743 | 0.21430872 | -0.339842146 | 3.10090457 | -0.1095945 | 0.91273096 | NA |
| TFB2M | 977.441407 | 0.468577283 | 0.24495705 | 1.91289565 | 0.05576141 | 0.15716495 |
| CNST | 1130.29411 | 0.013404387 | 0.14814187 | 0.09048344 | 0.92790305 | 0.96504253 |
| AL591848.3 | 32.8612934 | -0.54962532 | 0.40463306 | -1.3583302 | 0.17435892 | 0.34829853 |
| AL591848.2 | 2.544898 | 1.489496851 | 0.87031249 | 1.71145062 | 0.08699796 | 0.21499323 |
| SCCPDH | 3767.9565 | -0.262893659 | 0.26919551 | -0.9765901 | 0.32887214 | 0.52658294 |
| AL591848.4 | 39.337295 | 0.191940119 | 0.35834201 | 0.53563388 | 0.59221161 | 0.75166635 |
| AHCTF1 | 1880.62125 | -0.0421488 | 0.19575588 | -0.2153131 | 0.82952324 | 0.90939253 |
| ZNF695 | 8.07876354 | 0.636544997 | 0.53052245 | 1.19984554 | 0.23019933 | 0.4177809 |
| ZNF670-ZNF | 21.7914395 | -0.014613704 | 0.26541977 | -0.0550588 | 0.95609159 | 0.97900515 |
| ZNF670 | 148.718568 | -0.416104325 | 0.25769154 | -1.614738 | 0.10636744 | 0.24780542 |
| ZNF669 | 176.29597 | -0.193927301 | 0.30387212 | -0.6381872 | 0.52335182 | 0.69882538 |
| C1orf229 | 45.046683 | 0.490486713 | 0.26601476 | 1.84383273 | 0.06520752 | 0.17519331 |
| ZNF124 | 290.569641 | 0.172291726 | 0.15447332 | 1.11534938 | 0.26470073 | 0.45867777 |
| AL390728.6 | 705.740961 | -0.929846316 | 0.20690032 | -4.4941754 | 6.98E-06 | 0.00015386 |
| AL390728.5 | 29.3553932 | 0.738011173 | 0.3439714 | 2.14555971 | 0.03190812 | 0.10542754 |
| ZNF496 | 1711.668 | -0.024488749 | 0.14949011 | -0.1638152 | 0.86987664 | 0.93220007 |
| NLRP3 | 99.2167596 | 0.292107435 | 0.4199369 | 0.6955984 | 0.48668037 | 0.67073071 |
| OR2B11 | 1.7369699 | -0.780838201 | 1.16680902 | -0.6692082 | 0.50336265 | 0.68347012 |
| GCSAML | 7.60612877 | 0.223400928 | 0.89033313 | 0.25091836 | 0.80187723 | 0.89259192 |
| OR2C3 | 8.76401764 | -0.183540403 | 0.57362894 | -0.3199636 | 0.7489959 | 0.8597841 |
| GCSAML-AS1 | 7.96090999 | -0.164918907 | 0.57058946 | -0.2890325 | 0.7725565 | 0.87468614 |
| AL606804.1 | 0.81796288 | 0.561587264 | 1.38126273 | 0.40657527 | 0.68431996 | NA |
| OR2G2 | 0 | NA | NA | NA | NA | NA |
| OR2G3 | 0 | NA | NA | NA | NA | NA |
| AL390860.1 | 1.26647985 | 0.674497445 | 0.98206044 | 0.68681867 | 0.492197 | 0.67508764 |
| OR13G1 | 0.17515154 | 1.45420457 | 3.2783972 | 0.44357181 | 0.65735219 | NA |
| OR6F1 | 0.08666371 | -0.642905668 | 3.30558948 | -0.1944905 | 0.84579185 | NA |
| AC118470.1 | 0.08666371 | -0.642905668 | 3.30558948 | -0.1944905 | 0.84579185 | NA |
| OR14A2 | 0 | NA | NA | NA | NA | NA |
| OR14K1 | 0.06785815 | 0.795868262 | 3.28898349 | 0.24198001 | 0.80879565 | NA |
| OR1C1 | 0.24147734 | 0.0780252 | 3.07248537 | 0.02539482 | 0.97974005 | NA |
| OR14A16 | 0 | NA | NA | NA | NA | NA |
| OR11L1 | 0 | NA | NA | NA | NA | NA |
| TRIM58 | 11.7341894 | -0.260545572 | 0.64164302 | -0.40606 | 0.68469851 | 0.81783568 |
| OR2W3 | 5.39477747 | -0.685575717 | 0.63483527 | -1.0799269 | 0.28017471 | 0.47576838 |
| OR2T8 | 1.8061259 | 0.306608315 | 0.96573677 | 0.31748643 | 0.75087454 | 0.86108117 |
| OR2AJ1 | 0 | NA | NA | NA | NA | NA |
| OR2L8 | 0 | NA | NA | NA | NA | NA |
| OR2AK2 | 0 | NA | NA | NA | NA | NA |
| OR2L5 | 0.16421342 | 1.36803136 | 3.28033652 | 0.41703994 | 0.67664919 | NA |
| OR2L2 | 0.63065401 | -0.408618276 | 1.69978629 | -0.2403939 | 0.8100249 | NA |
| OR2L3 | 0.35086179 | -0.260527483 | 2.31348649 | -0.1126125 | 0.91033778 | NA |
| OR2L13 | 1.25748902 | 1.070564977 | 1.05954885 | 1.010397 | 0.31230513 | 0.50918254 |
| OR2M5 | 0 | NA | NA | NA | NA | NA |
| OR2M2 | 0 | NA | NA | NA | NA | NA |
| OR2M3 | 0.17332742 | -1.003574173 | 3.30558948 | -0.3035992 | 0.7614333 | NA |
| OR2M4 | 0 | NA | NA | NA | NA | NA |
| OR2T33 | 0.17423948 | 0.18657164 | 3.29377195 | 0.05664376 | 0.95482897 | NA |
| OR2T12 | 0 | NA | NA | NA | NA | NA |
| OR2M7 | 0 | NA | NA | NA | NA | NA |
| OR14C36 | 0 | NA | NA | NA | NA | NA |
| OR2T4 | 0.03392908 | 0.380740593 | 3.29738709 | 0.11546736 | 0.90807469 | NA |
| OR2T6 | 4.26823392 | 2.541823199 | 1.25382893 | 2.0272488 | 0.04263697 | 0.1297262 |
| OR2T1 | 0.09997307 | 0.547239394 | 3.29377202 | 0.16614368 | 0.86804388 | NA |
| OR2T2 | 19.939019 | 3.527801021 | 0.6118709 | 5.76559698 | 8.14E-09 | 5.77E-07 |
| OR2T3 | 0.2839041 | 2.085287735 | 2.91926552 | 0.71431931 | 0.47502975 | NA |
| AC138089.1 | 49.9531091 | 2.750503411 | 0.72905662 | 3.77268834 | 0.0001615 | 0.00185218 |
| OR2T5 | 2.07577841 | 2.548079593 | 1.13645663 | 2.24212656 | 0.02495319 | 0.08821769 |
| OR2G6 | 9.78848765 | 3.67126002 | 0.64185675 | 5.71974977 | 1.07E-08 | 7.33E-07 |
| AC098483.1 | 44.2596019 | 4.062469204 | 0.88158983 | 4.60811713 | 4.06E-06 | 9.99E-05 |
| OR2T29 | 18.6773747 | 4.554357004 | 1.16708058 | 3.90235007 | 9.53E-05 | 0.00121769 |
| OR2T34 | 0 | NA | NA | NA | NA | NA |
| OR2T10 | 117.244837 | 3.471693491 | 0.51546635 | 6.73505359 | 1.64E-11 | 3.04E-09 |
| OR2T11 | 5.17905231 | 3.821953409 | 0.84635291 | 4.51579165 | 6.31E-06 | 0.00014145 |
| OR2T35 | 11.0619543 | 3.53606974 | 0.69714875 | 5.07218832 | 3.93E-07 | 1.49E-05 |
| OR2T27 | 0 | NA | NA | NA | NA | NA |
| OR14I1 | 2.26561289 | 2.671786437 | 1.13660189 | 2.3506792 | 0.01873918 | 0.07160078 |
| LYPD8 | 118.584965 | 0.049236378 | 0.46453018 | 0.10599177 | 0.91558888 | 0.95882128 |
| SH3BP5L | 2635.8529 | -0.163996903 | 0.19387363 | -0.8458959 | 0.39761084 | 0.59247318 |
| ZNF672 | 2229.29296 | -0.287412596 | 0.11438922 | -2.5125846 | 0.01198504 | 0.05148699 |
| ZNF692 | 2827.27337 | -0.36599983 | 0.2521187 | -1.4516965 | 0.14658602 | 0.30955982 |
| AL672291.1 | 1.45807462 | 1.75196855 | 1.28426696 | 1.36417786 | 0.1725116 | 0.34559985 |
| PGBD2 | 423.385169 | 0.195395674 | 0.19976285 | 0.97813819 | 0.32800598 | 0.52568381 |
| FAM110C | 3763.81779 | -2.397100076 | 0.42098696 | -5.6940008 | 1.24E-08 | 8.40E-07 |
| AC079779.1 | 0.10721287 | -0.642905668 | 3.30558948 | -0.1944905 | 0.84579185 | NA |
| SH3YL1 | 5339.65901 | 0.507185577 | 0.27973577 | 1.81308804 | 0.06981824 | 0.1838144 |
| ACP1 | 4398.15544 | 0.056439806 | 0.13699344 | 0.41198911 | 0.6803474 | 0.8146636 |
| ALKAL2 | 491.435487 | -0.305396835 | 0.39240824 | -0.778263 | 0.43641396 | 0.62855623 |
| AC079779.3 | 0.91774093 | 3.296620246 | 1.32678289 | 2.48467195 | 0.01296709 | NA |
| AC079779.4 | 0.37509767 | 2.500457353 | 3.2139383 | 0.77800415 | 0.43656656 | NA |
| LINC01865 | 2.1451611 | 1.153588933 | 1.14808834 | 1.00479109 | 0.31499746 | 0.51209416 |
| AC105393.2 | 0.3350536 | -0.556688828 | 2.29550265 | -0.2425128 | 0.80838282 | NA |
| AC105393.1 | 0.07659028 | -0.642905668 | 3.30558948 | -0.1944905 | 0.84579185 | NA |

| | | | | | | | |
|---|---|---|---|---|---|---|---|
| LINC01874 | 172.36357 | | 3.023123924 | 0.61249456 | 4.93575638 | 7.98E-07 | 2.68E-05 |
| AC093326.2 | 0.09989236 | | -0.642905668 | 3.30558948 | -0.1944905 | 0.84579185 | NA |
| LINC01875 | 0 | NA | | NA | NA | NA | NA |
| AC093326.1 | 7.13938804 | | -0.604848963 | 1.11199174 | -0.543933 | 0.5864876 | 0.74768058 |
| TMEM18 | 1308.82813 | | 0.297328972 | 0.1560286 | 1.90560563 | 0.05670139 | 0.15905049 |
| AC092159.2 | 247.6229 | | 0.431932108 | 0.22212623 | 1.94453448 | 0.05183103 | 0.14953876 |
| AC092159.3 | 2.26926116 | | 2.139814871 | 0.95576524 | 2.23884986 | 0.02516569 | 0.08880872 |
| AC092159.1 | 0.85548273 | | 1.428745863 | 1.7019839 | 0.8394591 | 0.40121173 | NA |
| AC116609.3 | 2.1374904 | | 1.398716494 | 1.85990664 | 0.75203586 | 0.45202949 | 0.64175825 |
| AC116609.2 | 0.50529136 | | 0.831926949 | 2.12703301 | 0.39112085 | 0.69570791 | NA |
| AC116609.1 | 2.53531319 | | 1.627901365 | 1.43069527 | 1.13783934 | 0.25518757 | 0.44784349 |
| LINC01115 | 5.67631763 | | 0.220036575 | 0.7878012 | 0.2793047 | 0.780011 | 0.87953862 |
| LINC01939 | 2.69044534 | | -0.195190565 | 1.00307991 | -0.1945912 | 0.84571296 | 0.91863438 |
| AC113607.1 | 0.24089599 | | 0.077994266 | 3.07420509 | 0.02537055 | 0.9797594 | NA |
| AC116614.1 | 38.0400902 | | -1.56928212 | 0.6202623 | -2.5300298 | 0.01140528 | 0.04971854 |
| SNTG2 | 22.5605207 | | -0.661617587 | 0.4782152 | -1.3835143 | 0.16650721 | 0.33728646 |
| AC114808.1 | 0.16421342 | | 1.36803136 | 3.28033652 | 0.41703994 | 0.67664919 | NA |
| AC114808.2 | 0.12059279 | | 0.02007176 | 3.29738704 | 0.00608717 | 0.99514317 | NA |
| AC108462.1 | 0.10721287 | | -0.642905668 | 3.30558948 | -0.1944905 | 0.84579185 | NA |
| TPO | 3.14478991 | | -0.023680321 | 1.20801157 | -0.0196027 | 0.98436029 | 0.99326005 |
| AC141930.1 | 0.18028734 | | 0.18657164 | 3.29377195 | 0.05664376 | 0.95482897 | NA |
| AC141930.3 | 0.97976508 | | -2.413210334 | 1.47406839 | -1.6371088 | 0.10160774 | NA |
| AC141930.2 | 0.09635527 | | 0.547239394 | 3.29377202 | 0.16614368 | 0.86804388 | NA |
| AC144450.1 | 0.17391096 | | -1.003574173 | 3.30558948 | -0.3035992 | 0.7614333 | NA |
| PXDN | 2045.59154 | | -1.089130394 | 0.34879684 | -3.1225351 | 0.00179301 | 0.01240635 |
| MYT1L | 5.35589782 | | -1.935344743 | 1.06540026 | -1.8165424 | 0.06928721 | 0.18281726 |
| AC093390.2 | 1.18692323 | | 0.982807758 | 2.22437127 | 0.4418362 | 0.65860774 | 0.79946078 |
| AC093390.1 | 3.38254929 | | 0.203401647 | 1.13072559 | 0.17988595 | 0.85724211 | 0.92531618 |
| MYT1L-AS1 | 0 | NA | | NA | NA | NA | NA |
| AC018685.2 | 0.16786415 | | -1.003574173 | 3.30558948 | -0.3035992 | 0.7614333 | NA |
| AC018685.1 | 0 | NA | | NA | NA | NA | NA |
| AC011995.2 | 0 | NA | | NA | NA | NA | NA |
| AC011995.1 | 0 | NA | | NA | NA | NA | NA |
| LINC01250 | 1.77133115 | | 0.678474859 | 0.89877335 | 0.75488983 | 0.45031509 | 0.64018594 |
| AC019118.2 | 0.43480817 | | 2.041214287 | 2.05017235 | 0.99563058 | 0.31942967 | NA |
| AC019118.1 | 3.86456813 | | -0.918276158 | 0.8077918 | -1.1367733 | 0.25563306 | 0.44823751 |
| EIPR1 | 1303.32963 | | 0.049678832 | 0.13437472 | 0.3697037 | 0.71160327 | 0.83606132 |
| TRAPPC12 | 2210.39329 | | 0.082643362 | 0.10883039 | 0.75937762 | 0.4476267 | 0.63783587 |
| TRAPPC12-A! | 70.0876559 | | 0.398614665 | 0.37478734 | 1.06357558 | 0.28752101 | 0.48378822 |
| AC114810.1 | 45.0399553 | | 0.506991672 | 0.35641776 | 1.42246468 | 0.1548914 | 0.32100645 |
| ADI1 | 12616.3993 | | 0.904126656 | 0.38476277 | 2.34982884 | 0.01878205 | 0.07170733 |
| AC231981.1 | 1410.34703 | | 1.127628823 | 0.45597645 | 2.47299796 | 0.01339849 | 0.05594018 |
| AC108488.1 | 265.781639 | | 0.109297974 | 0.1774118 | 0.61606935 | 0.53784875 | 0.71139447 |
| AC108488.2 | 247.51615 | | -0.521503483 | 0.21219693 | -2.4576392 | 0.01398536 | 0.05785911 |
| RNASEH1 | 522.610626 | | -0.352727251 | 0.25075023 | -1.4066876 | 0.15952003 | 0.32726678 |
| RNASEH1-AS | 247.279385 | | 0.572942529 | 0.18031464 | 3.17745983 | 0.00148571 | 0.01072495 |
| RPS7 | 22219.9844 | | -0.291664467 | 0.21083642 | -1.3833686 | 0.16655189 | 0.33732514 |
| COLEC11 | 1065.31711 | | 1.803579055 | 0.38284505 | 4.71098965 | 2.47E-06 | 6.73E-05 |
| AC010907.2 | 13.895596 | | 1.34556368 | 0.64211577 | 2.09551571 | 0.03612517 | 0.11513691 |
| ALLC | 121.762869 | | 2.456384214 | 0.64027481 | 3.83645303 | 0.00012482 | 0.00150577 |
| AC010907.1 | 13.8293741 | | 2.285874324 | 0.56425353 | 4.05114756 | 5.10E-05 | 0.00075506 |
| DCDC2C | 4.59395889 | | 0.318434248 | 0.8768692 | 0.36314909 | 0.71649352 | 0.83868671 |
| LINC01304 | 0 | NA | | NA | NA | NA | NA |
| AC012445.1 | 0.03392908 | | 0.380740593 | 3.29738709 | 0.11546736 | 0.90807469 | NA |
| LINC01249 | 0 | NA | | NA | NA | NA | NA |
| AC107057.1 | 27.0249032 | | 2.547405881 | 0.73144531 | 3.48270178 | 0.00049638 | 0.00450123 |
| LINC01248 | 0 | NA | | NA | NA | NA | NA |
| SOX11 | 3.65034233 | | 0.168404359 | 0.67551703 | 0.24929698 | 0.80313106 | 0.89318485 |
| AC010729.1 | 0.06785815 | | 0.795868262 | 3.28898349 | 0.24198001 | 0.80879565 | NA |
| LINC01810 | 0 | NA | | NA | NA | NA | NA |
| AC017053.1 | 0 | NA | | NA | NA | NA | NA |
| LINC01247 | 0 | NA | | NA | NA | NA | NA |
| LINC01824 | 0 | NA | | NA | NA | NA | NA |
| MIR7515HG | 0.91908334 | | -3.106617151 | 3.25689602 | -0.9538583 | 0.3401554 | NA |
| LINC00487 | 14.1832568 | | -0.158937737 | 0.59554983 | -0.2668756 | 0.78956492 | 0.88561578 |
| NRIR | 23.3463523 | | -0.304183987 | 0.65716323 | -0.4628743 | 0.64345445 | 0.78927779 |
| CMPK2 | 210.277042 | | 0.287003012 | 0.36730523 | 0.7813747 | 0.43458215 | 0.62657677 |
| RSAD2 | 262.382187 | | -0.236899656 | 0.61476951 | -0.3853471 | 0.69998026 | 0.82792844 |
| AC017076.1 | 17.9386479 | | -0.316460204 | 0.6020678 | -0.5256222 | 0.59915073 | 0.7570849 |
| RNF144A | 961.206316 | | 0.626215278 | 0.23901184 | 2.62001784 | 0.00879252 | 0.04087776 |
| AC068481.1 | 3.08491417 | | 0.906391421 | 0.65760911 | 1.37831337 | 0.16810656 | 0.33943108 |
| AC010904.2 | 8.06582135 | | 0.057094667 | 0.57920458 | 0.09857427 | 0.9214763 | 0.96166416 |
| AC092580.2 | 0.15543392 | | 1.368031615 | 3.28033652 | 0.41704002 | 0.67664913 | NA |
| LINC01871 | 13.4787623 | | -1.599664642 | 0.8160637 | -1.9602203 | 0.04997005 | 0.14561942 |
| AC007463.1 | 0.62037336 | | -0.012776959 | 1.84111974 | -0.0069398 | 0.9944629 | NA |
| LINC00298 | 1.16345061 | | 1.636889385 | 1.26422189 | 1.29478013 | 0.19539609 | 0.37552619 |
| LINC00299 | 10.181101 | | 0.087178907 | 0.6187894 | 0.14088623 | 0.88795982 | 0.94202528 |
| U91324.1 | 0.13037664 | | 0.02007176 | 3.29738704 | 0.00608717 | 0.99514317 | NA |
| AC011747.2 | 4.09010299 | | 0.53926054 | 0.86168488 | 0.62582105 | 0.53143232 | 0.7057751 |
| LINC01814 | 44.077583 | | 0.953080777 | 0.48973019 | 1.94613441 | 0.0516386 | 0.14919577 |
| AC011747.1 | 3.71332686 | | 1.400534566 | 1.40883403 | 0.99410899 | 0.32016981 | 0.51758692 |
| ID2-AS1 | 42.1088528 | | 1.317879939 | 0.40778011 | 3.23183971 | 0.00122996 | 0.00923258 |
| ID2 | 5407.75434 | | 0.057609081 | 0.14960057 | 0.38508596 | 0.70017373 | 0.82803754 |
| KIDINS220 | 3665.66046 | | 0.432002493 | 0.09242641 | 4.67401558 | 2.95E-06 | 7.74E-05 |
| MBOAT2 | 1639.09376 | | 0.213676972 | 0.19222545 | 1.11159563 | 0.26631205 | 0.46047137 |
| AC093904.4 | 58.0309077 | | 1.725932407 | 0.53182307 | 3.24531315 | 0.00117322 | 0.00889781 |
| AC093904.3 | 8.68144574 | | 0.763034099 | 0.63508611 | 1.20146557 | 0.22957065 | 0.41704175 |
| AC093904.1 | 0 | NA | | NA | NA | NA | NA |
| ASAP2 | 3566.94902 | | 0.419427666 | 0.14016654 | 2.99235228 | 0.00276837 | 0.01727854 |
| ITGB1BP1 | 6448.109 | | 0.075203726 | 0.20979763 | 0.35845841 | 0.72000029 | 0.84070901 |
| CPSF3 | 1263.06029 | | -0.224866972 | 0.13052862 | -1.7227407 | 0.08493542 | 0.21125703 |
| IAH1 | 2633.94963 | | -0.443491107 | 0.10810467 | -4.1024231 | 4.09E-05 | 0.0006327 |
| ADAM17 | 1594.50097 | | -0.584765555 | 0.18240255 | -3.2059067 | 0.00134638 | 0.00994787 |
| AC080162.1 | 12.4295795 | | 0.643035102 | 0.4286317 | 1.50020427 | 0.1335615 | 0.29035587 |
| AC073195.1 | 43.1189053 | | 0.411795894 | 0.3108738 | 1.32464006 | 0.18529056 | 0.36268231 |
| YWHAQ | 11410.6653 | | -0.354233856 | 0.20090661 | -1.7631767 | 0.07787069 | 0.19867404 |
| AC082651.1 | 0.34423316 | | -0.279239303 | 2.90031402 | -0.096279 | 0.923299 | NA |
| AC082651.4 | 0 | NA | | NA | NA | NA | NA |
| AC082651.3 | 1.53555608 | | 0.615138991 | 1.03803142 | 0.59260151 | 0.55344787 | 0.72299723 |
| TAF1B | 650.543168 | | 0.182740191 | 0.19646606 | 0.93013618 | 0.35230058 | 0.54985388 |
| AC010969.2 | 331.287224 | | -0.023949904 | 0.21963088 | -0.1090462 | 0.91316588 | 0.95750066 |
| GRHL1 | 572.42 | | -0.286740466 | 0.25396657 | -1.1290481 | 0.25887755 | 0.45167971 |
| AC010969.1 | 1.13116507 | | 1.30663803 | 1.445938 | 0.90366117 | 0.36617509 | 0.56326414 |
| AC104794.3 | 0.50692592 | | 2.936710059 | 2.57104369 | 1.14222487 | 0.25336056 | NA |
| AC104794.2 | 381.407714 | | -0.561234771 | 0.30605867 | -1.833749 | 0.06669127 | 0.17796486 |
| KLF11 | 2437.334 | | 0.176402758 | 0.14157373 | 1.24601333 | 0.2127595 | 0.39661163 |
| AC104794.4 | 819.728807 | | -0.203553441 | 0.15157141 | -1.3429541 | 0.17928684 | 0.35488621 |
| CYS1 | 15302.1842 | | 0.478049099 | 0.29945209 | 1.59641264 | 0.1103967 | 0.25449125 |
| AC104794.1 | 5.0049749 | | 0.920137321 | 0.54826388 | 1.67827457 | 0.09329351 | 0.22634556 |
| RRM2 | 148.511278 | | -0.445726433 | 0.58839114 | -0.7575342 | 0.44872986 | 0.63886225 |
| AC007240.1 | 70.9785409 | | -0.463594057 | 0.61397296 | -0.7550724 | 0.45020552 | 0.64007469 |
| C2orf48 | 11.5631535 | | 0.476974798 | 0.70435566 | 0.67717891 | 0.49829245 | 0.67987397 |
| HPCAL1 | 3739.89475 | | 0.107767934 | 0.17230729 | 0.62544038 | 0.53168207 | 0.70599237 |

| | | | | | | |
|---|---|---|---|---|---|---|
| ODC1 | 6568.31976 | -0.145722475 | 0.1946418 | -0.74867 | 0.45405614 | 0.64372316 |
| AC007249.2 | 51.3967674 | -0.16146487 | 0.41040284 | -0.3934302 | 0.69400177 | 0.82380494 |
| AC007249.1 | 17.5961106 | 1.273528484 | 0.46860645 | 2.717693 | 0.00657388 | 0.03295689 |
| NOL10 | 1221.07553 | -0.204732215 | 0.18566306 | -1.1027084 | 0.27015381 | 0.46449207 |
| AC007314.1 | 285.506571 | -0.209251549 | 0.19612484 | -1.0669304 | 0.28600328 | 0.48203152 |
| RN7SL832P | 39.3638887 | 1.144062327 | 0.40640721 | 2.81506406 | 0.00487675 | 0.02638489 |
| AC092687.2 | 0 | NA | NA | NA | NA | NA |
| ATP6V1C2 | 8429.3022 | -0.206712465 | 0.20047638 | -1.0311063 | 0.30249096 | 0.49938563 |
| AC092687.3 | 158.315023 | 0.160768251 | 0.38129361 | 0.42163899 | 0.67328855 | 0.80949341 |
| PDIA6 | 14419.1415 | -0.266357899 | 0.20590287 | -1.2936094 | 0.19580037 | 0.37610915 |
| AC092687.1 | 0.39252305 | 1.874558999 | 2.15790216 | 0.86869509 | 0.38501393 | NA |
| LINC01954 | 20.4941118 | 3.046435586 | 0.70432761 | 4.32531047 | 1.52E-05 | 0.00028731 |
| KCNF1 | 25.7967194 | 1.418522494 | 0.64330578 | 2.20505169 | 0.02745047 | 0.09467288 |
| AC062028.1 | 5.30817126 | -0.144898633 | 0.59844584 | -0.2421249 | 0.80868339 | 0.89660916 |
| C2orf50 | 18.4561747 | -0.496126399 | 0.52263106 | -0.9492861 | 0.34247512 | 0.54018364 |
| PQLC3 | 1301.10799 | -0.026083477 | 0.18010761 | -0.1448216 | 0.88485171 | 0.94039898 |
| ROCK2 | 2211.36767 | -0.1372536 | 0.16558245 | -0.8289139 | 0.40715311 | 0.6017494 |
| AC099344.1 | 0 | NA | NA | NA | NA | NA |
| AC099344.2 | 0.35887416 | -0.278661391 | 2.87409007 | -0.0969564 | 0.92276103 | NA |
| LINC00570 | 3.60255569 | 0.9946035 | 0.86683505 | 1.14739649 | 0.25121779 | 0.44354643 |
| E2F6 | 740.911774 | -0.514856934 | 0.27018096 | -1.9056004 | 0.05670207 | 0.15905049 |
| GREB1 | 249.720515 | 2.437545958 | 0.3451424 | 7.06243562 | 1.64E-12 | 4.15E-10 |
| NTSR2 | 0.08666371 | -0.642905668 | 3.30558948 | -0.1944905 | 0.84579185 | NA |
| AC106875.2 | 0 | NA | NA | NA | NA | NA |
| LPIN1 | 2246.33782 | -0.639800853 | 0.32147009 | -1.9902345 | 0.04656511 | 0.13827951 |
| AC012456.1 | 0 | NA | NA | NA | NA | NA |
| AC012456.2 | 0.95513307 | 0.537672586 | 1.40834554 | 0.38177604 | 0.70262749 | NA |
| MIR3681HG | 11.0060596 | 0.11071802 | 0.49663856 | 0.2229348 | 0.82358625 | 0.90564302 |
| AC096559.1 | 0 | NA | NA | NA | NA | NA |
| AC009486.1 | 10.8703036 | 0.09797602 | 0.59473479 | 0.16473901 | 0.86914941 | 0.93191744 |
| TRIB2 | 5261.48692 | -0.502211551 | 0.24118132 | -2.0822987 | 0.0373152 | 0.11766133 |
| AC073062.1 | 0.10178723 | 1.087385526 | 3.28420134 | 0.33109588 | 0.74057207 | NA |
| LINC00276 | 1.00050947 | -0.552453416 | 1.35618242 | -0.4073592 | 0.68374417 | NA |
| AC016730.1 | 0.13390214 | 1.04981649 | 3.28477129 | 0.31960109 | 0.74927075 | NA |
| AC011897.1 | 0.80641031 | -2.90029717 | 1.78266856 | -1.6269413 | 0.10374956 | NA |
| FAM84A | 1235.76077 | -2.071852185 | 0.32527721 | -6.3694969 | 1.90E-10 | 2.49E-08 |
| AC068286.1 | 0.26351882 | -0.178709043 | 2.9796477 | -0.0599766 | 0.9521743 | NA |
| NBAS | 1656.88911 | 0.196289678 | 0.13945954 | 1.40750266 | 0.15927839 | 0.3269888 |
| AC008278.2 | 0.48875674 | 1.716208818 | 2.11232371 | 0.81247434 | 0.4165195 | NA |
| DDX1 | 5190.81827 | 0.229992568 | 0.13428345 | 1.7127395 | 0.08676048 | 0.2146223 |
| AC008278.1 | 1.17084131 | 0.889086282 | 1.14976979 | 0.7732733 | 0.43936065 | 0.63084261 |
| LINC01804 | 0 | NA | NA | NA | NA | NA |
| AC113608.2 | 0.09635527 | 0.547239394 | 3.29377202 | 0.16614368 | 0.86804388 | NA |
| MYCNUT | 4.0761098 | 5.544398296 | 1.11579504 | 4.96901143 | 6.73E-07 | 2.32E-05 |
| MYCNOS | 128.214435 | 5.241348995 | 0.66104531 | 7.92888006 | 2.21E-15 | 1.46E-12 |
| MYCN | 368.578644 | 4.098941908 | 0.43368155 | 9.4515017 | 3.34E-21 | 6.78E-18 |
| AC010145.1 | 0 | NA | NA | NA | NA | NA |
| GACAT3 | 0.68547691 | 2.845718449 | 1.46736565 | 1.93933833 | 0.05246015 | NA |
| AC130710.1 | 2.74067987 | 2.403710565 | 0.9643032 | 2.49269167 | 0.01267789 | 0.0537408 |
| AC010745.1 | 0 | NA | NA | NA | NA | NA |
| AC010745.2 | 0 | NA | NA | NA | NA | NA |
| AC010745.3 | 0 | NA | NA | NA | NA | NA |
| AC010745.5 | 0 | NA | NA | NA | NA | NA |
| AC010745.4 | 0 | NA | NA | NA | NA | NA |
| AC010880.1 | 0 | NA | NA | NA | NA | NA |
| AC104623.1 | 0 | NA | NA | NA | NA | NA |
| FAM49A | 288.537147 | -0.478187677 | 0.35114652 | -1.3617896 | 0.17326429 | 0.34689693 |
| AC008164.1 | 0.09635527 | 0.547239394 | 3.29377202 | 0.16614368 | 0.86804388 | NA |
| LINC01866 | 0.40449689 | 1.869563418 | 1.79216876 | 1.04318492 | 0.29686266 | NA |
| RAD51AP2 | 6.32252975 | 0.27312384 | 0.62524922 | 0.43682396 | 0.66223902 | 0.80128673 |
| VSNL1 | 62.4679932 | 2.069445576 | 0.46170783 | 4.48215396 | 7.39E-06 | 0.00016064 |
| SMC6 | 988.388862 | 0.041041821 | 0.11466555 | 0.35792634 | 0.72039844 | 0.84093782 |
| GEN1 | 414.053218 | 0.16320697 | 0.4456201 | 0.36624688 | 0.71418086 | 0.83763364 |
| MSGN1 | 0 | NA | NA | NA | NA | NA |
| KCNS3 | 523.684587 | -0.610327261 | 0.28891511 | -2.1124796 | 0.03464534 | 0.11185892 |
| AC079148.1 | 24.007367 | -0.545135675 | 0.43171996 | -1.2627067 | 0.20669461 | 0.38929725 |
| RDH14 | 1277.19093 | -0.374820695 | 0.14860556 | -2.5222521 | 0.01166061 | 0.05051591 |
| NT5C1B-RDH | 453.632478 | -0.202684843 | 0.15199159 | -1.3335267 | 0.18235902 | 0.35902698 |
| NT5C1B | 43.4394489 | 0.098468777 | 0.35173367 | 0.27995266 | 0.77951383 | 0.87916597 |
| AC106053.1 | 0.63591652 | -1.507667339 | 2.21581058 | -0.6804135 | 0.4962427 | NA |
| LINC01376 | 37.8066554 | 0.540750003 | 0.29577383 | 1.82825504 | 0.06751128 | 0.17942679 |
| OSR1 | 219.122906 | -0.895635196 | 0.4539859 | -1.972826 | 0.04851539 | 0.14257783 |
| AC010096.1 | 0.17648264 | -1.003574173 | 3.30558948 | -0.3035992 | 0.7614333 | NA |
| LINC01808 | 1.10773833 | 1.056746893 | 1.35243778 | 0.78136452 | 0.43458813 | 0.62657677 |
| AC019055.1 | 2.08409223 | 1.096189747 | 0.97972857 | 1.11887086 | 0.26319523 | 0.4569097 |
| LINC00954 | 51.7279824 | 0.234431989 | 0.40142167 | 0.58400432 | 0.55921741 | 0.72772184 |
| TTC32 | 559.015165 | -0.327244711 | 0.21408199 | -1.5285952 | 0.12636482 | 0.2795805 |
| AC013400.1 | 7.57274237 | 1.534786179 | 0.63639483 | 2.41168864 | 0.01587884 | 0.06345278 |
| WDR35 | 1135.37527 | 0.282841417 | 0.13084864 | 2.16159234 | 0.03064961 | 0.1024373 |
| AC079145.1 | 23.5443605 | 0.990799949 | 0.42321953 | 2.3411017 | 0.01922693 | 0.07277366 |
| MATN3 | 235.519087 | -0.838894682 | 0.36437418 | -2.3022891 | 0.02131888 | 0.07861268 |
| LAPTM4A | 24574.1591 | -0.546543052 | 0.17072429 | -3.2013197 | 0.001368 | 0.01007093 |
| AC098828.2 | 2.91169329 | 0.661487709 | 0.82809336 | 0.79880813 | 0.42440168 | 0.61688194 |
| AC098828.3 | 4.13960283 | 2.752085305 | 0.97288738 | 2.82878098 | 0.00467257 | 0.0255634 |
| SDC1 | 21426.2259 | 0.241709761 | 0.30860737 | 0.78322745 | 0.43349356 | 0.62557806 |
| PUM2 | 4954.20378 | 0.039681758 | 0.14275155 | 0.27797778 | 0.78102941 | 0.88016021 |
| RHOB | 22237.2195 | -0.826059711 | 0.32641107 | -2.5307343 | 0.0113824 | 0.04965025 |
| AC023137.1 | 2.16316915 | -1.790754147 | 1.07604933 | -1.6641934 | 0.0960738 | 0.23118938 |
| AC012065.1 | 0.88889771 | -3.045038281 | 2.19108886 | -1.3897375 | 0.16460861 | NA |
| HS1BP3 | 2674.65718 | 0.34708629 | 0.16717536 | 2.07618086 | 0.03787723 | 0.11894538 |
| AC012065.3 | 42.3843787 | -0.237773081 | 0.34895839 | -0.6813795 | 0.49563141 | 0.67769982 |
| GDF7 | 1177.56368 | 0.131126621 | 0.3921431 | 0.33438462 | 0.73808935 | 0.85298022 |
| AC012065.4 | 507.848307 | -0.386727585 | 0.42640076 | -0.906958 | 0.36442901 | 0.56156097 |
| LDAH | 1031.11329 | 0.611731712 | 0.19894368 | 3.07489891 | 0.00210574 | 0.01400463 |
| AC012361.1 | 0 | NA | NA | NA | NA | NA |
| AC115619.1 | 0.38679398 | 0.760854655 | 2.84844468 | 0.26711232 | 0.78938269 | NA |
| APOB | 110.569977 | -0.90214906 | 0.68127109 | -1.3242145 | 0.18543181 | 0.36278024 |
| TDRD15 | 1.50565042 | 0.115706369 | 1.20122951 | 0.09632328 | 0.92326382 | 0.96257727 |
| AC018742.1 | 8.21647664 | 0.363281408 | 0.6424682 | 0.56544652 | 0.57177009 | 0.73660547 |
| AC011752.1 | 0.11786115 | 0.02007176 | 3.29738704 | 0.00608717 | 0.99514317 | NA |
| AC009411.2 | 0 | NA | NA | NA | NA | NA |
| AC009411.1 | 0 | NA | NA | NA | NA | NA |
| LINC01822 | 0.31730519 | 0.328539062 | 2.2746369 | 0.14443583 | 0.88515633 | NA |
| LINC01830 | 0 | NA | NA | NA | NA | NA |
| AC068490.1 | 0 | NA | NA | NA | NA | NA |
| AC096570.1 | 0.09635527 | 0.547239394 | 3.29377202 | 0.16614368 | 0.86804388 | NA |
| LINC01884 | 0 | NA | NA | NA | NA | NA |
| AC018467.1 | 73.8563376 | 1.564006896 | 0.49392815 | 3.16646639 | 0.00154303 | 0.01104499 |
| AC012506.1 | 0.61607397 | 1.316884971 | 1.5113015 | 0.87135821 | 0.38355859 | NA |
| AC012506.3 | 1.68328552 | 1.758209384 | 1.08219023 | 1.62467683 | 0.10423144 | 0.2444706 |
| AC012506.2 | 10.1380284 | 1.637809874 | 0.86015506 | 1.90408678 | 0.05689888 | 0.15941602 |
| AC012506.4 | 1.01874137 | -1.709987276 | 1.63006319 | -1.0490313 | 0.29416372 | 0.49086233 |
| KLHL29 | 1094.96261 | -1.170536601 | 0.47839317 | -2.4468087 | 0.01441273 | 0.05915388 |

| | | | | | | |
|---|---|---|---|---|---|---|
| AC011239.1 | 4.16382438 | -1.235441054 | 1.08596155 | -1.1376471 | 0.25526785 | 0.44786515 |
| AC009242.1 | 77.2199722 | -0.795236369 | 0.48750959 | -1.631222 | 0.10284348 | 0.24207647 |
| ATAD2B | 629.886647 | -0.10854635 | 0.1237582 | -0.8770841 | 0.38044096 | 0.5765317 |
| UBXN2A | 904.73394 | -0.534889934 | 0.16792001 | -3.1853853 | 0.00144561 | 0.0105074 |
| MFSD2B | 310.032863 | -1.087121018 | 0.22976143 | -4.7315209 | 2.23E-06 | 6.20E-05 |
| WDCP | 364.15394 | 0.219493423 | 0.15646552 | 1.40282294 | 0.16066964 | 0.32905083 |
| FKBP1B | 390.697434 | -1.145605192 | 0.25201005 | -4.545871 | 5.47E-06 | 0.000126 |
| SF3B6 | 2626.21748 | -0.440728608 | 0.11273224 | -3.9095172 | 9.25E-05 | 0.00118904 |
| FAM228B | 1037.65779 | 0.266737863 | 0.20911511 | 1.27555519 | 0.2021128 | 0.38395754 |
| TP53I3 | 2646.79069 | 0.371124729 | 0.25324017 | 1.46550495 | 0.1427832 | 0.30408602 |
| PFN4 | 49.6972867 | 0.41845449 | 0.33621776 | 1.24459365 | 0.21328116 | 0.3973032 |
| AC008073.3 | 278.980446 | 0.057218812 | 0.30972023 | 0.18474354 | 0.85343019 | 0.9232762 |
| AC008073.2 | 11.0280942 | 0.408820227 | 0.45380611 | 0.90086981 | 0.36765755 | 0.5644908 |
| FAM228A | 28.6733942 | -0.448002351 | 0.52918698 | -0.8465861 | 0.39722585 | 0.59212807 |
| AC008073.1 | 2.61557996 | 0.333568628 | 0.7450953 | 0.44768586 | 0.65437993 | 0.79726477 |
| ITSN2 | 2563.33032 | -0.176947507 | 0.13125611 | -1.3481087 | 0.17762341 | 0.35275879 |
| AC009228.1 | 127.640467 | 0.225479054 | 0.22087614 | 1.02083937 | 0.30733055 | 0.50428141 |
| NCOA1 | 2585.56025 | 0.161016046 | 0.11905562 | 1.35244389 | 0.17623336 | 0.35087004 |
| PTRHD1 | 640.448697 | 0.121390113 | 0.18326728 | 0.66236655 | 0.50773634 | 0.68658055 |
| CENPO | 1990.27114 | -0.573064527 | 0.24043525 | -2.3834464 | 0.01715138 | 0.06705715 |
| ADCY3 | 2085.14922 | -0.546750513 | 0.30317115 | -1.8034385 | 0.07131938 | 0.18681754 |
| AC012073.1 | 152.165296 | -0.268031559 | 0.31812497 | -0.8425354 | 0.39948831 | 0.59426337 |
| DNAJC27 | 525.492925 | 0.234253666 | 0.08584988 | 2.72864275 | 0.00635956 | 0.03216369 |
| DNAJC27-AS | 97.4869509 | 1.081361998 | 0.4101091 | 2.63676662 | 0.00837004 | 0.03937173 |
| EFR3B | 120.526421 | 0.568010688 | 0.43071985 | 1.3187474 | 0.18725357 | 0.36520056 |
| POMC | 64.7128857 | -0.226018221 | 0.48567548 | -0.4653688 | 0.64166737 | 0.78769526 |
| LINC01381 | 0.17507102 | 0.435199075 | 3.28898345 | 0.13232024 | 0.894731 | NA |
| DNMT3A | 1769.35325 | 0.166201054 | 0.1622491 | 1.02435732 | 0.30566653 | 0.50270434 |
| AC012074.1 | 39.8196615 | -0.134479501 | 0.31308047 | -0.4295365 | 0.66753281 | 0.80542508 |
| DTNB | 774.642832 | -0.029400427 | 0.16100086 | -0.1826104 | 0.85510374 | 0.92410315 |
| AC104699.1 | 124.564334 | -0.178660411 | 0.21701976 | -0.8232449 | 0.41036875 | 0.60471773 |
| ASXL2 | 1564.68203 | -0.002499958 | 0.18269736 | -0.0136836 | 0.9890824 | 0.99508233 |
| KIF3C | 391.395023 | -0.713288576 | 0.16843106 | -4.2348994 | 2.29E-05 | 0.00039616 |
| RAB10 | 6816.20041 | -0.444517478 | 0.2527034 | -1.7590483 | 0.07856931 | 0.19992982 |
| GAREM2 | 289.386801 | -0.926061659 | 0.26943545 | -3.4370446 | 0.0005881 | 0.00514538 |
| HADHA | 16070.3593 | 0.301532999 | 0.22163302 | 1.36050574 | 0.17366994 | 0.34752438 |
| HADHB | 14291.1362 | 0.382340832 | 0.24406988 | 1.56652198 | 0.11722647 | 0.26561797 |
| ADGRF3 | 222.694367 | 1.59063775 | 0.36786826 | 4.32393309 | 1.53E-05 | 0.0002889 |
| SELENOI | 1701.74755 | -0.246767402 | 0.22820345 | -1.0813482 | 0.27954224 | 0.47514828 |
| DRC1 | 34.6561427 | -2.952599584 | 0.87918273 | -3.3583457 | 0.0007841 | 0.00646865 |
| OTOF | 5.69757095 | -1.712659925 | 0.74810844 | -2.2893204 | 0.02206074 | 0.08065185 |
| C2orf70 | 82.3025108 | -1.08132833 | 0.54513743 | -1.9835885 | 0.04730173 | 0.1398214 |
| CIB4 | 10.5668432 | -2.509355665 | 0.6777477 | -3.7024923 | 0.00021349 | 0.00230257 |
| AC015977.2 | 4.46898196 | -4.682818011 | 1.169042 | -4.0056884 | 6.18E-05 | 0.00086875 |
| KCNK3 | 1002.9912 | -1.90714715 | 0.44240638 | -4.3108491 | 1.63E-05 | 0.00030116 |
| SLC35F6 | 5255.22383 | -0.898026189 | 0.33613794 | -2.6716002 | 0.00754905 | 0.03661541 |
| CENPA | 223.328228 | -0.338794696 | 0.29442949 | -1.1506819 | 0.24986311 | 0.44194046 |
| DPYSL5 | 2.43031649 | 2.027061907 | 0.7860055 | 2.57894113 | 0.00991037 | 0.04478773 |
| AC013472.2 | 0.71247188 | 1.718469411 | 1.48113259 | 1.16024009 | 0.24595107 | NA |
| MAPRE3 | 2195.93135 | -0.461609166 | 0.19109279 | -2.4156284 | 0.01570808 | 0.06295467 |
| TMEM214 | 5904.39903 | -0.375050169 | 0.14858467 | -2.5241511 | 0.01159781 | 0.05033478 |
| AGBL5 | 3690.77503 | 0.17138709 | 0.12237038 | 1.40056025 | 0.16134561 | 0.32998604 |
| AGBL5-AS1 | 1.33811918 | 0.078188142 | 0.98756967 | 0.07917228 | 0.93689559 | 0.96969938 |
| AC013472.3 | 257.250469 | 0.825638817 | 0.20640921 | 4.00000961 | 6.33E-05 | 0.00088703 |
| AC013403.2 | 33.466783 | -0.091197928 | 0.22060151 | -0.4134057 | 0.67930938 | 0.81394894 |
| OST4 | 4586.93126 | -0.361491747 | 0.16486353 | -2.1926726 | 0.02833098 | 0.09697087 |
| EMILIN1 | 4463.63804 | -0.389457524 | 0.45654855 | -0.8530473 | 0.39363305 | 0.58894535 |
| KHK | 11093.2079 | 2.092919795 | 0.45427767 | 4.60713772 | 4.08E-06 | 0.00010024 |
| CGREF1 | 5840.15462 | 1.888069207 | 0.40464702 | 4.66596589 | 3.07E-06 | 7.98E-05 |
| ABHD1 | 207.260183 | 0.023085559 | 0.13614237 | 0.16956924 | 0.86534892 | 0.93001264 |
| PREB | 3738.09954 | -0.180745439 | 0.1258435 | -1.4362715 | 0.15092509 | 0.31566678 |
| PRR30 | 0 | NA | NA | NA | NA | NA |
| TCF23 | 7.04072579 | 0.94990849 | 0.53687677 | 1.76932313 | 0.07683996 | 0.19685902 |
| SLC5A6 | 2819.55897 | -0.015133016 | 0.10904739 | -0.1387747 | 0.88962821 | 0.94296756 |
| ATRAID | 4734.98137 | 0.064238689 | 0.12204503 | 0.52635234 | 0.59864342 | 0.75677743 |
| CAD | 1750.38933 | -0.103805984 | 0.1308677 | -0.7932132 | 0.42765363 | 0.62007072 |
| SLC30A3 | 10.8655177 | 0.034913658 | 0.61953595 | 0.05635453 | 0.95505938 | 0.97847297 |
| DNAJC5G | 0.48531344 | 2.176824213 | 3.2122032 | 0.67767326 | 0.49797889 | NA |
| TRIM54 | 72.7672275 | 0.317990065 | 0.53931266 | 0.58962099 | 0.55544478 | 0.72486671 |
| UCN | 92.7376428 | -1.509864326 | 0.38898446 | -3.8815544 | 0.00010379 | 0.00130518 |
| MPV17 | 2608.48619 | -0.262485715 | 0.16101275 | -1.6302169 | 0.10305566 | 0.24248931 |
| GTF3C2 | 3243.5051 | 0.130686809 | 0.15200258 | 0.85976706 | 0.38991746 | 0.58544231 |
| GTF3C2-AS1 | 697.833432 | 0.577823182 | 0.2315748 | 2.4951903 | 0.01258896 | 0.05346848 |
| AC074117.1 | 1216.67571 | -0.393169391 | 0.18643799 | -2.108848 | 0.0349577 | 0.1125836 |
| EIF2B4 | 2436.68774 | -0.393903038 | 0.1587834 | -2.480757 | 0.01311037 | 0.05505981 |
| SNX17 | 5831.69898 | 0.041563805 | 0.08023025 | 0.51805654 | 0.60441882 | 0.76144298 |
| ZNF513 | 2395.47999 | -0.456136672 | 0.15729403 | -2.8998982 | 0.00373284 | 0.02164357 |
| PPM1G | 4542.59989 | -0.513228142 | 0.1872561 | -2.7407819 | 0.00612932 | 0.03131759 |
| NRBP1 | 6647.87429 | -0.678523325 | 0.33903887 | -2.0013143 | 0.04535853 | 0.13562814 |
| KRTCAP3 | 803.965276 | -0.394791365 | 0.22096742 | -1.7866497 | 0.07399413 | 0.19195525 |
| IFT172 | 1847.58032 | 0.646541109 | 0.24881108 | 2.59852215 | 0.0093626 | 0.04287804 |
| FNDC4 | 930.159967 | -3.311119615 | 0.40685828 | -8.1382628 | 4.01E-16 | 3.11E-13 |
| GCKR | 89.7671836 | -3.201466647 | 0.33488108 | -9.5600105 | 1.18E-21 | 2.82E-18 |
| C2orf16 | 54.3527228 | 0.58748402 | 0.27966943 | 2.10063727 | 0.03567282 | 0.1141117 |
| ZNF512 | 1772.43316 | 0.030029779 | 0.18871826 | 0.15912492 | 0.87357045 | 0.93417196 |
| CCDC121 | 206.837978 | 0.720806216 | 0.23579043 | 3.05697823 | 0.00223581 | 0.01472279 |
| GPN1 | 2023.50293 | -0.378018331 | 0.21428602 | -1.764083 | 0.077718 | 0.19841454 |
| SUPT7L | 3556.49929 | -0.269448081 | 0.12277866 | -2.1945839 | 0.02819346 | 0.0965965 |
| SLC4A1AP | 1673.75878 | -0.013880482 | 0.13085937 | -0.1060718 | 0.91552542 | 0.95882128 |
| LINC01460 | 15.2219224 | 1.199251028 | 0.47061243 | 2.54827742 | 0.01082563 | 0.04777774 |
| MRPL33 | 3621.13073 | -0.341492769 | 0.1772501 | -1.9266154 | 0.05402758 | 0.1538958 |
| RBKS | 1938.56745 | 0.125376723 | 0.2757338 | 0.45470205 | 0.64932361 | 0.79367125 |
| BABAM2 | 4218.14554 | -0.178508103 | 0.12212344 | -1.4617022 | 0.14382284 | 0.30553519 |
| AC093690.1 | 35.4036471 | -0.257422452 | 0.23326281 | -1.1035726 | 0.26977856 | 0.46402671 |
| AC104695.2 | 57.115524 | 0.167037659 | 0.3689048 | 0.4527934 | 0.65069751 | 0.79476116 |
| FOSL2 | 12943.0968 | -1.083521351 | 0.30365048 | -3.5683176 | 0.00035928 | 0.0035012 |
| AC104695.3 | 64.4686788 | -1.132243964 | 0.40846347 | -2.7719589 | 0.00557201 | 0.02926386 |
| AC104695.1 | 0.26057467 | -1.364242679 | 3.30558948 | -0.4127078 | 0.67982068 | NA |
| PLB1 | 136.169215 | -0.246506797 | 0.47766456 | -0.5160667 | 0.60580779 | 0.76257346 |
| AC074011.1 | 6.19086288 | -0.503573389 | 0.68608438 | -0.7339817 | 0.46295989 | 0.65100676 |
| AC092164.1 | 50.3644515 | 0.091126983 | 0.30441307 | 0.29935306 | 0.76467068 | 0.86986609 |
| PPP1CB | 11656.557 | -0.561711086 | 0.18784988 | -2.9902126 | 0.00278783 | 0.0173555 |
| SPDYA | 1571.14773 | -0.010057109 | 0.22981472 | -0.0437618 | 0.96509426 | 0.98359669 |
| AC097724.1 | 2.09141498 | 0.090608227 | 1.05729461 | 0.08569818 | 0.93170635 | 0.96704554 |
| TRMT61B | 814.447235 | 0.061928153 | 0.19045734 | 0.32515499 | 0.74506377 | 0.85721343 |
| WDR43 | 1469.47229 | -0.670063885 | 0.39134271 | -1.7122176 | 0.08685657 | 0.21479358 |
| TOGARAM2 | 105.010029 | -0.425661617 | 0.36753507 | -1.1581524 | 0.24680185 | 0.43826026 |
| C2orf71 | 49.0076395 | 0.682357493 | 0.75001896 | 0.909787 | 0.36293485 | 0.56014234 |
| AC105398.1 | 0.83155549 | 2.623648607 | 1.50418034 | 1.74423807 | 0.08111759 | NA |
| CLIP4 | 1677.75851 | -0.72402244 | 0.18622552 | -3.8878798 | 0.00010112 | 0.00127651 |
| ALK | 14.5702117 | -1.82764409 | 0.70521805 | -2.5916014 | 0.00955304 | 0.04353095 |
| AC106870.2 | 0 | NA | NA | NA | NA | NA |
| AC106870.1 | 0 | NA | NA | NA | NA | NA |

| | | | | | | |
|---|---|---|---|---|---|---|
| AC016907.2 | 5.68499149 | 1.037903676 | 0.57574097 | 1.8027268 | 0.07143113 | 0.18701737 |
| YPEL5 | 6988.73729 | -0.415136121 | 0.12729467 | -3.2612216 | 0.00110933 | 0.00849393 |
| LBH | 3249.95726 | 0.538775094 | 0.27464191 | 1.96173666 | 0.04979315 | 0.14524035 |
| LINC01936 | 325.667219 | -0.192200456 | 0.59882564 | -0.3209623 | 0.74823896 | 0.85929193 |
| LCLAT1 | 601.494674 | 0.33807518 | 0.16433906 | 2.05718093 | 0.03966883 | 0.12308756 |
| CAPN13 | 534.200815 | 0.011116808 | 0.47599953 | 0.02335466 | 0.98136737 | 0.9915968 |
| GALNT14 | 4910.53861 | 0.423419943 | 0.30042155 | 1.40941935 | 0.15871121 | 0.3262066 |
| AC009305.1 | 1.32727914 | 1.077987081 | 1.22924036 | 0.87695386 | 0.3805117 | 0.57660576 |
| CAPN14 | 50.5497277 | -0.723043523 | 0.62522799 | -1.1564478 | 0.24749804 | 0.43905304 |
| EHD3 | 1739.2749 | 1.470136138 | 0.43499141 | 3.37969005 | 0.00072568 | 0.00608654 |
| XDH | 39.8408001 | -1.696520876 | 0.52552829 | -3.2282199 | 0.00124563 | 0.00933162 |
| SRD5A2 | 48.0110926 | 1.706621148 | 0.85838967 | 1.988166 | 0.04679333 | 0.1387697 |
| AL133247.1 | 0 | NA | NA | NA | NA | NA |
| LINC01946 | 0.03392908 | 0.380740593 | 3.29738709 | 0.11546736 | 0.90807469 | NA |
| AL121652.1 | 2.92532044 | -0.497734706 | 0.71700417 | -0.6941866 | 0.48756518 | 0.67138871 |
| MEMO1 | 955.084659 | -0.382605026 | 0.2016167 | -1.8976852 | 0.05773756 | 0.1611153 |
| DPY30 | 2385.18886 | -0.399782148 | 0.13714099 | -2.9151179 | 0.00355554 | 0.02081697 |
| AL121655.1 | 17.8765208 | 0.134075309 | 0.37290328 | 0.35954446 | 0.71918783 | 0.84021138 |
| SPAST | 680.777284 | -0.043272277 | 0.11994806 | -0.3607585 | 0.71828002 | 0.83985952 |
| AL121658.1 | 18.7351823 | -0.053253014 | 0.40492428 | -0.1315135 | 0.8953691 | 0.94625213 |
| SLC30A6 | 1412.34269 | 0.048692999 | 0.10988406 | 0.44313071 | 0.6576712 | 0.79917006 |
| NLRC4 | 112.117554 | -0.325790947 | 0.3009933 | -1.082386 | 0.27908103 | 0.47471035 |
| YIPF4 | 2787.63506 | 0.222156742 | 0.13147146 | 1.68977158 | 0.09107166 | 0.22249025 |
| AL133245.1 | 2.80863074 | 0.337621671 | 0.88468554 | 0.38162901 | 0.70273656 | 0.82987743 |
| BIRC6 | 3091.71616 | 0.128788767 | 0.08244365 | 1.5621429 | 0.11825433 | 0.26723512 |
| BIRC6-AS1 | 3.71234343 | 0.528984254 | 0.8044973 | 0.65753391 | 0.51083768 | 0.68920615 |
| TTC27 | 707.289779 | -0.103048663 | 0.19337904 | -0.5328843 | 0.59411364 | 0.75317421 |
| LINC00486 | 1.8100732 | 1.92952767 | 1.07485476 | 1.795152 | 0.07262949 | 0.18953385 |
| AL121656.1 | 0.17361919 | -1.003574173 | 3.30558948 | -0.3035992 | 0.7614333 | NA |
| LTBP1 | 2048.81278 | 0.707604851 | 0.5626895 | 1.25754052 | 0.20855798 | 0.39169675 |
| RASGRP3 | 933.53722 | 0.386299078 | 0.29920754 | 1.29107401 | 0.19667801 | 0.37724754 |
| AC020594.1 | 98.0933907 | 0.273889063 | 0.28794811 | 0.95117506 | 0.34151552 | 0.53949345 |
| FAM98A | 2543.9131 | 0.112136207 | 0.19284782 | 0.5814751 | 0.5609203 | 0.72882327 |
| LINC01317 | 9.80824391 | 1.511791294 | 0.50365683 | 3.0016297 | 0.00268539 | 0.01688451 |
| LINC01318 | 0 | NA | NA | NA | NA | NA |
| AC068599.1 | 3.89537212 | 0.619374415 | 0.71619269 | 0.86481532 | 0.38714016 | 0.5827655 |
| LINC01320 | 2753.71154 | -0.93381031 | 0.52026577 | -1.7948717 | 0.07267414 | 0.18963162 |
| AC073218.2 | 31.9825139 | -1.552113664 | 0.62647038 | -2.4775531 | 0.01322867 | 0.05543641 |
| AC073218.1 | 327.139489 | -1.164755667 | 0.5791141 | -2.0112715 | 0.04429679 | 0.13331504 |
| AC012593.1 | 5.2817912 | -0.475927123 | 0.98938735 | -0.4810321 | 0.63049365 | 0.78042234 |
| AC019064.1 | 0.80424259 | -2.178247293 | 1.90403256 | -1.1440179 | 0.25261622 | NA |
| AC009414.2 | 128.080908 | 0.04668036 | 0.23654637 | 0.19734126 | 0.84356049 | 0.9170421 |
| CRIM1 | 8683.68129 | 0.3628817 | 0.22647536 | 1.60230099 | 0.10908907 | 0.25245096 |
| AC007378.1 | 154.362592 | 2.050622034 | 0.36120376 | 5.67718907 | 1.37E-08 | 9.10E-07 |
| FEZ2 | 2432.33962 | -0.154720786 | 0.17334557 | -0.8925569 | 0.37209451 | 0.56863396 |
| VIT | 17.9067242 | 1.183001418 | 1.27171981 | 0.93023747 | 0.35224814 | 0.54985388 |
| STRN | 1597.08535 | -0.247094253 | 0.18798467 | -1.3144383 | 0.18869874 | 0.36687944 |
| HEATR5B | 1348.27997 | 0.121609015 | 0.16500837 | 0.73698695 | 0.46113028 | 0.64964235 |
| GPATCH11 | 503.961737 | 0.119473542 | 0.23761892 | 0.50279474 | 0.61510859 | 0.76990046 |
| EIF2AK2 | 2164.31255 | -0.013424536 | 0.10959099 | -0.1224967 | 0.90250565 | 0.95085997 |
| SULT6B1 | 143.797047 | -0.056105254 | 0.202987 | -0.2763983 | 0.78224219 | 0.88079261 |
| CEBPZOS | 3363.80978 | -0.240129111 | 0.1136649 | -2.1126056 | 0.03463454 | 0.11184335 |
| CEBPZ | 2933.29289 | -0.146455636 | 0.14243681 | -1.0282148 | 0.30384879 | 0.5007182 |
| NDUFAF7 | 1294.0375 | -0.264398037 | 0.12647625 | -2.0904956 | 0.03657331 | 0.1161274 |
| PRKD3 | 1660.31676 | -0.118712217 | 0.14107005 | -0.8415125 | 0.40006087 | 0.59481541 |
| AC007391.1 | 5.89780856 | -0.154971892 | 0.62536421 | -0.2478106 | 0.80428094 | 0.89390914 |
| QPCT | 852.427969 | -1.591484188 | 0.41532394 | -3.8319105 | 0.00012715 | 0.00152895 |
| AC006369.1 | 416.59537 | -0.157228558 | 0.26148209 | -0.6012976 | 0.54764177 | 0.71858058 |
| CDC42EP3 | 2858.04326 | -0.355542523 | 0.19840669 | -1.7919886 | 0.0731348 | 0.1902937 |
| AC006369.2 | 0 | NA | NA | NA | NA | NA |
| LINC00211 | 3.22305907 | 0.363078882 | 0.71661298 | 0.50665965 | 0.61239365 | 0.76779926 |
| RMDN2 | 363.491537 | 0.779888847 | 0.19097828 | 4.08365213 | 4.43E-05 | 0.00067382 |
| RMDN2-AS1 | 11.0451502 | 1.243969091 | 0.41040756 | 3.03105792 | 0.00243698 | 0.0156778 |
| CYP1B1 | 2663.93109 | 0.458512961 | 0.40630247 | 1.12850152 | 0.25910817 | 0.45187688 |
| CYP1B1-AS1 | 167.87496 | 1.505947355 | 0.36411097 | 4.13595714 | 3.53E-05 | 0.00056186 |
| AC009229.2 | 1.78715433 | -0.309243959 | 1.00649106 | -0.3072496 | 0.75865341 | 0.86635037 |
| AC009229.3 | 1.08989487 | 0.632987573 | 1.35304055 | 0.46782602 | 0.63990901 | 0.78705286 |
| AC009229.1 | 0.55244465 | -0.106760101 | 1.71426098 | -0.0622776 | 0.95034175 | NA |
| ATL2 | 2270.2382 | 0.314465822 | 0.15909925 | 1.97653866 | 0.0480938 | 0.14162295 |
| AC011247.1 | 19.0305879 | 0.781104268 | 0.55188473 | 1.41533952 | 0.15696897 | 0.32390157 |
| LINC01883 | 0 | NA | NA | NA | NA | NA |
| HNRNPLL | 1409.38956 | -0.042365946 | 0.1563305 | -0.2710024 | 0.78638917 | 0.88369898 |
| AC011247.2 | 6.6187014 | 0.025144908 | 0.54840408 | 0.04585106 | 0.96342896 | 0.98291994 |
| GALM | 5098.99091 | 1.638683727 | 0.47187612 | 3.47269896 | 0.00051525 | 0.00464049 |
| AC074366.1 | 0.15318056 | -1.003574173 | 3.30558948 | -0.3035992 | 0.7614333 | NA |
| SRSF7 | 5862.11241 | -0.121869393 | 0.11759491 | -1.0363492 | 0.30003925 | 0.4967375 |
| GEMIN6 | 539.89037 | 0.116247959 | 0.11667678 | 0.99632475 | 0.31909238 | 0.5164467 |
| DHX57 | 983.990256 | 0.187962716 | 0.14869028 | 1.26412239 | 0.2061861 | 0.38882079 |
| AC018693.1 | 7.06698794 | 1.076138826 | 0.63398688 | 1.69741497 | 0.08961823 | 0.21990879 |
| MORN2 | 1145.27852 | 0.221367222 | 0.12838605 | 1.72423107 | 0.08466614 | 0.21076114 |
| ARHGEF33 | 94.6246489 | 0.152227449 | 0.30298097 | 0.50243238 | 0.61536341 | 0.7700438 |
| AC019171.1 | 38.5115767 | 0.099837226 | 0.34451469 | 0.28979091 | 0.7719762 | 0.87421716 |
| SOS1 | 2160.80616 | 0.015722478 | 0.10291682 | 0.15276879 | 0.87858061 | 0.93684475 |
| CDKL4 | 10.9425681 | -0.076929507 | 0.60963979 | -0.1261885 | 0.89958275 | 0.94902072 |
| MAP4K3 | 2208.88645 | 0.421102712 | 0.13700853 | 3.07355105 | 0.00211527 | 0.01403972 |
| AC007388.1 | 623.11517 | 0.626328069 | 0.23409246 | 2.67555846 | 0.00746049 | 0.036311 |
| TMEM178A | 1083.50815 | 0.983059682 | 0.33304026 | 2.95177427 | 0.00315954 | 0.01912198 |
| THUMPD2 | 722.01956 | -0.419241234 | 0.12668406 | -3.3093448 | 0.00093515 | 0.00745685 |
| SLC8A1-AS1 | 248.923281 | 2.265769034 | 0.37196279 | 6.0913863 | 1.12E-09 | 1.14E-07 |
| SLC8A1 | 1310.07016 | 1.953521671 | 0.38914025 | 5.02009662 | 5.16E-07 | 1.86E-05 |
| LINC01794 | 0 | NA | NA | NA | NA | NA |
| LINC01913 | 0.15318056 | -1.003574173 | 3.30558948 | -0.3035992 | 0.7614333 | NA |
| LINC01914 | 21.1970464 | -0.971206713 | 0.70915553 | -1.3695257 | 0.17083501 | 0.34307316 |
| C2orf91 | 4.71328951 | 1.007970213 | 0.55761561 | 1.80764345 | 0.07066201 | 0.18551944 |
| AC013480.1 | 0.28738828 | 1.315010961 | 2.90677361 | 0.45239538 | 0.65098417 | NA |
| PKDCC | 2243.53426 | -0.346643307 | 0.31747171 | -1.0918872 | 0.27488266 | 0.47007807 |
| AC083949.1 | 16.1432111 | 1.352092285 | 0.4466377 | 3.02726861 | 0.00246775 | 0.01583705 |
| EML4 | 3167.58244 | 0.252582019 | 0.10054303 | 2.5121783 | 0.01199884 | 0.05152411 |
| COX7A2L | 5642.07277 | -0.080518676 | 0.22805106 | -0.353073 | 0.7240337 | 0.84369283 |
| KCNG3 | 18.4006007 | -0.892881296 | 0.75893925 | -1.1764859 | 0.23940078 | 0.42925128 |
| MTA3 | 1825.59805 | -0.05091447 | 0.17206372 | -0.2959047 | 0.76730284 | 0.87152635 |
| OXER1 | 684.615651 | 2.17543971 | 0.38344989 | 5.67333514 | 1.40E-08 | 9.23E-07 |
| HAAO | 1897.14686 | 2.350018566 | 0.31030531 | 7.57324647 | 3.64E-14 | 1.55E-11 |
| AC016735.1 | 2.54451854 | -0.674558191 | 0.82516619 | -0.8174816 | 0.41365325 | 0.60761193 |
| LINC01819 | 41.939505 | -0.679315229 | 0.62712975 | -1.0832132 | 0.2787138 | 0.4743261 |
| LINC02590 | 0 | NA | NA | NA | NA | NA |
| LINC02580 | 8.51073861 | 1.41238159 | 0.55626872 | 2.53902751 | 0.01111611 | 0.04872536 |
| ZFP36L2 | 14610.0203 | -1.105940329 | 0.23758034 | -4.6550162 | 3.24E-06 | 8.33E-05 |
| LINC01126 | 29.0159231 | 0.124251641 | 0.29054327 | 0.4276528 | 0.66890392 | 0.80661224 |
| AC010883.1 | 336.968107 | -0.436141496 | 0.16820881 | -2.5928576 | 0.00951822 | 0.04341729 |
| THADA | 1646.00389 | -0.162916021 | 0.14107527 | -1.1548163 | 0.24816567 | 0.43979958 |
| PLEKHH2 | 1652.81348 | 0.212798495 | 0.31557127 | 0.67432784 | 0.50010291 | 0.68130141 |
| C1GALT1C1L | 57.3304094 | 0.490992366 | 0.23945587 | 2.05045035 | 0.0403205 | 0.12451901 |

| | | | | | | |
|---|---|---|---|---|---|---|
| DYNC2LI1 | 1769.04541 | 0.222654307 | 0.26831436 | 0.82982629 | 0.406637 | 0.60140134 |
| ABCG5 | 32.1462739 | 0.596010141 | 0.32838957 | 1.81494844 | 0.06953183 | 0.1833236 |
| ABCG8 | 22.0254605 | 2.747490272 | 0.62863472 | 4.37056716 | 1.24E-05 | 0.00024369 |
| LRPPRC | 6581.70638 | 0.603613671 | 0.18243182 | 3.30870822 | 0.00093727 | 0.00747156 |
| AC019129.2 | 3.73191462 | 1.235206315 | 0.82154091 | 1.5035238 | 0.13270403 | 0.2890637 |
| PPM1B | 2598.39564 | -0.031573451 | 0.0902693 | -0.3497695 | 0.72651167 | 0.84540695 |
| SLC3A1 | 30357.1477 | 0.780227056 | 0.35614027 | 2.19078582 | 0.0284673 | 0.09723463 |
| PREPL | 16783.7885 | 0.526450411 | 0.28880421 | 1.82286268 | 0.06832419 | 0.1810217 |
| CAMKMT | 391.227754 | -0.009325009 | 0.2206894 | -0.042254 | 0.96629622 | 0.98410057 |
| LINC01833 | 87.4953432 | 0.022378039 | 0.66767812 | 0.03351621 | 0.97326294 | 0.98764503 |
| SIX3-AS1 | 1.73215344 | -1.903651293 | 1.34028802 | -1.42033 | 0.15551163 | 0.32176082 |
| SIX3 | 11.9414892 | -1.88237298 | 0.71123436 | -2.6466283 | 0.00812987 | 0.03858613 |
| AC012354.1 | 10.256092 | 2.203399955 | 0.6526119 | 3.37627916 | 0.00073473 | 0.00614102 |
| SIX2 | 21.5568478 | 0.045882812 | 0.52042157 | 0.0881647 | 0.92974577 | 0.96597439 |
| AC093702.1 | 0.84268756 | 0.257739847 | 1.90193579 | 0.13551448 | 0.89220511 | NA |
| AC093833.1 | 0.15543392 | 1.368031615 | 3.28033652 | 0.41704002 | 0.67664913 | NA |
| SRBD1 | 603.329858 | 0.447789796 | 0.14906496 | 3.00399097 | 0.00266463 | 0.01678203 |
| PRKCE | 1181.4611 | 1.143957658 | 0.14190123 | 8.06164726 | 7.53E-16 | 5.67E-13 |
| U51244.1 | 0.08666371 | -0.642905668 | 3.30558948 | -0.1944905 | 0.84579185 | NA |
| AC017006.2 | 0.08757577 | 0.547239394 | 3.29377202 | 0.16614368 | 0.86804388 | NA |
| AC017006.1 | 0.13390214 | 1.04981649 | 3.28477129 | 0.31960109 | 0.74927075 | NA |
| EPAS1 | 21033.5883 | 0.098744118 | 0.35091254 | 0.28139239 | 0.77840945 | 0.87861887 |
| LINC01820 | 0 | NA | NA | NA | NA | NA |
| LINC02583 | 0 | NA | NA | NA | NA | NA |
| TMEM247 | 0.15318056 | -1.003574173 | 3.30558948 | -0.3035992 | 0.7614333 | NA |
| ATP6V1E2 | 201.227322 | 0.45369818 | 0.2829405 | 1.60351091 | 0.1088219 | 0.25214248 |
| AC018682.2 | 0.62356192 | -0.006929622 | 1.4876035 | -0.0046582 | 0.99628327 | NA |
| RHOQ | 6266.47829 | -0.774595925 | 0.24493113 | -3.162505 | 0.00156418 | 0.01116304 |
| AC018682.1 | 77.627434 | -1.140169897 | 0.42115275 | -2.7072597 | 0.00678412 | 0.03374781 |
| PIGF | 2337.54685 | -0.539865877 | 0.21591224 | -2.500395 | 0.01240549 | 0.05289131 |
| CRIPT | 749.087034 | -0.064592469 | 0.17196426 | -0.3756157 | 0.70720265 | 0.83325953 |
| SOCS5 | 1252.11468 | -0.097772476 | 0.18582139 | -0.5261637 | 0.59877444 | 0.75677743 |
| AC016722.3 | 17.1408611 | 0.334060231 | 0.35775021 | 0.93378068 | 0.35041704 | 0.54813722 |
| LINC01118 | 7.68573163 | 0.333116299 | 0.64749609 | 0.51446843 | 0.60692452 | 0.76327544 |
| LINC01119 | 34.9052435 | -0.440909155 | 0.45336976 | -0.9725156 | 0.33079412 | 0.52867582 |
| AC016722.1 | 1.78677883 | -2.11253186 | 1.12406974 | -1.8793601 | 0.06019533 | 0.16590207 |
| AC016722.2 | 305.828842 | 0.304673638 | 0.36169974 | 0.84233856 | 0.39959847 | 0.59436019 |
| MCFD2 | 5713.75707 | 0.021438897 | 0.24882114 | 0.08616188 | 0.93133773 | 0.96690482 |
| TTC7A | 2385.44897 | 0.091530561 | 0.21525645 | 0.42521635 | 0.67067897 | 0.80768162 |
| AC093732.2 | 60.5552265 | 0.938070899 | 0.40461878 | 2.31840674 | 0.02042723 | 0.07608276 |
| AC093732.1 | 0.35996639 | -0.557452575 | 2.27507718 | -0.2450258 | 0.80643647 | NA |
| STPG4 | 2.87368279 | 0.571269529 | 0.70170099 | 0.81412103 | 0.41557562 | 0.60905514 |
| AC073283.3 | 2724.94457 | -0.250459561 | 0.18741914 | -1.3363606 | 0.18143143 | 0.35776837 |
| AC073283.1 | 2.24585869 | 0.438884333 | 0.74285362 | 0.59080864 | 0.55464865 | 0.72417116 |
| CALM2 | 17685.4066 | -0.368499926 | 0.16706902 | -2.2056748 | 0.02740678 | 0.09457705 |
| AC106869.1 | 117.60527 | 0.091018644 | 0.4653052 | 0.19561063 | 0.84491494 | 0.91817386 |
| AC073283.2 | 2.06393151 | 0.452800764 | 0.93419214 | 0.48469768 | 0.62789081 | 0.77859976 |
| EPCAM | 14956.4407 | -0.397668156 | 0.2039863 | -1.9494846 | 0.05123758 | 0.14826456 |
| MSH2 | 686.495285 | 0.258591796 | 0.13393055 | 1.93079021 | 0.053509 | 0.15284146 |
| KCNK12 | 6.12034212 | -0.769398176 | 0.73592956 | -1.045478 | 0.2958021 | 0.49266098 |
| AC138655.1 | 1.49855664 | 0.648535589 | 1.1924477 | 0.54386921 | 0.58653148 | 0.74768058 |
| MSH6 | 3591.70389 | -0.275085275 | 0.1026443 | -2.6799859 | 0.00736253 | 0.03595367 |
| FBXO11 | 3036.61672 | -0.194027884 | 0.10649987 | -1.8218603 | 0.06847619 | 0.18125325 |
| AC079807.1 | 83.098579 | 0.42850764 | 0.25834635 | 1.65865566 | 0.0971852 | 0.23283782 |
| AC092650.1 | 0.92803112 | 0.715307767 | 1.28858432 | 0.55511134 | 0.57881851 | NA |
| FOXN2 | 586.445306 | -0.042498668 | 0.18587345 | -0.228643 | 0.81914638 | 0.90289626 |
| AC093635.1 | 19.5409031 | 0.275038073 | 0.29943191 | 0.91853293 | 0.35833993 | 0.55597928 |
| PPP1R21 | 2679.83334 | 0.974864316 | 0.20679544 | 4.714148 | 2.43E-06 | 6.65E-05 |
| STON1 | 1292.13872 | 0.282605113 | 0.219332 | 1.28848098 | 0.19757859 | 0.37834197 |
| STON1-GTF2 | 191.085132 | 0.494914985 | 0.3613644 | 1.36957315 | 0.17082019 | 0.34307316 |
| GTF2A1L | 54.9347789 | -1.770840413 | 0.75935075 | -2.3320454 | 0.0196983 | 0.074038 |
| AC073082.1 | 20.5529751 | -0.675757574 | 0.64987444 | -1.0398279 | 0.29841986 | 0.49530098 |
| LHCGR | 11.3507105 | 2.78750774 | 0.55339318 | 5.03711978 | 4.73E-07 | 1.73E-05 |
| AC009975.1 | 4.07470981 | -1.775191498 | 0.98312587 | -1.8056605 | 0.07097139 | 0.18618341 |
| FSHR | 5.37007053 | -0.920243527 | 1.02873938 | -0.8945351 | 0.37103564 | 0.56783674 |
| AC009975.2 | 0.3490625 | 0.732866073 | 3.27839706 | 0.22354402 | 0.82311212 | NA |
| AC009971.1 | 0.10721287 | -0.642905668 | 3.30558948 | -0.1944905 | 0.84579185 | NA |
| NRXN1 | 35.8068528 | 0.769833348 | 0.70853958 | 1.08650719 | 0.27725466 | 0.4726665 |
| AC009234.1 | 0 | NA | NA | NA | NA | NA |
| AC007402.1 | 0.34847896 | 0.732866073 | 3.27839706 | 0.22354402 | 0.82311212 | NA |
| LINC01867 | 0 | NA | NA | NA | NA | NA |
| AC139712.3 | 0.09635527 | 0.547239394 | 3.29377202 | 0.16614368 | 0.86804388 | NA |
| AC010967.1 | 0.26413911 | 1.101280607 | 3.0105956 | 0.36580157 | 0.71451315 | NA |
| AC010967.2 | 0 | NA | NA | NA | NA | NA |
| ASB3 | 827.233654 | -0.1948506 | 0.15127078 | -1.2880915 | 0.19771412 | 0.3784641 |
| GPR75-ASB3 | 778.727551 | -0.163908186 | 0.15052358 | -1.0889203 | 0.27618902 | 0.47149878 |
| CHAC2 | 128.130847 | -0.34279408 | 0.28038938 | -1.2225644 | 0.22149427 | 0.40759273 |
| ERLEC1 | 2711.44689 | -0.123093057 | 0.1302398 | -0.9451263 | 0.34459442 | 0.54214873 |
| GPR75 | 16.1911336 | -0.221418586 | 0.39373035 | -0.562361 | 0.5738701 | 0.73844506 |
| PSME4 | 2665.59142 | -0.615177186 | 0.18672592 | -3.2945464 | 0.00098581 | 0.0077576 |
| ACYP2 | 605.797801 | -0.154220967 | 0.1453477 | -1.0610486 | 0.28866782 | 0.48485162 |
| AC008280.3 | 34.1644663 | 0.078728683 | 0.30821409 | 0.25543506 | 0.7983871 | 0.8908508 |
| TSPYL6 | 1.42555665 | 1.102209201 | 1.04960318 | 1.05011991 | 0.29366298 | 0.490306 |
| C2orf73 | 31.0988078 | 0.432576862 | 0.54926577 | 0.78755475 | 0.43095719 | 0.62301087 |
| SPTBN1 | 20161.5063 | 0.038089997 | 0.23175867 | 0.16435198 | 0.86945406 | 0.93205023 |
| AC092839.1 | 1.35379153 | 0.233122857 | 1.23615001 | 0.18858784 | 0.85041587 | 0.92153139 |
| AC092839.2 | 3.98222317 | 1.405420142 | 0.64093619 | 2.1927614 | 0.02832457 | 0.09696469 |
| AC093110.1 | 2387.31967 | 0.148260043 | 0.24545228 | 0.60402796 | 0.54582505 | 0.71715108 |
| EML6 | 291.190474 | 0.76195164 | 0.33227108 | 2.29316265 | 0.02183864 | 0.080073 |
| AC104781.1 | 0.18692855 | 0.18657164 | 3.29377195 | 0.05664376 | 0.95482897 | NA |
| RTN4 | 23760.0272 | -0.141004654 | 0.22508457 | -0.6264519 | 0.53101858 | 0.70536772 |
| CLHC1 | 366.423366 | -0.373248605 | 0.2520034 | -1.4811253 | 0.13857319 | 0.29778951 |
| AC012358.1 | 4.2834815 | 1.316415412 | 0.69894215 | 1.88343972 | 0.0596408 | 0.16494263 |
| RPS27A | 27707.0192 | -0.472116265 | 0.20555566 | -2.2967807 | 0.02163128 | 0.07948955 |
| MTIF2 | 1852.19893 | 0.358788331 | 0.17599191 | 2.03866373 | 0.0414836 | 0.12724181 |
| AC012358.3 | 116.196946 | 0.549426963 | 0.22612446 | 2.42975468 | 0.01510905 | 0.06120476 |
| CCDC88A | 907.257537 | -0.014379133 | 0.12306443 | -0.1168423 | 0.90698501 | 0.95356011 |
| CFAP36 | 2828.767 | -0.264835641 | 0.13205393 | -2.0055113 | 0.04490842 | 0.13466397 |
| PPP4R3B | 4239.08851 | 0.155325264 | 0.11141479 | 1.39411709 | 0.16328225 | 0.33251625 |
| AC015982.1 | 64.4807578 | -0.084423025 | 0.28333246 | -0.2979645 | 0.76573023 | 0.87077089 |
| PNPT1 | 1892.17914 | -0.517522611 | 0.40887197 | -1.2657327 | 0.20560881 | 0.38819391 |
| EFEMP1 | 6386.89381 | -0.747250249 | 0.33652335 | -2.2205004 | 0.02638482 | 0.09195976 |
| AC011306.1 | 2.9435015 | 0.692201648 | 1.19575662 | 0.57888172 | 0.56266899 | 0.73005221 |
| MIR217HG | 0.47508126 | 2.881667052 | 2.45460907 | 1.17398208 | 0.2404022 | NA |
| LINC01813 | 0 | NA | NA | NA | NA | NA |
| AC007743.1 | 61.0048416 | 0.889673512 | 0.47923996 | 1.85642597 | 0.06339287 | 0.17179132 |
| CCDC85A | 105.279741 | 1.002791802 | 0.45072772 | 2.22482835 | 0.02609276 | 0.0911829 |
| VRK2 | 995.007776 | -0.580294232 | 0.18989835 | -3.055815 | 0.0022445 | 0.01477468 |
| FANCL | 1105.50736 | -0.434642624 | 0.20495571 | -2.1206661 | 0.03394991 | 0.1101993 |
| AC007250.1 | 1.30391543 | -0.547858437 | 1.33941019 | -0.4090296 | 0.68251793 | 0.81630977 |
| LINC01795 | 0 | NA | NA | NA | NA | NA |
| LINC01122 | 6.22192765 | 0.990482854 | 0.46662833 | 2.12263762 | 0.03378423 | 0.10982373 |
| LINC01793 | 0.33489606 | -0.534763868 | 3.29377181 | -0.1623561 | 0.87102546 | NA |

| | | | | | | |
|---|---|---|---|---|---|---|
| AC007100.1 | 0.48761983 | -0.862897364 | 1.91359347 | -0.4509303 | 0.65203975 | NA |
| AC007179.2 | 0.29261638 | -0.695437658 | 2.34121901 | -0.2970408 | 0.76643533 | NA |
| AC007179.1 | 0 | NA | NA | NA | NA | NA |
| AC007381.1 | 0.43787885 | 2.733170053 | 3.20371799 | 0.85312442 | 0.39359031 | NA |
| MIR4432HG | 0.17515154 | 1.45420457 | 3.2783972 | 0.44357181 | 0.65735219 | NA |
| BCL11A | 88.370265 | 0.361373772 | 0.416645 | 0.86734216 | 0.38575456 | 0.58152559 |
| AC009970.1 | 0.18393104 | 1.469068379 | 3.27783996 | 0.44818185 | 0.65402196 | NA |
| PAPOLG | 772.856883 | -0.208268377 | 0.15936744 | -1.306844 | 0.19126571 | 0.37036838 |
| LINC01185 | 1.09158684 | 1.115115895 | 1.23257086 | 0.90470733 | 0.36562046 | 0.56262123 |
| REL | 777.296494 | -0.129982664 | 0.24902943 | -0.521957 | 0.60170024 | 0.75903919 |
| PUS10 | 492.813819 | 0.172376279 | 0.25201725 | 0.68398603 | 0.49398399 | 0.67665876 |
| PEX13 | 1668.05146 | -0.28322006 | 0.19711592 | -1.4368198 | 0.1507692 | 0.31551335 |
| KIAA1841 | 316.748986 | -0.072619621 | 0.12688647 | -0.5723197 | 0.56710543 | 0.73319386 |
| C2orf74 | 861.935918 | 0.939467224 | 0.31797712 | 2.95451204 | 0.00313164 | 0.01897549 |
| AC016747.2 | 26.5460645 | 1.281757696 | 0.34529684 | 3.71204581 | 0.00020559 | 0.00222921 |
| USP34 | 5918.80733 | -0.036218887 | 0.12338263 | -0.2935493 | 0.76910231 | 0.87240608 |
| AC016727.1 | 875.060425 | 0.031289142 | 0.20932595 | 0.1494757 | 0.88117829 | 0.93836253 |
| XPO1 | 5570.22935 | -0.162901886 | 0.22192357 | -0.734045 | 0.46292132 | 0.65100676 |
| FAM161A | 287.205362 | 0.492441568 | 0.21465454 | 2.29411204 | 0.02178406 | 0.07993957 |
| CCT4 | 6312.47944 | -0.150741553 | 0.16629641 | -0.9064631 | 0.3646908 | 0.56182657 |
| AC107081.2 | 1071.87771 | -0.478852413 | 0.15896712 | -3.0122732 | 0.00259299 | 0.0164374 |
| COMMD1 | 1185.25612 | -0.090317837 | 0.11019747 | -0.8195999 | 0.41244421 | 0.6062516 |
| AC018462.1 | 57.1132195 | -0.070312126 | 0.21164271 | -0.3322209 | 0.73972249 | 0.85396119 |
| B3GNT2 | 2233.0695 | 0.14114126 | 0.19296084 | 0.73145028 | 0.46450416 | 0.65190107 |
| AC093159.1 | 10.3739537 | -0.099338305 | 0.52493784 | -0.1892382 | 0.84990611 | 0.92113042 |
| TMEM17 | 195.324124 | 0.60932465 | 0.26234469 | 2.32261096 | 0.02020006 | 0.07541776 |
| AC092155.1 | 0.58933706 | -0.325877607 | 2.19311619 | -0.1485911 | 0.88187627 | NA |
| EHBP1 | 1482.77271 | 0.400122004 | 0.09544474 | 4.19218508 | 2.76E-05 | 0.00045994 |
| AC092567.1 | 21.5846992 | 1.946013455 | 0.47496928 | 4.09713538 | 4.18E-05 | 0.00064468 |
| AC009501.1 | 523.481582 | 0.164721349 | 0.09976816 | 1.6510412 | 0.09873016 | 0.23558262 |
| OTX1 | 1.94475095 | -0.67795763 | 1.07369405 | -0.6314253 | 0.52776245 | 0.7023701 |
| WDPCP | 261.543836 | 0.113233889 | 0.13986638 | 0.80958619 | 0.41817805 | 0.61099046 |
| MDH1 | 12628.3621 | 0.389774971 | 0.25905705 | 1.50459127 | 0.13242919 | 0.28871794 |
| UGP2 | 8211.71536 | -0.259743026 | 0.18132967 | -1.4324353 | 0.15201928 | 0.31707904 |
| VPS54 | 2125.67965 | -0.036451799 | 0.06599287 | -0.5523596 | 0.58070199 | 0.74352011 |
| AC012368.2 | 3.40158828 | 0.278319287 | 0.68118483 | 0.40858116 | 0.68284706 | 0.81659231 |
| PELI1 | 1625.2138 | -0.421486745 | 0.24887322 | -1.6935801 | 0.0903451 | 0.22126872 |
| AC012368.1 | 385.598354 | 0.432266609 | 0.608891 | 0.70992445 | 0.47775099 | 0.66285875 |
| LINC00309 | 0.08393208 | -0.642905668 | 3.30558948 | -0.1944905 | 0.84579185 | NA |
| AC114752.1 | 0 | NA | NA | NA | NA | NA |
| AC114752.2 | 0.44246721 | 2.092065913 | 2.05574777 | 1.01766664 | 0.30883641 | NA |
| AC008074.1 | 10.6167998 | 0.177498251 | 0.41153248 | 0.43131044 | 0.66624266 | 0.80439203 |
| LGALSL | 1011.69587 | -0.082183058 | 0.27647267 | -0.2972556 | 0.76627135 | 0.87113838 |
| LINC01805 | 0 | NA | NA | NA | NA | NA |
| AC008074.3 | 83.5160049 | 2.106429776 | 0.3546245 | 5.93988793 | 2.85E-09 | 2.40E-07 |
| AFTPH | 3660.30101 | 0.617761573 | 0.18426354 | 3.3525979 | 0.00080057 | 0.00658074 |
| LINC02579 | 0.14616582 | 0.435199075 | 3.28898345 | 0.13232024 | 0.894731 | NA |
| SERTAD2 | 1407.36528 | -0.482331354 | 0.22205317 | -2.1721435 | 0.02984484 | 0.10051192 |
| AC007365.1 | 3.10529453 | 0.156732874 | 0.90261678 | 0.17364277 | 0.86214622 | 0.92814027 |
| LINC01800 | 0.83578649 | 2.651769317 | 2.03246668 | 1.30470494 | 0.19199334 | NA |
| LINC02245 | 1.0981878 | 0.411702732 | 2.2280345 | 0.18478292 | 0.8533993 | 0.9232762 |
| SLC1A4 | 624.058542 | -0.856386638 | 0.5041603 | -1.6986396 | 0.08938712 | 0.21954805 |
| LINC02576 | 6.23681469 | -0.542214104 | 0.73295245 | -0.7397671 | 0.45944133 | 0.64811921 |
| CEP68 | 1404.35862 | 0.056423447 | 0.17132774 | 0.32933048 | 0.74190591 | 0.85503513 |
| RAB1A | 11007.4123 | -0.216148961 | 0.14620052 | -1.4784418 | 0.13928957 | 0.29889054 |
| ACTR2 | 7863.9432 | -0.081016091 | 0.17852123 | -0.4538177 | 0.64996006 | 0.7941547 |
| SPRED2 | 1905.88462 | -0.203227984 | 0.29156618 | -0.6970218 | 0.48578915 | 0.66985253 |
| AC012370.2 | 2.48055503 | 0.203530241 | 1.00133103 | 0.2032597 | 0.83893206 | 0.91454277 |
| AC007389.3 | 0.70525301 | 2.250517995 | 1.91559442 | 1.17484055 | 0.24005851 | NA |
| AC007389.5 | 9.21339943 | 0.028704244 | 0.76755435 | 0.03739702 | 0.97016845 | 0.98622863 |
| AC118345.1 | 1.06169633 | -0.254203097 | 1.41921563 | -0.1791152 | 0.85784724 | 0.92574201 |
| AC092669.1 | 0.7732681 | 0.674195443 | 1.76173177 | 0.38268904 | 0.70195034 | NA |
| LINC01873 | 0.07659028 | -0.642905668 | 3.30558948 | -0.1944905 | 0.84579185 | NA |
| MEIS1-AS3 | 2.37101367 | 1.178206053 | 1.15364701 | 1.02128818 | 0.30711792 | 0.50405799 |
| MEIS1 | 1262.20128 | -0.013314014 | 0.21902872 | -0.0607866 | 0.95152915 | 0.97675915 |
| MEIS1-AS2 | 9.61888869 | 0.050625715 | 0.52315413 | 0.09677017 | 0.92290891 | 0.96235944 |
| LINC01798 | 12.0592535 | 0.670979358 | 0.419092 | 1.60103117 | 0.10937002 | 0.25296793 |
| LINC01797 | 3.71088538 | 1.112982485 | 0.73513119 | 1.51399165 | 0.13002796 | 0.28514239 |
| LINC01799 | 2.66240113 | 2.77125847 | 1.04114888 | 2.66173121 | 0.00777399 | 0.03740879 |
| LINC01628 | 0 | NA | NA | NA | NA | NA |
| AC007403.1 | 0.34887651 | -0.556945191 | 2.28858666 | -0.2433577 | 0.80772831 | NA |
| LINC01828 | 8.99232916 | 1.698088782 | 0.61452425 | 2.76325756 | 0.00572276 | 0.02980039 |
| LINC01829 | 15.4909641 | -0.285968251 | 0.54048412 | -0.5290965 | 0.59673852 | 0.75537609 |
| AC023115.1 | 3.16521432 | 0.264027654 | 0.77613057 | 0.34018458 | 0.73371753 | 0.85001894 |
| ETAA1 | 823.140853 | -0.021746255 | 0.13934173 | -0.1560642 | 0.87598242 | 0.93550556 |
| AC007422.1 | 0.40443956 | -0.633798438 | 2.89858161 | -0.2186581 | 0.82691637 | NA |
| AC007422.2 | 1.12940392 | -2.154176288 | 1.63072168 | -1.3209957 | 0.18650281 | 0.36424818 |
| LINC01812 | 3.75567699 | -1.94641447 | 1.19920384 | -1.6230889 | 0.1045704 | 0.24502589 |
| C1D | 1295.10235 | 0.113598215 | 0.12541924 | 0.90574793 | 0.36506929 | 0.56205495 |
| WDR92 | 496.298871 | 0.372905459 | 0.11145552 | 3.34577823 | 0.00082052 | 0.00671126 |
| AC017083.3 | 851.500374 | 0.163516811 | 0.18525536 | 0.88265631 | 0.37742199 | 0.57343962 |
| PNO1 | 1036.40729 | -0.584819638 | 0.29378412 | -1.9906442 | 0.04652002 | 0.13817673 |
| PPP3R1 | 2671.78988 | -0.432126488 | 0.15883609 | -2.7205812 | 0.00651673 | 0.03273882 |
| AC017083.2 | 981.471567 | -0.340523572 | 0.1694633 | -2.0094237 | 0.04449222 | 0.13375056 |
| AC017083.1 | 32.8060338 | -0.152394998 | 0.36732213 | -0.4148811 | 0.67822897 | 0.81313471 |
| CNRIP1 | 527.940497 | -0.531092168 | 0.29287169 | -1.8133954 | 0.06977086 | 0.18376525 |
| AC015969.1 | 0.18331075 | 0.18657164 | 3.29377195 | 0.05664376 | 0.95482897 | NA |
| PLEK | 530.763872 | 0.082763479 | 0.51391448 | 0.16104524 | 0.87205777 | 0.93327291 |
| FBXO48 | 188.310741 | -0.015717404 | 0.18566079 | -0.0846566 | 0.93253443 | 0.96746494 |
| APLF | 168.255109 | -0.151990408 | 0.21461252 | -0.7082085 | 0.4788158 | 0.66367267 |
| PROKR1 | 3.29494469 | -0.664433445 | 0.89969444 | -0.7385101 | 0.46020452 | 0.64875405 |
| ARHGAP25 | 613.661309 | -0.319438033 | 0.25396392 | -1.2578087 | 0.20846094 | 0.39156431 |
| LINC01888 | 0.41781029 | -0.897576225 | 2.25933535 | -0.3972745 | 0.69116508 | NA |
| LINC01890 | 0.56322155 | -2.357939534 | 1.85891515 | -1.2684493 | 0.20463756 | NA |
| BMP10 | 0 | NA | NA | NA | NA | NA |
| GKN2 | 0.32447948 | -1.685682445 | 3.30319546 | -0.5103187 | 0.6098282 | NA |
| GKN1 | 0.08666371 | -0.642905668 | 3.30558948 | -0.1944905 | 0.84579185 | NA |
| ANTXR1 | 1390.6603 | 0.428155923 | 0.36555451 | 1.17125056 | 0.24149808 | 0.43169091 |
| GFPT1 | 4059.25675 | -0.591601969 | 0.21904767 | -2.7007909 | 0.00691748 | 0.03429478 |
| NFU1 | 1520.01777 | 0.072921387 | 0.14526616 | 0.50198469 | 0.61567829 | 0.77018414 |
| AAK1 | 3345.67321 | -0.310131728 | 0.14580696 | -2.1270023 | 0.0334199 | 0.10896175 |
| ANXA4 | 10018.2691 | -0.47263477 | 0.27871997 | -1.6957334 | 0.08993638 | 0.22053397 |
| AC092431.1 | 1.65679154 | 1.333356762 | 1.0604404 | 1.25736134 | 0.20862282 | 0.39176687 |
| GMCL1 | 1221.583 | 0.146674262 | 0.16187193 | 0.90611302 | 0.36487603 | 0.56195427 |
| SNRNP27 | 1271.06886 | -0.503760396 | 0.10301149 | -4.8903321 | 1.01E-06 | 3.27E-05 |
| MXD1 | 1866.48162 | -0.961954703 | 0.21896895 | -4.3931101 | 1.12E-05 | 0.00022306 |
| ASPRV1 | 158.30503 | 0.712909133 | 0.17383574 | 4.10105038 | 4.11E-05 | 0.00063535 |
| PCBP1 | 16587.6595 | -0.289832332 | 0.16433516 | -1.763666 | 0.07778823 | 0.19856097 |
| AC016700.3 | 1.73220037 | -0.537756919 | 0.92926355 | -0.5786915 | 0.56279735 | 0.73014691 |
| LINC01816 | 36.3550568 | 1.307667391 | 0.26925219 | 4.85666387 | 1.19E-06 | 3.76E-05 |
| C2orf42 | 647.894367 | 0.323838982 | 0.14154637 | 2.28786491 | 0.02214539 | 0.08084922 |
| TIA1 | 4687.39839 | 0.25885992 | 0.22887578 | 1.13100619 | 0.25805248 | 0.45056241 |
| PCYOX1 | 5923.24901 | 1.11166085 | 0.32466593 | 3.42401448 | 0.00061703 | 0.00532782 |

| | | | | | | |
|---|---|---|---|---|---|---|
| SNRPG | 1674.51668 | -0.6094849 | 0.16218967 | -3.7578527 | 0.00017138 | 0.00193958 |
| FAM136A | 2520.64667 | -0.164668231 | 0.16429965 | -1.0022433 | 0.31622608 | 0.5133515 |
| AC022201.2 | 221.240792 | 0.152711027 | 0.1513549 | 1.00895988 | 0.31299387 | 0.50986215 |
| AC022201.1 | 0.20908062 | 1.675937956 | 3.26598797 | 0.51314885 | 0.60784721 | NA |
| TGFA | 1490.31936 | -0.372612101 | 0.1993882 | -1.8687771 | 0.06165383 | 0.16856523 |
| ADD2 | 31.8526836 | -0.940584092 | 0.56270812 | -1.671531 | 0.09461683 | 0.22879834 |
| AC005234.1 | 0 | NA | NA | NA | NA | NA |
| FIGLA | 0.46140314 | -0.932448929 | 2.23907753 | -0.4164433 | 0.67708562 | NA |
| CLEC4F | 29.9420441 | 0.629123151 | 0.63769441 | 0.98655898 | 0.32385889 | 0.52162159 |
| CD207 | 9.24953794 | 1.502845253 | 1.23183394 | 1.22000637 | 0.22246246 | 0.4084406 |
| LINC01143 | 0.78036837 | -0.065005115 | 1.55940158 | -0.0416859 | 0.96674907 | NA |
| VAX2 | 4166.33874 | 1.761724422 | 0.66742117 | 2.63959926 | 0.00830041 | 0.03913135 |
| ATP6V1B1 | 4315.50193 | 1.780183093 | 0.67025109 | 2.65599434 | 0.00790749 | 0.03784403 |
| AC007040.2 | 481.199235 | 0.956192009 | 0.21162403 | 4.51835272 | 6.23E-06 | 0.00014035 |
| ATP6V1B1-A | 8.91192398 | -0.86896319 | 0.74951589 | -1.1593659 | 0.24630706 | 0.43769955 |
| ANKRD53 | 79.2182483 | -0.261112607 | 0.42123063 | -0.6198804 | 0.53533653 | 0.70904967 |
| TEX261 | 4353.00493 | 0.4994306 | 0.11027732 | 4.52886068 | 5.93E-06 | 0.00013493 |
| AC007040.1 | 77.6418874 | 1.000066349 | 0.21322629 | 4.69016428 | 2.73E-06 | 7.29E-05 |
| AC007881.2 | 0.93021793 | -1.119762 | 2.20269836 | -0.5083592 | 0.61120145 | NA |
| NAGK | 3496.40529 | -0.04964075 | 0.13776697 | -0.360324 | 0.71860482 | 0.84001549 |
| AC007881.3 | 16.0561107 | 0.908788468 | 0.56199811 | 1.61706677 | 0.10586387 | 0.24698135 |
| MCEE | 705.223749 | 0.55890975 | 0.13099741 | 4.26657102 | 1.99E-05 | 0.00035178 |
| MPHOSPH10 | 1200.19081 | -0.238317218 | 0.20270047 | -1.1757112 | 0.23971028 | 0.42963095 |
| PAIP2B | 1345.90111 | -0.014185519 | 0.4148769 | -0.0341921 | 0.97272395 | 0.98725004 |
| ZNF638 | 4615.10155 | -0.17177358 | 0.11825199 | -1.4526063 | 0.14633309 | 0.30917609 |
| AC007878.1 | 5.50141966 | 0.478420014 | 0.71711178 | 0.66714845 | 0.50467731 | 0.68403209 |
| DYSF | 1833.45113 | 0.023062182 | 0.24431076 | 0.09439691 | 0.92479387 | 0.96356293 |
| CYP26B1 | 1784.49308 | -0.865328332 | 0.66406421 | -1.3030793 | 0.19254769 | 0.37192225 |
| EXOC6B | 1143.05092 | -0.204195861 | 0.11166046 | -1.8287213 | 0.06744137 | 0.17929518 |
| SPR | 2518.19922 | -0.057218933 | 0.21107318 | -0.2710858 | 0.78632507 | 0.88367322 |
| EMX1 | 3754.67518 | 1.010775302 | 0.32540057 | 3.1062493 | 0.00189477 | 0.01292885 |
| AC012366.1 | 1045.84853 | 0.914357291 | 0.32179105 | 2.8414628 | 0.00449071 | 0.02484162 |
| SFXN5 | 4265.83838 | 0.151790506 | 0.26588941 | 0.57087835 | 0.56808211 | 0.73384869 |
| RAB11FIP5 | 8974.73696 | 0.298177409 | 0.25940148 | 1.14948229 | 0.25035717 | 0.44257682 |
| NOTO | 0.47864515 | -0.930299525 | 2.75094139 | -0.338175 | 0.73523134 | NA |
| SMYD5 | 1828.79985 | 0.009553901 | 0.15492054 | 0.06166969 | 0.95082588 | 0.97652119 |
| PRADC1 | 996.153474 | 0.292697017 | 0.28076893 | 1.04248363 | 0.29718751 | 0.49400117 |
| CCT7 | 8364.24471 | -0.108607541 | 0.15585573 | -0.6968466 | 0.48589877 | 0.66993363 |
| FBXO41 | 236.582464 | -0.823811222 | 0.39054629 | -2.1093818 | 0.03491164 | 0.112449 |
| EGR4 | 2.24320533 | 0.076886822 | 1.33739694 | 0.0574899 | 0.95415495 | 0.9780936 |
| ALMS1 | 574.98315 | 0.083832012 | 0.15487161 | 0.54130007 | 0.58830077 | 0.74904598 |
| NAT8 | 18398.6943 | 2.487440253 | 0.58141478 | 4.27825422 | 1.88E-05 | 0.0003379 |
| TPRKB | 927.733795 | 0.097021655 | 0.1245077 | 0.77924221 | 0.43583705 | 0.62793111 |
| AC092653.1 | 4.3246132 | 0.754151596 | 0.616959 | 1.22236907 | 0.22156809 | 0.40759381 |
| DUSP11 | 1061.20997 | -0.150173826 | 0.11372661 | -1.320481 | 0.18667446 | 0.36450245 |
| C2orf78 | 0 | NA | NA | NA | NA | NA |
| STAMBP | 1933.75002 | 0.011553122 | 0.1495127 | 0.07727184 | 0.93840729 | 0.97065193 |
| ACTG2 | 283.17802 | 1.981287298 | 0.98595855 | 2.00950363 | NA | NA |
| DGUOK | 1716.54785 | -0.754778942 | 0.11367065 | -6.6400512 | 3.14E-11 | 5.40E-09 |
| DGUOK-AS1 | 712.139564 | -0.761808016 | 0.12525681 | -6.081969 | 1.19E-09 | 1.19E-07 |
| AC073046.1 | 2.40807223 | -0.077917519 | 0.9768822 | -0.0797614 | 0.936427 | 0.96955436 |
| TET3 | 603.206292 | -0.079343343 | 0.20868331 | -0.3802093 | 0.70379003 | 0.83074815 |
| AC073263.1 | 14.9388189 | 0.825234362 | 0.60007209 | 1.37522537 | 0.16906159 | 0.34073326 |
| BOLA3 | 632.061293 | 0.035797265 | 0.20412355 | 0.17537058 | 0.86078846 | 0.92743582 |
| BOLA3-AS1 | 96.305668 | 0.062449828 | 0.25335978 | 0.24648674 | 0.80530546 | 0.8944712 |
| MOB1A | 6011.46885 | 0.00083256 | 0.12120808 | 0.00686885 | 0.9945195 | 0.99750786 |
| AC073263.2 | 0.29061591 | -1.364245083 | 2.89386475 | -0.4714267 | 0.63733606 | NA |
| MTHFD2 | 1641.06573 | -1.870688302 | 0.34593719 | -5.4075952 | 6.39E-08 | 3.25E-06 |
| AC006030.1 | 3439.59987 | -0.339780328 | 0.13179435 | -2.57811 | 0.00993424 | 0.0448436 |
| SLC4A5 | 324.533769 | -0.802588612 | 0.27367538 | -2.93263 | 0.00336104 | 0.01999339 |
| DCTN1 | 6940.86591 | 0.259664285 | 0.07393047 | 3.51227688 | 0.00044428 | 0.00414425 |
| DCTN1-AS1 | 100.413292 | 1.191967523 | 0.28258272 | 4.2181189 | 2.46E-05 | 0.00041992 |
| AC005041.1 | 431.551982 | -0.2710701 | 0.34838044 | -0.7780865 | 0.43651804 | 0.62867181 |
| C2orf81 | 448.175323 | -0.232608895 | 0.33185693 | -0.7009313 | 0.48334592 | 0.66791552 |
| WDR54 | 1082.74608 | -0.637094941 | 0.20180495 | -3.1569837 | 0.0015941 | 0.01132443 |
| RTKN | 6513.13682 | 0.64652031 | 0.13089987 | 4.93904459 | 7.85E-07 | 2.65E-05 |
| INO80B | 1538.97226 | -0.061787753 | 0.12448326 | -0.4963539 | 0.61964475 | 0.77271053 |
| INO80B-WBI | 5260.10122 | 0.150422826 | 0.13092671 | 1.14890865 | 0.25059365 | 0.44275742 |
| WBP1 | 4584.34953 | 0.219073561 | 0.16607976 | 1.31908646 | 0.18714021 | 0.36506328 |
| MOGS | 5100.01844 | -0.490022181 | 0.1376572 | -3.559728 | 0.00037124 | 0.00359911 |
| MRPL53 | 2048.11246 | -0.140903782 | 0.13098207 | -1.0757486 | 0.28203969 | 0.47782049 |
| CCDC142 | 2298.39687 | 0.002264572 | 0.12595017 | 0.0179799 | 0.98565488 | 0.99368571 |
| TTC31 | 1707.50359 | 0.348104252 | 0.14834074 | 2.34665302 | 0.01894289 | 0.07213342 |
| LBX2 | 253.078648 | 0.349395758 | 0.30988444 | 1.1275034 | 0.25952971 | 0.45227322 |
| AC005041.3 | 25.7269782 | -0.742094874 | 0.46474596 | -1.5967753 | 0.1103158 | 0.25435233 |
| LBX2-AS1 | 647.069553 | 0.155619177 | 0.35338278 | 0.44037001 | 0.65966914 | 0.79999826 |
| PCGF1 | 978.149643 | -0.577916284 | 0.20256376 | -2.8530092 | 0.00433074 | 0.02418499 |
| TLX2 | 0.48045953 | 1.448727339 | 2.1674194 | 0.66841117 | 0.50387117 | NA |
| DQX1 | 8.05199628 | -1.04189389 | 0.64575366 | -1.6134541 | 0.10664589 | 0.24830055 |
| AUP1 | 8678.21629 | -0.647904495 | 0.1641621 | -3.9467361 | 7.92E-05 | 0.00105246 |
| HTRA2 | 3202.64517 | -0.477108079 | 0.16626216 | -2.8696131 | 0.00410974 | 0.0232263 |
| LOXL3 | 1147.34262 | -0.605855616 | 0.18133545 | -3.3410765 | 0.00083454 | 0.00680064 |
| DOK1 | 563.958412 | -0.698441489 | 0.20382001 | -3.4267562 | 0.00061084 | 0.00528124 |
| M1AP | 83.4064211 | -0.527653459 | 0.41965834 | -1.2573406 | 0.20863034 | 0.39176687 |
| SEMA4F | 446.908684 | -0.357356615 | 0.26540299 | -1.3464679 | 0.17815166 | 0.35361395 |
| AC019069.1 | 14.5576708 | -1.183008961 | 0.60837163 | -1.9445498 | 0.05182918 | 0.14953876 |
| HK2 | 697.64241 | -2.65150899 | 0.48183887 | -5.5028955 | 3.74E-08 | 2.09E-06 |
| LINC01291 | 111.809792 | 0.749036834 | 0.82108464 | 0.91225288 | 0.36163561 | 0.55906728 |
| AC104135.1 | 8.40638708 | 0.920082716 | 0.66571291 | 1.38210135 | 0.16694058 | 0.3379307 |
| LINC01293 | 4.74477738 | 0.538688091 | 0.83037227 | 0.64873083 | 0.51651237 | 0.69389328 |
| POLE4 | 1071.00652 | -0.48722164 | 0.17341409 | -2.8095851 | 0.00496054 | 0.0267104 |
| TACR1 | 154.75064 | -0.701139032 | 0.5873965 | -1.1936384 | 0.23261944 | 0.42066727 |
| AC007681.1 | 54.5804948 | -0.51152671 | 0.53805937 | -0.9506882 | 0.34176266 | 0.53967632 |
| EVA1A | 2137.18647 | -0.708175618 | 0.36388105 | -1.9461734 | 0.05163392 | 0.14919577 |
| AC007099.2 | 0.30439593 | -0.695250385 | 2.32336129 | -0.2992433 | 0.76475439 | NA |
| AC007099.1 | 2.60746026 | -0.010920224 | 1.07658386 | -0.0101434 | 0.99190687 | 0.99613883 |
| MRPL19 | 2359.27471 | 0.187265617 | 0.17654557 | 1.06072111 | 0.28881666 | 0.48502327 |
| GCFC2 | 1429.12711 | 0.244934148 | 0.07747109 | 3.16162016 | 0.00156894 | 0.01118793 |
| AC005034.3 | 371.702286 | 0.003114693 | 0.23421634 | 0.01329836 | 0.98938976 | 0.99519138 |
| AC005034.2 | 3.40775012 | 1.563528915 | 0.82676998 | 1.89112928 | 0.05860709 | 0.16276661 |
| AC005034.4 | 22.1043254 | 1.380727445 | 0.61777485 | 2.23500104 | 0.02541728 | 0.08942678 |
| LRRTM4 | 2.89077628 | 1.139487275 | 0.93637404 | 1.21691464 | 0.2236367 | 0.40975107 |
| AC079117.1 | 0 | NA | NA | NA | NA | NA |
| AC084149.1 | 5.42721924 | 1.052147207 | 0.8852641 | 1.18851223 | 0.23463167 | 0.42334841 |
| AC084149.2 | 0 | NA | NA | NA | NA | NA |
| LINC01851 | 0 | NA | NA | NA | NA | NA |
| AC012494.1 | 0 | NA | NA | NA | NA | NA |
| AC092660.1 | 0 | NA | NA | NA | NA | NA |
| REG3G | 94.4399077 | -2.09885016 | 1.30146915 | -1.6126776 | 0.10681457 | 0.24862743 |
| REG1B | 8.33519361 | -2.889130207 | 1.30596184 | -2.2122623 | 0.02694854 | 0.09332017 |
| REG1A | 1004.99644 | -1.104538181 | 1.17856081 | -0.9371924 | 0.34865962 | 0.5461982 |
| REG3A | 0.17391096 | -1.003574173 | 3.30558948 | -0.3035992 | 0.7614333 | NA |
| CTNNA2 | 9.9256236 | 0.185005761 | 1.06775257 | 0.17326651 | 0.86244195 | 0.92830704 |

| | | | | | | |
|---|---|---|---|---|---|---|
| AC010975.1 | 0 | NA | NA | NA | NA | NA |
| AC016716.2 | 0 | NA | NA | NA | NA | NA |
| LRRTM1 | 16.5668559 | 1.189380978 | 0.56284459 | 2.11316055 | 0.03458703 | 0.11173097 |
| AC008067.1 | 1.10652374 | -0.410895531 | 1.38567136 | -0.2965317 | 0.76682404 | 0.87145103 |
| AC012355.1 | 0 | NA | NA | NA | NA | NA |
| LINC01815 | 0.03392908 | 0.380740593 | 3.29738709 | 0.11546736 | 0.90807469 | NA |
| AC079896.1 | 0.33023047 | 2.310906901 | 2.77720112 | 0.83209922 | 0.40535294 | NA |
| AC010105.1 | 0 | NA | NA | NA | NA | NA |
| LINC01809 | 0 | NA | NA | NA | NA | NA |
| AC106874.1 | 0.13390214 | 1.04981649 | 3.28477129 | 0.31960109 | 0.74927075 | NA |
| SUCLG1 | 13553.9633 | 1.231611359 | 0.25623085 | 4.80664738 | 1.53E-06 | 4.60E-05 |
| DNAH6 | 157.963865 | 0.777426917 | 0.56069757 | 1.38653519 | 0.16558354 | 0.33600908 |
| TRABD2A | 460.602761 | -0.986858711 | 0.30509172 | -3.2346296 | 0.00121801 | 0.00915849 |
| TMSB10 | 21337.6995 | -1.595274428 | 0.25485495 | -6.2595387 | 3.86E-10 | 4.49E-08 |
| KCMF1 | 2739.87725 | -0.677731619 | 0.25854065 | -2.6213735 | 0.00875763 | 0.04074428 |
| LINC01964 | 0.06785815 | 0.795868262 | 3.28898349 | 0.24198001 | 0.80879565 | NA |
| TCF7L1 | 1353.91271 | -0.175638178 | 0.25227156 | -0.6962266 | 0.48628692 | 0.67036364 |
| TGOLN2 | 15032.8691 | 0.348347653 | 0.16555827 | 2.10407886 | 0.03537157 | 0.11351131 |
| RETSAT | 7337.69242 | 0.74691389 | 0.21690829 | 3.44345487 | 0.00057433 | 0.00505344 |
| ELMOD3 | 1813.8402 | 0.474169376 | 0.12333279 | 3.84463344 | 0.00012073 | 0.00147326 |
| AC062037.2 | 34.3780164 | 0.517393747 | 0.26598401 | 1.94520624 | 0.05175016 | 0.14937081 |
| CAPG | 6227.20523 | -0.392991945 | 0.24016861 | -1.6363168 | 0.1017733 | 0.24039429 |
| SH2D6 | 83.5579677 | 2.229371083 | 0.4047036 | 5.50865151 | 3.62E-08 | 2.05E-06 |
| MAT2A | 16497.6054 | 0.353010546 | 0.23024881 | 1.53316991 | 0.125234 | 0.27765432 |
| GGCX | 7240.89014 | -0.018294626 | 0.12512991 | -0.1462051 | 0.88375951 | 0.93974509 |
| VAMP8 | 7135.58176 | -0.520601608 | 0.15746414 | -3.3061598 | 0.00094584 | 0.00752393 |
| VAMP5 | 1398.68059 | -0.690350458 | 0.18435679 | -3.7446435 | 0.00018065 | 0.00201342 |
| RNF181 | 3417.62988 | -0.251392968 | 0.15463584 | -1.6257096 | 0.10401145 | 0.24416075 |
| TMEM150A | 3365.35512 | 0.655159938 | 0.17000452 | 3.85377949 | 0.00011631 | 0.00142853 |
| USP39 | 2199.58654 | -0.271539035 | 0.13770186 | -1.9719344 | 0.04861709 | 0.14280853 |
| C2orf68 | 1737.32497 | -0.126073411 | 0.17559884 | -0.7179627 | 0.47278031 | 0.65885276 |
| SFTPB | 101.412452 | -1.736790074 | 0.67115628 | -2.5877581 | 0.00966028 | 0.04391194 |
| GNLY | 375.895421 | -0.946648374 | 0.75718277 | -1.2502244 | 0.21121758 | 0.39461916 |
| ATOH8 | 814.648753 | 0.173435647 | 0.31820468 | 0.54504429 | 0.58572306 | 0.74722314 |
| AC105053.1 | 2.36319546 | 1.447133922 | 0.89061913 | 1.62486284 | 0.10419179 | 0.24444081 |
| ST3GAL5 | 445.99729 | 0.030307536 | 0.30777814 | 0.09847202 | 0.92155749 | 0.96167277 |
| ST3GAL5-AS | 4.43015689 | 1.979774722 | 0.95850751 | 2.06547648 | 0.03887794 | 0.12126937 |
| AC012511.1 | 3.50656917 | 1.316781573 | 0.86221245 | 1.52721243 | 0.1267082 | 0.28005156 |
| POLR1A | 1839.40192 | -0.056557568 | 0.12238696 | -0.4621209 | 0.64399466 | 0.78968329 |
| PTCD3 | 3513.75506 | 0.100649299 | 0.11196378 | 0.89894519 | 0.36868186 | 0.56547087 |
| IMMT | 6228.46063 | 0.359663644 | 0.12925544 | 2.78258025 | 0.00539285 | 0.02848748 |
| AC009309.1 | 30.899228 | 0.46456565 | 0.40753357 | 1.13994449 | 0.25430943 | 0.44708166 |
| MRPL35 | 2131.72268 | 0.511282173 | 0.13311269 | 3.84097255 | 0.00012255 | 0.00148462 |
| REEP1 | 301.233715 | 0.032890494 | 0.48464562 | 0.06786504 | 0.94589307 | 0.97407741 |
| KDM3A | 3173.6002 | -0.702336203 | 0.21191878 | -3.3141763 | 0.00091914 | 0.0073581 |
| CHMP3 | 4727.89356 | 0.037159496 | 0.15492667 | 0.23985216 | 0.81044487 | 0.89747017 |
| RNF103-CHN | 2629.18332 | -0.047821853 | 0.15860605 | -0.3015134 | 0.76302302 | 0.86877586 |
| AC015971.1 | 1453.61664 | -0.229049015 | 0.17634702 | -1.2988539 | 0.19399407 | 0.37378516 |
| RNF103 | 2294.50779 | -0.291736576 | 0.2000747 | -1.4581383 | 0.14480244 | 0.30684991 |
| RMND5A | 2861.08187 | 0.180856583 | 0.2265207 | 0.79841085 | 0.42463211 | 0.61704682 |
| CD8A | 262.322041 | -1.045061295 | 0.72212348 | -1.4472058 | 0.14783929 | 0.3113825 |
| CD8B | 63.0796719 | -0.91374222 | 0.75001136 | -1.2183045 | 0.22310828 | 0.40919219 |
| RGPD1 | 20.7303626 | -0.097982596 | 0.58880573 | -0.166409 | 0.86783506 | 0.93122322 |
| PLGLB1 | 75.7630196 | 0.098532127 | 0.24617715 | 0.40024888 | 0.68897321 | 0.82056925 |
| LINC01955 | 0 | NA | NA | NA | NA | NA |
| IGKV3OR2-2 | 0.56403089 | 1.857774957 | 1.72861365 | 1.07471959 | 0.28250029 | NA |
| LINC01943 | 209.373968 | -1.279092964 | 0.36897807 | -3.4665826 | 0.00052712 | 0.00472111 |
| CYTOR | 388.075475 | -1.166117673 | 0.3600416 | -3.2388415 | 0.00120016 | 0.00905013 |
| AC133644.2 | 12.8674757 | -0.829812595 | 0.73591941 | -1.1275862 | 0.2594947 | 0.452272 |
| PLGLB2 | 34.709589 | 0.633920979 | 0.44302627 | 1.43088803 | 0.15246231 | 0.31766997 |
| RGPD2 | 59.9752521 | 0.198476918 | 0.53656707 | 0.36990141 | 0.71145595 | 0.8360186 |
| KRCC1 | 1621.77151 | -0.103435415 | 0.1830394 | -0.5650992 | 0.57200629 | 0.73683777 |
| SMYD1 | 1.34806238 | 0.561739525 | 2.65714959 | 0.21140681 | 0.83256984 | 0.9111315 |
| FABP1 | 3941.88355 | 2.308119622 | 0.7293938 | 3.1644355 | 0.00155384 | 0.01110729 |
| THNSL2 | 2832.05889 | 1.666689648 | 0.57569039 | 2.89511458 | 0.0037902 | 0.02192261 |
| FOXI3 | 0.44223139 | -1.131714835 | 2.84814584 | -0.3973514 | 0.69110832 | NA |
| TEX37 | 0 | NA | NA | NA | NA | NA |
| AC104134.1 | 353.942278 | -0.637420521 | 0.349084 | -1.8259803 | 0.06785323 | 0.18004722 |
| EIF2AK3 | 1407.95289 | -0.983303091 | 0.28214222 | -3.4851328 | 0.00049189 | 0.0044686 |
| AC062029.1 | 87.2818229 | 1.019514853 | 0.38051007 | 2.6793374 | 0.0073768 | 0.03599672 |
| RPIA | 769.02814 | -0.39130165 | 0.1789831 | -2.1862491 | 0.02879739 | 0.09799807 |
| IGKC | 19554.4406 | -0.390467674 | 0.94879098 | -0.4115424 | 0.68067489 | 0.8148446 |
| IGKJ5 | 379.657859 | -0.839805999 | 0.99488387 | -0.8441246 | 0.39859976 | 0.59351443 |
| IGKJ4 | 660.617397 | -0.684714154 | 0.84865329 | -0.8068244 | 0.41976768 | 0.6125435 |
| IGKJ3 | 624.283692 | 2.06969575 | 1.21734909 | 1.70016618 | NA | NA |
| IGKJ2 | 638.821909 | -0.819062776 | 0.89763381 | -0.9124687 | 0.36152202 | 0.55900878 |
| IGKJ1 | 1096.11267 | -0.952267118 | 0.935178 | -1.0182737 | 0.30854793 | 0.505524 |
| IGKV4-1 | 438.676128 | 0.365149973 | 0.84048185 | 0.43445313 | 0.66395942 | 0.80260972 |
| IGKV5-2 | 2.86296014 | 0.832885986 | 1.34158385 | 0.62082291 | 0.53471615 | 0.70842368 |
| IGKV7-3 | 3.94130122 | 0.833469401 | 1.66284246 | 0.50123173 | 0.61620805 | 0.77049015 |
| IGKV2-4 | 0.7549468 | -2.827197357 | 2.05337894 | -1.3768513 | 0.16855824 | NA |
| IGKV1-5 | 745.427149 | -0.008218736 | 0.85891298 | -0.0095688 | 0.99236535 | 0.99637048 |
| IGKV1-6 | 82.986519 | -0.01076657 | 0.88905493 | -0.0121101 | 0.99033775 | 0.99533983 |
| IGKV3-7 | 18.1114468 | -0.361851363 | 1.09454129 | -0.3305964 | 0.7409494 | 0.85460507 |
| IGKV1-8 | 32.3744662 | 0.711741641 | 1.11714615 | 0.63710701 | 0.52405515 | 0.69921748 |
| IGKV1-9 | 217.86915 | -0.474216111 | 0.95567555 | -0.4962104 | 0.619746 | 0.77271053 |
| IGKV2-10 | 0 | NA | NA | NA | NA | NA |
| IGKV3-11 | 733.919146 | -0.101566849 | 0.98270697 | -0.1033542 | 0.9176819 | 0.96005997 |
| IGKV1-12 | 177.690715 | -0.687720649 | 0.99338887 | -0.6922975 | 0.48875051 | 0.67221358 |
| IGKV1-13 | 11.4962681 | 0.06977171 | 0.91726704 | 0.07606477 | 0.93936757 | 0.97110795 |
| IGKV2-14 | 0.93765298 | -3.115404345 | 2.64778144 | -1.1766093 | 0.23935147 | NA |
| IGKV3-15 | 629.362997 | -0.475717601 | 1.16146538 | -0.409584 | 0.68211116 | 0.81589727 |
| IGKV1-16 | 226.065207 | -1.582102321 | 1.32307954 | -1.1957726 | 0.2317853 | 0.4197736 |
| IGKV1-17 | 135.419822 | 0.941107386 | 0.80841536 | 1.16413843 | NA | NA |
| IGKV2-18 | 2.44861492 | -0.594642765 | 1.9836509 | -0.2997719 | 0.76435116 | 0.86961516 |
| IGKV2-19 | 0 | NA | NA | NA | NA | NA |
| IGKV3-20 | 2554.20499 | 1.685240491 | 1.22195233 | 1.37913767 | 0.16785231 | 0.3390735 |
| IGKV6-21 | 18.7738679 | 0.973213226 | 0.99215514 | 0.98090832 | 0.32663796 | 0.52457021 |
| IGKV1-22 | 0.5710773 | -1.32647576 | 2.79974483 | -0.4737845 | 0.63565357 | NA |
| IGKV2-23 | 0.08757577 | 0.547239394 | 3.29377202 | 0.16614368 | 0.86804388 | NA |
| IGKV2-24 | 97.5157131 | -0.194636132 | 1.26962824 | -0.1533017 | NA | NA |
| IGKV3-25 | 0.91152058 | -3.088418688 | 3.25728697 | -0.9481568 | 0.34304965 | NA |
| IGKV2-26 | 13.2160858 | -2.527044977 | 1.52595799 | -1.6560384 | 0.09771405 | 0.23388577 |
| IGKV1-27 | 153.321078 | 0.191946956 | 0.90114695 | 0.21300295 | 0.83132466 | 0.91040231 |
| IGKV2-28 | 189.372326 | -0.778462492 | 0.85145555 | -0.9142726 | 0.36057362 | 0.55819692 |
| IGKV2-29 | 37.7362804 | -1.160664549 | 1.5266243 | -0.7602817 | 0.44708621 | 0.63731355 |
| IGKV2-30 | 162.306669 | 0.677840991 | 0.97557099 | 0.69481462 | 0.48717148 | 0.67117308 |
| IGKV3-31 | 0.35715186 | -1.747804551 | 2.8380943 | -0.6158374 | 0.53800186 | NA |
| IGKV1-32 | 0 | NA | NA | NA | NA | NA |
| IGKV1-33 | 149.789824 | 0.411496403 | 0.98762546 | 0.41665229 | 0.67693276 | 0.81209758 |
| IGKV3-34 | 2.6019398 | -1.708697569 | 1.37094576 | -1.2463641 | 0.21263076 | 0.39645571 |
| IGKV1-35 | 0.5193916 | -2.27975872 | 2.16660953 | -1.0522241 | 0.29269675 | NA |
| IGKV2-36 | 0 | NA | NA | NA | NA | NA |

| | | | | | | |
|---|---|---|---|---|---|---|
| IGKV1-37 | 8.98611166 | -1.12052436 | 1.19070958 | -0.941056 | 0.34667618 | 0.54425497 |
| IGKV2-38 | 0 | NA | NA | NA | NA | NA |
| IGKV1-39 | 348.311975 | -0.15976568 | 0.97567804 | -0.1637484 | 0.86992925 | 0.93221856 |
| IGKV2-40 | 19.0702691 | -2.585154758 | 1.04769475 | -2.4674694 | 0.01360718 | 0.05658752 |
| IGKV2D-40 | 17.4949497 | -2.516230224 | 1.03192014 | -2.4383963 | 0.01475259 | 0.06009754 |
| IGKV1D-39 | 26.0790986 | 0.28369985 | 1.05104405 | 0.26992194 | 0.78722031 | 0.88414299 |
| IGKV2D-38 | 0 | NA | NA | NA | NA | NA |
| IGKV1D-37 | 0.44563969 | 0.341158896 | 2.2746778 | 0.1499812 | 0.88077945 | NA |
| IGKV2D-36 | 0.51998226 | -2.27683341 | 3.28074098 | -0.6939997 | 0.48768237 | NA |
| IGKV1D-35 | 0 | NA | NA | NA | NA | NA |
| IGKV3D-34 | 0 | NA | NA | NA | NA | NA |
| IGKV1D-33 | 6.90029087 | 0.953878131 | 0.87833179 | 1.08601117 | 0.27747405 | 0.47297936 |
| IGKV1D-32 | 0 | NA | NA | NA | NA | NA |
| IGKV3D-31 | 0 | NA | NA | NA | NA | NA |
| IGKV2D-30 | 5.29954218 | 0.510973446 | 1.22937178 | 0.41563785 | 0.67767502 | 0.81265532 |
| IGKV2D-29 | 53.5028623 | -1.535250105 | 1.54558544 | -0.993313 | 0.32055745 | 0.51805478 |
| IGKV2D-28 | 7.54174822 | 0.093572886 | 0.88061354 | 0.10625874 | 0.91537707 | 0.95871369 |
| IGKV1D-27 | 3.60498628 | -0.519095249 | 1.09888103 | -0.4723853 | 0.63665179 | 0.78476604 |
| IGKV2D-26 | 0.69909435 | -1.702568133 | 2.27710353 | -0.7476903 | 0.45464701 | NA |
| IGKV3D-25 | 0 | NA | NA | NA | NA | NA |
| IGKV2D-24 | 4.89433418 | -0.390938497 | 1.34878466 | -0.289845 | 0.77193482 | 0.87421716 |
| IGKV2D-23 | 0.17332742 | -1.003574173 | 3.30558948 | -0.3035992 | 0.7614333 | NA |
| IGKV1D-22 | 0.08757577 | 0.547239394 | 3.29377202 | 0.16614368 | 0.86804388 | NA |
| IGKV6D-21 | 5.14597676 | -0.025371166 | 1.73162164 | -0.0146517 | 0.98831007 | 0.99471234 |
| IGKV3D-20 | 101.422954 | 2.937093946 | 1.41021853 | 2.08272255 | NA | NA |
| IGKV2D-19 | 0 | NA | NA | NA | NA | NA |
| IGKV2D-18 | 0.51559131 | 0.007087053 | 2.26713838 | 0.00312599 | 0.99750582 | NA |
| IGKV6D-41 | 0 | NA | NA | NA | NA | NA |
| IGKV1D-17 | 3.17392763 | -1.006714509 | 1.3774075 | -0.7308763 | 0.46485471 | 0.65205955 |
| IGKV1D-16 | 17.8381401 | -1.02151581 | 1.19439749 | -0.8552562 | 0.39240935 | 0.58811555 |
| IGKV3D-15 | 90.8927378 | 4.649463282 | 1.47722654 | 3.14742739 | NA | NA |
| IGKV2D-14 | 0.08757577 | 0.547239394 | 3.29377202 | 0.16614368 | 0.86804388 | NA |
| IGKV1D-13 | 8.21570931 | -0.283208483 | 1.27441186 | -0.2222268 | 0.82413732 | 0.90587524 |
| IGKV1D-12 | 10.2100613 | -0.511526171 | 1.17841535 | -0.4340797 | 0.66423057 | 0.8027705 |
| IGKV3D-11 | 22.0941971 | -0.140527848 | 1.05866427 | -0.1327407 | 0.89439845 | 0.94571903 |
| IGKV2D-10 | 0.08393208 | -0.642905668 | 3.30558948 | -0.1944905 | 0.84579185 | NA |
| IGKV1D-42 | 0.49399025 | -1.100247467 | 2.75200797 | -0.3997981 | 0.68930525 | NA |
| IGKV1D-43 | 2.49500475 | 0.725505468 | 1.10198009 | 0.65836531 | 0.51030342 | 0.68891634 |
| IGKV1D-8 | 7.51240078 | 1.877539828 | 1.22075924 | 1.53800993 | NA | NA |
| IGKV3D-7 | 10.0604686 | 2.641259723 | 1.31263562 | 2.01218044 | NA | NA |
| IGKV1OR2-1 | 0 | NA | NA | NA | NA | NA |
| AC233263.6 | 2.62192312 | -1.234901937 | 0.88562537 | -1.3943841 | 0.16320165 | 0.33240345 |
| AC233266.2 | 8.20914813 | -0.526556179 | 0.69373516 | -0.7590161 | 0.44784291 | 0.63796545 |
| IGKV1OR2-1 | 1.48062469 | -2.288217338 | 1.60841657 | -1.4226522 | 0.15483701 | 0.32092427 |
| AL845331.2 | 73.5793108 | 1.290897873 | 0.57541914 | 2.24340448 | 0.02487074 | 0.08798518 |
| ANKRD20A8I | 1.1459661 | -0.115669776 | 1.44855053 | -0.0798521 | 0.9363549 | 0.96951778 |
| TEKT4 | 5.00235696 | -2.747762916 | 1.31168351 | -2.0948368 | 0.0361855 | 0.11525343 |
| AC103563.7 | 81.5611296 | 0.788281827 | 0.50270451 | 1.56808186 | 0.11686203 | 0.26520955 |
| MAL | 9323.68989 | 0.496990177 | 0.43536112 | 1.14155848 | 0.25363759 | 0.44628166 |
| AC103563.2 | 199.723488 | 1.232504026 | 0.42271725 | 2.91567005 | 0.00354926 | 0.02078479 |
| MRPS5 | 4055.04855 | -0.382781314 | 0.13892569 | -2.7552954 | 0.00586392 | 0.03030208 |
| ZNF514 | 1107.46588 | 0.024526631 | 0.22367791 | 0.10965155 | 0.91268572 | 0.95711137 |
| ZNF2 | 314.711144 | 0.196830106 | 0.11962574 | 1.64538261 | 0.09989093 | 0.23737948 |
| AC092835.1 | 8.0817533 | -0.173782639 | 0.52190448 | -0.3329779 | 0.73915099 | 0.85357616 |
| PROM2 | 5702.84607 | 1.01924824 | 0.37236341 | 2.73724058 | 0.0061957 | 0.03160776 |
| KCNIP3 | 1341.08938 | -0.096683682 | 0.34353871 | -0.2814346 | 0.77837708 | 0.87861887 |
| FAHD2A | 1767.16136 | 0.104818048 | 0.27013727 | 0.38801772 | 0.69800292 | 0.82654528 |
| TRIM43B | 0.25307292 | -1.364242679 | 3.30558948 | -0.4127078 | 0.67982068 | NA |
| AC009237.14 | 303.470152 | -0.180085458 | 0.3032526 | -0.5938464 | 0.55261486 | 0.72258908 |
| AC009237.15 | 22.6114584 | 0.597779632 | 0.50523494 | 1.1831716 | 0.23674114 | 0.42601653 |
| TRIM43 | 0.15318056 | -1.003574173 | 3.30558948 | -0.3035992 | 0.7614333 | NA |
| AC008268.1 | 0.12088456 | 0.02007176 | 3.29738704 | 0.00608717 | 0.99514317 | NA |
| LINC00342 | 827.455952 | -0.261749886 | 0.29025741 | -0.9017854 | 0.36717089 | 0.56403916 |
| ANKRD36C | 61.9107656 | -0.449632686 | 0.34659125 | -1.2972996 | 0.19452813 | 0.37448581 |
| GPAT2 | 442.798332 | -1.083844306 | 0.36967389 | -2.931893 | 0.00336903 | 0.02002282 |
| ADRA2B | 446.369958 | -0.107378253 | 0.44260834 | -0.2426033 | 0.80831271 | 0.89635013 |
| ASTL | 4.59995784 | 0.754353475 | 0.86355081 | 0.87354845 | 0.38236421 | 0.57848217 |
| DUSP2 | 546.609574 | 1.808357812 | 0.77684788 | 2.32781455 | 0.01992195 | 0.07470754 |
| AC012307.1 | 2.92280971 | 2.258262546 | 1.01816293 | 2.21797757 | 0.02655636 | 0.09236587 |
| STARD7 | 7571.68981 | 0.297758775 | 0.17510048 | 1.70050234 | 0.08903648 | 0.21891766 |
| TMEM127 | 6217.67199 | -0.14963707 | 0.14138672 | -1.0583531 | 0.28989451 | 0.48604516 |
| CIAO1 | 4320.05668 | 0.362043978 | 0.16726861 | 2.16444657 | 0.03043009 | 0.10185854 |
| SNRNP200 | 7116.13807 | 0.094804004 | 0.10861985 | 0.87280548 | 0.38276911 | 0.5788687 |
| AC021188.1 | 4.82678687 | -0.974332057 | 0.90255518 | -1.0795263 | 0.28035318 | 0.47604078 |
| ITPRIPL1 | 52.0977619 | -0.600052388 | 0.48285177 | -1.2427259 | 0.21396886 | 0.39802193 |
| NCAPH | 32.6498464 | -0.80773402 | 0.52560934 | -1.5367574 | 0.12435274 | 0.27630449 |
| NEURL3 | 761.016822 | -1.083451911 | 0.49166159 | -2.2036538 | 0.0275487 | 0.09493718 |
| AC013270.1 | 0.5920687 | -0.332696691 | 2.19375193 | -0.1516565 | 0.87945789 | NA |
| ARID5A | 2114.14635 | -1.281464625 | 0.39979662 | -3.2052913 | 0.00134926 | 0.00994965 |
| KANSL3 | 4142.18134 | 0.090878891 | 0.14481833 | 0.62753721 | 0.53030715 | 0.70472412 |
| FER1L5 | 102.061089 | -0.17470589 | 0.35778527 | -0.4882982 | 0.62533866 | 0.77684932 |
| LMAN2L | 1907.00737 | 0.027457518 | 0.22169389 | 0.12385329 | 0.90143144 | 0.95006982 |
| CNNM4 | 1195.80732 | -0.315357036 | 0.21869166 | -1.4420167 | 0.14929765 | 0.3134045 |
| CNNM3-DT | 45.5556079 | 0.792704238 | 0.21913918 | 3.61735508 | 0.00029763 | 0.00301054 |
| CNNM3 | 1714.36289 | 0.177077309 | 0.09661575 | 1.83279969 | 0.06683236 | 0.17824241 |
| ANKRD23 | 1013.5626 | -0.187136881 | 0.10173171 | -1.8395137 | 0.06583966 | 0.17644225 |
| ANKRD39 | 629.221854 | -0.273045723 | 0.11627823 | -2.3482102 | 0.01886387 | 0.0719122 |
| SEMA4C | 3223.89712 | -0.46531039 | 0.23760461 | -1.9583391 | 0.05019024 | 0.14602601 |
| FAM178B | 5.8887732 | 0.100153319 | 0.66077478 | 0.15156953 | 0.87952647 | 0.93747425 |
| IGKV2OR2-1 | 0 | NA | NA | NA | NA | NA |
| IGKV2OR2-2 | 0 | NA | NA | NA | NA | NA |
| IGKV1OR2-3 | 0 | NA | NA | NA | NA | NA |
| FAHD2B | 476.85846 | 0.632971965 | 0.36196219 | 1.74872397 | 0.08033874 | 0.20300146 |
| ANKRD36 | 135.242022 | 0.255930333 | 0.30045064 | 0.85182157 | 0.39431313 | 0.58959159 |
| AC159540.2 | 4.77838628 | -1.280078123 | 0.6762193 | -1.8929926 | 0.05835886 | 0.16231148 |
| AC159540.1 | 0 | NA | NA | NA | NA | NA |
| IGKV1OR2-9 | 0 | NA | NA | NA | NA | NA |
| IGKV2OR2-1 | 0 | NA | NA | NA | NA | NA |
| IGKV2OR2-7I | 0 | NA | NA | NA | NA | NA |
| IGKV3OR2-5 | 0 | NA | NA | NA | NA | NA |
| IGKV1OR2-6 | 0 | NA | NA | NA | NA | NA |
| IGKV2OR2-7 | 0 | NA | NA | NA | NA | NA |
| IGKV2OR2-8 | 0 | NA | NA | NA | NA | NA |
| IGKV1OR2-1 | 0 | NA | NA | NA | NA | NA |
| AC092683.1 | 32.693522 | -0.9845375 | 0.51433215 | -1.9142056 | 0.05559389 | 0.15686509 |
| AC092683.2 | 29.2270938 | -0.936681043 | 0.50805329 | -1.8436669 | 0.06523169 | 0.1752404 |
| ANKRD36B | 40.5088279 | -0.215571402 | 0.43019145 | -0.5011057 | 0.61629671 | 0.77049015 |
| COX5B | 9088.25847 | 0.08329119 | 0.15264093 | 0.54566748 | 0.58529453 | 0.74691332 |
| ACTR1B | 5119.38627 | -0.044309292 | 0.12415867 | -0.3568763 | 0.72118438 | 0.84137646 |
| C2orf92 | 160.634343 | 0.060318002 | 0.24657872 | 0.24461966 | 0.80675094 | 0.89552207 |
| ZAP70 | 465.614112 | 0.121266824 | 0.602461 | 0.20128576 | 0.84047513 | 0.91507243 |
| TMEM131 | 2748.54501 | -0.017419677 | 0.12205528 | -0.1427196 | 0.88651166 | 0.94122211 |
| VWA3B | 17.7580684 | -0.093753593 | 0.70802105 | -0.1324164 | 0.89465495 | 0.94587648 |

| | | | | | | |
|---|---|---|---|---|---|---|
| AC092675.1 | 0.95647651 | -1.52441814 | 1.92187614 | -0.7931927 | 0.42766555 | NA |
| CNGA3 | 4.0920174 | -0.317022802 | 0.7771498 | -0.4079301 | 0.68332499 | 0.81705268 |
| INPP4A | 1124.75991 | 0.081345354 | 0.11797429 | 0.68951764 | 0.49049758 | 0.67369536 |
| COA5 | 1753.57231 | 0.230455917 | 0.12237056 | 1.88326274 | 0.05966477 | 0.16495701 |
| UNC50 | 2211.33094 | -0.188431811 | 0.09440537 | -1.9959862 | 0.04593543 | 0.13690293 |
| MGAT4A | 3807.47468 | 0.066961329 | 0.25646809 | 0.2610903 | 0.79402287 | 0.88844572 |
| AC109826.1 | 21.4891815 | -0.443128357 | 0.73037225 | -0.6067158 | 0.54403956 | 0.71615206 |
| KIAA1211L | 426.81649 | 0.279422936 | 0.27511498 | 1.01565874 | 0.30979192 | 0.50697377 |
| TSGA10 | 434.595681 | 0.131288825 | 0.26765074 | 0.49052293 | 0.62376391 | 0.77580536 |
| C2orf15 | 209.126112 | 0.404627477 | 0.19235276 | 2.10356994 | 0.03541598 | 0.11357404 |
| AC079447.1 | 1103.69145 | 0.109008803 | 0.12214954 | 0.89242091 | 0.37216737 | 0.56863396 |
| LIPT1 | 397.326701 | 0.011411858 | 0.16321544 | 0.06991899 | 0.94425814 | 0.97338209 |
| AC092587.1 | 428.352988 | 0.269437448 | 0.13417087 | 2.00816645 | 0.04462561 | 0.13398352 |
| MITD1 | 1421.22528 | -0.330546892 | 0.14384552 | -2.2979298 | 0.02156579 | 0.07927095 |
| MRPL30 | 1452.41262 | 0.178958812 | 0.08800074 | 2.03360584 | 0.04199135 | 0.12833936 |
| LYG2 | 7.00331339 | 0.271915116 | 0.60967362 | 0.44600112 | 0.65559644 | 0.79801642 |
| LYG1 | 1727.14094 | -0.769806233 | 0.38698785 | -1.9892259 | 0.04667628 | 0.13849967 |
| TXNDC9 | 1678.41734 | -0.442062054 | 0.18398791 | -2.402669 | 0.01627591 | 0.06458928 |
| EIF5B | 5309.82171 | -0.585128827 | 0.16249091 | -3.6009943 | 0.000317 | 0.00316044 |
| REV1 | 2215.6321 | 0.002203952 | 0.11666335 | 0.01889155 | 0.98492762 | 0.99347171 |
| AC018690.1 | 45.5396412 | -0.174870303 | 0.43589108 | -0.4011789 | 0.68828841 | 0.82004995 |
| AFF3 | 240.705124 | 0.802399426 | 0.35231223 | 2.27752364 | 0.02275497 | 0.08246958 |
| AC092667.1 | 4.30666977 | 0.61568894 | 0.61642856 | 0.99880016 | 0.31789151 | 0.51510153 |
| LINC01104 | 0.4732412 | 2.210871823 | 1.99778565 | 1.10666118 | 0.26844045 | NA |
| LONRF2 | 564.613066 | 0.133395766 | 0.32259576 | 0.4135075 | 0.67923483 | 0.8138966 |
| CHST10 | 396.326943 | 0.138402501 | 0.21529825 | 0.64284082 | 0.52032739 | 0.69695908 |
| NMS | 0 | NA | NA | NA | NA | NA |
| PDCL3 | 1047.19181 | -0.274651741 | 0.25412424 | -1.0807774 | 0.27979614 | 0.47540104 |
| LINC01849 | 0 | NA | NA | NA | NA | NA |
| LINC01868 | 1.33573254 | -0.616442976 | 1.38413902 | -0.445362 | 0.65605814 | 0.79815617 |
| AC092168.1 | 0.08393208 | -0.642905668 | 3.30558948 | -0.1944905 | 0.84579185 | NA |
| NPAS2 | 3262.5813 | -1.041294502 | 0.32869469 | -3.1679687 | 0.00153508 | 0.01100899 |
| AC016738.2 | 153.735948 | -0.810646705 | 0.29314874 | -2.7653085 | 0.0056869 | 0.02966637 |
| AC016738.1 | 1130.88855 | -0.940738641 | 0.35290859 | -2.6656723 | 0.00768346 | 0.03706104 |
| RPL31 | 19652.0052 | -0.427931625 | 0.15215138 | -2.8125386 | 0.00491521 | 0.02654396 |
| TBC1D8 | 2268.09223 | -0.569111131 | 0.21110219 | -2.6959035 | 0.0070198 | 0.03469768 |
| TBC1D8-AS1 | 49.5042551 | 1.101384286 | 0.35291774 | 3.12079607 | 0.00180363 | 0.01244745 |
| CNOT11 | 3251.7177 | -0.82367814 | 0.20437875 | -4.0301555 | 5.57E-05 | 0.00080717 |
| RNF149 | 3909.82023 | -1.037526673 | 0.18655318 | -5.5615599 | 2.67E-08 | 1.59E-06 |
| CREG2 | 24.3012525 | -2.095021538 | 0.49079158 | -4.2686582 | 1.97E-05 | 0.00034928 |
| RFX8 | 47.9054151 | -1.165032212 | 0.63501846 | -1.8346431 | 0.06655859 | 0.17770073 |
| LINC01870 | 0 | NA | NA | NA | NA | NA |
| MAP4K4 | 3373.24531 | -0.855761195 | 0.31425156 | -2.7231724 | 0.00646583 | 0.03258867 |
| LINC01127 | 122.253202 | -0.327617176 | 0.56710451 | -0.5777016 | 0.56346561 | 0.7305885 |
| IL1R2 | 1075.84498 | -1.713488853 | 0.530802 | -3.228113 | 0.0012461 | 0.00933246 |
| IL1R1 | 5034.39192 | -1.001396883 | 0.24461269 | -4.0938059 | 4.24E-05 | 0.00065097 |
| AC007271.1 | 39.7689028 | -0.236999606 | 0.29237001 | -0.8106153 | 0.41758663 | 0.61036996 |
| IL1RL2 | 262.990889 | -0.34888738 | 0.28159889 | -1.2389515 | 0.21536345 | 0.39983565 |
| IL1RL1 | 13570.2885 | -3.054408153 | 0.76084127 | -4.0145143 | 5.96E-05 | 0.00084635 |
| IL18R1 | 590.644414 | -1.303850561 | 0.31088545 | -4.1939902 | 2.74E-05 | 0.00045728 |
| IL18RAP | 131.319316 | -1.62443705 | 0.34953226 | -4.6474596 | 3.36E-06 | 8.57E-05 |
| SLC9A4 | 592.169059 | 2.598049246 | 0.41974735 | 6.18955484 | 6.03E-10 | 6.68E-08 |
| SLC9A2 | 368.617785 | 0.951213973 | 0.35961746 | 2.6450717 | 0.00816736 | 0.0387014 |
| MFSD9 | 726.891734 | -0.07836132 | 0.26475901 | -0.2959722 | 0.76725128 | 0.87152635 |
| TMEM182 | 167.098932 | 0.123359699 | 0.27855204 | 0.44286051 | 0.65786664 | 0.79937071 |
| LINC01796 | 0 | NA | NA | NA | NA | NA |
| LINC01935 | 0.15318056 | -1.003574173 | 3.30558948 | -0.3035992 | 0.7614333 | NA |
| AC108051.1 | 0.08695548 | -0.642905668 | 3.30558948 | -0.1944905 | 0.84579185 | NA |
| AC073987.1 | 0.08666371 | -0.642905668 | 3.30558948 | -0.1944905 | 0.84579185 | NA |
| AC013727.1 | 0 | NA | NA | NA | NA | NA |
| AC013727.2 | 0 | NA | NA | NA | NA | NA |
| LINC01965 | 4.67946386 | 0.420732891 | 0.95072706 | 0.44253804 | 0.65809991 | 0.79943711 |
| AC096554.1 | 0 | NA | NA | NA | NA | NA |
| AC068535.1 | 0.7890783 | 3.131847049 | 1.48897442 | 2.10335853 | 0.03543444 | NA |
| LINC01831 | 3.15439907 | 2.195627209 | 0.96732685 | 2.26978835 | 0.02322043 | 0.08375787 |
| LINC01102 | 8.52972704 | 2.244960785 | 0.99319601 | 2.26034012 | 0.02380015 | 0.08515739 |
| LINC01103 | 0.43468659 | 1.429435933 | 2.73259424 | 0.52310581 | 0.60090062 | NA |
| AC013402.2 | 0 | NA | NA | NA | NA | NA |
| AC013402.3 | 0 | NA | NA | NA | NA | NA |
| AC013402.1 | 0.24089599 | 0.077994266 | 3.07420509 | 0.02537055 | 0.9797594 | NA |
| AC068057.1 | 1.1425737 | -0.554490748 | 1.71792173 | -0.3227683 | 0.74687069 | 0.85831867 |
| LINC01114 | 30.6058599 | -0.311652589 | 0.65135416 | -0.4784687 | 0.63231664 | 0.78161486 |
| POU3F3 | 5800.34782 | -0.263468824 | 0.26158946 | -1.0071844 | 0.31384617 | 0.51084029 |
| AC018730.1 | 59.1645245 | 0.944005332 | 0.4483351 | 2.10557982 | 0.03524087 | 0.11320522 |
| AC010884.1 | 16.3117724 | 1.063318378 | 0.61916295 | 1.71734819 | 0.08591557 | 0.21309195 |
| MRPS9 | 1834.78566 | 0.032254461 | 0.14892321 | 0.21658452 | 0.82853215 | 0.90879717 |
| AC104655.1 | 755.414247 | -0.061547489 | 0.15126547 | -0.4068839 | 0.68409324 | 0.81752646 |
| LINC01918 | 1.73059538 | -0.292670782 | 1.23684158 | -0.2366275 | 0.81294575 | 0.89895498 |
| GPR45 | 4.76262193 | 0.910774912 | 0.73086764 | 1.24615575 | 0.21270722 | 0.3965422 |
| AC012360.2 | 20.8863005 | 0.27305277 | 0.49339602 | 0.55341503 | 0.57997925 | 0.74307904 |
| TGFBRAP1 | 1607.73811 | -0.223492284 | 0.18464474 | -1.2103907 | 0.22612899 | 0.41281666 |
| AC012360.1 | 2.33323632 | -1.035871228 | 0.94441625 | -1.0968376 | 0.2727124 | 0.46762862 |
| AC012360.3 | 242.668605 | 0.070787431 | 0.29149769 | 0.24284045 | 0.808129 | 0.89624087 |
| C2orf49 | 987.364296 | -0.083200652 | 0.13141062 | -0.6331349 | 0.52664554 | 0.7015024 |
| FHL2 | 2220.58556 | -0.969093431 | 0.32105888 | -3.018429 | 0.00254089 | 0.0161961 |
| AC108058.1 | 32.3728779 | -0.593283059 | 0.39622433 | -1.4973413 | 0.13430447 | 0.29151444 |
| NCK2 | 2585.21513 | -0.396243635 | 0.15161165 | -2.6135435 | 0.00896087 | 0.04152136 |
| AC010978.1 | 418.258816 | -0.097763206 | 0.1582784 | -0.6176661 | 0.53679544 | 0.71053163 |
| AC009505.1 | 0 | NA | NA | NA | NA | NA |
| C2orf40 | 534.462033 | 2.690386861 | 0.40306713 | 6.67478601 | 2.48E-11 | 4.41E-09 |
| UXS1 | 3674.64711 | 0.156085021 | 0.21724112 | 0.71848745 | 0.47245678 | 0.65870177 |
| AC018878.1 | 0.22673191 | 0.689158461 | 3.03348976 | 0.22718338 | 0.82028115 | NA |
| RGPD3 | 30.1244633 | 2.025100562 | 0.57808474 | 3.50312059 | 0.00045984 | 0.00425325 |
| CD8B2 | 36.9941157 | 2.700666392 | 0.73359239 | 3.6814264 | 0.00023193 | 0.00245921 |
| AC108868.1 | 0.59450169 | 3.168051306 | 2.44735881 | 1.29447766 | 0.19550048 | NA |
| ST6GAL2 | 54.406325 | 1.624530994 | 0.39050359 | 4.16009239 | 3.18E-05 | 0.00051402 |
| AC016994.1 | 0 | NA | NA | NA | NA | NA |
| LINC01789 | 11.3608609 | 6.317538989 | 0.91098687 | 6.93482991 | 4.07E-12 | 9.49E-10 |
| AC096669.1 | 0.66943403 | 2.120849991 | 1.89294049 | 1.12039972 | 0.26254346 | NA |
| LINC01885 | 0.99077615 | 3.892151455 | 1.70358529 | 2.28468247 | 0.02233145 | NA |
| LINC01886 | 15.8077581 | 4.499521246 | 0.9353612 | 4.81046384 | 1.51E-06 | 4.55E-05 |
| GACAT1 | 0 | NA | NA | NA | NA | NA |
| RGPD4-AS1 | 0 | NA | NA | NA | NA | NA |
| RGPD4 | 2.27894644 | 0.279574771 | 1.16024966 | 0.24096087 | 0.80958544 | 0.89703256 |
| SLC5A7 | 44.3220576 | 0.989147526 | 0.58140593 | 1.70130279 | 0.08888614 | 0.21862957 |
| LINC01593 | 9.32826368 | 0.981878719 | 0.55487478 | 1.76955008 | 0.07680212 | 0.19685769 |
| LINC01594 | 60.7472094 | 0.125984996 | 0.453074 | 0.27806715 | 0.78096081 | 0.88016021 |
| SULT1C3 | 1.1719759 | 1.728151289 | 1.29582896 | 1.33362607 | 0.18232643 | 0.359008 |
| SULT1C2 | 8060.25358 | 0.410226176 | 0.2967874 | 1.38222232 | 0.16690345 | 0.33788146 |
| SULT1C4 | 829.907135 | -0.155690775 | 0.40509749 | -0.3843292 | 0.7007345 | 0.8284778 |
| GCC2 | 2312.94753 | -0.290529822 | 0.11972665 | -2.4266095 | 0.01524065 | 0.06154609 |
| GCC2-AS1 | 174.644142 | -0.070106646 | 0.17557123 | -0.399306 | 0.68966775 | 0.82118262 |
| LIMS1 | 2353.4336 | -0.798782126 | 0.23469361 | -3.4035103 | 0.00066526 | 0.00565352 |

| | | | | | | |
|---|---|---|---|---|---|---|
| LIMS1-AS1 | 3.01178946 | -1.482221685 | 0.86156292 | -1.720387 | 0.08536211 | 0.2119424 |
| RANBP2 | 4211.45917 | 0.13104799 | 0.17239066 | 0.76018033 | 0.4471468 | 0.63733102 |
| CCDC138 | 133.946566 | 0.108159888 | 0.30457501 | 0.35511741 | 0.72250164 | 0.84246598 |
| EDAR | 111.272839 | 0.586283954 | 0.41862545 | 1.40049762 | 0.16136436 | 0.32998604 |
| SH3RF3-AS1 | 46.6902518 | -0.795659372 | 0.48112868 | -1.6537351 | 0.09818135 | 0.23469819 |
| SH3RF3 | 682.482382 | -0.84703781 | 0.26947079 | -3.1433381 | 0.00167033 | 0.01173323 |
| 10-Sep | 3476.48564 | -0.661150493 | 0.28873614 | -2.2898086 | 0.02203241 | 0.08057966 |
| SOWAHC | 2002.69878 | -0.48164122 | 0.49354344 | -0.9758842 | 0.32912187 | 0.52689812 |
| AC074387.1 | 0.912353 | -3.064141984 | 2.65003369 | -1.1562653 | 0.24757266 | NA |
| RGPD5 | 3.18225012 | -1.546002409 | 0.7785466 | -1.9857545 | 0.04706059 | 0.13931171 |
| LIMS3 | 0.90746626 | -2.112650763 | 1.88965986 | -1.1180058 | 0.2635645 | NA |
| AC013271.1 | 91.7150572 | -1.031443404 | 0.53955361 | -1.9116606 | 0.05591974 | 0.15749262 |
| LINC01123 | 190.546293 | -0.301278654 | 0.25272187 | -1.1921353 | 0.23320821 | 0.42150107 |
| MALL | 109.073516 | 0.667010466 | 0.32547709 | 2.04933152 | 0.04042971 | 0.1247521 |
| NPHP1 | 784.276815 | 0.17102977 | 0.31236416 | 0.54753328 | 0.58401241 | 0.74614377 |
| SMIM37 | 861.653615 | -0.415368457 | 0.15652535 | -2.6536817 | 0.00796189 | 0.03802844 |
| LINC01106 | 238.644847 | 0.202715698 | 0.23389033 | 0.8667126 | 0.3860995 | 0.58174897 |
| AC112229.3 | 97.9026874 | -1.156713 | 0.47710693 | -2.4244314 | 0.01533238 | 0.06180295 |
| LIMS4 | 36.9676611 | -1.511143373 | 0.50692114 | -2.9810226 | 0.00287288 | 0.01773543 |
| AC112229.2 | 0 | NA | NA | NA | NA | NA |
| RGPD6 | 1.64729324 | -2.157959739 | 1.41889026 | -1.5208785 | 0.12829032 | 0.28245446 |
| AC226101.1 | 0.28674021 | -1.364242679 | 3.30558948 | -0.4127078 | 0.67982068 | NA |
| BUB1 | 57.8910664 | -1.820677742 | 0.6339511 | -2.871953 | 0.00407944 | 0.02307976 |
| AC114776.1 | 0.16325399 | -1.003574173 | 3.30558948 | -0.3035992 | 0.7614333 | NA |
| ACOXL | 22.9780743 | -0.508183624 | 0.44866163 | -1.1326657 | 0.25735468 | 0.4499697 |
| MIR4435-2H[illegible] | 1095.81507 | -1.218466228 | 0.3285954 | -3.708105 | 0.00020882 | 0.00225768 |
| ACOXL-AS1 | 9.40694771 | 0.316160229 | 0.65224562 | 0.48472572 | 0.62787092 | 0.77859976 |
| BCL2L11 | 1704.90065 | -0.242065976 | 0.22344104 | -1.083355 | 0.27865086 | 0.47428021 |
| AC108463.2 | 6.14940691 | -1.363136923 | 0.62494674 | -2.1812049 | 0.02916826 | 0.09897648 |
| AC108463.3 | 3.20237142 | -0.748692722 | 0.92404328 | -0.8102356 | 0.4178048 | 0.61059148 |
| AC068491.2 | 0.88329345 | -1.206231687 | 1.31961786 | -0.9140765 | 0.36067664 | NA |
| AC068491.3 | 5.47027473 | -1.667089367 | 0.69589016 | -2.3956214 | 0.01659222 | 0.06550933 |
| AC017002.5 | 0.24357081 | -1.364242679 | 3.30558948 | -0.4127078 | 0.67982068 | NA |
| AC017002.3 | 6.28249584 | -0.262088608 | 0.54011757 | -0.4852436 | 0.62750354 | 0.77838442 |
| AC017002.1 | 5.37629888 | -1.331441012 | 0.69755292 | -1.9087312 | 0.05629678 | 0.1581997 |
| AC017002.2 | 0.75559553 | -0.803662209 | 1.85264042 | -0.4337929 | 0.66443886 | NA |
| ANAPC1 | 1495.87555 | -0.182101815 | 0.10847883 | -1.6786852 | 0.09321341 | 0.22621365 |
| MERTK | 1428.32953 | -0.321004582 | 0.24381402 | -1.3165961 | 0.18797405 | 0.36613054 |
| TMEM87B | 1722.56893 | -0.809362625 | 0.19952952 | -4.0563552 | 4.98E-05 | 0.00074248 |
| FBLN7 | 111.984723 | -0.253144513 | 0.5830902 | -0.434143 | 0.66418462 | 0.80275887 |
| ZC3H8 | 655.59063 | 0.19188111 | 0.0992549 | 1.93321541 | 0.05320967 | 0.15217051 |
| ZC3H6 | 651.854609 | 0.534944739 | 0.27645322 | 1.9350281 | 0.05298685 | 0.15167859 |
| RGPD8 | 64.3393583 | 0.213015734 | 0.36487161 | 0.5838101 | 0.55934808 | 0.72772184 |
| TTL | 1839.32239 | -0.415098787 | 0.25540355 | -1.6252663 | 0.10410583 | 0.24430636 |
| POLR1B | 959.572371 | 0.145138194 | 0.19023912 | 0.76292509 | 0.44550807 | 0.63595777 |
| CHCHD5 | 941.457102 | 0.002112369 | 0.19077518 | 0.01107256 | 0.99116556 | 0.99583588 |
| AC012442.1 | 14.8906003 | 0.298459636 | 0.351126 | 0.85000722 | 0.39532107 | 0.59059638 |
| AC079922.2 | 91.2876117 | 0.766068412 | 0.28811785 | 2.65887179 | 0.00784028 | 0.03763848 |
| SLC20A1 | 3408.76976 | -0.781452102 | 0.34551717 | -2.2616882 | 0.02371667 | 0.08495673 |
| NT5DC4 | 11.2907801 | 0.496844316 | 0.41323289 | 1.20233487 | 0.22923381 | 0.41665947 |
| CKAP2L | 27.6702291 | -0.815582017 | 0.56165405 | -1.4521074 | 0.14647174 | 0.30944238 |
| IL1A | 8.93404804 | -2.913260111 | 1.09677447 | -2.6562071 | 0.00790251 | 0.03782703 |
| IL1B | 164.194232 | 0.554137853 | 0.6856895 | 0.80814691 | 0.41900602 | 0.61189458 |
| IL37 | 0.85987558 | -2.223840476 | 1.38261623 | -1.6084293 | 0.10774118 | NA |
| IL36G | 2.13218625 | 0.412474782 | 1.63989162 | 0.25152564 | 0.80140774 | 0.89237067 |
| IL36A | 0.08695548 | -0.642905668 | 3.30558948 | -0.1944905 | 0.84579185 | NA |
| IL36B | 0.42885147 | -1.859940617 | 3.29773397 | -0.5640057 | 0.57275028 | NA |
| IL36RN | 0.29657586 | 1.235893319 | 2.82872083 | 0.4369089 | 0.66217741 | NA |
| IL1F10 | 0 | NA | NA | NA | NA | NA |
| IL1RN | 169.540537 | -1.167267268 | 0.67667901 | -1.7249941 | 0.08452854 | 0.21064238 |
| PSD4 | 2765.10798 | -0.359382278 | 0.25984883 | -1.3830437 | 0.16665148 | 0.337475 |
| AC016683.1 | 6.40078628 | 1.144855806 | 0.77706745 | 1.473303 | 0.14066935 | 0.30086598 |
| PAX8 | 42641.4946 | -1.158066188 | 0.29571746 | -3.9161238 | 9.00E-05 | 0.00116546 |
| IGKV1OR2-1[illegible] | 0.59374939 | 0.056860426 | 2.25109281 | 0.02525903 | 0.97984835 | NA |
| AC016745.1 | 0.31629348 | 1.314934116 | 2.84234801 | 0.46262249 | 0.64363498 | NA |
| AC016745.2 | 0.43020504 | -0.90630825 | 2.25788748 | -0.4013966 | 0.68812819 | NA |
| CBWD2 | 1040.83645 | -0.06987529 | 0.09179891 | -0.7611778 | 0.44655087 | 0.63689471 |
| FOXD4L1 | 11.2789977 | -0.289104155 | 0.45465381 | -0.6358776 | 0.52485623 | 0.69972119 |
| LINC01961 | 0.09997307 | 0.547239394 | 3.29377202 | 0.16614368 | 0.86804388 | NA |
| PGM5P4-AS1 | 20.4285871 | 1.231110244 | 0.42059803 | 2.92704708 | 0.00342197 | 0.02025985 |
| FAM138B | 9.80218766 | 1.83168159 | 0.67012894 | 2.7333271 | 0.0062698 | 0.031838 |
| RABL2A | 999.005779 | 0.400471038 | 0.15424061 | 2.59640463 | 0.00942051 | 0.04306307 |
| AC017074.2 | 1.3416182 | 0.003224725 | 1.49488494 | 0.00215717 | 0.99827883 | 0.99911237 |
| AC017074.1 | 68.8885795 | -0.837350942 | 0.39245424 | -2.133627 | 0.03287333 | 0.10768567 |
| SLC35F5 | 4062.20206 | -0.530529374 | 0.17229746 | -3.0791479 | 0.00207594 | 0.01385534 |
| AC104653.2 | 6.06103598 | -0.257874141 | 0.60698169 | -0.4248467 | 0.67094847 | 0.80780558 |
| AC104653.1 | 6.86748876 | 0.45567029 | 0.51011399 | 0.89327151 | 0.37171179 | 0.56836599 |
| AC110769.2 | 8.1165557 | -0.053134447 | 0.57450396 | -0.0924875 | 0.92631071 | 0.96430513 |
| ACTR3 | 8291.80094 | -0.622830777 | 0.17952944 | -3.4692403 | 0.00052193 | 0.00468939 |
| LINC01191 | 25.782654 | -2.955905084 | 0.50858957 | -5.8119656 | 6.17E-09 | 4.59E-07 |
| AC110769.1 | 0.87225068 | -0.603413983 | 1.49589311 | -0.4033804 | 0.68666839 | NA |
| AC010982.1 | 0 | NA | NA | NA | NA | NA |
| DPP10 | 23.8344181 | 1.155845552 | 0.70173673 | 1.64712134 | 0.09953311 | 0.23671429 |
| DPP10-AS3 | 0 | NA | NA | NA | NA | NA |
| DPP10-AS2 | 0 | NA | NA | NA | NA | NA |
| DPP10-AS1 | 5.58208311 | 0.370485645 | 0.99227709 | 0.37336914 | 0.70887372 | 0.83447565 |
| AC009312.1 | 0.80777368 | 2.559876964 | 1.81461355 | 1.4107009 | 0.15833283 | NA |
| DDX18 | 2508.52361 | -0.133794218 | 0.18694578 | -0.7156846 | 0.47418612 | 0.66003563 |
| AC009404.1 | 173.233964 | -0.175783113 | 0.34794427 | -0.5052048 | 0.613415 | 0.76841076 |
| CCDC93 | 2655.48215 | -0.342212536 | 0.23926267 | -1.4302797 | 0.15263677 | 0.31800835 |
| AC009303.2 | 28.171634 | 0.031173477 | 0.45700177 | 0.06821303 | 0.94561605 | 0.97395466 |
| AC009303.3 | 0.03392908 | 0.380740593 | 3.29738709 | 0.11546736 | 0.90807469 | NA |
| INSIG2 | 2190.21366 | -0.589941359 | 0.17908632 | -3.2941732 | 0.00098712 | 0.00776096 |
| AC093901.1 | 4.72544694 | 0.216822926 | 0.65409924 | 0.33148323 | 0.74027951 | 0.85420604 |
| LINC01956 | 1.07842342 | -1.100443598 | 1.84599104 | -0.5961262 | 0.55109093 | 0.72138337 |
| EN1 | 1.71829451 | 0.316634124 | 2.1744418 | 0.14561628 | 0.88422432 | 0.93997401 |
| MARCO | 399.490806 | -2.495734202 | 0.83243591 | -2.9981097 | 0.0027166 | 0.01703243 |
| AC013457.1 | 0.09644756 | -0.642905668 | 3.30558948 | -0.1944905 | 0.84579185 | NA |
| C1QL2 | 1.38618475 | -1.734345568 | 1.35916509 | -1.2760375 | 0.20194228 | 0.38380716 |
| STEAP3 | 1094.16669 | -0.709253713 | 0.25186072 | -2.8160553 | 0.00486173 | 0.02630901 |
| STEAP3-AS1 | 96.4876998 | 0.232644202 | 0.29037119 | 0.80119589 | 0.42301825 | 0.61559059 |
| C2orf76 | 292.17017 | 0.013299273 | 0.21327296 | 0.06235799 | 0.95027775 | 0.97619105 |
| DBI | 12610.3462 | -0.660651247 | 0.30298133 | -2.1805015 | 0.02922031 | 0.09906778 |
| TMEM37 | 10010.2661 | -0.059978093 | 0.44688515 | -0.1342137 | 0.89323362 | 0.94490415 |
| SCTR | 1203.65591 | -0.656704902 | 0.66551048 | -0.9867687 | 0.32375605 | 0.5215653 |
| AC013275.1 | 493.89876 | -0.140852827 | 0.59192493 | -0.2379572 | 0.81191425 | 0.89845484 |
| CFAP221 | 1218.65929 | -0.642226089 | 0.41324748 | -1.5540956 | 0.12016162 | 0.27015618 |
| TMEM177 | 919.697221 | 0.379585866 | 0.19806743 | 1.91644772 | 0.05530813 | 0.15635457 |
| PTPN4 | 1147.87245 | 0.657186051 | 0.14652432 | 4.48516697 | 7.29E-06 | 0.00015917 |
| EPB41L5 | 3792.12392 | 0.203104913 | 0.16426357 | 1.2364574 | 0.21628859 | 0.40109213 |
| AC012363.1 | 52.0890731 | 1.152483893 | 0.38610396 | 2.98490565 | 0.00283666 | 0.01756756 |
| TMEM185B | 737.636438 | -0.391849735 | 0.24982541 | -1.5684943 | 0.11676581 | 0.2650981 |
| RALB | 5461.02343 | -0.488088964 | 0.22883118 | -2.1329652 | 0.03292758 | 0.1078232 |

| | | | | | | |
|---|---|---|---|---|---|---|
| AC012363.2 | 0.35343227 | 1.682145278 | 2.83454133 | 0.59344532 | 0.55288317 | NA |
| INHBB | 587.139074 | -2.696960276 | 0.48626171 | -5.5463143 | 2.92E-08 | 1.71E-06 |
| AC073257.2 | 9.32520077 | -1.890655149 | 0.82252319 | -2.2986041 | 0.02152743 | 0.07918516 |
| AC073257.1 | 0 | NA | NA | NA | NA | NA |
| AC018866.1 | 0 | NA | NA | NA | NA | NA |
| GLI2 | 167.216539 | 0.146124185 | 0.32810951 | 0.44535187 | 0.65606549 | 0.79815617 |
| AC017033.1 | 1.89961288 | -0.801649625 | 1.0919078 | -0.7341734 | 0.46284308 | 0.65098743 |
| AC067960.1 | 0.12150485 | 1.04981648 | 3.28477129 | 0.31960109 | 0.74927075 | NA |
| AC079988.1 | 0.97420135 | 1.274853827 | 1.30916023 | 0.9737951 | 0.33015828 | NA |
| TFCP2L1 | 10948.6216 | 0.292961727 | 0.26110927 | 1.12198899 | 0.26186712 | 0.45541357 |
| CLASP1 | 2513.76665 | 0.180140969 | 0.14502177 | 1.24216505 | 0.21417567 | 0.39823808 |
| AC012447.1 | 13.1483248 | 0.27335905 | 0.49443551 | 0.552871 | 0.58035175 | 0.7433396 |
| NIFK-AS1 | 674.056153 | -0.265583901 | 0.14907857 | -1.7815029 | 0.07483034 | 0.19332576 |
| NIFK | 1220.93648 | -0.620082119 | 0.23381654 | -2.6520028 | 0.00800159 | 0.03813739 |
| TSN | 3122.83672 | 0.040408691 | 0.15673041 | 0.25782291 | 0.79654358 | 0.88965766 |
| LINC01823 | 0.40098181 | -1.795740071 | 2.80639654 | -0.639874 | 0.52225454 | NA |
| AC062020.1 | 0 | NA | NA | NA | NA | NA |
| LINC01826 | 0.08393208 | -0.642905668 | 3.30558948 | -0.1944905 | 0.84579185 | NA |
| AC073409.1 | 0 | NA | NA | NA | NA | NA |
| AC079154.1 | 0 | NA | NA | NA | NA | NA |
| CNTNAP5 | 11.6303667 | -0.892426285 | 0.97752882 | -0.9129412 | 0.36127349 | 0.55878832 |
| LINC01889 | 0.65621188 | 2.814113215 | 1.95705848 | 1.43793006 | 0.1504539 | NA |
| LINC01941 | 0.09989236 | -0.642905668 | 3.30558948 | -0.1944905 | 0.84579185 | NA |
| AC023347.1 | 1.28039989 | -1.677488259 | 1.61336314 | -1.0397462 | 0.29845781 | 0.49530098 |
| GYPC | 2118.32634 | -0.750035931 | 0.18848578 | -3.9792705 | 6.91E-05 | 0.00094802 |
| TEX51 | 9.22163454 | 4.733683253 | 0.92861388 | 5.09757968 | 3.44E-07 | 1.33E-05 |
| AC012508.2 | 12.2158363 | 1.209558229 | 0.76427825 | 1.58261501 | 0.11350924 | 0.25964943 |
| AC012508.1 | 7.47831027 | 1.096720624 | 0.93040961 | 1.17875032 | 0.23849761 | 0.42801021 |
| BIN1 | 8588.52793 | -0.018476797 | 0.37153029 | -0.0497316 | 0.96033627 | 0.98093621 |
| CYP27C1 | 29.5278284 | 0.411077544 | 0.45011862 | 0.91326492 | 0.36110323 | 0.55878717 |
| ERCC3 | 2396.79617 | -0.159743351 | 0.14076155 | -1.1348508 | 0.25643785 | 0.44921834 |
| MAP3K2 | 2408.65425 | -0.031341515 | 0.19732313 | -0.1588335 | 0.87380009 | 0.93424052 |
| AC068282.1 | 84.4596231 | 0.505013347 | 0.39823802 | 1.26811937 | 0.20475532 | 0.38711193 |
| PROC | 1870.60016 | 0.868900938 | 0.27896156 | 3.11476942 | 0.00184089 | 0.01263843 |
| IWS1 | 2274.16411 | -0.300277829 | 0.12231014 | -2.4550527 | 0.01408639 | 0.05818581 |
| AC010976.1 | 253.946814 | 0.402185053 | 0.17984372 | 2.23630303 | 0.02533193 | 0.08923363 |
| MYO7B | 3987.12706 | 0.748651258 | 0.43048289 | 1.73909645 | 0.0820178 | 0.20598185 |
| AC010976.2 | 253.472773 | 0.142559649 | 0.39307944 | 0.36267388 | 0.71684851 | 0.83899051 |
| LIMS2 | 3528.04216 | 0.413917089 | 0.34015702 | 1.21684124 | 0.22366463 | 0.40975658 |
| GPR17 | 363.240455 | 1.06010927 | 0.51406683 | 2.06220126 | 0.03918858 | 0.12196422 |
| WDR33 | 4538.30578 | -0.073630799 | 0.12318849 | -0.5977084 | 0.55003449 | 0.72039384 |
| SFT2D3 | 704.63614 | 0.07812914 | 0.18675726 | 0.41834593 | 0.67569421 | 0.81134996 |
| POLR2D | 1279.71964 | -0.320511115 | 0.19396961 | -1.652378 | 0.09845752 | 0.23508917 |
| AMMECR1L | 1503.15672 | -0.191969743 | 0.16234409 | -1.1824868 | 0.23701261 | 0.42624913 |
| AC012306.2 | 90.0018797 | 0.204869258 | 0.20773278 | 0.98621535 | 0.32402745 | 0.52163909 |
| SAP130 | 1085.21643 | 0.141008167 | 0.16521436 | 0.85348616 | 0.39338976 | 0.58884532 |
| UGGT1 | 2456.69279 | -0.034722151 | 0.15349232 | -0.2262143 | 0.82103478 | 0.90407079 |
| HS6ST1 | 6584.00377 | -0.476071727 | 0.30882821 | -1.5415422 | 0.12318487 | 0.2746124 |
| AC012451.1 | 0.81924651 | -0.973513029 | 1.84224268 | -0.5284391 | 0.59719462 | NA |
| AC068483.1 | 0 | NA | NA | NA | NA | NA |
| LINC01854 | 0.14114194 | 0.02007176 | 3.29738704 | 0.00608717 | 0.99514317 | NA |
| LINC02572 | 5.30288414 | 0.70473282 | 0.56290094 | 1.25196597 | 0.21058226 | 0.39397334 |
| LINC01856 | 0.35176226 | 2.413587081 | 2.07427612 | 1.16358042 | 0.24459409 | NA |
| RAB6C | 10.1479007 | -0.088748865 | 0.462751 | -0.1917854 | 0.84791034 | 0.92010671 |
| POTEF | 1.36890204 | -0.365110891 | 1.39175831 | -0.2623379 | 0.79306098 | 0.8878826 |
| AC018865.1 | 0 | NA | NA | NA | NA | NA |
| CCDC74B | 272.856301 | -0.294074387 | 0.46245794 | -0.6358943 | 0.52484529 | 0.69972119 |
| SMPD4 | 5341.15046 | -0.150297461 | 0.11072814 | -1.3573556 | 0.17466825 | 0.34859621 |
| MZT2B | 2815.00429 | -0.361975065 | 0.15460454 | -2.3412965 | 0.0192169 | 0.07276704 |
| TUBA3E | 0.58385584 | -0.015939823 | 2.72425271 | -0.0058511 | 0.99533154 | NA |
| CCDC115 | 2544.34575 | 0.147895682 | 0.1826732 | 0.80961895 | 0.41815921 | 0.61099046 |
| IMP4 | 3061.24111 | -0.272712915 | 0.18107727 | -1.5060583 | 0.13205222 | 0.28819142 |
| PTPN18 | 4186.04123 | -0.062408645 | 0.09719198 | -0.6421172 | 0.52079709 | 0.69741109 |
| POTEI | 2.49175447 | 0.052312119 | 0.76735002 | 0.06817243 | 0.94564837 | 0.97395466 |
| AC013269.2 | 0 | NA | NA | NA | NA | NA |
| CFC1B | 0 | NA | NA | NA | NA | NA |
| AC013269.1 | 0 | NA | NA | NA | NA | NA |
| PRSS40B | 0 | NA | NA | NA | NA | NA |
| PRSS40A | 0.2622387 | 1.101408102 | 3.0147524 | 0.36533949 | 0.714858 | NA |
| AC140481.2 | 0 | NA | NA | NA | NA | NA |
| CFC1 | 0 | NA | NA | NA | NA | NA |
| POTEJ | 18.3855459 | -0.873049869 | 0.54765742 | -1.5941533 | 0.11090171 | 0.25532373 |
| GPR148 | 0.08757577 | 0.547239394 | 3.29377202 | 0.16614368 | 0.86804388 | NA |
| AC140481.1 | 0.17423948 | 0.18657164 | 3.29377195 | 0.05664376 | 0.95482897 | NA |
| AMER3 | 1.12645236 | 0.087355527 | 1.37157846 | 0.06368978 | 0.94921724 | 0.97551965 |
| AC133785.1 | 0.08666371 | -0.642905668 | 3.30558948 | -0.1944905 | 0.84579185 | NA |
| ARHGEF4 | 115.392169 | 2.05139585 | 0.24811434 | 8.2679454 | 1.36E-16 | 1.16E-13 |
| AC009477.2 | 1.83535491 | 1.420224009 | 0.96993214 | 1.46425089 | 0.14312541 | 0.304618 |
| FAM168B | 6423.34452 | 0.327218648 | 0.28757045 | 1.13787301 | 0.25517351 | 0.44784349 |
| PLEKHB2 | 13989.6685 | 0.156272179 | 0.33411149 | 0.46772464 | 0.63998152 | 0.78705286 |
| POTEE | 27.0009938 | -0.796711876 | 0.54579405 | -1.4597299 | 0.14436433 | 0.3062311 |
| RAB6D | 10.8551474 | 0.121959726 | 0.42752287 | 0.28527065 | 0.7754368 | 0.87674575 |
| AC073869.5 | 3.47608088 | -0.937938685 | 0.87693461 | -1.0695651 | 0.28481509 | 0.48095192 |
| LINC01120 | 0.53613195 | -2.336985844 | 3.27852296 | -0.7128167 | 0.4759592 | NA |
| AC073869.3 | 10.9728733 | 0.272201094 | 0.55430382 | 0.49106841 | 0.62337807 | 0.77543656 |
| MZT2A | 1044.07548 | -0.407588167 | 0.18962933 | -2.1493941 | 0.03160318 | 0.10474256 |
| TUBA3D | 8.87687 | 0.22166922 | 0.49915244 | 0.44409123 | 0.65697663 | 0.7986941 |
| CCDC74A | 457.320535 | -0.520516659 | 0.40275654 | -1.2923854 | 0.19622372 | 0.37675788 |
| LINC01087 | 0 | NA | NA | NA | NA | NA |
| C2orf27B | 0.08695548 | -0.642905668 | 3.30558948 | -0.1944905 | 0.84579185 | NA |
| AC103564.1 | 0.35030308 | 2.408990581 | 3.21841175 | 0.74850292 | 0.45415687 | NA |
| AJ239322.2 | 0 | NA | NA | NA | NA | NA |
| AJ239322.1 | 0.09989236 | -0.642905668 | 3.30558948 | -0.1944905 | 0.84579185 | NA |
| LINC01945 | 0 | NA | NA | NA | NA | NA |
| ANKRD30BL | 0.87949852 | 1.394816501 | 1.53963417 | 0.90594021 | 0.3649675 | NA |
| AC097532.1 | 0 | NA | NA | NA | NA | NA |
| AC097532.2 | 6.75945283 | 1.017001931 | 0.57202194 | 1.77790721 | 0.0754191 | 0.19446578 |
| GPR39 | 903.561132 | -1.075888414 | 0.38119645 | -2.8223988 | 0.00476659 | 0.02594857 |
| LYPD1 | 922.115594 | -1.606143695 | 0.41421489 | -3.8775615 | 0.00010551 | 0.00132237 |
| AC010974.1 | 0.66595837 | 0.604700407 | 2.25037187 | 0.26871132 | 0.78815184 | NA |
| NCKAP5 | 332.465546 | 0.626424094 | 0.49263373 | 1.27158181 | 0.20352174 | 0.3857431 |
| AC016909.2 | 0.29434379 | -0.695409893 | 2.33854317 | -0.2973688 | 0.76618492 | NA |
| AC010890.1 | 11.8885507 | 1.316785488 | 0.43589889 | 3.02085073 | 0.00252066 | 0.01609436 |
| AC011243.1 | 0.3108733 | -1.364242679 | 3.30558948 | -0.4127078 | 0.67982068 | NA |
| MGAT5 | 2488.77635 | 0.351611862 | 0.17765381 | 1.97919682 | 0.04779385 | 0.14092853 |
| TMEM163 | 312.548163 | -1.043987554 | 0.40680737 | -2.5662946 | 0.01027915 | 0.04600494 |
| CCNT2-AS1 | 967.21815 | 1.026807298 | 0.54341446 | 1.88954723 | 0.05881854 | 0.16314767 |
| ACMSD | 4741.45928 | 1.016828316 | 0.56885737 | 1.78749256 | 0.07385791 | 0.19169618 |
| CCNT2 | 2254.327 | -0.237182677 | 0.14933045 | -1.5883075 | 0.11221682 | 0.25767262 |
| MAP3K19 | 8.1055876 | -1.087848537 | 0.68319358 | -1.5922991 | 0.11131751 | 0.25603512 |
| RAB3GAP1 | 2858.41812 | 0.03795537 | 0.10622248 | 0.35731955 | 0.7208526 | 0.84121268 |
| ZRANB3 | 387.667066 | 0.167559921 | 0.15081015 | 1.1110653 | 0.26654024 | 0.4606243 |
| R3HDM1 | 967.48473 | -0.064410301 | 0.08304451 | -0.7756118 | 0.43797826 | 0.6299491 |

| | | | | | | |
|---|---|---|---|---|---|---|
| UBXN4 | 7090.4336 | -0.178863746 | 0.17542524 | -1.019601 | 0.30791774 | 0.50478876 |
| LCT | 2.3623215 | 0.83201957 | 0.87186591 | 0.95429762 | 0.33993301 | 0.53809861 |
| AC011893.1 | 0.41843058 | 0.363671541 | 2.84353267 | 0.12789427 | 0.89823264 | NA |
| MCM6 | 821.906753 | -0.382248173 | 0.21638555 | -1.7665143 | 0.0773096 | 0.19783155 |
| DARS | 4547.27159 | -0.417457717 | 0.15464017 | -2.6995426 | 0.00694349 | 0.03440431 |
| DARS-AS1 | 421.527492 | -0.165488174 | 0.24116084 | -0.6862149 | 0.49257757 | 0.67542236 |
| AC068492.1 | 0.98317717 | -3.185623388 | 1.53149419 | -2.0800754 | 0.03751862 | NA |
| CXCR4 | 1785.0928 | -0.041266393 | 0.62134503 | -0.0664146 | 0.94704773 | 0.97446733 |
| THSD7B | 85.6740848 | 1.217734248 | 0.58317588 | 2.08810806 | 0.03678809 | 0.11657076 |
| HNMT | 3258.93332 | 0.764447581 | 0.32356243 | 2.36259689 | 0.0181474 | 0.06985066 |
| LINC01832 | 0.41348746 | 1.035701344 | 2.20827781 | 0.46900863 | 0.63906347 | NA |
| AC114763.1 | 9.27125569 | -0.01182166 | 0.38139995 | -0.0309954 | 0.97527318 | 0.98862634 |
| AC114763.2 | 0.54633891 | -0.670872773 | 1.90890189 | -0.3514443 | 0.72525502 | NA |
| SPOPL | 1242.12327 | -0.056855806 | 0.17177889 | -0.3309825 | 0.74065771 | 0.8544555 |
| AC092620.3 | 1.51593636 | 0.34842066 | 1.14274553 | 0.30489785 | 0.76044395 | 0.86715659 |
| AC092620.1 | 4.47062066 | 0.064435461 | 0.77262384 | 0.08339823 | 0.9335349 | 0.96800422 |
| NXPH2 | 65.4535834 | 0.512500698 | 0.79139724 | 0.6475897 | 0.51725036 | 0.69442481 |
| AC062021.1 | 0.6847025 | 2.167288715 | 1.692617 | 1.28043658 | 0.20039164 | NA |
| AC016710.1 | 0 | NA | NA | NA | NA | NA |
| AC078851.1 | 0 | NA | NA | NA | NA | NA |
| LINC01853 | 0.08393208 | -0.642905668 | 3.30558948 | -0.1944905 | 0.84579185 | NA |
| LRP1B | 87.2346364 | 0.02611022 | 0.32116841 | 0.0812976 | 0.93520528 | 0.96907653 |
| KYNU | 750.963676 | 1.030981964 | 0.35526262 | 2.90202767 | 0.00370756 | 0.02152065 |
| ARHGAP15 | 269.315668 | 0.229711001 | 0.50576658 | 0.45418383 | 0.64969652 | 0.79394306 |
| AC013437.1 | 0 | NA | NA | NA | NA | NA |
| AC096558.1 | 2.0122749 | -1.822471982 | 1.20729697 | -1.5095474 | 0.13115896 | 0.28676409 |
| AC096558.2 | 0.13028434 | 1.049816488 | 3.28477129 | 0.31960109 | 0.74927075 | NA |
| AC079793.1 | 5.21796886 | -0.531528366 | 0.82968679 | -0.6406374 | 0.52175831 | 0.69787989 |
| AC092652.1 | 0.98984784 | 0.300104504 | 1.32723479 | 0.2261126 | 0.82111385 | NA |
| AC092652.2 | 1.84051096 | 0.257440683 | 1.10733334 | 0.23248707 | 0.81615973 | 0.90104821 |
| AC079584.2 | 4.48798768 | 0.75939757 | 0.66925908 | 1.134684 | 0.25650774 | 0.44923353 |
| AC079584.1 | 0 | NA | NA | NA | NA | NA |
| AC016910.1 | 4.8177569 | 1.8576894 | 0.63981707 | 2.90346957 | 0.00369053 | 0.02146432 |
| GTDC1 | 900.857653 | 0.051503678 | 0.13801432 | 0.37317632 | 0.70901722 | 0.83449159 |
| ZEB2 | 2114.65334 | 0.547903667 | 0.20997447 | 2.60938233 | 0.00907058 | 0.04191191 |
| AC009951.2 | 0.60598168 | -0.030059045 | 1.84181101 | -0.0163204 | 0.9869788 | NA |
| ZEB2-AS1 | 89.7041738 | 1.993844565 | 0.30998086 | 6.43215382 | 1.26E-10 | 1.74E-08 |
| LINC01412 | 3.17287764 | -0.174436221 | 0.75402748 | -0.2313393 | 0.81705118 | 0.90160437 |
| TEX41 | 198.263833 | 1.85581053 | 0.32010614 | 5.797485 | 6.73E-09 | 4.92E-07 |
| AC023128.1 | 0.09635527 | 0.547239394 | 3.29377202 | 0.16614368 | 0.86804388 | NA |
| LINC01966 | 0 | NA | NA | NA | NA | NA |
| AC096666.1 | 0.03392908 | 0.380740593 | 3.29738709 | 0.11546736 | 0.90807469 | NA |
| AC092484.1 | 0.1857193 | 0.726716446 | 3.28420131 | 0.22127646 | 0.82487718 | NA |
| AC079163.1 | 0 | NA | NA | NA | NA | NA |
| AC062031.1 | 0.35856807 | 0.711047847 | 2.3196214 | 0.30653616 | 0.75919645 | NA |
| LINC01911 | 0.16416605 | 0.18657164 | 3.29377195 | 0.05664376 | 0.95482897 | NA |
| AC062032.1 | 0.11223674 | 0.02007176 | 3.29738704 | 0.00608717 | 0.99514317 | NA |
| ACVR2A | 897.254652 | 0.090450345 | 0.08311022 | 1.08831799 | 0.27645475 | 0.47179704 |
| AC009480.1 | 22.8833586 | -0.027796349 | 0.40288284 | -0.0689936 | 0.94499469 | 0.9737607 |
| ORC4 | 2148.72347 | 0.146289764 | 0.10990919 | 1.33100575 | 0.18318712 | 0.35993177 |
| MBD5 | 803.405801 | 0.137430234 | 0.1310905 | 1.04836154 | 0.29447207 | 0.49122143 |
| EPC2 | 1080.66597 | 0.070587185 | 0.18636254 | 0.37876274 | 0.70486406 | 0.83144403 |
| AC105402.3 | 0.34847896 | 0.732866073 | 3.27839706 | 0.22354402 | 0.82311212 | NA |
| KIF5C | 383.629128 | -1.508006009 | 0.50411098 | -2.9914167 | 0.00277686 | 0.01731517 |
| LYPD6B | 391.039753 | 0.126068391 | 0.46069189 | 0.27365012 | 0.78435351 | 0.88217189 |
| LYPD6 | 105.518986 | 1.868885854 | 0.38329805 | 4.87580325 | 1.08E-06 | 3.48E-05 |
| MMADHC | 4223.21358 | -0.135441806 | 0.15535838 | -0.8718024 | 0.38331619 | 0.57917139 |
| AC007364.1 | 478.767959 | 0.084341805 | 0.20129723 | 0.41899139 | 0.67522242 | 0.81102072 |
| LINC01931 | 2.67390222 | 0.433686143 | 0.75359075 | 0.57549293 | 0.56495797 | 0.73176196 |
| LINC01818 | 0 | NA | NA | NA | NA | NA |
| LINC01817 | 0 | NA | NA | NA | NA | NA |
| RND3 | 1789.19024 | -1.386684975 | 0.54233744 | -2.5568675 | 0.01056194 | 0.04688863 |
| LINC01920 | 0.21442573 | -1.003574173 | 3.30558948 | -0.3035992 | 0.7614333 | NA |
| AC104777.3 | 0 | NA | NA | NA | NA | NA |
| AC104777.2 | 0.3216386 | -1.364242679 | 3.30558948 | -0.4127078 | 0.67982068 | NA |
| AC104777.1 | 0.08666371 | -0.642905668 | 3.30558948 | -0.1944905 | 0.84579185 | NA |
| AC023469.2 | 0 | NA | NA | NA | NA | NA |
| AC023469.1 | 0 | NA | NA | NA | NA | NA |
| AC018731.1 | 0 | NA | NA | NA | NA | NA |
| RBM43 | 574.758816 | 0.315160194 | 0.30885701 | 1.02040811 | 0.30753495 | 0.50445985 |
| NMI | 784.559581 | -0.616197766 | 0.20430959 | -3.0160003 | 0.00256133 | 0.01629488 |
| TNFAIP6 | 99.1615281 | -3.118820386 | 0.72259915 | -4.3161141 | 1.59E-05 | 0.00029599 |
| RIF1 | 1776.2864 | -0.212496946 | 0.16509803 | -1.2870956 | 0.19806097 | 0.37901798 |
| NEB | 252.447142 | 0.942972665 | 0.38548405 | 2.44620408 | 0.01443693 | 0.05922322 |
| ARL5A | 2038.33507 | 0.211892847 | 0.15348455 | 1.38054834 | 0.16741788 | 0.33860415 |
| AC068547.1 | 896.82827 | 0.231648214 | 0.21621755 | 1.07136639 | 0.28400472 | 0.48004643 |
| CACNB4 | 208.352685 | 1.479342502 | 0.36683951 | 4.03266955 | 5.51E-05 | 0.00080167 |
| AC079790.1 | 2.17018624 | 0.833075364 | 1.02038508 | 0.81643232 | 0.41425292 | 0.60794725 |
| STAM2 | 1747.82686 | 0.325716056 | 0.11719548 | 2.77925449 | 0.00544838 | 0.02872328 |
| FMNL2 | 3605.63864 | -0.879436913 | 0.31072376 | -2.8302854 | 0.00465065 | 0.02548099 |
| PRPF40A | 2997.73011 | -0.157097054 | 0.10890436 | -1.442523 | 0.14915487 | 0.31330365 |
| ARL6IP6 | 615.883779 | -0.345666247 | 0.13001197 | -2.6587264 | 0.00784366 | 0.03764787 |
| AC079150.1 | 0 | NA | NA | NA | NA | NA |
| LINC01850 | 0 | NA | NA | NA | NA | NA |
| RPRM | 72.3933755 | 0.508854014 | 0.59008043 | 0.86234687 | 0.38849667 | 0.5841377 |
| GALNT13 | 27.4359372 | 0.37791739 | 0.45773865 | 0.82561827 | 0.40902067 | 0.60343935 |
| AC009227.1 | 9.23368237 | 0.505669871 | 0.59318116 | 0.85247123 | 0.3939526 | 0.58925298 |
| AC009227.2 | 0.23568937 | 1.881156793 | 3.2502805 | 0.57876752 | 0.56274605 | NA |
| AC061961.1 | 14.3505007 | 2.439389147 | 0.63875315 | 3.81898569 | 0.000134 | 0.00159244 |
| KCNJ3 | 739.692179 | 1.211778247 | 0.30785251 | 3.93622987 | 8.28E-05 | 0.00109298 |
| AC092625.1 | 0 | NA | NA | NA | NA | NA |
| LINC01876 | 10.985836 | -0.076135216 | 0.6382682 | -0.1192841 | 0.90505032 | 0.95236141 |
| AC074099.1 | 0.34103497 | 1.629530304 | 3.24275583 | 0.50251403 | 0.61530599 | NA |
| NR4A2 | 928.383313 | 1.881224998 | 0.71331845 | 2.63728634 | 0.00835723 | 0.03932549 |
| GPD2 | 1217.72759 | -0.295635159 | 0.17128176 | -1.7260166 | 0.08434442 | 0.21040227 |
| LINC01958 | 0.09997307 | 0.547239394 | 3.29377202 | 0.16614368 | 0.86804388 | NA |
| GALNT5 | 11.9871915 | -0.143299318 | 0.73645522 | -0.1945798 | 0.84572191 | 0.91863438 |
| ERMN | 40.9934206 | 0.263856559 | 0.51982796 | 0.50758439 | 0.61174484 | 0.76749276 |
| CYTIP | 518.382574 | -0.734357814 | 0.40983308 | -1.7918461 | 0.07315762 | 0.19030939 |
| ACVR1C | 68.1086595 | 0.138047448 | 0.38253869 | 0.36087186 | 0.71819524 | 0.83985952 |
| AC019186.1 | 27.5685571 | -0.809455487 | 0.37046399 | -2.1849775 | 0.02889049 | 0.09821353 |
| ACVR1 | 2101.76709 | -0.096071864 | 0.22356762 | -0.4297217 | 0.66739808 | 0.80538565 |
| UPP2 | 1693.29511 | 3.053354677 | 0.75634171 | 4.03700423 | 5.41E-05 | 0.00079312 |
| AC013731.1 | 31.365709 | 0.322406509 | 0.38949409 | 0.82775713 | 0.40780806 | 0.60242555 |
| AC091488.1 | 10.544517 | 0.665867993 | 0.49022762 | 1.35828331 | 0.1743738 | 0.34829853 |
| CCDC148-AS1 | 31.2476663 | 0.041083446 | 0.41063043 | 0.10004969 | 0.92030488 | 0.96099929 |
| CCDC148 | 290.192051 | 0.534154055 | 0.37382576 | 1.42888511 | 0.15303726 | 0.31857242 |
| PKP4 | 9290.65664 | 0.429309291 | 0.22660586 | 1.89451983 | 0.05815604 | 0.16199164 |
| PKP4-AS1 | 553.265649 | 0.981802156 | 0.22338054 | 4.39520012 | 1.11E-05 | 0.00022143 |
| AC005042.2 | 0 | NA | NA | NA | NA | NA |
| DAPL1 | 38.7578451 | -1.065046705 | 0.67805937 | -1.5707278 | 0.11624589 | 0.26428175 |
| TANC1 | 2174.14944 | -0.173767107 | 0.18626622 | -0.9328965 | 0.35087339 | 0.54857388 |
| WDSUB1 | 1148.66664 | 0.47854018 | 0.19228162 | 2.48874634 | 0.01281944 | 0.05416578 |

| | | | | | | |
|---|---|---|---|---|---|---|
| BAZ2B | 1460.9389 | -0.132203954 | 0.17304914 | -0.7639677 | 0.44488648 | 0.63551603 |
| AC008277.1 | 9.90013283 | 0.399221665 | 0.4614912 | 0.86506886 | 0.387001 | 0.58265579 |
| AC009506.1 | 163.618138 | 0.596380716 | 0.24660837 | 2.41833123 | 0.01559188 | 0.06256205 |
| AC009961.1 | 24.0417256 | -0.070304846 | 0.27509631 | -0.2555645 | 0.79828715 | 0.89081427 |
| 7-Mar | 4532.47555 | -0.130203226 | 0.13524217 | -0.9627413 | 0.33567735 | 0.53388013 |
| CD302 | 1719.6036 | 0.253600253 | 0.28241291 | 0.89797684 | 0.3691979 | 0.5660648 |
| LY75-CD302 | 503.915119 | 0.354431912 | 0.26518343 | 1.33655377 | 0.18136834 | 0.3577309 |
| LY75 | 702.090254 | -0.290135653 | 0.3405202 | -0.8520365 | 0.39419382 | 0.58948004 |
| PLA2R1 | 2636.29207 | 0.75780992 | 0.45932496 | 1.64983397 | 0.0989769 | 0.2359021 |
| ITGB6 | 11364.8351 | -2.877311696 | 0.4767315 | -6.0354973 | 1.58E-09 | 1.48E-07 |
| LINC02478 | 62.1156525 | -2.248077425 | 0.60237865 | -3.7320005 | 0.00018997 | 0.00209247 |
| RBMS1 | 4141.16526 | -0.645692187 | 0.20454522 | -3.156721 | 0.00159554 | 0.01132554 |
| AC009313.1 | 0.28145235 | -0.179698839 | 2.92317751 | -0.0614738 | 0.95098188 | NA |
| TANK | 1987.21142 | -0.504131487 | 0.08662322 | -5.8198192 | 5.89E-09 | 4.42E-07 |
| AC009299.3 | 25.3614175 | 0.007913694 | 0.53860509 | 0.01469294 | 0.98827715 | 0.99471234 |
| LINC01806 | 66.094724 | -0.165136844 | 0.68872111 | -0.2397732 | 0.81050611 | 0.8974992 |
| PSMD14 | 3010.16683 | -0.64673161 | 0.25697267 | -2.516733 | 0.01184486 | 0.05109583 |
| TBR1 | 1.10886226 | -0.112911686 | 1.29578211 | -0.0871379 | 0.93056193 | 0.96655578 |
| AC009487.1 | 0.08695548 | -0.642905668 | 3.30558948 | -0.1944905 | 0.84579185 | NA |
| SLC4A10 | 4.91315682 | 0.574333312 | 0.85564335 | 0.6712298 | 0.50207415 | 0.68251297 |
| DPP4 | 7694.13262 | 0.641974809 | 0.55625325 | 1.15410527 | 0.24845701 | 0.44013243 |
| AC008063.1 | 196.478546 | 1.395957117 | 0.75073691 | 1.85944916 | 0.0629635 | 0.17101616 |
| AC008063.2 | 0.69199782 | -2.653370068 | 2.23111937 | -1.1892551 | 0.2343393 | NA |
| GCG | 0.07659028 | -0.642905668 | 3.30558948 | -0.1944905 | 0.84579185 | NA |
| AC007750.1 | 47.9172328 | -2.070327946 | 0.58138351 | -3.5610366 | 0.00036939 | 0.00358385 |
| FAP | 103.281669 | -1.585092058 | 0.58947954 | -2.6889688 | 0.00716731 | 0.03525618 |
| IFIH1 | 819.855021 | 0.247256829 | 0.22144178 | 1.11657716 | 0.26417516 | 0.45803794 |
| GCA | 1586.39771 | -0.595027503 | 0.19196427 | -3.0996784 | 0.00193731 | 0.01315993 |
| KCNH7 | 11.0914074 | -0.746184706 | 0.47120958 | -1.5835516 | 0.11329579 | 0.25929612 |
| AC011900.1 | 0 | NA | NA | NA | NA | NA |
| FIGN | 429.280702 | 0.391645831 | 0.27254403 | 1.43700022 | 0.15071793 | 0.3154311 |
| AC016766.1 | 0.03392908 | 0.380740593 | 3.29738709 | 0.11546736 | 0.90807469 | NA |
| GRB14 | 2057.04382 | 0.195104181 | 0.21984845 | 0.88744853 | 0.37483747 | 0.5707954 |
| COBLL1 | 6316.15015 | 0.889290439 | 0.22661525 | 3.92423039 | 8.70E-05 | 0.0011364 |
| AC019197.1 | 477.462603 | 0.160609193 | 0.32784555 | 0.48989286 | 0.62420972 | 0.77611992 |
| SLC38A11 | 538.84645 | 0.971864201 | 0.41477243 | 2.34312631 | 0.01912291 | 0.07255699 |
| SCN3A | 360.040953 | -0.873264853 | 0.32065897 | -2.7233446 | 0.00646246 | 0.03257792 |
| SCN2A | 850.913137 | -0.143978467 | 0.41310575 | -0.3485269 | 0.72744452 | 0.84605971 |
| CSRNP3 | 229.202956 | 1.027795764 | 0.38683694 | 2.65692247 | 0.00788576 | 0.03778801 |
| GALNT3 | 1308.42256 | -0.809796374 | 0.21476541 | -3.7706089 | 0.00016285 | 0.00186225 |
| AC009495.3 | 16.0888001 | -0.523188108 | 0.51168677 | -1.0224773 | 0.30655505 | 0.50341616 |
| AC009495.1 | 7.97068253 | 0.227422639 | 0.48052326 | 0.47328123 | 0.63601255 | 0.7841886 |
| AC009495.2 | 3.83191243 | 0.86146918 | 0.65435007 | 1.31652645 | 0.18799741 | 0.36613054 |
| TTC21B | 844.942419 | 0.534414495 | 0.16296648 | 3.27929091 | 0.00104068 | 0.00808807 |
| AC010127.1 | 31.9394606 | 0.790961087 | 0.45342708 | 1.74440638 | 0.08108825 | 0.20436751 |
| SCN1A | 12.0996264 | 1.701189963 | 0.78409843 | 2.16961277 | 0.03003619 | 0.10091151 |
| SCN9A | 301.294079 | 0.964790576 | 0.37270122 | 2.58864348 | 0.00963548 | 0.04383088 |
| SCN7A | 233.476447 | 0.676440707 | 0.5376902 | 1.25804917 | 0.20837397 | 0.39152061 |
| XIRP2 | 2.62649924 | 2.128463625 | 1.13744182 | 1.87127252 | 0.06130732 | 0.16791379 |
| XIRP2-AS1 | 0 | NA | NA | NA | NA | NA |
| AC073050.1 | 0.25254714 | -0.178391175 | 2.99848453 | -0.0594938 | 0.95255882 | NA |
| AC016723.1 | 2.32743845 | 0.216092472 | 0.83262337 | 0.25953208 | 0.79522474 | 0.88893922 |
| B3GALT1 | 14.7077237 | 0.018221544 | 0.50965165 | 0.03575294 | 0.97147936 | 0.98670705 |
| STK39 | 1621.90492 | -1.365402335 | 0.19887299 | -6.8657001 | 6.62E-12 | 1.40E-09 |
| CERS6 | 1568.81436 | 0.174134885 | 0.17414456 | 0.99994444 | 0.3173374 | 0.51455372 |
| NOSTRIN | 2122.27062 | -0.468099955 | 0.39908802 | -1.1729241 | 0.24082624 | 0.43092814 |
| SPC25 | 68.9162648 | -1.02607808 | 0.66615213 | -1.5403059 | 0.1234858 | 0.27502924 |
| G6PC2 | 7.22312053 | -0.483953625 | 1.81410742 | -0.2667723 | 0.78964448 | 0.88562967 |
| ABCB11 | 0.19718306 | -0.340099831 | 3.16382928 | -0.1074963 | 0.91439529 | NA |
| DHRS9 | 32.8388939 | -1.082722017 | 0.49416657 | -2.1910062 | 0.02845134 | 0.09722197 |
| AC007556.1 | 0.08757577 | 0.547239394 | 3.29377202 | 0.16614368 | 0.86804388 | NA |
| LRP2 | 17443.4912 | 1.309792513 | 0.46227584 | 2.83335706 | 0.00460619 | 0.02533165 |
| BBS5 | 1017.36808 | -0.099261974 | 0.20586266 | -0.4821757 | 0.62968112 | 0.78003905 |
| AC093899.2 | 298.80495 | 0.285425099 | 0.22381104 | 1.27529502 | 0.20220484 | 0.38401466 |
| KLHL41 | 76.8142896 | 0.286251966 | 0.64034827 | 0.44702544 | 0.65485669 | 0.79747572 |
| FASTKD1 | 1041.1357 | 0.074808169 | 0.17018683 | 0.43956498 | 0.66025221 | 0.80009424 |
| PPIG | 2127.87039 | -0.011350857 | 0.0983111 | -0.1154585 | 0.90808167 | 0.95414248 |
| CCDC173 | 37.4923133 | 0.45072291 | 0.40265892 | 1.11936651 | 0.26298381 | 0.45663549 |
| PHOSPHO2 | 149.457456 | -0.039574938 | 0.22705667 | -0.1742954 | 0.8616333 | 0.92800482 |
| KLHL23 | 513.9342 | 0.313402959 | 0.23500564 | 1.33359762 | 0.18233576 | 0.359008 |
| SSB | 4231.83405 | -0.555354077 | 0.13146874 | -4.2242292 | 2.40E-05 | 0.00041241 |
| METTL5 | 2777.64194 | -0.656174104 | 0.12200943 | -5.3780604 | 7.53E-08 | 3.75E-06 |
| UBR3 | 1889.44676 | 0.367779851 | 0.10159072 | 3.62021111 | 0.00029436 | 0.00299124 |
| MYO3B | 425.237482 | 1.520694052 | 0.51796284 | 2.93591344 | 0.00332567 | 0.01984764 |
| AC012594.1 | 2.55826299 | 0.86447292 | 0.82823676 | 1.04375097 | 0.29660062 | 0.49341406 |
| AC007277.1 | 2.5304278 | 1.10986623 | 0.95022518 | 1.16800339 | 0.2428054 | 0.43329352 |
| LINC01124 | 20.0669201 | 0.620097828 | 0.61592505 | 1.00677482 | 0.314043 | 0.51094247 |
| SP5 | 30.6456726 | 0.551568838 | 0.49815879 | 1.1072149 | 0.26820103 | 0.46241125 |
| AC007405.1 | 39.8587002 | 0.295948806 | 0.44018791 | 0.6723238 | 0.50137758 | 0.68210661 |
| AC007405.3 | 197.062586 | -0.075995762 | 0.22979088 | -0.3307171 | 0.74085822 | 0.85453728 |
| ERICH2 | 55.2811401 | -0.833039502 | 0.26338459 | -3.1628255 | 0.00156246 | 0.0111568 |
| GAD1 | 576.456311 | 1.400139623 | 0.32881375 | 4.2581541 | 2.06E-05 | 0.00036333 |
| AC007405.2 | 30.1935952 | 1.06312015 | 0.45968579 | 2.31271049 | 0.02073856 | 0.07694891 |
| GORASP2 | 4038.23864 | -0.357246583 | 0.18102036 | -1.973516 | 0.0484368 | 0.14239447 |
| TLK1 | 2547.11626 | -0.021407999 | 0.15023809 | -0.1424938 | 0.88668996 | 0.94134282 |
| METTL8 | 581.797485 | 0.276443717 | 0.17178362 | 1.60925536 | 0.10756052 | 0.24985648 |
| DCAF17 | 809.685388 | 0.354298764 | 0.09514171 | 3.72390567 | 0.00019616 | 0.00214998 |
| CYBRD1 | 3014.83952 | 0.341988165 | 0.35231176 | 0.97069755 | 0.33169891 | 0.52963369 |
| DYNC1I2 | 5197.68289 | 0.011987079 | 0.11266134 | 0.10639923 | 0.91526561 | 0.95863503 |
| SLC25A12 | 2025.66693 | 0.078391252 | 0.2176183 | 0.36022361 | 0.71867992 | 0.84001549 |
| HAT1 | 1782.99357 | -0.331983381 | 0.17103896 | -1.940981 | 0.05226058 | 0.15021925 |
| METAP1D | 429.012476 | 0.481811698 | 0.26236178 | 1.83644011 | 0.06629259 | 0.17726497 |
| DLX1 | 4.03209065 | 0.224857926 | 0.60946453 | 0.36894342 | 0.7121699 | 0.83637304 |
| DLX2 | 1.81762328 | 0.076475729 | 0.90606146 | 0.08440457 | 0.93273477 | 0.96756079 |
| DLX2-DT | 0 | NA | NA | NA | NA | NA |
| AC104088.1 | 0.21775352 | -0.339791733 | 3.08904691 | -0.1099989 | 0.91241025 | NA |
| ITGA6 | 7093.64432 | -0.458427687 | 0.45404159 | -1.0096601 | 0.31265815 | 0.50944114 |
| AC078883.2 | 27.2052876 | 2.358701125 | 0.53270012 | 4.42782168 | 9.52E-06 | 0.00019624 |
| AC078883.3 | 65.4316851 | 0.296825491 | 0.34011648 | 0.87271716 | 0.38281726 | 0.5788687 |
| AC078883.1 | 415.023255 | -0.164178422 | 0.3879384 | -0.4232075 | 0.67214391 | 0.80870716 |
| PDK1 | 1269.36419 | -1.699448931 | 0.32321706 | -5.2579185 | 1.46E-07 | 6.58E-06 |
| RAPGEF4-AS | 3.06169306 | -1.812307839 | 0.92081195 | -1.9681628 | 0.04904931 | 0.14383657 |
| RAPGEF4 | 475.435841 | 0.691302086 | 0.27596641 | 2.50502255 | 0.01224436 | 0.05234634 |
| MAP3K20 | 957.595602 | -0.41694411 | 0.37443446 | -1.1135303 | 0.26548075 | 0.4595764 |
| MAP3K20-AS | 53.1883933 | -0.584080149 | 0.3870554 | -1.509035 | 0.13128983 | 0.28695506 |
| CDCA7 | 44.7404379 | 0.067782882 | 0.36660719 | 0.18489239 | 0.85331344 | 0.92322961 |
| AC106900.2 | 3.20164598 | -1.39389576 | 1.24039132 | -1.1237548 | 0.26111704 | 0.45431884 |
| AC106900.1 | 1.92555087 | -3.27235711 | 2.07045967 | -1.5804979 | 0.11399289 | 0.26039437 |
| SP3 | 2772.32298 | 0.092520125 | 0.14829506 | 0.62389216 | 0.53269841 | 0.7068228 |
| AC016737.1 | 39.6329726 | 0.189699439 | 0.36995211 | 0.51276756 | 0.60811393 | 0.76430886 |
| AC016737.2 | 4.87049093 | 0.915654954 | 0.68913891 | 1.32869431 | 0.18394884 | 0.36101253 |
| LINC01960 | 2.63795456 | 0.292498124 | 1.00640762 | 0.29063584 | 0.77132985 | 0.87389766 |
| OLA1 | 3063.09746 | -0.237493321 | 0.17935985 | -1.3241164 | 0.18546439 | 0.36280942 |

| | | | | | | | |
|---|---|---|---|---|---|---|---|
| LINC01305 | 1.17867327 | | -2.594422753 | 1.55899174 | -1.6641671 | 0.09607904 | 0.23118938 |
| SP9 | 0.16416605 | | 0.18657164 | 3.29377195 | 0.05664376 | 0.95482897 | NA |
| CIR1 | 1369.83978 | | -0.258180857 | 0.16623251 | -1.553131 | 0.12039185 | 0.27049642 |
| SCRN3 | 713.305146 | | 0.772475161 | 0.20538739 | 3.76106413 | 0.00016919 | 0.0019205 |
| GPR155 | 2106.40268 | | 1.055214907 | 0.39350066 | 2.68160897 | 0.0073269 | 0.0358328 |
| AC010894.2 | 35.4769661 | | 0.334809773 | 0.28565683 | 1.17206991 | 0.24116899 | 0.43133656 |
| AC010894.3 | 6.51077761 | | -0.327157624 | 0.47159939 | -0.6937193 | 0.48785823 | 0.67152554 |
| WIPF1 | 2216.33125 | | -0.096343868 | 0.23972722 | -0.4018896 | 0.68776531 | 0.81958773 |
| AC010894.4 | 85.4340517 | | 0.361599806 | 0.32456721 | 1.11409839 | 0.26523698 | 0.45930973 |
| CHRNA1 | 0.76172945 | | 1.1576278 | 1.58792046 | 0.72902128 | 0.46598865 | NA |
| CHN1 | 332.321922 | | -0.485410057 | 0.30346171 | -1.599576 | 0.10969269 | 0.25340308 |
| ATF2 | 2245.83059 | | -0.07577979 | 0.14397989 | -0.526322 | 0.59866447 | 0.75677743 |
| AC096649.1 | 45.111637 | | 0.657010629 | 0.33392397 | 1.96754559 | 0.04912035 | 0.1439546 |
| ATP5MC3 | 14369.9332 | | 0.257225684 | 0.19563908 | 1.31479702 | 0.18857813 | 0.3668072 |
| AC093459.1 | 0 | NA | | NA | NA | NA | NA |
| AC016751.2 | 0.15318056 | | -1.003574173 | 3.30558948 | -0.3035992 | 0.7614333 | NA |
| AC016751.1 | 0.18340304 | | -1.003574173 | 3.30558948 | -0.3035992 | 0.7614333 | NA |
| LNPK | 1172.42142 | | -0.17450543 | 0.22066335 | -0.790822 | 0.42904787 | 0.62112297 |
| EVX2 | 0.72618214 | | 0.747715259 | 2.24548878 | 0.33298552 | 0.73914521 | NA |
| HOXD13 | 2.10305598 | | 0.265329324 | 0.83070297 | 0.31940336 | 0.74942066 | 0.85998627 |
| HOXD12 | 0.40148714 | | -0.032128629 | 2.23936149 | -0.0143472 | 0.98855296 | NA |
| HOXD11 | 427.569674 | | -0.536614655 | 0.37432659 | -1.4335467 | 0.15170166 | 0.31677565 |
| HOXD10 | 1303.76159 | | 0.246884783 | 0.33237429 | 0.74279146 | 0.45760795 | 0.64682256 |
| HOXD-AS2 | 416.876469 | | 0.222473698 | 0.29104901 | 0.76438569 | 0.44463743 | 0.63537947 |
| HOXD9 | 1867.05307 | | -0.125203059 | 0.34975556 | -0.357973 | 0.7203635 | 0.84093782 |
| HOXD8 | 2893.79266 | | 0.547806336 | 0.23566683 | 2.32449491 | 0.02009898 | 0.07520009 |
| HOXD3 | 1198.79018 | | 0.242301628 | 0.28404374 | 0.85304336 | 0.39363526 | 0.58894535 |
| AC009336.2 | 1783.36583 | | 0.370548596 | 0.26969169 | 1.37397116 | 0.16945064 | 0.34129147 |
| HOXD4 | 901.939227 | | 0.520041668 | 0.24027859 | 2.16432795 | 0.03043919 | 0.10187605 |
| HAGLR | 1862.97878 | | -0.29797916 | 0.20215765 | -1.473994 | 0.14048321 | 0.300627 |
| AC009336.1 | 52.8785634 | | 0.296262132 | 0.22998519 | 1.28817919 | 0.19768359 | 0.37843313 |
| HAGLROS | 110.561419 | | -0.207557756 | 0.35162802 | -0.5902765 | 0.5550053 | 0.72445835 |
| HOXD1 | 449.259416 | | -0.977531323 | 0.32484281 | -3.0092442 | 0.00261899 | 0.01655781 |
| MTX2 | 2617.9306 | | -0.127970944 | 0.09233413 | -1.385955 | 0.16576065 | 0.33626497 |
| AC017048.1 | 0 | NA | | NA | NA | NA | NA |
| AC017048.2 | 0.58045948 | | 1.068351981 | 2.72667138 | 0.39181545 | 0.69519458 | NA |
| AC017048.3 | 52.5781526 | | -0.175113543 | 0.37869876 | -0.4624086 | 0.64378837 | 0.78950376 |
| LINC01116 | 563.800221 | | -0.065600917 | 0.28485395 | -0.2302967 | 0.81786124 | 0.90212076 |
| LINC01117 | 56.3929141 | | -0.105552929 | 0.31488808 | -0.3352078 | 0.73746837 | 0.85252908 |
| AC092162.2 | 8.64338735 | | 1.015172426 | 0.4720741 | 2.15045143 | 0.03151952 | 0.10456282 |
| AC073636.1 | 1.33827259 | | -0.602424834 | 1.1841992 | -0.5087192 | 0.61094908 | 0.76677585 |
| AC074286.1 | 1.98609529 | | 0.034752043 | 1.04210976 | 0.03334778 | 0.97339726 | 0.98766731 |
| AC079305.3 | 2.68188144 | | 0.405226897 | 1.14756189 | 0.35311986 | 0.72399858 | 0.84369283 |
| HNRNPA3 | 14506.7068 | | -0.18400541 | 0.16047314 | -1.1466431 | 0.25152917 | 0.4438156 |
| NFE2L2 | 8745.36015 | | -0.372338267 | 0.20584742 | -1.808807 | 0.07048099 | 0.18515822 |
| AC079305.1 | 276.826765 | | -0.40306521 | 0.266599 | -1.5118782 | 0.13056485 | 0.28589181 |
| AC019080.3 | 12.0418095 | | 1.277865 | 0.80686159 | 1.58374747 | 0.1132512 | 0.25927842 |
| AC019080.4 | 2.96763826 | | 2.192919027 | 0.8333755 | 2.6313697 | 0.00850415 | 0.03983997 |
| AGPS | 2146.77008 | | 0.368223684 | 0.14915097 | 2.46879838 | 0.01355676 | 0.05647578 |
| TTC30B | 353.10159 | | 0.645108869 | 0.18447578 | 3.4969841 | 0.00047055 | 0.00432952 |
| AC073834.1 | 53.4288447 | | 1.366179773 | 0.24804931 | 5.50769436 | 3.64E-08 | 2.05E-06 |
| TTC30A | 527.398347 | | 0.691476981 | 0.18655344 | 3.70658924 | 0.00021007 | 0.00226937 |
| PDE11A | 156.328266 | | 0.311699321 | 0.35052259 | 0.88924174 | 0.37387317 | 0.57001824 |
| AC012499.1 | 22.910748 | | 1.567289563 | 0.41366991 | 3.78874438 | 0.00015141 | 0.0017659 |
| RBM45 | 584.53486 | | 0.251739602 | 0.13783974 | 1.82632093 | 0.06780193 | 0.18000947 |
| OSBPL6 | 229.307443 | | 1.084923454 | 0.41878875 | 2.59062223 | 0.00958026 | 0.04363237 |
| AC009948.1 | 866.786095 | | -0.558692316 | 0.29198476 | -1.9134297 | 0.05569306 | 0.15705025 |
| PRKRA | 2246.99799 | | -0.232449213 | 0.12660992 | -1.8359479 | 0.06636536 | 0.1773644 |
| PJVK | 271.108149 | | 0.359075024 | 0.26885309 | 1.33558079 | 0.18168633 | 0.35814742 |
| FKBP7 | 331.202997 | | -0.070350177 | 0.28671249 | -0.2453684 | 0.80617121 | 0.89505795 |
| PLEKHA3 | 1343.81355 | | -0.136605585 | 0.13506281 | -1.0114227 | 0.31181418 | 0.50885203 |
| TTN-AS1 | 568.60247 | | -0.124977116 | 0.18810533 | -0.6643997 | 0.50643455 | 0.68566368 |
| TTN | 988.337158 | | -0.039255426 | 0.19120459 | -0.2053059 | 0.83733315 | 0.91386632 |
| AC009948.4 | 15.1948683 | | -0.893411594 | 0.52747564 | -1.6937495 | 0.0903129 | 0.22122233 |
| AC009948.3 | 25.0564195 | | -0.476603591 | 0.34404311 | -1.3853019 | 0.16596016 | 0.33652392 |
| AC010680.3 | 19.3380167 | | -0.477447742 | 0.458426 | -1.0414936 | 0.29764652 | 0.49454947 |
| AC010680.2 | 80.2561685 | | -0.294073211 | 0.40230257 | -0.7309752 | 0.46479429 | 0.65205955 |
| AC010680.4 | 21.1594173 | | 0.303836465 | 0.33578832 | 0.90484525 | 0.36554737 | 0.56260407 |
| AC010680.5 | 12.5187944 | | 0.33683701 | 0.38919202 | 0.86547769 | 0.38677666 | 0.58241586 |
| AC010680.1 | 8.1823576 | | 0.103605352 | 0.49633875 | 0.2087392 | 0.83465184 | 0.91223215 |
| CCDC141 | 12.5051536 | | 1.012162097 | 0.65524695 | 1.54470325 | 0.12241806 | 0.27355006 |
| SESTD1 | 2707.02165 | | -0.869252415 | 0.14527467 | -5.9835098 | 2.18E-09 | 1.92E-07 |
| AC093911.1 | 0 | NA | | NA | NA | NA | NA |
| ZNF385B | 757.793693 | | 1.590353368 | 0.35413965 | 4.49075212 | 7.10E-06 | 0.0001557 |
| CWC22 | 1017.46815 | | 0.131198819 | 0.14634034 | 0.89653216 | 0.36996861 | 0.56678512 |
| AC009478.1 | 7.9537649 | | 0.201645197 | 0.74632041 | 0.27018583 | 0.7870173 | 0.88407438 |
| SCHLAP1 | 11.9598338 | | 0.224014206 | 0.67199004 | 0.33335941 | 0.738863 | 0.85343045 |
| UBE2E3 | 2929.84104 | | 0.049832282 | 0.13792591 | 0.36129747 | 0.71787708 | 0.83983935 |
| AC104076.1 | 3.08346728 | | 1.293839553 | 0.93258959 | 1.38736221 | 0.16533134 | 0.33567812 |
| AC068196.1 | 44.9282442 | | -1.151195808 | 0.52297494 | -2.2012447 | 0.02771871 | 0.09538592 |
| LINC01934 | 16.38266 | | 0.866078835 | 0.7514016 | 1.15261776 | 0.24906731 | 0.44079899 |
| AC020595.1 | 0 | NA | | NA | NA | NA | NA |
| ITGA4 | 251.614604 | | 0.89481314 | 0.46186044 | 1.93741023 | 0.05269522 | 0.15112266 |
| CERKL | 615.133554 | | 0.715345645 | 0.24193939 | 2.95671426 | 0.00310936 | 0.01886739 |
| NEUROD1 | 3.64944424 | | 5.002444544 | 1.65894556 | 3.01543623 | 0.0025661 | 0.01631341 |
| AC013733.2 | 0 | NA | | NA | NA | NA | NA |
| AC013733.1 | 0 | NA | | NA | NA | NA | NA |
| AC009962.1 | 25.2290921 | | 0.288341596 | 0.42556876 | 0.67754409 | 0.49806081 | 0.67967569 |
| SSFA2 | 5350.95476 | | 0.570748785 | 0.12332954 | 4.62783528 | 3.70E-06 | 9.24E-05 |
| PPP1R1C | 59.2986285 | | -0.335657606 | 0.35513409 | -0.9451574 | 0.34457854 | 0.54214873 |
| PDE1A | 3175.05743 | | 0.348690206 | 0.39227669 | 0.88888841 | 0.37406305 | 0.57018461 |
| DNAJC10 | 4160.33189 | | -0.187164825 | 0.12477449 | -1.5000247 | 0.133608 | 0.29043301 |
| FRZB | 1012.03616 | | 1.434824135 | 0.46671624 | 3.07429658 | 0.00211 | 0.01401922 |
| NCKAP1 | 8322.96605 | | -0.040355725 | 0.1764859 | -0.2286626 | 0.81913117 | 0.90289626 |
| DUSP19 | 180.000572 | | 0.9064958 | 0.25817921 | 3.51111069 | 0.00044624 | 0.00415659 |
| AC064871.1 | 10.874924 | | 1.272662048 | 0.48306157 | 2.63457522 | 0.00842426 | 0.03957033 |
| NUP35 | 655.594594 | | -0.362042087 | 0.16076915 | -2.2519375 | 0.02432622 | 0.08647646 |
| AC021851.1 | 33.8678213 | | -0.498145087 | 0.30246937 | -1.6469274 | 0.09957297 | 0.23676639 |
| AC093639.1 | 0 | NA | | NA | NA | NA | NA |
| AC096667.1 | 8.88016681 | | -0.304321831 | 0.56239472 | -0.5411179 | 0.58842634 | 0.74908617 |
| ZNF804A | 37.4794333 | | 0.754918334 | 0.39397873 | 1.91613982 | 0.0553473 | 0.156415 |
| AC007966.1 | 39.0301218 | | 0.024124003 | 0.52174145 | 0.04623747 | 0.96312098 | 0.9827613 |
| FSIP2 | 98.3536631 | | 0.138731882 | 0.26176945 | 0.52997736 | 0.59612763 | 0.7549479 |
| FSIP2-AS1 | 37.1876028 | | 0.101824489 | 0.44895383 | 0.22680392 | 0.82057621 | 0.90366551 |
| LINC01473 | 45.4259718 | | 0.009932775 | 0.3831332 | 0.02592512 | 0.97931706 | 0.99065111 |
| AC017071.1 | 15.277093 | | -0.133498627 | 0.41135516 | -0.3245337 | 0.74553398 | 0.85740777 |
| ZC3H15 | 3133.77202 | | -0.249247895 | 0.16965442 | -1.4691506 | 0.14179195 | 0.30251455 |
| ITGAV | 7763.25746 | | -0.134009186 | 0.28513893 | -0.4699786 | 0.63837033 | 0.78603967 |
| AC017101.1 | 15.5834525 | | 0.888610019 | 0.52771582 | 1.68387983 | 0.0922049 | 0.22446853 |
| FAM171B | 221.132303 | | -0.842789139 | 0.25459161 | -3.3103571 | 0.00093177 | 0.00743695 |
| ZSWIM2 | 0.58836532 | | 1.281986106 | 2.15065229 | 0.59609176 | 0.55111393 | NA |
| AC007319.1 | 736.847242 | | -1.748533541 | 0.40589584 | -4.307838 | 1.65E-05 | 0.00030443 |
| CALCRL | 1140.85933 | | 0.043281672 | 0.27228059 | 0.15895982 | 0.87370053 | 0.93417196 |

| | | | | | | |
|---|---|---|---|---|---|---|
| TFPI | 11849.9735 | -1.977121242 | 0.33414578 | -5.9169421 | 3.28E-09 | 2.69E-07 |
| LINC01090 | 0.91818181 | -1.049505653 | 1.51188922 | -0.6941684 | 0.48757664 | NA |
| GULP1 | 1812.52873 | -0.487360389 | 0.46147125 | -1.0561013 | 0.29092193 | 0.48690769 |
| DIRC1 | 0.15318056 | -1.003574173 | 3.30558948 | -0.3035992 | 0.7614333 | NA |
| COL3A1 | 10287.4927 | -0.228940138 | 0.45416906 | -0.5040857 | 0.61420114 | 0.76913687 |
| COL5A2 | 1078.40684 | 0.173865904 | 0.36221142 | 0.48001221 | 0.63121871 | 0.78077009 |
| AC133106.1 | 0 | NA | NA | NA | NA | NA |
| WDR75 | 1758.25509 | -0.426749384 | 0.11310988 | -3.7728745 | 0.00016138 | 0.00185218 |
| SLC40A1 | 5077.8689 | 0.759283479 | 0.21973313 | 3.4554802 | 0.00054931 | 0.00488382 |
| ASNSD1 | 1927.71045 | -0.003943416 | 0.23434003 | -0.0168278 | 0.98657403 | 0.99400325 |
| ANKAR | 203.189875 | 0.285133279 | 0.21583228 | 1.32108728 | 0.18647226 | 0.36424249 |
| OSGEPL1 | 434.033195 | 0.75084033 | 0.24697123 | 3.04019346 | 0.00236426 | 0.01535984 |
| OSGEPL1-AS | 54.1855083 | 0.729148552 | 0.23692222 | 3.07758616 | 0.00208685 | 0.01390703 |
| AC013468.1 | 18.6482088 | 0.1329802 | 0.45313357 | 0.29346799 | 0.76916446 | 0.87240608 |
| ORMDL1 | 3049.53275 | -0.414993019 | 0.13275491 | -3.1260089 | 0.00177196 | 0.01229996 |
| PMS1 | 816.255908 | -0.090067162 | 0.13916378 | -0.6472026 | 0.51750082 | 0.69466488 |
| C2orf88 | 523.569608 | 0.689555587 | 0.2268221 | 3.04007236 | 0.00236521 | 0.01536093 |
| MSTN | 54.0199748 | 1.022900139 | 0.36597691 | 2.79498547 | 0.0051902 | 0.02765498 |
| HIBCH | 4574.35195 | 1.605152829 | 0.33141515 | 4.84332973 | 1.28E-06 | 3.97E-05 |
| INPP1 | 1628.91831 | -0.858033839 | 0.14627561 | -5.865871 | 4.47E-09 | 3.44E-07 |
| MFSD6 | 1329.0977 | -0.249100767 | 0.18182952 | -1.3699688 | 0.17069664 | 0.34290169 |
| AC093388.1 | 12.9505834 | 0.20202435 | 0.36208489 | 0.55794747 | 0.57688026 | 0.74083629 |
| NEMP2 | 320.889521 | -0.109661687 | 0.12778071 | -0.8582022 | 0.39078083 | 0.5864381 |
| AC108047.1 | 15.2155502 | 0.120865599 | 0.31880321 | 0.37912291 | 0.7045966 | 0.83125502 |
| NAB1 | 1637.93516 | 0.261020188 | 0.14831905 | 1.75985611 | 0.07843221 | 0.19967728 |
| AC006460.1 | 206.265373 | 0.695338746 | 0.24402731 | 2.84943006 | 0.00437976 | 0.02441741 |
| AC005540.1 | 198.254878 | 0.809873189 | 0.4586521 | 1.76576799 | 0.07743478 | 0.19802062 |
| GLS | 20625.5902 | 0.619864101 | 0.2594625 | 2.3890316 | 0.01689285 | 0.06638814 |
| STAT1 | 16213.9651 | -0.741336825 | 0.51215367 | -1.447489 | 0.14776 | 0.31128625 |
| AC067945.2 | 0.29967709 | -1.364242679 | 3.30558948 | -0.4127078 | 0.67982068 | NA |
| STAT4 | 255.195303 | -1.095842014 | 0.52643907 | -2.0816122 | 0.0373779 | 0.11778863 |
| AC092614.1 | 15.5898881 | 0.79213689 | 0.42615023 | 1.85882075 | 0.06305255 | 0.17120014 |
| MYO1B | 3162.72982 | -0.130824276 | 0.1870592 | -0.6993736 | 0.48431855 | 0.668712 |
| NABP1 | 2779.12787 | -0.877322227 | 0.31612678 | -2.7752227 | 0.00551639 | 0.029018 |
| AC098872.1 | 0.09635527 | 0.547239394 | 3.29377202 | 0.16614368 | 0.86804388 | NA |
| CAVIN2 | 1955.83613 | 0.033152209 | 0.29225865 | 0.11343448 | 0.9096861 | 0.95506719 |
| AC098617.1 | 262.318337 | 0.662402572 | 0.32555016 | 2.03471739 | 0.04187932 | 0.12808601 |
| TMEFF2 | 10.8259773 | 1.358684371 | 0.4966528 | 2.73568251 | 0.00622511 | 0.03170872 |
| AC062039.1 | 0.09989236 | -0.642905668 | 3.30558948 | -0.1944905 | 0.84579185 | NA |
| AC013401.1 | 0.08666371 | -0.642905668 | 3.30558948 | -0.1944905 | 0.84579185 | NA |
| PCGEM1 | 8.36944082 | -2.480729415 | 1.12914451 | -2.196999 | 0.02802051 | 0.09616003 |
| AC068135.2 | 0 | NA | NA | NA | NA | NA |
| LINC01821 | 0.07830766 | -0.642905668 | 3.30558948 | -0.1944905 | 0.84579185 | NA |
| LINC01790 | 0 | NA | NA | NA | NA | NA |
| AC010983.1 | 0.15481363 | 0.435199075 | 3.28898345 | 0.13232024 | 0.894731 | NA |
| AC104823.1 | 0 | NA | NA | NA | NA | NA |
| AC064834.1 | 0.18655607 | -1.003574173 | 3.30558948 | -0.3035992 | 0.7614333 | NA |
| LINC01825 | 0.21430872 | -0.339842146 | 3.10090457 | -0.1095945 | 0.91273096 | NA |
| LINC01827 | 0.37147987 | 2.480127684 | 2.736026 | 0.9064708 | 0.36468671 | NA |
| SLC39A10 | 1099.34578 | 0.190761588 | 0.22478188 | 0.84865198 | 0.39607498 | 0.59122987 |
| DNAH7 | 247.94722 | -0.009418205 | 0.40493296 | -0.0232587 | 0.98144393 | 0.9915968 |
| STK17B | 874.432441 | -0.197907682 | 0.28147164 | -0.7031177 | 0.48198243 | 0.66652035 |
| AC114760.2 | 0.97403783 | 0.328903445 | 1.34890749 | 0.24382951 | 0.80736287 | NA |
| HECW2 | 299.187075 | 0.912338112 | 0.42429352 | 2.15025228 | 0.03153526 | 0.10458872 |
| AC020571.1 | 10.1423711 | 0.231937842 | 0.77608875 | 0.29885479 | 0.76505084 | 0.87022347 |
| AC068544.1 | 0.26322705 | -0.178695909 | 2.98041902 | -0.0599566 | 0.95219017 | NA |
| CCDC150 | 216.543494 | 2.057726125 | 0.40734736 | 5.05152688 | 4.38E-07 | 1.62E-05 |
| GTF3C3 | 1390.05547 | 0.143506458 | 0.13764383 | 1.04259273 | 0.29713696 | 0.49394828 |
| C2orf66 | 21.7879687 | -0.591331106 | 0.3184377 | -1.8569758 | 0.0633146 | 0.17171715 |
| PGAP1 | 548.931765 | 0.386423744 | 0.20490504 | 1.88586746 | 0.05931282 | 0.16427675 |
| ANKRD44 | 704.359012 | 0.28456205 | 0.23171449 | 1.2280719 | 0.21941998 | 0.40496427 |
| AC013264.1 | 3.6253243 | -1.256676278 | 0.81490325 | -1.5421171 | 0.12304513 | 0.27453254 |
| AC010746.1 | 0.07659028 | -0.642905668 | 3.30558948 | -0.1944905 | 0.84579185 | NA |
| SF3B1 | 13204.3061 | 0.092466747 | 0.07907499 | 1.16935513 | 0.24226058 | 0.43264333 |
| COQ10B | 2222.14744 | -0.578215958 | 0.22796973 | -2.5363717 | 0.01120078 | 0.04901502 |
| HSPD1 | 22114.1957 | 0.197886426 | 0.28926278 | 0.68410607 | 0.49390819 | 0.67659007 |
| HSPE1 | 6254.05872 | -0.193062121 | 0.20213906 | -0.9550956 | 0.33952938 | 0.53765964 |
| HSPE1-MOB4 | 5809.96727 | -0.169113863 | 0.19978034 | -0.846499 | 0.39727442 | 0.59213918 |
| MOB4 | 1533.06453 | -0.156083082 | 0.11857665 | -1.3163054 | 0.18807156 | 0.36613054 |
| RFTN2 | 210.615752 | 0.362597587 | 0.34139401 | 1.06210882 | 0.28818629 | 0.48432046 |
| AC011997.1 | 0.2622128 | -0.178650184 | 2.98310902 | -0.0598872 | 0.95224544 | NA |
| MARS2 | 445.726134 | 0.028383251 | 0.30490553 | 0.09308867 | 0.92583312 | 0.9640363 |
| BOLL | 3.05052429 | -0.039885999 | 1.08313252 | -0.0368247 | 0.9706248 | 0.98639399 |
| PLCL1 | 1995.73229 | 2.137095733 | 0.29850076 | 7.15943147 | 8.10E-13 | 2.25E-10 |
| AC020719.1 | 0.10178723 | 1.087385526 | 3.28420134 | 0.33109588 | 0.74057207 | NA |
| LINC01923 | 0 | NA | NA | NA | NA | NA |
| AC019330.1 | 0.28413854 | -0.695575488 | 2.35465137 | -0.2954049 | 0.76768461 | NA |
| AC020718.1 | 0.09635527 | 0.547239394 | 3.29377202 | 0.16614368 | 0.86804388 | NA |
| SATB2 | 678.17572 | 0.49850643 | 0.26150807 | 1.90627553 | 0.05661447 | 0.15887235 |
| AC016746.1 | 15.1128508 | 1.281966192 | 0.46499776 | 2.75692982 | 0.00583469 | 0.03016877 |
| SATB2-AS1 | 43.9697862 | 0.751103588 | 0.38414837 | 1.95524348 | 0.05055434 | 0.14677902 |
| LINC01877 | 1.08951436 | 1.319820297 | 1.88574145 | 0.69989462 | 0.48399312 | 0.66849458 |
| FTCDNL1 | 132.228524 | 1.186215705 | 0.38515052 | 3.07987567 | 0.00207087 | 0.01382853 |
| C2orf69 | 1035.79468 | 0.177810251 | 0.17918603 | 0.99232204 | 0.32104045 | 0.51858112 |
| TYW5 | 677.803858 | -0.091657973 | 0.13443115 | -0.6818209 | 0.49535218 | 0.67754743 |
| MAIP1 | 1458.47011 | 0.26308584 | 0.2471204 | 1.06460592 | 0.2870543 | 0.48334532 |
| SPATS2L | 5926.75602 | -0.372751662 | 0.2773768 | -1.3438459 | 0.17899824 | 0.35470833 |
| KCTD18 | 855.006803 | 0.457344375 | 0.17528118 | 2.6092041 | 0.00907531 | 0.0419264 |
| SGO2 | 93.1209834 | 0.332089764 | 0.29835833 | 1.11305677 | 0.26568405 | 0.45983777 |
| AOX1 | 13161.3286 | 0.437150289 | 0.47654349 | 0.91733556 | 0.35896683 | 0.55662444 |
| LINC01792 | 1.08738845 | -0.669472294 | 1.27628935 | -0.5245459 | 0.59989893 | 0.75773985 |
| AC007163.1 | 247.984735 | 0.013337847 | 0.44841173 | 0.02974464 | 0.97627071 | 0.98894663 |
| BZW1 | 5760.77507 | -0.525958822 | 0.35174585 | -1.4952808 | 0.13484117 | 0.29227041 |
| CLK1 | 9432.38613 | -0.290804224 | 0.14834901 | -1.9602708 | 0.04996415 | 0.14561832 |
| PPIL3 | 832.599434 | 0.222670817 | 0.13051146 | 1.70614001 | 0.08798202 | 0.21687098 |
| NIF3L1 | 1009.79443 | 0.108031343 | 0.16095244 | 0.67120042 | 0.50209286 | 0.68251297 |
| ORC2 | 1327.60996 | -0.089357361 | 0.12356047 | -0.7231873 | 0.46956484 | 0.6562311 |
| AC005037.1 | 83.1186675 | 0.567617089 | 0.23397296 | 2.42599438 | 0.01526651 | 0.06161276 |
| FAM126B | 1292.79679 | 0.22209746 | 0.13442569 | 1.65219499 | 0.09849481 | 0.23515691 |
| NDUFB3 | 2339.71959 | 0.174990407 | 0.20175926 | 0.86732282 | 0.38576516 | 0.58152559 |
| AC007272.1 | 2.20635525 | 0.619234526 | 1.12360153 | 0.55111577 | 0.58155431 | 0.744196 |
| CFLAR | 11615.7431 | -0.359407767 | 0.25669891 | -1.4001141 | 0.16147914 | 0.330119 |
| CFLAR-AS1 | 556.084343 | -0.259629228 | 0.27633007 | -0.939562 | 0.34744229 | 0.54493863 |
| CASP10 | 1751.7931 | -0.724531013 | 0.28225951 | -2.5668968 | 0.01026132 | 0.04594073 |
| CASP8 | 1349.64522 | -0.279471934 | 0.14375093 | -1.94414 | 0.05187857 | 0.14959794 |
| ALS2CR12 | 48.1034102 | -0.080098505 | 0.31370686 | -0.2553291 | 0.79846889 | 0.89090398 |
| TRAK2 | 3466.66108 | -0.296225415 | 0.23146752 | -1.279771 | 0.20062569 | 0.38215108 |
| STRADB | 3988.16776 | 0.17647877 | 0.36605694 | 0.48210743 | 0.62972963 | 0.7800625 |
| C2CD6 | 22.4980078 | 0.32991465 | 0.38071458 | 0.86656688 | 0.38617937 | 0.58179847 |
| TMEM237 | 624.382633 | 0.280102974 | 0.25247253 | 1.10943943 | 0.26724066 | 0.46132078 |
| MPP4 | 10.751834 | 0.666119025 | 0.49860942 | 1.33595356 | 0.18156445 | 0.35797044 |
| ALS2 | 1754.30601 | -0.076304593 | 0.14550555 | -0.5244102 | 0.59999328 | 0.75775321 |
| CDK15 | 13.5351698 | 1.55239973 | 0.68968476 | 2.25088303 | 0.02439295 | 0.08666694 |

| | | | | | | |
|---|---|---|---|---|---|---|
| AC069148.1 | 11.1914202 | -0.591344185 | 0.7126436 | -0.8297895 | 0.4066578 | 0.60140134 |
| FZD7 | 774.256453 | 0.195018676 | 0.31805648 | 0.61315737 | 0.53977229 | 0.71283029 |
| KIAA2012 | 6.85347365 | -0.269217106 | 0.70399305 | -0.3824144 | 0.70215398 | 0.82947179 |
| AC079354.1 | 0.56503786 | 0.540605217 | 1.58315621 | 0.34147307 | 0.73274747 | NA |
| AC079354.3 | 3.54327539 | -3.120540203 | 1.31830805 | -2.3670797 | 0.01792907 | 0.06918198 |
| SUMO1 | 4860.48451 | -0.083144595 | 0.08541554 | -0.973413 | 0.33034809 | 0.52821242 |
| NOP58 | 1882.18582 | -0.597173368 | 0.19582917 | -3.0494607 | 0.00229253 | 0.01500469 |
| AC064836.2 | 3.33506137 | 0.335500215 | 0.63401839 | 0.5291648 | 0.59669113 | 0.75537609 |
| AC064836.3 | 42.1708637 | 0.302588627 | 0.39243903 | 0.7710462 | 0.44067955 | 0.63200738 |
| BMPR2 | 3577.30602 | -0.110490325 | 0.20221231 | -0.5464075 | 0.58478585 | 0.74662548 |
| FAM117B | 865.064602 | 0.105640726 | 0.15274001 | 0.69163754 | 0.48916498 | 0.67257742 |
| ICA1L | 399.617707 | -0.169175377 | 0.28812882 | -0.5871519 | 0.55710171 | 0.72602654 |
| WDR12 | 1504.82649 | 0.303752041 | 0.23458535 | 1.29484659 | 0.19537316 | 0.37552619 |
| CARF | 549.126226 | 0.636108676 | 0.25255069 | 2.51873666 | 0.01177767 | 0.05088926 |
| NBEAL1 | 470.351994 | 0.41141771 | 0.15328752 | 2.68396082 | 0.00727556 | 0.03563458 |
| CYP20A1 | 717.120293 | 0.066411506 | 0.14099896 | 0.47100707 | 0.63763569 | 0.78557527 |
| ABI2 | 4558.5335 | -0.036569869 | 0.15442673 | -0.2368105 | 0.81280382 | 0.89895498 |
| AC080075.1 | 49.0898318 | 0.146704092 | 0.32560389 | 0.45056001 | 0.65230669 | 0.79580511 |
| RAPH1 | 2053.71328 | -0.316387153 | 0.1764958 | -1.7926044 | 0.0730362 | 0.19013454 |
| CD28 | 76.9186327 | 0.568624606 | 0.65796386 | 0.86421859 | 0.38746782 | 0.5830724 |
| CTLA4 | 55.0542673 | -0.0387838 | 0.86874845 | -0.0446433 | 0.96439163 | 0.98336618 |
| ICOS | 26.1470295 | 0.216155884 | 0.94424931 | 0.22891823 | 0.81893247 | 0.90284917 |
| AC009965.1 | 0 | NA | NA | NA | NA | NA |
| AC009498.1 | 0 | NA | NA | NA | NA | NA |
| AC016903.1 | 0 | NA | NA | NA | NA | NA |
| AC016903.2 | 0 | NA | NA | NA | NA | NA |
| PARD3B | 895.069571 | 0.589841045 | 0.17182647 | 3.43277174 | 0.00059744 | 0.00520642 |
| NRP2 | 987.55116 | -0.736088386 | 0.31306626 | -2.3512224 | 0.01871185 | 0.07153254 |
| AC007362.1 | 19.7254858 | -0.488385415 | 0.47967977 | -1.0181489 | 0.30860722 | 0.50555832 |
| AC007679.1 | 33.0405757 | 0.831917038 | 0.42701059 | 1.94823512 | 0.05138684 | 0.14858233 |
| INO80D | 1803.27425 | -0.29436271 | 0.20057967 | -1.46756 | 0.14222377 | 0.30321294 |
| AC007383.2 | 256.31594 | -0.143764112 | 0.14587283 | -0.9855442 | 0.32435685 | 0.52189213 |
| NDUFS1 | 8093.54825 | 0.713009189 | 0.20772061 | 3.43253951 | 0.00059796 | 0.00520743 |
| AC007383.3 | 1086.90561 | 0.91173175 | 0.20043562 | 4.54875104 | 5.40E-06 | 0.0001245 |
| EEF1B2 | 11241.152 | -0.174390538 | 0.22084438 | -0.7896535 | 0.42973016 | 0.62180135 |
| GPR1 | 31.6283852 | -0.180024598 | 0.45641984 | -0.3944276 | 0.69326535 | 0.82352701 |
| ZDBF2 | 527.701419 | -0.058327318 | 0.23226719 | -0.2511216 | 0.80172007 | 0.89256768 |
| ADAM23 | 88.4286078 | 0.378413805 | 0.45584985 | 0.83012818 | 0.40646632 | 0.60140134 |
| AC010731.2 | 0.19633993 | -1.003574173 | 3.30558948 | -0.3035992 | 0.7614333 | NA |
| AC010731.3 | 0 | NA | NA | NA | NA | NA |
| FAM237A | 0 | NA | NA | NA | NA | NA |
| DYTN | 0.82784833 | -2.937461565 | 2.65157659 | -1.107817 | 0.26794087 | NA |
| MDH1B | 377.118649 | 0.825242625 | 0.47857623 | 1.72437029 | 0.08464102 | 0.21073676 |
| FASTKD2 | 1364.62533 | -0.066118372 | 0.11686493 | -0.5657674 | 0.57155188 | 0.73647771 |
| AC008269.2 | 0.70840047 | -0.100355605 | 1.65219651 | -0.0607407 | 0.9515657 | NA |
| AC008269.1 | 1.74266511 | 3.381358004 | 1.1360125 | 2.97651479 | 0.00291545 | 0.01795013 |
| CPO | 93.5895197 | 0.379543781 | 0.24251902 | 1.56500627 | 0.11758144 | 0.26615845 |
| KLF7 | 1400.75992 | -0.199975545 | 0.24234726 | -0.8251612 | 0.40928009 | 0.60378831 |
| AC007879.1 | 0.32869078 | 1.372149513 | 2.23627951 | 0.61358587 | 0.53948902 | NA |
| AC007879.3 | 7.26765601 | -0.284215166 | 0.47289055 | -0.6010168 | 0.5478288 | 0.71871868 |
| AC007879.2 | 1.01844109 | 0.605617825 | 1.28528643 | 0.47119289 | 0.637503 | 0.78549111 |
| AC007879.4 | 5.26397131 | -0.1889043 | 0.65977685 | -0.2863154 | 0.77463653 | 0.87617592 |
| MYOSLID | 5.60283807 | 0.047920823 | 0.53838496 | 0.08900847 | 0.92907518 | 0.96558201 |
| AC009226.1 | 8.58189232 | 0.456494147 | 0.47337418 | 0.96434104 | 0.33487497 | 0.53297581 |
| LINC01802 | 21.8076606 | 1.612877826 | 0.8360856 | 1.92908218 | 0.05372066 | 0.15330706 |
| CREB1 | 1552.2903 | 0.028398629 | 0.09851997 | 0.28825251 | 0.77315345 | 0.87515623 |
| METTL21A | 546.866855 | -0.055887953 | 0.15248475 | -0.366515 | 0.71398079 | 0.83749271 |
| LINC01857 | 7.05460738 | -0.891446378 | 0.97411993 | -0.91513 | 0.36012338 | 0.55779463 |
| CCNYL1 | 1380.74279 | 0.084359573 | 0.2781421 | 0.30329667 | 0.76166378 | 0.86791967 |
| AC096772.1 | 315.318132 | 0.043354759 | 0.30045078 | 0.14429904 | 0.88526434 | 0.94058667 |
| FZD5 | 1683.27648 | -0.451262283 | 0.2758312 | -1.6360088 | 0.10183774 | 0.24050341 |
| PLEKHM3 | 591.222376 | 0.308277579 | 0.20400051 | 1.5111608 | 0.13074749 | 0.28624419 |
| CRYGD | 0.50349154 | 1.394498274 | 3.2347309 | 0.43110179 | 0.66639436 | NA |
| CRYGC | 0 | NA | NA | NA | NA | NA |
| CRYGB | 0 | NA | NA | NA | NA | NA |
| CRYGA | 0 | NA | NA | NA | NA | NA |
| C2orf80 | 0.40797086 | -1.94656494 | 2.79501165 | -0.6964425 | 0.48615175 | NA |
| IDH1 | 5812.0478 | 1.152201584 | 0.40269619 | 2.86121801 | 0.00422017 | 0.02368791 |
| IDH1-AS1 | 25.9907272 | 0.641726589 | 0.45444455 | 1.41211198 | 0.15791699 | 0.32507244 |
| PIKFYVE | 1924.08993 | 0.109477192 | 0.0954804 | 1.14659341 | 0.2515497 | 0.44382214 |
| PTH2R | 944.42989 | 0.558196485 | 0.28282603 | 1.97363903 | 0.0484228 | 0.14238507 |
| MAP2 | 479.396363 | -0.211656765 | 0.33769831 | -0.6267629 | 0.53081468 | 0.70516792 |
| UNC80 | 14.6075821 | 1.541891709 | 0.53116546 | 2.90284634 | 0.00369788 | 0.02148812 |
| RPE | 2985.83012 | -0.086016347 | 0.20547986 | -0.4186121 | 0.67549967 | 0.81122718 |
| KANSL1L | 2889.94605 | 0.155471241 | 0.25751047 | 0.60374726 | 0.54601169 | 0.71722681 |
| AC007038.2 | 238.055342 | 0.010878414 | 0.2198056 | 0.04949107 | 0.96052796 | 0.98098634 |
| AC007038.1 | 121.53539 | 0.703220494 | 0.23647399 | 2.97377528 | 0.0029416 | 0.01807174 |
| ACADL | 1548.34905 | 1.011993916 | 0.29758752 | 3.40065978 | 0.00067223 | 0.00570912 |
| MYL1 | 6.1750918 | 19.4820941 | 3.14849116 | 6.1877557 | NA | NA |
| LANCL1-AS1 | 12.7818227 | 0.816682165 | 0.40661169 | 2.00850638 | 0.04458951 | 0.13392088 |
| LANCL1 | 2872.79785 | 0.579011771 | 0.17799856 | 3.25290137 | 0.00114233 | 0.00871623 |
| CPS1 | 180.273661 | 0.792674733 | 0.35284883 | 2.24649956 | 0.02467203 | 0.08739943 |
| ERBB4 | 1317.32505 | 0.470254405 | 0.24845492 | 1.89271519 | 0.05839576 | 0.16236779 |
| LINC01878 | 0.07830766 | -0.642905668 | 3.30558948 | -0.1944905 | 0.84579185 | NA |
| AC108066.1 | 0.09644756 | -0.642905668 | 3.30558948 | -0.1944905 | 0.84579185 | NA |
| IKZF2 | 254.745162 | -0.302748036 | 0.42058478 | -0.7198264 | 0.47163187 | 0.65803638 |
| AC079610.2 | 22.53516 | -0.114268593 | 0.465114 | -0.2456787 | 0.80593098 | 0.89487557 |
| LINC01953 | 0 | NA | NA | NA | NA | NA |
| AC079610.3 | 2.08039863 | 1.967331013 | 0.98127655 | 2.00486909 | 0.04497704 | 0.1347778 |
| AC079610.1 | 7.58071665 | 1.024281557 | 0.5059003 | 2.02467079 | 0.04290118 | 0.13038222 |
| SPAG16 | 976.601905 | 0.330128289 | 0.19526126 | 1.6907004 | 0.09089404 | 0.2222004 |
| AC068051.1 | 1.21813215 | 0.12482917 | 1.11724807 | 0.11172914 | 0.91103817 | 0.95600514 |
| VWC2L | 5.52738357 | -0.014989448 | 0.64367864 | -0.0232872 | 0.98142121 | 0.9915968 |
| BARD1 | 149.400789 | -0.186171627 | 0.24440802 | -0.7617247 | 0.44622433 | 0.6365667 |
| AC016708.1 | 1.83119689 | -0.552819095 | 0.98513044 | -0.5611633 | 0.57468619 | 0.73902637 |
| ABCA12 | 34.2070974 | -1.753529477 | 0.66884828 | -2.6217149 | 0.00874886 | 0.04071068 |
| AC072062.1 | 1.96077118 | 1.207145856 | 1.04945778 | 1.15025672 | 0.25003815 | 0.44213143 |
| AC073284.1 | 0.03392908 | 0.380740593 | 3.29738709 | 0.11546736 | 0.90807469 | NA |
| ATIC | 3291.2125 | -0.314421659 | 0.18659963 | -1.6850069 | 0.09198725 | 0.22416632 |
| FN1 | 8886.60444 | -0.07373785 | 0.45795452 | -0.1610157 | 0.87208106 | 0.93327291 |
| AC012462.3 | 1.00204518 | 1.308506918 | 1.16529885 | 1.12289386 | 0.26148258 | NA |
| AC012462.1 | 1.28780286 | 0.155667682 | 1.13228002 | 0.13748161 | 0.89065013 | 0.94323068 |
| AC012462.2 | 0 | NA | NA | NA | NA | NA |
| AC012668.2 | 0 | NA | NA | NA | NA | NA |
| AC012668.3 | 0.93636742 | -1.119197175 | 1.4148758 | -0.7910215 | 0.42893145 | NA |
| AC012668.1 | 0 | NA | NA | NA | NA | NA |
| LINC00607 | 5.93136446 | -0.539504826 | 0.8216703 | -0.6565953 | 0.5114412 | 0.68961008 |
| LINC01614 | 1.61180983 | 0.085508364 | 0.94707221 | 0.09028706 | 0.9280591 | 0.96509656 |
| AC122136.1 | 0 | NA | NA | NA | NA | NA |
| AC093382.1 | 5.25683278 | 0.401158485 | 0.5823297 | 0.6888855 | 0.49089533 | 0.67396833 |
| MREG | 476.872771 | -0.40967431 | 0.25462062 | -1.6089597 | 0.10762516 | 0.2499626 |
| PECR | 1684.07258 | 1.218112061 | 0.37342309 | 3.26201592 | 0.00110623 | 0.00847304 |
| TMEM169 | 156.147763 | 0.483583038 | 0.46403357 | 1.04212942 | 0.29735168 | 0.49419756 |

| | | | | | | | |
|---|---|---|---|---|---|---|---|
| XRCC5 | 11228.4926 | | -0.300195446 | 0.14972033 | -2.0050413 | 0.04495863 | 0.13476856 |
| LINC01963 | 483.569707 | | 0.412376764 | 0.41063315 | 1.00424617 | 0.31525998 | 0.51241405 |
| 4-Mar | 3.55420514 | | 1.080063475 | 1.20336605 | 0.89753527 | 0.36943336 | 0.56630227 |
| AC012513.1 | 0.58436591 | | 0.908193048 | 3.2427555 | 0.28006831 | 0.7794251 | NA |
| AC069155.1 | 0 | NA | | NA | NA | NA | NA |
| AC098820.2 | 13.0142481 | | 0.726393565 | 0.45651427 | 1.59117385 | 0.11157046 | 0.25650505 |
| SMARCAL1 | 812.102265 | | 0.090688115 | 0.14629155 | 0.61991354 | 0.53531471 | 0.70904967 |
| AC098820.3 | 24.6410513 | | 0.899804308 | 0.44426758 | 2.02536568 | 0.04282983 | 0.13020782 |
| RPL37A | 21859.4725 | | -0.463778062 | 0.14531878 | -3.191453 | 0.00141559 | 0.01034335 |
| LINC01280 | 0 | NA | | NA | NA | NA | NA |
| AC073321.1 | 0.37282257 | | -1.746732392 | 2.81866545 | -0.6197019 | 0.53545405 | NA |
| IGFBP2 | 5095.789 | | -0.470070246 | 0.35787694 | -1.3134969 | 0.18901557 | 0.3672788 |
| IGFBP5 | 47767.3303 | | 0.509170911 | 0.39108469 | 1.3019454 | 0.19293505 | 0.37249168 |
| AC007563.2 | 1030.29605 | | 0.961568071 | 0.41043692 | 2.34279134 | 0.01914008 | 0.07257773 |
| AC007563.1 | 0.1905444 | | -0.340597073 | 3.29738699 | -0.103293 | 0.91773041 | NA |
| AC007557.2 | 0 | NA | | NA | NA | NA | NA |
| TNP1 | 0 | NA | | NA | NA | NA | NA |
| AC007557.4 | 0 | NA | | NA | NA | NA | NA |
| AC007557.1 | 0.08757577 | | 0.547239394 | 3.29377202 | 0.16614368 | 0.86804388 | NA |
| AC007557.3 | 0.09635527 | | 0.547239394 | 3.29377202 | 0.16614368 | 0.86804388 | NA |
| LINC01921 | 0 | NA | | NA | NA | NA | NA |
| AC007749.1 | 0 | NA | | NA | NA | NA | NA |
| DIRC3-AS1 | 1.79122516 | | 0.274887526 | 0.99794541 | 0.27545347 | 0.78296787 | 0.88125233 |
| DIRC3 | 49.2984926 | | 0.110785937 | 0.26737038 | 0.41435381 | 0.67861501 | 0.8134496 |
| TNS1 | 23730.1032 | | 0.662883038 | 0.22188384 | 2.98752277 | 0.00281248 | 0.01746296 |
| AC010136.1 | 79.6470743 | | 0.052105938 | 0.48967636 | 0.10640893 | 0.91525791 | 0.95863503 |
| RUFY4 | 15.6044674 | | 0.206777753 | 0.76775956 | 0.26932618 | 0.78767868 | 0.88436526 |
| CXCR2 | 54.1250936 | | 0.053067742 | 0.75530852 | 0.07025969 | 0.94398697 | 0.97322897 |
| CXCR1 | 62.9450362 | | -0.706871999 | 0.65967581 | -1.0715445 | 0.28392466 | 0.47995946 |
| ARPC2 | 9977.53584 | | -0.640554359 | 0.17057682 | -3.7552252 | 0.00017319 | 0.00195669 |
| GPBAR1 | 130.867123 | | 1.156515585 | 0.63934271 | 1.80891339 | 0.07046446 | 0.18515822 |
| AAMP | 7248.24312 | | 0.014325687 | 0.13623725 | 0.10515249 | 0.91625481 | 0.95922174 |
| PNKD | 7662.76263 | | -0.074982904 | 0.0998811 | -0.7507216 | 0.45282021 | 0.64250035 |
| TMBIM1 | 18426.6693 | | -0.931267102 | 0.19947536 | -4.6685821 | 3.03E-06 | 7.91E-05 |
| CATIP-AS2 | 35.6627757 | | 0.676285234 | 0.30452368 | 2.22079685 | 0.02636472 | 0.09192422 |
| CATIP | 316.085642 | | 0.530163207 | 0.33348632 | 1.58975997 | 0.11188892 | 0.25709391 |
| SLC11A1 | 1362.20173 | | -2.64032829 | 0.49401581 | -5.344623 | 9.06E-08 | 4.42E-06 |
| CTDSP1 | 10201.0536 | | 0.11030613 | 0.10521233 | 1.04841443 | 0.29444771 | 0.49122143 |
| AC021016.3 | 119.234704 | | 0.511381225 | 0.18119604 | 2.82225394 | 0.00476874 | 0.02595494 |
| VIL1 | 2762.9145 | | 1.584126101 | 0.22850577 | 6.93254323 | 4.13E-12 | 9.56E-10 |
| USP37 | 897.289631 | | 0.13459139 | 0.14442871 | 0.93188803 | 0.3513944 | 0.54907977 |
| CNOT9 | 2143.93113 | | -0.607767757 | 0.14252515 | -4.2642844 | 2.01E-05 | 0.00035515 |
| PLCD4 | 1517.64889 | | -0.297457814 | 0.41912307 | -0.7097147 | 0.47788106 | 0.66293459 |
| AC012510.1 | 504.215723 | | -1.092407175 | 0.5540003 | -1.971853 | 0.04862638 | 0.14280853 |
| ZNF142 | 1237.18105 | | -0.401036474 | 0.27224593 | -1.4730669 | 0.140733 | 0.3009593 |
| BCS1L | 2134.38024 | | 0.022494475 | 0.13208232 | 0.17030648 | 0.86476912 | 0.92971142 |
| RNF25 | 1384.61425 | | -0.542514937 | 0.1964088 | -2.7621723 | 0.00574182 | 0.02985835 |
| STK36 | 2485.53816 | | 0.22002126 | 0.22223951 | 0.99001863 | 0.32216501 | 0.51998356 |
| TTLL4 | 1055.46646 | | -0.783071219 | 0.20601042 | -3.8011242 | 0.00014404 | 0.00169391 |
| CYP27A1 | 4252.63074 | | 0.923301497 | 0.34903185 | 2.64532159 | 0.00816133 | 0.03867978 |
| AC009974.1 | 20.3279712 | | 0.674524185 | 0.32677679 | 2.06417408 | 0.03900121 | 0.12156066 |
| PRKAG3 | 2.38013603 | | 1.431465907 | 0.73672671 | 1.94300801 | 0.05201519 | 0.14987331 |
| WNT6 | 45.1420608 | | -0.453727406 | 0.49559227 | -0.9155256 | 0.35991578 | 0.55760408 |
| WNT10A | 112.715485 | | 0.919076281 | 0.73779021 | 1.24571493 | 0.21286907 | 0.39675978 |
| LINC01494 | 0 | NA | | NA | NA | NA | NA |
| AC097468.2 | 0 | NA | | NA | NA | NA | NA |
| CDK5R2 | 0.18301898 | | 0.18657164 | 3.29377195 | 0.05664376 | 0.95482897 | NA |
| LINC00608 | 0.08393208 | | -0.642905668 | 3.30558948 | -0.1944905 | 0.84579185 | NA |
| FEV | 1.07819441 | | -1.027241153 | 1.51806242 | -0.6766791 | 0.49860956 | 0.6799953 |
| CRYBA2 | 0.39041598 | | -0.558914409 | 2.23742937 | -0.249802 | 0.80274045 | NA |
| AC097468.1 | 2.76575632 | | -0.284383391 | 1.10064197 | -0.2583796 | 0.79611399 | 0.88938094 |
| CFAP65 | 93.8973 | | 0.115861237 | 0.48756151 | 0.23763409 | 0.81216491 | 0.89861908 |
| IHH | 432.548561 | | 0.11576505 | 0.56840752 | 0.20366558 | 0.83861484 | 0.91450415 |
| AC097468.3 | 171.253369 | | 0.289162002 | 0.40472709 | 0.71446169 | 0.47494174 | 0.660316 |
| NHEJ1 | 1444.99417 | | 1.114482608 | 0.34270711 | 3.25199734 | 0.00114597 | 0.00873641 |
| AC068946.1 | 4608.66569 | | 2.623305623 | 0.41653711 | 6.29789169 | 3.02E-10 | 3.72E-08 |
| SLC23A3 | 4250.90539 | | 3.637230786 | 0.51301201 | 7.08995251 | 1.34E-12 | 3.47E-10 |
| CNPPD1 | 9957.80699 | | 0.043804165 | 0.147691 | 0.29659332 | 0.76677701 | 0.87143515 |
| RETREG2 | 10270.9912 | | -0.1284353 | 0.18496982 | -0.6943581 | 0.48745765 | 0.67137744 |
| ZFAND2B | 2717.21469 | | -0.053983824 | 0.08896271 | -0.606814 | 0.54397433 | 0.71612017 |
| ABCB6 | 2823.80757 | | -1.146442012 | 0.32283255 | -3.551197 | 0.00038348 | 0.00368933 |
| AC068946.2 | 6207.36432 | | -0.814513072 | 0.20546407 | -3.9642604 | 7.36E-05 | 0.0009946 |
| ATG9A | 4734.08461 | | -0.340635986 | 0.14476479 | -2.3530306 | 0.0186211 | 0.0712574 |
| ANKZF1 | 4956.49086 | | -0.541938007 | 0.29929343 | -1.8107247 | 0.07018347 | 0.18457547 |
| GLB1L | 2029.24397 | | 0.291415036 | 0.24732293 | 1.17827745 | 0.23868601 | 0.42826088 |
| STK16 | 2289.64819 | | -0.035926993 | 0.12185958 | -0.2948229 | 0.76812918 | 0.87195724 |
| TUBA4A | 5645.63091 | | -0.948778579 | 0.28705346 | -3.305233 | 0.00094897 | 0.00754509 |
| TUBA4B | 129.778739 | | 0.704412177 | 0.33264343 | 2.11761941 | 0.03420731 | 0.11074278 |
| DNAJB2 | 8205.32073 | | -0.366960801 | 0.17604242 | -2.0845022 | 0.03711451 | 0.11720142 |
| PTPRN | 12.7981851 | | -2.392582573 | 0.81550047 | -2.9338825 | 0.00334751 | 0.01993987 |
| AC114803.1 | 3.85449445 | | -4.356610617 | 1.52001512 | -2.8661627 | 0.00415481 | 0.02344079 |
| RESP18 | 0.58655806 | | -1.427616895 | 2.25569148 | -0.6328955 | 0.52680191 | NA |
| DNPEP | 3424.04971 | | -0.203165457 | 0.20280767 | -1.0017642 | 0.31645751 | 0.51363233 |
| AC053503.2 | 5.49543193 | | 0.831398942 | 0.71442142 | 1.16373742 | 0.24453044 | 0.43554662 |
| DES | 1026.45139 | | 1.456337053 | 0.77044576 | 1.89025253 | NA | NA |
| AC053503.4 | 284.073868 | | 1.372910934 | 0.82363419 | 1.66689406 | NA | NA |
| SPEG | 1089.73048 | | -0.551431486 | 0.51355976 | -1.0737436 | 0.28293763 | 0.47888737 |
| AC053503.1 | 1.7606191 | | 0.759858655 | 1.19174673 | 0.63760079 | 0.52373358 | 0.69892989 |
| AC053503.5 | 35.7210796 | | -0.416559155 | 0.45296626 | -0.9196251 | 0.35776869 | 0.55541979 |
| GMPPA | 2720.06248 | | -0.176927874 | 0.16486961 | -1.0731382 | 0.28320911 | 0.47907013 |
| ASIC4 | 7.19420266 | | 0.646586851 | 0.69391703 | 0.93179274 | 0.35144365 | 0.54912419 |
| CHPF | 3963.23034 | | -0.564284253 | 0.20427842 | -2.7623292 | 0.00573906 | 0.02985121 |
| TMEM198 | 437.257719 | | -0.175264135 | 0.2429799 | -0.7213112 | 0.47071805 | 0.65735502 |
| AC009955.3 | 49.530672 | | -0.054033022 | 0.33465845 | -0.1614572 | 0.8717333 | 0.93317628 |
| OBSL1 | 10209.7881 | | 0.32843487 | 0.24153842 | 1.35976244 | 0.17390511 | 0.34786299 |
| AC009955.4 | 396.898234 | | 0.942342995 | 0.26995167 | 3.49078412 | 0.00048161 | 0.00440053 |
| INHA | 165.230502 | | -0.281296209 | 0.29267799 | -0.9611116 | 0.33649606 | 0.53445331 |
| STK11IP | 1812.89145 | | 0.010665349 | 0.14414415 | 0.07399085 | 0.94101766 | 0.97201754 |
| AC009955.1 | 2.65042539 | | 1.366525968 | 0.73581841 | 1.8571511 | 0.06328966 | 0.17166718 |
| SLC4A3 | 1089.37529 | | -0.186053573 | 0.38620199 | -0.481752 | 0.62998215 | 0.7801554 |
| AC009955.2 | 1.97452796 | | -0.674412078 | 0.96643926 | -0.6978318 | 0.48528237 | 0.66953885 |
| AC009502.1 | 0.62632086 | | -0.254608819 | 2.80442186 | -0.0907883 | 0.92766077 | NA |
| LINC01803 | 0 | NA | | NA | NA | NA | NA |
| AC019211.1 | 0.08666371 | | -0.642905668 | 3.30558948 | -0.1944905 | 0.84579185 | NA |
| AC093083.1 | 0.29459006 | | 1.281678157 | 2.8378264 | 0.45164079 | 0.65152778 | NA |
| AC067956.1 | 0.2602829 | | -1.364242679 | 3.30558948 | -0.4127078 | 0.67982068 | NA |
| AC093843.1 | 0.08695548 | | -0.642905668 | 3.30558948 | -0.1944905 | 0.84579185 | NA |
| EPHA4 | 506.535736 | | 1.278008407 | 0.34366353 | 3.71877808 | 0.00020019 | 0.0021821 |
| AC079834.2 | 16.4359826 | | 0.351547025 | 0.42575123 | 0.82570994 | 0.40896865 | 0.60343013 |
| AC068489.1 | 0 | NA | | NA | NA | NA | NA |
| PAX3 | 1.26991802 | | 0.404300323 | 1.20261025 | 0.33618566 | 0.73673086 | 0.85219954 |
| CCDC140 | 0.06785815 | | 0.795868262 | 3.28898349 | 0.24198001 | 0.80879565 | NA |
| AC010980.2 | 24.2342982 | | 0.162260872 | 0.42078335 | 0.38561619 | 0.69978094 | 0.82783291 |

| | | | | | | |
|---|---|---|---|---|---|---|
| SGPP2 | 1398.80277 | -0.333275808 | 0.23884339 | -1.3953738 | 0.16290315 | 0.33192366 |
| AC097461.1 | 41.1447983 | 0.730392265 | 0.3929736 | 1.85862935 | 0.06307969 | 0.17122095 |
| FARSB | 1590.33239 | -0.469832467 | 0.14371136 | -3.2692786 | 0.00107822 | 0.00831122 |
| MOGAT1 | 47.486972 | 0.021062587 | 0.45523052 | 0.04626796 | 0.96309668 | 0.9827613 |
| ACSL3 | 3561.14944 | 0.016471341 | 0.16236619 | 0.10144563 | 0.91919671 | 0.96065382 |
| AC013476.1 | 2.4321959 | -0.775018063 | 1.05325288 | -0.7358328 | 0.46183243 | 0.65014385 |
| KCNE4 | 595.800468 | -1.242077241 | 0.54935247 | -2.2609842 | 0.02376024 | 0.08504775 |
| AC013448.1 | 0.16497137 | -1.003574173 | 3.30558948 | -0.3035992 | 0.7614333 | NA |
| AC013448.2 | 0 | NA | NA | NA | NA | NA |
| SCG2 | 59.5394576 | -1.622166933 | 0.41501752 | -3.908671 | 9.28E-05 | 0.00119205 |
| AP1S3 | 393.761856 | -0.248566296 | 0.23093678 | -1.0763392 | 0.2817756 | 0.47765319 |
| WDFY1 | 2133.45513 | -0.287175795 | 0.19097179 | -1.5037603 | 0.13264309 | 0.28900258 |
| MRPL44 | 2245.20945 | 0.074445022 | 0.20184876 | 0.36881585 | 0.71226499 | 0.83644747 |
| AC073641.1 | 291.655849 | -0.206233155 | 0.21820564 | -0.9451321 | 0.34459145 | 0.54214873 |
| SERPINE2 | 11491.8295 | -2.239544078 | 0.72636328 | -3.0832286 | 0.00204768 | 0.01373277 |
| FAM124B | 53.5932173 | -0.793569494 | 0.61960708 | -1.2807625 | 0.20027711 | 0.38170768 |
| AC073052.2 | 1230.16172 | -0.391440167 | 0.14966538 | -2.6154357 | 0.00891137 | 0.04132832 |
| CUL3 | 3761.34607 | 0.028560578 | 0.16727632 | 0.17073891 | 0.86442906 | 0.92964904 |
| CCDC195 | 1.79207194 | 3.867535971 | 1.21374709 | 3.18644303 | 0.00144034 | 0.0104835 |
| DOCK10 | 419.975603 | 0.411314623 | 0.34727468 | 1.18440718 | 0.23625192 | 0.425573 |
| NYAP2 | 4.26646616 | -0.443930102 | 0.61408875 | -0.7229087 | 0.46973598 | 0.65636592 |
| AC016717.2 | 4.36301107 | -0.633806138 | 0.74610359 | -0.8494881 | 0.39560974 | 0.5909272 |
| AC062015.1 | 6.61055842 | 4.843149745 | 0.98154669 | 4.93420213 | 8.05E-07 | 2.70E-05 |
| IRS1 | 1725.87301 | -0.091931092 | 0.26075427 | -0.3525583 | 0.72441957 | 0.84387647 |
| AC010735.2 | 22.2836752 | 0.53778408 | 0.3828441 | 1.40470776 | 0.1601082 | 0.32823238 |
| RHBDD1 | 1216.67666 | 0.057781029 | 0.17388579 | 0.332293 | 0.73966803 | 0.85394886 |
| COL4A4 | 4040.3493 | 0.248826116 | 0.20831659 | 1.19446135 | 0.23229755 | 0.42040261 |
| COL4A3 | 5337.55533 | 0.228537229 | 0.20675927 | 1.10533002 | 0.26901661 | 0.46314344 |
| AC097662.1 | 140.846319 | 0.253776523 | 0.14158027 | 1.79245678 | 0.07305983 | 0.19013454 |
| MFF | 4115.91121 | -0.027419641 | 0.10059018 | -0.2725876 | 0.78517021 | 0.88278922 |
| TM4SF20 | 6.73993247 | 0.05083337 | 0.72413916 | 0.07019834 | 0.94403579 | 0.97322897 |
| SCYGR1 | 0 | NA | NA | NA | NA | NA |
| AGFG1 | 3755.38399 | -0.714218474 | 0.19732344 | -3.6195319 | 0.00029514 | 0.00299278 |
| SCYGR2 | 0.36529963 | -1.01398843 | 2.89769059 | -0.3499298 | 0.72639135 | NA |
| SCYGR10 | 1.44172663 | -2.820841332 | 2.30137653 | -1.2257192 | 0.22030437 | 0.40616456 |
| C2orf83 | 1.43539594 | -1.589161451 | 1.21815789 | -1.3045611 | 0.19204233 | 0.37124523 |
| SCYGR3 | 0 | NA | NA | NA | NA | NA |
| AC064853.1 | 0.85326367 | -2.975808668 | 2.64639753 | -1.1244753 | 0.26081144 | NA |
| SCYGR4 | 1.50333509 | -2.606701024 | 1.98955427 | -1.3101935 | 0.19013039 | 0.36885418 |
| SCYGR5 | 0 | NA | NA | NA | NA | NA |
| SLC19A3 | 771.23721 | -0.57624217 | 0.3200672 | -1.8003787 | 0.07180086 | 0.18781754 |
| SCYGR6 | 0.40557068 | -1.788706826 | 3.29999086 | -0.5420339 | 0.58779517 | NA |
| SCYGR7 | 0 | NA | NA | NA | NA | NA |
| SCYGR8 | 0 | NA | NA | NA | NA | NA |
| CCL20 | 231.846053 | -2.61337643 | 0.72418286 | -3.6087245 | 0.00030771 | 0.00309349 |
| DAW1 | 27.2123959 | -1.21484388 | 0.57822585 | -2.1009851 | 0.03564228 | 0.11407778 |
| SPHKAP | 1.27049214 | 2.901973343 | 1.3757075 | 2.10944067 | 0.03490656 | 0.11244638 |
| LINC01807 | 0.76993867 | -0.435073952 | 1.56895896 | -0.277301 | 0.78154896 | NA |
| AC012070.1 | 0 | NA | NA | NA | NA | NA |
| PID1 | 387.133057 | 0.619596235 | 0.39398218 | 1.57265042 | 0.11579979 | 0.26349472 |
| DNER | 1443.30172 | -0.419603134 | 0.25124085 | -1.6701231 | 0.09489502 | 0.22918289 |
| TRIP12 | 6729.74618 | -0.282459487 | 0.13918999 | -2.0293089 | 0.04242684 | 0.12937046 |
| FBXO36 | 363.89591 | 0.370967506 | 0.204747 | 1.81183365 | 0.0700119 | 0.18426709 |
| SLC16A14 | 211.933939 | -0.118534766 | 0.35621303 | -0.3327637 | 0.73931266 | 0.85372547 |
| AC009950.1 | 103.767186 | 0.198267659 | 0.23488083 | 0.84412023 | 0.39860222 | 0.59351443 |
| SP110 | 1129.12478 | -0.93726861 | 0.5269519 | -1.7786607 | 0.07529541 | 0.19429886 |
| SP140 | 137.865078 | -0.548853578 | 0.66690485 | -0.8229863 | 0.41051576 | 0.60490057 |
| SP140L | 795.346324 | -1.124503337 | 0.32903213 | -3.4176095 | 0.00063174 | 0.00543698 |
| SP100 | 3272.53655 | -0.722473763 | 0.3675059 | -1.9658834 | 0.04931208 | 0.14427609 |
| AC010149.1 | 4.22332885 | 0.104966622 | 1.06098969 | 0.09893274 | 0.92119167 | 0.9615743 |
| LINC01907 | 2.89186179 | 0.802882836 | 1.05515995 | 0.76091103 | 0.44671021 | 0.63701862 |
| CAB39 | 4613.761 | 0.201665622 | 0.14839757 | 1.35895499 | 0.17416085 | 0.34811668 |
| ITM2C | 13752.7526 | -0.555209238 | 0.35211803 | -1.5767703 | 0.11484839 | 0.26205348 |
| AC012507.2 | 1.56566881 | -0.784071597 | 1.44182716 | -0.5438042 | 0.58657625 | 0.74768058 |
| GPR55 | 11.5686051 | 0.705752174 | 0.66370367 | 1.06335434 | 0.28762129 | 0.48386433 |
| AC012507.1 | 2.67536153 | 1.250425998 | 0.95535994 | 1.30885329 | 0.19058405 | 0.36937391 |
| AC012507.4 | 0.08666371 | -0.642905668 | 3.30558948 | -0.1944905 | 0.84579185 | NA |
| SPATA3-AS1 | 1.15909637 | -3.394554662 | 1.62489464 | -2.0890922 | 0.03669943 | 0.11637372 |
| SPATA3 | 0 | NA | NA | NA | NA | NA |
| C2orf72 | 336.855673 | 0.430981906 | 0.54733811 | 0.78741439 | 0.43103933 | 0.62306129 |
| PSMD1 | 3506.53024 | -0.182797723 | 0.19787102 | -0.9238226 | 0.35557868 | 0.55305686 |
| HTR2B | 174.660162 | 0.655862791 | 0.28109549 | 2.33323836 | 0.01963564 | 0.07389636 |
| ARMC9 | 508.021856 | -0.304237791 | 0.25480614 | -1.1939971 | 0.2324791 | 0.42058507 |
| B3GNT7 | 295.457382 | -1.18069103 | 0.2584485 | -4.5683803 | 4.92E-06 | 0.00011593 |
| AC017104.3 | 1.66486726 | -0.075902873 | 1.80297652 | -0.0420986 | 0.96642006 | 0.98410057 |
| NCL | 16958.3073 | -0.199633631 | 0.20684052 | -0.9651572 | 0.33446606 | 0.53253652 |
| LINC00471 | 13.690155 | 0.334076703 | 0.48851898 | 0.68385614 | 0.49406601 | 0.67670081 |
| NMUR1 | 82.9450742 | 0.86016233 | 0.41921375 | 2.05184665 | 0.04018457 | 0.12418458 |
| TEX44 | 0.50106323 | 1.497125818 | 1.78401894 | 0.83918718 | 0.40136428 | NA |
| PTMA | 30804.4121 | -0.576578152 | 0.24897237 | -2.3158319 | 0.02056745 | 0.0764862 |
| PDE6D | 860.363149 | -0.275117532 | 0.15214151 | -1.8083002 | 0.07055979 | 0.18532486 |
| COPS7B | 2199.08078 | -0.261118413 | 0.17678379 | -1.4770495 | 0.13966236 | 0.29948448 |
| NPPC | 3.75791186 | -0.483721414 | 0.75522287 | -0.6405015 | 0.52184659 | 0.69787989 |
| DIS3L2 | 1513.02075 | 0.129935512 | 0.16546386 | 0.78528029 | 0.43228925 | 0.62437624 |
| AC019130.1 | 65.7886524 | 0.813666403 | 0.20960287 | 3.88194302 | 0.00010363 | 0.00130372 |
| ALPP | 0.56905306 | 1.855270561 | 2.03888973 | 0.90994159 | 0.36285331 | NA |
| AC068134.1 | 0.09635527 | 0.547239394 | 3.29377202 | 0.16614368 | 0.86804388 | NA |
| ALPG | 0.5583431 | -2.349234845 | 1.85724931 | -1.2649001 | 0.20590714 | NA |
| AC068134.2 | 0.09997307 | 0.547239394 | 3.29377202 | 0.16614368 | 0.86804388 | NA |
| ALPI | 523.136483 | 0.41282484 | 0.81482334 | 0.50664336 | 0.61240508 | 0.76779926 |
| ECEL1 | 282.221866 | -2.410550875 | 0.47473369 | -5.0776908 | 3.82E-07 | 1.45E-05 |
| PRSS56 | 3.82540936 | -1.849562662 | 0.9626236 | -1.9213768 | 0.05468422 | 0.15510626 |
| CHRND | 2.32334659 | -2.585797811 | 1.4435001 | -1.7913389 | 0.07323893 | 0.19042701 |
| CHRNG | 5.29687562 | -0.48937232 | 0.59923314 | -0.8166643 | 0.41412029 | 0.60793588 |
| TIGD1 | 470.067912 | -0.195884055 | 0.22160043 | -0.8839516 | 0.37672235 | 0.57260755 |
| EIF4E2 | 4537.81994 | -0.071453528 | 0.17610742 | -0.4057383 | 0.68493489 | 0.81791843 |
| AC073254.1 | 927.479656 | 0.603831596 | 0.27925499 | 2.16229474 | 0.03059547 | 0.10228225 |
| EFHD1 | 8494.03661 | 0.838132039 | 0.26748192 | 3.13341564 | 0.00172785 | 0.01205697 |
| GIGYF2 | 2972.67413 | 0.203329487 | 0.12239522 | 1.66125356 | 0.09666253 | 0.23204668 |
| KCNJ13 | 109.400845 | 2.895479275 | 0.48258168 | 5.99997761 | 1.97E-09 | 1.79E-07 |
| SNORC | 620.921844 | 0.023515759 | 0.09167505 | 0.2565121 | 0.79755544 | 0.89022379 |
| NGEF | 1015.84201 | 1.322479367 | 0.43358544 | 3.05010098 | 0.00228764 | 0.01498018 |
| NEU2 | 0 | NA | NA | NA | NA | NA |
| INPP5D | 1126.79906 | -0.456142837 | 0.39498202 | -1.1548446 | 0.24815408 | 0.43979958 |
| ATG16L1 | 1762.86229 | -0.456598135 | 0.24913539 | -1.8327309 | 0.06684259 | 0.17824241 |
| SAG | 0.89802101 | -0.702155739 | 1.4654709 | -0.4791332 | 0.63184388 | NA |
| AC013726.1 | 1.66543995 | 0.441122154 | 1.19186725 | 0.37011014 | 0.71130043 | 0.8359476 |
| DGKD | 1525.34964 | -0.91921887 | 0.28371301 | -3.2399603 | 0.00119546 | 0.00902501 |
| USP40 | 2687.64421 | 0.21202456 | 0.2258866 | 0.93863275 | 0.34791933 | 0.54542145 |
| UGT1A8 | 33818.3688 | 0.063888633 | 0.50061514 | 0.12762026 | 0.8984495 | 0.94840532 |
| UGT1A10 | 34021.5966 | 0.063088966 | 0.49976131 | 0.1262382 | 0.89954338 | 0.94902072 |
| UGT1A9 | 41120.2974 | 0.226520995 | 0.52274902 | 0.43332649 | 0.6647776 | 0.80325262 |
| UGT1A7 | 33883.6502 | 0.061638924 | 0.50012597 | 0.1232468 | 0.90191167 | 0.95040422 |

| | | | | | | |
|---|---|---|---|---|---|---|
| UGT1A6 | 36559.2507 | -0.001447099 | 0.49268689 | -0.0029372 | 0.99765649 | 0.99867903 |
| AC114812.2 | 2090.68705 | -1.036130702 | 0.50966761 | -2.0329538 | 0.04205719 | 0.1285108 |
| UGT1A5 | 33806.2494 | 0.063619928 | 0.50053438 | 0.12710401 | 0.89885808 | 0.94857081 |
| UGT1A4 | 33915.5997 | 0.062639875 | 0.50013094 | 0.12524695 | 0.90032805 | 0.94951124 |
| UGT1A3 | 33816.6137 | 0.063172802 | 0.50056407 | 0.12620323 | 0.89957106 | 0.94902072 |
| UGT1A1 | 34358.0222 | 0.065634377 | 0.49870003 | 0.13161093 | 0.89529204 | 0.94620861 |
| MROH2A | 73.2886747 | 1.763747325 | 0.51319337 | 3.43680848 | 0.00058861 | 0.00514783 |
| HJURP | 36.5540288 | -1.746199381 | 0.70712497 | -2.4694353 | 0.01353265 | 0.05641099 |
| TRPM8 | 28.7948585 | -3.642065029 | 0.7166653 | -5.0819609 | 3.74E-07 | 1.43E-05 |
| AC005538.1 | 0 | NA | NA | NA | NA | NA |
| SPP2 | 0 | NA | NA | NA | NA | NA |
| AC006037.1 | 0 | NA | NA | NA | NA | NA |
| AC122134.1 | 2.21658148 | -2.942630248 | 1.38583987 | -2.1233552 | 0.0337241 | 0.10968236 |
| AC097713.1 | 0 | NA | NA | NA | NA | NA |
| LINC01891 | 0.25960926 | -0.178584176 | 2.98700511 | -0.059787 | 0.95232525 | NA |
| ARL4C | 1497.63551 | -1.232424015 | 0.46010656 | -2.6785622 | 0.0073939 | 0.03605347 |
| LINC01173 | 0.97858749 | -2.447802021 | 2.80580074 | -0.8724076 | 0.38298603 | NA |
| SH3BP4 | 3246.26897 | -0.151910484 | 0.32451167 | -0.4681202 | 0.6396986 | 0.78704498 |
| AC114814.1 | 0.08757577 | 0.547239394 | 3.29377202 | 0.16614368 | 0.86804388 | NA |
| AC114814.3 | 0 | NA | NA | NA | NA | NA |
| AGAP1 | 2667.94046 | 0.406526908 | 0.29882363 | 1.36042423 | 0.17369571 | 0.34754959 |
| AC064874.1 | 0.28759251 | -1.364245045 | 2.90093914 | -0.470277 | 0.63815711 | NA |
| GBX2 | 0.16416605 | 0.18657164 | 3.29377195 | 0.05664376 | 0.95482897 | NA |
| AC019068.1 | 0.17391096 | -1.003574173 | 3.30558948 | -0.3035992 | 0.7614333 | NA |
| ASB18 | 1.33855289 | 0.375624378 | 1.33558534 | 0.28124326 | 0.77852382 | 0.87861887 |
| IQCA1 | 655.072467 | -0.18283412 | 0.25406954 | -0.7196224 | 0.47175754 | 0.6580622 |
| AC093915.1 | 84.3833249 | 0.672528672 | 0.23487445 | 2.86335385 | 0.00419182 | 0.023604 |
| IQCA1-AS1 | 26.7341498 | 0.75229692 | 0.3547417 | 2.1206893 | 0.03394796 | 0.1101993 |
| ACKR3 | 853.646067 | 0.157479046 | 0.39186093 | 0.40187484 | 0.68777614 | 0.81958773 |
| AC084030.1 | 0.38007207 | -0.542633463 | 2.80507923 | -0.1934468 | 0.84660911 | NA |
| AC105760.1 | 0.19616881 | -0.340115482 | 3.16778127 | -0.1073671 | 0.91449776 | NA |
| AC105760.2 | 33.3555364 | 0.772526408 | 0.26858317 | 2.87630237 | 0.00402364 | 0.02285234 |
| COPS8 | 2537.19646 | 0.099641052 | 0.14227631 | 0.70033478 | 0.48371826 | 0.66827486 |
| AC107079.1 | 0.56980579 | 1.702533984 | 2.01346027 | 0.84557615 | 0.39778923 | NA |
| AC112715.1 | 1.52930217 | -1.549270315 | 1.28442271 | -1.2061997 | 0.22774049 | 0.41474563 |
| COL6A3 | 2679.95841 | -0.377860429 | 0.44448649 | -0.8501055 | 0.39526641 | 0.59054817 |
| AC112721.1 | 0.17423948 | 0.18657164 | 3.29377195 | 0.05664376 | 0.95482897 | NA |
| AC112721.2 | 0.16588343 | 0.18657164 | 3.29377195 | 0.05664376 | 0.95482897 | NA |
| MLPH | 392.806401 | -0.740241218 | 0.59343498 | -1.2473839 | 0.21225678 | 0.39601041 |
| PRLH | 0 | NA | NA | NA | NA | NA |
| RAB17 | 8966.95334 | -0.030425974 | 0.221257 | -0.1375142 | 0.89062439 | 0.94323068 |
| AC104667.2 | 134.988117 | -0.126419131 | 0.28569836 | -0.4424916 | 0.65813349 | 0.79943711 |
| AC104667.1 | 36.3970069 | 0.730711751 | 0.29177074 | 2.5044038 | 0.0122658 | 0.05241436 |
| LRRFIP1 | 6220.70621 | -0.551113197 | 0.21222436 | -2.5968422 | 0.00940851 | 0.04302106 |
| RBM44 | 55.5733266 | -0.278640473 | 0.42267056 | -0.659238 | 0.50974298 | 0.68848199 |
| RAMP1 | 554.217982 | -1.155358333 | 0.56055632 | -2.0610923 | 0.03929423 | 0.12222091 |
| UBE2F | 1622.76371 | -0.572678605 | 0.13931752 | -4.1105999 | 3.95E-05 | 0.00061723 |
| UBE2F-SCLY | 2486.35969 | -0.04951265 | 0.16382665 | -0.3022259 | 0.76247989 | 0.86838536 |
| SCLY | 1599.04771 | 0.330096354 | 0.18206138 | 1.81310479 | 0.06981566 | 0.1838144 |
| ESPNL | 80.6591549 | -0.855888336 | 0.68608701 | -1.2474924 | 0.212217 | 0.39596423 |
| KLHL30 | 175.265277 | -1.405004912 | 0.33769501 | -4.1605735 | 3.17E-05 | 0.00051325 |
| ERFE | 887.088995 | -3.929932971 | 0.50318313 | -7.8101445 | 5.71E-15 | 3.20E-12 |
| ILKAP | 1753.63724 | -0.417366474 | 0.16028011 | -2.6039816 | 0.00921477 | 0.0423395 |
| HES6 | 333.142504 | -1.506328963 | 0.25859155 | -5.8251284 | 5.71E-09 | 4.31E-07 |
| PER2 | 748.420562 | 0.54417686 | 0.33663751 | 1.6165069 | 0.10598476 | 0.24719779 |
| AC012485.3 | 13.3578032 | 1.113944931 | 0.75148709 | 1.48232079 | 0.13825496 | 0.29729946 |
| AC012485.2 | 11.1467858 | 1.066486304 | 0.79546305 | 1.34071131 | 0.1800142 | 0.35587184 |
| TRAF3IP1 | 937.68069 | 0.045751239 | 0.12985892 | 0.35231494 | 0.72460209 | 0.84389181 |
| ASB1 | 1563.21871 | 0.015822982 | 0.17221266 | 0.09188048 | 0.926793 | 0.96442648 |
| AC016999.1 | 22.9283594 | -0.286208772 | 0.38791201 | -0.7378188 | 0.46062457 | 0.64911904 |
| LINC01107 | 0.99706866 | 1.071234018 | 1.6241199 | 0.65957816 | 0.50952457 | NA |
| LINC01937 | 0 | NA | NA | NA | NA | NA |
| AC145625.1 | 0 | NA | NA | NA | NA | NA |
| TWIST2 | 49.7424406 | -1.121803359 | 0.5007847 | -2.2400911 | 0.02508501 | 0.08860055 |
| LINC01940 | 0.76614194 | 0.736972979 | 1.55276095 | 0.47462102 | 0.63505712 | NA |
| HDAC4 | 861.020804 | -0.429118884 | 0.19189661 | -2.2361983 | 0.02533878 | 0.08923394 |
| AC062017.1 | 49.2843962 | -0.097441078 | 0.36652421 | -0.2658517 | 0.79035344 | 0.88615293 |
| AC079612.1 | 5.13880248 | -1.355915601 | 1.34760395 | -1.0061677 | 0.3143349 | 0.51132016 |
| AC079612.2 | 0 | NA | NA | NA | NA | NA |
| AC093802.2 | 15.2649956 | -2.178713958 | 0.96077714 | -2.267658 | 0.02335006 | 0.08396972 |
| AC093802.1 | 2.29506388 | 0.73867244 | 1.09495113 | 0.67461681 | 0.49991925 | 0.68121489 |
| NDUFA10 | 6325.23451 | 0.366700005 | 0.17822381 | 2.05752536 | 0.03963572 | 0.1230074 |
| OR6B2 | 0 | NA | NA | NA | NA | NA |
| OR6B3 | 0 | NA | NA | NA | NA | NA |
| COPS9 | 1362.22703 | -0.282478648 | 0.19979778 | -1.4138227 | 0.15741396 | 0.32452554 |
| OTOS | 32.2967485 | -3.519709825 | 0.92125716 | -3.8205509 | 0.00013315 | 0.00158737 |
| AC124861.1 | 7.10237792 | -0.51164119 | 0.51385199 | -0.9956976 | 0.3193971 | 0.51681301 |
| GPC1 | 5699.70327 | -1.628252313 | 0.31956731 | -5.0951779 | 3.48E-07 | 1.35E-05 |
| ANKMY1 | 984.764604 | 0.05731422 | 0.11827658 | 0.48457793 | 0.62797577 | 0.77865807 |
| DUSP28 | 303.800436 | 0.081750555 | 0.15967611 | 0.51197738 | 0.60866684 | 0.76471221 |
| RNPEPL1 | 5753.16053 | -0.615832662 | 0.15195185 | -4.0528146 | 5.06E-05 | 0.00075096 |
| CAPN10-DT | 276.070016 | -0.047397744 | 0.26321193 | -0.1800744 | 0.85709412 | 0.92523219 |
| CAPN10 | 2627.77911 | -0.546789061 | 0.18127857 | -3.0162918 | 0.00255887 | 0.01628499 |
| GPR35 | 496.135913 | 0.57677245 | 0.2440982 | 2.36287054 | 0.018134 | 0.06980929 |
| AQP12B | 0.6216543 | -1.428682616 | 2.79937759 | -0.5103572 | 0.60980121 | NA |
| AC011298.1 | 13.8443677 | 1.171749723 | 0.69499191 | 1.68599044 | 0.09179764 | 0.22380336 |
| AQP12A | 1.06670303 | -1.777859438 | 3.23431971 | -0.5496857 | 0.58253494 | 0.74490659 |
| KIF1A | 104.226555 | 0.801880505 | 0.50308411 | 1.5939293 | 0.11095187 | 0.25539462 |
| AGXT | 1699.93197 | 3.163509301 | 0.96177508 | 3.28924026 | 0.00100458 | 0.00786783 |
| C2orf54 | 1030.77685 | 2.6238422 | 0.80107519 | 3.27540064 | 0.00105512 | 0.00817859 |
| CROCC2 | 122.524176 | 1.121446564 | 0.76908462 | 1.45815757 | 0.14479712 | 0.30684991 |
| AC104809.1 | 16.098173 | -0.386881662 | 0.91167807 | -0.4243621 | 0.67130172 | 0.80802567 |
| SNED1 | 2343.75129 | -0.001070129 | 0.17400958 | -0.0061498 | 0.99509318 | 0.99770729 |
| AC093585.1 | 4.24048638 | -0.050243251 | 0.57633246 | -0.0871775 | 0.93053038 | 0.96655578 |
| AC005237.1 | 24.1886624 | 0.16706686 | 0.30968511 | 0.53947333 | 0.58956029 | 0.74967234 |
| MTERF4 | 2602.27813 | 0.048647251 | 0.12261496 | 0.3967481 | 0.69155323 | 0.82241065 |
| PASK | 797.915745 | 0.19894702 | 0.21589605 | 0.9214945 | 0.35679231 | 0.55429559 |
| PPP1R7 | 2583.60535 | -0.182814019 | 0.13874581 | -1.3176183 | 0.18763146 | 0.36569414 |
| ANO7 | 145.186744 | -0.723130279 | 0.30190631 | -2.3952142 | 0.01661066 | 0.06557232 |
| HDLBP | 17346.0987 | 0.175528439 | 0.16137797 | 1.08768525 | 0.27673408 | 0.4721453 |
| 2-Sep | 17702.4619 | 0.047674167 | 0.16752897 | 0.28457268 | 0.77597154 | 0.87702653 |
| AC005104.1 | 1919.76758 | -0.152694352 | 0.17346815 | -0.8802443 | 0.37872697 | 0.57476003 |
| FARP2 | 2017.68303 | 0.111626508 | 0.13339194 | 0.83683098 | 0.40268758 | 0.5977748 |
| STK25 | 7796.19868 | -0.088052092 | 0.1087655 | -0.809559 | 0.41819366 | 0.61099046 |
| BOK-AS1 | 57.8168657 | 0.438533325 | 0.40030766 | 1.09549071 | 0.27330171 | 0.46825847 |
| BOK | 3241.77206 | 0.417339394 | 0.21262732 | 1.96277413 | 0.04967243 | 0.14504062 |
| AC133528.1 | 21.3131186 | -0.56806068 | 0.44988597 | -1.262677 | 0.20670527 | 0.38929725 |
| THAP4 | 4276.16886 | -0.377565421 | 0.10999971 | -3.432422 | 0.00059822 | 0.00520743 |
| ATG4B | 7936.11882 | -0.433694483 | 0.24310591 | -1.7839734 | 0.07442798 | 0.1927016 |
| DTYMK | 845.467571 | -0.287466763 | 0.17227369 | -1.6686631 | 0.09518416 | 0.22966477 |
| ING5 | 2623.25116 | 0.178043373 | 0.1401749 | 1.27015161 | 0.20403063 | 0.3863484 |
| AC114730.3 | 184.752928 | 0.444939639 | 0.18522352 | 2.40217681 | 0.01629783 | 0.06466653 |
| D2HGDH | 5666.92432 | 1.04066702 | 0.36536786 | 2.84827195 | 0.00439573 | 0.02447024 |

| | | | | | | |
|---|---|---|---|---|---|---|
| AC114730.1 | 379.321279 | 1.515742352 | 0.39443711 | 3.84279856 | 0.00012164 | 0.00148013 |
| AC114730.2 | 1035.1597 | 0.989435867 | 0.42537888 | 2.32601081 | 0.02001797 | 0.07497145 |
| GAL3ST2 | 36.6066663 | 1.988863824 | 0.78423038 | 2.53607086 | 0.0112104 | 0.04904901 |
| AC131097.1 | 19.0372566 | 1.723921856 | 0.80757725 | 2.13468352 | 0.03278687 | 0.10749593 |
| AC131097.3 | 113.350022 | 1.624956706 | 0.62284705 | 2.60891772 | 0.00908291 | 0.04194026 |
| NEU4 | 389.865186 | 1.736907225 | 0.5916275 | 2.93581216 | 0.00332676 | 0.01984764 |
| AC131097.4 | 18.0831098 | -1.024757353 | 0.71069222 | -1.4419144 | 0.14932652 | 0.31343949 |
| PDCD1 | 92.6382966 | -1.3468704 | 0.66880593 | -2.0138434 | 0.04402598 | 0.13290956 |
| RTP5 | 22.1426228 | -3.131109621 | 0.83614253 | -3.744708 | 0.0001806 | 0.00201342 |
| AC131097.2 | 1.68957747 | 1.595600865 | 1.09095001 | 1.46257927 | 0.14358254 | 0.30517221 |
| LINC01238 | 124.565681 | -0.455904355 | 0.4403511 | -1.0353201 | 0.30051944 | 0.49734515 |
| AC093642.1 | 0 | NA | NA | NA | NA | NA |
| LINC01880 | 0.73420087 | -1.174806365 | 1.50437331 | -0.7809274 | 0.43484519 | NA |
| LINC01986 | 0.08695548 | -0.642905668 | 3.30558948 | -0.1944905 | 0.84579185 | NA |
| AC066595.1 | 0.4752119 | -0.409426421 | 1.90086674 | -0.2153893 | 0.82946379 | NA |
| CHL1 | 1336.17631 | 0.574605691 | 0.42013882 | 1.36765676 | 0.17141954 | 0.34388234 |
| CHL1-AS1 | 25.1167254 | 1.020999286 | 0.33907178 | 3.01115975 | 0.00260252 | 0.01648533 |
| LINC01266 | 41.5711861 | 0.051063511 | 0.57364031 | 0.0890166 | 0.92906872 | 0.96558201 |
| AC090044.1 | 3.80839584 | 2.042667987 | 0.85974301 | 2.3759053 | 0.01750595 | 0.06803067 |
| AC087430.1 | 0 | NA | NA | NA | NA | NA |
| CNTN6 | 31.4383184 | -0.270147431 | 0.60074744 | -0.4496855 | 0.65293721 | 0.79602192 |
| AC018814.1 | 0 | NA | NA | NA | NA | NA |
| CNTN4 | 323.302879 | 0.769122901 | 0.23216038 | 3.31289477 | 0.00092336 | 0.00738518 |
| CNTN4-AS2 | 1.51751319 | 0.386403661 | 1.23015355 | 0.31411011 | 0.75343741 | 0.86263219 |
| CNTN4-AS1 | 33.1805474 | 1.170161431 | 0.25628147 | 4.56592297 | 4.97E-06 | 0.00011667 |
| IL5RA | 17.6220041 | 0.690914058 | 0.71412127 | 0.96750242 | 0.33329293 | 0.53127816 |
| TRNT1 | 1558.4903 | 0.022283155 | 0.14559224 | 0.1530518 | 0.87835742 | 0.93677394 |
| CRBN | 2347.71722 | 0.31844239 | 0.14473425 | 2.20018677 | 0.02779365 | 0.0955939 |
| AC024060.1 | 405.273831 | -0.248249595 | 0.13840718 | -1.7936179 | 0.07287418 | 0.1898155 |
| AC034195.1 | 8.82643767 | 0.894677741 | 0.80805106 | 1.10720446 | 0.26820554 | 0.46241125 |
| SUMF1 | 3244.25999 | 0.170574866 | 0.2563463 | 0.66540795 | 0.5057896 | 0.68500189 |
| LRRN1 | 82.9086411 | -0.282948663 | 0.4238455 | -0.667575 | 0.50440492 | 0.68392997 |
| SETMAR | 517.119852 | 0.517920366 | 0.15723386 | 3.29394922 | 0.0009879 | 0.00776483 |
| ITPR1-DT | 67.0576501 | 0.669589955 | 0.43778601 | 1.52949145 | 0.12614266 | 0.27915256 |
| ITPR1 | 2468.95054 | 0.406876852 | 0.14531531 | 2.79995856 | 0.00511092 | 0.02732662 |
| EGOT | 234.54382 | 2.022112445 | 0.54919638 | 3.6819479 | 0.00023146 | 0.00245518 |
| AC018816.1 | 360.78253 | 0.652653894 | 0.15734711 | 4.14786057 | 3.36E-05 | 0.00053765 |
| BHLHE40-AS | 807.048659 | -0.516847394 | 0.49008222 | -1.0546136 | 0.29160206 | 0.48779157 |
| BHLHE40 | 21202.1398 | -1.062321038 | 0.489611 | -2.1697246 | 0.03002772 | 0.10090874 |
| ARL8B | 5430.86293 | -0.407622159 | 0.2882551 | -1.4141021 | 0.15733191 | 0.32442096 |
| AC026202.2 | 12.1319177 | 1.193188157 | 0.37984398 | 3.14125858 | 0.00168223 | 0.01180176 |
| AC026202.3 | 17.1885259 | 0.662776892 | 0.44937961 | 1.47487085 | 0.14024727 | 0.30035901 |
| EDEM1 | 2641.06403 | -0.368056489 | 0.22759381 | -1.6171639 | 0.10584291 | 0.24695847 |
| AC026202.1 | 0 | NA | NA | NA | NA | NA |
| AC087857.1 | 3.21477797 | 0.553994825 | 0.75606184 | 0.73273745 | 0.46371858 | 0.65147889 |
| AC026167.1 | 1.79134461 | 1.850157868 | 1.16801111 | 1.5840242 | 0.11318821 | 0.2592074 |
| AC069277.1 | 2.2920128 | 1.729455054 | 1.05552931 | 1.63847185 | 0.1013233 | 0.23971788 |
| GRM7-AS3 | 2.6130438 | 4.88572152 | 1.16323513 | 4.20011517 | 2.67E-05 | 0.00044837 |
| GRM7 | 2.97706661 | 1.957434025 | 1.00879755 | 1.94036358 | 0.05233552 | 0.15036911 |
| GRM7-AS2 | 0.11223674 | 0.02007176 | 3.29738704 | 0.00608717 | 0.99514317 | NA |
| GRM7-AS1 | 0.23025741 | 1.79377171 | 3.01520403 | 0.5949089 | 0.55190437 | NA |
| AC068313.1 | 0.57166649 | 0.519277547 | 1.58752673 | 0.32709846 | 0.74359342 | NA |
| LMCD1-AS1 | 26.281375 | 0.77689802 | 0.3047334 | 2.54943509 | 0.01078976 | 0.04766727 |
| AC018832.1 | 0.72973537 | 1.134123985 | 1.74607795 | 0.64952655 | 0.51599809 | NA |
| AC023481.1 | 0.24111501 | 0.689160557 | 2.98884029 | 0.23057791 | 0.81764273 | NA |
| LMCD1 | 3275.77997 | -1.194170833 | 0.66648902 | -1.7917337 | 0.07317564 | 0.19033749 |
| AC034187.1 | 21.5671359 | -0.121275772 | 0.55705515 | -0.2177087 | 0.82765606 | 0.90825178 |
| SSUH2 | 68.8642617 | 2.159999297 | 0.50012896 | 4.31888467 | 1.57E-05 | 0.00029379 |
| CAV3 | 1.73339383 | 1.013702194 | 1.55748727 | 0.65085745 | 0.51513851 | 0.69281728 |
| OXTR | 60.6242736 | -0.721194747 | 0.4306299 | -1.6747438 | 0.09398451 | 0.22772872 |
| RAD18 | 344.016071 | -0.205467789 | 0.22708327 | -0.9048125 | 0.36556472 | 0.56260407 |
| SRGAP3 | 709.665244 | 0.516863865 | 0.40189719 | 1.28605993 | 0.19842215 | 0.37937877 |
| SRGAP3-AS1 | 3.86969224 | 1.219041231 | 0.80540688 | 1.51357191 | 0.13013445 | 0.2852813 |
| SRGAP3-AS2 | 4.90920292 | -1.933516766 | 1.29103945 | -1.4976434 | 0.13422591 | 0.29139189 |
| SRGAP3-AS3 | 0.13028434 | 1.049816488 | 3.28477129 | 0.31960109 | 0.74927075 | NA |
| SRGAP3-AS4 | 1.91577044 | 1.516247867 | 0.86133837 | 1.76033939 | 0.07835028 | 0.19950723 |
| AC026191.1 | 135.506733 | 0.411766767 | 0.22698042 | 1.814107 | 0.06966125 | 0.18362328 |
| THUMPD3-A! | 1531.50041 | -0.116804357 | 0.15184409 | -0.7692388 | 0.44175159 | 0.63299405 |
| THUMPD3 | 1817.43989 | -0.154272475 | 0.14301047 | -1.0787495 | 0.28069942 | 0.4763946 |
| SETD5 | 4174.2449 | -0.399476791 | 0.18959197 | -2.1070343 | 0.0351146 | 0.11292034 |
| LHFPL4 | 16.3213997 | 0.219620498 | 0.5454155 | 0.4026664 | 0.68719365 | 0.81934716 |
| MTMR14 | 2580.19683 | 0.15485181 | 0.14167359 | 1.09301817 | 0.27438582 | 0.46959723 |
| CPNE9 | 17.0828711 | 0.316821183 | 0.51141651 | 0.61949737 | 0.53558875 | 0.70933029 |
| BRPF1 | 1136.65727 | -0.014578832 | 0.21231567 | -0.0686658 | 0.94525562 | 0.97390119 |
| OGG1 | 4377.72049 | 0.274580132 | 0.25740857 | 1.06670936 | 0.2861031 | 0.48214077 |
| CAMK1 | 2352.50449 | 0.035831261 | 0.24066093 | 0.1488869 | 0.88164288 | 0.93857409 |
| TADA3 | 4445.53061 | -0.148562968 | 0.07513789 | -1.9772044 | 0.04801853 | 0.14143288 |
| ARPC4 | 8049.95519 | -0.400468639 | 0.12209967 | -3.2798502 | 0.00103862 | 0.00807681 |
| ARPC4-TTLL3 | 6195.38438 | -0.20909911 | 0.11139349 | -1.8771214 | 0.06050146 | 0.16648477 |
| TTLL3 | 3853.44564 | -0.056042496 | 0.27615771 | -0.2029366 | 0.83918462 | 0.91464269 |
| AC022382.1 | 124.478024 | 0.80367722 | 0.28691963 | 2.80105341 | 0.00509361 | 0.02725065 |
| RPUSD3 | 1558.01061 | -0.075469756 | 0.15193755 | -0.4967156 | 0.6193896 | 0.77261532 |
| CIDEC | 45.0992288 | -0.34151754 | 1.06763417 | -0.3198826 | NA | NA |
| JAGN1 | 1832.81721 | -0.297176383 | 0.14581831 | -2.0379909 | 0.04155085 | 0.12739168 |
| IL17RE | 540.317026 | 0.714994789 | 0.41540531 | 1.72119798 | 0.0852149 | 0.21181233 |
| IL17RC | 2453.67956 | 0.230475556 | 0.24523521 | 0.93981431 | 0.34731282 | 0.5447942 |
| CRELD1 | 3327.1754 | -0.017359446 | 0.32391745 | -0.0535922 | 0.95726008 | 0.97947184 |
| PRRT3 | 233.308231 | 0.033517258 | 0.23655907 | 0.14168663 | 0.88732754 | 0.94171639 |
| PRRT3-AS1 | 31.764694 | -0.333440663 | 0.31805177 | -1.0483849 | 0.29446133 | 0.49122143 |
| AC018809.2 | 139.726891 | 0.317532773 | 0.19752655 | 1.6075448 | 0.10793491 | 0.25049523 |
| EMC3 | 3316.27978 | -0.309003572 | 0.15910167 | -1.9421768 | 0.0521157 | 0.15003178 |
| AC022007.1 | 190.481695 | -0.028517369 | 0.28044912 | -0.1016846 | 0.91900699 | 0.96062334 |
| FANCD2 | 185.743705 | -0.26736663 | 0.32187779 | -0.8306464 | 0.4061734 | 0.60108667 |
| FANCD2OS | 49.6370936 | -0.008364553 | 0.38967581 | -0.0214654 | 0.98287439 | 0.99239475 |
| BRK1 | 6509.25005 | -0.260748192 | 0.1811441 | -1.4394518 | 0.15002257 | 0.31444997 |
| VHL | 2266.40922 | -0.397105026 | 0.14195117 | -2.7974762 | 0.00515036 | 0.02749289 |
| IRAK2 | 823.149288 | -1.541819601 | 0.24400061 | -6.318917 | 2.63E-10 | 3.34E-08 |
| TATDN2 | 2604.67034 | 0.01013595 | 0.15570198 | 0.0650984 | 0.94809565 | 0.97504763 |
| AC022384.1 | 1358.0768 | 0.392613305 | 0.19245191 | 2.04005923 | 0.04134443 | 0.12700945 |
| LINC00852 | 73.154818 | -0.188999455 | 0.47763011 | -0.3957026 | 0.69232447 | 0.82296233 |
| GHRL | 81.4282475 | 0.802008906 | 0.27098214 | 2.9596375 | 0.00308001 | 0.01873153 |
| GHRLOS | 79.4972627 | 0.21398987 | 0.26713682 | 0.80104971 | 0.42310287 | 0.61567281 |
| SEC13 | 5561.4768 | -0.150509314 | 0.21831286 | -0.6894203 | 0.49055882 | 0.67369536 |
| ATP2B2 | 32.3725223 | 2.064232624 | 0.38245215 | 5.39736175 | 6.76E-08 | 3.42E-06 |
| LINC00606 | 0 | NA | NA | NA | NA | NA |
| AC018495.1 | 0 | NA | NA | NA | NA | NA |
| SLC6A11 | 1.20114636 | -1.58851118 | 1.71102319 | -0.9283984 | 0.35320095 | 0.5507604 |
| SLC6A1 | 159.669836 | 0.964536096 | 0.55661468 | 1.73286139 | 0.08312031 | 0.2080967 |
| SLC6A1-AS1 | 0.53853128 | 1.113961452 | 1.53746828 | 0.72454272 | 0.46873261 | NA |
| HRH1 | 475.480689 | -1.522876614 | 0.34309197 | -4.4386833 | 9.05E-06 | 0.00018823 |
| ATG7 | 3980.75317 | -0.329771784 | 0.09557636 | -3.4503487 | 0.00055986 | 0.00495922 |
| VGLL4 | 4822.54638 | -0.172581416 | 0.09735944 | -1.7726214 | 0.07629147 | 0.19604425 |
| AC022001.2 | 2.7144932 | 0.400377088 | 1.06918084 | 0.37447088 | 0.70805402 | 0.83387899 |

| | | | | | | |
|---|---|---|---|---|---|---|
| AC022001.3 | 3.9554986 | -0.926799915 | 0.8915856 | -1.0394963 | 0.29857399 | 0.49546259 |
| TAMM41 | 578.91743 | 0.086970438 | 0.16487131 | 0.527505 | 0.59784295 | 0.75633844 |
| SYN2 | 273.914226 | 0.58426564 | 0.34458758 | 1.69555047 | 0.08997104 | 0.22059846 |
| TIMP4 | 121.28038 | -1.044215741 | 0.51883459 | -2.0126178 | 0.04415485 | 0.13310053 |
| PPARG | 1228.26229 | -0.582089023 | 0.2127234 | -2.7363658 | 0.00621219 | 0.03166741 |
| TSEN2 | 609.34258 | 0.244112279 | 0.1376481 | 1.77345181 | 0.07615387 | 0.19585319 |
| MKRN2OS | 83.5209386 | -0.047675842 | 0.28091782 | -0.1697146 | 0.86523463 | 0.92994649 |
| MKRN2 | 1855.14959 | 0.077400292 | 0.11397279 | 0.67911202 | 0.49706689 | 0.67890681 |
| RAF1 | 6203.13242 | 0.131561519 | 0.15244441 | 0.86301306 | 0.38813029 | 0.58378939 |
| TMEM40 | 9.97691427 | -2.244016265 | 0.80297301 | -2.7946347 | 0.00519584 | 0.02766288 |
| CAND2 | 327.069195 | 0.525051861 | 0.19360178 | 2.71201977 | 0.00668746 | 0.03341829 |
| AC034198.2 | 9.10418578 | 0.666471743 | 0.42746623 | 1.55912139 | 0.11896765 | 0.26824942 |
| RPL32 | 19208.6204 | -0.429331634 | 0.15080832 | -2.8468697 | 0.00441514 | 0.02453167 |
| LINC02022 | 0.08666371 | -0.642905668 | 3.30558948 | -0.1944905 | 0.84579185 | NA |
| IQSEC1 | 2551.01853 | 0.504130874 | 0.18020153 | 2.79759488 | 0.00514846 | 0.02748836 |
| NUP210 | 2080.60451 | 0.568167939 | 0.22020968 | 2.58012242 | 0.00987653 | 0.04466542 |
| HDAC11-AS1 | 4.00634597 | 0.564681714 | 0.80739327 | 0.69938868 | 0.48430916 | 0.668712 |
| HDAC11 | 2524.83354 | 0.181876613 | 0.30098543 | 0.60427049 | 0.54566383 | 0.71712709 |
| FBLN2 | 1478.26151 | -0.34848934 | 0.70064364 | -0.4973846 | 0.61891788 | 0.77231656 |
| LINC00620 | 0 | NA | NA | NA | NA | NA |
| WNT7A | 36.1026289 | -0.19297653 | 0.65465393 | -0.2947764 | 0.76816469 | 0.87196001 |
| CHCHD4 | 956.730895 | 0.163963302 | 0.23864876 | 0.68704862 | 0.49205209 | 0.6750144 |
| TMEM43 | 4619.78508 | -0.35825616 | 0.13336557 | -2.6862717 | 0.00722543 | 0.03548792 |
| AC090004.1 | 2317.20549 | 0.184419137 | 0.104455 | 1.76553676 | 0.0774736 | 0.19802062 |
| AC093495.1 | 2873.72899 | 0.485907531 | 0.1666125 | 2.91639298 | 0.00354104 | 0.02075512 |
| XPC | 3928.77052 | 0.665127269 | 0.19014433 | 3.49801267 | 0.00046874 | 0.00431737 |
| LSM3 | 2005.04743 | 0.350872255 | 0.18154746 | 1.93267511 | 0.05327623 | 0.15230857 |
| AC093496.1 | 79.3906134 | 3.225517655 | 0.84222205 | 3.82977108 | 0.00012826 | 0.0015402 |
| LINC01267 | 3.77900454 | -1.447370117 | 0.89888668 | -1.6101809 | 0.10735838 | 0.24951881 |
| SLC6A6 | 7791.52471 | -1.132115941 | 0.42707497 | -2.65086 | 0.00802871 | 0.03822299 |
| GRIP2 | 1201.87933 | -0.306417638 | 0.3744537 | -0.8183058 | 0.41318259 | 0.607085 |
| AC090952.2 | 1.82712332 | -0.349045976 | 1.00744116 | -0.3464679 | 0.72899114 | 0.84685006 |
| AC090952.1 | 2.05101461 | 0.897341763 | 0.92934611 | 0.96556252 | 0.33426314 | 0.53240549 |
| CCDC174 | 1256.30645 | -0.155693909 | 0.13206307 | -1.178936 | 0.23842367 | 0.42796489 |
| C3orf20 | 9.16487531 | 0.886848154 | 0.69708468 | 1.27222442 | 0.20329339 | 0.38549293 |
| AC090957.1 | 0.38371577 | 0.734535401 | 2.8327963 | 0.25929694 | 0.79540614 | NA |
| LINC02011 | 2.53334942 | 2.049098407 | 1.07583808 | 1.90465317 | 0.05682517 | 0.15927037 |
| FGD5 | 2485.23827 | 0.266497276 | 0.267757 | 0.99529527 | 0.31959268 | 0.51702829 |
| FGD5-AS1 | 6899.49808 | 0.396331286 | 0.14558228 | 2.72238694 | 0.00648122 | 0.0326226 |
| NR2C2 | 4017.33155 | 0.000619064 | 0.19734952 | 0.00313689 | 0.99749713 | 0.99867114 |
| MRPS25 | 4877.5374 | -0.072067889 | 0.15579509 | -0.4625813 | 0.64366454 | 0.78946204 |
| RBSN | 1333.8661 | -0.146865528 | 0.11138939 | -1.3184875 | 0.18734049 | 0.36526977 |
| CAPN7 | 1691.75809 | 0.023132778 | 0.11295107 | 0.20480354 | 0.83772562 | 0.91410871 |
| SH3BP5-AS1 | 6295.65843 | -0.426647922 | 0.30651405 | -1.3919359 | 0.1639418 | 0.33341303 |
| SH3BP5 | 6414.38113 | -0.414455138 | 0.31784338 | -1.3039603 | 0.19224713 | 0.37150497 |
| METTL6 | 656.476936 | 0.088431981 | 0.15919322 | 0.55550093 | 0.57855207 | 0.74204368 |
| EAF1 | 1603.42976 | 0.017797505 | 0.13019569 | 0.13669811 | 0.89126942 | 0.94362126 |
| EAF1-AS1 | 548.683142 | 0.436411184 | 0.18442703 | 2.36630815 | 0.01796648 | 0.06928578 |
| COLQ | 358.870761 | -0.134540181 | 0.27767665 | -0.484521 | 0.62801613 | 0.7786715 |
| HACL1 | 1400.54731 | 0.568923178 | 0.24662445 | 2.30684009 | 0.02106374 | 0.07783783 |
| BTD | 1650.42708 | 0.783778368 | 0.19442738 | 4.03121388 | 5.55E-05 | 0.00080487 |
| ANKRD28 | 2304.35467 | -0.201799142 | 0.14242885 | -1.4168418 | 0.15652918 | 0.32325479 |
| GALNT15 | 1217.58455 | -1.605761796 | 0.38015416 | -4.2239754 | 2.40E-05 | 0.00041241 |
| DPH3 | 1322.62085 | -0.37146746 | 0.25645738 | -1.4484569 | 0.14748932 | 0.31084505 |
| OXNAD1 | 880.726583 | 0.134769333 | 0.1296608 | 1.03939917 | 0.29861913 | 0.49550632 |
| RFTN1 | 1501.98597 | 0.108330125 | 0.27886619 | 0.38846632 | 0.69767097 | 0.82630063 |
| AC090948.1 | 22.965571 | -0.21026844 | 0.39180522 | -0.5366657 | 0.59149853 | 0.75108663 |
| AC090948.2 | 5.85662819 | 1.403904796 | 0.52620344 | 2.66798865 | 0.00763068 | 0.0368807 |
| AC090948.3 | 38.6088337 | -0.26676281 | 0.37721451 | -0.7071913 | 0.47944762 | 0.66418343 |
| AC010139.1 | 0 | NA | NA | NA | NA | NA |
| LINC00690 | 0.08695548 | -0.642905668 | 3.30558948 | -0.1944905 | 0.84579185 | NA |
| DAZL | 2.09162731 | 1.556010273 | 1.47699934 | 1.05349422 | 0.29211454 | 0.48833907 |
| AC091493.1 | 0.39115079 | -0.032171752 | 2.84325992 | -0.0113151 | 0.99097205 | NA |
| PLCL2 | 1163.49717 | 0.855764792 | 0.1739122 | 4.92067137 | 8.62E-07 | 2.86E-05 |
| PLCL2-AS1 | 117.123597 | 0.706326979 | 0.20686562 | 3.41442424 | 0.00063917 | 0.00548599 |
| TBC1D5 | 2173.72281 | 0.494920055 | 0.20879203 | 2.37039723 | 0.01776898 | 0.068705 |
| AC132807.1 | 0.21442573 | -1.003574173 | 3.30558948 | -0.3035992 | 0.7614333 | NA |
| AC132807.2 | 1.68619734 | -3.269169626 | 1.94291742 | -1.6826086 | 0.09245088 | 0.22483904 |
| SATB1 | 1381.09111 | 0.20104303 | 0.25194851 | 0.79795283 | 0.42489786 | 0.61726293 |
| KCNH8 | 27.8118065 | -0.979354481 | 0.49338643 | -1.9849644 | 0.04714844 | 0.139509 |
| EFHB | 153.185249 | -0.58512027 | 0.41399051 | -1.4133664 | 0.15754803 | 0.32471037 |
| RAB5A | 3540.6314 | -0.417210891 | 0.15796033 | -2.6412384 | 0.00826036 | 0.03899492 |
| PP2D1 | 610.009472 | -0.473010475 | 0.20046854 | -2.3595247 | 0.01829836 | 0.07032908 |
| KAT2B | 1568.63531 | 0.464981878 | 0.12396349 | 3.75095825 | 0.00017616 | 0.0019818 |
| SGO1 | 22.7847836 | -0.561212364 | 0.48099555 | -1.1667725 | 0.24330226 | 0.4338939 |
| AC116096.1 | 1.51419271 | 1.551567684 | 0.93729316 | 1.65537075 | 0.09784932 | 0.23406082 |
| AC099753.1 | 0 | NA | NA | NA | NA | NA |
| ZNF385D | 322.271602 | -0.47227375 | 0.52702085 | -0.8961197 | 0.37018885 | 0.56696095 |
| ZNF385D-AS | 0.35439893 | -1.716771674 | 2.26315439 | -0.7585747 | 0.44810701 | NA |
| ZNF385D-AS | 0 | NA | NA | NA | NA | NA |
| UBE2E2-AS1 | 1.85387252 | 1.316306162 | 1.06670162 | 1.23399659 | 0.21720417 | 0.40208312 |
| UBE2E2 | 1515.99405 | -0.046191395 | 0.18403505 | -0.2509924 | 0.80182 | 0.89258426 |
| UBE2E1-AS1 | 4.62550737 | -0.380477117 | 0.61243924 | -0.6212488 | 0.53443597 | 0.70812363 |
| UBE2E1 | 3357.42935 | -0.41133886 | 0.12999531 | -3.1642593 | 0.00155478 | 0.011111 |
| NKIRAS1 | 837.460635 | -0.0970773 | 0.18559394 | -0.5230629 | 0.6009305 | 0.75857048 |
| RPL15 | 30562.7271 | -0.21557131 | 0.20198441 | -1.0672671 | 0.28585126 | 0.48183954 |
| NR1D2 | 1641.17035 | 0.066336858 | 0.441643 | 0.15020471 | 0.88060311 | 0.93801519 |
| LINC00691 | 0.33403707 | -0.699367545 | 2.86233616 | -0.2443345 | 0.80697175 | NA |
| THRB | 864.551336 | 0.848792104 | 0.23100717 | 3.67431072 | 0.00023849 | 0.00252268 |
| AC012087.1 | 28.391493 | 1.20913816 | 0.36001633 | 3.35856479 | 0.00078348 | 0.0064665 |
| AC092422.1 | 3.13768367 | 0.399665085 | 0.717679 | 0.55688558 | 0.57760561 | 0.7412624 |
| RARB | 824.414764 | 0.22916232 | 0.28093909 | 0.81570108 | 0.41467112 | 0.60825557 |
| TOP2B | 4807.77813 | 0.052123621 | 0.14353088 | 0.36315265 | 0.71649086 | 0.83868671 |
| NGLY1 | 1852.22341 | -0.205783839 | 0.16061715 | -1.2812072 | 0.20012091 | 0.3814927 |
| OXSM | 986.211757 | -0.050193607 | 0.19682138 | -0.2550211 | 0.79870679 | 0.89101861 |
| AC099754.1 | 0 | NA | NA | NA | NA | NA |
| LRRC3B | 2.0968575 | 1.121529277 | 0.99340216 | 1.1289781 | 0.25890707 | 0.45167971 |
| AC114877.1 | 0 | NA | NA | NA | NA | NA |
| NEK10 | 123.02814 | -0.728643766 | 0.45040761 | -1.617743 | 0.10571799 | 0.24677201 |
| SLC4A7 | 1430.09662 | -0.80971398 | 0.33865192 | -2.3909919 | 0.01680293 | 0.06614318 |
| AC098614.4 | 11.0504796 | 0.066460298 | 0.5209663 | 0.1275712 | 0.89848832 | 0.94840833 |
| LINC02084 | 12.7341003 | -1.347814369 | 0.5109504 | -2.6378576 | 0.00834316 | 0.03928034 |
| EOMES | 42.5485288 | -0.083430166 | 0.70225612 | -0.118803 | 0.9054314 | 0.95257235 |
| LINC01980 | 1.22388705 | 0.685925812 | 1.55657238 | 0.44066426 | 0.65945608 | 0.79994664 |
| LINC01981 | 0.17515154 | 1.45420457 | 3.2783972 | 0.44357181 | 0.65735219 | NA |
| LINC01967 | 1.01997033 | 2.411592087 | 2.03115707 | 1.18729966 | 0.23510944 | 0.42393108 |
| CMC1 | 3037.60639 | -0.200983617 | 0.11648405 | -1.7254174 | 0.08445227 | 0.2105518 |
| AZI2 | 2667.12831 | -0.195418415 | 0.1601606 | -1.2201404 | 0.22241167 | 0.40839745 |
| ZCWPW2 | 76.2403766 | 0.64825338 | 0.25850931 | 2.50765966 | 0.01215337 | 0.05202667 |
| AC098650.1 | 159.769067 | -0.352306371 | 0.38778594 | -0.9085073 | 0.36361024 | 0.56082017 |
| RBMS3 | 981.234453 | -0.000330217 | 0.33214856 | -0.0009942 | 0.99920676 | 0.99973988 |
| RBMS3-AS3 | 23.7787515 | -0.045567132 | 0.65971683 | -0.0690707 | 0.94493331 | 0.9737607 |
| RBMS3-AS2 | 9.07860028 | -2.14288885 | 0.82696096 | -2.5912818 | 0.00956192 | 0.04356388 |

| | | | | | | |
|---|---|---|---|---|---|---|
| RBMS3-AS1 | 0.42728964 | -0.626487624 | 2.29408122 | -0.2730887 | 0.78478504 | NA |
| LINC01985 | 2.66497165 | -1.081361861 | 1.00441344 | -1.0766103 | 0.2816544 | 0.47757372 |
| AC116035.1 | 0 | NA | NA | NA | NA | NA |
| TGFBR2 | 10859.1158 | 0.167699165 | 0.26747047 | 0.62698198 | 0.53067105 | 0.70501262 |
| GADL1 | 21.8709183 | 3.712300145 | 0.95985915 | 3.86754675 | 0.00010994 | 0.00136552 |
| STT3B | 5923.64284 | 0.022779932 | 0.11035211 | 0.20642951 | 0.83645542 | 0.91319572 |
| OSBPL10 | 2353.66234 | -0.094313544 | 0.19736959 | -0.4778525 | 0.63275522 | 0.7819936 |
| OSBPL10-AS | 4.04007672 | 0.177580193 | 0.84047414 | 0.21128573 | 0.83266432 | 0.9111315 |
| ZNF860 | 92.1202691 | 0.213990508 | 0.20875154 | 1.02509669 | 0.30531757 | 0.50238514 |
| GPD1L | 2359.60666 | 0.978313071 | 0.28732241 | 3.4049313 | 0.00066181 | 0.00563618 |
| CMTM8 | 398.801765 | -0.38179408 | 0.25730275 | -1.4838321 | 0.13785347 | 0.2966536 |
| CMTM7 | 1695.6096 | -0.945513647 | 0.17937082 | -5.2712791 | 1.35E-07 | 6.20E-06 |
| CMTM6 | 4503.7584 | -0.579469237 | 0.15337486 | -3.7781239 | 0.00015801 | 0.00182676 |
| DYNC1LI1 | 1944.52447 | -0.063837465 | 0.1775389 | -0.3595689 | 0.71916955 | 0.84021138 |
| CNOT10 | 1145.83342 | -0.086005054 | 0.12304545 | -0.6989698 | 0.48457091 | 0.66894225 |
| CNOT10-AS1 | 90.6682157 | 0.141489623 | 0.16098843 | 0.87888072 | 0.37946595 | 0.57545098 |
| TRIM71 | 225.79238 | 0.432802002 | 0.63426216 | 0.68237084 | 0.49500449 | 0.67717723 |
| CCR4 | 44.6324131 | -0.250559677 | 0.7333052 | -0.3416854 | 0.73258766 | 0.8492009 |
| GLB1 | 5822.23608 | -0.102722394 | 0.19103316 | -0.5377202 | 0.59077022 | 0.7504724 |
| TMPPE | 82.6352629 | 0.023943477 | 0.32403949 | 0.07389061 | 0.94109742 | 0.97202378 |
| CRTAP | 8165.21384 | 0.060472397 | 0.21256068 | 0.28449474 | 0.77603126 | 0.87702653 |
| AC112211.1 | 537.490709 | 0.228632482 | 0.31150603 | 0.73395844 | 0.46297407 | 0.65100676 |
| SUSD5 | 125.23698 | 0.603344271 | 0.47848133 | 1.26095677 | 0.20732442 | 0.39014808 |
| FBXL2 | 750.366458 | -0.163738806 | 0.25424883 | -0.6440101 | 0.5195689 | 0.69631663 |
| UBP1 | 3117.67545 | -0.058735613 | 0.12164106 | -0.4828601 | 0.62919509 | 0.77972999 |
| CLASP2 | 1732.58627 | 0.522238705 | 0.10265611 | 5.08726373 | 3.63E-07 | 1.40E-05 |
| AC112220.3 | 54.3601113 | -0.067757912 | 0.42509763 | -0.1593938 | 0.87335865 | 0.9340817 |
| AC112220.4 | 80.4855697 | 0.114936525 | 0.46848809 | 0.245335 | 0.80619706 | 0.89505795 |
| AC112220.2 | 34.6189977 | -0.204066638 | 0.42566747 | -0.4794039 | 0.63165133 | 0.78108538 |
| PDCD6IP | 7704.07423 | -0.090354294 | 0.1660905 | -0.5440064 | 0.58643707 | 0.74768058 |
| LINC01811 | 0.97970509 | -1.282713712 | 1.63722571 | -0.7834679 | 0.43335242 | NA |
| AC123023.1 | 4.65101672 | -2.766710993 | 1.00116376 | -2.7634949 | 0.0057186 | 0.0297846 |
| AC018359.1 | 0 | NA | NA | NA | NA | NA |
| ARPP21 | 47.6163371 | 1.519157509 | 0.43717254 | 3.47496098 | 0.00051093 | 0.00461094 |
| ARPP21-AS1 | 0.21783422 | 0.689157102 | 3.06342475 | 0.22496296 | 0.82200809 | NA |
| STAC | 239.234871 | 0.77653362 | 0.28097382 | 2.76372232 | 0.00571461 | 0.02977562 |
| DCLK3 | 2.56785563 | 1.505454189 | 0.8388186 | 1.79473153 | 0.07269648 | 0.18964217 |
| LINC02033 | 6.40362995 | -0.391721744 | 0.68779008 | -0.5695368 | 0.56899192 | 0.7344976 |
| TRANK1 | 986.011096 | -1.184946065 | 0.65451019 | -1.8104318 | 0.07022885 | 0.18465797 |
| AC011816.2 | 4.53279415 | 0.27709633 | 0.62851091 | 0.44087752 | 0.65930167 | 0.7999349 |
| EPM2AIP1 | 1906.8444 | 0.182844894 | 0.14128513 | 1.29415528 | 0.19561179 | 0.37587495 |
| MLH1 | 1437.49695 | 0.111379415 | 0.10660183 | 1.04481712 | 0.2961075 | 0.49280217 |
| LRRFIP2 | 3166.65989 | -0.527454239 | 0.18251404 | -2.8899379 | 0.00385318 | 0.02214654 |
| AC126118.1 | 40.9441877 | -0.294180048 | 0.32557909 | -0.9035594 | 0.36622907 | 0.56331382 |
| AC097359.3 | 4.86916638 | 0.068951354 | 0.60211652 | 0.11451497 | 0.90882958 | 0.95447195 |
| AC097359.2 | 212.199167 | 1.092262087 | 0.38801758 | 2.81498093 | 0.00487801 | 0.0263863 |
| GOLGA4 | 4484.95042 | 0.125913958 | 0.21906558 | 0.57477745 | 0.56544182 | 0.73214268 |
| ITGA9 | 859.823193 | 0.183694967 | 0.34845545 | 0.52716916 | 0.59807612 | 0.75648826 |
| CTDSPL | 5017.70227 | 0.055768872 | 0.13853389 | 0.40256484 | 0.68726838 | 0.81935672 |
| VILL | 519.174055 | 0.121355683 | 0.28384872 | 0.42753649 | 0.66898861 | 0.80664072 |
| PLCD1 | 1652.85597 | -0.001757183 | 0.27539092 | -0.0063807 | 0.99490898 | 0.99770729 |
| DLEC1 | 4861.59826 | 1.174485256 | 0.3704454 | 3.17046791 | 0.00152194 | 0.01094748 |
| ACAA1 | 14097.0372 | 1.303183313 | 0.40405125 | 3.22529212 | 0.00125844 | 0.00940088 |
| MYD88 | 2407.62034 | -0.258499153 | 0.24897274 | -1.0382629 | 0.2991477 | 0.4959153 |
| OXSR1 | 3926.68205 | -0.480935728 | 0.33604935 | -1.4311461 | 0.15238834 | 0.3175783 |
| SLC22A13 | 789.500947 | 5.008685937 | 0.65583819 | 7.63707571 | 2.22E-14 | 9.77E-12 |
| SLC22A14 | 69.187053 | 2.534434517 | 0.48313021 | 5.24586217 | 1.56E-07 | 6.91E-06 |
| XYLB | 455.636408 | 2.238638352 | 0.44149591 | 5.07057551 | 3.97E-07 | 1.50E-05 |
| ACVR2B-AS1 | 136.495671 | 0.344475746 | 0.26766056 | 1.28698732 | 0.19809871 | 0.37906269 |
| ACVR2B | 789.780275 | 0.153241316 | 0.12313928 | 1.24445523 | 0.21333207 | 0.39731383 |
| EXOG | 510.761867 | -0.21230238 | 0.18444109 | -1.1510579 | 0.2497084 | 0.44175571 |
| SCN5A | 19.1663552 | -0.653002362 | 0.54149635 | -1.205922 | 0.22784756 | 0.41487185 |
| SCN10A | 0.08757577 | 0.547239394 | 3.29377202 | 0.16614368 | 0.86804388 | NA |
| AC116038.1 | 0 | NA | NA | NA | NA | NA |
| SCN11A | 54.978741 | 1.268904879 | 0.49581429 | 2.55923418 | 0.0104903 | 0.04668062 |
| WDR48 | 2494.89755 | 0.136690287 | 0.14380398 | 0.95053199 | 0.34184201 | 0.53968591 |
| GORASP1 | 4048.01866 | 0.016273483 | 0.09417583 | 0.17279893 | 0.86280948 | 0.92851314 |
| TTC21A | 581.686157 | -0.066628081 | 0.31912616 | -0.2087829 | 0.83461773 | 0.91223215 |
| CSRNP1 | 3539.38869 | -1.084841774 | 0.37095877 | -2.9244268 | 0.00345091 | 0.0203763 |
| AC092053.3 | 17.1400156 | 0.05213774 | 0.34326635 | 0.15188713 | 0.87927596 | 0.93728301 |
| AC092053.2 | 82.9288203 | -0.887222881 | 0.42283722 | -2.0982611 | 0.03588209 | 0.11459497 |
| XIRP1 | 3.79554231 | 0.040254706 | 1.14467888 | 0.03516681 | 0.97194673 | 0.98676504 |
| CX3CR1 | 150.572424 | 2.731248906 | 0.51801099 | 5.27256942 | 1.35E-07 | 6.17E-06 |
| CCR8 | 0.81959603 | 2.515423883 | 2.10540099 | 1.19474812 | 0.23218546 | NA |
| SLC25A38 | 2590.23075 | -0.021442664 | 0.20898909 | -0.1026018 | 0.91827899 | 0.96022749 |
| RPSA | 26265.3516 | -0.6232445 | 0.19098456 | -3.263324 | 0.00110114 | 0.00844588 |
| MOBP | 7.18066656 | -0.483175943 | 0.75673904 | -0.6384974 | 0.52314992 | 0.69865339 |
| MYRIP | 224.4888 | 0.637302915 | 0.37253475 | 1.71072071 | 0.08713268 | 0.2151797 |
| EIF1B | 3330.57731 | -0.44957117 | 0.25728893 | -1.7473397 | 0.08057842 | 0.20347151 |
| ENTPD3-AS1 | 333.238022 | 0.757052971 | 0.2308753 | 3.27905573 | 0.00104155 | 0.00809243 |
| ENTPD3 | 110.876502 | 1.666655355 | 0.41821382 | 3.9851752 | 6.74E-05 | 0.00093079 |
| RPL14 | 19249.4757 | -0.396141153 | 0.19642377 | -2.0167679 | 0.04371974 | 0.13228796 |
| ZNF619 | 206.904358 | 0.491295354 | 0.182933 | 2.68565727 | 0.00723873 | 0.03552654 |
| ZNF620 | 169.838308 | 0.103292866 | 0.23474969 | 0.44001279 | 0.65992784 | 0.79999826 |
| ZNF621 | 1273.35015 | -0.139154725 | 0.12725448 | -1.0935154 | 0.27416759 | 0.46931511 |
| AC122683.1 | 0.24118557 | 0.074529887 | 3.28898342 | 0.02266046 | 0.98192111 | NA |
| AC099560.1 | 1.39586575 | 0.719291071 | 0.98263445 | 0.7320027 | 0.46416692 | 0.65178001 |
| AC099541.1 | 0.19890045 | -0.340073434 | 3.15719848 | -0.1077137 | 0.91422283 | NA |
| AC009743.1 | 0.10721287 | -0.642905668 | 3.30558948 | -0.1944905 | 0.84579185 | NA |
| CTNNB1 | 12845.1097 | 0.140112788 | 0.13213114 | 1.06040704 | 0.28895946 | 0.4851505 |
| ULK4 | 308.253917 | 0.80107695 | 0.29848567 | 2.68380374 | 0.00727898 | 0.03564471 |
| TRAK1 | 3704.44964 | 0.12139163 | 0.14803267 | 0.82003271 | 0.41219746 | 0.60616749 |
| CCK | 7.16752607 | -3.326391282 | 1.22260203 | -2.7207474 | 0.00651345 | 0.0327286 |
| AC018358.1 | 0.71975802 | -2.012410025 | 2.22714733 | -0.9035819 | 0.36621714 | NA |
| LYZL4 | 0.21733212 | -0.339797876 | 3.09048411 | -0.1099497 | 0.91244925 | NA |
| VIPR1 | 161.429856 | 1.216980367 | 0.53857386 | 2.25963504 | 0.02384391 | 0.08529322 |
| VIPR1-AS1 | 26.7096116 | 1.646196501 | 0.46814323 | 3.5164377 | 0.00043738 | 0.00409141 |
| SEC22C | 2266.04538 | -0.125136495 | 0.16107926 | -0.7768629 | 0.43723968 | 0.62929868 |
| SS18L2 | 765.78835 | -0.441080869 | 0.1432372 | -3.0793738 | 0.00207436 | 0.01384834 |
| NKTR | 4462.30906 | -0.070626566 | 0.14077549 | -0.5016965 | 0.61588105 | 0.77026348 |
| AC006059.1 | 19.4025306 | -0.765283072 | 0.44252883 | -1.7293406 | 0.08374815 | 0.20921657 |
| ZBTB47 | 1460.5922 | -0.407212619 | 0.28264459 | -1.4407232 | 0.14966289 | 0.31392066 |
| KLHL40 | 1.43658401 | -1.384094361 | 2.78564597 | -0.4968666 | 0.61928315 | 0.77252929 |
| CCDC13 | 294.795713 | -0.505762413 | 0.27291411 | -1.8531926 | 0.06385476 | 0.17263173 |
| HHATL | 22.2480394 | -0.456950939 | 0.45745968 | -0.9988879 | 0.317849 | 0.51506655 |
| HHATL-AS1 | 14.1781609 | -0.536046307 | 0.44592037 | -1.2021122 | 0.22932007 | 0.41675881 |
| CCDC13-AS1 | 136.390013 | -0.660751039 | 0.37599591 | -1.7573357 | 0.07886061 | 0.20045828 |
| AC099329.3 | 3539.43472 | -0.352694917 | 0.18622867 | -1.8938808 | 0.05824083 | 0.16207752 |
| LINC02158 | 22.2647148 | -0.419640137 | 0.33611347 | -1.2485073 | 0.21184532 | 0.3954386 |
| HIGD1A | 8871.85605 | -0.464055618 | 0.16462767 | -2.8188191 | 0.00482007 | 0.02613182 |
| AC099329.1 | 4972.76697 | -0.375707101 | 0.18265112 | -2.0569658 | 0.03968951 | 0.12311104 |
| ACKR2 | 118.586746 | -0.099943877 | 0.34803294 | -0.2871679 | 0.77398379 | 0.8756706 |
| AC092042.3 | 134.536188 | 0.069596137 | 0.38889398 | 0.17895915 | 0.85796978 | 0.92579847 |

| | | | | | | |
|---|---|---|---|---|---|---|
| KRBOX1 | 113.998687 | -0.003676352 | 0.36495468 | -0.0100734 | 0.99196269 | 0.99613883 |
| CYP8B1 | 1363.1005 | 1.144525221 | 0.52757481 | 2.16940839 | 0.03005169 | 0.10095072 |
| ZNF662 | 476.922435 | 0.079883571 | 0.27558243 | 0.28987178 | 0.77191433 | 0.87421716 |
| KRBOX1-AS1 | 17.9247493 | 0.131585988 | 0.45516591 | 0.28909456 | 0.77250902 | 0.87466993 |
| FAM198A | 161.935252 | 0.384772144 | 0.36094753 | 1.06600577 | 0.28642103 | 0.48258403 |
| POMGNT2 | 1365.21847 | 0.08549768 | 0.09950962 | 0.85919009 | 0.39023565 | 0.58578665 |
| AC092042.1 | 0.10721287 | -0.642905668 | 3.30558948 | -0.1944905 | 0.84579185 | NA |
| SNRK | 3588.48493 | 0.037327733 | 0.19500497 | 0.19141939 | 0.84819703 | 0.92016931 |
| AC104184.1 | 216.946941 | 1.197233557 | 0.27440136 | 4.36307445 | 1.28E-05 | 0.00024959 |
| SNRK-AS1 | 679.361661 | 0.011497478 | 0.19786757 | 0.05810694 | 0.95366345 | 0.97769978 |
| ANO10 | 2737.71941 | -0.086024545 | 0.18832677 | -0.4567834 | 0.64782673 | 0.79277053 |
| ABHD5 | 2109.3188 | -0.963079078 | 0.40819929 | -2.3593355 | 0.0183077 | 0.07034445 |
| AC006055.1 | 0.18380315 | -1.003574173 | 3.30558948 | -0.3035992 | 0.7614333 | NA |
| AC006058.2 | 0.51960258 | -1.211711578 | 1.87845265 | -0.6450584 | 0.51888937 | NA |
| AC006058.3 | 25.1745978 | 0.919007588 | 0.43627491 | 2.10648738 | 0.03516204 | 0.11302084 |
| AC006058.1 | 412.379516 | -0.195083335 | 0.55090713 | -0.3541129 | 0.72325426 | 0.84311842 |
| TOPAZ1 | 0.58326623 | 2.537242408 | 1.9992702 | 1.26908429 | 0.204411 | NA |
| AC104187.1 | 48.8709608 | 0.753130091 | 0.31132283 | 2.41912903 | 0.01555772 | 0.06247253 |
| TCAIM | 2351.96683 | 0.470816767 | 0.18901643 | 2.49087753 | 0.0127428 | 0.05392838 |
| LINC01988 | 10.8959764 | -0.068476185 | 0.55842396 | -0.122624 | 0.90240484 | 0.95082973 |
| ZNF445 | 1291.48433 | 0.160021875 | 0.1536682 | 1.04134672 | 0.29771466 | 0.49454947 |
| ZNF852 | 56.5131285 | 0.31686094 | 0.25705224 | 1.23267137 | 0.2176984 | 0.40265881 |
| ZKSCAN7 | 255.009301 | 0.390549899 | 0.23560788 | 1.65762665 | 0.09739285 | 0.23324397 |
| AC099669.1 | 23.6895707 | 0.77285002 | 0.46825541 | 1.65048817 | 0.09884313 | 0.23568979 |
| ZNF660 | 118.700911 | -0.342479194 | 0.24731457 | -1.3847918 | 0.16611614 | 0.33677886 |
| ZNF197 | 657.194763 | 0.00882773 | 0.14353008 | 0.06150439 | 0.95095752 | 0.97657082 |
| ZNF197-AS1 | 15.893759 | 0.571049632 | 0.42882226 | 1.33166974 | 0.18296874 | 0.35970684 |
| ZNF35 | 267.023608 | -0.357111089 | 0.2866342 | -1.2458774 | 0.21280939 | 0.39667659 |
| AC124045.1 | 138.079033 | -0.300051571 | 0.37920244 | -0.7912702 | 0.42878636 | 0.62111967 |
| ZNF502 | 313.651457 | 0.271929568 | 0.21167343 | 1.28466555 | 0.19890918 | 0.37995184 |
| ZNF501 | 117.432631 | 0.480311637 | 0.23880931 | 2.01127685 | 0.04429622 | 0.13331504 |
| KIAA1143 | 1002.51003 | 0.1605428 | 0.15160521 | 1.05895305 | 0.28962116 | 0.48586488 |
| KIF15 | 30.6483071 | -0.395299985 | 0.45969845 | -0.8599115 | 0.38983783 | 0.58538941 |
| TMEM42 | 933.974252 | 0.074304764 | 0.2669376 | 0.27836005 | 0.78073598 | 0.88005505 |
| TGM4 | 6.13932103 | -0.369796685 | 0.63587978 | -0.5815513 | 0.56086898 | 0.7288024 |
| AC098649.1 | 0 | NA | NA | NA | NA | NA |
| ZDHHC3 | 4137.07378 | -0.139396891 | 0.09175667 | -1.5192017 | 0.12871174 | 0.28296194 |
| EXOSC7 | 3354.15379 | 0.023487424 | 0.2166354 | 0.10841914 | 0.91366322 | 0.95775549 |
| CLEC3B | 2462.66228 | 0.410651059 | 0.35111587 | 1.16955994 | 0.24217811 | 0.43255464 |
| CDCP1 | 1005.11665 | -2.139389339 | 0.27973065 | -7.6480334 | 2.04E-14 | 9.12E-12 |
| TMEM158 | 122.866041 | -0.719588796 | 0.51883973 | -1.3869192 | 0.1654664 | 0.33584108 |
| LARS2 | 1665.63774 | 0.626158985 | 0.14425562 | 4.34062117 | 1.42E-05 | 0.0002713 |
| LARS2-AS1 | 1.80700603 | -0.607958653 | 1.33174169 | -0.4565139 | 0.64802044 | 0.79277053 |
| LIMD1 | 1431.43226 | -0.230549278 | 0.1154103 | -1.9976491 | 0.04575472 | 0.13655571 |
| SACM1L | 3131.25075 | 0.382831497 | 0.08794552 | 4.35305264 | 1.34E-05 | 0.00025917 |
| SLC6A20 | 1669.172 | -1.985411943 | 0.37140476 | -5.3456825 | 9.01E-08 | 4.40E-06 |
| LZTFL1 | 1409.66637 | 0.165600853 | 0.1623211 | 1.02020534 | 0.30763109 | 0.5044676 |
| CCR9 | 52.7143186 | 0.261172376 | 0.62534787 | 0.41764335 | 0.6762079 | 0.81152334 |
| FYCO1 | 1945.48037 | 0.153787016 | 0.14076049 | 1.09254388 | 0.27459411 | 0.46986222 |
| CXCR6 | 61.4039676 | -1.066510339 | 0.5980844 | -1.7832104 | 0.07455205 | 0.19294707 |
| XCR1 | 31.811437 | 1.056599734 | 0.94437995 | 1.11882906 | 0.26321307 | 0.45691059 |
| CCR3 | 7.30015787 | 1.51569682 | 0.79775541 | 1.89995179 | 0.05743945 | 0.16057227 |
| CCR1 | 500.930228 | 0.263753279 | 0.45560566 | 0.57890694 | 0.56265197 | 0.73005221 |
| CCR2 | 122.36748 | 0.601185714 | 0.65994411 | 0.91096458 | 0.36231404 | 0.55963011 |
| AC098613.1 | 14.9842545 | -1.075325405 | 0.60196716 | -1.7863523 | 0.07404224 | 0.19202339 |
| CCR5 | 144.654286 | -0.37925261 | 0.52425246 | -0.723416 | 0.46942435 | 0.65606954 |
| CCRL2 | 141.22042 | -0.845926783 | 0.29600789 | -2.8577846 | 0.0042661 | 0.02389487 |
| LINC02009 | 548.057597 | -4.052914155 | 0.58954918 | -6.874599 | 6.22E-12 | 1.33E-09 |
| LTF | 68812.6916 | -5.739800546 | 0.8901023 | -6.4484729 | 1.13E-10 | 1.58E-08 |
| RTP3 | 10.2679242 | 0.508378409 | 0.49156148 | 1.03421124 | 0.30103742 | 0.49768474 |
| LRRC2 | 922.481783 | -0.205994633 | 0.38235044 | -0.5387587 | 0.59005335 | 0.74993767 |
| LRRC2-AS1 | 8.56239069 | 0.993992077 | 0.46142483 | 2.15417988 | 0.03122607 | 0.10390302 |
| TDGF1 | 1167.04079 | 0.583556849 | 0.46638147 | 1.25124364 | 0.2108456 | 0.39427021 |
| AC104304.1 | 178.617028 | 0.444301633 | 0.41884564 | 1.06077655 | 0.28879146 | 0.48502327 |
| FAM240A | 37.3637987 | -3.102909427 | 0.6753249 | -4.5946913 | 4.33E-06 | 0.00010514 |
| ALS2CL | 8851.41945 | -0.350190113 | 0.26121891 | -1.3406002 | 0.18005031 | 0.35591653 |
| TMIE | 70.8834006 | -0.250978074 | 0.24063253 | -1.0429931 | 0.29695149 | 0.49372822 |
| PRSS50 | 259.189638 | -0.16308666 | 0.39582325 | -0.4120189 | 0.68032557 | 0.8146636 |
| PRSS45 | 60.1429894 | 1.143409663 | 0.43516204 | 2.62754917 | 0.00860024 | 0.04018933 |
| AC109583.3 | 128.755998 | 0.542735107 | 0.41142311 | 1.31916534 | 0.18711384 | 0.36506304 |
| PRSS42 | 42.4412851 | 0.824621978 | 0.53476773 | 1.54201895 | 0.12306899 | 0.27456254 |
| MYL3 | 1098.12299 | 0.40724621 | 0.297744 | 1.36777304 | 0.17138313 | 0.34388234 |
| PTH1R | 17412.2481 | 1.534153745 | 0.44703373 | 3.43185231 | 0.00059947 | 0.00521033 |
| CCDC12 | 1898.77361 | -0.290375985 | 0.17090609 | -1.6990382 | 0.08931198 | 0.21943137 |
| NBEAL2 | 4899.19662 | -0.050912765 | 0.20393588 | -0.2496509 | 0.80285737 | 0.89311744 |
| SETD2 | 3222.75141 | -0.002435513 | 0.16158647 | -0.0150725 | 0.98797433 | 0.99454413 |
| KIF9-AS1 | 295.573266 | -0.232109556 | 0.25481827 | -0.9108827 | 0.36235718 | 0.55964384 |
| KIF9 | 552.400081 | 0.062141942 | 0.14098989 | 0.44075459 | 0.65939067 | 0.79994664 |
| KLHL18 | 935.821379 | -0.150233251 | 0.18662108 | -0.8050176 | 0.42080953 | 0.61352042 |
| AC099778.1 | 107.271052 | -0.065792219 | 0.26591281 | -0.2474203 | 0.80458299 | 0.89420103 |
| PTPN23 | 6217.87162 | -0.405571387 | 0.24309223 | -1.6683848 | 0.09523937 | 0.22975593 |
| SCAP | 5826.66202 | 0.16101706 | 0.08387625 | 1.91969794 | 0.05489606 | 0.1555578 |
| ELP6 | 1247.72314 | 0.151842963 | 0.13127427 | 1.15668485 | 0.24740114 | 0.43894019 |
| CSPG5 | 23.5470084 | -0.553027724 | 0.41767011 | -1.3240778 | 0.1854772 | 0.36280942 |
| SMARCC1 | 3712.46116 | 0.116189256 | 0.12762347 | 0.91040666 | 0.36260809 | 0.55991227 |
| DHX30 | 4506.54699 | 0.095186157 | 0.11335149 | 0.83974336 | 0.4010523 | 0.59591757 |
| MAP4 | 12546.6273 | -0.040960735 | 0.10996383 | -0.3724928 | 0.70952598 | 0.83497858 |
| CDC25A | 95.8251617 | -0.009549009 | 0.25579725 | -0.0373304 | 0.97022158 | 0.98622863 |
| CAMP | 6.19476465 | -1.704462557 | 1.0403556 | -1.6383461 | 0.10134951 | 0.23975838 |
| ZNF589 | 933.919554 | 0.434651477 | 0.20503732 | 2.11986514 | 0.03401742 | 0.11038441 |
| NME6 | 1204.11859 | -0.1448604 | 0.15380905 | -0.9418197 | 0.34628495 | 0.54387071 |
| SPINK8 | 0.30399583 | -0.695256667 | 2.32395324 | -0.2991698 | 0.76481048 | NA |
| FBXW12 | 34.3686576 | 1.320450108 | 0.29568096 | 4.46579345 | 7.98E-06 | 0.00017061 |
| PLXNB1 | 19754.239 | 0.158518828 | 0.15849032 | 1.00017984 | 0.31722348 | 0.51448733 |
| CCDC51 | 3081.67255 | -0.046767461 | 0.17644912 | -0.2650478 | 0.7909726 | 0.88641183 |
| TMA7 | 3443.41999 | -0.28065833 | 0.20235937 | -1.3869302 | 0.16546305 | 0.33584108 |
| AC134772.1 | 2929.50044 | -0.299942212 | 0.20036789 | -1.4969574 | 0.13440433 | 0.29163517 |
| ATRIP | 1283.49893 | -0.016903438 | 0.14849683 | -0.1138303 | 0.90937233 | 0.9548873 |
| TREX1 | 897.090196 | 8.24E-05 | 0.16502777 | 0.00049925 | 0.99960166 | 0.99979217 |
| SHISA5 | 11198.5357 | -1.294981183 | 0.25658911 | -5.0469063 | 4.49E-07 | 1.66E-05 |
| PFKFB4 | 1231.82752 | -2.756534701 | 0.44260261 | -6.2280127 | 4.72E-10 | 5.39E-08 |
| UCN2 | 93.6399589 | -2.849661311 | 0.87271445 | -3.2652849 | 0.00109354 | 0.00840964 |
| COL7A1 | 6215.90528 | -2.055763471 | 0.52895987 | -3.8864262 | 0.00010173 | 0.00128295 |
| UQCRC1 | 15165.3418 | 0.321579695 | 0.14886503 | 2.16020973 | 0.03075644 | 0.10270321 |
| TMEM89 | 15.2242027 | 1.595241955 | 0.7044565 | 2.26450029 | 0.02354336 | 0.08449689 |
| SLC26A6 | 4039.53353 | 0.880266268 | 0.34072596 | 2.58350222 | 0.00978028 | 0.04434424 |
| CELSR3 | 154.300756 | -0.591932668 | 0.32614511 | -1.8149365 | 0.06953366 | 0.1833236 |
| LINC02585 | 12.6310517 | -0.930359849 | 0.45855905 | -2.0288769 | 0.04247083 | 0.12944167 |
| AC141002.1 | 8.86774189 | 0.149967583 | 0.55269187 | 0.27134031 | 0.78612931 | 0.88350577 |
| NCKIPSD | 2268.80849 | -0.570277316 | 0.24142093 | -2.3621702 | 0.0181683 | 0.06991071 |
| IP6K2 | 6098.49784 | -0.266792301 | 0.15953451 | -1.6723171 | 0.0944618 | 0.22850727 |
| PRKAR2A | 2217.13909 | 0.107842486 | 0.20056415 | 0.53769573 | 0.59078713 | 0.7504724 |
| PRKAR2A-AS | 81.360001 | 0.637213324 | 0.25431497 | 2.50560682 | 0.01222415 | 0.05227027 |

| | | | | | | |
|---|---|---|---|---|---|---|
| SLC25A20 | 2015.30409 | 0.384431368 | 0.44309444 | 0.86760594 | 0.38561009 | 0.58142479 |
| ARIH2OS | 188.071255 | 0.394359557 | 0.18895565 | 2.0870482 | 0.03688378 | 0.11676671 |
| ARIH2 | 4755.18683 | -0.123626108 | 0.11482286 | -1.0766681 | 0.28162858 | 0.47756066 |
| AC137630.2 | 500.726179 | 0.048774658 | 0.12779734 | 0.38165628 | 0.70271634 | 0.82987743 |
| AC137630.1 | 957.28741 | -0.362374088 | 0.12111095 | -2.9920835 | 0.0027708 | 0.01728966 |
| AC137630.4 | 67.0274455 | -0.565646792 | 0.29354436 | -1.9269551 | 0.05398523 | 0.15380294 |
| P4HTM | 3844.14675 | -0.188824104 | 0.13246548 | -1.425459 | 0.15402456 | 0.31981321 |
| WDR6 | 14615.8935 | 0.03785098 | 0.17981588 | 0.21049854 | 0.83327859 | 0.91140944 |
| DALRD3 | 6263.7357 | -0.32606502 | 0.12337576 | -2.6428613 | 0.00822087 | 0.03884328 |
| NDUFAF3 | 3155.60451 | 0.053765947 | 0.18336194 | 0.29322305 | 0.76935167 | 0.87240608 |
| IMPDH2 | 5819.90983 | -0.555404518 | 0.18907872 | -2.9374248 | 0.0033095 | 0.01978051 |
| AC137630.3 | 556.909506 | -0.283964469 | 0.19618865 | -1.4474052 | 0.14778347 | 0.31129085 |
| QRICH1 | 3706.6857 | -0.037155584 | 0.11365329 | -0.3269204 | 0.74372805 | 0.85622495 |
| QARS | 8465.96285 | 0.210216425 | 0.11988517 | 1.75348154 | 0.07951939 | 0.20160813 |
| USP19 | 4144.18219 | 0.586482408 | 0.12753406 | 4.59863357 | 4.25E-06 | 0.00010349 |
| LAMB2 | 19130.4479 | 0.484721681 | 0.20157484 | 2.4046735 | 0.01618692 | 0.06436684 |
| CCDC71 | 1007.16244 | -0.11004663 | 0.15550449 | -0.707675 | 0.47914713 | 0.66392254 |
| KLHDC8B | 1637.71059 | 0.145149787 | 0.24158343 | 0.60082676 | 0.54795538 | 0.71876822 |
| C3orf84 | 0.22977084 | -1.364242679 | 3.30558948 | -0.4127078 | 0.67982068 | NA |
| CCDC36 | 19.238608 | -0.120342858 | 0.32876459 | -0.3660457 | 0.71433099 | 0.83764267 |
| AC121247.1 | 5.54679304 | -0.72499658 | 0.72529794 | -0.9995845 | 0.31751163 | 0.51474128 |
| C3orf62 | 828.185384 | -0.584753006 | 0.21141186 | -2.7659423 | 0.00567586 | 0.02961692 |
| USP4 | 2956.37735 | -0.134283944 | 0.10203754 | -1.3160249 | 0.18816568 | 0.36619403 |
| RHOA | 23997.1628 | -0.490576838 | 0.20709615 | -2.3688361 | 0.01784416 | 0.06894513 |
| TCTA | 3640.38567 | 0.579866823 | 0.16111444 | 3.59909893 | 0.00031932 | 0.00317995 |
| AMT | 2421.24478 | 1.392770581 | 0.31282126 | 4.45228878 | 8.50E-06 | 0.00017923 |
| AC104452.1 | 3725.39195 | 1.254088727 | 0.29594084 | 4.23763315 | 2.26E-05 | 0.00039214 |
| NICN1 | 1331.59841 | 0.997608503 | 0.31568085 | 3.16018066 | 0.00157671 | 0.01123424 |
| DAG1 | 6354.35547 | 0.249278223 | 0.1780003 | 1.40043709 | 0.16138247 | 0.32998604 |
| BSN-DT | 13.2553797 | 0.903878599 | 0.60516051 | 1.49361795 | 0.13527551 | 0.2927306 |
| BSN | 162.447065 | 0.78893887 | 0.29415695 | 2.68203376 | 0.00731761 | 0.03580201 |
| BSN-AS1 | 0.28750021 | -0.17981576 | 2.91671556 | -0.0616501 | 0.95084149 | NA |
| APEH | 8444.9217 | 0.526132087 | 0.19131097 | 2.75014068 | 0.00595697 | 0.03062091 |
| MST1 | 9415.0808 | 0.418337722 | 0.30106735 | 1.38951541 | 0.16467608 | 0.33465692 |
| RNF123 | 3780.66963 | 0.320818933 | 0.147147 | 2.18026149 | 0.02923808 | 0.09908858 |
| AMIGO3 | 336.956412 | 0.132390079 | 0.16100466 | 0.82227483 | 0.4109205 | 0.60529402 |
| GMPPB | 3630.24792 | -0.171844218 | 0.09189954 | -1.8699138 | 0.06149579 | 0.16825523 |
| IP6K1 | 3713.04105 | -0.114646607 | 0.2117295 | -0.5414768 | 0.58817901 | 0.74897126 |
| CDHR4 | 8.61869863 | 0.541853874 | 0.52703554 | 1.02811639 | 0.30389508 | 0.50076071 |
| INKA1 | 106.665678 | 0.491271818 | 0.31631732 | 1.55309806 | 0.12039971 | 0.27049642 |
| UBA7 | 4248.29611 | -0.149710245 | 0.61250157 | -0.2444243 | 0.80690225 | 0.89560375 |
| TRAIP | 90.3289001 | -0.059590056 | 0.35931778 | -0.1658422 | 0.86828112 | 0.93147444 |
| CAMKV | 2.29076528 | -1.664631132 | 1.50935084 | -1.1028789 | 0.27007978 | 0.46442802 |
| MST1R | 288.453531 | -1.2777749 | 0.28279644 | -4.5183557 | 6.23E-06 | 0.00014035 |
| AC105935.2 | 8.07903596 | 0.037609559 | 0.56401286 | 0.0666821 | 0.94683479 | 0.97438741 |
| AC105935.1 | 3.08676889 | 0.763841828 | 0.6455721 | 1.18320142 | 0.23672933 | 0.42601653 |
| MON1A | 819.495444 | 0.256498689 | 0.21086376 | 1.21641902 | 0.22382534 | 0.40990863 |
| RBM6 | 8299.39562 | -0.147036557 | 0.21475203 | -0.6846806 | 0.49354548 | 0.67619153 |
| RBM5 | 7819.9737 | -0.063875598 | 0.14134604 | -0.4519093 | 0.65133429 | 0.795097 |
| RBM5-AS1 | 146.180946 | 1.306736388 | 0.2746364 | 4.75805976 | 1.95E-06 | 5.57E-05 |
| SEMA3F-AS1 | 2851.40575 | -0.307661541 | 0.16003791 | -1.9224291 | 0.05455178 | 0.15479721 |
| SEMA3F | 4347.28181 | -1.046243762 | 0.26080961 | -4.011523 | 6.03E-05 | 0.00085346 |
| GNAT1 | 4.5832359 | -1.464391349 | 1.12877534 | -1.2973275 | 0.19451851 | 0.37448581 |
| SLC38A3 | 455.224543 | 0.824481612 | 0.59586127 | 1.38368049 | 0.16645631 | 0.33723516 |
| GNAI2 | 11556.1392 | -0.537233368 | 0.17733942 | -3.0294075 | 0.00245034 | 0.01574451 |
| U73166.1 | 72.100707 | 0.402595963 | 0.41058189 | 0.98054974 | 0.32681483 | 0.52471967 |
| SEMA3B-AS1 | 94.2727321 | 0.217931532 | 0.36275542 | 0.60076713 | 0.5479951 | 0.71876822 |
| SEMA3B | 9850.60034 | -0.053411869 | 0.2400691 | -0.2224854 | 0.82393604 | 0.90578475 |
| LSMEM2 | 671.393678 | -0.781819179 | 0.23657396 | -3.3047559 | 0.00095059 | 0.00755033 |
| IFRD2 | 3390.01861 | -0.578280686 | 0.21420138 | -2.6997057 | 0.00694008 | 0.03439391 |
| HYAL3 | 227.954802 | -0.368172828 | 0.21413401 | -1.7193571 | 0.08554937 | 0.21232347 |
| NAA80 | 688.394108 | -0.299323636 | 0.22294103 | -1.3426135 | 0.17939716 | 0.35502462 |
| HYAL1 | 5072.86514 | 0.576562484 | 0.34975711 | 1.64846535 | 0.09925722 | 0.23629395 |
| HYAL2 | 6070.93024 | -0.494640625 | 0.21475827 | -2.3032436 | 0.02126514 | 0.07843918 |
| TUSC2 | 3966.46966 | -0.288384103 | 0.13786102 | -2.0918466 | 0.03645224 | 0.11585459 |
| RASSF1 | 2226.79607 | -0.8787126 | 0.23172571 | -3.7920375 | 0.00014942 | 0.00174805 |
| RASSF1-AS1 | 52.2369461 | 0.49220396 | 0.29590105 | 1.66340726 | 0.09623094 | 0.23138597 |
| ZMYND10-AS | 43.7990978 | -0.455355944 | 0.54310687 | -0.8384279 | 0.40179043 | 0.59664454 |
| ZMYND10 | 223.406443 | -0.492067077 | 0.48042369 | -1.0242357 | 0.30572398 | 0.50274017 |
| NPRL2 | 1605.32126 | 0.035069419 | 0.0917956 | 0.38203812 | 0.70243309 | 0.829742 |
| CYB561D2 | 2511.98812 | -0.086253891 | 0.12981407 | -0.6644418 | 0.50640761 | 0.68566279 |
| Z84492.1 | 555.393534 | -0.018230771 | 0.20376334 | -0.0894703 | 0.92870814 | 0.96531469 |
| TMEM115 | 3328.04009 | -0.183494461 | 0.16582496 | -1.1065551 | 0.26848632 | 0.46271383 |
| CACNA2D2 | 1062.46801 | -0.235086105 | 0.38645925 | -0.6083076 | 0.54298347 | 0.71531565 |
| C3orf18 | 891.840773 | 0.784166714 | 0.24817779 | 3.15969734 | 0.00157933 | 0.01124985 |
| HEMK1 | 2310.4905 | 0.290518991 | 0.2640771 | 1.10012942 | 0.27127574 | 0.46596803 |
| AC096920.1 | 6.47960689 | 0.651414647 | 0.63680689 | 1.02293907 | 0.30633665 | 0.50318286 |
| CISH | 5946.41067 | -0.009492134 | 0.24710781 | -0.0384129 | 0.96935845 | 0.98580727 |
| MAPKAPK3 | 4188.15915 | 0.140429102 | 0.13946163 | 1.00693718 | 0.31396496 | 0.51093903 |
| LINC02019 | 144.608788 | -1.36192633 | 0.37799464 | -3.6030308 | 0.00031453 | 0.00314528 |
| DOCK3 | 221.466875 | -0.183130837 | 0.37344485 | -0.4903825 | 0.62386323 | 0.77584318 |
| MANF | 2268.90547 | -0.815425843 | 0.30870415 | -2.6414476 | 0.00825526 | 0.03897782 |
| RBM15B | 2862.50238 | -0.328034324 | 0.22956655 | -1.4289291 | 0.15302461 | 0.31857242 |
| DCAF1 | 2007.56035 | -0.034359036 | 0.17599941 | -0.1952225 | 0.8452188 | 0.91831485 |
| RAD54L2 | 1389.71609 | -0.19295673 | 0.09310996 | -2.0723533 | 0.0382325 | 0.11976611 |
| TEX264 | 5156.70573 | 0.239124972 | 0.13532444 | 1.76704938 | 0.07721996 | 0.19763444 |
| GRM2 | 30.2373778 | -0.175162369 | 0.52194887 | -0.335593 | 0.73717783 | 0.8524157 |
| IQCF6 | 3.14409685 | 1.383170363 | 0.7628666 | 1.8131222 | 0.06981298 | 0.1838144 |
| IQCF3 | 4.28684532 | 1.915076493 | 0.7947904 | 2.40953653 | 0.0159728 | 0.06377993 |
| IQCF2 | 0 | NA | NA | NA | NA | NA |
| IQCF5-AS1 | 0.26948541 | -1.364242679 | 3.30558948 | -0.4127078 | 0.67982068 | NA |
| IQCF5 | 0.26948541 | -1.364242679 | 3.30558948 | -0.4127078 | 0.67982068 | NA |
| IQCF1 | 0.97599533 | -3.178310062 | 1.3387184 | -2.3741439 | 0.0175897 | NA |
| RRP9 | 1100.5101 | -0.540970706 | 0.23104704 | -2.3413877 | 0.01921221 | 0.07275971 |
| PARP3 | 2617.6807 | 0.10150339 | 0.35682155 | 0.28446541 | 0.77605373 | 0.87702653 |
| GPR62 | 30.7341776 | 0.247313496 | 0.30662145 | 0.80657598 | 0.41991082 | 0.61265067 |
| PCBP4 | 2491.68049 | -0.035644505 | 0.2275523 | -0.1566431 | 0.87552611 | 0.93536537 |
| ABHD14B | 6802.2474 | 0.717866336 | 0.19307265 | 3.71811509 | 0.00020071 | 0.00218622 |
| ABHD14A | 2540.97177 | 1.313736359 | 0.23418769 | 5.60975831 | 2.03E-08 | 1.26E-06 |
| ABHD14A-AC | 26742.1414 | 2.198258851 | 0.39756781 | 5.52926771 | 3.22E-08 | 1.86E-06 |
| ACY1 | 25685.2328 | 2.223131929 | 0.40868622 | 5.43970373 | 5.34E-08 | 2.80E-06 |
| RPL29 | 19204.1888 | -0.41596628 | 0.13366212 | -3.1120729 | 0.00185779 | 0.01272532 |
| DUSP7 | 1117.06749 | -0.195869429 | 0.30664074 | -0.6387587 | 0.52297995 | 0.69862624 |
| POC1A | 183.14447 | -0.509208922 | 0.28484278 | -1.7876842 | 0.07382698 | 0.19163475 |
| ALAS1 | 4374.88827 | -0.419925413 | 0.21711573 | -1.9341086 | 0.05309977 | 0.15195236 |
| TLR9 | 68.1761024 | -0.603862894 | 0.39430501 | -1.5314614 | 0.12565541 | 0.27842568 |
| AC097637.1 | 1449.13918 | -0.885615855 | 0.17510319 | -5.0576797 | 4.24E-07 | 1.59E-05 |
| TWF2 | 2646.86829 | -0.967330096 | 0.18270424 | -5.2945137 | 1.19E-07 | 5.55E-06 |
| AC006252.1 | 14.7372856 | -0.034074675 | 0.42992114 | -0.079258 | 0.93682744 | 0.96969938 |
| PPM1M | 1247.9245 | -0.210486534 | 0.26172091 | -0.8042404 | 0.42125814 | 0.61391338 |
| WDR82 | 3882.13341 | 0.032422042 | 0.14197418 | 0.22836577 | 0.8193619 | 0.90302057 |
| GLYCTK | 2660.96999 | 1.654244533 | 0.34911828 | 4.73834985 | 2.15E-06 | 6.03E-05 |
| DNAH1 | 942.627133 | 0.618666957 | 0.17385563 | 3.55850976 | 0.00037296 | 0.00361053 |

| | | | | | | |
|---|---|---|---|---|---|---|
| BAP1 | 5474.8229 | 0.028599601 | 0.12756043 | 0.22420433 | 0.82259832 | 0.90522546 |
| PHF7 | 321.422099 | 0.410215568 | 0.22146937 | 1.85224514 | 0.06399063 | 0.17294588 |
| SEMA3G | 3114.39129 | 0.606448293 | 0.39525362 | 1.53432698 | 0.12494923 | 0.27727941 |
| TNNC1 | 210.607706 | 1.076385395 | 0.48451661 | 2.22156552 | 0.02631268 | 0.09180543 |
| NISCH | 7170.0536 | 0.113821847 | 0.19404252 | 0.586582 | 0.55748447 | 0.72629147 |
| STAB1 | 10234.888 | -0.899080257 | 0.37545774 | -2.3946244 | 0.0166374 | 0.06565765 |
| NT5DC2 | 2601.18413 | -0.79740049 | 0.32293168 | -2.4692544 | 0.01353949 | 0.05642168 |
| SMIM4 | 819.85277 | -0.182243861 | 0.26429303 | -0.6895523 | 0.49047579 | 0.67369536 |
| PBRM1 | 1560.63185 | -0.079775139 | 0.16422522 | -0.4857667 | 0.62713259 | 0.77806973 |
| GNL3 | 2707.26471 | -1.099621489 | 0.25698485 | -4.278935 | 1.88E-05 | 0.00033756 |
| GLT8D1 | 2580.90155 | 0.14025995 | 0.11226931 | 1.24931693 | 0.21154918 | 0.3950536 |
| SPCS1 | 5350.267 | -0.057036715 | 0.10280583 | -0.5548004 | 0.5790312 | 0.74245887 |
| NEK4 | 1133.40928 | 0.300285372 | 0.10980715 | 2.73466145 | 0.00624445 | 0.03177041 |
| ITIH1 | 4.42365249 | 0.1058348 | 0.72495072 | 0.14598896 | 0.88393011 | 0.93980604 |
| ITIH3 | 157.615286 | 0.425122664 | 0.39296096 | 1.08184454 | 0.27932161 | 0.47492119 |
| ITIH4 | 1204.51239 | 0.800031602 | 0.39723519 | 2.01399984 | 0.04400955 | 0.13287518 |
| AC006254.1 | 1852.09636 | 0.492503768 | 0.35919981 | 1.37111365 | 0.17033953 | 0.34236687 |
| ITIH4-AS1 | 113.041005 | 1.089988047 | 0.46152865 | 2.36169095 | 0.0181918 | 0.0699705 |
| MUSTN1 | 547.03443 | -0.246271834 | 0.43343471 | -0.5681867 | 0.5699082 | 0.73521253 |
| TMEM110-M | 1913.10075 | 0.226076629 | 0.19974846 | 1.13180662 | 0.25771575 | 0.45029234 |
| STIMATE | 1312.11701 | 0.406971822 | 0.18987688 | 2.14334585 | 0.03208534 | 0.10585392 |
| SFMBT1 | 604.67912 | -0.068438267 | 0.21917021 | -0.3122608 | 0.75484233 | 0.86367823 |
| AC096887.1 | 291.76599 | -0.037402036 | 0.19566884 | -0.1911497 | 0.84840833 | 0.92025018 |
| RFT1 | 880.792967 | -0.059181243 | 0.16142591 | -0.3666155 | 0.71390584 | 0.83749271 |
| PRKCD | 2879.47926 | 0.1977778 | 0.17048703 | 1.16007535 | 0.24601813 | 0.43731029 |
| TKT | 19731.6226 | -1.263638163 | 0.30294855 | -4.1711312 | 3.03E-05 | 0.00049245 |
| DCP1A | 1286.8458 | -0.463728189 | 0.17478058 | -2.653202 | 0.00797321 | 0.03806875 |
| CACNA1D | 1999.68007 | 0.519054298 | 0.27971629 | 1.85564557 | 0.06350409 | 0.1719658 |
| AC012467.1 | 49.22044 | -0.199555118 | 0.7600383 | -0.2625593 | 0.79289028 | 0.8878351 |
| CHDH | 7457.0336 | 1.41348577 | 0.37179933 | 3.80174375 | 0.00014368 | 0.00169146 |
| IL17RB | 12597.3785 | -0.673883073 | 0.54398354 | -1.2387931 | 0.21542211 | 0.39990716 |
| AC012467.2 | 2325.81175 | -0.602210795 | 0.46982574 | -1.2817748 | 0.19992165 | 0.38127822 |
| ACTR8 | 1306.81808 | -0.047343693 | 0.17914921 | -0.2642696 | 0.79157217 | 0.88684871 |
| SELENOK | 2690.51002 | -0.591061666 | 0.17520532 | -3.3735373 | 0.00074209 | 0.00619269 |
| CACNA2D3 | 537.718731 | -0.120709302 | 0.30797624 | -0.3919436 | 0.69509992 | 0.82446857 |
| CACNA2D3-A | 35.6870464 | -0.237201724 | 0.33447281 | -0.7091809 | 0.47821223 | 0.66311492 |
| LRTM1 | 0.36199939 | 0.724933013 | 2.86723138 | 0.2528338 | 0.80039665 | NA |
| AC092059.1 | 0 | NA | NA | NA | NA | NA |
| LINC02017 | 0.03392908 | 0.380740593 | 3.29738709 | 0.11546736 | 0.90807469 | NA |
| LINC02030 | 0.18874271 | 0.726716446 | 3.28420131 | 0.22127646 | 0.82487718 | NA |
| AC121764.2 | 0 | NA | NA | NA | NA | NA |
| AC121764.3 | 0 | NA | NA | NA | NA | NA |
| WNT5A | 354.374265 | -0.274707914 | 0.6570079 | -0.4181196 | 0.67585964 | 0.81147469 |
| WNT5A-AS1 | 3.68505161 | 0.636622305 | 0.99601213 | 0.63917124 | 0.52271154 | 0.69859758 |
| AC121764.1 | 0.19994613 | 1.506623855 | 3.27645372 | 0.45983371 | 0.64563559 | NA |
| ERC2 | 75.8382614 | -0.471566067 | 0.30782717 | -1.5319183 | 0.12554261 | 0.27822158 |
| AC025572.1 | 0 | NA | NA | NA | NA | NA |
| CCDC66 | 1219.3551 | 0.187209957 | 0.17138153 | 1.09235781 | 0.27467586 | 0.46994112 |
| FAM208A | 2970.64475 | -0.170158558 | 0.10323016 | -1.6483416 | 0.0992826 | 0.23633166 |
| ARHGEF3 | 2570.96251 | 0.059414733 | 0.16089519 | 0.36927601 | 0.71192201 | 0.83623089 |
| ARHGEF3-AS | 0 | NA | NA | NA | NA | NA |
| SPATA12 | 3.76509903 | -0.196815862 | 0.96626328 | -0.2036876 | 0.83859762 | 0.91450415 |
| AC097358.2 | 16.9650276 | -0.880870507 | 0.37905872 | -2.3238366 | 0.02013426 | 0.0753 |
| IL17RD | 1006.8185 | 0.651918002 | 0.20269011 | 3.21632857 | 0.00129842 | 0.00965577 |
| HESX1 | 85.3880794 | -0.030582933 | 0.5550824 | -0.0550962 | 0.95606182 | 0.97900515 |
| APPL1 | 2091.20233 | 0.022818772 | 0.0817601 | 0.27909421 | 0.78017252 | 0.87964552 |
| ASB14 | 166.232225 | -0.175997937 | 0.18780799 | -0.9371163 | 0.34869872 | 0.546227 |
| DNAH12 | 31.1495744 | -0.043606173 | 0.50423441 | -0.08648 | 0.93108488 | 0.96688001 |
| PDE12 | 1661.67785 | -0.100214204 | 0.24166689 | -0.4146791 | 0.67837685 | 0.81323805 |
| ARF4 | 8881.938 | -0.811560314 | 0.33260237 | -2.4400317 | 0.01468597 | 0.05995807 |
| ARF4-AS1 | 46.1268496 | -0.084689429 | 0.40405773 | -0.2095973 | 0.83398195 | 0.91200564 |
| DENND6A | 1504.47318 | -0.20930088 | 0.14347173 | -1.4588301 | 0.14461188 | 0.30664324 |
| DENND6A-A! | 68.7382336 | 0.854824279 | 0.28156772 | 3.03594556 | 0.00239783 | 0.01549012 |
| DENND6A-D' | 1.35236023 | 0.741842149 | 0.98020442 | 0.75682392 | 0.44915536 | 0.63932996 |
| SLMAP | 2402.06043 | -0.130735262 | 0.14221794 | -0.91926 | 0.35795959 | 0.55558532 |
| FLNB | 10820.3423 | 0.192383357 | 0.11873385 | 1.62029078 | 0.10516983 | 0.24597147 |
| FLNB-AS1 | 1951.68089 | 0.105507001 | 0.11199682 | 0.94205352 | 0.34616524 | 0.54374418 |
| AC137936.1 | 1.90189927 | -0.077210988 | 0.97519861 | -0.0791746 | 0.93689372 | 0.96969938 |
| DNASE1L3 | 1308.55519 | 0.480642193 | 0.39813858 | 1.20722336 | 0.22734614 | 0.41422771 |
| ABHD6 | 1939.7295 | 1.202619831 | 0.4242549 | 2.83466337 | 0.0045874 | 0.0252441 |
| RPP14 | 1776.27259 | 0.264265916 | 0.18059855 | 1.46327815 | 0.14339128 | 0.30496235 |
| HTD2 | 818.873401 | 0.409602807 | 0.22004844 | 1.86142105 | 0.06268474 | 0.17049991 |
| AC098479.1 | 6.08556613 | -0.011546148 | 0.63417314 | -0.0182066 | 0.98547402 | 0.99361283 |
| PXK | 1553.24323 | 0.121135192 | 0.16311792 | 0.74262345 | 0.45770969 | 0.64686234 |
| PDHB | 5205.45362 | 0.208938675 | 0.13610397 | 1.53514018 | 0.1247494 | 0.27692917 |
| AC135507.1 | 1725.39331 | 0.108597877 | 0.14134297 | 0.7683288 | 0.44229187 | 0.63338963 |
| AC116036.2 | 28.0964322 | 0.931892821 | 0.46323873 | 2.01169021 | 0.0442526 | 0.13327002 |
| KCTD6 | 1226.45953 | -0.04145888 | 0.17076573 | -0.2427822 | 0.80817413 | 0.89624087 |
| ACOX2 | 3322.3287 | 2.425280157 | 0.45751415 | 5.30099486 | 1.15E-07 | 5.38E-06 |
| FAM107A | 3714.06626 | -0.129537189 | 0.35128775 | -0.3687495 | 0.71231443 | 0.83646828 |
| AC119424.1 | 37.1092464 | 2.995524261 | 0.7431743 | 4.0307156 | 5.56E-05 | 0.00080614 |
| FAM3D-AS1 | 100.027532 | 0.070268626 | 0.81554487 | 0.08616157 | 0.93133798 | 0.96690482 |
| FAM3D | 58.2494694 | 0.384361568 | 0.69376512 | 0.55402262 | 0.57956337 | 0.74277814 |
| C3orf67 | 66.4340882 | -0.677339534 | 0.24163988 | -2.803095 | 0.00506148 | 0.02711174 |
| C3orf67-AS1 | 4.77378089 | -1.16993657 | 0.83347276 | -1.403689 | 0.16041147 | 0.32864972 |
| AC114400.1 | 0 | NA | NA | NA | NA | NA |
| AC138057.1 | 0 | NA | NA | NA | NA | NA |
| AC126121.2 | 0 | NA | NA | NA | NA | NA |
| FHIT | 902.890463 | 0.725602233 | 0.32450178 | 2.23605006 | 0.02534849 | 0.08925621 |
| PTPRG | 3329.5457 | 0.002954581 | 0.17869825 | 0.01653391 | 0.98680845 | 0.99400325 |
| C3orf14 | 315.694839 | -0.029185035 | 0.33178589 | -0.0879635 | 0.92990572 | 0.9661025 |
| FEZF2 | 0 | NA | NA | NA | NA | NA |
| CADPS | 15.8170012 | 1.183786276 | 0.61321652 | 1.930454 | 0.05355061 | 0.15291062 |
| LINC00698 | 0 | NA | NA | NA | NA | NA |
| SYNPR | 1.95058068 | 1.446588761 | 1.88854505 | 0.76598054 | 0.44368788 | 0.63473387 |
| SYNPR-AS1 | 0.31046549 | 1.449838875 | 2.87844687 | 0.50368791 | 0.61448071 | NA |
| SNTN | 0.65770508 | -1.858761331 | 1.87811683 | -0.9896942 | 0.32232361 | NA |
| C3orf49 | 10.6235075 | 0.770487629 | 0.48554733 | 1.58684351 | 0.11254809 | 0.25816747 |
| THOC7 | 1577.64742 | -0.352367525 | 0.1414083 | -2.4918447 | 0.01270816 | 0.05383359 |
| THOC7-AS1 | 0.58388377 | -0.72867751 | 1.66296165 | -0.4381806 | 0.66125538 | NA |
| ATXN7 | 1340.67329 | -0.046081057 | 0.22167571 | -0.207876 | 0.83532579 | 0.91268039 |
| SCAANT1 | 4.15075291 | 0.802135667 | 0.58593699 | 1.36897939 | 0.17100572 | 0.34328721 |
| PSMD6-AS2 | 170.601081 | -0.487869438 | 0.21513602 | -2.2677255 | 0.02334594 | 0.08396972 |
| AC012557.1 | 17.1962335 | 0.840428456 | 0.29923903 | 2.80855223 | 0.00497648 | 0.02677067 |
| PSMD6 | 3870.47316 | -0.255492251 | 0.17338158 | -1.4735836 | 0.14059374 | 0.30075907 |
| PSMD6-AS1 | 81.2833791 | -0.359084643 | 0.22806974 | -1.574451 | 0.11538321 | 0.26281898 |
| AC012557.2 | 20.4989842 | -0.557956861 | 0.34610914 | -1.6120836 | 0.10694375 | 0.24877441 |
| AC092040.1 | 162.538487 | 0.765534049 | 0.24443636 | 3.13183374 | 0.00173718 | 0.01211251 |
| LINC00994 | 2.29360552 | 0.709143071 | 1.21646796 | 0.58295252 | 0.55992526 | 0.72827885 |
| PRICKLE2 | 1018.5807 | 0.328798762 | 0.13594982 | 2.41853026 | 0.01558335 | 0.06253735 |
| PRICKLE2-AS | 10.2896851 | 0.088440404 | 0.47794785 | 0.18504195 | 0.85319613 | 0.92319951 |
| PRICKLE2-AS | 0.15489794 | -1.003574173 | 3.30558948 | -0.3035992 | 0.7614333 | NA |
| PRICKLE2-AS | 2.89261421 | 1.256538836 | 0.78334122 | 1.60407599 | 0.1086973 | 0.25198663 |

| | | | | | | |
|---|---|---|---|---|---|---|
| AC096888.1 | 0.91104688 | 2.232622741 | 1.24462772 | 1.79380766 | 0.07284387 | NA |
| ADAMTS9 | 4256.89628 | -1.495231792 | 0.31805243 | -4.7012116 | 2.59E-06 | 6.99E-05 |
| ADAMTS9-AS | 7489.9766 | 0.89081669 | 0.41094679 | 2.16771785 | 0.03018016 | 0.10124299 |
| ADAMTS9-AS | 246.561636 | 0.637271604 | 0.23147086 | 2.7531396 | 0.00590267 | 0.03042 |
| LINC02040 | 0.08666371 | -0.642905668 | 3.30558948 | -0.1944905 | 0.84579185 | NA |
| MAGI1 | 2106.23434 | 0.023789673 | 0.11702205 | 0.20329223 | 0.83890663 | 0.91454277 |
| MAGI1-AS1 | 0.94611489 | 0.641404173 | 1.80293628 | 0.35575532 | 0.72202381 | NA |
| SLC25A26 | 1588.81577 | -0.172898855 | 0.18119664 | -0.9542056 | 0.33997956 | 0.53809861 |
| LRIG1 | 1371.74786 | 3.11E-05 | 0.13002997 | 0.00023928 | 0.99980908 | 0.99992284 |
| KBTBD8 | 64.0461507 | -0.086924291 | 0.40713349 | -0.2135032 | 0.83093452 | 0.91040231 |
| AC114401.1 | 0 | NA | NA | NA | NA | NA |
| SUCLG2 | 7622.59181 | 0.988454003 | 0.24368597 | 4.05626143 | 4.99E-05 | 0.00074248 |
| SUCLG2-AS1 | 72.9255157 | 0.699760393 | 0.28230596 | 2.47873049 | 0.01318509 | 0.05529434 |
| FAM19A1 | 16.76321 | 0.307829885 | 0.53905717 | 0.57105239 | 0.56796413 | 0.73384869 |
| FAM19A4 | 3.30094039 | 1.589922413 | 0.92248071 | 1.72352918 | 0.08479287 | 0.21098208 |
| EOGT | 826.586027 | -0.604390078 | 0.22451783 | -2.6919469 | 0.00710363 | 0.03501141 |
| AC109587.1 | 897.131539 | -0.093009364 | 0.12511832 | -0.7433712 | 0.45725695 | 0.64655504 |
| TMF1 | 1975.72462 | -0.208298885 | 0.1471816 | -1.4152509 | 0.15699495 | 0.32391494 |
| UBA3 | 2778.75264 | -0.21815035 | 0.10084446 | -2.1632359 | 0.03052304 | 0.10207896 |
| ARL6IP5 | 6494.65293 | -0.381930878 | 0.21056733 | -1.8138183 | 0.0697057 | 0.18367289 |
| LMOD3 | 11.2553611 | 0.151351231 | 0.46348501 | 0.32655043 | 0.74400794 | 0.85638322 |
| FRMD4B | 2010.99477 | 0.450738909 | 0.13894823 | 3.24393411 | 0.00117891 | 0.00892816 |
| AC099328.2 | 0 | NA | NA | NA | NA | NA |
| AC104445.1 | 0 | NA | NA | NA | NA | NA |
| MITF | 1818.6519 | 0.418086655 | 0.18411802 | 2.27075362 | 0.0231619 | 0.08364547 |
| SAMMSON | 1.86915898 | 0.36106047 | 1.05663569 | 0.34170762 | 0.73257094 | 0.8492009 |
| MDFIC2 | 0 | NA | NA | NA | NA | NA |
| FOXP1 | 1436.81807 | -0.36190609 | 0.17566589 | -2.0601956 | 0.03937985 | 0.12242944 |
| FOXP1-AS1 | 3.23515692 | 1.193382484 | 0.80598017 | 1.48065987 | 0.13869723 | 0.2980075 |
| AC097634.2 | 0.36836239 | -1.044722622 | 2.30140842 | -0.4539492 | 0.64986536 | NA |
| AC097634.3 | 3.49397542 | -0.636989617 | 0.67507482 | -0.9435837 | 0.34538241 | 0.54312923 |
| AC097634.1 | 20.6137999 | 0.198401034 | 0.3953702 | 0.50181079 | 0.61580062 | 0.77023922 |
| EIF4E3 | 1596.44802 | -0.007961889 | 0.15061316 | -0.0528632 | 0.95784093 | 0.97980004 |
| GPR27 | 274.662592 | 0.174624464 | 0.40887073 | 0.42708967 | 0.66931402 | 0.80688537 |
| PROK2 | 43.0613627 | -0.988125802 | 0.79825312 | -1.2378602 | 0.21576788 | 0.40037992 |
| LINC00877 | 15.1215923 | -0.392996327 | 0.55491167 | -0.7082142 | 0.47881225 | 0.66367267 |
| AC105265.3 | 0.66683835 | 0.209278782 | 1.77188727 | 0.11811066 | 0.90597997 | NA |
| AC105265.4 | 0 | NA | NA | NA | NA | NA |
| LINC00870 | 21.2295814 | 3.215984952 | 0.51801243 | 6.20831619 | 5.36E-10 | 6.01E-08 |
| AC112219.2 | 3.33295456 | 4.440996697 | 1.15539905 | 3.84369079 | 0.0001212 | 0.0014762 |
| AC112219.1 | 0.09997307 | 0.547239394 | 3.29377202 | 0.16614368 | 0.86804388 | NA |
| AC134508.1 | 0 | NA | NA | NA | NA | NA |
| RYBP | 2162.63458 | -0.22239326 | 0.16280598 | -1.3660018 | 0.1719384 | 0.34476585 |
| AC104435.2 | 0.57995337 | -1.384128526 | 2.25181746 | -0.6146717 | 0.53877154 | NA |
| SHQ1 | 749.680474 | -0.328295153 | 0.19710311 | -1.6656011 | 0.0957929 | 0.23069029 |
| GXYLT2 | 106.81901 | 0.119726003 | 0.25470976 | 0.47004874 | 0.6383202 | 0.78602447 |
| PPP4R2 | 2531.50288 | -0.352253278 | 0.17449403 | -2.0187125 | 0.04351711 | 0.13181096 |
| EBLN2 | 9.31929178 | 0.716486105 | 0.57246439 | 1.25158196 | 0.21072223 | 0.3940954 |
| PDZRN3 | 863.025849 | 1.307001588 | 0.28942526 | 4.51585185 | 6.31E-06 | 0.00014145 |
| PDZRN3-AS1 | 11.7648545 | 1.792123311 | 0.43250588 | 4.14358133 | 3.42E-05 | 0.00054593 |
| LINC02005 | 0.24327904 | -1.364242679 | 3.30558948 | -0.4127078 | 0.67982068 | NA |
| LINC02047 | 0 | NA | NA | NA | NA | NA |
| CNTN3 | 422.59293 | 0.749990156 | 0.25174398 | 2.97917809 | 0.00289023 | 0.01781564 |
| LINC02018 | 27.044033 | 0.814665503 | 0.44623711 | 1.82563368 | 0.06790546 | 0.18011788 |
| FRG2C | 16.9056601 | 3.060821838 | 0.67566936 | 4.53005867 | 5.90E-06 | 0.00013428 |
| LINC00960 | 145.965811 | -0.163322177 | 0.53954455 | -0.3027038 | 0.76211561 | 0.86815969 |
| ZNF717 | 416.639652 | 0.394231041 | 0.2856987 | 1.37988389 | 0.1676224 | 0.33884663 |
| ROBO2 | 100.86909 | 1.011620541 | 0.52995046 | 1.90889643 | 0.05627545 | 0.15815663 |
| LINC02077 | 0 | NA | NA | NA | NA | NA |
| AC117462.1 | 0 | NA | NA | NA | NA | NA |
| AC108752.1 | 35.5946759 | -3.133951411 | 1.12325448 | -2.7900636 | 0.00526977 | 0.02796615 |
| ROBO1 | 449.373146 | 0.149206285 | 0.32532965 | 0.45863106 | 0.64649913 | 0.79168673 |
| AC068756.1 | 2.83501311 | -1.048382631 | 1.78714581 | -0.586624 | 0.55745626 | 0.72629087 |
| LINC02050 | 0.46297644 | -2.127129727 | 3.28667147 | -0.6471988 | 0.51750331 | NA |
| LINC02027 | 109.62019 | 0.725352848 | 0.5800363 | 1.2505301 | 0.21110597 | 0.39447735 |
| AC099542.2 | 0 | NA | NA | NA | NA | NA |
| AC099542.1 | 4.64432467 | -0.023588852 | 0.66218016 | -0.035623 | 0.97158296 | 0.98670705 |
| GBE1 | 4273.16826 | -0.335265073 | 0.35665185 | -0.9400346 | 0.34719982 | 0.54472033 |
| AC017015.2 | 1.89453444 | 0.771074501 | 0.95279736 | 0.80927439 | 0.41835733 | 0.61113582 |
| AC129807.1 | 0.17648264 | -1.003574173 | 3.30558948 | -0.3035992 | 0.7614333 | NA |
| LINC02008 | 0 | NA | NA | NA | NA | NA |
| LINC00971 | 0 | NA | NA | NA | NA | NA |
| LINC02025 | 0.03392908 | 0.380740593 | 3.29738709 | 0.11546736 | 0.90807469 | NA |
| CADM2 | 11.690448 | 0.161635089 | 0.79128711 | 0.20426857 | 0.83814363 | 0.9143814 |
| CADM2-AS2 | 0 | NA | NA | NA | NA | NA |
| CADM2-AS1 | 0 | NA | NA | NA | NA | NA |
| AC108733.1 | 0 | NA | NA | NA | NA | NA |
| LINC02070 | 0 | NA | NA | NA | NA | NA |
| VGLL3 | 289.793594 | -0.530721275 | 0.56472825 | -0.9397817 | 0.34732955 | 0.5447942 |
| AC107204.1 | 14.3691857 | 0.255335578 | 0.48158377 | 0.53019972 | 0.59597347 | 0.75484247 |
| CHMP2B | 2993.14897 | -0.462953915 | 0.26428629 | -1.7517137 | 0.07982305 | 0.2021448 |
| POU1F1 | 4.47464771 | 0.352664966 | 0.59477304 | 0.5929404 | 0.55322104 | 0.7229157 |
| AC117513.1 | 5.09872084 | 1.801940974 | 0.80293268 | 2.24419932 | 0.02481958 | 0.08782774 |
| HTR1F | 5.11823545 | 0.132060404 | 1.00883866 | 0.13090339 | 0.89585173 | 0.94661044 |
| CGGBP1 | 4232.60583 | 0.068986211 | 0.10707821 | 0.64426002 | 0.51940684 | 0.69628475 |
| ZNF654 | 776.87844 | -0.31979809 | 0.14093127 | -2.2691777 | 0.02325752 | 0.08379705 |
| C3orf38 | 636.126218 | -0.233164505 | 0.15988076 | -1.458365 | 0.14473995 | 0.30684078 |
| CSNKA2IP | 0.25082976 | -0.178310673 | 3.00331177 | -0.0593713 | 0.95265633 | NA |
| EPHA3 | 106.37437 | 0.879418595 | 0.46499008 | 1.89126315 | 0.05858923 | 0.16275128 |
| PROS1 | 4478.92921 | 0.009934456 | 0.25509168 | 0.03894465 | 0.96893452 | 0.9856042 |
| ARL13B | 585.147517 | 0.039796149 | 0.18186792 | 0.21881896 | 0.82679107 | 0.90760472 |
| STX19 | 10.7531459 | -0.116815798 | 0.70589673 | -0.1654857 | 0.86856172 | 0.93156219 |
| DHFR2 | 462.570416 | 0.82726954 | 0.1433454 | 5.77116219 | 7.87E-09 | 5.62E-07 |
| NSUN3 | 386.723617 | -0.059262984 | 0.14197156 | -0.4174286 | 0.67636496 | 0.81167489 |
| LINC00879 | 0.19632833 | 1.506623996 | 3.27645372 | 0.45983375 | 0.64563556 | NA |
| MTRNR2L12 | 386.112327 | -0.458734346 | 0.32551857 | -1.4092417 | 0.15876371 | 0.32623697 |
| EPHA6 | 12.9601162 | 0.085209513 | 0.54274546 | 0.15699719 | 0.87524706 | 0.93514302 |
| AC110491.1 | 1.97011883 | -3.024613116 | 1.88032118 | -1.608562 | 0.10771216 | 0.25009858 |
| ARL6 | 608.954454 | 0.250763497 | 0.13015896 | 1.92659418 | 0.05403022 | 0.1538958 |
| AC110491.3 | 83.617195 | -0.143176405 | 0.25349001 | -0.5648207 | 0.57219573 | 0.73695281 |
| CRYBG3 | 1870.8699 | -0.478711499 | 0.16114038 | -2.970773 | 0.00297051 | 0.01819568 |
| AC110491.2 | 1.75880581 | 0.849255061 | 0.86317568 | 0.98387279 | 0.32517807 | 0.5227881 |
| RIOX2 | 2899.5336 | -0.577085012 | 0.13567958 | -4.2532931 | 2.11E-05 | 0.00036914 |
| GABRR3 | 0.84524145 | -0.494846551 | 1.95031642 | -0.2537263 | 0.79970702 | NA |
| OR5AC2 | 0 | NA | NA | NA | NA | NA |
| AC117460.1 | 0 | NA | NA | NA | NA | NA |
| OR5H1 | 0 | NA | NA | NA | NA | NA |
| OR5H14 | 0 | NA | NA | NA | NA | NA |
| OR5H15 | 0 | NA | NA | NA | NA | NA |
| AC117473.1 | 0.10178723 | 1.087385526 | 3.28420134 | 0.33109588 | 0.74057207 | NA |
| OR5H6 | 0.19632833 | 1.506623996 | 3.27645372 | 0.45983375 | 0.64563556 | NA |
| OR5H2 | 0 | NA | NA | NA | NA | NA |
| OR5K4 | 0 | NA | NA | NA | NA | NA |

| | | | | | | |
|---|---|---|---|---|---|---|
| OR5K3 | 0 | NA | NA | NA | NA | NA |
| OR5K1 | 0.15489794 | -1.003574173 | 3.30558948 | -0.3035992 | 0.7614333 | NA |
| OR5K2 | 10.3611588 | -0.385676055 | 0.60630235 | -0.6361118 | 0.52470358 | 0.69962246 |
| CLDND1 | 3083.15163 | -0.459712859 | 0.13978171 | -3.2887911 | 0.00100619 | 0.00787801 |
| CPOX | 1336.9819 | -0.114867066 | 0.10275711 | -1.1178503 | 0.26363094 | 0.45742518 |
| AC021660.2 | 6.83806267 | 0.563148724 | 0.58496379 | 0.96270698 | 0.33569459 | 0.53388013 |
| GPR15 | 5.48364082 | 2.658861952 | 1.24934617 | 2.12820275 | 0.03332028 | 0.10871761 |
| ST3GAL6-AS | 130.412852 | 0.71729638 | 0.48254012 | 1.48650103 | 0.13714664 | 0.29564593 |
| ST3GAL6 | 1393.62292 | 0.14080852 | 0.30235034 | 0.46571312 | 0.64142086 | 0.78758115 |
| DCBLD2 | 1712.40406 | -0.086252592 | 0.21909018 | -0.3936853 | 0.69381336 | 0.82376229 |
| AC091212.1 | 2.24130907 | 1.273627652 | 0.84183204 | 1.51292372 | 0.13029904 | 0.28547571 |
| LINC00973 | 2.23756171 | 0.68889249 | 0.92804821 | 0.74230248 | 0.45790409 | 0.64706542 |
| AC078828.1 | 0 | NA | NA | NA | NA | NA |
| AC107029.1 | 0.2838234 | 1.099696404 | 2.96053436 | 0.371452 | 0.7103009 | NA |
| AC107029.2 | 0.27653641 | 1.145957499 | 3.27645355 | 0.34975545 | 0.72652223 | NA |
| COL8A1 | 1101.1808 | -0.762820714 | 0.52147221 | -1.4628214 | 0.14351626 | 0.3051051 |
| AC069222.1 | 17.0774028 | 0.639574806 | 0.42783347 | 1.49491533 | 0.13493654 | 0.29235696 |
| CMSS1 | 585.228247 | -0.397462376 | 0.18507779 | -2.1475423 | 0.03175013 | 0.10505031 |
| FILIP1L | 944.686381 | 1.104587758 | 0.31903568 | 3.46227024 | 0.00053564 | 0.00478513 |
| TBC1D23 | 1836.23295 | -0.145978416 | 0.1492298 | -0.9782122 | 0.32796936 | 0.52568381 |
| NIT2 | 5443.2174 | 0.518565096 | 0.2742015 | 1.89118259 | 0.05859997 | 0.16276399 |
| TOMM70 | 3903.54024 | 0.392526918 | 0.17568155 | 2.23430934 | 0.02546273 | 0.08947922 |
| LNP1 | 209.028474 | 1.412360093 | 0.23197488 | 6.08841823 | 1.14E-09 | 1.15E-07 |
| TMEM45A | 249.304408 | -1.502328224 | 0.35496592 | -4.2323168 | 2.31E-05 | 0.00040022 |
| ADGRG7 | 1.99659786 | -1.444494328 | 1.80558538 | -0.8000144 | 0.42370245 | 0.61621563 |
| TFG | 5423.72493 | -0.508838453 | 0.16527762 | -3.0786894 | 0.00207913 | 0.01386966 |
| ABI3BP | 1183.98471 | 0.611088815 | 0.43939124 | 1.39076242 | 0.16429748 | 0.33396475 |
| IMPG2 | 44.7858621 | 0.081623366 | 0.50320006 | 0.16220858 | 0.8711416 | 0.93306272 |
| SENP7 | 1107.78274 | 0.533194622 | 0.1966865 | 2.71088575 | 0.00671037 | 0.03348837 |
| TRMT10C | 878.087679 | -0.224048643 | 0.1986903 | -1.1276275 | 0.25947727 | 0.452272 |
| PCNP | 6266.67143 | -0.155272107 | 0.1070694 | -1.4502006 | 0.14700257 | 0.31019108 |
| ZBTB11 | 1104.06643 | 0.137340673 | 0.14207223 | 0.96669614 | 0.33369596 | 0.53170565 |
| ZBTB11-AS1 | 249.484281 | 0.189607999 | 0.23490233 | 0.80717803 | 0.41956392 | 0.61243738 |
| RPL24 | 15494.9395 | -0.400391143 | 0.16146065 | -2.4798063 | 0.01314538 | 0.05519802 |
| CEP97 | 378.855213 | -0.017270292 | 0.1724383 | -0.1001535 | 0.9202225 | 0.96099929 |
| NXPE3 | 496.482581 | -0.179004693 | 0.16654381 | -1.0748204 | 0.28245513 | 0.47843921 |
| AC020651.1 | 29.6791303 | 0.296669927 | 0.33252214 | 0.89218099 | 0.37229593 | 0.56878538 |
| NFKBIZ | 4958.50773 | -1.886987391 | 0.36346547 | -5.1916552 | 2.08E-07 | 8.81E-06 |
| LINC02085 | 4.58166521 | -1.654266564 | 1.06786221 | -1.5491386 | 0.12134841 | 0.27192025 |
| AC106712.1 | 4.95145322 | -1.681832844 | 1.03687282 | -1.6220242 | 0.10479816 | 0.24527624 |
| ZPLD1 | 182.865923 | -4.118246885 | 0.71702475 | -5.7435213 | 9.27E-09 | 6.49E-07 |
| AC108748.1 | 0.17332742 | -1.003574173 | 3.30558948 | -0.3035992 | 0.7614333 | NA |
| ALCAM | 2023.31895 | -0.006011114 | 0.20080552 | -0.029935 | 0.97611889 | 0.98894663 |
| CBLB | 1318.07385 | -0.379982135 | 0.14645845 | -2.5944705 | 0.00947368 | 0.0432815 |
| LINC00882 | 54.0982011 | 1.328527728 | 0.36564312 | 3.63340004 | 0.00027971 | 0.00287787 |
| DUBR | 197.080844 | 0.277363853 | 0.28692311 | 0.96668354 | 0.33370226 | 0.53170565 |
| AC063944.1 | 5.92591665 | -0.260822131 | 0.52781202 | -0.4941572 | 0.62119514 | 0.77367004 |
| AC063944.3 | 7.08134338 | -0.083496332 | 0.57414398 | -0.1454275 | 0.88437334 | 0.94009453 |
| CCDC54 | 2.58643393 | 0.261496517 | 0.73129625 | 0.35757946 | 0.72065805 | 0.8410601 |
| LINC01990 | 6.06818729 | 0.133987572 | 0.53729748 | 0.24937316 | 0.80307214 | 0.89318485 |
| BBX | 2577.92529 | -0.018748899 | 0.15876993 | -0.1180885 | 0.90599755 | 0.95297788 |
| LINC00635 | 8.28812994 | 0.570505308 | 0.69174057 | 0.82473883 | 0.40951988 | 0.6040069 |
| LINC00636 | 0.96004816 | -0.210551569 | 1.54422772 | -0.1363475 | 0.89154658 | NA |
| CD47 | 3602.59961 | -0.721169535 | 0.16192737 | -4.4536604 | 8.44E-06 | 0.00017823 |
| LINC01215 | 21.6658673 | 0.268350196 | 0.83732982 | 0.32048327 | 0.74860201 | 0.85955914 |
| IFT57 | 2589.52638 | -0.081888459 | 0.14147997 | -0.578799 | 0.56272484 | 0.73008875 |
| HHLA2 | 574.55392 | -0.279009946 | 0.5090246 | -0.5481266 | 0.58360495 | 0.74580385 |
| MYH15 | 12.6185959 | 0.422853624 | 0.42172955 | 1.0026654 | 0.31602233 | 0.5131471 |
| CIP2A | 103.100296 | 0.362453554 | 0.35677211 | 1.01592458 | 0.3096653 | 0.50691378 |
| DZIP3 | 1129.3644 | 0.704044492 | 0.35279952 | 1.99559365 | 0.04597817 | 0.13698389 |
| RETNLB | 0 | NA | NA | NA | NA | NA |
| TRAT1 | 28.6300965 | -0.261446063 | 0.80885216 | -0.323231 | 0.74652033 | 0.85800118 |
| GUCA1C | 10.5676921 | 1.485225942 | 0.71037622 | 2.09075965 | 0.03654961 | 0.11609216 |
| MORC1 | 1.2813747 | 1.781669775 | 1.38612399 | 1.28536104 | 0.19866615 | 0.37968468 |
| MORC1-AS1 | 0 | NA | NA | NA | NA | NA |
| C3orf85 | 206.150198 | 3.141438345 | 0.64176924 | 4.89496557 | 9.83E-07 | 3.21E-05 |
| AC063923.2 | 0 | NA | NA | NA | NA | NA |
| LINC00488 | 39.9194758 | -0.805592247 | 0.52605899 | -1.5313724 | 0.12567737 | 0.27842664 |
| DPPA2 | 5.23514019 | -0.192637041 | 0.75667869 | -0.2545824 | 0.79904569 | 0.89108294 |
| DPPA4 | 3.14959651 | -2.07280465 | 1.39313402 | -1.4878717 | 0.13678474 | 0.2951488 |
| LINC01205 | 0 | NA | NA | NA | NA | NA |
| AC078980.1 | 1.25705759 | 4.284951615 | 1.46618732 | 2.92251309 | 0.00347219 | 0.02046528 |
| AC117430.1 | 0 | NA | NA | NA | NA | NA |
| AC117402.1 | 4.18984606 | -0.112407464 | 0.85140391 | -0.132026 | 0.89496372 | 0.94598591 |
| NECTIN3-AS: | 10.8601209 | 0.521778183 | 0.45923363 | 1.13619332 | 0.25587567 | 0.44857341 |
| NECTIN3 | 684.030317 | 0.183078307 | 0.20906517 | 0.87569972 | 0.3811933 | 0.57724057 |
| CD96 | 147.136227 | -0.216474969 | 0.74990805 | -0.2886687 | 0.77283493 | 0.87488869 |
| AC092916.1 | 0.09635527 | 0.547239394 | 3.29377202 | 0.16614368 | 0.86804388 | NA |
| ZBED2 | 16.4493015 | 0.442181242 | 0.67118557 | 0.65880624 | 0.5100202 | 0.68868005 |
| PLCXD2 | 553.570771 | 0.216144989 | 0.29991898 | 0.72067792 | 0.4711077 | 0.65760449 |
| PLCXD2-AS1 | 0.30088013 | 0.328536705 | 2.29840135 | 0.1429414 | 0.88633646 | NA |
| PHLDB2 | 4697.41502 | 0.225928971 | 0.15613455 | 1.4470146 | 0.14789284 | 0.31139684 |
| ABHD10 | 2477.13344 | 1.286779257 | 0.23242546 | 5.53630934 | 3.09E-08 | 1.80E-06 |
| TAGLN3 | 154.935202 | 2.848135119 | 0.49445647 | 5.76013318 | 8.40E-09 | 5.94E-07 |
| TMPRSS7 | 1.3791588 | -1.088300647 | 1.23354134 | -0.8822571 | 0.37763777 | 0.57370135 |
| C3orf52 | 482.353909 | -0.450784535 | 0.34002793 | -1.325728 | 0.18492982 | 0.36222235 |
| GCSAM | 138.244378 | -0.157506394 | 0.42841952 | -0.3676452 | 0.71313778 | 0.83695061 |
| AC128688.2 | 160.928917 | -0.473104629 | 0.2887015 | -1.6387328 | 0.10126891 | 0.2396322 |
| SLC9C1 | 47.178519 | 0.016200651 | 0.31391261 | 0.05160879 | 0.95884041 | 0.98017606 |
| AC112487.1 | 15.5630763 | -0.141609478 | 0.43585037 | -0.3249039 | 0.74525382 | 0.85728246 |
| CD200 | 1009.67177 | -0.035484205 | 0.25578343 | -0.1387275 | 0.88966546 | 0.94296756 |
| AC092894.1 | 87.6060597 | 0.885764847 | 0.30745455 | 2.88096194 | 0.00396464 | 0.02261952 |
| BTLA | 24.9154555 | 0.975649399 | 0.67486041 | 1.44570548 | 0.14825983 | 0.31203872 |
| ATG3 | 2565.40446 | -0.241368958 | 0.14476495 | -1.6673163 | 0.09545153 | 0.23012039 |
| SLC35A5 | 1699.73752 | 0.207014364 | 0.16099584 | 1.28583669 | 0.19850006 | 0.37950023 |
| CCDC80 | 1690.99167 | 0.012606764 | 0.63334726 | 0.01990498 | 0.98411917 | 0.99314982 |
| LINC02042 | 0.07830766 | -0.642905668 | 3.30558948 | -0.1944905 | 0.84579185 | NA |
| CD200R1L | 0 | NA | NA | NA | NA | NA |
| CD200R1 | 49.4062793 | 0.812492398 | 0.57338637 | 1.41700682 | 0.15648093 | 0.32320845 |
| AC074044.1 | 89.6848516 | 0.046482372 | 0.20764148 | 0.22385879 | 0.82286718 | 0.90541456 |
| GTPBP8 | 461.49838 | -0.203401817 | 0.16311081 | -1.2470161 | 0.21239158 | 0.39620426 |
| NEPRO | 2259.76394 | -0.189140356 | 0.14558063 | -1.2992138 | 0.19387057 | 0.37365641 |
| AC078785.1 | 85.7092377 | 0.115394834 | 0.25203962 | 0.45784403 | 0.6470645 | 0.7920848 |
| AC078785.2 | 0.61108572 | -0.333252258 | 1.94408257 | -0.1714188 | 0.86389449 | NA |
| LINC02044 | 16.9338003 | 0.420108798 | 0.51341616 | 0.81826173 | 0.41320775 | 0.607085 |
| BOC | 1115.96116 | -0.282307874 | 0.25869697 | -1.0912686 | 0.27515472 | 0.47033318 |
| CFAP44 | 861.291698 | -0.070879063 | 0.25133725 | -0.2820078 | 0.77793754 | 0.87847694 |
| CFAP44-AS1 | 13.9095552 | -0.100086913 | 0.35355173 | -0.2830899 | 0.77710795 | 0.8778793 |
| SPICE1 | 589.663387 | -0.158436176 | 0.17322483 | -0.9146274 | 0.36038726 | 0.5580355 |
| SIDT1 | 151.795135 | -0.169547162 | 0.47011968 | -0.3606468 | 0.71836349 | 0.839869 |
| SIDT1-AS1 | 0.36311387 | 0.952165612 | 2.1947578 | 0.4338363 | 0.66440732 | NA |
| USF3 | 1161.6796 | 0.464450054 | 0.13972546 | 3.32401885 | 0.0008873 | 0.00714666 |
| NAA50 | 5189.5164 | -0.380236842 | 0.23521372 | -1.616559 | 0.10597351 | 0.24719341 |

| | | | | | | |
|---|---|---|---|---|---|---|
| AC108693.2 | 48.8290972 | 2.347974764 | 0.48069001 | 4.88459239 | 1.04E-06 | 3.35E-05 |
| ATP6V1A | 8793.85407 | 0.05529742 | 0.18420354 | 0.30019737 | 0.76402661 | 0.86935843 |
| GRAMD1C | 2195.04416 | 0.691434157 | 0.32589183 | 2.12166767 | 0.03386566 | 0.10999351 |
| AC128687.2 | 166.824432 | 0.108854372 | 0.31558231 | 0.34493179 | 0.73014565 | 0.84751607 |
| ZDHHC23 | 1194.0426 | -0.38198762 | 0.18962695 | -2.0144163 | 0.04396585 | 0.13282248 |
| CCDC191 | 635.884082 | 0.231532645 | 0.36808617 | 0.62901751 | 0.52933759 | 0.70387785 |
| QTRT2 | 1002.07562 | 0.07473127 | 0.18904877 | 0.39530154 | 0.69262037 | 0.82316563 |
| DRD3 | 0.22663961 | 1.777046332 | 3.25798548 | 0.54544329 | 0.58544867 | NA |
| AC093010.2 | 7.38852715 | -0.559725322 | 0.80687632 | -0.6936941 | 0.48787408 | 0.67152554 |
| ZNF80 | 10.9281014 | 0.249816362 | 0.74938014 | 0.333364 | 0.73885953 | 0.85343045 |
| TIGIT | 74.3671664 | 0.199797269 | 0.81805722 | 0.24423386 | 0.80704971 | 0.8957029 |
| ZBTB20 | 4174.51957 | 0.362167495 | 0.21198189 | 1.7084832 | 0.08754672 | 0.21600507 |
| ZBTB20-AS1 | 6.69321665 | 0.423458613 | 0.8536749 | 0.49604201 | 0.61986477 | 0.77274237 |
| ZBTB20-AS5 | 0.84779498 | 2.610603804 | 1.37806822 | 1.89439374 | 0.05817277 | NA |
| ZBTB20-AS2 | 0.34574733 | 0.725628444 | 2.89607834 | 0.25055553 | 0.80215777 | NA |
| ZBTB20-AS3 | 0.69900169 | 0.992494573 | 1.78748633 | 0.55524597 | 0.57872643 | NA |
| ZBTB20-AS4 | 3.03231969 | 2.051825619 | 0.98090946 | 2.09175841 | 0.03646013 | 0.11586571 |
| AC046136.1 | 1.45015611 | -0.047115672 | 1.16872814 | -0.0403136 | 0.96784309 | 0.98494992 |
| GAP43 | 3.59125523 | 0.278033951 | 1.01780384 | 0.27317047 | 0.78472218 | 0.88239836 |
| LSAMP | 165.021905 | 1.592881697 | 0.45323415 | 3.51447858 | 0.00044062 | 0.00411441 |
| LSAMP-AS1 | 0.29926206 | 0.605228892 | 2.92846486 | 0.20667104 | 0.83626677 | NA |
| LINC00903 | 0 | NA | NA | NA | NA | NA |
| TUSC7 | 0.14444843 | 0.435199075 | 3.28898345 | 0.13232024 | 0.894731 | NA |
| LINC00901 | 0 | NA | NA | NA | NA | NA |
| AC092691.1 | 2.84315898 | 0.64621823 | 0.89617466 | 0.72108514 | 0.47085713 | 0.65748881 |
| LINC02024 | 0.20816856 | 0.689155606 | 3.09692809 | 0.22252877 | 0.82390228 | NA |
| AC092691.3 | 0.09989236 | -0.642905668 | 3.30558948 | -0.1944905 | 0.84579185 | NA |
| AC068633.1 | 0.52134398 | -0.028836153 | 1.84778152 | -0.0156058 | 0.98754886 | NA |
| IGSF11 | 359.264207 | 2.080080232 | 0.45552411 | 4.56634495 | 4.96E-06 | 0.00011667 |
| IGSF11-AS1 | 0.03392908 | 0.380740593 | 3.29738709 | 0.11546736 | 0.90807469 | NA |
| C3orf30 | 6.13300393 | 0.11717915 | 0.8239924 | 0.14220902 | 0.88691491 | 0.94143003 |
| AC083800.1 | 61.5724729 | 0.843161285 | 0.5381814 | 1.56668604 | 0.11718809 | 0.26561337 |
| UPK1B | 354.084858 | 0.831609357 | 0.51010892 | 1.63025842 | 0.10304689 | 0.24248931 |
| B4GALT4 | 1912.00349 | 0.42424788 | 0.23053124 | 1.84030536 | 0.06572342 | 0.17620237 |
| B4GALT4-AS | 173.835506 | 0.860666274 | 0.29394805 | 2.9279537 | 0.00341201 | 0.02021903 |
| ARHGAP31 | 1109.38243 | 0.268876834 | 0.20276546 | 1.3260485 | 0.18482363 | 0.36207983 |
| ARHGAP31-A | 14.3404632 | 0.839081291 | 0.4005544 | 2.09479983 | 0.03618879 | 0.11525343 |
| TMEM39A | 1180.64437 | -0.179015204 | 0.15057387 | -1.1888863 | 0.23448443 | 0.42322754 |
| POGLUT1 | 826.71994 | -0.266751472 | 0.13903207 | -1.9186326 | 0.05503085 | 0.15577108 |
| AC073352.1 | 12.2876141 | 0.45238025 | 0.43045487 | 1.05093537 | 0.29328828 | 0.4897734 |
| TIMMDC1 | 3461.5901 | 0.027076653 | 0.1404346 | 0.19280614 | 0.8471108 | 0.91972627 |
| CD80 | 29.7127574 | -1.533244523 | 0.84239994 | -1.820091 | 0.06874515 | 0.18177175 |
| AC073352.2 | 8.66832822 | 0.710899719 | 0.56042336 | 1.26850479 | 0.20461774 | 0.38700964 |
| ADPRH | 573.126888 | -0.220191251 | 0.14350711 | -1.5343578 | 0.12494166 | 0.27727941 |
| PLA1A | 1378.05861 | -1.285777279 | 0.48477064 | -2.6523415 | 0.00799357 | 0.03812449 |
| POPDC2 | 201.010638 | 0.221550452 | 0.2420972 | 0.91513017 | 0.3601233 | 0.55779463 |
| COX17 | 1955.8978 | -0.52255763 | 0.22925187 | -2.279404 | 0.02264306 | 0.08219955 |
| MAATS1 | 625.612553 | -0.246346501 | 0.36340747 | -0.6778796 | 0.49784805 | 0.67955138 |
| AC069444.1 | 29.9155129 | -0.12676336 | 0.43177334 | -0.2935877 | 0.76907294 | 0.87240608 |
| NR1I2 | 24.3816708 | -0.30026435 | 0.59690551 | -0.503035 | 0.61493968 | 0.76976927 |
| GSK3B | 2195.97236 | -0.244068319 | 0.20104176 | -1.214018 | 0.22474086 | 0.41095738 |
| AC092910.3 | 63.4655485 | 0.774078454 | 0.25068668 | 3.08783243 | 0.00201622 | 0.01358052 |
| GPR156 | 15.7113275 | -0.071619847 | 0.50328669 | -0.1423043 | 0.88683967 | 0.94143003 |
| LRRC58 | 1962.59335 | 0.018036941 | 0.29340177 | 0.06147523 | 0.95098074 | 0.97657082 |
| AC063952.4 | 1.13766427 | 2.624642731 | 2.17263956 | 1.20804333 | 0.2270306 | 0.41385295 |
| FSTL1 | 6612.05989 | -0.307183427 | 0.34816798 | -0.8822851 | 0.37762264 | 0.57370135 |
| NDUFB4 | 5357.64827 | -0.39190528 | 0.22030173 | -1.7789478 | 0.07524831 | 0.19419633 |
| HGD | 7525.38169 | 0.726237982 | 0.39448408 | 1.84098175 | 0.06562423 | 0.17602594 |
| RABL3 | 1286.74312 | 0.221589414 | 0.13643088 | 1.62418807 | 0.10433568 | 0.24460631 |
| GTF2E1 | 563.341976 | -0.694869742 | 0.20388792 | -3.4080966 | 0.00065418 | 0.00558093 |
| LINC02049 | 0 | NA | NA | NA | NA | NA |
| STXBP5L | 0.94361717 | 1.307400013 | 1.4716183 | 0.88840973 | 0.3743204 | NA |
| POLQ | 24.2689894 | -0.272712014 | 0.5825394 | -0.4681435 | 0.639682 | 0.78704498 |
| ARGFX | 0.07830766 | -0.642905668 | 3.30558948 | -0.1944905 | 0.84579185 | NA |
| FBXO40 | 10.5036624 | 3.225049463 | 1.1787893 | 2.73589985 | 0.006221 | 0.03169391 |
| HCLS1 | 2139.25778 | -0.650175603 | 0.37053466 | -1.7546958 | 0.07931136 | 0.20115808 |
| GOLGB1 | 6574.12051 | 0.18945549 | 0.1364632 | 1.38832664 | 0.1650376 | 0.33521079 |
| IQCB1 | 1240.43921 | -0.459327436 | 0.13791819 | -3.3304341 | 0.00086711 | 0.00702908 |
| EAF2 | 370.240706 | 1.988825832 | 0.32841326 | 6.0558634 | 1.40E-09 | 1.35E-07 |
| SLC15A2 | 1020.45985 | 1.647354151 | 0.43650396 | 3.77397295 | 0.00016067 | 0.00184691 |
| ILDR1 | 968.35278 | -0.163824784 | 0.26599566 | -0.6158927 | 0.53796534 | 0.71150739 |
| CD86 | 275.154534 | -0.686565215 | 0.62894929 | -1.0916066 | 0.27500604 | 0.47019955 |
| CASR | 2535.84702 | 1.831113267 | 0.38551119 | 4.74983167 | 2.04E-06 | 5.74E-05 |
| CSTA | 58.513896 | -1.638183216 | 0.60159539 | -2.7230648 | 0.00646794 | 0.03258928 |
| CCDC58 | 923.78259 | -0.147672438 | 0.24679374 | -0.5983638 | 0.54959723 | 0.72007149 |
| FAM162A | 3684.93881 | -0.112699935 | 0.21055317 | -0.5352564 | 0.59247256 | 0.75183615 |
| WDR5B | 460.610274 | -0.101374567 | 0.21000057 | -0.4827347 | 0.62928411 | 0.77980148 |
| AC083798.2 | 1591.84811 | -0.424041993 | 0.20376337 | -2.0810511 | 0.03742923 | 0.11788273 |
| KPNA1 | 3117.7423 | -0.300000321 | 0.18532367 | -1.6187912 | 0.10549219 | 0.24646739 |
| PARP9 | 2477.3183 | -0.645846715 | 0.17936046 | -3.600831 | 0.0003172 | 0.00316123 |
| DTX3L | 2483.51183 | -1.228022897 | 0.39519392 | -3.1073932 | 0.00188745 | 0.01290101 |
| PARP15 | 156.224735 | -0.052236559 | 0.60462005 | -0.0863957 | 0.93115188 | 0.96688001 |
| PARP14 | 4079.65704 | -0.661274664 | 0.19418988 | -3.4052994 | 0.00066092 | 0.00563295 |
| HSPBAP1 | 1089.43136 | -0.850129584 | 0.18541435 | -4.5850259 | 4.54E-06 | 0.00010902 |
| DIRC2 | 1477.42666 | -1.034215433 | 0.20038687 | -5.1610939 | 2.46E-07 | 1.00E-05 |
| LINC02035 | 705.534552 | -0.501567022 | 0.17050075 | -2.9417291 | 0.00326385 | 0.01957866 |
| SEMA5B | 496.660484 | 1.158881783 | 0.33830697 | 3.42553331 | 0.00061359 | 0.00530159 |
| PDIA5 | 1823.84826 | -0.278707137 | 0.16281615 | -1.7117905 | 0.08693528 | 0.2149136 |
| SEC22A | 805.777626 | -0.20689012 | 0.10656223 | -1.9414958 | 0.05219817 | 0.15017389 |
| AC112503.2 | 4.19254501 | 0.400515523 | 0.71571271 | 0.55960376 | 0.57574974 | 0.73980078 |
| ADCY5 | 2074.18779 | 0.336768461 | 0.30550928 | 1.10231825 | 0.27032334 | 0.46463475 |
| AC112503.1 | 190.272793 | 0.368761701 | 0.32356018 | 1.1397005 | 0.2544111 | 0.44713547 |
| HACD2 | 2147.48827 | 0.176880957 | 0.15521122 | 1.13961451 | 0.25444694 | 0.44713886 |
| MYLK-AS1 | 323.093011 | 1.742976866 | 0.26709673 | 6.52563906 | 6.77E-11 | 1.04E-08 |
| MYLK | 5674.25549 | 0.883260716 | 0.42143435 | 2.09584417 | 0.03609602 | 0.11508306 |
| MYLK-AS2 | 8.37491658 | 1.507640739 | 0.60802217 | 2.47958186 | 0.01315365 | 0.05522398 |
| AC020634.1 | 8.17547769 | -0.285855932 | 0.53772127 | -0.5316061 | 0.59499881 | 0.75401092 |
| CCDC14 | 2198.64398 | 0.00062826 | 0.16730715 | 0.00375513 | 0.99700385 | 0.99850237 |
| ROPN1 | 0.42804161 | 0.310632805 | 2.19437699 | 0.14155854 | 0.88742872 | NA |
| KALRN | 1014.56787 | -0.339709162 | 0.28602745 | -1.1876803 | 0.23495939 | 0.42379475 |
| UMPS | 1103.33257 | 0.118669732 | 0.16942062 | 0.70044444 | 0.48364979 | 0.66823038 |
| ITGB5 | 7822.16667 | -0.004055052 | 0.20143203 | -0.0201311 | 0.98393878 | 0.99305611 |
| ITGB5-AS1 | 0.33725126 | 0.669857205 | 2.22872422 | 0.30055634 | 0.76375283 | NA |
| MUC13 | 1623.81522 | 3.399957998 | 0.91567911 | 3.71304529 | 0.00020478 | 0.00222499 |
| HEG1 | 2410.75306 | 0.590608004 | 0.30381713 | 1.94395885 | 0.05190041 | 0.14964071 |
| SLC12A8 | 517.897704 | -0.835437088 | 0.30612214 | -2.7290973 | 0.0063508 | 0.0321564 |
| ZNF148 | 1866.18936 | 0.031663498 | 0.10681205 | 0.29644126 | 0.76689312 | 0.87149199 |
| SNX4 | 3398.32658 | 0.211734585 | 0.23795415 | 0.88981254 | 0.37356655 | 0.56978136 |
| OSBPL11 | 895.549669 | -0.063481877 | 0.17402045 | -0.3647955 | 0.71526406 | 0.8380436 |
| AC092902.3 | 0.86158347 | 2.663629544 | 1.28320458 | 2.07576375 | 0.03791581 | NA |
| AC092903.2 | 24.2875164 | 0.318598179 | 0.44763976 | 0.71172895 | 0.47663264 | 0.66179458 |
| ALG1L | 68.6291111 | 0.609868723 | 0.38948518 | 1.56583293 | 0.11738774 | 0.26586916 |
| ROPN1B | 9.67590286 | -1.079552885 | 0.55330502 | -1.951099 | 0.05104527 | 0.14780539 |

| | | | | | | | |
|---|---|---|---|---|---|---|---|
| SLC41A3 | 2785.69808 | | -0.082368117 | 0.13836238 | -0.5953072 | 0.55163818 | 0.72166983 |
| AC117422.1 | 1.64480738 | | 2.409617614 | 1.10242213 | 2.18574858 | 0.028834 | 0.09809737 |
| AC079848.1 | 47.6616373 | | 0.481335583 | 0.37478221 | 1.28430746 | 0.1990344 | 0.3800809 |
| ALDH1L1 | 14391.3543 | | 1.217931732 | 0.44281292 | 2.75044308 | 0.00595147 | 0.03060459 |
| ALDH1L1-AS1 | 1828.92769 | | 0.675438352 | 0.39256542 | 1.72057525 | 0.08532793 | 0.21189889 |
| ALDH1L1-AS2 | 803.32229 | | 1.638972014 | 0.46078894 | 3.55688229 | 0.00037528 | 0.00363163 |
| AC016924.1 | 134.017494 | | 1.670125962 | 0.48400891 | 3.45060998 | 0.00055932 | 0.00495608 |
| AC063919.1 | 70.7383254 | | 2.029748351 | 0.5644332 | 3.59608252 | 0.00032305 | 0.00321348 |
| KLF15 | 2377.1033 | | 0.379473605 | 0.41945108 | 0.90469097 | 0.36562913 | 0.56262123 |
| CCDC37-DT | 0.43270659 | | -2.017828363 | 1.88018873 | -1.0732052 | 0.28317905 | NA |
| CFAP100 | 46.5677189 | | -0.147081483 | 0.51029239 | -0.2882298 | 0.77317082 | 0.87515623 |
| ZXDC | 2549.91359 | | 0.20739428 | 0.20838147 | 0.99526257 | 0.31960858 | 0.51702829 |
| UROC1 | 6.19224144 | | 0.733530368 | 0.60243173 | 1.21761577 | 0.22337002 | 0.40943055 |
| CHST13 | 283.613991 | | 0.233486209 | 0.43999563 | 0.53065575 | 0.59565736 | 0.75465958 |
| C3orf22 | 2.1742007 | | 1.853549454 | 0.93447173 | 1.98352652 | 0.04730865 | 0.13982616 |
| TXNRD3NB | 185.518512 | | -0.359734112 | 0.20320781 | -1.770277 | 0.07668101 | 0.19666195 |
| TXNRD3 | 454.058535 | | 0.017988299 | 0.14897285 | 0.12074884 | 0.90388997 | 0.95167363 |
| CHCHD6 | 492.198937 | | -0.018735769 | 0.20690618 | -0.090552 | 0.92784857 | 0.96504253 |
| PLXNA1 | 2341.07371 | | -0.534296078 | 0.175933 | -3.0369292 | 0.00239002 | 0.0154586 |
| C3orf56 | 0 | NA | | NA | NA | NA | NA |
| LINC02016 | 0 | NA | | NA | NA | NA | NA |
| LINC01471 | 0.38404007 | | -1.782947976 | 2.81648432 | -0.6330403 | 0.52670735 | NA |
| AC016968.1 | 0 | NA | | NA | NA | NA | NA |
| LINC02034 | 1.62009236 | | -3.905323252 | 2.53285767 | -1.5418645 | 0.12310653 | 0.2746124 |
| AC084035.1 | 6.59429329 | | 0.499339059 | 0.46777984 | 1.06746596 | 0.28576148 | 0.48176817 |
| TPRA1 | 2361.17608 | | -0.445525827 | 0.17820193 | -2.5001178 | 0.0124152 | 0.0529156 |
| MCM2 | 438.316353 | | 0.377782965 | 0.32126662 | 1.17591725 | 0.23962794 | 0.42951257 |
| PODXL2 | 519.501539 | | 0.921483065 | 0.33402778 | 2.7587019 | 0.00580314 | 0.03007287 |
| ABTB1 | 3833.95649 | | -0.231633264 | 0.17054654 | -1.3581821 | 0.17440591 | 0.34831081 |
| MGLL | 15578.1824 | | -0.802321553 | 0.19339128 | -4.1486955 | 3.34E-05 | 0.00053628 |
| KBTBD12 | 171.431719 | | -0.487889581 | 0.26826534 | -1.8186828 | 0.06895983 | 0.18216467 |
| SEC61A1 | 14490.5452 | | -0.398241362 | 0.15293526 | -2.6039866 | 0.00921464 | 0.0423395 |
| RUVBL1 | 2645.21665 | | 0.162698916 | 0.13309919 | 1.22238845 | 0.22156076 | 0.40759381 |
| RUVBL1-AS1 | 0.2892504 | | -0.174096114 | 3.29377188 | -0.0528562 | 0.95784652 | NA |
| EEFSEC | 1345.78768 | | 0.664935393 | 0.21285789 | 3.12384652 | 0.00178504 | 0.01235794 |
| DNAJB8 | 0.24605818 | | 0.078267493 | 3.05912219 | 0.02558495 | 0.97958839 | NA |
| DNAJB8-AS1 | 0 | NA | | NA | NA | NA | NA |
| GATA2 | 2030.25743 | | 0.70032066 | 0.33158683 | 2.11202798 | 0.03468405 | 0.11192875 |
| GATA2-AS1 | 183.269946 | | -0.237828422 | 0.35099772 | -0.6775782 | 0.49803914 | 0.67967569 |
| LINC01565 | 2.66811749 | | -1.20252144 | 0.82349077 | -1.4602731 | 0.14421502 | 0.30597469 |
| RPN1 | 14322.9277 | | -0.439798391 | 0.16355721 | -2.6889576 | 0.00716755 | 0.03525618 |
| RAB7A | 13646.6244 | | -0.278730771 | 0.18968468 | -1.4694427 | 0.14171275 | 0.30242894 |
| AC112484.3 | 44.4631058 | | 0.825109452 | 0.2628233 | 3.13940752 | 0.0016929 | 0.01185388 |
| AC112484.1 | 42.0689936 | | 0.776946886 | 0.25011546 | 3.10635287 | 0.00189411 | 0.01292885 |
| ACAD9 | 2777.28079 | | 0.151536938 | 0.0954858 | 1.5870102 | 0.11251034 | 0.25810331 |
| KIAA1257 | 497.999134 | | 0.017172163 | 0.10774238 | 0.1593817 | 0.87336816 | 0.9340817 |
| EFCC1 | 210.092718 | | 0.529596135 | 0.24337021 | 2.1760927 | 0.02954833 | 0.09980651 |
| GP9 | 4.18469112 | | -1.695297891 | 0.96584194 | -1.755254 | 0.07921588 | 0.20107062 |
| RAB43 | 1906.55243 | | -0.535223866 | 0.19151552 | -2.7946762 | 0.00519517 | 0.02766288 |
| ISY1-RAB43 | 2618.00606 | | -0.372135518 | 0.1326928 | -2.8044891 | 0.00503964 | 0.02701673 |
| AC108673.2 | 25.5732688 | | -1.028323878 | 0.49038311 | -2.0969806 | 0.03599529 | 0.11484522 |
| ISY1 | 1378.43694 | | -0.148598442 | 0.13551641 | -1.0965347 | 0.27284485 | 0.46768788 |
| AC108673.3 | 61.687194 | | -0.576340588 | 0.23966423 | -2.4047835 | 0.01618205 | 0.06436684 |
| CNBP | 14219.4719 | | -0.216648712 | 0.13326829 | -1.6256584 | 0.10402235 | 0.24416075 |
| COPG1 | 7980.62362 | | -0.049758142 | 0.13123625 | -0.3791494 | 0.70457694 | 0.83125502 |
| HMCES | 2117.65791 | | -0.080425084 | 0.13648089 | -0.5892772 | 0.55567531 | 0.72497317 |
| H1FX | 2912.17084 | | -0.788510186 | 0.23736178 | -3.3219761 | 0.00089382 | 0.00718601 |
| H1FX-AS1 | 1702.9696 | | -0.588117002 | 0.22758344 | -2.5841819 | 0.00976103 | 0.04427217 |
| EFCAB12 | 205.538543 | | 0.395576441 | 0.45772344 | 0.86422588 | 0.38746382 | 0.5830724 |
| MBD4 | 2420.7297 | | -0.433537512 | 0.17243228 | -2.514248 | 0.01192865 | 0.05134812 |
| IFT122 | 2592.82827 | | 0.069091819 | 0.15547908 | 0.44438016 | 0.65676776 | 0.79862426 |
| RHO | 4.66957489 | | -0.293779113 | 0.90396657 | -0.3249889 | 0.74518946 | 0.85726411 |
| H1FOO | 0.08695548 | | -0.642905668 | 3.30558948 | -0.1944905 | 0.84579185 | NA |
| PLXND1 | 4951.9096 | | 0.024648977 | 0.28037445 | 0.08791449 | 0.92994464 | 0.96610488 |
| TMCC1 | 1720.56116 | | -0.964040896 | 0.34193395 | -2.8193776 | 0.00481169 | 0.02610865 |
| TMCC1-AS1 | 90.3475367 | | 0.319595458 | 0.34238033 | 0.93345156 | 0.35058687 | 0.54831609 |
| AC083906.3 | 0.60676336 | | -1.210775863 | 2.2200862 | -0.5453734 | 0.58549676 | NA |
| TRH | 3.73806303 | | -1.117859799 | 1.40647386 | -0.794796 | 0.42673218 | 0.61900696 |
| ALG1L2 | 28.8874889 | | 0.533281278 | 0.40639056 | 1.31223835 | 0.18943973 | 0.36780486 |
| LINC02014 | 3.16978456 | | 0.112894682 | 0.89594469 | 0.12600631 | 0.89972693 | 0.9491079 |
| LINC02021 | 80.6567924 | | 1.01271397 | 0.53465275 | 1.89415275 | 0.05820474 | 0.1620588 |
| COL6A5 | 16.864527 | | -2.139826545 | 1.0366019 | -2.0642703 | 0.03899209 | 0.12155355 |
| COL6A6 | 17.0199009 | | 0.367202834 | 0.48288283 | 0.76043878 | 0.44699235 | 0.6372831 |
| PIK3R4 | 1261.04302 | | 0.236423628 | 0.10759301 | 2.19738837 | 0.02799272 | 0.09611563 |
| ATP2C1 | 4621.44292 | | 0.089464345 | 0.16010207 | 0.55879567 | 0.57630118 | 0.74034501 |
| AC055733.2 | 0.23387521 | | 1.816617118 | 3.25499482 | 0.55810139 | 0.57677515 | NA |
| ASTE1 | 599.861206 | | -0.024929988 | 0.10476033 | -0.2379717 | 0.81190308 | 0.89845484 |
| NEK11 | 681.936632 | | -0.217957389 | 0.23994754 | -0.9083543 | 0.36369105 | 0.56082876 |
| AC116424.1 | 0.14444843 | | 0.435199075 | 3.28898345 | 0.13232024 | 0.894731 | NA |
| NUDT16 | 3846.3918 | | 0.551701244 | 0.2221975 | 2.48293186 | 0.0130306 | 0.05481208 |
| MRPL3 | 3695.96547 | | -0.106887786 | 0.16222362 | -0.6588916 | 0.50996535 | 0.68864125 |
| AC107027.1 | 4.80213592 | | 0.810821303 | 0.76579307 | 1.05879948 | 0.28969111 | 0.48588949 |
| CPNE4 | 82.9120914 | | 2.316533842 | 0.76991891 | 3.00880239 | 0.0026228 | 0.01656602 |
| ACPP | 1135.19248 | | 1.125343844 | 0.44389148 | 2.53517782 | 0.01123903 | 0.04915792 |
| DNAJC13 | 2731.42739 | | 0.115544192 | 0.16144823 | 0.71567333 | 0.47419308 | 0.66003563 |
| ACAD11 | 5971.05457 | | 0.514311183 | 0.48313867 | 1.06452083 | 0.28709282 | 0.48334532 |
| NPHP3-ACAD | 6403.98854 | | 0.477400468 | 0.4409881 | 1.08256997 | 0.27899934 | 0.47466576 |
| ACKR4 | 711.844247 | | 1.064018022 | 0.51031998 | 2.08500169 | 0.03706915 | 0.11710955 |
| UBA5 | 3435.6581 | | 0.06918551 | 0.1806579 | 0.38296421 | 0.7017463 | 0.82913351 |
| NPHP3 | 1504.09149 | | -0.032980167 | 0.18144153 | -0.1817675 | 0.85576522 | 0.92451673 |
| NPHP3-AS1 | 27.9882693 | | -0.199380364 | 0.40905901 | -0.4874122 | 0.62596623 | 0.77729476 |
| AC079942.1 | 0 | NA | | NA | NA | NA | NA |
| TMEM108 | 36.4748263 | | 0.266141128 | 0.45403836 | 0.58616442 | 0.55776503 | 0.72649725 |
| TMEM108-AS | 0.15318056 | | -1.003574173 | 3.30558948 | -0.3035992 | 0.7614333 | NA |
| BFSP2 | 7.46415044 | | 0.300357307 | 0.62797917 | 0.47829183 | 0.63244251 | 0.78171075 |
| BFSP2-AS1 | 0.19114494 | | -1.003574173 | 3.30558948 | -0.3035992 | 0.7614333 | NA |
| AC022296.3 | 5.41532698 | | -0.126137985 | 0.80239995 | -0.1572009 | 0.87508653 | 0.93501854 |
| CDV3 | 8207.03442 | | -0.716656493 | 0.36051254 | -1.9878823 | 0.04682471 | 0.13884714 |
| TOPBP1 | 930.544964 | | -0.130675553 | 0.15913597 | -0.8211566 | 0.41155708 | 0.60570403 |
| TF | 95.4679889 | | 0.074539302 | 0.53318224 | 0.1398008 | 0.88881738 | 0.94259161 |
| SRPRB | 3258.70094 | | -0.383666246 | 0.1868169 | -2.053702 | 0.04000454 | 0.12382171 |
| RAB6B | 869.987822 | | -0.672988691 | 0.27710425 | -2.4286481 | 0.01515524 | 0.06131449 |
| C3orf36 | 68.6941028 | | -1.045658438 | 0.54747841 | -1.9099538 | 0.05613917 | 0.15789141 |
| SLCO2A1 | 6033.12889 | | -0.354094636 | 0.31100861 | -1.1385364 | 0.25489655 | 0.44760086 |
| LINC02000 | 2.74873406 | | 0.690150187 | 1.47981554 | 0.46637582 | 0.64094652 | 0.78748356 |
| RYK | 2586.94037 | | -0.258911392 | 0.11492012 | -2.2529683 | 0.02426115 | 0.08631496 |
| LINC02004 | 5.78732545 | | 0.69722786 | 0.4924385 | 1.41586789 | 0.15681419 | 0.32372702 |
| AC010207.1 | 25.2423588 | | 0.504096696 | 0.43375573 | 1.16216724 | 0.24516754 | 0.43618502 |
| AMOTL2 | 4938.69056 | | 1.050426793 | 0.26136383 | 4.0190213 | 5.84E-05 | 0.00083482 |
| ANAPC13 | 3080.13513 | | 0.136215175 | 0.16276773 | 0.83686846 | 0.40266651 | 0.5977748 |
| CEP63 | 1367.35117 | | 0.096789041 | 0.14207128 | 0.68127097 | 0.49570005 | 0.67770688 |
| EPHB1 | 115.45541 | | 0.431395422 | 0.4487611 | 0.96130306 | 0.33639981 | 0.53442067 |
| KY | 21.8307139 | | 2.229829385 | 0.42416312 | 5.2570091 | 1.46E-07 | 6.58E-06 |

| | | | | | | |
|---|---|---|---|---|---|---|
| PPP2R3A | 4161.09668 | -0.806650335 | 0.20838996 | -3.8708694 | 0.00010845 | 0.00135133 |
| AC072039.2 | 124.173056 | -0.667530235 | 0.28335617 | -2.3557992 | 0.01848291 | 0.07083968 |
| MSL2 | 1309.39726 | -0.027149681 | 0.19599839 | -0.1385199 | 0.88982953 | 0.94296756 |
| PCCB | 4760.47791 | 0.707664946 | 0.18378645 | 3.8504739 | 0.00011789 | 0.00144458 |
| STAG1 | 1756.09509 | 0.080930219 | 0.20555981 | 0.39370643 | 0.69379779 | 0.82376229 |
| AC117382.2 | 2.19354154 | 1.19063237 | 0.87735095 | 1.35707651 | 0.1747569 | 0.34864121 |
| SLC35G2 | 402.531037 | -1.178758492 | 0.20750805 | -5.6805435 | 1.34E-08 | 8.96E-07 |
| AC096992.2 | 74.410829 | 0.28751769 | 0.31713671 | 0.90660489 | 0.36461577 | 0.56178311 |
| NCK1-DT | 96.6165016 | -0.221183242 | 0.29827398 | -0.7415439 | 0.45836374 | 0.64730101 |
| NCK1 | 1151.26323 | -0.510508722 | 0.16230046 | -3.1454546 | 0.00165829 | 0.01167666 |
| IL20RB | 296.533157 | -0.82006644 | 0.18904079 | -4.3380396 | 1.44E-05 | 0.00027391 |
| IL20RB-AS1 | 2.98774219 | 0.369475452 | 0.70673806 | 0.5227898 | 0.60112054 | 0.75869641 |
| SOX14 | 0.07830766 | -0.642905668 | 3.30558948 | -0.1944905 | 0.84579185 | NA |
| LINC01210 | 0.18301898 | 0.18657164 | 3.29377195 | 0.05664376 | 0.95482897 | NA |
| AC007159.1 | 0 | NA | NA | NA | NA | NA |
| CLDN18 | 32.9074081 | 0.159299 | 0.4364134 | 0.36501858 | 0.71509753 | 0.83801965 |
| AC016252.1 | 7.62787638 | 0.84397791 | 0.52006822 | 1.62282153 | 0.10462756 | 0.24507504 |
| DZIP1L | 465.330464 | 0.414823686 | 0.26756365 | 1.55037387 | 0.12105181 | 0.27150941 |
| A4GNT | 37.3771554 | 2.668402385 | 0.66984283 | 3.9836246 | 6.79E-05 | 0.00093559 |
| DBR1 | 732.803375 | 0.018987601 | 0.15185432 | 0.12503827 | 0.90049325 | 0.94961238 |
| ARMC8 | 2310.21454 | -0.282491444 | 0.13854374 | -2.0390055 | 0.04144948 | 0.12716961 |
| NME9 | 66.2228702 | -0.539733434 | 0.37102865 | -1.4546948 | 0.14575379 | 0.30834556 |
| MRAS | 3481.47405 | -1.283061198 | 0.3284471 | -3.9064471 | 9.37E-05 | 0.0012019 |
| ESYT3 | 250.727028 | -0.419875148 | 0.245702 | -1.7088797 | 0.08747324 | 0.21589943 |
| CEP70 | 1339.86377 | -0.292373154 | 0.26890287 | -1.0872816 | 0.27691236 | 0.47229668 |
| FAIM | 685.936956 | -0.281963905 | 0.12704119 | -2.2194683 | 0.02645488 | 0.09207211 |
| PIK3CB | 2156.66942 | 0.622842353 | 0.19292086 | 3.22848625 | 0.00124447 | 0.00932559 |
| LINC01391 | 0 | NA | NA | NA | NA | NA |
| FOXL2 | 0.35640006 | -1.70666969 | 3.3025702 | -0.5167701 | 0.60531662 | NA |
| FOXL2NB | 0.18684785 | -1.003574173 | 3.30558948 | -0.3035992 | 0.7614333 | NA |
| PRR23A | 0 | NA | NA | NA | NA | NA |
| MRPS22 | 1831.33233 | 0.000504802 | 0.1560026 | 0.00323585 | 0.99741817 | 0.99866792 |
| PRR23B | 0 | NA | NA | NA | NA | NA |
| PRR23C | 0 | NA | NA | NA | NA | NA |
| BPESC1 | 0 | NA | NA | NA | NA | NA |
| PISRT1 | 0 | NA | NA | NA | NA | NA |
| AC024933.1 | 10.1211773 | 0.753312285 | 0.42740211 | 1.76253757 | 0.07797851 | 0.1987525 |
| COPB2 | 6319.83424 | -0.100645786 | 0.16628915 | -0.6052457 | 0.54501578 | 0.71682576 |
| AC097103.2 | 82.5697902 | 0.5108317 | 0.23396989 | 2.18332241 | 0.02901208 | 0.09855963 |
| RBP2 | 31.9094279 | 1.101730014 | 0.618115 | 1.78240296 | 0.07468354 | 0.19311679 |
| AC097103.1 | 0.18754884 | 1.469068384 | 3.27783996 | 0.44818185 | 0.65402196 | NA |
| RBP1 | 566.617659 | -0.304983432 | 0.35764796 | -0.8527476 | 0.3937993 | 0.58912394 |
| NMNAT3 | 496.03792 | 1.099527181 | 0.2470045 | 4.4514459 | 8.53E-06 | 0.00017965 |
| AC110716.2 | 0.09635527 | 0.547239394 | 3.29377202 | 0.16614368 | 0.86804388 | NA |
| AC110716.1 | 0 | NA | NA | NA | NA | NA |
| AC016933.1 | 0 | NA | NA | NA | NA | NA |
| CLSTN2 | 898.119018 | 1.264066184 | 0.55306737 | 2.28555553 | 0.02228027 | 0.08125167 |
| AC010181.1 | 3.01806188 | -0.770564974 | 0.77604477 | -0.9929388 | 0.32073977 | 0.51819065 |
| AC010181.2 | 0.38734656 | -1.053097982 | 2.28095921 | -0.4616908 | 0.64430304 | NA |
| CLSTN2-AS1 | 16.2257444 | 0.390886369 | 0.6683144 | 0.58488395 | 0.55862575 | 0.72720349 |
| TRIM42 | 1.6180888 | 0.286375698 | 0.99471545 | 0.2878971 | 0.7734255 | 0.87536172 |
| AC121333.1 | 0 | NA | NA | NA | NA | NA |
| SLC25A36 | 4338.15672 | 0.164515751 | 0.20648229 | 0.79675478 | 0.42559346 | 0.61793303 |
| SPSB4 | 29.9185438 | 1.784042418 | 0.53593214 | 3.32885879 | 0.00087203 | 0.00705722 |
| AC108727.2 | 0 | NA | NA | NA | NA | NA |
| PXYLP1 | 724.274418 | 0.22345959 | 0.2192953 | 1.01898941 | 0.30820799 | 0.50512901 |
| AC117383.1 | 6.97161509 | -2.281314953 | 1.04388676 | -2.1854046 | 0.02885919 | 0.09815773 |
| ZBTB38 | 4047.79011 | -0.534691054 | 0.32234007 | -1.6587794 | 0.09716026 | 0.23281381 |
| RASA2 | 1123.88277 | -0.571954131 | 0.14352333 | -3.9850952 | 6.75E-05 | 0.00093079 |
| RNF7 | 3176.73095 | -0.441760371 | 0.19018068 | -2.3228457 | 0.02018744 | 0.07540267 |
| GRK7 | 6.62319979 | 0.404869337 | 0.59676559 | 0.67843948 | 0.49749308 | 0.67924265 |
| ATP1B3 | 6587.06859 | -0.750259649 | 0.34764588 | -2.1581146 | 0.03091892 | 0.10308914 |
| ATP1B3-AS1 | 7.1790236 | 1.631455506 | 0.5368028 | 3.03920826 | 0.00237201 | 0.01538363 |
| TFDP2 | 3053.97739 | 0.707217126 | 0.13333642 | 5.30400552 | 1.13E-07 | 5.32E-06 |
| GK5 | 1325.05662 | 0.020700379 | 0.17844934 | 0.11600143 | 0.90765141 | 0.95385641 |
| XRN1 | 1628.91746 | -0.193905284 | 0.27770538 | -0.698241 | 0.48502649 | 0.66929087 |
| ATR | 1415.31812 | -0.23979192 | 0.12870129 | -1.8631664 | 0.06243886 | 0.17000648 |
| PLS1 | 1997.35397 | -0.105411681 | 0.29116233 | -0.3620375 | 0.71732401 | 0.83943526 |
| PLS1-AS1 | 0.2912362 | -0.180122781 | 2.89994773 | -0.0621124 | 0.9504733 | NA |
| TRPC1 | 477.274922 | -0.220922159 | 0.17944095 | -1.2311692 | 0.2182596 | 0.40332906 |
| PCOLCE2 | 884.104137 | 0.511945443 | 0.48091556 | 1.06452252 | 0.28709206 | 0.48334532 |
| AC021074.3 | 0.08695548 | -0.642905668 | 3.30558948 | -0.1944905 | 0.84579185 | NA |
| PAQR9 | 4.68218131 | -3.856668335 | 1.28455484 | -3.0023384 | 0.00267914 | 0.01685328 |
| PAQR9-AS1 | 94.2399285 | 0.352861386 | 0.21445735 | 1.6453686 | 0.09989382 | 0.23737948 |
| U2SURP | 3367.65375 | -0.062282708 | 0.11675572 | -0.5334446 | 0.59372586 | 0.75282738 |
| AC026304.1 | 92.1953062 | 0.376334864 | 0.22312777 | 1.68663394 | 0.09167376 | 0.22352631 |
| AC018450.2 | 0.08666371 | -0.642905668 | 3.30558948 | -0.1944905 | 0.84579185 | NA |
| CHST2 | 1111.75806 | -1.126743182 | 0.48870404 | -2.3055737 | 0.02113446 | 0.07803357 |
| SLC9A9 | 313.765452 | -0.328053488 | 0.2671654 | -1.2279041 | 0.21948297 | 0.40499641 |
| SLC9A9-AS1 | 0.13382144 | 0.02007176 | 3.29738704 | 0.00608717 | 0.99514317 | NA |
| SLC9A9-AS2 | 14.8735745 | 0.336242608 | 0.45481 | 0.73930346 | 0.45972275 | 0.64842153 |
| C3orf58 | 801.90345 | -0.567851415 | 0.18892876 | -3.0056377 | 0.00265025 | 0.0166983 |
| AC055758.2 | 0.26343812 | -1.364244712 | 2.96119245 | -0.4607079 | 0.6450082 | NA |
| AC055758.1 | 0.18380315 | -1.003574173 | 3.30558948 | -0.3035992 | 0.7614333 | NA |
| AC107021.1 | 3.22745062 | -1.157973253 | 0.85776408 | -1.3499904 | 0.17701907 | 0.35203566 |
| LNCSRLR | 13.3811877 | -2.090458677 | 0.74668948 | -2.7996359 | 0.00511603 | 0.02734521 |
| PLOD2 | 4739.86526 | -1.328384647 | 0.28649623 | -4.6366567 | 3.54E-06 | 8.94E-05 |
| PLSCR4 | 1771.64602 | 0.223621827 | 0.18481228 | 1.20999439 | 0.22628105 | 0.41305768 |
| PLSCR2 | 52.7614746 | -0.628968621 | 0.36353933 | -1.7301254 | 0.08360788 | 0.20899988 |
| PLSCR1 | 6506.5783 | -1.547313043 | 0.40636557 | -3.8076874 | 0.00014027 | 0.00165651 |
| PLSCR5 | 0.43384874 | 0.343328281 | 2.28792571 | 0.15006094 | 0.88071654 | NA |
| PLSCR5-AS1 | 0.26090319 | -0.174096114 | 3.29377188 | -0.0528562 | 0.95784652 | NA |
| AC092957.1 | 0 | NA | NA | NA | NA | NA |
| LINC02010 | 0 | NA | NA | NA | NA | NA |
| ZIC4 | 0.65800168 | -0.588264221 | 1.90656562 | -0.3085465 | 0.75766649 | NA |
| ZIC4-AS1 | 0 | NA | NA | NA | NA | NA |
| ZIC1 | 2.45110259 | -0.603292397 | 1.49209159 | -0.4043267 | 0.68597253 | 0.81840002 |
| AC092925.1 | 0 | NA | NA | NA | NA | NA |
| AC092958.1 | 0.58120394 | -1.142419552 | 1.88479534 | -0.6061239 | 0.54443246 | NA |
| LINC02032 | 0 | NA | NA | NA | NA | NA |
| AC092958.2 | 0 | NA | NA | NA | NA | NA |
| AC025566.1 | 0 | NA | NA | NA | NA | NA |
| LINC02045 | 0 | NA | NA | NA | NA | NA |
| LINC02046 | 0.5256099 | 1.788471952 | 2.04210082 | 0.87580003 | 0.38113875 | NA |
| AGTR1 | 541.64457 | 0.888201401 | 0.29651862 | 2.99543217 | 0.00274056 | 0.01716187 |
| CPB1 | 4.46729621 | 0.251269504 | 0.79094878 | 0.31768113 | 0.75072683 | 0.86094922 |
| AC092979.1 | 0 | NA | NA | NA | NA | NA |
| CPA3 | 214.931842 | -0.462812772 | 0.92197037 | -0.5019823 | 0.61568 | 0.77018414 |
| GYG1 | 2020.79234 | -0.788614057 | 0.19501934 | -4.0437736 | 5.26E-05 | 0.00077552 |
| HLTF | 1400.75488 | 0.798157873 | 0.20347217 | 3.92268808 | 8.76E-05 | 0.00114146 |
| HLTF-AS1 | 9.3521676 | 2.072676196 | 0.66341535 | 3.12425117 | 0.00178258 | 0.01234419 |
| HPS3 | 2568.79289 | -0.752810647 | 0.24800871 | -3.0354201 | 0.00240201 | 0.01550954 |
| CP | 2464.86898 | -1.600685605 | 0.40225825 | -3.9792486 | 6.91E-05 | 0.00094802 |

| | | | | | | |
|---|---|---|---|---|---|---|
| AC093001.1 | 76.823812 | 0.429557865 | 0.32712209 | 1.31314233 | 0.18913499 | 0.36740256 |
| TM4SF18 | 1431.67442 | 0.52351665 | 0.28138458 | 1.86050226 | 0.0628145 | 0.17071453 |
| TM4SF1 | 6364.96531 | -0.543161521 | 0.35114491 | -1.5468301 | 0.12190421 | 0.27277167 |
| TM4SF1-AS1 | 8.89388031 | 0.499048787 | 0.51395268 | 0.97100143 | 0.33154757 | 0.52952028 |
| AC108751.4 | 1.90829104 | 0.812312026 | 1.16005068 | 0.70023839 | 0.48377844 | 0.66830308 |
| TM4SF4 | 31.9881444 | 0.394405998 | 0.63524153 | 0.62087565 | 0.53468145 | 0.70841329 |
| WWTR1 | 5611.62629 | 0.166263758 | 0.21635991 | 0.76845918 | 0.44221443 | 0.63334752 |
| WWTR1-AS1 | 312.335598 | -0.119022029 | 0.22987379 | -0.5177712 | 0.60461791 | 0.76159798 |
| COMMD2 | 1396.14984 | -0.008281825 | 0.13360331 | -0.0619882 | 0.95057225 | 0.97632675 |
| ANKUB1 | 1104.11146 | 0.484941223 | 0.20513586 | 2.36400024 | 0.0180788 | 0.0696578 |
| RNF13 | 3903.73037 | 0.042420904 | 0.14302006 | 0.29660808 | 0.76676574 | 0.87143515 |
| PFN2 | 6280.29841 | 0.065562326 | 0.17440224 | 0.37592594 | 0.70697196 | 0.83325953 |
| AC117395.1 | 4.93047232 | -0.388296286 | 0.65411545 | -0.5936204 | 0.55276602 | 0.7226434 |
| AC117386.2 | 10.7215917 | 0.752510066 | 0.53221853 | 1.41391182 | 0.15738779 | 0.32449696 |
| LINC01998 | 0.13390214 | 1.04981649 | 3.28477129 | 0.31960109 | 0.74927075 | NA |
| AC117386.1 | 1.22032458 | 2.247217151 | 1.19632069 | 1.87844043 | 0.06032094 | 0.16617877 |
| LINC01213 | 8.46126045 | 1.988050587 | 0.49089102 | 4.04988174 | 5.12E-05 | 0.0007583 |
| LINC01214 | 2.27438979 | -1.238485344 | 0.88732806 | -1.3957468 | 0.16279074 | 0.33184848 |
| TSC22D2 | 2215.08984 | -0.001926491 | 0.17672022 | -0.0109014 | 0.99130215 | 0.99583588 |
| SERP1 | 6178.09672 | -0.187431419 | 0.15874887 | -1.1806787 | 0.23773037 | 0.42708476 |
| EIF2A | 3753.97669 | 0.054500451 | 0.20360473 | 0.26767773 | 0.78894739 | 0.88537504 |
| SELENOT | 5134.98196 | -0.127079367 | 0.12640073 | -1.0053689 | 0.31471924 | 0.51173087 |
| ERICH6 | 50.6218764 | 0.663360067 | 0.27893347 | 2.37820172 | 0.01739731 | 0.06772468 |
| ERICH6-AS1 | 112.742161 | 0.368927707 | 0.17040084 | 2.165058 | 0.03038324 | 0.10172955 |
| SIAH2 | 1545.67078 | -0.975544845 | 0.2048871 | -4.7613777 | 1.92E-06 | 5.51E-05 |
| SIAH2-AS1 | 175.47507 | -0.563180528 | 0.2171105 | -2.593981 | 0.00948717 | 0.04332815 |
| CLRN1-AS1 | 2.22909204 | 0.787068433 | 0.98461114 | 0.79936982 | 0.424076 | 0.61657589 |
| MINDY4B | 1.86345612 | -0.11492768 | 0.94426848 | -0.1217108 | 0.90312805 | 0.95128767 |
| AC020636.2 | 0.35072212 | -0.557433638 | 2.27557718 | -0.2449636 | 0.8064846 | NA |
| AC020636.1 | 0.91199697 | -0.4021566 | 1.90003538 | -0.2116574 | 0.8323743 | NA |
| CLRN1 | 0.28006796 | 1.300104138 | 2.87559618 | 0.45211638 | 0.65118514 | NA |
| MED12L | 74.9188168 | 0.042568628 | 0.27392433 | 0.15540287 | 0.87650372 | 0.93557592 |
| GPR171 | 73.2790609 | -1.142258957 | 0.66929509 | -1.7066597 | 0.08788532 | 0.21666185 |
| P2RY14 | 66.5363079 | 1.231800761 | 0.46109088 | 2.67149234 | 0.00755148 | 0.03661711 |
| GPR87 | 6.83847031 | -3.275593487 | 0.9080452 | -3.6073023 | 0.0003094 | 0.00310457 |
| P2RY13 | 67.6919824 | 1.347442737 | 0.50871529 | 2.64871681 | 0.0080798 | 0.03837614 |
| P2RY12 | 23.1357763 | 0.558738467 | 0.50582445 | 1.10460946 | 0.26932885 | 0.46352968 |
| IGSF10 | 95.6628526 | 0.638784079 | 0.33723871 | 1.89416 | 0.05820378 | 0.1620588 |
| LINC02066 | 0.08666371 | -0.642905668 | 3.30558948 | -0.1944905 | 0.84579185 | NA |
| AADACL2 | 1.10711901 | 0.418605148 | 1.76846129 | 0.23670586 | 0.81288499 | 0.89895498 |
| AADACL2-AS | 0.03392908 | 0.380740593 | 3.29738709 | 0.11546736 | 0.90807469 | NA |
| AADAC | 1.54074434 | -0.883204189 | 2.33558261 | -0.3781516 | 0.70531801 | 0.83188282 |
| SUCNR1 | 2094.53251 | 1.600017192 | 0.37610977 | 4.25412292 | 2.10E-05 | 0.00036822 |
| AC108718.1 | 0.27693877 | 2.099412187 | 3.23574276 | 0.64881925 | 0.51645521 | NA |
| MBNL1 | 4017.75082 | 0.006619791 | 0.14813434 | 0.04468775 | 0.96435619 | 0.98336618 |
| MBNL1-AS1 | 175.604439 | 0.331906095 | 0.49213052 | 0.67442697 | 0.5000399 | 0.68130141 |
| AC026347.1 | 3.40670801 | 1.556581374 | 0.75411858 | 2.06410692 | 0.03900758 | 0.12156066 |
| AC092924.1 | 0.03392908 | 0.380740593 | 3.29738709 | 0.11546736 | 0.90807469 | NA |
| P2RY1 | 90.7213021 | 0.145046741 | 0.3471869 | 0.41777712 | 0.67611008 | 0.81149917 |
| AC117394.2 | 0.957587 | -0.433176773 | 1.28453862 | -0.3372236 | 0.73594833 | NA |
| RAP2B | 1606.83333 | -1.575547974 | 0.28496037 | -5.5290074 | 3.22E-08 | 1.86E-06 |
| AC078788.1 | 0.7681323 | 1.249140217 | 1.72339781 | 0.72481247 | 0.46856709 | NA |
| AC078788.2 | 1.65307061 | -0.016499567 | 1.58124614 | -0.0104345 | 0.9916746 | 0.99605449 |
| LINC02083 | 0.68524979 | -2.68073762 | 2.1084846 | -1.2714049 | 0.20358465 | NA |
| C3orf79 | 1.3799566 | -1.419807678 | 1.3844611 | -1.0255309 | 0.30511274 | 0.5021421 |
| AC068985.1 | 0.03392908 | 0.380740593 | 3.29738709 | 0.11546736 | 0.90807469 | NA |
| ARHGEF26 | 345.045902 | -0.335204701 | 0.36463161 | -0.9192969 | 0.35794031 | 0.55558532 |
| DHX36 | 2185.15529 | -0.068718705 | 0.15975553 | -0.4301491 | 0.66708715 | 0.80515782 |
| GPR149 | 0.09635527 | 0.547239394 | 3.29377202 | 0.16614368 | 0.86804388 | NA |
| AC092994.1 | 0 | NA | NA | NA | NA | NA |
| AC073359.2 | 0.09997307 | 0.547239394 | 3.29377202 | 0.16614368 | 0.86804388 | NA |
| AC073359.1 | 0.39067777 | -0.032129813 | 2.25086966 | -0.0142744 | 0.98861106 | NA |
| MME | 12477.3407 | 2.061545394 | 0.64417957 | 3.20026511 | 0.00137301 | 0.01010503 |
| MME-AS1 | 2108.54342 | 1.692971468 | 0.64691598 | 2.61698816 | 0.00887094 | 0.04118428 |
| LINC01487 | 0.09635527 | 0.547239394 | 3.29377202 | 0.16614368 | 0.86804388 | NA |
| DWORF | 74.1523998 | 3.073814016 | 0.97728013 | 3.14527425 | 0.00165931 | 0.01168074 |
| AC104831.1 | 28.6270634 | 5.043676307 | 0.87363003 | 5.77324052 | 7.78E-09 | 5.62E-07 |
| PLCH1 | 221.650851 | -0.118350657 | 0.4259716 | -0.277837 | 0.78113751 | 0.88024427 |
| PLCH1-AS1 | 0.08666371 | -0.642905668 | 3.30558948 | -0.1944905 | 0.84579185 | NA |
| PLCH1-AS2 | 0.23148822 | -1.364242679 | 3.30558948 | -0.4127078 | 0.67982068 | NA |
| C3orf33 | 452.006141 | 0.3194565 | 0.2085302 | 1.53194356 | 0.12553636 | 0.27822158 |
| AC104472.3 | 1302.17928 | 0.547477886 | 0.20595925 | 2.65818549 | 0.00785626 | 0.03768092 |
| SLC33A1 | 2245.85601 | 0.351112797 | 0.18572603 | 1.89048781 | 0.05869275 | 0.16290157 |
| GMPS | 1852.25635 | -0.2711227 | 0.13833283 | -1.9599302 | 0.05000394 | 0.1457021 |
| KCNAB1 | 258.789219 | 0.441763516 | 0.3498662 | 1.26266416 | 0.2067099 | 0.38929725 |
| KCNAB1-AS2 | 2.13279508 | 3.269360179 | 1.3654424 | 2.39435964 | 0.01664941 | 0.06568598 |
| KCNAB1-AS1 | 0.25179623 | -1.364242679 | 3.30558948 | -0.4127078 | 0.67982068 | NA |
| AC084036.1 | 172.778854 | -0.517709742 | 0.42156203 | -1.2280749 | 0.21941885 | 0.40496427 |
| SSR3 | 5869.10027 | -0.273266432 | 0.20446485 | -1.3364959 | 0.18138724 | 0.3577309 |
| TIPARP-AS1 | 59.8332459 | 0.569919238 | 0.34249212 | 1.66403607 | 0.09610522 | 0.23121017 |
| TIPARP | 1372.821 | -0.676901416 | 0.26634614 | -2.541435 | 0.01103985 | 0.0484878 |
| LINC00886 | 221.545404 | 1.863627684 | 0.447417 | 4.16530374 | 3.11E-05 | 0.0005039 |
| LEKR1 | 72.7389858 | 1.019959592 | 0.43328538 | 2.35401341 | 0.01857194 | 0.07112142 |
| LINC00880 | 10.6579332 | -0.888591735 | 0.51590973 | -1.7223783 | 0.08500101 | 0.21138703 |
| LINC02029 | 10.2961067 | -1.117167487 | 0.5457082 | -2.0471884 | 0.04063959 | 0.1252678 |
| LINC00881 | 2.90564401 | -1.577572837 | 0.94902442 | -1.6623101 | 0.09645062 | 0.2317934 |
| CCNL1 | 7631.66103 | -0.769858735 | 0.1888346 | -4.0768945 | 4.56E-05 | 0.00069075 |
| AC104411.1 | 10.5685044 | -0.276351868 | 0.49720674 | -0.5558088 | 0.57834159 | 0.74196507 |
| VEPH1 | 1058.59603 | 1.444770709 | 0.47156314 | 3.06379058 | 0.00218552 | 0.01445138 |
| PTX3 | 845.404038 | -3.975293004 | 0.84624378 | -4.6975743 | 2.63E-06 | 7.08E-05 |
| PQLC2L | 97.5489103 | -1.02133121 | 0.4422706 | -2.3092903 | 0.02092748 | 0.07746457 |
| AC079943.1 | 0 | NA | NA | NA | NA | NA |
| AC079943.2 | 0 | NA | NA | NA | NA | NA |
| SHOX2 | 4.27919574 | -1.199769743 | 0.7687117 | -1.5607538 | 0.11858184 | 0.26767704 |
| RSRC1 | 895.773188 | 0.097979864 | 0.08844972 | 1.10774645 | 0.26797133 | 0.46209809 |
| AC106707.1 | 191.053038 | -0.347420754 | 0.18530473 | -1.8748618 | 0.06081174 | 0.16705259 |
| MLF1 | 767.793468 | -0.075251549 | 0.18846053 | -0.3992961 | 0.68967506 | 0.82118262 |
| GFM1 | 2977.4877 | 0.612167914 | 0.19647206 | 3.11580143 | 0.00183446 | 0.01260413 |
| LXN | 909.36409 | -0.297625706 | 0.21499362 | -1.3843467 | 0.16625232 | 0.33697723 |
| AC080013.4 | 64.4253114 | 0.037047422 | 0.36701539 | 0.10094242 | 0.91959617 | 0.96074349 |
| AC080013.6 | 11.673136 | 0.434124494 | 0.43996544 | 0.98672407 | 0.32377793 | 0.5215653 |
| RARRES1 | 2308.36254 | -2.667782224 | 0.42408109 | -6.290736 | 3.16E-10 | 3.86E-08 |
| MFSD1 | 3853.49894 | -0.130685309 | 0.15406305 | -0.8482586 | 0.39629396 | 0.59128615 |
| AC080013.1 | 165.286571 | -1.7964233 | 0.3998203 | -4.4930768 | 7.02E-06 | 0.0001544 |
| AC080013.5 | 10.0368976 | 1.0233847 | 0.57456009 | 1.78116217 | 0.07488596 | 0.19341262 |
| IQCJ-SCHIP1 | 1174.11177 | -0.194411506 | 0.25370639 | -0.7662854 | 0.44350649 | 0.63454322 |
| IQCJ | 2.2348078 | 1.84838342 | 1.14765593 | 1.61057279 | 0.10727287 | 0.24936403 |
| IQCJ-SCHIP1- | 1.10801001 | -1.365544352 | 1.46134832 | -0.9344414 | 0.35007626 | 0.54776662 |
| SCHIP1 | 1093.18522 | -0.159543479 | 0.24384184 | -0.6542908 | 0.51292446 | 0.6910094 |
| IL12A-AS1 | 6.23800862 | -0.345260425 | 0.56123347 | -0.6151815 | 0.53843489 | 0.71181271 |
| IL12A | 29.204679 | -0.584703046 | 0.52888807 | -1.1055327 | 0.26892884 | 0.46313275 |
| LINC01100 | 0 | NA | NA | NA | NA | NA |
| C3orf80 | 19.3970522 | 0.123107117 | 0.56415665 | 0.21821442 | 0.82726205 | 0.90800101 |

| | | | | | | | |
|---|---|---|---|---|---|---|---|
| AC079594.2 | 779.434889 | | 0.553046071 | 0.2388054 | 2.31588592 | 0.0205645 | 0.07648601 |
| IFT80 | 1633.38275 | | 0.322204608 | 0.25285048 | 1.27428909 | 0.20256098 | 0.38445871 |
| SMC4 | 837.399657 | | -0.326312611 | 0.14646254 | -2.2279595 | 0.02588322 | 0.0906414 |
| TRIM59 | 704.383517 | | -0.542082685 | 0.1386164 | -3.9106677 | 9.20E-05 | 0.00118512 |
| KPNA4 | 3860.79312 | | -0.321978388 | 0.24949158 | -1.2905381 | 0.19686389 | 0.37749424 |
| ARL14 | 14.4139031 | | -0.708024604 | 0.80284162 | -0.8818982 | 0.37783185 | 0.57382306 |
| AC069224.1 | 85.0714658 | | 0.883849591 | 0.34746326 | 2.54372101 | 0.01096786 | 0.04826812 |
| PPM1L | 897.859501 | | 0.61269979 | 0.21105241 | 2.90306943 | 0.00369525 | 0.02147755 |
| B3GALNT1 | 1270.71969 | | 0.818446306 | 0.19360882 | 4.22731929 | 2.36E-05 | 0.0004076 |
| NMD3 | 2347.81936 | | -0.127043847 | 0.14230469 | -0.8927594 | 0.37198606 | 0.56855459 |
| SPTSSB | 32.3841394 | | 0.205121749 | 0.6682443 | 0.30695623 | 0.75887669 | 0.8664871 |
| LINC02067 | 136.772789 | | -0.416449703 | 0.21815699 | -1.908945 | 0.05626919 | 0.15815587 |
| AC112491.1 | 144.016427 | | -0.411999307 | 0.24058923 | -1.7124595 | 0.08681203 | 0.21470954 |
| OTOL1 | 0.10721287 | | -0.642905668 | 3.30558948 | -0.1944905 | 0.84579185 | NA |
| AC131211.1 | 0.09989236 | | -0.642905668 | 3.30558948 | -0.1944905 | 0.84579185 | NA |
| AC128685.1 | 0.18393104 | | 1.469068379 | 3.27783996 | 0.44818185 | 0.65402196 | NA |
| LINC01192 | 0.03392908 | | 0.380740593 | 3.29738709 | 0.11546736 | 0.90807469 | NA |
| AC079910.1 | 0 | NA | | NA | NA | NA | NA |
| LINC01323 | 0.92825812 | | 1.044202355 | 1.31438736 | 0.79444035 | 0.42693912 | NA |
| LINC01324 | 0.26410578 | | 1.012519641 | 2.90828201 | 0.34815043 | 0.72772721 | NA |
| SI | 0 | NA | | NA | NA | NA | NA |
| LINC02023 | 0 | NA | | NA | NA | NA | NA |
| SLITRK3 | 2.9685304 | | 1.184289276 | 0.90753547 | 1.30495095 | 0.19190955 | 0.37117902 |
| LINC01322 | 43.3026919 | | -1.604205744 | 0.80152766 | -2.0014353 | 0.0453455 | 0.13560454 |
| BCHE | 163.698728 | | 0.252603623 | 0.5858426 | 0.43118002 | 0.66633748 | 0.80443709 |
| LINC01326 | 0 | NA | | NA | NA | NA | NA |
| AC069439.2 | 0 | NA | | NA | NA | NA | NA |
| ZBBX | 47.2095112 | | -1.988818443 | 0.81439027 | -2.442095 | 0.0146023 | 0.05969964 |
| LINC01327 | 0 | NA | | NA | NA | NA | NA |
| SERPINI2 | 7.19653896 | | -1.097589887 | 0.76402461 | -1.4365897 | 0.15083461 | 0.3156252 |
| WDR49 | 14.2627188 | | 0.196066254 | 0.72762701 | 0.26945983 | 0.78757585 | 0.88431742 |
| PDCD10 | 1779.57386 | | -0.531190087 | 0.12256614 | -4.3339058 | 1.46E-05 | 0.00027771 |
| SERPINI1 | 1188.02503 | | 0.216338104 | 0.55071603 | 0.39283059 | 0.69444461 | 0.8239763 |
| AC026353.1 | 0.20908062 | | 1.675937956 | 3.26598797 | 0.51314885 | 0.60784721 | NA |
| GOLIM4 | 2388.2382 | | 0.746405566 | 0.22778971 | 3.2767309 | 0.00105016 | 0.00814974 |
| AC124893.1 | 0 | NA | | NA | NA | NA | NA |
| LINC02082 | 6.22558686 | | 0.360356434 | 0.69279844 | 0.52014614 | 0.60296172 | 0.76022475 |
| AC092954.1 | 5.82281423 | | 0.724989352 | 0.7287852 | 0.99479155 | 0.31983766 | 0.51727194 |
| AC092954.2 | 1.77749811 | | 0.320155403 | 1.10462402 | 0.28983201 | 0.77194476 | 0.87421716 |
| LINC01997 | 0.03392908 | | 0.380740593 | 3.29738709 | 0.11546736 | 0.90807469 | NA |
| MECOM | 4422.95621 | | 0.137239805 | 0.26213231 | 0.52355167 | 0.60059041 | 0.75824464 |
| AC074033.1 | 12.7587851 | | -0.638333051 | 0.59298349 | -1.0764769 | 0.28171401 | 0.47759195 |
| TERC | 2.2929345 | | 0.577942986 | 0.76434771 | 0.75612575 | 0.44957381 | 0.63964936 |
| ACTRT3 | 160.577231 | | -0.244275151 | 0.1904776 | -1.2824351 | 0.19969007 | 0.38094677 |
| AC078802.1 | 21.899222 | | 0.074684726 | 0.42072354 | 0.17751497 | 0.85910391 | 0.92642086 |
| MYNN | 901.175212 | | 0.11496945 | 0.12927216 | 0.88935967 | 0.37380981 | 0.57001824 |
| AC078795.1 | 293.043447 | | 0.214032253 | 0.18794941 | 1.13877586 | 0.25479665 | 0.44753011 |
| LRRC34 | 221.347768 | | 0.100190435 | 0.23985474 | 0.41771297 | 0.67615699 | 0.81149917 |
| AC078795.3 | 37.4728554 | | 0.213977523 | 0.40105686 | 0.53353413 | 0.59366389 | 0.7528212 |
| AC078795.2 | 81.5110278 | | 0.095309153 | 0.28771626 | 0.33126092 | 0.74044741 | 0.85426378 |
| LRRIQ4 | 3.1434394 | | 1.084410041 | 0.84983378 | 1.27602604 | 0.20194631 | 0.38380716 |
| LRRC31 | 13.0619433 | | -0.441445254 | 0.50909281 | -0.8671214 | 0.38587551 | 0.58161074 |
| SAMD7 | 0.61785681 | | -0.914685647 | 1.62812818 | -0.561802 | 0.57425094 | NA |
| AC008040.1 | 3.64883765 | | 0.407685957 | 0.66822588 | 0.61010202 | 0.54179423 | 0.71442713 |
| SEC62 | 7326.70451 | | 0.324193216 | 0.12143021 | 2.66979054 | 0.00758986 | 0.036771 |
| SEC62-AS1 | 6.07663702 | | 0.961226479 | 0.5939147 | 1.6184588 | 0.10556376 | 0.24656965 |
| GPR160 | 2161.85795 | | 0.602671828 | 0.26677274 | 2.25912076 | 0.02387587 | 0.08534727 |
| PHC3 | 2500.2826 | | -0.008855444 | 0.11184112 | -0.0791788 | 0.93689042 | 0.96969938 |
| AC008040.5 | 1619.89851 | | -0.230574739 | 0.12853429 | -1.7938773 | 0.07283275 | 0.18975113 |
| PRKCI | 2441.37126 | | -0.419759217 | 0.16718783 | -2.5107044 | 0.01204905 | 0.0516892 |
| AC073288.1 | 0.07830766 | | -0.642905668 | 3.30558948 | -0.1944905 | 0.84579185 | NA |
| SKIL | 1887.09655 | | 0.028281433 | 0.21657194 | 0.13058678 | 0.8961022 | 0.94679922 |
| AC073288.2 | 6.17855324 | | 1.543832831 | 0.73639789 | 2.09646559 | 0.03604091 | 0.11495632 |
| CLDN11 | 399.357757 | | -0.227419184 | 0.39013968 | -0.5829173 | 0.55994895 | 0.72827885 |
| AC026316.5 | 101.461158 | | 0.175119418 | 0.45848872 | 0.38194924 | 0.70249901 | 0.82978271 |
| SLC7A14 | 9.6256375 | | 1.551719903 | 0.6278394 | 2.47152362 | 0.01345387 | 0.0561536 |
| AC026316.2 | 0.59000123 | | -2.422780352 | 2.22312769 | -1.0898071 | 0.27579812 | NA |
| RPL22L1 | 855.344298 | | -1.337812146 | 0.18372369 | -7.2816528 | 3.30E-13 | 1.06E-10 |
| EIF5A2 | 412.394804 | | -0.100059964 | 0.33047194 | -0.302779 | 0.76205829 | 0.86815969 |
| SLC2A2 | 1399.2173 | | 0.831674875 | 0.4381642 | 1.89808953 | 0.05768429 | 0.16100071 |
| TNIK | 904.544741 | | -0.501951298 | 0.32596306 | -1.5399024 | 0.12358414 | 0.27512991 |
| AC092919.1 | 0.96971995 | | 1.721614555 | 1.49092873 | 1.15472626 | 0.24820254 | NA |
| PLD1 | 1230.42568 | | 1.263409578 | 0.34267687 | 3.68688317 | 0.00022702 | 0.0024217 |
| TMEM212 | 0.84930141 | | -0.98875814 | 1.87268496 | -0.5279896 | 0.59750657 | NA |
| TMEM212-A: | 0.09989236 | | -0.642905668 | 3.30558948 | -0.1944905 | 0.84579185 | NA |
| FNDC3B | 3359.72634 | | -0.699684815 | 0.18642468 | -3.753177 | 0.00017461 | 0.00196881 |
| GHSR | 0.6728135 | | -1.587974953 | 2.22639714 | -0.7132487 | 0.47569185 | NA |
| TNFSF10 | 18039.3348 | | -0.771774529 | 0.36581446 | -2.1097431 | 0.03488049 | 0.11241733 |
| LINC02068 | 36.3897808 | | 1.569587118 | 0.52446852 | 2.99271939 | 0.00276504 | 0.01726594 |
| NCEH1 | 2252.52212 | | -0.158845705 | 0.29053448 | -0.5467362 | 0.58456001 | 0.74648301 |
| ECT2 | 625.350122 | | -1.031766295 | 0.2822938 | -3.6549379 | 0.00025724 | 0.00267595 |
| SPATA16 | 7.36008037 | | 2.135830096 | 0.6035489 | 3.53878553 | 0.00040197 | 0.00383225 |
| AC068759.1 | 0.65578014 | | -2.581288801 | 1.86510616 | -1.3839903 | 0.16636143 | NA |
| NLGN1 | 342.564104 | | -0.174702494 | 0.41917334 | -0.4167786 | 0.67684033 | 0.81209758 |
| NLGN1-AS1 | 0.09635527 | | 0.547239394 | 3.29377202 | 0.16614368 | 0.86804388 | NA |
| NAALADL2 | 409.534948 | | 0.73563608 | 0.39086276 | 1.88208281 | 0.05982477 | 0.16522615 |
| NAALADL2-A | 0.19416835 | | -1.003574173 | 3.30558948 | -0.3035992 | 0.7614333 | NA |
| NAALADL2-A | 0 | NA | | NA | NA | NA | NA |
| AC104640.1 | 0 | NA | | NA | NA | NA | NA |
| LINC01208 | 1.56791945 | | 0.320340413 | 1.13871414 | 0.28131767 | 0.77846675 | 0.87861887 |
| AC092920.1 | 0.99990598 | | -0.198054471 | 1.24934447 | -0.1585267 | 0.87404177 | NA |
| LINC01209 | 0.08695548 | | -0.642905668 | 3.30558948 | -0.1944905 | 0.84579185 | NA |
| TBL1XR1 | 5966.67777 | | -0.55426882 | 0.21010507 | -2.6380555 | 0.00833829 | 0.03927846 |
| TBL1XR1-AS: | 32.6651555 | | -0.275088913 | 0.32350236 | -0.8503459 | 0.39513279 | 0.59044889 |
| LINC00501 | 67.1925372 | | -0.080790792 | 0.57797251 | -0.1397831 | 0.88883136 | 0.94259161 |
| AC117465.1 | 0.84214915 | | 2.677252074 | 1.87585805 | 1.42721464 | 0.15351803 | NA |
| LINC00578 | 6.97348648 | | -0.831268554 | 0.97501669 | -0.8525685 | 0.39389862 | 0.58920565 |
| AC026355.2 | 0 | NA | | NA | NA | NA | NA |
| AC026355.1 | 2.4383136 | | 0.300592293 | 0.80527736 | 0.37327796 | 0.70894158 | 0.83447565 |
| LINC02015 | 32.6555578 | | 1.097865943 | 0.68746254 | 1.59698292 | 0.11026951 | 0.25431232 |
| AC007953.1 | 11.7603975 | | 2.914052349 | 0.7090691 | 4.10968742 | 3.96E-05 | 0.00061857 |
| AC007953.2 | 0.08695548 | | -0.642905668 | 3.30558948 | -0.1944905 | 0.84579185 | NA |
| AC109779.1 | 0.84794704 | | 2.679052234 | 1.33122331 | 2.01247396 | 0.04417 | NA |
| KCNMB2 | 201.567377 | | 0.876274217 | 0.49948853 | 1.75434303 | 0.07937175 | 0.2012818 |
| AC117457.1 | 142.326495 | | 0.841040647 | 0.45078004 | 1.86574511 | 0.06207704 | 0.16939516 |
| KCNMB2-AS: | 9.96938192 | | -0.587044069 | 0.46178828 | -1.2712407 | 0.20364302 | 0.38583607 |
| ZMAT3 | 1443.31113 | | -0.480427507 | 0.21939206 | -2.1898126 | 0.02853783 | 0.09742912 |
| AC076966.1 | 0.06785815 | | 0.795868262 | 3.28898349 | 0.24198001 | 0.80879565 | NA |
| PIK3CA | 1498.88631 | | -0.225325095 | 0.13380535 | -1.6839768 | 0.09218616 | 0.22444363 |
| KCNMB3 | 694.331865 | | -0.306679031 | 0.13959059 | -2.1969893 | 0.02802121 | 0.09616003 |
| ZNF639 | 1362.99888 | | -0.382873693 | 0.18296149 | -2.0926464 | 0.03638074 | 0.1156831 |
| AC007823.1 | 14.0741062 | | -0.09837155 | 0.534023 | -0.1842085 | 0.85384993 | 0.92346906 |
| MFN1 | 2456.04244 | | -0.378628922 | 0.14278515 | -2.6517389 | 0.00800785 | 0.0381512 |

| | | | | | | |
|---|---|---|---|---|---|---|
| GNB4 | 1093.16235 | -0.427438806 | 0.27079846 | -1.5784388 | 0.11446483 | 0.26135921 |
| AC007620.2 | 514.240856 | -0.150748586 | 0.25967865 | -0.5805197 | 0.56156417 | 0.7294083 |
| AC007620.3 | 5.77318714 | -0.800957416 | 0.67408222 | -1.1882192 | 0.23474707 | 0.42349868 |
| ACTL6A | 1487.13565 | -0.268844567 | 0.19548375 | -1.3752784 | 0.16904517 | 0.34072621 |
| AC090425.2 | 382.489589 | -0.521288026 | 0.2198478 | -2.3711314 | 0.01773372 | 0.0686274 |
| AC090425.1 | 8.93723414 | -0.106599266 | 0.47424326 | -0.2247776 | 0.82215229 | 0.90496121 |
| MRPL47 | 1656.53182 | -0.430723091 | 0.17111219 | -2.517197 | 0.01182927 | 0.05105227 |
| NDUFB5 | 4546.99313 | 0.283716813 | 0.21308011 | 1.33150307 | 0.18302354 | 0.35972297 |
| USP13 | 588.975784 | 0.518214169 | 0.15034317 | 3.4468754 | 0.00056711 | 0.00500995 |
| PEX5L | 17.3688457 | 0.931470047 | 0.5114088 | 1.82138055 | 0.06854903 | 0.18139869 |
| AC007687.1 | 4.712134 | 0.676824949 | 0.58888141 | 1.14933998 | 0.25041582 | 0.44262116 |
| PEX5L-AS1 | 0.53815482 | 0.99269848 | 1.88706575 | 0.526054 | 0.59885069 | NA |
| PEX5L-AS2 | 0.94788595 | -0.994872195 | 2.242666 | -0.4436114 | 0.65732356 | NA |
| LINC02053 | 0 | NA | NA | NA | NA | NA |
| AC125618.1 | 3.50182671 | 0.935099513 | 0.76459687 | 1.22299678 | 0.22133091 | 0.40744344 |
| TTC14 | 2621.42816 | 0.212872188 | 0.20032899 | 1.06261299 | 0.2879575 | 0.48416119 |
| CCDC39 | 847.573738 | 0.417460462 | 0.1784907 | 2.33883589 | 0.01934393 | 0.07313253 |
| CCDC39-AS1 | 1.40447794 | 0.672376577 | 1.22320391 | 0.54968478 | 0.58253559 | 0.74490659 |
| AC108734.4 | 8.68628772 | 0.273814764 | 0.80249161 | 0.34120577 | 0.73294868 | 0.84942677 |
| FXR1 | 4309.19195 | 0.085937047 | 0.08799706 | 0.97659 | 0.32877217 | 0.52658294 |
| DNAJC19 | 2238.48204 | 0.639818281 | 0.15089169 | 4.24024851 | 2.23E-05 | 0.00038786 |
| AC022294.1 | 1.2529895 | 1.467606658 | 1.1040583 | 1.32928366 | 0.1837544 | 0.36079376 |
| AC125613.1 | 0.08695548 | -0.642905668 | 3.30558948 | -0.1944905 | 0.84579185 | NA |
| SOX2 | 7.00457965 | -0.613861194 | 0.85831601 | -0.7151925 | 0.47449009 | 0.66007262 |
| LINC01206 | 0 | NA | NA | NA | NA | NA |
| AC012081.1 | 0 | NA | NA | NA | NA | NA |
| AC084211.1 | 0 | NA | NA | NA | NA | NA |
| LINC01994 | 0 | NA | NA | NA | NA | NA |
| LINC01995 | 0 | NA | NA | NA | NA | NA |
| LINC02031 | 0 | NA | NA | NA | NA | NA |
| AC083801.2 | 0.41317169 | -1.957544338 | 3.29391586 | -0.5942909 | 0.55231754 | NA |
| AC069431.1 | 1.57102309 | 0.845190075 | 0.9371978 | 0.90182678 | 0.36714889 | 0.56403824 |
| ATP11B | 2283.26152 | -0.262616619 | 0.13681809 | -1.9194584 | 0.05492635 | 0.1555755 |
| AC092953.2 | 287.186593 | -0.429054568 | 0.1788679 | -2.3987231 | 0.01645235 | 0.06510332 |
| DCUN1D1 | 1709.19817 | 0.152640301 | 0.12933565 | 1.18018734 | 0.23792571 | 0.42730359 |
| MCCC1 | 3676.54219 | 0.78087115 | 0.22590985 | 3.45656084 | 0.00054712 | 0.0048692 |
| MCCC1-AS1 | 861.367942 | 0.476640449 | 0.23165132 | 2.05757707 | 0.03963075 | 0.12300646 |
| LAMP3 | 92.5494519 | -0.296168596 | 0.6754318 | -0.4384878 | NA | NA |
| MCF2L2 | 461.302239 | -1.21064889 | 0.29484909 | -4.105995 | 4.03E-05 | 0.00062557 |
| B3GNT5 | 692.35534 | -0.92901957 | 0.31409277 | -2.9577872 | 0.00309856 | 0.01882262 |
| KLHL6 | 213.780826 | 0.015012147 | 0.43594461 | 0.03443591 | 0.97252955 | 0.98712873 |
| KLHL6-AS1 | 10.9880035 | 2.105451599 | 0.57990182 | 3.63070356 | 0.00028265 | 0.00289568 |
| KLHL24 | 3107.78155 | -0.270495096 | 0.1836733 | -1.4726969 | 0.14083279 | 0.30109945 |
| YEATS2 | 1684.48188 | -0.794078136 | 0.31105638 | -2.5528431 | 0.01068476 | 0.0472985 |
| YEATS2-AS1 | 324.482835 | -0.285163933 | 0.35840429 | -0.7956488 | 0.42623622 | 0.61866188 |
| AC131160.1 | 1771.96556 | -0.187275613 | 0.13518908 | -1.3852865 | 0.16596487 | 0.33652392 |
| MAP6D1 | 112.908686 | -0.547107285 | 0.23545927 | -2.323575 | 0.02014828 | 0.07532041 |
| PARL | 1951.74817 | -0.234496549 | 0.12085011 | -1.9403917 | 0.05233211 | 0.15036911 |
| ABCC5 | 4025.46447 | -0.318978312 | 0.29309516 | -1.0883097 | 0.2764584 | 0.47179704 |
| ABCC5-AS1 | 0.45150051 | -0.923257859 | 2.24522813 | -0.4112089 | 0.68091934 | NA |
| HTR3D | 2.22682493 | 0.330197408 | 0.90898049 | 0.36326127 | 0.71640972 | 0.83866307 |
| HTR3C | 0.86277734 | 2.185465663 | 1.29640188 | 1.68579335 | 0.09183561 | NA |
| HTR3E-AS1 | 0.06785815 | 0.795868262 | 3.28898349 | 0.24198001 | 0.80879565 | NA |
| HTR3E | 0.44160456 | 0.608096459 | 2.1810853 | 0.27880453 | 0.78039484 | NA |
| AC131235.4 | 9.81597428 | 0.768235511 | 0.49952126 | 1.53794357 | 0.12406241 | 0.27579884 |
| AC131235.3 | 21.5097716 | 1.037292361 | 0.38931704 | 2.66438984 | 0.00771281 | 0.03718224 |
| EIF2B5 | 3652.21709 | -0.149847595 | 0.10546061 | -1.4208868 | 0.15534967 | 0.32157711 |
| DVL3 | 6715.4414 | -0.096066884 | 0.17850462 | -0.5381759 | 0.59045561 | 0.75022224 |
| AP2M1 | 17304.6717 | -0.341625147 | 0.12416232 | -2.7514397 | 0.00593339 | 0.03054736 |
| ABCF3 | 3794.85132 | 0.080384205 | 0.12860569 | 0.62504391 | 0.53194224 | 0.70620333 |
| VWA5B2 | 45.0841374 | -0.850496121 | 0.40313931 | -2.109683 | 0.03488567 | 0.11242029 |
| ALG3 | 1922.03542 | -0.618288161 | 0.19187403 | -3.222365 | 0.00127137 | 0.00948003 |
| EEF1AKMT4 | 354.234347 | -1.096712656 | 0.34856111 | -3.1463999 | 0.00165294 | 0.01165454 |
| EEF1AKMT4- | 236.713964 | -0.998694571 | 0.34233999 | -2.9172595 | 0.00353122 | 0.02071135 |
| CAMK2N2 | 24.659005 | -1.016800525 | 0.41626075 | -2.4427009 | 0.01457781 | 0.05962725 |
| ECE2 | 1.04600494 | 0.790377954 | 1.66542959 | 0.47457903 | 0.63508706 | 0.78346022 |
| PSMD2 | 9144.51214 | -0.227429165 | 0.30139363 | -0.7545918 | 0.45049393 | 0.64037114 |
| EIF4G1 | 17041.4122 | 0.035977124 | 0.19097293 | 0.1883886 | 0.85057203 | 0.92161226 |
| FAM131A | 1650.22182 | -0.258988269 | 0.19074316 | -1.3577853 | 0.17453181 | 0.34848215 |
| CLCN2 | 916.737145 | 0.216022987 | 0.15182308 | 1.42285996 | 0.15477675 | 0.32088146 |
| POLR2H | 4191.36044 | -0.57172898 | 0.13539791 | -4.2225836 | 2.42E-05 | 0.00041464 |
| THPO | 502.725984 | 0.419763878 | 0.29749911 | 1.41097522 | 0.15825192 | 0.32542912 |
| CHRD | 3381.67807 | 0.004458815 | 0.36467033 | 0.01222698 | 0.99024453 | 0.99533983 |
| LINC02054 | 4.84179898 | 0.00079269 | 0.60154139 | 0.00131776 | 0.99894858 | 0.99955506 |
| LINC01839 | 0.46051482 | 2.798984569 | 2.49280695 | 1.12282444 | 0.26151206 | NA |
| LINC01840 | 0 | NA | NA | NA | NA | NA |
| EPHB3 | 422.472156 | 1.429627208 | 0.56431002 | 2.53340744 | NA | NA |
| MAGEF1 | 2372.05476 | -0.089219418 | 0.17328297 | -0.514877 | 0.60663896 | 0.76318255 |
| AC107294.2 | 11.8909247 | 0.108948489 | 0.60924267 | 0.1788261 | 0.85807425 | 0.92582473 |
| AC107294.3 | 10.3165517 | 1.158084629 | 0.66420628 | 1.74356169 | 0.08123555 | 0.20456278 |
| AC107294.1 | 3.87475424 | 0.989026497 | 0.63768018 | 1.55097576 | 0.12090749 | 0.27130065 |
| LINC02069 | 0.08393208 | -0.642905668 | 3.30558948 | -0.1944905 | 0.84579185 | NA |
| VPS8 | 1561.19544 | 0.067283954 | 0.10072709 | 0.66798272 | 0.50414463 | 0.68375733 |
| C3orf70 | 97.6174633 | 0.640929542 | 0.48091633 | 1.33272569 | 0.18262184 | 0.35938343 |
| EHHADH-AS1 | 1276.9859 | 1.976399478 | 0.62366246 | 3.16902106 | 0.00152953 | 0.01099013 |
| EHHADH | 7669.05492 | 2.176365693 | 0.61855431 | 3.51847147 | 0.00043404 | 0.00406451 |
| MAP3K13 | 2079.97752 | 0.059698351 | 0.18353922 | 0.32526209 | 0.74498272 | 0.85715757 |
| TMEM41A | 2019.66896 | -0.305381287 | 0.26983059 | -1.1317519 | 0.25773875 | 0.45029234 |
| LIPH | 814.933583 | -0.267448355 | 0.40465792 | -0.6609245 | 0.5086607 | 0.68744274 |
| SENP2 | 1897.59815 | 0.115435614 | 0.15846203 | 0.72847492 | 0.46632292 | 0.65353611 |
| IGF2BP2 | 1278.58464 | -0.48530462 | 0.26288022 | -1.8461055 | 0.06487689 | 0.1745718 |
| IGF2BP2-AS1 | 9.03424585 | 2.033258732 | 1.76068981 | 1.15480803 | 0.24816905 | 0.43979958 |
| TRA2B | 4684.41247 | -0.073869885 | 0.17035896 | -0.4336132 | 0.66456938 | 0.80303783 |
| ETV5 | 1342.41718 | -0.634654531 | 0.28108658 | -2.2578614 | 0.0239543 | 0.08553487 |
| ETV5-AS1 | 3.02025915 | 1.434941897 | 0.75203587 | 1.9080764 | 0.05638134 | 0.15838671 |
| DGKG | 331.607251 | -0.170340483 | 0.37656027 | -0.4523592 | 0.65101027 | 0.79488544 |
| LINC02020 | 1.01756938 | -2.000405145 | 1.93628256 | -1.0331163 | 0.30154946 | 0.49834165 |
| LINC02052 | 1.59761458 | -2.690476251 | 2.00652983 | -1.3408603 | 0.1799658 | 0.3558199 |
| LINC02051 | 0.23148822 | -1.364242679 | 3.30558948 | -0.4127078 | 0.67982068 | NA |
| CRYGS | 415.711408 | -0.237083559 | 0.15042691 | -1.5760714 | 0.11500934 | 0.26226188 |
| TBCCD1 | 490.652061 | 0.176832082 | 0.1233822 | 1.43320577 | 0.15179904 | 0.31683335 |
| DNAJB11 | 2997.61588 | -0.419374848 | 0.22292955 | -1.881199 | 0.05994485 | 0.16540189 |
| AC068631.1 | 7355.65193 | 2.305215918 | 0.53839182 | 4.28166966 | 1.85E-05 | 0.00033458 |
| AC068631.2 | 1554.43366 | -0.595426024 | 0.23872114 | -2.4942325 | 0.01262298 | 0.05358468 |
| AHSG | 7.74597641 | 1.726813289 | 0.62459065 | 2.76471202 | 0.00569731 | 0.0296913 |
| FETUB | 54.7323104 | 0.526895023 | 0.45412861 | 1.16023306 | 0.24595393 | 0.43722564 |
| HRG | 1101.02717 | 2.876213358 | 0.55018854 | 5.2276868 | 1.72E-07 | 7.53E-06 |
| KNG1 | 15696.9576 | 2.321909226 | 0.55897288 | 4.15388526 | 3.27E-05 | 0.00052585 |
| AC112907.3 | 682.345197 | 0.680912471 | 0.21842245 | 3.11741068 | 0.00182447 | 0.0125486 |
| EIF4A2 | 19485.3662 | 0.062223911 | 0.12502765 | 0.4976812 | 0.61870876 | 0.7722146 |
| RFC4 | 501.19003 | -0.443995818 | 0.19073514 | -2.3278135 | 0.01992201 | 0.07470754 |
| AC112907.1 | 0.39439756 | 1.069716182 | 2.23419263 | 0.47879317 | 0.63208577 | NA |
| LINC02043 | 9.33539783 | -0.713456487 | 0.7324254 | -0.9741012 | 0.33000628 | 0.52788993 |

| | | | | | | |
|---|---|---|---|---|---|---|
| ADIPOQ | 24.0172713 | 3.078931139 | 2.08578891 | 1.47614704 | NA | NA |
| ADIPOQ-AS1 | 4.58396528 | 2.931862146 | 2.21054047 | 1.3263101 | NA | NA |
| ST6GAL1 | 8543.54187 | -0.061150681 | 0.25372269 | -0.2410138 | 0.80954439 | 0.89703256 |
| RPL39L | 103.249481 | -0.849622035 | 0.22069615 | -3.8497366 | 0.00011824 | 0.00144827 |
| AC007920.1 | 0.2943225 | -0.699798508 | 2.94526751 | -0.237601 | 0.81219057 | NA |
| RTP1 | 0.49241762 | 0.002916002 | 2.22360234 | 0.00131139 | 0.99895367 | NA |
| MASP1 | 596.499579 | 0.806998861 | 0.43962463 | 1.83565433 | 0.0664088 | 0.17742654 |
| AC007920.2 | 15.6189259 | 0.28797013 | 0.50537812 | 0.56981123 | 0.56880574 | 0.73436511 |
| RTP4 | 230.75379 | -0.638447854 | 0.34098568 | -1.8723597 | 0.06115686 | 0.16762356 |
| LINC02041 | 5.39254923 | -0.700384923 | 0.6403929 | -1.09368 | 0.27409534 | 0.46922189 |
| SST | 49.1562614 | -0.764051285 | 0.92215547 | -0.8285493 | 0.40735948 | 0.60196523 |
| RTP2 | 0.18390514 | 0.18657164 | 3.29377195 | 0.05664376 | 0.95482897 | NA |
| AC072022.1 | 261.054507 | -1.466773152 | 0.38379406 | -3.8217714 | 0.0001325 | 0.00158025 |
| BCL6 | 8100.12337 | -2.506068331 | 0.37678393 | -6.6512081 | 2.91E-11 | 5.08E-09 |
| AC108681.1 | 0.60776075 | -0.263016415 | 2.27410188 | -0.1156573 | 0.90792417 | NA |
| AC068295.1 | 0 | NA | NA | NA | NA | NA |
| LINC01991 | 0.9045206 | -0.702726179 | 1.59746633 | -0.4399005 | 0.6600092 | NA |
| AC092941.1 | 0.35327105 | -0.260389715 | 2.31051322 | -0.1126978 | 0.91027016 | NA |
| AC092941.2 | 0.29835781 | -1.364242679 | 3.30558948 | -0.4127078 | 0.67982068 | NA |
| AC022498.1 | 0.06785815 | 0.795868262 | 3.28898349 | 0.24198001 | 0.80879565 | NA |
| LPP-AS2 | 122.450954 | 0.546704775 | 0.2396491 | 2.28127201 | 0.02253236 | 0.08186704 |
| LPP | 6224.95882 | -0.323798878 | 0.29013083 | -1.1160443 | 0.26440315 | 0.45837296 |
| LPP-AS1 | 0.34916828 | -0.556958688 | 2.28822425 | -0.2434021 | 0.80769388 | NA |
| TPRG1-AS1 | 13.7796878 | 0.105866244 | 0.47319462 | 0.22372664 | 0.82297001 | 0.90546648 |
| TPRG1 | 71.2390001 | -0.612520875 | 0.35176655 | -1.7412709 | 0.0816361 | 0.20531714 |
| TPRG1-AS2 | 4.20492696 | -1.250720826 | 0.69774863 | -1.7925092 | 0.07305144 | 0.19013454 |
| TP63 | 22.2903849 | 0.367964405 | 0.66418962 | 0.55400506 | 0.57957539 | 0.74277814 |
| P3H2 | 5843.64254 | -0.231903044 | 0.2249717 | -1.0308098 | 0.30263 | 0.49952135 |
| P3H2-AS1 | 49.6721599 | -0.159261523 | 0.51319213 | -0.3103351 | 0.75630615 | 0.86482777 |
| CLDN1 | 2816.34579 | -2.097964783 | 0.30331117 | -6.9168727 | 4.62E-12 | 1.05E-09 |
| CLDN16 | 2019.62915 | 2.277821938 | 0.37938004 | 6.00406368 | 1.92E-09 | 1.76E-07 |
| TMEM207 | 223.218903 | 3.705408663 | 0.6986645 | 5.30355939 | 1.14E-07 | 5.32E-06 |
| IL1RAP | 350.425718 | -0.630071212 | 0.25682958 | -2.4532658 | 0.01415657 | 0.05839336 |
| AC108747.1 | 0 | NA | NA | NA | NA | NA |
| LINC02013 | 0 | NA | NA | NA | NA | NA |
| GMNC | 150.264503 | 1.787649226 | 0.70245653 | 2.54485387 | 0.01093235 | 0.04813592 |
| OSTN | 10.1862934 | 0.432456321 | 0.66445475 | 0.6508439 | 0.51514726 | 0.69281728 |
| OSTN-AS1 | 7.1362821 | 0.06691603 | 0.5074536 | 0.1318663 | 0.89509004 | 0.94607096 |
| UTS2B | 131.870235 | -0.645130756 | 0.33395725 | -1.9317765 | 0.0533871 | 0.1525759 |
| CCDC50 | 5650.99399 | -0.413303842 | 0.21804776 | -1.895474 | 0.05802964 | 0.16172496 |
| AC073365.1 | 2.67674977 | -1.431148273 | 0.94165063 | -1.5198294 | 0.12855387 | 0.28279116 |
| PYDC2 | 0 | NA | NA | NA | NA | NA |
| AC026320.1 | 0 | NA | NA | NA | NA | NA |
| FGF12 | 95.5997117 | 0.869817155 | 0.4241318 | 2.05081806 | 0.04028467 | 0.12445018 |
| FGF12-AS1 | 0 | NA | NA | NA | NA | NA |
| FGF12-AS2 | 0.81710492 | -1.320156473 | 2.2646433 | -0.5829423 | 0.55993211 | NA |
| FGF12-AS3 | 0.25020947 | -1.364244509 | 2.99735315 | -0.4551497 | 0.64900151 | NA |
| MB21D2 | 340.028784 | 0.714755118 | 0.23883799 | 2.9926358 | 0.0027658 | 0.01726658 |
| AC092966.1 | 0 | NA | NA | NA | NA | NA |
| HRASLS | 19.4507948 | -0.52373579 | 0.51334266 | -1.020246 | 0.30761181 | 0.5044676 |
| ATP13A5 | 7.23734794 | 1.360128208 | 0.81650659 | 1.66578963 | 0.09575532 | 0.23064193 |
| ATP13A5-AS | 1.59844954 | 1.67925103 | 1.48191922 | 1.13315963 | 0.25714722 | 0.44990528 |
| ATP13A4 | 43.7862927 | 1.150839946 | 0.5196914 | 2.21446796 | 0.0267966 | 0.09292086 |
| ATP13A4-AS | 1.54646246 | 0.341770617 | 1.13780941 | 0.30037598 | 0.76389038 | 0.86935843 |
| OPA1 | 2771.12433 | 0.323604681 | 0.14014983 | 2.30899087 | 0.02094409 | 0.07750257 |
| OPA1-AS1 | 41.8133748 | 1.585632885 | 0.47309991 | 3.35158149 | 0.00080351 | 0.00660083 |
| LINC02038 | 31.7241278 | 1.807856006 | 0.57454444 | 3.1465904 | 0.00165186 | 0.01165052 |
| LINC02026 | 4.7330644 | 0.875034232 | 0.65991061 | 1.32598903 | 0.18484333 | 0.36207983 |
| AC024559.1 | 0.38290237 | 0.288792915 | 2.2860653 | 0.1263275 | 0.89947269 | NA |
| AC024559.2 | 0.47173475 | 1.488015062 | 1.73878428 | 0.85577899 | 0.39212004 | NA |
| AC080129.1 | 1.09709937 | -2.437511924 | 1.46173747 | -1.6675443 | 0.09540623 | 0.23007429 |
| AC080129.2 | 24.3961006 | 0.05378265 | 0.39116568 | 0.13749328 | 0.89064091 | 0.94323068 |
| HES1 | 3638.37912 | -0.03551809 | 0.44440842 | -0.0799222 | 0.93629915 | 0.96951778 |
| LINC02036 | 2.16880365 | -2.662605389 | 1.25263453 | -2.1256043 | 0.03353622 | 0.10928696 |
| LINC02037 | 0.35114197 | 1.644829455 | 2.15780252 | 0.76227062 | 0.44589851 | NA |
| LINC02048 | 14.1896799 | -3.378291929 | 0.84752325 | -3.9860758 | 6.72E-05 | 0.00092987 |
| LINC00887 | 55.8024834 | -1.669983287 | 0.52899793 | -3.1568806 | 0.00159467 | 0.01132443 |
| CPN2 | 2972.81491 | 0.181977368 | 0.56106591 | 0.32434223 | 0.74567895 | 0.85743487 |
| LRRC15 | 33.2693731 | -0.044603404 | 0.71219655 | -0.0626279 | 0.95006278 | 0.97604626 |
| GP5 | 14.7417831 | 1.143597957 | 0.45087103 | 2.53641926 | 0.01119926 | 0.04901502 |
| ATP13A3 | 5635.60164 | -0.51661679 | 0.29702599 | -1.7392983 | 0.08198231 | 0.20593194 |
| LINC00884 | 80.0892613 | 0.008829464 | 0.3048288 | 0.02896532 | 0.97689225 | 0.98937845 |
| TMEM44-AS | 330.839433 | 0.96581111 | 0.2482345 | 3.89072068 | 9.99E-05 | 0.00126499 |
| TMEM44 | 733.662847 | -0.320222138 | 0.19618123 | -1.6322772 | 0.10262111 | 0.24172566 |
| AC046143.1 | 230.485017 | 0.035344545 | 0.23949994 | 0.14757643 | 0.88267706 | 0.93902736 |
| AC046143.2 | 6.95132997 | -0.347541063 | 0.56476945 | -0.615368 | 0.5383117 | 0.71168553 |
| LSG1 | 2056.05028 | -0.034141577 | 0.21766092 | -0.1568567 | 0.87535777 | 0.93522342 |
| FAM43A | 1555.84332 | -1.008071591 | 0.64371773 | -1.566015 | 0.11734511 | 0.26580341 |
| AC106706.1 | 0.17332742 | -1.003574173 | 3.30558948 | -0.3035992 | 0.7614333 | NA |
| LINC01968 | 2.53978933 | 0.707243089 | 1.25434193 | 0.56383596 | 0.57286578 | 0.73753036 |
| LINC01972 | 0.62068332 | 0.726840343 | 1.78237918 | 0.40779221 | 0.68342623 | NA |
| AC090505.2 | 0.92598263 | 0.68745506 | 2.13718116 | 0.32166438 | 0.74770696 | NA |
| XXYLT1 | 553.912145 | 0.250540508 | 0.15001435 | 1.67011033 | 0.09489754 | 0.22918289 |
| XXYLT1-AS1 | 1.87883427 | 3.537534663 | 1.07506951 | 3.29051715 | 0.00100003 | 0.00783682 |
| XXYLT1-AS2 | 4.6431042 | -1.265253538 | 0.75568037 | -1.6743237 | 0.094067 | 0.22790765 |
| ACAP2 | 1821.92383 | -0.11825704 | 0.13585751 | -0.870449 | 0.38405507 | 0.57963524 |
| PPP1R2 | 2633.81925 | -0.205583746 | 0.12175881 | -1.6884506 | 0.09132476 | 0.22288144 |
| AC069213.1 | 2.80452365 | 0.074358229 | 0.79762864 | 0.09322412 | 0.92572551 | 0.9640363 |
| APOD | 1879.5712 | -1.664606484 | 0.63974849 | -2.6019702 | 0.00926899 | 0.04253799 |
| AC233280.1 | 25.6059631 | -0.762838834 | 0.58618756 | -1.3013562 | 0.19313655 | 0.37266019 |
| MUC20-OT1 | 6785.56571 | -0.746639412 | 0.34584184 | -2.1589042 | 0.0308576 | 0.10294151 |
| MUC20 | 4597.01829 | -0.877030963 | 0.36321781 | -2.4146144 | 0.01575188 | 0.06307927 |
| MUC4 | 455.140759 | 0.399828991 | 0.37760952 | 1.05884245 | 0.28967154 | 0.48588757 |
| LINC01983 | 35.9847859 | 1.209393076 | 0.70525203 | 1.71483812 | 0.08637491 | 0.2139897 |
| TNK2 | 2958.16677 | -0.155176389 | 0.24645877 | -0.6296241 | 0.52894054 | 0.7035288 |
| TNK2-AS1 | 95.5202768 | 0.466205855 | 0.3251043 | 1.43401934 | 0.15156674 | 0.3165885 |
| AC124944.1 | 33.8097315 | 0.192640715 | 0.56378559 | 0.34169145 | 0.73258311 | 0.8492009 |
| TFRC | 3986.98777 | -0.406797678 | 0.43130957 | -0.9431687 | 0.34559464 | 0.54326859 |
| LINC00885 | 32.5007707 | 1.595484803 | 0.67904077 | 2.34961563 | 0.01879281 | 0.07172765 |
| ZDHHC19 | 33.9141372 | -1.735231039 | 0.59927645 | -2.8955435 | 0.00378503 | 0.02190281 |
| SLC51A | 748.656341 | 0.29267028 | 0.46799715 | 0.62536765 | 0.53172979 | 0.70599237 |
| PCYT1A | 3068.35907 | -0.115508835 | 0.15064265 | -0.7667738 | 0.44321602 | 0.63434371 |
| AC069257.3 | 459.093813 | -0.230414944 | 0.16112459 | -1.4300421 | 0.15270495 | 0.31810013 |
| AC069257.1 | 0.79762483 | -0.068428739 | 1.40438046 | -0.0487252 | 0.96113828 | NA |
| TCTEX1D2 | 389.71722 | -0.287583165 | 0.32442288 | -0.8864454 | 0.37537757 | 0.57132094 |
| TM4SF19-TC | 220.979973 | -0.237890885 | 0.287677 | -0.8269375 | 0.40827252 | 0.60277881 |
| TM4SF19-AS | 16.7647672 | 1.060412888 | 0.33710688 | 3.14562812 | 0.00165731 | 0.01167597 |
| TM4SF19 | 21.4124144 | 0.428925991 | 0.43480749 | 0.98647333 | 0.32390089 | 0.52162159 |
| UBXN7 | 1578.20463 | -0.144039243 | 0.13598343 | -1.0592411 | 0.28948998 | 0.48571068 |
| UBXN7-AS1 | 13.9024889 | -0.24257564 | 0.50157841 | -0.4836246 | 0.62865234 | 0.77927443 |
| RNF168 | 853.21613 | -0.465288083 | 0.19646537 | -2.3682957 | 0.01787025 | 0.06899537 |
| AC117490.2 | 16.060713 | 0.316750686 | 0.37445576 | 0.84589616 | 0.39761067 | 0.59247318 |
| SMCO1 | 1.08252233 | 0.708193839 | 1.2059658 | 0.58724206 | 0.55704115 | 0.72600085 |

| | | | | | | |
|---|---|---|---|---|---|---|
| WDR53 | 323.391145 | -0.243929008 | 0.20792932 | -1.1731342 | 0.24074198 | 0.43089309 |
| FBXO45 | 450.78066 | -0.35464408 | 0.24982833 | -1.4195511 | 0.15573842 | 0.32211906 |
| LINC01063 | 5.41084772 | 0.201179533 | 0.53118484 | 0.37873734 | 0.70488293 | 0.83144403 |
| NRROS | 331.178958 | 0.385417438 | 0.3008876 | 1.28093493 | 0.20021652 | 0.38161978 |
| PIGX | 1070.46361 | -0.030105273 | 0.14919089 | -0.2017903 | 0.84008067 | 0.91492195 |
| CEP19 | 202.865315 | 0.04957745 | 0.27883265 | 0.17780361 | 0.85887722 | 0.92636068 |
| PAK2 | 3842.47742 | -0.573352999 | 0.22396871 | -2.5599692 | 0.01046814 | 0.04666075 |
| SENP5 | 1640.52298 | -0.323344573 | 0.13963566 | -2.3156304 | 0.02057846 | 0.07650556 |
| NCBP2 | 3817.82875 | -0.257852764 | 0.14499646 | -1.7783384 | 0.0753483 | 0.19435925 |
| NCBP2-AS1 | 255.476178 | -0.031790549 | 0.19594898 | -0.1622389 | 0.87111772 | 0.93306272 |
| NCBP2-AS2 | 736.224285 | -0.346320483 | 0.19054547 | -1.8175215 | 0.0691373 | 0.18251584 |
| PIGZ | 260.179237 | -0.435951132 | 0.38781599 | -1.1241185 | 0.26096275 | 0.45414028 |
| MELTF | 905.295388 | 1.825054661 | 0.67686504 | 2.69633466 | 0.00701072 | 0.03467229 |
| MELTF-AS1 | 264.287415 | 0.536292983 | 0.49930588 | 1.07407705 | 0.28278815 | 0.47875729 |
| DLG1 | 2749.57792 | 0.213955142 | 0.12117063 | 1.76573427 | 0.07744044 | 0.19802062 |
| DLG1-AS1 | 21.900478 | 1.011368878 | 0.44988874 | 2.24804223 | 0.0245735 | 0.0871089 |
| AC128709.2 | 0.16325399 | -1.003574173 | 3.30558948 | -0.3035992 | 0.7614333 | NA |
| AC128709.1 | 0.41710525 | 1.032177812 | 1.85718639 | 0.55577503 | 0.57836466 | NA |
| AC128709.3 | 0.07830766 | -0.642905668 | 3.30558948 | -0.1944905 | 0.84579185 | NA |
| LINC02012 | 68.8917599 | 0.95989832 | 0.43488149 | 2.20726412 | 0.02729561 | 0.09428678 |
| BDH1 | 2146.30814 | 0.777882268 | 0.29248328 | 2.65957859 | 0.00782385 | 0.03758697 |
| AC024560.3 | 7.75739566 | 0.309880971 | 0.47921851 | 0.64663816 | 0.51786616 | 0.69488619 |
| AC024560.1 | 71.7254402 | -1.759631904 | 1.55253043 | -1.1333961 | 0.25704796 | 0.44976144 |
| RUBCN | 1898.14153 | -0.748032182 | 0.18813938 | -3.9759469 | 7.01E-05 | 0.0009563 |
| FYTTD1 | 2951.26638 | -0.356282148 | 0.15142788 | -2.3528174 | 0.01863178 | 0.07127811 |
| AC055764.2 | 15.726829 | -1.343721606 | 0.44931146 | -2.9906239 | 0.00278408 | 0.01734788 |
| LRCH3 | 1785.9136 | -0.226730269 | 0.15051429 | -1.5063703 | 0.13197213 | 0.28813588 |
| AC055764.1 | 23.416136 | 0.842508121 | 0.43757822 | 1.92538861 | 0.05418076 | 0.15415911 |
| IQCG | 491.018161 | -0.059827902 | 0.19352433 | -0.3091493 | 0.75720799 | 0.8656338 |
| RPL35A | 11335.3378 | -0.407426135 | 0.16370377 | -2.4888011 | 0.01281746 | 0.05416578 |
| LMLN | 621.672334 | -0.138202616 | 0.19762489 | -0.6993179 | 0.4843534 | 0.668712 |
| LMLN-AS1 | 10.3984786 | 0.766747307 | 0.46553216 | 1.64703402 | 0.09955105 | 0.23673562 |
| ZNF595 | 416.96656 | -0.050609998 | 0.19666097 | -0.2573464 | 0.79691135 | 0.88983348 |
| ZNF718 | 296.94907 | 0.187639552 | 0.18486382 | 1.01501499 | 0.31009868 | 0.50718998 |
| ZNF732 | 23.351927 | -0.678728076 | 0.38484219 | -1.763653 | 0.07779042 | 0.19856097 |
| ZNF141 | 497.443586 | 0.062215975 | 0.16964951 | 0.36673242 | 0.71381862 | 0.8374651 |
| AC092574.2 | 1.39049968 | 2.487257232 | 1.27052164 | 1.95766617 | 0.0502692 | 0.14618375 |
| AC092574.1 | 10.2245796 | -0.010817884 | 0.57577581 | -0.0187884 | 0.98500994 | 0.99347171 |
| ZNF721 | 1247.72219 | 0.086936024 | 0.15833448 | 0.54906565 | 0.5829604 | 0.7452691 |
| PIGG | 2907.9246 | -0.256934722 | 0.13589937 | -1.8906249 | 0.05867444 | 0.16287786 |
| AC116565.1 | 0 | NA | NA | NA | NA | NA |
| AC116565.2 | 0.08393208 | -0.642905668 | 3.30558948 | -0.1944905 | 0.84579185 | NA |
| PDE6B | 507.391842 | -0.057550343 | 0.26481003 | -0.2173269 | 0.8279536 | 0.90837491 |
| AC107464.1 | 32.9689738 | 0.623983857 | 0.45236799 | 1.37937226 | 0.16778001 | 0.33895341 |
| AC107464.3 | 83.9208758 | -0.466338657 | 0.46292336 | -1.0073777 | 0.31375332 | 0.51080113 |
| ATP5ME | 4120.05419 | -0.021461196 | 0.14823848 | -0.1447748 | 0.88488868 | 0.94039898 |
| MYL5 | 5024.60501 | -0.009896599 | 0.1138549 | -0.0869229 | 0.93073279 | 0.96662347 |
| SLC49A3 | 1064.20528 | -0.387112733 | 0.24862315 | -1.5570261 | 0.11946428 | 0.26902418 |
| PCGF3 | 3500.22987 | -0.393255681 | 0.23658606 | -1.6622098 | 0.09647071 | 0.2317934 |
| AC107464.2 | 1.49356899 | 0.273633092 | 1.04122661 | 0.26279879 | 0.79270567 | 0.88770378 |
| AC139887.4 | 17.9345778 | 1.125199712 | 0.37676723 | 2.98645853 | 0.00282229 | 0.0175042 |
| AC139887.2 | 488.911755 | 0.091221364 | 0.22665839 | 0.40246188 | 0.68734413 | 0.81936914 |
| AC139887.1 | 168.822398 | -0.627164113 | 0.27805897 | -2.2555076 | 0.02410149 | 0.08588597 |
| CPLX1 | 418.709535 | -0.881783639 | 0.4098494 | -2.1514821 | 0.03143817 | 0.10437171 |
| AC139887.3 | 0.17150785 | 0.18657164 | 3.29377195 | 0.05664376 | 0.95482897 | NA |
| GAK | 9672.96305 | -0.268996641 | 0.17344611 | -1.5508946 | 0.12092694 | 0.27130065 |
| TMEM175 | 1758.07126 | 0.33290298 | 0.24754384 | 1.34482431 | 0.17868198 | 0.3542207 |
| DGKQ | 2182.00957 | -0.53071832 | 0.17780855 | -2.9847739 | 0.00283788 | 0.017571 |
| SLC26A1 | 1827.63066 | 1.029179392 | 0.40911517 | 2.51562268 | 0.01188223 | 0.05119844 |
| IDUA | 2155.70079 | -0.302770598 | 0.1360013 | -2.2262331 | 0.02599857 | 0.09098326 |
| FGFRL1 | 3519.02253 | -0.236220181 | 0.15794709 | -1.4955652 | 0.13476699 | 0.29218265 |
| AC019103.1 | 0.67304256 | -1.601119146 | 2.74353067 | -0.583598 | 0.55949077 | NA |
| RNF212 | 373.956239 | 0.592828145 | 0.30321024 | 1.95517189 | 0.05056279 | 0.14677902 |
| AC092535.3 | 6.88450048 | 1.208158289 | 0.62631138 | 1.9290058 | 0.05373014 | 0.15330706 |
| AC092535.1 | 9.9304992 | 0.845601003 | 0.60509142 | 1.39747645 | 0.16227034 | 0.33124857 |
| SPON2 | 4141.86043 | -1.982028841 | 0.47690084 | -4.1560607 | 3.24E-05 | 0.00052189 |
| AC092535.4 | 368.98612 | -2.357794836 | 0.544884 | -4.3271501 | 1.51E-05 | 0.00028533 |
| CTBP1-AS | 3434.17969 | -0.185942883 | 0.0992728 | -1.8730497 | 0.06106153 | 0.16745495 |
| CTBP1 | 7707.76239 | -0.282583136 | 0.11127274 | -2.5395541 | 0.01109939 | 0.04868444 |
| CTBP1-DT | 715.396353 | 0.043824705 | 0.15388553 | 0.28478769 | 0.77580681 | 0.87701057 |
| MAEA | 5715.13204 | -0.047757984 | 0.10615195 | -0.4499021 | 0.65278105 | 0.79594194 |
| UVSSA | 2022.03364 | 0.381453486 | 0.15756274 | 2.42096249 | 0.01547947 | 0.06226934 |
| NKX1-1 | 0.03392908 | 0.380740593 | 3.29738709 | 0.11546736 | 0.90807469 | NA |
| AC147067.2 | 13.5783849 | -1.086773203 | 0.60648234 | -1.7919288 | 0.07314437 | 0.1902937 |
| AC147067.1 | 22.3623727 | -0.30598355 | 0.39532801 | -0.7739992 | 0.43893129 | 0.63042582 |
| FAM53A | 99.1478547 | 0.105890501 | 0.30646904 | 0.34551778 | 0.72970515 | 0.84734355 |
| SLBP | 2021.0788 | -0.595275863 | 0.14907382 | -3.9931617 | 6.52E-05 | 0.00090919 |
| AC016773.1 | 74.6242788 | -0.673385561 | 0.38198504 | -1.7628585 | 0.07792436 | 0.19871861 |
| TMEM129 | 5439.53008 | 0.184455398 | 0.16705144 | 1.10418323 | 0.26951366 | 0.46378721 |
| TACC3 | 545.439974 | -0.499238835 | 0.30496918 | -1.637014 | 0.10162754 | 0.24022214 |
| FGFR3 | 5963.1535 | 1.543365969 | 0.52933335 | 2.91567867 | 0.00354916 | 0.02078479 |
| LETM1 | 3350.90385 | 0.31319667 | 0.20507455 | 1.52723324 | 0.12670302 | 0.28005156 |
| NSD2 | 1972.06917 | -0.41160019 | 0.13346362 | -3.0839879 | 0.00204246 | 0.01370823 |
| NELFA | 2451.92975 | -0.430663155 | 0.16101388 | -2.6746958 | 0.00747971 | 0.03639112 |
| C4orf48 | 142.018226 | -1.760595706 | 0.4954931 | -3.5532194 | 0.00038055 | 0.00366911 |
| NAT8L | 1542.67426 | 1.735302503 | 0.52413887 | 3.31076857 | 0.0009304 | 0.00743297 |
| POLN | 299.318709 | 0.091432791 | 0.15074918 | 0.60652266 | 0.54416774 | 0.71620188 |
| AL136360.2 | 0.73106255 | -0.321388706 | 1.82828768 | -0.1757867 | 0.8604615 | NA |
| HAUS3 | 664.950665 | -0.317340013 | 0.16198036 | -1.9591265 | 0.05009797 | 0.14583106 |
| MXD4 | 6482.02773 | -0.110108544 | 0.19980267 | -0.5510864 | 0.58157442 | 0.744196 |
| ZFYVE28 | 1525.53622 | 0.043370601 | 0.2490414 | 0.17415017 | 0.86174745 | 0.92800735 |
| AL158068.2 | 3.36119552 | 1.559957086 | 0.84424496 | 1.8477541 | 0.06463792 | 0.17408866 |
| CFAP99 | 103.775853 | -0.024777397 | 0.39177128 | -0.0632445 | 0.94957177 | 0.9757319 |
| RNF4 | 3884.6712 | -0.216746203 | 0.18081954 | -1.1986879 | 0.23064931 | 0.41825211 |
| FAM193A | 1540.29013 | -0.11972986 | 0.1433445 | -0.8352595 | 0.40357159 | 0.59858164 |
| TNIP2 | 3518.71074 | -1.074957282 | 0.17166742 | -6.2618598 | 3.80E-10 | 4.48E-08 |
| SH3BP2 | 9824.79232 | -0.916058167 | 0.20479091 | -4.4731388 | 7.71E-06 | 0.00016566 |
| ADD1 | 12328.8132 | 0.297050641 | 0.10867421 | 2.73340509 | 0.00626832 | 0.03183659 |
| MFSD10 | 5008.61271 | -1.070818922 | 0.169779 | -6.3071342 | 2.84E-10 | 3.54E-08 |
| NOP14-AS1 | 1488.8164 | -0.157923182 | 0.15854486 | -0.9960788 | 0.31921185 | 0.51660835 |
| NOP14 | 1950.4863 | -0.289237154 | 0.19018391 | -1.5208288 | 0.12830281 | 0.28245446 |
| GRK4 | 239.399288 | -0.255370479 | 0.23113193 | -1.1048689 | 0.26921641 | 0.46336641 |
| HTT-AS | 4.16086053 | -0.977975082 | 0.89717051 | -1.090066 | 0.27568406 | 0.47099377 |
| HTT | 3468.4744 | 0.054230529 | 0.09881174 | 0.54882678 | 0.58312433 | 0.7454064 |
| MSANTD1 | 61.2032228 | -0.09903022 | 0.18034583 | -0.5491129 | 0.58292798 | 0.74526378 |
| RGS12 | 2794.55689 | -0.010642052 | 0.16519249 | -0.0644221 | 0.94863411 | 0.97519151 |
| HGFAC | 104.184149 | 0.035245232 | 0.42129636 | 0.083659 | 0.93332756 | 0.96782855 |
| DOK7 | 230.500141 | 0.353199767 | 0.34407827 | 1.02650996 | 0.30465127 | 0.5016331 |
| LRPAP1 | 10485.4166 | 0.265378504 | 0.17886399 | 1.48368885 | 0.13789149 | 0.29668721 |
| AL590235.2 | 0.42353689 | -1.989192329 | 2.79004072 | -0.7129618 | 0.47586942 | NA |
| AL121796.1 | 36.5672311 | 5.00454298 | 0.64818126 | 7.7209004 | 1.16E-14 | 5.70E-12 |
| LINC02171 | 0.34707843 | -1.716757871 | 2.27232659 | -0.7555067 | 0.44994504 | NA |
| AC141928.1 | 352.232993 | 0.585640425 | 0.34844237 | 1.68073825 | 0.09281377 | 0.22549282 |

| | | | | | | |
|---|---|---|---|---|---|---|
| ADRA2C | 1170.67963 | -0.22541064 | 0.31340047 | -0.7192415 | 0.47199211 | 0.65821986 |
| OTOP1 | 0.10721287 | -0.642905668 | 3.30558948 | -0.1944905 | 0.84579185 | NA |
| TMEM128 | 1225.80397 | 0.036537238 | 0.1894376 | 0.19287215 | 0.84705909 | 0.91970801 |
| LYAR | 336.062581 | -0.700924807 | 0.24759512 | -2.8309314 | 0.00464127 | 0.02545024 |
| ZBTB49 | 280.461207 | -0.232610204 | 0.17424396 | -1.3349685 | 0.18188666 | 0.35836455 |
| AC105415.1 | 0.76300575 | -0.287434174 | 1.87542695 | -0.1532633 | 0.87819062 | NA |
| NSG1 | 179.812966 | 0.392934598 | 0.56894312 | 0.69063951 | 0.48979211 | 0.67311398 |
| STX18 | 2844.49587 | -0.027070765 | 0.10672779 | -0.2536431 | 0.79977132 | 0.8913389 |
| STX18-AS1 | 100.082389 | 0.684529604 | 0.30751436 | 2.22600857 | 0.02601361 | 0.09100276 |
| LINC01396 | 0.20754827 | -0.339942428 | 3.12492333 | -0.1087842 | 0.91337362 | NA |
| MSX1 | 128.229872 | -0.438928567 | 0.18294278 | -2.3992669 | 0.01642793 | 0.06502621 |
| CYTL1 | 20.6186577 | 0.883685156 | 0.65046416 | 1.35854549 | 0.17429066 | 0.34824908 |
| STK32B | 1248.3103 | 1.321293655 | 0.38153276 | 3.46311977 | 0.00053395 | 0.00477783 |
| EVC2 | 576.40615 | -0.874668217 | 0.29282531 | -2.9869967 | 0.00281733 | 0.01748892 |
| EVC | 3382.01794 | 0.293207075 | 0.22513918 | 1.30233695 | 0.19280123 | 0.37230289 |
| CRMP1 | 2898.5556 | -0.371829376 | 0.19812146 | -1.8767749 | 0.06054895 | 0.16654594 |
| C4orf50 | 3.88356686 | -1.589904636 | 0.98127372 | -1.6202458 | 0.10517948 | 0.24597224 |
| JAKMIP1 | 23.5445342 | -1.144295635 | 0.95002235 | -1.2044934 | 0.22839894 | 0.41554298 |
| AC092442.1 | 3.17588797 | -0.992672569 | 1.15184705 | -0.8618094 | 0.38879243 | 0.58437501 |
| AC113615.1 | 0 | NA | NA | NA | NA | NA |
| WFS1 | 2192.65074 | -0.070501406 | 0.13774092 | -0.5118407 | 0.60876252 | 0.76476423 |
| PPP2R2C | 32.0146256 | -0.964225742 | 0.40369274 | -2.388514 | 0.01691666 | 0.06645181 |
| MAN2B2 | 3535.50993 | 0.412586529 | 0.28337383 | 1.45597967 | 0.14539824 | 0.30774335 |
| MRFAP1 | 16324.1701 | -0.310115853 | 0.16869532 | -1.8383192 | 0.06601538 | 0.17678741 |
| LINC02482 | 111.541702 | 0.288207091 | 0.29093134 | 0.99063611 | 0.3218633 | 0.51955663 |
| AC093323.1 | 756.129001 | -0.022672594 | 0.20806621 | -0.1089682 | 0.91322774 | 0.95752745 |
| LINC002481 | 22.9122631 | 0.332289515 | 0.46727111 | 0.71112788 | 0.47700499 | 0.66212483 |
| S100P | 37.7181653 | -2.033162328 | 0.4374061 | -4.6482258 | 3.35E-06 | 8.55E-05 |
| MRFAP1L1 | 3430.42702 | -0.072236156 | 0.12821192 | -0.5634122 | 0.57315426 | 0.73777594 |
| BLOC1S4 | 690.434366 | -0.847493405 | 0.24277224 | -3.4908992 | 0.0004814 | 0.00440053 |
| KIAA0232 | 2377.44853 | 0.17438117 | 0.15899568 | 1.09676672 | 0.27274338 | 0.46762862 |
| TBC1D14 | 5533.427 | 0.236517338 | 0.18117168 | 1.30548733 | 0.19172696 | 0.3709347 |
| AC097382.2 | 4693.06035 | 0.058233626 | 0.17058686 | 0.34137229 | 0.73282334 | 0.84935616 |
| CCDC96 | 169.077141 | -0.922021245 | 0.29087956 | -3.1697698 | 0.0015256 | 0.01096783 |
| TADA2B | 2257.88333 | -0.20473841 | 0.18272572 | -1.1204685 | 0.26251417 | 0.45611772 |
| GRPEL1 | 3721.85643 | -0.622288929 | 0.29693197 | -2.095729 | 0.03610624 | 0.11510173 |
| LINC02447 | 69.8854933 | 1.562510765 | 0.39903918 | 3.91568255 | 9.01E-05 | 0.001167 |
| SORCS2 | 1269.34029 | 0.418851928 | 0.44070442 | 0.95041462 | 0.34190162 | 0.53971539 |
| PSAPL1 | 0.32549373 | -1.692331154 | 2.2973018 | -0.7366604 | 0.46132892 | NA |
| AFAP1-AS1 | 32.889432 | -0.173983026 | 0.34036486 | -0.5111662 | 0.60923466 | 0.76527975 |
| AFAP1 | 1056.28132 | -0.39386605 | 0.16851267 | -2.3373083 | 0.01942316 | 0.07332733 |
| AC097381.1 | 6.96143986 | 0.265442574 | 0.55551693 | 0.47782986 | 0.63277131 | 0.7819936 |
| ABLIM2 | 1020.44709 | -1.599129502 | 0.36252513 | -4.411086 | 1.03E-05 | 0.00020863 |
| AC097381.3 | 0.07659028 | -0.642905668 | 3.30558948 | -0.1944905 | 0.84579185 | NA |
| AC097381.2 | 0.09989236 | -0.642905668 | 3.30558948 | -0.1944905 | 0.84579185 | NA |
| SH3TC1 | 5660.81366 | -0.461138094 | 0.17909371 | -2.5748425 | 0.01002857 | 0.04516467 |
| HTRA3 | 364.981189 | -0.119360743 | 0.45452986 | -0.2626026 | 0.79285686 | 0.8878351 |
| LINC02517 | 0.39020491 | 0.70643347 | 2.28377771 | 0.30932672 | 0.75707301 | NA |
| AC104825.1 | 522.316402 | 0.298834181 | 0.15072149 | 1.98269132 | 0.04740192 | 0.14005475 |
| ACOX3 | 908.417319 | -0.037312798 | 0.2039834 | -0.1829208 | 0.8548602 | 0.92401312 |
| TRMT44 | 458.892835 | 0.172438416 | 0.17053382 | 1.01116843 | 0.31193582 | 0.50898754 |
| AC105345.2 | 35.4110832 | 0.229438207 | 0.30674211 | 0.74798405 | 0.45446978 | 0.64400924 |
| AC105345.1 | 148.556986 | 0.047604761 | 0.29694693 | 0.16031403 | 0.8726337 | 0.93366644 |
| GPR78 | 231.091248 | -0.713491099 | 0.46506614 | -1.5341713 | 0.12498753 | 0.27734106 |
| CPZ | 231.673393 | -0.723580329 | 0.46863491 | -1.5440171 | 0.12258418 | 0.27375888 |
| AC209005.1 | 0 | NA | NA | NA | NA | NA |
| HMX1 | 1.16473453 | -1.876319972 | 1.87771465 | -0.9992572 | 0.31767009 | 0.51490321 |
| FAM90A26 | 1.07682944 | 0.346405587 | 1.43156306 | 0.24197718 | 0.80879785 | 0.89660916 |
| USP17L10 | 0 | NA | NA | NA | NA | NA |
| USP17L11 | 0 | NA | NA | NA | NA | NA |
| USP17L12 | 0 | NA | NA | NA | NA | NA |
| USP17L13 | 0 | NA | NA | NA | NA | NA |
| USP17L15 | 0 | NA | NA | NA | NA | NA |
| USP17L17 | 0 | NA | NA | NA | NA | NA |
| USP17L18 | 0 | NA | NA | NA | NA | NA |
| USP17L19 | 0 | NA | NA | NA | NA | NA |
| USP17L20 | 0 | NA | NA | NA | NA | NA |
| USP17L21 | 0 | NA | NA | NA | NA | NA |
| USP17L22 | 0 | NA | NA | NA | NA | NA |
| USP17L23 | 0 | NA | NA | NA | NA | NA |
| USP17L24 | 0 | NA | NA | NA | NA | NA |
| USP17L25 | 0 | NA | NA | NA | NA | NA |
| USP17L26 | 0 | NA | NA | NA | NA | NA |
| USP17L5 | 0 | NA | NA | NA | NA | NA |
| USP17L27 | 0 | NA | NA | NA | NA | NA |
| USP17L28 | 0 | NA | NA | NA | NA | NA |
| USP17L29 | 0 | NA | NA | NA | NA | NA |
| USP17L30 | 0 | NA | NA | NA | NA | NA |
| DEFB131A | 0 | NA | NA | NA | NA | NA |
| SLC2A9 | 3157.59137 | 1.369608646 | 0.37923809 | 3.61147434 | 0.00030446 | 0.00307023 |
| DRD5 | 1.25459319 | 2.189940128 | 1.65039541 | 1.32691845 | 0.18453566 | 0.36171897 |
| AC108199.1 | 0.289369 | 0.328493799 | 2.93027205 | 0.11210352 | 0.91074133 | NA |
| AC005674.1 | 10.2767093 | 1.225709479 | 0.43049229 | 2.8472275 | 0.00441018 | 0.024522 |
| WDR1 | 18612.7549 | -0.2436369 | 0.15968198 | -1.5257632 | 0.12706884 | 0.28057117 |
| ZNF518B | 624.844754 | -0.28740153 | 0.22977071 | -1.2508188 | 0.21100059 | 0.39438912 |
| CLNK | 419.457241 | 0.343815994 | 0.5517071 | 0.62318574 | 0.53316247 | 0.70700465 |
| AC084048.1 | 0.57541034 | 1.857436751 | 1.72295962 | 1.07805008 | 0.28101141 | NA |
| LINC02498 | 0.19114494 | -1.003574173 | 3.30558948 | -0.3035992 | 0.7614333 | NA |
| HS3ST1 | 2026.6466 | -2.258208224 | 0.48450242 | -4.6608813 | 3.15E-06 | 8.12E-05 |
| AC025539.1 | 10.1641132 | -1.0149447 | 0.73217834 | -1.3861987 | 0.16568622 | 0.33619158 |
| AC005699.1 | 95.4349591 | 0.079130989 | 0.58023455 | 0.13637759 | 0.89152279 | 0.94377583 |
| LINC02360 | 7.25479133 | -0.526526923 | 0.96405658 | -0.5461577 | 0.58495754 | 0.74673623 |
| AC108037.1 | 0 | NA | NA | NA | NA | NA |
| LINC02270 | 0.08666371 | -0.642905668 | 3.30558948 | -0.1944905 | 0.84579185 | NA |
| AC007370.2 | 1.07658507 | -0.061658066 | 1.14219359 | -0.0539821 | 0.95694939 | 0.97944167 |
| AC007370.1 | 0.08666371 | -0.642905668 | 3.30558948 | -0.1944905 | 0.84579185 | NA |
| RAB28 | 1002.63072 | -0.073012913 | 0.11933549 | -0.611829 | 0.54065089 | 0.7136332 |
| LINC01097 | 0.08757577 | 0.547239394 | 3.29377202 | 0.16614368 | 0.86804388 | NA |
| NKX3-2 | 5.49224075 | 1.044906829 | 0.7347461 | 1.4221332 | 0.15498759 | 0.32113009 |
| LINC01096 | 0.39127983 | -0.032129707 | 2.24976281 | -0.0142814 | 0.9886055 | NA |
| BOD1L1 | 1847.54099 | -0.148868192 | 0.24285222 | -0.6129991 | 0.53987694 | 0.71287615 |
| LINC01182 | 6.88852906 | 0.760098265 | 0.5073331 | 1.49822329 | 0.13407525 | 0.29118467 |
| AC007126.1 | 3.08233279 | -1.298501585 | 1.02306563 | -1.2692261 | 0.20436044 | 0.38666749 |
| AC073848.1 | 3.45729977 | 0.971879733 | 0.85671349 | 1.13442796 | 0.25661508 | 0.44936509 |
| AC092546.1 | 1.53850033 | 2.3594155 | 1.11346275 | 2.11898916 | 0.03409138 | 0.11050249 |
| AC006296.2 | 0.08666371 | -0.642905668 | 3.30558948 | -0.1944905 | 0.84579185 | NA |
| AC006296.1 | 0 | NA | NA | NA | NA | NA |
| LINC00504 | 15.6189913 | 1.432237844 | 0.50634755 | 2.82856677 | 0.00467569 | 0.0255699 |
| CPEB2-DT | 17.8637707 | -0.054639612 | 0.37759723 | -0.1447034 | 0.88494504 | 0.94039898 |
| CPEB2 | 2238.91854 | -0.291364386 | 0.24231592 | -1.2024154 | 0.22920264 | 0.41665947 |
| AC098829.1 | 87.3706525 | -0.283450731 | 0.29984388 | -0.9453277 | 0.3444916 | 0.54210975 |
| C1QTNF7 | 169.044513 | 3.120300425 | 0.51523164 | 6.05611183 | 1.39E-09 | 1.35E-07 |
| AC099550.1 | 2.53691593 | 2.278442998 | 0.862469 | 2.641768 | 0.00824745 | 0.03895492 |

| | | | | | | | |
|---|---|---|---|---|---|---|---|
| CC2D2A | 1208.8284 | | 1.007944345 | 0.26112034 | 3.86007594 | 0.00011335 | 0.00139738 |
| AC116651.1 | 13.3935872 | | 0.214857139 | 0.44511936 | 0.48269556 | 0.62931192 | 0.77980148 |
| FBXL5 | 7772.8815 | | 0.91551662 | 0.16134712 | 5.67420487 | 1.39E-08 | 9.21E-07 |
| FAM200B | 1535.2581 | | 0.394174961 | 0.13137858 | 3.00029853 | 0.00269715 | 0.01694636 |
| BST1 | 417.886315 | | 1.267858801 | 0.26381608 | 4.8058435 | 1.54E-06 | 4.62E-05 |
| CD38 | 331.784321 | | -1.581789345 | 0.6184799 | -2.5575437 | 0.01054143 | 0.04684489 |
| FGFBP1 | 21.1753847 | | -1.013071982 | 0.85329557 | -1.1872463 | 0.2351305 | 0.42393108 |
| FGFBP2 | 710.252623 | | -0.773743879 | 0.41144375 | -1.8805581 | 0.06003205 | 0.16556445 |
| PROM1 | 5318.9857 | | -0.652540781 | 0.41260349 | -1.5815203 | 0.11375913 | 0.26005644 |
| TAPT1 | 2048.98815 | | 0.319017849 | 0.1275325 | 2.50146306 | 0.01236813 | 0.05278325 |
| AC108063.1 | 4.7214368 | | 0.670043121 | 0.71782269 | 0.9334382 | 0.35059377 | 0.54831609 |
| TAPT1-AS1 | 132.276553 | | 0.285205624 | 0.41048487 | 0.6948018 | 0.48717952 | 0.67117308 |
| AC097515.1 | 276.755134 | | 0.558004378 | 0.36540262 | 1.52709461 | 0.12673749 | 0.28009282 |
| LDB2 | 2032.9685 | | 0.160287869 | 0.38732499 | 0.41383302 | 0.6789964 | 0.8137958 |
| AC106894.1 | 0 | NA | NA | NA | NA | NA | |
| LINC02493 | 0 | NA | NA | NA | NA | NA | |
| QDPR | 4058.73238 | | 1.291261891 | 0.32437351 | 3.98078709 | 6.87E-05 | 0.00094436 |
| CLRN2 | 1.21163378 | | 3.703575303 | 1.38224489 | 2.67939156 | 0.00737561 | 0.03599672 |
| LAP3 | 9077.64146 | | -0.346174271 | 0.47183649 | -0.7336742 | 0.46314731 | 0.65105787 |
| AC006160.1 | 15.9656481 | | -1.013945927 | 0.6495486 | -1.5610009 | 0.11852355 | 0.26761418 |
| MED28 | 2987.3215 | | -0.183351243 | 0.13788639 | -1.3297269 | 0.18360826 | 0.36062489 |
| FAM184B | 901.321404 | | 0.213300214 | 0.11672717 | 1.82733992 | 0.06764868 | 0.17972676 |
| DCAF16 | 1346.34607 | | 0.09229558 | 0.18513956 | 0.49851895 | 0.61811832 | 0.77169681 |
| NCAPG | 202.297697 | | -0.58702727 | 0.17118188 | -3.4292606 | 0.00060523 | 0.00525341 |
| LCORL | 602.403012 | | -0.194561444 | 0.1192084 | -1.6321118 | 0.10265593 | 0.24176097 |
| AC093871.1 | 0.07659028 | | -0.642905668 | 3.30558948 | -0.1944905 | 0.84579185 | NA |
| LINC02438 | 0 | NA | NA | NA | NA | NA | |
| AC024230.1 | 0.2891581 | | 1.145957498 | 3.27645355 | 0.34975545 | 0.72652223 | NA |
| AC110767.1 | 0.09997307 | | 0.547239394 | 3.29377202 | 0.16614368 | 0.86804388 | NA |
| SLIT2 | 1191.9116 | | 1.482676019 | 0.43513123 | 3.40742266 | 0.00065579 | 0.00559111 |
| PACRGL | 573.798048 | | -0.118085536 | 0.13656324 | -0.8646949 | 0.38720626 | 0.58279846 |
| KCNIP4 | 855.961582 | | -0.997785465 | 0.38268058 | -2.6073585 | 0.00912438 | 0.04207936 |
| AC097505.1 | 7.91781886 | | -0.129434572 | 0.5039064 | -0.2568623 | 0.79728505 | 0.88999732 |
| AC110296.1 | 0.50851525 | | -1.153215071 | 2.2433008 | -0.5140706 | 0.60720261 | NA |
| AC096576.3 | 0 | NA | NA | NA | NA | NA | |
| AC096576.2 | 0 | NA | NA | NA | NA | NA | |
| KCNIP4-IT1 | 0 | NA | NA | NA | NA | NA | |
| AC096719.1 | 2.73310218 | | -0.835059379 | 0.98407845 | -0.8485699 | 0.39612065 | 0.5912411 |
| ADGRA3 | 1562.80339 | | 0.330661697 | 0.14874729 | 2.22297631 | 0.0262174 | 0.09152141 |
| AC097512.1 | 0.91759139 | | 2.23749758 | 1.70714113 | 1.31066936 | 0.18996949 | NA |
| AC114912.1 | 0.34069606 | | 0.670043202 | 2.22420583 | 0.30125054 | 0.76322345 | NA |
| AC092440.1 | 0 | NA | NA | NA | NA | NA | |
| AC093607.1 | 9.7869904 | | 0.600962924 | 0.36407337 | 1.65066435 | 0.09880713 | 0.23564656 |
| ERVH-1 | 81.854528 | | -2.196133915 | 0.82244867 | -2.6702383 | 0.00757974 | 0.03672875 |
| PPARGC1A | 3335.4736 | | 0.345499402 | 0.27420386 | 1.26000928 | 0.20766602 | 0.39062364 |
| AC092834.1 | 10.5591671 | | 0.19806683 | 0.60770322 | 0.3259269 | 0.74447966 | 0.85673198 |
| DHX15 | 5801.03661 | | -0.284386456 | 0.19316818 | -1.472222 | 0.14096093 | 0.30129785 |
| LINC02473 | 2.13487152 | | 0.407858505 | 1.2331383 | 0.33074839 | 0.74083455 | 0.85453728 |
| SOD3 | 6522.46626 | | -0.694296031 | 0.53597344 | -1.2953926 | 0.19518483 | 0.37531165 |
| CCDC149 | 832.710267 | | -0.345666428 | 0.21603658 | -1.6000366 | 0.10959047 | 0.2532779 |
| LGI2 | 467.311059 | | 0.958797189 | 0.5291963 | 1.81179874 | 0.0700173 | 0.18426709 |
| SEPSECS | 1505.42104 | | 0.489157101 | 0.27751516 | 1.76263203 | 0.07796257 | 0.19873108 |
| SEPSECS-AS1 | 92.2906165 | | 0.849703255 | 0.29416948 | 2.88848201 | 0.00387106 | 0.0222251 |
| AC104662.2 | 339.789469 | | 0.294165962 | 0.33918274 | 0.86727869 | 0.38578933 | 0.58152879 |
| AC104662.1 | 3.44192326 | | 1.013375444 | 1.19495744 | 0.84804313 | 0.39641395 | 0.59135803 |
| PI4K2B | 1245.95069 | | -0.346496817 | 0.26882698 | -1.2889213 | 0.19742546 | 0.3782685 |
| ZCCHC4 | 410.958354 | | -0.027903926 | 0.16851702 | -0.1655852 | 0.86848339 | 0.93156219 |
| ANAPC4 | 2368.48814 | | -0.712504704 | 0.24174503 | -2.9473396 | 0.00320521 | 0.01932801 |
| AC092436.2 | 3.79881257 | | -2.217045227 | 0.93911429 | -2.3607832 | 0.01823639 | 0.07012155 |
| SLC34A2 | 18946.5758 | | -4.154769228 | 0.49215549 | -8.4419849 | 3.12E-17 | 3.16E-14 |
| SEL1L3 | 3426.17735 | | -0.877934881 | 0.2350405 | -3.7352494 | 0.00018753 | 0.0020717 |
| AC092436.4 | 0.28759251 | | -1.364245045 | 2.90093914 | -0.470277 | 0.63815711 | NA |
| SMIM20 | 1407.19761 | | 0.0087258 | 0.1923692 | 0.04535965 | 0.96382064 | 0.98305633 |
| AC133961.1 | 3.90863106 | | -3.068038002 | 1.48455213 | -2.0666422 | 0.03876789 | 0.1210263 |
| LINC02357 | 0.09997307 | | 0.547239394 | 3.29377202 | 0.16614368 | 0.86804388 | NA |
| RBPJ | 2962.15958 | | -0.300049335 | 0.09965872 | -3.0107685 | 0.00260587 | 0.01650262 |
| CCKAR | 1.87202305 | | 1.250279018 | 1.05702321 | 1.18283025 | 0.23687643 | 0.4261158 |
| TBC1D19 | 591.454603 | | 0.535913828 | 0.27308853 | 1.96241794 | 0.04971385 | 0.14511336 |
| STIM2 | 1768.57348 | | -0.661789864 | 0.25882157 | -2.5569347 | 0.0105599 | 0.04688746 |
| AC106047.1 | 16.3044366 | | 0.685196978 | 0.6256714 | 1.09513872 | 0.27345587 | 0.46843129 |
| AC024132.2 | 7.59549042 | | -0.774716127 | 0.6239073 | -1.2417167 | 0.2143411 | 0.3984694 |
| AC024132.1 | 0 | NA | NA | NA | NA | NA | |
| LINC02261 | 0.6169507 | | -0.408347199 | 1.69289752 | -0.241212 | 0.80939081 | NA |
| AC024132.3 | 0 | NA | NA | NA | NA | NA | |
| AC007106.1 | 0 | NA | NA | NA | NA | NA | |
| AC109588.1 | 0 | NA | NA | NA | NA | NA | |
| AC097480.1 | 0.16416605 | | 0.18657164 | 3.29377195 | 0.05664376 | 0.95482897 | NA |
| LINC02364 | 0 | NA | NA | NA | NA | NA | |
| AC068944.1 | 0.09635527 | | 0.547239394 | 3.29377202 | 0.16614368 | 0.86804388 | NA |
| AC109349.1 | 0 | NA | NA | NA | NA | NA | |
| LINC02472 | 0 | NA | NA | NA | NA | NA | |
| PCDH7 | 205.259867 | | 0.20936301 | 0.36982932 | 0.56610712 | 0.57132096 | 0.73631464 |
| AC097716.1 | 0.38516334 | | 0.969430309 | 2.1710357 | 0.44652896 | 0.6552152 | NA |
| LINC02497 | 0.03392908 | | 0.380740593 | 3.29738709 | 0.11546736 | 0.90807469 | NA |
| AC104071.1 | 0 | NA | NA | NA | NA | NA | |
| LINC02501 | 0 | NA | NA | NA | NA | NA | |
| LINC02506 | 4.91095214 | | -0.389191722 | 2.03316646 | -0.1914215 | 0.8481954 | 0.92016931 |
| LINC02353 | 0 | NA | NA | NA | NA | NA | |
| AC097535.1 | 0 | NA | NA | NA | NA | NA | |
| AC016687.3 | 0.62778896 | | 2.708283985 | 1.52297594 | 1.77828415 | 0.0753572 | NA |
| AC016687.2 | 0 | NA | NA | NA | NA | NA | |
| LINC02484 | 2.12491439 | | 2.929223555 | 1.03042329 | 2.84273812 | 0.00447278 | 0.02477362 |
| AC093689.1 | 0.43331855 | | -2.014555905 | 3.29154549 | -0.6120395 | 0.54051161 | NA |
| ARAP2 | 813.898306 | | -0.509222971 | 0.21711987 | -2.345354 | 0.01900902 | 0.07230172 |
| AC104078.1 | 3.25719578 | | 0.038835302 | 0.7508831 | 0.0517195 | 0.9587522 | 0.98017606 |
| DTHD1 | 13.8664855 | | -0.576235685 | 0.74518335 | -0.7732804 | 0.43935645 | 0.63084261 |
| AC104078.2 | 0 | NA | NA | NA | NA | NA | |
| LINC02505 | 0.99803134 | | 1.447592979 | 1.29436009 | 1.11838505 | 0.26340257 | NA |
| AL136537.1 | 0 | NA | NA | NA | NA | NA | |
| NWD2 | 1.64633506 | | -1.415420136 | 1.33389455 | -1.0611185 | 0.28863607 | 0.48482917 |
| C4orf19 | 1391.79895 | | -1.190271837 | 0.32520024 | -3.6601198 | 0.0002521 | 0.00263892 |
| AC027607.1 | 105.696884 | | -0.730323652 | 0.32782241 | -2.2278027 | 0.02589368 | 0.09065537 |
| RELL1 | 663.314609 | | -0.932009435 | 0.29432436 | -3.1666065 | 0.00154229 | 0.01104267 |
| PGM2 | 1447.50612 | | -0.714499867 | 0.16871486 | -4.2349552 | 2.29E-05 | 0.00039616 |
| TBC1D1 | 8937.78794 | | -0.304294884 | 0.19998725 | -1.5215715 | 0.1281165 | 0.28219445 |
| PTTG2 | 7.7955177 | | -1.331340495 | 0.77244471 | -1.7235415 | 0.08479065 | 0.21098208 |
| AC098680.1 | 0 | NA | NA | NA | NA | NA | |
| LINC02513 | 0.17423948 | | 0.18657164 | 3.29377195 | 0.05664376 | 0.95482897 | NA |
| LINC01258 | 0.11786115 | | 0.02007176 | 3.29738704 | 0.00608717 | 0.99514317 | NA |
| LINC01259 | 0.43567286 | | 2.075329418 | 2.73531287 | 0.75871738 | 0.44802164 | NA |
| LINC02278 | 0.08757577 | | 0.547239394 | 3.29377202 | 0.16614368 | 0.86804388 | NA |
| AC021860.1 | 0 | NA | NA | NA | NA | NA | |
| KLF3 | 2903.14413 | | -0.44177365 | 0.17414426 | -2.5368258 | 0.01118626 | 0.04897587 |

| | | | | | | |
|---|---|---|---|---|---|---|
| TLR10 | 37.6512731 | 1.181132017 | 0.79034461 | 1.49445192 | 0.13505754 | 0.29242691 |
| TLR1 | 293.237063 | -0.402607444 | 0.35939333 | -1.1202418 | 0.26261071 | 0.4561951 |
| TLR6 | 71.7769307 | -0.642848267 | 0.38824249 | -1.6557906 | 0.09776424 | 0.23394219 |
| FAM114A1 | 1451.28772 | -0.610496517 | 0.21748009 | -2.8071375 | 0.00499839 | 0.02687208 |
| TMEM156 | 43.8684534 | 0.157395243 | 0.51191618 | 0.30746292 | 0.75849104 | 0.86635037 |
| KLHL5 | 1447.21472 | -0.114680769 | 0.1583778 | -0.7240962 | 0.46900665 | 0.65569182 |
| AC079921.1 | 288.751743 | -0.110645639 | 0.18349877 | -0.6029775 | 0.54652363 | 0.71768478 |
| AC079921.2 | 16.402826 | -0.308891476 | 0.42287702 | -0.7304523 | 0.46511378 | 0.65218816 |
| WDR19 | 2028.08557 | -0.386425107 | 0.18823183 | -2.0529212 | 0.04008022 | 0.1239803 |
| RFC1 | 1964.53381 | 0.363502937 | 0.12416061 | 2.92768321 | 0.00341498 | 0.02023208 |
| KLB | 38.8921457 | 0.688893233 | 0.59524225 | 1.15733256 | 0.24713652 | 0.43864777 |
| RPL9 | 22846.6172 | -0.453130265 | 0.17991992 | -2.5185108 | 0.01178523 | 0.05090616 |
| LIAS | 4234.57571 | -0.041038366 | 0.16177465 | -0.2536761 | 0.79974578 | 0.8913389 |
| UGDH | 3828.85005 | -0.823346141 | 0.40837925 | -2.0161312 | 0.04378626 | 0.13245884 |
| UGDH-AS1 | 187.183501 | 1.14679338 | 0.2789452 | 4.11117811 | 3.94E-05 | 0.00061605 |
| SMIM14 | 3069.76362 | 0.139959298 | 0.1640216 | 0.85329796 | 0.39349409 | 0.5889012 |
| AC108471.2 | 55.4228029 | 0.722224088 | 0.42004285 | 1.71940573 | 0.08554052 | 0.21232149 |
| UBE2K | 4008.18802 | -0.309636835 | 0.18584624 | -1.6660915 | 0.09569519 | 0.23057597 |
| PDS5A | 3748.27319 | -0.158275157 | 0.13087345 | -1.2093756 | 0.22651858 | 0.4133196 |
| N4BP2 | 321.286334 | -0.132222435 | 0.24235707 | -0.5455687 | 0.58536243 | 0.74695087 |
| RHOH | 122.936174 | -0.562500884 | 0.70038483 | -0.8031312 | 0.42189892 | 0.61446342 |
| AC095057.2 | 0.08393208 | -0.642905668 | 3.30558948 | -0.1944905 | 0.84579185 | NA |
| LINC02265 | 0.63376154 | 1.349220238 | 1.71818868 | 0.78525732 | 0.43230271 | NA |
| CHRNA9 | 1.29572117 | -0.015341926 | 0.99765594 | -0.015378 | 0.98773064 | 0.99451916 |
| RBM47 | 8451.93413 | 0.465904611 | 0.19567686 | 2.3809898 | 0.01726619 | 0.06738337 |
| AC098869.2 | 229.999395 | 1.465596122 | 0.33742867 | 4.34342503 | 1.40E-05 | 0.00026844 |
| NSUN7 | 672.568367 | 0.048678607 | 0.23453941 | 0.20754979 | 0.83558051 | 0.9127696 |
| APBB2 | 2266.79616 | 0.271162748 | 0.12622532 | 2.14824376 | 0.0316944 | 0.10494491 |
| AC131953.1 | 218.195625 | 0.164258998 | 0.14971013 | 1.09718023 | 0.27256262 | 0.46756936 |
| UCHL1-AS1 | 1.34904173 | 0.505930601 | 1.34698225 | 0.37560302 | 0.70721205 | 0.83325953 |
| UCHL1 | 5068.11206 | -0.92865381 | 0.33837113 | -2.744483 | 0.00606063 | 0.03101471 |
| LIMCH1 | 2790.09203 | 0.463600977 | 0.20407063 | 2.2717673 | 0.02310057 | 0.08348184 |
| AC105389.2 | 2.18874231 | 0.00344705 | 0.78947885 | 0.00436624 | 0.99651626 | 0.99824844 |
| PHOX2B | 0.24118557 | 0.074529887 | 3.28898342 | 0.02266046 | 0.98192111 | NA |
| AC105389.3 | 0.03392908 | 0.380740593 | 3.29738709 | 0.11546736 | 0.90807469 | NA |
| AC105389.1 | 0 | NA | NA | NA | NA | NA |
| LINC00682 | 0.2441911 | -0.174096114 | 3.29377188 | -0.0528562 | 0.95784652 | NA |
| AC108210.1 | 0 | NA | NA | NA | NA | NA |
| TMEM33 | 2993.39636 | -0.022324602 | 0.16420753 | -0.1359536 | 0.89185798 | 0.94397908 |
| DCAF4L1 | 119.676511 | -1.63237334 | 0.6114607 | -2.6696292 | 0.00759351 | 0.03677489 |
| AC106052.1 | 15.1079214 | 0.729036278 | 0.50563836 | 1.44181364 | 0.14935495 | 0.31347422 |
| SLC30A9 | 5238.43695 | 0.74660719 | 0.17178586 | 4.34615056 | 1.39E-05 | 0.00026571 |
| BEND4 | 9.93391048 | 0.109480425 | 0.53688783 | 0.20391676 | 0.83841855 | 0.91449053 |
| AC111006.1 | 0.65893903 | 2.223134004 | 1.61442537 | 1.37704353 | 0.16849879 | NA |
| AC024022.1 | 31.020014 | 1.171359069 | 0.77949828 | 1.50270899 | 0.1329141 | 0.28937788 |
| SHISA3 | 1520.46509 | 0.345224622 | 0.32326863 | 1.06791873 | 0.28555717 | 0.48163515 |
| ATP8A1 | 673.068709 | 0.479931036 | 0.22317622 | 2.15045777 | 0.03151902 | 0.10456282 |
| AC096734.2 | 1.1932704 | 0.715482018 | 1.05741174 | 0.67663521 | 0.49863743 | 0.6799953 |
| AC096734.1 | 2.30543639 | 0.707718314 | 1.3299026 | 0.53215801 | 0.59461657 | 0.75370069 |
| GRXCR1 | 0 | NA | NA | NA | NA | NA |
| AC111198.1 | 0 | NA | NA | NA | NA | NA |
| LINC02383 | 0 | NA | NA | NA | NA | NA |
| AC110799.1 | 0.7542462 | 1.876462955 | 2.13133903 | 0.88041505 | 0.37863451 | NA |
| LINC02475 | 0.97306469 | 0.145053895 | 1.30074213 | 0.11151626 | 0.91120697 | NA |
| KCTD8 | 75.6685194 | -0.134953168 | 0.19844178 | -0.6800643 | 0.49646376 | 0.67832899 |
| YIPF7 | 15.3625813 | 0.93882434 | 0.31256771 | 3.00358706 | 0.00266817 | 0.01680031 |
| GUF1 | 1319.55615 | 0.45829671 | 0.13438791 | 3.41025254 | 0.00064903 | 0.00554956 |
| GNPDA2 | 1095.08509 | -0.010018527 | 0.15442172 | -0.0648777 | 0.94827137 | 0.97510358 |
| AC096586.2 | 32.283027 | -0.128343767 | 0.22652658 | -0.5665727 | 0.57100455 | 0.73605074 |
| AC096586.1 | 56.0456664 | 0.238849472 | 0.30189619 | 0.79116425 | 0.42884815 | 0.62111967 |
| AC108467.1 | 0.10721287 | -0.642905668 | 3.30558948 | -0.1944905 | 0.84579185 | NA |
| GABRG1 | 4.21496259 | 2.909008932 | 1.05974979 | 2.74499599 | 0.00605116 | 0.03098278 |
| GABRA2 | 308.59896 | 0.480118793 | 0.37270903 | 1.28818663 | 0.19768101 | 0.37843313 |
| AC095060.1 | 3.29837002 | 0.606258328 | 0.78847406 | 0.76890079 | 0.44195221 | 0.63319617 |
| COX7B2 | 0.08666371 | -0.642905668 | 3.30558948 | -0.1944905 | 0.84579185 | NA |
| GABRA4 | 9.79597557 | 2.971607038 | 0.70071712 | 4.24080836 | 2.23E-05 | 0.00038737 |
| GABRB1 | 7.24431743 | 0.555359528 | 0.4844609 | 1.14634542 | 0.25165226 | 0.44391398 |
| COMMD8 | 1472.49113 | 0.496205996 | 0.18632169 | 2.66316813 | 0.00774087 | 0.03729025 |
| ATP10D | 1797.74328 | 0.212221628 | 0.22345508 | 0.94972836 | 0.3422503 | 0.54006337 |
| AC107398.2 | 43.7541308 | 0.70344056 | 0.2829382 | 2.48619863 | 0.01291159 | 0.05444173 |
| CORIN | 101.613087 | 0.895556438 | 0.37565354 | 2.38399571 | 0.0171258 | 0.06700406 |
| AC107068.2 | 65.789556 | 0.464800102 | 0.21926182 | 2.11984053 | 0.0340195 | 0.11038441 |
| AC107068.1 | 166.358859 | 0.209475556 | 0.22548803 | 0.92898746 | 0.35289559 | 0.5505121 |
| NFXL1 | 502.399252 | -0.085287454 | 0.14324308 | -0.5954036 | 0.55157369 | 0.72162126 |
| NIPAL1 | 268.853515 | 0.484688046 | 0.32235409 | 1.50358892 | 0.13268725 | 0.28905244 |
| CNGA1 | 233.35041 | -0.06664189 | 0.52356189 | -0.1272856 | 0.89871436 | 0.94852624 |
| TXK | 76.409359 | 0.101415336 | 0.21979079 | 0.46141758 | 0.64449904 | 0.78997117 |
| TEC | 260.812758 | -0.954651517 | 0.21523378 | -4.4354167 | 9.19E-06 | 0.00019078 |
| SLAIN2 | 2168.29463 | 0.039743296 | 0.149875 | 0.26517629 | 0.79087366 | 0.88640506 |
| SLC10A4 | 10.6439503 | -0.252368874 | 0.54270749 | -0.4650182 | 0.64191841 | 0.7878334 |
| ZAR1 | 0.64403822 | -0.830732385 | 1.66985061 | -0.497489 | 0.61884423 | NA |
| FRYL | 1666.18824 | -0.139471448 | 0.1102091 | -1.2655167 | 0.20568617 | 0.38823862 |
| OCIAD1 | 8012.24945 | -0.00262339 | 0.08663673 | -0.0302803 | 0.97584348 | 0.98881307 |
| OCIAD1-AS1 | 66.68513 | 0.179156145 | 0.21184603 | 0.84569034 | 0.39772551 | 0.59261056 |
| OCIAD2 | 5526.12742 | 0.200537859 | 0.36695256 | 0.54649533 | 0.5847255 | 0.74662073 |
| CWH43 | 397.730363 | 1.678523699 | 0.35396413 | 4.74207282 | 2.12E-06 | 5.94E-05 |
| DCUN1D4 | 2648.51668 | -0.291884901 | 0.15603475 | -1.8706404 | 0.06139494 | 0.16804905 |
| AC027271.1 | 23.1476638 | -0.834644125 | 0.29098725 | -2.8683185 | 0.0041266 | 0.02330159 |
| LRRC66 | 167.582275 | 0.186499548 | 0.40379294 | 0.46186927 | 0.64417507 | 0.78975762 |
| SGCB | 3573.01883 | 0.385632712 | 0.14968449 | 2.57630378 | 0.00998629 | 0.0450228 |
| LINC02480 | 0.14444843 | 0.435199075 | 3.28898345 | 0.13232024 | 0.894731 | NA |
| SPATA18 | 690.17487 | 0.424046403 | 0.34525336 | 1.22821802 | 0.21936513 | 0.40494635 |
| USP46 | 1109.21502 | 0.275649098 | 0.23753721 | 1.16044598 | 0.24586727 | 0.43713053 |
| USP46-AS1 | 96.217056 | 0.826963608 | 0.25518384 | 3.2406582 | 0.00119254 | 0.00900811 |
| AC104066.2 | 0.93690846 | -0.728933087 | 1.35896561 | -0.5363882 | 0.5916903 | NA |
| ERVMER34-1 | 232.19792 | 0.535396812 | 0.39639587 | 1.35066194 | 0.17680375 | 0.35174009 |
| LINC01618 | 0.18402333 | 0.18657164 | 3.29377195 | 0.05664376 | 0.95482897 | NA |
| RASL11B | 867.686961 | 0.222328586 | 0.39667502 | 0.56048044 | 0.57515178 | 0.73938042 |
| SCFD2 | 732.784405 | 0.396086229 | 0.18704568 | 2.11759086 | 0.03420973 | 0.11074278 |
| AC023154.1 | 357.449481 | -0.133762659 | 0.73347564 | -0.1823682 | 0.85529375 | 0.92411978 |
| FIP1L1 | 1886.73341 | -0.30941298 | 0.12467856 | -2.4816855 | 0.01307626 | 0.05493406 |
| AC058822.1 | 1005.15502 | 0.060940456 | 0.26110601 | 0.23339353 | 0.81545583 | 0.90052223 |
| LNX1 | 1605.89379 | -0.154015297 | 0.22553538 | -0.6828875 | 0.49467792 | 0.67694119 |
| LNX1-AS1 | 1.48847246 | -1.588651074 | 1.59990606 | -0.9929652 | 0.3207269 | 0.51819065 |
| AC110792.3 | 13.8638344 | -0.288921543 | 0.38051778 | -0.7592853 | 0.44768193 | 0.63783587 |
| CHIC2 | 660.483779 | -0.798321598 | 0.20491129 | -3.8959376 | 9.78E-05 | 0.00124134 |
| AC110792.2 | 0.07830766 | -0.642905668 | 3.30558948 | -0.1944905 | 0.84579185 | NA |
| GSX2 | 0.81548438 | 0.142745058 | 1.88100359 | 0.07588771 | 0.93950843 | NA |
| PDGFRA | 3450.04305 | -0.309813758 | 0.3723778 | -0.8319877 | 0.40541587 | 0.60018288 |
| LINC02283 | 0 | NA | NA | NA | NA | NA |
| AC098587.1 | 0 | NA | NA | NA | NA | NA |
| AC098868.1 | 0.15318056 | -1.003574173 | 3.30558948 | -0.3035992 | 0.7614333 | NA |
| LINC02260 | 0 | NA | NA | NA | NA | NA |

| | | | | | | |
|---|---|---|---|---|---|---|
| KIT | 1050.05491 | -0.309502907 | 0.45167097 | -0.6852398 | 0.49319263 | 0.67589075 |
| AC097494.1 | 0 | NA | NA | NA | NA | NA |
| LINC02358 | 0 | NA | NA | NA | NA | NA |
| AC111194.1 | 0 | NA | NA | NA | NA | NA |
| AC111194.2 | 0 | NA | NA | NA | NA | NA |
| KDR | 2565.89174 | 0.569408938 | 0.34655017 | 1.64307792 | 0.10036681 | 0.23816007 |
| SRD5A3 | 596.991719 | 0.153870465 | 0.22447781 | 0.68545958 | 0.49305396 | 0.67578034 |
| AC069200.1 | 4.16273991 | 0.24135819 | 0.79971273 | 0.30180611 | 0.76279987 | 0.86868167 |
| TMEM165 | 5050.15532 | -0.895375721 | 0.18834661 | -4.7538723 | 2.00E-06 | 5.65E-05 |
| CLOCK | 1389.69324 | -0.022742241 | 0.12813844 | -0.1774818 | 0.85912996 | 0.92642086 |
| AC024243.1 | 8.2745067 | 0.714790189 | 0.44789527 | 1.59588688 | 0.11051405 | 0.25463133 |
| PDCL2 | 0.08393208 | -0.642905668 | 3.30558948 | -0.1944905 | 0.84579185 | NA |
| NMU | 3.19862709 | -1.677325411 | 1.18616081 | -1.4140793 | 0.15733863 | 0.32442096 |
| EXOC1L | 0.16416605 | 0.18657164 | 3.29377195 | 0.05664376 | 0.95482897 | NA |
| EXOC1 | 2152.6061 | 0.002094559 | 0.10407147 | 0.02012615 | 0.98394274 | 0.99305611 |
| CEP135 | 538.540385 | 0.779978984 | 0.19138221 | 4.07550406 | 4.59E-05 | 0.00069306 |
| KIAA1211 | 62.7729947 | 0.731515114 | 0.56495803 | 1.2948132 | 0.19538468 | 0.37552619 |
| AASDH | 735.620947 | 0.012551798 | 0.17197294 | 0.07298705 | 0.94181642 | 0.97233647 |
| AC068620.1 | 10.3358276 | 0.111144032 | 0.48505092 | 0.2291389 | 0.81876096 | 0.90273557 |
| PPAT | 590.570668 | -0.025017995 | 0.15270156 | -0.1638359 | 0.86986034 | 0.93220007 |
| AC068620.2 | 78.6877334 | 0.120672423 | 0.20867173 | 0.57828831 | 0.56306949 | 0.73032032 |
| PAICS | 3922.19005 | -0.094392228 | 0.1474443 | -0.6401891 | 0.52204968 | 0.69798967 |
| SRP72 | 3040.06846 | -0.237678434 | 0.15706922 | -1.5132083 | 0.13022677 | 0.2854258 |
| ARL9 | 14.2070577 | -1.117103581 | 0.49880414 | -2.2395636 | 0.02511927 | 0.08870381 |
| THEGL | 0.91409985 | -0.788812766 | 1.81884964 | -0.4336877 | 0.66451522 | NA |
| HOPX | 536.489817 | -1.181481873 | 0.44477773 | -2.6563422 | 0.00789934 | 0.03781874 |
| SPINK2 | 4.81986439 | -0.037499494 | 0.99044931 | -0.0378611 | 0.96979843 | 0.98606457 |
| REST | 1958.6368 | 0.026706857 | 0.11685272 | 0.22855143 | 0.81921758 | 0.90290073 |
| AC069307.1 | 9.89899753 | -0.223654873 | 0.65876844 | -0.3395045 | 0.73422968 | 0.85046282 |
| NOA1 | 1192.32184 | -0.033285241 | 0.15622596 | -0.2130583 | 0.83128147 | 0.91040231 |
| POLR2B | 13584.5469 | -0.428748706 | 0.13239017 | -3.2385238 | 0.0012015 | 0.00905244 |
| IGFBP7 | 67084.7734 | -0.409620755 | 0.22038801 | -1.8586345 | 0.06307896 | 0.17122095 |
| IGFBP7-AS1 | 19318.1836 | -0.289408614 | 0.24280987 | -1.1919145 | 0.23329476 | 0.42159969 |
| AC105390.1 | 0.08757577 | 0.547239394 | 3.29377202 | 0.16614368 | 0.86804388 | NA |
| LINC02380 | 0 | NA | NA | NA | NA | NA |
| AC013724.1 | 3.1625904 | -0.92680641 | 0.94647581 | -0.9792183 | 0.32747214 | 0.52526221 |
| AC107396.1 | 4.23830448 | 4.117013945 | 1.02752598 | 4.00672491 | 6.16E-05 | 0.00086797 |
| AC093725.2 | 1.78179171 | 4.746803003 | 1.28397031 | 3.69697257 | 0.00021819 | 0.0023417 |
| AC093725.1 | 0.4907454 | 2.216505017 | 2.56323245 | 0.8647304 | 0.38718678 | NA |
| LINC02494 | 0 | NA | NA | NA | NA | NA |
| AC019133.1 | 0 | NA | NA | NA | NA | NA |
| AC017091.1 | 0 | NA | NA | NA | NA | NA |
| LINC02429 | 0 | NA | NA | NA | NA | NA |
| AC097501.2 | 0 | NA | NA | NA | NA | NA |
| AC108517.1 | 0 | NA | NA | NA | NA | NA |
| AC096588.1 | 0 | NA | NA | NA | NA | NA |
| AC093857.1 | 0 | NA | NA | NA | NA | NA |
| LINC02496 | 0 | NA | NA | NA | NA | NA |
| LINC02271 | 0 | NA | NA | NA | NA | NA |
| ADGRL3 | 157.180368 | 0.763868643 | 0.48628999 | 1.57080889 | 0.11622705 | 0.26426951 |
| AC020741.1 | 0.82631713 | -0.24835544 | 1.36667249 | -0.1817227 | 0.85580034 | NA |
| ADGRL3-AS1 | 8.55802846 | 0.271851913 | 0.67311261 | 0.40387286 | 0.68630622 | 0.81857592 |
| AC097491.1 | 0 | NA | NA | NA | NA | NA |
| AC093730.1 | 0 | NA | NA | NA | NA | NA |
| TECRL | 0.70261245 | 2.860669728 | 1.46652037 | 1.95065121 | 0.05109855 | NA |
| LINC02232 | 1.458304 | 3.085423862 | 1.47949186 | 2.08546187 | 0.0370274 | 0.11701971 |
| EPHA5 | 7.38175319 | -2.023941881 | 1.00779966 | -2.008278 | 0.04461376 | 0.1339632 |
| EPHA5-AS1 | 1.94627663 | -4.17027248 | 1.87076553 | -2.2291797 | 0.02580195 | 0.09040626 |
| AC104137.1 | 0.09644756 | -0.642905668 | 3.30558948 | -0.1944905 | 0.84579185 | NA |
| AC097110.1 | 0.09635527 | 0.547239394 | 3.29377202 | 0.16614368 | 0.86804388 | NA |
| AC116049.2 | 1.65544275 | 1.658748487 | 0.99155976 | 1.67286789 | 0.09435331 | 0.22834955 |
| AC096721.1 | 0.08393208 | -0.642905668 | 3.30558948 | -0.1944905 | 0.84579185 | NA |
| AC104806.2 | 10.3531295 | -0.571331972 | 0.51572569 | -1.1078214 | 0.26793894 | 0.46209809 |
| CENPC | 866.021818 | 0.59771409 | 0.15624115 | 3.82558676 | 0.00013046 | 0.0015606 |
| STAP1 | 347.124897 | -0.215699046 | 0.54797614 | -0.3936285 | 0.69385531 | 0.82376229 |
| UBA6 | 1849.12932 | -0.677952039 | 0.18094847 | -3.746658 | 0.00017921 | 0.00200513 |
| UBA6-AS1 | 672.859668 | 0.091402698 | 0.18418739 | 0.4962484 | 0.61971917 | 0.77271053 |
| GNRHR | 8.66803076 | 0.72514345 | 0.57242593 | 1.26679 | 0.20523039 | 0.38772928 |
| TMPRSS11D | 2.59676099 | 0.405988184 | 0.95017317 | 0.4272781 | 0.66917678 | 0.80675681 |
| TMPRSS11A | 0.96804569 | 1.378521771 | 1.77224079 | 0.77784113 | 0.43666268 | NA |
| TMPRSS11F | 0.58532195 | -0.726390675 | 1.87332962 | -0.3877538 | 0.69819824 | NA |
| TMPRSS11B | 0.03392908 | 0.380740593 | 3.29738709 | 0.11546736 | 0.90807469 | NA |
| YTHDC1 | 3838.82303 | -0.11630404 | 0.12119819 | -0.9596186 | 0.33724721 | 0.53527683 |
| TMPRSS11E | 7.44670656 | -3.343503243 | 1.40754317 | -2.3754179 | 0.01752909 | 0.06809696 |
| UGT2B17 | 2.94482936 | -0.528527191 | 1.19003026 | -0.4441292 | 0.65694918 | 0.7986941 |
| UGT2B15 | 0.13390214 | 1.04981649 | 3.28477129 | 0.31960109 | 0.74927075 | NA |
| UGT2B10 | 31.6973241 | 1.475693815 | 0.62992173 | 2.34266219 | 0.01914671 | 0.07257773 |
| AC021146.12 | 8.17216867 | 1.198363482 | 0.59770608 | 2.00493774 | 0.0449697 | 0.1347778 |
| UGT2A3 | 5118.22372 | 0.80225833 | 0.43324323 | 1.85175043 | 0.06406167 | 0.17308465 |
| UGT2B7 | 36413.5309 | 1.967725478 | 0.51882909 | 3.7926275 | 0.00014906 | 0.00174545 |
| AC111000.4 | 241.58347 | 0.759145599 | 0.57874181 | 1.31171722 | 0.18961556 | 0.36809205 |
| UGT2B11 | 63.7767481 | 1.211017023 | 0.53685968 | 2.25574217 | 0.02408678 | 0.08584518 |
| UGT2B28 | 147.71042 | -0.180711678 | 1.20141478 | -0.1504157 | 0.88043663 | 0.93798942 |
| UGT2B4 | 5.10208905 | 0.757994971 | 0.85955175 | 0.88184914 | 0.37785839 | 0.57382306 |
| UGT2A1 | 182.816609 | 3.174954758 | 1.20048825 | 2.64471957 | 0.00817586 | 0.03872778 |
| UGT2A2 | 108.016793 | 3.39094882 | 1.02712682 | 3.30139255 | 0.00096206 | 0.00762307 |
| SULT1B1 | 94.5535017 | 1.193119668 | 0.64623904 | 1.84625131 | 0.06485572 | 0.17455388 |
| AC108941.2 | 14.3567237 | 0.384736454 | 0.63635338 | 0.6045956 | 0.54544773 | 0.71702177 |
| SULT1E1 | 22.3947744 | 0.412466897 | 0.60953229 | 0.67669409 | 0.49860007 | 0.6799953 |
| CSN1S1 | 0.26795541 | -0.17909897 | 2.95701559 | -0.0605675 | 0.95170368 | NA |
| CSN2 | 0.19289513 | -1.003574173 | 3.30558948 | -0.3035992 | 0.7614333 | NA |
| STATH | 0 | NA | NA | NA | NA | NA |
| HTN3 | 0 | NA | NA | NA | NA | NA |
| HTN1 | 0 | NA | NA | NA | NA | NA |
| PRR27 | 0 | NA | NA | NA | NA | NA |
| ODAM | 18.7276176 | 1.805243476 | 0.63967697 | 2.82211735 | 0.00477077 | 0.0259573 |
| FDCSP | 4.02759002 | 0.870352412 | 2.39065483 | 0.36406444 | NA | NA |
| CSN3 | 0 | NA | NA | NA | NA | NA |
| CABS1 | 0 | NA | NA | NA | NA | NA |
| SMR3A | 0 | NA | NA | NA | NA | NA |
| SMR3B | 0 | NA | NA | NA | NA | NA |
| OPRPN | 0 | NA | NA | NA | NA | NA |
| MUC7 | 0 | NA | NA | NA | NA | NA |
| AMTN | 0.54277899 | -0.067023698 | 1.92712683 | -0.0347791 | 0.9722559 | NA |
| AMBN | 0.64767489 | -1.557343501 | 2.22069422 | -0.7012868 | 0.48312406 | NA |
| ENAM | 990.683262 | 0.762639397 | 0.45432754 | 1.67861143 | 0.0932278 | 0.22622775 |
| JCHAIN | 1879.33497 | 0.468999581 | 0.93290676 | 0.50272932 | 0.61515459 | 0.76990046 |
| UTP3 | 1981.75385 | -0.406694012 | 0.19735569 | -2.0607159 | 0.03933015 | 0.12228936 |
| AC009570.1 | 122.179655 | -0.560297553 | 0.5847107 | -0.9582475 | 0.33793798 | 0.53590196 |
| RUFY3 | 3123.90269 | -0.389423766 | 0.22891497 | -1.7011721 | 0.08891068 | 0.21866952 |
| GRSF1 | 4985.70953 | -0.20963026 | 0.13305087 | -1.5755647 | 0.11512615 | 0.2623921 |
| MOB1B | 3042.45992 | 0.084133925 | 0.13786958 | 0.61024284 | 0.54170095 | 0.71442107 |
| DCK | 738.381876 | 0.202366577 | 0.13523647 | 1.49639054 | 0.13455191 | 0.29188335 |

| | | | | | | |
|---|---|---|---|---|---|---|
| SLC4A4 | 10544.6526 | 1.553218292 | 0.52639953 | 2.95064528 | 0.00317111 | 0.0191662 |
| GC | 2284.89405 | 0.886147265 | 0.94890393 | 0.93386405 | 0.35037403 | 0.54813497 |
| AC068721.1 | 0.5853672 | 3.146319852 | 3.1890834 | 0.98659065 | 0.32384336 | NA |
| NPFFR2 | 3.73345729 | -0.710767648 | 1.00909674 | -0.7043603 | 0.48120844 | 0.66581781 |
| ADAMTS3 | 186.862076 | -0.772718144 | 0.37343827 | -2.0691991 | 0.03852741 | 0.12045282 |
| AC095056.1 | 0.66334371 | 2.086888816 | 1.73699814 | 1.20143411 | 0.22958285 | NA |
| COX18 | 1058.77932 | 0.432744882 | 0.15541384 | 2.78446805 | 0.00536156 | 0.02836765 |
| ANKRD17 | 3972.91244 | 0.150411472 | 0.12408354 | 1.21217905 | 0.22544385 | 0.41190011 |
| AC053527.1 | 20.5647339 | 0.565183211 | 0.26704318 | 2.1164488 | 0.03430666 | 0.11101565 |
| ALB | 8370.90728 | 2.308011225 | 1.18870788 | 1.94161346 | 0.05218391 | 0.15017389 |
| AFP | 84.3413119 | 3.361813004 | 0.66750867 | 5.03635856 | 4.74E-07 | 1.73E-05 |
| AFM | 1551.15886 | 4.320104648 | 1.1413558 | 3.78506392 | 0.00015367 | 0.00178592 |
| LINC02499 | 198.389554 | 6.699419013 | 0.93598993 | 7.15757594 | 8.21E-13 | 2.26E-10 |
| RASSF6 | 1399.05239 | -1.036365459 | 0.34065841 | -3.0422424 | 0.00234823 | 0.01529334 |
| AC112518.1 | 0.06785815 | 0.795868262 | 3.28898349 | 0.24198001 | 0.80879565 | NA |
| CXCL8 | 1466.4316 | -3.507908613 | 0.93242117 | -3.7621503 | 0.00016846 | 0.00191394 |
| CXCL6 | 319.567056 | -2.250486141 | 0.87947269 | -2.558904 | 0.01050027 | 0.04670922 |
| PF4V1 | 19.7782634 | -0.143610956 | 0.51633493 | -0.2781353 | 0.78090852 | 0.88016021 |
| CXCL1 | 1566.22669 | -3.589350486 | 0.78964578 | -4.5455197 | 5.48E-06 | 0.0001261 |
| PF4 | 16.1266988 | -0.817544219 | 0.79036477 | -1.0343885 | 0.30095459 | 0.49768474 |
| PPBP | 27.250471 | -1.08251036 | 0.94219556 | -1.1489232 | 0.25058764 | 0.44275742 |
| CXCL5 | 21.8645199 | -1.971206852 | 0.9209342 | -2.1404427 | 0.03231901 | 0.10628952 |
| CXCL3 | 122.731386 | -1.327919584 | 0.63371354 | -2.0954572 | 0.03613037 | 0.11513691 |
| CXCL2 | 1815.77016 | -2.658508023 | 0.73865489 | -3.5991206 | 0.0003193 | 0.00317995 |
| MTHFD2L | 771.148243 | 0.019072633 | 0.16397294 | 0.11631574 | 0.90740231 | 0.95380871 |
| AC093677.2 | 137.884343 | -0.009638142 | 0.21607848 | -0.0446048 | 0.9644223 | 0.98336618 |
| EPGN | 3.36448848 | 0.133124638 | 0.78987679 | 0.16853848 | 0.86615967 | 0.93052353 |
| EREG | 26.3478334 | -1.32231293 | 0.78777244 | -1.6785468 | 0.0932404 | 0.22623751 |
| AC021180.1 | 0 | NA | NA | NA | NA | NA |
| AREG | 99.9119854 | -1.969539565 | 0.68081025 | -2.8929347 | 0.00381661 | 0.02200829 |
| AC239584.1 | 0.47186847 | 0.002178752 | 2.25208566 | 0.00096744 | 0.9992281 | NA |
| BTC | 430.154778 | 0.120348517 | 0.45103332 | 0.26682844 | 0.78960126 | 0.88561887 |
| PARM1 | 5043.82413 | 1.091334586 | 0.27511623 | 3.9668128 | 7.28E-05 | 0.00098591 |
| AC110760.2 | 3.3789677 | 2.831523246 | 1.04103554 | 2.71991027 | 0.00652996 | 0.03278658 |
| AC110760.1 | 1.79413172 | 2.939281153 | 1.179939 | 2.49104501 | 0.0127368 | 0.05392027 |
| LINC02562 | 9.9764616 | 3.428889189 | 1.08911598 | 3.14832328 | 0.0016421 | 0.01160294 |
| AC025244.1 | 0 | NA | NA | NA | NA | NA |
| AC025244.2 | 0.17361919 | -1.003574173 | 3.30558948 | -0.3035992 | 0.7614333 | NA |
| LINC02483 | 0.10721287 | -0.642905668 | 3.30558948 | -0.1944905 | 0.84579185 | NA |
| AC096759.1 | 0 | NA | NA | NA | NA | NA |
| AC096759.2 | 0.03392908 | 0.380740593 | 3.29738709 | 0.11546736 | 0.90807469 | NA |
| RCHY1 | 1364.49296 | -0.034992594 | 0.13202806 | -0.2650391 | 0.79097937 | 0.88641183 |
| THAP6 | 723.897214 | 0.239522109 | 0.13166244 | 1.81921362 | 0.06887885 | 0.18201818 |
| ODAPH | 6.61968535 | -0.60837626 | 0.65186629 | -0.9332838 | 0.35067345 | 0.54834976 |
| CDKL2 | 420.758577 | 0.412451457 | 0.28308757 | 1.45697483 | 0.14512333 | 0.30735701 |
| G3BP2 | 5767.17743 | 0.167806243 | 0.22664779 | 0.74038332 | 0.45906743 | 0.64784375 |
| USO1 | 4051.62032 | 0.0934791 | 0.14296193 | 0.65387407 | 0.51319294 | 0.69122454 |
| AC110615.1 | 0 | NA | NA | NA | NA | NA |
| PPEF2 | 4.39011708 | -0.738038006 | 0.76188476 | -0.9687003 | 0.33269474 | 0.53064539 |
| NAAA | 1383.35138 | 0.506891627 | 0.28622775 | 1.77093812 | 0.07657099 | 0.19647534 |
| SDAD1 | 1657.52033 | -0.027405785 | 0.13234767 | -0.2070742 | 0.83595192 | 0.91287279 |
| AC112719.2 | 0.18754884 | 1.469068384 | 3.27783996 | 0.44818185 | 0.65402196 | NA |
| CXCL9 | 397.05524 | 1.696737371 | 0.95022972 | 1.78560755 | NA | NA |
| ART3 | 9.90663829 | 0.271563862 | 0.55984193 | 0.48507239 | 0.627625 | 0.77844275 |
| CXCL10 | 276.775529 | -0.421545922 | 0.81584994 | -0.5166954 | NA | NA |
| CXCL11 | 57.6913899 | -0.212392353 | 1.13271889 | -0.1875067 | NA | NA |
| NUP54 | 1441.96659 | -0.16983031 | 0.11519502 | -1.4742851 | 0.14040484 | 0.30057441 |
| SCARB2 | 9740.64318 | -0.391137395 | 0.17958358 | -2.1780242 | 0.02940423 | 0.09943583 |
| FAM47E | 549.887849 | 0.305939882 | 0.26878697 | 1.13822436 | 0.25502681 | 0.44771091 |
| FAM47E-STB | 1715.97384 | -1.027867086 | 0.26369805 | -3.8978942 | 9.70E-05 | 0.00123552 |
| STBD1 | 1586.07427 | -1.333378826 | 0.30169781 | -4.4195841 | 9.89E-06 | 0.00020137 |
| CCDC158 | 52.6048031 | 1.047060275 | 0.45783278 | 2.2869928 | 0.02219624 | 0.08097882 |
| SHROOM3 | 2546.32438 | -0.668259473 | 0.20795575 | -3.2134696 | 0.00131142 | 0.00974417 |
| AC107072.2 | 3.50016783 | -0.434084127 | 0.81165815 | -0.5348115 | 0.5927802 | 0.75209843 |
| AC107072.1 | 1.23346635 | 1.188223286 | 1.21444569 | 0.97840792 | 0.32787261 | 0.52559275 |
| SOWAHB | 544.60769 | 0.740147199 | 0.35487771 | 2.08564015 | 0.03701123 | 0.11700003 |
| 11-Sep | 5783.85144 | -0.069256178 | 0.21491522 | -0.3222488 | 0.74726419 | 0.85862121 |
| CCNI | 15311.4028 | -0.049289462 | 0.18581053 | -0.2652673 | 0.79080354 | 0.88640183 |
| CCNG2 | 4782.33514 | -0.087494449 | 0.25806228 | -0.3390439 | 0.73457663 | 0.85065526 |
| AC092674.1 | 0.19478864 | 0.18657164 | 3.29377195 | 0.05664376 | 0.95482897 | NA |
| CXCL13 | 41.781917 | -0.639038712 | 1.10375934 | -0.5789656 | 0.56261237 | 0.7300506 |
| CNOT6L | 1975.20902 | -0.133376083 | 0.09492142 | -1.4051211 | 0.15998528 | 0.32805691 |
| MRPL1 | 1046.01486 | 0.169560187 | 0.15714862 | 1.07897981 | 0.28059673 | 0.47633164 |
| FRAS1 | 1908.48386 | 1.383182976 | 0.40350381 | 3.42793041 | 0.0006082 | 0.00526881 |
| ANXA3 | 1465.6115 | -0.332817584 | 0.31314792 | -1.0628127 | 0.28786691 | 0.48409223 |
| LINC01094 | 55.1176215 | -1.694315619 | 0.49656355 | -3.4120822 | 0.00064469 | 0.00551961 |
| AC098818.2 | 3.61500126 | -0.383201701 | 0.99391301 | -0.3855485 | 0.69983106 | 0.82785506 |
| BMP2K | 1726.96251 | -0.497597607 | 0.26238554 | -1.8964368 | 0.05790231 | 0.16145541 |
| PAQR3 | 1894.24403 | -0.262275429 | 0.20154832 | -1.301303 | 0.19315477 | 0.37266019 |
| LINC01088 | 34.3118082 | 0.06972297 | 0.57231083 | 0.1218271 | 0.90303595 | 0.95126665 |
| NAA11 | 1.07482808 | 0.100703241 | 1.09232947 | 0.09219127 | 0.92654607 | 0.96437374 |
| GK2 | 0 | NA | NA | NA | NA | NA |
| LINC00989 | 41.1875441 | 0.369365708 | 0.51662479 | 0.71495931 | 0.47463419 | 0.66013204 |
| AC092542.1 | 9.87239075 | 0.441105418 | 0.72395329 | 0.60930094 | 0.54232498 | 0.71480531 |
| LINC02469 | 3.81666321 | 0.173608725 | 0.83462018 | 0.20800926 | 0.83522174 | 0.91264233 |
| PCAT4 | 0.40691305 | 1.069107225 | 2.22093617 | 0.48137684 | 0.6302487 | NA |
| ANTXR2 | 1381.83835 | 0.402197944 | 0.2294671 | 1.75274772 | 0.07964532 | 0.201811 |
| AC021127.1 | 1.10245961 | -0.180438454 | 1.61992768 | -0.1113867 | 0.91130968 | 0.95620071 |
| PRDM8 | 102.458017 | 0.045040278 | 0.35678187 | 0.12624038 | 0.89954165 | 0.94902072 |
| FGF5 | 100.505749 | -0.564889763 | 0.79404175 | -0.7114107 | 0.4768298 | 0.66203109 |
| CFAP299 | 10.8656909 | 0.099861531 | 0.58104692 | 0.17186483 | 0.8635438 | 0.92899031 |
| BMP3 | 13.5904614 | -0.941435877 | 0.73460206 | -1.281559 | 0.19999739 | 0.38139509 |
| PRKG2 | 102.130947 | -0.673691922 | 0.47669769 | -1.4132477 | 0.1575829 | 0.3247215 |
| AC139722.1 | 0 | NA | NA | NA | NA | NA |
| RASGEF1B | 881.631723 | -0.503636589 | 0.27480982 | -1.8326732 | 0.06685118 | 0.17824729 |
| AC124016.2 | 33.1847521 | 0.445993905 | 0.42507012 | 1.04922431 | 0.29407489 | 0.49080728 |
| HNRNPD | 9338.97521 | -0.465011133 | 0.11186813 | -4.1567794 | 3.23E-05 | 0.00052057 |
| AC124016.1 | 59.4737127 | -0.17021665 | 0.29028835 | -0.5863709 | 0.55762628 | 0.72636855 |
| HNRNPDL | 16824.4846 | -0.499471343 | 0.1640677 | -3.0443003 | 0.00233222 | 0.01521165 |
| ENOPH1 | 2145.97811 | -0.207673391 | 0.16132646 | -1.2872866 | 0.1979944 | 0.37891809 |
| TMEM150C | 1321.60303 | 0.633115331 | 0.31563699 | 2.00583379 | 0.04487398 | 0.13459133 |
| LINC00575 | 0.09997307 | 0.547239394 | 3.29377202 | 0.16614368 | 0.86804388 | NA |
| SCD5 | 2930.09961 | 1.050955895 | 0.33667339 | 3.12158884 | 0.00179878 | 0.01242549 |
| SEC31A | 7898.52849 | 0.240958978 | 0.18098882 | 1.33134729 | 0.18307476 | 0.35976463 |
| THAP9-AS1 | 2614.11245 | -0.559724181 | 0.2357693 | -2.3740334 | 0.01759496 | 0.06829179 |
| THAP9 | 303.02173 | 0.455080295 | 0.29840716 | 1.52503143 | 0.12725126 | 0.28080466 |
| LIN54 | 657.606398 | -0.114213962 | 0.25840553 | -0.441995 | 0.65849279 | 0.79946078 |
| COPS4 | 1742.87121 | 0.082964991 | 0.11030117 | 0.7521678 | 0.45195015 | 0.64173558 |
| PLAC8 | 121.823586 | -1.010331347 | 0.55454029 | -1.821926 | 0.06846622 | 0.18125325 |
| AC114781.2 | 1.69856484 | -1.110488952 | 0.99290716 | -1.1184217 | 0.26338692 | 0.45715217 |
| COQ2 | 421.527679 | -0.107937051 | 0.18921931 | -0.5704336 | 0.56838365 | 0.73404883 |
| HPSE | 376.159287 | -1.29216398 | 0.30889187 | -4.1832243 | 2.87E-05 | 0.00047486 |
| HELQ | 732.297575 | 0.015136272 | 0.09141799 | 0.16557214 | 0.86849368 | 0.93156219 |

| | | | | | | |
|---|---|---|---|---|---|---|
| MRPS18C | 1410.01029 | -0.121875009 | 0.14683702 | -0.8300019 | 0.4065377 | 0.60140134 |
| ABRAXAS1 | 1429.52193 | -0.016975953 | 0.14412283 | -0.1177881 | 0.90623557 | 0.95303816 |
| GPAT3 | 5390.83628 | 1.687836672 | 0.63439319 | 2.66055296 | 0.00780125 | 0.03752624 |
| AC021192.1 | 1.38055933 | -2.185301586 | 1.77761836 | -1.2293424 | 0.21894346 | 0.4043661 |
| AC104063.1 | 6.09509555 | -0.927809982 | 0.81170715 | -1.1430354 | 0.2530239 | 0.44550402 |
| NKX6-1 | 3.75375617 | 0.233042224 | 0.76324295 | 0.30533164 | 0.76011358 | 0.86700095 |
| CDS1 | 1910.76137 | -0.149471424 | 0.13400842 | -1.1153883 | 0.26468406 | 0.45867777 |
| WDFY3 | 3701.09601 | 0.559257858 | 0.18716497 | 2.98804765 | 0.00280766 | 0.01744943 |
| WDFY3-AS1 | 18.8516721 | 0.700874079 | 0.43287702 | 1.61910669 | 0.10542431 | 0.24637 |
| ARHGAP24 | 5101.92623 | 1.19559711 | 0.26649026 | 4.486457 | 7.24E-06 | 0.00015835 |
| MAPK10 | 3413.85049 | 0.743576403 | 0.28474498 | 2.61137665 | 0.00901785 | 0.04172675 |
| AC104827.1 | 1.18267571 | -0.791830393 | 1.49652075 | -0.5291142 | 0.59672622 | 0.75537609 |
| PTPN13 | 2674.80626 | 0.806345552 | 0.19914134 | 4.04911191 | 5.14E-05 | 0.00075952 |
| SLC10A6 | 9.17416534 | 0.688625446 | 0.42954585 | 1.60314772 | 0.10890205 | 0.25230439 |
| C4orf36 | 96.1645483 | -0.285825221 | 0.22273635 | -1.2832446 | 0.19940639 | 0.38057079 |
| AC093827.5 | 193.403516 | 0.128245175 | 0.22833813 | 0.56164589 | 0.5743573 | 0.73878358 |
| AC093827.4 | 285.611011 | 0.450973336 | 0.29267093 | 1.54088871 | 0.12334388 | 0.27480382 |
| AFF1 | 7350.76148 | 0.18387292 | 0.17708989 | 1.03830273 | 0.29912914 | 0.4959153 |
| KLHL8 | 998.271038 | 0.070790763 | 0.24565608 | 0.28817021 | 0.77321645 | 0.87517031 |
| AC108516.1 | 0.21220981 | 0.689156227 | 3.08302815 | 0.22353225 | 0.82312128 | NA |
| HSD17B13 | 36.2608499 | 0.798007068 | 0.49866671 | 1.60028141 | 0.10953617 | 0.25324117 |
| HSD17B11 | 3316.99852 | 0.359907714 | 0.23343387 | 1.54179733 | 0.12312285 | 0.2746124 |
| NUDT9 | 2903.37586 | -0.240220773 | 0.18070364 | -1.3293632 | 0.18372817 | 0.36077979 |
| AC112250.2 | 5.69448316 | -1.703807026 | 1.02185594 | -1.6673652 | 0.09544181 | 0.230118 |
| SPARCL1 | 10019.2676 | -0.349812273 | 0.45211645 | -0.7737216 | 0.43909543 | 0.63062723 |
| AC093895.1 | 3.98151962 | -1.456968608 | 0.93879684 | -1.551953 | 0.12067346 | 0.27098349 |
| DSPP | 0.03392908 | 0.380740593 | 3.29738709 | 0.11546736 | 0.90807469 | NA |
| DMP1 | 2.58052403 | -0.772111692 | 1.05684179 | -0.730584 | 0.46503329 | 0.65212372 |
| IBSP | 1.96278989 | -1.984592501 | 1.38942098 | -1.4283594 | 0.15318844 | 0.31872962 |
| MEPE | 1.0257924 | -1.713918897 | 1.26710631 | -1.3526244 | 0.17617566 | 0.35080813 |
| SPP1 | 629934.688 | -2.472723114 | 0.36687259 | -6.7400051 | 1.58E-11 | 2.96E-09 |
| PKD2 | 4077.12694 | 0.391738482 | 0.14771263 | 2.65203103 | 0.00800092 | 0.03813739 |
| ABCG2 | 290.177654 | 1.310169354 | 0.36095135 | 3.62976716 | 0.00028368 | 0.00289944 |
| PPM1K | 2405.23104 | 0.683900687 | 0.35898407 | 1.90510037 | 0.05676702 | 0.15916496 |
| AC107067.1 | 13.0500927 | 1.324893436 | 0.77312715 | 1.71368116 | 0.0865873 | 0.21439503 |
| HERC6 | 668.719438 | 0.067840815 | 0.29911251 | 0.22680701 | 0.82057381 | 0.90366551 |
| HERC5 | 545.643257 | -0.300238843 | 0.12648888 | -2.3736382 | 0.0176138 | 0.06833542 |
| PYURF | 3273.92601 | -0.010112148 | 0.11180494 | -0.0904446 | 0.92793395 | 0.96504253 |
| PIGY | 1250.90204 | 0.051717807 | 0.12737917 | 0.40601463 | 0.68473185 | 0.81783568 |
| AC098582.1 | 50.8516567 | 1.341626276 | 0.32240084 | 4.161361 | 3.16E-05 | 0.0005118 |
| AC083829.2 | 8.90330228 | 0.230826872 | 0.64232685 | 0.35936046 | 0.71932545 | 0.84021138 |
| HERC3 | 1803.47948 | -0.075317389 | 0.14765856 | -0.5100781 | 0.60999678 | 0.76596961 |
| NAP1L5 | 392.154392 | 0.467842722 | 0.25401727 | 1.84177524 | 0.06550803 | 0.17578578 |
| FAM13A-AS1 | 1710.16596 | 0.103735652 | 0.23547537 | 0.44053717 | 0.6595481 | 0.79999826 |
| FAM13A | 3478.46723 | 0.437244352 | 0.25809855 | 1.69409847 | 0.09024657 | 0.22110955 |
| TIGD2 | 380.105058 | 0.189980058 | 0.21028596 | 0.90343671 | 0.36629416 | 0.56331382 |
| AC104785.1 | 18.7613191 | 0.556364784 | 0.31688087 | 1.75575376 | 0.07913046 | 0.20093118 |
| GPRIN3 | 1189.34462 | -0.151993643 | 0.32492946 | -0.4677743 | 0.63994602 | 0.78705286 |
| AC097478.1 | 105.018627 | -0.233037589 | 0.31148055 | -0.748161 | 0.45436307 | 0.64389264 |
| SNCA | 285.422166 | -0.078135647 | 0.34167494 | -0.2286842 | 0.8191144 | 0.90289626 |
| AC097478.2 | 0.66283729 | -1.335591185 | 1.6369491 | -0.8159027 | 0.41455579 | NA |
| AC097478.4 | 0.34213586 | -1.677037466 | 3.30336094 | -0.5076761 | 0.6116805 | NA |
| AC097478.3 | 0.29934637 | -0.281107011 | 2.99073357 | -0.0939927 | 0.92511499 | NA |
| SNCA-AS1 | 0.79065327 | 0.601622367 | 1.37258077 | 0.43831473 | 0.66115815 | NA |
| MMRN1 | 263.963975 | -1.213395938 | 0.60659513 | -2.0003391 | 0.04546366 | 0.13586546 |
| CCSER1 | 240.536472 | 1.087776603 | 0.23787358 | 4.57291898 | 4.81E-06 | 0.00011396 |
| AC004054.1 | 0.08695548 | -0.642905668 | 3.30558948 | -0.1944905 | 0.84579185 | NA |
| AC074124.1 | 10.1111702 | -0.838295093 | 0.97567021 | -0.8591992 | 0.39023061 | 0.58578665 |
| AC093810.1 | 0 | NA | NA | NA | NA | NA |
| LNCPRESS2 | 0 | NA | NA | NA | NA | NA |
| AC112695.1 | 0.16416605 | 0.18657164 | 3.29377195 | 0.05664376 | 0.95482897 | NA |
| GRID2 | 5.33719972 | 1.319059826 | 0.71054209 | 1.85641336 | 0.06339466 | 0.17179132 |
| AC095059.1 | 0 | NA | NA | NA | NA | NA |
| AC110800.1 | 0 | NA | NA | NA | NA | NA |
| ATOH1 | 0 | NA | NA | NA | NA | NA |
| AC096746.1 | 0.09635527 | 0.547239394 | 3.29377202 | 0.16614368 | 0.86804388 | NA |
| SMARCAD1 | 1387.74363 | 0.089257882 | 0.1327547 | 0.67235194 | 0.50135967 | 0.68210661 |
| HPGDS | 55.8541464 | 0.702944866 | 0.59355013 | 1.18430581 | 0.23629203 | 0.42559533 |
| PDLIM5 | 4519.00694 | -0.060496969 | 0.15163858 | -0.398955 | 0.68992636 | 0.82140766 |
| AC108067.1 | 0.07659028 | -0.642905668 | 3.30558948 | -0.1944905 | 0.84579185 | NA |
| BMPR1B-DT | 15.8591019 | 1.030457149 | 0.41563815 | 2.47921698 | 0.01316712 | 0.05526292 |
| BMPR1B | 571.12721 | 0.658804311 | 0.24711079 | 2.6660281 | 0.00767533 | 0.03704762 |
| UNC5C | 167.981027 | 0.64208874 | 0.20757053 | 3.09335211 | 0.00197909 | 0.01337827 |
| AC106881.1 | 31.3460149 | 0.934709754 | 0.29412824 | 3.17789871 | 0.00148347 | 0.0107146 |
| PDHA2 | 0.03392908 | 0.380740593 | 3.29738709 | 0.11546736 | 0.90807469 | NA |
| LINC02267 | 0 | NA | NA | NA | NA | NA |
| AC108515.1 | 0.22329207 | 1.817076245 | 3.25496057 | 0.55824831 | 0.57667483 | NA |
| STPG2 | 38.3585453 | 1.555578106 | 0.4430205 | 3.51130054 | 0.00044592 | 0.00415509 |
| STPG2-AS1 | 1.21111969 | 1.848830644 | 1.28777027 | 1.43568359 | 0.1510924 | 0.31593898 |
| AC019077.1 | 41.7626539 | 0.271856999 | 0.29127893 | 0.93332187 | 0.35065381 | 0.54834976 |
| RAP1GDS1 | 1622.01299 | -0.064111522 | 0.14665898 | -0.4371469 | 0.66200481 | 0.80118715 |
| TSPAN5 | 566.268916 | -0.267713046 | 0.26903439 | -0.9950886 | 0.31969318 | 0.51710403 |
| AC108159.1 | 0.13390214 | 1.04981649 | 3.28477129 | 0.31960109 | 0.74927075 | NA |
| AC114811.2 | 25.6860819 | -0.535907988 | 0.36117506 | -1.4837901 | 0.13786461 | 0.2966536 |
| EIF4E | 2213.28515 | -0.42724745 | 0.3487937 | -1.2249288 | 0.22060205 | 0.40651824 |
| AC019131.1 | 2.93380228 | -0.219320477 | 0.84074618 | -0.2608641 | 0.79419733 | 0.88844572 |
| METAP1 | 2296.7228 | -0.06515673 | 0.15310849 | -0.4255592 | 0.67042905 | 0.80749128 |
| AC019131.2 | 64.6626976 | 0.538298996 | 0.39696894 | 1.35602296 | 0.17509186 | 0.34909817 |
| ADH5 | 8384.58898 | 0.831084946 | 0.20347939 | 4.08436922 | 4.42E-05 | 0.00067251 |
| AP002026.1 | 95.6083585 | 0.338789169 | 0.49566046 | 0.68351058 | 0.49428427 | 0.67673841 |
| ADH4 | 51.3714688 | -0.811116763 | 0.52539719 | -1.5438163 | 0.12263284 | 0.27378586 |
| ADH6 | 1933.43812 | 2.81622654 | 0.64552387 | 4.36269933 | 1.28E-05 | 0.00024983 |
| ADH1A | 19.6254386 | 0.269501634 | 0.45150409 | 0.59689743 | 0.55057586 | 0.72092385 |
| ADH1B | 5659.20303 | 0.492242246 | 0.34730639 | 1.41731409 | 0.15639112 | 0.32307354 |
| ADH1C | 724.082169 | 0.11643929 | 0.85094388 | 0.13683545 | 0.89116086 | 0.94360054 |
| ADH7 | 0.15489794 | -1.003574173 | 3.30558948 | -0.3035992 | 0.7614333 | NA |
| C4orf17 | 0.10721287 | -0.642905668 | 3.30558948 | -0.1944905 | 0.84579185 | NA |
| TRMT10A | 334.705204 | 0.548869128 | 0.23773484 | 2.30874504 | 0.02095773 | 0.07752215 |
| MTTP | 656.473726 | 2.291578254 | 0.72323294 | 3.1685203 | 0.00153217 | 0.0109941 |
| AC083902.1 | 30.7504808 | 2.837685488 | 0.87977183 | 3.2254789 | 0.00125762 | 0.00939741 |
| C4orf54 | 138.721253 | 1.357828418 | 0.57249225 | 2.37178481 | 0.0177024 | 0.06855812 |
| DAPP1 | 151.910586 | -0.224248307 | 0.60897911 | -0.3682365 | 0.71269693 | 0.83673114 |
| LAMTOR3 | 2788.93162 | -0.037857805 | 0.16027734 | -0.2362019 | 0.81327604 | 0.89913137 |
| DNAJB14 | 1850.00416 | -0.052033631 | 0.09756968 | -0.5332971 | 0.59382791 | 0.75288673 |
| H2AFZ | 5419.54877 | -0.969567199 | 0.34032989 | -2.848904 | 0.00438701 | 0.02444456 |
| AC097460.1 | 1535.09033 | -0.805760267 | 0.34845836 | -2.3123574 | 0.020758 | 0.07697769 |
| DDIT4L | 1942.63547 | -0.897303831 | 0.66790879 | -1.3434526 | 0.17912545 | 0.35485975 |
| AP001961.1 | 4.73934165 | 0.552481891 | 0.80470653 | 0.6865632 | 0.49235802 | 0.67523825 |
| EMCN | 3413.11589 | 1.195911411 | 0.50395652 | 2.3730448 | 0.01764212 | 0.06841512 |
| PPP3CA | 1362.60726 | -0.221383359 | 0.19099819 | -1.1590862 | 0.24642106 | 0.4377238 |
| AP001816.1 | 190.873257 | -0.064787697 | 0.138524 | -0.4677002 | 0.63999903 | 0.78705286 |
| BANK1 | 251.039307 | 0.300267451 | 0.32390781 | 0.92701516 | 0.35391866 | 0.55148822 |
| AP002075.1 | 0.09635527 | 0.547239394 | 3.29377202 | 0.16614368 | 0.86804388 | NA |

| | | | | | | |
|---|---|---|---|---|---|---|
| SLC39A8 | 1808.64673 | 0.017240164 | 0.31698098 | 0.05438864 | 0.95662553 | 0.97931865 |
| AC098487.1 | 27.1458333 | 0.026858386 | 0.39250787 | 0.06842764 | 0.94544522 | 0.97394598 |
| AF213884.3 | 5.45053133 | 0.703959354 | 0.4840444 | 1.45432807 | 0.14585537 | 0.30843912 |
| NFKB1 | 2493.4187 | -0.207260964 | 0.21726504 | -0.9539545 | 0.34010666 | 0.53820287 |
| MANBA | 1599.70245 | -0.048912039 | 0.16558052 | -0.2953973 | 0.76769039 | 0.87183444 |
| UBE2D3 | 16144.3201 | -0.482992251 | 0.2290989 | -2.1082259 | 0.03501145 | 0.11267414 |
| AC018797.2 | 73.3564722 | -0.622714954 | 0.3375071 | -1.8450425 | 0.06503135 | 0.17484469 |
| CISD2 | 2602.74798 | -0.407144502 | 0.25640328 | -1.5879067 | 0.11230743 | 0.25781727 |
| SLC9B1 | 863.24482 | -0.384677841 | 0.2683778 | -1.4333445 | 0.15175941 | 0.31680619 |
| SLC9B2 | 5845.22328 | 0.552008579 | 0.22693567 | 2.43244515 | 0.01499726 | 0.06088021 |
| BDH2 | 9866.13731 | 0.673913084 | 0.25488857 | 2.64395178 | 0.00819443 | 0.03878797 |
| CENPE | 33.7435338 | -1.186649489 | 0.48985396 | -2.4224556 | 0.01541601 | 0.06210207 |
| LINC02428 | 0.18311127 | -1.003574173 | 3.30558948 | -0.3035992 | 0.7614333 | NA |
| AC105460.2 | 0 | NA | NA | NA | NA | NA |
| AC105460.1 | 0 | NA | NA | NA | NA | NA |
| TACR3 | 14.2578642 | 1.596946979 | 0.79344527 | 2.01267438 | 0.0441489 | 0.13310053 |
| LINC02503 | 0 | NA | NA | NA | NA | NA |
| AC004052.1 | 0.21421642 | 0.68915655 | 3.07571484 | 0.22406386 | 0.82270761 | NA |
| CXXC4 | 67.7956404 | 0.401397386 | 0.47351435 | 0.84769847 | 0.39660592 | 0.59161094 |
| CXXC4-AS1 | 1.54525881 | 1.616445095 | 1.01317091 | 1.5954318 | 0.11061571 | 0.25479877 |
| AC004063.1 | 0.29992972 | 2.233572672 | 3.22779297 | 0.69198139 | 0.48894902 | NA |
| AC004053.1 | 5.04632527 | 1.36189472 | 0.67611861 | 2.01428374 | 0.04397975 | 0.13283084 |
| AC096577.1 | 108.619287 | 0.191755426 | 0.79589295 | 0.24093118 | 0.80960846 | 0.89703256 |
| AC004069.1 | 24.7075628 | -0.747683647 | 0.47644744 | -1.5692888 | 0.11658065 | 0.26479171 |
| TET2 | 998.414756 | 0.155884104 | 0.16967031 | 0.91874708 | 0.35822788 | 0.55590356 |
| TET2-AS1 | 4.15436772 | -0.857057724 | 0.58173796 | -1.4732711 | 0.14067795 | 0.30086598 |
| PPA2 | 4144.65369 | 0.064278027 | 0.19403893 | 0.33126356 | 0.74044541 | 0.85426378 |
| AC004066.2 | 2.12612885 | 0.945063515 | 0.77165094 | 1.22472931 | 0.22067723 | 0.40657154 |
| ARHGEF38 | 1500.09557 | -0.964409188 | 0.21045748 | -4.5824419 | 4.60E-06 | 0.00011002 |
| INTS12 | 1056.42767 | -0.331546609 | 0.13285492 | -2.4955538 | 0.01257607 | 0.05343714 |
| GSTCD | 392.507062 | 0.103755651 | 0.15340331 | 0.67635862 | 0.49881298 | 0.68014482 |
| AC008243.1 | 9.09031979 | -0.853910648 | 0.59326817 | -1.4393333 | 0.15005611 | 0.3144947 |
| NPNT | 5863.73504 | 0.219194329 | 0.35833176 | 0.61170778 | 0.54073111 | 0.71370336 |
| AC109361.2 | 125.528989 | 1.55915662 | 0.46033314 | 3.38701795 | 0.00070657 | 0.00595468 |
| AC109361.1 | 20.3299204 | 0.145663004 | 0.46955745 | 0.31021338 | 0.75639869 | 0.86489609 |
| TBCK | 1852.34134 | 0.402663223 | 0.20514585 | 1.96281433 | 0.04966775 | 0.14504062 |
| AIMP1 | 2540.48242 | -0.234951476 | 0.13091656 | -1.7946659 | 0.07270695 | 0.18964217 |
| GIMD1 | 0.20356813 | 0.18657164 | 3.29377195 | 0.05664376 | 0.95482897 | NA |
| LINC02173 | 0 | NA | NA | NA | NA | NA |
| DKK2 | 17.8100675 | 0.411072295 | 0.64421279 | 0.63810017 | 0.52340847 | 0.69882538 |
| AC104663.1 | 0 | NA | NA | NA | NA | NA |
| PAPSS1 | 2169.25312 | 0.220431551 | 0.13724962 | 1.60606315 | 0.10826004 | 0.2510833 |
| SGMS2 | 1688.21212 | -0.287604063 | 0.29084109 | -0.9888701 | 0.32272669 | 0.52050781 |
| AC096564.1 | 109.216741 | -0.584936122 | 0.39361066 | -1.486078 | 0.1372585 | 0.29583218 |
| CYP2U1 | 490.265417 | 0.474082637 | 0.18797986 | 2.52198627 | 0.01166943 | 0.05054579 |
| AC096564.2 | 3.33034375 | -0.891933084 | 0.92266665 | -0.9666905 | 0.33369878 | 0.53170565 |
| HADH | 6034.77003 | 1.65014808 | 0.38293669 | 4.30919289 | 1.64E-05 | 0.00030285 |
| LEF1 | 294.128231 | -0.35341888 | 0.319536 | -1.1060377 | 0.26871018 | 0.46288787 |
| RPL34-AS1 | 26.9187604 | 0.97595945 | 0.38201423 | 2.55477249 | 0.01062572 | 0.04710832 |
| RPL34 | 12612.689 | -0.397956337 | 0.16643195 | -2.3911055 | 0.01679773 | 0.06613494 |
| OSTC | 2820.04752 | -0.549731466 | 0.17425931 | -3.154675 | 0.00160677 | 0.01139605 |
| ETNPPL | 178.002839 | 5.415916243 | 0.65332197 | 8.2898119 | 1.13E-16 | 1.03E-13 |
| COL25A1 | 190.292538 | 0.335814119 | 0.52076374 | 0.64484927 | 0.51902487 | 0.695886 |
| ZCCHC23 | 1.04869283 | 0.937574336 | 1.08728215 | 0.86231006 | 0.38851692 | 0.5841377 |
| SEC24B-AS1 | 74.744226 | 0.310152675 | 0.3491215 | 0.88838033 | 0.3743362 | 0.57052628 |
| SEC24B | 2040.54672 | -0.145787723 | 0.12867309 | -1.1330086 | 0.25721064 | 0.44990535 |
| MCUB | 431.11303 | -1.834490136 | 0.3920352 | -4.6794016 | 2.88E-06 | 7.59E-05 |
| CASP6 | 1065.2679 | -0.511168749 | 0.24767923 | -2.0638337 | 0.03903349 | 0.12162508 |
| AC004067.1 | 24.2626603 | -0.143193273 | 0.38428816 | -0.3726195 | 0.70943164 | 0.83490482 |
| PLA2G12A | 2524.48033 | -0.125790474 | 0.20789495 | -0.6050675 | 0.54513415 | 0.7168597 |
| AC126283.2 | 16628.4455 | 0.043582865 | 0.21690296 | 0.20093255 | 0.84075132 | 0.91512686 |
| CFI | 18415.5354 | 0.020918859 | 0.21463923 | 0.09746056 | 0.92236066 | 0.96203347 |
| AC126283.1 | 5.29321089 | 1.191473357 | 0.54656464 | 2.1799313 | 0.02926255 | 0.09915333 |
| GAR1 | 740.144547 | -0.643627461 | 0.27283531 | -2.3590329 | 0.01832263 | 0.07037108 |
| RRH | 19.5706781 | -0.228714129 | 0.56351766 | -0.4058686 | 0.68483914 | 0.81786188 |
| LRIT3 | 58.9346045 | 0.760013944 | 0.34301641 | 2.21567807 | 0.02671356 | 0.09276225 |
| EGF | 5292.62854 | 3.186170189 | 0.3798978 | 8.38691411 | 4.99E-17 | 4.87E-14 |
| ELOVL6 | 950.762651 | -0.930941108 | 0.21319143 | -4.366691 | 1.26E-05 | 0.00024676 |
| ENPEP | 11445.7395 | 1.128195993 | 0.38732242 | 2.9128084 | 0.00358194 | 0.02092972 |
| AC098798.1 | 535.485926 | 1.662477378 | 0.50323107 | 3.30360637 | 0.0009545 | 0.00757451 |
| PANCR | 23.0810543 | 1.521977669 | 0.63402139 | 2.40051469 | 0.01637203 | 0.06487589 |
| PITX2 | 107.479801 | 0.37948354 | 0.53563914 | 0.70846866 | 0.47865427 | 0.6635883 |
| LINC01438 | 0 | NA | NA | NA | NA | NA |
| AC083795.2 | 0 | NA | NA | NA | NA | NA |
| AC139718.2 | 0 | NA | NA | NA | NA | NA |
| AC139718.1 | 0.08393208 | -0.642905668 | 3.30558948 | -0.1944905 | 0.84579185 | NA |
| AC004704.1 | 0 | NA | NA | NA | NA | NA |
| FAM241A | 776.458306 | -0.322297836 | 0.28091529 | -1.1473133 | 0.25125218 | 0.44354643 |
| AC109347.1 | 69.44057 | 0.523238906 | 0.46088629 | 1.13528849 | 0.25625446 | 0.44905696 |
| AP1AR | 1697.4282 | -0.173057893 | 0.38464938 | -0.4499108 | 0.65277479 | 0.79594194 |
| AC109347.2 | 10.2730248 | -1.183787105 | 0.55900217 | -2.1176789 | 0.03420227 | 0.11074278 |
| TIFA | 573.051854 | -1.030691801 | 0.26051582 | -3.9563501 | 7.61E-05 | 0.00102182 |
| ALPK1 | 1671.97559 | -1.021555557 | 0.28345366 | -3.6039597 | 0.00031341 | 0.00313763 |
| NEUROG2 | 0.08666371 | -0.642905668 | 3.30558948 | -0.1944905 | 0.84579185 | NA |
| AC023886.1 | 0.09989236 | -0.642905668 | 3.30558948 | -0.1944905 | 0.84579185 | NA |
| ZGRF1 | 142.033376 | -0.029587888 | 0.34571968 | -0.0855835 | 0.93179754 | 0.96704554 |
| LARP7 | 1839.15141 | -0.050670245 | 0.14097176 | -0.3594354 | 0.71926938 | 0.84021138 |
| AC106864.1 | 460.115395 | 0.192671987 | 0.17122183 | 1.12527702 | 0.26047166 | 0.45355506 |
| AC106864.2 | 2.01882133 | -0.062315802 | 0.84029461 | -0.0741595 | 0.9408835 | 0.97191702 |
| ANK2 | 6340.29335 | 2.119089501 | 0.41182459 | 5.14561194 | 2.67E-07 | 1.08E-05 |
| AC017007.5 | 0.13382144 | 0.02007176 | 3.29738704 | 0.00608717 | 0.99514317 | NA |
| AC093879.1 | 0 | NA | NA | NA | NA | NA |
| AC093879.2 | 0.09997307 | 0.547239394 | 3.29377202 | 0.16614368 | 0.86804388 | NA |
| CAMK2D | 1690.59133 | -0.143813222 | 0.11343474 | -1.2678058 | 0.20486733 | 0.38721589 |
| ARSJ | 505.401654 | 0.247599633 | 0.1883632 | 1.31447989 | 0.18868476 | 0.36687944 |
| AC104779.1 | 2.55515017 | 1.719109014 | 0.84374572 | 2.03747288 | 0.04160268 | 0.1274913 |
| UGT8 | 2508.81813 | 0.227975019 | 0.26703673 | 0.85372157 | 0.39325928 | 0.58868343 |
| NDST4 | 0.798702 | 2.017708206 | 1.42731226 | 1.41364176 | 0.15746711 | NA |
| MTRNR2L13 | 0 | NA | NA | NA | NA | NA |
| AC106892.1 | 5.64948225 | 1.630638838 | 1.49948535 | 1.08746567 | 0.27683106 | 0.4722191 |
| TRAM1L1 | 334.66718 | 0.32384834 | 0.4574804 | 0.70789555 | 0.47901013 | 0.66382897 |
| LINC02262 | 0.35047431 | -0.541570177 | 2.85870395 | -0.1894461 | 0.84974323 | NA |
| LINC02263 | 1.00025433 | -3.224942911 | 2.56640191 | -1.2566009 | 0.20889819 | NA |
| LINC01378 | 4.53841194 | 5.75221544 | 1.20279465 | 4.7823753 | 1.73E-06 | 5.06E-05 |
| LINC02264 | 0 | NA | NA | NA | NA | NA |
| NDST3 | 21.46821 | 1.711375896 | 0.70911146 | 2.41340888 | 0.01580408 | 0.0632403 |
| SNHG8 | 1960.77954 | -0.338698658 | 0.20310057 | -1.6676402 | 0.09538718 | 0.23004939 |
| PRSS12 | 106.546796 | 0.957900186 | 0.57639591 | 1.66187886 | 0.09653707 | 0.23186833 |
| AC110079.2 | 13.303122 | 0.964734385 | 0.57114397 | 1.68912646 | 0.0911952 | 0.22270159 |
| METTL14 | 1151.04822 | 0.067999622 | 0.13249676 | 0.51321724 | 0.60779937 | 0.76405917 |
| SEC24D | 1534.9923 | -1.056829427 | 0.24373674 | -4.3359463 | 1.45E-05 | 0.00027613 |
| SYNPO2 | 2275.16551 | -0.060232466 | 0.57633855 | -0.1045088 | 0.91676556 | 0.95955498 |
| MYOZ2 | 50.0541286 | 1.760887761 | 0.61962644 | 2.84185383 | 0.00448521 | 0.02482159 |

| | | | | | | |
|---|---|---|---|---|---|---|
| USP53 | 2695.17789 | -0.611059076 | 0.25777985 | -2.3704688 | 0.01776555 | 0.068705 |
| C4orf3 | 8483.07891 | -0.212832075 | 0.15559526 | -1.367857 | 0.17135683 | 0.34386116 |
| FABP2 | 10.295816 | 1.200654406 | 1.34902603 | 0.89001575 | 0.37345743 | 0.56968083 |
| AC093752.3 | 36.0321263 | 1.933415311 | 0.53213766 | 3.63329915 | 0.00027982 | 0.00287787 |
| AC093752.2 | 2.80430541 | 0.591056062 | 0.68196748 | 0.86669244 | 0.38611055 | 0.58174897 |
| PDE5A | 989.6198 | 0.441613202 | 0.21519674 | 2.05213705 | 0.04015635 | 0.12412646 |
| LINC01365 | 0 | NA | NA | NA | NA | NA |
| LINC02502 | 0.27048815 | -1.364244814 | 2.9428804 | -0.4635747 | 0.6429525 | NA |
| AC097173.2 | 0.13382144 | 0.02007176 | 3.29738704 | 0.00608717 | 0.99514317 | NA |
| MAD2L1 | 145.858244 | -0.213521766 | 0.38507586 | -0.5544927 | 0.57924168 | 0.74259293 |
| AC108866.1 | 18.1024476 | 0.697142066 | 0.32320525 | 2.15696396 | 0.03100847 | 0.10332239 |
| AC025741.1 | 0.3120295 | -0.695131526 | 2.31225482 | -0.3006293 | 0.76369719 | NA |
| PRDM5 | 273.004642 | 0.005571704 | 0.27058716 | 0.02059116 | 0.98357179 | 0.99290974 |
| NDNF | 746.663226 | 1.041959999 | 0.60373357 | 1.72586063 | 0.08437249 | 0.21041513 |
| AC105254.1 | 0.71395029 | -0.684960621 | 1.67561415 | -0.4087818 | 0.68269977 | NA |
| TNIP3 | 7.87896619 | -0.550848965 | 0.93980555 | -0.5861308 | 0.55778763 | 0.72649725 |
| QRFPR | 106.196237 | -1.230510208 | 0.56422308 | -2.1808931 | 0.02919132 | 0.0990192 |
| ANXA5 | 18099.4192 | -1.006471836 | 0.3052933 | -3.2967374 | 0.00097815 | 0.00771346 |
| AC079341.3 | 1.66006748 | -0.363704414 | 1.10852303 | -0.3280982 | 0.74283741 | 0.85562255 |
| TMEM155 | 1.66006748 | -0.363704414 | 1.10852303 | -0.3280982 | 0.74283741 | 0.85562255 |
| AC079341.1 | 0.34054637 | 0.737233786 | 2.90718031 | 0.25359066 | 0.79981181 | NA |
| EXOSC9 | 1002.71643 | -0.27104875 | 0.14799674 | -1.8314509 | 0.06703327 | 0.17860639 |
| CCNA2 | 317.608503 | -0.89897871 | 0.27733421 | -3.2414995 | 0.00118903 | 0.00898482 |
| BBS7 | 629.332686 | -0.107468278 | 0.14432667 | -0.7446183 | 0.4565025 | 0.64589761 |
| AC079341.2 | 0.15661533 | -1.003574173 | 3.30558948 | -0.3035992 | 0.7614333 | NA |
| TRPC3 | 23.8057067 | 0.127652878 | 0.40655263 | 0.31398857 | 0.75352972 | 0.86270042 |
| KIAA1109 | 3245.96062 | 0.413296043 | 0.14583646 | 2.8339692 | 0.00459738 | 0.02528845 |
| ADAD1 | 0 | NA | NA | NA | NA | NA |
| IL2 | 1.34637256 | 0.083754927 | 1.66451376 | 0.05031795 | 0.95986902 | 0.98061932 |
| IL21 | 0.18746814 | 0.18657164 | 3.29377195 | 0.05664376 | 0.95482897 | NA |
| IL21-AS1 | 0.89904022 | -2.105832129 | 2.16629893 | -0.9720875 | 0.33100702 | NA |
| BBS12 | 758.666325 | 0.204864819 | 0.19594868 | 1.0455024 | 0.29579083 | 0.49266098 |
| FGF2 | 1033.14478 | 0.055577724 | 0.30800403 | 0.18044479 | 0.85680339 | 0.92506983 |
| AC021205.3 | 40.5730129 | 1.145055061 | 0.3394065 | 3.37369804 | 0.00074166 | 0.00619104 |
| NUDT6 | 651.216616 | 0.753104838 | 0.19118316 | 3.93917968 | 8.18E-05 | 0.00108126 |
| SPATA5 | 360.303545 | 0.164667914 | 0.17845418 | 0.9227462 | 0.35613948 | 0.5535105 |
| SPRY1 | 2628.84061 | -1.075409403 | 0.31842471 | -3.37728 | 0.00073206 | 0.0061226 |
| AC093821.1 | 1.36407853 | -0.973755063 | 1.56089687 | -0.6238433 | 0.5327305 | 0.7068228 |
| LINC02435 | 5.3828593 | 0.444319678 | 0.73821219 | 0.60188613 | 0.54724994 | 0.71835237 |
| LINC01091 | 39.4281232 | 0.482717606 | 0.50836438 | 0.94955041 | 0.34234074 | 0.54014442 |
| AC096732.1 | 0 | NA | NA | NA | NA | NA |
| AC121154.1 | 0.38833669 | -1.77728758 | 3.30034214 | -0.5385162 | 0.59022075 | NA |
| LINC02516 | 6.23235046 | -0.041957149 | 0.70274166 | -0.0597049 | 0.95239064 | 0.97717271 |
| ANKRD50 | 703.469549 | -0.196602853 | 0.27776452 | -0.707804 | 0.47906696 | 0.66384633 |
| FAT4 | 511.811095 | 0.828630037 | 0.2879574 | 2.87761329 | 0.00400696 | 0.02277722 |
| AC105919.1 | 0.18754884 | 1.469068384 | 3.27783996 | 0.44818185 | 0.65402196 | NA |
| LINC02379 | 0 | NA | NA | NA | NA | NA |
| AC096773.1 | 2.07376805 | 0.817538023 | 0.9186342 | 0.88994947 | 0.37349302 | 0.56970216 |
| AC093772.1 | 5.92155025 | 1.120133705 | 0.57889456 | 1.93495289 | 0.05299608 | 0.15168854 |
| INTU | 804.58927 | 0.110879533 | 0.34945048 | 0.31729685 | 0.75101837 | 0.86117122 |
| SLC25A31 | 0.96689327 | -0.855058658 | 1.45132434 | -0.5891575 | 0.55575561 | NA |
| HSPA4L | 1801.72526 | 0.003791921 | 0.26477447 | 0.01432132 | 0.98857363 | 0.99491209 |
| AC093591.2 | 165.242317 | -0.459956113 | 0.34581956 | -1.3300466 | 0.18350293 | 0.36044486 |
| PLK4 | 67.1773473 | -0.135156035 | 0.36083928 | -0.3745602 | 0.70798757 | 0.83387039 |
| MFSD8 | 1301.20774 | 0.009946266 | 0.14060505 | 0.07073904 | 0.94360545 | 0.97315146 |
| ABHD18 | 1143.39607 | 0.055192968 | 0.13128248 | 0.4204138 | 0.67418319 | 0.81027171 |
| LARP1B | 1573.69966 | -0.233505031 | 0.24765767 | -0.942854 | 0.34575557 | 0.543392 |
| PGRMC2 | 5598.6358 | 0.083986618 | 0.17669062 | 0.47533151 | 0.6345507 | 0.78334825 |
| AC108062.1 | 242.425305 | 0.225116191 | 0.22182716 | 1.014827 | 0.3101883 | 0.50725613 |
| AC110609.1 | 5.78925478 | 0.988460305 | 0.56172574 | 1.75968491 | 0.07846125 | 0.19973193 |
| AC078850.1 | 40.607135 | -1.411423932 | 0.99156523 | -1.4234302 | 0.15461147 | 0.32065297 |
| AC078850.2 | 22.5341276 | 2.728921348 | 0.7101574 | 3.84269931 | 0.00012169 | 0.00148013 |
| JADRR | 0 | NA | NA | NA | NA | NA |
| JADE1 | 3239.68536 | -0.175337507 | 0.1700125 | -1.0313213 | 0.30239018 | 0.49928294 |
| SCLT1 | 734.37309 | -0.337228778 | 0.20027868 | -1.6837977 | 0.09222077 | 0.22448646 |
| C4orf33 | 767.213915 | 0.337292155 | 0.37860182 | 0.89088889 | 0.37298878 | 0.56921038 |
| AC082650.1 | 0 | NA | NA | NA | NA | NA |
| LINC02466 | 0 | NA | NA | NA | NA | NA |
| LINC02465 | 0.09989236 | -0.642905668 | 3.30558948 | -0.1944905 | 0.84579185 | NA |
| LINC02479 | 0 | NA | NA | NA | NA | NA |
| AC093831.1 | 0 | NA | NA | NA | NA | NA |
| LINC02377 | 3.35539443 | -4.968145446 | 2.26583398 | -2.1926344 | 0.02833373 | 0.09697087 |
| SNHG27 | 2.91687414 | -0.284022402 | 1.70051169 | -0.1670217 | 0.86735295 | 0.93105 |
| AC096711.2 | 3.84973077 | -0.310022674 | 0.93969613 | -0.329918 | 0.74146192 | 0.85474757 |
| AC096711.3 | 1.02757035 | -0.556909401 | 1.88982587 | -0.2946882 | 0.76823207 | 0.87196142 |
| AC096711.1 | 1.03687941 | -2.334107842 | 2.70366992 | -0.8633109 | 0.38796653 | 0.58360966 |
| AC114741.1 | 0 | NA | NA | NA | NA | NA |
| LINC01256 | 0 | NA | NA | NA | NA | NA |
| AC115622.1 | 0.19271054 | 1.506623853 | 3.27645372 | 0.4598337 | 0.64563559 | NA |
| AC110751.1 | 0.09989236 | -0.642905668 | 3.30558948 | -0.1944905 | 0.84579185 | NA |
| AC105383.1 | 5.23678867 | 0.983438367 | 0.65134663 | 1.50985409 | 0.13108066 | 0.28671176 |
| PCDH10 | 258.903274 | 0.756630132 | 0.34905431 | 2.16765732 | 0.03018477 | 0.10124299 |
| AC079380.1 | 3.04004906 | -0.007156261 | 0.82875196 | -0.008635 | 0.99311036 | 0.99685282 |
| PABPC4L | 86.5894398 | 0.278800196 | 0.29828207 | 0.9346864 | 0.34994994 | 0.54763396 |
| AC109830.1 | 0 | NA | NA | NA | NA | NA |
| LINC02462 | 0 | NA | NA | NA | NA | NA |
| AC093722.1 | 0.21694805 | 0.689156974 | 3.06622645 | 0.22475736 | 0.82216804 | NA |
| AC104619.2 | 0 | NA | NA | NA | NA | NA |
| AC107463.1 | 0 | NA | NA | NA | NA | NA |
| LINC02485 | 0.57868538 | -2.415981693 | 3.27574472 | -0.7375366 | 0.46079608 | NA |
| AC105362.1 | 0 | NA | NA | NA | NA | NA |
| LINC00613 | 0 | NA | NA | NA | NA | NA |
| AC073429.2 | 0.32742004 | -0.540950409 | 2.89140423 | -0.1870892 | 0.85159072 | NA |
| AC018680.1 | 0.28028631 | 2.074917451 | 3.237272 | 0.64094628 | 0.52155758 | NA |
| AC073429.1 | 0 | NA | NA | NA | NA | NA |
| LINC02511 | 1.91644071 | 0.673715315 | 1.14207354 | 0.58990537 | 0.55525409 | 0.72471047 |
| LINC02510 | 0.51416151 | 2.938899239 | 1.94766677 | 1.5089333 | 0.13131583 | NA |
| PCDH18 | 489.843012 | 0.672275784 | 0.35144442 | 1.91289362 | 0.05576167 | 0.15716495 |
| LINC02172 | 27.2713987 | 1.886251237 | 1.65747573 | 1.13802646 | 0.25510943 | 0.44782558 |
| AC116563.1 | 0.42641274 | 1.198768173 | 2.12454046 | 0.56424822 | 0.57258522 | NA |
| AC131956.2 | 1.41232584 | -1.845487443 | 1.94667147 | -0.948022 | 0.34311824 | 0.54076189 |
| AC131956.1 | 0 | NA | NA | NA | NA | NA |
| LINC00616 | 0 | NA | NA | NA | NA | NA |
| SLC7A11 | 75.0819585 | -2.715519687 | 0.5373537 | -5.0535052 | 4.34E-07 | 1.61E-05 |
| AC093903.1 | 0.19978473 | -1.003574173 | 3.30558948 | -0.3035992 | 0.7614333 | NA |
| LINC00498 | 0.10721287 | -0.642905668 | 3.30558948 | -0.1944905 | 0.84579185 | NA |
| LINC00499 | 0 | NA | NA | NA | NA | NA |
| LINC00500 | 0 | NA | NA | NA | NA | NA |
| AC098859.2 | 0.13390214 | 1.04981649 | 3.28477129 | 0.31960109 | 0.74927075 | NA |
| AC093766.1 | 0.17648264 | -1.003574173 | 3.30558948 | -0.3035992 | 0.7614333 | NA |
| AC109927.1 | 2.11463505 | -0.005020895 | 0.95849749 | -0.0052383 | 0.99582046 | 0.99795473 |
| AC109927.2 | 0 | NA | NA | NA | NA | NA |
| NOCT | 387.057801 | -0.607758455 | 0.45031933 | -1.3496166 | 0.17713899 | 0.35216954 |

| | | | | | | |
|---|---|---|---|---|---|---|
| ELF2 | 1798.03832 | -0.17790233 | 0.09943221 | -1.789182 | 0.07358551 | 0.19115849 |
| MGARP | 47.6014462 | 0.791836291 | 0.39942413 | 1.98244482 | 0.04742948 | 0.14007339 |
| NDUFC1 | 3409.46339 | 0.136169812 | 0.14650726 | 0.92944072 | 0.35266074 | 0.55030836 |
| NAA15 | 1348.26067 | -0.505156096 | 0.18143288 | -2.7842588 | 0.00536502 | 0.02838026 |
| AC097376.2 | 240.929452 | 0.374139623 | 0.23022023 | 1.62513792 | 0.10413317 | 0.2443488 |
| RAB33B | 582.62917 | -0.017635223 | 0.1963811 | -0.089801 | 0.92844534 | 0.96526981 |
| SETD7 | 1892.49524 | -0.527177548 | 0.25130375 | -2.0977703 | 0.03592544 | 0.11469176 |
| AC112236.2 | 5.0196129 | -1.456307999 | 0.64311131 | -2.2644727 | 0.02354506 | 0.08449689 |
| AC112236.1 | 1.33999391 | 0.446037685 | 2.28133803 | 0.19551582 | 0.84498915 | 0.91821667 |
| MGST2 | 1847.88484 | 0.196146954 | 0.18209883 | 1.07714562 | 0.28141521 | 0.47732162 |
| MAML3 | 443.124728 | 0.371164721 | 0.12920202 | 2.87274705 | 0.0040692 | 0.0230515 |
| AC131182.1 | 0.27086062 | -0.179036626 | 2.96059975 | -0.0604731 | 0.95177884 | NA |
| SCOC | 5186.44721 | -0.239101079 | 0.15252763 | -1.567592 | 0.11697639 | 0.26531355 |
| SCOC-AS1 | 40.4256023 | 0.381327878 | 0.34367363 | 1.10956396 | 0.26718697 | 0.46125829 |
| CLGN | 30.5767054 | -0.528305631 | 0.566394 | -0.9327529 | 0.35094758 | 0.54857388 |
| MGAT4D | 0.1981425 | 1.631193554 | 3.26970143 | 0.4988815 | 0.61786287 | NA |
| ELMOD2 | 1377.49332 | 0.159379912 | 0.14732606 | 1.08181752 | 0.27933362 | 0.47492119 |
| UCP1 | 1.9641262 | -2.375500606 | 1.80843045 | -1.3135703 | 0.18899083 | 0.3672788 |
| TBC1D9 | 2279.47286 | -0.035146775 | 0.27476869 | -0.127914 | 0.89821699 | 0.9482358 |
| AC096733.1 | 0.19280283 | 0.18657164 | 3.29377195 | 0.05664376 | 0.95482897 | NA |
| AC096733.2 | 32.7240385 | 0.700056794 | 0.40334716 | 1.73561851 | 0.08263131 | 0.20723636 |
| RNF150 | 1232.26232 | 0.642070964 | 0.33918291 | 1.89299326 | 0.05835877 | 0.16231148 |
| ZNF330 | 2163.94564 | -0.227829776 | 0.18448394 | -1.2349573 | 0.21684641 | 0.40161819 |
| AC107220.1 | 8.93923848 | 5.954696641 | 1.14768707 | 5.18843226 | 2.12E-07 | 8.93E-06 |
| LINC02432 | 288.016836 | 1.174267499 | 0.44670471 | 2.62873322 | 0.00857036 | 0.04007808 |
| AC097504.2 | 8.5066689 | 1.237534526 | 0.55779924 | 2.21860204 | 0.02651381 | 0.09225085 |
| LINC02276 | 0.08393208 | -0.642905668 | 3.30558948 | -0.1944905 | 0.84579185 | NA |
| IL15 | 451.523431 | -0.504964517 | 0.22948853 | -2.2003911 | 0.02777916 | 0.09555653 |
| INPP4B | 631.109472 | 0.672061179 | 0.23187423 | 2.89838665 | 0.00375088 | 0.02173861 |
| AC139720.2 | 0 | NA | NA | NA | NA | NA |
| AC139720.1 | 1.47634609 | 0.293198758 | 1.33234766 | 0.22006175 | 0.82582306 | 0.90695123 |
| AC104596.1 | 21.4097863 | 1.16670782 | 0.45744245 | 2.55050183 | 0.0107568 | 0.04756149 |
| USP38 | 1488.30186 | -0.193176858 | 0.24040471 | -0.8035486 | 0.42165774 | 0.61421392 |
| AC097658.2 | 0.43167188 | 1.250711029 | 2.71751393 | 0.46024089 | 0.64534332 | NA |
| GAB1 | 1490.57023 | 0.114341537 | 0.20576839 | 0.55568077 | 0.57842911 | 0.74199376 |
| SMARCA5 | 3517.12334 | -0.253831503 | 0.1126404 | -2.2534677 | 0.02422967 | 0.08623789 |
| SMARCA5-AS | 77.2847035 | 0.919575196 | 0.39430112 | 2.33216479 | 0.01969202 | 0.074038 |
| FREM3 | 2.9863068 | 1.307878997 | 0.77712143 | 1.68297893 | 0.09237917 | 0.22476829 |
| GYPE | 24.9228019 | 0.150108697 | 0.32496003 | 0.46192972 | 0.64413172 | 0.78974795 |
| GYPB | 3.07079522 | 0.770494356 | 1.12815073 | 0.68297111 | 0.4946251 | 0.67694119 |
| GYPA | 74.2254788 | 2.31478098 | 0.66544914 | 3.47852428 | 0.00050418 | 0.00455943 |
| AC106871.1 | 0 | NA | NA | NA | NA | NA |
| HHIP-AS1 | 81.0210519 | 0.146596488 | 0.25461031 | 0.57576806 | 0.56477196 | 0.73164184 |
| HHIP | 132.6125 | 0.764069161 | 0.42586982 | 1.79413785 | 0.07279117 | 0.18969885 |
| ANAPC10 | 586.371595 | -0.074770859 | 0.10386129 | -0.7199107 | 0.47157996 | 0.65803596 |
| AC109811.1 | 0.18037964 | -1.003574173 | 3.30558948 | -0.3035992 | 0.7614333 | NA |
| ABCE1 | 2510.22329 | -0.312330986 | 0.21776592 | -1.434251 | 0.15150065 | 0.31657328 |
| OTUD4 | 2961.70202 | -0.397726896 | 0.15402049 | -2.5822986 | 0.00981446 | 0.04445335 |
| LINC02266 | 0.62202363 | -2.530524878 | 2.16361076 | -1.1695842 | 0.24216835 | NA |
| SMAD1 | 741.272962 | -0.129657912 | 0.19858075 | -0.6529229 | 0.513806 | 0.69166697 |
| SMAD1-AS2 | 0.13037664 | 0.02007176 | 3.29738704 | 0.00608717 | 0.99514317 | NA |
| SMAD1-AS1 | 158.370426 | 0.314955304 | 0.26274307 | 1.19871976 | 0.23063693 | 0.41825211 |
| LINC02491 | 0.09635527 | 0.547239394 | 3.29377202 | 0.16614368 | 0.86804388 | NA |
| MMAA | 952.697502 | -0.134117362 | 0.20244017 | -0.6625037 | 0.50764846 | 0.68656733 |
| AC093864.1 | 101.672731 | -0.516702193 | 0.24450168 | -2.1132869 | 0.03457622 | 0.11172344 |
| C4orf51 | 6.7749192 | -0.711697414 | 0.61131585 | -1.1642057 | 0.24434065 | 0.4353556 |
| ZNF827 | 1049.46941 | 0.358584489 | 0.24930523 | 1.43833518 | 0.15033897 | 0.31488791 |
| AC104791.1 | 2.87697153 | 2.108761953 | 0.78989434 | 2.66967598 | 0.00759245 | 0.03677489 |
| AC108206.1 | 1.05969632 | -1.800819307 | 1.51121036 | -1.1916404 | 0.23340228 | 0.4217651 |
| AC097372.2 | 0.09997307 | 0.547239394 | 3.29377202 | 0.16614368 | 0.86804388 | NA |
| LINC01095 | 1.63799495 | 0.70386046 | 1.5903929 | 0.44257017 | 0.65807666 | 0.79943711 |
| LSM6 | 510.043904 | -0.243228181 | 0.09192845 | -2.6458424 | 0.00814878 | 0.03862953 |
| REELD1 | 75.3437103 | 0.94086595 | 0.45616159 | 2.06257162 | 0.03915335 | 0.12189773 |
| AC097372.1 | 0.10721287 | -0.642905668 | 3.30558948 | -0.1944905 | 0.84579185 | NA |
| SLC10A7 | 381.139279 | 0.051223034 | 0.12787978 | 0.40055617 | 0.68874692 | 0.82037384 |
| POU4F2 | 0.03392908 | 0.380740593 | 3.29738709 | 0.11546736 | 0.90807469 | NA |
| TTC29 | 14.9281547 | -0.315601245 | 0.9504081 | -0.3320692 | 0.73983702 | 0.85403179 |
| AC092435.2 | 0 | NA | NA | NA | NA | NA |
| AC092435.1 | 0 | NA | NA | NA | NA | NA |
| AC092435.3 | 0 | NA | NA | NA | NA | NA |
| EDNRA | 498.07353 | 0.175262105 | 0.29133052 | 0.60159198 | 0.54744577 | 0.71842513 |
| LINC02507 | 4.14413275 | 1.301268508 | 1.08831817 | 1.19566919 | 0.23182568 | 0.41977911 |
| AC093835.1 | 2.72146523 | 0.283827151 | 0.79351347 | 0.3576841 | 0.72057973 | 0.84102421 |
| TMEM184C | 1934.01686 | 0.111240448 | 0.18668675 | 0.59586687 | 0.55126416 | 0.72144992 |
| PRMT9 | 752.89886 | 0.550766159 | 0.1798049 | 3.06313214 | 0.00219033 | 0.01446871 |
| ARHGAP10 | 963.087341 | 0.264016004 | 0.23224327 | 1.13680798 | 0.25561857 | 0.44823751 |
| NR3C2 | 1108.49589 | 0.656429923 | 0.20011215 | 3.28031016 | 0.00103693 | 0.00806841 |
| AC069272.1 | 1.69034611 | 1.796222051 | 0.88487894 | 2.02990711 | 0.04236598 | 0.12927458 |
| AC108935.1 | 0 | NA | NA | NA | NA | NA |
| AC108156.1 | 5.66090069 | 0.194094069 | 0.64469944 | 0.30106133 | 0.76336773 | 0.86905872 |
| LINC02430 | 0.2121291 | -0.339874279 | 3.10853787 | -0.1093357 | 0.9129362 | NA |
| LINC02355 | 3.42311958 | -1.372530537 | 0.91023759 | -1.5078816 | 0.13158482 | 0.28743305 |
| IQCM | 0.27421255 | 1.100476972 | 2.98484026 | 0.36868873 | 0.71235975 | NA |
| AC093893.1 | 0 | NA | NA | NA | NA | NA |
| AC105343.1 | 0 | NA | NA | NA | NA | NA |
| DCLK2 | 541.545727 | 0.561969851 | 0.33585322 | 1.67326026 | 0.09427607 | 0.22824642 |
| LRBA | 3242.07036 | 0.526242625 | 0.11990302 | 4.38890234 | 1.14E-05 | 0.00022707 |
| AC110813.1 | 0.56246991 | 1.127102743 | 2.15222498 | 0.52369188 | 0.60049286 | NA |
| MAB21L2 | 2.63435928 | 0.207702118 | 1.48694915 | 0.1396834 | 0.88891014 | 0.94259304 |
| RPS3A | 28445.4725 | -0.278160987 | 0.24929425 | -1.1157938 | 0.2645104 | 0.45846048 |
| SH3D19 | 20596.8377 | -0.130751545 | 0.23527473 | -0.5557399 | 0.5783887 | 0.7419785 |
| AC095055.1 | 25.9476455 | -0.20905019 | 0.45310241 | -0.4613751 | 0.64452948 | 0.78997176 |
| PRSS48 | 0.56694594 | -0.074234186 | 1.92374516 | -0.0385884 | 0.96921858 | NA |
| AC104819.3 | 2.39281939 | 0.389425518 | 0.87173451 | 0.44672491 | 0.6550737 | 0.79755729 |
| FAM160A1-D | 3.47734914 | 0.487488651 | 0.81955766 | 0.59481922 | 0.55196432 | 0.72188162 |
| FAM160A1 | 157.089166 | -0.701916934 | 0.31677724 | -2.2158061 | 0.02670479 | 0.09274401 |
| AC092611.2 | 82.510647 | -0.215126509 | 0.24417615 | -0.88103 | 0.37830159 | 0.57427972 |
| GATB | 1576.98727 | 0.675797005 | 0.20206698 | 3.3444208 | 0.00082455 | 0.00674001 |
| AC092611.1 | 50.4925071 | 0.406668463 | 0.50458735 | 0.80594265 | 0.42027592 | 0.61304768 |
| AC107074.1 | 0 | NA | NA | NA | NA | NA |
| AC112242.1 | 0 | NA | NA | NA | NA | NA |
| AC097375.2 | 0 | NA | NA | NA | NA | NA |
| AC097375.1 | 0.72902276 | 2.875400689 | 2.54550498 | 1.12959932 | 0.2586451 | NA |
| LINC02273 | 8.90537998 | 0.855166637 | 1.08920928 | 0.78512611 | 0.43237964 | 0.62441681 |
| AC079340.1 | 1.53263158 | 3.125242726 | 1.21417945 | 2.57395456 | 0.01005435 | 0.04523685 |
| AC079340.2 | 24.0925805 | 3.023178438 | 0.61503 | 4.91549755 | 8.86E-07 | 2.93E-05 |
| AC080078.1 | 0.16416605 | 0.18657164 | 3.29377195 | 0.05664376 | 0.95482897 | NA |
| AC080078.2 | 0.88436292 | 0.582449405 | 1.38699597 | 0.4199359 | 0.67453228 | NA |
| FBXW7 | 865.344636 | -0.485115544 | 0.13310593 | -3.6445824 | 0.00026783 | 0.00277508 |
| FBXW7-AS1 | 14.5338638 | 0.992133247 | 0.53617921 | 1.8503762 | 0.06425935 | 0.17339145 |
| TMEM154 | 157.80121 | -1.006582935 | 0.40498051 | -2.4855096 | 0.01293661 | 0.05452108 |
| AC106882.1 | 1.31616915 | -3.63142282 | 1.74854402 | -2.0768267 | 0.03781756 | 0.1188333 |
| TIGD4 | 44.4980088 | 0.236794843 | 0.30138688 | 0.78568398 | 0.43205265 | 0.62420732 |

| | | | | | | |
|---|---|---|---|---|---|---|
| ARFIP1 | 2053.8749 | -0.220319097 | 0.10728958 | -2.0534995 | 0.04002416 | 0.12386305 |
| FHDC1 | 446.327167 | -1.045974765 | 0.26110717 | -4.0059213 | 6.18E-05 | 0.00086836 |
| AC093599.1 | 0 | NA | NA | NA | NA | NA |
| TRIM2 | 3691.37372 | 0.370333999 | 0.20796007 | 1.78079377 | 0.07494615 | 0.19353015 |
| MND1 | 25.5901724 | -0.942154655 | 0.44143509 | -2.1342994 | 0.03281828 | 0.10755971 |
| TMEM131L | 510.038101 | 0.052557785 | 0.23447948 | 0.22414663 | 0.82264321 | 0.90523708 |
| TLR2 | 1907.09825 | -2.287892793 | 0.48864057 | -4.6821589 | 2.84E-06 | 7.52E-05 |
| RNF175 | 106.073958 | -2.610096354 | 0.49077247 | -5.318343 | 1.05E-07 | 4.98E-06 |
| AC020703.1 | 2.45762026 | -1.066915367 | 0.91768357 | -1.1626179 | 0.24498456 | 0.43602949 |
| SFRP2 | 233.645049 | -0.247817796 | 1.16299222 | -0.2130864 | 0.83125959 | 0.91040231 |
| AC079298.3 | 7.84368673 | -0.302092607 | 0.67777399 | -0.4457129 | 0.65580465 | 0.79807876 |
| DCHS2 | 40.1684894 | 1.005507391 | 0.4620895 | 2.17600138 | 0.02955516 | 0.09981679 |
| AC079298.1 | 3.28787323 | 1.313767078 | 0.7157851 | 1.83542109 | 0.06644332 | 0.17748282 |
| AC079298.2 | 0 | NA | NA | NA | NA | NA |
| PLRG1 | 2599.92451 | -0.039298506 | 0.15109071 | -0.2600988 | 0.7947876 | 0.88867661 |
| FGB | 19327.5248 | -5.407225062 | 0.9148882 | -5.9102577 | 3.42E-09 | 2.77E-07 |
| FGA | 9233.89525 | -6.281159759 | 1.09701446 | -5.7256855 | 1.03E-08 | 7.11E-07 |
| FGG | 5124.35484 | -8.058196621 | 1.34286966 | -6.0007288 | 1.96E-09 | 1.79E-07 |
| LRAT | 59.7136441 | -1.569131235 | 0.47442821 | -3.3074155 | 0.00094161 | 0.00749707 |
| AC009567.1 | 0 | NA | NA | NA | NA | NA |
| RBM46 | 4.68989018 | -1.031092197 | 1.01549062 | -1.0153636 | 0.30993255 | 0.50703649 |
| AC097467.3 | 67.6235629 | 0.415417535 | 0.20632637 | 2.01340006 | 0.04407256 | 0.13298929 |
| AC104407.1 | 4.02991706 | 1.395351085 | 1.44840448 | 0.96337115 | 0.3353613 | 0.53359089 |
| NPY2R | 1.11614027 | 1.151909186 | 1.23995887 | 0.92898984 | 0.35289435 | 0.5505121 |
| AC097467.1 | 0 | NA | NA | NA | NA | NA |
| MAP9 | 693.390489 | 0.22473495 | 0.15340374 | 1.46499003 | 0.14292364 | 0.3042868 |
| AC107208.1 | 0 | NA | NA | NA | NA | NA |
| GUCY1A1 | 1026.417 | 0.335998805 | 0.23281257 | 1.44321592 | 0.14895966 | 0.31303205 |
| GUCY1B1 | 1333.17486 | 0.048307666 | 0.19097134 | 0.25295768 | 0.80030092 | 0.89162767 |
| ASIC5 | 6.05624647 | 1.707713263 | 1.15547757 | 1.47792852 | 0.13942691 | 0.29912852 |
| TDO2 | 52.0096861 | -0.999914332 | 0.97693149 | -1.0235255 | 0.30605943 | 0.5030244 |
| CTSO | 2696.70369 | 0.173992739 | 0.25623948 | 0.67902394 | 0.49712269 | 0.67893661 |
| AC096736.3 | 0 | NA | NA | NA | NA | NA |
| LINC02272 | 0 | NA | NA | NA | NA | NA |
| AC096736.2 | 0 | NA | NA | NA | NA | NA |
| AC096736.1 | 0 | NA | NA | NA | NA | NA |
| PDGFC | 1732.0023 | 0.357115942 | 0.2770018 | 1.28921883 | 0.19732202 | 0.37811436 |
| AC092608.1 | 0 | NA | NA | NA | NA | NA |
| GLRB | 295.920557 | 0.11507909 | 0.30785757 | 0.37380628 | 0.70854845 | 0.83427499 |
| GRIA2 | 15.4530372 | 0.677625887 | 0.64024984 | 1.05837728 | 0.28988347 | 0.48604516 |
| LINC02433 | 0.08695548 | -0.642905668 | 3.30558948 | -0.1944905 | 0.84579185 | NA |
| AC093817.1 | 0 | NA | NA | NA | NA | NA |
| FAM198B | 1370.91543 | -0.275427827 | 0.46444759 | -0.5930224 | 0.55316616 | 0.72290971 |
| FAM198B-AS | 135.880844 | 0.672371306 | 0.42660331 | 1.5761043 | 0.11500177 | 0.26226188 |
| AC098679.1 | 7.7151355 | 0.388146287 | 0.81561176 | 0.47589589 | 0.63414855 | 0.7831451 |
| TMEM144 | 469.40446 | -0.458818285 | 0.29660846 | -1.546882 | 0.1218917 | 0.27276682 |
| RXFP1 | 184.590835 | 0.579891705 | 0.4905869 | 1.18203667 | 0.23719115 | 0.42639107 |
| AC121161.2 | 0.13028434 | 1.049816488 | 3.28477129 | 0.31960109 | 0.74927075 | NA |
| C4orf46 | 226.700745 | 0.027627158 | 0.18587152 | 0.14863578 | 0.88184104 | 0.93863819 |
| ETFDH | 3456.16586 | 0.423511383 | 0.24482243 | 1.72987166 | 0.08365321 | 0.20905374 |
| PPID | 2574.06001 | 0.150286835 | 0.14530152 | 1.03431014 | 0.3009912 | 0.49768474 |
| FNIP2 | 6036.43607 | -0.70504286 | 0.45503693 | -1.549419 | 0.12128103 | 0.27183858 |
| C4orf45 | 1.54185325 | 0.595715338 | 1.02716134 | 0.57996277 | 0.56193972 | 0.72964972 |
| RAPGEF2 | 2078.19222 | 0.199136326 | 0.1870449 | 1.0646445 | 0.28703683 | 0.48334532 |
| AC074344.2 | 0.11223674 | 0.02007176 | 3.29738704 | 0.00608717 | 0.99514317 | NA |
| LINC02233 | 0.34108807 | 1.448580016 | 2.81680543 | 0.51426343 | 0.60706783 | NA |
| LINC02477 | 1.41360158 | 3.102209535 | 1.12498701 | 2.75755142 | 0.00582361 | 0.03012573 |
| AC104793.1 | 1.09677961 | 2.077054832 | 1.19779519 | 1.7340651 | 0.08290653 | 0.20771927 |
| FSTL5 | 5.09002877 | 1.270438442 | 0.83790064 | 1.5162161 | 0.12946472 | 0.28432185 |
| AC023136.1 | 0 | NA | NA | NA | NA | NA |
| AC005150.1 | 0 | NA | NA | NA | NA | NA |
| AC022272.1 | 2.12806034 | 0.317398076 | 1.07905997 | 0.29414313 | 0.76864854 | 0.87217134 |
| NAF1 | 391.104867 | -0.125031318 | 0.18453915 | -0.6775328 | 0.498068 | 0.67967569 |
| NPY1R | 2629.82179 | 0.525249294 | 0.33320803 | 1.57634044 | 0.11494737 | 0.26216589 |
| NPY5R | 202.565833 | 1.040325386 | 0.27919449 | 3.72616735 | 0.00019441 | 0.00213523 |
| TKTL2 | 1.51860145 | 0.907274561 | 1.02023283 | 0.88928187 | 0.37385161 | 0.57001824 |
| TMA16 | 538.952416 | -0.109419111 | 0.16649928 | -0.6571747 | 0.5110686 | 0.689262 |
| 1-Mar | 435.313306 | -0.246970937 | 0.32382478 | -0.7626684 | 0.44566116 | 0.63610743 |
| AC093788.1 | 79.9547435 | -0.54226314 | 0.30998663 | -1.7493114 | 0.08023721 | 0.20282355 |
| SMIM31 | 2.71149632 | 0.53211935 | 1.29524943 | 0.41082384 | 0.68120171 | 0.81527767 |
| APELA | 64.0373033 | 4.753593395 | 0.54868879 | 8.66355118 | 4.57E-18 | 5.74E-15 |
| TRIM61 | 15.3682449 | 0.758244408 | 0.47773263 | 1.58717316 | 0.11247343 | 0.25806355 |
| FAM218A | 72.8308977 | 0.292914662 | 0.39207834 | 0.74708199 | 0.45501407 | 0.64446885 |
| AC106872.1C | 0.27512461 | 2.050195067 | 3.23884076 | 0.63300274 | 0.52673186 | NA |
| TRIM60 | 0 | NA | NA | NA | NA | NA |
| TRIM75P | 0.65580907 | 0.75920356 | 1.54165136 | 0.49246125 | 0.62239331 | NA |
| TMEM192 | 3021.20939 | 0.738733524 | 0.34268937 | 2.1556943 | 0.03110754 | 0.10358706 |
| KLHL2 | 2121.56162 | -0.756720898 | 0.2363132 | -3.2021948 | 0.00136385 | 0.01004319 |
| GK3P | 2.29494182 | 0.56405198 | 0.88743671 | 0.63559685 | 0.52503922 | 0.69982229 |
| MSMO1 | 3855.09875 | -0.84067093 | 0.34949862 | -2.4053627 | 0.01615642 | 0.06431837 |
| CPE | 4078.11696 | -0.027520828 | 0.45067586 | -0.0610657 | 0.9513069 | 0.97675756 |
| LINC01179 | 3.54502188 | 3.022994221 | 0.96174421 | 3.14324141 | 0.00167088 | 0.01173398 |
| TLL1 | 249.081873 | -0.396501473 | 0.48894217 | -0.8109374 | 0.41740164 | 0.61027293 |
| AC097487.1 | 0 | NA | NA | NA | NA | NA |
| SPOCK3 | 10.431382 | 0.899465762 | 0.62675049 | 1.43512574 | 0.15125127 | 0.31617091 |
| AC079349.1 | 0 | NA | NA | NA | NA | NA |
| AC116634.1 | 0.07659028 | -0.642905668 | 3.30558948 | -0.1944905 | 0.84579185 | NA |
| ANXA10 | 3.2811307 | 0.053479271 | 1.36857081 | 0.03907673 | 0.96882921 | 0.98557312 |
| DDX60 | 761.084305 | 0.217766897 | 0.37167895 | 0.58590053 | 0.55794236 | 0.72652906 |
| DDX60L | 591.062182 | -0.482111428 | 0.25668366 | -1.8782319 | 0.06034945 | 0.16622259 |
| PALLD | 6026.95892 | 0.209572535 | 0.39663077 | 0.52838193 | 0.59723427 | 0.75582223 |
| AC079858.1 | 0.08757577 | 0.547239394 | 3.29377202 | 0.16614368 | 0.86804388 | NA |
| AC080188.1 | 3.46227024 | 0.053239835 | 1.02300587 | 0.05204255 | 0.95849479 | 0.98016472 |
| CBR4 | 2273.84439 | 0.968330789 | 0.15082796 | 6.42010155 | 1.36E-10 | 1.86E-08 |
| AC021151.1 | 0.90887459 | 2.074911215 | 1.45836451 | 1.42276584 | 0.15480405 | NA |
| SH3RF1 | 1587.24511 | 0.14795261 | 0.17844922 | 0.82910201 | 0.40704669 | 0.60168362 |
| AC096741.1 | 0.34589109 | 0.785291142 | 3.27645337 | 0.23967719 | 0.81058052 | NA |
| NEK1 | 688.744406 | 0.240132574 | 0.15956008 | 1.50496652 | 0.13233268 | 0.28859477 |
| CLCN3 | 3587.31037 | -0.040771766 | 0.14459734 | -0.2819676 | 0.77796835 | 0.87847694 |
| HPF1 | 668.43014 | -0.175490615 | 0.16614879 | -1.0562257 | 0.29086513 | 0.48690769 |
| LINC02275 | 201.537449 | 1.122697357 | 0.56893203 | 1.9733418 | 0.04845663 | 0.1424369 |
| AC084866.2 | 0.77623886 | 0.714582036 | 1.55747188 | 0.45880895 | 0.64637137 | NA |
| AC084866.1 | 21.5785388 | 2.215769333 | 0.55792992 | 3.97141156 | 7.14E-05 | 0.00097068 |
| MFAP3L | 3183.39739 | 0.277579112 | 0.2115059 | 1.31239418 | 0.18938717 | 0.36772989 |
| AADAT | 318.647583 | 0.629032088 | 0.43757986 | 1.43752523 | 0.15056881 | 0.31521907 |
| LINC01612 | 40.1727756 | -0.200896309 | 0.26831907 | -0.7487217 | 0.45402497 | 0.64372316 |
| LINC02512 | 16.4305257 | -0.932962702 | 0.6812497 | -1.3694871 | 0.17084705 | 0.34307316 |
| LINC02382 | 0 | NA | NA | NA | NA | NA |
| LINC02431 | 0 | NA | NA | NA | NA | NA |
| LINC02504 | 0.09989236 | -0.642905668 | 3.30558948 | -0.1944905 | 0.84579185 | NA |
| AC074254.1 | 0.03392908 | 0.380740593 | 3.29738709 | 0.11546736 | 0.90807469 | NA |
| LINC02174 | 0 | NA | NA | NA | NA | NA |
| GALNTL6 | 10.6194982 | -0.537004351 | 0.47447238 | -1.1317926 | 0.25772162 | 0.45029234 |

| | | | | | | |
|---|---|---|---|---|---|---|
| AC105285.1 | 84.4908868 | -0.067936003 | 0.2636068 | -0.2577172 | 0.79662518 | 0.88965766 |
| GALNT7 | 1052.6206 | -0.663352308 | 0.17275102 | -3.8399327 | 0.00012307 | 0.0014867 |
| AC097534.1 | 241.317204 | -0.703695776 | 0.1884585 | -3.7339563 | 0.0001885 | 0.0020802 |
| HMGB2 | 2676.03837 | -0.686256302 | 0.30652566 | -2.2388217 | 0.02516752 | 0.08880872 |
| AC097534.2 | 113.690315 | -1.330036155 | 0.36893642 | -3.6050551 | 0.00031209 | 0.00312561 |
| SAP30 | 428.921738 | -1.73993371 | 0.25269458 | -6.8855204 | 5.76E-12 | 1.27E-09 |
| SCRG1 | 28.0053369 | -0.766252014 | 0.7019131 | -1.0916622 | NA | NA |
| AC093849.1 | 0.44623614 | 0.296724979 | 1.93255515 | 0.15354024 | 0.87797226 | NA |
| HAND2 | 11.6872403 | -2.867343781 | 1.09180534 | -2.6262409 | 0.00863337 | 0.04030129 |
| HAND2-AS1 | 12.8065044 | -2.851163887 | 0.91839274 | -3.1045148 | 0.00190591 | 0.0129801 |
| AC106895.1 | 0 | NA | NA | NA | NA | NA |
| LINC02269 | 0 | NA | NA | NA | NA | NA |
| AC106895.2 | 0 | NA | NA | NA | NA | NA |
| LINC02268 | 3.31134994 | 0.234957644 | 1.01497345 | 0.23149142 | 0.81693305 | 0.90154947 |
| AC012055.1 | 0.07830766 | -0.642905668 | 3.30558948 | -0.1944905 | 0.84579185 | NA |
| AC012055.2 | 0 | NA | NA | NA | NA | NA |
| FBXO8 | 1356.38085 | -0.064903046 | 0.13792169 | -0.470579 | 0.63794142 | 0.78565828 |
| CEP44 | 717.719392 | -0.076935787 | 0.11149144 | -0.6900601 | 0.49015642 | 0.67347983 |
| AC116616.1 | 0.20167959 | 0.726716446 | 3.28420131 | 0.22127646 | 0.82487718 | NA |
| HPGD | 1536.96134 | 2.579320928 | 0.43217815 | 5.96818915 | 2.40E-09 | 2.07E-07 |
| AC096751.2 | 6.82479347 | 1.283024425 | 0.83482786 | 1.53687304 | 0.1243244 | 0.27626481 |
| GLRA3 | 3.57388122 | 2.289800774 | 1.59112013 | 1.43911244 | 0.15011867 | 0.3145264 |
| ADAM29 | 2.33755531 | 1.611247686 | 1.01567145 | 1.58638671 | 0.11265161 | 0.25832775 |
| AC105914.2 | 2.25991763 | 0.69799599 | 1.28387392 | 0.54366397 | 0.58667273 | 0.74773127 |
| AC095050.1 | 0.75057998 | 1.88816289 | 1.68878455 | 1.11806026 | 0.26354126 | NA |
| AC131094.1 | 0 | NA | NA | NA | NA | NA |
| GPM6A | 600.612785 | 0.361251297 | 0.44567016 | 0.81057996 | 0.41760693 | 0.61036996 |
| AC097537.1 | 2.72038692 | -0.19500451 | 0.80904046 | -0.2410318 | 0.80953044 | 0.89703256 |
| WDR17 | 216.936927 | 1.312159619 | 0.33860758 | 3.87516316 | 0.00010655 | 0.0013327 |
| SPATA4 | 56.5823333 | 0.880824693 | 0.44191755 | 1.99318784 | 0.04624088 | 0.13758006 |
| ASB5 | 31.8547794 | 0.771204059 | 0.5886673 | 1.31008476 | 0.19016716 | 0.36885418 |
| AC019163.1 | 267.579891 | -0.318252142 | 0.26433117 | -1.2039902 | 0.22859337 | 0.41583934 |
| SPCS3 | 4513.19157 | -0.079525461 | 0.17641064 | -0.4507974 | 0.65213557 | 0.79566996 |
| AC093801.1 | 1.21736066 | -0.951457008 | 1.3151935 | -0.723435 | 0.46941268 | 0.65606954 |
| VEGFC | 491.279147 | -0.967008897 | 0.44348395 | -2.1804823 | 0.02922173 | 0.09906778 |
| AC092673.1 | 0 | NA | NA | NA | NA | NA |
| LINC02509 | 0.09997307 | 0.547239394 | 3.29377202 | 0.16614368 | 0.86804388 | NA |
| AC097518.1 | 1.80649063 | 2.273212118 | 1.48357181 | 1.53225621 | 0.12545922 | 0.27808352 |
| NEIL3 | 5.69961521 | -2.224355411 | 0.83893221 | -2.6514126 | 0.00801559 | 0.03817428 |
| AC027627.1 | 0 | NA | NA | NA | NA | NA |
| AGA | 986.160616 | 0.33430831 | 0.20657838 | 1.61831219 | 0.10559534 | 0.24659536 |
| AC078881.1 | 27.3981353 | 0.441743079 | 0.30535891 | 1.44663563 | 0.147999 | 0.3115955 |
| AC103879.1 | 0.65638997 | 0.656737785 | 1.76613437 | 0.37185041 | 0.71000423 | NA |
| LINC01098 | 0.47693918 | -0.413442254 | 2.85073472 | -0.1450301 | 0.88468713 | NA |
| LINC01099 | 10.2345485 | 2.942085797 | 0.63533196 | 4.6307851 | 3.64E-06 | 9.14E-05 |
| AC020551.1 | 0 | NA | NA | NA | NA | NA |
| AC104803.1 | 0 | NA | NA | NA | NA | NA |
| LINC00290 | 0 | NA | NA | NA | NA | NA |
| LINC02500 | 0 | NA | NA | NA | NA | NA |
| TEMN3-AS1 | 0.29967709 | -1.364242679 | 3.30558948 | -0.4127078 | 0.67982068 | NA |
| AC098864.1 | 42.2979902 | -0.149402613 | 0.51252442 | -0.2915034 | 0.77066634 | 0.87337107 |
| TENM3 | 212.853934 | -0.328588696 | 0.28043253 | -1.171721 | 0.24130908 | 0.43141155 |
| AC079226.2 | 0 | NA | NA | NA | NA | NA |
| AC079226.1 | 1.48386103 | 1.805323594 | 1.17958694 | 1.53047099 | 0.12590019 | 0.27870953 |
| AC079766.1 | 4.91760146 | 0.805888722 | 0.56281335 | 1.43189341 | 0.15217433 | 0.31725782 |
| DCTD | 3985.02072 | -0.285448405 | 0.10115569 | -2.821872 | 0.00477442 | 0.02596392 |
| AC019193.3 | 2.37293575 | 1.850774329 | 0.87769086 | 2.10868589 | 0.0349717 | 0.11258812 |
| AC019193.2 | 16.9240821 | 0.781454908 | 0.48971114 | 1.59574665 | 0.11054537 | 0.25468123 |
| WWC2-AS2 | 66.0918843 | 0.839148874 | 0.30024122 | 2.79491565 | 0.00519133 | 0.02765498 |
| WWC2 | 2207.29443 | 0.634231327 | 0.1409999 | 4.49809761 | 6.86E-06 | 0.00015211 |
| WWC2-AS1 | 0.7707142 | -0.803632808 | 1.81734739 | -0.442201 | 0.65834377 | NA |
| CLDN22 | 303.5357 | 0.293202654 | 0.17435497 | 1.6816421 | 0.09263826 | 0.22521167 |
| CLDN24 | 3.51341636 | 0.60038828 | 0.98224338 | 0.61124187 | 0.54103947 | 0.71388779 |
| CDKN2AIP | 1403.08116 | 0.202009353 | 0.19373426 | 1.04271364 | 0.29708094 | 0.49388629 |
| AC107214.1 | 8.35797193 | -0.18703214 | 0.54030384 | -0.346161 | 0.72922169 | 0.84700595 |
| ING2 | 469.022355 | -0.681158044 | 0.22063867 | -3.0872105 | 0.00202044 | 0.01360202 |
| AC107214.2 | 23.5640734 | -1.207036734 | 0.44262815 | -2.7269769 | 0.00639175 | 0.03229555 |
| RWDD4 | 964.918496 | 0.000630848 | 0.18819397 | 0.00335212 | 0.9973254 | 0.9986244 |
| TRAPPC11 | 1593.75191 | 0.132482812 | 0.1317764 | 1.00536066 | 0.31472322 | 0.51173087 |
| AC108477.1 | 8.13370395 | 1.587118835 | 0.4924195 | 3.22310311 | 0.0012681 | 0.00946498 |
| STOX2 | 760.263442 | -0.355158167 | 0.27348142 | -1.2986556 | 0.19406216 | 0.37380709 |
| ENPP6 | 1716.99529 | 2.441574294 | 0.52560499 | 4.64526471 | 3.40E-06 | 8.65E-05 |
| LINC02363 | 6.59894055 | 0.200901456 | 0.59539356 | 0.33742632 | 0.73579555 | 0.85164064 |
| LINC02362 | 15.9087653 | -0.460169294 | 0.65631295 | -0.7011431 | 0.48321372 | 0.66777016 |
| IRF2 | 1675.85046 | -0.366507256 | 0.1689134 | -2.1697938 | 0.03002247 | 0.10090874 |
| LINC02427 | 10.491706 | 2.126225876 | 0.58119731 | 3.65835463 | 0.00025384 | 0.00265099 |
| LINC02365 | 18.2684982 | 0.217512889 | 0.87965416 | 0.24727092 | 0.80469857 | 0.89426024 |
| CASP3 | 1066.00562 | -0.890418296 | 0.21563207 | -4.1293408 | 3.64E-05 | 0.00057446 |
| PRIMPOL | 500.399237 | 0.004646635 | 0.14685129 | 0.03164177 | 0.97475773 | 0.98843899 |
| CENPU | 228.230494 | -0.425227787 | 0.25120542 | -1.6927492 | 0.09050321 | 0.2215325 |
| ACSL1 | 19355.4414 | 0.214400808 | 0.22587361 | 0.94920699 | 0.34251535 | 0.54018364 |
| AC084871.2 | 5.38306976 | 0.998608582 | 0.57344428 | 1.74142217 | 0.08160961 | 0.20530865 |
| MIR3945HG | 4.76426725 | -1.027166122 | 0.77402836 | -1.3270394 | 0.18449565 | 0.36166743 |
| LINC01093 | 5.86450253 | -1.665892169 | 0.87285649 | -1.9085522 | 0.05631988 | 0.15824777 |
| LINC02437 | 172.847511 | 2.234295292 | 0.70411919 | 3.17317765 | 0.0015078 | 0.01086359 |
| HELT | 32.1524193 | 1.719799171 | 0.61157105 | 2.81210035 | 0.00492191 | 0.0265584 |
| LINC02436 | 0.96854524 | 1.818021905 | 1.69995177 | 1.06945499 | 0.2848647 | NA |
| SLC25A4 | 5845.95947 | 0.488676692 | 0.1737878 | 2.81191599 | 0.00492474 | 0.02656819 |
| CFAP97 | 1542.2089 | -0.050699411 | 0.1574664 | -0.3219697 | 0.74747564 | 0.85882272 |
| SNX25 | 1168.97107 | -0.13711566 | 0.15463512 | -0.8867045 | 0.37523799 | 0.57129724 |
| LRP2BP | 389.907884 | -0.694920064 | 0.24855966 | -2.7957878 | 0.00517734 | 0.0276034 |
| AC112722.1 | 42.2580433 | -0.909494272 | 0.33629946 | -2.7044178 | 0.00684242 | 0.03398284 |
| ANKRD37 | 2059.20583 | -1.172820296 | 0.48303951 | -2.4280007 | 0.01518231 | 0.06136689 |
| UFSP2 | 3025.8226 | -0.276112603 | 0.30244216 | -0.9129435 | 0.36127226 | 0.55878832 |
| C4orf47 | 198.972276 | -1.140284368 | 0.37489669 | -3.0415963 | 0.00235327 | 0.01531486 |
| CCDC110 | 95.6847775 | -0.150302765 | 0.2911879 | -0.5161711 | 0.60573494 | 0.76251816 |
| AC106897.1 | 16.0950393 | -0.737929355 | 0.38016138 | -1.941095 | 0.05224676 | 0.15020333 |
| PDLIM3 | 656.588883 | 0.293827339 | 0.3405466 | 0.86281097 | 0.38824141 | 0.58388992 |
| SORBS2 | 1859.61848 | 1.079248532 | 0.29914484 | 3.60777922 | 0.00030883 | 0.00310208 |
| AC093797.1 | 94.9484583 | 1.568426609 | 0.2620094 | 5.98614642 | 2.15E-09 | 1.91E-07 |
| AC108472.1 | 2.38630763 | 0.84799952 | 0.89332377 | 0.94926335 | 0.34248669 | 0.54018364 |
| AC096659.1 | 0 | NA | NA | NA | NA | NA |
| TLR3 | 794.677784 | -0.169008174 | 0.4290802 | -0.3938848 | 0.69366608 | 0.82376229 |
| FAM149A | 2005.0751 | 0.729678749 | 0.22857472 | 3.19229859 | 0.00141145 | 0.01032169 |
| CYP4V2 | 4540.53151 | 1.017720834 | 0.25696578 | 3.96053065 | 7.48E-05 | 0.00100666 |
| KLKB1 | 1027.42854 | 1.851467734 | 0.30944724 | 5.98314516 | 2.19E-09 | 1.92E-07 |
| F11 | 872.031189 | 2.848444745 | 0.46802692 | 6.08607036 | 1.16E-09 | 1.16E-07 |
| F11-AS1 | 399.405287 | 2.801077615 | 0.4678662 | 5.98692022 | 2.14E-09 | 1.91E-07 |
| AC018709.1 | 7.8601927 | 1.330769289 | 0.66329038 | 2.00631478 | 0.04482267 | 0.13448332 |
| MTNR1A | 29.9876769 | 1.997066974 | 0.75120762 | 2.65847539 | 0.00784951 | 0.03766907 |
| FAT1 | 6750.15386 | 0.216420193 | 0.17021352 | 1.27146302 | 0.20356398 | 0.38576946 |
| AC108865.1 | 3.25477546 | -0.779374764 | 0.90950402 | -0.8569228 | 0.39148753 | 0.5872047 |
| AC108865.2 | 0.8014827 | -1.667833737 | 1.80269674 | -0.9251882 | 0.35486803 | NA |

| | | | | | | |
|---|---|---|---|---|---|---|
| AC110772.1 | 0.59647956 | -0.008150091 | 1.798955 | -0.0045305 | 0.99638523 | NA |
| AC108474.1 | 7.74953323 | -0.840407937 | 0.76955775 | -1.0920661 | 0.27480404 | 0.47005661 |
| LINC02374 | 0 | NA | NA | NA | NA | NA |
| AC097652.1 | 3.12057783 | -1.267001832 | 1.5183972 | -0.8344337 | 0.40403662 | 0.59897135 |
| LINC02514 | 0 | NA | NA | NA | NA | NA |
| LINC02515 | 0.23140391 | 0.077483112 | 3.10308039 | 0.02496974 | 0.9800791 | NA |
| AC093763.1 | 0.08666371 | -0.642905668 | 3.30558948 | -0.1944905 | 0.84579185 | NA |
| LINC02492 | 3.04322936 | -3.430550898 | 1.19117144 | -2.8799808 | 0.00397699 | 0.02264614 |
| AC097521.1 | 0 | NA | NA | NA | NA | NA |
| ZFP42 | 3.93685674 | -0.845620466 | 0.960206 | -0.8806657 | 0.3784988 | 0.57447987 |
| TRIML2 | 0.33501852 | -0.699357289 | 2.8604516 | -0.2444919 | 0.80684986 | NA |
| TRIML1 | 0.29363063 | -0.695421343 | 2.33964545 | -0.2972336 | 0.76628814 | NA |
| AC138781.1 | 0.09644756 | -0.642905668 | 3.30558948 | -0.1944905 | 0.84579185 | NA |
| LINC02434 | 0 | NA | NA | NA | NA | NA |
| LINC01060 | 89.1559469 | 1.487413633 | 0.81737048 | 1.81975452 | 0.0687964 | 0.18187077 |
| AC093909.2 | 0.03392908 | 0.380740593 | 3.29738709 | 0.11546736 | 0.90807469 | NA |
| LINC02508 | 0.5888236 | 1.817533675 | 2.64822528 | 0.6863214 | 0.49251046 | NA |
| AC093909.4 | 0.10721287 | -0.642905668 | 3.30558948 | -0.1944905 | 0.84579185 | NA |
| AC122138.1 | 0.03392908 | 0.380740593 | 3.29738709 | 0.11546736 | 0.90807469 | NA |
| AC107391.1 | 0 | NA | NA | NA | NA | NA |
| AC105924.1 | 0 | NA | NA | NA | NA | NA |
| LINC01262 | 0 | NA | NA | NA | NA | NA |
| AF250324.1 | 0.78584543 | 0.210813829 | 1.53798046 | 0.13707185 | 0.890974 | NA |
| FRG1-DT | 30.5674706 | 0.778576882 | 0.35679351 | 2.18214981 | 0.02909848 | 0.09878049 |
| LINC01596 | 0.18393104 | 1.469068379 | 3.27783996 | 0.44818185 | 0.65402196 | NA |
| FRG1 | 973.240663 | -0.31121583 | 0.16171451 | -1.9244768 | 0.05429484 | 0.15425069 |
| FRG2 | 0.07659028 | -0.642905668 | 3.30558948 | -0.1944905 | 0.84579185 | NA |
| DBET | 0 | NA | NA | NA | NA | NA |
| DUX4 | 0.27046685 | -1.364244813 | 2.94293477 | -0.4635661 | 0.64295864 | NA |
| PLEKHG4B | 179.835669 | 0.666292229 | 0.70102388 | 0.95045582 | 0.34188069 | 0.53971467 |
| LRRC14B | 5.80776066 | -0.104281105 | 0.96824416 | -0.1077012 | 0.91423268 | 0.9579717 |
| CCDC127 | 1007.07015 | -0.301549361 | 0.19531302 | -1.5439286 | 0.12260563 | 0.2737836 |
| SDHA | 14403.9356 | 0.532776699 | 0.19109912 | 2.78796004 | 0.00530411 | 0.0281314 |
| AC021087.1 | 14.6806412 | 1.386277664 | 0.50650178 | 2.73696505 | 0.00620089 | 0.03162813 |
| PDCD6 | 3974.54221 | -0.391013248 | 0.15003659 | -2.6061193 | 0.00915745 | 0.04218765 |
| AHRR | 765.426572 | -0.23879061 | 0.20202586 | -1.1819804 | 0.23721348 | 0.42640215 |
| EXOC3-AS1 | 266.899597 | 0.764203882 | 0.25336162 | 3.01625745 | 0.00255916 | 0.01628499 |
| EXOC3 | 11138.4232 | 0.830034903 | 0.2330312 | 3.56190459 | 0.00036817 | 0.00357548 |
| SLC9A3 | 17470.6705 | 1.366245679 | 0.31887854 | 4.2845331 | 1.83E-05 | 0.00033189 |
| AC106772.2 | 39.7951514 | 2.486658526 | 0.64343371 | 3.86466934 | 0.00011124 | 0.00137524 |
| AC106772.1 | 1.55645056 | 2.770496764 | 1.10032992 | 2.51787824 | 0.01180641 | 0.05098004 |
| CEP72 | 222.400435 | 0.686518962 | 0.20718597 | 3.31353985 | 0.00092123 | 0.0073704 |
| TPPP | 879.193784 | 0.832706532 | 0.31758132 | 2.62202617 | 0.00874087 | 0.04068787 |
| AC026740.1 | 19.996844 | -0.185345139 | 0.5326593 | -0.3479619 | 0.7278688 | 0.84611931 |
| ZDHHC11B | 576.01309 | 0.610512582 | 0.45358244 | 1.34597932 | 0.17830919 | 0.35377423 |
| ZDHHC11 | 556.505153 | 0.493800773 | 0.36055987 | 1.36953893 | 0.17083087 | 0.34307316 |
| BRD9 | 1485.2194 | -0.04475241 | 0.15253782 | -0.2933857 | 0.76922738 | 0.87240608 |
| TRIP13 | 140.248714 | 0.561485023 | 0.32096032 | 1.74939079 | 0.08022349 | 0.20280831 |
| AC122719.1 | 0.5356674 | 2.391263229 | 1.62612816 | 1.47052569 | 0.14141943 | NA |
| AC122719.2 | 0.29988235 | 1.471533266 | 2.82742967 | 0.52044911 | 0.60275059 | NA |
| AC116351.1 | 54.7946698 | 0.155418385 | 0.50023249 | 0.3106923 | 0.75603455 | 0.86462968 |
| AC116351.2 | 15.0770822 | -0.551480825 | 0.47263136 | -1.1668308 | 0.24327871 | 0.4338939 |
| NKD2 | 626.389572 | -1.149894123 | 0.31771454 | -3.6192682 | 0.00029544 | 0.00299296 |
| SLC12A7 | 11143.508 | -0.243523559 | 0.22034092 | -1.1052126 | 0.26906749 | 0.4632008 |
| AC114291.1 | 1120.53996 | 6.266275906 | 0.63601935 | 9.85233529 | 6.70E-23 | 2.94E-19 |
| SLC6A19 | 11301.9263 | 3.372345348 | 0.60243183 | 5.59788702 | 2.17E-08 | 1.34E-06 |
| SLC6A18 | 1173.72725 | 4.289146318 | 0.9024299 | 4.75288586 | 2.01E-06 | 5.67E-05 |
| TERT | 1.11622708 | -2.508711984 | 1.47212347 | -1.7041451 | 0.08835399 | 0.21762513 |
| CLPTM1L | 7458.28969 | 0.195397798 | 0.20488074 | 0.95371482 | 0.34022802 | 0.538298 |
| LINC01511 | 1.32582045 | -2.922044025 | 2.36491991 | -1.2355784 | 0.21661531 | 0.40150036 |
| SLC6A3 | 24.8367324 | 1.122384742 | 0.53269429 | 2.10699599 | 0.03511793 | 0.11292034 |
| LPCAT1 | 4057.65765 | -1.494664665 | 0.33173644 | -4.5055788 | 6.62E-06 | 0.00014768 |
| AC091849.2 | 44.2412292 | 0.581086713 | 0.47650457 | 1.21947774 | 0.22266292 | 0.40867689 |
| AC026412.3 | 16.0125778 | -0.513080399 | 0.61832702 | -0.8297881 | 0.4066586 | 0.60140134 |
| AC112176.1 | 0.10721287 | -0.642905668 | 3.30558948 | -0.1944905 | 0.84579185 | NA |
| MRPL36 | 1015.74701 | -0.668552362 | 0.22043437 | -3.0328863 | 0.00242227 | 0.01560978 |
| NDUFS6 | 2510.49884 | -0.073088092 | 0.15875333 | -0.4603878 | 0.64523791 | 0.79061957 |
| AC025183.3 | 0.03392908 | 0.380740593 | 3.29738709 | 0.11546736 | 0.90807469 | NA |
| LINC02116 | 0 | NA | NA | NA | NA | NA |
| IRX4 | 5.42904943 | -0.919900145 | 0.87689211 | -1.049046 | 0.29415695 | 0.49086233 |
| AC025183.1 | 0.20452486 | -0.339987873 | 3.13600186 | -0.1084144 | 0.91366695 | NA |
| AC025183.2 | 0 | NA | NA | NA | NA | NA |
| AC126768.2 | 8.36677961 | 0.377457255 | 0.59100727 | 0.6386677 | 0.52303913 | 0.69862624 |
| AC126768.1 | 0.3357283 | -1.716651648 | 3.30224936 | -0.5198431 | 0.60317291 | NA |
| AC126768.3 | 0.06785815 | 0.795868262 | 3.28898349 | 0.24198001 | 0.80879565 | NA |
| AC091891.2 | 3.40874041 | 2.197300925 | 0.70557287 | 3.11420833 | 0.00184439 | 0.01265919 |
| AC091891.1 | 16.2280749 | 0.241755843 | 0.56504186 | 0.42785475 | 0.66875687 | 0.80650868 |
| IRX2 | 1657.41271 | -0.588211429 | 0.38789775 | -1.5164085 | 0.1294161 | 0.28425007 |
| C5orf38 | 422.433195 | -1.073634161 | 0.40897383 | -2.6251904 | 0.00866005 | 0.04039223 |
| AC094105.1 | 0 | NA | NA | NA | NA | NA |
| AC112192.1 | 0 | NA | NA | NA | NA | NA |
| AC091832.1 | 0 | NA | NA | NA | NA | NA |
| LINC01019 | 0 | NA | NA | NA | NA | NA |
| LINC02162 | 0 | NA | NA | NA | NA | NA |
| LINC01017 | 0 | NA | NA | NA | NA | NA |
| IRX1 | 1561.25994 | -0.939618443 | 0.44895844 | -2.0928851 | 0.03635941 | 0.11562924 |
| AC016595.1 | 493.594046 | -0.98461739 | 0.46364746 | -2.1236337 | 0.03370078 | 0.1096289 |
| AC025773.1 | 0 | NA | NA | NA | NA | NA |
| AC025187.1 | 0.43491452 | -0.297985305 | 2.28996532 | -0.1301266 | 0.8964663 | NA |
| LINC02063 | 0 | NA | NA | NA | NA | NA |
| AC106799.3 | 0.338427 | 0.669960177 | 2.22621401 | 0.3009415 | 0.76345911 | NA |
| AC106799.2 | 0.19994613 | 1.506623855 | 3.27645372 | 0.45983371 | 0.64563559 | NA |
| AC106799.1 | 8.93498819 | 0.468774086 | 0.672206 | 0.69736671 | 0.48557333 | 0.66965999 |
| AC010634.1 | 0 | NA | NA | NA | NA | NA |
| LINC02114 | 0 | NA | NA | NA | NA | NA |
| AC026719.1 | 0 | NA | NA | NA | NA | NA |
| AC026719.2 | 0 | NA | NA | NA | NA | NA |
| AC026415.1 | 0 | NA | NA | NA | NA | NA |
| AC010451.3 | 0 | NA | NA | NA | NA | NA |
| AC010451.1 | 0.73714804 | 1.67766434 | 1.78648597 | 0.93908621 | 0.34768648 | NA |
| LINC01020 | 197.952568 | 4.01568938 | 0.9909398 | 4.05240496 | 5.07E-05 | 0.00075185 |
| AC010451.2 | 0.38387717 | 2.522961291 | 2.72442213 | 0.92605374 | 0.35441805 | NA |
| LINC02121 | 62.8095673 | 4.229224 | 1.03066202 | 4.10340529 | 4.07E-05 | 0.0006315 |
| AC022424.1 | 160.656459 | 1.333661909 | 0.47550359 | 2.80473575 | 0.00503578 | 0.02700751 |
| ADAMTS16 | 452.505676 | 1.412899768 | 0.43140879 | 3.27508341 | 0.00105631 | 0.00818325 |
| AC022424.2 | 4.17545566 | 1.365406442 | 0.84118425 | 1.62319544 | 0.10454763 | 0.24499438 |
| ICE1 | 1938.8997 | -0.092022477 | 0.1736418 | -0.5299558 | 0.59614259 | 0.7549479 |
| AC010266.2 | 7.80261113 | 0.354479101 | 0.88178231 | 0.40200296 | 0.68768185 | 0.81954943 |
| AC010266.1 | 0.09635527 | 0.547239394 | 3.29377202 | 0.16614368 | 0.86804388 | NA |
| LINC02145 | 50.4911086 | 0.983000714 | 0.46591216 | 2.10984128 | 0.03487203 | 0.11240379 |
| MED10 | 1193.07161 | -0.758392255 | 0.25205587 | -3.008826 | 0.00262259 | 0.01656602 |
| UBE2QL1 | 220.800894 | 1.017368186 | 0.4394345 | 2.31517596 | 0.0206033 | 0.07656582 |
| LINC01018 | 51.5447873 | 2.488096824 | 0.42189037 | 5.89749613 | 3.69E-09 | 2.95E-07 |

| | | | | | | |
|---|---|---|---|---|---|---|
| NSUN2 | 2804.9684 | -0.039147806 | 0.21932211 | -0.1784946 | 0.85833459 | 0.92596478 |
| SRD5A1 | 962.563027 | 0.525709126 | 0.27311602 | 1.92485643 | 0.05424732 | 0.1542496 |
| LINC02102 | 8.91125669 | 0.434013575 | 0.44612311 | 0.97285607 | 0.33062484 | 0.52855878 |
| TENT4A | 1842.33982 | -0.463777581 | 0.19185302 | -2.4173588 | 0.0156336 | 0.06270083 |
| AC122710.2 | 0.56989375 | -0.218416632 | 2.26772052 | -0.0963155 | 0.92327 | NA |
| AC122710.3 | 0.57821457 | 1.107249769 | 2.06619999 | 0.53588703 | 0.59203663 | NA |
| LINC02236 | 0 | NA | NA | NA | NA | NA |
| AC122710.1 | 1.73317406 | 2.504074166 | 1.39472866 | 1.79538447 | 0.07259247 | 0.1894935 |
| LINC02196 | 1.21194033 | -0.080327323 | 1.23473998 | -0.0650561 | 0.94812936 | 0.97504763 |
| AC091951.4 | 0 | NA | NA | NA | NA | NA |
| AC027343.3 | 0 | NA | NA | NA | NA | NA |
| LINC02123 | 0 | NA | NA | NA | NA | NA |
| AC027343.1 | 0 | NA | NA | NA | NA | NA |
| AC027343.2 | 0 | NA | NA | NA | NA | NA |
| LINC02142 | 0 | NA | NA | NA | NA | NA |
| ADCY2 | 60.7474936 | -0.295492534 | 0.45237898 | -0.6531969 | 0.51362937 | 0.69160572 |
| AC093305.1 | 1.71350783 | -0.47422789 | 1.31618211 | -0.3603057 | 0.71861855 | 0.84001549 |
| C5orf49 | 261.487922 | 0.08799801 | 0.34018102 | 0.25867995 | 0.79588219 | 0.88929717 |
| MTRR | 2622.83703 | -0.077146777 | 0.18972955 | -0.4066144 | 0.68429118 | 0.81757775 |
| FASTKD3 | 758.640083 | 0.54220397 | 0.19984931 | 2.71306402 | 0.00666642 | 0.03333843 |
| AC025174.1 | 1.70166371 | -0.518682711 | 1.11842787 | -0.4637605 | 0.64281932 | 0.78864546 |
| LINC02226 | 6.67469452 | 1.71065793 | 0.50731755 | 3.37196677 | 0.00074633 | 0.00622024 |
| AC091965.1 | 12.0354097 | 1.664794819 | 0.46541474 | 3.57701351 | 0.00034754 | 0.00341075 |
| AC091965.4 | 8.5634487 | 1.164066998 | 0.77984347 | 1.49269313 | 0.13551753 | 0.29314673 |
| MIR4458HG | 610.289986 | -0.020649129 | 0.28896654 | -0.0714585 | 0.94303281 | 0.97306994 |
| AC027328.1 | 0 | NA | NA | NA | NA | NA |
| AC091932.1 | 0 | NA | NA | NA | NA | NA |
| LINC02199 | 0.09989236 | -0.642905668 | 3.30558948 | -0.1944905 | 0.84579185 | NA |
| AC021088.1 | 47.7286645 | 1.380512494 | 0.43852672 | 3.1480693 | 0.00164353 | 0.01160992 |
| SEMA5A | 1876.61455 | 0.834380102 | 0.43582892 | 1.91446705 | 0.0555605 | 0.15679907 |
| AC091906.1 | 0.71425496 | -0.65533452 | 1.7122501 | -0.3827329 | 0.70191779 | NA |
| SEMA5A-AS1 | 0.54189314 | -0.034194371 | 1.84450527 | -0.0185385 | 0.98520926 | NA |
| SNHG18 | 786.223158 | 0.116524361 | 0.15104178 | 0.77147103 | 0.44042778 | 0.63178083 |
| TAS2R1 | 1.20146517 | 1.780766888 | 1.25731073 | 1.41632999 | 0.15667891 | 0.32351601 |
| LINC02112 | 1.31675785 | -0.01332274 | 1.28822203 | -0.010342 | 0.99174846 | 0.99605449 |
| LINC02221 | 0 | NA | NA | NA | NA | NA |
| AC034229.1 | 0.47583483 | 0.658734614 | 1.87262967 | 0.35176983 | 0.72501089 | NA |
| AC034229.2 | 15.3196386 | 0.478380417 | 0.38025366 | 1.25805603 | 0.20837149 | 0.39152061 |
| AC034229.3 | 15.1414385 | 0.505557305 | 0.39587889 | 1.27705041 | 0.20158445 | 0.3834234 |
| AC034229.4 | 12.3108045 | 0.246208182 | 0.35501545 | 0.693514 | 0.48798704 | 0.67152554 |
| FAM173B | 556.142801 | 0.183183953 | 0.21300134 | 0.86001316 | 0.38978179 | 0.58538532 |
| AC012640.1 | 26.1582109 | 0.254134587 | 0.31191232 | 0.81476291 | 0.41520804 | 0.6087871 |
| CCT5 | 7733.0964 | -0.460893065 | 0.18950185 | -2.4321296 | 0.01501034 | 0.06091231 |
| AC012640.4 | 2175.25519 | -0.560668971 | 0.18632353 | -3.0091153 | 0.0026201 | 0.01656087 |
| AC012640.3 | 24.6148429 | 0.468747221 | 0.94724138 | 0.49485509 | 0.62070241 | 0.77342166 |
| CMBL | 12965.3213 | 1.476059072 | 0.4544017 | 3.24835729 | 0.00116073 | 0.00882091 |
| AC012640.2 | 73.9167522 | 0.467156078 | 0.30825824 | 1.51546987 | 0.12965345 | 0.28460514 |
| 6-Mar | 7453.20648 | -0.145121489 | 0.12051287 | -1.2041991 | 0.22851264 | 0.41572117 |
| ROPN1L-AS1 | 1.17764115 | -0.345144206 | 1.17373541 | -0.2940562 | 0.76871494 | 0.87219868 |
| ROPN1L | 44.8927831 | -1.189856212 | 0.45211799 | -2.6317383 | 0.00849493 | 0.03982567 |
| LINC01513 | 3.43662367 | -0.714546102 | 0.81540251 | -0.8763109 | 0.38086104 | 0.57691402 |
| AC092336.1 | 229.309174 | -2.413217208 | 0.67580633 | -3.570871 | 0.0003558 | 0.0034801 |
| LINC02212 | 443.068918 | -2.427192235 | 0.69139397 | -3.5105777 | 0.00044713 | 0.00416199 |
| LINC02213 | 0.31764588 | -0.699541525 | 2.89491861 | -0.2416446 | 0.80905554 | NA |
| ANKRD33B | 1579.0874 | 0.190764657 | 0.34581475 | 0.55163829 | 0.5811962 | 0.74384312 |
| ANKRD33B-A | 0.09989236 | -0.642905668 | 3.30558948 | -0.1944905 | 0.84579185 | NA |
| DAP | 9752.41668 | -0.032217543 | 0.19452711 | -0.1656198 | 0.86845616 | 0.93156219 |
| AC012629.2 | 372.166992 | 0.330381461 | 0.28969126 | 1.14046056 | 0.25409447 | 0.44675765 |
| AC012629.1 | 0 | NA | NA | NA | NA | NA |
| CTNND2 | 17.7519129 | 1.808965035 | 0.40741973 | 4.44005259 | 8.99E-06 | 0.00018733 |
| AC106771.1 | 0 | NA | NA | NA | NA | NA |
| LINC01194 | 0 | NA | NA | NA | NA | NA |
| LINC02220 | 0.66711778 | 1.489403446 | 1.47014229 | 1.01310156 | 0.31101165 | NA |
| AC016553.1 | 0 | NA | NA | NA | NA | NA |
| DNAH5 | 900.223604 | -1.008253118 | 0.63055398 | -1.5989957 | 0.10982155 | 0.25360748 |
| AC016576.1 | 4.01482189 | -0.442264683 | 1.33865454 | -0.33038 | 0.74111285 | 0.8546067 |
| TRIO | 3465.88948 | -0.881477964 | 0.29112025 | -3.0278827 | 0.00246274 | 0.01580876 |
| OTULINL | 424.979936 | 1.037502441 | 0.3460884 | 2.99779604 | 0.0027194 | 0.01704551 |
| AC010491.1 | 52.3146391 | 0.155610124 | 0.42924348 | 0.36252182 | 0.71696212 | 0.839049 |
| OTULIN | 1121.38614 | -0.464535855 | 0.19535838 | -2.377865 | 0.0174132 | 0.06777654 |
| ANKH | 2004.35059 | 0.565085194 | 0.25907649 | 2.18115195 | 0.02917218 | 0.09897648 |
| AC016575.1 | 4.42908531 | -0.630903355 | 0.76605718 | -0.8235721 | 0.41018271 | 0.60457873 |
| AC016651.1 | 0.58047997 | 2.487706935 | 1.56343797 | 1.59117726 | 0.11156969 | NA |
| LINC02149 | 5.8478373 | 1.898080848 | 0.90190931 | 2.1045141 | 0.03533363 | 0.11344185 |
| FBXL7 | 599.181668 | 0.531405823 | 0.28184153 | 1.88547737 | 0.05936542 | 0.16436433 |
| AC010638.1 | 0.40682075 | 1.932042247 | 2.15525612 | 0.89643279 | 0.37002166 | NA |
| 11-Mar | 1.60882628 | 0.990381823 | 1.40548386 | 0.70465542 | 0.4810247 | 0.66565116 |
| AC016650.1 | 0 | NA | NA | NA | NA | NA |
| AC092335.1 | 0 | NA | NA | NA | NA | NA |
| LINC02150 | 0.41429116 | -0.543625002 | 2.75767873 | -0.1971314 | 0.84372473 | NA |
| AC020980.1 | 0 | NA | NA | NA | NA | NA |
| ZNF622 | 1481.39256 | -0.678420639 | 0.27926504 | -2.4293074 | 0.0151277 | 0.06124941 |
| RETREG1 | 6601.54742 | -1.380656759 | 0.35875352 | -3.8484828 | 0.00011885 | 0.001453 |
| AC022113.1 | 33.7057126 | -1.020122907 | 0.51554495 | -1.9787274 | 0.04784671 | 0.14103707 |
| AC022113.2 | 16.5090553 | -0.821368016 | 0.5978328 | -1.3739093 | 0.16946985 | 0.34129147 |
| MYO10 | 4229.37373 | -0.370698805 | 0.21420399 | -1.7305877 | 0.08352532 | 0.20883068 |
| BASP1 | 584.862887 | -0.322062142 | 0.42011975 | -0.766596 | 0.44332173 | 0.63444412 |
| AC026790.1 | 0 | NA | NA | NA | NA | NA |
| BASP1-AS1 | 1.72218906 | 2.213312402 | 1.09652112 | 2.01848588 | 0.04354068 | 0.1318407 |
| AC026785.2 | 99.0164359 | -1.326453016 | 0.43510643 | -3.0485714 | 0.00229932 | 0.01503798 |
| AC026785.3 | 0 | NA | NA | NA | NA | NA |
| LINC02111 | 0 | NA | NA | NA | NA | NA |
| LINC02217 | 0.4875428 | 1.513107275 | 2.13313304 | 0.70933563 | 0.47811622 | NA |
| LINC02218 | 0 | NA | NA | NA | NA | NA |
| AC106774.1C | 0.09989236 | -0.642905668 | 3.30558948 | -0.1944905 | 0.84579185 | NA |
| AC106774.8 | 0.16783122 | 1.368031398 | 3.28033652 | 0.41703996 | 0.67664918 | NA |
| AC106774.9 | 0 | NA | NA | NA | NA | NA |
| AC106774.7 | 0 | NA | NA | NA | NA | NA |
| AC106774.6 | 0 | NA | NA | NA | NA | NA |
| AC106774.5 | 0 | NA | NA | NA | NA | NA |
| AC233724.2C | 0 | NA | NA | NA | NA | NA |
| AC233724.17 | 0 | NA | NA | NA | NA | NA |
| AC233724.18 | 0 | NA | NA | NA | NA | NA |
| AC233724.19 | 0 | NA | NA | NA | NA | NA |
| AC233724.15 | 0 | NA | NA | NA | NA | NA |
| AC233724.6 | 0.07830766 | -0.642905668 | 3.30558948 | -0.1944905 | 0.84579185 | NA |
| AC233724.16 | 0.07830766 | -0.642905668 | 3.30558948 | -0.1944905 | 0.84579185 | NA |
| AC233724.12 | 0.07659028 | -0.642905668 | 3.30558948 | -0.1944905 | 0.84579185 | NA |
| H3.Y | 0.23492299 | -1.364242679 | 3.30558948 | -0.4127078 | 0.67982068 | NA |
| LINC02223 | 0 | NA | NA | NA | NA | NA |
| LINC02100 | 1.68649701 | -0.969146008 | 1.07101491 | -0.9048856 | 0.36552598 | 0.56260407 |
| AC106744.1 | 0 | NA | NA | NA | NA | NA |
| AC106744.2 | 0 | NA | NA | NA | NA | NA |

| | | | | | | |
|---|---|---|---|---|---|---|
| CDH18 | 6.51826242 | 5.497400671 | 1.36276941 | 4.03399183 | 5.48E-05 | 0.00079893 |
| CDH18-AS1 | 0 | NA | NA | NA | NA | NA |
| LINC02241 | 0.17828073 | 0.18657164 | 3.29377195 | 0.05664376 | 0.95482897 | NA |
| LINC02146 | 0 | NA | NA | NA | NA | NA |
| AC093274.1 | 3.73803746 | 0.608586363 | 0.99282792 | 0.61298273 | 0.53988776 | 0.71287615 |
| AC091946.2 | 3.62461628 | -0.666398567 | 1.03958081 | -0.6410262 | 0.52150564 | 0.69776958 |
| AC091946.1 | 1.48284266 | -0.041905005 | 1.13412254 | -0.0369493 | 0.97052545 | 0.98638542 |
| CDH12 | 14.8382056 | 1.292043304 | 0.97704245 | 1.32240242 | 0.18603418 | 0.36355441 |
| AC010445.1 | 0 | NA | NA | NA | NA | NA |
| PRDM9 | 0.89735565 | 2.228013106 | 1.21431802 | 1.83478552 | 0.06653748 | NA |
| C5orf17 | 9.52066879 | -1.173658111 | 1.06080363 | -1.1063858 | 0.26855955 | 0.46279822 |
| AC010387.1 | 0.21442573 | -1.003574173 | 3.30558948 | -0.3035992 | 0.7614333 | NA |
| CDH10 | 13.3604565 | 1.098722158 | 0.79860675 | 1.37579873 | 0.16888395 | 0.34047935 |
| AC091885.2 | 0 | NA | NA | NA | NA | NA |
| LINC02239 | 0 | NA | NA | NA | NA | NA |
| AC106821.1 | 0 | NA | NA | NA | NA | NA |
| LINC02228 | 2.61464756 | -3.903341393 | 2.01088507 | -1.9411062 | 0.0522454 | 0.15020333 |
| LINC02211 | 1.5441621 | -1.093078336 | 1.21593369 | -0.8989621 | 0.36867283 | 0.56547087 |
| AC099499.2 | 0.43372084 | -2.014539614 | 2.74681672 | -0.7334088 | 0.46330912 | NA |
| AC099499.1 | 0.08393208 | -0.642905668 | 3.30558948 | -0.1944905 | 0.84579185 | NA |
| AC106754.1 | 1.33567661 | -1.751271166 | 1.57975331 | -1.1085726 | 0.26761462 | 0.46178495 |
| AC022140.1 | 0.08393208 | -0.642905668 | 3.30558948 | -0.1944905 | 0.84579185 | NA |
| AC025475.1 | 0 | NA | NA | NA | NA | NA |
| AC113347.4 | 0 | NA | NA | NA | NA | NA |
| CDH9 | 301.832251 | 1.956954486 | 0.53429456 | 3.66268843 | 0.00024958 | 0.00261712 |
| AC113355.1 | 0 | NA | NA | NA | NA | NA |
| PURPL | 11.9253106 | -0.415669864 | 0.69563202 | -0.5975427 | 0.55014508 | 0.72047687 |
| LINC02103 | 0 | NA | NA | NA | NA | NA |
| AC008825.1 | 0.68765306 | -0.559128579 | 2.2434662 | -0.2492253 | 0.8031865 | NA |
| LINC02109 | 0.08666371 | -0.642905668 | 3.30558948 | -0.1944905 | 0.84579185 | NA |
| LINC02064 | 0 | NA | NA | NA | NA | NA |
| AC113386.1 | 149.439648 | -0.568639268 | 0.46757649 | -1.2161417 | 0.22393095 | 0.4100166 |
| CDH6 | 7330.41945 | -1.062237616 | 0.48933206 | -2.170791 | 0.02994698 | 0.10073177 |
| DROSHA | 1585.02741 | 0.010856197 | 0.12125443 | 0.08953237 | 0.92865883 | 0.96531469 |
| C5orf22 | 929.665229 | -0.051854735 | 0.14122759 | -0.3671714 | 0.71349116 | 0.83731181 |
| PDZD2 | 1328.7119 | 1.525982055 | 0.40723792 | 3.74715115 | 0.00017885 | 0.00200356 |
| AC022447.3 | 0.13028434 | 1.049816488 | 3.28477129 | 0.31960109 | 0.74927075 | NA |
| AC022447.1 | 0.09997307 | 0.547239394 | 3.29377202 | 0.16614368 | 0.86804388 | NA |
| AC022447.7 | 0 | NA | NA | NA | NA | NA |
| AC025178.1 | 41.0391637 | -0.988874493 | 0.40241558 | -2.4573464 | 0.01399676 | 0.05787449 |
| GOLPH3 | 5613.73549 | -0.420134029 | 0.16199544 | -2.5934929 | 0.00950065 | 0.04336717 |
| AC025181.2 | 158.951095 | -0.588072499 | 0.19727421 | -2.9809902 | 0.00287318 | 0.01773543 |
| MTMR12 | 2264.84666 | -0.015425466 | 0.09437094 | -0.1634557 | 0.87015967 | 0.9323196 |
| ZFR | 3838.61741 | -0.032334061 | 0.11888522 | -0.2719771 | 0.7856396 | 0.88309106 |
| SUB1 | 5486.19278 | -0.146356446 | 0.08575119 | -1.706757 | 0.08786723 | 0.21666185 |
| LINC02061 | 127.2166 | 7.693810468 | 1.69625346 | 4.53576699 | 5.74E-06 | 0.00013134 |
| NPR3 | 5712.47148 | 2.958155482 | 0.35643936 | 8.29918302 | 1.05E-16 | 9.87E-14 |
| AC010343.3 | 0.58574028 | 2.602513884 | 1.96939022 | 1.32148208 | 0.18634067 | NA |
| LINC02120 | 1.04601337 | -0.018686545 | 1.80150791 | -0.0103727 | 0.99172391 | 0.99605449 |
| AC034223.1 | 0.28228389 | 0.074529887 | 3.28898342 | 0.02266046 | 0.98192111 | NA |
| AC034223.2 | 0.27482507 | 0.374358866 | 2.98435262 | 0.12544056 | 0.90017478 | NA |
| LINC02160 | 0.31428178 | 0.32853863 | 2.27894242 | 0.14416276 | 0.88537195 | NA |
| AC034231.1 | 32.5861985 | 0.811317992 | 0.34345002 | 2.36225928 | 0.01816393 | 0.06990411 |
| TARS | 3379.00683 | -0.4738245 | 0.22312311 | -2.1236012 | 0.03370351 | 0.1096289 |
| AC034232.2 | 0 | NA | NA | NA | NA | NA |
| ADAMTS12 | 128.355068 | -0.072269329 | 0.31198156 | -0.2316462 | 0.81681286 | 0.90145455 |
| RXFP3 | 0.28174412 | -0.179711611 | 2.92246957 | -0.0614931 | 0.95096654 | NA |
| SLC45A2 | 55.2661085 | 0.495069234 | 0.36620081 | 1.35190643 | 0.17640525 | 0.35115924 |
| AMACR | 11564.5928 | -0.15423854 | 0.43892381 | -0.3514016 | 0.72528707 | 0.84425796 |
| C1QTNF3-AM | 11566.7599 | -0.181366105 | 0.42593946 | -0.4258025 | 0.67025173 | 0.80742522 |
| C1QTNF3 | 1018.61249 | -1.084665383 | 0.48104819 | -2.2547957 | 0.02414616 | 0.08599867 |
| AC139792.1 | 0.11051935 | 0.02007176 | 3.29738704 | 0.00608717 | 0.99514317 | NA |
| AC025754.2 | 7.64724457 | 0.865666865 | 0.45469172 | 1.90385446 | 0.05692914 | 0.15946691 |
| RAI14 | 2191.1943 | 0.177751743 | 0.11653605 | 1.52529406 | 0.12718577 | 0.28077762 |
| AC025754.1 | 15.3509762 | 0.665076232 | 0.31574421 | 2.10637668 | 0.03517165 | 0.11303794 |
| AC026801.2 | 125.568425 | 1.026727043 | 0.32816124 | 3.12872735 | 0.00175565 | 0.01221544 |
| TTC23L | 76.455863 | 0.857397354 | 0.35498149 | 2.41532977 | 0.01572097 | 0.06298419 |
| RAD1 | 1193.37358 | -0.067577054 | 0.18845928 | -0.3585764 | 0.71991198 | 0.84070812 |
| BRIX1 | 1106.7638 | -0.453138203 | 0.19148112 | -2.3664903 | 0.01795765 | 0.06926183 |
| DNAJC21 | 1953.26343 | -0.044817529 | 0.12891851 | -0.3476423 | 0.72810883 | 0.84627247 |
| AGXT2 | 7694.42368 | 2.372193342 | 0.6342863 | 3.73994099 | 0.00018406 | 0.00204627 |
| PRLR | 1823.72228 | 1.758600002 | 0.48014206 | 3.66266599 | 0.0002496 | 0.00261712 |
| SPEF2 | 951.908397 | 0.839272856 | 0.40744123 | 2.05986235 | 0.0394117 | 0.12248434 |
| AC137810.1 | 0.54205048 | 0.532554733 | 2.76217846 | 0.19280244 | 0.8471137 | NA |
| IL7R | 548.370023 | -0.33876754 | 0.75646813 | -0.4478279 | 0.6542774 | 0.79717669 |
| CAPSL | 30.0552397 | -0.994555315 | 0.69759605 | -1.4256894 | 0.15395801 | 0.31975054 |
| AC112204.2 | 0 | NA | NA | NA | NA | NA |
| UGT3A1 | 4231.04156 | 0.765727526 | 0.40205041 | 1.904556 | 0.05683781 | 0.15927875 |
| UGT3A2 | 145.254996 | 1.449413253 | 0.53432305 | 2.71261602 | 0.00667544 | 0.03337088 |
| LMBRD2 | 898.867244 | 0.458467662 | 0.17117621 | 2.67833756 | 0.00739886 | 0.03607099 |
| SKP2 | 369.698173 | 0.37120517 | 0.12033853 | 3.08467426 | 0.00203775 | 0.01368361 |
| NADK2 | 1967.29879 | 0.11970092 | 0.25681556 | 0.46609683 | 0.64114619 | 0.78758115 |
| NADK2-AS1 | 2.44494333 | 0.640135028 | 0.86674108 | 0.73855393 | 0.46017791 | 0.64875121 |
| RANBP3L | 821.119504 | 1.880722875 | 0.55041891 | 3.41689365 | 0.0006334 | 0.00544951 |
| SLC1A3 | 368.825951 | -1.788918394 | 0.51114594 | -3.4998192 | 0.00046557 | 0.00429723 |
| AC008957.2 | 0.48484069 | -2.1930953 | 2.7127443 | -0.8084416 | 0.41883642 | NA |
| AC008957.1 | 4.85495043 | 0.240369818 | 0.66852875 | 0.35955046 | 0.71918334 | 0.84021138 |
| AC026741.1 | 3.93713002 | 0.543353568 | 0.58601676 | 0.92719799 | 0.35382374 | 0.55143807 |
| NIPBL | 2485.73803 | -0.243513628 | 0.17456711 | -1.3949571 | 0.16302877 | 0.33212831 |
| CPLANE1 | 734.714192 | 0.035120701 | 0.22582449 | 0.15552211 | 0.87640973 | 0.93555133 |
| NUP155 | 747.97884 | 0.126146556 | 0.12014803 | 1.04992611 | 0.29375209 | 0.49039267 |
| WDR70 | 1021.73134 | -0.125813468 | 0.12756097 | -0.9863007 | 0.32398558 | 0.52163909 |
| GDNF-AS1 | 113.77171 | 2.067487575 | 0.47423738 | 4.35960486 | 1.30E-05 | 0.00025246 |
| GDNF | 70.1563195 | 1.216030828 | 0.41364577 | 2.93978786 | 0.00328437 | 0.01967936 |
| LINC02117 | 0.09635527 | 0.547239394 | 3.29377202 | 0.16614368 | 0.86804388 | NA |
| AC008869.1 | 0 | NA | NA | NA | NA | NA |
| LINC02110 | 4.08901983 | 1.933467846 | 0.67052078 | 2.88353156 | 0.00393243 | 0.02248444 |
| AC034226.1 | 0.52299198 | 0.878951938 | 1.77903951 | 0.49405982 | 0.62126393 | NA |
| LINC02107 | 1.38403217 | 3.926428424 | 1.24271739 | 3.15955056 | 0.00158013 | 0.01125248 |
| LINC02119 | 3.08154589 | 5.143736598 | 1.25596344 | 4.09545091 | 4.21E-05 | 0.00064781 |
| EGFLAM | 348.180346 | -0.619392563 | 0.45742519 | -1.3540849 | 0.17570928 | 0.34999137 |
| EGFLAM-AS4 | 0 | NA | NA | NA | NA | NA |
| EGFLAM-AS3 | 0 | NA | NA | NA | NA | NA |
| EGFLAM-AS2 | 0 | NA | NA | NA | NA | NA |
| EGFLAM-AS1 | 5.11244515 | 0.32295581 | 0.89484092 | 0.36090863 | 0.71816775 | 0.83985952 |
| AC010457.1 | 12.2849589 | -2.006200577 | 0.77997699 | -2.5721279 | 0.01010755 | 0.04542199 |
| LIFR | 3335.11412 | 0.33147828 | 0.22888905 | 1.44820509 | 0.1475597 | 0.31091884 |
| LIFR-AS1 | 231.323549 | 1.030654556 | 0.33939456 | 3.03674445 | 0.00239148 | 0.01546292 |
| AC010425.1 | 2.82292542 | 1.08840706 | 0.74276837 | 1.46533846 | 0.1428286 | 0.30413357 |
| OSMR-AS1 | 47.6995793 | -1.531674599 | 0.35069474 | -4.3675437 | 1.26E-05 | 0.00024647 |
| AC091435.2 | 9.93797202 | -0.837698182 | 0.47524613 | -1.7626618 | 0.07795755 | 0.19873108 |
| OSMR | 6442.22874 | -2.065009131 | 0.31256901 | -6.6065703 | 3.93E-11 | 6.69E-09 |
| RICTOR | 1936.05892 | -0.32771097 | 0.20306075 | -1.6138568 | 0.10655849 | 0.24814087 |

| | | | | | | |
|---|---|---|---|---|---|---|
| FYB1 | 744.976517 | -0.109391503 | 0.34499652 | -0.31708 | 0.75118289 | 0.86132241 |
| C9 | 1714.25617 | 0.763261899 | 0.35420802 | 2.1548408 | 0.03117429 | 0.10377003 |
| DAB2 | 22146.2578 | 0.22993889 | 0.24919003 | 0.92274514 | 0.35614004 | 0.5535105 |
| LINC02104 | 15.219748 | 0.370429218 | 0.38402281 | 0.96460212 | 0.33474414 | 0.53285059 |
| LINC00603 | 0.74925216 | 1.417786371 | 1.5124066 | 0.93743731 | 0.34853366 | NA |
| LINC00604 | 0.08393208 | -0.642905668 | 3.30558948 | -0.1944905 | 0.84579185 | NA |
| AC093277.1 | 0.23379451 | 0.689159497 | 3.01123709 | 0.22886258 | 0.81897572 | NA |
| TTC33 | 667.495196 | 0.038635117 | 0.15909627 | 0.24284112 | 0.80812848 | 0.89624087 |
| PTGER4 | 998.057112 | 0.393001313 | 0.25178742 | 1.56084568 | 0.11856017 | 0.26765102 |
| PRKAA1 | 3554.59161 | 0.092004112 | 0.10433348 | 0.88182729 | 0.37787021 | 0.57382306 |
| RPL37 | 19641.5978 | -0.527275187 | 0.13725993 | -3.8414357 | 0.00012232 | 0.0014836 |
| CARD6 | 549.397314 | -0.520444761 | 0.30415995 | -1.711089 | 0.08706468 | 0.21509233 |
| C7 | 42196.4943 | 0.930027525 | 0.31746794 | 2.92951632 | 0.0033949 | 0.02014027 |
| MROH2B | 11.7172886 | -0.382851046 | 0.73280756 | -0.5224442 | 0.6013611 | 0.75889606 |
| C6 | 593.514119 | -0.774869578 | 0.5486098 | -1.4124239 | 0.15782519 | 0.32494147 |
| PLCXD3 | 666.643745 | 0.923326268 | 0.4472762 | 2.06433133 | 0.03898631 | 0.1215499 |
| OXCT1 | 2856.13511 | 1.51452654 | 0.37598378 | 4.02816988 | 5.62E-05 | 0.00081224 |
| C5orf51 | 1459.18338 | -0.250287698 | 0.1960892 | -1.2763971 | 0.20181517 | 0.38371614 |
| FBXO4 | 402.363752 | -0.108438473 | 0.17831338 | -0.6081342 | 0.54309845 | 0.71536126 |
| AC108099.1 | 0 | NA | NA | NA | NA | NA |
| GHR | 731.258899 | 1.269919357 | 0.21044578 | 6.03442536 | 1.60E-09 | 1.48E-07 |
| CCDC152 | 18830.7918 | 1.52475911 | 0.31367736 | 4.86091544 | 1.17E-06 | 3.70E-05 |
| SELENOP | 23990.7023 | 1.57726786 | 0.3111035 | 5.06991361 | 3.98E-07 | 1.50E-05 |
| AC008945.1 | 3420.92914 | 1.992247236 | 0.3319723 | 6.00124534 | 1.96E-09 | 1.79E-07 |
| AC008875.1 | 29.5505731 | -1.149756158 | 0.35369983 | -3.2506551 | 0.00115139 | 0.00876289 |
| AC025171.3 | 10.3058348 | 0.466815084 | 0.55007407 | 0.84864041 | 0.39608141 | 0.59122987 |
| AC025171.5 | 13.9559315 | -0.668060375 | 0.45660629 | -1.4630994 | 0.14344019 | 0.30499973 |
| ANXA2R | 101.544765 | -0.360430639 | 0.4034922 | -0.8932778 | 0.3717084 | 0.56836599 |
| AC025171.2 | 40.3058168 | 0.440954576 | 0.29183683 | 1.51096274 | 0.13079794 | 0.28629404 |
| AC025171.4 | 34.3150813 | -0.192540866 | 0.53895452 | -0.3572488 | 0.72090554 | 0.84122492 |
| ZNF131 | 1208.79865 | -0.167912382 | 0.1557684 | -1.0779618 | 0.28105082 | 0.47686346 |
| NIM1K | 56.8719105 | -0.092267459 | 0.38399725 | -0.2402816 | 0.81011199 | 0.89738066 |
| AC114947.2 | 1556.57733 | -0.448969159 | 0.34532883 | -1.3001207 | 0.19355961 | 0.3731389 |
| HMGCS1 | 3791.28014 | -0.196791486 | 0.34906058 | -0.5637746 | 0.57290754 | 0.73753036 |
| CCL28 | 482.384439 | -0.86842362 | 0.40495864 | -2.1444749 | 0.03199486 | 0.10564794 |
| TMEM267 | 694.821791 | -0.165571984 | 0.15838297 | -1.0453901 | 0.29584272 | 0.49269752 |
| C5orf34 | 86.5465565 | -0.288199723 | 0.29432431 | -0.979191 | 0.32748561 | 0.52526221 |
| AC114956.2 | 24.4327803 | 0.066756808 | 0.28896754 | 0.23101836 | 0.81730054 | 0.9017286 |
| AC114956.1 | 11.8495686 | 0.176134929 | 0.4447608 | 0.3960217 | 0.69208902 | 0.82279372 |
| PAIP1 | 3881.0251 | -0.32003038 | 0.13925005 | -2.2982424 | 0.021548 | 0.07922767 |
| NNT-AS1 | 1012.35865 | 0.218022858 | 0.14084034 | 1.54801431 | 0.12161885 | 0.27234113 |
| NNT | 5666.90894 | 0.74134752 | 0.21977916 | 3.37314754 | 0.00074314 | 0.0061995 |
| AC104119.1 | 0 | NA | NA | NA | NA | NA |
| FGF10 | 24.9115192 | 0.27884373 | 0.62721548 | 0.44457405 | 0.6566276 | 0.79852748 |
| FGF10-AS1 | 3.10650195 | -0.300414812 | 0.77571009 | -0.3872772 | 0.69855102 | 0.82697151 |
| LINC02224 | 1.09619152 | 0.641219481 | 1.20933338 | 0.53022557 | 0.59595555 | 0.75484247 |
| AC093292.1 | 0 | NA | NA | NA | NA | NA |
| MRPS30-DT | 63.7498224 | 0.570641132 | 0.36744183 | 1.55301081 | 0.12042055 | 0.2705077 |
| AC093297.1 | 0.40759478 | -0.591312805 | 2.30004901 | -0.257087 | 0.79711158 | NA |
| MRPS30 | 1853.10401 | -0.162091527 | 0.20731111 | -0.7818758 | 0.4342876 | 0.62628043 |
| AC093297.2 | 639.939401 | 0.30385494 | 0.11000011 | 2.76231475 | 0.00573931 | 0.02985121 |
| AC114954.1 | 0 | NA | NA | NA | NA | NA |
| HCN1 | 2.01511214 | -2.351733422 | 1.40809376 | -1.670154 | 0.0948889 | 0.22918289 |
| AC122694.1 | 0.09989236 | -0.642905668 | 3.30558948 | -0.1944905 | 0.84579185 | NA |
| EMB | 466.159709 | -0.061235319 | 0.50906982 | -0.1202886 | 0.9042545 | 0.95182784 |
| AC112187.3 | 3.91752681 | -0.626274705 | 0.91967907 | -0.6809709 | 0.49588989 | 0.67772047 |
| PARP8 | 1228.25605 | -0.901591169 | 0.25438569 | -3.5441898 | 0.00039382 | 0.00377365 |
| AC022441.1 | 0.08757577 | 0.547239394 | 3.29377202 | 0.16614368 | 0.86804388 | NA |
| AC022441.2 | 0 | NA | NA | NA | NA | NA |
| AC008808.2 | 0.44419657 | -2.04182234 | 3.29033058 | -0.6205523 | 0.53489421 | NA |
| LINC02106 | 0 | NA | NA | NA | NA | NA |
| AC008808.1 | 0 | NA | NA | NA | NA | NA |
| AC010478.1 | 0.40615694 | -0.638495507 | 2.89867842 | -0.2202712 | 0.82565991 | NA |
| ISL1 | 0.23282952 | 0.077560627 | 3.09864508 | 0.0250305 | 0.98003064 | NA |
| AC091860.2 | 0 | NA | NA | NA | NA | NA |
| LINC02118 | 0 | NA | NA | NA | NA | NA |
| AC022126.1 | 27.700445 | 0.513904514 | 0.32943822 | 1.55994201 | 0.11877359 | 0.26797223 |
| ITGA1 | 4389.35435 | 0.388576177 | 0.23230368 | 1.67270777 | 0.09438484 | 0.22838394 |
| PELO | 2379.90149 | -0.681539529 | 0.24488415 | -2.78311 | 0.00538405 | 0.02845811 |
| AC025180.1 | 14.1933983 | 0.531463163 | 0.48037389 | 1.10635314 | 0.2685737 | 0.46279822 |
| ITGA2 | 2330.4682 | -0.706700979 | 0.46145581 | -1.5314597 | 0.12565582 | 0.27842568 |
| AC008966.2 | 17.3435905 | 0.441613924 | 0.37048754 | 1.19198051 | 0.23326889 | 0.42158183 |
| MOCS2 | 3364.32014 | 0.466791907 | 0.16971465 | 2.75045141 | 0.00595132 | 0.03060459 |
| AC008966.1 | 204.287916 | 2.039534761 | 0.29649805 | 6.8787459 | 6.04E-12 | 1.31E-09 |
| FST | 203.83843 | -0.633214989 | 0.44157126 | -1.4340041 | 0.1515711 | 0.3165885 |
| NDUFS4 | 2546.06659 | 0.027909252 | 0.15686395 | 0.17792012 | 0.85878571 | 0.92629987 |
| LINC02105 | 0.03392908 | 0.380740593 | 3.29738709 | 0.11546736 | 0.90807469 | NA |
| AC025175.1 | 69.8439532 | -0.073722707 | 0.42643571 | -0.1728812 | 0.86274483 | 0.92851314 |
| ARL15 | 1563.19603 | 0.369995566 | 0.24205133 | 1.52858305 | 0.12636784 | 0.2795805 |
| LINC01033 | 1.245296 | -1.206229934 | 1.42202095 | -0.8482505 | 0.3962985 | 0.59128615 |
| HSPB3 | 12.6217497 | -1.452290339 | 0.67052343 | -2.1659054 | 0.03031841 | 0.10154917 |
| SNX18 | 1774.29288 | 0.459587071 | 0.1738308 | 2.64387601 | 0.00819627 | 0.03878959 |
| AC112198.3 | 0 | NA | NA | NA | NA | NA |
| ESM1 | 1188.89758 | 0.014362755 | 0.54424753 | 0.02639012 | 0.97894618 | 0.99050094 |
| GZMK | 138.222244 | -0.162098525 | 0.7927714 | -0.2044707 | 0.83798568 | 0.91426076 |
| AC034238.3 | 0 | NA | NA | NA | NA | NA |
| AC034238.2 | 0 | NA | NA | NA | NA | NA |
| GZMA | 138.619214 | -0.4820585 | 0.69005976 | -0.698575 | 0.48481767 | 0.6691144 |
| CDC20B | 47.2378402 | 2.862349915 | 1.03255364 | 2.77210771 | 0.00556946 | 0.02925631 |
| GPX8 | 632.310278 | -0.438205876 | 0.29755185 | -1.4727043 | 0.1408308 | 0.30109945 |
| MCIDAS | 8.10939731 | 0.102340502 | 0.69042759 | 0.14822771 | 0.88216306 | 0.93886715 |
| CCNO | 217.558577 | -1.087341924 | 0.510419 | -2.1302928 | 0.03314745 | 0.1083548 |
| AC026704.1 | 36.2121761 | 1.120649549 | 0.46122223 | 2.42973878 | 0.01510971 | 0.06120476 |
| DHX29 | 1939.85645 | 0.325688723 | 0.13107511 | 2.48474872 | 0.01296429 | 0.0545969 |
| MTREX | 4941.27062 | -0.132554131 | 0.18894475 | -0.7015497 | 0.48296002 | 0.66755718 |
| PLPP1 | 6197.43274 | -0.32179931 | 0.31471051 | -1.0225248 | 0.30653257 | 0.50341059 |
| SLC38A9 | 652.155949 | -0.455541158 | 0.1551979 | -2.9352276 | 0.00333303 | 0.019877 |
| DDX4 | 3.02264103 | -0.129150646 | 0.82686077 | -0.1561939 | 0.87588015 | 0.93547831 |
| IL31RA | 8.38368943 | -1.968296561 | 0.69102664 | -2.8483657 | 0.00439444 | 0.0244682 |
| IL6ST | 11305.8682 | -0.263071487 | 0.1960448 | -1.3418947 | 0.17963014 | 0.355379 |
| AC008914.1 | 196.62103 | 0.590250913 | 0.33054913 | 1.7856677 | 0.07415308 | 0.19223522 |
| AC008892.1 | 9.46575375 | 1.148162314 | 0.44332312 | 2.5898995 | 0.0096004 | 0.04370898 |
| AC016596.1 | 24.9274629 | -0.083043886 | 0.55691324 | -0.1491146 | 0.88146322 | 0.93851436 |
| ANKRD55 | 26.4486721 | 0.509383056 | 0.35176508 | 1.44807738 | 0.14759541 | 0.31096924 |
| AC034245.1 | 0.85984127 | 0.896135599 | 1.54426223 | 0.58030015 | 0.56171223 | NA |
| LINC01948 | 34.9789901 | 0.922766928 | 0.49142945 | 1.87772006 | 0.06041947 | 0.16632858 |
| C5orf67 | 212.052879 | 1.015549257 | 0.70118008 | 1.448343 | 0.14752115 | 0.31087031 |
| AC008940.1 | 2.53143483 | 0.042691335 | 0.70898359 | 0.06021484 | 0.95198452 | 0.97699287 |
| MAP3K1 | 1847.92678 | -0.177466366 | 0.22909398 | -0.7746444 | 0.43854982 | 0.63025154 |
| AC008937.2 | 4.30186223 | 0.702711746 | 0.54200997 | 1.29649229 | 0.19480594 | 0.37480173 |
| AC008937.1 | 7.28343229 | 1.198786322 | 0.42964354 | 2.79018815 | 0.00526774 | 0.02796102 |
| SETD9 | 344.024559 | 0.458762689 | 0.30502814 | 1.50400121 | 0.13258106 | 0.28893905 |
| MIER3 | 1005.48691 | -0.079865278 | 0.13314241 | -0.5998485 | 0.54860718 | 0.7192389 |

| | | | | | | |
|---|---|---|---|---|---|---|
| AC008937.3 | 21.2279994 | 0.411935379 | 0.30060658 | 1.37034718 | 0.17057855 | 0.34269057 |
| AC016644.1 | 5.43418367 | 1.227185204 | 0.58441634 | 2.09984752 | 0.03574226 | 0.11424525 |
| GPBP1 | 4122.19764 | -0.242568232 | 0.15163314 | -1.5997045 | 0.10966414 | 0.25335933 |
| AC025470.2 | 1.5927728 | 2.73518013 | 1.34093137 | 2.03976146 | 0.04137409 | 0.12704736 |
| ACTBL2 | 1.8184232 | -1.304058983 | 1.19756844 | -1.0889223 | 0.27618815 | 0.47149878 |
| AC008780.1 | 0.03392908 | 0.380740593 | 3.29738709 | 0.11546736 | 0.90807469 | NA |
| AC008780.2 | 0.24176911 | 0.074529887 | 3.28898342 | 0.02266046 | 0.98192111 | NA |
| AC106822.1 | 0 | NA | NA | NA | NA | NA |
| LINC02225 | 0.08695548 | -0.642905668 | 3.30558948 | -0.1944905 | 0.84579185 | NA |
| LINC02101 | 0.24209763 | 1.011468467 | 2.97311454 | 0.34020501 | 0.73370214 | NA |
| PLK2 | 1276.49157 | -0.100130632 | 0.25581126 | -0.3914239 | 0.69548395 | 0.8247678 |
| GAPT | 52.9352773 | 0.877688469 | 0.63871511 | 1.37414703 | 0.16939604 | 0.34125085 |
| LINC02108 | 0.09997307 | 0.547239394 | 3.29377202 | 0.16614368 | 0.86804388 | NA |
| RAB3C | 40.5921383 | -1.449636133 | 0.75911029 | -1.9096515 | 0.05617809 | 0.15793351 |
| AC016642.1 | 0.08695548 | -0.642905668 | 3.30558948 | -0.1944905 | 0.84579185 | NA |
| AC008852.1 | 0.07830766 | -0.642905668 | 3.30558948 | -0.1944905 | 0.84579185 | NA |
| PDE4D | 1890.64679 | 0.106812271 | 0.21515524 | 0.49644281 | 0.61958203 | 0.77271053 |
| AC092343.1 | 1.16860058 | -1.168712701 | 1.46524766 | -0.7976213 | 0.4250903 | 0.61747446 |
| AC034234.1 | 0.86192236 | 0.813521195 | 1.7697924 | 0.45967041 | 0.64575281 | NA |
| PART1 | 78.8332647 | 0.99621979 | 0.50215097 | 1.98390496 | 0.04726644 | 0.13977697 |
| DEPDC1B | 131.92479 | 1.679413382 | 0.32200824 | 5.215436 | 1.83E-07 | 7.95E-06 |
| ELOVL7 | 1713.60287 | 0.048138643 | 0.18425018 | 0.26126783 | 0.79388598 | 0.88844572 |
| AC104113.1 | 27.2980265 | 0.249184763 | 0.25904483 | 0.96193684 | 0.33608133 | 0.53425171 |
| ERCC8 | 429.370294 | 0.181292244 | 0.12291827 | 1.47490073 | 0.14023923 | 0.30035901 |
| AC022445.1 | 44.8870879 | 0.528669314 | 0.20646759 | 2.56054377 | 0.01045085 | 0.04659941 |
| NDUFAF2 | 601.296244 | -0.734439971 | 0.1746127 | -4.2061084 | 2.60E-05 | 0.00043777 |
| SMIM15 | 2417.71069 | -0.112388812 | 0.1447633 | -0.7763626 | 0.43753493 | 0.62954394 |
| SMIM15-AS1 | 315.180173 | 0.224437675 | 0.2243066 | 1.00058435 | 0.3170278 | 0.514368 |
| LINC02057 | 0.69313557 | 0.2151135 | 1.42022874 | 0.15146398 | 0.87960972 | NA |
| ZSWIM6 | 1044.79771 | -0.521990996 | 0.17075579 | -3.0569446 | 0.00223606 | 0.01472279 |
| AC026746.1 | 0.19642063 | 0.18657164 | 3.29377195 | 0.05664376 | 0.95482897 | NA |
| C5orf64-AS1 | 8.75761231 | -0.964682099 | 0.62377501 | -1.5465225 | 0.12197842 | 0.27284511 |
| AC026434.1 | 0 | NA | NA | NA | NA | NA |
| KIF2A | 722.506822 | -0.598038224 | 0.19364188 | -3.0883724 | 0.00201256 | 0.01356627 |
| DIMT1 | 1151.74403 | -0.477947218 | 0.1387432 | -3.4448334 | 0.00057141 | 0.00503614 |
| IPO11 | 1106.04176 | -0.072035137 | 0.11390194 | -0.6324312 | 0.52710517 | 0.70183102 |
| LRRC70 | 211.417234 | -0.139147163 | 0.30894703 | -0.4503917 | 0.65242806 | 0.79583057 |
| HTR1A | 0 | NA | NA | NA | NA | NA |
| RNF180 | 681.087041 | 0.296565883 | 0.20940989 | 1.41619806 | 0.15671752 | 0.32357039 |
| RGS7BP | 46.2812991 | 1.149981584 | 0.46735837 | 2.46059909 | 0.01387053 | 0.05748083 |
| SHISAL2B | 1.61577691 | -0.200492527 | 1.17551319 | -0.1705574 | 0.86457176 | 0.92965085 |
| SREK1IP1 | 860.358516 | -0.189883174 | 0.12730266 | -1.4915885 | 0.13580707 | 0.29356769 |
| CWC27 | 709.902069 | -0.106988428 | 0.12148211 | -0.8806929 | 0.37848407 | 0.57447987 |
| AC092354.2 | 15.3867 | 0.385562418 | 0.2977492 | 1.29492344 | 0.19534664 | 0.37552619 |
| AC092354.1 | 4.54163487 | 0.200170935 | 0.59498282 | 0.33643145 | 0.73654553 | 0.85207763 |
| ADAMTS6 | 84.4859431 | 1.112620007 | 0.40681201 | 2.73497337 | 0.00623853 | 0.03174645 |
| AC025176.1 | 3.58416519 | 3.757242561 | 1.45081832 | 2.58974023 | 0.00960484 | 0.04372166 |
| CENPK | 56.6354607 | -0.318176096 | 0.42882341 | -0.7419746 | 0.4581027 | 0.64707098 |
| PPWD1 | 1358.74716 | 0.022646485 | 0.19163484 | 0.1181752 | 0.90592884 | 0.95294362 |
| TRIM23 | 686.053118 | 0.260357177 | 0.17632867 | 1.47654475 | 0.13979771 | 0.29968829 |
| TRAPPC13 | 1352.96486 | 0.265547903 | 0.08405568 | 3.15919045 | 0.00158208 | 0.01126336 |
| SHLD3 | 216.095088 | -0.248319604 | 0.23226142 | -1.0691384 | 0.28500731 | 0.48109992 |
| SGTB | 554.619799 | -0.578182533 | 0.22204134 | -2.6039409 | 0.00921586 | 0.0423395 |
| NLN | 1396.30668 | -0.317098044 | 0.2768622 | -1.145328 | 0.25207329 | 0.44432973 |
| AC010359.1 | 25.2200383 | 0.632301213 | 0.38134673 | 1.65807431 | 0.09730247 | 0.23306986 |
| ERBIN | 3657.83144 | 0.167979383 | 0.12008354 | 1.39885435 | 0.16185666 | 0.33063446 |
| SREK1 | 3535.01371 | -0.103906456 | 0.15249853 | -0.6813604 | 0.49564349 | 0.67769982 |
| AC025442.1 | 37.4646921 | 1.24223268 | 0.35753616 | 3.47442527 | 0.00051195 | 0.00461857 |
| LINC02065 | 0.4244122 | -1.988468613 | 3.29269062 | -0.6039039 | 0.54590756 | NA |
| AC113378.1 | 1.6001104 | -0.438144651 | 1.14070598 | -0.3840995 | 0.70090467 | 0.82856754 |
| LINC02229 | 0.16497137 | -1.003574173 | 3.30558948 | -0.3035992 | 0.7614333 | NA |
| MAST4 | 2211.28308 | 0.096599592 | 0.26469462 | 0.36494732 | 0.71515073 | 0.83801965 |
| MAST4-AS1 | 5.4516939 | -0.559253655 | 0.68112975 | -0.8210677 | 0.41160769 | 0.60570403 |
| CD180 | 97.5197407 | -0.345418127 | 0.49753981 | -0.6942522 | 0.48752404 | 0.67137744 |
| AC010420.1 | 0.59080159 | 0.277343652 | 1.64369659 | 0.16873166 | 0.86600772 | NA |
| AC112206.2 | 0.35815827 | -1.741051034 | 2.83291694 | -0.6145789 | 0.53883284 | NA |
| LINC02242 | 0.08666371 | -0.642905668 | 3.30558948 | -0.1944905 | 0.84579185 | NA |
| AC106798.1 | 0.72581425 | 2.362987006 | 1.37932466 | 1.7131478 | 0.08668535 | NA |
| AC024581.1 | 0.25174182 | 1.093535322 | 3.27839713 | 0.33355792 | 0.73871318 | NA |
| LINC02219 | 0 | NA | NA | NA | NA | NA |
| PIK3R1 | 3424.02847 | 0.076953004 | 0.30055105 | 0.25603971 | 0.79792018 | 0.89055551 |
| AC024579.1 | 0.25540699 | 1.967289346 | 2.88471024 | 0.68197122 | 0.49525715 | NA |
| AC010280.2 | 5.61182761 | 1.288772729 | 0.60209002 | 2.14049839 | 0.03231451 | 0.10628952 |
| AC010280.3 | 0.4538796 | 0.297968922 | 1.93957974 | 0.15362551 | 0.87790502 | NA |
| AC010280.1 | 1.14883108 | 3.644248219 | 1.73811618 | 2.09666549 | 0.0360232 | 0.11492036 |
| AC093523.1 | 1.18384924 | 3.661022119 | 1.26530231 | 2.89339717 | 0.00381099 | 0.02199405 |
| AC093248.1 | 1.06607505 | 0.636664414 | 1.26395561 | 0.50370789 | 0.61446666 | 0.76942073 |
| LINC02198 | 13.6890815 | 0.108586933 | 0.48784057 | 0.22258693 | 0.82385701 | 0.905779 |
| AC010273.2 | 13.0508899 | 0.032913286 | 0.50086417 | 0.065713 | 0.94760632 | 0.97478875 |
| SLC30A5 | 2492.94282 | 0.049072084 | 0.1391481 | 0.35266083 | 0.72434273 | 0.84387647 |
| AC010273.1 | 681.973497 | 0.210076503 | 0.15290512 | 1.373901 | 0.16947242 | 0.34129147 |
| CCNB1 | 310.811722 | -0.522475208 | 0.27026337 | -1.9332076 | 0.05321063 | 0.15217051 |
| CENPH | 90.8976916 | -0.798549158 | 0.25912085 | -3.0817634 | 0.00205778 | 0.01377665 |
| MRPS36 | 1702.35759 | 0.221893569 | 0.20363626 | 1.08965648 | 0.27586449 | 0.47121044 |
| CDK7 | 1421.91462 | -0.57490991 | 0.16765784 | -3.4290666 | 0.00060566 | 0.00525427 |
| CCDC125 | 815.89247 | 0.19957582 | 0.27491384 | 0.72595769 | 0.46786471 | 0.65469547 |
| AK6 | 1831.90921 | -0.380465145 | 0.20204619 | -1.8830603 | 0.0596922 | 0.16496368 |
| TAF9 | 2274.82826 | -0.472514452 | 0.21264844 | -2.2220453 | 0.02628025 | 0.09170439 |
| RAD17 | 2087.02459 | 0.133332524 | 0.13723969 | 0.97153034 | 0.33128425 | 0.52919589 |
| MARVELD2 | 1074.33285 | 0.554311908 | 0.15409911 | 3.59711291 | 0.00032177 | 0.00320311 |
| AC145146.1 | 0.03392908 | 0.380740593 | 3.29738709 | 0.11546736 | 0.90807469 | NA |
| OCLN | 1107.14712 | 0.087667375 | 0.21559404 | 0.40663173 | 0.68427849 | 0.81757775 |
| GTF2H2C | 635.831647 | -0.056654917 | 0.21747482 | -0.2605125 | 0.79446845 | 0.88857752 |
| SERF1B | 35.3431941 | 0.021397143 | 0.47160777 | 0.04537063 | 0.96381189 | 0.98305633 |
| SMN2 | 203.792688 | -0.016711797 | 0.72275316 | -0.0231224 | 0.98155263 | 0.99160566 |
| AC140134.2 | 0 | NA | NA | NA | NA | NA |
| AC146944.4 | 13.3550903 | 0.433117674 | 1.19032115 | 0.36386624 | 0.7159579 | 0.83832645 |
| SERF1A | 6.80135167 | 0.418947216 | 1.86591534 | 0.22452638 | 0.82234774 | 0.90506302 |
| SMN1 | 342.970777 | -0.594864663 | 0.11708193 | -5.0807556 | 3.76E-07 | 1.43E-05 |
| AC139834.1 | 0 | NA | NA | NA | NA | NA |
| NAIP | 371.289065 | 0.37283427 | 0.29565862 | 1.26102961 | 0.20729818 | 0.39012654 |
| GTF2H2 | 550.533779 | 0.028352289 | 0.42576674 | 0.06659113 | 0.94690721 | 0.97439878 |
| LINC02197 | 4.75699873 | 1.056767174 | 1.08363697 | 0.97520406 | 0.32945905 | 0.52723848 |
| AC145141.2 | 0.09635527 | 0.547239394 | 3.29377202 | 0.16614368 | 0.86804388 | NA |
| AC145141.1 | 1.39640987 | 1.091583082 | 1.24526287 | 0.87658847 | 0.3807102 | 0.57680713 |
| BDP1 | 1261.43225 | -0.118017871 | 0.13356748 | -0.8835823 | 0.37692171 | 0.57284454 |
| MCCC2 | 5624.75067 | 0.772154274 | 0.19067718 | 4.04953694 | 5.13E-05 | 0.00075899 |
| CARTPT | 1.65645374 | -1.186801109 | 1.01515133 | -1.1690879 | 0.24236823 | 0.43277697 |
| MAP1B | 1937.04795 | -1.012073889 | 0.5474319 | -1.8487667 | 0.0644915 | 0.17383634 |
| MRPS27 | 2753.44194 | 0.133348911 | 0.13199422 | 1.01026325 | 0.31236918 | 0.50922202 |
| PTCD2 | 560.686415 | 0.381415137 | 0.17647403 | 2.16131027 | 0.03067138 | 0.10248407 |
| ZNF366 | 46.2214858 | 1.055545178 | 0.28151151 | 3.74956317 | 0.00017714 | 0.00198946 |
| LINC02056 | 1.47394552 | 1.416262429 | 1.03697542 | 1.36576277 | 0.17201343 | 0.34489008 |

| | | | | | | |
|---|---|---|---|---|---|---|
| AC063979.2 | 2.47319645 | 1.856400593 | 0.8837402 | 2.10061802 | 0.03567451 | 0.1141117 |
| AC035140.1 | 3.17211802 | 0.428853148 | 0.66942124 | 0.64063272 | 0.52176134 | 0.69787989 |
| TNPO1 | 5102.66306 | -0.443164762 | 0.26038424 | -1.7019646 | 0.088762 | 0.21836494 |
| AC008972.1 | 18.1164132 | -0.833729304 | 0.38783487 | -2.1497018 | 0.03157881 | 0.10470681 |
| AC008972.2 | 22.3855453 | 0.013929261 | 0.43610806 | 0.03193993 | 0.97451996 | 0.98835011 |
| FCHO2 | 2135.58283 | -0.274026913 | 0.12504262 | -2.191468 | 0.02841794 | 0.09716141 |
| TMEM171 | 1000.43083 | -1.130979908 | 0.34867039 | -3.2436936 | 0.00117991 | 0.00893314 |
| AC116345.3 | 73.2508842 | 2.5452873 | 0.60619818 | 4.19877092 | 2.68E-05 | 0.00045047 |
| TMEM174 | 3795.1122 | 2.592320281 | 0.82611343 | 3.13797136 | 0.00170121 | 0.01190264 |
| AC116345.1 | 51.526617 | -1.990227198 | 0.56089437 | -3.5483102 | 0.00038771 | 0.00372729 |
| LINC02230 | 0.13390214 | 1.04981649 | 3.28477129 | 0.31960109 | 0.74927075 | NA |
| FOXD1 | 210.741686 | -0.967787216 | 0.27024536 | -3.5811427 | 0.00034209 | 0.0033648 |
| FOXD1-AS1 | 133.654244 | -1.020766649 | 0.27009052 | -3.7793501 | 0.00015724 | 0.00181938 |
| LINC01385 | 1.1672113 | -0.038215287 | 1.44015475 | -0.0265355 | 0.97883018 | 0.99044497 |
| LINC01386 | 0.38566543 | 1.848884861 | 2.6792717 | 0.69006994 | 0.49015021 | NA |
| AC099522.2 | 32.0246214 | 0.700156556 | 0.44634349 | 1.56864965 | 0.1167296 | 0.26504314 |
| BTF3 | 12406.0336 | -0.368381448 | 0.16236879 | -2.2687947 | 0.02328081 | 0.08383922 |
| ANKRA2 | 754.940507 | 0.011738811 | 0.18584911 | 0.06316313 | 0.94963661 | 0.97576049 |
| UTP15 | 551.261868 | -0.173230437 | 0.21694742 | -0.7984904 | 0.42458595 | 0.61701375 |
| ARHGEF28 | 4075.3284 | 0.141683622 | 0.19062997 | 0.74323898 | 0.45733701 | 0.64661289 |
| AC091868.2 | 3.00718894 | -0.792211524 | 0.90960834 | -0.870937 | 0.38378857 | 0.57948708 |
| AC093283.1 | 0.50828345 | 0.949863968 | 2.17585169 | 0.43654812 | 0.66243909 | NA |
| LINC02122 | 0.03392908 | 0.380740593 | 3.29738709 | 0.11546736 | 0.90807469 | NA |
| LINC01335 | 0 | NA | NA | NA | NA | NA |
| AC106732.1 | 0 | NA | NA | NA | NA | NA |
| LINC01333 | 0 | NA | NA | NA | NA | NA |
| LINC01332 | 0 | NA | NA | NA | NA | NA |
| LINC01331 | 0 | NA | NA | NA | NA | NA |
| ENC1 | 639.736718 | 0.635151006 | 0.3902884 | 1.62738888 | 0.10365453 | 0.24355074 |
| HEXB | 8260.90202 | -0.72694924 | 0.26757207 | -2.7168352 | 0.00659094 | 0.03301731 |
| GFM2 | 2352.87527 | 0.227659684 | 0.13646607 | 1.66825116 | 0.09526588 | 0.22978305 |
| NSA2 | 3119.4849 | -0.16466327 | 0.13903915 | -1.1842943 | 0.2362966 | 0.42559533 |
| FAM169A | 449.185067 | 0.174257559 | 0.26483924 | 0.65797485 | 0.51055429 | 0.68897788 |
| AC010501.1 | 40.3329447 | 0.447881636 | 0.62364724 | 0.71816503 | 0.47265554 | 0.65880472 |
| AC116337.3 | 6.44157237 | 0.370558766 | 0.61226199 | 0.60522909 | 0.54502679 | 0.71682576 |
| GCNT4 | 399.527171 | 0.456434439 | 0.27708416 | 1.64727725 | 0.09950107 | 0.23665944 |
| LINC01336 | 14.8605108 | 0.527418926 | 0.41926885 | 1.25794923 | 0.20841012 | 0.39152061 |
| ANKRD31 | 15.4663428 | 1.150020588 | 0.36429623 | 3.15682814 | 0.00159495 | 0.01132443 |
| AC008897.2 | 0.58923357 | 1.872067013 | 1.65853884 | 1.12874475 | 0.25900552 | NA |
| HMGCR | 2376.43801 | -0.230877798 | 0.34375809 | -0.6716287 | 0.5018201 | 0.68228905 |
| COL4A3BP | 2563.28789 | -0.028571282 | 0.23186269 | -0.123225 | 0.90192892 | 0.95040422 |
| AC008897.3 | 11.6532172 | -0.312997325 | 0.67998217 | -0.4603023 | 0.64529928 | 0.79065803 |
| POLK | 1048.11033 | 0.141535196 | 0.16824508 | 0.84124419 | 0.40021115 | 0.5949359 |
| AC010245.1 | 322.997038 | -0.07582239 | 0.14789991 | -0.5126601 | 0.60818908 | 0.76436689 |
| AC010245.2 | 115.499268 | -0.409470425 | 0.23933786 | -1.7108469 | 0.08710938 | 0.21514231 |
| ANKDD1B | 256.193223 | 0.655285252 | 0.49239028 | 1.33082492 | 0.18324662 | 0.35999505 |
| POC5 | 406.078965 | -0.509466487 | 0.14376986 | -3.5436252 | 0.00039467 | 0.00377899 |
| AC026774.1 | 0.2873605 | -0.174096114 | 3.29377188 | -0.0528562 | 0.95784652 | NA |
| SV2C | 6.35179187 | -1.061880969 | 0.66334804 | -1.6007901 | 0.10942342 | 0.25306925 |
| AC113404.1 | 1.25955687 | -0.58227432 | 1.70432932 | -0.3416443 | 0.73261863 | 0.8492009 |
| IQGAP2 | 2556.13049 | 1.431860003 | 0.35451159 | 4.03896533 | 5.37E-05 | 0.0007874 |
| AC026725.1 | 0.28374458 | 0.328534161 | 2.32489037 | 0.14131168 | 0.88762372 | NA |
| F2RL2 | 26.9758932 | -0.477820916 | 0.55110451 | -0.8670241 | 0.38592878 | 0.58163922 |
| AC025188.1 | 25.3193532 | 1.165299257 | 0.43521729 | 2.67751141 | 0.00741713 | 0.03612667 |
| F2R | 1723.05505 | 0.266370668 | 0.2910112 | 0.91532789 | 0.36001952 | 0.55773204 |
| F2RL1 | 2909.15483 | -0.253951798 | 0.17263637 | -1.4710214 | 0.14128532 | 0.3016983 |
| S100Z | 9.82098966 | 0.307770043 | 0.82451259 | 0.37327513 | 0.70894368 | 0.83447565 |
| CRHBP | 1257.69226 | 1.039430185 | 0.73720271 | 1.40996523 | 0.15854995 | 0.32595012 |
| AGGF1 | 1605.17975 | 0.12565621 | 0.14362573 | 0.8748865 | 0.38163566 | 0.57757877 |
| AC008581.2 | 782.46247 | 0.391551665 | 0.19711877 | 1.98637437 | 0.04699177 | 0.13921728 |
| ZBED3 | 1753.16574 | 0.198299053 | 0.22785979 | 0.87026787 | 0.38415403 | 0.57969338 |
| AC008581.1 | 773.009325 | -0.1939165 | 0.23832485 | -0.8136646 | 0.41583711 | 0.60913368 |
| ZBED3-AS1 | 219.634355 | 0.604358285 | 0.34063583 | 1.77420646 | 0.07602901 | 0.1956367 |
| AC022414.1 | 354.075151 | 0.843309948 | 0.23745497 | 3.55145203 | 0.00038311 | 0.00368796 |
| PDE8B | 666.406096 | 0.405728401 | 0.28508854 | 1.42316632 | 0.15468795 | 0.32076128 |
| WDR41 | 1428.90687 | -0.188635954 | 0.1647325 | -1.1451046 | 0.25216582 | 0.44443341 |
| OTP | 0.43457194 | 0.310906642 | 2.18808206 | 0.14209094 | 0.88700817 | NA |
| TBCA | 3391.041 | -0.553943628 | 0.14244875 | -3.8887222 | 0.00010077 | 0.00127331 |
| AC108482.1 | 0.37093407 | -1.019052299 | 2.28923705 | -0.4451493 | 0.65621185 | NA |
| AP3B1 | 2839.91196 | 0.157302785 | 0.1369585 | 1.14854344 | 0.25074429 | 0.44297323 |
| SCAMP1-AS1 | 579.245839 | -0.150855065 | 0.38697047 | -0.3898361 | 0.69665774 | 0.82558262 |
| SCAMP1 | 2200.22845 | 0.030804271 | 0.13509591 | 0.22801779 | 0.81963241 | 0.90320546 |
| LHFPL2 | 1123.52931 | -1.825133925 | 0.37145267 | -4.9135033 | 8.95E-07 | 2.95E-05 |
| ARSB | 1513.5746 | 0.625816937 | 0.21634315 | 2.89270506 | 0.0038194 | 0.02201476 |
| DMGDH | 9604.41432 | 3.128813671 | 0.47855607 | 6.53802941 | 6.23E-11 | 9.78E-09 |
| BHMT2 | 18117.6142 | 1.499278293 | 0.37904021 | 3.95545971 | 7.64E-05 | 0.00102407 |
| BHMT | 19172.4314 | 3.674372178 | 0.47860672 | 7.67722649 | 1.63E-14 | 7.39E-12 |
| JMY | 1457.16318 | -0.106036321 | 0.13482189 | -0.7864919 | 0.43157938 | 0.62363681 |
| HOMER1 | 518.15438 | -0.109229873 | 0.33849046 | -0.322697 | 0.74692469 | 0.8583325 |
| TENT2 | 2554.11227 | 0.014584838 | 0.13687871 | 0.106553 | 0.9151436 | 0.95863503 |
| CMYA5 | 913.75494 | 0.650277272 | 0.26357994 | 2.46709699 | 0.01362135 | 0.05663207 |
| AC008496.2 | 3.78758453 | 1.503101835 | 1.01313312 | 1.4836173 | 0.13791048 | 0.29670386 |
| LINC01455 | 0 | NA | NA | NA | NA | NA |
| MTX3 | 1226.41102 | 0.31174377 | 0.11354823 | 2.74547452 | 0.00604235 | 0.03094517 |
| THBS4 | 212.834334 | -0.626011964 | 0.70427228 | -0.8888778 | 0.37406878 | 0.57018461 |
| AC012636.1 | 54.2930968 | -0.932544948 | 0.62264181 | -1.497723 | 0.13420524 | 0.291371 |
| SERINC5 | 1139.9148 | -0.273756625 | 0.10486378 | -2.6105927 | 0.00903855 | 0.0417959 |
| AC010260.1 | 11.2240614 | 0.783999317 | 0.36677408 | 2.13755379 | 0.03255298 | 0.10687524 |
| SPZ1 | 0 | NA | NA | NA | NA | NA |
| ZFYVE16 | 1658.50545 | 0.179813198 | 0.1366159 | 1.31619528 | 0.18810851 | 0.36613054 |
| AC008771.1 | 113.857393 | -0.039837399 | 0.22603922 | -0.1762411 | 0.86010454 | 0.92705166 |
| FAM151B | 105.138622 | -0.369653052 | 0.26470555 | -1.3964688 | 0.16257336 | 0.33152742 |
| ANKRD34B | 89.3505566 | -0.622811143 | 0.45421195 | -1.3711906 | 0.17031555 | 0.34234478 |
| LINC01337 | 3.79585493 | 0.999594957 | 0.6601184 | 1.51426616 | 0.12995835 | 0.28507772 |
| DHFR | 739.862438 | 1.228216573 | 0.35082767 | 3.50091137 | 0.00046367 | 0.00428415 |
| AC008434.1 | 1.42330171 | 1.320312398 | 1.0476432 | 1.26026914 | 0.20757229 | 0.39047519 |
| MSH3 | 803.509963 | 1.214360553 | 0.21803067 | 5.56967758 | 2.55E-08 | 1.54E-06 |
| RASGRF2-AS | 21.2414459 | 0.259928303 | 0.35800897 | 0.72603851 | 0.46781516 | 0.65468799 |
| RASGRF2 | 984.364278 | 0.325307656 | 0.23639976 | 1.37609133 | 0.16879336 | 0.3404008 |
| AC026427.1 | 19.1867656 | 0.683068692 | 0.36292315 | 1.8821304 | 0.05981831 | 0.16522561 |
| CKMT2-AS1 | 366.522241 | 0.661618982 | 0.2138312 | 3.09411801 | 0.00197399 | 0.01335405 |
| CKMT2 | 754.482474 | 0.267538792 | 0.34500372 | 0.77546639 | 0.43806412 | 0.63000386 |
| ZCCHC9 | 1015.8707 | -0.015590099 | 0.10679359 | -0.1459835 | 0.88393443 | 0.93980604 |
| ACOT12 | 118.669997 | 2.574471349 | 0.46079263 | 5.58704971 | 2.31E-08 | 1.42E-06 |
| SSBP2 | 1114.02679 | -0.026777094 | 0.23329117 | -0.1147797 | 0.90861973 | 0.95432757 |
| AC026276.1 | 1.05243128 | 1.092530934 | 1.06960606 | 1.02143301 | 0.30704933 | 0.50401785 |
| ATG10 | 18982.8457 | -0.4924131 | 0.2175737 | -2.2632014 | 0.02362328 | 0.08470773 |
| ATG10-AS1 | 0.18692855 | 0.18657164 | 3.29377195 | 0.05664376 | 0.95482897 | NA |
| RPS23 | 23011.2529 | -0.442880221 | 0.2198559 | -2.0144113 | 0.04396636 | 0.13282248 |
| ATP6AP1L | 312.104712 | 0.46237952 | 0.33678925 | 1.37290463 | 0.169782 | 0.34157691 |
| AC026782.1 | 0 | NA | NA | NA | NA | NA |
| AC026782.2 | 0.4427946 | 1.207159214 | 2.11657443 | 0.57033629 | 0.56844963 | NA |
| AC008885.2 | 0 | NA | NA | NA | NA | NA |

| | | | | | | |
|---|---|---|---|---|---|---|
| LINC01338 | 9.78144587 | 1.88132253 | 0.49495472 | 3.80099929 | 0.00014411 | 0.00169391 |
| AC108174.1 | 11.1091928 | 2.153738337 | 0.63014098 | 3.41786742 | 0.00063114 | 0.0054336 |
| AC027338.2 | 0 | NA | NA | NA | NA | NA |
| AC027338.1 | 3.68953129 | 0.665932391 | 0.93418656 | 0.71284733 | 0.47594024 | 0.66132069 |
| AC104118.1 | 46.2838944 | -0.141153336 | 0.43745892 | -0.3226665 | 0.74694783 | 0.8583325 |
| TMEM167A | 3829.90659 | 0.009289062 | 0.19292565 | 0.0481484 | 0.96159797 | 0.98169256 |
| XRCC4 | 387.25088 | -0.1047828 | 0.13118708 | -0.7987281 | 0.4244481 | 0.61691542 |
| VCAN | 2062.52692 | -1.300473437 | 0.51628545 | -2.5189039 | 0.01177208 | 0.0508766 |
| VCAN-AS1 | 403.242573 | -1.451058477 | 0.50769348 | -2.8581389 | 0.00426134 | 0.02387327 |
| HAPLN1 | 45.0535334 | -2.295297478 | 0.94284697 | -2.4344327 | 0.01491515 | 0.06057487 |
| EDIL3 | 787.678968 | -0.220933283 | 0.38203293 | -0.5783095 | 0.56305517 | 0.73032032 |
| AC113383.1 | 23.1758076 | -0.077622771 | 0.34184052 | -0.2270731 | 0.82036694 | 0.90356178 |
| AC114928.1 | 0 | NA | NA | NA | NA | NA |
| AC117522.2 | 0 | NA | NA | NA | NA | NA |
| AC010486.1 | 0 | NA | NA | NA | NA | NA |
| AC010486.2 | 0 | NA | NA | NA | NA | NA |
| AC010486.3 | 0 | NA | NA | NA | NA | NA |
| AC010595.1 | 0 | NA | NA | NA | NA | NA |
| AC026414.1 | 0 | NA | NA | NA | NA | NA |
| AC016550.1 | 0 | NA | NA | NA | NA | NA |
| AC016550.2 | 0.47124435 | 0.293890545 | 1.91686907 | 0.15331801 | 0.8781475 | NA |
| AC016550.3 | 0 | NA | NA | NA | NA | NA |
| COX7C | 13759.8922 | 0.024181868 | 0.14161785 | 0.17075438 | 0.8644169 | 0.92964904 |
| LINC02059 | 0.19387658 | -1.003574173 | 3.30558948 | -0.3035992 | 0.7614333 | NA |
| AC008539.1 | 0 | NA | NA | NA | NA | NA |
| AC109492.1 | 0 | NA | NA | NA | NA | NA |
| LINC01949 | 0 | NA | NA | NA | NA | NA |
| RASA1 | 1752.99769 | 0.164394247 | 0.09898869 | 1.66073766 | 0.09676615 | 0.23219163 |
| CCNH | 2196.08507 | -0.133522696 | 0.09812388 | -1.3607564 | 0.17359068 | 0.34741851 |
| AC018754.1 | 328.10445 | 0.014858281 | 0.19047101 | 0.07800809 | 0.93782161 | 0.97027448 |
| LINC02488 | 0.15179023 | 0.435199075 | 3.28898345 | 0.13232024 | 0.894731 | NA |
| LINC02144 | 0.17150785 | 0.18657164 | 3.29377195 | 0.05664376 | 0.95482897 | NA |
| TMEM161B | 1144.49049 | -0.119539657 | 0.12738749 | -0.938394 | 0.34804198 | 0.54548987 |
| TMEM161B-. | 719.563369 | 0.258165801 | 0.26334908 | 0.98031783 | 0.32692926 | 0.52484625 |
| LINC02060 | 17.8919622 | -0.163995265 | 0.37792116 | -0.4339404 | 0.66433171 | 0.80282421 |
| AC091826.2 | 2.63382172 | 1.388007826 | 0.69572897 | 1.995041 | 0.04603841 | 0.13711688 |
| LINC00461 | 30.116686 | 0.701651405 | 0.41485608 | 1.69131284 | 0.09077707 | 0.22204556 |
| MEF2C-AS2 | 77.1577761 | 0.365341695 | 0.29034916 | 1.25828398 | 0.20828907 | 0.39152061 |
| AC008525.1 | 0 | NA | NA | NA | NA | NA |
| AC008525.2 | 0 | NA | NA | NA | NA | NA |
| MEF2C | 1176.27348 | 0.120706878 | 0.22043246 | 0.5475912 | 0.58397263 | 0.74612909 |
| MEF2C-AS1 | 43.198216 | 0.064744336 | 0.39241378 | 0.16498997 | 0.86895188 | 0.9318649 |
| AC074131.1 | 1.41093934 | 0.084167075 | 1.43761974 | 0.05854613 | 0.95331362 | 0.97744392 |
| LINC02161 | 0 | NA | NA | NA | NA | NA |
| AC008448.1 | 0.28647622 | 0.328534579 | 2.32045628 | 0.14158189 | 0.88741027 | NA |
| AC113167.1 | 0.20859201 | 0.689155664 | 3.09576263 | 0.22261257 | 0.82383706 | NA |
| AC099554.1 | 0.15318056 | -1.003574173 | 3.30558948 | -0.3035992 | 0.7614333 | NA |
| CETN3 | 606.520922 | 0.054292395 | 0.19311679 | 0.28113762 | 0.77860485 | 0.87862932 |
| MBLAC2 | 315.42786 | 0.97002718 | 0.20738342 | 4.6774578 | 2.90E-06 | 7.64E-05 |
| POLR3G | 122.263852 | 0.251833083 | 0.20694811 | 1.21688998 | 0.22364608 | 0.40975107 |
| LYSMD3 | 1222.81645 | -0.097446865 | 0.20443631 | -0.4766613 | 0.63360336 | 0.78276508 |
| ADGRV1 | 733.071004 | 3.006997077 | 0.37946064 | 7.92439787 | 2.29E-15 | 1.47E-12 |
| AC093281.1 | 0 | NA | NA | NA | NA | NA |
| AC093281.2 | 0.64159086 | -2.566420414 | 2.22100405 | -1.1555226 | 0.24787648 | NA |
| AC123595.1 | 19.6355193 | -0.335598409 | 0.39605581 | -0.8473513 | 0.39679934 | 0.59179903 |
| LUCAT1 | 79.6548131 | -3.854332773 | 0.80967814 | -4.7603271 | 1.93E-06 | 5.52E-05 |
| AC123595.2 | 23.8947574 | -3.298939206 | 0.83331714 | -3.958804 | 7.53E-05 | 0.00101293 |
| AC008799.2 | 0.13037664 | 0.02007176 | 3.29738704 | 0.00608717 | 0.99514317 | NA |
| ARRDC3 | 9232.13424 | -0.510096889 | 0.37924128 | -1.3450458 | 0.17861044 | 0.35416547 |
| ARRDC3-AS1 | 36.7830127 | -0.05669069 | 0.32601321 | -0.1738908 | 0.8619513 | 0.92808198 |
| AC114316.2 | 1.985757 | 1.754116305 | 0.81604742 | 2.14952742 | 0.03159261 | 0.10473941 |
| AC114316.1 | 0.09635527 | 0.547239394 | 3.29377202 | 0.16614368 | 0.86804388 | NA |
| AC124854.1 | 15.5925041 | 1.797921072 | 0.69978281 | 2.56925585 | 0.01019172 | 0.04569896 |
| AC026780.1 | 4.36242105 | 2.711632311 | 0.78574871 | 3.45101718 | 0.00055848 | 0.00495194 |
| AC026780.2 | 2.29245237 | -0.222125172 | 0.85115407 | -0.2609694 | 0.7941161 | 0.88844572 |
| AC008517.1 | 0.10721287 | -0.642905668 | 3.30558948 | -0.1944905 | 0.84579185 | NA |
| LINC02058 | 2.0427679 | 1.102561127 | 1.27383365 | 0.86554561 | 0.38673939 | 0.58239494 |
| AC012625.1 | 1.61936588 | -1.634634191 | 1.16623044 | -1.4016391 | 0.16102304 | 0.32959541 |
| NR2F1-AS1 | 763.586883 | 0.171864202 | 0.23450188 | 0.73289051 | 0.46362521 | 0.65138243 |
| NR2F1 | 3137.43321 | 0.053374802 | 0.31489007 | 0.16950297 | 0.86540104 | 0.93001264 |
| FAM172A | 993.033325 | 0.482846025 | 0.177221 | 2.72454185 | 0.00643908 | 0.03247245 |
| POU5F2 | 4.68134634 | -0.145976914 | 0.63090411 | -0.2313773 | 0.81702167 | 0.90160437 |
| AC108102.1 | 1.24808365 | -0.38621646 | 1.30455512 | -0.2960522 | 0.76719019 | 0.87152635 |
| KIAA0825 | 63.9452519 | 0.886864685 | 0.26225981 | 3.38162634 | 0.00072058 | 0.00604718 |
| SLF1 | 259.505915 | 0.21815073 | 0.1866151 | 1.16898757 | 0.24240864 | 0.43281981 |
| MCTP1 | 305.645766 | -0.568360813 | 0.5823838 | -0.9759214 | 0.32910342 | 0.52689812 |
| AC008534.1 | 0.21964838 | 0.952737031 | 3.28103697 | 0.2903768 | 0.77152799 | NA |
| AC012312.1 | 0 | NA | NA | NA | NA | NA |
| FAM81B | 57.1542497 | -1.180195528 | 0.70682972 | -1.6697027 | 0.09497819 | 0.22931467 |
| TTC37 | 2991.49541 | 0.176672176 | 0.09221553 | 1.91586144 | 0.05538273 | 0.15646496 |
| ARSK | 469.246462 | 0.160356475 | 0.17483879 | 0.9171676 | 0.35905482 | 0.55672815 |
| GPR150 | 4.34444078 | -1.566609196 | 0.83081259 | -1.8856349 | 0.05934417 | 0.16434634 |
| RFESD | 79.2709611 | 0.202000519 | 0.2781068 | 0.72634153 | 0.46762943 | 0.65456549 |
| SPATA9 | 64.1447496 | -0.036463932 | 0.27425948 | -0.1329541 | 0.89422966 | 0.94565432 |
| AC008840.1 | 60.8854716 | 0.563800402 | 0.43862433 | 1.28538333 | 0.19865836 | 0.37968468 |
| RHOBTB3 | 3033.17868 | -0.101382033 | 0.27637903 | -0.3668225 | 0.71375146 | 0.83745188 |
| GLRX | 9086.56748 | -0.45468513 | 0.3384799 | -1.343315 | 0.17916999 | 0.3548671 |
| AC008592.4 | 79.3899817 | -0.130687291 | 0.62424448 | -0.2093527 | 0.83417288 | 0.91206315 |
| AC008592.5 | 6.93457062 | -1.575290564 | 0.59593699 | -2.6433844 | 0.00820818 | 0.03879721 |
| LINC01554 | 345.305287 | -0.734928754 | 0.63531878 | -1.1567874 | 0.24735923 | 0.43894019 |
| ELL2 | 2997.7111 | -1.250924474 | 0.2791531 | -4.4811413 | 7.42E-06 | 0.00016101 |
| AC104123.1 | 7.12625017 | 1.028521634 | 0.49337358 | 2.08467109 | 0.03709917 | 0.11717631 |
| AC008592.2 | 0.17475523 | -1.003574173 | 3.30558948 | -0.3035992 | 0.7614333 | NA |
| MIR583HG | 22.6300007 | -0.901822216 | 0.42917704 | -2.1012825 | 0.03561617 | 0.11402191 |
| AC099509.1 | 42.1897175 | -1.469361601 | 0.69574856 | -2.1119147 | 0.03469377 | 0.11193866 |
| AC099509.2 | 0.18301898 | 0.18657164 | 3.29377195 | 0.05664376 | 0.95482897 | NA |
| PCSK1 | 38.3346901 | -1.577904052 | 0.50570088 | -3.120232 | 0.00180709 | 0.01246479 |
| CAST | 8980.49175 | -0.360679551 | 0.11707307 | -3.0808073 | 0.0020644 | 0.01379931 |
| AC020900.1 | 539.33782 | -0.055587341 | 0.21396259 | -0.2597993 | 0.79501856 | 0.88885946 |
| ERAP1 | 3756.1632 | -0.243704325 | 0.13458284 | -1.8108127 | 0.07016985 | 0.18455803 |
| AC008906.1 | 176.29516 | -0.101741022 | 0.30050496 | -0.3385669 | 0.73493605 | 0.85090725 |
| AC008906.2 | 10.5522166 | -1.035487431 | 0.67296689 | -1.53869 | 0.12387999 | 0.27548621 |
| AC009126.1 | 125.73089 | 0.451664823 | 0.26097965 | 1.73065152 | 0.08351394 | 0.20882444 |
| ERAP2 | 1324.11927 | 0.268664604 | 0.68992582 | 0.38941086 | 0.69697224 | 0.8258207 |
| LNPEP | 3295.025 | -0.250804161 | 0.15797961 | -1.5875729 | 0.11238294 | 0.25792325 |
| LIX1-AS1 | 49.8360016 | 0.103070208 | 0.2264933 | 0.45506957 | 0.64905919 | 0.79349874 |
| LIX1 | 170.669125 | 0.590840631 | 0.8703239 | 0.67887442 | 0.49721744 | 0.67893661 |
| RIOK2 | 789.058881 | -0.04781845 | 0.17877176 | -0.2674832 | 0.78909711 | 0.88545939 |
| AC008883.1 | 0.54639137 | 1.064221708 | 1.74082977 | 0.61133014 | 0.54098104 | NA |
| AC008883.2 | 0.69936807 | -1.727799137 | 1.62217485 | -1.0651128 | 0.2868249 | NA |
| LINC01340 | 0.71506591 | -1.747976592 | 1.66277618 | -1.0512399 | 0.29314844 | NA |
| LINC02234 | 1.67347451 | -0.016546476 | 1.00220091 | -0.0165101 | 0.98682741 | 0.99400325 |
| LINC01846 | 0 | NA | NA | NA | NA | NA |

| | | | | | | |
|---|---|---|---|---|---|---|
| RGMB | 689.743978 | -0.472815222 | 0.1861155 | -2.5404399 | 0.01107131 | 0.04858554 |
| RGMB-AS1 | 35.0064341 | -0.169853461 | 0.43906757 | -0.3868504 | 0.69886697 | 0.82715805 |
| AC008522.1 | 462.808538 | -0.380451854 | 0.19621914 | -1.9389131 | 0.05251192 | 0.15071173 |
| CHD1 | 1808.93232 | -0.294966684 | 0.13187867 | -2.2366519 | 0.0253091 | 0.08917704 |
| LINC02062 | 82.2695298 | -0.048793918 | 0.32373741 | -0.1507207 | 0.88019607 | 0.9378468 |
| LINC02113 | 0 | NA | NA | NA | NA | NA |
| AC113385.1 | 0 | NA | NA | NA | NA | NA |
| AC021086.1 | 2.74309435 | 0.458861538 | 0.72481401 | 0.63307487 | 0.52668475 | 0.70151919 |
| FAM174A | 1212.85649 | -0.211173212 | 0.13665182 | -1.5453377 | 0.1222646 | 0.27325345 |
| AC027315.1 | 0 | NA | NA | NA | NA | NA |
| ST8SIA4 | 241.089751 | -0.687121521 | 0.42033155 | -1.6347132 | 0.10210919 | 0.24094565 |
| AC008948.1 | 0.09635527 | 0.547239394 | 3.29377202 | 0.16614368 | 0.86804388 | NA |
| SLCO4C1 | 3054.6265 | 1.378008734 | 0.31166098 | 4.4214991 | 9.80E-06 | 0.00020037 |
| SLCO6A1 | 1.19004787 | 0.314553211 | 1.46423057 | 0.21482492 | 0.82990383 | 0.90970031 |
| AC094108.1 | 0.16783122 | 1.368031398 | 3.28033652 | 0.41703996 | 0.67664918 | NA |
| LINC00492 | 0.42232614 | 2.01143182 | 2.62380653 | 0.76660828 | 0.44331445 | NA |
| AC099487.1 | 0 | NA | NA | NA | NA | NA |
| LINC00491 | 0.32670497 | 1.41436792 | 2.2468522 | 0.62948863 | 0.52902921 | NA |
| PAM | 10411.4781 | -0.106803722 | 0.18650694 | -0.5726528 | 0.56687981 | 0.73302704 |
| GIN1 | 387.130004 | 0.255204891 | 0.11037214 | 2.31222196 | 0.02076545 | 0.07699452 |
| PPIP5K2 | 1662.85301 | 0.300042075 | 0.14618343 | 2.05250402 | 0.0401207 | 0.12407447 |
| AC011362.1 | 0 | NA | NA | NA | NA | NA |
| C5orf30 | 587.076388 | -0.265390582 | 0.25115567 | -1.0566777 | 0.29065873 | 0.48670756 |
| AC010406.1 | 0.15481363 | 0.435199075 | 3.28898345 | 0.13232024 | 0.894731 | NA |
| LINC02115 | 4.54327797 | -0.54306105 | 0.73506582 | -0.7387924 | 0.46003306 | 0.6486998 |
| NUDT12 | 1367.56291 | 1.05304073 | 0.20073831 | 5.24583847 | 1.56E-07 | 6.91E-06 |
| LINC02163 | 0 | NA | NA | NA | NA | NA |
| AC099520.1 | 0.10721287 | -0.642905668 | 3.30558948 | -0.1944905 | 0.84579185 | NA |
| AC099520.2 | 0.09635527 | 0.547239394 | 3.29377202 | 0.16614368 | 0.86804388 | NA |
| AC091987.1 | 0 | NA | NA | NA | NA | NA |
| LINC01950 | 1.09071404 | -2.445767689 | 1.86271148 | -1.3130148 | 0.18917797 | 0.36745898 |
| EFNA5 | 1307.95057 | -1.145845659 | 0.28336522 | -4.0437061 | 5.26E-05 | 0.00077552 |
| AC024587.2 | 0.30905368 | 2.226555015 | 2.80385542 | 0.79410479 | 0.42713443 | NA |
| FBXL17 | 1231.26876 | 0.660910966 | 0.15755935 | 4.19467947 | 2.73E-05 | 0.00045664 |
| LINC01023 | 135.733907 | -0.159237645 | 0.26159094 | -0.6087277 | 0.54270496 | 0.7151932 |
| FER | 810.169554 | -0.111264807 | 0.14992368 | -0.742143 | 0.45800071 | 0.64706542 |
| AC008871.1 | 1.01514599 | -1.702058637 | 1.3181917 | -1.2912072 | 0.19663183 | 0.37721878 |
| AC008467.1 | 80.838139 | -0.561193455 | 0.35410256 | -1.5848331 | 0.11300426 | 0.25889854 |
| PJA2 | 5385.38427 | 0.312457219 | 0.13339155 | 2.34240634 | 0.01915984 | 0.07260311 |
| AC091917.3 | 0 | NA | NA | NA | NA | NA |
| AC091917.2 | 0.20356813 | 0.18657164 | 3.29377195 | 0.05664376 | 0.95482897 | NA |
| AC091917.1 | 0 | NA | NA | NA | NA | NA |
| AC012603.1 | 9.05894286 | 0.198192473 | 0.44283732 | 0.44755142 | 0.65447697 | 0.79728574 |
| MAN2A1 | 3283.24506 | -0.360437697 | 0.19659205 | -1.8334297 | 0.06673869 | 0.17805538 |
| LINC01848 | 0.12150485 | 1.04981648 | 3.28477129 | 0.31960109 | 0.74927075 | NA |
| TMEM232 | 175.054879 | 0.737099206 | 0.40837763 | 1.80494513 | 0.07108327 | 0.18641151 |
| SLC25A46 | 2856.45239 | 0.163428913 | 0.18771163 | 0.87063818 | 0.38395174 | 0.57958713 |
| AC010395.1 | 0.66836975 | -0.555114824 | 1.83898159 | -0.3018599 | 0.76275886 | NA |
| TSLP | 49.3173204 | -0.172105482 | 0.39126497 | -0.4398694 | 0.66003171 | 0.79999826 |
| AC008086.1 | 0.86832277 | -0.254667054 | 1.30695418 | -0.1948554 | 0.84550617 | NA |
| WDR36 | 1250.93424 | -0.208425824 | 0.14301296 | -1.4573912 | 0.14500843 | 0.30721236 |
| CAMK4 | 118.969671 | 0.968539717 | 0.45343976 | 2.1359832 | 0.03268078 | 0.10724145 |
| AC010468.2 | 0 | NA | NA | NA | NA | NA |
| AC010275.1 | 0 | NA | NA | NA | NA | NA |
| STARD4 | 1097.19173 | -0.404682131 | 0.26087139 | -1.5512707 | 0.12083683 | 0.27121178 |
| STARD4-AS1 | 922.378458 | 1.301948103 | 0.22332143 | 5.8299291 | 5.55E-09 | 4.23E-07 |
| NREP | 2087.81272 | 1.372849831 | 0.24639451 | 5.57175502 | 2.52E-08 | 1.53E-06 |
| NREP-AS1 | 0 | NA | NA | NA | NA | NA |
| EPB41L4A | 1931.92036 | 0.39904244 | 0.12917338 | 3.08920027 | 0.00200696 | 0.01354236 |
| EPB41L4A-AS | 2387.56993 | -0.008224885 | 0.19372704 | -0.0424561 | 0.96613515 | 0.98410057 |
| AC010261.1 | 0.63140237 | 2.052218686 | 1.64380471 | 1.2484565 | 0.21186393 | NA |
| AC010261.2 | 105.431142 | 0.988736629 | 0.2214591 | 4.46464666 | 8.02E-06 | 0.00017111 |
| AC104126.1 | 58.0863288 | 3.352134529 | 0.95608897 | 3.50609058 | 0.00045474 | 0.00421237 |
| EPB41L4A-DT | 116.433292 | -0.285416178 | 0.29954238 | -0.9528407 | 0.34067077 | 0.53870762 |
| LINC02200 | 3.53993305 | 4.937340205 | 1.03338742 | 4.77782109 | 1.77E-06 | 5.16E-05 |
| APC | 1154.02821 | -0.033494278 | 0.26104313 | -0.1283094 | 0.89790416 | 0.94805736 |
| AC008575.1 | 1115.03212 | -0.70756946 | 0.22683748 | -3.1192793 | 0.00181294 | 0.01248231 |
| SRP19 | 1620.14098 | -0.452047901 | 0.21244249 | -2.1278601 | 0.03334868 | 0.10878337 |
| REEP5 | 11808.6543 | -0.019251112 | 0.13135513 | -0.1465577 | 0.88348111 | 0.93956271 |
| DCP2 | 1739.44385 | -0.372876777 | 0.12332705 | -3.0234793 | 0.00249886 | 0.01599781 |
| MCC | 889.174733 | 0.180536619 | 0.185361 | 0.97397304 | 0.33006993 | 0.52795972 |
| AC079465.1 | 4.41144388 | 1.254160812 | 0.71140523 | 1.76293449 | 0.07791154 | 0.19871861 |
| TSSK1B | 1.49710577 | 2.236334998 | 1.0498044 | 2.13023969 | 0.03315183 | 0.10835569 |
| YTHDC2 | 1826.81648 | 0.04260737 | 0.07844624 | 0.543141 | 0.58703273 | 0.74800933 |
| AC106789.1 | 0.2423894 | 1.011482133 | 2.97224145 | 0.34030954 | 0.73362343 | NA |
| KCNN2 | 87.2258909 | 2.817265607 | 0.44133279 | 6.38354025 | 1.73E-10 | 2.30E-08 |
| AC010230.1 | 16.4594085 | 2.547324829 | 0.50797218 | 5.0146936 | 5.31E-07 | 1.90E-05 |
| LINC01957 | 0 | NA | NA | NA | NA | NA |
| AC094104.2 | 0 | NA | NA | NA | NA | NA |
| AC094104.1 | 0 | NA | NA | NA | NA | NA |
| TRIM36 | 51.2909138 | -0.523445172 | 0.51230358 | -1.021748 | 0.30690017 | 0.50392015 |
| AC008494.2 | 2.41464856 | -0.38539636 | 1.0406189 | -0.370353 | 0.71111947 | 0.83587657 |
| PGGT1B | 1256.17012 | 0.049588282 | 0.12737457 | 0.3893107 | 0.69704632 | 0.8258207 |
| AC008494.3 | 64.5864106 | 0.338053974 | 0.3070298 | 1.10104614 | 0.27087658 | 0.46549465 |
| CCDC112 | 185.911363 | -0.149273907 | 0.20356759 | -0.7332892 | 0.46338208 | 0.65117961 |
| FEM1C | 1465.07801 | -0.826138369 | 0.27405891 | -3.0144555 | 0.00257441 | 0.01635427 |
| TMED7-TICA | 968.737053 | 0.144333955 | 0.19587457 | 0.73686928 | 0.46120185 | 0.64968454 |
| TICAM2 | 363.34321 | -0.572226025 | 0.26397941 | -2.1676919 | 0.03018214 | 0.10124299 |
| AC010226.1 | 413.136835 | 0.506749748 | 0.23459311 | 2.160122 | 0.03076323 | 0.10271287 |
| TMED7 | 4603.96554 | -0.030676708 | 0.17859954 | -0.1717625 | 0.86362423 | 0.92901081 |
| AC008549.2 | 0.03392908 | 0.380740593 | 3.29738709 | 0.11546736 | 0.90807469 | NA |
| AC008549.1 | 0.18754884 | 1.469068384 | 3.27783996 | 0.44818185 | 0.65402196 | NA |
| CDO1 | 178.840143 | 1.140178631 | 0.47129286 | 2.41925716 | 0.01555224 | 0.06246004 |
| ATG12 | 1798.12125 | -0.48392889 | 0.11682413 | -4.1423711 | 3.44E-05 | 0.00054769 |
| AP3S1 | 2207.03994 | -0.57646159 | 0.16321721 | -3.5318677 | 0.00041264 | 0.0039141 |
| LVRN | 19.8589881 | -1.261834049 | 0.80701693 | -1.5635782 | 0.11791667 | 0.26660915 |
| ARL14EPL | 7.03324005 | -1.832744354 | 0.72825566 | -2.5166222 | 0.01184858 | 0.05110355 |
| AC034236.3 | 4.51870524 | 0.157760257 | 0.68817705 | 0.22924371 | 0.8186795 | 0.9026835 |
| AC034236.2 | 263.319829 | 0.083402558 | 0.19126424 | 0.43605933 | 0.66279368 | 0.80177391 |
| COMMD10 | 1245.34342 | 0.075101708 | 0.18191798 | 0.4128328 | 0.67972912 | 0.81422989 |
| AC018752.1 | 1.33309125 | 3.391706169 | 1.30955708 | 2.58996437 | 0.00959859 | 0.04370829 |
| SEMA6A | 2346.4679 | -0.788903386 | 0.39829062 | -1.980723 | 0.04762235 | 0.14058001 |
| SEMA6A-AS1 | 199.065642 | -0.183816701 | 0.28768926 | -0.6389418 | 0.52286077 | 0.69862624 |
| SEMA6A-AS2 | 8.84758279 | -0.093422515 | 0.75272212 | -0.1241129 | 0.9012259 | 0.95000507 |
| LINC02214 | 0 | NA | NA | NA | NA | NA |
| AC010267.1 | 0.08666371 | -0.642905668 | 3.30558948 | -0.1944905 | 0.84579185 | NA |
| AC093534.2 | 0 | NA | NA | NA | NA | NA |
| LINC00992 | 29.5922952 | 0.483458885 | 0.54556309 | 0.88616494 | 0.37552865 | 0.57148491 |
| LINC02147 | 20.22335 | 0.525113224 | 0.55828962 | 0.94057494 | 0.34692273 | 0.54451237 |
| LINC02148 | 6.04590482 | 0.660758484 | 0.60996867 | 1.08326626 | 0.27869026 | 0.47431664 |
| LINC02208 | 9.28474705 | 2.572211656 | 0.67907657 | 3.78780798 | 0.00015198 | 0.001771 |
| LINC02216 | 0.47262264 | -0.925128525 | 2.79711416 | -0.3307439 | 0.74083792 | NA |
| LINC02215 | 0.51423212 | 0.657249855 | 1.85035281 | 0.35520245 | 0.72243793 | NA |

| | | | | | | |
|---|---|---|---|---|---|---|
| AC093206.1 | 6.24754456 | -1.467312143 | 0.99741983 | -1.4711078 | 0.14126195 | 0.30167458 |
| DTWD2 | 399.011635 | 1.142991329 | 0.16268532 | 7.02578056 | 2.13E-12 | 5.20E-10 |
| AC008629.1 | 12.4459284 | 0.539858162 | 0.50842 | 1.06183503 | 0.28831058 | 0.48440587 |
| DMXL1 | 2225.78513 | 0.715456069 | 0.19549478 | 3.6597195 | 0.00025249 | 0.00264005 |
| TNFAIP8 | 1241.52159 | -0.741641944 | 0.22358945 | -3.3169809 | 0.00090996 | 0.00728906 |
| HSD17B4 | 7750.64435 | 0.163754531 | 0.25668928 | 0.63794847 | 0.52350722 | 0.69887539 |
| FAM170A | 4.12022272 | -0.61250235 | 0.83551869 | -0.7330804 | 0.46350941 | 0.65128914 |
| AC008574.1 | 0 | NA | NA | NA | NA | NA |
| AC113418.1 | 0 | NA | NA | NA | NA | NA |
| PRR16 | 120.493627 | -0.911951167 | 0.5259514 | -1.7339077 | 0.08293446 | 0.20774977 |
| AC114284.1 | 6.05992956 | -0.712422201 | 0.81804841 | -0.8708803 | 0.38381954 | 0.57948708 |
| AC008568.1 | 0.37570419 | -1.067942514 | 2.86151185 | -0.3732092 | 0.70899276 | NA |
| FTMT | 0 | NA | NA | NA | NA | NA |
| SRFBP1 | 431.561651 | -0.060057983 | 0.20989488 | -0.2861336 | 0.77477577 | 0.87626409 |
| LOX | 732.551892 | -0.576641581 | 0.30784984 | -1.8731261 | 0.06105097 | 0.16745495 |
| AC010255.1 | 2.24058758 | -1.261844238 | 1.07496717 | -1.1738444 | 0.24045734 | 0.43058943 |
| ZNF474 | 11.0400136 | -0.571550236 | 0.42058372 | -1.3589452 | 0.17416395 | 0.34811668 |
| AC010255.3 | 1.34132857 | 1.191405807 | 1.31712807 | 0.90454819 | 0.3657048 | 0.562672 |
| AC010255.2 | 1.55203008 | -1.333553845 | 1.29189625 | -1.0322453 | 0.30195721 | 0.4987855 |
| AC113349.2 | 0.84423108 | -1.276025727 | 1.65984446 | -0.7687622 | 0.44203447 | NA |
| SNCAIP | 166.369118 | 0.139853072 | 0.35412557 | 0.39492509 | 0.69289818 | 0.8232818 |
| AC113349.1 | 0.51998226 | -2.27683341 | 3.28074098 | -0.6939997 | 0.48768237 | NA |
| AC119150.1 | 1.74268872 | -2.809087239 | 1.59732772 | -1.7586167 | 0.07864263 | 0.20009584 |
| AC022101.1 | 37.2462839 | -0.383681062 | 0.37950872 | -1.0109941 | 0.31201927 | 0.50905783 |
| LINC02201 | 1.17367801 | 0.534924145 | 1.43829358 | 0.37191583 | 0.70995552 | 0.8351859 |
| SNX2 | 4566.87203 | -0.009856265 | 0.12258008 | -0.0804067 | 0.93591376 | 0.96936551 |
| SNX24 | 1624.70856 | -0.348554022 | 0.15681967 | -2.2226423 | 0.02623993 | 0.09157584 |
| PPIC | 2217.22089 | -0.618294266 | 0.22254472 | -2.7782923 | 0.00546454 | 0.02879697 |
| AC106786.2 | 512.818865 | -0.494455966 | 0.21615004 | -2.287559 | 0.02216321 | 0.0808919 |
| AC106786.1 | 5.4749783 | -0.469476351 | 0.69423988 | -0.6762451 | 0.49888501 | 0.68019222 |
| PRDM6 | 131.748207 | 0.120875001 | 0.24216206 | 0.49914921 | 0.61767428 | 0.77143728 |
| CEP120 | 848.555079 | 0.408540659 | 0.14206536 | 2.87572328 | 0.00403103 | 0.02288444 |
| CSNK1G3 | 1110.88461 | -0.326017197 | 0.11074628 | -2.9438208 | 0.00324188 | 0.01949117 |
| LINC01170 | 0.62723577 | 0.567434 | 1.79172186 | 0.31669759 | 0.75147308 | NA |
| AC016556.1 | 0 | NA | NA | NA | NA | NA |
| AC025465.2 | 0 | NA | NA | NA | NA | NA |
| AC025465.4 | 0.09635527 | 0.547239394 | 3.29377202 | 0.16614368 | 0.86804388 | NA |
| AC025465.3 | 0 | NA | NA | NA | NA | NA |
| AC025465.1 | 0.18301898 | 0.18657164 | 3.29377195 | 0.05664376 | 0.95482897 | NA |
| ZNF608 | 835.556277 | -0.010692864 | 0.20506111 | -0.0521448 | 0.95841334 | 0.98011944 |
| AC112196.1 | 0.17507102 | 0.435199075 | 3.28898345 | 0.13232024 | 0.894731 | NA |
| LINC02240 | 10.7527072 | -0.927008105 | 0.42241599 | -2.1945384 | 0.02819673 | 0.0965965 |
| AC109464.1 | 0 | NA | NA | NA | NA | NA |
| AC109464.2 | 0 | NA | NA | NA | NA | NA |
| AC109471.1 | 0 | NA | NA | NA | NA | NA |
| AC010235.1 | 0.79326873 | 0.607754278 | 1.51435109 | 0.40132984 | 0.6881773 | NA |
| LINC02039 | 0 | NA | NA | NA | NA | NA |
| GRAMD2B | 3023.74451 | -0.707836542 | 0.19710428 | -3.591178 | 0.00032919 | 0.00326463 |
| AC093535.1 | 539.535735 | -0.377197664 | 0.23639638 | -1.5956152 | 0.11057472 | 0.25472661 |
| ALDH7A1 | 6552.53987 | 1.636875651 | 0.269286 | 6.07857694 | 1.21E-09 | 1.21E-07 |
| PHAX | 1255.49388 | 0.141077955 | 0.20881163 | 0.67562307 | 0.49927999 | 0.68062518 |
| TEX43 | 1.2676311 | 2.34259623 | 1.52547242 | 1.53565296 | 0.12462352 | 0.27671962 |
| LMNB1-DT | 0.36778754 | -1.757845562 | 2.82391078 | -0.6224862 | 0.5336222 | NA |
| LMNB1 | 304.877332 | -0.909015277 | 0.46762183 | -1.9439111 | 0.05190617 | 0.14964094 |
| 3-Mar | 221.932274 | -1.882735888 | 0.42059948 | -4.4763153 | 7.59E-06 | 0.00016401 |
| C5orf63 | 244.368018 | -0.065341472 | 0.22924475 | -0.2850293 | 0.7756217 | 0.8768389 |
| AC011416.3 | 9.853879 | -0.165102618 | 0.57454223 | -0.2873638 | 0.7738338 | 0.87556841 |
| MEGF10 | 6.88863807 | 0.188421676 | 0.55152785 | 0.34163583 | 0.73262497 | 0.8492009 |
| AC010424.2 | 0 | NA | NA | NA | NA | NA |
| PRRC1 | 2494.63168 | -0.042220636 | 0.25362745 | -0.1664671 | 0.86778934 | 0.93121205 |
| AC010424.3 | 0 | NA | NA | NA | NA | NA |
| CTXN3 | 1053.5319 | 5.45651039 | 0.72220743 | 7.555323 | 4.18E-14 | 1.72E-11 |
| AC022118.1 | 0.57143021 | 2.443389514 | 2.57273449 | 0.94972471 | 0.34225215 | NA |
| CCDC192 | 67.2330752 | 0.050965461 | 0.29323919 | 0.17380167 | 0.86202133 | 0.92808258 |
| AC068658.1 | 0.09997307 | 0.547239394 | 3.29377202 | 0.16614368 | 0.86804388 | NA |
| LINC01184 | 802.875539 | 0.170284435 | 0.16720925 | 1.01839123 | 0.30849207 | 0.50550464 |
| SLC12A2 | 873.726925 | 0.083917741 | 0.2137228 | 0.39264758 | 0.69457979 | 0.82407273 |
| FBN2 | 50.5005884 | 0.570009099 | 0.48458706 | 1.176278 | 0.2394838 | 0.42934176 |
| SLC27A6 | 3.67183637 | -3.595420124 | 1.82944764 | -1.9653036 | 0.0493791 | 0.14439199 |
| ISOC1 | 1686.56112 | 0.448053031 | 0.16435197 | 2.72617985 | 0.00640721 | 0.03233028 |
| ADAMTS19-/ | 2.75337242 | 0.446729294 | 1.06873903 | 0.41799661 | 0.6759496 | 0.81149917 |
| ADAMTS19 | 46.7702889 | 0.443316859 | 0.77991659 | 0.56841573 | 0.56975271 | 0.73511982 |
| AC008591.1 | 7.41571659 | 0.762939206 | 0.61138001 | 1.24789688 | 0.21206882 | 0.39574374 |
| KIAA1024L | 1.2138382 | -1.567202764 | 1.66705749 | -0.9401012 | 0.34716565 | 0.54472033 |
| CHSY3 | 48.9282775 | -0.412183475 | 0.33968306 | -1.2134355 | 0.22496337 | 0.41125019 |
| HINT1 | 12117.9221 | -0.586667209 | 0.16504275 | -3.5546378 | 0.0003785 | 0.00365338 |
| LYRM7 | 815.130658 | 0.714056564 | 0.16727827 | 4.26867501 | 1.97E-05 | 0.00034928 |
| CDC42SE2 | 5961.882 | -1.026332599 | 0.1763106 | -5.8211622 | 5.84E-09 | 4.40E-07 |
| RAPGEF6 | 623.861529 | -0.031394375 | 0.14172439 | -0.2215171 | 0.82468983 | 0.90622499 |
| AC008695.1 | 158.944524 | 0.688574772 | 0.21615828 | 3.18551196 | 0.00144498 | 0.0105074 |
| FNIP1 | 1360.25266 | -0.597783193 | 0.326679 | -1.8298795 | 0.06726797 | 0.17901466 |
| MEIKIN | 4.97987093 | -0.667597907 | 0.92759735 | -0.7197066 | 0.47170568 | 0.65803729 |
| AC026398.1 | 102.470254 | 1.081571178 | 0.38942698 | 2.77734013 | 0.00548058 | 0.02886417 |
| AC034228.3 | 33.0815547 | 1.173261028 | 0.48487703 | 2.41970843 | 0.01553296 | 0.06240159 |
| ACSL6 | 233.800011 | 1.466292886 | 0.4223499 | 3.47174913 | 0.00051708 | 0.00465213 |
| AC034228.1 | 0 | NA | NA | NA | NA | NA |
| AC034228.2 | 1.29661435 | 1.520920936 | 1.03038358 | 1.47607257 | 0.13992441 | 0.29988676 |
| IL3 | 0.27512461 | 2.050195067 | 3.23884076 | 0.63300274 | 0.52673186 | NA |
| CSF2 | 1.93553535 | -1.748545002 | 1.26608153 | -1.3810682 | 0.16725798 | 0.33839135 |
| AC063976.1 | 0.27351155 | -1.364244856 | 2.93521793 | -0.4647849 | 0.64208553 | NA |
| P4HA2-AS1 | 534.553484 | 0.411493211 | 0.26892891 | 1.53011891 | 0.1259873 | 0.27885555 |
| P4HA2 | 3541.67034 | 0.831452045 | 0.30939383 | 2.68735822 | 0.00720197 | 0.03539245 |
| PDLIM4 | 970.252221 | -1.314455255 | 0.4454932 | -2.9505619 | 0.00317196 | 0.01916698 |
| SLC22A4 | 826.910989 | 0.270807995 | 0.31093279 | 0.87095347 | 0.38377956 | 0.57948708 |
| SLC22A5 | 6614.35144 | -0.408666222 | 0.46713017 | -0.8748444 | 0.38165857 | 0.57758029 |
| AC116366.3 | 3051.42631 | -0.008339051 | 0.14354608 | -0.0580932 | 0.95367439 | 0.97769978 |
| AC116366.2 | 2.50387497 | -1.268463861 | 1.0283118 | -1.2335401 | 0.21737432 | 0.4023416 |
| AC116366.1 | 140.669908 | 0.631251522 | 0.45898428 | 1.37532274 | 0.16903141 | 0.34072453 |
| IRF1 | 3294.96787 | -0.684410495 | 0.40320187 | -1.6974388 | 0.08961373 | 0.21990879 |
| IL5 | 3.00360516 | 0.352011814 | 0.95287659 | 0.36942015 | 0.71181458 | 0.83618048 |
| RAD50 | 2350.4218 | 0.142296704 | 0.12557968 | 1.13311889 | 0.25716433 | 0.44990535 |
| TH2LCRR | 453.061023 | -0.183734593 | 0.15114293 | -1.2156347 | 0.2241241 | 0.41014244 |
| IL13 | 3.69681892 | 0.028772306 | 0.96308784 | 0.02987506 | 0.9761667 | 0.98894663 |
| IL4 | 8.11453944 | -0.911296694 | 0.57989263 | -1.5714921 | 0.11606839 | 0.26397902 |
| AC004039.1 | 14.6149664 | -0.167789389 | 0.41178932 | -0.4074642 | 0.68366712 | 0.81727648 |
| KIF3A | 761.63754 | -0.025172275 | 0.15145691 | -0.1662009 | 0.86799885 | 0.93135426 |
| CCNI2 | 314.133236 | 0.171816064 | 0.28305594 | 0.60700392 | 0.54384831 | 0.71604108 |
| 8-Sep | 3795.23256 | 0.05622022 | 0.26500113 | 0.21215087 | 0.83198933 | 0.91069523 |
| SOWAHA | 554.913254 | 0.220655401 | 0.33053431 | 0.66757186 | 0.50440693 | 0.68392997 |
| AC004775.1 | 1.53696224 | 0.156532922 | 1.39737523 | 0.11201925 | 0.91080815 | 0.95585446 |
| SHROOM1 | 1434.80836 | -0.993158059 | 0.45731633 | -2.1717092 | 0.02987761 | 0.10058375 |
| GDF9 | 2936.11835 | 0.178768215 | 0.19576283 | 0.91318771 | 0.36114382 | 0.55878832 |
| UQCRQ | 6276.0976 | 0.076507772 | 0.17300614 | 0.44222576 | 0.65832585 | 0.79946078 |

| | | | | | | |
|---|---|---|---|---|---|---|
| LEAP2 | 389.553327 | 1.498039065 | 0.43940369 | 3.40925459 | 0.00065141 | 0.00556196 |
| AFF4 | 4873.287 | -0.143722161 | 0.16346282 | -0.8792345 | 0.37927412 | 0.57529241 |
| AC010240.3 | 4.64393565 | 0.516700965 | 0.61071247 | 0.84606257 | 0.39751784 | 0.59240155 |
| ZCCHC10 | 729.942244 | -0.249509868 | 0.10602965 | -2.3532085 | 0.01861219 | 0.07124452 |
| HSPA4 | 5306.45839 | 0.041870583 | 0.23350863 | 0.17931064 | 0.85769379 | 0.92565217 |
| AC113410.2 | 0.09989236 | -0.642905668 | 3.30558948 | -0.1944905 | 0.84579185 | NA |
| AC010307.2 | 6.89755148 | 1.878786503 | 0.65035891 | 2.88884566 | 0.00386659 | 0.02220909 |
| FSTL4 | 213.527203 | 1.617615045 | 0.53868178 | 3.00291399 | 0.00267408 | 0.01683108 |
| AC010307.3 | 0.25087713 | 1.011917048 | 2.94486708 | 0.34362062 | 0.73113162 | NA |
| AC010307.4 | 16.7547998 | 2.373662208 | 1.14344169 | 2.07589265 | 0.03790388 | 0.11899111 |
| AC010608.1 | 0 | NA | NA | NA | NA | NA |
| WSPAR | 3.85249987 | 5.078439794 | 1.08030497 | 4.70093161 | 2.59E-06 | 6.99E-05 |
| C5orf15 | 4947.36387 | 0.045041938 | 0.16229276 | 0.2775351 | 0.78136926 | 0.88036187 |
| VDAC1 | 13680.3327 | -0.1656197 | 0.20583036 | -0.8046418 | 0.42102644 | 0.6136333 |
| AC008608.2 | 59.2972475 | 0.269320649 | 0.26753062 | 1.00669092 | 0.31408333 | 0.51094247 |
| TCF7 | 409.707433 | -0.18079947 | 0.4767087 | -0.3792661 | 0.70449024 | 0.83125502 |
| SKP1 | 14851.5229 | -0.082230001 | 0.13234453 | -0.6213328 | 0.53438065 | 0.70808591 |
| AC104109.3 | 6864.9033 | -0.030722505 | 0.15881891 | -0.1934436 | 0.84661156 | 0.9193833 |
| PPP2CA | 7083.00707 | -0.292121717 | 0.17463564 | -1.6727497 | 0.09437658 | 0.22838394 |
| AC104109.2 | 30.937567 | 1.104108711 | 0.34284121 | 3.22046672 | 0.00127982 | 0.0095282 |
| CDKL3 | 183.116787 | -0.101492887 | 0.17165185 | -0.5912717 | 0.55433838 | 0.72390942 |
| UBE2B | 3512.02087 | -0.613931847 | 0.13521962 | -4.5402573 | 5.62E-06 | 0.00012884 |
| AC109454.2 | 7.20847991 | 0.781287942 | 0.57896055 | 1.34946663 | 0.17718713 | 0.35221051 |
| CDKN2AIPNL | 730.844503 | 0.378940801 | 0.11746743 | 3.22592227 | 0.00125567 | 0.00938819 |
| AC109454.3 | 2.74891527 | 1.566590549 | 0.93780532 | 1.67048588 | 0.09482327 | 0.22913618 |
| AC005355.1 | 0.4482908 | -0.661119756 | 2.2924875 | -0.2883853 | 0.7730518 | NA |
| LINC01843 | 38.8479033 | 1.594124594 | 0.52101724 | 3.05963888 | 0.00221604 | 0.0146129 |
| JADE2 | 3065.75207 | -0.009764876 | 0.24703272 | -0.0395287 | 0.96846889 | 0.9853206 |
| SAR1B | 3912.60164 | 0.235873124 | 0.18841933 | 1.25185203 | 0.21062379 | 0.39399513 |
| SEC24A | 1449.0402 | -0.453028719 | 0.24540752 | -1.8460262 | 0.0648884 | 0.17457838 |
| CAMLG | 1845.03354 | -0.419205926 | 0.13119135 | -3.1953777 | 0.00139648 | 0.01023206 |
| DDX46 | 2563.622 | 0.153434309 | 0.13341904 | 1.1500181 | 0.25013642 | 0.44227553 |
| C5orf24 | 2846.9444 | 0.676672875 | 0.10286942 | 6.57797908 | 4.77E-11 | 7.83E-09 |
| TXNDC15 | 2758.22908 | 0.130179957 | 0.15898535 | 0.81881731 | 0.41289066 | 0.60677784 |
| PCBD2 | 760.110676 | 0.64856281 | 0.217361 | 2.98380481 | 0.00284688 | 0.01761021 |
| CATSPER3 | 128.771455 | 0.459957116 | 0.23681694 | 1.94224754 | 0.05210714 | 0.15002351 |
| PITX1 | 2.95911545 | -0.536230988 | 1.13717224 | -0.4715477 | 0.63724964 | 0.7853198 |
| C5orf66 | 1623.45639 | -0.453916262 | 0.13929305 | -3.2587144 | 0.00111918 | 0.00855941 |
| AC008406.3 | 0 | NA | NA | NA | NA | NA |
| C5orf66-AS1 | 0 | NA | NA | NA | NA | NA |
| AC008406.2 | 0 | NA | NA | NA | NA | NA |
| AC008406.1 | 0.09989236 | -0.642905668 | 3.30558948 | -0.1944905 | 0.84579185 | NA |
| C5orf66-AS2 | 0 | NA | NA | NA | NA | NA |
| H2AFY | 5913.39175 | -0.531712575 | 0.14893936 | -3.5699938 | 0.00035699 | 0.00348531 |
| AC026691.1 | 42.6866219 | 1.143096514 | 0.48523024 | 2.3557817 | 0.01848378 | 0.07083968 |
| DCANP1 | 11.8872397 | 1.559175072 | 1.13658001 | 1.37181286 | 0.1701217 | 0.3420855 |
| TIFAB | 17.3143275 | 1.633602925 | 1.08576387 | 1.50456556 | 0.13243581 | 0.28871794 |
| AC022092.1 | 0.08666371 | -0.642905668 | 3.30558948 | -0.1944905 | 0.84579185 | NA |
| NEUROG1 | 0.10721287 | -0.642905668 | 3.30558948 | -0.1944905 | 0.84579185 | NA |
| AC034206.1 | 36.3588868 | 1.343758857 | 0.68275834 | 1.96813247 | 0.0490528 | 0.14383657 |
| CXCL14 | 22905.1445 | 0.272909411 | 0.44642873 | 0.61131686 | 0.54098983 | 0.71386618 |
| SLC25A48 | 555.218809 | 2.20743517 | 0.62478361 | 3.53311954 | 0.00041069 | 0.00389842 |
| LINC01959 | 1.48608181 | 2.26629324 | 1.23299003 | 1.83804669 | 0.06605553 | 0.17684104 |
| IL9 | 3.97710996 | 3.929507586 | 0.97347464 | 4.0365793 | 5.42E-05 | 0.0007941 |
| AC002428.2 | 9.69978514 | 2.856205095 | 0.81819187 | 3.49087446 | 0.00048144 | 0.00440053 |
| AC002428.1 | 1.115847 | 3.09888598 | 1.54226445 | 2.00930909 | 0.04450436 | 0.13376225 |
| LECT2 | 90.4217001 | 0.850382492 | 0.4949522 | 1.71811034 | 0.08577649 | 0.21278702 |
| TGFBI | 5134.48283 | -1.04504435 | 0.44586252 | -2.3438713 | 0.01908476 | 0.07249568 |
| SMAD5-AS1 | 10.6292096 | 0.030734388 | 0.57779171 | 0.05319285 | 0.95757825 | 0.97960734 |
| SMAD5 | 2156.53368 | -0.067588666 | 0.12405277 | -0.544838 | 0.58586493 | 0.74726448 |
| SMIM32 | 2358.01639 | 0.981381486 | 0.4347367 | 2.25741579 | 0.02398211 | 0.08556462 |
| TRPC7 | 126.885646 | 0.690428426 | 0.6717023 | 1.0278786 | 0.30400694 | 0.50088085 |
| TRPC7-AS1 | 61.8828057 | 2.194098624 | 0.5452331 | 4.02414788 | 5.72E-05 | 0.00082306 |
| TRPC7-AS2 | 9.07045686 | 1.281867284 | 0.95430419 | 1.34324809 | 0.17919165 | 0.3548671 |
| AC063980.1 | 0 | NA | NA | NA | NA | NA |
| AC112178.1 | 0.03392908 | 0.380740593 | 3.29738709 | 0.11546736 | 0.90807469 | NA |
| AC109439.2 | 2.00422304 | 3.217006165 | 1.43919883 | 2.2352757 | 0.02539925 | 0.08937528 |
| SPOCK1 | 2278.83342 | -0.01637194 | 0.44114522 | -0.0371124 | 0.97039542 | 0.98629128 |
| KLHL3 | 873.119855 | 1.642715511 | 0.30099303 | 5.45765308 | 4.82E-08 | 2.58E-06 |
| HNRNPA0 | 6695.2133 | -0.447004624 | 0.18227479 | -2.4523667 | 0.014192 | 0.05848698 |
| AC106791.2 | 1.8033109 | -0.092662418 | 1.08939731 | -0.0850584 | 0.93221494 | 0.96732019 |
| AC106791.1 | 41.0169808 | 0.504848605 | 0.402268 | 1.25500564 | 0.20947671 | 0.39262925 |
| MYOT | 86.7430575 | 0.556412445 | 0.47725128 | 1.16586894 | 0.24366744 | 0.43439091 |
| PKD2L2 | 348.279039 | 0.361381824 | 0.25402028 | 1.4226495 | 0.15483779 | 0.32092427 |
| FAM13B | 1611.96246 | 0.158930508 | 0.21838635 | 0.72774928 | 0.46676708 | 0.65395005 |
| AC113382.1 | 15.6189057 | 1.379351309 | 0.55070679 | 2.50469276 | 0.01225578 | 0.05238005 |
| AC113382.2 | 0.50137837 | -0.026419023 | 1.85733029 | -0.0142242 | 0.98865112 | NA |
| WNT8A | 0.66674278 | -1.873298147 | 1.88095846 | -0.9959274 | 0.3192854 | NA |
| NME5 | 668.654083 | 0.417400625 | 0.40491078 | 1.03084592 | 0.30261308 | 0.49952135 |
| BRD8 | 3315.87679 | 0.128107045 | 0.16870055 | 0.75937539 | 0.44762803 | 0.63783587 |
| KIF20A | 72.8290536 | -0.593424144 | 0.45484105 | -1.3046847 | 0.19200025 | 0.37121831 |
| CDC23 | 1564.64567 | 0.249950331 | 0.12203107 | 2.04825164 | 0.04053535 | 0.12501955 |
| GFRA3 | 34.4637078 | 1.200662975 | 0.575788 | 2.08525184 | 0.03704645 | 0.11705186 |
| CDC25C | 48.5724925 | 0.130597572 | 0.45054414 | 0.28986632 | 0.77191851 | 0.87421716 |
| FAM53C | 2083.02659 | -0.36614238 | 0.26422362 | -1.3857292 | 0.16582961 | 0.3363406 |
| AC104116.1 | 609.556853 | -0.834320764 | 0.25593231 | -3.2599275 | 0.00111441 | 0.00853031 |
| KDM3B | 3576.08107 | 0.037832522 | 0.08117737 | 0.46604764 | 0.6411814 | 0.78758115 |
| REEP2 | 113.684169 | -0.357325656 | 0.23507656 | -1.5200395 | 0.12850105 | 0.28271089 |
| EGR1 | 7661.67972 | 1.387849451 | 0.65698561 | 2.11245029 | 0.03464785 | 0.11185892 |
| ETF1 | 5810.7531 | -0.486378109 | 0.23425973 | -2.0762344 | 0.03787228 | 0.11894538 |
| HSPA9 | 20608.1601 | 0.021878484 | 0.287498 | 0.07609961 | 0.93933985 | 0.97110795 |
| CTNNA1 | 19873.6059 | 0.010695224 | 0.15947995 | 0.06706313 | 0.94653145 | 0.97435431 |
| AC034243.1 | 21.4856155 | 2.213219387 | 0.70386254 | 3.1443915 | 0.00166433 | 0.01170354 |
| LRRTM2 | 13.8074738 | 2.637833789 | 0.61161047 | 4.31293103 | 1.61E-05 | 0.00029918 |
| SIL1 | 2756.43588 | -0.182340474 | 0.15983391 | -1.1408122 | 0.25394809 | 0.44656685 |
| AC011405.1 | 890.198784 | -0.098460469 | 0.16137157 | -0.6101476 | 0.54176407 | 0.71442309 |
| MATR3 | 11001.7175 | 0.15988558 | 0.09390512 | 1.702629 | 0.08863752 | 0.21813013 |
| SNHG4 | 153.423635 | -0.686320222 | 0.34354142 | -1.9977802 | 0.0457405 | 0.13654798 |
| PAIP2 | 5456.74585 | -0.372026811 | 0.15076764 | -2.4675509 | 0.01360409 | 0.05658357 |
| AC135457.1 | 3495.40055 | -0.446246559 | 0.15739385 | -2.8352223 | 0.00457938 | 0.02520524 |
| SLC23A1 | 2718.70009 | 2.467902119 | 0.41661967 | 5.92363328 | 3.15E-09 | 2.59E-07 |
| MZB1 | 326.492238 | -1.077264979 | 0.82242969 | -1.3098566 | 0.19024434 | 0.36890531 |
| PROB1 | 644.458358 | 1.122459361 | 0.3134305 | 3.58120657 | 0.00034201 | 0.0033648 |
| SPATA24 | 213.822229 | 0.60667264 | 0.19687427 | 3.08152317 | 0.00205944 | 0.01377665 |
| DNAJC18 | 455.545347 | 0.356587138 | 0.2105665 | 1.69346569 | 0.09036686 | 0.22129256 |
| AC142391.1 | 975.255973 | -0.634524139 | 0.34960304 | -1.8149846 | 0.06952627 | 0.1833236 |
| ECSCR | 1032.21994 | -0.653683829 | 0.3513493 | -1.8604956 | 0.06281544 | 0.17071453 |
| SMIM33 | 0.93272198 | -2.374735176 | 1.66131899 | -1.4294276 | 0.15288138 | NA |
| TMEM173 | 3068.33958 | -0.449143537 | 0.23017254 | -1.9513341 | 0.05101731 | 0.14777312 |
| UBE2D2 | 4434.44288 | -0.720053369 | 0.16685247 | -4.3155093 | 1.59E-05 | 0.00029654 |
| AC113361.1 | 7.93071501 | 0.674212199 | 0.55096171 | 1.22370064 | 0.22106518 | 0.40717251 |
| CXXC5 | 5807.33434 | -0.026902498 | 0.15769904 | -0.1705939 | 0.86454308 | 0.92965085 |
| CXXC5-AS1 | 89.6866436 | 0.790007786 | 0.30742423 | 2.5697642 | 0.01017678 | 0.04566302 |

| | | | | | | |
|---|---|---|---|---|---|---|
| AC008667.1 | 3.36389219 | 0.425377951 | 0.68909507 | 0.61729937 | 0.53703729 | 0.71074897 |
| PSD2-AS1 | 0.84413231 | 2.108525459 | 1.6137873 | 1.30656962 | 0.19135892 | NA |
| AC008667.3 | 2.37555477 | 0.298604637 | 0.85410092 | 0.34961283 | 0.72662928 | 0.84548446 |
| AC008667.4 | 0.37431598 | -0.542600628 | 2.80669061 | -0.193324 | 0.84670524 | NA |
| PSD2 | 16.3430642 | 0.233174724 | 0.43322155 | 0.53823436 | 0.59041526 | 0.75022132 |
| NRG2 | 101.844979 | 0.009484444 | 0.40543488 | 0.02339326 | 0.98133658 | 0.9915968 |
| AC008667.2 | 15.5883287 | 0.305681682 | 0.51292148 | 0.59596194 | 0.55120065 | 0.72143103 |
| MALINC1 | 188.952188 | 0.794610365 | 0.19742049 | 4.02496408 | 5.70E-05 | 0.00082083 |
| PURA | 1712.24317 | 0.225305514 | 0.15148242 | 1.48733773 | 0.13692564 | 0.29530784 |
| IGIP | 1501.05816 | 0.217653957 | 0.23374566 | 0.9311572 | 0.35177225 | 0.54940976 |
| CYSTM1 | 12219.414 | -0.400823005 | 0.13688324 | -2.928211 | 0.00340919 | 0.02020684 |
| PFDN1 | 2445.93271 | -0.184780067 | 0.08929667 | -2.0692827 | 0.03851956 | 0.12045282 |
| HBEGF | 944.139612 | 0.166684066 | 0.42305043 | 0.3940052 | 0.6935772 | 0.82370883 |
| AC008438.2 | 0.26413911 | 1.101280607 | 3.0105956 | 0.36580157 | 0.71451315 | NA |
| SLC4A9 | 1659.03872 | 2.106860153 | 0.59839559 | 3.52084838 | 0.00043017 | 0.00403685 |
| AC008438.1 | 224.472054 | 1.484014653 | 0.48878081 | 3.03615571 | 0.00239616 | 0.01548312 |
| ANKHD1 | 4619.41127 | 0.125728692 | 0.11334915 | 1.109216 | 0.26733701 | 0.4614569 |
| ANKHD1-EIF4 | 5983.49356 | 0.08350181 | 0.11413132 | 0.73162921 | 0.46439492 | 0.65183871 |
| SRA1 | 2904.63833 | -0.34341658 | 0.14279897 | -2.4048953 | 0.0161771 | 0.06436541 |
| EIF4EBP3 | 1518.22744 | -0.084420299 | 0.19104001 | -0.4418985 | 0.65856264 | 0.79946078 |
| APBB3 | 2340.31511 | 0.244963722 | 0.24717037 | 0.99107237 | 0.32165024 | 0.51937523 |
| SLC35A4 | 5823.85038 | 0.02896301 | 0.10180062 | 0.28450719 | 0.77602172 | 0.87702653 |
| CD14 | 3132.16459 | -0.43508841 | 0.50854959 | -0.8555477 | 0.39224803 | 0.58803756 |
| NDUFA2 | 3073.03181 | 0.045752084 | 0.12416867 | 0.36846723 | 0.71252488 | 0.83660363 |
| TMCO6 | 733.14282 | -0.093981259 | 0.17578717 | -0.5346309 | 0.59290508 | 0.7521845 |
| IK | 6891.46232 | -0.097179734 | 0.08437365 | -1.1517783 | 0.2494122 | 0.44131903 |
| WDR55 | 3006.30725 | -0.346035979 | 0.17349394 | -1.9945133 | 0.04609599 | 0.13725736 |
| DND1 | 905.075822 | -0.517211485 | 0.18730708 | -2.7613023 | 0.00575714 | 0.02991442 |
| HARS | 3405.46942 | -0.207523014 | 0.12282568 | -1.6895735 | 0.09110957 | 0.22255418 |
| HARS2 | 2520.80489 | 0.000516166 | 0.15096375 | 0.00341914 | 0.99727193 | 0.9986244 |
| ZMAT2 | 3858.48676 | -0.184995163 | 0.11748309 | -1.5746535 | 0.11533644 | 0.26275782 |
| PCDHA1 | 682.474584 | 0.277530176 | 0.31970434 | 0.86808385 | 0.38534843 | 0.58117277 |
| AC005609.5 | 8.14911281 | 0.932944909 | 0.56951873 | 1.63812858 | 0.10139487 | 0.23980115 |
| PCDHA2 | 692.368121 | 0.275691139 | 0.31367762 | 0.87889962 | 0.37945571 | 0.57545098 |
| PCDHA3 | 713.493482 | 0.307222067 | 0.31901103 | 0.9630453 | 0.33552479 | 0.53370665 |
| PCDHA4 | 705.763024 | 0.266833083 | 0.31439085 | 0.84873043 | 0.39603131 | 0.59122987 |
| PCDHA5 | 683.674354 | 0.264104135 | 0.31839095 | 0.82949637 | 0.40682359 | 0.60159114 |
| PCDHA6 | 691.960052 | 0.279088968 | 0.31786409 | 0.87801353 | 0.37993638 | 0.57591719 |
| PCDHA7 | 689.477574 | 0.257857664 | 0.31418021 | 0.82073173 | 0.4117991 | 0.60580988 |
| PCDHA8 | 674.539479 | 0.269279131 | 0.31698746 | 0.84949458 | 0.39560615 | 0.5909272 |
| PCDHA9 | 673.936692 | 0.26786256 | 0.31623268 | 0.84704263 | 0.39697136 | 0.59189006 |
| AC005609.2 | 0.23379451 | 0.689159497 | 3.01123709 | 0.22886258 | 0.81897572 | NA |
| PCDHA10 | 814.201271 | 0.293640134 | 0.30538621 | 0.96153699 | 0.33628223 | 0.53433132 |
| AC005609.4 | 20.4899328 | -0.311001639 | 0.42679083 | -0.728698 | 0.46618639 | 0.65341422 |
| PCDHA11 | 700.021569 | 0.319464282 | 0.31404536 | 1.01725522 | 0.30903203 | 0.50606562 |
| PCDHA12 | 689.361156 | 0.324196863 | 0.31649892 | 1.02432217 | 0.30568313 | 0.50270434 |
| AC005609.3 | 4.90597595 | 3.564116362 | 1.05817393 | 3.3681763 | 0.00075667 | 0.00628059 |
| AC005609.1 | 0.62688686 | 3.238515124 | 1.78560128 | 1.81368324 | 0.06972651 | NA |
| PCDHA13 | 674.161253 | 0.276018236 | 0.31677202 | 0.87134663 | 0.38356492 | 0.57933483 |
| PCDHAC1 | 683.296905 | 0.283959903 | 0.31782026 | 0.89346068 | 0.37161052 | 0.56836599 |
| AC010223.1 | 0.21220981 | 0.689156227 | 3.08302815 | 0.22353225 | 0.82312128 | NA |
| PCDHAC2 | 680.901536 | 0.285230543 | 0.31289544 | 0.91158421 | 0.36198764 | 0.55930239 |
| AC244517.5 | 41.4046696 | 0.535471416 | 0.49185636 | 1.08867437 | 0.27629751 | 0.47161404 |
| PCDHB1 | 29.3787353 | 1.306942791 | 0.64954294 | 2.01209606 | 0.04420981 | 0.13319008 |
| AC244517.9 | 0.03392908 | 0.380740593 | 3.29738709 | 0.11546736 | 0.90807469 | NA |
| PCDHB2 | 105.306898 | 0.229545839 | 0.43661532 | 0.52573931 | 0.59906935 | 0.7570307 |
| AC244517.2 | 65.4365132 | 0.243312675 | 0.48517105 | 0.50149875 | 0.61602016 | 0.77034489 |
| PCDHB3 | 120.524685 | 0.002204705 | 0.37204388 | 0.00592593 | 0.99527182 | 0.99776308 |
| AC244517.1 | 29.1977648 | -0.025234175 | 0.37800139 | -0.0667568 | 0.94677529 | 0.97438741 |
| PCDHB4 | 185.941961 | -0.074817742 | 0.3568017 | -0.20969 | 0.83390965 | 0.91196788 |
| PCDHB5 | 254.275808 | 0.178880541 | 0.29592041 | 0.6044887 | 0.54551878 | 0.71707942 |
| AC244517.11 | 61.3658983 | 0.247886926 | 0.3079385 | 0.80498843 | 0.42082636 | 0.61352042 |
| PCDHB6 | 67.875088 | -0.03499853 | 0.38952864 | -0.0898484 | 0.92840768 | 0.96526981 |
| AC244517.12 | 1.16173516 | 0.261152637 | 1.5548965 | 0.167955 | 0.86661868 | 0.9307387 |
| PCDHB7 | 108.73096 | 0.240285893 | 0.56280986 | 0.42693974 | 0.66942322 | 0.80698013 |
| PCDHB8 | 25.1683008 | 0.827614499 | 0.70526121 | 1.17348649 | 0.24060077 | 0.43072928 |
| PCDHB16 | 232.335137 | 0.080241575 | 0.39962294 | 0.20079321 | 0.84086027 | 0.91512686 |
| AC244517.6 | 26.5835471 | 0.835621367 | 0.43720923 | 1.91126194 | 0.05597093 | 0.15760313 |
| AC244517.4 | 57.9487824 | 0.447578984 | 0.41051143 | 1.09029604 | 0.27558276 | 0.47090048 |
| PCDHB9 | 54.2193971 | -0.2258883 | 0.27755715 | -0.8138443 | 0.41573417 | 0.60908136 |
| AC244517.7 | 3.59050873 | 0.704105031 | 0.7413984 | 0.94969861 | 0.34226541 | 0.54006337 |
| PCDHB10 | 64.8356134 | 0.52490148 | 0.43389379 | 1.20974646 | 0.2263762 | 0.41317416 |
| PCDHB11 | 87.8685795 | 0.0634958 | 0.41299858 | 0.15374339 | 0.87781207 | 0.9364039 |
| PCDHB12 | 61.1632543 | 0.204679121 | 0.4566212 | 0.44824708 | 0.65397489 | 0.79691637 |
| PCDHB13 | 102.958028 | 0.063302233 | 0.30430772 | 0.20802047 | 0.83521299 | 0.91264233 |
| PCDHB14 | 245.263801 | 0.160820166 | 0.28783905 | 0.5587156 | 0.57635583 | 0.74037915 |
| AC244517.1C | 44.2262612 | 0.599090029 | 0.32021698 | 1.87088775 | 0.06136064 | 0.16799005 |
| PCDHB15 | 251.623758 | -0.194209685 | 0.35242206 | -0.5510713 | 0.5815848 | 0.744196 |
| TAF7 | 4434.03607 | -0.25956187 | 0.1051322 | -2.4689093 | 0.01355256 | 0.0564672 |
| SLC25A2 | 3.64648983 | 0.824192835 | 0.69482069 | 1.186195 | 0.2355453 | 0.4244741 |
| AC005618.1 | 21.2692345 | 0.797057173 | 0.40958184 | 1.94602666 | 0.05165154 | 0.14920046 |
| PCDHGA1 | 4168.80735 | -0.197391954 | 0.23544658 | -0.8383726 | 0.4018215 | 0.59665707 |
| PCDHGA2 | 4242.31909 | -0.1794155 | 0.23471255 | -0.7644052 | 0.4446258 | 0.63537947 |
| PCDHGA3 | 4177.20835 | -0.188045853 | 0.23417593 | -0.803011 | 0.42196837 | 0.61449668 |
| PCDHGB1 | 4177.32636 | -0.18907843 | 0.23529522 | -0.8035796 | 0.42163983 | 0.61421392 |
| AC005618.3 | 7.28830513 | 1.401727824 | 0.65029894 | 2.15551301 | 0.03112171 | 0.10362115 |
| PCDHGA4 | 4175.79732 | -0.187849563 | 0.23563235 | -0.7972146 | 0.42532638 | 0.61768128 |
| PCDHGB2 | 4178.56239 | -0.185900295 | 0.2350671 | -0.7908393 | 0.42903779 | 0.62112297 |
| PCDHGA5 | 4178.1134 | -0.18979901 | 0.23561314 | -0.8055536 | 0.42050029 | 0.61323929 |
| PCDHGB3 | 4158.61078 | -0.191003492 | 0.23519812 | -0.8120962 | 0.41673645 | 0.60986792 |
| PCDHGA6 | 4271.79722 | -0.176329245 | 0.23444703 | -0.752107 | 0.45198673 | 0.64173558 |
| PCDHGA7 | 4212.74531 | -0.187441493 | 0.2344975 | -0.7993326 | 0.42409759 | 0.61657589 |
| PCDHGB4 | 4313.91967 | -0.175624328 | 0.23043752 | -0.7621343 | 0.44597988 | 0.63639012 |
| PCDHGA8 | 4262.24095 | -0.191140844 | 0.23303913 | -0.8202092 | 0.41209685 | 0.60607886 |
| PCDHGB5 | 4290.26838 | -0.155201659 | 0.23035197 | -0.6737588 | 0.5004647 | 0.68155397 |
| PCDHGA9 | 4197.80085 | -0.18766665 | 0.23502198 | -0.7985068 | 0.42457644 | 0.61701375 |
| PCDHGB6 | 4255.68145 | -0.176076675 | 0.2355076 | -0.7476475 | 0.45467279 | 0.64412386 |
| PCDHGA10 | 4206.42784 | -0.185368174 | 0.2338868 | -0.7925551 | 0.42803708 | 0.62042202 |
| PCDHGB7 | 4311.87642 | -0.149163522 | 0.23343289 | -0.6389996 | 0.52282319 | 0.69862624 |
| PCDHGA11 | 4187.1526 | -0.194539624 | 0.23345932 | -0.8332913 | 0.40468046 | 0.59945083 |
| AC005618.2 | 0.54993225 | 1.033829647 | 3.23473075 | 0.319603 | 0.7492693 | NA |
| PCDHGA12 | 4211.94254 | -0.174116775 | 0.23418456 | -0.7435024 | 0.4571776 | 0.64655504 |
| PCDHGC3 | 4477.55814 | -0.135070217 | 0.235221 | -0.5742269 | 0.5658143 | 0.73240345 |
| PCDHGC4 | 4147.63025 | -0.199003978 | 0.23560656 | -0.8446453 | 0.39830889 | 0.59327866 |
| PCDHGC5 | 4152.09695 | -0.19641199 | 0.23550039 | -0.8340198 | 0.40426982 | 0.59907812 |
| DIAPH1 | 9326.24658 | -0.402120529 | 0.13876158 | -2.8979242 | 0.00375641 | 0.02176112 |
| AC008781.2 | 0.90408626 | -0.341847193 | 1.54897051 | -0.2206932 | 0.82533136 | NA |
| AC008781.1 | 511.283222 | -0.024326003 | 0.1244947 | -0.1953979 | 0.84508146 | 0.91827913 |
| HDAC3 | 2789.27352 | -0.130701951 | 0.10816136 | -1.2083979 | 0.22689426 | 0.4137687 |
| RELL2 | 1374.46974 | -0.836740602 | 0.27123576 | -3.08492 | 0.00203607 | 0.01367927 |
| FCHSD1 | 2134.73215 | -0.740499627 | 0.204881 | -3.6142913 | 0.00030117 | 0.00304169 |
| ARAP3 | 2101.21539 | 0.170379008 | 0.26917868 | 0.63295878 | 0.52676056 | 0.70158472 |
| AC005753.1 | 15.5919566 | -0.123334382 | 0.52109971 | -0.236681 | 0.8129043 | 0.89895498 |

| | | | | | | |
|---|---|---|---|---|---|---|
| PCDH1 | 4380.93166 | -0.329534416 | 0.19765269 | -1.6672397 | 0.09546675 | 0.23013604 |
| DELE1 | 4497.66854 | 0.274863191 | 0.09131745 | 3.00997447 | 0.0026127 | 0.01654185 |
| PCDH12 | 1254.55906 | 0.051661349 | 0.23542294 | 0.21944059 | 0.82630686 | 0.90733757 |
| AC005740.4 | 27.2516073 | 0.110290403 | 0.39290065 | 0.28070812 | 0.77893428 | 0.87885063 |
| RNF14 | 3431.38548 | 0.045654318 | 0.20994126 | 0.21746234 | 0.82784805 | 0.90834907 |
| AC005740.3 | 175.83442 | 0.437577386 | 0.28675426 | 1.52596648 | 0.12701821 | 0.28057117 |
| GNPDA1 | 5790.475 | -0.026829889 | 0.47402799 | -0.0565998 | 0.95486399 | 0.97841508 |
| NDFIP1 | 9964.14282 | 0.008532211 | 0.10841643 | 0.07869851 | 0.93727243 | 0.96999796 |
| SPRY4 | 2071.12169 | -0.20949057 | 0.24130751 | -0.8681478 | 0.38531345 | 0.58117277 |
| SPRY4-AS1 | 17.3915483 | -0.236395501 | 0.3576664 | -0.6609385 | 0.50865174 | 0.68744274 |
| FGF1 | 3475.39404 | 0.692923651 | 0.59610682 | 1.16241523 | 0.24506684 | 0.43610474 |
| AC005592.1 | 4.72267557 | 0.505673322 | 0.75530607 | 0.66949459 | 0.50318003 | 0.68332771 |
| LINC01844 | 15.882791 | 0.021012436 | 0.57994282 | 0.03623191 | 0.97109744 | 0.9865502 |
| ARHGAP26 | 1634.33699 | -0.903700148 | 0.17372879 | -5.2017869 | 1.97E-07 | 8.45E-06 |
| ARHGAP26-A | 1.30362863 | -0.048375206 | 1.09379446 | -0.044227 | 0.96472349 | 0.98354657 |
| NR3C1 | 2052.51221 | 0.38805265 | 0.14325139 | 2.70889265 | 0.00675082 | 0.03363928 |
| AC016598.1 | 0 | NA | NA | NA | NA | NA |
| AC016598.2 | 0.26930081 | 1.145957498 | 3.27645355 | 0.34975545 | 0.72652223 | NA |
| HMHB1 | 0.51657952 | -1.1507074 | 2.74895712 | -0.4185978 | 0.6755101 | NA |
| YIPF5 | 2171.76536 | -0.156460954 | 0.24679211 | -0.6339787 | 0.52609469 | 0.70101652 |
| KCTD16 | 171.820184 | 1.359440695 | 0.52512149 | 2.58881179 | 0.00963077 | 0.04381702 |
| AC132803.1 | 1.61890878 | 0.105835539 | 1.05079563 | 0.10071943 | 0.91977319 | 0.96074349 |
| AC137770.1 | 1.34526754 | -0.111314293 | 1.00810044 | -0.1104198 | 0.91207642 | 0.95681217 |
| AC008700.1 | 0.25540699 | 1.967289346 | 2.88471024 | 0.68197122 | 0.49525715 | NA |
| PRELID2 | 84.955147 | -0.891127626 | 0.29316267 | -3.0397036 | 0.00236811 | 0.01536591 |
| GRXCR2 | 0.77493108 | 1.897532847 | 1.63811557 | 1.15836324 | 0.24671584 | NA |
| SH3RF2 | 216.826869 | -0.892453613 | 0.62498174 | -1.4279675 | 0.15330121 | 0.31886362 |
| PLAC8L1 | 24.3643298 | -0.615780435 | 0.35371734 | -1.7408828 | 0.08170413 | 0.20542885 |
| AC091887.1 | 654.655616 | -0.295800786 | 0.16824664 | -1.7581378 | 0.07872406 | 0.20022699 |
| LARS | 4387.12592 | 0.028170167 | 0.10241857 | 0.27504941 | 0.78327828 | 0.88142381 |
| AC091959.3 | 187.440796 | 0.482170808 | 0.20502607 | 2.35175362 | 0.01868515 | 0.07145118 |
| RBM27 | 1118.22272 | -0.033650782 | 0.15481345 | -0.2173634 | 0.82792513 | 0.90837491 |
| AC011396.2 | 0 | NA | NA | NA | NA | NA |
| POU4F3 | 9.55435123 | -1.005892098 | 0.69451875 | -1.4483297 | 0.14752488 | 0.31087031 |
| AC011396.1 | 0 | NA | NA | NA | NA | NA |
| TCERG1 | 2265.6572 | -0.444252604 | 0.1768077 | -2.5126315 | 0.01198344 | 0.05148699 |
| GPR151 | 0.35648076 | -0.541777668 | 2.84799994 | -0.1902309 | 0.84912818 | NA |
| AC008728.1 | 0 | NA | NA | NA | NA | NA |
| PPP2R2B | 258.788237 | 0.561134441 | 0.33604379 | 1.66982534 | 0.09495392 | 0.22927708 |
| STK32A-AS1 | 9.91559516 | 0.518061187 | 0.52038718 | 0.99553026 | 0.31947844 | 0.51688118 |
| STK32A | 392.325993 | 0.785583406 | 0.53779949 | 1.46073661 | 0.14408774 | 0.30587616 |
| DPYSL3 | 858.285696 | 0.747286554 | 0.32798907 | 2.27838861 | 0.02270343 | 0.08232997 |
| AC011373.1 | 0.64985408 | -0.033467476 | 1.59491687 | -0.0209838 | 0.98325855 | NA |
| JAKMIP2-AS1 | 5.25860673 | 0.964527646 | 0.7770898 | 1.24120488 | 0.21453007 | 0.39870026 |
| JAKMIP2 | 28.9226226 | 0.212399656 | 0.50269497 | 0.42252194 | 0.67264409 | 0.80897677 |
| SPINK1 | 1186.38977 | -2.969732198 | 0.8222996 | -3.6114966 | 0.00030444 | 0.00307023 |
| SCGB3A2 | 0.45428956 | 0.341779313 | 2.26925244 | 0.15061317 | 0.88028087 | NA |
| C5orf46 | 0.58927277 | 1.264524676 | 1.76136238 | 0.7179242 | 0.47280403 | NA |
| AC011352.1 | 0.2441911 | -0.174096114 | 3.29377188 | -0.0528562 | 0.95784652 | NA |
| AC011352.3 | 0.17515154 | 1.45420457 | 3.2783972 | 0.44357181 | 0.65735219 | NA |
| SPINK5 | 36.0198201 | 0.627299428 | 0.53252814 | 1.17796485 | 0.23881062 | 0.42839701 |
| SPINK14 | 0.03392908 | 0.380740593 | 3.29738709 | 0.11546736 | 0.90807469 | NA |
| SPINK6 | 0.61748072 | -0.015552686 | 1.59050352 | -0.0097785 | 0.99219804 | NA |
| MARCOL | 2.72953698 | 0.663538435 | 0.99272089 | 0.66840382 | 0.50387586 | 0.68370929 |
| SPINK13 | 31.8198765 | -0.955107793 | 0.52007695 | -1.836474 | 0.06628759 | 0.17726497 |
| AC091948.1 | 42.9067743 | 0.475346642 | 0.39553992 | 1.20176656 | 0.22945399 | 0.41691787 |
| SPINK7 | 3.31386358 | 0.600107298 | 0.92655877 | 0.64767322 | 0.51719633 | 0.69438762 |
| SPINK9 | 1.28400143 | 0.385015121 | 1.02698409 | 0.37489882 | 0.70773572 | 0.83365305 |
| FBXO38 | 1646.17534 | 0.11367065 | 0.08499546 | 1.33737317 | 0.18110086 | 0.35729684 |
| AC114939.1 | 3.23532414 | -0.119916509 | 1.13479728 | -0.1056722 | 0.91584245 | 0.9589345 |
| HTR4 | 2.12164154 | 1.487205016 | 0.98214887 | 1.51423584 | 0.12996603 | 0.28507772 |
| ADRB2 | 89.2848817 | -0.05887064 | 0.35140198 | -0.1675308 | 0.86695245 | 0.93095802 |
| SH3TC2 | 21.0474727 | -0.248941795 | 0.50328522 | -0.4946336 | 0.62085876 | 0.77350682 |
| AC116312.1 | 2.03953832 | -0.342182204 | 1.16461496 | -0.2938157 | 0.76889871 | 0.8723426 |
| SH3TC2-DT | 0.14114194 | 0.02007176 | 3.29738704 | 0.00608717 | 0.99514317 | NA |
| ABLIM3 | 4211.07398 | -1.223463187 | 0.47531128 | -2.5740252 | 0.0100523 | 0.04523533 |
| AC012613.1 | 1.38639489 | 0.789268645 | 1.12906312 | 0.69904741 | 0.48452241 | 0.66891031 |
| AC012613.2 | 143.804845 | -0.474252438 | 0.58099114 | -0.8162817 | 0.41433904 | 0.60798622 |
| AFAP1L1 | 952.952707 | -0.737202305 | 0.26888609 | -2.74169 | 0.0061124 | 0.03124932 |
| GRPEL2 | 962.33793 | -0.453653668 | 0.29425388 | -1.5417083 | 0.12314448 | 0.2746124 |
| GRPEL2-AS1 | 45.7982092 | -0.412978519 | 0.36230783 | -1.1398554 | 0.25434657 | 0.44708166 |
| PCYOX1L | 435.972919 | 0.189829008 | 0.31300383 | 0.60647503 | 0.54419936 | 0.71620188 |
| IL17B | 31.0379969 | 0.464249018 | 0.55360496 | 0.83859259 | 0.40169797 | 0.59661395 |
| AC131025.1 | 5.46808616 | 0.772756422 | 0.55046131 | 1.40383422 | 0.16036822 | 0.32858663 |
| CARMN | 2665.83981 | 0.344128248 | 0.51030695 | 0.6743554 | 0.50008539 | 0.68130141 |
| AC131025.2 | 40.0118499 | 0.015714569 | 0.5981706 | 0.02627105 | 0.97904115 | 0.99050094 |
| CSNK1A1 | 8482.09296 | -0.504750253 | 0.13225695 | -3.8164364 | 0.00013539 | 0.00160752 |
| ARHGEF37 | 1136.66935 | 0.229084915 | 0.25097379 | 0.91278424 | 0.36135603 | 0.55885556 |
| PPARGC1B | 281.621134 | 0.691030249 | 0.3488754 | 1.98073653 | 0.04762083 | 0.14058001 |
| PDE6A | 2.65055109 | -0.268682729 | 0.82437572 | -0.3259227 | 0.74448287 | 0.85673198 |
| SLC26A2 | 1028.23177 | -1.020056617 | 0.42957553 | -2.3745687 | 0.01756946 | 0.06823369 |
| TIGD6 | 387.52194 | 0.021894629 | 0.19516639 | 0.11218442 | 0.91067718 | 0.95582382 |
| HMGXB3 | 2902.80768 | -0.22276212 | 0.12730993 | -1.7497623 | 0.08015933 | 0.20272386 |
| CSF1R | 1862.90778 | 0.365523667 | 0.43354075 | 0.8431126 | 0.39916547 | 0.59400004 |
| PDGFRB | 7121.98776 | 0.336877477 | 0.28019857 | 1.20228123 | 0.22925458 | 0.41666851 |
| CDX1 | 6.84239655 | -0.805953827 | 0.74009795 | -1.0889826 | 0.27616154 | 0.47149878 |
| SLC6A7 | 13.6203795 | 1.749850302 | 0.47696811 | 3.66869452 | 0.00024379 | 0.00256844 |
| CAMK2A | 146.623118 | 2.538279978 | 0.52912744 | 4.79710521 | 1.61E-06 | 4.80E-05 |
| ARSI | 121.887704 | 0.099035748 | 0.38730066 | 0.25570767 | 0.79817658 | 0.89073665 |
| TCOF1 | 1680.71553 | -0.139266021 | 0.16530298 | -0.8424895 | 0.39951403 | 0.5942681 |
| CD74 | 86319.2861 | -0.623981343 | 0.33445297 | -1.8656774 | 0.06208652 | 0.16939516 |
| RPS14 | 19865.906 | -0.327512137 | 0.1940668 | -1.6876258 | 0.09148309 | 0.22314393 |
| NDST1-AS1 | 16.2984652 | 0.457340653 | 0.41884064 | 1.09192042 | 0.27486808 | 0.47007807 |
| NDST1 | 4631.95766 | 0.448461421 | 0.16288026 | 2.75331961 | 0.00589943 | 0.03041406 |
| SYNPO | 8683.03351 | -0.068069007 | 0.28687961 | -0.2372738 | 0.81244441 | 0.89882785 |
| AC011383.1 | 4.22057314 | -0.53532288 | 0.72881422 | -0.7345121 | 0.46263668 | 0.65082578 |
| MYOZ3 | 43.414201 | 0.091268921 | 0.36654683 | 0.24899661 | 0.8033634 | 0.89330352 |
| AC008453.1 | 2.77156466 | 0.530647351 | 0.8785101 | 0.60403102 | 0.54582302 | 0.71715108 |
| RBM22 | 2437.81444 | -0.303496185 | 0.16397157 | -1.8509073 | 0.06418289 | 0.17325241 |
| DCTN4 | 2491.76423 | 0.052336081 | 0.12283376 | 0.42607244 | 0.67005506 | 0.80733565 |
| SMIM3 | 2313.34661 | -1.524472971 | 0.28201233 | -5.4056962 | 6.46E-08 | 3.27E-06 |
| IRGM | 4.98667423 | -0.208313651 | 0.65406801 | -0.3184893 | 0.75011382 | 0.86047075 |
| ZNF300 | 518.319803 | -0.092977759 | 0.2346312 | -0.3962719 | 0.69190444 | 0.82267358 |
| GPX3 | 613948.37 | 0.564524006 | 0.34958283 | 1.61485051 | 0.10634307 | 0.24777053 |
| TNIP1 | 12140.2749 | -0.341090159 | 0.11070144 | -3.0811719 | 0.00206188 | 0.01378942 |
| ANXA6 | 7284.02206 | 0.363677213 | 0.29182362 | 1.24622266 | 0.21268266 | 0.39652445 |
| AC008641.1 | 1.2341342 | -0.459914573 | 1.57396349 | -0.2922016 | 0.77013253 | 0.87291382 |
| CCDC69 | 1239.45962 | -0.15126245 | 0.32790432 | -0.4613006 | 0.64458298 | 0.79000061 |
| GM2A | 1650.48976 | 0.060417548 | 0.23022242 | 0.2624312 | 0.79298902 | 0.8878826 |
| SLC36A3 | 0.73700806 | 1.722865714 | 1.7959574 | 0.95930211 | 0.33740657 | NA |
| SLC36A2 | 6318.76166 | 2.589607306 | 0.64794876 | 3.99662357 | 6.43E-05 | 0.00089742 |
| SLC36A1 | 992.032512 | -0.095378421 | 0.36202201 | -0.2634603 | 0.79219583 | 0.8873257 |
| FAT2 | 96.3739869 | 0.378999106 | 0.36780389 | 1.03043801 | 0.30280444 | 0.49971544 |

| | | | | | | |
|---|---|---|---|---|---|---|
| AC011337.1 | 68.8181444 | -0.430303742 | 0.35078856 | -1.2266755 | 0.21994457 | 0.4056752 |
| AC011374.1 | 9.98304471 | 0.427624227 | 0.59773958 | 0.71540222 | 0.47436054 | 0.66007262 |
| SPARC | 25153.9434 | 0.113283334 | 0.31663683 | 0.35777056 | 0.72051503 | 0.84100484 |
| CLMAT3 | 1208.78147 | 0.4297533 | 0.35144598 | 1.22281466 | 0.22139971 | 0.40753248 |
| AC011374.2 | 23.1097556 | 0.327834913 | 0.44191008 | 0.74185886 | 0.45817285 | 0.64713541 |
| ATOX1 | 2564.0366 | 0.129364667 | 0.15182947 | 0.85203926 | 0.3941923 | 0.58948004 |
| AC091982.1 | 8.75265402 | 0.994613726 | 0.63346773 | 1.57010953 | 0.11638963 | 0.26454013 |
| AC091982.3 | 77.4614231 | 0.870402916 | 0.30420365 | 2.8612507 | 0.00421973 | 0.02368791 |
| G3BP1 | 5988.15847 | -0.208261839 | 0.14743219 | -1.4125941 | 0.15777511 | 0.32488909 |
| GLRA1 | 0.82433814 | -0.440474842 | 1.30789283 | -0.3367821 | 0.7362812 | NA |
| LINC01933 | 43.1015904 | -0.439367296 | 0.45194444 | -0.972171 | 0.33096547 | 0.52884684 |
| NMUR2 | 17.4630863 | 1.115901756 | 0.94978722 | 1.17489657 | 0.2400361 | 0.42998111 |
| LINC01470 | 4.45653432 | -0.381711917 | 0.55369465 | -0.6893907 | 0.49057746 | 0.67369536 |
| GRIA1 | 8.54607443 | 0.993526646 | 0.54505106 | 1.82281391 | 0.06833158 | 0.18102308 |
| LINC01861 | 0.34844221 | -0.534763868 | 3.29377181 | -0.1623561 | 0.87102546 | NA |
| AC091962.1 | 0.08757577 | 0.547239394 | 3.29377202 | 0.16614368 | 0.86804388 | NA |
| FAM114A2 | 1137.92558 | 0.107140578 | 0.12950463 | 0.82731081 | 0.40806092 | 0.60264172 |
| MFAP3 | 358.168644 | 0.332972955 | 0.23179721 | 1.43648391 | 0.15086469 | 0.31565379 |
| GALNT10 | 1815.42772 | 0.079782775 | 0.11192086 | 0.71284992 | 0.47593863 | 0.66132069 |
| SAP30L-AS1 | 27.0343297 | 0.508506754 | 0.27010126 | 1.88265225 | 0.05974751 | 0.16508192 |
| SAP30L | 1979.57465 | 0.225942999 | 0.15761854 | 1.43347982 | 0.15172076 | 0.31677565 |
| HAND1 | 0.66329105 | -1.618589229 | 2.21164741 | -0.7318478 | 0.46426149 | NA |
| AC026688.2 | 0 | NA | NA | NA | NA | NA |
| LARP1 | 8711.54591 | 0.095316526 | 0.18934465 | 0.50340226 | 0.61468148 | 0.76958946 |
| FAXDC2 | 1973.19538 | 1.346762335 | 0.25649278 | 5.25068315 | 1.52E-07 | 6.76E-06 |
| CNOT8 | 2813.86835 | -0.431620159 | 0.15227454 | -2.8344866 | 0.00458994 | 0.0252528 |
| GEMIN5 | 686.824915 | 0.205418778 | 0.12138894 | 1.69223637 | 0.09060091 | 0.22168935 |
| MRPL22 | 1216.60447 | -0.304291285 | 0.1297353 | -2.345478 | 0.0190027 | 0.07229855 |
| KIF4B | 0.41444243 | -0.032127264 | 2.22635397 | -0.0144304 | 0.98848658 | NA |
| SGCD | 109.916298 | 1.327603663 | 0.38399797 | 3.45731949 | 0.00054558 | 0.00485715 |
| AC025434.1 | 1.28832917 | -0.13214836 | 1.38756846 | -0.0952374 | 0.92412629 | 0.96312021 |
| PPP1R2B | 5.4654277 | -0.395384755 | 0.5101451 | -0.7750437 | 0.43831382 | 0.63008807 |
| TIMD4 | 160.005064 | -1.061830091 | 0.42589465 | -2.4931754 | 0.01266063 | 0.05369262 |
| HAVCR1 | 362.847486 | -0.97635917 | 0.41063806 | -2.3776636 | 0.01742271 | 0.06780355 |
| HAVCR2 | 2768.90495 | -0.781314092 | 0.45939572 | -1.7007431 | 0.08899124 | 0.21882684 |
| AC011377.1 | 4.31601528 | 1.483613086 | 0.7197099 | 2.06140432 | 0.03926448 | 0.12217161 |
| MED7 | 698.622684 | -0.123735501 | 0.12204824 | -1.0138245 | 0.3106665 | 0.50760738 |
| ITK | 169.971956 | -0.171992641 | 0.69524678 | -0.2473836 | 0.80461138 | 0.89420103 |
| FAM71B | 0.10721287 | -0.642905668 | 3.30558948 | -0.1944905 | 0.84579185 | NA |
| AC010609.1 | 1.00701834 | 2.87879874 | 1.28439906 | 2.24135848 | 0.02500286 | 0.08834591 |
| AC009185.1 | 42.3990566 | 2.433473601 | 0.41960148 | 5.79948765 | 6.65E-09 | 4.89E-07 |
| CYFIP2 | 13640.9854 | 1.680152959 | 0.28607224 | 5.87317725 | 4.28E-09 | 3.32E-07 |
| FNDC9 | 4.23775379 | 0.359110272 | 0.62918865 | 0.57075135 | 0.5681682 | 0.73390132 |
| AC008676.1 | 6.82703926 | 1.088966382 | 0.56727406 | 1.91964778 | 0.0549024 | 0.1555578 |
| ADAM19 | 554.351701 | -0.073861051 | 0.39199841 | -0.1884218 | 0.850546 | 0.92161226 |
| NIPAL4 | 6.62000559 | -0.545031411 | 1.00931987 | -0.5399987 | 0.58919793 | 0.74953686 |
| AC106801.1 | 1.75519696 | -0.675345833 | 1.12734516 | -0.5990586 | 0.5491338 | 0.71964309 |
| SOX30 | 18.5150803 | 0.680825094 | 0.42134057 | 1.61585458 | 0.10612576 | 0.2474391 |
| C5orf52 | 0.70640431 | -1.430496164 | 1.90243206 | -0.7519302 | 0.45209302 | NA |
| THG1L | 441.660242 | -0.284219156 | 0.17586642 | -1.6161081 | 0.10607093 | 0.24734579 |
| LSM11 | 326.088391 | -0.438029978 | 0.21754024 | -2.0135584 | 0.04405591 | 0.13296949 |
| CLINT1 | 6997.88595 | -0.019945852 | 0.12128614 | -0.1644529 | 0.86937466 | 0.932003 |
| AC025437.5 | 0.19271054 | 1.506623853 | 3.27645372 | 0.4598337 | 0.64563559 | NA |
| AC025437.4 | 0.16354576 | -1.003574173 | 3.30558948 | -0.3035992 | 0.7614333 | NA |
| AC025437.3 | 0.15661533 | -1.003574173 | 3.30558948 | -0.3035992 | 0.7614333 | NA |
| AC025437.2 | 0 | NA | NA | NA | NA | NA |
| AC025437.1 | 0.08695548 | -0.642905668 | 3.30558948 | -0.1944905 | 0.84579185 | NA |
| LINC02227 | 1.06249072 | 0.622803784 | 1.07797372 | 0.57775414 | 0.56343012 | 0.7305885 |
| AC091979.1 | 0.16430572 | 0.435199075 | 3.28898345 | 0.13232024 | 0.894731 | NA |
| AC091979.2 | 0.09997307 | 0.547239394 | 3.29377202 | 0.16614368 | 0.86804388 | NA |
| AC091939.1 | 0.06785815 | 0.795868262 | 3.28898349 | 0.24198001 | 0.80879565 | NA |
| EBF1 | 534.724725 | 0.001862966 | 0.29644278 | 0.0062844 | 0.9949858 | 0.99770729 |
| AC136424.2 | 0.17294555 | 0.18657164 | 3.29377195 | 0.05664376 | 0.95482897 | NA |
| AC136424.1 | 0.32775918 | 0.669338683 | 2.24148455 | 0.29861401 | 0.76523458 | NA |
| LINC02202 | 89.7498817 | -0.111849382 | 0.44005102 | -0.2541737 | 0.79936139 | 0.89110801 |
| RNF145 | 3408.72678 | -0.888001644 | 0.26846569 | -3.3076913 | 0.00094068 | 0.00749386 |
| AC134043.1 | 3.4781935 | -1.491025884 | 1.0523866 | -1.4168043 | 0.15654015 | 0.32325479 |
| LINC01932 | 0.45449019 | -0.940708527 | 1.90037226 | -0.4950128 | 0.62059111 | NA |
| UBLCP1 | 1237.94028 | -0.162179263 | 0.18025893 | -0.8997017 | 0.36827902 | 0.56511596 |
| AC008691.1 | 39.9269361 | -1.155735244 | 0.56559063 | -2.043413 | 0.04101157 | 0.12621187 |
| IL12B | 17.4925553 | -0.419951303 | 0.57621435 | -0.7288109 | 0.46611734 | 0.6533869 |
| LINC01845 | 0.3143988 | -0.174096114 | 3.29377188 | -0.0528562 | 0.95784652 | NA |
| AC008703.1 | 0.07830766 | -0.642905668 | 3.30558948 | -0.1944905 | 0.84579185 | NA |
| LINC01847 | 1.19669982 | 2.672532301 | 1.20237633 | 2.22270868 | 0.02623545 | 0.09157231 |
| ADRA1B | 232.971786 | 0.959591854 | 0.49989196 | 1.91959849 | 0.05490864 | 0.15555874 |
| TTC1 | 2181.79251 | -0.242511299 | 0.18867274 | -1.2853542 | 0.19866853 | 0.37968468 |
| PWWP2A | 1015.77439 | 0.268909413 | 0.09894396 | 2.717795 | 0.00657185 | 0.03295299 |
| FABP6 | 3.11971031 | 0.750681788 | 0.79154961 | 0.94836985 | 0.3429412 | 0.54054749 |
| AC008609.1 | 2.09435628 | -2.282866498 | 2.11125676 | -1.0812832 | 0.27957115 | 0.47514828 |
| CCNJL | 372.646653 | -0.105625944 | 0.34573944 | -0.3055074 | 0.75997976 | 0.86700095 |
| C1QTNF2 | 56.2403571 | 0.686976365 | 0.54223733 | 1.2669293 | 0.20518057 | 0.3877185 |
| ZBED8 | 334.762233 | 0.259423071 | 0.25405902 | 1.02111342 | 0.30720071 | 0.50411861 |
| SLU7 | 1863.2053 | -0.047143199 | 0.11822492 | -0.3987586 | 0.6900711 | 0.8215058 |
| PTTG1 | 99.4521749 | -1.822053313 | 0.40957917 | -4.4485985 | 8.64E-06 | 0.00018175 |
| MIR3142HG | 102.460714 | -0.839366096 | 0.49238921 | -1.7046801 | 0.0882541 | 0.21739941 |
| ATP10B | 48.2750785 | -1.874540896 | 0.86250178 | -2.1733763 | 0.02975201 | 0.10031802 |
| AC008456.1 | 0.18382444 | -1.003574173 | 3.30558948 | -0.3035992 | 0.7614333 | NA |
| AC011363.1 | 0.13390214 | 1.04981649 | 3.28477129 | 0.31960109 | 0.74927075 | NA |
| LINC02159 | 18.0066391 | 0.002884434 | 0.90673958 | 0.00318111 | 0.99746185 | 0.99867114 |
| GABRB2 | 91.4389235 | 1.050334529 | 0.58799916 | 1.78628577 | 0.074053 | 0.19203241 |
| GABRA6 | 0.08695548 | -0.642905668 | 3.30558948 | -0.1944905 | 0.84579185 | NA |
| AC091944.1 | 0 | NA | NA | NA | NA | NA |
| GABRA1 | 0.28228389 | 0.074529887 | 3.28898342 | 0.02266046 | 0.98192111 | NA |
| LINC01202 | 0 | NA | NA | NA | NA | NA |
| GABRG2 | 0 | NA | NA | NA | NA | NA |
| AC113414.1 | 22.3589122 | 0.778309631 | 0.47188549 | 1.64936121 | 0.09907366 | 0.23602605 |
| CCNG1 | 6572.5605 | 0.278746354 | 0.21153694 | 1.31771953 | 0.18759756 | 0.36565508 |
| NUDCD2 | 1210.65445 | 0.036536435 | 0.099925 | 0.36563858 | 0.71463478 | 0.83781262 |
| HMMR | 61.397934 | 0.257571857 | 0.49388139 | 0.52152574 | 0.60200058 | 0.75930336 |
| HMMR-AS1 | 4.71583408 | -0.689886544 | 0.82766669 | -0.8335318 | 0.40454484 | 0.59935095 |
| MAT2B | 5331.75395 | 0.243958424 | 0.11479278 | 2.12520706 | 0.03356934 | 0.10936026 |
| AC008662.1 | 0.10721287 | -0.642905668 | 3.30558948 | -0.1944905 | 0.84579185 | NA |
| AC008662.2 | 0 | NA | NA | NA | NA | NA |
| AC109466.1 | 4.41824903 | -0.349000587 | 0.91414404 | -0.3817786 | 0.70262563 | 0.82987743 |
| LINC02143 | 0.40532843 | 1.840765306 | 2.07906074 | 0.88538313 | 0.37595003 | NA |
| LINC01938 | 0.189363 | 1.5839633 | 3.27302703 | 0.48394446 | 0.62842529 | NA |
| AC008415.1 | 0.99433644 | 1.03920531 | 1.5458237 | 0.67226639 | 0.50141413 | NA |
| AC114321.1 | 0 | NA | NA | NA | NA | NA |
| LINC01947 | 0 | NA | NA | NA | NA | NA |
| AC091819.2 | 0 | NA | NA | NA | NA | NA |
| AC091819.1 | 0 | NA | NA | NA | NA | NA |
| TENM2 | 129.21271 | 1.291265238 | 0.39213448 | 3.29291427 | 0.00099155 | 0.00778419 |
| AC091820.2 | 0 | NA | NA | NA | NA | NA |

| | | | | | | |
|---|---|---|---|---|---|---|
| AC091820.1 | 0 | NA | NA | NA | NA | NA |
| AC008601.1 | 0 | NA | NA | NA | NA | NA |
| AC008708.2 | 0 | NA | NA | NA | NA | NA |
| AC008464.1 | 0.48946982 | 0.295121083 | 2.70076391 | 0.10927319 | 0.91298581 | NA |
| AC008705.1 | 0 | NA | NA | NA | NA | NA |
| AC011369.1 | 0.771975 | -0.789009632 | 1.64101306 | -0.4808064 | 0.63065408 | NA |
| AC008637.1 | 137.375936 | 0.330682975 | 0.32930614 | 1.00418103 | 0.31529137 | 0.51243349 |
| WWC1 | 10998.8307 | -0.406830361 | 0.22969333 | -1.7711892 | 0.07652924 | 0.19640643 |
| RARS | 2821.88835 | -0.488908692 | 0.1978376 | -2.4712628 | 0.01346368 | 0.0561768 |
| FBLL1 | 35.461237 | -0.541047405 | 0.73715746 | -0.7339645 | 0.46297038 | 0.65100676 |
| PANK3 | 2994.23928 | -0.189273689 | 0.21324617 | -0.8875831 | 0.37476506 | 0.57075106 |
| AC011365.2 | 643.269026 | -0.091725539 | 0.39207079 | -0.2339515 | 0.81502265 | 0.9003455 |
| SLIT3 | 3539.66947 | 0.76070398 | 0.38492615 | 1.97623355 | 0.04812833 | 0.14167717 |
| AC011365.1 | 239.157501 | 0.979639295 | 0.36746759 | 2.66592025 | 0.00767779 | 0.03704762 |
| AC011389.2 | 5.20571004 | 0.790120803 | 0.82144804 | 0.96186339 | 0.33611823 | 0.5342602 |
| AC011389.1 | 1.26259066 | 0.555009296 | 1.35105829 | 0.41079596 | 0.68122215 | 0.81527767 |
| SPDL1 | 331.167874 | -0.103336641 | 0.19175335 | -0.538904 | 0.58995312 | 0.74984642 |
| DOCK2 | 431.793841 | -0.167079502 | 0.48978138 | -0.3411308 | 0.73300513 | 0.84942786 |
| AC008680.1 | 1.93207436 | -0.509849185 | 1.07099547 | -0.4760517 | 0.63403756 | 0.78307797 |
| FAM196B | 3.50273815 | 2.538646762 | 0.75337249 | 3.36970993 | 0.00075247 | 0.00625559 |
| FOXI1 | 1892.26444 | -0.29616799 | 0.46377585 | -0.6386016 | 0.52308216 | 0.69862624 |
| LINC01187 | 1011.06727 | 0.318138411 | 0.48473163 | 0.65631865 | 0.51161912 | 0.68975942 |
| C5orf58 | 106.518334 | -0.241985652 | 0.4295609 | -0.5633326 | 0.57320843 | 0.73780965 |
| LCP2 | 694.599229 | -0.253904025 | 0.42632313 | -0.5955671 | 0.55146447 | 0.72158577 |
| AC034199.1 | 4.30763078 | -3.22629154 | 1.51845059 | -2.1247261 | 0.03360948 | 0.10943103 |
| LINC01366 | 2.29975501 | 0.653773688 | 0.88823675 | 0.73603539 | 0.46170916 | 0.65011148 |
| KCNIP1 | 107.025505 | 0.47485076 | 0.35344042 | 1.34351008 | 0.17910687 | 0.35484959 |
| KCNMB1 | 249.909246 | -0.27427761 | 0.62038956 | -0.4421055 | 0.65841289 | 0.79946078 |
| AC027306.1 | 3.37337591 | 1.829859304 | 0.74564345 | 2.45406743 | 0.01412505 | 0.05832721 |
| AC008619.1 | 2.61946939 | 1.494654946 | 0.88827639 | 1.68264626 | 0.09244359 | 0.22483904 |
| AC027312.1 | 3.31582708 | 0.262759916 | 0.88400445 | 0.29723823 | 0.76628464 | 0.87113838 |
| AC008514.1 | 74.8117079 | -0.327277931 | 0.60195326 | -0.5436933 | 0.58665257 | 0.74773127 |
| GABRP | 342.911193 | -0.537880309 | 0.64318577 | -0.8362752 | 0.4030001 | 0.59801276 |
| RANBP17 | 168.108085 | 0.210416491 | 0.2714338 | 0.77520373 | 0.43821928 | 0.63008807 |
| AC091980.2 | 0 | NA | NA | NA | NA | NA |
| TLX3 | 0.08695548 | -0.642905668 | 3.30558948 | -0.1944905 | 0.84579185 | NA |
| NPM1 | 17732.1851 | -0.502649889 | 0.19974565 | -2.5164497 | 0.01185438 | 0.0511202 |
| FGF18 | 36.3752939 | -0.405872748 | 0.62450339 | -0.6499128 | 0.51574855 | 0.69339702 |
| AC011410.1 | 0 | NA | NA | NA | NA | NA |
| SMIM23 | 11.4209679 | -1.91750393 | 0.51883976 | -3.6957537 | 0.00021924 | 0.00235105 |
| FBXW11 | 2880.6937 | -0.059620215 | 0.16326025 | -0.3651851 | 0.71497322 | 0.83798586 |
| STK10 | 1499.38751 | -0.417581715 | 0.28596163 | -1.4602718 | 0.14421538 | 0.30597469 |
| EFCAB9 | 0.72532023 | 2.862608442 | 1.86428831 | 1.53549664 | 0.12466188 | NA |
| UBTD2 | 1545.60315 | -0.163545733 | 0.18876652 | -0.8663916 | 0.38627544 | 0.58184907 |
| SH3PXD2B | 644.795803 | -1.224783364 | 0.2723039 | -4.4978547 | 6.86E-06 | 0.00015211 |
| LINC01944 | 0.83265399 | -0.261010518 | 1.5489707 | -0.1685058 | 0.8661854 | NA |
| NEURL1B | 1872.31927 | -0.0552847 | 0.45114716 | -0.1225425 | 0.90246939 | 0.95085975 |
| AC027309.1 | 557.315198 | -0.31982105 | 0.45854385 | -0.697471 | 0.48550807 | 0.66964851 |
| AC022217.2 | 5.10094969 | -0.244791339 | 0.65446945 | -0.3740302 | 0.70838185 | 0.83417046 |
| AC022217.3 | 1.86458957 | 0.367645635 | 0.83190383 | 0.44193286 | 0.65853779 | 0.79946078 |
| DUSP1 | 18929.5728 | -0.325371876 | 0.4394401 | -0.7404237 | 0.45904293 | 0.64784375 |
| AC110011.1 | 0.16416605 | 0.18657164 | 3.29377195 | 0.05664376 | 0.95482897 | NA |
| ERGIC1 | 10482.3839 | -0.43217154 | 0.21224181 | -2.0362225 | 0.04172802 | 0.12780115 |
| AC008429.1 | 77.4129196 | -0.720833102 | 0.48954665 | -1.4724503 | 0.14089933 | 0.30119287 |
| AC008429.3 | 34.171308 | -0.703783116 | 0.59628005 | -1.1802896 | 0.23788507 | 0.42725986 |
| RPL26L1 | 791.55302 | -0.188391199 | 0.14722957 | -1.2795745 | 0.20069484 | 0.38222757 |
| ATP6V0E1 | 12168.2194 | -0.670393348 | 0.17535444 | -3.8230759 | 0.0001318 | 0.00157282 |
| CREBRF | 2295.39595 | -0.404658417 | 0.17837139 | -2.2686285 | 0.02329093 | 0.08384735 |
| AC008378.1 | 44.8882183 | 0.076993369 | 0.25553245 | 0.30130564 | 0.76318144 | 0.86888415 |
| BNIP1 | 560.843966 | -0.401224896 | 0.11238322 | -3.5701494 | 0.00035678 | 0.00348453 |
| NKX2-5 | 0.18331075 | 0.18657164 | 3.29377195 | 0.05664376 | 0.95482897 | NA |
| STC2 | 1134.70857 | -0.950625677 | 0.45942186 | -2.0691781 | 0.03852937 | 0.12045282 |
| AC016573.1 | 0.58669773 | -1.175456794 | 2.3039787 | -0.5101856 | 0.60992143 | NA |
| AC008632.1 | 0.66529716 | -1.365442182 | 1.94000938 | -0.7038328 | 0.48153692 | NA |
| AC008663.1 | 0.49720846 | 1.537423257 | 1.79460195 | 0.85669318 | 0.39161447 | NA |
| AC008663.3 | 5.5422515 | 0.40289612 | 0.52545097 | 0.76676254 | 0.44322271 | 0.63434371 |
| AC008663.2 | 17.2948411 | 0.700587712 | 0.46368331 | 1.51091855 | 0.1308092 | 0.28629404 |
| BOD1 | 1864.94524 | -0.399544141 | 0.14081802 | -2.8373084 | 0.00454956 | 0.02506205 |
| LINC01863 | 1.07299485 | 1.586115092 | 1.23505856 | 1.28424282 | 0.19905701 | 0.38009654 |
| LINC01942 | 16.4603169 | 1.789909119 | 0.6597783 | 2.71289481 | 0.00666983 | 0.03334914 |
| LINC01484 | 35.2050356 | 1.564540607 | 0.48930668 | 3.19746423 | 0.00138642 | 0.01018095 |
| LINC01485 | 108.556373 | 2.601069118 | 0.58561681 | 4.44158893 | 8.93E-06 | 0.00018615 |
| CPEB4 | 3829.35205 | 0.802472331 | 0.23147308 | 3.46680635 | 0.00052668 | 0.00471919 |
| C5orf47 | 9.75382154 | 0.862804325 | 0.54393897 | 1.58621533 | 0.11269047 | 0.25832775 |
| NSG2 | 35.8466328 | -0.120792188 | 0.50105667 | -0.2410749 | 0.80949707 | 0.89703256 |
| AC113423.1 | 0 | NA | NA | NA | NA | NA |
| AC113423.2 | 0.09989236 | -0.642905668 | 3.30558948 | -0.1944905 | 0.84579185 | NA |
| AC025752.1 | 0 | NA | NA | NA | NA | NA |
| LINC01411 | 0 | NA | NA | NA | NA | NA |
| AC108112.1 | 0 | NA | NA | NA | NA | NA |
| MSX2 | 9.12468219 | 0.052368704 | 0.56905942 | 0.09202678 | 0.92667677 | 0.96438573 |
| AC113346.2 | 0 | NA | NA | NA | NA | NA |
| AC113346.1 | 0.09989236 | -0.642905668 | 3.30558948 | -0.1944905 | 0.84579185 | NA |
| LINC01951 | 0.09635527 | 0.547239394 | 3.29377202 | 0.16614368 | 0.86804388 | NA |
| AC008413.2 | 0 | NA | NA | NA | NA | NA |
| DRD1 | 33.0686465 | 0.450978072 | 0.75784861 | 0.59507673 | 0.55179218 | 0.72178454 |
| SFXN1 | 3894.79561 | 0.590344815 | 0.30901122 | 1.91043163 | 0.05607766 | 0.15781942 |
| HRH2 | 377.987535 | -0.424606992 | 0.28362675 | -1.4970626 | 0.13437698 | 0.29159981 |
| CPLX2 | 0.57495301 | -1.340724299 | 1.8978335 | -0.7064499 | 0.47990839 | NA |
| THOC3 | 1044.93996 | 0.002596122 | 0.18485812 | 0.01404386 | 0.98879499 | 0.99494501 |
| AC138965.1 | 0.08666371 | -0.642905668 | 3.30558948 | -0.1944905 | 0.84579185 | NA |
| AC139491.3 | 0.08757577 | 0.547239394 | 3.29377202 | 0.16614368 | 0.86804388 | NA |
| AC139491.1 | 2.66204403 | 0.420749686 | 1.34313353 | 0.31325976 | 0.75408332 | 0.86318765 |
| FAM153B | 9.20806958 | 0.586230892 | 0.70856324 | 0.82735155 | 0.40803783 | 0.60264172 |
| AC139491.7 | 161.554462 | -0.233013177 | 0.46686235 | -0.4991047 | 0.61770565 | 0.77143728 |
| AC139491.4 | 31.9005719 | -0.30838479 | 0.51627686 | -0.5973245 | 0.55029078 | 0.72062214 |
| SIMC1 | 357.863757 | -0.056688015 | 0.14559087 | -0.3893652 | 0.69700602 | 0.8258207 |
| KIAA1191 | 15411.0817 | 0.667531153 | 0.35882303 | 1.86033533 | 0.0628381 | 0.17074092 |
| AC138956.1 | 1706.10161 | 1.092604524 | 0.39293072 | 2.78065439 | 0.00542495 | 0.0286398 |
| AC138956.2 | 20.9924071 | 1.695826543 | 0.51031289 | 3.32311132 | 0.00089019 | 0.00716343 |
| ARL10 | 982.487616 | -0.61201748 | 0.17500491 | -3.4971446 | 0.00047027 | 0.00432842 |
| NOP16 | 3080.03883 | -0.408538212 | 0.1902233 | -2.147677 | 0.03173942 | 0.10505031 |
| HIGD2A | 3326.76285 | -0.220403456 | 0.13828568 | -1.593827 | 0.11097479 | 0.25541422 |
| CLTB | 3396.64105 | -0.323038686 | 0.16723889 | -1.9316003 | 0.05340886 | 0.15262154 |
| FAF2 | 2342.36522 | -0.304233878 | 0.22974822 | -1.3242056 | 0.18543477 | 0.36278024 |
| RNF44 | 5239.2606 | -0.368076236 | 0.14819657 | -2.4837028 | 0.01300243 | 0.05473723 |
| CDHR2 | 3571.49303 | 0.328794961 | 0.57379075 | 0.57302241 | 0.56662952 | 0.73291904 |
| GPRIN1 | 292.888881 | -2.770060778 | 0.4697352 | -5.8970687 | 3.70E-09 | 2.95E-07 |
| SNCB | 0.59686431 | -2.479637097 | 2.74517745 | -0.9032702 | 0.36638249 | NA |
| EIF4E1B | 1.56553238 | 1.694956241 | 1.25935227 | 1.34589525 | 0.1783363 | 0.35377828 |
| TSPAN17 | 3163.31815 | -0.772760191 | 0.1461059 | -5.2890416 | 1.23E-07 | 5.70E-06 |
| AC113391.1 | 0 | NA | NA | NA | NA | NA |
| AC113391.2 | 0 | NA | NA | NA | NA | NA |

| | | | | | | |
|---|---|---|---|---|---|---|
| LINC01574 | 0.07659028 | -0.642905668 | 3.30558948 | -0.1944905 | 0.84579185 | NA |
| UNC5A | 21.3713211 | -1.30346607 | 0.43241523 | -3.0143852 | 0.00257501 | 0.01635427 |
| HK3 | 473.971571 | -1.714693905 | 0.56703067 | -3.0239879 | 0.00249466 | 0.01597481 |
| UIMC1 | 1629.8391 | -0.638878468 | 0.19575337 | -3.2636907 | 0.00109971 | 0.00843741 |
| ZNF346 | 772.345651 | -0.104964099 | 0.17641038 | -0.5949996 | 0.55184376 | 0.72178454 |
| FGFR4 | 5327.58966 | 0.301009943 | 0.27938598 | 1.07739818 | 0.28130242 | 0.47719168 |
| NSD1 | 2279.88954 | 0.023204256 | 0.10416624 | 0.22276176 | 0.82372093 | 0.90566715 |
| RAB24 | 3344.81023 | -0.622925334 | 0.26193046 | -2.3782088 | 0.01739697 | 0.06772468 |
| MXD3 | 8751.30609 | -0.466180549 | 0.22548072 | -2.0674963 | 0.03868741 | 0.12081798 |
| PRELID1 | 6931.00198 | -0.36557269 | 0.27516594 | -1.3285536 | 0.1839953 | 0.36102647 |
| LMAN2 | 8472.36561 | -0.379604296 | 0.12821641 | -2.960653 | 0.00306988 | 0.01869297 |
| RGS14 | 3343.71485 | -0.818418022 | 0.2715387 | -3.0140014 | 0.00257827 | 0.01637103 |
| SLC34A1 | 11217.3324 | 3.376503483 | 0.70814202 | 4.76811625 | 1.86E-06 | 5.36E-05 |
| PFN3 | 16.505482 | 4.43855989 | 0.67794446 | 6.54708479 | 5.87E-11 | 9.26E-09 |
| F12 | 690.505192 | 0.392384804 | 0.42519432 | 0.92283642 | 0.35609246 | 0.55350181 |
| GRK6 | 1419.22987 | -0.480562763 | 0.13835193 | -3.4734806 | 0.00051375 | 0.00463328 |
| PRR7-AS1 | 37.4844055 | -1.32926702 | 0.47374846 | -2.8058498 | 0.00501841 | 0.02695772 |
| PRR7 | 206.673567 | -2.54362742 | 0.32950598 | -7.7195181 | 1.17E-14 | 5.70E-12 |
| DBN1 | 5601.75104 | -0.423579077 | 0.25358855 | -1.6703399 | 0.09485212 | 0.22913618 |
| PDLIM7 | 4582.24127 | -0.945135028 | 0.40719781 | -2.321071 | 0.02028301 | 0.07566575 |
| AC145098.1 | 52.9052425 | -0.674431056 | 0.34284681 | -1.9671499 | 0.04916593 | 0.14404018 |
| DOK3 | 494.511789 | -0.778167297 | 0.30568038 | -2.5456894 | 0.01090622 | 0.04806099 |
| DDX41 | 4231.45473 | -0.473673537 | 0.14812504 | -3.1977953 | 0.00138483 | 0.0101721 |
| FAM193B | 5577.23095 | -0.442237602 | 0.21796921 | -2.0288994 | 0.04246854 | 0.12944167 |
| AC139795.3 | 17.3145938 | -0.093412826 | 0.61091204 | -0.1529072 | 0.87847149 | 0.93677394 |
| TMED9 | 8677.8185 | -0.770340613 | 0.19703094 | -3.9097443 | 9.24E-05 | 0.0011885 |
| B4GALT7 | 1798.8674 | -0.247786678 | 0.15794611 | -1.5688052 | 0.11669333 | 0.26500198 |
| AC139795.2 | 116.751342 | -0.247169028 | 0.27164745 | -0.909889 | 0.36288105 | 0.56012425 |
| AC138819.1 | 34.9108708 | -0.680446175 | 0.61359654 | -1.1089472 | 0.26745295 | 0.46156638 |
| FAM153A | 128.80423 | -0.271535131 | 0.45674695 | -0.594498 | 0.5521791 | 0.72212671 |
| AC140125.2 | 4.85393212 | -1.77210846 | 1.31597291 | -1.3466147 | 0.17810436 | 0.35355405 |
| AC140125.1 | 0 | NA | NA | NA | NA | NA |
| AC140125.3 | 0.08666371 | -0.642905668 | 3.30558948 | -0.1944905 | 0.84579185 | NA |
| AC106795.3 | 4.48010307 | 1.454093872 | 0.57203923 | 2.54194783 | 0.01102366 | 0.04844705 |
| AC106795.2 | 41.7208748 | 0.166448327 | 0.30381267 | 0.54786499 | 0.58378461 | 0.74595509 |
| AC106795.5 | 0.08666371 | -0.642905668 | 3.30558948 | -0.1944905 | 0.84579185 | NA |
| PROP1 | 0 | NA | NA | NA | NA | NA |
| FAM153C | 45.5605571 | -0.961590751 | 0.68070393 | -1.4126417 | 0.1577611 | 0.32488561 |
| N4BP3 | 691.975386 | 0.18278757 | 0.22839144 | 0.80032585 | 0.42352203 | 0.61607877 |
| RMND5B | 3588.09111 | -0.139755327 | 0.11703188 | -1.1941646 | 0.2324136 | 0.4205549 |
| NHP2 | 2475.48306 | -0.318171399 | 0.17620013 | -1.8057387 | 0.07095916 | 0.18616983 |
| HNRNPAB | 6260.92543 | -0.745051327 | 0.18960512 | -3.9294895 | 8.51E-05 | 0.00111681 |
| PHYKPL | 5039.54803 | -0.206496525 | 0.16298268 | -1.2669845 | 0.20516083 | 0.38770898 |
| COL23A1 | 424.488351 | -0.046448909 | 0.42860048 | -0.1083734 | 0.91369947 | 0.95775549 |
| AC136601.1 | 0.34741699 | -0.279112419 | 2.89448939 | -0.0964289 | 0.92317994 | NA |
| CLK4 | 2197.22309 | 0.242053118 | 0.16573459 | 1.46048641 | 0.14415644 | 0.3059685 |
| ZNF354A | 970.704786 | -0.288704234 | 0.21519958 | -1.341565 | 0.17973707 | 0.35556388 |
| ZNF354B | 681.479195 | -0.103264149 | 0.16216321 | -0.6367915 | 0.52426068 | 0.69941521 |
| ZFP2 | 223.503987 | 0.531842327 | 0.25639508 | 2.07430784 | 0.03805073 | 0.11935265 |
| AC104117.5 | 7.08764513 | -0.48495857 | 0.67544055 | -0.7179885 | 0.47276435 | 0.65885276 |
| ZNF454 | 55.7814034 | 0.203803481 | 0.3011423 | 0.67676804 | 0.49855314 | 0.6799953 |
| AC104117.3 | 20.0564634 | 0.074586341 | 0.44177656 | 0.16883273 | 0.86592822 | 0.93035068 |
| GRM6 | 39.4893418 | -0.524478657 | 0.36410729 | -1.4404509 | 0.14973988 | 0.31400722 |
| ZNF879 | 183.535484 | 0.103586459 | 0.18548507 | 0.55846251 | 0.5765286 | 0.74052894 |
| ZNF354C | 242.995757 | 0.1066683 | 0.16336768 | 0.65293393 | 0.51379887 | 0.69166697 |
| ADAMTS2 | 1094.29717 | -1.427699699 | 0.36643766 | -3.8961599 | 9.77E-05 | 0.0012408 |
| AC109479.3 | 10.763821 | -1.912013946 | 0.83888231 | -2.2792398 | 0.02265281 | 0.08221232 |
| AC109479.1 | 18.272231 | -2.150816946 | 0.8136221 | -2.6435085 | 0.00820517 | 0.03879721 |
| AC136628.1 | 0 | NA | NA | NA | NA | NA |
| AC136628.4 | 0.46772854 | 1.304365512 | 1.86566698 | 0.69914166 | 0.4844635 | NA |
| RUFY1 | 3073.04393 | -0.177930051 | 0.15727429 | -1.1313359 | 0.25791374 | 0.45046928 |
| AC136604.2 | 322.064717 | 0.08890764 | 0.18323365 | 0.4852146 | 0.62752413 | 0.77838442 |
| HNRNPH1 | 23528.2378 | -0.144945849 | 0.1580995 | -0.9168014 | 0.35924671 | 0.55689471 |
| C5orf60 | 5.11884174 | 0.69978411 | 0.60959105 | 1.14795666 | 0.25098645 | 0.44330293 |
| AC136604.3 | 11.6894967 | 0.479831356 | 0.48901428 | 0.98122157 | 0.32648349 | 0.52436398 |
| CBY3 | 26.0411058 | 1.554746313 | 0.35756943 | 4.34809624 | 1.37E-05 | 0.00026384 |
| CANX | 28630.2812 | -0.061383905 | 0.15608909 | -0.393262 | 0.69412599 | 0.82386304 |
| MAML1 | 2300.24454 | -0.242787499 | 0.17945086 | -1.352947 | 0.17607257 | 0.35062933 |
| LTC4S | 316.919281 | -0.725195011 | 0.45340619 | -1.5994378 | 0.10972335 | 0.25345172 |
| MGAT4B | 15756.7211 | -1.278018827 | 0.34960062 | -3.6556537 | 0.00025653 | 0.0026706 |
| SQSTM1 | 59609.9272 | -1.992412275 | 0.55587714 | -3.5842674 | 0.00033803 | 0.00333473 |
| MRNIP | 37036.9464 | -1.907251538 | 0.52240884 | -3.6508792 | 0.00026134 | 0.00271325 |
| AC008393.1 | 109.942699 | 0.35210325 | 0.3072855 | 1.14585052 | 0.25185701 | 0.44410074 |
| TBC1D9B | 9046.64961 | 0.19765053 | 0.15360106 | 1.28677844 | 0.19817152 | 0.37911949 |
| RNF130 | 4227.52676 | -0.185140956 | 0.17230487 | -1.0744964 | 0.28260027 | 0.47857314 |
| AC010285.1 | 1.68651157 | 0.115910327 | 0.8729431 | 0.13278108 | 0.89436652 | 0.94571903 |
| RASGEF1C | 15.5590138 | 0.44462673 | 0.54434532 | 0.81681006 | 0.41403698 | 0.60790934 |
| MAPK9 | 2067.53345 | -0.041610767 | 0.10000815 | -0.4160738 | 0.67735602 | 0.81242055 |
| AC008610.1 | 105.528768 | 0.087441509 | 0.36504535 | 0.23953602 | 0.81068997 | 0.89761671 |
| GFPT2 | 925.166206 | -1.79062456 | 0.38950508 | -4.5971789 | 4.28E-06 | 0.00010408 |
| AC034213.1 | 12.1828774 | -5.39715402 | 1.04497378 | -5.1648703 | 2.41E-07 | 9.91E-06 |
| CNOT6 | 1723.25978 | -0.186763973 | 0.14867008 | -1.256231 | 0.20903222 | 0.39223514 |
| SCGB3A1 | 14.3094265 | -1.033033963 | 0.59737622 | -1.7292854 | 0.08375803 | 0.20921657 |
| FLT4 | 3464.24534 | -0.270323797 | 0.31702263 | -0.8526956 | 0.39382814 | 0.58913366 |
| LINC02222 | 0.17453125 | 0.18657164 | 3.29377195 | 0.05664376 | 0.95482897 | NA |
| OR2Y1 | 0 | NA | NA | NA | NA | NA |
| MGAT1 | 8560.77793 | -0.716606554 | 0.18942237 | -3.7831146 | 0.00015488 | 0.0017968 |
| LINC00847 | 1514.04513 | -0.150365867 | 0.17860407 | -0.841895 | 0.39984675 | 0.59466239 |
| ZFP62 | 887.12985 | 0.072929392 | 0.15631682 | 0.4665486 | 0.64082287 | 0.78740502 |
| BTNL8 | 19.004466 | -0.08559276 | 0.63201212 | -0.135429 | 0.89227271 | 0.94422855 |
| BTNL3 | 5.54887338 | -2.606591183 | 1.39584874 | -1.867388 | 0.06184743 | 0.16895885 |
| BTNL9 | 1781.15922 | 0.05491045 | 0.46590047 | 0.11785876 | 0.90617957 | 0.95301728 |
| OR2V1 | 0.42659734 | 1.031665254 | 1.85056022 | 0.55748807 | 0.57719401 | NA |
| OR2V2 | 0 | NA | NA | NA | NA | NA |
| LINC01962 | 1.94306284 | -0.751539611 | 1.18743964 | -0.6329076 | 0.52679397 | 0.70159378 |
| AC008443.5 | 84.3214362 | -0.022950451 | 0.35379597 | -0.0648692 | 0.94827817 | 0.97510358 |
| TRIM7 | 239.10035 | -0.538174313 | 0.3546758 | -1.5173697 | 0.12917337 | 0.28378753 |
| AC008443.6 | 23.5174953 | -0.621014654 | 0.46783404 | -1.3274251 | 0.1843681 | 0.36149801 |
| AC008443.4 | 4.57674643 | -0.498251314 | 0.85339446 | -0.5838464 | 0.55932364 | 0.72772184 |
| TRIM41 | 4052.1807 | 0.083264019 | 0.11082913 | 0.75128278 | 0.45248249 | 0.64215937 |
| AC008443.2 | 515.845461 | 0.432641977 | 0.17987478 | 2.40523974 | 0.01616186 | 0.0643303 |
| RACK1 | 38324.0979 | -0.357103913 | 0.19068729 | -1.87272 | 0.06110705 | 0.16752133 |
| TRIM52 | 1961.88622 | 0.109572272 | 0.15855489 | 0.69106841 | 0.48952255 | 0.6728543 |
| AC008443.3 | 1.72198334 | 1.185196281 | 1.15555845 | 1.02564806 | 0.3050575 | 0.50211388 |
| AC138035.1 | 66.2176716 | 1.031331143 | 0.39320147 | 2.62290767 | 0.00871829 | 0.0406152 |
| AC138035.2 | 0.65131268 | -0.060718314 | 1.49170605 | -0.0407039 | 0.96753192 | NA |
| OR4F3 | 0 | NA | NA | NA | NA | NA |
| AL035696.3 | 0.3614598 | 0.735935719 | 2.87056228 | 0.25637337 | 0.79766255 | NA |
| AL035696.1 | 0 | NA | NA | NA | NA | NA |
| DUSP22 | 2306.96743 | -0.6925374 | 0.15079131 | -4.5926876 | 4.38E-06 | 0.00010596 |
| IRF4 | 105.927619 | -0.334026553 | 0.61673949 | -0.5416007 | 0.58809359 | 0.74892684 |
| EXOC2 | 1325.04631 | -0.110365583 | 0.12759514 | -0.864967 | 0.38705691 | 0.58268347 |
| AL031770.1 | 3.5334641 | -1.066984609 | 0.76109479 | -1.4019076 | 0.16094283 | 0.32946417 |

| | | | | | | | |
|---|---|---|---|---|---|---|---|
| HUS1B | 20.3287068 | | -1.006948658 | 0.61682048 | -1.6324825 | 0.10257788 | 0.24166699 |
| AL357054.4 | 34.1787877 | | -0.707157973 | 0.66766767 | -1.0591467 | 0.289533 | 0.48574788 |
| AL357054.2 | 0.92485079 | | -0.028189629 | 1.17630822 | -0.0239645 | 0.98088093 | NA |
| AL357054.3 | 1.75572582 | | -1.740342173 | 1.05276999 | -1.6531077 | 0.09830895 | 0.23485592 |
| AL392183.1 | 1.15441064 | | -0.835580104 | 1.28961762 | -0.6479286 | 0.51703115 | 0.69437797 |
| AL356130.1 | 2.88896008 | | 1.33664074 | 0.95645394 | 1.39749619 | 0.16226441 | 0.33124857 |
| AL356130.2 | 0.34823675 | | 1.594838211 | 2.1504546 | 0.7416284 | 0.45831251 | NA |
| AL033381.2 | 33.4857168 | | 0.23477999 | 0.83689425 | 0.28053722 | 0.77906537 | 0.87896093 |
| AL033381.1 | 23.8979523 | | 2.08365832 | 0.72439143 | 2.87642597 | 0.00402207 | 0.02284831 |
| FOXQ1 | 1060.48324 | | -0.971889725 | 0.36188348 | -2.6856427 | 0.00723905 | 0.03552654 |
| LINC01394 | 0 | NA | | NA | NA | NA | NA |
| AL034346.1 | 0 | NA | | NA | NA | NA | NA |
| FOXF2 | 11.0367846 | | 1.044487739 | 0.67536123 | 1.5465616 | 0.121969 | 0.27284511 |
| AL512329.2 | 12.5876768 | | -0.504240121 | 0.73550617 | -0.6855689 | 0.49298503 | 0.67578034 |
| FOXCUT | 78.9385628 | | -0.951878096 | 0.52229825 | -1.8224799 | 0.06838219 | 0.18112078 |
| FOXC1 | 2104.08896 | | 0.37985841 | 0.26341091 | 1.44207544 | 0.14928109 | 0.3134045 |
| GMDS | 1585.03243 | | -0.477190563 | 0.22919682 | -2.0820122 | 0.03734136 | 0.11770161 |
| GMDS-DT | 348.572597 | | 0.037461516 | 0.24334128 | 0.15394641 | 0.87765199 | 0.93634677 |
| AL031768.1 | 11.9357609 | | -0.513120854 | 0.57362575 | -0.894522 | 0.37104266 | 0.56783674 |
| LINC01600 | 11.9331134 | | 0.292451596 | 0.38221748 | 0.76514448 | 0.44418551 | 0.63510611 |
| LINC02521 | 0.29111801 | | -0.179913104 | 2.91136792 | -0.0617968 | 0.95072468 | NA |
| MYLK4 | 420.896622 | | 0.681769568 | 0.65209237 | 1.04551072 | 0.29578699 | 0.49266098 |
| WRNIP1 | 3294.72028 | | -0.139211984 | 0.0996264 | -1.3973403 | 0.16231127 | 0.33129854 |
| SERPINB1 | 3659.99955 | | -0.855259989 | 0.16824506 | -5.0834181 | 3.71E-07 | 1.42E-05 |
| SERPINB9P1 | 24.0247244 | | 1.3971731 | 0.60174566 | 2.32186653 | 0.02024012 | 0.07553525 |
| SERPINB9 | 910.597544 | | -0.258398118 | 0.31868113 | -0.8108359 | 0.41745989 | 0.61029426 |
| AL133351.2 | 0 | NA | | NA | NA | NA | NA |
| SERPINB6 | 6915.38669 | | -0.337713006 | 0.18462374 | -1.829196 | 0.06737026 | 0.1792356 |
| LINC01011 | 320.201226 | | -0.012578097 | 0.40700532 | -0.030904 | 0.97534609 | 0.98862634 |
| NQO2 | 7587.96533 | | 0.930905328 | 0.48106168 | 1.93510598 | 0.05297729 | 0.1516677 |
| AL133351.1 | 139.901059 | | 0.125891195 | 0.67369189 | 0.18686761 | 0.85176443 | 0.92231052 |
| AL031963.2 | 2.76683024 | | -0.938689454 | 1.21861754 | -0.7702905 | 0.44112762 | 0.63239724 |
| RIPK1 | 2459.95173 | | -0.222680349 | 0.23082319 | -0.9647226 | 0.33468377 | 0.53281883 |
| AL031963.3 | 41.0431979 | | 0.178794358 | 0.36526386 | 0.4894937 | 0.62449221 | 0.7763098 |
| BPHL | 3885.10732 | | 1.966365713 | 0.34828775 | 5.64580779 | 1.64E-08 | 1.06E-06 |
| AL031963.1 | 1725.86388 | | 1.872212617 | 0.34573736 | 5.41512954 | 6.12E-08 | 3.14E-06 |
| TUBB2A | 1234.50267 | | -0.448267206 | 0.33839503 | -1.3246861 | 0.18527528 | 0.36268231 |
| LINC02525 | 2.01697835 | | 0.170189825 | 0.81359002 | 0.20918377 | 0.83430478 | 0.91213172 |
| TUBB2B | 687.314953 | | 0.575037646 | 0.39912713 | 1.44073805 | 0.1496587 | 0.31392066 |
| PSMG4 | 1685.73687 | | 0.032754631 | 0.19473313 | 0.16820266 | 0.86642385 | 0.93065568 |
| SLC22A23 | 1777.04144 | | 0.184404047 | 0.30695568 | 0.60075137 | 0.5480056 | 0.71876822 |
| AL445309.1 | 32.9949448 | | 0.231474036 | 0.50519972 | 0.45818322 | 0.64682081 | 0.7919677 |
| AL138881.1 | 4.86167053 | | 1.357266924 | 0.95352258 | 1.423424 | 0.15461328 | 0.32065297 |
| PXDC1 | 2947.00489 | | -0.734768701 | 0.27965146 | -2.6274446 | 0.00860289 | 0.04019456 |
| AL391422.3 | 651.578158 | | -0.809780734 | 0.24906825 | -3.2512403 | 0.00114903 | 0.00875212 |
| FAM50B | 853.089607 | | 0.668161581 | 0.17765419 | 3.76102358 | 0.00016922 | 0.0019205 |
| AL590004.3 | 0.82820327 | | -0.483008388 | 1.40921032 | -0.3427511 | 0.73178572 | NA |
| AL590004.4 | 7.49004216 | | 0.130437435 | 0.6454119 | 0.20209952 | 0.83983892 | 0.91484357 |
| AL138831.2 | 4.32138098 | | -0.594154909 | 0.94129386 | -0.6312109 | 0.52790266 | 0.70250268 |
| AL138831.1 | 4.92027892 | | 0.466239984 | 0.65161258 | 0.71551716 | 0.47428954 | 0.66003563 |
| PRPF4B | 3668.1392 | | -0.06080401 | 0.16915205 | -0.3594636 | 0.71924828 | 0.84021138 |
| FAM217A | 29.9584568 | | 1.307275066 | 0.34820856 | 3.75428758 | 0.00017384 | 0.00196234 |
| C6orf201 | 4110.8486 | | 0.968590601 | 0.30798611 | 3.1449165 | 0.00166134 | 0.01169191 |
| ECI2 | 7445.78903 | | 0.798976715 | 0.27697644 | 2.88463777 | 0.00391864 | 0.02242018 |
| AL136309.4 | 619.520119 | | 1.156158831 | 0.35022299 | 3.30120774 | 0.0009627 | 0.0076258 |
| AL136309.2 | 21.3583395 | | 0.231551104 | 0.67684775 | 0.3421022 | 0.73227398 | 0.8490202 |
| AL136309.3 | 0.95794476 | | -0.078877499 | 1.257613 | -0.06272 | 0.94998946 | NA |
| AL162718.1 | 0.08695548 | | -0.642905668 | 3.30558948 | -0.1944905 | 0.84579185 | NA |
| AL159166.1 | 0.25283891 | | -0.178404819 | 2.99766869 | -0.0595145 | 0.9525423 | NA |
| LINC02533 | 0.3034605 | | -0.180487739 | 2.88038455 | -0.062661 | 0.95003646 | NA |
| AL034376.2 | 0 | NA | | NA | NA | NA | NA |
| AL034376.1 | 0.86827641 | | -2.264120928 | 1.81789999 | -1.2454596 | 0.21296287 | NA |
| CDYL | 1028.11938 | | -0.6100128 | 0.34093193 | -1.789251 | 0.0735744 | 0.19114847 |
| AL356747.1 | 0.18746814 | | 0.18657164 | 3.29377195 | 0.05664376 | 0.95482897 | NA |
| AL022725.1 | 0.10721287 | | -0.642905668 | 3.30558948 | -0.1944905 | 0.84579185 | NA |
| RPP40 | 191.427482 | | -0.274071268 | 0.25475867 | -1.0758074 | 0.28201338 | 0.47782049 |
| AL359643.3 | 46.4392493 | | 0.523226348 | 0.30821276 | 1.69761418 | 0.0895806 | 0.21988649 |
| AL359643.2 | 39.0799877 | | -0.258706338 | 0.26173466 | -0.9884298 | 0.32294219 | 0.52066432 |
| PPP1R3G | 224.301533 | | -1.584737587 | 0.47430105 | -3.3412062 | 0.00083415 | 0.00679957 |
| LYRM4 | 844.683769 | | -0.430239017 | 0.17976066 | -2.3933992 | 0.01669307 | 0.06581586 |
| FARS2 | 798.945167 | | 0.546958693 | 0.16147681 | 3.38722752 | 0.00070603 | 0.00595203 |
| AL021328.1 | 41.8225035 | | -1.368570239 | 0.60374901 | -2.2667867 | 0.02340326 | 0.08412539 |
| AL022097.1 | 0.66247599 | | 1.598354223 | 1.66100562 | 0.96228105 | 0.33590845 | NA |
| AL136307.1 | 0.93645037 | | 0.301733716 | 1.49136948 | 0.20231989 | 0.83966665 | NA |
| NRN1 | 409.30143 | | -0.226477111 | 0.47118143 | -0.480658 | 0.6307596 | 0.78054869 |
| F13A1 | 3922.45991 | | -0.142199832 | 0.63993538 | -0.2222097 | 0.82415066 | 0.90587524 |
| LY86-AS1 | 21.874351 | | 0.233251715 | 0.49566055 | 0.47058761 | 0.63793526 | 0.78565828 |
| LY86 | 134.922688 | | -0.502244665 | 0.58881482 | -0.8529756 | 0.39367284 | 0.58896817 |
| AL031123.2 | 112.985622 | | 0.875285437 | 0.41010717 | 2.13428464 | 0.03281949 | 0.10755971 |
| AL031123.1 | 128.081108 | | 0.349833669 | 0.50297085 | 0.69553468 | 0.48672028 | 0.67075066 |
| AL136361.1 | 1.75090267 | | 3.394484396 | 1.36533089 | 2.48619908 | 0.01291157 | 0.05444173 |
| AL158817.1 | 9.19793228 | | 2.890771932 | 0.97580229 | 2.96245659 | 0.00305195 | 0.01860376 |
| AL139390.1 | 0 | NA | | NA | NA | NA | NA |
| RREB1 | 2368.61485 | | -0.009473309 | 0.13358169 | -0.0709177 | 0.94346324 | 0.97315146 |
| SSR1 | 6646.2523 | | -0.043480594 | 0.1620131 | -0.268377 | 0.78840912 | 0.88492161 |
| AL139095.4 | 421.521693 | | 0.041308392 | 0.1974629 | 0.20919571 | 0.83429546 | 0.91213172 |
| CAGE1 | 3.95633005 | | 1.204899224 | 0.69295427 | 1.73878605 | 0.0820724 | 0.20609935 |
| RIOK1 | 648.2833 | | -0.453755982 | 0.21200685 | -2.1402893 | 0.0323314 | 0.1063066 |
| AL031058.1 | 75.3220363 | | -0.255764914 | 0.22856306 | -1.1190125 | 0.26313481 | 0.45686496 |
| DSP | 5018.69465 | | -0.019724404 | 0.18247095 | -0.1080961 | 0.91391944 | 0.9577987 |
| SNRNP48 | 771.1811 | | -0.043638973 | 0.13440289 | -0.3246878 | 0.74541739 | 0.85738944 |
| BMP6 | 889.042633 | | -0.131767838 | 0.50846785 | -0.2591468 | 0.79552194 | 0.88912069 |
| TXNDC5 | 9857.6523 | | -0.436607454 | 0.23929373 | -1.824567 | 0.06806639 | 0.18046559 |
| BLOC1S5-TXI | 10058.4565 | | -0.41208277 | 0.23484011 | -1.7547376 | 0.07930421 | 0.20115808 |
| BLOC1S5 | 1496.4515 | | 0.216816037 | 0.17625651 | 1.23011649 | 0.21865348 | 0.4039154 |
| EEF1E1-BLO( | 1424.73859 | | -0.291441475 | 0.19742215 | -1.4762349 | 0.13988084 | 0.29981775 |
| EEF1E1 | 1685.09634 | | -0.553780117 | 0.22092184 | -2.5066789 | 0.01218714 | 0.05213508 |
| AL359378.1 | 1.25141628 | | 0.561361817 | 1.07304658 | 0.52314767 | 0.60087149 | 0.7585323 |
| SLC35B3 | 1373.61065 | | 0.149493033 | 0.13088237 | 1.14219382 | 0.25337346 | 0.44593594 |
| AL137220.1 | 0.30341448 | | -0.699696944 | 2.92503995 | -0.2392094 | 0.81094324 | NA |
| TFAP2A | 938.27297 | | 0.370127307 | 0.29067726 | 1.27332736 | 0.2029019 | 0.38495415 |
| AL138885.3 | 260.095822 | | 0.557648881 | 0.3012128 | 1.85134523 | 0.0641199 | 0.17315329 |
| TFAP2A-AS1 | 87.5596313 | | -0.288841325 | 0.42120343 | -0.6857525 | 0.49286917 | 0.67565814 |
| AL138885.2 | 0.13028434 | | 1.049816488 | 3.28477129 | 0.31960109 | 0.74927075 | NA |
| LINC00518 | 0.54303581 | | -0.484034261 | 2.29042112 | -0.2113298 | 0.83262992 | NA |
| MIR5689HG | 0.65429116 | | 1.494060601 | 1.72515767 | 0.86604293 | 0.38646662 | NA |
| LINC02522 | 0.36739698 | | -0.032132436 | 2.27708452 | -0.0141112 | 0.98874125 | NA |
| GCNT2 | 1395.64796 | | 0.160404738 | 0.1806551 | 0.88790595 | 0.37459135 | 0.57064323 |
| AL139039.3 | 4.67707029 | | 1.218844308 | 0.64197028 | 1.89859927 | 0.05761718 | 0.16088429 |
| C6orf52 | 51.0512122 | | 0.567371286 | 0.32263128 | 1.75857495 | 0.07864973 | 0.20009584 |
| PAK1IP1 | 660.207162 | | -0.497279403 | 0.26191238 | -1.8986479 | 0.05761078 | 0.16088429 |
| TMEM14C | 4633.99992 | | -0.039301452 | 0.14753742 | -0.2663829 | 0.7899443 | 0.88585291 |
| AL024498.1 | 27.0911506 | | 0.120432753 | 0.3034068 | 0.39693492 | 0.69141545 | 0.82241065 |

| | | | | | | |
|---|---|---|---|---|---|---|
| TMEM14B | 2647.45469 | 0.071962075 | 0.1558515 | 0.4617349 | 0.64427144 | 0.78980233 |
| AL024498.2 | 966.268591 | 0.298303271 | 0.19317969 | 1.5441751 | 0.12254592 | 0.27373127 |
| MAK | 146.721939 | -0.515505826 | 0.27761458 | -1.8569119 | 0.06332369 | 0.17172415 |
| GCM2 | 0.87273312 | -0.659019303 | 1.6001923 | -0.4118376 | 0.68045848 | NA |
| AL357497.1 | 2.96949079 | -1.890521087 | 1.02136552 | -1.8509741 | 0.06417328 | 0.17324422 |
| SYCP2L | 68.7999823 | 0.053360683 | 0.28957015 | 0.18427549 | 0.85379734 | 0.92346684 |
| ELOVL2 | 30.4231612 | -0.296508418 | 0.55970342 | -0.5297599 | 0.59627842 | 0.75505948 |
| ELOVL2-AS1 | 0.18655607 | -1.003574173 | 3.30558948 | -0.3035992 | 0.7614333 | NA |
| SMIM13 | 583.488045 | -0.319999684 | 0.18299246 | -1.7487042 | 0.08034216 | 0.20300146 |
| ERVFRD-1 | 38.1936382 | 0.232007865 | 0.92190897 | 0.25166028 | 0.80130366 | 0.89233014 |
| AL139807.1 | 420.142383 | -1.052210259 | 0.37877835 | -2.777905 | 0.00547106 | 0.02882555 |
| NEDD9 | 5130.22243 | -0.838127541 | 0.3410487 | -2.4575011 | 0.01399074 | 0.0578632 |
| AL445430.1 | 0.45419896 | -0.711512978 | 2.86677272 | -0.248193 | 0.80398507 | NA |
| AL445430.2 | 0.5702106 | -2.432077345 | 2.14983664 | -1.1312847 | 0.25793527 | NA |
| TMEM170B | 604.29002 | 0.138054083 | 0.17988217 | 0.7674695 | 0.44280242 | 0.63398305 |
| AL357518.1 | 4.09476484 | -1.194717886 | 0.77888015 | -1.5338918 | 0.12505627 | 0.27744018 |
| ADTRP | 257.660815 | 0.686817565 | 0.53847423 | 1.27548828 | 0.20213647 | 0.38397484 |
| AL022724.1 | 0.26772358 | 1.012697542 | 2.89776332 | 0.34947559 | 0.72673229 | NA |
| AL157373.1 | 0 | NA | NA | NA | NA | NA |
| AL157373.2 | 183.325853 | -0.339499648 | 0.30297108 | -1.1205678 | 0.26247186 | 0.45610431 |
| HIVEP1 | 847.251478 | -0.450479619 | 0.18278883 | -2.4644811 | 0.01372118 | 0.05694484 |
| EDN1 | 681.621643 | -0.851656793 | 0.32397058 | -2.628809 | 0.00856845 | 0.04007808 |
| LINC02530 | 0.03392908 | 0.380740593 | 3.29738709 | 0.11546736 | 0.90807469 | NA |
| PHACTR1 | 951.02224 | 0.026239709 | 0.22251731 | 0.1179221 | 0.90612938 | 0.9530025 |
| AL008729.1 | 205.088075 | 0.156373248 | 0.29369656 | 0.53243133 | 0.59442729 | 0.75353561 |
| TBC1D7 | 846.254605 | -0.023526345 | 0.1811849 | -0.1298472 | 0.89668735 | 0.94711389 |
| AL008729.2 | 260.746475 | -0.580887175 | 0.32365747 | -1.7947591 | 0.0726921 | 0.18964217 |
| GFOD1 | 508.399764 | -1.360101877 | 0.38573796 | -3.5259736 | 0.00042193 | 0.00397367 |
| GFOD1-AS1 | 13.1095831 | -1.262858025 | 0.48017749 | -2.6299817 | 0.00853895 | 0.03996839 |
| SIRT5 | 1333.40919 | 0.348969571 | 0.28343406 | 1.23121961 | 0.21824074 | 0.40332906 |
| AL441883.1 | 103.928244 | -0.139364495 | 0.21144694 | -0.6590991 | 0.50983212 | 0.68853185 |
| NOL7 | 2942.95844 | -0.233363274 | 0.16245749 | -1.4364574 | 0.15087222 | 0.31565379 |
| RANBP9 | 2361.91599 | -0.064409789 | 0.09103265 | -0.707546 | 0.47922723 | 0.66399864 |
| MCUR1 | 2391.14078 | 0.652325721 | 0.25711611 | 2.53708618 | 0.01117794 | 0.04895447 |
| AL023583.1 | 25.4808036 | 1.715212088 | 0.40181965 | 4.26861172 | 1.97E-05 | 0.00034928 |
| RNF182 | 111.959718 | -0.191869359 | 0.27389553 | -0.7005202 | 0.48360248 | 0.66823038 |
| AL022396.1 | 0 | NA | NA | NA | NA | NA |
| CD83 | 976.750199 | 1.002092284 | 0.25176141 | 3.98032521 | 6.88E-05 | 0.00094521 |
| AL353152.1 | 29.1016991 | 1.426178361 | 0.68650873 | 2.07743659 | 0.03776128 | 0.11869889 |
| LINC01108 | 10.4849965 | 6.190843859 | 0.95231762 | 6.50081834 | 7.99E-11 | 1.19E-08 |
| AL080313.2 | 0.47923968 | 1.371760163 | 2.2054327 | 0.6219914 | 0.53394753 | NA |
| AL080313.1 | 0 | NA | NA | NA | NA | NA |
| AL109914.1 | 0.25999113 | -1.364242679 | 3.30558948 | -0.4127078 | 0.67982068 | NA |
| AL138720.1 | 1.05706499 | 0.277067691 | 1.25594715 | 0.22060458 | 0.82540034 | 0.90675212 |
| AL136162.1 | 16.1041448 | 0.359613439 | 0.49642704 | 0.72440341 | 0.46881811 | 0.65560447 |
| JARID2 | 611.71714 | 0.123303164 | 0.17978134 | 0.68585072 | 0.49280725 | 0.67562436 |
| JARID2-AS1 | 5.29627962 | 0.437957575 | 0.61636087 | 0.71055383 | 0.47736076 | 0.66249162 |
| DTNBP1 | 1539.1914 | -0.439472445 | 0.19919325 | -2.2062618 | 0.02736568 | 0.09447317 |
| LINC02543 | 0 | NA | NA | NA | NA | NA |
| MYLIP | 1829.96105 | 0.39533245 | 0.16773065 | 2.35694816 | 0.01842582 | 0.07070559 |
| GMPR | 2352.18173 | -1.150010847 | 0.51350376 | -2.2395374 | 0.02512097 | 0.08870381 |
| AL009031.1 | 6.97256208 | 0.192007713 | 0.66038216 | 0.29075242 | 0.77124068 | 0.87383418 |
| ATXN1 | 1549.47035 | 0.039801035 | 0.17840051 | 0.22309934 | 0.82345819 | 0.90564302 |
| AL137003.1 | 39.6495956 | 0.667493874 | 0.32074992 | 2.08104143 | 0.03743011 | 0.11788273 |
| AL137003.2 | 199.749907 | 0.82959719 | 0.3359696 | 2.46926269 | 0.01353918 | 0.05642168 |
| STMND1 | 1.08285134 | 2.543997595 | 2.00456953 | 1.2690992 | 0.20440568 | 0.3866976 |
| RBM24 | 102.720368 | -0.838296612 | 0.44762094 | -1.8727824 | 0.06109844 | 0.16751569 |
| CAP2 | 678.852301 | 0.526153035 | 0.26163207 | 2.01104184 | 0.04432104 | 0.1333606 |
| FAM8A1 | 2603.97952 | 0.207846928 | 0.179315 | 1.15911623 | 0.24640881 | 0.4377238 |
| NUP153 | 1946.34131 | -0.147293958 | 0.20551245 | -0.7167155 | 0.47354966 | 0.65952807 |
| AL138724.1 | 76.283194 | -1.390316667 | 0.46419141 | -2.9951366 | 0.00274322 | 0.01717036 |
| KIF13A | 1430.04785 | 0.363068065 | 0.12372063 | 2.93457988 | 0.00334 | 0.01989961 |
| NHLRC1 | 64.3329752 | 0.849200135 | 0.43288704 | 1.96171302 | 0.0497959 | 0.14524035 |
| TPMT | 4764.14191 | 1.000594887 | 0.26374658 | 3.79377395 | 0.00014837 | 0.00173895 |
| KDM1B | 979.833936 | 0.159920659 | 0.19658822 | 0.81348037 | 0.4159427 | 0.60920587 |
| DEK | 3879.90848 | -0.428336698 | 0.19956544 | -2.1463471 | 0.0318453 | 0.10524634 |
| RNF144B | 2894.52002 | -1.040280455 | 0.36650219 | -2.8384017 | 0.00453401 | 0.02499201 |
| MIR548A1HG | 6.30464942 | -0.890614246 | 0.66579 | -1.3376804 | 0.18100064 | 0.35717929 |
| AL357052.1 | 0.56664154 | -0.6869828 | 1.86620281 | -0.368118 | 0.71278527 | NA |
| AL022068.1 | 84.0433054 | 0.186542156 | 0.38604048 | 0.48321916 | 0.62894013 | 0.77952394 |
| ID4 | 1884.7804 | 0.201516157 | 0.24848433 | 0.81098136 | 0.41737638 | 0.61027293 |
| MBOAT1 | 222.307773 | 0.422648242 | 0.28719299 | 1.47165238 | 0.14111477 | 0.30148234 |
| AL158198.1 | 10.0055076 | 0.879347122 | 0.60874174 | 1.44453233 | 0.14858929 | 0.31253885 |
| E2F3 | 1125.55512 | -0.424316768 | 0.19117892 | -2.2194747 | 0.02645445 | 0.09207211 |
| CDKAL1 | 624.225593 | -0.116069029 | 0.14116636 | -0.8222145 | 0.41095481 | 0.60530474 |
| AL513188.1 | 0.49758283 | -1.407704121 | 1.90758217 | -0.737952 | 0.4605436 | NA |
| AL031767.1 | 0 | NA | NA | NA | NA | NA |
| LINC00581 | 0.30865737 | 0.328537818 | 2.28709442 | 0.14364856 | 0.88577799 | NA |
| AL512380.1 | 2.16997312 | 0.180387808 | 1.08676279 | 0.16598637 | 0.86816768 | 0.93146642 |
| AL512380.2 | 13.4515584 | -0.028981927 | 0.60238285 | -0.0481121 | 0.96162687 | 0.98169256 |
| SOX4 | 2323.33075 | 0.012131108 | 0.34269945 | 0.03539868 | 0.97176184 | 0.98670705 |
| CASC15 | 300.349435 | 0.880146034 | 0.46990374 | 1.87303474 | 0.06106359 | 0.16745495 |
| NBAT1 | 28.77592 | 1.949147402 | 0.68965138 | 2.82627926 | 0.00470922 | 0.02569994 |
| AL359694.2 | 11.938539 | -0.729031963 | 0.55341171 | -1.317341 | 0.18772433 | 0.36579404 |
| PRL | 2.85882878 | 0.932503117 | 1.31876241 | 0.70710471 | NA | NA |
| HDGFL1 | 0.38499583 | -0.570714416 | 2.3308409 | -0.2448534 | 0.80656991 | NA |
| AL033539.1 | 0.18552053 | -1.003574173 | 3.30558948 | -0.3035992 | 0.7614333 | NA |
| AL035401.1 | 0 | NA | NA | NA | NA | NA |
| AL139231.1 | 0.60150134 | -1.694693685 | 1.88320551 | -0.8998984 | 0.3681743 | NA |
| NRSN1 | 11.8440887 | -1.414824329 | 0.66069314 | -2.1414243 | 0.03223984 | 0.10621746 |
| DCDC2 | 4445.62337 | -0.928349764 | 0.22006351 | -4.2185538 | 2.46E-05 | 0.00041939 |
| KAAG1 | 587.350155 | -0.908272402 | 0.24644641 | -3.6854763 | 0.00022828 | 0.00243021 |
| MRS2 | 2102.07884 | 0.18857121 | 0.12631488 | 1.49286614 | 0.13547223 | 0.29308415 |
| GPLD1 | 389.805123 | 0.471841554 | 0.12628434 | 3.73634251 | 0.00018672 | 0.00206709 |
| ALDH5A1 | 1876.09197 | 1.63865987 | 0.41452873 | 3.9530671 | 7.72E-05 | 0.00103332 |
| KIAA0319 | 43.878173 | -2.453073981 | 0.71387863 | -3.436262 | 0.0005898 | 0.00515342 |
| TDP2 | 5766.61932 | -0.355844465 | 0.54160938 | -0.6570131 | 0.51117248 | 0.6893535 |
| ACOT13 | 2859.81591 | 0.702875839 | 0.2592709 | 2.71097083 | 0.00670865 | 0.03348612 |
| AL031775.1 | 808.942008 | 0.964655741 | 0.35367126 | 2.72754913 | 0.00638068 | 0.03224865 |
| C6orf62 | 7354.41356 | -0.279238692 | 0.16817796 | -1.6603763 | 0.09683878 | 0.23227035 |
| AL031775.2 | 53.5033314 | 0.698516199 | 0.21742728 | 3.21264294 | 0.0013152 | 0.00976675 |
| AL133264.2 | 1.36089834 | -1.430462855 | 1.33361536 | -1.0726203 | 0.28344153 | 0.47930955 |
| GMNN | 663.624842 | -0.161439169 | 0.15671303 | -1.0301579 | 0.30293587 | 0.49979568 |
| ARMH2 | 1.46976599 | 0.796676198 | 1.15742633 | 0.68831698 | 0.49125319 | 0.6742895 |
| AL512428.1 | 26.4684892 | 0.961843943 | 0.38506391 | 2.49788133 | 0.0124938 | 0.05314753 |
| RIPOR2 | 527.27281 | -0.002210873 | 0.41686712 | -0.0053035 | 0.9957684 | 0.99795473 |
| AL133268.3 | 7.19945086 | 0.323888415 | 0.78240449 | 0.41396544 | 0.67889942 | 0.81371654 |
| AL133268.2 | 0.09997307 | 0.547239394 | 3.29377202 | 0.16614368 | 0.86804388 | NA |
| CARMIL1 | 1695.81624 | 0.329620092 | 0.149968 | 2.19793619 | 0.02795365 | 0.09605616 |
| SCGN | 72.4904276 | 0.400669714 | 0.87277375 | 0.45907627 | 0.6461794 | 0.79136942 |
| HIST1H2AA | 5.62356165 | 2.697055621 | 0.94888455 | 2.84234326 | 0.00447832 | 0.02479391 |
| HIST1H2BA | 5.30308986 | 2.579664806 | 0.62978297 | 4.09611712 | 4.20E-05 | 0.00064638 |
| SLC17A4 | 411.807776 | 2.225773032 | 0.60372225 | 3.68675006 | 0.00022714 | 0.00242199 |

| | | | | | | |
|---|---|---|---|---|---|---|
| SLC17A1 | 6807.22036 | 0.796035965 | 0.55882726 | 1.42447591 | 0.15430875 | 0.32022679 |
| SLC17A3 | 9913.66071 | 0.242544337 | 0.74886206 | 0.32388386 | 0.74602596 | 0.8576469 |
| SLC17A2 | 81.4103334 | -0.872582658 | 0.4781942 | -1.8247454 | 0.06803946 | 0.18043047 |
| TRIM38 | 4930.87025 | -1.002206408 | 0.27598093 | -3.6314336 | 0.00028185 | 0.00289128 |
| U91328.2 | 467.254735 | -0.997280455 | 0.25593903 | -3.8965548 | 9.76E-05 | 0.00123937 |
| U91328.1 | 354.269306 | 0.379736485 | 0.17241259 | 2.20248702 | 0.02763092 | 0.09514592 |
| U91328.3 | 27.1485979 | -0.506552645 | 0.41098849 | -1.2325227 | 0.2177539 | 0.40273322 |
| HIST1H1A | 0.13382144 | 0.02007176 | 3.29738704 | 0.00608717 | 0.99514317 | NA |
| HIST1H3A | 5.41858952 | -0.837499537 | 0.72591164 | -1.153721 | 0.24861456 | 0.44038193 |
| HIST1H4A | 0.68197346 | -1.013542965 | 1.68300289 | -0.6022229 | 0.54702574 | NA |
| HIST1H4B | 2.88193099 | 0.499054907 | 0.77846111 | 0.64107879 | 0.52147149 | 0.69776958 |
| HIST1H3B | 2.43008184 | -3.673391348 | 1.40423585 | -2.6159362 | 0.00889832 | 0.04128231 |
| HIST1H2AB | 0.85878041 | -1.799308054 | 2.33455041 | -0.77073 | 0.44086698 | NA |
| HIST1H2BB | 0.36072882 | -1.025911653 | 2.30367303 | -0.4453374 | 0.65607598 | NA |
| HIST1H3C | 4.47489611 | -0.246308742 | 0.74459991 | -0.3307934 | 0.74080054 | 0.85453728 |
| HIST1H1C | 2220.2074 | -0.504791876 | 0.34651917 | -1.4567502 | 0.14518535 | 0.30741429 |
| HFE | 670.607686 | 0.229343171 | 0.23812849 | 0.96310683 | 0.33549392 | 0.53368973 |
| HIST1H4C | 4.60086194 | -0.68410805 | 0.56992475 | -1.200348 | 0.2300042 | 0.41757045 |
| HIST1H1T | 2.86698654 | -2.671169032 | 1.69748254 | -1.5736062 | 0.11557852 | 0.26308205 |
| HIST1H2BC | 296.114551 | -1.717890956 | 0.78826988 | -2.1793183 | 0.02930803 | 0.09923636 |
| HIST1H2AC | 2403.33826 | -0.310219687 | 0.34645529 | -0.8954104 | 0.37056772 | 0.56737332 |
| HIST1H1E | 31.3107047 | 1.531119635 | 0.73335585 | 2.0878263 | 0.03681351 | 0.11661207 |
| HIST1H2BD | 1110.20841 | 0.141493856 | 0.35152535 | 0.40251394 | 0.68730583 | 0.81936052 |
| AL353759.1 | 54.9504145 | 0.351053874 | 0.33951666 | 1.03398128 | 0.30114492 | 0.49781792 |
| HIST1H2BE | 86.3015277 | 0.512581304 | 0.37635543 | 1.3619607 | 0.17321029 | 0.34683567 |
| HIST1H4D | 3.92605233 | -0.141369077 | 0.80498346 | -0.1756174 | 0.86059456 | 0.92734057 |
| AL031777.3 | 63.5910817 | 0.221042227 | 0.59792463 | 0.36968243 | 0.71161913 | 0.83606132 |
| HIST1H3D | 30.6924181 | -0.209505758 | 0.57157545 | -0.3665409 | 0.71396153 | 0.83749271 |
| HIST1H2AD | 22.799304 | -0.57833081 | 0.69972696 | -0.8265093 | 0.40851527 | 0.60299735 |
| HIST1H2BF | 24.9203692 | 0.741364687 | 0.73784104 | 1.00477562 | 0.31500491 | 0.51209416 |
| HIST1H4E | 34.7313431 | 1.120012772 | 0.54336076 | 2.06126914 | 0.03927737 | 0.12219729 |
| HIST1H2BG | 97.4440823 | 0.324079192 | 0.40165594 | 0.80685771 | 0.41974847 | 0.6125435 |
| HIST1H2AE | 48.3189174 | 0.077917761 | 0.45138848 | 0.17261797 | 0.86295173 | 0.92862833 |
| HIST1H3E | 100.387078 | 0.486958247 | 0.40565313 | 1.20043015 | 0.22997232 | 0.41754132 |
| HIST1H1D | 6.44667187 | 0.491212366 | 1.02014522 | 0.4815122 | 0.63015251 | 0.7801882 |
| HIST1H4F | 0.27872071 | -0.179578873 | 2.92985236 | -0.0612928 | 0.95112602 | NA |
| HIST1H4G | 0 | NA | NA | NA | NA | NA |
| HIST1H3F | 0.88496736 | 1.645390289 | 1.67188391 | 0.98415343 | 0.32504008 | NA |
| HIST1H2BH | 47.9538824 | -0.67087948 | 0.69347179 | -0.9674214 | 0.33333339 | 0.53131054 |
| HIST1H3G | 5.56566049 | -1.408408508 | 1.07145256 | -1.3144852 | 0.18868298 | 0.36687944 |
| HIST1H2BI | 1.41856227 | -1.150216531 | 1.22048221 | -0.9424279 | 0.34597359 | 0.54354339 |
| HIST1H4H | 164.52703 | -0.265932833 | 0.33956861 | -0.783149 | 0.43353959 | 0.62557806 |
| BTN3A2 | 1839.49784 | -0.415764883 | 0.46754232 | -0.8892561 | 0.37386544 | 0.57001824 |
| BTN2A2 | 938.174724 | -0.220504608 | 0.25883453 | -0.8519134 | 0.39426216 | 0.5895488 |
| BTN3A1 | 1465.27194 | -0.185454912 | 0.53120617 | -0.3491204 | 0.72699892 | 0.84575518 |
| BTN3A3 | 1103.86282 | -0.238356966 | 0.51252701 | -0.4650623 | 0.64188688 | 0.78783137 |
| BTN2A1 | 2220.10955 | -0.351752513 | 0.16326178 | -2.1545306 | 0.03119858 | 0.10383777 |
| BTN1A1 | 2.40840818 | 0.180924399 | 0.8100102 | 0.22336064 | 0.82325483 | 0.90564302 |
| HCG11 | 464.617472 | 0.509154511 | 0.15520591 | 3.28050987 | 0.0010362 | 0.00806746 |
| HMGN4 | 2605.10363 | -0.400065441 | 0.12124789 | -3.2995663 | 0.00096834 | 0.00764299 |
| AL121936.1 | 0.71564022 | 0.930168216 | 1.52426904 | 0.61023887 | 0.54170359 | NA |
| ABT1 | 1184.24018 | -0.313880905 | 0.20881146 | -1.5031785 | 0.13279301 | 0.28918587 |
| AL513548.3 | 0.54794267 | 0.274222758 | 1.60893827 | 0.17043709 | 0.8646664 | NA |
| ZNF322 | 293.635612 | 0.301598782 | 0.22713611 | 1.32783284 | 0.18423333 | 0.3612875 |
| AL513548.1 | 5.53289632 | -0.443673924 | 0.791134 | -0.5608076 | 0.57492873 | 0.73915805 |
| LINC00240 | 166.071932 | -1.007995532 | 0.36512696 | -2.7606713 | 0.00576827 | 0.02994866 |
| AL133255.1 | 4.3419737 | -0.159684903 | 0.65992839 | -0.2419731 | 0.80880103 | 0.89660916 |
| AL021807.1 | 5.83579039 | -0.863243822 | 0.92220329 | -0.9360667 | 0.34923883 | 0.5468132 |
| HIST1H2BJ | 26.7522572 | -1.333940469 | 0.71306522 | -1.8707131 | 0.06138485 | 0.16803889 |
| HIST1H2AG | 12.6533679 | -0.751433742 | 0.48124279 | -1.5614441 | 0.11841899 | 0.2674468 |
| HIST1H4I | 198.613145 | -0.820696022 | 0.27599422 | -2.9735986 | 0.0029433 | 0.01807517 |
| HIST1H2BK | 368.860305 | -0.787253541 | 0.27502654 | -2.8624639 | 0.00420361 | 0.02364027 |
| HIST1H2AH | 0.8342339 | -0.206585689 | 1.87699589 | -0.1100619 | 0.9123603 | NA |
| PRSS16 | 327.52942 | -0.53434875 | 0.37649217 | -1.4192825 | 0.15581668 | 0.32222289 |
| POM121L2 | 1.05847561 | -0.453608981 | 1.90676345 | -0.2378947 | 0.81196274 | 0.89847079 |
| ZNF391 | 243.199908 | 0.328854347 | 0.20778709 | 1.58265053 | 0.11350114 | 0.25964943 |
| AL031118.1 | 42.1304513 | 0.227971656 | 0.60689897 | 0.37563362 | 0.70718929 | 0.83325953 |
| ZNF184 | 397.642398 | 0.18265744 | 0.17854657 | 1.02302406 | 0.30629647 | 0.5031515 |
| AL009179.1 | 8.22149353 | -0.66059821 | 0.66781996 | -0.9891861 | 0.3225721 | 0.5203279 |
| HIST1H2BL | 1.5997873 | -0.524147555 | 1.24194134 | -0.4220389 | 0.67299663 | 0.809264 |
| HIST1H2AI | 1.39189904 | -0.438928701 | 1.08066291 | -0.4061662 | 0.68462052 | 0.81782302 |
| HIST1H3H | 30.8558235 | -1.435484777 | 0.66992827 | -2.142744 | 0.03213366 | 0.10597357 |
| HIST1H2AJ | 0.39645959 | -1.917872495 | 2.76087028 | -0.6946623 | 0.48726696 | NA |
| HIST1H2BM | 2.05715932 | -1.903397974 | 1.59301252 | -1.1948418 | 0.23214884 | 0.42019115 |
| HIST1H4J | 1.84469978 | 0.641841967 | 1.0423101 | 0.61578792 | 0.5380345 | 0.71153309 |
| HIST1H4K | 4.0075866 | 1.922070094 | 0.84606937 | 2.27176417 | 0.02310076 | 0.08348184 |
| HIST1H2BN | 92.604374 | -0.401756857 | 0.27389282 | -1.4668397 | 0.14241967 | 0.30360605 |
| HIST1H2AK | 14.976143 | 0.745930316 | 0.50061622 | 1.49002426 | 0.13621786 | 0.29432633 |
| HIST1H2AL | 0.58681048 | -1.430127981 | 2.20314853 | -0.6491292 | 0.51625489 | NA |
| HIST1H1B | 0.24327904 | -1.364242679 | 3.30558948 | -0.4127078 | 0.67982068 | NA |
| HIST1H3I | 0.16416605 | 0.18657164 | 3.29377195 | 0.05664376 | 0.95482897 | NA |
| HIST1H4L | 0.42435055 | -1.993544365 | 2.21243756 | -0.9010624 | 0.36755513 | NA |
| HIST1H3J | 2.58528234 | -2.238265714 | 1.35113777 | -1.6565785 | 0.09760473 | 0.23370896 |
| HIST1H2AM | 2.11543636 | -1.916302494 | 1.29047597 | -1.4849579 | 0.13755497 | 0.29619521 |
| HIST1H2BO | 1.75147885 | -2.173096283 | 1.32517991 | -1.63985 | 0.10103636 | 0.23918922 |
| OR2B2 | 0 | NA | NA | NA | NA | NA |
| OR2B6 | 1.88112057 | -4.145110054 | 1.30818764 | -3.1685898 | 0.0015318 | 0.0109941 |
| AL121944.1 | 106.247368 | -0.082308292 | 0.46925187 | -0.1754032 | 0.86076281 | 0.92743582 |
| ZNF165 | 466.036802 | -1.693430854 | 0.65290291 | -2.5936947 | 0.00949507 | 0.04335673 |
| ZSCAN16-AS | 518.945895 | 0.279620726 | 0.15328794 | 1.82415346 | 0.06812887 | 0.18059494 |
| ZSCAN16 | 311.396638 | -0.410585714 | 0.20309147 | -2.0216788 | 0.04320955 | 0.13106002 |
| AL358933.1 | 11.1437147 | 0.795513683 | 0.59957906 | 1.32678697 | 0.18457916 | 0.36175046 |
| ZKSCAN8 | 1462.02074 | -0.068016601 | 0.15038995 | -0.4522683 | 0.65107574 | 0.79489177 |
| ZSCAN9 | 703.288592 | -0.451357633 | 0.18856733 | -2.3936152 | 0.01668324 | 0.0657899 |
| ZKSCAN4 | 264.950471 | 0.183810763 | 0.13263345 | 1.38585522 | 0.16579112 | 0.3363009 |
| NKAPL | 45.5183337 | 0.012022825 | 0.36766945 | 0.03270009 | 0.97391375 | 0.98796328 |
| ZSCAN26 | 791.251299 | 0.284587261 | 0.19284828 | 1.47570546 | 0.14002298 | 0.3000643 |
| AL021997.3 | 111.16346 | 0.951079995 | 0.20759574 | 4.58140422 | 4.62E-06 | 0.00011042 |
| PGBD1 | 344.330711 | 0.324536379 | 0.15074147 | 2.15293367 | 0.03132389 | 0.10412201 |
| ZSCAN31 | 366.118805 | 0.212756449 | 0.42032918 | 0.50616626 | 0.61273994 | 0.76799695 |
| ZKSCAN3 | 364.040759 | 0.301818303 | 0.20371045 | 1.48160445 | 0.13844557 | 0.29759525 |
| ZSCAN12 | 271.212917 | 0.25719929 | 0.12232377 | 2.10261093 | 0.03549979 | 0.113746 |
| ZSCAN23 | 10.7364584 | 0.889840929 | 0.59221904 | 1.50255372 | 0.13295417 | 0.28939342 |
| GPX6 | 0.03392908 | 0.380740593 | 3.29738709 | 0.11546736 | 0.90807469 | NA |
| GPX5 | 0 | NA | NA | NA | NA | NA |
| ZBED9 | 112.390363 | 0.070183787 | 0.25308504 | 0.27731305 | 0.78153974 | 0.88036187 |
| AL049543.1 | 4.60251883 | 0.673449004 | 0.92020228 | 0.73184888 | 0.46426081 | 0.65178001 |
| LINC00533 | 1.20607593 | 0.718324235 | 1.32078461 | 0.54386175 | 0.58653661 | 0.74768058 |
| AL662890.1 | 2.09409935 | -0.755129127 | 0.97297674 | -0.7761019 | 0.4376888 | 0.62967299 |
| LINC01623 | 0.07659028 | -0.642905668 | 3.30558948 | -0.1944905 | 0.84579185 | NA |
| HCG14 | 8.7087457 | 1.008027929 | 0.57800948 | 1.74396434 | 0.08116531 | 0.20448353 |
| TRIM27 | 3329.87285 | -0.475503611 | 0.1481126 | -3.2104197 | 0.00132541 | 0.00983431 |
| LINC01556 | 2.67631585 | -1.288764733 | 1.29807276 | -0.9928293 | 0.32079313 | 0.5182451 |

| | | | | | | |
|---|---|---|---|---|---|---|
| HCG15 | 16.1970823 | 0.447588877 | 0.39715272 | 1.12699438 | 0.25974486 | 0.45261825 |
| AL662791.1 | 10.7028496 | 0.580264202 | 0.4775948 | 1.21497178 | 0.22437686 | 0.41043408 |
| ZNF311 | 83.5918711 | 0.306397764 | 0.32188191 | 0.95189495 | 0.34115026 | 0.53918739 |
| AL662791.2 | 0.09644756 | -0.642905668 | 3.30558948 | -0.1944905 | 0.84579185 | NA |
| OR2W1 | 0 | NA | NA | NA | NA | NA |
| OR2B3 | 0 | NA | NA | NA | NA | NA |
| OR2J1 | 0 | NA | NA | NA | NA | NA |
| OR2J3 | 0.10721287 | -0.642905668 | 3.30558948 | -0.1944905 | 0.84579185 | NA |
| AL645937.2 | 0.08666371 | -0.642905668 | 3.30558948 | -0.1944905 | 0.84579185 | NA |
| AL645937.4 | 0 | NA | NA | NA | NA | NA |
| OR2J2 | 0 | NA | NA | NA | NA | NA |
| OR14J1 | 0.26210702 | 1.093535322 | 3.27839713 | 0.33355792 | 0.73871318 | NA |
| OR5V1 | 0 | NA | NA | NA | NA | NA |
| OR12D3 | 0 | NA | NA | NA | NA | NA |
| OR12D2 | 0.99285407 | 2.346969568 | 2.1173525 | 1.10844537 | 0.26766952 | NA |
| OR11A1 | 0 | NA | NA | NA | NA | NA |
| OR10C1 | 0.15661533 | -1.003574173 | 3.30558948 | -0.3035992 | 0.7614333 | NA |
| OR2H1 | 0.18746814 | 0.18657164 | 3.29377195 | 0.05664376 | 0.95482897 | NA |
| MAS1L | 1.32859721 | 4.319215753 | 1.42903987 | 3.02245993 | 0.00250729 | 0.0160401 |
| AL662860.1 | 1.00484738 | 0.879944653 | 1.79463348 | 0.49031998 | 0.62390749 | NA |
| LINC01015 | 0 | NA | NA | NA | NA | NA |
| OR2I1P | 1625.80134 | -1.048906507 | 0.40719055 | -2.5759599 | 0.00999623 | 0.04505991 |
| UBD | 1175.32191 | -1.064431398 | 0.39703414 | -2.6809569 | 0.0073412 | 0.03588939 |
| GABBR1 | 4341.91041 | -0.541621385 | 0.28568207 | -1.8958886 | 0.05797477 | 0.16162329 |
| OR2H2 | 5.09749062 | -2.003005044 | 0.9498452 | -2.10877 | 0.03496444 | 0.11258812 |
| MOG | 0.42557226 | -0.621461299 | 2.29374525 | -0.2709374 | 0.7864392 | NA |
| ZFP57 | 21.5811151 | -3.125457611 | 1.12201774 | -2.7855688 | 0.00534339 | 0.02828854 |
| HLA-F | 7437.16199 | -1.486827366 | 0.52826959 | -2.8145239 | 0.00488496 | 0.02641843 |
| AL645939.4 | 6.25242051 | -0.342998867 | 0.61261516 | -0.5598929 | 0.5755525 | 0.73970755 |
| HLA-G | 2169.76631 | -1.138210015 | 0.55521391 | -2.0500387 | 0.04036065 | 0.12458275 |
| AL645929.2 | 14.9223488 | 0.203796793 | 0.59758652 | 0.34103311 | 0.73307865 | 0.84942807 |
| HLA-A | 78310.9856 | -1.014935951 | 0.33758262 | -3.0064817 | 0.0026429 | 0.01666506 |
| HCG9 | 8.97063732 | -0.999179713 | 0.70224297 | -1.4228405 | 0.15478241 | 0.32088146 |
| ZNRD1 | 820.082183 | -0.518559882 | 0.12399506 | -4.1821011 | 2.89E-05 | 0.00047662 |
| PPP1R11 | 3707.37392 | -0.094327928 | 0.10559284 | -0.8933175 | 0.37168718 | 0.56836599 |
| RNF39 | 358.524514 | -0.649961727 | 0.12783937 | -5.0842062 | 3.69E-07 | 1.42E-05 |
| TRIM31 | 10.7505144 | -0.512179584 | 0.64784828 | -0.7905857 | 0.4291858 | 0.62123337 |
| TRIM31-AS1 | 2.12752686 | -0.11573295 | 1.02763041 | -0.1126212 | 0.91033089 | 0.95559196 |
| TRIM40 | 13.6329048 | 0.564781876 | 0.91617492 | 0.61645638 | 0.53759335 | 0.71119935 |
| TRIM10 | 272.751112 | 2.340424957 | 0.4813726 | 4.86198212 | 1.16E-06 | 3.68E-05 |
| TRIM15 | 576.399852 | 0.495817884 | 0.46193561 | 1.07334848 | 0.28311479 | 0.47900276 |
| TRIM26 | 3470.60244 | 0.270557816 | 0.20593403 | 1.31380822 | 0.18891075 | 0.36716397 |
| HCG17 | 24.9990291 | -0.029433878 | 0.35662416 | -0.0825347 | 0.9342215 | 0.96837347 |
| HCG18 | 1036.14415 | -0.284050739 | 0.25147839 | -1.1295235 | 0.25867708 | 0.45145198 |
| TRIM39 | 1187.38738 | 0.018353887 | 0.19649265 | 0.0934075 | 0.92557983 | 0.96400096 |
| TRIM39-RPP. | 1161.89482 | -0.15274186 | 0.16709073 | -0.9141253 | 0.36065101 | 0.55828396 |
| RPP21 | 941.635502 | -0.496810257 | 0.16464558 | -3.0174527 | 0.00254909 | 0.01623266 |
| HLA-E | 47074.2966 | -0.800266441 | 0.2542472 | -3.1475919 | 0.00164621 | 0.01162266 |
| LINC02569 | 5.38982544 | 0.843186435 | 0.58069204 | 1.45203719 | 0.14649126 | 0.30945884 |
| GNL1 | 4543.43837 | -0.507129078 | 0.13279122 | -3.8189956 | 0.000134 | 0.00159244 |
| PRR3 | 785.45145 | -0.256055499 | 0.16109866 | -1.5894328 | 0.11196271 | 0.25719417 |
| ABCF1 | 4460.81775 | -0.380245586 | 0.19946903 | -1.9062889 | 0.05661274 | 0.15887235 |
| PPP1R10 | 3632.79945 | 0.679928328 | 0.19326492 | 3.51811552 | 0.00043462 | 0.00406741 |
| MRPS18B | 4375.16033 | 0.102605325 | 0.18166579 | 0.56480269 | 0.57220798 | 0.73695281 |
| ATAT1 | 1028.48166 | -0.313577716 | 0.16079973 | -1.9501134 | 0.0511626 | 0.14809635 |
| C6orf136 | 2163.62324 | -0.019831844 | 0.12749231 | -0.1555533 | 0.87638517 | 0.93555133 |
| DHX16 | 3079.51526 | -0.029697411 | 0.14351036 | -0.2069357 | 0.83606011 | 0.91295155 |
| PPP1R18 | 2858.26654 | -1.06080897 | 0.33322862 | -3.183427 | 0.00145543 | 0.01056127 |
| NRM | 658.690061 | -0.528083903 | 0.31935592 | -1.6535905 | 0.09821075 | 0.23469819 |
| MDC1 | 2043.69956 | 0.141398423 | 0.12525122 | 1.12891853 | 0.2589322 | 0.45167971 |
| MDC1-AS1 | 277.212341 | 0.15776229 | 0.16645368 | 0.94778491 | 0.34323896 | 0.5408875 |
| TUBB | 16998.7848 | -0.565234411 | 0.34288213 | -1.6484802 | 0.09925416 | 0.23629395 |
| AL662797.1 | 6785.74584 | -0.474938277 | 0.36580012 | -1.2983546 | 0.1941655 | 0.37397884 |
| FLOT1 | 14174.0998 | -0.742293661 | 0.20750161 | -3.5772912 | 0.00034717 | 0.00340967 |
| IER3-AS1 | 2908.92729 | -2.042215345 | 0.44018656 | -4.6394314 | 3.49E-06 | 8.84E-05 |
| IER3 | 9477.51456 | -1.920982168 | 0.4556385 | -4.2160225 | 2.49E-05 | 0.00042302 |
| HCG20 | 6.90731577 | 0.016907203 | 0.68605491 | 0.0246441 | 0.98033885 | 0.99116384 |
| LINC00243 | 5.56667971 | -0.305476477 | 0.82034953 | -0.3723736 | 0.70961474 | 0.83500637 |
| LINC02570 | 0.76537478 | -0.898200381 | 2.29158478 | -0.391956 | 0.69509073 | NA |
| DDR1-DT | 0.31399703 | 1.532035203 | 2.80770895 | 0.54565314 | 0.58530439 | NA |
| DDR1 | 19618.4834 | -0.839652776 | 0.15568376 | -5.3933229 | 6.92E-08 | 3.48E-06 |
| GTF2H4 | 1721.68531 | -0.250778661 | 0.15929639 | -1.5742896 | 0.11542051 | 0.26288122 |
| VARS2 | 4809.20641 | 0.229560868 | 0.14148113 | 1.62255464 | 0.10468464 | 0.24516288 |
| SFTA2 | 79.1885669 | -1.353312071 | 0.43817525 | -3.0885178 | 0.00201158 | 0.01356309 |
| MUCL3 | 8.39978072 | 0.370421578 | 0.61407276 | 0.60322099 | 0.54636169 | 0.71750785 |
| HCG21 | 5.59606064 | 0.614604936 | 0.82332363 | 0.74649252 | 0.45536995 | 0.64476511 |
| MUC21 | 0 | NA | NA | NA | NA | NA |
| MUC22 | 0.45168795 | 2.130928619 | 2.68664233 | 0.79315679 | 0.42768648 | NA |
| HCG22 | 2.93183825 | 0.893685173 | 0.90413986 | 0.98843687 | 0.32293874 | 0.52066432 |
| C6orf15 | 0.07659028 | -0.642905668 | 3.30558948 | -0.1944905 | 0.84579185 | NA |
| PSORS1C1 | 208.101622 | -2.248165144 | 0.6611948 | -3.4001555 | 0.00067348 | 0.00571598 |
| CDSN | 36.0404935 | -1.622524686 | 0.75778702 | -2.1411355 | 0.03226311 | 0.10622638 |
| PSORS1C2 | 50.834095 | -2.195251873 | 0.77873772 | -2.8189875 | 0.00481754 | 0.02612886 |
| CCHCR1 | 2103.35699 | -0.502322577 | 0.29850991 | -1.6827669 | 0.09242023 | 0.22482671 |
| TCF19 | 858.685687 | -0.745123384 | 0.31791386 | -2.3437902 | 0.01908891 | 0.072501 |
| POU5F1 | 2760.66807 | -1.98276971 | 0.49811971 | -3.9805084 | 6.88E-05 | 0.00094498 |
| AL662844.4 | 230.092485 | 0.406642802 | 0.34852282 | 1.16676092 | 0.24330694 | 0.4338939 |
| HCG27 | 200.787569 | -0.091949859 | 0.37803124 | -0.2432335 | 0.80782452 | 0.89620864 |
| AL662844.3 | 30.0582133 | 0.548417139 | 0.50589559 | 1.08405202 | 0.27834173 | 0.47399881 |
| HLA-C | 68659.9039 | -1.193502837 | 0.4105809 | -2.906864 | 0.00365072 | 0.02127982 |
| HLA-B | 100489.981 | -1.451969111 | 0.4472361 | -3.2465383 | 0.00116818 | 0.00886725 |
| LINC02571 | 2.72070299 | 0.721859399 | 1.11446516 | 0.64771823 | 0.51716721 | 0.69438388 |
| AL645933.2 | 181.511663 | 0.74604202 | 0.35471021 | 2.10324374 | 0.03544447 | 0.11363777 |
| MICA | 2140.15446 | -0.344683512 | 0.16770743 | -2.0552668 | 0.03985324 | 0.12350806 |
| LINC01149 | 0.43740202 | 0.331067391 | 1.84802846 | 0.17914626 | 0.85782286 | NA |
| MICB | 271.327474 | -1.073916692 | 0.36626212 | -2.9320987 | 0.0033668 | 0.02001859 |
| MCCD1 | 1413.66951 | 1.022475261 | 0.66568908 | 1.5359652 | 0.12454691 | 0.2766194 |
| DDX39B | 18735.6994 | -0.373547091 | 0.17920348 | -2.0844857 | 0.03711601 | 0.11720142 |
| ATP6V1G2-D | 11778.739 | -0.442748877 | 0.17109876 | -2.5876802 | 0.00966246 | 0.04391194 |
| DDX39B-AS1 | 10.9747728 | -0.082427548 | 0.62415284 | -0.1320631 | 0.8949344 | 0.94598591 |
| ATP6V1G2 | 495.739229 | -0.016086788 | 0.13177928 | -0.1220737 | 0.90284063 | 0.95113689 |
| NFKBIL1 | 1336.39747 | -0.432229397 | 0.11980721 | -3.6077078 | 0.00030891 | 0.00310208 |
| LTA | 23.8715003 | -0.07991194 | 0.66041119 | -0.1210033 | 0.90368841 | 0.95167363 |
| TNF | 89.1635855 | -1.24859083 | 0.42918629 | -2.9092048 | 0.00362349 | 0.0211444 |
| LTB | 308.919688 | 0.164990545 | 0.59948261 | 0.27522157 | 0.78314602 | 0.88134007 |
| LST1 | 291.26434 | -0.190961767 | 0.46766616 | -0.4083292 | 0.68303198 | 0.8167764 |
| NCR3 | 20.0911353 | 0.167690933 | 0.79284812 | 0.21150448 | 0.83249363 | 0.91109596 |
| AIF1 | 865.21773 | -0.127832858 | 0.38287559 | -0.3338757 | 0.73847336 | 0.85320463 |
| PRRC2A | 15085.2087 | -0.029079611 | 0.13735059 | -0.2117181 | 0.83232694 | 0.91098914 |
| BAG6 | 14061.712 | -0.045056925 | 0.13649517 | -0.330099 | 0.74132512 | 0.85469576 |
| APOM | 5293.89975 | 1.079496876 | 0.61038984 | 1.76853676 | 0.07697121 | 0.19714377 |
| C6orf47 | 2198.60295 | -0.282783042 | 0.16840559 | -1.6791785 | 0.09311726 | 0.22603775 |
| C6orf47-AS1 | 687.34655 | -0.332228744 | 0.16701014 | -1.9892729 | 0.04667109 | 0.13849967 |

| | | | | | | |
|---|---|---|---|---|---|---|
| GPANK1 | 2293.20467 | 0.067555736 | 0.11195235 | 0.60343292 | 0.54622073 | 0.71739419 |
| CSNK2B | 6064.50372 | -0.283199223 | 0.097981 | -2.8903485 | 0.00384815 | 0.02213396 |
| AL662899.2 | 6760.22288 | -0.291808486 | 0.09592877 | -3.0419288 | 0.00235067 | 0.01530551 |
| LY6G5B | 1394.68788 | -0.257737646 | 0.20604662 | -1.2508705 | 0.21098171 | 0.39438489 |
| LY6G5C | 101.297252 | 0.097027395 | 0.26963468 | 0.3598476 | 0.7189611 | 0.84015795 |
| ABHD16A | 3312.03722 | -0.087386768 | 0.17521916 | -0.4987284 | 0.61797074 | 0.77162216 |
| AL662899.1 | 3235.69333 | -0.095117463 | 0.17675036 | -0.5381458 | 0.59047639 | 0.75022224 |
| LY6G6F | 2.21387172 | -0.057581423 | 0.96902198 | -0.0594222 | 0.95261583 | 0.97717271 |
| LY6G6F-LY6( | 1.98731741 | 0.029097575 | 1.03472551 | 0.02812106 | 0.9775656 | 0.98965006 |
| LY6G6E | 2.05124512 | -1.069279268 | 1.17855192 | -0.9072823 | 0.36425754 | 0.56142798 |
| LY6G6D | 0.54385493 | -1.274765892 | 1.88538323 | -0.6761309 | 0.49895753 | NA |
| MPIG6B | 16.5555345 | -0.078379079 | 0.45225719 | -0.1733064 | 0.86241057 | 0.92830704 |
| LY6G6C | 11.5684155 | -0.328197049 | 0.50473328 | -0.6502386 | 0.51553813 | 0.69322016 |
| DDAH2 | 4091.32926 | -0.074109511 | 0.16698652 | -0.4438054 | 0.65718331 | 0.79883488 |
| CLIC1 | 18461.5436 | -1.432961499 | 0.20196241 | -7.095189 | 1.29E-12 | 3.37E-10 |
| MSH5 | 1470.30423 | 0.227584297 | 0.25183064 | 0.90371963 | 0.36614409 | 0.56326414 |
| MSH5-SAPCI | 2251.39726 | 0.125931901 | 0.22347001 | 0.56352931 | 0.57307451 | 0.73770929 |
| SAPCD1 | 1076.8085 | 0.101330112 | 0.23150224 | 0.43770683 | 0.66159882 | 0.80084282 |
| SAPCD1-AS1 | 735.547055 | 0.077492838 | 0.22860784 | 0.33897717 | 0.73462693 | 0.85065526 |
| VWA7 | 1005.59317 | 0.821292204 | 0.24927822 | 3.29468092 | 0.00098533 | 0.0077576 |
| VARS | 3897.66636 | -0.170038592 | 0.1608523 | -1.0571101 | 0.29046135 | 0.48652584 |
| LSM2 | 969.402744 | -0.524220277 | 0.18205165 | -2.879514 | 0.00398289 | 0.0226697 |
| HSPA1L | 111.930337 | 0.589010573 | 0.2505733 | 2.35065181 | 0.01874056 | 0.07160078 |
| HSPA1A | 7522.09411 | -0.261998351 | 0.41975439 | -0.6241706 | 0.53251555 | 0.70675097 |
| HSPA1B | 8203.40488 | -0.343434472 | 0.35879403 | -0.9571912 | 0.33847078 | 0.53651565 |
| C6orf48 | 4791.52879 | -0.500436505 | 0.1255767 | -3.9851063 | 6.74E-05 | 0.00093079 |
| NEU1 | 7733.96446 | -0.639841177 | 0.25596546 | -2.4997169 | 0.01242926 | 0.05296199 |
| SLC44A4 | 7769.94773 | -0.147249904 | 0.2887203 | -0.5100088 | 0.61004527 | 0.76596961 |
| EHMT2-AS1 | 1172.17488 | 0.246626692 | 0.14092543 | 1.75005105 | 0.0801095 | 0.20261727 |
| EHMT2 | 2827.56488 | 0.346426603 | 0.12243574 | 2.8294565 | 0.00466271 | 0.02551479 |
| C2 | 2741.83838 | -1.350481164 | 0.50172761 | -2.691662 | 0.0071097 | 0.03503039 |
| ZBTB12 | 192.176921 | 0.211945586 | 0.22799422 | 0.92960947 | 0.35257332 | 0.55020467 |
| AL645922.1 | 14476.2518 | -1.908049907 | 0.45434903 | -4.1995245 | 2.67E-05 | 0.00044926 |
| C2-AS1 | 77.1388126 | -0.544719341 | 0.64475112 | -0.8448521 | 0.39819342 | 0.59314017 |
| CFB | 12670.1026 | -2.012075586 | 0.49937944 | -4.0291518 | 5.60E-05 | 0.00080929 |
| NELFE | 3450.04985 | -0.531781836 | 0.13031009 | -4.0808952 | 4.49E-05 | 0.00068068 |
| SKIV2L | 3884.21641 | -0.221244927 | 0.14226133 | -1.5552007 | 0.11989828 | 0.26977112 |
| DXO | 1983.37455 | -0.309190802 | 0.14913359 | -2.0732472 | 0.03814928 | 0.11959064 |
| STK19 | 2099.7711 | -0.161426163 | 0.146577 | -1.1013062 | 0.2707634 | 0.4653608 |
| C4A | 5808.4554 | 0.336424455 | 0.36089736 | 0.93218874 | 0.35123899 | 0.54890199 |
| C4A-AS1 | 48.5366424 | -1.122288038 | 0.61698117 | -1.8189988 | 0.06891161 | 0.18208407 |
| C4B | 6658.28461 | -0.492144934 | 0.51418113 | -0.9571431 | 0.33849502 | 0.53652182 |
| C4B-AS1 | 47.3075576 | -1.271267884 | 0.63682111 | -1.9962716 | 0.04590437 | 0.13686098 |
| CYP21A2 | 872.214272 | -0.935424473 | 0.41859986 | -2.2346507 | 0.02544029 | 0.08943613 |
| TNXB | 3637.55244 | -0.042803814 | 0.41639223 | -0.1027969 | 0.9181242 | 0.96017986 |
| ATF6B | 5659.18482 | 0.054493282 | 0.15404224 | 0.35375545 | 0.72352217 | 0.84328307 |
| FKBPL | 335.690261 | -0.48566762 | 0.18981321 | -2.5586608 | 0.01050762 | 0.04673402 |
| PRRT1 | 375.089472 | -0.611262702 | 0.38652165 | -1.5814449 | 0.11377634 | 0.2600574 |
| AL662884.4 | 369.77898 | 0.092079318 | 0.24669007 | 0.37325911 | 0.70895561 | 0.83447565 |
| PPT2 | 1944.3167 | 0.142458429 | 0.180057 | 0.79118519 | 0.42883593 | 0.62111967 |
| PPT2-EGFL8 | 6258.8346 | 0.183926102 | 0.17778086 | 1.03456639 | 0.30087146 | 0.49764664 |
| EGFL8 | 2193.78682 | -0.211995714 | 0.24365616 | -0.870061 | 0.38426708 | 0.57979762 |
| AGPAT1 | 5385.31527 | 0.318519033 | 0.18469531 | 1.72456484 | 0.08460592 | 0.21071574 |
| RNF5 | 4483.75293 | 0.128610901 | 0.11980764 | 1.07347831 | 0.28305657 | 0.4789542 |
| AGER | 893.480626 | -0.629543301 | 0.31080527 | -2.0255232 | 0.04281367 | 0.13017369 |
| PBX2 | 6060.56425 | -0.196907188 | 0.19646486 | -1.0022514 | 0.31622217 | 0.5133515 |
| GPSM3 | 1236.13005 | -0.390081846 | 0.34596225 | -1.1275272 | 0.25951966 | 0.45227322 |
| NOTCH4 | 3781.03128 | 0.228809405 | 0.33474588 | 0.68353164 | 0.49427096 | 0.67673841 |
| AL662796.1 | 54.1083139 | 0.576779078 | 0.31635024 | 1.82322949 | 0.06826864 | 0.18089269 |
| C6orf10 | 0.78163938 | 0.260861433 | 2.12577524 | 0.12271355 | 0.90233393 | NA |
| BTNL2 | 4.10385284 | 0.924979763 | 0.80685637 | 1.14639953 | 0.25162988 | 0.44390419 |
| HLA-DRA | 20419.1128 | -0.279848442 | 0.35370749 | -0.7911861 | 0.42883541 | 0.62111967 |
| HLA-DRB5 | 6393.05331 | 1.092632576 | 0.45560322 | 2.39821085 | 0.01647538 | 0.06515533 |
| HLA-DRB1 | 17128.0955 | -0.059677735 | 0.35667375 | -0.1673174 | 0.86712029 | 0.93098661 |
| HLA-DQA1 | 2480.1443 | 0.389661744 | 0.50422166 | 0.7727985 | 0.43964164 | 0.63101113 |
| HLA-DQB1 | 4351.7444 | 0.335191803 | 0.58543183 | 0.5725548 | 0.56694617 | 0.73307691 |
| HLA-DQB1-A | 1085.49498 | 0.239800495 | 0.60628708 | 0.39552302 | 0.69245694 | 0.82304273 |
| HLA-DQA2 | 606.271642 | -0.899788296 | 0.98054334 | -0.9176426 | 0.35880603 | 0.5564673 |
| HLA-DQB2 | 426.018026 | -1.819015093 | 0.69013478 | -2.6357389 | 0.00839543 | 0.03944895 |
| HLA-DOB | 136.713642 | 0.056133554 | 0.5633509 | 0.09964226 | 0.92062835 | 0.96119767 |
| AL669918.1 | 3250.58624 | -1.46064396 | 0.51245985 | -2.8502603 | 0.00436835 | 0.02436406 |
| TAP2 | 4717.59647 | -1.425740689 | 0.48800778 | -2.9215532 | 0.00348291 | 0.02049624 |
| PSMB8 | 3966.65604 | -1.285518571 | 0.35356706 | -3.635855 | 0.00027706 | 0.00285728 |
| PSMB8-AS1 | 3028.95284 | -1.746578124 | 0.46867309 | -3.7266448 | 0.00019405 | 0.00213208 |
| PSMB9 | 1536.52143 | -1.424978798 | 0.53956023 | -2.6410004 | 0.00826616 | 0.03901534 |
| TAP1 | 6043.72776 | -1.714628663 | 0.49608881 | -3.4562938 | 0.00054766 | 0.00487238 |
| HLA-DMB | 2402.54411 | -0.115455222 | 0.36505593 | -0.3162672 | 0.7517997 | 0.86150521 |
| AL645941.2 | 1686.5203 | -0.075074905 | 0.32848392 | -0.2285497 | 0.81921891 | 0.90290073 |
| HLA-DMA | 4235.64522 | -0.264100798 | 0.34179879 | -0.7726791 | 0.4397123 | 0.63103201 |
| BRD2 | 13700.4669 | -0.179491502 | 0.24448505 | -0.7341615 | 0.46285033 | 0.65098743 |
| AL645941.3 | 1207.37775 | 0.629843523 | 0.35386749 | 1.77988523 | 0.07509474 | 0.1938759 |
| HLA-DOA | 1220.65059 | 0.090647543 | 0.37459604 | 0.24198745 | 0.80878988 | 0.89660916 |
| HLA-DPA1 | 10556.4114 | 0.213388659 | 0.43981312 | 0.48518029 | 0.62754846 | 0.77838442 |
| HLA-DPB1 | 5875.05519 | -0.016363362 | 0.44468117 | -0.036798 | 0.9706461 | 0.98639399 |
| HCG24 | 0.13028434 | 1.049816488 | 3.28477129 | 0.31960109 | 0.74927075 | NA |
| COL11A2 | 228.84977 | 0.804148758 | 0.24079985 | 3.3394903 | 0.00083932 | 0.00683326 |
| RXRB | 4297.28805 | 0.091694124 | 0.1102434 | 0.8317425 | 0.40555431 | 0.60034058 |
| SLC39A7 | 9571.94027 | -0.460273708 | 0.2504274 | -1.8379527 | 0.06606938 | 0.17686016 |
| HSD17B8 | 1222.77596 | 0.920197814 | 0.26847374 | 3.42751517 | 0.00060913 | 0.00526995 |
| RING1 | 3878.07183 | -0.114227055 | 0.15809887 | -0.7225039 | 0.4699847 | 0.65659606 |
| AL645940.1 | 36.4348134 | -0.719130002 | 0.36763581 | -1.9560934 | 0.05045415 | 0.14660612 |
| HCG25 | 1505.25431 | 0.106235097 | 0.16065792 | 0.66125031 | 0.5084518 | 0.68730556 |
| VPS52 | 3442.29009 | 0.419442711 | 0.13999965 | 2.99602688 | 0.00273522 | 0.01713657 |
| RPS18 | 31835.7207 | -0.497503767 | 0.19521981 | -2.5484287 | 0.01082094 | 0.04776502 |
| B3GALT4 | 489.154111 | -0.554733088 | 0.22687612 | -2.4450925 | 0.0144815 | 0.05933455 |
| WDR46 | 2306.0886 | -0.435458218 | 0.18815957 | -2.3143028 | 0.02065112 | 0.0767108 |
| PFDN6 | 2009.34729 | -0.368009532 | 0.11567718 | -3.1813495 | 0.00146591 | 0.01060811 |
| RGL2 | 6431.57803 | -0.165304678 | 0.21544672 | -0.7672648 | 0.4429241 | 0.63408841 |
| TAPBP | 10360.0066 | -0.902273983 | 0.22440385 | -4.0207599 | 5.80E-05 | 0.00083138 |
| ZBTB22 | 1397.04336 | -0.053602198 | 0.12447409 | -0.4306293 | 0.6667379 | 0.8048468 |
| DAXX | 2704.13303 | -0.560726532 | 0.19063886 | -2.9413024 | 0.00326835 | 0.0196012 |
| AL662820.1 | 361.48514 | -0.151554131 | 0.24296839 | -0.6237607 | 0.53278476 | 0.70682367 |
| KIFC1 | 74.7416971 | -1.291017802 | 0.48929912 | -2.6385042 | 0.00832727 | 0.03924053 |
| PHF1 | 7198.15293 | -0.62081271 | 0.15442083 | -4.0202654 | 5.81E-05 | 0.00083177 |
| CUTA | 4753.4385 | -0.037936658 | 0.1592242 | -0.2382594 | 0.81167993 | 0.89834633 |
| SYNGAP1 | 3034.26581 | -0.438329788 | 0.2615836 | -1.6756777 | 0.09380135 | 0.22734757 |
| SYNGAP1-AS | 255.049123 | -0.41859489 | 0.29824903 | -1.403508 | 0.16046542 | 0.32873471 |
| ZBTB9 | 547.129427 | -0.059571775 | 0.2247673 | -0.2650376 | 0.79098053 | 0.88641183 |
| BAK1 | 1735.99958 | -0.739221257 | 0.19716848 | -3.7491857 | 0.00017741 | 0.00199161 |
| LINC00336 | 27.0193122 | -1.210635414 | 0.57980334 | -2.0880104 | 0.03679689 | 0.11658466 |
| ITPR3 | 4794.22075 | -1.138179692 | 0.22879109 | -4.9747552 | 6.53E-07 | 2.26E-05 |
| UQCC2 | 3890.01229 | -0.842092515 | 0.18725873 | -4.4969466 | 6.89E-06 | 0.0001525 |
| IP6K3 | 352.004323 | 0.85518796 | 0.57208358 | 1.49486543 | 0.13494957 | 0.29236116 |

| | | | | | | |
|---|---|---|---|---|---|---|
| LEMD2 | 3634.87665 | -0.618033105 | 0.14621354 | -4.2269213 | 2.37E-05 | 0.00040779 |
| MLN | 0.48585202 | 0.295012208 | 2.70585764 | 0.10902725 | 0.91318088 | NA |
| LINC01016 | 5.79487129 | 2.363949652 | 0.64510571 | 3.66443764 | 0.00024788 | 0.00260321 |
| AL138889.1 | 2.15216127 | 0.665757481 | 1.39372134 | 0.47768335 | 0.6328756 | 0.78205156 |
| GRM4 | 3.1288665 | -1.262582784 | 0.76075023 | -1.6596548 | 0.09698392 | 0.23251803 |
| HMGA1 | 5510.24374 | -2.352303299 | 0.60363239 | -3.8969136 | 9.74E-05 | 0.00123825 |
| SMIM29 | 1061.24428 | -0.930268253 | 0.24404964 | -3.8117993 | 0.00013796 | 0.00163284 |
| NUDT3 | 5407.79451 | 0.171078292 | 0.18488496 | 0.92532293 | 0.35479795 | 0.55230354 |
| RPS10-NUDT | 19137.8521 | -0.573256051 | 0.16339325 | -3.5084439 | 0.00045074 | 0.0041834 |
| RPS10 | 18107.5401 | -0.573655713 | 0.17141009 | -3.3466858 | 0.00081784 | 0.00669348 |
| PACSIN1 | 206.808846 | -0.061047555 | 0.53253718 | -0.1146353 | 0.9087342 | 0.95440979 |
| SPDEF | 10.2589858 | -2.32935824 | 0.79264935 | -2.9386995 | 0.00329592 | 0.01972618 |
| C6orf106 | 8668.63496 | -0.454910948 | 0.26416611 | -1.722064 | 0.08505793 | 0.21148194 |
| AL451165.2 | 321.758307 | 0.85196386 | 0.34434664 | 2.47414602 | 0.01335551 | 0.05579607 |
| SNRPC | 2405.53464 | -0.241712364 | 0.15224177 | -1.5876876 | 0.11235701 | 0.25788618 |
| UHRF1BP1 | 1390.69897 | -0.230218568 | 0.18729752 | -1.2291597 | 0.21901194 | 0.40446423 |
| TAF11 | 1399.9128 | -0.211649196 | 0.11693762 | -1.8099325 | 0.07030626 | 0.18482466 |
| ANKS1A | 2737.62959 | 0.179607277 | 0.15987231 | 1.12344207 | 0.26124979 | 0.45448984 |
| TCP11 | 22.2919062 | 0.154554079 | 0.44851163 | 0.34459324 | 0.73040018 | 0.84755348 |
| SCUBE3 | 364.000124 | 0.894918719 | 0.49955856 | 1.79141904 | 0.07322608 | 0.19041235 |
| Z97832.2 | 184.579315 | 0.718265799 | 0.31545915 | 2.27689002 | 0.02279279 | 0.08258395 |
| ZNF76 | 3609.8512 | 0.287755037 | 0.12579814 | 2.28743472 | 0.02217046 | 0.08090715 |
| DEF6 | 481.381162 | -0.487219641 | 0.49396889 | -0.9863367 | 0.32396792 | 0.52163909 |
| PPARD | 3077.1 | -0.287833384 | 0.17274452 | -1.6662375 | 0.09566613 | 0.23053237 |
| FANCE | 170.433893 | -0.361817213 | 0.23870626 | -1.5157424 | 0.12958449 | 0.28452482 |
| RPL10A | 19611.8183 | -0.35470278 | 0.21145401 | -1.6774465 | 0.09345521 | 0.2266336 |
| TEAD3 | 2878.77094 | -0.658438002 | 0.25782482 | -2.5538193 | 0.01065485 | 0.04718987 |
| TULP1 | 8.50824163 | -0.108312192 | 0.5356003 | -0.2022258 | 0.83974022 | 0.91484357 |
| AL033519.3 | 0.48607992 | 0.152981963 | 2.78621562 | 0.05490672 | 0.95621278 | NA |
| FKBP5 | 8969.11222 | -1.024063733 | 0.60411513 | -1.6951467 | 0.0900476 | 0.22072461 |
| AL157823.2 | 7.68887159 | 0.220384752 | 0.59107815 | 0.37285214 | 0.7092585 | 0.83473831 |
| ARMC12 | 79.9270434 | -0.764198274 | 0.48760087 | -1.5672619 | 0.11705348 | 0.26537355 |
| CLPSL2 | 2.14488911 | -4.321873092 | 1.91265123 | -2.2596242 | 0.02384458 | 0.08529322 |
| CLPSL1 | 0.77393049 | -2.854840992 | 2.09970314 | -1.3596403 | 0.17394378 | NA |
| CLPS | 0 | NA | NA | NA | NA | NA |
| LHFPL5 | 670.402365 | -0.633981877 | 0.20128503 | -3.1496723 | 0.00163454 | 0.01156189 |
| SRPK1 | 2682.87112 | -0.302365286 | 0.21368727 | -1.4149897 | 0.15707152 | 0.32402221 |
| SLC26A8 | 33.7767423 | -0.653759214 | 0.49402882 | -1.323322 | 0.18572831 | 0.36311207 |
| MAPK14 | 4034.65209 | -0.69988031 | 0.21662685 | -3.2308106 | 0.0012344 | 0.00925535 |
| MAPK13 | 2734.18641 | -1.002286334 | 0.14596194 | -6.8667647 | 6.57E-12 | 1.40E-09 |
| Z84485.1 | 57.9424729 | 0.682237751 | 0.52405894 | 1.30183401 | 0.19297313 | 0.37249168 |
| BRPF3 | 3175.85232 | 0.587172128 | 0.26667385 | 2.20183615 | 0.02767688 | 0.09529174 |
| PNPLA1 | 236.323112 | 4.373756281 | 0.60350603 | 7.24724538 | 4.25E-13 | 1.29E-10 |
| C6orf222 | 26.1586479 | -0.360584339 | 0.75104385 | -0.4801109 | 0.63114854 | 0.78075653 |
| ETV7 | 291.004408 | -2.805126782 | 0.84457514 | -3.3213466 | 0.00089584 | 0.00719785 |
| Z84484.1 | 6.22906127 | -0.789029348 | 0.77752248 | -1.0147994 | 0.31020146 | 0.50725613 |
| PXT1 | 6.55682802 | -1.651053217 | 0.70124605 | -2.3544564 | 0.01854982 | 0.07104703 |
| KCTD20 | 2704.18038 | -0.21474427 | 0.17090988 | -1.2564766 | 0.20894322 | 0.39217671 |
| STK38 | 3195.81784 | -0.225862296 | 0.1534439 | -1.4719536 | 0.14103342 | 0.30135737 |
| SRSF3 | 9489.44461 | -0.213672132 | 0.16580483 | -1.2886966 | 0.19750357 | 0.37830821 |
| PANDAR | 2.07923287 | -0.603106668 | 1.09535885 | -0.5506019 | 0.5819066 | 0.74445491 |
| CDKN1A | 12326.03 | -2.628553502 | 0.38885768 | -6.75968 | 1.38E-11 | 2.73E-09 |
| RAB44 | 25.7550291 | -1.593665278 | 0.67858828 | -2.348501 | 0.01884915 | 0.07188026 |
| CPNE5 | 304.889914 | 0.156508462 | 0.2198853 | 0.71177318 | 0.47660524 | 0.66179458 |
| PPIL1 | 1124.94539 | -0.274566882 | 0.17073749 | -1.6081229 | 0.10780826 | 0.25027764 |
| C6orf89 | 5678.86071 | -0.201196414 | 0.14155648 | -1.4213154 | 0.15522507 | 0.32144536 |
| AL122034.1 | 0.26049542 | -0.178572513 | 2.98769512 | -0.0597693 | 0.95233936 | NA |
| PI16 | 161.477056 | 1.680466897 | 0.84217598 | 1.99538688 | 0.0460007 | 0.13703553 |
| MTCH1 | 15050.893 | -0.465395676 | 0.16766932 | -2.7756758 | 0.00550871 | 0.02898338 |
| FGD2 | 628.515062 | -0.392562905 | 0.55320075 | -0.7096211 | 0.47793915 | 0.66294542 |
| PIM1 | 2828.12488 | -0.813382365 | 0.31187901 | -2.6080061 | 0.00910713 | 0.04202189 |
| TMEM217 | 177.86299 | -0.857275928 | 0.53920638 | -1.5898846 | 0.11186082 | 0.25706054 |
| TBC1D22B | 1465.28531 | -0.411362661 | 0.19925136 | -2.0645413 | 0.03896642 | 0.12151661 |
| RNF8 | 1209.0777 | -0.575260375 | 0.16678102 | -3.4491956 | 0.00056226 | 0.00497543 |
| CMTR1 | 2917.21481 | -0.052449514 | 0.33695274 | -0.1556584 | 0.87630231 | 0.93555133 |
| CCDC167 | 332.937569 | -0.227369367 | 0.24306377 | -0.9354309 | 0.34956627 | 0.54721682 |
| LINC02520 | 2.15680127 | 2.349329566 | 1.54491961 | 1.52068079 | 0.12833996 | 0.28245721 |
| AL353597.1 | 0.86019791 | 0.71497577 | 2.18370415 | 0.32741421 | 0.74335462 | NA |
| MDGA1 | 301.307677 | 0.97819974 | 0.40406061 | 2.42092329 | 0.01548114 | 0.06226934 |
| AL121574.1 | 9.83517394 | 2.224860421 | 0.54315746 | 4.09616105 | 4.20E-05 | 0.00064638 |
| ZFAND3 | 5056.28793 | -0.059652982 | 0.13185661 | -0.452408 | 0.65097509 | 0.79488544 |
| BTBD9 | 1274.37842 | 0.128831406 | 0.13697942 | 0.94051653 | 0.34695267 | 0.54452696 |
| BTBD9-AS1 | 0.73905284 | 0.940320547 | 1.52713573 | 0.6157413 | 0.53806527 | NA |
| GLO1 | 4844.09142 | -0.108626684 | 0.14186019 | -0.7657306 | 0.44383662 | 0.6348089 |
| AL391415.1 | 0.41728634 | 0.310172637 | 2.20509832 | 0.14066159 | 0.88813729 | NA |
| DNAH8 | 13.3428671 | 0.984190816 | 0.70584653 | 1.39434108 | 0.16321463 | 0.33240422 |
| AL034345.2 | 1.14246798 | 0.749394422 | 2.23955323 | 0.33461782 | 0.7379134 | 0.85283548 |
| GLP1R | 10.7522877 | 0.513978673 | 0.64380948 | 0.79833971 | 0.42467338 | 0.61707279 |
| SAYSD1 | 1003.83727 | -0.570949319 | 0.13951077 | -4.0925107 | 4.27E-05 | 0.00065385 |
| KCNK5 | 6278.99768 | -0.249658393 | 0.50218278 | -0.4971465 | 0.61908577 | 0.77236923 |
| KCNK17 | 5.90196898 | 0.690096126 | 0.69226978 | 0.99686011 | 0.31883241 | 0.5160893 |
| KCNK16 | 0.19809512 | 0.689153972 | 3.13478683 | 0.21984078 | 0.82599516 | NA |
| KIF6 | 118.111933 | 0.008329728 | 0.61680695 | 0.0135046 | 0.98922522 | 0.99515008 |
| DAAM2 | 1924.71787 | 0.590320933 | 0.29476142 | 2.0027076 | 0.04520868 | 0.13525675 |
| AL590999.1 | 464.876738 | 0.641559098 | 0.35758911 | 1.79412369 | 0.07279343 | 0.18969885 |
| MOCS1 | 4154.49855 | 0.247947774 | 0.24509654 | 1.01163311 | 0.31171351 | 0.50879963 |
| AL139275.2 | 16.2530561 | 1.050074482 | 0.74966853 | 1.40071837 | 0.16129831 | 0.32996583 |
| TDRG1 | 0.34416416 | 0.737030442 | 2.90133682 | 0.25403133 | 0.79947136 | NA |
| LINC00951 | 0.31777585 | -0.280321477 | 2.95159789 | -0.0949728 | 0.92433644 | NA |
| AL139275.1 | 0.22085762 | 0.689157565 | 3.05316003 | 0.22571944 | 0.82141964 | NA |
| LRFN2 | 2.46164529 | 1.971940121 | 0.91582667 | 2.1531805 | 0.0313045 | 0.1040983 |
| AL583854.1 | 0 | NA | NA | NA | NA | NA |
| UNC5CL | 9834.78133 | -0.348757629 | 0.43570887 | -0.8004373 | 0.42345748 | 0.61601885 |
| OARD1 | 3200.80263 | 0.204549857 | 0.20806379 | 0.98311127 | 0.32555268 | 0.52314589 |
| TSPO2 | 28.6439053 | -0.687041174 | 0.41551252 | -1.6534789 | 0.09823344 | 0.23472416 |
| APOBEC2 | 10.1161084 | 0.259394024 | 0.54563744 | 0.4753963 | 0.63450453 | 0.78332792 |
| NFYA | 929.029847 | -0.066036224 | 0.15070282 | -0.4381884 | 0.66124973 | 0.80071801 |
| TREML1 | 18.5903254 | -0.743191704 | 0.54117584 | -1.3732906 | 0.16966202 | 0.34142151 |
| TREM2 | 82.1945108 | -1.211919347 | 0.56146692 | -2.1584875 | 0.03088995 | 0.10301857 |
| TREML2 | 19.9008015 | 0.05388882 | 0.65501181 | 0.08227152 | 0.9344308 | 0.96855234 |
| TREML4 | 2.08974644 | -4.270940578 | 1.42362775 | -3.0000403 | 0.00269944 | 0.01695266 |
| TREM1 | 50.9483275 | -0.160550449 | 0.60432311 | -0.2656699 | 0.79049346 | 0.88620494 |
| NCR2 | 0.78682232 | -1.856838434 | 3.25533655 | -0.5703983 | 0.56840758 | NA |
| FOXP4-AS1 | 12.9625213 | 1.433742632 | 0.52115909 | 2.75106518 | 0.00594018 | 0.03057634 |
| LINC01276 | 0.52212323 | -2.285276025 | 2.20857281 | -1.0347298 | 0.30079513 | NA |
| FOXP4 | 3412.26646 | -0.07069363 | 0.16399222 | -0.4310792 | 0.66641081 | 0.80448879 |
| MDFI | 329.964512 | -1.905904872 | 0.39114947 | -4.8725743 | 1.10E-06 | 3.53E-05 |
| TFEB | 1806.37813 | -0.050687996 | 0.22540953 | -0.2248707 | 0.82207988 | 0.90491928 |
| AL035588.1 | 5.46930165 | 0.657982756 | 0.57774897 | 1.13887309 | 0.25475609 | 0.44753011 |
| PGC | 3.45241253 | -0.050419295 | 0.68966956 | -0.0731065 | 0.94172141 | 0.97233647 |
| AL365205.3 | 0.81322897 | 1.369217371 | 1.50465534 | 0.90998738 | 0.36282917 | NA |
| FRS3 | 538.232904 | 0.069637794 | 0.25556779 | 0.27248267 | 0.78525091 | 0.88280468 |
| AL365205.1 | 5339.17215 | -0.098128883 | 0.10937612 | -0.8971691 | 0.36962867 | 0.56639596 |

| | | | | | | | |
|---|---|---|---|---|---|---|---|
| PRICKLE4 | 1125.94334 | | 0.095982515 | 0.31539946 | 0.30432048 | 0.76088375 | 0.86746668 |
| TOMM6 | 4057.01275 | | -0.148029487 | 0.12884076 | -1.1489337 | 0.25058334 | 0.44275742 |
| USP49 | 421.387345 | | -0.360630881 | 0.17937638 | -2.0104703 | 0.04438144 | 0.13347274 |
| AL365205.2 | 17.0481611 | | -0.012557332 | 0.35450241 | -0.0354224 | 0.97174291 | 0.98670705 |
| AL160163.1 | 0.09635527 | | 0.547239394 | 3.29377202 | 0.16614368 | 0.86804388 | NA |
| MED20 | 1063.95657 | | 0.028833845 | 0.13559051 | 0.21265385 | 0.83159696 | 0.9104925 |
| BYSL | 670.925445 | | -0.657663308 | 0.28192905 | -2.3327263 | 0.01966251 | 0.07396257 |
| CCND3 | 3577.62549 | | -0.698762332 | 0.27530282 | -2.538159 | 0.01114373 | 0.04882517 |
| AL513008.1 | 2.64756933 | | 0.434877613 | 0.77541278 | 0.5608337 | 0.57491091 | 0.73915805 |
| TAF8 | 1565.28538 | | 0.069636554 | 0.12222314 | 0.56974936 | 0.56884771 | 0.73438334 |
| AL512274.1 | 62.3676402 | | -1.486941885 | 0.41438308 | -3.5883267 | 0.00033281 | 0.00329187 |
| C6orf132 | 612.059322 | | -0.299147162 | 0.29090538 | -1.0283315 | 0.30379392 | 0.50069036 |
| GUCA1A | 0.73551557 | | -0.319995878 | 1.37200489 | -0.2332323 | 0.81558101 | NA |
| AL096814.1 | 0.17453125 | | 0.18657164 | 3.29377195 | 0.05664376 | 0.95482897 | NA |
| GUCA1B | 91.6658778 | | 0.184418838 | 0.29009651 | 0.63571548 | 0.52496189 | 0.69978996 |
| MRPS10 | 2009.95916 | | 0.088567657 | 0.1568848 | 0.56453945 | 0.57238706 | 0.73707622 |
| TRERF1 | 601.618877 | | 0.685801591 | 0.3055181 | 2.24471673 | 0.02478632 | 0.08773115 |
| UBR2 | 2569.87539 | | 0.005089587 | 0.12053812 | 0.04222388 | 0.96632023 | 0.98410057 |
| PRPH2 | 96.810364 | | 0.117593515 | 0.40212408 | 0.29243092 | 0.76995717 | 0.87279244 |
| TBCC | 840.303974 | | -0.193921373 | 0.12947443 | -1.4977581 | 0.13419611 | 0.291371 |
| BICRAL | 864.321279 | | 0.431618856 | 0.07662005 | 5.63323673 | 1.77E-08 | 1.12E-06 |
| RPL7L1 | 5053.35179 | | 0.019332852 | 0.15207807 | 0.12712452 | 0.89884184 | 0.94857081 |
| C6orf226 | 276.078469 | | -0.272243862 | 0.2709306 | -1.0048472 | 0.31497041 | 0.51209416 |
| PTCRA | 3.9982786 | | 0.671295836 | 0.80471357 | 0.83420469 | 0.40416565 | 0.59899107 |
| CNPY3 | 2272.03117 | | -0.51600582 | 0.16468417 | -3.1333056 | 0.00172849 | 0.0120583 |
| AL035587.1 | 90.1050508 | | -0.347840026 | 0.20940545 | -1.6610839 | 0.09669659 | 0.23208238 |
| GNMT | 48.4388734 | | 0.71140586 | 0.36488312 | 1.94968148 | 0.0512141 | 0.14821287 |
| PEX6 | 2856.6467 | | 0.126999137 | 0.28985821 | 0.43814227 | 0.66128315 | 0.80071801 |
| PPP2R5D | 2650.79817 | | -0.05502844 | 0.1368004 | -0.4022535 | 0.68749747 | 0.81947556 |
| MEA1 | 3908.83364 | | -0.222653 | 0.11568437 | -1.9246594 | 0.05427199 | 0.15425001 |
| KLHDC3 | 4872.78306 | | -0.13605136 | 0.12612092 | -1.0787375 | 0.28070477 | 0.4763946 |
| RRP36 | 1751.03952 | | -0.331882227 | 0.17539043 | -1.8922482 | 0.05845792 | 0.1625235 |
| AL136304.1 | 94.5564925 | | -0.022902338 | 0.1917421 | -0.1194435 | 0.90492404 | 0.95226653 |
| CUL7 | 2716.30332 | | 0.389593951 | 0.14857227 | 2.62225206 | 0.00873508 | 0.04067124 |
| KLC4 | 3846.0071 | | 0.869616186 | 0.17392528 | 4.99994131 | 5.73E-07 | 2.04E-05 |
| MRPL2 | 2382.36158 | | 0.007724415 | 0.1704663 | 0.04531344 | 0.96385747 | 0.98305633 |
| AL355385.1 | 542.428519 | | 0.669158554 | 0.18987774 | 3.52415482 | 0.00042484 | 0.00399819 |
| PTK7 | 2709.51289 | | -0.075496686 | 0.28973592 | -0.2605707 | 0.7944236 | 0.88857752 |
| SRF | 2676.75932 | | -0.262869459 | 0.2060137 | -1.2759805 | 0.20196243 | 0.38381012 |
| CUL9 | 2434.38079 | | 0.047514603 | 0.1281536 | 0.37076292 | 0.71081412 | 0.83571128 |
| AL133375.1 | 671.305613 | | -0.239769948 | 0.14605172 | -1.6416784 | 0.10065668 | 0.23858837 |
| DNPH1 | 3106.26291 | | 0.615964465 | 0.17801232 | 3.46023498 | 0.0005397 | 0.00481204 |
| TTBK1 | 190.122266 | | 1.6694563 | 0.52472482 | 3.18158437 | 0.00146472 | 0.01060546 |
| SLC22A7 | 8051.85222 | | 1.52572261 | 0.71890706 | 2.12228075 | 0.03381417 | 0.10989397 |
| CRIP3 | 1253.23763 | | 0.55325494 | 0.51649268 | 1.07117673 | 0.28408997 | 0.48012899 |
| ZNF318 | 1354.92386 | | 0.02781431 | 0.18841843 | 0.1476199 | 0.88264275 | 0.93902736 |
| ABCC10 | 1665.17414 | | 0.149849785 | 0.17070703 | 0.8778185 | 0.38004223 | 0.5759962 |
| DLK2 | 106.226225 | | -0.332156609 | 0.30619659 | -1.0847822 | 0.27801814 | 0.47353948 |
| TJAP1 | 2690.01621 | | -0.20326685 | 0.14449064 | -1.4067821 | 0.159492 | 0.32726678 |
| LRRC73 | 219.958569 | | -0.581181899 | 0.2376091 | -2.4459581 | 0.01444678 | 0.0592383 |
| POLR1C | 2557.04152 | | -0.430013002 | 0.18008128 | -2.3878829 | 0.01694574 | 0.06651086 |
| YIPF3 | 6447.22012 | | -0.243982409 | 0.12198324 | -2.0001306 | 0.04548616 | 0.1359019 |
| AL355802.2 | 392.886104 | | -0.453407225 | 0.26672442 | -1.6999089 | 0.08914806 | 0.21908987 |
| XPO5 | 1896.50268 | | -0.654515565 | 0.1514507 | -4.3216411 | 1.55E-05 | 0.00029109 |
| POLH | 427.84606 | | 0.277982228 | 0.1518901 | 1.83015373 | 0.06722695 | 0.1789597 |
| POLH-AS1 | 20.2834284 | | -0.736621468 | 0.33611029 | -2.1916064 | 0.02840794 | 0.09715307 |
| GTPBP2 | 4622.39631 | | -1.116406595 | 0.26533588 | -4.2075221 | 2.58E-05 | 0.0004356 |
| MAD2L1BP | 1157.99063 | | -0.591787999 | 0.21983246 | -2.6919955 | 0.00710259 | 0.03501141 |
| RSPH9 | 1426.87529 | | -0.229912425 | 0.17185743 | -1.337809 | 0.1809587 | 0.35712326 |
| MRPS18A | 1984.46027 | | -0.192769087 | 0.16897673 | -1.1408026 | 0.25395208 | 0.44656685 |
| AL136131.2 | 2.65995382 | | 0.624388813 | 0.92943394 | 0.67179472 | 0.50171439 | 0.68226509 |
| VEGFA | 21120.025 | | -0.384322564 | 0.30303561 | -1.2682422 | 0.20471146 | 0.38708977 |
| AL136131.3 | 912.802366 | | 0.199719507 | 0.42328566 | 0.4718315 | 0.63704707 | 0.78513112 |
| AL157371.2 | 2.73530009 | | -1.189287887 | 1.19631614 | -0.9941251 | 0.32016197 | 0.51758692 |
| LINC02537 | 0.61251436 | | 1.875308487 | 1.96916113 | 0.95233877 | 0.3409252 | NA |
| AL157371.1 | 0 | NA | | NA | NA | NA | NA |
| AL109615.3 | 134.585574 | | -0.515333027 | 0.29028632 | -1.7752577 | 0.07585534 | 0.19534232 |
| C6orf223 | 451.207771 | | -1.032459509 | 0.38683592 | -2.6689856 | 0.00760807 | 0.03679839 |
| AL109615.2 | 1.4704666 | | -2.049783291 | 1.36125192 | -1.5058075 | 0.13211661 | 0.28824615 |
| MRPL14 | 1586.6003 | | -0.664282644 | 0.21534242 | -3.0847737 | 0.00203707 | 0.01368251 |
| TMEM63B | 3220.06786 | | -0.375545638 | 0.16249378 | -2.3111386 | 0.0208252 | 0.07715097 |
| CAPN11 | 28.5739219 | | -0.346171259 | 0.6237215 | -0.5550093 | 0.57888827 | 0.74240002 |
| MYMX | 17.3519077 | | -0.148396638 | 0.70106766 | -0.2116723 | 0.83236266 | 0.91099043 |
| SLC29A1 | 4076.59959 | | -0.633474742 | 0.24921273 | -2.5419036 | 0.01102506 | 0.04844705 |
| HSP90AB1 | 41981.5916 | | -0.366502704 | 0.14983108 | -2.446106 | 0.01444085 | 0.05922322 |
| SLC35B2 | 3885.05053 | | -0.431400286 | 0.14413113 | -2.9931097 | 0.0027615 | 0.01725204 |
| NFKBIE | 1389.82759 | | -1.180279519 | 0.22862199 | -5.1625809 | 2.44E-07 | 9.97E-06 |
| TMEM151B | 104.679011 | | 0.804369619 | 0.21187147 | 3.79649799 | 0.00014675 | 0.00172149 |
| AL353588.1 | 102.742965 | | 0.802817826 | 0.23205122 | 3.45965786 | 0.00054086 | 0.00482005 |
| TCTE1 | 14.6642394 | | -0.176866836 | 0.62736122 | -0.2819219 | 0.77800343 | 0.87847694 |
| AARS2 | 1165.10111 | | 0.450988835 | 0.17382524 | 2.59449565 | 0.00947298 | 0.0432815 |
| SPATS1 | 2.73268129 | | -3.521242063 | 2.33884605 | -1.5055467 | 0.13218357 | 0.28834008 |
| CDC5L | 1654.03909 | | -0.072399329 | 0.14224527 | -0.5089753 | 0.61076954 | 0.76665998 |
| AL136140.1 | 0.08757577 | | 0.547239394 | 3.29377202 | 0.16614368 | 0.86804388 | NA |
| SUPT3H | 403.740966 | | 0.140604701 | 0.20286235 | 0.69310396 | 0.48824431 | 0.67176263 |
| RUNX2 | 232.126274 | | -0.309792419 | 0.27426277 | -1.129546 | 0.25866756 | 0.45145198 |
| AL096865.1 | 30.0973503 | | -0.596102116 | 0.52560383 | -1.1341282 | 0.25674079 | 0.44945587 |
| RUNX2-AS1 | 1.27755575 | | -1.66930186 | 1.96561379 | -0.8492522 | 0.39574097 | 0.59105626 |
| CLIC5 | 3829.44058 | | 0.92101953 | 0.64690803 | 1.42372562 | 0.15452592 | 0.32057654 |
| AL035701.1 | 26.747774 | | 0.365520796 | 0.40292526 | 0.90716774 | 0.3643181 | 0.5614885 |
| ENPP4 | 2051.73113 | | -0.15170777 | 0.21289482 | -0.7125949 | 0.47609644 | 0.66150263 |
| ENPP5 | 1783.18645 | | 0.254909389 | 0.25942428 | 0.98259648 | 0.32580608 | 0.52348929 |
| RCAN2 | 1762.2055 | | 0.864676979 | 0.33728189 | 2.56366263 | 0.01035742 | 0.0462767 |
| AL359633.2 | 1.50466677 | | 2.163553666 | 1.30739409 | 1.65485961 | 0.09795298 | 0.23426629 |
| CYP39A1 | 335.383786 | | 0.225743291 | 0.38980947 | 0.57911187 | 0.56251369 | 0.72998405 |
| SLC25A27 | 1722.38517 | | -0.116734216 | 0.37061669 | -0.3149729 | 0.75278223 | 0.86218152 |
| AL591242.1 | 0.07659028 | | -0.642905668 | 3.30558948 | -0.1944905 | 0.84579185 | NA |
| TDRD6 | 281.951096 | | -0.075185415 | 0.38561772 | -0.194974 | 0.84541333 | 0.91848836 |
| PLA2G7 | 54.72853 | | -1.394705203 | 0.64070856 | -2.1768169 | 0.02949423 | 0.09970042 |
| ANKRD66 | 1.23630444 | | -3.522849739 | 2.71826051 | -1.2959942 | 0.1949775 | 0.37502236 |
| AL161618.1 | 0.7039133 | | -0.538136468 | 2.85534049 | -0.1884667 | 0.85051085 | NA |
| MEP1A | 0.69923639 | | -1.740564898 | 1.622976 | -1.0724526 | 0.28351677 | NA |
| ADGRF5 | 6356.72583 | | 0.343665893 | 0.3930067 | 0.87445302 | 0.38187159 | 0.57780319 |
| AL096772.1 | 0.5264487 | | 1.099975914 | 1.75955977 | 0.62514268 | 0.53187742 | NA |
| ADGRF1 | 3659.76947 | | -0.769219662 | 0.32652741 | -2.3557584 | 0.01848494 | 0.07083968 |
| TNFRSF21 | 3974.01319 | | 0.899099484 | 0.30124367 | 2.98462528 | 0.00283926 | 0.01757541 |
| AL355353.1 | 100.513143 | | 0.160616134 | 0.40305993 | 0.39849194 | 0.6902676 | 0.8215683 |
| CD2AP | 4139.39757 | | -0.300700007 | 0.169948 | -1.7693647 | 0.07683303 | 0.19685902 |
| ADGRF2 | 10.7960419 | | 3.640252329 | 1.14620587 | 3.17591492 | NA | NA |
| ADGRF4 | 19.0588145 | | 3.750562379 | 1.0832963 | 3.46217594 | 0.00053583 | 0.00478513 |
| AL356421.2 | 0 | NA | | NA | NA | NA | NA |
| OPN5 | 1.43109528 | | -2.86489824 | 1.59486814 | -1.796323 | 0.07244317 | 0.18914123 |
| PTCHD4 | 74.0892496 | | -1.065108901 | 0.54510678 | -1.9539454 | 0.05070767 | 0.14703894 |

| | | | | | | |
|---|---|---|---|---|---|---|
| MUT | 3452.71392 | 0.982108677 | 0.24874294 | 3.94828768 | 7.87E-05 | 0.00104936 |
| CENPQ | 133.445288 | -0.097952451 | 0.18331145 | -0.5343499 | 0.5930995 | 0.75237031 |
| GLYATL3 | 0 | NA | NA | NA | NA | NA |
| C6orf141 | 36.6778307 | -0.554338189 | 0.62935791 | -0.8807996 | 0.3784263 | 0.57443597 |
| RHAG | 11.6203793 | -3.665575968 | 1.37072035 | -2.6741968 | 0.00749085 | 0.03642516 |
| CRISP2 | 19.4023049 | 1.655616285 | 0.86896335 | 1.90527746 | 0.05674401 | 0.15912331 |
| CRISP3 | 2.82889966 | -4.720276551 | 1.83786296 | -2.5683507 | 0.01021837 | 0.04580291 |
| PGK2 | 0.03392908 | 0.380740593 | 3.29738709 | 0.11546736 | 0.90807469 | NA |
| AL121974.1 | 11.7655657 | 2.479188049 | 0.73111039 | 3.39099003 | 0.00069641 | 0.00588032 |
| AL359458.1 | 0 | NA | NA | NA | NA | NA |
| CRISP1 | 0.68569526 | 3.363073487 | 1.84237792 | 1.82539828 | 0.06794095 | NA |
| DEFB133 | 0 | NA | NA | NA | NA | NA |
| DEFB114 | 0 | NA | NA | NA | NA | NA |
| DEFB113 | 0 | NA | NA | NA | NA | NA |
| DEFB110 | 0 | NA | NA | NA | NA | NA |
| DEFB112 | 0 | NA | NA | NA | NA | NA |
| AL138826.1 | 460.415648 | 4.162072946 | 0.94250361 | 4.41597559 | 1.01E-05 | 0.00020429 |
| AL360175.1 | 0.19978473 | -1.003574173 | 3.30558948 | -0.3035992 | 0.7614333 | NA |
| AL135908.1 | 0 | NA | NA | NA | NA | NA |
| TFAP2D | 0 | NA | NA | NA | NA | NA |
| TFAP2B | 1298.23021 | 0.714726423 | 0.28098549 | 2.54364175 | 0.01097035 | 0.04827102 |
| AL355997.1 | 25.4438741 | 1.749966279 | 0.31440023 | 5.5660464 | 2.61E-08 | 1.55E-06 |
| PKHD1 | 3962.58985 | 0.97192863 | 0.3937503 | 2.46838827 | 0.0135723 | 0.05651375 |
| LINCMD1 | 2.22527321 | 2.161942577 | 0.93933997 | 2.30155497 | 0.02136028 | 0.07872406 |
| IL17A | 0.18340304 | -1.003574173 | 3.30558948 | -0.3035992 | 0.7614333 | NA |
| IL17F | 1.5218162 | 0.120439809 | 1.27545662 | 0.09442878 | 0.92476856 | 0.96356293 |
| MCM3 | 2207.36388 | -0.017287212 | 0.2048307 | -0.0843976 | 0.93274034 | 0.96756079 |
| PAQR8 | 633.548537 | 0.925579289 | 0.38015504 | 2.4347416 | 0.01490243 | 0.06055116 |
| EFHC1 | 1488.68235 | -0.100741143 | 0.21996762 | -0.4579817 | 0.64696559 | 0.79207403 |
| TRAM2 | 1381.82847 | 0.016160686 | 0.19091025 | 0.0846507 | 0.93253909 | 0.96746494 |
| TRAM2-AS1 | 435.419103 | 0.959474309 | 0.23213417 | 4.13327479 | 3.58E-05 | 0.00056675 |
| TMEM14A | 1069.68938 | 0.74949291 | 0.24123605 | 3.106886 | 0.00189069 | 0.01291312 |
| GSTA2 | 7055.72751 | 4.129919273 | 0.91362626 | 4.52035963 | 6.17E-06 | 0.00013955 |
| GSTA1 | 8552.90663 | 2.870042963 | 0.71806574 | 3.99690838 | 6.42E-05 | 0.00089682 |
| GSTA5 | 7.48465238 | 2.329910035 | 0.72631279 | 3.20786042 | 0.00133726 | 0.00989163 |
| GSTA3 | 2.01531098 | -0.281936316 | 1.23982675 | -0.2273998 | 0.8201129 | 0.90347066 |
| GSTA4 | 1027.34402 | 0.615095288 | 0.27566979 | 2.23127567 | 0.02566287 | 0.09001333 |
| ICK | 594.590522 | 0.201583274 | 0.16256781 | 1.239995 | 0.21497724 | 0.39924994 |
| FBXO9 | 3801.70299 | 0.363130959 | 0.12606504 | 2.88050476 | 0.00397039 | 0.02263601 |
| GCM1 | 118.706486 | 1.798381315 | 0.40866745 | 4.40059838 | 1.08E-05 | 0.00021714 |
| ELOVL5 | 3878.15176 | -0.214519607 | 0.25554995 | -0.839443 | 0.40122079 | 0.59613433 |
| AL034374.1 | 6.65742499 | 2.828527657 | 0.86287067 | 3.27804359 | 0.00104529 | 0.00811911 |
| GCLC | 4179.47118 | -1.154785575 | 0.52356561 | -2.2056177 | 0.02741078 | 0.09457705 |
| AL033397.2 | 233.842061 | -1.111145958 | 0.55755394 | -1.9928941 | 0.04627304 | 0.13764469 |
| AL033397.1 | 118.901064 | 0.580037538 | 0.77155521 | 0.7517771 | 0.45218511 | 0.64184097 |
| LINC01564 | 122.022734 | -2.157178662 | 0.75147133 | -2.8706068 | 0.00409685 | 0.02316336 |
| KLHL31 | 73.9811328 | -0.53409303 | 0.32341031 | -1.651441 | 0.09864857 | 0.23546006 |
| AL590652.1 | 25.4157641 | 1.153751927 | 0.54305254 | 2.12456778 | 0.0336227 | 0.10946055 |
| AL513211.1 | 15.6218972 | 0.810006655 | 0.51147968 | 1.58365364 | 0.11327256 | 0.25928797 |
| LRRC1 | 857.261328 | 0.021324786 | 0.16875618 | 0.12636447 | 0.89944342 | 0.94902072 |
| AL033384.2 | 169.580998 | 0.137290608 | 0.18711766 | 0.73371271 | 0.46312385 | 0.65105787 |
| MLIP | 23.4301802 | 0.926662732 | 0.37989277 | 2.43927448 | 0.01471679 | 0.06001881 |
| AL033384.1 | 22.2533767 | 0.320346089 | 0.54628743 | 0.58640575 | 0.55760288 | 0.72636855 |
| MLIP-AS1 | 0.28299204 | 1.12993107 | 2.97686151 | 0.37957126 | 0.7042637 | NA |
| TINAG | 3134.53949 | 0.868630842 | 0.39429007 | 2.2030249 | 0.02759299 | 0.09506496 |
| AL589946.1 | 0.19718306 | -0.340099831 | 3.16382928 | -0.1074963 | 0.91439529 | NA |
| AL512363.1 | 9.4306611 | 0.317648529 | 0.77543765 | 0.40963774 | 0.6820717 | 0.81588709 |
| FAM83B | 99.0424157 | -0.61172986 | 0.32718261 | -1.8696894 | 0.06152696 | 0.1683056 |
| HCRTR2 | 8.42118044 | 4.626460768 | 0.76596394 | 6.04005036 | 1.54E-09 | 1.46E-07 |
| GFRAL | 0.31692112 | 1.414901731 | 2.2605791 | 0.62590233 | 0.531379 | NA |
| HMGCLL1 | 36.9020638 | 1.23618322 | 0.28063133 | 4.4050079 | 1.06E-05 | 0.00021391 |
| BMP5 | 110.409934 | 1.770331478 | 0.68959548 | 2.56720283 | 0.01025226 | 0.045916 |
| COL21A1 | 117.317459 | 0.382673678 | 0.62655106 | 0.61076216 | 0.54135705 | 0.7142078 |
| DST | 5219.56645 | 0.128548179 | 0.23563376 | 0.54554228 | 0.58538061 | 0.74695087 |
| AL512422.1 | 5.80408058 | -0.88315535 | 0.87386589 | -1.0106303 | 0.31219341 | 0.50910117 |
| BEND6 | 51.4994662 | -0.434517209 | 0.29071235 | -1.4946638 | 0.13500222 | 0.29242691 |
| KIAA1586 | 270.668803 | -0.167830887 | 0.17641872 | -0.9513213 | 0.34144129 | 0.53944082 |
| ZNF451 | 1605.40466 | 0.007719475 | 0.10245605 | 0.07534426 | 0.9399408 | 0.97124761 |
| ZNF451-AS1 | 199.794493 | 0.518181409 | 0.18995699 | 2.72788811 | 0.00637412 | 0.032225 |
| BAG2 | 1012.6227 | -0.332836221 | 0.28359874 | -1.1736167 | 0.24054858 | 0.43069432 |
| RAB23 | 668.830686 | -0.031322959 | 0.24615625 | -0.1272483 | 0.8987439 | 0.94852624 |
| PRIM2 | 268.054245 | -0.091184238 | 0.1295339 | -0.7039411 | 0.48146944 | 0.66600139 |
| FO680682.1 | 1.35245026 | -0.517097082 | 1.24296254 | -0.4160198 | 0.67739548 | 0.81243093 |
| AL021368.3 | 24.5830785 | 0.525059594 | 0.37726525 | 1.39175182 | 0.16399756 | 0.33348363 |
| AL021368.1 | 6.5696797 | 0.614790361 | 0.71452618 | 0.86041685 | 0.3895593 | 0.58520444 |
| AL021368.2 | 199.426426 | 0.251538402 | 0.14340634 | 1.75402561 | 0.07942612 | 0.20139103 |
| AL445250.1 | 15.948713 | 0.25722735 | 0.38340424 | 0.67090376 | 0.50228184 | 0.68263189 |
| AL590227.1 | 0 | NA | NA | NA | NA | NA |
| KHDRBS2 | 42.284472 | 1.190351708 | 0.36284241 | 3.28063 | 0.00103576 | 0.00806641 |
| FKBP1C | 37.2172228 | -0.829583954 | 0.34006397 | -2.4394938 | 0.01470785 | 0.06001023 |
| LGSN | 149.890188 | 3.436634 | 0.59879838 | 5.73921727 | 9.51E-09 | 6.64E-07 |
| PTP4A1 | 9349.53086 | -1.011344194 | 0.25864845 | -3.9101111 | 9.23E-05 | 0.00118728 |
| AL135905.1 | 62.6881749 | 0.321098733 | 0.2509015 | 1.27978005 | 0.2006225 | 0.38215108 |
| PHF3 | 2632.61336 | 0.066503594 | 0.1138857 | 0.58395033 | 0.55925373 | 0.72772184 |
| EYS | 53.320094 | 0.130970411 | 0.20366585 | 0.64306516 | 0.52018182 | 0.69687028 |
| AL354719.2 | 14.3062461 | -2.035354099 | 0.57811851 | -3.520652 | 0.00043049 | 0.0040384 |
| AL357375.1 | 0 | NA | NA | NA | NA | NA |
| AL713998.1 | 4.19661726 | 4.795556013 | 0.93130512 | 5.14928556 | 2.61E-07 | 1.06E-05 |
| AL646090.1 | 0.08666371 | -0.642905668 | 3.30558948 | -0.1944905 | 0.84579185 | NA |
| LINC02549 | 0 | NA | NA | NA | NA | NA |
| AL391807.1 | 35.8184646 | -0.00669553 | 0.47458598 | -0.0141081 | 0.9887437 | 0.99493137 |
| ADGRB3 | 57.7388262 | 0.156529591 | 0.46629622 | 0.33568702 | 0.73710689 | 0.8524157 |
| LMBRD1 | 5907.54238 | 0.248683649 | 0.14127505 | 1.76028008 | 0.07836033 | 0.19951355 |
| COL19A1 | 69.7677431 | 2.403449109 | 0.37466643 | 6.41490381 | 1.41E-10 | 1.91E-08 |
| COL9A1 | 45.1668674 | 2.202702639 | 0.67512453 | 3.26266124 | 0.00110371 | 0.00846074 |
| AL160262.1 | 4.73516202 | 2.484211867 | 0.81486563 | 3.04861534 | 0.00229899 | 0.01503798 |
| AL365226.2 | 11.3751343 | -3.138432556 | 0.97689948 | -3.2126464 | 0.00131518 | 0.00976675 |
| AL365226.1 | 7.83441273 | -0.342414509 | 0.63383302 | -0.5402283 | 0.58903963 | 0.74948009 |
| FAM135A | 1422.76849 | -0.917697373 | 0.19884738 | -4.615084 | 3.93E-06 | 9.74E-05 |
| SDHAF4 | 642.309314 | 0.148118689 | 0.24311761 | 0.60924706 | 0.54236069 | 0.71481665 |
| AL583856.2 | 20.4795539 | 0.601563308 | 0.36851662 | 1.63239126 | 0.10259709 | 0.24169066 |
| SMAP1 | 1787.97036 | -0.387212366 | 0.17290239 | -2.2394853 | 0.02512435 | 0.08870387 |
| B3GAT2 | 1619.62966 | -0.440332853 | 0.16777058 | -2.6246131 | 0.00867475 | 0.04044433 |
| OGFRL1 | 2275.41192 | -0.96535555 | 0.49594362 | -1.9465026 | 0.05159439 | 0.14910931 |
| AL136164.2 | 246.298467 | -0.335109407 | 0.3322622 | -1.0085692 | 0.3131813 | 0.51007294 |
| AL136164.1 | 0.28015841 | -0.179498944 | 2.93432484 | -0.0611721 | 0.95122211 | NA |
| LINC00472 | 556.058076 | 0.677280212 | 0.3754467 | 1.80393172 | 0.07124201 | 0.18668904 |
| LINC01626 | 0.54082319 | -2.32594509 | 2.75817312 | -0.8432919 | 0.3990652 | NA |
| RIMS1 | 7.37944664 | 0.242286545 | 0.77084038 | 0.31431481 | 0.75328195 | 0.8625291 |
| KCNQ5 | 11.7937432 | 0.331340456 | 0.69668411 | 0.4755964 | 0.63436194 | 0.7832619 |
| KCNQ5-AS1 | 0 | NA | NA | NA | NA | NA |
| AL365232.1 | 13.1687869 | 0.387880353 | 0.34141342 | 1.13610167 | 0.25591402 | 0.44861082 |
| KHDC1L | 1.33130645 | -0.74858953 | 1.09613906 | -0.682933 | 0.49464918 | 0.67694119 |

| | | | | | | |
|---|---|---|---|---|---|---|
| KHDC1 | 62.9802611 | -0.148522899 | 0.31817366 | -0.4667982 | 0.64064425 | 0.78733983 |
| AC019205.1 | 73.9620863 | 0.336165789 | 0.29567842 | 1.1369304 | 0.25556738 | 0.44821175 |
| DPPA5 | 0.68584303 | 1.148162992 | 1.51919334 | 0.75577148 | 0.44978622 | NA |
| KHDC3L | 0.41194161 | 0.30983507 | 2.21308553 | 0.1400014 | 0.88865889 | NA |
| OOEP | 18.303865 | 0.89905632 | 0.71458529 | 1.25815116 | 0.20833709 | 0.39152061 |
| OOEP-AS1 | 1.01712642 | 1.467449157 | 2.13637109 | 0.6868887 | 0.49215287 | 0.67506222 |
| DDX43 | 85.0214721 | 0.485767973 | 0.65348744 | 0.74334707 | 0.45727158 | 0.64655504 |
| CGAS | 175.920633 | -0.513733606 | 0.32344621 | -1.5883124 | 0.11221573 | 0.25767262 |
| MTO1 | 1493.87701 | 0.410974202 | 0.13945656 | 2.94696939 | 0.00320905 | 0.01934674 |
| AL603910.1 | 208.080094 | 0.215069113 | 0.16626526 | 1.29353009 | 0.19582779 | 0.37610915 |
| EEF1A1 | 332682.26 | 0.097358944 | 0.25198711 | 0.38636477 | 0.69922653 | 0.82736264 |
| AL121972.1 | 16.4537597 | 0.448600047 | 0.41696777 | 1.07586265 | 0.28198869 | 0.47782049 |
| SLC17A5 | 3605.48321 | -0.147586339 | 0.39954666 | -0.3693845 | 0.71184116 | 0.83618048 |
| AL590428.1 | 2.8811912 | -0.583166962 | 0.83961004 | -0.6945688 | 0.48732555 | 0.67123399 |
| CD109 | 1011.88182 | 1.336382727 | 0.28910172 | 4.62253466 | 3.79E-06 | 9.44E-05 |
| AL357507.1 | 0.24261338 | 0.078085528 | 3.06914044 | 0.02544215 | 0.97970229 | NA |
| AL356277.3 | 0.24118557 | 0.074529887 | 3.28898342 | 0.02266046 | 0.98192111 | NA |
| AL356277.2 | 0.03392908 | 0.380740593 | 3.29738709 | 0.11546736 | 0.90807469 | NA |
| COL12A1 | 2492.15638 | 0.63334488 | 0.42947277 | 1.47470321 | 0.14029235 | 0.30042883 |
| COX7A2 | 5538.39211 | -0.227589006 | 0.16336831 | -1.3931037 | 0.16358843 | 0.33295977 |
| TMEM30A | 7999.43361 | -0.080933721 | 0.14057016 | -0.5757532 | 0.564782 | 0.73164184 |
| AL080250.1 | 55.5302582 | 0.490792258 | 0.19281989 | 2.54534041 | 0.01091713 | 0.04810102 |
| FILIP1 | 223.714561 | -0.010837622 | 0.39897227 | -0.0271638 | 0.97832905 | 0.99019487 |
| AL445465.1 | 4.10148556 | 0.650393997 | 0.67188328 | 0.96801634 | 0.33303621 | 0.53100478 |
| AL445465.2 | 1.0309561 | 2.032072585 | 1.82628358 | 1.11268185 | 0.2658451 | 0.45996556 |
| SENP6 | 3975.2119 | 0.009116297 | 0.16129725 | 0.05651861 | 0.95492867 | 0.97841508 |
| MYO6 | 8988.52175 | -0.112895416 | 0.20220793 | -0.5583135 | 0.57663034 | 0.74062355 |
| IMPG1 | 77.3166462 | 0.462869622 | 0.25152918 | 1.84022235 | 0.0657356 | 0.17621711 |
| LINC02540 | 0.06785815 | 0.795868262 | 3.28898349 | 0.24198001 | 0.80879565 | NA |
| AL591030.1 | 0.11223674 | 0.02007176 | 3.29738704 | 0.00608717 | 0.99514317 | NA |
| AL590426.2 | 6.24534544 | 2.889303396 | 0.77120566 | 3.74647586 | 0.00017934 | 0.00200555 |
| HTR1B | 18.8308282 | 0.423113592 | 0.62025065 | 0.6821655 | 0.49513431 | 0.67731969 |
| MEI4 | 260.181702 | 1.172758056 | 0.41615324 | 2.81809187 | 0.004831 | 0.02618032 |
| AL121718.1 | 0 | NA | NA | NA | NA | NA |
| AL450327.1 | 2.47632993 | 0.621151808 | 1.10920635 | 0.55999662 | 0.57548174 | 0.73968871 |
| IRAK1BP1 | 343.849361 | 0.58446128 | 0.27506534 | 2.12480893 | 0.03360256 | 0.10943103 |
| PHIP | 1936.96085 | -0.142093198 | 0.10923292 | -1.3008276 | 0.19331748 | 0.37286292 |
| HMGN3 | 5223.95523 | 0.038612661 | 0.15356654 | 0.25143929 | 0.80147449 | 0.89240731 |
| HMGN3-AS1 | 1294.90748 | 0.235386488 | 0.14258956 | 1.65079746 | 0.09877994 | 0.23562433 |
| AL078601.2 | 18.9049968 | 1.710009644 | 0.53349043 | 3.20532393 | 0.00134911 | 0.00994965 |
| LCA5 | 291.153062 | -0.022318476 | 0.1715451 | -0.1301027 | 0.8964852 | 0.94701417 |
| AL451064.1 | 12.0117558 | 0.524186717 | 0.52786284 | 0.99303583 | 0.32069249 | 0.51819065 |
| SH3BGRL2 | 1630.02542 | 0.311347464 | 0.18689652 | 1.66588151 | 0.09573702 | 0.23063997 |
| LINC01621 | 7.49159436 | 3.621745158 | 0.84859929 | 4.2679097 | 1.97E-05 | 0.00035014 |
| AL132875.2 | 0.28933606 | 2.158099797 | 2.78663725 | 0.7744459 | 0.43866715 | NA |
| ELOVL4 | 105.310857 | 0.014128517 | 0.49532969 | 0.02852346 | 0.97724466 | 0.98951899 |
| TTK | 24.7870743 | -0.733189524 | 0.55694933 | -1.3164385 | 0.18802691 | 0.36613054 |
| BCKDHB | 1436.38154 | 1.281311704 | 0.27350092 | 4.68485328 | 2.80E-06 | 7.44E-05 |
| AL359715.3 | 61.1645355 | 0.821182311 | 0.36745756 | 2.23476777 | 0.0254326 | 0.08943613 |
| AL359715.1 | 83.5013847 | 0.282680476 | 0.47029766 | 0.60106715 | 0.54779526 | 0.71871044 |
| AL359715.2 | 89.882449 | 0.606488573 | 0.34363693 | 1.76491092 | 0.07757874 | 0.19819331 |
| TENT5A | 797.856289 | -0.214378771 | 0.34416877 | -0.6228885 | 0.53335779 | 0.70715696 |
| AL359693.1 | 0.73957597 | 1.214382051 | 1.58761443 | 0.76490993 | 0.44432518 | NA |
| LINC01526 | 0.26775652 | -1.364242679 | 3.30558948 | -0.4127078 | 0.67982068 | NA |
| LINC02542 | 6.67431821 | 0.17524829 | 0.74207977 | 0.23615829 | 0.81330984 | 0.89913137 |
| AL360157.1 | 0 | NA | NA | NA | NA | NA |
| IBTK | 2768.592 | -0.037034603 | 0.18413129 | -0.2011315 | 0.84059576 | 0.91507243 |
| TPBG | 1302.84629 | -1.629623865 | 0.2926927 | -5.5676956 | 2.58E-08 | 1.54E-06 |
| UBE3D | 237.838025 | -0.27351498 | 0.15473054 | -1.7676858 | 0.07711345 | 0.19740624 |
| AL355613.1 | 0.49576106 | -0.412613629 | 1.68093788 | -0.2454663 | 0.80609539 | NA |
| DOPEY1 | 2004.9637 | -0.090601608 | 0.17118 | -0.5292768 | 0.59661342 | 0.75532651 |
| PGM3 | 2637.7137 | -0.499331643 | 0.22196021 | -2.2496449 | 0.0244715 | 0.0868641 |
| RWDD2A | 652.592886 | 0.060446456 | 0.15232643 | 0.39682186 | 0.69149883 | 0.82241065 |
| ME1 | 670.24097 | -1.04454426 | 0.31633187 | -3.3020519 | 0.0009598 | 0.00760746 |
| PRSS35 | 125.41361 | 2.505680031 | 0.62009674 | 4.04078892 | 5.33E-05 | 0.00078348 |
| SNAP91 | 6.14797528 | -0.563604335 | 0.82761296 | -0.6809999 | 0.49587156 | 0.67772047 |
| AL136972.1 | 0 | NA | NA | NA | NA | NA |
| RIPPLY2 | 2.37741952 | 0.222700498 | 1.01053395 | 0.22037904 | 0.82557597 | 0.9068371 |
| CYB5R4 | 973.518773 | -0.375274339 | 0.22964678 | -1.6341371 | 0.10223006 | 0.24114918 |
| AL034347.1 | 0.75442765 | -1.195953398 | 1.47826193 | -0.8090267 | 0.41849978 | NA |
| AL161621.1 | 3.13586274 | -0.659488722 | 0.98945341 | -0.6665182 | 0.50507993 | 0.68449777 |
| MRAP2 | 237.618603 | -0.114751318 | 0.39082996 | -0.2936093 | 0.76905646 | 0.87240608 |
| CEP162 | 550.36208 | -0.120443658 | 0.22012184 | -0.5471681 | 0.58426322 | 0.74635572 |
| LINC01611 | 0.07830766 | -0.642905668 | 3.30558948 | -0.1944905 | 0.84579185 | NA |
| TBX18 | 68.5526361 | 0.112888675 | 0.67148185 | 0.16811873 | 0.86648988 | 0.93068869 |
| TBX18-AS1 | 2.55238507 | 1.013202186 | 0.91078502 | 1.11244933 | 0.265945 | 0.46004787 |
| LINC02535 | 382.82752 | 2.054256086 | 0.85040842 | 2.41561117 | 0.01570883 | 0.06295467 |
| NT5E | 1678.95366 | 0.341958946 | 0.26471407 | 1.29180496 | 0.1964247 | 0.37700846 |
| SNX14 | 3745.26329 | 0.147136375 | 0.13574382 | 1.08392687 | 0.27839722 | 0.47406269 |
| AL589666.1 | 1308.06381 | -0.069713532 | 0.23530549 | -0.2962682 | 0.76702528 | 0.87152635 |
| SYNCRIP | 6290.14726 | -0.302302139 | 0.22069875 | -1.3697501 | 0.17076493 | 0.34301273 |
| HTR1E | 0 | NA | NA | NA | NA | NA |
| CGA | 17.2805749 | -1.352558005 | 0.83190778 | -1.6258509 | 0.10398138 | 0.24416075 |
| AL139274.2 | 28.5112498 | 0.27903004 | 0.43253676 | 0.64510133 | 0.51886152 | 0.69573774 |
| ZNF292 | 1898.58471 | -0.982911992 | 0.3406468 | -2.8854285 | 0.00390881 | 0.02237863 |
| GJB7 | 31.1670681 | 0.373570003 | 0.23379274 | 1.59786828 | 0.11007229 | 0.25403522 |
| SMIM8 | 351.527756 | 0.296866097 | 0.21985836 | 1.35026066 | 0.17693239 | 0.35191635 |
| C6orf163 | 59.2231889 | -0.121928754 | 0.28771482 | -0.4237834 | 0.67172381 | 0.80834924 |
| CFAP206 | 114.730076 | 0.0336799 | 0.27040849 | 0.12455193 | 0.90087828 | 0.9498184 |
| AL049697.1 | 1128.51948 | -0.025589355 | 0.12137362 | -0.2108313 | 0.83301894 | 0.91136194 |
| SLC35A1 | 1584.58197 | -0.192342408 | 0.15023842 | -1.2802478 | 0.200458 | 0.38199722 |
| RARS2 | 1633.11663 | -0.238024719 | 0.12435889 | -1.9140146 | 0.05561829 | 0.156895 |
| ORC3 | 1062.06466 | 0.186533758 | 0.18818248 | 0.99123872 | 0.32156903 | 0.51927589 |
| AKIRIN2 | 1848.93332 | -0.789013461 | 0.18258431 | -4.3213651 | 1.55E-05 | 0.00029125 |
| SPACA1 | 14.3194798 | 1.336555769 | 0.60452573 | 2.21091625 | 0.02704164 | 0.09359338 |
| CNR1 | 140.966202 | -2.373804763 | 0.84006712 | -2.8257323 | 0.00471727 | 0.02573853 |
| AL139042.1 | 1.17670394 | 0.117182074 | 1.58962409 | 0.07371685 | 0.94123569 | 0.97207934 |
| RNGTT | 676.046274 | -0.137819901 | 0.15796542 | -0.8724688 | 0.38295268 | 0.57895142 |
| AL353135.1 | 2.85918196 | -0.117842236 | 0.79516545 | -0.1481984 | 0.88218621 | 0.93886715 |
| PNRC1 | 9218.20097 | -1.048937187 | 0.23131391 | -4.5346914 | 5.77E-06 | 0.00013182 |
| SRSF12 | 204.238233 | -1.896976634 | 0.27398588 | -6.9236292 | 4.40E-12 | 1.01E-09 |
| PM20D2 | 1420.30286 | -0.554736493 | 0.17820787 | -3.1128618 | 0.00185283 | 0.01269925 |
| GABRR1 | 4.34799177 | 0.551154547 | 0.61508565 | 0.89606146 | 0.37021994 | 0.56696095 |
| GABRR2 | 28.4660802 | 1.068284613 | 0.57595151 | 1.85481694 | 0.06362237 | 0.17221919 |
| UBE2J1 | 3348.75022 | -0.316358679 | 0.18756782 | -1.6866362 | 0.09167332 | 0.22352631 |
| RRAGD | 3504.71103 | -0.109239065 | 0.27303807 | -0.4000873 | 0.68909222 | 0.82063171 |
| ANKRD6 | 1553.72625 | -0.008408385 | 0.12345192 | -0.0681106 | 0.94569759 | 0.97395466 |
| AL159174.1 | 23.6195863 | 0.77257433 | 0.3563839 | 2.16781492 | 0.03017277 | 0.10124138 |
| LYRM2 | 2295.91539 | 0.100947266 | 0.13320292 | 0.75784574 | 0.44854334 | 0.63866565 |
| MDN1 | 1735.39712 | 0.156939645 | 0.11141253 | 1.40863552 | 0.15894297 | 0.32649649 |
| AL096678.1 | 7.778495 | 0.208560359 | 0.54416726 | 0.38326518 | 0.70152316 | 0.82903853 |
| CASP8AP2 | 448.088752 | 0.103925435 | 0.11376704 | 0.91349336 | 0.36098312 | 0.55866688 |
| GJA10 | 0.09989236 | -0.642905668 | 3.30558948 | -0.1944905 | 0.84579185 | NA |

| | | | | | | |
|---|---|---|---|---|---|---|
| BACH2 | 176.16525 | -0.65374862 | 0.22896915 | -2.8551821 | 0.00430122 | 0.02405579 |
| AL121787.1 | 1.98657767 | 1.033999424 | 0.8507189 | 1.21544193 | 0.22419758 | 0.41021997 |
| AL132996.1 | 0.52408751 | 1.709765445 | 1.71515121 | 0.99685989 | 0.31883252 | NA |
| MAP3K7 | 2333.74554 | -0.389071605 | 0.13447569 | -2.8932486 | 0.00381279 | 0.02199593 |
| AL080284.1 | 0 | NA | NA | NA | NA | NA |
| CASC6 | 0.38044396 | 0.734766785 | 2.83891761 | 0.25881934 | 0.79577463 | NA |
| AL133457.1 | 0.31986932 | -1.67393407 | 2.89142001 | -0.5789315 | 0.56263541 | NA |
| LINC02531 | 1.1139693 | -0.542621482 | 2.27341325 | -0.2386814 | 0.81135263 | 0.89817258 |
| AL138731.1 | 0 | NA | NA | NA | NA | NA |
| EPHA7 | 737.743524 | 0.796434331 | 0.42258513 | 1.88467191 | 0.05947415 | 0.16456802 |
| AL391336.1 | 0.08695548 | -0.642905668 | 3.30558948 | -0.1944905 | 0.84579185 | NA |
| AL157378.1 | 0 | NA | NA | NA | NA | NA |
| MANEA-DT | 41.977134 | 0.179218486 | 0.29265794 | 0.61238211 | 0.54028497 | 0.71329302 |
| MANEA | 837.137314 | 0.562896167 | 0.12360788 | 4.55388576 | 5.27E-06 | 0.00012225 |
| FUT9 | 23.2494107 | -0.74037084 | 1.01165648 | -0.7318402 | 0.46426614 | 0.65178001 |
| UFL1-AS1 | 10.4408278 | 0.194478302 | 0.53218913 | 0.3654308 | 0.71478985 | 0.83791991 |
| UFL1 | 1692.05102 | 0.359587675 | 0.16708416 | 2.15213499 | 0.03138672 | 0.10427969 |
| FHL5 | 254.326762 | -0.339414389 | 0.60077439 | -0.5649615 | 0.57209997 | 0.73688644 |
| AL033379.1 | 8.09484262 | -0.021522758 | 0.43309439 | -0.0496953 | 0.9603652 | 0.98093621 |
| GPR63 | 51.7570729 | 0.117097258 | 0.27228967 | 0.43004664 | 0.66716172 | 0.80517412 |
| NDUFAF4 | 1093.03578 | -0.329202152 | 0.18873965 | -1.744213 | 0.08112195 | 0.20441336 |
| KLHL32 | 141.428539 | 1.269479249 | 0.32576021 | 3.89697461 | 9.74E-05 | 0.00123825 |
| MMS22L | 280.807149 | -0.69001763 | 0.28744758 | -2.400499 | 0.01637274 | 0.06487589 |
| AL589740.1 | 3.00374205 | -0.748197894 | 0.79928819 | -0.9360803 | 0.34923186 | 0.5468132 |
| AL080316.1 | 1.13150881 | 0.000782841 | 1.37807273 | 0.00056807 | 0.99954675 | 0.99979217 |
| AL590727.1 | 0 | NA | NA | NA | NA | NA |
| AL589826.2 | 0.26410578 | 1.012519641 | 2.90828201 | 0.34815043 | 0.72772721 | NA |
| POU3F2 | 4.25056162 | -0.786809267 | 0.61779579 | -1.273575 | 0.20281408 | 0.38484617 |
| FBXL4 | 1560.30279 | 0.32001841 | 0.12120466 | 2.64031431 | 0.00828292 | 0.03907883 |
| FAXC | 132.792428 | 0.08107273 | 0.27486898 | 0.29495045 | 0.76803173 | 0.87195714 |
| COQ3 | 442.913835 | 0.552454026 | 0.20890487 | 2.64452435 | 0.00818058 | 0.03874317 |
| PNISR | 6608.54128 | 0.138946817 | 0.15700058 | 0.88500832 | 0.37615214 | 0.57197501 |
| AL513550.1 | 233.285957 | 0.910544293 | 0.20710711 | 4.39648986 | 1.10E-05 | 0.00022028 |
| USP45 | 821.635894 | -0.014404079 | 0.1370548 | -0.1050972 | 0.91629866 | 0.95922174 |
| TSTD3 | 139.960496 | -0.377833575 | 0.1962642 | -1.9251273 | 0.05421343 | 0.15421878 |
| CCNC | 3855.38282 | -0.317413273 | 0.1429644 | -2.2202259 | 0.02640343 | 0.09197418 |
| AL137784.2 | 332.907654 | -0.223376691 | 0.26755282 | -0.8348882 | 0.40378067 | 0.5987907 |
| PRDM13 | 0 | NA | NA | NA | NA | NA |
| MCHR2 | 0.07659028 | -0.642905668 | 3.30558948 | -0.1944905 | 0.84579185 | NA |
| MCHR2-AS1 | 0.08757577 | 0.547239394 | 3.29377202 | 0.16614368 | 0.86804388 | NA |
| SIM1 | 1308.0867 | 0.553596566 | 0.31129989 | 1.77833844 | 0.07534829 | 0.19435925 |
| Z86062.2 | 0.31589717 | -0.695072024 | 2.30675949 | -0.3013197 | 0.76317074 | NA |
| ASCC3 | 1115.08996 | -0.666028935 | 0.1746866 | -3.8127077 | 0.00013745 | 0.00162831 |
| AL133338.1 | 67.0373494 | 0.020196332 | 0.24913812 | 0.0810648 | 0.93539042 | 0.9690899 |
| GRIK2 | 98.3873832 | 0.550187123 | 0.51524379 | 1.06781903 | 0.28560215 | 0.48163515 |
| HACE1 | 701.610974 | 0.299942658 | 0.1636687 | 1.83262071 | 0.066859 | 0.1782501 |
| LIN28B-AS1 | 0 | NA | NA | NA | NA | NA |
| LIN28B | 0 | NA | NA | NA | NA | NA |
| BVES | 151.07453 | 0.679344319 | 0.49227327 | 1.38001463 | 0.16758214 | 0.33881323 |
| BVES-AS1 | 7.12136779 | 1.963561987 | 0.88206588 | 2.22609449 | 0.02600785 | 0.0909947 |
| POPDC3 | 5.84146593 | 0.448350116 | 0.70782442 | 0.63341996 | 0.52645944 | 0.70136078 |
| AL133406.3 | 5.78278296 | 0.109895075 | 0.54388404 | 0.20205607 | 0.83987289 | 0.91484357 |
| PREP | 1323.24555 | -0.141676688 | 0.12946576 | -1.0943178 | 0.27381561 | 0.46886478 |
| AL133406.2 | 137.7123 | -0.199577849 | 0.1482915 | -1.3458482 | 0.17835148 | 0.35377828 |
| Z97206.2 | 0.25178919 | 1.950259806 | 2.89159949 | 0.6744571 | 0.50002075 | NA |
| AL591518.1 | 0.36792847 | 0.704561606 | 2.29485125 | 0.30701842 | 0.75882935 | NA |
| ATG5 | 1893.05928 | -0.34444936 | 0.14506403 | -2.3744642 | 0.01757444 | 0.0682417 |
| PRDM1 | 294.883224 | -0.628529967 | 0.41618072 | -1.5102333 | 0.13098391 | 0.28657145 |
| AL356859.1 | 0.70494132 | -0.599709208 | 1.60523179 | -0.3735966 | 0.70870443 | NA |
| CRYBG1 | 2156.18141 | 0.547051367 | 0.31932746 | 1.71313602 | 0.08668752 | 0.21450482 |
| AL109920.1 | 79.3915964 | 1.433117343 | 0.78632383 | 1.82255362 | 0.06837103 | 0.18110939 |
| RTN4IP1 | 499.985263 | 0.178922717 | 0.23489002 | 0.76172974 | 0.44622132 | 0.6365667 |
| QRSL1 | 1228.87594 | 0.079141579 | 0.16125229 | 0.49079353 | 0.62357249 | 0.77560523 |
| LINC02526 | 6.17732917 | -0.973526404 | 0.5918748 | -1.6448181 | 0.10000732 | 0.23748976 |
| LINC02532 | 5119.68184 | -0.60559852 | 0.46095307 | -1.3137965 | 0.18891471 | 0.36716397 |
| CD24 | 65871.4081 | -0.8590378 | 0.20115722 | -4.2704795 | 1.95E-05 | 0.00034754 |
| C6orf203 | 908.056164 | 0.256018489 | 0.17763365 | 1.44127249 | 0.14950772 | 0.31369498 |
| BEND3 | 182.909193 | -0.326404356 | 0.229272 | -1.4236556 | 0.1545462 | 0.32059341 |
| PDSS2 | 1638.45384 | 0.62172463 | 0.23113828 | 2.68983849 | 0.00714866 | 0.03518951 |
| SOBP | 538.580647 | 0.320520211 | 0.2257013 | 1.42010795 | 0.15557626 | 0.32184403 |
| AL121957.1 | 3.94238609 | 1.40298886 | 0.8134435 | 1.72475269 | 0.08457205 | 0.21069107 |
| SCML4 | 70.8640957 | -0.432412365 | 0.60745232 | -0.7118458 | 0.47656029 | 0.66179458 |
| SEC63 | 3893.1616 | -0.169105214 | 0.13878575 | -1.2184624 | 0.22304831 | 0.40912086 |
| AL024507.2 | 99.553837 | 0.454240477 | 0.33047274 | 1.37451724 | 0.16928116 | 0.34109759 |
| Z98200.1 | 24.3941813 | -1.590111633 | 0.56195185 | -2.8296226 | 0.00466029 | 0.02551214 |
| OSTM1 | 2067.2878 | -0.251175033 | 0.2027704 | -1.2387164 | 0.21545053 | 0.39991664 |
| OSTM1-AS1 | 0 | NA | NA | NA | NA | NA |
| NR2E1 | 6.83479893 | 1.747682371 | 0.80362269 | 2.17475488 | 0.02964849 | 0.10004231 |
| SNX3 | 7151.64791 | -0.311203793 | 0.11254501 | -2.7651496 | 0.00568967 | 0.02967496 |
| Z98742.1 | 0.08666371 | -0.642905668 | 3.30558948 | -0.1944905 | 0.84579185 | NA |
| AFG1L | 358.238464 | 0.809764319 | 0.22283166 | 3.63397344 | 0.00027909 | 0.0028726 |
| FOXO3 | 2728.66263 | -0.386898743 | 0.12493873 | -3.0967079 | 0.00195683 | 0.01326858 |
| LINC00222 | 21.4736577 | -1.029464812 | 0.57388159 | -1.7938628 | 0.07283508 | 0.18975113 |
| ARMC2 | 213.067047 | 0.109771402 | 0.32630133 | 0.33641114 | 0.73656085 | 0.85207763 |
| ARMC2-AS1 | 0.79565273 | 1.390222573 | 1.57372301 | 0.88339725 | 0.37702168 | NA |
| SESN1 | 1572.54556 | -0.340000108 | 0.22944468 | -1.4818391 | 0.13838311 | 0.29751492 |
| AL390208.1 | 212.658863 | 0.221688885 | 0.27026109 | 0.82027674 | 0.41205836 | 0.60605605 |
| CEP57L1 | 367.978815 | -0.080047706 | 0.19426325 | -0.4120579 | 0.68029699 | 0.8146636 |
| CD164 | 19950.8886 | 0.348441283 | 0.13396114 | 2.60106242 | 0.00929355 | 0.04260702 |
| AL359711.2 | 15.7133694 | 0.489222602 | 0.47888655 | 1.02158351 | 0.30697807 | 0.50401666 |
| PPIL6 | 390.906649 | 0.073327503 | 0.17514895 | 0.41865797 | 0.67546612 | 0.81122383 |
| SMPD2 | 912.436648 | -0.592225944 | 0.15002407 | -3.9475396 | 7.90E-05 | 0.00105158 |
| MICAL1 | 5396.2076 | -0.33259766 | 0.20303234 | -1.6381511 | 0.10139017 | 0.23980115 |
| ZBTB24 | 715.567815 | -0.062688113 | 0.17872473 | -0.3507523 | 0.72577418 | 0.84467583 |
| AL109947.1 | 22.2744756 | 0.638653564 | 0.29441766 | 2.16920941 | 0.03006679 | 0.10098857 |
| AK9 | 384.650043 | 0.341098361 | 0.142668 | 2.39085394 | 0.01680924 | 0.06614828 |
| FIG4 | 1021.30608 | 0.044379127 | 0.13354689 | 0.3323112 | 0.73965429 | 0.85394886 |
| AL512303.1 | 8.02242584 | 0.568959564 | 0.47575341 | 1.19591275 | 0.23173061 | 0.4197736 |
| GPR6 | 0.93431451 | -3.096565969 | 2.73103534 | -1.1338432 | 0.25686031 | NA |
| WASF1 | 1036.85148 | -0.590106638 | 0.22423315 | -2.6316655 | 0.00849675 | 0.03982567 |
| CDC40 | 1048.84036 | 0.38531721 | 0.11660181 | 3.30455603 | 0.00095127 | 0.00755344 |
| METTL24 | 78.0468795 | 0.71269136 | 0.28190048 | 2.52816649 | 0.011466 | 0.04992707 |
| DDO | 531.584706 | 1.375305773 | 0.49403932 | 2.78379818 | 0.00537265 | 0.0284092 |
| SLC22A16 | 9.42410348 | -1.951532002 | 0.90600034 | -2.154008 | 0.03123954 | 0.10393474 |
| AC002464.1 | 2.1356717 | -4.315320207 | 1.79931839 | -2.3983083 | 0.016471 | 0.06514777 |
| CDK19 | 883.879946 | 0.100265964 | 0.12753718 | 0.78617048 | 0.43176762 | 0.62387463 |
| AMD1 | 5343.87912 | -0.568808147 | 0.32524395 | -1.7488662 | 0.08031415 | 0.20297913 |
| GTF3C6 | 2261.35948 | -0.426089056 | 0.15499045 | -2.749131 | 0.00597535 | 0.03068708 |
| RPF2 | 1188.47784 | -0.561699446 | 0.16008023 | -3.5088621 | 0.00045003 | 0.00418323 |
| SLC16A10 | 845.866691 | 0.526953033 | 0.26293065 | 2.00415218 | 0.04505376 | 0.1349157 |
| AL360227.1 | 11.6321026 | 2.097726917 | 0.60359556 | 3.47538496 | 0.00051012 | 0.00460523 |
| MFSD4B | 348.091116 | 0.332332655 | 0.24191853 | 1.37373793 | 0.16952306 | 0.3413242 |
| AL080317.1 | 208.32407 | -0.435487039 | 0.28991215 | -1.5021345 | 0.13306238 | 0.28953338 |

| | | | | | | |
|---|---|---|---|---|---|---|
| AL080317.2 | 93.2830439 | -1.041156107 | 0.48801568 | -2.1334481 | 0.03288799 | 0.10771682 |
| REV3L | 1457.29795 | -0.243552128 | 0.15871925 | -1.5344838 | 0.12491067 | 0.27724049 |
| AL080317.3 | 798.071576 | 0.286636728 | 0.18172116 | 1.57734373 | 0.11471646 | 0.26184307 |
| TRAF3IP2-AS | 244.35241 | -0.098184243 | 0.17989719 | -0.5457797 | 0.58521736 | 0.74685098 |
| TRAF3IP2 | 1371.69831 | -0.959606703 | 0.16933503 | -5.6669121 | 1.45E-08 | 9.49E-07 |
| Z97989.1 | 145.81985 | 0.038751155 | 0.29974059 | 0.12928231 | 0.89713427 | 0.94743415 |
| FYN | 1854.17776 | 0.197692122 | 0.23574475 | 0.83858549 | 0.40170196 | 0.59661395 |
| LINC02527 | 0.09989236 | -0.642905668 | 3.30558948 | -0.1944905 | 0.84579185 | NA |
| WISP3 | 215.103275 | -0.386204076 | 0.34124832 | -1.1317391 | 0.25774413 | 0.45029234 |
| TUBE1 | 1181.95479 | 0.200122899 | 0.20013156 | 0.99995673 | 0.31733145 | 0.51455372 |
| FAM229B | 584.406209 | 0.039854418 | 0.28730313 | 0.13871905 | 0.88967216 | 0.94296756 |
| LAMA4 | 1083.68988 | -0.455749595 | 0.2727066 | -1.6712085 | 0.0946805 | 0.22888932 |
| Z99289.1 | 13.5166137 | 0.834113115 | 0.6128424 | 1.36105646 | 0.17349585 | 0.34725506 |
| Z99289.2 | 7.78849114 | -0.524988126 | 0.64049507 | -0.8196599 | 0.41241002 | 0.6062516 |
| Z99289.3 | 0.34982625 | -0.260586879 | 2.31477241 | -0.1125756 | 0.91036704 | NA |
| AL365214.2 | 12.6782484 | -0.584701467 | 0.61058038 | -0.9576159 | 0.33825649 | 0.53628886 |
| AL365214.3 | 19.2136031 | -2.791612812 | 1.16891784 | -2.3882028 | 0.016931 | 0.06648852 |
| RFPL4B | 45.2035735 | -3.166840397 | 1.00256535 | -3.1587371 | 0.00158454 | 0.01127549 |
| AL357514.1 | 0 | NA | NA | NA | NA | NA |
| AL589684.1 | 0.06785815 | 0.795868262 | 3.28898349 | 0.24198001 | 0.80879565 | NA |
| LINC02518 | 0.38027097 | 0.703328506 | 2.27976787 | 0.30850882 | 0.75769519 | NA |
| AL513123.1 | 0 | NA | NA | NA | NA | NA |
| LINC02541 | 1.83044002 | -2.259211895 | 1.47606532 | -1.5305636 | 0.12587728 | 0.27868221 |
| FO393415.3 | 0.25725949 | -1.364242679 | 3.30558948 | -0.4127078 | 0.67982068 | NA |
| MARCKS | 3541.88799 | -0.746682768 | 0.25898136 | -2.8831525 | 0.00393717 | 0.02250121 |
| LINC01268 | 141.819021 | 0.068122102 | 0.49544353 | 0.13749721 | 0.8906378 | 0.94323068 |
| FO393415.1 | 4.63864947 | 0.544850117 | 0.91149152 | 0.59775665 | 0.55000232 | 0.72038748 |
| HDAC2 | 3090.43502 | -0.302888753 | 0.13437661 | -2.2540289 | 0.02419435 | 0.08613542 |
| HDAC2-AS2 | 12.2380426 | -0.094369304 | 0.43015598 | -0.2193839 | 0.826351 | 0.90734826 |
| HS3ST5 | 8.56191905 | -0.369010108 | 0.96479039 | -0.382477 | 0.70210761 | 0.82947179 |
| AL138830.2 | 1.05598452 | 2.10338124 | 1.59526274 | 1.31851712 | 0.1873306 | 0.36526977 |
| AL138830.1 | 0.20357445 | 1.728110593 | 3.26179266 | 0.52980394 | 0.59624787 | NA |
| LINC02534 | 0 | NA | NA | NA | NA | NA |
| FRK | 2352.74598 | 0.332457685 | 0.23930598 | 1.38925773 | 0.16475439 | 0.33473868 |
| NT5DC1 | 2769.09629 | 0.771964757 | 0.26657965 | 2.8958128 | 0.00378178 | 0.02189844 |
| COL10A1 | 19.4973881 | 1.527632515 | 0.67935829 | 2.24864044 | 0.02453538 | 0.08702058 |
| AL050331.2 | 304.649323 | 1.160417903 | 0.29519483 | 3.93102375 | 8.46E-05 | 0.00111192 |
| TSPYL4 | 2077.53966 | 0.563155091 | 0.21408165 | 2.63056215 | 0.00852438 | 0.0399197 |
| DSE | 3020.23132 | 0.353471555 | 0.22122526 | 1.59779018 | 0.11008968 | 0.25405311 |
| TSPYL1 | 7340.61708 | 0.729891319 | 0.15225355 | 4.79392001 | 1.64E-06 | 4.86E-05 |
| Z84488.2 | 228.935678 | -0.288752139 | 0.39477485 | -0.731435 | 0.4645135 | 0.65190107 |
| Z84488.1 | 45.1770327 | -1.482181881 | 0.72456633 | -2.0456124 | 0.04079452 | 0.12573065 |
| CALHM6 | 129.063518 | -1.295787328 | 0.71031803 | -1.8242354 | 0.06811649 | 0.18058025 |
| AL445224.1 | 7.02465623 | -0.084371426 | 0.92428963 | -0.0912825 | 0.92726816 | 0.96473058 |
| TRAPPC3L | 36.1280788 | 0.659136267 | 0.50346292 | 1.30920518 | 0.19046486 | 0.3691901 |
| CALHM5 | 372.675597 | 0.691363903 | 0.2813766 | 2.45707671 | 0.01400728 | 0.05790436 |
| CALHM4 | 3.6569428 | 0.8537311 | 1.06224744 | 0.80370267 | 0.42156871 | 0.61418602 |
| RWDD1 | 1748.24196 | -0.239234984 | 0.14041273 | -1.7037984 | 0.08841875 | 0.21770331 |
| RSPH4A | 86.1533586 | -0.228715965 | 0.3839813 | -0.5956435 | 0.55141341 | 0.72155476 |
| ZUP1 | 484.175256 | -0.601974829 | 0.16978379 | -3.5455377 | 0.00039181 | 0.00375713 |
| KPNA5 | 732.141141 | -0.044617171 | 0.14511834 | -0.3074537 | 0.75849806 | 0.86635037 |
| FAM162B | 86.5316216 | 0.375642405 | 0.38815965 | 0.96775232 | 0.33316808 | 0.53114336 |
| GPRC6A | 36.0412464 | 0.396865489 | 0.82456863 | 0.48130074 | 0.63030277 | 0.78028359 |
| RFX6 | 1.05181024 | -0.961185377 | 1.50371544 | -0.639207 | 0.5226883 | 0.69859758 |
| VGLL2 | 0.30714886 | -0.699655751 | 2.91696002 | -0.2398578 | 0.81044046 | NA |
| ROS1 | 33.8551592 | -0.149081566 | 0.81326454 | -0.1833125 | 0.85455282 | 0.92399813 |
| AL132671.2 | 118.192321 | 0.774870121 | 0.28631038 | 2.70639896 | 0.00680173 | 0.03382266 |
| AL132671.1 | 0 | NA | NA | NA | NA | NA |
| DCBLD1 | 689.20848 | -0.427649339 | 0.20391367 | -2.0972078 | 0.03597518 | 0.11480678 |
| GOPC | 1730.22456 | 0.176376018 | 0.09211582 | 1.91472019 | 0.0555282 | 0.15672467 |
| NUS1 | 2438.51189 | -0.068233999 | 0.21448661 | -0.3181271 | 0.75038854 | 0.86064026 |
| SLC35F1 | 55.8035567 | 0.103610869 | 0.52151276 | 0.1986737 | 0.842518 | 0.91630899 |
| CEP85L | 308.570827 | -0.274229901 | 0.17503812 | -1.5666867 | 0.11718794 | 0.26561337 |
| PLN | 831.306267 | 0.225757439 | 0.58377975 | 0.38671681 | 0.69896587 | 0.82720268 |
| MCM9 | 919.425684 | 0.142899209 | 0.15655419 | 0.91277792 | 0.36135935 | 0.55885556 |
| ASF1A | 710.191154 | 0.313156625 | 0.16781886 | 1.86603958 | 0.06203583 | 0.16932348 |
| AL365275.1 | 43.3270298 | 1.193056491 | 0.21792059 | 5.47473043 | 4.38E-08 | 2.38E-06 |
| FAM184A | 471.617072 | 1.028353526 | 0.26113538 | 3.93800916 | 8.22E-05 | 0.001086 |
| MAN1A1 | 6513.76027 | 0.416531126 | 0.17965489 | 2.31850706 | 0.02042178 | 0.07608276 |
| TBC1D32 | 207.727172 | 0.112011591 | 0.24558571 | 0.4560998 | 0.64831822 | 0.79292015 |
| GJA1 | 2490.62594 | -0.895208731 | 0.2800659 | -3.1964217 | 0.00139144 | 0.01020077 |
| HSF2 | 693.58995 | -0.43559189 | 0.14649748 | -2.9733747 | 0.00294545 | 0.01808415 |
| SERINC1 | 14552.007 | 0.077346625 | 0.1332686 | 0.58038147 | 0.5616574 | 0.72942161 |
| PKIB | 185.725349 | -0.523471488 | 0.52794674 | -0.9915233 | 0.32143013 | 0.51908338 |
| FABP7 | 1.74327075 | 1.770103407 | 1.05111612 | 1.6840227 | 0.09217728 | 0.22444274 |
| SMPDL3A | 2232.96958 | 0.418900962 | 0.47250745 | 0.8865489 | 0.37532182 | 0.57129724 |
| CLVS2 | 0.55087932 | -0.069524214 | 1.92641741 | -0.0360899 | 0.97121067 | NA |
| TRDN | 11.2623948 | 0.965029499 | 1.02747613 | 0.93922328 | 0.34761612 | 0.545114 |
| TRDN-AS1 | 0.07830766 | -0.642905668 | 3.30558948 | -0.1944905 | 0.84579185 | NA |
| NKAIN2 | 14.41694 | 1.830102308 | 0.565359 | 3.23706231 | 0.00120767 | 0.00909114 |
| RNF217-AS1 | 40.4770964 | -0.63749676 | 0.38076011 | -1.674274 | 0.09407676 | 0.22791035 |
| RNF217 | 932.343007 | -1.084244799 | 0.17241942 | -6.2884147 | 3.21E-10 | 3.88E-08 |
| TPD52L1 | 2792.22855 | -0.525117293 | 0.30056879 | -1.7470786 | 0.08062371 | 0.20353903 |
| HDDC2 | 2850.07029 | 0.070394212 | 0.14077991 | 0.50003025 | 0.61705377 | 0.77106137 |
| AL450332.1 | 4.96876937 | 0.538328344 | 0.61516653 | 0.87509368 | 0.38152294 | 0.57756127 |
| AL365259.1 | 23.8731187 | 2.058404151 | 0.34267224 | 6.00691835 | 1.89E-09 | 1.74E-07 |
| LINC02523 | 0.68396063 | 1.071430796 | 1.56446616 | 0.68485393 | 0.4934361 | NA |
| HEY2 | 230.086024 | 0.552916962 | 0.2442049 | 2.26415178 | 0.02356478 | 0.08455617 |
| NCOA7 | 7331.54586 | -0.80954302 | 0.31859558 | -2.5409738 | 0.01105442 | 0.04852756 |
| NCOA7-AS1 | 2.09375502 | 1.361584483 | 1.02500002 | 1.32837509 | 0.18405423 | 0.36107052 |
| HINT3 | 2007.59039 | 0.083488576 | 0.1422378 | 0.58696476 | 0.55722737 | 0.72610001 |
| TRMT11 | 977.494465 | -0.418739531 | 0.12575327 | -3.3298499 | 0.00086893 | 0.00703815 |
| AC020559.1 | 10.2167337 | -0.402374517 | 0.39642603 | -1.0150053 | 0.3101033 | 0.50718998 |
| CENPW | 110.921468 | -1.265405253 | 0.3600927 | -3.5141097 | 0.00044123 | 0.00411867 |
| RSPO3 | 185.573786 | -0.227950481 | 0.52178806 | -0.4368641 | 0.66220989 | 0.80128673 |
| RNF146 | 2525.12023 | 0.238303661 | 0.11383202 | 2.09346765 | 0.03630743 | 0.11556149 |
| ECHDC1 | 4049.41781 | 0.505755159 | 0.16729812 | 3.02307727 | 0.00250218 | 0.01601519 |
| AL096711.2 | 209.249029 | -0.635763928 | 0.48127197 | -1.3210076 | 0.18649883 | 0.36424818 |
| KIAA0408 | 178.90849 | -0.652608092 | 0.48406447 | -1.3481843 | 0.17759913 | 0.35275879 |
| SOGA3 | 27.3346218 | -0.11720035 | 0.48334491 | -0.2424777 | 0.80841006 | 0.89642038 |
| C6orf58 | 4.48082544 | 0.095141514 | 0.91757261 | 0.10368827 | 0.91741674 | 0.9600034 |
| LINC02536 | 0 | NA | NA | NA | NA | NA |
| THEMIS | 41.1491159 | 0.149426634 | 0.79154171 | 0.18877923 | 0.85026585 | 0.92144456 |
| PTPRK | 2220.71901 | -0.042712813 | 0.10561168 | -0.4044327 | 0.68589459 | 0.81838108 |
| AL590006.1 | 39.8317165 | 0.917913671 | 0.28483883 | 3.22257215 | 0.00127045 | 0.00947717 |
| AL034349.1 | 0.46390404 | -1.336692482 | 1.89797943 | -0.7042713 | 0.48126382 | NA |
| LAMA2 | 1183.85943 | 0.072220762 | 0.33840692 | 0.21341396 | 0.83100409 | 0.91040231 |
| AL356124.2 | 79.2985249 | 0.430518193 | 0.38164492 | 1.12805955 | 0.25929477 | 0.45207293 |
| AL356124.1 | 10.1021474 | -0.35545808 | 0.56439082 | -0.6298084 | 0.52881995 | 0.70340389 |
| ARHGAP18 | 2285.96431 | 0.466701204 | 0.16842772 | 2.77092872 | 0.00558967 | 0.02929825 |
| TMEM244 | 1.24545099 | 0.320293519 | 1.62648468 | 0.19692378 | 0.84388718 | 0.9172838 |
| L3MBTL3 | 637.756 | -0.117528519 | 0.17576787 | -0.6686576 | 0.50371394 | 0.68364658 |
| AL355581.1 | 155.890788 | -0.245714569 | 0.23085161 | -1.0643832 | 0.28715516 | 0.48339437 |

| | | | | | | |
|---|---|---|---|---|---|---|
| SAMD3 | 97.1298899 | -0.214994156 | 0.4490289 | -0.478798 | 0.63208232 | 0.78149288 |
| TMEM200A | 1115.12156 | 0.357684077 | 0.26650555 | 1.34212619 | 0.17955508 | 0.35525717 |
| AL583803.1 | 0.19289513 | -1.003574173 | 3.30558948 | -0.3035992 | 0.7614333 | NA |
| SMLR1 | 173.838435 | 2.51665285 | 0.62268613 | 4.04160739 | 5.31E-05 | 0.00078118 |
| EPB41L2 | 2768.33564 | 0.348275396 | 0.21435235 | 1.62477991 | 0.10420946 | 0.24444081 |
| AKAP7 | 497.565458 | 0.154704619 | 0.28632683 | 0.5403078 | 0.58898478 | 0.74944647 |
| ARG1 | 31.3190149 | -2.165836208 | 0.49283268 | -4.3946684 | 1.11E-05 | 0.0002218 |
| MED23 | 1548.62026 | 0.226756918 | 0.10654612 | 2.12825133 | 0.03331625 | 0.10871761 |
| ENPP3 | 606.184012 | 0.80138524 | 0.3529422 | 2.27058491 | 0.02317212 | 0.08364841 |
| OR2A4 | 2.33295798 | -1.590760866 | 1.25185935 | -1.2707185 | 0.2038288 | 0.38607951 |
| CTAGE9 | 4.54135138 | -0.414041039 | 0.64514318 | -0.6417816 | 0.52101498 | 0.69753682 |
| ENPP1 | 1283.5817 | -0.145206138 | 0.44043249 | -0.3296899 | 0.74163429 | 0.85479681 |
| AL117378.1 | 0.26056869 | 1.992294186 | 3.24261664 | 0.61440941 | 0.53894482 | NA |
| CTGF | 13740.3637 | 0.06137413 | 0.49541068 | 0.12388536 | 0.90140605 | 0.95006982 |
| AL133346.1 | 196.065539 | -0.236114351 | 0.4727162 | -0.4994844 | 0.6174382 | 0.77133479 |
| AL021408.1 | 2.68491858 | 0.906528652 | 0.74171048 | 1.22221361 | 0.22162686 | 0.4076376 |
| LINC01013 | 9.66066662 | -1.551984602 | 1.10955116 | -1.3987499 | 0.16188799 | 0.33067283 |
| MOXD1 | 2203.81811 | 0.093888281 | 0.42658822 | 0.22009112 | 0.82580019 | 0.90695123 |
| STX7 | 4691.00663 | 0.514223872 | 0.13507233 | 3.80702599 | 0.00014065 | 0.0016602 |
| TAAR9 | 0.60173728 | 3.168409927 | 1.84491378 | 1.71737561 | 0.08591057 | NA |
| TAAR8 | 1.41570766 | 2.136880491 | 1.18241321 | 1.80721974 | 0.07072802 | 0.18566652 |
| TAAR6 | 4.91107211 | 1.615950217 | 0.73900344 | 2.1866613 | 0.02876726 | 0.09793346 |
| TAAR5 | 0 | NA | NA | NA | NA | NA |
| TAAR2 | 0.06785815 | 0.795868262 | 3.28898349 | 0.24198001 | 0.80879565 | NA |
| TAAR1 | 2.24244872 | 2.101324566 | 1.20672275 | 1.74134827 | 0.08162255 | 0.20531714 |
| VNN1 | 3652.16137 | -0.599858079 | 0.8061498 | -0.7441025 | 0.45681447 | 0.64622056 |
| VNN3 | 17.9449888 | -1.285198417 | 0.54278798 | -2.3677725 | 0.01789554 | 0.06906269 |
| VNN2 | 257.805054 | -1.076070785 | 0.4272658 | -2.5185044 | 0.01178544 | 0.05090616 |
| AL032821.1 | 29.3010365 | -0.870752636 | 0.42683698 | -2.0400122 | 0.04134911 | 0.12700945 |
| SLC18B1 | 1543.2426 | -0.135907826 | 0.23261759 | -0.5842543 | 0.55904924 | 0.7276111 |
| RPS12 | 24780.3256 | -0.628936551 | 0.18241628 | -3.4478093 | 0.00056515 | 0.0049964 |
| LINC00326 | 0.14114194 | 0.02007176 | 3.29738704 | 0.00608717 | 0.99514317 | NA |
| EYA4 | 394.688384 | 0.521434528 | 0.38134502 | 1.36735632 | 0.17151365 | 0.34399261 |
| AL024497.1 | 14.9531219 | 0.662449545 | 0.59016955 | 1.12247327 | 0.26166127 | 0.4551156 |
| AL024497.2 | 17.79746 | 0.600966287 | 0.54217408 | 1.10843788 | 0.26767275 | 0.46182482 |
| TARID | 1308.97653 | 1.040683745 | 0.34264173 | 3.03723584 | 0.00238759 | 0.01545046 |
| AL450270.1 | 5.78502008 | -0.097030101 | 0.58216114 | -0.1666722 | 0.86762795 | 0.93109788 |
| LINC01312 | 8.02806314 | 1.853135578 | 0.71759246 | 2.58243457 | 0.0098106 | 0.04444347 |
| TCF21 | 1834.50749 | 1.057495334 | 0.40156792 | 2.63341585 | 0.00845308 | 0.03967741 |
| TBPL1 | 875.101432 | 0.052548775 | 0.12052336 | 0.43600488 | 0.66283319 | 0.80178143 |
| SLC2A12 | 435.614087 | 1.43486866 | 0.28784099 | 4.98493515 | 6.20E-07 | 2.19E-05 |
| SGK1 | 9538.13855 | -0.147611648 | 0.41370504 | -0.3568041 | 0.72123848 | 0.84140235 |
| AL078590.2 | 37.1415586 | 1.336274418 | 0.43944976 | 3.04078993 | 0.00235958 | 0.01534079 |
| LINC01010 | 13.8250972 | 1.182062347 | 0.72039369 | 1.64085606 | 0.1008273 | 0.23888843 |
| AL078590.3 | 42.150364 | 0.968933195 | 0.4136944 | 2.34214722 | 0.01917315 | 0.0726431 |
| AL078590.1 | 0.17515154 | 1.45420457 | 3.2783972 | 0.44357181 | 0.65735219 | NA |
| AL596188.1 | 0.30481559 | 1.356039782 | 2.26732163 | 0.59808003 | 0.54978654 | NA |
| AL121970.1 | 2.9051417 | 2.988416028 | 1.19475428 | 2.50128086 | 0.0123745 | 0.05280187 |
| ALDH8A1 | 4522.17635 | 1.424186171 | 0.44309147 | 3.21420351 | 0.00130807 | 0.00972204 |
| AL445190.1 | 291.599229 | 1.944250081 | 0.53136378 | 3.6589812 | 0.00025322 | 0.00264568 |
| HBS1L | 1686.01539 | 0.125714008 | 0.19181589 | 0.65538891 | 0.51221739 | 0.69037427 |
| AL353596.1 | 6.6352954 | 1.115257707 | 0.93208117 | 1.19652423 | 0.23149205 | 0.41940538 |
| MYB | 13.9323822 | -0.694008104 | 0.74413429 | -0.9326383 | 0.35100678 | 0.54860412 |
| MYB-AS1 | 1.26905216 | -0.32908255 | 2.35890947 | -0.1395062 | 0.88905014 | 0.94263408 |
| AL023693.1 | 0 | NA | NA | NA | NA | NA |
| AHI1 | 1199.1322 | -0.600893687 | 0.17467732 | -3.4400212 | 0.00058167 | 0.00510367 |
| AL049552.1 | 77.1570963 | -0.852975031 | 0.34094418 | -2.5018026 | 0.01235628 | 0.05274119 |
| LINC00271 | 139.125578 | 0.451074421 | 0.43802667 | 1.02978758 | 0.30310973 | 0.49991508 |
| AL035604.1 | 20.0297722 | 0.717487935 | 0.78737428 | 0.91124126 | 0.36216827 | 0.55945037 |
| AL133319.1 | 0.46601102 | 1.398511074 | 2.74601299 | 0.50928786 | 0.61055047 | NA |
| PDE7B | 929.25417 | 0.083975487 | 0.19962801 | 0.42065984 | 0.67400349 | 0.81024216 |
| AL138828.1 | 598.572967 | -2.169759218 | 0.68859682 | -3.1509864 | 0.0016272 | 0.01151659 |
| MTFR2 | 53.134552 | -1.165767294 | 0.37919919 | -3.0742874 | 0.00211006 | 0.01401922 |
| BCLAF1 | 4626.73692 | -0.043695769 | 0.11960788 | -0.3653252 | 0.71486868 | 0.83793782 |
| AL023284.4 | 284.341796 | -0.457419695 | 0.29448098 | -1.5533081 | 0.12034954 | 0.2704634 |
| MAP7 | 2903.06556 | 0.819924669 | 0.20426789 | 4.01396752 | 5.97E-05 | 0.0008474 |
| AL024508.2 | 47.469134 | 0.040776818 | 0.38227687 | 0.10666828 | 0.91505215 | 0.95860182 |
| MAP3K5 | 1646.15857 | -0.415629722 | 0.23669363 | -1.7559819 | 0.0790915 | 0.20087094 |
| AL024508.1 | 9.55039592 | -0.371797388 | 0.44902356 | -0.8280131 | 0.40766308 | 0.60231261 |
| PEX7 | 650.401464 | 0.402841555 | 0.19792877 | 2.03528552 | 0.04182215 | 0.12797054 |
| SLC35D3 | 0.55153733 | 0.006963237 | 2.24628336 | 0.00309989 | 0.99752665 | NA |
| AL135902.1 | 0 | NA | NA | NA | NA | NA |
| AL135902.2 | 0.19978473 | -1.003574173 | 3.30558948 | -0.3035992 | 0.7614333 | NA |
| IL20RA | 36.383767 | 0.059126145 | 0.48958452 | 0.120768 | 0.9038748 | 0.95167363 |
| IL22RA2 | 0.42632748 | -0.902539273 | 2.83781619 | -0.3180401 | 0.75045454 | NA |
| IFNGR1 | 3530.13685 | -0.864901658 | 0.14027178 | -6.1658993 | 7.01E-10 | 7.70E-08 |
| OLIG3 | 0.8759643 | -3.028659172 | 1.73479123 | -1.745835 | 0.08083964 | NA |
| AL356234.2 | 0.34626307 | -0.557224145 | 2.28113027 | -0.2442755 | 0.80701749 | NA |
| AL356234.3 | 6.42925941 | -2.065609578 | 0.90110714 | -2.2923019 | 0.02188823 | 0.08018792 |
| LINC02539 | 0.19387658 | -1.003574173 | 3.30558948 | -0.3035992 | 0.7614333 | NA |
| AL357060.1 | 57.4572102 | -1.080891313 | 0.34797562 | -3.1062271 | 0.00189491 | 0.01292885 |
| TNFAIP3 | 2770.31766 | -1.812326136 | 0.63403157 | -2.8584162 | 0.00425762 | 0.02385748 |
| AL591468.1 | 0.27442362 | -0.179388694 | 2.94052751 | -0.0610056 | 0.95135474 | NA |
| LINC02528 | 0.19387658 | -1.003574173 | 3.30558948 | -0.3035992 | 0.7614333 | NA |
| PERP | 3990.56967 | -0.759698663 | 0.16886714 | -4.498795 | 6.83E-06 | 0.00015195 |
| ARFGEF3 | 647.521293 | 0.061967316 | 0.22931189 | 0.27023159 | 0.78698209 | 0.88407249 |
| PBOV1 | 0.40830231 | -1.802811494 | 3.29956038 | -0.5463793 | 0.58480524 | NA |
| SMIM28 | 0.38477927 | 1.84039758 | 2.10207876 | 0.87551314 | 0.38129476 | NA |
| HEBP2 | 3197.75395 | -0.44836197 | 0.12607716 | -3.5562506 | 0.00037619 | 0.00363903 |
| NHSL1 | 1309.82892 | 0.983817532 | 0.1892091 | 5.19963121 | 2.00E-07 | 8.53E-06 |
| AL590617.2 | 179.410977 | -0.644901827 | 0.32176981 | -2.0042335 | 0.04504505 | 0.13490495 |
| GVQW2 | 28.7538874 | 0.024573468 | 0.43845486 | 0.0560456 | 0.95530548 | 0.97856915 |
| CCDC28A | 1866.87123 | 0.144612984 | 0.17299088 | 0.83595729 | 0.40317893 | 0.59815921 |
| ECT2L | 23.9799714 | 0.38264421 | 0.36478352 | 1.04896244 | 0.2941954 | 0.49088413 |
| REPS1 | 1625.20351 | -0.074589348 | 0.09905402 | -0.7530169 | 0.45143979 | 0.64117486 |
| ABRACL | 1702.80124 | -0.685915939 | 0.39455741 | -1.7384439 | 0.08213262 | 0.20623093 |
| HECA | 1564.97149 | -0.667408042 | 0.2164943 | -3.0827973 | 0.00205065 | 0.0137457 |
| AL031772.1 | 22.629574 | 0.71178077 | 0.37073438 | 1.91992113 | 0.05486786 | 0.15552677 |
| TXLNB | 34.614526 | -1.05675928 | 0.4403267 | -2.3999437 | 0.0163976 | 0.06494512 |
| AL158850.1 | 3.10061404 | 0.169289265 | 0.90136511 | 0.18781431 | 0.85102222 | 0.92184442 |
| CITED2 | 6271.65633 | -0.380918838 | 0.28150428 | -1.3531547 | 0.17600621 | 0.35052366 |
| LINC01625 | 32.7004074 | 3.163493569 | 0.70119856 | 4.51155169 | 6.44E-06 | 0.00014406 |
| FILNC1 | 220.247129 | -1.94204378 | 0.83192395 | -2.3344006 | 0.01957475 | 0.07373037 |
| AL050338.1 | 43.8902745 | -1.976712488 | 0.85079559 | -2.3233695 | 0.02015931 | 0.07534325 |
| AL138737.1 | 2.67294344 | -0.628459529 | 0.94936475 | -0.661979 | 0.50798468 | 0.68682727 |
| AL035446.1 | 4.09706055 | -1.225497685 | 1.03592292 | -1.1830008 | 0.23680881 | 0.42602323 |
| AL355596.1 | 6.54580652 | -0.920759874 | 0.7715166 | -1.1934414 | 0.23269654 | 0.42074929 |
| NMBR | 2.58949133 | 0.805886027 | 0.79057243 | 1.01937027 | 0.30802722 | 0.50492184 |
| NMBR-AS1 | 0 | NA | NA | NA | NA | NA |
| GJE1 | 0.18380315 | -1.003574173 | 3.30558948 | -0.3035992 | 0.7614333 | NA |
| VTA1 | 1846.60506 | 0.080415056 | 0.21184333 | 0.37959682 | 0.70424472 | 0.83110017 |
| AL360007.1 | 3.17838731 | 1.754472553 | 0.99856181 | 1.75699946 | 0.0789179 | 0.20054592 |
| ADGRG6 | 924.969185 | 0.300229516 | 0.34305422 | 0.87516637 | 0.38148339 | 0.57754719 |

| | | | | | | |
|---|---|---|---|---|---|---|
| AL359313.1 | 0.5923131 | -1.469089681 | 1.8470773 | -0.7953591 | 0.42640466 | NA |
| AL023584.1 | 218.057662 | -0.351739718 | 0.34329577 | -1.0245967 | 0.30555352 | 0.50261658 |
| HIVEP2 | 1636.91402 | -0.624705329 | 0.25881795 | -2.4136862 | 0.01579206 | 0.06321497 |
| AL023584.2 | 10.1215169 | -1.420924483 | 0.64850956 | -2.1910617 | 0.02844733 | 0.09722084 |
| AL355304.1 | 11.0886946 | -1.467785115 | 0.58415687 | -2.5126557 | 0.01198262 | 0.05148699 |
| LINC01277 | 41.6448552 | 1.251168882 | 0.40686952 | 3.07511086 | 0.00210424 | 0.01399821 |
| AL023581.2 | 20.2498495 | 0.847099273 | 0.66900871 | 1.26620067 | 0.20544125 | 0.38801645 |
| AIG1 | 4313.31083 | 0.671364987 | 0.227571 | 2.95013424 | 0.00317636 | 0.01918914 |
| AL023581.1 | 0.12150485 | 1.04981648 | 3.28477129 | 0.31960109 | 0.74927075 | NA |
| ADAT2 | 610.725676 | -0.390085375 | 0.22911046 | -1.7026083 | 0.08864139 | 0.21813013 |
| PEX3 | 1370.16448 | 0.080491769 | 0.18227035 | 0.44160648 | 0.65887399 | 0.79954802 |
| FUCA2 | 4765.56862 | -0.083030312 | 0.20711907 | -0.400882 | 0.68850698 | 0.82016213 |
| PHACTR2 | 970.056854 | 0.027230047 | 0.1438276 | 0.18932421 | 0.84983872 | 0.92113042 |
| PHACTR2-AS | 1.31892734 | -0.273884135 | 1.04694087 | -0.2616042 | 0.7936266 | 0.88835782 |
| AL049844.1 | 153.709325 | 0.072535906 | 0.22420966 | 0.3235182 | 0.74630283 | 0.85787909 |
| LTV1 | 1088.99282 | -0.320095134 | 0.23484101 | -1.3630291 | 0.17287333 | 0.34627191 |
| AL049844.3 | 424.599133 | -0.171778004 | 0.26597238 | -0.645849 | 0.51837714 | 0.69530036 |
| ZC2HC1B | 1.59820358 | 0.017545667 | 1.03027727 | 0.01703004 | 0.98641265 | 0.99391317 |
| PLAGL1 | 1784.57767 | -0.374565249 | 0.20054135 | -1.8677707 | 0.06179404 | 0.16889603 |
| SF3B5 | 3755.54516 | -0.15129482 | 0.12609385 | -1.1998588 | 0.23019418 | 0.4177809 |
| STX11 | 251.813754 | -1.216336811 | 0.34085313 | -3.5685071 | 0.00035902 | 0.00349996 |
| UTRN | 2953.34365 | 0.430536841 | 0.27721578 | 1.55307479 | 0.12040527 | 0.27049642 |
| AL024474.2 | 0.17507102 | 0.435199075 | 3.28898345 | 0.13232024 | 0.894731 | NA |
| EPM2A | 803.301969 | 0.933401406 | 0.26437772 | 3.53055995 | 0.00041468 | 0.00392503 |
| AL023806.1 | 33.9895652 | 1.569385389 | 0.44952313 | 3.49122276 | 0.00048082 | 0.00439788 |
| AL023806.2 | 8.53773326 | 1.348174048 | 0.46283666 | 2.91285059 | 0.00358146 | 0.02092972 |
| FBXO30 | 952.950922 | -0.401240825 | 0.34706621 | -1.156093 | 0.24764311 | 0.43919542 |
| AL356599.1 | 932.349685 | 0.287824367 | 0.1139615 | 2.52562806 | 0.01154917 | 0.05016498 |
| SHPRH | 953.109938 | 0.340074418 | 0.14713906 | 2.31124495 | 0.02081933 | 0.07714005 |
| GRM1 | 85.5829124 | 3.272795678 | 0.62224775 | 5.25963441 | 1.44E-07 | 6.53E-06 |
| RAB32 | 1187.27109 | -0.910858955 | 0.33800872 | -2.6947795 | 0.00704352 | 0.0348019 |
| AL359547.1 | 0 | NA | NA | NA | NA | NA |
| ADGB | 3.71889719 | 0.951165485 | 1.04744652 | 0.90808024 | 0.36383583 | 0.56097473 |
| STXBP5-AS1 | 69.9704892 | -0.164966326 | 0.18006032 | -0.9161726 | 0.3595764 | 0.55720924 |
| AL138916.1 | 0.98184247 | 1.14879114 | 1.56962293 | 0.73188988 | 0.46423578 | NA |
| AL138916.2 | 0.08666371 | -0.642905668 | 3.30558948 | -0.1944905 | 0.84579185 | NA |
| LUADT1 | 0.2602829 | -1.364242679 | 3.30558948 | -0.4127078 | 0.67982068 | NA |
| STXBP5 | 738.318002 | -0.56113435 | 0.14936766 | -3.7567325 | 0.00017215 | 0.00194661 |
| SAMD5 | 2239.09967 | -0.695131825 | 0.50099956 | -1.3874899 | 0.16529243 | 0.33562496 |
| AL033504.1 | 18.9919014 | -1.669048819 | 0.68080109 | -2.4515954 | 0.01422245 | 0.05856433 |
| AL365271.1 | 1.05621835 | 0.805621057 | 1.2325398 | 0.65362681 | 0.51335227 | 0.6913032 |
| AL445123.1 | 0.21775352 | -0.339791733 | 3.08904691 | -0.1099989 | 0.91241025 | NA |
| AL359382.1 | 0 | NA | NA | NA | NA | NA |
| AL513164.1 | 0 | NA | NA | NA | NA | NA |
| SASH1 | 3885.25395 | -0.838715618 | 0.34818999 | -2.4087873 | 0.01600562 | 0.06387232 |
| UST | 327.800316 | 0.314134422 | 0.23480675 | 1.33784238 | 0.18094782 | 0.35712326 |
| UST-AS1 | 10.7661839 | 0.603457828 | 0.42535522 | 1.418715 | 0.15598212 | 0.32246944 |
| AL357992.1 | 0.64972926 | 0.683228233 | 2.24413038 | 0.30445122 | 0.76078415 | NA |
| AL031056.1 | 0.91833135 | 0.684045485 | 1.73695948 | 0.39381776 | 0.69371559 | NA |
| TAB2 | 3313.03172 | 0.01588159 | 0.16969824 | 0.09358724 | 0.92543705 | 0.96392839 |
| TAB2-AS1 | 0.48355709 | 0.837714183 | 2.13991563 | 0.39147066 | 0.69544937 | NA |
| AL031056.2 | 0.36569594 | 0.954230895 | 2.81736006 | 0.33869682 | 0.73483814 | NA |
| SUMO4 | 16.6151146 | -0.741586344 | 0.40612291 | -1.8260145 | 0.06784807 | 0.18004722 |
| ZC3H12D | 194.398437 | 0.726876508 | 0.35016339 | 2.07582097 | 0.03791051 | 0.11899681 |
| PPIL4 | 1361.78465 | -0.169713043 | 0.11160322 | -1.5206823 | 0.12833959 | 0.28245721 |
| AL078581.2 | 0 | NA | NA | NA | NA | NA |
| GINM1 | 5045.02223 | 0.157954379 | 0.14741132 | 1.07152138 | 0.28393506 | 0.47995946 |
| AL078581.3 | 1072.80418 | 0.166107013 | 0.14477165 | 1.14737253 | 0.25122769 | 0.44354643 |
| AL078581.1 | 25.1859557 | 0.815636922 | 0.30564562 | 2.66857062 | 0.00761748 | 0.03682362 |
| KATNA1 | 981.240108 | -0.118355287 | 0.13593015 | -0.8707067 | 0.38391435 | 0.57957258 |
| LATS1 | 1423.83798 | 0.053475359 | 0.13587481 | 0.39356346 | 0.69390336 | 0.82376229 |
| AL358852.1 | 31.689385 | 0.513856891 | 0.41359158 | 1.24242589 | 0.21407947 | 0.3981715 |
| NUP43 | 1585.80371 | 0.060073246 | 0.10591708 | 0.56717239 | 0.57059706 | 0.7357357 |
| PCMT1 | 3122.28327 | -0.321958808 | 0.20593576 | -1.5633943 | 0.11795987 | 0.2666611 |
| AL355312.2 | 897.996885 | -0.208417758 | 0.23902751 | -0.8719405 | 0.38324086 | 0.57911074 |
| LRP11 | 1998.86469 | 0.231737544 | 0.15506327 | 1.49447087 | 0.13505259 | 0.29242691 |
| AL355312.4 | 90.4876359 | 0.839408348 | 0.33866373 | 2.47858948 | 0.0131903 | 0.0553074 |
| AL355312.3 | 53.2352968 | 0.279713817 | 0.3813652 | 0.73345397 | 0.46328159 | 0.65111486 |
| RAET1E | 66.3985941 | 0.273668813 | 0.53734252 | 0.5093005 | 0.61054161 | 0.76644683 |
| RAET1E-AS1 | 32.6029304 | 0.418804533 | 0.40226471 | 1.04111677 | 0.29782136 | 0.49464944 |
| RAET1G | 41.1789071 | 0.271469184 | 0.31143282 | 0.87167816 | 0.38338398 | 0.5792028 |
| ULBP2 | 59.5515628 | -1.41382754 | 0.33027165 | -4.280802 | 1.86E-05 | 0.00033543 |
| ULBP1 | 22.4548665 | -1.310302322 | 0.3945613 | -3.3209094 | 0.00089725 | 0.00720475 |
| RAET1L | 8.63603141 | 1.102131585 | 0.67730192 | 1.62723824 | 0.10368651 | 0.24360417 |
| AL355497.2 | 0 | NA | NA | NA | NA | NA |
| ULBP3 | 31.5777575 | 0.399946633 | 0.42783382 | 0.93481771 | 0.34988225 | 0.54756054 |
| PPP1R14C | 742.465829 | -0.474223115 | 0.25435515 | -1.8644133 | 0.06226369 | 0.16966969 |
| IYD | 1715.27973 | 2.554195843 | 0.67242375 | 3.79849144 | 0.00014558 | 0.00170923 |
| PLEKHG1 | 922.352049 | -0.782694132 | 0.24879685 | -3.1459165 | 0.00165567 | 0.01167061 |
| AL450344.2 | 4.1838906 | 0.64935039 | 0.66332981 | 0.97892538 | 0.32761685 | 0.52534243 |
| MTHFD1L | 1031.26474 | -1.441131875 | 0.27124795 | -5.3129687 | 1.08E-07 | 5.11E-06 |
| AL133260.2 | 0.21773222 | -0.339792043 | 3.08911945 | -0.1099964 | 0.91241222 | NA |
| AL138733.1 | 0.78558509 | 1.295748204 | 1.34265472 | 0.96506435 | 0.33451258 | NA |
| AL138733.2 | 1.56562477 | 1.627797015 | 1.42008635 | 1.14626622 | 0.25168501 | 0.44394206 |
| AKAP12 | 6351.86056 | -0.77124297 | 0.23027461 | -3.3492314 | 0.00081036 | 0.00664672 |
| ZBTB2 | 842.331066 | -0.435989268 | 0.27614845 | -1.5788221 | 0.11437688 | 0.26118101 |
| RMND1 | 1635.299 | 0.854040943 | 0.29402433 | 2.90466077 | 0.00367651 | 0.02139697 |
| ARMT1 | 2098.91273 | 0.479081337 | 0.20975767 | 2.2839753 | 0.02237298 | 0.0814944 |
| CCDC170 | 79.1081072 | 0.948139768 | 0.54126043 | 1.75172562 | 0.079821 | 0.2021448 |
| ESR1 | 268.149679 | 1.70129311 | 0.32942381 | 5.16445094 | 2.41E-07 | 9.92E-06 |
| AL356311.1 | 0.09997307 | 0.547239394 | 3.29377202 | 0.16614368 | 0.86804388 | NA |
| SYNE1 | 6617.61792 | 0.669092011 | 0.21125197 | 3.16726998 | 0.00153877 | 0.01102948 |
| SYNE1-AS1 | 38.1108259 | -0.451743367 | 0.26404479 | -1.7108588 | 0.08710717 | 0.21514231 |
| AL049548.1 | 25.9513539 | 0.230344066 | 0.60058122 | 0.38353524 | 0.70132294 | 0.82893183 |
| NANOGP11 | 0.13028434 | 1.049816488 | 3.28477129 | 0.31960109 | 0.74927075 | NA |
| MYCT1 | 586.226987 | -0.129106683 | 0.27623157 | -0.4673857 | 0.64022396 | 0.78714964 |
| VIP | 69.1211165 | 0.329098701 | 0.49501775 | 0.66482202 | 0.50616434 | 0.68543899 |
| AL160162.1 | 0 | NA | NA | NA | NA | NA |
| FBXO5 | 237.943817 | -0.30104544 | 0.25652234 | -1.1735642 | 0.24056961 | 0.43070275 |
| AL080276.2 | 401.777578 | 0.190699539 | 0.28365621 | 0.67229108 | 0.50139841 | 0.68210661 |
| MTRF1L | 872.302788 | -0.036145021 | 0.23131009 | -0.1562622 | 0.87582636 | 0.93547831 |
| RGS17 | 91.4210966 | 0.218236938 | 0.31232789 | 0.69874305 | 0.48471263 | 0.66910286 |
| AL590867.1 | 0.09644756 | -0.642905668 | 3.30558948 | -0.1944905 | 0.84579185 | NA |
| AL590867.3 | 0 | NA | NA | NA | NA | NA |
| OPRM1 | 3.64335224 | 0.736185871 | 0.67155024 | 1.0962484 | 0.27297008 | 0.46784227 |
| IPCEF1 | 357.019288 | 0.179045098 | 0.38409482 | 0.46614817 | 0.64110944 | 0.78758115 |
| CNKSR3 | 3031.10376 | -0.071887346 | 0.15191458 | -0.473209 | 0.63606409 | 0.7841886 |
| SCAF8 | 1894.67609 | 0.245488567 | 0.23400793 | 1.04906089 | 0.2941501 | 0.49086233 |
| TIAM2 | 1203.35836 | 0.002212681 | 0.1703974 | 0.01298542 | 0.98963943 | 0.99519138 |
| AL596202.1 | 141.782212 | 0.850588157 | 0.32091916 | 2.65047481 | 0.00803787 | 0.03824588 |
| TFB1M | 516.367513 | -0.089107434 | 0.16391793 | -0.54361 | 0.58670985 | 0.74774244 |
| CLDN20 | 4.66388571 | 2.082147786 | 0.88090039 | 2.36365861 | 0.01809548 | 0.06971187 |
| AL031773.1 | 0 | NA | NA | NA | NA | NA |

| | | | | | | |
|---|---|---|---|---|---|---|
| NOX3 | 0.51868141 | 2.366373314 | 1.95949696 | 1.20764327 | 0.22718451 | NA |
| AL512658.1 | 0 | NA | NA | NA | NA | NA |
| AL355297.3 | 13.5518416 | 0.338715058 | 0.57543965 | 0.5886196 | 0.55611648 | 0.72529758 |
| AL355297.4 | 49.2489255 | 0.249339466 | 0.25995191 | 0.95917535 | 0.33747042 | 0.53547716 |
| ARID1B | 2506.66649 | 0.257108767 | 0.10087009 | 2.54890984 | 0.01080602 | 0.04772313 |
| AL355297.2 | 0.07659028 | -0.642905668 | 3.30558948 | -0.1944905 | 0.84579185 | NA |
| AL049820.1 | 0 | NA | NA | NA | NA | NA |
| TMEM242 | 1247.03927 | -0.026332273 | 0.17724391 | -0.1485652 | 0.88189675 | 0.93863819 |
| AL390955.2 | 13.2542285 | 0.252802246 | 0.3968257 | 0.63706118 | 0.52408499 | 0.69922192 |
| ZDHHC14 | 1488.30754 | -0.378352975 | 0.2001056 | -1.8907665 | 0.05865552 | 0.16287786 |
| SNX9 | 5121.00948 | -0.302672934 | 0.24673918 | -1.2266918 | 0.21993844 | 0.4056752 |
| AL391863.2 | 0.26148673 | -0.174096114 | 3.29377188 | -0.0528562 | 0.95784652 | NA |
| AL391863.1 | 216.622806 | 0.638011471 | 0.3983546 | 1.60161694 | 0.10924034 | 0.25271233 |
| AL035634.1 | 0.27140021 | -0.179253794 | 2.94817043 | -0.0608017 | 0.95151713 | NA |
| SYNJ2 | 1054.11633 | -0.81689595 | 0.33589039 | -2.4320313 | 0.01501441 | 0.06091231 |
| SERAC1 | 776.020987 | -0.358663131 | 0.24006373 | -1.494033 | 0.135167 | 0.29259183 |
| GTF2H5 | 1151.85639 | 0.109633002 | 0.15519903 | 0.70640264 | 0.47993778 | 0.66459903 |
| TULP4 | 1267.82525 | -0.01930471 | 0.16758686 | -0.1151923 | 0.90829273 | 0.95418579 |
| AL360169.1 | 0 | NA | NA | NA | NA | NA |
| TMEM181 | 1516.4566 | -0.426413494 | 0.14589156 | -2.9228113 | 0.00346887 | 0.02045026 |
| DYNLT1 | 4993.8757 | -1.031969394 | 0.22512473 | -4.5839894 | 4.56E-06 | 0.00010936 |
| SYTL3 | 738.686324 | -2.182369104 | 0.29012377 | -7.5222002 | 5.39E-14 | 2.15E-11 |
| EZR | 20998.0571 | 0.105405921 | 0.17441889 | 0.60432629 | 0.54562673 | 0.71712709 |
| EZR-AS1 | 823.305302 | 0.559757064 | 0.30232106 | 1.85153184 | 0.06409308 | 0.17311631 |
| AL627422.2 | 2.07807601 | 0.302069602 | 0.88571581 | 0.34104574 | 0.73306915 | 0.84942807 |
| C6orf99 | 20.502393 | 1.201353908 | 0.46785164 | 2.56780957 | 0.01023434 | 0.04585887 |
| RSPH3 | 1093.59476 | -0.03569264 | 0.15298805 | -0.2333035 | 0.81552577 | 0.90056175 |
| AL035530.2 | 119.902531 | 0.605632967 | 0.24408708 | 2.48121678 | 0.01309347 | 0.05499759 |
| TAGAP | 149.246078 | 0.596157348 | 0.48728671 | 1.22342213 | 0.2211703 | 0.40730645 |
| AL356417.3 | 3.73292954 | -0.513399181 | 0.99378535 | -0.5166097 | 0.60542862 | 0.76224172 |
| AL356417.1 | 2.71398978 | -0.148577274 | 1.05362652 | -0.1410151 | 0.88785799 | 0.94201397 |
| AL356417.2 | 1.56560137 | -0.890721071 | 1.11397944 | -0.7995848 | 0.42395138 | 0.61643128 |
| FNDC1 | 99.9770968 | 1.337871948 | 0.43212508 | 3.09602943 | 0.00196131 | 0.01328872 |
| AL357832.1 | 0 | NA | NA | NA | NA | NA |
| LINC02529 | 1.85985706 | 3.952702426 | 1.3459913 | 2.93664783 | 0.00331781 | 0.01982115 |
| AL078604.2 | 74.9842423 | -3.089916919 | 0.56751667 | -5.4446275 | 5.19E-08 | 2.75E-06 |
| SOD2 | 168798.075 | -3.13430198 | 0.40512049 | -7.7367155 | 1.02E-14 | 5.27E-12 |
| WTAP | 6780.88303 | -1.133080453 | 0.18483583 | -6.1301992 | 8.78E-10 | 9.26E-08 |
| SOD2-OT1 | 218.665076 | -0.052610466 | 0.33409271 | -0.1574727 | 0.87487236 | 0.93495454 |
| ACAT2 | 2688.73161 | -0.498354715 | 0.28875741 | -1.7258595 | 0.08437269 | 0.21041513 |
| TCP1 | 9841.53568 | -0.356728492 | 0.22629935 | -1.5763567 | 0.11494362 | 0.26216589 |
| MRPL18 | 2230.8115 | -0.303357533 | 0.165192 | -1.8363936 | 0.06629946 | 0.17726497 |
| PNLDC1 | 25.4271086 | -1.437617265 | 0.81418217 | -1.7657194 | 0.07744293 | 0.19802062 |
| AL035691.1 | 0.12150485 | 1.04981648 | 3.28477129 | 0.31960109 | 0.74927075 | NA |
| MAS1 | 0.76436996 | 3.052085487 | 1.82596272 | 1.67149387 | 0.09462417 | NA |
| IGF2R | 7359.24594 | -0.016287605 | 0.24057568 | -0.0677026 | 0.94602236 | 0.97409658 |
| AIRN | 0.63559725 | 1.289644281 | 1.70311873 | 0.75722512 | 0.448915 | NA |
| SLC22A1 | 102.836607 | 1.147473198 | 0.42714727 | 2.68636436 | 0.00722343 | 0.03548468 |
| SLC22A2 | 7145.28681 | 1.372622997 | 0.46900145 | 2.92669245 | 0.00342587 | 0.02027841 |
| AL162582.1 | 41.1951453 | -0.506142911 | 0.56074054 | -0.902633 | 0.36672071 | 0.56380744 |
| SLC22A3 | 772.752994 | 0.099180554 | 0.3701951 | 0.26791428 | 0.7887653 | 0.88524604 |
| LPA | 115.677284 | 1.994517198 | 0.89834375 | 2.22021603 | 0.02640411 | 0.09197418 |
| AL109933.1 | 6.54682339 | 6.639705588 | 1.68833792 | 3.9326876 | 8.40E-05 | 0.0011059 |
| PLG | 14763.7534 | 2.474873566 | 0.69758963 | 3.54774995 | 0.00038854 | 0.0037325 |
| AL391361.2 | 0.59662445 | 1.289790168 | 1.72121352 | 0.74934931 | 0.45364669 | NA |
| AL391361.3 | 0 | NA | NA | NA | NA | NA |
| AL139393.1 | 0.41960597 | 0.363629406 | 2.84187143 | 0.12795421 | 0.89818521 | NA |
| AL139393.2 | 34.0610228 | -0.378072663 | 0.50158128 | -0.7537615 | 0.45099244 | 0.64079343 |
| MAP3K4 | 1187.04701 | 0.06910885 | 0.08835407 | 0.78218069 | 0.43410839 | 0.62605624 |
| AGPAT4 | 884.940349 | 0.253373883 | 0.32833885 | 0.77168414 | 0.44030152 | 0.63167136 |
| PRKN | 640.273749 | 1.842776531 | 0.25363614 | 7.26543363 | 3.72E-13 | 1.17E-10 |
| PACRG | 313.486589 | -0.17112169 | 0.31341464 | -0.5459914 | 0.58507187 | 0.74684603 |
| PACRG-AS2 | 0.23025741 | 1.79377171 | 3.01520403 | 0.5949089 | 0.55190437 | NA |
| PACRG-AS3 | 0.12059279 | 0.02007176 | 3.29738704 | 0.00608717 | 0.99514317 | NA |
| PACRG-AS1 | 8.02874582 | -0.515319473 | 0.55193474 | -0.9336602 | 0.35047923 | 0.54820198 |
| CAHM | 68.3082963 | -0.146481554 | 0.29989101 | -0.4884493 | 0.62523163 | 0.77675295 |
| QKI | 4368.1197 | -0.048135618 | 0.14190249 | -0.3392161 | 0.7344469 | 0.85063971 |
| AL445307.1 | 3.89400702 | 0.015210573 | 0.87847051 | 0.01731484 | 0.98618545 | 0.99391317 |
| AL137005.1 | 0 | NA | NA | NA | NA | NA |
| Z97205.2 | 0.19624763 | 0.18657164 | 3.29377195 | 0.05664376 | 0.95482897 | NA |
| Z97205.1 | 0 | NA | NA | NA | NA | NA |
| AL358972.1 | 0 | NA | NA | NA | NA | NA |
| AL450345.1 | 0.18380315 | -1.003574173 | 3.30558948 | -0.3035992 | 0.7614333 | NA |
| C6orf118 | 9.18682233 | -0.832543887 | 0.985329 | -0.84494 | 0.39814434 | 0.59310056 |
| PDE10A | 841.605733 | 0.480733125 | 0.20413005 | 2.35503362 | 0.01852103 | 0.07094706 |
| AL590302.2 | 2.34383438 | -1.828817631 | 1.38425349 | -1.321158 | 0.18644867 | 0.36422339 |
| LINC00473 | 582.688503 | 0.160617224 | 0.96836515 | 0.16586432 | 0.86826373 | 0.93147444 |
| LINC00602 | 105.861733 | 0.494621889 | 0.84102399 | 0.58811864 | 0.55645266 | 0.72552072 |
| AL627443.1 | 0.22147791 | 1.745651581 | 3.03531355 | 0.57511409 | 0.56521414 | NA |
| TBXT | 0.69414459 | 1.05638876 | 1.77076729 | 0.59657119 | 0.55079371 | NA |
| AL121956.1 | 0.28652914 | -0.179717927 | 2.92211968 | -0.0615026 | 0.95095895 | NA |
| AL121956.4 | 1.31765909 | 4.309413898 | 1.35993151 | 3.16884627 | 0.00153045 | 0.01099075 |
| PRR18 | 11.6603695 | -0.903878245 | 0.74182767 | -1.2184477 | 0.22305387 | 0.40912086 |
| SFT2D1 | 2337.84538 | -0.289921966 | 0.09867589 | -2.9381235 | 0.00330205 | 0.01974822 |
| MPC1 | 6105.242 | 0.653375879 | 0.21087655 | 3.0983809 | 0.00194581 | 0.01321088 |
| AL022069.1 | 27.4671145 | 2.143412949 | 0.44750255 | 4.78972226 | 1.67E-06 | 4.93E-05 |
| RPS6KA2 | 2480.4028 | -0.285174563 | 0.22766918 | -1.2525831 | 0.21035748 | 0.3937409 |
| AL159163.1 | 1380.90265 | -0.751191555 | 0.34179392 | -2.1977909 | 0.02796401 | 0.09605616 |
| RPS6KA2-AS | 0.06785815 | 0.795868262 | 3.28898349 | 0.24198001 | 0.80879565 | NA |
| RNASET2 | 3130.66176 | -1.030643294 | 0.4222803 | -2.4406615 | 0.01466039 | 0.05989069 |
| Z94721.1 | 5.6171592 | 0.787327123 | 0.55178856 | 1.42686379 | 0.15361916 | 0.31929828 |
| FGFR1OP | 1565.43936 | 0.166104351 | 0.19435907 | 0.85462617 | 0.39275813 | 0.58833401 |
| CCR6 | 83.45535 | 0.43023125 | 0.33575219 | 1.28139523 | 0.20005488 | 0.38143769 |
| AL121935.1 | 1.13109503 | 1.042311466 | 1.36505889 | 0.7635652 | 0.44512639 | 0.63568822 |
| AL121935.2 | 3.10563688 | 1.962289282 | 1.33417206 | 1.47079177 | 0.14134744 | 0.30175939 |
| TCP10L2 | 2.03262761 | 0.880107023 | 1.27864778 | 0.68831076 | 0.49125711 | 0.6742895 |
| GPR31 | 0.94357561 | 1.736884769 | 1.5139782 | 1.14723235 | 0.2512856 | NA |
| AL353747.4 | 0 | NA | NA | NA | NA | NA |
| AL353747.3 | 0.84626876 | -0.990544002 | 1.90856716 | -0.5189988 | 0.6037616 | NA |
| AL353747.2 | 1.57337226 | 0.338697701 | 1.6753249 | 0.20216837 | 0.8397851 | 0.91484357 |
| UNC93A | 55.5314209 | 2.971693394 | 0.71675563 | 4.14603424 | 3.38E-05 | 0.00054097 |
| TTLL2 | 10.0121688 | -1.349797784 | 0.67308251 | -2.0053972 | 0.0449206 | 0.13468032 |
| TCP10 | 0.37809939 | 0.703529056 | 2.2821952 | 0.30826857 | 0.75787798 | NA |
| LINC02538 | 9.00930546 | 2.164414128 | 0.62635777 | 3.45555567 | 0.00054916 | 0.00488382 |
| LINC02487 | 4.45077955 | 1.838018499 | 0.86978948 | 2.11317628 | 0.03458568 | 0.11173097 |
| LINC01558 | 176.640229 | 0.43404766 | 0.42939685 | 1.01083102 | 0.31209731 | 0.50910117 |
| AL009178.2 | 107.344809 | 0.291661347 | 0.4360771 | 0.66882977 | 0.50360407 | 0.68362192 |
| AL009178.3 | 75.3138874 | 0.028881021 | 0.47409969 | 0.06091761 | 0.95142482 | 0.97675915 |
| AFDN-DT | 131.233777 | 0.068297806 | 0.49551648 | 0.13783155 | 0.89037355 | 0.94316 |
| AFDN | 5402.55313 | -0.137582702 | 0.11102349 | -1.2392216 | 0.21526346 | 0.39969705 |
| KIF25-AS1 | 36.9346741 | 1.820259608 | 1.2511209 | 1.45490305 | 0.1456961 | 0.30825717 |
| KIF25 | 33.4119006 | 1.037824775 | 0.63047033 | 1.646112 | 0.0997407 | 0.23707971 |
| FRMD1 | 1516.00965 | 2.709932434 | 0.53500793 | 5.06521916 | 4.08E-07 | 1.53E-05 |

| | | | | | | |
|---|---|---|---|---|---|---|
| AL611929.1 | 1.4340379 | 0.476933134 | 1.54974838 | 0.30774875 | 0.75827352 | 0.86631656 |
| AL606970.4 | 0.54115819 | 1.803940942 | 2.00047145 | 0.90175791 | 0.36718549 | NA |
| AL606970.2 | 1.0539286 | 2.076230243 | 1.37906245 | 1.50553751 | 0.13218594 | 0.28834008 |
| AL606970.3 | 26.3744575 | 2.996711139 | 0.60587718 | 4.94607033 | 7.57E-07 | 2.57E-05 |
| DACT2 | 422.092478 | 1.807745191 | 0.54089812 | 3.34211771 | 0.00083142 | 0.00678147 |
| AL138918.1 | 0 | NA | NA | NA | NA | NA |
| SMOC2 | 949.108335 | 0.301668 | 0.52838177 | 0.5709281 | 0.56804839 | 0.73384869 |
| AL136099.1 | 0.54045117 | 0.27284675 | 2.23922053 | 0.12184899 | 0.90301862 | NA |
| AL109924.1 | 0.21442573 | -1.003574173 | 3.30558948 | -0.3035992 | 0.7614333 | NA |
| AL109924.2 | 0.40098181 | -1.795740071 | 2.80639654 | -0.639874 | 0.52225454 | NA |
| AL109924.3 | 0 | NA | NA | NA | NA | NA |
| LINC01615 | 6.16100992 | -1.372371317 | 0.71754608 | -1.9125898 | 0.05580059 | 0.15724102 |
| LINC02544 | 1.98587622 | -1.990457077 | 1.16678073 | -1.7059393 | 0.08801939 | 0.21694282 |
| BX322234.1 | 70.745446 | -0.30472445 | 0.30925998 | -0.9853342 | 0.32445992 | 0.52196243 |
| THBS2 | 2878.44866 | -1.4580059 | 0.37155529 | -3.9240618 | 8.71E-05 | 0.00113663 |
| BX322234.2 | 3.09707455 | 0.053491265 | 0.98176451 | 0.05448482 | 0.9565489 | 0.97931415 |
| LINC02519 | 21.1939859 | 0.713646409 | 0.43987638 | 1.62237946 | 0.10472213 | 0.24520711 |
| AL009176.1 | 23.7185192 | -0.240493383 | 0.43270056 | -0.5557963 | 0.57835009 | 0.74196507 |
| WDR27 | 1321.53547 | -0.089830987 | 0.20461411 | -0.4390264 | 0.66064243 | 0.80027291 |
| C6orf120 | 1752.95804 | -0.110934036 | 0.13727814 | -0.8080969 | 0.41903481 | 0.61189458 |
| PHF10 | 3993.63865 | -0.316559568 | 0.08865607 | -3.5706474 | 0.0003561 | 0.00348178 |
| AL354892.2 | 68.5059172 | -0.073106281 | 0.20838703 | -0.3508197 | 0.7257236 | 0.84465425 |
| AL354892.1 | 22.6077046 | -0.183852141 | 0.31733957 | -0.5793546 | 0.56234992 | 0.72983653 |
| TCTE3 | 330.151569 | -0.211712255 | 0.21608078 | -0.9797829 | 0.32719328 | 0.52502871 |
| ERMARD | 1191.69468 | 0.399207427 | 0.20401282 | 1.95677616 | 0.05037379 | 0.14640768 |
| LINC00242 | 26.6384005 | 1.227020603 | 0.43396363 | 2.82747334 | 0.00469169 | 0.0256202 |
| LINC00574 | 13.8751932 | 2.260777181 | 0.46528653 | 4.8588924 | 1.18E-06 | 3.73E-05 |
| AL049612.1 | 12.7404248 | 2.305397591 | 0.52011071 | 4.43251324 | 9.31E-06 | 0.00019294 |
| AL596442.1 | 0 | NA | NA | NA | NA | NA |
| AL596442.2 | 65.6571117 | -1.756581601 | 0.47617956 | -3.688906 | 0.00022522 | 0.00240545 |
| AL109910.1 | 0.08393208 | -0.642905668 | 3.30558948 | -0.1944905 | 0.84579185 | NA |
| AL109910.2 | 1.22422993 | 0.971559514 | 1.30083052 | 0.74687632 | 0.45513822 | 0.64454083 |
| LINC01624 | 3.77201826 | 1.00767151 | 0.70344804 | 1.43247469 | 0.15200802 | 0.31707904 |
| DLL1 | 960.508302 | -0.379834269 | 0.32505971 | -1.1685061 | 0.24260266 | 0.43301964 |
| FAM120B | 1743.83939 | 0.10814041 | 0.08784334 | 1.23105981 | 0.21830049 | 0.40337636 |
| AL078605.1 | 2.92865059 | 0.677571181 | 0.76045768 | 0.89100445 | 0.37292678 | 0.56916768 |
| PSMB1 | 7056.39568 | -0.425228457 | 0.17821377 | -2.3860584 | 0.01703005 | 0.06674827 |
| TBP | 886.140124 | -0.022089066 | 0.10404344 | -0.2123062 | 0.83186817 | 0.91063821 |
| PDCD2 | 2291.72073 | -0.326634072 | 0.1513555 | -2.1580588 | 0.03092326 | 0.10309056 |
| AL731661.2 | 0 | NA | NA | NA | NA | NA |
| AL731661.1 | 1.90835833 | -1.970131743 | 1.1531543 | -1.7084719 | 0.08754881 | 0.21600507 |
| AL731684.2 | 2.72877197 | 0.330879189 | 0.97656803 | 0.33881837 | 0.73474656 | 0.85072522 |
| AL731684.1 | 2.85902353 | -0.473495614 | 0.86253651 | -0.5489572 | 0.58303483 | 0.74532812 |
| AC215522.3 | 0 | NA | NA | NA | NA | NA |
| AC093627.1 | 1.17076779 | -3.438941773 | 1.75669488 | -1.9576204 | 0.05027457 | 0.14618375 |
| AC093627.6 | 8.7284115 | -1.116735262 | 0.94585914 | -1.180657 | 0.23773899 | 0.42708476 |
| AC093627.5 | 76.6508861 | 0.936697799 | 0.35720722 | 2.62228127 | 0.00873433 | 0.04067124 |
| AC093627.4 | 382.223716 | -0.288413546 | 0.33738838 | -0.8548414 | 0.39263897 | 0.58819298 |
| AC093627.7 | 487.360668 | 4.077677226 | 0.60405774 | 6.75047593 | 1.47E-11 | 2.86E-09 |
| AC093627.2 | 77.8745696 | 7.201244367 | 0.99285404 | 7.25307456 | 4.07E-13 | 1.26E-10 |
| AC093627.3 | 18.5900018 | 2.099922932 | 0.55834583 | 3.7609718 | 0.00016925 | 0.0019205 |
| FAM20C | 8011.57724 | 0.308398088 | 0.21101606 | 1.4614911 | 0.14388072 | 0.30558429 |
| AC187653.1 | 0.94520228 | -3.145204834 | 1.72358841 | -1.8248004 | 0.06803115 | NA |
| AC226118.1 | 7.92468897 | 1.541161738 | 0.60920761 | 2.52978084 | 0.01141338 | 0.0497308 |
| AC147651.3 | 0.91974761 | -1.585377203 | 2.00903423 | -0.789124 | 0.43003951 | NA |
| PDGFA | 1957.46455 | -0.672854609 | 0.22956892 | -2.9309482 | 0.00337929 | 0.02007025 |
| AC147651.1 | 148.205554 | 0.321423586 | 0.26204881 | 1.22657905 | 0.21998085 | 0.40571373 |
| PRKAR1B | 1608.35324 | 0.041534292 | 0.20685791 | 0.20078658 | 0.84086546 | 0.91512686 |
| AC147651.2 | 49.2049625 | 1.979171317 | 0.57441189 | 3.44556118 | 0.00056987 | 0.00502595 |
| AC147651.4 | 49.9353088 | -1.058279475 | 0.31822158 | -3.3256057 | 0.00088227 | 0.00711479 |
| DNAAF5 | 1629.90903 | 0.074623895 | 0.18718126 | 0.39867183 | 0.69013503 | 0.82150773 |
| SUN1 | 11162.1685 | -0.142011409 | 0.13472226 | -1.054105 | 0.29183484 | 0.48802618 |
| GET4 | 6043.66589 | -0.195383002 | 0.20710008 | -0.9434231 | 0.34546453 | 0.54322597 |
| AC073957.3 | 2234.59025 | 0.568735037 | 0.23255837 | 2.44555824 | 0.01446281 | 0.05929482 |
| ADAP1 | 1488.33021 | 0.239779041 | 0.27106052 | 0.88459597 | 0.37637458 | 0.57222018 |
| COX19 | 1781.34581 | 0.079719828 | 0.1145339 | 0.69603699 | 0.48640567 | 0.67045724 |
| CYP2W1 | 23.4082011 | 0.782934369 | 0.49717658 | 1.57476116 | 0.11531159 | 0.26274661 |
| C7orf50 | 3259.4016 | 0.1324124 | 0.16741372 | 0.79092918 | 0.42898532 | 0.62112297 |
| AC073957.1 | 1.59935743 | 1.649311731 | 1.24492389 | 1.32482936 | 0.18522775 | 0.36264428 |
| GPR146 | 679.798066 | -0.619000649 | 0.27265883 | -2.2702387 | 0.0231931 | 0.08367794 |
| AC073957.2 | 592.423816 | -0.689673332 | 0.27829107 | -2.4782446 | 0.01320306 | 0.0553521 |
| AC091729.1 | 6.46802232 | -0.02815189 | 0.64562195 | -0.0436043 | 0.96521983 | 0.98359669 |
| AC091729.2 | 11.0082937 | -1.856599905 | 1.15369779 | -1.6092602 | 0.10755946 | 0.24985648 |
| GPER1 | 846.115862 | 0.489821996 | 0.43980709 | 1.1137201 | 0.26539928 | 0.45946554 |
| ZFAND2A | 854.332505 | -0.555628569 | 0.40723822 | -1.3643822 | 0.17244731 | 0.34552357 |
| AC091729.3 | 264.527734 | 0.486329818 | 0.38957882 | 1.24834769 | 0.21190376 | 0.39549169 |
| UNCX | 51.5645946 | 0.300163459 | 0.69511208 | 0.43182023 | 0.66587207 | 0.80416975 |
| AC073094.1 | 0.39027547 | -0.032171818 | 2.84461719 | -0.0113097 | 0.99097634 | NA |
| MICALL2 | 4401.09981 | -1.288202837 | 0.28789155 | -4.4746115 | 7.66E-06 | 0.00016508 |
| AC102953.1 | 6.31104552 | 0.003764864 | 0.61739063 | 0.00609803 | 0.99513451 | 0.99770729 |
| AC102953.2 | 248.298779 | 0.196181475 | 0.20806183 | 0.9428999 | 0.34573211 | 0.54338751 |
| INTS1 | 6635.58351 | 0.334225359 | 0.11925816 | 2.80253655 | 0.00507025 | 0.0271477 |
| AC093734.1 | 4.6954879 | 0.27335736 | 0.86643436 | 0.31549691 | 0.7523844 | 0.86198795 |
| MAFK | 5681.97093 | -0.36796726 | 0.19228919 | -1.9136138 | 0.05566952 | 0.1570059 |
| TMEM184A | 3555.80627 | 0.025977327 | 0.10821754 | 0.24004727 | 0.81029361 | 0.89745233 |
| PSMG3 | 1116.34598 | -0.422585012 | 0.14604919 | -2.893443 | 0.00381043 | 0.02199405 |
| PSMG3-AS1 | 504.318547 | 0.92704943 | 0.19874379 | 4.66454542 | 3.09E-06 | 8.01E-05 |
| AC074389.2 | 0.03392908 | 0.380740593 | 3.29738709 | 0.11546736 | 0.90807469 | NA |
| ELFN1 | 410.760646 | 0.77824311 | 0.47512858 | 1.63796316 | 0.10142938 | 0.23986124 |
| AC074389.3 | 0.5065795 | 0.883799204 | 1.57345855 | 0.56169208 | 0.57432583 | NA |
| AC074389.1 | 0.22780565 | -0.339647149 | 3.05581276 | -0.1111479 | 0.91149907 | NA |
| ELFN1-AS1 | 4.70412858 | 0.610338777 | 0.76575454 | 0.79704232 | 0.42542645 | 0.61775857 |
| MAD1L1 | 1066.72632 | -0.252014334 | 0.17841337 | -1.4125305 | 0.15779381 | 0.32490223 |
| AC104129.1 | 0.79990294 | -0.454258052 | 1.58163522 | -0.2872078 | 0.77395318 | NA |
| MRM2 | 1141.84605 | 0.010927249 | 0.15118228 | 0.07227864 | 0.94238017 | 0.97270099 |
| NUDT1 | 217.608694 | -0.142660281 | 0.21336069 | -0.6686343 | 0.50372876 | 0.68364658 |
| SNX8 | 2198.32057 | -0.032201383 | 0.30638572 | -0.1051008 | 0.91629583 | 0.95922174 |
| EIF3B | 7352.24928 | -0.414622909 | 0.19736441 | -2.1007987 | 0.03565864 | 0.11410245 |
| CHST12 | 1259.32345 | -0.091982978 | 0.1831879 | -0.5021237 | 0.61558054 | 0.77018414 |
| GRIFIN | 4.14668652 | 1.981962196 | 0.65927627 | 3.0062696 | 0.00264474 | 0.0166727 |
| LFNG | 533.487435 | 0.846792135 | 0.35211484 | 2.40487487 | 0.016178 | 0.06436541 |
| BRAT1 | 3014.54801 | 0.20255034 | 0.10738139 | 1.88627047 | 0.05925852 | 0.1641436 |
| IQCE | 1989.82449 | 0.095153766 | 0.16353268 | 0.58186391 | 0.56065835 | 0.72862652 |
| TTYH3 | 12302.4039 | -0.113467469 | 0.31571577 | -0.3593975 | 0.71929772 | 0.84021138 |
| AMZ1 | 19.6881737 | -0.926073296 | 0.85153706 | -1.0875314 | 0.27680203 | 0.47220012 |
| GNA12 | 2442.17466 | -0.217370609 | 0.21051204 | -1.0325804 | 0.3018003 | 0.49868249 |
| CARD11 | 279.317078 | -0.411441589 | 0.48788542 | -0.843316 | 0.39905173 | 0.59390477 |
| AC004906.1 | 11.4361769 | 0.357262911 | 0.43720555 | 0.81715091 | 0.41384219 | 0.60767896 |
| AC024028.1 | 0 | NA | NA | NA | NA | NA |
| AC073316.1 | 2.1331361 | 0.051833878 | 1.26091764 | 0.04110806 | 0.96720975 | 0.98450423 |
| AC073316.2 | 11.1954048 | 0.313539789 | 0.41759255 | 0.75082707 | 0.45275674 | 0.64247942 |
| AC073316.3 | 2.57076045 | -0.841993361 | 0.92581086 | -0.9094658 | 0.36310428 | 0.56027266 |
| SDK1 | 841.350522 | -0.078750001 | 0.32900946 | -0.2393548 | 0.81083046 | 0.89770758 |

| | | | | | | |
|---|---|---|---|---|---|---|
| AC011284.1 | 1.4308413 | -0.009051208 | 1.23265877 | -0.0073428 | 0.99414132 | 0.99728027 |
| AC017000.1 | 78.1540434 | 2.195974019 | 0.47808174 | 4.59330243 | 4.36E-06 | 0.00010574 |
| FOXK1 | 2575.81757 | -0.301499364 | 0.15522928 | -1.9422842 | 0.05210271 | 0.15002351 |
| AP5Z1 | 3887.91425 | -0.623569349 | 0.31372405 | -1.9876364 | 0.04685191 | 0.13891218 |
| RADIL | 90.4174727 | 0.047081812 | 0.45068693 | 0.10446678 | 0.91679893 | 0.95955498 |
| PAPOLB | 0.2516613 | 0.078560428 | 3.04321442 | 0.02581495 | 0.97940494 | NA |
| MMD2 | 0.03392908 | 0.380740593 | 3.29738709 | 0.11546736 | 0.90807469 | NA |
| RBAK-RBAKI | 130.708254 | -0.27061832 | 0.27981831 | -0.9671216 | 0.33348327 | 0.53148518 |
| RBAK | 898.955493 | -0.297269099 | 0.20636066 | -1.4405319 | 0.14971698 | 0.31399129 |
| RBAKDN | 15.833357 | -0.397924057 | 0.48549335 | -0.8196282 | 0.41242807 | 0.6062516 |
| WIPI2 | 5218.21777 | -0.290396491 | 0.16034848 | -1.8110336 | 0.07013565 | 0.18450489 |
| SLC29A4 | 702.460481 | -0.301152749 | 0.38268169 | -0.7869536 | 0.431309 | 0.62338276 |
| TNRC18 | 7970.91023 | 0.381405212 | 0.20303691 | 1.87850183 | 0.06031255 | 0.16617301 |
| AC093620.1 | 23.8619221 | -0.234786277 | 0.40761274 | -0.5760033 | 0.56461296 | 0.73156144 |
| AC092171.1 | 15.1265881 | -0.109677244 | 0.3730201 | -0.294025 | 0.76873878 | 0.87219868 |
| AC092171.4 | 57.0822735 | 0.154523671 | 0.4322426 | 0.35749292 | 0.72072283 | 0.84109846 |
| AC092171.3 | 108.919181 | 0.536097993 | 0.27654537 | 1.93855351 | 0.05255573 | 0.15080463 |
| AC092171.5 | 54.4282321 | 0.441304534 | 0.37181508 | 1.18689252 | 0.23527002 | 0.42412294 |
| FBXL18 | 184.25264 | 0.597579603 | 0.27183485 | 2.19831859 | 0.02792641 | 0.09597543 |
| AC092171.2 | 599.901457 | -0.140300489 | 0.28961607 | -0.4844361 | 0.62807638 | 0.77870958 |
| ACTB | 129016.807 | -0.217513596 | 0.27304595 | -0.796619 | 0.42567232 | 0.6180135 |
| AC006483.2 | 2.70121501 | -0.652367411 | 1.16172711 | -0.5615496 | 0.57442292 | 0.7387959 |
| FSCN1 | 1562.99695 | -0.145318861 | 0.22516718 | -0.6453821 | 0.51867962 | 0.69552919 |
| RNF216 | 2329.88124 | -0.051634154 | 0.19791016 | -0.2608969 | 0.79417199 | 0.88844572 |
| OCM | 9.69018553 | 1.648116274 | 0.51155158 | 3.22179884 | 0.00127389 | 0.00949473 |
| CCZ1 | 664.545942 | -0.479514898 | 0.29034562 | -1.6515314 | 0.09863012 | 0.23543735 |
| RSPH10B | 262.462508 | -0.731125071 | 0.34497218 | -2.119374 | 0.03405887 | 0.11043065 |
| PMS2 | 862.239873 | 0.601283463 | 0.22028247 | 2.72960191 | 0.00634108 | 0.03211339 |
| AIMP2 | 1657.61683 | -0.145021565 | 0.18504341 | -0.7837164 | 0.43320651 | 0.62533697 |
| EIF2AK1 | 6720.56394 | 0.083383837 | 0.09212199 | 0.90514582 | 0.36538814 | 0.56248017 |
| ANKRD61 | 11.7423878 | 0.168665552 | 0.4061062 | 0.41532375 | 0.67790491 | 0.81285707 |
| AC004895.1 | 16.4359889 | 0.963354465 | 0.35097768 | 2.74477413 | 0.00605526 | 0.03099323 |
| USP42 | 751.910732 | -0.194500146 | 0.1670786 | -1.1641236 | 0.24437392 | 0.43538547 |
| CYTH3 | 1584.2016 | 0.310517157 | 0.21759567 | 1.42703736 | 0.15356913 | 0.31921944 |
| FAM220A | 629.746211 | 0.015982957 | 0.09921648 | 0.16109176 | 0.87202113 | 0.93327291 |
| RAC1 | 13121.7868 | -0.463693387 | 0.12870561 | -3.6027442 | 0.00031488 | 0.00314637 |
| DAGLB | 1765.42679 | -0.531754173 | 0.21732077 | -2.4468631 | 0.01441056 | 0.05915388 |
| KDELR2 | 8601.49889 | -0.209433236 | 0.15904735 | -1.316798 | 0.18790633 | 0.36609456 |
| GRID2IP | 76.2071021 | -0.023232978 | 0.31092443 | -0.0747223 | 0.94043569 | 0.97160669 |
| ZDHHC4 | 1987.34189 | 0.016398426 | 0.09111773 | 0.17996966 | 0.85717639 | 0.92528312 |
| AC079742.1 | 835.944857 | -6.77E-06 | 0.08892157 | -7.61E-05 | 0.99993924 | 0.99993924 |
| C7orf26 | 1421.59533 | -0.168767579 | 0.15941652 | -1.058658 | 0.28975554 | 0.48593234 |
| ZNF853 | 472.336866 | -0.206735803 | 0.34180808 | -0.6048301 | 0.54529191 | 0.71697611 |
| ZNF316 | 3771.54998 | -0.466013293 | 0.19606455 | -2.376836 | 0.01746185 | 0.06788574 |
| AC073343.2 | 9.19540796 | 1.057899449 | 0.43528214 | 2.43037642 | 0.01508315 | 0.06113653 |
| ZNF12 | 1279.80101 | -0.071046175 | 0.11319821 | -0.6276263 | 0.53024876 | 0.70472412 |
| RSPH10B2 | 189.210207 | -0.56209549 | 0.38152785 | -1.4732751 | 0.14067686 | 0.30086598 |
| CCZ1B | 821.074468 | -0.256584194 | 0.27868901 | -0.9206829 | 0.35721603 | 0.55475776 |
| C1GALT1 | 2237.59248 | 0.124572869 | 0.23941368 | 0.52032477 | 0.60283723 | 0.76012907 |
| COL28A1 | 691.739242 | -0.074620099 | 0.39148314 | -0.1906087 | 0.84883216 | 0.92053414 |
| AC004982.1 | 214.566438 | -0.396818877 | 0.18615021 | -2.1317133 | 0.03303042 | 0.10806603 |
| AC004982.2 | 469.926865 | -0.520351579 | 0.22691044 | -2.2932024 | 0.02183635 | 0.080073 |
| MIOS | 1453.42298 | -0.40493063 | 0.15338829 | -2.6399057 | 0.00829291 | 0.03910659 |
| AC004948.1 | 10.3802671 | 0.234385505 | 0.49271409 | 0.47570287 | 0.63428608 | 0.7832049 |
| RPA3 | 1037.83329 | 0.259482577 | 0.16128547 | 1.60884042 | 0.10765124 | 0.25000116 |
| UMAD1 | 1134.84403 | 0.122387844 | 0.11390182 | 1.07450293 | 0.28259733 | 0.47857314 |
| AC006042.4 | 4.99084496 | 1.499676783 | 0.54543761 | 2.74949282 | 0.00596876 | 0.03066357 |
| AC006042.2 | 6.19155594 | 1.202173866 | 0.65864458 | 1.82522396 | 0.06796724 | 0.18025707 |
| GLCCI1 | 686.942335 | 0.230802732 | 0.18026228 | 1.28037177 | 0.20041442 | 0.38194177 |
| ICA1 | 1733.96873 | -0.572787286 | 0.18262303 | -3.136446 | 0.00171009 | 0.01194989 |
| AC006042.1 | 443.297437 | -0.586536739 | 0.24861912 | -2.3591779 | 0.01831547 | 0.07035383 |
| AC007009.1 | 49.1298671 | -0.63507619 | 0.31513838 | -2.0152296 | 0.0438806 | 0.13269855 |
| AC007128.1 | 49.1877639 | -0.613351681 | 0.31322706 | -1.9581695 | 0.05021012 | 0.14604466 |
| AC007128.2 | 0.08666371 | -0.642905668 | 3.30558948 | -0.1944905 | 0.84579185 | NA |
| NXPH1 | 0 | NA | NA | NA | NA | NA |
| AC004852.2 | 0 | NA | NA | NA | NA | NA |
| AC060834.2 | 0.15661533 | -1.003574173 | 3.30558948 | -0.3035992 | 0.7614333 | NA |
| AC004936.1 | 1.49036249 | 0.446579667 | 1.35969698 | 0.32844058 | 0.74257856 | 0.85547591 |
| AC004949.1 | 16.3684271 | 0.172205327 | 0.34289624 | 0.50220827 | 0.61552103 | 0.77018414 |
| AC004415.1 | 0 | NA | NA | NA | NA | NA |
| NDUFA4 | 10715.1996 | 0.108236354 | 0.17126945 | 0.63196532 | 0.52740954 | 0.70212993 |
| AC007029.1 | 1.40774148 | 0.792366969 | 1.06408243 | 0.74464811 | 0.45648448 | 0.64589761 |
| PHF14 | 1093.66467 | 0.021296139 | 0.12067518 | 0.17647489 | 0.85992088 | 0.92701929 |
| THSD7A | 1504.54633 | 0.141939417 | 0.30276863 | 0.46880491 | 0.63920909 | 0.78670511 |
| AC004160.2 | 20.6237115 | 0.864183275 | 0.43338052 | 1.99405194 | 0.04614638 | 0.13734537 |
| TMEM106B | 3445.78296 | 0.220559931 | 0.18652315 | 1.18248022 | 0.23701521 | 0.42624913 |
| VWDE | 30.8196707 | 0.156729046 | 0.55518426 | 0.28230095 | 0.77771275 | 0.87833472 |
| AC013470.2 | 47.876172 | 0.808019484 | 0.57442038 | 1.40666924 | 0.15952549 | 0.32726678 |
| AC005281.1 | 257.670596 | 0.532260359 | 0.27049787 | 1.96770632 | 0.04910184 | 0.14391636 |
| SCIN | 5920.35769 | 0.057948759 | 0.2202267 | 0.2631323 | 0.79244861 | 0.8874536 |
| ARL4A | 527.617687 | -1.172483814 | 0.38590961 | -3.0382343 | 0.00237969 | 0.01542586 |
| AC011287.1 | 1.70821601 | 0.598758363 | 1.3226353 | 0.45270103 | 0.65076404 | 0.7948056 |
| AC011287.2 | 0.10721287 | -0.642905668 | 3.30558948 | -0.1944905 | 0.84579185 | NA |
| AC005019.2 | 0.08695548 | -0.642905668 | 3.30558948 | -0.1944905 | 0.84579185 | NA |
| ETV1 | 315.763845 | -0.472294823 | 0.33427127 | -1.4129088 | 0.15768254 | 0.32479992 |
| DGKB | 56.9722374 | 3.150322718 | 0.52323925 | 6.02080734 | 1.74E-09 | 1.60E-07 |
| AC006150.1 | 0 | NA | NA | NA | NA | NA |
| AC006458.1 | 0.36186486 | -1.005075207 | 2.29764841 | -0.4374365 | 0.66179484 | NA |
| AGMO | 292.726327 | 1.402716816 | 0.37044217 | 3.78660131 | 0.00015272 | 0.00177883 |
| MEOX2 | 34.4082218 | 0.857426076 | 0.68399863 | 1.2535494 | 0.21000583 | 0.39331348 |
| AC005550.2 | 1.46058093 | 0.091771752 | 1.18874778 | 0.07720036 | 0.93846416 | 0.97067268 |
| LINC02587 | 2.00400267 | -0.460955285 | 1.32234191 | -0.3485901 | 0.72739707 | 0.84604913 |
| AC005550.1 | 0.43450912 | 0.512284934 | 2.22932558 | 0.22979368 | 0.81825209 | NA |
| AC006041.1 | 1.61747768 | 1.782076333 | 1.3978587 | 1.27486157 | 0.20235824 | 0.38417472 |
| ISPD | 170.659024 | 0.790950532 | 0.30183322 | 2.62048869 | 0.00878038 | 0.04082855 |
| ISPD-AS1 | 2.75114551 | 0.073493457 | 0.7889228 | 0.09315672 | 0.92577906 | 0.9640363 |
| SOSTDC1 | 654.516218 | 2.275431495 | 0.49199898 | 4.62487031 | 3.75E-06 | 9.35E-05 |
| AC005014.3 | 3.49476128 | 0.549105977 | 1.02930125 | 0.53347451 | 0.59370515 | 0.75282738 |
| LRRC72 | 0.19616881 | -0.340115482 | 3.16778127 | -0.1073671 | 0.91449776 | NA |
| AC005014.2 | 18.0667971 | -0.321171385 | 0.39334623 | -0.8165107 | 0.41420813 | 0.60794725 |
| ANKMY2 | 1112.71079 | 0.307309865 | 0.19973486 | 1.538589 | 0.12390467 | 0.27551784 |
| BZW2 | 1922.22349 | -1.055296322 | 0.27468296 | -3.8418704 | 0.0001221 | 0.00148241 |
| AC073333.1 | 391.067774 | -1.114290403 | 0.29875125 | -3.7298268 | 0.00019161 | 0.00210885 |
| TSPAN13 | 814.532635 | -0.981542965 | 0.33166389 | -2.9594508 | 0.00308188 | 0.01873856 |
| AGR2 | 701.159863 | -2.119698228 | 0.62417694 | -3.3959893 | 0.00068381 | 0.00578507 |
| AGR3 | 94.9185475 | -3.098883398 | 0.62549797 | -4.9542661 | 7.26E-07 | 2.48E-05 |
| AHR | 1696.19153 | -0.628576043 | 0.19796567 | -3.175177 | 0.00149745 | 0.01079491 |
| AC073332.1 | 34.8454569 | -0.017195169 | 0.2610431 | -0.065871 | 0.94748053 | 0.97472249 |
| AC019117.3 | 172.96512 | 0.257982899 | 0.3056806 | 0.84396229 | 0.39869048 | 0.59358697 |
| AC019117.2 | 28.9091665 | -3.237024463 | 0.92649935 | -3.4938227 | 0.00047616 | 0.0043674 |
| AC019117.1 | 23.3227452 | -0.921297983 | 0.56970274 | -1.6171556 | 0.10584469 | 0.24695847 |
| AC006482.1 | 0 | NA | NA | NA | NA | NA |
| SNX13 | 2053.49789 | 0.134450827 | 0.07836308 | 1.71574189 | 0.08620929 | 0.21365969 |

| | | | | | | |
|---|---|---|---|---|---|---|
| PRPS1L1 | 0.34715973 | -0.995035912 | 3.29552073 | -0.3019359 | 0.76270095 | NA |
| HDAC9 | 269.453854 | 0.393399594 | 0.27832581 | 1.41344991 | 0.15752348 | 0.32470055 |
| AC010082.1 | 0 | NA | NA | NA | NA | NA |
| AC004994.1 | 0.25163508 | -1.364242679 | 3.30558948 | -0.4127078 | 0.67982068 | NA |
| TWIST1 | 48.6188402 | -1.228492 | 0.42434175 | -2.8950533 | 0.00379094 | 0.02192261 |
| AC003986.2 | 2.03325029 | -1.253080623 | 1.11043867 | -1.1284555 | 0.2591276 | 0.45187688 |
| AC003986.3 | 0 | NA | NA | NA | NA | NA |
| AC003986.1 | 0 | NA | NA | NA | NA | NA |
| FERD3L | 0 | NA | NA | NA | NA | NA |
| AC007091.1 | 0.19642063 | 0.18657164 | 3.29377195 | 0.05664376 | 0.95482897 | NA |
| TWISTNB | 664.700968 | 0.034350442 | 0.16847224 | 0.20389377 | 0.83843651 | 0.91449053 |
| TMEM196 | 4.61577092 | -1.798137394 | 1.47604676 | -1.2182117 | 0.22314354 | 0.40922841 |
| AC004543.1 | 2.17518638 | -0.119260409 | 1.18951703 | -0.1002595 | 0.92013829 | 0.96099041 |
| AC007001.1 | 4.69293218 | -1.229162631 | 0.75343582 | -1.6314099 | 0.10280386 | 0.24201911 |
| MACC1 | 1679.73521 | -0.548825703 | 0.19690772 | -2.7872229 | 0.00531619 | 0.02817848 |
| MACC1-AS1 | 5.52856822 | -1.153084807 | 0.96359656 | -1.1966469 | 0.23144422 | 0.41934754 |
| AC099342.1 | 1.0368505 | -0.355531081 | 1.30465926 | -0.2725088 | 0.78523085 | 0.88280468 |
| AC004130.1 | 92.192364 | 0.410119729 | 0.3860424 | 1.06236965 | 0.28806791 | 0.48426636 |
| ITGB8 | 5280.73704 | 0.109935527 | 0.25127365 | 0.43751315 | 0.66173924 | 0.80097603 |
| ABCB5 | 6.12583085 | -0.751423326 | 0.78038141 | -0.9628924 | 0.33560152 | 0.5337965 |
| SP8 | 0.24357081 | -1.364242679 | 3.30558948 | -0.4127078 | 0.67982068 | NA |
| LINC01162 | 0 | NA | NA | NA | NA | NA |
| SP4 | 275.994107 | 0.317255024 | 0.16577944 | 1.9137175 | 0.05565626 | 0.15698531 |
| DNAH11 | 200.865615 | -0.292993332 | 0.43396 | -0.6751621 | 0.4995728 | 0.68095398 |
| CDCA7L | 278.146997 | -0.144806229 | 0.32272112 | -0.4487039 | 0.65364526 | 0.79659051 |
| RAPGEF5 | 2243.86418 | -1.246159611 | 0.25652605 | -4.8578287 | 1.19E-06 | 3.74E-05 |
| STEAP1B | 8.97203707 | -1.4579822 | 0.70150237 | -2.078371 | 0.0376752 | 0.11844782 |
| AC002480.2 | 60.4259709 | -1.060781568 | 0.59880488 | -1.7714979 | 0.07647795 | 0.19632326 |
| AC002480.3 | 5.11317935 | -0.245687836 | 0.95157579 | -0.2581905 | 0.79625988 | 0.8894177 |
| AC002480.4 | 0.42861074 | 2.048321891 | 3.21841163 | 0.63643875 | 0.52449049 | NA |
| AC073072.1 | 64.0903709 | -0.663987176 | 0.8422068 | -0.7883897 | NA | NA |
| IL6 | 237.324932 | -0.979780552 | 0.86144987 | -1.1373622 | NA | NA |
| TOMM7 | 5493.43477 | 0.056947075 | 0.1826387 | 0.31180179 | 0.75519116 | 0.86400238 |
| FAM126A | 627.10284 | 0.115824838 | 0.19295128 | 0.60028021 | 0.5483195 | 0.71900474 |
| KLHL7-DT | 22.5404785 | 1.491462157 | 0.72205624 | 2.06557617 | 0.03886852 | 0.12125432 |
| KLHL7 | 1205.09786 | 0.181075904 | 0.12054086 | 1.50219517 | 0.13304671 | 0.28952317 |
| NUPL2 | 1443.35187 | -0.16875474 | 0.13154036 | -1.2829122 | 0.19952283 | 0.3807379 |
| AC005082.1 | 503.901386 | -0.304383003 | 0.40563837 | -0.7503802 | 0.45302575 | 0.64266256 |
| GPNMB | 3216.24831 | -0.295299083 | 0.44263614 | -0.6671373 | 0.50468443 | 0.68403209 |
| MALSU1 | 1075.33667 | -0.251416064 | 0.13406496 | -1.8753301 | 0.06074732 | 0.16691996 |
| IGF2BP3 | 25.2628137 | -0.459361694 | 0.4579528 | -1.0030765 | 0.31582395 | 0.51295132 |
| TRA2A | 4282.93482 | -0.153931791 | 0.19441848 | -0.7917549 | 0.42850358 | 0.62089342 |
| CCDC126 | 412.230611 | 0.040380568 | 0.19374704 | 0.20841902 | 0.83490181 | 0.91244398 |
| FAM221A | 470.51907 | -0.202852014 | 0.1768887 | -1.1467777 | 0.25147352 | 0.44374835 |
| AC006026.3 | 36.1595817 | 0.10464894 | 0.35835302 | 0.29202751 | 0.77026559 | 0.87299194 |
| STK31 | 36.639158 | -0.124610989 | 0.53314981 | -0.233726 | 0.81519768 | 0.90043158 |
| AC003087.1 | 0 | NA | NA | NA | NA | NA |
| AC004485.1 | 111.163585 | 1.859994522 | 0.68775229 | 2.70445413 | 0.00684167 | 0.03398284 |
| NPY | 5.3203408 | -0.22612752 | 0.8358953 | -0.2705213 | 0.7867592 | 0.88392136 |
| MPP6 | 1046.37625 | -0.103095583 | 0.24205013 | -0.4259266 | 0.67016136 | 0.80739012 |
| GSDME | 1050.40217 | 0.343918376 | 0.2260254 | 1.52159169 | 0.12811142 | 0.28219445 |
| OSBPL3 | 2920.67928 | -0.172445752 | 0.30084963 | -0.5731958 | 0.56651212 | 0.73284279 |
| CYCS | 7981.96502 | 0.0689795 | 0.17241268 | 0.40008368 | 0.68909488 | 0.82063171 |
| C7orf31 | 203.911364 | -0.01231066 | 0.1813053 | -0.0679002 | 0.94586511 | 0.97407741 |
| NPVF | 0 | NA | NA | NA | NA | NA |
| AC003985.1 | 0.17515154 | 1.45420457 | 3.2783972 | 0.44357181 | 0.65735219 | NA |
| AC003985.2 | 0.36244621 | -1.02786398 | 1.94512821 | -0.5284299 | 0.59720096 | NA |
| AC091705.1 | 0 | NA | NA | NA | NA | NA |
| AC005165.1 | 20.5160767 | 0.152644314 | 0.63989175 | 0.23854709 | 0.8114568 | 0.89821247 |
| AC010719.1 | 61.5642232 | 0.2249212 | 0.39146301 | 0.57456566 | 0.56558508 | 0.73214268 |
| NFE2L3 | 1124.63513 | -0.817955782 | 0.21301214 | -3.8399492 | 0.00012306 | 0.0014867 |
| AC004520.1 | 202.966096 | 0.239589134 | 0.30130937 | 0.79515992 | 0.42652048 | 0.61883605 |
| HNRNPA2B1 | 40885.3517 | -0.170935699 | 0.13354176 | -1.2800168 | 0.20053921 | 0.38209372 |
| CBX3 | 7736.50689 | -0.952290829 | 0.21744385 | -4.3794793 | 1.19E-05 | 0.00023569 |
| SNX10 | 4572.804 | -1.34334717 | 0.26682312 | -5.0345981 | 4.79E-07 | 1.75E-05 |
| AC004540.2 | 562.349706 | -0.571224146 | 0.20137354 | -2.8366395 | 0.00455911 | 0.02510936 |
| AC004540.1 | 181.238705 | 0.515487354 | 0.25825777 | 1.99601873 | 0.04593188 | 0.13690293 |
| KIAA0087 | 17.7680065 | 2.04741115 | 0.71952304 | 2.84551158 | 0.00443401 | 0.02462097 |
| AC004947.1 | 29.8444608 | -0.014423118 | 0.54289048 | -0.0265673 | 0.97880488 | 0.99044497 |
| C7orf71 | 3.13102671 | -0.023445365 | 0.77021452 | -0.03044 | 0.97571611 | 0.98876465 |
| SKAP2 | 2609.93391 | 0.176036136 | 0.18976462 | 0.92765521 | 0.35358645 | 0.55115076 |
| HOXA1 | 90.0700844 | -0.497552538 | 0.19047379 | -2.6121837 | 0.00899659 | 0.04166454 |
| HOTAIRM1 | 1574.27793 | -0.848555672 | 0.25640955 | -3.3093763 | 0.00093504 | 0.00745685 |
| HOXA2 | 186.370722 | 0.091604595 | 0.35612068 | 0.25722908 | 0.79700194 | 0.88983348 |
| HOXA3 | 1223.09146 | 0.564426959 | 0.2570655 | 2.19565423 | 0.0281167 | 0.09641579 |
| HOXA-AS2 | 1408.13385 | 0.436349756 | 0.2879554 | 1.51533796 | 0.12968684 | 0.28465473 |
| HOXA4 | 573.910822 | -0.169612074 | 0.24599003 | -0.6895079 | 0.49050368 | 0.67369536 |
| HOXA-AS3 | 1156.52402 | 0.509888831 | 0.26132239 | 1.95118691 | 0.05103482 | 0.14780293 |
| HOXA5 | 1304.93349 | -0.073539183 | 0.22297733 | -0.3298056 | 0.74154682 | 0.85475107 |
| HOXA6 | 172.104714 | 0.938474155 | 0.31481445 | 2.98103897 | 0.00287272 | 0.01773543 |
| HOXA7 | 1104.33909 | 0.548440889 | 0.31676032 | 1.73140653 | 0.08337928 | 0.20856685 |
| HOXA9 | 2454.26849 | -0.038364195 | 0.17500175 | -0.2192218 | 0.82647729 | 0.90741548 |
| AC004080.3 | 902.833181 | 0.184576484 | 0.17021608 | 1.08436576 | 0.27820266 | 0.47379257 |
| HOXA10-AS | 793.78999 | 0.114618447 | 0.18984359 | 0.603752 | 0.54600854 | 0.71722681 |
| HOXA10 | 1192.95012 | 0.126592747 | 0.19075256 | 0.66364899 | 0.50691499 | 0.68596235 |
| HOXA11 | 250.446166 | 0.405095061 | 0.30170091 | 1.34270412 | 0.1793678 | 0.35499316 |
| HOXA11-AS | 211.958241 | 0.196249796 | 0.33755635 | 0.58138381 | 0.5609818 | 0.72886727 |
| AC004080.2 | 14.4173955 | -0.192121049 | 0.55871154 | -0.3438645 | 0.73094821 | 0.84798182 |
| HOXA13 | 9.89214813 | 0.613729939 | 0.60717693 | 1.01079258 | 0.31211572 | 0.50910117 |
| HOTTIP | 1.48282264 | -0.120593775 | 1.22845926 | -0.0981667 | 0.92179994 | 0.96181159 |
| AC004080.1 | 3.70074414 | 1.297302909 | 0.81322577 | 1.59525553 | 0.11065511 | 0.25486725 |
| EVX1-AS | 4.39990734 | 1.334609688 | 1.05158657 | 1.26913915 | 0.20439144 | 0.3866976 |
| EVX1 | 4.95967647 | 1.109323239 | 0.89843807 | 1.23472421 | 0.21693316 | 0.40175065 |
| AC004009.1 | 0.29111801 | -0.179913104 | 2.91136792 | -0.0617968 | 0.95072468 | NA |
| HIBADH | 6541.42383 | 1.012424381 | 0.32938467 | 3.07368394 | 0.00211433 | 0.014037 |
| AC005091.1 | 230.26287 | 0.563503613 | 0.29498583 | 1.91027348 | 0.05609801 | 0.15784299 |
| TAX1BP1 | 7225.25406 | -0.08992105 | 0.14580844 | -0.6167067 | 0.53742817 | 0.71112353 |
| JAZF1 | 619.685319 | -0.372675284 | 0.25632278 | -1.4539297 | 0.14596581 | 0.30859604 |
| JAZF1-AS1 | 30.293108 | 0.23968781 | 0.35555317 | 0.67412649 | 0.5002309 | 0.6813579 |
| CREB5 | 754.772775 | -0.631251903 | 0.3412497 | -1.849824 | 0.06433891 | 0.17352959 |
| TRIL | 676.416492 | 1.697116107 | 0.39991865 | 4.24365329 | 2.20E-05 | 0.00038353 |
| AC005013.1 | 66.7048476 | 2.037112092 | 0.25797237 | 7.89662906 | 2.87E-15 | 1.80E-12 |
| AC005162.3 | 179.954374 | 0.823727479 | 0.38637523 | 2.13193655 | 0.03301206 | 0.10803278 |
| AC005162.2 | 0.40577507 | 0.329533408 | 1.86763548 | 0.17644418 | 0.859945 | NA |
| CPVL | 5432.60238 | 0.098853441 | 0.29946979 | 0.33009487 | 0.74132828 | 0.85469576 |
| AC005162.1 | 0.65345429 | 2.20091634 | 1.61009681 | 1.36694659 | 0.17164205 | NA |
| AC005232.1 | 5.38497129 | 0.288004973 | 0.92528403 | 0.31126115 | 0.7556021 | 0.86427612 |
| AC004593.3 | 1193.70488 | 0.632040087 | 0.21211737 | 2.97967153 | 0.00288558 | 0.01779945 |
| CHN2 | 747.323778 | 0.770319642 | 0.22036141 | 3.49571022 | 0.0004728 | 0.00434242 |
| AC004593.1 | 0.74028969 | 1.756802888 | 1.43308835 | 1.22588596 | 0.22024159 | NA |
| AC007255.1 | 91.1919418 | 0.259524823 | 0.27296674 | 0.95075622 | 0.34172814 | 0.53967632 |
| PRR15 | 1122.73391 | -0.859394495 | 0.1793173 | -4.792591 | 1.65E-06 | 4.87E-05 |
| WIPF3 | 520.725533 | -0.308821286 | 0.30615723 | -1.0087016 | 0.31311777 | 0.51003247 |

| | | | | | | |
|---|---|---|---|---|---|---|
| SCRN1 | 3253.11778 | -0.106004553 | 0.17483212 | -0.606322 | 0.54430097 | 0.71629986 |
| AC007285.1 | 9.48222333 | -0.317648118 | 0.44938381 | -0.7068526 | 0.47965808 | 0.66428148 |
| FKBP14 | 570.274448 | -0.350755771 | 0.2095506 | -1.6738476 | 0.09416055 | 0.22800861 |
| PLEKHA8 | 937.696398 | -0.278304686 | 0.22836585 | -1.2186791 | 0.22296602 | 0.40907354 |
| AC007285.2 | 2.68540713 | -0.894312612 | 0.73639168 | -1.2144524 | 0.22457503 | 0.41071112 |
| MTURN | 4921.4133 | 0.947001014 | 0.32595486 | 2.90531342 | 0.00366886 | 0.02136222 |
| AC007036.1 | 8.45373152 | 0.072764294 | 0.7509332 | 0.09689849 | 0.92280701 | 0.96229124 |
| ZNRF2 | 1686.44232 | -0.009199926 | 0.13029229 | -0.0706099 | 0.94370823 | 0.97315146 |
| AC006978.2 | 668.781852 | 0.072811122 | 0.12994096 | 0.56034002 | 0.57524753 | 0.73945976 |
| LINC01176 | 257.867032 | -0.786828275 | 0.28911353 | -2.7215201 | 0.00649824 | 0.03268956 |
| NOD1 | 876.888053 | -0.602004027 | 0.22607664 | -2.6628316 | 0.00774862 | 0.03731393 |
| AC006027.1 | 456.293034 | -1.012915328 | 0.28577817 | -3.5444112 | 0.00039349 | 0.00377185 |
| GGCT | 1700.026 | -0.131137835 | 0.28023741 | -0.4679526 | 0.63981846 | 0.78705102 |
| AC005154.6 | 1191.02711 | 0.064011644 | 0.27395557 | 0.23365703 | 0.81525125 | 0.90043158 |
| AC005154.5 | 1.97713256 | 1.328051447 | 0.96800677 | 1.37194438 | 0.17008075 | 0.34205533 |
| AC005154.4 | 48.578169 | 0.556141712 | 0.30070994 | 1.8494291 | 0.06439588 | 0.17364958 |
| GARS | 5536.14189 | -0.897061934 | 0.31288056 | -2.8671067 | 0.00414243 | 0.02338099 |
| CRHR2 | 21.8650773 | -0.181894067 | 0.88821998 | -0.2047849 | 0.83774016 | 0.91410871 |
| INMT | 1103.63394 | -0.542407518 | 0.47815208 | -1.1343829 | 0.25663399 | 0.44936509 |
| INMT-MINDY | 630.014994 | 0.252453244 | 0.23108845 | 1.0924529 | 0.27463408 | 0.46990013 |
| MINDY4 | 564.513438 | 0.175460838 | 0.25235146 | 0.69530343 | 0.48686516 | 0.6708802 |
| AC004691.1 | 1.10180792 | 1.032599791 | 1.1470936 | 0.90018791 | 0.36802026 | 0.56485037 |
| AC004691.2 | 19614.266 | 1.577475871 | 0.48982163 | 3.22051084 | 0.00127962 | 0.0095282 |
| AQP1 | 62382.436 | 1.228440326 | 0.46956415 | 2.61612886 | 0.0088933 | 0.04126628 |
| GHRHR | 55.4834748 | -0.079026708 | 0.76752568 | -0.1029629 | 0.91799238 | 0.9601219 |
| ADCYAP1R1 | 5.10114179 | -0.855817519 | 1.32558227 | -0.6456163 | 0.51852788 | 0.69539644 |
| NEUROD6 | 0 | NA | NA | NA | NA | NA |
| AC005090.1 | 0.46268499 | 1.383927999 | 2.78404769 | 0.49709206 | 0.61912413 | NA |
| CCDC129 | 2.1743542 | 1.045678369 | 0.81006153 | 1.29086289 | 0.19675123 | 0.37733308 |
| PPP1R17 | 1.54459138 | 1.047179758 | 1.5910235 | 0.65817994 | 0.51042251 | 0.68892277 |
| PDE1C | 423.952327 | -0.02971869 | 0.42019684 | -0.0707256 | 0.94361612 | 0.97315146 |
| AC018637.1 | 0.09997307 | 0.547239394 | 3.29377202 | 0.16614368 | 0.86804388 | NA |
| LSM5 | 1070.29872 | 0.043549755 | 0.14279494 | 0.30498107 | 0.76038057 | 0.86715659 |
| AVL9 | 2147.41545 | -0.31138885 | 0.14973709 | -2.0795705 | 0.03756494 | 0.11822264 |
| AC018645.2 | 83.8581629 | -1.109261857 | 0.19221284 | -5.7710082 | 7.88E-09 | 5.62E-07 |
| LINC00997 | 263.193753 | -0.865997019 | 0.21891188 | -3.955916 | 7.62E-05 | 0.00102316 |
| AC018648.1 | 26.5348826 | -0.608080514 | 0.36600449 | -1.6614018 | 0.09663278 | 0.23201369 |
| KBTBD2 | 2517.8499 | -0.500205905 | 0.16466117 | -3.037789 | 0.00238321 | 0.01543359 |
| FKBP9 | 3796.17202 | 0.46751591 | 0.15635091 | 2.99017076 | 0.00278822 | 0.0173555 |
| NT5C3A | 1457.73703 | -0.380675415 | 0.17779673 | -2.141071 | 0.03226832 | 0.10622638 |
| RP9 | 405.714044 | -0.388211555 | 0.11472902 | -3.383726 | 0.00071509 | 0.00601116 |
| BBS9 | 690.918773 | 0.312989671 | 0.20306955 | 1.54129299 | 0.12324549 | 0.27470067 |
| AC008080.1 | 6.44454794 | 0.32664028 | 0.51543076 | 0.6337229 | 0.52626167 | 0.70114232 |
| AC008080.2 | 0.27946654 | -0.179468827 | 2.93601537 | -0.0611267 | 0.95125833 | NA |
| AC008080.4 | 5.87716305 | 0.716007959 | 0.57565139 | 1.24382216 | 0.21356502 | 0.39755147 |
| AC008080.3 | 0.09635527 | 0.547239394 | 3.29377202 | 0.16614368 | 0.86804388 | NA |
| BMPER | 40.4476903 | -0.361516555 | 0.35086952 | -1.0303447 | 0.30284822 | 0.49975641 |
| AC009262.1 | 0 | NA | NA | NA | NA | NA |
| NPSR1 | 0.73164797 | 1.621871089 | 1.68312226 | 0.9636086 | 0.33524219 | NA |
| DPY19L1 | 1187.08895 | -0.749155205 | 0.22154127 | -3.3815605 | 0.00072075 | 0.00604718 |
| TBX20 | 0.08757577 | 0.547239394 | 3.29377202 | 0.16614368 | 0.86804388 | NA |
| AC009531.1 | 0 | NA | NA | NA | NA | NA |
| AC007652.1 | 0 | NA | NA | NA | NA | NA |
| HERPUD2 | 1594.7917 | -0.042643835 | 0.10436927 | -0.4085861 | 0.68284342 | 0.81659231 |
| AC018647.2 | 462.221555 | 0.888636005 | 0.19827651 | 4.48180166 | 7.40E-06 | 0.00016077 |
| AC018647.1 | 31.6280193 | 1.303160814 | 0.43807857 | 2.97471942 | 0.00293257 | 0.01802606 |
| SEPT7-AS1 | 130.063903 | 0.891755816 | 0.26567161 | 3.35660942 | 0.00078904 | 0.00649816 |
| 7-Sep | 5945.53286 | -0.094380746 | 0.14732878 | -0.6406131 | 0.52177409 | 0.69787989 |
| AC083864.2 | 0 | NA | NA | NA | NA | NA |
| AC083864.1 | 0.69439003 | 3.393703775 | 2.27326563 | 1.49287603 | 0.13546964 | NA |
| EEPD1 | 505.747849 | -0.00926341 | 0.26665377 | -0.0347395 | 0.97228749 | 0.986997 |
| AC007327.2 | 0.03392908 | 0.380740593 | 3.29738709 | 0.11546736 | 0.90807469 | NA |
| AC006960.3 | 3.2631393 | -0.252784233 | 0.93167245 | -0.2713231 | 0.78614256 | 0.88350577 |
| KIAA0895 | 884.931906 | 0.290000785 | 0.13922982 | 2.08289277 | 0.037261 | 0.11754666 |
| ANLN | 62.7606546 | -1.409630469 | 0.52320324 | -2.6942312 | 0.00705512 | 0.03483962 |
| AC006960.2 | 32.329171 | 2.602798698 | 1.21582443 | 2.14076854 | 0.03229271 | 0.10627215 |
| AOAH | 308.308626 | -0.224197862 | 0.49372969 | -0.4540903 | 0.64976383 | 0.79398852 |
| AC007349.4 | 1.43459174 | 1.090300053 | 1.34169323 | 0.81262991 | 0.41643028 | 0.60966391 |
| AC007349.1 | 10.0775582 | 1.425680087 | 0.45137224 | 3.15854622 | 0.00158558 | 0.0112761 |
| AC007349.3 | 7.03362298 | 0.914290538 | 0.57523874 | 1.58941057 | 0.11196773 | 0.25719417 |
| AC007349.2 | 0.03392908 | 0.380740593 | 3.29738709 | 0.11546736 | 0.90807469 | NA |
| ELMO1 | 530.083914 | 0.042628355 | 0.30261641 | 0.14086598 | 0.88797582 | 0.94202528 |
| ELMO1-AS1 | 29.0376036 | 0.598897371 | 0.32288932 | 1.85480702 | 0.06362379 | 0.17221919 |
| GPR141 | 12.414792 | 0.207457896 | 0.7212508 | 0.28763628 | 0.77362517 | 0.87544504 |
| EPDR1 | 1605.78597 | -0.088319613 | 0.23962509 | -0.3685741 | 0.71244517 | 0.83654729 |
| NME8 | 19.78084 | 0.092205343 | 0.64807265 | 0.14227624 | 0.88686181 | 0.94143003 |
| SFRP4 | 245.105483 | 0.617400857 | 0.54331703 | 1.1363547 | 0.25580815 | 0.44848486 |
| STARD3NL | 2148.82606 | -0.447082946 | 0.1065479 | -4.1960745 | 2.72E-05 | 0.00045442 |
| TRGC2 | 68.1483019 | -0.723670076 | 0.50170798 | -1.4424129 | 0.14918592 | 0.3133439 |
| TRGJ2 | 0 | NA | NA | NA | NA | NA |
| TRGJP2 | 1.83975396 | -1.524639009 | 2.0366478 | -0.7486022 | 0.45409701 | 0.64372316 |
| TRGC1 | 61.252747 | 0.276110426 | 0.54573085 | 0.50594616 | 0.61289445 | 0.76804764 |
| TRGJ1 | 1.50519836 | -2.32969446 | 1.36125802 | -1.7114275 | 0.08700222 | 0.21499323 |
| TRGJP | 5.24648702 | 0.985299138 | 1.06847527 | 0.92215437 | 0.35644806 | 0.55382603 |
| TRGJP1 | 0.9092375 | -1.545376749 | 1.69378825 | -0.912379 | 0.36156927 | NA |
| TRGV11 | 0 | NA | NA | NA | NA | NA |
| TRGVB | 0.34136971 | -0.261076059 | 2.32545742 | -0.1122687 | 0.91061037 | NA |
| TRGV10 | 8.76572304 | -0.162183996 | 0.77466023 | -0.2093615 | 0.83416608 | 0.91206315 |
| TRGV9 | 4.80117046 | -1.068268751 | 0.88891735 | -1.2017639 | 0.22945503 | 0.41691787 |
| TRGVA | 0.6846766 | 1.490281817 | 1.39524673 | 1.06811346 | 0.28546934 | NA |
| AC006033.2 | 8.70688759 | -0.21534825 | 0.65244534 | -0.3300633 | 0.74135215 | 0.85469576 |
| TRGV8 | 2.3071014 | 0.601429792 | 0.97352592 | 0.61778508 | 0.53671702 | 0.71052979 |
| TRGV7 | 4.10287566 | 0.359088638 | 0.68705155 | 0.52265167 | 0.60121668 | 0.75875012 |
| TRGV6 | 0.62221669 | -1.53454141 | 1.65201567 | -0.9288903 | 0.35294591 | NA |
| TRG-AS1 | 51.5040969 | -0.649663707 | 0.56454996 | -1.1507639 | 0.24982939 | 0.44194009 |
| TRGV5P | 12.6418187 | -1.363785241 | 0.91827633 | -1.4851578 | 0.13750203 | 0.29616341 |
| TRGV5 | 10.108703 | -2.624041512 | 0.92675417 | -2.8314321 | 0.00463401 | 0.02544217 |
| TRGV4 | 18.3533826 | -0.358791782 | 0.69939323 | -0.5130044 | 0.60794827 | 0.76420992 |
| TRGV3 | 10.3462544 | 0.109188998 | 0.80040994 | 0.13641634 | 0.89149215 | 0.94377583 |
| TRGV2 | 6.91493832 | -0.839909747 | 0.77677799 | -1.0812739 | 0.2795753 | 0.47514828 |
| TRGV1 | 0.34807119 | 0.706453744 | 2.31872532 | 0.30467332 | 0.76061497 | NA |
| AMPH | 145.177798 | 1.441318417 | 0.4076053 | 3.53606397 | 0.00040614 | 0.00386776 |
| VPS41 | 2431.38505 | 0.26011992 | 0.15356779 | 1.69384432 | 0.09029487 | 0.22120733 |
| POU6F2 | 5.22950441 | 0.866638333 | 0.60167464 | 1.44037703 | 0.14976076 | 0.31402602 |
| POU6F2-AS2 | 0.17656334 | 0.18657164 | 3.29377195 | 0.05664376 | 0.95482897 | NA |
| POU6F2-AS1 | 6.12503038 | 1.261561071 | 0.78358735 | 1.60998141 | 0.10740191 | 0.249576 |
| AC011290.1 | 1.28066387 | 2.90951626 | 1.6483712 | 1.7650856 | 0.07754938 | 0.1981375 |
| YAE1D1 | 411.770935 | 0.002417212 | 0.14812582 | 0.01631864 | 0.98698019 | 0.99410333 |
| AC004837.3 | 6.39791746 | 0.858039062 | 0.6153397 | 1.39441526 | 0.16319224 | 0.33240345 |
| RALA | 2184.21901 | -0.627033695 | 0.30673449 | -2.044223 | 0.04093152 | 0.12606506 |
| AC004837.2 | 4.38475958 | -0.383473795 | 0.93119487 | -0.4118083 | 0.68047992 | 0.81472216 |
| LINC00265 | 253.313891 | 0.616604968 | 0.27169876 | 2.26944341 | 0.02324137 | 0.08377188 |
| AC072061.1 | 71.7873297 | 0.579866707 | 0.22494154 | 2.57785516 | 0.00994157 | 0.04486129 |

| | | | | | | |
|---|---|---|---|---|---|---|
| CDK13 | 2524.90517 | 0.006125181 | 0.11155551 | 0.05490703 | 0.95621253 | 0.9790458 |
| MPLKIP | 1356.44692 | 0.15847261 | 0.15972084 | 0.99218494 | 0.32110732 | 0.51865735 |
| SUGCT | 1028.99839 | 2.221451792 | 0.33876419 | 6.5575166 | 5.47E-11 | 8.74E-09 |
| AC004988.1 | 5.20917285 | -2.632391799 | 1.07726905 | -2.4435788 | 0.01454239 | 0.05950084 |
| LINC01450 | 1.24614789 | 1.880465489 | 1.29697798 | 1.44988235 | 0.14709133 | 0.31030387 |
| LINC01449 | 2.92665635 | 1.521793782 | 0.75265416 | 2.02190311 | 0.04318636 | 0.13104998 |
| INHBA | 389.998353 | -1.339662586 | 0.75608299 | -1.7718459 | 0.07642013 | 0.1962219 |
| INHBA-AS1 | 5.22560843 | 1.492364172 | 0.53864749 | 2.77057668 | 0.00559571 | 0.02931829 |
| GLI3 | 296.99691 | 1.057562501 | 0.3329719 | 3.1761314 | 0.00149253 | 0.01076535 |
| LINC01448 | 0 | NA | NA | NA | NA | NA |
| AC010132.1 | 0.10721287 | -0.642905668 | 3.30558948 | -0.1944905 | 0.84579185 | NA |
| AC010132.4 | 13.2026282 | 0.225360958 | 0.36063987 | 0.62489197 | 0.53204196 | 0.70623305 |
| C7orf25 | 727.520045 | 0.130204908 | 0.13008325 | 1.00093525 | 0.31685811 | 0.51415596 |
| AC010132.3 | 5295.14761 | -0.265537787 | 0.11394967 | -2.3303076 | 0.0197899 | 0.07431782 |
| PSMA2 | 4942.15771 | -0.296909361 | 0.11165409 | -2.6591892 | 0.0078329 | 0.03762358 |
| MRPL32 | 2663.91785 | -0.24519135 | 0.1477769 | -1.6591994 | 0.09707561 | 0.23267442 |
| AC005537.1 | 9.21401366 | -0.915276781 | 0.84960675 | -1.0772946 | 0.28134867 | 0.47723944 |
| HECW1 | 721.502451 | 3.495656341 | 0.86653754 | 4.03405065 | 5.48E-05 | 0.00079893 |
| AC004692.2 | 0.10721287 | -0.642905668 | 3.30558948 | -0.1944905 | 0.84579185 | NA |
| LUARIS | 91.2805332 | 3.539116774 | 0.80859528 | 4.37687044 | 1.20E-05 | 0.00023835 |
| STK17A | 1258.07452 | -0.461359329 | 0.31814753 | -1.4501427 | 0.14701872 | 0.31020032 |
| COA1 | 1891.52938 | 0.004217385 | 0.11821224 | 0.03567638 | 0.9715404 | 0.98670705 |
| BLVRA | 2347.02925 | -0.615727864 | 0.1254769 | -4.9071015 | 9.24E-07 | 3.04E-05 |
| MRPS24 | 2763.89835 | 0.002963361 | 0.1591648 | 0.01861819 | 0.98514569 | 0.99347171 |
| URGCP-MRP | 2890.50116 | 0.076170265 | 0.15564769 | 0.48937614 | 0.62457543 | 0.77634004 |
| URGCP | 4266.42956 | 0.287126547 | 0.13690315 | 2.09729687 | 0.0359673 | 0.1147976 |
| UBE2D4 | 1292.93908 | 0.420041226 | 0.14794261 | 2.83921742 | 0.00452243 | 0.02495951 |
| SPDYE1 | 36.1537497 | 0.969592387 | 0.2701271 | 3.58939323 | 0.00033145 | 0.00328212 |
| AC004951.1 | 27.3555699 | 0.681954118 | 0.23099388 | 2.95226056 | 0.00315457 | 0.01909686 |
| LINC00957 | 339.785343 | 0.44950761 | 0.30402786 | 1.47850797 | 0.13927187 | 0.29889054 |
| DBNL | 5284.0974 | -0.02144865 | 0.12153355 | -0.1764834 | 0.85991421 | 0.92701929 |
| PGAM2 | 364.327099 | 1.188919585 | 0.27202472 | 4.37063064 | 1.24E-05 | 0.00024369 |
| POLM | 2665.26809 | 0.072287634 | 0.13461803 | 0.53698328 | 0.59127916 | 0.75084424 |
| AEBP1 | 15537.7901 | -0.178082624 | 0.40656856 | -0.4380138 | 0.66137631 | 0.80075727 |
| POLD2 | 3631.04467 | -0.081216024 | 0.13578481 | -0.5981231 | 0.54975782 | 0.72018351 |
| MYL7 | 3.01128834 | 0.408333514 | 0.71189446 | 0.57358715 | 0.56624721 | 0.73281994 |
| GCK | 24.1232523 | -0.219916895 | 0.41453394 | -0.530516 | 0.5957542 | 0.75474602 |
| YKT6 | 5190.26962 | -0.447925085 | 0.28386581 | -1.5779466 | 0.11457788 | 0.26157206 |
| CAMK2B | 193.702718 | 1.875750548 | 0.2912558 | 6.44021697 | 1.19E-10 | 1.66E-08 |
| NUDCD3 | 4180.70629 | -0.114695653 | 0.11496293 | -0.9976751 | 0.31843694 | 0.51570239 |
| NPC1L1 | 173.609285 | 0.711736415 | 0.38878786 | 1.83065496 | 0.06715206 | 0.1788506 |
| DDX56 | 5521.61786 | -0.192149003 | 0.14738042 | -1.303762 | 0.19231473 | 0.37158114 |
| TMED4 | 7134.43907 | 0.442296072 | 0.18095454 | 2.44423866 | 0.01451582 | 0.05944746 |
| OGDH | 14016.602 | 0.884006971 | 0.17498419 | 5.05192476 | 4.37E-07 | 1.62E-05 |
| ZMIZ2 | 10510.5323 | -0.203918329 | 0.17130258 | -1.1903985 | 0.23388981 | 0.42244345 |
| AC013436.1 | 0.10721287 | -0.642905668 | 3.30558948 | -0.1944905 | 0.84579185 | NA |
| PPIA | 35257.3013 | -0.63760522 | 0.18372749 | -3.4703855 | 0.00051971 | 0.00467262 |
| H2AFV | 5336.14057 | 0.451468005 | 0.21064455 | 2.14326935 | 0.03209148 | 0.10586093 |
| LINC01952 | 4.20712449 | 0.234070878 | 0.81427481 | 0.28745931 | 0.77376065 | 0.87552321 |
| PURB | 2351.00862 | -0.215132579 | 0.11338578 | -1.8973507 | 0.05778167 | 0.16120007 |
| AC004854.2 | 46.6194453 | -0.504491859 | 0.25039824 | -2.014758 | 0.04393001 | 0.13280229 |
| MYO1G | 443.050763 | -0.654538287 | 0.55403311 | -1.1814064 | 0.2374413 | 0.42669532 |
| SNHG15 | 1218.6552 | -1.239352256 | 0.25086654 | -4.9402852 | 7.80E-07 | 2.63E-05 |
| CCM2 | 2213.81424 | -0.530860147 | 0.14890435 | -3.5651083 | 0.00036371 | 0.00353778 |
| NACAD | 363.507398 | -0.108996582 | 0.32712603 | -0.3331945 | 0.7389875 | 0.85348667 |
| TBRG4 | 4026.35872 | -0.292915268 | 0.17746241 | -1.6505764 | 0.0988251 | 0.23566811 |
| RAMP3 | 3112.06619 | -0.20956575 | 0.28811455 | -0.7273696 | 0.4669996 | 0.65413276 |
| AC073325.1 | 0 | NA | NA | NA | NA | NA |
| ADCY1 | 340.004245 | 0.873508989 | 0.44163422 | 1.9779015 | 0.04793982 | 0.14123243 |
| AC096582.3 | 0 | NA | NA | NA | NA | NA |
| IGFBP1 | 68.0749275 | -0.921252332 | 0.87842882 | -1.0487501 | 0.29429314 | 0.49101614 |
| IGFBP3 | 7125.25278 | -0.628609985 | 0.37233213 | -1.6883045 | 0.0913528 | 0.22292924 |
| AC073115.2 | 0.43414051 | 0.297942357 | 1.93942884 | 0.15362376 | 0.8779064 | NA |
| AC073115.1 | 2.29051618 | 1.424581348 | 0.95469761 | 1.4921807 | 0.13565178 | 0.29335226 |
| AC023669.1 | 3.97734039 | -2.791579816 | 1.18979593 | -2.3462678 | 0.01896248 | 0.07217676 |
| AC023669.2 | 0.95039116 | -1.217707647 | 1.66120943 | -0.7330248 | 0.46354333 | NA |
| AC004869.2 | 0 | NA | NA | NA | NA | NA |
| AC004869.1 | 0 | NA | NA | NA | NA | NA |
| AC011294.1 | 18.4250672 | 1.392493476 | 0.89778169 | 1.55103795 | 0.12089259 | 0.27130065 |
| AC004870.4 | 21.5865488 | 1.787767727 | 1.01088191 | 1.76852282 | 0.07697354 | 0.19714377 |
| AC004870.3 | 3.14169098 | -0.95205768 | 0.86090616 | -1.1058786 | 0.26877908 | 0.46297631 |
| TNS3 | 11136.0927 | -0.077212849 | 0.24778729 | -0.3116094 | 0.75533739 | 0.86405639 |
| LINC01447 | 8.79898923 | 4.94789947 | 0.92331816 | 5.35882398 | 8.38E-08 | 4.12E-06 |
| C7orf65 | 1.07496293 | -0.477175639 | 1.69527085 | -0.2814746 | 0.77834643 | 0.87861887 |
| HUS1 | 717.762124 | -0.209933156 | 0.14134365 | -1.4852677 | 0.13747292 | 0.29612489 |
| LINC00525 | 1.71797824 | 0.272159779 | 1.02569709 | 0.26534128 | 0.79074657 | 0.886377 |
| PKD1L1 | 18.7299674 | 0.596914251 | 0.40182068 | 1.48552396 | 0.13740508 | 0.29604946 |
| C7orf69 | 0.38049381 | -0.032083374 | 1.90704923 | -0.0168236 | 0.98657737 | NA |
| SUN3 | 2.33631784 | 1.002956742 | 0.88359656 | 1.13508448 | 0.25633992 | 0.44911858 |
| C7orf57 | 21.8567338 | -0.732231728 | 0.58573315 | -1.2501115 | 0.21125883 | 0.39465113 |
| UPP1 | 4937.37414 | -2.172665747 | 0.42755442 | -5.0816121 | 3.74E-07 | 1.43E-05 |
| ABCA13 | 97.5877727 | -0.175416432 | 0.63236816 | -0.2773961 | 0.78147601 | 0.88036187 |
| AC091770.1 | 0 | NA | NA | NA | NA | NA |
| AC091730.1 | 0.03392908 | 0.380740593 | 3.29738709 | 0.11546736 | 0.90807469 | NA |
| VWC2 | 7.90503857 | 1.907422627 | 0.834746 | 2.28503357 | 0.02231086 | 0.08132947 |
| ZPBP | 7.70188714 | 1.521725771 | 0.549061 | 2.77150585 | 0.00557977 | 0.02926964 |
| SPATA48 | 0 | NA | NA | NA | NA | NA |
| AC020743.1 | 0 | NA | NA | NA | NA | NA |
| AC020743.2 | 0.10178723 | 1.087385526 | 3.28420134 | 0.33109588 | 0.74057207 | NA |
| AC020743.3 | 0.08666371 | -0.642905668 | 3.30558948 | -0.1944905 | 0.84579185 | NA |
| IKZF1 | 316.733742 | 0.032729259 | 0.53601052 | 0.06106085 | 0.95131075 | 0.97675756 |
| AC124014.1 | 6.89564989 | 1.0360504 | 0.99455004 | 1.04172778 | 0.29753791 | 0.49442779 |
| FIGNL1 | 262.002667 | 0.024630777 | 0.21938885 | 0.11226996 | 0.91060936 | 0.95580818 |
| DDC | 11206.5276 | 2.4797466 | 0.5314994 | 4.66556801 | 3.08E-06 | 7.99E-05 |
| DDC-AS1 | 1519.96123 | 2.937060826 | 0.58209993 | 5.04562997 | 4.52E-07 | 1.66E-05 |
| GRB10 | 3846.8461 | 0.575893123 | 0.27140178 | 2.12192094 | 0.03384438 | 0.10993794 |
| AC004920.1 | 0 | NA | NA | NA | NA | NA |
| AC004830.1 | 272.311929 | 0.155079909 | 0.20021996 | 0.77454769 | 0.43860698 | 0.63025154 |
| COBL | 2109.45071 | 0.467559447 | 0.20871863 | 2.24014234 | 0.02508168 | 0.08860055 |
| AC005999.1 | 0 | NA | NA | NA | NA | NA |
| AC079763.1 | 0.18301898 | 0.18657164 | 3.29377195 | 0.05664376 | 0.95482897 | NA |
| POM121L12 | 0 | NA | NA | NA | NA | NA |
| AC009468.2 | 0 | NA | NA | NA | NA | NA |
| AC009468.1 | 0 | NA | NA | NA | NA | NA |
| LINC01446 | 0.08393208 | -0.642905668 | 3.30558948 | -0.1944905 | 0.84579185 | NA |
| AC073521.1 | 0 | NA | NA | NA | NA | NA |
| HPVC1 | 2.45088649 | 1.945809685 | 1.03005898 | 1.88902744 | 0.05888815 | 0.1633064 |
| LINC01445 | 0.08695548 | -0.642905668 | 3.30558948 | -0.1944905 | 0.84579185 | NA |
| VSTM2A | 2.52531496 | -2.387824188 | 1.87257868 | -1.2751529 | 0.20225512 | 0.38403424 |
| AC011228.1 | 0 | NA | NA | NA | NA | NA |
| SEC61G | 1998.00038 | -0.99326327 | 0.20883511 | -4.7562082 | 1.97E-06 | 5.60E-05 |
| AC074351.1 | 2.19318137 | 1.168381094 | 0.98187139 | 1.18995329 | 0.23406475 | 0.42267256 |
| EGFR | 2973.70756 | -0.108310654 | 0.12234588 | -0.8852824 | 0.37600435 | 0.57184583 |

| | | | | | | | |
|---|---|---|---|---|---|---|---|
| EGFR-AS1 | 42.2930317 | | -0.359107218 | 0.52252706 | -0.6872509 | 0.4919246 | 0.67488954 |
| ELDR | 0.69092408 | | -1.623837734 | 1.91812131 | -0.8465772 | 0.39723084 | NA |
| LANCL2 | 1151.25798 | | 0.495797671 | 0.29097569 | 1.7039144 | 0.08839708 | 0.21769061 |
| VOPP1 | 2713.36194 | | -0.805971966 | 0.19644032 | -4.1028846 | 4.08E-05 | 0.00063218 |
| AC099681.1 | 0.37673387 | | 1.705319083 | 2.7987996 | 0.60930375 | 0.54232312 | NA |
| AC099681.3 | 2.13153923 | | 0.2019326 | 0.77394678 | 0.26091277 | 0.79415978 | 0.88844572 |
| AC099681.2 | 2.76238948 | | -1.26976571 | 1.05412227 | -1.2045716 | 0.22836874 | 0.41551671 |
| 14-Sep | 0.07659028 | | -0.642905668 | 3.30558948 | -0.1944905 | 0.84579185 | NA |
| AC092647.5 | 569.192147 | | -0.331536407 | 0.17066708 | -1.9425914 | 0.05206555 | 0.14996926 |
| ZNF713 | 84.6976168 | | 0.877187596 | 0.23730172 | 3.6965075 | 0.00021859 | 0.00234504 |
| NIPSNAP2 | 4089.79142 | | -0.077161031 | 0.14713968 | -0.5244067 | 0.59999572 | 0.75775321 |
| MRPS17 | 847.089784 | | -0.362756706 | 0.1842166 | -1.9691857 | 0.04893177 | 0.14359276 |
| PSPH | 1007.86735 | | 0.442830958 | 0.3623093 | 1.22224563 | 0.22161475 | 0.4076376 |
| CCT6A | 8499.17162 | | -0.474306305 | 0.28377948 | -1.6713904 | 0.09464459 | 0.2288235 |
| SUMF2 | 8941.13495 | | -0.021957423 | 0.14224467 | -0.1543638 | 0.87732292 | 0.93618506 |
| PHKG1 | 1621.0051 | | -0.528690343 | 0.16164707 | -3.270646 | 0.00107302 | 0.0082881 |
| CHCHD2 | 11741.5847 | | -0.345188077 | 0.1925304 | -1.7929016 | 0.07298865 | 0.19005735 |
| NUPR2 | 186.827582 | | -0.552277509 | 0.26303317 | -2.0996496 | 0.03575967 | 0.11427322 |
| AC092447.4 | 0 | NA | | NA | NA | NA | NA |
| AC092447.8 | 0 | NA | | NA | NA | NA | NA |
| AC092447.5 | 1.39849353 | | 1.884485809 | 1.44430621 | 1.30476889 | 0.19197156 | 0.37121727 |
| AC118758.3 | 3.85860871 | | 1.008177352 | 1.06688094 | 0.94497644 | 0.3446709 | 0.54223671 |
| ZNF479 | 0 | NA | | NA | NA | NA | NA |
| AC099654.1 | 0 | NA | | NA | NA | NA | NA |
| AC237221.1 | 2.84172389 | | -0.137352586 | 1.36404553 | -0.100695 | 0.91979257 | 0.96074349 |
| ZNF716 | 0.08393208 | | -0.642905668 | 3.30558948 | -0.1944905 | 0.84579185 | NA |
| AC069285.1 | 0 | NA | | NA | NA | NA | NA |
| AC006455.1 | 0 | NA | | NA | NA | NA | NA |
| AC006455.4 | 0 | NA | | NA | NA | NA | NA |
| AC073188.3 | 0 | NA | | NA | NA | NA | NA |
| AC073188.4 | 0 | NA | | NA | NA | NA | NA |
| AC073188.2 | 0 | NA | | NA | NA | NA | NA |
| AC073188.5 | 0 | NA | | NA | NA | NA | NA |
| AC092634.5 | 1.85314816 | | -1.114349832 | 1.0950789 | -1.0175978 | 0.30886915 | 0.50589314 |
| AC092634.3 | 10.9561569 | | -0.42349428 | 0.52242707 | -0.8106285 | 0.41757904 | 0.61036996 |
| AC115220.1 | 0.65568698 | | -0.376906488 | 2.75140464 | -0.1369869 | 0.89104112 | NA |
| LINC01005 | 0.07659028 | | -0.642905668 | 3.30558948 | -0.1944905 | 0.84579185 | NA |
| ZNF727 | 14.380177 | | -0.34102092 | 0.56355724 | -0.6051221 | 0.54509789 | 0.71685262 |
| AC091685.1 | 3.52214099 | | -1.602717373 | 0.89606772 | -1.7886119 | 0.07367734 | 0.19137821 |
| ZNF735 | 0 | NA | | NA | NA | NA | NA |
| ZNF679 | 0.09644756 | | -0.642905668 | 3.30558948 | -0.1944905 | 0.84579185 | NA |
| ZNF736 | 892.162907 | | 0.122522354 | 0.22125694 | 0.55375597 | 0.57974586 | 0.74288831 |
| ZNF680 | 717.488679 | | 0.207405871 | 0.19101238 | 1.08582423 | 0.27755677 | 0.47303938 |
| ZNF107 | 309.419411 | | -0.185126936 | 0.37470623 | -0.4940589 | 0.6212646 | 0.77368586 |
| ZNF138 | 403.468949 | | 0.076258012 | 0.21159662 | 0.36039333 | 0.71855301 | 0.84001549 |
| ZNF273 | 203.41715 | | -0.547086277 | 0.33171024 | -1.6492897 | 0.0990883 | 0.23603962 |
| AC073349.1 | 13.1656667 | | -0.271128901 | 0.51504417 | -0.5264187 | 0.5985973 | 0.75677743 |
| ZNF117 | 1077.1491 | | -0.082857746 | 0.2043654 | -0.4054392 | 0.6851547 | 0.81810054 |
| ERV3-1 | 359.861321 | | 0.284945898 | 0.32217986 | 0.88443114 | 0.37646352 | 0.5723131 |
| AC104073.4 | 0 | NA | | NA | NA | NA | NA |
| ZNF92 | 326.592856 | | -0.035570521 | 0.19345896 | -0.183866 | 0.85411859 | 0.92360811 |
| AC114501.3 | 0 | NA | | NA | NA | NA | NA |
| VKORC1L1 | 1304.72022 | | 0.01129658 | 0.13453002 | 0.0839707 | 0.93307973 | 0.96776057 |
| GUSB | 3655.86834 | | 0.118253509 | 0.145072 | 0.81513668 | 0.41499408 | 0.60850722 |
| ASL | 6688.58984 | | 0.110139055 | 0.17338941 | 0.63521213 | 0.52529007 | 0.7000505 |
| AC068533.4 | 4004.18206 | | 0.076598099 | 0.15533673 | 0.49311002 | 0.62193485 | 0.77429876 |
| CRCP | 1905.63668 | | -0.274266861 | 0.13121399 | -2.0902257 | 0.03659753 | 0.11617633 |
| AC068533.3 | 5.05369503 | | 0.112545501 | 0.52914165 | 0.21269447 | 0.83156528 | 0.9104925 |
| TPST1 | 1789.86357 | | -0.910004683 | 0.27995301 | -3.2505622 | 0.00115177 | 0.00876289 |
| AC008267.5 | 98.8491079 | | 0.460008327 | 0.38164738 | 1.20532289 | 0.22807868 | 0.41516082 |
| KCTD7 | 846.068676 | | 0.701781427 | 0.15943838 | 4.40158413 | 1.07E-05 | 0.00021649 |
| AC027644.4 | 1968.85457 | | -0.326711662 | 0.1830588 | -1.7847362 | 0.07430412 | 0.1924754 |
| RABGEF1 | 2698.79663 | | -0.212571686 | 0.14080637 | -1.5096738 | 0.13112668 | 0.28674107 |
| AC027644.3 | 518.680533 | | 0.319493406 | 0.25157791 | 1.2699581 | 0.20409956 | 0.38645116 |
| AC073335.2 | 895.615251 | | 0.675324747 | 0.3215424 | 2.10026658 | 0.0357054 | 0.11416384 |
| TMEM248 | 5293.76394 | | -0.131917148 | 0.08503204 | -1.5513817 | 0.12081023 | 0.27117997 |
| SBDS | 6044.35653 | | -0.277439485 | 0.17784907 | -1.5599716 | 0.1187666 | 0.26797223 |
| TYW1 | 2149.73427 | | 0.027433751 | 0.11610043 | 0.23629327 | 0.81320511 | 0.89913137 |
| AC006480.2 | 79.7771185 | | 0.382427574 | 0.27324996 | 1.39955216 | 0.16164747 | 0.33033507 |
| LINC01372 | 107.04477 | | 0.062807537 | 0.28171387 | 0.22294797 | 0.823576 | 0.90564302 |
| AC005482.1 | 0 | NA | | NA | NA | NA | NA |
| AC092637.1 | 0 | NA | | NA | NA | NA | NA |
| AC006013.1 | 0 | NA | | NA | NA | NA | NA |
| AC093655.1 | 0 | NA | | NA | NA | NA | NA |
| AC104688.1 | 0 | NA | | NA | NA | NA | NA |
| AC069280.1 | 0 | NA | | NA | NA | NA | NA |
| AC092100.1 | 0.03392908 | | 0.380740593 | 3.29738709 | 0.11546736 | 0.90807469 | NA |
| AC006012.1 | 7.63869795 | | 0.397927877 | 0.58407523 | 0.68129559 | 0.49568447 | 0.67770688 |
| AUTS2 | 1845.41478 | | 0.871585178 | 0.18743401 | 4.65009086 | 3.32E-06 | 8.48E-05 |
| GALNT17 | 298.732744 | | 0.714119561 | 0.4750957 | 1.50310677 | 0.13281151 | 0.28920228 |
| CALN1 | 1.09642624 | | 0.556794571 | 1.31346579 | 0.4239125 | 0.67162963 | 0.80832284 |
| TYW1B | 226.778835 | | 0.212555598 | 0.4379998 | 0.48528697 | 0.6274728 | 0.77838442 |
| POM121 | 4786.99376 | | 0.049990397 | 0.08479372 | 0.58955307 | 0.55549032 | 0.72487529 |
| AC211476.2 | 7.14715829 | | 0.393773171 | 0.54302935 | 0.72514159 | 0.46836517 | 0.65515146 |
| TRIM74 | 58.4385753 | | -0.250824675 | 0.38092905 | -0.6584551 | 0.51024574 | 0.68887878 |
| NSUN5 | 1675.0592 | | -0.344789194 | 0.10043996 | -3.432789 | 0.00059741 | 0.00520642 |
| TRIM50 | 374.650623 | | 2.378695262 | 0.89521853 | 2.6571113 | 0.00788134 | 0.03777372 |
| FKBP6 | 6.43176005 | | -1.194406105 | 0.78812111 | -1.5155109 | 0.12964308 | 0.28460514 |
| FZD9 | 9.89709695 | | -0.407922672 | 0.54690986 | -0.7458682 | 0.45574703 | 0.64517131 |
| BAZ1B | 2647.06004 | | -0.058751619 | 0.12077313 | -0.4864627 | 0.62663918 | 0.77783715 |
| BCL7B | 3567.53884 | | -0.630135125 | 0.22007392 | -2.8632885 | 0.00419269 | 0.023604 |
| TBL2 | 1617.50829 | | -0.505694384 | 0.14984383 | -3.3748096 | 0.00073867 | 0.00616999 |
| MLXIPL | 8209.31998 | | 0.256706505 | 0.53414554 | 0.48059281 | 0.63080593 | 0.7805524 |
| VPS37D | 796.657438 | | -0.150489628 | 0.22491975 | -0.6690814 | 0.50344354 | 0.68350956 |
| DNAJC30 | 897.871385 | | 0.374536241 | 0.17833059 | 2.10023552 | 0.03570813 | 0.11416384 |
| BUD23 | 3982.99752 | | -0.232272198 | 0.11652148 | -1.9933853 | 0.04621927 | 0.13754081 |
| STX1A | 391.863719 | | -1.084593433 | 0.2569052 | -4.2217653 | 2.42E-05 | 0.00041545 |
| ABHD11 | 2991.88962 | | -0.560543068 | 0.17582446 | -3.1880836 | 0.00143219 | 0.01043612 |
| CLDN3 | 2889.594 | | -1.566764726 | 0.28546828 | -5.4884023 | 4.06E-08 | 2.24E-06 |
| CLDN4 | 13965.3677 | | -1.896383578 | 0.37123937 | -5.1082502 | 3.25E-07 | 1.27E-05 |
| METTL27 | 337.344406 | | 0.261361811 | 0.45773135 | 0.5709939 | 0.56800377 | 0.73384869 |
| TMEM270 | 3.08336686 | | -0.583695784 | 1.08146211 | -0.5397284 | 0.58938437 | 0.74961886 |
| AC099398.1 | 0 | NA | | NA | NA | NA | NA |
| ELN | 1881.11039 | | 1.186974236 | 0.62277271 | 1.90595094 | 0.05665657 | 0.15897359 |
| ELN-AS1 | 184.022288 | | 0.844561423 | 0.56881588 | 1.48477118 | 0.13760445 | 0.2962291 |
| LIMK1 | 1599.15241 | | -1.023243674 | 0.19853741 | -5.1539085 | 2.55E-07 | 1.03E-05 |
| EIF4H | 12636.5298 | | -0.011087968 | 0.13973301 | -0.0793511 | 0.93675336 | 0.96969938 |
| LAT2 | 340.39043 | | -0.247869925 | 0.36755178 | -0.674381 | 0.50006914 | 0.68130141 |
| RFC2 | 861.828351 | | -0.456084446 | 0.16489447 | -2.7659171 | 0.0056763 | 0.02961692 |
| CLIP2 | 1966.29151 | | -0.20277762 | 0.29404602 | -0.6896118 | 0.49043832 | 0.67369536 |
| GTF2IRD1 | 2946.89006 | | -0.881132746 | 0.29297865 | -3.0074982 | 0.00263408 | 0.01662533 |
| AC211433.2 | 8.22507805 | | -1.019346082 | 0.61912758 | -1.6464233 | 0.09967664 | 0.23696479 |
| GTF2I | 7738.85284 | | 0.606998289 | 0.22605933 | 2.68512827 | 0.0072502 | 0.03554392 |

| | | | | | | |
|---|---|---|---|---|---|---|
| NCF1 | 252.242808 | -0.063471432 | 0.55245872 | -0.114889 | 0.9085331 | 0.95432284 |
| GTF2IRD2 | 404.865415 | 0.113521906 | 0.27437661 | 0.41374484 | 0.67906099 | 0.81383622 |
| CASTOR2 | 86.1825063 | -0.323749309 | 0.46693742 | -0.6933462 | 0.48809229 | 0.6715885 |
| RCC1L | 3768.01702 | 0.065862792 | 0.13802462 | 0.47718148 | 0.6332329 | 0.78238071 |
| GTF2IRD2B | 781.233525 | 0.236383048 | 0.22442671 | 1.05327502 | 0.29221496 | 0.48841406 |
| AC211486.5 | 0 | NA | NA | NA | NA | NA |
| AC211486.2 | 0 | NA | NA | NA | NA | NA |
| TRIM73 | 148.494965 | -0.121663925 | 0.24595432 | -0.4946606 | 0.62083969 | 0.77350682 |
| POM121C | 2668.71257 | 0.00778929 | 0.15135876 | 0.05146243 | 0.95895704 | 0.98021928 |
| SPDYE5 | 48.6248202 | 0.578584218 | 0.28367683 | 2.03958929 | 0.04139125 | 0.12704736 |
| HIP1 | 2377.96386 | 0.593901286 | 0.30021554 | 1.97824966 | 0.04790055 | 0.14116422 |
| CCL26 | 20.118342 | -0.06774282 | 0.63473563 | -0.106726 | 0.91500632 | 0.9585919 |
| CCL24 | 8.60471042 | -1.265370946 | 1.02450782 | -1.2351013 | 0.2167928 | 0.40160353 |
| RHBDD2 | 6867.72454 | -0.279660564 | 0.12857905 | -2.1750088 | 0.02962946 | 0.10000369 |
| POR | 13372.0413 | -0.919869769 | 0.3724169 | -2.47 | 0.0135113 | 0.05634874 |
| TMEM120A | 3403.67743 | 0.312186905 | 0.16959458 | 1.84078351 | 0.06565329 | 0.17608597 |
| STYXL1 | 1035.62573 | 0.091030803 | 0.29555327 | 0.30800134 | 0.75808131 | 0.86618156 |
| MDH2 | 11207.3486 | 0.140020694 | 0.20308766 | 0.6894594 | 0.49053421 | 0.67369536 |
| SRRM3 | 149.910538 | -1.035959717 | 0.4562448 | -2.2706225 | 0.02316984 | 0.08364841 |
| HSPB1 | 22170.5025 | -0.630112506 | 0.24545738 | -2.5670954 | 0.01025544 | 0.04592222 |
| YWHAG | 7030.84626 | -0.318161241 | 0.1836253 | -1.7326656 | 0.08315512 | 0.20810483 |
| SSC4D | 385.41874 | 0.179044826 | 0.34293514 | 0.5220953 | 0.60160398 | 0.75900851 |
| ZP3 | 184.184805 | -0.718495251 | 0.37708947 | -1.9053708 | 0.05673189 | 0.15911721 |
| DTX2 | 956.765444 | -0.554600007 | 0.1417087 | -3.9136624 | 9.09E-05 | 0.00117395 |
| AC005522.1 | 0.80537598 | 2.51691401 | 1.54401328 | 1.63011164 | 0.1030779 | NA |
| UPK3B | 42.0405081 | 0.52466338 | 0.41416539 | 1.26679676 | 0.20522797 | 0.38772928 |
| SPDYE16 | 27.5292147 | 1.076168284 | 0.2972113 | 3.62088618 | 0.0002936 | 0.00298577 |
| POMZP3 | 547.635885 | 0.039718807 | 0.35992324 | 0.11035355 | 0.91212899 | 0.95681217 |
| AC114737.1 | 0.20784004 | -0.340597073 | 3.29738699 | -0.103293 | 0.91773041 | NA |
| CCDC146 | 811.810945 | 0.625886598 | 0.37804998 | 1.65556576 | 0.09780979 | 0.23402995 |
| FGL2 | 3042.41518 | 0.535055732 | 0.39098961 | 1.36846534 | 0.17116646 | 0.34353145 |
| AC098851.1 | 5.33996871 | 0.593852933 | 0.72715939 | 0.81667506 | 0.41411415 | 0.60793588 |
| GSAP | 1428.93218 | 0.502080506 | 0.30667979 | 1.63714899 | 0.10159934 | 0.24017702 |
| AC004921.1 | 7.43898779 | -0.093739996 | 0.58741774 | -0.1595798 | 0.8732121 | 0.93399056 |
| PTPN12 | 3674.50317 | -0.573184854 | 0.18377233 | -3.1189943 | 0.00181469 | 0.01249112 |
| APTR | 614.497865 | -0.215475487 | 0.20250174 | -1.0640674 | 0.28729819 | 0.48356758 |
| RSBN1L | 1116.74813 | 0.132195374 | 0.1340019 | 0.98651865 | 0.32387867 | 0.52162159 |
| TMEM60 | 747.770839 | -0.284510059 | 0.20200804 | -1.4084096 | 0.15900983 | 0.32654925 |
| PHTF2 | 810.167731 | -0.108737856 | 0.16238844 | -0.6696158 | 0.50310277 | 0.68332771 |
| AC004990.1 | 0.08757577 | 0.547239394 | 3.29377202 | 0.16614368 | 0.86804388 | NA |
| MAGI2 | 676.846576 | 0.549757637 | 0.23430271 | 2.34635634 | 0.01895797 | 0.07217151 |
| MAGI2-AS1 | 0 | NA | NA | NA | NA | NA |
| MAGI2-AS2 | 0.25999332 | -1.364244661 | 2.97037558 | -0.4592836 | 0.64603055 | NA |
| AC006355.2 | 0.20710523 | -1.003574173 | 3.30558948 | -0.3035992 | 0.7614333 | NA |
| GNAI1 | 3652.92696 | -0.303994411 | 0.20021269 | -1.5183574 | 0.12892434 | 0.28333484 |
| AC003988.1 | 0 | NA | NA | NA | NA | NA |
| AC004862.1 | 1.39940607 | 2.628705157 | 1.11554225 | 2.35643711 | 0.01845119 | 0.07077315 |
| CD36 | 722.04859 | -0.085660329 | 0.74596834 | -0.1148311 | 0.90857904 | 0.95432284 |
| GNAT3 | 0 | NA | NA | NA | NA | NA |
| SEMA3C | 556.780806 | 0.455081371 | 0.40290079 | 1.12951223 | 0.25868181 | 0.45145198 |
| AC005008.2 | 0 | NA | NA | NA | NA | NA |
| HGF | 1059.15872 | -0.058414674 | 0.36200327 | -0.161365 | 0.8718059 | 0.93317628 |
| CACNA2D1 | 874.471311 | 0.427307654 | 0.12121327 | 3.52525476 | 0.00042308 | 0.00398304 |
| AC006145.1 | 0.39628499 | 2.615556686 | 3.20868416 | 0.81514931 | 0.41498685 | NA |
| PCLO | 740.134402 | 0.786256754 | 0.40813914 | 1.92644292 | 0.05404909 | 0.15390072 |
| AC079799.1 | 0 | NA | NA | NA | NA | NA |
| SEMA3E | 30.5689469 | -0.766279059 | 0.57217686 | -1.3392346 | 0.18049431 | 0.35647355 |
| SEMA3A | 106.027912 | -1.709026129 | 0.41846712 | -4.0840153 | 4.43E-05 | 0.00067315 |
| AC003984.1 | 45.2394231 | 1.973756529 | 0.50191463 | 3.93245465 | 8.41E-05 | 0.00110642 |
| AC074183.1 | 1.54459359 | -1.091751549 | 1.02776238 | -1.0622607 | 0.28811737 | 0.48426636 |
| SEMA3D | 235.405226 | 0.793241449 | 0.37151496 | 2.13515346 | 0.03274848 | 0.10739676 |
| LINC00972 | 0 | NA | NA | NA | NA | NA |
| GRM3 | 14.1147415 | 2.270431689 | 0.62687778 | 3.62180918 | NA | NA |
| AC005009.1 | 5.69099729 | -0.023743339 | 0.88829713 | -0.0267291 | 0.97867584 | 0.99041113 |
| AC005009.2 | 5.15111626 | -0.123744376 | 0.84611416 | -0.1462502 | 0.88372387 | 0.93974508 |
| KIAA1324L | 296.023164 | 0.002357054 | 0.25800333 | 0.00913575 | 0.99271083 | 0.99664146 |
| AC004023.1 | 0.03392908 | 0.380740593 | 3.29738709 | 0.11546736 | 0.90807469 | NA |
| AC005076.1 | 39.570075 | 0.353277721 | 0.44604128 | 0.7920292 | 0.42834365 | 0.62074424 |
| DMTF1 | 3814.99627 | -0.018191617 | 0.17442621 | -0.1042941 | 0.916936 | 0.9596537 |
| TMEM243 | 1737.44734 | -0.736661589 | 0.1587231 | -4.6411743 | 3.46E-06 | 8.78E-05 |
| TP53TG1 | 620.174241 | 0.263596496 | 0.24478994 | 1.07682733 | 0.28155741 | 0.47750138 |
| CROT | 1197.53111 | 0.548975429 | 0.23423282 | 2.34371691 | 0.01909266 | 0.0725048 |
| ABCB4 | 68.7657729 | 0.983829881 | 0.27761806 | 3.54382524 | 0.00039437 | 0.0037775 |
| ABCB1 | 3277.83213 | 1.051029737 | 0.30782565 | 3.41436698 | 0.0006393 | 0.00548599 |
| RUNDC3B | 184.081956 | 1.157701289 | 0.33090157 | 3.49862734 | 0.00046766 | 0.00431131 |
| SLC25A40 | 896.388031 | 0.33254364 | 0.14030815 | 2.370095 | 0.01778352 | 0.06873095 |
| DBF4 | 343.1351 | -0.820745002 | 0.2391093 | -3.4325097 | 0.00059802 | 0.00520743 |
| ADAM22 | 585.590933 | 0.82299123 | 0.3322636 | 2.47692266 | 0.01325206 | 0.05550458 |
| SRI | 3395.06912 | -0.380452073 | 0.18379943 | -2.0699306 | 0.03845884 | 0.12033218 |
| AC003991.2 | 720.951927 | -0.52341412 | 0.21298621 | -2.4575023 | 0.01399069 | 0.0578632 |
| AC003991.1 | 797.726996 | -2.041821476 | 0.55906597 | -3.6522014 | 0.00026 | 0.00270144 |
| STEAP4 | 2332.58419 | -1.928503329 | 0.54800369 | -3.5191429 | 0.00043294 | 0.00405605 |
| AC002127.2 | 0.36036649 | -0.541910702 | 2.84119942 | -0.1907331 | 0.84873472 | NA |
| ZNF804B | 11.8732557 | 4.987494459 | 0.8099723 | 6.15761117 | 7.39E-10 | 8.05E-08 |
| TEX47 | 0.08757577 | 0.547239394 | 3.29377202 | 0.16614368 | 0.86804388 | NA |
| AC002383.1 | 1.01804303 | 2.93746895 | 1.49422359 | 1.96588313 | 0.04931211 | 0.14427609 |
| AC004969.1 | 10.6048792 | 1.832874573 | 0.49127762 | 3.73083266 | 0.00019085 | 0.00210132 |
| STEAP1 | 341.937593 | -0.112570788 | 0.27697636 | -0.4064274 | 0.68442857 | 0.81770485 |
| STEAP2 | 934.389961 | 0.457122064 | 0.29009757 | 1.57575279 | 0.11508279 | 0.26234531 |
| CFAP69 | 786.40034 | 0.425060152 | 0.2695184 | 1.57710995 | 0.11477023 | 0.26193616 |
| AC002064.2 | 1.91405304 | 1.968597394 | 1.45003501 | 1.3576206 | 0.1745841 | 0.34856018 |
| AC002064.1 | 8.73533104 | 0.293212367 | 0.56569884 | 0.51831884 | 0.60423583 | 0.76132156 |
| FAM237B | 43.4180835 | 1.267605891 | 0.26496766 | 4.78400229 | 1.72E-06 | 5.03E-05 |
| GTPBP10 | 1063.29542 | 0.07283208 | 0.11944827 | 0.60973741 | 0.54203577 | 0.7146027 |
| CLDN12 | 3194.98195 | -0.432172694 | 0.22371717 | -1.9317815 | 0.05338648 | 0.1525759 |
| CDK14 | 1431.29882 | 1.068263727 | 0.28009222 | 3.81397138 | 0.00013675 | 0.00162146 |
| AC002456.1 | 15.8773348 | -0.090029087 | 0.65321436 | -0.1378247 | 0.89037895 | 0.94316 |
| FZD1 | 2385.22299 | 0.66688169 | 0.19853829 | 3.3589576 | 0.00078237 | 0.00645933 |
| AC079760.2 | 14.5115388 | 1.489171096 | 0.60558443 | 2.45906439 | 0.01392996 | 0.05766611 |
| AC079760.1 | 2.42265188 | -0.375686048 | 1.1560712 | -0.3249679 | 0.74520535 | 0.85726411 |
| AC000058.1 | 0.09635527 | 0.547239394 | 3.29377202 | 0.16614368 | 0.86804388 | NA |
| MTERF1 | 492.695523 | 0.036502853 | 0.15707076 | 0.23239751 | 0.81622928 | 0.90107402 |
| AKAP9 | 2732.83907 | 0.602759335 | 0.11415666 | 5.28010678 | 1.29E-07 | 5.94E-06 |
| CYP51A1 | 3872.25388 | -0.534894731 | 0.28004392 | -1.9100387 | 0.05612823 | 0.15787748 |
| LRRD1 | 3775.58965 | -0.558561086 | 0.27988634 | -1.9956711 | 0.04596973 | 0.13697422 |
| CYP51A1-AS1 | 31.2139138 | 0.581892209 | 0.47447211 | 1.22639918 | 0.22004849 | 0.40575336 |
| KRIT1 | 1921.79238 | 0.332619215 | 0.10958625 | 3.03522765 | 0.00240354 | 0.01551564 |
| ANKIB1 | 2622.35443 | 0.607408627 | 0.19719068 | 3.08031106 | 0.00206785 | 0.01381813 |
| GATAD1 | 2742.48988 | 0.040902593 | 0.10351089 | 0.39515257 | 0.6927303 | 0.82325918 |
| AC007566.1 | 608.919712 | 0.221327608 | 0.11865059 | 1.86537302 | 0.06212914 | 0.16946063 |
| ERVW-1 | 2.55386657 | 1.257986021 | 0.77830563 | 1.61631366 | 0.10602652 | 0.2472733 |
| PEX1 | 1729.75953 | 0.549438346 | 0.13783704 | 3.9861443 | 6.72E-05 | 0.00092987 |
| RBM48 | 777.682348 | -0.113989604 | 0.12529854 | -0.9097441 | 0.36295748 | 0.56014449 |

| | | | | | | |
|---|---|---|---|---|---|---|
| FAM133B | 528.928672 | -0.176659212 | 0.1333933 | -1.3243485 | 0.18538734 | 0.36274128 |
| CDK6 | 1200.42873 | 0.168687294 | 0.17326441 | 0.97358306 | 0.3302636 | 0.52810934 |
| AC002454.1 | 7.78392841 | -0.933963745 | 0.58819189 | -1.5878555 | 0.11231901 | 0.25782141 |
| SAMD9 | 420.847119 | -0.121295079 | 0.17211068 | -0.7047504 | 0.48096556 | 0.66563776 |
| SAMD9L | 777.46908 | 0.149853518 | 0.30674994 | 0.48852013 | 0.62518147 | 0.77672723 |
| HEPACAM2 | 423.840837 | 1.366806743 | 0.56006297 | 2.44045191 | 0.0146689 | 0.05991618 |
| VPS50 | 1235.6975 | 0.41588629 | 0.14828378 | 2.80466467 | 0.0050369 | 0.02700751 |
| CALCR | 189.408276 | -0.395288678 | 0.39107773 | -1.0107675 | 0.3121277 | 0.50910117 |
| GNGT1 | 151.611499 | -1.332124454 | 0.58104687 | -2.2926282 | 0.02186942 | 0.08014127 |
| TFPI2 | 1109.96668 | -1.564441014 | 0.58936717 | -2.6544421 | 0.00794397 | 0.03797035 |
| AC002076.1 | 3.07471225 | -0.237132127 | 1.21417665 | -0.1953028 | 0.84515589 | 0.91828433 |
| GNG11 | 2282.50997 | -0.193176963 | 0.25976583 | -0.7436581 | 0.45708335 | 0.64649689 |
| BET1 | 1368.02046 | -0.238495551 | 0.13829142 | -1.7245868 | 0.08460196 | 0.21071574 |
| AC006378.1 | 17.3029955 | -0.002423253 | 0.36181262 | -0.0066975 | 0.99465618 | 0.99760701 |
| AC003092.1 | 0.88290652 | -0.301481214 | 1.67203265 | -0.1803082 | 0.85691061 | NA |
| AC003092.2 | 0.89047302 | -3.046035971 | 1.79417302 | -1.6977381 | 0.0895572 | NA |
| AC002074.1 | 6.89121972 | -1.09797025 | 0.88428134 | -1.2416526 | 0.21436475 | 0.39847727 |
| COL1A2 | 10490.2154 | 0.07511503 | 0.40134423 | 0.18715861 | 0.85153627 | 0.92224271 |
| CASD1 | 1164.14336 | 0.215402646 | 0.14362458 | 1.49976172 | 0.13367614 | 0.29048535 |
| SGCE | 1641.94622 | -0.33055405 | 0.22842815 | -1.4470811 | 0.14787421 | 0.3113825 |
| PEG10 | 706.134814 | 0.970185439 | 0.32148858 | 3.01779131 | 0.00254624 | 0.01621845 |
| PPP1R9A | 881.364194 | 0.074211635 | 0.18624965 | 0.39845249 | 0.69029668 | 0.8215683 |
| AC002429.2 | 1.91935449 | 0.777354938 | 0.99941583 | 0.77780931 | 0.43668144 | 0.62877147 |
| PON1 | 13.6542151 | -0.355612332 | 0.58340306 | -0.6095483 | 0.54216108 | 0.71466072 |
| PON3 | 125.455766 | 0.776231007 | 0.41912886 | 1.85201039 | 0.06402433 | 0.1730015 |
| PON2 | 5376.13532 | -1.305963056 | 0.18164336 | -7.1897097 | 6.49E-13 | 1.86E-10 |
| AC005021.1 | 392.398447 | -0.965548345 | 0.22036659 | -4.381555 | 1.18E-05 | 0.00023363 |
| AC004012.1 | 1.07545696 | 1.143040863 | 1.32918418 | 0.85995671 | 0.3898129 | 0.58538532 |
| ASB4 | 25.585706 | -0.641010274 | 0.58136756 | -1.1025904 | 0.2702051 | 0.46449207 |
| PDK4 | 16295.2381 | -1.607826708 | 0.6708221 | -2.3968004 | 0.01653893 | 0.06535764 |
| AC002451.1 | 18.9175241 | 0.263774029 | 0.39156843 | 0.67363457 | 0.50054367 | 0.6815873 |
| DYNC1I1 | 187.497794 | 0.192680012 | 0.40925252 | 0.47080959 | 0.63777672 | 0.78561835 |
| SLC25A13 | 2684.18504 | 0.446415351 | 0.22362601 | 1.99625864 | 0.04590578 | 0.13686098 |
| SEM1 | 2299.77107 | -0.603137795 | 0.1766213 | -3.4148645 | 0.00063814 | 0.00547955 |
| DLX6-AS1 | 7.03845196 | 0.941930504 | 0.84725412 | 1.11174497 | 0.26624782 | 0.46044605 |
| DLX6 | 6.97695574 | -0.384561525 | 1.41456574 | -0.2718584 | 0.78573093 | 0.88315607 |
| DLX5 | 22.3795199 | -0.766226146 | 0.76416573 | -1.0026963 | 0.31600742 | 0.5131471 |
| SDHAF3 | 695.27237 | 0.240141295 | 0.15479277 | 1.55137278 | 0.12081237 | 0.27117997 |
| TAC1 | 324.407284 | -2.454850966 | 0.86930544 | -2.8239222 | 0.00474399 | 0.02585051 |
| ASNS | 1552.75006 | -1.147204563 | 0.18136504 | -6.3253897 | 2.53E-10 | 3.23E-08 |
| OCM2 | 0 | NA | NA | NA | NA | NA |
| LMTK2 | 2143.92975 | -0.220848411 | 0.27362542 | -0.8071195 | 0.41959764 | 0.61243738 |
| BHLHA15 | 32.0116831 | -0.393275996 | 0.46207231 | -0.8511135 | 0.39470629 | 0.58994529 |
| TECPR1 | 2172.87512 | 0.127319624 | 0.13000274 | 0.97936106 | 0.32740161 | 0.52522085 |
| BRI3 | 5399.14761 | -0.426945221 | 0.28575122 | -1.4941151 | 0.13514554 | 0.29256939 |
| BAIAP2L1 | 1598.85155 | -0.942088421 | 0.21093163 | -4.4663213 | 7.96E-06 | 0.00017033 |
| AC093799.1 | 34.7997286 | -0.796188785 | 0.57489786 | -1.3849222 | 0.16607627 | 0.33672392 |
| NPTX2 | 99.2274028 | -2.585386615 | 0.68125558 | -3.7950318 | 0.00014762 | 0.00173093 |
| TMEM130 | 293.545552 | 1.438678851 | 0.45125489 | 3.18817344 | 0.00143175 | 0.01043612 |
| TRRAP | 2620.32904 | 0.007008191 | 0.12666107 | 0.05533027 | 0.95587535 | 0.97896664 |
| SMURF1 | 2432.40457 | -0.590033569 | 0.22679535 | -2.6016122 | 0.00927867 | 0.04256759 |
| KPNA7 | 4.57565033 | -1.518307103 | 0.71736537 | -2.1165046 | 0.03430191 | 0.11101393 |
| AC004834.1 | 15.0383616 | 0.453458673 | 0.48176301 | 0.94124842 | 0.34657757 | 0.54422977 |
| AC004922.1 | 9460.795 | -0.846863957 | 0.20836622 | -4.0643055 | 4.82E-05 | 0.00072222 |
| ARPC1A | 4078.44315 | -0.235250465 | 0.18092034 | -1.3002986 | 0.19349864 | 0.37310319 |
| ARPC1B | 6100.16876 | -1.329950901 | 0.27912912 | -4.764644 | 1.89E-06 | 5.43E-05 |
| PDAP1 | 6330.10803 | -0.57207848 | 0.15671506 | -3.6504371 | 0.00026179 | 0.00271683 |
| BUD31 | 2791.24902 | -0.473446016 | 0.12489668 | -3.7907015 | 0.00015022 | 0.00175592 |
| PTCD1 | 4425.21271 | -0.22292849 | 0.1061686 | -2.0997592 | 0.03575003 | 0.11425625 |
| ATP5MF-PTC | 4281.1641 | -0.085908297 | 0.11634619 | -0.7383851 | 0.46028044 | 0.64882638 |
| CPSF4 | 1602.11484 | -0.406770691 | 0.15610856 | -2.6056912 | 0.00916891 | 0.04221828 |
| ATP5MF | 5013.21895 | -0.19828715 | 0.11494934 | -1.7249959 | 0.08452821 | 0.21064238 |
| ZNF789 | 854.790254 | 0.505418657 | 0.24852393 | 2.03368205 | 0.04198366 | 0.12833073 |
| ZNF394 | 1184.09932 | -0.294274887 | 0.12579663 | -2.3392906 | 0.0193204 | 0.0730848 |
| ZKSCAN5 | 741.974342 | -0.020208528 | 0.12621687 | -0.1601096 | 0.87279477 | 0.93368745 |
| FAM200A | 543.660357 | -0.008399135 | 0.13855138 | -0.0606211 | 0.95166098 | 0.97678185 |
| ZNF655 | 4029.59532 | -0.067661992 | 0.1547578 | -0.4372122 | 0.66195749 | 0.80116666 |
| AC005020.2 | 384.301644 | 0.039992264 | 0.17988685 | 0.222319 | 0.82406556 | 0.90585722 |
| TMEM225B | 134.312902 | -0.14196408 | 0.40307225 | -0.352205 | 0.72468449 | 0.84389181 |
| ZSCAN25 | 739.502681 | 0.256570848 | 0.16230388 | 1.58080535 | 0.11392255 | 0.26029797 |
| CYP3A5 | 1302.90509 | 1.019487695 | 0.52156043 | 1.95468759 | 0.05061996 | 0.14686415 |
| CYP3A7 | 277.501726 | 4.266676024 | 0.59155309 | 7.21266799 | 5.49E-13 | 1.61E-10 |
| CYP3A7-CYP3 | 232.588469 | 4.232744432 | 0.58080163 | 7.28776263 | 3.15E-13 | 1.04E-10 |
| CYP3A4 | 129.670513 | 4.055331113 | 0.7539187 | 5.37900325 | 7.49E-08 | 3.74E-06 |
| AC069294.1 | 5.81739916 | 0.674464161 | 0.64916486 | 1.03897208 | 0.29881772 | 0.49563286 |
| CYP3A43 | 7.94693327 | 1.098583874 | 0.50469726 | 2.17671853 | 0.02950157 | 0.09970779 |
| OR2AE1 | 0.40087909 | 1.874276558 | 2.1488127 | 0.87223822 | 0.38307843 | NA |
| TRIM4 | 1795.4141 | 0.074996993 | 0.18441879 | 0.40666676 | 0.68425276 | 0.81757775 |
| GJC3 | 50.7220981 | 1.053310584 | 0.80024985 | 1.31622716 | 0.18809781 | 0.36613054 |
| AC004522.2 | 10.2315933 | 1.235179132 | 0.93315965 | 1.32365253 | 0.18561847 | 0.36296132 |
| AZGP1 | 10609.106 | 1.409171443 | 0.50308488 | 2.801061 | 0.00509349 | 0.02725065 |
| ZKSCAN1 | 3393.50792 | 0.079915045 | 0.17665534 | 0.45237831 | 0.65099647 | 0.79488544 |
| ZSCAN21 | 964.786135 | 0.270123546 | 0.16724577 | 1.61512935 | 0.10628269 | 0.24765435 |
| ZNF3 | 1911.30955 | 0.02070899 | 0.14577047 | 0.14206574 | 0.88702808 | 0.94150285 |
| COPS6 | 4025.06182 | -0.05127322 | 0.11943398 | -0.4293018 | 0.66770363 | 0.8055332 |
| MCM7 | 2888.52185 | -0.106071691 | 0.19731346 | -0.5375796 | 0.59086731 | 0.7505381 |
| AP4M1 | 3378.93541 | -0.346544821 | 0.12620926 | -2.7457955 | 0.00603644 | 0.03092692 |
| TAF6 | 4007.62892 | -0.211699011 | 0.13592535 | -1.5574653 | 0.11936005 | 0.26888131 |
| AC073842.2 | 52.5095697 | 0.289652071 | 0.25565123 | 1.132997 | 0.25721551 | 0.44990535 |
| CNPY4 | 703.307755 | 0.069309878 | 0.22401259 | 0.3094017 | 0.75701598 | 0.86548933 |
| MBLAC1 | 130.23624 | -0.227332852 | 0.27235154 | -0.8347037 | 0.40388453 | 0.59887737 |
| AC073842.1 | 12.6434157 | -0.895542379 | 0.64748774 | -1.3831032 | 0.16663322 | 0.33746394 |
| LAMTOR4 | 3893.93735 | -0.174994371 | 0.11629802 | -1.5047064 | 0.13239957 | 0.28868668 |
| C7orf43 | 2106.88866 | -0.228215569 | 0.17192059 | -1.3274476 | 0.18436067 | 0.36149801 |
| GAL3ST4 | 127.920023 | 0.126244767 | 0.41576111 | 0.30364737 | 0.76139656 | 0.8677512 |
| GPC2 | 65.2692415 | -1.024450357 | 0.26221287 | -3.9069416 | 9.35E-05 | 0.00120002 |
| STAG3 | 1153.3149 | -0.66932108 | 0.28627794 | -2.3380113 | 0.01938666 | 0.07324158 |
| CASTOR3 | 1451.80897 | -0.90553343 | 0.26669859 | -3.3953439 | 0.00068542 | 0.00579501 |
| PVRIG | 123.622462 | 0.203019705 | 0.41072492 | 0.49429604 | 0.62109712 | 0.77363364 |
| SPDYE3 | 417.825292 | 0.321028723 | 0.21571935 | 1.4881777 | 0.13670404 | 0.2950471 |
| PILRB | 11448.2065 | -0.042345784 | 0.32160014 | -0.1316722 | 0.89524361 | 0.94619535 |
| PILRA | 2030.17669 | -0.524329987 | 0.34902412 | -1.5022744 | 0.13302627 | 0.28950258 |
| ZCWPW1 | 400.460739 | 0.698794308 | 0.24681948 | 2.83119599 | 0.00463743 | 0.02545024 |
| MEPCE | 3755.65309 | -0.186308162 | 0.31245328 | -0.5962753 | 0.55099136 | 0.72132463 |
| AC092849.2 | 605.715186 | -1.205232252 | 0.18932084 | -6.3660833 | 1.94E-10 | 2.52E-08 |
| PPP1R35 | 1174.75605 | -1.250870006 | 0.19145784 | -6.5333965 | 6.43E-11 | 1.00E-08 |
| AC092849.1 | 230.372656 | -1.239491436 | 0.23688619 | -5.2324344 | 1.67E-07 | 7.36E-06 |
| C7orf61 | 49.5609052 | -1.620310863 | 0.41797238 | -3.8765979 | 0.00010593 | 0.00132698 |
| TSC22D4 | 5342.88418 | -0.587087506 | 0.2347331 | -2.5010853 | 0.01238133 | 0.05281394 |
| NYAP1 | 153.192983 | -0.207211633 | 0.64962276 | -0.3189723 | 0.74974756 | 0.86027513 |
| AC069281.1 | 1.46302875 | -0.988243599 | 1.20277816 | -0.8216341 | 0.41128515 | 0.60556055 |
| AGFG2 | 2438.1658 | -0.670632046 | 0.28336171 | -2.3666996 | 0.01794749 | 0.0692328 |
| SAP25 | 792.7321 | -0.800004433 | 0.27555306 | -2.9032681 | 0.0036929 | 0.02146866 |

| | | | | | | |
|---|---|---|---|---|---|---|
| LRCH4 | 5516.36392 | -0.409625712 | 0.16440747 | -2.4915274 | 0.01271951 | 0.05386439 |
| FBXO24 | 151.723452 | 0.471115158 | 0.45522414 | 1.03490811 | 0.30071183 | 0.4974536 |
| PCOLCE-AS1 | 774.638523 | -0.153872296 | 0.51740814 | -0.2973906 | 0.76616835 | 0.87108129 |
| PCOLCE | 2631.18612 | -0.56464677 | 0.42140536 | -1.3399136 | 0.18027343 | 0.35621088 |
| MOSPD3 | 1005.9503 | -0.226047452 | 0.11024473 | -2.0504151 | 0.04032394 | 0.12451901 |
| TFR2 | 31.2437672 | -1.567452967 | 0.58879472 | -2.6621383 | 0.0077646 | 0.0373704 |
| ACTL6B | 0.06785815 | 0.795868262 | 3.28898349 | 0.24198001 | 0.80879565 | NA |
| GNB2 | 10969.3156 | -0.752470609 | 0.15437196 | -4.8743995 | 1.09E-06 | 3.50E-05 |
| GIGYF1 | 8569.6635 | -0.149340435 | 0.20863465 | -0.7157988 | 0.47411558 | 0.66000253 |
| POP7 | 841.798159 | -0.509819326 | 0.2117263 | -2.4079168 | 0.01604384 | 0.06399282 |
| EPO | 120.541056 | 0.269013506 | 1.06628359 | 0.25229077 | 0.80081632 | 0.89197577 |
| ZAN | 0.39418504 | -0.032129452 | 2.2473689 | -0.0142965 | 0.98859345 | NA |
| EPHB4 | 4667.18383 | -0.099229808 | 0.24929716 | -0.3980383 | 0.69060199 | 0.82183882 |
| SLC12A9 | 3568.41844 | 0.218341043 | 0.13362553 | 1.63397699 | 0.10226368 | 0.24117165 |
| SLC12A9-AS1 | 7.26410481 | -0.045187829 | 0.59844707 | -0.0755085 | 0.93981015 | 0.97115066 |
| TRIP6 | 6498.56326 | -0.060542247 | 0.16298842 | -0.3714512 | 0.71030148 | 0.83536599 |
| SRRT | 7429.45263 | -0.302034392 | 0.21004957 | -1.4379196 | 0.15045688 | 0.31505979 |
| UFSP1 | 65.9737983 | 0.377856383 | 0.32372599 | 1.16721053 | 0.24312536 | 0.43368841 |
| ACHE | 445.50701 | -1.508052307 | 0.37877127 | -3.9814327 | 6.85E-05 | 0.00094278 |
| MUC3A | 305.019579 | -1.782535749 | 0.43551054 | -4.0929796 | 4.26E-05 | 0.00065291 |
| AC254629.1 | 138.211549 | -2.20267101 | 0.44241478 | -4.9787465 | 6.40E-07 | 2.24E-05 |
| MUC12 | 26.8981587 | -1.491285217 | 0.50897861 | -2.9299566 | 0.00339009 | 0.02012081 |
| AC105446.1 | 15.8381719 | -1.568197306 | 0.50486957 | -3.1061435 | 0.00189545 | 0.01292885 |
| MUC17 | 0 | NA | NA | NA | NA | NA |
| TRIM56 | 4457.25728 | -0.683624659 | 0.20951695 | -3.2628609 | 0.00110294 | 0.00845724 |
| SERPINE1 | 9612.41705 | -2.745902462 | 0.7351018 | -3.7354044 | 0.00018741 | 0.00207129 |
| AP1S1 | 3047.58212 | -0.503624407 | 0.16590961 | -3.0355349 | 0.00240109 | 0.01550743 |
| VGF | 232.645301 | -4.138271325 | 1.23807082 | -3.3425158 | NA | NA |
| NAT16 | 15.3920978 | -0.807228293 | 0.89114182 | -0.9058359 | 0.36502269 | 0.56205495 |
| MOGAT3 | 168.693811 | 0.626209199 | 0.65830742 | 0.95124128 | 0.34148191 | 0.53947267 |
| PLOD3 | 4465.46603 | -0.758792843 | 0.17565279 | -4.3198451 | 1.56E-05 | 0.00029305 |
| ZNHIT1 | 4190.10271 | -0.146234054 | 0.10577021 | -1.3825637 | 0.16679868 | 0.33772122 |
| CLDN15 | 1305.29717 | -0.154199648 | 0.30469014 | -0.5060868 | 0.61279574 | 0.76799695 |
| FIS1 | 4919.01479 | 0.028111057 | 0.09855794 | 0.28522365 | 0.7754728 | 0.87674575 |
| LNCPRESS1 | 0.23140391 | 0.077483112 | 3.10308039 | 0.02496974 | 0.9800791 | NA |
| AC006329.1 | 68.3536455 | -0.387206731 | 0.30832413 | -1.2558431 | 0.20917286 | 0.39235447 |
| IFT22 | 1751.6259 | 0.043887917 | 0.10268305 | 0.42741152 | 0.66907962 | 0.80667655 |
| COL26A1 | 343.987451 | 0.139711901 | 0.79540647 | 0.17564844 | 0.86057015 | 0.92734057 |
| LINC01007 | 0.45358783 | 0.345814902 | 2.28417101 | 0.15139624 | 0.87966316 | NA |
| MYL10 | 0.07830766 | -0.642905668 | 3.30558948 | -0.1944905 | 0.84579185 | NA |
| CUX1 | 6516.28102 | 0.606942617 | 0.14242209 | 4.26157626 | 2.03E-05 | 0.00035901 |
| AC005096.1 | 0.76694977 | -1.830255373 | 1.88597975 | -0.9704534 | 0.33182057 | NA |
| AC005086.1 | 1.07751733 | 1.138902973 | 1.24868262 | 0.91208363 | 0.3617247 | 0.55911494 |
| SH2B2 | 617.846914 | -1.472851945 | 0.32017627 | -4.6001284 | 4.22E-06 | 0.0001029 |
| SPDYE6 | 40.5143096 | 0.361014263 | 0.50634382 | 0.71298246 | 0.47585661 | 0.66130903 |
| PRKRIP1 | 3241.28137 | -0.463300459 | 0.15816 | -2.929315 | 0.0033971 | 0.0201446 |
| AC073517.1 | 9.73682823 | -0.44084029 | 0.60083748 | -0.7337097 | 0.46312568 | 0.65105787 |
| ORAI2 | 2306.38465 | -0.250898499 | 0.27954476 | -0.8975253 | 0.36943868 | 0.56630227 |
| ALKBH4 | 727.39965 | 0.122940098 | 0.10772884 | 1.14119947 | 0.25378692 | 0.44642527 |
| LRWD1 | 1498.16816 | -0.310872454 | 0.21176274 | -1.4680225 | 0.14209812 | 0.30304307 |
| POLR2J | 1499.85284 | -0.191505029 | 0.16185745 | -1.1831709 | 0.23674142 | 0.42601653 |
| RASA4B | 216.376067 | 0.977557992 | 0.46658163 | 2.09514891 | 0.03615776 | 0.11519633 |
| AC093668.2 | 628.982489 | 0.89156461 | 0.38946355 | 2.28921195 | 0.02206704 | 0.08066369 |
| POLR2J3 | 1106.47556 | 0.577822217 | 0.35463253 | 1.62935483 | 0.10323793 | 0.24283151 |
| AC093668.3 | 493.988797 | 0.736960629 | 0.37631998 | 1.95833508 | 0.05019071 | 0.14602601 |
| AC093668.1 | 556.559133 | 0.837203046 | 0.33078344 | 2.53097023 | 0.01137475 | 0.0496363 |
| SPDYE2 | 28.9633949 | 0.989332258 | 0.48398219 | 2.04415014 | 0.04093872 | 0.12606506 |
| RASA4 | 561.938087 | 0.560574492 | 0.48859517 | 1.14731893 | 0.25124983 | 0.44354643 |
| AC105052.1 | 732.106123 | 0.263119716 | 0.37899313 | 0.69425986 | 0.48751926 | 0.67137744 |
| UPK3BL1 | 1363.66424 | 0.053838189 | 0.39981193 | 0.13465879 | 0.89288166 | 0.94464551 |
| AC105052.3 | 1381.73949 | 0.06883404 | 0.40270392 | 0.17092965 | 0.86427908 | 0.92963749 |
| SPDYE2B | 7.50699148 | 2.299092044 | 0.63123485 | 3.64221343 | 0.0002703 | 0.00279636 |
| POLR2J2 | 43.7597984 | 0.580635357 | 0.59217208 | 0.98051795 | 0.32683051 | 0.52471967 |
| FAM185A | 375.039469 | 0.128099384 | 0.1464109 | 0.87493066 | 0.38161163 | 0.57757877 |
| FBXL13 | 56.0960598 | 0.074000366 | 0.22245733 | 0.3326497 | 0.73939872 | 0.85378746 |
| LRRC17 | 132.738029 | 0.359197646 | 0.3192597 | 1.12509548 | 0.26054858 | 0.45363767 |
| NFE4 | 3.11856561 | -0.201774377 | 0.72098828 | -0.2798581 | 0.77958641 | 0.87921023 |
| AC073127.1 | 0.54802852 | -2.342153202 | 2.70793381 | -0.8649226 | 0.38708126 | NA |
| ARMC10 | 1795.3792 | -0.250827509 | 0.12364393 | -2.0286278 | 0.04249622 | 0.12946225 |
| NAPEPLD | 2055.76334 | 0.78657272 | 0.21637505 | 3.63522839 | 0.00027773 | 0.00286312 |
| PMPCB | 4971.35236 | 0.086441248 | 0.09532494 | 0.90680617 | 0.3645093 | 0.56165189 |
| DNAJC2 | 1406.05892 | -0.422556908 | 0.2010542 | -2.1017065 | 0.035579 | 0.11394439 |
| PSMC2 | 4021.22473 | -0.113317276 | 0.16453557 | -0.6887099 | 0.49100584 | 0.67404987 |
| SLC26A5 | 12.7348254 | 0.25021791 | 0.45843528 | 0.54580858 | 0.58519752 | 0.74685098 |
| AC005064.1 | 5.29343798 | 0.606233083 | 0.63023301 | 0.96191896 | 0.33609032 | 0.53425171 |
| RELN | 166.368305 | 0.013945466 | 0.41097745 | 0.03393244 | 0.97293103 | 0.98741337 |
| ORC5 | 700.519783 | -0.209522616 | 0.17129665 | -1.2231565 | 0.22127059 | 0.40743702 |
| LHFPL3 | 98.0842078 | 0.729234695 | 0.27463689 | 2.65526851 | 0.00792453 | 0.03790493 |
| AC007384.1 | 17.6341802 | -0.866314466 | 0.77806883 | -1.1134162 | 0.26552971 | 0.45963098 |
| LINC01004 | 354.952267 | 0.226305806 | 0.17887131 | 1.26518781 | 0.20580401 | 0.38836781 |
| KMT2E-AS1 | 420.887904 | -0.246091012 | 0.25396727 | -0.9689871 | 0.33255162 | 0.53051338 |
| KMT2E | 2689.12343 | -0.020391046 | 0.1648849 | -0.1236684 | 0.90157786 | 0.95014819 |
| AC005070.3 | 31.7052033 | 0.557248617 | 0.26165491 | 2.12970823 | 0.03319571 | 0.1083932 |
| SRPK2 | 1531.02236 | -0.067058471 | 0.12049518 | -0.5565241 | 0.57785261 | 0.74147114 |
| AC004884.2 | 6.80844688 | 0.266242064 | 0.43682122 | 0.60949892 | 0.54219379 | 0.71466811 |
| PUS7 | 587.035816 | -0.645010617 | 0.32440168 | -1.9883085 | 0.04677758 | 0.13873859 |
| RINT1 | 1094.1735 | -0.114865681 | 0.1389632 | -0.8265907 | 0.4084691 | 0.60296296 |
| EFCAB10 | 517.071761 | -0.468326183 | 0.14081526 | -3.3258198 | 0.00088159 | 0.00711368 |
| AC073073.2 | 87.1895684 | 0.161664013 | 0.21544579 | 0.75036979 | 0.45303202 | 0.64266256 |
| ATXN7L1 | 691.802452 | 1.44153288 | 0.32772502 | 4.39860495 | 1.09E-05 | 0.00021864 |
| CDHR3 | 459.522008 | 1.419892781 | 0.37649349 | 3.77136078 | 0.00016236 | 0.00185826 |
| SYPL1 | 8778.69358 | -0.169860949 | 0.10651149 | -1.5947664 | 0.11076448 | 0.2550881 |
| NAMPT | 18106.3941 | -2.840508299 | 0.45311859 | -6.2687966 | 3.64E-10 | 4.31E-08 |
| AC007032.1 | 50.5126128 | -0.862285443 | 0.32350984 | -2.6654072 | 0.00768952 | 0.03708349 |
| AC005050.1 | 4.30920847 | 0.508117792 | 0.70906536 | 0.7166022 | 0.47361959 | 0.65956659 |
| CCDC71L | 1472.6088 | -0.900903741 | 0.3293934 | -2.7350388 | 0.00623729 | 0.03174626 |
| LINC02577 | 0.17648264 | -1.003574173 | 3.30558948 | -0.3035992 | 0.7614333 | NA |
| AC002384.1 | 0.18382444 | -1.003574173 | 3.30558948 | -0.3035992 | 0.7614333 | NA |
| PIK3CG | 50.2991677 | 0.596790501 | 0.46551592 | 1.28199806 | 0.19984333 | 0.38117169 |
| PRKAR2B | 732.631293 | 0.339880235 | 0.34098991 | 0.99674571 | 0.31888796 | 0.51614752 |
| HBP1 | 4566.80319 | -0.179108873 | 0.12047739 | -1.4866596 | 0.13710474 | 0.29562159 |
| AC004492.1 | 14.221466 | -0.25746024 | 0.43272723 | -0.5949712 | 0.55186272 | 0.72178454 |
| COG5 | 3529.34878 | -0.311309927 | 0.10336326 | -3.0118045 | 0.002597 | 0.01645827 |
| GPR22 | 4.88931818 | 1.083165091 | 0.83204994 | 1.30180297 | 0.19298375 | 0.37249168 |
| DUS4L | 492.95085 | 0.083549143 | 0.23524391 | 0.35515965 | 0.72246999 | 0.84246598 |
| AC004839.1 | 8.30256313 | -0.365797622 | 0.45902945 | -0.7968936 | 0.42551284 | 0.61784998 |
| BCAP29 | 1978.14681 | -0.359042286 | 0.1372283 | -2.6163866 | 0.00888659 | 0.0412424 |
| SLC26A4-AS1 | 13.0096807 | 2.888150482 | 0.52220223 | 5.53071263 | 3.19E-08 | 1.85E-06 |
| SLC26A4 | 282.523308 | 2.171459033 | 0.4240424 | 5.1208536 | 3.04E-07 | 1.20E-05 |
| AC002467.1 | 97.2032777 | 0.20743655 | 0.25431167 | 0.81567846 | 0.41468406 | 0.60825557 |
| CBLL1 | 1032.22329 | -0.09845228 | 0.1866555 | -0.5274545 | 0.59787803 | 0.75634654 |
| SLC26A3 | 7.12402483 | -0.220273335 | 0.57198077 | -0.3851062 | 0.70015874 | 0.82803754 |
| DLD | 9437.56166 | 0.314042149 | 0.11425706 | 2.74855785 | 0.00598581 | 0.03072124 |

| | | | | | | |
|---|---|---|---|---|---|---|
| LAMB1 | 17874.3434 | -0.201921793 | 0.18165033 | -1.111596 | 0.26631187 | 0.46047137 |
| AC005046.1 | 122.055945 | -0.663843331 | 0.51136946 | -1.2981677 | 0.1942297 | 0.37404785 |
| LAMB4 | 97.2171436 | 0.598073673 | 0.33637587 | 1.77799218 | 0.07540514 | 0.19444881 |
| NRCAM | 1404.26186 | -0.51132667 | 0.30360632 | -1.6841766 | 0.09214754 | 0.22442832 |
| PNPLA8 | 2721.78394 | -0.385389637 | 0.27339596 | -1.4096391 | 0.15864628 | 0.32612273 |
| THAP5 | 1075.36446 | 0.223485018 | 0.12748912 | 1.75297329 | 0.07960659 | 0.20177103 |
| DNAJB9 | 4180.79323 | -0.098205349 | 0.23770747 | -0.4131353 | 0.67950748 | 0.81406334 |
| AC005487.1 | 3.85233835 | 1.683340727 | 1.32570509 | 1.26977013 | 0.20416652 | 0.3864947 |
| C7orf66 | 0.03392908 | 0.380740593 | 3.29738709 | 0.11546736 | 0.90807469 | NA |
| AC004014.1 | 0 | NA | NA | NA | NA | NA |
| AC002386.1 | 0 | NA | NA | NA | NA | NA |
| AC073071.1 | 0 | NA | NA | NA | NA | NA |
| AC092167.1 | 1.00735423 | -1.262058749 | 1.42950807 | -0.8828623 | 0.37731069 | 0.57330353 |
| IMMP2L | 1276.16325 | 0.192083523 | 0.30538964 | 0.62897851 | 0.52936312 | 0.70387785 |
| LRRN3 | 20.6595098 | 1.791444057 | 0.47451106 | 3.77534734 | 0.00015978 | 0.00184156 |
| DOCK4 | 992.73403 | -0.020153257 | 0.20836229 | -0.0967222 | 0.92294702 | 0.96236112 |
| DOCK4-AS1 | 1.41807184 | -1.478902043 | 1.39896968 | -1.0571366 | 0.29044925 | 0.48652584 |
| ZNF277 | 1015.26739 | -0.374262564 | 0.13175178 | -2.8406641 | 0.00450197 | 0.02488303 |
| AC004112.1 | 5.16924134 | 0.482454317 | 0.64520042 | 0.74775884 | 0.45460563 | 0.64406332 |
| IFRD1 | 3492.61339 | -0.84838256 | 0.25368963 | -3.3441751 | 0.00082528 | 0.00674389 |
| LSMEM1 | 507.300442 | -0.868406349 | 0.36399962 | -2.3857342 | 0.01704506 | 0.06678727 |
| AC002463.1 | 0.08695548 | -0.642905668 | 3.30558948 | -0.1944905 | 0.84579185 | NA |
| TMEM168 | 1445.53506 | 0.2257657 | 0.1151218 | 1.96110293 | 0.04986702 | 0.1454156 |
| BMT2 | 572.076061 | -0.376449848 | 0.19393898 | -1.9410737 | 0.05224934 | 0.15020333 |
| AC018464.1 | 0.09989236 | -0.642905668 | 3.30558948 | -0.1944905 | 0.84579185 | NA |
| GPR85 | 43.8793964 | -1.493140123 | 0.56452523 | -2.6449484 | 0.00817034 | 0.03870855 |
| SMIM30 | 2054.1977 | -0.03678857 | 0.0931493 | -0.394942 | 0.69288568 | 0.8232818 |
| PPP1R3A | 0.18845094 | 0.726716446 | 3.28420131 | 0.22127646 | 0.82487718 | NA |
| FOXP2 | 280.746479 | 0.034181016 | 0.34982846 | 0.09770793 | 0.92216421 | 0.96203347 |
| AC073626.1 | 0 | NA | NA | NA | NA | NA |
| MDFIC | 1180.31609 | 0.277455047 | 0.21427404 | 1.29486078 | 0.19536826 | 0.37552619 |
| LINC01392 | 0.11223674 | 0.02007176 | 3.29738704 | 0.00608717 | 0.99514317 | NA |
| LINC01393 | 7.7411719 | -0.337475733 | 0.56393543 | -0.5984297 | 0.54955322 | 0.72004961 |
| AC092590.1 | 0.17332742 | -1.003574173 | 3.30558948 | -0.3035992 | 0.7614333 | NA |
| AC093714.1 | 0 | NA | NA | NA | NA | NA |
| AC093714.2 | 0 | NA | NA | NA | NA | NA |
| TFEC | 1443.05583 | 1.696800801 | 0.48025255 | 3.53314271 | 0.00041065 | 0.00389842 |
| TES | 2497.86811 | -0.775820228 | 0.24041941 | -3.2269451 | 0.00125119 | 0.00936266 |
| AC073130.2 | 7.9852633 | 0.901464552 | 0.43067852 | 2.09312636 | 0.03633788 | 0.11558876 |
| AC002066.1 | 4.48969474 | 0.5068802 | 0.73965492 | 0.68529282 | 0.49315916 | 0.67588 |
| CAV2 | 1468.28331 | -0.646237525 | 0.28162651 | -2.2946616 | 0.02175253 | 0.07985718 |
| CAV1 | 1237.63546 | 0.319841078 | 0.46871371 | 0.68238046 | 0.49499841 | 0.67717723 |
| AC006159.1 | 12.947666 | 2.314195928 | 0.62871563 | 3.68083093 | 0.00023248 | 0.00246397 |
| AC006159.2 | 228.385874 | 2.239240081 | 0.71635769 | 3.12586871 | 0.00177281 | 0.0123017 |
| LINC01510 | 420.561809 | 2.207007133 | 0.77443138 | 2.84984207 | 0.00437409 | 0.02439096 |
| MET | 5301.90524 | -1.191359354 | 0.31508042 | -3.7811279 | 0.00015612 | 0.00180906 |
| CAPZA2 | 8281.86521 | -0.214247144 | 0.15568987 | -1.3761149 | 0.16878607 | 0.3404008 |
| ST7-AS1 | 200.019873 | 0.423809839 | 0.18778138 | 2.25693223 | 0.02401231 | 0.08562605 |
| ST7 | 1798.33672 | 0.497391433 | 0.14052257 | 3.53958392 | 0.00040076 | 0.00382621 |
| AC106873.1 | 0.55295916 | 1.699879266 | 2.16009931 | 0.78694496 | 0.43131409 | NA |
| ST7-AS2 | 221.593198 | 0.913129665 | 0.24405136 | 3.74154708 | 0.00018289 | 0.00203495 |
| WNT2 | 12.127283 | 1.075606183 | 0.66305847 | 1.62218905 | 0.10476288 | 0.24525897 |
| AC002465.1 | 0.09635527 | 0.547239394 | 3.29377202 | 0.16614368 | 0.86804388 | NA |
| ASZ1 | 0.59679702 | -0.785202642 | 2.25179446 | -0.3487008 | 0.72731391 | NA |
| CFTR | 301.635016 | -1.996171943 | 0.55854539 | -3.5738759 | 0.00035174 | 0.00344549 |
| CFTR-AS1 | 0.19478864 | 0.18657164 | 3.29377195 | 0.05664376 | 0.95482897 | NA |
| AC000061.1 | 1.36437775 | -2.939142339 | 2.28823856 | -1.2844563 | 0.19898236 | 0.38006409 |
| CTTNBP2 | 629.742923 | 0.245820156 | 0.17068189 | 1.44022397 | 0.14980404 | 0.31408276 |
| AC003084.1 | 0.32377634 | -1.684644171 | 2.88199296 | -0.5845414 | 0.55885613 | NA |
| LSM8 | 1327.5642 | -0.651052099 | 0.20496683 | -3.1763778 | 0.00149127 | 0.01075915 |
| ANKRD7 | 30.0556823 | 0.414049499 | 0.32549405 | 1.27206471 | 0.20335013 | 0.38553044 |
| LINC02476 | 0 | NA | NA | NA | NA | NA |
| AC004872.1 | 0 | NA | NA | NA | NA | NA |
| AC004946.1 | 0 | NA | NA | NA | NA | NA |
| KCND2 | 5.70584245 | 0.246910058 | 0.77928126 | 0.31684331 | 0.7513625 | 0.86135902 |
| AC004888.1 | 0.09635527 | 0.547239394 | 3.29377202 | 0.16614368 | 0.86804388 | NA |
| TSPAN12 | 5566.95885 | 0.090828868 | 0.31057816 | 0.29245092 | 0.76994188 | 0.87279244 |
| ING3 | 668.555118 | 0.072785899 | 0.16817337 | 0.43280277 | 0.66515806 | 0.80360187 |
| CPED1 | 534.970686 | 0.603228755 | 0.38742231 | 1.55703154 | 0.119463 | 0.26902418 |
| WNT16 | 20.4559825 | -0.066225668 | 0.46993816 | -0.1409242 | 0.88792981 | 0.94202528 |
| FAM3C | 3990.2559 | -0.541407113 | 0.18519147 | -2.9234993 | 0.00346121 | 0.02040969 |
| AC004875.1 | 0 | NA | NA | NA | NA | NA |
| PTPRZ1 | 26.2733466 | -0.188059527 | 0.67166901 | -0.2799884 | 0.77948641 | 0.87916597 |
| AASS | 780.84237 | 1.090633898 | 0.29010498 | 3.75944566 | 0.00017029 | 0.00193059 |
| AC015983.2 | 6.09328484 | 1.722073874 | 0.92752115 | 1.85664109 | 0.06336223 | 0.17177569 |
| FEZF1 | 0.91312557 | -1.540711467 | 1.45483413 | -1.059029 | 0.28958659 | NA |
| FEZF1-AS1 | 2.87648344 | -1.694708785 | 0.90452293 | -1.8735941 | 0.0609864 | 0.16736033 |
| CADPS2 | 2022.32525 | 0.725893114 | 0.23656327 | 3.06849464 | 0.0021514 | 0.01425439 |
| AC004594.1 | 16.8963631 | 0.417676075 | 0.49931031 | 0.83650601 | 0.40287029 | 0.59794505 |
| RNF133 | 7.19831407 | -0.124655718 | 0.63645487 | -0.1958595 | 0.84472016 | 0.91800002 |
| RNF148 | 10.0444714 | -0.176577036 | 0.57384553 | -0.3077083 | 0.7583043 | 0.86631656 |
| TAS2R16 | 0 | NA | NA | NA | NA | NA |
| AC073311.1 | 1.67580507 | 2.853387249 | 1.21280544 | 2.35271639 | 0.01863684 | 0.07128714 |
| SLC13A1 | 3849.21479 | 1.577387202 | 0.5735395 | 2.75026775 | 0.00595466 | 0.030615 |
| IQUB | 182.171743 | 0.46011721 | 0.29845644 | 1.54165617 | 0.12315717 | 0.2746124 |
| AC073323.1 | 47.5881632 | 0.354528794 | 0.29356733 | 1.20765751 | 0.22717903 | 0.41406628 |
| NDUFA5 | 5905.81394 | 0.400442316 | 0.13738459 | 2.91475423 | 0.00355969 | 0.02082737 |
| ASB15 | 195.780041 | 4.304541267 | 1.2558714 | 3.42753348 | 0.00060909 | 0.00526995 |
| AC006333.1 | 14.4825166 | 4.739721541 | 0.99755756 | 4.75132636 | 2.02E-06 | 5.71E-05 |
| LMOD2 | 5.73191553 | -1.176649882 | 1.21005612 | -0.9723928 | 0.33085516 | 0.52870262 |
| WASL | 4590.59822 | 0.27622537 | 0.16292067 | 1.69545928 | 0.08998833 | 0.22062033 |
| AC006333.2 | 362.083805 | -0.082226421 | 0.19760366 | -0.4161179 | 0.67732372 | 0.81241988 |
| HYAL4 | 11.6704407 | 1.090167566 | 0.81102764 | 1.34418053 | 0.17889002 | 0.35455315 |
| SPAM1 | 1.54622014 | -0.824890539 | 1.34469219 | -0.6134419 | 0.53958419 | 0.71261757 |
| AC004690.2 | 0 | NA | NA | NA | NA | NA |
| TMEM229A | 1.27293198 | 0.98606826 | 1.10694222 | 0.89080373 | 0.37303447 | 0.56921038 |
| AC006148.1 | 0.40478866 | 1.869548399 | 1.79195237 | 1.0433025 | 0.29680821 | NA |
| AC006148.2 | 1.41031464 | -2.980776979 | 1.91654786 | -1.5552844 | 0.11987835 | 0.26974931 |
| AC004930.1 | 0.36210517 | -0.55754913 | 2.27253273 | -0.2453426 | 0.80619116 | NA |
| SSU72P8 | 0 | NA | NA | NA | NA | NA |
| GPR37 | 61.90938 | -1.261193882 | 0.43708067 | -2.8854945 | 0.00390799 | 0.02237863 |
| C7orf77 | 0.07659028 | -0.642905668 | 3.30558948 | -0.1944905 | 0.84579185 | NA |
| POT1 | 991.07951 | 0.084045115 | 0.16319226 | 0.51500674 | 0.60654831 | 0.76313157 |
| POT1-AS1 | 40.9566574 | 0.321480226 | 0.31525182 | 1.01975692 | 0.30784376 | 0.50474662 |
| AC019155.3 | 1.39713258 | 0.534853186 | 1.13852839 | 0.46977589 | 0.63851514 | 0.78614457 |
| AC010099.4 | 0.52277961 | 1.559535525 | 2.6660734 | 0.58495596 | 0.55857733 | NA |
| AC019155.2 | 0 | NA | NA | NA | NA | NA |
| AC073486.1 | 322.081653 | 0.582792047 | 0.71799188 | 0.81169727 | 0.41696536 | 0.61000703 |
| AC003975.1 | 0.03392908 | 0.380740593 | 3.29738709 | 0.11546736 | 0.90807469 | NA |
| AC000372.1 | 0 | NA | NA | NA | NA | NA |
| GRM8 | 48.0912238 | 0.185724253 | 0.50907844 | 0.36482443 | 0.71524246 | 0.8380436 |
| AC000099.1 | 0.09997307 | 0.547239394 | 3.29377202 | 0.16614368 | 0.86804388 | NA |
| ZNF800 | 809.49877 | -0.404229034 | 0.23117745 | -1.748566 | 0.08036607 | 0.2030325 |

| | | | | | | |
|---|---|---|---|---|---|---|
| AC000123.1 | 93.6996442 | -0.688023127 | 0.22996169 | -2.9919033 | 0.00277244 | 0.01729578 |
| AC000124.1 | 0.8987393 | -0.571203637 | 1.46986456 | -0.3886097 | 0.69756489 | NA |
| GCC1 | 2741.94904 | -0.419280323 | 0.09973011 | -4.2041497 | 2.62E-05 | 0.00044129 |
| ARF5 | 5777.32412 | -0.48311646 | 0.11853296 | -4.0757985 | 4.59E-05 | 0.00069258 |
| FSCN3 | 1982.99611 | -0.56155214 | 0.12527904 | -4.4824109 | 7.38E-06 | 0.00016064 |
| PAX4 | 0.12150485 | 1.04981648 | 3.28477129 | 0.31960109 | 0.74927075 | NA |
| AC073934.1 | 0.9273572 | 2.726669254 | 1.36167535 | 2.00243711 | 0.04523774 | NA |
| SND1 | 6778.12574 | -0.072282766 | 0.15870072 | -0.4554659 | 0.6487741 | 0.79325696 |
| SND1-IT1 | 8.75546386 | 0.011947183 | 0.40839028 | 0.02925433 | 0.97666175 | 0.98922875 |
| LRRC4 | 376.320888 | 0.26070567 | 0.3573753 | 0.7295011 | 0.4656952 | 0.65288037 |
| LEP | 5.92190885 | 1.506628572 | 1.05563144 | 1.42722972 | NA | NA |
| AC018635.2 | 2.70149406 | -0.506347717 | 0.93553447 | -0.541239 | 0.58834288 | 0.74906343 |
| RBM28 | 1766.011 | -0.137547773 | 0.19376885 | -0.7098549 | 0.4777941 | 0.66288369 |
| PRRT4 | 21.6324621 | -1.437080203 | 1.00563995 | -1.4290206 | 0.15299831 | 0.31857242 |
| IMPDH1 | 1719.5109 | -1.009788693 | 0.24427755 | -4.133776 | 3.57E-05 | 0.00056585 |
| HILPDA | 4665.09882 | -4.329822169 | 0.7752911 | -5.5847696 | 2.34E-08 | 1.43E-06 |
| METTL2B | 1125.3768 | -0.187805167 | 0.16133829 | -1.1640459 | 0.24440542 | 0.43541217 |
| AC090114.2 | 354.590985 | 0.091904329 | 0.31648785 | 0.29038817 | 0.77151929 | 0.87403719 |
| AC018638.7 | 157.942336 | 0.011091842 | 0.40288177 | 0.02753126 | 0.97803601 | 0.99001227 |
| FAM71F2 | 314.030446 | 0.151437128 | 0.25040519 | 0.60476833 | 0.54533294 | 0.71697611 |
| AC018638.6 | 90.5530722 | -0.543180177 | 0.32829828 | -1.6545325 | 0.09801937 | 0.23438255 |
| FAM71F1 | 15.8950176 | -0.536307295 | 0.68477852 | -0.7831836 | 0.43351932 | 0.62557806 |
| CALU | 4830.62482 | -0.662473725 | 0.27135982 | -2.4413111 | 0.01463404 | 0.0598016 |
| OPN1SW | 133.920265 | -0.74845159 | 0.57493365 | -1.3018052 | 0.192983 | 0.37249168 |
| CCDC136 | 58.4631356 | -0.927184262 | 0.32524663 | -2.8507113 | 0.00436216 | 0.02433468 |
| FLNC | 1272.48596 | -0.319466388 | 0.50063007 | -0.6381286 | 0.52338994 | 0.69882538 |
| FLNC-AS1 | 68.0493142 | -0.115067868 | 0.47892574 | -0.2402624 | 0.81012681 | 0.89738066 |
| KCP | 6508.1606 | -0.035959978 | 0.23805795 | -0.1510556 | 0.87993188 | 0.93771686 |
| ATP6V1F | 6603.1819 | -0.507279125 | 0.22287262 | -2.2760945 | 0.02284036 | 0.08273355 |
| ATP6V1FNB | 1570.68575 | 0.30936918 | 0.44341061 | 0.69770361 | 0.48536257 | 0.6696032 |
| IRF5 | 1868.90046 | -0.044304689 | 0.22574803 | -0.1962573 | 0.84440882 | 0.91773731 |
| TNPO3 | 2604.57758 | -0.178598649 | 0.16921882 | -1.0554301 | 0.29122864 | 0.48732148 |
| AC011005.1 | 6.08962778 | 0.114118816 | 0.46027903 | 0.24793399 | 0.80418548 | 0.8938784 |
| TSPAN33 | 7955.7424 | 1.088726066 | 0.25434967 | 4.2804304 | 1.87E-05 | 0.00033553 |
| SMO | 2358.68937 | 0.896857683 | 0.26926977 | 3.33070314 | 0.00086627 | 0.00702445 |
| AC011005.4 | 238.989922 | 0.443515171 | 0.32702846 | 1.35619748 | 0.17503634 | 0.34904026 |
| AHCYL2 | 2186.2722 | 0.95751077 | 0.18053012 | 5.30388384 | 1.13E-07 | 5.32E-06 |
| STRIP2 | 754.34189 | 0.87839413 | 0.73402313 | 1.19668454 | 0.23142953 | 0.41934754 |
| SMKR1 | 35.9329406 | 0.021029954 | 0.32040094 | 0.06563637 | 0.94766733 | 0.9747917 |
| AC078846.1 | 197.026938 | 0.26814571 | 0.20254684 | 1.32387009 | 0.18554619 | 0.36286362 |
| NRF1 | 1110.05236 | 0.022442939 | 0.10070256 | 0.22286363 | 0.82364164 | 0.90564302 |
| AC084864.1 | 1.25540588 | -0.660947065 | 1.2454062 | -0.530708 | 0.59562112 | 0.75464993 |
| UBE2H | 7663.00929 | -0.581306437 | 0.17186446 | -3.3823541 | 0.00071867 | 0.00603553 |
| AC073320.1 | 162.087386 | 0.134408423 | 0.27658649 | 0.48595441 | 0.62699949 | 0.77804072 |
| ZC3HC1 | 1234.43396 | -0.053485967 | 0.22269815 | -0.2401725 | 0.81019654 | 0.89742021 |
| KLHDC10 | 2550.86375 | 0.188104457 | 0.13836808 | 1.35944978 | 0.17400411 | 0.34788772 |
| AC087071.1 | 4.45389004 | -0.320992056 | 1.05042093 | -0.3055842 | 0.75992124 | 0.86700095 |
| TMEM209 | 751.40839 | 0.140565274 | 0.16507839 | 0.85150622 | 0.39448821 | 0.58975305 |
| AC087071.2 | 93.5641599 | 0.718210505 | 0.2377448 | 3.0209305 | 0.00251999 | 0.01609402 |
| SSMEM1 | 0.36538442 | -0.558109291 | 2.25793403 | -0.247177 | 0.80477127 | NA |
| CPA2 | 2.04041832 | 0.51493891 | 0.92308467 | 0.5578458 | 0.57694968 | 0.74088937 |
| CPA4 | 63.7386361 | -2.780209749 | 0.65301635 | -4.2574887 | 2.07E-05 | 0.00036393 |
| CPA5 | 4.80984857 | 0.685989237 | 0.70422679 | 0.97410273 | 0.33000553 | 0.52788993 |
| CPA1 | 15.7360717 | 1.6303489 | 0.74395729 | 2.19145497 | 0.02841888 | 0.09716141 |
| CEP41 | 398.110838 | 0.093684068 | 0.17132943 | 0.54680664 | 0.58451158 | 0.74648301 |
| AC007938.1 | 8.2264523 | 1.533866685 | 0.8064181 | 1.90207374 | 0.05716152 | 0.15988431 |
| MESTIT1 | 37.8195092 | 0.810971516 | 0.38216802 | 2.12202875 | 0.03383532 | 0.10993794 |
| MEST | 1520.20447 | 0.126083134 | 0.42696854 | 0.29529841 | 0.76776593 | 0.87183611 |
| AC007938.2 | 263.78758 | 0.677171622 | 0.39524097 | 1.71331334 | 0.08665491 | 0.21446175 |
| COPG2 | 1163.7309 | 0.193029549 | 0.17941869 | 1.07586089 | 0.28198948 | 0.47782049 |
| AC007938.3 | 314.207283 | -0.022472758 | 0.20643166 | -0.1088629 | 0.9133112 | 0.95756858 |
| TSGA13 | 21.2747169 | 1.208372952 | 0.34205894 | 3.5326454 | 0.00041142 | 0.00390401 |
| KLF14 | 1.61787657 | 0.037217838 | 1.24956842 | 0.02978455 | 0.97623888 | 0.98894663 |
| AC016831.6 | 1.66073987 | -1.427026455 | 1.36936598 | -1.0421074 | 0.29736188 | 0.49419756 |
| AC016831.7 | 796.217359 | -0.904484273 | 0.23139168 | -3.9088885 | 9.27E-05 | 0.00119156 |
| LINC00513 | 91.9502227 | -0.274409725 | 0.66432625 | -0.4130647 | 0.67955921 | 0.81406334 |
| AC016831.1 | 343.981405 | -2.094438759 | 0.43029504 | -4.867448 | 1.13E-06 | 3.60E-05 |
| AC016831.5 | 9.12471342 | -0.409000198 | 0.86874223 | -0.4707958 | 0.63778656 | 0.78561835 |
| AC058791.1 | 27.0060111 | -0.720428022 | 0.53447084 | -1.3479276 | 0.17768165 | 0.35284775 |
| MKLN1 | 4484.16936 | -0.398722624 | 0.19443599 | -2.0506627 | 0.04029981 | 0.12448236 |
| AC008264.2 | 2853.9627 | -0.307912569 | 0.15032686 | -2.0482871 | 0.04053188 | 0.12501955 |
| PODXL | 16483.8562 | 0.63511878 | 0.42984195 | 1.47756349 | 0.13952465 | 0.29929757 |
| AC009518.1 | 0.42413399 | 1.031797514 | 1.85226182 | 0.55704734 | 0.57749509 | NA |
| PLXNA4 | 205.964774 | 0.285438286 | 0.41149 | 0.69367004 | 0.48788916 | 0.67152554 |
| AC018643.1 | 0 | NA | NA | NA | NA | NA |
| AC011625.1 | 0.66392787 | 2.138135051 | 2.46479015 | 0.86747143 | 0.38568376 | NA |
| AC009365.1 | 0 | NA | NA | NA | NA | NA |
| AC009365.2 | 0.07659028 | -0.642905668 | 3.30558948 | -0.1944905 | 0.84579185 | NA |
| CHCHD3 | 3205.93951 | 0.06119289 | 0.17349502 | 0.35270689 | 0.72430819 | 0.84387647 |
| AC009365.3 | 0.40497678 | -1.934841972 | 3.29483137 | -0.5872355 | 0.55704555 | NA |
| EXOC4 | 2129.86018 | 0.092030828 | 0.1777052 | 0.51788482 | 0.60453863 | 0.76155753 |
| LRGUK | 80.6780438 | -0.168008409 | 0.23453249 | -0.7163545 | 0.47377247 | 0.65959453 |
| AC008154.2 | 1.29013772 | 0.373878382 | 1.33109199 | 0.28088095 | 0.77880171 | 0.87873866 |
| SLC35B4 | 1676.67796 | 0.615580872 | 0.16433764 | 3.74582998 | 0.0001798 | 0.00200902 |
| AC008154.1 | 2.48322573 | 0.000676667 | 0.91208655 | 0.00074189 | 0.99940806 | 0.99979217 |
| AC009275.1 | 17.7879727 | 0.401779576 | 0.7478483 | 0.53724743 | 0.59109671 | 0.75076202 |
| AKR1B1 | 12790.0461 | -1.117236781 | 0.48969853 | -2.2814787 | 0.02252014 | 0.081866 |
| AKR1B10 | 3476.2133 | -3.041962501 | 1.2253073 | -2.4826119 | 0.01304231 | 0.05481761 |
| AKR1B15 | 16.0823684 | -1.830673849 | 1.03999729 | -1.7602679 | NA | NA |
| BPGM | 2091.07261 | -0.206611021 | 0.22870407 | -0.903399 | 0.3663142 | 0.56331382 |
| AC009276.1 | 0 | NA | NA | NA | NA | NA |
| CALD1 | 12421.1647 | 0.430492286 | 0.29182527 | 1.47517139 | 0.14016647 | 0.30025912 |
| AGBL3 | 460.583863 | 0.370196584 | 0.27790442 | 1.33210037 | 0.18282721 | 0.35969104 |
| CYREN | 4921.95984 | -1.044242476 | 0.26777888 | -3.8996447 | 9.63E-05 | 0.00122899 |
| TMEM140 | 4881.61063 | -1.071362646 | 0.35825949 | -2.9904655 | 0.00278553 | 0.01735278 |
| AC083862.2 | 1454.61177 | -1.196553355 | 0.30313307 | -3.9472874 | 7.90E-05 | 0.00105163 |
| WDR91 | 6283.89991 | -0.566047027 | 0.29210698 | -1.9378072 | 0.05264674 | 0.15100007 |
| AC009542.1 | 276.363998 | 0.102162514 | 0.31590832 | 0.32339292 | 0.74639769 | 0.85792464 |
| STRA8 | 81.0370972 | -1.250634515 | 0.5537337 | -2.2585487 | 0.02391147 | 0.0854282 |
| CNOT4 | 1137.63605 | -0.006696953 | 0.12924135 | -0.0518174 | 0.95867418 | 0.98017606 |
| NUP205 | 1466.98262 | -0.165974136 | 0.097782 | -1.6973895 | 0.08962305 | 0.21990879 |
| STMP1 | 2247.48868 | 0.018350169 | 0.19516645 | 0.09402317 | 0.92509075 | 0.96368188 |
| SLC13A4 | 199.832145 | -0.308178115 | 0.35058605 | -0.879037 | 0.37938121 | 0.57542175 |
| AC091736.1 | 13.3290038 | -0.428537136 | 0.50814038 | -0.843344 | 0.39903609 | 0.59390477 |
| FAM180A | 176.882923 | 0.94030808 | 0.55627628 | 1.69036164 | 0.09095879 | 0.22229687 |
| AC015987.1 | 2.80055389 | -0.29826553 | 0.83516168 | -0.3571351 | 0.72099069 | 0.84122492 |
| MTPN | 5331.49575 | 0.007619039 | 0.11927409 | 0.06387841 | 0.94906704 | 0.9754033 |
| LUZP6 | 1319.38883 | -0.426059238 | 0.11618678 | -3.6670199 | 0.00024539 | 0.00258325 |
| AC024084.1 | 0.27406511 | -0.700031684 | 2.99343044 | -0.233856 | 0.81509677 | NA |
| AC078845.1 | 0.56865298 | 0.472297715 | 2.18050223 | 0.21660043 | 0.82851975 | NA |
| AC009264.1 | 0.08393208 | -0.642905668 | 3.30558948 | -0.1944905 | 0.84579185 | NA |
| CHRM2 | 0.68316298 | 1.471778236 | 1.6641003 | 0.8844288 | 0.37646478 | NA |
| PTN | 977.651877 | 1.10337409 | 0.39100855 | 2.8218669 | 0.0047745 | 0.02596392 |

| | | | | | | |
|---|---|---|---|---|---|---|
| AC078842.2 | 0.88370565 | 3.240585424 | 1.42612091 | 2.27230763 | 0.02306794 | NA |
| AC078842.1 | 2.19015204 | 0.934604891 | 0.85189939 | 1.09708365 | 0.27260483 | 0.46759554 |
| DGKI | 157.549909 | 0.796725267 | 0.55922273 | 1.42470116 | 0.1542436 | 0.32016145 |
| CREB3L2 | 3600.02698 | 0.499984301 | 0.18457132 | 2.7088948 | 0.00675077 | 0.03363928 |
| AC022173.1 | 1.0508738 | 0.806002552 | 1.3177817 | 0.61163587 | 0.5407787 | 0.71373045 |
| AKR1D1 | 2.03988663 | -1.061230495 | 1.21775617 | -0.8714639 | 0.38350093 | 0.57930123 |
| AC083867.1 | 0.06785815 | 0.795868262 | 3.28898349 | 0.24198001 | 0.80879565 | NA |
| TRIM24 | 1081.30385 | -0.115258879 | 0.24697677 | -0.466679 | 0.64072953 | 0.7873637 |
| SVOPL | 157.466139 | 0.80220012 | 0.48771431 | 1.64481562 | 0.10000784 | 0.23748976 |
| ATP6V0A4 | 5002.96058 | 0.781235673 | 0.34243318 | 2.28142516 | 0.0225233 | 0.08186622 |
| TMEM213 | 5452.7223 | 1.037742854 | 0.35921626 | 2.88890833 | 0.00386582 | 0.02220909 |
| KIAA1549 | 1351.79345 | 0.66907915 | 0.29229564 | 2.28904938 | 0.02207648 | 0.08068702 |
| ZC3HAV1L | 46.4962608 | -0.131541667 | 0.26917381 | -0.4886867 | 0.62506351 | 0.77665385 |
| ZC3HAV1 | 2596.29685 | -0.495566126 | 0.23055981 | -2.1494037 | 0.03160241 | 0.10474256 |
| TTC26 | 507.507171 | -0.008718103 | 0.14907068 | -0.058483 | 0.95336389 | 0.97745746 |
| UBN2 | 1588.38117 | -0.0832511 | 0.16636995 | -0.5003974 | 0.61679525 | 0.77088415 |
| FMC1 | 1101.38943 | 0.359823589 | 0.25009919 | 1.43872353 | 0.15022887 | 0.31468229 |
| LUC7L2 | 5260.19282 | 0.045292744 | 0.07788343 | 0.58154528 | 0.56087301 | 0.7288024 |
| C7orf55-LUC | 4713.13672 | 0.142931746 | 0.07632489 | 1.87267553 | 0.0611132 | 0.16752133 |
| AC083880.1 | 39.6100619 | -0.195341867 | 0.33131036 | -0.5896039 | 0.55545626 | 0.72486671 |
| KLRG2 | 146.523315 | 0.957366806 | 0.58658154 | 1.63211207 | 0.10265588 | 0.24176097 |
| CLEC2L | 2.55457636 | 3.338170252 | 1.03213185 | 3.23424788 | 0.00121964 | 0.00916813 |
| HIPK2 | 9372.70516 | 0.068196344 | 0.20927108 | 0.32587562 | 0.74451846 | 0.85673555 |
| TBXAS1 | 851.021513 | -1.147642556 | 0.25798794 | -4.4484349 | 8.65E-06 | 0.00018175 |
| PARP12 | 1676.64807 | -0.408785218 | 0.21192727 | -1.9288939 | 0.05374404 | 0.15333011 |
| KDM7A | 1332.207 | -0.656239984 | 0.2378509 | -2.7590393 | 0.00579716 | 0.03006904 |
| SLC37A3 | 2014.26062 | -0.229481539 | 0.18784387 | -1.221661 | 0.22183584 | 0.40779443 |
| RAB19 | 41.3021327 | 0.669273195 | 0.35922374 | 1.86310958 | 0.06244685 | 0.17001068 |
| MKRN1 | 5455.65292 | -0.329619679 | 0.11601928 | -2.8410768 | 0.00449615 | 0.02486624 |
| DENND2A | 1275.62485 | 0.51635583 | 0.28977833 | 1.78189942 | 0.07476564 | 0.19324739 |
| ADCK2 | 1019.58181 | -0.209766929 | 0.13213564 | -1.5875121 | 0.11239672 | 0.25793242 |
| NDUFB2 | 5439.94332 | 0.072610817 | 0.12579798 | 0.57720178 | 0.56380315 | 0.7309124 |
| NDUFB2-AS1 | 2476.95339 | 0.145671721 | 0.14725724 | 0.98923302 | 0.32254914 | 0.5203279 |
| BRAF | 2058.25819 | -0.104873817 | 0.09863764 | -1.0632231 | 0.28768078 | 0.4838718 |
| MRPS33 | 1469.51483 | 0.371050776 | 0.1907386 | 1.9453366 | 0.05173448 | 0.14937081 |
| TMEM178B | 443.335235 | -0.078033231 | 0.45347627 | -0.1720779 | 0.86337631 | 0.92889569 |
| AC005692.1 | 0.21442573 | -1.003574173 | 3.30558948 | -0.3035992 | 0.7614333 | NA |
| AC005692.2 | 0.17361919 | -1.003574173 | 3.30558948 | -0.3035992 | 0.7614333 | NA |
| AC073878.2 | 1.54635289 | -0.616011707 | 1.38440035 | -0.4449664 | 0.656344 | 0.79832986 |
| AC073878.1 | 1.90075583 | 0.709770839 | 0.80155155 | 0.88549618 | 0.37588908 | 0.57173646 |
| AGK | 2023.97652 | 0.406529373 | 0.14790112 | 2.74865649 | 0.00598401 | 0.03071798 |
| AC004918.1 | 561.702802 | 0.378638617 | 0.13478478 | 2.80920904 | 0.00496634 | 0.02672701 |
| KIAA1147 | 2424.84111 | 0.684423643 | 0.19362695 | 3.53475396 | 0.00040816 | 0.00388272 |
| WEE2-AS1 | 129.373346 | 1.002053401 | 0.26650862 | 3.75992866 | 0.00016996 | 0.0019277 |
| WEE2 | 44.663186 | 2.095618044 | 0.38276004 | 5.47501781 | 4.37E-08 | 2.38E-06 |
| SSBP1 | 2397.82641 | -0.519997551 | 0.17800453 | -2.9212602 | 0.00348619 | 0.02050652 |
| TAS2R3 | 0.51765562 | 0.674034934 | 2.18093844 | 0.30905729 | 0.75727794 | NA |
| TAS2R4 | 41.5823015 | -0.438295222 | 0.32917718 | -1.3314873 | 0.18302874 | 0.35972297 |
| TAS2R5 | 43.6965663 | 0.29098484 | 0.38970239 | 0.74668478 | 0.45525386 | 0.64465663 |
| PRSS37 | 5.37562726 | 0.419031744 | 0.78289424 | 0.53523416 | 0.59248794 | 0.75183615 |
| MGAM | 2526.79954 | 2.560401617 | 0.49555731 | 5.16671147 | 2.38E-07 | 9.86E-06 |
| OR9A4 | 0.44479748 | 0.374347945 | 2.84560722 | 0.13155292 | 0.89533792 | NA |
| CLEC5A | 76.8907757 | 0.250395384 | 0.64306118 | 0.38938035 | 0.69699481 | 0.8258207 |
| TAS2R38 | 0.513301 | -0.005919815 | 1.6411117 | -0.0036072 | 0.99712188 | NA |
| MGAM2 | 21.3565147 | 2.402028634 | 0.60416041 | 3.97581273 | 7.01E-05 | 0.00095635 |
| PRSS58 | 0.51441671 | -1.461804462 | 2.28493023 | -0.6397589 | 0.52232935 | NA |
| AC245088.2 | 0 | NA | NA | NA | NA | NA |
| TRBV1 | 0.34939102 | 1.629530304 | 3.24275583 | 0.50251403 | 0.61530599 | NA |
| TRBV2 | 5.395804 | 0.685419105 | 1.16643717 | 0.58761768 | 0.55678894 | 0.72574388 |
| TRBV3-1 | 3.93879368 | 0.660226379 | 1.27242707 | 0.51887169 | 0.60385022 | 0.76105388 |
| TRBV4-1 | 2.6880794 | 0.803314542 | 1.30809392 | 0.61411075 | 0.53914214 | 0.71231917 |
| TRBV5-1 | 9.89775059 | 1.373437583 | 1.4150987 | 0.97055956 | 0.33176765 | 0.5296793 |
| TRBV6-1 | 3.56542364 | 0.604490802 | 1.39855898 | 0.43222403 | 0.66557859 | 0.80385214 |
| TRBV7-1 | 0.43605473 | 1.268861676 | 3.24275566 | 0.39129118 | 0.69558202 | NA |
| TRBV4-2 | 5.3046116 | -0.394403062 | 0.83791433 | -0.4706962 | 0.63785772 | 0.78562859 |
| TRBV6-2 | 5.12237152 | 1.098242949 | 1.25053423 | 0.87821902 | 0.37982487 | 0.5757966 |
| TRBV7-2 | 17.2263934 | -0.186938481 | 1.18177238 | -0.1581848 | 0.87431114 | 0.93458499 |
| TRBV8-1 | 0 | NA | NA | NA | NA | NA |
| TRBV5-2 | 0 | NA | NA | NA | NA | NA |
| TRBV6-4 | 1.23591374 | 2.198245422 | 1.40668884 | 1.56270908 | 0.11812104 | 0.26697966 |
| TRBV7-3 | 6.48456825 | 1.63692832 | 0.92391792 | 1.77172483 | 0.07644025 | 0.19625443 |
| TRBV8-2 | 3.29857466 | 1.498963966 | 1.19123167 | 1.25833119 | NA | NA |
| TRBV5-3 | 0.30452382 | 1.356048466 | 2.26776054 | 0.5979681 | 0.54986122 | NA |
| TRBV9 | 5.49759768 | 0.042706156 | 0.79665022 | 0.05360716 | 0.95724815 | 0.97947184 |
| TRBV10-1 | 0.62004132 | 0.507136613 | 2.76893316 | 0.18315235 | 0.85467848 | NA |
| TRBV11-1 | 2.47515728 | -0.918749833 | 1.8013202 | -0.5100425 | 0.6100217 | 0.76596961 |
| TRBV12-1 | 0 | NA | NA | NA | NA | NA |
| TRBV10-2 | 0.86446864 | 1.575354861 | 1.85488248 | 0.84930171 | 0.39571343 | NA |
| TRBV11-2 | 3.8972702 | 0.94930106 | 0.94570349 | 1.00380412 | 0.31547304 | 0.51256204 |
| TRBV12-2 | 0.17361919 | -1.003574173 | 3.30558948 | -0.3035992 | 0.7614333 | NA |
| TRBV6-5 | 10.451724 | 0.452370062 | 1.26349045 | 0.35803204 | 0.72031934 | 0.84093782 |
| TRBV7-4 | 1.17811315 | -0.591469216 | 1.28839218 | -0.4590754 | 0.64617999 | 0.79136942 |
| TRBV5-4 | 4.08817476 | 2.231360099 | 1.11710487 | 1.99744908 | NA | NA |
| TRBV6-6 | 5.92116628 | -0.945314469 | 1.043471 | -0.9059327 | 0.36497149 | 0.56205495 |
| TRBV7-5 | 0.35835569 | -0.55778788 | 2.26627675 | -0.2461252 | 0.80558529 | NA |
| TRBV5-5 | 1.60881185 | 0.639201481 | 1.38507821 | 0.46149126 | 0.64444619 | 0.78997117 |
| TRBV6-7 | 0.18382444 | -1.003574173 | 3.30558948 | -0.3035992 | 0.7614333 | NA |
| TRBV7-6 | 3.52900251 | 1.348505936 | 1.08360235 | 1.24446569 | 0.21332822 | 0.39731383 |
| TRBV5-6 | 2.84618491 | 0.994444673 | 1.40481529 | 0.70788286 | 0.479018 | 0.66382897 |
| TRBV6-8 | 0.11051935 | 0.02007176 | 3.29738704 | 0.00608717 | 0.99514317 | NA |
| TRBV7-7 | 0.78972315 | -0.061612806 | 1.88225957 | -0.0327334 | 0.97388717 | NA |
| TRBV5-7 | 0.43059146 | 1.268861676 | 3.24275566 | 0.39129118 | 0.69558202 | NA |
| TRBV7-9 | 15.2780797 | -0.689341011 | 1.08326989 | -0.636352 | 0.52454699 | 0.69951978 |
| TRBV13 | 4.04588775 | -1.228022675 | 1.35925029 | -0.9034559 | 0.36628399 | 0.56331382 |
| TRBV10-3 | 4.10363487 | 1.684002829 | 1.33331108 | 1.2630232 | 0.20658084 | 0.38924304 |
| TRBV11-3 | 1.02447868 | -0.069734291 | 2.20001536 | -0.0316972 | 0.97471354 | 0.98843899 |
| TRBV12-3 | 2.29500164 | 1.047988632 | 1.13949462 | 0.91969599 | 0.35773165 | 0.55541979 |
| TRBV12-4 | 2.90925623 | 0.767307016 | 1.2454697 | 0.61607843 | 0.53784276 | 0.71139447 |
| TRBV12-5 | 0.62392042 | 0.524739997 | 1.66218599 | 0.31569271 | 0.75223576 | NA |
| TRBV14 | 1.87594343 | 1.134352137 | 1.18608954 | 0.95637985 | 0.33888036 | 0.53700349 |
| TRBV15 | 1.66384757 | 1.462259915 | 1.78032648 | 0.82134369 | 0.41145053 | 0.60566878 |
| TRBV16 | 0.08757577 | 0.547239394 | 3.29377202 | 0.16614368 | 0.86804388 | NA |
| TRBV17 | 0 | NA | NA | NA | NA | NA |
| TRBV18 | 4.79777677 | -0.202664757 | 1.30178357 | -0.1556824 | 0.87628339 | 0.93555133 |
| TRBV19 | 10.9410521 | 0.333226316 | 1.03700123 | 0.32133647 | 0.74795542 | 0.85907855 |
| TRBV20-1 | 11.8659893 | 0.44599472 | 0.73870297 | 0.60375379 | 0.54600734 | 0.71722681 |
| TRBV21-1 | 1.95857937 | -2.000271352 | 1.62747213 | -1.2290664 | 0.2190469 | 0.40450048 |
| TRBV22-1 | 0 | NA | NA | NA | NA | NA |
| TRBV23-1 | 2.67747712 | 1.268006279 | 1.17580053 | 1.07841955 | 0.28084657 | 0.47657168 |
| TRBV24-1 | 0.83904971 | 3.146232628 | 1.93634989 | 1.6248265 | 0.10419953 | NA |
| MTRNR2L6 | 15.3236285 | -1.150978456 | 0.55579558 | -2.0708665 | 0.03837127 | 0.12015799 |
| TRBV25-1 | 2.20108085 | 1.143051245 | 1.42042864 | 0.80472275 | 0.42097969 | 0.61359907 |
| TRBVA | 0 | NA | NA | NA | NA | NA |

| | | | | | | |
|---|---|---|---|---|---|---|
| TRBV26 | 0.08666371 | -0.642905668 | 3.30558948 | -0.1944905 | 0.84579185 | NA |
| TRBVB | 0 | NA | NA | NA | NA | NA |
| TRBV27 | 5.0874907 | 0.61140038 | 0.91452995 | 0.66854057 | 0.50378859 | 0.68364658 |
| TRBV28 | 7.88088549 | 0.580294884 | 1.22622545 | 0.4732367 | 0.63604432 | 0.7841886 |
| TRBV29-1 | 12.4024436 | 1.528946762 | 0.98899452 | 1.5459608 | 0.12211404 | 0.27305581 |
| PRSS1 | 0.73592708 | -1.791117162 | 1.67078859 | -1.072019 | 0.28371148 | NA |
| PRSS2 | 0.25908362 | 1.093535322 | 3.27839713 | 0.33355792 | 0.73871318 | NA |
| TRBD1 | 5.52727431 | 0.495609361 | 1.25779529 | 0.39403022 | 0.69355873 | 0.82370883 |
| TRBJ1-1 | 9.21776896 | 0.443042716 | 1.26676185 | 0.34974428 | 0.72653062 | 0.84540695 |
| TRBJ1-2 | 8.17726005 | 0.218724193 | 1.4042381 | 0.15576005 | 0.87622216 | 0.93555133 |
| TRBJ1-3 | 2.27328825 | 1.250303691 | 1.2490107 | 1.00103521 | 0.31680978 | 0.51410917 |
| TRBJ1-4 | 2.91960363 | 0.7212849 | 1.61965803 | 0.4453316 | 0.65608014 | 0.79815617 |
| TRBJ1-5 | 10.0029786 | 0.645738198 | 0.78037886 | 0.82746757 | 0.4079721 | 0.60260036 |
| TRBJ1-6 | 4.18547502 | 2.14932028 | 1.29754444 | 1.65645215 | 0.09763029 | 0.23372771 |
| TRBC1 | 273.621984 | -0.065841722 | 0.87112738 | -0.0755822 | 0.9397515 | 0.97115066 |
| TRBJ2-1 | 18.9376485 | 0.895037251 | 0.8367223 | 1.06969451 | NA | NA |
| TRBJ2-2 | 8.33461018 | 0.141101567 | 0.89764571 | 0.15719071 | 0.87509455 | 0.93501854 |
| TRBJ2-2P | 5.35069714 | 0.406312726 | 0.89412021 | 0.45442741 | 0.64952123 | 0.79382745 |
| TRBJ2-3 | 23.7693537 | 0.180477606 | 0.93766353 | 0.19247587 | 0.84736947 | 0.91981776 |
| TRBJ2-4 | 7.46850862 | -0.054302716 | 1.13067244 | -0.0480269 | 0.96169479 | 0.98172389 |
| TRBJ2-5 | 7.03466211 | 0.232050987 | 0.93289924 | 0.24874175 | 0.80356055 | 0.89344739 |
| TRBJ2-6 | 4.40165069 | -0.357137693 | 0.85202488 | -0.4191635 | 0.67509668 | 0.8109648 |
| TRBJ2-7 | 22.1442093 | 0.464933634 | 0.88160341 | 0.52737277 | 0.59793475 | 0.75638201 |
| TRBC2 | 457.358786 | 0.042508006 | 0.78851534 | 0.05390891 | 0.95700773 | 0.97944167 |
| TRBV30 | 3.08688846 | 1.016296469 | 1.47827414 | 0.6874885 | 0.49177495 | 0.67481148 |
| EPHB6 | 729.285742 | 0.137774453 | 0.31218509 | 0.44132297 | 0.6589792 | 0.79965402 |
| TRPV6 | 975.226406 | -0.789310348 | 0.32361342 | -2.4390532 | 0.0147258 | 0.06002772 |
| AC245427.1 | 67.2810929 | -0.521431654 | 0.39143124 | -1.3321156 | 0.18282222 | 0.35969104 |
| TRPV5 | 139.84379 | -1.024395289 | 0.57510838 | -1.7812213 | 0.07487631 | 0.19340662 |
| LLCFC1 | 4.73213229 | -0.527854291 | 0.71407492 | -0.7392142 | 0.45977697 | 0.64846332 |
| KEL | 48.4227036 | -0.985547058 | 0.34338121 | -2.8701252 | 0.00410309 | 0.02319369 |
| OR9A2 | 0 | NA | NA | NA | NA | NA |
| OR6V1 | 0.09644756 | -0.642905668 | 3.30558948 | -0.1944905 | 0.84579185 | NA |
| PIP | 2.45155866 | -0.713576261 | 1.11833135 | -0.6380723 | 0.52342661 | 0.69882538 |
| TAS2R39 | 0.08757577 | 0.547239394 | 3.29377202 | 0.16614368 | 0.86804388 | NA |
| AC073342.2 | 1.07655821 | 0.200221291 | 1.29992928 | 0.15402476 | 0.87759022 | 0.93634106 |
| TAS2R40 | 0.11051935 | 0.02007176 | 3.29738704 | 0.00608717 | 0.99514317 | NA |
| GSTK1 | 11084.1547 | 0.518030575 | 0.16571659 | 3.12600319 | 0.001772 | 0.01229996 |
| AC073342.1 | 1865.98891 | 1.038943415 | 0.39298403 | 2.64372934 | 0.00819982 | 0.03879247 |
| TMEM139 | 3052.92162 | 1.012258806 | 0.38419033 | 2.63478473 | 0.00841907 | 0.03955296 |
| CASP2 | 1178.08726 | -0.103042069 | 0.14639764 | -0.7038506 | 0.48152581 | 0.66600139 |
| CLCN1 | 9.42458225 | -0.908364476 | 0.7536881 | -1.205226 | 0.22811608 | 0.41517157 |
| FAM131B | 59.0786507 | -0.223406706 | 0.44728724 | -0.4994703 | 0.61744808 | 0.77133479 |
| AC093673.2 | 0.67709901 | -1.386688817 | 1.94367359 | -0.7134371 | 0.47557533 | NA |
| AC093673.1 | 325.304539 | -1.752733522 | 0.33534417 | -5.2266705 | 1.73E-07 | 7.56E-06 |
| ZYX | 9095.6195 | -1.102862334 | 0.26478063 | -4.1651927 | 3.11E-05 | 0.0005039 |
| EPHA1 | 2818.35104 | -0.897826203 | 0.21492626 | -4.1773686 | 2.95E-05 | 0.00048422 |
| EPHA1-AS1 | 10.2089132 | -0.94016511 | 0.60264112 | -1.5600746 | 0.11874225 | 0.26797223 |
| TAS2R60 | 0.08666371 | -0.642905668 | 3.30558948 | -0.1944905 | 0.84579185 | NA |
| TAS2R41 | 0 | NA | NA | NA | NA | NA |
| CTAGE15 | 21.5541235 | -0.353773846 | 1.31089466 | -0.2698721 | 0.78725868 | 0.88414299 |
| TCAF2 | 376.049122 | -1.626056129 | 0.35949454 | -4.5231733 | 6.09E-06 | 0.00013837 |
| TCAF2C | 2.63094511 | -0.687292439 | 1.45148524 | -0.4735098 | 0.63584953 | 0.78407071 |
| CTAGE6 | 5.88004039 | -0.954327113 | 0.70297913 | -1.3575469 | 0.17460751 | 0.34858052 |
| TCAF1 | 896.034663 | 0.001295462 | 0.2215395 | 0.00584755 | 0.99533436 | 0.99776308 |
| OR2F2 | 0 | NA | NA | NA | NA | NA |
| OR2F1 | 1.54896528 | 0.10088185 | 1.12276204 | 0.0898515 | 0.92840522 | 0.96526981 |
| OR6B1 | 0.31835939 | -0.280297048 | 2.95040738 | -0.0950028 | 0.92431258 | NA |
| OR2A5 | 3.16959413 | 0.283827249 | 0.72721521 | 0.39029334 | 0.69631965 | 0.82529324 |
| OR2A25 | 2.09368308 | -0.294105816 | 1.02526365 | -0.2868587 | 0.7742205 | 0.87581805 |
| OR2A12 | 5.55996764 | -0.704240571 | 0.78630498 | -0.8956329 | 0.37044888 | 0.56722431 |
| OR2A2 | 0.33747028 | 1.4137913 | 2.23234043 | 0.63332245 | 0.5265231 | NA |
| OR2A14 | 1.11914651 | -0.773535555 | 1.59135398 | -0.4860864 | 0.6269059 | 0.77804072 |
| CTAGE4 | 67.1238316 | -0.615016957 | 0.70650271 | -0.870509 | 0.38402231 | 0.57962729 |
| ARHGEF35 | 560.761965 | -0.026928543 | 0.7593671 | -0.0354618 | 0.97171149 | 0.98670705 |
| AC004889.1 | 155.122989 | 0.589646751 | 0.38147506 | 1.54570198 | 0.12217656 | 0.27307983 |
| OR2A42 | 36.7140144 | 0.544032108 | 0.33864148 | 1.6065135 | 0.10816113 | 0.25089805 |
| AC074386.1 | 297.04131 | 0.17213478 | 0.32752063 | 0.52556927 | 0.59918751 | 0.7570951 |
| OR2A7 | 72.4605168 | 0.338795117 | 0.36067149 | 0.93934542 | 0.34755342 | 0.54508052 |
| CTAGE8 | 45.8191006 | -0.45209092 | 0.43760536 | -1.0331019 | 0.30155623 | 0.49834165 |
| OR2A1-AS1 | 163.242369 | 0.206533349 | 0.35377305 | 0.58380182 | 0.55935366 | 0.72772184 |
| OR2A1 | 20.2756037 | 0.230408568 | 0.33951101 | 0.67864829 | 0.49736074 | 0.67909712 |
| ARHGEF5 | 2323.51462 | -0.692067154 | 0.22823452 | -3.0322633 | 0.00242727 | 0.01563439 |
| NOBOX | 0.21733212 | -0.339797876 | 3.09048411 | -0.1099497 | 0.91244925 | NA |
| TPK1 | 406.720441 | -0.438522854 | 0.1591806 | -2.7548763 | 0.00587143 | 0.03032903 |
| AC006007.1 | 0.89470167 | 3.795304667 | 1.57607879 | 2.40806786 | 0.0160372 | NA |
| CNTNAP2 | 11.7913722 | 2.681575892 | 0.47690064 | 5.62292363 | 1.88E-08 | 1.18E-06 |
| AC006004.1 | 0 | NA | NA | NA | NA | NA |
| AC006974.1 | 0 | NA | NA | NA | NA | NA |
| AC006974.2 | 0.56487181 | -0.261436567 | 1.86226021 | -0.1403867 | 0.88835446 | NA |
| C7orf33 | 0.10721287 | -0.642905668 | 3.30558948 | -0.1944905 | 0.84579185 | NA |
| AC005229.4 | 134.227537 | -0.263778802 | 0.23415556 | -1.1265109 | 0.25994931 | 0.45291466 |
| CUL1 | 2597.20221 | -0.309382814 | 0.17877729 | -1.7305488 | 0.08353227 | 0.20883068 |
| EZH2 | 318.593698 | -0.882730519 | 0.23139372 | -3.8148422 | 0.00013627 | 0.00161648 |
| GHET1 | 8.33403558 | -0.517052322 | 0.579724 | -0.8918939 | 0.37244978 | 0.56883475 |
| PDIA4 | 7681.93944 | -0.695975599 | 0.25120142 | -2.7705878 | 0.00559552 | 0.02931829 |
| ZNF786 | 359.013074 | 0.236475831 | 0.18863518 | 1.25361468 | 0.20998209 | 0.39329695 |
| ZNF425 | 187.58801 | 0.874263575 | 0.20544662 | 4.25542936 | 2.09E-05 | 0.00036656 |
| ZNF398 | 602.246516 | -0.109097805 | 0.20822824 | -0.5239338 | 0.60032462 | 0.75798708 |
| ZNF282 | 1702.20062 | -0.21713852 | 0.15929631 | -1.3631108 | 0.17284761 | 0.34624669 |
| ZNF212 | 894.376101 | 0.063073524 | 0.18170132 | 0.3471275 | 0.72849553 | 0.84642348 |
| ZNF783 | 1581.95421 | -0.210364232 | 0.16367001 | -1.2852949 | 0.19868925 | 0.37969676 |
| AC004941.1 | 0.30322989 | 1.411981162 | 2.88745696 | 0.48900509 | 0.62483809 | NA |
| AC073314.1 | 2.01729653 | -0.013303664 | 1.0399358 | -0.0127928 | 0.98979312 | 0.99521246 |
| ZNF777 | 1007.2179 | -0.255554832 | 0.21696877 | -1.1778415 | 0.23885979 | 0.42842692 |
| ZNF746 | 1991.18608 | -0.842127663 | 0.31853212 | -2.6437763 | 0.00819868 | 0.03879247 |
| KRBA1 | 1098.03411 | -0.211641986 | 0.20303445 | -1.0423945 | 0.29722884 | 0.49403873 |
| ZNF467 | 1299.80939 | -0.132834701 | 0.3638037 | -0.3651274 | 0.7150163 | 0.83799911 |
| SSPO | 463.572937 | 0.607122319 | 0.38567006 | 1.57420133 | 0.11544091 | 0.26290198 |
| ZNF862 | 1446.28282 | 0.320400952 | 0.20512135 | 1.56200685 | 0.11828637 | 0.26728464 |
| AC004877.1 | 140.06906 | 1.100595314 | 0.33003292 | 3.33480465 | 0.00085359 | 0.00693874 |
| ATP6V0E2-A! | 1183.39596 | 0.938927632 | 0.31072128 | 3.02176804 | 0.00251303 | 0.01605733 |
| ATP6V0E2 | 3543.37491 | 1.183779612 | 0.38140645 | 3.10372205 | 0.00191103 | 0.01300821 |
| AC093458.1 | 26.0884368 | -1.409056578 | 0.51960103 | -2.7118048 | 0.0066918 | 0.03343363 |
| AC092681.3 | 29.7604909 | -0.393201555 | 0.46051355 | -0.8538328 | 0.39319767 | 0.58865802 |
| AC092681.2 | 12.883537 | -0.255360841 | 0.38620729 | -0.6612015 | 0.50848309 | 0.68730838 |
| AC092666.1 | 2.74658789 | -0.784949444 | 1.16897854 | -0.6714832 | 0.50191276 | 0.68234467 |
| ACTR3C | 717.86039 | -0.598538374 | 0.26752843 | -2.2372888 | 0.02526747 | 0.08906605 |
| LRRC61 | 1539.39009 | -1.12511618 | 0.32449932 | -3.4672374 | 0.00052584 | 0.00471393 |
| ZBED6CL | 529.782383 | -0.443876075 | 0.25737642 | -1.7246182 | 0.0845963 | 0.21071574 |
| RARRES2 | 5803.24783 | -0.96254315 | 0.36057936 | -2.6694349 | 0.0075979 | 0.03678293 |
| AC005586.1 | 458.883288 | 0.027947551 | 0.16742772 | 0.16692308 | 0.86743057 | 0.93105439 |
| REPIN1 | 6961.67713 | -0.048433468 | 0.10690311 | -0.4530595 | 0.65050591 | 0.79460075 |

| | | | | | | |
|---|---|---|---|---|---|---|
| AC073111.3 | 227.969051 | 0.200868052 | 0.14935347 | 1.34491718 | 0.17865198 | 0.35418786 |
| ZNF775 | 255.165244 | -0.295057988 | 0.16795703 | -1.7567469 | 0.07896096 | 0.20061667 |
| AC073111.1 | 5.13305957 | 0.962253816 | 0.77200872 | 1.24642869 | 0.21260705 | 0.39645571 |
| AC073111.5 | 752.422384 | 0.078092497 | 0.37358851 | 0.20903345 | 0.83442212 | 0.9121801 |
| LINC00996 | 17.6374162 | 2.005564717 | 0.91591638 | 2.18968103 | 0.02854738 | 0.09744911 |
| GIMAP8 | 1078.26224 | 0.2286171 | 0.3331699 | 0.68618775 | 0.49259472 | 0.67542236 |
| GIMAP7 | 847.886491 | -0.117565184 | 0.26718716 | -0.4400106 | 0.65992943 | 0.79999826 |
| GIMAP4 | 1639.9402 | -0.760868443 | 0.2965617 | -2.5656329 | 0.01029878 | 0.04606153 |
| GIMAP6 | 1446.82686 | 0.320910472 | 0.24598773 | 1.30457917 | 0.19203618 | 0.37124523 |
| GIMAP2 | 421.75308 | -0.47156783 | 0.31729328 | -1.4862207 | 0.13722076 | 0.29577501 |
| GIMAP1 | 621.108335 | 0.084565521 | 0.35054776 | 0.24123823 | 0.80937048 | 0.89703256 |
| GIMAP1-GIM | 1827.21204 | -0.71458571 | 0.2418879 | -2.954202 | 0.00313479 | 0.0189902 |
| GIMAP5 | 1747.70337 | -0.73647361 | 0.24252455 | -3.0366971 | 0.00239186 | 0.01546292 |
| TMEM176B | 20796.4773 | 0.014696759 | 0.19774385 | 0.0743222 | 0.94075401 | 0.97182134 |
| TMEM176A | 17854.8155 | 0.031326413 | 0.17764045 | 0.17634729 | 0.86002111 | 0.92704165 |
| AOC1 | 13526.1141 | 2.517969212 | 0.46814707 | 5.37858589 | 7.51E-08 | 3.74E-06 |
| KCNH2 | 136.636454 | -0.081337954 | 0.42509235 | -0.1913418 | 0.84825778 | 0.92017597 |
| NOS3 | 1943.84628 | -0.139150629 | 0.3130364 | -0.444519 | 0.65666739 | 0.79853904 |
| ATG9B | 796.826494 | -0.40902664 | 0.31854905 | -1.2840303 | 0.19913135 | 0.38018344 |
| ABCB8 | 3239.5326 | 0.327037557 | 0.06376021 | 5.12917951 | 2.91E-07 | 1.16E-05 |
| AC010973.1 | 13.6387493 | 1.44857934 | 0.36284521 | 3.99227907 | 6.54E-05 | 0.00091065 |
| ASIC3 | 333.591331 | -0.448049846 | 0.44130385 | -1.0152865 | 0.30996927 | 0.50706512 |
| CDK5 | 607.409089 | 0.54645929 | 0.19097172 | 2.86146711 | 0.00421685 | 0.02368443 |
| SLC4A2 | 10799.1622 | -0.344126109 | 0.26688499 | -1.2894173 | 0.19725306 | 0.37804806 |
| AC010973.2 | 4028.20919 | -0.659180662 | 0.27620416 | -2.3865704 | 0.01700635 | 0.0666951 |
| FASTK | 5514.92646 | -0.068880681 | 0.13496624 | -0.5103549 | 0.60980286 | 0.76577463 |
| TMUB1 | 2652.34879 | -0.825809187 | 0.17306574 | -4.7716502 | 1.83E-06 | 5.28E-05 |
| AGAP3 | 3779.95862 | -0.497274989 | 0.16452525 | -3.0224843 | 0.00250709 | 0.0160401 |
| GBX1 | 0.81902602 | 1.34132952 | 1.52344961 | 0.88045546 | 0.37861262 | NA |
| ASB10 | 5.06538445 | 1.454116809 | 0.91012784 | 1.59770611 | 0.1101084 | 0.25407407 |
| IQCA1L | 0.41466729 | -0.871780633 | 2.80041163 | -0.3113045 | 0.75556918 | NA |
| AC021097.2 | 13.2350772 | -0.178767247 | 0.48748029 | -0.3667169 | 0.71383022 | 0.8374651 |
| ABCF2 | 2814.93241 | -0.254795766 | 0.19078804 | -1.3354913 | 0.18171559 | 0.35814742 |
| CHPF2 | 3432.70818 | -0.919326592 | 0.29838002 | -3.0810595 | 0.00206265 | 0.01379113 |
| SMARCD3 | 3656.87952 | -1.165998083 | 0.24129693 | -4.8322126 | 1.35E-06 | 4.15E-05 |
| AC021097.1 | 1037.98387 | -1.254149319 | 0.27835505 | -4.5055742 | 6.62E-06 | 0.00014768 |
| AC005486.1 | 5.32490841 | 0.246962268 | 0.7145163 | 0.3456356 | 0.72961659 | 0.84732662 |
| NUB1 | 3589.15294 | -0.363692574 | 0.30741849 | -1.1830537 | 0.23678788 | 0.42601653 |
| WDR86 | 853.969281 | -0.350974626 | 0.30790667 | -1.1398734 | 0.25433905 | 0.44708166 |
| AC005996.1 | 0.92919039 | 1.097063968 | 1.13509133 | 0.96649841 | 0.33379484 | NA |
| CRYGN | 9.07004786 | 0.190198769 | 0.55133909 | 0.34497603 | 0.73011239 | 0.84751607 |
| RHEB | 4574.13794 | -0.750944356 | 0.19631083 | -3.8252823 | 0.00013062 | 0.00156143 |
| PRKAG2 | 2688.59002 | -0.255908055 | 0.26920045 | -0.9506227 | 0.34179595 | 0.53967782 |
| PRKAG2-AS1 | 194.985803 | 0.007341224 | 0.30161541 | 0.02433968 | 0.98058166 | 0.99131724 |
| AC074257.1 | 0 | NA | NA | NA | NA | NA |
| GALNTL5 | 0 | NA | NA | NA | NA | NA |
| GALNT11 | 9459.5176 | 1.500736474 | 0.31352599 | 4.7866413 | 1.70E-06 | 4.99E-05 |
| AC006017.1 | 2301.66466 | 1.316239945 | 0.31120096 | 4.22954983 | 2.34E-05 | 0.00040437 |
| KMT2C | 2446.94726 | -0.031266728 | 0.1124793 | -0.2779776 | 0.78102954 | 0.88016021 |
| LINC01003 | 248.057484 | 0.093100266 | 0.30591191 | 0.30433684 | 0.76087128 | 0.86746668 |
| XRCC2 | 24.9497693 | 0.565539707 | 0.42468831 | 1.33165827 | 0.18297251 | 0.35970684 |
| ACTR3B | 536.616853 | -0.134423975 | 0.18955941 | -0.709139 | 0.4782382 | 0.66311607 |
| LINC01287 | 1.81299114 | 1.0151552 | 0.86844633 | 1.16893257 | 0.2424308 | 0.43283008 |
| DPP6 | 355.33143 | 0.94001938 | 0.72372497 | 1.29886272 | 0.19399105 | 0.37378516 |
| AC006019.3 | 0 | NA | NA | NA | NA | NA |
| AC006019.1 | 0 | NA | NA | NA | NA | NA |
| AC006019.2 | 2.14987368 | -1.578010177 | 1.12681609 | -1.400415 | 0.16138907 | 0.32998604 |
| AC073336.1 | 9.64513649 | 6.831641401 | 3.14699109 | 2.17084866 | 0.02994261 | 0.10073177 |
| PAXIP1-AS2 | 1060.65607 | 0.076860383 | 0.15585913 | 0.49314006 | 0.62191362 | 0.77429876 |
| PAXIP1 | 910.423036 | -0.019703741 | 0.19317441 | -0.1019997 | 0.91875688 | 0.96046034 |
| AC093726.1 | 207.071321 | 0.17652975 | 0.29402615 | 0.60038793 | 0.54824773 | 0.71894637 |
| AC093726.2 | 66.9475874 | -0.182815597 | 0.32618171 | -0.5604716 | 0.57515778 | 0.73938042 |
| PAXIP1-AS1 | 725.747244 | -0.040861868 | 0.2057218 | -0.1986268 | 0.84255467 | 0.91630899 |
| HTR5A-AS1 | 0.08695548 | -0.642905668 | 3.30558948 | -0.1944905 | 0.84579185 | NA |
| HTR5A | 0.17648264 | -1.003574173 | 3.30558948 | -0.3035992 | 0.7614333 | NA |
| AC099552.1 | 5.71888175 | 0.365027921 | 1.60254215 | 0.22778054 | NA | NA |
| AC099552.3 | 36.0860778 | 1.009611794 | 0.84334537 | 1.19715105 | 0.23124769 | 0.41907783 |
| AC099552.2 | 0 | NA | NA | NA | NA | NA |
| AC144652.1 | 46.3528424 | -0.268216775 | 0.52918305 | -0.5068507 | 0.61225961 | 0.76779926 |
| INSIG1 | 3504.86337 | -1.125738396 | 0.44099338 | -2.5527331 | 0.01068814 | 0.04730479 |
| BLACE | 0 | NA | NA | NA | NA | NA |
| AC008060.1 | 0 | NA | NA | NA | NA | NA |
| AC008060.3 | 0 | NA | NA | NA | NA | NA |
| EN2 | 6.35448134 | -1.549935787 | 0.72413252 | -2.1404035 | 0.03232217 | 0.10628952 |
| CNPY1 | 0.62895418 | -1.236022954 | 1.93766911 | -0.6378917 | 0.52354421 | NA |
| AC008060.2 | 0 | NA | NA | NA | NA | NA |
| AC009403.1 | 484.467221 | 0.592932338 | 0.26379892 | 2.24766782 | 0.02459738 | 0.08718184 |
| RBM33 | 3757.15729 | -0.614136729 | 0.12682034 | -4.8425726 | 1.28E-06 | 3.98E-05 |
| SHH | 626.118747 | 0.193371618 | 0.33607889 | 0.57537568 | 0.56503725 | 0.7318287 |
| AC021218.1 | 215.493094 | 0.87193089 | 0.60432872 | 1.44280895 | 0.1490743 | 0.31318429 |
| AC073133.1 | 9.73010764 | 0.113383586 | 0.6509716 | 0.17417593 | 0.8617272 | 0.92800735 |
| AC073133.2 | 11.673574 | 0.837931053 | 0.55502942 | 1.50970566 | 0.13111855 | 0.28674107 |
| LINC01006 | 522.799569 | 0.004652141 | 0.14893704 | 0.03123563 | 0.97508163 | 0.98857757 |
| LINC00244 | 13.812506 | -0.285032596 | 0.37445424 | -0.7611947 | 0.44654076 | 0.63689471 |
| RNF32 | 257.697602 | -0.084713533 | 0.21978961 | -0.3854301 | 0.69991878 | 0.82792169 |
| AC005534.1 | 50.1923922 | 0.031633859 | 0.33472051 | 0.09450828 | 0.92470541 | 0.96356293 |
| LMBR1 | 1980.62473 | 0.062006167 | 0.12435777 | 0.49861113 | 0.61805337 | 0.77165225 |
| AC006357.1 | 1.35628046 | -0.74243223 | 1.581197 | -0.4695381 | 0.63868506 | 0.78618782 |
| NOM1 | 1084.94515 | -0.225721782 | 0.19296919 | -1.1697296 | 0.24210979 | 0.43253528 |
| MNX1 | 14.3753 | -1.718887032 | 0.68110559 | -2.5236719 | 0.01161363 | 0.05037857 |
| MNX1-AS2 | 4.06296893 | -1.543414711 | 0.82848169 | -1.8629437 | 0.06247019 | 0.17005667 |
| MNX1-AS1 | 0.39027547 | -0.032171818 | 2.84461719 | -0.0113097 | 0.99097634 | NA |
| AC006967.2 | 0 | NA | NA | NA | NA | NA |
| AC006967.3 | 0 | NA | NA | NA | NA | NA |
| UBE3C | 3598.19299 | -0.093687567 | 0.15367121 | -0.6096624 | 0.54208543 | 0.71463245 |
| AC004975.2 | 1.96784362 | -0.555389537 | 1.03423259 | -0.5370064 | 0.59126318 | 0.75084424 |
| DNAJB6 | 4773.31947 | -0.245812854 | 0.18817273 | -1.306315 | 0.19144547 | 0.37060762 |
| AC006372.3 | 1.14401378 | 1.33149791 | 1.45980335 | 0.91210772 | 0.36171201 | 0.55911494 |
| AC006372.1 | 0.60978931 | 1.187909683 | 1.79413128 | 0.66210856 | 0.50790165 | NA |
| AC006372.2 | 1.42744659 | 2.111573097 | 1.26490471 | 1.6693535 | 0.09504734 | 0.22943962 |
| PTPRN2 | 655.384183 | 0.015580402 | 0.30150713 | 0.05167507 | 0.9587876 | 0.98017606 |
| AC005481.1 | 2.48517667 | 0.372203551 | 0.86657938 | 0.42950889 | 0.66755292 | 0.80542508 |
| AC011899.2 | 22.065747 | 0.016912794 | 0.54656261 | 0.03094393 | 0.97531426 | 0.98862634 |
| AC011899.3 | 2.89758688 | -0.848772134 | 0.90262236 | -0.9403402 | 0.34704306 | 0.54463641 |
| AC011899.1 | 0.39111697 | 1.696948813 | 2.21948353 | 0.76456923 | 0.4445281 | NA |
| AC078942.1 | 0.85314962 | 2.58833879 | 1.75741333 | 1.47281163 | 0.14080184 | NA |
| LINC01022 | 0 | NA | NA | NA | NA | NA |
| NCAPG2 | 265.756344 | 0.448569085 | 0.17921151 | 2.50301495 | 0.01231403 | 0.05258641 |
| ESYT2 | 4593.74191 | 0.085016882 | 0.17072888 | 0.49796428 | 0.61850922 | 0.77200209 |
| WDR60 | 1582.04629 | -0.084318299 | 0.19258274 | -0.4378289 | 0.66151029 | 0.80084282 |
| VIPR2 | 149.979014 | 0.384009257 | 0.28444724 | 1.35001927 | 0.1770098 | 0.35203566 |
| PLCXD1 | 697.84182 | -0.121695201 | 0.28941483 | -0.4204871 | 0.67412966 | 0.81024608 |
| GTPBP6 | 4231.80931 | -0.007545214 | 0.15955502 | -0.0472891 | 0.96228281 | 0.98209605 |

| | | | | | | |
|---|---|---|---|---|---|---|
| LINC00685 | 81.8860232 | -0.262839714 | 0.28280562 | -0.9294006 | 0.35268151 | 0.55030836 |
| PPP2R3B | 1014.65614 | -0.311513618 | 0.19689867 | -1.5821012 | 0.11362648 | 0.25984127 |
| AL732314.6 | 6.90903759 | 0.266241288 | 0.449943 | 0.59172225 | 0.55403659 | 0.72358699 |
| AL732314.4 | 9.73805576 | -0.320214904 | 0.58043843 | -0.5516776 | 0.58116923 | 0.74384312 |
| SHOX | 1.06472492 | -2.613522013 | 1.95486504 | -1.3369322 | 0.18124477 | 0.35752727 |
| AL672277.1 | 3.30795444 | -0.422269302 | 1.17020101 | -0.3608519 | 0.71821014 | 0.83985952 |
| CRLF2 | 4.12054563 | 1.236621477 | 1.10817315 | 1.11590998 | 0.26446067 | 0.4584124 |
| CSF2RA | 281.602007 | -0.197887563 | 0.52781181 | -0.3749207 | 0.70771946 | 0.83365305 |
| IL3RA | 892.900352 | -0.230970547 | 0.27857594 | -0.8291116 | 0.40704125 | 0.60168362 |
| SLC25A6 | 28544.9124 | 0.101426617 | 0.20750865 | 0.48878259 | 0.62499563 | 0.7766061 |
| LINC00106 | 389.213642 | 0.671579126 | 0.21624499 | 3.10564011 | 0.00189868 | 0.01294727 |
| ASMTL-AS1 | 1649.49682 | -0.007152182 | 0.16339538 | -0.0437722 | 0.96508595 | 0.98359669 |
| ASMTL | 2706.50655 | 0.186462697 | 0.1602972 | 1.16323119 | 0.24473572 | 0.43576508 |
| P2RY8 | 397.758727 | -0.253945744 | 0.4082716 | -0.622002 | 0.53394057 | 0.70771615 |
| AKAP17A | 5081.81698 | -0.398118419 | 0.14018343 | -2.839982 | 0.00451161 | 0.0249102 |
| ASMT | 12.3263757 | -0.281020043 | 0.46219836 | -0.6080074 | 0.54318254 | 0.71539924 |
| AL683807.1 | 9.83890081 | 0.390328299 | 0.85180425 | 0.45823709 | 0.64678212 | 0.79195971 |
| AL683807.2 | 0 | NA | NA | NA | NA | NA |
| DHRSX | 1425.46597 | 0.161789525 | 0.17537716 | 0.92252335 | 0.35625566 | 0.55359232 |
| ZBED1 | 2786.24413 | 0.686195818 | 0.15193648 | 4.5163336 | 6.29E-06 | 0.00014145 |
| LINC00102 | 0.15489794 | -1.003574173 | 3.30558948 | -0.3035992 | 0.7614333 | NA |
| CD99 | 4316.74818 | -0.638678538 | 0.23140852 | -2.7599612 | 0.00578082 | 0.02999613 |
| XG | 22.9205066 | -0.255434168 | 0.60063592 | -0.4252729 | 0.67063776 | 0.80766888 |
| GYG2 | 181.945556 | 0.588297531 | 0.40530089 | 1.45150813 | 0.14663842 | 0.30961048 |
| GYG2-AS1 | 0.07830766 | -0.642905668 | 3.30558948 | -0.1944905 | 0.84579185 | NA |
| ARSD | 5199.21617 | 0.557098518 | 0.31417013 | 1.77323831 | 0.07618923 | 0.19589605 |
| ARSD-AS1 | 1410.11974 | 0.643505463 | 0.30600709 | 2.10291033 | 0.03547361 | 0.11369468 |
| ARSE | 2356.61269 | 0.675328401 | 0.4382537 | 1.54095311 | 0.1233282 | 0.27479211 |
| ARSH | 1.85371054 | 0.050407854 | 0.89554431 | 0.05628739 | 0.95511286 | 0.97848026 |
| ARSF | 364.178784 | 3.558113438 | 0.53242875 | 6.68279735 | 2.34E-11 | 4.26E-09 |
| LINC01546 | 0.45117825 | 2.129723975 | 2.5939157 | 0.82104595 | 0.4116201 | NA |
| MXRA5 | 1015.1057 | 0.585973675 | 0.50039234 | 1.17102846 | 0.24158734 | 0.4318212 |
| PRKX | 2813.83586 | -0.912265005 | 0.43013383 | -2.1208864 | 0.03393136 | 0.11017978 |
| PRKX-AS1 | 0.63601428 | -2.58264421 | 2.67769503 | -0.9645027 | 0.33479393 | NA |
| BX890604.1 | 548.368567 | -0.828848793 | 0.33502421 | -2.4739967 | 0.01336109 | 0.05581055 |
| FAM239A | 105.303473 | -2.046567385 | 1.67340997 | -1.2229922 | 0.22133263 | 0.40744344 |
| FAM239B | 53.2213459 | -1.007300982 | 1.82984067 | -0.5504856 | 0.58198634 | 0.74445693 |
| FAM239C | 6.03198075 | 0.786870293 | 3.07939241 | 0.25552778 | NA | NA |
| AC074035.1 | 0 | NA | NA | NA | NA | NA |
| AC110995.1 | 14.7235123 | 0.121674913 | 0.53479275 | 0.22751788 | 0.82002107 | 0.90344498 |
| NLGN4X | 56.6957808 | -0.342079702 | 0.44004855 | -0.7773681 | 0.43694163 | 0.62901194 |
| VCX3A | 2.57175725 | 0.693379038 | 0.88686806 | 0.78182885 | 0.43431516 | 0.62628593 |
| AC097347.1 | 5.26811961 | -0.359846899 | 0.87267905 | -0.4123474 | 0.68008484 | 0.81450801 |
| PUDP | 1130.30872 | -0.526883208 | 0.28781031 | -1.8306613 | 0.06715111 | 0.1788506 |
| STS | 708.277956 | -0.295908219 | 0.26403161 | -1.1207303 | 0.2624027 | 0.45603853 |
| VCX | 1.89104801 | -0.775374237 | 0.98260427 | -0.7891012 | 0.43005284 | 0.62207863 |
| PNPLA4 | 1383.17319 | 0.98656566 | 0.32690351 | 3.01791084 | 0.00254524 | 0.01621845 |
| AC078960.1 | 1.39059182 | 0.602074955 | 1.00337781 | 0.6000481 | 0.54847418 | 0.71910029 |
| VCX2 | 0.99279223 | -0.028259341 | 1.80959988 | -0.0156163 | 0.98754046 | NA |
| VCX3B | 10.1321476 | 0.183108772 | 0.5244896 | 0.34911802 | 0.72700071 | 0.84575518 |
| ANOS1 | 85.6916234 | -0.299487039 | 0.47697483 | -0.6278886 | 0.53007695 | 0.70461383 |
| FAM9A | 0.12150485 | 1.04981648 | 3.28477129 | 0.31960109 | 0.74927075 | NA |
| AC003685.1 | 0 | NA | NA | NA | NA | NA |
| FAM9B | 4.1445721 | -0.673138959 | 1.10366288 | -0.6099136 | 0.54191907 | 0.71449304 |
| AC003684.1 | 0.22447348 | 0.074529887 | 3.28898342 | 0.02266046 | 0.98192111 | NA |
| TBL1X | 2683.9093 | 0.367670869 | 0.16017854 | 2.29538155 | 0.02171127 | 0.07972791 |
| GPR143 | 421.396867 | -0.607115741 | 0.36377135 | -1.6689487 | 0.09512754 | 0.22957017 |
| SHROOM2 | 707.220508 | -0.047012976 | 0.27664458 | -0.16994 | 0.86505733 | 0.92986964 |
| CLDN34 | 2.8374515 | -0.111162786 | 0.70091794 | -0.158596 | 0.87398718 | 0.93436479 |
| WWC3 | 1512.13907 | -0.073147121 | 0.12730799 | -0.5745682 | 0.56558338 | 0.73214268 |
| WWC3-AS1 | 0.37988765 | 0.049486319 | 2.28544026 | 0.02165286 | 0.98272487 | NA |
| CLCN4 | 481.189082 | 0.582731542 | 0.27783304 | 2.09741625 | 0.03595674 | 0.11477779 |
| AC003666.1 | 0.60545969 | -0.794082327 | 1.87119336 | -0.4243721 | 0.67129444 | NA |
| MID1 | 1262.07513 | -1.134629949 | 0.30248003 | -3.7510905 | 0.00017607 | 0.0019818 |
| AC073529.1 | 4.21742632 | 0.676355145 | 0.58170765 | 1.16270629 | 0.24494869 | 0.4359971 |
| HCCS | 1235.21901 | -0.217644988 | 0.1568362 | -1.3877216 | 0.16522182 | 0.33554038 |
| ARHGAP6 | 1063.38019 | 0.288753237 | 0.1967161 | 1.46786784 | 0.14214013 | 0.30308362 |
| AMELX | 1.27903817 | -0.17394955 | 1.18548708 | -0.1467326 | 0.88334312 | 0.9394864 |
| AC004554.2 | 79.5760145 | 1.156907676 | 0.37081177 | 3.11993244 | 0.00180893 | 0.01247019 |
| MSL3 | 1317.21778 | -0.440244563 | 0.14166808 | -3.1075776 | 0.00188627 | 0.01289631 |
| FRMPD4 | 3.14842304 | 2.4414657 | 1.30136734 | 1.87607728 | 0.06064467 | 0.16676532 |
| FRMPD4-AS1 | 0 | NA | NA | NA | NA | NA |
| PRPS2 | 1753.38776 | 0.632308234 | 0.18761574 | 3.37023018 | 0.00075105 | 0.00624773 |
| TLR7 | 68.4795369 | 0.819059732 | 0.55057892 | 1.48763367 | 0.13684753 | 0.29526013 |
| TLR8-AS1 | 0.29148034 | -0.695455975 | 2.34298984 | -0.2968242 | 0.76660076 | NA |
| TLR8 | 73.9565389 | -0.458221404 | 0.60429836 | -0.7582701 | 0.44828928 | 0.63837284 |
| TMSB4X | 39811.4222 | -0.647009487 | 0.2945101 | -2.1969008 | 0.02802753 | 0.09616003 |
| FAM9C | 1.50692097 | 0.229316856 | 0.95746332 | 0.23950458 | 0.81071434 | 0.89761671 |
| AC079171.1 | 0.98045425 | 1.819247828 | 1.54543707 | 1.17717367 | 0.2391262 | NA |
| LINC02154 | 1.13318164 | -3.388853765 | 2.5503472 | -1.3287813 | 0.18392012 | 0.36099543 |
| AC004674.1 | 0 | NA | NA | NA | NA | NA |
| ATXN3L | 0 | NA | NA | NA | NA | NA |
| LINC01203 | 0.57344844 | -1.62954015 | 1.53186878 | -1.0637596 | 0.28743761 | NA |
| AC135371.1 | 0.62347545 | -1.247276879 | 2.22514837 | -0.5605365 | 0.57511355 | NA |
| EGFL6 | 22.3148729 | 2.91036478 | 0.60355484 | 4.82203868 | 1.42E-06 | 4.33E-05 |
| TCEANC | 170.726309 | 0.593870299 | 0.24518011 | 2.42217977 | 0.01542772 | 0.06213974 |
| RAB9A | 2136.38326 | -1.050383733 | 0.31521844 | -3.3322407 | 0.0008615 | 0.00699221 |
| TRAPPC2 | 834.777367 | -0.226891628 | 0.12205551 | -1.8589216 | 0.06303825 | 0.17117894 |
| OFD1 | 1812.3409 | -0.406211151 | 0.18508424 | -2.1947366 | 0.0281825 | 0.09659118 |
| GPM6B | 581.80903 | 0.380347469 | 0.32255545 | 1.17916927 | 0.23833078 | 0.42788399 |
| AC003035.2 | 0.37176169 | 0.952917273 | 2.18438705 | 0.43624012 | 0.66266252 | NA |
| AC003035.1 | 4.29366197 | -0.101297348 | 0.66248254 | -0.1529057 | 0.87847265 | 0.93677394 |
| GEMIN8 | 998.337003 | -0.244298548 | 0.16188721 | -1.5090664 | 0.13128181 | 0.28695506 |
| GLRA2 | 10.502743 | -1.13539348 | 1.0928255 | -1.0389522 | 0.29882696 | 0.49563286 |
| FANCB | 16.8575207 | -0.330766573 | 0.34864359 | -0.9487241 | 0.34276095 | 0.54048703 |
| MOSPD2 | 720.83873 | 0.281991868 | 0.11512425 | 2.44945669 | 0.01430719 | 0.05882143 |
| ASB9 | 484.084133 | 2.644020092 | 0.35551094 | 7.43723984 | 1.03E-13 | 3.99E-11 |
| ASB11 | 5.07896222 | -2.274958029 | 0.94049775 | -2.4188873 | 0.01556806 | 0.06249503 |
| PIGA | 773.367646 | -0.77145647 | 0.22700841 | -3.3983608 | 0.00067791 | 0.00574621 |
| VEGFD | 131.691171 | 0.551144259 | 0.50160559 | 1.09876021 | 0.27187267 | 0.46667822 |
| PIR | 900.491447 | -0.766997901 | 0.50012007 | -1.5336275 | 0.12512132 | 0.27747452 |
| BMX | 77.7652841 | -1.08288142 | 0.52414678 | -2.0659889 | 0.03882953 | 0.12119004 |
| ACE2 | 3553.93604 | 1.246156388 | 0.64127434 | 1.94325003 | 0.05198596 | 0.1498382 |
| AC097625.1 | 2.40006343 | -1.141682169 | 1.15086201 | -0.9920235 | 0.32118606 | 0.51875276 |
| CLTRN | 14611.511 | 0.015548127 | 0.46362169 | 0.03353624 | 0.97324697 | 0.98764503 |
| CA5B | 782.458063 | -0.417801456 | 0.31355508 | -1.3324659 | 0.18270714 | 0.35952445 |
| INE2 | 254.957704 | -0.495278138 | 0.33040097 | -1.4990214 | 0.13386807 | 0.29081145 |
| ZRSR2 | 834.068313 | -0.537687232 | 0.22140718 | -2.4284995 | 0.01516145 | 0.06132578 |
| AP1S2 | 1217.99121 | -0.439220126 | 0.21500154 | -2.0428697 | 0.04106534 | 0.12631437 |
| GRPR | 5.08970094 | -0.485993823 | 0.63719179 | -0.762712 | 0.44563518 | 0.63610477 |
| AC078993.1 | 5.67664237 | -0.910081342 | 0.56873554 | -1.6001837 | 0.10955783 | 0.25324685 |
| MAGEB17 | 4.91583746 | -0.489986142 | 0.61042685 | -0.8026943 | 0.42215146 | 0.61460404 |
| CTPS2 | 821.87535 | 0.156195613 | 0.16573944 | 0.94241667 | 0.34597936 | 0.54354339 |

| | | | | | | |
|---|---|---|---|---|---|---|
| S100G | 0.14114194 | 0.02007176 | 3.29738704 | 0.00608717 | 0.99514317 | NA |
| SYAP1 | 3227.00145 | -0.011301181 | 0.15981086 | -0.070716 | 0.9436238 | 0.97315146 |
| TXLNG | 1073.0608 | -0.243395857 | 0.14619438 | -1.6648784 | 0.09593703 | 0.23093194 |
| RBBP7 | 4643.66766 | -0.119102784 | 0.10669293 | -1.1163137 | 0.26428786 | 0.45820322 |
| REPS2 | 1780.13113 | 0.038903017 | 0.42314817 | 0.0919371 | 0.92674801 | 0.96441772 |
| NHS | 1273.51256 | 0.107606534 | 0.19571381 | 0.54981573 | 0.58244577 | 0.74490659 |
| NHS-AS1 | 9.63778855 | 0.209876189 | 0.83523051 | 0.25127936 | 0.80159813 | 0.89249269 |
| SCML1 | 1217.82466 | -0.134644803 | 0.21668412 | -0.6213875 | 0.53434469 | 0.70808591 |
| RAI2 | 1387.10148 | -0.509622704 | 0.22945816 | -2.2209831 | 0.0263521 | 0.09192422 |
| LINC01456 | 0.13028434 | 1.049816488 | 3.28477129 | 0.31960109 | 0.74927075 | NA |
| BEND2 | 0.53570507 | 1.66552361 | 1.69707905 | 0.98140603 | 0.32639255 | NA |
| SCML2 | 84.1849394 | 0.195024312 | 0.21965276 | 0.88787556 | 0.37460769 | 0.57064323 |
| CDKL5 | 679.043587 | 0.529549817 | 0.17016072 | 3.11205679 | 0.00185789 | 0.01272532 |
| RS1 | 18.1860497 | 1.196811017 | 0.51399417 | 2.32845252 | 0.01988809 | 0.07462278 |
| PPEF1 | 10.4923021 | 1.205706689 | 0.47615213 | 2.53218795 | 0.01133532 | 0.0495052 |
| PPEF1-AS1 | 0 | NA | NA | NA | NA | NA |
| PHKA2-AS1 | 846.788975 | -0.065578914 | 0.21676379 | -0.3025363 | 0.76224327 | 0.86819087 |
| PHKA2 | 1951.11236 | 0.169667666 | 0.2183478 | 0.77705233 | 0.4371279 | 0.62917928 |
| ADGRG2 | 46.1240546 | 0.277450184 | 0.35609179 | 0.77915356 | 0.43588926 | 0.62797202 |
| PDHA1 | 9420.75417 | 0.219119207 | 0.22690371 | 0.96569248 | 0.33419808 | 0.53236701 |
| MAP3K15 | 1496.38089 | 0.539163605 | 0.24387811 | 2.21079132 | 0.02705029 | 0.09360189 |
| SH3KBP1 | 1330.00557 | 0.421148229 | 0.21551996 | 1.95410311 | 0.05068902 | 0.14701601 |
| BCLAF3 | 927.727239 | -0.017134725 | 0.18173675 | -0.0942832 | 0.92488418 | 0.96358089 |
| MAP7D2 | 421.978986 | 1.040982047 | 0.47118935 | 2.20926481 | 0.02715623 | 0.09387909 |
| EIF1AX | 3631.10237 | 0.038368235 | 0.21470201 | 0.17870459 | 0.85816966 | 0.92582473 |
| EIF1AX-AS1 | 0.5328827 | -2.306080255 | 2.70803855 | -0.8515685 | 0.39445364 | NA |
| RPS6KA3 | 2965.10169 | -0.169465023 | 0.27188506 | -0.6232966 | 0.53308966 | 0.70697919 |
| AL928596.1 | 0.09997307 | 0.547239394 | 3.29377202 | 0.16614368 | 0.86804388 | NA |
| AL807740.1 | 0 | NA | NA | NA | NA | NA |
| BX088723.1 | 0.09635527 | 0.547239394 | 3.29377202 | 0.16614368 | 0.86804388 | NA |
| CNKSR2 | 21.9130946 | 0.712279396 | 0.73451891 | 0.96972234 | 0.33218491 | 0.53009359 |
| KLHL34 | 4.84755287 | 0.288219407 | 0.79252 | 0.36367461 | 0.716101 | 0.83845056 |
| SMPX | 4.96855442 | -0.113066891 | 1.42272109 | -0.0794723 | 0.93665698 | 0.96969938 |
| MBTPS2 | 876.41553 | 0.287987079 | 0.15607293 | 1.84520844 | 0.06500722 | 0.17480267 |
| YY2 | 72.7910662 | 0.661258262 | 0.30241564 | 2.18658751 | 0.02877265 | 0.09793917 |
| SMS | 7233.28367 | -0.174746162 | 0.25144324 | -0.6949726 | 0.48707247 | 0.67109572 |
| PHEX | 41.3408412 | -0.603754003 | 0.55215204 | -1.0934561 | 0.2741936 | 0.46932917 |
| PHEX-AS1 | 2.80296641 | -0.339778454 | 1.00927378 | -0.3366564 | 0.73637595 | 0.85207763 |
| PTCHD1-AS | 1.04472838 | 0.78496626 | 1.10564249 | 0.7099639 | 0.47772652 | 0.66285875 |
| CBLL2 | 0.09989236 | -0.642905668 | 3.30558948 | -0.1944905 | 0.84579185 | NA |
| AC092832.2 | 0 | NA | NA | NA | NA | NA |
| AC004470.1 | 0.08757577 | 0.547239394 | 3.29377202 | 0.16614368 | 0.86804388 | NA |
| DDX53 | 0 | NA | NA | NA | NA | NA |
| PTCHD1 | 16.3008216 | 0.933744299 | 0.40333569 | 2.31505498 | 0.02060992 | 0.07657935 |
| PRDX4 | 2240.03651 | -0.856236707 | 0.23318305 | -3.6719509 | 0.00024071 | 0.00253998 |
| ACOT9 | 1919.2145 | -0.423492518 | 0.28779917 | -1.4714863 | 0.14115966 | 0.30155381 |
| AC131011.1 | 22.8962271 | -0.548910122 | 0.71934122 | -0.7630734 | 0.44541965 | 0.63590039 |
| SAT1 | 40237.7121 | -1.261663918 | 0.24322859 | -5.187153 | 2.14E-07 | 8.95E-06 |
| APOO | 1123.56667 | 0.265783341 | 0.16626601 | 1.59854284 | 0.10992222 | 0.25379993 |
| CXorf58 | 127.281093 | 0.335317351 | 0.19891257 | 1.68575245 | 0.09184349 | 0.22383738 |
| KLHL15 | 459.970491 | 0.308210302 | 0.14691217 | 2.09792213 | 0.03591202 | 0.1146628 |
| EIF2S3 | 6309.53076 | -0.032000238 | 0.15766093 | -0.2029687 | 0.83915948 | 0.91464269 |
| ZFX-AS1 | 3.19851195 | -0.242784008 | 0.8278358 | -0.2932756 | 0.76931153 | 0.87240608 |
| ZFX | 1241.93801 | -0.067191565 | 0.21705755 | -0.3095565 | 0.75689827 | 0.86539227 |
| PDK3 | 363.893415 | -0.530525069 | 0.28212969 | -1.8804298 | 0.06004953 | 0.16558436 |
| PCYT1B | 23.0968086 | 2.880503081 | 0.50676082 | 5.68414718 | 1.31E-08 | 8.84E-07 |
| PCYT1B-AS1 | 0 | NA | NA | NA | NA | NA |
| POLA1 | 386.337531 | 0.189532678 | 0.16737279 | 1.13239836 | 0.25746699 | 0.45010638 |
| ARX | 4.70459215 | 0.527980413 | 0.78064285 | 0.67634055 | 0.49882445 | 0.68014482 |
| AC121342.1 | 0 | NA | NA | NA | NA | NA |
| MAGEB18 | 0.08666371 | -0.642905668 | 3.30558948 | -0.1944905 | 0.84579185 | NA |
| MAGEB6B | 0 | NA | NA | NA | NA | NA |
| MAGEB6 | 0 | NA | NA | NA | NA | NA |
| MAGEB5 | 0.08695548 | -0.642905668 | 3.30558948 | -0.1944905 | 0.84579185 | NA |
| VENTXP1 | 0 | NA | NA | NA | NA | NA |
| AC112493.1 | 0.19387658 | -1.003574173 | 3.30558948 | -0.3035992 | 0.7614333 | NA |
| AC107419.1 | 1.09547997 | -0.289189598 | 1.50341109 | -0.1923556 | 0.84746364 | 0.91988212 |
| PPP4R3C | 0 | NA | NA | NA | NA | NA |
| DCAF8L2 | 0 | NA | NA | NA | NA | NA |
| MAGEB10 | 0 | NA | NA | NA | NA | NA |
| DCAF8L1 | 0.18754884 | 1.469068384 | 3.27783996 | 0.44818185 | 0.65402196 | NA |
| AC112495.1 | 0 | NA | NA | NA | NA | NA |
| IL1RAPL1 | 0.77668041 | 1.787339248 | 1.41277167 | 1.26512959 | 0.20582488 | NA |
| MAGEB2 | 0.03392908 | 0.380740593 | 3.29738709 | 0.11546736 | 0.90807469 | NA |
| MAGEB3 | 0 | NA | NA | NA | NA | NA |
| MAGEB4 | 0 | NA | NA | NA | NA | NA |
| MAGEB1 | 0 | NA | NA | NA | NA | NA |
| NR0B1 | 2.2615656 | -1.938121199 | 1.22374916 | -1.5837569 | 0.11324905 | 0.25927842 |
| CXorf21 | 51.7360432 | 1.127890123 | 0.58505695 | 1.92782964 | 0.05387632 | 0.15355799 |
| GK | 3212.86976 | 1.867748088 | 0.47874658 | 3.90132939 | 9.57E-05 | 0.00122165 |
| GK-AS1 | 394.116208 | 1.673415765 | 0.46781548 | 3.57708509 | 0.00034745 | 0.00341075 |
| TAB3 | 2158.05683 | -0.183323959 | 0.14772976 | -1.2409413 | 0.21462745 | 0.39877895 |
| TAB3-AS1 | 4.19821871 | -0.249997309 | 0.54764307 | -0.4564968 | 0.64803277 | 0.79277053 |
| TAB3-AS2 | 24.3857793 | 0.031258172 | 0.41912311 | 0.07457993 | 0.94054894 | 0.97168563 |
| FTHL17 | 0 | NA | NA | NA | NA | NA |
| DMD | 644.903578 | 0.202424053 | 0.22610417 | 0.89526899 | 0.37064331 | 0.56745612 |
| DMD-AS3 | 0 | NA | NA | NA | NA | NA |
| AL591501.1 | 1.11342439 | 0.552509528 | 1.36686758 | 0.40421584 | 0.68605401 | 0.8184602 |
| FAM47A | 0 | NA | NA | NA | NA | NA |
| AL592043.1 | 0 | NA | NA | NA | NA | NA |
| TMEM47 | 2443.38567 | 0.034782556 | 0.22833545 | 0.15233095 | 0.87892592 | 0.93705294 |
| FAM47B | 0.18331075 | 0.18657164 | 3.29377195 | 0.05664376 | 0.95482897 | NA |
| MAGEB16 | 0.07659028 | -0.642905668 | 3.30558948 | -0.1944905 | 0.84579185 | NA |
| CFAP47 | 25.13275 | -0.608346953 | 0.63362968 | -0.9600986 | 0.33700561 | 0.53499716 |
| AL606516.1 | 3.55043756 | -1.104574799 | 0.85615245 | -1.2901614 | 0.19699464 | 0.37769003 |
| FAM47C | 1.12016852 | 1.25726541 | 1.31222478 | 0.95811741 | 0.33800356 | 0.53593613 |
| PRRG1 | 1244.94507 | -0.254143525 | 0.2830171 | -0.8979794 | 0.36919653 | 0.5660648 |
| AF241726.2 | 1089.76369 | 1.217221695 | 0.3941564 | 3.08816932 | 0.00201394 | 0.01357207 |
| LANCL3 | 34.0852037 | 0.006477268 | 0.38618596 | 0.01677241 | 0.98661818 | 0.99400325 |
| XK | 78.6275744 | 0.624156846 | 0.19481334 | 3.20387121 | 0.00135593 | 0.00999606 |
| CYBB | 970.612373 | 0.333387065 | 0.50549576 | 0.65952495 | 0.50955873 | 0.68830364 |
| DYNLT3 | 2949.1682 | -0.087651715 | 0.23640222 | -0.3707736 | 0.71080613 | 0.83571128 |
| AL121578.3 | 2.05108967 | 0.216342518 | 1.20783413 | 0.17911608 | 0.85784655 | 0.92574201 |
| HYPM | 1.2809364 | -1.614212829 | 1.34289477 | -1.2020397 | 0.22934815 | 0.41678111 |
| SYTL5 | 131.418695 | -0.43872612 | 0.41790726 | -1.0498169 | 0.29380228 | 0.49041437 |
| SRPX | 473.269536 | -1.014708837 | 0.50450768 | -2.0112852 | 0.04429534 | 0.13331504 |
| AL606748.1 | 0 | NA | NA | NA | NA | NA |
| RPGR | 629.547861 | -0.763887169 | 0.20523301 | -3.7220482 | 0.00019761 | 0.00215868 |
| OTC | 0.95129214 | 1.898833634 | 1.65155932 | 1.14972173 | 0.2502585 | NA |
| TSPAN7 | 2313.38778 | 1.041858707 | 0.40727391 | 2.55812781 | 0.01052374 | 0.04678972 |
| AF241728.1 | 5.03304235 | 0.669951933 | 0.65436107 | 1.02382609 | 0.30591743 | 0.50289283 |
| MID1IP1 | 3078.25909 | -0.341658733 | 0.2348351 | -1.4548878 | 0.14570031 | 0.30825717 |
| MID1IP1-AS1 | 93.4728342 | 0.463913989 | 0.21345446 | 2.17336279 | 0.02975302 | 0.10031802 |

| | | | | | | |
|---|---|---|---|---|---|---|
| AC108879.1 | 0.08757577 | 0.547239394 | 3.29377202 | 0.16614368 | 0.86804388 | NA |
| LINC01281 | 2.08905418 | 1.252131849 | 2.7466258 | 0.45588003 | 0.64847625 | 0.7930031 |
| LINC01282 | 0.60276243 | -2.468103995 | 3.2739918 | -0.7538516 | 0.45093833 | NA |
| AC091808.1 | 0.3216386 | -1.364242679 | 3.30558948 | -0.4127078 | 0.67982068 | NA |
| AL592164.1 | 0.76202333 | 2.379788275 | 1.76612318 | 1.34746449 | 0.17783068 | NA |
| AC092198.1 | 4.83295679 | -2.065604958 | 0.72997597 | -2.829689 | 0.00465933 | 0.02551214 |
| BCOR | 1401.90026 | 0.099543289 | 0.20016448 | 0.49730747 | 0.61897225 | 0.77232413 |
| AC091806.1 | 8.95881708 | -0.822062147 | 0.72033517 | -1.1412217 | 0.25377766 | 0.44642527 |
| ATP6AP2 | 9955.65611 | 0.074556915 | 0.16003756 | 0.46587136 | 0.64130758 | 0.78758115 |
| MPC1L | 0.62274934 | 1.427850304 | 2.04096935 | 0.69959419 | 0.48418077 | NA |
| CXorf38 | 1406.13404 | -0.687308812 | 0.18346874 | -3.7461904 | 0.00017954 | 0.00200699 |
| MED14 | 2172.7148 | -0.169744995 | 0.1334695 | -1.2717887 | 0.20344821 | 0.38566095 |
| MED14OS | 42.0097618 | -0.102259862 | 0.28365549 | -0.3605073 | 0.71846783 | 0.83995375 |
| USP9X | 5016.31809 | 0.065590566 | 0.11791737 | 0.55624178 | 0.57804558 | 0.74165288 |
| AL109797.1 | 6.05210681 | 0.443200126 | 0.50380143 | 0.87971193 | 0.37901539 | 0.57504694 |
| DDX3X | 13280.1716 | -0.168516633 | 0.25959515 | -0.6491517 | 0.51624034 | 0.69363383 |
| NYX | 49.8843904 | 1.293826706 | 0.60912979 | 2.1240575 | 0.03366534 | 0.10955879 |
| CASK | 1426.51131 | 0.330322204 | 0.17825902 | 1.85304619 | 0.06387574 | 0.17267076 |
| CASK-AS1 | 240.393962 | 0.22518061 | 0.22044943 | 1.02146151 | 0.30703584 | 0.50401785 |
| GPR34 | 236.212736 | 0.392810585 | 0.40736883 | 0.96426274 | 0.33491422 | 0.53299263 |
| GPR82 | 13.1611769 | 0.466816236 | 0.49486877 | 0.94331318 | 0.34552074 | 0.54324657 |
| AL133329.1 | 0 | NA | NA | NA | NA | NA |
| Z93403.1 | 37.0085653 | 1.353497724 | 0.71544644 | 1.89182258 | 0.05851462 | 0.16261257 |
| PPP1R2C | 0.16416605 | 0.18657164 | 3.29377195 | 0.05664376 | 0.95482897 | NA |
| PINCR | 0.16416605 | 0.18657164 | 3.29377195 | 0.05664376 | 0.95482897 | NA |
| MAOA | 12341.237 | -0.427439205 | 0.42657607 | -1.0020234 | 0.31633229 | 0.51346069 |
| MAOB | 6068.97553 | 1.522265266 | 0.4114101 | 3.70011643 | 0.0002155 | 0.00232044 |
| NDP | 3.8824189 | 0.533321979 | 1.41319966 | 0.37738615 | 0.70588666 | 0.8324418 |
| NDP-AS1 | 1.25829035 | 0.67276895 | 2.71202429 | 0.24806892 | 0.80408108 | 0.89380003 |
| EFHC2 | 191.995733 | -0.661608788 | 0.4165699 | -1.5882299 | 0.11223436 | 0.25767262 |
| FUNDC1 | 716.020014 | -0.096257769 | 0.1284067 | -0.749632 | 0.45347637 | 0.64311992 |
| DUSP21 | 0 | NA | NA | NA | NA | NA |
| KDM6A | 1181.99287 | -0.075061076 | 0.26908949 | -0.2789447 | 0.7802873 | 0.8796997 |
| CXorf36 | 903.199517 | -0.973455172 | 0.41555979 | -2.3425153 | 0.01915425 | 0.07259234 |
| AC136489.1 | 0.3020015 | -0.174096114 | 3.29377188 | -0.0528562 | 0.95784652 | NA |
| LINC01204 | 0.47434806 | -0.961631994 | 1.90439687 | -0.5049536 | 0.61359145 | NA |
| MIR222HG | 397.438678 | -0.446631437 | 0.42055101 | -1.0620149 | 0.28822893 | 0.48436125 |
| AC234772.3 | 4.44473699 | -0.422513535 | 0.97343581 | -0.4340436 | 0.66425682 | 0.8027705 |
| AC234772.2 | 0.87763155 | -0.124447759 | 1.63671617 | -0.076035 | 0.93939123 | NA |
| AL031584.2 | 0.07830766 | -0.642905668 | 3.30558948 | -0.1944905 | 0.84579185 | NA |
| LINC01186 | 22.8507577 | 2.197825083 | 0.42370727 | 5.18713097 | 2.14E-07 | 8.95E-06 |
| KRBOX4 | 800.850027 | 0.001547391 | 0.14025551 | 0.01103266 | 0.99119739 | 0.99583588 |
| ZNF674 | 178.545092 | 0.354483396 | 0.18643522 | 1.90137572 | 0.05725282 | 0.16010146 |
| ZNF674-AS1 | 252.007998 | 0.126332586 | 0.24556219 | 0.5144627 | 0.60692853 | 0.76327544 |
| CHST7 | 261.637916 | -0.208109104 | 0.33259495 | -0.6257134 | 0.53150297 | 0.70583338 |
| SLC9A7 | 847.171148 | -0.093453001 | 0.12533563 | -0.745622 | 0.4558958 | 0.64526715 |
| RP2 | 584.625769 | -0.119472155 | 0.28902556 | -0.4133619 | 0.67934149 | 0.81395043 |
| LINC01545 | 3.88216806 | 0.890357767 | 0.60914504 | 1.4616515 | 0.14383673 | 0.30554009 |
| JADE3 | 604.113736 | -0.242848755 | 0.15412281 | -1.5756834 | 0.11509879 | 0.26235243 |
| RGN | 2679.07144 | 1.186360181 | 0.44265544 | 2.68009852 | 0.00736005 | 0.03595367 |
| NDUFB11 | 3337.90003 | -0.049166829 | 0.13243232 | -0.37126 | 0.71044386 | 0.8353877 |
| RBM10 | 3568.81053 | -0.104979464 | 0.12114652 | -0.8665495 | 0.38618887 | 0.58179847 |
| UBA1 | 19082.439 | 0.155826709 | 0.08642453 | 1.80303803 | 0.07138224 | 0.18692649 |
| CDK16 | 6809.56477 | -0.260154576 | 0.13364203 | -1.9466524 | 0.05157642 | 0.14908146 |
| USP11 | 4314.59507 | 0.192150437 | 0.18826379 | 1.02064469 | 0.30742281 | 0.50440141 |
| AL591503.1 | 0.1981425 | 1.631193554 | 3.26970143 | 0.4988815 | 0.61786287 | NA |
| ZNF157 | 14.5051172 | 0.87205273 | 0.47334564 | 1.842317 | 0.06542879 | 0.17560889 |
| ZNF41 | 506.236049 | 0.297559593 | 0.10741808 | 2.77010717 | 0.00560379 | 0.02935475 |
| LINC01560 | 103.999644 | 0.471539656 | 0.25399001 | 1.85652833 | 0.06337829 | 0.17178779 |
| ARAF | 5393.75734 | 0.044959371 | 0.08824222 | 0.50949957 | 0.6104021 | 0.76634467 |
| SYN1 | 119.650677 | -1.124965864 | 0.24653464 | -4.5631148 | 5.04E-06 | 0.00011772 |
| TIMP1 | 15781.4647 | -2.301094016 | 0.48600924 | -4.7346713 | 2.19E-06 | 6.12E-05 |
| CFP | 160.094556 | -0.042917219 | 0.38264375 | -0.1121597 | 0.91069676 | 0.95582382 |
| ELK1 | 1593.07257 | -0.447840029 | 0.14266715 | -3.1390549 | 0.00169494 | 0.01186501 |
| UXT | 1995.25343 | -0.276128856 | 0.16338688 | -1.6900308 | 0.09102206 | 0.22238967 |
| UXT-AS1 | 883.721886 | -0.148761453 | 0.22798552 | -0.6525039 | 0.51407613 | 0.69195996 |
| ZNF81 | 151.882113 | -0.104027288 | 0.1358495 | -0.7657539 | 0.44382272 | 0.6348089 |
| ZNF182 | 665.946939 | -0.007912696 | 0.12988923 | -0.0609188 | 0.95142388 | 0.97675915 |
| ZNF630 | 223.873674 | 0.293889586 | 0.16947824 | 1.73408452 | 0.08290309 | 0.20771927 |
| SPACA5 | 0.18028734 | 0.18657164 | 3.29377195 | 0.05664376 | 0.95482897 | NA |
| ZNF630-AS1 | 0.48055917 | -0.391595429 | 1.67830038 | -0.2333286 | 0.81550627 | NA |
| SPACA5B | 0 | NA | NA | NA | NA | NA |
| SSX5 | 0.12150485 | 1.04981648 | 3.28477129 | 0.31960109 | 0.74927075 | NA |
| SSX1 | 0 | NA | NA | NA | NA | NA |
| AL606490.3 | 0 | NA | NA | NA | NA | NA |
| SSX3 | 0 | NA | NA | NA | NA | NA |
| SSX4 | 0 | NA | NA | NA | NA | NA |
| SSX4B | 0.03392908 | 0.380740593 | 3.29738709 | 0.11546736 | 0.90807469 | NA |
| SLC38A5 | 297.276588 | -1.400787476 | 0.478298 | -2.9286919 | 0.00340392 | 0.02018014 |
| FTSJ1 | 1553.63222 | -0.409124675 | 0.20541967 | -1.9916529 | 0.04640916 | 0.1379562 |
| AF196972.1 | 13.4849274 | 0.824105577 | 0.45627301 | 1.80616772 | 0.07089215 | 0.18601252 |
| PORCN | 511.937853 | -0.05827652 | 0.21909723 | -0.2659847 | 0.79025096 | 0.88610295 |
| EBP | 2685.8717 | 0.221037072 | 0.24573769 | 0.89948382 | 0.36839501 | 0.56515711 |
| TBC1D25 | 1183.87687 | -0.316282346 | 0.17027417 | -1.8574886 | 0.06324167 | 0.17159398 |
| AC115618.1 | 1106.41255 | 0.112161097 | 0.44425201 | 0.25247179 | 0.80067642 | 0.89197065 |
| RBM3 | 8947.38096 | -0.23913951 | 0.36278977 | -0.6591683 | 0.50978771 | 0.68850714 |
| AC115618.2 | 337.812328 | -0.485399797 | 0.20126347 | -2.411763 | 0.0158756 | 0.06344945 |
| WDR13 | 8724.86801 | -0.406496453 | 0.15194335 | -2.6753158 | 0.00746589 | 0.03633059 |
| WAS | 520.12848 | -0.471326971 | 0.43730841 | -1.0777908 | 0.28112713 | 0.47692502 |
| SUV39H1 | 302.936825 | 0.41045381 | 0.30559397 | 1.34313452 | 0.17922842 | 0.3548671 |
| AC231533.1 | 65.9881303 | 0.770205799 | 0.33329446 | 2.31088689 | 0.0208391 | 0.07719162 |
| GLOD5 | 178.082242 | 1.187766102 | 0.41990841 | 2.82863139 | 0.00467475 | 0.0255699 |
| GATA1 | 13.320996 | -1.122621398 | 0.49797903 | -2.2543548 | 0.02417386 | 0.08607408 |
| HDAC6 | 7951.28935 | 1.237461057 | 0.26526766 | 4.66495252 | 3.09E-06 | 8.00E-05 |
| ERAS | 23.7081504 | 0.820266948 | 0.4021066 | 2.0399241 | 0.04135789 | 0.1270216 |
| PCSK1N | 2870.17309 | 0.548729805 | 0.3059989 | 1.7932411 | 0.07293438 | 0.18995354 |
| TIMM17B | 2308.26679 | -0.242301538 | 0.0958564 | -2.5277553 | 0.01147943 | 0.04997424 |
| PQBP1 | 2920.5302 | -0.397800371 | 0.17141311 | -2.3207114 | 0.02030242 | 0.07571417 |
| SLC35A2 | 2091.77186 | -0.438241699 | 0.13802512 | -3.1750866 | 0.00149792 | 0.01079532 |
| PIM2 | 1033.84563 | -0.619090124 | 0.38599154 | -1.6038956 | 0.10873707 | 0.25202218 |
| OTUD5 | 4723.20304 | -0.381847271 | 0.08082955 | -4.7241048 | 2.31E-06 | 6.38E-05 |
| KCND1 | 252.488869 | -0.866073993 | 0.37822292 | -2.2898506 | 0.02202998 | 0.08057966 |
| GRIPAP1 | 3757.04495 | -0.225096575 | 0.16069235 | -1.4007921 | 0.16127625 | 0.32996001 |
| TFE3 | 6651.26631 | -1.113808558 | 0.24382395 | -4.5680852 | 4.92E-06 | 0.00011599 |
| CCDC120 | 2261.89971 | -0.895217858 | 0.23264709 | -3.8479649 | 0.0001191 | 0.0014554 |
| PRAF2 | 1407.49211 | -0.764058796 | 0.32603037 | -2.3435203 | 0.01910272 | 0.07253232 |
| WDR45 | 6402.93987 | -0.509976976 | 0.16723196 | -3.0495186 | 0.00229208 | 0.01500469 |
| GPKOW | 1354.87515 | -0.319060478 | 0.12519085 | -2.5485927 | 0.01081585 | 0.04775855 |
| MAGIX | 535.927557 | 0.004959735 | 0.26493272 | 0.01872073 | 0.98506389 | 0.99347171 |
| PLP2 | 2966.83072 | -1.614555561 | 0.31700152 | -5.0932107 | 3.52E-07 | 1.36E-05 |
| PRICKLE3 | 610.743508 | -0.879668922 | 0.18517427 | -4.7504921 | 2.03E-06 | 5.72E-05 |
| SYP | 322.840042 | 0.901917465 | 0.41039781 | 2.19766638 | 0.02797289 | 0.09606004 |
| SYP-AS1 | 22.5311108 | 1.36267572 | 0.60782146 | 2.24190129 | 0.02496775 | 0.08825732 |

| | | | | | | |
|---|---|---|---|---|---|---|
| CACNA1F | 33.2829207 | -0.864866301 | 0.47416978 | -1.8239591 | 0.06815825 | 0.18061834 |
| CCDC22 | 1442.73471 | -0.168839601 | 0.10550353 | -1.6003218 | 0.10952722 | 0.25324117 |
| FOXP3 | 133.147266 | -0.183172412 | 0.27850668 | -0.6576949 | 0.51073423 | 0.68915017 |
| PPP1R3F | 451.359725 | -0.231815105 | 0.20230926 | -1.1458453 | 0.25185918 | 0.44410074 |
| AC232271.1 | 232.831395 | 0.186489869 | 0.26091614 | 0.71475021 | 0.47476341 | 0.66021715 |
| GAGE10 | 2.78627148 | -0.630914607 | 0.78332086 | -0.8054357 | 0.42056828 | 0.61330335 |
| GAGE12J | 0.08666371 | -0.642905668 | 3.30558948 | -0.1944905 | 0.84579185 | NA |
| GAGE13 | 0.08757577 | 0.547239394 | 3.29377202 | 0.16614368 | 0.86804388 | NA |
| GAGE2E | 0 | NA | NA | NA | NA | NA |
| GAGE12B | 0 | NA | NA | NA | NA | NA |
| GAGE12C | 0 | NA | NA | NA | NA | NA |
| GAGE12D | 0 | NA | NA | NA | NA | NA |
| GAGE12F | 0 | NA | NA | NA | NA | NA |
| GAGE12E | 0 | NA | NA | NA | NA | NA |
| GAGE12G | 0 | NA | NA | NA | NA | NA |
| GAGE12H | 0 | NA | NA | NA | NA | NA |
| GAGE1 | 0 | NA | NA | NA | NA | NA |
| GAGE2A | 0 | NA | NA | NA | NA | NA |
| PAGE1 | 0.08393208 | -0.642905668 | 3.30558948 | -0.1944905 | 0.84579185 | NA |
| PAGE4 | 5.51910155 | 2.21370872 | 1.41011793 | 1.56987488 | 0.11644422 | 0.26459579 |
| USP27X-AS1 | 115.216658 | 0.693736069 | 0.2909235 | 2.38459959 | 0.01709772 | 0.06693392 |
| USP27X | 276.877791 | -0.086746806 | 0.17074224 | -0.5080571 | 0.6114133 | 0.76723122 |
| CLCN5 | 5462.87111 | 1.445076117 | 0.27960624 | 5.16825413 | 2.36E-07 | 9.80E-06 |
| AKAP4 | 2.28468014 | 3.059024392 | 0.96224699 | 3.17904282 | 0.00147762 | 0.0106841 |
| CCNB3 | 25.3808726 | 0.376023197 | 0.31952691 | 1.17681228 | 0.23927044 | 0.42907227 |
| DGKK | 2.15438868 | 0.367346412 | 0.86504754 | 0.42465459 | 0.67108849 | 0.80785905 |
| SHROOM4 | 1056.27178 | 0.407702003 | 0.12747213 | 3.19836179 | 0.00138211 | 0.0101578 |
| BMP15 | 0.21442573 | -1.003574173 | 3.30558948 | -0.3035992 | 0.7614333 | NA |
| LINC01284 | 2.53687176 | -0.240100192 | 1.19691969 | -0.2005984 | 0.8410126 | 0.91512686 |
| NUDT10 | 236.278036 | -0.906242943 | 0.59484767 | -1.5234874 | 0.1276368 | 0.28139641 |
| AC233976.1 | 7.39233763 | 0.688897375 | 0.53829375 | 1.27977964 | 0.20062265 | 0.38215108 |
| AL158055.1 | 8.20292826 | 1.016167311 | 0.5481979 | 1.85365048 | 0.06378918 | 0.17250909 |
| CXorf67 | 14.4528698 | -0.662374983 | 0.44810806 | -1.4781591 | 0.13936521 | 0.29902853 |
| NUDT11 | 42.0363009 | -0.501846636 | 0.41707876 | -1.2032419 | 0.22888273 | 0.41621113 |
| LINC01496 | 0 | NA | NA | NA | NA | NA |
| CENPVL3 | 0.47371947 | 0.70848424 | 2.22197847 | 0.31885288 | 0.74983807 | NA |
| CENPVL2 | 0 | NA | NA | NA | NA | NA |
| CENPVL1 | 0 | NA | NA | NA | NA | NA |
| GSPT2 | 520.60391 | 0.305001708 | 0.11075027 | 2.75395909 | 0.00588791 | 0.03037251 |
| MAGED1 | 6703.04598 | 0.089225451 | 0.20362745 | 0.43817987 | 0.6612559 | 0.80071801 |
| AC239585.1 | 0 | NA | NA | NA | NA | NA |
| AC239585.2 | 0 | NA | NA | NA | NA | NA |
| MAGED4B | 1.3918499 | -0.00609053 | 1.12596914 | -0.0054091 | 0.99568415 | 0.99795473 |
| MAGED4 | 0.90746187 | -1.096277348 | 1.45396021 | -0.753994 | 0.4508528 | NA |
| AC245177.1 | 0 | NA | NA | NA | NA | NA |
| XAGE2 | 2.84466616 | 0.609136773 | 0.77647495 | 0.78448992 | 0.43275269 | 0.62475027 |
| XAGE1A | 0 | NA | NA | NA | NA | NA |
| XAGE1B | 0 | NA | NA | NA | NA | NA |
| SSX7 | 0 | NA | NA | NA | NA | NA |
| SSX2 | 0 | NA | NA | NA | NA | NA |
| SSX2B | 0 | NA | NA | NA | NA | NA |
| SPANXN5 | 0.10721287 | -0.642905668 | 3.30558948 | -0.1944905 | 0.84579185 | NA |
| XAGE5 | 0 | NA | NA | NA | NA | NA |
| XAGE3 | 2.66916516 | 0.416893227 | 1.03871416 | 0.4013551 | 0.6881587 | 0.81993245 |
| FAM156B | 3.37131345 | -0.40631933 | 0.79112271 | -0.5135984 | 0.60753282 | 0.76386977 |
| FAM156A | 17.3535573 | -0.130241875 | 0.34712113 | -0.3752058 | 0.70750739 | 0.83353303 |
| GPR173 | 376.428528 | -0.419071466 | 0.30781847 | -1.3614241 | 0.17337972 | 0.34710167 |
| TSPYL2 | 3270.52465 | -0.711827655 | 0.26545796 | -2.6815081 | 0.00732911 | 0.03583697 |
| AL591212.1 | 3.40970633 | -0.117583568 | 0.70406416 | -0.1670069 | 0.86736463 | 0.93105 |
| KANTR | 934.293194 | 0.245740702 | 0.21464445 | 1.14487333 | 0.25226163 | 0.44448345 |
| KDM5C | 6492.23497 | -0.434529684 | 0.23301322 | -1.8648284 | 0.06220546 | 0.16952851 |
| IQSEC2 | 1772.27852 | 0.457211148 | 0.19780336 | 2.31144275 | 0.02080841 | 0.07711043 |
| SMC1A | 2744.14315 | 0.154201256 | 0.1621577 | 0.95093389 | 0.34163793 | 0.53958986 |
| RIBC1 | 157.876646 | 0.394060497 | 0.36785862 | 1.07122811 | 0.28406688 | 0.48012072 |
| HSD17B10 | 4110.60396 | 0.344471907 | 0.16346077 | 2.10736749 | 0.03508574 | 0.11287189 |
| AC233728.1 | 40.135826 | 0.081019116 | 0.26438056 | 0.30644883 | 0.75926293 | 0.86663046 |
| HUWE1 | 8587.4964 | 0.222590433 | 0.08627658 | 2.57996361 | 0.00988107 | 0.04467831 |
| PHF8 | 2363.06657 | -0.226236788 | 0.22309002 | -1.0141054 | 0.31053248 | 0.50745129 |
| FAM120C | 668.441685 | 0.245615628 | 0.10655114 | 2.30514317 | 0.02115856 | 0.07810066 |
| WNK3 | 243.780213 | 0.139581023 | 0.22518154 | 0.61985998 | 0.53534998 | 0.70904967 |
| TSR2 | 2662.9466 | 0.185657093 | 0.12286841 | 1.51102382 | 0.13078238 | 0.28629404 |
| FGD1 | 888.603223 | -0.049780373 | 0.152449 | -0.3265379 | 0.74401743 | 0.85638322 |
| GNL3L | 197.536371 | 0.773394765 | 0.26220095 | 2.94962612 | 0.00318159 | 0.01921632 |
| ITIH6 | 0.89008252 | 0.324368888 | 1.35341329 | 0.23966728 | 0.8105882 | NA |
| MAGED2 | 12876.0904 | 0.160198918 | 0.10913056 | 1.46795646 | 0.14211605 | 0.30305679 |
| TRO | 1382.13118 | -0.088307015 | 0.28012059 | -0.3152464 | 0.75257455 | 0.86206078 |
| PFKFB1 | 40.7149562 | 0.111228673 | 0.7543212 | 0.14745532 | 0.88277264 | 0.93903649 |
| APEX2 | 1668.80527 | -0.436694316 | 0.22478411 | -1.9427277 | 0.05204907 | 0.14993816 |
| ALAS2 | 67.8087477 | -2.409698845 | 0.74271546 | -3.2444441 | 0.0011768 | 0.00891475 |
| PAGE2B | 8.15448456 | -1.940750812 | 0.62347117 | -3.1128156 | 0.00185312 | 0.01269925 |
| PAGE2 | 2.63982558 | -0.469632181 | 0.90579711 | -0.5184739 | 0.60412765 | 0.761258 |
| FAM104B | 627.649141 | 0.609549396 | 0.17413615 | 3.50041846 | 0.00046453 | 0.00429058 |
| MTRNR2L10 | 44.0591694 | -0.912117766 | 0.53931497 | -1.6912525 | 0.0907886 | 0.22204556 |
| PAGE5 | 3.1527751 | 0.633151348 | 1.13569842 | 0.55749954 | 0.57718617 | 0.74104871 |
| PAGE3 | 0.08666371 | -0.642905668 | 3.30558948 | -0.1944905 | 0.84579185 | NA |
| MAGEH1 | 1276.47514 | 0.220932385 | 0.26798583 | 0.82441815 | 0.409702 | 0.60414035 |
| USP51 | 253.668898 | 1.080752067 | 0.3261501 | 3.3136647 | 0.00092082 | 0.00736934 |
| FOXR2 | 0 | NA | NA | NA | NA | NA |
| RRAGB | 1080.31922 | -0.114275896 | 0.12957857 | -0.8819043 | 0.37782859 | 0.57382306 |
| AL445472.1 | 72.1254068 | -0.132552106 | 0.2433902 | -0.5446074 | 0.58602355 | 0.74736507 |
| KLF8 | 922.33939 | 0.087826129 | 0.36830768 | 0.23845859 | 0.81152543 | 0.89825074 |
| UBQLN2 | 2553.67412 | -0.411512961 | 0.20130994 | -2.044176 | 0.04093616 | 0.12606506 |
| AL354793.1 | 1.65486804 | 1.227628225 | 1.10908197 | 1.10688683 | 0.26834286 | 0.46255662 |
| NBDY | 1621.56324 | -0.221490395 | 0.13513238 | -1.6390623 | 0.10120028 | 0.23951278 |
| SPIN3 | 746.542165 | 0.303858726 | 0.15910485 | 1.90980178 | 0.05615874 | 0.15791287 |
| AL139397.1 | 4.76005608 | 0.384458935 | 0.65099718 | 0.59056928 | 0.55480906 | 0.72430887 |
| SPIN2B | 197.811137 | -0.139479941 | 0.18545327 | -0.7521029 | 0.45198919 | 0.64173558 |
| AL022157.1 | 19.7781018 | 0.060208434 | 0.35003987 | 0.1720045 | 0.86343399 | 0.92891984 |
| SPIN2A | 8.86250294 | -0.051355788 | 0.43296588 | -0.1186139 | 0.90558122 | 0.95262374 |
| FAAH2 | 908.541796 | -0.994309517 | 0.20769597 | -4.7873319 | 1.69E-06 | 4.97E-05 |
| ZXDB | 536.639027 | -0.2387853 | 0.14796809 | -1.6137621 | 0.10657904 | 0.24816683 |
| NLRP2B | 2.14979034 | 0.477190703 | 0.9590916 | 0.49754445 | 0.61880516 | 0.77226182 |
| ZXDA | 161.76473 | 0.486885752 | 0.19515802 | 2.49482826 | 0.01260181 | 0.05350342 |
| SPIN4 | 176.266723 | 0.365166325 | 0.11878801 | 3.07410098 | 0.00211138 | 0.01402445 |
| SPIN4-AS1 | 10.8598927 | 0.973070884 | 0.42983594 | 2.26381929 | 0.02358523 | 0.08460652 |
| ARHGEF9 | 1293.82495 | 0.751677024 | 0.15470832 | 4.85867229 | 1.18E-06 | 3.73E-05 |
| AMER1 | 343.023071 | 0.65219894 | 0.17415608 | 3.74491062 | 0.00018046 | 0.00201298 |
| ASB12 | 27.4195906 | 0.115421492 | 0.25745534 | 0.44831656 | 0.65392475 | 0.79689429 |
| MTMR8 | 46.1254796 | 0.290981708 | 0.19146796 | 1.51974097 | 0.12857609 | 0.2828052 |
| ZC4H2 | 581.647263 | 0.044043115 | 0.17455123 | 0.252322 | 0.80079219 | 0.89197577 |
| ZC3H12B | 160.698457 | 0.316491706 | 0.22155356 | 1.42851102 | 0.15314483 | 0.31866401 |
| LAS1L | 1897.95855 | 0.258750931 | 0.18686541 | 1.38469141 | 0.16614685 | 0.33681522 |
| MSN | 25989.9684 | -0.223501266 | 0.21358755 | -1.0464153 | 0.29536935 | 0.49218893 |

| | | | | | | |
|---|---|---|---|---|---|---|
| AL034397.2 | 0.44524338 | -2.045962029 | 1.88409541 | -1.0859121 | 0.27751788 | NA |
| AL034397.3 | 194.871631 | -0.754466225 | 0.55124633 | -1.3686553 | 0.17110704 | 0.34343832 |
| VSIG4 | 1665.96718 | -1.563361239 | 0.42451728 | -3.6826798 | 0.00023079 | 0.00244962 |
| HEPH | 672.574257 | 0.281117242 | 0.40193513 | 0.69940948 | 0.48429616 | 0.668712 |
| EDA2R | 170.274609 | -0.908903143 | 0.45847294 | -1.9824575 | 0.04742805 | 0.14007339 |
| AR | 689.427542 | 1.11528338 | 0.31944428 | 3.49132368 | 0.00048063 | 0.00439775 |
| AL157700.1 | 39.6720665 | 0.732806118 | 0.29490423 | 2.48489528 | 0.01295896 | 0.0545969 |
| OPHN1 | 532.479012 | 0.682604819 | 0.22279337 | 3.06384714 | 0.00218511 | 0.01445138 |
| YIPF6 | 2669.47262 | 0.156519093 | 0.09485421 | 1.65010175 | 0.09892213 | 0.23581661 |
| STARD8 | 2638.05997 | 1.074234184 | 0.2015943 | 5.32869323 | 9.89E-08 | 4.75E-06 |
| EFNB1 | 3978.32893 | -0.271093382 | 0.16472079 | -1.6457751 | 0.09981007 | 0.23722321 |
| PJA1 | 1280.47073 | -0.030109973 | 0.16662868 | -0.180701 | 0.85660225 | 0.9249284 |
| LINC00269 | 0.3373959 | 1.578568919 | 3.24620529 | 0.4862813 | 0.62676774 | NA |
| FAM155B | 221.35706 | -0.979461123 | 0.37322674 | -2.6243059 | 0.00868258 | 0.04045937 |
| AL158069.1 | 0.03392908 | 0.380740593 | 3.29738709 | 0.11546736 | 0.90807469 | NA |
| EDA | 389.545 | 0.906089082 | 0.18381454 | 4.92936572 | 8.25E-07 | 2.75E-05 |
| AWAT2 | 1.61044067 | -3.929648746 | 1.26937337 | -3.0957391 | 0.00196323 | 0.01329833 |
| OTUD6A | 0 | NA | NA | NA | NA | NA |
| IGBP1 | 2725.29069 | 0.188408446 | 0.14820073 | 1.27130581 | 0.20361988 | 0.38583607 |
| IGBP1-AS2 | 720.513534 | 0.230747444 | 0.16513674 | 1.39731139 | 0.16231995 | 0.33129854 |
| IGBP1-AS1 | 29.6039157 | 0.023989196 | 0.29877325 | 0.08029231 | 0.93600477 | 0.96939601 |
| DGAT2L6 | 1.25731015 | 0.673895956 | 1.05448798 | 0.6390741 | 0.52277473 | 0.69862624 |
| AWAT1 | 0.49554303 | 0.890428204 | 1.79711257 | 0.49547714 | 0.62026335 | NA |
| P2RY4 | 0.41702455 | -0.032126964 | 2.22348912 | -0.0144489 | 0.98847185 | NA |
| ARR3 | 32.5165134 | 0.023907208 | 0.35535573 | 0.06727683 | 0.94636132 | 0.97427563 |
| RAB41 | 36.4774035 | 0.216903816 | 0.23603086 | 0.91896378 | 0.35811452 | 0.55576035 |
| PDZD11 | 1206.39426 | -0.012172339 | 0.11308767 | -0.1076363 | 0.9142842 | 0.95798761 |
| KIF4A | 34.3425858 | -0.428412049 | 0.71696972 | -0.5975316 | 0.55015253 | 0.72047687 |
| GDPD2 | 52.7240918 | -0.256762739 | 0.73791666 | -0.3479563 | 0.72787302 | 0.84611931 |
| AL139398.1 | 9.92954328 | 0.241354662 | 0.9898794 | 0.24382229 | 0.80736846 | 0.89590613 |
| DLG3 | 2298.55682 | -0.039308171 | 0.17559963 | -0.2238511 | 0.82287316 | 0.90541456 |
| DLG3-AS1 | 21.1567247 | -0.722894415 | 0.36099114 | -2.0025268 | 0.04522811 | 0.13529951 |
| TEX11 | 14.516847 | -0.728146307 | 1.01627621 | -0.7164847 | 0.47369214 | 0.65958716 |
| SLC7A3 | 2.13826694 | 0.912139999 | 0.88452699 | 1.03121783 | 0.30243868 | 0.49933058 |
| SNX12 | 2833.23406 | 0.093929308 | 0.17005831 | 0.55233589 | 0.58071825 | 0.74352011 |
| FOXO4 | 1516.38335 | 0.511958431 | 0.19132335 | 2.6758806 | 0.00745332 | 0.03628282 |
| CXorf65 | 21.3223727 | -0.294886122 | 0.69264867 | -0.4257369 | 0.67029955 | 0.80744594 |
| AL590764.2 | 474.255718 | 0.050867168 | 0.63156084 | 0.08054199 | 0.9358062 | 0.96929217 |
| IL2RG | 584.027588 | -0.06648469 | 0.64390551 | -0.1032522 | 0.91776277 | 0.96010649 |
| MED12 | 1842.18375 | 0.099286484 | 0.18221138 | 0.54489726 | 0.58582418 | 0.74726375 |
| NLGN3 | 75.2468994 | 1.00987641 | 0.27737464 | 3.64083898 | 0.00027175 | 0.00280912 |
| AL590764.1 | 9.68627839 | -0.789529969 | 0.69310918 | -1.1391134 | 0.25465585 | 0.44747469 |
| GJB1 | 2530.6123 | 1.527219969 | 0.57615951 | 2.65068898 | 0.00803278 | 0.03823544 |
| ZMYM3 | 2650.5382 | 0.394148406 | 0.29651746 | 1.32925867 | 0.18376264 | 0.36079376 |
| NONO | 11622.4769 | -0.082950269 | 0.14675344 | -0.5652356 | 0.57191352 | 0.73675425 |
| ITGB1BP2 | 59.4560581 | 0.017666659 | 0.3699655 | 0.04775218 | 0.96191375 | 0.98183339 |
| TAF1 | 1428.09132 | 0.116831083 | 0.11641892 | 1.00354031 | 0.31560024 | 0.5126827 |
| OGT | 9282.64636 | -0.166197892 | 0.16530736 | -1.0053871 | 0.31471049 | 0.51173087 |
| GCNA | 227.801259 | -0.50859008 | 0.28177926 | -1.8049237 | 0.07108662 | 0.18641151 |
| CXCR3 | 49.7041188 | 0.474289168 | 0.66092838 | 0.71761054 | 0.47299746 | 0.65907229 |
| BX276092.9 | 6.89737437 | -0.376517471 | 0.55087942 | -0.6834844 | 0.49430082 | 0.67673841 |
| LINC00891 | 4.88322473 | 0.091524711 | 0.53099031 | 0.17236607 | 0.86314975 | 0.92871584 |
| CXorf49 | 0 | NA | NA | NA | NA | NA |
| CXorf49B | 0.08666371 | -0.642905668 | 3.30558948 | -0.1944905 | 0.84579185 | NA |
| NHSL2 | 361.917153 | 0.489226562 | 0.34753875 | 1.40768926 | 0.1592231 | 0.32690075 |
| RTL5 | 682.425395 | 0.792323491 | 0.21824682 | 3.63040108 | 0.00028298 | 0.00289795 |
| PIN4 | 927.859384 | 0.008688049 | 0.17570427 | 0.049447 | 0.96056307 | 0.98098634 |
| ERCC6L | 10.9149898 | -0.407431153 | 0.61461394 | -0.6629058 | 0.50739088 | 0.6863686 |
| RPS4X | 35321.001 | -0.329386343 | 0.24236548 | -1.3590481 | 0.17413135 | 0.3481043 |
| CITED1 | 110.390306 | -3.827510328 | 0.74221354 | -5.1568856 | 2.51E-07 | 1.02E-05 |
| HDAC8 | 1024.51196 | -0.016910571 | 0.14394947 | -0.1174757 | 0.90648306 | 0.95326042 |
| PHKA1 | 547.23494 | -0.420605954 | 0.2994522 | -1.4045846 | 0.16014484 | 0.32825646 |
| PHKA1-AS1 | 1.11689286 | -2.672590425 | 1.5382701 | -1.7374 | 0.0823166 | 0.20659452 |
| DMRTC1B | 3.18210006 | -0.586439181 | 0.94135436 | -0.6229739 | 0.53330169 | 0.70711813 |
| FAM236B | 0 | NA | NA | NA | NA | NA |
| FAM236D | 0 | NA | NA | NA | NA | NA |
| DMRTC1 | 0.37171431 | -0.032084529 | 1.91353623 | -0.0167671 | 0.98662239 | NA |
| FAM236C | 0 | NA | NA | NA | NA | NA |
| FAM236A | 0 | NA | NA | NA | NA | NA |
| PABPC1L2B-/ | 0 | NA | NA | NA | NA | NA |
| PABPC1L2B | 0.08666371 | -0.642905668 | 3.30558948 | -0.1944905 | 0.84579185 | NA |
| PABPC1L2A | 0.11223674 | 0.02007176 | 3.29738704 | 0.00608717 | 0.99514317 | NA |
| AL662864.1 | 0 | NA | NA | NA | NA | NA |
| NAP1L2 | 455.853208 | 0.657937607 | 0.30600939 | 2.15005692 | 0.03155071 | 0.1046268 |
| CDX4 | 0 | NA | NA | NA | NA | NA |
| CHIC1 | 935.714465 | 0.083754203 | 0.10377594 | 0.80706766 | 0.4196275 | 0.61244062 |
| TSIX | 7163.52884 | -0.755010161 | 2.33590858 | -0.3232191 | 0.74652935 | 0.85800118 |
| XIST | 7315.58192 | -0.741389696 | 2.63450598 | -0.2814151 | 0.77839206 | 0.87861887 |
| JPX | 1371.35535 | 0.007840781 | 0.24016688 | 0.03264722 | 0.97395591 | 0.98796805 |
| FTX | 4282.69711 | -0.067332026 | 0.18522563 | -0.3635135 | 0.71622129 | 0.83854488 |
| AL353804.1 | 13.8815448 | 0.778326774 | 0.387122 | 2.01054647 | 0.04437338 | 0.13346939 |
| ZCCHC13 | 0 | NA | NA | NA | NA | NA |
| SLC16A2 | 1808.17495 | 0.734556164 | 0.27515898 | 2.66956998 | 0.00759484 | 0.03677489 |
| RLIM | 2214.01005 | -0.26800024 | 0.13047233 | -2.0540772 | 0.03996822 | 0.12377708 |
| NEXMIF | 205.863782 | -0.427986039 | 0.29450249 | -1.453251 | 0.14615408 | 0.30887709 |
| ABCB7 | 1283.64591 | -0.091123891 | 0.11609778 | -0.7848892 | 0.43251855 | 0.62454901 |
| UPRT | 690.842273 | 0.161527925 | 0.11590201 | 1.39365934 | 0.1634205 | 0.33267806 |
| ZDHHC15 | 195.928056 | 0.45470541 | 0.20617785 | 2.20540375 | 0.02742578 | 0.09461247 |
| MAGEE2 | 6.64586127 | 0.795950397 | 0.48316161 | 1.64737923 | 0.09948012 | 0.23663096 |
| AL451105.2 | 0 | NA | NA | NA | NA | NA |
| PBDC1 | 1101.03227 | -0.801424993 | 0.30424679 | -2.634128 | 0.00843537 | 0.03961543 |
| MAGEE1 | 350.348234 | 0.373475977 | 0.13757674 | 2.71467388 | 0.00663411 | 0.03321462 |
| AC233296.1 | 0.74005056 | 0.402983554 | 1.87242187 | 0.21522049 | 0.82959542 | NA |
| FGF16 | 2.77571842 | -0.334171384 | 0.93249953 | -0.3583609 | 0.72007324 | 0.84074967 |
| ATRX | 2156.47734 | 0.258068546 | 0.10270744 | 2.51265682 | 0.01198258 | 0.05148699 |
| MAGT1 | 4428.06759 | 0.197798685 | 0.13799365 | 1.43338978 | 0.15174648 | 0.31680426 |
| COX7B | 7248.52095 | 0.05911233 | 0.14498548 | 0.40771206 | 0.68348508 | 0.81717 |
| ATP7A | 624.094158 | 0.192043143 | 0.11471395 | 1.67410451 | 0.09411006 | 0.22794914 |
| PGK1 | 29243.1295 | -0.439130444 | 0.19415257 | -2.2617802 | 0.02371099 | 0.08495363 |
| PGAM4 | 3.18694399 | -0.848660418 | 0.86236342 | -0.9841099 | 0.32506146 | 0.52264317 |
| TAF9B | 1413.72167 | 0.040088491 | 0.16176529 | 0.24781887 | 0.80427456 | 0.89390914 |
| CYSLTR1 | 86.3128193 | 0.631789716 | 0.42360726 | 1.49145158 | 0.13584297 | 0.29362125 |
| RTL3 | 1.07993639 | -0.774424531 | 1.53020001 | -0.5060937 | 0.6127909 | 0.76799695 |
| LPAR4 | 3.1737182 | 0.194439179 | 0.66913702 | 0.29058201 | 0.77137102 | 0.87390676 |
| P2RY10 | 31.1364662 | 1.007144255 | 0.78813735 | 1.27787911 | 0.20129206 | 0.38306066 |
| GPR174 | 81.357902 | 0.637492658 | 0.75109634 | 0.84874951 | 0.39602069 | 0.59122987 |
| ITM2A | 839.029546 | 0.562613034 | 0.47715872 | 1.17908991 | 0.23836237 | 0.42788399 |
| TBX22 | 2.04994275 | 0.968059164 | 1.19892547 | 0.80743898 | 0.41941361 | 0.61235293 |
| TENT5D | 1.704594 | 0.506611675 | 0.91960244 | 0.55090292 | 0.58170022 | 0.74428759 |
| BRWD3 | 778.063715 | -0.154949714 | 0.13769635 | -1.1253001 | 0.26046189 | 0.45355506 |
| AL391294.1 | 0.97013515 | 1.207084022 | 1.30548346 | 0.92462605 | 0.35516046 | NA |
| HMGN5 | 430.076513 | 0.59004766 | 0.27909257 | 2.11416468 | 0.0345012 | 0.11152203 |
| SH3BGRL | 5543.05375 | 0.13689721 | 0.21333911 | 0.64168829 | 0.52107559 | 0.69754901 |

| | | | | | | |
|---|---|---|---|---|---|---|
| POU3F4 | 100.253301 | -4.46E-05 | 0.57066728 | -7.81E-05 | 0.9999377 | 0.99993924 |
| CYLC1 | 0.08757577 | 0.547239394 | 3.29377202 | 0.16614368 | 0.86804388 | NA |
| RPS6KA6 | 836.705532 | 0.436971274 | 0.20875132 | 2.09326229 | 0.03632575 | 0.11558876 |
| HDX | 230.443073 | -0.689567234 | 0.26421777 | -2.6098443 | 0.00905834 | 0.04187738 |
| APOOL | 1725.57394 | 0.592081198 | 0.23375615 | 2.532901 | 0.01131229 | 0.04942099 |
| SATL1 | 588.087883 | 0.311677573 | 0.23222146 | 1.34215662 | 0.17954522 | 0.35525717 |
| AC003001.1 | 0.64905605 | 0.036779453 | 1.63211093 | 0.0225349 | 0.98202127 | NA |
| ZNF711 | 541.300314 | 0.740891954 | 0.28757568 | 2.57633732 | 0.00998532 | 0.0450228 |
| POF1B | 154.505535 | -0.434842093 | 0.39402747 | -1.1035832 | 0.26977399 | 0.46402671 |
| CHM | 1006.2475 | 0.297629959 | 0.13860619 | 2.14730645 | 0.03176889 | 0.10505979 |
| DACH2 | 62.7745601 | 0.128763087 | 0.59416221 | 0.21671369 | 0.82843148 | 0.90876235 |
| KLHL4 | 107.556104 | 1.628873444 | 0.3347371 | 4.86612758 | 1.14E-06 | 3.62E-05 |
| Z83818.2 | 0 | NA | NA | NA | NA | NA |
| CPXCR1 | 0 | NA | NA | NA | NA | NA |
| AL121872.1 | 0 | NA | NA | NA | NA | NA |
| TGIF2LX | 0 | NA | NA | NA | NA | NA |
| AL121823.1 | 0 | NA | NA | NA | NA | NA |
| PABPC5-AS1 | 1.3060349 | -1.373527284 | 1.4443332 | -0.9509767 | 0.34161618 | 0.53958782 |
| PABPC5 | 45.5176391 | 0.034289252 | 0.23292069 | 0.14721428 | 0.88296289 | 0.93920098 |
| PCDH11X | 2.13738515 | 0.948803645 | 0.99857191 | 0.95016056 | 0.34203068 | 0.5398868 |
| NAP1L3 | 140.025481 | 1.329504125 | 0.3162768 | 4.20360936 | 2.63E-05 | 0.00044207 |
| FAM133A | 3.427785 | 1.493440944 | 0.73797887 | 2.02369066 | 0.04300199 | 0.13062579 |
| DIAPH2 | 599.092922 | -0.22856127 | 0.17000508 | -1.3444379 | 0.17880683 | 0.35441492 |
| RPA4 | 9.78934207 | -1.971707105 | 0.86278982 | -2.2852693 | 0.02229704 | 0.08130157 |
| DIAPH2-AS1 | 26.9232836 | 0.404186141 | 0.25545752 | 1.58220492 | 0.1136028 | 0.25981837 |
| AL157778.1 | 0.09644756 | -0.642905668 | 3.30558948 | -0.1944905 | 0.84579185 | NA |
| PCDH19 | 35.8700463 | 1.007064141 | 0.36634182 | 2.748974 | 0.00597821 | 0.03069421 |
| TNMD | 29.111167 | 2.207562952 | 0.64196498 | 3.43875917 | 0.00058439 | 0.00511969 |
| TSPAN6 | 2893.23261 | 0.309610587 | 0.19766041 | 1.56637633 | 0.11726054 | 0.26567235 |
| SRPX2 | 1505.44064 | -0.717148781 | 0.43893382 | -1.6338426 | 0.1022919 | 0.24118718 |
| SYTL4 | 1220.93366 | -0.27529479 | 0.35635057 | -0.7725392 | 0.43979513 | 0.63111654 |
| CSTF2 | 529.769186 | -0.057493452 | 0.13797829 | -0.4166848 | 0.67690899 | 0.81209758 |
| NOX1 | 27.426553 | 0.022327886 | 0.29242775 | 0.07635352 | 0.93913785 | 0.97102671 |
| XKRX | 18.254578 | -0.662379345 | 0.4283953 | -1.5461872 | 0.12205936 | 0.27295668 |
| ARL13A | 12.4055782 | 0.754681682 | 0.41294496 | 1.82755998 | 0.06761562 | 0.17966786 |
| TRMT2B | 1077.25018 | 0.266621558 | 0.1036801 | 2.57157891 | 0.01012359 | 0.04547856 |
| TMEM35A | 29.2310314 | 0.588469035 | 0.74796779 | 0.78675719 | 0.43142402 | 0.62348065 |
| CENPI | 15.8113484 | 0.049631277 | 0.60597197 | 0.08190358 | 0.93472338 | 0.96870331 |
| DRP2 | 9.01320441 | 0.36795106 | 0.7343656 | 0.50104616 | 0.61633864 | 0.77049635 |
| TAF7L | 27.3616662 | 0.423491292 | 0.69149977 | 0.61242434 | 0.54025704 | 0.71329185 |
| TIMM8A | 307.26605 | 0.210663965 | 0.19998153 | 1.05341713 | 0.29214986 | 0.48836714 |
| BTK | 178.612897 | -0.223434224 | 0.45727891 | -0.4886169 | 0.62511291 | 0.77667864 |
| RPL36A | 8535.8076 | -0.620955704 | 0.16385121 | -3.7897536 | 0.0001508 | 0.00176072 |
| RPL36A-HNR | 9564.60264 | -0.488623895 | 0.15618073 | -3.1285799 | 0.00175653 | 0.01221835 |
| GLA | 2309.16746 | -1.407197228 | 0.48983057 | -2.8728245 | 0.0040682 | 0.02305081 |
| HNRNPH2 | 6591.13668 | -0.159268502 | 0.15328594 | -1.0390288 | 0.29879134 | 0.49563286 |
| ARMCX4 | 982.23447 | 0.335197981 | 0.22097543 | 1.5169016 | 0.12929153 | 0.28402346 |
| ARMCX1 | 1373.31692 | -0.339875851 | 0.30830374 | -1.1024059 | 0.27028527 | 0.46459961 |
| ARMCX6 | 770.421836 | -0.289101921 | 0.12104112 | -2.3884604 | 0.01691913 | 0.06645181 |
| ARMCX3 | 4093.49637 | -0.056341086 | 0.12469241 | -0.4518406 | 0.65138385 | 0.79512068 |
| ARMCX3-AS1 | 349.470807 | 0.566060795 | 0.23427156 | 2.41625914 | 0.0156809 | 0.06287141 |
| ARMCX2 | 2889.48769 | -0.222750064 | 0.18721748 | -1.1897931 | 0.23412774 | 0.42272673 |
| NXF5 | 13.6896221 | 1.067116126 | 0.53640992 | 1.98936687 | 0.04666072 | 0.13848546 |
| ZMAT1 | 1264.82109 | 0.766066488 | 0.40713375 | 1.8816089 | 0.05988914 | 0.16530004 |
| TCEAL2 | 392.582228 | 0.03078103 | 0.32874902 | 0.09363079 | 0.92540245 | 0.96392839 |
| TCEAL6 | 14.4989331 | -0.921451849 | 0.60870364 | -1.5137939 | 0.13007813 | 0.28521479 |
| BEX5 | 290.743449 | 0.044120692 | 0.28852928 | 0.15291582 | 0.87846466 | 0.93677394 |
| TCP11X1 | 0.09635527 | 0.547239394 | 3.29377202 | 0.16614368 | 0.86804388 | NA |
| NXF2 | 0.54884854 | 1.754246905 | 1.80915707 | 0.96964876 | 0.3322216 | NA |
| AC235565.2 | 1.88624112 | 1.683756758 | 0.94312143 | 1.7853022 | 0.07421231 | 0.19231316 |
| NXF2B | 0.40501264 | 1.032842498 | 1.86590893 | 0.55353318 | 0.57989837 | NA |
| TCP11X2 | 1.6773973 | 1.439950208 | 0.94433569 | 1.52482875 | 0.12730182 | 0.28088759 |
| TMSB15A | 23.6333445 | 0.508344223 | 0.5061184 | 1.00439784 | 0.31518689 | 0.51235842 |
| ARMCX5 | 739.855879 | 0.175693229 | 0.18042182 | 0.97379146 | 0.3301601 | 0.52807192 |
| AL035427.1 | 86.9417354 | 0.328421405 | 0.20367939 | 1.61244303 | 0.10686557 | 0.24863641 |
| GPRASP1 | 1029.09036 | 0.664914743 | 0.3854232 | 1.72515495 | 0.08449955 | 0.21062985 |
| GPRASP2 | 964.239459 | 0.915205783 | 0.23442972 | 3.90396652 | 9.46E-05 | 0.00121252 |
| BHLHB9 | 382.181837 | 0.633681087 | 0.19128297 | 3.31279404 | 0.00092369 | 0.0073856 |
| LINC00630 | 61.3168857 | -0.108560256 | 0.23032589 | -0.4713333 | 0.63740276 | 0.78543508 |
| Z68871.1 | 21.5575214 | 0.079551142 | 0.29545121 | 0.26925306 | 0.78773495 | 0.88439078 |
| RAB40AL | 7.40504566 | -1.451812894 | 0.75005699 | -1.9356035 | 0.05291629 | 0.15157729 |
| Z95624.1 | 5.74838148 | -1.301000425 | 0.85423854 | -1.5229943 | 0.12776013 | 0.28157415 |
| BEX1 | 235.633328 | -0.04552296 | 0.53626346 | -0.0848892 | 0.9323495 | 0.96734562 |
| NXF3 | 66.4441236 | 0.755006847 | 0.47266697 | 1.59733362 | 0.11019136 | 0.25419877 |
| BEX4 | 3488.46849 | 0.04249305 | 0.17387175 | 0.24439307 | 0.80692641 | 0.89560375 |
| TCEAL8 | 2134.82802 | -0.20486816 | 0.08230771 | -2.4890518 | 0.01280843 | 0.05414529 |
| TCEAL5 | 15.9317854 | -1.984003491 | 0.81405843 | -2.4371758 | 0.01480248 | 0.06024717 |
| BEX2 | 1549.97204 | 0.043703764 | 0.28785319 | 0.15182658 | 0.87932372 | 0.93729603 |
| TCEAL7 | 122.317476 | -0.159760588 | 0.23981868 | -0.6661724 | 0.50530091 | 0.68458616 |
| TCEAL9 | 3087.43617 | -0.850578592 | 0.18758657 | -4.5343256 | 5.78E-06 | 0.00013194 |
| BEX3 | 5312.93362 | 0.136899216 | 0.20666668 | 0.66241553 | 0.50770495 | 0.68658055 |
| AL606763.1 | 17.5332003 | 0.350190877 | 0.44374759 | 0.78916682 | 0.43001451 | 0.62207863 |
| Z69733.1 | 46.7634732 | -0.257487695 | 0.38016588 | -0.6773035 | 0.49821339 | 0.67983892 |
| RAB40A | 98.2241488 | -0.165985963 | 0.30149897 | -0.5505357 | 0.58195196 | 0.74445491 |
| TCEAL4 | 4535.7829 | -0.415633121 | 0.16631854 | -2.4990185 | 0.01245378 | 0.05303658 |
| TCEAL3 | 1876.68222 | -0.612037514 | 0.22444907 | -2.7268436 | 0.00639433 | 0.03230241 |
| TCEAL3-AS1 | 0.69155369 | 1.044153673 | 1.78440157 | 0.58515622 | 0.55844268 | NA |
| TCEAL1 | 1176.12912 | -0.228165348 | 0.18050894 | -1.2640113 | 0.20622596 | 0.38883016 |
| MORF4L2 | 9703.06602 | -0.587464129 | 0.16897421 | -3.4766497 | 0.00050772 | 0.00458828 |
| MORF4L2-AS | 219.653204 | 0.33383108 | 0.16832329 | 1.98327326 | 0.04733691 | 0.13989403 |
| GLRA4 | 7.84556697 | -0.198436324 | 0.55481345 | -0.3576631 | 0.72059541 | 0.84102421 |
| TMEM31 | 13.5597997 | -0.120655345 | 0.5240775 | -0.2302242 | 0.81791752 | 0.9021451 |
| PLP1 | 80.3519405 | -0.514991172 | 0.84959355 | -0.6061618 | 0.54440731 | 0.7163683 |
| RAB9B | 44.7130947 | 0.1646568 | 0.4800143 | 0.34302478 | 0.73157982 | 0.84851161 |
| TMSB15B-AS | 6.45727795 | -0.473003824 | 0.57363692 | -0.8245701 | 0.40961573 | 0.60405105 |
| TMSB15B | 10.978306 | -0.285933856 | 0.47313479 | -0.6043391 | 0.54561821 | 0.71712709 |
| H2BFWT | 0.03392908 | 0.380740593 | 3.29738709 | 0.11546736 | 0.90807469 | NA |
| H2BFM | 0.38432268 | -1.056481192 | 2.28623022 | -0.4621062 | 0.64400515 | NA |
| SLC25A53 | 127.105366 | -0.091477258 | 0.20044188 | -0.456378 | 0.6481182 | 0.79279496 |
| ZCCHC18 | 42.1466318 | 0.162465717 | 0.44959522 | 0.36135997 | 0.71783037 | 0.83983935 |
| FAM199X | 3471.39539 | -0.381657283 | 0.12509191 | -3.0510148 | 0.00228069 | 0.01493837 |
| ESX1 | 0.17332742 | -1.003574173 | 3.30558948 | -0.3035992 | 0.7614333 | NA |
| IL1RAPL2 | 8.60266142 | 0.857550082 | 0.738482 | 1.16123357 | 0.24554693 | 0.4367082 |
| TEX13A | 0 | NA | NA | NA | NA | NA |
| NRK | 805.410116 | 1.443007539 | 0.30102618 | 4.79362802 | 1.64E-06 | 4.86E-05 |
| SERPINA7 | 7.30281421 | 2.486162036 | 0.67042037 | 3.70836291 | 0.0002086 | 0.00225723 |
| MUM1L1 | 114.49789 | 1.014603255 | 0.46579181 | 2.17823337 | 0.02938867 | 0.09942003 |
| CXorf57 | 269.542822 | 0.118320364 | 0.20124608 | 0.58793874 | 0.55657341 | 0.72557054 |
| RNF128 | 2340.20488 | 0.259939737 | 0.232512 | 1.11796265 | 0.26358294 | 0.457372 |
| TBC1D8B | 745.287132 | 0.231494115 | 0.30955912 | 0.74781874 | 0.4545695 | 0.64404672 |
| MORC4 | 924.800487 | -0.053426162 | 0.1255283 | -0.4256105 | 0.6703917 | 0.80748317 |
| RIPPLY1 | 66.2667773 | 0.225125145 | 0.51003017 | 0.44139574 | 0.65892652 | 0.79962689 |
| CLDN2 | 14908.1946 | 0.571194061 | 0.43203349 | 1.32210597 | 0.18613287 | 0.36366704 |

| | | | | | | |
|---|---|---|---|---|---|---|
| RBM41 | 368.872126 | -0.001030715 | 0.22701302 | -0.0045403 | 0.99637735 | 0.99823988 |
| NUP62CL | 207.935516 | -0.091605524 | 0.3313173 | -0.2764888 | 0.78217266 | 0.8807709 |
| PIH1D3 | 1.69283999 | -0.791126763 | 1.02422746 | -0.7724132 | 0.43986977 | 0.63115495 |
| FRMPD3-AS1 | 0.16497137 | -1.003574173 | 3.30558948 | -0.3035992 | 0.7614333 | NA |
| FRMPD3 | 10.4946386 | -0.2613641 | 0.59112972 | -0.4421434 | 0.65838545 | 0.79946078 |
| PRPS1 | 1618.32022 | -0.020546173 | 0.20009454 | -0.1026823 | 0.9182151 | 0.96022749 |
| TSC22D3 | 7789.44878 | -0.450384344 | 0.45050939 | -0.9997224 | 0.31744485 | 0.51466467 |
| NCBP2L | 78.7784171 | 0.227224583 | 0.24844161 | 0.91459955 | 0.36040189 | 0.5580355 |
| MID2 | 342.299396 | -1.204570959 | 0.29484409 | -4.0854506 | 4.40E-05 | 0.00066978 |
| AL034399.2 | 1.96680787 | -4.189722889 | 1.79198918 | -2.3380291 | 0.01938574 | 0.07324158 |
| TEX13B | 0.17303784 | -1.003574173 | 3.30558948 | -0.3035992 | 0.7614333 | NA |
| VSIG1 | 11.1682722 | -0.939737387 | 0.62983561 | -1.492036 | 0.13568972 | 0.29339127 |
| PSMD10 | 1434.78912 | -0.165797711 | 0.0932284 | -1.7784034 | 0.07533762 | 0.19435925 |
| ATG4A | 1110.80656 | -0.002160242 | 0.13391107 | -0.0161319 | 0.98712915 | 0.99410333 |
| COL4A6 | 486.92506 | 1.527727139 | 0.30507455 | 5.00771736 | 5.51E-07 | 1.97E-05 |
| COL4A5 | 1632.13614 | 0.585580476 | 0.2573053 | 2.27581968 | 0.0228568 | 0.08274766 |
| AL035425.2 | 0.85295111 | 2.026629719 | 1.33460927 | 1.51851914 | 0.12888358 | NA |
| IRS4 | 0.53181971 | -0.022190739 | 2.16432368 | -0.010253 | 0.99181946 | NA |
| AL035425.1 | 4.48529697 | -0.299042796 | 0.85036586 | -0.3516637 | 0.7250905 | 0.84414094 |
| GUCY2F | 0.77404102 | -2.855773329 | 1.8241669 | -1.565522 | 0.11746057 | NA |
| NXT2 | 387.289413 | 0.302586053 | 0.17164463 | 1.76286354 | 0.07792351 | 0.19871861 |
| KCNE5 | 45.789313 | -2.627012284 | 0.4827982 | -5.4412222 | 5.29E-08 | 2.78E-06 |
| ACSL4 | 9554.03356 | -1.450745014 | 0.27888444 | -5.2019575 | 1.97E-07 | 8.45E-06 |
| TMEM164 | 2323.26043 | 0.135647506 | 0.12702291 | 1.06789791 | 0.28556656 | 0.48163515 |
| AMMECR1 | 575.560955 | -0.158507116 | 0.18896714 | -0.8388078 | 0.40157715 | 0.59649578 |
| RTL9 | 5.40923503 | 1.242912309 | 0.56821814 | 2.18738585 | 0.02871437 | 0.09780826 |
| CHRDL1 | 1223.98013 | 0.908453784 | 0.47758691 | 1.90217479 | 0.05714831 | 0.15987703 |
| PAK3 | 58.9617937 | -0.115300555 | 0.38755545 | -0.2975072 | 0.76607928 | 0.8710362 |
| CAPN6 | 570.996159 | 1.031193531 | 0.58114198 | 1.77442616 | 0.07599269 | 0.19559832 |
| DCX | 3.74903746 | 1.508439681 | 0.69231601 | 2.17883114 | 0.02934421 | 0.09929513 |
| SERTM2 | 34.6611894 | -1.093571106 | 0.78917881 | -1.3857076 | 0.1658362 | 0.3363406 |
| ALG13 | 1905.68699 | -0.029529941 | 0.13773557 | -0.2143959 | 0.83023835 | 0.91002515 |
| ALG13-AS1 | 1.13121439 | 0.255962764 | 1.1897551 | 0.21513904 | 0.82965892 | 0.90950346 |
| TRPC5 | 4.13379849 | 1.139378291 | 0.8823008 | 1.2913717 | 0.19657482 | 0.37715934 |
| TRPC5OS | 9.71831342 | 2.249575473 | 0.44492877 | 5.05603512 | 4.28E-07 | 1.60E-05 |
| RTL4 | 33.2137215 | 3.221800495 | 0.59915433 | 5.37724646 | 7.56E-08 | 3.76E-06 |
| LHFPL1 | 3.548451 | 0.853464458 | 0.96867871 | 0.88106041 | 0.37828512 | 0.57427972 |
| AMOT | 1407.15213 | 0.789090759 | 0.26985337 | 2.92414641 | 0.00345402 | 0.02038554 |
| XACT | 0.03392908 | 0.380740593 | 3.29738709 | 0.11546736 | 0.90807469 | NA |
| HTR2C | 0.16783122 | 1.368031398 | 3.28033652 | 0.41703996 | 0.67664918 | NA |
| IL13RA2 | 62.314676 | 1.272888442 | 0.75702166 | 1.68144256 | 0.09267698 | 0.22528504 |
| LRCH2 | 165.800355 | -0.479656912 | 0.33937591 | -1.4133499 | 0.15755287 | 0.32471037 |
| RBMXL3 | 0 | NA | NA | NA | NA | NA |
| LUZP4 | 0.21442573 | -1.003574173 | 3.30558948 | -0.3035992 | 0.7614333 | NA |
| PLS3-AS1 | 42.7831679 | 0.942833933 | 0.34535529 | 2.7300405 | 0.00633265 | 0.03207685 |
| PLS3 | 7552.56388 | 0.446957639 | 0.18586517 | 2.40474118 | 0.01618392 | 0.06436684 |
| DANT1 | 0.18301898 | 0.18657164 | 3.29377195 | 0.05664376 | 0.95482897 | NA |
| DANT2 | 9.49173169 | -0.066700187 | 0.47115885 | -0.1415662 | 0.88742264 | 0.94177941 |
| AGTR2 | 26.7518759 | 5.40976934 | 1.08686289 | 4.97741657 | 6.44E-07 | 2.25E-05 |
| SLC6A14 | 5.2122855 | -3.489276504 | 1.5430683 | -2.2612586 | 0.02374325 | 0.08501146 |
| CT83 | 0 | NA | NA | NA | NA | NA |
| BX119904.2 | 0.23568937 | 1.881156793 | 3.2502805 | 0.57876752 | 0.56274605 | NA |
| KLHL13 | 1463.493 | -0.001612925 | 0.20596348 | -0.0078311 | 0.99375173 | 0.99711705 |
| WDR44 | 743.601317 | -0.057120319 | 0.17657619 | -0.3234882 | 0.74632553 | 0.85787909 |
| DOCK11 | 854.151696 | 0.084822227 | 0.30745728 | 0.27588297 | 0.78263795 | 0.88106052 |
| IL13RA1 | 4643.76056 | -0.354993133 | 0.19944121 | -1.7799387 | 0.07508599 | 0.19387228 |
| ZCCHC12 | 4.56044048 | -0.774460412 | 0.63526712 | -1.2191099 | 0.2228025 | 0.40880196 |
| LINC01285 | 9.34017146 | 1.734597198 | 0.58412176 | 2.96958157 | 0.00298206 | 0.01824944 |
| LONRF3 | 127.573316 | -0.467634422 | 0.42276955 | -1.1061214 | 0.268674 | 0.46285577 |
| AL772284.2 | 0.03392908 | 0.380740593 | 3.29738709 | 0.11546736 | 0.90807469 | NA |
| KIAA1210 | 13.8953827 | 2.197877163 | 1.26977981 | 1.73091205 | 0.08346745 | 0.20874779 |
| PGRMC1 | 10803.235 | 0.408176501 | 0.20160224 | 2.02466252 | 0.04290203 | 0.13038222 |
| AC004835.1 | 0.28934269 | -1.364242679 | 3.30558948 | -0.4127078 | 0.67982068 | NA |
| AC004973.1 | 0.99633267 | -0.401454225 | 2.2788637 | -0.1761642 | 0.86016494 | NA |
| SLC25A43 | 1034.86352 | -0.256901526 | 0.20193443 | -1.2722027 | 0.20330111 | 0.38549293 |
| SLC25A5 | 22516.9079 | 0.4751573 | 0.25871875 | 1.83657853 | 0.06627214 | 0.17726497 |
| CXorf56 | 960.092589 | -0.042418048 | 0.12474923 | -0.3400265 | 0.73383654 | 0.85011947 |
| UBE2A | 3719.01241 | -0.479241872 | 0.14017367 | -3.4189152 | 0.00062871 | 0.00541626 |
| NKRF | 699.108464 | -0.10126007 | 0.1159316 | -0.8734467 | 0.38241966 | 0.57853286 |
| 6-Sep | 2234.37086 | 0.116864498 | 0.24970137 | 0.46801705 | 0.6397724 | 0.78704634 |
| SOWAHD | 43.4629742 | -0.466733986 | 0.30604408 | -1.5250548 | 0.12724544 | 0.28080466 |
| RPL39 | 5856.94298 | -0.626767942 | 0.15536111 | -4.0342652 | 5.48E-05 | 0.00079888 |
| UPF3B | 792.55363 | -0.296074523 | 0.12174735 | -2.4318764 | 0.01502083 | 0.06092898 |
| RNF113A | 641.577708 | -0.588458701 | 0.10359187 | -5.6805492 | 1.34E-08 | 8.96E-07 |
| NDUFA1 | 5299.16936 | 0.012227983 | 0.14056073 | 0.08699431 | 0.93067604 | 0.96662347 |
| AKAP14 | 4.23368019 | -1.289896689 | 0.91199013 | -1.4143757 | 0.15725162 | 0.32431763 |
| NKAP | 864.185882 | -0.173108821 | 0.10744414 | -1.6111518 | 0.10714664 | 0.24913947 |
| RHOXF1-AS1 | 32.6546614 | -0.778323165 | 0.58075809 | -1.3401848 | 0.18018527 | 0.35610325 |
| RHOXF2B | 0 | NA | NA | NA | NA | NA |
| RHOXF1 | 43.9998394 | 0.160010652 | 0.40572161 | 0.39438533 | 0.69329657 | 0.82352701 |
| LINC01402 | 8.47467715 | 0.808395824 | 0.48639823 | 1.66200405 | 0.09651196 | 0.23185027 |
| RHOXF2 | 0.57543764 | -1.130176205 | 1.91985215 | -0.5886788 | 0.55607678 | NA |
| ZBTB33 | 778.405228 | -0.272675733 | 0.17631009 | -1.5465691 | 0.12196719 | 0.27284511 |
| TMEM255A | 210.301084 | -0.729066279 | 0.84777183 | -0.8599794 | 0.38980042 | 0.58538532 |
| ATP1B4 | 0.98764676 | -3.17493279 | 2.18825754 | -1.4508954 | 0.14680899 | NA |
| LAMP2 | 12314.0264 | -0.233217387 | 0.17415929 | -1.3391039 | 0.18053684 | 0.35650414 |
| CUL4B | 2340.4762 | 0.014487445 | 0.11128289 | 0.13018574 | 0.89641948 | 0.94698268 |
| MCTS1 | 1186.70833 | -0.49234408 | 0.14105367 | -3.4904733 | 0.00048217 | 0.0044026 |
| C1GALT1C1 | 1606.17006 | -0.410039129 | 0.13513544 | -3.0342828 | 0.00241108 | 0.0155529 |
| CT47B1 | 0 | NA | NA | NA | NA | NA |
| AC008162.2 | 0 | NA | NA | NA | NA | NA |
| CT47A12 | 0 | NA | NA | NA | NA | NA |
| CT47A11 | 0 | NA | NA | NA | NA | NA |
| CT47A10 | 0 | NA | NA | NA | NA | NA |
| CT47A9 | 0 | NA | NA | NA | NA | NA |
| CT47A8 | 0 | NA | NA | NA | NA | NA |
| CT47A7 | 0 | NA | NA | NA | NA | NA |
| CT47A6 | 0 | NA | NA | NA | NA | NA |
| CT47A5 | 0 | NA | NA | NA | NA | NA |
| CT47A4 | 0 | NA | NA | NA | NA | NA |
| CT47A3 | 0 | NA | NA | NA | NA | NA |
| CT47A2 | 0 | NA | NA | NA | NA | NA |
| CT47A1 | 0 | NA | NA | NA | NA | NA |
| GLUD2 | 156.281776 | 1.394408785 | 0.24671977 | 5.6517919 | 1.59E-08 | 1.03E-06 |
| AL359851.1 | 1.00695846 | 0.875317663 | 1.13044025 | 0.77431573 | 0.4387441 | 0.63036297 |
| GRIA3 | 140.484611 | 1.310364275 | 0.49278305 | 2.65910986 | 0.00783474 | 0.03762559 |
| THOC2 | 2250.20787 | -0.125074621 | 0.16055143 | -0.7790315 | 0.43596117 | 0.6280413 |
| XIAP | 2426.71289 | -0.043037752 | 0.14613879 | -0.2944992 | 0.7683765 | 0.87201276 |
| XIAP-AS1 | 0.31722449 | -0.695051713 | 2.30489341 | -0.3015548 | 0.76299145 | NA |
| AL121601.1 | 21.8776465 | 0.756155504 | 0.34289798 | 2.2051909 | 0.0274407 | 0.09465158 |
| STAG2 | 3033.54926 | 0.146644839 | 0.0927099 | 1.58176032 | 0.1137043 | 0.25996032 |
| SH2D1A | 54.0621257 | -0.062928313 | 0.81278256 | -0.0774233 | 0.9382868 | 0.97060692 |
| TEX13D | 0.12150485 | 1.04981648 | 3.28477129 | 0.31960109 | 0.74927075 | NA |

| | | | | | | |
|---|---|---|---|---|---|---|
| TENM1 | 336.595323 | 1.036753945 | 0.4234219 | 2.44851279 | 0.01434473 | 0.05893902 |
| TEX13C | 0.39442346 | 1.870085995 | 1.79976333 | 1.03907329 | 0.29877065 | NA |
| AL445072.1 | 0.14114194 | 0.02007176 | 3.29738704 | 0.00608717 | 0.99514317 | NA |
| DCAF12L2 | 22.5044764 | -0.023421204 | 0.3946093 | -0.0593529 | 0.95267103 | 0.97717271 |
| DCAF12L1 | 255.885768 | -0.279659882 | 0.33490338 | -0.8350465 | 0.40369155 | 0.59870627 |
| PRR32 | 0.63167585 | -0.886327943 | 2.240135 | -0.3956583 | 0.69235714 | NA |
| AL591643.1 | 0 | NA | NA | NA | NA | NA |
| ACTRT1 | 0.60137097 | 0.302515464 | 1.87020958 | 0.16175485 | 0.87149891 | NA |
| AL442647.1 | 2.5508534 | -1.321971556 | 0.88796988 | -1.4887572 | 0.13655132 | 0.29487545 |
| SMARCA1 | 2678.58905 | -0.186979728 | 0.18500427 | -1.0106779 | 0.31217062 | 0.50910117 |
| OCRL | 1571.00211 | 0.011792498 | 0.12803194 | 0.09210591 | 0.9266139 | 0.96438573 |
| APLN | 780.466731 | -0.175342417 | 0.68812682 | -0.2548112 | 0.79886893 | 0.89107318 |
| XPNPEP2 | 13473.5432 | 2.656258041 | 0.67789982 | 3.91836369 | 8.92E-05 | 0.00115639 |
| SASH3 | 537.811827 | -0.410132921 | 0.52261399 | -0.7847722 | 0.43258715 | 0.62461387 |
| ZDHHC9 | 4486.96614 | -0.031127066 | 0.21157906 | -0.1471179 | 0.88303897 | 0.93924401 |
| AL034405.1 | 1.56637024 | -0.710650595 | 1.49359299 | -0.4757994 | 0.63421732 | 0.78318369 |
| UTP14A | 723.431677 | -0.332287547 | 0.18518608 | -1.7943441 | 0.07275826 | 0.18969885 |
| BCORL1 | 1000.40564 | -0.051578232 | 0.25343134 | -0.2035195 | 0.83872897 | 0.91450415 |
| ELF4 | 1061.02616 | -0.745190807 | 0.23467361 | -3.1754351 | 0.00149612 | 0.01078826 |
| AIFM1 | 9888.58363 | 0.586894622 | 0.3576602 | 1.64092794 | 0.10081237 | 0.23888843 |
| RAB33A | 44.308418 | -1.557387051 | 0.47871449 | -3.2532691 | 0.00114085 | 0.00870747 |
| ZNF280C | 183.984851 | -0.008495551 | 0.16339112 | -0.0519952 | 0.95853253 | 0.98016531 |
| SLC25A14 | 544.611112 | -0.168749666 | 0.14706465 | -1.1474523 | 0.25119475 | 0.44354643 |
| GPR119 | 0.21723982 | 0.689157019 | 3.06522228 | 0.22483101 | 0.82211074 | NA |
| RBMX2 | 727.850635 | -0.360642236 | 0.11902668 | -3.0299278 | 0.00244612 | 0.01572507 |
| ENOX2 | 438.796034 | -0.234161859 | 0.19115978 | -1.2249536 | 0.2205927 | 0.40651824 |
| LINC01201 | 0 | NA | NA | NA | NA | NA |
| ARHGAP36 | 0.51444143 | -1.018018245 | 2.80187185 | -0.363335 | 0.71635461 | NA |
| IGSF1 | 69.7625332 | 0.603416241 | 0.47274477 | 1.27641019 | 0.20181056 | 0.38371614 |
| OR13H1 | 0.07659028 | -0.642905668 | 3.30558948 | -0.1944905 | 0.84579185 | NA |
| FIRRE | 16.0488829 | -0.989929904 | 0.71345597 | -1.3875137 | 0.16528519 | 0.33562496 |
| STK26 | 372.434241 | -0.330368528 | 0.20963074 | -1.5759546 | 0.11503627 | 0.2623006 |
| FRMD7 | 77.5489349 | 2.798447488 | 0.7287775 | 3.83992024 | 0.00012307 | 0.0014867 |
| RAP2C | 2954.22916 | 0.32220361 | 0.17214031 | 1.87174992 | 0.06124121 | 0.16778501 |
| RAP2C-AS1 | 252.284398 | 0.85819625 | 0.1998577 | 4.29403656 | 1.75E-05 | 0.00032041 |
| MBNL3 | 920.356459 | 0.354849394 | 0.20365107 | 1.74243814 | 0.08143181 | 0.20493957 |
| HS6ST2 | 1049.80498 | 1.199229231 | 0.36553447 | 3.28075556 | 0.00103529 | 0.0080652 |
| HS6ST2-AS1 | 0.07659028 | -0.642905668 | 3.30558948 | -0.1944905 | 0.84579185 | NA |
| USP26 | 0.09997307 | 0.547239394 | 3.29377202 | 0.16614368 | 0.86804388 | NA |
| TFDP3 | 0.08666371 | -0.642905668 | 3.30558948 | -0.1944905 | 0.84579185 | NA |
| GPC4 | 5426.60832 | -0.657809009 | 0.19964817 | -3.2948411 | 0.00098477 | 0.00775641 |
| GPC3 | 4156.48137 | 1.02119977 | 0.53031061 | 1.92566347 | 0.05414641 | 0.15407799 |
| CCDC160 | 266.725268 | 0.262958153 | 0.38024383 | 0.6915514 | 0.48921909 | 0.67257742 |
| PHF6 | 990.027295 | 0.149951927 | 0.17389424 | 0.8623168 | 0.38851321 | 0.5841377 |
| HPRT1 | 1854.58368 | -0.605095717 | 0.19018712 | -3.1815809 | 0.00146474 | 0.01060546 |
| MIR503HG | 823.666401 | 0.604152178 | 0.33691701 | 1.79317804 | 0.07294446 | 0.18996103 |
| LINC00629 | 24.1539357 | 1.974014103 | 0.52220546 | 3.78014832 | 0.00015673 | 0.00181436 |
| PLAC1 | 1.36917304 | 0.739034207 | 1.07757287 | 0.68583223 | 0.49281891 | 0.67562436 |
| AL672032.1 | 0.10721287 | -0.642905668 | 3.30558948 | -0.1944905 | 0.84579185 | NA |
| FAM122B | 1710.29216 | -0.196745965 | 0.12718116 | -1.5469741 | 0.12186949 | 0.27274026 |
| FAM122C | 278.121323 | -0.155765094 | 0.32811638 | -0.4747251 | 0.6349829 | 0.78344233 |
| MOSPD1 | 644.842708 | -0.310796729 | 0.22027488 | -1.4109495 | 0.1582595 | 0.32542912 |
| LINC02243 | 0.37137328 | 0.704239854 | 2.29087628 | 0.30741069 | 0.75853079 | NA |
| SMIM10 | 249.874934 | -0.394347731 | 0.23131405 | -1.7048153 | 0.08822887 | 0.21737788 |
| RTL8B | 888.50509 | -0.168420909 | 0.16719292 | -1.0073447 | 0.31376915 | 0.51080113 |
| RTL8C | 5878.1953 | -0.412436751 | 0.19745099 | -2.0888057 | 0.03672522 | 0.11642751 |
| RTL8A | 2296.33262 | -0.754103253 | 0.14069159 | -5.3599739 | 8.32E-08 | 4.10E-06 |
| SMIM10L2B- | 0.53065351 | -0.614686635 | 1.85754992 | -0.3309126 | 0.74071052 | NA |
| SMIM10L2B | 380.352947 | 0.865141827 | 0.31613647 | 2.7366087 | 0.00620761 | 0.03165016 |
| ETDB | 0.5194031 | -2.276098206 | 2.67045699 | -0.8523254 | 0.39403354 | NA |
| AC234771.1 | 0.32016109 | -1.673938555 | 2.89062307 | -0.5790926 | 0.56252666 | NA |
| CT55 | 1.33002493 | -0.537775448 | 1.12028852 | -0.480033 | 0.63120394 | 0.78077009 |
| ZNF75D | 1116.63173 | -0.069507816 | 0.15532418 | -0.4475016 | 0.65451294 | 0.79728574 |
| ETDA | 2.12512372 | 0.101768783 | 0.87832684 | 0.11586664 | 0.90775823 | 0.95386955 |
| ETDC | 0 | NA | NA | NA | NA | NA |
| AL590282.2 | 2.61161543 | -0.187166933 | 0.79367078 | -0.2358244 | 0.81356894 | 0.8993047 |
| ZNF449 | 484.883104 | -0.279407768 | 0.13299743 | -2.1008509 | 0.03565406 | 0.11410165 |
| SMIM10L2A | 1864.33033 | -0.744130575 | 0.49071927 | -1.5164079 | 0.12941624 | 0.28425007 |
| INTS6L-AS1 | 4.44531949 | 1.054909804 | 0.84260029 | 1.25196944 | 0.210581 | 0.39397334 |
| INTS6L | 677.284989 | -0.184983497 | 0.15295247 | -1.2094182 | 0.22650222 | 0.4133196 |
| CT45A1 | 0 | NA | NA | NA | NA | NA |
| CT45A3 | 0 | NA | NA | NA | NA | NA |
| CT45A5 | 0 | NA | NA | NA | NA | NA |
| CT45A6 | 0 | NA | NA | NA | NA | NA |
| CT45A2 | 0 | NA | NA | NA | NA | NA |
| CT45A7 | 0 | NA | NA | NA | NA | NA |
| CT45A8 | 0 | NA | NA | NA | NA | NA |
| CT45A9 | 0 | NA | NA | NA | NA | NA |
| CT45A10 | 0 | NA | NA | NA | NA | NA |
| SAGE1 | 0.51452118 | -2.274619551 | 1.84488543 | -1.2329327 | 0.21760088 | NA |
| MMGT1 | 1534.52433 | -0.042913744 | 0.19036813 | -0.225425 | 0.82164864 | 0.90455786 |
| SLC9A6 | 996.0396 | -0.073164448 | 0.09461861 | -0.7732564 | 0.43937064 | 0.63084261 |
| FHL1 | 3487.4419 | 0.328262646 | 0.36955635 | 0.8882614 | 0.37440016 | 0.57055999 |
| MAP7D3 | 572.02006 | -0.669598945 | 0.14394997 | -4.6516088 | 3.29E-06 | 8.44E-05 |
| ADGRG4 | 4.80792501 | -0.733539296 | 0.99798376 | -0.7350213 | 0.46232654 | 0.6505283 |
| BRS3 | 1.34199328 | 0.007645245 | 1.52124576 | 0.00502565 | 0.99599013 | 0.99803389 |
| HTATSF1 | 2990.46655 | 0.287189345 | 0.15836344 | 1.81348259 | 0.06975742 | 0.1837482 |
| VGLL1 | 126.679333 | -0.442953256 | 0.89988792 | -0.4922316 | 0.62255564 | 0.77481557 |
| CD40LG | 34.6923155 | 0.807342538 | 0.7451812 | 1.08341775 | 0.27862304 | 0.47426347 |
| ARHGEF6 | 783.605964 | 0.436083374 | 0.25937023 | 1.68131618 | 0.09270152 | 0.22528843 |
| AL683813.1 | 189.618355 | 0.053390542 | 0.23407599 | 0.22809063 | 0.81957578 | 0.9031808 |
| RBMX | 7366.8243 | 0.019546612 | 0.15216545 | 0.12845631 | 0.89778787 | 0.94799743 |
| GPR101 | 0 | NA | NA | NA | NA | NA |
| AL035443.1 | 0.2072565 | -0.340597073 | 3.29738699 | -0.103293 | 0.91773041 | NA |
| ZIC3 | 1.46972455 | 1.478649633 | 1.27696142 | 1.15794386 | 0.24688695 | 0.43838185 |
| Z96074.1 | 0 | NA | NA | NA | NA | NA |
| FGF13 | 449.944133 | -0.054376 | 0.29495509 | -0.1843535 | 0.85373615 | 0.92346684 |
| AL031386.1 | 0 | NA | NA | NA | NA | NA |
| FGF13-AS1 | 8.46866829 | 0.543791359 | 0.54891886 | 0.9906589 | 0.32185216 | 0.51955663 |
| F9 | 0 | NA | NA | NA | NA | NA |
| MCF2 | 63.1335854 | 1.499606026 | 0.38200383 | 3.92563091 | 8.65E-05 | 0.00113133 |
| ATP11C | 1270.4967 | 0.037167547 | 0.17176926 | 0.21638067 | 0.82869103 | 0.90885803 |
| CXorf66 | 0.09635527 | 0.547239394 | 3.29377202 | 0.16614368 | 0.86804388 | NA |
| AL589987.2 | 0 | NA | NA | NA | NA | NA |
| FO393408.1 | 0 | NA | NA | NA | NA | NA |
| SOX3 | 0.75131164 | 1.142110613 | 1.53626281 | 0.7434344 | 0.45721873 | NA |
| AL844175.1 | 0.19994613 | 1.506623855 | 3.27645372 | 0.45983371 | 0.64563559 | NA |
| LINC00632 | 6.12737352 | -1.480918397 | 0.90821428 | -1.6305826 | 0.10297843 | 0.24235083 |
| CDR1 | 3.86314182 | -0.700114443 | 0.73234081 | -0.9559954 | 0.33907456 | 0.53721438 |
| AL078639.1 | 3.61423837 | -0.560340399 | 0.7555695 | -0.7416133 | 0.45832165 | 0.64727624 |
| AL451048.1 | 0.39021123 | 1.903032154 | 2.69570202 | 0.70595049 | 0.48021893 | NA |
| SPANXB1 | 1.7667084 | 2.981268896 | 1.15455353 | 2.58218335 | 0.00981774 | 0.04446056 |
| AC234778.2 | 0.19986543 | 0.18657164 | 3.29377195 | 0.05664376 | 0.95482897 | NA |

| | | | | | | |
|---|---|---|---|---|---|---|
| LDOC1 | 1844.95334 | -0.158447868 | 0.1974426 | -0.8025009 | 0.42226326 | 0.61472244 |
| SPANXC | 0 | NA | NA | NA | NA | NA |
| SPANXA2-OT | 1.49613323 | -0.581410549 | 1.36896339 | -0.4247086 | 0.6710491 | 0.80785905 |
| SPANXA1 | 0 | NA | NA | NA | NA | NA |
| SPANXA2 | 0 | NA | NA | NA | NA | NA |
| SPANXD | 0.07659028 | -0.642905668 | 3.30558948 | -0.1944905 | 0.84579185 | NA |
| MAGEC3 | 8.59434298 | -0.716480789 | 0.80214802 | -0.8932027 | 0.37174863 | 0.56838936 |
| MAGEC1 | 0.53551403 | -2.313163024 | 2.21130245 | -1.0460636 | 0.29553167 | NA |
| MAGEC2 | 0.07659028 | -0.642905668 | 3.30558948 | -0.1944905 | 0.84579185 | NA |
| SPANXN4 | 0 | NA | NA | NA | NA | NA |
| AC239727.1 | 0 | NA | NA | NA | NA | NA |
| SPANXN3 | 0 | NA | NA | NA | NA | NA |
| SLITRK4 | 454.247818 | 0.365243586 | 0.38193114 | 0.95630743 | 0.33891694 | 0.53701303 |
| SPANXN2 | 0 | NA | NA | NA | NA | NA |
| AL500522.1 | 0 | NA | NA | NA | NA | NA |
| AL713923.1 | 0 | NA | NA | NA | NA | NA |
| SPANXN1 | 0.07659028 | -0.642905668 | 3.30558948 | -0.1944905 | 0.84579185 | NA |
| SLITRK2 | 156.662085 | -0.727930984 | 0.40332385 | -1.80483 | 0.07110129 | 0.1864129 |
| CXorf51B | 0 | NA | NA | NA | NA | NA |
| CXorf51A | 0 | NA | NA | NA | NA | NA |
| AL589669.1 | 260.918294 | 1.165312409 | 0.55871197 | 2.08571225 | 0.0370047 | 0.11700003 |
| L29074.1 | 0.17423948 | 0.18657164 | 3.29377195 | 0.05664376 | 0.95482897 | NA |
| FMR1 | 3755.21879 | 0.033295437 | 0.2361062 | 0.1410189 | 0.88785501 | 0.94201397 |
| FMR1NB | 1.56304191 | -0.89763702 | 1.16536265 | -0.7702641 | 0.44114324 | 0.63239724 |
| AC002368.1 | 0.45606984 | -0.368183447 | 1.68567207 | -0.2184194 | 0.82710237 | NA |
| AFF2 | 69.667082 | -0.665791967 | 0.54004167 | -1.232853 | 0.21763063 | 0.40265393 |
| IDS | 5779.07715 | 0.009807381 | 0.1659236 | 0.05910781 | 0.95286624 | 0.97723075 |
| AC244197.3 | 1458.29038 | -0.029208545 | 0.21631355 | -0.1350287 | 0.89258915 | 0.94443203 |
| AC244197.2 | 52.6202634 | 0.594307329 | 0.55373117 | 1.07327773 | 0.28314652 | 0.47902572 |
| LINC00893 | 563.306908 | 0.604802193 | 0.29033136 | 2.08314458 | 0.03723805 | 0.11749468 |
| CXorf40A | 1027.94732 | 0.080859495 | 0.10446005 | 0.77407103 | 0.43888879 | 0.63042582 |
| HSFX3 | 27.7325955 | -0.473074289 | 0.42087531 | -1.1240248 | 0.26100251 | 0.4541795 |
| MAGEA9B | 0 | NA | NA | NA | NA | NA |
| HSFX2 | 0.9002478 | 1.160076601 | 2.21271435 | 0.52427761 | 0.60008546 | NA |
| TMEM185A | 647.246687 | -0.086983716 | 0.1247736 | -0.6971324 | 0.48571997 | 0.66982719 |
| MAGEA11 | 0 | NA | NA | NA | NA | NA |
| HSFX1 | 4.15334013 | 0.909550198 | 0.62324541 | 1.45937728 | 0.1444613 | 0.3063732 |
| MAGEA9 | 0.07659028 | -0.642905668 | 3.30558948 | -0.1944905 | 0.84579185 | NA |
| LINC00850 | 0.60473989 | -0.247480268 | 2.27229508 | -0.108912 | 0.91327227 | NA |
| MAGEA8-AS | 1.92472838 | 0.578761358 | 0.95463723 | 0.60626313 | 0.54434004 | 0.71631552 |
| MAGEA8 | 2.00833375 | 0.582873316 | 0.97879199 | 0.59550274 | 0.55150747 | 0.72160623 |
| CXorf40B | 1175.21624 | -0.200976803 | 0.16999266 | -1.1822675 | 0.23709957 | 0.42637178 |
| HSFX4 | 58.6801157 | -0.575577639 | 0.24369276 | -2.3618988 | 0.0181816 | 0.06994148 |
| LINC00894 | 341.361707 | 0.311513944 | 0.32640066 | 0.95439128 | 0.33988562 | 0.53807911 |
| MAMLD1 | 497.561467 | -0.544489533 | 0.38634103 | -1.4093495 | 0.15873185 | 0.32622234 |
| MTM1 | 988.493281 | 0.485977818 | 0.16096194 | 3.01920943 | 0.00253435 | 0.01616224 |
| MTMR1 | 1395.46669 | -0.164271687 | 0.11454492 | -1.4341246 | 0.15153672 | 0.31657328 |
| CD99L2 | 3555.30877 | 0.363716648 | 0.15965103 | 2.27819796 | 0.02271479 | 0.08234656 |
| AF274573.1 | 0.03392908 | 0.380740593 | 3.29738709 | 0.11546736 | 0.90807469 | NA |
| HMGB3 | 1172.89278 | -0.672585423 | 0.37081023 | -1.8138265 | 0.06970444 | 0.18367289 |
| GPR50-AS1 | 0.18663678 | 0.18657164 | 3.29377195 | 0.05664376 | 0.95482897 | NA |
| GPR50 | 0.84243671 | -2.946190695 | 1.84627668 | -1.5957471 | 0.11054526 | NA |
| KC877982.1 | 0.16354576 | -1.003574173 | 3.30558948 | -0.3035992 | 0.7614333 | NA |
| VMA21 | 1856.36728 | -0.076661179 | 0.14843111 | -0.5164765 | 0.60552164 | 0.76232244 |
| BX546450.2 | 2.66487544 | -1.883898739 | 1.04347242 | -1.8054131 | 0.07101007 | 0.18624782 |
| BX546450.1 | 0.18882702 | -0.340230161 | 3.19721851 | -0.1064144 | 0.91525355 | NA |
| PASD1 | 0.09635527 | 0.547239394 | 3.29377202 | 0.16614368 | 0.86804388 | NA |
| PRRG3 | 33.3237184 | 0.590992862 | 0.61718925 | 0.95755534 | 0.33828703 | 0.53628886 |
| FATE1 | 14.2412008 | -0.912346461 | 0.57209723 | -1.5947402 | 0.11077034 | 0.2550881 |
| CNGA2 | 0.08393208 | -0.642905668 | 3.30558948 | -0.1944905 | 0.84579185 | NA |
| MAGEA4-AS | 0.03392908 | 0.380740593 | 3.29738709 | 0.11546736 | 0.90807469 | NA |
| MAGEA4 | 0.22977084 | -1.364242679 | 3.30558948 | -0.4127078 | 0.67982068 | NA |
| GABRE | 1404.39536 | -0.673333647 | 0.40449025 | -1.6646474 | 0.09598312 | 0.2310218 |
| MAGEA10 | 0.22048515 | -0.339752076 | 3.07981855 | -0.1103156 | 0.91215907 | NA |
| AC116666.1 | 0.03392908 | 0.380740593 | 3.29738709 | 0.11546736 | 0.90807469 | NA |
| GABRA3 | 3.33696825 | 5.663325356 | 1.5503577 | 3.6529153 | 0.00025928 | 0.002695 |
| GABRQ | 7.47059829 | 0.527995676 | 0.68811125 | 0.7673115 | 0.44289633 | 0.63408308 |
| MAGEA3 | 0 | NA | NA | NA | NA | NA |
| CSAG2 | 0.78229184 | -1.263620992 | 2.31129804 | -0.5467149 | 0.58457464 | NA |
| MAGEA2B | 0 | NA | NA | NA | NA | NA |
| CSAG1 | 0.88869953 | -0.308668023 | 1.67259863 | -0.184544 | 0.85358672 | NA |
| MAGEA12 | 0.13028434 | 1.049816488 | 3.28477129 | 0.31960109 | 0.74927075 | NA |
| AC244102.1 | 0 | NA | NA | NA | NA | NA |
| MAGEA2 | 0 | NA | NA | NA | NA | NA |
| CSAG3 | 1.50199574 | 0.570382678 | 1.04226247 | 0.54725436 | 0.584204 | 0.74631622 |
| MAGEA6 | 0 | NA | NA | NA | NA | NA |
| CETN2 | 2715.25915 | -0.083096673 | 0.16272587 | -0.5106544 | 0.60959311 | 0.7656206 |
| NSDHL | 1154.49914 | -0.110044797 | 0.16720158 | -0.6581565 | 0.5104376 | 0.68892277 |
| ZNF185 | 1044.02615 | 0.284290331 | 0.25904409 | 1.09745926 | 0.27244068 | 0.46741265 |
| PNMA5 | 7.07718015 | 2.630035024 | 0.93714214 | 2.80644195 | 0.00500919 | 0.02691918 |
| PNMA3 | 476.456882 | 0.666572666 | 0.58121086 | 1.1468689 | 0.25143581 | 0.44374835 |
| PNMA6A | 402.286712 | 0.791410305 | 0.28046173 | 2.82181213 | 0.00477531 | 0.02596392 |
| MAGEA1 | 0 | NA | NA | NA | NA | NA |
| AC236972.3 | 0.28112383 | -1.364242679 | 3.30558948 | -0.4127078 | 0.67982068 | NA |
| PNMA6F | 0.9243696 | 0.378947621 | 1.43986964 | 0.2631819 | 0.79241038 | NA |
| ZNF275 | 968.856829 | 0.389981606 | 0.14131885 | 2.75958665 | 0.00578745 | 0.03002462 |
| PNMA6E | 2.69451368 | 1.246371716 | 1.61291641 | 0.77274415 | 0.43967381 | 0.63101113 |
| ZFP92 | 196.579959 | 0.00921964 | 0.38070288 | 0.02421742 | 0.98067919 | 0.9913179 |
| TREX2 | 1203.92606 | 0.395154038 | 0.24637001 | 1.6039048 | 0.10873504 | 0.25202218 |
| HAUS7 | 1206.83579 | 0.361745045 | 0.2424912 | 1.49178628 | 0.13575518 | 0.2934796 |
| BGN | 16348.4303 | 0.1139326 | 0.42386124 | 0.26879693 | 0.78808596 | 0.88470952 |
| ATP2B3 | 41.091864 | 2.067663005 | 0.84464494 | 2.44796707 | 0.01436648 | 0.05899722 |
| CCNQ | 950.115108 | -0.34257878 | 0.21811584 | -1.5706277 | 0.11626916 | 0.26431187 |
| DUSP9 | 3651.34295 | 1.795210787 | 0.5124971 | 3.50287011 | 0.00046027 | 0.00425576 |
| PNCK | 423.459587 | -0.303453296 | 0.28668674 | -1.0584839 | 0.28983489 | 0.486007 |
| SLC6A8 | 16160.8914 | -0.592550212 | 0.30541609 | -1.9401408 | 0.05236258 | 0.15041042 |
| BCAP31 | 12080.4515 | -0.485783675 | 0.11981413 | -4.0544773 | 5.02E-05 | 0.0007469 |
| ABCD1 | 1132.4337 | -0.236717919 | 0.18171402 | -1.3026948 | 0.19267897 | 0.37209686 |
| U52111.1 | 13.4649138 | 0.760489796 | 0.47068659 | 1.61570311 | 0.10615852 | 0.24748969 |
| PLXNB3 | 311.603797 | 0.343609212 | 0.40746338 | 0.84328857 | 0.39906707 | 0.59390477 |
| SRPK3 | 286.502698 | -0.100677647 | 0.37496029 | -0.2685022 | 0.78831281 | 0.88485118 |
| IDH3G | 4731.3442 | 0.002587893 | 0.14571454 | 0.01776002 | 0.9858303 | 0.99369329 |
| SSR4 | 6434.59469 | -0.402017743 | 0.15687135 | -2.5627225 | 0.0103855 | 0.04638646 |
| PDZD4 | 399.791905 | -0.164104146 | 0.48562321 | -0.3379248 | 0.73541983 | 0.85135523 |
| L1CAM | 3567.46346 | 0.396609847 | 0.30665831 | 1.29332822 | 0.19589757 | 0.37618838 |
| U52112.2 | 581.638728 | 0.050835831 | 0.314586 | 0.16159598 | 0.87162403 | 0.93317628 |
| AVPR2 | 457.336076 | 0.667240378 | 0.40802475 | 1.63529389 | 0.10198745 | 0.24072764 |
| ARHGAP4 | 5409.54712 | -0.27800855 | 0.25738333 | -1.0801343 | 0.2800824 | 0.47564225 |
| NAA10 | 3581.19456 | -0.484212811 | 0.17169723 | -2.820155 | 0.00480005 | 0.02607152 |
| RENBP | 3659.0494 | 0.770835818 | 0.35619776 | 2.16406699 | 0.03045921 | 0.10191718 |
| HCFC1 | 3840.84835 | -0.029308088 | 0.18977719 | -0.1544342 | 0.87726739 | 0.9361637 |
| HCFC1-AS1 | 1.51828725 | -0.135244067 | 1.03659159 | -0.13047 | 0.89619462 | 0.94685893 |
| TMEM187 | 513.803476 | 0.107715655 | 0.21083118 | 0.51090951 | 0.60941442 | 0.7654691 |

| | | | | | | |
|---|---|---|---|---|---|---|
| IRAK1 | 6848.80186 | -0.717892757 | 0.26639446 | -2.6948486 | 0.00704206 | 0.03480119 |
| MECP2 | 3541.77719 | -0.127141242 | 0.12868687 | -0.9879892 | 0.32315794 | 0.52085294 |
| OPN1LW | 0.71475495 | -1.135895844 | 1.84548824 | -0.6154988 | 0.53822535 | NA |
| OPN1MW | 0 | NA | NA | NA | NA | NA |
| OPN1MW2 | 0 | NA | NA | NA | NA | NA |
| OPN1MW3 | 0 | NA | NA | NA | NA | NA |
| TEX28 | 0.03392908 | 0.380740593 | 3.29738709 | 0.11546736 | 0.90807469 | NA |
| TKTL1 | 16.9999381 | -0.203488491 | 0.88068571 | -0.2310569 | 0.81727061 | 0.9017286 |
| AC245140.1 | 5.26672963 | 0.010454319 | 0.58968332 | 0.0177287 | 0.98585528 | 0.99369329 |
| FLNA | 40225.1609 | -0.068376819 | 0.51426197 | -0.1329611 | 0.89422417 | 0.94565432 |
| AC245140.3 | 20.0934025 | -0.612941236 | 0.50184715 | -1.2213704 | 0.22194582 | 0.40784067 |
| EMD | 3471.63674 | -0.533207189 | 0.13134179 | -4.0596917 | 4.91E-05 | 0.00073372 |
| RPL10 | 48902.2454 | -0.425893164 | 0.201453 | -2.1141068 | 0.03450614 | 0.11152433 |
| AC245140.2 | 13526.9385 | -0.299472854 | 0.22611962 | -1.3244001 | 0.1853702 | 0.36273464 |
| DNASE1L1 | 2131.44621 | 0.133526782 | 0.09709963 | 1.37515232 | 0.16908423 | 0.34073806 |
| TAZ | 2749.35713 | -0.427836342 | 0.16542856 | -2.5862303 | 0.00970321 | 0.04406295 |
| AC244090.1 | 166.749521 | 0.502892001 | 0.27633371 | 1.81987208 | 0.06877849 | 0.18184166 |
| ATP6AP1 | 11236.2719 | -0.012043163 | 0.11577653 | -0.1040208 | 0.91715287 | 0.95977306 |
| GDI1 | 9854.6506 | -0.494432864 | 0.14091513 | -3.5087281 | 0.00045025 | 0.0041834 |
| FAM50A | 4660.51227 | -0.94307472 | 0.20965047 | -4.4983191 | 6.85E-06 | 0.00015211 |
| PLXNA3 | 2745.72667 | -1.05721131 | 0.23663625 | -4.4676642 | 7.91E-06 | 0.00016954 |
| LAGE3 | 389.052524 | -0.240286898 | 0.22320162 | -1.0765464 | 0.28168296 | 0.47759144 |
| UBL4A | 2419.41271 | -0.19505775 | 0.20429077 | -0.9548045 | 0.33967656 | 0.53784501 |
| SLC10A3 | 1797.0692 | -0.560983058 | 0.20478519 | -2.7393732 | 0.00615565 | 0.03142776 |
| FAM3A | 3195.0687 | -0.105383983 | 0.13695316 | -0.7694892 | 0.44160294 | 0.63288422 |
| G6PD | 3188.89954 | -1.003283497 | 0.29539493 | -3.3964141 | 0.00068275 | 0.00577796 |
| IKBKG | 1489.64239 | -0.426708537 | 0.19975124 | -2.1361997 | 0.03266314 | 0.10721024 |
| FAM223A | 0 | NA | NA | NA | NA | NA |
| CTAG1A | 0.19978473 | -1.003574173 | 3.30558948 | -0.3035992 | 0.7614333 | NA |
| CTAG1B | 0 | NA | NA | NA | NA | NA |
| FAM223B | 0 | NA | NA | NA | NA | NA |
| CTAG2 | 13.7180294 | 6.42509899 | 3.14603362 | 2.04228555 | NA | NA |
| GAB3 | 304.460354 | -0.179439602 | 0.26749037 | -0.6708264 | 0.50233113 | 0.68263189 |
| DKC1 | 2011.29018 | -0.514799286 | 0.22038037 | -2.335958 | 0.01949343 | 0.07350805 |
| MPP1 | 3270.60941 | 0.396592937 | 0.29053256 | 1.36505503 | 0.17223575 | 0.34520459 |
| SMIM9 | 1.69476747 | -1.5230523 | 1.41949461 | -1.0729539 | 0.28329179 | 0.47917925 |
| F8 | 1691.95275 | -0.236517658 | 0.32529437 | -0.7270881 | 0.46717199 | 0.65430877 |
| H2AFB1 | 0.72654834 | 2.324909912 | 2.05519622 | 1.13123501 | 0.25795619 | NA |
| F8A1 | 739.552147 | 0.106363087 | 0.26492871 | 0.40147814 | 0.68806813 | 0.81986158 |
| FUNDC2 | 4474.62926 | -0.330975717 | 0.22025105 | -1.5027203 | 0.13291119 | 0.28937788 |
| CMC4 | 750.176011 | -0.29516594 | 0.13643358 | -2.1634405 | 0.03050732 | 0.10205226 |
| MTCP1 | 837.421602 | -0.366088047 | 0.13714896 | -2.6692731 | 0.00760156 | 0.0367939 |
| BRCC3 | 1145.24792 | 0.126675352 | 0.10930071 | 1.15896186 | 0.24647173 | 0.43775299 |
| VBP1 | 1739.69291 | -0.17547622 | 0.14467227 | -1.2129223 | 0.22515954 | 0.41151291 |
| RAB39B | 64.5600042 | -0.078698757 | 0.26752245 | -0.2941763 | 0.76862321 | 0.87217134 |
| CLIC2 | 515.528835 | -0.468389284 | 0.26785282 | -1.7486815 | 0.08034608 | 0.20300146 |
| AC234781.1 | 5.09778395 | -0.615571343 | 0.69926469 | -0.8803124 | 0.37869012 | 0.57473718 |
| H2AFB2 | 0 | NA | NA | NA | NA | NA |
| F8A2 | 5.29194008 | -0.598302871 | 0.63823521 | -0.9374332 | 0.34853577 | 0.54606905 |
| F8A3 | 3.0972807 | 0.570397807 | 0.71963601 | 0.79261988 | 0.42799932 | 0.62042202 |
| H2AFB3 | 0.46564926 | 0.294514906 | 1.92028687 | 0.15337026 | 0.87810629 | NA |
| TMLHE-AS1 | 9.76972568 | -0.63181345 | 0.4714651 | -1.3401065 | 0.18021072 | 0.35612685 |
| BX571846.1 | 0.3432521 | -0.541317636 | 2.87189509 | -0.188488 | 0.85049416 | NA |
| TMLHE | 614.457579 | 0.153943721 | 0.20908894 | 0.73625951 | 0.46157278 | 0.64998795 |
| SPRY3 | 147.793959 | 0.209283803 | 0.22048765 | 0.94918604 | 0.342526 | 0.54018364 |
| VAMP7 | 2765.23139 | -0.010872447 | 0.13423159 | -0.0809977 | 0.9354438 | 0.96910714 |
| IL9R | 8.41073175 | -0.963229813 | 0.82881432 | -1.162178 | 0.24516315 | 0.43618502 |
| WASIR1 | 1.23431792 | -0.757636486 | 1.96444185 | -0.3856752 | 0.69973724 | 0.82781835 |
| AC144568.2 | 0.18746814 | 0.18657164 | 3.29377195 | 0.05664376 | 0.95482897 | NA |
| OR4F21 | 0.07659028 | -0.642905668 | 3.30558948 | -0.1944905 | 0.84579185 | NA |
| ZNF596 | 407.437212 | 0.28613003 | 0.15947535 | 1.79419596 | 0.0727819 | 0.18969885 |
| AC004908.1 | 80.9902308 | -0.01295613 | 0.3080479 | -0.0420588 | 0.96645181 | 0.98410057 |
| AC004908.3 | 39.3616331 | -0.123618268 | 0.22801363 | -0.542153 | 0.58771313 | 0.74865926 |
| AC136777.1 | 0 | NA | NA | NA | NA | NA |
| FAM87A | 1.59054631 | 1.699055056 | 1.06178274 | 1.60019088 | 0.10955624 | 0.25324685 |
| FBXO25 | 2008.26531 | 0.008057603 | 0.13074386 | 0.06162892 | 0.95085835 | 0.97652119 |
| AC083964.1 | 335.780198 | -0.148272084 | 0.17194252 | -0.8623352 | 0.38850311 | 0.5841377 |
| TDRP | 2220.01541 | -0.154705606 | 0.18219139 | -0.8491378 | 0.3958046 | 0.59110713 |
| ERICH1 | 1318.43252 | -0.260288394 | 0.20826603 | -1.249788 | 0.21137699 | 0.39481592 |
| DLGAP2 | 178.247349 | 1.78208628 | 0.78457341 | 2.27140795 | 0.02312229 | 0.08354822 |
| AC100797.4 | 0 | NA | NA | NA | NA | NA |
| AC026950.2 | 0 | NA | NA | NA | NA | NA |
| AC129915.3 | 0.4529328 | 0.627479411 | 1.82399782 | 0.34401325 | 0.73083632 | NA |
| AC110288.1 | 0.08666371 | -0.642905668 | 3.30558948 | -0.1944905 | 0.84579185 | NA |
| AF067845.2 | 0.66243165 | 1.447333212 | 2.1343295 | 0.67812079 | 0.49769511 | NA |
| AF067845.1 | 1.23954416 | 0.386192949 | 1.25978366 | 0.30655497 | 0.75918213 | 0.86663046 |
| AF067845.5 | 0.32101177 | 0.652944617 | 2.84542999 | 0.22947133 | 0.8185026 | NA |
| AF067845.4 | 0.43126934 | 0.330999332 | 1.84888258 | 0.17902669 | 0.85791674 | NA |
| DLGAP2-AS1 | 2.63697131 | 1.877219489 | 1.04758965 | 1.79194162 | 0.07314232 | 0.1902937 |
| CLN8 | 1878.89912 | 0.237069853 | 0.22410886 | 1.05783348 | 0.29013137 | 0.48622587 |
| AC100810.3 | 10.1502299 | 0.00528603 | 0.64571303 | 0.00818634 | 0.99346831 | 0.99694729 |
| AC100810.1 | 774.750885 | 0.636281521 | 0.27329155 | 2.32821512 | 0.01990068 | 0.07465941 |
| AC019257.8 | 1465.85012 | -0.027185311 | 0.14292106 | -0.1902121 | 0.84914295 | 0.92060599 |
| ARHGEF10 | 1301.66073 | -0.018388377 | 0.1820091 | -0.10103 | 0.91952666 | 0.96074349 |
| KBTBD11-OT | 67.6921466 | -0.909801825 | 0.43888832 | -2.0729689 | 0.03817518 | 0.11962916 |
| AC019257.1 | 119.921423 | -0.371683607 | 0.35571683 | -1.0448862 | 0.29607556 | 0.49280217 |
| KBTBD11 | 3152.98459 | -0.005289652 | 0.28003891 | -0.018889 | 0.98492966 | 0.99347171 |
| AC019257.2 | 19.4413547 | -0.719308671 | 0.48652061 | -1.4784752 | 0.13928063 | 0.29889054 |
| AC245164.1 | 0 | NA | NA | NA | NA | NA |
| MYOM2 | 675.356683 | 1.906664315 | 0.36268078 | 5.25714188 | 1.46E-07 | 6.58E-06 |
| AC245519.1 | 6.82143669 | 3.899163181 | 1.17529746 | 3.31759686 | 0.00090795 | 0.00727522 |
| AC245519.2 | 0.63048279 | 2.022555474 | 2.14129992 | 0.94454562 | 0.3448909 | NA |
| AC245187.2 | 7.39419768 | 1.593583732 | 0.71752565 | 2.22094323 | 0.02635481 | 0.09192422 |
| AC245187.1 | 3.69092083 | 2.066449724 | 0.67709091 | 3.05195313 | 0.00227358 | 0.01490286 |
| AC245123.1 | 6.86797885 | 0.655406077 | 0.5385015 | 1.21709238 | 0.22356907 | 0.4096669 |
| AC246817.2 | 25.4570425 | 1.264419822 | 0.34843726 | 3.6288307 | 0.00028471 | 0.00290773 |
| AC246817.1 | 20.8259849 | 0.794239126 | 0.4774393 | 1.6635395 | 0.09620449 | 0.23136456 |
| CSMD1 | 333.349506 | 2.54343378 | 0.56259578 | 4.52089022 | 6.16E-06 | 0.00013939 |
| AC004944.1 | 0 | NA | NA | NA | NA | NA |
| AC084768.1 | 0 | NA | NA | NA | NA | NA |
| AC079054.1 | 0.09989236 | -0.642905668 | 3.30558948 | -0.1944905 | 0.84579185 | NA |
| AC009435.1 | 0.12059279 | 0.02007176 | 3.29738704 | 0.00608717 | 0.99514317 | NA |
| AC016065.1 | 198.927397 | 0.125848859 | 0.232364 | 0.54160223 | 0.58809256 | 0.74892684 |
| MCPH1 | 818.586327 | -0.11730172 | 0.14389556 | -0.8151865 | 0.41496556 | 0.60849924 |
| ANGPT2 | 393.13132 | -0.733834955 | 0.45955584 | -1.5968352 | 0.11030244 | 0.25434376 |
| AF287957.1 | 43.6058218 | 0.255512559 | 0.35898608 | 0.71176174 | 0.47661233 | 0.66179458 |
| MCPH1-AS1 | 143.018018 | -0.003684714 | 0.19728612 | -0.018677 | 0.98509877 | 0.99347171 |
| AGPAT5 | 1790.41259 | -0.3144871 | 0.26080497 | -1.2058325 | 0.2278821 | 0.41488891 |
| XKR5 | 3.45758829 | 0.438657406 | 0.87399851 | 0.5018972 | 0.61573983 | 0.77022245 |
| AF233439.1 | 14.3356319 | 2.086641833 | 0.50325884 | 4.14625971 | 3.38E-05 | 0.00054076 |
| DEFB1 | 16977.363 | 0.277386499 | 0.3616431 | 0.76701726 | 0.44307126 | 0.63422761 |
| DEFA6 | 0 | NA | NA | NA | NA | NA |
| AF233439.2 | 0.08666371 | -0.642905668 | 3.30558948 | -0.1944905 | 0.84579185 | NA |

| | | | | | | |
|---|---|---|---|---|---|---|
| DEFA4 | 23.5173566 | -5.036594181 | 1.04565306 | -4.8166972 | 1.46E-06 | 4.43E-05 |
| DEFA1 | 3.14516692 | -4.884113179 | 1.58801295 | -3.0756129 | 0.0021007 | 0.01398171 |
| DEFA1B | 3.14516692 | -4.884113179 | 1.58801295 | -3.0756129 | 0.0021007 | 0.01398171 |
| DEFA3 | 203.472078 | -10.20383241 | 1.5273654 | -6.6806754 | 2.38E-11 | 4.30E-09 |
| DEFA5 | 0.09644756 | -0.642905668 | 3.30558948 | -0.1944905 | 0.84579185 | NA |
| FAM66B | 23.5167722 | 0.589613114 | 0.30818491 | 1.91317968 | 0.05572505 | 0.15711212 |
| USP17L1 | 0.09997307 | 0.547239394 | 3.29377202 | 0.16614368 | 0.86804388 | NA |
| USP17L4 | 0 | NA | NA | NA | NA | NA |
| ZNF705G | 0 | NA | NA | NA | NA | NA |
| DEFB4B | 0.53606433 | -2.287253828 | 3.28035022 | -0.697259 | 0.4856407 | NA |
| DEFB103B | 1.08235868 | 0.659778118 | 1.56893355 | 0.42052649 | 0.67410088 | 0.81024608 |
| SPAG11B | 0.13382144 | 0.02007176 | 3.29738704 | 0.00608717 | 0.99514317 | NA |
| DEFB104B | 0 | NA | NA | NA | NA | NA |
| DEFB106B | 0 | NA | NA | NA | NA | NA |
| DEFB105B | 0 | NA | NA | NA | NA | NA |
| DEFB107B | 0 | NA | NA | NA | NA | NA |
| PRR23D1 | 0 | NA | NA | NA | NA | NA |
| PRR23D2 | 0 | NA | NA | NA | NA | NA |
| DEFB107A | 0 | NA | NA | NA | NA | NA |
| DEFB105A | 0 | NA | NA | NA | NA | NA |
| DEFB106A | 0 | NA | NA | NA | NA | NA |
| DEFB104A | 0 | NA | NA | NA | NA | NA |
| SPAG11A | 0.59849685 | -0.067911444 | 3.26841084 | -0.0207781 | 0.98342265 | NA |
| DEFB103A | 0.85770293 | -2.975577396 | 3.25982302 | -0.9128034 | 0.36134597 | NA |
| DEFB4A | 0.24823779 | 0.078381887 | 3.05287807 | 0.02567475 | 0.97951676 | NA |
| ZNF705B | 0 | NA | NA | NA | NA | NA |
| FAM66E | 0.63745462 | 2.73078873 | 1.52674749 | 1.78863155 | 0.07367417 | NA |
| USP17L8 | 0 | NA | NA | NA | NA | NA |
| USP17L3 | 0 | NA | NA | NA | NA | NA |
| FAM85B | 13.3903428 | 0.551074636 | 0.41476755 | 1.32863489 | 0.18396845 | 0.36101253 |
| AC068020.1 | 1.05767463 | 0.535317599 | 1.24124907 | 0.43127331 | 0.66626965 | 0.80439203 |
| PRAG1 | 874.21934 | -0.427635728 | 0.17162702 | -2.4916573 | 0.01271486 | 0.05385335 |
| AC103957.1 | 0.27291861 | 1.131425716 | 3.00149036 | 0.37695464 | 0.70620731 | NA |
| AC103957.2 | 3.56159149 | 0.978453505 | 0.84603205 | 1.15652061 | 0.24746827 | 0.43902976 |
| AC114550.2 | 0.07830766 | -0.642905668 | 3.30558948 | -0.1944905 | 0.84579185 | NA |
| AC114550.3 | 0.25125321 | -0.178199516 | 3.01001542 | -0.0592022 | 0.95279106 | NA |
| AC114550.1 | 0.08695548 | -0.642905668 | 3.30558948 | -0.1944905 | 0.84579185 | NA |
| CLDN23 | 603.334308 | -0.654334676 | 0.2747677 | -2.3814105 | 0.01724648 | 0.0673563 |
| AC087269.1 | 2.65770698 | -3.84756111 | 1.15604776 | -3.3282026 | 0.00087408 | 0.00706608 |
| MFHAS1 | 3418.78827 | -1.049552559 | 0.13339418 | -7.8680534 | 3.60E-15 | 2.13E-12 |
| ERI1 | 1045.25395 | -1.174927181 | 0.23587011 | -4.9812467 | 6.32E-07 | 2.22E-05 |
| PPP1R3B | 2100.13261 | -1.065102748 | 0.35208954 | -3.0250906 | 0.00248559 | 0.01592442 |
| AC022784.7 | 30.6244846 | 0.155053631 | 0.52385268 | 0.29598709 | 0.76723994 | 0.87152635 |
| AC022784.3 | 46.086171 | 1.213664095 | 0.25242604 | 4.80799888 | 1.52E-06 | 4.58E-05 |
| AC022784.8 | 12.2041074 | 1.102237487 | 0.6639095 | 1.66022251 | 0.0968697 | 0.23230756 |
| AC022784.2 | 0.2320242 | 1.01099135 | 3.00413042 | 0.33653377 | 0.73646839 | NA |
| AC022784.1 | 26.0515912 | 1.05910014 | 0.49970955 | 2.11943148 | 0.03405402 | 0.11042857 |
| AC022784.6 | 2.1789921 | 0.647004259 | 0.87144992 | 0.74244571 | 0.45781733 | 0.6469798 |
| AC022784.4 | 0 | NA | NA | NA | NA | NA |
| AC021736.1 | 0 | NA | NA | NA | NA | NA |
| AC021242.2 | 0 | NA | NA | NA | NA | NA |
| AC021242.3 | 40.7176456 | -0.337003242 | 0.26921945 | -1.251779 | 0.21065042 | 0.39401699 |
| TNKS | 1810.22303 | -0.245893905 | 0.1214221 | -2.0251166 | 0.04285539 | 0.13027051 |
| LINC00599 | 19.3462184 | -0.165076415 | 0.54961336 | -0.3003501 | 0.76391014 | 0.86935843 |
| AC034111.1 | 6.77842609 | 1.46128574 | 0.60155697 | 2.42917266 | 0.01513332 | 0.06126278 |
| MSRA | 4333.83258 | 1.500096332 | 0.35978999 | 4.16936655 | 3.05E-05 | 0.00049567 |
| AC104964.1 | 33.4106132 | 0.650334251 | 0.35760696 | 1.81857269 | 0.06897665 | 0.18216467 |
| AC104964.2 | 9.32487953 | 0.850687678 | 0.47690612 | 1.7837634 | 0.07446212 | 0.19275214 |
| PRSS51 | 29.5812948 | 0.675881215 | 0.48990263 | 1.37962357 | 0.16770258 | 0.33887486 |
| AC104964.4 | 220.094403 | 0.631144768 | 0.30625257 | 2.06086361 | 0.03931605 | 0.12225995 |
| PRSS55 | 32.6497997 | 1.365271622 | 0.57391556 | 2.37887195 | 0.01736571 | 0.0676516 |
| RP1L1 | 8.74638568 | 0.662275767 | 0.52294186 | 1.26644245 | 0.20535472 | 0.3879086 |
| C8orf74 | 0.15452186 | 0.435199075 | 3.28898345 | 0.13232024 | 0.894731 | NA |
| SOX7 | 713.373008 | -0.784991002 | 0.38836113 | -2.0212914 | 0.04324961 | 0.13113568 |
| PINX1 | 767.658964 | -0.497579847 | 0.25173426 | -1.9766076 | 0.048086 | 0.1416158 |
| AC105001.1 | 20.497499 | 0.236337174 | 0.2619149 | 0.90234338 | 0.36687449 | 0.56387888 |
| AC011008.1 | 1.08096349 | 1.663430882 | 1.59379669 | 1.04369076 | 0.29662848 | 0.49341406 |
| XKR6 | 72.5192125 | -1.007503125 | 0.29552459 | -3.4092024 | 0.00065153 | 0.00556196 |
| AF131215.3 | 0.33501852 | -0.699357289 | 2.8604516 | -0.2444919 | 0.80684986 | NA |
| AF131215.7 | 20.5398481 | -0.399471118 | 0.41921753 | -0.952897 | 0.34064227 | 0.53869486 |
| LINC00529 | 0 | NA | NA | NA | NA | NA |
| AF131216.4 | 24.159342 | 0.408964093 | 0.363426 | 1.12530226 | 0.26046097 | 0.45355506 |
| MTMR9 | 846.30616 | -0.074813449 | 0.13326625 | -0.5613833 | 0.57453626 | 0.73890564 |
| AF131216.1 | 90.8223807 | 0.169944506 | 0.1770879 | 0.95966185 | 0.33722543 | 0.53527683 |
| SLC35G5 | 13.5426188 | -0.773667017 | 0.5610986 | -1.3788432 | 0.16794309 | 0.33920491 |
| AF131216.3 | 3.28953412 | -0.788728634 | 1.29637558 | -0.6084106 | 0.54291519 | 0.7152972 |
| FAM167A-AS | 6.4250858 | 2.126177699 | 0.7374532 | 2.88313576 | 0.00393738 | 0.02250121 |
| FAM167A | 850.777848 | 0.434703218 | 0.38376376 | 1.13273649 | 0.25732493 | 0.4499697 |
| AF131216.5 | 0.21442573 | -1.003574173 | 3.30558948 | -0.3035992 | 0.7614333 | NA |
| BLK | 98.1916043 | 0.348088478 | 0.86798578 | 0.40103016 | 0.68839792 | 0.8201273 |
| AC022239.1 | 36.1073751 | 0.261956637 | 0.87600778 | 0.2990346 | 0.76491364 | 0.87010494 |
| LINC00208 | 0 | NA | NA | NA | NA | NA |
| GATA4 | 0 | NA | NA | NA | NA | NA |
| C8orf49 | 0 | NA | NA | NA | NA | NA |
| NEIL2 | 1064.44989 | -0.045358301 | 0.22364528 | -0.2028136 | 0.83928074 | 0.91465195 |
| FDFT1 | 5577.81322 | -0.358575203 | 0.22375952 | -1.6025025 | 0.10904452 | 0.25239218 |
| AC069185.1 | 52.6822696 | 1.141024176 | 0.25379141 | 4.49591338 | 6.93E-06 | 0.00015299 |
| CTSB | 98971.3911 | -0.367876586 | 0.46677998 | -0.7881156 | 0.43062909 | 0.62264716 |
| DEFB136 | 0 | NA | NA | NA | NA | NA |
| DEFB135 | 0 | NA | NA | NA | NA | NA |
| DEFB134 | 0 | NA | NA | NA | NA | NA |
| DEFB130B | 0 | NA | NA | NA | NA | NA |
| ZNF705D | 0.37500701 | 0.018400558 | 2.84942556 | 0.00645764 | 0.99484759 | NA |
| FAM66D | 16.5509272 | 0.364055894 | 0.57018359 | 0.63848891 | 0.52315547 | 0.69865339 |
| USP17L7 | 0.19978473 | -1.003574173 | 3.30558948 | -0.3035992 | 0.7614333 | NA |
| USP17L2 | 0.3774791 | -0.542547472 | 2.80930516 | -0.1931251 | 0.84686096 | NA |
| FAM86B1 | 277.088292 | 0.359999684 | 0.38292412 | 0.94013321 | 0.34714924 | 0.54472033 |
| AC145124.1 | 79.2597911 | 1.061580782 | 0.40657599 | 2.61102674 | 0.00902708 | 0.04175481 |
| DEFB130A | 0 | NA | NA | NA | NA | NA |
| FAM66A | 11.1387289 | 1.383820385 | 0.55607195 | 2.48856356 | 0.01282603 | 0.05416759 |
| AC087203.3 | 4.94014839 | -0.327891734 | 0.92488662 | -0.354521 | 0.72294846 | 0.84283792 |
| FAM86B2 | 32.2754194 | 0.904469686 | 0.59100417 | 1.53039475 | 0.12591905 | 0.27872788 |
| AC130352.1 | 8.60092879 | 0.436201708 | 0.49024591 | 0.88976104 | 0.37359421 | 0.56979059 |
| LONRF1 | 1413.57775 | -0.626789684 | 0.21868539 | -2.8661708 | 0.0041547 | 0.02344079 |
| AC123777.1 | 1.06016931 | -0.452210287 | 1.91060944 | -0.2366838 | 0.81290211 | 0.89895498 |
| LINC00681 | 0.17820003 | -1.003574173 | 3.30558948 | -0.3035992 | 0.7614333 | NA |
| TRMT9B | 1802.96127 | 0.030448622 | 0.32592396 | 0.09342247 | 0.92556794 | 0.96400096 |
| DLC1 | 2010.70047 | 0.227292097 | 0.28333873 | 0.80219211 | 0.42244184 | 0.61491451 |
| AC019270.1 | 0.25668517 | 0.372875031 | 3.03140481 | 0.12300404 | 0.9021039 | NA |
| C8orf48 | 58.2102682 | 0.379697919 | 0.44063827 | 0.86169982 | 0.38885272 | 0.58440934 |
| AC022690.2 | 0 | NA | NA | NA | NA | NA |
| SGCZ | 1.17373407 | 0.141540683 | 1.70114209 | 0.08320333 | 0.93368987 | 0.96806738 |
| AC022039.1 | 0.73657393 | -2.76428257 | 2.17561515 | -1.2705752 | 0.20387983 | NA |

| | | | | | | |
|---|---|---|---|---|---|---|
| TUSC3 | 2648.82796 | -0.505861961 | 0.21544303 | -2.348008 | 0.01887412 | 0.07192347 |
| AC018437.2 | 0 | NA | NA | NA | NA | NA |
| MSR1 | 853.487848 | -1.013294203 | 0.37102235 | -2.7310867 | 0.00631259 | 0.03200594 |
| AC011586.1 | 0.94268841 | -0.080334576 | 1.54426428 | -0.0520213 | 0.95851175 | NA |
| AC068992.1 | 4.02573148 | 0.309127406 | 0.71150509 | 0.4344697 | 0.66394739 | 0.80260972 |
| FGF20 | 3.56667567 | -1.367041492 | 1.25099726 | -1.0927614 | 0.27449858 | 0.46975972 |
| MICU3 | 380.656447 | 0.515546994 | 0.20055613 | 2.57058701 | 0.01015263 | 0.04558571 |
| AC079193.2 | 9.8498834 | -0.456091131 | 0.47855601 | -0.953057 | 0.34056122 | 0.53859897 |
| ZDHHC2 | 2617.878 | 0.115478028 | 0.14870689 | 0.77654791 | 0.43742555 | 0.62949747 |
| CNOT7 | 3327.28122 | -0.078663156 | 0.14475035 | -0.5434402 | 0.58682676 | 0.74778302 |
| VPS37A | 2598.7738 | -0.473619598 | 0.19415368 | -2.4394057 | 0.01471144 | 0.06001557 |
| MTMR7 | 180.422066 | -0.160955085 | 0.25665167 | -0.6271344 | 0.53057116 | 0.70495094 |
| SLC7A2 | 2080.93991 | -1.340094931 | 0.28570702 | -4.6904516 | 2.73E-06 | 7.29E-05 |
| PDGFRL | 944.547937 | -1.025523004 | 0.22231141 | -4.6130021 | 3.97E-06 | 9.82E-05 |
| MTUS1 | 3586.78008 | -0.549313928 | 0.21897154 | -2.5086088 | 0.01212076 | 0.05192238 |
| AC124069.1 | 0.29512404 | 0.328535886 | 2.30676753 | 0.14242263 | 0.88674619 | NA |
| AC027117.2 | 6.24087056 | -0.404465365 | 0.66757165 | -0.6058756 | 0.54459737 | 0.71643964 |
| FGL1 | 186.347025 | -5.900661211 | 1.00282736 | -5.8840249 | 4.00E-09 | 3.17E-07 |
| AC087273.1 | 51.3556443 | -6.392214498 | 1.26316245 | -5.0604849 | 4.18E-07 | 1.57E-05 |
| AC087273.2 | 6.92590732 | -1.877403554 | 0.86136837 | -2.1795594 | 0.02929013 | 0.09920124 |
| PCM1 | 4011.84097 | -0.068852635 | 0.1169527 | -0.5887221 | 0.55604774 | 0.72524378 |
| ASAH1 | 20181.3075 | 0.202503411 | 0.27759867 | 0.72948265 | 0.46570648 | 0.65288037 |
| AC124242.1 | 64.879242 | 1.054808472 | 0.27757468 | 3.80008901 | 0.00014464 | 0.001699 |
| NAT1 | 364.320994 | 1.143706396 | 0.28339267 | 4.03576568 | 5.44E-05 | 0.0007952 |
| NAT2 | 45.6617514 | 1.286829974 | 0.32965599 | 3.90355409 | 9.48E-05 | 0.001214 |
| PSD3 | 476.304462 | -0.824149774 | 0.28476231 | -2.8941674 | 0.00380165 | 0.02196049 |
| AC100800.1 | 0.42315254 | 1.051852187 | 2.20153287 | 0.47778173 | 0.63280556 | NA |
| AC009884.1 | 0.09989236 | -0.642905668 | 3.30558948 | -0.1944905 | 0.84579185 | NA |
| AC100849.1 | 5.8731262 | -0.664413377 | 0.73198781 | -0.9076837 | 0.36404538 | 0.56116658 |
| AC100849.2 | 0.49969646 | -2.237634025 | 2.20419336 | -1.0151714 | 0.31002414 | NA |
| AC068880.3 | 0.08757577 | 0.547239394 | 3.29377202 | 0.16614368 | 0.86804388 | NA |
| AC068880.2 | 0 | NA | NA | NA | NA | NA |
| SH2D4A | 972.701298 | -0.344431635 | 0.18745827 | -1.8373777 | 0.06615416 | 0.17702015 |
| CSGALNACT1 | 1289.51031 | -0.904415052 | 0.24328039 | -3.717583 | 0.00020114 | 0.00218993 |
| AC090541.1 | 11.6213476 | -2.542584651 | 1.26407128 | -2.0114251 | 0.04428058 | 0.13331504 |
| INTS10 | 3346.83839 | -0.306905868 | 0.14444357 | -2.1247458 | 0.03360783 | 0.10943103 |
| LPL | 1395.75987 | 1.507645854 | 0.70336261 | 2.14348308 | 0.03207433 | 0.10583083 |
| AC100802.1 | 0 | NA | NA | NA | NA | NA |
| SLC18A1 | 1.46089219 | 1.828307011 | 1.23139394 | 1.48474582 | 0.13761117 | 0.2962291 |
| ATP6V1B2 | 9822.657 | 0.366465234 | 0.41062768 | 0.89245137 | 0.37215105 | 0.56863396 |
| LZTS1 | 411.455608 | -0.134205482 | 0.33683267 | -0.3984337 | 0.69031053 | 0.8215683 |
| LZTS1-AS1 | 0.39655556 | -1.848090578 | 3.29816102 | -0.5603397 | 0.57524775 | NA |
| AC023403.1 | 0.2939202 | -0.701265545 | 3.29738693 | -0.2126731 | 0.83158194 | NA |
| AC018541.1 | 39.4961304 | 5.806361992 | 0.62150864 | 9.34236737 | 9.42E-21 | 1.77E-17 |
| AC015468.4 | 0 | NA | NA | NA | NA | NA |
| AC015468.3 | 0 | NA | NA | NA | NA | NA |
| AC015468.1 | 0.11051935 | 0.02007176 | 3.29738704 | 0.00608717 | 0.99514317 | NA |
| AC015468.2 | 0.11051935 | 0.02007176 | 3.29738704 | 0.00608717 | 0.99514317 | NA |
| LINC02153 | 0.17648264 | -1.003574173 | 3.30558948 | -0.3035992 | 0.7614333 | NA |
| AC103719.1 | 0.47507074 | 2.84394444 | 3.19938614 | 0.88890316 | 0.37405513 | NA |
| AC021355.1 | 0 | NA | NA | NA | NA | NA |
| GFRA2 | 97.1032644 | 0.88119498 | 0.57813029 | 1.52421521 | 0.12745496 | 0.28106601 |
| DOK2 | 280.184233 | -0.278267439 | 0.51498453 | -0.5403414 | 0.58896164 | 0.74944647 |
| XPO7 | 3197.92833 | 0.749442072 | 0.16297353 | 4.59855096 | 4.25E-06 | 0.00010349 |
| NPM2 | 395.215754 | -0.73448102 | 0.28520304 | -2.5752917 | 0.01001556 | 0.04511621 |
| FGF17 | 194.23445 | -0.990372658 | 0.34515938 | -2.8693199 | 0.00411356 | 0.02324287 |
| DMTN | 6462.81569 | 0.111112666 | 0.2119923 | 0.52413539 | 0.60018438 | 0.75785859 |
| FAM160B2 | 5764.08898 | -0.500818706 | 0.18094014 | -2.7678695 | 0.0056424 | 0.02951015 |
| NUDT18 | 485.583386 | -0.160045565 | 0.19677175 | -0.8133564 | 0.41601375 | 0.60925706 |
| HR | 174.035291 | 0.12409309 | 0.5030336 | 0.24668947 | 0.80514856 | 0.89442108 |
| REEP4 | 892.013616 | -1.047905326 | 0.23514117 | -4.4564945 | 8.33E-06 | 0.00017674 |
| LGI3 | 67.8862457 | -1.259278056 | 0.7957167 | -1.5825709 | 0.11351931 | 0.25964994 |
| SFTPC | 0.70189702 | -0.713525175 | 1.52639666 | -0.4674572 | 0.64017278 | NA |
| BMP1 | 3044.26775 | -1.246709274 | 0.2039976 | -6.1113918 | 9.88E-10 | 1.02E-07 |
| PHYHIP | 683.623529 | -1.052972213 | 0.30043883 | -3.5047807 | 0.00045698 | 0.0042283 |
| POLR3D | 932.812545 | -0.603526013 | 0.17903563 | -3.3709828 | 0.00074901 | 0.00623856 |
| AC105206.2 | 39.8640693 | 0.10098153 | 0.25484731 | 0.39624327 | 0.69192557 | 0.82267358 |
| PIWIL2 | 55.8570422 | -0.052158357 | 0.395099 | -0.1320134 | 0.8949737 | 0.94598591 |
| SLC39A14 | 11086.0094 | -1.648019781 | 0.39965133 | -4.1236439 | 3.73E-05 | 0.00058501 |
| PPP3CC | 1195.51738 | -0.628203759 | 0.15550862 | -4.0396716 | 5.35E-05 | 0.0007859 |
| AC037459.2 | 230.596344 | -0.494049157 | 0.17521484 | -2.8196764 | 0.00480721 | 0.02609431 |
| SORBS3 | 6191.47922 | -0.046268332 | 0.18736393 | -0.2469436 | 0.80495184 | 0.89435325 |
| AC037459.4 | 1173.76169 | 0.200615245 | 0.18824806 | 1.06569624 | 0.28656098 | 0.48273999 |
| PDLIM2 | 3029.55134 | 0.91202295 | 0.32412383 | 2.81381021 | 0.00489581 | 0.02646629 |
| AC037459.1 | 1612.43933 | 0.586248619 | 0.25655449 | 2.28508423 | 0.02230789 | 0.08132947 |
| C8orf58 | 1006.78091 | -0.454883208 | 0.17015051 | -2.6734167 | 0.00750829 | 0.03648817 |
| CCAR2 | 6464.02878 | -0.131737107 | 0.14993319 | -0.8786387 | 0.3795972 | 0.57561692 |
| AC037459.3 | 1741.62028 | 0.328919253 | 0.1928239 | 1.70580128 | 0.08804509 | 0.21698588 |
| BIN3 | 1940.98717 | -0.651646466 | 0.15585314 | -4.1811571 | 2.90E-05 | 0.000478 |
| EGR3 | 256.834251 | 0.921802634 | 0.77757598 | 1.18548239 | 0.23582677 | 0.42492045 |
| PEBP4 | 81.8120639 | 0.802301486 | 0.40029419 | 2.00427964 | 0.04504011 | 0.13490495 |
| AC037441.1 | 0.47648675 | 1.371928337 | 2.20774611 | 0.62141581 | 0.53432607 | NA |
| AC107959.1 | 736.955721 | 0.037940408 | 0.32683559 | 0.11608408 | 0.90758591 | 0.95385641 |
| RHOBTB2 | 2577.96567 | -0.227589729 | 0.30141912 | -0.7550607 | 0.45021256 | 0.64007469 |
| TNFRSF10B | 4458.9551 | -1.068253301 | 0.16377128 | -6.5228366 | 6.90E-11 | 1.05E-08 |
| AC107959.2 | 2.46359749 | -0.160744872 | 0.86291645 | -0.1862809 | 0.85222447 | 0.92254348 |
| AC107959.3 | 13.081461 | -0.282974972 | 0.50176597 | -0.5639581 | 0.57278266 | 0.73747761 |
| AC107959.4 | 95.1719106 | -0.388222045 | 0.28735712 | -1.351009 | 0.17669256 | 0.35157194 |
| AC107959.5 | 78.9013067 | -0.747752324 | 0.33756147 | -2.215159 | 0.02674916 | 0.09282984 |
| TNFRSF10C | 173.670613 | -0.971272324 | 0.32071484 | -3.0284608 | 0.00245803 | 0.01578622 |
| TNFRSF10D | 1793.9281 | -1.070044916 | 0.24479448 | -4.3711971 | 1.24E-05 | 0.00024353 |
| TNFRSF10A- | 126.628755 | -1.198512886 | 0.29716862 | -4.0331072 | 5.50E-05 | 0.00080061 |
| TNFRSF10A | 596.902358 | -1.359153088 | 0.37545692 | -3.6199975 | 0.00029461 | 0.00299124 |
| AC100861.2 | 58.1133927 | -0.582125606 | 0.28038738 | -2.0761477 | 0.03788029 | 0.11894538 |
| CHMP7 | 2320.37529 | 0.012348601 | 0.14595406 | 0.08460608 | 0.93257456 | 0.96746494 |
| R3HCC1 | 1572.68452 | 0.361708512 | 0.14495439 | 2.49532643 | 0.01258413 | 0.05346279 |
| LOXL2 | 638.67514 | -0.818265655 | 0.36217749 | -2.2592946 | 0.02386506 | 0.08533949 |
| ENTPD4 | 2362.8542 | -0.442711554 | 0.15094011 | -2.9330279 | 0.00335674 | 0.01997679 |
| AC104561.1 | 1.54243965 | 0.747110935 | 0.90785257 | 0.82294302 | 0.4105404 | 0.60490308 |
| AC104561.3 | 8.62673514 | 0.6690294 | 0.51455582 | 1.30020761 | 0.19352982 | 0.37310874 |
| SLC25A37 | 4173.84357 | -1.05386178 | 0.30859778 | -3.4150012 | 0.00063782 | 0.00547858 |
| NKX3-1 | 135.299375 | -0.574893419 | 0.3923471 | -1.4652674 | 0.14284797 | 0.30415026 |
| NKX2-6 | 0.08695548 | -0.642905668 | 3.30558948 | -0.1944905 | 0.84579185 | NA |
| AC012574.1 | 0 | NA | NA | NA | NA | NA |
| AC012574.2 | 0 | NA | NA | NA | NA | NA |
| STC1 | 2277.5855 | -0.062067886 | 0.46741882 | -0.1327886 | 0.89436058 | 0.94571903 |
| AC023202.1 | 0.64261388 | 0.772902391 | 1.59240654 | 0.48536751 | 0.62741568 | NA |
| ADAM28 | 517.963923 | 0.73610727 | 0.31539338 | 2.33393382 | 0.01959919 | 0.07379077 |
| AC120193.1 | 0.90177617 | 1.5813492 | 1.54813834 | 1.02145213 | 0.30704028 | NA |
| ADAMDEC1 | 33.7310332 | -1.569191523 | 1.17875313 | -1.33123 | 0.18311335 | 0.35981363 |
| ADAM7 | 24.702511 | -0.245414762 | 0.8026955 | -0.3057383 | 0.75980391 | 0.86698524 |
| AC024958.1 | 2.7217719 | 0.081284412 | 0.94594838 | 0.08592901 | 0.93152285 | 0.96698254 |
| AF106564.1 | 56.1130093 | 0.294552349 | 0.93199984 | 0.31604335 | 0.75196961 | 0.86163275 |

| | | | | | | |
|---|---|---|---|---|---|---|
| NEFM | 202.1462 | -0.746234674 | 0.95603457 | -0.780552 | 0.43506605 | 0.62716294 |
| NEFL | 344.5378 | 2.463427758 | 0.96252 | 2.55935228 | 0.01048674 | 0.04668062 |
| AC107373.2 | 8.09197215 | 3.894670747 | 0.80389501 | 4.84475047 | 1.27E-06 | 3.95E-05 |
| AC107373.1 | 0.38178775 | -0.032130764 | 2.26018777 | -0.014216 | 0.98865768 | NA |
| DOCK5 | 1296.72504 | -0.838784889 | 0.18339201 | -4.5737263 | 4.79E-06 | 0.00011372 |
| GNRH1 | 279.330095 | -0.868290012 | 0.26076196 | -3.3298185 | 0.00086903 | 0.00703815 |
| AC091185.1 | 17.1487153 | 0.003466392 | 0.44555045 | 0.00778002 | 0.9937925 | 0.99712002 |
| KCTD9 | 1536.17417 | -0.355051419 | 0.14770074 | -2.4038567 | 0.01622313 | 0.06442831 |
| CDCA2 | 34.2250277 | -1.509634387 | 0.66391979 | -2.2738204 | 0.02297679 | 0.08307937 |
| AC009623.1 | 0 | NA | NA | NA | NA | NA |
| AC009623.2 | 0 | NA | NA | NA | NA | NA |
| AC090103.1 | 0.03392908 | 0.380740593 | 3.29738709 | 0.11546736 | 0.90807469 | NA |
| EBF2 | 64.0291607 | -0.062401743 | 0.66290787 | -0.0941334 | 0.92500322 | 0.96362877 |
| PPP2R2A | 2448.91781 | -0.388120055 | 0.14931291 | -2.5993737 | 0.0093394 | 0.04278667 |
| BNIP3L | 6992.81197 | -0.348405118 | 0.21639697 | -1.6100277 | 0.10739181 | 0.24957452 |
| PNMA2 | 844.600537 | -1.692362116 | 0.35783242 | -4.7294823 | 2.25E-06 | 6.25E-05 |
| DPYSL2 | 3712.19527 | -0.083953836 | 0.24184396 | -0.3471405 | 0.72848576 | 0.84642348 |
| ADRA1A | 26.0500419 | 0.952544428 | 0.39394561 | 2.41795924 | 0.01560782 | 0.06261651 |
| AC090150.2 | 0.64572959 | 1.447912 | 2.14384826 | 0.67537989 | 0.49943444 | NA |
| AC090150.1 | 178.552063 | 2.19548574 | 0.67994991 | 3.22889334 | 0.0012427 | 0.00931497 |
| STMN4 | 4.20978971 | 1.241064196 | 0.67590568 | 1.83614996 | 0.06633548 | 0.17733847 |
| TRIM35 | 1216.84025 | -0.506252024 | 0.28174052 | -1.7968733 | 0.07235573 | 0.18893164 |
| PTK2B | 2606.49363 | 0.127073527 | 0.23987628 | 0.52974612 | 0.59628797 | 0.75505948 |
| CHRNA2 | 1.88970643 | 0.389595348 | 0.98161752 | 0.39689119 | 0.6914477 | 0.82241065 |
| EPHX2 | 7184.76262 | 2.086640372 | 0.44350659 | 4.7048689 | 2.54E-06 | 6.89E-05 |
| CLU | 81214.2177 | -2.531018076 | 0.37286917 | -6.7879521 | 1.14E-11 | 2.32E-09 |
| SCARA3 | 1225.53966 | -0.773953997 | 0.54673192 | -1.4156005 | 0.1568925 | 0.3238049 |
| CCDC25 | 2109.46145 | 0.179649594 | 0.09782804 | 1.83638143 | 0.06630126 | 0.17726497 |
| AC013643.2 | 191.128934 | -0.337289923 | 0.17588514 | -1.9176715 | 0.05515268 | 0.15603833 |
| ESCO2 | 14.3497328 | -0.963736706 | 0.68395854 | -1.4090572 | 0.15881826 | 0.32631944 |
| PBK | 26.5461197 | -1.051740673 | 0.9151454 | -1.1492607 | 0.25044848 | 0.44264923 |
| SCARA5 | 178.883514 | -0.000483532 | 0.99283528 | -0.000487 | 0.99961141 | 0.99979217 |
| AC069113.2 | 0.25532629 | 1.012079767 | 2.9348562 | 0.34484816 | 0.73020852 | NA |
| NUGGC | 354.772564 | 3.279845019 | 0.41691389 | 7.86696033 | 3.63E-15 | 2.13E-12 |
| ELP3 | 1700.32041 | 0.108179598 | 0.14456602 | 0.74830587 | 0.45427569 | 0.64388479 |
| AC021678.2 | 17.5239633 | -0.36689162 | 0.57297843 | -0.6403236 | 0.52196223 | 0.69790813 |
| PNOC | 24.5299839 | -0.450665194 | 0.62411747 | -0.7220839 | 0.47024288 | 0.65686538 |
| ZNF395 | 5077.6903 | -0.761165158 | 0.26699169 | -2.8508946 | 0.00435964 | 0.02432581 |
| FBXO16 | 888.216237 | 0.179597179 | 0.39026413 | 0.46019392 | 0.64537704 | 0.7907115 |
| AC025871.1 | 16.1015672 | -0.241867204 | 0.51404505 | -0.4705175 | 0.63798532 | 0.78567566 |
| FZD3 | 679.083429 | -0.157933349 | 0.12856126 | -1.2284676 | 0.21927148 | 0.4048018 |
| EXTL3 | 2121.16011 | -0.127297595 | 0.17521617 | -0.7265174 | 0.46752165 | 0.65455658 |
| EXTL3-AS1 | 69.1243414 | 0.482825945 | 0.29441988 | 1.63992302 | 0.10102117 | 0.23918837 |
| INTS9 | 1054.3094 | -0.184846757 | 0.11058999 | -1.6714601 | 0.09463083 | 0.2288112 |
| INTS9-AS1 | 2.07550334 | 1.194349511 | 0.84992513 | 1.40524085 | 0.15994967 | 0.32800939 |
| HMBOX1 | 1498.78952 | -0.626866553 | 0.15374981 | -4.0771858 | 4.56E-05 | 0.00069044 |
| KIF13B | 6241.03715 | -0.041237485 | 0.11277087 | -0.365675 | 0.71460762 | 0.83781262 |
| AC108449.3 | 2090.33601 | -0.34285425 | 0.13584385 | -2.5238849 | 0.01160659 | 0.05036462 |
| AC108449.1 | 15.4627127 | 1.97183567 | 0.48784294 | 4.04194776 | 5.30E-05 | 0.00078048 |
| DUSP4 | 173.209154 | -0.933868057 | 0.46764284 | -1.9969686 | 0.04582859 | 0.13667723 |
| AC084026.2 | 0.28585593 | -0.695547354 | 2.35188919 | -0.2957398 | 0.76742876 | NA |
| AC084026.1 | 0 | NA | NA | NA | NA | NA |
| LINC00589 | 0.10721287 | -0.642905668 | 3.30558948 | -0.1944905 | 0.84579185 | NA |
| LINC02099 | 1.36854366 | -3.675951175 | 1.74720722 | -2.1039011 | 0.03538708 | 0.11352277 |
| AC131254.1 | 0 | NA | NA | NA | NA | NA |
| AC131254.3 | 0 | NA | NA | NA | NA | NA |
| AC131254.2 | 0 | NA | NA | NA | NA | NA |
| LINC02209 | 0 | NA | NA | NA | NA | NA |
| SARAF | 16866.9897 | -0.05711419 | 0.20150435 | -0.283439 | 0.77684034 | 0.8776898 |
| AC044849.1 | 1349.2966 | 0.218507118 | 0.26416032 | 0.82717617 | 0.40813722 | 0.60264172 |
| LEPROTL1 | 3416.12268 | -0.225632178 | 0.17575732 | -1.2837712 | 0.19922204 | 0.38032905 |
| MBOAT4 | 21.4654501 | -1.005504606 | 0.5472419 | -1.8374043 | 0.06615024 | 0.17702015 |
| AC026979.2 | 40.4828633 | -0.31453494 | 0.34060534 | -0.9234586 | 0.35576825 | 0.55319445 |
| DCTN6 | 1711.17787 | -0.322366429 | 0.1368767 | -2.3551593 | 0.01851477 | 0.07093814 |
| AC026979.1 | 10.734765 | 0.088766951 | 0.39265866 | 0.22606645 | 0.82114974 | 0.90408983 |
| AC026979.3 | 22.1177266 | 0.17831922 | 0.3829256 | 0.46567589 | 0.64144751 | 0.78758115 |
| RBPMS-AS1 | 1786.25222 | -1.116672723 | 0.27863085 | -4.0077139 | 6.13E-05 | 0.0008652 |
| RBPMS | 15045.8181 | -0.918319615 | 0.22912214 | -4.0079916 | 6.12E-05 | 0.00086516 |
| GTF2E2 | 1274.85631 | -0.788520597 | 0.18433064 | -4.277751 | 1.89E-05 | 0.00033844 |
| AC102945.1 | 2.4534072 | -1.835137103 | 1.63822053 | -1.1202015 | 0.2626279 | 0.4561951 |
| SMIM18 | 5.43260148 | 2.026242584 | 0.72108862 | 2.8099772 | 0.0049545 | 0.02669054 |
| GSR | 6392.03659 | -0.391322851 | 0.32745423 | -1.195046 | 0.23206907 | 0.42010445 |
| UBXN8 | 705.757358 | -0.058687483 | 0.16797245 | -0.3493876 | 0.72679837 | 0.84564389 |
| PPP2CB | 7389.41514 | -0.307221842 | 0.17350124 | -1.7707185 | 0.07660753 | 0.19651173 |
| TEX15 | 0.95133558 | 0.112313814 | 1.55477503 | 0.07223798 | 0.94241252 | NA |
| PURG | 30.2916845 | -0.059684945 | 0.5669575 | -0.1052723 | 0.91615971 | 0.95919051 |
| WRN | 419.66935 | -0.369482233 | 0.24237737 | -1.524409 | 0.12740658 | 0.28102982 |
| AC009563.1 | 95.631316 | -0.508196627 | 0.29535382 | -1.7206367 | 0.08531677 | 0.21189889 |
| AC068672.2 | 0.03392908 | 0.380740593 | 3.29738709 | 0.11546736 | 0.90807469 | NA |
| AC068672.1 | 0 | NA | NA | NA | NA | NA |
| NRG1 | 872.124163 | -0.091810965 | 0.41273035 | -0.2224478 | 0.82396529 | 0.90578475 |
| AC090204.1 | 59.6654537 | 0.509526507 | 0.36511544 | 1.39552167 | 0.16285859 | 0.33188415 |
| AC067838.1 | 74.5109763 | 0.050024403 | 0.33277253 | 0.15032612 | 0.88050733 | 0.93801192 |
| FUT10 | 290.369282 | 0.194481205 | 0.22447887 | 0.86636752 | 0.38628865 | 0.58184907 |
| TTI2 | 787.636164 | -0.16419194 | 0.14777062 | -1.1111271 | 0.26651363 | 0.4606085 |
| MAK16 | 942.710386 | -0.327642346 | 0.19722691 | -1.6612457 | 0.09666412 | 0.23204668 |
| RNF122 | 705.461517 | -1.007661736 | 0.28199119 | -3.5733802 | 0.0003524 | 0.00345074 |
| DUSP26 | 123.917921 | 1.039768027 | 0.44492833 | 2.33693374 | 0.01944263 | 0.07335844 |
| AF279873.3 | 2.19632864 | 4.618057942 | 1.35784971 | 3.40100815 | 0.00067138 | 0.00570369 |
| AF279873.4 | 0 | NA | NA | NA | NA | NA |
| AC087855.2 | 0.2072565 | -0.340597073 | 3.29738699 | -0.103293 | 0.91773041 | NA |
| AC090993.1 | 0.03392908 | 0.380740593 | 3.29738709 | 0.11546736 | 0.90807469 | NA |
| AC087855.1 | 0 | NA | NA | NA | NA | NA |
| LINC01288 | 0 | NA | NA | NA | NA | NA |
| UNC5D | 237.714477 | 0.13028859 | 0.46539504 | 0.27995268 | 0.77951381 | 0.87916597 |
| AC100818.1 | 0 | NA | NA | NA | NA | NA |
| AC124290.1 | 0 | NA | NA | NA | NA | NA |
| AC090809.1 | 0 | NA | NA | NA | NA | NA |
| KCNU1 | 0.17332742 | -1.003574173 | 3.30558948 | -0.3035992 | 0.7614333 | NA |
| AC092818.1 | 0.11051935 | 0.02007176 | 3.29738704 | 0.00608717 | 0.99514317 | NA |
| AC091182.2 | 2.59447279 | 1.476430888 | 0.84131796 | 1.75490239 | 0.07927601 | 0.20115808 |
| LINC01605 | 31.1879474 | 0.527595655 | 0.42874424 | 1.23056032 | 0.21848735 | 0.40366507 |
| AC124067.1 | 0 | NA | NA | NA | NA | NA |
| AC124067.3 | 5.71965213 | -0.608525726 | 0.84809515 | -0.7175206 | 0.47305294 | 0.65911477 |
| AC124067.4 | 134.602238 | -0.460098292 | 0.6727352 | -0.6839218 | 0.49402452 | 0.67667913 |
| AC137579.2 | 0.09997307 | 0.547239394 | 3.29377202 | 0.16614368 | 0.86804388 | NA |
| AC137579.1 | 0 | NA | NA | NA | NA | NA |
| ZNF703 | 564.699777 | -0.868770813 | 0.33692756 | -2.5785092 | 0.00992276 | 0.04482839 |
| AC138356.3 | 0.03392908 | 0.380740593 | 3.29738709 | 0.11546736 | 0.90807469 | NA |
| AC138356.1 | 87.6441586 | 1.558142064 | 0.32403127 | 4.80861635 | 1.52E-06 | 4.57E-05 |
| ERLIN2 | 3598.23118 | 0.654763035 | 0.12958164 | 5.05289992 | 4.35E-07 | 1.62E-05 |
| PLPBP | 3350.04626 | 0.34290809 | 0.20496547 | 1.67300419 | 0.09432647 | 0.22832651 |
| ADGRA2 | 2097.06933 | 0.405207533 | 0.36381457 | 1.11377491 | 0.26537576 | 0.45945899 |

| | | | | | | |
|---|---|---|---|---|---|---|
| BRF2 | 1211.41077 | 0.001857678 | 0.232892 | 0.00797656 | 0.99363569 | 0.99703855 |
| RAB11FIP1 | 2579.15782 | -0.34140271 | 0.15143097 | -2.2545105 | 0.02416407 | 0.08605086 |
| GOT1L1 | 0.08393208 | -0.642905668 | 3.30558948 | -0.1944905 | 0.84579185 | NA |
| ADRB3 | 3.24024036 | -2.33472817 | 1.38598634 | -1.6845247 | 0.09208032 | 0.22433093 |
| EIF4EBP1 | 1149.16871 | -1.156865648 | 0.25100101 | -4.6090079 | 4.05E-06 | 9.95E-05 |
| AP006545.2 | 12.5099786 | 0.554185079 | 0.53242032 | 1.0408789 | 0.29793176 | 0.49480164 |
| ASH2L | 1646.96115 | 0.391920209 | 0.15406105 | 2.54392794 | 0.01096137 | 0.04824759 |
| AC084024.1 | 0.51801375 | 0.932266777 | 1.79225183 | 0.52016506 | 0.60294854 | NA |
| STAR | 9.73669792 | 0.280334044 | 0.61133355 | 0.45856152 | 0.64654907 | 0.79171112 |
| AC084024.3 | 865.680798 | -0.609309915 | 0.14467329 | -4.2116268 | 2.54E-05 | 0.00042968 |
| LSM1 | 1520.80503 | -0.519003037 | 0.13366368 | -3.8829025 | 0.00010322 | 0.0012992 |
| BAG4 | 1252.60908 | -0.104068194 | 0.13988574 | -0.7439514 | 0.45690588 | 0.64628054 |
| DDHD2 | 2528.65392 | 0.195952307 | 0.09791607 | 2.00122718 | 0.04536791 | 0.1356408 |
| PLPP5 | 2552.15034 | -0.195538878 | 0.20524815 | -0.952695 | 0.34074465 | 0.53879213 |
| NSD3 | 3142.88459 | -0.275216737 | 0.16486964 | -1.6692991 | 0.09505812 | 0.22944463 |
| AC087362.2 | 21.3003087 | -0.882214236 | 0.35673151 | -2.4730482 | 0.01339661 | 0.05594018 |
| AC087362.1 | 78.7898833 | 0.564994915 | 0.30947726 | 1.82564272 | 0.0679041 | 0.18011788 |
| AC087623.3 | 34.8018791 | -0.569648931 | 0.44217203 | -1.2882971 | 0.19764257 | 0.37843313 |
| LETM2 | 391.454256 | -0.739287444 | 0.30128223 | -2.4538037 | 0.01413541 | 0.05835619 |
| AC087623.1 | 21.0867123 | -0.985463296 | 0.63264748 | -1.5576815 | 0.11930876 | 0.26881169 |
| AC087623.4 | 107.503847 | -1.156484377 | 0.34177279 | -3.3837813 | 0.00071495 | 0.00601116 |
| FGFR1 | 5886.47885 | -0.931674954 | 0.35753145 | -2.6058545 | 0.00916454 | 0.04220553 |
| AC087623.2 | 212.713024 | -0.519947011 | 0.41086221 | -1.2655021 | 0.20569138 | 0.38823862 |
| C8orf86 | 0.87194008 | -0.234019024 | 1.21103787 | -0.1932384 | 0.84677226 | NA |
| AC069120.1 | 24.0103537 | 0.469867974 | 0.69207056 | 0.67893073 | 0.49718175 | 0.67893661 |
| AC016813.1 | 1.22525662 | -2.659667369 | 1.35067437 | -1.9691403 | 0.04893698 | 0.14359276 |
| TACC1 | 5432.79544 | 0.342040672 | 0.25393859 | 1.34694247 | 0.17799876 | 0.35337103 |
| AC067817.2 | 6.3017099 | -0.770031236 | 0.62352312 | -1.2349682 | 0.21684233 | 0.40161819 |
| PLEKHA2 | 2816.45576 | -0.077721752 | 0.21574201 | -0.3602532 | 0.71865778 | 0.84001549 |
| AC108863.1 | 732.476391 | -0.168636714 | 0.19457559 | -0.86669 | 0.38611189 | 0.58174897 |
| HTRA4 | 13.148452 | -1.08299936 | 0.48790849 | -2.2196772 | 0.02644069 | 0.09206931 |
| TM2D2 | 2592.40231 | -0.091178794 | 0.18266012 | -0.4991719 | 0.6176583 | 0.77143728 |
| ADAM9 | 6685.72818 | -0.520771791 | 0.25676669 | -2.0281906 | 0.04254079 | 0.12957693 |
| ADAM32 | 45.5022116 | -1.112904605 | 0.25913018 | -4.2947703 | 1.75E-05 | 0.00031979 |
| ADAM18 | 0.17648264 | -1.003574173 | 3.30558948 | -0.3035992 | 0.7614333 | NA |
| ADAM2 | 1.45297559 | -0.17414128 | 1.22670822 | -0.1419582 | 0.88711303 | 0.94152663 |
| IDO1 | 194.436482 | -0.74701113 | 0.99204327 | -0.7530026 | 0.45144837 | 0.64117486 |
| AC007991.3 | 2.78739056 | 3.688296769 | 1.37004133 | 2.69210622 | 0.00710023 | 0.03501141 |
| AC007991.4 | 28.6891688 | -0.417512164 | 0.92553443 | -0.4511039 | 0.65191469 | 0.7955477 |
| IDO2 | 79.9200082 | 4.931723033 | 0.57344624 | 8.60014888 | 7.96E-18 | 9.54E-15 |
| AC087518.1 | 0 | NA | NA | NA | NA | NA |
| AC022733.1 | 0.19271054 | 1.506623853 | 3.27645372 | 0.4598337 | 0.64563559 | NA |
| TCIM | 12838.8919 | -1.55852933 | 0.38536333 | -4.0443115 | 5.25E-05 | 0.00077438 |
| AC022733.2 | 10.2571929 | -2.343586403 | 0.6165322 | -3.8012393 | 0.00014397 | 0.00169391 |
| AC105999.2 | 31.8557092 | 1.511733922 | 0.56784459 | 2.66223182 | 0.00776244 | 0.0373704 |
| AC105999.1 | 0.09635527 | 0.547239394 | 3.29377202 | 0.16614368 | 0.86804388 | NA |
| AC010857.1 | 0.75049007 | -2.780432901 | 3.26469898 | -0.8516659 | 0.39439953 | NA |
| ZMAT4 | 11.1854848 | 1.297288038 | 0.72761579 | 1.78293003 | 0.07459769 | 0.19300836 |
| AC048387.1 | 0.17648264 | -1.003574173 | 3.30558948 | -0.3035992 | 0.7614333 | NA |
| SFRP1 | 9116.25772 | 0.471459897 | 0.28096028 | 1.6780304 | 0.09334116 | 0.22641951 |
| AC016868.1 | 0.16325399 | -1.003574173 | 3.30558948 | -0.3035992 | 0.7614333 | NA |
| GOLGA7 | 2717.04018 | -0.084288106 | 0.17554734 | -0.4801446 | 0.63112458 | 0.78075653 |
| GINS4 | 110.424197 | -0.429833854 | 0.29379405 | -1.4630448 | 0.14345513 | 0.30499973 |
| AC009630.1 | 12.7201367 | 0.327072675 | 0.34826468 | 0.93914972 | 0.34765388 | 0.5451408 |
| AC009630.2 | 43.9638105 | -0.538141484 | 0.31206783 | -1.7244376 | 0.08462888 | 0.2107331 |
| GPAT4 | 7391.11649 | -0.700616711 | 0.28552593 | -2.4537762 | 0.01413649 | 0.05835619 |
| AC009630.3 | 3178.52743 | -0.870846266 | 0.27483496 | -3.1686153 | 0.00153167 | 0.0109941 |
| NKX6-3 | 0.54767118 | 0.228380064 | 1.87325654 | 0.12191606 | 0.90296549 | NA |
| ANK1 | 116.381932 | -0.632160637 | 0.47569261 | -1.3289268 | 0.18387213 | 0.3609281 |
| AC113133.1 | 3.19776616 | 0.886341412 | 0.70214106 | 1.26234095 | 0.20682613 | 0.38946051 |
| AC027702.1 | 15.2882135 | 0.655761464 | 0.34636786 | 1.89325148 | 0.05832443 | 0.1622555 |
| KAT6A | 1597.79576 | 0.008797136 | 0.1286196 | 0.06839654 | 0.94546997 | 0.97394598 |
| AC103724.4 | 1.69420191 | -1.413465201 | 1.21493221 | -1.1634108 | 0.24466289 | 0.43572367 |
| AC103724.3 | 0.29052362 | -0.179944783 | 2.90963392 | -0.0618445 | 0.95068669 | NA |
| AP3M2 | 1150.78642 | 0.213763118 | 0.10516269 | 2.03268973 | 0.04208388 | 0.12856255 |
| PLAT | 6972.14348 | -0.164232319 | 0.3716566 | -0.4418926 | 0.65856689 | 0.79946078 |
| AC083973.1 | 1.8695806 | 0.693574434 | 1.13718648 | 0.60990387 | 0.54192549 | 0.71449304 |
| IKBKB | 3072.43177 | -0.215133033 | 0.12473264 | -1.7247533 | 0.08457194 | 0.21069107 |
| POLB | 662.071466 | -0.275837952 | 0.13999952 | -1.9702778 | 0.04880654 | 0.14325785 |
| DKK4 | 0.7439111 | 0.41401149 | 1.57135112 | 0.26347484 | 0.79218461 | NA |
| VDAC3 | 4971.12273 | -0.115657292 | 0.1458964 | -0.7927358 | 0.42793179 | 0.62042202 |
| SLC20A2 | 3282.3032 | -0.232606227 | 0.21952732 | -1.0595776 | 0.28933682 | 0.48560412 |
| AC090739.1 | 15.8594173 | -0.956126352 | 0.44900922 | -2.1294136 | 0.03322005 | 0.10844427 |
| SMIM19 | 1497.97412 | 0.852890349 | 0.21608993 | 3.94692316 | 7.92E-05 | 0.00105246 |
| CHRNB3 | 0.1357163 | 1.313406857 | 3.28103699 | 0.40030236 | 0.68893383 | NA |
| AC103843.1 | 0.18301898 | 0.18657164 | 3.29377195 | 0.05664376 | 0.95482897 | NA |
| CHRNA6 | 2.36339023 | 0.151255682 | 0.89814313 | 0.16840933 | 0.86626127 | 0.93055685 |
| THAP1 | 441.926846 | -0.144646584 | 0.22685779 | -0.6376091 | 0.5237282 | 0.69892989 |
| RNF170 | 1358.17572 | 0.466199515 | 0.1892568 | 2.46331712 | 0.01376581 | 0.0571031 |
| HOOK3 | 1814.64167 | -0.07798349 | 0.12505997 | -0.6235687 | 0.53291084 | 0.70688425 |
| AC110275.1 | 1040.49679 | -0.318790382 | 0.14910395 | -2.1380412 | 0.0325134 | 0.10677944 |
| FNTA | 3313.8876 | -0.56704221 | 0.14996775 | -3.7810943 | 0.00015614 | 0.00180906 |
| POMK | 9.29574764 | 0.278757274 | 0.47346143 | 0.58876448 | 0.55601927 | 0.72524253 |
| HGSNAT | 4792.30178 | 0.145248789 | 0.1213014 | 1.19742053 | 0.23114269 | 0.41900272 |
| LINC00293 | 0 | NA | NA | NA | NA | NA |
| AC091163.1 | 0.81865461 | 0.561101795 | 1.56813792 | 0.35781406 | 0.72048247 | NA |
| AC091163.2 | 0.06785815 | 0.795868262 | 3.28898349 | 0.24198001 | 0.80879565 | NA |
| AC120036.4 | 42.9842261 | -0.303919638 | 0.30306738 | -1.0028121 | 0.31595153 | 0.51312692 |
| IGLV8OR8-1 | 0 | NA | NA | NA | NA | NA |
| SPIDR | 2833.42672 | -0.873247398 | 0.22991961 | -3.7980552 | 0.00014584 | 0.00171148 |
| CEBPD | 10399.2082 | -2.415630711 | 0.38256915 | -6.3142329 | 2.72E-10 | 3.43E-08 |
| PRKDC | 3229.55363 | 0.026287733 | 0.15508825 | 0.16950177 | 0.86540198 | 0.93001264 |
| MCM4 | 1136.54204 | 0.00887253 | 0.31736249 | 0.02795709 | 0.97769638 | 0.9897433 |
| UBE2V2 | 1898.746 | -0.192862322 | 0.15176628 | -1.2707851 | 0.20380512 | 0.38607951 |
| AC026904.2 | 0.07830766 | -0.642905668 | 3.30558948 | -0.1944905 | 0.84579185 | NA |
| AC026904.3 | 14.4684633 | 1.50805184 | 0.43771901 | 3.44525093 | 0.00057053 | 0.00503004 |
| AC022915.2 | 10.0530483 | 1.563754188 | 0.50379404 | 3.1039553 | 0.00190952 | 0.01300131 |
| AC026904.1 | 2.0404421 | 2.585262891 | 1.28259746 | 2.01564636 | 0.04383697 | 0.13258182 |
| AC026904.4 | 0.74027313 | 2.326155286 | 1.61483879 | 1.44048762 | 0.14972949 | NA |
| AC022915.1 | 0.27359226 | -0.179157115 | 2.95368447 | -0.0606555 | 0.9516336 | NA |
| AC022915.3 | 0.08757577 | 0.547239394 | 3.29377202 | 0.16614368 | 0.86804388 | NA |
| EFCAB1 | 17.974976 | -0.805363542 | 0.72633656 | -1.1088022 | 0.26751554 | 0.46164418 |
| SNAI2 | 468.42518 | 0.455302531 | 0.40370439 | 1.12781169 | 0.25939946 | 0.45219565 |
| PPDPFL | 1.97124682 | 0.285598347 | 1.05870064 | 0.26976308 | 0.78734253 | 0.88417607 |
| AC022568.1 | 0.08695548 | -0.642905668 | 3.30558948 | -0.1944905 | 0.84579185 | NA |
| AC090155.2 | 0 | NA | NA | NA | NA | NA |
| AC090155.1 | 0 | NA | NA | NA | NA | NA |
| AC113145.1 | 0.31567319 | 0.328538905 | 2.27609247 | 0.14434339 | 0.88522932 | NA |
| AC023762.1 | 0.07830766 | -0.642905668 | 3.30558948 | -0.1944905 | 0.84579185 | NA |
| SNTG1 | 16.7690979 | 2.045549845 | 0.55508175 | 3.68513262 | 0.00022858 | 0.00243152 |
| AC012413.1 | 17.5742233 | 2.945344684 | 0.98083835 | 3.00288491 | 0.00267434 | 0.01683108 |
| PXDNL | 91.7415905 | 3.348227959 | 0.66767265 | 5.01477475 | 5.31E-07 | 1.90E-05 |
| AC090186.1 | 2.41993819 | -0.904687676 | 0.90750264 | -0.9968981 | 0.31881397 | 0.5160893 |

| | | | | | | | |
|---|---|---|---|---|---|---|---|
| PCMTD1 | 3833.85708 | | 0.256980507 | 0.20342706 | 1.26325626 | 0.2064971 | 0.38915008 |
| AC064807.1 | 113.400054 | | 0.851024547 | 0.27645673 | 3.07832817 | 0.00208166 | 0.01388011 |
| AC064807.4 | 14.6225247 | | 0.227970805 | 0.48923055 | 0.46597827 | 0.64123105 | 0.78758115 |
| AC064807.2 | 96.0424605 | | 1.690690735 | 0.56445123 | 2.99528222 | 0.00274191 | 0.01716623 |
| AC064807.3 | 0.09997307 | | 0.547239394 | 3.29377202 | 0.16614368 | 0.86804388 | NA |
| ST18 | 47.0854657 | | 1.286604931 | 0.68244226 | 1.88529493 | 0.05939003 | 0.16438702 |
| AC021915.2 | 0.19280283 | | 0.18657164 | 3.29377195 | 0.05664376 | 0.95482897 | NA |
| AC021915.1 | 0 | NA | NA | NA | NA | NA | |
| AC103831.1 | 0.19978473 | | -1.003574173 | 3.30558948 | -0.3035992 | 0.7614333 | NA |
| ALKAL1 | 4.47621844 | | -0.452418746 | 0.64589019 | -0.7004577 | 0.48364153 | 0.66823038 |
| RB1CC1 | 2226.12169 | | -0.319126753 | 0.26159326 | -1.2199349 | 0.22248956 | 0.40845514 |
| NPBWR1 | 2.97371148 | | -1.101907165 | 0.83721147 | -1.3161635 | 0.18811918 | 0.36613054 |
| AC009646.2 | 0.07830766 | | -0.642905668 | 3.30558948 | -0.1944905 | 0.84579185 | NA |
| OPRK1 | 1.09463561 | | -0.3475889 | 1.47729312 | -0.2352877 | 0.81398544 | 0.89962526 |
| AC131902.2 | 0.72941629 | | 1.314674523 | 1.47829917 | 0.88931561 | 0.37383349 | NA |
| AC131902.1 | 1.27393871 | | -2.709902177 | 1.48166831 | -1.8289533 | 0.0674066 | 0.17927504 |
| AC022034.4 | 3.3476566 | | -0.415895543 | 0.73664727 | -0.564579 | 0.57236019 | 0.73707622 |
| AC022034.2 | 9.6188892 | | 0.420637241 | 0.76451973 | 0.55019802 | 0.58218356 | 0.74463698 |
| ATP6V1H | 4526.3673 | | 0.120827317 | 0.1467201 | 0.8235226 | 0.41021087 | 0.60458644 |
| RGS20 | 15.1551779 | | -1.681081547 | 0.47757807 | -3.5200141 | 0.00043152 | 0.00404578 |
| AC113194.1 | 0.17648264 | | -1.003574173 | 3.30558948 | -0.3035992 | 0.7614333 | NA |
| TCEA1 | 3117.64451 | | 0.057629054 | 0.1423587 | 0.40481581 | 0.6856129 | 0.81834121 |
| AC100821.2 | 5.99135393 | | -1.236584403 | 0.61526437 | -2.0098424 | 0.04444787 | 0.13363248 |
| LYPLA1 | 3751.98568 | | 0.134815481 | 0.14303496 | 0.94253521 | 0.3459187 | 0.54354339 |
| MRPL15 | 2479.44805 | | -0.158573654 | 0.15641911 | -1.0137742 | 0.31069052 | 0.50761518 |
| SOX17 | 239.738095 | | 0.07645723 | 0.38286137 | 0.19969951 | 0.84171559 | 0.915716 |
| RP1 | 106.099821 | | -0.090589908 | 0.57630828 | -0.15719 | 0.87509509 | 0.93501854 |
| XKR4 | 123.56085 | | -1.845561995 | 0.4384163 | -4.2096108 | 2.56E-05 | 0.00043214 |
| AC022679.2 | 0.09644756 | | -0.642905668 | 3.30558948 | -0.1944905 | 0.84579185 | NA |
| AC022679.1 | 0.48920812 | | -1.018123215 | 2.80745741 | -0.3626496 | 0.71686668 | NA |
| AC090200.1 | 0 | NA | NA | NA | NA | NA | |
| TMEM68 | 1069.89159 | | 0.027274706 | 0.14309279 | 0.19060852 | 0.84883231 | 0.92053414 |
| TGS1 | 909.817421 | | -0.11859608 | 0.16671436 | -0.7113729 | 0.47685319 | 0.66203109 |
| LYN | 1416.29247 | | -0.822860147 | 0.23194625 | -3.5476329 | 0.00038871 | 0.0037328 |
| RPS20 | 22100.5677 | | -0.575840962 | 0.13440724 | -4.2843001 | 1.83E-05 | 0.00033201 |
| CERNA3 | 0.35085019 | | 1.644831918 | 2.15817781 | 0.7621392 | 0.44597693 | NA |
| MOS | 0 | NA | NA | NA | NA | NA | |
| PLAG1 | 300.590826 | | -0.247360358 | 0.26038355 | -0.9499846 | 0.34212008 | 0.53994876 |
| CHCHD7 | 1552.85212 | | -0.208312236 | 0.20162551 | -1.0331641 | 0.30152712 | 0.49834165 |
| AC107952.2 | 59.7263938 | | 0.011399103 | 0.31090985 | 0.03666369 | 0.97075316 | 0.98642676 |
| SDR16C5 | 1.73374684 | | -1.257443953 | 1.24831241 | -1.0073151 | 0.31378337 | 0.51080113 |
| PENK | 1.66427587 | | 0.962135904 | 1.57002995 | 0.61281372 | 0.53999951 | 0.71298754 |
| AC012349.1 | 0.83656804 | | 0.551252134 | 1.46791462 | 0.37553419 | 0.70726322 | NA |
| LINC00968 | 2.24229331 | | -0.73670207 | 1.18138169 | -0.6235936 | 0.53289451 | 0.70688425 |
| AC013644.1 | 0.08393208 | | -0.642905668 | 3.30558948 | -0.1944905 | 0.84579185 | NA |
| AC009597.1 | 0 | NA | NA | NA | NA | NA | |
| IMPAD1 | 4327.96914 | | -0.245175823 | 0.18803216 | -1.3039037 | 0.19226643 | 0.37151504 |
| LINC01606 | 761.97305 | | -0.044814846 | 0.44102699 | -0.1016147 | 0.91906247 | 0.96062334 |
| AC025674.2 | 0.09997307 | | 0.547239394 | 3.29377202 | 0.16614368 | 0.86804388 | NA |
| LINC00588 | 1.07683965 | | 0.321903351 | 1.60429401 | 0.2006511 | 0.8409714 | 0.91512686 |
| AC068075.1 | 0 | NA | NA | NA | NA | NA | |
| AC090796.1 | 0.2320242 | | 1.01099135 | 3.00413042 | 0.33653377 | 0.73646839 | NA |
| AC104051.1 | 0 | NA | NA | NA | NA | NA | |
| AC104051.2 | 1.13157523 | | 0.285030443 | 1.11471387 | 0.2556983 | 0.79818381 | 0.89073665 |
| AC012103.1 | 0.18874271 | | 0.726716446 | 3.28420131 | 0.22127646 | 0.82487718 | NA |
| LINC01602 | 0.10721287 | | -0.642905668 | 3.30558948 | -0.1944905 | 0.84579185 | NA |
| FAM110B | 770.345906 | | -0.074661757 | 0.20458996 | -0.3649336 | 0.71516095 | 0.83801965 |
| AC104350.1 | 29.2926668 | | -0.061890491 | 0.27515027 | -0.2249334 | 0.82203107 | 0.90490334 |
| AC027698.1 | 0.40358483 | | 1.052679206 | 2.22146157 | 0.47386784 | 0.63559415 | NA |
| UBXN2B | 1877.48913 | | 0.000735878 | 0.14022743 | 0.00524775 | 0.99581292 | 0.99795473 |
| CYP7A1 | 5.18574444 | | -0.312208745 | 0.63004727 | -0.4955323 | 0.62022446 | 0.77304528 |
| SDCBP | 11662.6537 | | -1.039649088 | 0.26475732 | -3.9268001 | 8.61E-05 | 0.00112712 |
| NSMAF | 1293.36452 | | -0.422499058 | 0.1900403 | -2.2232077 | 0.0262018 | 0.09147907 |
| TOX | 451.39946 | | -0.187778914 | 0.280457 | -0.6695462 | 0.50314713 | 0.68332771 |
| AC090152.1 | 123.505798 | | -0.211997363 | 0.25225199 | -0.840419 | 0.40067351 | 0.59547525 |
| AC087664.1 | 0 | NA | NA | NA | NA | NA | |
| AC087664.2 | 0 | NA | NA | NA | NA | NA | |
| AC021393.1 | 0.03392908 | | 0.380740593 | 3.29738709 | 0.11546736 | 0.90807469 | NA |
| CA8 | 810.601953 | | -0.516538414 | 0.47571714 | -1.08581 | 0.27756307 | 0.47303938 |
| LINC01301 | 39.204306 | | 0.427479072 | 0.41248167 | 1.03635896 | 0.30003471 | 0.4967375 |
| RAB2A | 6468.58917 | | -0.112275227 | 0.14349291 | -0.7824444 | 0.43395344 | 0.62592658 |
| AC068389.4 | 0.34595857 | | -0.260809688 | 2.31961822 | -0.1124365 | 0.91047734 | NA |
| AC068389.1 | 0.52460164 | | -0.547203839 | 1.90342767 | -0.2874834 | 0.77374222 | NA |
| AC068389.2 | 4.90586894 | | -0.537236091 | 0.69829957 | -0.769349 | 0.44168614 | 0.63293465 |
| AC068389.3 | 0.29614 | | -0.174096114 | 3.29377188 | -0.0528562 | 0.95784652 | NA |
| CHD7 | 938.102026 | | -0.393749973 | 0.11632674 | -3.3848622 | 0.00071214 | 0.00599206 |
| AC113143.1 | 2.4260382 | | -1.72967776 | 0.97972695 | -1.7654692 | 0.07748494 | 0.19803042 |
| AC022182.1 | 8.46954144 | | 0.088961988 | 0.80664941 | 0.11028582 | 0.91218271 | 0.95681217 |
| AC022182.2 | 7.72327599 | | 0.028030736 | 0.838622 | 0.03342476 | 0.97333587 | 0.98766731 |
| CLVS1 | 830.53171 | | -0.284978753 | 0.21395576 | -1.3319518 | 0.18287604 | 0.35969577 |
| AC023866.2 | 0 | NA | NA | NA | NA | NA | |
| AC023866.1 | 0.08666371 | | -0.642905668 | 3.30558948 | -0.1944905 | 0.84579185 | NA |
| ASPH | 5478.28381 | | -0.161827068 | 0.17282268 | -0.9363763 | 0.34907947 | 0.54672599 |
| AC091173.1 | 0.2854129 | | -1.364245016 | 2.9060998 | -0.4694419 | 0.6387538 | NA |
| AC025524.2 | 0.08757577 | | 0.547239394 | 3.29377202 | 0.16614368 | 0.86804388 | NA |
| AC025524.1 | 0 | NA | NA | NA | NA | NA | |
| LINC02155 | 0 | NA | NA | NA | NA | NA | |
| NKAIN3 | 3.5609574 | | 0.599862497 | 0.8280442 | 0.72443294 | 0.46879998 | 0.65560447 |
| AC018861.2 | 0 | NA | NA | NA | NA | NA | |
| AC120042.3 | 15.9574964 | | 1.572411268 | 0.73011987 | 2.15363441 | 0.03126885 | 0.1039929 |
| AC120042.1 | 2.85403091 | | 3.765850059 | 1.040064 | 3.62078687 | 0.00029371 | 0.00298577 |
| GGH | 2461.98474 | | 1.098082165 | 0.42647244 | 2.57480217 | 0.01002974 | 0.04516467 |
| AC120042.2 | 1.22467707 | | 0.443270569 | 1.50192133 | 0.29513568 | 0.76789024 | 0.87183611 |
| TTPA | 52.4214173 | | 2.6043948 | 0.69702199 | 3.73646003 | 0.00018663 | 0.00206709 |
| YTHDF3-AS1 | 48.5597311 | | 0.995249294 | 0.33243518 | 2.99381456 | 0.00275513 | 0.01722448 |
| YTHDF3 | 2874.39185 | | 0.079068786 | 0.21281185 | 0.37154315 | 0.71023302 | 0.83536329 |
| AC011978.2 | 58.4188099 | | -0.414230147 | 0.2177873 | -1.901994 | 0.05717194 | 0.15989224 |
| AC011124.1 | 0 | NA | NA | NA | NA | NA | |
| AC011124.2 | 0 | NA | NA | NA | NA | NA | |
| AC018953.1 | 0 | NA | NA | NA | NA | NA | |
| AC022639.1 | 0 | NA | NA | NA | NA | NA | |
| AC069133.1 | 0 | NA | NA | NA | NA | NA | |
| LINC01289 | 0 | NA | NA | NA | NA | NA | |
| MIR124-2HG | 0 | NA | NA | NA | NA | NA | |
| AC090136.3 | 4.85636083 | | 0.855946453 | 0.70386078 | 1.21607351 | 0.22395692 | 0.4100272 |
| BHLHE22 | 13.9635973 | | 0.173209456 | 0.50238606 | 0.34477361 | 0.73026457 | 0.8475453 |
| CYP7B1 | 147.152052 | | -0.449063681 | 0.27623469 | -1.62566 | 0.10402201 | 0.24416075 |
| AC104232.1 | 3.96318046 | | 0.966936473 | 0.83921434 | 1.1521925 | 0.24924198 | 0.44107851 |
| LINC00251 | 0 | NA | NA | NA | NA | NA | |
| LINC01299 | 0.27275909 | | -0.695764746 | 2.37351103 | -0.2931374 | 0.76941716 | NA |
| AC100814.1 | 59.8169894 | | 0.435233872 | 0.30975956 | 1.40507001 | 0.16000046 | 0.32806253 |
| ARMC1 | 1479.63474 | | 0.070377834 | 0.20216931 | 0.34811334 | 0.72775506 | 0.84611931 |
| MTFR1 | 2257.41638 | | 0.444724223 | 0.26049113 | 1.70725283 | 0.08777507 | 0.21648228 |

| | | | | | | |
|---|---|---|---|---|---|---|
| AC055822.1 | 44.0963697 | -0.37525863 | 0.36769747 | -1.0205635 | 0.30746127 | 0.50442087 |
| PDE7A | 1690.33891 | 0.288555419 | 0.32664499 | 0.88339152 | 0.37702477 | 0.57295187 |
| AC100812.1 | 2.83990331 | 0.796071325 | 0.90161426 | 0.88294003 | 0.37726867 | 0.57327273 |
| DNAJC5B | 7.9955502 | -1.146685042 | 1.03514211 | -1.1077562 | 0.26796714 | 0.46209809 |
| AC084082.1 | 1.71291026 | 0.153051996 | 0.94436566 | 0.16206857 | 0.87125185 | 0.93306711 |
| TRIM55 | 417.283755 | 0.536000045 | 0.36979784 | 1.44944069 | 0.14721455 | 0.31046447 |
| CRH | 1.2121817 | 1.114972974 | 1.87906851 | 0.59336473 | 0.55293709 | 0.72282076 |
| LINC00967 | 2.076957 | 1.523602909 | 1.03366445 | 1.47398211 | 0.14048641 | 0.300627 |
| RRS1-AS1 | 17.6304275 | 2.105438252 | 0.50415422 | 4.17617896 | 2.96E-05 | 0.00048585 |
| RRS1 | 863.273506 | -0.778979955 | 0.34045645 | -2.2880458 | 0.02213485 | 0.08082194 |
| ADHFE1 | 1622.60698 | 0.475765817 | 0.2992739 | 1.58973374 | 0.11189484 | 0.25709391 |
| VXN | 675.184297 | 1.2979948 | 0.30202032 | 4.29770687 | 1.73E-05 | 0.00031603 |
| MYBL1 | 137.419433 | 0.127242573 | 0.26744631 | 0.47576866 | 0.6342392 | 0.78318369 |
| VCPIP1 | 1231.46224 | -0.319589765 | 0.17092285 | -1.8697896 | 0.06151304 | 0.16828497 |
| SGK3 | 1075.72771 | 0.781641613 | 0.22814308 | 3.42610271 | 0.00061231 | 0.00529223 |
| C8orf44 | 441.211852 | 0.011846068 | 0.27794441 | 0.04262028 | 0.96600423 | 0.98402795 |
| MCMDC2 | 681.277324 | -0.305855073 | 0.12913291 | -2.3685292 | 0.01785897 | 0.06897204 |
| TCF24 | 27.5322503 | 2.473558689 | 0.55871855 | 4.4271999 | 9.55E-06 | 0.00019624 |
| PPP1R42 | 950.976017 | -0.005627525 | 0.11624327 | -0.0484116 | 0.9613882 | 0.98156291 |
| COPS5 | 3278.50571 | -0.032235603 | 0.09277395 | -0.347464 | 0.72824279 | 0.84631627 |
| CSPP1 | 988.227088 | 0.564547011 | 0.18319978 | 3.08159224 | 0.00205897 | 0.01377665 |
| ARFGEF1 | 3926.42524 | -0.116205915 | 0.10273606 | -1.1311113 | 0.25800826 | 0.45051502 |
| AC021321.1 | 38.928067 | 0.020573651 | 0.37854665 | 0.05434905 | 0.95665707 | 0.97931865 |
| CPA6 | 3.42661487 | -0.181752887 | 0.73163757 | -0.2484193 | 0.80381001 | 0.89361172 |
| AC022874.1 | 0 | NA | NA | NA | NA | NA |
| PREX2 | 399.636296 | 0.3425833 | 0.39464838 | 0.86807223 | 0.38535479 | 0.58117277 |
| AC011853.2 | 0 | NA | NA | NA | NA | NA |
| C8orf34-AS1 | 17.1375808 | 0.564880941 | 1.00539588 | 0.56184927 | 0.57421872 | 0.73864963 |
| C8orf34 | 19.1043233 | 0.425598886 | 0.94731573 | 0.44926826 | 0.65323815 | 0.79625709 |
| AC083967.1 | 0.15452186 | 0.435199075 | 3.28898345 | 0.13232024 | 0.894731 | NA |
| LINC01592 | 1.09316604 | -0.407181119 | 2.26616321 | -0.1796786 | 0.85740486 | 0.9254517 |
| AC021785.1 | 1.36134641 | 0.187944101 | 2.26099969 | 0.08312434 | 0.93375267 | 0.96807777 |
| LINC01603 | 1.54813185 | 1.427080404 | 1.08732317 | 1.31247125 | 0.18936118 | 0.36772989 |
| SULF1 | 1373.76245 | -0.15603712 | 0.33749434 | -0.46234 | 0.64383754 | 0.78952734 |
| SLCO5A1 | 44.167127 | -2.169893631 | 0.68409277 | -3.1719289 | 0.0015143 | 0.01090148 |
| AC091047.1 | 0.18380315 | -1.003574173 | 3.30558948 | -0.3035992 | 0.7614333 | NA |
| AC079089.1 | 3.42864018 | -4.981040395 | 1.38291686 | -3.6018365 | 0.00031598 | 0.00315457 |
| PRDM14 | 4.38386209 | 2.61442591 | 1.22019119 | 2.14263628 | 0.03214232 | 0.10598802 |
| NCOA2 | 2323.66567 | 0.106713831 | 0.13677156 | 0.78023407 | 0.43525311 | 0.62732972 |
| AC022730.3 | 1.80928597 | 0.539095779 | 1.26093555 | 0.42753635 | 0.66898872 | 0.80664072 |
| AC022730.4 | 13.1668633 | -0.014822077 | 1.22355826 | -0.0121139 | 0.99033473 | 0.99533983 |
| TRAM1 | 13050.5403 | -0.623784237 | 0.18746023 | -3.3275551 | 0.00087612 | 0.00707776 |
| LACTB2-AS1 | 1352.73709 | -0.011963441 | 0.29575868 | -0.04045 | 0.96773436 | 0.98487728 |
| LACTB2 | 3240.64963 | -0.107908316 | 0.28980371 | -0.3723497 | 0.70963252 | 0.83500637 |
| XKR9 | 49.3906153 | -0.362894271 | 0.65774919 | -0.5517213 | 0.58113928 | 0.74384312 |
| AC022858.1 | 3.67568253 | 0.621598627 | 0.79087544 | 0.78596274 | 0.43188931 | 0.62401629 |
| EYA1 | 17.7211668 | 0.324569095 | 0.48875925 | 0.66406742 | 0.50664716 | 0.68574068 |
| AC009446.1 | 0.03392908 | 0.380740593 | 3.29738709 | 0.11546736 | 0.90807469 | NA |
| AC104012.1 | 0 | NA | NA | NA | NA | NA |
| MSC-AS1 | 449.253536 | -0.867383 | 0.4032654 | -2.1508986 | 0.0314842 | 0.10449823 |
| MSC | 587.154606 | -1.436805752 | 0.52651597 | -2.728893 | 0.00635473 | 0.03216192 |
| TRPA1 | 119.721464 | -0.527204826 | 0.42757831 | -1.2330018 | 0.2175751 | 0.40262844 |
| AC078906.1 | 9.5763157 | -1.030547505 | 0.50625469 | -2.0356305 | 0.04178747 | 0.12793865 |
| AC011131.1 | 0.08666371 | -0.642905668 | 3.30558948 | -0.1944905 | 0.84579185 | NA |
| KCNB2 | 1.80970191 | 1.948904123 | 1.19244948 | 1.63437039 | 0.1021811 | 0.24107682 |
| AC090735.1 | 0 | NA | NA | NA | NA | NA |
| AC022893.2 | 2.02290238 | 0.866467954 | 1.27431833 | 0.67994624 | 0.4965385 | 0.67836081 |
| TERF1 | 1535.4132 | 0.084934446 | 0.14741433 | 0.57616139 | 0.5645061 | 0.73150827 |
| AC022893.3 | 2.79979911 | 0.027126198 | 1.02268876 | 0.02652439 | 0.97883908 | 0.99044497 |
| AC022893.1 | 114.558459 | 0.047767473 | 0.33584511 | 0.14223066 | 0.88689781 | 0.94143003 |
| SBSPON | 1190.04052 | -0.425959369 | 0.27728985 | -1.536152 | 0.1245011 | 0.27654093 |
| C8orf89 | 2.97883131 | 0.649327199 | 0.86701024 | 0.74892679 | 0.45390133 | 0.64358565 |
| RPL7 | 34263.0576 | -0.284416046 | 0.25087828 | -1.1336814 | 0.2569282 | 0.44958173 |
| RDH10 | 5784.61545 | -0.457039572 | 0.41311916 | -1.1063141 | 0.26859057 | 0.46279822 |
| RDH10-AS1 | 144.054935 | 0.25669747 | 0.54991405 | 0.46679562 | 0.64064611 | 0.78733983 |
| AC111149.2 | 0 | NA | NA | NA | NA | NA |
| AC111149.1 | 0.07659028 | -0.642905668 | 3.30558948 | -0.1944905 | 0.84579185 | NA |
| STAU2-AS1 | 4.34691725 | 0.327859601 | 0.6372114 | 0.5145225 | 0.60688673 | 0.76327544 |
| STAU2 | 1865.41495 | 0.02525884 | 0.15269144 | 0.16542407 | 0.86861021 | 0.93156219 |
| AC027018.1 | 0.48918151 | 0.837831966 | 2.13514625 | 0.39240027 | 0.69476249 | NA |
| AC022826.2 | 430.788568 | 0.163033988 | 0.19944919 | 0.81742117 | 0.41368778 | 0.60761193 |
| UBE2W | 1999.36648 | -0.164289098 | 0.11438951 | -1.4362252 | 0.15093828 | 0.31566678 |
| ELOC | 2199.27516 | -0.724441057 | 0.23343769 | -3.1033594 | 0.00191337 | 0.01301808 |
| TMEM70 | 1400.98035 | -0.474915657 | 0.20883998 | -2.2740649 | 0.02296209 | 0.08305342 |
| LY96 | 148.495436 | -1.496461812 | 0.43141089 | -3.4687623 | 0.00052286 | 0.00469614 |
| AC087672.2 | 6.77415022 | -0.694737437 | 0.59219476 | -1.173157 | 0.24073284 | 0.43089309 |
| AC087672.1 | 0.9893134 | -2.28666119 | 1.48525902 | -1.5395706 | 0.12366505 | NA |
| AC087627.1 | 2.42543148 | -0.444455104 | 0.91848537 | -0.4839 | 0.62845682 | 0.77907137 |
| JPH1 | 157.797266 | -0.692730008 | 0.43724221 | -1.5843164 | 0.11312173 | 0.25910016 |
| GDAP1 | 471.762381 | 0.597724 | 0.21306145 | 2.80540659 | 0.00502531 | 0.02698382 |
| AC103952.1 | 0.97393196 | 1.742366677 | 1.23477723 | 1.41107775 | 0.15822169 | NA |
| MIR2052HG | 2.23037428 | 0.008592683 | 0.94811401 | 0.00906292 | 0.99276893 | 0.99666185 |
| AC115837.2 | 0.34145402 | -1.700893945 | 2.27276716 | -0.7483802 | 0.45423086 | NA |
| AC011632.1 | 1.04239804 | -1.29957574 | 1.44082046 | -0.9019692 | 0.36707321 | 0.56395482 |
| PI15 | 665.496087 | -4.138457775 | 0.67387506 | -6.1412835 | 8.19E-10 | 8.74E-08 |
| AC011632.2 | 0.03392908 | 0.380740593 | 3.29738709 | 0.11546736 | 0.90807469 | NA |
| CRISPLD1 | 106.661738 | 0.435740545 | 0.43041187 | 1.01238041 | 0.3113562 | 0.50845077 |
| AC100782.1 | 14.0660208 | 0.510718756 | 0.54692606 | 0.93379854 | 0.35040782 | 0.54813722 |
| AC022274.1 | 0.33973663 | -1.692524189 | 2.84715844 | -0.5944608 | 0.55220393 | NA |
| CASC9 | 19.3992683 | 1.268548766 | 0.59130626 | 2.1453329 | 0.03192624 | 0.10547418 |
| HNF4G | 811.04761 | 1.908686643 | 0.4867061 | 3.9216411 | 8.79E-05 | 0.00114528 |
| AC067773.1 | 0 | NA | NA | NA | NA | NA |
| LINC01109 | 0.9876691 | 1.916113221 | 1.253495 | 1.52861657 | 0.12635953 | NA |
| LINC01111 | 7.04740842 | 0.855401304 | 1.05926767 | 0.80754028 | 0.41935528 | 0.61231444 |
| ZFHX4-AS1 | 2.82904515 | -1.188380763 | 1.17932134 | -1.0076819 | 0.3136072 | 0.51067166 |
| ZFHX4 | 51.9592863 | -0.504594283 | 0.48454315 | -1.0413815 | 0.29769851 | 0.49454947 |
| AC023200.1 | 0 | NA | NA | NA | NA | NA |
| PEX2 | 2175.34865 | 0.033209684 | 0.08601842 | 0.38607645 | 0.69944005 | 0.82757815 |
| AC062004.1 | 0.38469857 | 0.954021709 | 2.16942743 | 0.43975737 | 0.66011284 | NA |
| AC084706.1 | 0 | NA | NA | NA | NA | NA |
| PKIA-AS1 | 6.16821668 | 0.614084132 | 0.59963679 | 1.02409349 | 0.30579112 | 0.50275652 |
| PKIA | 337.477392 | 0.008072049 | 0.3240532 | 0.02490964 | 0.98012704 | 0.99106512 |
| ZC2HC1A | 524.994514 | 0.078551026 | 0.16430367 | 0.47808444 | 0.63259011 | 0.78184295 |
| IL7 | 308.693198 | -0.269796069 | 0.21357279 | -1.2632511 | 0.20649894 | 0.38915008 |
| AC083837.1 | 45.0138456 | -2.066101111 | 0.69729852 | -2.963008 | 0.00304649 | 0.01858984 |
| AC138646.1 | 0 | NA | NA | NA | NA | NA |
| STMN2 | 1.38917736 | -0.909839242 | 1.18376896 | -0.7685953 | 0.4421336 | 0.63331779 |
| HEY1 | 220.273387 | 0.141733959 | 0.51087062 | 0.27743611 | 0.78144526 | 0.88036187 |
| LINC01607 | 13.658652 | 0.79733195 | 0.51125985 | 1.55954345 | 0.11886781 | 0.268116 |
| AC036214.1 | 2.47111744 | 1.041109028 | 0.80562129 | 1.29230576 | 0.19625128 | 0.37675788 |
| MRPS28 | 1474.92719 | 0.160435408 | 0.14222171 | 1.12806554 | 0.25929224 | 0.45207293 |
| AC036214.3 | 2212.40301 | 0.311966148 | 0.13764022 | 2.26653331 | 0.02341875 | 0.08414667 |
| AC009686.2 | 30.2831504 | -0.887540979 | 0.54843654 | -1.6183112 | 0.10559555 | 0.24659536 |

| | | | | | | | |
|---|---|---|---|---|---|---|---|
| TPD52 | 3739.91562 | | 0.310852442 | 0.14862423 | 2.09153276 | 0.03648033 | 0.11591594 |
| AC009686.1 | 0.84292407 | | 1.363787107 | 1.68954812 | 0.80719045 | 0.41955677 | NA |
| AC034114.2 | 7.05255245 | | 1.405589813 | 0.60409264 | 2.32677859 | 0.01997705 | 0.07486072 |
| AC009812.1 | 197.590961 | | 0.107973806 | 0.27736738 | 0.38928084 | 0.69706841 | 0.8258207 |
| ZBTB10 | 1693.17963 | | -0.189860507 | 0.12212945 | -1.5545841 | 0.12004515 | 0.26998642 |
| AC009812.3 | 3.43948436 | | 1.597105371 | 0.75843198 | 2.10579909 | 0.03522181 | 0.11315778 |
| AC009812.4 | 158.454407 | | -0.512905965 | 0.23238137 | -2.2071734 | 0.02730195 | 0.09429632 |
| ZNF704 | 1806.55072 | | 0.942370464 | 0.16805485 | 5.60751725 | 2.05E-08 | 1.28E-06 |
| PAG1 | 549.972432 | | -0.457705391 | 0.21360026 | -2.1428129 | 0.03212813 | 0.10596857 |
| AC022778.1 | 0.08393208 | | -0.642905668 | 3.30558948 | -0.1944905 | 0.84579185 | NA |
| AC079209.2 | 0.13390214 | | 1.04981649 | 3.28477129 | 0.31960109 | 0.74927075 | NA |
| AC079209.1 | 3.04349694 | | -1.446250988 | 0.97237216 | -1.4873431 | 0.13692422 | 0.29530784 |
| AC009902.3 | 2.35106767 | | -0.124241566 | 0.99777579 | -0.1245185 | 0.90090474 | 0.9498184 |
| AC009902.2 | 100.047504 | | 0.32062479 | 0.52517654 | 0.6105086 | 0.54152495 | 0.71428636 |
| FABP5 | 216.903328 | | 0.124213234 | 0.4954831 | 0.25069116 | 0.80205289 | 0.89266577 |
| AC018616.1 | 71.7254352 | | -1.507722553 | 0.39166829 | -3.8494884 | 0.00011836 | 0.00144906 |
| PMP2 | 26.4507757 | | 0.190387729 | 0.94541233 | 0.20138063 | 0.84040096 | 0.91505077 |
| FABP9 | 0.20718593 | | 0.18657164 | 3.29377195 | 0.05664376 | 0.95482897 | NA |
| FABP4 | 211.054408 | | 0.447822709 | 0.60784152 | 0.73674254 | 0.46127893 | 0.64971293 |
| AC023644.1 | 71.9969868 | | -1.529638727 | 0.3946968 | -3.8754779 | 0.00010642 | 0.00133184 |
| FABP12 | 0.27872071 | | -0.179578873 | 2.92985236 | -0.0612928 | 0.95112602 | NA |
| IMPA1 | 1461.54456 | | 0.406759776 | 0.14722848 | 2.76277912 | 0.00573115 | 0.02982643 |
| SLC10A5 | 42.5732423 | | 1.048186652 | 0.22445476 | 4.6699239 | 3.01E-06 | 7.87E-05 |
| ZFAND1 | 1985.32251 | | -0.128143428 | 0.1154428 | -1.1100166 | 0.26699187 | 0.4611392 |
| CHMP4C | 1061.69358 | | -0.636750212 | 0.18033216 | -3.5309853 | 0.00041401 | 0.00392295 |
| AC132219.1 | 55.3514807 | | -0.140455509 | 0.25885425 | -0.5426046 | 0.58740207 | 0.74833531 |
| SNX16 | 614.038677 | | -0.397443294 | 0.23664321 | -1.6795043 | 0.0930538 | 0.22593027 |
| LINC02235 | 4.53544208 | | -0.07047153 | 0.69970646 | -0.1007158 | 0.91977603 | 0.96074349 |
| AC027701.1 | 0 | NA | | NA | NA | NA | NA |
| AC060765.1 | 0 | NA | | NA | NA | NA | NA |
| AC105031.2 | 0 | NA | | NA | NA | NA | NA |
| AC105031.3 | 0 | NA | | NA | NA | NA | NA |
| LINC01419 | 0 | NA | | NA | NA | NA | NA |
| AC015522.1 | 3.01551567 | | -0.266512058 | 1.34193447 | -0.1986029 | 0.8425734 | 0.91630899 |
| RALYL | 385.354636 | | 2.84244759 | 0.49157872 | 5.78228366 | 7.37E-09 | 5.34E-07 |
| LRRCC1 | 388.587952 | | 0.663969372 | 0.21924105 | 3.02849017 | 0.00245779 | 0.01578622 |
| AC011773.4 | 5.03861186 | | 1.179714785 | 0.58075477 | 2.03134757 | 0.04221975 | 0.12890295 |
| E2F5 | 1865.88698 | | -0.442083429 | 0.12229339 | -3.6149413 | 0.00030042 | 0.0030364 |
| AC011773.1 | 31.9155809 | | -0.290273085 | 0.34396819 | -0.8438951 | 0.398728 | 0.59358697 |
| C8orf59 | 1700.22397 | | -0.509762792 | 0.1364631 | -3.7355358 | 0.00018732 | 0.00207108 |
| CA13 | 653.483978 | | -0.462395851 | 0.23793164 | -1.9433979 | 0.0519681 | 0.14980309 |
| AC011773.3 | 10.1945813 | | -0.643858471 | 0.48610894 | -1.3245148 | 0.18533214 | 0.36268759 |
| AC011773.2 | 0.29445039 | | 1.356350883 | 2.28325128 | 0.59404363 | 0.55248293 | NA |
| CA1 | 70.7185589 | | -4.10911831 | 0.90008902 | -4.5652354 | 4.99E-06 | 0.00011687 |
| CA3 | 248.58798 | | -0.006591863 | 0.37887005 | -0.0173987 | 0.98611851 | 0.99391317 |
| CA3-AS1 | 118.105757 | | 0.456655641 | 0.26802053 | 1.70380846 | 0.08841688 | 0.21770331 |
| CA2 | 9927.63522 | | 0.911218488 | 0.22626422 | 4.02723195 | 5.64E-05 | 0.00081459 |
| AC093331.2 | 0.19416835 | | -1.003574173 | 3.30558948 | -0.3035992 | 0.7614333 | NA |
| AC100801.1 | 0.42382268 | | 1.269910657 | 2.14203385 | 0.59285275 | 0.5532797 | NA |
| ATP6V0D2 | 2220.03718 | | 0.865422303 | 0.51596799 | 1.67727906 | 0.09348792 | 0.2266921 |
| PSKH2 | 6.34394464 | | 1.866279336 | 0.67652834 | 2.7586122 | 0.00580474 | 0.03007287 |
| AC023194.3 | 885.111364 | | 0.810175132 | 0.51620561 | 1.56948144 | 0.1165358 | 0.26472695 |
| AC084128.1 | 5.62789425 | | 6.410717139 | 1.07883208 | 5.9422752 | 2.81E-09 | 2.38E-07 |
| SLC7A13 | 927.516729 | | 6.24435652 | 1.70770973 | 3.65656787 | 0.00025561 | 0.00266425 |
| WWP1 | 3560.43969 | | 0.095930428 | 0.14776682 | 0.64920139 | 0.51620821 | 0.69363383 |
| RMDN1 | 2615.79344 | | 0.666343147 | 0.1695233 | 3.93068757 | 8.47E-05 | 0.00111292 |
| CPNE3 | 4179.62442 | | 0.35620646 | 0.12508987 | 2.84760426 | 0.00440497 | 0.02451128 |
| CNGB3 | 3.33136376 | | 2.441497278 | 0.93965265 | 2.59829765 | 0.00936872 | 0.04289864 |
| AC090572.3 | 0.42885147 | | -1.859940617 | 3.29773397 | -0.5640057 | 0.57275028 | NA |
| AC090572.2 | 1.11466255 | | -1.494508055 | 1.68676137 | -0.8860222 | 0.37560555 | 0.57152566 |
| CNBD1 | 0.15661533 | | -1.003574173 | 3.30558948 | -0.3035992 | 0.7614333 | NA |
| AF121898.1 | 0 | NA | | NA | NA | NA | NA |
| DCAF4L2 | 0 | NA | | NA | NA | NA | NA |
| AC037450.1 | 0.24176911 | | 0.074529887 | 3.28898342 | 0.02266046 | 0.98192111 | NA |
| MMP16 | 12.7540178 | | 0.893421352 | 0.65335612 | 1.36743398 | 0.17148932 | 0.34398835 |
| AC090568.2 | 0.31911136 | | 0.668858826 | 2.25352608 | 0.29680545 | 0.76661504 | NA |
| AC090578.1 | 5.8769161 | | -0.118850184 | 0.69982412 | -0.1698286 | 0.8651449 | 0.9299135 |
| AF117829.1 | 304.387825 | | -0.378607876 | 0.25196875 | -1.5025985 | 0.1329426 | 0.28939214 |
| RIPK2 | 758.037111 | | -1.197143086 | 0.28976343 | -4.1314499 | 3.60E-05 | 0.00057024 |
| OSGIN2 | 1652.72787 | | -0.563223094 | 0.35950647 | -1.5666564 | 0.11719504 | 0.26561337 |
| NBN | 2141.59006 | | -0.11596615 | 0.16681084 | -0.6951955 | 0.48693278 | 0.67093832 |
| DECR1 | 6521.53218 | | 0.302940798 | 0.20806783 | 1.45597136 | 0.14540054 | 0.30774335 |
| CALB1 | 7006.39855 | | 3.072259242 | 0.70963693 | 4.32933955 | 1.50E-05 | 0.00028297 |
| LINC00534 | 1.35047717 | | 0.39404819 | 1.57782965 | 0.24974064 | 0.80278793 | 0.89307786 |
| AB015752.1 | 2.13608272 | | -0.434471217 | 1.01379218 | -0.4285604 | 0.66824314 | 0.80603658 |
| LINC01030 | 1.28742695 | | -1.657761684 | 1.62114778 | -1.0225852 | 0.30650402 | 0.50339506 |
| TMEM64 | 1577.90385 | | 0.133710324 | 0.23069103 | 0.57960782 | 0.56217912 | 0.72981162 |
| AC106038.1 | 6.06874491 | | 0.739861044 | 0.59695776 | 1.23938593 | 0.21520261 | 0.3996122 |
| NECAB1 | 463.954657 | | -0.381920834 | 0.20466753 | -1.8660548 | 0.0620337 | 0.16932348 |
| AC103770.1 | 4.71575795 | | -1.923651361 | 0.88707536 | -2.1685321 | 0.03011823 | 0.10110982 |
| C8orf88 | 219.293387 | | -0.559324827 | 0.24282422 | -2.3034145 | 0.02125554 | 0.07841473 |
| PIP4P2 | 895.679464 | | -0.071907359 | 0.1624893 | -0.442536 | 0.65810142 | 0.79943711 |
| AC087439.1 | 195.421269 | | 0.089363465 | 0.18965639 | 0.47118616 | 0.6375078 | 0.78549111 |
| OTUD6B-AS1 | 1435.33149 | | 0.249422669 | 0.17951621 | 1.38941585 | 0.16470633 | 0.33469261 |
| OTUD6B | 485.729475 | | 0.027071296 | 0.16453615 | 0.16453099 | 0.86931315 | 0.932003 |
| LRRC69 | 76.001593 | | -0.48583137 | 0.27164789 | -1.7884599 | 0.07370184 | 0.19141645 |
| SLC26A7 | 1485.34605 | | 1.810418743 | 0.715145 | 2.53154078 | 0.01135626 | 0.04957203 |
| AC103409.1 | 0.19994613 | | 1.506623855 | 3.27645372 | 0.45983371 | 0.64563559 | NA |
| RUNX1T1 | 215.218085 | | 0.622021965 | 0.36996415 | 1.68130333 | 0.09270401 | 0.22528843 |
| AF181450.1 | 2.88601012 | | 1.513180154 | 0.77406695 | 1.95484403 | 0.05060148 | 0.14684286 |
| AC022695.2 | 0.15661533 | | -1.003574173 | 3.30558948 | -0.3035992 | 0.7614333 | NA |
| AC022695.3 | 0 | NA | | NA | NA | NA | NA |
| AC091096.1 | 0 | NA | | NA | NA | NA | NA |
| AC091096.2 | 0 | NA | | NA | NA | NA | NA |
| AC104211.2 | 12.3131882 | | 1.779897521 | 0.7499823 | 2.3732527 | 0.0176322 | 0.06838668 |
| AC104211.4 | 0 | NA | | NA | NA | NA | NA |
| AC104211.3 | 0 | NA | | NA | NA | NA | NA |
| AC117834.1 | 2.81838865 | | 0.158528353 | 0.71887593 | 0.22052255 | 0.82546421 | 0.90675212 |
| TRIQK | 1195.97428 | | 0.083961167 | 0.13442924 | 0.62457517 | 0.53224992 | 0.70646955 |
| C8orf87 | 2.54371753 | | -0.134622969 | 1.06576758 | -0.1263155 | 0.89948218 | 0.94902072 |
| LINC00535 | 39.3897453 | | 0.496513722 | 0.44110984 | 1.12560111 | 0.2603344 | 0.45340586 |
| AC016885.1 | 0 | NA | | NA | NA | NA | NA |
| AC016885.2 | 0 | NA | | NA | NA | NA | NA |
| FAM92A | 1759.92479 | | 0.145344399 | 0.15723493 | 0.92437727 | 0.35528992 | 0.55280831 |
| AC120053.1 | 213.68093 | | -0.197876493 | 0.29924819 | -0.6612454 | 0.50845494 | 0.68730556 |
| AC010834.3 | 30.9598508 | | 0.247719562 | 0.27116917 | 0.91352406 | 0.36096699 | 0.55866688 |
| RBM12B | 856.023493 | | 0.0352853 | 0.14281663 | 0.24706716 | 0.80485625 | 0.89428472 |
| AC010834.1 | 247.16339 | | -0.366277863 | 0.14473395 | -2.5306976 | 0.0113836 | 0.04965025 |
| AC010834.2 | 4.06466273 | | 0.646705557 | 0.689011 | 0.93859976 | 0.34793628 | 0.54542145 |
| TMEM67 | 644.934944 | | -0.04466869 | 0.18742401 | -0.2383296 | 0.81162546 | 0.89832375 |
| AC084346.1 | 6.35397269 | | 0.006858669 | 0.61842337 | 0.01109057 | 0.99115118 | 0.99583588 |
| PDP1 | 1401.98525 | | -0.212202653 | 0.35414594 | -0.5991955 | 0.54904252 | 0.71963077 |
| AC084346.2 | 6.62677837 | | -0.488363893 | 0.89465383 | -0.5458691 | 0.58515591 | 0.74685098 |

| | | | | | | |
|---|---|---|---|---|---|---|
| AP003351.1 | 0.63413805 | -0.922363544 | 1.83772295 | -0.5019057 | 0.61573389 | NA |
| CDH17 | 22.7990368 | 0.620215555 | 0.77949709 | 0.79566115 | 0.42622901 | 0.61866188 |
| GEM | 2665.69056 | -0.596689762 | 0.438748 | -1.3599829 | 0.17383535 | 0.34774982 |
| RAD54B | 288.293624 | -0.412934727 | 0.51158873 | -0.8071615 | 0.41957345 | 0.61243738 |
| FSBP | 266.12036 | -0.458510845 | 0.52402685 | -0.8749759 | 0.38158704 | 0.57757877 |
| VIRMA | 1619.62625 | 0.06959215 | 0.11135498 | 0.62495767 | 0.53199884 | 0.70623305 |
| AC023632.5 | 0.09997307 | 0.547239394 | 3.29377202 | 0.16614368 | 0.86804388 | NA |
| AC023632.2 | 25.8741724 | 0.344041982 | 0.29643573 | 1.16059554 | 0.24580642 | 0.43705606 |
| AC108860.2 | 84.4719806 | 0.466638274 | 0.42440613 | 1.0995088 | 0.2715462 | 0.46622001 |
| ESRP1 | 1205.05861 | -0.498174883 | 0.24455564 | -2.0370615 | 0.04164388 | 0.12758789 |
| DPY19L4 | 1421.55983 | 0.413533002 | 0.14801146 | 2.79392551 | 0.00520725 | 0.02770125 |
| AP003692.1 | 520.307806 | 0.405402009 | 0.16832334 | 2.40847171 | 0.01601947 | 0.06390896 |
| INTS8 | 1219.63161 | -0.428181943 | 0.17616958 | -2.4305101 | 0.01507759 | 0.06113099 |
| CCNE2 | 314.870268 | -0.722908998 | 0.20664628 | -3.4982918 | 0.00046825 | 0.00431436 |
| AC087752.4 | 12.6596686 | -0.092104984 | 0.47629555 | -0.1933778 | 0.8466631 | 0.91939162 |
| NDUFAF6 | 592.539371 | 0.172108683 | 0.14854271 | 1.15864777 | 0.24659979 | 0.43793093 |
| TP53INP1 | 2071.19489 | -0.242801266 | 0.19885656 | -1.220987 | 0.22209095 | 0.40805956 |
| MIR3150BHG | 1.37524238 | 0.231173535 | 1.16563026 | 0.19832493 | 0.84279085 | 0.91639431 |
| AC068189.1 | 0.59342838 | -1.142260835 | 1.66978969 | -0.6840747 | 0.49392801 | NA |
| PLEKHF2 | 1357.76982 | 0.468221493 | 0.26628353 | 1.75835695 | 0.07868679 | 0.20015151 |
| C8orf37-AS1 | 24.1640765 | 1.530167614 | 0.37252896 | 4.10751313 | 4.00E-05 | 0.00062331 |
| LINC01298 | 2.61919479 | 2.710301084 | 0.80734089 | 3.3570715 | 0.00078773 | 0.00649137 |
| C8orf37 | 185.913985 | 0.692342274 | 0.18800185 | 3.68263538 | 0.00023084 | 0.00244962 |
| AC083836.1 | 0.42538609 | -2.004632472 | 1.87973344 | -1.0664451 | 0.28622249 | NA |
| AC012339.1 | 0 | NA | NA | NA | NA | NA |
| AP003465.2 | 0 | NA | NA | NA | NA | NA |
| GDF6 | 22.8539325 | 2.616530295 | 0.42164321 | 6.20555538 | 5.45E-10 | 6.09E-08 |
| UQCRB | 10532.2044 | 0.074798412 | 0.16608752 | 0.45035539 | 0.6524542 | 0.79583057 |
| AP003465.1 | 2991.97929 | 0.157962966 | 0.16950499 | 0.93190748 | 0.35138434 | 0.54907977 |
| MTERF3 | 846.529681 | -0.221270591 | 0.10460395 | -2.1153177 | 0.03440288 | 0.11125875 |
| PTDSS1 | 3093.70014 | 0.230205855 | 0.19882084 | 1.15785578 | 0.2469229 | 0.43841519 |
| AP003548.1 | 2.10225798 | 1.227765278 | 0.95235111 | 1.28919394 | 0.19733067 | 0.37811436 |
| SDC2 | 5470.75654 | -0.152157554 | 0.17224647 | -0.8833711 | 0.37703579 | 0.57295187 |
| CPQ | 3788.90896 | 0.535732498 | 0.21787415 | 2.45890809 | 0.01393603 | 0.05768217 |
| AP003117.2 | 0.20908062 | 1.675937956 | 3.26598797 | 0.51314885 | 0.60784721 | NA |
| AP003115.1 | 0.22809742 | -0.339643007 | 3.05487708 | -0.1111806 | 0.91147315 | NA |
| TSPYL5 | 946.444181 | 0.196250728 | 0.20560926 | 0.9544839 | 0.33983876 | 0.5380566 |
| MTDH | 3853.6379 | -0.168046117 | 0.22840373 | -0.7357416 | 0.46188799 | 0.65014385 |
| LAPTM4B | 8242.03953 | -0.964441115 | 0.24447683 | -3.9449183 | 7.98E-05 | 0.00105887 |
| MATN2 | 2555.31412 | 1.042904546 | 0.4220033 | 2.47131846 | 0.01346159 | 0.0561768 |
| RPL30 | 16854.2016 | -0.244331224 | 0.16375791 | -1.492027 | 0.13569207 | 0.29339127 |
| AP003352.1 | 11597.2534 | -0.276839016 | 0.16357191 | -1.6924606 | 0.09055819 | 0.22162594 |
| ERICH5 | 346.723887 | -0.436473325 | 0.29657805 | -1.471698 | 0.14110244 | 0.30148043 |
| RIDA | 6737.0762 | 1.437965672 | 0.34985121 | 4.11022068 | 3.95E-05 | 0.00061788 |
| POP1 | 354.401609 | 0.358585403 | 0.19974203 | 1.79524258 | 0.07261506 | 0.18951495 |
| NIPAL2 | 854.886973 | 0.022402048 | 0.21187471 | 0.10573253 | 0.91579457 | 0.95892245 |
| AP003438.1 | 4.53758689 | 0.481512457 | 0.64358284 | 0.7481748 | 0.45435473 | 0.64389264 |
| STK3 | 1620.76381 | -0.446854943 | 0.2058807 | -2.1704557 | 0.02997234 | 0.10078687 |
| KCNS2 | 12.1385217 | -0.168607978 | 0.66048173 | -0.2552803 | 0.79850661 | 0.89090837 |
| AP003355.2 | 12.717647 | -0.773500115 | 0.53040691 | -1.4583145 | 0.14475386 | 0.30684561 |
| AP003467.1 | 0.19280283 | 0.18657164 | 3.29377195 | 0.05664376 | 0.95482897 | NA |
| AC016877.3 | 0.68227872 | -0.25287281 | 1.62295825 | -0.1558098 | 0.87618294 | NA |
| OSR2 | 105.261804 | 0.633644895 | 0.67520301 | 0.93845093 | 0.34801272 | 0.54548839 |
| AC016877.1 | 0.41816123 | 2.676048776 | 3.20608098 | 0.8346791 | 0.40389841 | NA |
| AC104986.2 | 83.4964676 | 0.392873399 | 0.28024441 | 1.40189558 | 0.16094643 | 0.32946417 |
| VPS13B | 1218.81429 | 0.171307932 | 0.10364714 | 1.65279941 | 0.09837169 | 0.23496935 |
| AC107909.2 | 1.25489997 | 0.279455579 | 1.07320047 | 0.26039457 | 0.79455943 | 0.88857752 |
| AC018442.2 | 0.81809255 | -1.966079532 | 1.47888865 | -1.3294304 | 0.18370601 | NA |
| COX6C | 6099.35628 | -0.110641842 | 0.16945433 | -0.6529301 | 0.51380132 | 0.69166697 |
| RGS22 | 17.1060268 | -0.44972403 | 0.59188781 | -0.759813 | 0.4473664 | 0.63760955 |
| FBXO43 | 14.4385986 | 0.305481368 | 0.58057767 | 0.52616796 | 0.59877152 | 0.75677743 |
| POLR2K | 1796.81551 | -0.129298042 | 0.07237745 | -1.7864411 | 0.07402787 | 0.19200541 |
| SPAG1 | 1029.34827 | -0.528558483 | 0.25528477 | -2.0704662 | 0.03840871 | 0.12021811 |
| RNF19A | 4575.92309 | -0.330038813 | 0.19653015 | -1.6793292 | 0.09308791 | 0.22599228 |
| AP001574.1 | 9.76894165 | 2.906920665 | 0.73186795 | 3.97191963 | 7.13E-05 | 0.00096911 |
| AP003472.1 | 6.87123523 | 2.103744668 | 0.79418949 | 2.6489203 | 0.00807494 | 0.03836686 |
| AP003472.2 | 0.79015022 | -2.881086252 | 1.45074405 | -1.985937 | 0.04704032 | NA |
| AP000424.1 | 92.8169087 | -2.546074133 | 0.59084386 | -4.3092165 | 1.64E-05 | 0.00030285 |
| AP000424.2 | 73.0637192 | -1.260224195 | 0.80526212 | -1.5649863 | 0.11758612 | 0.26615845 |
| ANKRD46 | 1418.85806 | 0.966902097 | 0.18909643 | 5.1132753 | 3.17E-07 | 1.25E-05 |
| SNX31 | 8.15004178 | 0.134097034 | 0.88520899 | 0.1514863 | 0.87959212 | 0.93749138 |
| AP001205.1 | 0 | NA | NA | NA | NA | NA |
| PABPC1 | 19714.3565 | -0.18178973 | 0.26406197 | -0.6884359 | 0.49117834 | 0.67425158 |
| YWHAZ | 21394.3879 | -0.50315454 | 0.19657188 | -2.5596466 | 0.01047786 | 0.04668062 |
| AC018781.1 | 2.33978375 | -0.777437995 | 1.02896355 | -0.7555545 | 0.44991637 | 0.63993921 |
| AP003469.2 | 1.65490278 | -0.192258114 | 0.92763988 | -0.2072551 | 0.83581062 | 0.91280681 |
| AP003469.3 | 0.50546304 | -2.201222026 | 2.77479786 | -0.7932909 | 0.42760833 | NA |
| AP003469.4 | 404.608766 | -0.580245758 | 0.25326895 | -2.291026 | 0.02196191 | 0.08039083 |
| ZNF706 | 2350.81199 | -0.261867174 | 0.16860672 | -1.5531241 | 0.12039348 | 0.27049642 |
| AP001330.5 | 14.9144964 | -0.136200742 | 0.42635238 | -0.3194558 | 0.7493809 | 0.85998627 |
| AP001330.2 | 0 | NA | NA | NA | NA | NA |
| AP001330.1 | 0.08393208 | -0.642905668 | 3.30558948 | -0.1944905 | 0.84579185 | NA |
| AP001208.1 | 0 | NA | NA | NA | NA | NA |
| AP001208.2 | 0 | NA | NA | NA | NA | NA |
| AP001207.3 | 13.9705129 | 0.62958234 | 0.60834513 | 1.03490981 | 0.30071103 | 0.4974536 |
| GRHL2 | 678.687358 | 0.027704926 | 0.35504462 | 0.07803224 | 0.93780241 | 0.97027448 |
| NCALD | 891.632154 | 1.568855442 | 0.30387343 | 5.16285822 | 2.43E-07 | 9.97E-06 |
| AP000426.1 | 128.101708 | 1.661631612 | 0.30105853 | 5.51929755 | 3.40E-08 | 1.95E-06 |
| RRM2B | 2019.489 | 0.105119004 | 0.18480286 | 0.56881696 | 0.56948036 | 0.73494823 |
| UBR5-AS1 | 420.633439 | -0.076418008 | 0.21471533 | -0.3559038 | 0.72191259 | 0.84200252 |
| UBR5 | 3192.04979 | -0.203041461 | 0.11343354 | -1.7899597 | 0.07346039 | 0.19094632 |
| AP002907.1 | 491.647615 | -0.359501616 | 0.13823944 | -2.600572 | 0.00930685 | 0.04265235 |
| AP002852.1 | 5.11400397 | -4.049593364 | 1.54657393 | -2.6184286 | 0.00883358 | 0.04103249 |
| ODF1 | 0 | NA | NA | NA | NA | NA |
| KLF10 | 4331.29537 | 0.067868791 | 0.32160933 | 0.21102868 | 0.83286489 | 0.91131316 |
| GASAL1 | 29.9443208 | 0.084297892 | 0.43774226 | 0.19257426 | 0.8472924 | 0.91980983 |
| AZIN1 | 3895.98212 | -0.127272882 | 0.16219171 | -0.7847065 | 0.43262569 | 0.62463532 |
| AP003696.1 | 0 | NA | NA | NA | NA | NA |
| AZIN1-AS1 | 244.594228 | -0.675452929 | 0.31129719 | -2.1698009 | 0.03002193 | 0.10090874 |
| AP003354.1 | 0.89094253 | -0.264431574 | 1.28487894 | -0.2058027 | 0.83694502 | NA |
| AP003354.2 | 57.7115315 | 0.147849803 | 0.32667555 | 0.45258913 | 0.65084463 | 0.79486596 |
| AP003550.1 | 269.503951 | 0.006092607 | 0.31373838 | 0.01941939 | 0.98450654 | 0.99328299 |
| ATP6V1C1 | 5899.23196 | -0.294436341 | 0.29860478 | -0.9860403 | 0.32411334 | 0.52169132 |
| LINC01181 | 0.13390214 | 1.04981649 | 3.28477129 | 0.31960109 | 0.74927075 | NA |
| BAALC-AS2 | 6.21743717 | 0.050344157 | 0.59021602 | 0.08529785 | 0.9320246 | 0.96716074 |
| BAALC | 199.997792 | -1.050233177 | 0.29320731 | -3.5818792 | 0.00034113 | 0.00335783 |
| BAALC-AS1 | 87.9266665 | 0.702228894 | 0.21362622 | 3.28718493 | 0.00101194 | 0.00791603 |
| AC025370.1 | 66.7779532 | -0.952012207 | 0.35298776 | -2.697012 | 0.00699648 | 0.03462133 |
| FZD6 | 1602.99219 | 0.049426572 | 0.13701331 | 0.36074285 | 0.71829169 | 0.83985952 |
| CTHRC1 | 152.234309 | -1.609104811 | 0.33387066 | -4.8195454 | 1.44E-06 | 4.38E-05 |
| AC012213.3 | 12.2565031 | -0.318490652 | 0.41588621 | -0.765812 | 0.44378817 | 0.6348089 |
| SLC25A32 | 1156.63123 | -0.153779106 | 0.25610887 | -0.6004443 | 0.54821019 | 0.71893289 |
| DCAF13 | 2018.76255 | 0.069117632 | 0.2411391 | 0.28662972 | 0.77439586 | 0.87594127 |

| | | | | | | |
|---|---|---|---|---|---|---|
| AC012213.2 | 7.74260245 | 0.99861786 | 0.546813 | 1.82625113 | 0.06781244 | 0.18000947 |
| AC012213.4 | 8.38080707 | -1.61244972 | 0.8494723 | -1.8981781 | 0.05767262 | 0.16098519 |
| AC012213.1 | 6.61046319 | -2.004727605 | 1.13624006 | -1.7643522 | 0.07767271 | 0.19836392 |
| RIMS2 | 54.8437493 | -0.666877573 | 0.50742763 | -1.3142319 | 0.18876818 | 0.36698739 |
| DPYS | 8857.09487 | 1.581090611 | 0.61259089 | 2.58098942 | 0.00985176 | 0.04459188 |
| DCSTAMP | 21.1539181 | 0.692201843 | 0.45768313 | 1.51240412 | 0.13043109 | 0.28569381 |
| LRP12 | 507.108783 | -0.012152667 | 0.18740119 | -0.0648484 | 0.94829471 | 0.97510358 |
| ZFPM2 | 150.733626 | 0.201882966 | 0.27718292 | 0.72833841 | 0.46640646 | 0.65361844 |
| AC012564.1 | 0 | NA | NA | NA | NA | NA |
| AC021546.1 | 0.46088298 | -0.820674226 | 2.86622347 | -0.286326 | 0.77462847 | NA |
| AC103853.1 | 1.17885197 | -0.823432013 | 1.29508761 | -0.6358118 | 0.52489908 | 0.69974161 |
| AC090802.1 | 0.12059279 | 0.02007176 | 3.29738704 | 0.00608717 | 0.99514317 | NA |
| AC027031.1 | 0.14444843 | 0.435199075 | 3.28898345 | 0.13232024 | 0.894731 | NA |
| AC027031.2 | 545.14654 | -0.960778469 | 0.38999369 | -2.4635744 | 0.01375594 | 0.05707318 |
| OXR1 | 2068.92642 | 0.491572408 | 0.17862987 | 2.75190492 | 0.00592497 | 0.03052188 |
| AC090579.1 | 4.08579623 | 0.397070725 | 0.61953778 | 0.64091447 | 0.52157826 | 0.69778391 |
| ABRA | 4.64794758 | -0.265253546 | 0.87928696 | -0.3016689 | 0.76290446 | 0.86875635 |
| ANGPT1 | 430.466532 | 1.614375845 | 0.42034759 | 3.84057359 | 0.00012275 | 0.00148547 |
| RSPO2 | 1.06534657 | 1.105959229 | 1.48759245 | 0.7434558 | 0.45720577 | 0.64655504 |
| EIF3E | 8835.22775 | -0.102568383 | 0.19161494 | -0.5352839 | 0.59245359 | 0.75183615 |
| AC087620.1 | 0 | NA | NA | NA | NA | NA |
| EMC2 | 2146.8869 | -0.193169386 | 0.20700809 | -0.933149 | 0.35074305 | 0.54841946 |
| AC022634.2 | 0 | NA | NA | NA | NA | NA |
| TMEM74 | 4.86511235 | -0.042555354 | 0.752114 | -0.056581 | 0.95487898 | 0.97841508 |
| AC104248.1 | 2.99084829 | -1.26011487 | 0.88745047 | -1.419927 | 0.15562894 | 0.32192776 |
| TRHR | 0 | NA | NA | NA | NA | NA |
| NUDCD1 | 1220.49909 | -0.437313257 | 0.136538 | -3.2028686 | 0.00136066 | 0.01002252 |
| AC021237.1 | 0.33280259 | 0.66965421 | 2.23368702 | 0.29979769 | 0.76433148 | NA |
| ENY2 | 2702.85305 | -0.151817412 | 0.12256214 | -1.2386975 | 0.21545756 | 0.39991664 |
| PKHD1L1 | 104.368333 | -0.868287789 | 0.70776089 | -1.2268095 | 0.2198942 | 0.40563903 |
| EBAG9 | 1237.21336 | 0.20501105 | 0.13898949 | 1.47501112 | 0.14020956 | 0.30032702 |
| SYBU | 329.855922 | -0.869130963 | 0.40248424 | -2.1594161 | 0.0308179 | 0.10284333 |
| AC079061.1 | 1.46964617 | -0.363245747 | 1.28444026 | -0.2828047 | 0.77732655 | 0.87808863 |
| KCNV1 | 46.4166181 | -1.766706422 | 0.52389189 | -3.3722729 | 0.00074551 | 0.0062153 |
| AC027451.1 | 7.63090162 | -1.830104527 | 0.78500375 | -2.3313322 | 0.01973585 | 0.07414651 |
| AC073023.1 | 0 | NA | NA | NA | NA | NA |
| AC025366.1 | 0.2745553 | -0.179253712 | 2.94817509 | -0.0608016 | 0.95151723 | NA |
| AP005357.1 | 0 | NA | NA | NA | NA | NA |
| LINC01608 | 0 | NA | NA | NA | NA | NA |
| LINC01609 | 0 | NA | NA | NA | NA | NA |
| AC009930.1 | 0 | NA | NA | NA | NA | NA |
| LINC02237 | 0.08666371 | -0.642905668 | 3.30558948 | -0.1944905 | 0.84579185 | NA |
| CSMD3 | 7.97973001 | 0.402594798 | 0.90075451 | 0.44695285 | 0.6549091 | 0.79747572 |
| AC103993.1 | 0 | NA | NA | NA | NA | NA |
| AC064802.1 | 0 | NA | NA | NA | NA | NA |
| TRPS1 | 916.325465 | -0.08928215 | 0.19402503 | -0.4601579 | 0.64540289 | 0.7907115 |
| AF178030.1 | 0.29115023 | 2.197172564 | 3.22988048 | 0.68026435 | 0.49633709 | NA |
| LINC00536 | 0.26369991 | -0.701265545 | 3.29738693 | -0.2126731 | 0.83158194 | NA |
| AC105177.1 | 0.10721287 | -0.642905668 | 3.30558948 | -0.1944905 | 0.84579185 | NA |
| EIF3H | 7029.70068 | -0.112073497 | 0.15226667 | -0.7360343 | 0.46170981 | 0.65011148 |
| UTP23 | 887.590284 | 0.293261317 | 0.13648966 | 2.14859732 | 0.03166633 | 0.1048915 |
| RAD21 | 5775.14206 | 0.10522578 | 0.11876571 | 0.88599459 | 0.37562044 | 0.57152566 |
| RAD21-AS1 | 66.272542 | 1.261711558 | 0.34198615 | 3.6893645 | 0.00022481 | 0.00240209 |
| AARD | 55.563914 | 2.2711106 | 0.34644331 | 6.55550426 | 5.55E-11 | 8.81E-09 |
| SLC30A8 | 146.541351 | 4.891308337 | 0.71306475 | 6.85955704 | 6.91E-12 | 1.45E-09 |
| MED30 | 304.665953 | -0.499053564 | 0.20362916 | -2.4507962 | 0.01425406 | 0.05864382 |
| EXT1 | 1853.99172 | -0.352217182 | 0.11509298 | -3.0602837 | 0.00221127 | 0.01458877 |
| SAMD12 | 933.11527 | -0.526917786 | 0.20384208 | -2.5849314 | 0.00973984 | 0.04419124 |
| AC023590.1 | 1.49110027 | -0.9868117 | 1.27195551 | -0.7758225 | 0.4378538 | 0.62980443 |
| SAMD12-AS1 | 18.3589742 | -1.330364023 | 0.37051064 | -3.590623 | 0.00032989 | 0.00327036 |
| TNFRSF11B | 3565.52819 | -0.867066914 | 0.36415361 | -2.3810471 | 0.0172635 | 0.06738286 |
| COLEC10 | 40.9661088 | -1.356659997 | 0.42544503 | -3.1888021 | 0.00142864 | 0.01042287 |
| AC107953.2 | 2.19565692 | -3.127536646 | 1.41467676 | -2.2107783 | 0.0270512 | 0.09360189 |
| MAL2 | 6495.95727 | -0.109431814 | 0.22057506 | -0.4961205 | 0.61980937 | 0.77271053 |
| MAL2-AS1 | 2001.93977 | -0.081655039 | 0.21609984 | -0.3778579 | 0.70553613 | 0.83210286 |
| NOV | 1177.33997 | -0.810936947 | 0.59751305 | -1.357187 | 0.1747218 | 0.34862393 |
| AC021733.1 | 0.07830766 | -0.642905668 | 3.30558948 | -0.1944905 | 0.84579185 | NA |
| ENPP2 | 7154.49998 | 0.091449827 | 0.47900145 | 0.19091764 | 0.84859012 | 0.92038504 |
| TAF2 | 1118.86656 | 0.214425713 | 0.12174475 | 1.7612727 | 0.07819226 | 0.19914331 |
| DSCC1 | 71.0261728 | -0.07755867 | 0.29443243 | -0.2634176 | 0.79222876 | 0.8873257 |
| AP005717.1 | 677.993244 | -1.203565747 | 0.33134591 | -3.6323544 | 0.00028085 | 0.00288429 |
| DEPTOR | 6485.39626 | -1.414476643 | 0.3378519 | -4.1866766 | 2.83E-05 | 0.00046888 |
| AC091563.1 | 645.192312 | -1.125485546 | 0.26208239 | -4.2943959 | 1.75E-05 | 0.00032011 |
| COL14A1 | 2196.39168 | 0.67617056 | 0.72286395 | 0.93540501 | 0.34957962 | 0.54721682 |
| MRPL13 | 1772.91225 | -0.141382785 | 0.17557817 | -0.8052413 | 0.42068047 | 0.61340028 |
| MTBP | 312.398399 | -0.031827583 | 0.23777286 | -0.1338571 | 0.89351558 | 0.94512659 |
| SNTB1 | 2147.08935 | -0.781899159 | 0.19179459 | -4.076753 | 4.57E-05 | 0.00069075 |
| AC104958.1 | 1.78134961 | -0.853209886 | 1.31877622 | -0.6469709 | 0.51765075 | 0.69475012 |
| AC104958.2 | 165.050465 | -1.737859217 | 0.44301364 | -3.922812 | 8.75E-05 | 0.00114146 |
| AC068413.1 | 0.16325399 | -1.003574173 | 3.30558948 | -0.3035992 | 0.7614333 | NA |
| HAS2 | 94.815315 | -0.380786734 | 0.77374648 | -0.4921337 | 0.62262482 | 0.77486509 |
| HAS2-AS1 | 5.90362304 | 1.119496848 | 0.54078703 | 2.07012518 | 0.03844062 | 0.12028946 |
| AC104233.2 | 0 | NA | NA | NA | NA | NA |
| AC104233.1 | 0.09997307 | 0.547239394 | 3.29377202 | 0.16614368 | 0.86804388 | NA |
| AC104036.1 | 0.09635527 | 0.547239394 | 3.29377202 | 0.16614368 | 0.86804388 | NA |
| AC037486.1 | 0 | NA | NA | NA | NA | NA |
| SMILR | 0.14114194 | 0.02007176 | 3.29738704 | 0.00608717 | 0.99514317 | NA |
| AC108136.1 | 0.69790895 | -1.066518158 | 2.24442103 | -0.4751863 | 0.63465418 | NA |
| LINC01151 | 1.48703506 | 0.050187196 | 1.58822513 | 0.03159955 | 0.9747914 | 0.98843899 |
| AC100872.1 | 0.74239049 | 1.225604591 | 1.50916427 | 0.81210814 | 0.41672958 | NA |
| AC016405.3 | 24.9646094 | -0.329755151 | 0.41261631 | -0.7991811 | 0.42418542 | 0.61664005 |
| AC016405.2 | 8.37411875 | -0.05280382 | 0.51947652 | -0.1016481 | 0.91903597 | 0.96062334 |
| ZHX2 | 1791.34806 | -0.467893941 | 0.14694728 | -3.184094 | 0.00145208 | 0.01054276 |
| AC104316.2 | 1.43575466 | -3.017255573 | 1.27049114 | -2.3748734 | 0.01755497 | 0.06818745 |
| AC104316.1 | 386.34209 | -0.384022735 | 0.23109144 | -1.6617783 | 0.09655724 | 0.23189566 |
| DERL1 | 3875.91895 | -0.579136966 | 0.14945129 | -3.8750885 | 0.00010659 | 0.0013327 |
| TBC1D31 | 601.573208 | -0.236547819 | 0.16541856 | -1.4299956 | 0.15271827 | 0.31810274 |
| FAM83A | 12.0878799 | -4.671340365 | 1.05016806 | -4.4481836 | 8.66E-06 | 0.00018182 |
| AC068228.1 | 0.74778079 | -2.039413613 | 1.48224607 | -1.3758941 | 0.16885442 | NA |
| FAM83A-AS1 | 0.76224017 | -2.839483033 | 1.82012372 | -1.5600495 | 0.11874819 | NA |
| C8orf76 | 686.468915 | -0.675559159 | 0.16288898 | -4.1473595 | 3.36E-05 | 0.0005385 |
| ZHX1-C8orf7 | 636.325556 | -0.527962385 | 0.15854504 | -3.3300467 | 0.00086831 | 0.00703671 |
| ZHX1 | 1469.54647 | 0.319668381 | 0.10468215 | 3.0537049 | 0.00226034 | 0.01483456 |
| ATAD2 | 249.779688 | -0.228245706 | 0.25547171 | -0.8934285 | 0.37162775 | 0.56836599 |
| WDYHV1 | 475.95375 | -0.470007458 | 0.24203029 | -1.9419365 | 0.05214479 | 0.15008516 |
| FBXO32 | 904.193497 | -1.025224682 | 0.25206035 | -4.0673778 | 4.75E-05 | 0.00071358 |
| AC090193.2 | 0 | NA | NA | NA | NA | NA |
| AC090193.1 | 0.3216386 | -1.364242679 | 3.30558948 | -0.4127078 | 0.67982068 | NA |
| KLHL38 | 3.86614704 | 1.114686624 | 0.76646374 | 1.45432401 | 0.1458565 | 0.30843912 |
| ANXA13 | 142.811413 | 1.891385412 | 0.62270832 | 3.03735369 | 0.00238665 | 0.01544821 |
| FAM91A1 | 2498.22886 | -0.046259914 | 0.19720599 | -0.2345766 | 0.81453736 | 0.8999602 |
| FER1L6 | 86.0609075 | 2.565316311 | 0.57981561 | 4.42436576 | 9.67E-06 | 0.00019834 |
| AC090753.1 | 0.39257597 | 1.760399417 | 2.78348343 | 0.63244473 | 0.52709632 | NA |

| | | | | | | |
|---|---|---|---|---|---|---|
| FER1L6-AS1 | 0.23749721 | 0.689160045 | 2.99954805 | 0.22975463 | 0.81828244 | NA |
| AC100871.2 | 0 | NA | NA | NA | NA | NA |
| FER1L6-AS2 | 5.24938673 | 2.129125292 | 0.74001292 | 2.87714613 | 0.0040129 | 0.02280606 |
| AC090921.1 | 2.70882181 | -0.46028422 | 0.82462438 | -0.5581744 | 0.5767253 | 0.74070944 |
| AC090192.2 | 0.85038243 | -2.963982637 | 3.26009488 | -0.9091707 | 0.36326005 | NA |
| TMEM65 | 1423.46141 | 0.026427358 | 0.13487581 | 0.19593845 | 0.84465834 | 0.91797067 |
| TRMT12 | 531.615389 | 0.024069192 | 0.15524898 | 0.15503608 | 0.87679288 | 0.93577093 |
| RNF139-AS1 | 313.402019 | -0.177453483 | 0.26117299 | -0.6794481 | 0.496854 | 0.67865838 |
| RNF139 | 1385.38323 | -0.25187682 | 0.18779821 | -1.3412099 | 0.17985231 | 0.35569525 |
| TATDN1 | 3205.65467 | -0.01296707 | 0.19455054 | -0.0666514 | 0.94685921 | 0.97438741 |
| AC090198.1 | 425.275079 | -0.189714184 | 0.10493724 | -1.8078824 | 0.07062481 | 0.18547158 |
| NDUFB9 | 7330.70483 | 0.050077751 | 0.19363849 | 0.25861466 | 0.79593257 | 0.88931579 |
| MTSS1 | 3435.70594 | 0.554405933 | 0.28785337 | 1.92600119 | 0.05410423 | 0.15399119 |
| AC090922.1 | 30.7674039 | -0.065940847 | 0.45179863 | -0.1459519 | 0.8839594 | 0.93980604 |
| AC100858.3 | 5.86420529 | -0.391476008 | 0.56290484 | -0.6954568 | 0.48676906 | 0.67078282 |
| LINC00964 | 34.035017 | 2.164910565 | 0.59975394 | 3.6096646 | 0.00030659 | 0.00308465 |
| AC100858.2 | 0.79454012 | 3.007508207 | 1.5352816 | 1.95892936 | 0.05012106 | NA |
| ZNF572 | 112.660777 | 0.76101447 | 0.25671722 | 2.96440757 | 0.00303266 | 0.01851617 |
| AC009908.1 | 0.58337333 | -1.142479351 | 2.85072535 | -0.4007679 | 0.68859098 | NA |
| SQLE | 1778.48736 | -1.209432123 | 0.41678783 | -2.9017933 | 0.00371033 | 0.02153201 |
| WASHC5 | 2084.04633 | 0.392219867 | 0.19820882 | 1.97882147 | 0.04783611 | 0.1410216 |
| WASHC5-AS | 317.090779 | 0.353437415 | 0.22994645 | 1.53704228 | 0.12428296 | 0.27620135 |
| NSMCE2 | 749.379552 | -0.327293504 | 0.1391649 | -2.3518395 | 0.01868083 | 0.07144505 |
| AC084083.1 | 1.71931869 | -1.184515968 | 1.14239647 | -1.0368694 | 0.29979671 | 0.49649183 |
| TRIB1 | 2900.74318 | -0.557122718 | 0.27770314 | -2.0061809 | 0.04483695 | 0.13451086 |
| AC091114.1 | 9.36043256 | 0.468884511 | 0.45188132 | 1.03762756 | 0.29944348 | 0.49613236 |
| AC016074.1 | 0 | NA | NA | NA | NA | NA |
| LINC00861 | 93.7019466 | 0.542673404 | 0.72653937 | 0.74692911 | 0.45510635 | 0.64453917 |
| AC024681.2 | 0.18028734 | 0.18657164 | 3.29377195 | 0.05664376 | 0.95482897 | NA |
| AC087667.1 | 0.67634663 | 2.166377968 | 1.63658615 | 1.32371764 | 0.18559684 | NA |
| AC084116.2 | 0.61051678 | 3.188596849 | 1.89330964 | 1.68413913 | 0.09215478 | NA |
| AC084116.1 | 1.68035643 | -0.7292903 | 1.07739861 | -0.6768992 | 0.49846989 | 0.6799953 |
| AC084116.3 | 1.86056749 | 3.939313986 | 1.32881091 | 2.96454066 | 0.00303135 | 0.01851245 |
| FAM84B | 5327.12253 | 0.144808384 | 0.41621308 | 0.34791887 | 0.72790111 | 0.84611931 |
| PCAT1 | 547.796696 | 0.631433383 | 0.38992624 | 1.61936624 | 0.10536849 | 0.24626137 |
| AC024382.1 | 0.11051935 | 0.02007176 | 3.29738704 | 0.00608717 | 0.99514317 | NA |
| PRNCR1 | 8.14282296 | -0.447133221 | 0.47627305 | -0.938817 | 0.34782473 | 0.54534385 |
| CASC8 | 8.02472756 | -1.795456668 | 0.58816089 | -3.0526625 | 0.00226821 | 0.01488248 |
| POU5F1B | 84.000018 | 0.198973477 | 0.40801459 | 0.48766265 | 0.62578881 | 0.77729476 |
| CCAT2 | 0.09989236 | -0.642905668 | 3.30558948 | -0.1944905 | 0.84579185 | NA |
| CASC11 | 2.01419755 | -0.758378321 | 1.30905381 | -0.5793332 | 0.56236437 | 0.72983653 |
| MYC | 2299.40126 | -1.714153167 | 0.46717912 | -3.6691562 | 0.00024335 | 0.00256585 |
| PVT1 | 170.42073 | -1.822303966 | 0.45380641 | -4.0155977 | 5.93E-05 | 0.00084383 |
| CCDC26 | 4.44575141 | 0.069661434 | 0.77565233 | 0.08981013 | 0.9284381 | 0.96526981 |
| AC011257.1 | 0.29992972 | 2.233572672 | 3.22779297 | 0.69198139 | 0.48894902 | NA |
| AC103718.1 | 104.46026 | 0.666387718 | 0.70949468 | 0.93924273 | 0.34760613 | 0.545114 |
| GSDMC | 11.7388918 | -2.945467393 | 1.08151701 | -2.7234591 | 0.00646022 | 0.03257285 |
| AC022973.4 | 84.1331443 | 0.025628569 | 0.23291173 | 0.11003554 | 0.91238119 | 0.95694423 |
| FAM49B | 2098.77654 | -0.494938943 | 0.13296334 | -3.7223714 | 0.00019736 | 0.00215682 |
| AC022973.3 | 0.29449776 | 2.177142669 | 3.23105086 | 0.67381875 | 0.50042655 | NA |
| AC022973.2 | 2.29078059 | -1.925236691 | 0.94180037 | -2.0442089 | 0.04093292 | 0.12606506 |
| ASAP1 | 1406.51915 | -0.353738558 | 0.1824242 | -1.9390988 | 0.0524893 | 0.15067219 |
| AC090987.1 | 0 | NA | NA | NA | NA | NA |
| ADCY8 | 38.7690461 | 1.109171864 | 0.85906965 | 1.29113147 | 0.19665809 | 0.37723677 |
| AC103726.2 | 2.00344575 | 0.808823506 | 1.28289559 | 0.63046714 | 0.528389 | 0.7029725 |
| AC087341.1 | 0 | NA | NA | NA | NA | NA |
| AC104257.1 | 0.51158304 | -0.035299358 | 1.85753482 | -0.0190033 | 0.98483844 | NA |
| EFR3A | 2947.16706 | 0.043016827 | 0.14329304 | 0.30020178 | 0.76402324 | 0.86935843 |
| OC90 | 0.45449073 | -0.712910449 | 2.86691215 | -0.2486684 | 0.8036173 | NA |
| AC100868.1 | 0.45449073 | -0.712910449 | 2.86691215 | -0.2486684 | 0.8036173 | NA |
| HHLA1 | 0.12088456 | 0.02007176 | 3.29738704 | 0.00608717 | 0.99514317 | NA |
| KCNQ3 | 176.328452 | -0.086548106 | 0.36819185 | -0.2350625 | 0.8141602 | 0.89973198 |
| HPYR1 | 11.5290643 | 0.387462093 | 0.44436307 | 0.87194936 | 0.383236 | 0.57911074 |
| LRRC6 | 615.811383 | 0.009436829 | 0.25934548 | 0.0363871 | 0.9709737 | 0.9865502 |
| TMEM71 | 104.032949 | -0.247657776 | 0.49354363 | -0.5017951 | 0.61581166 | 0.77023922 |
| PHF20L1 | 1613.16729 | 0.076546507 | 0.11744001 | 0.65179242 | 0.5145351 | 0.69239713 |
| AF230666.1 | 139.773655 | 0.404777128 | 0.20108063 | 2.01300901 | 0.04411368 | 0.13306769 |
| TG | 313.24767 | 0.7475306 | 0.45681812 | 1.6363856 | 0.10175891 | 0.24039429 |
| SLA | 960.912757 | -1.164794924 | 0.38860721 | -2.997358 | 0.00272331 | 0.01706597 |
| WISP1 | 1566.77493 | -0.883045847 | 0.42249775 | -2.0900605 | 0.03661237 | 0.11620946 |
| NDRG1 | 52390.7734 | -0.369213946 | 0.47423453 | -0.7785472 | 0.43624651 | 0.62838371 |
| ST3GAL1 | 5527.20866 | 0.766765038 | 0.41410384 | 1.85162503 | 0.06407969 | 0.17309786 |
| AC103706.1 | 50.1793919 | -0.623752387 | 0.27298168 | -2.2849606 | 0.02231514 | 0.08133382 |
| AC133634.1 | 0 | NA | NA | NA | NA | NA |
| AC090821.1 | 2.59646126 | -1.106704154 | 1.74134862 | -0.6355443 | 0.5250735 | 0.69983261 |
| AC110741.1 | 0.97849826 | 0.543609291 | 1.76694558 | 0.3076548 | 0.75834502 | NA |
| AC105180.1 | 0 | NA | NA | NA | NA | NA |
| ZFAT | 1008.18186 | 1.29497777 | 0.38088672 | 3.39990267 | 0.0006741 | 0.00571943 |
| ZFAT-AS1 | 92.580028 | 1.677445717 | 0.45770032 | 3.66494329 | 0.00024739 | 0.00260015 |
| AC087045.2 | 3.69248319 | -0.524060075 | 0.81352189 | -0.6441868 | 0.5194543 | 0.69628475 |
| AC083843.3 | 1006.78398 | -0.231137592 | 0.35484056 | -0.6513844 | 0.5147984 | 0.69256857 |
| AC083843.2 | 2.9335096 | 1.432149697 | 0.73980803 | 1.93583963 | 0.05288735 | 0.15154181 |
| AC083843.1 | 1.21743389 | 1.529921367 | 1.20161375 | 1.27322225 | 0.20293919 | 0.38495415 |
| AC103764.1 | 0.99757497 | 0.806759314 | 1.82286421 | 0.44257785 | 0.65807111 | NA |
| LINC01591 | 0.20687462 | 0.68915539 | 3.1020265 | 0.22216296 | 0.82418703 | NA |
| AC040914.1 | 3.67855062 | -0.078034337 | 1.11831027 | -0.0697788 | 0.94436973 | 0.97345853 |
| KHDRBS3 | 1351.74436 | -0.530612911 | 0.3065197 | -1.7310891 | 0.08343587 | 0.2086886 |
| AC079098.1 | 0 | NA | NA | NA | NA | NA |
| AC013546.1 | 0.03392908 | 0.380740593 | 3.29738709 | 0.11546736 | 0.90807469 | NA |
| LINC02055 | 4.19540001 | -0.859512697 | 1.08076676 | -0.7952805 | 0.42645037 | 0.61876837 |
| AC046195.1 | 53.5057494 | 2.085962813 | 0.8731083 | 2.38912266 | 0.01688866 | 0.06638158 |
| AC046195.2 | 0 | NA | NA | NA | NA | NA |
| AC079015.1 | 5.15647693 | 1.290420905 | 0.6731404 | 1.91701599 | 0.0552359 | 0.15620061 |
| FAM135B | 93.578971 | -0.298334069 | 0.77421471 | -0.3853376 | 0.69998728 | 0.82792844 |
| COL22A1 | 194.881006 | 0.991159614 | 0.55902177 | 1.77302507 | 0.07622455 | 0.19594867 |
| AC027541.1 | 0 | NA | NA | NA | NA | NA |
| AC100807.1 | 0 | NA | NA | NA | NA | NA |
| AC087354.1 | 0.239366 | 1.011340114 | 2.98135597 | 0.33922152 | 0.73444286 | NA |
| KCNK9 | 5.94267364 | -0.703413475 | 0.56434354 | -1.2464278 | 0.21260738 | 0.39645571 |
| TRAPPC9 | 1850.64319 | 0.324766672 | 0.2168094 | 1.49793633 | 0.13414979 | 0.29129859 |
| AC021744.1 | 41.083235 | 4.345272411 | 0.64055108 | 6.78364704 | 1.17E-11 | 2.38E-09 |
| AC107375.1 | 229.61611 | -0.047358276 | 0.24483152 | -0.1934321 | 0.84662058 | 0.9193833 |
| CHRAC1 | 1349.96789 | -0.154324332 | 0.2167632 | -0.711949 | 0.47649638 | 0.66179458 |
| AGO2 | 2062.58166 | -0.748628154 | 0.20610182 | -3.6323219 | 0.00028088 | 0.00288429 |
| ERICD | 66.1588462 | -0.556969376 | 0.24829699 | -2.243158 | 0.02488663 | 0.08802957 |
| PTK2 | 4651.09448 | 0.202271643 | 0.09276835 | 2.18039507 | 0.02922819 | 0.09906778 |
| AC100860.1 | 1.97583832 | 4.498671116 | 1.25538237 | 3.5835067 | 0.00033901 | 0.00334071 |
| DENND3 | 3098.02232 | -1.008240346 | 0.32029807 | -3.147819 | 0.00164494 | 0.01161675 |
| AC040970.1 | 50.6432837 | 0.205384883 | 0.46512489 | 0.44156932 | 0.65880089 | 0.79954802 |
| SLC45A4 | 3434.62338 | -0.497665324 | 0.31423916 | -1.5837151 | 0.11325856 | 0.25927842 |
| AC011676.1 | 12.6740014 | 0.996980634 | 0.66782965 | 1.49286668 | 0.13547209 | 0.29308415 |
| AC011676.2 | 21.1901548 | 0.825192613 | 0.40816891 | 2.02169393 | 0.04320798 | 0.13106002 |
| AC011676.3 | 0.18684785 | -1.003574173 | 3.30558948 | -0.3035992 | 0.7614333 | NA |

| | | | | | | |
|---|---|---|---|---|---|---|
| AC011676.4 | 0.09989236 | -0.642905668 | 3.30558948 | -0.1944905 | 0.84579185 | NA |
| LINC01300 | 4.77816725 | 0.735786779 | 0.96413337 | 0.76315871 | 0.44536875 | 0.63586215 |
| GPR20 | 24.0397958 | 1.686865094 | 0.62522233 | 2.69802437 | 0.00697523 | 0.03453566 |
| AC100803.1 | 7.92347112 | -2.291659341 | 0.91214407 | -2.5123875 | 0.01199173 | 0.05150196 |
| PTP4A3 | 4763.23774 | -2.065666629 | 0.34964378 | -5.9079175 | 3.46E-09 | 2.80E-07 |
| AC100803.3 | 587.942009 | -2.486265768 | 0.47521391 | -5.2318877 | 1.68E-07 | 7.37E-06 |
| AC138647.1 | 0 | NA | NA | NA | NA | NA |
| AC104417.2 | 0.10721287 | -0.642905668 | 3.30558948 | -0.1944905 | 0.84579185 | NA |
| LINC00051 | 2.78941969 | 2.110027841 | 0.89018379 | 2.37032831 | 0.0177723 | 0.06870773 |
| TSNARE1 | 1224.62046 | 0.370019328 | 0.15390534 | 2.40420076 | 0.01620787 | 0.06438709 |
| AC134682.1 | 18.7920154 | 1.079122178 | 0.4059593 | 2.65820286 | 0.00785586 | 0.03768092 |
| ADGRB1 | 130.909271 | 1.008821126 | 0.48976261 | 2.05981653 | 0.03941608 | 0.12248434 |
| ARC | 150.858873 | -0.313865388 | 0.47934324 | -0.6547821 | 0.51260804 | 0.69079487 |
| AP006547.1 | 1.70335754 | -0.244287919 | 1.00489236 | -0.2430986 | 0.80792902 | 0.89620864 |
| AC108002.3 | 0.26704335 | -1.364244764 | 2.95174887 | -0.4621819 | 0.64395091 | NA |
| JRK | 1109.62198 | 0.565891065 | 0.1428392 | 3.9617351 | 7.44E-05 | 0.00100252 |
| PSCA | 87.9609828 | -0.77178153 | 0.84659395 | -0.9116313 | 0.36196285 | 0.55929686 |
| LY6K | 107.44998 | -1.385224363 | 0.39192407 | -3.5344202 | 0.00040867 | 0.00388348 |
| AC108002.1 | 81.6940033 | -0.961501804 | 0.34384821 | -2.7962973 | 0.00516918 | 0.02757107 |
| THEM6 | 2292.09724 | 0.693442687 | 0.12296211 | 5.63948272 | 1.71E-08 | 1.08E-06 |
| AC108002.2 | 101.575902 | 0.495421933 | 0.31773554 | 1.55922734 | 0.11894258 | 0.26823425 |
| SLURP1 | 0.56722069 | -0.684153933 | 2.82741016 | -0.2419719 | 0.8088019 | NA |
| LYPD2 | 0.94703395 | 2.856609833 | 1.26668449 | 2.25518656 | 0.02412162 | NA |
| AC083841.1 | 20.1628708 | 4.655283963 | 0.8743273 | 5.32441795 | 1.01E-07 | 4.84E-06 |
| LYNX1-SLURI | 236.605829 | 0.838787785 | 0.44370965 | 1.89039788 | 0.05870477 | 0.16291778 |
| SLURP2 | 10.9715964 | 0.679487414 | 0.65565342 | 1.03635152 | 0.30003818 | 0.4967375 |
| LYNX1 | 2127.95711 | 0.707192831 | 0.46139975 | 1.53271179 | 0.12534688 | 0.27788124 |
| LY6D | 0.84531658 | -0.24751349 | 2.66535996 | -0.0928631 | 0.92601235 | NA |
| AC083841.2 | 0.36207013 | 0.017594066 | 2.8711013 | 0.00612799 | 0.99511061 | NA |
| AC083841.3 | 0 | NA | NA | NA | NA | NA |
| GML | 0 | NA | NA | NA | NA | NA |
| CYP11B1 | 0.09989236 | -0.642905668 | 3.30558948 | -0.1944905 | 0.84579185 | NA |
| CYP11B2 | 0.08393208 | -0.642905668 | 3.30558948 | -0.1944905 | 0.84579185 | NA |
| LY6E-DT | 123.099935 | -0.654700054 | 0.60604416 | -1.0802844 | 0.28001555 | 0.47558998 |
| LY6E | 15253.1712 | -1.780938289 | 0.590513 | -3.0159172 | 0.00256203 | 0.01629542 |
| LY6L | 21.4822627 | 1.655704524 | 0.83012221 | 1.99453105 | 0.04609405 | 0.13725736 |
| LY6H | 14.2402401 | -2.338956408 | 0.81083065 | -2.8846423 | 0.00391859 | 0.02242018 |
| GPIHBP1 | 540.161514 | 0.084332502 | 0.70049804 | 0.12038935 | 0.90417473 | 0.95178646 |
| ZFP41 | 802.889548 | 0.444457702 | 0.19175224 | 2.31787493 | 0.02045612 | 0.07615811 |
| AC138696.1 | 790.807982 | 0.315714695 | 0.1551036 | 2.03550852 | 0.04179973 | 0.12794648 |
| GLI4 | 1469.60325 | 0.025452102 | 0.13813467 | 0.18425572 | 0.85381285 | 0.92346684 |
| MINCR | 274.786436 | 0.039283996 | 0.23246561 | 0.16898842 | 0.86580575 | 0.93029491 |
| ZNF696 | 397.2231 | 0.354973659 | 0.18496028 | 1.91918858 | 0.05496047 | 0.155622 |
| AC138696.2 | 33.1116745 | 0.727215505 | 0.30282108 | 2.40146924 | 0.01632938 | 0.06476249 |
| TOP1MT | 1208.38135 | -0.463154273 | 0.17293251 | -2.6782372 | 0.00740108 | 0.03607513 |
| RHPN1-AS1 | 58.6755088 | 0.057124533 | 0.33601445 | 0.17000618 | 0.86500528 | 0.92986964 |
| RHPN1 | 5284.59753 | -0.798734534 | 0.30493701 | -2.6193427 | 0.00880994 | 0.04094432 |
| AC105118.1 | 11.6770253 | -2.129561992 | 0.5277804 | -4.0349395 | 5.46E-05 | 0.00079703 |
| MAFA-AS1 | 9.12938431 | -0.867046859 | 0.46046466 | -1.8829824 | 0.05970275 | 0.16497554 |
| MAFA | 14.1649312 | -0.985151543 | 0.47993383 | -2.052682 | 0.04010343 | 0.12403558 |
| ZC3H3 | 1942.36089 | 0.087770898 | 0.22431608 | 0.39128224 | 0.69558863 | 0.82479752 |
| AC067930.4 | 72.5972276 | 1.108673458 | 0.80752705 | 1.37292423 | 0.1697759 | 0.34157691 |
| AC067930.1 | 55.8992112 | 1.12522741 | 0.83264195 | 1.35139409 | 0.17656923 | 0.35137956 |
| GSDMD | 6893.33764 | -0.440422913 | 0.31248582 | -1.4094173 | 0.15871182 | 0.3262066 |
| MROH6 | 712.81038 | -1.118835883 | 0.36198789 | -3.0908102 | 0.00199611 | 0.0134795 |
| AC067930.5 | 2886.18338 | 0.468124955 | 0.16467356 | 2.84274501 | 0.00447268 | 0.02477362 |
| NAPRT | 4624.91319 | 0.451132763 | 0.16833236 | 2.68001217 | 0.00736195 | 0.03595367 |
| AC067930.2 | 8612.62385 | -0.517845794 | 0.16078651 | -3.2207042 | 0.00127876 | 0.00952568 |
| EEF1D | 17738.9829 | -0.446396806 | 0.15297286 | -2.9181437 | 0.00352122 | 0.0206665 |
| TIGD5 | 667.246135 | -0.447816549 | 0.23355212 | -1.9174159 | 0.05518512 | 0.15607375 |
| PYCR3 | 853.618998 | 0.176282943 | 0.16988273 | 1.03767429 | 0.29942172 | 0.49613236 |
| TSTA3 | 2599.11117 | -0.507253358 | 0.19463279 | -2.6062071 | 0.00915511 | 0.04218421 |
| AC067930.3 | 10.6356526 | 1.024226908 | 0.6193793 | 1.65363439 | 0.09820182 | 0.23469819 |
| ZNF623 | 903.519618 | -0.175637802 | 0.13822125 | -1.2707004 | 0.20383524 | 0.38607951 |
| ZNF707 | 746.22653 | -0.198594561 | 0.16013798 | -1.2401465 | 0.2149212 | 0.39917397 |
| CCDC166 | 0.17515154 | 1.45420457 | 3.2783972 | 0.44357181 | 0.65735219 | NA |
| AC105219.3 | 0.89988812 | 0.36551792 | 1.28215423 | 0.28508109 | 0.77558202 | NA |
| MAPK15 | 1481.4652 | -0.978174039 | 0.49587749 | -1.9726123 | 0.04853975 | 0.14263353 |
| AC105219.2 | 206.752904 | -0.815703002 | 0.47972771 | -1.7003458 | 0.0890659 | 0.21892876 |
| FAM83H | 2996.22816 | -0.298361463 | 0.1584437 | -1.8830756 | 0.05969012 | 0.16496368 |
| IQANK1 | 288.036388 | -0.329913716 | 0.36008713 | -0.9162053 | 0.35955925 | 0.55720924 |
| AC105219.4 | 12.9126969 | 0.270609422 | 0.36752924 | 0.7362936 | 0.46155204 | 0.64998795 |
| SCRIB | 6242.9089 | -0.541307715 | 0.12928078 | -4.1870702 | 2.83E-05 | 0.00046836 |
| PUF60 | 6749.85321 | -0.488770441 | 0.16968811 | -2.8804048 | 0.00397165 | 0.02263601 |
| NRBP2 | 10860.0319 | -0.832415839 | 0.36949274 | -2.2528611 | 0.02426791 | 0.08631571 |
| EPPK1 | 355.722646 | -0.076731935 | 0.45752285 | -0.1677117 | 0.86681009 | 0.9308678 |
| PLEC | 17154.7261 | -0.107252527 | 0.22946848 | -0.4673955 | 0.64021696 | 0.78714964 |
| PARP10 | 6770.25689 | 0.316979598 | 0.24870524 | 1.27451917 | 0.20247948 | 0.38437722 |
| GRINA | 14511.6263 | 0.240837832 | 0.1596991 | 1.50807257 | 0.13153595 | 0.28735011 |
| SPATC1 | 61.1768276 | 0.6361781 | 0.50372408 | 1.26294955 | 0.20660731 | 0.38924304 |
| OPLAH | 3586.10518 | 0.777328782 | 0.24388424 | 3.18728581 | 0.00143615 | 0.01045877 |
| AC109322.1 | 393.388345 | -0.367200923 | 0.24066038 | -1.5258055 | 0.12705832 | 0.28057117 |
| EXOSC4 | 959.814685 | -0.505923121 | 0.24257946 | -2.0855975 | 0.03701509 | 0.11700003 |
| GPAA1 | 7119.68937 | 0.189680585 | 0.13425766 | 1.41281016 | 0.15771155 | 0.3248343 |
| CYC1 | 8098.40675 | 0.233665678 | 0.16631175 | 1.40498598 | 0.16002545 | 0.32808825 |
| SHARPIN | 4004.65008 | -0.232230447 | 0.10831747 | -2.1439796 | 0.03203452 | 0.10572943 |
| MAF1 | 6292.77493 | -0.404885335 | 0.16573076 | -2.4430308 | 0.0145645 | 0.05958203 |
| WDR97 | 336.951833 | 0.183841467 | 0.225476 | 0.81534826 | 0.41487299 | 0.60846501 |
| HGH1 | 1243.63824 | 0.023111299 | 0.10690774 | 0.21617986 | 0.82884755 | 0.90895409 |
| MROH1 | 4841.04789 | -0.532209341 | 0.24536506 | -2.1690511 | 0.03007881 | 0.10101606 |
| BOP1 | 2717.73658 | -0.49250633 | 0.23121626 | -2.1300679 | 0.03316601 | 0.10836174 |
| SCX | 197.49276 | -1.35223688 | 0.33588425 | -4.0259015 | 5.68E-05 | 0.00081871 |
| HSF1 | 5493.10574 | -0.583347179 | 0.13508958 | -4.3182248 | 1.57E-05 | 0.00029437 |
| DGAT1 | 3088.97401 | -0.100311451 | 0.15342862 | -0.6537988 | 0.51324143 | 0.69122454 |
| AC233992.1 | 623.408066 | -0.492082322 | 0.11973382 | -4.1098023 | 3.96E-05 | 0.00061857 |
| SCRT1 | 5.98486871 | 0.218868428 | 0.6623129 | 0.33046076 | 0.74105183 | 0.8546067 |
| TMEM249 | 29.5813166 | -0.147846937 | 0.3163479 | -0.4673555 | 0.64024554 | 0.78714964 |
| AC233992.2 | 232.118544 | -0.780196382 | 0.17488126 | -4.461292 | 8.15E-06 | 0.00017353 |
| SLC52A2 | 2776.06196 | -0.940348505 | 0.22550406 | -4.1699848 | 3.05E-05 | 0.00049463 |
| FBXL6 | 1924.86756 | -0.710711819 | 0.16678593 | -4.2612216 | 2.03E-05 | 0.00035934 |
| ADCK5 | 710.373367 | -0.152440129 | 0.14811294 | -1.0292155 | 0.30337842 | 0.50024347 |
| CPSF1 | 6610.18022 | -0.162763758 | 0.27354481 | -0.5950168 | 0.55183224 | 0.72178454 |
| AC233992.3 | 9.72251476 | -0.289156733 | 0.49896254 | -0.5795159 | 0.56224111 | 0.72982024 |
| SLC39A4 | 3568.71674 | 0.639242834 | 0.26554641 | 2.40727349 | 0.01607213 | 0.06406985 |
| VPS28 | 7452.96205 | -0.221666266 | 0.13224334 | -1.6761999 | 0.09369905 | 0.22714138 |
| TONSL | 547.509943 | -0.036973507 | 0.1791954 | -0.2063307 | 0.83653262 | 0.9132422 |
| TONSL-AS1 | 37.1755183 | -0.054767415 | 0.25494534 | -0.2148202 | 0.82990748 | 0.90970031 |
| CYHR1 | 4491.29207 | -0.088801345 | 0.09519581 | -0.9328283 | 0.35090862 | 0.54857388 |
| AC084125.1 | 713.525761 | 0.109089655 | 0.24782872 | 0.44018165 | 0.65980555 | 0.79999826 |
| KIFC2 | 4405.62723 | -0.294293993 | 0.20346777 | -1.4463912 | 0.14806751 | 0.31168998 |
| FOXH1 | 1629.09621 | -0.419860586 | 0.21473489 | -1.9552509 | 0.05055346 | 0.14677902 |
| PPP1R16A | 6917.27819 | 0.185741025 | 0.17899886 | 1.03766599 | 0.29942558 | 0.49613236 |
| GPT | 2933.53354 | 2.215620891 | 0.46762818 | 4.73799694 | 2.16E-06 | 6.04E-05 |

| | | | | | | |
|---|---|---|---|---|---|---|
| MFSD3 | 1726.78297 | 0.554330114 | 0.17445318 | 3.17752937 | 0.00148536 | 0.01072495 |
| RECQL4 | 554.537283 | 0.522152825 | 0.24021598 | 2.17368063 | 0.02972913 | 0.10026309 |
| AC084125.4 | 141.40188 | 0.621111264 | 0.25162073 | 2.46844236 | 0.01357025 | 0.05651375 |
| LRRC14 | 2000.18732 | 0.141996251 | 0.11896488 | 1.19359809 | 0.23263522 | 0.42066727 |
| LRRC24 | 973.202196 | -0.063356183 | 0.15443927 | -0.4102336 | 0.68163457 | 0.81559291 |
| C8orf82 | 1591.4735 | 0.378552637 | 0.20923735 | 1.80920201 | 0.07041963 | 0.18508577 |
| ARHGAP39 | 841.27833 | -0.150016443 | 0.2286569 | -0.6560766 | 0.51177484 | 0.68990587 |
| AC084125.3 | 0.25178919 | 1.950259806 | 2.89159949 | 0.6744571 | 0.50002075 | NA |
| AF186192.1 | 23.6571638 | 0.140502875 | 0.40682984 | 0.3453603 | 0.72982352 | 0.8473873 |
| AF186192.3 | 4.48971984 | -1.572170314 | 1.04371194 | -1.5063259 | 0.13198354 | 0.28813692 |
| ZNF251 | 1514.2459 | 0.491057517 | 0.13031277 | 3.76829937 | 0.00016436 | 0.00187468 |
| ZNF34 | 397.895157 | 0.249580606 | 0.20922772 | 1.19286588 | 0.2329219 | 0.42109904 |
| RPL8 | 41176.5279 | -0.485373743 | 0.17803922 | -2.7262182 | 0.00640646 | 0.03233028 |
| ZNF517 | 872.849885 | 0.193317981 | 0.12978175 | 1.48956213 | 0.13633941 | 0.29450116 |
| ZNF7 | 1270.6532 | -0.060150214 | 0.13222325 | -0.454914 | 0.64917107 | 0.79355837 |
| COMMD5 | 1401.84193 | -0.310762404 | 0.23837241 | -1.3036845 | 0.19234119 | 0.37159853 |
| ZNF250 | 978.078117 | -0.294096827 | 0.23171152 | -1.269237 | 0.20435656 | 0.38666749 |
| ZNF16 | 462.127256 | 0.446155158 | 0.17417915 | 2.56147285 | 0.01042294 | 0.04650641 |
| AF235103.1 | 18.1534109 | 0.984291487 | 0.32568133 | 3.02225337 | 0.002509 | 0.01604605 |
| ZNF252P-AS | 95.0612082 | 0.53401575 | 0.28058653 | 1.90321235 | 0.05701284 | 0.15959964 |
| C8orf33 | 2283.93988 | -0.136138929 | 0.22087261 | -0.6163685 | 0.53765131 | 0.71124035 |
| WASHC1 | 4555.42746 | 0.077903713 | 0.18329735 | 0.42501277 | 0.67082737 | 0.80778655 |
| AL928970.1 | 7.34099265 | 0.975554409 | 0.64389555 | 1.51508177 | 0.1297517 | 0.28474969 |
| FAM138C | 0.32045822 | 0.328539521 | 2.2700964 | 0.14472492 | 0.88492806 | NA |
| PGM5P3-AS1 | 11.2531377 | 0.906859776 | 0.58060272 | 1.56192823 | 0.1183049 | 0.26730358 |
| AL449043.1 | 0.42455947 | -0.842919892 | 2.8121666 | -0.2997404 | 0.76437519 | NA |
| LINC01388 | 0.59722944 | -1.368596979 | 2.22788639 | -0.6143029 | 0.53901521 | NA |
| FOXD4 | 41.2445979 | 0.409228851 | 0.39455817 | 1.03718257 | 0.29965078 | 0.49633563 |
| CBWD1 | 1373.92824 | 0.208921991 | 0.22522035 | 0.92763373 | 0.3535976 | 0.55115076 |
| C9orf66 | 946.308831 | 3.309882833 | 0.7548299 | 4.38493868 | 1.16E-05 | 0.00023055 |
| DOCK8 | 1664.0947 | 0.801431559 | 0.25755404 | 3.11170258 | 0.00186012 | 0.01273399 |
| AL158832.1 | 0.17515154 | 1.45420457 | 3.2783972 | 0.44357181 | 0.65735219 | NA |
| AL161725.1 | 20.5482768 | 1.649132363 | 0.38724909 | 4.25858288 | 2.06E-05 | 0.00036288 |
| KANK1 | 3166.43272 | 0.783869121 | 0.15880814 | 4.93595063 | 7.98E-07 | 2.68E-05 |
| AL136979.1 | 0.5136765 | -1.194611853 | 2.20472316 | -0.5418421 | 0.58792728 | NA |
| DMRT1 | 0.10721287 | -0.642905668 | 3.30558948 | -0.1944905 | 0.84579185 | NA |
| DMRT3 | 0.61896276 | 0.686883642 | 2.70772856 | 0.25367522 | 0.79974648 | NA |
| LINC01230 | 431.377963 | -0.253981073 | 0.5011536 | -0.5067929 | 0.61230016 | 0.76779926 |
| DMRT2 | 1590.18557 | -0.18557293 | 0.57312811 | -0.3237896 | 0.74609732 | 0.85769154 |
| SMARCA2 | 4206.64474 | 0.576907835 | 0.16578438 | 3.47986846 | 0.00050166 | 0.004542 |
| AL359076.1 | 40.9672045 | -0.715915007 | 0.21419585 | -3.3423384 | 0.00083076 | 0.00677818 |
| VLDLR-AS1 | 275.533131 | -0.113254254 | 0.28739894 | -0.3940664 | 0.69353205 | 0.82370883 |
| AL353614.1 | 0.09644756 | -0.642905668 | 3.30558948 | -0.1944905 | 0.84579185 | NA |
| VLDLR | 3076.42712 | -0.956829215 | 0.24879858 | -3.8457985 | 0.00012016 | 0.00146696 |
| KCNV2 | 10.6402134 | -0.72549064 | 0.48856812 | -1.4849324 | 0.13756173 | 0.29619521 |
| PUM3 | 2213.5899 | -0.903664176 | 0.26059523 | -3.4676927 | 0.00052495 | 0.00471166 |
| LINC01231 | 0.07659028 | -0.642905668 | 3.30558948 | -0.1944905 | 0.84579185 | NA |
| RFX3 | 585.866791 | -0.042420512 | 0.21425643 | -0.1979894 | 0.84305333 | 0.91660411 |
| RFX3-AS1 | 46.9152054 | 0.592516232 | 0.36464445 | 1.62491501 | 0.10418067 | 0.2444385 |
| AL354977.2 | 3.10310995 | -1.523237944 | 0.98932677 | -1.5396712 | 0.12364052 | 0.27519739 |
| AL354977.1 | 5.89990903 | 0.618395851 | 0.74996607 | 0.82456511 | 0.40961854 | 0.60405105 |
| GLIS3 | 1769.11462 | -0.464505232 | 0.21223213 | -2.188666 | 0.02862113 | 0.09757455 |
| AL137071.1 | 0.68347334 | -0.58769701 | 1.60835518 | -0.3654025 | 0.71481097 | NA |
| GLIS3-AS1 | 37.7925296 | -0.176930863 | 0.36827606 | -0.48043 | 0.63092168 | 0.78062237 |
| AL162419.1 | 17.3464091 | 0.207960952 | 0.48186546 | 0.43157472 | 0.66605054 | 0.80431161 |
| SLC1A1 | 4476.69253 | 0.798257446 | 0.33302925 | 2.39695898 | 0.01653177 | 0.06533916 |
| SPATA6L | 388.343477 | -0.84588955 | 0.30669387 | -2.7580908 | 0.005814 | 0.03009723 |
| PLPP6 | 571.107085 | 0.262621035 | 0.15091776 | 1.74015989 | 0.08183095 | 0.20566928 |
| CDC37L1-DT | 65.4067539 | 0.904168747 | 0.31850679 | 2.83877388 | 0.00452872 | 0.02497333 |
| CDC37L1 | 1578.90805 | -0.138075224 | 0.14153001 | -0.9755898 | 0.3292678 | 0.52702834 |
| AK3 | 7710.19775 | 0.232371682 | 0.1773832 | 1.30999824 | 0.19019643 | 0.36886665 |
| RCL1 | 938.788774 | -0.275667852 | 0.21390208 | -1.2887572 | 0.19748251 | 0.3783034 |
| AL158147.1 | 0 | NA | NA | NA | NA | NA |
| JAK2 | 825.390382 | -0.022602987 | 0.22328982 | -0.1012271 | 0.91937016 | 0.96074349 |
| IGHEP2 | 2.74687698 | 0.110554479 | 0.73472365 | 0.15047083 | 0.88039316 | 0.937981 |
| INSL6 | 4.63740037 | -0.604620614 | 0.76071886 | -0.7948017 | 0.42672888 | 0.61900696 |
| INSL4 | 3.38754314 | 0.363896077 | 1.16687294 | 0.31185578 | 0.75515013 | 0.86399292 |
| RLN2 | 26.070085 | -0.22692019 | 0.42727689 | -0.5310846 | 0.59536013 | 0.7543555 |
| RLN1 | 9.07216104 | -0.688790505 | 0.4965955 | -1.3870253 | 0.16543406 | 0.33583496 |
| AL135786.2 | 3.6469459 | 0.63353902 | 0.6440254 | 0.98371744 | 0.32525446 | 0.52282598 |
| PLGRKT | 1087.14898 | 0.006205419 | 0.20978558 | 0.02957982 | 0.97640216 | 0.9890038 |
| CD274 | 482.562745 | -1.816389502 | 0.49310181 | -3.6835993 | 0.00022996 | 0.00244423 |
| PDCD1LG2 | 114.125279 | -0.627325381 | 0.43972011 | -1.426647 | 0.15368167 | 0.31937786 |
| AL162253.1 | 0.03392908 | 0.380740593 | 3.29738709 | 0.11546736 | 0.90807469 | NA |
| RIC1 | 1178.39878 | -0.062048588 | 0.15335423 | -0.4046096 | 0.68576453 | 0.81835792 |
| AL136980.1 | 32.6006108 | 0.031422672 | 0.4751931 | 0.06612611 | 0.94727742 | 0.97458958 |
| ERMP1 | 4966.51294 | -0.39128549 | 0.38750804 | -1.0097481 | 0.31261601 | 0.50941908 |
| KIAA2026 | 1322.76437 | -0.161850467 | 0.12818705 | -1.2626117 | 0.20672877 | 0.38930498 |
| MLANA | 26.0520492 | -0.251894428 | 0.37679592 | -0.6685169 | 0.50380372 | 0.68364658 |
| RANBP6 | 1343.99743 | 0.028682046 | 0.12266879 | 0.23381698 | 0.81512706 | 0.90042312 |
| AL162384.1 | 0.08757577 | 0.547239394 | 3.29377202 | 0.16614368 | 0.86804388 | NA |
| IL33 | 856.682278 | -1.217669055 | 0.68173635 | -1.786129 | 0.07407838 | 0.19207933 |
| TPD52L3 | 0.09989236 | -0.642905668 | 3.30558948 | -0.1944905 | 0.84579185 | NA |
| UHRF2 | 1925.91119 | -0.260534187 | 0.12851392 | -2.0272838 | 0.0426334 | 0.1297262 |
| AL133480.1 | 0.03392908 | 0.380740593 | 3.29738709 | 0.11546736 | 0.90807469 | NA |
| GLDC | 4688.5501 | 1.267804046 | 0.36850116 | 3.44043435 | 0.00058078 | 0.00509997 |
| AL162411.1 | 67.1793868 | 0.216318759 | 0.23873013 | 0.90612257 | 0.36487098 | 0.56195427 |
| AL354707.2 | 1.84748825 | -0.931147367 | 1.01493317 | -0.917447 | 0.35890847 | 0.55656667 |
| AL354707.1 | 64.1809792 | 0.544961083 | 0.5216166 | 1.0447541 | 0.29613664 | 0.49281352 |
| KDM4C | 1320.49222 | -0.059419863 | 0.12519819 | -0.4746064 | 0.63506754 | 0.78346022 |
| AL354694.1 | 4.76969961 | -0.634776875 | 0.67700081 | -0.9376309 | 0.34843413 | 0.54600713 |
| DMAC1 | 1797.01381 | 0.210596344 | 0.17224345 | 1.22266681 | 0.22145557 | 0.4075784 |
| AL135923.1 | 0 | NA | NA | NA | NA | NA |
| AL135923.2 | 0 | NA | NA | NA | NA | NA |
| PTPRD | 1719.24297 | 1.125415571 | 0.34482759 | 3.26370516 | 0.00109966 | 0.00843741 |
| PTPRD-AS1 | 81.9367683 | 1.361735875 | 0.47965178 | 2.8390093 | 0.00452538 | 0.02496103 |
| AL596451.1 | 0 | NA | NA | NA | NA | NA |
| AL513422.1 | 0 | NA | NA | NA | NA | NA |
| PTPRD-AS2 | 0 | NA | NA | NA | NA | NA |
| AL135790.1 | 0 | NA | NA | NA | NA | NA |
| AL162413.1 | 0.08393208 | -0.642905668 | 3.30558948 | -0.1944905 | 0.84579185 | NA |
| AL451129.1 | 0 | NA | NA | NA | NA | NA |
| AL589678.1 | 0 | NA | NA | NA | NA | NA |
| TYRP1 | 430.203176 | 1.23765267 | 0.34823331 | 3.5540904 | 0.00037929 | 0.00365966 |
| LURAP1L-AS | 0.75444719 | -1.789507187 | 1.88035516 | -0.9516857 | 0.34125639 | NA |
| LURAP1L | 374.975717 | -0.807669103 | 0.29552868 | -2.7329635 | 0.00627673 | 0.03186703 |
| MPDZ | 1824.6552 | 0.702286815 | 0.21426952 | 3.27758614 | 0.00104699 | 0.00812988 |
| LINC01235 | 42.4782526 | -0.598016487 | 0.64280725 | -0.9303201 | 0.35220538 | 0.54982572 |
| AL583785.1 | 7.91831227 | -1.016261411 | 1.00280909 | -1.0134146 | 0.31086215 | 0.50786413 |
| LINC00583 | 0 | NA | NA | NA | NA | NA |
| AL360089.1 | 0.8547947 | 1.627765337 | 1.69539932 | 0.96010734 | 0.33700119 | NA |
| NFIB | 2650.79018 | 0.221747164 | 0.19828941 | 1.11830059 | 0.26343864 | 0.45721185 |
| AL136366.1 | 15.4666735 | -0.116940672 | 0.60389317 | -0.1936446 | 0.84645415 | 0.919354 |

| | | | | | | | |
|---|---|---|---|---|---|---|---|
| AL159169.3 | 19.2568972 | | -0.492125491 | 0.39310734 | -1.2518858 | 0.21061149 | 0.39399513 |
| AL159169.2 | 88.2969136 | | -0.332336422 | 0.31064265 | -1.0698351 | 0.28469353 | 0.4808388 |
| ZDHHC21 | 1446.83139 | | -0.048332751 | 0.17296331 | -0.2794393 | 0.77990769 | 0.87952228 |
| CER1 | 11.5184308 | | 1.157876508 | 0.4545607 | 2.54724289 | 0.01085778 | 0.04790359 |
| FREM1 | 1192.60602 | | 1.701866155 | 0.50901369 | 3.34345854 | 0.00082741 | 0.00675505 |
| AL512643.1 | 1.87781333 | | 1.566934664 | 1.03588517 | 1.51265286 | 0.13036786 | 0.28557904 |
| TTC39B | 1922.84939 | | 0.146356614 | 0.18489914 | 0.79154838 | 0.42862405 | 0.62103386 |
| SNAPC3 | 2160.79437 | | -0.016470349 | 0.15815777 | -0.1041387 | 0.91705926 | 0.95971317 |
| PSIP1 | 2705.9214 | | 0.213311418 | 0.13090111 | 1.62956158 | 0.10319419 | 0.24277195 |
| CCDC171 | 128.925134 | | 0.499918449 | 0.31303286 | 1.59701588 | 0.11026217 | 0.25431232 |
| C9orf92 | 8.10714528 | | 0.344256047 | 0.99216266 | 0.34697541 | 0.72860979 | 0.84651893 |
| BNC2 | 932.711422 | | 0.555803166 | 0.25892198 | 2.14660481 | 0.03182476 | 0.10519163 |
| AL449983.1 | 0.46349046 | | 0.34438482 | 2.27337599 | 0.15148608 | 0.87959229 | NA |
| AL450003.1 | 2.16270278 | | 2.594921132 | 0.95460378 | 2.71832271 | 0.00656138 | 0.03292552 |
| BNC2-AS1 | 48.5809971 | | 0.099153257 | 0.29506406 | 0.33603977 | 0.73684088 | 0.85226566 |
| AL358117.1 | 0 | NA | | NA | NA | NA | NA |
| AL162725.2 | 15.7713877 | | 3.33985905 | 1.21589704 | 2.7468272 | 0.00601748 | 0.03084179 |
| CNTLN | 374.156767 | | 0.113667195 | 0.10859242 | 1.04673235 | 0.29522304 | 0.49206952 |
| SH3GL2 | 766.62824 | | 0.973570938 | 0.27444451 | 3.54742366 | 0.00038902 | 0.00373441 |
| ADAMTSL1 | 405.985498 | | 0.684547268 | 0.33493543 | 2.0438186 | 0.04097147 | 0.12613508 |
| AL442638.1 | 0 | NA | | NA | NA | NA | NA |
| SAXO1 | 7.47986798 | | -0.211800091 | 0.48062658 | -0.4406749 | 0.65944834 | 0.79994664 |
| RRAGA | 4245.18407 | | -0.440697922 | 0.15647026 | -2.8164964 | 0.00485506 | 0.0262945 |
| HAUS6 | 737.787563 | | -0.580428543 | 0.18741565 | -3.0970122 | 0.00195482 | 0.01326179 |
| PLIN2 | 35966.3773 | | -2.250493314 | 0.6051833 | -3.718697 | 0.00020025 | 0.0021821 |
| DENND4C | 1562.30464 | | -0.028167483 | 0.13962122 | -0.2017421 | 0.84011832 | 0.91492195 |
| AL161909.2 | 0.15543392 | | 1.368031615 | 3.28033652 | 0.41704002 | 0.67664913 | NA |
| AL391834.1 | 137.074092 | | 0.403339787 | 0.2082964 | 1.93637422 | 0.05282188 | 0.1514036 |
| AL391834.2 | 103.566606 | | -0.22450731 | 0.21360285 | -1.0510502 | 0.29323554 | 0.48977138 |
| RPS6 | 45215.9718 | | -0.302667083 | 0.22722668 | -1.332005 | 0.18285854 | 0.35969577 |
| ACER2 | 57.8937925 | | -0.097761502 | 0.35244532 | -0.2773806 | 0.78148786 | 0.88036187 |
| SLC24A2 | 20.6488459 | | -0.126186006 | 0.59231487 | -0.2130387 | 0.83129676 | 0.91040231 |
| AL591222.1 | 0.47101374 | | 1.509729963 | 1.80441945 | 0.8366846 | 0.40276988 | NA |
| MLLT3 | 789.746789 | | -0.111395468 | 0.18695367 | -0.5958453 | 0.55127857 | 0.72144992 |
| FOCAD | 1180.8997 | | 0.052707096 | 0.21135851 | 0.24937295 | 0.80307231 | 0.89318485 |
| FOCAD-AS1 | 3.04824676 | | -0.486241478 | 0.86632896 | -0.5612666 | 0.57461583 | 0.73897193 |
| HACD4 | 316.814759 | | 0.612857297 | 0.33022119 | 1.85589936 | 0.0634679 | 0.17190855 |
| IFNB1 | 0.27380333 | | -1.364242679 | 3.30558948 | -0.4127078 | 0.67982068 | NA |
| IFNW1 | 1.22664988 | | 0.123343788 | 1.06344948 | 0.11598462 | 0.90766473 | 0.95385641 |
| IFNA21 | 1.61332671 | | 0.291614646 | 1.13935765 | 0.25594654 | 0.79799212 | 0.89059811 |
| IFNA4 | 0.18390514 | | 0.18657164 | 3.29377195 | 0.05664376 | 0.95482897 | NA |
| IFNA7 | 0.2331213 | | 0.077576459 | 3.09774174 | 0.02504291 | 0.98002074 | NA |
| IFNA10 | 0.22977084 | | -1.364242679 | 3.30558948 | -0.4127078 | 0.67982068 | NA |
| IFNA16 | 0.19890045 | | -0.340073434 | 3.15719848 | -0.1077137 | 0.91422283 | NA |
| IFNA17 | 0.92668564 | | -1.459246783 | 1.52791544 | -0.9550573 | 0.33954872 | NA |
| IFNA14 | 0.63209959 | | 1.494196963 | 2.08820351 | 0.71554183 | 0.4742743 | NA |
| IFNA5 | 1.58255061 | | -1.609540463 | 1.40189301 | -1.1481193 | 0.2509193 | 0.44324371 |
| KLHL9 | 1718.82333 | | 0.150048819 | 0.10484461 | 1.43115428 | 0.152386 | 0.3175783 |
| IFNA6 | 0 | NA | | NA | NA | NA | NA |
| IFNA13 | 0.46125906 | | -2.122368367 | 3.28687016 | -0.645711 | 0.51846651 | NA |
| IFNA2 | 0.70945937 | | -2.733505325 | 3.26597188 | -0.8369654 | 0.40261204 | NA |
| AL353732.1 | 0.22077692 | | -0.339747857 | 3.07884176 | -0.1103492 | 0.91213241 | NA |
| IFNA8 | 0.6039901 | | -1.258892392 | 3.26177852 | -0.3859528 | 0.69953166 | NA |
| IFNA1 | 1.75462693 | | -2.509770026 | 1.82357769 | -1.3762891 | 0.16873216 | 0.3403815 |
| IFNE | 20.1126757 | | -0.80961852 | 0.44372455 | -1.8245971 | 0.06806185 | 0.18046559 |
| MTAP | 1423.58353 | | -0.329530262 | 0.12806449 | -2.5731588 | 0.0100775 | 0.04531007 |
| AL359922.1 | 162.662439 | | -0.080784292 | 0.18130987 | -0.4455593 | 0.65591565 | 0.79814017 |
| AL359922.2 | 138.179259 | | -0.289515202 | 0.22831835 | -1.268033 | 0.20478618 | 0.38711193 |
| CDKN2A-DT | 0.03392908 | | 0.380740593 | 3.29738709 | 0.11546736 | 0.90807469 | NA |
| CDKN2A | 22.1601083 | | -1.223866994 | 0.53820195 | -2.2739921 | 0.02296646 | 0.08305342 |
| CDKN2B-AS1 | 9.93445441 | | -1.624575607 | 0.60111906 | -2.7025854 | 0.00688025 | 0.03412946 |
| CDKN2B | 104.731848 | | -0.965688419 | 0.3172062 | -3.0443555 | 0.00233179 | 0.01521165 |
| AL157937.1 | 0 | NA | | NA | NA | NA | NA |
| DMRTA1 | 499.947614 | | 0.681520926 | 0.25072091 | 2.71824531 | 0.00656292 | 0.03292696 |
| LINC01239 | 49.3570876 | | 1.626613502 | 0.64900513 | 2.50631842 | 0.01219957 | 0.05217364 |
| AL391117.1 | 10.1698861 | | 2.580996295 | 0.54880191 | 4.70296526 | 2.56E-06 | 6.94E-05 |
| AC017067.1 | 4.16742614 | | 2.12699283 | 1.01627367 | 2.09293312 | 0.03635512 | 0.11562924 |
| AL445623.2 | 1.27439253 | | 2.393815133 | 1.65800809 | 1.44378978 | 0.14879812 | 0.31286357 |
| AL445623.1 | 0 | NA | | NA | NA | NA | NA |
| ELAVL2 | 12.6385266 | | -0.270413162 | 0.95383177 | -0.2835019 | 0.77679209 | 0.87767289 |
| AL365204.1 | 0 | NA | | NA | NA | NA | NA |
| AL365204.2 | 0 | NA | | NA | NA | NA | NA |
| IZUMO3 | 0 | NA | | NA | NA | NA | NA |
| AL353811.1 | 0 | NA | | NA | NA | NA | NA |
| TUSC1 | 934.010846 | | -0.364023636 | 0.18576472 | -1.9595951 | 0.05004314 | 0.14571984 |
| LINC01241 | 0 | NA | | NA | NA | NA | NA |
| AL353753.1 | 0.29483227 | | 0.3284944 | 2.91766247 | 0.11258821 | 0.91035703 | NA |
| AL451137.1 | 2.05888487 | | -2.370437414 | 1.33886725 | -1.7704798 | 0.07664725 | 0.19659449 |
| AL356133.2 | 5.33061584 | | -1.953493741 | 0.80231004 | -2.4348365 | 0.01489852 | 0.06054462 |
| CAAP1 | 1681.08951 | | -0.113974423 | 0.13211493 | -0.8626915 | 0.38830711 | 0.58395543 |
| PLAA | 1680.79736 | | -0.227491757 | 0.17578733 | -1.2941306 | 0.19562032 | 0.37587495 |
| IFT74 | 1125.37204 | | -0.015822244 | 0.18258707 | -0.0866559 | 0.93094504 | 0.96680138 |
| IFT74-AS1 | 2.79556149 | | -0.023905762 | 1.04461725 | -0.0228847 | 0.98174224 | 0.99170821 |
| LRRC19 | 2843.1784 | | 0.787866871 | 0.4777763 | 1.64902878 | 0.09914174 | 0.23612424 |
| AL355432.1 | 8.1697848 | | 2.324736778 | 0.49740506 | 4.67372963 | 2.96E-06 | 7.75E-05 |
| TEK | 1967.85668 | | 0.657792313 | 0.29979399 | 2.19414775 | 0.02822479 | 0.0966733 |
| EQTN | 5.63169565 | | -0.11296648 | 0.68679438 | -0.1644837 | 0.86935038 | 0.932003 |
| MOB3B | 1051.38319 | | 0.187682383 | 0.1245335 | 1.50708353 | 0.13178925 | 0.28783191 |
| AL163192.1 | 0.17361919 | | -1.003574173 | 3.30558948 | -0.3035992 | 0.7614333 | NA |
| IFNK | 0.18380315 | | -1.003574173 | 3.30558948 | -0.3035992 | 0.7614333 | NA |
| C9orf72 | 897.155277 | | -0.571832485 | 0.23523379 | -2.4309113 | 0.0150609 | 0.06107272 |
| AL360014.1 | 1.52696997 | | -0.169968026 | 1.79961897 | -0.0944467 | 0.92475434 | 0.96356293 |
| LINGO2 | 28.6628917 | | 1.554581692 | 0.39648368 | 3.92092226 | 8.82E-05 | 0.00114813 |
| LINC01242 | 0 | NA | | NA | NA | NA | NA |
| LINC01243 | 0 | NA | | NA | NA | NA | NA |
| ACO1 | 8431.5268 | | 0.971038929 | 0.41801343 | 2.32298501 | 0.02017996 | 0.07538539 |
| DDX58 | 1008.55897 | | -0.368160127 | 0.23279322 | -1.5814899 | 0.11376606 | 0.26005644 |
| TOPORS | 1257.50907 | | 0.104837927 | 0.18289633 | 0.57320956 | 0.56650281 | 0.73284279 |
| SMIM27 | 339.790277 | | 0.198181607 | 0.14977553 | 1.32319086 | 0.18577191 | 0.36314347 |
| NDUFB6 | 2292.46778 | | 0.153489116 | 0.17727701 | 0.86581511 | 0.38659156 | 0.58227207 |
| TAF1L | 0.67741986 | | -0.535598172 | 1.45802522 | -0.3673449 | 0.71336175 | NA |
| AL589642.1 | 0 | NA | | NA | NA | NA | NA |
| AL589642.2 | 0.31417979 | | -0.695098386 | 2.30918892 | -0.3010141 | 0.76340376 | NA |
| TMEM215 | 0.58104723 | | -0.01523461 | 2.7567254 | -0.0055263 | 0.99559064 | NA |
| AL157884.2 | 7.39358388 | | 0.090082723 | 0.469829 | 0.19173512 | 0.84794969 | 0.92010671 |
| APTX | 1428.61885 | | -0.221007804 | 0.06220683 | -3.5527901 | 0.00038117 | 0.00367376 |
| DNAJA1 | 8660.2034 | | -0.240903234 | 0.16702101 | -1.4423528 | 0.14920286 | 0.31335453 |
| SMU1 | 2776.37517 | | 0.039153684 | 0.08715332 | 0.44925061 | 0.65325088 | 0.79625709 |
| B4GALT1 | 9538.4995 | | -0.372648459 | 0.17257522 | -2.1593394 | 0.03082385 | 0.10285017 |
| B4GALT1-AS | 147.025633 | | 0.646093769 | 0.32024449 | 2.01750161 | 0.04364319 | 0.13209578 |
| SPINK4 | 1.50770609 | | -0.32697888 | 1.50967298 | -0.2165892 | 0.82852849 | 0.90879717 |
| BAG1 | 5219.39434 | | -0.144842489 | 0.1695412 | -0.8543203 | 0.39292755 | 0.58844158 |
| CHMP5 | 4347.31406 | | -0.18300893 | 0.17380071 | -1.0529815 | 0.29234947 | 0.48857694 |

| | | | | | | |
|---|---|---|---|---|---|---|
| NFX1 | 2322.63859 | -0.186939857 | 0.19922951 | -0.9383141 | 0.34808302 | 0.54549255 |
| AQP7 | 3406.49518 | 1.135454833 | 0.35841386 | 3.16799922 | 0.00153492 | 0.01100899 |
| AL356218.1 | 9.29037271 | 0.824888379 | 0.6936457 | 1.18920708 | 0.23435819 | 0.4230576 |
| AQP3 | 17846.3134 | -0.215618115 | 0.35890067 | -0.6007738 | 0.54799067 | 0.71876822 |
| NOL6 | 2149.15033 | -0.236531411 | 0.20485131 | -1.1546493 | 0.24823408 | 0.43985572 |
| AL139008.2 | 0.2926952 | -0.180036946 | 2.90460663 | -0.0619832 | 0.95057617 | NA |
| ANKRD18B | 3.71484846 | -1.791155277 | 0.89073273 | -2.0108785 | 0.04433829 | 0.13338227 |
| CYP4F26P | 2.51475552 | -2.493510281 | 1.34705979 | -1.8510762 | 0.0641586 | 0.1732223 |
| TRBV20OR9- | 0.28054029 | -1.364242679 | 3.30558948 | -0.4127078 | 0.67982068 | NA |
| TRBV21OR9- | 0 | NA | NA | NA | NA | NA |
| TRBV22OR9- | 0 | NA | NA | NA | NA | NA |
| TRBV23OR9- | 0.23371381 | -0.340597073 | 3.29738699 | -0.103293 | 0.91773041 | NA |
| TRBV24OR9- | 0 | NA | NA | NA | NA | NA |
| TRBV25OR9- | 0 | NA | NA | NA | NA | NA |
| AL356489.3 | 1.4980884 | 2.211652898 | 1.13667711 | 1.94571781 | 0.05168865 | 0.1492913 |
| TRBV26OR9- | 0.32661268 | 2.296964916 | 2.78156108 | 0.82578266 | 0.40892739 | NA |
| AL356489.2 | 7.40684406 | 0.615986106 | 0.87499136 | 0.70399107 | 0.48143833 | 0.66600139 |
| AL356489.1 | 0 | NA | NA | NA | NA | NA |
| LINC01251 | 4.64754579 | 0.187494535 | 0.62609886 | 0.29946475 | 0.76458547 | 0.86980668 |
| PRSS3 | 76.5246822 | 0.275958227 | 0.56086004 | 0.4920269 | 0.62270033 | 0.77488006 |
| UBE2R2-AS1 | 37.6020021 | -0.011161481 | 0.5166843 | -0.0216021 | 0.98276533 | 0.99232988 |
| TRBV29OR9- | 0.07659028 | -0.642905668 | 3.30558948 | -0.1944905 | 0.84579185 | NA |
| UBE2R2 | 3703.25469 | -0.420923016 | 0.18140408 | -2.3203613 | 0.02032134 | 0.07577286 |
| UBAP2 | 1780.0956 | -0.023687536 | 0.16968095 | -0.1396004 | 0.88897569 | 0.94259304 |
| AL354989.1 | 7.15825565 | 0.791065267 | 0.47017603 | 1.68248744 | 0.09247436 | 0.22485526 |
| DCAF12 | 1963.06017 | -0.100798285 | 0.113575 | -0.8875042 | 0.37480751 | 0.57078275 |
| UBAP1 | 5370.73644 | -0.898644447 | 0.22809872 | -3.9397171 | 8.16E-05 | 0.00107992 |
| KIF24 | 363.91801 | -0.673794177 | 0.18095348 | -3.7235768 | 0.00019642 | 0.0021501 |
| NUDT2 | 636.209408 | 0.193589267 | 0.24199676 | 0.79996638 | 0.42373028 | 0.61621563 |
| MYORG | 2089.06512 | 1.176944962 | 0.39208238 | 3.0017798 | 0.00268406 | 0.01688021 |
| C9orf24 | 101.730428 | 0.468361695 | 0.27835403 | 1.68261152 | 0.09245032 | 0.22483904 |
| FAM219A | 2820.36 | -1.241508229 | 0.20950494 | -5.9259139 | 3.11E-09 | 2.56E-07 |
| DNAI1 | 175.551858 | -0.607219167 | 0.43511795 | -1.3955277 | 0.16285676 | 0.33188415 |
| ENHO | 21.1009282 | 0.574991066 | 0.56669999 | 1.01463046 | 0.31028201 | 0.50726204 |
| AL160270.1 | 12.1159905 | 0.616097248 | 0.54102703 | 1.13875502 | 0.25480535 | 0.44753011 |
| CNTFR | 202.539843 | 0.710842431 | 0.53330256 | 1.33290647 | 0.1825625 | 0.35929346 |
| CNTFR-AS1 | 2.13820722 | 0.952869851 | 1.17087857 | 0.81380758 | 0.41575519 | 0.60908136 |
| RPP25L | 944.909103 | 0.027806159 | 0.15332224 | 0.18135764 | 0.85608687 | 0.92463693 |
| DCTN3 | 2293.2592 | 0.219866868 | 0.13588085 | 1.61808577 | 0.10564412 | 0.24665673 |
| ARID3C | 2.61830097 | 0.849756977 | 0.96004819 | 0.88511909 | 0.37609241 | 0.57194676 |
| SIGMAR1 | 2652.81137 | -0.341226006 | 0.19881273 | -1.7163187 | 0.08610372 | 0.21343816 |
| GALT | 2863.14995 | 0.479294495 | 0.32860335 | 1.45858068 | 0.14468055 | 0.30673953 |
| AL162231.3 | 1595.8061 | 0.677263347 | 0.28140807 | 2.40669486 | 0.01609761 | 0.06414235 |
| IL11RA | 1177.37813 | 0.435827224 | 0.250053 | 1.74293943 | 0.0813442 | 0.20475816 |
| CCL27 | 24.426142 | 0.037394139 | 0.39174822 | 0.09545452 | 0.92395381 | 0.96310558 |
| AL162231.1 | 197.993495 | 0.162140051 | 0.16558654 | 0.9791862 | 0.32748798 | 0.52526221 |
| AL162231.2 | 226.92642 | -0.008812343 | 0.22710626 | -0.0388027 | 0.96904767 | 0.98564326 |
| CCL19 | 818.623111 | -0.7693423 | 1.14092648 | -0.6743137 | 0.50011193 | 0.68130141 |
| CCL21 | 1951.04242 | -1.153480933 | 0.70495839 | -1.6362397 | 0.10178942 | 0.24041085 |
| FAM205A | 0.73820835 | -1.06672632 | 1.6816633 | -0.6343281 | 0.52586671 | NA |
| FAM205C | 16.1941421 | 3.186456292 | 0.71305056 | 4.46876626 | 7.87E-06 | 0.0001688 |
| PHF24 | 70.9627006 | -1.798270917 | 0.79512869 | -2.2616099 | 0.02372152 | 0.08495673 |
| DNAJB5-DT | 14.9103305 | 0.156164717 | 0.85339934 | 0.18299137 | 0.85480479 | 0.92400911 |
| DNAJB5 | 968.075234 | -0.138037771 | 0.36044469 | -0.3829652 | 0.70174557 | 0.82913351 |
| C9orf131 | 9.51117483 | 0.445918775 | 0.66775501 | 0.66778799 | 0.50426893 | 0.68389075 |
| VCP | 15763.6195 | -0.335304407 | 0.23106614 | -1.4511187 | 0.1467468 | 0.30980011 |
| FANCG | 581.173986 | 0.044065033 | 0.2261693 | 0.19483207 | 0.84552441 | 0.9185712 |
| PIGO | 2621.29771 | 0.505450242 | 0.11951842 | 4.22905733 | 2.35E-05 | 0.00040499 |
| AL353795.2 | 12.3960188 | 1.051248008 | 0.53023996 | 1.9825892 | 0.04741333 | 0.14005709 |
| STOML2 | 5082.79679 | -0.119825479 | 0.14447134 | -0.8294066 | 0.40687437 | 0.60161922 |
| FAM214B | 2380.96909 | -0.628231502 | 0.24313888 | -2.5838381 | 0.00977076 | 0.0443087 |
| UNC13B | 3407.99635 | 0.366110617 | 0.16599044 | 2.20561264 | 0.02741113 | 0.09457705 |
| RUSC2 | 4476.34405 | -0.01122861 | 0.28795928 | -0.0389937 | 0.96889538 | 0.9856024 |
| FAM166B | 139.909889 | -0.043388017 | 0.28779572 | -0.1507598 | 0.88016522 | 0.9378468 |
| TESK1 | 3520.45705 | -0.764551146 | 0.18299732 | -4.1779362 | 2.94E-05 | 0.00048361 |
| CD72 | 1424.53833 | -1.029687131 | 0.20103268 | -5.1219889 | 3.02E-07 | 1.20E-05 |
| AL357874.2 | 4.84863732 | 0.071713994 | 0.62711684 | 0.11435508 | 0.90895632 | 0.95452902 |
| AL357874.1 | 16.2450331 | -0.090501336 | 0.45184786 | -0.2002916 | 0.84125252 | 0.91528774 |
| SIT1 | 53.7190155 | -0.009971097 | 0.68840392 | -0.0144844 | NA | NA |
| RMRP | 37.5081106 | -0.449841085 | 0.56550444 | -0.7954687 | 0.42634091 | 0.61867047 |
| CCDC107 | 1610.0776 | -0.861956235 | 0.16045835 | -5.3718379 | 7.79E-08 | 3.86E-06 |
| ARHGEF39 | 1689.12117 | -0.964740837 | 0.15965586 | -6.0426271 | 1.52E-09 | 1.44E-07 |
| CA9 | 84.180301 | -3.545979376 | 0.80098916 | -4.4270005 | 9.56E-06 | 0.00019624 |
| TPM2 | 7909.04217 | -0.611954857 | 0.48350011 | -1.2656768 | 0.20562883 | 0.38820392 |
| TLN1 | 16090.131 | 0.162896281 | 0.16682106 | 0.9764731 | 0.32883007 | 0.52661539 |
| CREB3 | 3250.59639 | -0.341490008 | 0.14023127 | -2.4351916 | 0.01488391 | 0.06052089 |
| GBA2 | 7999.61077 | 0.411758212 | 0.11327641 | 3.63498629 | 0.000278 | 0.00286469 |
| RGP1 | 2936.61956 | 0.044724707 | 0.19654891 | 0.22755001 | 0.81999609 | 0.90344498 |
| MSMP | 644.464144 | 0.358036253 | 0.17801178 | 2.01130651 | 0.04429309 | 0.13331504 |
| AL133410.2 | 529.391773 | -0.503229893 | 0.20207515 | -2.4903107 | 0.01276315 | 0.05397982 |
| AL133410.1 | 75.0874234 | -0.666780538 | 0.36046415 | -1.8497832 | 0.0643448 | 0.17352959 |
| NPR2 | 1311.34167 | -0.361055494 | 0.25091275 | -1.4389683 | 0.1501595 | 0.31458696 |
| SPAG8 | 445.190435 | -0.124732749 | 0.16455964 | -0.757979 | 0.44846355 | 0.63858653 |
| HINT2 | 1819.96783 | 0.706110732 | 0.19000999 | 3.71617697 | 0.00020226 | 0.00220033 |
| TMEM8B | 2477.49829 | 0.357732522 | 0.23366027 | 1.53099424 | 0.12577082 | 0.27856345 |
| FAM221B | 7.30180922 | 0.507430702 | 0.56833953 | 0.89283021 | 0.37194811 | 0.56855459 |
| OR13J1 | 3.65930107 | -0.93455308 | 0.84340693 | -1.108069 | 0.26783202 | 0.46200891 |
| HRCT1 | 257.805101 | -1.373392009 | 0.42471837 | -3.2336534 | 0.00122218 | 0.00918461 |
| SPAAR | 172.614322 | 0.114111253 | 0.30467081 | 0.3745395 | 0.70800297 | 0.83387039 |
| OR2S2 | 0.47463758 | 0.839972316 | 1.80359276 | 0.46572172 | 0.64141471 | NA |
| RECK | 644.391701 | 0.02886498 | 0.26715271 | 0.10804674 | 0.91395861 | 0.9577987 |
| GLIPR2 | 578.120506 | -0.5973382 | 0.45303764 | -1.3185178 | 0.18733037 | 0.36526977 |
| CCIN | 3.33965144 | 0.469860142 | 0.98162801 | 0.47865397 | 0.63218482 | 0.78153143 |
| CLTA | 7130.5984 | -0.545856485 | 0.17353452 | -3.1455211 | 0.00165791 | 0.01167666 |
| GNE | 2161.69016 | -0.3331467 | 0.26124579 | -1.2752232 | 0.20223024 | 0.38401466 |
| RNF38 | 2551.60214 | -0.071075768 | 0.21631397 | -0.3285769 | 0.74247553 | 0.85544394 |
| MELK | 51.17659 | -2.3189238 | 0.57178625 | -4.0555781 | 5.00E-05 | 0.00074381 |
| PAX5 | 19.8495858 | 1.18418191 | 1.07116679 | 1.10550656 | 0.26894015 | 0.46313275 |
| AL450267.1 | 0 | NA | NA | NA | NA | NA |
| AL161781.2 | 1.28609102 | 1.222333884 | 1.93492866 | 0.63172039 | 0.5275696 | 0.70223445 |
| LINC01400 | 0 | NA | NA | NA | NA | NA |
| AL512604.3 | 1.29692281 | 0.369503161 | 1.12773598 | 0.32765041 | 0.743176 | 0.85590038 |
| EBLN3P | 2952.59557 | 0.206287988 | 0.10633564 | 1.93997027 | 0.0523833 | 0.15042445 |
| AL512604.2 | 0.84027588 | -0.255443219 | 1.61393209 | -0.1582738 | 0.87424103 | NA |
| ZCCHC7 | 634.911392 | -0.315615769 | 0.10100121 | -3.1248712 | 0.00177883 | 0.01232312 |
| LINC01627 | 0 | NA | NA | NA | NA | NA |
| GRHPR | 8647.79182 | 1.091788228 | 0.28581122 | 3.8199628 | 0.00013347 | 0.00158973 |
| ZBTB5 | 1360.13368 | -0.324337555 | 0.18166543 | -1.7853565 | 0.07420351 | 0.19230924 |
| POLR1E | 465.751813 | -0.481306501 | 0.23314234 | -2.064432 | 0.03897678 | 0.12153455 |
| AL513165.1 | 81.8846142 | -0.567798666 | 0.3606129 | -1.5745379 | 0.11536315 | 0.26279599 |
| FBXO10 | 790.991822 | -0.334535251 | 0.29781506 | -1.1232986 | 0.26131069 | 0.45456579 |
| AL513165.2 | 2121.25313 | -0.329633251 | 0.2631925 | -1.2524416 | 0.21040898 | 0.3937609 |
| TOMM5 | 2361.58529 | -0.386506116 | 0.23813765 | -1.6230366 | 0.10458159 | 0.24503034 |

| | | | | | | |
|---|---|---|---|---|---|---|
| AL138752.2 | 3868.38449 | -0.42218566 | 0.18705514 | -2.2570118 | 0.02400734 | 0.0856199 |
| FRMPD1 | 63.1286256 | 1.510354854 | 0.36165184 | 4.17626753 | 2.96E-05 | 0.00048585 |
| TRMT10B | 631.157404 | 0.031804414 | 0.18710373 | 0.16998279 | 0.86502367 | 0.92986964 |
| EXOSC3 | 767.04945 | -0.528430869 | 0.14013719 | -3.7708112 | 0.00016272 | 0.00186155 |
| DCAF10 | 1694.25484 | -0.189827807 | 0.18922767 | -1.0031715 | 0.31577811 | 0.51290845 |
| SLC25A51 | 532.904754 | -0.052501207 | 0.25038026 | -0.2096859 | 0.83391284 | 0.91196788 |
| SHB | 2232.35844 | -0.520285017 | 0.22084076 | -2.3559284 | 0.01847648 | 0.07083968 |
| AL135785.1 | 30.6569952 | 3.617828238 | 0.85081787 | 4.25217708 | 2.12E-05 | 0.00037044 |
| ALDH1B1 | 6652.15758 | 0.52032233 | 0.59222918 | 0.87858272 | 0.37962757 | 0.57562986 |
| IGFBPL1 | 8.81101843 | 0.517709941 | 0.48298673 | 1.07189268 | 0.28376824 | 0.47976973 |
| FAM95C | 864.603656 | -0.661709316 | 0.19776965 | -3.3458588 | 0.00082028 | 0.00671126 |
| ANKRD18A | 112.942055 | -1.051514572 | 0.29266736 | -3.5928659 | 0.00032706 | 0.00324966 |
| FAM201A | 204.968149 | -0.094866264 | 0.24447604 | -0.3880391 | 0.6979871 | 0.82654528 |
| AL591543.1 | 0.09989236 | -0.642905668 | 3.30558948 | -0.1944905 | 0.84579185 | NA |
| FAM240B | 0 | NA | NA | NA | NA | NA |
| AL845311.1 | 0 | NA | NA | NA | NA | NA |
| CNTNAP3 | 320.777396 | -0.517259467 | 0.32366665 | -1.5981241 | 0.11001537 | 0.25394829 |
| SPATA31A1 | 0 | NA | NA | NA | NA | NA |
| FAM74A1 | 0.11051935 | 0.02007176 | 3.29738704 | 0.00608717 | 0.99514317 | NA |
| BX005214.1 | 0 | NA | NA | NA | NA | NA |
| BX005214.2 | 0 | NA | NA | NA | NA | NA |
| GLIDR | 247.998089 | 0.114086515 | 0.23441952 | 0.48667669 | 0.62648746 | 0.77772688 |
| AL773545.1 | 0 | NA | NA | NA | NA | NA |
| ANKRD20A2 | 1.43357674 | -0.696012194 | 1.38449323 | -0.5027198 | 0.61516127 | 0.76990046 |
| FAM95B1 | 64.3082553 | -1.136414668 | 0.59103275 | -1.9227609 | 0.05451009 | 0.15472136 |
| BMS1P14 | 0.26948541 | -1.364242679 | 3.30558948 | -0.4127078 | 0.67982068 | NA |
| FP325318.1 | 0.6645712 | -1.861945673 | 1.67327762 | -1.1127536 | 0.26581427 | NA |
| IGKV1OR9-2 | 0 | NA | NA | NA | NA | NA |
| AL353626.1 | 0.48399843 | 0.868413299 | 1.80203907 | 0.48190592 | 0.62987278 | NA |
| BX255923.1 | 7.3583201 | 0.083024533 | 0.65514553 | 0.12672685 | 0.89915659 | 0.94884786 |
| BX255923.2 | 5.86327556 | -0.092336651 | 0.99550806 | -0.0927533 | 0.92609956 | 0.9641707 |
| FOXD4L6 | 1.13180124 | -1.100078614 | 1.18224453 | -0.9305001 | 0.35211223 | 0.54974538 |
| CBWD6 | 365.677386 | -0.645605698 | 0.21839765 | -2.9561019 | 0.00311554 | 0.01889966 |
| AL845472.1 | 8.59539164 | -0.636790958 | 0.79317866 | -0.8028342 | 0.42207057 | 0.6145437 |
| AL354718.2 | 0 | NA | NA | NA | NA | NA |
| AL591926.2 | 5.34042909 | -0.458175359 | 0.63262078 | -0.7242496 | 0.4689125 | 0.65563216 |
| CNTNAP3B | 412.731583 | 0.320157545 | 0.44893337 | 0.71315159 | 0.47575196 | 0.66123329 |
| SPATA31A6 | 0 | NA | NA | NA | NA | NA |
| FAM74A7 | 0.08695548 | -0.642905668 | 3.30558948 | -0.1944905 | 0.84579185 | NA |
| AL445584.2 | 0 | NA | NA | NA | NA | NA |
| BX088651.4 | 144.146881 | -1.527644299 | 0.49429309 | -3.0905637 | 0.00199777 | 0.01348724 |
| BX088651.2 | 0.06785815 | 0.795868262 | 3.28898349 | 0.24198001 | 0.80879565 | NA |
| BX664718.1 | 0.20357445 | 1.728110593 | 3.26179266 | 0.52980394 | 0.59624787 | NA |
| AL591441.2 | 1.53781669 | -0.103300508 | 1.80034648 | -0.0573781 | 0.95424399 | 0.9780936 |
| AL591441.1 | 8.34871161 | -0.583722997 | 1.5086564 | -0.3869158 | 0.69881854 | 0.82713968 |
| SPATA31A5 | 0 | NA | NA | NA | NA | NA |
| AL590491.2 | 0.08666371 | -0.642905668 | 3.30558948 | -0.1944905 | 0.84579185 | NA |
| SPATA31A7 | 0 | NA | NA | NA | NA | NA |
| BX005040.3 | 0 | NA | NA | NA | NA | NA |
| BX005040.1 | 0.08393208 | -0.642905668 | 3.30558948 | -0.1944905 | 0.84579185 | NA |
| BX088645.1 | 1.60422816 | -1.489468382 | 1.14685464 | -1.2987421 | 0.19403246 | 0.37380449 |
| AL935212.2 | 0.17332742 | -1.003574173 | 3.30558948 | -0.3035992 | 0.7614333 | NA |
| AL935212.1 | 0.07659028 | -0.642905668 | 3.30558948 | -0.1944905 | 0.84579185 | NA |
| FAM27C | 61.2320765 | -0.991991494 | 0.42209048 | -2.3501868 | 0.01876399 | 0.07166052 |
| AL391987.4 | 20.2300729 | -0.076396835 | 0.51331433 | -0.1488305 | 0.88168737 | 0.93857409 |
| AL391987.2 | 0.19289513 | -1.003574173 | 3.30558948 | -0.3035992 | 0.7614333 | NA |
| AL391987.3 | 0 | NA | NA | NA | NA | NA |
| CR769775.2 | 0.19978473 | -1.003574173 | 3.30558948 | -0.3035992 | 0.7614333 | NA |
| AL590399.3 | 1.07845576 | -1.286713096 | 1.23090467 | -1.0453394 | 0.29586615 | 0.49270442 |
| AL590399.1 | 57.7484674 | -0.032335728 | 0.3352608 | -0.0964495 | 0.9231636 | 0.96251084 |
| AL590399.5 | 0.23568937 | 1.881156793 | 3.2502805 | 0.57876752 | 0.56274605 | NA |
| LINC01189 | 0.09635527 | 0.547239394 | 3.29377202 | 0.16614368 | 0.86804388 | NA |
| BX005266.2 | 43.0825413 | -0.371307363 | 0.64466649 | -0.5759681 | 0.56463671 | 0.73156144 |
| LINC01410 | 85.9885804 | -0.278131557 | 0.43009972 | -0.6466676 | 0.51784709 | 0.69488619 |
| AL512625.2 | 6.30088249 | 0.976503871 | 0.56107266 | 1.74042318 | 0.08178474 | 0.20558795 |
| AL512625.3 | 147.412927 | 0.839042583 | 0.66792098 | 1.25620037 | 0.20904333 | 0.39223514 |
| AL512625.1 | 156.666182 | 0.54210133 | 0.41613974 | 1.30269059 | 0.19268042 | 0.37209686 |
| AL772155.1 | 0 | NA | NA | NA | NA | NA |
| CR786580.1 | 3.07520883 | -1.215981279 | 0.80411963 | -1.5121895 | 0.13048565 | 0.28576586 |
| ANKRD20A4 | 6.85270498 | -2.160349935 | 1.24130314 | -1.7403887 | 0.08179079 | 0.20558795 |
| IGKV1OR-2 | 0.09644756 | -0.642905668 | 3.30558948 | -0.1944905 | 0.84579185 | NA |
| IGKV1OR9-1 | 0 | NA | NA | NA | NA | NA |
| FOXD4L5 | 0.52922195 | -1.248244638 | 1.88160055 | -0.6633951 | 0.50707752 | NA |
| CBWD5 | 724.510519 | -0.214777136 | 0.26433768 | -0.8125105 | 0.41649878 | 0.60973035 |
| FOXD4L4 | 0.99727717 | 0.536282865 | 1.61151459 | 0.33278189 | 0.73929893 | NA |
| IGKV1OR-3 | 0.08695548 | -0.642905668 | 3.30558948 | -0.1944905 | 0.84579185 | NA |
| AL136317.2 | 2.13516379 | -0.831643094 | 0.91685186 | -0.9070638 | 0.36437308 | 0.5615216 |
| ANKRD20A3 | 0.63414507 | -1.791710997 | 2.77378123 | -0.6459453 | 0.51831477 | NA |
| AL513478.2 | 0.49300312 | -2.179047281 | 3.28454609 | -0.6634242 | 0.50705893 | NA |
| ZNF658 | 137.645196 | 0.557328675 | 0.16761124 | 3.32512713 | 0.00088378 | 0.00712484 |
| AL353770.4 | 0.25999113 | -1.364242679 | 3.30558948 | -0.4127078 | 0.67982068 | NA |
| AL353770.3 | 0.09635527 | 0.547239394 | 3.29377202 | 0.16614368 | 0.86804388 | NA |
| SPATA31A3 | 0.2803786 | 1.099964891 | 2.9688179 | 0.37050601 | 0.7110055 | NA |
| BX664730.1 | 0 | NA | NA | NA | NA | NA |
| ANKRD20A1 | 5.1290236 | 0.121997998 | 0.81787321 | 0.14916493 | 0.8814235 | 0.93850996 |
| AL353608.2 | 0.07659028 | -0.642905668 | 3.30558948 | -0.1944905 | 0.84579185 | NA |
| CBWD3 | 178.024668 | -0.175890653 | 0.19321905 | -0.9103173 | 0.36265518 | 0.55991227 |
| FOXD4L3 | 2.08549738 | -0.92816599 | 0.95744327 | -0.9694214 | 0.33233498 | 0.53024413 |
| AL353608.3 | 1.2948257 | 0.089332814 | 1.40487727 | 0.06358763 | 0.94929858 | 0.97552721 |
| AL353608.1 | 0 | NA | NA | NA | NA | NA |
| PGM5 | 1311.40913 | 0.267611507 | 0.39668568 | 0.67461852 | 0.49991817 | 0.68121489 |
| PGM5-AS1 | 24.728244 | 0.11426498 | 0.51590285 | 0.22148546 | 0.82471446 | 0.90622499 |
| AL161457.2 | 17.8333093 | 0.583099186 | 0.37400661 | 1.55906119 | 0.1189819 | 0.26825861 |
| TMEM252 | 3436.41647 | -1.860367452 | 0.61367244 | -3.0315317 | 0.00243316 | 0.01566086 |
| AL353616.2 | 279.006409 | -2.045753717 | 0.58787429 | -3.4799169 | 0.00050157 | 0.004542 |
| LINC01506 | 85.2177912 | -0.806516668 | 0.56781458 | -1.4203874 | 0.15549494 | 0.32175397 |
| PIP5K1B | 568.50527 | 0.25367775 | 0.29948719 | 0.84704039 | 0.39697261 | 0.59189006 |
| FAM122A | 1325.46929 | 0.393180419 | 0.10124415 | 3.88348795 | 0.00010297 | 0.0012967 |
| AL354794.1 | 20.8178839 | 1.005674069 | 0.52254519 | 1.9245686 | 0.05428335 | 0.15425001 |
| PRKACG | 0.56347454 | -1.313695776 | 2.25830876 | -0.5817166 | 0.56075756 | NA |
| FXN | 517.115204 | -0.118143629 | 0.14564464 | -0.8111773 | 0.41726385 | 0.61024058 |
| AL358113.1 | 8376.40921 | -0.584463506 | 0.18281784 | -3.196972 | 0.00138878 | 0.01018983 |
| TJP2 | 7442.57459 | -0.5383382 | 0.19469216 | -2.7650738 | 0.00569099 | 0.02967496 |
| BANCR | 8.14664675 | -1.818163598 | 1.06302053 | -1.7103749 | 0.08719658 | 0.21527701 |
| FAM189A2 | 386.940808 | -0.167481116 | 0.20221944 | -0.8282147 | 0.40754891 | 0.60217767 |
| APBA1 | 283.650759 | 0.350219349 | 0.21496361 | 1.62920295 | 0.10327006 | 0.24288544 |
| AL353693.1 | 0.42296795 | 2.671030466 | 2.65059627 | 1.00770928 | 0.31359405 | NA |
| AL162412.1 | 4.59592286 | 1.613165771 | 0.57397779 | 2.81050208 | 0.00494643 | 0.02666884 |
| PTAR1 | 2404.6729 | 0.270724102 | 0.16173488 | 1.67387578 | 0.09415502 | 0.22800861 |
| C9orf135-DT | 0.45955539 | 1.387663068 | 2.22499817 | 0.62366931 | 0.53284478 | NA |
| C9orf135 | 48.8448224 | 0.020144594 | 0.69776141 | 0.02887032 | 0.97696802 | 0.98938694 |
| MAMDC2-AS | 142.888841 | 0.639410303 | 0.31531843 | 2.02782409 | 0.0425782 | 0.12963714 |
| MAMDC2 | 197.445404 | 1.388731945 | 0.50022496 | 2.77621483 | 0.00549959 | 0.02894757 |

| | | | | | | |
|---|---|---|---|---|---|---|
| SMC5-AS1 | 88.7882641 | 0.437279561 | 0.28966465 | 1.50960623 | 0.13114393 | 0.28675501 |
| SMC5 | 1667.14376 | -0.386662712 | 0.14934568 | -2.5890453 | 0.00962424 | 0.04379517 |
| KLF9 | 5240.62278 | 0.029590853 | 0.30386786 | 0.09738066 | 0.9224241 | 0.96204412 |
| TRPM3 | 1459.03913 | 1.59760665 | 0.31851385 | 5.01581525 | 5.28E-07 | 1.90E-05 |
| AL159990.2 | 0 | NA | NA | NA | NA | NA |
| AL159990.1 | 0.08393208 | -0.642905668 | 3.30558948 | -0.1944905 | 0.84579185 | NA |
| CEMIP2 | 4003.4574 | -1.826446475 | 0.26980132 | -6.7695979 | 1.29E-11 | 2.60E-09 |
| ABHD17B | 852.197661 | 0.028078595 | 0.18784289 | 0.14947914 | 0.88117557 | 0.93836253 |
| C9orf85 | 438.685288 | -0.091980173 | 0.14124193 | -0.6512243 | 0.51490172 | 0.69264696 |
| C9orf57 | 0.16421342 | 1.36803136 | 3.28033652 | 0.41703994 | 0.67664919 | NA |
| GDA | 5102.25974 | 0.349743028 | 0.52625307 | 0.66459095 | 0.50631216 | 0.68556876 |
| AL135924.2 | 121.228275 | -2.051939763 | 0.74035711 | -2.7715541 | 0.00557894 | 0.02926964 |
| LINC01504 | 89.0927343 | 0.193590391 | 0.74216073 | 0.26084699 | 0.79421051 | 0.88844572 |
| ZFAND5 | 13321.3715 | -0.950124828 | 0.21391544 | -4.4415906 | 8.93E-06 | 0.00018615 |
| TMC1 | 64.4867313 | 2.210160702 | 0.68484016 | 3.22726504 | 0.0012498 | 0.00935485 |
| LINC01474 | 32.6733314 | -1.02001864 | 0.36830421 | -2.7695004 | 0.00561423 | 0.02940364 |
| ALDH1A1 | 21910.6021 | 0.175590069 | 0.46652681 | 0.37637723 | 0.70663647 | 0.83308876 |
| ANXA1 | 4049.00384 | -0.682918345 | 0.38728457 | -1.7633503 | 0.07784143 | 0.19865272 |
| AL451127.1 | 0 | NA | NA | NA | NA | NA |
| AL355674.1 | 0 | NA | NA | NA | NA | NA |
| RORB-AS1 | 1.30895436 | 3.385046217 | 1.34879399 | 2.50968364 | 0.01208394 | 0.05182142 |
| RORB | 50.6056802 | 1.902341802 | 0.37200343 | 5.11377495 | 3.16E-07 | 1.24E-05 |
| TRPM6 | 267.269742 | 2.308727794 | 0.47720989 | 4.83797142 | 1.31E-06 | 4.06E-05 |
| C9orf40 | 266.903687 | -0.600115234 | 0.1409035 | -4.2590512 | 2.05E-05 | 0.00036236 |
| C9orf41-AS1 | 3.7719058 | -0.501192056 | 0.90381913 | -0.5545269 | 0.57921827 | 0.74259293 |
| CARNMT1 | 919.346078 | 0.44836041 | 0.12968253 | 3.45736953 | 0.00054548 | 0.00485715 |
| NMRK1 | 1965.49465 | 0.245747249 | 0.26628014 | 0.92288987 | 0.3560646 | 0.55349113 |
| OSTF1 | 1804.45093 | -0.37389428 | 0.15333385 | -2.4384327 | 0.01475111 | 0.06009754 |
| AL353780.1 | 0 | NA | NA | NA | NA | NA |
| PCSK5 | 884.354587 | -0.251916412 | 0.36307617 | -0.693839 | 0.48778317 | 0.67152554 |
| RFK | 2393.65222 | -0.317406405 | 0.17235536 | -1.8415813 | 0.06553641 | 0.17584404 |
| GCNT1 | 247.13503 | -0.968953711 | 0.25649892 | -3.7776132 | 0.00015834 | 0.0018289 |
| PRUNE2 | 3406.32009 | 0.227377809 | 0.3170419 | 0.71718536 | 0.47325973 | 0.65929839 |
| PCA3 | 75.1213672 | 0.270324224 | 0.54500712 | 0.49600127 | 0.61989352 | 0.77274237 |
| FOXB2 | 0.12150485 | 1.04981648 | 3.28477129 | 0.31960109 | 0.74927075 | NA |
| VPS13A-AS1 | 4.63778114 | -0.152936191 | 0.64617663 | -0.2366786 | 0.81290612 | 0.89895498 |
| VPS13A | 2539.83811 | 0.201074435 | 0.20467784 | 0.98239477 | 0.3259054 | 0.52358508 |
| GNA14 | 662.756162 | 0.255509225 | 0.4019534 | 0.63566877 | 0.52499233 | 0.69979517 |
| GNA14-AS1 | 5.78328493 | -0.603463996 | 0.69055954 | -0.8738769 | 0.38218532 | 0.5782447 |
| GNAQ | 3526.40386 | -0.364999977 | 0.16092197 | -2.2681799 | 0.02331825 | 0.08391132 |
| CEP78 | 635.735371 | 0.224147357 | 0.17484531 | 1.28197522 | 0.19985134 | 0.38117169 |
| PSAT1 | 5760.66121 | 0.947484842 | 0.5486453 | 1.72695337 | 0.08417604 | 0.21006172 |
| AL512634.1 | 0 | NA | NA | NA | NA | NA |
| AL358975.1 | 0.08393208 | -0.642905668 | 3.30558948 | -0.1944905 | 0.84579185 | NA |
| TLE4 | 977.673482 | -0.010594017 | 0.20656873 | -0.0512857 | 0.95909788 | 0.98028725 |
| AL161912.1 | 0 | NA | NA | NA | NA | NA |
| AL161912.2 | 0 | NA | NA | NA | NA | NA |
| AL161782.1 | 33.6859358 | 1.775654581 | 0.43669576 | 4.06611359 | 4.78E-05 | 0.00071705 |
| LINC01507 | 72.0990164 | 1.404810016 | 0.34051859 | 4.1255017 | 3.70E-05 | 0.000581 |
| AL138749.1 | 0.64803858 | 1.531785873 | 1.75780113 | 0.8714216 | 0.383524 | NA |
| TLE1 | 1818.73967 | -0.964531155 | 0.1720781 | -5.6051942 | 2.08E-08 | 1.29E-06 |
| AL591368.1 | 2.88702833 | -0.436556321 | 0.70499975 | -0.619229 | 0.53576548 | 0.70949308 |
| AL158154.1 | 0 | NA | NA | NA | NA | NA |
| SPATA31D4 | 0 | NA | NA | NA | NA | NA |
| AL158154.2 | 0.09635527 | 0.547239394 | 3.29377202 | 0.16614368 | 0.86804388 | NA |
| SPATA31D3 | 0 | NA | NA | NA | NA | NA |
| SPATA31D1 | 0.07830766 | -0.642905668 | 3.30558948 | -0.1944905 | 0.84579185 | NA |
| AL162726.1 | 0.03392908 | 0.380740593 | 3.29738709 | 0.11546736 | 0.90807469 | NA |
| AL162726.4 | 0.1981425 | 1.631193554 | 3.26970143 | 0.4988815 | 0.61786287 | NA |
| RASEF | 691.75421 | -0.360461244 | 0.22762159 | -1.5835986 | 0.11328508 | 0.25929412 |
| AL137847.1 | 27.6490973 | 0.0258113 | 0.32409095 | 0.07964215 | 0.93652187 | 0.96961451 |
| FRMD3 | 1194.35058 | 1.172343789 | 0.25158395 | 4.65985119 | 3.16E-06 | 8.15E-05 |
| AL137847.2 | 36.5436476 | 0.556627539 | 0.35156268 | 1.58329528 | 0.11335418 | 0.25938885 |
| IDNK | 888.163234 | 1.649631079 | 0.36563355 | 4.51170598 | 6.43E-06 | 0.00014406 |
| UBQLN1 | 5815.95224 | -0.078568293 | 0.16736132 | -0.4694531 | 0.63874578 | 0.78620829 |
| AL354920.1 | 257.926028 | 0.739825588 | 0.34217365 | 2.1621349 | 0.03060778 | 0.10231045 |
| GKAP1 | 365.889897 | -0.09289337 | 0.13949445 | -0.6659288 | 0.50545661 | 0.68472675 |
| AL354733.1 | 97.9059972 | 0.046914271 | 0.23454967 | 0.20001849 | 0.84146612 | 0.91548236 |
| AL354733.2 | 90.4201808 | 0.142850146 | 0.23650781 | 0.60399758 | 0.54584525 | 0.71715108 |
| KIF27 | 162.724544 | 0.002512876 | 0.197595 | 0.01271731 | 0.98985333 | 0.99521246 |
| C9orf64 | 907.948788 | 0.527147996 | 0.17628481 | 2.9903201 | 0.00278685 | 0.0173555 |
| HNRNPK | 24870.3668 | -0.096233934 | 0.13369928 | -0.719779 | 0.47166109 | 0.65803638 |
| AL354733.3 | 2239.54728 | 0.255532904 | 0.18161993 | 1.40696513 | 0.15943773 | 0.32718855 |
| RMI1 | 330.568218 | 0.465368506 | 0.11789212 | 3.94740973 | 7.90E-05 | 0.00105162 |
| AL390838.1 | 0.08666371 | -0.642905668 | 3.30558948 | -0.1944905 | 0.84579185 | NA |
| SLC28A3 | 24.9391515 | -3.380578708 | 0.71219286 | -4.746718 | 2.07E-06 | 5.82E-05 |
| AL356134.1 | 1.01858091 | -3.228826386 | 1.82402475 | -1.7701659 | 0.07669951 | 0.19669027 |
| NTRK2 | 2492.3987 | 0.868373922 | 0.39685666 | 2.18812988 | 0.02866014 | 0.09767369 |
| AGTPBP1 | 963.539888 | -0.092697437 | 0.09230436 | -1.0042585 | 0.31525401 | 0.51241405 |
| AL157882.1 | 0.20718593 | 0.18657164 | 3.29377195 | 0.05664376 | 0.95482897 | NA |
| AL353743.2 | 4.89733153 | 1.744103462 | 0.73073905 | 2.38676648 | 0.01699728 | 0.06667932 |
| NAA35 | 957.408362 | -0.385824475 | 0.1486248 | -2.595963 | 0.00943262 | 0.04310888 |
| GOLM1 | 7981.25971 | 0.092733477 | 0.21325428 | 0.43484931 | 0.66367181 | 0.80246755 |
| C9orf153 | 5.39957283 | -0.095321897 | 0.61181136 | -0.1558028 | 0.8761885 | 0.93555133 |
| ISCA1 | 3539.40512 | -0.012060438 | 0.24822452 | -0.0485868 | 0.96124858 | 0.98149636 |
| TUT7 | 951.966596 | -0.601179686 | 0.1545 | -3.8911307 | 9.98E-05 | 0.00126376 |
| AL158828.1 | 0.25543992 | -0.174096114 | 3.29377188 | -0.0528562 | 0.95784652 | NA |
| AL353613.1 | 0.55277203 | -1.001703492 | 1.93215216 | -0.5184392 | 0.60415185 | NA |
| GAS1 | 666.799203 | 0.353472603 | 0.45980831 | 0.76873905 | 0.44204824 | 0.63328147 |
| GAS1RR | 23.2365431 | 1.093100657 | 0.31244178 | 3.49857386 | 0.00046775 | 0.00431131 |
| AL513318.2 | 168.869684 | 0.238429282 | 0.35092041 | 0.67943977 | 0.49685926 | 0.67865838 |
| C9orf170 | 9.73845035 | -0.105146196 | 0.5473611 | -0.1920966 | 0.84766655 | 0.92002663 |
| DAPK1 | 4884.38021 | -0.370723028 | 0.23799513 | -1.5576916 | 0.11930636 | 0.26881169 |
| CTSL | 25315.3242 | -1.26376947 | 0.592007 | -2.1347205 | 0.03278385 | 0.10749593 |
| AL772337.2 | 50.5439598 | 3.826082463 | 0.65413948 | 5.84903159 | 4.94E-09 | 3.80E-07 |
| AL772337.1 | 165.638325 | 1.34760047 | 0.65445325 | 2.05912413 | 0.03948235 | 0.12262363 |
| SPATA31E1 | 0.73115034 | -1.163929472 | 1.64163536 | -0.7090061 | 0.4783207 | NA |
| AL353572.3 | 0.08393208 | -0.642905668 | 3.30558948 | -0.1944905 | 0.84579185 | NA |
| CDK20 | 1305.33473 | 0.114693715 | 0.15907735 | 0.72099339 | 0.47091358 | 0.65748881 |
| SPIN1 | 4480.272 | 0.584187256 | 0.12266374 | 4.76250961 | 1.91E-06 | 5.49E-05 |
| NXNL2 | 259.559864 | 1.571145262 | 0.40531006 | 3.87640333 | 0.00010601 | 0.00132741 |
| AL592486.1 | 0.06785815 | 0.795868262 | 3.28898349 | 0.24198001 | 0.80879565 | NA |
| AL772202.1 | 0 | NA | NA | NA | NA | NA |
| C9orf47 | 248.075126 | -1.028319007 | 0.72843184 | -1.4116887 | 0.15804164 | 0.32516038 |
| S1PR3 | 2265.22754 | -0.636008057 | 0.42990784 | -1.4794056 | 0.13903195 | 0.298485 |
| SHC3 | 261.708461 | 0.235213274 | 0.36252531 | 0.64881891 | 0.51645543 | 0.69385214 |
| CKS2 | 676.677261 | -1.25590446 | 0.41763122 | -3.0072093 | 0.00263658 | 0.0166292 |
| SECISBP2 | 3487.96982 | 0.039128853 | 0.08544658 | 0.45793352 | 0.6470002 | 0.79207964 |
| SEMA4D | 5511.73344 | -0.61123708 | 0.27762178 | -2.20169 | 0.02768722 | 0.09531487 |
| GADD45G | 666.502277 | -0.853626162 | 0.3487037 | -2.4479986 | 0.01436522 | 0.05899722 |
| AL606807.1 | 3.41125196 | -1.075737606 | 0.76595306 | -1.4044432 | 0.1601869 | 0.32831716 |
| BX470209.1 | 0 | NA | NA | NA | NA | NA |
| BX470209.2 | 0 | NA | NA | NA | NA | NA |

| | | | | | | |
|---|---|---|---|---|---|---|
| AL161629.1 | 0.03392908 | 0.380740593 | 3.29738709 | 0.11546736 | 0.90807469 | NA |
| MIR4290HG | 0.56346809 | 1.810064128 | 2.01884013 | 0.89658617 | 0.36993978 | NA |
| LINC01508 | 60.8638882 | 0.326390633 | 0.45461022 | 0.7179571 | 0.47278374 | 0.65885276 |
| LINC01501 | 0.31959997 | 1.548086548 | 2.78922242 | 0.55502442 | 0.57887796 | NA |
| DIRAS2 | 87.4985129 | -0.162597968 | 0.41977581 | -0.3873448 | 0.69850098 | 0.82697006 |
| SYK | 1555.55171 | 0.467413518 | 0.19519705 | 2.39457266 | 0.01663975 | 0.06565765 |
| AL355607.1 | 12.8494024 | -0.462958034 | 0.62954391 | -0.7353864 | 0.46210419 | 0.65031954 |
| AL355607.2 | 1.0921352 | -1.840505585 | 1.43150772 | -1.2857113 | 0.19854385 | 0.37955643 |
| AL158071.3 | 88.3249674 | -0.518446868 | 0.47665131 | -1.0876858 | 0.27673383 | 0.4721453 |
| AL158071.1 | 13.457655 | -0.499943418 | 0.42902718 | -1.1652954 | 0.24389942 | 0.43471631 |
| AL158071.4 | 12.6917796 | -1.686295132 | 0.59386424 | -2.8395297 | 0.00451801 | 0.02494032 |
| LINC00484 | 9.79791813 | 0.009446026 | 0.51059747 | 0.01849995 | 0.98524002 | 0.99348815 |
| AL158071.5 | 73.0117618 | 0.466716168 | 0.28221101 | 1.65378443 | 0.09817132 | 0.23469819 |
| AUH | 2634.3113 | 0.369586721 | 0.1586866 | 2.32903554 | 0.01985718 | 0.07454135 |
| NFIL3 | 4089.23896 | -1.900864721 | 0.39934565 | -4.7599484 | 1.94E-06 | 5.52E-05 |
| AL353764.1 | 12.5917642 | -1.02010809 | 0.66189571 | -1.5411916 | 0.12327017 | 0.27473245 |
| ROR2 | 260.342549 | -0.61392541 | 0.27285819 | -2.2499798 | 0.02445023 | 0.08680779 |
| SPTLC1 | 4045.30342 | -0.186011896 | 0.14401347 | -1.2916284 | 0.19648585 | 0.37704351 |
| AL136097.2 | 0 | NA | NA | NA | NA | NA |
| IARS | 4719.36432 | -0.310105089 | 0.23350538 | -1.3280426 | 0.18416404 | 0.36120533 |
| NOL8 | 1154.27695 | -0.237317931 | 0.10841918 | -2.1888924 | 0.02860466 | 0.09755624 |
| CENPP | 940.054658 | -0.33627548 | 0.14717275 | -2.2849031 | 0.02231851 | 0.08133486 |
| OGN | 300.622093 | 1.426018245 | 0.7093592 | 2.01029076 | 0.04440042 | 0.13350504 |
| OMD | 76.9692523 | 1.455618585 | 0.52381454 | 2.77888159 | 0.00545464 | 0.02875052 |
| ASPN | 352.289056 | 1.806959743 | 0.41372368 | 4.36755219 | 1.26E-05 | 0.00024647 |
| ECM2 | 774.783755 | 1.886955129 | 0.36271989 | 5.20223778 | 1.97E-07 | 8.45E-06 |
| IPPK | 1002.84372 | -0.341844086 | 0.15887827 | -2.1516101 | 0.03142808 | 0.10436449 |
| BICD2 | 1905.62983 | 0.005117681 | 0.19599385 | 0.02611144 | 0.97916845 | 0.99058424 |
| ZNF484 | 186.168609 | 0.215942808 | 0.23349135 | 0.92484285 | 0.35504766 | 0.55256178 |
| FGD3 | 897.862113 | 0.455463863 | 0.34628384 | 1.31529055 | 0.18841227 | 0.36651163 |
| SUSD3 | 1078.01426 | 0.801952497 | 0.48843331 | 1.64188738 | 0.10061335 | 0.23850898 |
| AL451065.1 | 0.39398426 | 0.292662794 | 2.8241804 | 0.10362751 | 0.91746495 | NA |
| CARD19 | 2591.74717 | -1.157103399 | 0.18840269 | -6.1416502 | 8.17E-10 | 8.74E-08 |
| NINJ1 | 6023.988 | 0.031632755 | 0.15197081 | 0.2081502 | 0.83511169 | 0.9125977 |
| AL390760.1 | 15.2399645 | 0.752874183 | 0.6531557 | 1.15267184 | 0.2490451 | 0.44078927 |
| WNK2 | 200.711734 | -0.000789805 | 0.31267588 | -0.002526 | 0.99798458 | 0.99887805 |
| C9orf129 | 18.918693 | -0.189905171 | 0.35452899 | -0.5356548 | 0.59219712 | 0.75166635 |
| AL583839.2 | 0.37139917 | 1.664841297 | 2.80034203 | 0.59451356 | 0.55216868 | NA |
| AL583839.1 | 25.9554082 | 0.636782439 | 0.45203263 | 1.40870902 | 0.15892123 | 0.32648432 |
| FAM120AOS | 1870.96582 | 0.276294919 | 0.11741096 | 2.35322939 | 0.01861115 | 0.07124452 |
| FAM120A | 7411.21655 | 0.232675873 | 0.13225042 | 1.75935825 | 0.07851668 | 0.19983446 |
| PHF2 | 2149.61578 | 0.157709509 | 0.12972228 | 1.21574724 | 0.22408122 | 0.41012089 |
| AL442224.1 | 0.82112469 | -0.102734782 | 1.59798159 | -0.0642903 | 0.94873904 | NA |
| BARX1 | 1.55931982 | -2.238820574 | 1.43941635 | -1.5553669 | 0.11985871 | 0.26972812 |
| BARX1-DT | 0 | NA | NA | NA | NA | NA |
| PTPDC1 | 589.923552 | -0.443190456 | 0.21829096 | -2.030274 | 0.0423287 | 0.12917576 |
| AL158152.2 | 63.5050189 | -0.839883378 | 0.27251326 | -3.0819909 | 0.00205621 | 0.01377599 |
| AL158152.1 | 42.8133515 | 0.547315891 | 0.53760786 | 1.01805784 | 0.30865047 | 0.50559777 |
| MIRLET7DHG | 64.4284716 | 0.136431507 | 0.32114906 | 0.42482301 | 0.6709657 | 0.80780558 |
| ZNF169 | 263.079485 | 0.494704066 | 0.2200517 | 2.24812652 | 0.02456812 | 0.08710156 |
| NUTM2F | 12.5961839 | 0.229863207 | 0.76237287 | 0.30151021 | 0.76302546 | 0.86877586 |
| AL691447.2 | 11.2161953 | 0.890475296 | 0.51620589 | 1.72503899 | 0.08452045 | 0.21064238 |
| MFSD14B | 2370.86952 | -0.308066946 | 0.19955101 | -1.5438005 | 0.12263668 | 0.27378586 |
| PCAT7 | 5306.73199 | 2.4813489 | 0.45426587 | 5.46232741 | 4.70E-08 | 2.53E-06 |
| FBP2 | 22.4177905 | 1.178720674 | 0.62972984 | 1.87178786 | 0.06123596 | 0.16778501 |
| FBP1 | 11618.6142 | 2.668214082 | 0.4826935 | 5.52776051 | 3.24E-08 | 1.87E-06 |
| C9orf3 | 2911.81314 | 0.261473297 | 0.23141844 | 1.12987234 | 0.25853002 | 0.45133642 |
| AL353768.1 | 16.7128516 | -0.07749519 | 0.40710679 | -0.1903559 | 0.84903023 | 0.92059733 |
| FANCC | 1071.93677 | 1.009803719 | 0.16704949 | 6.04493733 | 1.49E-09 | 1.43E-07 |
| AL354893.2 | 3.090717 | 0.355484316 | 0.78124354 | 0.45502369 | 0.6490922 | 0.79349874 |
| AL161729.3 | 27.4840902 | -0.116640526 | 0.36870519 | -0.3163517 | 0.75173555 | 0.86146914 |
| PTCH1 | 908.711279 | 0.275255724 | 0.17171168 | 1.60301106 | 0.10893221 | 0.25230939 |
| AL161729.2 | 0.17303784 | -1.003574173 | 3.30558948 | -0.3035992 | 0.7614333 | NA |
| AL161729.1 | 32.3966557 | 0.000488258 | 0.29191525 | 0.0016726 | 0.99866546 | 0.99938553 |
| AL161729.4 | 32.5308098 | -0.909554991 | 0.29274678 | -3.1069684 | 0.00189017 | 0.01291287 |
| AL392185.1 | 0 | NA | NA | NA | NA | NA |
| AL354861.2 | 0.17332742 | -1.003574173 | 3.30558948 | -0.3035992 | 0.7614333 | NA |
| LINC00476 | 223.26165 | 0.183125095 | 0.21733266 | 0.84260273 | 0.39945067 | 0.59426337 |
| AL354861.3 | 7.01527143 | 1.063705318 | 0.83726175 | 1.27045732 | 0.20392178 | 0.38619773 |
| ERCC6L2 | 455.604376 | 0.000690385 | 0.15863994 | 0.0043519 | 0.9965277 | 0.99824844 |
| LINC00092 | 39.9672679 | -0.043673004 | 0.5383272 | -0.0811272 | 0.93534076 | 0.96907653 |
| AL449403.1 | 7.20308426 | -0.073368224 | 0.53020354 | -0.1383775 | 0.8899421 | 0.94297317 |
| AL449403.2 | 0.66377986 | -0.200284728 | 2.18927292 | -0.0914846 | 0.92710755 | NA |
| HSD17B3 | 248.970369 | 0.00665237 | 0.26415154 | 0.02518392 | 0.97990827 | 0.99099448 |
| AL160269.1 | 2752.92862 | -0.464567652 | 0.19152438 | -2.4256319 | 0.01528176 | 0.06166489 |
| HSD17B3-AS | 64.8401603 | 0.012120223 | 0.30610346 | 0.03959519 | 0.96841587 | 0.98530467 |
| SLC35D2 | 2627.45758 | -0.55154611 | 0.19681185 | -2.802403 | 0.00507235 | 0.02715343 |
| ZNF367 | 211.550089 | 0.645085376 | 0.44791464 | 1.44019711 | 0.14981164 | 0.31408276 |
| HABP4 | 2403.8615 | 0.301650196 | 0.20032683 | 1.50579028 | 0.13212102 | 0.28824615 |
| CDC14B | 2495.00296 | 0.350747646 | 0.2138198 | 1.64038896 | 0.10092432 | 0.23902534 |
| AAED1 | 771.327939 | -0.966656937 | 0.25615341 | -3.7737422 | 0.00016082 | 0.00184782 |
| AL589843.1 | 35.2290633 | -0.565566094 | 0.48371027 | -1.1692249 | 0.24231303 | 0.43270771 |
| ZNF510 | 435.061211 | 0.17062154 | 0.11052079 | 1.54379587 | 0.12263779 | 0.27378586 |
| ZNF782 | 264.002447 | 0.405185158 | 0.22279367 | 1.81865654 | 0.06896385 | 0.18216467 |
| MFSD14C | 1400.53402 | -0.14043767 | 0.16423551 | -0.8550993 | 0.39249619 | 0.58817551 |
| NUTM2G | 34.534709 | -0.64990326 | 0.44620178 | -1.4565232 | 0.14524803 | 0.30749764 |
| CTSV | 634.594689 | 2.222657071 | 0.47234413 | 4.70558842 | 2.53E-06 | 6.88E-05 |
| AL590705.3 | 16.6015173 | 0.468902021 | 0.50394679 | 0.93045938 | 0.35213328 | 0.54974571 |
| AL590705.1 | 39.9542597 | -0.622467575 | 0.46555364 | -1.337048 | 0.18120696 | 0.35747943 |
| AL590705.2 | 0.583636 | 1.068243309 | 2.72375675 | 0.39219483 | 0.69491426 | NA |
| CCDC180 | 560.160208 | 0.40128646 | 0.31635413 | 1.26847231 | 0.20462933 | 0.38700964 |
| TDRD7 | 1030.85525 | -0.503438287 | 0.32823846 | -1.5337578 | 0.12508925 | 0.27747338 |
| TMOD1 | 1593.09219 | -0.470124376 | 0.23628736 | -1.9896298 | 0.04663173 | 0.13842219 |
| AL162385.1 | 0.16223974 | -1.003574173 | 3.30558948 | -0.3035992 | 0.7614333 | NA |
| TSTD2 | 1614.18736 | -0.138746043 | 0.1420451 | -0.9767746 | 0.32868075 | 0.52650416 |
| NCBP1 | 1477.81179 | -0.005249509 | 0.1040055 | -0.0504734 | 0.95974516 | 0.98056877 |
| AL162385.2 | 5.91966651 | 0.379996796 | 0.64114214 | 0.59268728 | 0.55339045 | 0.72299723 |
| XPA | 790.204109 | 0.097119561 | 0.15657623 | 0.62027013 | 0.53507996 | 0.70879884 |
| AL445531.1 | 1.05710552 | -1.046917712 | 1.66381095 | -0.6292288 | 0.5291993 | 0.70376648 |
| PTCSC2 | 0.08695548 | -0.642905668 | 3.30558948 | -0.1944905 | 0.84579185 | NA |
| FOXE1 | 5.59698386 | 0.062150807 | 0.9661735 | 0.06432676 | 0.94871005 | 0.97522644 |
| TRMO | 600.074396 | 0.036987666 | 0.2691404 | 0.13742889 | 0.8906918 | 0.94323693 |
| HEMGN | 15.134801 | -3.226976549 | 1.25927268 | -2.5625717 | 0.01039001 | 0.04639875 |
| ANP32B | 4406.80679 | -0.580043075 | 0.14189967 | -4.0876987 | 4.36E-05 | 0.00066486 |
| AL354726.1 | 0.65111999 | 2.087383911 | 1.74201074 | 1.19826121 | 0.23081534 | NA |
| NANS | 1934.31226 | -0.730559031 | 0.17163693 | -4.256421 | 2.08E-05 | 0.00036542 |
| TRIM14 | 2723.45386 | -0.334590777 | 0.36784346 | -0.909601 | 0.36303298 | 0.56022821 |
| CORO2A | 247.516139 | -1.167370569 | 0.47460176 | -2.4596844 | 0.01390592 | 0.05759372 |
| TBC1D2 | 756.244283 | -0.557875046 | 0.13635706 | -4.0912811 | 4.29E-05 | 0.00065619 |
| GABBR2 | 1.67524594 | 1.43633723 | 0.96626121 | 1.4864896 | 0.13714967 | 0.29564593 |
| ANKS6 | 1951.65238 | 0.132470933 | 0.12930662 | 1.02447142 | 0.30561266 | 0.5026825 |
| GALNT12 | 474.43681 | -0.252856709 | 0.29962349 | -0.8439148 | 0.398717 | 0.59358697 |
| AL136084.2 | 236.057643 | -0.347986659 | 0.3106029 | -1.1203587 | 0.26256095 | 0.45616489 |

| | | | | | | |
|---|---|---|---|---|---|---|
| COL15A1 | 2420.84535 | -0.49718279 | 0.61988476 | -0.8020568 | 0.4225201 | 0.61499448 |
| AL136084.3 | 10.6644367 | 0.388939866 | 0.90385111 | 0.43031409 | 0.66696718 | 0.80504987 |
| TGFBR1 | 2755.16735 | -0.315013678 | 0.38357493 | -0.8212572 | 0.41149978 | 0.60567365 |
| AL162427.1 | 2.1261621 | 1.998135537 | 0.98831409 | 2.02176166 | 0.04320098 | 0.13106002 |
| ALG2 | 2581.22002 | -0.392385823 | 0.19503918 | -2.0118308 | 0.04423778 | 0.13324671 |
| SEC61B | 3552.57957 | -0.429247206 | 0.19812789 | -2.1665159 | 0.03027179 | 0.10144454 |
| AL137067.1 | 17.2215455 | -0.071846292 | 0.50145083 | -0.1432768 | 0.88607154 | 0.94110305 |
| AL162394.1 | 0 | NA | NA | NA | NA | NA |
| AL359710.1 | 11.2181004 | 0.8448761 | 0.5031585 | 1.67914503 | 0.09312378 | 0.22603775 |
| NR4A3 | 825.74869 | 1.242487314 | 0.84401728 | 1.47211122 | 0.14099085 | 0.30131523 |
| STX17-AS1 | 31.2965966 | 0.018376003 | 0.35955007 | 0.05110833 | 0.9592392 | 0.9803557 |
| STX17 | 479.863007 | 0.043388424 | 0.23093166 | 0.18788426 | 0.85096738 | 0.92182598 |
| AL358937.1 | 38.4159351 | -0.052133391 | 0.22726321 | -0.2293965 | 0.81856074 | 0.9026318 |
| ERP44 | 4003.89198 | -0.465198739 | 0.10803324 | -4.3060706 | 1.66E-05 | 0.00030587 |
| INVS | 666.367724 | 0.252946307 | 0.22285564 | 1.13502313 | 0.25636563 | 0.44913378 |
| TEX10 | 1146.89267 | -0.102247619 | 0.15244068 | -0.6707371 | 0.50238802 | 0.68265066 |
| MSANTD3 | 1042.30481 | -1.372880538 | 0.39002515 | -3.5199795 | 0.00043158 | 0.00404578 |
| MSANTD3-TI | 299.604255 | -0.989265699 | 0.3248639 | -3.0451696 | 0.00232549 | 0.01518026 |
| TMEFF1 | 88.3115798 | -0.672035725 | 0.32953398 | -2.0393518 | 0.04141493 | 0.12707838 |
| CAVIN4 | 24.5917409 | -0.818046417 | 0.4024314 | -2.0327599 | 0.04207678 | 0.12855576 |
| PLPPR1 | 723.278205 | 3.069830056 | 0.39850827 | 7.70330327 | 1.33E-14 | 6.24E-12 |
| BAAT | 12.4942431 | 0.879252183 | 0.61483671 | 1.43005804 | 0.15270036 | 0.31810013 |
| MRPL50 | 1461.75357 | -0.239607314 | 0.15544121 | -1.5414658 | 0.12320346 | 0.27463019 |
| ZNF189 | 1390.39294 | 0.165310188 | 0.21444752 | 0.77086547 | 0.44078668 | 0.63212664 |
| ALDOB | 198110.464 | 2.213598226 | 0.59133466 | 3.74339334 | 0.00018155 | 0.00202176 |
| TMEM246-A! | 200.342929 | 3.433045566 | 0.39342297 | 8.72609326 | 2.64E-18 | 3.66E-15 |
| TMEM246 | 976.059761 | -0.134115876 | 0.22159047 | -0.6052421 | 0.54501816 | 0.71682576 |
| RNF20 | 1489.91282 | 0.612484107 | 0.18620847 | 3.28923876 | 0.00100459 | 0.00786783 |
| GRIN3A | 37.9413019 | 1.299475906 | 0.29254362 | 4.44199026 | 8.91E-06 | 0.00018609 |
| PPP3R2 | 1.76349797 | 3.407566158 | 1.28462284 | 2.65258101 | 0.0079879 | 0.03811124 |
| LINC00587 | 0 | NA | NA | NA | NA | NA |
| CYLC2 | 0 | NA | NA | NA | NA | NA |
| LINC01492 | 0 | NA | NA | NA | NA | NA |
| AL390962.1 | 0 | NA | NA | NA | NA | NA |
| AL590381.1 | 0.17466293 | 0.18657164 | 3.29377195 | 0.05664376 | 0.95482897 | NA |
| SMC2-AS1 | 25.9790806 | 0.725908047 | 0.42774768 | 1.69704731 | 0.08968772 | 0.22002652 |
| SMC2 | 513.366856 | 0.46345976 | 0.15146527 | 3.05984173 | 0.00221454 | 0.01460666 |
| OR13F1 | 0 | NA | NA | NA | NA | NA |
| AL450426.1 | 0 | NA | NA | NA | NA | NA |
| OR13C4 | 0.12059279 | 0.02007176 | 3.29738704 | 0.00608717 | 0.99514317 | NA |
| OR13C3 | 0 | NA | NA | NA | NA | NA |
| OR13C8 | 0 | NA | NA | NA | NA | NA |
| OR13C5 | 1.13644407 | 0.022276103 | 1.34890604 | 0.0165142 | 0.98682417 | 0.99400325 |
| OR13C2 | 0.98549692 | 1.477386292 | 1.23728067 | 1.19405914 | 0.23245483 | NA |
| OR13C9 | 0.07659028 | -0.642905668 | 3.30558948 | -0.1944905 | 0.84579185 | NA |
| OR13D1 | 0.33923707 | 0.669854558 | 2.22878938 | 0.30054637 | 0.76376043 | NA |
| NIPSNAP3A | 1412.32992 | 0.259237781 | 0.21992559 | 1.17875222 | 0.23849685 | 0.42801021 |
| NIPSNAP3B | 219.184319 | 0.237150847 | 0.32256304 | 0.73520774 | 0.46221299 | 0.65043793 |
| ABCA1 | 1822.75744 | -0.283754053 | 0.35932828 | -0.7896791 | 0.42971517 | 0.62180135 |
| AL359182.1 | 3.0311849 | 0.954355317 | 0.7665248 | 1.24504167 | 0.21311643 | 0.3970244 |
| AL359182.2 | 0.65890576 | 0.267850895 | 1.55361604 | 0.17240482 | 0.86311928 | NA |
| SLC44A1 | 3461.95096 | -0.199001819 | 0.15400952 | -1.2921397 | 0.19630876 | 0.37684079 |
| FSD1L | 483.025308 | -0.535799914 | 0.19071972 | -2.8093578 | 0.00496405 | 0.02672012 |
| AL158070.1 | 3.32162892 | 1.093662435 | 0.67713832 | 1.61512412 | 0.10628382 | 0.24765435 |
| FKTN | 770.836554 | 0.15737846 | 0.12510657 | 1.2579552 | 0.20840796 | 0.39152061 |
| TAL2 | 17.3590968 | 2.143287902 | 0.37691024 | 5.68646764 | 1.30E-08 | 8.75E-07 |
| TMEM38B | 1860.18261 | 0.110437355 | 0.32011358 | 0.34499428 | 0.73009867 | 0.84751607 |
| LINC01505 | 6.04422018 | 0.698606893 | 0.53962586 | 1.29461343 | 0.19545362 | 0.37560938 |
| AL732323.1 | 0.28699197 | -0.699881757 | 2.96218094 | -0.2362725 | 0.81322126 | NA |
| AL451140.1 | 0 | NA | NA | NA | NA | NA |
| AL512649.2 | 0.09989236 | -0.642905668 | 3.30558948 | -0.1944905 | 0.84579185 | NA |
| ZNF462 | 714.110944 | -0.010066223 | 0.13144315 | -0.0765823 | 0.93895582 | 0.9709527 |
| AL807761.4 | 4.02935209 | -0.33422081 | 0.72475219 | -0.4611518 | 0.64468967 | 0.79005793 |
| AL807761.3 | 1.1670468 | -1.767284259 | 1.43310181 | -1.2331882 | 0.21750555 | 0.40252799 |
| RAD23B | 7160.74502 | -0.255189638 | 0.17675512 | -1.4437468 | 0.14881021 | 0.31286357 |
| LINC01509 | 0.36221177 | 1.698992863 | 2.25240544 | 0.75430153 | 0.45066819 | NA |
| KLF4 | 1242.3983 | -0.072711074 | 0.34102549 | -0.213213 | 0.83116082 | 0.91040231 |
| AL353742.1 | 0 | NA | NA | NA | NA | NA |
| AL358779.1 | 0 | NA | NA | NA | NA | NA |
| ACTL7B | 0.73681295 | -2.027601256 | 1.63778475 | -1.2380145 | 0.21571069 | NA |
| ACTL7A | 0.18746814 | 0.18657164 | 3.29377195 | 0.05664376 | 0.95482897 | NA |
| ELP1 | 2019.30019 | 0.324143411 | 0.15293887 | 2.11943118 | 0.03405404 | 0.11042857 |
| FAM206A | 1059.86989 | -0.360973735 | 0.1278079 | -2.824346 | 0.00473772 | 0.02582875 |
| CTNNAL1 | 3233.20309 | -0.666192261 | 0.2935839 | -2.2691716 | 0.02325789 | 0.08379705 |
| TMEM245 | 5587.30321 | 0.144992068 | 0.24351283 | 0.5954186 | 0.55156369 | 0.72162126 |
| FRRS1L | 8.44726015 | 0.821510137 | 0.89201616 | 0.9209588 | 0.35707194 | 0.5545993 |
| EPB41L4B | 731.510908 | 0.274037307 | 0.18905592 | 1.44950399 | 0.14719689 | 0.31045204 |
| PTPN3 | 5768.96987 | -0.246492648 | 0.1987027 | -1.2405098 | 0.21478689 | 0.39900882 |
| PALM2 | 11.1904167 | -0.02861297 | 0.4040013 | -0.070824 | 0.94353787 | 0.97315146 |
| AL158829.1 | 0.57498079 | 0.220068063 | 1.64861085 | 0.13348697 | 0.89380827 | NA |
| PALM2-AKAF | 5280.50005 | -0.056482999 | 0.24336458 | -0.2320921 | 0.81646647 | 0.90122314 |
| AKAP2 | 5277.04823 | -0.056833263 | 0.24348983 | -0.2334112 | 0.81544209 | 0.90052223 |
| C9orf152 | 25.0105843 | -1.59525562 | 0.50166119 | -3.1799462 | 0.00147302 | 0.01065377 |
| TXN | 11677.6813 | -0.631922118 | 0.4719238 | -1.3390342 | 0.18055953 | 0.35652226 |
| TXNDC8 | 0.89421968 | 1.478486776 | 1.58616933 | 0.93211156 | 0.35127887 | NA |
| SVEP1 | 1581.04952 | -0.308946727 | 0.43680943 | -0.7072804 | 0.47939227 | 0.66418343 |
| AL592463.1 | 0 | NA | NA | NA | NA | NA |
| MUSK | 5.66491141 | -0.735213556 | 0.77111722 | -0.9534394 | 0.34036748 | 0.53838944 |
| LPAR1 | 1085.62851 | -0.947864481 | 0.35301415 | -2.6850608 | 0.00725166 | 0.03554392 |
| AL162414.1 | 1.64895591 | 0.293002885 | 0.96568128 | 0.30341573 | 0.76157306 | 0.86789099 |
| OR2K2 | 1.18716136 | -2.714579131 | 1.66008317 | -1.6352067 | 0.10200572 | 0.24074921 |
| ECPAS | 4842.46474 | -0.317355667 | 0.14886931 | -2.1317736 | 0.03302546 | 0.10806321 |
| ZNF483 | 67.7021089 | -0.764186365 | 0.34788138 | -2.1966866 | 0.02804284 | 0.09620003 |
| PTGR1 | 14047.7341 | -0.386274216 | 0.4990491 | -0.7740205 | 0.43891869 | 0.63042582 |
| AL135787.1 | 9.96341302 | -1.144820333 | 0.86215171 | -1.3278641 | 0.184223 | 0.3612875 |
| DNAJC25 | 1262.3363 | -0.716284374 | 0.34278619 | -2.0895952 | 0.03665418 | 0.11628571 |
| DNAJC25-GN | 1728.17188 | -0.731354241 | 0.25996861 | -2.8132406 | 0.00490449 | 0.02649693 |
| GNG10 | 1648.29573 | -0.752820773 | 0.26503691 | -2.8404375 | 0.00450517 | 0.02488507 |
| C9orf84 | 143.580484 | 0.314507557 | 0.57698991 | 0.54508329 | 0.58569624 | 0.74722314 |
| UGCG | 8756.63099 | -1.65713706 | 0.34071074 | -4.8637653 | 1.15E-06 | 3.65E-05 |
| SUSD1 | 1104.39613 | -0.203377389 | 0.21599411 | -0.9415876 | 0.3464038 | 0.54402169 |
| PTBP3 | 5080.30817 | -0.573831753 | 0.28299577 | -2.0277043 | 0.04259043 | 0.12964444 |
| HSDL2 | 5156.65088 | 0.304214146 | 0.34871783 | 0.87237911 | 0.3830016 | 0.57895142 |
| C9orf147 | 47.4771251 | 0.842346806 | 0.24789082 | 3.39805571 | 0.00067867 | 0.00575078 |
| KIAA1958 | 940.334969 | 0.442026013 | 0.21459218 | 2.05984214 | 0.03941363 | 0.12248434 |
| INIP | 1269.09581 | 0.181919102 | 0.09607012 | 1.89360752 | 0.05827712 | 0.16215762 |
| AL390067.1 | 14.3272112 | 1.547763956 | 0.4894421 | 3.16230243 | 0.00156527 | 0.01116779 |
| SNX30 | 2628.58126 | 0.91259264 | 0.22842028 | 3.99523483 | 6.46E-05 | 0.00090174 |
| SLC46A2 | 31.5207361 | 0.857012486 | 0.37819388 | 2.26606654 | 0.02344731 | 0.08422634 |
| AL139041.1 | 15.0906879 | -0.018039449 | 0.49844799 | -0.0361912 | 0.97112987 | 0.9865502 |
| ZNF883 | 114.128882 | 0.348189703 | 0.36797437 | 0.94623357 | 0.34402947 | 0.54158301 |
| ZFP37 | 157.804497 | 0.545896883 | 0.24257474 | 2.25042758 | 0.02442182 | 0.08674501 |
| FAM225B | 5.25429077 | -1.362326489 | 0.94269686 | -1.4451374 | 0.14841929 | 0.31228092 |

| | | | | | | |
|---|---|---|---|---|---|---|
| FAM225A | 8.08763891 | -1.846118509 | 0.95686785 | -1.9293349 | 0.0536893 | 0.15327346 |
| SLC31A2 | 1617.03124 | -0.183655113 | 0.29029243 | -0.6326555 | 0.52695861 | 0.70174218 |
| FKBP15 | 3644.71419 | -0.137520623 | 0.171909 | -0.7999618 | 0.42373296 | 0.61621563 |
| SLC31A1 | 2793.03238 | 0.329491061 | 0.28759695 | 1.14566951 | 0.25193192 | 0.44419931 |
| CDC26 | 718.632359 | -0.382383932 | 0.14297008 | -2.6745732 | 0.00748244 | 0.03639675 |
| PRPF4 | 1919.60454 | 0.047159136 | 0.23447313 | 0.20112811 | 0.8405984 | 0.91507243 |
| RNF183 | 805.038574 | 0.750815427 | 0.34501464 | 2.17618427 | 0.02954148 | 0.09979617 |
| WDR31 | 264.558908 | 1.216732248 | 0.30531604 | 3.9851566 | 6.74E-05 | 0.00093079 |
| BSPRY | 1771.17049 | -0.734117763 | 0.22253728 | -3.298853 | 0.00097081 | 0.00766014 |
| HDHD3 | 4737.84282 | 0.699445169 | 0.26519334 | 2.63749143 | 0.00835217 | 0.03931575 |
| ALAD | 5958.558 | 2.17368223 | 0.26569143 | 8.18122834 | 2.81E-16 | 2.25E-13 |
| POLE3 | 1571.97941 | -0.377875946 | 0.26596402 | -1.4207785 | 0.15538118 | 0.32161708 |
| C9orf43 | 225.63922 | 0.312169781 | 0.26584834 | 1.17424012 | 0.24029886 | 0.43039333 |
| RGS3 | 5172.90334 | -0.307839527 | 0.22824063 | -1.3487499 | 0.17741731 | 0.35252117 |
| AL162727.2 | 4.03185819 | 0.640430688 | 0.71546627 | 0.89512352 | 0.37072106 | 0.56749134 |
| AL162727.1 | 15.4170853 | -0.323101894 | 0.33880133 | -0.9536618 | 0.34025485 | 0.53830815 |
| AL157702.2 | 3.28974611 | -0.298562122 | 0.76475587 | -0.3904019 | 0.6962394 | 0.82523522 |
| ZNF618 | 1631.91997 | 0.588011739 | 0.2521447 | 2.33204081 | 0.01969854 | 0.074038 |
| AMBP | 992.050972 | -0.652627903 | 0.54334406 | -1.2011319 | 0.22970003 | 0.41719056 |
| KIF12 | 24591.246 | 0.471758294 | 0.33174603 | 1.42204655 | 0.15501274 | 0.32115697 |
| COL27A1 | 8833.23484 | -0.116830666 | 0.33700812 | -0.3466702 | 0.72883911 | 0.84674806 |
| ORM1 | 4.84940708 | -2.224533856 | 1.07333055 | -2.0725524 | 0.03821395 | 0.11972221 |
| ORM2 | 8.79571965 | -1.832186364 | 0.98006148 | -1.8694606 | 0.06155875 | 0.16834018 |
| AKNA | 1633.68688 | -0.499198635 | 0.32148246 | -1.5528021 | 0.12047042 | 0.27059667 |
| AL138895.1 | 5.3039548 | -0.139858293 | 0.57785138 | -0.2420316 | 0.80875568 | 0.89660916 |
| WHRN | 1345.8139 | -0.688090946 | 0.23645956 | -2.9099731 | 0.0036146 | 0.02109716 |
| ATP6V1G1 | 6327.11725 | -0.246052275 | 0.17171281 | -1.432929 | 0.15187812 | 0.31692854 |
| TMEM268 | 834.849334 | -0.496118441 | 0.31653223 | -1.5673552 | 0.11703169 | 0.26535904 |
| TEX53 | 3.14892788 | 0.504708058 | 0.86064385 | 0.5864308 | 0.55758605 | 0.72636855 |
| TEX48 | 0.08695548 | -0.642905668 | 3.30558948 | -0.1944905 | 0.84579185 | NA |
| TNFSF15 | 1492.12907 | -1.695639143 | 0.43863421 | -3.8657248 | 0.00011076 | 0.00137124 |
| TNFSF8 | 50.3972062 | 0.129034473 | 0.50099438 | 0.25755673 | 0.79674903 | 0.88970027 |
| TNC | 8180.71585 | -0.991224924 | 0.57059956 | -1.7371638 | 0.08235826 | 0.20667941 |
| AL162425.1 | 0.06785815 | 0.795868262 | 3.28898349 | 0.24198001 | 0.80879565 | NA |
| 1-Dec | 3.52290693 | 0.142331879 | 0.71594231 | 0.19880356 | 0.84241641 | 0.91630899 |
| AL355601.1 | 0.96661236 | -2.219629992 | 1.97961098 | -1.1212455 | 0.26218336 | NA |
| AL691420.1 | 0.26885207 | -0.701265545 | 3.29738693 | -0.2126731 | 0.83158194 | NA |
| LINC00474 | 0 | NA | NA | NA | NA | NA |
| AL691426.1 | 0 | NA | NA | NA | NA | NA |
| PAPPA | 2203.35051 | -1.139722012 | 0.38247444 | -2.9798645 | 0.00288376 | 0.0177924 |
| PAPPA-AS2 | 13.6792684 | 0.534508797 | 0.65809236 | 0.8122094 | 0.41667149 | 0.60984776 |
| AL137024.1 | 12.7256553 | -0.190356481 | 0.5958195 | -0.3194868 | 0.74935738 | 0.85998627 |
| PAPPA-AS1 | 736.714788 | -1.140186804 | 0.41973569 | -2.71644 | 0.00659882 | 0.03305048 |
| ASTN2 | 581.956912 | 0.213350741 | 0.25872696 | 0.82461736 | 0.40958886 | 0.60405105 |
| ASTN2-AS1 | 0.51028404 | 0.998653935 | 2.12720774 | 0.46946705 | 0.63873583 | NA |
| AL133284.1 | 9.38692373 | 0.026606045 | 0.60291352 | 0.04412912 | 0.96480148 | 0.98354657 |
| TRIM32 | 592.236537 | -0.058338303 | 0.21091761 | -0.2765929 | 0.78209275 | 0.8807709 |
| AL445644.1 | 0 | NA | NA | NA | NA | NA |
| AL161630.1 | 19.084393 | 0.230585905 | 0.82289356 | 0.28021353 | 0.77931369 | 0.87909064 |
| AL160272.2 | 28.6448302 | 0.151167752 | 0.29876289 | 0.50597901 | 0.61287138 | 0.76804764 |
| TLR4 | 1349.04043 | -0.537497152 | 0.29956191 | -1.7942774 | 0.07276891 | 0.18969885 |
| AL160272.1 | 41.1979365 | -0.502620858 | 0.31597973 | -1.5906744 | 0.11168288 | 0.25674113 |
| AL354754.1 | 2.00929543 | 0.839333779 | 0.99580972 | 0.84286563 | 0.3993036 | 0.59408914 |
| AL355592.1 | 0.12150485 | 1.04981648 | 3.28477129 | 0.31960109 | 0.74927075 | NA |
| LINC02578 | 0 | NA | NA | NA | NA | NA |
| BRINP1 | 147.69277 | 2.326647074 | 0.43653196 | 5.32984362 | 9.83E-08 | 4.73E-06 |
| AC006288.1 | 0 | NA | NA | NA | NA | NA |
| LINC01613 | 0.06785815 | 0.795868262 | 3.28898349 | 0.24198001 | 0.80879565 | NA |
| AL441989.1 | 2.76545498 | 0.92034073 | 1.31571195 | 0.69950017 | 0.48423951 | 0.668712 |
| CDK5RAP2 | 1347.24626 | 0.159308721 | 0.11943028 | 1.33390565 | 0.18223478 | 0.35888956 |
| MEGF9 | 1805.4731 | 0.069771659 | 0.20941993 | 0.33316628 | 0.73900877 | 0.85348667 |
| FBXW2 | 3469.10379 | 0.050296971 | 0.10805071 | 0.46549414 | 0.64157763 | 0.78767178 |
| B3GNT10 | 143.35817 | 0.7017877 | 0.17284087 | 4.06031108 | 4.90E-05 | 0.00073261 |
| PSMD5 | 1503.25949 | 0.018052496 | 0.18155255 | 0.099434 | 0.92079369 | 0.96133224 |
| PHF19 | 649.392383 | -0.067967675 | 0.29205099 | -0.2327254 | 0.81597466 | 0.90094427 |
| TRAF1 | 1035.76569 | 0.173779861 | 0.32914997 | 0.5279656 | 0.59752321 | 0.75607904 |
| C5 | 416.719224 | 1.344639314 | 0.38041521 | 3.53466234 | 0.0004083 | 0.00388272 |
| CNTRL | 550.66611 | 0.147445151 | 0.18703269 | 0.78833893 | 0.43049848 | 0.62251837 |
| RAB14 | 6129.61217 | 0.04673578 | 0.10530656 | 0.44380691 | 0.65718219 | 0.79883488 |
| GSN | 19804.4128 | -0.166536469 | 0.33282807 | -0.5003679 | 0.61681609 | 0.77088415 |
| AL513122.2 | 19.5845138 | 0.253651464 | 0.42163989 | 0.60158317 | 0.54745163 | 0.71842513 |
| GSN-AS1 | 629.364032 | -0.378408375 | 0.29179714 | -1.29682 | 0.19469312 | 0.37469402 |
| STOM | 16522.4792 | -0.638772764 | 0.22454741 | -2.8447122 | 0.00444516 | 0.02467244 |
| AL359644.1 | 21.8405181 | 0.362471588 | 0.42129894 | 0.86036671 | 0.38958693 | 0.58521261 |
| DAB2IP | 8330.77281 | -0.319347024 | 0.15246115 | -2.0946124 | 0.03620546 | 0.11526758 |
| AL357936.1 | 0.86981206 | -0.119645588 | 1.88599972 | -0.0634388 | 0.94941708 | NA |
| TTLL11 | 180.55337 | -0.168104603 | 0.24690828 | -0.6808383 | 0.49597384 | 0.67780007 |
| TTLL11-IT1 | 5.13356333 | -1.465714773 | 0.66749511 | -2.1958435 | 0.02810315 | 0.0963899 |
| NDUFA8 | 2225.10263 | 0.197635641 | 0.15480422 | 1.27668125 | 0.2017148 | 0.38358833 |
| MORN5 | 29.0686797 | -0.046635525 | 0.46250794 | -0.1008318 | 0.91968395 | 0.96074349 |
| LHX6 | 130.025443 | -0.237613231 | 0.48890651 | -0.4860095 | 0.6269604 | 0.77804072 |
| RBM18 | 1439.57984 | 0.023077309 | 0.12340642 | 0.18700251 | 0.85165867 | 0.92224978 |
| MRRF | 2672.6797 | -0.158421771 | 0.11115849 | -1.4251883 | 0.15410279 | 0.31995044 |
| PTGS1 | 598.938162 | 0.801282927 | 0.58540772 | 1.36876043 | 0.17107417 | 0.34339849 |
| AL359636.3 | 0 | NA | NA | NA | NA | NA |
| AL359636.1 | 0 | NA | NA | NA | NA | NA |
| AL359636.2 | 0.41038523 | 0.309875973 | 2.21210731 | 0.1400818 | 0.88859536 | NA |
| OR1J1 | 0 | NA | NA | NA | NA | NA |
| OR1J2 | 0.07659028 | -0.642905668 | 3.30558948 | -0.1944905 | 0.84579185 | NA |
| OR1J4 | 0.07830766 | -0.642905668 | 3.30558948 | -0.1944905 | 0.84579185 | NA |
| OR1N1 | 1.09049787 | -2.40570391 | 1.87212168 | -1.2850147 | 0.19878714 | 0.37982877 |
| OR1N2 | 0 | NA | NA | NA | NA | NA |
| OR1L8 | 11.6280276 | 0.443316963 | 0.5015647 | 0.88386795 | 0.3767675 | 0.57264317 |
| AL162254.1 | 1.74275156 | -2.174893502 | 1.38185543 | -1.5738937 | 0.11551204 | 0.26299882 |
| OR1Q1 | 0.52694218 | -1.186973624 | 2.79800618 | -0.4242212 | 0.67140448 | NA |
| OR1B1 | 0 | NA | NA | NA | NA | NA |
| OR1L1 | 0 | NA | NA | NA | NA | NA |
| OR1L3 | 0 | NA | NA | NA | NA | NA |
| OR1L4 | 0.07659028 | -0.642905668 | 3.30558948 | -0.1944905 | 0.84579185 | NA |
| OR1L6 | 0.34741699 | -0.279112419 | 2.89448939 | -0.0964289 | 0.92317994 | NA |
| OR5C1 | 0 | NA | NA | NA | NA | NA |
| PDCL | 689.404802 | -0.065473768 | 0.17671896 | -0.3704966 | 0.71101253 | 0.83587002 |
| OR1K1 | 0.17294555 | 0.18657164 | 3.29377195 | 0.05664376 | 0.95482897 | NA |
| RC3H2 | 2072.36593 | -0.275940679 | 0.11330846 | -2.4353052 | 0.01487924 | 0.06051287 |
| ZBTB6 | 396.058424 | 0.105603403 | 0.16152097 | 0.65380615 | 0.5132367 | 0.69122454 |
| ZBTB26 | 498.160457 | 0.16050988 | 0.19378904 | 0.8282712 | 0.40751693 | 0.60216415 |
| RABGAP1 | 2411.01396 | 0.137882858 | 0.10128524 | 1.36133223 | 0.17340872 | 0.34713338 |
| GPR21 | 6.91513506 | 1.560012648 | 0.72456271 | 2.15304021 | 0.03131552 | 0.10412183 |
| STRBP | 1357.9734 | 0.032875494 | 0.10211958 | 0.32193135 | 0.7475047 | 0.85882272 |
| CRB2 | 1193.98654 | 0.730113078 | 0.30181816 | 2.41904954 | 0.01556112 | 0.06247667 |
| DENND1A | 2712.09286 | 0.608988931 | 0.15420195 | 3.94929455 | 7.84E-05 | 0.00104602 |
| AL445489.1 | 0.82438868 | 0.438232891 | 1.51294333 | 0.28965585 | 0.77207953 | NA |
| AL390774.2 | 0.61273061 | -0.340481159 | 1.9445318 | -0.1750967 | 0.86100363 | NA |

| | | | | | | |
|---|---|---|---|---|---|---|
| LHX2 | 0.67225073 | 0.237588756 | 1.42704509 | 0.16649001 | 0.86777134 | NA |
| AC006450.3 | 0 | NA | NA | NA | NA | NA |
| AC006450.2 | 0.42627686 | -0.869589478 | 2.23745239 | -0.3886516 | 0.69753388 | NA |
| NEK6 | 2132.32542 | -1.056511686 | 0.40488151 | -2.6094343 | 0.00906921 | 0.04191191 |
| AL162724.2 | 10.7133492 | -0.530106699 | 0.56933383 | -0.9311 | 0.35180186 | 0.54942346 |
| PSMB7 | 5185.97627 | -0.294960035 | 0.2484394 | -1.1872514 | 0.23512846 | 0.42393108 |
| ADGRD2 | 1.99650238 | 1.39880966 | 1.22583328 | 1.14110922 | 0.25382448 | 0.44646154 |
| NR5A1 | 1.24302305 | -2.662625542 | 1.41762033 | -1.878236 | 0.06034889 | 0.16622259 |
| AL354979.1 | 1.68024923 | -2.088612684 | 1.63429669 | -1.2779887 | 0.20125342 | 0.38301476 |
| NR6A1 | 503.765467 | -0.290458998 | 0.24021508 | -1.2091622 | 0.22660053 | 0.41341192 |
| MIR181A2HC | 14.9898404 | -0.481931517 | 0.63930948 | -0.7538313 | 0.45095051 | 0.64079343 |
| AL354928.1 | 6.60905272 | 0.445941984 | 0.67743345 | 0.65828162 | 0.51035719 | 0.68891634 |
| OLFML2A | 514.191952 | 0.893197435 | 0.43388119 | 2.05862214 | 0.03953045 | 0.12272404 |
| WDR38 | 532.285485 | -0.898555436 | 0.14456978 | -6.2153753 | 5.12E-10 | 5.79E-08 |
| RPL35 | 11888.4568 | -0.722739319 | 0.13725357 | -5.2657233 | 1.40E-07 | 6.35E-06 |
| ARPC5L | 2533.86761 | -0.977450271 | 0.22771518 | -4.2924248 | 1.77E-05 | 0.0003223 |
| GOLGA1 | 1264.26576 | 0.342898148 | 0.17398936 | 1.97079948 | 0.04874682 | 0.1431144 |
| SCAI | 430.604813 | 0.517608926 | 0.22033507 | 2.34918994 | 0.01881431 | 0.07177186 |
| PPP6C | 3422.73664 | -0.000471857 | 0.12705022 | -0.0037139 | 0.99703671 | 0.99850237 |
| RABEPK | 814.494377 | -0.078959122 | 0.19142302 | -0.412485 | 0.67998399 | 0.8144612 |
| HSPA5 | 27235.803 | -0.585234326 | 0.30322233 | -1.9300502 | 0.05360062 | 0.15303685 |
| AL354710.2 | 2.81857249 | 1.59321283 | 0.89833215 | 1.77352311 | 0.07614207 | 0.19585118 |
| GAPVD1 | 2020.29746 | -0.130761442 | 0.13293198 | -0.9836718 | 0.3252769 | 0.52283017 |
| MAPKAP1 | 4976.81031 | 0.089354781 | 0.30472407 | 0.29323178 | 0.769345 | 0.87240608 |
| AL162584.1 | 1.46153131 | 1.04851372 | 1.03334113 | 1.01468304 | 0.31025694 | 0.50726204 |
| PBX3 | 620.237848 | -0.366455492 | 0.15972625 | -2.2942722 | 0.02177487 | 0.07991695 |
| AL589923.1 | 0 | NA | NA | NA | NA | NA |
| AL162391.1 | 0 | NA | NA | NA | NA | NA |
| MVB12B | 640.81437 | 0.161828266 | 0.17964937 | 0.90080065 | 0.36769432 | 0.5645144 |
| AL356309.2 | 0.18301898 | 0.18657164 | 3.29377195 | 0.05664376 | 0.95482897 | NA |
| AL356309.1 | 0 | NA | NA | NA | NA | NA |
| AL161908.1 | 7.49115827 | 0.432787938 | 0.79202775 | 0.54643027 | 0.58477021 | 0.74662548 |
| LMX1B | 338.364208 | 0.971147678 | 0.72345833 | 1.3423685 | 0.17947654 | 0.35515507 |
| ZBTB43 | 1131.39034 | -0.724573512 | 0.3316617 | -2.1846765 | 0.02891257 | 0.09826326 |
| ZBTB34 | 816.78042 | -0.518400311 | 0.22532681 | -2.3006597 | 0.02141087 | 0.07884531 |
| RALGPS1 | 1373.15983 | 0.428554222 | 0.14191321 | 3.01983318 | 0.00252914 | 0.0161368 |
| ANGPTL2 | 1364.95243 | -0.470018625 | 0.57043015 | -0.8239723 | 0.40995532 | 0.60441248 |
| AL450263.1 | 132.113521 | 0.429471211 | 0.17836434 | 2.40783119 | 0.0160476 | 0.06399282 |
| GARNL3 | 348.604204 | 0.606908968 | 0.19949019 | 3.04229981 | 0.00234778 | 0.01529334 |
| SLC2A8 | 1265.67958 | 0.015454454 | 0.16240194 | 0.09516176 | 0.92418634 | 0.96312021 |
| ZNF79 | 205.463291 | 0.059293783 | 0.2483879 | 0.23871446 | 0.811327 | 0.89817258 |
| RPL12 | 23774.337 | -0.568872642 | 0.15167709 | -3.7505509 | 0.00017645 | 0.00198418 |
| LRSAM1 | 2047.89583 | -0.403884232 | 0.16467789 | -2.4525711 | 0.01418394 | 0.05846965 |
| FAM129B | 12622.296 | -1.396492769 | 0.28026243 | -4.9828041 | 6.27E-07 | 2.20E-05 |
| STXBP1 | 2225.5827 | -0.500524994 | 0.23084968 | -2.1681858 | 0.03014455 | 0.10117243 |
| AL162426.1 | 191.867424 | -0.230512731 | 0.24843918 | -0.9278437 | 0.35348865 | 0.55107867 |
| PTRH1 | 859.0071 | -0.361408757 | 0.19577966 | -1.8459974 | 0.06489258 | 0.17457838 |
| CFAP157 | 842.782127 | -0.39738404 | 0.23025958 | -1.7258089 | 0.0843818 | 0.21041589 |
| TTC16 | 93.3633893 | 0.186252008 | 0.28010301 | 0.66494111 | 0.50608816 | 0.68537104 |
| TOR2A | 853.521459 | -0.385841789 | 0.14846848 | -2.5988129 | 0.00935467 | 0.04284919 |
| SH2D3C | 1382.2879 | -0.104921776 | 0.31910356 | -0.3288016 | 0.74230562 | 0.85534626 |
| CDK9 | 4179.96325 | -0.325297864 | 0.11899446 | -2.7337228 | 0.00626227 | 0.03181814 |
| FPGS | 3430.56652 | -0.618868824 | 0.16428191 | -3.7671149 | 0.00016515 | 0.00188034 |
| ENG | 19751.0341 | -0.205572777 | 0.46357028 | -0.4434555 | 0.65743632 | 0.79902876 |
| AL162586.1 | 4947.34123 | -0.120720495 | 0.42953663 | -0.2810482 | 0.77867344 | 0.87865683 |
| AK1 | 2278.20378 | -0.692432285 | 0.2353615 | -2.9419947 | 0.00326106 | 0.01956891 |
| AL157935.2 | 7086.03316 | -0.623791004 | 0.18843675 | -3.3103468 | 0.0009318 | 0.00743695 |
| ST6GALNAC6 | 5203.10002 | -0.621822369 | 0.18536702 | -3.3545469 | 0.00079495 | 0.00654067 |
| ST6GALNAC4 | 1975.54687 | -1.489833832 | 0.42666706 | -3.4917949 | 0.00047979 | 0.00439152 |
| PIP5KL1 | 124.374748 | -1.646307924 | 0.31842944 | -5.170087 | 2.34E-07 | 9.72E-06 |
| AL157935.1 | 886.611547 | -2.008783865 | 0.73557418 | -2.7309059 | 0.00631605 | 0.03200855 |
| DPM2 | 2399.31933 | -0.615922462 | 0.14345637 | -4.2934481 | 1.76E-05 | 0.00032104 |
| FAM102A | 9453.99954 | -0.691047499 | 0.44858256 | -1.5405135 | 0.12343523 | 0.27497254 |
| AL360268.2 | 0.17332742 | -1.003574173 | 3.30558948 | -0.3035992 | 0.7614333 | NA |
| NAIF1 | 470.52673 | -0.184496545 | 0.17308463 | -1.0659326 | 0.28645411 | 0.48260893 |
| SLC25A25 | 4495.03149 | -0.201370014 | 0.36202354 | -0.5562346 | 0.5780505 | 0.74165288 |
| AL360268.1 | 1.02943844 | 0.780722332 | 1.2937965 | 0.60343519 | 0.54621922 | 0.71739419 |
| SLC25A25-AS | 2543.39126 | -0.195524603 | 0.19814756 | -0.9867626 | 0.32375904 | 0.5215653 |
| PTGES2 | 4098.00842 | -0.353840329 | 0.13014868 | -2.7187393 | 0.00655312 | 0.03289034 |
| AL590708.1 | 63.6356791 | -0.500390188 | 0.39301504 | -1.2732088 | 0.20294397 | 0.38495415 |
| LCN2 | 7543.33108 | -7.19494262 | 0.62864795 | -11.445106 | 2.49E-30 | 3.28E-26 |
| C9orf16 | 2006.30734 | -0.754711944 | 0.20352201 | -3.7082572 | 0.00020869 | 0.00225725 |
| CIZ1 | 4229.62676 | -0.045620144 | 0.10489954 | -0.4348936 | 0.66363963 | 0.80246755 |
| DNM1 | 2327.7345 | -0.056246074 | 0.29562008 | -0.1902647 | 0.84910169 | 0.92060599 |
| GOLGA2 | 4771.3241 | -0.152927504 | 0.22279971 | -0.68639 | 0.49246718 | 0.67535283 |
| SWI5 | 807.849974 | 0.03863399 | 0.17418314 | 0.22180097 | 0.82446882 | 0.90611165 |
| TRUB2 | 2758.14632 | 0.476450226 | 0.1349643 | 3.53019443 | 0.00041525 | 0.00392905 |
| AL359091.4 | 39.6101631 | -0.217695951 | 0.28893172 | -0.7534512 | 0.45117882 | 0.6409301 |
| COQ4 | 4892.11824 | 0.254415278 | 0.14900306 | 1.70744999 | 0.08773845 | 0.21645269 |
| SLC27A4 | 3546.8429 | -0.350270335 | 0.24344926 | -1.4387817 | 0.15021239 | 0.31467276 |
| URM1 | 2828.55057 | -0.304879001 | 0.11523034 | -2.6458223 | 0.00814926 | 0.03862953 |
| AL359091.3 | 99.4236561 | -0.770666974 | 0.18472964 | -4.1718641 | 3.02E-05 | 0.00049117 |
| AL359091.1 | 0.61813478 | -1.209717537 | 1.89623066 | -0.6379591 | 0.52350033 | NA |
| CERCAM | 1727.95038 | -0.33872824 | 0.38037295 | -0.8905161 | 0.37318882 | 0.56940284 |
| ODF2 | 1760.86953 | -0.309766579 | 0.16526621 | -1.8743492 | 0.06088232 | 0.16718387 |
| ODF2-AS1 | 98.7500825 | 0.609829349 | 0.28723939 | 2.12307003 | 0.03374798 | 0.1097465 |
| GLE1 | 1717.30991 | 0.230606857 | 0.14408811 | 1.60045725 | 0.10949718 | 0.25319543 |
| AL356481.1 | 270.257553 | 0.140163316 | 0.16309785 | 0.85938176 | 0.39012993 | 0.58572798 |
| SPTAN1 | 11231.8561 | 0.230490062 | 0.0844749 | 2.72850336 | 0.00636224 | 0.03217112 |
| AL356481.3 | 1220.49208 | 0.04075358 | 0.11584491 | 0.35179432 | 0.72499252 | 0.84406414 |
| WDR34 | 3560.95443 | 0.187183508 | 0.22809324 | 0.82064469 | 0.41184869 | 0.60584904 |
| HMGA1P4 | 18.0725562 | -0.01466531 | 0.47369022 | -0.0309597 | 0.97530167 | 0.98862634 |
| SET | 10726.7131 | -0.41405076 | 0.17609827 | -2.3512483 | 0.01871054 | 0.07153254 |
| PKN3 | 977.36721 | -0.750149526 | 0.28433462 | -2.638263 | 0.00833319 | 0.03926144 |
| ZDHHC12 | 1385.17909 | -0.322027047 | 0.15743734 | -2.0454299 | 0.04081249 | 0.12577134 |
| AL441992.1 | 109.80059 | 0.25747163 | 0.29325057 | 0.87799193 | 0.3799481 | 0.57591719 |
| ZER1 | 4614.34524 | 0.358082447 | 0.14568185 | 2.45797578 | 0.01397226 | 0.05782305 |
| TBC1D13 | 4317.15869 | 1.156338852 | 0.22254846 | 5.19589685 | 2.04E-07 | 8.68E-06 |
| ENDOG | 1749.37891 | -0.106430772 | 0.16162028 | -0.6585236 | 0.51020173 | 0.68885463 |
| SPOUT1 | 3262.85161 | -0.114616651 | 0.13964511 | -0.820771 | 0.41177674 | 0.60580988 |
| KYAT1 | 775.531687 | 0.832075479 | 0.27601623 | 3.01458892 | 0.00257328 | 0.01635116 |
| LRRC8A | 3705.95232 | -0.336141496 | 0.29787769 | -1.1284548 | 0.25912791 | 0.45187688 |
| PHYHD1 | 2797.77444 | 0.565126365 | 0.23925068 | 2.36206794 | 0.01817331 | 0.06991979 |
| AL672142.1 | 789.584209 | 0.51516827 | 0.23712742 | 2.17253777 | 0.02981512 | 0.10046318 |
| DOLK | 938.308344 | 0.209801088 | 0.19347422 | 1.08438785 | 0.27819287 | 0.47379257 |
| NUP188 | 2151.66644 | -0.177786177 | 0.12901531 | -1.3780239 | 0.16819592 | 0.33950753 |
| AL592211.1 | 44.4668598 | -0.455371374 | 0.30443513 | -1.4957912 | 0.13470809 | 0.29215003 |
| SH3GLB2 | 9687.22876 | -0.238480349 | 0.23850691 | -0.9998886 | 0.3173644 | 0.51456586 |
| MIGA2 | 3044.45773 | -0.023217825 | 0.15453606 | -0.1502421 | 0.8805736 | 0.93801519 |
| DOLPP1 | 753.774546 | 0.170008028 | 0.19437882 | 0.8746222 | 0.38177951 | 0.57773015 |
| CRAT | 5894.15365 | 0.945758065 | 0.16030958 | 5.89957287 | 3.64E-09 | 2.93E-07 |
| AL158151.2 | 537.547916 | 1.036254954 | 0.16498513 | 6.28089905 | 3.37E-10 | 4.03E-08 |
| PTPA | 9099.61672 | -0.18016822 | 0.11514888 | -1.5646546 | 0.11766391 | 0.26626598 |

| | | | | | | |
|---|---|---|---|---|---|---|
| AL158151.4 | 17.9610246 | -2.809566216 | 0.5996862 | -4.6850606 | 2.80E-06 | 7.44E-05 |
| IER5L | 2088.90905 | -1.550102865 | 0.37317928 | -4.1537753 | 3.27E-05 | 0.00052585 |
| AL158151.3 | 276.783961 | -1.209758026 | 0.35046774 | -3.4518385 | 0.00055678 | 0.00494188 |
| AL158151.1 | 1046.51948 | -1.62461606 | 0.37751716 | -4.3034231 | 1.68E-05 | 0.00030905 |
| AL161785.2 | 2.21321479 | 0.766227914 | 0.8710188 | 0.87969159 | 0.3790264 | 0.57504694 |
| AL161785.3 | 1.2644851 | -1.611558469 | 1.61379483 | -0.9986142 | 0.31798161 | 0.51515482 |
| AL161785.1 | 91.1768561 | 0.029982169 | 0.3660177 | 0.08191453 | 0.93471468 | 0.96870331 |
| C9orf106 | 30.5558684 | 0.20292421 | 0.58712643 | 0.34562268 | 0.7296263 | 0.84732662 |
| AL353803.5 | 5.78089376 | -0.038662458 | 0.76195413 | -0.0507412 | 0.95953176 | 0.98050555 |
| LINC01503 | 2449.78539 | -2.250465265 | 0.42740634 | -5.2653998 | 1.40E-07 | 6.35E-06 |
| AL353803.2 | 44.307361 | -0.236574188 | 0.48030717 | -0.4925477 | 0.62233221 | 0.77470996 |
| AL353803.1 | 67.8969616 | 0.353816079 | 0.52798326 | 0.67012746 | 0.50277654 | 0.68306118 |
| AL353803.4 | 1.93751912 | -0.988142511 | 0.91293472 | -1.0823802 | 0.2790836 | 0.47471035 |
| AL391056.1 | 27.2145127 | -1.362775519 | 0.33569722 | -4.0595377 | 4.92E-05 | 0.00073379 |
| NTMT1 | 2220.88809 | -0.577926626 | 0.14241071 | -4.0581682 | 4.95E-05 | 0.00073769 |
| C9orf50 | 26.5376894 | -0.264737767 | 0.38327721 | -0.6907214 | 0.48974065 | 0.67311398 |
| ASB6 | 3447.71956 | -0.347858649 | 0.13949939 | -2.4936212 | 0.01264474 | 0.05363387 |
| AL590369.1 | 172.779254 | 0.117540978 | 0.20578416 | 0.57118573 | 0.56787375 | 0.73380854 |
| PRRX2 | 52.8394897 | -0.663964614 | 0.60930554 | -1.0897072 | 0.27584216 | 0.4712028 |
| PRRX2-AS1 | 0.29934637 | -0.281107011 | 2.99073357 | -0.0939927 | 0.92511499 | NA |
| PTGES | 679.230288 | -1.39098466 | 0.47923474 | -2.9025121 | 0.00370183 | 0.02149685 |
| TOR1B | 1283.26567 | -0.4319444 | 0.19369628 | -2.2300088 | 0.02574686 | 0.09023721 |
| TOR1A | 1933.28985 | -0.292929047 | 0.11576379 | -2.5304031 | 0.01139316 | 0.04968373 |
| C9orf78 | 3027.41373 | 0.250922479 | 0.15258348 | 1.64449308 | 0.10007439 | 0.23760371 |
| USP20 | 2942.70547 | 0.283769175 | 0.09873255 | 2.87411992 | 0.00405155 | 0.0229861 |
| FNBP1 | 2597.66602 | -0.128376644 | 0.18373743 | -0.6986962 | 0.48474191 | 0.66910826 |
| AL158207.2 | 0.03392908 | 0.380740593 | 3.29738709 | 0.11546736 | 0.90807469 | NA |
| AL136141.1 | 3.26950953 | 0.215370359 | 0.77672009 | 0.27728182 | 0.78156372 | 0.88036187 |
| GPR107 | 4449.55411 | 0.112835974 | 0.11903658 | 0.94791004 | 0.34317525 | 0.54081942 |
| AL360004.1 | 3.21105539 | -0.560791598 | 0.92525495 | -0.6060941 | 0.54445225 | 0.71639168 |
| NCS1 | 4092.06279 | -0.560061229 | 0.32359117 | -1.730768 | 0.08349315 | 0.20879227 |
| HMCN2 | 519.498371 | 2.311619453 | 0.6367873 | 3.63012806 | 0.00028328 | 0.00289806 |
| ASS1 | 39007.5705 | 2.334935872 | 0.42080751 | 5.54870291 | 2.88E-08 | 1.69E-06 |
| FUBP3 | 2847.35014 | 0.170667044 | 0.20563111 | 0.82996702 | 0.40655743 | 0.60140134 |
| PRDM12 | 3.9716031 | -1.97418916 | 0.9085038 | -2.1730115 | 0.02977945 | 0.1003943 |
| EXOSC2 | 1068.0255 | -0.013123363 | 0.16366099 | -0.0801863 | 0.93608912 | 0.96942144 |
| ABL1 | 5127.82825 | -0.205871219 | 0.15878744 | -1.2965208 | 0.19479611 | 0.37480173 |
| QRFP | 27.3348709 | -0.1553646 | 0.34217868 | -0.4540452 | 0.64979627 | 0.79399136 |
| FIBCD1 | 7.67571987 | -1.149230917 | 0.72932773 | -1.57574 | 0.11508572 | 0.26234531 |
| AL161733.1 | 0 | NA | NA | NA | NA | NA |
| LAMC3 | 2624.46531 | 0.341513117 | 0.51173447 | 0.66736391 | 0.50453971 | 0.68396433 |
| AIF1L | 27896.8796 | 0.453156157 | 0.28835975 | 1.57149586 | 0.11606751 | 0.26397902 |
| NUP214 | 3810.0678 | 0.139560572 | 0.16010753 | 0.87166778 | 0.38338964 | 0.5792028 |
| AL157938.2 | 45.6781343 | 1.697091114 | 0.37300872 | 4.54973583 | 5.37E-06 | 0.00012414 |
| AL157938.3 | 74.0145177 | 1.286834783 | 0.24131421 | 5.33261085 | 9.68E-08 | 4.67E-06 |
| FAM78A | 294.713709 | -0.405676982 | 0.37473827 | -1.0825609 | 0.27900338 | 0.47466576 |
| PLPP7 | 279.460533 | 0.046874316 | 0.23306457 | 0.20112158 | 0.84060351 | 0.91507243 |
| PRRC2B | 10388.5201 | -0.125936362 | 0.10931304 | -1.1520708 | 0.24929196 | 0.44113737 |
| AL358781.1 | 6428.95718 | -0.465404767 | 0.12883842 | -3.6123135 | 0.00030348 | 0.00306265 |
| POMT1 | 2742.32634 | 0.240408268 | 0.06943877 | 3.46216186 | 0.00053585 | 0.00478513 |
| AL358781.2 | 344.874888 | 0.161317238 | 0.1034102 | 1.55997415 | 0.11876599 | 0.26797223 |
| UCK1 | 2684.92036 | 0.107723836 | 0.15532181 | 0.69355253 | 0.48796287 | 0.67152554 |
| PRRT1B | 38.1927157 | 0.182744657 | 0.49774189 | 0.36714743 | 0.71350905 | 0.83731181 |
| RAPGEF1 | 5379.37649 | -0.070183336 | 0.2664716 | -0.2633802 | 0.79225757 | 0.8873257 |
| AL160271.1 | 0 | NA | NA | NA | NA | NA |
| MED27 | 739.679274 | -0.477506647 | 0.16049467 | -2.9752181 | 0.0029278 | 0.01800097 |
| NTNG2 | 83.8348126 | -1.114878984 | 0.39194449 | -2.8444818 | 0.00444837 | 0.0246851 |
| SETX | 3349.36221 | -0.309821082 | 0.17858118 | -1.7349033 | 0.08275794 | 0.20750469 |
| TTF1 | 537.655098 | -0.212558383 | 0.12185735 | -1.7443214 | 0.08110306 | 0.20438528 |
| CFAP77 | 13.9453315 | -1.655712086 | 0.84116621 | -1.9683531 | 0.04902743 | 0.14379416 |
| BARHL1 | 0.44582964 | -0.627625667 | 2.28357788 | -0.2748431 | 0.78343677 | NA |
| DDX31 | 1038.17192 | -0.571618413 | 0.22005509 | -2.5976151 | 0.00938737 | 0.04296907 |
| GTF3C4 | 1213.35662 | -0.157337336 | 0.20110243 | -0.7823741 | 0.43399475 | 0.62592658 |
| AK8 | 245.50301 | 0.483738663 | 0.31197675 | 1.55055996 | 0.12100717 | 0.27143239 |
| SPACA9 | 711.321864 | 0.514481956 | 0.26132011 | 1.96878058 | 0.04897829 | 0.14368201 |
| TSC1 | 2931.85323 | 0.076110073 | 0.13486904 | 0.56432576 | 0.57253245 | 0.73719144 |
| GFI1B | 19.2091452 | -0.715020122 | 0.48891653 | -1.4624585 | 0.14361561 | 0.30519331 |
| GTF3C5 | 3389.71902 | -0.515443609 | 0.24326012 | -2.1188989 | 0.03409901 | 0.11050249 |
| CEL | 562.638192 | 1.417107062 | 0.51747748 | 2.73849029 | 0.0061722 | 0.03149398 |
| RALGDS | 3401.48202 | -0.405942239 | 0.219702 | -1.8476948 | 0.0646465 | 0.17409398 |
| AL162417.1 | 3003.10412 | -0.359786209 | 0.21928869 | -1.6406966 | 0.10086042 | 0.23892285 |
| GBGT1 | 643.020364 | -0.463821571 | 0.32585732 | -1.4233885 | 0.15462355 | 0.32065297 |
| OBP2B | 4.93197809 | -1.914602014 | 0.80776867 | -2.3702356 | 0.01777676 | 0.0687149 |
| ABO | 2680.117 | 0.092034239 | 0.35708273 | 0.25773927 | 0.79660814 | 0.88965766 |
| SURF6 | 2219.35692 | -0.216799852 | 0.16185118 | -1.3395012 | 0.18040756 | 0.35632892 |
| MED22 | 2468.90552 | -0.267438129 | 0.18843366 | -1.4192694 | 0.15582049 | 0.32222289 |
| RPL7A | 41457.3562 | -0.369703847 | 0.19554414 | -1.8906414 | 0.05867223 | 0.16287786 |
| SURF1 | 2820.02356 | 0.239298622 | 0.13469856 | 1.77654918 | 0.07564244 | 0.19490827 |
| SURF2 | 734.008887 | -0.160377498 | 0.19169162 | -0.8366433 | 0.40279312 | 0.59789781 |
| SURF4 | 12514.6825 | -0.281221395 | 0.22630439 | -1.2426687 | 0.21398993 | 0.39803304 |
| STKLD1 | 113.558277 | 0.586000347 | 0.23046293 | 2.54270977 | 0.01099966 | 0.04836767 |
| REXO4 | 1369.14874 | -0.20794649 | 0.16228428 | -1.2813717 | 0.20006313 | 0.38143769 |
| ADAMTS13 | 667.32788 | 0.575866744 | 0.36305042 | 1.58618946 | 0.11269634 | 0.25832775 |
| CACFD1 | 4032.01381 | -0.175169862 | 0.23545134 | -0.7439748 | 0.45689172 | 0.64628054 |
| SLC2A6 | 462.50079 | -0.925352316 | 0.37827959 | -2.4462126 | 0.01443658 | 0.05922322 |
| MYMK | 8.8188285 | 3.190893229 | 0.60343269 | 5.28790251 | 1.24E-07 | 5.71E-06 |
| ADAMTSL2 | 3453.38475 | 0.826324461 | 0.51300105 | 1.61076564 | 0.10723081 | 0.24928824 |
| FAM163B | 30.091929 | -0.002555496 | 0.74712128 | -0.0034205 | 0.99727088 | 0.9986244 |
| AC002101.1 | 0 | NA | NA | NA | NA | NA |
| DBH | 324.165762 | 2.862728973 | 0.37134517 | 7.70907823 | 1.27E-14 | 6.08E-12 |
| DBH-AS1 | 777.871424 | 1.992731702 | 0.34677818 | 5.74641605 | 9.12E-09 | 6.41E-07 |
| SARDH | 1266.26702 | 2.868397098 | 0.48367235 | 5.93045494 | 3.02E-09 | 2.51E-07 |
| VAV2 | 5043.57032 | -0.259890548 | 0.17557756 | -1.4802036 | 0.13881891 | 0.29817176 |
| BRD3OS | 2164.52477 | -0.183656643 | 0.15566531 | -1.1798174 | 0.23807283 | 0.42748063 |
| BRD3 | 1873.66077 | -0.017400409 | 0.16442122 | -0.1058282 | 0.91571863 | 0.958881 |
| AL445931.1 | 3.62981965 | 2.036840243 | 0.72104731 | 2.82483578 | 0.00473049 | 0.02579998 |
| BX649632.1 | 41.0893879 | 0.782421718 | 0.30121675 | 2.59753724 | 0.00938949 | 0.04297136 |
| WDR5 | 2540.25025 | -0.243353825 | 0.13309111 | -1.8284755 | 0.06747822 | 0.17936893 |
| BX649601.1 | 16.3557089 | 0.96032394 | 0.43623823 | 2.20137501 | 0.02770949 | 0.09536663 |
| LINC02247 | 0.09989236 | -0.642905668 | 3.30558948 | -0.1944905 | 0.84579185 | NA |
| RXRA | 5752.52901 | 0.476276983 | 0.26316045 | 1.80983495 | 0.07032139 | 0.18484599 |
| AL669970.1 | 0.91432736 | -0.801101141 | 1.87055847 | -0.4282684 | 0.6684557 | NA |
| AL669970.3 | 0.36665097 | 0.048303582 | 2.30045981 | 0.02099736 | 0.98324776 | NA |
| AL669970.2 | 0 | NA | NA | NA | NA | NA |
| COL5A1 | 2014.2987 | -0.034444208 | 0.40882279 | -0.0842522 | 0.93285594 | 0.96760457 |
| COL5A1-AS1 | 3.15786506 | -0.444398028 | 0.97799088 | -0.4543989 | 0.64954171 | 0.79382745 |
| FCN2 | 164.853188 | -2.30240042 | 0.52305988 | -4.4017913 | 1.07E-05 | 0.00021645 |
| FCN1 | 390.987317 | 0.773731126 | 0.92490832 | 0.83654899 | 0.40284613 | 0.59794283 |
| AL353611.1 | 0.50494281 | 1.685182043 | 1.7198993 | 0.97981437 | 0.32717776 | NA |
| OLFM1 | 534.175831 | -0.556091531 | 0.5080287 | -1.0946065 | 0.27368906 | 0.4687394 |
| AL390778.1 | 0 | NA | NA | NA | NA | NA |
| AL390778.2 | 0 | NA | NA | NA | NA | NA |
| C9orf62 | 0.16223974 | -1.003574173 | 3.30558948 | -0.3035992 | 0.7614333 | NA |

| | | | | | | |
|---|---|---|---|---|---|---|
| AL353615.1 | 0 | NA | NA | NA | NA | NA |
| PPP1R26-AS: | 273.351922 | 0.754931694 | 0.32300281 | 2.33722953 | 0.01942725 | 0.0733319 |
| PPP1R26 | 2133.34774 | 0.258881199 | 0.16088204 | 1.6091367 | 0.10758645 | 0.24989472 |
| C9orf116 | 278.877917 | -0.253340313 | 0.16098083 | -1.5737297 | 0.11554996 | 0.26303975 |
| MRPS2 | 3067.0993 | 0.032173024 | 0.12320038 | 0.26114387 | 0.79398156 | 0.88844572 |
| AL161452.1 | 1530.86207 | 0.064987979 | 0.13132355 | 0.49486919 | 0.62069246 | 0.77342166 |
| LCN1 | 0.18393104 | 1.469068379 | 3.27783996 | 0.44818185 | 0.65402196 | NA |
| OBP2A | 0.67406757 | 0.920666716 | 1.52329295 | 0.60439242 | 0.54558277 | NA |
| PAEP | 0.53332363 | -2.305007193 | 2.20936514 | -1.0432894 | 0.29681429 | NA |
| LINC01502 | 1.01077419 | -0.539575385 | 2.79390164 | -0.1931261 | 0.8468602 | 0.91956777 |
| AL354761.1 | 0.40986573 | -0.266347271 | 2.89257372 | -0.0920797 | 0.92663473 | NA |
| GLT6D1 | 0 | NA | NA | NA | NA | NA |
| LCN9 | 0.34782192 | -1.749510989 | 3.30120699 | -0.529961 | 0.59613896 | NA |
| SOHLH1 | 0.34968658 | -0.557385158 | 2.27685874 | -0.2448045 | 0.80660785 | NA |
| KCNT1 | 34.661906 | -0.074607544 | 0.4715517 | -0.1582171 | 0.87428573 | 0.93458499 |
| CAMSAP1 | 1219.53184 | -0.424295878 | 0.25829315 | -1.6426912 | 0.10044685 | 0.2383071 |
| AL355574.1 | 32.1070417 | 0.105460262 | 0.28939763 | 0.36441301 | 0.71554962 | 0.8381399 |
| UBAC1 | 2234.32349 | -0.123361371 | 0.13423365 | -0.9190048 | 0.35809306 | 0.55575975 |
| NACC2 | 2707.7101 | -0.050863599 | 0.23093797 | -0.2202479 | 0.82567811 | 0.90691152 |
| AL138781.1 | 30.6179065 | -0.20726028 | 0.33580736 | -0.6171999 | 0.53710287 | 0.71080009 |
| TMEM250 | 2247.57203 | -0.564403956 | 0.23579272 | -2.3936445 | 0.01668191 | 0.0657899 |
| AL138781.2 | 14.7863689 | 0.548287365 | 0.37196312 | 1.47403691 | 0.14047165 | 0.300627 |
| LHX3 | 0.77816196 | -0.820306506 | 2.30551861 | -0.3558013 | 0.72198938 | NA |
| QSOX2 | 1207.47742 | -0.252287541 | 0.12944889 | -1.9489356 | 0.05130312 | 0.14842163 |
| CCDC187 | 75.169677 | 0.387348274 | 0.46634303 | 0.83060805 | 0.40619509 | 0.60108667 |
| CR392000.2 | 1.948014 | 1.576351881 | 0.90074456 | 1.75005428 | 0.08010895 | 0.20261727 |
| AC174065.1 | 8.11785897 | 0.128803944 | 0.66297828 | 0.19428079 | 0.84595603 | 0.91885083 |
| GPSM1 | 1354.52202 | -0.521033806 | 0.27525323 | -1.8929253 | 0.05836781 | 0.16231148 |
| DNLZ | 435.622269 | -0.3966546 | 0.27548757 | -1.4398276 | 0.14991619 | 0.31425199 |
| CARD9 | 650.446366 | -0.569124356 | 0.2909174 | -1.9563091 | 0.05042875 | 0.14655126 |
| SNAPC4 | 2039.78586 | -0.548573954 | 0.27963948 | -1.9617185 | 0.04979526 | 0.14524035 |
| ENTR1 | 2153.16078 | -0.409572715 | 0.22785289 | -1.7975314 | 0.0722513 | 0.18869632 |
| PMPCA | 3395.80488 | 0.111640204 | 0.17181621 | 0.64976526 | 0.51584386 | 0.69342133 |
| INPP5E | 2109.31624 | -0.247519771 | 0.15003966 | -1.6496957 | 0.0990052 | 0.23594822 |
| SEC16A | 4905.23921 | -0.208961204 | 0.17058508 | -1.2249677 | 0.22058741 | 0.40651824 |
| C9orf163 | 59.5060043 | 0.243407693 | 0.31312001 | 0.77736231 | 0.43694504 | 0.62901194 |
| NOTCH1 | 4599.71164 | 0.213977507 | 0.15355697 | 1.39347312 | 0.16347677 | 0.33275819 |
| AL592301.1 | 6.46515423 | -0.409581436 | 0.60806511 | -0.6735815 | 0.50057739 | 0.6815873 |
| NALT1 | 151.364023 | 0.504145705 | 0.47815664 | 1.05435261 | 0.29172151 | 0.48792134 |
| AL590226.2 | 47.3949308 | 1.739302525 | 0.59046471 | 2.94565027 | 0.00322277 | 0.01941434 |
| AL590226.1 | 72.2524925 | -0.491745051 | 0.38343587 | -1.2824701 | 0.19967777 | 0.38094677 |
| EGFL7 | 6981.7859 | -0.831878656 | 0.26545083 | -3.1338333 | 0.00172539 | 0.01204301 |
| AGPAT2 | 3979.71362 | -0.405929756 | 0.21147992 | -1.9194719 | 0.05492464 | 0.1555755 |
| FAM69B | 1124.39712 | 0.671279751 | 0.31292209 | 2.14519769 | 0.03193704 | 0.10548776 |
| SNHG7 | 2642.31803 | 0.062911302 | 0.2010361 | 0.31293535 | 0.75432978 | 0.86329626 |
| LCN10 | 174.110473 | -0.039398786 | 0.81501553 | -0.0483411 | 0.96144436 | 0.98158225 |
| AL355987.1 | 208.159412 | -0.061697244 | 0.80215586 | -0.0769143 | 0.93869173 | 0.97076919 |
| LCN6 | 74.6274757 | 0.122014923 | 0.90264283 | 0.1351752 | 0.89247336 | 0.94440299 |
| LCN8 | 22.1787561 | 2.691045883 | 0.9517995 | 2.82732432 | 0.00469388 | 0.02562682 |
| LCN15 | 1.39056039 | 3.116361877 | 1.32993336 | 2.34324663 | 0.01911674 | 0.07254403 |
| TMEM141 | 3447.37029 | 0.429485388 | 0.26503345 | 1.62049504 | 0.10512598 | 0.24589073 |
| AL355987.3 | 2510.2797 | 0.485364109 | 0.26401547 | 1.8383927 | 0.06600456 | 0.17677638 |
| CCDC183 | 728.477813 | 1.17005514 | 0.24400089 | 4.79529048 | 1.62E-06 | 4.83E-05 |
| CCDC183-AS: | 659.818251 | 1.655765939 | 0.30510166 | 5.42693184 | 5.73E-08 | 2.96E-06 |
| RABL6 | 6416.44563 | 0.100957561 | 0.0995447 | 1.01419322 | 0.31049056 | 0.50743407 |
| AJM1 | 779.126751 | -0.146869291 | 0.36581809 | -0.4014818 | 0.68806546 | 0.81986158 |
| PHPT1 | 4614.56419 | -0.002485055 | 0.19006552 | -0.0130747 | 0.98956817 | 0.99519138 |
| MAMDC4 | 2140.98922 | -0.008888259 | 0.34908308 | -0.0254617 | 0.97968667 | 0.99084638 |
| EDF1 | 9574.48401 | -0.200632885 | 0.11605255 | -1.7288106 | 0.083843 | 0.20940897 |
| TRAF2 | 1055.83234 | -0.354837777 | 0.15212661 | -2.3325162 | 0.01967355 | 0.07398625 |
| FBXW5 | 7575.00651 | -0.20439259 | 0.1364965 | -1.49742 | 0.134284 | 0.291494 |
| C8G | 163.680951 | -0.92200359 | 0.55409258 | -1.6639883 | 0.09611476 | 0.23121203 |
| LCN12 | 1983.94441 | 0.339452918 | 0.51902582 | 0.65401933 | 0.51309936 | 0.6911744 |
| PRR31 | 1.41826518 | 2.19686587 | 1.13935041 | 1.92817403 | 0.05383348 | 0.15348567 |
| PTGDS | 6635.69942 | -0.183169665 | 0.45686847 | -0.4009243 | 0.68847587 | 0.82016212 |
| AL807752.7 | 6349.59106 | -0.178618172 | 0.46144694 | -0.3870828 | 0.69869492 | 0.82710473 |
| LCNL1 | 197.622947 | 0.30415993 | 0.67483323 | 0.45071866 | 0.65219233 | 0.7957024 |
| PAXX | 1729.43978 | -0.523787629 | 0.17801278 | -2.9424158 | 0.00325662 | 0.01955308 |
| CLIC3 | 25.950378 | 0.697588051 | 0.5733321 | 1.21672596 | 0.2237085 | 0.40978002 |
| ABCA2 | 4743.88909 | 0.856564543 | 0.29465906 | 2.90696831 | 0.0036495 | 0.02127982 |
| C9orf139 | 90.6523166 | -0.135944895 | 0.40629177 | -0.3345992 | 0.73792746 | 0.85283548 |
| FUT7 | 26.0506465 | 0.216156227 | 0.54734839 | 0.39491525 | 0.69290544 | 0.8232818 |
| AL807752.6 | 1176.93995 | -0.877990854 | 0.38193224 | -2.2988131 | 0.02151555 | 0.0791525 |
| NPDC1 | 4200.97197 | -0.871083249 | 0.39136103 | -2.2257792 | 0.02602897 | 0.09100949 |
| ENTPD2 | 814.404312 | 0.413817883 | 0.41905778 | 0.98749601 | 0.32339954 | 0.52121048 |
| AL807752.4 | 8.39192603 | 0.875140664 | 0.84768176 | 1.03239294 | 0.30188808 | 0.49870254 |
| AL807752.2 | 20.426663 | 0.792835382 | 0.46078682 | 1.72061212 | 0.08532123 | 0.21189889 |
| SAPCD2 | 135.084792 | 0.756967517 | 0.52782947 | 1.43411377 | 0.1515398 | 0.31657328 |
| AL807752.3 | 43.1516109 | 0.685394075 | 0.62481571 | 1.09695397 | 0.27266151 | 0.46759554 |
| UAP1L1 | 3371.8485 | -0.132130812 | 0.32150472 | -0.4109763 | 0.68108994 | 0.81526744 |
| MAN1B1-DT | 141.922949 | 0.579965382 | 0.2921192 | 1.98537234 | 0.04710306 | 0.13939038 |
| MAN1B1 | 4494.35412 | -0.182172986 | 0.15311811 | -1.1897547 | 0.23414284 | 0.42272673 |
| DPP7 | 7696.66347 | -0.485230836 | 0.08926817 | -5.4356533 | 5.46E-08 | 2.83E-06 |
| GRIN1 | 8.93241923 | -0.721831972 | 0.81265522 | -0.8882389 | 0.37441227 | 0.57055999 |
| LRRC26 | 1.36449709 | 1.145965787 | 2.17512579 | 0.52685035 | 0.59829751 | 0.75662313 |
| AL929554.1 | 1.12338226 | 1.065514346 | 1.80761576 | 0.58945843 | 0.55555379 | 0.72490182 |
| TMEM210 | 0.31739398 | 0.652794139 | 2.8538039 | 0.22874527 | 0.81906691 | NA |
| ANAPC2 | 3513.4001 | 0.155001623 | 0.11359317 | 1.36453297 | 0.17239988 | 0.34545478 |
| SSNA1 | 2361.57714 | -0.332631563 | 0.19335776 | -1.7202907 | 0.08537961 | 0.21196199 |
| TPRN | 3619.86397 | -0.593588741 | 0.27043199 | -2.1949649 | 0.02816611 | 0.09656012 |
| TMEM203 | 1811.27648 | 0.090807935 | 0.09489059 | 0.95697515 | 0.3385798 | 0.5366066 |
| NDOR1 | 1596.16867 | -0.236992973 | 0.16614332 | -1.426437 | 0.15374224 | 0.31942832 |
| BX255925.3 | 1458.36727 | -0.64835103 | 0.14701813 | -4.4100075 | 1.03E-05 | 0.00020951 |
| RNF208 | 452.770795 | -0.124703295 | 0.25507464 | -0.4888894 | 0.62491998 | 0.77654869 |
| CYSRT1 | 234.136681 | 1.897576732 | 0.41018129 | 4.62619038 | 3.72E-06 | 9.30E-05 |
| RNF224 | 213.219152 | 2.710467386 | 0.52104086 | 5.20202465 | 1.97E-07 | 8.45E-06 |
| SLC34A3 | 1584.45495 | 2.949931511 | 0.59787666 | 4.93401354 | 8.06E-07 | 2.70E-05 |
| TUBB4B | 13635.8666 | -0.445972495 | 0.27048932 | -1.6487619 | 0.09919642 | 0.23621181 |
| FAM166A | 1344.83733 | -0.664627342 | 0.27926133 | -2.3799476 | 0.0173151 | 0.06753427 |
| STPG3-AS1 | 768.565843 | 1.755461835 | 0.642394 | 2.73268716 | 0.006282 | 0.03188596 |
| STPG3 | 722.869875 | 1.87250684 | 0.64816689 | 2.88892702 | 0.00386559 | 0.02220909 |
| NELFB | 4109.33909 | -0.341779098 | 0.12651208 | -2.7015531 | 0.00690165 | 0.0342245 |
| TOR4A | 1355.76021 | -0.942701748 | 0.25392004 | -3.712593 | 0.00020515 | 0.00222554 |
| NRARP | 271.178599 | 0.264017464 | 0.38610872 | 0.68379047 | 0.49410749 | 0.67670324 |
| EXD3 | 2388.00445 | -0.489439845 | 0.26372312 | -1.8558853 | 0.06346991 | 0.17190855 |
| NOXA1 | 2037.71808 | -0.204236574 | 0.2388213 | -0.8551858 | 0.39244831 | 0.58813719 |
| ENTPD8 | 387.257892 | 0.976698476 | 0.29764787 | 3.28138911 | 0.00103297 | 0.00805186 |
| NSMF | 6267.02056 | 0.172096781 | 0.19938964 | 0.86311796 | 0.38807262 | 0.58373595 |
| PNPLA7 | 2016.19958 | 0.178639568 | 0.35935453 | 0.49711233 | 0.61910984 | 0.77236923 |
| MRPL41 | 2510.11726 | -0.205414569 | 0.13009289 | -1.5789838 | 0.11433977 | 0.26111889 |
| DPH7 | 1799.72377 | -0.419410949 | 0.1441082 | -2.9103893 | 0.00360979 | 0.02107375 |
| ZMYND19 | 1102.7605 | -0.807927446 | 0.26133625 | -3.0915247 | 0.00199131 | 0.01345055 |
| ARRDC1 | 3880.65314 | -0.681358217 | 0.12018937 | -5.6690387 | 1.44E-08 | 9.42E-07 |

| | | | | | | |
|---|---|---|---|---|---|---|
| ARRDC1-AS1 | 1068.62297 | -0.263073385 | 0.1621091 | -1.6228169 | 0.10462855 | 0.24507504 |
| EHMT1 | 2557.7208 | -0.18551665 | 0.13301576 | -1.3946968 | 0.1631073 | 0.33226261 |
| AL772363.1 | 0.35804263 | -0.557773459 | 2.26665319 | -0.246078 | 0.80562186 | NA |
| CACNA1B | 4.16411678 | 0.10980436 | 0.68183865 | 0.16104156 | 0.87206067 | 0.93327291 |
| AL954642.1 | 0.61279331 | -0.280226725 | 2.23889786 | -0.1251628 | 0.90039467 | NA |
| AC069287.1 | 0 | NA | NA | NA | NA | NA |
| LINC01001 | 73.1778759 | 0.737338605 | 0.40057512 | 1.84069996 | 0.06566554 | 0.17610092 |
| AC069287.3 | 0.07830766 | -0.642905668 | 3.30558948 | -0.1944905 | 0.84579185 | NA |
| BET1L | 4396.87417 | 0.080496736 | 0.11439132 | 0.7036962 | 0.48162198 | 0.66609177 |
| SCGB1C1 | 2.02845426 | -1.501137669 | 1.07811952 | -1.3923667 | 0.1638114 | 0.33331069 |
| ODF3 | 0.72212678 | 0.244406453 | 1.65104769 | 0.14803113 | 0.8823182 | NA |
| AC069287.2 | 1682.77537 | 0.13370022 | 0.12079375 | 1.10684715 | 0.26836002 | 0.46255662 |
| RIC8A | 4980.76909 | -0.287156747 | 0.21084573 | -1.3619282 | 0.17322054 | 0.34683567 |
| SIRT3 | 1836.61461 | 0.405141242 | 0.16890954 | 2.39856933 | 0.01645926 | 0.06511347 |
| PSMD13 | 5220.15937 | -0.384688689 | 0.13750368 | -2.797661 | 0.00514741 | 0.0274883 |
| NLRP6 | 245.189067 | 0.619776751 | 0.5533308 | 1.1200836 | 0.26267814 | 0.45625231 |
| AC136475.3 | 585.527191 | 0.237270177 | 0.78839819 | 0.30095221 | 0.76345094 | 0.8690784 |
| PGGHG | 11393.6265 | -0.67467816 | 0.54667196 | -1.2341554 | 0.21714499 | 0.40203001 |
| IFITM5 | 4.56246749 | -1.494441131 | 0.91489075 | -1.633464 | 0.10237144 | 0.24128842 |
| IFITM2 | 24341.4337 | -2.266123813 | 0.46686592 | -4.8539071 | 1.21E-06 | 3.80E-05 |
| AC136475.2 | 149.90836 | -0.998491179 | 0.47703349 | -2.0931259 | 0.03633791 | 0.11558876 |
| IFITM1 | 15690.0932 | -2.744878125 | 0.68610997 | -4.0006388 | 6.32E-05 | 0.00088514 |
| AC136475.1 | 91.3073665 | -0.536685308 | 0.31677031 | -1.6942412 | 0.09021946 | 0.22109924 |
| IFITM3 | 38680.451 | -2.225705909 | 0.40393739 | -5.510027 | 3.59E-08 | 2.04E-06 |
| AC136475.4 | 1.50561125 | 0.115725896 | 1.03339827 | 0.11198577 | 0.91083469 | 0.95585446 |
| AC136475.8 | 2.55149752 | 0.126722878 | 0.79694866 | 0.15901009 | 0.87366093 | 0.93417196 |
| AC136475.7 | 10.899028 | -0.844426106 | 0.48496335 | -1.7412163 | 0.08164567 | 0.20531714 |
| AC136475.5 | 59.5110404 | -1.262621884 | 0.53404378 | -2.3642666 | 0.01806581 | 0.0696179 |
| AC136475.9 | 47.2781825 | -2.30791842 | 0.61769548 | -3.7363369 | 0.00018672 | 0.00206709 |
| B4GALNT4 | 85.8277417 | -1.288068037 | 0.58564097 | -2.1994158 | 0.02784837 | 0.09574465 |
| PKP3 | 305.713451 | -1.030086915 | 0.38451427 | -2.6789302 | 0.00738578 | 0.03603385 |
| SIGIRR | 5544.09738 | 0.370272398 | 0.16773462 | 2.20748936 | 0.02727989 | 0.09425717 |
| ANO9 | 2737.65654 | -0.499924911 | 0.3118882 | -1.6028978 | 0.10895722 | 0.25232299 |
| PTDSS2 | 2959.79357 | -0.195034661 | 0.10907828 | -1.7880248 | 0.07377201 | 0.19156749 |
| AC138230.1 | 48.8033989 | -0.581706431 | 0.30975312 | -1.8779679 | 0.06038555 | 0.16628728 |
| RNH1 | 12193.6273 | -0.179858344 | 0.11728624 | -1.5334991 | 0.12515294 | 0.27749793 |
| AC137894.1 | 14.3037006 | 0.379033141 | 0.40451193 | 0.9370135 | 0.34875161 | 0.5462774 |
| HRAS | 1923.57967 | -0.850321559 | 0.20750097 | -4.097916 | 4.17E-05 | 0.00064288 |
| LRRC56 | 471.749432 | -0.694993599 | 0.1961885 | -3.5424787 | 0.00039639 | 0.00379408 |
| LMNTD2 | 3662.53803 | -0.623122235 | 0.35966652 | -1.7325 | 0.08318458 | 0.20814161 |
| AP006284.1 | 2839.9556 | -0.730878846 | 0.3723806 | -1.96272 | 0.04967872 | 0.14504294 |
| RASSF7 | 6624.52943 | -1.006437172 | 0.26068678 | -3.8607142 | 0.00011306 | 0.00139508 |
| MIR210HG | 1597.67447 | -3.031714985 | 0.45028768 | -6.7328401 | 1.66E-11 | 3.07E-09 |
| PHRF1 | 4013.0176 | -0.244695383 | 0.11485176 | -2.1305324 | 0.03312769 | 0.10833158 |
| IRF7 | 1884.52465 | -1.132644229 | 0.25430988 | -4.4537956 | 8.44E-06 | 0.00017823 |
| CDHR5 | 10768.3206 | 1.588803041 | 0.49554043 | 3.20620263 | 0.00134499 | 0.00994043 |
| SCT | 7.14754813 | 0.186443643 | 0.65857031 | 0.28310363 | 0.7770974 | 0.8778793 |
| DRD4 | 95.7849926 | -1.115895194 | 0.36750887 | -3.0363762 | 0.0023944 | 0.0154756 |
| DEAF1 | 2482.52255 | -0.091871723 | 0.12148638 | -0.7562306 | 0.44951093 | 0.63959456 |
| AC131934.1 | 10.4293557 | 0.665340254 | 0.41689 | 1.59596118 | 0.11049746 | 0.25461535 |
| EPS8L2 | 21249.782 | 0.281447585 | 0.20107905 | 1.39968627 | 0.16160729 | 0.3303041 |
| TMEM80 | 1799.26965 | 0.579961411 | 0.17575393 | 3.29984891 | 0.00096737 | 0.00764032 |
| AP006621.4 | 1642.80983 | 0.340306712 | 0.18727644 | 1.81713575 | 0.06919633 | 0.18261682 |
| TALDO1 | 10689.8593 | -1.055352022 | 0.25396213 | -4.1555488 | 3.25E-05 | 0.00052242 |
| GATD1 | 5174.35747 | -0.009153046 | 0.15439248 | -0.0592843 | 0.95272569 | 0.97717271 |
| AP006621.3 | 598.188473 | 0.187716772 | 0.39004206 | 0.48127316 | 0.63032237 | 0.78028359 |
| AP006621.2 | 130.953522 | 0.98412774 | 0.4247692 | 2.31685287 | 0.02051175 | 0.07633288 |
| CEND1 | 48.6810726 | 0.392457683 | 0.44785344 | 0.87630829 | 0.38086246 | 0.57691402 |
| SLC25A22 | 2376.39329 | -1.09891349 | 0.18154081 | -6.0532587 | 1.42E-09 | 1.37E-07 |
| PANO1 | 74.6031053 | -0.113166943 | 0.19785938 | -0.5719564 | 0.5673515 | 0.73338525 |
| PIDD1 | 2221.55063 | -0.130472654 | 0.18829245 | -0.6929256 | 0.48835625 | 0.67184654 |
| RPLP2 | 21007.0924 | -0.781078769 | 0.1942108 | -4.0218091 | 5.78E-05 | 0.00082859 |
| PNPLA2 | 12047.1899 | -0.767980335 | 0.16023164 | -4.792938 | 1.64E-06 | 4.87E-05 |
| AP006621.1 | 4188.69449 | -0.702339394 | 0.17185854 | -4.0867297 | 4.37E-05 | 0.00066687 |
| CRACR2B | 1387.98756 | -0.401788678 | 0.38915653 | -1.0324603 | 0.30185652 | 0.49870254 |
| CD151 | 17602.6303 | -0.718443318 | 0.20568798 | -3.4928794 | 0.00047784 | 0.00437828 |
| POLR2L | 9000.87495 | -0.398350052 | 0.17566627 | -2.2676525 | 0.0233504 | 0.08396972 |
| TSPAN4 | 3701.15931 | -0.197616279 | 0.17673272 | -1.1181646 | 0.26349668 | 0.4572825 |
| AP006623.1 | 233.457799 | -0.157730836 | 0.15236472 | -1.0352189 | 0.30056669 | 0.49739112 |
| CHID1 | 3943.54572 | 0.047212218 | 0.14757337 | 0.3199237 | 0.74902617 | 0.8597841 |
| AP2A2 | 7838.92966 | 0.115660631 | 0.14929544 | 0.77470972 | 0.4385112 | 0.63025154 |
| MUC6 | 371.690797 | 0.333521019 | 0.78499307 | 0.42487129 | 0.67093051 | 0.80780558 |
| AC139749.1 | 98.0242526 | 1.555952012 | 0.40407366 | 3.85066425 | 0.0001178 | 0.00144413 |
| MUC2 | 0.91672958 | 0.096204845 | 1.89374391 | 0.0508014 | 0.95948377 | NA |
| MUC5AC | 1.06294871 | -0.341919694 | 1.25956475 | -0.2714586 | 0.78603833 | 0.88346393 |
| MUC5B | 26.0064266 | 0.124751566 | 0.4664495 | 0.26744924 | 0.78912329 | 0.88545939 |
| AC061979.1 | 0.43005373 | -1.177085761 | 2.86395127 | -0.4110006 | 0.68107208 | NA |
| TOLLIP | 6752.34802 | 0.292705343 | 0.20202902 | 1.44882818 | 0.14738557 | 0.31072574 |
| TOLLIP-AS1 | 94.1386501 | 0.550002628 | 0.32871996 | 1.67316467 | 0.09429488 | 0.22827101 |
| BRSK2 | 213.461373 | -0.17689944 | 0.45152577 | -0.3917815 | 0.69521967 | 0.82450833 |
| MOB2 | 2097.98078 | -0.47155614 | 0.16809976 | -2.805216 | 0.00502829 | 0.02698879 |
| DUSP8 | 1217.25005 | -0.870869141 | 0.3363672 | -2.589043 | 0.00962431 | 0.04379517 |
| KRTAP5-AS1 | 40.0310727 | -0.200193283 | 0.51739992 | -0.3869217 | 0.69881414 | 0.82713968 |
| KRTAP5-1 | 77.6271284 | -1.369465258 | 0.53174206 | -2.5754315 | 0.01001151 | 0.04511339 |
| KRTAP5-2 | 16.3147325 | -0.108523407 | 0.59760295 | -0.1815978 | 0.85589834 | 0.92454691 |
| KRTAP5-3 | 0.30157196 | 0.366047064 | 3.28420129 | 0.11145695 | 0.911254 | NA |
| KRTAP5-4 | 2.60337876 | -1.567151454 | 1.20299206 | -1.3027114 | 0.19267331 | 0.37209686 |
| KRTAP5-5 | 0.46293108 | -2.094014416 | 2.7808484 | -0.7530128 | 0.45144223 | NA |
| AP006285.2 | 0.6983163 | -2.705057914 | 1.58957062 | -1.7017538 | 0.08880152 | NA |
| FAM99A | 10.7422124 | -1.761491785 | 0.99770317 | -1.7655469 | 0.07747189 | 0.19802062 |
| FAM99B | 2.80267443 | -0.475816675 | 1.12594245 | -0.4225941 | 0.67259141 | 0.80897677 |
| AP006285.1 | 2.08884099 | -2.440826397 | 1.35012597 | -1.8078509 | 0.07062972 | 0.18547158 |
| KRTAP5-6 | 0.92604048 | 0.042535498 | 1.86512112 | 0.02280576 | 0.98180522 | NA |
| IFITM10 | 1078.51101 | -0.808960375 | 0.56495739 | -1.4318963 | 0.15217351 | 0.31725782 |
| AC068580.4 | 84498.9692 | -0.852191632 | 0.24012941 | -3.5488848 | 0.00038687 | 0.00372052 |
| CTSD | 136009.342 | -0.987558853 | 0.22590896 | -4.3714904 | 1.23E-05 | 0.00024342 |
| AC068580.3 | 17679.8697 | -0.776043009 | 0.23788096 | -3.2623166 | 0.00110506 | 0.00846857 |
| AC068580.2 | 0 | NA | NA | NA | NA | NA |
| SYT8 | 36.2185905 | -1.100672877 | 0.77681608 | -1.4169028 | 0.15651135 | 0.32324595 |
| TNNI2 | 18.3210772 | 0.76690713 | 0.58944506 | 1.30106633 | 0.19323575 | 0.37275982 |
| LSP1 | 655.990852 | 0.335723602 | 0.65407924 | 0.51327665 | 0.60775781 | 0.76405917 |
| PRR33 | 243.379371 | 0.033958215 | 0.66749054 | 0.05087445 | 0.95942556 | 0.98047016 |
| LINC01150 | 4.5943264 | 0.363991498 | 0.82856617 | 0.43930287 | 0.66044209 | 0.80018573 |
| TNNT3 | 56.8627148 | -0.393505052 | 0.80843803 | -0.4867473 | 0.6264374 | 0.77770135 |
| MRPL23 | 491.183694 | -0.227379206 | 0.50549215 | -0.4498175 | 0.65284205 | 0.79597951 |
| MRPL23-AS1 | 35.271786 | 0.064551374 | 0.52908038 | 0.12200674 | 0.90289368 | 0.95115477 |
| LINC01219 | 2.25470484 | -0.117937193 | 1.26790031 | -0.0930177 | 0.92588949 | 0.96405693 |
| IGF2 | 4535.47972 | -0.940702623 | 0.62683967 | -1.5007069 | 0.13343139 | 0.29014478 |
| INS-IGF2 | 413.513512 | -0.356032214 | 0.65863254 | -0.5405627 | 0.588809 | 0.74929512 |
| IGF2-AS | 23.4039121 | -0.879761444 | 0.85996522 | -1.0230198 | 0.30629848 | 0.5031515 |
| INS | 0 | NA | NA | NA | NA | NA |
| TH | 4.74024402 | -1.554148562 | 0.75050884 | -2.0707931 | 0.03837813 | 0.1201652 |
| ASCL2 | 105.024911 | -1.591091392 | 0.54658757 | -2.9109542 | 0.00360327 | 0.02104456 |

| | | | | | | |
|---|---|---|---|---|---|---|
| C11orf21 | 117.317829 | 0.164401705 | 0.60754274 | 0.27060105 | 0.78669789 | 0.88390384 |
| TSPAN32 | 134.573052 | 0.264188907 | 0.47648671 | 0.55445178 | 0.57926969 | 0.74259293 |
| CD81-AS1 | 1466.66336 | -0.433285108 | 0.19472922 | -2.2250647 | 0.02607689 | 0.09113952 |
| CD81 | 27266.4612 | -0.727926781 | 0.18573943 | -3.9190751 | 8.89E-05 | 0.00115448 |
| TSSC4 | 1944.10644 | -0.600751665 | 0.2313367 | -2.5968714 | 0.00940771 | 0.04302106 |
| TRPM5 | 170.312921 | -0.751102244 | 0.44808863 | -1.6762359 | 0.093692 | 0.22714138 |
| AC124057.1 | 50.8301751 | -1.368751641 | 0.51222887 | -2.6721486 | 0.00753673 | 0.03658754 |
| KCNQ1 | 3778.94572 | -0.288034234 | 0.29236316 | -0.9851933 | 0.32452912 | 0.5220419 |
| KCNQ1OT1 | 1351.78469 | -0.090262566 | 0.25376152 | -0.3556984 | 0.72206645 | 0.84214472 |
| KCNQ1-AS1 | 2.23731872 | 0.446197392 | 0.80610629 | 0.55352178 | 0.57990617 | 0.74302151 |
| KCNQ1DN | 2.09604003 | -0.413017399 | 1.05355509 | -0.3920226 | 0.69504152 | 0.82444536 |
| CDKN1C | 3959.16921 | -0.394813359 | 0.28748174 | -1.3733511 | 0.16964323 | 0.34140975 |
| SLC22A18AS | 1018.60902 | 0.627806052 | 0.42131394 | 1.4901146 | 0.13619411 | 0.29430784 |
| SLC22A18 | 4918.73224 | 0.336220778 | 0.27945681 | 1.20312251 | 0.22892891 | 0.41624874 |
| PHLDA2 | 257.483304 | -1.559528863 | 0.3537933 | -4.4080226 | 1.04E-05 | 0.00021128 |
| NAP1L4 | 6378.78396 | -0.176895726 | 0.11300133 | -1.5654305 | 0.11748199 | 0.26601409 |
| AC131971.1 | 6.62635674 | 0.264938922 | 0.86304947 | 0.30698 | 0.7588586 | 0.8664871 |
| CARS | 2457.27015 | -0.809288074 | 0.2228635 | -3.6313173 | 0.00028198 | 0.00289128 |
| CARS-AS1 | 363.433793 | -0.333079999 | 0.26553127 | -1.2543908 | 0.20969998 | 0.39288017 |
| OSBPL5 | 2258.7967 | -0.449632244 | 0.2843692 | -1.5811566 | 0.11384224 | 0.26014036 |
| AC108448.1 | 0.12088456 | 0.02007176 | 3.29738704 | 0.00608717 | 0.99514317 | NA |
| MRGPRG | 0.08666371 | -0.642905668 | 3.30558948 | -0.1944905 | 0.84579185 | NA |
| MRGPRG-AS | 0.36995912 | -1.765571688 | 2.26742542 | -0.778668 | 0.43617529 | NA |
| AC109309.1 | 2.00152708 | -2.319731743 | 1.21558706 | -1.9083222 | 0.05634959 | 0.15831437 |
| MRGPRE | 3.13234212 | -1.716046104 | 0.99871647 | -1.7182515 | 0.08575075 | 0.21274317 |
| AC123788.1 | 0.83511453 | 0.936629655 | 1.86389018 | 0.50251333 | 0.61530648 | NA |
| ZNF195 | 1199.29225 | -0.046979271 | 0.17908493 | -0.2623296 | 0.79306738 | 0.8878826 |
| AC127526.1 | 2.76857972 | 0.281033196 | 0.76737583 | 0.36622628 | 0.71419623 | 0.83763364 |
| ART5 | 54.6415274 | 0.090784735 | 0.7109427 | 0.12769628 | 0.89838934 | 0.94837978 |
| ART1 | 3.02728315 | 2.615660808 | 0.84704403 | 3.08798682 | 0.00201517 | 0.01357694 |
| CHRNA10 | 105.934622 | -0.500315665 | 0.23229594 | -2.1537856 | 0.03125699 | 0.10397292 |
| NUP98 | 3527.08658 | -0.318624203 | 0.17828766 | -1.7871354 | 0.0739156 | 0.19179651 |
| PGAP2 | 1102.53191 | 0.345054544 | 0.20155374 | 1.71197289 | 0.08690166 | 0.21485064 |
| RHOG | 1905.63678 | -1.065064293 | 0.22262531 | -4.7841114 | 1.72E-06 | 5.03E-05 |
| AC090587.1 | 26.6221854 | 0.452630388 | 0.35480486 | 1.27571642 | 0.20205578 | 0.38389726 |
| STIM1 | 3023.09231 | -0.58983758 | 0.18135829 | -3.2523332 | 0.00114462 | 0.00872862 |
| RRM1 | 1176.82232 | 0.182790963 | 0.20617632 | 0.88657592 | 0.37530726 | 0.57129724 |
| RRM1-AS1 | 177.057002 | 0.012568001 | 0.23548971 | 0.05336964 | 0.95743739 | 0.97953925 |
| AC015689.1 | 3.15184751 | 0.388612294 | 0.81882025 | 0.47460025 | 0.63507193 | 0.78346022 |
| SSU72P5 | 0.20452486 | -0.339987873 | 3.13600186 | -0.1084144 | 0.91366695 | NA |
| SSU72P2 | 0.07659028 | -0.642905668 | 3.30558948 | -0.1944905 | 0.84579185 | NA |
| SSU72P4 | 0 | NA | NA | NA | NA | NA |
| SSU72P3 | 0 | NA | NA | NA | NA | NA |
| SSU72P7 | 0 | NA | NA | NA | NA | NA |
| OR52B4 | 0.07659028 | -0.642905668 | 3.30558948 | -0.1944905 | 0.84579185 | NA |
| TRIM21 | 980.476497 | -0.565385411 | 0.1440162 | -3.9258458 | 8.64E-05 | 0.00113104 |
| OR52K2 | 0 | NA | NA | NA | NA | NA |
| OR52K1 | 1.85236241 | -0.819986763 | 1.80746376 | -0.4536671 | 0.65006848 | 0.79421358 |
| OR52M1 | 0.15318056 | -1.003574173 | 3.30558948 | -0.3035992 | 0.7614333 | NA |
| C11orf40 | 0.03392908 | 0.380740593 | 3.29738709 | 0.11546736 | 0.90807469 | NA |
| OR52I2 | 1.20507999 | -0.096353568 | 1.78672856 | -0.0539274 | 0.95699303 | 0.97944167 |
| OR52I1 | 1.40776432 | -0.99147169 | 1.64573671 | -0.6024485 | 0.5468756 | 0.71796822 |
| TRIM68 | 1113.1444 | -0.048871476 | 0.21303933 | -0.2294012 | 0.81855711 | 0.9026318 |
| OR51D1 | 0 | NA | NA | NA | NA | NA |
| OR51E1 | 47.3330519 | 0.526186787 | 0.42366765 | 1.24198009 | 0.21424391 | 0.39833687 |
| OR51E2 | 47.9385221 | -0.644289444 | 0.50196919 | -1.2835239 | 0.1993086 | 0.38045043 |
| MMP26 | 0.71347005 | 2.266142975 | 1.37126341 | 1.65259494 | 0.09841333 | NA |
| OR51F1 | 0 | NA | NA | NA | NA | NA |
| OR52R1 | 0.09635527 | 0.547239394 | 3.29377202 | 0.16614368 | 0.86804388 | NA |
| OR51F2 | 0 | NA | NA | NA | NA | NA |
| OR51S1 | 0 | NA | NA | NA | NA | NA |
| OR51T1 | 0.03392908 | 0.380740593 | 3.29738709 | 0.11546736 | 0.90807469 | NA |
| OR51A7 | 0 | NA | NA | NA | NA | NA |
| OR51G2 | 0 | NA | NA | NA | NA | NA |
| OR51G1 | 0 | NA | NA | NA | NA | NA |
| OR51A4 | 0.2999192 | 2.164538337 | 3.23179536 | 0.66976343 | 0.50300861 | NA |
| OR51A2 | 0 | NA | NA | NA | NA | NA |
| OR51L1 | 0.03392908 | 0.380740593 | 3.29738709 | 0.11546736 | 0.90807469 | NA |
| OR52J3 | 0 | NA | NA | NA | NA | NA |
| OR52E2 | 0 | NA | NA | NA | NA | NA |
| AC113331.1 | 0 | NA | NA | NA | NA | NA |
| OR52A5 | 0.07659028 | -0.642905668 | 3.30558948 | -0.1944905 | 0.84579185 | NA |
| OR52A1 | 0.09997307 | 0.547239394 | 3.29377202 | 0.16614368 | 0.86804388 | NA |
| OR51V1 | 0 | NA | NA | NA | NA | NA |
| AC104389.1 | 0.08393208 | -0.642905668 | 3.30558948 | -0.1944905 | 0.84579185 | NA |
| HBB | 7415.06566 | -0.621687273 | 0.69661997 | -0.8924339 | 0.37216041 | 0.56863396 |
| HBD | 75.4838314 | -2.036595324 | 0.75655021 | -2.69195 | 0.00710356 | 0.03501141 |
| BGLT3 | 0 | NA | NA | NA | NA | NA |
| HBG1 | 3.31352475 | -1.389529254 | 1.37657662 | -1.0094093 | 0.31277838 | 0.50960555 |
| AC104389.5 | 20.1259615 | -0.776151459 | 0.91135442 | -0.8516461 | NA | NA |
| HBG2 | 18.3693739 | -0.715900183 | 0.92675085 | -0.772484 | 0.43982783 | 0.63112912 |
| AC104389.4 | 329.859906 | -0.110432441 | 0.43904888 | -0.2515265 | 0.80140706 | 0.89237067 |
| HBE1 | 15.6543719 | 0.938273454 | 0.91618135 | 1.02411324 | 0.30578179 | 0.50275652 |
| OR51B4 | 3.89208925 | 3.778122026 | 1.40565648 | 2.68779895 | 0.00719247 | 0.03535236 |
| OR51B5 | 14.2761287 | 2.051055906 | 1.07348789 | 1.91064653 | 0.05605002 | 0.1577753 |
| OR51B2 | 0.38749497 | 2.544415503 | 3.21188376 | 0.79218792 | 0.42825111 | NA |
| OR51B6 | 0.09644756 | -0.642905668 | 3.30558948 | -0.1944905 | 0.84579185 | NA |
| OR51M1 | 0.35486015 | 0.017137201 | 2.88362477 | 0.00594294 | 0.99525825 | NA |
| OR51Q1 | 0.09989236 | -0.642905668 | 3.30558948 | -0.1944905 | 0.84579185 | NA |
| OR51I1 | 1.08396781 | -1.463336982 | 1.66964038 | -0.8764384 | 0.38079174 | 0.57689751 |
| OR51I2 | 0.19747483 | -0.340095337 | 3.16269738 | -0.1075333 | 0.9143659 | NA |
| OR52D1 | 0.88981057 | -0.41220475 | 1.56265667 | -0.2637846 | 0.79194591 | NA |
| UBQLN3 | 0.27664261 | 1.145957498 | 3.27645355 | 0.34975545 | 0.72652223 | NA |
| UBQLNL | 61.2673421 | -0.425381482 | 0.56881365 | -0.7478398 | 0.4545568 | 0.64404672 |
| OR52H1 | 0.09644756 | -0.642905668 | 3.30558948 | -0.1944905 | 0.84579185 | NA |
| OR52B6 | 0.34298276 | 2.387254166 | 2.67736855 | 0.89164197 | 0.37258487 | NA |
| TRIM6 | 1512.04803 | 1.408746756 | 0.62252865 | 2.26294284 | 0.02363921 | 0.08474253 |
| TRIM6-TRIM | 870.711259 | 0.975805582 | 0.46826083 | 2.08389323 | 0.03716988 | 0.11731535 |
| TRIM34 | 449.691639 | -0.03831284 | 0.41150569 | -0.093104 | 0.92582091 | 0.9640363 |
| TRIM5 | 1340.51778 | -0.112292526 | 0.23736217 | -0.4730852 | 0.63615241 | 0.78426083 |
| TRIM22 | 4571.55141 | -1.523084471 | 0.74395225 | -2.0472879 | 0.04062982 | 0.12526385 |
| OR56B1 | 4.86409003 | -0.995595422 | 1.31253564 | -0.7585283 | 0.44813478 | 0.63822176 |
| OR52N4 | 34.4937249 | -1.787858426 | 1.06435617 | -1.6797558 | 0.09300484 | 0.2258322 |
| OR52N5 | 0.34782192 | -1.749510989 | 3.30120699 | -0.529961 | 0.59613896 | NA |
| OR52N1 | 0.80700747 | -2.182368791 | 1.96139127 | -1.1126637 | 0.26585291 | NA |
| OR52N2 | 3.98836632 | -2.586810275 | 1.09930939 | -2.3531231 | 0.01861647 | 0.07125056 |
| OR52E6 | 0 | NA | NA | NA | NA | NA |
| OR52E8 | 0 | NA | NA | NA | NA | NA |
| OR52E4 | 0 | NA | NA | NA | NA | NA |
| OR52E5 | 0 | NA | NA | NA | NA | NA |
| OR56A3 | 0.10178723 | 1.087385526 | 3.28420134 | 0.33109588 | 0.74057207 | NA |
| OR56A5 | 0 | NA | NA | NA | NA | NA |
| OR52L1 | 0 | NA | NA | NA | NA | NA |

| | | | | | | |
|---|---|---|---|---|---|---|
| OR56A4 | 0.07830766 | -0.642905668 | 3.30558948 | -0.1944905 | 0.84579185 | NA |
| OR56A1 | 0.03392908 | 0.380740593 | 3.29738709 | 0.11546736 | 0.90807469 | NA |
| OR56B4 | 2.49524442 | 1.348161675 | 0.91689901 | 1.47034914 | 0.14146722 | 0.30199065 |
| OR52B2 | 0.10178723 | 1.087385526 | 3.28420134 | 0.33109588 | 0.74057207 | NA |
| OR52W1 | 0 | NA | NA | NA | NA | NA |
| AC022762.2 | 162.377749 | 0.396555557 | 0.26980845 | 1.46976702 | 0.14162486 | 0.30227824 |
| C11orf42 | 4.28904114 | 1.881167565 | 0.58850365 | 3.19652661 | 0.00139093 | 0.0101999 |
| FAM160A2 | 2974.34601 | 0.222508097 | 0.15327158 | 1.45172447 | 0.14657823 | 0.30955982 |
| CNGA4 | 21.7786669 | -0.057221686 | 0.68223831 | -0.0838735 | 0.93315704 | 0.96776463 |
| CCKBR | 2.33991002 | 0.327769697 | 0.88158521 | 0.37179582 | 0.71004487 | 0.83521649 |
| CAVIN3 | 856.638274 | -0.781109974 | 0.23963293 | -3.2596103 | 0.00111565 | 0.00853737 |
| AC068733.3 | 4.78546132 | 0.753954206 | 0.69075885 | 1.09148685 | 0.27505871 | 0.47019955 |
| AC068733.1 | 0.51898464 | 0.899037133 | 1.88198268 | 0.47770744 | 0.63285845 | NA |
| SMPD1 | 7613.62632 | 0.123293419 | 0.2430584 | 0.50725841 | 0.61197352 | 0.76758665 |
| APBB1 | 3737.53749 | -0.184652365 | 0.18721042 | -0.9863359 | 0.32396829 | 0.52163909 |
| HPX | 52.2790389 | -0.169157597 | 0.48674455 | -0.3475285 | 0.72819432 | 0.84629817 |
| TRIM3 | 1434.13944 | -0.140154076 | 0.15017437 | -0.9332756 | 0.35067768 | 0.54834976 |
| ARFIP2 | 4164.2 | -0.365636775 | 0.17044563 | -2.1451813 | 0.03193835 | 0.10548776 |
| TIMM10B | 1932.0701 | 0.036336176 | 0.14348373 | 0.25324248 | 0.80008084 | 0.89152759 |
| AC084337.2 | 431.308991 | 0.159446669 | 0.18153426 | 0.87832827 | 0.3797656 | 0.57573985 |
| DNHD1 | 1314.09317 | -0.253824875 | 0.35295132 | -0.7191498 | 0.47204862 | 0.65823712 |
| RRP8 | 1053.59997 | -0.211486172 | 0.12856107 | -1.645025 | 0.09996465 | 0.23748358 |
| AC091564.2 | 1121.1636 | 0.0991998 | 0.20351276 | 0.48743775 | 0.62594815 | 0.77729476 |
| ILK | 8594.08519 | -0.163026807 | 0.14625897 | -1.1146449 | 0.26500264 | 0.45901994 |
| TAF10 | 10315.4231 | -0.238546357 | 0.13252604 | -1.7999961 | 0.07186125 | 0.18791499 |
| TPP1 | 13346.682 | 0.096060158 | 0.21235107 | 0.45236485 | 0.65100617 | 0.79488544 |
| AC091564.6 | 649.470738 | 0.875716757 | 0.29277576 | 2.99108358 | 0.00277989 | 0.01732998 |
| DCHS1 | 1539.41322 | 0.036354121 | 0.28419755 | 0.12791849 | 0.89821348 | 0.9482358 |
| AC091564.4 | 582.434728 | -0.560586327 | 0.30544425 | -1.8353147 | 0.06645907 | 0.1775069 |
| AC091564.5 | 6.89674381 | 1.883691967 | 0.64073864 | 2.9398757 | 0.00328344 | 0.01967825 |
| MRPL17 | 2437.08556 | -0.539551547 | 0.12851753 | -4.198272 | 2.69E-05 | 0.00045117 |
| OR2AG2 | 2.57798058 | 0.370312777 | 0.72081832 | 0.51373941 | 0.6074342 | 0.76378217 |
| OR2AG1 | 0.50832025 | -0.537686628 | 2.23216458 | -0.2408813 | 0.80964714 | NA |
| OR6A2 | 2.2879991 | -1.614399959 | 1.07757957 | -1.4981724 | 0.13408847 | 0.2911894 |
| AC087280.2 | 15.190166 | 0.284277417 | 0.39524997 | 0.7192345 | 0.47199644 | 0.65821986 |
| OR10A5 | 0 | NA | NA | NA | NA | NA |
| OR10A2 | 1.22152573 | 0.424288342 | 1.13999403 | 0.37218471 | 0.70975533 | 0.8350644 |
| OR10A4 | 0 | NA | NA | NA | NA | NA |
| OR2D2 | 0.62427208 | -0.048867747 | 2.22081013 | -0.0220045 | 0.98244439 | NA |
| OR2D3 | 0.09635527 | 0.547239394 | 3.29377202 | 0.16614368 | 0.86804388 | NA |
| ZNF215 | 244.946491 | 0.057051118 | 0.27933311 | 0.20424044 | 0.83816561 | 0.9143814 |
| ZNF214 | 199.003165 | 0.391870249 | 0.29880596 | 1.3114539 | 0.18970446 | 0.3682104 |
| NLRP14 | 12.0561803 | 1.650454419 | 0.54296442 | 3.03971005 | 0.00236806 | 0.01536591 |
| RBMXL2 | 6.57418525 | 0.451776133 | 0.8204284 | 0.55065882 | 0.58186758 | 0.74444947 |
| AC027804.1 | 0 | NA | NA | NA | NA | NA |
| SYT9 | 177.319844 | 0.12478489 | 0.38196011 | 0.32669613 | 0.74389773 | 0.85633247 |
| AC107884.1 | 166.535326 | 0.837412686 | 0.25533603 | 3.27964953 | 0.00103936 | 0.00808017 |
| OLFML1 | 316.203435 | 1.274282798 | 0.40524817 | 3.1444505 | 0.00166399 | 0.01170354 |
| PPFIBP2 | 2469.30785 | 0.480349708 | 0.27545739 | 1.74382582 | 0.08118947 | 0.20452486 |
| AC107884.2 | 2.08355057 | 0.578345406 | 0.88379111 | 0.65439152 | 0.5128596 | 0.69095732 |
| CYB5R2 | 336.085359 | -0.510335034 | 0.28284308 | -1.8043045 | 0.07118359 | 0.18657302 |
| OVCH2 | 167.92196 | 1.324668415 | 0.46770662 | 2.83226358 | 0.00462197 | 0.02539846 |
| AC044810.3 | 25.515101 | 1.788969792 | 0.43826514 | 4.08193494 | 4.47E-05 | 0.00067803 |
| AC104237.1 | 0.59733386 | -0.011625701 | 1.83901983 | -0.0063217 | 0.99495606 | NA |
| OR5P2 | 0 | NA | NA | NA | NA | NA |
| OR5P3 | 0.17820003 | -1.003574173 | 3.30558948 | -0.3035992 | 0.7614333 | NA |
| OR10A6 | 2.3672203 | -1.117708873 | 1.34247056 | -0.8325761 | 0.40508385 | 0.59984621 |
| OR10A3 | 2.90106848 | -1.448080322 | 1.27975147 | -1.1315325 | 0.25783105 | 0.45038453 |
| NLRP10 | 0.61276399 | -0.329318246 | 1.91279887 | -0.1721656 | 0.86330731 | NA |
| EIF3F | 6159.39363 | -0.091160265 | 0.17406262 | -0.5237211 | 0.60047255 | 0.75813757 |
| CASC23 | 0.19978473 | -1.003574173 | 3.30558948 | -0.3035992 | 0.7614333 | NA |
| TUB | 1894.68524 | 1.073843762 | 0.41776899 | 2.57042477 | 0.01015739 | 0.04559155 |
| AC116456.1 | 0.57729161 | 0.247465683 | 1.64997536 | 0.14998144 | 0.88077926 | NA |
| RIC3 | 1160.44995 | 0.161278867 | 0.24665763 | 0.65385721 | 0.51320381 | 0.69122454 |
| AC091013.1 | 1.92204163 | 0.300925673 | 0.95085664 | 0.31647849 | 0.75163935 | 0.86143377 |
| LMO1 | 54.4765729 | 1.219950047 | 0.47408803 | 2.57325638 | 0.01007465 | 0.04530782 |
| STK33 | 764.810602 | 0.16487428 | 0.23112302 | 0.71336156 | 0.47562205 | 0.66108758 |
| TRIM66 | 1116.68267 | 0.839621524 | 0.26154706 | 3.21021209 | 0.00132637 | 0.00983865 |
| AC091053.2 | 0.14114194 | 0.02007176 | 3.29738704 | 0.00608717 | 0.99514317 | NA |
| RPL27A | 22026.2645 | -0.522989556 | 0.16715525 | -3.1287654 | 0.00175542 | 0.01221544 |
| ST5 | 3997.50907 | -1.066238131 | 0.27238272 | -3.9144852 | 9.06E-05 | 0.0011711 |
| AC091053.1 | 1870.76097 | -1.510880554 | 0.28378904 | -5.3239567 | 1.02E-07 | 4.84E-06 |
| AC026894.1 | 0.7394485 | 0.42254247 | 1.43077381 | 0.29532444 | 0.76774605 | NA |
| AKIP1 | 883.022648 | -0.477650577 | 0.16154855 | -2.9566999 | 0.00310951 | 0.01886739 |
| C11orf16 | 37.5988483 | -1.381636397 | 0.37644458 | -3.6702252 | 0.00024234 | 0.00255617 |
| ASCL3 | 5.90569648 | -2.113992309 | 0.87924492 | -2.404327 | 0.01620227 | 0.06437456 |
| TMEM9B | 3382.65722 | -0.071191243 | 0.08239846 | -0.8639875 | 0.38759476 | 0.58321677 |
| TMEM9B-AS | 220.369251 | 0.544883358 | 0.26715864 | 2.03954985 | 0.04139518 | 0.12704736 |
| NRIP3 | 91.949504 | -0.556965677 | 0.5462688 | -1.0195817 | 0.30792689 | 0.50478876 |
| AC079296.1 | 1.6011081 | -0.027614041 | 1.1722484 | -0.0235565 | 0.98120639 | 0.99154675 |
| SCUBE2 | 104.282139 | 0.055239449 | 0.30957031 | 0.1784391 | 0.85837814 | 0.92597389 |
| DENND5A | 3313.30325 | -0.44917325 | 0.18831196 | -2.3852613 | 0.01706699 | 0.06685332 |
| TMEM41B | 2271.62343 | -0.340857845 | 0.17806891 | -1.9141907 | 0.05559579 | 0.15686509 |
| IPO7 | 5464.67544 | -0.167438981 | 0.21048396 | -0.7954952 | 0.42632551 | 0.61867047 |
| AC132192.1 | 3.38379604 | 0.875645996 | 0.76909511 | 1.13854058 | 0.25489483 | 0.44760086 |
| AC132192.2 | 164.193515 | 0.391633337 | 0.26748278 | 1.4641441 | 0.14315458 | 0.3046555 |
| ZNF143 | 928.387319 | -0.344968524 | 0.23318091 | -1.4794029 | 0.13903266 | 0.298485 |
| WEE1 | 1193.40681 | -0.056342711 | 0.22242509 | -0.253311 | 0.80002793 | 0.89152759 |
| SWAP70 | 2991.20041 | -0.525463486 | 0.24321581 | -2.1604824 | 0.03073534 | 0.10265876 |
| AC026250.1 | 16.7831641 | -0.515739929 | 0.41475571 | -1.2434788 | 0.21369145 | 0.39767421 |
| SBF2-AS1 | 1075.76953 | 0.118341739 | 0.16170684 | 0.73182889 | 0.46427301 | 0.65178001 |
| SBF2 | 1940.64749 | 0.178980534 | 0.12548074 | 1.42635865 | 0.15376484 | 0.31942832 |
| AC011092.2 | 3.11796012 | -2.033565041 | 1.03036232 | -1.9736407 | 0.04842261 | 0.14238507 |
| AC100763.1 | 0.03392908 | 0.380740593 | 3.29738709 | 0.11546736 | 0.90807469 | NA |
| AC080023.1 | 20.0174016 | 0.396208563 | 0.43213219 | 0.91686889 | 0.35921135 | 0.55687261 |
| ADM | 9641.77196 | -2.122485881 | 0.58338313 | -3.6382366 | 0.00027451 | 0.00283543 |
| AMPD3 | 1002.1303 | -0.114852521 | 0.15935198 | -0.7207474 | 0.47106497 | 0.65759572 |
| MTRNR2L8 | 177.543658 | -0.147677823 | 0.40586539 | -0.3638591 | 0.71596322 | 0.83832645 |
| RNF141 | 1890.05216 | 0.336541957 | 0.15225978 | 2.21031417 | 0.02708337 | 0.09370091 |
| MRVI1-AS1 | 21.6263409 | 1.252930339 | 0.4197408 | 2.98500964 | 0.00283569 | 0.01756571 |
| LYVE1 | 2103.9325 | -1.491365866 | 0.41954329 | -3.5547366 | 0.00037836 | 0.00365335 |
| MRVI1 | 1143.98577 | 0.310171802 | 0.50479941 | 0.61444566 | 0.53892087 | 0.71213387 |
| CTR9 | 2209.75327 | 0.202445683 | 0.16589586 | 1.2203179 | 0.22234439 | 0.4083592 |
| EIF4G2 | 28203.8222 | 0.139555545 | 0.22969764 | 0.60756195 | 0.54347804 | 0.71573402 |
| AC116535.1 | 25.27938 | 0.220924986 | 0.32817484 | 0.67319294 | 0.50082455 | 0.68181432 |
| ZBED5 | 2546.79922 | -0.002286014 | 0.17740984 | -0.0128855 | 0.98971915 | 0.99519138 |
| ZBED5-AS1 | 637.388748 | 1.029547557 | 0.36505986 | 2.8202157 | 0.00479914 | 0.02607152 |
| AC069360.1 | 1.3765908 | 0.654630949 | 1.13317785 | 0.5776948 | 0.56347019 | 0.7305885 |
| AC111188.1 | 3.22922385 | 0.643160838 | 1.00930049 | 0.63723425 | 0.52397227 | 0.69914611 |
| KC877392.1 | 2.73722757 | -1.110372722 | 1.141763 | -0.9725072 | 0.33079829 | 0.52867582 |
| GALNT18 | 2496.38979 | 0.402137155 | 0.32332901 | 1.24373979 | 0.21359535 | 0.39755147 |
| CSNK2A3 | 9.88979001 | 0.061420064 | 0.42949519 | 0.14300524 | 0.88628604 | 0.94110344 |
| AC023946.1 | 5.88528208 | 0.121422714 | 0.49785703 | 0.24389073 | 0.80731545 | 0.89590613 |

| | | | | | | |
|---|---|---|---|---|---|---|
| AC104031.1 | 12.6962615 | 0.366889277 | 0.62429991 | 0.58768113 | 0.55674635 | 0.72574388 |
| USP47 | 3088.26465 | -0.195708404 | 0.12590414 | -1.5544239 | 0.12008334 | 0.27002624 |
| DKK3 | 1978.38653 | 0.65303989 | 0.41569511 | 1.57095878 | 0.11619222 | 0.26422809 |
| LINC02547 | 0.09989236 | -0.642905668 | 3.30558948 | -0.1944905 | 0.84579185 | NA |
| AC124276.1 | 0.30010799 | -1.364242679 | 3.30558948 | -0.4127078 | 0.67982068 | NA |
| AC124276.2 | 2.63113765 | 1.267262933 | 1.70193455 | 0.74460145 | 0.45651269 | 0.64589761 |
| MICAL2 | 985.557017 | -0.453514989 | 0.42423448 | -1.0690196 | 0.28506083 | 0.48111504 |
| AC079329.1 | 325.236718 | -0.633803652 | 0.43687662 | -1.4507612 | 0.14684637 | 0.30988626 |
| PARVA | 3186.51706 | -0.142301645 | 0.15326893 | -0.9284442 | 0.35317722 | 0.55075596 |
| AC009806.1 | 14.6627346 | 0.532858806 | 0.42030302 | 1.26779676 | 0.20487054 | 0.38721589 |
| TEAD1 | 2828.46481 | -0.003839157 | 0.1898072 | -0.0202266 | 0.9838626 | 0.99305611 |
| AC013549.2 | 0.10721287 | -0.642905668 | 3.30558948 | -0.1944905 | 0.84579185 | NA |
| AC013549.1 | 1.49406077 | -1.303720941 | 1.62368694 | -0.8029386 | 0.42201022 | 0.61452369 |
| LINC00958 | 580.429603 | -0.909186957 | 0.29147252 | -3.1192888 | 0.00181288 | 0.01248231 |
| AC084859.1 | 11.6151975 | 1.472221709 | 0.51260812 | 2.87202185 | 0.00407855 | 0.02307968 |
| RASSF10 | 247.240038 | 0.978832518 | 0.27899924 | 3.50836987 | 0.00045086 | 0.0041834 |
| AC013762.1 | 0.18754884 | 1.469068384 | 3.27783996 | 0.44818185 | 0.65402196 | NA |
| ARNTL | 972.1401 | -0.075470292 | 0.30059469 | -0.2510699 | 0.80176004 | 0.8925745 |
| BTBD10 | 1106.40735 | -0.446296783 | 0.16569354 | -2.6935075 | 0.00707046 | 0.03488306 |
| PTH | 0 | NA | NA | NA | NA | NA |
| FAR1 | 2438.9779 | -0.216215893 | 0.10148701 | -2.1304785 | 0.03313213 | 0.10833158 |
| LINC02548 | 0 | NA | NA | NA | NA | NA |
| AC027779.1 | 4.94896954 | 2.237277453 | 1.40477566 | 1.59262259 | 0.11124488 | 0.25595738 |
| LINC02545 | 0.18754884 | 1.469068384 | 3.27783996 | 0.44818185 | 0.65402196 | NA |
| AC022240.1 | 3.04495642 | -0.911743206 | 1.04431752 | -0.8730517 | 0.38263489 | 0.57875886 |
| SPON1 | 2258.47153 | -0.381497515 | 0.3719227 | -1.0257441 | 0.30501222 | 0.50208732 |
| SPON1-AS1 | 144.67046 | -0.076305461 | 0.42116786 | -0.1811759 | 0.85622952 | 0.92471767 |
| RRAS2 | 2836.31207 | -0.971315533 | 0.46540546 | -2.0870308 | 0.03688535 | 0.11676671 |
| COPB1 | 6515.49965 | -0.215860785 | 0.18914238 | -1.1412608 | 0.25376141 | 0.44642527 |
| AC018523.2 | 4767.56096 | -0.381846263 | 0.1950993 | -1.9571893 | 0.05032522 | 0.14629875 |
| PSMA1 | 4952.04032 | -0.543788803 | 0.18728834 | -2.9034845 | 0.00369035 | 0.02146432 |
| PDE3B | 354.43539 | -0.070218698 | 0.32087955 | -0.2188319 | 0.82678096 | 0.90760472 |
| CYP2R1 | 606.339636 | -0.112096079 | 0.12634921 | -0.8871925 | 0.37497524 | 0.57097224 |
| CALCB | 34.8647795 | -1.543484061 | 0.75546603 | -2.0430886 | 0.04104366 | 0.12627715 |
| CALCA | 237.591497 | 0.076646589 | 0.70918773 | 0.10807659 | 0.91393494 | 0.9577987 |
| INSC | 112.667362 | -0.567922392 | 0.69757465 | -0.8141385 | 0.4155656 | 0.60905514 |
| AC073172.2 | 1.70735159 | 2.529313678 | 1.22857609 | 2.05873588 | 0.03951955 | 0.12270463 |
| AC073172.1 | 56.2791226 | 0.425760731 | 0.76158346 | 0.55904672 | 0.57612984 | 0.74016096 |
| AC087379.2 | 84.614721 | -0.491754332 | 0.78128533 | -0.6294171 | 0.52907604 | 0.70367354 |
| AC087379.1 | 10.4121636 | 0.32758684 | 0.6857863 | 0.47768064 | 0.63287752 | 0.78205156 |
| AC009869.1 | 0.03392908 | 0.380740593 | 3.29738709 | 0.11546736 | 0.90807469 | NA |
| SOX6 | 1158.32942 | 0.745350568 | 0.27990739 | 2.66284701 | 0.00774826 | 0.03731393 |
| AC103794.1 | 0.44092752 | 1.251349279 | 1.77304496 | 0.70576286 | 0.48033562 | NA |
| C11orf58 | 5963.1577 | -0.19571334 | 0.08208008 | -2.3844194 | 0.0171061 | 0.06695677 |
| PLEKHA7 | 4197.41257 | 0.732941595 | 0.16760443 | 4.37304428 | 1.23E-05 | 0.0002422 |
| RPS13 | 13213.2118 | -0.492804178 | 0.19555932 | -2.5199728 | 0.01173639 | 0.0507608 |
| PIK3C2A | 3329.47813 | 0.097746983 | 0.11563229 | 0.84532603 | 0.39792883 | 0.59281302 |
| NUCB2 | 1849.43446 | -0.486199241 | 0.19558276 | -2.4859003 | 0.01292242 | 0.05446996 |
| NCR3LG1 | 51.0059282 | 0.350430429 | 0.35710025 | 0.98132226 | 0.32643385 | 0.52433822 |
| AC124798.1 | 552.661399 | -0.915737818 | 0.32858336 | -2.786927 | 0.00532105 | 0.02819771 |
| KCNJ11 | 429.160606 | -1.484647291 | 0.31047564 | -4.7818479 | 1.74E-06 | 5.07E-05 |
| ABCC8 | 83.382952 | -0.350955686 | 0.59502997 | -0.5898118 | 0.55531684 | 0.7247565 |
| USH1C | 5554.78081 | 0.564628405 | 0.40901001 | 1.38047577 | 0.1674402 | 0.33860415 |
| OTOG | 12.7970115 | 1.012461508 | 0.5535419 | 1.8290603 | 0.06739058 | 0.17926858 |
| AC124301.1 | 0 | NA | NA | NA | NA | NA |
| AC124301.2 | 0 | NA | NA | NA | NA | NA |
| MYOD1 | 0.435637 | -0.299274917 | 2.28700398 | -0.1308589 | 0.8958869 | NA |
| KCNC1 | 7.38557114 | 0.686896191 | 0.51795831 | 1.32616114 | 0.18478633 | 0.36205213 |
| SERGEF | 1163.56484 | -0.314730653 | 0.17857267 | -1.7624794 | 0.07798834 | 0.19875833 |
| TPH1 | 55.2501839 | 0.760814683 | 0.44954517 | 1.69240986 | 0.09056785 | 0.22162902 |
| SAAL1 | 368.617574 | -0.076492194 | 0.14045162 | -0.544616 | 0.58601766 | 0.74736507 |
| MRGPRX3 | 0.61770743 | -2.521239388 | 2.16190703 | -1.1662108 | 0.24352922 | NA |
| AC090099.4 | 0.55704427 | 1.86899668 | 2.64217904 | 0.70736943 | 0.47933693 | NA |
| MRGPRX4 | 0 | NA | NA | NA | NA | NA |
| SAA4 | 88.669313 | -5.26712269 | 0.88797212 | -5.9316307 | 3.00E-09 | 2.50E-07 |
| SAA2-SAA4 | 2382.59949 | -7.733618863 | 1.17164109 | -6.6006723 | 4.09E-11 | 6.83E-09 |
| SAA2 | 3181.77347 | -7.849987852 | 1.16353485 | -6.7466719 | 1.51E-11 | 2.91E-09 |
| SAA1 | 7144.22913 | -7.211201938 | 1.19504605 | -6.0342461 | 1.60E-09 | 1.48E-07 |
| HPS5 | 971.952192 | 0.193472365 | 0.13590988 | 1.42353427 | 0.15458133 | 0.32064106 |
| GTF2H1 | 1975.34158 | -0.312900483 | 0.2764625 | -1.1318008 | 0.25771819 | 0.45029234 |
| LDHA | 54518.4753 | -1.987601758 | 0.30807997 | -6.4515774 | 1.11E-10 | 1.57E-08 |
| LDHC | 71.5968912 | 1.2561701 | 0.93415866 | 1.34470744 | 0.17871973 | 0.35426891 |
| LDHAL6A | 8.96332695 | -0.028688763 | 0.58282743 | -0.0492234 | 0.96074124 | 0.9810923 |
| TSG101 | 3910.0501 | -0.392577734 | 0.14386193 | -2.7288507 | 0.00635555 | 0.03216192 |
| UEVLD | 1246.10089 | -0.056513472 | 0.13702845 | -0.4124214 | 0.68003055 | 0.81447997 |
| SPTY2D1OS | 192.2993 | 0.036504087 | 0.23342652 | 0.15638363 | 0.87573064 | 0.93547018 |
| SPTY2D1 | 1217.87566 | -0.184566908 | 0.31220218 | -0.5911775 | 0.55440149 | 0.72395599 |
| TMEM86A | 1496.45283 | 0.881205676 | 0.32886314 | 2.67955139 | 0.00737209 | 0.0359937 |
| IGSF22 | 289.689344 | 0.138893781 | 0.26266299 | 0.52879083 | 0.59695056 | 0.7555357 |
| AC103974.1 | 110.332059 | 0.000250829 | 0.2645495 | 0.00094814 | 0.9992435 | 0.99973988 |
| PTPN5 | 34.7942514 | -0.691921537 | 0.70918082 | -0.9756631 | 0.32923145 | 0.52700215 |
| MRGPRX1 | 0 | NA | NA | NA | NA | NA |
| AC023078.5 | 0 | NA | NA | NA | NA | NA |
| MRGPRX2 | 0.55718922 | 1.203574063 | 2.11033827 | 0.57032281 | 0.56845877 | NA |
| ZDHHC13 | 1080.79013 | -0.707566176 | 0.14752991 | -4.7960862 | 1.62E-06 | 4.82E-05 |
| CSRP3 | 0.82461767 | 2.10685111 | 1.69320699 | 1.24429625 | 0.21339055 | NA |
| AC009652.2 | 2.35585831 | -1.382631818 | 1.0851219 | -1.2741719 | 0.20260251 | 0.38450005 |
| E2F8 | 15.9352331 | -1.005343609 | 0.64800274 | -1.5514496 | 0.12079397 | 0.27117997 |
| AC009652.1 | 1.62930311 | -1.349486257 | 1.17295893 | -1.1504974 | 0.24993904 | 0.44201546 |
| NAV2 | 2379.63957 | 0.214587813 | 0.26261101 | 0.81713181 | 0.41385311 | 0.60767896 |
| NAV2-AS5 | 0.57361788 | -0.269708234 | 1.67878026 | -0.1606573 | 0.87236336 | NA |
| NAV2-AS4 | 1.02572511 | 1.08159942 | 1.50508813 | 0.71862863 | 0.47236977 | 0.65865011 |
| AC009549.1 | 50.2779404 | -2.543993581 | 0.64144198 | -3.9660541 | 7.31E-05 | 0.00098816 |
| NAV2-AS3 | 2.10977284 | 3.70847831 | 1.16145645 | 3.19295511 | 0.00140825 | 0.01030683 |
| NAV2-AS2 | 15.0491716 | 1.24638863 | 0.48771108 | 2.55558809 | 0.01060085 | 0.04705344 |
| NAV2-AS1 | 747.745894 | 0.265371495 | 0.26909016 | 0.98618061 | 0.32404449 | 0.52163909 |
| DBX1 | 0.385634 | 0.2490405 | 2.83387848 | 0.08787974 | 0.92997226 | NA |
| HTATIP2 | 2756.91593 | -1.58833864 | 0.35929648 | -4.4206908 | 9.84E-06 | 0.00020065 |
| PRMT3 | 754.482756 | -0.317837291 | 0.16925112 | -1.8779037 | 0.06039434 | 0.16629413 |
| SLC6A5 | 0.30684769 | 1.415461554 | 2.2752879 | 0.62210218 | 0.53387468 | NA |
| NELL1 | 608.684985 | 3.036606425 | 0.42928219 | 7.07368374 | 1.51E-12 | 3.86E-10 |
| AC090707.1 | 0.57456572 | 2.584977715 | 1.92103885 | 1.34561449 | 0.17842688 | NA |
| AC090857.2 | 0.06785815 | 0.795868262 | 3.28898349 | 0.24198001 | 0.80879565 | NA |
| AC107882.1 | 0 | NA | NA | NA | NA | NA |
| ANO5 | 460.369753 | 0.583857748 | 0.23891076 | 2.44383198 | 0.01453219 | 0.05947409 |
| AC107886.1 | 2.6534754 | 0.964918304 | 0.73893455 | 1.30582378 | 0.19161249 | 0.37079485 |
| AC104009.1 | 1.77447057 | -0.703664586 | 1.11388132 | -0.6317231 | 0.52756782 | 0.70223445 |
| SLC17A6 | 0 | NA | NA | NA | NA | NA |
| AC040936.1 | 0 | NA | NA | NA | NA | NA |
| LINC01495 | 0 | NA | NA | NA | NA | NA |
| AC055878.1 | 0 | NA | NA | NA | NA | NA |
| FANCF | 727.705094 | 0.260846612 | 0.1416249 | 1.84181323 | 0.06550247 | 0.17578578 |
| GAS2 | 284.55092 | 1.366996322 | 0.25375735 | 5.38702153 | 7.16E-08 | 3.58E-06 |

| | | | | | | |
|---|---|---|---|---|---|---|
| SVIP | 1601.30272 | 0.188452221 | 0.17540006 | 1.07441364 | 0.28263733 | 0.47859413 |
| AC006299.1 | 10.3945163 | 0.465118456 | 0.34892355 | 1.33300964 | 0.18252864 | 0.35928046 |
| CCDC179 | 0.2408037 | 1.011447186 | 2.97445912 | 0.34004407 | 0.73382334 | NA |
| AC090092.1 | 7.15418184 | 0.219726029 | 0.68909412 | 0.31886214 | 0.74983105 | 0.86029607 |
| AC100768.2 | 0 | NA | NA | NA | NA | NA |
| AC100768.1 | 0 | NA | NA | NA | NA | NA |
| AC015771.1 | 0 | NA | NA | NA | NA | NA |
| LUZP2 | 8.22479977 | 1.361563378 | 0.91380268 | 1.48999714 | 0.13622499 | 0.29432633 |
| AC090592.1 | 0 | NA | NA | NA | NA | NA |
| AC013799.1 | 0.07830766 | -0.642905668 | 3.30558948 | -0.1944905 | 0.84579185 | NA |
| AC099687.1 | 3.96895512 | 0.839697411 | 0.67371189 | 1.24637464 | 0.21262689 | 0.39645571 |
| AC021698.1 | 1.28598963 | 1.511130071 | 2.6018912 | 0.58078142 | NA | NA |
| ANO3 | 347.657129 | 1.565864456 | 0.42174641 | 3.71281036 | 0.00020497 | 0.00222522 |
| MUC15 | 1068.5481 | 1.248880675 | 0.54003459 | 2.31259388 | 0.02074498 | 0.07696189 |
| SLC5A12 | 14786.6174 | 2.521549071 | 0.53813199 | 4.6857446 | 2.79E-06 | 7.42E-05 |
| FIBIN | 1234.86692 | -0.579260093 | 0.42352138 | -1.3677234 | 0.17139868 | 0.34388234 |
| BBOX1 | 11393.0876 | 1.287489083 | 0.4878807 | 2.63894243 | 0.00831651 | 0.03919685 |
| BBOX1-AS1 | 45.501591 | 1.071741571 | 0.52896281 | 2.02611896 | 0.0427526 | 0.13001798 |
| CCDC34 | 166.605641 | 0.118209902 | 0.24526249 | 0.48197302 | 0.62982511 | 0.78011215 |
| LGR4 | 6093.10706 | 0.021694111 | 0.17855558 | 0.1214978 | 0.90329676 | 0.95141786 |
| AC100771.2 | 24.7357501 | 1.448785562 | 0.47615021 | 3.04270696 | 0.00234461 | 0.01528727 |
| LIN7C | 1448.55133 | -0.08336242 | 0.11194457 | -0.7446759 | 0.45646768 | 0.64589761 |
| BDNF-AS | 278.532476 | 0.764636836 | 0.33025741 | 2.31527534 | 0.02059787 | 0.07656582 |
| LINC00678 | 0.2320242 | 1.01099135 | 3.00413042 | 0.33653377 | 0.73646839 | NA |
| BDNF | 123.924599 | 0.380471772 | 0.39127017 | 0.97240167 | 0.33085076 | 0.52870262 |
| AC090159.1 | 0.1357163 | 1.313406857 | 3.28103699 | 0.40030236 | 0.68893383 | NA |
| KIF18A | 38.0833117 | -0.399403005 | 0.34109316 | -1.1709499 | 0.24161891 | 0.43184836 |
| METTL15 | 678.469582 | 0.184792396 | 0.17796222 | 1.0383799 | 0.29909322 | 0.49588735 |
| AC013714.1 | 0 | NA | NA | NA | NA | NA |
| AC100773.1 | 0.40660282 | 1.970115679 | 2.06762612 | 0.95283942 | 0.34067144 | NA |
| AC090833.1 | 0.2121291 | -0.339874279 | 3.10853787 | -0.1093357 | 0.9129362 | NA |
| AC090791.1 | 0 | NA | NA | NA | NA | NA |
| AC090124.1 | 0 | NA | NA | NA | NA | NA |
| AC090124.2 | 0.26066098 | 1.012364238 | 2.91757497 | 0.34698825 | 0.72860015 | NA |
| AC110058.1 | 0.03392908 | 0.380740593 | 3.29738709 | 0.11546736 | 0.90807469 | NA |
| LINC02546 | 0.08695548 | -0.642905668 | 3.30558948 | -0.1944905 | 0.84579185 | NA |
| AC107973.1 | 0 | NA | NA | NA | NA | NA |
| LINC01616 | 3.25498685 | 0.70380976 | 0.93333767 | 0.75407838 | 0.45080216 | 0.64064962 |
| KCNA4 | 11.1658431 | 0.794885489 | 0.7202616 | 1.10360665 | 0.2697638 | 0.46402671 |
| AC124657.1 | 1.00303789 | -0.005620451 | 1.50790548 | -0.0037273 | 0.99702603 | NA |
| FSHB | 0.75535704 | 3.50923531 | 2.29894933 | 1.52645179 | 0.12689739 | NA |
| ARL14EP | 1136.95352 | 0.169898662 | 0.11412616 | 1.48869161 | 0.1365686 | 0.29487545 |
| MPPED2 | 605.247615 | 0.751273928 | 0.3033268 | 2.47678054 | 0.01325734 | 0.05551787 |
| AL353699.1 | 4.35772859 | 0.449511032 | 0.80716493 | 0.5569011 | 0.57759501 | 0.7412624 |
| AL136088.1 | 32.6973356 | 0.311432877 | 0.46680417 | 0.66715959 | 0.5046702 | 0.68403209 |
| DCDC1 | 169.882854 | -0.039008395 | 0.35384983 | -0.11024 | 0.91221906 | 0.95681225 |
| AL137804.1 | 4.17363206 | -0.263462579 | 0.59222472 | -0.4448693 | 0.65641423 | 0.79835608 |
| DNAJC24 | 439.908542 | 0.319792756 | 0.1983217 | 1.61249504 | 0.10685426 | 0.24863608 |
| IMMP1L | 567.962768 | -0.11467575 | 0.11304927 | -1.0143874 | 0.31039794 | 0.50741242 |
| ELP4 | 889.979823 | 0.430526201 | 0.18149636 | 2.37209276 | 0.01768765 | 0.06853119 |
| AC131571.1 | 0.93306605 | 1.020291817 | 1.50339199 | 0.67865987 | 0.4973534 | NA |
| PAX6 | 130.55706 | 1.150239576 | 0.26671465 | 4.31262241 | 1.61E-05 | 0.00029938 |
| AL035078.4 | 2703.08472 | -0.976918126 | 0.25344295 | -3.8545879 | 0.00011592 | 0.00142448 |
| AL035078.1 | 0 | NA | NA | NA | NA | NA |
| AL035078.3 | 0 | NA | NA | NA | NA | NA |
| RCN1 | 5584.36071 | -1.309069977 | 0.22255137 | -5.8821026 | 4.05E-09 | 3.20E-07 |
| AL078612.1 | 1372.1165 | -1.310551983 | 0.22608218 | -5.7967946 | 6.76E-09 | 4.92E-07 |
| AL078612.2 | 0.08666371 | -0.642905668 | 3.30558948 | -0.1944905 | 0.84579185 | NA |
| WT1 | 772.541193 | 0.536492858 | 0.56923091 | 0.94248722 | 0.34594325 | 0.54354339 |
| EIF3M | 5604.3271 | -0.242057058 | 0.16600116 | -1.4581648 | 0.14479511 | 0.30684991 |
| CCDC73 | 1360.14746 | -0.410258361 | 0.1425394 | -2.8782103 | 0.00399938 | 0.02274887 |
| PRRG4 | 2126.75329 | -0.530435056 | 0.21773356 | -2.4361658 | 0.01484388 | 0.06038097 |
| QSER1 | 1033.41781 | -0.168528991 | 0.1185347 | -1.4217692 | 0.15509326 | 0.32122284 |
| DEPDC7 | 820.637855 | 1.307822699 | 0.44289477 | 2.95289714 | 0.00314807 | 0.0190619 |
| TCP11L1 | 725.589086 | -0.225757822 | 0.1484238 | -1.5210352 | 0.12825101 | 0.28239638 |
| LINC00294 | 1317.34877 | 0.049766172 | 0.21300374 | 0.2336399 | 0.81526455 | 0.90043158 |
| CSTF3 | 1460.86494 | -0.113950161 | 0.14917114 | -0.7638888 | 0.44493352 | 0.63551603 |
| CSTF3-DT | 1.1350066 | 0.427189605 | 1.18244284 | 0.36127717 | 0.71789226 | 0.83983935 |
| HIPK3 | 5307.8255 | 0.055068971 | 0.21455885 | 0.25666138 | 0.79744018 | 0.89013282 |
| KIAA1549L | 216.160388 | 0.785767754 | 0.23127371 | 3.39756621 | 0.00067988 | 0.00575738 |
| AL049629.1 | 192.938095 | -0.200565326 | 0.27335633 | -0.7337138 | 0.46312316 | 0.65105787 |
| AL049629.2 | 17667.8413 | -0.446415224 | 0.22746134 | -1.9625982 | 0.04969288 | 0.14506822 |
| C11orf91 | 96.7656762 | -1.909712093 | 0.55578236 | -3.436079 | 0.0005902 | 0.0051552 |
| CD59 | 45225.5878 | -0.659405029 | 0.23586837 | -2.7956483 | 0.00517957 | 0.02760973 |
| FBXO3 | 3003.76438 | 0.487316941 | 0.12087058 | 4.03172496 | 5.54E-05 | 0.00080401 |
| FBXO3-DT | 0.76668088 | 1.727541134 | 1.75054372 | 0.98685975 | 0.3237114 | NA |
| AC113192.5 | 0 | NA | NA | NA | NA | NA |
| AC113192.4 | 0.36407885 | 1.644721722 | 2.14154708 | 0.76800633 | 0.44248342 | NA |
| AC113199.1 | 0.03392908 | 0.380740593 | 3.29738709 | 0.11546736 | 0.90807469 | NA |
| AC113192.2 | 1.09931482 | -1.064254844 | 1.5319416 | -0.6947098 | 0.48723718 | 0.67119354 |
| LMO2 | 2235.39301 | 0.277752095 | 0.26631153 | 1.04295933 | 0.29696713 | 0.49372822 |
| AC132216.1 | 0.34346317 | -1.70467338 | 2.84472907 | -0.5992393 | 0.54901334 | NA |
| CAPRIN1 | 7550.84238 | 0.054721857 | 0.11907181 | 0.45957023 | 0.64582473 | 0.7910813 |
| NAT10 | 1850.46335 | -0.336373622 | 0.14779289 | -2.2759798 | 0.02284722 | 0.08274379 |
| ABTB2 | 2241.75979 | -0.541753105 | 0.26057905 | -2.0790356 | 0.03761408 | 0.11833493 |
| CAT | 15637.8494 | 0.950795289 | 0.30508571 | 3.11648584 | 0.0018302 | 0.01258167 |
| ELF5 | 514.334511 | 1.325523284 | 0.55717496 | 2.37900729 | 0.01735933 | 0.0676516 |
| AL137224.1 | 0 | NA | NA | NA | NA | NA |
| AC087783.1 | 0.2818758 | -0.179573506 | 2.93015205 | -0.0612847 | 0.95113247 | NA |
| EHF | 483.488795 | 0.499254224 | 0.3599811 | 1.38689009 | 0.16547528 | 0.33584108 |
| APIP | 1546.51674 | -0.135162556 | 0.20104907 | -0.6722864 | 0.50140138 | 0.68210661 |
| PDHX | 2105.14476 | -0.045297647 | 0.15245043 | -0.2971303 | 0.76636701 | 0.8711864 |
| AL356215.1 | 2.80984906 | -0.857741474 | 0.96168678 | -0.8919136 | 0.37243927 | 0.56883475 |
| CD44 | 5211.0319 | -1.402386261 | 0.47579451 | -2.947462 | 0.00320394 | 0.01932478 |
| CD44-AS1 | 444.057171 | -1.315194811 | 0.50155669 | -2.6222257 | 0.00873576 | 0.04067124 |
| SLC1A2 | 29.6015506 | 0.803633789 | 0.54221313 | 1.48213636 | 0.13830402 | 0.2973807 |
| AL133330.2 | 0.20543692 | 0.689155168 | 3.10695182 | 0.2218107 | 0.82446125 | NA |
| AC090625.2 | 0.22267178 | 0.952737031 | 3.28103697 | 0.2903768 | 0.77152799 | NA |
| PAMR1 | 368.394694 | -0.5441569 | 0.50138437 | -1.0853089 | 0.27778488 | 0.47329505 |
| AL135934.1 | 0 | NA | NA | NA | NA | NA |
| FJX1 | 441.035995 | -1.231075688 | 0.38646656 | -3.185465 | 0.00144522 | 0.0105074 |
| AL138812.1 | 0 | NA | NA | NA | NA | NA |
| TRIM44 | 5260.6304 | 0.114544141 | 0.09291814 | 1.23274245 | 0.21767187 | 0.40265881 |
| AC090692.1 | 0 | NA | NA | NA | NA | NA |
| LDLRAD3 | 514.81433 | -1.134049343 | 0.32488098 | -3.4906609 | 0.00048183 | 0.00440103 |
| COMMD9 | 2308.61788 | -0.054776725 | 0.15305469 | -0.3578899 | 0.72042573 | 0.84093782 |
| PRR5L | 545.926401 | -0.04394445 | 0.43553414 | -0.1008978 | 0.91963157 | 0.96074349 |
| AC087277.1 | 0 | NA | NA | NA | NA | NA |
| AC009656.1 | 0.63346037 | 2.700713519 | 1.90980009 | 1.41413414 | 0.15732252 | NA |
| TRAF6 | 701.470109 | -0.338584349 | 0.14620799 | -2.3157719 | 0.02057072 | 0.07648759 |
| RAG1 | 161.93611 | 0.643702834 | 0.31261289 | 2.05910524 | 0.03948416 | 0.12262363 |
| RAG2 | 82.1312705 | 0.654458178 | 0.18079383 | 3.6199144 | 0.0002947 | 0.00299124 |
| C11orf74 | 493.265841 | 0.29887621 | 0.18990903 | 1.57378621 | 0.11553688 | 0.26303269 |

| | | | | | | |
|---|---|---|---|---|---|---|
| AC103798.1 | 0 | NA | NA | NA | NA | NA |
| AC021713.1 | 0.16786415 | -1.003574173 | 3.30558948 | -0.3035992 | 0.7614333 | NA |
| AC022796.1 | 0.31739398 | 0.652794139 | 2.8538039 | 0.22874527 | 0.81906691 | NA |
| LINC01493 | 0 | NA | NA | NA | NA | NA |
| LRRC4C | 34.3019891 | -0.984162407 | 0.60326663 | -1.6313888 | 0.10280831 | 0.24201911 |
| AC090138.1 | 0 | NA | NA | NA | NA | NA |
| AC021006.2 | 0.17648264 | -1.003574173 | 3.30558948 | -0.3035992 | 0.7614333 | NA |
| LINC01499 | 0 | NA | NA | NA | NA | NA |
| AC023442.3 | 0.07659028 | -0.642905668 | 3.30558948 | -0.1944905 | 0.84579185 | NA |
| AC023442.2 | 0 | NA | NA | NA | NA | NA |
| AC079917.1 | 0.09997307 | 0.547239394 | 3.29377202 | 0.16614368 | 0.86804388 | NA |
| AC090458.1 | 0.03392908 | 0.380740593 | 3.29738709 | 0.11546736 | 0.90807469 | NA |
| API5 | 4157.89514 | 0.082644282 | 0.12367486 | 0.66823835 | 0.50398146 | 0.68375733 |
| AC087276.2 | 376.520742 | 0.838729647 | 0.27589575 | 3.04002377 | 0.00236559 | 0.01536093 |
| TTC17 | 3720.7942 | -0.187150405 | 0.16450165 | -1.137681 | 0.25525371 | 0.44786515 |
| AC087276.3 | 28.9672972 | 0.359841435 | 0.42183023 | 0.85304801 | 0.39363268 | 0.58894535 |
| AC068205.1 | 0.77839779 | 1.787321259 | 1.41232735 | 1.26551487 | 0.20568682 | NA |
| MIR670HG | 1.36423523 | 3.453334591 | 1.83470797 | 1.88222576 | 0.05980537 | 0.16520717 |
| HSD17B12 | 5435.93805 | 0.052223994 | 0.22068705 | 0.23664277 | 0.81293394 | 0.89895498 |
| AC087521.2 | 406.567338 | 0.439239616 | 0.2654552 | 1.65466571 | 0.09799233 | 0.23433913 |
| AC087521.4 | 2156.43167 | -0.280410875 | 0.24700369 | -1.1352497 | 0.25627069 | 0.44905696 |
| ALKBH3 | 780.783877 | -0.110163013 | 0.16401377 | -0.6716693 | 0.50179425 | 0.68228905 |
| ALKBH3-AS1 | 20.2231659 | 0.242597939 | 0.35805809 | 0.67753793 | 0.49806471 | 0.67967569 |
| C11orf96 | 5312.39466 | -1.544595709 | 0.63198585 | -2.4440353 | 0.01452401 | 0.05946252 |
| AC087521.1 | 1304.4814 | -1.451607511 | 0.5835318 | -2.4876237 | 0.01285997 | 0.05427617 |
| ACCSL | 1.05463273 | 0.123411542 | 1.37302 | 0.08988328 | 0.92837997 | 0.96526981 |
| ACCS | 1863.08827 | 0.325436496 | 0.44856903 | 0.72549925 | 0.4681458 | 0.65490742 |
| EXT2 | 3225.84579 | -0.399721384 | 0.16512022 | -2.42079 | 0.01548682 | 0.06227317 |
| ALX4 | 9.93185677 | 1.627818942 | 0.83676642 | 1.9453684 | 0.05173066 | 0.14937081 |
| AC010768.4 | 0 | NA | NA | NA | NA | NA |
| CD82 | 2472.02579 | -0.19050502 | 0.40064563 | -0.4754951 | 0.63443415 | 0.7832777 |
| AC010768.1 | 373.590206 | 0.049630219 | 0.41523403 | 0.11952349 | 0.90486063 | 0.9522378 |
| AC010768.2 | 0.70405072 | 1.755961933 | 1.4434888 | 1.21647077 | 0.22380564 | NA |
| AC104241.2 | 0.22663961 | 1.777046332 | 3.25798548 | 0.54544329 | 0.58544867 | NA |
| AC104241.1 | 0.09635527 | 0.547239394 | 3.29377202 | 0.16614368 | 0.86804388 | NA |
| TSPAN18 | 2513.11122 | 0.856534561 | 0.31097135 | 2.75438413 | 0.00588027 | 0.03035684 |
| TP53I11 | 3442.56239 | -0.425240776 | 0.26974445 | -1.5764579 | 0.11492031 | 0.26214953 |
| AC068858.1 | 0.77310998 | 2.376534307 | 1.88506939 | 1.26071449 | 0.20741173 | NA |
| PRDM11 | 737.223834 | 0.226507544 | 0.20304224 | 1.11556857 | 0.26460685 | 0.45857536 |
| AC103681.2 | 0.84828119 | -1.331724946 | 1.91798527 | -0.6943353 | 0.48747194 | NA |
| SYT13 | 1298.27287 | 0.669826742 | 0.35105907 | 1.9080172 | 0.05638899 | 0.15839132 |
| AC103681.1 | 29.8892283 | 2.62283585 | 0.59195971 | 4.43076749 | 9.39E-06 | 0.0001942 |
| AC103736.1 | 26.49616 | 0.675996537 | 0.40045143 | 1.6880862 | 0.09139468 | 0.22299018 |
| AC018716.2 | 0 | NA | NA | NA | NA | NA |
| AC103855.3 | 0.08393208 | -0.642905668 | 3.30558948 | -0.1944905 | 0.84579185 | NA |
| AC018716.1 | 0 | NA | NA | NA | NA | NA |
| AC103855.1 | 0 | NA | NA | NA | NA | NA |
| CHST1 | 290.791233 | -0.450096134 | 0.40284046 | -1.1173062 | 0.26386344 | 0.45761781 |
| AC087442.1 | 22.8765699 | -0.105942291 | 0.48327596 | -0.219217 | 0.82648104 | 0.90741548 |
| AC044839.4 | 0 | NA | NA | NA | NA | NA |
| AC044839.3 | 1.06864172 | -0.471326971 | 2.79805652 | -0.168448 | 0.86623086 | 0.93055685 |
| AC044839.1 | 14.9925922 | -0.273250847 | 0.47115722 | -0.5799568 | 0.56194373 | 0.72964972 |
| SLC35C1 | 1699.80666 | -1.137606476 | 0.2564348 | -4.4362406 | 9.15E-06 | 0.00019023 |
| AC044839.5 | 0.09997307 | 0.547239394 | 3.29377202 | 0.16614368 | 0.86804388 | NA |
| CRY2 | 3440.62566 | 0.685789667 | 0.20812418 | 3.29509847 | 0.00098387 | 0.00775353 |
| MAPK8IP1 | 1625.2972 | 0.426274034 | 0.20614741 | 2.06781177 | 0.03865773 | 0.12076819 |
| AC068385.1 | 395.216873 | 0.010698544 | 0.19901528 | 0.0537574 | 0.95712845 | 0.97945117 |
| C11orf94 | 23.0491909 | -0.345495119 | 0.42579157 | -0.8114184 | 0.41712542 | 0.61013964 |
| PEX16 | 1644.66567 | -0.141847639 | 0.17107601 | -0.8291498 | 0.40701964 | 0.60168362 |
| LARGE2 | 2000.77951 | 0.679518969 | 0.35999444 | 1.88758185 | 0.05908211 | 0.16375817 |
| PHF21A | 2047.55407 | -0.553336067 | 0.21755595 | -2.5434196 | 0.01097733 | 0.04829271 |
| AC024475.1 | 0.63851098 | -1.526825289 | 1.90027866 | -0.8034744 | 0.42170058 | NA |
| AC024475.4 | 0.99214913 | -0.866795202 | 1.58190711 | -0.5479432 | 0.58373092 | NA |
| AC024475.3 | 0.07659028 | -0.642905668 | 3.30558948 | -0.1944905 | 0.84579185 | NA |
| AC116021.1 | 1.97204891 | -1.443472109 | 1.1537486 | -1.2511149 | 0.21089255 | 0.39427412 |
| LINC02489 | 0.5632237 | 0.263388988 | 1.51053978 | 0.17436746 | 0.86157668 | NA |
| CREB3L1 | 339.061723 | -1.044955473 | 0.37174216 | -2.8109684 | 0.00493926 | 0.02664111 |
| DGKZ | 5631.42544 | 0.02782106 | 0.15987982 | 0.17401233 | 0.86185577 | 0.92801702 |
| MDK | 2142.36995 | -2.568640922 | 0.72937477 | -3.5217024 | 0.00042879 | 0.00402673 |
| CHRM4 | 7.40418117 | -1.083270888 | 0.78284756 | -1.3837571 | 0.16643284 | 0.33721351 |
| AMBRA1 | 1799.10699 | -0.122619872 | 0.16928782 | -0.7243278 | 0.46886452 | 0.6556107 |
| HARBI1 | 294.911452 | 0.479629368 | 0.26786825 | 1.79054206 | 0.07336681 | 0.19074069 |
| ATG13 | 4042.96553 | -0.009469828 | 0.11590988 | -0.0816999 | 0.93488534 | 0.96883307 |
| ARHGAP1 | 6055.77646 | -0.301957212 | 0.12083575 | -2.4989063 | 0.01245772 | 0.05303684 |
| ZNF408 | 654.644426 | -0.164872546 | 0.18633892 | -0.8847993 | 0.37626488 | 0.57211007 |
| F2 | 13.6115747 | 0.300613342 | 0.89962925 | 0.33415248 | 0.7382645 | 0.85304361 |
| CKAP5 | 1981.80759 | -0.090870157 | 0.11566146 | -0.7856563 | 0.43206887 | 0.62420732 |
| LRP4-AS1 | 7.34365356 | 0.886257904 | 0.45034693 | 1.96794482 | 0.04907439 | 0.14386793 |
| LRP4 | 222.254913 | 0.069418705 | 0.26794867 | 0.25907463 | 0.79557766 | 0.88912284 |
| C11orf49 | 2256.22662 | -0.059249576 | 0.19966682 | -0.2967422 | 0.76666332 | 0.87141861 |
| AC090589.2 | 1.41340458 | 0.392460391 | 1.30180684 | 0.3014736 | 0.76305338 | 0.86877586 |
| ARFGAP2 | 4952.76035 | 0.284641687 | 0.18263753 | 1.55850604 | 0.11911334 | 0.26848609 |
| PACSIN3 | 1464.32287 | -0.099151991 | 0.1898653 | -0.5222228 | 0.60151521 | 0.75894526 |
| DDB2 | 1030.6807 | -0.475195616 | 0.2194848 | -2.1650502 | 0.03038384 | 0.10172955 |
| AC018410.2 | 3.55978606 | -0.146464914 | 0.90606159 | -0.1616501 | 0.87158143 | 0.93317628 |
| ACP2 | 3505.05304 | 0.175688878 | 0.17752317 | 0.9896673 | 0.32233676 | 0.5201016 |
| NR1H3 | 2067.2514 | 0.259667607 | 0.26400049 | 0.98358758 | 0.32531833 | 0.52283304 |
| MADD | 3071.88411 | 0.237083035 | 0.14624577 | 1.62112745 | 0.10499031 | 0.24563876 |
| AC018410.1 | 0.501603 | 1.468143246 | 2.05598938 | 0.71408114 | 0.47517701 | NA |
| MYBPC3 | 24.658054 | 0.957855188 | 0.43722681 | 2.19075125 | 0.0284698 | 0.09723463 |
| SPI1 | 750.827914 | -0.604084872 | 0.42924108 | -1.4073324 | 0.15932885 | 0.32704148 |
| AC090559.1 | 84.1494314 | 1.141539453 | 0.24918087 | 4.58116808 | 4.62E-06 | 0.00011044 |
| SLC39A13 | 8301.39191 | -0.479918922 | 0.21880417 | -2.1933719 | 0.02828059 | 0.09683927 |
| PSMC3 | 4998.97105 | -0.190106467 | 0.17993121 | -1.0565508 | 0.29071665 | 0.48677364 |
| RAPSN | 64.7875746 | -1.044169776 | 0.55916411 | -1.8673763 | 0.06184906 | 0.16895885 |
| CELF1 | 5421.45735 | -0.346511244 | 0.15818671 | -2.1905205 | 0.02848651 | 0.0972665 |
| NDUFS3 | 5222.6518 | 0.144769123 | 0.11571176 | 1.25111852 | 0.21089124 | 0.39427412 |
| PTPMT1 | 2379.23312 | 0.028793018 | 0.10534021 | 0.27333359 | 0.78459679 | 0.88229499 |
| KBTBD4 | 1533.56939 | 0.161596385 | 0.18193315 | 0.8882185 | 0.37442323 | 0.57055999 |
| KF459542.1 | 156.551371 | 0.492477935 | 0.24515507 | 2.00884257 | 0.04455383 | 0.13385914 |
| FAM180B | 7.73950014 | 0.318627328 | 0.98866228 | 0.32228126 | 0.74723963 | 0.85862121 |
| C1QTNF4 | 31.7401129 | -0.533411646 | 0.5771225 | -0.9242607 | 0.3553506 | 0.55283748 |
| MTCH2 | 7131.60061 | 0.508919988 | 0.24387303 | 2.08682361 | 0.03690408 | 0.11679796 |
| AGBL2 | 199.010155 | 0.230519629 | 0.5246667 | 0.43936394 | 0.66039785 | 0.80018573 |
| FNBP4 | 4457.49053 | -0.589868911 | 0.20408781 | -2.8902702 | 0.00384911 | 0.02213396 |
| NUP160 | 1942.0139 | 0.14838514 | 0.15831011 | 0.93730676 | 0.34860079 | 0.54613847 |
| PTPRJ | 3458.04274 | -0.115224199 | 0.13367661 | -0.8619623 | 0.38870828 | 0.58435877 |
| PTPRJ-AS1 | 0.85595371 | -1.096592352 | 1.50464474 | -0.7288048 | 0.46612106 | NA |
| OR4B1 | 0 | NA | NA | NA | NA | NA |
| OR4X2 | 0.19289513 | -1.003574173 | 3.30558948 | -0.3035992 | 0.7614333 | NA |
| OR4X1 | 0 | NA | NA | NA | NA | NA |
| OR4S1 | 0 | NA | NA | NA | NA | NA |
| OR4C3 | 0 | NA | NA | NA | NA | NA |

| | | | | | | |
|---|---|---|---|---|---|---|
| OR4C5 | 0 | NA | NA | NA | NA | NA |
| OR4A47 | 0 | NA | NA | NA | NA | NA |
| TRIM49B | 0.16052235 | -1.003574173 | 3.30558948 | -0.3035992 | 0.7614333 | NA |
| TRIM64C | 0 | NA | NA | NA | NA | NA |
| FOLH1 | 826.991616 | 2.452582136 | 0.40685015 | 6.02821987 | 1.66E-09 | 1.53E-07 |
| OR4C13 | 0 | NA | NA | NA | NA | NA |
| OR4C12 | 0 | NA | NA | NA | NA | NA |
| AC109635.2 | 0 | NA | NA | NA | NA | NA |
| OR4C46 | 0 | NA | NA | NA | NA | NA |
| OR4A5 | 0 | NA | NA | NA | NA | NA |
| TRIM48 | 0.03392908 | 0.380740593 | 3.29738709 | 0.11546736 | 0.90807469 | NA |
| OR4A16 | 0 | NA | NA | NA | NA | NA |
| OR4A15 | 0 | NA | NA | NA | NA | NA |
| OR4C15 | 0 | NA | NA | NA | NA | NA |
| OR4C16 | 0 | NA | NA | NA | NA | NA |
| OR4C11 | 0 | NA | NA | NA | NA | NA |
| OR4P4 | 0 | NA | NA | NA | NA | NA |
| OR4S2 | 0 | NA | NA | NA | NA | NA |
| OR4C6 | 0.36593297 | -1.749352747 | 3.30121196 | -0.5299123 | 0.59617275 | NA |
| AP006437.1 | 0 | NA | NA | NA | NA | NA |
| OR5D3P | 0 | NA | NA | NA | NA | NA |
| OR5D13 | 0 | NA | NA | NA | NA | NA |
| OR5D14 | 0 | NA | NA | NA | NA | NA |
| OR5L1 | 0 | NA | NA | NA | NA | NA |
| OR5D18 | 0 | NA | NA | NA | NA | NA |
| OR5L2 | 0 | NA | NA | NA | NA | NA |
| OR5D16 | 0 | NA | NA | NA | NA | NA |
| TRIM51 | 0 | NA | NA | NA | NA | NA |
| OR5W2 | 0 | NA | NA | NA | NA | NA |
| OR5I1 | 0 | NA | NA | NA | NA | NA |
| OR10AG1 | 0.08666371 | -0.642905668 | 3.30558948 | -0.1944905 | 0.84579185 | NA |
| OR5F1 | 0 | NA | NA | NA | NA | NA |
| OR5AS1 | 0 | NA | NA | NA | NA | NA |
| OR8I2 | 0 | NA | NA | NA | NA | NA |
| OR8H2 | 0 | NA | NA | NA | NA | NA |
| OR8H3 | 0 | NA | NA | NA | NA | NA |
| OR8J3 | 0 | NA | NA | NA | NA | NA |
| OR8K5 | 0 | NA | NA | NA | NA | NA |
| OR5J2 | 0 | NA | NA | NA | NA | NA |
| OR5T2 | 0 | NA | NA | NA | NA | NA |
| OR5T3 | 0 | NA | NA | NA | NA | NA |
| OR5T1 | 0.03392908 | 0.380740593 | 3.29738709 | 0.11546736 | 0.90807469 | NA |
| OR8H1 | 0 | NA | NA | NA | NA | NA |
| OR8K3 | 0 | NA | NA | NA | NA | NA |
| OR8K1 | 0 | NA | NA | NA | NA | NA |
| OR8J1 | 0 | NA | NA | NA | NA | NA |
| OR8U1 | 0 | NA | NA | NA | NA | NA |
| OR5R1 | 0 | NA | NA | NA | NA | NA |
| AP002512.3 | 0 | NA | NA | NA | NA | NA |
| OR5M9 | 0 | NA | NA | NA | NA | NA |
| OR5M3 | 0 | NA | NA | NA | NA | NA |
| OR5M8 | 0.03392908 | 0.380740593 | 3.29738709 | 0.11546736 | 0.90807469 | NA |
| OR5M11 | 0.23374159 | 1.011073437 | 2.99871927 | 0.33716842 | 0.73598994 | NA |
| OR5M10 | 0.03392908 | 0.380740593 | 3.29738709 | 0.11546736 | 0.90807469 | NA |
| OR5M1 | 0 | NA | NA | NA | NA | NA |
| OR5AP2 | 0 | NA | NA | NA | NA | NA |
| OR5AR1 | 0 | NA | NA | NA | NA | NA |
| OR9G1 | 0 | NA | NA | NA | NA | NA |
| OR9G4 | 0 | NA | NA | NA | NA | NA |
| AP001803.2 | 0 | NA | NA | NA | NA | NA |
| OR5AK2 | 0.29114391 | 1.099131448 | 2.94336048 | 0.3734274 | 0.70883037 | NA |
| LRRC55 | 21.438051 | 2.546693509 | 0.62276007 | 4.08936543 | 4.33E-05 | 0.00066124 |
| APLNR | 500.872607 | 2.349165199 | 0.61163 | 3.84082728 | 0.00012262 | 0.00148462 |
| TNKS1BP1 | 10580.0596 | -1.28061316 | 0.23570045 | -5.4332232 | 5.53E-08 | 2.87E-06 |
| AP000781.1 | 772.297516 | -0.460391809 | 0.16126845 | -2.8548163 | 0.00430617 | 0.02407235 |
| SSRP1 | 4875.69194 | -0.302107046 | 0.14987 | -2.015794 | 0.04382153 | 0.13255031 |
| P2RX3 | 0.98507211 | -0.22634118 | 1.36222302 | -0.1661557 | 0.86803439 | NA |
| PRG3 | 2.42442824 | -3.671332265 | 1.47137304 | -2.4951744 | 0.01258953 | 0.05346848 |
| PRG2 | 58.0833873 | -2.33729616 | 0.68172411 | -3.4285074 | 0.00060691 | 0.00526108 |
| AP000781.2 | 1107.06417 | -1.262794557 | 0.35515502 | -3.5556151 | 0.0003771 | 0.0036425 |
| SLC43A3 | 1670.46619 | -1.334729196 | 0.35384442 | -3.7720792 | 0.00016189 | 0.00185452 |
| RTN4RL2 | 227.515649 | -0.645764651 | 0.41588674 | -1.5527416 | 0.12048487 | 0.27060609 |
| AP002893.1 | 118.403984 | -0.756793652 | 0.37576497 | -2.014008 | 0.04400869 | 0.13287518 |
| SLC43A1 | 728.562765 | 1.169345697 | 0.28978811 | 4.03517492 | 5.46E-05 | 0.00079667 |
| TIMM10 | 1191.04797 | -0.295294881 | 0.1993092 | -1.4815918 | 0.13844894 | 0.29759525 |
| SMTNL1 | 17.5457021 | -0.161885803 | 0.54417998 | -0.2974858 | 0.76609567 | 0.8710362 |
| UBE2L6 | 2199.85648 | -0.822967738 | 0.23122581 | -3.5591517 | 0.00037205 | 0.00360436 |
| SERPING1 | 38815.952 | -1.879579006 | 0.30035238 | -6.2579127 | 3.90E-10 | 4.51E-08 |
| YPEL4 | 90.1576257 | -1.565600107 | 0.36536094 | -4.285078 | 1.83E-05 | 0.0003313 |
| CLP1 | 570.518913 | -0.402300406 | 0.28769249 | -1.3983695 | 0.16200214 | 0.3308035 |
| ZDHHC5 | 6239.94965 | -0.452765112 | 0.19923897 | -2.2724726 | 0.02305798 | 0.0833501 |
| MED19 | 556.360188 | -0.676928112 | 0.21176863 | -3.1965458 | 0.00139084 | 0.0101999 |
| TMX2 | 5493.76516 | 0.012753894 | 0.12871112 | 0.09908929 | 0.92106737 | 0.96154183 |
| TMX2-CTNNI | 1103.38533 | 0.363000975 | 0.14805874 | 2.45173624 | 0.01421688 | 0.05855055 |
| SELENOH | 3175.00681 | -0.217019035 | 0.12750931 | -1.7019858 | 0.08875803 | 0.21836494 |
| AP001931.1 | 1624.22941 | -0.143734315 | 0.13989869 | -1.0274171 | 0.30422409 | 0.50114876 |
| BTBD18 | 6.89721796 | 0.007691687 | 0.52620112 | 0.01461739 | 0.98833743 | 0.99471234 |
| CTNND1 | 15180.3091 | 0.142559536 | 0.11776954 | 1.21049586 | 0.22608868 | 0.41281666 |
| OR9Q1 | 1.8907619 | 0.091051944 | 1.38593266 | 0.06569724 | 0.94761887 | 0.97478875 |
| OR6Q1 | 0 | NA | NA | NA | NA | NA |
| AP004247.2 | 2.69953714 | 0.380108241 | 0.81599099 | 0.46582407 | 0.64134144 | 0.78758115 |
| OR9I1 | 0.39627446 | 2.574957812 | 3.21049141 | 0.80204476 | 0.42252707 | NA |
| OR9Q2 | 0.03392908 | 0.380740593 | 3.29738709 | 0.11546736 | 0.90807469 | NA |
| OR1S2 | 0 | NA | NA | NA | NA | NA |
| OR1S1 | 0 | NA | NA | NA | NA | NA |
| OR10Q1 | 0.230269 | -0.33961227 | 3.04796217 | -0.1114227 | 0.91128114 | NA |
| OR10W1 | 0 | NA | NA | NA | NA | NA |
| OR5B17 | 0 | NA | NA | NA | NA | NA |
| OR5B3 | 0 | NA | NA | NA | NA | NA |
| OR5B2 | 1.74141148 | 1.147803803 | 1.16392801 | 0.98614673 | 0.32406112 | 0.52163909 |
| OR5B12 | 0.98108136 | 0.348919458 | 1.21776869 | 0.28652359 | 0.77447713 | NA |
| AP003557.1 | 2.60335655 | 1.713122552 | 0.94619456 | 1.81053942 | 0.07021218 | 0.18463254 |
| OR5B21 | 0 | NA | NA | NA | NA | NA |
| LPXN | 770.371014 | 0.317304982 | 0.17239055 | 1.84061706 | 0.0656777 | 0.1761156 |
| ZFP91 | 3411.35468 | 0.032289436 | 0.20183969 | 0.15997565 | 0.87290026 | 0.93369487 |
| ZFP91-CNTF | 267.534701 | 1.254653041 | 0.44308737 | 2.83161544 | 0.00463135 | 0.02543819 |
| AP001350.1 | 58.7132321 | 1.526109469 | 0.49151794 | 3.10489066 | 0.00190349 | 0.01296697 |
| CNTF | 42.4961939 | -0.449957711 | 0.5999911 | -0.7499406 | 0.45329046 | 0.64296 |
| GLYAT | 7471.55442 | 2.452918494 | 0.6693598 | 3.66457399 | 0.00024775 | 0.00260286 |
| GLYATL2 | 17.3563177 | 1.10408174 | 0.54260916 | 2.03476429 | 0.0418746 | 0.12808601 |
| GLYATL1 | 8716.63354 | 2.314776297 | 0.53349989 | 4.33885058 | 1.43E-05 | 0.0002731 |
| AP001652.1 | 5.671967 | 4.530367941 | 1.04118757 | 4.35115445 | 1.35E-05 | 0.00026085 |
| AP001636.3 | 16.9121269 | 0.577311802 | 0.38375028 | 1.50439448 | 0.13247983 | 0.28877269 |

| | | | | | | |
|---|---|---|---|---|---|---|
| GLYATL1B | 0.66753917 | 1.383313281 | 2.62989358 | 0.52599592 | 0.59889105 | NA |
| FAM111B | 24.9049319 | -0.636395814 | 0.70789995 | -0.8989912 | 0.36865736 | 0.56547087 |
| FAM111A-D1 | 221.258657 | 0.183594053 | 0.26538673 | 0.69179817 | 0.48906408 | 0.67250456 |
| FAM111A | 2018.17776 | 0.26674929 | 0.28644869 | 0.93122887 | 0.35173519 | 0.5493844 |
| DTX4 | 2072.36434 | -0.212690838 | 0.34016251 | -0.6252624 | 0.53179883 | 0.70604849 |
| MPEG1 | 950.777002 | 0.967114425 | 0.47089308 | 2.05378773 | 0.03999624 | 0.12382171 |
| AP002358.1 | 0.9692694 | 0.555883906 | 1.71780659 | 0.32360099 | 0.74624014 | NA |
| OR5AN1 | 0.26780428 | 2.026828142 | 3.24034724 | 0.6254972 | 0.53164478 | NA |
| OR5A2 | 0.09644756 | -0.642905668 | 3.30558948 | -0.1944905 | 0.84579185 | NA |
| OR5A1 | 0.07659028 | -0.642905668 | 3.30558948 | -0.1944905 | 0.84579185 | NA |
| OR4D6 | 0 | NA | NA | NA | NA | NA |
| OR4D10 | 0 | NA | NA | NA | NA | NA |
| OR4D11 | 0 | NA | NA | NA | NA | NA |
| OR4D9 | 0.19747483 | -0.340095337 | 3.16269738 | -0.1075333 | 0.9143659 | NA |
| AP000442.1 | 7.77394348 | -0.53417671 | 0.53634863 | -0.9959505 | 0.31927418 | 0.51667751 |
| OSBP | 5254.01774 | 0.178943257 | 0.15123722 | 1.18319587 | 0.23673153 | 0.42601653 |
| AP000442.2 | 296.357995 | -0.550986038 | 0.32748325 | -1.6824862 | 0.0924746 | 0.22485526 |
| PATL1 | 3495.35648 | -0.815196013 | 0.28897993 | -2.8209434 | 0.00478827 | 0.02601825 |
| OR10V1 | 0 | NA | NA | NA | NA | NA |
| STX3 | 7759.44059 | -0.675576384 | 0.54248698 | -1.245332 | 0.21300975 | 0.39688173 |
| AP000640.1 | 28.8384531 | 1.344133218 | 0.58439785 | 2.30003109 | 0.02144646 | 0.07893135 |
| MRPL16 | 2231.14381 | 0.426842855 | 0.13630982 | 3.13141677 | 0.00173965 | 0.01212331 |
| GIF | 9.22854274 | 1.349144698 | 0.61916178 | 2.17898576 | 0.02933273 | 0.09926899 |
| TCN1 | 52.1442184 | -2.223802828 | 0.53682923 | -4.1424772 | 3.44E-05 | 0.00054769 |
| AP002347.1 | 7.82763884 | 1.010974375 | 0.57677543 | 1.75280417 | 0.07963563 | 0.201811 |
| AP000790.3 | 9.3906617 | 1.147670973 | 0.616861 | 1.86050176 | 0.06281457 | 0.17071453 |
| AP000790.4 | 15.3591347 | 1.310060066 | 0.71301015 | 1.83736525 | 0.06615599 | 0.17702015 |
| AP000790.2 | 0.2999234 | 1.145957498 | 3.27645355 | 0.34975545 | 0.72652223 | NA |
| OOSP2 | 0 | NA | NA | NA | NA | NA |
| MS4A3 | 16.4089019 | -3.827583416 | 0.93798422 | -4.080648 | 4.49E-05 | 0.00068102 |
| MS4A2 | 32.4547994 | 1.000985891 | 0.89551635 | 1.11777511 | 0.26366305 | 0.4574508 |
| AP001257.1 | 2.42538639 | 1.041940495 | 1.17850816 | 0.88411818 | 0.37663242 | 0.57253686 |
| MS4A6A | 1312.08453 | -0.692853127 | 0.53005778 | -1.3071275 | 0.1911694 | 0.37023627 |
| MS4A4E | 54.4511413 | -0.22162415 | 0.61729447 | -0.359025 | 0.71957639 | 0.840362 |
| MS4A4A | 755.56933 | -1.393130159 | 0.47695907 | -2.920859 | 0.00349068 | 0.02052836 |
| MS4A6E | 4.97815795 | -2.347705105 | 0.8526266 | -2.7534974 | 0.00589622 | 0.03040349 |
| MS4A7 | 911.481728 | -0.616602504 | 0.43156719 | -1.428752 | 0.15307551 | 0.31857242 |
| MS4A14 | 191.646287 | -0.714125425 | 0.54341485 | -1.3141441 | 0.18879771 | 0.36701774 |
| MS4A5 | 0.15452186 | 0.435199075 | 3.28898345 | 0.13232024 | 0.894731 | NA |
| MS4A1 | 249.449584 | 0.795180503 | 0.94334344 | 0.8429385 | 0.39926284 | 0.59406202 |
| MS4A12 | 0.03392908 | 0.380740593 | 3.29738709 | 0.11546736 | 0.90807469 | NA |
| MS4A13 | 0 | NA | NA | NA | NA | NA |
| MS4A8 | 0.98112266 | 1.728094881 | 1.45800411 | 1.18524692 | 0.23591983 | NA |
| MS4A18 | 0 | NA | NA | NA | NA | NA |
| MS4A15 | 0 | NA | NA | NA | NA | NA |
| MS4A10 | 30.8585168 | 4.355210603 | 1.03397647 | 4.21209837 | 2.53E-05 | 0.00042933 |
| AP000777.1 | 0 | NA | NA | NA | NA | NA |
| AP000777.2 | 346.492668 | -0.568041713 | 0.25019614 | -2.2703856 | 0.0231842 | 0.08366872 |
| AP000777.3 | 335.816109 | -0.903945627 | 0.22979143 | -3.9337657 | 8.36E-05 | 0.00110209 |
| CCDC86 | 1332.51671 | -0.678707461 | 0.23761504 | -2.856332 | 0.00428567 | 0.02398918 |
| PTGDR2 | 166.67839 | -0.464193653 | 0.16339018 | -2.8410132 | 0.00449705 | 0.02486624 |
| ZP1 | 7.22217263 | -2.856793473 | 0.96507053 | -2.9601914 | 0.00307448 | 0.01871435 |
| PRPF19 | 4781.0128 | -0.386775182 | 0.20421892 | -1.8939243 | 0.05823506 | 0.16207752 |
| AP003721.4 | 17.7075765 | 1.789450068 | 0.60241513 | 2.97046003 | 0.00297354 | 0.01820578 |
| AP003721.3 | 38.3563469 | 0.683610263 | 0.4057559 | 1.68478206 | 0.09203063 | 0.22423059 |
| TMEM109 | 5605.58192 | 0.042198333 | 0.13769003 | 0.3064734 | 0.75924423 | 0.86663046 |
| AP003721.1 | 13.4191986 | -0.817533633 | 0.5403664 | -1.5129246 | 0.13029881 | 0.28547571 |
| TMEM132A | 1539.93306 | -1.505586612 | 0.37582355 | -4.0060997 | 6.17E-05 | 0.00086836 |
| SLC15A3 | 1290.25062 | -1.325702548 | 0.58446385 | -2.268237 | 0.02331476 | 0.08391024 |
| CD6 | 213.305983 | 0.044822715 | 0.68611595 | 0.06532819 | 0.9479127 | 0.97493887 |
| AP003721.2 | 0.45481756 | 2.105662906 | 2.67749569 | 0.78642999 | 0.43161562 | NA |
| CD5 | 149.745039 | 0.127473676 | 0.72571412 | 0.17565274 | 0.86056676 | 0.92734057 |
| VPS37C | 2129.23465 | -0.331870166 | 0.14548225 | -2.281173 | 0.02253821 | 0.08187529 |
| PGA3 | 0.39675137 | -1.918645508 | 2.76069497 | -0.6949864 | 0.48706381 | NA |
| PGA4 | 0.08666371 | -0.642905668 | 3.30558948 | -0.1944905 | 0.84579185 | NA |
| PGA5 | 4.49714434 | 1.474774019 | 0.70398623 | 2.09489044 | 0.03618073 | 0.11525343 |
| VWCE | 133.798465 | -0.062352137 | 0.30564506 | -0.2040018 | 0.83835211 | 0.91449053 |
| DDB1 | 11103.2096 | 0.396120098 | 0.15784003 | 2.50963011 | 0.01208577 | 0.05182142 |
| TKFC | 6478.71513 | 1.715869825 | 0.27184134 | 6.31202688 | 2.75E-10 | 3.46E-08 |
| CYB561A3 | 2389.12696 | 0.315061754 | 0.15353747 | 2.05201867 | 0.04016785 | 0.12414746 |
| TMEM138 | 1734.19156 | 0.045313861 | 0.13420425 | 0.33764849 | 0.7356281 | 0.85152157 |
| TMEM216 | 555.658456 | -0.247507485 | 0.13568955 | -1.8240718 | 0.06814122 | 0.18060951 |
| CPSF7 | 5787.20543 | 0.062508704 | 0.1625983 | 0.38443639 | 0.70065503 | 0.82842894 |
| AP003108.2 | 1609.71008 | -0.141744639 | 0.17484948 | -0.8106666 | 0.41755714 | 0.61036996 |
| SDHAF2 | 1682.49957 | -0.400770025 | 0.19546447 | -2.0503472 | 0.04033056 | 0.12451901 |
| PPP1R32 | 492.649549 | 0.123679245 | 0.29176959 | 0.42389354 | 0.67164345 | 0.80832284 |
| AP003108.1 | 2.10720104 | -2.437715666 | 2.04818429 | -1.1901838 | 0.23397418 | 0.42254485 |
| LRRC10B | 131.089575 | -1.515643484 | 0.57122995 | -2.6532983 | 0.00797094 | 0.03806477 |
| SYT7 | 1251.61297 | 1.21513536 | 0.34927949 | 3.47897709 | 0.00050333 | 0.00455329 |
| AP003559.1 | 14.4047296 | 0.092557029 | 0.44616804 | 0.20744881 | 0.83565937 | 0.91280681 |
| AP002754.1 | 3.06205073 | 1.817933044 | 1.03563534 | 1.7553795 | 0.07919442 | 0.20107062 |
| AP002754.2 | 0.15318056 | -1.003574173 | 3.30558948 | -0.3035992 | 0.7614333 | NA |
| DAGLA | 598.508004 | 0.386863858 | 0.28646623 | 1.35046932 | 0.17686549 | 0.35183637 |
| DKFZP434K0 | 107.493023 | -0.404249019 | 0.2971971 | -1.3602051 | 0.17376502 | 0.3476363 |
| MYRF | 3239.86983 | -0.720645934 | 0.34154639 | -2.1099504 | 0.03486263 | 0.11239245 |
| TMEM258 | 2014.22878 | -0.510310401 | 0.13559312 | -3.7635419 | 0.00016752 | 0.00190496 |
| FEN1 | 615.045412 | -0.355439264 | 0.33820093 | -1.0509707 | 0.29327205 | 0.4897734 |
| FADS2 | 2300.46375 | -0.352649858 | 0.42891979 | -0.8221814 | 0.41097369 | 0.60530474 |
| FADS1 | 2216.26643 | -1.277323799 | 0.525198 | -2.4320805 | 0.01501237 | 0.06091231 |
| FADS3 | 5360.7333 | -1.69898637 | 0.33647596 | -5.0493544 | 4.43E-07 | 1.64E-05 |
| RAB3IL1 | 1903.02293 | -0.177665989 | 0.34265509 | -0.518498 | 0.60411083 | 0.761258 |
| BEST1 | 172960.444 | -1.123429606 | 0.37226546 | -3.0178186 | 0.00254601 | 0.01621845 |
| FTH1 | 195105.032 | -1.125867533 | 0.37018066 | -3.0414002 | 0.00235481 | 0.01532105 |
| AP003733.3 | 15.4557702 | -3.321814026 | 0.96696148 | -3.4353116 | 0.00059187 | 0.00516701 |
| INCENP | 428.708485 | 0.006988098 | 0.23475967 | 0.02976703 | 0.97625285 | 0.98894663 |
| SCGB1D1 | 0 | NA | NA | NA | NA | NA |
| SCGB2A1 | 24.707388 | -0.247017622 | 0.54016344 | -0.4573016 | 0.64745426 | 0.79245155 |
| AP003306.1 | 0.09989236 | -0.642905668 | 3.30558948 | -0.1944905 | 0.84579185 | NA |
| SCGB1D2 | 67.7196891 | 1.363748574 | 0.49978286 | 2.72868215 | 0.0063588 | 0.03216369 |
| SCGB2A2 | 0.68206825 | -0.130839325 | 2.7285063 | -0.0479527 | 0.96175392 | NA |
| SCGB1D4 | 0.03392908 | 0.380740593 | 3.29738709 | 0.11546736 | 0.90807469 | NA |
| AP003306.2 | 549.873046 | 0.578678423 | 0.39418653 | 1.46803196 | 0.14209554 | 0.30304307 |
| ASRGL1 | 4281.36088 | 0.210986671 | 0.39386743 | 0.53567941 | 0.59218014 | 0.75166635 |
| AP003064.1 | 1989.42559 | 0.104601119 | 0.41516204 | 0.25195251 | 0.80107777 | 0.89215393 |
| SCGB1A1 | 1.95599501 | 0.451679902 | 0.90623333 | 0.49841458 | 0.61819186 | 0.77175209 |
| AP003064.2 | 4.14615494 | 0.910796292 | 0.66005589 | 1.37987752 | 0.16762436 | 0.33884663 |
| AHNAK | 21530.0953 | 0.243821432 | 0.19123921 | 1.2749552 | 0.2023251 | 0.38413945 |
| AP001363.1 | 1.11037917 | -0.495388445 | 1.2025178 | -0.4119593 | 0.68036922 | 0.8146636 |
| AP001363.2 | 9.23665154 | 0.25593156 | 0.43896584 | 0.58303298 | 0.5598711 | 0.72824944 |
| EEF1G | 52108.2526 | -0.080612257 | 0.23170887 | -0.3479032 | 0.72791288 | 0.84611931 |
| AP002990.1 | 53485.4093 | -0.069064478 | 0.22356638 | -0.3089216 | 0.75738118 | 0.86579425 |
| TUT1 | 2242.16973 | 0.188256155 | 0.20645209 | 0.91186364 | 0.3618405 | 0.55919679 |
| MTA2 | 4524.6145 | -0.614913855 | 0.1960559 | -3.136421 | 0.00171023 | 0.01194989 |

| | | | | | | |
|---|---|---|---|---|---|---|
| EML3 | 4800.97294 | -0.682720717 | 0.21582002 | -3.1633799 | 0.00155949 | 0.01113859 |
| AP001458.1 | 384.607841 | -0.518169748 | 0.21795469 | -2.3774195 | 0.01743425 | 0.06782683 |
| ROM1 | 462.631617 | -0.664603159 | 0.17367071 | -3.8268005 | 0.00012982 | 0.00155395 |
| B3GAT3 | 3169.17738 | -0.745078967 | 0.13380419 | -5.5684278 | 2.57E-08 | 1.54E-06 |
| GANAB | 18182.3329 | 0.059470994 | 0.12103516 | 0.49135304 | 0.62317677 | 0.77531577 |
| INTS5 | 1471.94374 | -0.088996285 | 0.13496179 | -0.6594184 | 0.50962714 | 0.6883608 |
| AP001458.2 | 1270.34406 | -0.386141355 | 0.14458125 | -2.6707568 | 0.00756805 | 0.03668555 |
| C11orf98 | 1535.05007 | -0.565278912 | 0.1480105 | -3.8191812 | 0.0001339 | 0.00159244 |
| LBHD1 | 2476.40069 | -0.347667898 | 0.12805075 | -2.715079 | 0.006626 | 0.03318031 |
| CSKMT | 328.980016 | -0.358496485 | 0.20143375 | -1.779724 | 0.07512114 | 0.19392506 |
| UQCC3 | 992.430637 | -0.072225744 | 0.1587296 | -0.4550238 | 0.64909212 | 0.79349874 |
| UBXN1 | 4366.31403 | -0.319381751 | 0.12229367 | -2.6115967 | 0.00901205 | 0.04171454 |
| LRRN4CL | 63.6056464 | -0.609440515 | 0.66948586 | -0.9103113 | 0.36265839 | 0.55991227 |
| BSCL2 | 4769.21934 | 0.197923223 | 0.12476872 | 1.58632086 | 0.11266654 | 0.25832775 |
| HNRNPUL2-E | 9969.65748 | 0.175065466 | 0.06794713 | 2.5764955 | 0.00998075 | 0.04501324 |
| GNG3 | 122.800884 | 0.680845653 | 0.14459138 | 4.70875693 | 2.49E-06 | 6.78E-05 |
| HNRNPUL2 | 7712.93766 | 0.120887735 | 0.09894251 | 1.22179771 | 0.22178413 | 0.40772969 |
| TTC9C | 931.975669 | -0.372880749 | 0.13270256 | -2.8098987 | 0.00495571 | 0.02669159 |
| ZBTB3 | 226.941888 | 0.516165794 | 0.21315728 | 2.42152551 | 0.01545552 | 0.0622137 |
| POLR2G | 2007.50163 | -0.264285106 | 0.16656449 | -1.5866834 | 0.11258438 | 0.25822824 |
| AP001160.2 | 128.579156 | 0.440638584 | 0.24779462 | 1.7782411 | 0.07536426 | 0.19438142 |
| TAF6L | 1648.28875 | 0.196426216 | 0.16439168 | 1.19486713 | 0.23213895 | 0.42019115 |
| TMEM223 | 2482.73253 | -0.321627847 | 0.10991009 | -2.9262813 | 0.00343041 | 0.02030067 |
| AP001160.3 | 589.3271 | 0.123627404 | 0.17902362 | 0.69056475 | 0.48983911 | 0.67311398 |
| TMEM179B | 2643.83633 | -0.334763133 | 0.13297115 | -2.5175621 | 0.01181701 | 0.05100911 |
| NXF1 | 7981.2424 | -0.272176018 | 0.23227032 | -1.1718071 | 0.2412745 | 0.43140822 |
| STX5 | 2417.02837 | -0.397444828 | 0.09414321 | -4.2217047 | 2.42E-05 | 0.00041545 |
| AP001160.1 | 829.723308 | -0.796299392 | 0.1973116 | -4.0357455 | 5.44E-05 | 0.0007952 |
| TEX54 | 5.62946731 | 0.68547855 | 0.48873442 | 1.40255837 | 0.16074857 | 0.32917177 |
| WDR74 | 1948.5978 | -0.598123506 | 0.18199094 | -3.2865564 | 0.0010142 | 0.00792902 |
| SLC3A2 | 20935.1 | -0.161509614 | 0.48385241 | -0.3337993 | 0.73853098 | 0.85323382 |
| CHRM1 | 25.4987932 | 1.366070019 | 0.6528899 | 2.09234361 | 0.03640779 | 0.11575517 |
| AP000438.1 | 16.2048404 | 1.902356026 | 0.75326613 | 2.52547666 | 0.01155415 | 0.05017833 |
| SLC22A6 | 19921.8232 | 1.925116995 | 0.62401781 | 3.08503536 | 0.00203528 | 0.0136775 |
| SLC22A8 | 20018.8764 | 2.434739379 | 0.87939952 | 2.76863852 | 0.00562911 | 0.02945813 |
| SLC22A24 | 105.072094 | 1.465464314 | 0.54486201 | 2.68960632 | 0.00715364 | 0.03520742 |
| SLC22A10 | 7.69312049 | 2.021885258 | 0.63420303 | 3.18807255 | 0.00143225 | 0.01043612 |
| SLC22A25 | 1.05253139 | 1.431590635 | 1.13364744 | 1.26281821 | 0.20665452 | 0.38929725 |
| SLC22A9 | 8.3682419 | 2.490767122 | 0.7123077 | 3.49675725 | 0.00047095 | 0.00433018 |
| HRASLS5 | 112.619471 | 2.243107965 | 0.64694911 | 3.4672093 | 0.00052589 | 0.00471393 |
| LGALS12 | 22.2035985 | 0.329971929 | 0.73759679 | 0.44736085 | 0.65461454 | 0.79736641 |
| RARRES3 | 5947.38241 | -1.162771457 | 0.4271945 | -2.7218784 | 0.0064912 | 0.03266543 |
| HRASLS2 | 508.643352 | 1.857027838 | 0.73825281 | 2.5154362 | 0.01188852 | 0.05120299 |
| PLA2G16 | 4505.91139 | -0.313066943 | 0.30476599 | -1.0272371 | 0.30430881 | 0.5012257 |
| AP000753.1 | 0.97670292 | -0.280873773 | 1.71568596 | -0.1637093 | 0.86995998 | NA |
| ATL3 | 3893.72609 | -0.478747684 | 0.18743077 | -2.5542641 | 0.01064125 | 0.04714546 |
| AP000753.2 | 77.2307152 | -1.340651797 | 0.39852024 | -3.3640745 | 0.00076801 | 0.00636067 |
| RTN3 | 7792.35389 | -0.163993612 | 0.16141027 | -1.0160048 | 0.30962708 | 0.50688268 |
| C11orf95 | 1819.54726 | 0.456417914 | 0.28408442 | 1.60662775 | 0.10813605 | 0.25086194 |
| SPINDOC | 1024.69368 | -0.881755566 | 0.16745876 | -5.2655088 | 1.40E-07 | 6.35E-06 |
| MARK2 | 3160.13525 | -0.443243057 | 0.13556799 | -3.2695259 | 0.00107728 | 0.00830882 |
| RCOR2 | 51.0680355 | 0.653884433 | 0.49571169 | 1.31908213 | 0.18714166 | 0.36506328 |
| NAA40 | 2484.62741 | 0.006455742 | 0.20983656 | 0.03076557 | 0.9754565 | 0.98865354 |
| COX8A | 9199.08964 | 0.310170437 | 0.12557206 | 2.47005923 | 0.01350907 | 0.05634833 |
| AP000721.1 | 7107.62907 | 0.339254578 | 0.12698288 | 2.67165601 | 0.0075478 | 0.03661541 |
| OTUB1 | 5152.41876 | -0.250121532 | 0.13001584 | -1.9237774 | 0.0543825 | 0.15446118 |
| MACROD1 | 2017.89336 | 0.788687729 | 0.19900149 | 3.96322518 | 7.39E-05 | 0.00099739 |
| AP000721.2 | 19.716866 | 2.436577572 | 0.3868001 | 6.2993199 | 2.99E-10 | 3.70E-08 |
| AP006333.2 | 7.89739899 | 2.006985081 | 0.83760431 | 2.39610167 | 0.01657049 | 0.06544867 |
| FLRT1 | 239.635142 | 2.2048401 | 0.39704532 | 5.55311942 | 2.81E-08 | 1.65E-06 |
| AP006333.1 | 56.1139231 | 2.061277828 | 0.52644713 | 3.9154508 | 9.02E-05 | 0.001167 |
| STIP1 | 5106.12608 | -0.46940089 | 0.15192284 | -3.0897321 | 0.00200337 | 0.0135216 |
| FERMT3 | 986.63221 | -0.736628293 | 0.33243045 | -2.215887 | 0.02669924 | 0.09273697 |
| TRPT1 | 2789.2457 | -0.287791888 | 0.12468844 | -2.3080879 | 0.02099425 | 0.07762455 |
| NUDT22 | 4191.92674 | -0.019220531 | 0.10452954 | -0.1838766 | 0.8541103 | 0.92360811 |
| AP001453.1 | 1555.40742 | -0.157770833 | 0.12957117 | -1.2176384 | 0.22336143 | 0.40943055 |
| DNAJC4 | 2440.17865 | 0.327677244 | 0.14776464 | 2.21756197 | 0.02658471 | 0.09242438 |
| VEGFB | 4107.92089 | 0.00115323 | 0.21903321 | 0.00526509 | 0.99579909 | 0.99795473 |
| FKBP2 | 5331.7678 | -0.191499326 | 0.15564137 | -1.2303883 | 0.21855173 | 0.40375572 |
| PPP1R14B | 2738.55649 | -1.409272602 | 0.23038027 | -6.1171583 | 9.53E-10 | 9.85E-08 |
| AP001453.2 | 216.749097 | -1.338744594 | 0.21986257 | -6.0890065 | 1.14E-09 | 1.15E-07 |
| AP001453.4 | 155.121364 | -1.317883387 | 0.24240509 | -5.4366985 | 5.43E-08 | 2.83E-06 |
| PLCB3 | 2187.79605 | -0.567516438 | 0.25282637 | -2.2446885 | 0.02478814 | 0.08773115 |
| BAD | 2481.61156 | -0.235159591 | 0.12302909 | -1.9114146 | 0.05595132 | 0.15756476 |
| GPR137 | 4225.81558 | -0.299425976 | 0.11940306 | -2.5076909 | 0.01215229 | 0.05202667 |
| KCNK4 | 4.17448322 | 0.454054197 | 0.6255551 | 0.72584205 | 0.46793561 | 0.65475231 |
| CATSPERZ | 14.2928188 | 0.353307195 | 0.46138495 | 0.76575362 | 0.44382292 | 0.6348089 |
| ESRRA | 7590.0108 | 0.039305437 | 0.18769062 | 0.20941609 | 0.83412343 | 0.91206315 |
| TRMT112 | 5596.54301 | -0.319938086 | 0.12165379 | -2.6299064 | 0.00854084 | 0.03996839 |
| PRDX5 | 12796.8453 | -0.131992548 | 0.15217327 | -0.8673832 | 0.38573206 | 0.58152559 |
| AP003774.4 | 4.72712997 | -0.132707197 | 0.75801824 | -0.1750712 | 0.86102365 | 0.92761342 |
| CCDC88B | 1126.24689 | -0.770354757 | 0.40330837 | -1.9100887 | 0.0561218 | 0.15787623 |
| RPS6KA4 | 2081.04566 | -0.596776925 | 0.17594352 | -3.3918664 | 0.00069418 | 0.00586342 |
| AP003774.3 | 7.13619142 | 2.645943375 | 0.98226902 | 2.6937054 | NA | NA |
| AP003774.2 | 0.10721287 | -0.642905668 | 3.30558948 | -0.1944905 | 0.84579185 | NA |
| AP003774.1 | 9.06621192 | 1.432426836 | 0.66354137 | 2.15876039 | 0.03086876 | 0.10296094 |
| AP005273.1 | 0.29665639 | 2.176798933 | 3.23107108 | 0.67370815 | 0.50049688 | NA |
| SLC22A11 | 8419.28321 | 0.857714813 | 0.42026844 | 2.0408737 | 0.04126338 | 0.1268052 |
| SLC22A12 | 26913.3994 | 1.18054444 | 0.49243955 | 2.39733878 | 0.01651465 | 0.06528125 |
| NRXN2 | 831.195551 | 0.351761969 | 0.31990868 | 1.09956995 | 0.27151954 | 0.4662046 |
| AP001092.1 | 2.96296091 | 0.032902077 | 0.75463189 | 0.04360017 | 0.96522312 | 0.98359669 |
| RASGRP2 | 1293.63668 | 0.133921626 | 0.41854034 | 0.31997304 | 0.74898876 | 0.8597841 |
| PYGM | 262.947125 | 0.084438445 | 0.41424621 | 0.20383637 | 0.83848137 | 0.91449887 |
| SF1 | 13793.7326 | -0.099434537 | 0.16261014 | -0.6114904 | 0.54087499 | 0.7138218 |
| AP001462.1 | 23.0749126 | 0.150172728 | 0.31485862 | 0.47695288 | 0.63339567 | 0.78254516 |
| MAP4K2 | 1315.72332 | 0.328260979 | 0.23951171 | 1.37054253 | 0.17051761 | 0.34262034 |
| MEN1 | 2887.29961 | -0.384264067 | 0.16623086 | -2.3116289 | 0.02079814 | 0.07708321 |
| CDC42BPG | 1670.67877 | 0.307602383 | 0.19038719 | 1.61566747 | 0.10616623 | 0.24748969 |
| EHD1 | 5549.05508 | 0.524871628 | 0.17350503 | 3.02510905 | 0.00248543 | 0.01592442 |
| MIR194-2HG | 198.38737 | 1.759791192 | 0.51164824 | 3.43945517 | 0.00058289 | 0.00511164 |
| ATG2A | 4476.41755 | -0.538080543 | 0.25507539 | -2.1094961 | 0.03490178 | 0.11244638 |
| PPP2R5B | 2377.02211 | -0.424947648 | 0.20582909 | -2.0645656 | 0.03896412 | 0.12151661 |
| GPHA2 | 23.9672312 | 2.265406917 | 0.43448171 | 5.21404439 | 1.85E-07 | 8.00E-06 |
| MAJIN | 63.8899981 | 2.383767356 | 0.56460911 | 4.22197824 | 2.42E-05 | 0.00041545 |
| BATF2 | 518.18815 | -0.282878104 | 0.17681181 | -1.5998825 | 0.10962466 | 0.25333471 |
| ARL2 | 3878.2074 | 0.137020896 | 0.1546697 | 0.8858936 | 0.37567487 | 0.57155421 |
| ARL2-SNX15 | 3811.80021 | 0.160859601 | 0.10448838 | 1.53949746 | 0.1236829 | 0.27523358 |
| SNX15 | 1788.99488 | 0.087609666 | 0.15038215 | 0.58258024 | 0.56017591 | 0.72838393 |
| SAC3D1 | 301.503308 | -0.353771496 | 0.17663679 | -2.0028189 | 0.04519673 | 0.13523633 |
| NAALADL1 | 220.938394 | -0.200659048 | 0.20929914 | -0.9587189 | 0.33770035 | 0.53568084 |
| CDCA5 | 97.5076666 | -1.025013831 | 0.46473213 | -2.2056014 | 0.02741192 | 0.09457705 |
| ZFPL1 | 1803.07703 | -0.104129459 | 0.15538089 | -0.6701561 | 0.50275827 | 0.68306118 |
| TMEM262 | 1664.71512 | -0.12368763 | 0.15147901 | -0.8165331 | 0.41419529 | 0.60794725 |

| | | | | | | |
|---|---|---|---|---|---|---|
| VPS51 | 5192.16814 | 0.207134829 | 0.1398746 | 1.48086087 | 0.13864365 | 0.29791665 |
| AP003068.1 | 1641.48255 | -0.036955536 | 0.13638299 | -0.2709688 | 0.78641502 | 0.88369898 |
| TM7SF2 | 1801.29346 | 0.933800728 | 0.24536473 | 3.80576596 | 0.00014137 | 0.00166644 |
| ZNHIT2 | 588.936794 | -0.364682366 | 0.15745324 | -2.3161312 | 0.02055111 | 0.07644699 |
| AP003068.3 | 199.813854 | -0.267534618 | 0.17096901 | -1.5648135 | 0.11762663 | 0.26621481 |
| FAU | 12989.5103 | -0.300814623 | 0.15560218 | -1.9332289 | 0.05320801 | 0.15217051 |
| SYVN1 | 12009.5524 | -0.234842264 | 0.1178125 | -1.993356 | 0.04622248 | 0.13754081 |
| MRPL49 | 5019.65854 | 0.109049544 | 0.09566338 | 1.13992989 | 0.25431551 | 0.44708166 |
| SPDYC | 3.28688246 | 0.692293976 | 0.77644458 | 0.89162059 | 0.37259633 | 0.56886089 |
| AP003068.2 | 344.57452 | -0.242767058 | 0.3035341 | -0.7998016 | 0.42382576 | 0.6163166 |
| CAPN1 | 11060.4869 | -0.404501553 | 0.13040965 | -3.1017763 | 0.00192363 | 0.01307713 |
| POLA2 | 557.832235 | -0.009349136 | 0.14480544 | -0.0645634 | 0.9485216 | 0.97514677 |
| CDC42EP2 | 1084.2155 | -0.966710097 | 0.20126327 | -4.8032118 | 1.56E-06 | 4.67E-05 |
| DPF2 | 2324.82247 | -0.188413318 | 0.11121064 | -1.6942023 | 0.09022685 | 0.22109924 |
| TIGD3 | 40.6826727 | -0.946439987 | 0.39006979 | -2.426335 | 0.01525218 | 0.06158323 |
| AP000944.1 | 94.0422252 | -0.181078076 | 0.30900932 | -0.5859955 | 0.55787853 | 0.7265177 |
| SLC25A45 | 1255.74869 | 0.188052305 | 0.33071494 | 0.56862356 | 0.56961163 | 0.73500743 |
| FRMD8 | 2637.85743 | -0.917738795 | 0.17532576 | -5.2344778 | 1.65E-07 | 7.30E-06 |
| NEAT1 | 157433.974 | 0.254633717 | 0.32541962 | 0.78247808 | 0.43393366 | 0.62592658 |
| AP000769.3 | 0.58168218 | 1.168974639 | 1.84166035 | 0.63473954 | 0.5255983 | NA |
| MALAT1 | 46457.5929 | -0.062840462 | 0.12664551 | -0.4961918 | 0.61975911 | 0.77271053 |
| AP000769.2 | 5.32303399 | 0.483195104 | 0.741374 | 0.6517562 | 0.51455846 | 0.69239713 |
| SCYL1 | 6218.24862 | -0.289896662 | 0.15698575 | -1.8466431 | 0.06479888 | 0.17445089 |
| LTBP3 | 8605.85367 | 0.131521761 | 0.23797608 | 0.55266798 | 0.58049078 | 0.74348156 |
| SSSCA1 | 2454.88828 | -0.536894981 | 0.21048135 | -2.550796 | 0.01074772 | 0.0475373 |
| FAM89B | 1551.44097 | -0.678155979 | 0.2401245 | -2.8241848 | 0.0047401 | 0.02583641 |
| EHBP1L1 | 4508.08829 | -0.743871342 | 0.31131439 | -2.3894538 | 0.01687345 | 0.06634155 |
| KCNK7 | 25.9407036 | -0.62530356 | 0.38833183 | -1.61023 | 0.10734765 | 0.24951587 |
| MAP3K11 | 5897.25142 | -0.231090668 | 0.10921841 | -2.1158582 | 0.03435687 | 0.11115086 |
| PCNX3 | 4351.5071 | -0.10633061 | 0.12413083 | -0.8566011 | 0.39166536 | 0.58736439 |
| SIPA1 | 3191.61241 | -0.981432383 | 0.2809438 | -3.4933406 | 0.00047702 | 0.00437309 |
| RELA | 6181.78608 | -0.790947551 | 0.19834479 | -3.9877405 | 6.67E-05 | 0.00092629 |
| KAT5 | 4058.51085 | 0.085120334 | 0.17482733 | 0.4868823 | 0.62634174 | 0.7776192 |
| RNASEH2C | 5063.42997 | 0.039152009 | 0.11019729 | 0.35529012 | 0.72237226 | 0.84246413 |
| AP001266.2 | 3.18481985 | 0.666734122 | 0.8449176 | 0.78911141 | 0.43004689 | 0.62207863 |
| AP5B1 | 1335.33333 | 0.101169619 | 0.25816137 | 0.3918852 | 0.69514304 | 0.82446857 |
| OVOL1 | 817.855974 | 0.292226189 | 0.27449936 | 1.06457877 | 0.28706659 | 0.48334532 |
| OVOL1-AS1 | 50.7701962 | 2.191691512 | 0.55083759 | 3.97883431 | 6.93E-05 | 0.00094852 |
| AP001266.1 | 385.636485 | -0.082842953 | 0.25878631 | -0.3201211 | 0.74887654 | 0.8597841 |
| CFL1 | 27999.5888 | -0.666567732 | 0.20229491 | -3.2950297 | 0.00098411 | 0.00775353 |
| SNX32 | 8074.3231 | -0.472576122 | 0.20845885 | -2.2669996 | 0.02339025 | 0.08409011 |
| MUS81 | 3146.80355 | -0.4518506 | 0.13726892 | -3.2917183 | 0.00099577 | 0.00781039 |
| EFEMP2 | 3664.05145 | -0.189517866 | 0.24202111 | -0.7830634 | 0.43358989 | 0.62558268 |
| CTSW | 178.980353 | -0.321019768 | 0.64976372 | -0.4940562 | 0.62126652 | 0.77368586 |
| FIBP | 2463.51308 | 0.004925339 | 0.11465332 | 0.04295854 | 0.96573459 | 0.98387257 |
| CCDC85B | 815.062434 | -1.170264021 | 0.36823183 | -3.1780632 | 0.00148262 | 0.01071145 |
| FOSL1 | 355.97028 | -1.899502377 | 0.68386261 | -2.7776082 | NA | NA |
| C11orf68 | 1861.90808 | -0.111650648 | 0.12964426 | -0.8612078 | 0.3891236 | 0.58468663 |
| DRAP1 | 2511.71886 | -0.606050975 | 0.13708418 | -4.4210133 | 9.82E-06 | 0.00020051 |
| TSGA10IP | 2.10903426 | 0.19370112 | 1.02904088 | 0.18823462 | 0.85069273 | 0.92161226 |
| SART1 | 3212.26415 | -0.211494381 | 0.14251074 | -1.4840592 | 0.13779321 | 0.29654836 |
| EIF1AD | 1147.9158 | -0.391623448 | 0.18582671 | -2.1074659 | 0.03507722 | 0.11285825 |
| BANF1 | 3275.41599 | -0.404754117 | 0.15371491 | -2.633148 | 0.00845975 | 0.03969013 |
| CST6 | 69.7974289 | -2.590181445 | 0.744284 | -3.4800983 | 0.00050123 | 0.004542 |
| CATSPER1 | 19.4787569 | -0.455681726 | 0.47673865 | -0.9558313 | 0.33915746 | 0.53727066 |
| GAL3ST3 | 151.367404 | 1.899629911 | 0.45936507 | 4.13533818 | 3.54E-05 | 0.0005627 |
| SF3B2 | 10639.0385 | -0.220649286 | 0.10316224 | -2.1388571 | 0.03244724 | 0.10660774 |
| AP006287.2 | 63.9593026 | -0.184664285 | 0.23674432 | -0.7800157 | 0.43538164 | 0.62748068 |
| PACS1 | 4933.45202 | -0.536728548 | 0.11195552 | -4.7941231 | 1.63E-06 | 4.86E-05 |
| AP000759.1 | 70.7653373 | 0.265469968 | 0.27960415 | 0.94944933 | 0.34239213 | 0.54016635 |
| KLC2 | 3283.26706 | -0.627870075 | 0.19073046 | -3.2919235 | 0.00099505 | 0.00780702 |
| AP001107.4 | 385.303245 | -0.198349374 | 0.18283091 | -1.0848788 | 0.27797534 | 0.47353948 |
| AP001107.8 | 421.294798 | -0.426382321 | 0.1877981 | -2.2704293 | 0.02318155 | 0.08366872 |
| AP001107.7 | 103.686018 | -1.157863431 | 0.24642682 | -4.6986096 | 2.62E-06 | 7.05E-05 |
| RAB1B | 11876.1203 | -0.212041956 | 0.12768538 | -1.6606597 | 0.09678182 | 0.23220239 |
| AP001107.1 | 2506.79838 | 0.023894348 | 0.16447549 | 0.14527604 | 0.88449293 | 0.94018376 |
| AP001107.2 | 5805.37561 | -0.359318899 | 0.11726187 | -3.0642434 | 0.00218221 | 0.01443677 |
| CNIH2 | 56.1287595 | -0.330689466 | 0.26766448 | -1.2354626 | 0.21665838 | 0.40152375 |
| YIF1A | 3206.64185 | -0.272969362 | 0.16870174 | -1.6180589 | 0.1056499 | 0.24665673 |
| TMEM151A | 9.41820909 | -1.612140951 | 0.49684391 | -3.2447634 | 0.00117548 | 0.00890732 |
| AP001107.3 | 1.14083627 | 2.215751341 | 1.39946251 | 1.58328739 | 0.11335598 | 0.25938885 |
| CD248 | 1073.43628 | 0.833549716 | 0.51869741 | 1.60700573 | 0.10805311 | 0.2507357 |
| RIN1 | 405.228849 | -0.543489768 | 0.46130684 | -1.1781524 | 0.23873585 | 0.42832116 |
| AP001107.6 | 56.1714397 | -0.413006349 | 0.40669255 | -1.0155247 | 0.30985576 | 0.50697377 |
| BRMS1 | 2399.64429 | -0.261341612 | 0.18898847 | -1.3828442 | 0.16671264 | 0.33757294 |
| B4GAT1 | 2095.44571 | 0.670023698 | 0.220567 | 3.03773314 | 0.00238365 | 0.01543359 |
| AP001107.9 | 472.028152 | 0.149078093 | 0.26855407 | 0.55511389 | 0.57881676 | 0.74234707 |
| SLC29A2 | 1680.76777 | 0.038894452 | 0.26856974 | 0.14482068 | 0.88485245 | 0.94039898 |
| NPAS4 | 12.4267169 | 1.122107083 | 0.70025615 | 1.60242374 | 0.10906194 | 0.25241033 |
| MRPL11 | 1392.34217 | -0.007080714 | 0.13782392 | -0.0513751 | 0.95902665 | 0.98025244 |
| PELI3 | 718.927994 | -0.163248689 | 0.18422951 | -0.8861158 | 0.37555511 | 0.57149219 |
| AP002748.3 | 266.953391 | 0.261064753 | 0.20289564 | 1.28669477 | 0.1982007 | 0.37914779 |
| DPP3 | 3067.60739 | -0.402604434 | 0.19073645 | -2.1107892 | 0.03479043 | 0.11219564 |
| AP002748.4 | 4600.77834 | 0.643927044 | 0.19408482 | 3.31776091 | 0.00090742 | 0.00727316 |
| BBS1 | 4192.41858 | 0.773058595 | 0.24276706 | 3.18436356 | 0.00145073 | 0.01053875 |
| ZDHHC24 | 1817.15645 | 0.111303803 | 0.13047491 | 0.85306672 | 0.3936223 | 0.58894535 |
| ACTN3 | 10.8944044 | -0.179540914 | 0.53360979 | -0.3364648 | 0.73652039 | 0.85207763 |
| AP002748.1 | 1.83883886 | 1.325893241 | 0.95770858 | 1.38444332 | 0.16622275 | 0.33694319 |
| CTSF | 4286.62519 | 0.630895329 | 0.2555695 | 2.46858617 | 0.0135648 | 0.05650035 |
| CCDC87 | 40.1926971 | 0.74833878 | 0.26597372 | 2.81358164 | 0.00489929 | 0.02647426 |
| CCS | 2901.13721 | 0.093448885 | 0.20153227 | 0.46369192 | 0.64286848 | 0.78866908 |
| RBM14 | 3227.55372 | -0.11527944 | 0.09737687 | -1.1838483 | 0.23647313 | 0.42576789 |
| RBM14-RBM | 4571.44565 | -0.291098749 | 0.14658446 | -1.9858773 | 0.04704695 | 0.13931171 |
| RBM4 | 4912.91678 | -0.256952899 | 0.14038025 | -1.8304064 | 0.06718919 | 0.17889529 |
| RBM4B | 1213.87003 | 0.143928539 | 0.13553775 | 1.06190742 | 0.28827772 | 0.48438151 |
| AP001157.1 | 802.711259 | 0.197021038 | 0.16992893 | 1.15943198 | 0.24628015 | 0.43768753 |
| SPTBN2 | 5766.83179 | 0.429168869 | 0.31450709 | 1.36457612 | 0.17238632 | 0.34545384 |
| C11orf80 | 1386.53403 | -0.454884536 | 0.29019047 | -1.5675378 | 0.11698904 | 0.26531355 |
| RCE1 | 1955.6617 | -0.733157066 | 0.18145499 | -4.0404349 | 5.34E-05 | 0.00078422 |
| PC | 8803.6187 | 0.588406103 | 0.52257483 | 1.12597482 | 0.26017618 | 0.45319016 |
| LRFN4 | 715.166048 | -0.863237882 | 0.26208758 | -3.2937001 | 0.00098878 | 0.0077694 |
| C11orf86 | 273.936365 | -2.632553957 | 1.4589 | -1.8044787 | 0.0711563 | 0.18653858 |
| SYT12 | 80.3444471 | -1.373351235 | 0.63055072 | -2.1780187 | 0.02940465 | 0.09943583 |
| RHOD | 987.742279 | -0.816121956 | 0.2616004 | -3.1197275 | 0.00181018 | 0.01247019 |
| KDM2A | 5436.64024 | -0.159246614 | 0.14550552 | -1.0944369 | 0.27376339 | 0.46881316 |
| GRK2 | 6941.56109 | -0.76944404 | 0.21038045 | -3.6573932 | 0.00025479 | 0.00265751 |
| ANKRD13D | 2889.37888 | -0.969341531 | 0.25880945 | -3.7453869 | 0.00018012 | 0.00201171 |
| SSH3 | 3384.68417 | -0.791135326 | 0.19696968 | -4.0165336 | 5.91E-05 | 0.0008414 |
| RAD9A | 3088.35094 | -0.565467637 | 0.15441888 | -3.6619073 | 0.00025034 | 0.0026228 |
| POLD4 | 5184.40695 | -0.649287507 | 0.19991966 | -3.2477421 | 0.00116325 | 0.00883746 |
| AP003419.1 | 4489.39741 | -0.648084556 | 0.21018112 | -3.0834575 | 0.0020461 | 0.0137257 |
| AP003419.2 | 10.0277916 | -0.09935186 | 0.48548093 | -0.2046463 | 0.8378485 | 0.91418674 |
| CLCF1 | 1247.61218 | -0.640793422 | 0.28658161 | -2.2359893 | 0.02535247 | 0.08925829 |

| | | | | | | |
|---|---|---|---|---|---|---|
| AP003419.3 | 59.3930046 | -0.261861743 | 0.38314858 | -0.683447 | 0.49432443 | 0.67673841 |
| PPP1CA | 8043.78004 | -0.241462952 | 0.13052315 | -1.8499626 | 0.06431894 | 0.17349533 |
| TBC1D10C | 505.14462 | 0.163675222 | 0.48809863 | 0.33533227 | 0.73737446 | 0.85252584 |
| CARNS1 | 134.944888 | 0.596793262 | 0.24790535 | 2.40734321 | 0.01606906 | 0.0640673 |
| RPS6KB2 | 2463.89314 | -0.267599814 | 0.11966505 | -2.2362404 | 0.02533603 | 0.08923394 |
| RPS6KB2-AS | 752.911212 | -0.292497694 | 0.12709679 | -2.3013776 | 0.0213703 | 0.07873896 |
| CORO1B | 6756.78197 | -0.229502046 | 0.098679 | -2.3257435 | 0.02003224 | 0.07500358 |
| PTPRCAP | 761.697522 | -0.489172403 | 0.30578189 | -1.5997429 | 0.10965563 | 0.25335933 |
| GPR152 | 6.49017396 | 0.274168301 | 0.43976003 | 0.62344979 | 0.53298899 | 0.70689184 |
| CABP4 | 73.5970718 | -0.451191865 | 0.25476981 | -1.7709785 | 0.07656427 | 0.19647534 |
| TMEM134 | 2156.06686 | -0.210547066 | 0.15713727 | -1.3398926 | 0.18028026 | 0.35621088 |
| AIP | 2908.09564 | -0.050573409 | 0.10197332 | -0.4959475 | 0.61993149 | 0.77275317 |
| PITPNM1 | 7649.53787 | -0.915673663 | 0.26928127 | -3.4004358 | 0.00067279 | 0.00571196 |
| CDK2AP2 | 5073.10538 | -0.641299308 | 0.24634497 | -2.6032571 | 0.00923427 | 0.04240815 |
| CABP2 | 0.41416255 | 1.931698142 | 2.14762388 | 0.89945831 | 0.36840859 | NA |
| GSTP1 | 22145.2312 | -0.804784416 | 0.28714439 | -2.8027168 | 0.00506741 | 0.02713804 |
| C11orf72 | 6.41943681 | 1.116155645 | 0.57087198 | 1.95517677 | 0.05056221 | 0.14677902 |
| AP003385.3 | 2.81714901 | 0.106389988 | 0.8024819 | 0.13257618 | 0.89452857 | 0.94578078 |
| NDUFV1 | 15078.3806 | 0.635665765 | 0.16504206 | 3.85153805 | 0.00011738 | 0.001441 |
| NUDT8 | 665.964983 | 0.193029197 | 0.28526604 | 0.67666378 | 0.4986193 | 0.6799953 |
| TBX10 | 11.1376897 | 0.728015236 | 0.63531252 | 1.14591672 | 0.25182961 | 0.44410074 |
| ACY3 | 2927.3076 | 2.265519883 | 0.57891292 | 3.91340355 | 9.10E-05 | 0.00117464 |
| ALDH3B2 | 94.8936541 | -0.481150674 | 0.74447676 | -0.6462937 | 0.51808914 | 0.69505548 |
| AP003716.1 | 266.476431 | 4.411622969 | 0.63589672 | 6.93764071 | 3.99E-12 | 9.39E-10 |
| UNC93B1 | 1913.38866 | -0.557879906 | 0.14777308 | -3.7752472 | 0.00015985 | 0.00184156 |
| ALDH3B1 | 2113.2281 | 1.192059292 | 0.43349878 | 2.74985613 | 0.00596214 | 0.03063557 |
| AC004923.4 | 40.9914125 | 0.458845815 | 0.37311223 | 1.2297796 | 0.21877965 | 0.40412015 |
| NDUFS8 | 6038.14178 | -0.036028659 | 0.14299521 | -0.2519571 | 0.80107422 | 0.89215393 |
| TCIRG1 | 5071.41772 | -0.854849365 | 0.39372675 | -2.1711742 | 0.02991801 | 0.10066835 |
| AP002807.1 | 2126.47323 | -0.77466566 | 0.23884434 | -3.2433913 | 0.00118116 | 0.00893749 |
| CHKA | 2998.36841 | -0.580359612 | 0.27812141 | -2.0867131 | 0.03691407 | 0.11679995 |
| AP002992.1 | 12.2924831 | 0.580596714 | 0.40876058 | 1.42038333 | 0.15549611 | 0.32175397 |
| KMT5B | 2282.89034 | 0.095157143 | 0.12505401 | 0.76092836 | 0.44669986 | 0.63701862 |
| C11orf24 | 1699.64726 | -0.406073211 | 0.20624374 | -1.9688995 | 0.04896463 | 0.14365791 |
| LRP5 | 4877.46537 | 0.461534181 | 0.25077125 | 1.84045891 | 0.06570089 | 0.17615989 |
| PPP6R3 | 4356.78915 | -0.120827983 | 0.09863979 | -1.2249416 | 0.22059722 | 0.40651824 |
| AP003096.1 | 374.440793 | 0.23263717 | 0.15103348 | 1.54030194 | 0.12348677 | 0.27502924 |
| GAL | 18.3404484 | -2.235464444 | 0.81623758 | -2.7387424 | 0.00616747 | 0.03147593 |
| TESMIN | 418.14065 | 0.418035443 | 0.16121133 | 2.5930898 | 0.00951179 | 0.04340301 |
| CPT1A | 7142.35695 | 0.268008966 | 0.28812743 | 0.93017512 | 0.35228042 | 0.54985388 |
| AP000808.1 | 49.7289136 | -0.316036274 | 0.63298044 | -0.4992828 | 0.61758014 | 0.77142666 |
| MRPL21 | 1475.13958 | -0.189481227 | 0.12227295 | -1.5496578 | 0.12122368 | 0.27175623 |
| IGHMBP2 | 1551.81176 | -0.117852492 | 0.1598591 | -0.7372273 | 0.46098413 | 0.64950585 |
| AP000808.2 | 2.32936877 | 0.072788096 | 0.94079746 | 0.07736851 | 0.93833039 | 0.97061046 |
| MRGPRD | 0.8432554 | -2.957014482 | 2.13341546 | -1.3860472 | 0.1657325 | NA |
| AP003071.3 | 368.164793 | -0.215045516 | 0.27132833 | -0.7925656 | 0.42803093 | 0.62042202 |
| MRGPRF | 894.704245 | -0.069147543 | 0.27521132 | -0.2512525 | 0.80161886 | 0.89249269 |
| MRGPRF-AS | 105.353344 | 0.190885247 | 0.26997511 | 0.70704758 | 0.47953691 | 0.66418343 |
| TPCN2 | 2413.59542 | 0.722882529 | 0.25053549 | 2.88534985 | 0.00390979 | 0.02237924 |
| AP003071.1 | 9.61386477 | 0.465683481 | 0.76088603 | 0.6120279 | 0.5405193 | 0.71356665 |
| AP003071.2 | 0.2963014 | 2.148440699 | 3.23275528 | 0.66458501 | 0.50631596 | NA |
| AP003071.5 | 108.594098 | 3.263908251 | 0.61112071 | 5.34085685 | 9.25E-08 | 4.48E-06 |
| MYEOV | 20.8704523 | -1.605574458 | 0.60972778 | -2.6332644 | 0.00845685 | 0.03968804 |
| AP005233.2 | 2.99112788 | -2.161861363 | 1.28016659 | -1.6887344 | 0.09127034 | 0.22285175 |
| AP005233.1 | 0 | NA | NA | NA | NA | NA |
| AP000439.3 | 0.34077676 | 1.644917858 | 2.17138189 | 0.75754425 | 0.44872387 | NA |
| AP000439.1 | 0.48209152 | 1.480407517 | 1.79164657 | 0.82628323 | 0.40864344 | NA |
| AP000439.4 | 0.4782967 | 2.250690012 | 2.58623104 | 0.87025868 | 0.38415905 | NA |
| AP000439.2 | 7.04210857 | 3.271638194 | 0.80073779 | 4.08577967 | 4.39E-05 | 0.00066921 |
| LINC01488 | 1.76170567 | 1.701489538 | 1.4807494 | 1.14907326 | 0.25052577 | 0.44272649 |
| CCND1 | 3354.93819 | 1.158642835 | 0.30622254 | 3.78366275 | 0.00015454 | 0.00179443 |
| ORAOV1 | 2224.68518 | 0.251782027 | 0.18590927 | 1.35432746 | 0.17563193 | 0.34990597 |
| FGF19 | 0.26272731 | 1.990198932 | 3.242756 | 0.61373687 | 0.53938922 | NA |
| FGF4 | 2.83808034 | -1.794554971 | 1.72059271 | -1.0429865 | 0.29695454 | 0.49372822 |
| FGF3 | 0 | NA | NA | NA | NA | NA |
| AP007216.1 | 0 | NA | NA | NA | NA | NA |
| AP007216.2 | 0.28581056 | 1.235001422 | 2.85411953 | 0.43270837 | 0.66522664 | NA |
| AP003555.3 | 1.39532838 | 1.657024908 | 1.06007496 | 1.5631205 | 0.11802425 | 0.26678378 |
| AP003555.2 | 0.34792611 | 1.729835917 | 2.75381143 | 0.62816063 | 0.52989872 | NA |
| AP003555.1 | 0 | NA | NA | NA | NA | NA |
| LINC02584 | 0.31777585 | -0.280321477 | 2.95159789 | -0.0949728 | 0.92433644 | NA |
| ANO1 | 728.502642 | 0.284841388 | 0.38387207 | 0.74202164 | 0.45807422 | 0.64706542 |
| ANO1-AS1 | 239.72931 | 0.245753681 | 0.39961931 | 0.61496949 | 0.53857487 | 0.71192638 |
| FADD | 1461.36351 | -0.511585422 | 0.14141575 | -3.6175986 | 0.00029735 | 0.00300886 |
| AP000879.1 | 1250.74458 | -0.498538516 | 0.13520619 | -3.6872463 | 0.00022669 | 0.00241923 |
| PPFIA1 | 2818.92082 | -0.115004612 | 0.25944417 | -0.4432731 | 0.65756823 | 0.79911859 |
| AP002336.2 | 10.5916025 | 0.226829429 | 0.44692719 | 0.50753106 | 0.61178225 | 0.76749276 |
| AP002336.3 | 27.309658 | 1.976373219 | 0.54732471 | 3.61097023 | 0.00030505 | 0.00307385 |
| AP000487.2 | 0.717051 | 1.783305784 | 1.69046691 | 1.05491907 | 0.29146234 | NA |
| AP000487.1 | 521.071424 | 0.179537286 | 0.30553087 | 0.58762404 | 0.55678467 | 0.72574388 |
| CTTN | 17972.9226 | -0.804310437 | 0.20264357 | -3.9690894 | 7.21E-05 | 0.00097817 |
| SHANK2 | 1430.50254 | 0.38427962 | 0.18406568 | 2.08773092 | 0.03682211 | 0.11662149 |
| SHANK2-AS1 | 0.66433152 | 1.457756624 | 1.69674539 | 0.85914872 | 0.39025847 | NA |
| SHANK2-AS2 | 0.93097878 | 0.541303298 | 1.21910122 | 0.44401834 | 0.65702933 | NA |
| SHANK2-AS3 | 0.09997307 | 0.547239394 | 3.29377202 | 0.16614368 | 0.86804388 | NA |
| DHCR7 | 2179.43559 | -0.383032539 | 0.31179402 | -1.2284794 | 0.21926705 | 0.4048018 |
| AP002387.2 | 116.64549 | -0.492748831 | 0.26740921 | -1.8426771 | 0.06537617 | 0.17552123 |
| NADSYN1 | 4405.14335 | -0.198203651 | 0.13904341 | -1.4254803 | 0.1540184 | 0.31981321 |
| KRTAP5-7 | 18.8287833 | 1.240267709 | 0.61640069 | 2.01211278 | 0.04420805 | 0.13319008 |
| KRTAP5-8 | 116.588719 | 0.246941127 | 0.74863339 | 0.32985588 | 0.74150886 | 0.85475107 |
| KRTAP5-9 | 19.8560052 | 0.017019176 | 0.44276306 | 0.03843856 | 0.96933801 | 0.98580727 |
| KRTAP5-10 | 24.6302009 | -0.181891943 | 0.44033015 | -0.4130808 | 0.6795474 | 0.81406334 |
| KRTAP5-11 | 2.37980782 | 0.537589082 | 1.10627446 | 0.48594549 | 0.62700582 | 0.77804072 |
| AP003498.1 | 1.31095904 | 1.535233128 | 1.2787601 | 1.20056383 | 0.22992044 | 0.41750461 |
| FAM86C1 | 338.669205 | 0.496437135 | 0.17382343 | 2.85598512 | 0.00429035 | 0.02401031 |
| ZNF705E | 12.9130796 | -0.430818136 | 0.42009197 | -1.0255329 | 0.30511181 | 0.5021421 |
| DEFB108B | 0.55344888 | 0.241030446 | 2.19555924 | 0.10978089 | 0.91258315 | NA |
| AP002495.2 | 99.9994845 | 0.544269849 | 0.32446077 | 1.67745966 | 0.09345263 | 0.2266336 |
| DEFB131B | 4.4684097 | 0.959009764 | 0.61163397 | 1.56794719 | 0.11689346 | 0.26521478 |
| RNF121 | 1747.4166 | 0.152450601 | 0.13202573 | 1.15470368 | 0.24821179 | 0.43984577 |
| IL18BP | 5867.94391 | -0.436515098 | 0.29847056 | -1.4625064 | 0.1436025 | 0.30519003 |
| NUMA1 | 13934.0192 | 0.507202371 | 0.18934406 | 2.67873407 | 0.00739011 | 0.03604163 |
| AP002490.1 | 1015.97581 | 2.119414979 | 0.31437374 | 6.74170488 | 1.57E-11 | 2.95E-09 |
| LRTOMT | 1473.99476 | 0.431993695 | 0.18563905 | 2.3270626 | 0.01996193 | 0.07481469 |
| AP000812.5 | 1161.6444 | 0.407727627 | 0.22034962 | 1.85036681 | 0.0642607 | 0.17339145 |
| LAMTOR1 | 5936.62783 | -0.118399396 | 0.171777 | -0.6892622 | 0.49065828 | 0.67371303 |
| AP000812.4 | 15.0093376 | 0.412813263 | 0.36138749 | 1.14230091 | 0.25332896 | 0.44588738 |
| ANAPC15 | 1006.57214 | 0.320789348 | 0.26273688 | 1.22095288 | 0.22210386 | 0.40805956 |
| FOLR3 | 256.210188 | 0.061089777 | 0.27414915 | 0.22283409 | 0.82366464 | 0.90564302 |
| AP000812.1 | 2.28146001 | 0.992855245 | 0.90448172 | 1.09770626 | 0.27233278 | 0.46731527 |
| FOLR1 | 5244.10026 | 1.124751056 | 0.19607062 | 5.73645891 | 9.67E-09 | 6.73E-07 |
| FOLR2 | 643.294348 | -0.350224876 | 0.55925354 | -0.6262363 | 0.53115995 | 0.70547921 |
| INPPL1 | 10406.7843 | 0.076179996 | 0.1604054 | 0.47492165 | 0.63484282 | 0.78341547 |

| | | | | | | |
|---|---|---|---|---|---|---|
| PHOX2A | 6.76306733 | -0.123698871 | 0.44603349 | -0.2773309 | 0.78152603 | 0.88036187 |
| CLPB | 1418.77653 | 0.427844444 | 0.15787197 | 2.71007226 | 0.00672686 | 0.03355792 |
| AP000593.3 | 77.3466797 | 0.688361735 | 0.20619561 | 3.33839178 | 0.00084265 | 0.00685611 |
| AP002892.2 | 0.13028434 | 1.049816488 | 3.28477129 | 0.31960109 | 0.74927075 | NA |
| AP002892.1 | 0 | NA | NA | NA | NA | NA |
| AP003785.1 | 0.12088456 | 0.02007176 | 3.29738704 | 0.00608717 | 0.99514317 | NA |
| LINC01537 | 68.6263126 | -0.27576691 | 0.39252095 | -0.7025533 | 0.48233414 | 0.66686685 |
| PDE2A | 2235.34333 | -0.316518684 | 0.34308431 | -0.9225682 | 0.35623226 | 0.55358858 |
| AP005019.1 | 123.715436 | 0.362101718 | 0.32556283 | 1.112233 | 0.26603798 | 0.46017852 |
| AP003065.1 | 0.2854129 | -1.364245016 | 2.9060998 | -0.4694419 | 0.6387538 | NA |
| ARAP1 | 7554.64821 | -0.256600943 | 0.20270338 | -1.2658937 | 0.20555113 | 0.38814058 |
| ARAP1-AS1 | 1695.47848 | -0.713925476 | 0.21190799 | -3.3690352 | 0.00075432 | 0.006265 |
| ARAP1-AS2 | 323.963121 | 0.367088421 | 0.24745657 | 1.48344587 | 0.13795599 | 0.29675336 |
| STARD10 | 2416.32494 | -0.217742348 | 0.19498093 | -1.1167366 | 0.26410695 | 0.4579799 |
| ATG16L2 | 3587.78206 | -0.18176008 | 0.26365021 | -0.6893986 | 0.49057248 | 0.67369536 |
| FCHSD2 | 1522.39885 | -0.154200653 | 0.13631755 | -1.1311871 | 0.25797636 | 0.45051502 |
| AP002761.2 | 1.98470032 | 2.794771904 | 1.04789518 | 2.66703384 | 0.0076524 | 0.03696532 |
| AP002761.1 | 3.04341939 | -0.321420099 | 0.86889772 | -0.3699171 | 0.71144428 | 0.8360186 |
| P2RY2 | 133.578771 | -0.568301021 | 0.28672733 | -1.982026 | 0.04747634 | 0.14019608 |
| P2RY6 | 577.418864 | -1.230138311 | 0.44493057 | -2.7647871 | 0.005696 | 0.02969034 |
| AP002761.3 | 156.699295 | 0.942233811 | 0.2990171 | 3.15110342 | 0.00162655 | 0.01151659 |
| ARHGEF17 | 6736.65046 | 0.50189907 | 0.27606804 | 1.81802672 | 0.06906005 | 0.18233014 |
| RELT | 844.100825 | -1.318638509 | 0.44810009 | -2.9427321 | 0.0032533 | 0.01954647 |
| AP000763.4 | 374.674001 | -1.546755903 | 0.45159701 | -3.4250801 | 0.00061462 | 0.00530871 |
| FAM168A | 2419.96263 | 0.457554037 | 0.12990234 | 3.52229251 | 0.00042783 | 0.00402208 |
| AP000763.3 | 55.3179049 | 0.443344092 | 0.33284357 | 1.33198936 | 0.18286368 | 0.35969577 |
| PLEKHB1 | 848.124193 | 0.135828707 | 0.28658208 | 0.47396091 | 0.63552778 | 0.78382057 |
| RAB6A | 6531.01053 | -0.177213318 | 0.12105961 | -1.4638517 | 0.14323448 | 0.30477634 |
| AP002993.1 | 1.73402103 | -1.599505894 | 1.97267413 | -0.8108313 | 0.41746257 | 0.61029426 |
| AP002770.1 | 560.234659 | 0.659729119 | 0.28077066 | 2.34970822 | 0.01878813 | 0.07172019 |
| MRPL48 | 959.280592 | 0.235447177 | 0.15823401 | 1.48796822 | 0.13675928 | 0.29514216 |
| COA4 | 2546.03736 | -0.035845143 | 0.17608404 | -0.2035684 | 0.83869081 | 0.91450415 |
| PAAF1 | 1333.67065 | 0.397545398 | 0.30356252 | 1.30959975 | 0.19033127 | 0.3690196 |
| DNAJB13 | 102.308312 | -1.10068673 | 0.55979292 | -1.9662391 | 0.049271 | 0.14425776 |
| AP003717.1 | 79.5912169 | -1.154810128 | 0.600608 | -1.9227352 | 0.05451332 | 0.15472136 |
| UCP2 | 1901.76911 | -0.307756935 | 0.38570866 | -0.7979 | 0.42492853 | 0.61727347 |
| UCP3 | 134.17876 | -0.30497616 | 0.27657958 | -1.1026706 | 0.27017026 | 0.46449207 |
| C2CD3 | 1044.52611 | -0.254792184 | 0.21120657 | -1.2063649 | 0.22767683 | 0.41468697 |
| PPME1 | 2921.16066 | -0.499702518 | 0.14113734 | -3.5405407 | 0.00039931 | 0.00381651 |
| P4HA3 | 281.053341 | -0.181825452 | 0.28516957 | -0.6376047 | 0.52373105 | 0.69892989 |
| P4HA3-AS1 | 3.06641869 | -1.039702453 | 1.08103742 | -0.9617636 | 0.33616836 | 0.5342602 |
| PGM2L1 | 483.472075 | -1.180437192 | 0.27466094 | -4.2977978 | 1.73E-05 | 0.00031603 |
| AP001372.1 | 8.6033535 | 0.893171891 | 0.43902824 | 2.03442926 | 0.04190833 | 0.12815989 |
| KCNE3 | 1021.67609 | 1.528862146 | 0.36847846 | 4.14912217 | 3.34E-05 | 0.000536 |
| AP001372.3 | 417.407089 | 1.246582774 | 0.36082199 | 3.45484148 | 0.00055062 | 0.00489375 |
| LIPT2 | 110.410118 | 1.107068368 | 0.32763814 | 3.3789362 | 0.00072767 | 0.00610131 |
| POLD3 | 1240.59367 | 0.816429349 | 0.28101304 | 2.90530765 | 0.00366892 | 0.02136222 |
| CHRDL2 | 72.6599079 | -1.004574174 | 0.3613944 | -2.779717 | 0.00544063 | 0.02870536 |
| RNF169 | 1343.39956 | -0.202575359 | 0.20918393 | -0.9684078 | 0.33284073 | 0.53079553 |
| XRRA1 | 1618.1035 | -0.066089004 | 0.24297882 | -0.2719949 | 0.78562593 | 0.88309106 |
| SPCS2 | 2976.23123 | -0.297228218 | 0.14903944 | -1.9942923 | 0.04612012 | 0.1372982 |
| NEU3 | 729.203875 | 0.151418474 | 0.13005141 | 1.16429706 | 0.24430365 | 0.43534851 |
| OR2AT4 | 1.3060152 | 0.938477063 | 1.14739179 | 0.81792207 | 0.41340169 | 0.60732048 |
| AP001972.4 | 0.48902353 | 1.371166986 | 2.19734465 | 0.62401089 | 0.53262044 | NA |
| SLCO2B1 | 3580.44382 | 0.863568381 | 0.38952898 | 2.21695543 | 0.02662614 | 0.09254399 |
| AP001972.3 | 3.41957803 | 0.461912462 | 0.86542451 | 0.5337409 | 0.59352081 | 0.75271214 |
| TPBGL | 18.946759 | -1.13761027 | 0.46415369 | -2.4509345 | 0.01424859 | 0.05864103 |
| AP001972.1 | 258.484468 | -0.31336244 | 0.41576973 | -0.7536923 | 0.45103401 | 0.64079343 |
| ARRB1 | 812.164548 | 0.195762851 | 0.2519328 | 0.77704391 | 0.43713287 | 0.62917928 |
| AP001972.2 | 41.1312462 | 0.484445657 | 0.30347422 | 1.59633217 | 0.11041465 | 0.25449125 |
| RPS3 | 27292.4526 | -0.545025291 | 0.19667495 | -2.7711984 | 0.00558504 | 0.02927982 |
| KLHL35 | 29.1785949 | -0.95458203 | 0.46106854 | -2.070369 | 0.0384178 | 0.12023231 |
| GDPD5 | 690.334689 | -0.380560439 | 0.34706997 | -1.0964949 | 0.27286227 | 0.46768788 |
| SERPINH1 | 6320.30915 | -0.79013337 | 0.25601539 | -3.0862729 | 0.00202683 | 0.01363803 |
| AP001922.5 | 15.3095265 | 0.090242476 | 0.39645126 | 0.22762565 | 0.81993728 | 0.90344498 |
| MAP6 | 644.320031 | 0.554593031 | 0.28302098 | 1.95954742 | 0.05004871 | 0.14571984 |
| MOGAT2 | 85.536388 | 2.929584177 | 0.65345447 | 4.48322616 | 7.35E-06 | 0.00016036 |
| AP001922.1 | 0 | NA | NA | NA | NA | NA |
| DGAT2 | 303.542609 | 0.504088546 | 0.292594 | 1.722826 | 0.08491999 | 0.21123859 |
| AP003031.1 | 104.852622 | 0.486718687 | 0.2690894 | 1.80876202 | 0.07048798 | 0.18515822 |
| AP003031.2 | 15.8010174 | 0.351209324 | 0.38021884 | 0.9237031 | 0.35564092 | 0.55309336 |
| UVRAG | 1197.86438 | -0.004499154 | 0.14868571 | -0.0302595 | 0.9758601 | 0.98881307 |
| AP002340.1 | 2.88336636 | 0.948987238 | 0.75110077 | 1.26346194 | 0.20642322 | 0.38909985 |
| WNT11 | 290.727262 | -1.832933742 | 0.83079678 | -2.206236 | 0.02736748 | 0.09447317 |
| AP000785.1 | 44.714735 | -1.695450565 | 0.8809707 | -1.9245255 | 0.05428875 | 0.15425001 |
| AP000785.2 | 5.25122181 | -0.499231641 | 0.81084959 | -0.6156896 | 0.53809942 | 0.71154757 |
| THAP12 | 1810.98748 | 0.041230179 | 0.15232322 | 0.2706756 | 0.78664054 | 0.88387707 |
| GVQW3 | 76.5542389 | 1.353177558 | 0.45022101 | 3.00558511 | 0.0026507 | 0.0166983 |
| AP002360.3 | 43.4268035 | 0.416786001 | 0.45330541 | 0.91943751 | 0.35786678 | 0.55553935 |
| AP002360.2 | 13.6013633 | 1.66827447 | 0.42748368 | 3.90254541 | 9.52E-05 | 0.0012173 |
| AP002360.1 | 271.714163 | 0.898509486 | 0.38485642 | 2.33466154 | 0.01956111 | 0.07372109 |
| EMSY | 922.16374 | -0.13534535 | 0.16007581 | -0.8455078 | 0.39782736 | 0.59270617 |
| AP001189.5 | 27.8475649 | -0.346819674 | 0.34473393 | -1.0060503 | 0.31439137 | 0.51138049 |
| AP001189.6 | 8.49852372 | 0.078195681 | 0.46533926 | 0.16804015 | 0.86655169 | 0.93071718 |
| AP001189.3 | 68.8387157 | 0.723424465 | 0.40993914 | 1.76471187 | 0.0776122 | 0.19825959 |
| AP001189.1 | 474.268004 | -0.015367099 | 0.37392154 | -0.0410971 | 0.96721847 | 0.98450423 |
| LRRC32 | 6749.56193 | -0.334920018 | 0.35504622 | -0.9433139 | 0.34552039 | 0.54324657 |
| AP001189.4 | 1.91880283 | 0.059188827 | 0.84081259 | 0.07039479 | 0.94387944 | 0.97318192 |
| AP003119.1 | 22.6786489 | 1.371791132 | 0.66845507 | 2.05218149 | 0.04015203 | 0.12412646 |
| TSKU | 8169.64808 | -0.589474444 | 0.6460066 | -0.9124898 | 0.36151093 | 0.55900878 |
| AP003119.2 | 32.5947563 | -0.383145077 | 0.45110473 | -0.8493484 | 0.39568746 | 0.59100981 |
| ACER3 | 1346.05811 | -0.273017058 | 0.16437794 | -1.6609106 | 0.09673141 | 0.23214482 |
| AP002498.1 | 19.5927041 | -0.512028049 | 0.29500089 | -1.7356831 | 0.08261988 | 0.20723636 |
| B3GNT6 | 0.23371381 | -0.340597073 | 3.29738699 | -0.103293 | 0.91773041 | NA |
| CAPN5 | 2983.14509 | 0.292049477 | 0.37006575 | 0.78918267 | 0.43000525 | 0.62207863 |
| OMP | 8.16259991 | 0.287197162 | 0.60458157 | 0.4750346 | 0.63476231 | 0.78341547 |
| MYO7A | 1162.83071 | 1.570968598 | 0.2975508 | 5.27966513 | 1.29E-07 | 5.95E-06 |
| GDPD4 | 18.6846404 | 1.442727487 | 0.74886571 | 1.92655034 | 0.05403569 | 0.1538958 |
| PAK1 | 2442.40686 | 0.207440392 | 0.10384883 | 1.99752265 | 0.04576844 | 0.13656075 |
| AP003680.1 | 45.726393 | 0.790548548 | 0.31626631 | 2.49962935 | 0.01243233 | 0.05296293 |
| CLNS1A | 2422.97447 | -0.082706047 | 0.11274139 | -0.7335908 | 0.46319818 | 0.65109466 |
| AQP11 | 360.328121 | 0.887533194 | 0.46862142 | 1.89392367 | 0.05823514 | 0.16207752 |
| RSF1 | 1863.6946 | -0.282753759 | 0.11699341 | -2.4168349 | 0.01565612 | 0.0627816 |
| AAMDC | 795.555225 | 0.192255973 | 0.27091452 | 0.70965549 | 0.4779178 | 0.66294542 |
| AP002812.2 | 324.480583 | 0.143834353 | 0.28905476 | 0.49760244 | 0.61876428 | 0.77224735 |
| AP002812.3 | 6.46866666 | -0.836192568 | 0.56845868 | -1.4709822 | 0.14129594 | 0.3016983 |
| AP002812.5 | 486.192652 | 0.214292019 | 0.28069826 | 0.76342483 | 0.44521008 | 0.63573887 |
| INTS4 | 1124.83418 | -0.10004755 | 0.11942116 | -0.8377707 | 0.40215949 | 0.5970917 |
| KCTD14 | 794.827729 | -0.404157644 | 0.23340766 | -1.7315526 | 0.08335325 | 0.20852151 |
| AP003032.2 | 488.4992 | -0.5226304 | 0.2469889 | -2.1160077 | 0.03434416 | 0.11112336 |
| NDUFC2-KCT | 3200.5864 | 0.064638857 | 0.15371051 | 0.42052333 | 0.67410319 | 0.81024608 |
| AP003032.1 | 3.77357188 | 0.7181406 | 0.75736274 | 0.94821221 | 0.34302143 | 0.54064163 |
| THRSP | 63.6107968 | 2.281216533 | 0.53188263 | 4.28894725 | 1.80E-05 | 0.00032671 |

| | | | | | | |
|---|---|---|---|---|---|---|
| NDUFC2 | 3706.48812 | 0.228572548 | 0.15251079 | 1.49873039 | 0.13394359 | 0.2909227 |
| ALG8 | 1297.78124 | -0.136039316 | 0.13454306 | -1.011121 | 0.31195851 | 0.50899307 |
| KCTD21-AS1 | 254.676252 | 0.460523852 | 0.30915339 | 1.48962901 | 0.13632181 | 0.29448727 |
| KCTD21 | 726.342402 | 0.027478143 | 0.19208364 | 0.14305301 | 0.88624831 | 0.94110344 |
| USP35 | 1078.07885 | -0.919214193 | 0.30300228 | -3.0336874 | 0.00241585 | 0.01557601 |
| GAB2 | 1688.07207 | -0.813050129 | 0.31612584 | -2.5719192 | 0.01011365 | 0.04544163 |
| AP003086.2 | 207.19111 | 0.453704855 | 0.20860596 | 2.17493714 | 0.02963483 | 0.10000901 |
| AP003086.3 | 0.08393208 | -0.642905668 | 3.30558948 | -0.1944905 | 0.84579185 | NA |
| AP003086.1 | 16.245245 | 0.252624746 | 0.51655745 | 0.4890545 | 0.62480311 | 0.77650129 |
| NARS2 | 1090.77143 | 0.507447624 | 0.2171229 | 2.33714461 | 0.01943167 | 0.07333806 |
| AP003110.1 | 24.22835 | 1.927453301 | 0.50455216 | 3.82012693 | 0.00013338 | 0.00158939 |
| TENM4 | 525.840406 | 0.033859475 | 0.34905297 | 0.09700383 | 0.92272335 | 0.96224205 |
| AP002768.1 | 0.27026623 | -0.174096114 | 3.29377188 | -0.0528562 | 0.95784652 | NA |
| AP002957.1 | 6.52184369 | -2.173154041 | 0.84951627 | -2.5581076 | 0.01052435 | 0.04678972 |
| AP001547.1 | 0.64396736 | -0.108212875 | 1.81675639 | -0.0595638 | 0.95250307 | NA |
| AP001978.1 | 0.13390214 | 1.04981649 | 3.28477129 | 0.31960109 | 0.74927075 | NA |
| AP006295.1 | 0 | NA | NA | NA | NA | NA |
| AP002759.1 | 0.03392908 | 0.380740593 | 3.29738709 | 0.11546736 | 0.90807469 | NA |
| MIR4300HG | 0.34301569 | 0.732866073 | 3.27839706 | 0.22354402 | 0.82311212 | NA |
| AP002802.1 | 0 | NA | NA | NA | NA | NA |
| AP000793.1 | 0 | NA | NA | NA | NA | NA |
| FAM181B | 70.0927332 | -0.263640002 | 0.36442697 | -0.7234371 | 0.46941136 | 0.65606954 |
| AP001646.3 | 5.63219037 | -0.140453051 | 0.59452 | -0.2362461 | 0.81324168 | 0.89913137 |
| PRCP | 9969.57413 | -0.199885255 | 0.23376889 | -0.855055 | 0.39252071 | 0.58817884 |
| DDIAS | 35.0258191 | -0.592505514 | 0.34550986 | -1.7148729 | 0.08636853 | 0.2139897 |
| RAB30 | 395.042746 | 0.152163378 | 0.17421847 | 0.87340554 | 0.38244207 | 0.57853359 |
| RAB30-AS1 | 378.511892 | 0.147407648 | 0.22976738 | 0.64155167 | 0.52116432 | 0.69758404 |
| AP001767.3 | 210.671943 | 0.285921888 | 0.23823472 | 1.20016884 | 0.23007378 | 0.41761051 |
| PCF11 | 3933.01767 | -0.22110798 | 0.16109984 | -1.3724904 | 0.16991083 | 0.34176571 |
| PCF11-AS1 | 317.846959 | -0.007340198 | 0.1614684 | -0.045459 | 0.96374142 | 0.98305633 |
| ANKRD42 | 1301.04127 | -0.181074457 | 0.14789915 | -1.2243103 | 0.22083519 | 0.4067773 |
| AP000873.3 | 63.4987365 | -0.670517223 | 0.22994871 | -2.9159426 | 0.00354616 | 0.02077587 |
| CCDC90B | 1524.54164 | -0.295661633 | 0.11437048 | -2.5851219 | 0.00973446 | 0.04417443 |
| AP000446.1 | 54.2484574 | 0.338081627 | 0.2081051 | 1.62457152 | 0.10425389 | 0.24447977 |
| DLG2 | 176.106045 | 1.029289528 | 0.30018198 | 3.42888516 | 0.00060607 | 0.00525549 |
| AP002370.2 | 0.93471965 | 1.189973603 | 1.43774907 | 0.82766432 | 0.40786063 | NA |
| AP001825.1 | 0.75342373 | -0.819414671 | 1.88053701 | -0.4357344 | 0.66302944 | NA |
| AP000857.2 | 0.42288725 | 1.92122963 | 2.72405655 | 0.70528258 | 0.4806344 | NA |
| TMEM126B | 2008.26505 | -0.19908549 | 0.12335755 | -1.6138898 | 0.10655134 | 0.24814087 |
| TMEM126A | 715.055791 | -0.105076971 | 0.13282799 | -0.7910755 | 0.42889993 | 0.62112297 |
| CREBZF | 4854.05767 | -0.05073602 | 0.12267839 | -0.4135693 | 0.67918954 | 0.8138966 |
| CCDC89 | 67.3458688 | 0.685643732 | 0.25064729 | 2.73549233 | 0.0062287 | 0.03171479 |
| SYTL2 | 2219.9844 | -0.084553714 | 0.25146524 | -0.3362441 | 0.73668677 | 0.85218591 |
| CCDC83 | 1.11799465 | 0.755196975 | 1.16047084 | 0.65076773 | 0.51519644 | 0.69283135 |
| AP003128.1 | 0.30843339 | 1.372906539 | 2.26508559 | 0.60611685 | 0.54443716 | NA |
| PICALM | 12351.9552 | 0.243915314 | 0.11360287 | 2.14708766 | 0.0317863 | 0.1050777 |
| AP003097.2 | 0.43885145 | 0.593550961 | 2.81016466 | 0.21121572 | 0.83271894 | NA |
| EED | 1042.73555 | -0.268170299 | 0.12528708 | -2.1404465 | 0.0323187 | 0.10628952 |
| HIKESHI | 1149.97813 | -0.436663744 | 0.18461191 | -2.3653064 | 0.01801516 | 0.06945317 |
| CCDC81 | 36.6773517 | -0.119523419 | 0.39414307 | -0.3032488 | 0.76170025 | 0.86791967 |
| AP001831.1 | 1.75164508 | -0.696171036 | 1.39720398 | -0.4982601 | 0.61830071 | 0.77185143 |
| ME3 | 1795.22696 | 0.361953809 | 0.18767806 | 1.92858884 | 0.05378193 | 0.15338003 |
| AP003059.1 | 3.49180932 | 0.719837301 | 0.62674603 | 1.14853109 | 0.25074939 | 0.44297323 |
| AP003059.2 | 10.4717266 | 0.679511766 | 0.5754326 | 1.18087117 | 0.23765391 | 0.42701918 |
| PRSS23 | 4334.82085 | 0.17461164 | 0.30491433 | 0.57265804 | 0.56687626 | 0.73302704 |
| AP000654.1 | 6.66062238 | 0.663999509 | 0.68318509 | 0.97191744 | 0.33109162 | 0.52898431 |
| AP001528.2 | 102.976765 | 0.874933676 | 0.52735285 | 1.65910487 | 0.09709466 | 0.23269892 |
| FZD4 | 4796.92747 | 0.390220151 | 0.26412297 | 1.47741846 | 0.1395635 | 0.29935656 |
| AP001528.1 | 36.5844881 | 1.115808798 | 0.31076828 | 3.59048487 | 0.00033006 | 0.00327086 |
| TMEM135 | 1037.54094 | 0.087529929 | 0.25354365 | 0.34522627 | 0.72992427 | 0.84744873 |
| AP000756.1 | 0.08393208 | -0.642905668 | 3.30558948 | -0.1944905 | 0.84579185 | NA |
| AP005436.3 | 0.57258401 | -2.385733165 | 2.17902427 | -1.0948631 | 0.27357664 | NA |
| AP005436.1 | 0.67979687 | -2.631540038 | 2.17771745 | -1.2083937 | 0.22689586 | NA |
| AP005436.2 | 2.20499414 | 0.042596642 | 0.84016589 | 0.05070028 | 0.95956436 | 0.98050555 |
| RAB38 | 279.528793 | -0.182338043 | 0.2104399 | -0.8664614 | 0.3862372 | 0.58183805 |
| CTSC | 12951.7146 | 0.363043633 | 0.17326421 | 2.09531813 | 0.03614272 | 0.11516235 |
| GRM5-AS1 | 47.9094734 | 0.115159391 | 0.33495746 | 0.34380303 | 0.73099442 | 0.84798182 |
| GRM5 | 53.942478 | 0.008977779 | 0.30281741 | 0.0296475 | 0.97634818 | 0.98898712 |
| TYR | 2.14610892 | 1.023566734 | 0.77337395 | 1.32350815 | 0.18566645 | 0.36301803 |
| NOX4 | 4588.14084 | 1.854351468 | 0.55460755 | 3.34353808 | 0.00082717 | 0.00675505 |
| AP003400.1 | 34.985882 | 0.149761588 | 0.89841077 | 0.16669612 | 0.86760916 | 0.93109788 |
| TRIM77 | 0 | NA | NA | NA | NA | NA |
| AP003122.2 | 0 | NA | NA | NA | NA | NA |
| TRIM49 | 0.07659028 | -0.642905668 | 3.30558948 | -0.1944905 | 0.84579185 | NA |
| TRIM64B | 0 | NA | NA | NA | NA | NA |
| TRIM49D1 | 0 | NA | NA | NA | NA | NA |
| TRIM49D2 | 0 | NA | NA | NA | NA | NA |
| TRIM64 | 0 | NA | NA | NA | NA | NA |
| TRIM49C | 0.07830766 | -0.642905668 | 3.30558948 | -0.1944905 | 0.84579185 | NA |
| UBTFL1 | 0 | NA | NA | NA | NA | NA |
| NAALAD2 | 166.269512 | 0.694277644 | 0.33150325 | 2.09433134 | 0.03623047 | 0.11533047 |
| CHORDC1 | 1385.98241 | -0.253053583 | 0.14656206 | -1.7265967 | 0.08424011 | 0.21018182 |
| AP003386.1 | 0 | NA | NA | NA | NA | NA |
| AP003028.1 | 0 | NA | NA | NA | NA | NA |
| FAT3 | 48.9693242 | 0.741748427 | 0.57166556 | 1.29752162 | 0.19445177 | 0.37444817 |
| AP000722.1 | 0.10721287 | -0.642905668 | 3.30558948 | -0.1944905 | 0.84579185 | NA |
| AP003718.1 | 0 | NA | NA | NA | NA | NA |
| AP003171.3 | 0 | NA | NA | NA | NA | NA |
| AP003171.1 | 0.07659028 | -0.642905668 | 3.30558948 | -0.1944905 | 0.84579185 | NA |
| MTNR1B | 0.07659028 | -0.642905668 | 3.30558948 | -0.1944905 | 0.84579185 | NA |
| SLC36A4 | 1252.62874 | -0.811827798 | 0.27278707 | -2.9760494 | 0.00291988 | 0.0179732 |
| AP003969.1 | 0.16052235 | -1.003574173 | 3.30558948 | -0.3035992 | 0.7614333 | NA |
| DEUP1 | 14.5028964 | -0.292524154 | 0.61692719 | -0.4741632 | 0.63538355 | 0.78367934 |
| SMCO4 | 931.770046 | -0.748828476 | 0.24337417 | -3.076861 | 0.00209193 | 0.01393034 |
| CEP295 | 945.801547 | -0.503778157 | 0.1405345 | -3.5847294 | 0.00033743 | 0.00333008 |
| SCARNA9 | 25.6819802 | 0.346654548 | 0.33676479 | 1.02936696 | 0.30330727 | 0.50016963 |
| TAF1D | 5111.34962 | -0.853831705 | 0.18361008 | -4.6502441 | 3.32E-06 | 8.48E-05 |
| C11orf54 | 15203.6482 | 1.556922329 | 0.49183259 | 3.16555338 | 0.00154788 | 0.0110707 |
| AP001273.2 | 8851.00955 | 1.50149442 | 0.42447376 | 3.53730796 | 0.00040423 | 0.00385097 |
| MED17 | 1588.31447 | -0.269604236 | 0.16236022 | -1.6605314 | 0.0968076 | 0.23224313 |
| VSTM5 | 114.96598 | 2.131064544 | 0.46806798 | 4.55289534 | 5.29E-06 | 0.00012261 |
| HEPHL1 | 38.5706789 | -1.180032197 | 0.7128358 | -1.6554053 | 0.0978423 | 0.23406082 |
| PANX1 | 704.080644 | -1.103839546 | 0.24469874 | -4.5110144 | 6.45E-06 | 0.00014418 |
| AP002784.1 | 2.19842664 | -1.425495345 | 1.19744657 | -1.1904459 | 0.23387119 | 0.42244345 |
| IZUMO1R | 3.66895356 | 2.489366785 | 1.11342842 | 2.23576724 | 0.02536702 | 0.08929689 |
| GPR83 | 22.0011896 | 0.638042509 | 0.40761247 | 1.56531644 | 0.11750873 | 0.26605179 |
| MRE11 | 1017.1395 | -0.245767762 | 0.09193591 | -2.6732511 | 0.007512 | 0.03649437 |
| AP000786.1 | 1.50427602 | 0.66686407 | 1.21903988 | 0.5470404 | 0.58435097 | 0.74635935 |
| ANKRD49 | 722.155598 | -0.304985472 | 0.13753159 | -2.2175667 | 0.02658439 | 0.09242438 |
| C11orf97 | 23.9928679 | 0.510882756 | 0.57596837 | 0.88699794 | 0.37508001 | 0.57109878 |
| FUT4 | 666.852751 | -1.259807947 | 0.27492645 | -4.5823454 | 4.60E-06 | 0.00011002 |
| PIWIL4 | 401.277628 | -0.989452946 | 0.30834244 | -3.2089418 | 0.00133224 | 0.0098628 |
| AP000943.3 | 224.376541 | -1.22550538 | 0.27656859 | -4.4311084 | 9.37E-06 | 0.00019405 |

| | | | | | | |
|---|---|---|---|---|---|---|
| AP000943.1 | 1.17118805 | -1.473732957 | 1.42474171 | -1.0343861 | 0.30095572 | 0.49768474 |
| AP000943.2 | 0.5339141 | -0.061938093 | 1.93139355 | -0.0320691 | 0.97441693 | NA |
| AMOTL1 | 1556.82923 | -0.435380018 | 0.23145406 | -1.8810645 | 0.05996314 | 0.16543503 |
| AP002383.2 | 170.384839 | 0.037971077 | 0.28552425 | 0.13298722 | 0.89420349 | 0.94565432 |
| CWC15 | 2224.5789 | -0.252553766 | 0.09523351 | -2.6519421 | 0.00800303 | 0.03813739 |
| KDM4D | 58.8989303 | 0.699527746 | 0.23848718 | 2.93318809 | 0.00335501 | 0.01997568 |
| KDM4E | 6.86221628 | -0.523260554 | 0.53695492 | -0.9744962 | 0.3298102 | 0.52767231 |
| KDM4F | 0 | NA | NA | NA | NA | NA |
| SRSF8 | 3029.5959 | 0.012513255 | 0.19494847 | 0.0641875 | 0.94882093 | 0.97530241 |
| ENDOD1 | 4537.625 | -0.88788661 | 0.28856532 | -3.0768999 | 0.00209166 | 0.01393034 |
| AP000787.1 | 110.410969 | 0.551080274 | 0.2803747 | 1.96551358 | 0.04935483 | 0.14434327 |
| SESN3 | 1544.65851 | 0.54714065 | 0.24014769 | 2.27835066 | 0.02270569 | 0.08232997 |
| AP001790.1 | 0.70428516 | -0.127469032 | 2.16448164 | -0.0588913 | 0.95303873 | NA |
| AP000820.1 | 0.35640006 | -1.70666969 | 3.3025702 | -0.5167701 | 0.60531662 | NA |
| FAM76B | 842.164279 | -0.016682196 | 0.11792206 | -0.141468 | 0.88750025 | 0.94182388 |
| CEP57 | 1740.58466 | -0.054599906 | 0.13542297 | -0.4031805 | 0.68681543 | 0.81899805 |
| MTMR2 | 2071.51966 | -0.214118303 | 0.16617375 | -1.2885206 | 0.19756481 | 0.37834197 |
| MAML2 | 1343.40649 | 0.383693525 | 0.22925995 | 1.67361775 | 0.09420575 | 0.22809711 |
| CCDC82 | 1875.1172 | -0.234713734 | 0.24741 | -0.9486833 | 0.34278171 | 0.54048703 |
| JRKL | 764.041683 | -0.334235751 | 0.18754883 | -1.7821266 | 0.0747286 | 0.19317646 |
| JRKL-AS1 | 0.18655607 | -1.003574173 | 3.30558948 | -0.3035992 | 0.7614333 | NA |
| AP003781.1 | 0 | NA | NA | NA | NA | NA |
| AP003066.1 | 0.09644756 | -0.642905668 | 3.30558948 | -0.1944905 | 0.84579185 | NA |
| LINC02553 | 0.07830766 | -0.642905668 | 3.30558948 | -0.1944905 | 0.84579185 | NA |
| AP003730.2 | 0.8098013 | 1.916027117 | 1.28800838 | 1.48758902 | 0.13685931 | NA |
| AP003715.1 | 0 | NA | NA | NA | NA | NA |
| AP002428.1 | 0 | NA | NA | NA | NA | NA |
| CNTN5 | 31.8831582 | 1.571373113 | 0.33852947 | 4.64176167 | 3.45E-06 | 8.77E-05 |
| AP001351.1 | 14.7912951 | 0.706979525 | 0.48207168 | 1.46654442 | 0.14250003 | 0.30372824 |
| ARHGAP42 | 925.899409 | 0.277100215 | 0.27604756 | 1.00381329 | 0.31546862 | 0.51256204 |
| PGR | 224.338889 | 0.805854071 | 0.22244136 | 3.62277075 | 0.00029146 | 0.00296868 |
| TRPC6 | 150.127452 | 0.210469101 | 0.44283852 | 0.47527279 | 0.63459255 | 0.78336324 |
| ANGPTL5 | 5.90246323 | -0.483241577 | 0.70731073 | -0.6832097 | 0.49447432 | 0.67689496 |
| CEP126 | 355.569551 | -0.122298778 | 0.29142352 | -0.4196599 | 0.67473389 | 0.81067472 |
| CFAP300 | 122.305651 | -1.136219036 | 0.39669161 | -2.8642377 | 0.00418014 | 0.02356239 |
| AP001527.1 | 5.23177177 | -2.096459491 | 0.87068665 | -2.4078232 | 0.01604795 | 0.06399282 |
| AP001527.2 | 29.8751455 | -0.413986127 | 0.35698839 | -1.1596627 | 0.24618616 | 0.43754998 |
| YAP1 | 5785.80893 | -0.421852335 | 0.15313531 | -2.7547685 | 0.00587337 | 0.03033308 |
| AP000942.2 | 718.519047 | -0.027648069 | 0.18395405 | -0.1502988 | 0.8805289 | 0.93801192 |
| BIRC3 | 2365.32527 | -1.922630646 | 0.46021078 | -4.1777176 | 2.94E-05 | 0.00048378 |
| BIRC2 | 3097.7775 | 0.02934956 | 0.14749196 | 0.19899092 | 0.84226985 | 0.91624339 |
| TMEM123 | 8235.66147 | -0.330461403 | 0.12590817 | -2.6246224 | 0.00867451 | 0.04044433 |
| AP001830.1 | 37.1807842 | -0.126330486 | 0.41478865 | -0.3045659 | 0.76069678 | 0.86736603 |
| AP001830.2 | 0 | NA | NA | NA | NA | NA |
| MMP7 | 10859.5392 | -2.107791653 | 0.49265871 | -4.2784013 | 1.88E-05 | 0.0003379 |
| MMP20 | 7.89980923 | 0.709335724 | 1.03481672 | 0.68546991 | 0.49304744 | 0.67578034 |
| AP000851.1 | 1.4737677 | -0.514726031 | 1.4797075 | -0.3478566 | 0.72794787 | 0.84612269 |
| AP000851.2 | 125.848898 | -0.179824654 | 1.74430232 | -0.1030926 | 0.91788948 | 0.9601219 |
| MMP27 | 5.3902762 | 0.942329477 | 1.2357911 | 0.76253137 | 0.44574293 | 0.63615528 |
| MMP8 | 98.745824 | -5.571964311 | 0.90230119 | -6.175282 | 6.60E-10 | 7.29E-08 |
| MMP10 | 6.42234399 | -2.27038898 | 0.83842677 | -2.7079156 | 0.00677072 | 0.03370026 |
| MMP1 | 121.847964 | -2.856095959 | 0.62156104 | -4.5950369 | 4.33E-06 | 0.00010506 |
| MMP3 | 12.9541256 | 1.116807821 | 1.24565569 | 0.89656221 | NA | NA |
| MMP12 | 19.9139218 | -0.600966809 | 0.4727347 | -1.271256 | 0.2036376 | 0.38583607 |
| MMP13 | 0.80186741 | -0.479502929 | 1.40624721 | -0.3409805 | 0.73311823 | NA |
| DCUN1D5 | 840.53976 | -0.877792453 | 0.22704301 | -3.8661945 | 0.00011055 | 0.00137118 |
| DYNC2H1 | 855.593071 | 0.550991147 | 0.24696508 | 2.23104878 | 0.0256779 | 0.0900314 |
| AP002989.1 | 1.92988468 | -1.209596185 | 1.05363047 | -1.148027 | 0.25095744 | 0.44328137 |
| PDGFD | 575.516101 | 0.529131674 | 0.3466881 | 1.52624702 | 0.12694836 | 0.28044133 |
| AP003043.1 | 0 | NA | NA | NA | NA | NA |
| DDI1 | 0.16416605 | 0.18657164 | 3.29377195 | 0.05664376 | 0.95482897 | NA |
| LINC02552 | 0.37832336 | -0.55838662 | 2.25080741 | -0.2480828 | 0.80407034 | NA |
| CASP4 | 3407.29607 | -1.600064236 | 0.29086872 | -5.5009842 | 3.78E-08 | 2.11E-06 |
| CASP5 | 9.07301091 | -1.745954734 | 0.83214215 | -2.0981448 | 0.03589235 | 0.11461388 |
| CASP1 | 903.820078 | -1.038274283 | 0.44315848 | -2.3428961 | 0.01913471 | 0.07257773 |
| CARD16 | 206.863628 | -0.81825812 | 0.48406653 | -1.6903836 | 0.09095459 | 0.22229687 |
| CARD17 | 2.57632879 | -0.386430919 | 1.17176512 | -0.3297853 | 0.74156219 | 0.85475107 |
| CARD18 | 0.08666371 | -0.642905668 | 3.30558948 | -0.1944905 | 0.84579185 | NA |
| AP003181.1 | 0 | NA | NA | NA | NA | NA |
| GRIA4 | 84.7246941 | 1.232225399 | 0.81941575 | 1.50378534 | 0.13263665 | 0.28900258 |
| MSANTD4 | 841.965103 | 0.022104308 | 0.14677599 | 0.15059893 | 0.8802921 | 0.93791122 |
| KBTBD3 | 510.883329 | 0.280888703 | 0.14844547 | 1.89220125 | 0.05846417 | 0.16252374 |
| AASDHPPT | 2375.65089 | 0.096754715 | 0.19308668 | 0.50109472 | 0.61630446 | 0.77049015 |
| AP001001.1 | 372.565577 | 0.153992659 | 0.20141864 | 0.76454025 | 0.44454536 | 0.63537357 |
| AP001001.2 | 0 | NA | NA | NA | NA | NA |
| GUCY1A2 | 490.726697 | 0.391089266 | 0.5057894 | 0.7732255 | 0.43938894 | 0.63084261 |
| AP000766.1 | 137.333267 | 0.855331858 | 0.23838193 | 3.58807334 | 0.00033313 | 0.00329383 |
| CWF19L2 | 656.393212 | 0.462349564 | 0.1545927 | 2.99075944 | 0.00278285 | 0.01734428 |
| ALKBH8 | 396.609546 | 0.437872225 | 0.19419363 | 2.25482277 | 0.02414446 | 0.08599867 |
| ELMOD1 | 189.177062 | 0.603745274 | 0.47560729 | 1.26941972 | 0.20429141 | 0.38664081 |
| SLN | 8.64287472 | -1.497125057 | 1.41549237 | -1.0576709 | NA | NA |
| SLC35F2 | 1298.41893 | -0.269988546 | 0.20693727 | -1.3046879 | 0.19199915 | 0.37121831 |
| RAB39A | 11.1175644 | 0.037176239 | 0.61674316 | 0.06027832 | 0.95193397 | 0.97699287 |
| CUL5 | 2069.06146 | 0.30416973 | 0.1173551 | 2.59187485 | 0.00954545 | 0.04351141 |
| AP002433.1 | 2300.31407 | 1.264042533 | 0.34497737 | 3.66413173 | 0.00024818 | 0.00260529 |
| ACAT1 | 20109.5859 | 1.102001697 | 0.31831604 | 3.46197349 | 0.00053623 | 0.00478686 |
| NPAT | 660.781325 | -0.003297091 | 0.1234514 | -0.0267076 | 0.97869295 | 0.99041113 |
| ATM | 1996.3995 | -0.038660822 | 0.18907604 | -0.2044723 | 0.8379844 | 0.91426076 |
| C11orf65 | 66.9567131 | 0.587613095 | 0.34391855 | 1.70858217 | 0.08752837 | 0.21599506 |
| KDELC2 | 1403.77952 | 0.432693114 | 0.13382207 | 3.23334656 | 0.00122349 | 0.00918925 |
| EXPH5 | 513.861343 | 0.258340251 | 0.27935085 | 0.92478777 | 0.35507632 | 0.55257376 |
| DDX10 | 1141.97401 | 0.288960354 | 0.24981581 | 1.15669361 | 0.24739756 | 0.43894019 |
| AP003027.1 | 0.17391096 | -1.003574173 | 3.30558948 | -0.3035992 | 0.7614333 | NA |
| AP003123.1 | 0 | NA | NA | NA | NA | NA |
| AP003049.2 | 1.1712644 | 4.146202233 | 2.27018241 | 1.82637405 | 0.06779394 | 0.18000947 |
| C11orf87 | 1.51925966 | 1.894839964 | 0.99238661 | 1.90937679 | 0.0562135 | 0.15801621 |
| AP002444.1 | 0.18390514 | 0.18657164 | 3.29377195 | 0.05664376 | 0.95482897 | NA |
| RDX | 3653.87543 | 0.696650675 | 0.16972113 | 4.10467859 | 4.05E-05 | 0.00062841 |
| ZC3H12C | 738.899791 | -0.144311961 | 0.18875259 | -0.7645562 | 0.44453587 | 0.63537357 |
| AP000924.1 | 66.4452207 | 4.086286695 | 0.6916735 | 5.90782599 | 3.47E-09 | 2.80E-07 |
| FDX1 | 2126.17677 | 0.556298237 | 0.16947827 | 3.28241624 | 0.00102922 | 0.00803059 |
| ARHGAP20 | 52.5211046 | 0.454473157 | 0.54805817 | 0.82924255 | 0.40696717 | 0.60168362 |
| AP003973.2 | 1.99022839 | -0.130337834 | 1.08514005 | -0.1201115 | 0.9043948 | 0.95193753 |
| LINC02550 | 4.24617729 | 0.066910251 | 0.78279169 | 0.08547645 | 0.93188262 | 0.96708952 |
| C11orf53 | 4.61392605 | 0.493127706 | 0.97904938 | 0.50368012 | 0.61448618 | 0.76942073 |
| COLCA1 | 921.953354 | 1.501429696 | 0.3655687 | 4.10710683 | 4.01E-05 | 0.00062353 |
| COLCA2 | 524.820753 | 0.30012227 | 0.24896773 | 1.20546653 | 0.22802325 | 0.41511724 |
| POU2AF1 | 178.636672 | -0.294311144 | 0.75363411 | -0.3905226 | 0.69615015 | 0.82520361 |
| BTG4 | 0.16421342 | 1.36803136 | 3.28033652 | 0.41703994 | 0.67664919 | NA |
| AP002008.1 | 1.53021525 | -1.570046623 | 1.69450518 | -0.9265517 | 0.35415935 | 0.55176546 |
| C11orf88 | 5.00686998 | -1.914185087 | 1.19861104 | -1.5970027 | 0.1102651 | 0.25431232 |
| LAYN | 259.459114 | -0.540409724 | 0.25024148 | -2.159553 | 0.03080729 | 0.10282094 |
| SIK2 | 3707.9518 | -0.055509843 | 0.20603067 | -0.2694251 | 0.78760254 | 0.88431742 |

| | | | | | | |
|---|---|---|---|---|---|---|
| PPP2R1B | 2320.95535 | -0.366522399 | 0.24063932 | -1.5231193 | 0.12772886 | 0.28152876 |
| AP001781.1 | 5.13694002 | -0.162912872 | 0.59377392 | -0.2743685 | 0.78380143 | 0.88185185 |
| ALG9 | 1189.86757 | -0.3087145 | 0.10786121 | -2.8621456 | 0.00420784 | 0.02365394 |
| AP001781.2 | 943.488415 | -0.189095198 | 0.12208541 | -1.5488763 | 0.12141146 | 0.2719958 |
| FDXACB1 | 173.83116 | 0.545177165 | 0.17615444 | 3.0948818 | 0.00196891 | 0.01332997 |
| C11orf1 | 922.818109 | 0.750936865 | 0.28283284 | 2.65505537 | 0.00792954 | 0.03791681 |
| CRYAB | 100975.149 | -1.989566052 | 0.60228493 | -3.3033635 | 0.00095532 | 0.00757662 |
| HSPB2 | 815.792072 | -1.103206504 | 0.37573604 | -2.9361211 | 0.00332345 | 0.01984382 |
| HSPB2-C11or | 3032.04891 | -0.749330254 | 0.36074045 | -2.0772005 | 0.03778306 | 0.11875318 |
| C11orf52 | 2259.99091 | -0.621211051 | 0.40147573 | -1.5473191 | 0.12178632 | 0.27262355 |
| DIXDC1 | 1238.48231 | 0.289305067 | 0.33493859 | 0.86375555 | 0.38772219 | 0.58332428 |
| DLAT | 2911.7949 | 0.424788796 | 0.13503189 | 3.14584053 | 0.0016561 | 0.01167061 |
| PIH1D2 | 340.147804 | 0.098001662 | 0.16999866 | 0.57648492 | 0.56428746 | 0.73132527 |
| NKAPD1 | 1546.99575 | -0.011412644 | 0.10774958 | -0.1059182 | 0.91564724 | 0.95884432 |
| TIMM8B | 1605.7086 | -0.203103413 | 0.158677 | -1.2799802 | 0.20055211 | 0.38209372 |
| SDHD | 9847.33271 | 0.603102287 | 0.17174072 | 3.51170231 | 0.00044525 | 0.00415028 |
| AP002884.3 | 4256.49176 | 0.793349027 | 0.20281848 | 3.91162101 | 9.17E-05 | 0.00118167 |
| IL18 | 1199.47139 | -0.770794154 | 0.39319873 | -1.960317 | 0.04995875 | 0.14561832 |
| AP002884.2 | 4.72716495 | 0.195005505 | 0.63307689 | 0.30802815 | 0.75806091 | 0.86618156 |
| TEX12 | 9.52330827 | 0.307756194 | 0.59761221 | 0.51497642 | 0.6065695 | 0.76313157 |
| BCO2 | 250.507938 | -0.280125829 | 0.28150927 | -0.9950856 | 0.31969461 | 0.51710403 |
| PTS | 1380.99889 | -0.343239228 | 0.14777912 | -2.3226504 | 0.02019794 | 0.07541776 |
| PLET1 | 1.97509485 | 0.408501584 | 0.82872081 | 0.49293028 | 0.62206185 | 0.77442031 |
| AP002884.4 | 0.72964263 | 1.757741351 | 1.41679234 | 1.24064854 | 0.21473561 | NA |
| AP002884.1 | 209.864707 | -1.63313286 | 0.23697201 | -6.89167 | 5.51E-12 | 1.23E-09 |
| AP003063.1 | 3.92851681 | 0.209685883 | 0.77561662 | 0.27034733 | 0.78689306 | 0.88401013 |
| AP003063.2 | 0 | NA | NA | NA | NA | NA |
| AP003100.1 | 0.07830766 | -0.642905668 | 3.30558948 | -0.1944905 | 0.84579185 | NA |
| AP000802.1 | 4.04146226 | 0.535145881 | 1.03634499 | 0.51637812 | 0.60559034 | 0.76237253 |
| NCAM1 | 239.601726 | 0.000993675 | 0.72016022 | 0.0013798 | 0.99889908 | 0.99954346 |
| NCAM1-AS1 | 9.81804738 | 0.846451576 | 0.80812055 | 1.04743231 | 0.29490023 | 0.4916937 |
| AP000880.1 | 4.4660705 | -0.096178856 | 0.62257381 | -0.1544859 | 0.87722665 | 0.93615811 |
| TTC12 | 819.727767 | 0.41629371 | 0.22065215 | 1.88665152 | 0.05920721 | 0.16405319 |
| ANKK1 | 68.3681044 | 0.2841821 | 0.38870956 | 0.73109111 | 0.46472351 | 0.65205955 |
| AP002840.1 | 0.85055061 | 1.549190542 | 1.40659579 | 1.10137578 | 0.27073314 | NA |
| DRD2 | 23.7145203 | 1.310209327 | 0.65949488 | 1.98668612 | 0.04695719 | 0.13914345 |
| TMPRSS5 | 81.1742786 | -0.288198925 | 0.41808744 | -0.6893269 | 0.49061756 | 0.67369536 |
| ZW10 | 847.707682 | -0.043301235 | 0.16720739 | -0.2589672 | 0.79566052 | 0.88916252 |
| AP003170.4 | 0.21442573 | -1.003574173 | 3.30558948 | -0.3035992 | 0.7614333 | NA |
| CLDN25 | 0.10721287 | -0.642905668 | 3.30558948 | -0.1944905 | 0.84579185 | NA |
| USP28 | 958.81929 | 0.139518315 | 0.21477011 | 0.64961702 | 0.51593964 | 0.69347715 |
| AP003170.3 | 1.07768173 | 0.307604381 | 1.20691463 | 0.25486838 | 0.79882476 | 0.89107318 |
| HTR3B | 12.1570863 | -0.159957468 | 0.89413214 | -0.1788969 | 0.85801866 | 0.92581333 |
| HTR3A | 27.689508 | -2.845117659 | 0.69742183 | -4.0794789 | 4.51E-05 | 0.00068406 |
| ZBTB16 | 356.338486 | -0.547100189 | 0.51505963 | -1.0622075 | 0.28814151 | 0.48427607 |
| AP002518.2 | 1.14537062 | -3.427247643 | 2.08735673 | -1.641908 | 0.10060908 | 0.23850898 |
| NNMT | 17941.876 | -4.991846225 | 0.6264238 | -7.9688004 | 1.60E-15 | 1.11E-12 |
| AP002518.1 | 0.88766976 | 0.611426411 | 1.3056163 | 0.46830482 | 0.63956662 | NA |
| C11orf71 | 900.887803 | 0.782956418 | 0.26875012 | 2.91332488 | 0.00357602 | 0.02090903 |
| RBM7 | 1872.56755 | -0.28805035 | 0.07013837 | -4.1068867 | 4.01E-05 | 0.00062353 |
| AP002373.1 | 1703.08367 | 0.02159966 | 0.17353022 | 0.12447203 | 0.90094154 | 0.94981923 |
| REXO2 | 3625.57422 | -0.148341741 | 0.17220471 | -0.8614267 | 0.38900305 | 0.58456863 |
| NXPE1 | 2.01556636 | -0.460434822 | 0.99871253 | -0.4610284 | 0.64477824 | 0.79009304 |
| NXPE4 | 6.54978852 | 1.015941488 | 0.91114308 | 1.1150186 | 0.26484245 | 0.45883645 |
| NXPE2 | 3.28602133 | -0.616978804 | 0.78829028 | -0.7826797 | 0.43381522 | 0.62580456 |
| CADM1 | 2237.13317 | 0.417793997 | 0.28177422 | 1.48272611 | 0.1381472 | 0.29711618 |
| AP000462.3 | 1.32860774 | 0.269611086 | 1.05677271 | 0.25512684 | 0.79862513 | 0.89098462 |
| AP000462.1 | 0.83138047 | -0.053630666 | 1.84420775 | -0.0290806 | 0.97680031 | NA |
| AP003174.2 | 0 | NA | NA | NA | NA | NA |
| AP003174.1 | 0.78844523 | -0.10396473 | 1.5508919 | -0.0670354 | 0.94655348 | NA |
| AP000997.1 | 0.18393104 | 1.469068379 | 3.27783996 | 0.44818185 | 0.65402196 | NA |
| AP000997.2 | 0.33384977 | -1.711569964 | 2.86877804 | -0.5966199 | 0.55076121 | NA |
| AP002991.1 | 0.43437247 | -0.662636422 | 2.3015859 | -0.2879043 | 0.77342001 | NA |
| LINC00900 | 13.2161243 | 0.517237321 | 0.52811827 | 0.97939676 | 0.32738398 | 0.52522085 |
| AP000797.3 | 0.47954304 | -0.97807301 | 2.24788683 | -0.4351078 | 0.66348421 | NA |
| LINC02151 | 0 | NA | NA | NA | NA | NA |
| AP000770.1 | 0.12150485 | 1.04981648 | 3.28477129 | 0.31960109 | 0.74927075 | NA |
| BUD13 | 825.872199 | -0.055145866 | 0.12269322 | -0.4494614 | 0.65309886 | 0.79614538 |
| AP006216.1 | 23.6067505 | -0.344022632 | 0.38356218 | -0.8969149 | 0.36976433 | 0.56653799 |
| ZPR1 | 1850.48579 | -0.6439827 | 0.22108228 | -2.9128644 | 0.0035813 | 0.02092972 |
| APOA5 | 0.45751595 | 2.048321891 | 3.21841163 | 0.63643875 | 0.52449049 | NA |
| AP006216.2 | 0 | NA | NA | NA | NA | NA |
| AP006216.3 | 0.3216386 | -1.364242679 | 3.30558948 | -0.4127078 | 0.67982068 | NA |
| APOA4 | 0.6250991 | -1.436928384 | 2.79951535 | -0.5132776 | 0.60775718 | NA |
| APOC3 | 130.994371 | 4.782199097 | 1.01138663 | 4.72835902 | 2.26E-06 | 6.28E-05 |
| APOA1 | 36.1602438 | 2.134969139 | 0.56507132 | 3.77822954 | 0.00015795 | 0.00182676 |
| APOA1-AS | 10.4098859 | 2.153349809 | 0.55914424 | 3.85115261 | 0.00011756 | 0.0014426 |
| SIK3 | 2410.59496 | 0.084846896 | 0.16741178 | 0.50681557 | 0.61228423 | 0.76779926 |
| AP000936.1 | 0.54301362 | 3.042736168 | 3.19237857 | 0.95312511 | 0.34052669 | NA |
| PAFAH1B2 | 4164.46764 | -0.557215196 | 0.18259115 | -3.0517097 | 0.00227542 | 0.01491124 |
| SIDT2 | 2687.65519 | 0.037810955 | 0.20652982 | 0.18307746 | 0.85473724 | 0.92400911 |
| TAGLN | 24824.7264 | -0.477519186 | 0.59183591 | -0.8068439 | 0.41975643 | 0.6125435 |
| PCSK7 | 13976.5134 | -0.558058996 | 0.51001297 | -1.0942055 | 0.27386486 | 0.46888822 |
| RNF214 | 1431.15153 | -0.046176386 | 0.10358918 | -0.4457646 | 0.65576732 | 0.79807876 |
| BACE1 | 2422.19464 | 0.355295151 | 0.10209154 | 3.48016261 | 0.00050111 | 0.004542 |
| BACE1-AS | 489.809908 | 0.412444853 | 0.10387212 | 3.97069846 | 7.17E-05 | 0.00097259 |
| CEP164 | 2121.53361 | 0.209507397 | 0.21374663 | 0.98016702 | 0.32700368 | 0.52487577 |
| AP000892.1 | 1.0245398 | 3.440806742 | 1.72838502 | 1.99076404 | 0.04650683 | 0.13816108 |
| DSCAML1 | 623.827247 | -0.126603865 | 0.37350479 | -0.3389618 | 0.73463849 | 0.85065526 |
| AP001554.1 | 0.71053654 | -2.721533217 | 1.8185746 | -1.49652 | 0.13451821 | NA |
| FXYD2 | 60877.7724 | -0.174911945 | 0.21912182 | -0.7982407 | 0.42473084 | 0.61708828 |
| AP000757.2 | 27231.0077 | -0.10745 | 0.20411432 | -0.5264207 | 0.59859595 | 0.75677743 |
| FXYD6-FXYD2 | 55617.213 | -0.105617052 | 0.20740581 | -0.509229 | 0.61059171 | 0.76647325 |
| AP000757.1 | 2684.66105 | 0.314624994 | 0.2461576 | 1.27814453 | 0.20119847 | 0.38296548 |
| FXYD6 | 3712.02563 | 0.258506811 | 0.24591528 | 1.05120273 | 0.29316549 | 0.48970402 |
| TMPRSS13 | 137.793999 | -0.570541825 | 0.63752144 | -0.8949375 | 0.37082052 | 0.5675956 |
| IL10RA | 1004.36431 | -0.038800879 | 0.45694372 | -0.0849139 | 0.93232983 | 0.96734562 |
| SMIM35 | 116.235183 | 4.247679029 | 0.82013075 | 5.17927051 | 2.23E-07 | 9.29E-06 |
| TMPRSS4 | 2349.06575 | -0.879013573 | 0.44761821 | -1.9637574 | 0.04955823 | 0.14477131 |
| SCN4B | 294.245237 | 2.036231746 | 0.62302384 | 3.26830472 | 0.00108194 | 0.00833257 |
| SCN2B | 34.2911704 | 1.883674491 | 0.73585905 | 2.55983057 | 0.01047232 | 0.04667147 |
| JAML | 446.970687 | 0.354416169 | 0.71423751 | 0.49621612 | 0.61974194 | 0.77271053 |
| MPZL3 | 545.80564 | -0.8367507 | 0.20168513 | -4.1487971 | 3.34E-05 | 0.00053628 |
| MPZL2 | 2871.61549 | -1.724312434 | 0.39491649 | -4.366271 | 1.26E-05 | 0.00024706 |
| CD3E | 311.021963 | -0.202776237 | 0.7610582 | -0.2664399 | 0.78990047 | 0.88584143 |
| CD3D | 107.150589 | -0.381029958 | 0.76610863 | -0.4973576 | 0.61893689 | 0.77231656 |
| CD3G | 63.2842277 | 0.231878872 | 0.79912859 | 0.29016466 | 0.77169027 | 0.87415579 |
| UBE4A | 2917.53008 | 0.07466699 | 0.10299541 | 0.72495454 | 0.46847992 | 0.65527054 |
| AP001267.1 | 6774.24955 | -0.03669171 | 0.12467332 | -0.2943028 | 0.76852652 | 0.87214547 |
| ATP5MG | 8515.05432 | 0.021364373 | 0.14962935 | 0.14278196 | 0.88646238 | 0.94121489 |
| AP001267.2 | 4.33940662 | -0.006322741 | 0.90627408 | -0.0069766 | 0.9944335 | 0.99749764 |
| KMT2A | 2444.8449 | -0.101442141 | 0.11052755 | -0.9177996 | 0.35872381 | 0.5564112 |
| AP001267.3 | 279.162729 | 2.491393693 | 0.39257654 | 6.34626224 | 2.21E-10 | 2.84E-08 |

| | | | | | | |
|---|---|---|---|---|---|---|
| TTC36 | 291.236262 | 2.925259132 | 0.48223351 | 6.06606361 | 1.31E-09 | 1.29E-07 |
| TMEM25 | 2210.48999 | 0.451103106 | 0.19615468 | 2.29973149 | 0.02146344 | 0.07898281 |
| IFT46 | 1348.59468 | 0.362567513 | 0.16847902 | 2.15200398 | 0.03139704 | 0.10430083 |
| ARCN1 | 5865.03916 | -0.042621959 | 0.20617809 | -0.206724 | 0.83622542 | 0.91299005 |
| PHLDB1 | 6941.04134 | 0.463250099 | 0.23173313 | 1.99906719 | 0.04560108 | 0.13616811 |
| AP000941.1 | 9.30153528 | 1.335511874 | 0.52018644 | 2.56737158 | 0.01024727 | 0.04590904 |
| TREH | 3609.52125 | 0.888389675 | 0.19828374 | 4.48039606 | 7.45E-06 | 0.00016144 |
| AP002954.1 | 127.997201 | 0.872859334 | 0.28788989 | 3.03192072 | 0.00243003 | 0.01564833 |
| DDX6 | 4347.3817 | 0.167225813 | 0.11606959 | 1.44073749 | 0.14965886 | 0.31392066 |
| AP004609.3 | 136.045126 | 0.452301821 | 0.22080176 | 2.04845206 | 0.04051572 | 0.12498826 |
| CXCR5 | 2872.12049 | -1.068309963 | 0.29392443 | -3.6346416 | 0.00027837 | 0.0028674 |
| AP004609.1 | 7.80337693 | 0.428885046 | 0.69440433 | 0.61763014 | 0.53681917 | 0.71053163 |
| BCL9L | 6391.89288 | -0.66478081 | 0.32461839 | -2.047884 | 0.04057137 | 0.12510138 |
| UPK2 | 88.1581304 | 0.277886555 | 0.37501791 | 0.74099543 | 0.45869621 | 0.64757615 |
| FOXR1 | 0.54790329 | 0.920269447 | 1.77360551 | 0.5188693 | 0.60385189 | NA |
| AP003392.4 | 57.5732741 | 1.189932466 | 0.27798041 | 4.28063432 | 1.86E-05 | 0.00033545 |
| CCDC84 | 3331.39352 | -0.42385777 | 0.1220531 | -3.4727325 | 0.00051519 | 0.00464049 |
| AP003392.1 | 4.26618223 | 0.581173 | 0.72082615 | 0.80625959 | 0.42009318 | 0.61281503 |
| RPS25 | 15596.966 | -0.326268485 | 0.16306586 | -2.0008387 | 0.04540978 | 0.13573519 |
| TRAPPC4 | 6280.65462 | 0.563527271 | 0.20958662 | 2.6887559 | 0.00717188 | 0.03526433 |
| SLC37A4 | 8474.39647 | 1.150720313 | 0.34767883 | 3.30972209 | 0.00093389 | 0.00745131 |
| AP003392.3 | 2470.35389 | -0.76773772 | 0.2793981 | -2.7478272 | 0.00599916 | 0.03077679 |
| HYOU1 | 8136.36795 | -0.301994902 | 0.24624423 | -1.226404 | 0.22004668 | 0.40575336 |
| AP003392.5 | 6.63260419 | -0.868783698 | 0.5045302 | -1.7219657 | 0.08507573 | 0.21150627 |
| AP003392.6 | 11.7667377 | -0.38468006 | 0.46671655 | -0.8242263 | 0.40981098 | 0.60426726 |
| VPS11 | 2765.8828 | 0.279802253 | 0.10792856 | 2.5924765 | 0.00952877 | 0.04345791 |
| HMBS | 988.201558 | -0.097794679 | 0.18227332 | -0.5365277 | 0.59159392 | 0.75117159 |
| H2AFX | 1076.18854 | -1.438084169 | 0.24166177 | -5.9508137 | 2.67E-09 | 2.28E-07 |
| DPAGT1 | 2023.933 | -0.182657906 | 0.15371601 | -1.1882816 | 0.23472249 | 0.42348332 |
| C2CD2L | 1773.6516 | 0.147065537 | 0.23350314 | 0.62982251 | 0.52881071 | 0.70340389 |
| HINFP | 1736.73092 | -0.413654529 | 0.14994332 | -2.7587393 | 0.00580248 | 0.03007287 |
| ABCG4 | 46.9653243 | 0.626400362 | 0.53868565 | 1.16283099 | 0.24489808 | 0.43599208 |
| NLRX1 | 1438.90443 | 0.305472673 | 0.17194604 | 1.77656126 | 0.07564045 | 0.19490827 |
| PDZD3 | 3900.8195 | 2.177398319 | 0.71178995 | 3.05904618 | 0.00222043 | 0.01463452 |
| CCDC153 | 868.571975 | 1.690508581 | 0.60720354 | 2.78408881 | 0.00536783 | 0.02838944 |
| CBL | 1268.51075 | -0.411399092 | 0.23886346 | -1.7223191 | 0.08501173 | 0.21138703 |
| MCAM | 5944.14037 | -0.413352176 | 0.39377914 | -1.0497056 | 0.29385348 | 0.49046878 |
| RNF26 | 1884.56878 | -0.076991499 | 0.0982978 | -0.7832474 | 0.43348184 | 0.62557806 |
| AP003396.1 | 728.192099 | -0.340805656 | 0.11611151 | -2.9351582 | 0.00333378 | 0.019877 |
| C1QTNF5 | 1558.62038 | -0.773721197 | 0.31607127 | -2.4479327 | 0.01436785 | 0.05899722 |
| MFRP | 1559.83136 | -0.774990184 | 0.31620642 | -2.4508996 | 0.01424997 | 0.05864103 |
| USP2 | 2457.10331 | 1.666092717 | 0.43098767 | 3.86575492 | 0.00011075 | 0.00137124 |
| AP003396.3 | 81.066732 | 2.624030119 | 0.43061162 | 6.09372805 | 1.10E-09 | 1.13E-07 |
| USP2-AS1 | 79.761321 | 0.57156703 | 0.40649039 | 1.40610221 | 0.15969378 | 0.32758654 |
| THY1 | 8730.98214 | 3.070037599 | 0.50731219 | 6.05157471 | 1.43E-09 | 1.38E-07 |
| AP003393.1 | 11.5653815 | 0.496271688 | 0.73976553 | 0.67084997 | 0.50231611 | 0.68263189 |
| NECTIN1 | 909.966167 | 0.825542328 | 0.21719406 | 3.80094341 | 0.00014415 | 0.00169391 |
| AP003390.2 | 0.11051935 | 0.02007176 | 3.29738704 | 0.00608717 | 0.99514317 | NA |
| AP003390.1 | 3.93308799 | 0.482149866 | 0.65653481 | 0.73438584 | 0.46271362 | 0.65089929 |
| AP001360.1 | 0.08757577 | 0.547239394 | 3.29377202 | 0.16614368 | 0.86804388 | NA |
| AP001360.2 | 0.03392908 | 0.380740593 | 3.29738709 | 0.11546736 | 0.90807469 | NA |
| TRIM29 | 104.005846 | 0.854911331 | 0.3752364 | 2.27832727 | 0.02270709 | 0.08232997 |
| AP000679.1 | 1.48779701 | 0.818106507 | 1.25423504 | 0.65227527 | 0.51422361 | 0.69208781 |
| OAF | 6321.3343 | -0.08889213 | 0.26152149 | -0.3399267 | 0.73391174 | 0.8501319 |
| POU2F3 | 40.0688353 | 0.906914183 | 0.55156026 | 1.64427035 | 0.10012037 | 0.23766083 |
| AP001150.1 | 0 | NA | NA | NA | NA | NA |
| TMEM136 | 589.819574 | -0.507916432 | 0.19711966 | -2.5766909 | 0.00997511 | 0.04499549 |
| ARHGEF12 | 9974.07267 | 0.123469508 | 0.1356771 | 0.91002466 | 0.3628095 | 0.56004722 |
| GRIK4 | 18.2490846 | 0.324353685 | 0.72011279 | 0.45042067 | 0.65240714 | 0.79583057 |
| AP004147.1 | 0 | NA | NA | NA | NA | NA |
| TBCEL | 1025.23084 | 0.062369542 | 0.16597583 | 0.37577484 | 0.7070843 | 0.83325953 |
| AP000646.1 | 118.51646 | 0.699671078 | 0.28656985 | 2.44153766 | 0.01462486 | 0.05977335 |
| TECTA | 124.13566 | 0.087465994 | 0.26961034 | 0.32441632 | 0.74562286 | 0.85740777 |
| SC5D | 2694.40143 | -0.164453359 | 0.27337936 | -0.6015573 | 0.54746883 | 0.71842513 |
| AP000977.1 | 1.66800726 | 0.610574317 | 1.01307986 | 0.6026912 | 0.54671413 | 0.71786345 |
| SORL1 | 3552.57797 | 0.901293787 | 0.34304985 | 2.62729684 | 0.00860662 | 0.04020489 |
| BLID | 1.48548246 | 0.987478432 | 1.0039393 | 0.98360372 | 0.3253104 | 0.52283304 |
| AP000755.1 | 5.73344143 | 0.846177518 | 0.78619056 | 1.07630078 | 0.28179276 | 0.47765319 |
| AP000755.2 | 0.07659028 | -0.642905668 | 3.30558948 | -0.1944905 | 0.84579185 | NA |
| UBASH3B | 376.297996 | -1.770205088 | 0.38402674 | -4.6095881 | 4.03E-06 | 9.94E-05 |
| CRTAM | 19.4791683 | 0.044782431 | 0.71960938 | 0.06223158 | 0.95037841 | 0.9761964 |
| JHY | 528.733498 | 0.228682977 | 0.24030544 | 0.95163462 | 0.34128232 | 0.53931892 |
| BSX | 0.17453125 | 0.18657164 | 3.29377195 | 0.05664376 | 0.95482897 | NA |
| HSPA8 | 49647.0873 | 0.572533039 | 0.23607902 | 2.42517539 | 0.01530099 | 0.06171416 |
| CLMP | 590.514919 | -0.105175039 | 0.41543947 | -0.2531657 | 0.80014015 | 0.89152759 |
| AP001970.1 | 0.45189822 | -0.669510801 | 2.29731454 | -0.2914319 | 0.770721 | NA |
| AP001782.1 | 0 | NA | NA | NA | NA | NA |
| GRAMD1B | 1501.67209 | 2.341784507 | 0.49471005 | 4.7336506 | 2.21E-06 | 6.15E-05 |
| SCN3B | 146.412443 | -0.092515717 | 0.30881946 | -0.2995787 | 0.76449857 | 0.86974534 |
| AP002765.1 | 0.13382144 | 0.02007176 | 3.29738704 | 0.00608717 | 0.99514317 | NA |
| ZNF202 | 708.423377 | -0.197121647 | 0.18348836 | -1.0743005 | 0.282688 | 0.4786492 |
| OR6X1 | 0 | NA | NA | NA | NA | NA |
| OR6M1 | 0 | NA | NA | NA | NA | NA |
| TMEM225 | 0 | NA | NA | NA | NA | NA |
| OR8D4 | 0.12059279 | 0.02007176 | 3.29738704 | 0.00608717 | 0.99514317 | NA |
| OR4D5 | 0 | NA | NA | NA | NA | NA |
| OR6T1 | 0 | NA | NA | NA | NA | NA |
| OR10S1 | 0 | NA | NA | NA | NA | NA |
| OR10G6 | 0 | NA | NA | NA | NA | NA |
| OR10G4 | 0.24613888 | 1.011711844 | 2.95767469 | 0.34206326 | 0.73230329 | NA |
| OR10G9 | 0 | NA | NA | NA | NA | NA |
| OR10G8 | 0 | NA | NA | NA | NA | NA |
| OR10G7 | 0 | NA | NA | NA | NA | NA |
| VWA5A | 2678.45644 | -0.743068509 | 0.20742112 | -3.5824149 | 0.00034043 | 0.00335346 |
| OR10D3 | 0 | NA | NA | NA | NA | NA |
| OR8G1 | 0 | NA | NA | NA | NA | NA |
| OR8G5 | 0.1357163 | 1.313406857 | 3.28103699 | 0.40030236 | 0.68893383 | NA |
| OR8D1 | 0.03392908 | 0.380740593 | 3.29738709 | 0.11546736 | 0.90807469 | NA |
| OR8D2 | 0 | NA | NA | NA | NA | NA |
| OR8B2 | 0 | NA | NA | NA | NA | NA |
| OR8B3 | 0 | NA | NA | NA | NA | NA |
| OR8B4 | 0 | NA | NA | NA | NA | NA |
| OR8B8 | 0 | NA | NA | NA | NA | NA |
| OR8B12 | 0 | NA | NA | NA | NA | NA |
| OR8A1 | 0 | NA | NA | NA | NA | NA |
| PANX3 | 0.53668769 | 1.788076691 | 1.69259535 | 1.0564112 | 0.29078039 | NA |
| TBRG1 | 3262.96753 | -0.146105787 | 0.11804342 | -1.2377292 | 0.2158165 | 0.40041379 |
| SIAE | 7209.56175 | 0.318504541 | 0.2058809 | 1.54703296 | 0.1218553 | 0.27274026 |
| SPA17 | 507.085889 | -0.040071321 | 0.17953665 | -0.223193 | 0.82338531 | 0.90564302 |
| NRGN | 502.172918 | 0.36028784 | 0.42448756 | 0.84875949 | 0.39601514 | 0.59122987 |
| AP000866.5 | 110.71859 | 0.451576672 | 0.43141562 | 1.04673232 | 0.29522305 | 0.49206952 |
| VSIG2 | 345.831481 | 0.057914256 | 0.29176702 | 0.19849487 | 0.8426579 | 0.9163631 |
| ESAM | 4878.51211 | -0.043904163 | 0.36654007 | -0.11978 | 0.90465744 | 0.95213796 |

| | | | | | | |
|---|---|---|---|---|---|---|
| AP000866.2 | 105.806759 | 0.781475087 | 0.25094603 | 3.11411615 | 0.00184497 | 0.01265985 |
| MSANTD2 | 1306.50798 | 0.136332852 | 0.17772334 | 0.7671072 | 0.44301778 | 0.6341881 |
| AP000866.1 | 53.7571199 | 1.586776936 | 0.24565264 | 6.45943356 | 1.05E-10 | 1.51E-08 |
| ROBO3 | 1097.39041 | -0.617921722 | 0.36381312 | -1.6984591 | 0.08942114 | 0.21957681 |
| AP003501.1 | 311.707567 | -0.063031707 | 0.34706094 | -0.1816157 | 0.85588435 | 0.92454691 |
| ROBO4 | 2560.5584 | 0.120346863 | 0.36193022 | 0.332514 | 0.73950117 | 0.85383099 |
| AP003501.2 | 258.875261 | 0.3323383 | 0.34833949 | 0.95406438 | 0.34005106 | 0.53817947 |
| HEPACAM | 18.7826695 | 1.668735403 | 0.84892552 | 1.96570296 | 0.04933293 | 0.14432099 |
| HEPN1 | 11.9909957 | 1.373323026 | 0.92270032 | 1.48837385 | 0.13665233 | 0.29498449 |
| CCDC15 | 38.3104865 | 0.467043586 | 0.40166537 | 1.16276786 | 0.2449237 | 0.43599208 |
| SLC37A2 | 189.361035 | 0.660055368 | 0.33946004 | 1.94442729 | 0.05184395 | 0.14955966 |
| TMEM218 | 714.150424 | 0.403271484 | 0.18586893 | 2.16965522 | 0.03003297 | 0.10091151 |
| AP001007.2 | 0.06785815 | 0.795868262 | 3.28898349 | 0.24198001 | 0.80879565 | NA |
| PKNOX2-AS1 | 0.68591745 | -0.57312448 | 1.60060987 | -0.3580663 | 0.72029369 | NA |
| PKNOX2 | 300.279174 | 0.095312681 | 0.47502139 | 0.20064924 | 0.84097285 | 0.91512686 |
| AP001007.1 | 1.44093309 | -0.49060441 | 1.24397456 | -0.3943846 | 0.69329711 | 0.82352701 |
| FEZ1 | 308.847379 | -0.193247898 | 0.38172049 | -0.506255 | 0.61267767 | 0.76799695 |
| EI24 | 4603.16467 | -0.268556364 | 0.146312 | -1.8355047 | 0.06643095 | 0.17746774 |
| STT3A-AS1 | 6.52768636 | 0.46717525 | 0.8294618 | 0.56322696 | 0.57328034 | 0.7378662 |
| STT3A | 5655.50655 | -0.172178639 | 0.15792278 | -1.0902711 | 0.27559375 | 0.47090048 |
| CHEK1 | 286.944983 | 0.07182078 | 0.21425359 | 0.33521389 | 0.73746375 | 0.85252908 |
| ACRV1 | 6.81449888 | -0.779901415 | 0.74888282 | -1.0414198 | 0.29768074 | 0.49454947 |
| PATE1 | 0.90977748 | 1.629104189 | 1.46571218 | 1.11147619 | 0.26636343 | NA |
| PATE2 | 0.49088144 | -0.936262384 | 2.2361138 | -0.4187007 | 0.67543489 | NA |
| PATE3 | 0.17423948 | 0.18657164 | 3.29377195 | 0.05664376 | 0.95482897 | NA |
| PATE4 | 0.24835481 | -0.340597073 | 3.29738699 | -0.103293 | 0.91773041 | NA |
| AP000842.3 | 3.02799938 | -0.410264614 | 1.0064393 | -0.4076397 | 0.68353821 | 0.81719648 |
| HYLS1 | 271.06066 | -0.208630923 | 0.24745881 | -0.8430935 | 0.39917614 | 0.59400004 |
| PUS3 | 520.94476 | -0.131314112 | 0.20688146 | -0.6347312 | 0.52560376 | 0.70039776 |
| AP000842.2 | 2.11852776 | 2.554827057 | 0.92143511 | 2.77266084 | 0.00556 | 0.02922992 |
| DDX25 | 146.896427 | 1.469647492 | 0.38094796 | 3.85786945 | 0.00011438 | 0.00140746 |
| AP000842.1 | 0 | NA | NA | NA | NA | NA |
| VSIG10L2 | 128.240204 | -0.298509491 | 0.46513548 | -0.6417689 | 0.52102324 | 0.69753682 |
| CDON | 202.989941 | -0.061056156 | 0.41879447 | -0.1457903 | 0.88408696 | 0.93990377 |
| AP000821.1 | 0.6466237 | -2.579672194 | 2.67658635 | -0.9637919 | 0.33515028 | NA |
| AP001893.3 | 1.82594175 | -0.20296978 | 1.0216094 | -0.1986765 | 0.84251581 | 0.91630899 |
| RPUSD4 | 1348.78106 | -0.018192996 | 0.18399117 | -0.0988797 | 0.92123377 | 0.9615743 |
| AP001893.1 | 1.68517659 | 2.790003634 | 1.13538549 | 2.45731838 | 0.01399786 | 0.05787449 |
| FAM118B | 1126.47697 | -0.475239193 | 0.17913119 | -2.6530231 | 0.00797744 | 0.03807513 |
| SRPRA | 8096.50991 | -0.176440147 | 0.24339973 | -0.7248987 | 0.46851418 | 0.6552837 |
| FOXRED1 | 2005.0843 | 0.690208187 | 0.12292709 | 5.61477669 | 1.97E-08 | 1.24E-06 |
| TIRAP | 367.12215 | 0.733600246 | 0.23357425 | 3.14075826 | 0.00168511 | 0.01181612 |
| AP001318.1 | 118.244717 | 0.716002518 | 0.28219746 | 2.53723941 | 0.01117305 | 0.04894242 |
| AP001318.2 | 261.590248 | 0.207095685 | 0.12466769 | 1.66118163 | 0.09667697 | 0.23205642 |
| DCPS | 1731.30944 | -0.430897651 | 0.35598501 | -1.2104376 | 0.22611102 | 0.41281666 |
| GSEC | 172.005124 | 0.464687691 | 0.37535961 | 1.23797999 | 0.21572348 | 0.4003257 |
| ST3GAL4 | 2510.27568 | 0.529535394 | 0.34000818 | 1.55741959 | 0.1193709 | 0.26888277 |
| KIRREL3 | 24.3856654 | 0.939095963 | 0.52332925 | 1.79446488 | 0.072739 | 0.18968823 |
| KIRREL3-AS1 | 0.81706512 | 1.899450287 | 1.38595086 | 1.37050335 | 0.17052983 | NA |
| AP001783.1 | 0.91400308 | -1.118520501 | 1.66856623 | -0.6703483 | 0.5026358 | NA |
| AP002833.1 | 0.27330048 | -0.174096114 | 3.29377188 | -0.0528562 | 0.95784652 | NA |
| KIRREL3-AS2 | 0 | NA | NA | NA | NA | NA |
| AP002833.2 | 0.08757577 | 0.547239394 | 3.29377202 | 0.16614368 | 0.86804388 | NA |
| KIRREL3-AS3 | 0 | NA | NA | NA | NA | NA |
| AP001993.1 | 0.37395748 | -1.027134238 | 2.28569092 | -0.4493758 | 0.65316057 | NA |
| AP003121.1 | 0 | NA | NA | NA | NA | NA |
| AP003481.1 | 0.36673167 | 0.968025711 | 2.19253091 | 0.44151063 | 0.65884337 | NA |
| AP000904.1 | 0.60134296 | -1.484148951 | 1.84974744 | -0.8023522 | 0.42234925 | NA |
| LINC02098 | 0 | NA | NA | NA | NA | NA |
| ETS1 | 5085.12953 | -0.393251078 | 0.32335717 | -1.2161508 | 0.22392748 | 0.4100166 |
| AP003397.1 | 0.64673218 | -0.020077177 | 1.66847462 | -0.0120333 | 0.99039908 | NA |
| FLI1 | 970.860566 | 0.080621801 | 0.26638483 | 0.30265163 | 0.76215537 | 0.86816102 |
| SENCR | 30.7473884 | 0.255581494 | 0.38227909 | 0.66857304 | 0.50376788 | 0.68364658 |
| KCNJ1 | 9856.18004 | 1.158147405 | 0.39117577 | 2.96068288 | 0.00306958 | 0.01869297 |
| KCNJ5 | 571.2635 | 1.584801065 | 0.43490161 | 3.64404504 | 0.00026839 | 0.0027787 |
| C11orf45 | 138.281132 | 0.424058277 | 0.33022786 | 1.28413839 | 0.19909354 | 0.38013878 |
| TP53AIP1 | 43.5394428 | 0.175740493 | 0.39709247 | 0.44256818 | 0.6580781 | 0.79943711 |
| ARHGAP32 | 1326.42141 | 0.249599234 | 0.16407161 | 1.52128232 | 0.128189 | 0.282307 |
| BARX2 | 288.970019 | 0.064151649 | 0.30084332 | 0.2132394 | 0.83114024 | 0.91040231 |
| AP003500.2 | 0 | NA | NA | NA | NA | NA |
| AP003500.1 | 0.85772874 | 2.632011917 | 1.43928368 | 1.82869573 | 0.0674452 | NA |
| TMEM45B | 1610.11442 | -0.281638717 | 0.35607461 | -0.7909542 | 0.42897071 | 0.62112297 |
| NFRKB | 1736.51475 | 0.235554591 | 0.09588988 | 2.4565115 | 0.01402933 | 0.05797735 |
| PRDM10 | 516.555338 | -0.134607053 | 0.24751694 | -0.5438297 | 0.5865587 | 0.74768058 |
| LINC00167 | 4.46391574 | 0.66530969 | 0.95371927 | 0.69759489 | 0.48543058 | 0.6696032 |
| APLP2 | 40992.13 | 0.208023777 | 0.15704589 | 1.32460502 | 0.18530219 | 0.36268231 |
| ST14 | 8659.44101 | -0.72113411 | 0.17180693 | -4.1973518 | 2.70E-05 | 0.00045244 |
| ZBTB44 | 3526.43295 | 0.523242743 | 0.18099345 | 2.89094851 | 0.00384081 | 0.02210918 |
| AP002986.1 | 59.5541673 | 1.38554909 | 0.52441188 | 2.64210086 | 0.00823935 | 0.03892363 |
| ADAMTS8 | 143.414373 | 0.721062302 | 0.6670738 | 1.08093333 | 0.27972677 | 0.47534443 |
| ADAMTS15 | 1068.30443 | 1.451809267 | 0.41223243 | 3.52182209 | 0.00042859 | 0.00402635 |
| C11orf44 | 0.34228177 | 0.67001768 | 2.22482457 | 0.30115529 | 0.76329608 | NA |
| LINC02551 | 9.15420805 | 0.205208803 | 0.74135236 | 0.27680333 | 0.78193112 | 0.88068696 |
| AP003486.1 | 567.985647 | 0.396615757 | 0.1292778 | 3.06793397 | 0.00215544 | 0.01427757 |
| SNX19 | 3748.55916 | 0.194816363 | 0.13524032 | 1.44051983 | 0.14972038 | 0.31399129 |
| AP002856.3 | 0 | NA | NA | NA | NA | NA |
| AP002856.1 | 0 | NA | NA | NA | NA | NA |
| AP002856.4 | 0 | NA | NA | NA | NA | NA |
| AP002856.2 | 0 | NA | NA | NA | NA | NA |
| NTM | 95.8972183 | 0.00696834 | 0.4612251 | 0.01510833 | 0.98794576 | 0.99454413 |
| AP003025.2 | 1.70146233 | -0.917988449 | 1.61822737 | -0.5672803 | 0.57052379 | 0.73568275 |
| AP003025.1 | 0.52343183 | -1.496309406 | 2.79115999 | -0.5360887 | 0.59189723 | NA |
| NTM-AS1 | 0 | NA | NA | NA | NA | NA |
| AP000844.2 | 0.2873605 | -0.174096114 | 3.29377188 | -0.0528562 | 0.95784652 | NA |
| AP000844.1 | 0 | NA | NA | NA | NA | NA |
| OPCML | 94.6137388 | 2.601749317 | 0.61635521 | 4.22118491 | 2.43E-05 | 0.00041576 |
| AP001979.2 | 0.03392908 | 0.380740593 | 3.29738709 | 0.11546736 | 0.90807469 | NA |
| SPATA19 | 0 | NA | NA | NA | NA | NA |
| IGSF9B | 44.2609437 | -0.373334699 | 0.60666956 | -0.6153839 | 0.5383012 | 0.71168553 |
| AP000911.3 | 0.8167703 | -2.190693127 | 1.60524119 | -1.3647128 | 0.17234335 | NA |
| AP000911.1 | 13.6932747 | 0.342227971 | 0.49268611 | 0.69461664 | 0.48729558 | 0.67122777 |
| AP000911.2 | 10.5543232 | -0.458946508 | 0.47592555 | -0.9643242 | 0.33488343 | 0.53297581 |
| JAM3 | 1174.54751 | 0.113262752 | 0.28962083 | 0.39107253 | 0.69574362 | 0.82494422 |
| NCAPD3 | 935.596317 | 0.233191462 | 0.13702334 | 1.70183749 | 0.08878584 | 0.21840322 |
| AP001775.2 | 14.2934025 | 0.72474241 | 0.32720693 | 2.21493599 | 0.02676446 | 0.09285154 |
| VPS26B | 2210.12489 | 0.179628033 | 0.10008982 | 1.79466838 | 0.07270655 | 0.18964217 |
| THYN1 | 1452.50736 | -0.044764019 | 0.19400202 | -0.23074 | 0.81751682 | 0.90189176 |
| ACAD8 | 3190.57131 | 0.739856096 | 0.15613718 | 4.73850057 | 2.15E-06 | 6.03E-05 |
| GLB1L3 | 71.5627074 | 0.449070344 | 0.52124944 | 0.86152676 | 0.38894799 | 0.5845192 |
| GLB1L2 | 2198.68771 | 1.050324351 | 0.33474288 | 3.13770481 | 0.00170276 | 0.01191031 |
| B3GAT1 | 417.567403 | -1.449935939 | 0.61816976 | -2.3455304 | 0.01900003 | 0.07229855 |
| AP004550.1 | 21.0825151 | -0.951673182 | 0.66042078 | -1.4410103 | 0.14958176 | 0.31382536 |
| AP001999.2 | 0.13028434 | 1.049816488 | 3.28477129 | 0.31960109 | 0.74927075 | NA |

| | | | | | | |
|---|---|---|---|---|---|---|
| AP001999.3 | 0.32299488 | 2.288366988 | 3.2247442 | 0.70962744 | 0.4779352 | NA |
| AP001999.1 | 42.5398031 | 1.601948581 | 0.74572686 | 2.1481707 | 0.0317002 | 0.10495094 |
| AP003062.1 | 0.53043597 | -0.08909288 | 2.18722399 | -0.0407333 | 0.9675085 | NA |
| AP005135.1 | 0 | NA | NA | NA | NA | NA |
| AC215217.1 | 0 | NA | NA | NA | NA | NA |
| TUBB8 | 21.6344321 | 0.272327802 | 0.66464361 | 0.40973508 | 0.68200029 | 0.81588709 |
| ZMYND11 | 4019.11812 | 0.492129239 | 0.12719395 | 3.86912468 | 0.00010923 | 0.00135928 |
| DIP2C | 2568.30752 | 1.2068921 | 0.22791756 | 5.29530113 | 1.19E-07 | 5.54E-06 |
| AL669841.1 | 5.38230786 | 1.308714002 | 0.59377072 | 2.204073 | 0.02751921 | 0.09484794 |
| AL358216.1 | 9.70603808 | -1.280157114 | 0.67670538 | -1.8917496 | 0.05852436 | 0.16262247 |
| PRR26 | 546.784741 | -0.903015539 | 0.469337 | -1.9240237 | 0.05435161 | 0.15439533 |
| AL157709.1 | 0 | NA | NA | NA | NA | NA |
| LARP4B | 3320.66945 | -0.218426826 | 0.12259727 | -1.7816614 | 0.07480447 | 0.19327787 |
| AL359878.2 | 4.48220821 | 0.430040326 | 0.63714605 | 0.6749478 | 0.49970893 | 0.68105131 |
| GTPBP4 | 2028.84006 | -0.656942149 | 0.23359074 | -2.8123638 | 0.00491788 | 0.02654975 |
| IDI2 | 163.138915 | -0.436756298 | 0.33812668 | -1.2916943 | 0.19646304 | 0.37702736 |
| IDI2-AS1 | 422.443113 | -0.121976208 | 0.45618851 | -0.2673811 | 0.78917571 | 0.88546295 |
| IDI1 | 3639.3747 | -0.411482759 | 0.33837794 | -1.2160449 | 0.22396783 | 0.4100272 |
| WDR37 | 1693.53818 | -0.038420955 | 0.1890308 | -0.2032524 | 0.8389378 | 0.91454277 |
| LINC00200 | 0.33754084 | 0.653531307 | 2.81352574 | 0.23228197 | 0.81631901 | NA |
| ADARB2 | 59.9085975 | 0.445068934 | 0.4393247 | 1.01307513 | 0.31102427 | 0.50806604 |
| ADARB2-AS1 | 0.47137242 | 2.091372371 | 2.02886544 | 1.03080881 | 0.30263049 | NA |
| LINC00700 | 0 | NA | NA | NA | NA | NA |
| AL441943.1 | 0.36778787 | 2.525748504 | 3.21274886 | 0.78616432 | 0.43177123 | NA |
| AL441943.2 | 0 | NA | NA | NA | NA | NA |
| AL441943.3 | 0 | NA | NA | NA | NA | NA |
| LINC00701 | 0 | NA | NA | NA | NA | NA |
| AC024908.1 | 0.06785815 | 0.795868262 | 3.28898349 | 0.24198001 | 0.80879565 | NA |
| AL713851.2 | 0.03392908 | 0.380740593 | 3.29738709 | 0.11546736 | 0.90807469 | NA |
| AL713851.1 | 0.17515154 | 1.45420457 | 3.2783972 | 0.44357181 | 0.65735219 | NA |
| AL731533.1 | 0 | NA | NA | NA | NA | NA |
| AL731533.2 | 74.5196799 | -1.673620906 | 0.5433495 | -3.0801922 | 0.00206867 | 0.01381813 |
| PFKP | 8639.07704 | -1.385713089 | 0.36403947 | -3.8064913 | 0.00014095 | 0.0016623 |
| AL451164.3 | 1920.37776 | -1.762252123 | 0.39383595 | -4.4745842 | 7.66E-06 | 0.00016508 |
| PITRM1 | 5343.01059 | -0.349988353 | 0.23736904 | -1.4744482 | 0.14036096 | 0.30050486 |
| PITRM1-AS1 | 1457.7033 | -0.283057084 | 0.2500515 | -1.1319951 | 0.25763648 | 0.45029234 |
| AL451164.2 | 0 | NA | NA | NA | NA | NA |
| AL451164.1 | 0.43866041 | -0.306274467 | 2.28425996 | -0.1340804 | 0.89333901 | NA |
| AL139280.1 | 82.9881269 | -0.592788842 | 0.66760813 | -0.8879293 | 0.37457877 | 0.57064323 |
| AL357833.1 | 1.07863546 | 1.066975652 | 1.07404578 | 0.99341729 | 0.32050664 | 0.51800442 |
| AL357833.2 | 0.27151858 | 0.079570021 | 2.99037415 | 0.02660872 | 0.97877182 | NA |
| AL450322.2 | 1.21716696 | -0.05942791 | 1.11548043 | -0.0532756 | 0.95751231 | 0.97957788 |
| AL450322.1 | 0.5728494 | 1.042412246 | 2.23792028 | 0.46579508 | 0.64136219 | NA |
| KLF6 | 11556.6425 | -1.436453414 | 0.40648457 | -3.5338449 | 0.00040956 | 0.00389054 |
| AL513303.2 | 0.19616881 | -0.340115482 | 3.16778127 | -0.1073671 | 0.91449776 | NA |
| AL513303.1 | 0.17059578 | -1.003574173 | 3.30558948 | -0.3035992 | 0.7614333 | NA |
| AC025822.1 | 0 | NA | NA | NA | NA | NA |
| AC025822.2 | 1.64371515 | -0.74862402 | 1.1745899 | -0.6373493 | 0.52389736 | 0.69911307 |
| LINC00702 | 46.5194374 | -0.078103354 | 0.90422371 | -0.0863761 | 0.93116741 | 0.96688001 |
| LINC00703 | 0.12150485 | 1.04981648 | 3.28477129 | 0.31960109 | 0.74927075 | NA |
| AC022535.1 | 0 | NA | NA | NA | NA | NA |
| MANCR | 5.04276199 | -1.418912466 | 0.82871371 | -1.7121865 | 0.0868623 | 0.21479358 |
| LINC00705 | 0.9471058 | -1.219062756 | 1.66984838 | -0.730044 | 0.4653633 | NA |
| AKR1E2 | 73.4404416 | -0.31121069 | 0.28060743 | -1.1090608 | 0.26740397 | 0.46151205 |
| AKR1C1 | 25943.4916 | -3.917195539 | 0.87452678 | -4.4792174 | 7.49E-06 | 0.00016206 |
| AKR1C2 | 4168.26742 | -2.19317956 | 0.874265 | -2.5085981 | 0.01212113 | 0.05192238 |
| AL391427.1 | 88.8071879 | -1.810833389 | 0.53433597 | -3.3889416 | 0.00070163 | 0.00591874 |
| AKR1C3 | 11289.9978 | -0.919635784 | 0.49530386 | -1.8567103 | 0.06335238 | 0.17177501 |
| AKR1C4 | 47.4817631 | -1.00634899 | 0.77125802 | -1.3048149 | 0.19195587 | 0.37121416 |
| AL355303.1 | 2.8327937 | -3.303346298 | 1.84619279 | -1.7892748 | 0.07357056 | 0.19114847 |
| LINC02561 | 39.818623 | -3.830816917 | 0.9282668 | -4.126849 | 3.68E-05 | 0.00057864 |
| UCN3 | 30.3735735 | 0.902981036 | 0.55499206 | 1.62701613 | 0.10373367 | 0.24368787 |
| TUBAL3 | 502.323277 | -0.219964025 | 0.61750279 | -0.3562154 | 0.72167923 | 0.84180481 |
| NET1 | 3055.27094 | 0.681601708 | 0.28505871 | 2.39109241 | 0.01679832 | 0.06613494 |
| CALML5 | 1.22505803 | 3.773883313 | 2.27919537 | 1.65579632 | 0.09776307 | 0.23394219 |
| CALML3-AS1 | 150.104362 | 2.89705235 | 0.78887645 | 3.67237778 | 0.0002403 | 0.00253777 |
| CALML3 | 728.063116 | 3.002532985 | 0.95413296 | 3.14687064 | 0.00165028 | 0.01164514 |
| AL732437.1 | 528.000135 | 3.080329337 | 1.16102512 | 2.65311171 | 0.00797535 | 0.03807203 |
| AL365356.5 | 7.79809198 | -0.530746726 | 0.72478268 | -0.7322839 | 0.46399527 | 0.65164735 |
| AL365356.4 | 8.83412807 | -0.012269765 | 0.68517617 | -0.0179075 | 0.98571268 | 0.99368571 |
| AL365356.3 | 19.0326983 | 0.554001134 | 0.35155259 | 1.57586986 | 0.1150558 | 0.26232245 |
| ASB13 | 3424.4731 | 0.927543299 | 0.28279241 | 3.27994412 | 0.00103828 | 0.0080765 |
| FAM208B | 1589.19537 | -0.2527501 | 0.17145993 | -1.4741059 | 0.14045307 | 0.300627 |
| AL365356.1 | 1.10145889 | 1.414789725 | 1.09703368 | 1.28965022 | 0.19717213 | 0.37794789 |
| GDI2 | 9692.00819 | -0.158371056 | 0.18786615 | -0.8429994 | 0.39922876 | 0.59404483 |
| AL596094.1 | 38.6475864 | 0.216005351 | 0.30152561 | 0.7163748 | 0.47375995 | 0.65959453 |
| ANKRD16 | 401.042162 | 0.253785614 | 0.13391638 | 1.89510505 | 0.05807849 | 0.16184399 |
| FBH1 | 3972.45968 | -0.030127226 | 0.11763313 | -0.2561117 | 0.79786456 | 0.89053113 |
| AL137186.2 | 1538.43567 | -0.501785924 | 0.15231484 | -3.2943994 | 0.00098632 | 0.00775849 |
| IL15RA | 929.109231 | -1.896237033 | 0.41752363 | -4.541628 | 5.58E-06 | 0.00012822 |
| IL2RA | 124.774412 | -1.735914268 | 0.56059388 | -3.096563 | 0.00195778 | 0.01327165 |
| AL137186.1 | 11.6418711 | -1.717988723 | 0.62229554 | -2.760728 | 0.00576727 | 0.02994866 |
| RBM17 | 5035.1497 | -0.473322705 | 0.15581207 | -3.0377795 | 0.00238328 | 0.01543359 |
| PFKFB3 | 17950.0716 | -0.774590066 | 0.49605153 | -1.5615113 | 0.11840316 | 0.26743395 |
| AL157395.1 | 5.71834175 | 0.170958744 | 0.6363452 | 0.26865724 | 0.78819346 | 0.88475488 |
| AL137145.2 | 75.1401436 | -0.481865274 | 0.50721904 | -0.9500142 | 0.34210505 | 0.53994876 |
| AL137145.1 | 83.6448235 | -0.46697219 | 0.47679218 | -0.979404 | 0.32738038 | 0.52522085 |
| PRKCQ | 735.872176 | 0.191616399 | 0.29525029 | 0.64899647 | 0.51634066 | 0.69373328 |
| AL158210.1 | 12.8172866 | 0.576150667 | 0.47216221 | 1.22023885 | 0.22237434 | 0.40838578 |
| LINP1 | 23.535419 | -1.670179555 | 0.79559618 | -2.0992805 | 0.03579218 | 0.11434938 |
| AL590637.1 | 5.59890919 | -1.25570376 | 0.85742802 | -1.4645005 | 0.14305725 | 0.30449863 |
| LINC00707 | 27.0408438 | -1.378479705 | 0.54281691 | -2.5394929 | 0.01110133 | 0.04868476 |
| AL392086.1 | 0.37456774 | -0.03213158 | 2.26833961 | -0.0141652 | 0.98869815 | NA |
| AL590095.1 | 0 | NA | NA | NA | NA | NA |
| SFMBT2 | 537.977358 | -0.307575058 | 0.20679522 | -1.4873412 | 0.13692472 | 0.29530784 |
| AL139125.1 | 0.72921447 | -1.819045589 | 1.9175127 | -0.9486485 | 0.3427994 | NA |
| AL445070.1 | 4.14333184 | 2.55702819 | 0.93970802 | 2.72108797 | 0.00650674 | 0.03271918 |
| ITIH5 | 2181.40467 | 0.730004224 | 0.31216254 | 2.33853884 | 0.01935931 | 0.0731802 |
| ITIH2 | 61.9276939 | -0.143054251 | 0.31379074 | -0.4558906 | 0.64846865 | 0.7930031 |
| KIN | 621.814794 | -0.18436385 | 0.11788954 | -1.5638694 | 0.11784824 | 0.26654346 |
| ATP5F1C | 11914.3165 | 0.101199284 | 0.16153421 | 0.62648827 | 0.53099473 | 0.70536772 |
| TAF3 | 539.032591 | 0.059078433 | 0.13640575 | 0.43310808 | 0.66493625 | 0.80340752 |
| GATA3 | 1982.52159 | 0.507186357 | 0.23104418 | 2.19519212 | 0.02814982 | 0.0965168 |
| GATA3-AS1 | 381.234101 | 0.108249464 | 0.28438075 | 0.38064976 | 0.70346315 | 0.83054951 |
| AL390294.1 | 152.489995 | 0.341952364 | 0.30560005 | 1.11895389 | 0.26315981 | 0.45687828 |
| LINC00708 | 0 | NA | NA | NA | NA | NA |
| AL390835.1 | 0 | NA | NA | NA | NA | NA |
| AC025946.1 | 0 | NA | NA | NA | NA | NA |
| AC044784.2 | 0.07830766 | -0.642905668 | 3.30558948 | -0.1944905 | 0.84579185 | NA |
| AC044784.1 | 0.08757577 | 0.547239394 | 3.29377202 | 0.16614368 | 0.86804388 | NA |
| LINC00709 | 0 | NA | NA | NA | NA | NA |
| AL157401.1 | 0.08695548 | -0.642905668 | 3.30558948 | -0.1944905 | 0.84579185 | NA |
| AL138775.1 | 0 | NA | NA | NA | NA | NA |

| | | | | | | |
|---|---|---|---|---|---|---|
| AL583859.1 | 0.08757577 | 0.547239394 | 3.29377202 | 0.16614368 | 0.86804388 | NA |
| AL583859.2 | 0.08757577 | 0.547239394 | 3.29377202 | 0.16614368 | 0.86804388 | NA |
| SFTA1P | 0.80251589 | -2.174799733 | 1.60091403 | -1.3584738 | 0.1743134 | NA |
| CELF2 | 679.494265 | -0.409657798 | 0.33715631 | -1.2150382 | 0.22435152 | 0.41041621 |
| AL136369.2 | 0.47481916 | -2.127330852 | 3.28666309 | -0.6472616 | 0.51746264 | NA |
| AL136369.1 | 0 | NA | NA | NA | NA | NA |
| AL162408.1 | 0.07659028 | -0.642905668 | 3.30558948 | -0.1944905 | 0.84579185 | NA |
| CELF2-AS2 | 0.17453125 | 0.18657164 | 3.29377195 | 0.05664376 | 0.95482897 | NA |
| AL136320.1 | 0.58506411 | 0.2405096 | 1.90651845 | 0.12615121 | 0.89961224 | NA |
| CELF2-AS1 | 1.0930533 | -1.554590797 | 1.44443483 | -1.0762623 | 0.28180995 | 0.47765319 |
| USP6NL | 803.322534 | -0.082252963 | 0.13611304 | -0.6042989 | 0.54564492 | 0.71712709 |
| AL512631.2 | 12.6359432 | 0.55360651 | 0.44061632 | 1.25643669 | 0.20895769 | 0.39217671 |
| AL512631.1 | 1.41787482 | -0.423625185 | 1.47205959 | -0.2877772 | 0.77351729 | 0.8753901 |
| ECHDC3 | 4949.10385 | 0.828937872 | 0.37371842 | 2.21808139 | 0.02654928 | 0.09236209 |
| PROSER2 | 1160.51145 | -1.369605501 | 0.33706411 | -4.0633383 | 4.84E-05 | 0.00072399 |
| PROSER2-AS | 183.199278 | -1.161642535 | 0.30794434 | -3.7722484 | 0.00016178 | 0.00185407 |
| UPF2 | 1502.49484 | -0.107790137 | 0.15186523 | -0.709775 | 0.47784369 | 0.66291762 |
| DHTKD1 | 3286.46648 | 1.248263049 | 0.30377045 | 4.10923134 | 3.97E-05 | 0.00061943 |
| SEC61A2 | 1126.97451 | -0.342861111 | 0.16289023 | -2.1048599 | 0.03530351 | 0.11339263 |
| NUDT5 | 2820.68919 | -0.117107595 | 0.18024262 | -0.649722 | 0.51587181 | 0.69342133 |
| CDC123 | 2285.17636 | -0.450999306 | 0.1594588 | -2.8283124 | 0.00467941 | 0.02557962 |
| AL512770.1 | 839.118994 | -0.501909275 | 0.15496394 | -3.2388778 | 0.00120001 | 0.00905013 |
| CAMK1D | 366.618887 | 0.044204597 | 0.24264335 | 0.18217931 | 0.85544201 | 0.92420542 |
| CCDC3 | 1157.87059 | -0.10537219 | 0.66419026 | -0.1586476 | 0.87394653 | 0.93435921 |
| OPTN | 6690.5287 | 0.17138496 | 0.2188649 | 0.78306278 | 0.43359024 | 0.62558268 |
| MCM10 | 14.8748886 | -1.02230063 | 0.60146767 | -1.6996768 | 0.08919174 | 0.21915637 |
| UCMA | 0.40557068 | -1.788706826 | 3.29999086 | -0.5420339 | 0.58779517 | NA |
| PHYH | 6133.38316 | 1.551369711 | 0.33545172 | 4.62471825 | 3.75E-06 | 9.35E-05 |
| SEPHS1 | 3057.93365 | 0.401881363 | 0.16252117 | 2.47279399 | 0.01340614 | 0.05596326 |
| AL355870.1 | 5.38920392 | -0.28893912 | 0.70159713 | -0.4118305 | 0.68046364 | 0.81472216 |
| BEND7 | 2546.08432 | -0.149232017 | 0.26322122 | -0.5669452 | 0.57075139 | 0.73576038 |
| AL590677.1 | 34.2220941 | -0.203218102 | 0.4018076 | -0.5057597 | 0.61302533 | 0.76813865 |
| PRPF18 | 900.863776 | -0.064209043 | 0.14405159 | -0.4457364 | 0.65578765 | 0.79807876 |
| AL157392.5 | 1218.26584 | -0.130753745 | 0.11149579 | -1.1727236 | 0.24090668 | 0.43104283 |
| FRMD4A | 2622.83032 | 0.201500234 | 0.21409757 | 0.94116078 | 0.34662247 | 0.54425278 |
| AL157392.4 | 30.5290981 | -0.625850495 | 0.44846394 | -1.3955425 | 0.1628523 | 0.33188415 |
| AL157392.1 | 0.30722956 | 0.328495715 | 2.89077905 | 0.11363571 | 0.90952658 | NA |
| AL157392.2 | 30.2065815 | 1.524295991 | 0.68553866 | 2.22350115 | 0.02618203 | 0.09142214 |
| AC044781.1 | 1.61533104 | 0.209898318 | 0.981458 | 0.21386378 | 0.83065329 | 0.91033489 |
| AL157896.1 | 6.06198393 | -0.276665821 | 0.75988207 | -0.3640905 | 0.71579045 | 0.83823582 |
| FAM107B | 7585.87393 | 0.584702699 | 0.22175705 | 2.63668142 | 0.00837214 | 0.0393746 |
| AL158168.1 | 0.44433078 | 1.496009765 | 2.67591356 | 0.55906506 | 0.57611732 | NA |
| CDNF | 202.636875 | 1.175635952 | 0.33438876 | 3.51577592 | 0.00043847 | 0.00409872 |
| HSPA14 | 1294.86989 | -0.238836327 | 0.17154599 | -1.3922583 | 0.16384419 | 0.3333517 |
| AC069544.1 | 68.331563 | -0.349696451 | 0.26394528 | -1.3248824 | 0.18521016 | 0.36264428 |
| SUV39H2 | 240.668491 | -0.160937661 | 0.16912663 | -0.9515809 | 0.34130959 | 0.5393297 |
| DCLRE1C | 282.049549 | -0.149772033 | 0.16499699 | -0.9077258 | 0.36402308 | 0.56116501 |
| MEIG1 | 9.16662446 | 0.203129326 | 0.45984806 | 0.4417314 | 0.65868358 | 0.79950001 |
| OLAH | 50.9151504 | -1.14687112 | 0.5507385 | -2.0824241 | 0.03730375 | 0.11763931 |
| ACBD7 | 16.7364985 | 0.127761891 | 0.40612646 | 0.31458647 | 0.75307565 | 0.86236777 |
| RPP38 | 792.458332 | -0.547724993 | 0.15829303 | -3.4601966 | 0.00053978 | 0.00481204 |
| NMT2 | 1286.86117 | 0.053189386 | 0.16833729 | 0.31596912 | 0.75202595 | 0.86165215 |
| FAM171A1 | 3215.61605 | 0.740979049 | 0.27714276 | 2.67363664 | 0.00750337 | 0.0364726 |
| AL607028.1 | 12.4845506 | 0.116063755 | 0.69448786 | 0.16712136 | 0.86727456 | 0.93105 |
| ITGA8 | 1092.89645 | 0.779666529 | 0.436372 | 1.78670155 | 0.07398574 | 0.19195237 |
| MINDY3 | 1357.78496 | 0.210349728 | 0.13142579 | 1.60052089 | 0.10948308 | 0.25318502 |
| AL355339.1 | 0 | NA | NA | NA | NA | NA |
| PTER | 2334.94366 | 1.128979627 | 0.37687486 | 2.99563528 | 0.00273874 | 0.01715452 |
| C1QL3 | 103.388696 | 2.656677536 | 0.65180763 | 4.07586139 | 4.58E-05 | 0.00069258 |
| RSU1 | 3122.11947 | -0.281817306 | 0.21545966 | -1.3079818 | 0.1908795 | 0.36981062 |
| CUBN | 17721.6652 | 2.961251168 | 0.60872275 | 4.86469609 | 1.15E-06 | 3.65E-05 |
| AC067747.1 | 7.68721787 | 0.392288608 | 0.46926147 | 0.83597021 | 0.40317166 | 0.59815921 |
| TRDMT1 | 248.375449 | 0.817577915 | 0.17660906 | 4.62930896 | 3.67E-06 | 9.20E-05 |
| VIM-AS1 | 9083.73041 | -0.468203545 | 0.35633496 | -1.3139422 | 0.18886566 | 0.36712276 |
| VIM | 32612.2876 | -0.719398789 | 0.35738468 | -2.0129536 | 0.04411951 | 0.13307004 |
| AL133415.1 | 12888.2069 | -0.620572445 | 0.36782041 | -1.6871615 | 0.0915723 | 0.22332021 |
| ST8SIA6 | 20.6763002 | 1.046636815 | 0.77061679 | 1.35818065 | 0.17440637 | 0.34831081 |
| ST8SIA6-AS1 | 7.06363595 | 0.876259657 | 0.56535289 | 1.54993398 | 0.12115736 | 0.27165374 |
| HACD1 | 255.332739 | -0.88847359 | 0.3649986 | -2.4341836 | 0.01492542 | 0.06060724 |
| STAM-AS1 | 59.4353766 | 0.704866929 | 0.31452057 | 2.24108369 | 0.02502065 | 0.08839693 |
| STAM | 1267.91084 | -0.21957359 | 0.22865191 | -0.9602963 | 0.3369061 | 0.5349358 |
| AC069542.1 | 99.2488439 | 0.202514292 | 0.33421843 | 0.60593394 | 0.54455862 | 0.71643964 |
| TMEM236 | 45.8623348 | -0.484808856 | 0.32440534 | -1.494454 | 0.13505699 | 0.29242691 |
| MRC1 | 1342.25343 | -0.635883193 | 0.47075993 | -1.350759 | 0.17677266 | 0.35170477 |
| AC069023.1 | 0 | NA | NA | NA | NA | NA |
| SLC39A12 | 3.30702658 | -2.483867652 | 1.14632342 | -2.1668123 | 0.03024917 | 0.10141057 |
| SLC39A12-AS | 1.09138638 | -2.409974993 | 1.91644176 | -1.2575258 | 0.2085633 | 0.39169675 |
| CACNB2 | 406.700735 | 1.045611287 | 0.2357577 | 4.43510983 | 9.20E-06 | 0.00019078 |
| AL450384.2 | 207.723575 | 0.367957102 | 0.22836051 | 1.61129918 | 0.10711453 | 0.24912773 |
| AL450384.1 | 6.48507569 | 0.872182648 | 0.44668368 | 1.95257336 | 0.05087017 | 0.14741169 |
| NSUN6 | 686.473475 | 0.281192048 | 0.25090368 | 1.12071713 | 0.26240829 | 0.45603853 |
| ARL5B | 1346.79801 | -0.965662122 | 0.25768883 | -3.7473962 | 0.00017868 | 0.00200246 |
| AL512641.1 | 0 | NA | NA | NA | NA | NA |
| MALRD1 | 64.0523217 | 1.102751403 | 0.47046435 | 2.34396379 | 0.01908002 | 0.07248815 |
| AL590378.1 | 0 | NA | NA | NA | NA | NA |
| AL157895.1 | 6.18000674 | -0.10030523 | 0.98864201 | -0.1014576 | 0.91918723 | 0.96065382 |
| PLXDC2 | 2441.15657 | 0.934226726 | 0.21861828 | 4.27332401 | 1.93E-05 | 0.00034406 |
| AC069549.1 | 16.8580801 | 0.058584745 | 0.36204147 | 0.16181777 | 0.87144936 | 0.93316493 |
| NEBL | 2335.4529 | 0.830894481 | 0.22902031 | 3.62803843 | 0.00028558 | 0.00291553 |
| C10orf113 | 0.87803786 | 0.945211361 | 1.50223878 | 0.62920181 | 0.52921694 | NA |
| NEBL-AS1 | 41.6902889 | -0.048218599 | 0.59617583 | -0.0808798 | 0.93553752 | 0.96914261 |
| AL158209.1 | 0.16786415 | -1.003574173 | 3.30558948 | -0.3035992 | 0.7614333 | NA |
| CASC10 | 316.717967 | 1.266144604 | 0.40630655 | 3.1162299 | 0.00183179 | 0.01258911 |
| SKIDA1 | 152.811977 | 1.795391373 | 0.27669619 | 6.48867399 | 8.66E-11 | 1.28E-08 |
| MLLT10 | 1828.52847 | -0.007599173 | 0.11761589 | -0.0646101 | 0.94848445 | 0.97514677 |
| DNAJC1 | 2310.74163 | -0.616525644 | 0.16102186 | -3.828832 | 0.00012875 | 0.00154429 |
| EBLN1 | 0.08666371 | -0.642905668 | 3.30558948 | -0.1944905 | 0.84579185 | NA |
| AL157831.2 | 0 | NA | NA | NA | NA | NA |
| AL158211.1 | 9.31436742 | 0.141353764 | 0.56856757 | 0.24861384 | 0.8036595 | 0.89348207 |
| COMMD3 | 1600.88761 | 0.096518055 | 0.20075596 | 0.48077305 | 0.63067781 | 0.78054869 |
| COMMD3-BN | 1928.85529 | 0.275131755 | 0.15831 | 1.73793041 | 0.08222308 | 0.2063991 |
| BMI1 | 2231.84541 | 0.039525321 | 0.12963652 | 0.30489341 | 0.76044733 | 0.86715659 |
| AL158211.4 | 2.68445372 | 0.421091932 | 0.73412652 | 0.57359586 | 0.56624131 | 0.73281994 |
| AL158211.2 | 1.86130745 | -0.257298271 | 0.95396513 | -0.2697145 | 0.78737987 | 0.88418035 |
| AL158211.3 | 0.17423948 | 0.18657164 | 3.29377195 | 0.05664376 | 0.95482897 | NA |
| SPAG6 | 50.9175034 | -0.545294913 | 0.75918475 | -0.7182638 | 0.47259465 | 0.65875468 |
| AL513128.2 | 0.43705876 | 0.399535775 | 2.83916406 | 0.14072303 | 0.88808875 | NA |
| PIP4K2A | 1973.39191 | -0.336598108 | 0.21462243 | -1.5683268 | 0.11680489 | 0.26514117 |
| ARMC3 | 82.006537 | -0.865364908 | 0.62184841 | -1.3916011 | 0.16404324 | 0.33355079 |
| MSRB2 | 1486.41932 | 0.100412102 | 0.24452626 | 0.41063934 | 0.68133701 | 0.81534112 |
| PTF1A | 0.18028734 | 0.18657164 | 3.29377195 | 0.05664376 | 0.95482897 | NA |
| C10orf67 | 166.715468 | 1.503961437 | 0.38711374 | 3.88506349 | 0.0001023 | 0.00128893 |
| AL606469.1 | 31.8747187 | 0.719137415 | 0.42018058 | 1.71149608 | 0.08698957 | 0.21499323 |

| | | | | | | | |
|---|---|---|---|---|---|---|---|
| OTUD1 | 1482.74363 | | 0.472467131 | 0.19946403 | 2.36868341 | 0.01785153 | 0.06896349 |
| KIAA1217 | 3750.39279 | | 0.121051929 | 0.14945642 | 0.80994798 | 0.41797007 | 0.61076531 |
| ARHGAP21 | 2458.83689 | | -0.238468443 | 0.23574082 | -1.0115704 | 0.31174348 | 0.50879963 |
| PRTFDC1 | 217.402785 | | 0.385398205 | 0.24548234 | 1.56996303 | 0.11642371 | 0.26457198 |
| AL512598.1 | 8.37966008 | | 0.27167915 | 0.50035719 | 0.54297042 | 0.58715017 | 0.74812283 |
| ENKUR | 34.6399749 | | -0.785527848 | 0.57821987 | -1.3585279 | 0.17429622 | 0.34824908 |
| THNSL1 | 560.939136 | | 1.294001418 | 0.31191396 | 4.14858447 | 3.35E-05 | 0.00053628 |
| LINC01516 | 0.51546328 | | -2.265703084 | 2.70931325 | -0.8362647 | 0.403006 | NA |
| GPR158-AS1 | 0.08666371 | | -0.642905668 | 3.30558948 | -0.1944905 | 0.84579185 | NA |
| GPR158 | 4.15740148 | | 0.715915204 | 0.62067266 | 1.15345053 | 0.24872551 | 0.44046175 |
| LINC00836 | 0.54366959 | | 1.001265414 | 2.10170469 | 0.47640633 | 0.63378493 | NA |
| AL358612.1 | 0 | NA | | NA | NA | NA | NA |
| MYO3A | 50.6191934 | | 0.058586063 | 0.50473416 | 0.11607311 | 0.9075946 | 0.95385641 |
| GAD2 | 0 | NA | | NA | NA | NA | NA |
| APBB1IP | 602.458164 | | -0.511781528 | 0.40641473 | -1.2592593 | 0.20793669 | 0.391077 |
| AL390961.1 | 0 | NA | | NA | NA | NA | NA |
| PDSS1 | 414.693629 | | 0.355112527 | 0.27134427 | 1.30871577 | 0.19063065 | 0.36943707 |
| AL390961.2 | 1.26796328 | | -0.048062135 | 1.3379147 | -0.0359232 | 0.97134362 | 0.98667343 |
| ABI1 | 3388.35725 | | -0.162104489 | 0.15828799 | -1.0241111 | 0.30578278 | 0.50275652 |
| ANKRD26 | 310.26143 | | 0.261172991 | 0.25738764 | 1.0147068 | 0.31024561 | 0.50726204 |
| YME1L1 | 5475.5647 | | -0.060172723 | 0.14332103 | -0.4198457 | 0.67459815 | 0.81055056 |
| MASTL | 516.089926 | | -0.243575597 | 0.23672091 | -1.0289568 | 0.30349996 | 0.50039351 |
| ACBD5 | 2784.75066 | | 0.314358203 | 0.1675326 | 1.87640015 | 0.06060035 | 0.16666994 |
| AL160291.1 | 10.3580796 | | 0.581723691 | 0.5998836 | 0.96972761 | 0.33218228 | 0.53009359 |
| AL355493.2 | 2.96543496 | | 0.248912766 | 0.95069156 | 0.26182284 | 0.79345803 | 0.88828224 |
| RAB18 | 5270.70927 | | -0.345143922 | 0.23914744 | -1.4432265 | 0.14895668 | 0.31303205 |
| MKX | 27.6263853 | | -0.063545824 | 0.41306122 | -0.1538412 | 0.87773497 | 0.93635953 |
| MKX-AS1 | 0.3310045 | | -0.261686027 | 2.339021 | -0.1118784 | 0.91091979 | NA |
| ARMC4 | 9.17535426 | | 0.939830519 | 0.47047275 | 1.99763008 | 0.04575678 | 0.13655571 |
| AL390866.1 | 0.03392908 | | 0.380740593 | 3.29738709 | 0.11546736 | 0.90807469 | NA |
| MPP7 | 936.697661 | | 0.600658974 | 0.17108875 | 3.51080337 | 0.00044675 | 0.00415993 |
| AC022021.1 | 0.8239087 | | 1.417807916 | 1.22798982 | 1.15457628 | 0.24826399 | NA |
| AL358234.1 | 2.11906392 | | 0.97676861 | 1.14353101 | 0.85416889 | 0.39301142 | 0.58847942 |
| WAC-AS1 | 1285.93481 | | -0.064392664 | 0.13754126 | -0.4681698 | 0.63966316 | 0.78704498 |
| WAC | 6743.06862 | | -0.231527959 | 0.16852202 | -1.3738736 | 0.16948092 | 0.34129147 |
| BAMBI | 1243.7606 | | 0.252741378 | 0.21046041 | 1.20089746 | 0.22979098 | 0.41729826 |
| AL355376.1 | 0 | NA | | NA | NA | NA | NA |
| LINC00837 | 0.19271054 | | 1.506623853 | 3.27645372 | 0.4598337 | 0.64563559 | NA |
| LYZL1 | 0.27442362 | | -0.179388694 | 2.94052751 | -0.0610056 | 0.95135474 | NA |
| SVIL | 2942.49624 | | 0.155249527 | 0.21381342 | 0.72609814 | 0.46777861 | 0.65467158 |
| JCAD | 1715.19951 | | -0.421253168 | 0.38628855 | -1.0905142 | 0.27548668 | 0.4708151 |
| MTPAP | 1169.36666 | | -0.12272076 | 0.1579857 | -0.776784 | 0.43728624 | 0.62933135 |
| MAP3K8 | 1130.72288 | | -1.388459164 | 0.24967906 | -5.5609756 | 2.68E-08 | 1.59E-06 |
| AL590068.1 | 1.13210504 | | -1.920198476 | 1.5909105 | -1.2069808 | 0.22743953 | 0.41434061 |
| AL590068.2 | 0 | NA | | NA | NA | NA | NA |
| LYZL2 | 0 | NA | | NA | NA | NA | NA |
| AL451107.1 | 0 | NA | | NA | NA | NA | NA |
| ZNF438 | 454.415503 | | -0.265897091 | 0.10775122 | -2.4676945 | 0.01359863 | 0.05657872 |
| AL356258.1 | 3.19213311 | | 0.729854283 | 0.84303202 | 0.86574919 | 0.38662772 | 0.58229328 |
| ZEB1-AS1 | 466.696883 | | -0.445203613 | 0.23599394 | -1.8865044 | 0.05922701 | 0.16407356 |
| ZEB1 | 1304.48652 | | -0.306671413 | 0.27034162 | -1.1343848 | 0.25663319 | 0.44936509 |
| AL161935.1 | 2.26173617 | | 0.159515774 | 0.8731138 | 0.18269758 | 0.85503532 | 0.92406891 |
| AL161935.3 | 3.77104079 | | 0.4588804 | 0.70585859 | 0.65010245 | 0.51562605 | 0.69330302 |
| ARHGAP12 | 2198.42352 | | -0.04192452 | 0.13402926 | -0.3128013 | 0.75443166 | 0.86329626 |
| KIF5B | 4903.59137 | | -0.139911696 | 0.13466193 | -1.0389848 | 0.29881182 | 0.49563286 |
| AL158834.2 | 446.028782 | | 0.137759826 | 0.16031139 | 0.8593265 | 0.39016041 | 0.58574039 |
| EPC1 | 1427.27225 | | 0.247267055 | 0.17776045 | 1.39101273 | 0.16422157 | 0.33383617 |
| AL158834.1 | 0.73098585 | | -1.179756707 | 1.386266 | -0.851032 | 0.39475159 | NA |
| AL391839.2 | 64.5702538 | | 0.64275366 | 0.4478274 | 1.43527096 | 0.1512099 | 0.31610949 |
| AL391839.1 | 0.44106535 | | 1.048626814 | 2.71913916 | 0.38564662 | 0.6997584 | NA |
| CCDC7 | 176.663848 | | 0.261700091 | 0.19625739 | 1.33345345 | 0.18238304 | 0.35904747 |
| AL365203.2 | 362.173687 | | -1.207849696 | 0.49393262 | -2.4453734 | 0.01447022 | 0.05930678 |
| ITGB1 | 11060.5811 | | -0.168817756 | 0.28919101 | -0.5837587 | 0.5593827 | 0.72772184 |
| ITGB1-DT | 11.8421131 | | -1.389727962 | 0.62948437 | -2.2077243 | 0.0272635 | 0.09421287 |
| AL121748.1 | 1.11556443 | | 2.059198274 | 1.36892102 | 1.50424915 | 0.13251723 | 0.28882381 |
| NRP1 | 10963.6823 | | -0.447789223 | 0.22926544 | -1.9531475 | 0.05080211 | 0.14727922 |
| AL121748.2 | 693.123618 | | -0.4330373 | 0.21533085 | -2.0110323 | 0.04432205 | 0.1333606 |
| AL353600.1 | 6.29372689 | | -0.586014784 | 0.6093032 | -0.9617786 | 0.33616083 | 0.5342602 |
| AC018980.1 | 0 | NA | | NA | NA | NA | NA |
| AL450313.1 | 2.90988782 | | 3.233648416 | 2.03205936 | 1.59131592 | NA | NA |
| LINC00838 | 0 | NA | | NA | NA | NA | NA |
| AL117338.1 | 0.17332742 | | -1.003574173 | 3.30558948 | -0.3035992 | 0.7614333 | NA |
| PARD3 | 2727.34315 | | -0.179761513 | 0.2118676 | -0.8484616 | 0.39618096 | 0.5912411 |
| PARD3-AS1 | 33.5225115 | | -0.07811659 | 0.35275398 | -0.2214478 | 0.82474378 | 0.90622499 |
| AL392046.2 | 30.0966411 | | -0.464414183 | 0.39269601 | -1.1826302 | 0.23695572 | 0.42620028 |
| CUL2 | 1459.67489 | | -0.232404919 | 0.16212293 | -1.4335105 | 0.151712 | 0.31677565 |
| AL392046.1 | 38.2036589 | | -0.77045862 | 0.27163726 | -2.836351 | 0.00456323 | 0.02512681 |
| CREM | 2376.31072 | | -1.136391799 | 0.2962026 | -3.8365355 | 0.00012478 | 0.00150577 |
| AL117336.1 | 1.02778418 | | 0.657157108 | 1.88715127 | 0.34822704 | 0.72766968 | 0.84611931 |
| CCNY | 3640.40577 | | -0.046722311 | 0.16731648 | -0.2792451 | 0.78005671 | 0.87955255 |
| AL117336.3 | 193.671061 | | -0.050058498 | 0.30257505 | -0.1654416 | 0.86859642 | 0.93156219 |
| AL121749.1 | 5.18926268 | | -1.045500696 | 1.12986637 | -0.9253313 | 0.35479361 | 0.55230354 |
| GJD4 | 2.75567745 | | -0.838540847 | 1.00009446 | -0.8384616 | 0.40177148 | 0.59664454 |
| FZD8 | 1304.50481 | | -0.028876243 | 0.20947703 | -0.1378492 | 0.89035959 | 0.94316 |
| PCAT5 | 1.39378011 | | 2.498054714 | 1.21360741 | 2.05837134 | 0.0395545 | 0.12278425 |
| AL365500.1 | 0.03392908 | | 0.380740593 | 3.29738709 | 0.11546736 | 0.90807469 | NA |
| AL355300.1 | 0.59842268 | | 1.987823594 | 2.00718374 | 0.99035457 | 0.32200084 | NA |
| AL590730.1 | 0.07659028 | | -0.642905668 | 3.30558948 | -0.1944905 | 0.84579185 | NA |
| AL390061.1 | 0 | NA | | NA | NA | NA | NA |
| ANKRD30A | 0.37819168 | | -0.558687541 | 2.24315333 | -0.2490635 | 0.80331169 | NA |
| AL157387.1 | 0.2268242 | | -0.340597073 | 3.29738699 | -0.103293 | 0.91773041 | NA |
| MTRNR2L7 | 0 | NA | | NA | NA | NA | NA |
| AL132657.1 | 16.9765257 | | 0.455949115 | 0.35655309 | 1.2787692 | 0.20097834 | 0.38268459 |
| ZNF248 | 1002.02051 | | 0.212575385 | 0.22863662 | 0.92975213 | 0.35249944 | 0.55012193 |
| AL132657.2 | 0.44095431 | | -2.038225231 | 2.21739984 | -0.9191961 | 0.35799302 | NA |
| AL135791.1 | 21.2710019 | | 1.219406784 | 0.40928981 | 2.97932361 | 0.00288886 | 0.01781134 |
| ZNF25 | 719.923607 | | 0.508591409 | 0.14318097 | 3.55208794 | 0.00038219 | 0.00368089 |
| AL117337.1 | 0.17391096 | | -1.003574173 | 3.30558948 | -0.3035992 | 0.7614333 | NA |
| ZNF33A | 2281.10047 | | 0.284166708 | 0.16281534 | 1.7453313 | 0.08092722 | 0.20407868 |
| ZNF37A | 1213.4118 | | -0.116145079 | 0.17007402 | -0.682909 | 0.49466434 | 0.67694119 |
| AL117339.5 | 179.017088 | | -0.140780482 | 0.3131916 | -0.4495027 | 0.65306903 | 0.79614538 |
| AL117339.4 | 49.9634479 | | 0.646373163 | 0.49678779 | 1.30110516 | 0.19322246 | 0.37275982 |
| AL133216.1 | 0.25668517 | | 0.372875031 | 3.03140481 | 0.12300404 | 0.9021039 | NA |
| IGKV1OR10- | 0 | NA | | NA | NA | NA | NA |
| LINC00839 | 652.364771 | | 0.370683963 | 0.44995843 | 0.82381823 | 0.41004285 | 0.6044565 |
| ZNF33B | 1987.7256 | | 0.627581938 | 0.19300629 | 3.25161398 | 0.00114752 | 0.00874315 |
| LINC01518 | 0.08393208 | | -0.642905668 | 3.30558948 | -0.1944905 | 0.84579185 | NA |
| AL022344.1 | 0 | NA | | NA | NA | NA | NA |
| AL022344.2 | 14.5594864 | | -0.854931304 | 0.5154405 | -1.6586421 | 0.09718794 | 0.23283782 |
| BMS1 | 2238.98507 | | -0.296897891 | 0.1486723 | -1.9969953 | 0.04582569 | 0.13667723 |
| AL365199.1 | 0.89998316 | | 2.687087583 | 1.35477963 | 1.98341304 | 0.04732131 | NA |
| LINC01264 | 0 | NA | | NA | NA | NA | NA |
| RET | 111.32142 | | 0.298180599 | 0.41370757 | 0.7207521 | 0.47106205 | 0.65759572 |

| | | | | | | |
|---|---|---|---|---|---|---|
| AC010864.1 | 41.9368357 | 0.201196387 | 0.34497609 | 0.58321836 | 0.55974632 | 0.72812304 |
| CSGALNACT2 | 1352.33939 | -0.628603478 | 0.31217021 | -2.0136562 | 0.04404564 | 0.13295371 |
| RASGEF1A | 213.020338 | -0.349899372 | 0.30078286 | -1.1632956 | 0.2447096 | 0.435748 |
| AC068707.1 | 1.0362137 | 0.114751463 | 1.5580333 | 0.07365148 | 0.9412877 | 0.97207934 |
| FXYD4 | 6821.40171 | -0.218281999 | 0.63171472 | -0.3455389 | 0.72968928 | 0.84734355 |
| HNRNPF | 10578.4319 | -0.468782055 | 0.2561639 | -1.8300082 | 0.06724871 | 0.17899953 |
| AL450326.1 | 67.1149545 | 1.373452844 | 0.35745844 | 3.8422728 | 0.0001219 | 0.00148134 |
| ZNF487 | 304.419674 | -0.064982245 | 0.1480667 | -0.4388714 | 0.6607547 | 0.80033538 |
| ZNF239 | 228.603658 | 0.039379441 | 0.24584771 | 0.16017819 | 0.87274071 | 0.93368745 |
| ZNF485 | 114.755514 | -0.079780449 | 0.20215254 | -0.3946547 | 0.69309774 | 0.82343609 |
| ZNF32-AS3 | 0.08666371 | -0.642905668 | 3.30558948 | -0.1944905 | 0.84579185 | NA |
| AL645634.2 | 1.45264064 | 1.690670224 | 1.3726845 | 1.23165245 | 0.21807894 | 0.40322126 |
| ZNF32 | 1538.20699 | -0.030691574 | 0.10289379 | -0.298284 | 0.7654864 | 0.87056869 |
| ZNF32-AS1 | 380.000034 | -0.277227518 | 0.1486604 | -1.8648377 | 0.06220416 | 0.16952851 |
| ZNF32-AS2 | 322.748258 | 0.371267739 | 0.12311626 | 3.01558667 | 0.00256483 | 0.01630925 |
| AL645634.1 | 0 | NA | NA | NA | NA | NA |
| AL355989.1 | 0 | NA | NA | NA | NA | NA |
| LINC00619 | 4.55016046 | 0.398404232 | 0.97118241 | 0.41022595 | 0.6816402 | 0.81559291 |
| LINC00840 | 3.51208787 | 0.431851651 | 0.73399378 | 0.58835874 | 0.55629153 | 0.7253727 |
| AL139237.2 | 0.07830766 | -0.642905668 | 3.30558948 | -0.1944905 | 0.84579185 | NA |
| AL139237.1 | 0.08666371 | -0.642905668 | 3.30558948 | -0.1944905 | 0.84579185 | NA |
| LINC00841 | 0.03392908 | 0.380740593 | 3.29738709 | 0.11546736 | 0.90807469 | NA |
| AL137026.2 | 0.15179023 | 0.435199075 | 3.28898345 | 0.13232024 | 0.894731 | NA |
| AL137026.1 | 15.9151253 | 0.488504671 | 0.41622561 | 1.17365358 | 0.24053381 | 0.43069432 |
| C10orf142 | 16.4397722 | 0.900643092 | 0.50332295 | 1.78939406 | 0.07355137 | 0.1911302 |
| CXCL12 | 6349.55552 | 1.885720668 | 0.4687412 | 4.02294629 | 5.75E-05 | 0.00082594 |
| AL356157.1 | 2.92110763 | 1.756814673 | 0.78696486 | 2.23239279 | 0.02558901 | 0.08979601 |
| TMEM72-AS | 77.8850407 | 2.152801678 | 0.34656087 | 6.21190069 | 5.23E-10 | 5.90E-08 |
| AL353801.2 | 4.33634448 | 1.559632657 | 0.68853914 | 2.26513288 | 0.02350453 | 0.08439738 |
| TMEM72 | 5134.57305 | 2.249876653 | 0.48781173 | 4.6121824 | 3.98E-06 | 9.84E-05 |
| AL353801.1 | 37.0265054 | 1.991744463 | 0.57093958 | 3.48853808 | 0.00048567 | 0.00442999 |
| RASSF4 | 13783.194 | 0.737809244 | 0.29656336 | 2.48786377 | 0.0128513 | 0.05424822 |
| DEPP1 | 11669.5578 | -1.760057331 | 0.29961324 | -5.8744311 | 4.24E-09 | 3.32E-07 |
| C10orf25 | 121.537708 | 0.297116445 | 0.21912001 | 1.35595303 | 0.17511411 | 0.34911614 |
| ZNF22 | 1730.24354 | -0.3365972 | 0.15157241 | -2.2207023 | 0.02637113 | 0.09192422 |
| AL358394.1 | 0.60974681 | -0.762970987 | 1.66763159 | -0.4575177 | 0.64729898 | NA |
| OR13A1 | 3.06732586 | -0.196902682 | 0.96813852 | -0.2033828 | 0.83883588 | 0.91454277 |
| ALOX5 | 1016.82283 | -1.412940494 | 0.48093283 | -2.9379165 | 0.00330426 | 0.01975365 |
| AL731567.1 | 29.2031403 | -1.336023052 | 0.50608233 | -2.6399322 | 0.00829226 | 0.03910659 |
| 8-Mar | 2628.25671 | -0.016489436 | 0.16022763 | -0.1029126 | 0.91803237 | 0.9601219 |
| ZFAND4 | 353.054831 | 0.104453091 | 0.13452032 | 0.77648559 | 0.43746233 | 0.62951605 |
| WASHC2C | 2888.14312 | -0.12217518 | 0.22036496 | -0.554422 | 0.57929007 | 0.74259293 |
| AGAP4 | 582.29882 | 0.667042531 | 0.30712416 | 2.17189856 | 0.02986331 | 0.10054847 |
| TIMM23 | 2629.48553 | -0.207556793 | 0.17049173 | -1.217401 | 0.22345169 | 0.40950873 |
| NCOA4 | 9471.06468 | 0.926805513 | 0.16841316 | 5.50316569 | 3.73E-08 | 2.09E-06 |
| MSMB | 0.6804246 | -0.03709498 | 1.4956399 | -0.0248021 | 0.98021283 | NA |
| ANTXRL | 3.7282782 | 0.148974237 | 0.93016077 | 0.16015966 | 0.8727553 | 0.93368745 |
| ANXA8L1 | 1.53009509 | 0.308053224 | 3.13675425 | 0.09820764 | 0.92176742 | 0.96181159 |
| LINC00842 | 0.76355521 | -0.085408218 | 1.59392379 | -0.0535836 | 0.9572669 | NA |
| NPY4R | 0.19823479 | 0.726716446 | 3.28420131 | 0.22127646 | 0.82487718 | NA |
| GPRIN2 | 267.551737 | 0.250389806 | 0.46885044 | 0.53405048 | 0.59330661 | 0.75251288 |
| SYT15 | 52.2243035 | 0.352374237 | 0.42145326 | 0.83609328 | 0.40310243 | 0.59812126 |
| AL356056.2 | 14.9733527 | 0.233688099 | 0.43701384 | 0.53473844 | 0.59283074 | 0.75212637 |
| AL356056.1 | 11.4112031 | -0.031250441 | 0.47325741 | -0.0660326 | 0.94735183 | 0.97462811 |
| AL356056.3 | 38.1314557 | 0.665769208 | 0.36196779 | 1.8393051 | 0.06587032 | 0.1765065 |
| PTPN20 | 263.975972 | 0.021774654 | 0.30427378 | 0.0715627 | 0.94294992 | 0.97302248 |
| GDF10 | 11.0306466 | 1.066624225 | 0.80103653 | 1.33155503 | 0.18300645 | 0.35972297 |
| GDF2 | 0 | NA | NA | NA | NA | NA |
| RBP3 | 5.64189253 | 1.642161133 | 0.93505911 | 1.75621104 | 0.07905238 | 0.20081026 |
| ZNF488 | 70.9292544 | 2.123601002 | 0.76976628 | 2.75876074 | 0.0058021 | 0.03007287 |
| ANXA8 | 2.27285848 | -1.707955594 | 1.25085472 | -1.3654308 | 0.17211768 | 0.34506776 |
| FAM25G | 2.57184255 | 0.140586806 | 0.87063885 | 0.16147546 | 0.87171894 | 0.93317628 |
| AL591684.1 | 174.468127 | -0.116809684 | 0.60993216 | -0.1915126 | 0.84812402 | 0.92016931 |
| AGAP9 | 1248.785 | 0.024272559 | 0.52421256 | 0.0463029 | 0.96306883 | 0.9827613 |
| BX547991.1 | 162.869526 | 2.062151205 | 0.59938002 | 3.4404737 | 0.0005807 | 0.00509997 |
| FO681492.1 | 25.2617126 | -0.196632336 | 0.52414712 | -0.3751472 | 0.70755098 | 0.83354715 |
| AC245041.2 | 1.10283746 | -1.510669705 | 2.00432238 | -0.753706 | 0.45102581 | 0.64079343 |
| NPY4R2 | 0.03392908 | 0.380740593 | 3.29738709 | 0.11546736 | 0.90807469 | NA |
| AC245041.1 | 0.08666371 | -0.642905668 | 3.30558948 | -0.1944905 | 0.84579185 | NA |
| FAM25C | 0.46146185 | 0.64021426 | 1.81966587 | 0.35183067 | 0.72496526 | NA |
| FRMPD2 | 17.3080621 | 1.588935784 | 0.82894819 | 1.91680953 | 0.05526213 | 0.15625804 |
| MAPK8 | 2023.65029 | 0.009583207 | 0.23009085 | 0.04164967 | 0.96677798 | 0.98424582 |
| ARHGAP22 | 202.331359 | -1.522953938 | 0.36201683 | -4.2068595 | 2.59E-05 | 0.0004366 |
| AC068898.1 | 0 | NA | NA | NA | NA | NA |
| WDFY4 | 346.52357 | 1.537843781 | 0.47188057 | 3.25896824 | 0.00111818 | 0.00855424 |
| LRRC18 | 0.87487935 | 1.714999909 | 2.01582348 | 0.85076889 | 0.39489774 | NA |
| AC060234.1 | 54.4929956 | 1.378895437 | 0.3994206 | 3.45223917 | 0.00055595 | 0.0049362 |
| VSTM4 | 1201.8842 | 1.40187395 | 0.33426257 | 4.1939304 | 2.74E-05 | 0.00045728 |
| FAM170B-AS | 0.56493854 | 0.245041462 | 1.51166177 | 0.16210072 | 0.87122654 | NA |
| FAM170B | 2.1201901 | 1.522402123 | 1.18536128 | 1.28433597 | 0.19902443 | 0.3800809 |
| TMEM273 | 403.32227 | -0.375533808 | 0.50564669 | -0.7426802 | 0.45767529 | 0.64686234 |
| C10orf71-AS | 0.85090911 | 2.014145863 | 1.58416104 | 1.27142747 | 0.20357661 | NA |
| C10orf71 | 6.08961011 | 0.332476023 | 0.7909849 | 0.4203317 | 0.67424316 | 0.81027171 |
| DRGX | 0.6339508 | -1.8174024 | 1.86957282 | -0.972095 | 0.33100329 | NA |
| AL138760.1 | 0.48417503 | -0.856448443 | 1.91321877 | -0.4476479 | 0.6544073 | NA |
| ERCC6 | 665.752706 | 0.022608194 | 0.16678269 | 0.1355548 | 0.89217324 | 0.94419715 |
| CHAT | 0 | NA | NA | NA | NA | NA |
| SLC18A3 | 0.08695548 | -0.642905668 | 3.30558948 | -0.1944905 | 0.84579185 | NA |
| C10orf53 | 0.21694805 | 0.689156974 | 3.06622645 | 0.22475736 | 0.82216804 | NA |
| OGDHL | 14922.3661 | 1.915567042 | 0.37755388 | 5.07362561 | 3.90E-07 | 1.48E-05 |
| PARG | 1122.95244 | 0.686592439 | 0.19644954 | 3.49500662 | 0.00047405 | 0.0043511 |
| TIMM23B | 391.307789 | 0.102393617 | 0.22478274 | 0.45552259 | 0.64873332 | 0.79324389 |
| LINC00843 | 102.661566 | 0.243398484 | 0.22369972 | 1.08805893 | 0.27656909 | 0.47192487 |
| AGAP6 | 1435.46759 | -0.468447774 | 0.12684835 | -3.692975 | 0.00022165 | 0.00237208 |
| WASHC2A | 3080.66501 | -0.068453513 | 0.24339741 | -0.2812417 | 0.77852498 | 0.87861887 |
| ASAH2 | 60.1350722 | -0.170906784 | 0.48432738 | -0.3528745 | 0.72418253 | 0.84382896 |
| SGMS1 | 2424.51966 | -0.390574236 | 0.15941111 | -2.4501068 | 0.01428139 | 0.05874281 |
| AL117341.1 | 0.03392908 | 0.380740593 | 3.29738709 | 0.11546736 | 0.90807469 | NA |
| SGMS1-AS1 | 396.213584 | 1.095805644 | 0.26536578 | 4.12941577 | 3.64E-05 | 0.00057446 |
| ASAH2B | 413.916768 | -0.037940572 | 0.25169811 | -0.1507384 | 0.88018207 | 0.9378468 |
| A1CF | 1092.90712 | 1.901846746 | 0.56616114 | 3.35919691 | 0.00078169 | 0.00645779 |
| AL731537.2 | 1.48251675 | -1.549958517 | 1.65884056 | -0.9343626 | 0.3501169 | 0.54779772 |
| PRKG1 | 499.080553 | 0.379087567 | 0.33341978 | 1.13696782 | 0.25555174 | 0.44821175 |
| AL731537.1 | 4.24524358 | -0.41535236 | 0.66622922 | -0.6234376 | 0.53299698 | 0.70689184 |
| AC022537.1 | 0 | NA | NA | NA | NA | NA |
| CSTF2T | 1591.22431 | 0.202303767 | 0.13342875 | 1.51619328 | 0.12947049 | 0.28432185 |
| PRKG1-AS1 | 96.9247393 | 0.035101486 | 0.30672987 | 0.11443778 | 0.90889076 | 0.95449819 |
| DKK1 | 16.5109314 | -0.63859799 | 0.69570634 | -0.9179131 | 0.35866434 | 0.55635169 |
| LINC01468 | 0.42310618 | -1.988210508 | 2.79024851 | -0.7125568 | 0.47612006 | NA |
| AC074327.1 | 3.53207669 | -1.644638314 | 1.13493195 | -1.4491074 | 0.14730758 | 0.31063583 |
| MBL2 | 0.26086644 | -1.364242679 | 3.30558948 | -0.4127078 | 0.67982068 | NA |
| AC073174.1 | 66.0005433 | 0.945852776 | 1.74822214 | 0.54103695 | 0.58848211 | 0.74909603 |
| AC036101.1 | 0 | NA | NA | NA | NA | NA |

| | | | | | | |
|---|---|---|---|---|---|---|
| PCDH15 | 69.2870529 | 5.054087209 | 0.71612526 | 7.0575463 | 1.69E-12 | 4.26E-10 |
| AL353784.1 | 0 | NA | NA | NA | NA | NA |
| AL355314.1 | 0 | NA | NA | NA | NA | NA |
| AL355314.2 | 0 | NA | NA | NA | NA | NA |
| MTRNR2L5 | 0 | NA | NA | NA | NA | NA |
| ZWINT | 331.019034 | 0.140655738 | 0.35772543 | 0.39319469 | 0.69417569 | 0.82386304 |
| IPMK | 667.270834 | -1.055627699 | 0.19811136 | -5.3284561 | 9.91E-08 | 4.75E-06 |
| CISD1 | 2383.76746 | -0.04177656 | 0.18688703 | -0.2235391 | 0.82311594 | 0.90556834 |
| AC016396.1 | 10.1000689 | -0.515264896 | 0.5213365 | -0.9883538 | 0.32297942 | 0.5206925 |
| AC016396.2 | 0.43541621 | 0.310941864 | 2.18727693 | 0.14215935 | 0.88695414 | NA |
| UBE2D1 | 1120.03589 | -1.057410207 | 0.31232252 | -3.3856355 | 0.00071014 | 0.00597902 |
| TFAM | 1732.7183 | 0.137569577 | 0.16516076 | 0.83294346 | 0.40487662 | 0.59970771 |
| BICC1 | 5118.78626 | -0.15022993 | 0.23625663 | -0.635876 | 0.52485723 | 0.69972119 |
| LINC00844 | 53.0238849 | 1.35989379 | 0.58655796 | 2.31843035 | 0.02042594 | 0.07608276 |
| PHYHIPL | 1569.13412 | 0.795575722 | 0.33069263 | 2.40578603 | 0.01613771 | 0.06426329 |
| FAM13C | 565.117659 | 0.921634977 | 0.34858249 | 2.64395089 | 0.00819445 | 0.03878797 |
| AC025038.1 | 0 | NA | NA | NA | NA | NA |
| AC026391.1 | 12.3079741 | 3.328680941 | 0.8834909 | 3.76764596 | 0.00016479 | 0.00187878 |
| SLC16A9 | 7264.95801 | 2.897227561 | 0.71966722 | 4.02578784 | 5.68E-05 | 0.00081871 |
| MRLN | 82.706786 | 3.409277502 | 0.73265445 | 4.65332259 | 3.27E-06 | 8.39E-05 |
| CCDC6 | 2818.41071 | -0.023403621 | 0.14950744 | -0.1565382 | 0.87560884 | 0.93541586 |
| LINC01553 | 0 | NA | NA | NA | NA | NA |
| ANK3 | 5649.64281 | 0.684823909 | 0.28274868 | 2.42202334 | 0.01543436 | 0.062157 |
| AL592430.1 | 1129.47408 | 1.022200369 | 0.33824718 | 3.02205143 | 0.00251068 | 0.0160462 |
| AL592430.2 | 105.786122 | 1.063562365 | 0.41882801 | 2.53937733 | 0.011105 | 0.04868476 |
| AC022390.1 | 11.2383653 | 0.70141694 | 0.54154566 | 1.29521293 | 0.19524679 | 0.37540341 |
| CDK1 | 176.172298 | -0.158795136 | 0.51738958 | -0.306916 | 0.75890732 | 0.8664871 |
| RHOBTB1 | 2506.99421 | 1.231930643 | 0.42943976 | 2.86869256 | 0.00412172 | 0.02328402 |
| LINC00845 | 15.61049 | 5.567461052 | 0.87054058 | 6.39540666 | 1.60E-10 | 2.14E-08 |
| TMEM26 | 242.434502 | 1.139759752 | 0.254331 | 4.48140317 | 7.42E-06 | 0.00016094 |
| TMEM26-AS | 25.723941 | 1.471677987 | 0.31631849 | 4.65251961 | 3.28E-06 | 8.41E-05 |
| CABCOCO1 | 71.8539674 | -0.563201905 | 0.37257881 | -1.5116316 | 0.13062761 | 0.28600549 |
| AL451049.1 | 3.09023095 | 0.874264264 | 0.79093178 | 1.10535988 | 0.26900368 | 0.46314344 |
| AC022398.1 | 4.55975287 | 1.672641283 | 0.62666186 | 2.66912889 | 0.00760483 | 0.03679619 |
| ARID5B | 4981.72953 | -0.485202193 | 0.26223094 | -1.8502858 | 0.06427237 | 0.17339232 |
| RTKN2 | 70.5014804 | 0.107860782 | 0.3093229 | 0.34869963 | 0.72731482 | 0.84602081 |
| AC024597.1 | 3.06761046 | -2.178132358 | 1.27917404 | -1.7027647 | 0.08861212 | 0.21811832 |
| ZNF365 | 46.5351435 | -1.026838812 | 0.70813576 | -1.4500592 | 0.147042 | 0.31022462 |
| AL133417.1 | 120.391453 | 0.292106859 | 0.25267713 | 1.15604786 | 0.24766158 | 0.43919542 |
| ADO | 1182.57171 | -0.049626965 | 0.14340845 | -0.3460533 | 0.72930268 | 0.8470627 |
| EGR2 | 154.936908 | 1.662477119 | 0.67774999 | 2.45293565 | 0.01416957 | 0.0584287 |
| AL590502.1 | 0.08666371 | -0.642905668 | 3.30558948 | -0.1944905 | 0.84579185 | NA |
| NRBF2 | 929.090283 | -0.462318468 | 0.24907079 | -1.8561729 | 0.06342891 | 0.17183279 |
| JMJD1C | 2373.03695 | 0.05901129 | 0.16754185 | 0.35221821 | 0.72467462 | 0.84389181 |
| JMJD1C-AS1 | 37.5435564 | -0.212052599 | 0.29439109 | -0.7203092 | 0.47133466 | 0.65783283 |
| REEP3 | 2166.35693 | -0.553951366 | 0.20971245 | -2.6414806 | 0.00825445 | 0.03897782 |
| AC022387.1 | 1.79684187 | 0.083950785 | 1.10661422 | 0.07586274 | 0.9395283 | 0.97110795 |
| AC012558.1 | 0 | NA | NA | NA | NA | NA |
| AC024600.1 | 8.73243744 | 3.255835689 | 1.03613173 | 3.142299 | 0.00167627 | 0.01176868 |
| AL513321.2 | 0 | NA | NA | NA | NA | NA |
| AL592466.1 | 0 | NA | NA | NA | NA | NA |
| LINC01515 | 92.5761763 | 1.069407741 | 0.43436383 | 2.46200922 | 0.01381611 | 0.05730275 |
| CTNNA3 | 18.6745732 | 1.011697783 | 0.683299 | 1.48060773 | 0.13871113 | 0.29801309 |
| AC022017.1 | 0 | NA | NA | NA | NA | NA |
| LRRTM3 | 1.15801902 | 1.401321141 | 1.19455906 | 1.17308653 | 0.24076111 | 0.43089309 |
| AL139240.1 | 0 | NA | NA | NA | NA | NA |
| DNAJC12 | 624.524118 | 1.182844576 | 0.36127767 | 3.27405951 | 0.00106014 | 0.00820064 |
| AL133551.1 | 27.8006688 | 1.270179698 | 0.66177438 | 1.9193546 | 0.05493947 | 0.15558773 |
| SIRT1 | 1069.29867 | 0.092692011 | 0.25288845 | 0.36653319 | 0.71396725 | 0.83749271 |
| HERC4 | 2293.46392 | -0.238078314 | 0.11392808 | -2.0897246 | 0.03664255 | 0.11626327 |
| MYPN | 5.45220054 | -0.335211112 | 0.87022662 | -0.3851998 | 0.7000894 | 0.82801208 |
| ATOH7 | 91.1432485 | 1.400135948 | 0.37842008 | 3.69995155 | 0.00021564 | 0.002321 |
| AC016395.1 | 81.3937744 | 1.995050734 | 0.4980114 | 4.00603424 | 6.17E-05 | 0.00086836 |
| PBLD | 8390.78637 | 2.118483732 | 0.31718499 | 6.67901633 | 2.41E-11 | 4.32E-09 |
| HNRNPH3 | 8263.54654 | -0.234929847 | 0.10337726 | -2.2725485 | 0.0230534 | 0.08334497 |
| RUFY2 | 2347.45134 | -0.245834809 | 0.11293993 | -2.1766864 | 0.02950398 | 0.09970779 |
| DNA2 | 473.315624 | -0.768885198 | 0.3420036 | -2.2481786 | 0.0245648 | 0.08710149 |
| SLC25A16 | 2535.84544 | 0.071479156 | 0.20722673 | 0.34493212 | 0.7301454 | 0.84751607 |
| TET1 | 56.7520623 | 0.791212104 | 0.39124049 | 2.02231651 | 0.04314366 | 0.13098124 |
| CCAR1 | 3067.90742 | -0.341610136 | 0.19623827 | -1.7407926 | 0.08171993 | 0.205449 |
| STOX1 | 164.56437 | 0.651668829 | 0.32067211 | 2.03219678 | 0.04213374 | 0.12868505 |
| DDX50 | 1801.94211 | -0.193095366 | 0.13254761 | -1.4568 | 0.14517159 | 0.30740985 |
| DDX21 | 3685.65038 | -0.868800976 | 0.32047268 | -2.7109986 | 0.00670809 | 0.03348612 |
| KIF1BP | 1766.95781 | -0.315765804 | 0.1285173 | -2.4569906 | 0.01401064 | 0.05790916 |
| SRGN | 6036.36016 | -1.243907011 | 0.37143818 | -3.3488937 | 0.00081135 | 0.00665069 |
| VPS26A | 3099.26584 | -0.324031757 | 0.2071656 | -1.5641195 | 0.11778951 | 0.2664992 |
| SUPV3L1 | 1460.91411 | -0.128567424 | 0.15973899 | -0.8048594 | 0.42090085 | 0.61354889 |
| AL596223.1 | 4.71010158 | -1.582430529 | 1.15006646 | -1.375947 | 0.16883803 | 0.34041279 |
| HKDC1 | 4672.63653 | -2.134470202 | 0.60465989 | -3.5300344 | 0.00041551 | 0.00393002 |
| AL596223.2 | 665.368564 | -2.178097847 | 0.58784511 | -3.7052241 | 0.0002112 | 0.00228069 |
| HK1 | 6036.82604 | -0.843166504 | 0.27466231 | -3.0698297 | 0.00214181 | 0.01420293 |
| TACR2 | 18.4870861 | -0.751347909 | 0.43096911 | -1.7433915 | 0.08126524 | 0.20461802 |
| TSPAN15 | 1907.61303 | -0.872784216 | 0.27677127 | -3.1534495 | 0.00161353 | 0.01143786 |
| NEUROG3 | 5.75601758 | -0.914334629 | 0.76671683 | -1.1925324 | 0.23305257 | 0.42124864 |
| AL450311.2 | 7.27009441 | -1.010187523 | 0.64985667 | -1.5544774 | 0.12007058 | 0.27002056 |
| FAM241B | 452.507156 | -0.639635545 | 0.2283946 | -2.8005721 | 0.00510121 | 0.02728025 |
| AL450311.1 | 4.41667173 | 4.877321008 | 1.2022837 | 4.05671389 | 4.98E-05 | 0.00074188 |
| COL13A1 | 228.316498 | -0.883685542 | 0.49574935 | -1.7825249 | 0.07466368 | 0.1931033 |
| AC024601.1 | 4.35585936 | -0.355333359 | 0.72600267 | -0.4894381 | 0.62453158 | 0.77632213 |
| H2AFY2 | 988.305635 | -0.234585787 | 0.2630557 | -0.8917723 | 0.37251499 | 0.56884364 |
| AIFM2 | 2007.28561 | -0.8731175 | 0.23450902 | -3.7231723 | 0.00019674 | 0.00215266 |
| TYSND1 | 1439.91794 | 0.093107178 | 0.16716977 | 0.55696182 | 0.57755351 | 0.7412624 |
| SAR1A | 5705.38003 | -0.374910125 | 0.20883487 | -1.7952468 | 0.07261439 | 0.18951495 |
| PPA1 | 7436.14666 | -0.846184539 | 0.18165246 | -4.6582607 | 3.19E-06 | 8.20E-05 |
| NPFFR1 | 7.80718501 | -1.228788949 | 0.8523937 | -1.4415744 | 0.14942247 | 0.31354106 |
| LRRC20 | 300.590123 | 0.074928554 | 0.30308061 | 0.24722319 | 0.8047355 | 0.89426338 |
| EIF4EBP2 | 9495.05128 | 0.288988875 | 0.27695382 | 1.04345509 | 0.29673757 | 0.49351962 |
| NODAL | 35.3275409 | -0.070719337 | 0.40106595 | -0.1763285 | 0.86003591 | 0.92704165 |
| PALD1 | 678.749032 | -0.065805049 | 0.29054079 | -0.2264916 | 0.82081909 | 0.90387102 |
| PRF1 | 413.731031 | -1.540242975 | 0.59597551 | -2.5844065 | 0.00975468 | 0.04425095 |
| ADAMTS14 | 51.0066444 | -1.238438692 | 0.5258463 | -2.3551343 | 0.01851601 | 0.07093814 |
| TBATA | 18.0843956 | -0.350712662 | 0.84341087 | -0.4158266 | 0.6775369 | 0.81256359 |
| SGPL1 | 3625.9624 | -0.2605642 | 0.12074452 | -2.1579796 | 0.03092942 | 0.10309807 |
| PCBD1 | 5667.68026 | -0.114457775 | 0.36581625 | -0.3128832 | 0.75436937 | 0.86329626 |
| AC073176.3 | 0.08666371 | -0.642905668 | 3.30558948 | -0.1944905 | 0.84579185 | NA |
| AC073176.2 | 0.17515154 | 1.45420457 | 3.2783972 | 0.44357181 | 0.65735219 | NA |
| AC073176.1 | 0.08757577 | 0.547239394 | 3.29377202 | 0.16614368 | 0.86804388 | NA |
| UNC5B | 1391.12072 | 0.408967783 | 0.32242493 | 1.26841241 | 0.20465071 | 0.38702232 |
| UNC5B-AS1 | 8.98418877 | -2.244452491 | 0.78106456 | -2.8735813 | 0.00405847 | 0.02301543 |
| SLC29A3 | 370.361003 | 0.531691019 | 0.27577005 | 1.92802308 | 0.05385226 | 0.153506 |
| CDH23 | 200.90428 | 0.549192551 | 0.3983549 | 1.37865144 | 0.16800225 | 0.33929841 |
| CDH23-AS1 | 0 | NA | NA | NA | NA | NA |
| C10orf105 | 31.9801547 | 0.496103608 | 0.71459928 | 0.69424028 | 0.48753153 | 0.67137744 |

| | | | | | | |
|---|---|---|---|---|---|---|
| VSIR | 2291.91395 | 0.325514404 | 0.28202519 | 1.1542033 | 0.24841683 | 0.44009082 |
| PSAP | 85520.2228 | 0.101571703 | 0.22664374 | 0.44815579 | 0.65404077 | 0.79692519 |
| AC073370.1 | 0.72876416 | -0.235739503 | 1.42590467 | -0.1653263 | 0.86868718 | NA |
| CHST3 | 1750.97543 | -0.018316689 | 0.19120529 | -0.0957959 | 0.92368265 | 0.96286166 |
| AC022392.1 | 1.85845049 | 1.049929654 | 1.03791105 | 1.01157961 | 0.3117391 | 0.50879963 |
| SPOCK2 | 6513.44727 | 0.97875935 | 0.3754401 | 2.60696536 | 0.00913486 | 0.04211298 |
| ASCC1 | 1948.42172 | -0.016375296 | 0.1262437 | -0.1297118 | 0.89679445 | 0.94715114 |
| ANAPC16 | 6853.88057 | 0.145428436 | 0.15873107 | 0.91619387 | 0.35956522 | 0.55720924 |
| DDIT4 | 16774.0481 | -2.008581474 | 0.57307717 | -3.5049058 | 0.00045677 | 0.00422785 |
| DDIT4-AS1 | 11970.5016 | -2.070484407 | 0.57604012 | -3.5943406 | 0.00032521 | 0.00323375 |
| DNAJB12 | 3438.89274 | 0.137617795 | 0.13188729 | 1.04345004 | 0.29673991 | 0.49351962 |
| MICU1 | 4379.44481 | 0.879648087 | 0.1916832 | 4.58907248 | 4.45E-06 | 0.00010761 |
| AL513185.1 | 0.17150785 | 0.18657164 | 3.29377195 | 0.05664376 | 0.95482897 | NA |
| MCU | 1337.56705 | -0.390747904 | 0.14649289 | -2.6673507 | 0.00764519 | 0.03693726 |
| OIT3 | 171.199642 | 2.228267645 | 0.70234783 | 3.17259847 | 0.00151081 | 0.01087934 |
| PLA2G12B | 274.234503 | 1.760966929 | 0.68921849 | 2.55501984 | 0.01061817 | 0.04709861 |
| P4HA1 | 3737.90717 | -0.891204292 | 0.32388891 | -2.7515739 | 0.00593096 | 0.03054081 |
| AL731563.2 | 165.235946 | -0.270881606 | 0.39412014 | -0.6873072 | 0.49188917 | 0.67487604 |
| AL731563.3 | 87.0415793 | 0.092944757 | 0.25404769 | 0.36585554 | 0.71447287 | 0.83777178 |
| NUDT13 | 457.902073 | 1.000280641 | 0.24923742 | 4.01336466 | 5.99E-05 | 0.00084828 |
| AC016394.2 | 102.91657 | 0.895496244 | 0.23959454 | 3.73754858 | 0.00018582 | 0.00206062 |
| ECD | 1574.36123 | 0.036960081 | 0.08500733 | 0.434787 | 0.66371705 | 0.80246755 |
| FAM149B1 | 2297.53154 | 0.077438418 | 0.09631722 | 0.80399346 | 0.42140075 | 0.61404309 |
| DNAJC9 | 1519.06828 | -0.388800043 | 0.21116444 | -1.8412193 | 0.06558942 | 0.17595047 |
| DNAJC9-AS1 | 1102.40612 | -0.103973588 | 0.12388333 | -0.8392864 | 0.40130864 | 0.59616409 |
| MRPS16 | 3765.21096 | -0.175986181 | 0.18456534 | -0.9535169 | 0.34032822 | 0.53835964 |
| AC016394.1 | 289.5063 | 0.377797325 | 0.36612431 | 1.03188265 | 0.3021271 | 0.49897235 |
| CFAP70 | 796.439368 | 0.740117275 | 0.48619012 | 1.52227953 | 0.12793905 | 0.2818978 |
| ANXA7 | 9566.75401 | -0.131546944 | 0.15089355 | -0.8717864 | 0.38332492 | 0.57917139 |
| AL512656.1 | 2016.40453 | -0.027592813 | 0.18425246 | -0.1497555 | 0.88095755 | 0.93831693 |
| MSS51 | 229.265291 | -0.400036665 | 0.29661541 | -1.3486712 | 0.17744259 | 0.35252117 |
| PPP3CB | 2457.59252 | 0.227786435 | 0.12626345 | 1.80405682 | 0.0712224 | 0.1866562 |
| USP54 | 1185.09997 | 0.125880133 | 0.16031052 | 0.78522689 | 0.43232055 | 0.62437624 |
| AC073389.1 | 20.4524321 | 1.134491983 | 0.34437543 | 3.29434649 | 0.00098651 | 0.00775849 |
| AC073389.3 | 39.881531 | 0.302526649 | 0.34796887 | 0.86940722 | 0.38462443 | 0.5803036 |
| MYOZ1 | 124.063618 | 0.383282791 | 0.44887422 | 0.85387571 | 0.39317386 | 0.5886558 |
| SYNPO2L | 18.8719862 | 0.078387441 | 0.59719929 | 0.13125843 | 0.89557088 | 0.94638952 |
| AC073389.2 | 68.6119033 | -0.037531035 | 0.22496463 | -0.1668308 | 0.86750317 | 0.93109441 |
| AGAP5 | 340.907374 | -0.02909175 | 0.2073908 | -0.140275 | 0.8884427 | 0.94229312 |
| AC022400.6 | 57.3652169 | 0.30495306 | 0.23697416 | 1.2868621 | 0.19814236 | 0.37909258 |
| SEC24C | 6417.3098 | 0.024591372 | 0.13600918 | 0.18080671 | 0.8565193 | 0.92488775 |
| FUT11 | 1459.39203 | -1.479581315 | 0.29188807 | -5.0690024 | 4.00E-07 | 1.51E-05 |
| CHCHD1 | 1470.66495 | -0.282879064 | 0.13088023 | -2.1613583 | 0.03066767 | 0.10248407 |
| ZSWIM8 | 9369.36311 | 0.065045889 | 0.20527883 | 0.31686604 | 0.75134525 | 0.86135902 |
| ZSWIM8-AS1 | 3229.10316 | -0.017111899 | 0.19687514 | -0.0869175 | 0.93073707 | 0.96662347 |
| AC022400.7 | 5847.48219 | -0.147006148 | 0.15126584 | -0.9718397 | 0.33113031 | 0.52901408 |
| NDST2 | 1900.18142 | -0.158059683 | 0.12524669 | -1.2619869 | 0.20695351 | 0.38961691 |
| CAMK2G | 2531.20556 | 1.135607981 | 0.17016026 | 6.67375575 | 2.49E-11 | 4.41E-09 |
| AC022400.1 | 386.98622 | 1.065898395 | 0.20280649 | 5.25574105 | 1.47E-07 | 6.61E-06 |
| AC022400.2 | 0.40540895 | 2.603510162 | 2.06081207 | 1.26334186 | 0.20646635 | NA |
| PLAU | 4367.71596 | 1.475158071 | 0.51641234 | 2.85655075 | 0.00428272 | 0.02397775 |
| C10orf55 | 1536.85396 | 1.607068384 | 0.5241348 | 3.06613562 | 0.00216845 | 0.01435652 |
| VCL | 6385.79843 | 0.038303943 | 0.2469135 | 0.15513102 | 0.87671803 | 0.93572892 |
| AP3M1 | 2063.52915 | -0.05674641 | 0.12745717 | -0.4452194 | 0.65616117 | 0.79821793 |
| ADK | 1487.78045 | -0.14141085 | 0.17809814 | -0.7940052 | 0.42719241 | 0.61950416 |
| AC022540.1 | 4.11283602 | 0.121706557 | 0.64825187 | 0.18774579 | 0.85107594 | 0.92184442 |
| AC063962.1 | 0.6881356 | -0.094661509 | 2.2080771 | -0.0428706 | 0.96580471 | NA |
| KAT6B | 1049.42579 | 0.515400481 | 0.20124241 | 2.56109274 | 0.01043435 | 0.04654945 |
| AC018511.2 | 6.63982066 | 2.488869549 | 0.58259637 | 4.27203062 | 1.94E-05 | 0.00034583 |
| AC018511.1 | 0.11223674 | 0.02007176 | 3.29738704 | 0.00608717 | 0.99514317 | NA |
| DUPD1 | 0.3429728 | 0.788469417 | 3.27633747 | 0.24065574 | 0.80982195 | NA |
| DUSP13 | 6.02739347 | -3.794798656 | 1.11008526 | -3.418475 | 0.00062973 | 0.00542326 |
| SAMD8 | 1460.57586 | -0.477068752 | 0.14356462 | -3.3230246 | 0.00089047 | 0.00716343 |
| VDAC2 | 6552.18608 | -0.055873808 | 0.17453286 | -0.3201335 | 0.74886716 | 0.8597841 |
| COMTD1 | 1176.43252 | -0.525784676 | 0.20123309 | -2.6128143 | 0.00898001 | 0.04160275 |
| ZNF503-AS1 | 69.6626704 | 0.179587756 | 0.3319944 | 0.54093609 | 0.58855163 | 0.74914836 |
| AC010997.3 | 393.801416 | 0.869774824 | 0.26017503 | 3.34303721 | 0.00082867 | 0.00676322 |
| ZNF503 | 2819.18222 | 0.530184864 | 0.22048351 | 2.40464628 | 0.01618813 | 0.06436684 |
| ZNF503-AS2 | 476.55776 | 0.298938791 | 0.24183901 | 1.2361066 | 0.21641894 | 0.40127742 |
| AC010997.5 | 35.3068803 | 0.314464379 | 0.27762411 | 1.13269839 | 0.25734094 | 0.4499697 |
| AC010997.4 | 142.662938 | 0.851934985 | 0.26771315 | 3.18226803 | 0.00146127 | 0.01059196 |
| AC010997.2 | 1.52227905 | 1.267320686 | 1.16931032 | 1.08381895 | 0.27844508 | 0.47411356 |
| LRMDA | 498.559026 | 0.207815899 | 0.28080526 | 0.74007124 | 0.45925677 | 0.6480069 |
| AC013286.1 | 0.20718593 | 0.18657164 | 3.29377195 | 0.05664376 | 0.95482897 | NA |
| AC024603.1 | 3.12631964 | 2.231804402 | 0.85453741 | 2.61171059 | 0.00900905 | 0.04170797 |
| KCNMA1 | 536.200268 | -0.676298046 | 0.38866699 | -1.740045 | 0.08185112 | 0.20568077 |
| KCNMA1-AS: | 15.0339266 | 0.736594762 | 0.5466248 | 1.34753264 | 0.17780875 | 0.35304696 |
| KCNMA1-AS: | 0 | NA | NA | NA | NA | NA |
| AL731575.1 | 0.07830766 | -0.642905668 | 3.30558948 | -0.1944905 | 0.84579185 | NA |
| KCNMA1-AS: | 0.16783122 | 1.368031398 | 3.28033652 | 0.41703996 | 0.67664918 | NA |
| AL450306.1 | 1520.07742 | -0.890666889 | 0.23816976 | -3.7396305 | 0.00018429 | 0.00204794 |
| DLG5 | 2154.32455 | -0.61249994 | 0.26882244 | -2.2784554 | 0.02269946 | 0.08232997 |
| AL391421.1 | 11.0847505 | -0.641502573 | 1.08731006 | -0.5899905 | 0.55519704 | 0.72467188 |
| POLR3A | 1066.20404 | 0.086481228 | 0.12129478 | 0.71298393 | 0.4758557 | 0.66130903 |
| RPS24 | 23095.5767 | -0.563256537 | 0.17238813 | -3.2673742 | 0.0010855 | 0.00835561 |
| AL731555.1 | 0 | NA | NA | NA | NA | NA |
| AC012560.1 | 0.70313844 | 0.741494922 | 1.45407152 | 0.50994392 | 0.61009075 | NA |
| LINC00595 | 2.21224191 | 1.223150816 | 0.89008488 | 1.3741957 | 0.16938093 | 0.34124649 |
| AC010163.1 | 0.12150485 | 1.04981648 | 3.28477129 | 0.31960109 | 0.74927075 | NA |
| AC010163.2 | 0.76729713 | -1.2320393 | 2.23418088 | -0.5514501 | 0.58132516 | NA |
| AL583852.1 | 0.20718593 | 0.18657164 | 3.29377195 | 0.05664376 | 0.95482897 | NA |
| AC016820.1 | 0.56542701 | 1.121043418 | 2.17110981 | 0.51634579 | 0.60561292 | NA |
| ZMIZ1-AS1 | 178.901033 | -1.941344807 | 0.41261645 | -4.7049623 | 2.54E-06 | 6.89E-05 |
| AL356753.1 | 0 | NA | NA | NA | NA | NA |
| ZMIZ1 | 6102.09914 | 0.170896489 | 0.17426611 | 0.98066391 | 0.32675851 | 0.52469456 |
| PPIF | 6248.73914 | -0.677213119 | 0.32582406 | -2.0784626 | 0.03766677 | 0.11844417 |
| ZCCHC24 | 1455.79307 | -0.268581099 | 0.28795593 | -0.932716 | 0.35096662 | 0.54857388 |
| AL133481.1 | 310.789496 | -0.926336112 | 0.36193783 | -2.5593791 | 0.01048593 | 0.04668062 |
| AL133481.3 | 0 | NA | NA | NA | NA | NA |
| EIF5AL1 | 61.7193755 | -0.348678977 | 0.5741746 | -0.6072699 | 0.54367178 | 0.71593632 |
| SFTPA2 | 4.58897983 | 0.100684323 | 0.68145987 | 0.14774798 | 0.88254166 | 0.93901806 |
| SFTPA1 | 0.08757577 | 0.547239394 | 3.29377202 | 0.16614368 | 0.86804388 | NA |
| BX248123.1 | 2.21924034 | -0.409186498 | 1.04656734 | -0.3909796 | 0.6958123 | 0.82498855 |
| AL132656.2 | 134.065447 | -0.718229286 | 0.24787266 | -2.8975737 | 0.00376061 | 0.02178067 |
| NUTM2B-AS | 743.209211 | -0.500591 | 0.19952608 | -2.5089001 | 0.01211077 | 0.05191175 |
| NUTM2B | 27.403286 | 0.194855442 | 0.43747958 | 0.44540465 | 0.65602735 | 0.79815617 |
| AL135925.1 | 239.724808 | -0.280670609 | 0.13666677 | -2.0536859 | 0.0400061 | 0.12382171 |
| NUTM2E | 6.0448569 | -0.774792336 | 1.37902711 | -0.5618398 | 0.57422516 | 0.73864963 |
| SFTPD | 106.951864 | -0.463306974 | 0.34077088 | -1.359585 | 0.17396128 | 0.34788772 |
| SFTPD-AS1 | 47.1381264 | 0.540968374 | 0.29693631 | 1.82183305 | 0.06848032 | 0.18125325 |
| TMEM254-A: | 193.308787 | 1.407223303 | 0.31456574 | 4.4735428 | 7.69E-06 | 0.00016552 |
| TMEM254 | 1356.72358 | 0.550567409 | 0.27159574 | 2.02715772 | 0.04264629 | 0.12973956 |
| PLAC9 | 366.983785 | -0.106269324 | 0.44843304 | -0.2369792 | 0.81267289 | 0.89895498 |

| | | | | | | |
|---|---|---|---|---|---|---|
| ANXA11 | 16744.504 | -0.494150225 | 0.15268454 | -3.2364129 | 0.00121042 | 0.00910665 |
| LINC00857 | 93.5462676 | -0.156161439 | 0.2464234 | -0.6337119 | 0.52626886 | 0.70114232 |
| MAT1A | 100.318014 | 0.439128982 | 0.60846022 | 0.72170532 | 0.47047567 | 0.65712094 |
| AL359195.1 | 0 | NA | NA | NA | NA | NA |
| DYDC1 | 0.40473133 | -0.616137061 | 2.3120305 | -0.2664918 | 0.78986051 | NA |
| DYDC2 | 10.1935089 | -2.862833229 | 0.79698688 | -3.5920707 | 0.00032806 | 0.00325591 |
| FAM213A | 6899.98227 | 0.333851489 | 0.21843885 | 1.52835219 | 0.12642512 | 0.2796603 |
| TSPAN14 | 6891.64486 | -0.697824865 | 0.13250755 | -5.2663025 | 1.39E-07 | 6.35E-06 |
| AC021028.1 | 12.8873239 | -0.553396921 | 0.48333502 | -1.1449551 | 0.25222774 | 0.44445342 |
| SH2D4B | 7.19365228 | -0.44043707 | 0.6868953 | -0.6411997 | 0.52139293 | 0.69771296 |
| AL731544.1 | 0.03392908 | 0.380740593 | 3.29738709 | 0.11546736 | 0.90807469 | NA |
| NRG3 | 87.8858211 | -0.526739122 | 0.41871367 | -1.2579936 | 0.20839406 | 0.39152061 |
| AC010157.1 | 0.03392908 | 0.380740593 | 3.29738709 | 0.11546736 | 0.90807469 | NA |
| AC069540.2 | 0 | NA | NA | NA | NA | NA |
| AL390786.1 | 2.89441036 | 0.198805338 | 1.065635 | 0.18656044 | 0.85200528 | 0.92250934 |
| AL603756.1 | 1434.79296 | 0.547652493 | 0.33739162 | 1.62319532 | 0.10454766 | 0.24499438 |
| GHITM | 19696.8037 | 0.356600341 | 0.23919196 | 1.49085419 | 0.13599978 | 0.29393609 |
| C10orf99 | 37.7829766 | -2.127350047 | 1.05289037 | -2.0204858 | 0.04333302 | 0.13131564 |
| CDHR1 | 309.761972 | -0.394881634 | 0.79896709 | -0.4942402 | 0.62113657 | 0.77363364 |
| LRIT2 | 4.72966454 | 0.303619077 | 0.74908163 | 0.40532175 | 0.68524102 | 0.81810054 |
| LRIT1 | 0 | NA | NA | NA | NA | NA |
| RGR | 0.5657757 | 2.509559899 | 2.64168999 | 0.94998274 | 0.34212102 | NA |
| LINC00858 | 2.96874823 | -1.764540589 | 0.95668785 | -1.8444267 | 0.06512098 | 0.17503214 |
| CCSER2 | 1710.07081 | -0.084201327 | 0.15119766 | -0.5568957 | 0.57759869 | 0.7412624 |
| LINC01519 | 0 | NA | NA | NA | NA | NA |
| AC025428.1 | 0 | NA | NA | NA | NA | NA |
| AC025428.2 | 0 | NA | NA | NA | NA | NA |
| LINC01520 | 0 | NA | NA | NA | NA | NA |
| GRID1-AS1 | 0.12150485 | 1.04981648 | 3.28477129 | 0.31960109 | 0.74927075 | NA |
| GRID1 | 67.6865071 | 0.006547249 | 0.48994176 | 0.01336332 | 0.98933793 | 0.99519138 |
| AC022028.2 | 10.841311 | -0.034897916 | 0.70024236 | -0.0498369 | 0.96025235 | 0.98093494 |
| WAPL | 2430.93792 | -0.118048835 | 0.16695898 | -0.7070529 | 0.47953358 | 0.66418343 |
| AL731569.1 | 110.909578 | 0.349237739 | 0.24264828 | 1.43927557 | 0.15007246 | 0.3144947 |
| OPN4 | 2.66789446 | -0.156259323 | 1.13402063 | -0.1377923 | 0.89040456 | 0.94316 |
| LDB3 | 252.227714 | 0.133491801 | 0.60808179 | 0.21952935 | 0.82623772 | 0.90729943 |
| AC067750.1 | 116.744547 | 0.289451845 | 0.22918116 | 1.26298274 | 0.20659538 | 0.38924304 |
| BMPR1A | 1484.39435 | 0.048000414 | 0.1389607 | 0.34542437 | 0.72977536 | 0.8473873 |
| MMRN2 | 3317.47442 | 0.103683651 | 0.32274176 | 0.32125886 | 0.74801423 | 0.85910867 |
| SNCG | 542.875782 | -0.263710629 | 0.4397513 | -0.5996813 | 0.54871866 | 0.71931354 |
| ADIRF | 4974.33859 | 0.439540765 | 0.47435936 | 0.9265987 | 0.35413493 | 0.55176 |
| AL136982.7 | 4495.03904 | 0.432479634 | 0.47745416 | 0.90580347 | 0.36503988 | 0.56205495 |
| AL136982.2 | 38.1320249 | 0.725333487 | 0.47921987 | 1.51357141 | 0.13013458 | 0.2852813 |
| FAM25A | 1.61507306 | -0.934189952 | 1.14746258 | -0.8141354 | 0.41556737 | 0.60905514 |
| GLUD1 | 13951.8791 | 0.862604607 | 0.30571773 | 2.8215721 | 0.00477889 | 0.02597265 |
| SHLD2 | 3161.60298 | 0.933334507 | 0.24946586 | 3.74133162 | 0.00018305 | 0.00203584 |
| NUTM2A-AS | 781.367776 | -0.200741664 | 0.16664948 | -1.2045742 | 0.22836772 | 0.41551671 |
| NUTM2A | 52.8842854 | 0.249285483 | 0.68028262 | 0.366444 | 0.71403379 | 0.83751762 |
| NUTM2D | 159.902032 | 0.752095043 | 0.31924697 | 2.35584079 | 0.01848084 | 0.07083968 |
| LINC00864 | 118.136273 | 1.598005643 | 0.55360418 | 2.88654912 | 0.00389492 | 0.02233396 |
| MINPP1 | 1356.34696 | 0.114448925 | 0.23559369 | 0.48578943 | 0.62711647 | 0.77806973 |
| AL138767.2 | 0.09997307 | 0.547239394 | 3.29377202 | 0.16614368 | 0.86804388 | NA |
| AL138767.3 | 7.84803607 | 0.483341024 | 0.56471725 | 0.85589916 | 0.39205356 | 0.58779376 |
| AL138767.1 | 42.3718426 | 1.406004414 | 0.50069147 | 2.80812536 | 0.00498308 | 0.02679525 |
| PAPSS2 | 2415.5131 | 0.331382342 | 0.29909391 | 1.10795416 | 0.26788161 | 0.46206424 |
| ATAD1 | 2760.35285 | -0.136570755 | 0.10952868 | -1.246895 | 0.21243601 | 0.39623268 |
| KLLN | 68.8446399 | 1.524725412 | 0.35636937 | 4.27849741 | 1.88E-05 | 0.0003379 |
| PTEN | 4578.98252 | -0.179700118 | 0.15619203 | -1.1505076 | 0.24993485 | 0.44201546 |
| RNLS | 748.21714 | 0.89934438 | 0.23187043 | 3.87865052 | 0.00010504 | 0.00131709 |
| LIPJ | 8.73852935 | 0.829276496 | 0.47809999 | 1.73452522 | 0.08282494 | 0.20761346 |
| LIPF | 4.38632876 | 1.859252858 | 0.73952674 | 2.51411175 | 0.01193326 | 0.05135957 |
| LIPK | 0.13037664 | 0.02007176 | 3.29738704 | 0.00608717 | 0.99514317 | NA |
| LIPN | 2.74611561 | -1.078051837 | 0.96359567 | -1.1187803 | 0.26323388 | 0.45691664 |
| LIPM | 20.261267 | -0.615647531 | 0.47841064 | -1.28686 | 0.19814308 | 0.37909258 |
| ANKRD22 | 182.695393 | -0.798929868 | 0.35014997 | -2.2816791 | 0.02250829 | 0.08183423 |
| STAMBPL1 | 186.700752 | -0.053790054 | 0.35318152 | -0.1523014 | 0.87894919 | 0.93705294 |
| ACTA2-AS1 | 8507.87887 | 0.078285193 | 0.5424073 | 0.14432917 | 0.88524055 | 0.94058667 |
| ACTA2 | 13647.1281 | 0.180805317 | 0.54511507 | 0.33168284 | 0.74012876 | 0.8541816 |
| FAS | 1521.35254 | -1.283450453 | 0.25685566 | -4.996777 | 5.83E-07 | 2.07E-05 |
| CH25H | 184.389509 | -0.584020255 | 0.56076595 | -1.0414688 | 0.29765802 | 0.49454947 |
| LIPA | 4402.04847 | -0.389301489 | 0.17470716 | -2.2283087 | 0.02585994 | 0.09058535 |
| IFIT2 | 1320.70167 | -0.285469191 | 0.3077144 | -0.9277082 | 0.35355893 | 0.55115076 |
| AL353751.1 | 415.147882 | -2.23060098 | 0.61910713 | -3.6029321 | 0.00031465 | 0.00314528 |
| IFIT3 | 2238.21725 | -0.933254665 | 0.5706901 | -1.635309 | 0.10198429 | 0.24072764 |
| IFIT1B | 10.9523037 | -1.996236322 | 1.03069421 | -1.9367881 | 0.05277125 | 0.15129134 |
| IFIT1 | 1775.36296 | 1.063586223 | 0.22231332 | 4.78417688 | 1.72E-06 | 5.03E-05 |
| IFIT5 | 1291.18466 | -0.617122945 | 0.30098004 | -2.0503783 | 0.04032753 | 0.12451901 |
| SLC16A12 | 4620.43418 | 2.140189978 | 0.46405545 | 4.61192728 | 3.99E-06 | 9.84E-05 |
| SLC16A12-AS | 109.282174 | 3.487222637 | 0.4372994 | 7.97445099 | 1.53E-15 | 1.11E-12 |
| PANK1 | 2153.03848 | 1.777785375 | 0.46965398 | 3.78530884 | 0.00015352 | 0.00178495 |
| AL157400.2 | 6.11331549 | 2.052308729 | 0.93674648 | 2.19089024 | 0.02845974 | 0.09723463 |
| AL157400.1 | 0.69466431 | 2.82470359 | 1.50411115 | 1.8779886 | 0.06038273 | NA |
| AL157400.3 | 14.7313924 | 1.116185796 | 0.39429347 | 2.83085033 | 0.00464244 | 0.0254514 |
| AL157400.4 | 5.259799 | 1.426751876 | 0.5484242 | 2.60154801 | 0.00928041 | 0.04256805 |
| KIF20B | 154.721539 | 0.163157094 | 0.21396761 | 0.76253172 | 0.44574272 | 0.63615528 |
| LINC00865 | 216.621398 | 0.865975047 | 0.4667129 | 1.855477 | 0.06352814 | 0.17201325 |
| LINC01374 | 5.05120589 | 0.121096686 | 0.69762303 | 0.1735847 | 0.86219185 | 0.9281515 |
| LINC01375 | 0 | NA | NA | NA | NA | NA |
| AL391704.2 | 0.03392908 | 0.380740593 | 3.29738709 | 0.11546736 | 0.90807469 | NA |
| AL391704.1 | 0.19890045 | -0.340073434 | 3.15719848 | -0.1077137 | 0.91422283 | NA |
| AL391704.3 | 0 | NA | NA | NA | NA | NA |
| AL390862.1 | 0 | NA | NA | NA | NA | NA |
| HTR7 | 11.2133714 | 0.769516729 | 0.64827431 | 1.18702333 | 0.23521842 | 0.42405891 |
| RPP30 | 921.07996 | -0.114623183 | 0.12398201 | -0.9245146 | 0.35521843 | 0.55272969 |
| ANKRD1 | 37.8484563 | -0.970751446 | 0.44892273 | -2.1624021 | 0.0305872 | 0.10226758 |
| AL365434.1 | 0.07830766 | -0.642905668 | 3.30558948 | -0.1944905 | 0.84579185 | NA |
| AL365434.2 | 1.04907879 | 1.499109974 | 1.2505366 | 1.19877337 | 0.23061608 | 0.41824938 |
| LINC00502 | 0.68983927 | 1.632899349 | 1.46006441 | 1.11837487 | 0.26340692 | NA |
| PCGF5 | 3559.57445 | -0.116850907 | 0.25873733 | -0.4516198 | 0.65154289 | 0.7952412 |
| HECTD2 | 695.103134 | 0.210272913 | 0.19869988 | 1.05824379 | 0.28994431 | 0.48609775 |
| PPP1R3C | 1537.99407 | -1.122784746 | 0.59551482 | -1.8854019 | 0.0593756 | 0.16436433 |
| TNKS2-AS1 | 24.6140015 | 0.834795659 | 0.6223024 | 1.34146302 | 0.17977017 | 0.35557601 |
| TNKS2 | 3027.66886 | -0.238669067 | 0.19055011 | -1.2525265 | 0.21037807 | 0.3937409 |
| FGFBP3 | 80.0793863 | 0.168190849 | 0.37763418 | 0.44538037 | 0.6560449 | 0.79815617 |
| AL359198.1 | 19.6752735 | -0.172216856 | 0.45385662 | -0.3794521 | 0.70435216 | 0.83115257 |
| BTAF1 | 1921.00381 | -0.224864726 | 0.16061404 | -1.4000316 | 0.16150386 | 0.33014395 |
| CPEB3 | 565.592309 | 0.85375615 | 0.24581306 | 3.47319276 | 0.00051431 | 0.00463666 |
| 5-Mar | 2655.85578 | -0.250709597 | 0.15208693 | -1.6484625 | 0.09925779 | 0.23629395 |
| IDE | 1568.71624 | -0.218254491 | 0.15495106 | -1.4085382 | 0.15897177 | 0.32651184 |
| KIF11 | 108.348154 | 0.142525032 | 0.42464102 | 0.33563651 | 0.73714499 | 0.8524157 |
| HHEX | 342.596911 | -0.094135038 | 0.28746538 | -0.3274657 | 0.74331572 | 0.85602391 |
| EXOC6 | 637.321408 | -0.220689099 | 0.1903786 | -1.1592117 | 0.24636991 | 0.43769955 |
| AL358613.1 | 0.50834428 | -2.250444995 | 2.76168203 | -0.814882 | 0.41513985 | NA |
| CYP26C1 | 9.13660372 | -0.408058481 | 0.63057841 | -0.6471177 | 0.51755574 | 0.69466488 |

| | | | | | | |
|---|---|---|---|---|---|---|
| AL358613.2 | 0.97501211 | 0.780630319 | 2.68498132 | 0.29073957 | 0.7712505 | NA |
| CYP26A1 | 15.0887462 | -2.84657282 | 1.07012781 | -2.6600307 | 0.00781335 | 0.03755708 |
| MYOF | 3489.35142 | -0.919329443 | 0.26649805 | -3.4496666 | 0.00056128 | 0.00496842 |
| CEP55 | 41.3786016 | -1.579311428 | 0.7245939 | -2.1795815 | 0.0292885 | 0.09920124 |
| FFAR4 | 16.0960586 | -0.245860378 | 0.60543815 | -0.4060867 | 0.6846789 | 0.81783568 |
| RBP4 | 7578.1447 | 0.768119129 | 0.44357188 | 1.73166778 | 0.08333273 | 0.20848995 |
| PDE6C | 45.1884055 | 0.271115554 | 0.25202899 | 1.07573164 | 0.28204729 | 0.47782049 |
| FRA10AC1 | 925.11655 | -0.029530518 | 0.11430628 | -0.2583455 | 0.79614024 | 0.88938094 |
| LGI1 | 2.04319334 | 3.004745696 | 1.14944366 | 2.61408696 | 0.00894663 | 0.04146995 |
| AL358154.1 | 0.07659028 | -0.642905668 | 3.30558948 | -0.1944905 | 0.84579185 | NA |
| SLC35G1 | 143.42449 | -0.057408944 | 0.23866525 | -0.2405417 | 0.80991034 | 0.89729165 |
| PLCE1 | 841.353077 | 0.412691669 | 0.29381436 | 1.40460008 | 0.16014023 | 0.32825646 |
| PLCE1-AS2 | 0.26272731 | 1.990198932 | 3.242756 | 0.61373687 | 0.53938922 | NA |
| AL389885.1 | 0.03392908 | 0.380740593 | 3.29738709 | 0.11546736 | 0.90807469 | NA |
| PLCE1-AS1 | 28.6689003 | 1.125048995 | 0.55711727 | 2.01941145 | 0.04344447 | 0.13160606 |
| AL139124.1 | 4.25250995 | -0.307347596 | 0.67998323 | -0.4519929 | 0.6512741 | 0.795097 |
| NOC3L | 685.693394 | -0.142737896 | 0.1947407 | -0.7329639 | 0.46358047 | 0.65135428 |
| TBC1D12 | 758.595666 | 0.007088642 | 0.13045107 | 0.05433947 | 0.95666471 | 0.97931865 |
| HELLS | 368.629274 | 0.020741307 | 0.2818802 | 0.07358199 | 0.941343 | 0.97208705 |
| CYP2C18 | 16.4458217 | -0.501042371 | 0.51666169 | -0.9697688 | 0.33216176 | 0.53009359 |
| AL583836.1 | 9.0743079 | -0.64448547 | 0.64276654 | -1.0026743 | 0.31601805 | 0.5131471 |
| CYP2C19 | 1.41331242 | 0.621543921 | 1.1493287 | 0.54078865 | 0.58865326 | 0.74924156 |
| CYP2C9 | 163.751904 | 0.371819865 | 0.59769169 | 0.62209309 | 0.53388066 | 0.70767231 |
| CYP2C8 | 132.389963 | 0.662634165 | 0.42434974 | 1.56152838 | 0.11839913 | 0.26743395 |
| AL157834.1 | 7.01567304 | 0.353644993 | 0.50438542 | 0.7011404 | 0.4832154 | 0.66777016 |
| ACSM6 | 8.98369361 | 1.067952853 | 0.78046463 | 1.36835523 | NA | NA |
| AL157834.2 | 7.42075778 | -0.309061828 | 0.660269 | -0.4680847 | 0.63972401 | 0.78704498 |
| PDLIM1 | 3852.09853 | -0.699573315 | 0.32736453 | -2.1369857 | 0.03259915 | 0.10701353 |
| SORBS1 | 2968.38519 | -0.549221551 | 0.37111985 | -1.4799035 | 0.13889901 | 0.29824664 |
| ALDH18A1 | 3290.58263 | -0.8306816 | 0.19002377 | -4.3714615 | 1.23E-05 | 0.00024342 |
| TCTN3 | 2842.41075 | 0.138682757 | 0.11429985 | 1.213324 | 0.22500597 | 0.41129956 |
| ENTPD1 | 1407.82384 | -0.094683932 | 0.29909506 | -0.316568 | 0.7515714 | 0.86139333 |
| ENTPD1-AS1 | 321.178467 | 0.350375754 | 0.22286763 | 1.57212495 | 0.11592158 | 0.26372634 |
| AL365273.2 | 81.0489458 | -0.012969253 | 0.48105081 | -0.0269603 | 0.97849143 | 0.99032121 |
| CC2D2B | 83.9563692 | 0.296336563 | 0.23946621 | 1.23748799 | 0.21590597 | 0.40049525 |
| CCNJ | 435.192463 | -0.464167935 | 0.26293699 | -1.76532 | 0.07751 | 0.19805608 |
| ZNF518A | 1718.18374 | -0.011238436 | 0.20877053 | -0.0538315 | 0.95706939 | 0.97945117 |
| BLNK | 2179.05766 | -0.842691802 | 0.24893259 | -3.3852209 | 0.00071121 | 0.00598615 |
| AL136181.1 | 0.27293021 | -1.364244848 | 2.93668264 | -0.464553 | 0.64225157 | NA |
| DNTT | 5.88187757 | -2.653554286 | 1.01884981 | -2.6044607 | 0.0092019 | 0.04233322 |
| OPALIN | 0.08393208 | -0.642905668 | 3.30558948 | -0.1944905 | 0.84579185 | NA |
| TLL2 | 89.7321838 | -2.124113307 | 0.45343295 | -4.6845147 | 2.81E-06 | 7.44E-05 |
| TM9SF3 | 10366.4279 | -0.201896605 | 0.13793564 | -1.4637015 | 0.14327552 | 0.30479475 |
| PIK3AP1 | 2031.35421 | -0.404993698 | 0.32874145 | -1.2319521 | 0.217967 | 0.40307081 |
| LCOR | 1722.07967 | -0.22324261 | 0.12095052 | -1.845735 | 0.06493069 | 0.1746631 |
| SLIT1 | 20.3166847 | -0.151334299 | 0.35757025 | -0.4232296 | 0.67212778 | 0.80870716 |
| SLIT1-AS1 | 0 | NA | NA | NA | NA | NA |
| ARHGAP19-S | 18.4491395 | 1.564376652 | 0.40640783 | 3.84927788 | 0.00011847 | 0.00144963 |
| ARHGAP19 | 585.370083 | 0.142755816 | 0.14509976 | 0.98384599 | 0.32519125 | 0.5227881 |
| FRAT1 | 202.3881 | -0.612774968 | 0.2710268 | -2.2609386 | 0.02376306 | 0.08504775 |
| FRAT2 | 572.946698 | -0.731736153 | 0.26242921 | -2.7883183 | 0.00529825 | 0.02810596 |
| RRP12 | 1733.07288 | -0.811453938 | 0.29281322 | -2.771234 | 0.00558443 | 0.02927982 |
| AL355490.2 | 47.567826 | 0.231737016 | 0.35726102 | 0.64864904 | 0.51656525 | 0.69389362 |
| PGAM1 | 5206.5084 | -0.476291621 | 0.19073065 | -2.497195 | 0.01251801 | 0.0532098 |
| EXOSC1 | 904.024225 | -0.350229182 | 0.11681695 | -2.9981025 | 0.00271666 | 0.01703243 |
| ZDHHC16 | 2313.49971 | -0.253709882 | 0.18270048 | -1.3886657 | 0.16493443 | 0.33505285 |
| MMS19 | 3615.73181 | 0.055834763 | 0.11930512 | 0.4679997 | 0.6397848 | 0.78704634 |
| UBTD1 | 1383.64344 | -0.515229983 | 0.24231829 | -2.126253 | 0.0334822 | 0.1091379 |
| ANKRD2 | 380.972075 | 1.894411114 | 0.6513509 | 2.90843403 | 0.00363244 | 0.02118721 |
| HOGA1 | 3881.53547 | 2.047731246 | 0.45945229 | 4.45689633 | 8.32E-06 | 0.00017655 |
| AL355315.1 | 1820.21875 | 1.70805925 | 0.28202454 | 6.05642056 | 1.39E-09 | 1.35E-07 |
| C10orf62 | 16.7767473 | 0.724034122 | 0.38839516 | 1.86416873 | 0.06229801 | 0.16974567 |
| MORN4 | 879.609188 | 0.835128744 | 0.25227402 | 3.31040323 | 0.00093162 | 0.00743695 |
| PI4K2A | 2439.39792 | -0.22464417 | 0.19645875 | -1.1434674 | 0.25284459 | 0.4452726 |
| AVPI1 | 580.666479 | -0.05308831 | 0.25220082 | -0.2105001 | 0.83327734 | 0.91140944 |
| MARVELD1 | 1067.29512 | -0.236781954 | 0.22797975 | -1.0386096 | 0.29898634 | 0.49579957 |
| ZFYVE27 | 3617.62346 | -0.543105696 | 0.13496859 | -4.0239413 | 5.72E-05 | 0.00082306 |
| SFRP5 | 9.14473971 | 0.103449293 | 0.7239041 | 0.14290469 | 0.88636545 | 0.94114987 |
| LINC00866 | 0.15318056 | -1.003574173 | 3.30558948 | -0.3035992 | 0.7614333 | NA |
| GOLGA7B | 121.180766 | -0.246352394 | 0.23691187 | -1.0398483 | 0.2984104 | 0.49530098 |
| CRTAC1 | 483.213515 | 0.705585329 | 0.4921532 | 1.43367012 | 0.15166642 | 0.31673744 |
| R3HCC1L | 1047.25841 | -0.426421608 | 0.21048228 | -2.0259264 | 0.04277233 | 0.130063 |
| LOXL4 | 834.819191 | -0.310208471 | 0.40021994 | -0.775095 | 0.43828354 | 0.63008807 |
| AL139241.1 | 59.7931615 | 0.19851244 | 0.49344432 | 0.40229958 | 0.68746357 | 0.81947447 |
| PYROXD2 | 771.181635 | 0.605607024 | 0.30591158 | 1.97967995 | 0.0477395 | 0.14086278 |
| HPS1 | 4447.0997 | -0.039118578 | 0.11242552 | -0.3479511 | 0.72787694 | 0.84611931 |
| HPSE2 | 87.9381558 | -0.437888488 | 0.81309719 | -0.5385438 | 0.59020164 | 0.75008999 |
| CNNM1 | 402.925615 | -0.700987597 | 0.78416153 | -0.8939327 | 0.37135792 | 0.56825326 |
| GOT1 | 11109.8438 | -0.033413427 | 0.23789147 | -0.1404566 | 0.88829924 | 0.94222589 |
| AL391684.1 | 226.785281 | 0.736103802 | 0.36895188 | 1.99512141 | 0.04602964 | 0.13710625 |
| LINC01475 | 0.66300758 | -2.611177414 | 2.74246609 | -0.9521275 | 0.34103232 | NA |
| AL513542.1 | 0.19978473 | -1.003574173 | 3.30558948 | -0.3035992 | 0.7614333 | NA |
| NKX2-3 | 2.97826622 | -3.003297677 | 2.04637692 | -1.4676171 | 0.14220827 | 0.30320441 |
| SLC25A28 | 3125.03819 | -0.931797538 | 0.35693254 | -2.6105704 | 0.00903914 | 0.0417959 |
| AL353719.1 | 13.128873 | 0.819274756 | 0.49029195 | 1.67099369 | 0.09472292 | 0.2289709 |
| AL133353.1 | 2.8067349 | 0.382674042 | 0.80978966 | 0.4725598 | 0.63652727 | 0.78464959 |
| ENTPD7 | 329.026831 | -0.279173129 | 0.2512494 | -1.1111395 | 0.26650831 | 0.4606085 |
| CUTC | 956.61026 | -0.570865248 | 0.19359937 | -2.9486937 | 0.0031912 | 0.01926115 |
| COX15 | 2443.06291 | 0.345349223 | 0.18501676 | 1.86658342 | 0.06195978 | 0.16919121 |
| ABCC2 | 7496.87688 | -1.457847225 | 0.85198066 | -1.7111271 | 0.08705765 | 0.21509233 |
| DNMBP | 2103.99794 | -0.084243198 | 0.20697506 | -0.407021 | 0.68399256 | 0.81748139 |
| DNMBP-AS1 | 34.0757552 | 0.379235885 | 0.53783678 | 0.70511333 | 0.48073971 | 0.66542234 |
| CPN1 | 6.7945713 | 0.255064433 | 0.82659112 | 0.30857388 | 0.75764569 | 0.865984 |
| ERLIN1 | 2312.28177 | -0.013415228 | 0.20084243 | -0.0667948 | 0.94674507 | 0.97438741 |
| CHUK | 1798.82206 | 0.196173183 | 0.22797294 | 0.86051081 | 0.38950753 | 0.58516 |
| AL138921.2 | 29.4704387 | 1.018020297 | 0.40338295 | 2.52370683 | 0.01161247 | 0.05037857 |
| AL138921.1 | 225.732996 | -0.314978394 | 0.15127748 | -2.0821235 | 0.03733119 | 0.1176977 |
| CWF19L1 | 1142.64725 | -0.169863562 | 0.13438328 | -1.264023 | 0.20622177 | 0.38883016 |
| BLOC1S2 | 2033.36658 | -0.824900703 | 0.29526502 | -2.7937637 | 0.00520985 | 0.02770953 |
| PKD2L1 | 15.1513373 | -0.587190293 | 0.71008319 | -0.8269317 | 0.40827578 | 0.60277881 |
| AL139819.1 | 0.53579601 | 0.635181477 | 1.83690957 | 0.3457881 | 0.72950197 | NA |
| SCD | 4732.37384 | -1.669263407 | 0.67236433 | -2.4826769 | 0.01303993 | 0.05481745 |
| OLMALINC | 372.107359 | 0.171486367 | 0.41833374 | 0.40992717 | 0.68185937 | 0.81578112 |
| WNT8B | 32.6767894 | 1.369900363 | 0.67718689 | 2.02292806 | 0.04308057 | 0.13083433 |
| SEC31B | 1338.83349 | 0.367909131 | 0.19316613 | 1.90462543 | 0.05682878 | 0.15927037 |
| AL133352.1 | 6113.54841 | 0.100466507 | 0.12367428 | 0.81234761 | 0.4165922 | 0.6097994 |
| NDUFB8 | 6371.1367 | 0.078247513 | 0.12190818 | 0.64185613 | 0.5209666 | 0.69753183 |
| HIF1AN | 5366.488 | 0.052096546 | 0.11787109 | 0.441979 | 0.6585044 | 0.79946078 |
| PAX2 | 11607.1333 | 0.344276963 | 0.36630288 | 0.93986967 | 0.34728442 | 0.54478988 |
| AL138762.1 | 31.8537285 | 0.335019683 | 0.40993764 | 0.81724548 | 0.41378816 | 0.60767896 |
| SLF2 | 1500.45728 | -0.084829603 | 0.11450445 | -0.7408411 | 0.45878977 | 0.6476252 |
| AL133215.3 | 5.55733675 | -1.230631085 | 0.61417277 | -2.0037213 | 0.04509992 | 0.13499261 |
| MRPL43 | 3398.88026 | 0.236866578 | 0.12324824 | 1.92186585 | 0.05462264 | 0.15496493 |

| | | | | | | |
|---|---|---|---|---|---|---|
| SEMA4G | 1955.03769 | 0.998693788 | 0.29301436 | 3.40834416 | 0.00065358 | 0.00557767 |
| AL133215.1 | 234.920761 | 0.8994231 | 0.35996476 | 2.49864208 | 0.01246702 | 0.0530507 |
| TWNK | 1095.67927 | -0.11260678 | 0.19663698 | -0.5726633 | 0.56687269 | 0.73302704 |
| LZTS2 | 7906.04059 | -0.951454365 | 0.17724618 | -5.3679822 | 7.96E-08 | 3.93E-06 |
| PDZD7 | 740.13556 | -0.420274352 | 0.19321781 | -2.1751325 | 0.02962019 | 0.09998519 |
| SFXN3 | 3249.27432 | 0.129815501 | 0.20110846 | 0.64549995 | 0.51860325 | 0.69546215 |
| AL133215.2 | 5.03977342 | 0.881092785 | 0.5678176 | 1.551718 | 0.12072971 | 0.27108673 |
| KAZALD1 | 172.663498 | 0.537489806 | 0.3601641 | 1.49234698 | 0.13560821 | 0.29330613 |
| TLX1NB | 0 | NA | NA | NA | NA | NA |
| TLX1 | 0.16354576 | -1.003574173 | 3.30558948 | -0.3035992 | 0.7614333 | NA |
| LINC01514 | 0 | NA | NA | NA | NA | NA |
| LBX1 | 0.43648996 | -0.608162902 | 1.94123158 | -0.3132871 | 0.75406251 | NA |
| AL133387.2 | 0 | NA | NA | NA | NA | NA |
| AL133387.1 | 4.70209177 | 1.547513328 | 0.73607175 | 2.10239468 | 0.03551872 | 0.11377898 |
| AL133387.3 | 0.23387521 | 1.816617118 | 3.25499482 | 0.55810139 | 0.57677515 | NA |
| BTRC | 1339.32016 | 0.152900087 | 0.17377775 | 0.87985997 | 0.37893517 | 0.57500981 |
| DPCD | 1357.53869 | -0.259865903 | 0.15999105 | -1.6242528 | 0.10432187 | 0.24460631 |
| POLL | 2071.44323 | 0.071537142 | 0.17680823 | 0.40460302 | 0.68576934 | 0.81835792 |
| FBXW4 | 4083.54098 | -0.043611797 | 0.10181395 | -0.4283479 | 0.66839783 | 0.806131 |
| AC010789.1 | 13.6652831 | 0.209100575 | 0.74271838 | 0.28153413 | 0.77830075 | 0.87861887 |
| FGF8 | 2.26568929 | 2.761277797 | 1.17629218 | 2.34744212 | 0.01890281 | 0.07201201 |
| NPM3 | 658.023397 | -0.705671742 | 0.20688439 | -3.4109473 | 0.00064738 | 0.00553724 |
| OGA | 10178.4572 | -0.07063121 | 0.15972183 | -0.4422139 | 0.65833445 | 0.79946078 |
| KCNIP2-AS1 | 59.2493809 | 0.156466241 | 0.40024578 | 0.3909254 | 0.69585238 | 0.82499898 |
| KCNIP2 | 272.377927 | -0.211631009 | 0.39702274 | -0.5330451 | 0.59400239 | 0.75306938 |
| ARMH3 | 1576.84036 | -0.007177964 | 0.10418702 | -0.068895 | 0.9450732 | 0.97380355 |
| HPS6 | 1115.11919 | -0.374422818 | 0.14547106 | -2.5738646 | 0.01005696 | 0.04524089 |
| LDB1 | 4123.38857 | 0.183079103 | 0.12698288 | 1.44176205 | 0.14936951 | 0.31347983 |
| PPRC1 | 2017.74769 | -0.715050828 | 0.35139503 | -2.0348917 | 0.04186177 | 0.12807692 |
| NOLC1 | 3271.45718 | -0.349213955 | 0.22863424 | -1.5273913 | 0.12666374 | 0.28002371 |
| ELOVL3 | 31.3332904 | -1.303655762 | 0.75220767 | -1.7331062 | 0.0830768 | 0.20800752 |
| PITX3 | 11.5004491 | -0.145603562 | 0.82154754 | -0.1772308 | 0.85932707 | 0.9265427 |
| GBF1 | 5215.26891 | 0.060165924 | 0.11148174 | 0.53969306 | 0.58940873 | 0.74961886 |
| NFKB2 | 5903.53349 | -1.238257774 | 0.27270127 | -4.5407114 | 5.61E-06 | 0.00012867 |
| PSD | 557.71709 | -0.301773926 | 0.15676962 | -1.9249516 | 0.05423542 | 0.1542496 |
| FBXL15 | 1603.86032 | -0.459575033 | 0.17578248 | -2.614453 | 0.00893705 | 0.04143283 |
| CUEDC2 | 2637.98298 | -0.011541595 | 0.14777575 | -0.0781021 | 0.93774684 | 0.97027326 |
| C10orf95 | 252.496604 | 0.195006322 | 0.19435643 | 1.00334381 | 0.31569501 | 0.51280506 |
| MFSD13A | 357.679411 | -0.218226609 | 0.20189477 | -1.0808928 | 0.27974479 | 0.47534443 |
| ACTR1A | 6786.62551 | -0.438452988 | 0.1680047 | -2.6097661 | 0.00906041 | 0.0418796 |
| AL121928.1 | 593.161539 | -0.203063733 | 0.17097082 | -1.1877099 | 0.23494772 | 0.42379475 |
| SUFU | 1379.85344 | 0.601564623 | 0.1765295 | 3.40772859 | 0.00065506 | 0.00558665 |
| TRIM8 | 9426.08663 | -0.082797684 | 0.09602328 | -0.8622668 | 0.38854071 | 0.58414017 |
| AL391121.1 | 278.652254 | 0.136639753 | 0.21475342 | 0.63626345 | 0.52460473 | 0.69953025 |
| ARL3 | 1870.46517 | 0.118107081 | 0.20605167 | 0.57319158 | 0.56651499 | 0.73284279 |
| SFXN2 | 3160.45151 | 2.626094464 | 0.30088526 | 8.72789328 | 2.59E-18 | 3.66E-15 |
| WBP1L | 3725.27732 | -0.481093608 | 0.1152447 | -4.1745399 | 2.99E-05 | 0.00048865 |
| CYP17A1 | 2765.47243 | 2.373664999 | 0.63532308 | 3.73615421 | 0.00018686 | 0.0020673 |
| CYP17A1-AS | 73.8323594 | 0.812660796 | 0.33504207 | 2.4255485 | 0.01528527 | 0.06166962 |
| AL358790.1 | 338.121987 | 0.958256029 | 0.23601335 | 4.06017723 | 4.90E-05 | 0.00073262 |
| BORCS7 | 1971.55017 | 0.73445903 | 0.25204211 | 2.91403299 | 0.00356792 | 0.02087092 |
| BORCS7-ASN | 1481.56901 | 1.074957177 | 0.26979317 | 3.98437512 | 6.77E-05 | 0.00093313 |
| AS3MT | 946.415483 | 1.181559206 | 0.37333062 | 3.16491373 | 0.00155129 | 0.01109206 |
| AL356608.1 | 3.20698777 | 1.530446945 | 0.82102006 | 1.86407984 | 0.06231049 | 0.16976213 |
| CNNM2 | 5023.55534 | -0.049311418 | 0.15884424 | -0.3104388 | 0.75622729 | 0.86477509 |
| NT5C2 | 4882.78895 | -0.244345355 | 0.20060053 | -1.2180693 | 0.22319763 | 0.40924076 |
| RPEL1 | 1.85001722 | -0.418428149 | 1.0104354 | -0.4141068 | 0.67879591 | 0.81362946 |
| INA | 12.3718964 | -1.976462789 | 0.71856164 | -2.7505821 | 0.00594895 | 0.03060459 |
| PCGF6 | 452.120388 | -0.349003884 | 0.17435469 | -2.0016891 | 0.04531818 | 0.13555359 |
| TAF5 | 459.68319 | -0.332388064 | 0.19106923 | -1.7396211 | 0.08192557 | 0.20580902 |
| ATP5MD | 4044.15457 | -0.39636174 | 0.17543876 | -2.2592599 | 0.02386722 | 0.08533949 |
| PDCD11 | 1655.63628 | -0.488465058 | 0.14705064 | -3.3217473 | 0.00089456 | 0.00718972 |
| CALHM2 | 658.947836 | 0.052576396 | 0.35390392 | 0.14856122 | 0.88189988 | 0.93863819 |
| AL139339.2 | 68.1495605 | 0.496942664 | 0.32306121 | 1.53823067 | 0.12399223 | 0.27568929 |
| AL139339.1 | 5.40193328 | 0.511181554 | 0.73449983 | 0.69595871 | 0.48645469 | 0.67048976 |
| CALHM1 | 0.90454695 | -0.705308374 | 1.32520975 | -0.532224 | 0.59457089 | NA |
| CALHM3 | 1.13274358 | 0.652217037 | 1.31929961 | 0.49436612 | 0.62104763 | 0.77363364 |
| NEURL1-AS1 | 0.25441423 | 0.078703007 | 3.03556796 | 0.02592695 | 0.97931561 | NA |
| NEURL1 | 138.104264 | -0.509571094 | 0.36516794 | -1.3954431 | 0.16288225 | 0.33190673 |
| SH3PXD2A | 1821.01982 | -0.087039729 | 0.29548147 | -0.2945692 | 0.76832302 | 0.87199765 |
| AL121929.2 | 22.3676293 | -0.469966707 | 0.57260273 | -0.8207553 | 0.41178568 | 0.60580988 |
| SH3PXD2A-A | 1.60792238 | 0.361004741 | 1.08257306 | 0.33346917 | 0.73878016 | 0.85340953 |
| STN1 | 946.047972 | -0.104120713 | 0.18808137 | -0.553594 | 0.57985675 | 0.7429943 |
| SLK | 3432.96128 | -0.165740681 | 0.15339738 | -1.0804662 | 0.27993462 | 0.47554347 |
| COL17A1 | 52.2263791 | -1.367437431 | 0.53273928 | -2.5668042 | 0.01026406 | 0.04594519 |
| SFR1 | 246.93774 | -0.395473466 | 0.20095761 | -1.9679447 | 0.0490744 | 0.14386793 |
| CFAP43 | 375.59358 | 0.087895379 | 0.40766881 | 0.21560487 | 0.82929576 | 0.90929435 |
| GSTO1 | 5891.03514 | -0.768588662 | 0.2975347 | -2.58319 | 0.00978914 | 0.04436737 |
| GSTO2 | 1152.45323 | 0.373770238 | 0.23319615 | 1.6028148 | 0.10897555 | 0.25234328 |
| ITPRIP | 2670.6034 | -0.968606166 | 0.33866973 | -2.8600317 | 0.00423599 | 0.02375648 |
| AL162742.2 | 304.360273 | -1.08236651 | 0.3358512 | -3.2227562 | 0.00126964 | 0.00947377 |
| AL162742.1 | 4.74034041 | -0.776702692 | 0.67877657 | -1.1442686 | 0.25251227 | 0.4448359 |
| CFAP58-DT | 14.6802092 | -1.234770358 | 0.51135484 | -2.4147036 | 0.01574802 | 0.06307927 |
| CFAP58 | 43.8706293 | -0.614507219 | 0.25003886 | -2.4576469 | 0.01398506 | 0.05785911 |
| AL161646.1 | 0.72308802 | 1.706666703 | 1.76121579 | 0.9690276 | 0.33253142 | NA |
| AL161646.2 | 0 | NA | NA | NA | NA | NA |
| SORCS3 | 86.7931983 | 0.995790534 | 0.35615061 | 2.79598155 | 0.00517424 | 0.02759243 |
| SORCS3-AS1 | 0 | NA | NA | NA | NA | NA |
| AL731574.1 | 0.20543692 | 0.689155168 | 3.10695182 | 0.2218107 | 0.82446125 | NA |
| AL353148.1 | 0 | NA | NA | NA | NA | NA |
| SORCS1 | 331.099149 | 1.910798302 | 0.49125257 | 3.88964544 | 0.00010039 | 0.00126969 |
| LINC01435 | 4.40804255 | 0.480256891 | 0.82524411 | 0.58195737 | 0.5605954 | 0.72862652 |
| AL353740.1 | 2.42017666 | 0.409564895 | 0.95619699 | 0.4283269 | 0.66841313 | 0.806131 |
| AL356476.1 | 0 | NA | NA | NA | NA | NA |
| XPNPEP1 | 3494.83421 | -0.470972289 | 0.22296978 | -2.1122696 | 0.03466333 | 0.11188153 |
| ADD3 | 5763.93354 | 1.171887669 | 0.24529099 | 4.77754066 | 1.77E-06 | 5.16E-05 |
| MXI1 | 3141.80278 | -0.387403357 | 0.18979408 | -2.0411772 | 0.04123322 | 0.12672726 |
| AL360182.2 | 25.1208988 | -2.160389195 | 0.58536423 | -3.6906751 | 0.00022366 | 0.00239072 |
| SMNDC1 | 1572.0629 | -0.505183268 | 0.14006465 | -3.6067864 | 0.00031001 | 0.00310956 |
| AL355512.1 | 9.37070481 | -3.514823924 | 0.83363791 | -4.2162477 | 2.48E-05 | 0.00042302 |
| DUSP5 | 2382.00875 | -1.624484181 | 0.37647584 | -4.3149759 | 1.60E-05 | 0.00029705 |
| SMC3 | 1956.46292 | -0.008792844 | 0.14829425 | -0.0592932 | 0.95271856 | 0.97717271 |
| RBM20 | 176.052385 | 0.454024659 | 0.30416932 | 1.49267077 | 0.13552339 | 0.29314673 |
| AL136368.1 | 18.3808372 | 0.114349885 | 0.35641659 | 0.32083211 | 0.74833762 | 0.85936782 |
| PDCD4-AS1 | 58.2667121 | 0.044558037 | 0.25542018 | 0.17444995 | 0.86151186 | 0.92794984 |
| PDCD4 | 3492.21188 | -0.077077292 | 0.15714573 | -0.4904829 | 0.62379223 | 0.77580536 |
| BBIP1 | 1803.64466 | 0.051417784 | 0.08968923 | 0.57328825 | 0.56644954 | 0.73284279 |
| AL158163.2 | 31.8274548 | -0.431336452 | 0.40297714 | -1.0703745 | 0.28445078 | 0.48060335 |
| AL158163.1 | 42.264056 | -0.335445555 | 0.30955121 | -1.0836512 | 0.27851946 | 0.47414837 |
| SHOC2 | 2037.81002 | -0.182396216 | 0.14612363 | -1.2482321 | 0.21194606 | 0.39554265 |
| ADRA2A | 377.855239 | 0.645553002 | 0.30389888 | 2.12423617 | 0.0336504 | 0.1095237 |
| AC021035.1 | 0 | NA | NA | NA | NA | NA |
| GPAM | 532.787761 | 0.458219835 | 0.20298404 | 2.257418 | 0.02398197 | 0.08556462 |

| | | | | | | |
|---|---|---|---|---|---|---|
| TECTB | 1.21568825 | 0.070561601 | 1.24481745 | 0.0566843 | 0.95479668 | 0.97841508 |
| ACSL5 | 1032.13447 | -0.448139208 | 0.25673223 | -1.745551 | 0.080889 | 0.2040018 |
| AL157786.1 | 210.249119 | -0.21836865 | 0.18961363 | -1.1516506 | 0.24946468 | 0.44135415 |
| ZDHHC6 | 3746.71256 | -0.002256097 | 0.2035952 | -0.0110813 | 0.99115859 | 0.99583588 |
| VTI1A | 1023.82746 | -0.106611232 | 0.20578824 | -0.5180628 | 0.60441446 | 0.76144298 |
| AL139120.1 | 0.46384489 | -0.371171541 | 2.23702324 | -0.1659221 | 0.86821827 | NA |
| TCF7L2 | 1471.20367 | -0.071265781 | 0.16045112 | -0.4441588 | 0.65692776 | 0.7986941 |
| AL158212.2 | 82.2034845 | 0.179401532 | 0.26832687 | 0.66859325 | 0.50375498 | 0.68364658 |
| HABP2 | 2563.97745 | -1.685937471 | 0.58925782 | -2.8611202 | 0.00422147 | 0.02369018 |
| NRAP | 679.618613 | -1.724254098 | 0.61954054 | -2.7831175 | 0.00538393 | 0.02845811 |
| CASP7 | 1960.55562 | -0.747623893 | 0.23943347 | -3.1224703 | 0.0017934 | 0.01240635 |
| AL592546.1 | 13.7873377 | 1.062424671 | 0.63916037 | 1.66221926 | 0.09646882 | 0.2317934 |
| PLEKHS1 | 91.5263134 | -2.283185922 | 0.69078522 | -3.3052038 | 0.00094907 | 0.00754509 |
| DCLRE1A | 473.230979 | 0.645581516 | 0.24303331 | 2.65634988 | 0.00789916 | 0.03781874 |
| NHLRC2 | 1706.91986 | 0.150023274 | 0.10006101 | 1.49931804 | 0.13379114 | 0.29071131 |
| ADRB1 | 70.6634857 | 0.512236808 | 0.48138128 | 1.0640979 | 0.28728435 | 0.48356758 |
| CCDC186 | 1468.85546 | 0.23588296 | 0.11247701 | 2.09716598 | 0.03597888 | 0.11480678 |
| TDRD1 | 35.0785297 | 1.320040257 | 0.52469508 | 2.51582356 | 0.01187546 | 0.051186 |
| VWA2 | 57.0777921 | 0.386107644 | 0.56285147 | 0.68598496 | 0.49272259 | 0.67556256 |
| AC005383.1 | 6.11822649 | 0.519319576 | 0.81305801 | 0.63872389 | 0.52300257 | 0.69862624 |
| AFAP1L2 | 1886.36928 | 0.784476865 | 0.37042529 | 2.11777349 | 0.03419426 | 0.11074278 |
| ABLIM1 | 8810.88285 | -0.154456526 | 0.12006908 | -1.2863972 | 0.19830447 | 0.37923626 |
| AL137025.1 | 14.866919 | 1.552663463 | 0.5911926 | 2.62632425 | 0.00863126 | 0.04029855 |
| FAM160B1 | 1410.66448 | -0.441667721 | 0.1414823 | -3.1217172 | 0.001798 | 0.01242483 |
| TRUB1 | 1010.24708 | 0.504586386 | 0.15687983 | 3.21638799 | 0.00129815 | 0.00965577 |
| AL355340.1 | 6.17744279 | 0.104484399 | 0.54452037 | 0.19188336 | 0.84783357 | 0.9200943 |
| ATRNL1 | 341.617992 | 0.423966887 | 0.24115985 | 1.75803264 | 0.07874196 | 0.20022821 |
| GFRA1 | 933.968845 | 1.136998805 | 0.57335075 | 1.98307723 | 0.0473588 | 0.13994304 |
| CCDC172 | 0 | NA | NA | NA | NA | NA |
| PNLIPRP3 | 3.0636024 | -0.271797339 | 2.34003896 | -0.1161508 | 0.90753305 | 0.95385641 |
| PNLIP | 0 | NA | NA | NA | NA | NA |
| PNLIPRP1 | 3.59518243 | 0.985479327 | 0.93621346 | 1.05262248 | 0.29251405 | 0.48882101 |
| C10orf82 | 62.0476496 | 0.205448258 | 0.38990386 | 0.5269203 | 0.59824893 | 0.75659797 |
| AC016825.1 | 744.831386 | 0.817186344 | 0.09099272 | 8.98078851 | 2.69E-19 | 4.73E-16 |
| HSPA12A | 3207.77653 | 0.800093095 | 0.10918265 | 7.32802384 | 2.34E-13 | 7.90E-11 |
| ENO4 | 137.650781 | -0.2847992 | 0.23587298 | -1.2074261 | 0.22726808 | 0.41416927 |
| SHTN1 | 2857.86589 | -0.365481333 | 0.18673092 | -1.957262 | 0.05031667 | 0.14629004 |
| VAX1 | 0.22147791 | 1.745651581 | 3.03531355 | 0.57511409 | 0.56521414 | NA |
| MIR3663HG | 0 | NA | NA | NA | NA | NA |
| KCNK18 | 0 | NA | NA | NA | NA | NA |
| AL731557.1 | 2.37930479 | 0.967327586 | 0.74851746 | 1.29232467 | 0.19624473 | 0.37675788 |
| SLC18A2 | 1146.49556 | -0.305112427 | 0.36884645 | -0.8272072 | 0.40811966 | 0.60264172 |
| AL391988.1 | 125.641327 | 0.247281876 | 0.26565725 | 0.9308305 | 0.35194125 | 0.5496086 |
| PDZD8 | 2994.90429 | -0.260765234 | 0.36230067 | -0.7197481 | 0.47168008 | 0.65803638 |
| AC005871.2 | 48.3539143 | 1.590849117 | 0.38071462 | 4.17858683 | 2.93E-05 | 0.00048262 |
| EMX2OS | 5169.43152 | 0.22844778 | 0.2317633 | 0.98569438 | 0.32428311 | 0.52186901 |
| EMX2 | 3581.81532 | -0.141172017 | 0.20686068 | -0.6824497 | 0.49495462 | 0.67717723 |
| AC005871.1 | 1.95073149 | 2.117302823 | 1.53210871 | 1.38195339 | 0.16698601 | 0.33798385 |
| AL139121.1 | 4.42946376 | -0.45239613 | 0.67013644 | -0.6750806 | 0.49962454 | 0.68098926 |
| AL513324.1 | 0.75537086 | -0.332084717 | 1.62004214 | -0.2049852 | 0.83758366 | NA |
| RAB11FIP2 | 1600.00189 | -0.110634038 | 0.13295818 | -0.8320965 | 0.40535446 | 0.60014582 |
| AC022395.1 | 11.2283002 | 1.780684522 | 0.52821305 | 3.3711483 | 0.00074856 | 0.00623678 |
| CASC2 | 220.902524 | 0.736289249 | 0.36338386 | 2.02620237 | 0.04274405 | 0.13000699 |
| FAM204A | 1770.52488 | -0.072306879 | 0.13533127 | -0.5342954 | 0.59313714 | 0.75237031 |
| LINC00867 | 3.38454575 | -0.571430026 | 0.70075769 | -0.815446 | 0.41481709 | 0.60841685 |
| PRLHR | 0.10721287 | -0.642905668 | 3.30558948 | -0.1944905 | 0.84579185 | NA |
| CACUL1 | 2420.63754 | -0.620701572 | 0.15998141 | -3.8798356 | 0.00010453 | 0.00131131 |
| AL157788.1 | 1.4393016 | 3.048368391 | 1.11413327 | 2.73608954 | 0.00621741 | 0.03168788 |
| NANOS1 | 94.2347729 | 1.752205711 | 0.36313551 | 4.82521171 | 1.40E-06 | 4.27E-05 |
| EIF3A | 7124.58727 | 0.050420204 | 0.24906697 | 0.20243633 | 0.83957563 | 0.91484357 |
| FAM45A | 1507.74843 | -0.090946364 | 0.18763982 | -0.4846858 | 0.62789922 | 0.77859976 |
| SFXN4 | 1653.87636 | 0.446676415 | 0.2381251 | 1.87580571 | 0.06068196 | 0.16679962 |
| PRDX3 | 12553.964 | 0.575780151 | 0.16057269 | 3.58579124 | 0.00033606 | 0.00332029 |
| GRK5 | 900.684654 | -0.755568595 | 0.25077434 | -3.0129423 | 0.00258728 | 0.01642036 |
| AL583824.1 | 2.99534598 | -0.643665642 | 0.78500161 | -0.8199545 | 0.41224202 | 0.60616749 |
| RGS10 | 381.729845 | -0.399224497 | 0.35530366 | -1.1236149 | 0.26117642 | 0.45439218 |
| TIAL1 | 4459.10653 | -0.438256798 | 0.15006167 | -2.9205112 | 0.00349458 | 0.02054213 |
| BAG3 | 5036.19414 | -0.257770564 | 0.25022182 | -1.0301682 | 0.30293106 | 0.49979568 |
| INPP5F | 1382.59438 | -0.001674419 | 0.13136368 | -0.0127464 | 0.98983009 | 0.99521246 |
| MCMBP | 2635.88405 | -0.424111191 | 0.16878787 | -2.5126877 | 0.01198154 | 0.05148699 |
| SEC23IP | 2097.52321 | -0.024503677 | 0.1556723 | -0.1574055 | 0.87492527 | 0.93495454 |
| PLPP4 | 274.248542 | 0.91872029 | 0.29377553 | 3.12728668 | 0.00176428 | 0.01226252 |
| LINC01561 | 54.2607196 | 2.134361673 | 0.48074575 | 4.43968915 | 9.01E-06 | 0.0001875 |
| WDR11-AS1 | 15.3000726 | 0.737383111 | 0.37587339 | 1.96178587 | 0.04978742 | 0.14524035 |
| WDR11 | 3645.00518 | 0.386016383 | 0.10847162 | 3.55868537 | 0.00037272 | 0.00360944 |
| AC010998.2 | 6.84434427 | 0.666333398 | 0.69482064 | 0.95900058 | 0.33755845 | 0.53552018 |
| AC010998.1 | 0.09989236 | -0.642905668 | 3.30558948 | -0.1944905 | 0.84579185 | NA |
| AC010998.3 | 0.79289396 | 2.507735958 | 1.54175231 | 1.62654918 | 0.10383288 | NA |
| LINC01153 | 0 | NA | NA | NA | NA | NA |
| FGFR2 | 2789.85306 | 0.306588421 | 0.35475126 | 0.8642349 | 0.38745887 | 0.5830724 |
| AC009988.1 | 0 | NA | NA | NA | NA | NA |
| AC025947.1 | 0 | NA | NA | NA | NA | NA |
| ATE1 | 1817.54809 | 0.154173612 | 0.16669631 | 0.9248772 | 0.35502979 | 0.55256178 |
| AL731566.2 | 11.8409881 | 0.885875957 | 0.413953 | 2.14003994 | 0.03235154 | 0.10635957 |
| NSMCE4A | 1449.29831 | -0.057055614 | 0.1526962 | -0.3736544 | 0.70866143 | 0.83437076 |
| AL731566.1 | 263.188136 | 0.11431669 | 0.17329279 | 0.65967368 | 0.50946326 | 0.68820994 |
| TACC2 | 3175.74128 | -0.280420254 | 0.25849464 | -1.0848204 | 0.27800121 | 0.47353948 |
| BTBD16 | 53.0323344 | -2.164145347 | 0.42243957 | -5.1229702 | 3.01E-07 | 1.19E-05 |
| PLEKHA1 | 9257.93919 | -1.521709727 | 0.23316258 | -6.5263892 | 6.74E-11 | 1.04E-08 |
| ARMS2 | 4.88530332 | -0.960183746 | 0.64663912 | -1.4848835 | 0.13757469 | 0.29619894 |
| HTRA1 | 5472.21861 | 0.327731935 | 0.28066667 | 1.16769099 | 0.24293143 | 0.4334891 |
| DMBT1 | 5.64179206 | 0.044177688 | 0.78276958 | 0.05643767 | 0.95499315 | 0.97844314 |
| C10orf120 | 0 | NA | NA | NA | NA | NA |
| CUZD1 | 301.791302 | -0.490643689 | 0.20969504 | -2.3397964 | 0.01929426 | 0.07300753 |
| FAM24B | 272.425478 | -0.33416811 | 0.28496863 | -1.1726487 | 0.2409367 | 0.43106436 |
| FAM24A | 0.08695548 | -0.642905668 | 3.30558948 | -0.1944905 | 0.84579185 | NA |
| C10orf88 | 482.39418 | -0.364461571 | 0.18106468 | -2.0128805 | 0.0441272 | 0.13307458 |
| PSTK | 288.1787 | -0.227714291 | 0.26836832 | -0.848514 | 0.39615176 | 0.5912411 |
| IKZF5 | 1250.61219 | -0.103629739 | 0.16934354 | -0.6119498 | 0.540571 | 0.71359918 |
| ACADSB | 7407.03742 | 1.290821631 | 0.60537803 | 2.13225715 | 0.03298571 | 0.10797334 |
| HMX3 | 5.22686543 | 0.042813927 | 1.23997729 | 0.03452799 | 0.97245612 | 0.98710875 |
| HMX2 | 60.1153975 | 0.869677416 | 0.59936199 | 1.4510053 | 0.14677838 | 0.30983482 |
| BUB3 | 3275.46003 | -0.193436646 | 0.13711723 | -1.4107392 | 0.15832154 | 0.32553131 |
| AL160290.2 | 3.51063226 | -0.84056861 | 0.7951002 | -1.0571858 | 0.29042682 | 0.48652584 |
| AL160290.1 | 0.57428354 | -1.127694543 | 1.88197262 | -0.5992088 | 0.54903366 | NA |
| AL357127.1 | 0.16775052 | 0.435199075 | 3.28898345 | 0.13232024 | 0.894731 | NA |
| GPR26 | 7.28088698 | -0.476707694 | 0.89743083 | -0.5311916 | 0.59528603 | 0.75429785 |
| CPXM2 | 1437.7328 | 0.091525096 | 0.47726938 | 0.19176821 | 0.84792377 | 0.92010671 |
| AC009987.1 | 2.04024716 | -0.12844303 | 0.95520027 | -0.1344671 | 0.89303321 | 0.94473004 |
| CHST15 | 1451.48325 | -0.525714554 | 0.28346461 | -1.8546038 | 0.06365282 | 0.17226241 |
| OAT | 5523.56465 | 0.465679757 | 0.2623 | 1.7753708 | 0.07583668 | 0.19531334 |
| NKX1-2 | 0.4841247 | 2.175206133 | 1.97806485 | 1.09966371 | 0.27147867 | NA |
| LHPP | 1874.89541 | 0.884475061 | 0.3045546 | 2.90415923 | 0.00368241 | 0.02142655 |

| | | | | | | |
|---|---|---|---|---|---|---|
| AC068896.1 | 829.261531 | 0.463797499 | 0.17931278 | 2.58652788 | 0.00969483 | 0.04404009 |
| FAM53B | 2085.48818 | 0.248902571 | 0.17769883 | 1.40069896 | 0.16130412 | 0.32996583 |
| AL513190.1 | 113.477587 | 1.065127304 | 0.24818776 | 4.29161891 | 1.77E-05 | 0.00032325 |
| FAM53B-AS1 | 3.96571204 | 0.554280515 | 0.76202718 | 0.72737631 | 0.46699546 | 0.65413276 |
| EEF1AKMT2 | 887.404186 | 0.234683168 | 0.22026694 | 1.06544892 | 0.28667283 | 0.48288486 |
| ABRAXAS2 | 1064.56551 | -0.142120778 | 0.2152967 | -0.6601159 | 0.50917946 | 0.68800961 |
| ZRANB1 | 2662.27759 | -0.324990743 | 0.24141829 | -1.3461728 | 0.17824679 | 0.35372853 |
| AL731577.1 | 0.63659424 | -0.048392718 | 1.59613122 | -0.0303188 | 0.97581284 | NA |
| CTBP2 | 6563.32711 | -0.488067538 | 0.17850248 | -2.7342339 | 0.00625256 | 0.03179943 |
| AL731571.1 | 616.486505 | 0.166476995 | 0.25726237 | 0.64710978 | 0.5175609 | 0.69466488 |
| AL157888.1 | 6.50358074 | -0.623060488 | 0.54400038 | -1.145331 | 0.25207208 | 0.44432973 |
| TEX36-AS1 | 0 | NA | NA | NA | NA | NA |
| TEX36 | 0.03392908 | 0.380740593 | 3.29738709 | 0.11546736 | 0.90807469 | NA |
| AL158835.1 | 108.781167 | 1.136173895 | 0.42766376 | 2.65669901 | 0.00789099 | 0.0378062 |
| AL158835.4 | 151.486089 | 0.496823827 | 0.21830144 | 2.27586146 | 0.0228543 | 0.08274766 |
| EDRF1 | 715.440645 | 0.082629608 | 0.09735787 | 0.84872042 | 0.39603688 | 0.59122987 |
| EDRF1-AS1 | 150.16327 | 0.217701708 | 0.14416652 | 1.51007117 | 0.13102526 | 0.28663815 |
| MMP21 | 7.97546856 | 1.27197101 | 0.5330784 | 2.38608619 | 0.01702876 | 0.06674827 |
| UROS | 2007.35489 | 0.323538153 | 0.18130868 | 1.78446041 | 0.07434888 | 0.19253563 |
| BCCIP | 3268.30365 | -0.148216047 | 0.1630061 | -0.9092669 | 0.36320927 | 0.56030352 |
| DHX32 | 2422.30165 | -0.707227481 | 0.21707069 | -3.2580514 | 0.0011218 | 0.00857695 |
| FANK1 | 510.855088 | -0.609392883 | 0.30579441 | -1.9928189 | 0.04628128 | 0.13765367 |
| FANK1-AS1 | 0.10721287 | -0.642905668 | 3.30558948 | -0.1944905 | 0.84579185 | NA |
| ADAM12 | 148.697384 | -1.39489383 | 0.38867783 | -3.5888176 | 0.00033218 | 0.00328691 |
| LINC00601 | 2.36574946 | -3.039309471 | 1.14767398 | -2.6482342 | 0.00809135 | 0.03842405 |
| C10orf90 | 4.32339513 | 0.83589419 | 0.66422696 | 1.25844664 | 0.20823027 | 0.39146163 |
| DOCK1 | 3435.77476 | 0.183791082 | 0.12697532 | 1.44745517 | 0.14776948 | 0.31128625 |
| AL359094.1 | 69.8060472 | 0.745782562 | 0.29875481 | 2.4963031 | 0.01254953 | 0.05333298 |
| AL359094.2 | 17.6884296 | 0.676045229 | 0.31622035 | 2.13789284 | 0.03252544 | 0.10679814 |
| FAM196A | 43.236482 | -0.847671049 | 0.44670321 | -1.8976158 | 0.05774672 | 0.16112379 |
| NPS | 0 | NA | NA | NA | NA | NA |
| FOXI2 | 138.669539 | 0.816565711 | 0.68115711 | 1.19879202 | 0.23060883 | 0.41824938 |
| CLRN3 | 973.940722 | 0.954201908 | 0.49531231 | 1.92646516 | 0.05404632 | 0.15390072 |
| PTPRE | 790.034176 | -0.487360705 | 0.29794798 | -1.6357241 | 0.10189734 | 0.24057953 |
| AL158166.2 | 2.57844711 | -0.481075197 | 0.82816185 | -0.5808951 | 0.56131113 | 0.72922329 |
| MKI67 | 119.06456 | -1.727248883 | 0.69811558 | -2.4741589 | 0.01335503 | 0.05579607 |
| AL390763.2 | 1.20062029 | -0.063046098 | 1.46531461 | -0.0430256 | 0.9656811 | 0.98387257 |
| AL390763.1 | 0.17423948 | 0.18657164 | 3.29377195 | 0.05664376 | 0.95482897 | NA |
| AL355537.1 | 0.26369991 | -0.701265545 | 3.29738693 | -0.2126731 | 0.83158194 | NA |
| AL355537.2 | 0 | NA | NA | NA | NA | NA |
| AL391869.1 | 0 | NA | NA | NA | NA | NA |
| MGMT | 1565.91381 | 0.335814996 | 0.25076696 | 1.33915169 | 0.1805213 | 0.35650016 |
| AL157832.2 | 1.47992307 | -1.846212488 | 1.33456252 | -1.383384 | 0.16654714 | 0.33732514 |
| AL157832.1 | 3.35952884 | 0.149658918 | 0.87743763 | 0.1705636 | 0.86456692 | 0.92965085 |
| AL157832.3 | 13.747933 | 0.727609019 | 0.47909378 | 1.51871941 | 0.12883314 | 0.28315801 |
| EBF3 | 91.0068252 | 0.265786934 | 0.3170734 | 0.8382505 | 0.40189003 | 0.59672523 |
| AL354950.2 | 10.9209179 | 0.520822674 | 0.47715168 | 1.09152434 | 0.27504223 | 0.47019955 |
| AL354950.1 | 11.762668 | 0.527456827 | 0.42503844 | 1.24096264 | 0.21461955 | 0.39877895 |
| C10orf143 | 222.26726 | 0.553349888 | 0.24493118 | 2.25920557 | 0.0238706 | 0.08533999 |
| GLRX3 | 2658.50713 | -0.799719146 | 0.30062531 | -2.6601857 | 0.00780976 | 0.03755348 |
| AL355500.1 | 0.64012707 | 2.050229968 | 2.05503574 | 0.99766147 | 0.31844354 | NA |
| TCERG1L | 3.70901109 | 1.240761066 | 0.95965892 | 1.29291881 | 0.19603914 | 0.37643284 |
| TCERG1L-AS | 0.72327109 | -1.165951701 | 1.855238 | -0.6284648 | 0.52969953 | NA |
| LINC01164 | 1.28962956 | 0.460592586 | 1.91590502 | 0.24040471 | 0.81001653 | 0.89733389 |
| FP565171.1 | 0.09997307 | 0.547239394 | 3.29377202 | 0.16614368 | 0.86804388 | NA |
| PPP2R2D | 2265.90438 | -0.35669401 | 0.26478556 | -1.3471052 | 0.17794635 | 0.35329357 |
| BNIP3 | 8904.94979 | -0.780791816 | 0.36467297 | -2.141074 | 0.03226807 | 0.10622638 |
| AL162274.2 | 60.1976559 | -0.983595704 | 0.38005225 | -2.5880539 | 0.00965199 | 0.04389083 |
| AL162274.1 | 11.6031338 | -0.379853211 | 0.41372583 | -0.9181279 | 0.35855192 | 0.55624649 |
| JAKMIP3 | 38.1811665 | -0.137164974 | 0.3694386 | -0.3712795 | 0.71042937 | 0.8353877 |
| AL512622.1 | 9.51940958 | -0.514457421 | 0.46108087 | -1.115764 | 0.26452318 | 0.45846048 |
| DPYSL4 | 72.069681 | -2.724295624 | 0.66656614 | -4.0870598 | 4.37E-05 | 0.0006663 |
| STK32C | 757.264037 | -0.880555391 | 0.25215139 | -3.4921695 | 0.00047911 | 0.00438689 |
| LRRC27 | 762.821325 | 0.033996676 | 0.2538648 | 0.13391646 | 0.89346863 | 0.94511484 |
| PWWP2B | 990.136949 | -0.703384178 | 0.18197008 | -3.8653836 | 0.00011091 | 0.00137251 |
| AL451069.2 | 4.89119679 | -0.484113409 | 0.51441285 | -0.941099 | 0.34665414 | 0.54425278 |
| AL451069.3 | 3.45822828 | -0.777635006 | 0.71525839 | -1.0872085 | 0.27694467 | 0.47232124 |
| C10orf91 | 3.78311845 | -1.58928768 | 0.78378138 | -2.0277181 | 0.04258903 | 0.12964444 |
| AL451069.1 | 14.9512121 | -0.329968665 | 0.52151824 | -0.6327078 | 0.52692447 | 0.70173215 |
| LINC01165 | 3.2713663 | 2.275604836 | 0.91296693 | 2.49253808 | 0.01268337 | 0.05374587 |
| INPP5A | 1465.30187 | 0.083416809 | 0.19904478 | 0.41908565 | 0.67515354 | 0.81099616 |
| NKX6-2 | 330.47957 | -0.54149434 | 0.26552193 | -2.0393582 | 0.04141429 | 0.12707838 |
| CFAP46 | 378.078249 | 0.411776501 | 0.35981934 | 1.1443979 | 0.25245866 | 0.44477117 |
| LINC01166 | 15.0019246 | 0.190701232 | 0.80335831 | 0.23738004 | 0.81236197 | 0.89879941 |
| LINC01167 | 11.3738082 | -0.060415924 | 0.79746256 | -0.0757602 | 0.93960988 | 0.97112068 |
| LINC01168 | 1.73535894 | 0.19602421 | 1.41582059 | 0.13845272 | 0.88988263 | 0.94297317 |
| ADGRA1 | 20.5626574 | 1.801234535 | 0.85957454 | 2.09549545 | 0.03612697 | 0.11513691 |
| ADGRA1-AS1 | 3.35296194 | 2.158391135 | 0.85606424 | 2.52129575 | 0.01169235 | 0.05062847 |
| KNDC1 | 315.524708 | -0.551310195 | 0.42188108 | -1.3067905 | 0.19128387 | 0.37037636 |
| UTF1 | 0.46212209 | 0.838647407 | 1.8115028 | 0.46295673 | 0.64339538 | NA |
| VENTX | 168.53089 | 0.152745661 | 0.43334193 | 0.352483 | 0.72447607 | 0.84387647 |
| MIR202HG | 0.25254714 | -0.178391175 | 2.99848453 | -0.0594938 | 0.95255882 | NA |
| AL592071.1 | 1.37322216 | -1.096812599 | 1.1139832 | -0.9845863 | 0.32482732 | 0.52236227 |
| ADAM8 | 779.544335 | 0.192642082 | 0.30072133 | 0.6406 | 0.5217826 | 0.69787989 |
| TUBGCP2 | 5326.17017 | 0.307355316 | 0.08297352 | 3.70425793 | 0.00021201 | 0.00228846 |
| AL360181.2 | 114.718503 | 0.958395478 | 0.28466459 | 3.36675342 | 0.00076059 | 0.00630911 |
| ZNF511 | 1385.38034 | -0.253131589 | 0.14143443 | -1.7897452 | 0.07349488 | 0.19101715 |
| ZNF511-PRA | 6015.87579 | 1.547950724 | 0.53338278 | 2.90213854 | 0.00370625 | 0.02151777 |
| CALY | 8.39970472 | -0.963897421 | 0.54072506 | -1.7826017 | 0.07465116 | 0.19309295 |
| AL360181.1 | 31.2180263 | 2.132006083 | 0.38620203 | 5.52044249 | 3.38E-08 | 1.94E-06 |
| PRAP1 | 5276.60207 | 1.801829272 | 0.65940765 | 2.73249678 | 0.00628563 | 0.03189379 |
| FUOM | 1114.41984 | 1.003691764 | 0.23252549 | 4.31648056 | 1.59E-05 | 0.00029587 |
| ECHS1 | 19428.1337 | 0.86332244 | 0.31576143 | 2.73409721 | 0.00625516 | 0.03180425 |
| AL360181.4 | 2.69090084 | 0.491912986 | 1.28418616 | 0.38305427 | 0.70167953 | 0.82913351 |
| PAOX | 547.218159 | 0.105038745 | 0.17964215 | 0.58471103 | 0.55874203 | 0.72731896 |
| AL360181.3 | 3234.39998 | -0.391792088 | 0.21941553 | -1.7856169 | 0.0741613 | 0.19223764 |
| MTG1 | 3972.19403 | -0.449177935 | 0.24650107 | -1.8222149 | 0.06842238 | 0.18117317 |
| SPRN | 933.301614 | -0.521044623 | 0.33862467 | -1.5387084 | 0.1238755 | 0.27548621 |
| SCART1 | 313.100073 | 0.904498097 | 0.28427638 | 3.1817561 | 0.00146385 | 0.01060546 |
| CYP2E1 | 325.486019 | 0.68353609 | 0.34664399 | 1.97186771 | 0.04862471 | 0.14280853 |
| AL161645.1 | 31.8704091 | 1.446648587 | 0.37337732 | 3.87449508 | 0.00010685 | 0.00133532 |
| SYCE1 | 57.327033 | 0.213189883 | 0.67646083 | 0.31515481 | 0.75264412 | 0.86206078 |
| FRG2B | 2.99879854 | 3.89297845 | 1.19973412 | 3.24486766 | 0.00117505 | 0.00890732 |
| FAM138D | 0.19978473 | -1.003574173 | 3.30558948 | -0.3035992 | 0.7614333 | NA |
| IQSEC3 | 767.092694 | 2.423955768 | 0.61368313 | 3.94984909 | 7.82E-05 | 0.00104465 |
| AC026369.3 | 1.04353621 | -3.286039417 | 1.51895024 | -2.1633621 | 0.03051334 | 0.10205945 |
| AC026369.1 | 187.143619 | 2.68061454 | 0.75271202 | 3.56127504 | 0.00036906 | 0.00358192 |
| AC026369.2 | 70.4636182 | 1.986498838 | 0.46453538 | 4.27631335 | 1.90E-05 | 0.00034017 |
| AC007406.3 | 29.4012999 | 1.452544756 | 0.42766668 | 3.39644128 | 0.00068268 | 0.00577796 |
| SLC6A12 | 9040.13846 | -0.33499236 | 0.50550821 | -0.6626843 | 0.50753276 | 0.68648128 |
| AC007406.1 | 49.5302113 | 0.867486086 | 0.35768144 | 2.4253036 | 0.01529559 | 0.0617018 |
| AC007406.4 | 423.29707 | 0.029196233 | 0.4712137 | 0.06195964 | 0.95059497 | 0.97632675 |
| SLC6A13 | 8995.48245 | 2.275651969 | 0.38764116 | 5.87051173 | 4.34E-09 | 3.36E-07 |

| | | | | | | |
|---|---|---|---|---|---|---|
| KDM5A | 1569.79164 | -0.269812892 | 0.12564402 | -2.1474392 | 0.03175834 | 0.10505979 |
| CCDC77 | 293.348147 | -0.002827285 | 0.20923884 | -0.0135122 | 0.98921912 | 0.99515008 |
| B4GALNT3 | 1457.58774 | 0.550448657 | 0.29376334 | 1.87378266 | 0.06096039 | 0.16736033 |
| NINJ2 | 108.889791 | 0.005034235 | 0.23312825 | 0.02159427 | 0.9827716 | 0.99232988 |
| AC006205.1 | 0.09635527 | 0.547239394 | 3.29377202 | 0.16614368 | 0.86804388 | NA |
| AC021054.1 | 386.804382 | 0.282920139 | 0.28710557 | 0.98542197 | 0.32441684 | 0.52192497 |
| LINC02455 | 0.25020947 | -1.364244509 | 2.99735315 | -0.4551497 | 0.64900151 | NA |
| WNK1 | 9121.1539 | 0.631337091 | 0.20506562 | 3.07870762 | 0.00207901 | 0.01386966 |
| RAD52 | 848.796191 | 0.215306885 | 0.28368644 | 0.7589608 | 0.44787601 | 0.63796545 |
| AC004803.1 | 105.484756 | 0.554958203 | 0.20016127 | 2.77255538 | 0.00556181 | 0.02923357 |
| ERC1 | 2090.70055 | 0.002639614 | 0.15332341 | 0.01721599 | 0.98626431 | 0.99391317 |
| AC004672.1 | 0.40378873 | 0.880139507 | 2.70057155 | 0.32590861 | 0.7444935 | NA |
| LINC00942 | 9.59834214 | -0.961723975 | 0.65311533 | -1.4725178 | 0.14088109 | 0.30117831 |
| WNT5B | 141.282471 | -0.273800201 | 0.36323437 | -0.7537838 | 0.45097903 | 0.64079343 |
| FBXL14 | 618.766903 | 0.136975098 | 0.14562222 | 0.94061949 | 0.34689989 | 0.54450894 |
| ADIPOR2 | 3740.99302 | -0.42270789 | 0.17171237 | -2.4617207 | 0.01382723 | 0.05733985 |
| CACNA2D4 | 166.62242 | 0.439156408 | 0.28468036 | 1.54262979 | 0.12292063 | 0.27427796 |
| LRTM2 | 0.66139313 | 0.964954807 | 1.37964101 | 0.69942456 | 0.48428674 | NA |
| AC005342.2 | 1.62820543 | -0.126362606 | 1.07133085 | -0.1179492 | 0.90610792 | 0.9530025 |
| LINC00940 | 3.29046667 | 4.063545933 | 0.9651024 | 4.21048163 | 2.55E-05 | 0.00043103 |
| DCP1B | 776.925277 | -0.031705175 | 0.16684465 | -0.1900281 | 0.8492871 | 0.92068657 |
| CACNA1C | 662.996687 | -0.040107746 | 0.37120033 | -0.1080488 | 0.91395699 | 0.9577987 |
| AC005342.1 | 1.72903452 | 0.100746781 | 1.00773604 | 0.09997338 | 0.92036546 | 0.96099929 |
| AC005344.1 | 0.84625297 | -1.402860975 | 1.46023136 | -0.9607114 | 0.33669728 | NA |
| CACNA1C-AS | 1.28029322 | -0.023852211 | 1.15729372 | -0.0206103 | 0.9835565 | 0.99290974 |
| CACNA1C-AS | 0 | NA | NA | NA | NA | NA |
| CACNA1C-AS | 38.2474916 | 0.184809713 | 0.23908167 | 0.77299826 | 0.43952341 | 0.63096697 |
| CACNA1C-AS | 126.49896 | 0.03737852 | 0.41303918 | 0.09049631 | 0.92789283 | 0.96504253 |
| LINC02371 | 3.83147043 | 0.114007357 | 0.72048699 | 0.15823652 | 0.87427042 | 0.93458499 |
| FKBP4 | 7993.81083 | -0.307713018 | 0.1247769 | -2.4661056 | 0.0136591 | 0.05675868 |
| ITFG2-AS1 | 193.022784 | 0.416284237 | 0.17420994 | 2.38955505 | 0.0168688 | 0.06633316 |
| AC005841.1 | 2011.95766 | -0.375473335 | 0.13999643 | -2.6820208 | 0.00731789 | 0.03580201 |
| ITFG2 | 1740.86138 | 0.469033346 | 0.17872602 | 2.62431486 | 0.00868235 | 0.04045937 |
| NRIP2 | 754.123505 | 0.72028508 | 0.27320268 | 2.63644958 | 0.00837786 | 0.03938747 |
| TEX52 | 61.7194522 | 1.482538768 | 0.30579119 | 4.84820633 | 1.25E-06 | 3.90E-05 |
| FOXM1 | 485.393056 | 0.266572899 | 0.18790142 | 1.41868487 | 0.15599091 | 0.32246944 |
| RHNO1 | 744.15818 | -0.125006256 | 0.21068386 | -0.5933357 | 0.55295651 | 0.72282076 |
| TULP3 | 2466.79622 | -0.160949255 | 0.18735402 | -0.8590649 | 0.39030473 | 0.5858507 |
| TEAD4 | 1415.90897 | -1.466372055 | 0.32656868 | -4.490241 | 7.11E-06 | 0.00015582 |
| AC125807.2 | 398.639241 | -0.911154977 | 0.43739867 | -2.0831224 | 0.03724007 | 0.11749468 |
| TSPAN9 | 9827.46863 | 0.155183894 | 0.1260157 | 1.23146473 | 0.2181491 | 0.40326616 |
| AC005865.1 | 7.83585242 | 0.938088272 | 0.65271159 | 1.43721712 | 0.15065631 | 0.31532716 |
| LINC02417 | 25.8401369 | 2.72069796 | 0.80846291 | 3.36527245 | 0.00076468 | 0.00633511 |
| PRMT8 | 13.0691074 | 2.518363552 | 0.84486963 | 2.98077179 | 0.00287523 | 0.01774393 |
| AC005908.2 | 0.16325399 | -1.003574173 | 3.30558948 | -0.3035992 | 0.7614333 | NA |
| CRACR2A | 61.1431496 | -0.759921783 | 0.29744261 | -2.5548518 | 0.0106233 | 0.04710572 |
| AC006207.1 | 0.82050192 | 0.737790213 | 1.51580494 | 0.48673163 | 0.62644853 | NA |
| PARP11 | 385.7193 | 0.056488099 | 0.20345962 | 0.27763789 | 0.78129034 | 0.88034125 |
| AC005842.1 | 1.68212186 | 0.673575324 | 1.14924812 | 0.58610087 | 0.55780773 | 0.72649725 |
| AC007207.2 | 20.9550101 | 1.437185867 | 0.64021783 | 2.24483886 | 0.02477848 | 0.08772939 |
| CCND2 | 1522.85477 | -0.226300939 | 0.4394345 | -0.5149822 | 0.60656547 | 0.76313157 |
| TIGAR | 464.494529 | -0.80214302 | 0.27852531 | -2.8799646 | 0.0039772 | 0.02264614 |
| FGF23 | 1.05859328 | -2.381360271 | 1.69588799 | -1.4041967 | 0.16026029 | 0.32839101 |
| FGF6 | 0 | NA | NA | NA | NA | NA |
| C12orf4 | 683.828878 | -0.055452528 | 0.14450711 | -0.3837356 | 0.70117441 | 0.82884925 |
| RAD51AP1 | 66.707342 | -0.153652914 | 0.34115984 | -0.4503839 | 0.65243362 | 0.79583057 |
| DYRK4 | 866.275641 | -0.427814906 | 0.12950904 | -3.3033595 | 0.00095534 | 0.00757662 |
| AC005832.4 | 2832.36874 | 0.192438392 | 0.137312 | 1.40146812 | 0.16107413 | 0.3296744 |
| AKAP3 | 132.888557 | 1.280508318 | 0.52663376 | 2.43149683 | 0.01503658 | 0.06098348 |
| NDUFA9 | 4792.02585 | 0.146611651 | 0.14173601 | 1.03439948 | 0.30094945 | 0.49768474 |
| AC005833.2 | 3509.72116 | 0.114667427 | 0.15001812 | 0.76435719 | 0.44465441 | 0.63537947 |
| AC005833.1 | 5.16999584 | 0.214828772 | 0.58912349 | 0.3646583 | 0.71536649 | 0.8380436 |
| GALNT8 | 26.6017356 | 0.399604779 | 0.48698444 | 0.82056991 | 0.41189129 | 0.60587791 |
| AC005833.3 | 0.56751246 | -0.68817541 | 2.284075 | -0.3012928 | 0.76319121 | NA |
| KCNA6 | 3.16170908 | 2.352480342 | 0.91206543 | 2.57928901 | 0.00990039 | 0.04475799 |
| KCNA1 | 2.86772951 | 3.135334252 | 1.0248843 | 3.05920802 | 0.00221923 | 0.01463027 |
| AC006063.2 | 0 | NA | NA | NA | NA | NA |
| KCNA5 | 102.903118 | 0.309517095 | 0.45155398 | 0.68544871 | 0.49306082 | 0.67578034 |
| LINC02443 | 0.18390514 | 0.18657164 | 3.29377195 | 0.05664376 | 0.95482897 | NA |
| AC006206.2 | 2.67934544 | -1.66894517 | 2.03732054 | -0.8191863 | 0.4126801 | 0.60653073 |
| AC006206.1 | 0.08666371 | -0.642905668 | 3.30558948 | -0.1944905 | 0.84579185 | NA |
| AC007848.2 | 0.07659028 | -0.642905668 | 3.30558948 | -0.1944905 | 0.84579185 | NA |
| AC007848.1 | 1.78094322 | -4.054188364 | 2.171739 | -1.8667936 | 0.06193042 | 0.16914607 |
| NTF3 | 154.186209 | 0.241182324 | 0.46419502 | 0.51957112 | 0.60336253 | 0.76063835 |
| ANO2 | 75.413161 | -0.269090641 | 0.37365152 | -0.7201647 | 0.47142359 | 0.65787603 |
| VWF | 4278.22657 | -1.096445262 | 0.64895825 | -1.6895467 | 0.09111471 | 0.22255418 |
| AC005845.1 | 4.14346467 | -0.061518168 | 0.6151017 | -0.100013 | 0.920334 | 0.96099929 |
| CD9 | 12091.1402 | -0.648185072 | 0.26702668 | -2.4274169 | 0.01520677 | 0.06144691 |
| PLEKHG6 | 263.718623 | -1.260680552 | 0.32649653 | -3.8612372 | 0.00011281 | 0.0013934 |
| TNFRSF1A | 10672.481 | -0.966494233 | 0.23061552 | -4.1909332 | 2.78E-05 | 0.00046175 |
| SCNN1A | 20843.459 | -0.306011725 | 0.22319521 | -1.3710497 | 0.17035946 | 0.34238083 |
| LTBR | 10625.8839 | -0.833119912 | 0.22903585 | -3.6375088 | 0.00027529 | 0.00284234 |
| AC005840.2 | 216.314567 | -0.665297796 | 0.48172516 | -1.3810734 | 0.16725639 | 0.33839135 |
| CD27 | 602.500362 | 0.200440695 | 0.33604557 | 0.5964688 | 0.55086209 | 0.72122702 |
| TAPBPL | 2472.23815 | 0.007018691 | 0.32286513 | 0.02173877 | 0.98265634 | 0.99232748 |
| VAMP1 | 1003.96021 | 0.073660492 | 0.21593786 | 0.34111893 | 0.73301405 | 0.84942786 |
| AC005840.4 | 72.1208434 | -0.039098433 | 0.39480884 | -0.0990313 | 0.92111342 | 0.96155183 |
| MRPL51 | 3086.75337 | -0.246193875 | 0.1479891 | -1.6635946 | 0.09619347 | 0.23135915 |
| NCAPD2 | 1367.30106 | 0.060029614 | 0.11639251 | 0.51575152 | 0.60602796 | 0.76277779 |
| AC006064.3 | 19.5576198 | -0.074880145 | 0.62049031 | -0.120679 | 0.9039453 | 0.95167363 |
| GAPDH | 134900.226 | -0.666920681 | 0.29537116 | -2.2579072 | 0.02395144 | 0.08553487 |
| AC006064.4 | 67388.4597 | -0.748965938 | 0.29535638 | -2.5358041 | 0.01121895 | 0.04907824 |
| IFFO1 | 1687.11793 | 0.046873619 | 0.16011796 | 0.2927443 | 0.76971761 | 0.87270139 |
| AC006064.1 | 4.9572068 | 1.53803927 | 0.69390933 | 2.21648449 | 0.02665834 | 0.09261929 |
| NOP2 | 2079.90403 | -0.733385144 | 0.31874181 | -2.3008753 | 0.02139868 | 0.07883253 |
| AC006064.6 | 5697.40634 | -0.444946648 | 0.16267505 | -2.7351868 | 0.00623449 | 0.03173812 |
| CHD4 | 7583.4696 | -0.022140835 | 0.08899474 | -0.2487881 | 0.80352467 | 0.89344517 |
| AC006064.2 | 936.687184 | 0.482927157 | 0.16704594 | 2.89098419 | 0.00384037 | 0.02210918 |
| LPAR5 | 79.7922419 | 0.403378274 | 0.41546933 | 0.97089785 | 0.33159915 | 0.52953852 |
| ACRBP | 139.053322 | 0.534108002 | 0.30659683 | 1.74205322 | 0.08149913 | 0.20508059 |
| ING4 | 1798.38391 | 0.018201397 | 0.23151201 | 0.07861967 | 0.93733514 | 0.96999796 |
| ZNF384 | 3467.05296 | -0.05561282 | 0.1902617 | -0.2922965 | 0.77005997 | 0.87287145 |
| PIANP | 42.118667 | -0.022171419 | 0.44982549 | -0.0492889 | 0.96068904 | 0.98107699 |
| AC125494.3 | 1.4545015 | -0.436397349 | 1.07882951 | -0.40451 | 0.68583771 | 0.81835792 |
| COPS7A | 3775.30511 | 0.268959214 | 0.11661844 | 2.30631809 | 0.02109287 | 0.07791271 |
| AC125494.2 | 52.0261752 | 0.333652695 | 0.34095125 | 0.97859355 | 0.32778085 | 0.52554149 |
| MLF2 | 11016.9408 | -0.258602071 | 0.10204674 | -2.5341532 | 0.01127194 | 0.04927738 |
| PTMS | 7479.64957 | -0.047553999 | 0.26054589 | -0.1825168 | 0.85517718 | 0.92410315 |
| LAG3 | 179.208483 | -1.339789924 | 0.61726898 | -2.1705123 | 0.02996806 | 0.10078531 |
| CD4 | 1416.21021 | 0.490469168 | 0.4593057 | 1.06784907 | 0.2855886 | 0.48163515 |
| GPR162 | 552.371683 | -0.396282491 | 0.42816435 | -0.9255383 | 0.35468599 | 0.55227628 |
| P3H3 | 926.864702 | -0.51139064 | 0.31577813 | -1.6194619 | 0.10534792 | 0.24625695 |
| GNB3 | 261.537571 | -0.096045796 | 0.38985264 | -0.2463644 | 0.80540019 | 0.89451219 |

| | | | | | | |
|---|---|---|---|---|---|---|
| CDCA3 | 272.36998 | -0.355376607 | 0.31797612 | -1.1176204 | 0.26372914 | 0.45750525 |
| USP5 | 4086.3614 | 0.047416934 | 0.11024556 | 0.4301029 | 0.66712079 | 0.80516157 |
| TPI1 | 37700.617 | -0.465275675 | 0.16865074 | -2.7588119 | 0.00580119 | 0.03007287 |
| SPSB2 | 1212.85592 | -0.223243341 | 0.27875591 | -0.800856 | 0.42321502 | 0.6157794 |
| LRRC23 | 763.782526 | 0.378941023 | 0.15170934 | 2.49780946 | 0.01249633 | 0.05314973 |
| U47924.3 | 39.9526627 | -1.631644436 | 0.39372716 | -4.1440992 | 3.41E-05 | 0.00054522 |
| ENO2 | 3147.5242 | -2.379361119 | 0.28736026 | -8.2800631 | 1.23E-16 | 1.08E-13 |
| ATN1 | 19698.4566 | -0.850247093 | 0.28991097 | -2.9327869 | 0.00335934 | 0.01998779 |
| C12orf57 | 3581.10345 | -0.406244808 | 0.26328195 | -1.5430029 | 0.12283007 | 0.2740991 |
| U47924.2 | 2970.91769 | -0.373930513 | 0.26826057 | -1.3939079 | 0.16334543 | 0.33259355 |
| PTPN6 | 2070.46185 | -0.116170179 | 0.22397453 | -0.5186759 | 0.6039868 | 0.76115326 |
| MIR200CHG | 77.1667469 | -1.167327191 | 0.29630889 | -3.9395618 | 8.16E-05 | 0.00108008 |
| U47924.1 | 75.908061 | -0.367203385 | 0.20593617 | -1.7830932 | 0.07457113 | 0.19297749 |
| PHB2 | 7156.60402 | -0.017426897 | 0.0939127 | -0.1855649 | 0.852786 | 0.92285671 |
| EMG1 | 6618.96573 | -0.513849432 | 0.19375264 | -2.65209 | 0.00799952 | 0.03813739 |
| LPCAT3 | 6624.40428 | -0.413634842 | 0.21632848 | -1.9120684 | 0.05586743 | 0.15737892 |
| C1S | 15024.6451 | -1.257802628 | 0.38415095 | -3.2742406 | 0.00105946 | 0.00819779 |
| C1R | 23160.0368 | -1.551294939 | 0.35582086 | -4.3597639 | 1.30E-05 | 0.00025246 |
| C1RL | 9981.78246 | -1.27882063 | 0.30241944 | -4.2286323 | 2.35E-05 | 0.00040549 |
| C1RL-AS1 | 1772.55094 | 0.550215008 | 0.25401528 | 2.16607048 | 0.0303058 | 0.10151982 |
| AC233309.1 | 328.685071 | 0.990980457 | 0.45588761 | 2.17373854 | 0.02972478 | 0.10026124 |
| RBP5 | 10708.3729 | 1.428431434 | 0.44899972 | 3.18136372 | 0.00146583 | 0.01060811 |
| AC018653.3 | 452.595378 | 1.154801099 | 0.28518115 | 4.04935986 | 5.14E-05 | 0.00075914 |
| CLSTN3 | 8546.50473 | -0.02379705 | 0.20023128 | -0.1188478 | 0.90539593 | 0.95257235 |
| AC018653.1 | 10.0368969 | 1.609176593 | 0.6601648 | 2.4375377 | 0.01478767 | 0.06019619 |
| PEX5 | 3735.09774 | 0.345561512 | 0.20564604 | 1.68037041 | 0.09288527 | 0.22562496 |
| ACSM4 | 4.53196818 | -0.721140773 | 0.66899085 | -1.0779531 | 0.28105468 | 0.47686346 |
| CD163L1 | 837.261778 | -1.913241519 | 0.47546977 | -4.0238973 | 5.72E-05 | 0.00082306 |
| CD163 | 4891.40313 | -2.290351424 | 0.51023934 | -4.4887786 | 7.16E-06 | 0.00015676 |
| APOBEC1 | 1.98696328 | -4.186372564 | 1.81606259 | -2.3051918 | 0.02115583 | 0.07810066 |
| GDF3 | 61.3972763 | 0.182801587 | 0.88358304 | 0.20688671 | 0.83609834 | 0.91295155 |
| DPPA3 | 0.09635527 | 0.547239394 | 3.29377202 | 0.16614368 | 0.86804388 | NA |
| CLEC4C | 1.57882673 | 2.37150632 | 1.44900998 | 1.63663905 | 0.10170591 | 0.24034276 |
| NANOGNB | 0.08393208 | -0.642905668 | 3.30558948 | -0.1944905 | 0.84579185 | NA |
| NANOG | 2.01389266 | 0.765041575 | 0.84871941 | 0.90140695 | 0.36737199 | 0.56428234 |
| SLC2A14 | 28.0916346 | -0.825389232 | 0.64692287 | -1.2758696 | 0.20200162 | 0.38385693 |
| SLC2A3 | 3729.34948 | -1.434938125 | 0.59968603 | -2.3928157 | 0.01671964 | 0.06588409 |
| AC006511.4 | 0.35817052 | 1.665335538 | 2.8231258 | 0.58989066 | 0.55526396 | NA |
| FOXJ2 | 1280.13826 | -0.020688788 | 0.13363894 | -0.1548111 | 0.87697026 | 0.93592236 |
| C3AR1 | 482.697768 | -0.436742977 | 0.33901864 | -1.2882565 | 0.19765669 | 0.37843313 |
| NECAP1 | 1813.38733 | 0.074557956 | 0.13293975 | 0.56084018 | 0.57490649 | 0.73915805 |
| CLEC4A | 112.346179 | -0.216830569 | 0.25796219 | -0.8405518 | 0.4005991 | 0.59544547 |
| AC092111.3 | 47.3376647 | -0.36788646 | 0.26053539 | -1.4120402 | 0.15793811 | 0.32507244 |
| ZNF705A | 9.40014642 | 0.559953799 | 0.41842154 | 1.3382528 | 0.18081404 | 0.35691791 |
| FAM66C | 122.711045 | 0.275392812 | 0.27709276 | 0.99386505 | 0.32028857 | 0.51771544 |
| AC092111.2 | 8.05599014 | 0.005908667 | 0.46853816 | 0.01261085 | 0.98993826 | 0.9952599 |
| AC092111.1 | 65.6155692 | 0.519321049 | 0.56485408 | 0.91938974 | 0.35789175 | 0.55554541 |
| FAM90A1 | 53.2257827 | -0.462717566 | 0.40984435 | -1.129008 | 0.25889445 | 0.45167971 |
| LINC02449 | 172.17263 | 0.260990913 | 0.2851665 | 0.9152229 | 0.36007462 | 0.55778464 |
| LINC00937 | 108.719646 | -1.072531783 | 0.58458207 | -1.8346984 | 0.06655039 | 0.17769682 |
| AC092745.1 | 3.7256128 | 2.136929745 | 0.83898077 | 2.54705451 | 0.01086365 | 0.04792144 |
| CLEC6A | 2.02088372 | 1.48522055 | 0.96175949 | 1.5442744 | 0.12252187 | 0.27373127 |
| CLEC4D | 20.8064077 | -2.087162198 | 0.6281969 | -3.322465 | 0.00089226 | 0.00717562 |
| CLEC4E | 242.973271 | -2.641871817 | 0.65590635 | -4.0278187 | 5.63E-05 | 0.000813 |
| AC092746.1 | 0.32750288 | -1.694569198 | 2.87850449 | -0.5886978 | 0.55606403 | NA |
| AICDA | 3.53211201 | -3.643879052 | 1.55236995 | -2.3473007 | 0.01890999 | 0.07202894 |
| MFAP5 | 139.727636 | 0.199652806 | 0.99492163 | 0.20067189 | 0.84095514 | 0.91512686 |
| AC092490.2 | 0.10721287 | -0.642905668 | 3.30558948 | -0.1944905 | 0.84579185 | NA |
| RIMKLB | 2963.25703 | -0.877497295 | 0.27428564 | -3.1992097 | 0.00137805 | 0.01013644 |
| A2ML1-AS1 | 84.7273627 | -0.916667821 | 0.33847704 | -2.7082127 | 0.00676467 | 0.03368387 |
| AC092490.1 | 12.6367827 | 0.292745849 | 0.81779598 | 0.35796929 | 0.72036629 | 0.84093782 |
| A2ML1-AS2 | 0 | NA | NA | NA | NA | NA |
| A2ML1 | 8.4074529 | 0.880063379 | 0.57878711 | 1.52053036 | 0.12837773 | 0.2825103 |
| AC006581.2 | 0.5986314 | -1.152323256 | 1.67187941 | -0.6892383 | 0.49067334 | NA |
| PHC1 | 2720.98359 | -0.087337218 | 0.11837513 | -0.7378004 | 0.46063575 | 0.64911904 |
| M6PR | 4793.98321 | -0.16896317 | 0.17422907 | -0.969776 | 0.33215815 | 0.53009359 |
| KLRG1 | 79.7573533 | 0.229479962 | 0.39700097 | 0.57803376 | 0.56324132 | 0.73050726 |
| AC006581.1 | 0 | NA | NA | NA | NA | NA |
| A2M-AS1 | 4282.03903 | -0.820691223 | 0.39853716 | -2.059259 | 0.03946944 | 0.12262123 |
| A2M | 47238.6831 | 0.166179454 | 0.41196313 | 0.40338429 | 0.68666554 | 0.81889338 |
| PZP | 48.8402045 | 0.282704972 | 0.41360794 | 0.68350953 | 0.49428493 | 0.67673841 |
| LINC00987 | 113.284124 | -0.09809843 | 0.29908146 | -0.327999 | 0.74291239 | 0.85567153 |
| AC010175.1 | 7.24947776 | 0.755942572 | 0.56468381 | 1.33870063 | 0.18066815 | 0.35665662 |
| LINC02367 | 13.5483872 | -0.459391705 | 0.59442404 | -0.772835 | 0.43962003 | 0.63101113 |
| KLRB1 | 88.2948023 | 0.22807429 | 0.70251896 | 0.32465215 | 0.74544436 | 0.85738944 |
| AC010186.1 | 1.53759987 | -0.18652452 | 1.18252871 | -0.1577336 | 0.87466672 | 0.93482598 |
| CLEC2D | 726.697412 | -0.394945962 | 0.33487831 | -1.1793716 | 0.23825025 | 0.42777007 |
| LINC02390 | 1.84379797 | -0.752028712 | 0.98561711 | -0.7630029 | 0.44546167 | 0.63592595 |
| CLECL1 | 12.4624462 | 0.357526053 | 0.79002779 | 0.4525487 | 0.65087374 | 0.79486596 |
| CD69 | 235.652702 | 0.951226543 | 0.65113338 | 1.46087817 | 0.14404888 | 0.30583361 |
| KLRF1 | 68.5265816 | -0.0732022 | 0.50983277 | -0.1435808 | 0.8858315 | 0.94097225 |
| CLEC2B | 519.371935 | -1.219194014 | 0.29190714 | -4.1766503 | 2.96E-05 | 0.00048545 |
| AC091814.1 | 51.8976905 | -0.65770829 | 0.24932241 | -2.637983 | 0.00834008 | 0.03927978 |
| KLRF2 | 0.53006243 | 0.887040677 | 2.24255374 | 0.39554935 | 0.69243751 | NA |
| CLEC2A | 0.17515154 | 1.45420457 | 3.2783972 | 0.44357181 | 0.65735219 | NA |
| LINC02470 | 0.17820003 | -1.003574173 | 3.30558948 | -0.3035992 | 0.7614333 | NA |
| CLEC12A-AS1 | 1.56653339 | 1.616844517 | 0.97549241 | 1.65746499 | 0.0974255 | 0.23330099 |
| CLEC12A | 57.512536 | 0.572205845 | 0.69117943 | 0.82786874 | 0.40774484 | 0.60239965 |
| CLEC1B | 8.61739784 | -0.209388363 | 0.66081661 | -0.316863 | 0.75134752 | 0.86135902 |
| CLEC12B | 7.82363468 | -0.041661246 | 0.68341463 | -0.0609604 | 0.95139073 | 0.97675915 |
| AC024224.2 | 2.3338066 | -0.006432581 | 1.09941294 | -0.0058509 | 0.99533167 | 0.99776308 |
| CLEC9A | 25.4546142 | 1.343671443 | 0.5029493 | 2.6715843 | 0.00754941 | 0.03661541 |
| CLEC1A | 349.783859 | -1.155647945 | 0.3257056 | -3.5481365 | 0.00038797 | 0.00372839 |
| CLEC7A | 551.203469 | -0.416490368 | 0.47030303 | -0.8855787 | 0.37584457 | 0.57170174 |
| OLR1 | 452.221562 | 0.633771901 | 0.69606707 | 0.91050407 | 0.36255673 | 0.55991227 |
| TMEM52B | 4600.1942 | 2.00447994 | 0.62680504 | 3.19793207 | 0.00138417 | 0.01017011 |
| GABARAPL1 | 27003.3239 | -0.697058584 | 0.24904554 | -2.7989202 | 0.00512738 | 0.02739798 |
| AC115676.1 | 234.797852 | 0.00604451 | 0.31373365 | 0.01926637 | 0.98462861 | 0.99336814 |
| KLRD1 | 199.025868 | -1.56420733 | 0.41558607 | -3.7638589 | 0.00016731 | 0.00190336 |
| AC022075.3 | 0.2926836 | 2.132206094 | 3.2337338 | 0.65936352 | 0.50966236 | NA |
| AC022075.2 | 1.16032347 | 0.00474101 | 1.5694863 | 0.00302074 | 0.9975898 | 0.99867691 |
| AC022075.1 | 60.6011156 | -0.475125333 | 0.34201692 | -1.3891866 | 0.16477601 | 0.33475681 |
| KLRK1 | 224.355972 | -0.750398361 | 0.60095474 | -1.248677 | 0.21178324 | 0.39540668 |
| KLRC4-KLRK1 | 220.673564 | -0.757753764 | 0.5928496 | -1.2781552 | 0.20119473 | 0.38296548 |
| KLRC4 | 13.291284 | -0.179281277 | 0.65554072 | -0.2734861 | 0.78447957 | 0.88222274 |
| AC068775.1 | 5.92966071 | -2.092272593 | 0.8853577 | -2.3631947 | 0.01811815 | 0.06976863 |
| KLRC3 | 5.15336102 | -2.089470908 | 1.00380928 | -2.0815417 | 0.03738435 | 0.11779487 |
| KLRC2 | 3.75424634 | -0.640124535 | 0.86718709 | -0.738162 | 0.46041603 | 0.64891345 |
| KLRC1 | 8.59329883 | 0.362610243 | 0.72924496 | 0.49724065 | 0.61901936 | 0.77234636 |
| EIF2S3B | 25.7530588 | 1.194303996 | 0.90986263 | 1.31262013 | NA | NA |
| LINC02446 | 13.4262968 | -0.910941036 | 1.0677427 | -0.8531466 | 0.39357802 | 0.58894535 |
| MAGOHB | 443.757611 | -0.315689515 | 0.18442989 | -1.7117047 | 0.0869511 | 0.21493256 |
| STYK1 | 73.0832562 | -1.573936407 | 0.50997419 | -3.086306 | 0.0020266 | 0.01363803 |

| | | | | | | |
|---|---|---|---|---|---|---|
| YBX3 | 12764.8966 | -1.575188816 | 0.2706652 | -5.8196947 | 5.90E-09 | 4.42E-07 |
| LINC02366 | 0.19416835 | -1.003574173 | 3.30558948 | -0.3035992 | 0.7614333 | NA |
| TAS2R7 | 0 | NA | NA | NA | NA | NA |
| TAS2R8 | 0.28475883 | 0.328534317 | 2.32323766 | 0.14141227 | 0.88754426 | NA |
| TAS2R9 | 0.17332742 | -1.003574173 | 3.30558948 | -0.3035992 | 0.7614333 | NA |
| PRH1 | 485.24187 | -0.112003115 | 0.13095917 | -0.8552522 | 0.39241157 | 0.58811555 |
| TAS2R10 | 5.21823733 | 0.083233132 | 0.63997898 | 0.13005604 | 0.89652209 | 0.9470152 |
| PRR4 | 193.394566 | -0.225036315 | 0.24668659 | -0.9122357 | 0.36164466 | 0.55906728 |
| AC018630.2 | 577.791619 | -0.139622183 | 0.13101704 | -1.0656796 | 0.28656852 | 0.48273999 |
| AC006518.7 | 71.7234591 | -0.024563756 | 0.34772345 | -0.0706416 | 0.94368297 | 0.97315146 |
| TAS2R13 | 0.91720406 | -1.524320867 | 2.18806906 | -0.6966512 | 0.48602112 | NA |
| PRH2 | 1.37057672 | -0.009085946 | 1.3180477 | -0.0068935 | 0.99449984 | 0.99750786 |
| TAS2R14 | 473.598273 | -0.124625013 | 0.11328995 | -1.1000535 | 0.27130879 | 0.46598666 |
| TAS2R50 | 0.74098454 | 2.249046576 | 1.99122279 | 1.12948013 | 0.25869535 | NA |
| TAS2R20 | 69.3616312 | -0.016187302 | 0.32308421 | -0.0501024 | 0.96004077 | 0.98075679 |
| TAS2R19 | 15.1536052 | 0.212906493 | 0.38069645 | 0.55925527 | 0.57598752 | 0.74001417 |
| TAS2R31 | 8.07227173 | 0.267132862 | 0.55810551 | 0.47864222 | 0.63219318 | 0.78153143 |
| TAS2R46 | 2.34973126 | 0.091825937 | 1.06502763 | 0.0862193 | 0.93129209 | 0.96690482 |
| TAS2R43 | 2.90350791 | 0.56125844 | 1.33576758 | 0.42017672 | 0.67435636 | 0.81034701 |
| TAS2R30 | 2.77371075 | 0.003331031 | 0.87254711 | 0.00381759 | 0.99695401 | 0.99850237 |
| SMIM10L1 | 1386.19178 | -0.141160734 | 0.16100056 | -0.8767717 | 0.38061067 | 0.57668946 |
| TAS2R42 | 0.80727822 | -2.17946918 | 1.60205259 | -1.360423 | 0.17369611 | NA |
| AC244131.2 | 0 | NA | NA | NA | NA | NA |
| PRB3 | 2.17308211 | 0.046873235 | 1.27798226 | 0.03667753 | 0.97074212 | 0.98642676 |
| PRB4 | 0.37996032 | 0.969037958 | 2.17697013 | 0.44513149 | 0.65622473 | NA |
| PRB1 | 0.52061348 | -0.034165911 | 2.17312726 | -0.015722 | 0.98745617 | NA |
| PRB2 | 1.41511884 | 1.345051925 | 1.05698499 | 1.27253644 | 0.20318258 | 0.38532652 |
| AC007450.1 | 4.13077794 | -1.004916208 | 0.87035785 | -1.1546012 | 0.24825378 | 0.43986106 |
| LINC01252 | 77.1061534 | -0.131449814 | 0.5046584 | -0.2604729 | 0.79449905 | 0.88857752 |
| ETV6 | 2180.31492 | -1.177793673 | 0.21440294 | -5.4933654 | 3.94E-08 | 2.18E-06 |
| BCL2L14 | 34.9679584 | 0.269450751 | 0.66170915 | 0.40720421 | 0.68385801 | 0.81746762 |
| LRP6 | 2036.07582 | 0.41103276 | 0.19220112 | 2.13855553 | 0.03247168 | 0.10667476 |
| MANSC1 | 1405.20299 | -0.418281158 | 0.19749091 | -2.1179767 | 0.03417704 | 0.11071849 |
| LOH12CR2 | 75.6140308 | 1.142151299 | 0.37979807 | 3.00725935 | 0.00263615 | 0.0166292 |
| BORCS5 | 346.184278 | 0.106772516 | 0.14490269 | 0.73685669 | 0.46120951 | 0.64968454 |
| DUSP16 | 1847.95698 | -0.143110469 | 0.16628173 | -0.8606506 | 0.38943051 | 0.58507763 |
| AC007619.1 | 2.99678363 | 1.331273754 | 0.83727241 | 1.5900127 | 0.11183194 | 0.25701659 |
| CREBL2 | 4034.85299 | 0.472124452 | 0.15957615 | 2.95861541 | 0.00309025 | 0.01878077 |
| AC008115.4 | 12.3642141 | 0.638158203 | 0.55361168 | 1.15271809 | 0.24902611 | 0.44078927 |
| GPR19 | 6.29378464 | -1.007344306 | 0.61337398 | -1.6423004 | 0.10052777 | 0.23843473 |
| AC008115.2 | 0.2883433 | 0.375429037 | 2.95186421 | 0.12718371 | 0.898795 | NA |
| CDKN1B | 3010.60051 | -0.116909626 | 0.16132154 | -0.7246994 | 0.46863645 | 0.65541995 |
| AC008115.1 | 0.14444843 | 0.435199075 | 3.28898345 | 0.13232024 | 0.894731 | NA |
| APOLD1 | 4481.91322 | -0.821884901 | 0.23248466 | -3.5352221 | 0.00040743 | 0.0038773 |
| DDX47 | 2607.46355 | -0.363513939 | 0.13398053 | -2.7131848 | 0.00666399 | 0.03333843 |
| GPRC5A | 1978.95456 | -1.388159116 | 0.48238467 | -2.8777016 | 0.00400584 | 0.02277576 |
| GPRC5D-AS1 | 222.643305 | 0.021679643 | 0.26702998 | 0.08118805 | 0.9352924 | 0.96907653 |
| GPRC5D | 26.6397152 | -0.262753524 | 0.48024612 | -0.5471226 | 0.58429448 | 0.74635935 |
| AC007688.2 | 0.51046302 | 2.259002619 | 2.08084274 | 1.0856191 | 0.27764755 | NA |
| HEBP1 | 2725.3766 | -0.181247201 | 0.14247539 | -1.2721299 | 0.20332696 | 0.38551424 |
| AC023790.2 | 23.3538177 | 0.546578858 | 0.31099352 | 1.75752489 | 0.07882838 | 0.20039569 |
| FAM234B | 716.962053 | 0.16977531 | 0.25411586 | 0.66810198 | 0.5040685 | 0.68375733 |
| GSG1 | 14.9723938 | 0.977808503 | 0.50395528 | 1.9402684 | 0.05234708 | 0.15038594 |
| EMP1 | 6817.09235 | -0.656717647 | 0.47268934 | -1.3893219 | 0.16473489 | 0.33472485 |
| LINC01559 | 98.4249483 | -5.042053101 | 0.94375203 | -5.3425613 | 9.16E-08 | 4.46E-06 |
| GRIN2B | 18.1965031 | 0.082771551 | 0.65610387 | 0.12615617 | 0.89960831 | 0.94902072 |
| AC007527.2 | 0.68223657 | 0.150812065 | 2.20815537 | 0.06829776 | 0.94554861 | NA |
| AC092112.1 | 8.48015872 | 0.037828671 | 0.67910437 | 0.05570377 | 0.9555778 | 0.97873792 |
| ATF7IP | 2174.48171 | 0.071637873 | 0.14978342 | 0.47827639 | 0.6324535 | 0.78171075 |
| PLBD1 | 1819.99059 | -0.196094971 | 0.15731567 | -1.2465063 | 0.21257859 | 0.39645571 |
| AC008114.1 | 0.88778013 | 0.317097183 | 1.35852444 | 0.23341294 | 0.81544077 | NA |
| PLBD1-AS1 | 88.4098042 | 0.638527752 | 0.31600488 | 2.02062624 | 0.04331847 | 0.13129978 |
| GUCY2C | 88.4203653 | 0.956680863 | 0.27430452 | 3.4876598 | 0.00048727 | 0.0044415 |
| AC010168.1 | 8.60874348 | 0.79462372 | 0.50158827 | 1.58421512 | 0.11314477 | 0.25913042 |
| HIST4H4 | 49.8914158 | 0.004391777 | 0.33669081 | 0.01304395 | 0.98959273 | 0.99519138 |
| H2AFJ | 3126.62887 | -0.791577531 | 0.16943653 | -4.6718234 | 2.99E-06 | 7.80E-05 |
| WBP11 | 3542.89585 | -0.031107768 | 0.15190663 | -0.2047822 | 0.83774232 | 0.91410871 |
| C12orf60 | 11741.3111 | -0.740266082 | 0.43130567 | -1.7163375 | 0.08610029 | 0.21343816 |
| SMCO3 | 135.08846 | 0.775226091 | 0.62310936 | 1.24412525 | 0.21345347 | 0.39748379 |
| ART4 | 201.816121 | 0.06039466 | 0.51864188 | 0.11644771 | 0.90729772 | 0.953776 |
| AC007655.1 | 17.1470047 | 0.217965729 | 0.5927711 | 0.3677064 | 0.71309217 | 0.83695061 |
| MGP | 27835.6384 | -0.917570115 | 0.43717292 | -2.0988723 | 0.03582816 | 0.11445047 |
| ERP27 | 1523.39543 | -0.443512941 | 0.30925543 | -1.4341315 | 0.15153475 | 0.31657328 |
| ARHGDIB | 3350.39871 | -0.316186607 | 0.35184679 | -0.8986486 | 0.36883988 | 0.56564744 |
| PDE6H | 1.62460003 | -0.501140342 | 1.49196209 | -0.3358935 | 0.73695119 | 0.85234223 |
| LINC01489 | 7.21903179 | -0.60072049 | 0.52796819 | -1.1377967 | 0.25520536 | 0.44784491 |
| RERG | 2064.79957 | 0.2840666 | 0.2067533 | 1.37393987 | 0.16946035 | 0.34129147 |
| RERG-AS1 | 17.5863828 | 2.928312585 | 0.55630426 | 5.26386871 | 1.41E-07 | 6.39E-06 |
| PTPRO | 1550.66102 | 0.714626234 | 0.56881317 | 1.25634614 | 0.2089905 | 0.39219182 |
| EPS8 | 4982.01698 | 0.210261719 | 0.25335506 | 0.82990931 | 0.40659006 | 0.60140134 |
| AC073651.1 | 0.69954059 | -1.701565365 | 1.86058127 | -0.9145343 | 0.36043617 | NA |
| STRAP | 6955.6538 | -0.233003547 | 0.23175187 | -1.0054009 | 0.31470383 | 0.51173087 |
| DERA | 3238.34767 | 0.093814848 | 0.27179426 | 0.34516861 | 0.72996762 | 0.84746174 |
| SLC15A5 | 3.74788187 | 1.390020363 | 0.75080786 | 1.85136629 | 0.06411687 | 0.17315329 |
| MGST1 | 14472.8898 | -0.785473433 | 0.41763997 | -1.880743 | 0.06000689 | 0.1655211 |
| LMO3 | 286.982219 | 0.75739059 | 0.23725446 | 3.19231345 | 0.00141138 | 0.01032169 |
| AC007552.2 | 0.37374641 | -0.032131735 | 2.27000489 | -0.0141549 | 0.98870639 | NA |
| AC007529.2 | 3.18895141 | -0.953121403 | 1.00692033 | -0.9465708 | 0.34385752 | 0.54151789 |
| AC007529.1 | 0.57066768 | -0.698346993 | 1.65673317 | -0.4215205 | 0.67337505 | NA |
| LINC02378 | 0 | NA | NA | NA | NA | NA |
| AC087321.1 | 0 | NA | NA | NA | NA | NA |
| RERGL | 364.110303 | 0.565110577 | 0.48956278 | 1.15431687 | 0.24837028 | 0.44003791 |
| PIK3C2G | 292.541298 | 1.276349413 | 0.44389027 | 2.87537144 | 0.00403552 | 0.02290503 |
| PLCZ1 | 19.6226934 | 1.667118527 | 0.54577579 | 3.05458495 | 0.00225372 | 0.01480249 |
| AC087242.1 | 8.51794733 | 1.224830127 | 0.58249045 | 2.10274712 | 0.03548788 | 0.11372164 |
| CAPZA3 | 22.1413704 | 0.499271972 | 0.58638165 | 0.85144542 | 0.39452197 | 0.58977008 |
| PLEKHA5 | 2467.49052 | 0.932558431 | 0.20312987 | 4.5909469 | 4.41E-06 | 0.00010675 |
| AEBP2 | 1694.5337 | 0.088913294 | 0.11979364 | 0.74222046 | 0.45795377 | 0.64706542 |
| AC024901.1 | 2.61116638 | -0.326234853 | 0.9612291 | -0.3393934 | 0.73431336 | 0.85052239 |
| LINC02398 | 2.55964214 | 0.647400922 | 0.80531634 | 0.80390883 | 0.42144963 | 0.61408038 |
| LINC02468 | 1.19080767 | -0.024847073 | 1.12714687 | -0.0220442 | 0.98241268 | 0.99215742 |
| AC129102.1 | 6.28391498 | 1.269192492 | 0.51427727 | 2.46791481 | 0.01359027 | 0.05656816 |
| PDE3A | 724.848342 | 0.85109412 | 0.22252281 | 3.82475009 | 0.0001309 | 0.0015641 |
| SLCO1C1 | 24.8609595 | 0.562161187 | 0.46734565 | 1.20288098 | 0.22902238 | 0.41638998 |
| SLCO1B3 | 0.38696503 | -0.03213025 | 2.25519186 | -0.0142472 | 0.98863274 | NA |
| AC011604.2 | 0.25658839 | -0.174096114 | 3.29377188 | -0.0528562 | 0.95784652 | NA |
| SLCO1B7 | 0 | NA | NA | NA | NA | NA |
| AC022335.1 | 0 | NA | NA | NA | NA | NA |
| SLCO1B1 | 0.16783122 | 1.368031398 | 3.28033652 | 0.41703996 | 0.67664918 | NA |
| SLCO1A2 | 30.8416784 | 2.449160808 | 0.66551472 | 3.68010011 | 0.00023314 | 0.00246906 |
| IAPP | 0.31713219 | 0.328539074 | 2.27448921 | 0.14444521 | 0.88514892 | NA |
| PYROXD1 | 2107.46072 | -0.075107644 | 0.17694619 | -0.424466 | 0.67122598 | 0.80797138 |
| RECQL | 2019.00346 | -0.177313773 | 0.1934391 | -0.9166388 | 0.35933197 | 0.55696138 |

| | | | | | | |
|---|---|---|---|---|---|---|
| GOLT1B | 1855.274 | -0.569679738 | 0.16860666 | -3.3787498 | 0.00072816 | 0.00610157 |
| SPX | 2070.55427 | -0.130468933 | 0.30008764 | -0.4347694 | 0.6637298 | 0.80246755 |
| GYS2 | 70.3157775 | -0.166511193 | 0.41854721 | -0.3978313 | 0.69075452 | 0.82194832 |
| LDHB | 58109.0878 | 0.150765514 | 0.23560178 | 0.63991671 | 0.52222675 | 0.69814982 |
| AC010197.1 | 8.30084679 | -1.127763876 | 0.58469094 | -1.9288205 | 0.05375315 | 0.15333952 |
| AC010185.1 | 186.751881 | 0.521793964 | 0.26095707 | 1.99953948 | 0.04555001 | 0.13606184 |
| KCNJ8 | 791.247908 | 0.241121033 | 0.26095041 | 0.92401092 | 0.35548063 | 0.55300715 |
| ABCC9 | 656.092856 | 0.654501398 | 0.28507737 | 2.29587287 | 0.02168315 | 0.07964685 |
| AC008250.1 | 4.81682071 | 0.572484745 | 0.61963319 | 0.9239091 | 0.35553365 | 0.55305686 |
| CMAS | 2000.03064 | -0.424200854 | 0.1696036 | -2.5011312 | 0.01237973 | 0.05281394 |
| ST8SIA1 | 77.6328999 | 0.889046536 | 0.51948653 | 1.71139479 | 0.08700826 | 0.21499323 |
| AC087318.1 | 0.17423948 | 0.18657164 | 3.29377195 | 0.05664376 | 0.95482897 | NA |
| C2CD5 | 2397.7117 | 0.452746868 | 0.15597358 | 2.90271509 | 0.00369943 | 0.02149239 |
| AC053513.1 | 3.31058621 | 2.463133791 | 0.94479732 | 2.60704993 | 0.0091326 | 0.04210994 |
| AC087241.1 | 0.4552195 | 2.174927851 | 2.00446871 | 1.08503956 | 0.27790413 | NA |
| AC087241.3 | 2.98378357 | -0.185391568 | 0.7206727 | -0.2572479 | 0.79698738 | 0.88983348 |
| ETNK1 | 4036.2841 | -0.083609851 | 0.1254803 | -0.6663185 | 0.50520752 | 0.68452998 |
| AC084816.1 | 4.73866058 | 0.148709685 | 0.77613812 | 0.19160209 | 0.84805391 | 0.92016931 |
| AC084819.1 | 0.93918967 | -0.162332978 | 1.3342321 | -0.1216677 | 0.90316218 | NA |
| AC087258.1 | 0.908534 | 1.201610873 | 1.34798277 | 0.89141412 | 0.37270705 | NA |
| AC087235.1 | 0 | NA | NA | NA | NA | NA |
| SOX5 | 123.050724 | 0.53781421 | 0.33082008 | 1.6257 | 0.10401349 | 0.24416075 |
| AC087260.1 | 0 | NA | NA | NA | NA | NA |
| SOX5-AS1 | 0.38058391 | -1.083407007 | 3.29309194 | -0.3289939 | 0.74216033 | NA |
| LINC00477 | 0.15489794 | -1.003574173 | 3.30558948 | -0.3035992 | 0.7614333 | NA |
| AC023796.1 | 0 | NA | NA | NA | NA | NA |
| BCAT1 | 402.928193 | -0.444001 | 0.50193162 | -0.8845846 | 0.3763807 | 0.57222018 |
| AC023796.2 | 8.22052052 | -0.962055842 | 0.64659858 | -1.4878719 | 0.13678468 | 0.2951488 |
| AC026310.2 | 0.4116088 | -1.061841662 | 2.26633542 | -0.468528 | 0.63940703 | NA |
| C12orf77 | 0.22809742 | -0.339643007 | 3.05487708 | -0.1111806 | 0.91147315 | NA |
| LRMP | 778.92675 | 0.891496503 | 0.31654233 | 2.81635794 | 0.00485715 | 0.02630043 |
| AC023510.1 | 100.458374 | 1.067474 | 0.32839315 | 3.25059764 | 0.00115163 | 0.00876289 |
| AC023510.2 | 120.123212 | 0.853048002 | 0.33657603 | 2.53448824 | 0.01126117 | 0.04924661 |
| CASC1 | 159.889822 | 0.503007248 | 0.47711252 | 1.05427384 | 0.29175756 | 0.48792789 |
| ETFRF1 | 1427.0088 | 0.315708241 | 0.1737873 | 1.81663589 | 0.06927288 | 0.18280056 |
| KRAS | 2342.98823 | -0.36663568 | 0.1435347 | -2.5543349 | 0.01063909 | 0.0471438 |
| AC092794.1 | 26.6279309 | 0.587447928 | 0.32659362 | 1.79871221 | 0.07206422 | 0.1882596 |
| AC092794.2 | 42.1663033 | -0.175410458 | 0.26336025 | -0.6660476 | 0.50538069 | 0.68465907 |
| AC087239.1 | 15.0570261 | -0.696706835 | 0.40592805 | -1.7163308 | 0.08610151 | 0.21343816 |
| LMNTD1 | 1.047502 | 0.027147423 | 2.25512514 | 0.0120381 | 0.99039522 | 0.99533983 |
| AC019209.3 | 0 | NA | NA | NA | NA | NA |
| AC019209.2 | 0 | NA | NA | NA | NA | NA |
| AC019209.1 | 0 | NA | NA | NA | NA | NA |
| RASSF8-AS1 | 1387.2338 | 0.522814753 | 0.27010504 | 1.93559788 | 0.05291697 | 0.15157729 |
| RASSF8 | 1809.18263 | 0.23165367 | 0.28195715 | 0.82159176 | 0.41130928 | 0.60556227 |
| BHLHE41 | 1285.51307 | -1.934655017 | 0.5065403 | -3.8193506 | 0.0001338 | 0.00159244 |
| SSPN | 531.509153 | 0.315440445 | 0.34823889 | 0.90581624 | 0.36503312 | 0.56205495 |
| AC022509.3 | 23.1991643 | -0.899138787 | 0.56094529 | -1.6028993 | 0.1089569 | 0.25232299 |
| AC022509.2 | 106.469313 | 0.366132978 | 0.19249265 | 1.9020621 | 0.05716304 | 0.15988431 |
| AC022509.1 | 57.1946124 | 2.291193165 | 0.42460213 | 5.39609435 | 6.81E-08 | 3.43E-06 |
| AC022509.4 | 1.63292937 | -0.297855656 | 1.34318922 | -0.2217526 | 0.82450651 | 0.9061153 |
| AC055720.1 | 0 | NA | NA | NA | NA | NA |
| AC024145.1 | 9.68071462 | -0.046331708 | 0.59452852 | -0.0779302 | 0.9378836 | 0.97030054 |
| ITPR2 | 1981.6609 | 0.681096862 | 0.14094663 | 4.83230322 | 1.35E-06 | 4.15E-05 |
| AC023051.1 | 9.96931656 | -0.558669789 | 0.4712741 | -1.1854456 | 0.23584133 | 0.42492045 |
| INTS13 | 994.491477 | -0.273538977 | 0.16587189 | -1.6490978 | 0.0991276 | 0.23611189 |
| FGFR1OP2 | 2261.71589 | 0.078399236 | 0.10631776 | 0.73740489 | 0.46087616 | 0.64938843 |
| AC024896.1 | 831.352587 | 0.788535949 | 0.33144352 | 2.379096 | 0.01735516 | 0.0676516 |
| TM7SF3 | 6530.25927 | 1.22480889 | 0.33770788 | 3.62682946 | 0.00028692 | 0.00292808 |
| MED21 | 1497.01303 | -0.160111991 | 0.15519919 | -1.0316548 | 0.30223385 | 0.4991174 |
| AC092747.4 | 561.05129 | 0.373216503 | 0.13965089 | 2.67249634 | 0.00752892 | 0.03655636 |
| C12orf71 | 5.76974918 | 1.140432757 | 0.61733908 | 1.84733608 | 0.06469845 | 0.17419829 |
| AC092747.1 | 1.03505031 | 1.356127899 | 1.28010233 | 1.05939023 | 0.2894221 | 0.48568542 |
| STK38L | 1522.73504 | 0.225929146 | 0.31077449 | 0.72698742 | 0.46723366 | 0.65433492 |
| ARNTL2 | 227.911238 | -2.306399557 | 0.39988028 | -5.7677252 | 8.03E-09 | 5.71E-07 |
| ARNTL2-AS1 | 8.61199874 | -2.046262261 | 0.65475466 | -3.1252351 | 0.00177663 | 0.01231314 |
| SMCO2 | 9.69103632 | -1.988047807 | 0.63239734 | -3.1436688 | 0.00166844 | 0.0117231 |
| PPFIBP1 | 3693.4239 | 0.201931053 | 0.12147875 | 1.66227471 | 0.09645771 | 0.2317934 |
| AC087257.1 | 0.82149501 | -1.292285212 | 1.6730285 | -0.7724227 | 0.43986411 | NA |
| AC009509.1 | 99.1897948 | -0.334339561 | 0.50045544 | -0.6680706 | 0.50408854 | 0.68375733 |
| REP15 | 93.6119684 | -0.490462699 | 0.4589731 | -1.0686088 | 0.28524598 | 0.48132743 |
| AC009509.4 | 17.5583004 | 0.24575338 | 0.37344051 | 0.65807906 | 0.51048733 | 0.68892277 |
| AC009509.2 | 1.09279795 | -0.776233224 | 1.40296579 | -0.5532802 | 0.58007154 | 0.74312508 |
| MRPS35 | 3662.04548 | 0.135523164 | 0.20615452 | 0.65738633 | 0.51093254 | 0.68920615 |
| MANSC4 | 28.417536 | -1.766618774 | 0.81568247 | -2.1658168 | 0.03032519 | 0.10155896 |
| AC009511.2 | 84.8506821 | -1.359840836 | 0.51315611 | -2.6499555 | 0.00805024 | 0.038284 |
| KLHL42 | 1827.36104 | -0.595698174 | 0.20798168 | -2.8641858 | 0.00418083 | 0.02356239 |
| AC009511.1 | 41.1496654 | 0.159834736 | 0.26976489 | 0.59249643 | 0.55351821 | 0.7230174 |
| PTHLH | 34.4045282 | 0.256725237 | 0.36830194 | 0.69705099 | 0.48577088 | 0.66985253 |
| AC008011.2 | 21.1359618 | 0.721026038 | 0.42766869 | 1.68594534 | 0.09180633 | 0.22380336 |
| CCDC91 | 1083.93842 | 0.001026897 | 0.09869984 | 0.01040424 | 0.99169877 | 0.99605449 |
| AC022364.1 | 196.193978 | 0.503261338 | 0.30167197 | 1.6682403 | 0.09526803 | 0.22978305 |
| AC022079.2 | 21.5437136 | 0.494746108 | 0.35412201 | 1.39710635 | 0.16238159 | 0.33138509 |
| AC022081.1 | 0.36177836 | 0.005377682 | 3.28420126 | 0.00163744 | 0.99869351 | NA |
| AC012150.1 | 0.72829185 | 0.019315664 | 1.3975145 | 0.01382144 | 0.98897244 | NA |
| FAR2 | 572.732775 | -0.569332818 | 0.19429434 | -2.9302594 | 0.00338679 | 0.02010574 |
| AC012150.2 | 0.24613888 | 1.011711844 | 2.95767469 | 0.34206326 | 0.73230329 | NA |
| AC009318.4 | 31.9014218 | -0.297728365 | 0.46528588 | -0.6398827 | 0.52224889 | 0.69814982 |
| AC009318.1 | 79.4413449 | 0.044995047 | 0.29347731 | 0.15331695 | 0.87814833 | 0.93672471 |
| AC009318.3 | 16.7648135 | 0.100555986 | 0.31942609 | 0.31480205 | 0.75291196 | 0.86228066 |
| AC009318.2 | 50.9693109 | -0.025869035 | 0.21639012 | -0.1195481 | 0.90484111 | 0.9522378 |
| ERGIC2 | 1624.81284 | -0.542821224 | 0.10658095 | -5.0930418 | 3.52E-07 | 1.36E-05 |
| OVCH1-AS1 | 44.0875497 | -0.275022618 | 0.45714313 | -0.6016116 | 0.54743269 | 0.71842513 |
| OVCH1 | 23.9535642 | -0.552142189 | 0.5007645 | -1.1025985 | 0.27020156 | 0.46449207 |
| TMTC1 | 2237.29875 | 0.216617262 | 0.30629212 | 0.70722441 | 0.47942703 | 0.66418343 |
| AC009320.1 | 3.97168334 | -0.93879783 | 0.77711203 | -1.2080598 | 0.22702425 | 0.41385295 |
| AC023511.1 | 0.4786891 | 1.564576618 | 3.22487806 | 0.48515838 | 0.627564 | NA |
| LINC02386 | 0.17507102 | 0.435199075 | 3.28898345 | 0.13232024 | 0.894731 | NA |
| AC078776.1 | 0.92108585 | -3.077449715 | 2.73055671 | -1.1270411 | 0.25972509 | NA |
| IPO8 | 2411.92793 | 0.198211729 | 0.1136993 | 1.74329766 | 0.08128163 | 0.20463974 |
| CAPRIN2 | 1481.92525 | 0.101023952 | 0.0862438 | 1.17137635 | 0.24144753 | 0.43162982 |
| AC010198.1 | 5.20044646 | 2.034542705 | 0.99237778 | 2.05016955 | 0.04034789 | 0.12455793 |
| LINC00941 | 8.9264987 | -1.420830522 | 0.59847944 | -2.3740674 | 0.01759334 | 0.06829179 |
| TSPAN11 | 126.620443 | 0.585981657 | 0.48491502 | 1.20842134 | 0.22688524 | 0.4137687 |
| DDX11-AS1 | 21.6960126 | -0.153560104 | 0.34887803 | -0.4401541 | 0.65982548 | 0.79999826 |
| DDX11 | 1302.63886 | 0.048365951 | 0.3370967 | 0.14347797 | 0.88591271 | 0.94097225 |
| SINHCAF | 2924.12434 | -0.556714388 | 0.15921349 | -3.4966533 | 0.00047113 | 0.00433036 |
| AC024940.1 | 10.6005706 | -0.604191884 | 0.52682521 | -1.1468546 | 0.25144174 | 0.44374835 |
| LINC02387 | 1.10976806 | 0.261772663 | 1.48206986 | 0.1766264 | 0.85980186 | 0.92697882 |
| DENND5B | 1004.82693 | 0.467592489 | 0.24348311 | 1.92043088 | 0.0548035 | 0.15541114 |
| DENND5B-A[illegible] | 12.810195 | 0.220084489 | 0.50879193 | 0.43256286 | 0.66533238 | 0.80373362 |
| ETFBKMT | 769.771973 | 0.534122634 | 0.14764475 | 3.61762015 | 0.00029732 | 0.00300886 |
| AMN1 | 602.322332 | 0.09423302 | 0.16176566 | 0.58252797 | 0.56021111 | 0.72838393 |

| | | | | | | |
|---|---|---|---|---|---|---|
| AC023157.3 | 116.724066 | 0.110187849 | 0.27206688 | 0.4050028 | 0.68547545 | 0.81825122 |
| H3F3C | 16.8941514 | -0.729640435 | 0.51869603 | -1.4066821 | 0.15952167 | 0.32726678 |
| LINC02422 | 10.61716 | 1.484587239 | 1.00975753 | 1.47024131 | NA | NA |
| KIAA1551 | 1230.75946 | -0.379336097 | 0.21416278 | -1.7712513 | 0.07651893 | 0.19639908 |
| AC016957.2 | 30.0655221 | -0.515263969 | 0.38489131 | -1.3387259 | 0.18065992 | 0.35665662 |
| AC048344.1 | 32.9361214 | -0.261045744 | 0.28966849 | -0.9011879 | 0.36748842 | 0.56438321 |
| BICD1 | 736.518787 | -0.43473671 | 0.19110313 | -2.2748802 | 0.02291312 | 0.08292876 |
| FGD4 | 1918.96412 | -0.27865871 | 0.19183403 | -1.4526031 | 0.14633397 | 0.30917609 |
| DNM1L | 3463.81482 | -0.112394226 | 0.09100885 | -1.2349813 | 0.21683746 | 0.40161819 |
| AC084824.5 | 66.2174939 | -0.512784567 | 0.24383852 | -2.1029679 | 0.03546858 | 0.11369468 |
| YARS2 | 692.40859 | -0.5108905 | 0.16246412 | -3.1446359 | 0.00166294 | 0.01170001 |
| PKP2 | 612.091355 | 0.754369769 | 0.3348471 | 2.25287829 | 0.02426682 | 0.08631571 |
| AC087311.2 | 0 | NA | NA | NA | NA | NA |
| SYT10 | 24.850472 | 1.863896515 | 0.52903332 | 3.52321193 | 0.00042635 | 0.00400958 |
| ALG10 | 130.328036 | 0.062373705 | 0.22568365 | 0.27637672 | 0.78225874 | 0.88079261 |
| AC046130.1 | 27.6926745 | 0.28720233 | 0.35711545 | 0.80422824 | 0.42126517 | 0.61391338 |
| AC046130.2 | 2.11642048 | 0.306659427 | 1.18691996 | 0.25836572 | 0.79612467 | 0.88938094 |
| AC140847.1 | 0.17391096 | -1.003574173 | 3.30558948 | -0.3035992 | 0.7614333 | NA |
| AC140847.2 | 9.17063908 | 1.787769509 | 0.65666161 | 2.72251262 | 0.00647876 | 0.03261642 |
| ALG10B | 460.037222 | 0.30479098 | 0.17254936 | 1.76639881 | 0.07732896 | 0.19784368 |
| AC087897.2 | 0.16416605 | 0.18657164 | 3.29377195 | 0.05664376 | 0.95482897 | NA |
| CPNE8 | 1415.28011 | -0.274281661 | 0.15501252 | -1.7694162 | 0.07682445 | 0.19685902 |
| AC067735.1 | 33.8863712 | 0.023609514 | 0.25075866 | 0.09415234 | 0.92498815 | 0.96362877 |
| LINC02406 | 3.48564694 | 0.270141985 | 0.9143753 | 0.29543885 | 0.76765866 | 0.87183444 |
| KIF21A | 1888.7689 | 0.399731476 | 0.24863915 | 1.60767714 | 0.10790591 | 0.25048229 |
| ABCD2 | 38.8510588 | 1.542695295 | 0.77560419 | 1.98902394 | 0.04669856 | 0.13853538 |
| C12orf40 | 7.61784635 | 0.418527445 | 0.53376444 | 0.78410514 | 0.43297842 | 0.62504193 |
| SLC2A13 | 1157.41291 | 0.461813057 | 0.20629358 | 2.23862057 | 0.02518061 | 0.08883115 |
| LINC02555 | 16.7713913 | -1.616238945 | 0.813864 | -1.9858833 | 0.04704628 | 0.13931171 |
| LINC02471 | 186.440482 | -1.4134788 | 0.58019269 | -2.436223 | 0.01484153 | 0.06038097 |
| AC079630.1 | 469.039086 | 0.02934967 | 0.43960066 | 0.06676439 | 0.94676927 | 0.97438741 |
| LRRK2 | 3392.5764 | -0.203223542 | 0.37948436 | -0.5355255 | 0.59228654 | 0.75170299 |
| MUC19 | 59.2795059 | 0.24907011 | 0.67499508 | 0.36899544 | 0.71213112 | 0.83636475 |
| AC107023.1 | 0.03392908 | 0.380740593 | 3.29738709 | 0.11546736 | 0.90807469 | NA |
| CNTN1 | 630.50472 | 1.864591197 | 0.44075974 | 4.23040269 | 2.33E-05 | 0.00040311 |
| PDZRN4 | 51.7141765 | 1.552282757 | 0.63826237 | 2.43204491 | 0.01501385 | 0.06091231 |
| AC090531.1 | 0 | NA | NA | NA | NA | NA |
| LINC02400 | 0 | NA | NA | NA | NA | NA |
| AC090630.1 | 0.66865319 | -0.474386347 | 2.23501101 | -0.2122524 | 0.83191016 | NA |
| GXYLT1 | 1093.99572 | -0.555916876 | 0.23120146 | -2.4044696 | 0.01619595 | 0.0643689 |
| YAF2 | 813.997089 | -0.488735775 | 0.15197556 | -3.2158839 | 0.00130043 | 0.00966801 |
| PPHLN1 | 2184.28307 | -0.268365943 | 0.1297419 | -2.0684601 | 0.03859677 | 0.12063494 |
| ZCRB1 | 1631.9881 | -0.092402698 | 0.12290277 | -0.7518358 | 0.45214984 | 0.64182544 |
| PRICKLE1 | 475.54714 | 0.353635316 | 0.25316403 | 1.39686242 | 0.16245495 | 0.33138509 |
| AC079601.1 | 274.711974 | 0.391996757 | 0.27696227 | 1.41534355 | 0.15696779 | 0.32390157 |
| LINC02402 | 0 | NA | NA | NA | NA | NA |
| LINC02451 | 0 | NA | NA | NA | NA | NA |
| LINC02450 | 2.34817707 | -0.741208501 | 0.80085824 | -0.9255177 | 0.35469667 | 0.55227628 |
| AC012038.2 | 0.19280283 | 0.18657164 | 3.29377195 | 0.05664376 | 0.95482897 | NA |
| AC012038.1 | 0 | NA | NA | NA | NA | NA |
| LINC02461 | 0.81162295 | -1.950410809 | 1.83712204 | -1.0616664 | 0.28838714 | NA |
| ADAMTS20 | 0.09635527 | 0.547239394 | 3.29377202 | 0.16614368 | 0.86804388 | NA |
| PUS7L | 715.243353 | 0.062868974 | 0.15964695 | 0.39380002 | 0.69372869 | 0.82376229 |
| IRAK4 | 1386.41474 | -0.289459253 | 0.13871626 | -2.0867003 | 0.03691523 | 0.11679995 |
| TWF1 | 3417.1002 | -0.308117723 | 0.20243574 | -1.522052 | 0.12799606 | 0.28199983 |
| TMEM117 | 232.354712 | 0.054597789 | 0.22738863 | 0.24010782 | 0.81024667 | 0.89743804 |
| AC025030.2 | 0.09635527 | 0.547239394 | 3.29377202 | 0.16614368 | 0.86804388 | NA |
| AC025253.1 | 0.2892504 | -0.174096114 | 3.29377188 | -0.0528562 | 0.95784652 | NA |
| NELL2 | 71.812645 | 0.874308708 | 0.79235794 | 1.10342645 | 0.26984202 | 0.46404943 |
| DBX2 | 5.12183242 | -1.090114485 | 0.78776102 | -1.3838137 | 0.16641551 | 0.33721351 |
| AC008127.1 | 0.2804709 | -0.174096114 | 3.29377188 | -0.0528562 | 0.95784652 | NA |
| ANO6 | 4658.77021 | 0.005380894 | 0.29093261 | 0.01849533 | 0.9852437 | 0.99348815 |
| AC079950.1 | 0.17828073 | 0.18657164 | 3.29377195 | 0.05664376 | 0.95482897 | NA |
| AC008124.1 | 1098.1694 | 0.342801022 | 0.17631799 | 1.94422037 | 0.05186888 | 0.14959794 |
| ARID2 | 1041.71099 | 0.001283253 | 0.10639227 | 0.01206153 | 0.99037653 | 0.99533983 |
| SCAF11 | 3919.94011 | 0.027407778 | 0.08202333 | 0.33414612 | 0.7382693 | 0.85304361 |
| SLC38A1 | 4240.16943 | -0.931375484 | 0.2125495 | -4.3819227 | 1.18E-05 | 0.00023341 |
| SLC38A2 | 10683.05 | -1.101173846 | 0.34469473 | -3.194635 | 0.00140008 | 0.01025557 |
| AC025031.2 | 1.18964559 | -1.613011022 | 1.63249739 | -0.9880635 | 0.32312157 | 0.52085294 |
| AC008014.1 | 188.596069 | 0.898888419 | 0.25800966 | 3.48393317 | 0.0004941 | 0.00448365 |
| AC025031.1 | 4.26418563 | -0.124595919 | 0.56119821 | -0.2220177 | 0.82430013 | 0.90600177 |
| AC025031.4 | 16.7223498 | 0.661470815 | 0.40516134 | 1.63261088 | 0.10255086 | 0.24162493 |
| AC008035.1 | 67.3200257 | 0.857965439 | 0.28509371 | 3.0094155 | 0.00261751 | 0.01655574 |
| SLC38A4 | 562.176239 | -0.648179095 | 0.66474932 | -0.975073 | 0.32952406 | 0.5272929 |
| AC079906.1 | 26.8355342 | 0.262250205 | 0.39904078 | 0.65720151 | 0.51105135 | 0.689262 |
| AMIGO2 | 844.025181 | -0.290255603 | 0.38710157 | -0.7498177 | 0.45336451 | 0.64299587 |
| PCED1B | 375.888471 | -0.134024099 | 0.2652482 | -0.5052781 | 0.61336355 | 0.76841076 |
| AC008083.1 | 0.08666371 | -0.642905668 | 3.30558948 | -0.1944905 | 0.84579185 | NA |
| AC008083.2 | 4.19803279 | -0.32580463 | 0.77433852 | -0.4207522 | 0.67393605 | 0.81019802 |
| AC008083.3 | 0.93893775 | 0.760862003 | 1.47764914 | 0.51491385 | 0.60661322 | NA |
| LINC02416 | 1.0733298 | -0.719995477 | 1.35841434 | -0.5300264 | 0.59609362 | 0.7549479 |
| LINC02156 | 0.08757577 | 0.547239394 | 3.29377202 | 0.16614368 | 0.86804388 | NA |
| RPAP3 | 987.400706 | -0.028777661 | 0.11082949 | -0.2596571 | 0.7951283 | 0.88891824 |
| AC004241.1 | 4008.6013 | -0.84239208 | 0.17022355 | -4.9487401 | 7.47E-07 | 2.54E-05 |
| ENDOU | 77.31214 | -1.037881457 | 0.58918356 | -1.7615588 | 0.07814388 | 0.19903932 |
| AC004241.2 | 19.3195017 | -0.844290128 | 0.51611324 | -1.6358622 | 0.10186844 | 0.24055437 |
| RAPGEF3 | 15471.9626 | -0.719125329 | 0.14734241 | -4.8806405 | 1.06E-06 | 3.41E-05 |
| SLC48A1 | 5509.1375 | -0.659458735 | 0.15578285 | -4.2331921 | 2.30E-05 | 0.00039892 |
| HDAC7 | 10774.7665 | -0.847608713 | 0.2428485 | -3.4902777 | 0.00048252 | 0.0044043 |
| AC004466.1 | 1479.83753 | -0.737904187 | 0.21981403 | -3.3569476 | 0.00078808 | 0.00649225 |
| AC004466.3 | 37.8726266 | -0.072078139 | 0.33835359 | -0.2130261 | 0.83130662 | 0.91040231 |
| LINC02354 | 6.93451781 | -0.671640274 | 0.5200941 | -1.2913822 | 0.19657117 | 0.37715934 |
| VDR | 2127.67195 | 0.918004628 | 0.3793697 | 2.41981539 | 0.01552839 | 0.06239275 |
| AC121338.1 | 35.9190708 | 2.688827652 | 0.68332103 | 3.9349406 | 8.32E-05 | 0.00109832 |
| AC121338.2 | 77.2099295 | 1.403764796 | 0.32780891 | 4.2822655 | 1.85E-05 | 0.00033437 |
| TMEM106C | 3880.25958 | -0.259483061 | 0.27164416 | -0.9552315 | 0.33946063 | 0.53763225 |
| COL2A1 | 4.85641262 | 0.429680144 | 0.73034015 | 0.5883288 | 0.55631162 | 0.7253727 |
| AC004801.5 | 0 | NA | NA | NA | NA | NA |
| AC004801.4 | 0 | NA | NA | NA | NA | NA |
| AC004801.3 | 0 | NA | NA | NA | NA | NA |
| AC004801.6 | 13.9754414 | -0.594274375 | 0.36491354 | -1.6285347 | 0.10341155 | 0.2431098 |
| SENP1 | 561.334687 | 0.000148942 | 0.15885691 | 0.00093759 | 0.99925191 | 0.99973988 |
| PFKM | 4999.38393 | 0.496636216 | 0.19060946 | 2.60551713 | 0.00917357 | 0.04223237 |
| ASB8 | 1832.20338 | 0.504478679 | 0.17736965 | 2.84422206 | 0.004452 | 0.02469484 |
| AC074029.3 | 4.405882 | 1.6740157 | 0.6236184 | 2.68435905 | 0.0072669 | 0.03559877 |
| CCDC184 | 56.3298178 | -0.45366388 | 0.46157162 | -0.9828678 | 0.32567252 | 0.52330658 |
| AC024257.3 | 42.2145803 | 0.451156692 | 0.3860747 | 1.16857358 | 0.24257548 | 0.43301964 |
| OR10AD1 | 4.78322622 | 0.812297941 | 0.78284878 | 1.03761795 | 0.29944796 | 0.49613236 |
| AC024257.1 | 0 | NA | NA | NA | NA | NA |
| AC024257.5 | 3.03148388 | 0.341649805 | 1.2493561 | 0.27346071 | 0.78449909 | 0.88222274 |
| H1FNT | 7.68661342 | 0.517196322 | 0.68146261 | 0.7589504 | 0.44788223 | 0.63796545 |
| ZNF641 | 1217.48857 | 0.461432874 | 0.09899463 | 4.66119087 | 3.14E-06 | 8.12E-05 |
| AC090115.1 | 4.48973017 | 0.440887812 | 0.65621739 | 0.67186243 | 0.50167128 | 0.68226259 |

| | | | | | | |
|---|---|---|---|---|---|---|
| AC024257.4 | 9.05446461 | 0.451899937 | 0.6078797 | 0.74340357 | 0.45723739 | 0.64655504 |
| ANP32D | 1.08449772 | -0.19969915 | 1.2420017 | -0.1607881 | 0.87226026 | 0.93338892 |
| C12orf54 | 9.25289708 | 0.707094563 | 0.48959573 | 1.44424169 | 0.148671 | 0.31268578 |
| AC089987.2 | 0.17466293 | 0.18657164 | 3.29377195 | 0.05664376 | 0.95482897 | NA |
| OR8S1 | 0.17150785 | 0.18657164 | 3.29377195 | 0.05664376 | 0.95482897 | NA |
| OR5BS1P | 0 | NA | NA | NA | NA | NA |
| LALBA | 0.03392908 | 0.380740593 | 3.29738709 | 0.11546736 | 0.90807469 | NA |
| KANSL2 | 1558.3839 | -0.115138738 | 0.13571516 | -0.8483852 | 0.39622346 | 0.5912411 |
| CCNT1 | 1509.47513 | -0.190487912 | 0.12010206 | -1.5860503 | 0.1127279 | 0.25835519 |
| TEX49 | 1.19750389 | -3.454205208 | 1.41855036 | -2.4350247 | 0.01489077 | 0.06052245 |
| AC117498.1 | 2060.21357 | -0.250197526 | 0.25921899 | -0.9651975 | 0.3344459 | 0.53253652 |
| ADCY6 | 5055.36132 | 0.135726933 | 0.22082644 | 0.61463171 | 0.53879797 | 0.71207851 |
| AC117498.2 | 19.5761676 | 0.83335661 | 0.34320192 | 2.42818168 | 0.01517474 | 0.06134745 |
| CACNB3 | 1020.11334 | -0.787538644 | 0.24546284 | -3.2083824 | 0.00133484 | 0.00987923 |
| DDX23 | 4352.72893 | -0.289919254 | 0.13634671 | -2.1263384 | 0.03347509 | 0.10912822 |
| RND1 | 352.850865 | -1.52007026 | 0.66028475 | -2.3021435 | 0.02132708 | 0.07861268 |
| AC073610.3 | 2590.35311 | 0.069295518 | 0.14762987 | 0.46938685 | 0.63879315 | 0.78622989 |
| CCDC65 | 263.880276 | -0.277788674 | 0.19316946 | -1.4380569 | 0.15041791 | 0.31500319 |
| FKBP11 | 1516.8188 | -1.199420908 | 0.26342234 | -4.5532239 | 5.28E-06 | 0.00012253 |
| AC073610.2 | 1767.96224 | -0.290450813 | 0.16258637 | -1.7864401 | 0.07402802 | 0.19200541 |
| ARF3 | 12000.5499 | -0.171028793 | 0.10893235 | -1.570046 | 0.11640441 | 0.26455092 |
| WNT10B | 28.8425907 | -0.573385322 | 0.68565141 | -0.8362636 | 0.40300662 | 0.59801276 |
| WNT1 | 4.64331385 | 0.2036789 | 0.81650733 | 0.2494514 | 0.80301163 | 0.89318485 |
| DDN | 1080.39993 | 1.680752669 | 0.95319184 | 1.76328899 | 0.07785176 | 0.19865986 |
| AC011603.3 | 163.989755 | 1.478278492 | 1.2060814 | 1.22568717 | 0.22031642 | 0.40616456 |
| DDN-AS1 | 478.871356 | -0.122415049 | 0.11187465 | -1.0942162 | 0.27386018 | 0.46888822 |
| PRKAG1 | 3156.7724 | 0.003311624 | 0.11652697 | 0.02841938 | 0.97732767 | 0.98956115 |
| KMT2D | 4602.77876 | 0.065529283 | 0.18975327 | 0.34533941 | 0.72983922 | 0.8473873 |
| RHEBL1 | 52.7833992 | -0.353240796 | 0.24842023 | -1.4219486 | 0.15504118 | 0.32116543 |
| DHH | 32.1085391 | 0.294885819 | 0.62544649 | 0.47148049 | 0.63729765 | 0.78534225 |
| AC011603.1 | 9.69231016 | 0.177669799 | 0.70232163 | 0.25297498 | 0.80028755 | 0.89162767 |
| LMBR1L | 3025.04556 | -0.597318155 | 0.18205229 | -3.2810252 | 0.0010343 | 0.00805988 |
| AC011603.2 | 16907.2611 | -0.800028353 | 0.37885208 | -2.1117169 | 0.03471073 | 0.11196974 |
| TUBA1B | 19566.0623 | -0.782632622 | 0.37839875 | -2.0682749 | 0.03861418 | 0.12067504 |
| TUBA1A | 4970.0221 | -0.513095368 | 0.41086059 | -1.2488308 | 0.21172696 | 0.39532959 |
| TUBA1C | 10083.4823 | -1.072096628 | 0.44549344 | -2.4065374 | 0.01610455 | 0.06416031 |
| AC010173.1 | 25.5297178 | 0.265666204 | 0.6249284 | 0.42511463 | 0.67075312 | 0.80773402 |
| AC125611.3 | 6467.89071 | -1.187246851 | 0.45093434 | -2.6328597 | 0.00846693 | 0.03971416 |
| AC125611.4 | 30.2415953 | -0.316093628 | 0.48941022 | -0.6458664 | 0.51836587 | 0.69530036 |
| PRPH | 41.1873808 | -0.256784651 | 0.51151715 | -0.5020059 | 0.61566334 | 0.77018414 |
| TROAP | 72.3940078 | -1.516836578 | 0.63212939 | -2.3995666 | 0.01641449 | 0.06498276 |
| C1QL4 | 16.7011977 | -0.128162957 | 0.58979869 | -0.2172995 | 0.82797495 | 0.90837491 |
| DNAJC22 | 1239.19476 | 1.315732081 | 0.3453148 | 3.8102395 | 0.00013883 | 0.00164171 |
| SPATS2 | 1223.42069 | -0.546709203 | 0.14433376 | -3.7878123 | 0.00015198 | 0.001771 |
| KCNH3 | 262.926458 | 0.66573097 | 0.40672897 | 1.6367926 | 0.10167381 | 0.24028844 |
| MCRS1 | 3498.89651 | 0.050980975 | 0.14062233 | 0.36253826 | 0.71694983 | 0.839049 |
| PRPF40B | 3257.35983 | -0.136703994 | 0.22704036 | -0.6021132 | 0.54709879 | 0.71822077 |
| AC020612.3 | 5.79478805 | 2.623745111 | 0.61017971 | 4.29995468 | 1.71E-05 | 0.00031349 |
| FAM186B | 55.266725 | 1.006796894 | 0.25377161 | 3.9673346 | 7.27E-05 | 0.00098438 |
| FMNL3 | 3618.76959 | -0.123904845 | 0.20871117 | -0.5936666 | 0.55273514 | 0.72263885 |
| TMBIM6 | 75084.7156 | 0.503054609 | 0.25953409 | 1.93829875 | 0.05258678 | 0.1508582 |
| NCKAP5L | 3198.15821 | -0.466697959 | 0.24655005 | -1.8929137 | 0.05836935 | 0.16231148 |
| BCDIN3D-AS | 282.015994 | 0.032831985 | 0.23254324 | 0.14118658 | 0.88772254 | 0.94201397 |
| BCDIN3D | 427.93278 | -0.10274835 | 0.2449462 | -0.4194731 | 0.67487039 | 0.81076684 |
| AC131157.1 | 0 | NA | NA | NA | NA | NA |
| FAIM2 | 168.960695 | 0.214683602 | 0.45942469 | 0.46728791 | 0.6402939 | 0.78715852 |
| LINC02396 | 0.38225813 | 0.87876065 | 2.7346317 | 0.32134516 | 0.74794884 | NA |
| LINC02395 | 1.5912552 | -1.607357382 | 2.30081008 | -0.698605 | 0.48479893 | 0.6691144 |
| AQP2 | 52565.086 | 0.891306081 | 0.43738142 | 2.03782339 | 0.0415676 | 0.12742822 |
| AC025154.2 | 4172.86732 | 1.217525013 | 0.39456215 | 3.08576234 | 0.00203031 | 0.01365798 |
| AC025154.1 | 9073.4934 | 1.202850398 | 0.41656692 | 2.88753219 | 0.00388277 | 0.02228261 |
| AQP5 | 117.949595 | 0.271080952 | 0.47676407 | 0.56858512 | 0.56963773 | 0.73500743 |
| AQP6 | 3310.24736 | 0.327399061 | 0.5327918 | 0.61449719 | 0.53888683 | 0.71212457 |
| RACGAP1 | 818.529146 | -0.25243676 | 0.27334027 | -0.9235257 | 0.35573332 | 0.55319445 |
| ASIC1 | 202.717022 | 0.910922682 | 0.38502558 | 2.36587576 | 0.01798748 | 0.06935661 |
| SMARCD1 | 3660.86605 | -0.16544166 | 0.15490917 | -1.0679914 | 0.28552439 | 0.4816272 |
| GPD1 | 9781.63766 | 2.144462217 | 0.49271753 | 4.35231567 | 1.35E-05 | 0.00025981 |
| COX14 | 2402.41609 | -0.088164095 | 0.18058674 | -0.4882091 | 0.62540172 | 0.77689106 |
| AC074032.1 | 1071.99863 | 0.051651862 | 0.18332052 | 0.28175712 | 0.77812976 | 0.87857682 |
| CERS5 | 2363.21651 | -0.506410293 | 0.15573181 | -3.2518102 | 0.00114673 | 0.00873964 |
| LIMA1 | 4884.38133 | -0.137992345 | 0.1741783 | -0.7922476 | 0.42821631 | 0.62061358 |
| AC008147.1 | 2.41311709 | 0.048518432 | 0.72262447 | 0.06714197 | 0.94646868 | 0.97432773 |
| FAM186A | 60.6329573 | 0.427090748 | 0.39953258 | 1.06897602 | 0.28508047 | 0.48111504 |
| LARP4 | 3654.11649 | -0.020269863 | 0.1778394 | -0.1139785 | 0.90925486 | 0.9548045 |
| DIP2B | 1542.28107 | -0.302109118 | 0.23782726 | -1.270288 | 0.20398205 | 0.38628415 |
| ATF1 | 1038.7005 | 0.115720913 | 0.21991644 | 0.52620401 | 0.59874647 | 0.75677743 |
| TMPRSS12 | 8.05214908 | 0.454314146 | 0.61219818 | 0.74210307 | 0.45802489 | 0.64706542 |
| METTL7A | 10607.4354 | 1.422247138 | 0.35784129 | 3.97451933 | 7.05E-05 | 0.00096007 |
| AC008121.2 | 14.181611 | -0.769190171 | 0.46156486 | -1.6664834 | 0.09561717 | 0.23045652 |
| HIGD1C | 4.48134277 | -0.109920225 | 0.59511691 | -0.1847036 | 0.85346154 | 0.9232762 |
| AC008121.3 | 4.36872255 | -1.316207419 | 0.66243349 | -1.9869276 | 0.04693042 | 0.13909802 |
| SLC11A2 | 4951.20147 | -0.796734547 | 0.22640177 | -3.519118 | 0.00043298 | 0.00405605 |
| LETMD1 | 3952.2802 | 0.331214562 | 0.12907011 | 2.56616014 | 0.01028313 | 0.04601498 |
| CSRNP2 | 1950.19355 | -0.36676671 | 0.15544699 | -2.3594326 | 0.0183029 | 0.07033628 |
| TFCP2 | 1640.94545 | 0.265293367 | 0.13586785 | 1.9525839 | 0.05086892 | 0.14741169 |
| POU6F1 | 1250.80955 | 0.004024536 | 0.18451422 | 0.02181152 | 0.9825983 | 0.99230688 |
| DAZAP2 | 21655.0821 | -0.076832382 | 0.18882068 | -0.4069066 | 0.68407659 | 0.81752646 |
| SMAGP | 1192.15103 | -0.589849249 | 0.2121241 | -2.78068 | 0.00542452 | 0.0286398 |
| BIN2 | 382.839809 | -0.346465103 | 0.45002012 | -0.769888 | 0.44136631 | 0.63264824 |
| CELA1 | 1.01390338 | -1.269463286 | 2.05194503 | -0.6186634 | 0.53613813 | 0.70988757 |
| GALNT6 | 456.969726 | -0.371001708 | 0.35293578 | -1.0511876 | 0.29317245 | 0.48970402 |
| SLC4A8 | 437.695954 | 0.874025294 | 0.49599202 | 1.76217612 | 0.07803955 | 0.1988504 |
| AC107031.1 | 1.14683518 | -1.445159996 | 1.37198494 | -1.0533352 | 0.2921874 | 0.48839895 |
| SCN8A | 391.890452 | 0.792695455 | 0.35712086 | 2.21968397 | 0.02644023 | 0.09206931 |
| AC068987.2 | 56.3889378 | 0.661793074 | 0.30268584 | 2.1864025 | 0.02878617 | 0.09797255 |
| AC068987.5 | 14.7456238 | 0.544587584 | 0.38706139 | 1.40697989 | 0.15943335 | 0.32718855 |
| AC068987.3 | 76.6954613 | 0.163852841 | 0.23917479 | 0.68507571 | 0.49329614 | 0.67596234 |
| FIGNL2 | 1225.75789 | 0.212733755 | 0.19428535 | 1.09495518 | 0.27353627 | 0.46853858 |
| AC068987.4 | 924.036468 | 0.146208723 | 0.19187653 | 0.76199377 | 0.44606372 | 0.63647532 |
| AC068987.1 | 65.0716224 | 0.766490247 | 0.39140221 | 1.95831865 | 0.05019263 | 0.14602601 |
| ANKRD33 | 35.4578326 | -2.182561548 | 0.51671196 | -4.2239423 | 2.40E-05 | 0.00041241 |
| ACVRL1 | 2352.73807 | -0.495128093 | 0.3057503 | -1.6193871 | 0.105364 | 0.24626137 |
| ACVR1B | 7496.18031 | -0.125052892 | 0.15584425 | -0.8024222 | 0.42230877 | 0.61475475 |
| GRASP | 1344.58318 | -0.183602323 | 0.27815695 | -0.6600674 | 0.5092106 | 0.68800961 |
| NR4A1 | 12055.3695 | 1.873136724 | 0.63934627 | 2.92976874 | 0.00339214 | 0.02012845 |
| AC025259.3 | 3502.86146 | 1.491351059 | 0.63372931 | 2.3532935 | 0.01860794 | 0.07124452 |
| ATG101 | 2229.29423 | -0.634972158 | 0.25905677 | -2.4510927 | 0.01424233 | 0.05863702 |
| AC025259.1 | 1277.25134 | -0.736252329 | 0.25473712 | -2.8902436 | 0.00384943 | 0.02213396 |
| AC078864.2 | 53.5150176 | -0.715026276 | 0.37103023 | -1.9271375 | 0.0539625 | 0.15377037 |
| AC078864.1 | 147.981187 | -0.573729781 | 0.44988922 | -1.275269 | 0.20221404 | 0.38401466 |
| LINC00592 | 0.80365575 | -0.872240833 | 1.8864118 | -0.4623809 | 0.64380818 | NA |
| KRT80 | 469.841665 | -1.145282734 | 0.40810483 | -2.8063445 | 0.00501071 | 0.02692184 |
| C12orf80 | 0.27421255 | 1.100476972 | 2.98484026 | 0.36868873 | 0.71235975 | NA |

| | | | | | | |
|---|---|---|---|---|---|---|
| KRT7 | 9857.99436 | -1.382984149 | 0.4265696 | -3.2421067 | 0.0011865 | 0.00897273 |
| KRT7-AS | 3673.30324 | -1.416927836 | 0.43629538 | -3.2476343 | 0.00116369 | 0.00883826 |
| KRT86 | 23.8442397 | -0.088186737 | 0.46513654 | -0.1895932 | 0.8496279 | 0.92098029 |
| AC021066.1 | 0.03392908 | 0.380740593 | 3.29738709 | 0.11546736 | 0.90807469 | NA |
| KRT81 | 3.10989308 | 0.051259411 | 0.68476147 | 0.07485732 | 0.94032823 | 0.97153373 |
| AC121757.1 | 5.15478833 | 0.325541329 | 0.53510815 | 0.60836548 | 0.5429451 | 0.71530087 |
| KRT83 | 0.10178723 | 1.087385526 | 3.28420134 | 0.33109588 | 0.74057207 | NA |
| KRT85 | 13.5444107 | 1.429139166 | 0.89531416 | 1.59624323 | 0.1104345 | 0.25451475 |
| KRT84 | 0.21442573 | -1.003574173 | 3.30558948 | -0.3035992 | 0.7614333 | NA |
| AC078865.1 | 0 | NA | NA | NA | NA | NA |
| KRT82 | 0 | NA | NA | NA | NA | NA |
| AC055736.1 | 0 | NA | NA | NA | NA | NA |
| KRT75 | 0.09997307 | 0.547239394 | 3.29377202 | 0.16614368 | 0.86804388 | NA |
| KRT6B | 1.90002825 | 4.870836143 | 3.15792795 | 1.54241522 | NA | NA |
| KRT6C | 0 | NA | NA | NA | NA | NA |
| KRT6A | 5.56065812 | 3.372431192 | 1.72234568 | 1.95804549 | NA | NA |
| KRT5 | 12.2346843 | 3.119098123 | 1.01214203 | 3.08168026 | 0.00205836 | 0.01377665 |
| KRT71 | 0.61282717 | -1.730578315 | 2.24447948 | -0.7710377 | 0.44068458 | NA |
| KRT74 | 0.4431024 | -2.042642583 | 3.29029438 | -0.6208085 | 0.53472566 | NA |
| KRT72 | 10.4283518 | -0.020101241 | 0.78048888 | -0.0257547 | 0.97945301 | 0.99069759 |
| KRT73-AS1 | 1.67776319 | 0.853904753 | 1.01454365 | 0.84166389 | 0.39997612 | 0.5947877 |
| KRT73 | 4.11849905 | 0.882155191 | 0.94392462 | 0.93456106 | 0.35001456 | 0.54770258 |
| KRT2 | 5.92602784 | 0.89185276 | 1.03131874 | 0.86476927 | 0.38716544 | 0.58277028 |
| KRT1 | 3.49811311 | 1.250212919 | 1.20551714 | 1.03707602 | 0.29970043 | 0.49636354 |
| KRT77 | 4.49098161 | 5.332596579 | 1.96146131 | 2.71868557 | NA | NA |
| AC055716.3 | 0.57679429 | 3.175255401 | 3.18820343 | 0.99593877 | 0.3192799 | NA |
| KRT76 | 0 | NA | NA | NA | NA | NA |
| KRT3 | 0 | NA | NA | NA | NA | NA |
| KRT4 | 1.53934904 | 3.191981107 | 1.3305126 | 2.39906117 | 0.01643717 | 0.065053 |
| KRT79 | 0.71472202 | 0.502261574 | 1.7800322 | 0.28216432 | 0.77781752 | NA |
| KRT78 | 0.38371765 | 0.997977569 | 1.87624253 | 0.53190222 | 0.59479372 | NA |
| KRT8 | 32757.1004 | -0.931080182 | 0.27462703 | -3.3903443 | 0.00069805 | 0.0058923 |
| KRT18 | 20194.111 | -1.353229311 | 0.26352042 | -5.1351972 | 2.82E-07 | 1.13E-05 |
| EIF4B | 16889.0072 | 0.27472872 | 0.24638684 | 1.11503003 | 0.26483755 | 0.45883645 |
| AC068888.1 | 4557.16304 | 0.357919712 | 0.22737131 | 1.5741639 | 0.11544956 | 0.26290198 |
| TNS2 | 15126.2508 | -0.097306756 | 0.22275523 | -0.4368326 | 0.66223272 | 0.80128673 |
| SPRYD3 | 4131.97587 | -0.233877777 | 0.13054259 | -1.7915822 | 0.07319991 | 0.19036309 |
| IGFBP6 | 1482.00909 | -0.46364889 | 0.42446936 | -1.0923024 | 0.27470022 | 0.4699523 |
| SOAT2 | 68.777872 | 1.16052834 | 0.66387936 | 1.74810126 | 0.0804465 | 0.20315783 |
| CSAD | 2971.4343 | 0.244108573 | 0.1951711 | 1.2507414 | 0.21102884 | 0.39438912 |
| AC073573.1 | 836.545059 | 0.253729667 | 0.19807268 | 1.28099273 | 0.20019622 | 0.38160867 |
| ZNF740 | 2615.03592 | 0.214139862 | 0.10275292 | 2.08402704 | 0.03715771 | 0.11729097 |
| ITGB7 | 827.424159 | 0.596860087 | 0.31894568 | 1.87135343 | 0.06129611 | 0.16790053 |
| RARG | 1183.56362 | -0.010333189 | 0.30947383 | -0.0333895 | 0.97336395 | 0.98766731 |
| AC021072.1 | 0.5462191 | 1.026381184 | 2.21675669 | 0.4630103 | 0.64335698 | NA |
| MFSD5 | 1850.58232 | -0.68890654 | 0.25317844 | -2.7210316 | 0.00650785 | 0.03271918 |
| ESPL1 | 562.238013 | 2.928326957 | 0.61929479 | 4.72848634 | 2.26E-06 | 6.28E-05 |
| PFDN5 | 10194.6236 | -0.250449906 | 0.1017435 | -2.4615814 | 0.0138326 | 0.0573531 |
| AC073611.1 | 690.495533 | 0.134562221 | 0.12657489 | 1.06310363 | 0.28773496 | 0.48393206 |
| C12orf10 | 2073.21838 | 0.081147418 | 0.13135861 | 0.61775484 | 0.53673695 | 0.71052979 |
| AAAS | 3374.143 | 0.093207867 | 0.19341925 | 0.48189549 | 0.62988019 | 0.78011215 |
| SP7 | 0.95203744 | 1.251575334 | 1.47977549 | 0.84578731 | 0.3976714 | NA |
| SP1 | 5008.79351 | -0.286892534 | 0.16747118 | -1.713086 | 0.08669672 | 0.21450482 |
| AMHR2 | 6.07644289 | -0.111117352 | 1.09835165 | -0.1011674 | 0.9194176 | 0.96074349 |
| PRR13 | 11746.847 | -0.002461485 | 0.25019541 | -0.0098382 | 0.99215034 | 0.99623046 |
| PCBP2 | 29949.4757 | 0.174188465 | 0.12171431 | 1.43112558 | 0.15239423 | 0.3175783 |
| MAP3K12 | 5564.25848 | 0.028180951 | 0.10028132 | 0.28101895 | 0.77869587 | 0.87865683 |
| AC023509.4 | 93.7407099 | 0.175927299 | 0.24068757 | 0.73093636 | 0.46481803 | 0.65205955 |
| TARBP2 | 1664.4706 | 0.005239271 | 0.12406896 | 0.0422287 | 0.96631638 | 0.98410057 |
| AC023509.3 | 709.567083 | -0.430223242 | 0.18997607 | -2.2646181 | 0.02353613 | 0.08448783 |
| NPFF | 326.624145 | -0.594126132 | 0.26710361 | -2.2243283 | 0.02612636 | 0.09125866 |
| ATF7 | 2038.21453 | -0.050160113 | 0.10996533 | -0.4561448 | 0.64828586 | 0.79291736 |
| AC023509.2 | 234.866632 | 0.95074083 | 0.22269856 | 4.26918265 | 1.96E-05 | 0.00034918 |
| ATP5MC2 | 9638.84179 | 0.083000635 | 0.1023336 | 0.81107902 | 0.41732029 | 0.6102566 |
| CALCOCO1 | 7727.47194 | 0.105315052 | 0.22171534 | 0.4750012 | 0.63478612 | 0.78341547 |
| CISTR | 1.11071694 | 1.659224841 | 1.4468605 | 1.14677596 | 0.25147423 | 0.44374835 |
| AC076968.1 | 0.19271054 | 1.506623853 | 3.27645372 | 0.4598337 | 0.64563559 | NA |
| HOXC13-AS | 1.4787478 | -0.492122505 | 0.98364912 | -0.5003029 | 0.61686182 | 0.77089456 |
| HOXC13 | 3.12791803 | -1.354768113 | 0.88929402 | -1.5234198 | 0.12765371 | 0.28141016 |
| HOXC12 | 1.48222285 | -0.62484538 | 1.26184268 | -0.4951849 | 0.62046964 | 0.77327778 |
| HOTAIR | 42.9383441 | 0.384690204 | 0.55912414 | 0.68802288 | 0.49143837 | 0.67443299 |
| HOXC11 | 83.820684 | -0.482153181 | 0.4106911 | -1.1740045 | 0.24039324 | 0.43053313 |
| HOXC10 | 2146.09005 | 0.41569721 | 0.20451177 | 2.03263222 | 0.04208969 | 0.12856541 |
| AC012531.3 | 505.332266 | 0.338309999 | 0.22730843 | 1.48833016 | 0.13666384 | 0.29498449 |
| HOXC6 | 1330.66767 | -0.201292608 | 0.24509512 | -0.8212836 | 0.41148473 | 0.60567365 |
| HOXC9 | 434.951148 | -0.200309899 | 0.16889489 | -1.1860033 | 0.235621 | 0.42458151 |
| HOXC-AS1 | 114.245947 | 0.083971411 | 0.21262141 | 0.39493393 | 0.69289165 | 0.8232818 |
| HOXC8 | 295.615103 | -0.362591417 | 0.31442412 | -1.1531921 | 0.24883154 | 0.44058871 |
| HOXC4 | 1330.02239 | 0.413394146 | 0.3345096 | 1.23582146 | 0.21652494 | 0.40139957 |
| AC012531.1 | 20.6006796 | 0.045211004 | 0.27984523 | 0.16155717 | 0.87165459 | 0.93317628 |
| HOXC5 | 157.586778 | 0.119070032 | 0.34788203 | 0.34227129 | 0.73214674 | 0.84898243 |
| AC023794.1 | 211.43577 | 0.550601711 | 0.4045086 | 1.36116196 | 0.17346251 | 0.3472147 |
| AC023794.3 | 50.9211031 | -0.204058504 | 0.41869378 | -0.4873693 | 0.62599663 | 0.77729476 |
| AC023794.4 | 97.1023337 | 0.636932067 | 0.41691246 | 1.52773574 | 0.12657816 | 0.27988463 |
| SMUG1 | 1630.77428 | 0.027704748 | 0.14777651 | 0.18747735 | 0.85128638 | 0.92202004 |
| LINC02381 | 2213.82947 | -0.223667987 | 0.29390359 | -0.761025 | 0.44664212 | 0.63699041 |
| SMUG1-AS1 | 51.0682074 | 0.349698954 | 0.44338993 | 0.78869395 | 0.4302909 | 0.62235471 |
| AC023794.7 | 47.6003985 | 0.285256308 | 0.41367502 | 0.6895662 | 0.49046703 | 0.67369536 |
| AC023794.6 | 1.76689807 | -0.054341101 | 1.01671455 | -0.0534477 | 0.95737516 | 0.97951358 |
| AC023794.2 | 4.74005425 | 0.012767784 | 0.69179261 | 0.01845609 | 0.98527501 | 0.99348815 |
| CBX5 | 4520.6057 | 0.112963552 | 0.17314377 | 0.6524263 | 0.5141262 | 0.69199204 |
| AC078778.1 | 91.4195218 | -0.85249527 | 0.33900762 | -2.5146788 | 0.01191409 | 0.05130217 |
| HNRNPA1 | 30047.7927 | -0.225690428 | 0.18233457 | -1.2377819 | 0.21579694 | 0.40040567 |
| NFE2 | 124.440407 | -2.456988809 | 0.40142782 | -6.1206241 | 9.32E-10 | 9.72E-08 |
| COPZ1 | 6233.88037 | -0.148271203 | 0.09272927 | -1.5989687 | 0.10982756 | 0.25360748 |
| AC079313.2 | 4.6024998 | -1.042665123 | 1.02202391 | -1.0201964 | 0.30763532 | 0.5044676 |
| AC079313.1 | 63.860236 | -0.267137885 | 0.52626049 | -0.5076153 | 0.61172315 | 0.76749276 |
| GPR84 | 80.6618458 | -1.806593398 | 0.62879564 | -2.873101 | 0.00406464 | 0.02303559 |
| ZNF385A | 1119.04532 | -0.977678142 | 0.33309703 | -2.9351152 | 0.00333424 | 0.019877 |
| ITGA5 | 4344.20172 | -1.310246266 | 0.51545338 | -2.5419297 | 0.01102423 | 0.04844705 |
| LINC01154 | 2.74402141 | -0.800385779 | 1.21300515 | -0.6598371 | 0.50935838 | 0.68813877 |
| GTSF1 | 8.14105131 | -0.850482728 | 0.92153768 | -0.9228952 | 0.35606181 | 0.55349113 |
| NCKAP1L | 605.454504 | -0.359750326 | 0.47131258 | -0.7632946 | 0.44528775 | 0.63581536 |
| AC068789.1 | 12.1660431 | 0.699219298 | 0.71816653 | 0.97361722 | 0.33024663 | 0.52810934 |
| PDE1B | 495.178076 | -0.907667493 | 0.42240881 | -2.1487892 | 0.03165111 | 0.10486743 |
| PPP1R1A | 10909.8773 | -0.544763712 | 0.44159254 | -1.2336343 | 0.2173392 | 0.40230483 |
| LACRT | 0.11223674 | 0.02007176 | 3.29738704 | 0.00608717 | 0.99514317 | NA |
| DCD | 1.26752969 | 1.674574515 | 1.43875696 | 1.16390368 | 0.24446305 | 0.435456 |
| AC079310.1 | 91.7267861 | 2.922670756 | 1.39958029 | 2.08824801 | 0.03677547 | 0.11654478 |
| MUCL1 | 2.53914911 | 0.069871633 | 0.86615117 | 0.0806691 | 0.93570511 | 0.96922554 |
| TESPA1 | 63.7573098 | 0.5439177 | 0.72252399 | 0.75280227 | 0.45156874 | 0.64131127 |
| AC027287.2 | 0.09997307 | 0.547239394 | 3.29377202 | 0.16614368 | 0.86804388 | NA |
| NEUROD4 | 0.46398281 | -0.756972994 | 2.87116932 | -0.2636462 | 0.79205252 | NA |

| | | | | | | |
|---|---|---|---|---|---|---|
| OR9K2 | 0 | NA | NA | NA | NA | NA |
| OR10A7 | 0 | NA | NA | NA | NA | NA |
| OR6C74 | 0.09635527 | 0.547239394 | 3.29377202 | 0.16614368 | 0.86804388 | NA |
| OR6C6 | 0 | NA | NA | NA | NA | NA |
| OR6C1 | 0 | NA | NA | NA | NA | NA |
| OR6C3 | 0 | NA | NA | NA | NA | NA |
| OR6C75 | 0 | NA | NA | NA | NA | NA |
| OR6C65 | 0 | NA | NA | NA | NA | NA |
| OR6C76 | 0 | NA | NA | NA | NA | NA |
| AC122685.1 | 0.46127146 | -0.952361494 | 2.76886933 | -0.3439532 | 0.73088147 | NA |
| OR6C2 | 0.23492299 | -1.364242679 | 3.30558948 | -0.4127078 | 0.67982068 | NA |
| OR6C70 | 0 | NA | NA | NA | NA | NA |
| OR6C68 | 0.28674021 | -1.364242679 | 3.30558948 | -0.4127078 | 0.67982068 | NA |
| OR6C4 | 0.08695548 | -0.642905668 | 3.30558948 | -0.1944905 | 0.84579185 | NA |
| OR2AP1 | 0 | NA | NA | NA | NA | NA |
| OR10P1 | 0 | NA | NA | NA | NA | NA |
| AC009779.5 | 0 | NA | NA | NA | NA | NA |
| METTL7B | 1452.02123 | 0.955151714 | 0.36826893 | 2.59362558 | 0.00949698 | 0.04335795 |
| ITGA7 | 3195.05197 | 0.005950429 | 0.45096269 | 0.01319495 | 0.98947226 | 0.99519138 |
| AC009779.3 | 3757.28736 | 0.325034169 | 0.19331526 | 1.68136837 | 0.09269138 | 0.22528843 |
| BLOC1S1 | 2867.92214 | 0.096180021 | 0.12705554 | 0.75699196 | 0.44905468 | 0.63923742 |
| RDH5 | 1242.08647 | 0.806928818 | 0.36894399 | 2.18713092 | 0.02873297 | 0.09785462 |
| CD63 | 33349.7068 | -1.023098165 | 0.29053396 | -3.5214409 | 0.00042921 | 0.00402927 |
| AC009779.2 | 602.407622 | -0.18068182 | 0.20738056 | -0.8712573 | 0.38361369 | 0.57937532 |
| GDF11 | 732.097294 | -0.243034951 | 0.232419 | -1.0456759 | 0.29571069 | 0.49263319 |
| SARNP | 2278.84719 | -0.651162415 | 0.12764069 | -5.101527 | 3.37E-07 | 1.31E-05 |
| AC023055.1 | 4004.46917 | -0.534259297 | 0.13013273 | -4.1054952 | 4.03E-05 | 0.00062656 |
| ORMDL2 | 2283.5558 | -0.721336924 | 0.19376054 | -3.7228267 | 0.000197 | 0.00215382 |
| DNAJC14 | 2467.84737 | 0.101469409 | 0.14693188 | 0.6905881 | 0.48982443 | 0.67311398 |
| MMP19 | 1451.1421 | 0.173449493 | 0.39623558 | 0.43774336 | 0.66157234 | 0.80084282 |
| PYM1 | 1095.23056 | -0.352870539 | 0.12985469 | -2.7174262 | 0.00657918 | 0.03297093 |
| DGKA | 1084.83889 | -0.578076565 | 0.4577901 | -1.2627546 | 0.20667737 | 0.38929725 |
| PMEL | 573.120098 | -0.301897027 | 0.15637616 | -1.9305821 | 0.05353475 | 0.15288189 |
| CDK2 | 1727.70085 | -1.182071669 | 0.22765989 | -5.1922701 | 2.08E-07 | 8.79E-06 |
| AC025162.1 | 264.889491 | -0.89162332 | 0.25173942 | -3.5418503 | 0.00039733 | 0.00380175 |
| RAB5B | 8675.36421 | 0.002850915 | 0.09535682 | 0.02989734 | 0.97614893 | 0.98894663 |
| SUOX | 3994.59975 | 0.72872792 | 0.37957551 | 1.91984969 | 0.05487689 | 0.15553563 |
| IKZF4 | 636.38665 | 0.20937864 | 0.16686235 | 1.25479857 | 0.20955189 | 0.39265852 |
| AC034102.8 | 1.7618078 | 0.488954366 | 0.95489235 | 0.51205182 | 0.60861474 | 0.76468319 |
| AC034102.3 | 1.04953771 | 2.496218778 | 1.29388211 | 1.92924746 | 0.05370015 | 0.15328782 |
| RPS26 | 7507.37519 | -0.718771024 | 0.22147249 | -3.245419 | 0.00117278 | 0.00889706 |
| ERBB3 | 5815.38681 | 0.54346852 | 0.31045609 | 1.75054875 | 0.08002367 | 0.20249728 |
| AC034102.2 | 2095.41696 | -0.027181074 | 0.17852818 | -0.1522509 | 0.87898906 | 0.93705294 |
| PA2G4 | 5287.67006 | -0.326940372 | 0.17656331 | -1.8516892 | 0.06407046 | 0.17309068 |
| AC034102.4 | 1208.84223 | -0.168307938 | 0.20828128 | -0.80808 | 0.41904453 | 0.61189458 |
| RPL41 | 16734.0834 | -0.474429792 | 0.14439793 | -3.285572 | 0.00101776 | 0.00795206 |
| ZC3H10 | 573.042467 | 0.192752525 | 0.15618834 | 1.2341032 | 0.21716445 | 0.40203781 |
| ESYT1 | 6022.37496 | -0.552430308 | 0.16897523 | -3.2692976 | 0.00107815 | 0.00831122 |
| AC034102.7 | 36.0059381 | 0.381597092 | 0.30177254 | 1.26451892 | 0.20604383 | 0.38870557 |
| AC034102.6 | 393.582415 | 0.442333076 | 0.19013006 | 2.32647629 | 0.01999316 | 0.07489978 |
| AC034102.5 | 379.242011 | 0.11184847 | 0.16148689 | 0.69261639 | 0.48855032 | 0.67197329 |
| MYL6B | 21615.9037 | -0.385334461 | 0.21722863 | -1.7738659 | 0.07608533 | 0.19572433 |
| MYL6 | 45781.4435 | -0.532367367 | 0.21338477 | -2.4948705 | 0.01260031 | 0.05350342 |
| SMARCC2 | 5162.36985 | 0.301759473 | 0.13168059 | 2.29160181 | 0.02192863 | 0.08031361 |
| RNF41 | 2078.74886 | -0.117066152 | 0.1474751 | -0.7938028 | 0.42731022 | 0.61960687 |
| NABP2 | 1226.76155 | 0.056820084 | 0.16317625 | 0.34821296 | 0.72768025 | 0.84611931 |
| SLC39A5 | 10018.2444 | 2.006846985 | 0.47183698 | 4.25326349 | 2.11E-05 | 0.00036914 |
| ANKRD52 | 5341.15737 | -0.182389418 | 0.15276749 | -1.193902 | 0.2325163 | 0.42059644 |
| COQ10A | 983.626606 | 0.730976761 | 0.16022134 | 4.56229337 | 5.06E-06 | 0.00011784 |
| AC073896.5 | 156.392163 | 1.257667319 | 0.19171049 | 6.56024258 | 5.37E-11 | 8.64E-09 |
| CS | 10062.6948 | 0.128363294 | 0.1762902 | 0.72813629 | 0.46653016 | 0.65372232 |
| AC073896.1 | 4602.28569 | -0.104670263 | 0.12888581 | -0.8121163 | 0.41672493 | 0.60986792 |
| AC073896.2 | 758.703609 | 0.494339159 | 0.20838003 | 2.37229622 | 0.01767791 | 0.06852367 |
| AC073896.3 | 36.3550021 | 0.498141718 | 0.25953926 | 1.91933089 | 0.05494247 | 0.15558773 |
| CNPY2 | 4211.02213 | -0.291467089 | 0.11635516 | -2.5049779 | 0.01224591 | 0.05234634 |
| PAN2 | 3466.55869 | 0.603627518 | 0.1790923 | 3.37048275 | 0.00075037 | 0.00624668 |
| IL23A | 70.8139631 | -0.97847086 | 0.43938306 | -2.2269198 | 0.02595263 | 0.09084971 |
| STAT2 | 4900.39915 | -0.198791303 | 0.16231312 | -1.2247396 | 0.22067336 | 0.40657154 |
| APOF | 2.09430735 | 0.608193365 | 1.12756998 | 0.53938414 | 0.58962182 | 0.74967828 |
| AC024884.2 | 4.78820598 | -0.449166546 | 0.76748585 | -0.5852441 | 0.55838361 | 0.72699595 |
| TIMELESS | 913.132732 | -0.275854813 | 0.21148313 | -1.3043821 | 0.19210333 | 0.37130871 |
| MIP | 301.930502 | 0.570431199 | 0.14731733 | 3.87212567 | 0.00010789 | 0.00134582 |
| SPRYD4 | 1840.27281 | 0.341485359 | 0.12175581 | 2.80467415 | 0.00503675 | 0.02700751 |
| GLS2 | 384.224777 | 0.352881196 | 0.17912891 | 1.96998465 | 0.04884013 | 0.14334048 |
| RBMS2 | 3487.06525 | -0.416144012 | 0.22786738 | -1.8262553 | 0.06781181 | 0.18000947 |
| BAZ2A | 5884.68836 | -0.405398606 | 0.18532116 | -2.1875463 | 0.02870267 | 0.09778933 |
| ATP5F1B | 52414.0416 | 0.41981252 | 0.19492064 | 2.15376121 | 0.0312589 | 0.10397292 |
| PTGES3 | 12640.2815 | -0.422274428 | 0.10003449 | -4.2212884 | 2.43E-05 | 0.00041576 |
| NACA | 21245.9096 | -0.197571325 | 0.16457661 | -1.2004824 | 0.22995203 | 0.41753322 |
| PRIM1 | 127.053576 | 0.389146805 | 0.22756853 | 1.71002027 | 0.08726212 | 0.21541867 |
| HSD17B6 | 92.6870338 | 0.396065498 | 0.38136977 | 1.03853408 | 0.29902148 | 0.49579957 |
| SDR9C7 | 3.571892 | -0.581676399 | 0.75565337 | -0.7697662 | 0.44143861 | 0.63269844 |
| RDH16 | 55.3866025 | -0.093022667 | 0.304196 | -0.3057985 | 0.7597581 | 0.86698524 |
| GPR182 | 157.337752 | -0.339200012 | 0.47525542 | -0.7137215 | 0.47539941 | 0.66084778 |
| ZBTB39 | 294.422437 | 0.188557429 | 0.13156643 | 1.43317277 | 0.15180847 | 0.31683335 |
| TAC3 | 2.58698135 | -1.686516971 | 1.11811743 | -1.5083541 | 0.13146393 | 0.28726416 |
| MYO1A | 5.24713302 | 0.012201614 | 0.53267487 | 0.02290631 | 0.98172501 | 0.99170821 |
| NEMP1 | 651.89659 | -0.009330572 | 0.1499803 | -0.062212 | 0.95039402 | 0.9761964 |
| NAB2 | 2665.87897 | -0.601952165 | 0.20393504 | -2.9516858 | 0.00316044 | 0.01912198 |
| STAT6 | 16949.3741 | -0.313050494 | 0.13447307 | -2.3279791 | 0.01991321 | 0.0746958 |
| LRP1 | 7105.48627 | 0.13395681 | 0.3238015 | 0.4137004 | 0.67909353 | 0.81383823 |
| LRP1-AS | 7.97660704 | 0.446936923 | 0.48743572 | 0.91691459 | 0.3591874 | 0.55686823 |
| NXPH4 | 39.7104147 | 0.174180947 | 0.43509117 | 0.40033207 | 0.68891195 | 0.82053334 |
| SHMT2 | 4215.99128 | 0.024093008 | 0.16173919 | 0.14896209 | 0.88158355 | 0.93857409 |
| AC137834.2 | 343.101915 | 0.486905308 | 0.20908616 | 2.32873047 | 0.01987335 | 0.0745887 |
| NDUFA4L2 | 942.7481 | -1.050534988 | 0.5054653 | -2.0783524 | 0.03767692 | 0.11844782 |
| STAC3 | 238.776926 | -0.438598928 | 0.34950574 | -1.2549119 | 0.20951073 | 0.39264177 |
| AC137834.1 | 1667.51579 | 0.741716279 | 0.18737059 | 3.95855231 | 7.54E-05 | 0.00101348 |
| R3HDM2 | 3250.42907 | 0.463535863 | 0.17793692 | 2.6050573 | 0.00918589 | 0.04228171 |
| AC126614.1 | 1.54866517 | 1.095517433 | 1.08231326 | 1.01219996 | 0.31144246 | 0.50856013 |
| INHBC | 16.070169 | 1.833455144 | 0.45869524 | 3.99710957 | 6.41E-05 | 0.00089682 |
| INHBE | 54.6231254 | -1.192528406 | 0.69585303 | -1.7137648 | 0.08657193 | 0.21437712 |
| GLI1 | 157.115715 | 0.734403675 | 0.51158558 | 1.43554414 | 0.1511321 | 0.31597189 |
| ARHGAP9 | 761.818522 | -0.903156132 | 0.51043764 | -1.769376 | 0.07683114 | 0.19685902 |
| MARS | 6687.59009 | -0.635967422 | 0.2097666 | -3.0317859 | 0.00243112 | 0.0156515 |
| DDIT3 | 3758.1011 | -1.293014873 | 0.43110371 | -2.9993128 | 0.00270589 | 0.01697702 |
| AC022506.1 | 726.380914 | -1.27494428 | 0.47827515 | -2.665713 | 0.00768252 | 0.03706104 |
| MBD6 | 8144.10196 | -0.271785213 | 0.2318462 | -1.1722651 | 0.24109063 | 0.43122567 |
| DCTN2 | 7598.36425 | -0.444979793 | 0.14772278 | -3.0122626 | 0.00259308 | 0.0164374 |
| KIF5A | 81.5416716 | -0.230279051 | 0.30284939 | -0.7603748 | 0.44703056 | 0.63728592 |
| PIP4K2C | 3970.27449 | -0.017468417 | 0.1729889 | -0.10098 | 0.91956635 | 0.96074349 |
| DTX3 | 3196.58834 | -0.285070782 | 0.24258376 | -1.1751437 | 0.23993721 | 0.42986802 |
| ARHGEF25 | 1116.42558 | -0.003097058 | 0.4207984 | -0.00736 | 0.99412766 | 0.99728027 |

| | | | | | | |
|---|---|---|---|---|---|---|
| AC025165.1 | 660.473223 | 0.097201354 | 0.41620005 | 0.23354479 | 0.81533839 | 0.90046796 |
| SLC26A10 | 291.078921 | -0.16873889 | 0.386176 | -0.4369482 | 0.66214895 | 0.80128673 |
| B4GALNT1 | 291.462573 | -0.331031542 | 0.40428321 | -0.818811 | 0.41289426 | 0.60677784 |
| OS9 | 16477.2171 | 5.73E-05 | 0.11827684 | 0.00048404 | 0.99961379 | 0.99979217 |
| AC025165.2 | 6239.11222 | -0.180218021 | 0.13276704 | -1.3574003 | 0.17465404 | 0.34859424 |
| AGAP2 | 351.164537 | -0.625690552 | 0.32653696 | -1.91614 | 0.05534727 | 0.156415 |
| AGAP2-AS1 | 233.966039 | -0.816543575 | 0.2820963 | -2.8945562 | 0.00379695 | 0.02193983 |
| TSPAN31 | 4822.47155 | -0.303232413 | 0.12213796 | -2.4827042 | 0.01303893 | 0.05481745 |
| CDK4 | 5030.54628 | -0.387269852 | 0.22329147 | -1.7343692 | 0.08285259 | 0.20765672 |
| 9-Mar | 2287.3004 | -0.36295647 | 0.14801157 | -2.4522169 | 0.0141979 | 0.05849612 |
| CYP27B1 | 1896.05378 | 1.386083857 | 0.54137246 | 2.56031467 | 0.01045774 | 0.04662226 |
| METTL1 | 1204.37233 | 0.174176289 | 0.23148771 | 0.75242132 | 0.45179773 | 0.64156737 |
| EEF1AKMT3 | 1099.4693 | 0.235652685 | 0.2266031 | 1.03993583 | 0.29836971 | 0.49530098 |
| AC025165.3 | 927.566542 | 0.061234987 | 0.16010736 | 0.38246204 | 0.70211868 | 0.82947179 |
| TSFM | 2391.6789 | -0.107761874 | 0.14030077 | -0.7680776 | 0.44244109 | 0.63356891 |
| AVIL | 723.591882 | 0.201166397 | 0.23294458 | 0.86358049 | 0.38781838 | 0.58342009 |
| AC025165.5 | 47.4867684 | 0.591488919 | 0.28454579 | 2.07871261 | 0.03764377 | 0.11839778 |
| AC025165.4 | 14.1095431 | -0.212502685 | 0.35498823 | -0.598619 | 0.54942701 | 0.71992001 |
| CTDSP2 | 10628.8062 | -0.092255025 | 0.16493317 | -0.5593479 | 0.57592429 | 0.73996899 |
| AC083805.2 | 2.13372345 | 2.059801192 | 0.84275765 | 2.44412044 | 0.01452058 | 0.05945771 |
| AC083805.1 | 33.2597115 | -0.118927275 | 0.55706633 | -0.2134885 | 0.83094593 | 0.91040231 |
| AC083805.3 | 26.3803978 | -0.232929421 | 0.56586578 | -0.4116337 | 0.68060793 | 0.81480144 |
| AC084033.3 | 210.11137 | -0.014443552 | 0.19030584 | -0.0758965 | 0.93950142 | 0.97110795 |
| ATP23 | 217.218195 | 0.776427784 | 0.20963566 | 3.70370099 | 0.00021248 | 0.00229256 |
| LINC02403 | 0 | NA | NA | NA | NA | NA |
| AC020637.1 | 0 | NA | NA | NA | NA | NA |
| LINC02388 | 1.39085815 | 0.544948167 | 1.83008084 | 0.29777273 | 0.76587663 | 0.87089982 |
| LRIG3 | 755.57531 | 0.549799144 | 0.26969119 | 2.03862481 | 0.04148749 | 0.12724181 |
| AC068305.2 | 3.55886043 | 1.613547878 | 0.77212429 | 2.08975148 | 0.03664013 | 0.11626327 |
| AC108721.1 | 0 | NA | NA | NA | NA | NA |
| LINC02448 | 0.14114194 | 0.02007176 | 3.29738704 | 0.00608717 | 0.99514317 | NA |
| SLC16A7 | 1877.06749 | 0.524810675 | 0.28130546 | 1.86562566 | 0.06209376 | 0.16939516 |
| AC080011.1 | 0 | NA | NA | NA | NA | NA |
| FAM19A2 | 63.3717315 | 0.526160362 | 0.46215819 | 1.13848541 | 0.25491785 | 0.44760844 |
| USP15 | 2275.36664 | -0.259260223 | 0.17923026 | -1.4465204 | 0.14803129 | 0.31163861 |
| MON2 | 2455.91936 | 0.15313873 | 0.09260908 | 1.65360377 | 0.09820805 | 0.23469819 |
| AC079035.1 | 1.63937822 | 0.375998709 | 1.05852617 | 0.35520965 | 0.72243254 | 0.84246598 |
| LINC01465 | 60.2604059 | -1.391036994 | 0.52770465 | -2.6360143 | 0.00838862 | 0.039431 |
| AC048341.1 | 883.960281 | -0.963768009 | 0.35810856 | -2.6912733 | 0.00711799 | 0.03506469 |
| AC048341.2 | 384.40436 | -1.49683146 | 0.33242044 | -4.5028262 | 6.71E-06 | 0.00014935 |
| PPM1H | 1809.26815 | 0.202382118 | 0.24894659 | 0.81295395 | 0.41624446 | 0.60949339 |
| AC078814.1 | 0 | NA | NA | NA | NA | NA |
| AVPR1A | 457.561721 | -0.179512156 | 0.53602149 | -0.3348973 | 0.73770255 | 0.85272503 |
| AC026116.1 | 0.37516828 | -0.558553231 | 2.2465623 | -0.2486257 | 0.80365029 | NA |
| DPY19L2 | 1797.06927 | 0.22155693 | 0.26397864 | 0.83929871 | 0.40130171 | 0.59616409 |
| AC084357.2 | 3.86442202 | 0.142722653 | 0.89634589 | 0.15922721 | 0.87348987 | 0.9341361 |
| RXYLT1 | 1046.47687 | -0.001950085 | 0.17785189 | -0.0109647 | 0.99125164 | 0.99583588 |
| RXYLT1-AS1 | 224.964097 | -0.108372014 | 0.18430045 | -0.5880182 | 0.5565201 | 0.72557054 |
| SRGAP1 | 1027.19769 | -0.181673916 | 0.12597366 | -1.4421579 | 0.14925783 | 0.31340282 |
| AC020611.2 | 10.1083089 | -0.807650206 | 0.53606084 | -1.5066391 | 0.13190319 | 0.28800921 |
| AC025576.2 | 2.5945226 | -0.803732282 | 0.94918565 | -0.8467598 | 0.397129 | 0.59205629 |
| AC025576.1 | 288.773148 | -0.359269828 | 0.14664347 | -2.4499544 | 0.01428743 | 0.0587585 |
| C12orf66 | 421.792357 | 0.185834099 | 0.19582267 | 0.94899175 | 0.34262481 | 0.54030715 |
| C12orf56 | 69.9434407 | 0.690648943 | 0.33373314 | 2.06946465 | 0.03850251 | 0.12043456 |
| XPOT | 2706.28321 | -0.356794191 | 0.22918476 | -1.5567972 | 0.11951865 | 0.26912363 |
| AC135279.3 | 26.284315 | 0.172460659 | 0.33566417 | 0.5137893 | 0.60739931 | 0.76377473 |
| TBK1 | 1538.69291 | -0.660288476 | 0.19354498 | -3.4115505 | 0.00064595 | 0.00552679 |
| RASSF3 | 974.158778 | -0.245832539 | 0.25116104 | -0.9787845 | 0.32768645 | 0.52542209 |
| AC078962.2 | 0.47800863 | -0.856632517 | 2.29277883 | -0.373622 | 0.70868559 | NA |
| AC078962.4 | 0 | NA | NA | NA | NA | NA |
| AC078962.3 | 0.28559037 | 0.375213434 | 2.95831617 | 0.12683345 | 0.89907222 | NA |
| GNS | 15744.2068 | 0.11676495 | 0.21734148 | 0.5372419 | 0.59110053 | 0.75076202 |
| AC025262.1 | 3.72803241 | 0.075347148 | 0.7107248 | 0.10601452 | 0.91557083 | 0.95882128 |
| TBC1D30 | 491.05516 | -0.60300854 | 0.17960167 | -3.3574774 | 0.00078657 | 0.00648387 |
| LINC02389 | 19.7520205 | 0.374094681 | 0.50171334 | 0.74563431 | 0.45588835 | 0.64526715 |
| LINC02231 | 0.14444843 | 0.435199075 | 3.28898345 | 0.13232024 | 0.894731 | NA |
| AC135895.1 | 0.13390214 | 1.04981649 | 3.28477129 | 0.31960109 | 0.74927075 | NA |
| WIF1 | 24.5837387 | 3.208021884 | 0.69277035 | 4.63071475 | 3.64E-06 | 9.14E-05 |
| LEMD3 | 1189.72525 | 0.088206004 | 0.11877435 | 0.74263511 | 0.45770262 | 0.64686234 |
| MSRB3 | 1062.46416 | 0.415299436 | 0.40388066 | 1.02827266 | 0.30382159 | 0.50070466 |
| AC025419.1 | 26.7728242 | 0.023964107 | 0.46179352 | 0.05189355 | 0.95861351 | 0.98017606 |
| AC090023.1 | 0 | NA | NA | NA | NA | NA |
| LINC02454 | 0.95070244 | 0.070493951 | 1.5203932 | 0.04636561 | 0.96301885 | NA |
| AC090023.2 | 0.51010802 | 0.671326405 | 1.85485773 | 0.36192879 | 0.71740525 | NA |
| HMGA2 | 27.9966835 | -0.656245236 | 0.459185 | -1.4291522 | 0.15296051 | 0.31855698 |
| AC090673.1 | 75.4969992 | 1.158733334 | 0.45533604 | 2.5447872 | 0.01093444 | 0.04813707 |
| AC090673.2 | 0.66277322 | -1.575824435 | 1.66394063 | -0.9470437 | 0.34361653 | NA |
| LINC02425 | 0 | NA | NA | NA | NA | NA |
| LLPH | 1097.42474 | -0.474780381 | 0.21803693 | -2.1775228 | 0.02944158 | 0.09954795 |
| AC078927.1 | 3802.41371 | -0.050823722 | 0.12925715 | -0.3931985 | 0.69417285 | 0.82386304 |
| LLPH-DT | 29.6305876 | -0.213031322 | 0.85413628 | -0.2494114 | 0.80304257 | 0.89318485 |
| TMBIM4 | 3991.74166 | 0.028823587 | 0.17588614 | 0.16387639 | 0.86982845 | 0.93220007 |
| IRAK3 | 897.752493 | -1.270063215 | 0.36679055 | -3.4626388 | 0.00053491 | 0.00478314 |
| AC078889.1 | 120.390137 | -1.076597679 | 0.37613634 | -2.8622538 | 0.0042064 | 0.0236509 |
| HELB | 476.591312 | -0.112219745 | 0.30157084 | -0.3721174 | 0.70980547 | 0.83508389 |
| GRIP1 | 296.406325 | -0.583512762 | 0.18108446 | -3.2223238 | 0.00127155 | 0.00948003 |
| AC073530.1 | 0.51580041 | 0.890548444 | 1.78534275 | 0.49881091 | 0.6179126 | NA |
| CAND1 | 4000.25215 | -0.011934334 | 0.10788282 | -0.1106231 | 0.91191521 | 0.95675993 |
| LINC02420 | 0 | NA | NA | NA | NA | NA |
| LINC02408 | 1.24173155 | 0.242401198 | 1.25237492 | 0.19355322 | 0.84652573 | 0.91937416 |
| LINC02442 | 0.7868244 | 1.245172833 | 1.50758983 | 0.82593608 | 0.40884035 | NA |
| DYRK2 | 1200.53027 | -0.408628882 | 0.12004654 | -3.4039205 | 0.00066426 | 0.00564868 |
| AC078777.1 | 20.0485866 | 0.94709207 | 0.40925848 | 2.31416604 | 0.02065862 | 0.07671704 |
| LINC02421 | 3.43740876 | 0.461267059 | 0.66398319 | 0.69469689 | 0.48724528 | 0.67119354 |
| LINC01479 | 2.20364613 | 0.464050836 | 0.79548489 | 0.58335594 | 0.55965371 | 0.72803849 |
| IFNG-AS1 | 7.3394964 | 0.923497655 | 0.8881725 | 1.03977286 | 0.29844544 | 0.49530098 |
| IFNG | 4.39715378 | 1.567673229 | 1.33791641 | 1.17172734 | 0.24130654 | 0.43141155 |
| IL26 | 0.44837132 | 0.362616802 | 2.80300273 | 0.12936727 | 0.89706705 | NA |
| IL22 | 0.25817155 | -0.178652511 | 2.98297197 | -0.0598908 | 0.95224263 | NA |
| MDM1 | 555.466128 | 0.670787216 | 0.15477665 | 4.33390459 | 1.46E-05 | 0.00027771 |
| LINC02384 | 91.2597166 | 0.795614653 | 0.43118175 | 1.84519558 | 0.06500909 | 0.17480267 |
| AC022511.1 | 0 | NA | NA | NA | NA | NA |
| AC008033.3 | 19.3765361 | 1.87079581 | 0.63295101 | 2.95567237 | 0.00311988 | 0.0189173 |
| RAP1B | 4380.5107 | -0.632379375 | 0.2312802 | -2.7342564 | 0.00625213 | 0.03179943 |
| AC090061.1 | 39.5896745 | 0.483974864 | 0.33991535 | 1.42380999 | 0.15450148 | 0.32055107 |
| NUP107 | 1266.6009 | -0.177669905 | 0.19886283 | -0.8934294 | 0.37162726 | 0.56836599 |
| SLC35E3 | 962.672394 | 0.057129895 | 0.26877506 | 0.21255653 | 0.83167288 | 0.91050001 |
| MDM2 | 4319.65234 | -0.788502276 | 0.14746137 | -5.3471786 | 8.93E-08 | 4.37E-06 |
| AC025423.1 | 0.80051783 | 1.259575829 | 1.33412474 | 0.94412148 | 0.34510757 | NA |
| AC025423.4 | 1110.09538 | -0.659939635 | 0.17468384 | -3.7779088 | 0.00015815 | 0.00182754 |
| CPM | 5257.25913 | -0.347387474 | 0.31962289 | -1.0868667 | 0.27709572 | 0.47251774 |
| AC127894.1 | 0 | NA | NA | NA | NA | NA |
| CPSF6 | 3761.366 | -0.204890842 | 0.1713926 | -1.1954474 | 0.23191227 | 0.4198911 |

| | | | | | | | |
|---|---|---|---|---|---|---|---|
| AC020656.2 | 37.8788863 | | 1.921562276 | 0.62695715 | 3.0649021 | 0.00217741 | 0.01440862 |
| LYZ | 1831.51279 | | 0.64835406 | 0.7958226 | 0.81469672 | 0.41524593 | 0.60880881 |
| AC020656.1 | 859.693407 | | 0.365220597 | 0.78925028 | 0.4627437 | 0.64354809 | 0.78935593 |
| YEATS4 | 557.82838 | | 0.008304593 | 0.24499927 | 0.0338964 | 0.97295977 | 0.98741337 |
| LINC02373 | 3.46002996 | | -0.906941648 | 0.75750237 | -1.197279 | 0.23119785 | 0.4190163 |
| FRS2 | 907.959304 | | 0.043842923 | 0.18372364 | 0.23863518 | 0.81138848 | 0.89817456 |
| CCT2 | 6476.46632 | | -0.41563698 | 0.21848193 | -1.9023861 | 0.0571207 | 0.15983371 |
| LRRC10 | 1.25626384 | | -2.602448872 | 2.30598705 | -1.1285618 | 0.25908274 | 0.45187688 |
| BEST3 | 51.9913346 | | -0.694511077 | 0.70523976 | -0.9847872 | 0.32472862 | 0.52233096 |
| AC025263.1 | 11.371979 | | 1.203699387 | 0.64391377 | 1.86934873 | 0.06157431 | 0.16836527 |
| RAB3IP | 3184.34099 | | 0.233065633 | 0.13136693 | 1.77415752 | 0.0760371 | 0.19563843 |
| AC025263.2 | 519.575831 | | -0.487774912 | 0.18820043 | -2.5917842 | 0.00954796 | 0.04351535 |
| MYRFL | 112.26332 | | 0.023689667 | 0.44046241 | 0.05378363 | 0.95710755 | 0.97945117 |
| AC025159.1 | 276.819509 | | -0.264476001 | 0.26722549 | -0.989711 | 0.3223154 | 0.52009896 |
| AC078922.1 | 3.14525743 | | 0.203064207 | 1.15086823 | 0.17644436 | 0.85994486 | 0.92701929 |
| AC084032.1 | 2.71051107 | | 2.228212256 | 1.01683476 | 2.19132188 | 0.02842851 | 0.09716912 |
| LINC01481 | 5.39339367 | | 0.715841268 | 0.5302916 | 1.34990119 | 0.17704768 | 0.352066 |
| CNOT2 | 2525.85149 | | -0.17224108 | 0.09234494 | -1.8651923 | 0.06215445 | 0.16949462 |
| KCNMB4 | 500.359572 | | 0.066602371 | 0.23207301 | 0.28698887 | 0.77412084 | 0.87574864 |
| AC025569.1 | 872.440125 | | 0.854148017 | 0.36588528 | 2.33446946 | 0.01957115 | 0.07372733 |
| PTPRB | 2906.90261 | | 0.845865254 | 0.35648458 | 2.37279617 | 0.017654 | 0.06844106 |
| AC083809.1 | 17.2423708 | | 1.991603842 | 0.3868678 | 5.14802176 | 2.63E-07 | 1.06E-05 |
| PTPRR | 12.8285015 | | 1.690690058 | 0.47905498 | 3.52921926 | 0.00041679 | 0.0039379 |
| AC123905.1 | 0.21786011 | | 1.725828273 | 3.04405575 | 0.56695029 | 0.57074796 | NA |
| AC025575.1 | 0.09997307 | | 0.547239394 | 3.29377202 | 0.16614368 | 0.86804388 | NA |
| AC025575.2 | 2.06967166 | | 0.728186493 | 0.77477559 | 0.93986762 | 0.34728547 | 0.54478988 |
| TSPAN8 | 2240.72751 | | -0.490323018 | 0.40594603 | -1.2078527 | 0.22710392 | 0.413958 |
| LGR5 | 43.0825046 | | 2.765206147 | 0.97243052 | 2.84360281 | 0.00446066 | 0.02472727 |
| AC078860.1 | 3.63631825 | | 2.520190284 | 1.32858011 | 1.89690503 | 0.05784047 | 0.16131711 |
| ZFC3H1 | 2205.41548 | | -0.388639064 | 0.1406537 | -2.7630917 | 0.00572567 | 0.02980964 |
| THAP2 | 397.980465 | | 0.385059542 | 0.18166525 | 2.11961035 | 0.03403892 | 0.11042857 |
| AC073612.1 | 514.569414 | | 1.283592745 | 0.28881848 | 4.44428883 | 8.82E-06 | 0.00018455 |
| TMEM19 | 1881.26428 | | 1.204136178 | 0.22546036 | 5.34078888 | 9.25E-08 | 4.48E-06 |
| AC011601.1 | 0.49946182 | | -2.196059689 | 3.28386579 | -0.6687422 | 0.50365993 | NA |
| RAB21 | 4789.5451 | | -0.048550335 | 0.12801921 | -0.3792426 | 0.70450774 | 0.83125502 |
| AC089984.1 | 174.897015 | | 0.426306729 | 0.30356692 | 1.40432538 | 0.16022198 | 0.32835907 |
| TBC1D15 | 2910.35364 | | -0.522754807 | 0.20692038 | -2.5263573 | 0.01152522 | 0.05010222 |
| TPH2 | 1.01732941 | | -1.291120906 | 1.89587811 | -0.6810147 | 0.49586218 | 0.67772047 |
| AC090109.1 | 1.54563802 | | -1.926724257 | 2.82927291 | -0.6809963 | 0.49587386 | 0.67772047 |
| TRHDE | 916.526599 | | 1.029288077 | 0.23154797 | 4.44524777 | 8.78E-06 | 0.00018402 |
| AC133480.1 | 0 | NA | | NA | NA | NA | NA |
| LINC02444 | 0 | NA | | NA | NA | NA | NA |
| AC090503.1 | 0 | NA | | NA | NA | NA | NA |
| LINC02445 | 0.07659028 | | -0.642905668 | 3.30558948 | -0.1944905 | 0.84579185 | NA |
| AC136188.1 | 0 | NA | | NA | NA | NA | NA |
| LINC02394 | 0 | NA | | NA | NA | NA | NA |
| AC090502.1 | 3.85723889 | | -1.114938493 | 1.54711585 | -0.7206561 | 0.47112112 | 0.65760449 |
| AC090502.3 | 0.18754884 | | 1.469068384 | 3.27783996 | 0.44818185 | 0.65402196 | NA |
| ATXN7L3B | 4890.1848 | | 0.477316599 | 0.13204923 | 3.61468666 | 0.00030071 | 0.00303822 |
| AC025257.1 | 441.21936 | | 1.497411856 | 0.32731056 | 4.5748963 | 4.76E-06 | 0.00011329 |
| AC073525.1 | 0.12150485 | | 1.04981648 | 3.28477129 | 0.31960109 | 0.74927075 | NA |
| KCNC2 | 0.03392908 | | 0.380740593 | 3.29738709 | 0.11546736 | 0.90807469 | NA |
| AC091534.1 | 0.33725126 | | 0.669857205 | 2.22872422 | 0.30055634 | 0.76375283 | NA |
| CAPS2 | 302.510995 | | 0.584013609 | 0.32102637 | 1.8192076 | 0.06887976 | 0.18201818 |
| GLIPR1L1 | 38.8148719 | | 0.478848531 | 0.27462162 | 1.74366656 | 0.08121725 | 0.20454547 |
| GLIPR1L2 | 70.0351284 | | 1.534507183 | 0.59519436 | 2.57816149 | 0.00993276 | 0.0448436 |
| GLIPR1 | 1328.52829 | | -0.221000833 | 0.23846032 | -0.9267824 | 0.35403951 | 0.55164393 |
| AC121761.1 | 118.608661 | | -0.601801932 | 0.67146726 | -0.8962491 | 0.37011973 | 0.56691783 |
| KRR1 | 1989.08964 | | 0.013330854 | 0.13326534 | 0.10003242 | 0.92031859 | 0.96099929 |
| AC078923.1 | 5.86446953 | | 0.956890594 | 0.79129262 | 1.20927527 | 0.22655711 | 0.41336131 |
| AC078820.1 | 1.91952188 | | 1.335588738 | 1.21903015 | 1.09561583 | 0.27324693 | 0.46819503 |
| AC011611.2 | 0.03392908 | | 0.380740593 | 3.29738709 | 0.11546736 | 0.90807469 | NA |
| PHLDA1 | 2769.28464 | | -0.215531106 | 0.35815378 | -0.6017837 | 0.54731813 | 0.71840613 |
| AC011611.3 | 767.125211 | | -0.156351205 | 0.43304136 | -0.3610538 | 0.71805927 | 0.83985952 |
| AC011611.4 | 20.1621105 | | 1.301633336 | 0.4110293 | 3.16676531 | 0.00154145 | 0.01103964 |
| NAP1L1 | 9459.52228 | | -0.450018531 | 0.20321235 | -2.2145235 | 0.02679279 | 0.09292086 |
| LINC02407 | 6.20987781 | | 2.197918835 | 1.20325593 | 1.82664284 | 0.06775349 | 0.17994354 |
| BBS10 | 702.417011 | | 0.730474582 | 0.1083019 | 6.74479958 | 1.53E-11 | 2.91E-09 |
| OSBPL8 | 2678.31771 | | 0.607104588 | 0.2145423 | 2.82976644 | 0.0046582 | 0.02551214 |
| AC107032.2 | 7.61035136 | | 1.028657898 | 0.47895641 | 2.14770673 | 0.03173706 | 0.10505031 |
| ZDHHC17 | 1797.7923 | | 0.182313285 | 0.13226601 | 1.37838344 | 0.16808493 | 0.33941341 |
| CSRP2 | 2295.64122 | | -0.235720256 | 0.32009132 | -0.7364156 | 0.46147779 | 0.64992359 |
| AC124784.1 | 1.31340357 | | 1.139259511 | 1.0061926 | 1.13224795 | 0.2575302 | 0.4501572 |
| E2F7 | 17.4417637 | | -0.683881885 | 0.60804452 | -1.1247234 | 0.26070627 | 0.45387366 |
| LINC02464 | 0.17515154 | | 1.45420457 | 3.2783972 | 0.44357181 | 0.65735219 | NA |
| NAV3 | 103.391395 | | -1.253033471 | 0.53443868 | -2.3445786 | 0.0190486 | 0.07240804 |
| AC073591.1 | 0.3216386 | | -1.364242679 | 3.30558948 | -0.4127078 | 0.67982068 | NA |
| AC138331.1 | 0.4926956 | | -0.327436422 | 2.2653007 | -0.1445444 | 0.88507064 | NA |
| AC073571.1 | 0.12088456 | | 0.02007176 | 3.29738704 | 0.00608717 | 0.99514317 | NA |
| LINC02424 | 0.39627446 | | 2.574957812 | 3.21049141 | 0.80204476 | 0.42252707 | NA |
| AC128707.1 | 15.4923234 | | 3.720689817 | 0.85539071 | 4.34969633 | 1.36E-05 | 0.0002624 |
| AC130415.1 | 0.58933999 | | 3.144576678 | 1.89354037 | 1.66068637 | 0.09677645 | NA |
| AC079362.1 | 0 | NA | | NA | NA | NA | NA |
| AC068993.2 | 0.69430933 | | 2.801604572 | 1.84325992 | 1.51991835 | 0.1285315 | NA |
| SYT1 | 162.678926 | | 0.677486423 | 0.49422792 | 1.37079757 | 0.17043807 | 0.34251272 |
| AC090709.1 | 25.6933005 | | 1.956266235 | 0.62334332 | 3.13834476 | 0.00169905 | 0.01189064 |
| AC027288.3 | 231.179861 | | 3.638299654 | 0.73688774 | 4.93738662 | 7.92E-07 | 2.67E-05 |
| AC027288.1 | 7.52241526 | | 0.948387573 | 0.58842831 | 1.61173003 | 0.1070207 | 0.24893145 |
| PAWR | 3298.84079 | | -0.103428666 | 0.19150814 | -0.5400745 | 0.58914564 | 0.74950649 |
| AC073569.2 | 112.604926 | | -0.071805313 | 0.3551398 | -0.2021889 | 0.83976908 | 0.91484357 |
| PPP1R12A | 2172.5209 | | 0.02941499 | 0.12970964 | 0.22677566 | 0.82059819 | 0.90366551 |
| AC073569.1 | 1.50520124 | | 0.23463042 | 1.25386388 | 0.18712591 | 0.85156191 | 0.92224271 |
| OTOGL | 371.299609 | | 3.512209893 | 0.8476382 | 4.14352479 | 3.42E-05 | 0.00054593 |
| PTPRQ | 435.449787 | | 0.554483912 | 0.75698844 | 0.73248663 | 0.4638716 | 0.65155396 |
| AC074031.1 | 1.82726205 | | 0.740987312 | 1.2646623 | 0.58591713 | 0.5579312 | 0.72652906 |
| MYF6 | 5.56623478 | | 0.510314027 | 0.70495455 | 0.72389635 | 0.46912937 | 0.65576154 |
| MYF5 | 0.75003833 | | 1.633314411 | 1.63822889 | 0.99700012 | 0.31876445 | NA |
| LINC01490 | 0 | NA | | NA | NA | NA | NA |
| LIN7A | 1014.00678 | | 0.263862076 | 0.24274768 | 1.08698085 | 0.27704527 | 0.47246225 |
| ACSS3 | 2943.34501 | | 0.953223772 | 0.27393579 | 3.47973436 | 0.00050191 | 0.004542 |
| AC078955.1 | 0.47097593 | | 0.296078276 | 1.92893743 | 0.15349294 | 0.87800956 | NA |
| PPFIA2 | 167.459476 | | 0.502340418 | 0.43551186 | 1.1534483 | 0.24872643 | 0.44046175 |
| PPFIA2-AS1 | 6.59793905 | | 0.76064688 | 0.48827514 | 1.55782429 | 0.11927491 | 0.26878134 |
| AC069228.1 | 0.23560867 | | 0.952737031 | 3.28103697 | 0.2903768 | 0.77152799 | NA |
| LINC02426 | 0 | NA | | NA | NA | NA | NA |
| CCDC59 | 700.687606 | | -0.327978268 | 0.12904981 | -2.5414859 | 0.01103824 | 0.0484878 |
| METTL25 | 382.289976 | | 0.17638739 | 0.16492092 | 1.06952707 | 0.28483224 | 0.48095192 |
| AC089998.4 | 0.41451299 | | -0.668280548 | 3.28467738 | -0.2034539 | 0.83878025 | NA |
| AC089998.2 | 0.17423948 | | 0.18657164 | 3.29377195 | 0.05664376 | 0.95482897 | NA |
| AC089998.1 | 12.1611808 | | 1.613167712 | 0.68924301 | 2.34049196 | 0.01925835 | 0.07288214 |
| TMTC2 | 1092.28616 | | 0.114527157 | 0.37948264 | 0.30179815 | 0.76280594 | 0.86868167 |
| AC090680.1 | 0 | NA | | NA | NA | NA | NA |
| AC093025.1 | 0 | NA | | NA | NA | NA | NA |

| | | | | | | |
|---|---|---|---|---|---|---|
| AC090679.2 | 0 | NA | NA | NA | NA | NA |
| AC090679.1 | 0 | NA | NA | NA | NA | NA |
| SLC6A15 | 10.57509 | -0.551621807 | 0.60085871 | -0.9180558 | 0.35858967 | 0.55626858 |
| TSPAN19 | 10.9642631 | -0.1026084 | 0.49501941 | -0.2072816 | 0.83578997 | 0.91280681 |
| LRRIQ1 | 105.614888 | 0.290802125 | 0.45467446 | 0.63958316 | 0.52244363 | 0.69833935 |
| AC133794.1 | 28.1248613 | 0.080124942 | 0.44945928 | 0.17826963 | 0.85851123 | 0.92607016 |
| ALX1 | 78.145669 | 0.717641552 | 0.45236393 | 1.58642525 | 0.11264288 | 0.25832775 |
| AC131532.1 | 0.42451639 | 1.401902864 | 2.14576484 | 0.65333481 | 0.51354046 | NA |
| RASSF9 | 210.5668 | 0.380561313 | 0.30076167 | 1.26532516 | 0.20575479 | 0.38833051 |
| NTS | 29.5567315 | -0.138813235 | 0.74466886 | -0.1864093 | 0.85212376 | 0.92251225 |
| MGAT4C | 14.6228779 | 1.747255909 | 0.44406095 | 3.93472093 | 8.33E-05 | 0.00109877 |
| AC016993.1 | 0.48475234 | 2.23618348 | 1.99461361 | 1.12111111 | 0.26224057 | NA |
| AC010196.1 | 0.35958232 | 0.710895303 | 2.31841065 | 0.30663045 | 0.75912467 | NA |
| LINC02258 | 0 | NA | NA | NA | NA | NA |
| AC079598.1 | 0.65445655 | 2.211423412 | 1.61286631 | 1.37111389 | 0.17033945 | NA |
| C12orf50 | 2.73007358 | 0.263320275 | 0.69356734 | 0.37966072 | 0.70419728 | 0.83108138 |
| C12orf29 | 1059.82696 | -0.213303403 | 0.17149645 | -1.2437773 | 0.21358152 | 0.39755147 |
| CEP290 | 910.736593 | -0.092072613 | 0.14405164 | -0.6391639 | 0.5227163 | 0.69859758 |
| TMTC3 | 1141.06086 | -0.067176811 | 0.12719243 | -0.528151 | 0.59739453 | 0.75595249 |
| KITLG | 2688.73875 | -0.514936338 | 0.34469537 | -1.4938882 | 0.13520485 | 0.29264974 |
| AC024941.2 | 16.3250617 | 1.02760797 | 0.67498597 | 1.52241383 | 0.12790542 | 0.28184724 |
| AC024941.1 | 2.42259714 | 0.952517744 | 1.13386694 | 0.84006131 | 0.40087401 | 0.59568622 |
| LINC02458 | 6.05973047 | -1.587522742 | 0.76458702 | -2.076314 | 0.03786492 | 0.11894538 |
| AC024909.1 | 0.65612486 | 1.448899666 | 1.69819098 | 0.85320184 | 0.39354738 | NA |
| DUSP6 | 5116.20214 | -0.978216432 | 0.28457868 | -3.4374199 | 0.00058728 | 0.00513996 |
| AC024909.2 | 992.380104 | -0.371288058 | 0.30051076 | -1.2355233 | 0.2166358 | 0.40151012 |
| AC010201.2 | 129.555271 | 0.234966015 | 0.42976241 | 0.54673468 | 0.58456102 | 0.74648301 |
| AC010201.1 | 45.1221583 | 0.327028357 | 0.46344725 | 0.7056431 | 0.48041011 | 0.66514831 |
| POC1B | 1195.1029 | -0.110634407 | 0.13430444 | -0.8237584 | 0.41007686 | 0.6044565 |
| GALNT4 | 653.437466 | -0.004557133 | 0.15753428 | -0.0289279 | 0.97692211 | 0.98937845 |
| POC1B-GALN | 734.575517 | 0.040622816 | 0.14655605 | 0.27718279 | 0.78163975 | 0.88039639 |
| POC1B-AS1 | 159.825686 | 0.068911295 | 0.19549172 | 0.35250237 | 0.72446154 | 0.84387647 |
| AC025034.1 | 69.7475877 | 0.45944028 | 0.32758677 | 1.40249948 | 0.16076614 | 0.32917177 |
| ATP2B1 | 1711.67859 | 0.078193284 | 0.18469878 | 0.42335572 | 0.67203575 | 0.80865082 |
| ATP2B1-AS1 | 103.905793 | 0.986985201 | 0.37052991 | 2.66371259 | 0.00772836 | 0.03723675 |
| AC084200.1 | 0.23749721 | 0.689160045 | 2.99954805 | 0.22975463 | 0.81828244 | NA |
| LINC02399 | 0 | NA | NA | NA | NA | NA |
| AC126178.1 | 0.94597637 | -0.453310533 | 1.43278315 | -0.3163846 | 0.7517106 | NA |
| LINC02392 | 0.14114194 | 0.02007176 | 3.29738704 | 0.00608717 | 0.99514317 | NA |
| AC084365.1 | 0 | NA | NA | NA | NA | NA |
| AC112481.1 | 0 | NA | NA | NA | NA | NA |
| CCER1 | 0 | NA | NA | NA | NA | NA |
| LINC00615 | 0.29479552 | -0.701265545 | 3.29738693 | -0.2126731 | 0.83158194 | NA |
| EPYC | 0.33489606 | -0.534763868 | 3.29377181 | -0.1623561 | 0.87102546 | NA |
| KERA | 0.17332742 | -1.003574173 | 3.30558948 | -0.3035992 | 0.7614333 | NA |
| LUM | 8613.68203 | -0.017651406 | 0.49704406 | -0.0355128 | 0.97167087 | 0.98670705 |
| DCN | 16337.3265 | 0.674269596 | 0.38463683 | 1.75300322 | 0.07960145 | 0.20177103 |
| AC025035.1 | 0.98456616 | 0.154071689 | 1.36901091 | 0.11254234 | 0.9103934 | NA |
| AC126175.1 | 3.92226495 | 0.186769862 | 0.75892891 | 0.24609665 | 0.80560742 | 0.89470466 |
| AC025254.1 | 0 | NA | NA | NA | NA | NA |
| LINC02404 | 1.39681073 | 3.084811881 | 1.35304389 | 2.27990526 | 0.02261331 | 0.08211415 |
| AC090049.1 | 0 | NA | NA | NA | NA | NA |
| BTG1 | 9637.92808 | -0.95323223 | 0.17320001 | -5.50365 | 3.72E-08 | 2.09E-06 |
| AC025164.1 | 553.861184 | -0.602771189 | 0.20285146 | -2.9714905 | 0.00296358 | 0.01816614 |
| LINC02391 | 12.4723564 | 0.825746855 | 0.48162143 | 1.71451435 | 0.0864343 | 0.21407651 |
| CLLU1OS | 0.9925964 | -1.685506055 | 1.46892119 | -1.1474448 | 0.25119782 | NA |
| AC063949.2 | 14.5423999 | 0.045869991 | 0.62875141 | 0.0729541 | 0.94184264 | 0.97233647 |
| LINC02397 | 23.0703738 | 0.27876451 | 0.64177968 | 0.4343617 | 0.66402581 | 0.80263845 |
| C12orf74 | 8.50870644 | -0.013425431 | 0.80860827 | -0.0166031 | 0.98675322 | 0.99400325 |
| PLEKHG7 | 66.9035864 | -0.192853662 | 0.40847389 | -0.4721322 | 0.63683246 | 0.78491572 |
| EEA1 | 1192.8418 | -0.173960505 | 0.17606624 | -0.9880401 | 0.32313302 | 0.52085294 |
| LINC02413 | 14.705691 | 1.034570368 | 0.52792224 | 1.95970217 | 0.05003061 | 0.14571984 |
| AC138123.1 | 70.4613014 | 1.050162559 | 0.32759788 | 3.2056452 | 0.0013476 | 0.00994965 |
| AC138123.2 | 2.16068114 | 2.378441387 | 0.83734109 | 2.84046897 | 0.00450473 | 0.02488507 |
| LINC02412 | 3.68373814 | 1.375983591 | 0.89555982 | 1.53645079 | 0.12442786 | 0.27641117 |
| AC124947.1 | 22.6552433 | 0.465270409 | 0.39486896 | 1.17829066 | 0.23868075 | 0.42826088 |
| NUDT4 | 6156.46596 | 0.583616756 | 0.35020795 | 1.66648631 | 0.09561659 | 0.23045652 |
| UBE2N | 3894.22105 | -0.206868624 | 0.1628023 | -1.2706738 | 0.20384471 | 0.38607951 |
| MRPL42 | 2844.08848 | 0.077582472 | 0.09751447 | 0.79559962 | 0.42626479 | 0.61866451 |
| SOCS2 | 1583.1773 | -0.065769774 | 0.25355242 | -0.2593932 | 0.79533188 | 0.8889978 |
| CRADD | 842.554278 | 0.43759367 | 0.18258767 | 2.39662224 | 0.01654697 | 0.06537963 |
| AC012085.2 | 12.0139932 | 0.402349234 | 0.48224538 | 0.83432471 | 0.40409803 | 0.59897135 |
| AC012464.1 | 4.86173032 | 0.944379889 | 0.60344101 | 1.56499123 | 0.11758497 | 0.26615845 |
| AC012464.3 | 14.4337234 | 0.32732794 | 0.3558531 | 0.91984006 | 0.35765635 | 0.55534345 |
| PLXNC1 | 434.594625 | 0.523455658 | 0.2539916 | 2.06091721 | 0.03931094 | 0.12225846 |
| AC123567.2 | 0.23140391 | 0.077483112 | 3.10308039 | 0.02496974 | 0.9800791 | NA |
| AC073655.1 | 1.85288156 | -0.531996768 | 0.94825769 | -0.5610255 | 0.57478014 | 0.73911114 |
| AC073655.2 | 47.2714667 | -0.235021271 | 0.28751309 | -0.8174281 | 0.41368383 | 0.60761193 |
| CEP83 | 527.003663 | -0.130671794 | 0.18272683 | -0.715121 | 0.47453427 | 0.66007262 |
| CEP83-DT | 29.7599184 | 0.263621252 | 0.4047273 | 0.65135526 | 0.51481719 | 0.69256857 |
| AC023161.1 | 0.16416605 | 0.18657164 | 3.29377195 | 0.05664376 | 0.95482897 | NA |
| TMCC3 | 935.137184 | 0.143189277 | 0.27711683 | 0.51671086 | 0.60535802 | 0.76218921 |
| NDUFA12 | 1868.47877 | -0.237253071 | 0.15664355 | -1.5146048 | 0.12987253 | 0.28492001 |
| NR2C1 | 1558.39426 | -0.232749586 | 0.14312473 | -1.626201 | 0.1039069 | 0.244035 |
| FGD6 | 712.137201 | 0.004238942 | 0.18345428 | 0.02310626 | 0.98156552 | 0.99160566 |
| VEZT | 2467.84917 | -0.252752909 | 0.14004004 | -1.8048617 | 0.07109633 | 0.1864129 |
| AC084879.2 | 0.32843282 | -0.27987943 | 2.93029431 | -0.0955124 | 0.92390784 | NA |
| METAP2 | 3694.39133 | -0.532267824 | 0.23785103 | -2.2378202 | 0.02523278 | 0.08895569 |
| USP44 | 141.755732 | -0.333810461 | 0.35356936 | -0.9441159 | 0.34511042 | 0.54275134 |
| NTN4 | 7148.81056 | 0.929783443 | 0.3309379 | 2.80954056 | 0.00496123 | 0.0267104 |
| AC090001.1 | 4.86327963 | 0.024319679 | 1.22405548 | 0.01986812 | 0.98414858 | 0.99314982 |
| LINC02410 | 9.34531508 | 3.875661935 | 0.89521743 | 4.32929675 | 1.50E-05 | 0.00028297 |
| SNRPF | 1127.7318 | -0.533058411 | 0.14243322 | -3.7425145 | 0.00018219 | 0.00202799 |
| CCDC38 | 18.0855927 | 1.044077034 | 0.33282297 | 3.13703418 | 0.00170666 | 0.01193442 |
| AMDHD1 | 510.637107 | 1.526245316 | 0.41119549 | 3.71172681 | 0.00020585 | 0.0022311 |
| HAL | 71.592386 | -1.526919563 | 0.46624637 | -3.27492 | 0.00105692 | 0.0081853 |
| AC007298.2 | 42.4552398 | 0.115447571 | 0.37329527 | 0.3092661 | 0.75711912 | 0.86556972 |
| LTA4H | 4056.05016 | -0.003251005 | 0.1893365 | -0.0171705 | 0.98630058 | 0.99391317 |
| AC007298.1 | 401.987849 | 0.270570431 | 0.19335801 | 1.39932364 | 0.16171595 | 0.33042381 |
| LINC02452 | 1.58916661 | 0.971903716 | 1.09024932 | 0.89145088 | 0.37268733 | 0.56890101 |
| ELK3 | 1869.04294 | -0.329769138 | 0.33881034 | -0.9733149 | 0.33039684 | 0.52823623 |
| AC008149.1 | 83.5316294 | 0.688126418 | 0.38861297 | 1.77072425 | 0.07660657 | 0.19651173 |
| CDK17 | 1483.54315 | -0.41405057 | 0.21934069 | -1.8877052 | 0.05906554 | 0.16372944 |
| AC007513.1 | 0.09997307 | 0.547239394 | 3.29377202 | 0.16614368 | 0.86804388 | NA |
| CFAP54 | 63.84718 | -1.428764525 | 0.54562382 | -2.618589 | 0.00882943 | 0.04102043 |
| NEDD1 | 686.762531 | 0.081889471 | 0.10610219 | 0.77179815 | 0.44023399 | 0.63160883 |
| AC007656.1 | 0.31734661 | -0.534763868 | 3.29377181 | -0.1623561 | 0.87102546 | NA |
| AC013417.1 | 0 | NA | NA | NA | NA | NA |
| LINC02409 | 0.94009952 | 1.703971597 | 1.68016775 | 1.01416754 | 0.31050281 | NA |
| AC016152.1 | 0.57471739 | -1.377448786 | 2.73981063 | -0.5027533 | 0.61513774 | NA |
| LINC02453 | 31.4412919 | 1.274688789 | 0.3824426 | 3.33301985 | 0.00085909 | 0.00697695 |
| TMPO-AS1 | 174.180468 | 1.076282644 | 0.21510005 | 5.00363741 | 5.63E-07 | 2.00E-05 |
| TMPO | 2359.30673 | 0.360692484 | 0.18944097 | 1.90398353 | 0.05691233 | 0.15943675 |

| | | | | | | |
|---|---|---|---|---|---|---|
| SLC25A3 | 26379.7359 | 0.036783946 | 0.23258208 | 0.15815468 | 0.87433491 | 0.93458499 |
| IKBIP | 898.749286 | -1.139830413 | 0.35002514 | -3.2564244 | 0.00112825 | 0.00862375 |
| APAF1 | 829.330663 | -0.409695683 | 0.20832509 | -1.9666171 | 0.04922737 | 0.14417211 |
| ANKS1B | 499.823744 | 0.248010787 | 0.27547863 | 0.90029049 | 0.36796568 | 0.56479948 |
| AC069437.1 | 0.29449776 | 2.177142669 | 3.23105086 | 0.67381875 | 0.50042655 | NA |
| AC117377.1 | 0.20908062 | 1.675937956 | 3.26598797 | 0.51314885 | 0.60784721 | NA |
| FAM71C | 0.72633607 | -0.243306801 | 1.64450352 | -0.1479515 | 0.88238103 | NA |
| UHRF1BP1L | 972.287899 | 0.028921512 | 0.2128171 | 0.13589844 | 0.89190157 | 0.94398733 |
| AC010203.2 | 6.40557848 | 1.122357627 | 0.79533032 | 1.41118426 | 0.15819029 | 0.32536292 |
| ACTR6 | 1278.25392 | -0.115573255 | 0.20175075 | -0.5728517 | 0.56674513 | 0.73296071 |
| DEPDC4 | 57.1737174 | 0.968133325 | 0.32193365 | 3.00724489 | 0.00263627 | 0.0166292 |
| SCYL2 | 2556.22092 | -0.001256843 | 0.18015539 | -0.0069764 | 0.99443365 | 0.99749764 |
| SLC17A8 | 8.33798175 | 0.268903489 | 0.74680447 | 0.36007215 | 0.71879318 | 0.84003617 |
| NR1H4 | 3078.06908 | -1.159075472 | 0.35262249 | -3.2870151 | 0.00101255 | 0.00791846 |
| GAS2L3 | 363.775553 | -1.364631605 | 0.29766745 | -4.5844166 | 4.55E-06 | 0.00010924 |
| ANO4 | 30.2644345 | -0.456646857 | 0.56676669 | -0.8057052 | 0.42041287 | 0.61314569 |
| AC138360.1 | 0.08757577 | 0.547239394 | 3.29377202 | 0.16614368 | 0.86804388 | NA |
| AC063947.1 | 0 | NA | NA | NA | NA | NA |
| SLC5A8 | 1138.27565 | 0.812925532 | 0.69676052 | 1.16672158 | 0.24332283 | 0.4338939 |
| UTP20 | 755.603816 | -0.130212799 | 0.26360919 | -0.4939615 | 0.62133335 | 0.77373255 |
| ARL1 | 4528.90435 | 0.178583814 | 0.13371474 | 1.33555819 | 0.18169372 | 0.35814742 |
| AC063948.1 | 34.6073083 | 0.307267325 | 0.38820576 | 0.79150635 | 0.42864857 | 0.62103527 |
| SPIC | 0.2923924 | 0.328535476 | 2.3110355 | 0.14215942 | 0.88695408 | NA |
| MYBPC1 | 9.90168316 | 2.770897829 | 1.03739246 | 2.67102176 | NA | NA |
| AC117505.1 | 1.57625346 | 0.705503375 | 1.91181761 | 0.36902232 | 0.71211109 | 0.83636475 |
| AC010205.1 | 11.515554 | 1.055409875 | 0.45209467 | 2.33448864 | 0.01957015 | 0.07372733 |
| CHPT1 | 4173.49005 | 0.045801057 | 0.28747191 | 0.15932359 | 0.87341393 | 0.93409277 |
| SYCP3 | 789.991165 | -0.229981529 | 0.29362218 | -0.7832567 | 0.43347641 | 0.62557806 |
| GNPTAB | 1565.40292 | 0.256984145 | 0.08782514 | 2.92608861 | 0.00343253 | 0.02030414 |
| DRAM1 | 3648.57732 | -0.64514397 | 0.31436363 | -2.052222 | 0.04014809 | 0.12412646 |
| AC084398.2 | 595.103772 | -1.147676851 | 0.30167992 | -3.8042866 | 0.00014221 | 0.00167568 |
| AC079907.2 | 2.68304754 | -0.258877305 | 0.90052379 | -0.2874741 | 0.7737493 | 0.87552321 |
| WASHC3 | 1321.41221 | 0.040853303 | 0.17855683 | 0.2287972 | 0.81902654 | 0.90287738 |
| AC079907.1 | 150.165151 | -0.234111368 | 0.18067514 | -1.2957586 | 0.19505865 | 0.37512374 |
| NUP37 | 578.02821 | -0.057042864 | 0.14521683 | -0.3928117 | 0.6944586 | 0.8239763 |
| PARPBP | 81.2093627 | -0.19814077 | 0.31708236 | -0.6248874 | 0.53204497 | 0.70623305 |
| PMCH | 39.7391158 | -0.322728114 | 0.39276186 | -0.821689 | 0.4112539 | 0.60554835 |
| LINC02456 | 0.8512769 | -1.005524014 | 1.52340707 | -0.6600495 | 0.50922209 | NA |
| IGF1 | 316.072574 | -0.795039968 | 0.56551277 | -1.4058745 | 0.1597614 | 0.32767428 |
| LINC00485 | 0.78590276 | 1.896072807 | 1.28706472 | 1.47317596 | 0.1407036 | NA |
| PAH | 13690.8095 | 3.504798523 | 0.71429231 | 4.90667261 | 9.26E-07 | 3.04E-05 |
| ASCL1 | 2.26745402 | -1.228705481 | 1.25495676 | -0.9790819 | 0.3275395 | 0.52526221 |
| AC068643.1 | 0 | NA | NA | NA | NA | NA |
| AC068643.2 | 0 | NA | NA | NA | NA | NA |
| C12orf42 | 5.48627465 | 1.624065978 | 0.72421013 | 2.2425342 | 0.02492687 | 0.08816007 |
| LINC02401 | 0.310781 | -0.174096114 | 3.29377188 | -0.0528562 | 0.95784652 | NA |
| STAB2 | 46.7818978 | -1.027447141 | 0.28024755 | -3.6662127 | 0.00024617 | 0.00258831 |
| AC025265.2 | 0 | NA | NA | NA | NA | NA |
| AC025265.3 | 0 | NA | NA | NA | NA | NA |
| AC025265.1 | 35.5553214 | -1.096376288 | 0.38170105 | -2.8723429 | 0.00407441 | 0.02307109 |
| NT5DC3 | 471.107801 | -0.649481327 | 0.21764088 | -2.9841881 | 0.00284332 | 0.01759229 |
| AC012555.2 | 0 | NA | NA | NA | NA | NA |
| AC012555.1 | 25.2565985 | 0.581472437 | 0.57472004 | 1.01174903 | 0.31165807 | 0.50875469 |
| HSP90B1 | 37487.2644 | -0.441190379 | 0.31306761 | -1.4092495 | 0.1587614 | 0.32623697 |
| C12orf73 | 521.78123 | -0.328970573 | 0.17991817 | -1.8284455 | 0.06748271 | 0.17936893 |
| TDG | 1083.86864 | -0.449976986 | 0.17692168 | -2.5433682 | 0.01097895 | 0.04829271 |
| GLT8D2 | 235.803253 | 0.330359152 | 0.44799015 | 0.73742504 | 0.46086392 | 0.64938843 |
| HCFC2 | 687.681983 | 0.15425294 | 0.09625358 | 1.60256834 | 0.10902999 | 0.25239218 |
| NFYB | 1675.84545 | 0.161707167 | 0.10641265 | 1.51962349 | 0.12860563 | 0.28282299 |
| LINC02385 | 0.54921022 | -0.63784723 | 2.24082026 | -0.284649 | 0.77591307 | NA |
| TXNRD1 | 14147.6073 | -2.560652545 | 0.63170528 | -4.0535557 | 5.04E-05 | 0.00074943 |
| AC089983.1 | 0.67805956 | -0.09225208 | 1.6783195 | -0.0549669 | 0.95616481 | NA |
| EID3 | 93.3093869 | -0.686961772 | 0.31851378 | -2.1567725 | 0.03102339 | 0.10335905 |
| CHST11 | 894.171954 | -1.40844273 | 0.27787407 | -5.0686368 | 4.01E-07 | 1.51E-05 |
| SLC41A2 | 842.680894 | -0.899302933 | 0.3032379 | -2.965668 | 0.00302026 | 0.01845755 |
| C12orf45 | 387.499292 | -0.159505551 | 0.18551513 | -0.8597981 | 0.38990036 | 0.58544231 |
| ALDH1L2 | 121.798719 | -0.727177515 | 0.40740183 | -1.7849147 | 0.07427515 | 0.19243818 |
| AC016257.1 | 2.84325856 | 0.185880666 | 0.76457999 | 0.24311474 | 0.80791651 | 0.89620864 |
| WASHC4 | 2909.40305 | 0.034328243 | 0.17820445 | 0.19263404 | 0.84724558 | 0.91979687 |
| APPL2 | 2149.26358 | 0.19642816 | 0.10345933 | 1.8986027 | 0.05761673 | 0.16088429 |
| C12orf75 | 3054.83882 | -0.777191098 | 0.20864688 | -3.7249113 | 0.00019538 | 0.00214499 |
| AC011313.1 | 37.3816285 | -0.701022625 | 0.54544005 | -1.2852423 | 0.19870762 | 0.37970435 |
| CASC18 | 16.8262766 | -0.011829808 | 0.49740181 | -0.0237832 | 0.98102554 | 0.99143998 |
| AC011595.2 | 2.14105479 | 0.599417532 | 0.90858509 | 0.65972636 | 0.50942945 | 0.68819951 |
| NUAK1 | 748.022735 | 0.088665843 | 0.25154953 | 0.35247867 | 0.72447931 | 0.84387647 |
| AC011595.1 | 1.17359878 | 0.09451078 | 1.24360739 | 0.07599728 | 0.93942126 | 0.97110795 |
| CKAP4 | 3041.3258 | -0.55586033 | 0.19945727 | -2.7868643 | 0.00532208 | 0.02819771 |
| AC079174.2 | 107.826802 | 0.486974921 | 0.4577638 | 1.06381266 | 0.28741358 | 0.48370005 |
| TCP11L2 | 233.951017 | 0.271738957 | 0.20882117 | 1.30129985 | 0.19315584 | 0.37266019 |
| POLR3B | 523.676791 | 0.764113942 | 0.17415757 | 4.38748634 | 1.15E-05 | 0.00022838 |
| AC079385.1 | 84.4155095 | 0.751265589 | 0.20826014 | 3.60734214 | 0.00030935 | 0.00310457 |
| RFX4 | 5.89587382 | 0.897683789 | 0.64736567 | 1.3866719 | 0.16554183 | 0.33595029 |
| AC079385.2 | 4.62695143 | 1.106791056 | 0.67037036 | 1.65101431 | 0.09873565 | 0.23558262 |
| AC079385.3 | 8.57926407 | 1.475309755 | 0.58650665 | 2.51541863 | 0.01188911 | 0.05120299 |
| RIC8B | 1047.76538 | 0.147346495 | 0.13275096 | 1.10994676 | 0.26702197 | 0.4611392 |
| AC007541.1 | 121.552269 | 0.811178927 | 0.31509364 | 2.57440593 | 0.01004124 | 0.045196 |
| TMEM263 | 3419.02389 | -0.001385608 | 0.15428548 | -0.0089808 | 0.99283445 | 0.9966781 |
| MTERF2 | 1527.97845 | 0.328246121 | 0.247124 | 1.32826486 | 0.18409062 | 0.3611083 |
| CRY1 | 1666.93868 | -0.337207753 | 0.29640147 | -1.1376723 | 0.25525733 | 0.44786515 |
| AC078929.1 | 60.1318781 | 0.400952552 | 0.47310946 | 0.84748369 | 0.39672558 | 0.59175595 |
| BTBD11 | 547.607009 | 0.354120594 | 0.52912365 | 0.66925868 | 0.50333048 | 0.68346162 |
| AC007540.1 | 7.7557839 | 1.75003376 | 0.78642578 | 2.2253006 | 0.02606107 | 0.09109627 |
| PWP1 | 2375.95589 | -0.198804616 | 0.20170868 | -0.9856027 | 0.32432812 | 0.52189213 |
| PRDM4 | 2961.23264 | -0.523674467 | 0.38583737 | -1.3572414 | 0.17470451 | 0.34861686 |
| AC007622.2 | 304.352754 | -0.400180454 | 0.47537313 | -0.8418239 | 0.39988654 | 0.59468803 |
| ASCL4 | 3.98383837 | 1.702848723 | 0.85484524 | 1.9919965 | 0.04637144 | 0.13787517 |
| AC126177.2 | 1.4103557 | 1.655422506 | 1.1134215 | 1.4867887 | 0.13707063 | 0.2955722 |
| AC126177.8 | 7.97149414 | 1.312021264 | 0.58388275 | 2.2470629 | 0.02463601 | 0.08729529 |
| AC126177.4 | 4.75843295 | 0.977983801 | 0.77740747 | 1.25800669 | 0.20838934 | 0.39152061 |
| WSCD2 | 105.16605 | 0.933197758 | 0.50904918 | 1.8332173 | 0.06677026 | 0.17812158 |
| CMKLR1 | 555.796044 | 0.209035656 | 0.36947979 | 0.56575667 | 0.57155921 | 0.73647771 |
| LINC01498 | 0.37171431 | -0.032084529 | 1.91353623 | -0.0167671 | 0.98662239 | NA |
| FICD | 692.896856 | -0.318690369 | 0.24988278 | -1.2753594 | 0.20218205 | 0.38401466 |
| SART3 | 2263.23145 | -0.112813538 | 0.1175009 | -0.9601078 | 0.33700095 | 0.53499716 |
| ISCU | 11151.1819 | -0.021683701 | 0.12450257 | -0.1741627 | 0.86173761 | 0.92800735 |
| TMEM119 | 342.043595 | 1.649006965 | 0.53832672 | 3.06320844 | 0.00218977 | 0.01446871 |
| SELPLG | 553.357754 | -0.409237043 | 0.45853643 | -0.8924853 | 0.37213284 | 0.56863396 |
| AC007569.1 | 9.3506729 | -0.143847155 | 0.61957458 | -0.2321708 | 0.81640532 | 0.90119337 |
| CORO1C | 4447.19136 | -0.664789298 | 0.27841012 | -2.3878058 | 0.0169493 | 0.06651086 |
| SSH1 | 2277.87684 | -0.28923513 | 0.1763562 | -1.6400622 | 0.10099224 | 0.2391492 |
| DAO | 2057.18211 | 4.407738466 | 0.67635087 | 6.51694064 | 7.18E-11 | 1.08E-08 |
| SVOP | 29.5915767 | 3.609629306 | 0.79862195 | 4.51982232 | 6.19E-06 | 0.00013973 |
| USP30 | 1088.46741 | 0.54785845 | 0.2423592 | 2.26052256 | 0.02378884 | 0.08512846 |

| | | | | | | | |
|---|---|---|---|---|---|---|---|
| USP30-AS1 | 55.1515644 | | 0.203875023 | 0.38450391 | 0.53022873 | 0.59595335 | 0.75484247 |
| ALKBH2 | 488.558708 | | -0.244170144 | 0.14337802 | -1.7029817 | 0.08857148 | 0.21803865 |
| UNG | 2003.75812 | | 0.006359874 | 0.17720467 | 0.03588999 | 0.97137008 | 0.98667343 |
| ACACB | 4051.37578 | | 0.495437029 | 0.35377804 | 1.40041771 | 0.16138827 | 0.32998604 |
| FOXN4 | 34.0548341 | | 1.668677929 | 0.57027073 | 2.92611536 | 0.00343224 | 0.02030414 |
| MYO1H | 27.7689119 | | 0.773340705 | 0.35687914 | 2.16695408 | 0.03023836 | 0.10140982 |
| LINC01486 | 0.08757577 | | 0.547239394 | 3.29377202 | 0.16614368 | 0.86804388 | NA |
| AC007570.1 | 16.1358918 | | 0.33488531 | 0.30958563 | 1.08172112 | 0.27937646 | 0.4749634 |
| KCTD10 | 3978.90713 | | -0.244421626 | 0.14549581 | -1.6799221 | 0.09297247 | 0.22577439 |
| UBE3B | 2408.31989 | | 0.292532256 | 0.16361581 | 1.78792166 | 0.07378865 | 0.19159137 |
| MMAB | 2086.33422 | | 0.05342414 | 0.13595614 | 0.39295128 | 0.69435546 | 0.82396716 |
| MVK | 4807.00266 | | -0.708322278 | 0.24963827 | -2.8373946 | 0.00454834 | 0.02506052 |
| FAM222A | 792.509562 | | 0.545916678 | 0.21367854 | 2.55485027 | 0.01062335 | 0.04710572 |
| FAM222A-AS | 19.4627945 | | 0.996269897 | 0.59960447 | 1.66154514 | 0.09660401 | 0.23198687 |
| TRPV4 | 1947.5139 | | 1.884634708 | 0.31502419 | 5.98250798 | 2.20E-09 | 1.92E-07 |
| GLTP | 3145.98753 | | 0.218819473 | 0.2421437 | 0.90367609 | 0.36616718 | 0.56326414 |
| AC007834.1 | 1.48597504 | | 1.443202496 | 1.04550759 | 1.38038453 | 0.16746828 | 0.33863496 |
| TCHP | 3094.17425 | | -0.292441077 | 0.12911544 | -2.2649583 | 0.02351524 | 0.08442435 |
| GIT2 | 2676.94606 | | -0.378302916 | 0.14150789 | -2.6733698 | 0.00750934 | 0.03648817 |
| AC084876.1 | 169.070384 | | -0.759797739 | 0.30252276 | -2.5115391 | 0.0120206 | 0.05158145 |
| ANKRD13A | 3948.74585 | | -0.510219667 | 0.17491768 | -2.916913 | 0.00353514 | 0.02072976 |
| C12orf76 | 905.785695 | | 0.144656988 | 0.15923714 | 0.90843748 | 0.36364713 | 0.56082017 |
| IFT81 | 991.896185 | | -0.019782546 | 0.20296986 | -0.0974654 | 0.92235678 | 0.96203347 |
| ATP2A2 | 9569.73058 | | -0.298780246 | 0.17362214 | -1.7208649 | 0.08527534 | 0.21186992 |
| ANAPC7 | 2736.21957 | | -0.156542628 | 0.13309694 | -1.176155 | 0.23953295 | 0.42937149 |
| ARPC3 | 9189.71153 | | -0.679322103 | 0.13102889 | -5.1845217 | 2.17E-07 | 9.07E-06 |
| GPN3 | 1213.72546 | | 0.036244383 | 0.13565651 | 0.26717761 | 0.78933241 | 0.88550565 |
| FAM216A | 279.81831 | | -0.144682747 | 0.20608236 | -0.7020627 | 0.48264003 | 0.66718482 |
| VPS29 | 3630.37408 | | -0.254071504 | 0.10820081 | -2.3481478 | 0.01886703 | 0.0719122 |
| RAD9B | 38.3529928 | | 0.201428309 | 0.29113474 | 0.69187316 | 0.48901699 | 0.67250456 |
| PPTC7 | 2364.60511 | | -0.160565933 | 0.16266336 | -0.9871057 | 0.32359082 | 0.52148689 |
| TCTN1 | 2482.95276 | | 0.429490163 | 0.24364776 | 1.76275032 | 0.07794261 | 0.19871861 |
| HVCN1 | 358.027882 | | 0.35350902 | 0.23363429 | 1.51308708 | 0.13025754 | 0.2854427 |
| PPP1CC | 4144.34714 | | 0.077502643 | 0.15696317 | 0.49376324 | 0.62147339 | 0.7738704 |
| AC002375.1 | 0 | NA | | NA | NA | NA | NA |
| CCDC63 | 0.48183801 | | 0.629882423 | 1.80581221 | 0.34880837 | 0.72723318 | NA |
| MYL2 | 9.53413448 | | 3.559238956 | 1.60145612 | 2.22250171 | NA | NA |
| LINC01405 | 0.44844207 | | -0.376763532 | 3.27988897 | -0.1148708 | 0.90854752 | NA |
| AC002351.1 | 0 | NA | | NA | NA | NA | NA |
| CUX2 | 5.87388025 | | 1.449500536 | 0.98251093 | 1.47530221 | 0.14013131 | 0.3002082 |
| PHETA1 | 1176.17877 | | -0.527187366 | 0.24487441 | -2.1528888 | 0.03132742 | 0.10412201 |
| LINC02356 | 4.7194746 | | -0.874152394 | 0.77353383 | -1.1300765 | 0.258444 | 0.45121612 |
| SH2B3 | 2040.59155 | | -0.199866191 | 0.26531458 | -0.7533178 | 0.45125897 | 0.64097491 |
| ATXN2 | 2797.99654 | | -0.072842332 | 0.12543888 | -0.5806998 | 0.56144279 | 0.72933859 |
| ATXN2-AS | 18.9281208 | | 0.142229383 | 0.31090668 | 0.45746647 | 0.6473358 | 0.79234334 |
| BRAP | 1033.31561 | | -0.198995001 | 0.15127041 | -1.3154919 | 0.18834462 | 0.36646113 |
| ACAD10 | 4109.71582 | | 0.679928909 | 0.12494874 | 5.44166281 | 5.28E-08 | 2.78E-06 |
| AC002996.1 | 10068.5588 | | 1.322216334 | 0.41335021 | 3.19877986 | 0.00138011 | 0.01014874 |
| ALDH2 | 24523.531 | | 1.132245292 | 0.39548048 | 2.86296128 | 0.00419702 | 0.02361308 |
| MAPKAPK5-A | 835.546012 | | -0.385120834 | 0.14519691 | -2.6524037 | 0.00799209 | 0.03812436 |
| MAPKAPK5 | 2189.67503 | | -0.040426659 | 0.10630323 | -0.3802957 | 0.70372594 | 0.830711 |
| TMEM116 | 1524.93737 | | 0.224148871 | 0.19229527 | 1.1656494 | 0.24375623 | 0.43449045 |
| ERP29 | 8466.48545 | | -0.071284084 | 0.12948286 | -0.5505291 | 0.58195652 | 0.74445491 |
| AC073575.2 | 92.5443829 | | -0.163779596 | 0.28878279 | -0.5671377 | 0.57062065 | 0.7357357 |
| NAA25 | 1140.95181 | | -0.467077275 | 0.22142179 | -2.1094459 | 0.03490611 | 0.11244638 |
| AC073575.1 | 0.03392908 | | 0.380740593 | 3.29738709 | 0.11546736 | 0.90807469 | NA |
| TRAFD1 | 2231.49179 | | -0.677600935 | 0.27374558 | -2.4752945 | 0.01331264 | 0.05568753 |
| HECTD4 | 2973.3646 | | 0.10126879 | 0.13407331 | 0.75532403 | 0.45005458 | 0.63999865 |
| AC004217.1 | 0.53009893 | | -0.453373289 | 2.27740731 | -0.1990743 | 0.84220462 | NA |
| RPL6 | 24723.513 | | -0.311946785 | 0.22946228 | -1.3594687 | 0.17399813 | 0.34788772 |
| PTPN11 | 4625.24674 | | -0.211973596 | 0.22954842 | -0.9234374 | 0.35577931 | 0.55319445 |
| RPH3A | 4.66058303 | | -0.248948784 | 0.97880726 | -0.2543389 | 0.79923373 | 0.89108294 |
| OAS1 | 1287.85836 | | -0.025718078 | 0.24949083 | -0.1030823 | 0.91789768 | 0.9601219 |
| AC004551.1 | 220.792808 | | -0.053440602 | 0.31640309 | -0.1689004 | 0.865875 | 0.93033141 |
| OAS3 | 849.05487 | | -0.162435401 | 0.46005287 | -0.3530799 | 0.72402858 | 0.84369283 |
| OAS2 | 665.029847 | | -0.159443108 | 0.41088453 | -0.3880485 | 0.69798017 | 0.82654528 |
| DTX1 | 1346.61224 | | -0.291842013 | 0.318487 | -0.9163388 | 0.3594892 | 0.55717234 |
| RASAL1 | 3757.04167 | | -1.531808834 | 0.27071376 | -5.6584077 | 1.53E-08 | 9.92E-07 |
| CFAP73 | 1359.65238 | | -0.508419443 | 0.14596262 | -3.4832167 | 0.00049543 | 0.00449412 |
| DDX54 | 4380.05301 | | -0.207012803 | 0.13625346 | -1.5193215 | 0.1286816 | 0.28291928 |
| RITA1 | 1451.28688 | | 0.071295298 | 0.20515014 | 0.34752742 | 0.72819512 | 0.84629817 |
| AC089999.1 | 267.809028 | | -0.05989423 | 0.25479205 | -0.235071 | 0.8141536 | 0.89973198 |
| IQCD | 235.200245 | | -0.238802466 | 0.29614527 | -0.8063693 | 0.42002992 | 0.61275665 |
| TPCN1 | 11520.9018 | | 0.853246987 | 0.28850431 | 2.95748433 | 0.00310161 | 0.01883393 |
| SLC8B1 | 3411.33318 | | -0.440056956 | 0.170189 | -2.5856957 | 0.00971827 | 0.04410856 |
| PLBD2 | 3085.66517 | | -0.332180408 | 0.11109682 | -2.9900083 | 0.0027897 | 0.01735827 |
| SDS | 87.2946671 | | -1.015174167 | 0.51948669 | -1.9541871 | 0.0506791 | 0.1470034 |
| SDSL | 1620.9413 | | 0.607575741 | 0.35250364 | 1.72360135 | 0.08477984 | 0.21098208 |
| LHX5 | 0.86496397 | | -2.073369151 | 1.64565364 | -1.2599062 | 0.20770322 | NA |
| LHX5-AS1 | 0 | NA | | NA | NA | NA | NA |
| LINC01234 | 1.80835561 | | -0.949810955 | 1.11030422 | -0.8554511 | 0.39230147 | 0.58808424 |
| RBM19 | 1962.4272 | | -0.061225562 | 0.13106887 | -0.4671251 | 0.64041036 | 0.78719159 |
| AC073863.1 | 122.741368 | | 0.092726075 | 0.22197928 | 0.417724 | 0.67614892 | 0.81149917 |
| AC010183.1 | 0 | NA | | NA | NA | NA | NA |
| AC010183.2 | 0 | NA | | NA | NA | NA | NA |
| LINC02459 | 0 | NA | | NA | NA | NA | NA |
| TBX5 | 0.83670857 | | -2.967799807 | 2.64246526 | -1.1231178 | 0.26138746 | NA |
| TBX5-AS1 | 0.28015841 | | -0.179498944 | 2.93432484 | -0.0611721 | 0.95122211 | NA |
| AC010177.1 | 0 | NA | | NA | NA | NA | NA |
| TBX3 | 1214.17203 | | 0.444171551 | 0.29100634 | 1.52632949 | 0.12692783 | 0.28041947 |
| AC026765.2 | 1.97214476 | | 1.046392469 | 1.06024678 | 0.98693294 | 0.32367552 | 0.5215653 |
| AC026765.1 | 0.08695548 | | -0.642905668 | 3.30558948 | -0.1944905 | 0.84579185 | NA |
| AC009804.2 | 0.08666371 | | -0.642905668 | 3.30558948 | -0.1944905 | 0.84579185 | NA |
| AC008125.1 | 3.18985994 | | 1.503515255 | 0.77018368 | 1.95215154 | 0.05092022 | 0.1475405 |
| AC078880.1 | 0.08757577 | | 0.547239394 | 3.29377202 | 0.16614368 | 0.86804388 | NA |
| AC078880.3 | 10.852946 | | 0.112546022 | 0.42291299 | 0.26612099 | 0.79014603 | 0.88604146 |
| AC078880.4 | 0.20718593 | | 0.18657164 | 3.29377195 | 0.05664376 | 0.95482897 | NA |
| AC078880.5 | 0.46136955 | | 1.487797521 | 1.74545852 | 0.85238205 | 0.39400208 | NA |
| AC078880.2 | 0.31873528 | | 1.515503097 | 2.79436541 | 0.54234249 | 0.58758259 | NA |
| AC009803.2 | 0 | NA | | NA | NA | NA | NA |
| AC009803.3 | 0 | NA | | NA | NA | NA | NA |
| AC009803.1 | 0 | NA | | NA | NA | NA | NA |
| AC009387.1 | 0.08393208 | | -0.642905668 | 3.30558948 | -0.1944905 | 0.84579185 | NA |
| LINC02463 | 0.08393208 | | -0.642905668 | 3.30558948 | -0.1944905 | 0.84579185 | NA |
| MED13L | 2527.00673 | | -0.13197901 | 0.11988208 | -1.1009069 | 0.27093719 | 0.46553813 |
| AC012157.1 | 184.430152 | | 0.619644322 | 0.2699388 | 2.29549932 | 0.02170453 | 0.07971425 |
| AC079384.1 | 1.61290719 | | -2.4585104 | 1.25569833 | -1.957883 | 0.05024374 | 0.14612634 |
| LINC02457 | 0 | NA | | NA | NA | NA | NA |
| MAP1LC3B2 | 44.2973408 | | -1.626296991 | 0.22341957 | -7.279116 | 3.36E-13 | 1.07E-10 |
| AC125603.4 | 4.00581087 | | -1.737845096 | 0.9952465 | -1.7461454 | 0.08078569 | 0.20383337 |
| AC125603.3 | 3.33294844 | | -1.067503636 | 0.80855975 | -1.3202532 | 0.18675048 | 0.36459689 |
| AC125603.2 | 2.94279819 | | 1.222687712 | 0.77285806 | 1.58203398 | 0.11364181 | 0.25984127 |
| AC125603.1 | 0.9430378 | | -0.802411338 | 1.9088371 | -0.4203666 | 0.67421767 | NA |

| | | | | | | | |
|---|---|---|---|---|---|---|---|
| C12orf49 | 3933.82167 | | -0.810100895 | 0.39216008 | -2.0657403 | 0.03885301 | 0.12122438 |
| RNFT2 | 154.0254 | | -0.350477664 | 0.30642698 | -1.1437559 | 0.25272489 | 0.44512124 |
| AC083806.3 | 1.08699853 | | 1.617836188 | 1.14853764 | 1.40860529 | 0.15895191 | 0.32649649 |
| HRK | 30.14734 | | -2.024408631 | 0.53704101 | -3.7695606 | 0.00016354 | 0.00186928 |
| FBXW8 | 681.887029 | | 0.148909881 | 0.14859426 | 1.00212405 | 0.31628368 | 0.51341341 |
| AC083806.2 | 28.2170845 | | 0.202734876 | 0.28216585 | 0.71849544 | 0.47245186 | 0.65870177 |
| AC127164.1 | 51.3797924 | | -0.085163375 | 0.25260568 | -0.3371396 | 0.73601167 | 0.85182775 |
| TESC | 1051.49777 | | -1.738191591 | 0.3552911 | -4.8923026 | 9.97E-07 | 3.24E-05 |
| TESC-AS1 | 12.9875411 | | 0.249903845 | 0.35220131 | 0.7095483 | 0.47798429 | 0.66297316 |
| FBXO21 | 3874.26265 | | 0.916824194 | 0.23206442 | 3.95073144 | 7.79E-05 | 0.00104189 |
| NOS1 | 270.541208 | | 1.752143715 | 0.63898265 | 2.7420834 | 0.00610508 | 0.03122402 |
| AC073864.1 | 57.10564 | | -0.232176218 | 0.53813304 | -0.4314476 | 0.66614293 | 0.80436322 |
| KSR2 | 307.221003 | | 0.2145576 | 0.45306341 | 0.4735708 | 0.635806 | 0.7840537 |
| AC084291.1 | 0.08393208 | | -0.642905668 | 3.30558948 | -0.1944905 | 0.84579185 | NA |
| RFC5 | 1133.95788 | | -0.199658451 | 0.15435087 | -1.2935362 | 0.19582566 | 0.37610915 |
| AC131159.2 | 98.1337106 | | 0.338715268 | 0.16884666 | 2.00605247 | 0.04485065 | 0.13453664 |
| WSB2 | 4077.77202 | | -0.177698172 | 0.26689321 | -0.6658025 | 0.50553734 | 0.68476972 |
| AC131159.1 | 5.51087141 | | 1.366916711 | 0.58735587 | 2.32723766 | 0.01995262 | 0.07479042 |
| VSIG10 | 1266.04822 | | -0.099339214 | 0.17964581 | -0.5529726 | 0.58028216 | 0.74328657 |
| AC131238.1 | 77.3116116 | | -0.040255449 | 0.24414045 | -0.1648864 | 0.86903337 | 0.9318649 |
| PEBP1 | 41073.8718 | | 0.88073442 | 0.25694345 | 3.42773636 | 0.00060864 | 0.00526995 |
| TAOK3 | 1574.00408 | | 0.277881742 | 0.08491002 | 3.27266153 | 0.0010654 | 0.00823405 |
| AC026367.1 | 24.9458771 | | 0.31084716 | 0.35488268 | 0.87591527 | 0.3810761 | 0.57712937 |
| SUDS3 | 2529.8689 | | 0.018375027 | 0.14675965 | 0.1252049 | 0.90036134 | 0.94951124 |
| AC026367.3 | 79.1159245 | | -0.0358287 | 0.31789531 | -0.112706 | 0.91026367 | 0.95555943 |
| AC026367.2 | 51.3893654 | | -0.235602255 | 0.28793297 | -0.8182538 | 0.41321227 | 0.607085 |
| LINC02460 | 0 | NA | | NA | NA | NA | NA |
| LINC02423 | 0 | NA | | NA | NA | NA | NA |
| LINC02440 | 0 | NA | | NA | NA | NA | NA |
| LINC02439 | 0 | NA | | NA | NA | NA | NA |
| AC087863.1 | 0.03392908 | | 0.380740593 | 3.29738709 | 0.11546736 | 0.90807469 | NA |
| SRRM4 | 20.9023735 | | 3.162384577 | 0.44462916 | 7.1124093 | 1.14E-12 | 3.01E-10 |
| AC084361.1 | 0.39808863 | | 2.60001047 | 2.57960067 | 1.007912 | 0.31349671 | NA |
| AC084880.4 | 0.42658979 | | -0.897626148 | 2.25818527 | -0.3974989 | 0.6909996 | NA |
| HSPB8 | 7322.54061 | | -1.694260971 | 0.34172403 | -4.9579802 | 7.12E-07 | 2.44E-05 |
| AC084880.3 | 7.27545226 | | -1.409723601 | 0.55808578 | -2.5259981 | 0.01153701 | 0.05012043 |
| AC084880.2 | 0.08695548 | | -0.642905668 | 3.30558948 | -0.1944905 | 0.84579185 | NA |
| LINC00934 | 0.504905 | | 0.650785088 | 1.64291037 | 0.39611722 | 0.69201856 | NA |
| CCDC60 | 5.10738582 | | -1.174492025 | 0.85556599 | -1.3727661 | 0.16982506 | 0.34161925 |
| AC002070.1 | 385.133545 | | 0.028919054 | 0.26706061 | 0.10828648 | 0.91376844 | 0.95778972 |
| TMEM233 | 85.0665563 | | -0.271498898 | 0.48938063 | -0.5547806 | 0.57904471 | 0.74245887 |
| PRKAB1 | 7713.48399 | | -0.321654535 | 0.19019508 | -1.6911822 | 0.09080202 | 0.2220493 |
| CIT | 3137.77124 | | -0.660258135 | 0.31161663 | -2.1188154 | 0.03410607 | 0.11050249 |
| AC002563.1 | 83.3916796 | | -0.184707241 | 0.28439636 | -0.6494712 | 0.51603387 | 0.69349777 |
| BICDL1 | 14253.7775 | | 0.42358627 | 0.43324063 | 0.97771594 | 0.32821483 | 0.5258856 |
| RAB35 | 3366.11475 | | -0.62644955 | 0.2141106 | -2.9258222 | 0.00343547 | 0.02031698 |
| AC004812.2 | 294.034639 | | -0.28933758 | 0.37281035 | -0.7760986 | 0.43769078 | 0.62967299 |
| GCN1 | 4105.84324 | | -0.126666208 | 0.13522722 | -0.9366916 | 0.34891721 | 0.54650432 |
| RPLP0 | 47290.1833 | | -0.8168702 | 0.24208892 | -3.3742569 | 0.00074015 | 0.00618044 |
| PXN-AS1 | 5482.44808 | | -0.577511866 | 0.15467476 | -3.7337175 | 0.00018867 | 0.00208086 |
| PXN | 12018.2982 | | -0.383868546 | 0.16515042 | -2.3243571 | 0.02010636 | 0.07521702 |
| AC004263.1 | 25.9902912 | | 0.90189255 | 0.35256749 | 2.55807068 | 0.01052547 | 0.04678972 |
| SIRT4 | 177.441656 | | 0.410042345 | 0.39145621 | 1.04747948 | 0.29487849 | 0.4916937 |
| PLA2G1B | 11.3231655 | | 0.358954502 | 0.63312361 | 0.56695801 | 0.57074271 | 0.73576038 |
| MSI1 | 112.173014 | | -0.802456345 | 0.45262011 | -1.7729136 | 0.07624303 | 0.19595796 |
| AC003982.1 | 0 | NA | | NA | NA | NA | NA |
| COX6A1 | 9990.89193 | | -0.093104478 | 0.16150399 | -0.5764841 | 0.56428803 | 0.73132527 |
| AL021546.1 | 8288.06271 | | -0.036012925 | 0.16457213 | -0.2188276 | 0.82678434 | 0.90760472 |
| TRIAP1 | 787.778218 | | -0.41955531 | 0.16603807 | -2.5268622 | 0.01150867 | 0.05005623 |
| GATC | 5174.01114 | | -0.313660094 | 0.09825994 | -3.1921461 | 0.0014122 | 0.01032428 |
| SRSF9 | 6798.57288 | | -0.355954815 | 0.10341884 | -3.4418759 | 0.0005777 | 0.00507963 |
| DYNLL1 | 6415.44869 | | -0.119037493 | 0.22691053 | -0.524601 | 0.59986059 | 0.75773985 |
| NRAV | 527.980648 | | 0.023689929 | 0.1920931 | 0.12332525 | 0.90184955 | 0.95039654 |
| AC063943.1 | 6.42456568 | | 0.560517547 | 0.48855486 | 1.14729704 | 0.25125888 | 0.44354643 |
| COQ5 | 1843.74598 | | 0.518278494 | 0.19092283 | 2.71459675 | 0.00663565 | 0.03321605 |
| RNF10 | 11496.3802 | | -0.289593708 | 0.15242696 | -1.899885 | 0.05744821 | 0.16057973 |
| POP5 | 1443.53993 | | 0.287732912 | 0.10187568 | 2.82435315 | 0.00473762 | 0.02582875 |
| AC125616.1 | 36.8819248 | | 1.347570688 | 0.378148 | 3.56360656 | 0.00036579 | 0.00355547 |
| CABP1 | 704.39357 | | 0.613435735 | 0.45150343 | 1.35865133 | 0.1742571 | 0.34824908 |
| MLEC | 10360.1531 | | 0.394924141 | 0.21009403 | 1.8797495 | 0.06014223 | 0.16577302 |
| AC069234.2 | 1072.02914 | | 1.255975351 | 0.32800603 | 3.82912276 | 0.0001286 | 0.00154356 |
| AC069234.5 | 6.89796761 | | 0.256253981 | 0.53400918 | 0.47986812 | 0.63132117 | 0.78082357 |
| AC069234.4 | 6.69473949 | | 1.035565017 | 0.48495894 | 2.1353664 | 0.03273109 | 0.10736645 |
| UNC119B | 3746.73044 | | 0.242589406 | 0.17650315 | 1.37441968 | 0.16931143 | 0.34113252 |
| AC069234.3 | 791.623272 | | 0.434863757 | 0.19844182 | 2.19139168 | 0.02842346 | 0.09716446 |
| ACADS | 4500.54712 | | 0.262031047 | 0.21424332 | 1.2230535 | 0.22130949 | 0.40744344 |
| AC069234.1 | 4.02672145 | | -0.220915249 | 0.83401452 | -0.2648818 | 0.79110054 | 0.88647096 |
| SPPL3 | 2904.22623 | | -0.247367177 | 0.1705072 | -1.4507726 | 0.14684319 | 0.30988626 |
| HNF1A | 2484.15658 | | 1.248682318 | 0.51591869 | 2.42030835 | 0.01550735 | 0.06233671 |
| C12orf43 | 2151.67441 | | 0.738312737 | 0.29326373 | 2.51757266 | 0.01181666 | 0.05100911 |
| OASL | 105.348289 | | -0.758659883 | 0.39992117 | -1.8970236 | 0.05782483 | 0.16129054 |
| P2RX7 | 115.774952 | | 0.650482039 | 0.2705304 | 2.40446932 | 0.01619597 | 0.0643689 |
| AC069209.1 | 2.826217 | | 1.111677225 | 0.71328607 | 1.55852929 | 0.11910783 | 0.26848609 |
| P2RX4 | 2213.39259 | | -0.558565411 | 0.29022689 | -1.9245819 | 0.05428169 | 0.15425001 |
| CAMKK2 | 2297.04994 | | 0.385952723 | 0.17484526 | 2.20739601 | 0.0272864 | 0.09426733 |
| ANAPC5 | 5641.23571 | | 0.191049055 | 0.08970693 | 2.12970242 | 0.03319619 | 0.1083932 |
| AC048337.1 | 4.05231739 | | 0.266196205 | 0.59798202 | 0.44515754 | 0.65620591 | 0.79823553 |
| RNF34 | 1569.45138 | | -0.348716616 | 0.14084258 | -2.4759318 | 0.0132889 | 0.05560587 |
| KDM2B | 1365.88378 | | -0.011767588 | 0.18382949 | -0.0640136 | 0.9489594 | 0.9753687 |
| AC145422.1 | 10.3504639 | | 0.575733906 | 0.41557502 | 1.38539103 | 0.16593293 | 0.33651091 |
| ORAI1 | 1460.03354 | | -1.404914427 | 0.28403704 | -4.9462366 | 7.57E-07 | 2.57E-05 |
| MORN3 | 125.474755 | | -0.243897221 | 0.35925661 | -0.6788942 | 0.49720491 | 0.67893661 |
| TMEM120B | 4743.78365 | | -0.451938418 | 0.31956593 | -1.4142259 | 0.15729558 | 0.32438293 |
| RHOF | 3773.36335 | | -0.587420299 | 0.35229776 | -1.6673972 | 0.09543545 | 0.230118 |
| LINC01089 | 2284.00241 | | 0.387259367 | 0.33026727 | 1.17256358 | 0.24097086 | 0.43106879 |
| AC084018.1 | 419.701859 | | 0.724597126 | 0.32554841 | 2.22577384 | 0.02602933 | 0.09100949 |
| AC084018.2 | 443.40549 | | 0.942563055 | 0.37362644 | 2.52274186 | 0.01164438 | 0.05047514 |
| SETD1B | 2608.04513 | | 0.045653335 | 0.19842759 | 0.23007554 | 0.81803307 | 0.90223481 |
| HPD | 14465.8408 | | 2.305377965 | 0.90402027 | 2.55013969 | 0.01076798 | 0.04760294 |
| AC079360.1 | 4157.64856 | | 2.485202377 | 0.93909908 | 2.64636867 | 0.00813611 | 0.03859434 |
| AC069503.3 | 4.02211934 | | 2.623592573 | 1.25254918 | 2.09460244 | 0.03620635 | 0.11526758 |
| PSMD9 | 3255.52447 | | -0.248596641 | 0.21907703 | -1.1347453 | 0.25648204 | 0.44921834 |
| AC069503.2 | 2922.59347 | | -0.273033256 | 0.21960727 | -1.2432797 | 0.21376479 | 0.39775456 |
| WDR66 | 634.571429 | | -0.383858435 | 0.20451384 | -1.8769314 | 0.0605275 | 0.16652683 |
| AC069503.1 | 57.7665883 | | 1.080880557 | 0.3579607 | 3.01955093 | 0.0025315 | 0.01614794 |
| BCL7A | 1831.40977 | | 0.511570849 | 0.12958434 | 3.94778311 | 7.89E-05 | 0.00105105 |
| MLXIP | 6503.817 | | -0.762974395 | 0.19249162 | -3.9636759 | 7.38E-05 | 0.00099653 |
| LRRC43 | 289.914807 | | 0.59075813 | 0.35245965 | 1.67610144 | 0.09371833 | 0.22716723 |
| IL31 | 0.42885147 | | -1.859940617 | 3.29773397 | -0.5640057 | 0.57275028 | NA |
| B3GNT4 | 2190.70792 | | -0.173597656 | 0.14910735 | -1.1642461 | 0.24432428 | 0.4353556 |
| DIABLO | 2749.97659 | | -0.237960478 | 0.13866741 | -1.716052 | 0.08615253 | 0.21353907 |
| AC048338.1 | 3205.53284 | | -0.211856488 | 0.14032841 | -1.5097191 | 0.1311151 | 0.28674107 |
| VPS33A | 1220.17583 | | -0.080718145 | 0.19914951 | -0.4053143 | 0.6852465 | 0.81810054 |

| | | | | | | |
|---|---|---|---|---|---|---|
| CLIP1 | 3200.27728 | -0.219088375 | 0.26250304 | -0.8346127 | 0.40393579 | 0.5989197 |
| CLIP1-AS1 | 1.49241409 | 0.787808826 | 0.89017524 | 0.8850042 | 0.37615437 | 0.57197501 |
| ZCCHC8 | 1522.24671 | -0.224185042 | 0.13936883 | -1.6085737 | 0.10770959 | 0.25009858 |
| AC127002.1 | 0.84773263 | 0.513690826 | 1.57439047 | 0.32627918 | 0.74421314 | NA |
| RSRC2 | 4088.21021 | -0.221603507 | 0.09459547 | -2.3426441 | 0.01914764 | 0.07257773 |
| KNTC1 | 446.538107 | -0.181217533 | 0.12727073 | -1.4238744 | 0.15448284 | 0.32053762 |
| AC026333.4 | 9.89111688 | 0.52631103 | 0.49271653 | 1.06818222 | 0.28543833 | 0.4815227 |
| AC026333.3 | 37.3602764 | 0.052892875 | 0.39301542 | 0.13458219 | 0.89294222 | 0.94467169 |
| HCAR2 | 89.1840044 | 0.035575153 | 0.51549149 | 0.06901211 | 0.94497998 | 0.9737607 |
| HCAR3 | 37.3816836 | -0.472646598 | 0.37549578 | -1.2587268 | 0.20812903 | 0.39135499 |
| HCAR1 | 95.9742715 | 0.371550065 | 0.41685297 | 0.89132162 | 0.37275666 | 0.56894393 |
| DENR | 2889.19432 | -0.315669468 | 0.11488188 | -2.7477742 | 0.00600013 | 0.03077679 |
| CCDC62 | 14.3952947 | 0.250459443 | 0.38864081 | 0.64444967 | 0.51928389 | 0.69619788 |
| HIP1R | 10853.2534 | 0.295719252 | 0.14036509 | 2.10678633 | 0.03513611 | 0.11296502 |
| VPS37B | 3296.40316 | -0.488739518 | 0.20919089 | -2.3363327 | 0.01947391 | 0.07345545 |
| AC027290.1 | 961.789588 | -0.568509733 | 0.20215135 | -2.8122974 | 0.0049189 | 0.02654975 |
| ABCB9 | 3372.6351 | 0.041693222 | 0.09074191 | 0.45947039 | 0.64589641 | 0.7910956 |
| OGFOD2 | 1739.63344 | 0.256731958 | 0.10008862 | 2.56504633 | 0.0103162 | 0.04610818 |
| ARL6IP4 | 6657.19715 | -0.081256746 | 0.0855368 | -0.9499624 | 0.34213135 | 0.53994876 |
| PITPNM2 | 4424.78306 | -0.413567841 | 0.21437276 | -1.9291996 | 0.05370609 | 0.1532882 |
| PITPNM2-AS | 7.29958542 | -0.518769151 | 0.58840586 | -0.8816519 | 0.37796508 | 0.57390102 |
| MPHOSPH9 | 619.942604 | 0.415426524 | 0.142197 | 2.92148576 | 0.00348366 | 0.02049624 |
| C12orf65 | 1057.55072 | -0.5318027 | 0.21020379 | -2.5299387 | 0.01140825 | 0.04971854 |
| AC068768.1 | 1757.64356 | -0.892653531 | 0.27125576 | -3.2908187 | 0.00099896 | 0.00783308 |
| CDK2AP1 | 2979.13533 | -0.839747367 | 0.26563671 | -3.1612625 | 0.00157087 | 0.01119865 |
| AC068768.2 | 7.70544227 | -0.553392925 | 0.46136331 | -1.1994732 | 0.230344 | 0.4179284 |
| SBNO1 | 2294.68951 | -0.123015539 | 0.17476236 | -0.7039018 | 0.48149394 | 0.66600139 |
| AC137767.1 | 255.444711 | 0.766513345 | 0.3033481 | 2.52684404 | 0.01150926 | 0.05005623 |
| KMT5A | 1768.00679 | -0.585501635 | 0.16541106 | -3.5396765 | 0.00040062 | 0.00382621 |
| RILPL2 | 2086.81266 | -0.13271609 | 0.14905364 | -0.8903915 | 0.37325571 | 0.56947195 |
| SNRNP35 | 1797.74274 | -0.319642497 | 0.15449038 | -2.0690122 | 0.03854494 | 0.1204872 |
| RILPL1 | 1517.49745 | -0.43295136 | 0.18317568 | -2.3635854 | 0.01809905 | 0.06971546 |
| AC055713.1 | 1296.98509 | 0.009838606 | 0.1527357 | 0.06441589 | 0.94863908 | 0.97519151 |
| TMED2 | 9241.11787 | -0.400484271 | 0.18445063 | -2.1712274 | 0.02991399 | 0.10066835 |
| DDX55 | 1036.07952 | -0.100796502 | 0.10471619 | -0.9625685 | 0.33576413 | 0.53393559 |
| EIF2B1 | 2505.47625 | -0.093181788 | 0.10739663 | -0.8676417 | 0.38559053 | 0.58142479 |
| GTF2H3 | 829.845016 | 0.27583066 | 0.13177611 | 2.09317652 | 0.0363334 | 0.11558876 |
| TCTN2 | 1216.22206 | 0.098305774 | 0.17654801 | 0.55682175 | 0.57764922 | 0.7412823 |
| AC117503.1 | 376.776867 | -0.364075201 | 0.24023638 | -1.5154874 | 0.12964902 | 0.28460514 |
| ATP6V0A2 | 938.035806 | -0.416316569 | 0.2058726 | -2.0222048 | 0.0431552 | 0.13100063 |
| DNAH10 | 238.576496 | 0.754924982 | 0.26994287 | 2.7966102 | 0.00516418 | 0.02754995 |
| CCDC92 | 2556.52494 | -0.048103052 | 0.14082004 | -0.3415924 | 0.73265767 | 0.84920148 |
| AC079315.1 | 158.627585 | -0.303180507 | 0.32419241 | -0.9351869 | 0.34969196 | 0.5473277 |
| DNAH10OS | 345.099523 | 0.722124749 | 0.25576352 | 2.82340793 | 0.00475161 | 0.02588306 |
| ZNF664 | 6786.05801 | 0.146059978 | 0.19030068 | 0.76752208 | 0.44277117 | 0.63397273 |
| RFLNA | 232.518042 | 0.763908391 | 0.35545604 | 2.14909384 | 0.03162696 | 0.10480058 |
| AC068790.8 | 197.743366 | 1.54100489 | 0.39521492 | 3.8991567 | 9.65E-05 | 0.00123028 |
| AC068790.9 | 9.77543043 | -0.188085249 | 0.47827206 | -0.39326 | 0.69412749 | 0.82386304 |
| AC026358.1 | 5.21576839 | 0.890064422 | 0.66134061 | 1.34584873 | 0.17835131 | 0.35377828 |
| NCOR2 | 7645.38036 | 0.228339931 | 0.1507176 | 1.51501838 | 0.12976775 | 0.28476122 |
| AC073592.1 | 30.7243553 | 0.620412848 | 0.31933823 | 1.94280792 | 0.05203937 | 0.1499266 |
| SCARB1 | 1414.9706 | -0.273952334 | 0.25755033 | -1.0636847 | 0.28747158 | 0.48376678 |
| UBC | 44365.0428 | -0.93401447 | 0.25471604 | -3.6668852 | 0.00024552 | 0.00258358 |
| DHX37 | 1407.82903 | -0.516705511 | 0.20511056 | -2.5191561 | 0.01176365 | 0.05085368 |
| BRI3BP | 1016.014 | 0.453333751 | 0.28860164 | 1.57079412 | 0.11623048 | 0.26426951 |
| THRIL | 60.117895 | 0.740133352 | 0.30819766 | 2.40148918 | 0.01632849 | 0.06476249 |
| AACS | 1794.98618 | 0.088679133 | 0.15694086 | 0.5650481 | 0.57204105 | 0.73684655 |
| AC122688.2 | 0.07659028 | -0.642905668 | 3.30558948 | -0.1944905 | 0.84579185 | NA |
| TMEM132B | 28.023548 | 0.069963365 | 0.69916778 | 0.10006663 | 0.92029143 | 0.96099929 |
| LINC00939 | 78.5002238 | 1.93378025 | 0.67544391 | 2.8629768 | 0.00419681 | 0.02361308 |
| AC005186.1 | 5.46412704 | 0.874783503 | 0.55257655 | 1.58309923 | 0.11339885 | 0.25944193 |
| LINC02359 | 0 | NA | NA | NA | NA | NA |
| AC007368.2 | 0.08666371 | -0.642905668 | 3.30558948 | -0.1944905 | 0.84579185 | NA |
| LINC02347 | 1.04176476 | -0.847009542 | 1.89717411 | -0.4464585 | 0.65526607 | 0.79771785 |
| AC006065.4 | 0.3069976 | -1.364242679 | 3.30558948 | -0.4127078 | 0.67982068 | NA |
| AC006065.3 | 0 | NA | NA | NA | NA | NA |
| AC069235.1 | 0 | NA | NA | NA | NA | NA |
| LINC00943 | 5.47211945 | -0.256066409 | 0.70567008 | -0.3628699 | 0.7167021 | 0.83885895 |
| LINC00944 | 14.8539129 | -1.369654055 | 0.68039351 | -2.0130322 | 0.04411124 | 0.13306769 |
| LINC02372 | 0 | NA | NA | NA | NA | NA |
| AC078878.2 | 0.18028734 | 0.18657164 | 3.29377195 | 0.05664376 | 0.95482897 | NA |
| LINC02405 | 1.44000257 | -1.528929074 | 1.61499776 | -0.9467066 | 0.34378829 | 0.54151789 |
| AC079949.1 | 11.6764942 | -0.002443614 | 0.56430747 | -0.0043303 | 0.99654494 | 0.99824844 |
| AC079949.2 | 34.6133611 | -0.198402015 | 0.46508764 | -0.4265906 | 0.66967754 | 0.80710223 |
| LINC02376 | 0 | NA | NA | NA | NA | NA |
| LINC02375 | 1.80563752 | 0.149711834 | 0.95774124 | 0.15631762 | 0.87578267 | 0.93547831 |
| AC025252.1 | 0 | NA | NA | NA | NA | NA |
| LINC02411 | 0 | NA | NA | NA | NA | NA |
| AC087894.2 | 0 | NA | NA | NA | NA | NA |
| LINC02393 | 0.17332742 | -1.003574173 | 3.30558948 | -0.3035992 | 0.7614333 | NA |
| LINC00507 | 0.84530462 | 3.190975701 | 2.32385079 | 1.37314139 | 0.1697084 | NA |
| LINC00508 | 0.03392908 | 0.380740593 | 3.29738709 | 0.11546736 | 0.90807469 | NA |
| LINC02441 | 0 | NA | NA | NA | NA | NA |
| LINC02369 | 0 | NA | NA | NA | NA | NA |
| LINC02368 | 0 | NA | NA | NA | NA | NA |
| AC061709.1 | 0 | NA | NA | NA | NA | NA |
| TMEM132C | 85.1802155 | 1.197813861 | 0.54673505 | 2.19084887 | 0.02846273 | 0.09723463 |
| SLC15A4 | 1818.31706 | -0.998663287 | 0.23228237 | -4.2993504 | 1.71E-05 | 0.00031413 |
| AC108704.1 | 7.00068116 | 0.061907541 | 0.55892128 | 0.11076254 | 0.91180465 | 0.956682 |
| GLT1D1 | 105.467062 | -0.131594587 | 0.49505638 | -0.2658174 | 0.79037986 | 0.88615293 |
| TMEM132D | 12.7777948 | 0.233927057 | 0.67882189 | 0.34460742 | 0.73038953 | 0.84755348 |
| TMEM132D-. | 0 | NA | NA | NA | NA | NA |
| TMEM132D-. | 0.27059475 | 1.100737003 | 2.99309129 | 0.36775925 | 0.71305276 | NA |
| AC130404.1 | 0 | NA | NA | NA | NA | NA |
| AC117373.1 | 0.03392908 | 0.380740593 | 3.29738709 | 0.11546736 | 0.90807469 | NA |
| AC055717.1 | 0.19114494 | -1.003574173 | 3.30558948 | -0.3035992 | 0.7614333 | NA |
| AC055717.2 | 0 | NA | NA | NA | NA | NA |
| LINC02418 | 0.34803786 | 0.670435996 | 2.2147669 | 0.30271176 | 0.76210953 | NA |
| AC135388.1 | 0 | NA | NA | NA | NA | NA |
| LINC02419 | 2.02889021 | 0.860717497 | 0.96383778 | 0.89301075 | 0.37185142 | 0.56851356 |
| AC026336.3 | 0.07830766 | -0.642905668 | 3.30558948 | -0.1944905 | 0.84579185 | NA |
| FZD10 | 16.4799219 | -0.168134356 | 0.88842075 | -0.1892508 | 0.84989624 | 0.92113042 |
| AC026336.2 | 0.08695548 | -0.642905668 | 3.30558948 | -0.1944905 | 0.84579185 | NA |
| PIWIL1 | 26.9896015 | 1.174968099 | 0.61103192 | 1.92292427 | 0.05448956 | 0.15468724 |
| RIMBP2 | 601.771913 | -0.734001915 | 0.32845197 | -2.2347314 | 0.02543499 | 0.08943613 |
| AC063926.1 | 2.34681279 | -1.166668002 | 0.98729004 | -1.1816872 | 0.23732984 | 0.42655315 |
| AC095350.1 | 0.12059279 | 0.02007176 | 3.29738704 | 0.00608717 | 0.99514317 | NA |
| STX2 | 1635.95284 | -0.375976078 | 0.20057701 | -1.8744724 | 0.06086534 | 0.16715466 |
| AC073912.1 | 3.65935391 | -0.256419672 | 0.87448334 | -0.2932242 | 0.76935079 | 0.87240608 |
| AC073912.2 | 2.65601241 | -0.373310344 | 1.07262429 | -0.3480346 | 0.72781421 | 0.84611931 |
| RAN | 9233.88827 | -0.734650871 | 0.21084348 | -3.4843424 | 0.00049335 | 0.00447834 |
| ADGRD1 | 216.75558 | 0.285142963 | 0.59386476 | 0.48014798 | 0.63112217 | 0.78075653 |
| AC073862.1 | 1.64910059 | -0.223463663 | 1.0538512 | -0.2120448 | 0.83207209 | 0.910748 |

| | | | | | | |
|---|---|---|---|---|---|---|
| ADGRD1-AS1 | 4.61549067 | 2.100431403 | 0.86427315 | 2.43028655 | 0.01508689 | 0.06113894 |
| AC078925.1 | 1.2080867 | -0.925359853 | 1.68873279 | -0.5479611 | 0.58371862 | 0.74591297 |
| LINC01257 | 2.004273 | 0.045652723 | 1.00962552 | 0.04521748 | 0.96393396 | 0.98309632 |
| LINC02415 | 3.78151436 | -1.699267247 | 1.08424073 | -1.5672417 | 0.11705821 | 0.26537355 |
| LINC02370 | 0.08757577 | 0.547239394 | 3.29377202 | 0.16614368 | 0.86804388 | NA |
| AC140118.1 | 0 | NA | NA | NA | NA | NA |
| AC073578.2 | 0.78517777 | -1.318429486 | 1.85030626 | -0.7125466 | 0.47612635 | NA |
| AC073578.1 | 0 | NA | NA | NA | NA | NA |
| AC117500.3 | 0 | NA | NA | NA | NA | NA |
| LINC02414 | 0.34968279 | 1.629530304 | 3.24275583 | 0.50251403 | 0.61530599 | NA |
| AC117500.1 | 1.34156105 | -3.649687723 | 2.03590877 | -1.7926578 | 0.07302766 | 0.19013454 |
| SFSWAP | 2519.55462 | 0.024249593 | 0.1448762 | 0.16738148 | 0.8670699 | 0.93098661 |
| AC117500.2 | 1.59902515 | 0.062558718 | 0.94469956 | 0.06622075 | 0.94720207 | 0.97455008 |
| MMP17 | 81.1590329 | -0.626047018 | 0.4809552 | -1.3016743 | 0.19302775 | 0.37251871 |
| AC131009.2 | 0.08393208 | -0.642905668 | 3.30558948 | -0.1944905 | 0.84579185 | NA |
| ULK1 | 5560.37035 | -0.331610257 | 0.18639381 | -1.7790841 | 0.07522598 | 0.19415768 |
| AC131009.1 | 62.9601525 | -0.373915813 | 0.28313244 | -1.3206392 | 0.18662169 | 0.36445337 |
| PUS1 | 1483.8046 | -0.728225563 | 0.21494306 | -3.3879929 | 0.00070406 | 0.00593735 |
| EP400 | 2404.79434 | -0.094506673 | 0.13456552 | -0.7023097 | 0.48248604 | 0.66704188 |
| AC137590.1 | 319.336701 | -0.216518716 | 0.15384343 | -1.4073965 | 0.15930984 | 0.3270279 |
| AC137590.2 | 0.03392908 | 0.380740593 | 3.29738709 | 0.11546736 | 0.90807469 | NA |
| DDX51 | 1730.33589 | 0.02704583 | 0.1144354 | 0.23634146 | 0.81316772 | 0.89913137 |
| NOC4L | 1121.99194 | -0.561705425 | 0.18443706 | -3.0455127 | 0.00232284 | 0.01516921 |
| AC138466.3 | 0.4799803 | -2.179794672 | 2.76515889 | -0.7883072 | 0.43051704 | NA |
| LINC02361 | 153.076093 | -0.073326317 | 0.44511996 | -0.1647338 | 0.86915349 | 0.93191744 |
| AC138466.1 | 16.5990822 | -0.501577373 | 0.57476355 | -0.8726673 | 0.38284443 | 0.57887659 |
| GALNT9 | 843.018963 | 0.040516505 | 0.36719867 | 0.11033946 | 0.91214016 | 0.95681217 |
| AC148477.3 | 96.4177264 | 0.688197104 | 0.42834801 | 1.60663081 | 0.10813538 | 0.25086194 |
| AC148477.2 | 561.013269 | 0.334774348 | 0.45050207 | 0.74311389 | 0.45741273 | 0.64668528 |
| AC148477.4 | 269.620559 | 1.052295037 | 0.32117273 | 3.27641461 | 0.00105134 | 0.00815168 |
| AC148477.1 | 19.9064179 | -1.798172653 | 0.87507032 | -2.0548893 | 0.0398897 | 0.12359198 |
| AC148476.1 | 9.39680835 | -1.103893436 | 0.68762857 | -1.605363 | 0.10841395 | 0.25139605 |
| AC079031.1 | 0 | NA | NA | NA | NA | NA |
| AC079031.2 | 0.17648264 | -1.003574173 | 3.30558948 | -0.3035992 | 0.7614333 | NA |
| FBRSL1 | 4702.8735 | -0.139487063 | 0.17804343 | -0.783444 | 0.43336642 | 0.62553355 |
| AC131212.1 | 9.6498751 | -1.933612732 | 0.89584811 | -2.1584158 | 0.03089552 | 0.10302412 |
| LRCOL1 | 85.116755 | -0.862946759 | 0.6167534 | -1.3991763 | 0.16176011 | 0.33046283 |
| P2RX2 | 94.6747783 | -1.975217727 | 0.76615362 | -2.5780962 | 0.00993463 | 0.0448436 |
| POLE | 943.887752 | 0.029541701 | 0.16940735 | 0.17438264 | 0.86156476 | 0.9279689 |
| PXMP2 | 1622.93217 | 1.291027769 | 0.32181131 | 4.01175387 | 6.03E-05 | 0.00085309 |
| AC135586.2 | 1583.95611 | 1.244790116 | 0.31933857 | 3.89802624 | 9.70E-05 | 0.00123544 |
| PGAM5 | 1750.54685 | -0.314766875 | 0.23583392 | -1.3346972 | 0.18197547 | 0.3584592 |
| ANKLE2 | 3896.5781 | -0.701642943 | 0.27656 | -2.5370369 | 0.01117952 | 0.04895447 |
| GOLGA3 | 3745.52428 | -0.779016428 | 0.25104693 | -3.1030709 | 0.00191524 | 0.01302678 |
| CHFR | 1767.71255 | -0.51195717 | 0.17017821 | -3.0083591 | 0.00262663 | 0.01658624 |
| AC127070.1 | 19.5697496 | 0.623170161 | 0.35553858 | 1.75274977 | 0.07964497 | 0.201811 |
| AC127070.2 | 47.8300411 | -0.096188751 | 0.28078729 | -0.342568 | 0.73192345 | 0.84883549 |
| ZNF605 | 944.971528 | 0.307449852 | 0.19846219 | 1.54916085 | 0.12134306 | 0.27192025 |
| ZNF26 | 1069.10931 | -0.077011739 | 0.19491726 | -0.3950996 | 0.69276936 | 0.8232685 |
| AC073911.1 | 2.41984752 | 1.174994947 | 0.76318083 | 1.53960228 | 0.12365733 | 0.27519992 |
| ZNF84 | 1775.33949 | -0.024113638 | 0.17180933 | -0.1403512 | 0.88838255 | 0.94226723 |
| ZNF140 | 1207.42452 | -0.096248493 | 0.15972814 | -0.6025769 | 0.54679017 | 0.71792754 |
| ZNF891 | 641.527709 | -0.13539534 | 0.16859338 | -0.8030881 | 0.42192381 | 0.61446572 |
| AC026786.2 | 15.7096654 | -0.045126939 | 0.40152734 | -0.1123882 | 0.9105156 | 0.95574781 |
| ZNF10 | 961.758659 | 0.484431884 | 0.19885139 | 2.43615031 | 0.01484451 | 0.06038097 |
| AC026786.1 | 170.463567 | 0.925136658 | 0.27621429 | 3.34934396 | 0.00081003 | 0.00664672 |
| ZNF268 | 945.524072 | 0.580397275 | 0.16618265 | 3.49252632 | 0.00047847 | 0.00438255 |
| ANHX | 0.07659028 | -0.642905668 | 3.30558948 | -0.1944905 | 0.84579185 | NA |
| FAM230C | 7.49065245 | 5.329482218 | 1.01641331 | 5.24342036 | 1.58E-07 | 6.98E-06 |
| LINC00349 | 0.09997307 | 0.547239394 | 3.29377202 | 0.16614368 | 0.86804388 | NA |
| LINC00388 | 0 | NA | NA | NA | NA | NA |
| LINC00387 | 0.17391096 | -1.003574173 | 3.30558948 | -0.3035992 | 0.7614333 | NA |
| LINC00408 | 0 | NA | NA | NA | NA | NA |
| AL355516.1 | 0 | NA | NA | NA | NA | NA |
| LINC00442 | 2.61111858 | -2.242474981 | 1.65808591 | -1.352448 | 0.17623204 | 0.35087004 |
| AL137001.1 | 0 | NA | NA | NA | NA | NA |
| TUBA3C | 0 | NA | NA | NA | NA | NA |
| AL139327.2 | 0.12088456 | 0.02007176 | 3.29738704 | 0.00608717 | 0.99514317 | NA |
| LINC00421 | 0 | NA | NA | NA | NA | NA |
| AL356259.1 | 1.9300276 | -0.396937907 | 1.29434726 | -0.3066703 | 0.75909431 | 0.86658803 |
| TPTE2 | 10.8289367 | -1.574204135 | 0.53811435 | -2.9254082 | 0.00344005 | 0.02032657 |
| LINC00350 | 0 | NA | NA | NA | NA | NA |
| MPHOSPH8 | 2804.56991 | 0.030155231 | 0.1101343 | 0.27380417 | 0.78423512 | 0.88211398 |
| PSPC1-AS2 | 184.078854 | -0.808283221 | 0.27692476 | -2.9187828 | 0.00351401 | 0.02063337 |
| PSPC1 | 1899.14902 | -0.338972489 | 0.24284504 | -1.3958387 | 0.16276309 | 0.33181775 |
| ZMYM5 | 848.624332 | -0.186225867 | 0.14775044 | -1.2604082 | 0.20752213 | 0.39041044 |
| AL355001.1 | 1.11914261 | 0.016304487 | 1.14686868 | 0.01421652 | 0.98865724 | 0.99493137 |
| AL355001.2 | 148.659936 | 0.191405645 | 0.24982808 | 0.76614945 | 0.44358738 | 0.63462453 |
| ZMYM2 | 3393.41938 | -0.172148072 | 0.13514906 | -1.2737645 | 0.20274689 | 0.38474636 |
| LINC01072 | 0.10721287 | -0.642905668 | 3.30558948 | -0.1944905 | 0.84579185 | NA |
| GJA3 | 81.8739537 | 0.477333904 | 0.68557456 | 0.69625382 | 0.48626989 | 0.67036364 |
| LINC00556 | 0 | NA | NA | NA | NA | NA |
| GJB2 | 966.978485 | 0.283294735 | 0.43756912 | 0.64742853 | 0.51735464 | 0.69452944 |
| GJB6 | 5.14015142 | 0.664891501 | 1.01897031 | 0.65251312 | 0.51407021 | 0.69195996 |
| CRYL1 | 14444.589 | 1.122758849 | 0.33730406 | 3.32862535 | 0.00087276 | 0.0070597 |
| AL590096.1 | 30.9952912 | 0.730333131 | 0.33715879 | 2.16614 | 0.03030049 | 0.10151492 |
| IFT88 | 1659.6873 | 0.418454931 | 0.13240114 | 3.16050859 | 0.00157494 | 0.01122464 |
| AL161772.1 | 79.6234786 | -1.574779985 | 0.32179955 | -4.8936675 | 9.90E-07 | 3.23E-05 |
| IL17D | 109.665459 | -1.177634972 | 0.28496575 | -4.1325492 | 3.59E-05 | 0.0005682 |
| EEF1AKMT1 | 237.136611 | -0.051414048 | 0.24102543 | -0.2133138 | 0.83108222 | 0.91040231 |
| XPO4 | 1105.97952 | 0.04472344 | 0.13472019 | 0.3319728 | 0.73990979 | 0.85407841 |
| LATS2 | 1484.73775 | 0.171735673 | 0.25606094 | 0.67068283 | 0.5024226 | 0.68265066 |
| LATS2-AS1 | 0.10721287 | -0.642905668 | 3.30558948 | -0.1944905 | 0.84579185 | NA |
| SAP18 | 7021.26389 | 0.159025558 | 0.11716836 | 1.35723977 | 0.17470504 | 0.34861686 |
| SKA3 | 26.6429804 | -0.128339368 | 0.57128674 | -0.2246497 | 0.82225183 | 0.90503301 |
| MRPL57 | 2086.50015 | 0.243651845 | 0.13144462 | 1.85364639 | 0.06378977 | 0.17250909 |
| LINC01046 | 0 | NA | NA | NA | NA | NA |
| LINC00539 | 36.9141763 | 0.271715639 | 0.30122559 | 0.90203371 | 0.36703896 | 0.56395482 |
| ZDHHC20 | 1553.76166 | -0.355224702 | 0.12547681 | -2.8309988 | 0.00464029 | 0.02545024 |
| MICU2 | 3906.07337 | 0.447780686 | 0.12410386 | 3.60811245 | 0.00030843 | 0.00309961 |
| FGF9 | 1067.25369 | 1.571939847 | 0.39152062 | 4.01496058 | 5.95E-05 | 0.00084566 |
| LINC00424 | 0.36695002 | 1.705742188 | 2.8152169 | 0.60590081 | 0.54458062 | NA |
| AL136962.1 | 42.0045532 | 0.031996507 | 0.36054645 | 0.08874448 | 0.92928498 | 0.965762 |
| LINC00621 | 1106.28352 | -2.608272077 | 0.69616376 | -3.7466358 | 0.00017922 | 0.00200513 |
| AL157931.1 | 934.985639 | -2.826362755 | 0.66648779 | -4.2406819 | 2.23E-05 | 0.00038737 |
| AL157931.2 | 24.3303337 | -0.893463823 | 0.49538285 | -1.8035825 | 0.07129679 | 0.1867769 |
| LINC00362 | 0 | NA | NA | NA | NA | NA |
| SGCG | 12.6844143 | 0.783272946 | 0.74081193 | 1.05731686 | 0.290367 | 0.4864971 |
| SACS | 494.664819 | -0.954761816 | 0.3580283 | -2.6667216 | 0.00765951 | 0.03698613 |
| SACS-AS1 | 1.02754045 | -0.400000447 | 1.79174839 | -0.2232459 | 0.82334415 | 0.90564302 |
| LINC00327 | 2.78159351 | 0.500227024 | 0.87005935 | 0.57493437 | 0.56533568 | 0.73208851 |
| LINC00352 | 0 | NA | NA | NA | NA | NA |
| TNFRSF19 | 1300.21542 | 1.634823567 | 0.31871089 | 5.12948767 | 2.91E-07 | 1.16E-05 |

| | | | | | | |
|---|---|---|---|---|---|---|
| MIPEP | 1899.77976 | 0.243398765 | 0.22626643 | 1.07571751 | 0.28205361 | 0.47782049 |
| C1QTNF9B | 69.7753284 | 0.116564236 | 0.5475583 | 0.21288004 | 0.83142053 | 0.91040231 |
| AL359736.1 | 81.9083793 | 0.342321629 | 0.29722138 | 1.1517396 | 0.24942809 | 0.44131903 |
| SPATA13 | 3557.70485 | -0.465788941 | 0.14746337 | -3.1586756 | 0.00158488 | 0.01127549 |
| AL445985.1 | 9.86495856 | 1.196611021 | 0.43474336 | 2.75245385 | 0.00591505 | 0.03047671 |
| SPATA13-AS | 0.79729163 | -0.482182536 | 1.51385684 | -0.3185126 | 0.75009611 | NA |
| C1QTNF9 | 14.3607998 | -0.473799529 | 0.48481055 | -0.977288 | 0.3284266 | 0.52616096 |
| C1QTNF9-AS | 7.84393825 | -0.641668986 | 0.47844896 | -1.3411441 | 0.17987368 | 0.3557007 |
| LINC00566 | 1.36666457 | -1.463592185 | 1.15073622 | -1.2718746 | 0.20341767 | 0.38563077 |
| PARP4 | 4067.61216 | -0.054910051 | 0.19971137 | -0.274947 | 0.78335693 | 0.88146457 |
| ATP12A | 260.946387 | 4.299345087 | 0.67419283 | 6.37702582 | 1.81E-10 | 2.39E-08 |
| RNF17 | 0.21786011 | 1.725828273 | 3.04405575 | 0.56695029 | 0.57074796 | NA |
| CENPJ | 294.332515 | -0.108768246 | 0.21236486 | -0.5121763 | 0.60852763 | 0.76464661 |
| PABPC3 | 14.3437925 | -0.388508487 | 0.35081652 | -1.1074407 | 0.26810345 | 0.46229568 |
| AMER2 | 2.67108468 | 0.661901808 | 0.99453487 | 0.66553907 | 0.50570576 | 0.68492352 |
| LINC00463 | 0 | NA | NA | NA | NA | NA |
| LINC01053 | 0 | NA | NA | NA | NA | NA |
| LINC01076 | 0.6098622 | -1.48130013 | 1.89093042 | -0.783371 | 0.43340926 | NA |
| MTMR6 | 1721.82162 | -0.023179082 | 0.15330675 | -0.1511941 | 0.87982258 | 0.93767616 |
| AL590787.1 | 7.534501 | 0.241465416 | 0.72139018 | 0.33472235 | 0.73783454 | 0.85283548 |
| NUP58 | 2675.82442 | -0.271684427 | 0.17521093 | -1.5506135 | 0.12099433 | 0.27142668 |
| ATP8A2 | 70.372928 | 0.17816532 | 0.37200614 | 0.47893112 | 0.63198763 | 0.78146459 |
| SHISA2 | 397.496407 | -1.232420497 | 0.49635464 | -2.4829435 | 0.01303018 | 0.05481208 |
| LINC00415 | 0.09635527 | 0.547239394 | 3.29377202 | 0.16614368 | 0.86804388 | NA |
| RNF6 | 1730.83723 | -0.117739152 | 0.33802574 | -0.3483142 | 0.72760426 | 0.84611931 |
| AL138966.2 | 35.1130927 | 0.473580218 | 0.41511363 | 1.14084478 | 0.25393452 | 0.44656685 |
| CDK8 | 956.281435 | -0.484975888 | 0.17449825 | -2.7792593 | 0.0054483 | 0.02872328 |
| WASF3 | 722.408027 | 0.209079562 | 0.12382721 | 1.6884784 | 0.09131943 | 0.22288144 |
| WASF3-AS1 | 0.68697706 | -1.077865945 | 1.8846249 | -0.571926 | 0.56737212 | NA |
| GPR12 | 2.74633208 | 2.54549817 | 1.01462148 | 2.50881559 | 0.01211367 | 0.05191425 |
| AL160035.1 | 0 | NA | NA | NA | NA | NA |
| USP12 | 2690.3328 | -0.251182669 | 0.2571221 | -0.9769004 | 0.32861848 | 0.52643639 |
| USP12-AS1 | 0.09997307 | 0.547239394 | 3.29377202 | 0.16614368 | 0.86804388 | NA |
| USP12-AS2 | 5.08709623 | 1.344112627 | 0.77157886 | 1.74202885 | 0.0815034 | 0.20508059 |
| LINC02340 | 24.7647171 | 0.301421499 | 0.39559372 | 0.76194713 | 0.44609156 | 0.6364806 |
| LINC00412 | 7.60806984 | 0.742505386 | 0.53715296 | 1.38229785 | 0.16688027 | 0.33786047 |
| RPL21 | 20119.937 | -0.419787683 | 0.22534995 | -1.8628257 | 0.06248679 | 0.17007145 |
| RASL11A | 1647.86767 | -0.649055465 | 0.28995014 | -2.2385071 | 0.025188 | 0.08883344 |
| LINC01079 | 0 | NA | NA | NA | NA | NA |
| GTF3A | 3807.37871 | -0.371976671 | 0.1709218 | -2.1762974 | 0.02953303 | 0.09978039 |
| MTIF3 | 2311.71416 | -0.119262597 | 0.12804549 | -0.931408 | 0.35164255 | 0.54930479 |
| LNX2 | 946.999633 | -0.214887907 | 0.2148444 | -1.0002025 | 0.31721251 | 0.51448733 |
| POLR1D | 3621.80555 | -0.357125428 | 0.19262045 | -1.8540369 | 0.06373388 | 0.17242868 |
| AL136439.1 | 0.08666371 | -0.642905668 | 3.30558948 | -0.1944905 | 0.84579185 | NA |
| GSX1 | 0 | NA | NA | NA | NA | NA |
| PLUT | 0 | NA | NA | NA | NA | NA |
| PDX1 | 0.62245633 | -0.357507842 | 1.94745384 | -0.1835771 | 0.85434526 | NA |
| LINC00543 | 74.2397422 | 0.15046037 | 0.41620687 | 0.36150381 | 0.71772286 | 0.8397902 |
| CDX2 | 0.35080282 | 0.706189817 | 2.31533966 | 0.30500485 | 0.76036246 | NA |
| URAD | 1.23526415 | 0.856274461 | 1.20254897 | 0.71204956 | 0.47643409 | 0.66179458 |
| FLT3 | 114.053432 | 1.28957556 | 0.84515476 | 1.52584547 | 0.12704835 | 0.28057117 |
| PAN3-AS1 | 139.623231 | -0.398714468 | 0.25068527 | -1.5904982 | 0.11172256 | 0.25680995 |
| PAN3 | 2514.95726 | -0.17611416 | 0.12121016 | -1.4529654 | 0.14623336 | 0.30901305 |
| FLT1 | 5969.73523 | -0.937826827 | 0.41364409 | -2.2672313 | 0.0233761 | 0.08405069 |
| POMP | 4184.06008 | -0.725187521 | 0.19363118 | -3.7452002 | 0.00018025 | 0.00201217 |
| SLC46A3 | 1613.08389 | 0.185563426 | 0.16991352 | 1.09210512 | 0.2747869 | 0.47005661 |
| MTUS2 | 10.2860496 | 0.11658688 | 0.57850495 | 0.20153134 | 0.84028312 | 0.91502585 |
| MTUS2-AS2 | 0.09644756 | -0.642905668 | 3.30558948 | -0.1944905 | 0.84579185 | NA |
| MTUS2-AS1 | 2.50118798 | -2.115247284 | 1.47590192 | -1.4331896 | 0.15180366 | 0.31683335 |
| SLC7A1 | 1396.69852 | -0.94308918 | 0.24973766 | -3.7763195 | 0.00015916 | 0.00183601 |
| UBL3 | 2600.558 | 0.373902488 | 0.1967725 | 1.90017653 | 0.05740996 | 0.16050685 |
| LINC00297 | 0 | NA | NA | NA | NA | NA |
| LINC00572 | 0.08695548 | -0.642905668 | 3.30558948 | -0.1944905 | 0.84579185 | NA |
| LINC00544 | 0.86751801 | -0.215780692 | 1.90146723 | -0.1134812 | 0.9096491 | NA |
| AL354674.1 | 0 | NA | NA | NA | NA | NA |
| LINC00365 | 2.68058983 | -0.170669461 | 0.84146282 | -0.2028247 | 0.83927204 | 0.91465195 |
| LINC00384 | 1.08511413 | -1.701194186 | 1.63594923 | -1.039882 | 0.29839472 | 0.49530098 |
| LINC00385 | 0 | NA | NA | NA | NA | NA |
| KATNAL1 | 693.017534 | -0.043477288 | 0.13409756 | -0.3242213 | 0.74577049 | 0.85746535 |
| LINC00427 | 3.90007641 | -0.961587546 | 0.70722245 | -1.3596677 | 0.17393509 | 0.34788772 |
| LINC00426 | 47.3407714 | -0.637775205 | 0.73427986 | -0.8685724 | 0.38508107 | 0.58085962 |
| LINC01058 | 3.20587024 | 3.095806435 | 0.84769963 | 3.6520087 | 0.0002602 | 0.0027024 |
| UBE2L5 | 2.58014332 | 0.192141139 | 0.77733406 | 0.24717962 | 0.80476922 | 0.89426338 |
| HMGB1 | 11749.5198 | -0.128559855 | 0.12345768 | -1.0413273 | 0.29772366 | 0.49454947 |
| USPL1 | 1141.67398 | -0.010308562 | 0.10214483 | -0.100921 | 0.91961314 | 0.96074349 |
| ALOX5AP | 501.915361 | -1.132625397 | 0.39145894 | -2.893344 | 0.00381164 | 0.02199405 |
| LINC00398 | 10.9213501 | 1.193412276 | 0.78412127 | 1.52197411 | 0.12801557 | 0.28201926 |
| TEX26-AS1 | 11.6985272 | -0.12293818 | 0.55098965 | -0.2231225 | 0.82344018 | 0.90564302 |
| LINC00545 | 7.6020277 | -0.436222241 | 0.75875976 | -0.5749148 | 0.5653489 | 0.73208851 |
| MEDAG | 780.190023 | -2.065188684 | 0.8933775 | -2.3116641 | 0.0207962 | 0.07708321 |
| TEX26 | 1.54089362 | -1.119282273 | 1.34130645 | -0.8344717 | 0.40401523 | 0.59897135 |
| AL353680.1 | 0.55085573 | 0.224040538 | 2.20899035 | 0.10142214 | 0.91921536 | NA |
| LINC01066 | 0 | NA | NA | NA | NA | NA |
| HSPH1 | 3772.23321 | -0.419043351 | 0.26748713 | -1.5665926 | 0.11720995 | 0.26561337 |
| B3GLCT | 661.933821 | 0.418763387 | 0.15181694 | 2.75834428 | 0.0058095 | 0.03007981 |
| AL161616.2 | 0.03392908 | 0.380740593 | 3.29738709 | 0.11546736 | 0.90807469 | NA |
| RXFP2 | 3.12570891 | -0.287538894 | 0.95760782 | -0.3002679 | 0.76397281 | 0.86935843 |
| FRY | 1570.3828 | 0.54467301 | 0.24240406 | 2.2469632 | 0.02464238 | 0.08730613 |
| FRY-AS1 | 15.7436625 | 0.553520554 | 0.59949702 | 0.92330827 | 0.35584658 | 0.55325006 |
| ZAR1L | 0.62676641 | -0.497455634 | 1.88902096 | -0.2633405 | 0.79228818 | NA |
| BRCA2 | 57.7596248 | -1.166115211 | 0.47252114 | -2.4678583 | 0.01359241 | 0.05656816 |
| N4BP2L1 | 1288.21566 | -0.313183299 | 0.22074901 | -1.4187302 | 0.15597768 | 0.32246944 |
| N4BP2L2 | 5706.7883 | 0.037364024 | 0.13078255 | 0.2856958 | 0.77511112 | 0.87656232 |
| PDS5B | 1191.31523 | 0.004259185 | 0.13157395 | 0.03237104 | 0.97417616 | 0.98807744 |
| AL138820.1 | 23.1000931 | 0.245381479 | 0.29748961 | 0.8248405 | 0.40946215 | 0.60395553 |
| LINC00423 | 0.55983075 | 1.923187641 | 2.02549128 | 0.94949194 | 0.34237047 | NA |
| KL | 6259.28659 | 2.852919012 | 0.42253463 | 6.75191759 | 1.46E-11 | 2.85E-09 |
| STARD13 | 1418.03297 | -0.309386601 | 0.21355059 | -1.4487743 | 0.14740063 | 0.31073263 |
| LINC02344 | 0.58959401 | -1.370403727 | 2.26440014 | -0.605195 | 0.54504941 | NA |
| AL139383.1 | 9.27792379 | 1.205579745 | 0.70068895 | 1.72056338 | 0.08533008 | 0.21189889 |
| AL161719.1 | 8.44441129 | 0.919368549 | 0.55022617 | 1.67089208 | 0.094743 | 0.22897744 |
| AL139081.1 | 0.20718593 | 0.18657164 | 3.29377195 | 0.05664376 | 0.95482897 | NA |
| RFC3 | 261.290388 | 0.320229768 | 0.28563156 | 1.12112881 | 0.26223304 | 0.45590135 |
| LINC02343 | 28.9580089 | 1.239710447 | 0.48012917 | 2.58203527 | 0.00982195 | 0.044472 |
| LINC00457 | 0 | NA | NA | NA | NA | NA |
| AL161716.1 | 2.04549014 | 0.082291015 | 1.07552894 | 0.07651213 | 0.93901166 | 0.97097237 |
| AL138690.1 | 3.22706417 | -1.301895231 | 0.98903291 | -1.3163316 | 0.18806278 | 0.36613054 |
| NBEA | 728.345825 | -0.076562744 | 0.16227662 | -0.4718039 | 0.63706676 | 0.78513112 |
| MAB21L1 | 14.483529 | 0.097339303 | 0.53901078 | 0.18058879 | 0.85669036 | 0.92498566 |
| LINC00445 | 0.09635527 | 0.547239394 | 3.29377202 | 0.16614368 | 0.86804388 | NA |
| DCLK1 | 296.111863 | -0.88724394 | 0.70676822 | -1.2553535 | 0.20935047 | 0.3925052 |
| SOHLH2 | 44.195171 | 0.154080179 | 0.60670328 | 0.25396299 | 0.79952415 | 0.8912141 |
| CCDC169-SOI | 45.5821686 | 0.414681601 | 0.49679318 | 0.83471677 | 0.40387719 | 0.59887737 |

| | | | | | | | |
|---|---|---|---|---|---|---|---|
| CCDC169 | 60.2615789 | | 0.388252354 | 0.43442434 | 0.89371685 | 0.37147341 | 0.56829807 |
| SPART | 4837.58704 | | -0.048374883 | 0.17353038 | -0.278769 | 0.78042214 | 0.8797765 |
| SPART-AS1 | 51.1847267 | | 0.184998513 | 0.25143215 | 0.73577906 | 0.46186516 | 0.65014385 |
| CCNA1 | 35.4841648 | | -0.897514023 | 0.51160174 | -1.7543217 | 0.0793754 | 0.2012818 |
| SERTM1 | 8.56811008 | | 1.492565549 | 0.85538204 | 1.74491102 | 0.08100036 | 0.20420454 |
| RFXAP | 269.732383 | | 0.186834629 | 0.17388347 | 1.07448183 | 0.28260679 | 0.47857314 |
| SMAD9 | 793.654868 | | -0.584261998 | 0.35128095 | -1.6632328 | 0.09626585 | 0.2314488 |
| ALG5 | 1711.33991 | | -0.290852746 | 0.12269458 | -2.3705427 | 0.01776199 | 0.068705 |
| EXOSC8 | 1497.82739 | | -0.350759008 | 0.16634424 | -2.1086333 | 0.03497625 | 0.11258833 |
| SUPT20H | 2185.79723 | | 0.003338596 | 0.1340378 | 0.02490787 | 0.98012845 | 0.99106512 |
| CSNK1A1L | 2.30902667 | | -0.991495361 | 0.99204775 | -0.9994432 | 0.31758005 | 0.51482055 |
| LINC01048 | 0 | NA | | NA | NA | NA | NA |
| LINC00547 | 0 | NA | | NA | NA | NA | NA |
| POSTN | 1891.32757 | | -0.423349395 | 0.64397623 | -0.6573991 | 0.51092433 | 0.68920615 |
| TRPC4 | 164.535378 | | -1.637181034 | 0.42761296 | -3.8286516 | 0.00012885 | 0.00154429 |
| LINC02334 | 35.3550637 | | 1.744092848 | 0.47687738 | 3.65731929 | 0.00025487 | 0.00265751 |
| LINC00571 | 26.8452236 | | 0.982886518 | 0.3007055 | 3.26860174 | 0.0010808 | 0.00832626 |
| UFM1 | 4572.0181 | | -0.363118984 | 0.1980944 | -1.8330603 | 0.06679361 | 0.17816583 |
| LINC00437 | 0.26061361 | | -0.1787649 | 2.9763739 | -0.0600613 | 0.95210681 | NA |
| LINC00366 | 0 | NA | | NA | NA | NA | NA |
| FREM2 | 1957.59405 | | 1.369024811 | 0.37009847 | 3.69908258 | 0.00021638 | 0.00232792 |
| FREM2-AS1 | 3.12572213 | | 1.576893172 | 0.81410706 | 1.93696044 | 0.05275018 | 0.15126383 |
| STOML3 | 1.30448539 | | 0.099053975 | 1.18205578 | 0.08379806 | 0.93321699 | 0.96778874 |
| PROSER1 | 1707.59535 | | -0.12952797 | 0.16598099 | -0.7803784 | 0.4351682 | 0.62724162 |
| NHLRC3 | 2095.96929 | | 0.243790824 | 0.2368762 | 1.02919089 | 0.30338998 | 0.50024347 |
| AL354809.1 | 9.52410344 | | 0.735062195 | 0.45316766 | 1.62205353 | 0.10479189 | 0.24527624 |
| LHFPL6 | 2490.22193 | | 0.105188873 | 0.37007644 | 0.28423553 | 0.77622989 | 0.87711283 |
| COG6 | 1222.40048 | | 0.367583686 | 0.15341103 | 2.39607075 | 0.01657189 | 0.06544867 |
| LINC00598 | 63.4180121 | | -0.245574916 | 0.32699519 | -0.7510047 | 0.45264984 | 0.6423623 |
| LINC00332 | 0.56524985 | | -0.131521741 | 1.71102377 | -0.0768673 | 0.93872913 | NA |
| FOXO1 | 1923.54241 | | -0.198070691 | 0.183283 | -1.0806823 | 0.27983848 | 0.47541172 |
| MRPS31 | 1141.51758 | | -0.177757698 | 0.15498738 | -1.1469172 | 0.25141583 | 0.44374835 |
| SLC25A15 | 262.822581 | | 0.760647285 | 0.2506602 | 3.03457549 | 0.00240874 | 0.01554162 |
| ELF1 | 2562.94354 | | -0.286140765 | 0.15577724 | -1.8368586 | 0.06623078 | 0.17720227 |
| WBP4 | 934.028469 | | -0.184674058 | 0.14735506 | -1.253259 | 0.21011145 | 0.39345181 |
| KBTBD6 | 509.999602 | | 0.570543144 | 0.12186764 | 4.68166236 | 2.85E-06 | 7.53E-05 |
| AL354696.2 | 59.800994 | | 1.302344026 | 0.30925502 | 4.21123001 | 2.54E-05 | 0.00043016 |
| KBTBD7 | 432.941491 | | 1.113670816 | 0.22097839 | 5.03972736 | 4.66E-07 | 1.71E-05 |
| MTRF1 | 732.684259 | | 0.097006481 | 0.19141908 | 0.50677541 | 0.61231241 | 0.76779926 |
| NAA16 | 952.551564 | | -0.295123151 | 0.14026982 | -2.1039676 | 0.03538127 | 0.11351797 |
| RGCC | 1063.77132 | | 0.005566507 | 0.48984114 | 0.0113639 | 0.99093311 | 0.99580451 |
| VWA8 | 1890.66975 | | 1.032003275 | 0.23113815 | 4.46487639 | 8.01E-06 | 0.00017106 |
| VWA8-AS1 | 7.82443245 | | 1.195245339 | 0.6780552 | 1.76275521 | 0.07794179 | 0.19871861 |
| DGKH | 1620.55974 | | -0.605293262 | 0.16476753 | -3.6736197 | 0.00023914 | 0.00252648 |
| AL157932.1 | 11.8913457 | | -0.094567932 | 0.41185379 | -0.2296153 | 0.81839071 | 0.90255377 |
| AKAP11 | 2148.81997 | | 0.317956142 | 0.11182013 | 2.84346069 | 0.00446265 | 0.02472791 |
| LINC02341 | 10.9769461 | | 0.955877235 | 0.53773425 | 1.77760156 | 0.07546932 | 0.19455723 |
| TNFSF11 | 8.55843175 | | -1.407005589 | 0.62873208 | -2.237846 | 0.0252311 | 0.08895569 |
| FAM216B | 0.66564087 | | -2.620903833 | 3.26919793 | -0.8016963 | 0.42272867 | NA |
| LINC01050 | 0.27406511 | | -0.700031684 | 2.99343044 | -0.233856 | 0.81509677 | NA |
| LINC00428 | 0 | NA | | NA | NA | NA | NA |
| EPSTI1 | 1015.51365 | | -0.387864711 | 0.33736841 | -1.149677 | 0.25027692 | 0.4424943 |
| DNAJC15 | 2782.62324 | | -0.231788194 | 0.13497298 | -1.7172933 | 0.08592559 | 0.21309676 |
| LINC00400 | 0 | NA | | NA | NA | NA | NA |
| ENOX1 | 210.397734 | | 1.376181404 | 0.31510014 | 4.36744149 | 1.26E-05 | 0.00024647 |
| ENOX1-AS2 | 3.56095597 | | 0.752590738 | 0.98205131 | 0.76634564 | 0.44347067 | 0.63454322 |
| ENOX1-AS1 | 0.96357292 | | 0.182022048 | 1.22593933 | 0.14847558 | 0.88196746 | NA |
| AL162713.1 | 3.81310605 | | 0.851242689 | 0.75891244 | 1.12166126 | 0.26200649 | 0.45561491 |
| CCDC122 | 211.725125 | | -0.071440384 | 0.23607768 | -0.3026139 | 0.76218414 | 0.86816102 |
| LACC1 | 688.90623 | | -0.40732497 | 0.17412702 | -2.3392405 | 0.01932299 | 0.0730848 |
| AL512506.3 | 131.063983 | | -0.064963337 | 0.40583225 | -0.1600744 | 0.8728225 | 0.93368745 |
| LINC00284 | 263.806798 | | -0.991432613 | 0.6690655 | -1.481817 | 0.138389 | 0.29751492 |
| LINC00390 | 4.04583789 | | 2.283082603 | 0.90586174 | 2.52034333 | 0.01172404 | 0.05073236 |
| SMIM2-AS1 | 2053.65818 | | 0.062520586 | 0.41822323 | 0.14949094 | 0.88116626 | 0.93836253 |
| SMIM2 | 102.816802 | | -0.412389247 | 0.42731212 | -0.9650773 | 0.33450607 | 0.53256808 |
| AL589745.2 | 0.79551886 | | -0.390839185 | 1.3853322 | -0.2821267 | 0.77784637 | NA |
| AL589745.1 | 349.262361 | | 0.171745589 | 0.51024986 | 0.33659116 | 0.73642512 | 0.85207763 |
| AL138960.1 | 27.5740013 | | -0.56025587 | 0.55216574 | -1.0146516 | 0.31027192 | 0.50726204 |
| SERP2 | 237.888523 | | -0.354542853 | 0.23888245 | -1.4841728 | 0.13776307 | 0.29650769 |
| TUSC8 | 0.89389878 | | 1.080293289 | 1.4952594 | 0.72247885 | 0.47000013 | NA |
| TSC22D1 | 22108.7987 | | -1.024088478 | 0.29516734 | -3.4695183 | 0.00052139 | 0.00468614 |
| TSC22D1-AS1 | 163.672497 | | 0.711651903 | 0.38914475 | 1.82875886 | 0.06743574 | 0.17929518 |
| LINC00407 | 1.04086112 | | -0.34580544 | 1.27715315 | -0.2707627 | 0.78657355 | 0.88383946 |
| AL356515.1 | 0.85350734 | | 0.024438254 | 1.60523366 | 0.01522411 | 0.98785339 | NA |
| LINC00330 | 11.8815572 | | 1.098697198 | 0.8180256 | 1.34310858 | 0.17923682 | 0.3548671 |
| NUFIP1 | 384.757277 | | 0.091027376 | 0.20577443 | 0.44236485 | 0.65822521 | 0.79946078 |
| AL359706.1 | 35.910047 | | 0.201296292 | 0.46554091 | 0.43239227 | 0.66545633 | 0.8037781 |
| GPALPP1 | 944.102033 | | 0.20515462 | 0.11792681 | 1.73967747 | 0.08191567 | 0.20580902 |
| GTF2F2 | 1234.94605 | | -0.713362313 | 0.15453318 | -4.6162405 | 3.91E-06 | 9.70E-05 |
| KCTD4 | 15.4019175 | | 0.513889613 | 0.62995809 | 0.81575207 | 0.41464195 | 0.60825557 |
| TPT1 | 127613.081 | | -0.270135171 | 0.24625493 | -1.0969737 | 0.2726529 | 0.46759554 |
| AL138963.3 | 71139.8188 | | -0.154776013 | 0.25296659 | -0.6118437 | 0.54064118 | 0.7136332 |
| AL138963.1 | 1.70988655 | | 1.801491028 | 1.31774118 | 1.36710536 | 0.17159228 | 0.34412415 |
| SLC25A30 | 3243.46487 | | 0.412209487 | 0.24474936 | 1.6842107 | 0.09214096 | 0.22442832 |
| SLC25A30-AS | 58.7527538 | | 0.989382574 | 0.27242872 | 3.63171168 | 0.00028155 | 0.00289 |
| COG3 | 1924.67985 | | -0.050210025 | 0.1357554 | -0.3698566 | 0.71148937 | 0.83602062 |
| ERICH6B | 186.685228 | | 0.278489492 | 0.17048626 | 1.63350114 | 0.10236364 | 0.24128842 |
| LINC01055 | 546.829142 | | 0.688742836 | 0.84663072 | 0.81351033 | 0.41592553 | 0.60920587 |
| SPERT | 4.0954675 | | 2.032712793 | 0.84552692 | 2.40407816 | 0.01621331 | 0.06439899 |
| SIAH3 | 63.7855138 | | 2.343475796 | 0.56561865 | 4.14320815 | 3.42E-05 | 0.00054635 |
| ZC3H13 | 1948.20177 | | 0.26450069 | 0.10492936 | 2.52075014 | 0.0117105 | 0.05068206 |
| CPB2-AS1 | 9.96338597 | | 1.047531884 | 0.51632096 | 2.02883857 | 0.04247474 | 0.12944167 |
| CPB2 | 2.20871245 | | 1.218407895 | 0.88812206 | 1.37189239 | 0.17009694 | 0.3420618 |
| LCP1 | 2075.37313 | | -0.151587174 | 0.51094441 | -0.2966804 | 0.76671055 | 0.87143473 |
| LRRC63 | 7.20501042 | | -0.04094253 | 0.78047967 | -0.0524582 | 0.95816363 | 0.97997808 |
| LINC00563 | 0.47556657 | | -0.851217142 | 2.25411837 | -0.3776275 | 0.70570731 | NA |
| RUBCNL | 166.195472 | | 0.21634611 | 0.50463176 | 0.42872076 | 0.66812645 | 0.80593269 |
| LINC01198 | 2.08516303 | | -0.755647592 | 0.96505766 | -0.7830077 | 0.43362258 | 0.62559511 |
| AL138686.1 | 0 | NA | | NA | NA | NA | NA |
| LRCH1 | 2096.00612 | | -0.014952168 | 0.13414567 | -0.1114622 | 0.91124986 | 0.95617599 |
| ESD | 5211.81799 | | 0.215589158 | 0.11640392 | 1.85207816 | 0.0640146 | 0.17299293 |
| HTR2A | 12.9287706 | | -0.730652586 | 0.89414462 | -0.8171526 | 0.41384124 | 0.60767896 |
| HTR2A-AS1 | 1.19574039 | | 1.455674134 | 2.18427076 | 0.66643484 | 0.5051332 | 0.68452748 |
| SUCLA2 | 3619.11275 | | 0.540022208 | 0.17578575 | 3.07204777 | 0.00212596 | 0.01410707 |
| AL138962.1 | 1.75371504 | | -0.049793729 | 1.2805419 | -0.0388849 | 0.96898216 | 0.98561465 |
| LINC00562 | 18.9850793 | | 0.685734945 | 0.34565591 | 1.98386581 | 0.0472708 | 0.13977697 |
| SUCLA2-AS1 | 31.9447993 | | -0.020774929 | 0.3172848 | -0.0654772 | 0.94779404 | 0.97485485 |
| NUDT15 | 1097.29268 | | 0.060371518 | 0.18359458 | 0.32883062 | 0.74228372 | 0.85534626 |
| AL158196.1 | 16.3664945 | | -0.760668816 | 0.5864723 | -1.2970243 | 0.19462283 | 0.37461344 |
| MED4 | 2032.57365 | | -0.327918016 | 0.13427657 | -2.442109 | 0.01460174 | 0.05969964 |
| MED4-AS1 | 833.679267 | | -0.300427354 | 0.14031138 | -2.1411474 | 0.03226215 | 0.10622638 |
| ITM2B | 79988.1582 | | 0.257873361 | 0.15746677 | 1.63763671 | 0.1014975 | 0.2399793 |
| RB1-DT | 7.68761931 | | 1.152556942 | 0.66084752 | 1.74405882 | 0.08114883 | 0.20446156 |

| | | | | | | |
|---|---|---|---|---|---|---|
| RB1 | 2287.93602 | -0.324995555 | 0.14940119 | -2.175321 | 0.02960607 | 0.09995034 |
| AL392048.1 | 4.08609829 | 0.493650695 | 0.76174174 | 0.6480552 | 0.51694925 | 0.69430334 |
| LPAR6 | 1867.05708 | -0.154884018 | 0.25637357 | -0.6041341 | 0.54575449 | 0.71715108 |
| RCBTB2 | 1646.95518 | 0.414439019 | 0.18428898 | 2.24885403 | 0.02452178 | 0.08699576 |
| LINC01077 | 0.38903887 | 2.550078411 | 3.21162345 | 0.79401538 | 0.42718648 | NA |
| LINC00462 | 4.10002526 | 0.798018761 | 1.09197262 | 0.73080474 | 0.46489843 | 0.65205955 |
| CYSLTR2 | 120.790357 | 0.63903323 | 0.341816 | 1.86952406 | 0.06154994 | 0.16833353 |
| AL161421.1 | 107.735558 | 0.346551897 | 0.29359999 | 1.1803539 | 0.23785949 | 0.42725597 |
| FNDC3A | 3498.4103 | 0.071724939 | 0.1386123 | 0.51745004 | 0.60484204 | 0.76179417 |
| MLNR | 3.88323261 | 1.655689032 | 0.69907483 | 2.36840028 | 0.0178652 | 0.06898597 |
| CDADC1 | 835.771919 | 0.75856278 | 0.22065643 | 3.4377552 | 0.00058656 | 0.0051353 |
| CAB39L | 909.625427 | 0.551801062 | 0.19509855 | 2.82831967 | 0.00467931 | 0.02557962 |
| SETDB2 | 1066.2452 | 0.17824644 | 0.1722845 | 1.03460519 | 0.30085333 | 0.49764664 |
| PHF11 | 2216.68944 | -0.743664427 | 0.32148879 | -2.3131893 | 0.02071223 | 0.07688368 |
| RCBTB1 | 1924.32913 | 0.530803397 | 0.14447145 | 3.67410587 | 0.00023868 | 0.00252307 |
| ARL11 | 40.0216641 | -0.265195309 | 0.40035176 | -0.6624057 | 0.50771122 | 0.68658055 |
| EBPL | 685.236733 | -0.541506955 | 0.22447895 | -2.4122839 | 0.01585293 | 0.06338446 |
| KPNA3 | 1622.77603 | -0.197244693 | 0.2387007 | -0.8263264 | 0.40861896 | 0.60308289 |
| AL136301.1 | 1.3966406 | -1.158596501 | 1.40602537 | -0.8240225 | 0.40992679 | 0.60440422 |
| SPRYD7 | 858.931795 | -0.00142271 | 0.17498567 | -0.0081304 | 0.99351292 | 0.9969533 |
| DLEU2 | 286.112248 | -0.202342544 | 0.13280741 | -1.5235787 | 0.12761398 | 0.28136963 |
| TRIM13 | 1842.59948 | -0.047083701 | 0.11484363 | -0.4099809 | 0.68181993 | 0.81577094 |
| KCNRG | 107.812922 | 1.204760986 | 0.20510957 | 5.87374336 | 4.26E-09 | 3.32E-07 |
| DLEU1-AS1 | 0.36399815 | 0.654464221 | 2.76512861 | 0.23668491 | 0.81290124 | NA |
| DLEU7 | 11.9130781 | 1.601294683 | 0.71714323 | 2.23287987 | 0.02555687 | 0.08972635 |
| DLEU7-AS1 | 3.61074668 | 0.008490551 | 0.95828842 | 0.00886012 | 0.99293074 | 0.99671045 |
| RNASEH2B-A | 14.8570214 | 0.316394493 | 0.37853917 | 0.83583027 | 0.40325039 | 0.59817937 |
| RNASEH2B | 867.796879 | -0.077636722 | 0.16347244 | -0.4749224 | 0.63484229 | 0.78341547 |
| C13orf42 | 7.56503942 | 2.917081901 | 0.98168036 | 2.97151905 | 0.0029633 | 0.01816614 |
| AL157817.1 | 0 | NA | NA | NA | NA | NA |
| FAM124A | 723.7092 | 0.096786766 | 0.46126417 | 0.20982936 | 0.83380086 | 0.91192105 |
| SERPINE3 | 78.3853719 | 0.534333091 | 0.1868369 | 2.85989064 | 0.00423787 | 0.023762 |
| INTS6 | 1520.3539 | 0.063206631 | 0.14364294 | 0.44002602 | 0.65991826 | 0.79999826 |
| INTS6-AS1 | 295.657406 | 0.45302053 | 0.23763068 | 1.90640592 | 0.05659756 | 0.15886407 |
| WDFY2 | 1265.39344 | 0.443849111 | 0.16080215 | 2.76021879 | 0.00577627 | 0.02997838 |
| AL162377.3 | 1309.80736 | 0.776660935 | 0.17141622 | 4.5308485 | 5.87E-06 | 0.0001339 |
| DHRS12 | 1040.40094 | 0.92558619 | 0.25837771 | 3.58229891 | 0.00034058 | 0.00335369 |
| AL162377.1 | 212.05725 | 0.77665504 | 0.28196602 | 2.75442783 | 0.00587949 | 0.03035684 |
| TMEM272 | 15.1982423 | 1.406943368 | 0.44344915 | 3.17272761 | 0.00151014 | 0.01087747 |
| CCDC70 | 0.26272731 | 1.990198932 | 3.242756 | 0.61373687 | 0.53938922 | NA |
| ATP7B | 605.495213 | 0.251818606 | 0.1650766 | 1.52546519 | 0.12714311 | 0.28070695 |
| ALG11 | 253.496032 | 0.005887408 | 0.15815367 | 0.03722587 | 0.97030491 | 0.98627531 |
| UTP14C | 1327.03255 | 0.463769206 | 0.09267231 | 5.00439902 | 5.60E-07 | 2.00E-05 |
| NEK5 | 125.965456 | 0.388767377 | 0.5055223 | 0.769041 | 0.44186897 | 0.63312784 |
| AL139082.1 | 37.4682446 | 0.021648553 | 0.34665046 | 0.06245067 | 0.95020395 | 0.97615326 |
| NEK3 | 979.452605 | -0.390330083 | 0.25351876 | -1.5396497 | 0.12364576 | 0.27519739 |
| LINC02333 | 0.45631946 | 0.399882992 | 2.84410371 | 0.14060071 | 0.88818539 | NA |
| THSD1 | 276.002155 | -0.129340092 | 0.26529319 | -0.4875364 | 0.62587824 | 0.77729476 |
| VPS36 | 2993.2125 | 0.355811218 | 0.07678517 | 4.63385333 | 3.59E-06 | 9.03E-05 |
| AL359513.1 | 25.624897 | 0.380356653 | 0.33624536 | 1.13118782 | 0.25797605 | 0.45051502 |
| CKAP2 | 432.443909 | 0.412424908 | 0.19866308 | 2.07600182 | 0.03789378 | 0.11897357 |
| LINC00345 | 0.81425937 | 0.592209437 | 1.52315642 | 0.38880408 | 0.69742108 | NA |
| AL137058.1 | 0.274636 | 1.131168378 | 2.99720785 | 0.37740739 | 0.70587088 | NA |
| HNRNPA1L2 | 439.320452 | 0.182237696 | 0.21353913 | 0.85341595 | 0.39342868 | 0.58885619 |
| AL139089.1 | 103.362174 | -0.042494726 | 0.30434648 | -0.1396261 | 0.88895538 | 0.94259304 |
| SUGT1 | 2728.43441 | -0.001842911 | 0.10017096 | -0.0183977 | 0.98532162 | 0.99349716 |
| CNMD | 1.90815733 | 1.18306211 | 1.05227834 | 1.12428629 | 0.26089159 | 0.45407638 |
| PCDH8 | 3.17628719 | -3.697594683 | 1.70234249 | -2.1720627 | 0.02985093 | 0.10051961 |
| OLFM4 | 7127.39595 | -5.081352022 | 0.65008332 | -7.8164627 | 5.43E-15 | 3.11E-12 |
| LINC01065 | 0.03392908 | 0.380740593 | 3.29738709 | 0.11546736 | 0.90807469 | NA |
| AL136359.1 | 0 | NA | NA | NA | NA | NA |
| AL450423.1 | 0.09997307 | 0.547239394 | 3.29377202 | 0.16614368 | 0.86804388 | NA |
| LINC00558 | 0 | NA | NA | NA | NA | NA |
| LINC00458 | 0 | NA | NA | NA | NA | NA |
| AL390964.1 | 0 | NA | NA | NA | NA | NA |
| LINC02335 | 0 | NA | NA | NA | NA | NA |
| AL354821.1 | 0.09635527 | 0.547239394 | 3.29377202 | 0.16614368 | 0.86804388 | NA |
| PRR20A | 0 | NA | NA | NA | NA | NA |
| PRR20C | 0 | NA | NA | NA | NA | NA |
| PRR20B | 0 | NA | NA | NA | NA | NA |
| PRR20D | 0 | NA | NA | NA | NA | NA |
| PRR20E | 0 | NA | NA | NA | NA | NA |
| PCDH17 | 466.595589 | -1.068061626 | 0.41953086 | -2.5458476 | 0.01090128 | 0.04804725 |
| AL445288.1 | 5.36982546 | 0.261059021 | 0.55926481 | 0.46678964 | 0.64065039 | 0.78733983 |
| LINC02338 | 0.18874271 | 0.726716446 | 3.28420131 | 0.22127646 | 0.82487718 | NA |
| LINC00374 | 0 | NA | NA | NA | NA | NA |
| DIAPH3 | 148.023722 | -0.623746072 | 0.51149664 | -1.2194529 | 0.22267234 | 0.40867689 |
| DIAPH3-AS1 | 1.85600325 | -0.039634251 | 1.06068544 | -0.0373666 | 0.97019267 | 0.98622863 |
| DIAPH3-AS2 | 0 | NA | NA | NA | NA | NA |
| LINC00434 | 1.05378201 | -0.933326914 | 1.37733675 | -0.6776316 | 0.4980053 | 0.67967569 |
| TDRD3 | 1064.9367 | 0.164453242 | 0.14816311 | 1.10994725 | 0.26702176 | 0.4611392 |
| LINC00378 | 0 | NA | NA | NA | NA | NA |
| LINC01442 | 0 | NA | NA | NA | NA | NA |
| PCDH20 | 107.709497 | 0.01789014 | 0.58338227 | 0.03066624 | 0.97553572 | 0.98866399 |
| AL592490.1 | 106.281838 | 0.007081414 | 0.58471678 | 0.01211084 | 0.99033718 | 0.99533983 |
| LINC02339 | 0.16588343 | 0.18657164 | 3.29377195 | 0.05664376 | 0.95482897 | NA |
| LINC00358 | 0 | NA | NA | NA | NA | NA |
| LINC01075 | 0 | NA | NA | NA | NA | NA |
| LINC01074 | 0 | NA | NA | NA | NA | NA |
| LINC00459 | 0 | NA | NA | NA | NA | NA |
| LINC00448 | 0 | NA | NA | NA | NA | NA |
| AL354810.1 | 0 | NA | NA | NA | NA | NA |
| LINC00376 | 0 | NA | NA | NA | NA | NA |
| LINC00395 | 0 | NA | NA | NA | NA | NA |
| AL445989.1 | 0 | NA | NA | NA | NA | NA |
| LINC00355 | 0.18301898 | 0.18657164 | 3.29377195 | 0.05664376 | 0.95482897 | NA |
| LINC01052 | 0 | NA | NA | NA | NA | NA |
| PCDH9 | 493.113346 | 1.07762019 | 0.36988229 | 2.91341388 | 0.003575 | 0.02090771 |
| PCDH9-AS1 | 7.36547243 | 1.777662427 | 0.6242832 | 2.84752565 | 0.00440605 | 0.02451216 |
| PCDH9-AS2 | 0.20718593 | 0.18657164 | 3.29377195 | 0.05664376 | 0.95482897 | NA |
| PCDH9-AS3 | 0.03392908 | 0.380740593 | 3.29738709 | 0.11546736 | 0.90807469 | NA |
| PCDH9-AS4 | 0.07659028 | -0.642905668 | 3.30558948 | -0.1944905 | 0.84579185 | NA |
| LINC00364 | 0 | NA | NA | NA | NA | NA |
| LINC00550 | 0 | NA | NA | NA | NA | NA |
| LINC02342 | 0 | NA | NA | NA | NA | NA |
| LINC00383 | 0 | NA | NA | NA | NA | NA |
| AL355478.1 | 0 | NA | NA | NA | NA | NA |
| KLHL1 | 7.42291545 | 3.273697904 | 0.77444621 | 4.22714691 | 2.37E-05 | 0.00040765 |
| ATXN8OS | 0.32318444 | 1.473795704 | 2.77691295 | 0.53073169 | 0.59560472 | NA |
| LINC00348 | 0.40653303 | -1.944918198 | 2.7564801 | -0.7055804 | 0.48044915 | NA |
| DACH1 | 746.143587 | 0.876980226 | 0.28057582 | 3.1256443 | 0.00177416 | 0.01230202 |
| MZT1 | 570.697455 | 0.014909702 | 0.17822322 | 0.08365746 | 0.93332878 | 0.96782855 |
| BORA | 302.321554 | -0.303684067 | 0.17522418 | -1.7331174 | 0.08307481 | 0.20800752 |

| | | | | | | |
|---|---|---|---|---|---|---|
| DIS3 | 1861.13582 | -0.101548675 | 0.16116304 | -0.630099 | 0.52862979 | 0.70325737 |
| PIBF1 | 731.859318 | -0.186955001 | 0.14783234 | -1.2646421 | 0.20599967 | 0.38865359 |
| KLF5 | 1456.07517 | -1.413662011 | 0.28435124 | -4.9715345 | 6.64E-07 | 2.29E-05 |
| LINC00393 | 0.36445536 | -1.037791048 | 2.86726535 | -0.3619445 | 0.71739347 | NA |
| LINC00392 | 0 | NA | NA | NA | NA | NA |
| KLF12 | 978.170096 | 0.779074852 | 0.17448898 | 4.46489423 | 8.01E-06 | 0.00017106 |
| LINC00402 | 4.54947885 | 3.096490226 | 1.30832531 | 2.36675863 | NA | NA |
| AL355390.1 | 0.10178723 | 1.087385526 | 3.28420134 | 0.33109588 | 0.74057207 | NA |
| LINC00381 | 0.60942373 | 1.347280264 | 1.82335863 | 0.73890031 | 0.45996753 | NA |
| AL355390.2 | 0.82948984 | 2.597205037 | 1.682309 | 1.54383352 | 0.12262867 | NA |
| LINC00347 | 0.54845261 | 1.092843893 | 2.72020152 | 0.40175108 | 0.68786723 | NA |
| LINC01078 | 0 | NA | NA | NA | NA | NA |
| TBC1D4 | 3859.87277 | 0.638527949 | 0.2286696 | 2.79236047 | 0.0052325 | 0.02781316 |
| AL139230.1 | 0.16052235 | -1.003574173 | 3.30558948 | -0.3035992 | 0.7614333 | NA |
| COMMD6 | 2403.76824 | -0.138169324 | 0.18187532 | -0.7596925 | 0.4474384 | 0.63765248 |
| UCHL3 | 1236.12008 | -0.793433705 | 0.24044616 | -3.2998394 | 0.0009674 | 0.00764032 |
| LMO7-AS1 | 426.212688 | -0.761267503 | 0.22079882 | -3.447788 | 0.0005652 | 0.0049964 |
| LMO7 | 4104.37529 | 0.807100681 | 0.27457955 | 2.93940568 | 0.00328842 | 0.01969469 |
| LMO7DN | 5.38137338 | -0.90627189 | 0.88719327 | -1.0215045 | 0.30701549 | 0.50401785 |
| LINC00561 | 0 | NA | NA | NA | NA | NA |
| LINC01034 | 0.17332742 | -1.003574173 | 3.30558948 | -0.3035992 | 0.7614333 | NA |
| AL365394.1 | 6.43259315 | 1.454374557 | 0.74645911 | 1.94836467 | 0.05137134 | 0.14855383 |
| KCTD12 | 3740.91902 | 0.21729932 | 0.32919902 | 0.66008496 | 0.50919931 | 0.68800961 |
| AC000403.1 | 16.3936093 | -0.166002167 | 0.54070451 | -0.3070109 | 0.75883509 | 0.8664871 |
| ACOD1 | 0.85842908 | 0.823332955 | 1.42200818 | 0.57899312 | 0.56259381 | NA |
| CLN5 | 4727.4611 | -0.09711494 | 0.1088929 | -0.8918391 | 0.3724792 | 0.56884364 |
| AC001226.2 | 1171.66785 | 0.048437354 | 0.15051972 | 0.32180071 | 0.74760368 | 0.85884766 |
| FBXL3 | 5558.13694 | 0.017923944 | 0.10805005 | 0.16588557 | 0.868247 | 0.93147444 |
| AC001226.1 | 12.3163815 | -0.216110544 | 0.44850393 | -0.4818476 | 0.62991421 | 0.78011215 |
| MYCBP2 | 1883.00612 | 0.245807895 | 0.10881236 | 2.25900705 | 0.02388295 | 0.08535593 |
| SCEL | 23.6791132 | -1.196907509 | 0.98539434 | -1.2146483 | 0.22450028 | 0.41060288 |
| SCEL-AS1 | 0.77850772 | -0.80164517 | 1.61304334 | -0.4969768 | 0.61920539 | NA |
| SLAIN1 | 434.495671 | -0.227546206 | 0.18612949 | -1.2225156 | 0.22151272 | 0.40759381 |
| EDNRB-AS1 | 0.64718716 | -0.468701249 | 2.23008264 | -0.2101721 | 0.83353332 | NA |
| EDNRB | 3479.37572 | -0.430818392 | 0.36363041 | -1.18477 | 0.23610841 | 0.42537259 |
| RNF219-AS1 | 190.456655 | 0.236604741 | 0.21350287 | 1.10820402 | 0.26777372 | 0.46196879 |
| AL139002.1 | 1.16614244 | 0.865561531 | 1.55784631 | 0.5556142 | 0.57847462 | 0.74199376 |
| LINC01069 | 0.3439254 | -1.717541376 | 2.27648342 | -0.7544713 | 0.45056628 | NA |
| LINC00446 | 0 | NA | NA | NA | NA | NA |
| AL138957.1 | 0 | NA | NA | NA | NA | NA |
| AL445209.1 | 0.26322705 | -0.178695909 | 2.98041902 | -0.0599566 | 0.95219017 | NA |
| POU4F1 | 1.5538482 | 0.496360122 | 0.91733476 | 0.54108941 | 0.58844596 | 0.74908617 |
| RNF219 | 531.831413 | -0.167977634 | 0.15547921 | -1.0803864 | 0.27997013 | 0.47554347 |
| LINC00331 | 0 | NA | NA | NA | NA | NA |
| RBM26 | 2477.97773 | -0.112810486 | 0.09070169 | -1.2437528 | 0.21359056 | 0.39755147 |
| RBM26-AS1 | 110.58584 | 0.930142394 | 0.28127862 | 3.3068364 | 0.00094356 | 0.00750909 |
| NDFIP2-AS1 | 2.97010675 | -0.111773758 | 0.9708261 | -0.1151326 | 0.90834 | 0.95418579 |
| NDFIP2 | 3522.66748 | -0.083089743 | 0.24202487 | -0.3433108 | 0.73136469 | 0.8482994 |
| LINC01068 | 16.2115706 | -1.371756195 | 0.83484545 | -1.6431259 | 0.10035688 | 0.23815794 |
| LINC01038 | 0 | NA | NA | NA | NA | NA |
| LINC00382 | 0 | NA | NA | NA | NA | NA |
| LINC01080 | 0 | NA | NA | NA | NA | NA |
| SPRY2 | 1384.89513 | -0.054890342 | 0.2785858 | -0.1970321 | 0.84380243 | 0.91722948 |
| LINC00377 | 4.20277689 | -1.123640212 | 0.75697886 | -1.4843746 | 0.13770956 | 0.29641672 |
| LINC00564 | 0.08695548 | -0.642905668 | 3.30558948 | -0.1944905 | 0.84579185 | NA |
| AL353633.1 | 0 | NA | NA | NA | NA | NA |
| SLITRK1 | 2.61923296 | 1.979193535 | 0.81185371 | 2.43786967 | 0.0147741 | 0.06015022 |
| LINC00333 | 0.09635527 | 0.547239394 | 3.29377202 | 0.16614368 | 0.86804388 | NA |
| LINC00375 | 0 | NA | NA | NA | NA | NA |
| LINC00351 | 0 | NA | NA | NA | NA | NA |
| SLITRK6 | 6.22545893 | 2.890843735 | 0.99080171 | 2.91768142 | 0.00352644 | 0.02069255 |
| AL354994.1 | 0 | NA | NA | NA | NA | NA |
| LINC00430 | 0 | NA | NA | NA | NA | NA |
| AL360267.1 | 38.8793715 | 1.460987052 | 0.79804407 | 1.83070975 | 0.06714388 | 0.1788506 |
| MIR4500HG | 50.891032 | 1.532291045 | 0.80489774 | 1.90370896 | 0.0569481 | 0.15948612 |
| AL355578.1 | 0.09644756 | -0.642905668 | 3.30558948 | -0.1944905 | 0.84579185 | NA |
| SLITRK5 | 18.4981825 | 0.739828555 | 0.65240073 | 1.13400939 | 0.25679061 | 0.44946026 |
| LINC00397 | 0 | NA | NA | NA | NA | NA |
| AL445647.1 | 0.33820869 | -0.556556868 | 2.29908657 | -0.2420774 | 0.8087202 | NA |
| AL354896.1 | 0.10721287 | -0.642905668 | 3.30558948 | -0.1944905 | 0.84579185 | NA |
| LINC00433 | 0 | NA | NA | NA | NA | NA |
| LINC00560 | 0.21442573 | -1.003574173 | 3.30558948 | -0.3035992 | 0.7614333 | NA |
| LINC01047 | 0 | NA | NA | NA | NA | NA |
| LINC00440 | 0 | NA | NA | NA | NA | NA |
| LINC01040 | 0 | NA | NA | NA | NA | NA |
| LINC00353 | 0.09989236 | -0.642905668 | 3.30558948 | -0.1944905 | 0.84579185 | NA |
| LINC02336 | 0.43725726 | -0.903320577 | 2.24951237 | -0.4015628 | 0.68800578 | NA |
| LINC00559 | 0.65684914 | 1.461211954 | 2.17671788 | 0.67129138 | 0.50203492 | NA |
| LINC01049 | 0 | NA | NA | NA | NA | NA |
| LINC00410 | 7.0243371 | 1.717573586 | 0.92009539 | 1.86673427 | 0.06193871 | 0.16915117 |
| LINC00380 | 7.76426294 | 2.459085969 | 0.85961561 | 2.86068092 | 0.00422732 | 0.02371798 |
| LINC00379 | 77.2087693 | 2.686190415 | 0.97970675 | 2.74183108 | 0.00610978 | 0.03124196 |
| GPC5 | 180.579681 | 2.199446287 | 0.8391562 | 2.62102131 | 0.00876668 | 0.04077201 |
| GPC5-AS2 | 0.12088456 | 0.02007176 | 3.29738704 | 0.00608717 | 0.99514317 | NA |
| GPC5-AS1 | 6.80132106 | 0.359845656 | 0.6338543 | 0.56771036 | 0.57023165 | 0.73541129 |
| LINC00363 | 0 | NA | NA | NA | NA | NA |
| AL354811.1 | 20.8765991 | -0.438260829 | 0.52740464 | -0.8309764 | 0.40598694 | 0.60084608 |
| GPC6 | 895.299068 | 0.676360098 | 0.33665076 | 2.00908529 | 0.04452809 | 0.13381262 |
| GPC6-AS2 | 0.79411377 | 1.174484601 | 1.73526971 | 0.67683116 | 0.49851309 | NA |
| GPC6-AS1 | 0.13028434 | 1.049816488 | 3.28477129 | 0.31960109 | 0.74927075 | NA |
| DCT | 37.0844455 | 1.415446916 | 0.52893175 | 2.67604829 | 0.00744959 | 0.03627137 |
| TGDS | 798.122682 | -0.233973552 | 0.10288967 | -2.2740238 | 0.02296455 | 0.08305342 |
| GPR180 | 836.930598 | -0.76139396 | 0.28922491 | -2.6325325 | 0.00847509 | 0.03974536 |
| LINC00391 | 0 | NA | NA | NA | NA | NA |
| SOX21 | 0.28329541 | -1.364244989 | 2.91116299 | -0.4686254 | 0.6393374 | NA |
| SOX21-AS1 | 0.72103591 | -2.009601099 | 1.63375804 | -1.2300482 | 0.21867906 | NA |
| LINC00557 | 0.17837751 | 0.726716446 | 3.28420131 | 0.22127646 | 0.82487718 | NA |
| ABCC4 | 2524.30781 | 0.159063529 | 0.24204714 | 0.6571593 | 0.51107849 | 0.689262 |
| CLDN10 | 6291.89848 | 0.455834235 | 0.3455045 | 1.31932937 | 0.18705902 | 0.36501786 |
| CLDN10-AS1 | 109.133362 | -0.117399291 | 0.46373295 | -0.2531614 | 0.80014348 | 0.89152759 |
| DZIP1 | 3476.0245 | 0.725245034 | 0.17293752 | 4.19368241 | 2.74E-05 | 0.00045749 |
| DNAJC3-DT | 321.58355 | 0.734597602 | 0.33840365 | 2.17077325 | 0.02994832 | 0.10073177 |
| DNAJC3 | 4177.962 | -0.384911567 | 0.22690615 | -1.696347 | 0.08982019 | 0.22026954 |
| UGGT2 | 1245.21718 | -0.347383127 | 0.15654647 | -2.2190415 | 0.0264839 | 0.09215893 |
| HS6ST3 | 97.5255344 | 0.010975437 | 0.44439635 | 0.02469741 | 0.98029632 | 0.99116093 |
| AL353581.1 | 0.56420008 | -1.123832195 | 2.29902378 | -0.4888302 | 0.62496193 | NA |
| OXGR1 | 96.4095252 | 1.526121155 | 0.62264408 | 2.45103297 | 0.01424469 | 0.0586376 |
| LINC00456 | 0 | NA | NA | NA | NA | NA |
| MBNL2 | 2927.93163 | 0.177604122 | 0.15971468 | 1.11200876 | 0.26613438 | 0.46028489 |
| RAP2A | 1660.36373 | 0.329532468 | 0.21878632 | 1.50618403 | 0.13201994 | 0.28815531 |
| AL359502.1 | 0.22077692 | -0.339747857 | 3.07884176 | -0.1103492 | 0.91213241 | NA |
| AL356580.1 | 0 | NA | NA | NA | NA | NA |
| IPO5 | 4841.16027 | -0.017776074 | 0.17988248 | -0.0988205 | 0.9212808 | 0.9615743 |

| | | | | | | |
|---|---|---|---|---|---|---|
| FARP1 | 17459.4729 | 0.047039097 | 0.20537139 | 0.22904406 | 0.81883467 | 0.90277909 |
| RNF113B | 5.79691143 | 0.026362091 | 0.52078186 | 0.05062022 | 0.95962815 | 0.98050555 |
| AL161896.1 | 1.75639555 | 1.328107456 | 0.84698827 | 1.56803524 | 0.11687291 | 0.26520955 |
| FARP1-AS1 | 762.127473 | 0.303612752 | 0.19604438 | 1.54869395 | 0.12145531 | 0.27204421 |
| STK24 | 13746.8203 | -0.391491909 | 0.1978032 | -1.9791991 | 0.0477936 | 0.14092853 |
| STK24-AS1 | 34.0402141 | 0.615933146 | 0.45796263 | 1.34494193 | 0.17864399 | 0.35418786 |
| SLC15A1 | 1685.10334 | -1.143394911 | 0.59561568 | -1.9196857 | 0.05489761 | 0.1555578 |
| DOCK9 | 1903.85142 | -0.268523881 | 0.26213894 | -1.0243571 | 0.30566664 | 0.50270434 |
| DOCK9-AS1 | 0.79744533 | 3.032145572 | 1.58248032 | 1.91607158 | 0.05535598 | NA |
| DOCK9-DT | 61.5876936 | -0.507210088 | 0.22170619 | -2.2877579 | 0.02215162 | 0.08086078 |
| UBAC2-AS1 | 52.2193126 | 0.654424395 | 0.36852226 | 1.77580696 | 0.07576474 | 0.19516618 |
| UBAC2 | 3497.89365 | -0.250510427 | 0.10104293 | -2.4792474 | 0.013166 | 0.05526292 |
| GPR18 | 22.7259256 | -0.293841013 | 0.63679047 | -0.4614407 | 0.64448249 | 0.78997117 |
| GPR183 | 373.190517 | -0.319649038 | 0.76477609 | -0.4179642 | 0.67597328 | 0.81149917 |
| AL136961.1 | 0.58142041 | -0.726274073 | 1.91314116 | -0.3796239 | 0.70422463 | NA |
| TM9SF2 | 12133.6424 | -0.021980477 | 0.15828625 | -0.1388654 | 0.88955655 | 0.94296756 |
| AL139035.1 | 8.92066612 | 1.559922478 | 0.629903 | 2.47644873 | 0.01326967 | 0.05554304 |
| LINC00449 | 7.99119002 | 1.615062897 | 0.68929306 | 2.34307147 | 0.01912572 | 0.07255722 |
| LINC01039 | 0.03392908 | 0.380740593 | 3.29738709 | 0.11546736 | 0.90807469 | NA |
| CLYBL | 1687.49885 | 1.10500264 | 0.37510267 | 2.94586716 | 0.00322051 | 0.01940694 |
| CLYBL-AS2 | 1.77663689 | -1.180856866 | 0.99004618 | -1.1927291 | 0.23297549 | 0.42116705 |
| CLYBL-AS1 | 0.09644756 | -0.642905668 | 3.30558948 | -0.1944905 | 0.84579185 | NA |
| ZIC5 | 1.56945313 | -3.874182769 | 2.2342878 | -1.7339677 | 0.08292382 | 0.20774285 |
| ZIC2 | 7.32409255 | -3.336635257 | 1.13457946 | -2.9408564 | 0.00327306 | 0.01962498 |
| LINC00554 | 0 | NA | NA | NA | NA | NA |
| AL355338.1 | 278.892162 | -0.267443977 | 0.18177287 | -1.4713086 | 0.14120769 | 0.30159593 |
| PCCA | 7069.34066 | 0.52972696 | 0.27786897 | 1.90639123 | 0.05659947 | 0.15886407 |
| PCCA-AS1 | 1.70279282 | 0.526139501 | 1.00260398 | 0.524773 | 0.599741 | 0.75764924 |
| GGACT | 3132.4812 | 1.833551455 | 0.36700973 | 4.99592053 | 5.86E-07 | 2.07E-05 |
| AL136526.1 | 14.335574 | 0.697629338 | 0.61211354 | 1.13970579 | 0.2544089 | 0.44713547 |
| TMTC4 | 1257.49081 | 0.410270856 | 0.13615625 | 3.01323558 | 0.00258478 | 0.01640845 |
| NALCN-AS1 | 10.8413703 | 0.299934395 | 0.51262262 | 0.58509785 | 0.55848192 | 0.72705215 |
| LINC00411 | 0 | NA | NA | NA | NA | NA |
| NALCN | 43.393491 | 0.563074197 | 0.42005153 | 1.3404884 | 0.18008661 | 0.35596162 |
| ITGBL1 | 250.306606 | 0.577508327 | 0.59583269 | 0.96924579 | 0.33242257 | 0.53033959 |
| FGF14 | 208.51489 | 0.486737346 | 0.5394935 | 0.9022117 | 0.36694442 | 0.56392123 |
| AL160153.1 | 11.2206832 | 0.824310724 | 0.62230532 | 1.32460819 | 0.18530114 | 0.36268231 |
| FGF14-AS1 | 0.16783122 | 1.368031398 | 3.28033652 | 0.41703996 | 0.67664918 | NA |
| FGF14-AS2 | 225.540487 | 0.217743088 | 0.27598261 | 0.78897395 | 0.43012723 | 0.62215211 |
| LINC00555 | 0.08757577 | 0.547239394 | 3.29377202 | 0.16614368 | 0.86804388 | NA |
| AL158063.1 | 23.5825557 | -0.08438857 | 0.58302829 | -0.1447418 | 0.88491473 | 0.94039898 |
| TPP2 | 1967.51168 | -0.044459537 | 0.11332342 | -0.3923244 | 0.69481857 | 0.82425506 |
| METTL21C | 0.30159974 | 1.471703365 | 2.82353541 | 0.52122717 | 0.60220853 | NA |
| CCDC168 | 11.4473696 | -0.494526079 | 0.43606848 | -1.134056 | 0.25677105 | 0.44945587 |
| LINC00283 | 0.16354576 | -1.003574173 | 3.30558948 | -0.3035992 | 0.7614333 | NA |
| TEX30 | 941.332455 | -0.223896251 | 0.20284745 | -1.1037666 | 0.26969438 | 0.4639947 |
| KDELC1 | 348.736617 | -0.134168954 | 0.21584549 | -0.6215972 | 0.53420676 | 0.7079978 |
| BIVM | 2525.09276 | 0.416603702 | 0.19126281 | 2.17817411 | 0.02939308 | 0.0994222 |
| BIVM-ERCC5 | 3677.27827 | 0.257498251 | 0.13139209 | 1.95976985 | 0.0500227 | 0.14571984 |
| ERCC5 | 3738.47818 | 0.258870757 | 0.13202687 | 1.96074293 | 0.04990902 | 0.14552198 |
| AL137246.1 | 4.46702982 | 0.361378957 | 0.62233601 | 0.58068142 | 0.56145519 | 0.72933859 |
| AL137246.2 | 4.9436329 | 1.338141115 | 0.78780664 | 1.69856542 | 0.0894011 | 0.21954805 |
| SLC10A2 | 465.607526 | 2.833114476 | 0.64893971 | 4.36575914 | 1.27E-05 | 0.00024745 |
| LINC01309 | 0.30897298 | 1.314953842 | 2.85791828 | 0.46010897 | 0.645438 | NA |
| DAOA-AS1 | 0 | NA | NA | NA | NA | NA |
| DAOA | 0 | NA | NA | NA | NA | NA |
| AL359745.1 | 0.03392908 | 0.380740593 | 3.29738709 | 0.11546736 | 0.90807469 | NA |
| LINC00460 | 0.19890045 | -0.340073434 | 3.15719848 | -0.1077137 | 0.91422283 | NA |
| EFNB2 | 1698.19039 | 0.31362103 | 0.3224607 | 0.97258683 | 0.33075869 | 0.52867582 |
| AL138689.2 | 270.084195 | 1.066777285 | 0.30911096 | 3.45111442 | 0.00055828 | 0.00495182 |
| ARGLU1 | 6474.97138 | -0.192049005 | 0.13712607 | -1.4005288 | 0.16135502 | 0.32998604 |
| LINC00551 | 125.68281 | 0.348379033 | 0.51862366 | 0.67173763 | 0.50175074 | 0.68226509 |
| LINC00443 | 8.74396272 | -0.198504088 | 0.60364995 | -0.3288397 | 0.74227683 | 0.85534626 |
| AL162574.2 | 0.96148313 | 1.797836665 | 1.69954456 | 1.05783438 | 0.29013096 | NA |
| AL162574.1 | 4.1398612 | 0.771135875 | 0.96242033 | 0.80124645 | 0.42298898 | 0.61559059 |
| FAM155A | 69.1347836 | -0.700517888 | 0.31027678 | -2.2577193 | 0.02396316 | 0.08554775 |
| AL136964.1 | 5.6093578 | 0.076959509 | 0.87147096 | 0.08830989 | 0.92963037 | 0.96596867 |
| LIG4 | 1038.69646 | -0.014350215 | 0.22689902 | -0.0632449 | 0.94957146 | 0.9757319 |
| ABHD13 | 941.340814 | -0.348883786 | 0.21267283 | -1.6404719 | 0.10090709 | 0.23901195 |
| TNFSF13B | 279.690225 | -1.169695121 | 0.53442009 | -2.1887185 | 0.02861731 | 0.09757414 |
| AL138694.1 | 0.32406811 | -1.684647081 | 2.88132138 | -0.5846786 | 0.55876381 | NA |
| MYO16 | 21.1744196 | 0.080774616 | 0.54639158 | 0.14783283 | 0.8824747 | 0.9389847 |
| MYO16-AS2 | 0 | NA | NA | NA | NA | NA |
| MYO16-AS1 | 0.39395308 | -1.797208921 | 2.81205183 | -0.6391095 | 0.52275173 | NA |
| AL161431.1 | 3.83081216 | -1.229162294 | 1.070965 | -1.1477147 | 0.25108635 | 0.44341998 |
| LINC00370 | 0.18382444 | -1.003574173 | 3.30558948 | -0.3035992 | 0.7614333 | NA |
| LINC01067 | 0.07830766 | -0.642905668 | 3.30558948 | -0.1944905 | 0.84579185 | NA |
| LINC00399 | 0.17515154 | 1.45420457 | 3.2783972 | 0.44357181 | 0.65735219 | NA |
| LINC00676 | 1.11597772 | -1.505528369 | 1.48344352 | -1.0148876 | 0.31015943 | 0.50725032 |
| IRS2 | 1274.86683 | -0.585384857 | 0.31952176 | -1.8320657 | 0.06694164 | 0.17839828 |
| AL162497.1 | 2.72111645 | 0.89788781 | 0.97253594 | 0.92324383 | 0.35588015 | 0.55326964 |
| AL355810.1 | 1.88004035 | 0.258968602 | 0.94870236 | 0.27297139 | 0.78487521 | 0.88249517 |
| AL355974.1 | 1.71467783 | 1.900517259 | 1.28972101 | 1.47358789 | 0.14059258 | 0.30075907 |
| AL355974.2 | 20.5652001 | -0.028918566 | 0.45213687 | -0.0639598 | 0.94900227 | 0.97537474 |
| AL355974.3 | 7.7040147 | -0.476910744 | 0.66103837 | -0.721457 | 0.4706284 | 0.65729945 |
| LINC00396 | 0.20710523 | -1.003574173 | 3.30558948 | -0.3035992 | 0.7614333 | NA |
| COL4A1 | 16114.5834 | -0.756186567 | 0.34453367 | -2.1948118 | 0.0281771 | 0.09658522 |
| COL4A2 | 24219.2455 | -0.555951354 | 0.27501454 | -2.0215344 | 0.04322447 | 0.13108371 |
| COL4A2-AS2 | 424.735989 | 0.086954994 | 0.31444338 | 0.27653625 | 0.78213622 | 0.8807709 |
| COL4A2-AS1 | 1542.14168 | -0.74619337 | 0.3174624 | -2.3504937 | 0.01874853 | 0.07162083 |
| RAB20 | 2523.01541 | -1.150885784 | 0.3210062 | -3.5852447 | 0.00033676 | 0.00332544 |
| AL139385.1 | 200.812204 | 0.411622342 | 0.20046546 | 2.05333303 | 0.04004029 | 0.12388389 |
| NAXD | 4515.59594 | 0.021034694 | 0.08641081 | 0.24342665 | 0.8076749 | 0.89613276 |
| CARS2 | 2813.90724 | -0.073932444 | 0.08189843 | -0.9027333 | 0.36666746 | 0.56375843 |
| ING1 | 916.326979 | -0.553445486 | 0.11570304 | -4.783327 | 1.72E-06 | 5.05E-05 |
| LINC00567 | 2.50231766 | -2.103250395 | 1.15111962 | -1.8271345 | 0.06767955 | 0.17976527 |
| LINC00346 | 89.0301517 | -0.237097129 | 0.30203898 | -0.7849885 | 0.43246031 | 0.62449911 |
| ANKRD10 | 7802.85999 | -0.639338343 | 0.13674543 | -4.675391 | 2.93E-06 | 7.70E-05 |
| AL442128.2 | 38.3676515 | 0.479858 | 0.31180042 | 1.53899088 | 0.12380652 | 0.27542002 |
| LINC00368 | 0.19271054 | 1.506623853 | 3.27645372 | 0.4598337 | 0.64563559 | NA |
| ARHGEF7-AS | 43.4837023 | 1.958274942 | 0.39101067 | 5.00823916 | 5.49E-07 | 1.97E-05 |
| ARHGEF7 | 3882.27396 | -0.272653454 | 0.13403156 | -2.0342481 | 0.04192658 | 0.12820084 |
| ARHGEF7-AS | 0.38874767 | 0.983752813 | 1.87003298 | 0.52606174 | 0.59884531 | NA |
| TEX29 | 35.9656287 | -0.262080078 | 0.34281195 | -0.764501 | 0.44456874 | 0.63537357 |
| AL359649.1 | 0 | NA | NA | NA | NA | NA |
| LINC02337 | 0 | NA | NA | NA | NA | NA |
| LINC00354 | 0 | NA | NA | NA | NA | NA |
| SOX1 | 0.27780865 | -1.364244915 | 2.92451635 | -0.4664857 | 0.64086792 | NA |
| AL356961.1 | 0 | NA | NA | NA | NA | NA |
| LINC00404 | 0.08393208 | -0.642905668 | 3.30558948 | -0.1944905 | 0.84579185 | NA |
| SOX1-OT | 0.16052235 | -1.003574173 | 3.30558948 | -0.3035992 | 0.7614333 | NA |
| LINC01070 | 0.12059279 | 0.02007176 | 3.29738704 | 0.00608717 | 0.99514317 | NA |
| LINC01043 | 0 | NA | NA | NA | NA | NA |

| | | | | | | |
|---|---|---|---|---|---|---|
| LINC01044 | 0 | NA | NA | NA | NA | NA |
| SPACA7 | 0 | NA | NA | NA | NA | NA |
| TUBGCP3 | 1059.70117 | 0.405967805 | 0.18754769 | 2.16461107 | 0.03041748 | 0.10182925 |
| AL139384.2 | 19.8444451 | 0.407981419 | 0.34341287 | 1.18802016 | 0.23482547 | 0.42361113 |
| ATP11AUN | 4.04998075 | 3.673611281 | 0.81286082 | 4.51936073 | 6.20E-06 | 0.00013992 |
| AL139384.1 | 70.3155919 | 0.116881844 | 0.43314219 | 0.26984636 | 0.78727845 | 0.88414299 |
| ATP11A | 4017.75722 | 0.189570415 | 0.2046842 | 0.92616047 | 0.35436259 | 0.55198426 |
| ATP11A-AS1 | 0.36038778 | -0.557471631 | 2.27457433 | -0.2450883 | 0.80638804 | NA |
| MCF2L | 3276.96626 | 0.491105792 | 0.28253502 | 1.73821211 | 0.08217345 | 0.20629415 |
| AL356740.1 | 62.9685229 | 0.423054072 | 0.40483169 | 1.04501223 | 0.29601732 | 0.49277045 |
| MCF2L-AS1 | 84.2958503 | 0.138869607 | 0.25512886 | 0.54431164 | 0.58622704 | 0.74756066 |
| F7 | 43.490154 | 1.997414848 | 0.30974852 | 6.44850499 | 1.13E-10 | 1.58E-08 |
| F10 | 288.077491 | 0.796360549 | 0.26927178 | 2.95746008 | 0.00310185 | 0.01883393 |
| F10-AS1 | 5.58072367 | 1.500476962 | 0.69256966 | 2.16653579 | 0.03027027 | 0.10144454 |
| AL137002.2 | 131.905716 | 1.306618771 | 0.22891333 | 5.70791911 | 1.14E-08 | 7.79E-07 |
| PROZ | 1078.03742 | 2.870519125 | 0.45073129 | 6.36858191 | 1.91E-10 | 2.49E-08 |
| AL137002.1 | 172.021922 | 2.970679372 | 0.49176286 | 6.04087789 | 1.53E-09 | 1.45E-07 |
| PCID2 | 1873.31654 | -0.236137831 | 0.13342561 | -1.7698089 | 0.07675898 | 0.19678536 |
| CUL4A | 4271.37594 | -0.098327055 | 0.17309833 | -0.5680416 | 0.57000671 | 0.73526766 |
| LAMP1 | 22387.4388 | 0.074462995 | 0.18992132 | 0.39207286 | 0.69500438 | 0.82443839 |
| GRTP1 | 1655.06626 | 0.202919655 | 0.29295249 | 0.69267087 | 0.48851612 | 0.6719613 |
| GRTP1-AS1 | 276.16865 | 0.738132764 | 0.3118609 | 2.36686539 | 0.01793946 | 0.06921192 |
| ADPRHL1 | 142.673806 | -0.158623201 | 0.21629924 | -0.7333507 | 0.46334456 | 0.65116158 |
| DCUN1D2 | 1027.69365 | -0.173099795 | 0.1897907 | -0.9120563 | 0.3617391 | 0.55911494 |
| DCUN1D2-AS | 5.29283775 | 1.142801607 | 0.5199818 | 2.19777232 | 0.02796534 | 0.09605616 |
| TMCO3 | 3557.8582 | -0.254804999 | 0.21214923 | -1.2010649 | 0.22972602 | 0.41720904 |
| TFDP1 | 2524.69749 | 0.038025094 | 0.19754353 | 0.19248969 | 0.84735864 | 0.91981776 |
| ATP4B | 25.5902926 | 1.182759948 | 0.58546094 | 2.02022007 | 0.04336057 | 0.13138207 |
| GRK1 | 9.24410529 | -0.733028037 | 0.7337908 | -0.9989605 | 0.31781382 | 0.5150412 |
| TMEM255B | 1520.2396 | -0.801608708 | 0.4539121 | -1.7659999 | 0.07739587 | 0.19799467 |
| GAS6-AS1 | 2363.14662 | 0.483231137 | 0.32854913 | 1.47080328 | 0.14134433 | 0.30175939 |
| GAS6 | 7971.37808 | 0.279606753 | 0.3004797 | 0.9305346 | 0.35209436 | 0.54974538 |
| GAS6-DT | 98.3251605 | 0.783166021 | 0.36745033 | 2.13135209 | 0.03306015 | 0.10814986 |
| BX072579.1 | 0.38025937 | 2.510697003 | 2.72775744 | 0.92042532 | 0.35735054 | NA |
| LINC00454 | 0.81386969 | 0.735250545 | 1.90019442 | 0.38693438 | 0.69880479 | NA |
| BX072579.2 | 0.33207757 | 0.65333381 | 2.82413371 | 0.23133955 | 0.81705103 | NA |
| LINC00452 | 4.51763149 | -0.805591455 | 0.72099444 | -1.1173338 | 0.26385163 | 0.45761781 |
| LINC00565 | 24.9991596 | -1.862608135 | 0.64325074 | -2.8956175 | 0.00378413 | 0.02190245 |
| C13orf46 | 18.7562843 | -0.880915153 | 0.51257635 | -1.7186028 | 0.08568672 | 0.21260434 |
| RASA3 | 1329.90049 | -0.840412448 | 0.31294041 | -2.6855351 | 0.00724138 | 0.03552654 |
| CFAP97D2 | 3.34133021 | -0.391332432 | 0.92736219 | -0.4219845 | 0.67303636 | 0.809264 |
| AL160396.1 | 0.25176329 | 1.011972812 | 2.94142038 | 0.34404222 | 0.73081453 | NA |
| CDC16 | 3546.66134 | -0.202399153 | 0.10789678 | -1.875859 | 0.06067465 | 0.16679962 |
| UPF3A | 2577.39952 | 0.051401271 | 0.21134816 | 0.24320662 | 0.80784534 | 0.89620864 |
| CHAMP1 | 799.242752 | 0.44025725 | 0.22248186 | 1.97884561 | 0.04783339 | 0.1410216 |
| LINC01054 | 0.08666371 | -0.642905668 | 3.30558948 | -0.1944905 | 0.84579185 | NA |
| CR383656.10 | 0 | NA | NA | NA | NA | NA |
| OR11H12 | 0 | NA | NA | NA | NA | NA |
| LINC02297 | 0 | NA | NA | NA | NA | NA |
| CR383656.12 | 0 | NA | NA | NA | NA | NA |
| AL929601.1 | 0 | NA | NA | NA | NA | NA |
| AL929601.2 | 0 | NA | NA | NA | NA | NA |
| POTEM | 0 | NA | NA | NA | NA | NA |
| AL929601.3 | 0 | NA | NA | NA | NA | NA |
| AL589182.1 | 0 | NA | NA | NA | NA | NA |
| AL589743.5 | 1.63226916 | 1.425388156 | 1.32491515 | 1.07583354 | 0.28200171 | 0.47782049 |
| AL589743.2 | 0 | NA | NA | NA | NA | NA |
| LINC01297 | 0.08757577 | 0.547239394 | 3.29377202 | 0.16614368 | 0.86804388 | NA |
| POTEG | 1.11824834 | -0.095660666 | 1.89982167 | -0.0503524 | 0.95984153 | 0.98061932 |
| AL512624.2 | 0 | NA | NA | NA | NA | NA |
| AL512310.10 | 0 | NA | NA | NA | NA | NA |
| AL512310.9 | 0 | NA | NA | NA | NA | NA |
| AL512310.2 | 0 | NA | NA | NA | NA | NA |
| OR11H2 | 0.09635527 | 0.547239394 | 3.29377202 | 0.16614368 | 0.86804388 | NA |
| OR4N2 | 0.03392908 | 0.380740593 | 3.29738709 | 0.11546736 | 0.90807469 | NA |
| OR4Q3 | 0 | NA | NA | NA | NA | NA |
| OR4M1 | 0 | NA | NA | NA | NA | NA |
| OR4K2 | 0 | NA | NA | NA | NA | NA |
| OR4K5 | 0 | NA | NA | NA | NA | NA |
| OR4K1 | 0 | NA | NA | NA | NA | NA |
| OR4K15 | 0 | NA | NA | NA | NA | NA |
| OR4K14 | 0 | NA | NA | NA | NA | NA |
| OR4K13 | 0.03392908 | 0.380740593 | 3.29738709 | 0.11546736 | 0.90807469 | NA |
| OR4L1 | 0 | NA | NA | NA | NA | NA |
| OR4K17 | 0 | NA | NA | NA | NA | NA |
| OR4N5 | 0 | NA | NA | NA | NA | NA |
| OR11G2 | 0 | NA | NA | NA | NA | NA |
| OR11H6 | 0.24013604 | -1.364242679 | 3.30558948 | -0.4127078 | 0.67982068 | NA |
| OR11H4 | 0.36348995 | -0.032132887 | 2.28169024 | -0.0140829 | 0.98876382 | NA |
| TTC5 | 694.077989 | 0.25020905 | 0.1403623 | 1.78259438 | 0.07465236 | 0.19309295 |
| AL356019.2 | 130.66796 | 0.578223193 | 0.22261011 | 2.59747049 | 0.00939132 | 0.04297225 |
| AL356019.1 | 34.6303257 | 0.47957914 | 0.31266133 | 1.53386137 | 0.12506376 | 0.27744018 |
| CCNB1IP1 | 2592.94123 | 0.277771889 | 0.19630593 | 1.4149949 | 0.15706999 | 0.32402221 |
| AL355075.4 | 97.2184512 | 0.267960609 | 0.40388798 | 0.66345279 | 0.50704061 | 0.68603166 |
| PARP2 | 1125.07284 | 0.350587468 | 0.13374694 | 2.62127469 | 0.00876016 | 0.0407489 |
| AL355075.1 | 97.3437555 | 0.627969587 | 0.22906224 | 2.7414802 | 0.0061163 | 0.03126322 |
| TEP1 | 2072.10272 | -0.560017275 | 0.14550863 | -3.8486878 | 0.00011875 | 0.00145246 |
| KLHL33 | 54.6756695 | 2.06435803 | 0.44836891 | 4.60415073 | 4.14E-06 | 0.0001014 |
| OSGEP | 2015.79849 | 0.143741038 | 0.17590723 | 0.81714118 | 0.41384775 | 0.60767896 |
| AL355075.2 | 79.583872 | 0.96411503 | 0.2591143 | 3.72080983 | 0.00019858 | 0.00216661 |
| APEX1 | 5334.79403 | -0.056058198 | 0.19935835 | -0.2811931 | 0.77856227 | 0.87861887 |
| PIP4P1 | 3005.02106 | -0.387288386 | 0.22443698 | -1.7255997 | 0.08441945 | 0.21048987 |
| PNP | 9702.63817 | 0.15263715 | 0.35107883 | 0.43476603 | 0.66373226 | 0.80246755 |
| AL355075.3 | 4247.58743 | 0.163525814 | 0.36156259 | 0.45227525 | 0.6510707 | 0.79489177 |
| RNASE10 | 0.23034971 | 0.68915899 | 3.0220972 | 0.22803998 | 0.81961516 | NA |
| RNASE9 | 0 | NA | NA | NA | NA | NA |
| AL163195.3 | 0.3069976 | -1.364242679 | 3.30558948 | -0.4127078 | 0.67982068 | NA |
| RNASE11 | 0.09989236 | -0.642905668 | 3.30558948 | -0.1944905 | 0.84579185 | NA |
| AL163195.2 | 1.19870838 | -3.469249132 | 3.25004244 | -1.0674473 | 0.28576988 | 0.48176817 |
| RNASE12 | 0.09989236 | -0.642905668 | 3.30558948 | -0.1944905 | 0.84579185 | NA |
| OR6S1 | 0 | NA | NA | NA | NA | NA |
| RNASE4 | 4599.64354 | -0.16181717 | 0.36623078 | -0.4418448 | 0.65860151 | 0.79946078 |
| ANG | 2088.71274 | -0.298607711 | 0.37388152 | -0.7986694 | 0.42448216 | 0.61693091 |
| AL163636.2 | 4689.76374 | -0.172013872 | 0.37191867 | -0.462504 | 0.64371993 | 0.78949326 |
| AL163636.1 | 1792.30919 | -0.151548747 | 0.36264704 | -0.417896 | 0.67602316 | 0.81149917 |
| EDDM3A | 17.8209336 | 1.014200988 | 0.43246353 | 2.34517116 | 0.01901835 | 0.07232677 |
| EDDM3B | 0.35900838 | 2.495453554 | 3.21417603 | 0.77638982 | 0.43751885 | NA |
| RNASE6 | 432.308845 | -0.724572435 | 0.51064 | -1.4189496 | 0.15591371 | 0.32239038 |
| AL133371.2 | 8.30170203 | 0.523936968 | 0.68587237 | 0.76389863 | 0.44492765 | 0.63551603 |
| RNASE1 | 7199.24394 | -0.271806717 | 0.36722148 | -0.7401711 | 0.4591962 | 0.64795611 |
| AL133371.3 | 1.07775971 | -0.116685438 | 1.38925482 | -0.0839914 | 0.93306328 | 0.96776057 |
| AL133371.1 | 0.29603988 | -0.695382734 | 2.33593535 | -0.2976892 | 0.76594038 | NA |
| RNASE3 | 17.3542232 | -3.133947079 | 0.77915807 | -4.0222224 | 5.77E-05 | 0.00082758 |

| | | | | | | |
|---|---|---|---|---|---|---|
| AL355922.5 | 0.57715502 | -1.113304423 | 1.66430895 | -0.6689289 | 0.50354081 | NA |
| AL355922.4 | 0.07659028 | -0.642905668 | 3.30558948 | -0.1944905 | 0.84579185 | NA |
| AL355922.1 | 4.91703726 | -2.564741482 | 0.98316794 | -2.6086505 | 0.00909 | 0.04195889 |
| RNASE2 | 97.9804648 | -2.804459959 | 0.7514444 | -3.7320924 | 0.0001899 | 0.00209247 |
| METTL17 | 4085.57514 | -0.00258988 | 0.10079992 | -0.0256933 | 0.97950199 | 0.99069759 |
| AL355922.2 | 1716.79556 | -0.230770334 | 0.12704091 | -1.8165041 | 0.06929308 | 0.18281726 |
| SLC39A2 | 3.14759049 | -0.490466914 | 1.06351899 | -0.4611736 | 0.64467404 | 0.79005793 |
| NDRG2 | 8995.6485 | 0.836048498 | 0.24302391 | 3.44019025 | 0.00058131 | 0.00510287 |
| TPPP2 | 120.66083 | 1.851567521 | 0.28607172 | 6.47238931 | 9.65E-11 | 1.40E-08 |
| AL161668.3 | 95.8160614 | 1.947330997 | 0.33426346 | 5.82573698 | 5.69E-09 | 4.31E-07 |
| AL161668.2 | 3.68187214 | 2.154326394 | 0.98826744 | 2.17990224 | 0.02926471 | 0.09915333 |
| RNASE13 | 8.29121753 | 2.133384044 | 0.75324751 | 2.83224837 | 0.00462219 | 0.02539846 |
| RNASE7 | 4.16592738 | -0.25770778 | 0.77967281 | -0.3305332 | 0.74099708 | 0.8546067 |
| RNASE8 | 0.3830574 | 0.761266088 | 2.85435907 | 0.26670299 | 0.78969785 | NA |
| ARHGEF40 | 3552.82445 | 0.316606147 | 0.09299094 | 3.40469895 | 0.00066237 | 0.00563806 |
| ZNF219 | 1843.55632 | -0.344173226 | 0.27006388 | -1.2744141 | 0.20251669 | 0.38442017 |
| TMEM253 | 51.1942056 | 0.436240538 | 0.29993449 | 1.45445274 | 0.14582083 | 0.30843789 |
| AL161668.1 | 0.34145041 | 0.67012343 | 2.22226185 | 0.30155017 | 0.762995 | NA |
| OR5AU1 | 0.25283891 | -0.178404819 | 2.99766869 | -0.0595145 | 0.9525423 | NA |
| HNRNPC | 17992.3639 | -0.439226324 | 0.13097848 | -3.3534237 | 0.00079818 | 0.00656318 |
| RPGRIP1 | 23.4802603 | 0.189963786 | 0.40938116 | 0.46402669 | 0.64262862 | 0.78844818 |
| SUPT16H | 3492.03483 | -0.057757123 | 0.12038718 | -0.4797614 | 0.63139707 | 0.78088081 |
| AL135744.1 | 4.2111435 | 0.321442119 | 0.79367738 | 0.4050035 | 0.68547494 | 0.81825122 |
| CHD8 | 2315.55709 | -0.125475484 | 0.14006607 | -0.8958307 | 0.37034319 | 0.56709541 |
| RAB2B | 1385.39387 | -0.138786019 | 0.16288298 | -0.8520597 | 0.39418098 | 0.58948004 |
| TOX4 | 4196.1122 | -0.168914403 | 0.08667649 | -1.9487915 | 0.05132033 | 0.14843887 |
| METTL3 | 3238.68897 | -0.263388564 | 0.12620977 | -2.0869111 | 0.03689617 | 0.11678694 |
| SALL2 | 641.759615 | 0.619040051 | 0.27691783 | 2.23546478 | 0.02538685 | 0.08934356 |
| AL161747.2 | 122.91861 | 0.292120521 | 0.30437559 | 0.95973702 | 0.33718759 | 0.53525383 |
| OR10G3 | 0.07830766 | -0.642905668 | 3.30558948 | -0.1944905 | 0.84579185 | NA |
| TRAV1-1 | 1.59060207 | 1.280236908 | 1.75477109 | 0.72957488 | 0.46565008 | 0.65288037 |
| OR10G2 | 0 | NA | NA | NA | NA | NA |
| TRAV1-2 | 2.82903682 | 1.191048303 | 1.41724796 | 0.84039515 | 0.40068687 | 0.59547525 |
| OR4E2 | 0 | NA | NA | NA | NA | NA |
| OR4E1 | 0 | NA | NA | NA | NA | NA |
| TRAV2 | 2.02459666 | 0.936841482 | 1.92596704 | 0.48642654 | 0.62666478 | 0.77783715 |
| AC243972.1 | 0.24953583 | -0.174096114 | 3.29377188 | -0.0528562 | 0.95784652 | NA |
| TRAV3 | 3.08743187 | 2.511251161 | 1.1707708 | 2.14495542 | 0.03195641 | 0.1055342 |
| TRAV4 | 3.90335679 | 0.980258387 | 1.17753905 | 0.83246359 | 0.40514732 | 0.59987898 |
| TRAV5 | 2.2459018 | 1.095371322 | 1.68912028 | 0.64848628 | 0.51667048 | 0.69396428 |
| TRAV6 | 1.00639685 | -1.30598051 | 1.87056837 | -0.6981731 | 0.48506895 | NA |
| TRAV7 | 0 | NA | NA | NA | NA | NA |
| TRAV8-1 | 1.58672322 | 1.990581833 | 1.91373302 | 1.0401565 | 0.2982672 | 0.49523398 |
| TRAV9-1 | 0 | NA | NA | NA | NA | NA |
| TRAV10 | 0.7699611 | -1.907184776 | 1.67350169 | -1.1396372 | 0.25443748 | NA |
| TRAV11 | 0.17332742 | -1.003574173 | 3.30558948 | -0.3035992 | 0.7614333 | NA |
| TRAV12-1 | 3.20124052 | -0.153758454 | 1.90492938 | -0.0807161 | 0.93566774 | 0.9692249 |
| TRAV8-2 | 42.0327464 | 0.668564324 | 0.67590122 | 0.98914502 | 0.3225922 | 0.5203279 |
| TRAV8-3 | 8.51996017 | 0.517413197 | 0.8391826 | 0.61656807 | 0.53751966 | 0.71113754 |
| TRAV13-1 | 5.5674509 | 0.055236177 | 1.04024251 | 0.05309933 | 0.95765277 | 0.97964557 |
| TRAV12-2 | 5.78136152 | -0.621936113 | 1.0489762 | -0.5928982 | 0.55324927 | 0.72291677 |
| TRAV8-4 | 4.28181869 | 1.392996401 | 1.19332766 | 1.16732097 | NA | NA |
| TRAV8-5 | 0.25543992 | -0.174096114 | 3.29377188 | -0.0528562 | 0.95784652 | NA |
| TRAV13-2 | 3.32637393 | -0.062399625 | 1.61328575 | -0.0386786 | 0.96914664 | 0.9857059 |
| TRAV14DV4 | 4.07800681 | 0.514997556 | 1.57380489 | 0.32723088 | 0.74349327 | 0.85611622 |
| TRAV9-2 | 5.20416096 | 1.723456715 | 1.50389657 | 1.14599418 | NA | NA |
| TRAV15 | 0.08393208 | -0.642905668 | 3.30558948 | -0.1944905 | 0.84579185 | NA |
| TRAV12-3 | 5.08020658 | 0.262391941 | 1.11188555 | 0.23598827 | 0.81344178 | 0.89922052 |
| TRAV8-6 | 6.44251249 | 0.337763711 | 1.22444963 | 0.27584941 | 0.78266373 | 0.88106052 |
| TRAV16 | 2.19396643 | 0.800285725 | 1.60449881 | 0.49877614 | 0.6179371 | 0.77161669 |
| TRAV17 | 6.03658723 | 0.871152844 | 1.26679767 | 0.68768112 | 0.49165361 | 0.67469326 |
| TRAV18 | 1.3001942 | 1.041469817 | 1.14215312 | 0.91184781 | 0.36184884 | 0.55919679 |
| TRAV19 | 6.52026297 | 0.18206945 | 0.92576843 | 0.19666846 | 0.84408699 | 0.91746317 |
| TRAV20 | 2.51598325 | -0.936063515 | 1.5696256 | -0.596361 | 0.55093408 | 0.72128546 |
| TRAV21 | 4.83186213 | 0.583889848 | 0.88723407 | 0.65810125 | 0.51047307 | 0.68892277 |
| TRAV22 | 3.54227093 | 1.052863818 | 1.14245991 | 0.92157616 | 0.3567497 | 0.55426204 |
| TRAV23DV6 | 2.59364361 | 2.395570378 | 1.30636052 | 1.83377432 | 0.0666875 | 0.17796486 |
| TRDV1 | 3.09014195 | 0.209519067 | 1.094699 | 0.19139422 | 0.84821674 | 0.92016931 |
| TRAV24 | 1.40714942 | -0.954768933 | 1.38149313 | -0.6911138 | 0.48949404 | 0.6728543 |
| TRAV25 | 2.04784063 | 0.640818348 | 1.37744837 | 0.46522132 | 0.64177297 | 0.78772823 |
| TRAV26-1 | 3.82538998 | 1.537214835 | 1.41341725 | 1.08758743 | 0.27677728 | 0.47218845 |
| TRAV8-7 | 0 | NA | NA | NA | NA | NA |
| TRAV27 | 4.26678188 | 2.568578628 | 0.89174638 | 2.88039141 | 0.00397182 | 0.02263601 |
| TRAV28 | 0.75690095 | 3.513603594 | 2.38017987 | 1.47619247 | 0.13989223 | NA |
| TRAV29DV5 | 6.13889476 | -0.18050508 | 0.98538145 | -0.1831829 | 0.85465447 | 0.92399813 |
| TRAV30 | 115.014678 | 0.626820679 | 0.74499856 | 0.84137167 | 0.40013975 | 0.59486331 |
| TRAV31 | 0 | NA | NA | NA | NA | NA |
| TRAV32 | 0 | NA | NA | NA | NA | NA |
| TRAV33 | 5.05177528 | 4.312736849 | 1.21835166 | 3.53981284 | 0.00040041 | 0.00382566 |
| TRAV26-2 | 1.9337594 | 1.767033573 | 1.1389293 | 1.55148662 | 0.12078511 | 0.27117997 |
| TRAV34 | 0.95964953 | 1.278585354 | 1.87456345 | 0.68207099 | 0.49519406 | NA |
| TRAV35 | 1.3571865 | 1.277287368 | 1.41864055 | 0.90036011 | 0.36792864 | 0.5647755 |
| TRAV36DV7 | 1.4374192 | 0.662747965 | 1.56849346 | 0.42253792 | 0.67263243 | 0.80897677 |
| TRAV37 | 0 | NA | NA | NA | NA | NA |
| TRAV38-1 | 0.71903698 | 1.709031628 | 2.10925694 | 0.81025294 | 0.41779482 | NA |
| TRAV38-2DV | 4.53054574 | 2.102100877 | 1.13677784 | 1.84917474 | 0.06443258 | 0.17371304 |
| TRAV39 | 6.24674009 | 2.349536421 | 0.71088037 | 3.30510803 | 0.0009494 | 0.0075454 |
| TRAV40 | 0.26090319 | -0.174096114 | 3.29377188 | -0.0528562 | 0.95784652 | NA |
| TRAV41 | 1.39775678 | 1.356137355 | 1.50976152 | 0.89824607 | 0.36905438 | 0.56591056 |
| AC244502.3 | 2.9185501 | -3.260551023 | 1.90331501 | -1.7130906 | 0.08669588 | 0.21450482 |
| TRDV2 | 1.50401835 | 1.706745381 | 1.40113045 | 1.21812025 | 0.22317828 | 0.40924076 |
| TRDD1 | 0 | NA | NA | NA | NA | NA |
| TRDD2 | 0 | NA | NA | NA | NA | NA |
| TRDD3 | 0 | NA | NA | NA | NA | NA |
| TRDJ1 | 3.41094195 | 0.179995639 | 0.95880693 | 0.18772876 | 0.85108928 | 0.92184442 |
| TRDJ4 | 0.25817155 | -0.178652511 | 2.98297197 | -0.0598908 | 0.95224263 | NA |
| TRDJ2 | 0.97236434 | 0.796600011 | 2.16239286 | 0.3683882 | 0.7125838 | NA |
| TRDJ3 | 0.26272731 | 1.990198932 | 3.242756 | 0.61373687 | 0.53938922 | NA |
| TRDC | 32.8275595 | 1.106300203 | 0.58838015 | 1.88024734 | 0.06007438 | 0.16560332 |
| TRDV3 | 0.58123986 | 1.318933024 | 1.51725331 | 0.86928993 | 0.38468857 | NA |
| TRAJ61 | 0.08666371 | -0.642905668 | 3.30558948 | -0.1944905 | 0.84579185 | NA |
| TRAJ60 | 0.09635527 | 0.547239394 | 3.29377202 | 0.16614368 | 0.86804388 | NA |
| TRAJ59 | 0 | NA | NA | NA | NA | NA |
| TRAJ58 | 0.46504618 | 0.61111714 | 1.65468539 | 0.36932528 | 0.71188529 | NA |
| TRAJ57 | 0.39108093 | -1.075706472 | 2.28686427 | -0.4703849 | 0.63808003 | NA |
| TRAJ56 | 0.6264177 | 1.322257035 | 1.84766601 | 0.71563639 | 0.4742159 | NA |
| TRAJ55 | 0.08393208 | -0.642905668 | 3.30558948 | -0.1944905 | 0.84579185 | NA |
| TRAJ54 | 1.15825716 | 0.87357285 | 1.87701662 | 0.46540496 | 0.64164148 | 0.78769526 |
| TRAJ53 | 0.72997948 | 2.363006201 | 2.11451731 | 1.11751565 | 0.26377391 | NA |
| TRAJ52 | 1.49275176 | 2.239406478 | 1.34482942 | 1.66519742 | 0.09587338 | 0.23084194 |
| TRAJ51 | 0 | NA | NA | NA | NA | NA |
| TRAJ50 | 0.66020778 | -2.625858405 | 3.2690507 | -0.803248 | 0.42183141 | NA |
| TRAJ49 | 2.73198739 | 1.278286644 | 1.2806454 | 0.99815815 | 0.31820268 | 0.51541797 |

| | | | | | | |
|---|---|---|---|---|---|---|
| TRAJ48 | 1.18733601 | -1.975695446 | 1.68486707 | -1.1726121 | 0.24095141 | 0.43106436 |
| TRAJ47 | 0.73616331 | 0.133077173 | 1.5062742 | 0.08834857 | 0.92959964 | NA |
| TRAJ46 | 0.08757577 | 0.547239394 | 3.29377202 | 0.16614368 | 0.86804388 | NA |
| TRAJ45 | 2.13165045 | 0.701149183 | 1.01454997 | 0.69109379 | 0.4895066 | 0.6728543 |
| TRAJ44 | 1.94568828 | 0.527898623 | 1.58913196 | 0.33219307 | 0.73974348 | 0.85396119 |
| TRAJ43 | 1.13065252 | 1.137214064 | 1.39916253 | 0.81278196 | 0.41634308 | 0.60960394 |
| TRAJ42 | 1.53158356 | 1.612099564 | 1.60134037 | 1.00671887 | 0.31406989 | 0.51094247 |
| TRAJ41 | 1.28783421 | -1.766941931 | 1.61821718 | -1.0919065 | 0.27487418 | 0.47007807 |
| TRAJ40 | 0.88930669 | 0.934437163 | 1.9032336 | 0.49097345 | 0.62344523 | NA |
| TRAJ39 | 5.68847478 | 1.573553408 | 0.62775476 | 2.50663714 | 0.01218858 | 0.05213508 |
| TRAJ38 | 0.45055796 | 1.404469946 | 2.08683053 | 0.67301581 | 0.50093723 | NA |
| TRAJ37 | 2.1027301 | 1.135954573 | 1.32597063 | 0.85669663 | 0.39161257 | 0.58731862 |
| TRAJ36 | 1.0560405 | 1.965707899 | 2.6438699 | 0.74349645 | 0.45718117 | 0.64655504 |
| TRAJ35 | 2.85179669 | 2.428021957 | 0.94510601 | 2.56904721 | 0.01019786 | 0.04571871 |
| TRAJ34 | 1.43104686 | 1.007249203 | 1.41433719 | 0.71217048 | 0.47635921 | 0.66176225 |
| TRAJ33 | 0.6903508 | -1.02888887 | 1.65253395 | -0.6226128 | 0.53353897 | NA |
| TRAJ32 | 1.00658624 | 0.94673363 | 1.71326723 | 0.55258959 | 0.58054447 | NA |
| TRAJ31 | 1.41088431 | 0.984419314 | 1.62797683 | 0.60468877 | 0.54538581 | 0.71697611 |
| TRAJ30 | 1.19239687 | 1.262883923 | 1.41627825 | 0.89169196 | 0.37255807 | 0.56884873 |
| TRAJ29 | 1.53330374 | 0.245385774 | 1.28892122 | 0.19038074 | 0.84901079 | 0.92059733 |
| TRAJ28 | 1.06929772 | -0.611181932 | 2.00606827 | -0.3046666 | 0.76062012 | 0.86731611 |
| TRAJ27 | 4.68109031 | 2.562206677 | 0.90528406 | 2.83027923 | 0.00465074 | 0.02548099 |
| TRAJ26 | 0.88409754 | 0.366566651 | 2.2282359 | 0.1645098 | 0.86932983 | NA |
| TRAJ25 | 0 | NA | NA | NA | NA | NA |
| TRAJ24 | 0.87389334 | 0.417113461 | 1.86520824 | 0.22362836 | 0.82304649 | NA |
| TRAJ23 | 1.68640406 | 0.04509384 | 1.90806097 | 0.02363333 | 0.98114508 | 0.9915228 |
| TRAJ22 | 2.26665634 | 1.405883513 | 0.93975841 | 1.49600525 | 0.13465229 | 0.29205304 |
| TRAJ21 | 0.46725217 | 1.448882221 | 2.17876694 | 0.66500101 | 0.50604985 | NA |
| TRAJ20 | 1.50722123 | 0.225116555 | 2.29776074 | 0.09797215 | 0.92195441 | 0.96185859 |
| TRAJ19 | 0.08757577 | 0.547239394 | 3.29377202 | 0.16614368 | 0.86804388 | NA |
| TRAJ18 | 1.52302483 | 1.826364965 | 1.50447503 | 1.21395499 | 0.22476491 | 0.41097287 |
| TRAJ17 | 0.86375341 | 1.553070666 | 2.68948202 | 0.57746089 | 0.56362815 | NA |
| TRAJ16 | 0.53660138 | 0.173153764 | 2.17327298 | 0.07967419 | 0.93649639 | NA |
| TRAJ14 | 0 | NA | NA | NA | NA | NA |
| TRAJ13 | 1.25448472 | 3.304035059 | 1.68917559 | 1.9560045 | 0.05046462 | 0.14660704 |
| TRAJ12 | 1.49130835 | -1.926492934 | 1.60810614 | -1.1979887 | 0.23092143 | 0.4186592 |
| TRAJ11 | 0.60300682 | -0.275062622 | 2.76134467 | -0.0996118 | 0.92065249 | NA |
| TRAJ10 | 1.39740751 | 0.052786788 | 1.67568095 | 0.03150169 | 0.97486944 | 0.98843899 |
| TRAJ9 | 1.49452343 | 0.824059692 | 1.73036372 | 0.47623496 | 0.633907 | 0.78295679 |
| TRAJ8 | 0.3833201 | 1.814406794 | 2.76357843 | 0.65654254 | 0.51147511 | NA |
| TRAJ7 | 0.12150485 | 1.04981648 | 3.28477129 | 0.31960109 | 0.74927075 | NA |
| TRAJ6 | 1.86820665 | -0.564314945 | 1.26226392 | -0.4470657 | 0.6548276 | 0.79747572 |
| TRAJ5 | 2.02838539 | -1.328855283 | 1.28374522 | -1.0351394 | 0.3006038 | 0.49739112 |
| TRAJ4 | 1.72746659 | 0.442045056 | 1.69380824 | 0.26097704 | 0.79411022 | 0.88844572 |
| TRAJ3 | 1.16023438 | -1.145407497 | 1.99226556 | -0.5749271 | 0.56534059 | 0.73208851 |
| TRAJ2 | 0.08666371 | -0.642905668 | 3.30558948 | -0.1944905 | 0.84579185 | NA |
| TRAJ1 | 0.27068704 | -0.17922184 | 2.94998951 | -0.0607534 | 0.95155562 | NA |
| TRAC | 403.595497 | 0.493540391 | 0.6793593 | 0.72647919 | 0.46754506 | 0.65455658 |
| LINC02332 | 5.46623612 | 0.938897418 | 0.79978039 | 1.17394403 | 0.24041745 | 0.43054724 |
| AC243965.1 | 4.98587642 | 1.165049757 | 0.8094839 | 1.43925006 | 0.15007969 | 0.3144947 |
| DAD1 | 6308.95587 | -0.50865075 | 0.21383309 | -2.378728 | 0.01737249 | 0.06766803 |
| AC243965.2 | 153.241683 | -0.367639528 | 0.26806058 | -1.3714793 | 0.1702256 | 0.34219004 |
| ABHD4 | 4953.40795 | -0.918030917 | 0.25493645 | -3.6010187 | 0.00031697 | 0.00316044 |
| OR6J1 | 0.81352629 | 0.589656151 | 1.5277524 | 0.38596317 | 0.69952395 | NA |
| OXA1L | 6840.53546 | 0.129395383 | 0.17032842 | 0.75968169 | 0.44744488 | 0.63765248 |
| SLC7A7 | 9608.1271 | 1.813439196 | 0.50816375 | 3.56861185 | 0.00035888 | 0.00349985 |
| MRPL52 | 2343.03217 | -0.422013159 | 0.13398991 | -3.149589 | 0.001635 | 0.01156209 |
| MMP14 | 9521.01476 | -1.226199811 | 0.2056547 | -5.9624204 | 2.49E-09 | 2.14E-07 |
| LRP10 | 22514.0072 | -0.750743884 | 0.15632617 | -4.8024196 | 1.57E-06 | 4.69E-05 |
| REM2 | 83.9419 | -0.15058935 | 0.24219141 | -0.6217782 | 0.53408769 | 0.70787558 |
| RBM23 | 4590.72871 | 0.061949407 | 0.08309341 | 0.74553933 | 0.45594574 | 0.6453032 |
| PRMT5-AS1 | 2655.26289 | -0.008959905 | 0.13059551 | -0.0686081 | 0.9453016 | 0.97390119 |
| PRMT5 | 3298.32189 | 0.019363647 | 0.13429074 | 0.14419197 | 0.88534888 | 0.9406386 |
| AL132780.1 | 259.66694 | 0.621531141 | 0.29706109 | 2.09226709 | 0.03641463 | 0.11576297 |
| HAUS4 | 1822.25807 | 0.552590317 | 0.24426528 | 2.26225483 | 0.02368166 | 0.08487163 |
| AL132780.3 | 1265.43358 | 0.520834732 | 0.23811021 | 2.18736834 | 0.02871565 | 0.09780826 |
| AJUBA | 2092.84187 | 0.672411403 | 0.28366994 | 2.37040059 | 0.01776882 | 0.068705 |
| AL132780.2 | 206.199331 | 0.504055876 | 0.18952283 | 2.6596051 | 0.00782323 | 0.03758697 |
| C14orf93 | 669.253909 | 0.253971204 | 0.18866711 | 1.34613393 | 0.17825933 | 0.35372853 |
| PSMB5 | 5501.79717 | -0.326287448 | 0.16614058 | -1.9639238 | 0.04953893 | 0.14473095 |
| PSMB11 | 0.80666941 | 0.418677592 | 1.6353978 | 0.25600963 | 0.7979434 | NA |
| CDH24 | 547.815247 | -0.021173472 | 0.26089989 | -0.0811555 | 0.93531826 | 0.96907653 |
| ACIN1 | 9650.20723 | -0.343497916 | 0.13729762 | -2.501849 | 0.01235466 | 0.05274119 |
| C14orf119 | 2671.60114 | -0.543704353 | 0.14059645 | -3.8671271 | 0.00011013 | 0.00136723 |
| LMLN2 | 23.7787776 | -0.101942497 | 0.28247527 | -0.36089 | 0.71818168 | 0.83985952 |
| CEBPE | 8.8496002 | -0.646329721 | 0.60827575 | -1.0625604 | 0.28798135 | 0.48416119 |
| SLC7A8 | 14895.1277 | 2.036227293 | 0.51495069 | 3.954218 | 7.68E-05 | 0.00102888 |
| RNF212B | 393.733646 | 3.430391859 | 0.52921179 | 6.48207749 | 9.05E-11 | 1.33E-08 |
| HOMEZ | 1357.43815 | 0.547379905 | 0.21389229 | 2.55913807 | 0.01049321 | 0.04668565 |
| PPP1R3E | 1453.8799 | 0.524443616 | 0.31014378 | 1.69096933 | 0.09084266 | 0.22211599 |
| BCL2L2 | 4911.19757 | 0.225738176 | 0.12849639 | 1.75676666 | 0.07895759 | 0.20061667 |
| BCL2L2-PABI | 6860.49896 | -0.047580278 | 0.12179803 | -0.390649 | 0.69605669 | 0.82515362 |
| PABPN1 | 13234.9041 | -0.232582698 | 0.14781284 | -1.5734946 | 0.11560435 | 0.26311813 |
| SLC22A17 | 3456.40244 | -0.015410336 | 0.26925565 | -0.0572331 | 0.95435951 | 0.97813596 |
| EFS | 644.001286 | 0.212206192 | 0.45570382 | 0.46566691 | 0.64145394 | 0.78758115 |
| AL049829.2 | 132.205524 | -0.108096778 | 0.42051215 | -0.2570598 | 0.79713259 | 0.88994016 |
| IL25 | 0.57104577 | 1.136382101 | 1.52919917 | 0.74312236 | 0.4574076 | NA |
| CMTM5 | 16.7693474 | 0.604487673 | 0.6583949 | 0.91812326 | 0.35855434 | 0.55624649 |
| MYH6 | 1.61451272 | -0.795345171 | 1.30558139 | -0.6091885 | 0.5423995 | 0.71483207 |
| MYH7 | 26.7805106 | 2.496273054 | 1.1339772 | 2.20134324 | NA | NA |
| AL132855.1 | 5.28087137 | 2.728285154 | 1.45008423 | 1.88146667 | NA | NA |
| NGDN | 1461.22193 | -0.359810154 | 0.14428177 | -2.4938018 | 0.01263831 | 0.05362384 |
| ZFHX2 | 447.717319 | 0.577086336 | 0.27784234 | 2.07702808 | 0.03779897 | 0.11878903 |
| AL135999.2 | 8.24752647 | 1.334456802 | 0.54700795 | 2.43955649 | 0.0147053 | 0.06001023 |
| THTPA | 2794.69869 | -0.256780506 | 0.15850278 | -1.6200379 | 0.10522415 | 0.24603712 |
| AP1G2 | 8508.40091 | -0.555815145 | 0.20355625 | -2.7305236 | 0.00632338 | 0.03203603 |
| AL135999.1 | 4161.38403 | -0.395287051 | 0.20425529 | -1.9352598 | 0.05295842 | 0.15163517 |
| JPH4 | 220.849108 | 0.572942187 | 0.32943754 | 1.7391527 | 0.08200791 | 0.20597662 |
| AL135999.3 | 20.0693281 | 1.11710804 | 0.80214977 | 1.39264273 | 0.16372785 | 0.33321784 |
| DHRS2 | 26.2631267 | 0.554774697 | 0.57886402 | 0.95838518 | 0.33786857 | 0.53585097 |
| LINC00596 | 0.51932321 | 2.968620637 | 2.56450659 | 1.15757965 | 0.24703562 | NA |
| DHRS4-AS1 | 3075.29389 | 1.64832804 | 0.386104 | 4.26912961 | 1.96E-05 | 0.00034918 |
| DHRS4 | 3229.17302 | 1.826319645 | 0.43668126 | 4.18227158 | 2.89E-05 | 0.00047656 |
| DHRS4L2 | 2320.42724 | 1.486238901 | 0.36237822 | 4.10134721 | 4.11E-05 | 0.00063491 |
| CARMIL3 | 210.655818 | 0.947313003 | 0.31011365 | 3.05472852 | 0.00225264 | 0.01480244 |
| CPNE6 | 266.604089 | 7.531920763 | 0.76051354 | 9.90373005 | 4.01E-23 | 2.12E-19 |
| NRL | 251.558168 | 0.862516145 | 0.23475787 | 3.67406704 | 0.00023872 | 0.00252307 |
| PCK2 | 12487.8729 | 3.016388728 | 0.40957809 | 7.36462431 | 1.78E-13 | 6.25E-11 |
| DCAF11 | 6126.12703 | 0.547970993 | 0.15712686 | 3.48744309 | 0.00048766 | 0.00444228 |
| AL136295.3 | 1326.21778 | 0.593353935 | 0.15106464 | 3.92781486 | 8.57E-05 | 0.00112349 |
| FITM1 | 93.739206 | -0.042899318 | 0.16610978 | -0.2582588 | 0.79620716 | 0.88939648 |
| PSME1 | 8877.22007 | -0.439955919 | 0.16643132 | -2.6434683 | 0.00820615 | 0.03879721 |
| EMC9 | 1116.47901 | -0.551518873 | 0.23176542 | -2.3796426 | 0.01732944 | 0.06758019 |

| | | | | | | |
|---|---|---|---|---|---|---|
| AL136295.2 | 450.558725 | -0.655557632 | 0.23555472 | -2.7830376 | 0.00538526 | 0.02845876 |
| PSME2 | 6320.29684 | -0.850270412 | 0.28546296 | -2.9785665 | 0.002896 | 0.01784706 |
| RNF31 | 3300.61343 | -0.184517498 | 0.19384548 | -0.9518793 | 0.3411582 | 0.53918739 |
| AL136295.5 | 7698.76015 | -0.740162134 | 0.29628301 | -2.4981593 | 0.01248401 | 0.05311445 |
| IRF9 | 6411.81782 | -0.891124785 | 0.34991104 | -2.5467181 | 0.01087412 | 0.04795161 |
| REC8 | 2215.49625 | -0.170346609 | 0.36501058 | -0.4666895 | 0.64072204 | 0.7873637 |
| IPO4 | 4212.54671 | -0.450840342 | 0.18462963 | -2.4418635 | 0.01461167 | 0.05972869 |
| AL136295.4 | 5673.69093 | -0.451337649 | 0.16516536 | -2.7326411 | 0.00628288 | 0.03188596 |
| TM9SF1 | 4458.62785 | -0.186411057 | 0.10746509 | -1.7346196 | 0.08280821 | 0.20759126 |
| AL136295.1 | 6093.73319 | -0.196065041 | 0.10178617 | -1.9262443 | 0.05407387 | 0.15395466 |
| AL136295.6 | 60.5876552 | -0.014809257 | 0.26326423 | -0.0562524 | 0.9551407 | 0.97848026 |
| TSSK4 | 44.3519495 | 1.192473419 | 0.30090063 | 3.96301399 | 7.40E-05 | 0.00099777 |
| CHMP4A | 2199.58713 | -0.292397507 | 0.15491861 | -1.8874266 | 0.05910297 | 0.16379876 |
| MDP1 | 574.300348 | 0.088058809 | 0.17173029 | 0.51277388 | 0.6081095 | 0.76430886 |
| NEDD8-MDP | 3981.10387 | -0.235015092 | 0.10079625 | -2.3315858 | 0.01972249 | 0.07411745 |
| NEDD8 | 4581.01414 | -0.354691876 | 0.10843222 | -3.2710931 | 0.00107133 | 0.00827743 |
| GMPR2 | 4552.8495 | 0.258771017 | 0.17079604 | 1.51508793 | 0.12975014 | 0.28474969 |
| TINF2 | 2825.36605 | -0.412799238 | 0.18592171 | -2.2202853 | 0.0263994 | 0.09197418 |
| TGM1 | 174.810348 | 0.132997469 | 0.37447213 | 0.35515986 | 0.72246984 | 0.84246598 |
| RABGGTA | 2032.37749 | 0.047577256 | 0.13973782 | 0.34047516 | 0.73349873 | 0.84980279 |
| DHRS1 | 2381.42148 | 0.055569221 | 0.12875191 | 0.43159919 | 0.66603275 | 0.80431161 |
| NOP9 | 5050.70187 | -0.224888337 | 0.32847286 | -0.6846482 | 0.49356597 | 0.67619153 |
| CIDEB | 4426.78774 | -0.33833582 | 0.35783427 | -0.9455098 | 0.34439866 | 0.54203477 |
| LTB4R2 | 1098.58971 | -0.405568895 | 0.38719647 | -1.0474499 | 0.29489213 | 0.4916937 |
| LTB4R | 1243.27634 | -1.720708147 | 0.45358017 | -3.7936142 | 0.00014847 | 0.0017393 |
| ADCY4 | 1988.59401 | 0.052579244 | 0.28212898 | 0.18636598 | 0.85215777 | 0.92251225 |
| AL096870.1 | 269.563813 | -0.29595145 | 0.2504338 | -1.1817552 | 0.23730285 | 0.42653371 |
| RIPK3 | 589.060758 | -0.705943925 | 0.28096581 | -2.5125616 | 0.01198582 | 0.05148699 |
| NFATC4 | 3756.21512 | -1.074545952 | 0.38447252 | -2.7948577 | 0.00519226 | 0.02765498 |
| NYNRIN | 1592.70014 | 0.595721393 | 0.46056751 | 1.29345074 | 0.19585521 | 0.37613443 |
| CBLN3 | 488.275189 | -0.611911569 | 0.34869485 | -1.7548626 | 0.07928281 | 0.20115808 |
| KHNYN | 7396.87954 | -0.331926432 | 0.10869262 | -3.0538083 | 0.00225956 | 0.01483313 |
| SDR39U1 | 5926.213 | -0.374068804 | 0.11913994 | -3.1397432 | 0.00169096 | 0.01184574 |
| AL132800.1 | 15.8870848 | 1.530738405 | 0.54068872 | 2.83108996 | 0.00463897 | 0.02545024 |
| CMA1 | 13.0698687 | 1.09109631 | 0.72214945 | 1.51090097 | 0.13081368 | 0.28629404 |
| CTSG | 84.8537541 | -1.395653886 | 0.94801614 | -1.4721837 | 0.14097129 | 0.30129785 |
| AL136018.1 | 1.15399751 | -0.448546371 | 1.28812483 | -0.3482165 | 0.72767757 | 0.84611931 |
| GZMH | 87.4905604 | -1.455800241 | 0.71584011 | -2.0336947 | 0.04198239 | 0.12833073 |
| GZMB | 125.91176 | -1.37501042 | 0.70218889 | -1.9581774 | 0.0502092 | 0.14604466 |
| STXBP6 | 138.38612 | 1.517946075 | 0.4140247 | 3.66631766 | 0.00024607 | 0.00258828 |
| LINC02286 | 0 | NA | NA | NA | NA | NA |
| LINC02306 | 0.46321216 | -1.338803912 | 1.89915447 | -0.7049474 | 0.480843 | NA |
| NOVA1 | 508.740294 | -0.4171127 | 0.22213667 | -1.8777301 | 0.0604181 | 0.16632858 |
| LINC02588 | 0.90881274 | 1.111126299 | 1.88138538 | 0.59058942 | 0.55479556 | NA |
| LINC02294 | 195.873893 | 2.1526012 | 0.68751039 | 3.13100897 | 0.00174207 | 0.01213695 |
| LINC02293 | 99.1597826 | 1.928335458 | 0.66078566 | 2.91824652 | 0.00352006 | 0.02066428 |
| AL110292.1 | 6.10074107 | 1.599500296 | 0.76588895 | 2.08842325 | 0.03675967 | 0.11652273 |
| MIR4307HG | 0.38379647 | 1.697458365 | 2.22752156 | 0.76203903 | 0.44603671 | NA |
| LINC00645 | 464.004428 | 1.694007767 | 0.35586251 | 4.76028721 | 1.93E-06 | 5.52E-05 |
| AL139023.1 | 0 | NA | NA | NA | NA | NA |
| LINC02300 | 0 | NA | NA | NA | NA | NA |
| FOXG1-AS1 | 0.03392908 | 0.380740593 | 3.29738709 | 0.11546736 | 0.90807469 | NA |
| FOXG1 | 2.95233074 | -0.228844724 | 1.10705405 | -0.206715 | 0.83623241 | 0.91299005 |
| LINC01551 | 0.08666371 | -0.642905668 | 3.30558948 | -0.1944905 | 0.84579185 | NA |
| LINC02282 | 0.03392908 | 0.380740593 | 3.29738709 | 0.11546736 | 0.90807469 | NA |
| LINC02281 | 0 | NA | NA | NA | NA | NA |
| AL049777.1 | 0 | NA | NA | NA | NA | NA |
| LINC02327 | 0.03392908 | 0.380740593 | 3.29738709 | 0.11546736 | 0.90807469 | NA |
| AL138974.1 | 0.06785815 | 0.795868262 | 3.28898349 | 0.24198001 | 0.80879565 | NA |
| LINC02326 | 0 | NA | NA | NA | NA | NA |
| AL158058.1 | 0 | NA | NA | NA | NA | NA |
| PRKD1 | 2297.88278 | -0.6005646 | 0.29594295 | -2.0293256 | 0.04242514 | 0.12937046 |
| AL356756.1 | 0.61663303 | -1.486623312 | 1.89265375 | -0.7854703 | 0.43217788 | NA |
| AL133372.3 | 3.0642501 | 1.163500498 | 0.7886435 | 1.4753187 | 0.14012688 | 0.3002082 |
| AL133372.2 | 1.00379869 | 3.930106208 | 1.55229574 | 2.53180248 | 0.01134779 | NA |
| G2E3-AS1 | 0.03392908 | 0.380740593 | 3.29738709 | 0.11546736 | 0.90807469 | NA |
| G2E3 | 518.413191 | 0.197217082 | 0.11371311 | 1.73433903 | 0.08285795 | 0.20765672 |
| SCFD1 | 2446.78175 | -0.079833952 | 0.10514743 | -0.7592573 | 0.44769867 | 0.63783587 |
| AL121852.1 | 18.4984128 | -0.478547957 | 0.32969573 | -1.4514836 | 0.14664524 | 0.30961048 |
| COCH | 488.978556 | 0.87696096 | 0.28844703 | 3.04028422 | 0.00236355 | 0.015359 |
| AL049830.3 | 218.161567 | 1.385857387 | 0.49156875 | 2.81925448 | 0.00481353 | 0.02611251 |
| STRN3 | 1963.78174 | -0.485135129 | 0.19673814 | -2.4658926 | 0.01366723 | 0.0567835 |
| AP4S1 | 458.775649 | 0.444232635 | 0.20460168 | 2.17120712 | 0.02991552 | 0.10066835 |
| HECTD1 | 5578.26565 | -0.141404137 | 0.25565841 | -0.5530979 | 0.58019635 | 0.74321276 |
| AL136418.1 | 81.2217851 | 0.797692733 | 0.24172472 | 3.30000474 | 0.00096683 | 0.00764032 |
| HEATR5A | 1146.34584 | 0.061934095 | 0.16456497 | 0.37635041 | 0.70665641 | 0.83308876 |
| AL139353.1 | 290.588198 | -0.110021252 | 0.28263621 | -0.3892681 | 0.69707785 | 0.8258207 |
| AL139353.2 | 52.0465275 | -0.943785404 | 0.3994052 | -2.3629773 | 0.01812878 | 0.06979938 |
| DTD2 | 485.623013 | 0.252647055 | 0.15334116 | 1.64761406 | 0.09943189 | 0.23657651 |
| GPR33 | 0.29665639 | 2.176798933 | 3.23107108 | 0.67370815 | 0.50049688 | NA |
| NUBPL | 778.712081 | 0.864305827 | 0.20706602 | 4.17405912 | 2.99E-05 | 0.00048918 |
| AL163973.3 | 0.2423894 | 1.011482133 | 2.97224145 | 0.34030954 | 0.73362343 | NA |
| AL163973.2 | 0.08757577 | 0.547239394 | 3.29377202 | 0.16614368 | 0.86804388 | NA |
| AL355112.1 | 0.16354576 | -1.003574173 | 3.30558948 | -0.3035992 | 0.7614333 | NA |
| LINC02313 | 9.49781878 | 0.276990189 | 0.70462306 | 0.39310406 | 0.69424262 | 0.8239054 |
| AL352984.1 | 0.17453125 | 0.18657164 | 3.29377195 | 0.05664376 | 0.95482897 | NA |
| ARHGAP5-AS | 241.100806 | -0.215525827 | 0.29928431 | -0.7201374 | 0.4714404 | 0.65787603 |
| ARHGAP5 | 4028.62381 | -0.137377098 | 0.23591713 | -0.5823108 | 0.56035733 | 0.72845072 |
| AKAP6 | 394.192358 | 0.76844295 | 0.30049725 | 2.55723788 | 0.0105507 | 0.0468624 |
| AL136298.1 | 1.26681561 | -0.257865096 | 1.72794828 | -0.149232 | 0.88137061 | 0.93849154 |
| NPAS3 | 83.5638993 | 0.016367011 | 0.24336853 | 0.06725196 | 0.94638112 | 0.97427563 |
| EGLN3 | 2052.68476 | -1.553160374 | 0.40430156 | -3.8415889 | 0.00012224 | 0.00148342 |
| EGLN3-AS1 | 0.17423948 | 0.18657164 | 3.29377195 | 0.05664376 | 0.95482897 | NA |
| SPTSSA | 2439.71679 | -0.440268664 | 0.28236961 | -1.5591928 | 0.11895075 | 0.26823425 |
| EAPP | 1521.61155 | -0.013335869 | 0.14465923 | -0.0921882 | 0.92654854 | 0.96437374 |
| AL445363.1 | 3.75381396 | -0.154533879 | 0.73131648 | -0.2113092 | 0.83264603 | 0.9111315 |
| AL445363.2 | 1.6869999 | -0.758352216 | 0.9968707 | -0.7607328 | 0.44681669 | 0.63710155 |
| SNX6 | 3047.59113 | -0.108239673 | 0.10579181 | -1.0231385 | 0.30624239 | 0.50312351 |
| CFL2 | 2181.71395 | 0.015684151 | 0.22216044 | 0.07059831 | 0.94371746 | 0.97315146 |
| BAZ1A | 1228.29273 | -1.220779308 | 0.25978378 | -4.6992129 | 2.61E-06 | 7.03E-05 |
| AL121603.2 | 124.010925 | -1.085232778 | 0.1643693 | -6.6024056 | 4.05E-11 | 6.79E-09 |
| SRP54 | 3264.34955 | -0.36198564 | 0.158579 | -2.2826833 | 0.02244903 | 0.08167508 |
| FAM177A1 | 3272.60098 | 0.157097847 | 0.17811924 | 0.88198134 | 0.3777869 | 0.57382306 |
| PPP2R3C | 962.658784 | -0.346311671 | 0.1739198 | -1.9912148 | 0.04645728 | 0.13808369 |
| KIAA0391 | 1876.71469 | 0.239793425 | 0.15566028 | 1.54049205 | 0.12344046 | 0.27497254 |
| AL121594.1 | 5569.9175 | -0.423418736 | 0.12902013 | -3.2818036 | 0.00103145 | 0.00804241 |
| PSMA6 | 4299.02269 | -0.727546146 | 0.17280093 | -4.2103139 | 2.55E-05 | 0.00043108 |
| AL133163.2 | 0.65626422 | -0.543634904 | 1.59878152 | -0.3400308 | 0.73383336 | NA |
| NFKBIA | 10050.2968 | -1.17739529 | 0.39739636 | -2.9627732 | 0.00304881 | 0.01858984 |
| AL133163.3 | 0 | NA | NA | NA | NA | NA |
| INSM2 | 1.44022776 | -1.728194361 | 1.29925912 | -1.3301383 | 0.1834727 | 0.36041233 |
| RALGAPA1 | 1137.19099 | 0.097921162 | 0.09263389 | 1.05707708 | 0.29047641 | 0.48652584 |
| AL162311.3 | 64.4959472 | 0.852412424 | 0.2321402 | 3.67197246 | 0.00024069 | 0.00253998 |

| | | | | | | |
|---|---|---|---|---|---|---|
| BRMS1L | 604.278743 | 0.35866246 | 0.14695112 | 2.44069224 | 0.01465914 | 0.05989069 |
| AL162311.1 | 1.17644427 | 0.55106087 | 1.156909 | 0.47632171 | 0.6338452 | 0.78295381 |
| AL133304.1 | 0 | NA | NA | NA | NA | NA |
| AL133304.2 | 0.36047453 | 0.04774254 | 2.30770608 | 0.02068831 | 0.98349429 | NA |
| LINC00609 | 0.70500674 | 0.945498087 | 1.38292418 | 0.68369481 | 0.4941679 | NA |
| PTCSC3 | 55.9698247 | 2.592410705 | 0.54487491 | 4.75780892 | 1.96E-06 | 5.57E-05 |
| AL162511.1 | 2.10499819 | -0.797208899 | 0.99643246 | -0.8000632 | 0.42367421 | 0.61621563 |
| MBIP | 1143.48619 | -0.119337865 | 0.13046699 | -0.9146978 | 0.3603503 | 0.5580355 |
| SFTA3 | 0.16783122 | 1.368031398 | 3.28033652 | 0.41703996 | 0.67664918 | NA |
| NKX2-1 | 0.17475523 | -1.003574173 | 3.30558948 | -0.3035992 | 0.7614333 | NA |
| NKX2-1-AS1 | 0 | NA | NA | NA | NA | NA |
| NKX2-8 | 0.3760064 | -1.802811494 | 3.29956038 | -0.5463793 | 0.58480524 | NA |
| AL079303.1 | 1.33895163 | -1.738685621 | 2.21353137 | -0.7854805 | 0.43217193 | 0.624322 |
| PAX9 | 25.8146836 | 1.164883599 | 0.39901417 | 2.91940412 | 0.00350701 | 0.02060604 |
| SLC25A21 | 144.364821 | 1.053606658 | 0.36987515 | 2.84854678 | 0.00439194 | 0.02446461 |
| AL162464.2 | 0 | NA | NA | NA | NA | NA |
| AL162464.1 | 0 | NA | NA | NA | NA | NA |
| SLC25A21-AS | 186.009068 | 0.958374322 | 0.47612959 | 2.01284343 | 0.0441311 | 0.13307458 |
| MIPOL1 | 304.681756 | 0.260840568 | 0.20803517 | 1.25382919 | 0.2099041 | 0.39317879 |
| AL121790.2 | 0.40531487 | -0.636076481 | 2.89743754 | -0.2195307 | 0.82623667 | NA |
| AL121790.1 | 3.32044868 | -0.286708454 | 0.9331561 | -0.307246 | 0.75865616 | 0.86635037 |
| FOXA1 | 4.82370263 | -2.658851356 | 1.05629484 | -2.5171489 | 0.01183088 | 0.05105227 |
| TTC6 | 1.32983837 | -1.673572755 | 1.44406187 | -1.1589342 | 0.24648299 | 0.43775299 |
| LINC00517 | 0.61575331 | -0.047932135 | 1.80492756 | -0.0265563 | 0.97881366 | NA |
| AL359233.1 | 3.74393956 | -0.504971435 | 1.34289399 | -0.3760322 | 0.70689293 | 0.83325953 |
| AL392023.2 | 1.96336202 | 0.034432271 | 1.37860898 | 0.0249761 | 0.98007403 | 0.99106512 |
| AL392023.1 | 0.23371381 | -0.340597073 | 3.29738699 | -0.103293 | 0.91773041 | NA |
| SSTR1 | 348.394623 | 0.998253923 | 0.3743685 | 2.66650088 | 0.00766454 | 0.03700365 |
| CLEC14A | 1517.94357 | -0.558592651 | 0.28964248 | -1.9285591 | 0.05378562 | 0.15338003 |
| AL161751.1 | 0.20176029 | 1.652197309 | 3.26794479 | 0.50557687 | 0.61315372 | NA |
| LINC00639 | 17.4574511 | 0.504136172 | 0.92150459 | 0.54707939 | 0.58432419 | 0.74635935 |
| SEC23A | 2431.49621 | -0.31628967 | 0.27512188 | -1.1496347 | 0.25029435 | 0.44249543 |
| SEC23A-AS1 | 40.2133424 | 0.918829345 | 0.32921358 | 2.79098253 | 0.00525483 | 0.02790372 |
| GEMIN2 | 360.003616 | -0.509497124 | 0.10691667 | -4.7653666 | 1.89E-06 | 5.42E-05 |
| TRAPPC6B | 1616.00921 | -0.064756535 | 0.07955602 | -0.813974 | 0.41565984 | 0.60908136 |
| AL132639.3 | 297.162072 | 0.260934914 | 0.18362503 | 1.4210204 | 0.15531083 | 0.32152195 |
| PNN | 6600.65454 | -0.19283538 | 0.14612565 | -1.3196545 | 0.18695039 | 0.36490612 |
| MIA2 | 6321.97847 | 0.54318223 | 0.23353978 | 2.32586601 | 0.0200257 | 0.07498973 |
| AL132639.2 | 114.262709 | 0.993439187 | 0.28674734 | 3.46451054 | 0.0005312 | 0.00475481 |
| FBXO33 | 851.099381 | -0.372974086 | 0.14934377 | -2.4974197 | 0.01251008 | 0.05319961 |
| AL049875.1 | 4.12722693 | 1.792967713 | 0.73412975 | 2.44230359 | 0.01459387 | 0.05968367 |
| AL352977.1 | 0.47590972 | 0.01802996 | 2.25210222 | 0.00800584 | 0.99361234 | NA |
| AL352977.2 | 1.04573103 | 0.620605852 | 1.22698246 | 0.50579847 | 0.61299813 | 0.76813865 |
| LINC02315 | 3.62173419 | 0.636010276 | 1.44674933 | 0.43961332 | 0.66021719 | 0.80008857 |
| AL121821.1 | 5.00407635 | 0.729119596 | 0.70228954 | 1.0382037 | 0.29917523 | 0.49592977 |
| LRFN5 | 28.3059279 | 0.235422094 | 0.39694328 | 0.59308749 | 0.5531226 | 0.72290971 |
| AL445074.1 | 0 | NA | NA | NA | NA | NA |
| AL450442.1 | 0 | NA | NA | NA | NA | NA |
| LINC02307 | 0.22056585 | 0.689157521 | 3.05414254 | 0.22564681 | 0.82147613 | NA |
| AL161752.1 | 1.87490885 | -2.253887974 | 1.54233317 | -1.4613496 | 0.14391952 | 0.30561747 |
| LINC02277 | 0.15452186 | 0.435199075 | 3.28898345 | 0.13232024 | 0.894731 | NA |
| FSCB | 0.08695548 | -0.642905668 | 3.30558948 | -0.1944905 | 0.84579185 | NA |
| AL356022.1 | 0 | NA | NA | NA | NA | NA |
| LINC02302 | 3.94383368 | -0.18547348 | 0.7396702 | -0.2507516 | 0.80200617 | 0.89266009 |
| AL049870.2 | 1.84123545 | 0.542277134 | 0.9650943 | 0.56189031 | 0.57419076 | 0.73864963 |
| C14orf28 | 357.471995 | -0.165352042 | 0.20174097 | -0.8196255 | 0.41242964 | 0.6062516 |
| AL049870.3 | 5.84538885 | -0.164447227 | 0.62351106 | -0.2637439 | 0.79197728 | 0.8871895 |
| KLHL28 | 567.674121 | -0.190392835 | 0.11942154 | -1.5942922 | 0.11087059 | 0.25527439 |
| TOGARAM1 | 964.58991 | 0.34726188 | 0.1307724 | 2.65546758 | 0.00791985 | 0.03788944 |
| AL121809.2 | 1.39164223 | -0.119377915 | 1.03938222 | -0.1148547 | 0.90856031 | 0.95432284 |
| AL121809.1 | 0.63831477 | 0.776361435 | 2.13352984 | 0.3638859 | 0.71594321 | NA |
| PRPF39 | 3336.74105 | -0.3903035 | 0.14095917 | -2.7689117 | 0.00562439 | 0.02943928 |
| FKBP3 | 2751.32844 | -0.334045694 | 0.19666266 | -1.698572 | 0.08939986 | 0.21954805 |
| FANCM | 210.292263 | 0.16024708 | 0.2343602 | 0.68376405 | 0.49412417 | 0.67670324 |
| MIS18BP1 | 412.867772 | -0.194676389 | 0.16941224 | -1.1491283 | 0.25050309 | 0.44271607 |
| AL162632.3 | 1.51537397 | 0.216760486 | 1.33741418 | 0.16207431 | 0.87124733 | 0.93306711 |
| AL162632.1 | 1.58429807 | 0.294453644 | 0.93544825 | 0.31477278 | 0.75293418 | 0.86228066 |
| AL139354.1 | 0.52662468 | -0.085126846 | 2.24298359 | -0.0379525 | 0.96972555 | NA |
| LINC02303 | 243.152422 | 1.39821481 | 0.81119563 | 1.72364688 | 0.08477161 | 0.21098208 |
| LINC00871 | 49.9494166 | 1.804781923 | 0.54844449 | 3.29072853 | 0.00099928 | 0.00783326 |
| RPL10L | 0.39277703 | -1.059762887 | 2.27846186 | -0.4651221 | 0.64184405 | NA |
| MDGA2 | 3.39976196 | 0.622597941 | 0.62392441 | 0.997874 | 0.31834046 | 0.51560948 |
| LINC00648 | 8.92271618 | 0.005022643 | 1.09675235 | 0.00457956 | 0.99634605 | 0.99823988 |
| AL358335.2 | 1.46106514 | 1.011209376 | 1.03848251 | 0.97373751 | 0.33018689 | 0.52808274 |
| AL110505.1 | 3.30283006 | 1.638946971 | 1.26845441 | 1.29208189 | 0.19632879 | 0.37685181 |
| RPS29 | 9012.51595 | -0.340448762 | 0.15529246 | -2.1923071 | 0.02835734 | 0.09700252 |
| LRR1 | 316.546707 | -0.906407032 | 0.30372621 | -2.9842898 | 0.00284237 | 0.01759057 |
| RPL36AL | 5565.25545 | -0.618882286 | 0.21929909 | -2.8220924 | 0.00477114 | 0.0259573 |
| MGAT2 | 1500.05537 | -0.241638258 | 0.19751609 | -1.2233852 | 0.22118424 | 0.40730645 |
| AL139099.1 | 1495.93388 | -0.243078053 | 0.19743127 | -1.2312034 | 0.21824681 | 0.40332906 |
| DNAAF2 | 757.353261 | -0.03871373 | 0.17473613 | -0.2215554 | 0.82466002 | 0.90622499 |
| POLE2 | 108.472318 | -0.199656257 | 0.19957724 | -1.0003959 | 0.31711893 | 0.51445256 |
| KLHDC1 | 344.274232 | 0.515149685 | 0.2149629 | 2.39645856 | 0.01655437 | 0.06539904 |
| AL591767.1 | 1.66292539 | 0.59602846 | 1.21741757 | 0.48958424 | 0.62442813 | 0.77626672 |
| KLHDC2 | 6157.59204 | 0.325444926 | 0.14376354 | 2.26375153 | 0.0235894 | 0.08460997 |
| NEMF | 3793.62643 | 0.067380377 | 0.11077439 | 0.60826676 | 0.54301057 | 0.71531565 |
| AL627171.2 | 32.5817805 | 0.978588795 | 0.38032072 | 2.57306203 | 0.01008031 | 0.04531501 |
| AL627171.1 | 11.1287455 | 1.243925598 | 0.40123614 | 3.10023322 | 0.00193368 | 0.01313868 |
| ARF6 | 5784.06105 | -0.482032246 | 0.15576108 | -3.09469 | 0.00197019 | 0.01333517 |
| LINC01588 | 109.174365 | -0.234203317 | 0.30592777 | -0.765551 | 0.44394349 | 0.63489288 |
| VCPKMT | 647.685006 | -0.381982652 | 0.20523598 | -1.8611875 | 0.0627177 | 0.17055436 |
| SOS2 | 2043.93708 | 0.378430982 | 0.12732889 | 2.97207488 | 0.00295795 | 0.01814399 |
| L2HGDH | 1478.36021 | 1.21807751 | 0.31057073 | 3.92206155 | 8.78E-05 | 0.00114385 |
| ATP5S | 2507.66252 | 1.05059529 | 0.16256367 | 6.46266961 | 1.03E-10 | 1.48E-08 |
| AL359397.2 | 513.317543 | 0.984033072 | 0.19263765 | 5.10820732 | 3.25E-07 | 1.27E-05 |
| CDKL1 | 710.913004 | 1.498061722 | 0.20362356 | 7.35701577 | 1.88E-13 | 6.53E-11 |
| AL359397.1 | 23.3258174 | 2.824544197 | 0.47256335 | 5.97706989 | 2.27E-09 | 1.98E-07 |
| MAP4K5 | 1907.35706 | -0.333929343 | 0.13966886 | -2.3908647 | 0.01680875 | 0.06614828 |
| AL118556.1 | 59.5464794 | 0.616400628 | 0.28705766 | 2.14730595 | 0.03176893 | 0.10505979 |
| ATL1 | 360.929284 | 0.795108172 | 0.18698125 | 4.25234178 | 2.12E-05 | 0.00037042 |
| SAV1 | 2084.09684 | -0.327030934 | 0.1188981 | -2.7505144 | 0.00595018 | 0.03060459 |
| NIN | 1296.37412 | 0.136666193 | 0.1330635 | 1.02707501 | 0.30438515 | 0.50132011 |
| AL133485.1 | 3.73282863 | -0.16746846 | 0.66106189 | -0.2533325 | 0.80001129 | 0.89152759 |
| AL358334.1 | 0 | NA | NA | NA | NA | NA |
| PYGL | 1712.06866 | -1.100936329 | 0.27510353 | -4.0018982 | 6.28E-05 | 0.00088091 |
| ABHD12B | 30.7268196 | 0.025121532 | 0.53369832 | 0.04707066 | 0.96245691 | 0.98223572 |
| AL358334.3 | 282.081051 | -0.995931393 | 0.31859013 | -3.1260585 | 0.00177166 | 0.01229996 |
| AL358334.2 | 61.3132838 | 3.310952422 | 0.76235967 | 4.34303198 | 1.41E-05 | 0.00026873 |
| TRIM9 | 166.461718 | -0.542330532 | 0.47282819 | -1.1469928 | 0.2513846 | 0.44373866 |
| AL591770.1 | 0.36760629 | -1.691981139 | 3.30304428 | -0.512249 | 0.60847675 | NA |
| TMX1 | 2104.76649 | -0.270428604 | 0.1710237 | -1.5812347 | 0.11382439 | 0.26012213 |
| LINC00519 | 1.12502564 | -1.470859631 | 1.66051944 | -0.8857828 | 0.37573456 | 0.57159919 |
| LINC00640 | 3.8016525 | 0.019337629 | 0.60109756 | 0.03217053 | 0.97433606 | 0.98820161 |

| | | | | | | |
|---|---|---|---|---|---|---|
| AL358332.1 | 0.96222765 | 0.417493613 | 1.63955312 | 0.25463866 | 0.7990022 | NA |
| LINC02310 | 0.18393104 | 1.469068379 | 3.27783996 | 0.44818185 | 0.65402196 | NA |
| FRMD6-AS2 | 0.48585202 | 0.295012208 | 2.70585764 | 0.10902725 | 0.91318088 | NA |
| FRMD6 | 416.545993 | 0.263543868 | 0.35267264 | 0.7472762 | 0.45489686 | 0.64433744 |
| AL122125.1 | 17.539728 | -0.602675441 | 0.46910212 | -1.2847425 | 0.19888228 | 0.37994704 |
| FRMD6-AS1 | 11.1561589 | -0.349895673 | 0.60895221 | -0.5745864 | 0.56557103 | 0.73214268 |
| AL079307.2 | 0 | NA | NA | NA | NA | NA |
| GNG2 | 538.543292 | 0.09306493 | 0.2913831 | 0.31939028 | 0.74943058 | 0.85998627 |
| AL358333.2 | 0.26695105 | -0.178915937 | 2.96757513 | -0.0602903 | 0.95192444 | NA |
| AL358333.3 | 86.2573407 | 0.178490699 | 0.24122226 | 0.7399429 | 0.45933464 | 0.64801707 |
| AL358333.1 | 0.07830766 | -0.642905668 | 3.30558948 | -0.1944905 | 0.84579185 | NA |
| RTRAF | 4751.9662 | -0.412317233 | 0.14189588 | -2.9057732 | 0.00366347 | 0.02134303 |
| NID2 | 1400.80449 | -0.423869101 | 0.28952569 | -1.4640121 | 0.14319065 | 0.30470767 |
| LINC02319 | 0.6028402 | -0.309203489 | 1.93748508 | -0.1595901 | 0.87320396 | NA |
| PTGDR | 45.2939304 | 0.298027677 | 0.4745051 | 0.62808109 | 0.52995082 | 0.70455272 |
| PTGER2 | 321.156889 | -0.63028129 | 0.25092185 | -2.5118629 | 0.01200957 | 0.05155338 |
| TXNDC16 | 378.943855 | 0.470087736 | 0.17620743 | 2.66780874 | 0.00763477 | 0.0368937 |
| GPR137C | 51.7454108 | 0.131231851 | 0.29649475 | 0.44261105 | 0.65804709 | 0.79943711 |
| ERO1A | 5067.66895 | -2.130152888 | 0.34027656 | -6.2600635 | 3.85E-10 | 4.49E-08 |
| AL133453.1 | 987.810173 | -1.94953876 | 0.34040513 | -5.7271134 | 1.02E-08 | 7.07E-07 |
| PSMC6 | 3988.39412 | -0.362036596 | 0.15065206 | -2.4031307 | 0.01625538 | 0.06452722 |
| STYX | 713.474789 | -0.078078699 | 0.12536094 | -0.6228312 | 0.53339547 | 0.70717137 |
| GNPNAT1 | 1685.71951 | -0.446146841 | 0.30043791 | -1.4849885 | 0.13754687 | 0.29619521 |
| AL139317.3 | 1121.39069 | -0.5590481 | 0.27525153 | -2.0310445 | 0.04225048 | 0.12896691 |
| AL139317.1 | 0.51897318 | 2.36637478 | 1.95926159 | 1.20778909 | 0.2271284 | NA |
| FERMT2 | 4771.37117 | -0.037588031 | 0.21178373 | -0.1774831 | 0.85912895 | 0.92642086 |
| DDHD1 | 835.486508 | -0.33002678 | 0.13046754 | -2.5295701 | 0.01142024 | 0.04975246 |
| AL352979.2 | 122.774869 | -0.89329254 | 0.2229521 | -4.0066567 | 6.16E-05 | 0.00086797 |
| AL356020.1 | 1.75996486 | -0.423403822 | 1.04925526 | -0.4035279 | 0.68655988 | 0.81884143 |
| AL163953.1 | 1.72403265 | 2.101451108 | 1.01332663 | 2.07381415 | 0.03809657 | 0.11946803 |
| LINC02331 | 4.05599013 | 2.282158209 | 0.83567763 | 2.73090738 | 0.00631602 | 0.03200855 |
| BMP4 | 616.658938 | 0.557345191 | 0.27440338 | 2.03111638 | 0.04224319 | 0.12895959 |
| CDKN3 | 68.5062444 | -1.930614537 | 0.49104503 | -3.9316446 | 8.44E-05 | 0.0011096 |
| CNIH1 | 3022.74437 | 0.014899432 | 0.17020808 | 0.08753658 | 0.93024501 | 0.96634079 |
| AL359792.1 | 0.55016869 | 2.380259907 | 1.93390255 | 1.23080654 | 0.21839523 | NA |
| GMFB | 3427.01448 | -0.181018383 | 0.1928502 | -0.9386476 | 0.3479117 | 0.54542145 |
| CGRRF1 | 948.291788 | -0.158169333 | 0.22435115 | -0.7050079 | 0.48080533 | 0.66545092 |
| SAMD4A | 1806.31364 | -1.259577882 | 0.28668135 | -4.3936513 | 1.11E-05 | 0.00022267 |
| GCH1 | 1674.02097 | 0.149176655 | 0.32833107 | 0.45434827 | 0.64957818 | 0.79383523 |
| WDHD1 | 165.03894 | -0.181147654 | 0.25375998 | -0.7138543 | 0.47531728 | 0.66076844 |
| SOCS4 | 1289.74665 | -0.376994338 | 0.18086982 | -2.0843408 | 0.03712918 | 0.11722899 |
| MAPK1IP1L | 6791.0197 | -0.384387403 | 0.16724248 | -2.2983837 | 0.02153996 | 0.07922018 |
| LGALS3 | 5518.30087 | -0.928818351 | 0.31791822 | -2.9215638 | 0.00348279 | 0.02049624 |
| DLGAP5 | 23.3865916 | -2.138347688 | 0.78930029 | -2.7091688 | 0.0067452 | 0.0336301 |
| AL139316.1 | 0.21723982 | 0.689157019 | 3.06522228 | 0.22483101 | 0.82211074 | NA |
| AL158801.2 | 7.24097546 | -0.061671894 | 0.50983634 | -0.1209641 | 0.90371947 | 0.95167363 |
| FBXO34 | 1663.28358 | 0.121813668 | 0.10427474 | 1.16819923 | 0.24272641 | 0.43318188 |
| AL158801.3 | 0.18028734 | 0.18657164 | 3.29377195 | 0.05664376 | 0.95482897 | NA |
| ATG14 | 1425.03254 | 0.225888993 | 0.13604956 | 1.66034348 | 0.09684537 | 0.23227035 |
| TBPL2 | 0.62377299 | -0.931183301 | 1.49500406 | -0.6228634 | 0.5333743 | NA |
| KTN1-AS1 | 110.267857 | -0.065233139 | 0.16067953 | -0.4059829 | 0.68475517 | 0.81783568 |
| KTN1 | 9740.41671 | -0.001458708 | 0.17110031 | -0.0085255 | 0.99319775 | 0.99686464 |
| AL355773.1 | 1.0831115 | 0.460432316 | 1.36179677 | 0.33810648 | 0.73528295 | 0.85123414 |
| LINC00520 | 1.35982069 | -0.994328582 | 1.17027012 | -0.8496573 | 0.39551563 | 0.59085357 |
| AL138995.1 | 33.7139876 | -0.624404595 | 0.30786718 | -2.0281623 | 0.04254369 | 0.12957693 |
| PELI2 | 1366.01512 | 0.231190542 | 0.1447328 | 1.5973611 | 0.11018524 | 0.25419877 |
| AL355073.1 | 0.88225624 | 0.228501406 | 1.33869246 | 0.17068999 | 0.86446753 | NA |
| AL355073.2 | 14.6448142 | 0.500669485 | 0.33678674 | 1.48660688 | 0.13711867 | 0.29562745 |
| LINC02284 | 3.13457095 | 0.765193373 | 0.72804912 | 1.05101889 | 0.29324992 | 0.48977138 |
| TMEM260 | 1082.22326 | 0.126043813 | 0.22282581 | 0.56566075 | 0.57162442 | 0.73652575 |
| AL355103.1 | 0.54186533 | 0.922819311 | 2.20225726 | 0.41903338 | 0.67519173 | NA |
| AL161757.2 | 179.555021 | 0.157314128 | 0.23780367 | 0.66152945 | 0.50827283 | 0.68712984 |
| AL161757.5 | 255.114925 | -0.17356093 | 0.26034982 | -0.6666451 | 0.50499886 | 0.68442308 |
| AL161757.3 | 0.10721287 | -0.642905668 | 3.30558948 | -0.1944905 | 0.84579185 | NA |
| OTX2 | 0.29525572 | 0.328535869 | 2.3070254 | 0.1424067 | 0.88675877 | NA |
| OTX2-AS1 | 0.18663678 | 0.18657164 | 3.29377195 | 0.05664376 | 0.95482897 | NA |
| AL161757.4 | 0.09989236 | -0.642905668 | 3.30558948 | -0.1944905 | 0.84579185 | NA |
| AL391152.1 | 4.30741152 | -0.261666875 | 0.85097633 | -0.3074902 | 0.75847029 | 0.86635037 |
| EXOC5 | 2347.1535 | -0.059054995 | 0.14374427 | -0.4108337 | 0.68119445 | 0.81527767 |
| AP5M1 | 2897.52994 | 0.392508938 | 0.10651141 | 3.68513517 | 0.00022858 | 0.00243152 |
| AL355834.1 | 0 | NA | NA | NA | NA | NA |
| NAA30 | 1416.34732 | -0.016424877 | 0.2164541 | -0.0758816 | 0.93951332 | 0.97110795 |
| CCDC198 | 5371.70477 | 0.808530346 | 0.34825332 | 2.32167301 | 0.02025055 | 0.07556346 |
| SLC35F4 | 18.5718084 | 0.646323407 | 0.36753718 | 1.75852522 | 0.07865818 | 0.20009804 |
| AL161804.1 | 15.8707663 | 0.913889748 | 0.58658757 | 1.55797668 | 0.11923878 | 0.26872289 |
| ARMH4 | 1138.65913 | 0.333261042 | 0.18238703 | 1.82721896 | 0.06766685 | 0.17974967 |
| ACTR10 | 3362.67162 | -0.140830919 | 0.11423303 | -1.2328389 | 0.21763589 | 0.40265393 |
| PSMA3 | 4439.18131 | -0.685785587 | 0.17175755 | -3.9927536 | 6.53E-05 | 0.00091027 |
| PSMA3-AS1 | 3766.80251 | 0.069364816 | 0.12727936 | 0.54498087 | 0.58576668 | 0.74722655 |
| AL132989.2 | 13.4503978 | 0.22549012 | 0.42927124 | 0.52528588 | 0.59938447 | 0.75727139 |
| ARID4A | 1002.51739 | 0.111484488 | 0.16993268 | 0.65605089 | 0.51179138 | 0.68990587 |
| AL139021.2 | 6.09423516 | 1.442936568 | 0.59255425 | 2.43511301 | 0.01488714 | 0.06052089 |
| TOMM20L | 75.3524462 | 0.190360326 | 0.20848273 | 0.91307481 | 0.3612032 | 0.55878832 |
| AL139021.1 | 260.390146 | -0.17369218 | 0.15336029 | -1.1325759 | 0.25739238 | 0.45000576 |
| TIMM9 | 1081.55108 | -0.420892898 | 0.11931269 | -3.5276458 | 0.00041927 | 0.00395855 |
| KIAA0586 | 517.117963 | 0.007204978 | 0.1207528 | 0.05966717 | 0.95242072 | 0.97717271 |
| DACT1 | 682.511208 | 0.335986946 | 0.42041682 | 0.79917579 | 0.42418849 | 0.61664005 |
| LINC01500 | 0.27654246 | 0.374496445 | 2.9801101 | 0.12566531 | 0.89999687 | NA |
| DAAM1 | 941.846121 | 0.30057081 | 0.19671664 | 1.52793789 | 0.12652796 | 0.27981737 |
| GPR135 | 174.121108 | -0.504066134 | 0.23413071 | -2.1529262 | 0.03132448 | 0.10412201 |
| L3HYPDH | 1500.18457 | 0.057492906 | 0.30467363 | 0.18870326 | 0.8503254 | 0.92147122 |
| JKAMP | 2619.38127 | 0.004215568 | 0.10628343 | 0.03966346 | 0.96836144 | 0.98528731 |
| CCDC175 | 200.082712 | -0.1131982 | 0.14984 | -0.7554605 | 0.44997272 | 0.6399407 |
| RTN1 | 246.00381 | 0.756853893 | 0.44157701 | 1.71397939 | 0.08653251 | 0.21429961 |
| AL133299.1 | 9.20686767 | 0.930203511 | 0.67934479 | 1.36926569 | 0.17091624 | 0.34315983 |
| LRRC9 | 10.3941449 | 1.442791223 | 0.76610732 | 1.8832756 | 0.05966303 | 0.16495701 |
| AL157911.1 | 4.65832108 | 0.908133164 | 0.93653267 | 0.9696759 | 0.33220807 | 0.53009359 |
| PCNX4 | 3474.39213 | 0.130442829 | 0.07716837 | 1.69036658 | 0.09095784 | 0.22229687 |
| DHRS7 | 4236.73209 | -0.300511043 | 0.14203831 | -2.1157041 | 0.03436998 | 0.11117964 |
| AL157756.1 | 114.427652 | 1.225188358 | 0.30302139 | 4.04324055 | 5.27E-05 | 0.00077663 |
| PPM1A | 4601.52976 | 0.065419908 | 0.25132406 | 0.26030101 | 0.79463159 | 0.88857752 |
| LINC02322 | 0.5411423 | -1.520939273 | 2.25245648 | -0.6752358 | 0.49952596 | NA |
| C14orf39 | 4.01823807 | -0.172182381 | 0.6879938 | -0.2502673 | 0.80238061 | 0.89288841 |
| SIX6 | 0.43270659 | -2.017828363 | 1.88018873 | -1.0732052 | 0.28317905 | NA |
| AL049874.3 | 0.81662807 | -0.413380666 | 1.52420886 | -0.27121 | 0.78622953 | NA |
| SALRNA1 | 0.32666247 | -0.69944518 | 2.87673143 | -0.2431389 | 0.80789783 | NA |
| SIX1 | 14.0632818 | -0.136491705 | 0.74663959 | -0.182808 | 0.85494866 | 0.92401312 |
| SIX4 | 194.57519 | -0.526185396 | 0.20334483 | -2.5876507 | 0.00966329 | 0.04391194 |
| MNAT1 | 815.958304 | -0.095813831 | 0.10023629 | -0.9558797 | 0.33913302 | 0.53727066 |
| TRMT5 | 812.316876 | 0.14903818 | 0.12357574 | 1.20604729 | 0.22779926 | 0.41482401 |
| SLC38A6 | 608.572747 | -0.067195568 | 0.18659192 | -0.3601205 | 0.71875705 | 0.84003117 |
| PRKCH | 1250.11256 | -0.265591338 | 0.26184648 | -1.0143017 | 0.3104388 | 0.50741242 |
| TMEM30B | 1914.85936 | -0.246350717 | 0.26817631 | -0.9186148 | 0.35829711 | 0.55594556 |

| | | | | | | |
|---|---|---|---|---|---|---|
| AL359220.1 | 29.1610088 | 0.050017254 | 0.27277995 | 0.18336118 | 0.85451463 | 0.92399813 |
| AL355916.3 | 283.744796 | -0.320846937 | 0.31809938 | -1.0086374 | 0.31314855 | 0.5100511 |
| AL355916.2 | 2.28998305 | -0.626561745 | 0.96613437 | -0.6485244 | 0.51664581 | 0.69396428 |
| LINC01303 | 115.341935 | -0.883018001 | 0.42674349 | -2.0692008 | 0.03852724 | 0.12045282 |
| AL355916.1 | 116.083037 | -1.898640582 | 0.63028667 | -3.0123445 | 0.00259238 | 0.0164374 |
| HIF1A-AS1 | 459.253647 | -0.52369023 | 0.45791746 | -1.1436346 | 0.25277523 | 0.44518017 |
| HIF1A | 23977.4744 | -1.814590054 | 0.34906345 | -5.1984533 | 2.01E-07 | 8.57E-06 |
| HIF1A-AS2 | 24.139557 | -2.133471488 | 0.53654051 | -3.9763475 | 7.00E-05 | 0.00095568 |
| SNAPC1 | 970.148025 | -1.3132752 | 0.29649376 | -4.429352 | 9.45E-06 | 0.00019518 |
| SYT16 | 82.9959791 | -2.229892754 | 0.51137279 | -4.3606011 | 1.30E-05 | 0.00025187 |
| AL390816.1 | 1.24821104 | -1.330788416 | 1.2246052 | -1.0867081 | 0.27716582 | 0.47260671 |
| AL390816.2 | 0 | NA | NA | NA | NA | NA |
| LINC00644 | 0 | NA | NA | NA | NA | NA |
| KCNH5 | 1.97427806 | 0.7078617 | 1.3389467 | 0.52867056 | 0.597034 | 0.75560504 |
| AL137191.1 | 0.87847841 | 0.410916498 | 1.34789538 | 0.30485786 | 0.76047441 | NA |
| RHOJ | 929.401162 | -0.581718956 | 0.41776549 | -1.3924533 | 0.16378516 | 0.33328303 |
| AL049871.1 | 5.41256812 | 0.21457416 | 0.70333545 | 0.30508083 | 0.76030459 | 0.86715659 |
| GPHB5 | 0.09997307 | 0.547239394 | 3.29377202 | 0.16614368 | 0.86804388 | NA |
| PPP2R5E | 1958.13747 | -0.523032695 | 0.18903571 | -2.7668459 | 0.00566015 | 0.02956192 |
| AL136038.3 | 9.17828034 | 0.707742698 | 0.46989709 | 1.50616531 | 0.13202474 | 0.28815531 |
| WDR89 | 620.731252 | 0.132813711 | 0.17184899 | 0.77285129 | 0.4396104 | 0.63101113 |
| AL136038.2 | 144.291928 | 0.458400658 | 0.20605489 | 2.22465311 | 0.02610453 | 0.09121195 |
| AL136038.5 | 53.8499559 | 0.024643175 | 0.20777589 | 0.1186046 | 0.90558862 | 0.95262374 |
| SGPP1 | 2617.43953 | 0.393376302 | 0.21469644 | 1.83224421 | 0.06691505 | 0.17834544 |
| SYNE2 | 14283.5291 | -0.189807692 | 0.28891937 | -0.6569573 | 0.51120836 | 0.68936664 |
| ESR2 | 268.077105 | -0.61533848 | 0.48069545 | -1.2801005 | 0.2005098 | 0.38206831 |
| AL161756.1 | 22.9694741 | 0.4379999 | 0.33854248 | 1.29378119 | 0.19574101 | 0.37605205 |
| AL161756.3 | 29.9276563 | -0.042449497 | 0.39006346 | -0.1088272 | 0.91333958 | 0.95756858 |
| MTHFD1 | 4648.27081 | 0.912345243 | 0.29637822 | 3.07831406 | 0.00208175 | 0.01388011 |
| AL122035.1 | 574.422975 | 0.758071426 | 0.29546099 | 2.56572427 | 0.01029606 | 0.0460572 |
| AL122035.2 | 787.387683 | 0.73142625 | 0.26949825 | 2.71402963 | 0.00664702 | 0.03326034 |
| ZBTB25 | 1990.37865 | -0.036740901 | 0.24108118 | -0.1524005 | 0.87887103 | 0.93705294 |
| AKAP5 | 158.102291 | 0.709757585 | 0.31272216 | 2.26961076 | 0.02323121 | 0.08375814 |
| ZBTB1 | 2077.94279 | -0.312025 | 0.13872212 | -2.2492808 | 0.02449463 | 0.08692282 |
| AL049869.3 | 90.5884872 | -0.698806734 | 0.28905658 | -2.4175431 | 0.01562568 | 0.06267862 |
| HSPA2 | 4061.45969 | 0.837984954 | 0.38960959 | 2.15083248 | 0.03148942 | 0.10450241 |
| PPP1R36 | 104.335556 | 0.426468343 | 0.48672906 | 0.87619248 | 0.38092541 | 0.57696743 |
| AL049869.2 | 10.287263 | 0.606633967 | 0.59865238 | 1.01333259 | 0.31090132 | 0.50789666 |
| PLEKHG3 | 6220.86137 | 0.114487637 | 0.13639314 | 0.83939438 | 0.40124804 | 0.59614123 |
| SPTB | 1587.63618 | 0.197256398 | 0.1772967 | 1.11257793 | 0.26588974 | 0.45998893 |
| CHURC1 | 2281.42082 | 0.203766607 | 0.09685634 | 2.10380255 | 0.03539567 | 0.11353376 |
| CHURC1-FNT | 1477.82895 | -0.000651886 | 0.10626086 | -0.0061348 | 0.99510519 | 0.99770729 |
| GPX2 | 673.788558 | -1.479330946 | 0.80077208 | -1.8473808 | 0.06469197 | 0.17419829 |
| RAB15 | 1973.30149 | -0.352401719 | 0.25460673 | -1.3841021 | 0.16632719 | 0.33710307 |
| FNTB | 1335.52239 | 0.089059953 | 0.14514879 | 0.61357698 | 0.5394949 | 0.71254659 |
| AL139022.2 | 6.17622138 | -0.553385866 | 0.50973117 | -1.0856426 | 0.27763715 | 0.47313504 |
| MAX | 3454.96982 | -0.48545533 | 0.14392555 | -3.3729615 | 0.00074364 | 0.00620173 |
| AL139022.1 | 489.56251 | -0.088395181 | 0.15174873 | -0.5825102 | 0.56022309 | 0.72838393 |
| LINC02324 | 0.54511066 | -0.096511863 | 1.93602681 | -0.0498505 | 0.96024154 | NA |
| AL355076.2 | 0.17423948 | 0.18657164 | 3.29377195 | 0.05664376 | 0.95482897 | NA |
| FUT8 | 914.831398 | -0.37946308 | 0.24624035 | -1.5410272 | 0.12331016 | 0.27477513 |
| FUT8-AS1 | 51.9065598 | -0.042457763 | 0.4460813 | -0.0951794 | 0.92417231 | 0.96312021 |
| AL391261.2 | 19.6205148 | 1.79080901 | 0.572044 | 3.13054416 | 0.00174483 | 0.01215102 |
| LINC02290 | 6.28102717 | 1.436347761 | 0.67270043 | 2.13519674 | 0.03274494 | 0.10739676 |
| AL359232.1 | 23.9583295 | 0.367736465 | 0.3680642 | 0.99910957 | 0.31774161 | 0.51495583 |
| CCDC196 | 8.84730492 | 1.563961357 | 0.73279334 | 2.13424613 | 0.03282264 | 0.10755971 |
| GPHN | 1655.10528 | 1.08881729 | 0.16913817 | 6.43744275 | 1.22E-10 | 1.69E-08 |
| AL049835.1 | 0.33009685 | 1.594575331 | 2.17484609 | 0.73318997 | 0.46344257 | NA |
| FAM71D | 16.7815363 | 0.022353956 | 0.37969299 | 0.05887376 | 0.95305265 | 0.97729034 |
| MPP5 | 2676.95937 | 0.634661084 | 0.17581835 | 3.60975462 | 0.00030649 | 0.00308465 |
| ATP6V1D | 4376.05791 | -0.137334627 | 0.17771682 | -0.7727722 | 0.43965719 | 0.63101113 |
| EIF2S1 | 2662.78066 | -0.469133795 | 0.21910682 | -2.141119 | 0.03226444 | 0.10622638 |
| PLEK2 | 364.00405 | -0.934605593 | 0.35674185 | -2.6198373 | 0.00879717 | 0.0408922 |
| TMEM229B | 278.416701 | -0.514478353 | 0.48121217 | -1.06913 | 0.28501111 | 0.48109992 |
| PLEKHH1 | 2189.92577 | -0.222479283 | 0.16555814 | -1.3438136 | 0.17900866 | 0.35470833 |
| PIGH | 2064.03648 | -0.203486507 | 0.14560824 | -1.397493 | 0.16226536 | 0.33124857 |
| ARG2 | 9199.17671 | -0.821410081 | 0.4352588 | -1.8871763 | 0.05913662 | 0.16387482 |
| VTI1B | 9993.39051 | -0.359074558 | 0.31784985 | -1.1296987 | 0.25860321 | 0.45140442 |
| AL049779.1 | 2850.03932 | 0.252618309 | 0.18474289 | 1.36740473 | 0.17149848 | 0.34398835 |
| RDH11 | 7221.1109 | 0.001243977 | 0.20241953 | 0.00614554 | 0.9950966 | 0.99770729 |
| RDH12 | 461.889853 | 3.701973985 | 0.83791765 | 4.41806421 | 9.96E-06 | 0.00020264 |
| ZFYVE26 | 1781.52602 | -0.016420527 | 0.24092251 | -0.0681569 | 0.94566075 | 0.97395466 |
| RAD51B | 162.1519 | 0.383837478 | 0.29556341 | 1.29866372 | 0.19405936 | 0.37380709 |
| AL133370.1 | 2.11968404 | -0.94901249 | 1.22582135 | -0.774185 | 0.4388214 | 0.63037483 |
| AL121820.2 | 3.73462305 | 0.071552412 | 0.73595759 | 0.09722355 | 0.92254886 | 0.96213619 |
| AL121820.1 | 3.50057931 | 0.994420824 | 0.75743199 | 1.31288465 | 0.18922182 | 0.36749025 |
| ZFP36L1 | 22300.2195 | -0.398074791 | 0.25455105 | -1.5638309 | 0.11785729 | 0.26654346 |
| ACTN1 | 6642.77152 | -0.601176559 | 0.40612007 | -1.4802927 | 0.13879516 | 0.29815117 |
| ACTN1-AS1 | 7.99711607 | -0.772114786 | 0.63660742 | -1.2128586 | 0.22518389 | 0.41151291 |
| DCAF5 | 4028.04086 | -0.08292547 | 0.12946495 | -0.6405245 | 0.52183169 | 0.69787989 |
| AL391262.1 | 1.74992627 | -0.184147295 | 1.46339328 | -0.1258358 | 0.89986189 | 0.94921229 |
| AL359317.2 | 18.513211 | 1.208267364 | 0.30512114 | 3.95995958 | 7.50E-05 | 0.00100855 |
| EXD2 | 1643.97773 | 0.079279653 | 0.13377492 | 0.59263465 | 0.55342569 | 0.72299723 |
| AL359317.1 | 54.2606194 | 0.715972583 | 0.3687843 | 1.94143999 | 0.05220493 | 0.15017389 |
| GALNT16 | 1333.7932 | -0.075783437 | 0.45961146 | -0.1648859 | 0.86903381 | 0.9318649 |
| ERH | 3194.59942 | -0.586192176 | 0.13481823 | -4.3480185 | 1.37E-05 | 0.00026384 |
| SLC39A9 | 4406.62929 | 0.132649163 | 0.09304884 | 1.42558637 | 0.15398777 | 0.31978718 |
| PLEKHD1 | 162.862471 | 1.333663444 | 0.61672551 | 2.16249113 | 0.03058034 | 0.10225762 |
| CCDC177 | 0.59306489 | -0.18264977 | 2.79196945 | -0.0654197 | 0.94783985 | NA |
| SUSD6 | 5118.28013 | -0.238250446 | 0.1690892 | -1.4090222 | 0.15882859 | 0.32631944 |
| SRSF5 | 19117.5475 | 0.148559654 | 0.13630255 | 1.08992575 | 0.27574585 | 0.47106881 |
| SLC10A1 | 5.66505566 | 0.681430092 | 0.5288587 | 1.2884918 | 0.19757482 | 0.37834197 |
| SMOC1 | 212.435019 | 1.030500156 | 0.41214837 | 2.50031353 | 0.01240834 | 0.05289493 |
| SLC8A3 | 9.25159864 | -1.401251628 | 0.80092305 | -1.7495459 | 0.0801967 | 0.20277947 |
| AL160191.1 | 0.22977084 | -1.364242679 | 3.30558948 | -0.4127078 | 0.67982068 | NA |
| AL356804.1 | 1.33073925 | 0.800128643 | 1.86488701 | 0.4290494 | 0.66788728 | 0.8057179 |
| COX16 | 1888.81707 | -0.214017562 | 0.13146241 | -1.6279754 | 0.1035301 | 0.24334512 |
| SYNJ2BP-CO | 2004.7724 | -0.232820119 | 0.10955554 | -2.1251331 | 0.0335755 | 0.10936093 |
| SYNJ2BP | 2999.12285 | 0.434302378 | 0.10434087 | 4.162342 | 3.15E-05 | 0.00050992 |
| AL357153.2 | 1.23972628 | -0.981517306 | 1.41411323 | -0.6940868 | 0.48762779 | 0.67143986 |
| ADAM21 | 10.2089215 | -0.476527566 | 0.39621119 | -1.202711 | 0.22908817 | 0.41648087 |
| ADAM20 | 6.72423591 | 0.019479409 | 0.60168996 | 0.0323745 | 0.9741734 | 0.98807744 |
| MED6 | 709.960928 | -0.354543302 | 0.17282513 | -2.0514569 | 0.04022248 | 0.12428717 |
| AL357153.3 | 181.533182 | -0.32003954 | 0.22128441 | -1.4462815 | 0.14809828 | 0.31172985 |
| AL357153.1 | 6.05159759 | -2.443451064 | 1.06613215 | -2.2918839 | 0.02191235 | 0.08026511 |
| TTC9 | 656.262168 | -1.564521735 | 0.43141948 | -3.6264513 | 0.00028734 | 0.00293123 |
| LINC01269 | 0.86688902 | -1.43936747 | 1.73687956 | -0.8287089 | 0.40726918 | NA |
| MAP3K9 | 795.063076 | -0.822451592 | 0.25621786 | -3.2099698 | 0.00132749 | 0.00984091 |
| AC004816.1 | 75.67609 | -0.253185809 | 0.22752966 | -1.1127596 | 0.26581171 | 0.45996556 |
| AC004816.2 | 9.05662208 | 0.891393498 | 0.47485154 | 1.87720459 | 0.06049006 | 0.16648477 |
| AC004825.2 | 24.8819434 | -0.554703353 | 0.48867168 | -1.1351248 | 0.25632303 | 0.44911858 |
| PCNX1 | 2816.18744 | -0.066956936 | 0.16021639 | -0.4179157 | 0.67600879 | 0.81149917 |
| AC004817.3 | 1.42164959 | 0.710304054 | 1.004487 | 0.70713116 | 0.47948497 | 0.66418343 |

| | | | | | | |
|---|---|---|---|---|---|---|
| AC004817.2 | 4.38579563 | 1.938219824 | 0.85420689 | 2.26902854 | 0.02326659 | 0.08381694 |
| AC004817.1 | 0.49056642 | 0.890751681 | 2.70866041 | 0.32885321 | 0.74226664 | NA |
| AC004817.4 | 0.62409696 | 2.724183794 | 2.5203331 | 1.08088244 | 0.27974941 | NA |
| AC005476.2 | 79.7338178 | 0.719955013 | 0.40368924 | 1.78343868 | 0.07451492 | 0.19286989 |
| SIPA1L1 | 3014.80396 | 0.175468222 | 0.16426736 | 1.06818676 | 0.28543628 | 0.4815227 |
| AC005993.1 | 0.15543392 | 1.368031615 | 3.28033652 | 0.41704002 | 0.67664913 | NA |
| RGS6 | 184.655962 | -0.175121599 | 0.49048936 | -0.3570345 | 0.72106601 | 0.84127559 |
| AC004828.1 | 1.3678392 | 0.497084514 | 0.99625001 | 0.49895559 | 0.61781067 | 0.77149535 |
| DPF3 | 437.404452 | 2.163627732 | 0.51843008 | 4.17342247 | 3.00E-05 | 0.00048933 |
| DCAF4 | 1058.37364 | -0.445038564 | 0.22960731 | -1.9382596 | 0.05259156 | 0.1508582 |
| ZFYVE1 | 2032.24523 | 0.115098526 | 0.2436752 | 0.47234403 | 0.63668125 | 0.78476604 |
| AL442663.3 | 84.1637836 | 0.728682896 | 0.25905635 | 2.81283548 | 0.00491068 | 0.0265249 |
| RBM25 | 4611.49373 | -0.069260562 | 0.14409742 | -0.480651 | 0.63076459 | 0.78054869 |
| PSEN1 | 4211.6028 | -0.393805633 | 0.20419863 | -1.928542 | 0.05378775 | 0.15338003 |
| PAPLN | 5388.56098 | 0.451946644 | 0.30030566 | 1.50495545 | 0.13233553 | 0.28859477 |
| AC004846.1 | 122.794073 | 0.971237787 | 0.20423777 | 4.75542682 | 1.98E-06 | 5.62E-05 |
| AC004846.2 | 1772.54661 | 0.269895943 | 0.30532116 | 0.88397393 | 0.37671029 | 0.57260755 |
| NUMB | 5516.22388 | 0.24487502 | 0.13053022 | 1.87600248 | 0.06065494 | 0.16676788 |
| AC005280.2 | 28.1386424 | 2.859608434 | 0.49171019 | 5.8156379 | 6.04E-09 | 4.50E-07 |
| HEATR4 | 198.860972 | -0.307520537 | 0.21701328 | -1.4170586 | 0.15646579 | 0.32320249 |
| RIOX1 | 605.429458 | -0.433486139 | 0.23899285 | -1.8138038 | 0.06970794 | 0.18367289 |
| AC005225.3 | 9.62499758 | -1.124974225 | 0.76816342 | -1.4644986 | 0.14305778 | 0.30449863 |
| ACOT1 | 826.570051 | 0.178897399 | 0.37255905 | 0.48018536 | 0.6310956 | 0.78075653 |
| ACOT2 | 1764.97585 | 0.807085201 | 0.6168146 | 1.30847291 | 0.19071296 | 0.36951511 |
| ACOT4 | 706.774467 | 0.532192998 | 0.44520507 | 1.19538844 | 0.2319353 | 0.4198911 |
| ACOT6 | 103.516064 | 1.858219015 | 0.61937809 | 3.00013681 | 0.00269858 | 0.01695132 |
| AC005225.2 | 76.9369155 | 1.560942512 | 0.54148006 | 2.88273314 | 0.00394241 | 0.02252197 |
| DNAL1 | 1227.86325 | 0.061414224 | 0.1474318 | 0.41656022 | 0.67700011 | 0.81214143 |
| AC006146.1 | 98.0150579 | 0.536083804 | 0.19690336 | 2.72257322 | 0.00647757 | 0.03261642 |
| PNMA1 | 3261.10986 | -0.177406187 | 0.1767586 | -1.0036637 | 0.31554074 | 0.51261762 |
| ELMSAN1 | 2228.09344 | -0.422347861 | 0.25571567 | -1.6516308 | 0.09860985 | 0.23541025 |
| AC005520.2 | 163.818646 | 0.363903496 | 0.7004187 | 0.51955137 | 0.6033763 | 0.76063835 |
| LINC02274 | 826.852298 | -1.246845337 | 0.42423935 | -2.9390139 | 0.00329258 | 0.01971065 |
| PTGR2 | 1462.47815 | 0.840638722 | 0.24390967 | 3.44651652 | 0.00056786 | 0.00501451 |
| AC005520.1 | 3312.42239 | -0.11558934 | 0.10305428 | -1.1216355 | 0.26201745 | 0.45561491 |
| AC005520.5 | 45.2609264 | 0.302766357 | 0.2660566 | 1.13797724 | 0.25512999 | 0.44783064 |
| ZNF410 | 2697.20562 | -0.441782206 | 0.11196148 | -3.945841 | 7.95E-05 | 0.00105587 |
| AC005480.1 | 420.338547 | 0.070555277 | 0.15786179 | 0.44694334 | 0.65491597 | 0.79747572 |
| FAM161B | 1996.21778 | -0.281580805 | 0.09386885 | -2.9997256 | 0.00270223 | 0.0169621 |
| COQ6 | 3553.01997 | 0.613713514 | 0.31215768 | 1.96603687 | 0.04929435 | 0.14426256 |
| ENTPD5 | 7731.59325 | 1.138337632 | 0.44727321 | 2.54506106 | 0.01092586 | 0.04812344 |
| BBOF1 | 3211.84587 | 2.167146284 | 0.31739536 | 6.82790797 | 8.62E-12 | 1.78E-09 |
| ALDH6A1 | 18879.7211 | 2.431051579 | 0.4507614 | 5.39321147 | 6.92E-08 | 3.48E-06 |
| LIN52 | 872.528374 | 1.134705052 | 0.25088539 | 4.52280244 | 6.10E-06 | 0.00013849 |
| VSX2 | 0.59399959 | -1.691657885 | 2.23981066 | -0.7552683 | 0.45008804 | NA |
| ABCD4 | 5362.62369 | -0.044164369 | 0.13314997 | -0.3316889 | 0.74012417 | 0.8541816 |
| VRTN | 0.12059279 | 0.02007176 | 3.29738704 | 0.00608717 | 0.99514317 | NA |
| SYNDIG1L | 32.4018321 | 0.282464452 | 0.58433109 | 0.48339795 | 0.6288132 | 0.77940325 |
| AC005479.2 | 19.340069 | -1.768584477 | 0.53591653 | -3.3001118 | 0.00096646 | 0.00764032 |
| AC005479.1 | 13.5704936 | -0.287605595 | 0.53078248 | -0.5418521 | 0.58792041 | 0.74877861 |
| NPC2 | 8899.40682 | -0.14531023 | 0.21748605 | -0.6681359 | 0.50404687 | 0.68375733 |
| ISCA2 | 1138.48971 | -0.021855843 | 0.14051365 | -0.1555425 | 0.87639366 | 0.93555133 |
| LTBP2 | 1148.71993 | 1.828965166 | 0.39138318 | 4.67308062 | 2.97E-06 | 7.76E-05 |
| AC013451.1 | 17.0409883 | 4.127886754 | 0.61964431 | 6.66170362 | 2.71E-11 | 4.76E-09 |
| AC013451.2 | 1.7788612 | 1.753095405 | 1.11329945 | 1.57468451 | 0.11532928 | 0.26275782 |
| AREL1 | 2252.59406 | 0.113044156 | 0.10575049 | 1.06897047 | 0.28508298 | 0.48111504 |
| FCF1 | 1452.85951 | -0.112360466 | 0.12872932 | -0.8728428 | 0.38274875 | 0.5788687 |
| YLPM1 | 2576.93745 | 0.181650001 | 0.08306222 | 2.18691471 | 0.02874875 | 0.09788469 |
| PROX2 | 84.1031834 | 0.497902029 | 0.25898987 | 1.92247684 | 0.05454579 | 0.15479685 |
| DLST | 9848.51816 | 0.466835555 | 0.20734808 | 2.25145828 | 0.02435653 | 0.0865492 |
| RPS6KL1 | 460.255653 | -0.032463676 | 0.37903717 | -0.0856477 | 0.93174645 | 0.96704554 |
| PGF | 561.307316 | -0.787140833 | 0.37220876 | -2.1147832 | 0.03444843 | 0.1113924 |
| EIF2B2 | 2162.05917 | -0.276579782 | 0.19255692 | -1.4363534 | 0.15090181 | 0.31566678 |
| AL049780.1 | 545.964087 | -0.123445829 | 0.2136378 | -0.5778277 | 0.56338048 | 0.73057995 |
| MLH3 | 1301.78617 | -0.052134178 | 0.16947604 | -0.3076198 | 0.75837169 | 0.86632565 |
| ACYP1 | 567.031093 | -0.518510112 | 0.25500695 | -2.0333176 | 0.04202045 | 0.1284134 |
| ZC2HC1C | 428.069647 | 0.141725057 | 0.22073679 | 0.64205455 | 0.52083777 | 0.69743015 |
| NEK9 | 4173.92002 | 0.021479847 | 0.18090041 | 0.11873852 | 0.90548251 | 0.95258812 |
| AL049780.2 | 18.4764121 | 1.822741793 | 0.31581272 | 5.77159076 | 7.85E-09 | 5.62E-07 |
| TMED10 | 10859.0144 | -0.209973576 | 0.11470638 | -1.8305309 | 0.06717059 | 0.17886383 |
| AL691403.2 | 3.4921257 | -0.613777572 | 0.74669009 | -0.8219977 | 0.41107818 | 0.60535722 |
| AL691403.1 | 3.76481445 | 0.021586443 | 0.81981278 | 0.02633094 | 0.97899338 | 0.99050094 |
| AF111167.1 | 3.23831585 | -0.154548289 | 0.73798165 | -0.2094202 | 0.8341202 | 0.91206315 |
| FOS | 8762.2788 | 1.692649625 | 0.578619 | 2.92532671 | 0.00344095 | 0.02032657 |
| LINC01220 | 16.9376411 | 0.036084348 | 0.39707094 | 0.09087633 | 0.92759086 | 0.96487598 |
| AF111167.2 | 87.688053 | 2.991343068 | 0.45787986 | 6.53303042 | 6.45E-11 | 1.00E-08 |
| JDP2 | 1115.12102 | 0.562735232 | 0.29906606 | 1.88164191 | 0.05988465 | 0.16530004 |
| BATF | 70.3229482 | -1.281393418 | 0.69465567 | -1.8446455 | 0.06508913 | 0.17496436 |
| AC007182.1 | 120.742219 | 1.745920679 | 0.37856886 | 4.61189722 | 3.99E-06 | 9.84E-05 |
| FLVCR2 | 868.087748 | 0.828086243 | 0.27171123 | 3.04767034 | 0.00230623 | 0.0150794 |
| TTLL5 | 927.985087 | 0.103671457 | 0.11478064 | 0.90321382 | 0.36641243 | 0.56339916 |
| ERG28 | 1852.46881 | -0.129024224 | 0.11524237 | -1.1195902 | 0.26288845 | 0.45652739 |
| AF107885.2 | 0.3915716 | -0.032081771 | 1.89806175 | -0.0169024 | 0.98651449 | NA |
| IFT43 | 1943.71787 | -0.571358168 | 0.17889246 | -3.1938639 | 0.00140382 | 0.01028015 |
| TGFB3 | 1302.98502 | -1.479078405 | 0.38016109 | -3.8906623 | 1.00E-04 | 0.00126499 |
| TGFB3-AS1 | 82.3118414 | -1.102758053 | 0.4506684 | -2.4469389 | 0.01440752 | 0.05915093 |
| AC016526.4 | 1.40446342 | -1.184847742 | 1.22460391 | -0.9675355 | 0.33327641 | 0.53127816 |
| GPATCH2L | 1953.21445 | -0.454235432 | 0.19058046 | -2.3834313 | 0.01715208 | 0.06705715 |
| AC016526.1 | 17.2145257 | 2.762944669 | 0.57137608 | 4.83559737 | 1.33E-06 | 4.09E-05 |
| ESRRB | 2214.59703 | 0.957631703 | 0.54228331 | 1.76592509 | 0.07740842 | 0.19800756 |
| AC008050.1 | 2.51245838 | -1.103818585 | 1.04324004 | -1.0580677 | 0.29002458 | 0.48615433 |
| AC007376.2 | 0.98705585 | -0.020530165 | 1.37045694 | -0.0149805 | 0.98804772 | NA |
| VASH1 | 1756.49126 | 0.040241446 | 0.23583205 | 0.17063603 | 0.86450996 | 0.92965085 |
| AF111169.3 | 206.582555 | 0.096441908 | 0.26859212 | 0.35906454 | 0.71954681 | 0.840362 |
| AF111169.1 | 707.248798 | -0.096141796 | 0.24114229 | -0.3986932 | 0.69011926 | 0.82150773 |
| VASH1-AS1 | 1261.61619 | -0.131053466 | 0.21876452 | -0.5990618 | 0.54913168 | 0.71964309 |
| ANGEL1 | 1235.39681 | 0.414300854 | 0.16952848 | 2.44384214 | 0.01453178 | 0.05947409 |
| LRRC74A | 1.07195937 | 0.608904936 | 1.45476852 | 0.41855795 | 0.67553922 | 0.81123773 |
| AF111169.2 | 0.13382144 | 0.02007176 | 3.29738704 | 0.00608717 | 0.99514317 | NA |
| LINC01629 | 9.36105901 | -3.474592283 | 0.8866925 | -3.918599 | 8.91E-05 | 0.00115583 |
| IRF2BPL | 4496.0252 | -0.538865727 | 0.23828368 | -2.2614462 | 0.02373164 | 0.08498145 |
| AC007686.3 | 636.479022 | 0.02174832 | 0.18546062 | 0.11726651 | 0.90664886 | 0.95339676 |
| LINC02288 | 32.7403352 | -2.206235362 | 0.50166257 | -4.3978472 | 1.09E-05 | 0.00021908 |
| AC007686.2 | 0.53617371 | 0.001855062 | 2.2668606 | 0.00081834 | 0.99934706 | NA |
| LINC02289 | 98.8179992 | -2.228568369 | 0.42391709 | -5.2570855 | 1.46E-07 | 6.58E-06 |
| CIPC | 1774.68932 | -0.083970934 | 0.15373948 | -0.5461898 | 0.58493549 | 0.74673623 |
| AC007375.2 | 36.4405498 | 1.535256899 | 0.3663347 | 4.19085848 | 2.78E-05 | 0.00046175 |
| TMEM63C | 770.972772 | -0.337224495 | 0.15043255 | -2.2416989 | 0.02498084 | 0.08829174 |
| ZDHHC22 | 0.64237202 | 1.48936252 | 1.68638145 | 0.8831706 | 0.3771441 | NA |
| AC007375.3 | 0 | NA | NA | NA | NA | NA |
| NGB | 1.39350769 | -1.153523289 | 1.13057452 | -1.0202983 | 0.307587 | 0.5044676 |
| POMT2 | 2151.55589 | -0.578415345 | 0.2640328 | -2.190695 | 0.02847387 | 0.09723594 |
| GSTZ1 | 1109.88911 | 0.574685666 | 0.24768151 | 2.32026065 | 0.02032678 | 0.07577286 |

| | | | | | | |
|---|---|---|---|---|---|---|
| TMED8 | 1740.57773 | 0.075128003 | 0.16228352 | 0.46294288 | 0.64340531 | 0.78925422 |
| SAMD15 | 94.2773685 | 0.421154849 | 0.32203294 | 1.30780051 | 0.19094098 | 0.36990256 |
| NOXRED1 | 82.974447 | 0.143678497 | 0.31613051 | 0.45449108 | 0.64947541 | 0.79382 |
| VIPAS39 | 1014.15726 | 0.124867845 | 0.17796622 | 0.70163791 | 0.48290501 | 0.66751612 |
| AHSA1 | 4092.3144 | -0.182392988 | 0.12570372 | -1.4509753 | 0.14678674 | 0.30983482 |
| ISM2 | 76.1933433 | 0.736134682 | 0.41394327 | 1.77834676 | 0.07534692 | 0.19435925 |
| SPTLC2 | 3515.66942 | -0.355093057 | 0.17211828 | -2.0630758 | 0.03910542 | 0.12177728 |
| ALKBH1 | 736.578235 | -0.431933559 | 0.18504516 | -2.3342062 | 0.01958493 | 0.07374761 |
| SLIRP | 2204.63114 | -0.331168322 | 0.14540201 | -2.2776049 | 0.02275013 | 0.08246335 |
| SNW1 | 2539.84374 | -0.449021357 | 0.15338744 | -2.9273672 | 0.00341845 | 0.0202481 |
| C14orf178 | 99.1665244 | 0.008306021 | 0.21670494 | 0.03832871 | 0.9694256 | 0.98583753 |
| ADCK1 | 767.829893 | 0.172846384 | 0.21924842 | 0.78835863 | 0.43048696 | 0.62251837 |
| NRXN3 | 136.086358 | 0.190292334 | 0.33014934 | 0.57638259 | 0.56435661 | 0.73137821 |
| AF099810.1 | 0 | NA | NA | NA | NA | NA |
| AC009396.2 | 0 | NA | NA | NA | NA | NA |
| AC009396.3 | 0 | NA | NA | NA | NA | NA |
| AC009396.1 | 0 | NA | NA | NA | NA | NA |
| AC026888.1 | 2.1941773 | 1.569439293 | 1.50259899 | 1.04448313 | 0.29626192 | 0.4929598 |
| AC022469.2 | 0 | NA | NA | NA | NA | NA |
| AC008056.2 | 0.62972836 | -2.541699683 | 2.16437319 | -1.1743352 | 0.24026078 | NA |
| AC008056.1 | 0 | NA | NA | NA | NA | NA |
| AF123462.1 | 0.69381272 | 1.021415841 | 1.52574321 | 0.66945462 | 0.50320552 | NA |
| DIO2 | 167.542061 | 1.620196065 | 0.54204951 | 2.98901859 | 0.00279875 | 0.01740737 |
| DIO2-AS1 | 1.25501614 | 1.916475259 | 1.49315516 | 1.2835071 | 0.19931447 | 0.38045043 |
| CEP128 | 110.051587 | 0.103252976 | 0.24079606 | 0.42879844 | 0.66806991 | 0.80590136 |
| TSHR | 10.2022155 | 0.178618799 | 0.55039293 | 0.32452961 | 0.74553711 | 0.85740777 |
| AC007262.2 | 34.9565313 | 0.830648034 | 0.38890198 | 2.13588021 | 0.03268918 | 0.10725565 |
| GTF2A1 | 1497.60238 | 0.348528876 | 0.16955157 | 2.05559215 | 0.03982184 | 0.12342525 |
| AL136040.1 | 33.7762637 | 0.770531237 | 0.33948431 | 2.26971088 | 0.02322513 | 0.08375814 |
| STON2 | 1656.24508 | -0.015205603 | 0.31141499 | -0.0488275 | 0.9610568 | 0.98133854 |
| LINC02308 | 5.14741196 | -0.621655568 | 0.66399429 | -0.9362363 | 0.34915153 | 0.5467739 |
| SEL1L | 4294.30509 | 0.020647593 | 0.16954919 | 0.12177937 | 0.90307375 | 0.95126847 |
| LINC01467 | 0.08757577 | 0.547239394 | 3.29377202 | 0.16614368 | 0.86804388 | NA |
| LINC02311 | 0 | NA | NA | NA | NA | NA |
| AL355838.1 | 0.03392908 | 0.380740593 | 3.29738709 | 0.11546736 | 0.90807469 | NA |
| LINC02301 | 2.24800188 | -0.337848002 | 1.13144429 | -0.2985989 | 0.76524612 | 0.87040805 |
| AL355095.1 | 0 | NA | NA | NA | NA | NA |
| AL359238.1 | 0 | NA | NA | NA | NA | NA |
| AL162872.1 | 0 | NA | NA | NA | NA | NA |
| LINC02305 | 0 | NA | NA | NA | NA | NA |
| AL357172.1 | 0 | NA | NA | NA | NA | NA |
| AL049775.2 | 0.08757577 | 0.547239394 | 3.29377202 | 0.16614368 | 0.86804388 | NA |
| LINC02329 | 0 | NA | NA | NA | NA | NA |
| LINC00911 | 0.09635527 | 0.547239394 | 3.29377202 | 0.16614368 | 0.86804388 | NA |
| AL049775.1 | 4.8162946 | -0.973444017 | 0.68984632 | -1.4111027 | 0.15821433 | 0.32538698 |
| AL049775.3 | 2.30339517 | -0.837801194 | 0.81215953 | -1.0315722 | 0.30227257 | 0.49915008 |
| FLRT2 | 455.438996 | 0.047697819 | 0.29643546 | 0.16090457 | 0.87216856 | 0.93332867 |
| LINC02328 | 27.653683 | 0.355165873 | 0.37688028 | 0.94238379 | 0.34599618 | 0.54354339 |
| LINC02316 | 0.16416605 | 0.18657164 | 3.29377195 | 0.05664376 | 0.95482897 | NA |
| LINC02309 | 0 | NA | NA | NA | NA | NA |
| LINC01148 | 0.08695548 | -0.642905668 | 3.30558948 | -0.1944905 | 0.84579185 | NA |
| AL358292.1 | 0 | NA | NA | NA | NA | NA |
| AL352955.1 | 0.17391096 | -1.003574173 | 3.30558948 | -0.3035992 | 0.7614333 | NA |
| AL157688.1 | 0.17391096 | -1.003574173 | 3.30558948 | -0.3035992 | 0.7614333 | NA |
| LINC02296 | 0.7193018 | 1.001829358 | 2.24056255 | 0.44713296 | 0.65477906 | NA |
| AL135746.1 | 0.08695548 | -0.642905668 | 3.30558948 | -0.1944905 | 0.84579185 | NA |
| LINC02330 | 0.27507702 | -1.364242679 | 3.30558948 | -0.4127078 | 0.67982068 | NA |
| AL359237.1 | 0.73511564 | 1.645989942 | 1.48744333 | 1.10659002 | 0.26847122 | NA |
| AL136501.1 | 0.06785815 | 0.795868262 | 3.28898349 | 0.24198001 | 0.80879565 | NA |
| GALC | 1747.59673 | 0.44208064 | 0.14663391 | 3.01485954 | 0.00257098 | 0.01634052 |
| GPR65 | 156.432149 | -0.247148317 | 0.47114619 | -0.5245682 | 0.59988338 | 0.75773985 |
| AL157955.1 | 35.5989602 | -0.267473401 | 0.57435851 | -0.4656907 | 0.64143693 | 0.78758115 |
| AL157955.2 | 0.23320561 | -1.364242679 | 3.30558948 | -0.4127078 | 0.67982068 | NA |
| LINC01146 | 11.0453538 | -0.926126863 | 0.85720235 | -1.0804063 | 0.27996127 | 0.47554347 |
| AL133279.2 | 0.09997307 | 0.547239394 | 3.29377202 | 0.16614368 | 0.86804388 | NA |
| AL133279.3 | 0.19632833 | 1.506623996 | 3.27645372 | 0.45983375 | 0.64563556 | NA |
| AL133279.1 | 1.42651437 | 2.967752869 | 1.2951756 | 2.29139034 | 0.02194085 | 0.08032489 |
| KCNK10 | 268.046169 | 3.29839809 | 0.58091216 | 5.67796362 | 1.36E-08 | 9.08E-07 |
| SPATA7 | 767.501877 | 0.375413527 | 0.1856323 | 2.02235026 | 0.04314018 | 0.13098124 |
| PTPN21 | 1368.99958 | 0.419471647 | 0.12541063 | 3.34478531 | 0.00082346 | 0.00673325 |
| AL162171.2 | 18.3106107 | 0.397581345 | 0.63969584 | 0.62151623 | 0.53426002 | 0.7080328 |
| AL162171.1 | 41.3614152 | 0.730845309 | 0.38568793 | 1.8949136 | 0.05810385 | 0.16188045 |
| ZC3H14 | 3228.31012 | 0.070860041 | 0.20449634 | 0.34651007 | 0.72895942 | 0.84685006 |
| EML5 | 1242.81084 | -0.081983814 | 0.31776325 | -0.2580028 | 0.79640473 | 0.88954182 |
| AL121768.1 | 6.78231297 | -0.100873825 | 0.68111516 | -0.148101 | 0.88226307 | 0.93888264 |
| TTC8 | 1045.30933 | 0.066060485 | 0.13364974 | 0.49428069 | 0.62110796 | 0.77363364 |
| AL137785.1 | 0.33747028 | 1.4137913 | 2.23234043 | 0.63332245 | 0.5265231 | NA |
| FOXN3 | 1827.71145 | 0.265688895 | 0.19307183 | 1.37611422 | 0.16878627 | 0.3404008 |
| AL138478.1 | 835.282021 | 0.114603691 | 0.20480271 | 0.55958092 | 0.57576532 | 0.73980078 |
| AL357093.2 | 1.51218365 | -1.652191497 | 1.74541125 | -0.9465915 | 0.34384696 | 0.54151789 |
| AL357093.1 | 0.15318056 | -1.003574173 | 3.30558948 | -0.3035992 | 0.7614333 | NA |
| AL356805.1 | 0.32784928 | -0.2861393 | 3.28898338 | -0.0869993 | 0.93067206 | NA |
| FOXN3-AS1 | 61.3341969 | 0.431711855 | 0.28717595 | 1.50330084 | 0.13276148 | 0.28914109 |
| FOXN3-AS2 | 0 | NA | NA | NA | NA | NA |
| AL137230.1 | 0.48852126 | 0.621755594 | 1.64592393 | 0.37775476 | 0.70561279 | NA |
| EFCAB11 | 348.383637 | 0.797017576 | 0.27983495 | 2.84817027 | 0.00439714 | 0.02447289 |
| TDP1 | 762.806168 | -0.415534931 | 0.16483088 | -2.5209774 | 0.01170294 | 0.05066597 |
| KCNK13 | 266.210607 | 1.014454139 | 0.35396056 | 2.86600897 | 0.00415682 | 0.02344716 |
| PSMC1 | 3151.8555 | -0.306026152 | 0.2122776 | -1.4416319 | 0.14940625 | 0.31353198 |
| NRDE2 | 2965.79266 | -0.158927231 | 0.16719066 | -0.9505748 | 0.34182025 | 0.53968388 |
| AL512791.2 | 21.8688499 | 1.540835723 | 0.44078885 | 3.49563223 | 0.00047294 | 0.00434242 |
| CALM1 | 18131.1545 | 0.606273564 | 0.25509446 | 2.376663 | 0.01747004 | 0.06790757 |
| AL512791.1 | 2269.90727 | 0.287185195 | 0.25376838 | 1.13168235 | 0.257768 | 0.45030423 |
| LINC02317 | 0.06785815 | 0.795868262 | 3.28898349 | 0.24198001 | 0.80879565 | NA |
| LINC00642 | 5.29899775 | 0.375018767 | 1.11846674 | 0.3352972 | 0.7374009 | 0.85252584 |
| AL096869.3 | 0.10178723 | 1.087385526 | 3.28420134 | 0.33109588 | 0.74057207 | NA |
| TTC7B | 1706.48515 | -0.141477304 | 0.23891845 | -0.5921573 | 0.55374526 | 0.72327815 |
| AL096869.1 | 0.2621321 | -1.364244692 | 2.96465539 | -0.4601697 | 0.64539439 | NA |
| AL096869.2 | 89.1786611 | 0.122330554 | 0.26348642 | 0.4642765 | 0.64244966 | 0.78833864 |
| AL139193.1 | 0.26773522 | -1.364244774 | 2.94995568 | -0.4624628 | 0.64374947 | NA |
| AL139193.2 | 0.35533229 | -0.541949506 | 2.83922491 | -0.1908794 | 0.84862009 | NA |
| LINC02321 | 4.85943843 | -0.240601642 | 0.90823663 | -0.2649107 | 0.79107822 | 0.88647096 |
| RPS6KA5 | 355.082008 | -0.107890902 | 0.22330542 | -0.483154 | 0.62898642 | 0.77954466 |
| DGLUCY | 4164.12157 | 0.459665323 | 0.28180576 | 1.63114242 | 0.10286027 | 0.24209438 |
| GPR68 | 174.808706 | -0.112083691 | 0.36386857 | -0.3080335 | 0.75805688 | 0.86618156 |
| AL135818.2 | 10.1007236 | 0.221149176 | 0.45602187 | 0.484953 | 0.62770969 | 0.77851117 |
| CCDC88C | 1433.49096 | 0.140167917 | 0.24683896 | 0.56785169 | 0.57013568 | 0.73533109 |
| AL133153.2 | 1.30362091 | 2.299315233 | 1.21762048 | 1.88836774 | 0.0589766 | 0.16353448 |
| PPP4R3A | 3780.49006 | -0.174836991 | 0.18146629 | -0.9634682 | 0.33531263 | 0.53356228 |
| CATSPERB | 170.228148 | -0.53732218 | 0.31599542 | -1.7004113 | 0.08905359 | 0.21892876 |
| TC2N | 1243.74721 | -0.47369753 | 0.2576213 | -1.8387359 | 0.06595405 | 0.17666047 |
| FBLN5 | 8330.00438 | 0.927674079 | 0.28904781 | 3.20941401 | 0.00133006 | 0.00985214 |
| TRIP11 | 1852.0926 | 0.410713077 | 0.10362477 | 3.96346434 | 7.39E-05 | 0.0009969 |

| | | | | | | |
|---|---|---|---|---|---|---|
| ATXN3 | 1027.32703 | -0.124998174 | 0.12669266 | -0.9866252 | 0.32382642 | 0.52161155 |
| NDUFB1 | 2803.47122 | 0.000271508 | 0.10868693 | 0.00249808 | 0.99800682 | 0.99887805 |
| CPSF2 | 1700.93786 | -0.153465048 | 0.158151 | -0.9703704 | 0.33186191 | 0.52979772 |
| AL133240.1 | 0.08757577 | 0.547239394 | 3.29377202 | 0.16614368 | 0.86804388 | NA |
| SLC24A4 | 32.7143759 | 0.080795044 | 0.48978407 | 0.16496054 | 0.86897504 | 0.9318649 |
| RIN3 | 1188.67954 | -0.145833737 | 0.17544661 | -0.8312143 | 0.40585256 | 0.60071464 |
| LGMN | 15673.5591 | 0.785986263 | 0.31187079 | 2.52023042 | 0.0117278 | 0.05074031 |
| GOLGA5 | 2137.11643 | -0.133908556 | 0.22006193 | -0.608504 | 0.54285328 | 0.71525137 |
| AL117192.1 | 0.15661533 | -1.003574173 | 3.30558948 | -0.3035992 | 0.7614333 | NA |
| LINC02287 | 1.49494323 | -1.494866733 | 1.27545939 | -1.1720222 | 0.24118814 | 0.43134156 |
| CHGA | 36.6951313 | -0.061589097 | 0.66928789 | -0.0920218 | 0.9266807 | 0.96438573 |
| ITPK1 | 6010.71893 | -0.083704128 | 0.14951764 | -0.5598278 | 0.5755969 | 0.73972494 |
| ITPK1-AS1 | 1.26691717 | 1.219383918 | 1.10980756 | 1.09873456 | 0.27188386 | 0.46667822 |
| MOAP1 | 3617.25514 | -0.448843284 | 0.17521841 | -2.5616216 | 0.01041847 | 0.04650223 |
| TMEM251 | 1221.88133 | -0.601668416 | 0.27840248 | -2.161146 | 0.03068406 | 0.10251345 |
| AL110118.2 | 828.482337 | -0.272228063 | 0.26322727 | -1.034194 | 0.30104548 | 0.49768474 |
| GON7 | 712.392 | -0.186673141 | 0.13489931 | -1.3837961 | 0.1664209 | 0.33721351 |
| UBR7 | 1597.31354 | 0.118414884 | 0.1669579 | 0.70924998 | 0.47816937 | 0.66311492 |
| BTBD7 | 2170.62108 | -0.279265042 | 0.133993 | -2.0841764 | 0.03714413 | 0.11726214 |
| UNC79 | 121.349682 | 1.072625612 | 0.40403851 | 2.65476086 | 0.00793646 | 0.03794137 |
| COX8C | 4.14081599 | 0.152917425 | 0.64919557 | 0.23554909 | 0.81378258 | 0.89946545 |
| PRIMA1 | 129.046024 | 0.931797314 | 0.78963471 | 1.18003592 | 0.23798593 | 0.42735369 |
| FAM181A-AS | 0.66343346 | -0.984837397 | 1.64232697 | -0.5996598 | 0.54873301 | NA |
| FAM181A | 0.07830766 | -0.642905668 | 3.30558948 | -0.1944905 | 0.84579185 | NA |
| ASB2 | 111.490664 | 0.189945571 | 0.65246114 | 0.29112166 | 0.77095827 | 0.87355577 |
| AL132642.1 | 6.96709676 | 0.363343081 | 0.92998619 | 0.39069728 | 0.69602101 | 0.82515362 |
| CCDC197 | 1.75918348 | -2.576098818 | 1.72668978 | -1.4919291 | 0.13571772 | 0.29342267 |
| OTUB2 | 132.779831 | -0.082891097 | 0.32575448 | -0.2544588 | 0.79914112 | 0.89108294 |
| DDX24 | 7423.93441 | -0.060642452 | 0.09507852 | -0.6378144 | 0.52359448 | 0.69892989 |
| IFI27L1 | 591.163045 | -0.447964108 | 0.25133345 | -1.7823498 | 0.07469221 | 0.19312027 |
| IFI27 | 4411.33732 | -1.750701077 | 0.73835409 | -2.371086 | 0.0177359 | 0.0686274 |
| IFI27L2 | 715.75542 | -0.856375041 | 0.35831398 | -2.3900129 | 0.01684778 | 0.06627029 |
| PPP4R4 | 27.8121373 | -0.069230834 | 0.32892974 | -0.210473 | 0.83329851 | 0.91140944 |
| SERPINA10 | 9.59535103 | 2.045495454 | 0.65701399 | 3.11332102 | 0.00184995 | 0.01268741 |
| SERPINA6 | 1000.71373 | 2.499227579 | 0.51208106 | 4.88053116 | 1.06E-06 | 3.41E-05 |
| SERPINA1 | 93488.5869 | -2.108143893 | 0.56542614 | -3.728416 | 0.00019269 | 0.00211803 |
| AL132708.1 | 0.32299488 | 2.288366988 | 3.2247442 | 0.70962744 | 0.4779352 | NA |
| SERPINA11 | 0.22139721 | 0.689157666 | 3.05072504 | 0.22589963 | 0.82127948 | NA |
| SERPINA9 | 0.36695002 | 1.705742188 | 2.8152169 | 0.60590081 | 0.54458062 | NA |
| SERPINA12 | 0.25755127 | -1.364244623 | 2.97698289 | -0.4582642 | 0.64676266 | NA |
| SERPINA4 | 256.917505 | 0.899301088 | 0.34990925 | 2.57009804 | 0.01016697 | 0.0456268 |
| SERPINA5 | 8817.24972 | 1.571513322 | 0.33339886 | 4.71361335 | 2.43E-06 | 6.66E-05 |
| AL049839.2 | 41238.4118 | -3.962608478 | 0.62926032 | -6.2972483 | 3.03E-10 | 3.72E-08 |
| SERPINA3 | 40392.2268 | -5.206186005 | 0.89209146 | -5.835933 | 5.35E-09 | 4.09E-07 |
| GSC | 6.3026449 | -1.44339113 | 0.76179265 | -1.8947297 | 0.05812823 | 0.16193127 |
| AL121612.1 | 0.07830766 | -0.642905668 | 3.30558948 | -0.1944905 | 0.84579185 | NA |
| LINC02279 | 5.34301276 | 0.102097219 | 0.9477052 | 0.10773099 | 0.91420909 | 0.9579717 |
| AL390254.1 | 0 | NA | NA | NA | NA | NA |
| DICER1 | 2994.39909 | -0.254651883 | 0.11158218 | -2.2821914 | 0.02247804 | 0.0817468 |
| DICER1-AS1 | 631.197361 | 0.249378398 | 0.29890859 | 0.83429651 | 0.40411392 | 0.59897135 |
| CLMN | 6208.21364 | 0.240564908 | 0.19907309 | 1.20842505 | 0.22688381 | 0.4137687 |
| LINC02292 | 4.42114729 | 1.746698959 | 0.69519042 | 2.51254751 | 0.0119863 | 0.05148699 |
| SYNE3 | 344.205304 | 0.392574555 | 0.27679323 | 1.41829537 | 0.15610455 | 0.32262241 |
| AL133467.1 | 7.63566264 | -1.625257968 | 0.91595492 | -1.7743864 | 0.07599926 | 0.19559832 |
| SNHG10 | 344.895824 | 0.017274832 | 0.19129678 | 0.09030383 | 0.92804578 | 0.96509656 |
| GLRX5 | 2880.88932 | 0.266639397 | 0.20021287 | 1.3317795 | 0.18293266 | 0.35970684 |
| LINC02318 | 0.2745553 | -0.179253712 | 2.94817509 | -0.0608016 | 0.95151723 | NA |
| AL133467.2 | 0.63019012 | -2.543163844 | 2.16475838 | -1.1748026 | 0.24007369 | NA |
| AL133467.4 | 0.38502367 | 0.997924502 | 1.87521946 | 0.53216411 | 0.59461234 | NA |
| TCL1B | 27.5547743 | 4.165209074 | 0.70630244 | 5.89720332 | 3.70E-09 | 2.95E-07 |
| TCL1A | 24.5580493 | 2.110972323 | 1.01315749 | 2.08355794 | NA | NA |
| AL139020.1 | 7.45098336 | 2.276726136 | 1.08595686 | 2.09651619 | NA | NA |
| AL133167.1 | 0 | NA | NA | NA | NA | NA |
| TUNAR | 18.330269 | 0.603642659 | 0.81516981 | 0.74051154 | 0.45898966 | 0.64780332 |
| C14orf132 | 687.643626 | 0.109246124 | 0.42930964 | 0.2544693 | 0.79913302 | 0.89108294 |
| BDKRB2 | 1696.23697 | -0.210835937 | 0.38992743 | -0.5407056 | 0.58871053 | 0.74924212 |
| AL355102.2 | 306.160351 | 0.003135552 | 0.41321832 | 0.00758812 | 0.99394561 | 0.9972357 |
| AL355102.5 | 0.44171524 | 1.302787166 | 2.67676359 | 0.48670236 | 0.62646927 | NA |
| BDKRB1 | 154.520968 | 0.342512829 | 0.27467443 | 1.24697749 | 0.21240575 | 0.39620426 |
| AL355102.4 | 40.4554614 | 3.521803306 | 0.51200275 | 6.87848508 | 6.05E-12 | 1.31E-09 |
| AL355102.3 | 0.88470264 | -1.786906406 | 1.86817165 | -0.9565001 | 0.33881962 | NA |
| ATG2B | 2650.37673 | -0.017869947 | 0.1480068 | -0.1207373 | 0.90389909 | 0.95167363 |
| GSKIP | 1175.25521 | -0.581729112 | 0.19833149 | -2.9331152 | 0.00335579 | 0.01997568 |
| AK7 | 416.067152 | -0.365488298 | 0.21366957 | -1.7105304 | 0.08716783 | 0.2152407 |
| AL163051.1 | 47.80218 | 0.499334771 | 0.34192667 | 1.46035631 | 0.14419217 | 0.30597469 |
| PAPOLA | 7225.8946 | -0.238874847 | 0.13613014 | -1.7547535 | 0.07930148 | 0.20115808 |
| AL137786.1 | 4.2568667 | -1.477123373 | 1.03197081 | -1.4313616 | 0.15232661 | 0.31751267 |
| LINC02299 | 0 | NA | NA | NA | NA | NA |
| VRK1 | 278.353875 | 0.021749123 | 0.23193336 | 0.09377316 | 0.92528936 | 0.96385069 |
| AL049833.2 | 0.07830766 | -0.642905668 | 3.30558948 | -0.1944905 | 0.84579185 | NA |
| AL049833.1 | 0 | NA | NA | NA | NA | NA |
| LINC02304 | 0 | NA | NA | NA | NA | NA |
| LINC02325 | 1.868721 | 1.471880172 | 0.99592665 | 1.47790018 | 0.1394345 | 0.29912852 |
| LINC02291 | 0 | NA | NA | NA | NA | NA |
| LINC02312 | 0.08757577 | 0.547239394 | 3.29377202 | 0.16614368 | 0.86804388 | NA |
| AL355097.1 | 0.27447648 | -0.700026849 | 2.99240644 | -0.2339344 | 0.8150359 | NA |
| LINC01550 | 116.85902 | 1.66052992 | 0.37065997 | 4.47992786 | 7.47E-06 | 0.00016166 |
| AL163932.1 | 3.98234306 | 3.996841644 | 0.86708656 | 4.60950708 | 4.04E-06 | 9.94E-05 |
| LINC02295 | 0.239366 | 1.011340114 | 2.98135597 | 0.33922152 | 0.73444286 | NA |
| AL132719.1 | 0 | NA | NA | NA | NA | NA |
| C14orf177 | 0 | NA | NA | NA | NA | NA |
| AL132796.1 | 0 | NA | NA | NA | NA | NA |
| AL132796.2 | 0 | NA | NA | NA | NA | NA |
| AL162151.1 | 0 | NA | NA | NA | NA | NA |
| BCL11B | 133.259734 | 0.466254323 | 0.51742513 | 0.90110491 | 0.36753254 | 0.56438321 |
| AL109767.1 | 0.83944854 | -2.032887059 | 1.86802091 | -1.0882571 | 0.27648161 | NA |
| AL132819.1 | 0 | NA | NA | NA | NA | NA |
| SETD3 | 6647.86502 | 0.469580141 | 0.20742787 | 2.26382378 | 0.02358496 | 0.08460652 |
| CCNK | 2566.87357 | -0.639371579 | 0.15764231 | -4.0558375 | 5.00E-05 | 0.0007434 |
| CCDC85C | 4536.24378 | -0.410998588 | 0.16532125 | -2.4860603 | 0.01291661 | 0.0544542 |
| AL110504.1 | 979.755195 | -0.49734949 | 0.27648938 | -1.7988014 | 0.0720501 | 0.1882596 |
| AL160313.1 | 6.84880954 | 0.362577787 | 0.49080868 | 0.73873548 | 0.46006764 | 0.6486998 |
| HHIPL1 | 84.5305223 | 0.02543788 | 0.34533281 | 0.07366193 | 0.9412794 | 0.97207934 |
| CYP46A1 | 180.694301 | 1.200653913 | 0.34438584 | 3.48636257 | 0.00048964 | 0.00445539 |
| AL160313.2 | 8.74054864 | 1.041127009 | 0.5658319 | 1.8399935 | 0.06576919 | 0.17628924 |
| EML1 | 964.000779 | 0.374066326 | 0.26322555 | 1.42108667 | 0.15529156 | 0.32150731 |
| EVL | 4163.3678 | -0.326370802 | 0.22428352 | -1.4551707 | 0.14562201 | 0.30816567 |
| AL133368.2 | 0 | NA | NA | NA | NA | NA |
| AL157912.1 | 0.4697416 | 2.164786889 | 1.98805247 | 1.08889826 | 0.27619875 | NA |
| DEGS2 | 1245.85145 | -0.185916506 | 0.33936855 | -0.5478307 | 0.58380816 | 0.74595509 |
| AL133523.1 | 14.0964324 | -0.00530343 | 0.38445891 | -0.0137945 | 0.98899391 | 0.99503126 |
| YY1 | 5497.81944 | -0.293940475 | 0.15690053 | -1.8734192 | 0.06101052 | 0.16737904 |
| SLC25A29 | 11981.3812 | -0.833289273 | 0.33808804 | -2.4647109 | 0.01371238 | 0.05691865 |

| | | | | | | |
|---|---|---|---|---|---|---|
| AL157871.1 | 4810.3942 | -1.128444937 | 0.34400209 | -3.2803432 | 0.00103681 | 0.00806841 |
| SLC25A47 | 167.66177 | -0.485331003 | 0.75509811 | -0.642739 | 0.52039347 | 0.69699386 |
| WARS | 7846.03796 | -0.793439849 | 0.34103741 | -2.3265478 | 0.01998934 | 0.07489614 |
| AL157871.4 | 1396.15937 | -0.852134516 | 0.36143878 | -2.3576178 | 0.01839262 | 0.07060901 |
| AL157871.2 | 9.84843103 | 0.535562739 | 0.76003673 | 0.70465376 | 0.48102573 | 0.66565116 |
| WDR25 | 523.596677 | -0.021231409 | 0.14054783 | -0.1510618 | 0.87992696 | 0.93771686 |
| BEGAIN | 129.295476 | -0.365812068 | 0.34721831 | -1.0535506 | 0.2920887 | 0.48832683 |
| AL845552.2 | 20.0657897 | -0.207315445 | 0.40866253 | -0.5073023 | 0.61194272 | 0.76758665 |
| LINC00523 | 0.18754884 | 1.469068384 | 3.27783996 | 0.44818185 | 0.65402196 | NA |
| AL132711.1 | 0 | NA | NA | NA | NA | NA |
| DLK1 | 30.7000325 | 1.271379351 | 0.91465884 | 1.3900039 | 0.16452769 | 0.33440691 |
| MEG3 | 4358.97485 | -0.019325119 | 0.34819311 | -0.0555012 | 0.95573921 | 0.97886523 |
| AL117190.2 | 626.643809 | -0.24381062 | 0.36024213 | -0.6767965 | 0.49853507 | 0.6799953 |
| AL117190.1 | 838.093875 | -0.465570401 | 0.35277182 | -1.3197494 | 0.1869187 | 0.36489828 |
| RTL1 | 0.48266291 | -0.132168728 | 2.81440656 | -0.0469615 | 0.96254392 | NA |
| MEG8 | 32.9783277 | -0.241490074 | 0.4625643 | -0.5220681 | 0.6016229 | 0.75900851 |
| MIR381HG | 0 | NA | NA | NA | NA | NA |
| MEG9 | 58.228955 | 0.498285467 | 0.36586177 | 1.36195009 | 0.17321364 | 0.34683567 |
| AL132709.1 | 1.0977174 | -0.100379155 | 1.66943255 | -0.0601277 | 0.95205392 | 0.97702607 |
| LINC02285 | 8.83953302 | 0.023092229 | 0.62190503 | 0.03713144 | 0.9703802 | 0.98629128 |
| AL359682.1 | 0 | NA | NA | NA | NA | NA |
| AL355096.1 | 0.11223674 | 0.02007176 | 3.29738704 | 0.00608717 | 0.99514317 | NA |
| LINC00524 | 0.22780565 | -0.339647149 | 3.05581276 | -0.1111479 | 0.91149907 | NA |
| LINC02314 | 0 | NA | NA | NA | NA | NA |
| AL049836.2 | 0 | NA | NA | NA | NA | NA |
| DIO3OS | 306.798548 | -0.055944273 | 0.32947463 | -0.1697984 | 0.86516867 | 0.9299135 |
| DIO3 | 38.3965616 | -1.641241343 | 0.78514518 | -2.0903667 | 0.03658487 | 0.11615014 |
| AL049836.1 | 4.42479494 | -2.426409588 | 0.97954777 | -2.4770712 | 0.01324655 | 0.05549912 |
| LINC02320 | 0.79262289 | -2.882405982 | 2.11967336 | -1.359835 | 0.17388215 | NA |
| LINC00239 | 11.7407174 | -1.639164176 | 0.60456795 | -2.7112985 | 0.00670203 | 0.03346571 |
| PPP2R5C | 4024.40787 | -0.070398413 | 0.12279273 | -0.5733109 | 0.56643418 | 0.73284279 |
| AL137779.2 | 15.6535642 | 0.801107761 | 0.64908808 | 1.234205 | 0.21712652 | 0.40202404 |
| AL137779.1 | 4.5271089 | -0.454326488 | 0.66026504 | -0.6880971 | 0.49139161 | 0.67440391 |
| AL118558.3 | 186.919802 | 0.610430963 | 0.34629124 | 1.76276754 | 0.0779397 | 0.19871861 |
| AL118558.4 | 46.0262703 | 0.499434297 | 0.3795951 | 1.31570269 | 0.18827385 | 0.36635705 |
| DYNC1H1 | 9880.68157 | 0.03636368 | 0.11460388 | 0.31729886 | 0.75101684 | 0.86117122 |
| HSP90AA1 | 39088.3722 | -0.024501191 | 0.18080483 | -0.1355118 | 0.89220722 | 0.94419715 |
| WDR20 | 1508.64971 | -0.187695401 | 0.16167463 | -1.1609453 | 0.24566414 | 0.4368872 |
| MOK | 1214.56878 | 0.268933789 | 0.21761803 | 1.23580653 | 0.21653049 | 0.40139957 |
| ZNF839 | 1246.79793 | 0.414906283 | 0.21120968 | 1.96442837 | 0.04948044 | 0.1445921 |
| CINP | 1817.5395 | 0.027350099 | 0.15298088 | 0.17878115 | 0.85810954 | 0.92582473 |
| TECPR2 | 1548.18727 | -0.04289841 | 0.12122094 | -0.3538861 | 0.72342421 | 0.84320617 |
| ANKRD9 | 2650.0524 | -0.322474906 | 0.24706787 | -1.3052078 | 0.1918221 | 0.3710371 |
| LINC02323 | 1.37023567 | -0.010716534 | 1.27352424 | -0.0084149 | 0.99328599 | 0.99691526 |
| RCOR1 | 1369.8434 | -0.19273861 | 0.13436797 | -1.4344089 | 0.1514556 | 0.31652276 |
| AL132801.1 | 0.44339417 | -2.043787703 | 2.78508174 | -0.733834 | 0.46304991 | NA |
| TRAF3 | 1531.77973 | -0.480505928 | 0.14020363 | -3.4272003 | 0.00060984 | 0.00527434 |
| AMN | 10168.1198 | 0.971612136 | 0.1654362 | 5.8730321 | 4.28E-09 | 3.32E-07 |
| CDC42BPB | 8282.42387 | 0.204379746 | 0.11233087 | 1.81944424 | 0.06884368 | 0.1819593 |
| AL117209.1 | 1365.82217 | -0.297237722 | 0.1616986 | -1.8382208 | 0.06602988 | 0.17680827 |
| LBHD2 | 2.94494584 | -0.42567981 | 0.74631015 | -0.5703792 | 0.56842052 | 0.73404883 |
| EXOC3L4 | 1101.15451 | 0.53456277 | 0.58934434 | 0.90704659 | 0.36438216 | 0.5615216 |
| LINC00677 | 7.67879295 | 0.223351628 | 0.50336927 | 0.44371327 | 0.6572499 | 0.79887899 |
| TNFAIP2 | 12509.0642 | -1.209011174 | 0.35239862 | -3.4308057 | 0.00060179 | 0.00522875 |
| LINC00605 | 43.2705537 | 1.187022251 | 0.52818544 | 2.2473589 | 0.0246171 | 0.08724001 |
| AL161669.3 | 196.569725 | -0.19421207 | 0.31642324 | -0.6137731 | 0.53936528 | 0.71247053 |
| AL138976.2 | 47.5090293 | -0.359883059 | 0.32378779 | -1.1114782 | 0.26636259 | 0.46052855 |
| EIF5 | 11768.4525 | -0.143478751 | 0.31947435 | -0.4491088 | 0.65335315 | 0.79630813 |
| MARK3 | 4554.51942 | -0.459109063 | 0.13972743 | -3.2857475 | 0.00101712 | 0.00794946 |
| CKB | 15675.7801 | -0.334663495 | 0.28869427 | -1.1592315 | 0.24636183 | 0.43769955 |
| AL133367.1 | 465.915873 | 0.885729215 | 0.3345746 | 2.64732952 | 0.00811303 | 0.03851314 |
| TRMT61A | 1640.76037 | -0.455711325 | 0.21272857 | -2.1422197 | 0.03217581 | 0.10607277 |
| AL139300.2 | 7.09226254 | -0.602943642 | 0.69401895 | -0.8687712 | 0.38497231 | 0.58076201 |
| BAG5 | 2824.67026 | -0.763178065 | 0.14606408 | -5.2249538 | 1.74E-07 | 7.62E-06 |
| KLC1 | 5263.93445 | -0.513839204 | 0.19740098 | -2.6030226 | 0.00924059 | 0.04242978 |
| AL139300.1 | 2456.94739 | -0.342040092 | 0.16994463 | -2.0126561 | 0.04415082 | 0.13310053 |
| APOPT1 | 1457.75461 | -0.53818405 | 0.17048119 | -3.156853 | 0.00159482 | 0.01132443 |
| AL049840.1 | 78.0839218 | 0.373413439 | 0.29443663 | 1.26823026 | 0.20471573 | 0.38708977 |
| AL049840.2 | 199.751152 | 0.16700983 | 0.24942902 | 0.66956857 | 0.50313286 | 0.68332771 |
| XRCC3 | 3015.16271 | -0.53347624 | 0.20308821 | -2.6268204 | 0.00861868 | 0.04024697 |
| ZFYVE21 | 2977.26868 | -0.130420776 | 0.11739013 | -1.1110029 | 0.26656709 | 0.46064051 |
| PPP1R13B | 2981.54042 | -0.102125922 | 0.14906594 | -0.6851057 | 0.49327725 | 0.67596234 |
| LINC00637 | 8.65620797 | 1.146404929 | 0.68819664 | 1.66581011 | 0.09575124 | 0.23064193 |
| AL132712.1 | 37.2562824 | -0.031618656 | 0.46750199 | -0.0676332 | 0.94607763 | 0.97409658 |
| AL132712.2 | 9.93146498 | 0.295692804 | 0.47444836 | 0.62323495 | 0.53313014 | 0.70699732 |
| ATP5MPL | 3383.6335 | -0.189297768 | 0.1574377 | -1.2023662 | 0.22922168 | 0.41665947 |
| TDRD9 | 373.495063 | -0.403504629 | 0.51382095 | -0.785302 | 0.43227651 | 0.62437624 |
| RD3L | 0.90858407 | -3.066618497 | 1.7468149 | -1.7555486 | 0.07916551 | NA |
| ASPG | 2152.86678 | 1.567357347 | 0.41928502 | 3.73816684 | 0.00018537 | 0.00205846 |
| AL359399.1 | 1.76077512 | -0.097497021 | 0.95214218 | -0.1023975 | 0.91844113 | 0.96032088 |
| KIF26A | 650.589432 | 1.245752384 | 0.45564243 | 2.73405704 | 0.00625592 | 0.03180425 |
| AL590079.1 | 7.01437373 | 0.828824665 | 0.76476758 | 1.0837602 | 0.27847114 | 0.47412258 |
| AL512357.1 | 0 | NA | NA | NA | NA | NA |
| TMEM179 | 13.299004 | -0.715593949 | 0.68122611 | -1.05045 | 0.29351128 | 0.49008373 |
| C14orf180 | 104.979351 | 0.377201252 | 0.70730393 | 0.53329444 | 0.59382978 | 0.75288673 |
| BX927359.1 | 57.179413 | 0.301482974 | 0.72451555 | 0.41611664 | 0.67732465 | 0.81241988 |
| AL583722.3 | 1.19100536 | 0.230700932 | 1.28007645 | 0.18022434 | 0.85697645 | 0.92514304 |
| LINC02280 | 2.34165875 | 0.233414472 | 1.09105105 | 0.21393543 | 0.83059741 | 0.91033489 |
| AL583722.1 | 4.4282404 | -0.50708256 | 0.95784727 | -0.5293981 | 0.59652929 | 0.75525626 |
| INF2 | 8225.88604 | -0.691558058 | 0.1677663 | -4.1221513 | 3.75E-05 | 0.00058847 |
| AL583722.4 | 2.15381627 | 0.603402515 | 0.86261803 | 0.6995014 | 0.48423874 | 0.668712 |
| ADSSL1 | 1200.00012 | 0.721917664 | 0.37345615 | 1.93307208 | 0.05322732 | 0.15220173 |
| SIVA1 | 2553.26474 | -0.00765386 | 0.11851767 | -0.0645799 | 0.94850848 | 0.97514677 |
| AKT1 | 9032.56761 | -0.49776927 | 0.14642388 | -3.3995088 | 0.00067507 | 0.00572583 |
| AL583722.2 | 3336.95211 | -0.752154889 | 0.14565364 | -5.1639965 | 2.42E-07 | 9.93E-06 |
| ZBTB42 | 769.539984 | -0.233222858 | 0.1313546 | -1.7755211 | 0.07581189 | 0.19526857 |
| LINC00638 | 53.604851 | 0.920344385 | 0.31617268 | 2.91089157 | 0.00360399 | 0.02104456 |
| AL583810.2 | 21.6829998 | -0.370845033 | 0.4266467 | -0.8692087 | 0.38473298 | 0.58043416 |
| AL583810.1 | 34.0919809 | 0.292377053 | 0.2308167 | 1.26670668 | 0.20526019 | 0.38775781 |
| CEP170B | 8374.80587 | -0.613423712 | 0.18761116 | -3.2696547 | 0.00107679 | 0.00830746 |
| AL583810.3 | 0.15661533 | -1.003574173 | 3.30558948 | -0.3035992 | 0.7614333 | NA |
| PLD4 | 227.998777 | -0.425752573 | 0.51754486 | -0.822639 | 0.41071333 | 0.60505645 |
| AHNAK2 | 477.983579 | -1.812293026 | 0.5848303 | -3.0988357 | 0.00194283 | 0.01319401 |
| CLBA1 | 1450.48688 | 0.089584135 | 0.2603659 | 0.34407015 | 0.73079353 | 0.84789798 |
| CDCA4 | 280.989353 | -0.560735577 | 0.20922671 | -2.6800382 | 0.00736138 | 0.03595367 |
| GPR132 | 557.22136 | 0.976736136 | 0.42794692 | 2.28237684 | 0.0224671 | 0.08171827 |
| LINC02298 | 84.850416 | -0.078381185 | 0.28372836 | -0.2762543 | 0.78235274 | 0.88085486 |
| JAG2 | 1505.97859 | -0.271200113 | 0.32799592 | -0.8268399 | 0.40832781 | 0.60278813 |
| NUDT14 | 1550.23131 | 0.307278207 | 0.22628126 | 1.35794809 | 0.17448016 | 0.3484054 |
| BRF1 | 2627.40314 | 0.04031483 | 0.12569641 | 0.32073176 | 0.74841368 | 0.85941773 |
| BTBD6 | 2258.51324 | 0.119371852 | 0.11266609 | 1.05951887 | 0.28936354 | 0.48561806 |
| PACS2 | 5603.20114 | -0.093868307 | 0.09928455 | -0.9454473 | 0.34443059 | 0.54205267 |
| TEX22 | 501.472336 | 0.092125886 | 0.2322089 | 0.3967371 | 0.69156134 | 0.82241065 |

| | | | | | | |
|---|---|---|---|---|---|---|
| AL928654.1 | 65.3406253 | 1.102326906 | 0.21864813 | 5.04155655 | 4.62E-07 | 1.70E-05 |
| MTA1 | 5457.12845 | -0.365818813 | 0.23993458 | -1.5246606 | 0.12734376 | 0.28093826 |
| AL928654.2 | 1065.00901 | -0.584786283 | 0.24914118 | -2.3472084 | 0.01891467 | 0.07203638 |
| CRIP2 | 7946.18264 | -0.177800945 | 0.37846149 | -0.4697993 | 0.63849841 | 0.78614457 |
| CRIP1 | 1320.26338 | 0.456841623 | 0.55055155 | 0.829789 | 0.40665809 | 0.60140134 |
| AL928654.3 | 1357.6773 | 0.438104936 | 0.53375316 | 0.82080064 | 0.41175983 | 0.60580988 |
| TEDC1 | 390.821199 | 0.3072584 | 0.42796806 | 0.71794704 | 0.47278994 | 0.65885276 |
| TMEM121 | 95.2457071 | -0.80079112 | 0.43321699 | -1.8484758 | 0.06453354 | 0.17391172 |
| IGHA2 | 1242.23674 | 1.776331229 | 0.92676251 | 1.91670597 | NA | NA |
| IGHE | 5.49421945 | 2.163073434 | 0.70115051 | 3.08503439 | 0.00203529 | 0.0136775 |
| IGHG4 | 1514.04826 | 0.120396836 | 1.26189762 | 0.09540935 | 0.92398969 | 0.96310558 |
| IGHG2 | 3212.42948 | 0.084321303 | 0.85615825 | 0.09848799 | 0.92154481 | 0.96167277 |
| AL928742.1 | 11.3814191 | 0.04876682 | 1.21256087 | 0.04021804 | 0.9679193 | 0.98498946 |
| IGHGP | 1153.67305 | -3.406203639 | 0.90287677 | -3.7726119 | 0.00016155 | 0.00185218 |
| IGHA1 | 12001.305 | 1.119913303 | 1.18169173 | 0.94772035 | NA | NA |
| IGHEP1 | 0.63888795 | -1.539108367 | 2.21801782 | -0.6939116 | 0.48773763 | NA |
| IGHG1 | 16575.1417 | -1.759993557 | 0.91649243 | -1.920358 | 0.0548127 | 0.1554164 |
| IGHG3 | 4084.00908 | -1.478309293 | 0.95507178 | -1.5478515 | 0.12165805 | 0.27240578 |
| IGHD | 235.771481 | -0.54672169 | 0.94729903 | -0.5771374 | 0.56384665 | 0.73093285 |
| IGHM | 4653.47638 | -0.438976437 | 0.94573549 | -0.4641641 | 0.6425302 | 0.78840079 |
| IGHJ6 | 381.713234 | -0.671831872 | 0.91411118 | -0.7349564 | 0.46236604 | 0.65054917 |
| IGHJ3P | 12.0559402 | 0.317249061 | 0.96912226 | 0.32735711 | 0.7433978 | 0.85604367 |
| IGHJ5 | 158.993304 | -0.188425296 | 0.92667775 | -0.2033342 | 0.83887381 | 0.91454277 |
| IGHJ4 | 754.864594 | 1.111831948 | 1.13852545 | 0.97655432 | 0.32878984 | 0.52658294 |
| IGHJ3 | 280.381667 | 0.661749102 | 1.0083011 | 0.65630108 | 0.51163042 | 0.68975942 |
| IGHJ2P | 1.27863537 | 0.425550337 | 1.62831029 | 0.26134474 | 0.79382666 | 0.88844572 |
| IGHJ2 | 70.1608095 | -0.24617236 | 1.01053839 | -0.2436052 | 0.80753664 | 0.89601937 |
| IGHJ1 | 29.602047 | 0.159244814 | 1.06297994 | 0.1498098 | 0.88091468 | 0.93830917 |
| IGHD7-27 | 0.26181525 | 1.093535322 | 3.27839713 | 0.33355792 | 0.73871318 | NA |
| IGHJ1P | 0.54925645 | 1.711921605 | 2.16509816 | 0.79069007 | 0.42912487 | NA |
| IGHD1-26 | 0.2441911 | -0.174096114 | 3.29377188 | -0.0528562 | 0.95784652 | NA |
| IGHD6-25 | 1.88191168 | 0.230764258 | 2.31176841 | 0.09982153 | 0.92048602 | 0.96108712 |
| IGHD5-24 | 0.08757577 | 0.547239394 | 3.29377202 | 0.16614368 | 0.86804388 | NA |
| IGHD4-23 | 0.23148822 | -1.364242679 | 3.30558948 | -0.4127078 | 0.67982068 | NA |
| IGHD3-22 | 1.15321275 | -1.206537417 | 1.97401345 | -0.6112103 | 0.54106034 | 0.71388779 |
| IGHD2-21 | 0.17515154 | 1.45420457 | 3.2783972 | 0.44357181 | 0.65735219 | NA |
| AC246787.2 | 3.37859095 | 1.11432 | 1.16673146 | 0.95507839 | 0.33953806 | 0.53765964 |
| IGHD1-20 | 0.2534592 | 1.093535322 | 3.27839713 | 0.33355792 | 0.73871318 | NA |
| IGHD6-19 | 0.90110549 | -0.319611899 | 1.90067337 | -0.1681572 | 0.86645962 | NA |
| IGHD5-18 | 0.16588343 | 0.18657164 | 3.29377195 | 0.05664376 | 0.95482897 | NA |
| IGHD4-17 | 0.09989236 | -0.642905668 | 3.30558948 | -0.1944905 | 0.84579185 | NA |
| IGHD3-16 | 0.08757577 | 0.547239394 | 3.29377202 | 0.16614368 | 0.86804388 | NA |
| IGHD2-15 | 0.18655607 | -1.003574173 | 3.30558948 | -0.3035992 | 0.7614333 | NA |
| IGHD1-14 | 0.29967709 | -1.364242679 | 3.30558948 | -0.4127078 | 0.67982068 | NA |
| IGHD6-13 | 0.07830766 | -0.642905668 | 3.30558948 | -0.1944905 | 0.84579185 | NA |
| IGHD5-12 | 0.95690442 | -1.132729361 | 2.04252825 | -0.5545722 | 0.57918732 | NA |
| IGHD4-11 | 0.08393208 | -0.642905668 | 3.30558948 | -0.1944905 | 0.84579185 | NA |
| IGHD3-10 | 0.51390703 | -2.265764565 | 2.20599444 | -1.0270944 | 0.30437601 | NA |
| IGHD3-9 | 0.59389986 | -1.423544711 | 2.79377692 | -0.5095413 | 0.61037286 | NA |
| IGHD2-8 | 0 | NA | NA | NA | NA | NA |
| IGHD1-7 | 0 | NA | NA | NA | NA | NA |
| IGHD6-6 | 0.42496705 | 1.263083491 | 2.81353046 | 0.44893187 | 0.65348081 | NA |
| IGHD5-5 | 0.07659028 | -0.642905668 | 3.30558948 | -0.1944905 | 0.84579185 | NA |
| IGHD4-4 | 0.16588343 | 0.18657164 | 3.29377195 | 0.05664376 | 0.95482897 | NA |
| IGHD3-3 | 0.60106341 | -1.458798675 | 2.20647809 | -0.6611435 | 0.50852028 | NA |
| IGHD2-2 | 0.79867441 | 1.911780163 | 2.64294816 | 0.72335137 | 0.46946404 | NA |
| FAM30A | 86.702295 | 0.344489808 | 0.94508382 | 0.36450715 | 0.71547934 | 0.83812505 |
| IGHD1-1 | 0.72746947 | -0.345905232 | 1.47019005 | -0.2352793 | 0.81399199 | NA |
| IGHV6-1 | 83.5967561 | -1.025596774 | 1.02865425 | -0.9970277 | 0.31875107 | 0.51608434 |
| IGHVII-1-1 | 0 | NA | NA | NA | NA | NA |
| IGHV1-2 | 281.551171 | 0.121701114 | 1.31912797 | 0.09225876 | NA | NA |
| IGHVIII-2-1 | 0 | NA | NA | NA | NA | NA |
| IGHV1-3 | 69.2205207 | -0.494962564 | 1.05897077 | -0.4673996 | 0.64021398 | 0.78714964 |
| IGHV4-4 | 63.8678818 | 0.197597549 | 0.95393719 | 0.20713895 | 0.83590134 | 0.91285536 |
| IGHV7-4-1 | 10.8616418 | -1.650835573 | 1.73639999 | -0.9507231 | 0.34174496 | 0.53967632 |
| IGHV2-5 | 130.982669 | 1.184450489 | 1.25134976 | 0.94653831 | 0.34387409 | 0.54151789 |
| IGHVIII-5-1 | 0 | NA | NA | NA | NA | NA |
| IGHVIII-5-2 | 0 | NA | NA | NA | NA | NA |
| IGHV3-6 | 1.89299339 | 0.020029535 | 1.55382325 | 0.01289048 | 0.98971517 | 0.99519138 |
| IGHV3-7 | 267.010852 | 0.319747353 | 1.14080914 | 0.2802812 | 0.77926177 | 0.87906969 |
| IGHV3-64D | 25.4081062 | 0.848730796 | 1.1376559 | 0.74603472 | 0.45564644 | 0.64508734 |
| IGHV5-10-1 | 77.9159516 | -1.47787753 | 1.92804416 | -0.7665164 | 0.44336908 | 0.63444989 |
| IGHV3-11 | 110.270476 | 0.665253917 | 1.06101138 | 0.6269998 | 0.53065937 | 0.70501262 |
| IGHVIII-11-1 | 0 | NA | NA | NA | NA | NA |
| IGHV1-12 | 7.08318988 | 0.350799465 | 1.30000382 | 0.26984495 | 0.78727954 | 0.88414299 |
| IGHV3-13 | 23.489196 | 0.018295802 | 1.21072927 | 0.01511139 | 0.98794331 | 0.99454413 |
| IGHVIII-13-1 | 0 | NA | NA | NA | NA | NA |
| IGHV1-14 | 3.28389307 | 0.521344262 | 1.54971069 | 0.33641393 | 0.73655875 | 0.85207763 |
| IGHV3-15 | 176.440023 | 1.397589927 | 0.95363994 | 1.46553208 | 0.14277581 | 0.30408602 |
| IGHVII-15-1 | 0 | NA | NA | NA | NA | NA |
| IGHV3-16 | 0.35908258 | 2.442028688 | 3.21676432 | 0.75915686 | 0.44775873 | NA |
| IGHVIII-16-1 | 0 | NA | NA | NA | NA | NA |
| IGHV1-17 | 1.63567517 | -0.88125513 | 2.81254134 | -0.3133306 | 0.75402954 | 0.86318765 |
| IGHV1-18 | 245.085392 | -0.715780614 | 0.83332652 | -0.8589438 | 0.39037154 | 0.5858814 |
| IGHV3-19 | 1.6951387 | -0.547323123 | 1.88527736 | -0.2903144 | 0.77157574 | 0.87406359 |
| AC247036.1 | 0 | NA | NA | NA | NA | NA |
| IGHV3-20 | 32.8545344 | -0.140716353 | 1.01370173 | -0.1388144 | 0.88959685 | 0.94296756 |
| IGHV3-21 | 169.476691 | 0.677906803 | 1.01053426 | 0.67084 | 0.50232246 | 0.68263189 |
| IGHV3-22 | 1.13203571 | 0.297695169 | 1.8381744 | 0.16195154 | 0.87134402 | 0.93311835 |
| IGHVII-22-1 | 0 | NA | NA | NA | NA | NA |
| IGHVIII-22-2 | 0.08757577 | 0.547239394 | 3.29377202 | 0.16614368 | 0.86804388 | NA |
| IGHV3-23 | 499.440832 | 0.585379173 | 1.19663513 | 0.48918769 | 0.62470882 | 0.77643779 |
| IGHV1-24 | 96.6135466 | -0.210139902 | 1.43928603 | -0.1460029 | NA | NA |
| LINC00226 | 0 | NA | NA | NA | NA | NA |
| IGHV3-25 | 2.1150199 | 3.346002418 | 2.03086125 | 1.64757805 | 0.09943929 | 0.23657651 |
| IGHVIII-25-1 | 0 | NA | NA | NA | NA | NA |
| IGHV2-26 | 37.9216238 | 0.528591028 | 1.01894269 | 0.51876424 | 0.60392516 | 0.76111195 |
| IGHVIII-26-1 | 0 | NA | NA | NA | NA | NA |
| IGHVII-26-2 | 0 | NA | NA | NA | NA | NA |
| IGHV7-27 | 1.30101505 | 0.15046502 | 1.65152156 | 0.09110691 | 0.92740764 | 0.96476151 |
| IGHV4-28 | 22.25375 | 0.779522946 | 1.1435827 | 0.68164982 | 0.4954604 | 0.67766031 |
| IGHVII-28-1 | 0 | NA | NA | NA | NA | NA |
| IGHV3-32 | 0.5919942 | -1.461412912 | 1.88444357 | -0.7755143 | 0.43803583 | NA |
| IGHV3-30 | 553.461893 | -1.447427382 | 1.28331979 | -1.1278774 | 0.2593717 | 0.45217717 |
| IGHVII-30-1 | 0.07830766 | -0.642905668 | 3.30558948 | -0.1944905 | 0.84579185 | NA |
| IGHV3-30-2 | 0.08757577 | 0.547239394 | 3.29377202 | 0.16614368 | 0.86804388 | NA |
| IGHV4-31 | 69.6921918 | 0.574206692 | 1.15974868 | 0.49511304 | 0.62052033 | 0.7733044 |
| IGHVII-30-21 | 0 | NA | NA | NA | NA | NA |
| IGHV3-29 | 0 | NA | NA | NA | NA | NA |
| IGHV3-33 | 135.060015 | 0.824889476 | 1.01262981 | 0.81460122 | 0.41530061 | 0.60885513 |
| IGHVII-33-1 | 0 | NA | NA | NA | NA | NA |
| IGHV3-33-2 | 0 | NA | NA | NA | NA | NA |
| IGHV4-34 | 236.556728 | -0.465158732 | 1.28793119 | -0.3611674 | 0.71797433 | 0.83985952 |

| | | | | | | |
|---|---|---|---|---|---|---|
| IGHV7-34-1 | 0.07830766 | -0.642905668 | 3.30558948 | -0.1944905 | 0.84579185 | NA |
| IGHV3-35 | 1.70753089 | -0.219269177 | 1.08060578 | -0.2029132 | 0.83920288 | 0.91464269 |
| IGHV3-36 | 0.33925836 | 0.669855702 | 2.22876149 | 0.30055064 | 0.76375717 | NA |
| IGHV3-37 | 0.44665835 | 2.760059159 | 3.20263664 | 0.8618084 | 0.38879296 | NA |
| IGHV3-38 | 1.35994776 | 1.647677719 | 1.29204359 | 1.27524932 | 0.20222101 | 0.38401466 |
| IGHVIII-38-1 | 0.67306063 | -1.676685695 | 1.90483642 | -0.8802256 | 0.37873713 | NA |
| IGHV4-39 | 176.77621 | -0.00065879 | 1.00098466 | -0.0006581 | 0.99947488 | 0.99979217 |
| IGHV7-40 | 0.46984598 | -2.146049573 | 3.28588824 | -0.6531109 | 0.51368476 | NA |
| IGHVII-40-1 | 0 | NA | NA | NA | NA | NA |
| IGHV3-41 | 3.07072798 | -0.460133373 | 1.35756028 | -0.3389414 | 0.73465388 | 0.85065526 |
| IGHV3-42 | 0.60153039 | 1.334097754 | 1.86086879 | 0.71692199 | 0.47342223 | NA |
| IGHV3-43 | 23.9741705 | 0.377601663 | 1.04853302 | 0.36012377 | 0.71875458 | 0.84003117 |
| IGHVII-43-1 | 0 | NA | NA | NA | NA | NA |
| IGHVIII-44 | 0 | NA | NA | NA | NA | NA |
| AC244452.3 | 0 | NA | NA | NA | NA | NA |
| LINC00221 | 0.08757577 | 0.547239394 | 3.29377202 | 0.16614368 | 0.86804388 | NA |
| IGHVIV-44-1 | 0.08757577 | 0.547239394 | 3.29377202 | 0.16614368 | 0.86804388 | NA |
| IGHVII-44-2 | 0 | NA | NA | NA | NA | NA |
| IGHV1-45 | 14.9109044 | 2.244719064 | 2.13840479 | 1.04971663 | NA | NA |
| IGHV1-46 | 80.89159 | -0.310416839 | 1.02473437 | -0.3029242 | 0.76194763 | 0.86807911 |
| IGHVII-46-1 | 0.03392908 | 0.380740593 | 3.29738709 | 0.11546736 | 0.90807469 | NA |
| IGHV3-47 | 1.05083968 | 1.98622345 | 1.64544948 | 1.20710084 | 0.22739331 | 0.41428504 |
| IGHVII-47-1 | 0.08757577 | 0.547239394 | 3.29377202 | 0.16614368 | 0.86804388 | NA |
| IGHV3-48 | 137.677395 | 1.730843497 | 1.03587702 | 1.6708967 | 0.09474208 | 0.22897744 |
| IGHV3-49 | 47.847303 | 0.461381298 | 0.81694752 | 0.56476247 | 0.57223534 | 0.73695281 |
| IGHVII-49-1 | 0 | NA | NA | NA | NA | NA |
| IGHV3-50 | 0.08393208 | -0.642905668 | 3.30558948 | -0.1944905 | 0.84579185 | NA |
| IGHV5-51 | 938.149776 | 3.997727197 | 1.31734818 | 3.03467775 | NA | NA |
| IGHVIII-51-1 | 0.18882702 | -0.340230161 | 3.19721851 | -0.1064144 | 0.91525355 | NA |
| IGHVII-51-2 | 0.12150485 | 1.04981648 | 3.28477129 | 0.31960109 | 0.74927075 | NA |
| IGHV3-52 | 0.86435963 | 0.289351694 | 1.50394242 | 0.19239546 | 0.84743245 | NA |
| IGHV3-53 | 43.2615652 | 0.410068799 | 0.88774462 | 0.46192203 | 0.64413723 | 0.78974795 |
| IGHVII-53-1 | 0 | NA | NA | NA | NA | NA |
| IGHV3-54 | 0.03392908 | 0.380740593 | 3.29738709 | 0.11546736 | 0.90807469 | NA |
| IGHV4-55 | 24.0369096 | 1.267442863 | 1.19993223 | 1.05626204 | 0.29084852 | 0.48690769 |
| IGHV7-56 | 1.84154317 | 0.326396079 | 1.14764445 | 0.28440523 | 0.77609985 | 0.87704106 |
| IGHV3-57 | 0 | NA | NA | NA | NA | NA |
| IGHV1-58 | 6.82743799 | 0.810278398 | 1.52434133 | 0.53155969 | 0.59503099 | 0.75401092 |
| IGHV4-59 | 181.869336 | 1.424932179 | 0.91713707 | 1.55367417 | 0.12026216 | 0.27033612 |
| IGHV3-60 | 0.80049853 | 3.053946315 | 3.18154973 | 0.95989269 | 0.33710923 | NA |
| IGHVII-60-1 | 0.67196573 | -0.571194928 | 1.91694386 | -0.2979717 | 0.7657248 | NA |
| IGHV4-61 | 59.6159733 | 0.909528646 | 1.01351264 | 0.89740237 | 0.36950425 | 0.56635208 |
| IGHV3-62 | 3.07265956 | 0.271618288 | 1.95965882 | 0.13860489 | 0.88976238 | 0.94296756 |
| IGHVII-62-1 | 0 | NA | NA | NA | NA | NA |
| IGHV3-63 | 2.93806123 | -1.874544672 | 1.79074732 | -1.0467946 | 0.29519431 | 0.49206952 |
| IGHV3-64 | 8.74801224 | 2.119067999 | 1.01745584 | 2.08271251 | NA | NA |
| IGHV3-65 | 5.35048762 | 0.168085197 | 1.25813109 | 0.13359911 | 0.89371959 | 0.94530446 |
| IGHVII-65-1 | 0 | NA | NA | NA | NA | NA |
| IGHV3-66 | 14.6583181 | 0.406486709 | 0.83121613 | 0.48902649 | 0.62482294 | 0.77650129 |
| IGHV1-67 | 3.758191 | 0.606268358 | 1.98797136 | 0.30496836 | 0.76039025 | 0.86715659 |
| IGHVII-67-1 | 0 | NA | NA | NA | NA | NA |
| IGHVIII-67-2 | 0 | NA | NA | NA | NA | NA |
| IGHVIII-67-3 | 0 | NA | NA | NA | NA | NA |
| IGHVIII-67-4 | 0 | NA | NA | NA | NA | NA |
| IGHV1-68 | 0 | NA | NA | NA | NA | NA |
| IGHV1-69 | 27.1681026 | 0.658089812 | 1.06798748 | 0.61619618 | 0.53776505 | 0.71135512 |
| IGHV2-70D | 9.07588148 | 0.433129782 | 1.86845787 | 0.23181137 | 0.81668452 | 0.90138837 |
| IGHV3-69-1 | 7.71476857 | -0.046160811 | 1.05140413 | -0.043904 | 0.96498095 | 0.98359669 |
| IGHV1-69-2 | 4.46587277 | 2.124298594 | 1.29709184 | 1.63773954 | 0.10147604 | 0.23995007 |
| IGHV1-69D | 120.188185 | -0.21788265 | 0.97919088 | -0.2225129 | 0.82391459 | 0.90578475 |
| IGHV2-70 | 17.8637343 | 1.825976015 | 1.14666767 | 1.59241955 | 0.11129047 | 0.25599526 |
| IGHV3-71 | 2.88299022 | 1.123449526 | 1.18358093 | 0.94919536 | 0.34252126 | 0.54018364 |
| IGHV3-72 | 28.8059943 | 1.827680356 | 1.07585745 | 1.69881275 | 0.08935447 | 0.21949487 |
| IGHV3-73 | 45.945073 | 1.397279122 | 0.94065134 | 1.48543787 | 0.13742787 | 0.29605203 |
| IGHV3-74 | 137.631598 | 1.218471843 | 0.80861307 | 1.50686637 | 0.13184491 | 0.28790579 |
| IGHVII-74-1 | 0.09989236 | -0.642905668 | 3.30558948 | -0.1944905 | 0.84579185 | NA |
| IGHV3-75 | 0.68536047 | 0.239531541 | 1.8933984 | 0.12650879 | 0.89932919 | NA |
| IGHV3-76 | 0 | NA | NA | NA | NA | NA |
| IGHVIII-76-1 | 1.6112409 | 1.887696596 | 1.33438675 | 1.41465478 | 0.15716974 | 0.32417411 |
| IGHV5-78 | 8.20714379 | 0.169714391 | 0.67119641 | 0.25285354 | 0.80038139 | 0.89167965 |
| IGHVII-78-1 | 1.17006991 | 0.300684227 | 1.19986982 | 0.25059738 | 0.80212541 | 0.89266577 |
| IGHV3-79 | 1.3489495 | 2.55809478 | 1.14908845 | 2.22619485 | 0.02600113 | 0.09098326 |
| IGHV4-80 | 5.65917153 | 0.304750966 | 0.8856641 | 0.34409317 | 0.73077622 | 0.84789798 |
| IGHV7-81 | 5.0226425 | 1.164032339 | 0.77787642 | 1.49642323 | 0.1345434 | 0.29188335 |
| IGHVIII-82 | 0 | NA | NA | NA | NA | NA |
| IGHV1OR15- | 0.64752809 | 2.718032491 | 2.53374378 | 1.07273376 | 0.28339059 | NA |
| IGHV1OR15- | 0.81341153 | 2.559927062 | 2.03577553 | 1.2574702 | 0.20858342 | NA |
| IGHV3OR15- | 0.712323 | -1.728946005 | 2.22555152 | -0.7768618 | 0.4372403 | NA |
| IGHD5OR15- | 0 | NA | NA | NA | NA | NA |
| IGHD4OR15- | 0 | NA | NA | NA | NA | NA |
| IGHD3OR15- | 0 | NA | NA | NA | NA | NA |
| IGHD2OR15- | 0 | NA | NA | NA | NA | NA |
| IGHD1OR15- | 0 | NA | NA | NA | NA | NA |
| AC087386.1 | 0.46267344 | 2.807834585 | 3.20076292 | 0.87723916 | 0.38035675 | NA |
| AC026495.1 | 76.73267 | 3.277003233 | 0.99302189 | 3.30003121 | 0.00096674 | 0.00764032 |
| GOLGA6L6 | 0.08393208 | -0.642905668 | 3.30558948 | -0.1944905 | 0.84579185 | NA |
| IGHV1OR15- | 0.18301898 | 0.18657164 | 3.29377195 | 0.05664376 | 0.95482897 | NA |
| AC023310.4 | 2.47812507 | -0.319206114 | 1.36628211 | -0.2336312 | 0.81527131 | 0.90043158 |
| AC012414.5 | 8.20286552 | -0.054527605 | 0.69395332 | -0.0785753 | 0.93737042 | 0.96999796 |
| AC012414.3 | 0 | 0 | 0 | 0 | 1 | NA |
| AC012414.4 | 0 | NA | NA | NA | NA | NA |
| POTEB2 | 0 | NA | NA | NA | NA | NA |
| LINC01193 | 0 | NA | NA | NA | NA | NA |
| IGHD5OR15- | 0 | NA | NA | NA | NA | NA |
| IGHD4OR15- | 0 | NA | NA | NA | NA | NA |
| IGHD3OR15- | 0 | NA | NA | NA | NA | NA |
| IGHD2OR15- | 0 | NA | NA | NA | NA | NA |
| IGHD1OR15- | 0 | NA | NA | NA | NA | NA |
| AC126335.1 | 0 | NA | NA | NA | NA | NA |
| AC068446.2 | 0.99571089 | -0.797494972 | 1.7248634 | -0.4623525 | 0.64382853 | NA |
| AC060814.5 | 0.09644756 | -0.642905668 | 3.30558948 | -0.1944905 | 0.84579185 | NA |
| AC060814.4 | 0 | NA | NA | NA | NA | NA |
| AC060814.3 | 0.17361919 | -1.003574173 | 3.30558948 | -0.3035992 | 0.7614333 | NA |
| POTEB3 | 0 | NA | NA | NA | NA | NA |
| LINC02203 | 0 | NA | NA | NA | NA | NA |
| AC135068.1 | 0 | NA | NA | NA | NA | NA |
| AC135068.3 | 0 | NA | NA | NA | NA | NA |
| AC135068.7 | 0 | NA | NA | NA | NA | NA |
| AC135068.8 | 1.86967805 | -3.219574579 | 2.79243372 | -1.1529637 | 0.2489253 | 0.44066597 |
| AC135068.9 | 1.04231882 | 0.034864843 | 2.31456296 | 0.01506325 | 0.98798172 | 0.99454413 |
| AC135068.2 | 0.27504391 | 1.093535322 | 3.27839713 | 0.33355792 | 0.73871318 | NA |
| AC134981.1 | 0.5323841 | 0.938199091 | 2.21170305 | 0.42419758 | 0.67142172 | NA |
| POTEB | 0 | NA | NA | NA | NA | NA |

| | | | | | | |
|---|---|---|---|---|---|---|
| AC134980.2 | 0 | NA | NA | NA | NA | NA |
| AC134980.3 | 0 | NA | NA | NA | NA | NA |
| OR4M2 | 0 | NA | NA | NA | NA | NA |
| OR4N4 | 0 | NA | NA | NA | NA | NA |
| AC010760.1 | 0 | NA | NA | NA | NA | NA |
| IGHV1OR15- | 1.59527338 | 0.363993728 | 1.72889787 | 0.21053512 | 0.83325004 | 0.91140944 |
| IGHV1OR15- | 0.36303872 | -1.73318485 | 2.82351077 | -0.6138404 | 0.53932083 | NA |
| IGHV4OR15- | 0 | NA | NA | NA | NA | NA |
| IGHV1OR15- | 0.09989236 | -0.642905668 | 3.30558948 | -0.1944905 | 0.84579185 | NA |
| GOLGA6L22 | 0.18340304 | -1.003574173 | 3.30558948 | -0.3035992 | 0.7614333 | NA |
| NIPA1 | 1500.75811 | -0.350389517 | 0.16029274 | -2.185935 | 0.02882036 | 0.0980636 |
| NIPA2 | 3173.79329 | -0.501452762 | 0.23145223 | -2.1665497 | 0.0302692 | 0.10144454 |
| CYFIP1 | 6058.72898 | -0.128163882 | 0.10757883 | -1.1913485 | 0.2335168 | 0.42191421 |
| TUBGCP5 | 820.694074 | 0.242388567 | 0.15692887 | 1.54457602 | 0.12244885 | 0.27359568 |
| GOLGA6L1 | 0.22977084 | -1.364242679 | 3.30558948 | -0.4127078 | 0.67982068 | NA |
| GOLGA8S | 9.08600635 | -0.312995085 | 0.78519487 | -0.3986209 | 0.69017256 | 0.82151532 |
| GOLGA6L2 | 0 | NA | NA | NA | NA | NA |
| MKRN3 | 6.66799532 | 2.389250326 | 1.21804725 | 1.96154158 | 0.04981588 | 0.14528254 |
| AC126407.1 | 0.92436967 | 2.799104761 | 1.92463743 | 1.45435432 | 0.1458481 | NA |
| MAGEL2 | 15.6348697 | -0.209939573 | 0.45658803 | -0.4598009 | 0.64565916 | 0.79091523 |
| NDN | 757.904492 | -0.140229499 | 0.19805579 | -0.7080303 | 0.47892645 | 0.66379116 |
| AC021439.1 | 0.09644756 | -0.642905668 | 3.30558948 | -0.1944905 | 0.84579185 | NA |
| PWRN4 | 0.33213693 | 1.533287534 | 2.77023021 | 0.55348741 | 0.5799297 | NA |
| PWRN2 | 0.15318056 | -1.003574173 | 3.30558948 | -0.3035992 | 0.7614333 | NA |
| AC087463.1 | 1.04890417 | 1.523420671 | 1.23977402 | 1.22878899 | 0.21915093 | 0.40463591 |
| AC087463.4 | 2.51677661 | 1.142273376 | 0.99958505 | 1.14274756 | 0.25314342 | 0.44565004 |
| AC087463.2 | 0 | NA | NA | NA | NA | NA |
| AC087463.3 | 0.68242637 | 0.804496198 | 1.84765877 | 0.43541384 | 0.66326207 | NA |
| PWRN3 | 4.9858667 | 3.105075262 | 0.74361459 | 4.17565134 | 2.97E-05 | 0.00048667 |
| PWRN1 | 293.772364 | 1.837367083 | 0.58744117 | 3.12774655 | 0.00176152 | 0.01224981 |
| NPAP1 | 92.0386385 | 2.195624663 | 0.46915156 | 4.67999016 | 2.87E-06 | 7.57E-05 |
| SNRPN | 10645.7048 | 0.165317994 | 0.17925798 | 0.92223505 | 0.35640599 | 0.55379329 |
| SNURF | 9024.82485 | 0.22027819 | 0.18305582 | 1.20333891 | 0.2288452 | 0.41621113 |
| AC124312.1 | 7710.98813 | 0.191104182 | 0.17356252 | 1.10106826 | 0.27086695 | 0.46549465 |
| AC124312.2 | 6.37637723 | 2.435157749 | 0.59016435 | 4.12623664 | 3.69E-05 | 0.00057949 |
| PWAR6 | 958.681854 | 0.938066511 | 0.16959685 | 5.53115543 | 3.18E-08 | 1.85E-06 |
| AC124312.3 | 15.2544131 | 1.392965435 | 0.49691598 | 2.80322123 | 0.00505949 | 0.02711174 |
| UBE3A | 3026.13097 | 0.1248841 | 0.1150001 | 1.08594775 | 0.27750211 | 0.47299661 |
| LINC02250 | 0.5441145 | 0.96008739 | 2.20698199 | 0.43502276 | 0.66354591 | NA |
| ATP10A | 634.475448 | 0.079854833 | 0.34040992 | 0.23458433 | 0.81453137 | 0.8999602 |
| AC016266.1 | 0 | NA | NA | NA | NA | NA |
| LINC02346 | 0.34236247 | 1.6090754 | 2.16135584 | 0.744475 | 0.45658916 | NA |
| AC100836.1 | 0.12088456 | 0.02007176 | 3.29738704 | 0.00608717 | 0.99514317 | NA |
| LINC00929 | 0 | NA | NA | NA | NA | NA |
| LINC02248 | 0 | NA | NA | NA | NA | NA |
| GABRB3 | 657.190388 | 0.945035237 | 0.34394765 | 2.74761359 | 0.00600307 | 0.03078588 |
| AC011196.1 | 0.07659028 | -0.642905668 | 3.30558948 | -0.1944905 | 0.84579185 | NA |
| GABRA5 | 4.09160888 | 1.889943524 | 1.53135999 | 1.23416018 | NA | NA |
| GABRG3 | 11.6781092 | 1.093427341 | 0.60706773 | 1.80116203 | 0.07167734 | 0.18753166 |
| GABRG3-AS1 | 0 | NA | NA | NA | NA | NA |
| AC104002.1 | 0 | NA | NA | NA | NA | NA |
| AC104002.3 | 0.60905363 | -0.291907463 | 2.76208472 | -0.1056837 | 0.91583328 | NA |
| AC021979.1 | 0 | NA | NA | NA | NA | NA |
| AC021979.2 | 0.03392908 | 0.380740593 | 3.29738709 | 0.11546736 | 0.90807469 | NA |
| OCA2 | 29.5379777 | 1.904578731 | 0.51215154 | 3.71877963 | 0.00020019 | 0.0021821 |
| AC090696.1 | 0 | NA | NA | NA | NA | NA |
| HERC2 | 3028.17811 | 0.20854164 | 0.11776924 | 1.77076489 | 0.07659981 | 0.19651173 |
| GOLGA8F | 0.07830766 | -0.642905668 | 3.30558948 | -0.1944905 | 0.84579185 | NA |
| GOLGA8G | 0.50195508 | -2.238999393 | 2.20312437 | -1.0162837 | 0.30949431 | NA |
| GOLGA8M | 45.5294161 | -0.134783077 | 0.60394692 | -0.2231704 | 0.82340289 | 0.90564302 |
| AC055876.5 | 14.109619 | -1.548807513 | 0.78992932 | -1.9606912 | 0.04991505 | 0.14552348 |
| GOLGA6L7 | 2.89869803 | 1.152762547 | 0.68611833 | 1.68012208 | 0.09293357 | 0.22572148 |
| APBA2 | 141.41399 | -0.65135977 | 0.43792665 | -1.4873718 | 0.13691663 | 0.29530784 |
| AC127522.1 | 0.30068768 | 0.605370132 | 2.92532295 | 0.2069413 | 0.8360557 | NA |
| FAM189A1 | 402.350015 | 0.019853107 | 0.47404634 | 0.0418801 | 0.96659428 | 0.98417279 |
| NSMCE3 | 1009.68752 | -0.3513189 | 0.13079782 | -2.6859691 | 0.00723198 | 0.03551346 |
| AC102941.1 | 1.03378314 | 2.423010799 | 1.84210542 | 1.31534861 | 0.18839277 | 0.36650072 |
| AC022613.1 | 153.745866 | 1.288015765 | 0.33299848 | 3.8679329 | 0.00010976 | 0.00136465 |
| TJP1 | 3908.42289 | -0.18210867 | 0.16780532 | -1.0852377 | 0.27781638 | 0.47331812 |
| AC022613.2 | 27.0303417 | 0.312475475 | 0.40004097 | 0.78110869 | 0.43473857 | 0.6267594 |
| AC111152.2 | 2.39634057 | -2.407850843 | 1.09522196 | -2.1985049 | 0.02791315 | 0.09594234 |
| GOLGA8J | 1.21848557 | -1.582783755 | 1.67437964 | -0.9452956 | 0.34450797 | 0.54210975 |
| GOLGA8T | 0.80928555 | 2.50474054 | 1.32937852 | 1.88414398 | 0.0595455 | NA |
| AC120045.1 | 0 | NA | NA | NA | NA | NA |
| LINC02249 | 0.41499581 | 0.363795023 | 2.84842212 | 0.12771809 | 0.89837208 | NA |
| CHRFAM7A | 12.2776033 | 0.52072568 | 0.59024965 | 0.88221261 | 0.37766185 | 0.57370487 |
| GOLGA8R | 8.36981678 | 2.015976617 | 0.55975844 | 3.6015118 | 0.00031637 | 0.00315654 |
| AC026150.1 | 1.74778591 | 1.171464716 | 0.89417049 | 1.31011337 | 0.19015749 | 0.36885418 |
| GOLGA8Q | 2.09486095 | 1.015642499 | 1.00375567 | 1.01184235 | 0.31161344 | 0.50871332 |
| GOLGA8H | 35.8435239 | 1.352164617 | 0.41572662 | 3.25253318 | 0.00114381 | 0.008725 |
| AC026150.3 | 10.5937825 | 1.092414052 | 0.63546214 | 1.71908598 | 0.08559872 | 0.21240596 |
| AC091057.3 | 17.846447 | -0.248578809 | 0.4733379 | -0.5251614 | 0.59947097 | 0.75734439 |
| AC091057.2 | 44.8034708 | 0.734455414 | 0.29190781 | 2.51605264 | 0.01186775 | 0.0511611 |
| ARHGAP11B | 43.2207383 | 0.548775274 | 0.30284007 | 1.81209601 | 0.06997136 | 0.18418296 |
| AC091057.6 | 22.4501916 | 0.236680013 | 0.41519214 | 0.57004935 | 0.56864422 | 0.73422848 |
| AC091057.4 | 8.1437546 | -1.054399219 | 0.53126337 | -1.9847015 | 0.04717769 | 0.13957991 |
| FAN1 | 2358.24909 | 0.46136299 | 0.11443804 | 4.03155271 | 5.54E-05 | 0.00080416 |
| MTMR10 | 6309.55333 | -0.035259002 | 0.19649538 | -0.1794393 | 0.85759274 | 0.925581 |
| TRPM1 | 13.7708896 | 0.809713667 | 0.47764702 | 1.69521348 | 0.09003493 | 0.22071406 |
| AC009562.1 | 2.84107326 | 0.151620697 | 0.89014221 | 0.17033312 | 0.86474817 | 0.92971142 |
| LINC02352 | 63.2142335 | -0.445447499 | 0.29924319 | -1.4885802 | 0.13659794 | 0.29491465 |
| AC012236.1 | 34.7393802 | -0.757576388 | 0.6458523 | -1.1729871 | 0.24080099 | 0.4309122 |
| KLF13 | 4998.98654 | 0.349021541 | 0.21710441 | 1.6076207 | 0.10791828 | 0.25048895 |
| OTUD7A | 271.577803 | -0.079502126 | 0.17940395 | -0.4431459 | 0.65766021 | 0.79917006 |
| CHRNA7 | 16.7484148 | 0.355908646 | 0.43009794 | 0.82750605 | 0.40795029 | 0.60260036 |
| AC139426.2 | 0 | NA | NA | NA | NA | NA |
| GOLGA8K | 3.8513468 | 0.762215977 | 0.68022984 | 1.1205271 | 0.26248921 | 0.45610441 |
| GOLGA8O | 5.41579946 | 0.949363597 | 0.72736275 | 1.30521338 | 0.1918202 | 0.3710371 |
| LINC02256 | 19.8822276 | 0.281354653 | 0.3699035 | 0.76061637 | 0.44688624 | 0.63716627 |
| AC123768.4 | 0.8854637 | 0.568919283 | 1.79083061 | 0.31768459 | 0.7507242 | NA |
| AC123768.3 | 173.823529 | -0.085139979 | 0.27821071 | -0.306027 | 0.75958412 | 0.86685293 |
| GOLGA8N | 363.269693 | 0.240964464 | 0.31859359 | 0.75633807 | 0.44944653 | 0.63959456 |
| AC123768.2 | 25.9869691 | 1.035983152 | 0.34783557 | 2.97837039 | 0.00289786 | 0.01785431 |
| ARHGAP11A | 122.266897 | -0.701803027 | 0.25901903 | -2.7094651 | 0.00673918 | 0.03360669 |
| SCG5 | 42.9792824 | -1.096842284 | 0.49109087 | -2.2334813 | 0.02551722 | 0.08961101 |
| AC123768.1 | 0.52753937 | -1.552131318 | 2.27839669 | -0.6812384 | 0.49572065 | NA |
| AC090877.2 | 1.92987085 | 0.149239698 | 0.91173415 | 0.16368773 | 0.86997697 | 0.93223182 |
| GREM1 | 130.183871 | -0.035420585 | 0.74333874 | -0.0476507 | 0.96199465 | 0.98187796 |
| FMN1 | 464.034152 | 0.377692515 | 0.24935495 | 1.51467824 | 0.12985391 | 0.28492001 |
| AC055874.1 | 1.22818711 | 0.511576092 | 1.07354669 | 0.47652896 | 0.63369758 | 0.7828448 |
| RYR3 | 309.581135 | 0.372891481 | 0.27025909 | 1.37975557 | 0.16766192 | 0.33885767 |
| AC010809.3 | 10.2713722 | 0.290132128 | 0.41192136 | 0.70433864 | 0.48122191 | 0.66581781 |
| AC010809.2 | 77.3069038 | 0.419200682 | 0.29005568 | 1.4452421 | 0.1483899 | 0.31224398 |

| | | | | | | |
|---|---|---|---|---|---|---|
| AC010809.1 | 28.6177821 | 0.122372902 | 0.40021881 | 0.30576499 | 0.75978358 | 0.86698524 |
| AVEN | 325.338408 | -0.413163097 | 0.23081512 | -1.7900175 | 0.0734511 | 0.190941 |
| CHRM5 | 15.650874 | 1.163031841 | 0.42281723 | 2.75067278 | 0.0059473 | 0.03060459 |
| EMC7 | 3551.71562 | -0.357298694 | 0.13262032 | -2.6941474 | 0.0070569 | 0.03484185 |
| PGBD4 | 151.37555 | 0.939611179 | 0.17342027 | 5.41811633 | 6.02E-08 | 3.09E-06 |
| KATNBL1 | 1315.97693 | -0.454296623 | 0.12412712 | -3.6599305 | 0.00025228 | 0.00263892 |
| EMC4 | 3240.36436 | -0.183346963 | 0.13915615 | -1.3175628 | 0.18765006 | 0.36570265 |
| SLC12A6 | 2962.29058 | 1.056951559 | 0.20349552 | 5.19397951 | 2.06E-07 | 8.76E-06 |
| AC079203.2 | 16.5131257 | 3.271486481 | 0.72760009 | 4.49627006 | 6.92E-06 | 0.00015286 |
| NOP10 | 3065.60612 | -0.837149346 | 0.23851725 | -3.5098064 | 0.00044843 | 0.00417114 |
| NUTM1 | 1.20479435 | -0.879209688 | 1.29501127 | -0.6789205 | 0.49718824 | 0.67893661 |
| LPCAT4 | 1636.73658 | -0.758387491 | 0.24494306 | -3.0961787 | 0.00196032 | 0.01328545 |
| GOLGA8A | 4089.73532 | -0.251768103 | 0.19930022 | -1.2632605 | 0.20649558 | 0.38915008 |
| GOLGA8B | 4049.34067 | -0.020948236 | 0.25760926 | -0.0813179 | 0.93518916 | 0.96907653 |
| LINC02252 | 0.11786115 | 0.02007176 | 3.29738704 | 0.00608717 | 0.99514317 | NA |
| GJD2 | 4.57283881 | 2.966937832 | 0.98590482 | 3.00935524 | 0.00261803 | 0.01655574 |
| AC087457.1 | 0.9470219 | 2.808837999 | 1.414596 | 1.98561144 | 0.04707649 | NA |
| ACTC1 | 364.410471 | -1.105474238 | 0.89538105 | -1.2346411 | 0.21696411 | 0.40177975 |
| AQR | 1752.21919 | 0.029616225 | 0.07676267 | 0.38581547 | 0.69963334 | 0.82773257 |
| ZNF770 | 2641.41269 | 0.087226452 | 0.1460957 | 0.59705008 | 0.55047395 | 0.7208262 |
| AC114546.1 | 1.46160051 | -0.252411821 | 1.24015134 | -0.2035331 | 0.83871839 | 0.91450415 |
| AC114546.2 | 1.64050453 | 1.313938055 | 1.21905419 | 1.077834 | 0.28110784 | 0.47692298 |
| NANOGP8 | 1.78099656 | 2.052479171 | 1.12874099 | 1.81837924 | 0.06900619 | 0.18222444 |
| AC021231.1 | 2.02560772 | 1.971529365 | 1.09436108 | 1.80153462 | 0.07161866 | 0.1874339 |
| DPH6 | 423.52953 | 0.213940329 | 0.17868331 | 1.19731568 | 0.23118353 | 0.4190163 |
| DPH6-DT | 55.046354 | 1.884881946 | 0.34504705 | 5.46268088 | 4.69E-08 | 2.53E-06 |
| AC021351.1 | 0.25846333 | -0.178665973 | 2.98217935 | -0.0599112 | 0.95222635 | NA |
| AC040918.1 | 4.45836722 | 0.655089505 | 0.56061653 | 1.16851622 | 0.2425986 | 0.43301964 |
| AC087516.1 | 0.31600365 | 1.515292019 | 2.80013962 | 0.54114874 | 0.58840507 | NA |
| AC087516.2 | 0.84006587 | 1.44968493 | 1.47091608 | 0.98556603 | 0.32434611 | NA |
| C15orf41 | 322.195435 | 0.150914952 | 0.16777586 | 0.89950337 | 0.3683846 | 0.56515711 |
| AC013640.1 | 0.07659028 | -0.642905668 | 3.30558948 | -0.1944905 | 0.84579185 | NA |
| MEIS2 | 1323.53709 | 0.504073526 | 0.27231202 | 1.85108808 | 0.06415688 | 0.1732223 |
| AC078909.1 | 29.8732939 | 1.578996042 | 0.32570014 | 4.84800542 | 1.25E-06 | 3.90E-05 |
| AC078909.2 | 15.1553188 | 0.245469251 | 0.5141264 | 0.47744922 | 0.63304227 | 0.78218269 |
| AC068875.1 | 0.99345656 | 1.820180662 | 1.46763136 | 1.24021652 | 0.21489532 | NA |
| TMCO5A | 0 | NA | NA | NA | NA | NA |
| LINC02345 | 8.21927683 | -0.142226921 | 0.55192262 | -0.2576936 | 0.7966434 | 0.88965766 |
| LINC01852 | 69.4767978 | 1.835784982 | 0.3889257 | 4.72014315 | 2.36E-06 | 6.48E-05 |
| AC087473.1 | 0.18655607 | -1.003574173 | 3.30558948 | -0.3035992 | 0.7614333 | NA |
| SPRED1 | 1641.65788 | -0.207966722 | 0.16921973 | -1.2289744 | 0.2190814 | 0.40453584 |
| FAM98B | 1618.8311 | 0.242782413 | 0.10714758 | 2.2658693 | 0.02345939 | 0.08424677 |
| RASGRP1 | 635.086954 | 0.364270977 | 0.15549477 | 2.34265745 | 0.01914695 | 0.07257773 |
| AC116158.1 | 25.204298 | 0.789341399 | 0.32299794 | 2.44379703 | 0.0145336 | 0.05947409 |
| AC116158.2 | 0.98573838 | -1.660645278 | 2.3622206 | -0.7030018 | 0.48205464 | NA |
| C15orf53 | 15.3175202 | 1.070922581 | 0.55676466 | 1.92347443 | 0.05442049 | 0.15454108 |
| AC022929.1 | 0.09635527 | 0.547239394 | 3.29377202 | 0.16614368 | 0.86804388 | NA |
| AC022929.2 | 3.32747741 | 1.675097354 | 0.80172493 | 2.08936667 | 0.03667473 | 0.11632338 |
| AC013652.1 | 23.4659504 | 0.978239998 | 0.49006333 | 1.99615015 | 0.04591758 | 0.1368807 |
| AC087878.1 | 1.25405553 | 3.316174405 | 1.14163685 | 2.90475416 | 0.00367542 | 0.02139531 |
| AC113146.1 | 2.35159834 | 2.015056455 | 1.06937073 | 1.88433852 | 0.0595192 | 0.16464086 |
| C15orf54 | 1.25665634 | -0.959956222 | 1.15812537 | -0.828888 | 0.40716779 | 0.6017494 |
| AC013652.2 | 0.09989236 | -0.642905668 | 3.30558948 | -0.1944905 | 0.84579185 | NA |
| AC109630.1 | 0.44075387 | -1.292214284 | 1.89582595 | -0.6816102 | 0.49548547 | NA |
| AC022196.1 | 0 | NA | NA | NA | NA | NA |
| THBS1 | 15638.5949 | 0.008628698 | 0.43870291 | 0.01966866 | 0.98430769 | 0.99326005 |
| AC037198.2 | 226.174213 | 1.485257932 | 0.49004079 | 3.03088635 | 0.00243837 | 0.01568288 |
| FSIP1 | 1321.11381 | 0.545968102 | 0.42726936 | 1.27780776 | 0.20131722 | 0.38307233 |
| AC023908.3 | 122.348197 | 0.78697372 | 0.25443124 | 3.09307037 | 0.00198097 | 0.01338755 |
| GPR176 | 490.0576 | -0.189211821 | 0.37947129 | -0.4986196 | 0.6180474 | 0.77165225 |
| AC012377.1 | 1.96510089 | 1.180604877 | 0.98731011 | 1.19577919 | 0.23178274 | 0.4197736 |
| EIF2AK4 | 1652.04351 | -0.036730492 | 0.14701443 | -0.2498428 | 0.80270895 | 0.89302768 |
| SRP14 | 11065.7056 | -0.235264421 | 0.14276421 | -1.647923 | 0.09936848 | 0.23645071 |
| SRP14-AS1 | 478.837491 | 0.565816578 | 0.26680326 | 2.12072591 | 0.03394487 | 0.1101993 |
| AC021755.2 | 11.9328091 | 0.8304114 | 0.58781601 | 1.41270634 | 0.15774209 | 0.32487182 |
| AC021755.3 | 7.32997473 | 0.718266274 | 0.57220544 | 1.25525943 | 0.2093846 | 0.3925403 |
| BMF | 1360.54869 | 0.269230797 | 0.36336436 | 0.74093893 | 0.45873047 | 0.64757615 |
| BMF-AS1 | 340.925117 | 0.16060975 | 0.40872926 | 0.39294899 | 0.69435716 | 0.82396716 |
| BUB1B | 49.4654255 | -0.444174469 | 0.55938535 | -0.7940402 | 0.42717202 | 0.61950416 |
| BUB1B-PAK6 | 405.365452 | 0.318143149 | 0.21827036 | 1.4575646 | 0.14496059 | 0.30713569 |
| PAK6 | 663.704119 | 0.17445045 | 0.20513345 | 0.85042422 | 0.39508928 | 0.59041733 |
| PLCB2 | 985.84248 | 0.027246798 | 0.45735398 | 0.05957486 | 0.95249424 | 0.97717271 |
| ANKRD63 | 2.98424681 | 5.495179838 | 1.20734172 | 4.5514702 | 5.33E-06 | 0.00012334 |
| PLCB2-AS1 | 3.48136909 | -0.512949002 | 0.91560911 | -0.5602271 | 0.57532457 | 0.73952274 |
| AC020658.2 | 8.42847009 | -1.466382149 | 0.51243278 | -2.8616088 | 0.00421497 | 0.02367889 |
| INAFM2 | 809.392942 | -0.598092599 | 0.29476487 | -2.0290498 | 0.04245322 | 0.12942096 |
| CCDC9B | 1536.54094 | -0.175428236 | 0.27750418 | -0.6321643 | 0.52727952 | 0.70199228 |
| PHGR1 | 0.09989236 | -0.642905668 | 3.30558948 | -0.1944905 | 0.84579185 | NA |
| DISP2 | 51.9988373 | -0.905405037 | 0.74907297 | -1.2087007 | 0.22677785 | 0.41364957 |
| AC013356.4 | 11.5256624 | -0.868062069 | 1.00846697 | -0.8607739 | 0.38936257 | 0.58500889 |
| KNSTRN | 400.602928 | -0.098789235 | 0.24566527 | -0.4021294 | 0.68758878 | 0.81947556 |
| IVD | 6137.88965 | 1.326318065 | 0.25446058 | 5.2122732 | 1.87E-07 | 8.06E-06 |
| BAHD1 | 2874.81965 | -0.156237008 | 0.09014568 | -1.7331613 | 0.08306701 | 0.20800752 |
| AC013356.3 | 1.11644469 | 1.198099687 | 1.40722056 | 0.85139439 | 0.39455031 | 0.58977902 |
| AC013356.2 | 608.563007 | 0.041925447 | 0.11842747 | 0.35401792 | 0.72332545 | 0.84312833 |
| CHST14 | 758.852085 | -0.176513661 | 0.16171243 | -1.0915281 | 0.27504057 | 0.47019955 |
| AC091045.1 | 796.29865 | -0.336985451 | 0.16162862 | -2.0849367 | 0.03707505 | 0.11711415 |
| CCDC32 | 1171.76502 | -0.318396333 | 0.13701925 | -2.3237343 | 0.02013974 | 0.07530983 |
| RPUSD2 | 440.775527 | -0.021649172 | 0.15643521 | -0.1383907 | 0.88993168 | 0.94297317 |
| KNL1 | 233.451146 | 2.158941527 | 0.61947596 | 3.48510943 | 0.00049194 | 0.0044686 |
| RAD51 | 55.2171538 | -0.255147011 | 0.41244984 | -0.6186134 | 0.53617106 | 0.70988757 |
| AC012476.1 | 39.3384569 | -0.673088221 | 0.47656725 | -1.4123678 | 0.15784171 | 0.3249501 |
| RMDN3 | 2846.58363 | 0.008929917 | 0.12598644 | 0.07087999 | 0.94349327 | 0.97315146 |
| GCHFR | 1854.30143 | 0.356173399 | 0.32717906 | 1.08861916 | 0.27632186 | 0.47162508 |
| DNAJC17 | 2693.02903 | 0.158832014 | 0.21655592 | 0.73344572 | 0.46328662 | 0.65111486 |
| C15orf62 | 279.269103 | -0.410920024 | 0.31138053 | -1.3196715 | 0.18694471 | 0.36490612 |
| ZFYVE19 | 2367.03097 | -0.235295361 | 0.19474339 | -1.2082329 | 0.22695771 | 0.41379753 |
| PPP1R14D | 116.504092 | 0.136248199 | 0.52719812 | 0.25843833 | 0.79606864 | 0.88938094 |
| SPINT1-AS1 | 944.297564 | -0.426373709 | 0.17874675 | -2.3853508 | 0.01706284 | 0.066847 |
| SPINT1 | 12435.4763 | -0.983352687 | 0.18463496 | -5.3259291 | 1.00E-07 | 4.81E-06 |
| RHOV | 316.266713 | -1.927739042 | 0.40621222 | -4.7456451 | 2.08E-06 | 5.84E-05 |
| AC025166.1 | 1.62201881 | -1.393489097 | 1.17219524 | -1.1887858 | 0.23452395 | 0.42324094 |
| VPS18 | 2512.86351 | -0.454803714 | 0.26852295 | -1.6937238 | 0.09031777 | 0.22122233 |
| AC020661.1 | 16.7461518 | -0.946502791 | 0.40754505 | -2.3224495 | 0.02020875 | 0.07542882 |
| DLL4 | 1491.42885 | -0.887071837 | 0.35203059 | -2.5198714 | 0.01173977 | 0.05076397 |
| CHAC1 | 255.179432 | 0.70761136 | 0.44694403 | 1.58322143 | 0.113371 | 0.25940073 |
| INO80 | 1547.38803 | -0.104536145 | 0.18731916 | -0.5580644 | 0.57680044 | 0.74076986 |
| AC020661.3 | 1.76772453 | 2.296745832 | 0.94140321 | 2.4397047 | 0.01469927 | 0.06000307 |
| AC020661.2 | 0.3261261 | -0.534763868 | 3.29377181 | -0.1623561 | 0.87102546 | NA |
| INO80-AS1 | 33.5298585 | 0.875444644 | 0.41487262 | 2.11015283 | 0.03484519 | 0.11235849 |
| EXD1 | 1.88520352 | 1.663266395 | 0.86023449 | 1.93350349 | 0.0531742 | 0.15213234 |
| CHP1 | 6765.81905 | 0.093934684 | 0.21241932 | 0.44221346 | 0.65833475 | 0.79946078 |
| OIP5 | 31.8732065 | -0.109216316 | 0.39497141 | -0.276517 | 0.78215099 | 0.8807709 |

| | | | | | | |
|---|---|---|---|---|---|---|
| NUSAP1 | 383.407325 | -0.845254913 | 0.36057629 | -2.3441777 | 0.01906908 | 0.07245702 |
| NDUFAF1 | 1072.1793 | 0.173545674 | 0.22354586 | 0.77633141 | 0.43755333 | 0.62954394 |
| RTF1 | 2168.89814 | 0.098710472 | 0.14108836 | 0.69963584 | 0.48415475 | 0.66868281 |
| ITPKA | 15.7953578 | -1.273137451 | 0.62782739 | -2.0278463 | 0.04257594 | 0.12963714 |
| LTK | 274.527714 | 0.559601013 | 0.34866672 | 1.60497396 | 0.10849953 | 0.25155027 |
| RPAP1 | 1312.58166 | 0.226308929 | 0.13731758 | 1.6480696 | 0.09933839 | 0.23640044 |
| TYRO3 | 1805.13852 | -0.048843799 | 0.50127329 | -0.0974395 | 0.92237741 | 0.96203347 |
| AC016134.1 | 0.5577484 | -1.617486405 | 2.23210355 | -0.7246467 | 0.46866882 | NA |
| MGA | 1270.73164 | -0.073919143 | 0.18631639 | -0.3967399 | 0.69155929 | 0.82241065 |
| AC073657.1 | 19.1665238 | -0.203280359 | 0.3852538 | -0.5276531 | 0.59774014 | 0.75631723 |
| MAPKBP1 | 1718.29988 | -0.03952622 | 0.27467297 | -0.1439028 | 0.88557719 | 0.94076746 |
| AC020659.2 | 698.902645 | -0.16503532 | 0.29061207 | -0.5678887 | 0.57011052 | 0.73533109 |
| JMJD7 | 600.292284 | 0.446713604 | 0.23193387 | 1.92603866 | 0.05409955 | 0.15399119 |
| JMJD7-PLA2( | 2891.73193 | 0.202715104 | 0.22466216 | 0.90231086 | 0.36689176 | 0.56387888 |
| PLA2G4B | 2031.34436 | -0.002291896 | 0.2380378 | -0.0096283 | 0.99231786 | 0.99636073 |
| SPTBN5 | 761.53689 | 0.439709742 | 0.43413962 | 1.01283024 | 0.31114125 | 0.50819417 |
| AC020659.1 | 1084.60443 | 0.174879178 | 0.39438972 | 0.44341718 | 0.65746401 | 0.79902876 |
| EHD4 | 3190.14051 | -0.264350891 | 0.15359479 | -1.7210928 | 0.08523398 | 0.21183979 |
| EHD4-AS1 | 6.88265999 | 0.401939918 | 0.51906917 | 0.7743475 | 0.43872532 | 0.63036297 |
| PLA2G4E-AS | 2.66499656 | 0.621216861 | 1.15011122 | 0.54013634 | 0.58910301 | 0.74948842 |
| PLA2G4E | 2.09169962 | -1.375510525 | 1.11570694 | -1.2328601 | 0.21762798 | 0.40265393 |
| AC039056.1 | 0 | NA | NA | NA | NA | NA |
| PLA2G4D | 1.848602 | -0.524385608 | 1.01857796 | -0.5148213 | 0.60667792 | 0.76318279 |
| PLA2G4F | 1711.58118 | 0.802181525 | 0.48591541 | 1.65086661 | 0.09876581 | 0.23562219 |
| VPS39 | 4159.7734 | 0.169498269 | 0.12559405 | 1.34957245 | 0.17715317 | 0.35216954 |
| AC036103.1 | 621.982415 | -0.372331205 | 0.16972363 | -2.19375 | 0.02825339 | 0.09675869 |
| TMEM87A | 2505.21126 | -0.293563645 | 0.16698821 | -1.7579902 | 0.07874917 | 0.20022821 |
| GANC | 781.537754 | 0.289043353 | 0.15531743 | 1.8609846 | 0.06274635 | 0.17061469 |
| AC012651.1 | 4560.31235 | 0.957432383 | 0.50857325 | 1.88258503 | 0.05975663 | 0.16508981 |
| CAPN3 | 4694.38511 | 0.992485141 | 0.5112843 | 1.94116099 | 0.05223875 | 0.15020333 |
| ZNF106 | 2579.63279 | 0.348301324 | 0.14424702 | 2.41461706 | 0.01575176 | 0.06307927 |
| SNAP23 | 3336.14017 | -0.120292707 | 0.14245024 | -0.8444542 | 0.39841563 | 0.59337062 |
| AC018362.1 | 1885.58073 | -0.338590384 | 0.12129458 | -2.7914716 | 0.0052469 | 0.0278672 |
| LRRC57 | 1125.50248 | -0.120505215 | 0.08335574 | -1.4456739 | 0.14826868 | 0.31203872 |
| HAUS2 | 947.801634 | -0.457630196 | 0.1795676 | -2.5485121 | 0.01081835 | 0.04776159 |
| AC018362.2 | 211.38803 | -0.65427272 | 0.21470151 | -3.0473597 | 0.00230861 | 0.01508777 |
| STARD9 | 564.96461 | 0.228280379 | 0.26944183 | 0.84723438 | 0.39686449 | 0.59183908 |
| CDAN1 | 1024.60146 | 0.310172157 | 0.12527105 | 2.47600821 | 0.01328605 | 0.05560279 |
| AC090510.3 | 325.672792 | -0.059609878 | 0.1728644 | -0.3448361 | 0.73021762 | 0.84752811 |
| AC090510.2 | 23.6022746 | 0.721910083 | 0.35469488 | 2.03529884 | 0.04182081 | 0.12797054 |
| TTBK2 | 937.687294 | -0.351011178 | 0.14993199 | -2.3411359 | 0.01922517 | 0.07277366 |
| AC090510.1 | 356.985123 | -0.128857572 | 0.16703626 | -0.7714347 | 0.44044929 | 0.63178083 |
| UBR1 | 1512.15487 | -0.152021499 | 0.11791523 | -1.289244 | 0.19731328 | 0.37811436 |
| AC068724.3 | 10.3982522 | -0.248380665 | 0.48953811 | -0.5073776 | 0.61188991 | 0.76758665 |
| TMEM62 | 1270.35606 | -0.765711904 | 0.29420146 | -2.6026788 | 0.00924986 | 0.04246496 |
| AC068724.1 | 444.603721 | -0.906063142 | 0.32073471 | -2.8249613 | 0.00472863 | 0.02579521 |
| CCNDBP1 | 2950.60367 | -0.090253083 | 0.15800891 | -0.5711898 | 0.56787097 | 0.73380854 |
| EPB42 | 14.1660585 | -4.343624212 | 1.25230604 | -3.4685006 | 0.00052337 | 0.00469912 |
| TGM5 | 1.89700793 | -0.005564994 | 1.22826528 | -0.0045308 | 0.99638498 | 0.99823988 |
| TGM7 | 3.7530895 | 2.266462203 | 0.73455756 | 3.08547938 | 0.00203224 | 0.0136675 |
| AC009852.1 | 0 | NA | NA | NA | NA | NA |
| LCMT2 | 616.397633 | 0.142650737 | 0.19493699 | 0.73177869 | 0.46430366 | 0.65178001 |
| ADAL | 604.987489 | 0.857470541 | 0.20897695 | 4.10318239 | 4.08E-05 | 0.00063174 |
| ZSCAN29 | 901.14689 | -0.218966454 | 0.18198245 | -1.2032284 | 0.22888793 | 0.41621113 |
| TUBGCP4 | 2516.34638 | -0.162697143 | 0.12869128 | -1.2642437 | 0.20614257 | 0.38879021 |
| TP53BP1 | 3194.06962 | -0.080467316 | 0.15938991 | -0.5048457 | 0.61366718 | 0.76859525 |
| MAP1A | 421.162859 | -0.095885404 | 0.30574355 | -0.3136138 | 0.75381436 | 0.86298883 |
| PPIP5K1 | 1203.1867 | 0.13326263 | 0.1688411 | 0.78927837 | 0.42994933 | 0.62206539 |
| CKMT1B | 211.670643 | -0.534294439 | 0.31725166 | -1.6841344 | 0.09215569 | 0.22442832 |
| STRC | 48.8940457 | -0.29109154 | 0.3935065 | -0.7397376 | 0.45945925 | 0.64811921 |
| AC011330.3 | 56.7295644 | -0.15469481 | 0.36010404 | -0.4295837 | 0.66749852 | 0.80542508 |
| CATSPER2 | 380.049584 | -0.045809844 | 0.36428015 | -0.1257544 | 0.89992631 | 0.94924227 |
| CKMT1A | 132.684038 | -0.625377355 | 0.54661109 | -1.1440993 | 0.25258247 | 0.44492984 |
| PDIA3 | 19797.0067 | -0.385947076 | 0.18725093 | -2.0611224 | 0.03929137 | 0.12222091 |
| ELL3 | 375.456545 | -0.354835362 | 0.19904331 | -1.7827043 | 0.07463445 | 0.1930845 |
| AC018512.1 | 1038.97965 | 0.044645149 | 0.09915845 | 0.45024048 | 0.65253705 | 0.79586523 |
| SERF2 | 21191.0365 | -0.376617824 | 0.12895074 | -2.9206334 | 0.00349321 | 0.02053865 |
| SERINC4 | 1866.0987 | 0.294064999 | 0.10677863 | 2.7539686 | 0.00588774 | 0.03037251 |
| HYPK | 2177.75861 | -0.545666932 | 0.13394188 | -4.0739083 | 4.62E-05 | 0.00069743 |
| MFAP1 | 1643.25037 | -0.175866938 | 0.10195555 | -1.7249373 | 0.08453878 | 0.21064797 |
| WDR76 | 140.804889 | 0.287199822 | 0.18628036 | 1.54176118 | 0.12313164 | 0.2746124 |
| FRMD5 | 52.9446441 | -0.229767263 | 0.48007681 | -0.4786052 | 0.63221951 | 0.78153143 |
| CASC4 | 4807.63566 | -0.047364355 | 0.18413424 | -0.2572273 | 0.7970033 | 0.88983348 |
| AC090519.2 | 0.21704035 | -0.339802133 | 3.09148137 | -0.1099156 | 0.91247628 | NA |
| AC025043.1 | 958.587785 | 0.159457611 | 0.23547443 | 0.6771759 | 0.49829436 | 0.67987397 |
| CTDSPL2 | 1329.47636 | -0.02255393 | 0.14635319 | -0.1541062 | 0.87752602 | 0.93632602 |
| AC025430.1 | 1.29328324 | 0.638017492 | 1.04483644 | 0.61063863 | 0.54143884 | 0.71427997 |
| EIF3J-DT | 940.532354 | -0.222536836 | 0.14385068 | -1.5469989 | 0.12186352 | 0.27274026 |
| EIF3J | 3280.03167 | -0.420271355 | 0.12548208 | -3.3492539 | 0.0008103 | 0.00664672 |
| SPG11 | 3089.29619 | 0.082641376 | 0.09880143 | 0.83643904 | 0.40290796 | 0.59796729 |
| PATL2 | 125.312747 | -0.593427911 | 0.6054401 | -0.9801596 | 0.32700736 | 0.52487577 |
| B2M | 105376.062 | -1.132480179 | 0.29771023 | -3.8039679 | 0.0001424 | 0.00167709 |
| TRIM69 | 2074.99469 | -0.640407455 | 0.33375157 | -1.9188148 | 0.05500778 | 0.15572249 |
| AC122108.2 | 1.13144528 | -0.774004033 | 1.16891699 | -0.6621548 | 0.507872 | 0.68672877 |
| AC122108.1 | 27.1952588 | 1.228823519 | 0.73001041 | 1.68329588 | 0.09231783 | 0.22466049 |
| TERB2 | 0.08393208 | -0.642905668 | 3.30558948 | -0.1944905 | 0.84579185 | NA |
| SORD | 7956.13772 | 1.367595518 | 0.76057918 | 1.79809749 | 0.07216156 | 0.18848062 |
| AC091117.2 | 0.09997307 | 0.547239394 | 3.29377202 | 0.16614368 | 0.86804388 | NA |
| AC091117.1 | 1508.72226 | 1.357600646 | 0.76999815 | 1.76312195 | 0.07787992 | 0.19867404 |
| DUOX2 | 82.6749737 | 0.838917376 | 0.57449139 | 1.46027842 | 0.14421357 | 0.30597469 |
| DUOXA2 | 19.8612373 | -0.305866802 | 0.90551749 | -0.3377812 | 0.73552806 | 0.85144315 |
| DUOXA1 | 29.9414799 | -0.468841453 | 0.64893472 | -0.7224786 | 0.47000027 | 0.65659606 |
| DUOX1 | 229.658576 | 0.068145036 | 0.18227882 | 0.37385055 | 0.70851551 | 0.83427345 |
| SHF | 246.243736 | 0.004580202 | 0.33037606 | 0.0138636 | 0.9889388 | 0.99503126 |
| AC051619.4 | 3.40183377 | 0.489729391 | 0.73786418 | 0.6637121 | 0.50687459 | 0.68594288 |
| AC051619.5 | 57.5097895 | 1.458468054 | 0.55997414 | 2.60452752 | 0.0092001 | 0.04233235 |
| SLC28A2 | 208.155748 | 2.584871706 | 0.63812209 | 4.0507479 | 5.11E-05 | 0.00075592 |
| GATM | 53307.1657 | 3.690935339 | 0.42520049 | 8.68045887 | 3.94E-18 | 5.20E-15 |
| AC025580.3 | 13.4337573 | 2.493425519 | 0.45991718 | 5.42146635 | 5.91E-08 | 3.04E-06 |
| SPATA5L1 | 618.521442 | -0.292429636 | 0.18948657 | -1.5432737 | 0.12276439 | 0.27398222 |
| C15orf48 | 128.339594 | -0.809505436 | 0.36375761 | -2.2253979 | 0.02605454 | 0.09108553 |
| AC025580.1 | 66.408553 | -0.000890447 | 0.47220028 | -0.0018857 | 0.9984954 | 0.99925327 |
| AC025580.2 | 9.35414936 | -0.11097386 | 0.51902497 | -0.2138122 | 0.83069353 | 0.91033489 |
| SLC30A4 | 823.908092 | 0.266165387 | 0.26066117 | 1.02111635 | 0.30719932 | 0.50411861 |
| AC090527.1 | 0.03392908 | 0.380740593 | 3.29738709 | 0.11546736 | 0.90807469 | NA |
| AC090527.3 | 13.9712042 | 0.927068919 | 0.42180685 | 2.19785174 | 0.02795967 | 0.09605616 |
| BLOC1S6 | 2410.40962 | -0.143352974 | 0.1870681 | -0.7663144 | 0.44348927 | 0.63454322 |
| AC090527.2 | 2726.1856 | -1.193384591 | 0.30216513 | -3.9494451 | 7.83E-05 | 0.00104589 |
| SQOR | 4603.98296 | -1.422211612 | 0.27789054 | -5.1178843 | 3.09E-07 | 1.22E-05 |
| AC068722.2 | 13.6718325 | -1.208695082 | 0.61935573 | -1.9515361 | 0.0509933 | 0.14772532 |
| AC068722.1 | 0.57435436 | -1.65493398 | 2.23677298 | -0.7398757 | 0.45937542 | NA |
| AC073941.1 | 0.88046973 | 2.695586823 | 1.46381831 | 1.84147637 | 0.06555178 | NA |
| SEMA6D | 1147.56121 | 2.034181273 | 0.37260982 | 5.45927982 | 4.78E-08 | 2.56E-06 |

| | | | | | | |
|---|---|---|---|---|---|---|
| AC023905.1 | 0 | NA | NA | NA | NA | NA |
| AC009558.1 | 0 | NA | NA | NA | NA | NA |
| AC012050.1 | 0 | NA | NA | NA | NA | NA |
| LINC01491 | 0.10178723 | 1.087385526 | 3.28420134 | 0.33109588 | 0.74057207 | NA |
| AC092078.2 | 32.8471055 | 4.377540594 | 2.05321566 | 2.1320413 | 0.03300345 | 0.108018 |
| AC092078.1 | 16.3193128 | 4.713434354 | 1.29256637 | 3.64657046 | 0.00026576 | 0.00275479 |
| SLC24A5 | 906.459933 | -0.354433 | 0.16819514 | -2.1072725 | 0.03509396 | 0.11288458 |
| MYEF2 | 2684.29404 | -0.159504149 | 0.14230822 | -1.1208358 | 0.26235775 | 0.45603853 |
| CTXN2 | 36.0601641 | -0.020921537 | 0.29085102 | -0.0719321 | 0.94265591 | 0.97275716 |
| SLC12A1 | 33945.0096 | 3.074659866 | 0.64446255 | 4.77088989 | 1.83E-06 | 5.30E-05 |
| AC066612.1 | 1.91595099 | 1.313489772 | 1.02105147 | 1.28640898 | 0.19830037 | 0.37923626 |
| AC066612.2 | 6.0198938 | 0.496168016 | 0.65594182 | 0.75642077 | 0.44939696 | 0.63959456 |
| AC023355.1 | 498.575807 | -0.120566444 | 0.29283953 | -0.4117151 | 0.68054829 | 0.81476702 |
| DUT | 2282.33778 | -0.173953525 | 0.20049144 | -0.8676357 | 0.38559381 | 0.58142479 |
| FBN1 | 1531.68727 | -0.172259973 | 0.34590289 | -0.498001 | 0.61848337 | 0.77200209 |
| AC084757.3 | 4.13758508 | -0.202202964 | 0.70099779 | -0.2884502 | 0.77300213 | 0.87504041 |
| AC084757.4 | 6.3924966 | -1.168021492 | 0.65432393 | -1.7850814 | 0.07424811 | 0.19238702 |
| CEP152 | 289.227863 | -0.009214445 | 0.21147385 | -0.0435725 | 0.96524517 | 0.98359669 |
| AC084757.2 | 4.29597588 | -1.616899043 | 0.75150831 | -2.1515385 | 0.03143372 | 0.10437009 |
| AC012379.1 | 13.6597579 | 2.268129173 | 0.6378773 | 3.55574525 | 0.00037691 | 0.00364236 |
| SHC4 | 23.4126589 | 0.282926684 | 0.4866884 | 0.58133025 | 0.5610179 | 0.72887826 |
| EID1 | 7818.54861 | 0.373186308 | 0.18587878 | 2.0076865 | 0.04467662 | 0.13409087 |
| AC091073.1 | 0 | NA | NA | NA | NA | NA |
| SECISBP2L | 3471.02392 | -0.0021806 | 0.1227934 | -0.0177583 | 0.98583168 | 0.99369329 |
| AC013452.2 | 5.2454873 | -0.031355464 | 0.60752556 | -0.0516118 | 0.95883805 | 0.98017606 |
| COPS2 | 3221.83559 | -0.079369346 | 0.15770995 | -0.5032615 | 0.61478044 | 0.769643 |
| GALK2 | 1907.15952 | 0.232747837 | 0.25535659 | 0.91146205 | 0.36205197 | 0.55933625 |
| FAM227B | 1042.14297 | 0.177978336 | 0.2550169 | 0.69790801 | 0.48523472 | 0.66950814 |
| AC022306.2 | 82.9534755 | 0.483235167 | 0.19219692 | 2.51427113 | 0.01192787 | 0.05134812 |
| FGF7 | 814.474622 | -1.447385198 | 0.51927891 | -2.7872983 | 0.00531495 | 0.02817848 |
| DTWD1 | 909.586612 | 0.004268089 | 0.12888137 | 0.03311642 | 0.97358175 | 0.98779582 |
| ATP8B4 | 1310.79652 | 0.485485321 | 0.50732223 | 0.95695653 | 0.3385892 | 0.5366066 |
| AC025040.1 | 3.16384016 | -0.147141669 | 1.05603035 | -0.1393347 | 0.88918567 | 0.94273988 |
| SLC27A2 | 8126.68216 | 0.258854082 | 0.52453131 | 0.49349596 | 0.6216622 | 0.7740324 |
| HDC | 105.933319 | -0.341818775 | 0.8544382 | -0.4000509 | 0.68911902 | 0.82063171 |
| GABPB1 | 734.133528 | -0.620151413 | 0.22697978 | -2.7321879 | 0.00629152 | 0.03191756 |
| GABPB1-AS1 | 1444.03705 | 0.306266316 | 0.34143577 | 0.8969954 | 0.36972137 | 0.56650509 |
| AC022087.1 | 9.20698939 | 0.882965969 | 0.47707764 | 1.85078047 | 0.06420114 | 0.17327417 |
| USP8 | 2598.76859 | 0.068915576 | 0.11225449 | 0.61392266 | 0.53926644 | 0.71237631 |
| AC012170.3 | 173.906052 | 0.298484128 | 0.20527939 | 1.45403847 | 0.14593564 | 0.30855699 |
| AC012170.2 | 503.941413 | 0.014680449 | 0.1217433 | 0.12058527 | 0.90401953 | 0.95169448 |
| USP50 | 224.932969 | -0.295773153 | 0.14075199 | -2.1013782 | 0.03560778 | 0.11400889 |
| TRPM7 | 3130.20844 | 0.489360659 | 0.18136755 | 2.698171 | 0.00697216 | 0.03452693 |
| AC084756.1 | 384.389205 | 0.229086627 | 0.18910837 | 1.21140393 | 0.22574063 | 0.41232808 |
| SPPL2A | 4743.79349 | -0.371981686 | 0.17530042 | -2.1219669 | 0.03384052 | 0.10993794 |
| AC012100.2 | 260.381106 | -0.181874165 | 0.25511205 | -0.7129188 | 0.47589602 | 0.66132069 |
| AC021752.1 | 12.0013328 | 0.439227146 | 0.57995851 | 0.75734236 | 0.44884478 | 0.63899135 |
| AP4E1 | 658.309745 | -0.442619266 | 0.17769409 | -2.490906 | 0.01274178 | 0.05392838 |
| MIR4713HG | 0.49462849 | 1.488580369 | 2.59844671 | 0.57287316 | 0.56673058 | NA |
| TNFAIP8L3 | 127.575768 | -0.666914834 | 0.33144534 | -2.0121412 | 0.04420505 | 0.13319008 |
| CYP19A1 | 34.195448 | -3.640985818 | 0.92989257 | -3.9154908 | 9.02E-05 | 0.001167 |
| AC020891.2 | 0.09644756 | -0.642905668 | 3.30558948 | -0.1944905 | 0.84579185 | NA |
| GLDN | 96.4239564 | -1.500577248 | 0.55418094 | -2.7077388 | 0.00677433 | 0.03371185 |
| AC066613.2 | 412.737034 | 0.11743384 | 0.18713274 | 0.62754301 | 0.53030335 | 0.70472412 |
| DMXL2 | 1809.0705 | 0.384572241 | 0.21260238 | 1.80888022 | 0.07046961 | 0.18515822 |
| AC066613.1 | 174.518771 | 1.047380961 | 0.30899685 | 3.38961694 | 0.0006999 | 0.00590607 |
| SCG3 | 7.1663505 | -0.288944664 | 0.52794899 | -0.5472966 | 0.58417501 | 0.74631534 |
| AC020892.1 | 0.26431171 | -1.364242679 | 3.30558948 | -0.4127078 | 0.67982068 | NA |
| AC020892.2 | 1.04170165 | -0.939549069 | 1.25718774 | -0.7473419 | 0.45485721 | 0.64431589 |
| LYSMD2 | 515.617667 | -0.435274209 | 0.20257942 | -2.1486596 | 0.03166139 | 0.10488831 |
| TMOD2 | 523.566079 | 0.182043398 | 0.21071763 | 0.86392106 | 0.38763126 | 0.5832384 |
| TMOD3 | 2984.28316 | -0.066476164 | 0.13052267 | -0.5093074 | 0.61053675 | 0.76644683 |
| AC090971.2 | 80.5340928 | 0.643941729 | 0.3384682 | 1.90251766 | 0.05710351 | 0.15980257 |
| AC090971.1 | 235.120285 | 0.398997201 | 0.25662699 | 1.55477489 | 0.11999969 | 0.26991662 |
| LEO1 | 854.995843 | 0.250401892 | 0.15373415 | 1.62879807 | 0.10335577 | 0.24301562 |
| MAPK6 | 2143.81063 | -0.242785113 | 0.38442786 | -0.6315492 | 0.52768148 | 0.70231468 |
| AC090970.2 | 5.09831264 | 1.152929665 | 0.62702424 | 1.83873221 | 0.06595458 | 0.17666047 |
| AC023906.5 | 4.83706222 | 0.779705499 | 0.59646757 | 1.30720518 | 0.19114304 | 0.37023627 |
| AC023906.2 | 0.51453373 | -2.233923824 | 3.2823796 | -0.6805806 | 0.49613692 | NA |
| BCL2L10 | 131.036652 | 0.921943422 | 0.31969476 | 2.88382399 | 0.00392878 | 0.02246845 |
| GNB5 | 3071.45099 | 0.780349287 | 0.21106901 | 3.69712868 | 0.00021805 | 0.00234122 |
| AC023906.3 | 588.367094 | 0.764547486 | 0.22188396 | 3.44570873 | 0.00056956 | 0.00502489 |
| AC023906.4 | 413.087452 | 1.094615415 | 0.24751943 | 4.42234139 | 9.76E-06 | 0.00019974 |
| CERNA1 | 134.311496 | 0.473858761 | 0.20639426 | 2.2958912 | 0.0216821 | 0.07964685 |
| MYO5C | 1121.93728 | 0.019097333 | 0.28612653 | 0.06674436 | 0.94678522 | 0.97438741 |
| MYO5A | 1229.57594 | -0.144875077 | 0.12463505 | -1.1623943 | 0.24507532 | 0.43610474 |
| ARPP19 | 6610.7798 | -0.25559797 | 0.20991872 | -1.2176045 | 0.22337431 | 0.40943055 |
| AC025917.1 | 315.654998 | 0.141217926 | 0.14683479 | 0.96174702 | 0.3361767 | 0.5342602 |
| FAM214A | 1649.80321 | 0.120993517 | 0.11045196 | 1.09544022 | 0.27332382 | 0.46826593 |
| ONECUT1 | 0.849997 | 0.342115638 | 1.53374676 | 0.22305875 | 0.82348978 | NA |
| AC016044.1 | 0.29479552 | -0.701265545 | 3.29738693 | -0.2126731 | 0.83158194 | NA |
| LINC02490 | 0.03392908 | 0.380740593 | 3.29738709 | 0.11546736 | 0.90807469 | NA |
| WDR72 | 7043.74228 | 1.190533317 | 0.44881693 | 2.65260343 | 0.00798737 | 0.03811124 |
| AC084759.2 | 0.42296795 | 2.671030466 | 2.65059627 | 1.00770928 | 0.31359405 | NA |
| UNC13C | 41.2392826 | 1.01270326 | 0.57354677 | 1.76568558 | 0.07744861 | 0.19802062 |
| AC025272.1 | 0.17303784 | -1.003574173 | 3.30558948 | -0.3035992 | 0.7614333 | NA |
| RSL24D1 | 2878.36466 | -0.329597336 | 0.17426843 | -1.8913198 | 0.05858166 | 0.16274742 |
| RAB27A | 1658.3458 | -0.95983192 | 0.24615202 | -3.8993462 | 9.65E-05 | 0.00122991 |
| PIGBOS1 | 465.154256 | -0.467338043 | 0.11088138 | -4.2147566 | 2.50E-05 | 0.00042486 |
| PIGB | 1367.19435 | -0.019369627 | 0.0850024 | -0.2278715 | 0.81974612 | 0.90329302 |
| CCPG1 | 5248.74298 | 0.016489779 | 0.11728584 | 0.14059481 | 0.88819005 | 0.94217674 |
| C15orf65 | 139.997026 | 0.245706433 | 0.30610686 | 0.80268188 | 0.42215863 | 0.61460404 |
| DNAAF4 | 221.948692 | 0.234075494 | 0.24129215 | 0.97009163 | 0.33200082 | 0.5299553 |
| PYGO1 | 161.142856 | 0.072194184 | 0.32714795 | 0.22067748 | 0.82534357 | 0.90673291 |
| PRTG | 199.827881 | -0.1571878 | 0.4478594 | -0.3509758 | 0.72560653 | 0.84459254 |
| AC012378.1 | 2.65582133 | -0.224388007 | 0.8642689 | -0.2596275 | 0.79515109 | 0.88891824 |
| NEDD4 | 973.727496 | -0.032547739 | 0.26113752 | -0.1246383 | 0.9008099 | 0.94979436 |
| RFX7 | 684.83387 | -0.116881162 | 0.11505714 | -1.0158532 | 0.30969932 | 0.50692551 |
| TEX9 | 557.400886 | 0.279974723 | 0.33299258 | 0.84078368 | 0.40046913 | 0.59528585 |
| AC084782.1 | 73.1668281 | 0.247838767 | 0.35823278 | 0.69183721 | 0.48903956 | 0.67250456 |
| AC084782.2 | 2.73941163 | 2.1425708 | 0.84870106 | 2.52452942 | 0.01158533 | 0.0502972 |
| MNS1 | 427.800478 | 0.395996894 | 0.36502175 | 1.08485836 | 0.27798439 | 0.47353948 |
| AC084782.3 | 16.5908839 | 0.224833035 | 0.36943116 | 0.60859251 | 0.54279457 | 0.71520975 |
| AC090518.1 | 1.39940688 | -0.890137365 | 1.12548953 | -0.7908891 | 0.42900873 | 0.62112297 |
| ZNF280D | 1537.55311 | 0.259663092 | 0.15377641 | 1.68857559 | 0.09130079 | 0.22288144 |
| AC090517.4 | 511.941163 | 0.491327001 | 0.13997362 | 3.51014 | 0.00044787 | 0.00416738 |
| AC010999.2 | 18.3282898 | 0.379302189 | 0.3677833 | 1.03131977 | 0.30239089 | 0.49928294 |
| TCF12 | 2462.67919 | 0.238699126 | 0.11230817 | 2.12539406 | 0.03355375 | 0.10933057 |
| LINC00926 | 280.943661 | 0.192804031 | 0.35729033 | 0.53962846 | 0.58945329 | 0.74961886 |
| LINC01413 | 0.48885242 | 1.537826555 | 1.79952166 | 0.85457518 | 0.39278637 | NA |
| CGNL1 | 10672.8051 | 0.846144567 | 0.21390855 | 3.95563699 | 7.63E-05 | 0.00102383 |
| AC025271.3 | 13.137573 | -0.328503167 | 0.62886004 | -0.5223788 | 0.6014066 | 0.75890644 |
| AC025271.2 | 128.233906 | 1.870837588 | 0.38246895 | 4.89147573 | 1.00E-06 | 3.25E-05 |

| | | | | | | |
|---|---|---|---|---|---|---|
| AC025271.4 | 181.193449 | 0.610817725 | 0.37135124 | 1.64485173 | 0.10000039 | 0.23748976 |
| GCOM1 | 830.479816 | 0.172739521 | 0.23596431 | 0.73205784 | 0.46413327 | 0.65178001 |
| MYZAP | 341.549425 | 0.3229995 | 0.49068775 | 0.65825874 | 0.51037189 | 0.68891634 |
| POLR2M | 1719.67089 | -0.035997942 | 0.11822963 | -0.3044748 | 0.76076619 | 0.86740767 |
| AC090651.1 | 0.35897598 | 0.725061207 | 2.87246869 | 0.25241744 | 0.80071842 | NA |
| ALDH1A2 | 1891.65426 | -0.121308528 | 0.22196127 | -0.5465302 | 0.58470157 | 0.74662073 |
| AC025431.1 | 89.7793521 | 0.116567503 | 0.62492635 | 0.18652998 | 0.85202917 | 0.92250934 |
| AQP9 | 264.684634 | -2.210404107 | 0.52745169 | -4.1907234 | 2.78E-05 | 0.00046175 |
| LIPC | 846.310026 | -0.122272055 | 0.27914595 | -0.4380219 | 0.66137037 | 0.80075727 |
| LIPC-AS1 | 10.9380768 | 0.679767325 | 0.43166308 | 1.57476364 | 0.11531101 | 0.26274661 |
| AC018904.1 | 193.741263 | -1.139712044 | 0.30743466 | -3.7071683 | 0.00020959 | 0.00226511 |
| ADAM10 | 4572.42632 | -0.593178217 | 0.18577536 | -3.1929865 | 0.0014081 | 0.01030683 |
| AC090515.2 | 37.4071747 | -0.140073712 | 0.33909218 | -0.4130845 | 0.67954473 | 0.81406334 |
| MINDY2 | 1961.32304 | -0.134634286 | 0.1325585 | -1.0156594 | 0.3097916 | 0.50697377 |
| AC090515.4 | 48.6685427 | -0.26940483 | 0.30339526 | -0.8879665 | 0.37455877 | 0.57064323 |
| RNF111 | 1251.74024 | -0.107288768 | 0.15211523 | -0.7053125 | 0.4806158 | 0.66539815 |
| SLTM | 4779.73093 | -0.229213916 | 0.18765746 | -1.2214485 | 0.22191626 | 0.40782854 |
| AC025918.1 | 42.80375 | -0.174234999 | 0.39757751 | -0.4382416 | 0.66121117 | 0.80071801 |
| CCNB2 | 73.5400909 | -0.814304637 | 0.43840427 | -1.8574286 | 0.0632502 | 0.17159398 |
| AC092757.3 | 9.63945605 | -0.162197701 | 0.47174845 | -0.3438224 | 0.73097983 | 0.84798182 |
| MYO1E | 3058.29213 | -0.301028262 | 0.237923 | -1.265234 | 0.20578746 | 0.38836437 |
| LDHAL6B | 4.53431532 | -0.460034929 | 0.75987196 | -0.6054111 | 0.54490587 | 0.71677397 |
| FAM81A | 480.070313 | 0.795694123 | 0.27792548 | 2.86297654 | 0.00419682 | 0.02361308 |
| GCNT3 | 2852.8922 | -3.810478101 | 0.48353793 | -7.8804119 | 3.26E-15 | 2.00E-12 |
| GTF2A2 | 1618.95113 | -0.434377473 | 0.13223643 | -3.2848548 | 0.00102035 | 0.00796761 |
| BNIP2 | 3196.90214 | -0.453367649 | 0.1645361 | -2.7554297 | 0.00586151 | 0.03029557 |
| AC092755.2 | 136.558205 | -0.086495795 | 0.22543832 | -0.3836783 | 0.70121688 | 0.82886231 |
| FOXB1 | 0.34968279 | 1.629530304 | 3.24275583 | 0.50251403 | 0.61530599 | NA |
| AC009654.1 | 0.15318056 | -1.003574173 | 3.30558948 | -0.3035992 | 0.7614333 | NA |
| ANXA2 | 20348.4962 | -1.46435895 | 0.297149 | -4.9280293 | 8.31E-07 | 2.77E-05 |
| ICE2 | 1423.89034 | 0.118603657 | 0.14871081 | 0.79754562 | 0.42513422 | 0.61749944 |
| RORA-AS1 | 211.698779 | -0.122165231 | 0.23595135 | -0.517756 | 0.60462851 | 0.76159798 |
| RORA | 2187.7603 | 0.168669223 | 0.22837407 | 0.73856557 | 0.46017084 | 0.64875121 |
| AC107241.1 | 3.22622348 | 1.143401159 | 0.70198567 | 1.62880983 | 0.10335328 | 0.24301562 |
| AC009560.1 | 35.2174297 | 0.999023191 | 0.38303079 | 2.60820599 | 0.00910182 | 0.04200471 |
| RORA-AS2 | 0.72629062 | 2.28977183 | 1.38610819 | 1.65194308 | 0.09854615 | NA |
| AC022898.1 | 4.02673983 | 1.103919923 | 0.63707548 | 1.73279299 | 0.08313247 | 0.20809952 |
| AC012404.1 | 0.79596044 | 0.882696695 | 1.50552582 | 0.58630459 | 0.55767085 | NA |
| AC012404.2 | 5.33265299 | 1.886159166 | 0.75670821 | 2.49258451 | 0.01268171 | 0.05374587 |
| AC107905.1 | 0.15452186 | 0.435199075 | 3.28898345 | 0.13232024 | 0.894731 | NA |
| AC104574.2 | 0.60934919 | -2.508655005 | 1.81093803 | -1.3852793 | 0.16596707 | NA |
| AC104574.1 | 0.03392908 | 0.380740593 | 3.29738709 | 0.11546736 | 0.90807469 | NA |
| AC018618.1 | 0.84023552 | -0.122054501 | 1.87603379 | -0.0650599 | 0.94812634 | NA |
| LINC02349 | 1.46817131 | -3.027054067 | 1.5302926 | -1.9780884 | 0.04791873 | 0.14119304 |
| AC009554.1 | 2.82622457 | -0.379202678 | 0.80757447 | -0.4695575 | 0.63867117 | 0.78618782 |
| VPS13C | 3288.09008 | -0.274114427 | 0.13891272 | -1.9732852 | 0.04846307 | 0.14243995 |
| AC104590.1 | 20.7770439 | -0.551654768 | 0.32583165 | -1.6930668 | 0.09044275 | 0.22142561 |
| C2CD4A | 1412.06572 | -6.492042521 | 0.48950023 | -13.262593 | 3.81E-40 | 1.01E-35 |
| C2CD4B | 162.170342 | -1.778399149 | 0.63439155 | -2.8033147 | 0.00505803 | 0.0271098 |
| AC126323.2 | 0 | NA | NA | NA | NA | NA |
| AC126323.3 | 1.01839785 | -2.502861661 | 2.23596972 | -1.1193629 | 0.26298533 | 0.45663549 |
| AC126323.6 | 1.39540333 | -2.247281405 | 1.35425072 | -1.6594279 | 0.0970296 | 0.23260642 |
| AC126323.5 | 0.07830766 | -0.642905668 | 3.30558948 | -0.1944905 | 0.84579185 | NA |
| AC032011.1 | 5.7444926 | 0.859104306 | 0.5565762 | 1.543552 | 0.1226969 | 0.27389463 |
| TLN2 | 5836.83175 | 2.00374951 | 0.41032358 | 4.88333991 | 1.04E-06 | 3.37E-05 |
| AC100839.1 | 0.11223674 | 0.02007176 | 3.29738704 | 0.00608717 | 0.99514317 | NA |
| AC100839.2 | 0.08757577 | 0.547239394 | 3.29377202 | 0.16614368 | 0.86804388 | NA |
| AC103740.1 | 920.475552 | 2.030984651 | 0.37933246 | 5.35410186 | 8.60E-08 | 4.22E-06 |
| AC103740.2 | 1.32628111 | -0.518189235 | 1.07616609 | -0.4815142 | 0.6301511 | 0.7801882 |
| TPM1 | 13842.3801 | -0.281589378 | 0.27798837 | -1.0129538 | 0.31108222 | 0.50812922 |
| TPM1-AS | 650.95055 | -0.390823451 | 0.33062861 | -1.1820618 | 0.23718117 | 0.42639107 |
| AC079328.2 | 3361.10492 | -0.536118451 | 0.28508822 | -1.8805353 | 0.06003516 | 0.16556445 |
| AC087612.1 | 0.35442022 | -1.739419877 | 2.83942826 | -0.6125951 | 0.54014409 | NA |
| LACTB | 1555.25657 | -0.214120807 | 0.2033256 | -1.0530932 | 0.29229828 | 0.48852236 |
| RPS27L | 4189.82405 | -0.119823196 | 0.20515426 | -0.5840639 | 0.55917733 | 0.72772184 |
| RAB8B | 1704.77164 | -0.614465343 | 0.16292875 | -3.7713745 | 0.00016235 | 0.00185826 |
| APH1B | 1019.5633 | -0.180851043 | 0.14033112 | -1.2887451 | 0.19748671 | 0.3783034 |
| CA12 | 33993.682 | 0.33003254 | 0.40062671 | 0.82379065 | 0.41005852 | 0.6044565 |
| LINC02568 | 185.816259 | 0.189203504 | 0.3420689 | 0.55311518 | 0.58018454 | 0.74321276 |
| AC007950.1 | 0.78609562 | 0.180153673 | 1.57418809 | 0.11444228 | 0.9088872 | NA |
| AC007950.2 | 6.47143426 | 0.138794797 | 0.47016356 | 0.29520535 | 0.76783702 | 0.87183611 |
| USP3 | 2460.96757 | -0.320745623 | 0.1300105 | -2.4670747 | 0.01362219 | 0.05663207 |
| USP3-AS1 | 1292.83638 | -0.375069045 | 0.13638945 | -2.7499858 | 0.00595979 | 0.03062942 |
| FBXL22 | 75.6544661 | 0.81503976 | 0.35160649 | 2.31804526 | 0.02044686 | 0.07613439 |
| HERC1 | 2198.50493 | 0.349690167 | 0.08747719 | 3.99750103 | 6.40E-05 | 0.000896 |
| AC073167.1 | 2.9334408 | 1.226772626 | 0.69811724 | 1.75725876 | 0.07887371 | 0.20046337 |
| DAPK2 | 348.565978 | 0.409489122 | 0.30227698 | 1.35468178 | 0.17551896 | 0.34973818 |
| AC015914.1 | 13.3558004 | 1.079193913 | 0.47096887 | 2.29143365 | 0.02193835 | 0.08032489 |
| FAM96A | 2767.5801 | -0.2962008 | 0.19884087 | -1.4896374 | 0.1363196 | 0.29448727 |
| SNX1 | 7634.94464 | -0.138487788 | 0.11984993 | -1.15551 | 0.24788164 | 0.43948736 |
| SNX22 | 13795.7729 | -0.681977391 | 0.21720306 | -3.1398148 | 0.00169055 | 0.01184574 |
| PPIB | 17704.6916 | -0.645637838 | 0.21363167 | -3.0222009 | 0.00250944 | 0.01604605 |
| CSNK1G1 | 1038.7392 | -0.396601187 | 0.1694047 | -2.3411464 | 0.01922463 | 0.07277366 |
| AC087632.1 | 104.505941 | -0.747501905 | 0.26184282 | -2.8547734 | 0.00430675 | 0.02407235 |
| PCLAF | 55.917481 | -0.418467819 | 0.50977073 | -0.8208942 | 0.41170656 | 0.60580893 |
| TRIP4 | 1717.21734 | 0.203961446 | 0.16469775 | 1.23839853 | 0.21556833 | 0.40009408 |
| ZNF609 | 1733.58606 | -0.379303435 | 0.15555717 | -2.4383539 | 0.01475432 | 0.06009754 |
| OAZ2 | 5334.07799 | -0.142020651 | 0.14278316 | -0.9946597 | 0.31990182 | 0.51734398 |
| AC100830.3 | 32.5864424 | 0.218280156 | 0.22780218 | 0.95820046 | 0.33796168 | 0.53590196 |
| AC100830.2 | 3.26058647 | 1.473984517 | 0.76964191 | 1.91515626 | 0.05547258 | 0.15661798 |
| AC100830.1 | 6.26366415 | -0.294675132 | 0.60883603 | -0.4839975 | 0.62838763 | 0.77902222 |
| RBPMS2 | 965.658689 | 0.187549901 | 0.29866423 | 0.62796237 | 0.53002859 | 0.70458506 |
| PIF1 | 73.10493 | -0.606146065 | 0.40411953 | -1.4999178 | 0.1336357 | 0.29046929 |
| AC069368.1 | 264.969908 | 0.049617622 | 0.31973832 | 0.15518197 | 0.87667786 | 0.93572392 |
| PLEKHO2 | 2360.91523 | -0.560834467 | 0.3244765 | -1.7284286 | 0.08391141 | 0.20953107 |
| ANKDD1A | 540.496857 | -0.347929609 | 0.18504036 | -1.8802904 | 0.06006851 | 0.16560332 |
| AC103691.2 | 17.8942786 | 0.038604761 | 0.32960085 | 0.11712579 | 0.90676037 | 0.95343797 |
| AC103691.1 | 166.000456 | -0.65388721 | 0.29281137 | -2.2331346 | 0.02554007 | 0.08967931 |
| SPG21 | 4650.23699 | -0.124536089 | 0.17007146 | -0.7322574 | 0.46401147 | 0.65164735 |
| MTFMT | 740.485918 | -0.105404128 | 0.14511716 | -0.7263382 | 0.46763149 | 0.65456549 |
| SLC51B | 589.706306 | 0.589122149 | 0.44749496 | 1.3164889 | 0.18801 | 0.36613054 |
| AC013553.3 | 13.5179426 | -0.009955122 | 0.53375837 | -0.018651 | 0.98511953 | 0.99347171 |
| RASL12 | 510.894276 | -0.219368839 | 0.59918225 | -0.3661137 | 0.71428022 | 0.83764267 |
| KBTBD13 | 1.23497215 | 0.28049703 | 1.36443169 | 0.20557792 | 0.83712063 | 0.91377063 |
| AC013553.4 | 8.55841404 | -1.583293981 | 0.73783324 | -2.1458697 | 0.03188338 | 0.10535898 |
| UBAP1L | 564.893068 | -0.02099991 | 0.201426 | -0.1042562 | 0.91696604 | 0.9596537 |
| PDCD7 | 1139.05386 | -0.525298616 | 0.15322267 | -3.4283349 | 0.0006073 | 0.0052627 |
| CLPX | 2824.40054 | -0.104834063 | 0.18080696 | -0.5798121 | 0.56204133 | 0.72974052 |
| CILP | 220.671173 | -0.150421537 | 0.65441447 | -0.2298567 | 0.81820314 | 0.90238464 |
| PARP16 | 692.443622 | 0.178526681 | 0.12370531 | 1.44316105 | 0.14897511 | 0.31303205 |
| IGDCC3 | 32.6584703 | 1.440215294 | 0.26516711 | 5.43134958 | 5.59E-08 | 2.89E-06 |
| IGDCC4 | 163.655756 | -1.407763961 | 0.52718889 | -2.6703217 | 0.00757786 | 0.03672637 |
| DPP8 | 1956.58424 | -0.029721412 | 0.13301817 | -0.2234387 | 0.82319405 | 0.90561649 |

| | | | | | | |
|---|---|---|---|---|---|---|
| HACD3 | 5540.72316 | 0.271263748 | 0.19718923 | 1.37565198 | 0.16892941 | 0.34054495 |
| INTS14 | 1975.96437 | -0.132728969 | 0.16001457 | -0.8294805 | 0.40683257 | 0.60159114 |
| SLC24A1 | 695.604504 | 0.259048097 | 0.22603993 | 1.14602803 | 0.25178355 | 0.44405647 |
| AC011939.3 | 76.9801507 | -0.341113554 | 0.3187749 | -1.0700766 | 0.28458483 | 0.48071886 |
| DENND4A | 1656.68264 | -0.53287983 | 0.21367564 | -2.4938726 | 0.01263579 | 0.05362178 |
| RAB11A | 7069.19593 | -0.368822674 | 0.11263644 | -3.2744526 | 0.00105867 | 0.00819645 |
| MEGF11 | 1861.1208 | -4.382119418 | 0.56170006 | -7.8015293 | 6.12E-15 | 3.29E-12 |
| AC055855.2 | 7.32585706 | 1.127918979 | 0.58016677 | 1.9441289 | 0.05187991 | 0.14959794 |
| DIS3L | 998.787215 | 0.719011009 | 0.17181268 | 4.1848541 | 2.85E-05 | 0.00047206 |
| AC055855.1 | 60.2178073 | 1.224932392 | 0.26281398 | 4.66083421 | 3.15E-06 | 8.12E-05 |
| TIPIN | 229.964442 | 0.309516297 | 0.185947 | 1.66454045 | 0.09600447 | 0.23105211 |
| MAP2K1 | 3782.21376 | -0.591723016 | 0.20095946 | -2.9444894 | 0.00323488 | 0.01946242 |
| AC116913.1 | 1268.44607 | -0.473516777 | 0.17634277 | -2.6852066 | 0.0072485 | 0.03554392 |
| SNAPC5 | 1871.86135 | -0.368659166 | 0.15722827 | -2.3447385 | 0.01904043 | 0.07238987 |
| RPL4 | 43731.2694 | -0.10623348 | 0.23650781 | -0.4491754 | 0.65330516 | 0.79628643 |
| ZWILCH | 414.29285 | -0.038028946 | 0.20767601 | -0.1831167 | 0.85470645 | 0.92400911 |
| LCTL | 118.106286 | -0.087159946 | 0.21524947 | -0.4049253 | 0.68553245 | 0.81828221 |
| LINC01169 | 1.72828725 | 2.071905703 | 1.1349069 | 1.82561732 | 0.06790792 | 0.18011788 |
| SMAD6 | 974.267193 | 0.373088617 | 0.29831785 | 1.25064127 | 0.21106538 | 0.39442947 |
| AC110048.2 | 25.8554116 | 0.602793077 | 0.34842686 | 1.73004192 | 0.08362279 | 0.20901734 |
| AC093334.1 | 1.70815574 | 0.720373975 | 1.16785406 | 0.61683562 | 0.53734316 | 0.71104672 |
| LINC02206 | 1.56588837 | 1.298404223 | 1.25088403 | 1.03798928 | 0.29927504 | 0.49606405 |
| AC087482.1 | 676.162506 | -0.125938599 | 0.74996958 | -0.1679249 | 0.86664233 | 0.9307387 |
| SMAD3 | 6028.52803 | -0.996783748 | 0.25053032 | -3.9786951 | 6.93E-05 | 0.00094852 |
| AC012568.1 | 0 | NA | NA | NA | NA | NA |
| AAGAB | 2904.41732 | -0.427561186 | 0.12692908 | -3.3685044 | 0.00075577 | 0.00627509 |
| IQCH | 181.093148 | 0.381362511 | 0.26398658 | 1.44462841 | 0.14856229 | 0.31253191 |
| IQCH-AS1 | 650.140657 | 0.544521142 | 0.1744395 | 3.12154724 | 0.00179903 | 0.01242549 |
| C15orf61 | 366.222916 | 0.408048483 | 0.21331432 | 1.91289773 | 0.05576115 | 0.15716495 |
| AC016355.1 | 67.9921403 | 0.979264642 | 0.32965748 | 2.97055186 | 0.00297265 | 0.01820456 |
| MAP2K5 | 973.645451 | -0.01100562 | 0.15149485 | -0.0726468 | 0.94208716 | 0.97252858 |
| SKOR1 | 70.7949709 | 0.019951113 | 0.28302647 | 0.07049204 | 0.94380204 | 0.97317822 |
| AC009292.1 | 10.6841192 | -0.881446834 | 0.83755435 | -1.0524055 | 0.29261353 | 0.48895626 |
| PIAS1 | 2292.96014 | 0.128708327 | 0.12097159 | 1.06395498 | 0.2873491 | 0.48362239 |
| AC107871.1 | 2970.58516 | 0.536713012 | 0.21768133 | 2.46559046 | 0.01367876 | 0.05681677 |
| CALML4 | 1769.08866 | 0.766137951 | 0.29025105 | 2.63956998 | 0.00830113 | 0.03913135 |
| CLN6 | 1979.5766 | 0.199681975 | 0.20792062 | 0.96037602 | 0.336866 | 0.53490435 |
| AC107871.2 | 14.7389181 | -0.210866636 | 0.5217956 | -0.4041173 | 0.68612645 | 0.81847636 |
| FEM1B | 2439.617 | -0.287412416 | 0.13646055 | -2.1061941 | 0.03518749 | 0.11306676 |
| ITGA11 | 748.971959 | 0.50484458 | 0.35497053 | 1.42221548 | 0.15496371 | 0.32110584 |
| CORO2B | 971.365089 | -0.409286452 | 0.32513123 | -1.2588346 | 0.20809007 | 0.39130965 |
| ANP32A | 5574.39077 | -0.053707645 | 0.16486554 | -0.3257663 | 0.74460115 | 0.85675592 |
| SPESP1 | 140.700236 | -0.560716556 | 0.41357218 | -1.3557889 | 0.17516636 | 0.34919389 |
| NOX5 | 75.5114867 | 0.150201133 | 0.43572463 | 0.34471573 | 0.73030809 | 0.84755348 |
| AC027088.3 | 0 | NA | NA | NA | NA | NA |
| EWSAT1 | 80.658546 | 1.382418103 | 0.50684076 | 2.72751961 | 0.00638125 | 0.03224865 |
| AC027088.2 | 10.7720429 | 1.511699184 | 0.50984701 | 2.96500548 | 0.00302678 | 0.01849306 |
| GLCE | 1492.64419 | 0.444457651 | 0.17963339 | 2.47424855 | 0.01335168 | 0.05579607 |
| AC026992.2 | 458.855151 | 1.369936319 | 0.3897444 | 3.51496087 | 0.00043982 | 0.0041084 |
| AC026992.1 | 166.153719 | 1.855659589 | 0.45661093 | 4.06398417 | 4.82E-05 | 0.00072246 |
| PAQR5 | 6963.0282 | -0.15935559 | 0.34918253 | -0.4563676 | 0.64812566 | 0.79279496 |
| AC027237.3 | 948.089409 | 0.039835856 | 0.36941791 | 0.10783412 | 0.91412728 | 0.95793738 |
| KIF23 | 743.519495 | -0.772373394 | 0.47994304 | -1.6093022 | 0.10755028 | 0.24985648 |
| RPLP1 | 40110.8508 | -0.524946003 | 0.17625876 | -2.9782691 | 0.00289881 | 0.01785604 |
| AC027237.2 | 5.04944471 | 1.251015185 | 0.71507145 | 1.74949676 | 0.08020519 | 0.20278148 |
| DRAIC | 578.281729 | 2.08962593 | 0.4151077 | 5.0339368 | 4.81E-07 | 1.75E-05 |
| AC100826.1 | 152.022571 | 2.110967493 | 0.3984791 | 5.29756138 | 1.17E-07 | 5.48E-06 |
| AC021818.1 | 0.37653498 | 0.761405404 | 2.85636983 | 0.26656401 | 0.78980487 | NA |
| TLE3 | 3727.849 | -0.42402879 | 0.16558669 | -2.5607662 | 0.01044416 | 0.04657747 |
| AC026583.1 | 0 | NA | NA | NA | NA | NA |
| AC048383.1 | 0 | NA | NA | NA | NA | NA |
| LINC02205 | 0 | NA | NA | NA | NA | NA |
| LINC02204 | 21.4235532 | 0.852032344 | 0.34119777 | 2.49718031 | 0.01251853 | 0.0532098 |
| SALRNA3 | 0.21049242 | 0.689155957 | 3.08915071 | 0.22308913 | 0.82346614 | NA |
| SALRNA2 | 1.05760067 | 3.061979069 | 1.45679511 | 2.10185979 | 0.03556556 | 0.1139152 |
| UACA | 3572.16122 | 0.560833504 | 0.16502125 | 3.39855319 | 0.00067743 | 0.00574402 |
| AC009269.2 | 2.66672327 | 0.884045202 | 0.72692137 | 1.21614969 | 0.2239279 | 0.4100166 |
| AC009269.3 | 0.08666371 | -0.642905668 | 3.30558948 | -0.1944905 | 0.84579185 | NA |
| LARP6 | 1069.09112 | -0.222859397 | 0.23857197 | -0.934139 | 0.35023217 | 0.54794556 |
| LRRC49 | 363.592925 | -0.013412276 | 0.26322874 | -0.0509529 | 0.95936303 | 0.98044425 |
| THAP10 | 155.394655 | 0.471087475 | 0.25273092 | 1.86398831 | 0.06232335 | 0.16977961 |
| THSD4 | 1728.09044 | -0.22073543 | 0.3655812 | -0.6037932 | 0.54598117 | 0.71722681 |
| CT62 | 4.04808632 | 1.146686754 | 0.7200385 | 1.59253532 | 0.11126447 | 0.25595779 |
| THSD4-AS1 | 0.39571739 | 1.854937698 | 2.15012479 | 0.86271165 | 0.38829603 | NA |
| AC108861.1 | 2.24326098 | -2.488815406 | 1.41721905 | -1.7561261 | 0.07906688 | 0.20082773 |
| NR2E3 | 35.3855626 | -0.317354806 | 0.29841215 | -1.0634782 | 0.28756516 | 0.48380077 |
| AC104938.1 | 910.39069 | -0.114319612 | 0.26314467 | -0.4344364 | 0.6639716 | 0.80260972 |
| MYO9A | 2837.02888 | 0.120695387 | 0.22896931 | 0.52712474 | 0.59810696 | 0.75649099 |
| AC022872.1 | 1.94100069 | 0.242842641 | 0.82005086 | 0.29613119 | 0.76712989 | 0.87152635 |
| SENP8 | 265.409796 | 0.435739319 | 0.23433948 | 1.85943621 | 0.06296533 | 0.17101616 |
| AC020779.2 | 69.4774746 | 0.46172496 | 0.31382399 | 1.47128637 | 0.14121369 | 0.30159593 |
| GRAMD2A | 673.5948 | 0.412611866 | 0.43943967 | 0.93894997 | 0.34775643 | 0.54526918 |
| PKM | 71302.8991 | -1.073554428 | 0.22251336 | -4.8246741 | 1.40E-06 | 4.28E-05 |
| PARP6 | 4055.38372 | -0.124153421 | 0.21418544 | -0.5796539 | 0.56214805 | 0.72980721 |
| AC009690.3 | 834.106367 | 0.755538727 | 0.23139239 | 3.26518403 | 0.00109393 | 0.00841018 |
| AC009690.2 | 6043.60044 | 0.145402304 | 0.18254113 | 0.79654545 | 0.42571507 | 0.61804153 |
| CELF6 | 1305.23175 | 0.618302211 | 0.19980553 | 3.09452003 | 0.00197132 | 0.01333939 |
| AC009690.1 | 6305.79977 | 0.311172491 | 0.16044677 | 1.93941261 | 0.05245112 | 0.15058638 |
| HEXA | 9413.64385 | 0.180120372 | 0.17512832 | 1.02850512 | 0.30371229 | 0.5005871 |
| HEXA-AS1 | 60.4912927 | 0.924612618 | 0.25643265 | 3.60567427 | 0.00031134 | 0.00312053 |
| TMEM202 | 0.51016396 | 0.832262939 | 2.12313303 | 0.39199755 | 0.69506003 | NA |
| TMEM202-A: | 119.170654 | -0.012296113 | 0.28207976 | -0.0435909 | 0.9652305 | 0.98359669 |
| AC079322.1 | 29.9571684 | -0.189118315 | 0.27972839 | -0.6760784 | 0.49899088 | 0.68030139 |
| ARIH1 | 3159.39426 | -0.245590761 | 0.12659069 | -1.940038 | 0.05237507 | 0.15041718 |
| LINC02259 | 6.29753164 | 0.298765486 | 0.50382599 | 0.5929934 | 0.55318557 | 0.72290971 |
| AC100827.4 | 7.87326714 | 0.279282643 | 0.48294444 | 0.57829146 | 0.56306736 | 0.73032032 |
| GOLGA6B | 0.09997307 | 0.547239394 | 3.29377202 | 0.16614368 | 0.86804388 | NA |
| HIGD2B | 3.08105526 | -1.082207579 | 0.78393563 | -1.3804802 | 0.16743886 | 0.33860415 |
| BBS4 | 1518.44503 | 0.117910398 | 0.1386835 | 0.85021216 | 0.39520714 | 0.59052654 |
| ADPGK | 3086.679 | -0.394906946 | 0.15930876 | -2.4788779 | 0.01317964 | 0.05529066 |
| ADPGK-AS1 | 80.7851165 | -0.060341221 | 0.27068389 | -0.2229214 | 0.82359671 | 0.90564302 |
| NEO1 | 3002.50914 | 0.384051465 | 0.13388924 | 2.86842674 | 0.00412519 | 0.0232986 |
| AC068397.1 | 0.7811522 | -1.307003282 | 1.6514249 | -0.7914397 | 0.42868743 | NA |
| HCN4 | 22.7036266 | 0.781082008 | 0.64095966 | 1.21861337 | 0.22299097 | 0.40909085 |
| AC068397.2 | 0.10721287 | -0.642905668 | 3.30558948 | -0.1944905 | 0.84579185 | NA |
| REC114 | 4.24062335 | -0.26519257 | 0.65535252 | -0.4046564 | 0.68573012 | 0.81835792 |
| NPTN | 6843.91411 | 0.184449042 | 0.15163096 | 1.21643396 | 0.22381966 | 0.40990863 |
| CD276 | 2712.92222 | -1.113944686 | 0.33872946 | -3.288597 | 0.00100688 | 0.00788111 |
| AC022188.1 | 0.3108733 | -1.364242679 | 3.30558948 | -0.4127078 | 0.67982068 | NA |
| C15orf59 | 761.758877 | 0.913392472 | 0.47948491 | 1.90494518 | 0.0567872 | 0.15920459 |
| C15orf59-AS | 129.323492 | 1.056488024 | 0.53764004 | 1.96504714 | 0.04940878 | 0.14444674 |
| AC018943.1 | 0.18663678 | 0.18657164 | 3.29377195 | 0.05664376 | 0.95482897 | NA |
| TBC1D21 | 0.82300791 | 0.815127473 | 1.59718885 | 0.51035134 | 0.60980534 | NA |

| | | | | | | |
|---|---|---|---|---|---|---|
| LOXL1-AS1 | 432.417105 | 0.012035161 | 0.2678184 | 0.04493777 | 0.96415691 | 0.98324765 |
| LOXL1 | 934.614375 | -0.218953194 | 0.25349531 | -0.8637367 | 0.38773258 | 0.58332428 |
| STOML1 | 891.067982 | -0.576192828 | 0.16295668 | -3.5358651 | 0.00040644 | 0.00386927 |
| PML | 5110.68711 | -1.11748488 | 0.37699413 | -2.9641971 | 0.00303474 | 0.01852455 |
| GOLGA6A | 0.79522928 | 1.354235528 | 1.56803474 | 0.86365148 | 0.38777937 | NA |
| ISLR2 | 23.842272 | 0.639540174 | 0.78010542 | 0.8198125 | 0.41232301 | 0.60620861 |
| AC010931.2 | 2.06185375 | -0.41979678 | 1.10506223 | -0.3798852 | 0.70403064 | 0.8309219 |
| AC010931.3 | 0.74674525 | -2.047686372 | 1.80378147 | -1.1352187 | 0.25628371 | NA |
| ISLR | 1430.5825 | 0.354129609 | 0.45609116 | 0.7764448 | 0.4374864 | 0.62951634 |
| STRA6 | 1161.16609 | 3.843881038 | 0.64704372 | 5.9406821 | 2.84E-09 | 2.39E-07 |
| CCDC33 | 61.8990282 | 3.109507953 | 0.55800557 | 5.57253929 | 2.51E-08 | 1.53E-06 |
| AC023300.2 | 0.08695548 | -0.642905668 | 3.30558948 | -0.1944905 | 0.84579185 | NA |
| CYP11A1 | 55.4436554 | 0.112868355 | 0.66269629 | 0.17031687 | 0.86476095 | 0.92971142 |
| AC090826.1 | 16.5054503 | 0.387277971 | 0.70381549 | 0.55025497 | 0.5821445 | 0.74462314 |
| AC090826.2 | 0 | NA | NA | NA | NA | NA |
| LINC02255 | 0.55449885 | 2.437265331 | 2.04790097 | 1.19012851 | 0.23399589 | NA |
| SEMA7A | 300.561369 | 0.008425676 | 0.39998404 | 0.02106503 | 0.98319378 | 0.99260413 |
| UBL7 | 1860.18803 | 0.133714923 | 0.14280939 | 0.93631743 | 0.34910978 | 0.54674099 |
| UBL7-AS1 | 89.9718422 | 0.264565341 | 0.20843977 | 1.26926518 | 0.2043465 | 0.38666749 |
| AC012435.2 | 6.38537336 | 0.200581886 | 0.54014847 | 0.37134584 | 0.71037996 | 0.83538707 |
| ARID3B | 578.122054 | -0.170350967 | 0.279031 | -0.6105091 | 0.5415246 | 0.71428636 |
| CLK3 | 4593.48243 | -0.321367704 | 0.15131532 | -2.1238279 | 0.03368454 | 0.10960776 |
| AC100835.2 | 39.7955937 | 1.123610425 | 0.42940466 | 2.61667029 | 0.00887921 | 0.0412154 |
| AC100835.1 | 4.47233489 | 0.318324432 | 0.6507282 | 0.48918186 | 0.62471295 | 0.77643779 |
| EDC3 | 1531.2267 | 0.023813213 | 0.13760979 | 0.17304882 | 0.86261305 | 0.92841542 |
| CYP1A1 | 25.441961 | 0.702775019 | 0.92625092 | 0.75873071 | 0.44801366 | 0.6381182 |
| CYP1A2 | 2.72848503 | 1.132254983 | 1.35756417 | 0.83403423 | 0.4042617 | 0.59907812 |
| CSK | 2439.12461 | -0.464338644 | 0.19272824 | -2.4092922 | 0.01598349 | 0.06380332 |
| LMAN1L | 1.27278061 | 1.317814486 | 1.13479687 | 1.16127787 | 0.24552892 | 0.43670562 |
| CPLX3 | 2.70738735 | -1.023195572 | 1.1327551 | -0.9032805 | 0.36637705 | 0.56337763 |
| ULK3 | 5151.06256 | 0.650247067 | 0.14992626 | 4.33711271 | 1.44E-05 | 0.00027487 |
| SCAMP2 | 7384.19546 | 0.095672425 | 0.16719703 | 0.57221366 | 0.56717724 | 0.73323187 |
| MPI | 5866.25832 | 0.010871544 | 0.10050961 | 0.10816422 | 0.91386542 | 0.9577987 |
| FAM219B | 4116.99849 | -0.074614313 | 0.07903143 | -0.9441094 | 0.34511376 | 0.54275134 |
| COX5A | 5256.35763 | -0.040172936 | 0.1958523 | -0.2051185 | 0.83747952 | 0.91397322 |
| RPP25 | 671.280697 | -1.072412721 | 0.33861592 | -3.1670475 | 0.00153995 | 0.01103492 |
| SCAMP5 | 2532.40607 | 0.736595189 | 0.19505459 | 3.77635409 | 0.00015914 | 0.00183601 |
| PPCDC | 591.392753 | 0.184234975 | 0.20623608 | 0.89332076 | 0.37168542 | 0.56836599 |
| C15orf39 | 1588.10842 | -0.573315967 | 0.17407225 | -3.2935518 | 0.0009893 | 0.00777119 |
| AC113208.3 | 172.771852 | -0.985893199 | 0.24494716 | -4.0249219 | 5.70E-05 | 0.00082083 |
| GOLGA6C | 0.40921437 | -0.534763868 | 3.29377181 | -0.1623561 | 0.87102546 | NA |
| GOLGA6D | 0 | NA | NA | NA | NA | NA |
| COMMD4 | 3197.08816 | -0.106570158 | 0.12796517 | -0.832806 | 0.40495418 | 0.59972246 |
| AC068338.3 | 120.632422 | 0.099383243 | 0.20934935 | 0.47472439 | 0.63498343 | 0.78344233 |
| NEIL1 | 5986.95731 | -0.142415616 | 0.1454593 | -0.9790753 | 0.32754275 | 0.52526221 |
| MAN2C1 | 9475.01821 | 0.165008197 | 0.14105643 | 1.16980271 | 0.24208037 | 0.43252656 |
| AC068338.2 | 515.65509 | 0.686325944 | 0.20465727 | 3.35353809 | 0.00079785 | 0.00656251 |
| SIN3A | 2506.04368 | -0.216773558 | 0.11848318 | -1.8295724 | 0.0673139 | 0.17911882 |
| PTPN9 | 2356.72732 | -0.354441749 | 0.13243504 | -2.6763442 | 0.00744302 | 0.03624605 |
| SNUPN | 1063.21351 | 0.053619447 | 0.14722519 | 0.36420023 | 0.71570849 | 0.83823283 |
| AC105020.3 | 70.6852638 | 0.482154276 | 0.2286518 | 2.1086835 | 0.03497191 | 0.11258812 |
| AC105020.2 | 552.04773 | -0.245231714 | 0.21958498 | -1.1167964 | 0.26408138 | 0.45796568 |
| IMP3 | 2434.52192 | -0.129919673 | 0.19329523 | -0.6721308 | 0.50150044 | 0.68220617 |
| AC105020.6 | 941.037391 | -0.061207548 | 0.18403095 | -0.3325938 | 0.73944095 | 0.85379883 |
| AC105020.5 | 14.8223023 | 0.264550679 | 0.49080814 | 0.53901037 | 0.5898797 | 0.74983272 |
| SNX33 | 2384.76316 | 0.021155216 | 0.13558727 | 0.15602656 | 0.87601208 | 0.93550556 |
| CSPG4 | 1351.22567 | 0.655190218 | 0.32245341 | 2.03189111 | 0.04216468 | 0.12876464 |
| AC105020.4 | 317.138834 | -0.281316483 | 0.64536387 | -0.4359037 | 0.66290662 | 0.80180018 |
| AC105020.1 | 123.893809 | -0.497121801 | 0.21677794 | -2.2932306 | 0.02183473 | 0.080073 |
| ODF3L1 | 14.2625739 | 0.51424346 | 0.66104466 | 0.77792544 | 0.43661297 | 0.62877147 |
| DNM1P35 | 30.1003025 | 0.158839005 | 0.37241716 | 0.42650828 | 0.66973751 | 0.80713762 |
| AC019294.3 | 2.73128799 | 1.060750433 | 0.97181793 | 1.09151148 | 0.27504788 | 0.47019955 |
| UBE2Q2 | 1981.42447 | -0.360294253 | 0.14970272 | -2.4067315 | 0.016096 | 0.06414235 |
| FBXO22 | 1559.05267 | 0.100376246 | 0.14002466 | 0.71684694 | 0.47346855 | 0.65944993 |
| NRG4 | 307.317192 | 0.009742206 | 0.22073856 | 0.04413459 | 0.96479712 | 0.98354657 |
| TMEM266 | 160.856934 | 0.76818675 | 0.3898616 | 1.97040888 | 0.04879153 | 0.14322973 |
| AC091100.1 | 20.07979 | 1.128515608 | 0.39288942 | 2.87234919 | 0.00407432 | 0.02307109 |
| ETFA | 7928.59493 | 0.540249403 | 0.28949842 | 1.86615668 | 0.06201945 | 0.16931906 |
| ISL2 | 1.87545293 | -2.698628778 | 1.27032306 | -2.1243642 | 0.0336397 | 0.10950241 |
| AC027243.1 | 0.60173911 | -2.496483468 | 1.80827922 | -1.3805852 | 0.16740655 | NA |
| AC027243.2 | 0 | NA | NA | NA | NA | NA |
| SCAPER | 728.819899 | 0.124788479 | 0.23730927 | 0.52584747 | 0.59899419 | 0.75699592 |
| RCN2 | 2829.02738 | 0.149678301 | 0.15254476 | 0.98120907 | 0.32648965 | 0.52436398 |
| PSTPIP1 | 363.018536 | -0.553138242 | 0.59593207 | -0.9281901 | 0.35330897 | 0.55089626 |
| TSPAN3 | 13916.3859 | -0.349598955 | 0.15574586 | -2.2446757 | 0.02478896 | 0.08773115 |
| AC090181.1 | 70.5094008 | -0.716992123 | 0.43383174 | -1.6526963 | 0.09839268 | 0.23497691 |
| PEAK1 | 1281.55539 | -0.116306331 | 0.18534365 | -0.6275172 | 0.53032027 | 0.70472412 |
| HMG20A | 1758.05158 | 0.029418456 | 0.09572725 | 0.30731539 | 0.75860332 | 0.86635037 |
| AC046168.1 | 2.73397362 | -1.217155065 | 1.25628319 | -0.9688541 | 0.33261801 | 0.53055511 |
| AC046168.2 | 1.8503385 | 0.01321919 | 0.88892543 | 0.01487098 | 0.98813511 | 0.99466057 |
| LINGO1 | 235.527977 | 0.669427908 | 0.3777249 | 1.77226313 | 0.07635089 | 0.1961526 |
| LINGO1-AS1 | 0.86497178 | -0.992763586 | 1.588043 | -0.6251491 | 0.53187323 | NA |
| LINGO1-AS2 | 0.19994613 | 1.506623855 | 3.27645372 | 0.45983371 | 0.64563559 | NA |
| AC105133.1 | 0 | NA | NA | NA | NA | NA |
| AC104758.4 | 1.77701351 | 1.731938224 | 1.34463038 | 1.28804038 | 0.19773191 | 0.37847067 |
| AC104758.5 | 0.18382444 | -1.003574173 | 3.30558948 | -0.3035992 | 0.7614333 | NA |
| TBC1D2B | 2083.81817 | -0.324708375 | 0.1871864 | -1.7346793 | 0.08279762 | 0.20758444 |
| AC104758.2 | 420.447744 | -1.042379278 | 0.27569067 | -3.780974 | 0.00015622 | 0.00180914 |
| SH2D7 | 152.274199 | -1.961667276 | 0.33087892 | -5.928656 | 3.05E-09 | 2.52E-07 |
| CIB2 | 574.182321 | -1.564305468 | 0.27633907 | -5.6608191 | 1.51E-08 | 9.81E-07 |
| IDH3A | 4351.72356 | -0.588706307 | 0.17345375 | -3.3940247 | 0.00068873 | 0.00582113 |
| AC090260.1 | 10.7035241 | 0.813234705 | 0.50632291 | 1.60615824 | 0.10823915 | 0.25105694 |
| ACSBG1 | 3085.79944 | -0.692636369 | 0.1737631 | -3.9860959 | 6.72E-05 | 0.00092987 |
| AC090607.3 | 0.2768272 | -1.364242679 | 3.30558948 | -0.4127078 | 0.67982068 | NA |
| DNAJA4 | 2385.28091 | -0.858526251 | 0.29381949 | -2.9219513 | 0.00347846 | 0.02048391 |
| WDR61 | 2607.04961 | 0.209518044 | 0.1335395 | 1.56895928 | 0.11665743 | 0.26494328 |
| AC090607.1 | 424.118053 | 0.425857138 | 0.19801273 | 2.15065532 | 0.03150342 | 0.10453568 |
| AC090607.5 | 4.59383552 | 0.9646002 | 0.8274531 | 1.16574608 | 0.24371712 | 0.43445012 |
| CRABP1 | 126.645621 | 2.534719976 | 0.79987338 | 3.16890153 | 0.00153016 | 0.01099075 |
| IREB2 | 2332.58527 | -0.050623661 | 0.13168591 | -0.3844273 | 0.70066176 | 0.82842894 |
| AC027228.1 | 0.16052235 | -1.003574173 | 3.30558948 | -0.3035992 | 0.7614333 | NA |
| HYKK | 537.067521 | 1.23243758 | 0.39409137 | 3.12728892 | 0.00176426 | 0.01226252 |
| PSMA4 | 4229.00742 | -0.530299609 | 0.16332249 | -3.2469479 | 0.0011665 | 0.00885705 |
| CHRNA5 | 32.1394528 | -0.916385645 | 0.58429157 | -1.5683705 | 0.1167947 | 0.26514086 |
| AC027228.2 | 21.3954781 | -0.736040598 | 0.5869908 | -1.2539219 | 0.20987041 | 0.39314362 |
| CHRNA3 | 17.9073577 | 1.274837727 | 0.64429281 | 1.97866203 | 0.04785407 | 0.14104301 |
| CHRNB4 | 27.3295463 | 2.44403167 | 0.46613734 | 5.24315794 | 1.58E-07 | 6.98E-06 |
| AC067863.1 | 0.84285229 | 3.086799068 | 1.43869351 | 2.1455571 | 0.03190833 | NA |
| ADAMTS7 | 500.001377 | -0.293755817 | 0.28516315 | -1.0301325 | 0.30294782 | 0.49979568 |
| MORF4L1 | 15508.6953 | -0.284724373 | 0.16943094 | -1.6804745 | 0.09286503 | 0.22559658 |
| CTSH | 25541.0587 | 0.502639145 | 0.2920076 | 1.72132215 | 0.08519238 | 0.21177631 |
| RASGRF1 | 246.58192 | -0.442086701 | 0.60245796 | -0.7338051 | 0.46306755 | 0.65105787 |
| AC011944.1 | 44.5789716 | -0.386080199 | 0.54982723 | -0.7021846 | 0.48256406 | 0.66711478 |

| | | | | | | |
|---|---|---|---|---|---|---|
| ANKRD34C-A | 1.58286885 | -0.852822089 | 1.00476142 | -0.8487807 | 0.39600334 | 0.59122987 |
| ANKRD34C | 2.29664853 | 1.412597755 | 1.0493474 | 1.34616788 | 0.17824838 | 0.35372853 |
| TMED3 | 2781.56137 | -0.794996813 | 0.22517037 | -3.5306458 | 0.00041455 | 0.00392503 |
| KIAA1024 | 38.3548514 | -0.803000059 | 0.26530878 | -3.0266622 | 0.0024727 | 0.01585728 |
| AC021483.1 | 1.90978075 | 0.41573426 | 1.26777642 | 0.32792396 | 0.74296916 | 0.85569954 |
| MTHFS | 2281.13322 | 0.355755779 | 0.27289146 | 1.30365302 | 0.19235191 | 0.37159853 |
| AC021483.2 | 19.7524265 | 1.087474817 | 0.33877307 | 3.21003914 | 0.00132717 | 0.00984091 |
| ST20-MTHFS | 2477.31406 | 0.351754323 | 0.26964828 | 1.30449311 | 0.19206551 | 0.37126282 |
| ST20 | 399.114176 | 0.147155565 | 0.25179382 | 0.58442882 | 0.55893184 | 0.72753011 |
| ST20-AS1 | 393.991468 | 0.359239524 | 0.30336135 | 1.18419674 | 0.23633519 | 0.42563578 |
| BCL2A1 | 116.956002 | -1.245266736 | 0.63333158 | -1.9662161 | 0.04927366 | 0.14425776 |
| ZFAND6 | 4232.73965 | -0.439197715 | 0.15012913 | -2.9254664 | 0.0034394 | 0.02032657 |
| FAH | 4118.333 | 0.866793724 | 0.35796662 | 2.42143729 | 0.01545927 | 0.06221931 |
| AC087761.1 | 1.77988491 | 0.323394052 | 1.22775723 | 0.26340228 | 0.79224053 | 0.8873257 |
| CTXND1 | 387.164679 | 0.974859561 | 0.36008686 | 2.70729002 | 0.0067835 | 0.03374781 |
| LINC00927 | 2.50854755 | 1.385803013 | 0.98397005 | 1.40837926 | 0.1590188 | 0.32654925 |
| AC016705.2 | 507.201949 | 0.73744949 | 0.33683189 | 2.18936959 | 0.02856999 | 0.09748843 |
| ARNT2 | 2352.31194 | 0.33755409 | 0.32341243 | 1.0437264 | 0.29661199 | 0.49341406 |
| AC016705.1 | 10.9373738 | 0.456311868 | 0.55225364 | 0.82627226 | 0.40864966 | 0.60309445 |
| AC108451.1 | 8.31293664 | 2.325071057 | 0.72866846 | 3.19084905 | 0.00141855 | 0.01035925 |
| AC108451.2 | 17.0149602 | 1.535024013 | 0.504089 | 3.04514486 | 0.00232568 | 0.01518026 |
| ABHD17C | 1080.50904 | 0.251586442 | 0.18279022 | 1.37636711 | 0.16870801 | 0.34035882 |
| AC023302.1 | 8.81803449 | 0.89714135 | 0.58635928 | 1.53001986 | 0.12601181 | 0.27888641 |
| CEMIP | 122.959806 | 0.009670086 | 0.31589605 | 0.03061161 | 0.97557928 | 0.98866399 |
| AC027808.1 | 43.3844703 | 0.139594987 | 0.25954085 | 0.53785363 | 0.5906781 | 0.7504062 |
| AC027808.2 | 20.9660523 | -0.112314237 | 0.38580445 | -0.291117 | 0.77096183 | 0.87355577 |
| MESD | 3127.89034 | -0.481215644 | 0.15464005 | -3.1118435 | 0.00185923 | 0.01273121 |
| AC068870.1 | 7.24201751 | 0.092018297 | 0.62220217 | 0.14789131 | 0.88242854 | 0.93897348 |
| AC068870.2 | 64.4469841 | 0.445932243 | 0.36842354 | 1.21037936 | 0.22613336 | 0.41281666 |
| TLNRD1 | 1682.04175 | -0.926313739 | 0.31809674 | -2.9120504 | 0.00359065 | 0.02097594 |
| CFAP161 | 8.38576211 | -0.288747215 | 0.4847129 | -0.5957077 | 0.5513705 | 0.72153443 |
| IL16 | 584.384516 | 0.316763052 | 0.40097076 | 0.78999041 | 0.42953337 | 0.6216341 |
| STARD5 | 329.357782 | -0.039500379 | 0.37700894 | -0.1047731 | 0.91655589 | 0.95937676 |
| TMC3-AS1 | 10.7785607 | 0.343727831 | 0.54463293 | 0.63111834 | 0.52796315 | 0.70254772 |
| TMC3 | 12.9094616 | -0.786997225 | 0.73791257 | -1.0665183 | 0.28618943 | 0.48225543 |
| AC103858.2 | 0.94533254 | 0.16035455 | 1.55921526 | 0.10284311 | 0.91808749 | NA |
| AC109809.1 | 1.41692248 | 1.08234253 | 1.18474727 | 0.91356407 | 0.36094595 | 0.55866688 |
| AC060809.1 | 0.20718593 | 0.18657164 | 3.29377195 | 0.05664376 | 0.95482897 | NA |
| AC023034.1 | 0.74142307 | 0.016343685 | 1.38830405 | 0.01177241 | 0.99060719 | NA |
| AC023034.2 | 1.14070967 | -0.508910017 | 1.32310923 | -0.3846319 | 0.70051016 | 0.82832397 |
| AC104041.1 | 0.62819056 | -0.527543257 | 1.89621196 | -0.278209 | 0.78085192 | NA |
| AC025219.1 | 0.57821645 | 1.302649754 | 1.73353541 | 0.7514411 | 0.45238723 | NA |
| MEX3B | 115.573082 | 0.282417604 | 0.38824353 | 0.72742386 | 0.46696634 | 0.65413276 |
| LINC01583 | 0 | NA | NA | NA | NA | NA |
| EFL1 | 1230.6142 | -0.598149499 | 0.30857917 | -1.9383988 | 0.05257458 | 0.15084232 |
| SAXO2 | 109.552899 | 1.031921508 | 0.46183294 | 2.23440432 | 0.02545648 | 0.0894692 |
| GOLGA6L10 | 64.9771051 | -0.483051319 | 0.59348856 | -0.8139185 | 0.41569164 | 0.60908136 |
| GOLGA6L9 | 498.886552 | -0.115463567 | 0.22084407 | -0.5228285 | 0.60109363 | 0.75869641 |
| RPS17 | 20043.3822 | -0.475232539 | 0.18585431 | -2.5570166 | 0.01055742 | 0.04688433 |
| AC245033.1 | 19686.2336 | -0.467228393 | 0.18848892 | -2.478811 | 0.01318211 | 0.05529066 |
| AC245033.2 | 1.1747674 | 0.506176622 | 1.47961745 | 0.34209966 | 0.7322759 | 0.8490202 |
| CPEB1 | 119.296417 | 0.326628371 | 0.44142676 | 0.73993786 | 0.4593377 | 0.64801707 |
| CPEB1-AS1 | 5.35367869 | 0.244388018 | 0.77094612 | 0.31699753 | 0.75124547 | 0.86135672 |
| AP3B2 | 264.463528 | -1.041467582 | 0.43809101 | -2.3772859 | 0.01744057 | 0.06783299 |
| AC105339.3 | 7.58897968 | 1.590984082 | 0.67806806 | 2.34634866 | 0.01895836 | 0.07217151 |
| SNHG21 | 241.076768 | 0.01825509 | 0.21666338 | 0.08425554 | 0.93285326 | 0.96760457 |
| FSD2 | 308.199602 | -0.111913228 | 0.19572773 | -0.5717801 | 0.56747093 | 0.73345276 |
| WHAMM | 1835.08518 | -0.941562816 | 0.2439566 | -3.8595505 | 0.0001136 | 0.00139912 |
| HOMER2 | 813.852929 | -0.204100772 | 0.37659441 | -0.5419644 | 0.58784302 | 0.74875494 |
| AC022558.2 | 0.77403962 | -0.394322135 | 1.59734328 | -0.2468612 | 0.80501562 | NA |
| RAMMET | 596.565019 | -0.566805076 | 0.26750832 | -2.1188316 | 0.0341047 | 0.11050249 |
| C15orf40 | 1699.26838 | 0.081483535 | 0.16774924 | 0.48574606 | 0.62714722 | 0.77806973 |
| AC024270.4 | 1.52991582 | 0.117353096 | 1.25815493 | 0.09327396 | 0.92568592 | 0.9640363 |
| BTBD1 | 3543.7851 | -0.142421551 | 0.19207535 | -0.7414879 | 0.45839766 | 0.64731425 |
| AC022558.3 | 279.730856 | 0.252093534 | 0.25464943 | 0.98996308 | 0.32219217 | 0.51999556 |
| AC024270.2 | 3.16641883 | 0.718371116 | 0.68107877 | 1.05475482 | 0.29153747 | 0.48774871 |
| TM6SF1 | 267.456451 | -0.045065794 | 0.32939605 | -0.1368134 | 0.89117829 | 0.94360054 |
| HDGFL3 | 1601.86488 | -0.403254264 | 0.24835302 | -1.623714 | 0.10443687 | 0.24480002 |
| AC024270.3 | 44.4538807 | -1.10631517 | 0.43885404 | -2.5209183 | 0.0117049 | 0.05066617 |
| AC103876.1 | 1.73911437 | -0.260914045 | 0.89137644 | -0.2927092 | 0.76974447 | 0.87270139 |
| BNC1 | 18.5684694 | -1.298790322 | 0.68882347 | -1.8855198 | 0.05935969 | 0.16436433 |
| SH3GL3 | 15.0003518 | 0.633293309 | 0.73993074 | 0.85588188 | 0.39206312 | 0.58779376 |
| ADAMTSL3 | 485.154859 | 1.186196272 | 0.3926897 | 3.02069616 | 0.00252194 | 0.01609868 |
| AC136698.1 | 0.38280371 | -0.542723645 | 2.80066761 | -0.1937837 | 0.84634527 | NA |
| UBE2Q2L | 6.55556407 | -0.65787017 | 0.62247429 | -1.0568632 | 0.29057403 | 0.48665844 |
| GOLGA6L4 | 138.088507 | 0.112846835 | 0.30212685 | 0.37350813 | 0.70877029 | 0.83446168 |
| AC243562.3 | 2.84483495 | -0.481649915 | 0.91651131 | -0.5255253 | 0.59921804 | 0.7570974 |
| AC048382.4 | 0.65856662 | -0.554569896 | 1.87540486 | -0.2957068 | 0.76745403 | NA |
| AC048382.5 | 232.439131 | 0.937640107 | 0.22439393 | 4.17854481 | 2.93E-05 | 0.00048262 |
| ZSCAN2 | 728.049928 | 0.731805175 | 0.16289007 | 4.49263224 | 7.03E-06 | 0.00015459 |
| AC048382.1 | 10.5410336 | 1.004566548 | 0.41735213 | 2.40699994 | 0.01608417 | 0.06410817 |
| AC048382.6 | 352.218834 | 0.628702051 | 0.20726161 | 3.03337442 | 0.00241835 | 0.01558836 |
| WDR73 | 3473.16778 | -0.057521004 | 0.17409943 | -0.3303917 | 0.74110402 | 0.8546067 |
| NMB | 427.087544 | -1.15488418 | 0.30911486 | -3.7361005 | 0.0001869 | 0.0020673 |
| SEC11A | 5539.99338 | -0.357636764 | 0.13917823 | -2.5696315 | 0.01018068 | 0.04567275 |
| AC115102.1 | 2.67514899 | -0.478532217 | 0.93601088 | -0.5112464 | 0.60917851 | 0.76524568 |
| ZNF592 | 2835.56474 | -0.004719199 | 0.18887441 | -0.0249859 | 0.9800662 | 0.99106512 |
| ALPK3 | 3765.24722 | -1.095114528 | 0.37104412 | -2.9514402 | 0.00316296 | 0.01913009 |
| SLC28A1 | 5321.4578 | 1.270122555 | 0.45367869 | 2.79960814 | 0.00511647 | 0.02734521 |
| PDE8A | 3253.35378 | 0.250618502 | 0.16546356 | 1.51464469 | 0.12986241 | 0.28492001 |
| AKAP13 | 4085.63901 | -0.359663801 | 0.16884324 | -2.1301641 | 0.03315807 | 0.10836174 |
| AC087286.3 | 9.19952642 | 1.959973089 | 0.66153664 | 2.96275816 | 0.00304896 | 0.01858984 |
| AC087286.2 | 2.39668996 | -0.907775296 | 1.24891243 | -0.7268526 | 0.46731623 | 0.65440651 |
| AC087286.1 | 0.35669117 | -1.016996084 | 2.3123762 | -0.4398056 | 0.66007789 | NA |
| AC021739.4 | 30.4681208 | -0.03928549 | 0.30589857 | -0.1284265 | 0.89781144 | 0.94799743 |
| AC021739.2 | 1023.70323 | -0.862554355 | 0.16788269 | -5.1378398 | 2.78E-07 | 1.12E-05 |
| AC021739.5 | 20.7923132 | -1.326297314 | 0.48476572 | -2.7359552 | 0.00621995 | 0.03169391 |
| AC021739.3 | 20.7816488 | -1.64937501 | 0.51567873 | -3.1984546 | 0.00138166 | 0.01015736 |
| KLHL25 | 787.867956 | -0.415724828 | 0.20696627 | -2.0086598 | 0.04457323 | 0.13388723 |
| LINC01584 | 0 | NA | NA | NA | NA | NA |
| AGBL1 | 0.62810719 | -1.503364216 | 2.79388645 | -0.5380907 | 0.59051446 | NA |
| AGBL1-AS1 | 0 | NA | NA | NA | NA | NA |
| AC012229.1 | 0 | NA | NA | NA | NA | NA |
| AC016987.1 | 0 | NA | NA | NA | NA | NA |
| AC078905.1 | 0 | NA | NA | NA | NA | NA |
| AC020687.1 | 0.13028434 | 1.049816488 | 3.28477129 | 0.31960109 | 0.74927075 | NA |
| LINC00052 | 1.94412978 | 0.812938296 | 1.56305435 | 0.52009599 | 0.60299667 | 0.76023247 |
| NTRK3 | 907.890502 | -0.238932115 | 0.44376615 | -0.538419 | 0.59028783 | 0.75016338 |
| NTRK3-AS1 | 8.76730983 | -0.663136211 | 0.84189976 | -0.7876665 | 0.43089182 | 0.62295052 |
| MRPL46 | 1590.48095 | 0.000865766 | 0.18340821 | 0.00472043 | 0.99623365 | 0.99823988 |
| MRPS11 | 1643.83186 | 0.00851602 | 0.09971893 | 0.08540023 | 0.93194321 | 0.96711434 |
| DET1 | 402.191351 | 0.536536509 | 0.26950807 | 1.99079944 | 0.04650294 | 0.13816108 |
| LINC01586 | 1.35270741 | 1.044768658 | 1.12566056 | 0.92813828 | 0.35333584 | 0.55090559 |

| | | | | | | |
|---|---|---|---|---|---|---|
| AEN | 1393.11094 | -1.678508284 | 0.39950482 | -4.201472 | 2.65E-05 | 0.00044598 |
| ISG20 | 648.855509 | -2.154042691 | 0.49384792 | -4.3617531 | NA | NA |
| AC103982.1 | 1.62691349 | 0.969388965 | 1.42017777 | 0.68258283 | 0.49487049 | 0.67712083 |
| ACAN | 284.596404 | -0.018610265 | 0.71876021 | -0.0258922 | 0.97934334 | 0.99065111 |
| HAPLN3 | 532.547052 | -1.245693214 | 0.60562973 | -2.0568561 | 0.03970007 | 0.12311104 |
| MFGE8 | 4887.63795 | -0.182533964 | 0.33631801 | -0.5427422 | 0.58730734 | 0.74828692 |
| AC013565.1 | 8.91225736 | 0.986709641 | 0.69798729 | 1.41364987 | 0.15746473 | 0.32460483 |
| AC013565.3 | 2.4052124 | 0.544499375 | 0.80675219 | 0.67492767 | 0.49972172 | 0.68105131 |
| ABHD2 | 5033.68619 | -0.183691437 | 0.30608135 | -0.6001393 | 0.54841343 | 0.71905639 |
| RLBP1 | 0.47853322 | 0.587471938 | 1.86025764 | 0.31580138 | 0.75215327 | NA |
| FANCI | 1244.57315 | -0.75672428 | 0.16586902 | -4.5621798 | 5.06E-06 | 0.00011784 |
| POLG | 3576.15309 | -0.315486791 | 0.16499221 | -1.9121314 | 0.05585934 | 0.15737295 |
| AC124068.2 | 7.10323607 | 0.487346797 | 0.57442737 | 0.84840456 | 0.39621271 | 0.5912411 |
| MIR9-3HG | 114.096015 | 0.035009574 | 0.40669062 | 0.08608405 | 0.93139961 | 0.96693074 |
| RHCG | 3488.17872 | 1.137023956 | 0.58573949 | 1.94117689 | 0.05223682 | 0.15020333 |
| LINC00928 | 0.31684042 | 0.328539033 | 2.27490269 | 0.14441894 | 0.88516966 | NA |
| AC013391.3 | 0.94363764 | -1.578724625 | 1.64884391 | -0.9574737 | 0.33832823 | NA |
| TICRR | 135.044849 | -0.502005165 | 0.34690652 | -1.4470906 | 0.14787155 | 0.3113825 |
| AC013391.1 | 0 | NA | NA | NA | NA | NA |
| KIF7 | 324.192789 | -0.061733588 | 0.35194834 | -0.1754053 | 0.86076121 | 0.92743582 |
| PLIN1 | 356.756773 | -1.432842406 | 0.80818293 | -1.7729184 | 0.07624223 | 0.19595796 |
| PEX11A | 928.772239 | 0.044222659 | 0.33351999 | 0.13259373 | 0.89451469 | 0.94578078 |
| WDR93 | 97.9623658 | 0.402966531 | 0.42484476 | 0.948503 | 0.34287345 | 0.54049181 |
| MESP1 | 92.569787 | -1.06932131 | 0.33779454 | -3.1655968 | 0.00154765 | 0.0110707 |
| MESP2 | 14.355725 | -0.812423598 | 0.70924019 | -1.1454844 | 0.25200854 | 0.44427498 |
| ANPEP | 54638.239 | 0.443014771 | 0.47003561 | 0.9425132 | 0.34592996 | 0.54354339 |
| AP3S2 | 3658.67627 | 0.290060518 | 0.13973598 | 2.07577547 | 0.03791472 | 0.11899681 |
| C15orf38-AP | 2464.46476 | 0.265551693 | 0.13403539 | 1.98120586 | 0.04756819 | 0.1404516 |
| ARPIN | 1637.77152 | 0.554791089 | 0.18567519 | 2.98796572 | 0.00280841 | 0.01745 |
| ZNF710 | 975.01765 | 0.287317907 | 0.23093336 | 1.24415938 | 0.21344091 | 0.39748379 |
| ZNF710-AS1 | 2537.17338 | 0.787975473 | 0.25257896 | 3.11971932 | 0.00181023 | 0.01247019 |
| IDH2 | 16834.0853 | 0.152995538 | 0.21405799 | 0.71473875 | 0.47477049 | 0.66021715 |
| IDH2-DT | 29.5455038 | -0.095948708 | 0.4050061 | -0.2369068 | 0.81272908 | 0.89895498 |
| SEMA4B | 11367.0156 | -2.332799244 | 0.31074819 | -7.5070404 | 6.05E-14 | 2.38E-11 |
| CIB1 | 10501.0396 | -0.986591625 | 0.14939239 | -6.6040287 | 4.00E-11 | 6.76E-09 |
| GDPGP1 | 2357.89644 | -0.731061276 | 0.15937673 | -4.5870014 | 4.50E-06 | 0.00010829 |
| AC091167.7 | 208.566043 | -0.124834664 | 0.19905355 | -0.6271411 | 0.53056674 | 0.70495094 |
| TTLL13P | 54.9467873 | 0.0445057 | 0.28489015 | 0.15622057 | 0.87585917 | 0.93547831 |
| AC091167.2 | 5845.49755 | -0.502257752 | 0.14867498 | -3.3782264 | 0.00072955 | 0.00610544 |
| NGRN | 7165.07497 | -0.447201339 | 0.14932904 | -2.9947379 | 0.00274681 | 0.01718465 |
| AC091167.6 | 1112.85639 | -0.295313576 | 0.18196628 | -1.6229027 | 0.1046102 | 0.24507504 |
| AC091167.5 | 2.87332984 | -0.222671601 | 0.84867466 | -0.2623757 | 0.79303181 | 0.8878826 |
| ZNF774 | 179.673609 | 0.317796101 | 0.22683851 | 1.40097947 | 0.16122021 | 0.32994158 |
| IQGAP1 | 7278.92633 | -0.502572089 | 0.17611248 | -2.8536995 | 0.00432134 | 0.02414273 |
| AC018946.1 | 1.7667676 | -0.395188685 | 1.0528265 | -0.3753597 | 0.70739294 | 0.83343543 |
| CRTC3 | 2241.41287 | -0.230326326 | 0.1153062 | -1.9975191 | 0.04576882 | 0.13656075 |
| AC103739.1 | 2.09463254 | 0.470017418 | 0.80013573 | 0.58742211 | 0.55692025 | 0.72587915 |
| AC103739.2 | 39.7767744 | 1.661117976 | 0.4099356 | 4.0521437 | 5.08E-05 | 0.00075227 |
| CRTC3-AS1 | 71.2060679 | 0.658809414 | 0.30176332 | 2.1831991 | 0.02902115 | 0.09856875 |
| AC021422.1 | 1.91170514 | -1.675999514 | 1.37198402 | -1.2215882 | 0.22186339 | 0.40780891 |
| BLM | 43.8737978 | -0.844153315 | 0.46711108 | -1.807179 | 0.07073438 | 0.18566652 |
| FURIN | 10371.4426 | -0.714427061 | 0.20445319 | -3.4943307 | 0.00047525 | 0.00436061 |
| FES | 1710.68081 | -0.869339968 | 0.36426405 | -2.3865654 | 0.01700658 | 0.0666951 |
| MAN2A2 | 4273.04129 | 0.088127276 | 0.10703905 | 0.82331893 | 0.41032665 | 0.6046895 |
| AC068831.4 | 1748.29879 | -0.370944126 | 0.17609195 | -2.1065365 | 0.03515778 | 0.11302084 |
| HDDC3 | 880.873516 | 0.054549392 | 0.22447523 | 0.24300852 | 0.8079988 | 0.89622816 |
| UNC45A | 5242.01956 | -0.082392317 | 0.12834089 | -0.6419803 | 0.52088599 | 0.69745931 |
| AC068831.1 | 869.589871 | -0.302269682 | 0.14485753 | -2.0866687 | 0.03691809 | 0.11679995 |
| RCCD1 | 942.105778 | -0.240131515 | 0.27338274 | -0.8783712 | 0.37974232 | 0.57573766 |
| PRC1 | 234.553734 | -0.467762695 | 0.33125858 | -1.4120772 | 0.15792723 | 0.32507244 |
| AC068831.7 | 1127.90247 | -0.072613133 | 0.13538047 | -0.5363634 | 0.59170739 | 0.7512795 |
| PRC1-AS1 | 94.9816553 | -0.773241591 | 0.30807922 | -2.5098791 | 0.01207725 | 0.05180175 |
| VPS33B | 1020.30157 | -0.085798402 | 0.11880088 | -0.7222035 | 0.4701694 | 0.65679754 |
| VPS33B-DT | 8.53470852 | 0.024098275 | 0.57624869 | 0.04181923 | 0.96664281 | 0.9841842 |
| AC068831.6 | 3.2455675 | -0.984276498 | 1.34632425 | -0.7310843 | 0.46472767 | 0.65205955 |
| SV2B | 90.8138136 | -0.948174616 | 0.53648592 | -1.7673803 | 0.07716457 | 0.19751792 |
| AC104035.1 | 10.9245266 | -0.436078302 | 0.93591354 | -0.4659387 | 0.64125941 | 0.78758115 |
| AC107958.3 | 0.03392908 | 0.380740593 | 3.29738709 | 0.11546736 | 0.90807469 | NA |
| AC107958.2 | 0.07659028 | -0.642905668 | 3.30558948 | -0.1944905 | 0.84579185 | NA |
| SLCO3A1 | 1760.99097 | -0.708961919 | 0.32304082 | -2.1946512 | 0.02818863 | 0.0965965 |
| AC116903.2 | 1.15936589 | 2.154564933 | 1.21836396 | 1.7684083 | 0.07699267 | 0.19717362 |
| AC116903.1 | 340.725795 | -0.729352508 | 0.32963404 | -2.2126128 | 0.02692435 | 0.09324863 |
| ST8SIA2 | 20.796546 | 0.408990849 | 0.4584974 | 0.89202437 | 0.37237987 | 0.56879386 |
| AC090985.1 | 0.07659028 | -0.642905668 | 3.30558948 | -0.1944905 | 0.84579185 | NA |
| C15orf32 | 0.25687762 | -0.178459274 | 2.99441912 | -0.0595973 | 0.95247637 | NA |
| LINC00930 | 7.45489823 | -0.646261358 | 0.54938221 | -1.176342 | 0.23945824 | 0.42932512 |
| AC091544.5 | 1.38749367 | 0.605356155 | 1.37341221 | 0.440768 | 0.65938096 | 0.79994664 |
| FAM174B | 432.464517 | -0.506334123 | 0.26453648 | -1.9140427 | 0.0556147 | 0.156895 |
| AC091544.7 | 0.09635527 | 0.547239394 | 3.29377202 | 0.16614368 | 0.86804388 | NA |
| AC106028.4 | 7.41674717 | -0.155220095 | 0.58526569 | -0.265213 | 0.79084535 | 0.88640506 |
| AC106028.5 | 0.50799168 | 0.949829785 | 2.17607822 | 0.43648697 | 0.66248345 | NA |
| AC013394.1 | 1697.68481 | -0.343041087 | 0.14719584 | -2.330508 | 0.01977932 | 0.07429923 |
| CHD2 | 5221.83311 | -0.338272702 | 0.18237441 | -1.8548255 | 0.06362115 | 0.17221919 |
| RGMA | 328.202571 | 0.066257004 | 0.39874444 | 0.16616408 | 0.86802782 | 0.93135426 |
| AC112693.1 | 0.03392908 | 0.380740593 | 3.29738709 | 0.11546736 | 0.90807469 | NA |
| AC091078.1 | 2.51810368 | 2.224949711 | 0.93230058 | 2.38651541 | 0.01700889 | 0.0666951 |
| AC091078.3 | 0.03392908 | 0.380740593 | 3.29738709 | 0.11546736 | 0.90807469 | NA |
| AC110023.1 | 0.16421342 | 1.36803136 | 3.28033652 | 0.41703994 | 0.67664919 | NA |
| LINC01579 | 2.72913351 | 0.616959818 | 1.05987733 | 0.58210493 | 0.560496 | 0.72859507 |
| AC103996.3 | 0.32876134 | 0.653167071 | 2.83320692 | 0.23053984 | 0.81767231 | NA |
| LINC02207 | 16.5429639 | -2.175746244 | 0.81524921 | -2.6688112 | 0.00761202 | 0.03680476 |
| AC103996.2 | 2.43966468 | -2.095010458 | 1.37106219 | -1.52802 | 0.12650757 | 0.27979574 |
| LINC01580 | 0.07659028 | -0.642905668 | 3.30558948 | -0.1944905 | 0.84579185 | NA |
| LINC01581 | 0.25087713 | 1.011917048 | 2.94486708 | 0.34362062 | 0.73113162 | NA |
| MCTP2 | 605.724046 | -0.72001871 | 0.29046374 | -2.4788592 | 0.01318033 | 0.05529066 |
| AC009432.1 | 0 | NA | NA | NA | NA | NA |
| AC107976.1 | 1.49641569 | -1.141001076 | 1.08673668 | -1.0499333 | 0.29374876 | 0.49039267 |
| AC087633.1 | 1.03041429 | -1.344274549 | 1.83001387 | -0.7345707 | 0.46260099 | 0.6508103 |
| AC087636.1 | 3.22008015 | -1.533990288 | 0.92931256 | -1.6506721 | 0.09880555 | 0.23564656 |
| LINC01197 | 137.427211 | 0.756466918 | 0.39860124 | 1.89780373 | 0.05772194 | 0.16108875 |
| AC104260.1 | 9.07971272 | 0.349171504 | 0.54265378 | 0.64345171 | 0.51993104 | 0.69660509 |
| LINC00924 | 524.435164 | 0.278175141 | 0.38752179 | 0.71783097 | 0.47286151 | 0.65891768 |
| AC027013.1 | 6.15136433 | 0.002667358 | 0.67089521 | 0.00397582 | 0.99682776 | 0.99845589 |
| AC015574.1 | 6.84836855 | -0.071412914 | 0.49595547 | -0.1439906 | 0.88550791 | 0.94073175 |
| AC012409.2 | 4.52135297 | 0.973717337 | 0.80592131 | 1.20820398 | 0.22696881 | 0.41379753 |
| AC024337.1 | 14.8387907 | 2.270153764 | 0.38631727 | 5.8763973 | 4.19E-09 | 3.30E-07 |
| AC024337.2 | 258.208316 | 0.490695034 | 0.34588854 | 1.41865076 | 0.15600086 | 0.32246944 |
| NR2F2-AS1 | 377.62693 | 0.599856181 | 0.32639471 | 1.83782446 | 0.06608828 | 0.1768928 |
| AC012409.3 | 118.621927 | 0.250695645 | 0.36507277 | 0.68670047 | 0.4922715 | 0.6751547 |
| AC012409.1 | 70.2089005 | 1.668483283 | 0.45584918 | 3.66016512 | 0.00025205 | 0.00263892 |
| AC012409.4 | 9.11955273 | 0.209356747 | 0.47873057 | 0.43731643 | 0.66188188 | 0.80111191 |
| LINC02157 | 2.83473404 | 2.826165732 | 0.8631496 | 3.27424785 | 0.00105944 | 0.00819779 |
| NR2F2 | 9181.65279 | 0.086440911 | 0.2658886 | 0.32510198 | 0.74510389 | 0.85722218 |

| | | | | | | | |
|---|---|---|---|---|---|---|---|
| AC103746.1 | 57.7530258 | | 0.506977546 | 0.32737191 | 1.54862871 | 0.121471 | 0.27205625 |
| AC087477.2 | 201.304623 | | 0.350406218 | 0.27840957 | 1.25859977 | 0.20817493 | 0.39140559 |
| AC087477.5 | 25.0556945 | | 0.560002579 | 0.30265939 | 1.85027328 | 0.06427417 | 0.17339232 |
| AC087477.3 | 0.17423948 | | 0.18657164 | 3.29377195 | 0.05664376 | 0.95482897 | NA |
| AC087477.4 | 0.40142484 | | -1.083777078 | 2.27936009 | -0.4754743 | 0.63444897 | NA |
| SPATA8-AS1 | 0.45848006 | | -2.07761151 | 2.21954539 | -0.9360527 | 0.34924604 | NA |
| SPATA8 | 4.6025968 | | -1.089927432 | 0.87879776 | -1.2402483 | 0.21488356 | 0.39913218 |
| AC020704.1 | 0.08393208 | | -0.642905668 | 3.30558948 | -0.1944905 | 0.84579185 | NA |
| LINC02253 | 5.38096384 | | -0.853625901 | 1.04883161 | -0.8138827 | 0.41571215 | 0.60908136 |
| LINC02254 | 1.49336049 | | -0.133154918 | 1.21255442 | -0.1098136 | 0.91255724 | 0.95701469 |
| AC026523.1 | 6.19737322 | | -1.437726799 | 0.75031051 | -1.9161757 | 0.05534273 | 0.156415 |
| LINC00923 | 20.0075712 | | -0.412553315 | 0.56412575 | -0.7313145 | 0.4645871 | 0.65193497 |
| AC024651.2 | 0 | NA | | NA | NA | NA | NA |
| ARRDC4 | 2504.58142 | | -0.151000852 | 0.21235757 | -0.7110688 | 0.47704158 | 0.66212483 |
| AC024651.1 | 0.15661533 | | -1.003574173 | 3.30558948 | -0.3035992 | 0.7614333 | NA |
| LINC02251 | 0.18874271 | | 0.726716446 | 3.28420131 | 0.22127646 | 0.82487718 | NA |
| AC022523.1 | 0.10721287 | | -0.642905668 | 3.30558948 | -0.1944905 | 0.84579185 | NA |
| LINC01582 | 0 | NA | | NA | NA | NA | NA |
| AC022523.3 | 0 | NA | | NA | NA | NA | NA |
| AC015722.1 | 0 | NA | | NA | NA | NA | NA |
| AC015722.2 | 42.0781738 | | -1.400976505 | 0.76003167 | -1.8433133 | 0.06528328 | 0.17534325 |
| LINC02351 | 14.7846002 | | -2.053701387 | 0.86492879 | -2.3744167 | 0.0175767 | 0.0682417 |
| FAM169B | 181.584151 | | -2.778484828 | 0.50946428 | -5.4537383 | 4.93E-08 | 2.63E-06 |
| AC118658.1 | 12.8100986 | | 0.092910765 | 0.40503031 | 0.22939213 | 0.81856415 | 0.9026318 |
| IGF1R | 5009.61194 | | 0.337532922 | 0.17308898 | 1.95005434 | 0.05116964 | 0.14810048 |
| AC069029.1 | 0.52521305 | | 0.646104361 | 3.26179209 | 0.19808263 | 0.84298041 | NA |
| AC036108.1 | 3.54730713 | | 0.574320817 | 0.70515591 | 0.81445934 | 0.41538186 | 0.60894038 |
| PGPEP1L | 22.7643548 | | 0.340579051 | 0.65150902 | 0.52275416 | 0.60114534 | 0.75869641 |
| AC036108.4 | 11.60602 | | 0.154923665 | 0.62767108 | 0.24682301 | 0.8050452 | 0.89438161 |
| SYNM | 890.820061 | | -0.271343071 | 0.4085312 | -0.6641918 | 0.50656756 | 0.68569375 |
| AC036108.2 | 89.7493341 | | 0.608638899 | 0.33745655 | 1.80360671 | 0.07129298 | 0.1867769 |
| TTC23 | 1176.31785 | | -0.102801606 | 0.19099966 | -0.5382293 | 0.59041877 | 0.75022132 |
| AC036108.3 | 126.462647 | | 0.165444105 | 0.25463604 | 0.64972775 | 0.51586809 | 0.69342133 |
| LRRC28 | 1193.17094 | | 0.740921179 | 0.21538607 | 3.43996799 | 0.00058178 | 0.00510367 |
| AC015660.2 | 19.2347651 | | -1.064652336 | 0.74666594 | -1.4258751 | 0.15390441 | 0.3196644 |
| AC015660.3 | 18.9879241 | | -0.668613452 | 0.69043766 | -0.9683908 | 0.33284925 | 0.53079553 |
| AC015660.4 | 33.8350242 | | 0.662550551 | 0.42211474 | 1.56959823 | 0.11650861 | 0.26469648 |
| AC015660.1 | 5.21457357 | | 0.926648486 | 0.49038843 | 1.88962143 | 0.05880861 | 0.16314767 |
| MEF2A | 3540.0265 | | 0.008942561 | 0.12169176 | 0.07348535 | 0.9414199 | 0.97209033 |
| LYSMD4 | 1690.90539 | | -0.197131871 | 0.12691079 | -1.5533105 | 0.12034897 | 0.2704634 |
| AC090825.1 | 35.0815497 | | 0.055941879 | 0.31522914 | 0.17746417 | 0.85914381 | 0.92642086 |
| AC084855.1 | 48.922972 | | 1.931930715 | 0.34061383 | 5.6719092 | 1.41E-08 | 9.29E-07 |
| ADAMTS17 | 210.156529 | | 1.693716654 | 0.42720174 | 3.96467635 | 7.35E-05 | 0.00099338 |
| AC084855.2 | 4.8946177 | | 2.24730003 | 0.82419786 | 2.72665115 | 0.00639806 | 0.03231506 |
| AC022710.1 | 0.23371381 | | -0.340597073 | 3.29738699 | -0.103293 | 0.91773041 | NA |
| SPATA41 | 25.6617782 | | 0.766954053 | 0.47859593 | 1.6025085 | 0.10904321 | 0.25239218 |
| CERS3-AS1 | 1.9116402 | | 0.797180121 | 0.99145594 | 0.80404997 | 0.42136811 | 0.61402946 |
| CERS3 | 2.98509102 | | 0.115642849 | 0.78089723 | 0.14808972 | 0.88227197 | 0.93888264 |
| AC027020.2 | 261.734794 | | -0.019242333 | 0.32435518 | -0.0593249 | 0.95269334 | 0.97717271 |
| AC027020.1 | 0.09635527 | | 0.547239394 | 3.29377202 | 0.16614368 | 0.86804388 | NA |
| LINS1 | 649.911375 | | -0.023020439 | 0.09798944 | -0.2349278 | 0.81426481 | 0.89980987 |
| ASB7 | 807.91812 | | 0.089692542 | 0.13389683 | 0.66986308 | 0.50294508 | 0.68322963 |
| AC087762.1 | 85.2990108 | | 0.917122014 | 0.50173941 | 1.82788515 | 0.06756679 | 0.17955621 |
| AC015712.1 | 50.7682747 | | -0.577505416 | 0.44743135 | -1.2907129 | 0.19680325 | 0.3774054 |
| AC015712.4 | 5.17795737 | | -0.222477085 | 0.63182577 | -0.3521178 | 0.72474994 | 0.84389349 |
| ALDH1A3 | 855.012807 | | -1.172942011 | 0.47529523 | -2.4678178 | 0.01359395 | 0.05656816 |
| AC015712.7 | 1.79186447 | | 0.565565203 | 0.92177038 | 0.61356409 | 0.53950342 | 0.71254659 |
| AC015712.2 | 576.949937 | | -1.027714667 | 0.46153925 | -2.2267113 | 0.02596657 | 0.09088645 |
| LRRK1 | 539.63973 | | -0.424750358 | 0.31690018 | -1.3403286 | 0.18013855 | 0.3560376 |
| AC090907.1 | 24.0360272 | | -1.043511274 | 0.44758625 | -2.3314194 | 0.01973126 | 0.07413981 |
| AC090907.2 | 55.3095804 | | -0.368713018 | 0.43161949 | -0.8542548 | 0.39296383 | 0.58844158 |
| AC019254.2 | 0.32504708 | | 0.328540168 | 2.26374193 | 0.14513146 | 0.88460708 | NA |
| AC019254.1 | 16.4701671 | | -0.856594546 | 0.4889899 | -1.7517633 | 0.07981452 | 0.2021448 |
| CHSY1 | 2263.17762 | | -0.843094392 | 0.32938836 | -2.5595755 | 0.01048001 | 0.04668062 |
| SELENOS | 2867.39495 | | -0.833630743 | 0.21212773 | -3.9298528 | 8.50E-05 | 0.00111568 |
| SNRPA1 | 2010.94087 | | -0.915950106 | 0.15416066 | -5.9415297 | 2.82E-09 | 2.39E-07 |
| AC023024.1 | 375.44799 | | -0.541001042 | 0.25868543 | -2.0913472 | 0.03649695 | 0.11594798 |
| PCSK6 | 651.751064 | | -0.388773204 | 0.28687146 | -1.3552174 | 0.17534829 | 0.34950373 |
| PCSK6-AS1 | 0 | NA | | NA | NA | NA | NA |
| AC090164.3 | 2.18770434 | | -1.763985295 | 0.99490082 | -1.7730263 | 0.07622435 | 0.19594867 |
| AC090164.4 | 3.35300935 | | -2.385779401 | 1.02397089 | -2.3299289 | 0.01980991 | 0.07438235 |
| LINC02348 | 3.52226273 | | -3.180469004 | 1.19843418 | -2.6538537 | 0.00795783 | 0.03801595 |
| TM2D3 | 2533.37473 | | -0.483435966 | 0.11088771 | -4.3596894 | 1.30E-05 | 0.00025246 |
| TARSL2 | 940.812766 | | -0.214494905 | 0.18603417 | -1.1529866 | 0.2489159 | 0.44066597 |
| OR4F6 | 0.09989236 | | -0.642905668 | 3.30558948 | -0.1944905 | 0.84579185 | NA |
| OR4F15 | 0.23560867 | | 0.952737031 | 3.28103697 | 0.2903768 | 0.77152799 | NA |
| OR4F4 | 0 | NA | | NA | NA | NA | NA |
| FAM138E | 4.81297168 | | -1.669859145 | 1.51820983 | -1.0998869 | 0.27138139 | 0.46605844 |
| AC140725.1 | 7.77126181 | | 0.959452358 | 0.45061709 | 2.12919658 | 0.033238 | 0.10848942 |
| WASIR2 | 3.79507734 | | -0.518155465 | 0.77562864 | -0.6680458 | 0.50410435 | 0.68375733 |
| POLR3K | 439.902682 | | -0.134589914 | 0.13181015 | -1.0210891 | 0.30721221 | 0.50411861 |
| SNRNP25 | 1100.79849 | | 0.05836267 | 0.13769474 | 0.42385547 | 0.67167122 | 0.80832284 |
| RHBDF1 | 2088.47824 | | -0.680273124 | 0.23606734 | -2.881691 | 0.00395547 | 0.02258191 |
| MPG | 1713.78215 | | -0.712844571 | 0.13327913 | -5.3485088 | 8.87E-08 | 4.35E-06 |
| NPRL3 | 3640.17091 | | -0.118801145 | 0.06851835 | -1.7338589 | 0.08294312 | 0.20775173 |
| Z69720.1 | 39.6494588 | | -0.099069982 | 0.24515416 | -0.404113 | 0.68612964 | 0.81847636 |
| Z69666.1 | 1.95503341 | | -0.357412405 | 0.91546118 | -0.3904179 | 0.69622758 | 0.82523522 |
| HBZ | 0.33725126 | | 0.669857205 | 2.22872422 | 0.30055634 | 0.76375283 | NA |
| HBM | 11.7858434 | | -3.041415346 | 1.15352907 | -2.6366178 | 0.00837371 | 0.03937496 |
| HBA2 | 1118.5181 | | -0.211124126 | 0.66735716 | -0.3163585 | 0.7517304 | 0.86146914 |
| HBA1 | 201.634453 | | -1.231363775 | 0.72737418 | -1.692889 | 0.09047659 | 0.2214879 |
| HBQ1 | 5.6411074 | | -1.874631823 | 1.08930362 | -1.7209452 | 0.08526077 | 0.21186992 |
| Z69706.1 | 21.4080335 | | -0.214257715 | 0.39684791 | -0.5398988 | 0.58926682 | 0.74958833 |
| LUC7L | 4613.42947 | | -0.231010129 | 0.16788275 | -1.3760206 | 0.16881524 | 0.34041279 |
| FAM234A | 5696.17093 | | -0.065569219 | 0.23968984 | -0.2735586 | 0.78442384 | 0.88221337 |
| AC004754.1 | 2858.04945 | | 0.194728188 | 0.236675 | 0.8227662 | 0.41064096 | 0.60498364 |
| RGS11 | 371.455869 | | 0.129437589 | 0.47125587 | 0.2746652 | 0.78357347 | 0.88163299 |
| ARHGDIG | 112.356329 | | 0.027107665 | 0.47237058 | 0.05738644 | 0.95423737 | 0.9780936 |
| PDIA2 | 16.9794904 | | -0.148031067 | 0.73789359 | -0.200613 | 0.84100117 | 0.91512686 |
| AXIN1 | 2629.32782 | | -0.833260831 | 0.20995756 | -3.9687108 | 7.23E-05 | 0.00097922 |
| MRPL28 | 3355.01029 | | -0.094972586 | 0.10456084 | -0.9082997 | 0.36371989 | 0.56082876 |
| TMEM8A | 9172.89808 | | -0.770610015 | 0.16471653 | -4.6784012 | 2.89E-06 | 7.62E-05 |
| NME4 | 3843.02174 | | 0.139438841 | 0.23977517 | 0.58153996 | 0.5608766 | 0.7288024 |
| DECR2 | 3212.02503 | | 0.941571332 | 0.24032509 | 3.91790689 | 8.93E-05 | 0.0011575 |
| AL023881.1 | 2.15694316 | | 1.1308773 | 1.00942027 | 1.12032354 | 0.26257591 | 0.45616489 |
| RAB11FIP3 | 16296.3606 | | 1.352470486 | 0.32966208 | 4.10259645 | 4.09E-05 | 0.0006326 |
| LINC00235 | 117.252654 | | -0.160403948 | 0.36401283 | -0.4406547 | 0.65946303 | 0.79994664 |
| CAPN15 | 5248.08878 | | -0.456401403 | 0.22802921 | -2.0015041 | 0.04533809 | 0.13559775 |
| Z97986.1 | 955.228845 | | -0.489495721 | 0.24300819 | -2.0143178 | 0.04397618 | 0.13283084 |
| PRR35 | 510.059763 | | 1.567550219 | 0.72999119 | 2.14735498 | 0.03176503 | 0.10505979 |
| PIGQ | 7661.7648 | | 0.327266158 | 0.18463208 | 1.77253142 | 0.07630638 | 0.19606347 |
| NHLRC4 | 470.156258 | | 1.208399553 | 0.55195534 | 2.18930675 | 0.02857455 | 0.0974908 |
| RAB40C | 2539.69792 | | -0.335182695 | 0.12428855 | -2.6968108 | 0.0070007 | 0.03463575 |

| | | | | | | |
|---|---|---|---|---|---|---|
| WFIKKN1 | 331.181205 | 0.118966901 | 0.34517986 | 0.34465192 | 0.73035607 | 0.84755348 |
| METTL26 | 1770.68607 | -0.099271718 | 0.14175362 | -0.7003117 | 0.48373267 | 0.66827486 |
| MCRIP2 | 4302.85181 | 0.161107315 | 0.25263774 | 0.63770089 | 0.52366841 | 0.69892989 |
| AL022341.1 | 40.4349799 | 0.375547743 | 0.29109953 | 1.2901008 | 0.19701566 | 0.37770287 |
| WDR90 | 3354.66145 | 0.012623577 | 0.20661593 | 0.06109682 | 0.9512821 | 0.97675756 |
| AL022341.2 | 323.932049 | 0.162843593 | 0.2176205 | 0.74829162 | 0.45428428 | 0.64388479 |
| RHOT2 | 8423.76637 | -0.265350396 | 0.12898873 | -2.0571595 | 0.03967088 | 0.12308756 |
| RHBDL1 | 280.020018 | 1.01579317 | 0.28909908 | 3.5136506 | 0.00044199 | 0.00412434 |
| Z92544.1 | 15.5897216 | 0.79995408 | 0.47326865 | 1.69027482 | 0.09097539 | 0.22230891 |
| STUB1 | 8518.34827 | -0.020135003 | 0.10626691 | -0.1894758 | 0.84971996 | 0.92104222 |
| JMJD8 | 8110.67867 | 0.039293004 | 0.10741444 | 0.36580746 | 0.71450875 | 0.83777661 |
| WDR24 | 981.752995 | 0.06854435 | 0.11927985 | 0.57465157 | 0.56552697 | 0.73214268 |
| FBXL16 | 567.943735 | -0.590028252 | 0.48255783 | -1.2227099 | 0.22143927 | 0.40757685 |
| Z97653.1 | 18.0936879 | 0.54400521 | 0.38900028 | 1.39846996 | 0.16197199 | 0.33076754 |
| METRN | 972.796548 | 0.178903605 | 0.25643649 | 0.69765269 | 0.48539442 | 0.6696032 |
| FAM173A | 806.10429 | -0.324437245 | 0.16134186 | -2.0108684 | 0.04433936 | 0.13338227 |
| CCDC78 | 350.63524 | -0.474381445 | 0.25400504 | -1.8676064 | 0.06181695 | 0.16894113 |
| HAGHL | 1107.84823 | -0.507924059 | 0.2044151 | -2.4847678 | 0.01296359 | 0.0545969 |
| NARFL | 2742.79395 | -0.314011552 | 0.11478806 | -2.7355769 | 0.0062271 | 0.03171277 |
| MSLN | 442.245431 | -1.333478278 | 0.47330818 | -2.8173573 | 0.00484206 | 0.02623488 |
| MSLNL | 8.43580115 | -2.842771517 | 1.02064031 | -2.7852824 | 0.00534811 | 0.02830785 |
| RPUSD1 | 1242.31484 | -0.507769321 | 0.28763178 | -1.765345 | 0.0775058 | 0.19805608 |
| CHTF18 | 897.427103 | -0.174166669 | 0.32353212 | -0.5383288 | 0.59035004 | 0.75020628 |
| GNG13 | 9.58607653 | -0.82214662 | 0.61124537 | -1.3450353 | 0.17861383 | 0.35416547 |
| PRR25 | 14.6290591 | -0.476988034 | 0.53488244 | -0.8917624 | 0.37252032 | 0.56884364 |
| AL031008.1 | 3.65698521 | -1.108269953 | 0.99477395 | -1.1140923 | 0.26523961 | 0.45930973 |
| LMF1 | 1515.81654 | 0.046999869 | 0.17599287 | 0.26705553 | 0.7894264 | 0.88557334 |
| AL031716.1 | 18.1407561 | 0.530955668 | 0.36773356 | 1.44385969 | 0.14877845 | 0.31286186 |
| LMF1-AS1 | 91.8109035 | 0.232921011 | 0.22485828 | 1.03585693 | 0.30026889 | 0.49704988 |
| AC009041.2 | 263.152546 | -0.828390716 | 0.27617896 | -2.9994707 | 0.00270449 | 0.01697226 |
| SOX8 | 80.100243 | -0.692005813 | 0.28602598 | -2.419381 | 0.01554695 | 0.06244829 |
| AC009041.1 | 1.19261865 | 0.3524818 | 1.64967228 | 0.21366777 | 0.83080615 | 0.91040231 |
| AC009041.3 | 381.694749 | -1.403349638 | 0.57873998 | -2.4248362 | 0.0153153 | 0.0617624 |
| SSTR5 | 161.20728 | -1.234402643 | 0.52899416 | -2.3334901 | 0.01962244 | 0.07386776 |
| C1QTNF8 | 5.83455165 | 1.073104161 | 0.82424541 | 1.30192312 | 0.19294267 | 0.37249168 |
| AL031598.1 | 8.04608329 | 0.167929363 | 0.67961902 | 0.24709338 | 0.80483596 | 0.89428472 |
| AC120498.8 | 4.79336902 | -0.037661674 | 0.87874596 | -0.0428584 | 0.96581438 | 0.98387257 |
| CACNA1H | 4611.41237 | 0.463374417 | 0.46830378 | 0.98947401 | 0.32243128 | 0.52022228 |
| AC120498.3 | 14.0232426 | 0.575893795 | 0.58758678 | 0.98009998 | 0.32703677 | 0.52489103 |
| AC120498.1 | 13.5246044 | 0.251614555 | 0.56437123 | 0.44583165 | 0.65571886 | 0.79807876 |
| AC120498.6 | 123.594089 | 1.238585707 | 0.48551503 | 2.55107593 | 0.01073909 | 0.04750711 |
| TPSG1 | 265.784591 | -0.067687078 | 0.48194478 | -0.1404457 | 0.88830784 | 0.94222589 |
| AC120498.1( | 40.1637084 | -0.221099548 | 0.61143499 | -0.3616076 | 0.71764528 | 0.83973668 |
| TPSB2 | 227.792112 | -0.130145583 | 0.9837549 | -0.1322947 | 0.89475118 | 0.94590237 |
| TPSAB1 | 141.701611 | 0.584325335 | 0.78499105 | 0.74437197 | 0.45665147 | 0.64604305 |
| TPSD1 | 44.0791803 | 0.473708803 | 1.16108481 | 0.40798811 | 0.6832824 | 0.81703881 |
| AC120498.4 | 81.4060122 | -0.734541013 | 0.86253368 | -0.8516085 | 0.39443142 | 0.58973502 |
| AC120498.9 | 60.1417367 | -1.266971382 | 0.43446515 | -2.9161635 | 0.00354365 | 0.02076578 |
| UBE2I | 4636.88498 | -0.765450269 | 0.16315565 | -4.6915339 | 2.71E-06 | 7.26E-05 |
| AL031714.1 | 2136.52259 | -0.776408126 | 0.17650951 | -4.3986759 | 1.09E-05 | 0.00021864 |
| BAIAP3 | 2661.93846 | 0.059176159 | 0.16792724 | 0.35239165 | 0.72454456 | 0.84389181 |
| TSR3 | 2355.06945 | -0.164119177 | 0.12375555 | -1.3261561 | 0.184788 | 0.36205213 |
| GNPTG | 5898.57688 | 0.062834005 | 0.08000118 | 0.78541351 | 0.43221116 | 0.62434447 |
| AL031709.1 | 157.610447 | 0.18532956 | 0.25367875 | 0.73056794 | 0.4650431 | 0.65212372 |
| UNKL | 2376.14416 | -0.271372748 | 0.11129951 | -2.4382204 | 0.01475977 | 0.06011045 |
| AL032819.1 | 0.83307744 | -0.22560972 | 1.39999465 | -0.1611504 | 0.87197493 | NA |
| C16orf91 | 564.56662 | -0.326882003 | 0.1711602 | -1.9098015 | 0.05615877 | 0.15791287 |
| AL032819.3 | 6.28297273 | 0.869191227 | 0.54680277 | 1.5895882 | 0.11192766 | 0.25714692 |
| CCDC154 | 402.687544 | 0.191910672 | 0.38125637 | 0.50336385 | 0.61470849 | 0.76958946 |
| CLCN7 | 11839.3908 | -0.280173988 | 0.22638391 | -1.2376056 | 0.21586235 | 0.40047068 |
| AL031600.3 | 4667.43558 | -0.507534937 | 0.21987196 | -2.3083204 | 0.02098132 | 0.07759852 |
| AL031600.1 | 498.027745 | -0.21458531 | 0.29723153 | -0.7219466 | 0.47032728 | 0.65694848 |
| PTX4 | 2.11562325 | -0.878864409 | 1.2601641 | -0.6974206 | 0.48553961 | 0.66964851 |
| TELO2 | 2989.74591 | -0.30120152 | 0.1975357 | -1.5247954 | 0.12731014 | 0.28088759 |
| IFT140 | 2363.04172 | 0.208775087 | 0.21788001 | 0.95821131 | 0.33795621 | 0.53590196 |
| AL031705.1 | 18.8445059 | 0.180077238 | 0.34831014 | 0.51700257 | 0.60515437 | 0.76207835 |
| TMEM204 | 1706.3438 | 0.089337863 | 0.34316567 | 0.2603345 | 0.79460577 | 0.88857752 |
| AL133297.2 | 4.7296801 | 1.441460225 | 0.6894734 | 2.09066835 | 0.0365578 | 0.11609216 |
| CRAMP1 | 1790.67901 | -0.139998818 | 0.16087941 | -0.8702097 | 0.38418583 | 0.57970819 |
| Z97652.1 | 4.26998127 | 1.333097118 | 0.67994119 | 1.96060651 | 0.04992494 | 0.14553623 |
| AL031708.1 | 854.019951 | 0.107825864 | 0.25449403 | 0.42368721 | 0.67179394 | 0.80839675 |
| JPT2 | 6112.93643 | -1.005204924 | 0.23949595 | -4.1971688 | 2.70E-05 | 0.00045251 |
| MAPK8IP3 | 10812.5231 | -0.554385028 | 0.19663451 | -2.8193679 | 0.00481183 | 0.02610865 |
| AL031710.1 | 67.6682393 | 1.792333632 | 0.61402339 | 2.91899896 | 0.00351157 | 0.02062539 |
| AL031717.1 | 14.2673764 | 0.46311815 | 0.44811097 | 1.03348987 | 0.30137471 | 0.49813533 |
| NME3 | 2109.2028 | -0.218899393 | 0.20242501 | -1.0813851 | 0.27952583 | 0.47514828 |
| MRPS34 | 2604.33079 | -0.10228178 | 0.2024574 | -0.5052015 | 0.61341732 | 0.76841076 |
| EME2 | 6613.53187 | -0.148339572 | 0.13586061 | -1.0918512 | 0.27489852 | 0.47007807 |
| SPSB3 | 4899.14016 | -0.195655509 | 0.1458499 | -1.3414853 | 0.17976292 | 0.35557601 |
| NUBP2 | 2395.53122 | -0.085447096 | 0.12248164 | -0.6976319 | 0.48540745 | 0.6696032 |
| IGFALS | 64.6484982 | 1.200641274 | 0.34544113 | 3.47567551 | 0.00050957 | 0.00460182 |
| HAGH | 5562.25717 | 0.878018128 | 0.27407307 | 3.20359142 | 0.00135725 | 0.01000019 |
| FAHD1 | 3532.69626 | 0.777857487 | 0.32415492 | 2.39964736 | 0.01641087 | 0.06497819 |
| MEIOB | 72.6393108 | 1.259902281 | 0.40561511 | 3.10615221 | 0.00189539 | 0.01292885 |
| LINC00254 | 0 | NA | NA | NA | NA | NA |
| LINC02124 | 0.83987373 | 1.951209941 | 1.44190261 | 1.35321896 | 0.17598569 | NA |
| HS3ST6 | 12.1587411 | 1.268917686 | 0.84463776 | 1.50232177 | 0.13301403 | 0.28949983 |
| MSRB1 | 5679.03064 | 0.095264995 | 0.30997951 | 0.30732675 | 0.75859468 | 0.86635037 |
| RPL3L | 14.029701 | 1.094797803 | 0.54615586 | 2.00455196 | 0.04501097 | 0.13484881 |
| NDUFB10 | 3985.57154 | 0.210677162 | 0.13898558 | 1.51582033 | 0.12956479 | 0.28450525 |
| RPS2 | 29934.9939 | -0.996800225 | 0.16972313 | -5.8730958 | 4.28E-09 | 3.32E-07 |
| SNHG9 | 209.919759 | -0.323599168 | 0.31980421 | -1.0118665 | 0.31160188 | 0.50871332 |
| RNF151 | 5.04222928 | 0.363215512 | 0.79493855 | 0.45691018 | 0.64773561 | 0.79272233 |
| AC005363.2 | 3.30446423 | 0.048480446 | 0.71706408 | 0.06760964 | 0.94609638 | 0.97409658 |
| TBL3 | 2305.12737 | 0.030093485 | 0.11057881 | 0.27214512 | 0.78551044 | 0.88302115 |
| NOXO1 | 80.5108449 | 0.099155464 | 0.42915929 | 0.23104583 | 0.8172792 | 0.9017286 |
| GFER | 1391.10062 | -0.117971048 | 0.10226645 | -1.1535655 | 0.24867836 | 0.44046175 |
| AC005606.2 | 758.082708 | -0.054337695 | 0.11302324 | -0.4807657 | 0.63068306 | 0.78054869 |
| SYNGR3 | 197.837431 | -0.15196239 | 0.36749114 | -0.4135131 | 0.67923075 | 0.8138966 |
| AC005606.1 | 795.388367 | -1.09825357 | 0.23795308 | -4.6154206 | 3.92E-06 | 9.73E-05 |
| ZNF598 | 2850.56338 | -0.890470989 | 0.2304086 | -3.8647472 | 0.0001112 | 0.00137524 |
| NPW | 14.4947512 | -0.377682102 | 0.6293235 | -0.6001398 | 0.54841304 | 0.71905639 |
| SLC9A3R2 | 5654.16684 | 0.858932649 | 0.30526701 | 2.81370934 | 0.00489735 | 0.02646917 |
| NTHL1 | 435.225961 | 0.43321242 | 0.28796687 | 1.50438283 | 0.13248282 | 0.28877269 |
| TSC2 | 7845.23768 | 0.235000175 | 0.17493771 | 1.34333628 | 0.17916311 | 0.3548671 |
| PKD1 | 5558.77563 | 0.063321521 | 0.22517476 | 0.28121057 | 0.7785489 | 0.87861887 |
| AC009065.6 | 110.702018 | 0.737955774 | 0.37073499 | 1.990521 | 0.04653357 | 0.13820141 |
| AC009065.3 | 62.2734861 | 0.874599307 | 0.32422405 | 2.69751525 | 0.00698591 | 0.03457834 |
| RAB26 | 81.15807 | 0.131826913 | 0.57189138 | 0.2305104 | 0.81769518 | 0.90201305 |
| SNHG19 | 471.147996 | -0.265487376 | 0.19599627 | -1.3545532 | 0.17555994 | 0.3497934 |
| TRAF7 | 5254.10923 | -0.328326482 | 0.10472577 | -3.1351069 | 0.00171792 | 0.0119972 |
| CASKIN1 | 1244.7783 | -0.729534845 | 0.12576752 | -5.8006617 | 6.61E-09 | 4.87E-07 |
| MLST8 | 2330.64296 | 0.183397832 | 0.14447725 | 1.26938901 | 0.20430235 | 0.38664081 |

| | | | | | | |
|---|---|---|---|---|---|---|
| BRICD5 | 948.087411 | -0.169307584 | 0.22123615 | -0.7652799 | 0.44410489 | 0.63508925 |
| AC009065.7 | 486.678927 | 0.152649379 | 0.17174513 | 0.88881342 | 0.37410336 | 0.57020436 |
| PGP | 1325.31773 | 0.010846406 | 0.16747997 | 0.06476241 | 0.94836317 | 0.97513596 |
| E4F1 | 2383.33249 | -0.209678016 | 0.17483755 | -1.1992734 | 0.23042166 | 0.41798301 |
| AC009065.1 | 169.340264 | -0.228190139 | 0.22299413 | -1.0233011 | 0.30616551 | 0.50305841 |
| DNASE1L2 | 223.436433 | -0.109641657 | 0.32042452 | -0.3421762 | 0.73221826 | 0.8490202 |
| ECI1 | 4173.0592 | 0.252304373 | 0.11382687 | 2.21656246 | 0.02665301 | 0.09261296 |
| AC009065.8 | 143.373064 | -0.118454533 | 0.22282979 | -0.531592 | 0.5950086 | 0.75401092 |
| RNPS1 | 5303.69822 | -0.461422437 | 0.15564395 | -2.9646024 | 0.00303074 | 0.01851245 |
| AC009065.4 | 105.121913 | -0.193907493 | 0.30819871 | -0.6291639 | 0.52924178 | 0.70378748 |
| ABCA3 | 1939.74321 | -0.381638102 | 0.17585664 | -2.170166 | 0.02999427 | 0.10084776 |
| CCNF | 181.913968 | -0.157319496 | 0.19017148 | -0.8272507 | 0.40809496 | 0.60264172 |
| AC106820.3 | 14.5793675 | 0.567047641 | 0.47704984 | 1.18865493 | 0.23457548 | 0.42330496 |
| AC106820.5 | 39.6123888 | 0.487876114 | 0.27963123 | 1.74471256 | 0.08103492 | 0.2042526 |
| TEDC2 | 65.625439 | 0.195917491 | 0.28090308 | 0.69745582 | 0.48551758 | 0.66964851 |
| AC106820.2 | 18.7531066 | 0.684214642 | 0.49926346 | 1.37044806 | 0.17054708 | 0.34265344 |
| NTN3 | 11.1244199 | 1.122451197 | 0.40792463 | 2.75161422 | 0.00593023 | 0.03054081 |
| TBC1D24 | 2589.49343 | 0.653427521 | 0.18789263 | 3.47766454 | 0.0005058 | 0.00457251 |
| AC093525.2 | 12056.1422 | -0.353806649 | 0.16741297 | -2.1133766 | 0.03456855 | 0.11171233 |
| ATP6V0C | 12321.7268 | -0.402942172 | 0.16986311 | -2.3721582 | 0.01768452 | 0.06852912 |
| AC093525.1 | 4980.91698 | -0.113087236 | 0.19152487 | -0.5904572 | 0.55488419 | 0.72437109 |
| AMDHD2 | 4188.13189 | 0.239511509 | 0.35221898 | 0.68000739 | 0.49649978 | 0.67834305 |
| CEMP1 | 1888.19467 | 0.133201784 | 0.38169199 | 0.34897715 | 0.72710646 | 0.84581576 |
| PDPK1 | 2916.90526 | 0.34697917 | 0.1345529 | 2.57875658 | 0.00991566 | 0.04480398 |
| AC093525.6 | 63.4169645 | -0.261114761 | 1.00485723 | -0.2598526 | 0.79497748 | 0.88885122 |
| AC093525.7 | 18.3652055 | 0.599040772 | 0.44874853 | 1.33491418 | 0.18190444 | 0.3583728 |
| AC093525.4 | 3.71768658 | -0.464586573 | 0.61728938 | -0.7526236 | 0.45167612 | 0.64142922 |
| AC093525.3 | 0.07659028 | -0.642905668 | 3.30558948 | -0.1944905 | 0.84579185 | NA |
| AC141586.3 | 182.466954 | 1.622445538 | 0.34755337 | 4.66819107 | 3.04E-06 | 7.92E-05 |
| AC141586.2 | 35.7971847 | 0.277115937 | 0.45826305 | 0.60470932 | 0.54537215 | 0.71697611 |
| KCTD5 | 1153.03154 | -0.400169231 | 0.16786668 | -2.3838514 | 0.01713252 | 0.0670204 |
| PRSS27 | 98.3707434 | -0.476365478 | 0.2848317 | -1.6724454 | 0.09443652 | 0.22846706 |
| SRRM2-AS1 | 102.823495 | 0.856311376 | 0.40200668 | 2.13009242 | 0.03316398 | 0.10836174 |
| SRRM2 | 39103.4042 | 0.118842376 | 0.16618145 | 0.71513622 | 0.47452488 | 0.66007262 |
| ELOB | 5129.24376 | -0.224222597 | 0.12381448 | -1.8109562 | 0.07014764 | 0.18451803 |
| AC092117.1 | 118.33817 | 0.699134286 | 0.45663757 | 1.53104855 | 0.12575739 | 0.27855712 |
| PRSS33 | 1.60409307 | -0.24558897 | 1.13444757 | -0.2164833 | 0.82861103 | 0.90880809 |
| PRSS41 | 0 | NA | NA | NA | NA | NA |
| PRSS21 | 10.8671029 | -1.964477586 | 0.78483619 | -2.5030415 | 0.01231311 | 0.05258641 |
| ZG16B | 15.4316948 | -0.82288834 | 0.44627435 | -1.8439069 | 0.06519671 | 0.17518212 |
| PRSS22 | 2091.4644 | -1.876787592 | 0.40210532 | -4.6674031 | 3.05E-06 | 7.93E-05 |
| AC003965.2 | 156.312393 | -1.677870069 | 0.41396783 | -4.0531412 | 5.05E-05 | 0.00075033 |
| AC003965.1 | 8.29360618 | 0.845757754 | 0.70514633 | 1.19940744 | 0.23036955 | 0.417946 |
| FLYWCH2 | 1017.47577 | -0.131862496 | 0.19699621 | -0.6693656 | 0.50326226 | 0.68340418 |
| FLYWCH1 | 3696.57143 | 0.155201422 | 0.27852616 | 0.55722387 | 0.57737448 | 0.7411933 |
| AC004034.1 | 30.0814333 | -0.11484979 | 0.34268077 | -0.335151 | 0.73751121 | 0.85254123 |
| KREMEN2 | 7.53312738 | -0.61099671 | 0.52197163 | -1.1705554 | 0.24177753 | 0.43204402 |
| PKMYT1 | 259.905816 | -1.255812969 | 0.34831541 | -3.6053902 | 0.00031168 | 0.00312276 |
| PAQR4 | 293.237958 | -1.171484222 | 0.32733977 | -3.578802 | 0.00034517 | 0.00339128 |
| AC004233.4 | 1.4603736 | -1.236607909 | 1.42887806 | -0.8654398 | 0.38679746 | 0.58241586 |
| AC004233.2 | 13.9314786 | 0.535591934 | 0.7969781 | 0.67202842 | 0.5015656 | 0.68222386 |
| AC004233.3 | 86.7071435 | -0.007182809 | 0.4162223 | -0.0172571 | 0.98623147 | 0.99391317 |
| CLDN9 | 37.751207 | 0.063328618 | 0.66517051 | 0.09520659 | 0.92415073 | 0.96312021 |
| CLDN6 | 11.7512076 | -0.766373587 | 0.61171246 | -1.252833 | 0.21026647 | 0.39360594 |
| TNFRSF12A | 5202.29164 | -1.537178426 | 0.5079882 | -3.0260121 | 0.00247802 | 0.01588755 |
| HCFC1R1 | 2988.72863 | 0.080538604 | 0.22433048 | 0.35901766 | 0.71958188 | 0.840362 |
| THOC6 | 1689.48041 | -0.602430097 | 0.25317987 | -2.379455 | 0.01733826 | 0.0676046 |
| BICDL2 | 1000.15008 | -1.077611412 | 0.39067164 | -2.7583559 | 0.00580929 | 0.03007981 |
| AC108134.1 | 206.523824 | -0.525274578 | 0.36178074 | -1.4519142 | 0.14652548 | 0.30948152 |
| MMP25 | 468.189064 | -0.516522739 | 0.33403528 | -1.5463119 | 0.12202925 | 0.2729125 |
| MMP25-AS1 | 909.367863 | -0.589937764 | 0.29360491 | -2.0092912 | 0.04450626 | 0.13376225 |
| IL32 | 8054.6405 | -1.052375413 | 0.46507056 | -2.2628296 | 0.0236462 | 0.08475605 |
| AC108134.3 | 157.407507 | -0.377623037 | 0.33221646 | -1.1366777 | 0.25567306 | 0.44827783 |
| ZSCAN10 | 1.14905886 | -0.760590399 | 1.17141303 | -0.6492931 | 0.51614894 | 0.69361706 |
| ZNF213-AS1 | 1018.62362 | 0.130250944 | 0.12353991 | 1.05432282 | 0.29173514 | 0.48792134 |
| ZNF205 | 1098.75495 | -0.198941102 | 0.14833132 | -1.3411941 | 0.17985743 | 0.35569525 |
| ZNF213 | 946.687749 | 0.143579336 | 0.17941079 | 0.8002826 | 0.42354708 | 0.61608123 |
| AC108134.2 | 10.8212412 | -0.910405185 | 0.64696301 | -1.4071982 | 0.15936862 | 0.32709765 |
| AJ003147.1 | 0 | NA | NA | NA | NA | NA |
| AJ003147.2 | 24.8757924 | 0.490726045 | 0.40163596 | 1.221818 | 0.22177646 | 0.40772969 |
| OR1F1 | 0 | NA | NA | NA | NA | NA |
| ZNF200 | 375.297277 | -0.15790592 | 0.12745263 | -1.2389381 | 0.21536844 | 0.39983565 |
| MEFV | 65.1098013 | -1.183489323 | 0.53433261 | -2.2148926 | 0.02676744 | 0.09285154 |
| LINC00921 | 78.0820514 | -0.214904404 | 0.26577369 | -0.8085992 | 0.41874571 | 0.61158634 |
| ZNF263 | 2334.41493 | -0.274793118 | 0.17327689 | -1.5858613 | 0.11277077 | 0.25843098 |
| TIGD7 | 572.85833 | 0.145912291 | 0.24174476 | 0.60357994 | 0.54612295 | 0.71733723 |
| ZNF75A | 1002.97866 | 0.065381782 | 0.26528798 | 0.24645587 | 0.80532936 | 0.8944712 |
| AC004232.2 | 33.9742181 | 0.719580351 | 0.28261788 | 2.54612469 | 0.01089263 | 0.04801715 |
| OR2C1 | 3.66268637 | 3.07833163 | 0.83009025 | 3.70843007 | 0.00020855 | 0.00225723 |
| AC025283.3 | 847.783015 | 0.496813149 | 0.17764094 | 2.79672671 | 0.00516232 | 0.02754559 |
| MTRNR2L4 | 1.45485181 | -2.02784422 | 1.29444545 | -1.5665737 | 0.11721436 | 0.26561337 |
| ZSCAN32 | 804.980097 | 0.077760098 | 0.17056642 | 0.45589336 | 0.64846666 | 0.7930031 |
| AC025283.2 | 765.141642 | 0.061931156 | 0.16119663 | 0.38419635 | 0.70083293 | 0.82855702 |
| ZNF174 | 552.326403 | 0.131013191 | 0.13660384 | 0.95907397 | 0.33752148 | 0.53552018 |
| ZNF597 | 220.921522 | 0.087844698 | 0.17605172 | 0.49897097 | 0.61779984 | 0.77149535 |
| NAA60 | 5538.90965 | 0.055906507 | 0.16493457 | 0.33896174 | 0.73463855 | 0.85065526 |
| C16orf90 | 2.54025765 | 0.565078288 | 0.92834652 | 0.60869328 | 0.54272776 | 0.7151932 |
| CLUAP1 | 1163.92733 | 0.186020494 | 0.24133164 | 0.77080857 | 0.44082041 | 0.63214063 |
| NLRC3 | 274.604197 | 0.185210347 | 0.37305564 | 0.49646843 | 0.61956395 | 0.77271053 |
| AC004494.1 | 13.5361753 | 0.989377678 | 0.52879745 | 1.87099557 | 0.0613457 | 0.16798401 |
| AC006111.1 | 109.319056 | -0.084629718 | 0.22171152 | -0.381711 | 0.70267576 | 0.82987743 |
| SLX4 | 459.03428 | 0.152380669 | 0.19640314 | 0.77585659 | 0.43783367 | 0.62980443 |
| DNASE1 | 6698.42674 | 0.93825531 | 0.19653043 | 4.77409676 | 1.81E-06 | 5.24E-05 |
| TRAP1 | 6535.75993 | 0.487542727 | 0.20545939 | 2.37293966 | 0.01764715 | 0.06842453 |
| AC006111.3 | 9.05898559 | 0.140031567 | 0.44944252 | 0.31156724 | 0.75536943 | 0.86405639 |
| CREBBP | 3339.31381 | 0.136135108 | 0.16765741 | 0.81198386 | 0.4168009 | 0.60986792 |
| ADCY9 | 2978.48097 | -0.014959471 | 0.28673104 | -0.0521725 | 0.95839126 | 0.98011944 |
| AC005736.3 | 0 | NA | NA | NA | NA | NA |
| AC009171.2 | 10.1532739 | 0.067100315 | 0.40155966 | 0.16709925 | 0.86729196 | 0.93105 |
| SRL | 185.035735 | 0.311816814 | 0.40798134 | 0.76429184 | 0.44469335 | 0.63537947 |
| LINC01569 | 294.978511 | 0.324098645 | 0.43786098 | 0.74018617 | 0.45918704 | 0.64795611 |
| TFAP4 | 453.786276 | -0.336260203 | 0.20384507 | -1.6495871 | 0.09902741 | 0.23597983 |
| GLIS2 | 6352.13069 | 0.226117095 | 0.26715291 | 0.84639579 | 0.39733199 | 0.59219152 |
| GLIS2-AS1 | 39.5609524 | 0.800981026 | 0.47655946 | 1.68075778 | 0.09280997 | 0.22549282 |
| PAM16 | 1179.8841 | -0.358828539 | 0.15172978 | -2.3649183 | 0.01803405 | 0.06951584 |
| CORO7-PAM16 | 2976.64287 | -0.621628978 | 0.13551436 | -4.5871816 | 4.49E-06 | 0.00010829 |
| CORO7 | 2878.02325 | -0.763313701 | 0.16973746 | -4.4970256 | 6.89E-06 | 0.0001525 |
| VASN | 5686.28321 | -0.272314143 | 0.27679916 | -0.9837968 | 0.32521541 | 0.52279507 |
| DNAJA3 | 3931.4415 | 0.076651793 | 0.20162589 | 0.3801684 | 0.70382042 | 0.83074815 |
| NMRAL1 | 2171.78504 | 0.02958822 | 0.18319329 | 0.16151367 | 0.87168885 | 0.93317628 |
| HMOX2 | 3461.81718 | -0.144447402 | 0.20073388 | -0.7195965 | 0.47177345 | 0.6580622 |
| CDIP1 | 4748.44339 | 0.178057164 | 0.18568842 | 0.95890292 | 0.33760765 | 0.53556601 |
| C16orf96 | 22.6014122 | 2.006423445 | 0.45288528 | 4.43031276 | 9.41E-06 | 0.00019446 |

| | | | | | | |
|---|---|---|---|---|---|---|
| AC023830.2 | 0.27291861 | 1.131425716 | 3.00149036 | 0.37695464 | 0.70620731 | NA |
| UBALD1 | 2468.30289 | -0.348762023 | 0.1814619 | -1.9219573 | 0.05461113 | 0.15494894 |
| MGRN1 | 5853.0332 | -0.196356612 | 0.20742127 | -0.9466561 | 0.34381403 | 0.54151789 |
| AC023830.1 | 11.0162653 | 0.32577688 | 0.53033989 | 0.61427942 | 0.5390307 | 0.7122433 |
| NUDT16L1 | 2009.35664 | 0.372953579 | 0.23834036 | 1.56479405 | 0.11763121 | 0.26621481 |
| ANKS3 | 2052.4572 | -0.238452605 | 0.11978091 | -1.9907397 | 0.04650951 | 0.13816108 |
| C16orf71 | 120.946217 | 1.074301516 | 0.26633649 | 4.03362492 | 5.49E-05 | 0.00079973 |
| ZNF500 | 1029.84497 | 0.200835833 | 0.21151369 | 0.94951696 | 0.34235775 | 0.54014442 |
| 12-Sep | 9.5252257 | 1.674881118 | 0.79022698 | 2.11949373 | 0.03404876 | 0.11042857 |
| SMIM22 | 247.090365 | -0.578617337 | 0.31608828 | -1.8305561 | 0.06716683 | 0.17886383 |
| AC020663.2 | 183.216009 | -0.501329235 | 0.28932599 | -1.7327487 | 0.08314034 | 0.20809952 |
| ROGDI | 2268.77097 | 0.890185954 | 0.26976703 | 3.29983234 | 0.00096743 | 0.00764032 |
| GLYR1 | 8254.72183 | 0.319207023 | 0.0772053 | 4.13452216 | 3.56E-05 | 0.00056436 |
| UBN1 | 2252.0091 | 0.143353137 | 0.16039265 | 0.89376375 | 0.37144831 | 0.56829264 |
| PPL | 1885.5373 | 0.068598323 | 0.28485699 | 0.24081671 | 0.80969718 | 0.89709317 |
| SEC14L5 | 28.8534655 | 2.163767756 | 0.42182511 | 5.12953761 | 2.90E-07 | 1.16E-05 |
| NAGPA-AS1 | 61.1786445 | 1.032110378 | 0.28646287 | 3.60294646 | 0.00031463 | 0.00314528 |
| NAGPA | 1395.66132 | 0.003846839 | 0.17052563 | 0.02255872 | 0.98200227 | 0.99187603 |
| ALG1 | 1849.24816 | -0.085690848 | 0.17080625 | -0.5016845 | 0.61588948 | 0.77026348 |
| C16orf89 | 906.1525 | 1.890610823 | 0.61943552 | 3.05215113 | 0.00227208 | 0.01489673 |
| EEF2KMT | 1311.79859 | -0.20551835 | 0.16184997 | -1.2698078 | 0.2041531 | 0.3864947 |
| AC026458.2 | 3.25549046 | 1.814330948 | 0.70371655 | 2.57821268 | 0.00993129 | 0.0448436 |
| LINC02164 | 0 | NA | NA | NA | NA | NA |
| RBFOX1 | 31.405685 | 0.672067134 | 0.43763539 | 1.53567822 | 0.12461732 | 0.27671962 |
| LINC01570 | 0.38861175 | -1.033955399 | 2.27349217 | -0.4547873 | 0.64926226 | NA |
| AC005774.1 | 0 | NA | NA | NA | NA | NA |
| AC093515.1 | 0.07830766 | -0.642905668 | 3.30558948 | -0.1944905 | 0.84579185 | NA |
| AC018767.2 | 0 | NA | NA | NA | NA | NA |
| LINC02152 | 0 | NA | NA | NA | NA | NA |
| AC018767.3 | 0 | NA | NA | NA | NA | NA |
| AC018767.1 | 0 | NA | NA | NA | NA | NA |
| AC074052.2 | 0 | NA | NA | NA | NA | NA |
| TMEM114 | 0.60625718 | 2.62693962 | 1.54567584 | 1.6995411 | 0.08921728 | NA |
| METTL22 | 771.926444 | -0.522851874 | 0.17355932 | -3.0125255 | 0.00259084 | 0.01643502 |
| ABAT | 8104.22736 | 2.822936486 | 0.47108824 | 5.99237302 | 2.07E-09 | 1.86E-07 |
| TMEM186 | 1072.49384 | 0.913000585 | 0.2320938 | 3.93375689 | 8.36E-05 | 0.00110209 |
| PMM2 | 2182.04622 | -0.819141182 | 0.27842073 | -2.9420983 | 0.00325996 | 0.01956869 |
| AC012173.1 | 705.817402 | -0.916211677 | 0.30666603 | -2.987653 | 0.00281129 | 0.01746084 |
| AC022167.2 | 3386.92391 | -1.171614039 | 0.25455186 | -4.6026537 | 4.17E-06 | 0.00010195 |
| CARHSP1 | 7438.0683 | -1.224966837 | 0.30392987 | -4.030426 | 5.57E-05 | 0.00080669 |
| AC022167.1 | 1569.37924 | -1.234875094 | 0.32254309 | -3.8285585 | 0.0001289 | 0.00154429 |
| AC022167.4 | 14.4577073 | -1.535223765 | 0.95551959 | -1.60669 | 0.10812239 | 0.25086194 |
| AC022167.5 | 0.9260414 | -1.113751525 | 1.30391418 | -0.8541601 | 0.39301626 | NA |
| USP7 | 4199.63993 | -0.107577581 | 0.15042758 | -0.7151453 | 0.47451924 | 0.66007262 |
| AC022167.3 | 3.01700143 | -0.130170343 | 0.69608639 | -0.1870031 | 0.85165817 | 0.92224978 |
| AC087190.2 | 0 | NA | NA | NA | NA | NA |
| C16orf72 | 5333.42535 | -0.197873039 | 0.22496922 | -0.879556 | 0.37909987 | 0.57509426 |
| AC087190.1 | 74.6626999 | -0.558039385 | 0.2502573 | -2.2298626 | 0.02575657 | 0.09025922 |
| LINC02177 | 3.42289554 | 0.78033036 | 0.95827226 | 0.81430966 | 0.41546758 | 0.60903218 |
| LINC01177 | 0 | NA | NA | NA | NA | NA |
| LINC01195 | 0.19271054 | 1.506623853 | 3.27645372 | 0.4598337 | 0.64563559 | NA |
| AC007221.1 | 1.91054931 | 1.331076104 | 0.95870736 | 1.3884071 | 0.16501311 | 0.33518686 |
| AC007221.2 | 0.74517688 | 2.368398571 | 1.71887406 | 1.37787789 | 0.16824099 | NA |
| AC007218.1 | 0.37402421 | -0.542590783 | 2.80717432 | -0.1932872 | 0.84673407 | NA |
| GRIN2A | 81.8606911 | 1.306503598 | 0.51175135 | 2.55300471 | 0.01067981 | 0.0472845 |
| AC007218.2 | 0 | NA | NA | NA | NA | NA |
| AC133565.1 | 1.72051032 | 0.096872131 | 0.98762053 | 0.09808639 | 0.9218637 | 0.96183415 |
| ATF7IP2 | 474.735928 | 0.307733899 | 0.23923399 | 1.28633016 | 0.19832786 | 0.3792535 |
| AC131649.1 | 0 | NA | NA | NA | NA | NA |
| LINC01290 | 1.33339179 | -0.517913042 | 1.08174848 | -0.478774 | 0.63209942 | 0.78149288 |
| EMP2 | 1247.30961 | -0.147979425 | 0.33681939 | -0.4393436 | 0.66041262 | 0.80018573 |
| AC027277.1 | 275.776029 | -0.022747859 | 0.34127946 | -0.0666546 | 0.94685665 | 0.97438741 |
| TEKT5 | 3.80430532 | -0.293424831 | 0.72228234 | -0.4062467 | 0.68456133 | 0.81778936 |
| AC007595.1 | 0.19994613 | 1.506623855 | 3.27645372 | 0.45983371 | 0.64563559 | NA |
| NUBP1 | 1494.17498 | -0.095256256 | 0.17662434 | -0.5393156 | 0.58966912 | 0.74970227 |
| TVP23A | 1008.70506 | -0.113442129 | 0.16505289 | -0.6873078 | 0.49188881 | 0.67487604 |
| AC133065.1 | 21.9480075 | 2.422841986 | 0.42959159 | 5.63987289 | 1.70E-08 | 1.08E-06 |
| CIITA | 4408.66968 | 0.066242531 | 0.17118531 | 0.38696388 | 0.69878295 | 0.82713968 |
| DEXI | 2440.1009 | -0.108399273 | 0.14517892 | -0.7466599 | 0.45526889 | 0.64465663 |
| CLEC16A | 2407.85174 | -0.54732632 | 0.19813268 | -2.7624233 | 0.0057374 | 0.02985121 |
| AC007014.1 | 0.44368594 | -2.044607519 | 2.74620012 | -0.7445224 | 0.45656049 | NA |
| AC007220.1 | 91.5940886 | -0.364172371 | 0.15588653 | -2.3361375 | 0.01948407 | 0.07348327 |
| RMI2 | 181.932324 | -0.267439771 | 0.5615265 | -0.4762728 | 0.63388007 | 0.78295679 |
| SOCS1 | 1096.844 | -2.707163631 | 0.38457088 | -7.0394401 | 1.93E-12 | 4.76E-10 |
| TNP2 | 0.07659028 | -0.642905668 | 3.30558948 | -0.1944905 | 0.84579185 | NA |
| PRM3 | 0 | NA | NA | NA | NA | NA |
| PRM2 | 0.09997307 | 0.547239394 | 3.29377202 | 0.16614368 | 0.86804388 | NA |
| PRM1 | 0 | NA | NA | NA | NA | NA |
| AC009121.2 | 4.32972371 | 0.256463517 | 0.68277431 | 0.37561975 | 0.7071996 | 0.83325953 |
| AC009121.1 | 3.27746121 | 0.226030812 | 0.79517148 | 0.28425417 | 0.7762156 | 0.87711283 |
| AC099489.1 | 226.114799 | -0.192158341 | 0.41593886 | -0.461987 | 0.64409065 | 0.78974795 |
| AC099489.3 | 1.76342952 | -0.295500465 | 1.14002154 | -0.259206 | 0.79547627 | 0.88910734 |
| LITAF | 12469.3771 | -0.921260258 | 0.29395368 | -3.134032 | 0.00172422 | 0.01203804 |
| SNN | 2462.91356 | -0.36460281 | 0.33807799 | -1.0784577 | 0.28082955 | 0.47657168 |
| TXNDC11 | 3820.70138 | 0.085300362 | 0.12693725 | 0.67198844 | 0.50159106 | 0.68222386 |
| AC007613.1 | 108.337412 | 0.650737674 | 0.25159786 | 2.58641971 | 0.00969788 | 0.04404633 |
| ZC3H7A | 2914.55945 | 0.051720993 | 0.10505546 | 0.49232084 | 0.62249255 | 0.77481557 |
| AC010654.1 | 19.868452 | 0.223054933 | 0.44908669 | 0.4966857 | 0.61941071 | 0.77261532 |
| BCAR4 | 0.99413899 | -1.950841626 | 2.01778763 | -0.9668221 | 0.33363299 | NA |
| RSL1D1 | 3907.29911 | -0.22255103 | 0.17550786 | -1.2680402 | 0.20478359 | 0.38711193 |
| AC007216.4 | 7.80893175 | 1.198901623 | 0.53006892 | 2.26178443 | 0.02371073 | 0.08495363 |
| GSPT1 | 6370.56867 | -0.469913092 | 0.19345209 | -2.4290929 | 0.01513666 | 0.06126686 |
| AC007216.2 | 3.85143199 | -0.194560962 | 0.66313931 | -0.2933938 | 0.76922116 | 0.87240608 |
| NPIPB2 | 146.967031 | -0.240052092 | 0.96323776 | -0.2492137 | 0.80319545 | 0.89318485 |
| TNFRSF17 | 13.4629893 | 0.063336367 | 0.76444854 | 0.08285236 | 0.93396893 | 0.96826391 |
| SNX29 | 3724.94972 | 0.430112274 | 0.27837492 | 1.54508273 | 0.12232625 | 0.27336808 |
| AC007216.1 | 29.5539762 | 0.278620119 | 0.62262248 | 0.44749448 | 0.65451808 | 0.79728574 |
| AC007601.1 | 11.8381705 | 0.476998625 | 0.55147228 | 0.86495486 | 0.38706357 | 0.58268347 |
| AC007601.2 | 0.56759961 | 0.907632257 | 1.64885896 | 0.55046082 | 0.58200334 | NA |
| AC007598.2 | 1.51836381 | -0.554283244 | 1.09504441 | -0.5061742 | 0.61273436 | 0.76799695 |
| AC010333.1 | 1.4322056 | 0.413831439 | 1.30690275 | 0.31665052 | 0.75150879 | 0.86135902 |
| AC010333.2 | 0.8912136 | 1.416427812 | 1.57050919 | 0.90189081 | 0.36711487 | NA |
| AC010333.3 | 3.42413804 | 0.332873439 | 0.91390815 | 0.36423074 | 0.71568572 | 0.83823283 |
| CPPED1 | 3665.29287 | 0.008033099 | 0.33080448 | 0.02428353 | 0.98062645 | 0.99131724 |
| AC109597.1 | 0.15543392 | 1.368031615 | 3.28033652 | 0.41704002 | 0.67664913 | NA |
| AC109597.2 | 0.99044742 | 0.829651259 | 1.16809962 | 0.71025728 | 0.47754461 | NA |
| SHISA9 | 274.166904 | -0.378563637 | 0.29923373 | -1.2651102 | 0.20583184 | 0.38839253 |
| AC009134.1 | 8.72643349 | 0.124955228 | 0.63357108 | 0.19722369 | 0.84365249 | 0.9171043 |
| U91319.1 | 17.3064331 | -0.433025067 | 0.51394615 | -0.8425495 | 0.39948045 | 0.59426337 |
| AC003009.1 | 0.09989236 | -0.642905668 | 3.30558948 | -0.1944905 | 0.84579185 | NA |
| U95743.1 | 1.66502093 | 1.372449345 | 1.0279015 | 1.33519539 | 0.18181241 | 0.35827179 |
| ERCC4 | 357.577609 | 0.129064138 | 0.15739937 | 0.81997874 | 0.41222823 | 0.60616749 |
| AC010401.1 | 45.4070869 | 0.226854083 | 0.23494948 | 0.96554408 | 0.33427237 | 0.53240549 |

| | | | | | | |
|---|---|---|---|---|---|---|
| AC010401.2 | 0 | NA | NA | NA | NA | NA |
| LINC02185 | 10.8863838 | 0.206163406 | 0.54179921 | 0.38051626 | 0.70356223 | 0.83059446 |
| LINC02186 | 0.38022378 | -0.558351028 | 2.25171835 | -0.2479666 | 0.80416022 | NA |
| MKL2 | 1478.10933 | -0.04102911 | 0.15883021 | -0.2583206 | 0.79615952 | 0.88938094 |
| AC130650.2 | 75.1836642 | -0.563838114 | 0.34208214 | -1.6482536 | 0.09930065 | 0.23635329 |
| AC130650.1 | 1.05159037 | -1.483532198 | 1.36594815 | -1.0860824 | 0.27744255 | 0.47295624 |
| MIR193BHG | 178.775467 | -0.233010142 | 0.34606853 | -0.6733064 | 0.50075241 | 0.68176568 |
| LINC02130 | 0.14444843 | 0.435199075 | 3.28898345 | 0.13232024 | 0.894731 | NA |
| AC040173.1 | 0.69183758 | 0.216649408 | 1.42001498 | 0.1525684 | 0.87873865 | NA |
| PARN | 1552.8165 | 0.030308956 | 0.11794526 | 0.25697476 | 0.79719826 | 0.88994647 |
| BFAR | 2643.12414 | -0.515140613 | 0.19338029 | -2.6638735 | 0.00772466 | 0.03722575 |
| PLA2G10 | 4.21530676 | -0.747538957 | 0.71309523 | -1.0483017 | 0.29449962 | 0.49123632 |
| AC009167.1 | 0 | NA | NA | NA | NA | NA |
| NPIPA3 | 9.07087585 | -1.526626676 | 0.66744119 | -2.2872827 | 0.02217933 | 0.08092831 |
| AC136443.4 | 0.09635527 | 0.547239394 | 3.29377202 | 0.16614368 | 0.86804388 | NA |
| NPIPA2 | 11.188928 | -0.930626285 | 0.55710696 | -1.6704625 | 0.09482789 | 0.22913618 |
| NOMO1 | 3177.41152 | 0.1197589 | 0.17692821 | 0.67687851 | 0.49848304 | 0.6799953 |
| AC136443.3 | 2.28504593 | -0.921214734 | 0.84253066 | -1.0933902 | 0.27422253 | 0.46934822 |
| AC138932.2 | 7.35226294 | -0.416985524 | 0.47048798 | -0.886283 | 0.37546502 | 0.57142106 |
| NPIPA1 | 1602.29696 | 0.188669315 | 0.27186275 | 0.69398737 | 0.48769013 | 0.67149064 |
| PDXDC1 | 8980.87802 | 0.374412155 | 0.1571003 | 2.38326826 | 0.01715968 | 0.06706138 |
| AC138932.5 | 498.372166 | 1.156852609 | 0.22655751 | 5.10622044 | 3.29E-07 | 1.28E-05 |
| NTAN1 | 1370.74489 | -0.584876167 | 0.22664071 | -2.5806315 | 0.00986198 | 0.04463024 |
| RRN3 | 2206.39512 | -0.183109675 | 0.17125838 | -1.0692013 | 0.28497898 | 0.48109992 |
| AC126763.1 | 0.23387521 | 1.816617118 | 3.25499482 | 0.55810139 | 0.57677515 | NA |
| NPIPA5 | 248.519602 | -0.640663977 | 0.70301326 | -0.9113114 | 0.36213133 | 0.55942609 |
| MPV17L | 860.656998 | 1.172506259 | 0.34817587 | 3.367569 | 0.00075834 | 0.00629246 |
| AC140504.1 | 270.697309 | 0.522396764 | 0.20560012 | 2.54083886 | 0.01105869 | 0.04853821 |
| C16orf45 | 1494.59425 | -0.03461288 | 0.19144962 | -0.1807937 | 0.85652953 | 0.92488775 |
| MARF1 | 3022.94138 | 0.323461753 | 0.11238221 | 2.87822917 | 0.00399914 | 0.02274887 |
| AC026401.1 | 23.2031792 | 1.918169693 | 0.53433998 | 3.58979254 | 0.00033094 | 0.00327833 |
| NDE1 | 1479.84383 | 0.516674792 | 0.52063659 | 0.99239047 | 0.32100709 | 0.51855899 |
| AC026401.3 | 90.2536545 | -0.918159855 | 0.34157881 | -2.6879883 | 0.00718839 | 0.0353389 |
| MYH11 | 7301.1862 | 0.225948538 | 0.80142554 | 0.28193329 | 0.77799466 | 0.87847694 |
| AF001548.2 | 461.508623 | 0.807744932 | 0.59652887 | 1.35407518 | 0.17571239 | 0.34999137 |
| AF001548.1 | 105.417754 | 1.447482156 | 0.7245564 | 1.99774945 | 0.04574383 | 0.13654798 |
| FOPNL | 1647.3631 | -0.195219216 | 0.15163923 | -1.2873925 | 0.19795751 | 0.37887498 |
| AC130651.1 | 1.1047933 | 2.120052839 | 1.35704725 | 1.56225425 | 0.11822811 | 0.26719876 |
| ABCC1 | 1747.4307 | -0.456617544 | 0.19206805 | -2.3773738 | 0.01743641 | 0.06782683 |
| ABCC6 | 3467.63722 | 2.222192134 | 0.38584092 | 5.75934802 | 8.44E-09 | 5.95E-07 |
| AC136624.1 | 2.90909257 | 0.525886603 | 0.6932716 | 0.75855783 | 0.44811711 | 0.63822176 |
| NOMO3 | 126.417092 | 0.540658506 | 0.46185686 | 1.1706192 | 0.24175187 | 0.43204402 |
| AC136624.2 | 0.38334918 | 0.761233862 | 2.85389455 | 0.26673511 | 0.78967312 | NA |
| AC138969.3 | 0.07830766 | -0.642905668 | 3.30558948 | -0.1944905 | 0.84579185 | NA |
| AC138969.1 | 1116.12525 | 0.482338805 | 0.2789217 | 1.72929822 | 0.08375574 | 0.20921657 |
| NPIPA7 | 2.50035377 | -1.088190815 | 0.9348835 | -1.1639855 | 0.2444299 | 0.43542636 |
| AC092326.1 | 0 | NA | NA | NA | NA | NA |
| XYLT1 | 253.855925 | 0.038565301 | 0.37825782 | 0.10195507 | 0.91879234 | 0.96046034 |
| AC109446.3 | 10.8394943 | -0.68004814 | 0.69522288 | -0.9781728 | 0.32798885 | 0.52568381 |
| AC109495.1 | 0 | NA | NA | NA | NA | NA |
| AC025277.1 | 0 | NA | NA | NA | NA | NA |
| AC010601.1 | 0 | NA | NA | NA | NA | NA |
| AC091489.1 | 0 | NA | NA | NA | NA | NA |
| NPIPA8 | 37.4815866 | -0.09277127 | 1.02025513 | -0.0909295 | 0.92754862 | 0.96487011 |
| AC126755.2 | 113.053467 | -0.05822903 | 0.65071193 | -0.0894851 | 0.92869639 | 0.96531469 |
| AC126755.4 | 0.08695548 | -0.642905668 | 3.30558948 | -0.1944905 | 0.84579185 | NA |
| AC126755.3 | 0.28283252 | -0.695596956 | 2.3567663 | -0.2951489 | 0.76788014 | NA |
| NOMO2 | 409.930656 | 0.361066093 | 0.24830548 | 1.45412053 | 0.14591289 | 0.30853364 |
| AC136618.1 | 0.9550908 | 0.661428029 | 1.23539429 | 0.53539832 | 0.59237445 | NA |
| RPS15A | 16347.1982 | -0.388979442 | 0.18994004 | -2.0479065 | 0.04056917 | 0.12510138 |
| AC138811.2 | 16532.8475 | -0.307348483 | 0.19228435 | -1.5984061 | 0.10995262 | 0.25384617 |
| ARL6IP1 | 9192.60522 | -0.3310141 | 0.21224973 | -1.5595502 | 0.11886622 | 0.268116 |
| AC138811.1 | 45.6756362 | 0.444689938 | 0.38268886 | 1.16201434 | 0.24522964 | 0.43623217 |
| SMG1 | 4979.40832 | -0.114868346 | 0.18683 | -0.6148282 | 0.53866821 | 0.71200553 |
| AC092287.1 | 24.0123932 | 0.279563805 | 0.42808933 | 0.65305016 | 0.51372394 | 0.69166243 |
| TMC7 | 79.5254155 | 0.376200888 | 0.30994455 | 1.21376838 | 0.22483618 | 0.41104619 |
| AC099518.4 | 72.2536527 | -0.036628225 | 0.22621473 | -0.1619179 | 0.87137047 | 0.93311835 |
| AC099518.2 | 2.65959841 | 0.219100725 | 0.72971474 | 0.30025531 | 0.76398242 | 0.86935843 |
| COQ7 | 1010.77998 | 0.333695048 | 0.17804737 | 1.87419255 | 0.06090389 | 0.1672257 |
| AC099518.1 | 3.34321162 | 0.74378318 | 0.74493358 | 0.9984557 | 0.31805844 | 0.51524764 |
| AC099518.6 | 6.33485502 | -0.914704159 | 0.65218738 | -1.4025174 | 0.16076079 | 0.32917177 |
| ITPRIPL2 | 1416.2618 | -0.463339637 | 0.20310655 | -2.2812638 | 0.02253284 | 0.08186704 |
| AC099518.3 | 164.634801 | -0.107819468 | 0.21302698 | -0.5061306 | 0.61276499 | 0.76799695 |
| AC099518.5 | 775.869588 | -0.913994471 | 0.2359832 | -3.8731336 | 0.00010744 | 0.00134153 |
| SYT17 | 427.118644 | 0.020083686 | 0.3313453 | 0.06061256 | 0.95166777 | 0.97678185 |
| CLEC19A | 0.61770576 | -0.538696707 | 2.20963903 | -0.243794 | 0.80739038 | NA |
| AC130456.2 | 0.41568007 | -0.671407531 | 3.2845786 | -0.2044121 | 0.83803149 | NA |
| TMC5 | 26.5549037 | -0.489400546 | 0.47122972 | -1.0385604 | 0.29900921 | 0.49579957 |
| AC130456.4 | 0.3041953 | 0.328495401 | 2.89713094 | 0.11338645 | 0.90972418 | NA |
| AC130456.3 | 6.75323845 | -2.744513957 | 1.03245502 | -2.6582407 | 0.00785498 | 0.03768092 |
| AC130456.5 | 936.370472 | -0.752597418 | 0.2133622 | -3.5273232 | 0.00041978 | 0.00396054 |
| GDE1 | 3869.82577 | -0.328067363 | 0.18619466 | -1.761959 | 0.07807622 | 0.19892464 |
| CCP110 | 712.218697 | -0.274074145 | 0.2052969 | -1.3350135 | 0.18187192 | 0.35836228 |
| VPS35L | 2820.69105 | 0.321290883 | 0.10958897 | 2.93178112 | 0.00337024 | 0.02002552 |
| AC002550.2 | 12.3387119 | -0.894903937 | 0.4941344 | -1.8110537 | 0.07013254 | 0.18450489 |
| KNOP1 | 595.551617 | -0.330442291 | 0.23750251 | -1.3913212 | 0.16412804 | 0.33369748 |
| AC002550.1 | 280.693032 | -0.355812027 | 0.28048507 | -1.2685596 | 0.20459818 | 0.38700625 |
| IQCK | 2608.39824 | -0.036642566 | 0.20040652 | -0.1828412 | 0.85492263 | 0.92401312 |
| AC027130.1 | 3.27370729 | 0.440199498 | 0.89819321 | 0.49009444 | 0.62406707 | 0.77600079 |
| GPRC5B | 6094.00156 | 0.003915729 | 0.24068373 | 0.01626919 | 0.98701964 | 0.99410333 |
| GPR139 | 0 | NA | NA | NA | NA | NA |
| AC092132.1 | 0 | NA | NA | NA | NA | NA |
| GP2 | 950.231119 | 2.783828246 | 0.8111409 | 3.43199098 | 0.00059917 | 0.00521033 |
| UMOD | 294924.126 | 1.021093272 | 0.5054509 | 2.02016314 | 0.04336647 | 0.13138485 |
| PDILT | 13.334952 | 1.757101513 | 0.66105096 | 2.65804244 | 0.0078596 | 0.03769006 |
| AC106796.1 | 1.01165038 | 2.491667356 | 1.29362539 | 1.92611198 | 0.05409039 | 0.15398507 |
| ACSM5 | 1658.50663 | 1.510379962 | 0.62190253 | 2.42864419 | 0.0151554 | 0.06131449 |
| AC137056.1 | 576.888229 | 1.775152521 | 0.64451681 | 2.75423775 | 0.0058829 | 0.03036448 |
| ACSM2A | 15122.4083 | 1.888650899 | 0.53557287 | 3.52641257 | 0.00042123 | 0.00397276 |
| ACSM2B | 20136.2908 | 1.547749196 | 0.51057663 | 3.03137493 | 0.00243443 | 0.01566517 |
| AC141273.1 | 52.5134629 | 1.90228437 | 0.74803051 | 2.54305721 | 0.01098872 | 0.04832766 |
| ACSM3 | 4242.52747 | 0.763916786 | 0.37386513 | 2.04329509 | 0.04102324 | 0.12622902 |
| ACSM1 | 82.0323443 | -0.493991033 | 0.44108577 | -1.1199432 | 0.26273795 | 0.45630758 |
| THUMPD1 | 2605.65753 | 0.04207788 | 0.11517746 | 0.36533085 | 0.71486445 | 0.83793782 |
| AC004381.1 | 22.1429082 | 1.1686328 | 0.47916805 | 2.43887882 | 0.01473291 | 0.06004742 |
| ERI2 | 2016.79532 | 0.662491218 | 0.2912287 | 2.27481435 | 0.02291707 | 0.08293168 |
| REXO5 | 107.044194 | 0.032766993 | 0.27304598 | 0.1200054 | 0.90447887 | 0.95198801 |
| DCUN1D3 | 1403.69428 | -0.530732192 | 0.22252824 | -2.3850106 | 0.01707863 | 0.06688898 |
| LYRM1 | 2062.58135 | 0.185453838 | 0.25816625 | 0.71835043 | 0.47254125 | 0.65875468 |
| DNAH3 | 18.0155064 | -0.922175831 | 0.44278878 | -2.082654 | 0.03728277 | 0.11758721 |
| AC008551.1 | 1.10413098 | -0.092594028 | 1.58135695 | -0.0585535 | 0.95330773 | 0.97744392 |
| TMEM159 | 1394.03782 | -0.634854582 | 0.22028523 | -2.8819662 | 0.00395202 | 0.02256709 |
| AF001550.1 | 1.79544362 | 3.420646037 | 1.4158769 | 2.41592051 | 0.01569549 | 0.06292034 |

| | | | | | | |
|---|---|---|---|---|---|---|
| ZP2 | 6.93885741 | 3.698732566 | 0.90255459 | 4.09807076 | 4.17E-05 | 0.00064283 |
| ANKS4B | 914.146652 | 1.261080053 | 0.65826693 | 1.91575788 | 0.05539592 | 0.15646856 |
| CRYM | 2501.11307 | 0.663377112 | 0.61613149 | 1.07668107 | 0.28162277 | 0.47756066 |
| NPIPB3 | 358.428274 | -0.186181621 | 0.30787642 | -0.6047284 | 0.54535946 | 0.71697611 |
| AC005632.4 | 0.08666371 | -0.642905668 | 3.30558948 | -0.1944905 | 0.84579185 | NA |
| METTL9 | 8264.10461 | 0.250151817 | 0.14862947 | 1.68305663 | 0.09236413 | 0.22475243 |
| IGSF6 | 171.03166 | -0.166688284 | 0.46669164 | -0.3571701 | 0.72096449 | 0.84122492 |
| OTOA | 23.5861034 | 0.139557266 | 0.4190796 | 0.33300897 | 0.73912751 | 0.85357616 |
| AC092375.2 | 53.435929 | 0.46543084 | 0.35435798 | 1.31344817 | 0.18903198 | 0.36728361 |
| NPIPB4 | 911.760251 | -0.259188787 | 0.244836 | -1.058622 | 0.28977194 | 0.48593234 |
| AC092119.2 | 155.870378 | 0.150139493 | 0.26715112 | 0.56200212 | 0.57411458 | 0.73864963 |
| UQCRC2 | 11723.5945 | 0.414170057 | 0.14625593 | 2.83181727 | 0.00462843 | 0.02542743 |
| PDZD9 | 10.5727652 | 0.733299747 | 0.35879279 | 2.04379735 | 0.04097357 | 0.12613508 |
| AC092119.3 | 6.99318388 | 0.539342107 | 0.63292123 | 0.85214729 | 0.39413235 | 0.58948004 |
| MOSMO | 926.756311 | -0.021797624 | 0.16004176 | -0.1361996 | 0.89166349 | 0.94385074 |
| AC009019.1 | 172.484812 | 0.007334067 | 0.16807119 | 0.04363667 | 0.96519402 | 0.98359669 |
| VWA3A | 70.6040709 | -1.321329784 | 0.70544701 | -1.873039 | 0.061063 | 0.16745495 |
| SDR42E2 | 14.3348897 | 0.476411467 | 0.40952904 | 1.16331546 | 0.24470154 | 0.435748 |
| EEF2K | 1284.15198 | -0.045860941 | 0.18589597 | -0.2467022 | 0.80513872 | 0.89442108 |
| AC009034.1 | 148.465695 | -0.280085792 | 0.25119491 | -1.1150138 | 0.26484451 | 0.45883645 |
| POLR3E | 2281.29635 | -0.161054029 | 0.12156427 | -1.3248467 | 0.18522199 | 0.36264428 |
| CDR2 | 2239.60475 | 0.117728172 | 0.18872972 | 0.62379245 | 0.5327639 | 0.70682367 |
| AC092338.1 | 6.65612624 | 0.406860648 | 0.56644876 | 0.71826558 | 0.47259355 | 0.65875468 |
| AC092338.2 | 3.34820864 | -0.837191855 | 0.82209357 | -1.0183657 | 0.30850422 | 0.50550464 |
| NPIPB5 | 1096.59162 | -0.014048022 | 0.26707352 | -0.0525998 | 0.95805075 | 0.97993865 |
| AC009021.2 | 0.21442573 | -1.003574173 | 3.30558948 | -0.3035992 | 0.7614333 | NA |
| AC009021.1 | 6.31721194 | 0.233336234 | 0.62974418 | 0.37052543 | 0.71099103 | 0.83587002 |
| AC130466.1 | 4.22534378 | -1.708876076 | 0.85204728 | -2.0056118 | 0.04489768 | 0.13464709 |
| HS3ST2 | 54.2841946 | -4.065730356 | 0.85403963 | -4.7605875 | 1.93E-06 | 5.52E-05 |
| AC127459.2 | 0.75033858 | 2.401610864 | 2.07501479 | 1.15739458 | 0.24711119 | NA |
| USP31 | 1451.68499 | -0.737226938 | 0.25550015 | -2.8854266 | 0.00390884 | 0.02237863 |
| SCNN1G | 3957.17598 | 0.984066177 | 0.34649992 | 2.8400185 | 0.00451109 | 0.0249102 |
| AC099482.1 | 166.631991 | 1.448160053 | 0.3374596 | 4.29135832 | 1.78E-05 | 0.0003234 |
| SCNN1B | 1856.30945 | 1.098745513 | 0.26870688 | 4.08901149 | 4.33E-05 | 0.00066187 |
| COG7 | 2254.11674 | 0.050010986 | 0.10544801 | 0.47427152 | 0.6353063 | 0.78365737 |
| AC008915.2 | 238.98979 | 0.237959275 | 0.33031779 | 0.72039497 | 0.47128185 | 0.65779401 |
| GGA2 | 7070.17043 | 0.194236702 | 0.262278 | 0.74057564 | 0.45895078 | 0.64778312 |
| EARS2 | 1436.41506 | 0.535825826 | 0.17147353 | 3.12483106 | 0.00177907 | 0.01232312 |
| UBFD1 | 2655.93721 | -0.122602275 | 0.13144163 | -0.9327507 | 0.3509487 | 0.54857388 |
| NDUFAB1 | 2924.21731 | 0.124237439 | 0.15954849 | 0.7786814 | 0.43616742 | 0.6283041 |
| PALB2 | 508.141926 | -0.125460867 | 0.19493433 | -0.6436058 | 0.51983107 | 0.69650655 |
| AC008870.4 | 71.2038041 | -0.186470745 | 0.28129729 | -0.6628956 | 0.5073974 | 0.6863686 |
| DCTN5 | 3196.65174 | -0.038993822 | 0.13288124 | -0.2934487 | 0.76917923 | 0.87240608 |
| PLK1 | 210.251017 | -0.630114823 | 0.15885028 | -3.9667216 | 7.29E-05 | 0.00098591 |
| AC008870.3 | 32.8260726 | -0.371002992 | 0.23296199 | -1.5925473 | 0.11126178 | 0.25595779 |
| ERN2 | 5.13967925 | -1.821521247 | 0.90779783 | -2.0065274 | 0.04480001 | 0.1344306 |
| AC012317.1 | 0 | NA | NA | NA | NA | NA |
| CHP2 | 75.9656642 | 4.415819554 | 0.9956287 | 4.43520716 | 9.20E-06 | 0.00019078 |
| PRKCB | 244.404504 | 0.315894097 | 0.49283018 | 0.64097961 | 0.52153593 | 0.69776958 |
| LINC2194 | 0 | NA | NA | NA | NA | NA |
| CACNG3 | 0.16783122 | 1.368031398 | 3.28033652 | 0.41703996 | 0.67664918 | NA |
| AC008938.1 | 0 | NA | NA | NA | NA | NA |
| RBBP6 | 2513.9132 | -0.33951256 | 0.18191762 | -1.8662984 | 0.06199963 | 0.16928249 |
| TNRC6A | 3240.99398 | 0.286975116 | 0.15828322 | 1.81304827 | 0.06982438 | 0.1838144 |
| LINC01567 | 3.75960934 | 1.559565654 | 0.99695835 | 1.56432377 | 0.11774155 | 0.2664188 |
| AC008731.1 | 2.85678674 | 1.739055121 | 0.79245684 | 2.19451084 | 0.02819871 | 0.0965965 |
| SLC5A11 | 774.170884 | 1.089773317 | 0.89894964 | 1.21227404 | 0.2254075 | 0.41186223 |
| ARHGAP17 | 2777.77331 | -0.350610849 | 0.14526075 | -2.4136655 | 0.01579296 | 0.06321497 |
| LINC02175 | 10.8345268 | 0.314267463 | 0.50631004 | 0.62070162 | 0.53479596 | 0.70849382 |
| LCMT1 | 1642.65939 | -0.211593024 | 0.11935512 | -1.7728023 | 0.07626147 | 0.19598627 |
| LCMT1-AS2 | 13.7250814 | 0.562664201 | 0.62342915 | 0.90253111 | 0.3667748 | 0.56385773 |
| AQP8 | 5.00664752 | -0.889802418 | 0.79007692 | -1.1262225 | 0.26007134 | 0.45303747 |
| ZKSCAN2 | 233.639173 | 0.146467484 | 0.17271222 | 0.84804353 | 0.39641373 | 0.59135803 |
| AC008741.1 | 19.311428 | 0.893780607 | 0.42559348 | 2.10008059 | 0.03572175 | 0.11419355 |
| AC008741.2 | 398.678627 | -0.037291468 | 0.23692015 | -0.157401 | 0.87492883 | 0.93495454 |
| LINC02191 | 0 | NA | NA | NA | NA | NA |
| HS3ST4 | 1.21281903 | -0.309049529 | 1.62463756 | -0.1902268 | 0.84913145 | 0.92060599 |
| AC009158.1 | 0 | NA | NA | NA | NA | NA |
| LINC02195 | 1.97908529 | -3.343856204 | 1.94320616 | -1.7207933 | 0.08528833 | 0.21187502 |
| AC009035.1 | 9.84131822 | 3.22909326 | 1.00122712 | 3.22513564 | 0.00125913 | 0.00940336 |
| C16orf82 | 0 | NA | NA | NA | NA | NA |
| AC092725.1 | 0 | NA | NA | NA | NA | NA |
| LINC02129 | 0 | NA | NA | NA | NA | NA |
| KDM8 | 314.561576 | 0.806591849 | 0.19136251 | 4.21499403 | 2.50E-05 | 0.00042468 |
| AC109449.1 | 80.4978863 | 0.94352398 | 0.20374507 | 4.63090469 | 3.64E-06 | 9.14E-05 |
| NSMCE1 | 2066.05106 | 0.023912693 | 0.23454351 | 0.10195419 | 0.91879304 | 0.96046034 |
| AC106739.1 | 42.1112849 | 0.467254432 | 0.39462873 | 1.18403551 | 0.23639901 | 0.42570859 |
| AC106739.2 | 51.9090463 | -0.979048964 | 0.35880397 | -2.7286459 | 0.0063595 | 0.03216369 |
| IL4R | 5439.03783 | -1.607357062 | 0.28474921 | -5.6448165 | 1.65E-08 | 1.06E-06 |
| IL21R | 145.783149 | -0.370226316 | 0.49320149 | -0.7506594 | 0.45285768 | 0.64251894 |
| IL21R-AS1 | 85.2246206 | -0.362017375 | 0.47030628 | -0.7697481 | 0.44144932 | 0.63269844 |
| GTF3C1 | 4271.12542 | -0.138847835 | 0.12921058 | -1.0745856 | 0.2825603 | 0.47857314 |
| KIAA0556 | 1870.17527 | -0.142442777 | 0.11346301 | -1.2554116 | 0.20932939 | 0.3925052 |
| AC016597.1 | 12.9244466 | 0.933529048 | 0.58072564 | 1.60752167 | 0.10793998 | 0.25049523 |
| GSG1L | 7.72669029 | 2.048892419 | 0.58789345 | 3.48514247 | 0.00049188 | 0.0044686 |
| XPO6 | 3760.60577 | -0.481952314 | 0.22184921 | -2.172432 | 0.0298231 | 0.1004772 |
| AC138904.3 | 0.6645014 | 0.966364914 | 1.84240624 | 0.5245124 | 0.5999222 | NA |
| AC138904.1 | 3.36226749 | 1.291284682 | 0.79776082 | 1.61863637 | 0.10552552 | 0.24651914 |
| SBK1 | 395.143883 | -0.514424026 | 0.35897259 | -1.4330454 | 0.15184485 | 0.3168842 |
| NPIPB6 | 26.253691 | -0.80283549 | 0.33732779 | -2.3799862 | 0.01731329 | 0.06753427 |
| EIF3CL | 26.1611566 | 0.877793633 | 0.27114237 | 3.23739013 | 0.00120628 | 0.0090833 |
| AC138894.2 | 0 | NA | NA | NA | NA | NA |
| AC138894.1 | 3368.15037 | -0.349847018 | 0.18263857 | -1.9155155 | 0.05542679 | 0.15653899 |
| NPIPB7 | 17.787043 | -0.020250041 | 0.36841174 | -0.0549658 | 0.95616572 | 0.97903588 |
| CLN3 | 3465.13391 | -0.349586595 | 0.17620644 | -1.9839604 | 0.04726025 | 0.13977697 |
| APOBR | 330.651327 | -0.348411198 | 0.47970515 | -0.7263028 | 0.46765316 | 0.65456549 |
| IL27 | 7.76498758 | -1.036878393 | 0.51221278 | -2.0243118 | 0.04293808 | 0.13047673 |
| NUPR1 | 5213.75634 | -1.453526099 | 0.49367969 | -2.9442696 | 0.00323718 | 0.01946737 |
| AC020765.2 | 187.653391 | -0.163070956 | 0.19679678 | -0.8286261 | 0.407316 | 0.6019347 |
| SGF29 | 767.517522 | -0.263067317 | 0.17785689 | -1.4790954 | 0.13911481 | 0.29863706 |
| SULT1A2 | 360.313339 | 1.232120665 | 0.48232081 | 2.55456668 | 0.01063201 | 0.04712552 |
| SULT1A1 | 1964.15941 | 1.132393201 | 0.40527097 | 2.7941631 | 0.00520342 | 0.02769208 |
| NPIPB8 | 1.59110374 | 0.527669644 | 1.11279122 | 0.47418567 | 0.63536751 | 0.78367934 |
| EIF3C | 433.270194 | 1.010915443 | 0.20314634 | 4.97629168 | 6.48E-07 | 2.26E-05 |
| NPIPB9 | 5.43812413 | 0.04496018 | 0.59445938 | 0.07563205 | 0.93971184 | 0.97115066 |
| AC145285.2 | 35.8421841 | 0.34309046 | 0.24866409 | 1.37973462 | 0.16766837 | 0.33885767 |
| AC145285.3 | 6.92564924 | 1.021393781 | 0.48061399 | 2.1251853 | 0.03357115 | 0.10936026 |
| AC145285.6 | 239.548425 | -0.469596254 | 0.2165245 | -2.1687903 | 0.0300986 | 0.10105681 |
| ATXN2L | 9937.50964 | -0.506183184 | 0.15698971 | -3.224308 | 0.00126278 | 0.00942791 |
| AC133550.1 | 1216.88398 | -0.76074963 | 0.13694712 | -5.5550611 | 2.78E-08 | 1.63E-06 |
| TUFM | 13534.7096 | 0.215046953 | 0.16600309 | 1.29543946 | 0.19516867 | 0.37530795 |
| SH2B1 | 6485.19775 | 0.16812188 | 0.17140989 | 0.98081787 | 0.32668257 | 0.5246099 |
| AC133550.2 | 45.9242767 | 0.93031529 | 0.20553696 | 4.52626755 | 6.00E-06 | 0.00013648 |

| | | | | | | |
|---|---|---|---|---|---|---|
| ATP2A1 | 320.325564 | 0.135736609 | 0.21784308 | 0.6230935 | 0.53322308 | 0.70704946 |
| ATP2A1-AS1 | 29.4841264 | -0.009086181 | 0.28790217 | -0.03156 | 0.97482297 | 0.98843899 |
| RABEP2 | 2106.10882 | -0.128333191 | 0.20425213 | -0.6283077 | 0.52980238 | 0.70439088 |
| CD19 | 108.054098 | -0.362260007 | 0.82603711 | -0.4385517 | 0.66098638 | 0.80054245 |
| NFATC2IP | 2676.13355 | -0.070979508 | 0.18676036 | -0.3800566 | 0.7039034 | 0.83080891 |
| AC109460.2 | 376.144827 | -0.293886158 | 0.15381995 | -1.9105854 | 0.05605788 | 0.15778058 |
| AC109460.1 | 451.458896 | -0.221324564 | 0.24421899 | -0.9062545 | 0.36480116 | 0.56191044 |
| SPNS1 | 3683.4646 | -0.458133322 | 0.2568198 | -1.7838707 | 0.07444468 | 0.19272592 |
| LAT | 1040.13814 | -0.064914205 | 0.19695791 | -0.3295841 | 0.7417142 | 0.85485155 |
| AC109460.4 | 179.18485 | -0.153488429 | 0.24267147 | -0.6324947 | 0.52706364 | 0.70181116 |
| AC009093.2 | 27.4512154 | -1.165419429 | 0.70215245 | -1.6597812 | 0.09695848 | 0.23248405 |
| AC009093.1 | 23.9032117 | -0.698046925 | 0.65214787 | -1.0703814 | 0.28444768 | 0.48060335 |
| AC009093.5 | 0.61935239 | 2.07824295 | 1.67590118 | 1.24007488 | 0.2149477 | NA |
| AC009093.4 | 7.76546253 | 1.279126831 | 1.00487849 | 1.27291692 | 0.20304752 | 0.38512287 |
| AC009093.6 | 30.0651989 | 0.246118287 | 0.65312742 | 0.37683043 | 0.70629963 | 0.83289156 |
| NPIPB11 | 257.090245 | -0.177473554 | 0.33979043 | -0.522303 | 0.6014594 | 0.75891116 |
| BOLA2 | 457.939578 | -0.181344083 | 0.31909521 | -0.5683071 | 0.56982644 | 0.73517898 |
| BOLA2-SMG | 460.685158 | -0.197072621 | 0.31960445 | -0.6166141 | 0.53748927 | 0.71113301 |
| SLX1B | 20.2354839 | -0.583464161 | 0.58644933 | -0.9949098 | 0.31978016 | 0.51721067 |
| SULT1A4 | 25.8623662 | -0.196610208 | 0.32729388 | -0.6007146 | 0.54803011 | 0.71876822 |
| AC133555.3 | 0.15661533 | -1.003574173 | 3.30558948 | -0.3035992 | 0.7614333 | NA |
| NPIPB12 | 69.1299195 | 0.082339278 | 0.42545588 | 0.19353188 | 0.84654244 | 0.91937416 |
| SPN | 225.919847 | 0.218932824 | 0.54959698 | 0.39835158 | 0.69037105 | 0.82160324 |
| AC009086.2 | 3065.66551 | 1.361417432 | 0.36960315 | 3.68345734 | 0.00023009 | 0.00244461 |
| QPRT | 10183.9619 | 1.240594314 | 0.34360303 | 3.61054528 | 0.00030555 | 0.00307771 |
| C16orf54 | 97.5223602 | -0.205917604 | 0.51139849 | -0.4026559 | 0.6872014 | 0.81934716 |
| AC009133.3 | 10.3147457 | 0.920810469 | 0.69889548 | 1.31752243 | 0.18766357 | 0.36570265 |
| ZG16 | 3.583109 | 1.787954999 | 1.2194256 | 1.46622721 | 0.1425864 | 0.30378955 |
| KIF22 | 1761.27538 | 0.112070714 | 0.18968933 | 0.59081191 | 0.55464646 | 0.72417116 |
| AC009133.4 | 807.589021 | -0.022103123 | 0.1836054 | -0.1203838 | 0.90417909 | 0.95178646 |
| MAZ | 6071.39855 | -0.567268267 | 0.2114026 | -2.6833552 | 0.00728875 | 0.03567931 |
| AC009133.2 | 1066.15106 | -0.368632238 | 0.20923805 | -1.761784 | 0.0781058 | 0.19898077 |
| AC009133.1 | 5408.86811 | -0.568272344 | 0.20801248 | -2.7319146 | 0.00629675 | 0.03193176 |
| PRRT2 | 881.690795 | 0.204797719 | 0.2132141 | 0.96052616 | 0.33679047 | 0.53481661 |
| AC009133.6 | 1761.47248 | 0.086640134 | 0.17395167 | 0.49807016 | 0.61843459 | 0.77198202 |
| PAGR1 | 1617.46739 | 0.06804688 | 0.19314512 | 0.3523096 | 0.72460609 | 0.84389181 |
| AC120114.5 | 2206.95372 | -0.080672181 | 0.17670924 | -0.456525 | 0.64801252 | 0.79277053 |
| MVP | 25862.9381 | -0.489181599 | 0.14664904 | -3.3357299 | 0.00085076 | 0.00691995 |
| CDIPT | 5907.7284 | -0.305429377 | 0.11016752 | -2.7724087 | 0.00556431 | 0.02924092 |
| AC120114.3 | 685.616147 | -0.110703278 | 0.15699758 | -0.7051273 | 0.48073102 | 0.66542234 |
| SEZ6L2 | 1021.39228 | -0.118952173 | 0.54063074 | -0.2200248 | 0.82585184 | 0.90695123 |
| ASPHD1 | 428.431467 | -1.926483251 | 0.38271459 | -5.0337334 | 4.81E-07 | 1.75E-05 |
| KCTD13 | 1026.20462 | -0.469368667 | 0.11960474 | -3.9243316 | 8.70E-05 | 0.0011364 |
| AC120114.1 | 22.1724931 | 1.739351615 | 0.4464715 | 3.89577299 | 9.79E-05 | 0.00124158 |
| TMEM219 | 5277.61073 | -0.115631597 | 0.20974212 | -0.5513036 | 0.58142554 | 0.74410053 |
| TAOK2 | 5376.01182 | 0.020251827 | 0.14147538 | 0.14314736 | 0.88617381 | 0.94110344 |
| HIRIP3 | 1553.47421 | 0.444226634 | 0.13902049 | 3.19540407 | 0.00139635 | 0.01023206 |
| INO80E | 5536.87929 | 0.250886771 | 0.24189958 | 1.03715257 | 0.29966476 | 0.49633563 |
| DOC2A | 1012.99338 | -0.245849857 | 0.23150332 | -1.0619712 | 0.28824876 | 0.48436371 |
| C16orf92 | 1.55031652 | 0.446800969 | 1.10580703 | 0.40404967 | 0.68617619 | 0.81849487 |
| FAM57B | 5.65179026 | -0.647242633 | 0.66063735 | -0.9797246 | 0.3272221 | 0.52502871 |
| AC093512.2 | 67738.6383 | -0.870100057 | 0.17798052 | -4.8887377 | 1.01E-06 | 3.29E-05 |
| ALDOA | 68470.695 | -0.87093718 | 0.17724771 | -4.9136724 | 8.94E-07 | 2.95E-05 |
| AC093512.1 | 1570.76578 | -0.936179436 | 0.31382052 | -2.9831683 | 0.00285281 | 0.0176386 |
| PPP4C | 4635.58198 | -0.677644978 | 0.16236624 | -4.1735585 | 3.00E-05 | 0.00048933 |
| TBX6 | 211.602509 | 0.123352677 | 0.33824853 | 0.3646806 | 0.71534984 | 0.8380436 |
| YPEL3 | 4556.70697 | -0.78518565 | 0.24826345 | -3.1627115 | 0.00156307 | 0.01115815 |
| AC012645.1 | 67.1652846 | 0.15747805 | 0.27581307 | 0.57095935 | 0.5680272 | 0.73384869 |
| GDPD3 | 885.501302 | 0.288037225 | 0.32177718 | 0.89514496 | 0.3707096 | 0.56749134 |
| AC012645.2 | 9.45768145 | -0.246566494 | 0.81987921 | -0.3007351 | 0.76361647 | 0.86922931 |
| AC012645.4 | 335.398242 | 0.425841622 | 0.35820384 | 1.18882483 | 0.2345086 | 0.42324094 |
| MAPK3 | 3655.82307 | -0.170303013 | 0.16558258 | -1.028508 | 0.30371091 | 0.5005871 |
| CORO1A | 1674.03092 | -0.221258071 | 0.51127153 | -0.4327604 | 0.66518884 | 0.80360225 |
| AC012645.3 | 219.688948 | 0.123256444 | 0.53763893 | 0.22925506 | 0.81867068 | 0.9026835 |
| BOLA2B | 29.5081702 | -0.52097943 | 0.25879742 | -2.0130781 | 0.04410641 | 0.13306769 |
| SLX1A | 125.40855 | -0.423110046 | 0.23508316 | -1.7998314 | 0.07188727 | 0.18795036 |
| SULT1A3 | 31.9364297 | -0.486087928 | 0.38530886 | -1.261554 | 0.20710932 | 0.38985458 |
| AC106782.1 | 30.4298603 | -1.598453787 | 0.40149374 | -3.981267 | 6.85E-05 | 0.00094295 |
| NPIPB13 | 34.190347 | -0.111421607 | 0.32863873 | -0.3390398 | 0.73457975 | 0.85065526 |
| CD2BP2 | 5180.45349 | -0.399675868 | 0.14786846 | -2.702915 | 0.00687343 | 0.03411489 |
| AC106782.2 | 116.171285 | 0.576690485 | 0.26463612 | 2.17918279 | 0.02931809 | 0.0992381 |
| TBC1D10B | 3346.31588 | -0.533560842 | 0.16866773 | -3.1633842 | 0.00155946 | 0.01113859 |
| AC106782.6 | 267.70889 | -0.298755064 | 0.15204247 | -1.9649448 | 0.04942062 | 0.14446534 |
| MYLPF | 13.9199637 | 0.748427229 | 0.57850141 | 1.29373449 | 0.19575715 | 0.37605565 |
| ZNF48 | 716.2251 | -0.356735185 | 0.16572249 | -2.1526058 | 0.03134967 | 0.10416972 |
| 1-Sep | 414.421061 | 0.083532931 | 0.47500512 | 0.17585691 | 0.86040636 | 0.92732725 |
| ZNF771 | 862.276126 | -0.468456914 | 0.21461142 | -2.1828145 | 0.02904948 | 0.09865224 |
| DCTPP1 | 1536.97383 | -0.531114153 | 0.18030715 | -2.9456078 | 0.00322321 | 0.01941434 |
| SEPHS2 | 14297.1684 | -0.29461901 | 0.34301316 | -0.8589146 | 0.39038764 | 0.5858814 |
| ITGAL | 595.014292 | 0.142165897 | 0.57276991 | 0.24820769 | 0.80397372 | 0.89372191 |
| AC116348.3 | 42.9202718 | 0.750498266 | 0.63043892 | 1.19043771 | 0.2338744 | 0.42244345 |
| AC116348.1 | 24.8662491 | -0.015568312 | 0.63042807 | -0.0246948 | 0.98029838 | 0.99116093 |
| AC116348.2 | 38.3626644 | 0.631235117 | 0.64830193 | 0.97367459 | 0.33021814 | 0.52810069 |
| ZNF768 | 3346.54903 | -0.130755322 | 0.16060306 | -0.8141521 | 0.41555781 | 0.60905514 |
| AC002310.5 | 927.334187 | 0.275881517 | 0.14380292 | 1.91846949 | 0.05505151 | 0.15579612 |
| ZNF747 | 981.18493 | 0.332894347 | 0.12332073 | 2.69941929 | 0.00694606 | 0.03440914 |
| AC002310.4 | 144.914871 | 0.621513163 | 0.17384122 | 3.57517711 | 0.00034999 | 0.00342968 |
| AC002310.1 | 148.126035 | 0.282670887 | 0.18275913 | 1.54668542 | 0.12193912 | 0.27282663 |
| ZNF764 | 461.973452 | 0.115342775 | 0.10965949 | 1.05182663 | 0.2928791 | 0.48930701 |
| ZNF688 | 753.22596 | -0.050356166 | 0.12679305 | -0.3971524 | 0.69125508 | 0.82241065 |
| AC002310.2 | 340.346502 | 0.342537238 | 0.18202902 | 1.88177272 | 0.05986688 | 0.16529051 |
| ZNF785 | 852.044368 | 0.002627182 | 0.19928956 | 0.01318274 | 0.989482 | 0.99519138 |
| AC093249.2 | 16.087292 | 1.073529327 | 0.45772117 | 2.34537835 | 0.01900778 | 0.07230172 |
| ZNF689 | 455.020721 | 0.413272437 | 0.18614346 | 2.22018236 | 0.02640639 | 0.09197418 |
| PRR14 | 3104.70406 | -0.597024919 | 0.19979457 | -2.9881939 | 0.00280631 | 0.01744639 |
| FBRS | 6536.19225 | -0.399339045 | 0.18086634 | -2.2079236 | 0.0272496 | 0.0941772 |
| AC093249.6 | 141.01337 | 0.103820842 | 0.18736518 | 0.55410958 | 0.57950386 | 0.74277814 |
| SRCAP | 6546.72204 | -0.058823307 | 0.23118604 | -0.2544414 | 0.79915455 | 0.89108294 |
| AC106886.5 | 6981.01569 | -0.155250109 | 0.21377199 | -0.7262416 | 0.46769068 | 0.65458327 |
| TMEM265 | 1100.39753 | -0.922778002 | 0.16821206 | -5.4858016 | 4.12E-08 | 2.26E-06 |
| PHKG2 | 3288.86297 | -0.32581326 | 0.14275118 | -2.2823858 | 0.02246658 | 0.08171827 |
| CCDC189 | 672.609299 | -0.045151835 | 0.13551954 | -0.3331758 | 0.73900157 | 0.85348667 |
| RNF40 | 5072.71935 | 0.041141809 | 0.12413928 | 0.33141653 | 0.74032988 | 0.85422678 |
| ZNF629 | 1518.13146 | -0.134359821 | 0.15691779 | -0.8562434 | 0.39186315 | 0.58760908 |
| AC106886.3 | 18.8491251 | -1.016996295 | 0.34672646 | -2.9331372 | 0.00335556 | 0.01997568 |
| BCL7C | 1920.99839 | 0.060267172 | 0.22462581 | 0.2683003 | 0.78846818 | 0.88495024 |
| MIR762HG | 1147.7268 | 0.011537729 | 0.23290742 | 0.04953783 | 0.96049069 | 0.98098634 |
| CTF1 | 461.905027 | 0.12365485 | 0.31967844 | 0.3868101 | 0.6988968 | 0.82715805 |
| FBXL19-AS1 | 111.372318 | -0.488436926 | 0.42827052 | -1.1404869 | 0.25408349 | 0.44675765 |
| FBXL19 | 1898.62379 | -0.816231071 | 0.21724328 | -3.7572212 | 0.00017181 | 0.00194365 |
| ORAI3 | 2111.83828 | -0.177119691 | 0.23176623 | -0.764217 | 0.44473794 | 0.63538066 |
| AC135048.3 | 10.746026 | 0.819216665 | 0.52452731 | 1.56181891 | 0.11833066 | 0.26733888 |
| SETD1A | 2821.10991 | -0.276581767 | 0.31354317 | -0.882117 | 0.37771353 | 0.57375034 |

| | | | | | | |
|---|---|---|---|---|---|---|
| HSD3B7 | 3982.10665 | -1.077258725 | 0.36137065 | -2.9810355 | 0.00287275 | 0.01773543 |
| STX1B | 221.581763 | 0.564857017 | 0.35884978 | 1.57407655 | 0.11546975 | 0.26292526 |
| STX4 | 3243.98681 | -0.558929232 | 0.1429038 | -3.9112272 | 9.18E-05 | 0.00118296 |
| AC135050.3 | 70.8109268 | -0.868063532 | 0.2703319 | -3.2111028 | 0.00132227 | 0.00981372 |
| AC135050.1 | 408.473652 | -0.823364895 | 0.26373614 | -3.1219267 | 0.00179672 | 0.0124225 |
| ZNF668 | 548.99198 | -0.716301856 | 0.24815879 | -2.8864658 | 0.00389595 | 0.02233396 |
| AC135050.4 | 3.20350938 | -0.240398085 | 0.94508161 | -0.2543675 | 0.79921163 | 0.89108294 |
| ZNF646 | 1333.55736 | -0.049018235 | 0.14930726 | -0.3283044 | 0.74268149 | 0.85555507 |
| PRSS53 | 724.976824 | -0.956764358 | 0.19107405 | -5.0072963 | 5.52E-07 | 1.97E-05 |
| AC135050.2 | 4333.78425 | -0.62007009 | 0.22970911 | -2.6993709 | 0.00694707 | 0.03440914 |
| VKORC1 | 4105.90086 | -0.587593962 | 0.24432932 | -2.4049262 | 0.01617573 | 0.06436541 |
| BCKDK | 4308.62355 | -0.329135032 | 0.1755084 | -1.8753235 | 0.06074823 | 0.16691996 |
| KAT8 | 2449.92841 | -0.09332792 | 0.13872188 | -0.67277 | 0.50109363 | 0.6819342 |
| AC009088.3 | 3628.71034 | -0.911020689 | 0.16735983 | -5.4434849 | 5.22E-08 | 2.76E-06 |
| PRSS8 | 10864.4508 | -0.627095586 | 0.13621359 | -4.6037667 | 4.15E-06 | 0.0001015 |
| PRSS36 | 124.571386 | -0.502234603 | 0.20836217 | -2.4103925 | 0.01593537 | 0.06364011 |
| FUS | 14740.2413 | -0.071140407 | 0.18071404 | -0.3936629 | 0.69382996 | 0.82376229 |
| AC009088.2 | 2150.51862 | 0.205329473 | 0.27366341 | 0.75029932 | 0.45307445 | 0.64268818 |
| AC009088.1 | 19.0955311 | -0.116525602 | 0.39337571 | -0.2962196 | 0.76706237 | 0.87152635 |
| PYCARD | 301.722763 | -0.867266084 | 0.39348084 | -2.2040872 | 0.02751821 | 0.09484794 |
| PYCARD-AS1 | 142.555996 | -0.746051168 | 0.33680786 | -2.2150646 | 0.02675563 | 0.092835 |
| TRIM72 | 15.3978249 | -1.065033556 | 0.67715883 | -1.5727973 | 0.11576577 | 0.26344878 |
| PYDC1 | 2.82536525 | -0.165631836 | 1.79473696 | -0.0922875 | 0.9264696 | 0.96437374 |
| ITGAM | 475.74872 | -0.274523059 | 0.48102804 | -0.5707007 | 0.56820251 | 0.73390969 |
| ITGAX | 756.973263 | -0.333567245 | 0.48229133 | -0.6916302 | 0.4891696 | 0.67257742 |
| ITGAD | 6.85252009 | 0.472327538 | 0.71192948 | 0.66344708 | 0.50704426 | 0.68603166 |
| COX6A2 | 43.1988693 | 0.775418536 | 0.41578032 | 1.86497173 | 0.06218537 | 0.16952851 |
| AC026471.5 | 32.7568143 | 0.806262581 | 0.39465816 | 2.04293911 | 0.04105847 | 0.12630795 |
| ZNF843 | 64.3122416 | 0.937166364 | 0.31897324 | 2.93807204 | 0.0033026 | 0.01974822 |
| AC026471.4 | 120.948333 | 0.04976581 | 0.24272292 | 0.20503136 | 0.83754762 | 0.91400972 |
| AC026471.1 | 286.617484 | 0.479171738 | 0.22576162 | 2.12246762 | 0.03379849 | 0.10985655 |
| ARMC5 | 1172.16795 | -0.085540351 | 0.17495183 | -0.4889366 | 0.6248866 | 0.77654381 |
| TGFB1I1 | 1765.53995 | -0.36124336 | 0.33763412 | -1.0699255 | 0.28465284 | 0.48080294 |
| SLC5A2 | 9466.79503 | 1.15924658 | 0.18840109 | 6.15307788 | 7.60E-10 | 8.21E-08 |
| AC026471.3 | 1588.53488 | 2.031654076 | 0.51283202 | 3.96163656 | 7.44E-05 | 0.00100252 |
| C16orf58 | 8402.66869 | 0.543893371 | 0.11560899 | 4.70459424 | 2.54E-06 | 6.89E-05 |
| AC026471.2 | 28.7066018 | 0.843302597 | 0.5160435 | 1.63416961 | 0.10222324 | 0.24114918 |
| AHSP | 11.3990181 | -3.4223636 | 1.04793934 | -3.2658032 | 0.00109154 | 0.00839671 |
| AC106730.1 | 0 | NA | NA | NA | NA | NA |
| LINC02190 | 0 | NA | NA | NA | NA | NA |
| AC074050.2 | 3.67525668 | 3.255807405 | 1.11075936 | 2.93115459 | 0.00337705 | 0.02006144 |
| AC074050.4 | 5.59911077 | -0.972797013 | 0.69493975 | -1.3998293 | 0.16156445 | 0.33024219 |
| AC074050.3 | 298.146092 | 0.881505544 | 0.23916102 | 3.68582447 | 0.00022796 | 0.00242787 |
| ZNF720 | 726.200856 | 0.105072166 | 0.11313452 | 0.92873655 | 0.35302563 | 0.55058469 |
| ZNF267 | 693.434414 | -0.700165603 | 0.2280096 | -3.0707725 | 0.00213506 | 0.01416389 |
| IGHV1OR16- | 0.03392908 | 0.380740593 | 3.29738709 | 0.11546736 | 0.90807469 | NA |
| IGHV1OR16- | 0.08393208 | -0.642905668 | 3.30558948 | -0.1944905 | 0.84579185 | NA |
| IGHV3OR16- | 0.29967709 | -1.364242679 | 3.30558948 | -0.4127078 | 0.67982068 | NA |
| AC133485.5 | 0 | NA | NA | NA | NA | NA |
| AC133485.3 | 0.75056297 | -0.787493129 | 1.88756622 | -0.4172003 | 0.67653193 | NA |
| TP53TG3D | 5.51821556 | -0.643046795 | 1.23501444 | -0.5206796 | 0.60259 | 0.75990141 |
| AC133485.2 | 0 | NA | NA | NA | NA | NA |
| AC138915.3 | 0 | NA | NA | NA | NA | NA |
| AC138907.7 | 0 | NA | NA | NA | NA | NA |
| AC138907.1 | 0 | NA | NA | NA | NA | NA |
| TP53TG3 | 0 | NA | NA | NA | NA | NA |
| AC138907.3 | 0 | NA | NA | NA | NA | NA |
| AC138907.2 | 0 | NA | NA | NA | NA | NA |
| AC137761.1 | 0.15543392 | 1.368031615 | 3.28033652 | 0.41704002 | 0.67664913 | NA |
| IGHV2OR16- | 0 | NA | NA | NA | NA | NA |
| AC142086.1 | 0 | NA | NA | NA | NA | NA |
| IGHV3OR16- | 0 | NA | NA | NA | NA | NA |
| IGHV3OR16- | 0.08757577 | 0.547239394 | 3.29377202 | 0.16614368 | 0.86804388 | NA |
| IGHV1OR16- | 0 | NA | NA | NA | NA | NA |
| IGHV3OR16- | 0 | NA | NA | NA | NA | NA |
| AC142086.4 | 0 | NA | NA | NA | NA | NA |
| IGHV1OR16- | 0.08757577 | 0.547239394 | 3.29377202 | 0.16614368 | 0.86804388 | NA |
| IGHV3OR16- | 0 | NA | NA | NA | NA | NA |
| AC145350.2 | 0.26039342 | -1.364242679 | 3.30558948 | -0.4127078 | 0.67982068 | NA |
| AC145350.4 | 0 | NA | NA | NA | NA | NA |
| AC138869.2 | 0 | NA | NA | NA | NA | NA |
| TP53TG3C | 0 | NA | NA | NA | NA | NA |
| AC138869.3 | 0 | NA | NA | NA | NA | NA |
| AC141257.2 | 0 | NA | NA | NA | NA | NA |
| TP53TG3E | 0 | NA | NA | NA | NA | NA |
| TP53TG3B | 0 | NA | NA | NA | NA | NA |
| AC141257.1 | 0 | NA | NA | NA | NA | NA |
| TP53TG3F | 0 | NA | NA | NA | NA | NA |
| AC136944.1 | 0 | NA | NA | NA | NA | NA |
| AC136944.2 | 0 | NA | NA | NA | NA | NA |
| AC136944.4 | 0 | NA | NA | NA | NA | NA |
| IGHV3OR16- | 0 | NA | NA | NA | NA | NA |
| IGHV3OR16- | 0.94834187 | 3.319417282 | 1.88987846 | 1.75641839 | 0.07901699 | NA |
| AC136428.1 | 0 | NA | NA | NA | NA | NA |
| AC140658.1 | 0 | NA | NA | NA | NA | NA |
| IGHV3OR16- | 0 | NA | NA | NA | NA | NA |
| IGHV3OR16- | 0.08757577 | 0.547239394 | 3.29377202 | 0.16614368 | 0.86804388 | NA |
| AC133561.2 | 0 | NA | NA | NA | NA | NA |
| LINC00273 | 0 | NA | NA | NA | NA | NA |
| LINC02184 | 0 | NA | NA | NA | NA | NA |
| AC135776.1 | 0 | NA | NA | NA | NA | NA |
| AC135776.2 | 0 | NA | NA | NA | NA | NA |
| AC023824.5 | 0 | NA | NA | NA | NA | NA |
| AC023824.1 | 0 | NA | NA | NA | NA | NA |
| AC023824.4 | 0 | NA | NA | NA | NA | NA |
| AC023824.3 | 0.07830766 | -0.642905668 | 3.30558948 | -0.1944905 | 0.84579185 | NA |
| LINC01566 | 0 | NA | NA | NA | NA | NA |
| AC018558.2 | 0 | NA | NA | NA | NA | NA |
| AC018558.3 | 0 | NA | NA | NA | NA | NA |
| AC018558.1 | 0 | NA | NA | NA | NA | NA |
| AC092325.1 | 30.5675461 | 3.713614257 | 1.16625087 | 3.18423279 | 0.00145138 | 0.01054061 |
| LINC02167 | 0.82499316 | -0.867565275 | 1.91668338 | -0.4526388 | 0.65080885 | NA |
| AC106785.2 | 0 | NA | NA | NA | NA | NA |
| SHCBP1 | 332.560619 | 1.348012591 | 0.37329009 | 3.61116633 | 0.00030482 | 0.0030727 |
| AC092368.3 | 328.220468 | -0.358294516 | 0.2312189 | -1.5495901 | 0.12123994 | 0.27176956 |
| VPS35 | 4841.32074 | -0.102376502 | 0.15855171 | -0.6456979 | 0.51847503 | 0.69536155 |
| ORC6 | 116.98041 | -0.840982528 | 0.28085507 | -2.9943648 | 0.00275017 | 0.0172016 |
| MYLK3 | 152.311836 | 1.114957348 | 0.65839765 | 1.69344066 | 0.09037162 | 0.22129256 |
| C16orf87 | 752.15815 | 0.019031138 | 0.30346221 | 0.06271337 | 0.94999475 | 0.9760144 |
| GPT2 | 3382.04874 | 0.694390367 | 0.53876014 | 1.28886738 | 0.1974442 | 0.37827691 |
| DNAJA2 | 3872.49729 | -0.135539808 | 0.10624856 | -1.2756861 | 0.20206651 | 0.38389726 |
| AC018845.3 | 10.6530279 | 0.080618349 | 0.57602915 | 0.13995533 | 0.88869529 | 0.94252311 |
| AC018845.1 | 0 | NA | NA | NA | NA | NA |

| | | | | | | |
|---|---|---|---|---|---|---|
| NETO2 | 245.820315 | -0.82144742 | 0.34631976 | -2.3719334 | 0.01769528 | 0.06855069 |
| ITFG1-AS1 | 865.454762 | -0.020702052 | 0.1798054 | -0.1151359 | 0.90833743 | 0.95418579 |
| ITFG1 | 4555.52687 | 0.210987902 | 0.16063049 | 1.31349845 | 0.18901504 | 0.3672788 |
| AC007494.2 | 0.93924253 | 1.091412426 | 1.19936831 | 0.90998938 | 0.36282811 | NA |
| AC007533.1 | 0.32721111 | -1.693792801 | 2.87919697 | -0.5882865 | 0.55633998 | NA |
| PHKB | 3825.13608 | 0.55382914 | 0.21464032 | 2.58026608 | 0.00987242 | 0.04466216 |
| LINC02133 | 2.39552304 | -0.191807005 | 0.92542517 | -0.2072637 | 0.83580395 | 0.91280681 |
| LINC02192 | 0.41120703 | 0.5088399 | 2.25188424 | 0.22596184 | 0.8212311 | NA |
| LINC02134 | 0 | NA | NA | NA | NA | NA |
| ABCC12 | 0.25999113 | -1.364242679 | 3.30558948 | -0.4127078 | 0.67982068 | NA |
| ABCC11 | 61.1481147 | 1.208750211 | 0.40287052 | 3.00034414 | 0.00269675 | 0.01694636 |
| LONP2 | 4797.58148 | 0.177339683 | 0.0738855 | 2.40019607 | 0.01638629 | 0.0649101 |
| SIAH1 | 2038.0096 | -0.102091879 | 0.14719902 | -0.6935636 | 0.48795595 | 0.67152554 |
| N4BP1 | 2194.98197 | -0.211952245 | 0.26061891 | -0.813265 | 0.41606613 | 0.60929993 |
| AC026470.2 | 3.64410771 | -0.801754582 | 0.74317924 | -1.0788173 | 0.2806692 | 0.4763946 |
| AC023813.1 | 0 | NA | NA | NA | NA | NA |
| AC007611.1 | 18.4184568 | 0.447440247 | 0.48610496 | 0.92046016 | 0.35733235 | 0.55490571 |
| AC023813.3 | 3.8691262 | -0.738026119 | 0.87407874 | -0.8443474 | 0.3984753 | 0.59342597 |
| AC023813.4 | 0 | NA | NA | NA | NA | NA |
| AC044798.1 | 0 | NA | NA | NA | NA | NA |
| AC044798.2 | 0 | NA | NA | NA | NA | NA |
| CBLN1 | 13.8019003 | 1.936069122 | 0.71012541 | 2.72637634 | 0.00640339 | 0.03232341 |
| AC007614.1 | 3.08439421 | 0.38716665 | 0.6493949 | 0.59619601 | 0.55104429 | 0.72135812 |
| C16orf78 | 0.27512461 | 2.050195067 | 3.23884076 | 0.63300274 | 0.52673186 | NA |
| AC007861.1 | 0.40655545 | 1.198259274 | 2.14506936 | 0.55861097 | 0.57642725 | NA |
| AC007861.2 | 0 | NA | NA | NA | NA | NA |
| LINC02179 | 0 | NA | NA | NA | NA | NA |
| ZNF423 | 204.730888 | 0.397888893 | 0.43614 | 0.91229626 | 0.36161278 | 0.55906728 |
| AC007603.1 | 0 | NA | NA | NA | NA | NA |
| CNEP1R1 | 783.091274 | -0.284495866 | 0.12131809 | -2.3450408 | 0.019025 | 0.07234163 |
| AC007610.1 | 20.3804568 | 0.240583207 | 0.40563585 | 0.59310144 | 0.55311327 | 0.72290971 |
| HEATR3 | 776.487785 | -0.142644302 | 0.1496495 | -0.9531893 | 0.34049417 | 0.53855754 |
| AC007610.2 | 74.5897493 | 0.229081163 | 0.19466835 | 1.17677663 | 0.23928467 | 0.42907227 |
| TENT4B | 952.770139 | 0.367425819 | 0.1177718 | 3.11981161 | 0.00180967 | 0.01247019 |
| ADCY7 | 756.282668 | 0.137496619 | 0.33330211 | 0.4125285 | 0.6799521 | 0.81445999 |
| BRD7 | 3366.81535 | -0.054760692 | 0.07957254 | -0.6881858 | 0.49133578 | 0.67436238 |
| AC007493.2 | 43.6860368 | 0.199926128 | 0.25291075 | 0.7905007 | 0.42923541 | 0.62127108 |
| LINC02178 | 0.21442573 | -1.003574173 | 3.30558948 | -0.3035992 | 0.7614333 | NA |
| AC007493.1 | 0.46952845 | -1.352447239 | 2.27634417 | -0.5941313 | 0.55242432 | NA |
| NKD1 | 517.351749 | 0.092061032 | 0.41715284 | 0.22068897 | 0.82533462 | 0.90673291 |
| AC007608.2 | 0.35467906 | 0.670828026 | 2.2054785 | 0.30416439 | 0.76100265 | NA |
| AC007608.1 | 17.1516048 | 0.225027461 | 0.45773662 | 0.49160904 | 0.62299575 | 0.77518039 |
| AC007608.3 | 53.5147704 | 0.135887884 | 0.59834396 | 0.22710664 | 0.82034082 | 0.90356178 |
| SNX20 | 170.946044 | -0.001596129 | 0.57679591 | -0.0027672 | 0.99779207 | 0.99873892 |
| NOD2 | 144.284515 | -0.981203239 | 0.59338257 | -1.6535761 | 0.09821367 | 0.23469819 |
| AC007728.2 | 78.7942697 | -0.742491081 | 0.51245756 | -1.4488831 | 0.14737024 | 0.31071827 |
| CYLD | 2328.80747 | -0.45180541 | 0.3742006 | -1.2073883 | 0.22728265 | 0.41416927 |
| AC007728.1 | 19.8379137 | 0.559965888 | 0.46286391 | 1.20978516 | 0.22636134 | 0.41317416 |
| LINC02168 | 0.90843402 | -0.331170914 | 1.90304835 | -0.1740213 | 0.86184874 | NA |
| LINC02128 | 41.3490793 | 0.589100279 | 0.42167015 | 1.39706422 | 0.16239426 | 0.33138509 |
| AC025810.1 | 12.9284387 | 1.224666882 | 0.60028377 | 2.04014657 | 0.04133573 | 0.12699792 |
| LINC02127 | 0.52379574 | 1.709772272 | 1.71531392 | 0.99676931 | 0.3188765 | NA |
| AC027688.1 | 0.30722956 | 0.328495715 | 2.89077905 | 0.11363571 | 0.90952658 | NA |
| AC027688.2 | 0.33566243 | 2.367239012 | 3.22054628 | 0.73504269 | 0.46231349 | NA |
| SALL1 | 4822.98071 | 0.013674487 | 0.26717888 | 0.05118102 | 0.95918128 | 0.98033449 |
| AC009166.1 | 21.4045643 | 0.639306748 | 0.44796158 | 1.42714639 | 0.1535377 | 0.31917929 |
| AC087564.1 | 108.670886 | -0.064507467 | 0.23353847 | -0.2762177 | 0.78238083 | 0.88085486 |
| AC007344.1 | 161.776998 | -2.625771231 | 0.60021841 | -4.3746929 | 1.22E-05 | 0.00024056 |
| HNRNPA1P4: | 162.28942 | -0.46355456 | 0.22571676 | -2.0537002 | 0.04000471 | 0.12382171 |
| LINC01571 | 57.1689022 | 0.550506787 | 0.51520792 | 1.06851383 | 0.2852888 | 0.48135297 |
| AC009039.1 | 0.09997307 | 0.547239394 | 3.29377202 | 0.16614368 | 0.86804388 | NA |
| LINC00919 | 1.12459581 | 1.108172153 | 1.29854923 | 0.85339249 | 0.39344169 | 0.58885619 |
| LINC02180 | 0.12150485 | 1.04981648 | 3.28477129 | 0.31960109 | 0.74927075 | NA |
| AC007333.2 | 0.75134293 | 2.360605917 | 1.71491444 | 1.37651527 | 0.16866216 | NA |
| AC007333.1 | 0.94816034 | 0.932017188 | 1.48969739 | 0.62564195 | 0.53154981 | NA |
| CASC22 | 10.1755472 | 2.526476837 | 0.98599809 | 2.56235469 | 0.01039651 | 0.0464199 |
| TOX3 | 906.99812 | 0.254802274 | 0.22379752 | 1.13853934 | 0.25489534 | 0.44760086 |
| CASC16 | 28.2988016 | -1.693281469 | 0.57509847 | -2.9443331 | 0.00323651 | 0.01946737 |
| AC026462.3 | 148.277143 | 1.482266156 | 0.46980178 | 3.1550884 | 0.00160449 | 0.01138604 |
| AC026462.2 | 0 | NA | NA | NA | NA | NA |
| AC026462.4 | 0.34785867 | -0.541687062 | 2.85265938 | -0.1898884 | 0.84939654 | NA |
| AC007906.2 | 1103.85407 | -0.98994722 | 0.56068526 | -1.7656024 | 0.07746258 | 0.19802062 |
| CHD9 | 2263.63176 | 0.209634807 | 0.12940183 | 1.62002974 | 0.10522589 | 0.24603712 |
| AC007342.4 | 50.955393 | 1.632880132 | 0.62588566 | 2.60891122 | 0.00908308 | 0.04194026 |
| AC007342.5 | 70.1860453 | 0.432869067 | 0.58979099 | 0.73393638 | 0.46298751 | 0.65100676 |
| RBL2 | 4846.77514 | 0.484823402 | 0.20346439 | 2.38284158 | 0.01717959 | 0.06712486 |
| AKTIP | 3231.6592 | 0.233320873 | 0.13389684 | 1.74254207 | 0.08141364 | 0.20491339 |
| RPGRIP1L | 328.831908 | -0.036502307 | 0.26335161 | -0.1386067 | 0.88976092 | 0.94296756 |
| FTO | 2116.94205 | 0.417342689 | 0.16807873 | 2.48301907 | 0.01302741 | 0.05481208 |
| AC007496.2 | 0.5618249 | 1.169646318 | 1.85145498 | 0.6317444 | 0.52755391 | NA |
| AC007496.1 | 0.69327712 | 0.472515957 | 1.55443951 | 0.30397835 | 0.76114439 | NA |
| AC007347.1 | 0.4762153 | 0.837559011 | 2.14624612 | 0.39024369 | 0.69635636 | NA |
| AC007907.1 | 0.21783422 | 0.689157102 | 3.06342475 | 0.22496296 | 0.82200809 | NA |
| LINC02169 | 0 | NA | NA | NA | NA | NA |
| IRX3 | 1672.80261 | -0.91800805 | 0.49998817 | -1.8360595 | 0.06634885 | 0.17734848 |
| AC018553.2 | 106.337377 | -0.702234922 | 0.55489887 | -1.2655187 | 0.20568544 | 0.38823862 |
| AC018553.1 | 43.3020436 | -0.98508117 | 0.38371794 | -2.5672012 | 0.01025231 | 0.045916 |
| LINC02140 | 0 | NA | NA | NA | NA | NA |
| LINC02183 | 0.38553722 | -0.032130408 | 2.2567608 | -0.0142374 | 0.98864058 | NA |
| AC007491.1 | 0 | NA | NA | NA | NA | NA |
| CRNDE | 463.93722 | -0.473872846 | 0.35551085 | -1.332935 | 0.18255314 | 0.35929346 |
| IRX5 | 574.550937 | -0.235515701 | 0.47650381 | -0.4942577 | 0.62112416 | 0.77363364 |
| AC106738.1 | 8.81357909 | -0.203412694 | 0.4654867 | -0.4369893 | 0.66211913 | 0.80128673 |
| AC106738.2 | 1.95858962 | 1.159462208 | 1.07441442 | 1.07915734 | 0.28051759 | 0.47625862 |
| AC109462.1 | 0.50905572 | -2.249318492 | 3.28178621 | -0.6853946 | 0.49309496 | NA |
| AC109462.3 | 4.91053148 | 0.341250057 | 0.76870474 | 0.44392865 | 0.65709417 | 0.79880017 |
| IRX6 | 174.663772 | -0.327312811 | 0.55531695 | -0.5894162 | 0.5555821 | 0.72490182 |
| AC109462.2 | 1.15051676 | -1.524100558 | 1.55869948 | -0.9778027 | 0.32817192 | 0.52584878 |
| MMP2 | 4438.90834 | -0.252093041 | 0.37470539 | -0.6727767 | 0.50108938 | 0.6819342 |
| AC007336.2 | 0.80402783 | -0.228432196 | 1.55287734 | -0.1471025 | 0.88305109 | NA |
| MMP2-AS1 | 6.86121832 | 0.648522638 | 1.134608 | 0.57158299 | 0.56760452 | 0.73353253 |
| LPCAT2 | 1145.3363 | 0.255496676 | 0.21706174 | 1.17706916 | 0.2391679 | 0.42892122 |
| AC007336.1 | 34.3071268 | -0.053584754 | 0.33209238 | -0.161355 | 0.87181383 | 0.93317628 |
| CAPNS2 | 2.70403303 | 0.764661996 | 0.84512412 | 0.90479254 | 0.3655753 | 0.56260407 |
| SLC6A2 | 25.1604611 | 0.720458253 | 1.05647343 | 0.6819464 | 0.49527284 | 0.67747406 |
| CES1 | 449.493479 | 0.217963121 | 0.61558995 | 0.35407193 | 0.72328498 | 0.84311842 |
| CES5A | 3.50394853 | -1.159385842 | 0.80723466 | -1.4362439 | 0.15093295 | 0.31566678 |
| AC007495.1 | 7.56349612 | 0.9358577 | 0.46720839 | 2.00308411 | 0.04516826 | 0.13516648 |
| AC007495.2 | 0.09635527 | 0.547239394 | 3.29377202 | 0.16614368 | 0.86804388 | NA |
| AC079411.1 | 0.16783122 | 1.368031398 | 3.28033652 | 0.41703996 | 0.67664918 | NA |
| GNAO1 | 245.526296 | 0.481292805 | 0.37011362 | 1.30039204 | 0.19346664 | 0.37306875 |
| DKFZP434H1 | 4.8535176 | 1.144409646 | 0.53460736 | 2.14065451 | 0.03230191 | 0.10628916 |
| AC009102.2 | 1.96563663 | 2.221771356 | 1.13991468 | 1.94906811 | 0.05128729 | 0.14839213 |

| | | | | | | |
|---|---|---|---|---|---|---|
| AMFR | 10500.0675 | -0.085766478 | 0.17818277 | -0.4813399 | 0.63027493 | 0.78028359 |
| AC092140.1 | 399.060272 | 0.716136411 | 0.20138913 | 3.55598339 | 0.00037657 | 0.0036414 |
| NUDT21 | 4149.58255 | -0.011071436 | 0.12142004 | -0.0911829 | 0.92734723 | 0.96473672 |
| OGFOD1 | 1791.03164 | -0.347242782 | 0.14072331 | -2.4675569 | 0.01360386 | 0.05658357 |
| AC092140.2 | 89.5542092 | 0.111222412 | 0.15813118 | 0.70335535 | 0.48183432 | 0.66635049 |
| BBS2 | 3794.33636 | 0.407227131 | 0.18228068 | 2.2340663 | 0.02547871 | 0.08951154 |
| MT4 | 0 | NA | NA | NA | NA | NA |
| MT3 | 54.4928609 | 1.099991408 | 0.50448632 | 2.1804187 | 0.02922644 | 0.09906778 |
| MT2A | 28310.1508 | -2.495557327 | 0.47618486 | -5.2407321 | 1.60E-07 | 7.06E-06 |
| AC026461.3 | 0.79139164 | -1.244541392 | 1.6265052 | -0.7651629 | 0.44417457 | NA |
| MT1E | 9209.77079 | -1.387940267 | 0.57865658 | -2.3985561 | 0.01645986 | 0.06511347 |
| MT1M | 1251.88836 | -2.796779291 | 0.69838965 | -4.0046116 | 6.21E-05 | 0.00087225 |
| MT1A | 373.733495 | -3.509592005 | 0.67935198 | -5.1660879 | 2.39E-07 | 9.87E-06 |
| MT1B | 2.53814061 | -3.013196385 | 1.16831423 | -2.5790976 | 0.00990588 | 0.04477512 |
| MT1F | 6302.21071 | -0.106257563 | 0.50465392 | -0.2105553 | 0.83323429 | 0.91140944 |
| MT1G | 24316.4144 | 0.080310962 | 0.60787494 | 0.13211757 | 0.8948913 | 0.94598591 |
| MT1H | 6921.54243 | -0.307528481 | 0.68838333 | -0.4467402 | 0.65506268 | 0.79755729 |
| AC026461.4 | 3.24326526 | -0.751669398 | 0.81223134 | -0.9254376 | 0.35473834 | 0.55230354 |
| MT1X | 11774.5804 | -1.109681783 | 0.53622445 | -2.0694352 | 0.03850526 | 0.12043456 |
| AC106779.1 | 7.17857272 | 0.446151358 | 0.80445213 | 0.55460274 | 0.57916641 | 0.7425788 |
| NUP93 | 1462.13817 | -0.37096054 | 0.15530712 | -2.3885611 | 0.0169145 | 0.06645181 |
| SLC12A3 | 10417.7726 | 3.810823142 | 0.88282308 | 4.31663289 | 1.58E-05 | 0.00029587 |
| HERPUD1 | 10814.7227 | 0.250945179 | 0.20798996 | 1.20652545 | 0.22761495 | 0.41460291 |
| CETP | 216.097217 | 0.145507135 | 0.41588587 | 0.34987276 | 0.72643419 | 0.84536936 |
| NLRC5 | 1918.6358 | -1.196514289 | 0.50716752 | -2.3592092 | 0.01831393 | 0.07035383 |
| AC023825.2 | 10.5970477 | -1.116306997 | 0.66334286 | -1.6828507 | 0.092404 | 0.22480796 |
| AC009090.2 | 0.56999341 | -1.624273453 | 3.27208431 | -0.4964033 | 0.6196099 | NA |
| CPNE2 | 1499.33214 | -0.286609807 | 0.20491778 | -1.3986576 | 0.16191569 | 0.33068846 |
| FAM192A | 3541.58141 | -0.244119281 | 0.10172996 | -2.3996793 | 0.01640944 | 0.06497819 |
| RSPRY1 | 1879.53014 | 0.036423988 | 0.13037017 | 0.27938897 | 0.77994633 | 0.87952228 |
| ARL2BP | 3319.04192 | -0.292307112 | 0.23253791 | -1.2570299 | 0.20874281 | 0.39192224 |
| AC009090.3 | 279.854627 | 0.025744796 | 0.29537688 | 0.08715914 | 0.93054501 | 0.96655578 |
| PLLP | 5538.17245 | -0.10172619 | 0.17290339 | -0.5883412 | 0.55630329 | 0.7253727 |
| CCL22 | 14.9970458 | 2.620162056 | 0.88142538 | 2.97264195 | NA | NA |
| CX3CL1 | 5596.03288 | 0.092971905 | 0.33326641 | 0.27897172 | 0.78026653 | 0.8796997 |
| CCL17 | 18.6955397 | 1.003353815 | 0.685482 | 1.46372014 | 0.14327043 | 0.30479475 |
| CIAPIN1 | 2817.6363 | -0.148945825 | 0.15386136 | -0.9680522 | 0.33301832 | 0.53100478 |
| COQ9 | 5958.90776 | 0.31151213 | 0.15966559 | 1.95102858 | 0.05105365 | 0.14781342 |
| POLR2C | 4592.30159 | 0.060379958 | 0.13224232 | 0.45658576 | 0.64796882 | 0.79277053 |
| DOK4 | 5969.67717 | -0.577896949 | 0.23029418 | -2.5093858 | 0.01209413 | 0.05184885 |
| CCDC102A | 495.491149 | -0.045097533 | 0.30958249 | -0.1456721 | 0.88418024 | 0.93996505 |
| ADGRG5 | 1077.63613 | 2.847388776 | 0.45389448 | 6.27323944 | 3.54E-10 | 4.22E-08 |
| ADGRG1 | 42176.2433 | -0.412299176 | 0.38473815 | -1.0716358 | 0.28388365 | 0.47993409 |
| AC018552.3 | 6.30050174 | 1.509242992 | 0.68286821 | 2.21015267 | 0.02709457 | 0.09372737 |
| ADGRG3 | 140.932737 | -1.408574885 | 0.41644828 | -3.3823525 | 0.00071868 | 0.00603553 |
| AC018552.2 | 25.2119981 | -0.966467344 | 0.34934157 | -2.7665398 | 0.00566547 | 0.02957798 |
| DRC7 | 91.0617791 | -0.641335507 | 0.63227908 | -1.0143235 | 0.31042843 | 0.50741242 |
| KATNB1 | 1144.60165 | -0.240365893 | 0.14638081 | -1.6420588 | 0.10057783 | 0.23848911 |
| KIFC3 | 11917.1027 | 0.394921741 | 0.22620664 | 1.74584507 | 0.08083788 | 0.20389238 |
| AC092118.2 | 2002.97495 | 0.139606439 | 0.2506092 | 0.55706829 | 0.57748077 | 0.74124649 |
| AC092118.1 | 179.613524 | 0.126609961 | 0.3098619 | 0.40860125 | 0.68283231 | 0.81659231 |
| CNGB1 | 9.32817522 | -1.714159405 | 0.64780972 | -2.6460847 | 0.00814294 | 0.03861345 |
| TEPP | 38.4203916 | -0.630517001 | 0.46520534 | -1.355352 | 0.17530543 | 0.34944472 |
| ZNF319 | 613.129273 | 0.088781628 | 0.10238927 | 0.86709892 | 0.38588781 | 0.58161074 |
| USB1 | 2302.49236 | -0.834526366 | 0.18292345 | -4.5621617 | 5.06E-06 | 0.00011784 |
| MMP15 | 2825.34105 | -0.534530792 | 0.15953263 | -3.3506047 | 0.00080635 | 0.00662209 |
| CFAP20 | 1110.36946 | 0.0569432 | 0.12743836 | 0.44682935 | 0.65499828 | 0.79753911 |
| AC026771.1 | 186.162065 | 0.291578096 | 0.21836152 | 1.33529983 | 0.18177823 | 0.35823123 |
| AC009107.2 | 39.755443 | -0.23298904 | 0.24139591 | -0.965174 | 0.33445769 | 0.53253652 |
| CSNK2A2 | 3136.03484 | -0.500163716 | 0.19453296 | -2.5711001 | 0.0101376 | 0.04553374 |
| AC009107.1 | 0.46797215 | -2.1071007 | 2.74195262 | -0.7684672 | 0.44220966 | NA |
| CCDC113 | 303.834229 | -0.51820037 | 0.26834687 | -1.9310841 | 0.05347265 | 0.15275419 |
| PRSS54 | 29.5416054 | -0.03390498 | 0.38351456 | -0.088406 | 0.92955401 | 0.96592738 |
| GINS3 | 172.308431 | -0.329697167 | 0.25628016 | -1.2864717 | 0.1982785 | 0.37923626 |
| AC009118.1 | 0.07830766 | -0.642905668 | 3.30558948 | -0.1944905 | 0.84579185 | NA |
| AC009118.2 | 36.5393556 | -0.425713758 | 0.34466975 | -1.2351352 | 0.21678018 | 0.40160353 |
| LINC02137 | 1.4478325 | -0.60362165 | 1.94890033 | -0.3097242 | 0.75677068 | 0.86528391 |
| NDRG4 | 336.632791 | -0.243347182 | 0.43934916 | -0.5538811 | 0.57966024 | 0.74281468 |
| SETD6 | 3217.71372 | -0.315453716 | 0.14879963 | -2.1199899 | 0.0340069 | 0.1103707 |
| CNOT1 | 5757.59225 | -0.04624501 | 0.10937222 | -0.4228223 | 0.67242493 | 0.80893455 |
| AC009118.3 | 1250.44172 | -0.299153949 | 0.10775634 | -2.7762075 | 0.00549971 | 0.02894757 |
| SLC38A7 | 2025.54097 | -0.310176881 | 0.30433817 | -1.0191849 | 0.30811517 | 0.50503462 |
| GOT2 | 8355.86777 | 0.178212977 | 0.17719698 | 1.00573372 | 0.31454368 | 0.51159667 |
| AC092378.1 | 393.41925 | 0.655714786 | 0.31518095 | 2.08043914 | 0.03748527 | 0.11801416 |
| AC092378.2 | 0.08757577 | 0.547239394 | 3.29377202 | 0.16614368 | 0.86804388 | NA |
| LINC02141 | 0 | NA | NA | NA | NA | NA |
| AC018554.3 | 0 | NA | NA | NA | NA | NA |
| AC018554.1 | 0 | NA | NA | NA | NA | NA |
| AC009081.1 | 0 | NA | NA | NA | NA | NA |
| CDH8 | 3.76821017 | 0.479011246 | 0.7215665 | 0.66384906 | 0.50678692 | 0.68589464 |
| AC012174.1 | 0.36210517 | -0.55754913 | 2.27253273 | -0.2453426 | 0.80619116 | NA |
| AC009110.1 | 0 | NA | NA | NA | NA | NA |
| AC138305.1 | 46.7105283 | 0.599520764 | 0.40911575 | 1.46540623 | 0.14281012 | 0.30411878 |
| AC138305.2 | 0.86995682 | 0.372624844 | 1.2472724 | 0.29875178 | 0.76512945 | NA |
| LINC02165 | 0 | NA | NA | NA | NA | NA |
| AC092337.1 | 0 | NA | NA | NA | NA | NA |
| AC093536.2 | 0 | NA | NA | NA | NA | NA |
| AC012322.1 | 0 | NA | NA | NA | NA | NA |
| AC092131.1 | 0.93600027 | 0.821396506 | 1.83147346 | 0.44848944 | 0.65380001 | NA |
| CDH11 | 1042.51365 | 0.274906934 | 0.36419899 | 0.75482619 | 0.45035328 | 0.64020571 |
| LINC02126 | 1.14495219 | 2.635550541 | 1.17092847 | 2.25082114 | 0.02439687 | 0.08666919 |
| AC009055.2 | 0 | NA | NA | NA | NA | NA |
| AC009055.1 | 0 | NA | NA | NA | NA | NA |
| LINC00922 | 0.85250844 | -1.031331127 | 1.69499897 | -0.6084553 | 0.54288553 | NA |
| AC092138.2 | 0.56954185 | 2.47640886 | 1.91924308 | 1.29030496 | 0.1969448 | NA |
| AC092138.1 | 0 | NA | NA | NA | NA | NA |
| AC022164.1 | 2.58359415 | -1.167000887 | 1.29746049 | -0.89945 | 0.368413 | 0.56515711 |
| CDH5 | 2639.19129 | -0.143399025 | 0.34293814 | -0.4181484 | 0.67583864 | 0.81147469 |
| LINC00920 | 21.5172523 | 0.145280508 | 0.56949451 | 0.25510432 | 0.79864253 | 0.89098462 |
| BEAN1 | 72.2780955 | -0.841482034 | 0.49598719 | -1.6965802 | 0.08977606 | 0.22020228 |
| BEAN1-AS1 | 33.8021388 | -0.606436351 | 0.50231965 | -1.2072718 | 0.22732748 | 0.41422234 |
| AC010542.3 | 346.682694 | 0.393083126 | 0.29573896 | 1.32915569 | 0.18379661 | 0.36082334 |
| TK2 | 2898.79909 | 0.092753944 | 0.23745195 | 0.39062196 | 0.69607669 | 0.82515362 |
| AC010542.2 | 632.989622 | -0.281925026 | 0.24270365 | -1.1616019 | 0.2453972 | 0.43650078 |
| AC010542.4 | 95.8074548 | 0.354667339 | 0.28404754 | 1.24861964 | 0.21180423 | 0.39541787 |
| CKLF | 642.254965 | -1.095758445 | 0.23262319 | -4.7104438 | 2.47E-06 | 6.73E-05 |
| CKLF-CMTM1 | 621.686428 | -1.185533849 | 0.24073024 | -4.9247401 | 8.45E-07 | 2.81E-05 |
| AC010542.5 | 360.544288 | -0.932359686 | 0.24087095 | -3.8707852 | 0.00010849 | 0.00135133 |
| CMTM1 | 69.7171307 | -0.288532613 | 0.25588112 | -1.1276042 | 0.25948712 | 0.452272 |
| CMTM2 | 36.6604117 | 0.28390867 | 0.24701019 | 1.14938038 | 0.25039917 | 0.44262116 |
| AC010542.1 | 11.7853013 | 0.1115126 | 0.48997432 | 0.22758866 | 0.81996603 | 0.90344498 |
| CMTM3 | 1012.16982 | -0.830956744 | 0.24242737 | -3.4276523 | 0.00060882 | 0.00526995 |
| CMTM4 | 4802.14144 | 0.489803519 | 0.2607094 | 1.87873363 | 0.06028087 | 0.1661031 |

| | | | | | | |
|---|---|---|---|---|---|---|
| DYNC1LI2 | 6316.14726 | 0.138289553 | 0.12452636 | 1.11052435 | 0.26677314 | 0.46090597 |
| AC018557.1 | 3182.64888 | -0.29984045 | 0.12852326 | -2.3329664 | 0.01964991 | 0.07393952 |
| AC018557.2 | 0.19624763 | 0.18657164 | 3.29377195 | 0.05664376 | 0.95482897 | NA |
| AC044802.1 | 13.466194 | 0.26132626 | 0.36917542 | 0.70786474 | 0.47902926 | 0.66382897 |
| TERB1 | 5.25294217 | 0.342151423 | 0.6380451 | 0.53624959 | 0.59178607 | 0.75134322 |
| NAE1 | 1837.45089 | -0.414203334 | 0.14966239 | -2.7675846 | 0.00564734 | 0.02951255 |
| AC044802.2 | 2.82968085 | 1.088883901 | 1.01437504 | 1.07345297 | 0.28306793 | 0.4789542 |
| CA7 | 4.80212923 | 1.952068144 | 0.77254688 | 2.52679571 | 0.01151084 | 0.05005623 |
| PDP2 | 1309.8671 | 0.160615188 | 0.2467872 | 0.65082464 | 0.5151597 | 0.69281728 |
| CDH16 | 69044.9027 | -0.125686078 | 0.2498025 | -0.5031418 | 0.61486459 | 0.76971181 |
| RRAD | 4441.87449 | -2.601879285 | 0.42522418 | -6.1188413 | 9.43E-10 | 9.79E-08 |
| FAM96B | 4098.97254 | -0.757440772 | 0.17951267 | -4.2194279 | 2.45E-05 | 0.00041803 |
| CES2 | 12200.4932 | 0.819249242 | 0.37502455 | 2.18452164 | 0.02892394 | 0.09827655 |
| CES3 | 1594.69847 | 2.326206343 | 0.61830819 | 3.76221174 | 0.00016842 | 0.00191394 |
| CES4A | 894.972394 | -0.393750823 | 0.41742879 | -0.9432766 | 0.34553943 | 0.54324657 |
| CBFB | 1986.50347 | -0.682520539 | 0.12864667 | -5.3053885 | 1.12E-07 | 5.30E-06 |
| C16orf70 | 2209.20655 | -0.097286807 | 0.21833082 | -0.4455936 | 0.65589086 | 0.79814017 |
| B3GNT9 | 1176.86633 | -0.232844956 | 0.15923288 | -1.4622919 | 0.14366124 | 0.30524106 |
| TRADD | 1765.97192 | -0.254355409 | 0.20687434 | -1.2295165 | 0.21887822 | 0.40427392 |
| FBXL8 | 653.093163 | -0.251795549 | 0.21223737 | -1.1863865 | 0.23546971 | 0.42436688 |
| AC074143.1 | 798.712325 | -0.207800506 | 0.28331718 | -0.7334554 | 0.46328074 | 0.65111486 |
| HSF4 | 2377.88192 | -0.225305218 | 0.3359145 | -0.6707219 | 0.5023977 | 0.68265066 |
| NOL3 | 2279.78775 | -0.61857798 | 0.25936959 | -2.3849287 | 0.01708243 | 0.06688945 |
| KIAA0895L | 3571.43201 | -0.107437848 | 0.25844731 | -0.415705 | 0.67762586 | 0.81263331 |
| EXOC3L1 | 556.184935 | -0.032742642 | 0.2811824 | -0.1164463 | 0.90729886 | 0.953776 |
| E2F4 | 4247.35543 | -0.341678891 | 0.17761128 | -1.9237454 | 0.0543865 | 0.15446118 |
| ELMO3 | 2770.52273 | 0.123264819 | 0.23376837 | 0.52729468 | 0.59798897 | 0.75641431 |
| LRRC29 | 431.857102 | 0.507593574 | 0.26971881 | 1.88193614 | 0.05984469 | 0.16526384 |
| TMEM208 | 2152.02993 | -0.32634475 | 0.14935791 | -2.1849847 | 0.02888997 | 0.09821353 |
| FHOD1 | 2460.32963 | -0.197863492 | 0.30897843 | -0.6403796 | 0.52192583 | 0.69790813 |
| SLC9A5 | 475.881822 | -0.014778345 | 0.26525524 | -0.0557137 | 0.9555699 | 0.97873792 |
| PLEKHG4 | 476.719121 | 0.470186794 | 0.25778151 | 1.82397405 | 0.06815599 | 0.18061834 |
| KCTD19 | 25.0843408 | 0.082782616 | 0.59263156 | 0.13968648 | 0.88890771 | 0.94259304 |
| LRRC36 | 120.803986 | 0.376146613 | 0.49926077 | 0.75340711 | 0.45120531 | 0.64093321 |
| TPPP3 | 773.958161 | 1.056981638 | 0.48106351 | 2.19717689 | 0.02800782 | 0.09615494 |
| ZDHHC1 | 1439.29623 | -0.228910284 | 0.1969695 | -1.162161 | 0.24517005 | 0.43618502 |
| HSD11B2 | 17951.1256 | 1.475042883 | 0.56709385 | 2.60105602 | 0.00929373 | 0.04260702 |
| AC009061.1 | 272.657422 | 1.558942001 | 0.52935812 | 2.94496663 | 0.00322989 | 0.01943804 |
| ATP6V0D1 | 7777.27527 | 0.085629221 | 0.17257623 | 0.49618201 | 0.61976601 | 0.77271053 |
| AC009061.2 | 148.202834 | 0.724133813 | 0.35676205 | 2.02973888 | 0.04238309 | 0.12931181 |
| AGRP | 21.0380174 | 1.245991213 | 0.35882368 | 3.47243308 | 0.00051576 | 0.00464346 |
| AC027682.6 | 158.945855 | 0.405257176 | 0.41804687 | 0.96940607 | 0.33234263 | 0.53024413 |
| RIPOR1 | 9882.89327 | -0.193430219 | 0.2043689 | -0.9464758 | 0.34390596 | 0.54151789 |
| AC027682.5 | 690.168226 | 0.51086486 | 0.29358874 | 1.74006969 | 0.08184678 | 0.20568077 |
| AC027682.2 | 1525.97245 | 0.100583892 | 0.2614819 | 0.38466867 | 0.70048291 | 0.82832397 |
| AC027682.3 | 866.930691 | 0.178607006 | 0.28060133 | 0.63651517 | 0.5244407 | 0.69951978 |
| AC027682.4 | 29.6823667 | -0.49401696 | 0.35785152 | -1.3805082 | 0.16743022 | 0.33860415 |
| AC027682.1 | 8.64427383 | 0.034584703 | 0.58656739 | 0.05896118 | 0.95298303 | 0.97725695 |
| CTCF | 2150.73836 | 0.136711926 | 0.2074548 | 0.6589962 | 0.50989821 | 0.68858585 |
| AC009095.1 | 2.28375528 | 1.639383579 | 0.93364676 | 1.75589275 | 0.07910672 | 0.20089024 |
| CARMIL2 | 208.173315 | -0.524286324 | 0.60484795 | -0.8668068 | 0.38604787 | 0.58174897 |
| ACD | 1201.34786 | -0.360470075 | 0.14816445 | -2.4329054 | 0.01497822 | 0.06081225 |
| PARD6A | 416.965862 | 0.057162064 | 0.14409098 | 0.39670812 | 0.69158271 | 0.82241065 |
| ENKD1 | 2359.82432 | -0.446067008 | 0.25814445 | -1.7279744 | 0.08399281 | 0.20968383 |
| C16orf86 | 505.283515 | 0.211543064 | 0.26253224 | 0.80577937 | 0.42037007 | 0.61311719 |
| GFOD2 | 1982.69028 | -0.346777795 | 0.13081647 | -2.6508727 | 0.00802841 | 0.03822299 |
| RANBP10 | 2401.33805 | 0.01016691 | 0.12573131 | 0.0808622 | 0.93555154 | 0.96914261 |
| TSNAXIP1 | 1561.05064 | -0.203460561 | 0.18672706 | -1.0896147 | 0.27588289 | 0.47121133 |
| CENPT | 4612.00104 | -0.429163329 | 0.16426587 | -2.6126141 | 0.00898527 | 0.04161981 |
| THAP11 | 1324.95582 | -0.111598307 | 0.14404503 | -0.774746 | 0.43848978 | 0.63025154 |
| NUTF2 | 3148.42967 | -0.839543116 | 0.20880888 | -4.0206294 | 5.80E-05 | 0.00083138 |
| EDC4 | 4107.59903 | -0.104520457 | 0.11789612 | -0.8865471 | 0.37532281 | 0.57129724 |
| AC040162.3 | 2323.50843 | -0.559532523 | 0.12233316 | -4.5738419 | 4.79E-06 | 0.00011372 |
| NRN1L | 79.35853 | -0.348293772 | 0.24445956 | -1.4247501 | 0.15422946 | 0.32016145 |
| PSKH1 | 2824.11651 | 0.366670878 | 0.21326731 | 1.71930183 | 0.08555943 | 0.21232845 |
| CTRL | 752.010598 | 0.101711901 | 0.14111989 | 0.72074815 | 0.47106448 | 0.65759572 |
| AC040162.1 | 2122.41094 | -1.091015371 | 0.33637066 | -3.2434914 | 0.00118074 | 0.00893691 |
| PSMB10 | 2566.61481 | -1.080835397 | 0.3489493 | -3.0973995 | 0.00195227 | 0.01324788 |
| LCAT | 1561.61068 | -0.182410603 | 0.2535773 | -0.7193491 | 0.47192585 | 0.65821986 |
| SLC12A4 | 4118.34767 | 0.128140815 | 0.19682997 | 0.6510229 | 0.51503171 | 0.6927644 |
| DPEP3 | 3.90800817 | 0.766538394 | 0.69265461 | 1.10666757 | 0.26843768 | 0.46266024 |
| DPEP2 | 141.085925 | 0.806033109 | 0.4853499 | 1.66072582 | 0.09676852 | 0.23219163 |
| DUS2 | 776.909893 | -0.127415928 | 0.17832657 | -0.7145089 | 0.47491253 | 0.66031021 |
| DPEP2NB | 0.91240889 | -1.414732173 | 1.62461397 | -0.8708113 | 0.3838572 | NA |
| DDX28 | 719.555439 | 0.018953379 | 0.27450052 | 0.06904678 | 0.94495238 | 0.9737607 |
| NFATC3 | 2836.85413 | 0.083618642 | 0.12423779 | 0.67305317 | 0.50091346 | 0.68184512 |
| AC020978.3 | 21.8827553 | -0.373729907 | 0.37896615 | -0.9861828 | 0.32404341 | 0.52163909 |
| AC020978.5 | 2592.95234 | 0.001621441 | 0.12434365 | 0.01304 | 0.98959588 | 0.99519138 |
| ESRP2 | 2581.93208 | -0.093505534 | 0.20093927 | -0.4653423 | 0.64168637 | 0.78769526 |
| PLA2G15 | 4630.67256 | -0.3186682 | 0.34340163 | -0.9279752 | 0.35342043 | 0.55100489 |
| AC020978.2 | 1312.74028 | -0.43248234 | 0.34615476 | -1.2493901 | 0.21152242 | 0.3950316 |
| SLC7A6 | 2393.64287 | -0.25749183 | 0.17167587 | -1.499872 | 0.13364756 | 0.29047112 |
| SLC7A6OS | 2299.02293 | -0.230741423 | 0.15681909 | -1.4713861 | 0.14118673 | 0.30158721 |
| PRMT7 | 1352.16031 | 0.439552473 | 0.27880996 | 1.57653076 | 0.11490353 | 0.26214953 |
| SMPD3 | 266.766976 | 0.056734758 | 0.53768303 | 0.10551711 | 0.9159655 | 0.95902526 |
| AC099521.1 | 10.9097662 | 0.030048797 | 0.71517684 | 0.0420159 | 0.96648602 | 0.98410057 |
| ZFP90 | 1112.18405 | 0.217185191 | 0.15731021 | 1.38061725 | 0.16739668 | 0.33860415 |
| AC126773.1 | 4.1281677 | -1.146179561 | 0.78574615 | -1.4587148 | 0.14464361 | 0.30668586 |
| AC126773.4 | 15.3170271 | 0.164286285 | 0.36984402 | 0.44420425 | 0.65689492 | 0.7986941 |
| CDH3 | 1667.73607 | -1.349757019 | 0.3125863 | -4.31803 | 1.57E-05 | 0.00029442 |
| AC126773.2 | 99.5701897 | -0.981723291 | 0.34082795 | -2.8804072 | 0.00397162 | 0.02263601 |
| CDH1 | 5421.97071 | 0.072601282 | 0.19325509 | 0.37567592 | 0.70715784 | 0.83325953 |
| AC099314.1 | 263.881589 | 1.368084647 | 0.25048254 | 5.46179654 | 4.71E-08 | 2.53E-06 |
| TANGO6 | 653.102324 | 0.486770565 | 0.16940221 | 2.87346053 | 0.00406002 | 0.02301928 |
| AC009137.1 | 52.9079083 | 0.692586219 | 0.23827535 | 2.90666335 | 0.00365306 | 0.02128867 |
| HAS3 | 2292.34428 | 0.014424503 | 0.19020114 | 0.07583815 | 0.93954786 | 0.97110795 |
| CHTF8 | 6763.84921 | 0.315202694 | 0.20874792 | 1.5099681 | 0.13105156 | 0.2866719 |
| UTP4 | 4045.98251 | 0.12292323 | 0.3167716 | 0.38805003 | 0.69797901 | 0.82654528 |
| SNTB2 | 2173.25903 | -0.307091756 | 0.23444089 | -1.3098899 | 0.19023309 | 0.36890531 |
| AC026464.4 | 3563.96335 | 0.219382577 | 0.08918966 | 2.45973111 | 0.01390411 | 0.05759372 |
| VPS4A | 7001.26719 | 0.151119529 | 0.07292507 | 2.07225761 | 0.03824142 | 0.11977982 |
| COG8 | 5363.05986 | 0.276231507 | 0.10394697 | 2.65742712 | 0.00787396 | 0.0377452 |
| PDF | 1171.37493 | 0.259839018 | 0.21313518 | 1.21912775 | 0.22279571 | 0.40880196 |
| AC026464.3 | 464.764398 | 0.354849017 | 0.08210413 | 4.32193851 | 1.55E-05 | 0.00029091 |
| AC026464.1 | 442.960403 | 1.101521003 | 0.31607938 | 3.48495053 | 0.00049223 | 0.00446972 |
| NIP7 | 1158.17922 | -0.51580597 | 0.25340228 | -2.0355223 | 0.04179835 | 0.12794648 |
| TMED6 | 319.676701 | 1.367554102 | 0.57458028 | 2.38009232 | 0.0173083 | 0.06752773 |
| TERF2 | 1469.52735 | -0.112605018 | 0.09705055 | -1.1602719 | 0.24593814 | 0.43722564 |
| CYB5B | 4236.23462 | -0.360494464 | 0.0996419 | -3.6179003 | 0.000297 | 0.00300767 |
| NFAT5 | 5608.75929 | -0.380836227 | 0.18066941 | -2.1079175 | 0.03503812 | 0.11274622 |
| NQO1 | 9404.86087 | -1.620646727 | 0.59772046 | -2.711379 | 0.0067004 | 0.03346392 |
| AC092115.2 | 1633.22422 | -1.585783271 | 0.65042983 | -2.4380543 | 0.01476655 | 0.06012879 |
| AC092115.3 | 952.245965 | -0.55763128 | 0.10986899 | -5.0754201 | 3.87E-07 | 1.47E-05 |

| | | | | | | |
|---|---|---|---|---|---|---|
| NOB1 | 2122.71043 | -0.463403577 | 0.12169065 | -3.8080459 | 0.00014007 | 0.00165485 |
| WWP2 | 5142.15222 | -0.102468028 | 0.14490855 | -0.707122 | 0.47949065 | 0.66418343 |
| CLEC18A | 971.66546 | 0.710383042 | 0.68952117 | 1.03025559 | 0.30289004 | 0.49979415 |
| PDPR | 3505.52443 | 0.383926644 | 0.26174957 | 1.46677087 | 0.1424384 | 0.30362141 |
| AC009060.1 | 1793.84043 | 0.529387064 | 0.26060851 | 2.03134988 | 0.04221951 | 0.12890295 |
| CLEC18C | 905.256248 | 0.819931728 | 0.80857353 | 1.01404721 | 0.31056023 | 0.50746519 |
| EXOSC6 | 1805.45872 | -0.459399186 | 0.27951219 | -1.6435748 | 0.10026407 | 0.23795909 |
| AARS | 7204.83927 | 0.125148404 | 0.16057693 | 0.7793673 | 0.43576338 | 0.62789358 |
| DDX19B | 1318.10812 | 0.156484777 | 0.12950098 | 1.20836753 | 0.22690592 | 0.4137687 |
| AC012184.2 | 2013.10945 | 0.084472135 | 0.12375819 | 0.68255793 | 0.49488623 | 0.67712083 |
| DDX19A | 2329.73634 | -0.105457553 | 0.10972566 | -0.961102 | 0.33650088 | 0.53445331 |
| ST3GAL2 | 1767.01726 | -0.0465704 | 0.14773378 | -0.3152319 | 0.75258558 | 0.86206078 |
| AC012184.1 | 212.22316 | -0.025216085 | 0.15065833 | -0.1673727 | 0.86707684 | 0.93098661 |
| FUK | 2544.95272 | 0.242708161 | 0.1633865 | 1.4854848 | 0.13741545 | 0.29604946 |
| COG4 | 3385.96973 | 0.158839744 | 0.15904294 | 0.99872241 | 0.31792918 | 0.51510153 |
| SF3B3 | 4552.50236 | -0.107375478 | 0.1245896 | -0.8618334 | 0.38877922 | 0.58437501 |
| IL34 | 274.388259 | 0.062572486 | 0.34552265 | 0.18109518 | 0.85629287 | 0.9247458 |
| MTSS1L | 5926.03072 | -0.955672242 | 0.43140276 | -2.2152669 | 0.02674175 | 0.09282348 |
| VAC14 | 3413.96983 | -0.086306117 | 0.21068563 | -0.4096441 | 0.68206707 | 0.81588709 |
| AC020763.1 | 4.23469079 | -2.273930744 | 1.51327767 | -1.5026527 | 0.13292863 | 0.28938562 |
| VAC14-AS1 | 245.400239 | -0.154382867 | 0.32017775 | -0.4821786 | 0.62967906 | 0.78003905 |
| AC027281.1 | 25.7927261 | -0.175738509 | 0.57923041 | -0.3034 | 0.76158504 | 0.86789099 |
| HYDIN | 386.2675 | -0.208543893 | 0.39030801 | -0.534306 | 0.59312987 | 0.75237031 |
| AC138625.2 | 0.08666371 | -0.642905668 | 3.30558948 | -0.1944905 | 0.84579185 | NA |
| AC138625.1 | 3.8660632 | -1.37580504 | 0.74606308 | -1.8440868 | 0.0651705 | 0.17512952 |
| CMTR2 | 1560.87765 | -0.129920718 | 0.31920009 | -0.4070197 | 0.68399353 | 0.81748139 |
| AC106736.1 | 0 | NA | NA | NA | NA | NA |
| CALB2 | 10.8497461 | -0.902897211 | 1.11228518 | -0.8117497 | 0.41693524 | 0.6099968 |
| LINC02136 | 0.09635527 | 0.547239394 | 3.29377202 | 0.16614368 | 0.86804388 | NA |
| TLE7 | 0.09644756 | -0.642905668 | 3.30558948 | -0.1944905 | 0.84579185 | NA |
| ZNF23 | 949.29888 | -0.11663031 | 0.13598181 | -0.8576905 | 0.39106339 | 0.5866952 |
| AC010547.4 | 975.49958 | -0.088208473 | 0.13524788 | -0.6521986 | 0.51427308 | 0.69211908 |
| AC010547.1 | 39.5481409 | 0.809995919 | 0.27065149 | 2.99276359 | 0.00276464 | 0.01726594 |
| ZNF19 | 328.550019 | 0.175385139 | 0.27004101 | 0.64947595 | 0.51603079 | 0.69349777 |
| AC010547.3 | 0.38697506 | -0.032130248 | 2.25518086 | -0.0142473 | 0.98863268 | NA |
| CHST4 | 3.96995927 | -0.045334204 | 0.66094133 | -0.0685904 | 0.94531569 | 0.97390119 |
| TAT | 31.4217717 | -0.131277618 | 0.66404696 | -0.1976933 | 0.84328505 | 0.91678046 |
| TAT-AS1 | 11.6036426 | 0.338700135 | 0.46614623 | 0.72659631 | 0.46747329 | 0.65455658 |
| AC009097.2 | 135.101731 | 0.258310961 | 0.4393378 | 0.58795524 | 0.55656234 | 0.72557054 |
| MARVELD3 | 1703.52642 | 0.005482791 | 0.36070254 | 0.01520031 | 0.98787237 | 0.99454413 |
| PHLPP2 | 1277.89824 | -0.434949797 | 0.22496718 | -1.9333922 | 0.0531879 | 0.15215501 |
| AC009097.1 | 8.92718467 | 0.050459249 | 0.5117333 | 0.09860458 | 0.92145223 | 0.96166416 |
| AC009097.4 | 4.03204735 | -0.677544905 | 0.63094763 | -1.0738528 | 0.28288864 | 0.47886593 |
| AC009097.3 | 0.61332334 | 0.031469644 | 1.58443801 | 0.01986171 | 0.98415369 | NA |
| AP1G1 | 5975.53226 | 0.018197618 | 0.14702741 | 0.12377024 | 0.90149719 | 0.95010115 |
| ATXN1L | 2736.66906 | 0.246594399 | 0.2032663 | 1.2131593 | 0.22506892 | 0.41138611 |
| IST1 | 9400.67615 | -0.29730009 | 0.10484403 | -2.8356416 | 0.00457337 | 0.02517743 |
| ZNF821 | 643.984124 | 0.053710606 | 0.21415593 | 0.25080139 | 0.80196767 | 0.89265491 |
| PKD1L3 | 109.461649 | 0.10487977 | 0.18930125 | 0.55403633 | 0.57955398 | 0.74277814 |
| DHODH | 1003.3796 | 0.385788522 | 0.11911855 | 3.23869381 | 0.00120078 | 0.00905222 |
| TXNL4B | 765.116821 | -0.144810185 | 0.1426451 | -1.0151781 | 0.31002092 | 0.50711814 |
| HP | 6107.1254 | -8.110351237 | 1.30356092 | -6.2216895 | 4.92E-10 | 5.59E-08 |
| HPR | 273.08862 | -2.24261478 | 0.90251218 | -2.4848582 | 0.01296031 | 0.0545969 |
| AC009087.1 | 125.820414 | -0.262388558 | 0.15565084 | -1.6857509 | 0.09184378 | 0.22383738 |
| DHX38 | 3366.78166 | 0.013643478 | 0.18689218 | 0.07300187 | 0.94180463 | 0.97233647 |
| PMFBP1 | 899.500363 | -0.555931962 | 0.19999232 | -2.7797665 | 0.0054398 | 0.02870536 |
| AC009075.1 | 1.02503966 | 0.186768971 | 1.42615964 | 0.13095937 | 0.89580745 | 0.94660158 |
| LINC01572 | 31.5486255 | -0.263767556 | 0.60421657 | -0.4365447 | 0.66244156 | 0.80142609 |
| AC004158.1 | 8.13064538 | 0.516685851 | 0.63842619 | 0.80931181 | 0.41833582 | 0.61113582 |
| AC004943.2 | 210.369199 | 0.260310469 | 0.20898118 | 1.24561679 | 0.21290512 | 0.39676223 |
| AC004943.3 | 2.83768815 | -0.280836249 | 0.88243364 | -0.318252 | 0.75029381 | 0.86062162 |
| ZFHX3 | 1253.91998 | -0.461489567 | 0.20267516 | -2.2769913 | 0.02278675 | 0.08257338 |
| AC004943.1 | 10.8188957 | 1.008117683 | 0.44797651 | 2.2503807 | 0.02442479 | 0.08674501 |
| AC116667.1 | 107.10707 | 0.119800234 | 0.38890409 | 0.3080457 | 0.75804756 | 0.86618156 |
| HCCAT5 | 2.98504358 | 0.229285406 | 1.10382067 | 0.2077198 | 0.83544776 | 0.91276071 |
| AC116667.2 | 1.87037295 | 1.522540868 | 1.09001369 | 1.39680894 | 0.16247103 | 0.33138509 |
| AC140912.1 | 1.33302833 | -1.879521091 | 1.47497347 | -1.2742745 | 0.20256615 | 0.38445871 |
| LINC01568 | 0 | NA | NA | NA | NA | NA |
| AC092114.1 | 0 | NA | NA | NA | NA | NA |
| AC087565.2 | 0 | NA | NA | NA | NA | NA |
| AC087565.1 | 0 | NA | NA | NA | NA | NA |
| AC138627.1 | 9.55750398 | 1.09978822 | 0.57306991 | 1.91911702 | 0.05496953 | 0.15563092 |
| AC009120.4 | 0.22673191 | 0.689158461 | 3.03348976 | 0.22718338 | 0.82028115 | NA |
| AC009120.5 | 8.28843355 | 0.180347191 | 0.5360158 | 0.33645872 | 0.73652497 | 0.85207763 |
| PSMD7 | 4488.67305 | -0.532771899 | 0.1836264 | -2.9013905 | 0.00371511 | 0.02155497 |
| AC009120.3 | 29.6632325 | 0.609385269 | 0.26694183 | 2.28283919 | 0.02243984 | 0.08165291 |
| AC009053.2 | 18.6877527 | 0.754522847 | 0.33440298 | 2.25632809 | 0.02405009 | 0.08573758 |
| NPIPB15 | 740.205345 | -1.286142476 | 1.11547203 | -1.1530029 | 0.2489092 | 0.44066597 |
| CLEC18B | 1996.52719 | 1.558625451 | 0.73926643 | 2.10834062 | 0.03500154 | 0.11265599 |
| AC009053.3 | 55.1545496 | 1.571344517 | 0.57812921 | 2.71798154 | 0.00656815 | 0.03294695 |
| GLG1 | 9047.98566 | 0.4206806 | 0.06734314 | 6.24682194 | 4.19E-10 | 4.82E-08 |
| RFWD3 | 1053.00883 | 0.056906047 | 0.18621672 | 0.30559043 | 0.7599165 | 0.86700095 |
| MLKL | 933.035926 | -1.23920051 | 0.3825401 | -3.2394003 | 0.00119781 | 0.00903758 |
| FA2H | 486.851172 | 0.231579879 | 0.33871676 | 0.68369773 | 0.49416606 | 0.67670324 |
| WDR59 | 3569.03924 | 0.028016438 | 0.15094097 | 0.18561189 | 0.85274913 | 0.92285671 |
| ZNRF1 | 1505.11285 | -0.047178325 | 0.16583592 | -0.284488 | 0.77603645 | 0.87702653 |
| AC099508.1 | 293.767207 | 0.509201202 | 0.1946727 | 2.61567854 | 0.00890504 | 0.0413062 |
| LDHD | 4746.64567 | 1.775324017 | 0.42866358 | 4.14153219 | 3.45E-05 | 0.00054936 |
| AC099508.2 | 0.78015163 | 3.554487843 | 2.37510333 | 1.49656135 | 0.13450743 | NA |
| ZFP1 | 452.966504 | -0.050723274 | 0.20576814 | -0.2465069 | 0.80528984 | 0.8944712 |
| CTRB2 | 18.6973775 | -2.705407288 | 0.81270865 | -3.3288772 | 0.00087197 | 0.00705722 |
| CTRB1 | 15.0648057 | -2.828730189 | 0.89304823 | -3.1674999 | 0.00153756 | 0.01102376 |
| BCAR1 | 12304.8855 | -1.072032854 | 0.25452293 | -4.2119303 | 2.53E-05 | 0.00042938 |
| CFDP1 | 3183.53729 | 0.174409801 | 0.18277993 | 0.95420652 | 0.33997912 | 0.53809861 |
| AC009163.4 | 788.534737 | 0.216151921 | 0.20683557 | 1.04504231 | 0.29600341 | 0.49277045 |
| AC009163.7 | 72.457693 | 1.580685422 | 0.34033517 | 4.64449631 | 3.41E-06 | 8.66E-05 |
| TMEM170A | 1568.83353 | -0.302639055 | 0.16799556 | -1.8014706 | 0.07162874 | 0.18744169 |
| CHST6 | 46.5975807 | 0.590023151 | 0.66268603 | 0.89035097 | 0.37327746 | 0.56947218 |
| AC009163.1 | 2.40655001 | 1.273005305 | 1.22492275 | 1.03925354 | 0.29868684 | 0.49558749 |
| AC009163.6 | 1.20486383 | -0.585953877 | 1.34749465 | -0.4348469 | 0.66367356 | 0.80246755 |
| AC009163.2 | 101.143606 | 0.353405568 | 0.24610017 | 1.43602328 | 0.15099572 | 0.31576186 |
| CHST5 | 41.3133695 | -0.064190131 | 0.29475782 | -0.2177724 | 0.82760642 | 0.9082351 |
| TMEM231 | 715.055 | -0.014494932 | 0.26991943 | -0.053701 | 0.95717342 | 0.97945918 |
| AC025287.2 | 2.93702253 | 0.168535785 | 0.94835522 | 0.17771377 | 0.85894777 | 0.92639888 |
| GABARAPL2 | 4869.19626 | -0.046828806 | 0.16982589 | -0.275746 | 0.78274318 | 0.88107472 |
| AC025287.3 | 10.2548948 | -0.313820505 | 0.48147817 | -0.6517855 | 0.51453954 | 0.69239713 |
| ADAT1 | 619.444079 | 0.239115771 | 0.13592059 | 1.75923144 | 0.07853821 | 0.19986996 |
| KARS | 5543.62572 | -0.200543247 | 0.1085062 | -1.8482193 | 0.06457062 | 0.17394293 |
| TERF2IP | 3552.2135 | -0.275167839 | 0.10956321 | -2.5114985 | 0.01202198 | 0.05158145 |
| AC025287.4 | 3.0090614 | -0.148251183 | 0.83719481 | -0.1770809 | 0.85944487 | 0.92663183 |
| DUXB | 6.99806027 | -0.550501992 | 0.54207896 | -1.0155384 | 0.30984926 | 0.50697377 |
| CPHXL | 0.23017671 | 0.689158974 | 3.02240241 | 0.22801695 | 0.81963307 | NA |
| AC105430.1 | 0 | NA | NA | NA | NA | NA |

| | | | | | | |
|---|---|---|---|---|---|---|
| AC099511.1 | 0 | NA | NA | NA | NA | NA |
| AC010528.1 | 0 | NA | NA | NA | NA | NA |
| CNTNAP4 | 5.47471499 | 2.179574207 | 0.7038144 | 3.09680251 | 0.0019562 | 0.01326776 |
| AC104151.1 | 0.47856485 | 0.340082806 | 2.24943679 | 0.15118576 | 0.87982918 | NA |
| AC108125.1 | 0 | NA | NA | NA | NA | NA |
| LINC02125 | 0.54526294 | -0.064264064 | 1.92583355 | -0.0333695 | 0.97337995 | NA |
| AC106729.1 | 0.25112153 | -0.178324373 | 3.00248861 | -0.0593922 | 0.95263973 | NA |
| MON1B | 2753.31702 | -0.16667157 | 0.2001328 | -0.8328048 | 0.4049548 | 0.59972246 |
| SYCE1L | 270.196582 | -0.952255765 | 0.17012889 | -5.5972607 | 2.18E-08 | 1.34E-06 |
| AC009139.2 | 522.381954 | -0.07094946 | 0.25540601 | -0.2777909 | 0.7811729 | 0.88024653 |
| AC009139.1 | 1.60752124 | 0.263572833 | 0.99796606 | 0.26411002 | 0.79169514 | 0.8869488 |
| ADAMTS18 | 26.1181957 | -0.61326988 | 0.37052406 | -1.6551419 | 0.09789572 | 0.23415059 |
| AC025284.1 | 0.16783122 | 1.368031398 | 3.28033652 | 0.41703996 | 0.67664918 | NA |
| LINC02131 | 0.03392908 | 0.380740593 | 3.29738709 | 0.11546736 | 0.90807469 | NA |
| NUDT7 | 458.738569 | 0.845068027 | 0.3500165 | 2.41436626 | 0.01576261 | 0.06311266 |
| AC092134.1 | 367.127734 | 0.814402136 | 0.36672598 | 2.22073747 | 0.02636875 | 0.09192422 |
| VAT1L | 291.406587 | 1.564750485 | 0.55927056 | 2.79784167 | 0.00514453 | 0.02747849 |
| AC079414.1 | 0 | NA | NA | NA | NA | NA |
| CLEC3A | 4.07871837 | 3.67033958 | 1.2501886 | 2.93582872 | 0.00332658 | 0.01984764 |
| WWOX | 770.04742 | 0.496126281 | 0.14492709 | 3.42328182 | 0.0006187 | 0.00533871 |
| WWOX-AS1 | 0.09635527 | 0.547239394 | 3.29377202 | 0.16614368 | 0.86804388 | NA |
| AC046158.1 | 8.70335453 | 4.688267788 | 0.79538035 | 5.89437213 | 3.76E-09 | 2.99E-07 |
| AC009145.3 | 0.11786115 | 0.02007176 | 3.29738704 | 0.00608717 | 0.99514317 | NA |
| AC092376.2 | 22.1169843 | 0.36987524 | 0.39113619 | 0.94564311 | 0.34433065 | 0.54199243 |
| AC092376.1 | 0.7056569 | 0.176138771 | 1.87803095 | 0.09378907 | 0.92527672 | NA |
| AC084064.1 | 1.85436811 | -0.551149164 | 1.23577912 | -0.4459933 | 0.65560211 | 0.79801642 |
| MAF | 8911.62299 | 0.879781363 | 0.34927455 | 2.51888197 | 0.01177281 | 0.0508766 |
| AC009159.2 | 7.94112466 | 1.748554613 | 0.52791647 | 3.31218048 | 0.00092572 | 0.00739957 |
| AC009159.3 | 149.702901 | 1.723015096 | 0.37556867 | 4.58774983 | 4.48E-06 | 0.00010812 |
| AC009159.1 | 3.94752949 | 0.93391959 | 0.89193842 | 1.04706734 | 0.29506852 | 0.49190523 |
| LINC01229 | 21.5943406 | 1.217124129 | 0.58745464 | 2.07186062 | 0.03827844 | 0.11988153 |
| MAFTRR | 51.1992728 | 1.958205149 | 0.55757236 | 3.51201974 | 0.00044471 | 0.00414679 |
| LINC01228 | 1.62065445 | 0.384339157 | 0.91396387 | 0.42051898 | 0.67410636 | 0.81024608 |
| AC092130.1 | 0 | NA | NA | NA | NA | NA |
| AC022166.1 | 0.35296529 | -1.691981139 | 3.30304428 | -0.512249 | 0.60847675 | NA |
| AC105411.1 | 5.99647305 | 0.62024342 | 0.72606186 | 0.85425699 | 0.39296261 | 0.58844158 |
| AC022166.2 | 0 | NA | NA | NA | NA | NA |
| DYNLRB2 | 90.9693634 | 0.64660886 | 0.31301492 | 2.06574454 | 0.03885261 | 0.12122438 |
| AC108097.1 | 6.29740431 | 1.097532474 | 0.60223507 | 1.82243202 | 0.06838946 | 0.18112183 |
| LINC01227 | 0.2891581 | 1.145957498 | 3.27645355 | 0.34975545 | 0.72652223 | NA |
| CDYL2 | 774.626797 | -0.248186407 | 0.17032369 | -1.4571456 | 0.1450762 | 0.30730656 |
| AC099313.1 | 0.16775052 | 0.435199075 | 3.28898345 | 0.13232024 | 0.894731 | NA |
| LINC02170 | 7.34763603 | 0.382327124 | 0.43808579 | 0.87272204 | 0.3828146 | 0.5788687 |
| CMC2 | 1627.50648 | -0.029968518 | 0.12208681 | -0.2454689 | 0.80609336 | 0.89501819 |
| CENPN | 611.352268 | -0.078740864 | 0.17113961 | -0.4600972 | 0.64544642 | 0.79072809 |
| AC092718.2 | 47.9206868 | -0.012726307 | 0.33686739 | -0.0377784 | 0.96986438 | 0.9860936 |
| AC092718.8 | 1086.17061 | 0.559717782 | 0.35483202 | 1.57741623 | 0.11469978 | 0.26182767 |
| ATMIN | 2652.74039 | -0.104496576 | 0.24976176 | -0.418385 | 0.67566565 | 0.81134996 |
| C16orf46 | 222.757808 | -0.357637461 | 0.16437124 | -2.175791 | 0.02957089 | 0.09984435 |
| AC092718.3 | 1277.72446 | 0.397881599 | 0.29883258 | 1.33145323 | 0.18303992 | 0.35972297 |
| AC092718.5 | 21.1970058 | -0.152644631 | 0.34689275 | -0.4400341 | 0.65991242 | 0.79999826 |
| AC092718.6 | 0.25082976 | -0.178310673 | 3.00331177 | -0.0593713 | 0.95265633 | NA |
| AC092718.1 | 5.31806491 | 0.453504375 | 0.66331794 | 0.6836908 | 0.49417044 | 0.67670324 |
| GCSH | 1160.42371 | 0.552760283 | 0.34508046 | 1.60183013 | 0.10919318 | 0.25262538 |
| BCO1 | 239.837103 | 0.177390743 | 0.38850262 | 0.45660114 | 0.64795776 | 0.79277053 |
| AC009148.1 | 31.843472 | -0.126295842 | 0.37857567 | -0.3336079 | 0.73867544 | 0.85332595 |
| GAN | 537.13378 | -0.720052011 | 0.20112595 | -3.580105 | 0.00034346 | 0.00337693 |
| AC092139.1 | 11.3646464 | 1.369003468 | 0.49438919 | 2.7690805 | 0.00562147 | 0.02943572 |
| CMIP | 4310.45489 | 0.116133801 | 0.13538883 | 0.85777976 | 0.39101409 | 0.58667712 |
| AC099524.1 | 12.6136625 | -0.095168831 | 0.49339317 | -0.1928864 | 0.84704794 | 0.91970801 |
| PLCG2 | 3151.94807 | 0.655940497 | 0.2290739 | 2.86344493 | 0.00419062 | 0.023604 |
| AC092142.2 | 19.6671466 | -0.37912067 | 0.33438007 | -1.1338016 | 0.25687777 | 0.44955314 |
| SDR42E1 | 572.293321 | -0.031655615 | 0.15689225 | -0.2017666 | 0.8400992 | 0.91492195 |
| HSD17B2 | 107.146571 | 0.095853329 | 0.58869436 | 0.16282359 | 0.87065733 | 0.93269556 |
| AC092142.1 | 46.8290859 | 0.356786221 | 0.59418297 | 0.60046524 | 0.54819622 | 0.71893289 |
| MPHOSPH6 | 664.878975 | -0.49263998 | 0.12105265 | -4.069634 | 4.71E-05 | 0.00070751 |
| AC138304.1 | 0.08695548 | -0.642905668 | 3.30558948 | -0.1944905 | 0.84579185 | NA |
| AC009117.3 | 0 | NA | NA | NA | NA | NA |
| AC009117.2 | 0 | NA | NA | NA | NA | NA |
| CDH13 | 528.523485 | 0.516754087 | 0.50199803 | 1.02939465 | 0.30329426 | 0.50016963 |
| AC125793.1 | 0 | NA | NA | NA | NA | NA |
| AC009142.1 | 0.59396971 | -0.80412446 | 2.23038 | -0.3605325 | 0.71844896 | NA |
| AC009063.3 | 0.16421342 | 1.36803136 | 3.28033652 | 0.41703994 | 0.67664919 | NA |
| AC009063.2 | 1.25759883 | 0.064324867 | 1.4003226 | 0.04593575 | 0.96336146 | 0.98289258 |
| AC009063.1 | 0 | NA | NA | NA | NA | NA |
| AC009119.1 | 4.79173587 | 0.750423812 | 0.59868844 | 1.25344631 | 0.21004333 | 0.39335577 |
| HSBP1 | 5895.81147 | -0.271516648 | 0.12322494 | -2.2034229 | 0.02756496 | 0.09498078 |
| MLYCD | 1567.85747 | 0.356593732 | 0.2399749 | 1.48596262 | 0.13728901 | 0.29587374 |
| AC009119.2 | 434.181515 | 0.540822551 | 0.23727049 | 2.27935022 | 0.02264625 | 0.08219983 |
| AC009119.3 | 5.09018296 | 0.335130592 | 0.65125632 | 0.51459092 | 0.60683891 | 0.76327544 |
| OSGIN1 | 4555.29027 | -2.658225776 | 0.75965649 | -3.4992471 | 0.00046657 | 0.00430413 |
| NECAB2 | 373.197736 | 2.673287637 | 0.62887788 | 4.25088515 | 2.13E-05 | 0.00037234 |
| SLC38A8 | 2.30255072 | -0.076079107 | 0.86289914 | -0.0881669 | 0.92974406 | 0.96597439 |
| MBTPS1 | 5582.08522 | 0.202307994 | 0.09695484 | 2.08662078 | 0.03692243 | 0.11679995 |
| AC040169.1 | 46.0695518 | 0.165441617 | 0.33064765 | 0.50035625 | 0.61682425 | 0.77088415 |
| HSDL1 | 1296.73453 | -0.053776383 | 0.28021373 | -0.191912 | 0.84781112 | 0.9200943 |
| DNAAF1 | 1409.2284 | -0.195285034 | 0.23472562 | -0.8319715 | 0.40542501 | 0.60018288 |
| TAF1C | 6701.54366 | -0.118357654 | 0.18762704 | -0.6308134 | 0.52816253 | 0.70274211 |
| ADAD2 | 36.7891658 | 1.471600243 | 0.76911209 | 1.91337551 | NA | NA |
| AC009123.1 | 25.5035997 | 1.719526378 | 0.84861769 | 2.02626743 | 0.04273739 | 0.13000172 |
| KCNG4 | 0.32679975 | -1.692381882 | 2.87486798 | -0.5886816 | 0.55607488 | NA |
| AC010551.3 | 0.23374159 | 1.011073437 | 2.99871927 | 0.33716842 | 0.73598994 | NA |
| WFDC1 | 646.377931 | -0.105795462 | 0.44108847 | -0.2398509 | 0.81044586 | 0.89747017 |
| AC010551.2 | 0.27685825 | 1.149138791 | 3.27633754 | 0.35073883 | 0.72578429 | NA |
| AC010551.1 | 0 | NA | NA | NA | NA | NA |
| ATP2C2 | 556.401747 | 0.788918456 | 0.27696159 | 2.84847604 | 0.00439292 | 0.02446488 |
| ATP2C2-AS1 | 312.402416 | 0.454132289 | 0.30206031 | 1.50344905 | 0.13272329 | 0.28908178 |
| TLDC1 | 949.802819 | -0.463651219 | 0.19111288 | -2.4260596 | 0.01526377 | 0.06161113 |
| AC022165.1 | 66.4715181 | -0.255309064 | 0.25823227 | -0.9886799 | 0.32281977 | 0.52056242 |
| COTL1 | 8175.87309 | 0.965359965 | 0.35202306 | 2.74232022 | 0.00610068 | 0.03120757 |
| AC092145.1 | 2.74741248 | 1.738783997 | 1.5257405 | 1.13963285 | 0.25443929 | 0.44713886 |
| KLHL36 | 3241.49071 | -0.13671953 | 0.18088265 | -0.7558466 | 0.44974118 | 0.63978395 |
| USP10 | 3385.81247 | -0.133605269 | 0.17233737 | -0.7752542 | 0.43818947 | 0.63008807 |
| CRISPLD2 | 3393.25118 | -0.528023229 | 0.62976103 | -0.8384502 | 0.40177793 | 0.59664454 |
| LINC02176 | 0.12150485 | 1.04981648 | 3.28477129 | 0.31960109 | 0.74927075 | NA |
| ZDHHC7 | 4653.83989 | -0.411266655 | 0.18004794 | -2.2842064 | 0.02235941 | 0.08146139 |
| KIAA0513 | 870.573697 | -0.510349118 | 0.21181667 | -2.4093907 | 0.01597918 | 0.06379576 |
| FAM92B | 8.01490322 | -3.968098844 | 1.34589112 | -2.948306 | 0.00319521 | 0.0192765 |
| LINC02139 | 43.4161842 | 3.307684585 | 0.45413545 | 7.28347583 | 3.25E-13 | 1.06E-10 |
| AC026469.1 | 41.1392453 | 3.273713276 | 0.45763782 | 7.15350254 | 8.46E-13 | 2.30E-10 |
| GSE1 | 4074.25976 | -0.420046722 | 0.14275057 | -2.9425222 | 0.0032555 | 0.01955082 |
| LINC00311 | 2.36135415 | 0.220355177 | 0.80718768 | 0.27299125 | 0.78485994 | 0.88249517 |
| AC133540.1 | 1.92494411 | -0.043172194 | 1.12047773 | -0.0385302 | 0.96926498 | 0.98578824 |

| | | | | | | |
|---|---|---|---|---|---|---|
| AC092127.2 | 5.88643118 | 0.215196656 | 0.58781787 | 0.3660941 | 0.71429486 | 0.83764267 |
| AC092127.1 | 1.45075958 | -0.417589133 | 1.00216326 | -0.4166877 | 0.67690683 | 0.81209758 |
| GINS2 | 50.82272 | -0.485247985 | 0.48194325 | -1.0068571 | 0.31400345 | 0.51094247 |
| C16orf74 | 44.4462913 | 0.245385461 | 0.38658729 | 0.63474787 | 0.52559287 | 0.70039776 |
| AC018695.2 | 0.11786115 | 0.02007176 | 3.29738704 | 0.00608717 | 0.99514317 | NA |
| EMC8 | 1295.08961 | -0.432583874 | 0.19636532 | -2.2029546 | 0.02759795 | 0.09506962 |
| AC018695.4 | 4.25413083 | 1.633981056 | 0.78356667 | 2.0853121 | 0.03704098 | 0.11704861 |
| AC018695.3 | 1.0435481 | -2.523885272 | 1.46539263 | -1.722327 | 0.0850103 | 0.21138703 |
| COX4I1 | 17079.7781 | 0.020020001 | 0.14517005 | 0.13790724 | 0.89031373 | 0.94316 |
| IRF8 | 968.60627 | -0.490938085 | 0.45427722 | -1.0807015 | 0.2798299 | 0.47541172 |
| AC092723.5 | 1.25577203 | 0.085986011 | 1.25037938 | 0.06876794 | 0.94517434 | 0.9738697 |
| LINC02132 | 1.05220389 | -0.078062965 | 1.30642088 | -0.0597533 | 0.95235212 | 0.97717271 |
| AC092723.4 | 0.08757577 | 0.547239394 | 3.29377202 | 0.16614368 | 0.86804388 | NA |
| AC092723.1 | 0 | NA | NA | NA | NA | NA |
| AC092723.3 | 0.49723436 | 2.259354852 | 2.09142231 | 1.08029585 | 0.28001046 | NA |
| AC135012.1 | 0.15179023 | 0.435199075 | 3.28898345 | 0.13232024 | 0.894731 | NA |
| AC135012.2 | 0 | NA | NA | NA | NA | NA |
| AC135012.3 | 0 | NA | NA | NA | NA | NA |
| LINC01082 | 0.08695548 | -0.642905668 | 3.30558948 | -0.1944905 | 0.84579185 | NA |
| AC040170.1 | 1.61137745 | 1.771511507 | 1.20927221 | 1.46494023 | 0.14293723 | 0.30429117 |
| LINC01081 | 2.17975364 | 1.129453684 | 1.06292284 | 1.06259235 | 0.28796686 | 0.48416119 |
| LINC02135 | 0 | NA | NA | NA | NA | NA |
| LINC00917 | 0.52674588 | 0.01501143 | 2.28026447 | 0.0065832 | 0.99474741 | NA |
| AC092327.2 | 0.3933166 | -1.901396697 | 3.2962053 | -0.5768441 | 0.56404475 | NA |
| FENDRR | 19.7160845 | -0.138683731 | 0.66784799 | -0.2076576 | 0.8354963 | 0.91276071 |
| FOXF1 | 193.063847 | -0.309258556 | 0.459911 | -0.6724313 | 0.50130916 | 0.68210661 |
| AC009108.3 | 23.1687447 | -0.175650359 | 0.56073147 | -0.3132522 | 0.75408907 | 0.86318765 |
| MTHFSD | 922.401306 | 0.021074997 | 0.18528778 | 0.11374197 | 0.90944234 | 0.9548873 |
| FOXC2-AS1 | 67.6239185 | 0.26440875 | 0.36131442 | 0.73179683 | 0.46429258 | 0.65178001 |
| FOXC2 | 376.819112 | -0.09003472 | 0.34482851 | -0.2611 | 0.7940154 | 0.88844572 |
| FOXL1 | 143.350021 | -0.953141397 | 0.54312195 | -1.7549307 | 0.07927116 | 0.20115808 |
| AC009108.2 | 0 | NA | NA | NA | NA | NA |
| AC009154.1 | 2.28840728 | -0.632459003 | 0.98055586 | -0.6450005 | 0.51892687 | 0.69578998 |
| AC009154.2 | 7.31772661 | 0.786216654 | 0.64523561 | 1.21849544 | 0.22303576 | 0.40912086 |
| LINC02189 | 0.03392908 | 0.380740593 | 3.29738709 | 0.11546736 | 0.90807469 | NA |
| LINC02188 | 78.7431252 | -0.270341706 | 0.40464257 | -0.6681 | 0.50406976 | 0.68375733 |
| LINC02181 | 4.14731402 | 0.945181444 | 0.61084345 | 1.54733827 | 0.1217817 | 0.27262355 |
| AC106745.1 | 3.46971866 | 0.73447688 | 0.73444395 | 1.00004483 | 0.31728881 | 0.51453822 |
| C16orf95 | 118.234544 | 0.616001734 | 0.25243004 | 2.44028697 | 0.0146756 | 0.05993426 |
| AC136285.1 | 0.08666371 | -0.642905668 | 3.30558948 | -0.1944905 | 0.84579185 | NA |
| AC136285.2 | 0.34004239 | -0.26115352 | 2.32716496 | -0.1122196 | 0.91064929 | NA |
| AC010531.1 | 276.697455 | 0.321244342 | 0.22232716 | 1.44491721 | 0.14848114 | 0.31238613 |
| AC010531.6 | 20.0613103 | 0.852299727 | 0.33299197 | 2.55952037 | 0.01048167 | 0.04668062 |
| FBXO31 | 2306.21133 | -0.361221148 | 0.20449607 | -1.7663965 | 0.07732934 | 0.19784368 |
| AC010531.7 | 143.675767 | -0.720270033 | 0.30201284 | -2.3848987 | 0.01708382 | 0.06688945 |
| AC010531.5 | 6.52305119 | 0.624483312 | 0.6489055 | 0.96236403 | 0.33586677 | 0.53405733 |
| MAP1LC3B | 6888.74819 | -0.90658011 | 0.28269346 | -3.2069369 | 0.00134156 | 0.00992065 |
| AC010531.3 | 5.22735305 | -0.234695085 | 0.59275253 | -0.3959411 | 0.69214849 | 0.82282732 |
| ZCCHC14 | 3134.56099 | -0.157263629 | 0.33831602 | -0.4648424 | 0.64204433 | 0.78795127 |
| AC092720.1 | 17.5574141 | 0.938135428 | 0.43955977 | 2.13426137 | 0.03282139 | 0.10755971 |
| AC092720.2 | 0 | NA | NA | NA | NA | NA |
| JPH3 | 30.6912765 | 0.074429676 | 0.30620032 | 0.24307511 | 0.80794721 | 0.89620864 |
| AC010536.3 | 8.7037848 | 0.365682101 | 0.54235833 | 0.67424447 | 0.5001559 | 0.68132613 |
| AC010536.1 | 37.2905999 | 0.013759251 | 0.32081536 | 0.04288838 | 0.96579051 | 0.98387257 |
| KLHDC4 | 2304.97366 | -0.088988338 | 0.11479173 | -0.7752156 | 0.43821226 | 0.63008807 |
| AC126696.3 | 0.47271494 | -2.119560063 | 2.77932556 | -0.7626167 | 0.44569203 | NA |
| AC126696.1 | 0.08695548 | -0.642905668 | 3.30558948 | -0.1944905 | 0.84579185 | NA |
| SLC7A5 | 2404.36209 | -0.483100835 | 0.39833033 | -1.2128146 | 0.22520071 | 0.41151291 |
| AC126696.2 | 168.479926 | -0.021868918 | 0.38603489 | -0.0566501 | 0.95482392 | 0.97841508 |
| CA5A | 7.13498276 | 0.607209226 | 0.60492564 | 1.00377498 | 0.31548708 | 0.51256204 |
| BANP | 877.675626 | -0.17889132 | 0.19590649 | -0.9131465 | 0.36116551 | 0.55878832 |
| AC134312.4 | 0.48866611 | 1.467906965 | 2.06687212 | 0.71020696 | 0.47757581 | NA |
| AC134312.3 | 0.23560867 | 0.952737031 | 3.28103697 | 0.2903768 | 0.77152799 | NA |
| AC134312.1 | 3.24349319 | 1.09887135 | 0.71788332 | 1.53071025 | 0.12584102 | 0.27862533 |
| AC134312.2 | 1.1994212 | -1.260021028 | 1.54973751 | -0.8130545 | 0.41618682 | 0.60944283 |
| AC134312.5 | 2.15827711 | -1.110973944 | 1.54879965 | -0.7173129 | 0.47318106 | 0.65922361 |
| AC134312.6 | 0.20754827 | -0.339942428 | 3.12492333 | -0.1087842 | 0.91337362 | NA |
| LINC02182 | 5.30521783 | 1.094124936 | 0.7480689 | 1.46259915 | 0.1435771 | 0.30517221 |
| AC138512.1 | 3.58829929 | 3.280575404 | 0.87283152 | 3.75854368 | 0.00017091 | 0.0019359 |
| ZNF469 | 62.2030302 | 0.774858038 | 0.29122947 | 2.66064437 | 0.00779913 | 0.0375229 |
| ZFPM1 | 306.810836 | -0.492160985 | 0.20675604 | -2.3803947 | 0.0172941 | 0.06748232 |
| AC116552.1 | 23.2400256 | 0.377459316 | 0.36382166 | 1.03748445 | 0.29951014 | 0.49614701 |
| ZC3H18 | 2656.01297 | -0.342096806 | 0.15084358 | -2.267891 | 0.02333585 | 0.08395176 |
| IL17C | 5.16274721 | -3.476931744 | 1.31238726 | -2.6493184 | 0.00806543 | 0.03833551 |
| CYBA | 8405.29536 | -0.733388169 | 0.27713201 | -2.6463495 | 0.00813657 | 0.03859434 |
| MVD | 2949.52177 | -0.611489999 | 0.17004371 | -3.5960755 | 0.00032305 | 0.00321348 |
| SNAI3-AS1 | 186.021553 | 0.052560327 | 0.21364152 | 0.24602113 | 0.80566587 | 0.89473191 |
| SNAI3 | 78.1693129 | -0.345264717 | 0.38897022 | -0.8876379 | 0.37473555 | 0.57073908 |
| RNF166 | 1552.8542 | -0.177625305 | 0.24533269 | -0.7240181 | 0.46905463 | 0.65569182 |
| AC138028.3 | 316.404384 | -0.359694622 | 0.23122773 | -1.555586 | 0.11980657 | 0.26965681 |
| CTU2 | 774.314574 | -0.36723518 | 0.14557173 | -2.5227094 | 0.01164546 | 0.05047514 |
| AC138028.6 | 153.236828 | -0.032581759 | 0.18751473 | -0.1737557 | 0.86205744 | 0.92808258 |
| PIEZO1 | 7715.44917 | -0.099329935 | 0.17744117 | -0.5597908 | 0.57562214 | 0.73972494 |
| AC138028.4 | 1126.88514 | 0.068537153 | 0.18299842 | 0.3745232 | 0.7080151 | 0.83387039 |
| AC138028.2 | 297.201273 | 0.977823363 | 0.33651589 | 2.90572718 | 0.00366401 | 0.02134303 |
| AC138028.5 | 85.3766724 | 1.267342772 | 0.36363059 | 3.48524794 | 0.00049168 | 0.0044686 |
| AC138028.1 | 10.4381463 | 0.123240186 | 0.40991003 | 0.3006518 | 0.76368003 | 0.86926414 |
| CDT1 | 83.3263581 | -0.800775893 | 0.30455816 | -2.6293037 | 0.00855599 | 0.0400322 |
| APRT | 4053.9058 | -0.128809993 | 0.10020675 | -1.2854423 | 0.19863776 | 0.37968468 |
| GALNS | 2031.48269 | -0.043647836 | 0.13954522 | -0.3127863 | 0.75444302 | 0.86329626 |
| TRAPPC2L | 1866.93919 | 0.00913722 | 0.1542136 | 0.05925041 | 0.95275266 | 0.97717271 |
| PABPN1L | 1.52708775 | -1.686706961 | 1.60442552 | -1.0512841 | 0.29312815 | 0.48969207 |
| CBFA2T3 | 167.332274 | 0.31975131 | 0.51254703 | 0.62384775 | 0.53272758 | 0.7068228 |
| AC092384.2 | 47.3423386 | 0.76548225 | 0.47313003 | 1.61791093 | 0.10568179 | 0.24670936 |
| AC092384.1 | 5.99256175 | -0.343490972 | 0.54401139 | -0.631404 | 0.5277764 | 0.7023701 |
| AC135782.2 | 0.08757577 | 0.547239394 | 3.29377202 | 0.16614368 | 0.86804388 | NA |
| AC135782.1 | 0.26181525 | 1.093535322 | 3.27839713 | 0.33355792 | 0.73871318 | NA |
| ACSF3 | 1987.38321 | 0.202552298 | 0.17120103 | 1.18312549 | 0.23675942 | 0.42601653 |
| AC135782.3 | 249.193535 | 0.521551527 | 0.21650242 | 2.408987 | 0.01599687 | 0.06384704 |
| LINC00304 | 44.2702973 | -0.634132189 | 0.47964168 | -1.3220957 | 0.18613629 | 0.36366704 |
| LINC02138 | 5.74537166 | 0.592991826 | 0.59969217 | 0.98882703 | 0.32274778 | 0.52050999 |
| CDH15 | 11.3468561 | 1.493446054 | 0.43312293 | 3.44808821 | 0.00056457 | 0.0049942 |
| SLC22A31 | 43.9410218 | -0.843573074 | 0.73008952 | -1.1554379 | 0.24791117 | 0.43949025 |
| AC009113.1 | 229.077698 | 0.791977825 | 0.29391064 | 2.69462117 | 0.00704687 | 0.03481191 |
| ZNF778 | 608.327254 | 0.08520936 | 0.23888522 | 0.35669582 | 0.72131954 | 0.84145968 |
| AC009113.2 | 28.4048351 | -0.490967523 | 0.26824004 | -1.830329 | 0.06720076 | 0.17890803 |
| ANKRD11 | 4923.21406 | -0.083731222 | 0.14691548 | -0.5699279 | 0.56872663 | 0.73429893 |
| AC137932.3 | 484.783971 | -0.344883104 | 0.19504647 | -1.7682099 | 0.07702581 | 0.19723903 |
| AC137932.1 | 43.6585019 | 0.561399248 | 0.36085923 | 1.5557292 | 0.1197725 | 0.26960314 |
| SPG7 | 7957.79127 | -0.077105993 | 0.18068488 | -0.4267429 | 0.66956659 | 0.80707917 |
| AC092123.1 | 35.5401303 | 0.981971134 | 0.43265581 | 2.26963584 | 0.02322969 | 0.08375814 |
| RPL13 | 35243.8771 | -0.58822644 | 0.16693499 | -3.5236858 | 0.00042559 | 0.00400384 |
| CPNE7 | 840.769242 | -2.853157305 | 0.56325047 | -5.0655214 | 4.07E-07 | 1.53E-05 |

| | | | | | | |
|---|---|---|---|---|---|---|
| DPEP1 | 17990.6357 | 1.602255905 | 0.50739905 | 3.1577826 | 0.00158974 | 0.01129959 |
| CHMP1A | 5588.81498 | -0.233763722 | 0.12236808 | -1.9103324 | 0.05609042 | 0.15783849 |
| SPATA33 | 618.531971 | 0.117187735 | 0.1055558 | 1.110197 | 0.26691415 | 0.46111937 |
| CDK10 | 7983.35234 | -0.183574864 | 0.18567281 | -0.9887008 | 0.32280954 | 0.52056242 |
| LINC02166 | 158.092811 | 0.77437597 | 0.31911319 | 2.42664986 | 0.01523896 | 0.06154609 |
| AC010538.1 | 75.7966741 | 0.665179599 | 0.3109019 | 2.13951603 | 0.0323939 | 0.10645901 |
| SPATA2L | 1002.90753 | -0.382117411 | 0.20009015 | -1.9097263 | 0.05616846 | 0.15792327 |
| VPS9D1 | 2875.89621 | -0.195197118 | 0.16406122 | -1.1897822 | 0.234132 | 0.42272673 |
| VPS9D1-AS1 | 958.277664 | -0.047173545 | 0.18076555 | -0.2609653 | 0.79411923 | 0.88844572 |
| ZNF276 | 2879.80898 | -0.103521857 | 0.15388547 | -0.6727202 | 0.50112534 | 0.68194216 |
| FANCA | 1847.1593 | -0.212748262 | 0.15907835 | -1.3373804 | 0.18109851 | 0.35729684 |
| SPIRE2 | 475.156045 | 0.219543801 | 0.30046626 | 0.73067706 | 0.46497643 | 0.65209958 |
| TCF25 | 7864.29465 | -0.30994059 | 0.10414807 | -2.975961 | 0.00292072 | 0.01797419 |
| MC1R | 1101.05108 | -1.041221333 | 0.3051026 | -3.4126925 | 0.00064324 | 0.00550905 |
| AC092143.1 | 1075.79859 | -2.326014451 | 0.39186438 | -5.9357639 | 2.92E-09 | 2.44E-07 |
| AC092143.2 | 415.25177 | -1.682156862 | 0.27477244 | -6.122 | 9.24E-10 | 9.67E-08 |
| TUBB3 | 586.794658 | -3.829505442 | 0.61482419 | -6.2286187 | 4.71E-10 | 5.39E-08 |
| DEF8 | 4232.73646 | -0.085649395 | 0.11365151 | -0.7536143 | 0.45108088 | 0.6408255 |
| CENPBD1 | 500.022985 | 0.181411901 | 0.21190389 | 0.85610463 | 0.39193991 | 0.58767586 |
| DBNDD1 | 1601.46449 | -0.702892507 | 0.20587297 | -3.4142049 | 0.00063968 | 0.00548747 |
| GAS8 | 3309.37829 | 0.249263218 | 0.1487776 | 1.67540822 | 0.09385416 | 0.22745467 |
| GAS8-AS1 | 8.70392805 | 2.29095768 | 0.71452762 | 3.20625488 | 0.00134475 | 0.00994043 |
| PRDM7 | 16.1899687 | 2.308828335 | 0.65087704 | 3.54725729 | 0.00038926 | 0.00373541 |
| FAM157C | 56.8224889 | 0.727240947 | 0.36872524 | 1.97231128 | 0.04857408 | 0.1427185 |
| AC133919.1 | 0.7669961 | 0.257767434 | 1.49089971 | 0.17289388 | 0.86273484 | NA |
| AC133919.2 | 96.7689754 | 0.904761043 | 0.91718788 | 0.98645115 | 0.32391177 | 0.52162159 |
| LINC02193 | 28.52783 | -0.785881881 | 0.48648969 | -1.6154132 | 0.10622124 | 0.24757416 |
| AC240565.2 | 27.1274522 | 0.890502609 | 0.51347 | 1.73428363 | 0.08286777 | 0.20766161 |
| AC240565.1 | 40.2331734 | 0.452425583 | 0.51670595 | 0.87559585 | 0.38124978 | 0.57725981 |
| SCGB1C2 | 1.9295351 | -1.536979255 | 1.12831208 | -1.3621934 | 0.17313686 | 0.34677344 |
| DOC2B | 1582.92801 | -0.901886136 | 0.31239004 | -2.8870515 | 0.00388871 | 0.02230698 |
| LINC02091 | 10.8743345 | 0.139924614 | 0.53156964 | 0.26322913 | 0.79237398 | 0.8874077 |
| RPH3AL | 1428.17285 | -0.343292453 | 0.18631449 | -1.8425429 | 0.06539577 | 0.17553812 |
| AC129507.4 | 137.728273 | -0.289627453 | 0.28336458 | -1.0221018 | 0.30673271 | 0.50367654 |
| AC129507.2 | 221.290238 | 0.523588051 | 0.37259381 | 1.40525162 | 0.15994647 | 0.32800939 |
| AC129507.3 | 96.079705 | 0.695680134 | 0.49764114 | 1.39795542 | 0.16212645 | 0.33103172 |
| AC141424.1 | 213.583081 | 0.512832299 | 0.31692846 | 1.61813269 | 0.105634 | 0.24665673 |
| C17orf97 | 446.479022 | 0.515423807 | 0.31878451 | 1.6168408 | 0.10591265 | 0.2470733 |
| RFLNB | 986.715713 | -0.597506726 | 0.5004031 | -1.1940508 | 0.2324581 | 0.42057771 |
| AC015853.2 | 5.72635707 | -0.320678997 | 0.9311205 | -0.3444012 | 0.7305446 | 0.84768376 |
| VPS53 | 2042.86473 | -0.189372034 | 0.1331128 | -1.4226434 | 0.15483957 | 0.32092427 |
| AC015853.3 | 2.24734583 | 1.622266569 | 0.82150503 | 1.97474941 | 0.04829659 | 0.14206148 |
| AC015853.1 | 38.1786422 | 0.353203169 | 0.28433177 | 1.24222196 | 0.21415468 | 0.39823808 |
| FAM57A | 988.882652 | -2.540677784 | 0.28327371 | -8.9689855 | 2.99E-19 | 4.93E-16 |
| GEMIN4 | 822.765839 | -0.361337161 | 0.17219582 | -2.0984085 | 0.03586908 | 0.11456729 |
| GLOD4 | 3906.27206 | 0.093999757 | 0.15458152 | 0.60809182 | 0.54312658 | 0.71536126 |
| AC087392.3 | 21.643258 | -0.23198747 | 0.39224102 | -0.5914411 | 0.55422489 | 0.72379706 |
| MRM3 | 1003.17992 | -0.171973215 | 0.18236607 | -0.9430111 | 0.34567525 | 0.54333196 |
| AC087392.4 | 7.99266037 | 0.206948403 | 0.40840619 | 0.50672201 | 0.61234989 | 0.76779926 |
| AC087392.5 | 6.16965503 | -0.006614 | 0.46756984 | -0.0141455 | 0.98871392 | 0.99493137 |
| NXN | 996.510642 | -0.856838003 | 0.33964695 | -2.5227313 | 0.01164473 | 0.05047514 |
| AC087392.2 | 2.57070862 | -0.992301618 | 0.92432675 | -1.0735399 | 0.28302896 | 0.4789542 |
| AC087392.1 | 3.49349228 | -3.790317188 | 1.30552023 | -2.9033002 | 0.00369252 | 0.02146866 |
| TIMM22 | 1375.7159 | -0.269847311 | 0.15261452 | -1.7681627 | 0.0770337 | 0.19723903 |
| ABR | 3670.7537 | 0.429796002 | 0.30004794 | 1.43242446 | 0.15202238 | 0.31707904 |
| AC016292.1 | 0 | NA | NA | NA | NA | NA |
| AC144836.1 | 0.08666371 | -0.642905668 | 3.30558948 | -0.1944905 | 0.84579185 | NA |
| BHLHA9 | 0 | NA | NA | NA | NA | NA |
| TRARG1 | 13.6038818 | 2.668658874 | 1.32068048 | 2.02066958 | NA | NA |
| YWHAE | 19898.9226 | -0.14097866 | 0.12657795 | -1.1137695 | 0.26537808 | 0.45945899 |
| CRK | 3877.78354 | -0.317307521 | 0.25529238 | -1.2429181 | 0.21389801 | 0.39794628 |
| AC032044.1 | 12.9125489 | 1.396767246 | 0.63121572 | 2.21282076 | 0.02691001 | 0.0932112 |
| MYO1C | 10144.2191 | 0.02312087 | 0.16161082 | 0.14306511 | 0.88623876 | 0.94110344 |
| INPP5K | 2147.78485 | -0.084092038 | 0.15656818 | -0.5370953 | 0.59120176 | 0.75081891 |
| PITPNA-AS1 | 370.971851 | -0.184624737 | 0.16678597 | -1.106956 | 0.26831294 | 0.46253594 |
| PITPNA | 4838.80037 | -0.041449823 | 0.12997986 | -0.3188942 | 0.74980674 | 0.86029607 |
| SLC43A2 | 6994.7034 | 0.972982117 | 0.31847656 | 3.05511377 | 0.00224975 | 0.0147945 |
| SCARF1 | 1825.26606 | 0.104454252 | 0.19357829 | 0.53959695 | 0.58947502 | 0.74961886 |
| RILP | 1049.07243 | 0.217051015 | 0.25404802 | 0.85437002 | 0.3929 | 0.58844158 |
| PRPF8 | 9145.65965 | 0.248493417 | 0.10603905 | 2.34341426 | 0.01910815 | 0.07253232 |
| AC130343.2 | 9.46066501 | 0.327252293 | 0.55522975 | 0.58939978 | 0.55559312 | 0.72490182 |
| TLCD2 | 392.724829 | -0.947145298 | 0.14525189 | -6.5207089 | 7.00E-11 | 1.06E-08 |
| MIR22HG | 3788.04134 | -0.441882374 | 0.61253324 | -0.7214015 | 0.47066255 | 0.65731233 |
| WDR81 | 3589.63646 | 0.29919468 | 0.30995354 | 0.96528878 | 0.33440019 | 0.53252811 |
| AC130343.1 | 499.629603 | 0.089420908 | 0.28337983 | 0.31555142 | 0.75234301 | 0.86197798 |
| SERPINF2 | 12495.4881 | 0.74076524 | 0.41852286 | 1.76995168 | 0.07673519 | 0.19674351 |
| SERPINF1 | 2699.12477 | -0.70353877 | 0.4068169 | -1.7293745 | 0.08374209 | 0.20921657 |
| SMYD4 | 555.699555 | -0.043371116 | 0.18131753 | -0.2391998 | 0.81095066 | 0.89780297 |
| RPA1 | 2409.72001 | 0.078050901 | 0.12221808 | 0.63861993 | 0.52307021 | 0.69862624 |
| RTN4RL1 | 231.397989 | -0.434468518 | 0.42900167 | -1.0127432 | 0.31118284 | 0.50823061 |
| AC099684.2 | 113.897129 | 3.752285713 | 0.48549791 | 7.72873707 | 1.09E-14 | 5.51E-12 |
| AC099684.1 | 47.0196064 | 3.01101272 | 0.55366607 | 5.43831902 | 5.38E-08 | 2.81E-06 |
| DPH1 | 4895.86917 | 0.00114443 | 0.1145653 | 0.00998933 | 0.9920298 | 0.99614736 |
| OVCA2 | 2924.04155 | -0.255431111 | 0.12380567 | -2.0631617 | 0.03909726 | 0.12176625 |
| AC090617.3 | 1943.13005 | -0.259397617 | 0.1363318 | -1.9026933 | 0.05708057 | 0.15977228 |
| AC090617.5 | 79.6645974 | -0.120134188 | 0.29265305 | -0.4105004 | 0.68143892 | 0.81542607 |
| HIC1 | 3422.61832 | -0.80329735 | 0.27506342 | -2.9204077 | 0.00349574 | 0.02054437 |
| SMG6 | 2951.36217 | -0.237602811 | 0.16960264 | -1.4009381 | 0.16123258 | 0.32994158 |
| AC090617.4 | 3.24457166 | 0.210754714 | 0.70614859 | 0.29845661 | 0.76535469 | 0.87045645 |
| AC090617.2 | 6.0233878 | -1.756100514 | 0.77757324 | -2.2584374 | 0.0239184 | 0.08544137 |
| AL450226.1 | 0.17507102 | 0.435199075 | 3.28898345 | 0.13232024 | 0.894731 | NA |
| SRR | 1568.10734 | -0.213048705 | 0.17104261 | -1.2455885 | 0.21291552 | 0.39676223 |
| TSR1 | 2315.35954 | -0.251064737 | 0.19138411 | -1.3118369 | 0.18957517 | 0.36804073 |
| SGSM2 | 5865.16849 | 0.004106408 | 0.15096291 | 0.02720144 | 0.97829907 | 0.99019487 |
| AC006435.3 | 20.5883392 | -0.903907379 | 0.68557002 | -1.3184757 | 0.18734447 | 0.36526977 |
| AC006435.2 | 1379.90619 | 0.159682208 | 0.16118738 | 0.99066195 | 0.32185068 | 0.51955663 |
| MNT | 2462.00227 | -0.709229136 | 0.1794561 | -3.9521039 | 7.75E-05 | 0.00103696 |
| AC006435.1 | 811.865525 | -0.766878179 | 0.18576912 | -4.1281251 | 3.66E-05 | 0.00057624 |
| METTL16 | 1233.45609 | -0.037064687 | 0.11895074 | -0.311597 | 0.75534685 | 0.86405639 |
| PAFAH1B1 | 6941.15629 | -0.085534718 | 0.109671 | -0.779921 | 0.43543737 | 0.6275267 |
| AC005696.2 | 1468.32186 | 0.239750332 | 0.09289093 | 2.58098761 | 0.00985181 | 0.04459188 |
| AC005696.3 | 13.5688032 | -0.624739058 | 0.63733899 | -0.9802304 | 0.32697239 | 0.52487577 |
| CLUH | 15604.8811 | -0.031762957 | 0.23354447 | -0.1360039 | 0.8918182 | 0.94397487 |
| AC005696.1 | 188.973834 | 1.121110987 | 0.25989344 | 4.31373338 | 1.61E-05 | 0.0002983 |
| AC005696.4 | 32.1153163 | -0.708550913 | 0.65076555 | -1.088796 | 0.27624386 | 0.471553 |
| hsa-mir-125: | 0 | NA | NA | NA | NA | NA |
| RAP1GAP2 | 1738.87191 | 0.392404684 | 0.18597981 | 2.10993158 | 0.03486425 | 0.11239245 |
| AC015921.1 | 27.4225426 | 2.023697637 | 0.39009679 | 5.18768081 | 2.13E-07 | 8.95E-06 |
| OR1D5 | 0 | NA | NA | NA | NA | NA |
| OR1D2 | 0.22673191 | 0.689158461 | 3.03348976 | 0.22718338 | 0.82028115 | NA |
| OR1G1 | 0.03392908 | 0.380740593 | 3.29738709 | 0.11546736 | 0.90807469 | NA |
| AC090282.1 | 0.14114194 | 0.02007176 | 3.29738704 | 0.00608717 | 0.99514317 | NA |
| OR1A2 | 0 | NA | NA | NA | NA | NA |

| | | | | | | |
|---|---|---|---|---|---|---|
| OR1A1 | 0.03392908 | 0.380740593 | 3.29738709 | 0.11546736 | 0.90807469 | NA |
| OR3A2 | 4.42229704 | 0.253141371 | 0.70045331 | 0.3613965 | 0.71780307 | 0.83983935 |
| OR3A1 | 0.41739998 | -0.857276453 | 2.80618186 | -0.3054957 | 0.75998865 | NA |
| AC087498.1 | 1.13338484 | -0.77107365 | 1.63777688 | -0.4708051 | 0.63777995 | 0.78561835 |
| OR1E1 | 0.87219032 | -0.058363719 | 1.77081885 | -0.0329586 | 0.9737076 | NA |
| OR3A3 | 1.59701034 | -0.402557135 | 1.6109969 | -0.2498808 | 0.80267956 | 0.89302768 |
| OR1E2 | 0.19978473 | -1.003574173 | 3.30558948 | -0.3035992 | 0.7614333 | NA |
| SPATA22 | 19.168274 | 0.203574811 | 0.6991717 | 0.29116569 | 0.7709246 | 0.87355577 |
| ASPA | 2258.63658 | 1.686523575 | 0.61935322 | 2.72303997 | 0.00646842 | 0.03258928 |
| TRPV3 | 49.0832575 | 0.337751445 | 0.2874148 | 1.17513587 | 0.23994036 | 0.42986802 |
| TRPV1 | 1435.70368 | 0.922962228 | 0.22850604 | 4.03911515 | 5.37E-05 | 0.00078733 |
| AC027796.3 | 1395.19062 | 1.162947127 | 0.23696488 | 4.90767709 | 9.22E-07 | 3.03E-05 |
| SHPK | 1552.64439 | 0.876982461 | 0.20978772 | 4.18033258 | 2.91E-05 | 0.00047944 |
| CTNS | 2547.36847 | -0.106288761 | 0.28349662 | -0.3749207 | 0.70771942 | 0.83365305 |
| AC027796.4 | 331.967984 | 0.167367219 | 0.22195385 | 0.75406313 | 0.45081131 | 0.64064962 |
| TAX1BP3 | 6903.13552 | -0.772572011 | 0.22388058 | -3.4508219 | 0.00055888 | 0.00495386 |
| P2RX5-TAX1I | 7871.62402 | -0.725896409 | 0.21151301 | -3.4319232 | 0.00059932 | 0.00521033 |
| EMC6 | 1015.59937 | -0.509060886 | 0.18989565 | -2.6807401 | 0.00734595 | 0.03590599 |
| P2RX5 | 129.88721 | -0.496227069 | 0.68388465 | -0.7256005 | 0.46808369 | 0.65488634 |
| ITGAE | 769.241392 | -0.457701963 | 0.20318431 | -2.2526442 | 0.02428159 | 0.08635273 |
| AC116914.2 | 40.8238574 | -0.488767435 | 0.27879536 | -1.7531405 | 0.0795779 | 0.20173708 |
| HASPIN | 8.99392364 | -0.901473242 | 0.50022355 | -1.8021407 | 0.07152326 | 0.18722141 |
| AC116914.1 | 1.95175334 | -1.781185314 | 1.21744299 | -1.4630544 | 0.1434525 | 0.30499973 |
| NCBP3 | 2447.39735 | -0.054155126 | 0.09758187 | -0.5549712 | 0.57891436 | 0.74240002 |
| CAMKK1 | 458.164723 | -0.328455995 | 0.25254217 | -1.3005986 | 0.19339588 | 0.37298685 |
| P2RX1 | 301.242902 | -0.064857093 | 0.63527658 | -0.1020927 | 0.91868311 | 0.96046034 |
| ATP2A3 | 1465.93145 | 0.432703931 | 0.36241109 | 1.19395884 | 0.23249407 | 0.42058507 |
| LINC01975 | 3.72815698 | 1.272388216 | 0.71686329 | 1.77493844 | 0.07590805 | 0.19545897 |
| ZZEF1 | 2825.10105 | 0.01846221 | 0.11988587 | 0.15399822 | 0.87761114 | 0.93634106 |
| CYB5D2 | 1813.88783 | 0.704627375 | 0.15915841 | 4.42720801 | 9.55E-06 | 0.00019624 |
| ANKFY1 | 3338.48688 | -0.122054213 | 0.22106628 | -0.5521159 | 0.58086893 | 0.74364083 |
| AC087292.1 | 1485.91188 | -0.486515679 | 0.23685059 | -2.0541038 | 0.03996565 | 0.12377708 |
| AC087742.1 | 11.0199323 | 0.345903082 | 0.54631961 | 0.63315151 | 0.52663471 | 0.7015024 |
| UBE2G1 | 2463.54024 | -0.233081498 | 0.17544368 | -1.328526 | 0.18400439 | 0.36102647 |
| SPNS3 | 472.915655 | 1.761385415 | 0.29002063 | 6.07331069 | 1.25E-09 | 1.24E-07 |
| AC127521.1 | 219.144991 | 1.798039507 | 0.30855782 | 5.82723691 | 5.64E-09 | 4.28E-07 |
| AC118754.1 | 64.5454866 | 1.695720542 | 0.32067996 | 5.28789053 | 1.24E-07 | 5.71E-06 |
| SPNS2 | 10946.7555 | -0.036010249 | 0.22072221 | -0.1631474 | 0.8704024 | 0.93249822 |
| MYBBP1A | 3693.91781 | -0.401891505 | 0.15433438 | -2.604031 | 0.00921344 | 0.0423395 |
| GGT6 | 3294.78727 | 2.04202158 | 0.2712721 | 7.52757692 | 5.17E-14 | 2.10E-11 |
| AC118754.2 | 1.0992078 | 1.997658671 | 1.33264951 | 1.49901279 | 0.13387031 | 0.29081145 |
| SMTNL2 | 3211.19767 | 0.329884172 | 0.29645504 | 1.11276289 | 0.26581028 | 0.45996556 |
| LINC01996 | 12.9603504 | 1.009918367 | 0.79101214 | 1.27674193 | 0.20169337 | 0.3835795 |
| ALOX15 | 4.90594425 | 1.918354615 | 0.92285575 | 2.07871556 | 0.0376435 | 0.11839778 |
| PELP1 | 3320.88441 | -0.296469961 | 0.11904853 | -2.4903286 | 0.0127625 | 0.05397982 |
| AC091153.2 | 54.9980303 | -0.498453002 | 0.27354258 | -1.8222136 | 0.06842258 | 0.18117317 |
| AC091153.3 | 13.7248679 | 0.596417777 | 0.49081872 | 1.21514879 | 0.22430935 | 0.41036753 |
| ARRB2 | 2271.07591 | -0.350759467 | 0.24698807 | -1.4201474 | 0.15556477 | 0.32184403 |
| MED11 | 1148.90619 | 0.225088778 | 0.10299974 | 2.18533352 | 0.0288644 | 0.09816279 |
| AC091153.4 | 488.064524 | 0.419808613 | 0.14045114 | 2.9890012 | 0.00279891 | 0.01740737 |
| CXCL16 | 3252.83164 | -1.236217544 | 0.21885965 | -5.6484489 | 1.62E-08 | 1.05E-06 |
| ZMYND15 | 354.278994 | -0.904466662 | 0.45403706 | -1.9920547 | 0.04636505 | 0.13787173 |
| TM4SF5 | 607.570556 | 1.011419805 | 0.40641469 | 2.4886399 | 0.01282328 | 0.05416759 |
| VMO1 | 180.468511 | -1.193896559 | 0.47775636 | -2.4989653 | 0.01245565 | 0.05303658 |
| GLTPD2 | 851.467117 | 1.528035189 | 0.54040904 | 2.82755297 | 0.00469053 | 0.02561913 |
| PSMB6 | 5073.56365 | -0.140460292 | 0.14726801 | -0.9537733 | 0.34019843 | 0.53828349 |
| AC233723.1 | 15.9539384 | 0.371433559 | 0.46791374 | 0.79380775 | 0.42730736 | 0.61960687 |
| PLD2 | 1668.87017 | -0.331746107 | 0.22197091 | -1.4945477 | 0.13503253 | 0.29242691 |
| MINK1 | 10958.7997 | -0.551717131 | 0.1559372 | -3.5380726 | 0.00040306 | 0.00384123 |
| CHRNE | 1121.66652 | -0.839118194 | 0.14951481 | -5.6122747 | 2.00E-08 | 1.25E-06 |
| C17orf107 | 246.37195 | 0.130221459 | 0.22453534 | 0.57995974 | 0.56194177 | 0.72964972 |
| GP1BA | 73.8622584 | 1.667056473 | 0.3379186 | 4.93330788 | 8.08E-07 | 2.70E-05 |
| SLC25A11 | 4753.73824 | 0.700199379 | 0.20514179 | 3.41324599 | 0.00064194 | 0.00550145 |
| RNF167 | 5611.65354 | -0.163786377 | 0.08849682 | -1.85076 | 0.06420409 | 0.17327417 |
| PFN1 | 23113.5629 | -1.007513407 | 0.20833723 | -4.835974 | 1.32E-06 | 4.09E-05 |
| ENO3 | 10391.1092 | -0.777225547 | 0.19550911 | -3.975393 | 7.03E-05 | 0.00095704 |
| SPAG7 | 3108.35099 | 0.04477857 | 0.1477806 | 0.30300709 | 0.76188446 | 0.86804464 |
| CAMTA2 | 5276.43582 | -0.336169311 | 0.1835096 | -1.8318895 | 0.06696789 | 0.1784502 |
| AC004771.4 | 1344.06925 | -0.808373473 | 0.21217199 | -3.8099915 | 0.00013897 | 0.00164262 |
| AC004771.2 | 757.314936 | -0.058168427 | 0.18260535 | -0.3185472 | 0.75006988 | 0.86045777 |
| AC004771.5 | 41.1728237 | 0.110557194 | 0.33316901 | 0.33183517 | 0.74001373 | 0.85412805 |
| AC004771.3 | 59.5542282 | 0.724917841 | 0.23359242 | 3.10334483 | 0.00191347 | 0.01301808 |
| INCA1 | 262.462093 | 0.730574085 | 0.15651251 | 4.66783203 | 3.04E-06 | 7.92E-05 |
| KIF1C | 14459.6429 | 0.16662636 | 0.19516127 | 0.85378806 | 0.39322243 | 0.58866168 |
| AC004771.1 | 1229.12889 | 0.31237641 | 0.19371802 | 1.61253147 | 0.10684634 | 0.24863608 |
| SLC52A1 | 31.521819 | -0.091954361 | 0.53057669 | -0.1733102 | 0.86240759 | 0.92830704 |
| AC012146.3 | 2.81568605 | 0.28909417 | 0.68370473 | 0.42283483 | 0.67241578 | 0.80893455 |
| ZFP3 | 629.436003 | 0.321669546 | 0.17616314 | 1.82597537 | 0.06785397 | 0.18004722 |
| ZNF232 | 642.480864 | -1.074972411 | 0.53771926 | -1.9991332 | 0.04559395 | 0.13616222 |
| USP6 | 208.198098 | -1.076448264 | 0.64357545 | -1.6726062 | 0.09440486 | 0.22841142 |
| ZNF594 | 356.269909 | 0.566588357 | 0.21561467 | 2.62778201 | 0.00859436 | 0.04016895 |
| AC087500.1 | 302.04558 | 0.525898571 | 0.26134804 | 2.01225375 | 0.0441932 | 0.13318566 |
| SCIMP | 278.813158 | 0.375449544 | 0.29504131 | 1.27253212 | 0.20318412 | 0.38532652 |
| AC087500.2 | 22.8900099 | 0.809958234 | 0.35701145 | 2.26871781 | 0.02328549 | 0.08383922 |
| RABEP1 | 2936.54538 | 0.043343525 | 0.14783978 | 0.29317903 | 0.76938531 | 0.87240672 |
| NUP88 | 2114.92692 | -0.306542142 | 0.18118517 | -1.6918721 | 0.09067036 | 0.2218181 |
| AC015727.1 | 3.2405948 | 0.115098688 | 0.74055302 | 0.15542262 | 0.87648816 | 0.93557592 |
| RPAIN | 2107.09157 | -0.170476838 | 0.11107882 | -1.5347375 | 0.12484832 | 0.27712544 |
| AC004148.2 | 1482.74155 | -0.465884685 | 0.15107837 | -3.0837285 | 0.00204424 | 0.0137167 |
| C1QBP | 3998.48857 | -0.056405235 | 0.17622676 | -0.3200719 | 0.74891383 | 0.8597841 |
| DHX33 | 836.710999 | -0.069416079 | 0.16423727 | -0.4226573 | 0.67254532 | 0.80897677 |
| AC004148.1 | 0.21230658 | 0.952737031 | 3.28103697 | 0.2903768 | 0.77152799 | NA |
| DERL2 | 2332.47446 | -0.335097815 | 0.12410592 | -2.7000953 | 0.00693196 | 0.03436012 |
| MIS12 | 1065.33699 | -0.006742763 | 0.15184741 | -0.0444049 | 0.96458169 | 0.98349067 |
| NLRP1 | 1254.54215 | 0.375476943 | 0.31348275 | 1.1977595 | 0.23101066 | 0.41879218 |
| AC007846.1 | 0.42120972 | 1.1986364 | 2.12980188 | 0.56279244 | 0.57357623 | NA |
| AC007846.2 | 42.676549 | 0.503697904 | 0.38240593 | 1.31718119 | 0.1877779 | 0.36587137 |
| WSCD1 | 904.48988 | 0.406580938 | 0.33733848 | 1.20526106 | 0.22810254 | 0.41517157 |
| AIPL1 | 0.35036362 | -0.999448737 | 2.92586748 | -0.3415906 | 0.73265904 | NA |
| PIMREG | 58.997958 | -0.781375756 | 0.35367533 | -2.2093024 | 0.02715361 | 0.09387909 |
| PITPNM3 | 417.625806 | 0.25411163 | 0.20278723 | 1.25309481 | 0.21017121 | 0.3934588 |
| KIAA0753 | 1099.20288 | -0.387121192 | 0.18596999 | -2.0816326 | 0.03737605 | 0.11778863 |
| TXNDC17 | 2553.72294 | -0.100876773 | 0.18855746 | -0.5349922 | 0.59265524 | 0.75199593 |
| MED31 | 562.43801 | 0.04063052 | 0.11635876 | 0.34918316 | 0.72695181 | 0.84575518 |
| C17orf100 | 161.21184 | -0.156539425 | 0.27849937 | -0.5620818 | 0.5740603 | 0.73863954 |
| AC004706.1 | 10.8556371 | 0.323533998 | 0.56298326 | 0.57467783 | 0.5655092 | 0.73214268 |
| SLC13A5 | 10.8233746 | 1.771679112 | 0.6625676 | 2.67395979 | 0.00749614 | 0.0364442 |
| XAF1 | 1385.31372 | 0.122964569 | 0.41929415 | 0.29326564 | 0.76931911 | 0.87240608 |
| FBXO39 | 8.37162566 | -1.657145463 | 0.9433626 | -1.7566368 | 0.07897974 | 0.20064505 |
| TEKT1 | 18.1214029 | -3.22597812 | 1.08746137 | -2.9665221 | 0.00301189 | 0.01841063 |
| ALOX12-AS1 | 307.151804 | 0.498242586 | 0.20822447 | 2.39281475 | 0.01671968 | 0.06588409 |
| AC027763.2 | 29.7530917 | 0.414481229 | 0.39083576 | 1.06049976 | 0.2889173 | 0.48514705 |
| AC040977.2 | 865.116379 | -0.203190039 | 0.16355525 | -1.2423327 | 0.21411384 | 0.39820734 |

| | | | | | | |
|---|---|---|---|---|---|---|
| AC040977.1 | 164.716588 | -0.388604296 | 0.20236405 | -1.9203228 | 0.05481714 | 0.1554164 |
| ALOX12 | 290.66015 | 0.403510549 | 0.17351483 | 2.32551043 | 0.02004468 | 0.07503951 |
| RNASEK | 8551.45434 | -0.497245506 | 0.18700369 | -2.6590144 | 0.00783696 | 0.0376294 |
| RNASEK-C17 | 9817.22021 | -0.448693695 | 0.16068011 | -2.7924658 | 0.0052308 | 0.02780972 |
| C17orf49 | 2439.63025 | -0.3667057 | 0.22058441 | -1.662428 | 0.09642699 | 0.2317934 |
| MIR497HG | 1536.68636 | -0.28490673 | 0.19604942 | -1.4532393 | 0.14615731 | 0.30887709 |
| BCL6B | 1089.99482 | -0.411335759 | 0.3630308 | -1.1330602 | 0.25718897 | 0.44990535 |
| SLC16A13 | 909.918444 | 1.081480466 | 0.53582683 | 2.0183395 | 0.04355592 | 0.13186791 |
| SLC16A11 | 219.696782 | 0.492765076 | 0.4212936 | 1.16964767 | 0.24214279 | 0.43253528 |
| CLEC10A | 251.403078 | 1.181956598 | 0.73176286 | 1.61521808 | 0.10626348 | 0.24765072 |
| ASGR2 | 156.852033 | 0.770932651 | 0.38747971 | 1.9896078 | 0.04663416 | 0.13842219 |
| ASGR1 | 140.094339 | -0.015387021 | 0.37433048 | -0.0411054 | 0.96721183 | 0.98450423 |
| DLG4 | 1657.96632 | -0.688304827 | 0.36856638 | -1.8675193 | 0.0618291 | 0.16895683 |
| ACADVL | 58242.8112 | -0.344800288 | 0.2905273 | -1.1868086 | 0.23530314 | 0.42415365 |
| DVL2 | 2758.40222 | -0.362186925 | 0.15891487 | -2.2791254 | 0.02265961 | 0.08222566 |
| PHF23 | 2536.50628 | -0.596677054 | 0.22186163 | -2.6894108 | 0.00715783 | 0.03522148 |
| GABARAP | 13463.3042 | -0.187849457 | 0.12165213 | -1.5441527 | 0.12255135 | 0.27373127 |
| AC120057.2 | 14516.1163 | -0.221165496 | 0.11572764 | -1.9110862 | 0.05599351 | 0.15764988 |
| CTDNEP1 | 5977.75079 | -0.506429565 | 0.13501264 | -3.7509789 | 0.00017615 | 0.0019818 |
| AC003688.1 | 3943.77943 | -0.80604014 | 0.13444554 | -5.995291 | 2.03E-09 | 1.83E-07 |
| ELP5 | 2502.20095 | -0.234604874 | 0.09852302 | -2.3812189 | 0.01725545 | 0.06737139 |
| CLDN7 | 6955.25759 | -1.141668634 | 0.17070345 | -6.6880233 | 2.26E-11 | 4.14E-09 |
| SLC2A4 | 328.848263 | -0.199908363 | 0.40051551 | -0.4991276 | 0.61768947 | 0.77143728 |
| AC003688.2 | 45.1101218 | 0.373120526 | 0.45073943 | 0.8277965 | 0.40778576 | 0.60242555 |
| YBX2 | 306.292563 | 0.665420229 | 0.35270974 | 1.88659441 | 0.0592149 | 0.16405725 |
| EIF5A | 14027.6227 | -0.666692579 | 0.19114579 | -3.4878748 | 0.00048688 | 0.00443946 |
| GPS2 | 9813.32559 | -0.420838706 | 0.17388381 | -2.4202294 | 0.01551072 | 0.06234076 |
| AC026954.2 | 5690.441 | -0.116275364 | 0.13491005 | -0.8618732 | 0.38875727 | 0.58437501 |
| NEURL4 | 3436.21873 | 0.438720633 | 0.10695918 | 4.10175773 | 4.10E-05 | 0.00063415 |
| ACAP1 | 863.87101 | -0.277308691 | 0.45963114 | -0.6033288 | 0.54629 | 0.71744944 |
| KCTD11 | 1109.01904 | -0.543761007 | 0.15811753 | -3.4389672 | 0.00058394 | 0.00511746 |
| AC026954.3 | 277.777062 | -0.54820571 | 0.17221084 | -3.1833404 | 0.00145586 | 0.01056152 |
| TMEM95 | 0.75163162 | -2.81489201 | 2.59317108 | -1.0855019 | 0.27769944 | NA |
| TNK1 | 977.641555 | -0.389096439 | 0.1325679 | -2.9350729 | 0.00333469 | 0.019877 |
| PLSCR3 | 1711.95944 | -0.414505815 | 0.21307775 | -1.9453266 | 0.05173568 | 0.14937081 |
| TMEM256-PI | 3247.85275 | -0.07229304 | 0.19570393 | -0.3694001 | 0.71182956 | 0.83618048 |
| TMEM256 | 1799.83096 | 0.228636028 | 0.30172819 | 0.75775494 | 0.4485977 | 0.63870857 |
| NLGN2 | 3139.42326 | -1.1652999 | 0.348812 | -3.3407678 | 0.00083547 | 0.00680611 |
| SPEM1 | 1.06804298 | -0.97120857 | 1.66896097 | -0.5819241 | 0.56061782 | 0.72862652 |
| SPEM2 | 0.73852648 | -2.801839386 | 2.04784875 | -1.3681867 | 0.17125365 | NA |
| SPEM3 | 0.03392908 | 0.380740593 | 3.29738709 | 0.11546736 | 0.90807469 | NA |
| TMEM102 | 566.459328 | -0.703310344 | 0.20476917 | -3.4346495 | 0.00059332 | 0.00517732 |
| AC113189.1 | 148.46564 | -0.852930489 | 0.25271592 | -3.3750564 | 0.00073801 | 0.00616642 |
| FGF11 | 687.319657 | -0.413450955 | 0.44057445 | -0.9384361 | 0.34802035 | 0.54548839 |
| AC113189.3 | 85.5207977 | 0.274722173 | 0.41277432 | 0.66555054 | 0.50569843 | 0.68492352 |
| CHRNB1 | 795.239537 | -0.314956307 | 0.23817309 | -1.3223841 | 0.18604028 | 0.36355441 |
| ZBTB4 | 6659.23408 | 0.264427277 | 0.15963042 | 1.65649677 | 0.09762126 | 0.23372731 |
| SLC35G6 | 1.90561907 | -0.105905883 | 1.39846501 | -0.0757301 | 0.93963384 | 0.97112068 |
| POLR2A | 8181.73502 | 0.103474919 | 0.17047321 | 0.60698639 | 0.54385995 | 0.71604108 |
| TNFSF12 | 1947.75723 | 0.08858022 | 0.26298015 | 0.33683234 | 0.73624329 | 0.85204668 |
| TNFSF12-TN | 5365.75296 | -0.169532985 | 0.26325519 | -0.6439872 | 0.5195837 | 0.69631663 |
| AC016876.3 | 172.027723 | -0.103572865 | 0.25674954 | -0.4034004 | 0.6866537 | 0.81889338 |
| TNFSF13 | 5594.58855 | -0.153942641 | 0.2689903 | -0.5722981 | 0.56712003 | 0.73319386 |
| SENP3 | 2816.16899 | -0.258741628 | 0.13918376 | -1.858993 | 0.06302814 | 0.1711691 |
| SENP3-EIF4A | 26490.783 | -1.079250517 | 0.21737303 | -4.9649697 | 6.87E-07 | 2.36E-05 |
| EIF4A1 | 25691.0511 | -1.16675564 | 0.22747589 | -5.1291398 | 2.91E-07 | 1.16E-05 |
| CD68 | 7345.59954 | -1.315685316 | 0.35326794 | -3.7243269 | 0.00019584 | 0.00214729 |
| AC016876.1 | 1198.12745 | -0.491280201 | 0.21644503 | -2.2697689 | 0.02322161 | 0.08375787 |
| MPDU1 | 5132.1235 | -0.166560475 | 0.15902538 | -1.047383 | 0.29492299 | 0.4916937 |
| SOX15 | 90.1717517 | -0.250927585 | 0.29174023 | -0.8601062 | 0.38973049 | 0.58536157 |
| FXR2 | 3542.66343 | -0.378840219 | 0.16801284 | -2.254829 | 0.02414407 | 0.08599867 |
| SHBG | 179.080714 | 1.242022043 | 0.55195109 | 2.25023933 | 0.02443376 | 0.08676517 |
| SAT2 | 9373.02656 | 0.526644507 | 0.35075578 | 1.50145639 | 0.13323756 | 0.28986671 |
| ATP1B2 | 669.052438 | 1.22359483 | 0.37126168 | 3.2957746 | 0.00098151 | 0.00773762 |
| TP53 | 1601.60162 | -0.330979192 | 0.27225988 | -1.2156738 | 0.22410922 | 0.41014244 |
| WRAP53 | 812.10429 | -0.241349527 | 0.18334442 | -1.3163724 | 0.18804909 | 0.36613054 |
| EFNB3 | 146.565313 | -0.119156023 | 0.6417119 | -0.1856846 | 0.8526921 | 0.92285671 |
| DNAH2 | 37.4112164 | -0.850608025 | 0.35785657 | -2.3769524 | 0.01745634 | 0.06787434 |
| KDM6B | 4904.48996 | -0.303655026 | 0.31220748 | -0.9726065 | 0.33074891 | 0.52867582 |
| TMEM88 | 521.303896 | -0.438653336 | 0.35002009 | -1.2532233 | 0.21012445 | 0.39345181 |
| NAA38 | 2168.7109 | -0.155860513 | 0.15191652 | -1.0259616 | 0.30490969 | 0.50199592 |
| CYB5D1 | 1977.02891 | 0.447788453 | 0.15127306 | 2.96013342 | 0.00307506 | 0.01871435 |
| CHD3 | 7330.08635 | 0.044503745 | 0.24968997 | 0.17823601 | 0.85853763 | 0.92607016 |
| RNF227 | 281.56593 | -0.354067551 | 0.32196242 | -1.099717 | 0.27145543 | 0.46613251 |
| KCNAB3 | 254.983112 | -0.211831124 | 0.35238943 | -0.6011279 | 0.54775479 | 0.7186931 |
| AC104581.2 | 1442.31081 | -0.512890133 | 0.10112459 | -5.0718638 | 3.94E-07 | 1.49E-05 |
| TRAPPC1 | 3693.96664 | -0.527563914 | 0.10017516 | -5.2664143 | 1.39E-07 | 6.35E-06 |
| CNTROB | 2242.92447 | -0.298742002 | 0.12060977 | -2.4769304 | 0.01325177 | 0.05550458 |
| GUCY2D | 16.1473323 | 0.033687224 | 0.5172395 | 0.06512887 | 0.94807139 | 0.97504763 |
| ALOX15B | 58.378572 | -4.912533827 | 1.01163289 | -4.856044 | 1.20E-06 | 3.76E-05 |
| AC129492.2 | 0.3357283 | -1.716651648 | 3.30224936 | -0.5198431 | 0.60317291 | NA |
| ALOX12B | 88.8580645 | 0.204815721 | 0.5288007 | 0.3873212 | 0.69851844 | 0.82697006 |
| AC129492.1 | 94.693106 | 0.125659391 | 0.52996184 | 0.23711026 | 0.81257125 | 0.89891781 |
| ALOXE3 | 7.30081835 | -2.187726465 | 0.75571078 | -2.8949256 | 0.00379248 | 0.02192673 |
| HES7 | 30.5935786 | -1.939776128 | 0.7175151 | -2.7034638 | 0.00686209 | 0.03406501 |
| PER1 | 10733.7522 | -0.739674653 | 0.33858607 | -2.1845986 | 0.02891829 | 0.09827001 |
| AC129492.3 | 1688.58312 | 0.320590696 | 0.18889579 | 1.69718286 | 0.08966209 | 0.21998413 |
| VAMP2 | 7488.35837 | 0.130241639 | 0.19141732 | 0.68040676 | 0.49624694 | 0.67810301 |
| TMEM107 | 627.248875 | -0.164818442 | 0.22729383 | -0.7251338 | 0.46836994 | 0.65515146 |
| AC129492.4 | 3.46281038 | 0.320188486 | 0.71102158 | 0.45032176 | 0.65247845 | 0.79583057 |
| BORCS6 | 752.522771 | -0.279035456 | 0.13530043 | -2.0623397 | 0.03917541 | 0.12195201 |
| AURKB | 48.29334 | -1.837732639 | 0.5608952 | -3.2764278 | 0.00105129 | 0.00815168 |
| LINC00324 | 169.000629 | 0.121551795 | 0.32726905 | 0.37141244 | 0.71033036 | 0.83536599 |
| CTC1 | 2190.40721 | -0.179831404 | 0.14734476 | -1.2204805 | 0.22228277 | 0.40830292 |
| PFAS | 1280.00361 | 0.617474981 | 0.17744752 | 3.47976117 | 0.00050186 | 0.004542 |
| AC135178.5 | 16.9564008 | -0.397231323 | 0.36691904 | -1.082613 | 0.27898023 | 0.47466576 |
| SLC25A35 | 2414.50563 | 0.063645766 | 0.14757123 | 0.43128844 | 0.66625865 | 0.80439203 |
| RANGRF | 1698.49027 | -0.152790472 | 0.13426957 | -1.1379382 | 0.25514628 | 0.44783064 |
| ARHGEF15 | 1370.15543 | 0.70710837 | 0.23864997 | 2.96295183 | 0.00304704 | 0.01858984 |
| AC135178.1 | 89.3989756 | 1.033429167 | 0.21893289 | 4.72030115 | 2.35E-06 | 6.48E-05 |
| AC135178.4 | 0 | NA | NA | NA | NA | NA |
| ODF4 | 0.83874792 | 1.48246385 | 1.52215893 | 0.97392185 | 0.33009534 | NA |
| AC135178.3 | 77.0474262 | 0.504112794 | 0.21098285 | 2.38935437 | 0.01687801 | 0.06634962 |
| KRBA2 | 176.071362 | 1.068909707 | 0.20913221 | 5.11116731 | 3.20E-07 | 1.26E-05 |
| AC135178.2 | 19774.1651 | -0.360696524 | 0.17512158 | -2.0596921 | 0.03942799 | 0.1225069 |
| RPL26 | 20261.1793 | -0.386248506 | 0.17336778 | -2.227914 | 0.02588625 | 0.0906414 |
| RNF222 | 3.20705802 | 0.208731022 | 0.73574604 | 0.28369982 | 0.77664043 | 0.87753913 |
| NDEL1 | 2597.44283 | -0.408480566 | 0.2517264 | -1.6227164 | 0.10465004 | 0.24510361 |
| MYH10 | 4575.98314 | 0.614079759 | 0.16382389 | 3.74841399 | 0.00017796 | 0.00199605 |
| CCDC42 | 6.57413411 | 0.771169712 | 0.78895598 | 0.97745594 | 0.32834348 | 0.52605976 |
| SPDYE4 | 1.19313357 | 1.264956702 | 1.11716971 | 1.13228697 | 0.25751381 | 0.4501572 |
| MFSD6L | 157.434596 | -0.616108926 | 0.23426993 | -2.6299104 | 0.00854074 | 0.03996839 |
| PIK3R6 | 63.8364224 | -0.306370379 | 0.6525209 | -0.4695181 | 0.63869934 | 0.78618782 |

| | | | | | | |
|---|---|---|---|---|---|---|
| PIK3R5 | 359.274335 | -0.929926231 | 0.41222883 | -2.2558496 | 0.02408005 | 0.08583279 |
| AC002091.2 | 3.26955003 | 0.692238567 | 0.76839019 | 0.90089459 | 0.36764437 | 0.5644908 |
| AC002091.1 | 4.35325146 | 0.032121855 | 0.78695085 | 0.04081812 | 0.96744089 | 0.98467589 |
| NTN1 | 361.781657 | -0.198278831 | 0.47814236 | -0.4146858 | 0.67837195 | 0.81323805 |
| AC005695.1 | 1.26736382 | 0.497582387 | 1.01492509 | 0.49026514 | 0.62394629 | 0.77588719 |
| AC005695.3 | 2.06659929 | -0.618422698 | 0.95468201 | -0.6477787 | 0.51712808 | 0.69438193 |
| STX8 | 1066.77339 | -0.111238187 | 0.14056946 | -0.7913396 | 0.42874582 | 0.62111903 |
| AC005695.2 | 0 | NA | NA | NA | NA | NA |
| AC087501.1 | 7.52537673 | 0.446858355 | 0.80112052 | 0.55779168 | 0.57698665 | 0.74090075 |
| AC087501.2 | 0.22977084 | -1.364242679 | 3.30558948 | -0.4127078 | 0.67982068 | NA |
| AC087501.3 | 0.2320242 | 1.01099135 | 3.00413042 | 0.33653377 | 0.73646839 | NA |
| CFAP52 | 51.3771105 | -0.583124893 | 0.44431181 | -1.3124227 | 0.18937756 | 0.36772989 |
| AC118755.2 | 21.9589174 | 0.33362467 | 0.38149574 | 0.87451743 | 0.38183653 | 0.57778329 |
| USP43 | 690.733765 | -0.747204095 | 0.2757059 | -2.7101491 | 0.0067253 | 0.0335565 |
| AC118755.1 | 1.47630439 | -1.60093363 | 1.19114872 | -1.344025 | 0.17894031 | 0.35462618 |
| AC027045.2 | 0.48623317 | -2.161135024 | 3.28527092 | -0.6578255 | 0.51065026 | NA |
| DHRS7C | 0.93624556 | 1.611091273 | 1.45044818 | 1.11075411 | 0.26667421 | NA |
| GSG1L2 | 0.56731998 | -1.11795736 | 3.26714905 | -0.3421813 | 0.73221445 | NA |
| AC027045.3 | 1.76610905 | -2.228468093 | 2.81851425 | -0.7906535 | 0.42914623 | 0.62121019 |
| GLP2R | 7.40304702 | 2.060141133 | 1.021185 | 2.01740246 | 0.04365353 | 0.13210278 |
| RCVRN | 1.46149631 | -0.315554659 | 1.3087482 | -0.2411118 | 0.80946845 | 0.89703256 |
| GAS7 | 403.384727 | -0.361026399 | 0.32684935 | -1.1045652 | 0.26934806 | 0.46353248 |
| AC005291.2 | 0.33921085 | -0.541378846 | 2.86868125 | -0.1887205 | 0.85031192 | NA |
| MYH13 | 1.10067936 | 1.212178023 | 1.15849009 | 1.04634302 | 0.29540269 | 0.49221338 |
| AC005291.1 | 0.07830766 | -0.642905668 | 3.30558948 | -0.1944905 | 0.84579185 | NA |
| AC005323.2 | 1.75166107 | 1.920338401 | 1.21813303 | 1.57646034 | 0.11491975 | 0.26214953 |
| MYHAS | 7.63892098 | 2.265820663 | 0.96060609 | 2.35874068 | 0.01833706 | 0.07041625 |
| MYH8 | 263.899397 | 1.697168127 | 0.55197157 | 3.07473828 | 0.00210687 | 0.01400864 |
| MYH4 | 3.16414863 | 2.521173302 | 0.85425204 | 2.95132253 | 0.00316416 | 0.01913299 |
| MYH1 | 15.2422624 | 3.547568831 | 0.95121142 | 3.72952718 | 0.00019184 | 0.00211047 |
| MYH2 | 7.76293286 | 5.412920023 | 1.28236612 | 4.22104105 | NA | NA |
| AC005323.1 | 4.08685703 | 1.967263051 | 1.20224186 | 1.63632886 | 0.10177078 | 0.24039429 |
| MYH3 | 308.349982 | -0.324029902 | 0.31054231 | -1.0434324 | 0.29674809 | 0.49351962 |
| AC002347.1 | 2.59484517 | -1.484385805 | 0.93271 | -1.5914762 | 0.11150244 | 0.25639338 |
| SCO1 | 2545.21026 | 0.149821893 | 0.162867 | 0.91990332 | 0.35762328 | 0.55532482 |
| AC002347.2 | 318.191101 | -0.070783254 | 0.15774147 | -0.4487295 | 0.6536268 | 0.79659051 |
| ADPRM | 339.645575 | -0.326942041 | 0.16714364 | -1.9560544 | 0.05045875 | 0.14660612 |
| TMEM220 | 931.679092 | 0.203283046 | 0.26854733 | 0.75697289 | 0.4490661 | 0.63923742 |
| TMEM220-A! | 309.15439 | 0.237847739 | 0.21912043 | 1.08546585 | 0.27771538 | 0.4732072 |
| AC015908.3 | 90.9856958 | 0.426171482 | 0.29375333 | 1.4507801 | 0.1468411 | 0.30988626 |
| AC015908.4 | 5.31530096 | -0.888416637 | 1.28243391 | -0.6927582 | 0.48846129 | 0.67195599 |
| TMEM238L | 121.780956 | 0.277456453 | 0.90569431 | 0.30634669 | 0.75934069 | 0.8666817 |
| PIRT | 0.1981425 | 1.631193554 | 3.26970143 | 0.4988815 | 0.61786287 | NA |
| AC015908.2 | 14.8465442 | 0.126092402 | 0.55472902 | 0.2273045 | 0.82018697 | 0.90351452 |
| SHISA6 | 113.356555 | 2.151556246 | 0.64100784 | 3.35652094 | 0.0007893 | 0.00649821 |
| AC005725.1 | 21.6910442 | 2.112555563 | 0.57304883 | 3.6865193 | 0.00022734 | 0.00242223 |
| DNAH9 | 113.305726 | -0.771918808 | 0.43818695 | -1.7616198 | 0.07813356 | 0.19903225 |
| AC005209.1 | 0.75982153 | -0.359994616 | 1.37767605 | -0.2613057 | 0.79385676 | NA |
| AC005410.2 | 59.3975832 | -0.182530636 | 0.21548611 | -0.8470645 | 0.39695916 | 0.59189006 |
| ZNF18 | 449.215698 | 0.15916885 | 0.18845665 | 0.84459132 | 0.39833906 | 0.59329008 |
| MAP2K4 | 1951.72111 | -0.131800925 | 0.21814945 | -0.6041772 | 0.54572583 | 0.71715108 |
| AC005244.2 | 0.32066929 | -0.699509041 | 2.8887459 | -0.2421497 | 0.80866415 | NA |
| LINC00670 | 4.18387125 | 0.346952182 | 0.80252592 | 0.4323252 | 0.66550507 | 0.80380015 |
| MYOCD | 108.859066 | 0.265044036 | 0.32424726 | 0.81741335 | 0.41369225 | 0.60761193 |
| AC005358.1 | 0.16223974 | -1.003574173 | 3.30558948 | -0.3035992 | 0.7614333 | NA |
| AC005358.2 | 4.47024559 | -0.053544357 | 0.67632788 | -0.0791692 | 0.93689802 | 0.96969938 |
| ARHGAP44 | 783.844844 | -0.027481712 | 0.18498164 | -0.1485645 | 0.88189726 | 0.93863819 |
| AC005277.2 | 1.20917015 | 0.410658365 | 1.16812242 | 0.35155422 | 0.7251726 | 0.84416976 |
| AC005277.1 | 191.743578 | -0.164299756 | 0.21949568 | -0.748533 | 0.45413875 | 0.64374771 |
| ELAC2 | 3698.45382 | -0.158379432 | 0.16176568 | -0.979067 | 0.3275469 | 0.52526221 |
| AC005303.1 | 0 | NA | NA | NA | NA | NA |
| LINC02093 | 0 | NA | NA | NA | NA | NA |
| HS3ST3A1 | 64.85602 | 0.169299799 | 0.34882009 | 0.48534991 | 0.62742816 | 0.77838166 |
| AC005304.2 | 0 | NA | NA | NA | NA | NA |
| AC005304.1 | 0.39465542 | 0.943937467 | 2.72130833 | 0.34686899 | 0.72868975 | NA |
| AC005304.3 | 0.38683783 | 1.154201553 | 2.16538742 | 0.53302312 | 0.59401758 | NA |
| COX10 | 900.114126 | 0.002895997 | 0.19067117 | 0.01518843 | 0.98788185 | 0.99454413 |
| AC005224.1 | 33.6225983 | 0.268365044 | 0.31619839 | 0.84872364 | 0.39603509 | 0.59122987 |
| CDRT15 | 5.40282271 | -0.738485137 | 0.87484996 | -0.8441278 | 0.39859802 | 0.59351443 |
| AC005224.2 | 0.4574167 | -0.952771099 | 1.90985129 | -0.4988719 | 0.61786965 | NA |
| HS3ST3B1 | 498.516815 | -0.373245367 | 0.35340016 | -1.0561551 | 0.29089738 | 0.48690769 |
| AC005224.4 | 16.5759849 | -0.93372447 | 0.46347669 | -2.0146093 | 0.0439456 | 0.13282248 |
| AC005224.3 | 16.7245675 | -0.831175229 | 0.47889888 | -1.7355965 | 0.08263521 | 0.20723636 |
| AC022816.1 | 0.42382866 | -1.990251205 | 2.75055779 | -0.7235809 | 0.46932304 | NA |
| AC013248.1 | 0 | NA | NA | NA | NA | NA |
| AC005863.1 | 0.21442573 | -1.003574173 | 3.30558948 | -0.3035992 | 0.7614333 | NA |
| LINC02096 | 0 | NA | NA | NA | NA | NA |
| AC005772.1 | 0 | NA | NA | NA | NA | NA |
| CDRT7 | 0 | NA | NA | NA | NA | NA |
| CDRT8 | 0 | NA | NA | NA | NA | NA |
| PMP22 | 2565.89853 | -1.014396217 | 0.32217967 | -3.148542 | 0.00164087 | 0.01159737 |
| AC005703.3 | 1.05397001 | 0.803003068 | 1.36885552 | 0.58662369 | 0.55745647 | 0.72629087 |
| AC005703.1 | 0.27631848 | 1.234200164 | 2.87756746 | 0.42890399 | 0.6679931 | NA |
| AC005703.2 | 0 | NA | NA | NA | NA | NA |
| TEKT3 | 24.2572287 | -0.793499435 | 0.36491495 | -2.1744777 | 0.02966928 | 0.10009964 |
| AC005703.4 | 2.02394698 | -0.116056725 | 0.88406204 | -0.1312767 | 0.89555644 | 0.94638952 |
| CDRT4 | 1101.47721 | -0.34484978 | 0.20962344 | -1.6450917 | 0.0999509 | 0.23748358 |
| TVP23C-CDR' | 1109.08531 | -0.3527863 | 0.19934466 | -1.7697303 | 0.07677207 | 0.1967998 |
| TVP23C | 295.068229 | -0.174869239 | 0.1941478 | -0.9007016 | 0.36774699 | 0.56452953 |
| AC005838.2 | 1.09797743 | -0.184908646 | 1.24251573 | -0.148818 | 0.88169728 | 0.93857409 |
| CDRT1 | 19.7010256 | -0.751710511 | 0.34400691 | -2.1851611 | 0.02887704 | 0.09819309 |
| AC005324.3 | 512.450413 | -1.246720417 | 0.42017099 | -2.9671739 | 0.00300551 | 0.01838016 |
| TRIM16 | 1106.87541 | -1.436992189 | 0.41860149 | -3.4328406 | 0.00059729 | 0.00520642 |
| AC005324.5 | 37.6417282 | -0.748618748 | 0.29785622 | -2.5133561 | 0.01195885 | 0.05145293 |
| ZNF286A | 358.519391 | -0.118159382 | 0.19364932 | -0.6101719 | 0.54174791 | 0.71442309 |
| AC005324.4 | 90.1451864 | 0.336956193 | 0.30283108 | 1.11268695 | 0.2658429 | 0.45996556 |
| TBC1D26 | 7.96391783 | -0.527498657 | 0.58480454 | -0.9020085 | 0.36705236 | 0.56395482 |
| AC005324.1 | 1.48307133 | -0.12834585 | 1.00329561 | -0.1279243 | 0.89820891 | 0.9482358 |
| AC005324.2 | 0.03392908 | 0.380740593 | 3.29738709 | 0.11546736 | 0.90807469 | NA |
| AC015922.3 | 156.884655 | -0.001334479 | 0.17940007 | -0.0074386 | 0.99406494 | 0.99728027 |
| AC015922.4 | 296.429308 | -0.682124957 | 0.22812559 | -2.9901291 | 0.0027886 | 0.0173555 |
| LINC02087 | 0.17294555 | 0.18657164 | 3.29377195 | 0.05664376 | 0.95482897 | NA |
| ADORA2B | 45.8415646 | 0.231754591 | 0.53108417 | 0.43638015 | 0.66256094 | 0.80152913 |
| ZSWIM7 | 1329.20379 | -0.198825154 | 0.12469461 | -1.5944968 | 0.11082482 | 0.25519127 |
| TTC19 | 3634.78319 | 0.040689443 | 0.11367462 | 0.35794661 | 0.72038327 | 0.84093782 |
| AC002553.2 | 45.6876096 | -0.008823861 | 0.28643829 | -0.0308055 | 0.97542469 | 0.98865354 |
| NCOR1 | 4371.29441 | 0.236031533 | 0.10765466 | 2.19248774 | 0.02834431 | 0.09698188 |
| AC002553.1 | 333.341982 | 0.335962054 | 0.11805258 | 2.84586802 | 0.00442905 | 0.02460379 |
| PIGL | 1073.94308 | 0.18148557 | 0.17271257 | 1.05079538 | 0.29335258 | 0.48984977 |
| CENPV | 724.431008 | 0.568009598 | 0.20900951 | 2.71762563 | 0.00657722 | 0.03295733 |
| UBB | 34497.8851 | 0.199932206 | 0.15901974 | 1.25727916 | 0.20865257 | 0.39178071 |
| AC093484.3 | 25270.8136 | 0.180084079 | 0.16066066 | 1.12089716 | 0.26233164 | 0.45603853 |
| AC093484.2 | 21.2013844 | 0.537811221 | 0.55461554 | 0.96970097 | 0.33219557 | 0.53009359 |

| | | | | | | |
|---|---|---|---|---|---|---|
| TRPV2 | 454.383228 | -0.336387301 | 0.34175221 | -0.9843018 | 0.32496716 | 0.52255528 |
| AC093484.4 | 42.7977976 | -1.155439121 | 0.42159839 | -2.7406156 | 0.00613242 | 0.03132131 |
| LRRC75A | 5819.52745 | -0.402849055 | 0.20353234 | -1.9792877 | 0.04778363 | 0.14092853 |
| ZNF287 | 248.522327 | -0.039086559 | 0.20615611 | -0.1895969 | 0.84962502 | 0.92098029 |
| ZNF624 | 135.836732 | 0.610543117 | 0.25419673 | 2.40185276 | 0.01631227 | 0.0647141 |
| AC098850.2 | 0 | NA | NA | NA | NA | NA |
| CCDC144A | 8.81894047 | 0.317080719 | 0.58559493 | 0.54146766 | 0.58818528 | 0.74897126 |
| AC098850.3 | 1035.22114 | -0.249990511 | 0.77051445 | -0.3244462 | 0.74560022 | 0.85740777 |
| AC098850.1 | 0.09635527 | 0.547239394 | 3.29377202 | 0.16614368 | 0.86804388 | NA |
| AC022596.1 | 0.09635527 | 0.547239394 | 3.29377202 | 0.16614368 | 0.86804388 | NA |
| AC022596.2 | 1.56136755 | 3.17376301 | 1.17326738 | 2.70506372 | 0.00682913 | 0.03393972 |
| TNFRSF13B | 20.5050724 | 0.13972979 | 1.07227226 | 0.13031186 | 0.8963197 | 0.94695314 |
| AC104024.1 | 0.57338042 | -0.020033981 | 2.79368677 | -0.0071712 | 0.99427829 | NA |
| LINC02090 | 0.49918457 | -0.397297004 | 1.88755959 | -0.2104818 | 0.83329163 | NA |
| AC104024.2 | 1.25255217 | 1.946021068 | 1.21809946 | 1.59758799 | 0.1101347 | 0.25411253 |
| MPRIP | 9956.27573 | -0.122530322 | 0.2653284 | -0.4618063 | 0.64422025 | 0.78977629 |
| MPRIP-AS1 | 1.8665113 | 0.521978603 | 0.96312858 | 0.54196149 | 0.58784503 | 0.74875494 |
| AC055811.1 | 312.58902 | -0.909195404 | 0.37139956 | -2.448025 | 0.01436417 | 0.05899722 |
| PLD6 | 292.157163 | 0.323887747 | 0.33659877 | 0.96223688 | 0.33593063 | 0.53409445 |
| AC055811.2 | 980.021546 | 0.090155177 | 0.40419237 | 0.22305017 | 0.82349646 | 0.90564302 |
| FLCN | 3407.48069 | -0.318329143 | 0.38197962 | -0.8333668 | 0.40463788 | 0.59945083 |
| AC055811.3 | 1.21607367 | -0.311059748 | 1.51504692 | -0.2053136 | 0.83732711 | 0.91386632 |
| COPS3 | 1814.66662 | -0.195955981 | 0.18905693 | -1.0364919 | 0.2999727 | 0.49672088 |
| NT5M | 211.035778 | 0.135764841 | 0.27934265 | 0.48601545 | 0.62695621 | 0.77804072 |
| MED9 | 1781.50308 | -0.846304629 | 0.19785839 | -4.277325 | 1.89E-05 | 0.00033886 |
| RASD1 | 13686.8827 | -2.279840571 | 0.37152623 | -6.1364188 | 8.44E-10 | 8.97E-08 |
| PEMT | 1102.93283 | 0.305415798 | 0.17435948 | 1.75164439 | 0.07983497 | 0.20215559 |
| AC020558.2 | 96.4204306 | -0.068581441 | 0.22410763 | -0.3060201 | 0.75958933 | 0.86685293 |
| AC020558.1 | 179.365119 | 0.047990392 | 0.15657075 | 0.30650932 | 0.75921689 | 0.86663046 |
| SMCR2 | 3.72522418 | -0.810952748 | 0.79046969 | -1.0259125 | 0.30493283 | 0.50200266 |
| RAI1 | 2910.01709 | -0.443380568 | 0.1846969 | -2.4005847 | 0.0163689 | 0.06487589 |
| RAI1-AS1 | 6.04718064 | 0.012165433 | 0.59367592 | 0.02049171 | 0.98365113 | 0.99295181 |
| SMCR5 | 4.13419065 | 2.880159159 | 0.76823911 | 3.74903999 | 0.00017751 | 0.00199192 |
| SREBF1 | 8073.71247 | -0.109840156 | 0.30122009 | -0.3646508 | 0.71537207 | 0.8380436 |
| TOM1L2 | 5978.97386 | 0.83677503 | 0.15817589 | 5.29015547 | 1.22E-07 | 5.67E-06 |
| AC122129.1 | 5.14751091 | 2.472639622 | 0.83677454 | 2.95496518 | 0.00312704 | 0.01895635 |
| DRC3 | 826.930489 | 0.687065345 | 0.27672613 | 2.48283505 | 0.01303414 | 0.05481745 |
| ATPAF2 | 1223.20512 | 0.233876072 | 0.12332275 | 1.89645528 | 0.05789987 | 0.16145541 |
| GID4 | 820.8413 | -0.133106213 | 0.18092139 | -0.735713 | 0.46190539 | 0.65014385 |
| DRG2 | 2324.89714 | -0.109172246 | 0.08685508 | -1.2569472 | 0.20877276 | 0.39195056 |
| MYO15A | 123.031306 | -0.403903664 | 0.30938556 | -1.3055026 | 0.19172175 | 0.3709347 |
| AC087164.1 | 487.080082 | 0.345785094 | 0.32234766 | 1.07270857 | 0.2834019 | 0.479304 |
| ALKBH5 | 5876.61919 | -0.368436618 | 0.23307435 | -1.5807686 | 0.11393095 | 0.26029797 |
| LLGL1 | 1678.06274 | 0.107384103 | 0.10634946 | 1.00972868 | 0.3126253 | 0.50941908 |
| FLII | 10068.4619 | -0.45174766 | 0.2040076 | -2.2143668 | 0.02680356 | 0.09292792 |
| MIEF2 | 1794.74567 | -0.404702057 | 0.21801237 | -1.8563261 | 0.06340709 | 0.17179132 |
| AC127537.1 | 23.1874368 | -0.200090277 | 0.35142378 | -0.5693703 | 0.56910484 | 0.73454422 |
| TOP3A | 1147.90576 | -0.15776671 | 0.15102531 | -1.0446375 | 0.29619053 | 0.49287211 |
| SMCR8 | 3736.25891 | 0.470939418 | 0.19175758 | 2.45591036 | 0.01405282 | 0.05805622 |
| SHMT1 | 19409.1378 | 1.376401235 | 0.3112294 | 4.42246535 | 9.76E-06 | 0.00019974 |
| EVPLL | 99.5367057 | 1.481229071 | 0.31518235 | 4.69959393 | 2.61E-06 | 7.03E-05 |
| AL353997.2 | 80.4832219 | 1.466737441 | 0.3198579 | 4.58559083 | 4.53E-06 | 0.00010882 |
| LINC02076 | 9.79863852 | -0.94465394 | 0.63167337 | -1.4954785 | 0.13478961 | 0.29218265 |
| LGALS9C | 2.45085722 | -1.856392707 | 1.34846737 | -1.3766686 | 0.16861473 | 0.34024521 |
| TBC1D28 | 1.70284139 | 0.478310989 | 1.16834252 | 0.40939278 | 0.68225144 | 0.81602805 |
| ZNF286B | 87.1969472 | 0.203970392 | 0.30547318 | 0.66771948 | 0.50431267 | 0.6839149 |
| TRIM16L | 1217.83421 | -1.524072795 | 0.55754724 | -2.7335312 | 0.00626592 | 0.03183053 |
| FBXW10 | 19.6237296 | 0.642748759 | 0.54557589 | 1.17811064 | 0.2387525 | 0.42832189 |
| TVP23B | 1568.56452 | -0.446814025 | 0.13427974 | -3.3274866 | 0.00087633 | 0.00707776 |
| AC107982.3 | 17.5768776 | 0.457201854 | 0.32587862 | 1.40298204 | 0.16062219 | 0.3289792 |
| PRPSAP2 | 1940.37782 | 0.269735629 | 0.16563823 | 1.62846242 | 0.10342687 | 0.24312414 |
| AC107982.2 | 16.3613221 | 0.388005577 | 0.6007146 | 0.64590669 | 0.51833979 | 0.69530036 |
| SLC5A10 | 8173.76197 | 0.995883283 | 0.30138579 | 3.30434722 | 0.00095198 | 0.00755679 |
| AC090286.1 | 601.51341 | 1.575740463 | 0.4111119 | 3.83287481 | 0.00012665 | 0.00152436 |
| FAM83G | 3708.05252 | 0.342589198 | 0.39267281 | 0.87245459 | 0.38296043 | 0.57895142 |
| GRAP | 1285.31608 | -0.063357676 | 0.30060987 | -0.2107638 | 0.83307159 | 0.91136194 |
| AC007952.3 | 0.495767 | 0.013132816 | 1.85104088 | 0.00709483 | 0.99433919 | NA |
| AC007952.7 | 0.43581956 | -0.90895699 | 2.8211851 | -0.3221898 | 0.74730893 | NA |
| AC007952.4 | 4.66364936 | -0.713519383 | 0.93793289 | -0.7607361 | 0.44681473 | 0.63710155 |
| GRAPL | 6.70224685 | -1.189970239 | 0.71982566 | -1.6531367 | 0.09830304 | 0.23485592 |
| AC007952.6 | 25.2124327 | 1.473250121 | 0.75736785 | 1.945224 | 0.05174802 | 0.14937081 |
| AC007952.1 | 4.65528675 | -1.32072456 | 0.92926293 | -1.4212604 | 0.15524108 | 0.32145327 |
| AC007952.5 | 0 | NA | NA | NA | NA | NA |
| AC007952.8 | 0.08757577 | 0.547239394 | 3.29377202 | 0.16614368 | 0.86804388 | NA |
| AC106017.1 | 0.34258644 | 0.670184554 | 2.22078833 | 0.30177777 | 0.76282148 | NA |
| EPN2 | 2595.90005 | -0.187480919 | 0.18026337 | -1.0400389 | 0.29832181 | 0.49529347 |
| AC106017.2 | 0.796636 | 1.966139234 | 1.71493902 | 1.14647764 | 0.25159758 | NA |
| EPN2-AS1 | 0.31629348 | 1.314934116 | 2.84234801 | 0.46262249 | 0.64363498 | NA |
| AC124066.1 | 1475.52014 | -0.314013265 | 0.18745211 | -1.6751653 | 0.0939018 | 0.22754921 |
| B9D1 | 1117.64412 | 0.049929449 | 0.16610285 | 0.30059357 | 0.76372443 | 0.86927716 |
| MAPK7 | 2019.97639 | -0.722813216 | 0.18396996 | -3.928974 | 8.53E-05 | 0.00111865 |
| MFAP4 | 3284.80497 | 0.480401696 | 0.59368623 | 0.8091845 | 0.41840903 | 0.61113582 |
| RNF112 | 239.709704 | -0.849047408 | 0.38322165 | -2.2155518 | 0.02672222 | 0.0927801 |
| AC004448.3 | 27.9204926 | -0.384010584 | 0.40558189 | -0.9468139 | 0.3437336 | 0.54151789 |
| AC004448.2 | 8.34603781 | -0.187527455 | 0.75258319 | -0.2491784 | 0.80322281 | 0.89318485 |
| LINC02094 | 3.69623754 | -0.374940517 | 1.02426196 | -0.3660592 | 0.7143209 | 0.83764267 |
| AC004448.1 | 0.10721287 | -0.642905668 | 3.30558948 | -0.1944905 | 0.84579185 | NA |
| SLC47A1 | 8018.91551 | 1.524712163 | 0.49925272 | 3.05398871 | 0.00225821 | 0.01482791 |
| AC025627.1 | 676.527149 | 1.956018328 | 0.57268874 | 3.41549986 | 0.00063665 | 0.00547034 |
| ALDH3A2 | 9428.88517 | 1.939854193 | 0.29190237 | 6.64555815 | 3.02E-11 | 5.24E-09 |
| AC115989.2 | 1.92553685 | 2.635395287 | 1.17131465 | 2.2499465 | 0.02445234 | 0.08680779 |
| SLC47A2 | 10912.4091 | -0.030075393 | 0.33205519 | -0.0905735 | 0.92783151 | 0.96504253 |
| AC005722.2 | 367.007034 | -0.190400381 | 0.39063628 | -0.4874109 | 0.62596717 | 0.77729476 |
| AC005722.3 | 31.4288824 | -0.537531407 | 0.43334324 | -1.2404287 | 0.21481685 | 0.39903638 |
| ALDH3A1 | 132.989052 | -0.622548016 | 0.33338074 | -1.8673785 | 0.06184875 | 0.16895885 |
| ULK2 | 1159.89664 | 0.160134613 | 0.19945058 | 0.80287864 | 0.42204488 | 0.61454023 |
| AC015726.2 | 1.36983094 | 0.594553815 | 1.17584804 | 0.50563831 | 0.61311058 | 0.76817248 |
| AC015726.1 | 104.801431 | 0.170518529 | 0.26522395 | 0.64292282 | 0.52027418 | 0.6969232 |
| AKAP10 | 1585.78903 | -0.133696112 | 0.15995762 | -0.8358221 | 0.40325501 | 0.59817937 |
| AC005730.2 | 2.4702512 | -0.143886493 | 0.77543877 | -0.1855549 | 0.85279379 | 0.92285671 |
| SPECC1 | 1007.41649 | -0.774070338 | 0.18291935 | -4.2317575 | 2.32E-05 | 0.00040095 |
| AC004702.1 | 0.09997307 | 0.547239394 | 3.29377202 | 0.16614368 | 0.86804388 | NA |
| LINC01984 | 1.17362289 | -2.558335576 | 1.86471186 | -1.3719737 | 0.17007164 | 0.34205533 |
| LGALS9B | 1.9751513 | -0.208428769 | 1.17389382 | -0.1775533 | 0.85907377 | 0.92642086 |
| AC015818.8 | 0 | NA | NA | NA | NA | NA |
| AC015818.2 | 1.48709988 | -1.234459293 | 1.23421351 | -1.0001991 | 0.31721414 | 0.51448733 |
| AC015818.9 | 0.03392908 | 0.380740593 | 3.29738709 | 0.11546736 | 0.90807469 | NA |
| AC015818.4 | 0 | NA | NA | NA | NA | NA |
| CDRT15L2 | 0 | NA | NA | NA | NA | NA |
| LINC02088 | 0 | NA | NA | NA | NA | NA |
| AC126365.1 | 1.64785144 | -1.638502175 | 1.13105371 | -1.4486511 | 0.14743505 | 0.31075551 |
| CCDC144NL | 0.71110193 | -0.603842289 | 1.92979581 | -0.3129048 | 0.75435302 | NA |
| AC090774.3 | 0.15489794 | -1.003574173 | 3.30558948 | -0.3035992 | 0.7614333 | NA |

| | | | | | | |
|---|---|---|---|---|---|---|
| CCDC144NL-/ | 964.062535 | 0.563256869 | 0.22248866 | 2.53162053 | 0.01135368 | 0.04957203 |
| AC107926.1 | 0.42437184 | -1.990508063 | 2.74987466 | -0.7238541 | 0.46915531 | NA |
| USP22 | 9635.34117 | 0.044399714 | 0.17745288 | 0.25020566 | 0.80242831 | 0.89289776 |
| AC087393.2 | 3265.35497 | -0.49565095 | 0.20137449 | -2.4613393 | 0.01384194 | 0.0573828 |
| LINC01563 | 2.97881569 | -0.065924607 | 0.76776347 | -0.0858658 | 0.93157312 | 0.96699666 |
| DHRS7B | 1743.08976 | -0.130411258 | 0.14625414 | -0.8916757 | 0.37256679 | 0.56884873 |
| TMEM11 | 1232.04097 | -0.743193564 | 0.21124896 | -3.5180934 | 0.00043466 | 0.00406741 |
| AC087294.1 | 35.0848768 | -0.597438903 | 0.33751437 | -1.770114 | 0.07670816 | 0.19669331 |
| NATD1 | 2149.39094 | -0.272346836 | 0.2052993 | -1.3265843 | 0.18464622 | 0.361855 |
| MAP2K3 | 4854.64817 | -0.523485881 | 0.35931631 | -1.4568943 | 0.14514556 | 0.30737941 |
| KCNJ12 | 297.254198 | 1.201450285 | 0.40712392 | 2.95106775 | 0.00316677 | 0.01914439 |
| AC068418.2 | 2.03247665 | 0.186113581 | 1.01115481 | 0.18406042 | 0.85396605 | 0.9235189 |
| AC068418.1 | 0.85819924 | -0.978875132 | 1.42213652 | -0.6883131 | 0.49125566 | NA |
| AC233702.5 | 0 | NA | NA | NA | NA | NA |
| C17orf51 | 504.316636 | -0.043294754 | 0.2667249 | -0.1623199 | 0.87105396 | 0.93304464 |
| AC233702.7 | 32.6538748 | -0.281373518 | 0.39504309 | -0.7122603 | 0.47630359 | 0.66172097 |
| KCNJ18 | 0.44609554 | 2.182452166 | 2.64182147 | 0.82611645 | 0.40873804 | NA |
| LINC02002 | 0 | NA | NA | NA | NA | NA |
| AC138761.3 | 0.25999113 | -1.364242679 | 3.30558948 | -0.4127078 | 0.67982068 | NA |
| FAM27E5 | 0 | NA | NA | NA | NA | NA |
| AC209154.2 | 0 | NA | NA | NA | NA | NA |
| AC132825.3 | 0 | NA | NA | NA | NA | NA |
| MTRNR2L1 | 5.2766056 | -1.022776645 | 0.61263969 | -1.6694587 | 0.09502651 | 0.22941034 |
| AC132825.2 | 0 | NA | NA | NA | NA | NA |
| AC131274.1 | 0 | NA | NA | NA | NA | NA |
| AC131274.4 | 0 | NA | NA | NA | NA | NA |
| AC069061.2 | 0.09644756 | -0.642905668 | 3.30558948 | -0.1944905 | 0.84579185 | NA |
| AC129926.1 | 0.68175792 | 0.804308374 | 1.59767608 | 0.50342393 | 0.61466625 | NA |
| WSB1 | 15632.2823 | -0.415739454 | 0.25459981 | -1.6329134 | 0.10248721 | 0.24151811 |
| AC026254.2 | 5.74254531 | 0.018061328 | 0.78098821 | 0.02312625 | 0.98154957 | 0.99160566 |
| KSR1 | 3267.66953 | -0.389272727 | 0.28348784 | -1.3731549 | 0.16970419 | 0.34148031 |
| AC015688.4 | 555.183898 | -0.715510535 | 0.49618778 | -1.4420156 | 0.14929796 | 0.3134045 |
| AC015688.6 | 1162.81316 | -0.832130725 | 0.31738621 | -2.6218238 | 0.00874606 | 0.04070485 |
| LGALS9 | 2289.37909 | -1.736253027 | 0.63370024 | -2.7398649 | 0.00614645 | 0.03138686 |
| NOS2 | 191.432103 | -0.79157698 | 0.6399356 | -1.2369635 | 0.21610063 | 0.40079995 |
| AC005697.1 | 258.930209 | 0.986682867 | 0.39283206 | 2.51171676 | 0.01201455 | 0.05156634 |
| AC005697.2 | 613.188123 | 0.859924683 | 0.38421345 | 2.23814313 | 0.02521172 | 0.08890521 |
| LYRM9 | 1139.94483 | 0.774907144 | 0.38433785 | 2.01621346 | 0.04377766 | 0.13244801 |
| LINC01992 | 0.13037664 | 0.02007176 | 3.29738704 | 0.00608717 | 0.99514317 | NA |
| NLK | 1674.14467 | 0.127418537 | 0.1187318 | 1.07316271 | 0.28319812 | 0.47907013 |
| AC061975.7 | 5.92958647 | 0.049391484 | 0.48847787 | 0.10111304 | 0.91946072 | 0.96074349 |
| ERVE-1 | 162.123835 | 1.377555598 | 0.26380087 | 5.22195242 | 1.77E-07 | 7.73E-06 |
| AC061975.6 | 5.58020126 | 0.595624364 | 0.7279308 | 0.81824312 | 0.41321838 | 0.607085 |
| AC061975.4 | 1.66288685 | 0.948154764 | 1.32506973 | 0.71555085 | 0.47426873 | 0.66003563 |
| AC061975.1 | 3.39705283 | 0.910066049 | 0.91062117 | 0.99939039 | 0.31760561 | 0.51483034 |
| TMEM97 | 1000.33971 | -0.213199254 | 0.17146026 | -1.2434325 | 0.21370851 | 0.3976779 |
| IFT20 | 1485.64992 | -0.297550311 | 0.14800257 | -2.0104402 | 0.04438462 | 0.13347274 |
| TNFAIP1 | 5214.36985 | -0.730049427 | 0.19952321 | -3.65897 | 0.00025323 | 0.00264568 |
| POLDIP2 | 9146.7315 | 0.249515846 | 0.15282493 | 1.63269073 | 0.10253406 | 0.24160692 |
| TMEM199 | 1560.5203 | 0.071374979 | 0.16606691 | 0.42979651 | 0.66734367 | 0.80535685 |
| AC002094.2 | 584.789474 | -0.006814889 | 0.18844503 | -0.0361638 | 0.97115175 | 0.9865502 |
| SEBOX | 1.16121214 | -1.555135553 | 1.6225763 | -0.958436 | 0.33784294 | 0.53584256 |
| AC002094.3 | 73.7029893 | 1.742200235 | 0.34501411 | 5.0496492 | 4.43E-07 | 1.64E-05 |
| SARM1 | 2197.29324 | 0.675060062 | 0.18276761 | 3.69354317 | 0.00022115 | 0.00236774 |
| VTN | 229.937794 | 1.967485803 | 0.42902515 | 4.58594511 | 4.52E-06 | 0.00010874 |
| SLC46A1 | 2202.37677 | 0.749438136 | 0.19310473 | 3.88099315 | 0.00010403 | 0.00130633 |
| AC015917.2 | 47.3349863 | 1.140686719 | 0.44497915 | 2.56346105 | 0.01036343 | 0.04629573 |
| AC005726.1 | 2611.73735 | 1.248691247 | 0.45725038 | 2.73086979 | 0.00631674 | 0.03200855 |
| SLC13A2 | 3643.69338 | 3.087116089 | 0.69151677 | 4.46426784 | 8.03E-06 | 0.00017127 |
| FOXN1 | 10.6992503 | 3.040462775 | 0.60770826 | 5.00316184 | 5.64E-07 | 2.00E-05 |
| UNC119 | 2055.05977 | -0.234215712 | 0.2218613 | -1.0556853 | 0.29111202 | 0.48715725 |
| PIGS | 4191.19819 | 0.084193703 | 0.14983707 | 0.56190169 | 0.574183 | 0.73864963 |
| ALDOC | 884.319295 | -0.441389942 | 0.24240539 | -1.8208751 | 0.06862584 | 0.18156552 |
| AC005726.4 | 319.517931 | -0.559537944 | 0.25319718 | -2.2098902 | 0.02711279 | 0.09376579 |
| SPAG5 | 3523.56578 | 1.495681805 | 0.46849372 | 3.1925333 | 0.00141031 | 0.01031903 |
| AC005726.5 | 280.756399 | 0.873879735 | 0.25670604 | 3.40420404 | 0.00066357 | 0.00564464 |
| SGK494 | 779.408618 | 1.597431961 | 0.30409179 | 5.25312426 | 1.50E-07 | 6.69E-06 |
| KIAA0100 | 10042.1078 | 0.037817286 | 0.27766501 | 0.13619752 | 0.89166514 | 0.94385074 |
| AC005726.2 | 215.277103 | 1.715260095 | 0.41314592 | 4.15170522 | 3.30E-05 | 0.0005303 |
| SDF2 | 2760.58367 | -0.401081828 | 0.15360894 | -2.6110579 | 0.00902626 | 0.04175481 |
| SUPT6H | 6401.00353 | -0.001966506 | 0.17242669 | -0.0114049 | 0.99090042 | 0.99580451 |
| AC010761.3 | 2.08870874 | 0.738385235 | 0.96271789 | 0.76697986 | 0.44309349 | 0.63422761 |
| PROCA1 | 305.391657 | 0.144639672 | 0.31020962 | 0.4662643 | 0.64102633 | 0.78754492 |
| RAB34 | 4021.7573 | -0.905053687 | 0.2169411 | -4.1718867 | 3.02E-05 | 0.00049117 |
| RPL23A | 13721.8203 | -0.436137903 | 0.17626753 | -2.4742952 | 0.01334994 | 0.05579607 |
| AC010761.1 | 9613.01935 | -0.441521968 | 0.18134915 | -2.4346514 | 0.01490614 | 0.06055692 |
| TLCD1 | 604.804784 | -0.802183204 | 0.18907293 | -4.2427184 | 2.21E-05 | 0.00038463 |
| NEK8 | 1085.54299 | -0.065267425 | 0.35051059 | -0.1862067 | 0.85228266 | 0.9225686 |
| AC010761.6 | 16.7380723 | 0.102794372 | 0.4195473 | 0.24501259 | 0.80644668 | 0.8952597 |
| AC010761.2 | 231.173571 | -0.561366226 | 0.38988109 | -1.4398396 | 0.14991279 | 0.31425199 |
| TRAF4 | 6534.66101 | -0.777388779 | 0.13008073 | -5.9762024 | 2.28E-09 | 1.98E-07 |
| AC010761.4 | 764.034915 | -0.479145466 | 0.16438479 | -2.9147799 | 0.0035594 | 0.02082737 |
| AC010761.5 | 849.027789 | -1.112835481 | 0.1265044 | -8.7968124 | 1.41E-18 | 2.18E-15 |
| FAM222B | 2237.7192 | -0.383105667 | 0.15247627 | -2.5125592 | 0.0119859 | 0.05148699 |
| ERAL1 | 2784.70622 | -0.037604991 | 0.13409795 | -0.2804293 | 0.77914818 | 0.87898785 |
| AC024267.1 | 0.3370857 | -0.541099294 | 2.88344717 | -0.1876571 | 0.85114547 | NA |
| FLOT2 | 11033.1838 | -0.848910582 | 0.21300257 | -3.9854477 | 6.74E-05 | 0.00093079 |
| AC024267.3 | 3.06552798 | 0.407410359 | 0.72407829 | 0.56266065 | 0.57366599 | 0.73821843 |
| AC024267.6 | 1230.85378 | -1.685805452 | 0.34631925 | -4.8677787 | 1.13E-06 | 3.60E-05 |
| DHRS13 | 1556.90811 | -1.623551605 | 0.33754936 | -4.8098198 | 1.51E-06 | 4.56E-05 |
| PHF12 | 2726.12595 | -0.108212057 | 0.1139313 | -0.9498009 | 0.34221342 | 0.54004595 |
| AC024267.4 | 11.0743979 | 0.674858873 | 0.61578698 | 1.0959291 | 0.2731098 | 0.46802089 |
| AC024267.5 | 0.75760184 | 0.443428595 | 1.39015325 | 0.31897821 | 0.74974304 | NA |
| PIPOX | 4609.34747 | 3.246746905 | 0.70511462 | 4.60456614 | 4.13E-06 | 0.0001013 |
| SEZ6 | 7.20127565 | 1.554663307 | 0.67151017 | 2.31517461 | 0.02060338 | 0.07656582 |
| AC024619.2 | 0.83559062 | 2.550353938 | 1.68200026 | 1.51626251 | 0.12945299 | NA |
| MYO18A | 4921.52136 | -0.366442884 | 0.25084192 | -1.4608519 | 0.1440561 | 0.30583361 |
| TIAF1 | 3220.2617 | -0.733460044 | 0.22561491 | -3.2509378 | 0.00115025 | 0.00875891 |
| AC024619.3 | 0.09997307 | 0.547239394 | 3.29377202 | 0.16614368 | 0.86804388 | NA |
| CRYBA1 | 7.31204566 | -1.126025575 | 0.55952528 | -2.012466 | 0.04417084 | 0.13313351 |
| NUFIP2 | 4209.95365 | -0.542654452 | 0.18244564 | -2.9743351 | 0.00293624 | 0.01804445 |
| AC068025.2 | 0.42907716 | 0.330812837 | 1.85122911 | 0.17869902 | 0.85817403 | NA |
| AC068025.1 | 4.3344258 | 1.776130624 | 0.65286654 | 2.72051105 | 0.00651811 | 0.03273953 |
| TAOK1 | 2590.02202 | -0.27573036 | 0.13940881 | -1.9778546 | 0.04794511 | 0.14123243 |
| ABHD15-AS1 | 30.0069776 | -0.068117088 | 0.35781154 | -0.1903714 | 0.8490181 | 0.92059733 |
| ABHD15 | 1354.60729 | 0.313969877 | 0.33176588 | 0.94635975 | 0.34396513 | 0.54155816 |
| TP53I13 | 3506.44792 | -0.395910534 | 0.19131131 | -2.069457 | 0.03850322 | 0.12043456 |
| GIT1 | 3955.80671 | -0.564476216 | 0.19071583 | -2.9597764 | 0.00307862 | 0.01872741 |
| ANKRD13B | 346.382721 | -1.036873313 | 0.3619219 | -2.864909 | 0.00417129 | 0.02352154 |
| AC104564.4 | 0.16421342 | 1.36803136 | 3.28033652 | 0.41703994 | 0.67664919 | NA |
| CORO6 | 400.761642 | -1.176608313 | 0.36213901 | -3.2490516 | 0.0011579 | 0.00880702 |
| SSH2 | 1479.751 | -0.086976434 | 0.15011046 | -0.5794162 | 0.56230837 | 0.72983653 |
| AC104564.1 | 0.46921604 | -2.098896459 | 2.22011143 | -0.9454014 | 0.34445399 | NA |

| | | | | | | |
|---|---|---|---|---|---|---|
| AC023389.1 | 4.90189402 | -0.586349733 | 0.59945884 | -0.9781318 | 0.32800916 | 0.52568381 |
| AC104982.1 | 0.14114194 | 0.02007176 | 3.29738704 | 0.00608717 | 0.99514317 | NA |
| EFCAB5 | 138.39661 | -0.41924677 | 0.27797799 | -1.5082013 | 0.13150301 | 0.28732574 |
| AC104996.1 | 2.41123318 | -1.727957897 | 0.92829735 | -1.8614271 | 0.06268388 | 0.17049991 |
| AC104984.2 | 6.64764625 | -0.726033197 | 0.49602828 | -1.4636932 | 0.1432778 | 0.30479475 |
| NSRP1 | 1571.10682 | -0.351849491 | 0.12762038 | -2.7570007 | 0.00583342 | 0.03016814 |
| AC104984.3 | 0.19718306 | -0.340099831 | 3.16382928 | -0.1074963 | 0.91439529 | NA |
| AC104984.6 | 1.55551557 | 0.589361749 | 1.09328085 | 0.53907626 | 0.58983424 | 0.74983272 |
| SLC6A4 | 67.2191121 | 0.5569939 | 0.39411339 | 1.41328336 | 0.15757243 | 0.3247215 |
| AC104984.1 | 1.19347668 | -1.504223443 | 1.48320966 | -1.0141678 | 0.3105027 | 0.50743407 |
| AC104984.4 | 13.3500405 | 0.719592401 | 0.41999352 | 1.71334167 | 0.0866497 | 0.21446175 |
| BLMH | 2395.06755 | -0.15514231 | 0.13109614 | -1.1834239 | 0.23664117 | 0.42598318 |
| TMIGD1 | 93.012505 | 2.073183921 | 0.74454766 | 2.78448787 | 0.00536123 | 0.02836765 |
| CPD | 7970.9376 | -1.534688952 | 0.29707414 | -5.1660133 | 2.39E-07 | 9.87E-06 |
| GOSR1 | 2480.26893 | 0.112604382 | 0.18005294 | 0.62539597 | 0.5317112 | 0.70599237 |
| AC011840.3 | 0.99982591 | 3.911206408 | 1.49538611 | 2.61551608 | 0.00890927 | NA |
| TBC1D29 | 41.3788707 | 1.290151601 | 0.51054695 | 2.52699894 | 0.01150418 | 0.05005203 |
| AC011840.5 | 6.62057744 | 1.678558082 | 0.58422947 | 2.87311438 | 0.00406447 | 0.02303559 |
| AC011840.2 | 2.27701248 | 2.09408438 | 1.07648462 | 1.94529893 | 0.05173901 | 0.14937081 |
| AC127024.3 | 1.26937377 | 0.299629825 | 1.13320867 | 0.26440834 | 0.79146528 | 0.88679396 |
| AC127024.2 | 3.08073751 | -0.527097128 | 0.70095897 | -0.7519657 | 0.45207166 | 0.6417624 |
| CRLF3 | 1024.67443 | -0.106616326 | 0.20037718 | -0.5320782 | 0.59467184 | 0.75370069 |
| AC127024.5 | 180.29746 | 0.242944653 | 0.23729772 | 1.02379685 | 0.30593124 | 0.50289283 |
| AC127024.6 | 97.8024046 | 0.105129897 | 0.18683236 | 0.5626964 | 0.57364163 | 0.73821843 |
| ATAD5 | 126.093509 | 0.150762518 | 0.19594094 | 0.76942839 | 0.44163904 | 0.63290156 |
| AC130324.2 | 2.03316648 | 0.658333671 | 0.95762762 | 0.68746312 | 0.49179093 | 0.67481148 |
| AC130324.1 | 6.10955658 | 0.270191791 | 0.5599607 | 0.4825192 | 0.62943716 | 0.77986909 |
| TEFM | 583.846742 | 0.026922017 | 0.14747042 | 0.18255876 | 0.85514425 | 0.92410315 |
| AC130324.3 | 259.800814 | -0.250285165 | 0.20560269 | -1.2173244 | 0.22348082 | 0.40953365 |
| ADAP2 | 756.741107 | -0.441582325 | 0.3271962 | -1.3495949 | 0.17714596 | 0.35216954 |
| AC138207.1 | 131.941099 | -0.097978892 | 0.33904708 | -0.2889831 | 0.77259428 | 0.87469136 |
| AC138207.5 | 47.2743883 | -0.393062544 | 0.33234355 | -1.1826995 | 0.23692827 | 0.42617998 |
| RNF135 | 1907.46328 | 0.140960372 | 0.20024601 | 0.70393598 | 0.48147264 | 0.66600139 |
| AC138207.7 | 3.11671694 | 1.739759432 | 0.8543901 | 2.03625888 | 0.04172437 | 0.12780115 |
| AC138207.2 | 43.5195018 | 0.021699636 | 0.3186083 | 0.06810757 | 0.94570001 | 0.97395466 |
| NF1 | 2752.96359 | 0.059318744 | 0.09316379 | 0.63671459 | 0.52431077 | 0.69941699 |
| OMG | 22.5229468 | -0.63935191 | 0.3853255 | -1.6592515 | 0.09706513 | 0.23267043 |
| EVI2B | 419.423688 | -0.299328299 | 0.49466226 | -0.6051165 | 0.54510158 | 0.71685262 |
| AC134669.1 | 273.408775 | -0.310072499 | 0.51567227 | -0.6012976 | 0.54764178 | 0.71858058 |
| EVI2A | 256.218321 | -0.505960019 | 0.51890418 | -0.9750548 | 0.32953307 | 0.5272929 |
| RAB11FIP4 | 1946.6092 | 0.537235987 | 0.18570513 | 2.89295182 | 0.0038164 | 0.02200829 |
| AC003101.2 | 12.2301683 | -0.641755504 | 0.54083945 | -1.1865915 | 0.23538879 | 0.42427904 |
| AC003101.1 | 0.10721287 | -0.642905668 | 3.30558948 | -0.1944905 | 0.84579185 | NA |
| AC007923.4 | 0.96475299 | -0.794072961 | 1.50477635 | -0.5277016 | 0.59770643 | NA |
| AC007923.1 | 0.26056869 | 1.992294186 | 3.24261664 | 0.61440941 | 0.53894482 | NA |
| AC004253.2 | 0.2875219 | 2.108261977 | 3.23519631 | 0.65166431 | 0.51461775 | NA |
| COPRS | 1842.29818 | -0.331372592 | 0.20066961 | -1.6513342 | 0.09867035 | 0.23549074 |
| UTP6 | 2053.15572 | -0.383918115 | 0.12957984 | -2.9627919 | 0.00304863 | 0.01858984 |
| AC004253.1 | 100.655214 | -0.084765878 | 0.34670422 | -0.2444905 | 0.80685098 | 0.89559542 |
| SUZ12 | 1612.3175 | 0.01166254 | 0.11271569 | 0.10346865 | 0.91759103 | 0.96005997 |
| LRRC37B | 383.558932 | -0.477740722 | 0.16553957 | -2.8859609 | 0.00390221 | 0.0223601 |
| AC116407.2 | 37.864293 | -0.457855759 | 0.40056367 | -1.1430287 | 0.25302668 | 0.44550402 |
| AC116407.1 | 88.6899116 | 0.462841879 | 0.19237429 | 2.40594453 | 0.01613071 | 0.06424512 |
| RHOT1 | 1600.23141 | 0.308859526 | 0.11053218 | 2.79429519 | 0.0052013 | 0.02768636 |
| RHBDL3 | 47.9901134 | 0.371181134 | 0.44936314 | 0.82601598 | 0.40879503 | 0.6032077 |
| AC026620.1 | 0 | NA | NA | NA | NA | NA |
| C17orf75 | 1016.15531 | 0.785198894 | 0.2092071 | 3.75321341 | 0.00017458 | 0.00196881 |
| AC005899.3 | 28.7046157 | 0.757290904 | 0.39343171 | 1.92483446 | 0.05425007 | 0.1542496 |
| ZNF207 | 7467.9949 | -0.261463312 | 0.16142116 | -1.6197586 | 0.10528414 | 0.24612969 |
| AC005899.6 | 43.6033126 | -0.218227548 | 0.34726461 | -0.6284186 | 0.52972974 | 0.70432981 |
| AC005899.7 | 22.9855018 | 0.205833649 | 0.40106319 | 0.51321999 | 0.60779744 | 0.76405917 |
| AC005899.5 | 2.67701968 | 0.757957879 | 0.79300545 | 0.95580412 | 0.3391712 | 0.53727066 |
| AC005899.8 | 248.792446 | -0.153054831 | 0.26302567 | -0.5819007 | 0.56063355 | 0.72862652 |
| AC005899.1 | 10.2559542 | -0.335384435 | 0.39907122 | -0.8404125 | 0.40067716 | 0.59547525 |
| PSMD11 | 4557.3578 | -0.360463951 | 0.23717714 | -1.519809 | 0.12855898 | 0.28279116 |
| CDK5R1 | 155.076359 | -0.024949619 | 0.41090625 | -0.0607185 | 0.95158338 | 0.97677127 |
| MYO1D | 4107.93818 | -0.211657969 | 0.21255376 | -0.9957856 | 0.31935434 | 0.51677553 |
| AC079336.5 | 30.2667961 | 0.093838066 | 0.31245528 | 0.30032479 | 0.76392942 | 0.86935843 |
| AC079336.4 | 0.63810679 | 2.108329567 | 1.63607904 | 1.28864775 | 0.19752058 | NA |
| AC025211.1 | 0.98374943 | -0.526579311 | 1.400978 | -0.3758655 | 0.70701688 | NA |
| AC084809.1 | 25.3603477 | 1.30898172 | 0.37372472 | 3.50252912 | 0.00046086 | 0.00425972 |
| AC084809.2 | 2.63099421 | -1.01268138 | 0.91159349 | -1.1108914 | 0.26661509 | 0.46069327 |
| TMEM98 | 1789.74042 | 0.200200176 | 0.25443412 | 0.78684484 | 0.4313727 | 0.62344065 |
| SPACA3 | 0.62402702 | 2.037824023 | 1.77956382 | 1.14512557 | 0.25215715 | NA |
| ASIC2 | 116.926329 | -0.079573618 | 0.55428062 | -0.143562 | 0.88584637 | 0.94097225 |
| AC008133.1 | 8.80855108 | 0.252763909 | 0.80711468 | 0.31316976 | 0.75415169 | 0.86318765 |
| AC011824.4 | 0 | NA | NA | NA | NA | NA |
| AC011824.2 | 1.7421185 | -0.204492591 | 1.16732559 | -0.1751804 | 0.86093787 | 0.9275589 |
| AC011824.1 | 0.3184437 | -1.669100907 | 2.89273496 | -0.5769975 | 0.56394113 | NA |
| AC024614.1 | 0 | NA | NA | NA | NA | NA |
| AC024614.2 | 0 | NA | NA | NA | NA | NA |
| AC024614.4 | 0.08666371 | -0.642905668 | 3.30558948 | -0.1944905 | 0.84579185 | NA |
| AC024610.2 | 1.88683767 | 0.992104647 | 1.32310301 | 0.74983175 | 0.45335604 | 0.64299587 |
| AC024610.1 | 0.09997307 | 0.547239394 | 3.29377202 | 0.16614368 | 0.86804388 | NA |
| AC004147.3 | 0.08695548 | -0.642905668 | 3.30558948 | -0.1944905 | 0.84579185 | NA |
| AC004147.2 | 0.22329207 | 1.817076245 | 3.25496057 | 0.55824831 | 0.57667483 | NA |
| LINC01989 | 0.78150354 | -0.088628434 | 1.54630725 | -0.0573162 | 0.95429333 | NA |
| AC005549.1 | 0 | NA | NA | NA | NA | NA |
| CCL2 | 4979.49703 | -1.009922278 | 0.70641278 | -1.4296489 | 0.1528178 | 0.31828493 |
| CCL7 | 24.2842308 | -1.975044238 | 1.23499974 | -1.5992264 | 0.1097703 | 0.25353794 |
| CCL11 | 275.067808 | -0.804816278 | 0.91987258 | -0.8749215 | 0.38161663 | 0.57757877 |
| CCL8 | 135.363513 | 0.379135103 | 0.68715587 | 0.55174542 | 0.58112279 | 0.74384312 |
| CCL13 | 68.1816053 | -2.186518753 | 0.52918184 | -4.1318854 | 3.60E-05 | 0.0005695 |
| CCL1 | 0.53848582 | -2.315429574 | 2.1696691 | -1.067181 | 0.28589012 | NA |
| AC011193.1 | 0 | NA | NA | NA | NA | NA |
| TMEM132E | 523.010181 | 2.245811577 | 0.34162736 | 6.57386342 | 4.90E-11 | 7.98E-09 |
| AC005552.1 | 1.08843796 | -1.499688822 | 2.23197267 | -0.6719118 | 0.50163983 | 0.68225501 |
| AC022903.1 | 0.38658711 | 0.760280982 | 2.84025691 | 0.26768036 | 0.78894537 | NA |
| CCT6B | 186.727932 | 0.811945732 | 0.47630322 | 1.70468243 | 0.08825367 | 0.21739941 |
| ZNF830 | 683.406098 | -0.247040291 | 0.10413957 | -2.3722039 | 0.01768233 | 0.06852912 |
| LIG3 | 2383.43847 | -0.638295032 | 0.20834208 | -3.0636875 | 0.00218627 | 0.01445274 |
| RFFL | 3744.39793 | -0.798601127 | 0.32118376 | -2.4864306 | 0.01290317 | 0.0544407 |
| AC004223.3 | 893.035168 | -0.073135374 | 0.35927579 | -0.2035633 | 0.83869476 | 0.91450415 |
| AC004223.4 | 4.39055465 | 0.449050292 | 0.65192797 | 0.68880354 | 0.49094691 | 0.67400406 |
| AC004223.2 | 8.74640558 | -0.964735809 | 0.50160707 | -1.9232899 | 0.05444365 | 0.15457355 |
| RAD51D | 730.10133 | 0.436977269 | 0.09826828 | 4.44677829 | 8.72E-06 | 0.00018286 |
| FNDC8 | 53.336583 | 0.167232194 | 0.31571284 | 0.52969716 | 0.59632192 | 0.75506621 |
| NLE1 | 647.310566 | 0.093427891 | 0.20467073 | 0.45647901 | 0.64804556 | 0.79277053 |
| UNC45B | 19.9855688 | 0.665896547 | 0.51157608 | 1.30165692 | 0.19303369 | 0.37251871 |
| AC022916.4 | 0.10721287 | -0.642905668 | 3.30558948 | -0.1944905 | 0.84579185 | NA |
| SLC35G3 | 0.08393208 | -0.642905668 | 3.30558948 | -0.1944905 | 0.84579185 | NA |
| AC022916.1 | 90.2669429 | 0.081046275 | 0.32427182 | 0.24993315 | 0.80263905 | 0.89302526 |
| SLFN5 | 1536.36979 | -1.028569267 | 0.34943494 | -2.9435215 | 0.00324501 | 0.01950558 |

| | | | | | | |
|---|---|---|---|---|---|---|
| AC022706.1 | 16.3041624 | -0.785937779 | 0.7664197 | -1.0254666 | 0.30514309 | 0.50216071 |
| SLFN11 | 843.789051 | -0.27534821 | 0.3696848 | -0.7448189 | 0.45638123 | 0.64586168 |
| AC060766.6 | 4.65192408 | 0.413791621 | 0.55751465 | 0.74220762 | 0.45796155 | 0.64706542 |
| AC060766.4 | 37.6258426 | -0.255510627 | 0.4400838 | -0.5805954 | 0.56151318 | 0.72937798 |
| SLFN12 | 181.062231 | -0.225952372 | 0.34012639 | -0.6643188 | 0.50648626 | 0.68566368 |
| SLFN13 | 564.063727 | -0.409713413 | 0.43580018 | -0.9401405 | 0.34714548 | 0.54472033 |
| SLFN12L | 48.7000229 | -0.989524195 | 0.75766737 | -1.306014 | 0.19154779 | 0.37072406 |
| AC015911.5 | 0.98777779 | -2.273229024 | 2.22066726 | -1.0236694 | 0.30599147 | NA |
| AC015911.2 | 0 | NA | NA | NA | NA | NA |
| SLFN14 | 6.36030319 | 0.070152527 | 0.81538905 | 0.08603565 | 0.93143808 | 0.96693261 |
| AC015911.6 | 4.29136556 | -0.76652029 | 0.85213482 | -0.8995294 | 0.36837076 | 0.56515711 |
| AC015911.9 | 0 | NA | NA | NA | NA | NA |
| LINC02001 | 362.144273 | -0.017102101 | 0.23135358 | -0.0739219 | 0.94107252 | 0.97202378 |
| PEX12 | 423.176613 | 0.861780932 | 0.136551 | 6.31105549 | 2.77E-10 | 3.46E-08 |
| AP2B1 | 10402.2067 | -0.082921492 | 0.14496888 | -0.5719951 | 0.56732529 | 0.73338525 |
| TAF15 | 6100.39611 | -0.133822023 | 0.12059566 | -1.1096753 | 0.26713896 | 0.46124006 |
| RASL10B | 260.545733 | -0.348137759 | 0.4453419 | -0.7817314 | 0.43437242 | 0.62633425 |
| GAS2L2 | 30.8765439 | -0.584507678 | 0.87578218 | -0.6674122 | 0.5045089 | 0.68396433 |
| MMP28 | 130.931557 | 0.727012304 | 0.4845352 | 1.5004324 | 0.13350243 | 0.29027533 |
| C17orf50 | 6.1556801 | 0.124840145 | 0.87005384 | 0.14348554 | 0.88590673 | 0.94097225 |
| AC015849.2 | 0.21694805 | 0.689156974 | 3.06622645 | 0.22475736 | 0.82216804 | NA |
| AC015849.3 | 7.67372897 | 0.436076165 | 0.51505869 | 0.84665335 | 0.39718837 | 0.59211133 |
| HEATR9 | 16.3605217 | -1.158452227 | 0.53106666 | -2.1813688 | 0.02915615 | 0.0989508 |
| AC015849.1 | 12.8064336 | -0.467723394 | 0.49886381 | -0.9375773 | 0.34846167 | 0.54600902 |
| CCL5 | 617.724076 | -0.997821685 | 0.73656448 | -1.354697 | 0.17551412 | 0.34973818 |
| AC015849.5 | 1.01304484 | -0.469035827 | 1.52076232 | -0.3084215 | 0.75776161 | 0.86607895 |
| RDM1 | 5.20653541 | -0.767849012 | 0.77021569 | -0.9969273 | 0.31879982 | 0.5160893 |
| LYZL6 | 0.2963014 | 2.148440699 | 3.23275528 | 0.66458501 | 0.50631596 | NA |
| CCL16 | 31.4483064 | 0.553508061 | 0.54359958 | 1.01822753 | 0.30856984 | 0.50552849 |
| CCL14 | 1567.43276 | -0.116827059 | 0.50279761 | -0.232354 | 0.81626304 | 0.90107402 |
| AC244100.2 | 1186.18442 | -0.160937111 | 0.50824042 | -0.3166555 | 0.75150504 | 0.86135902 |
| CCL15-CCL14 | 1948.0492 | -0.008818376 | 0.43536596 | -0.0202551 | 0.98383988 | 0.99305611 |
| CCL15 | 526.982383 | 1.334713142 | 0.34483378 | 3.87059857 | 0.00010857 | 0.00135172 |
| AC244100.3 | 241.915695 | 1.459094444 | 0.32343363 | 4.51126385 | 6.44E-06 | 0.00014413 |
| AC244100.4 | 2.9713097 | 5.085475076 | 0.99497514 | 5.11115792 | 3.20E-07 | 1.26E-05 |
| CCL23 | 147.740583 | -2.336044315 | 0.67830495 | -3.4439441 | 0.00057329 | 0.00504767 |
| CCL18 | 47.0557819 | -2.968462623 | 0.70721345 | -4.1974069 | 2.70E-05 | 0.00045244 |
| AC243829.4 | 189.704823 | 0.826429778 | 0.63758718 | 1.29618317 | 0.19491239 | 0.37495183 |
| CCL3 | 217.848028 | 0.813732667 | 0.63995672 | 1.27154328 | 0.20353544 | 0.3857431 |
| CCL4 | 229.171702 | 0.120348782 | 0.55715281 | 0.21600678 | 0.82898247 | 0.90906423 |
| AC243829.2 | 1.28422541 | -0.033321149 | 1.01604099 | -0.0327951 | 0.973838 | 0.98792445 |
| TBC1D3B | 7.21106865 | -0.354168528 | 0.47976432 | -0.7382136 | 0.46038468 | 0.64890395 |
| AC243829.1 | 18.3377426 | 1.058596376 | 0.7409235 | 1.42875261 | 0.15307535 | 0.31857242 |
| CCL3L1 | 54.2065621 | 1.442038807 | 0.89396839 | 1.61307583 | 0.10672803 | 0.24846988 |
| CCL4L2 | 72.6563622 | 1.330880534 | 0.89566682 | 1.48591027 | 0.13730286 | 0.29587941 |
| AC243829.5 | 0 | NA | NA | NA | NA | NA |
| TBC1D3I | 0 | NA | NA | NA | NA | NA |
| TBC1D3G | 0.48292446 | -1.043692077 | 3.28541311 | -0.3176745 | 0.75073183 | NA |
| TBC1D3H | 0.13390214 | 1.04981649 | 3.28477129 | 0.31960109 | 0.74927075 | NA |
| TBC1D3F | 1.22568121 | -1.919945558 | 1.99525034 | -0.962258 | 0.33592003 | 0.53409445 |
| ZNHIT3 | 2133.91964 | -0.148149719 | 0.16037191 | -0.9237884 | 0.35559647 | 0.55305686 |
| MYO19 | 2867.34433 | -0.12201525 | 0.13096168 | -0.9316866 | 0.35149849 | 0.54914481 |
| PIGW | 508.042547 | -0.281778712 | 0.42996801 | -0.6553481 | 0.51224367 | 0.6903744 |
| GGNBP2 | 3546.96821 | -0.288844374 | 0.16784993 | -1.7208489 | 0.08527825 | 0.21186992 |
| DHRS11 | 1072.71256 | 0.980651824 | 0.23684096 | 4.1405499 | 3.46E-05 | 0.00055139 |
| MRM1 | 470.22411 | 0.066034509 | 0.19074603 | 0.34619073 | 0.72919938 | 0.84700595 |
| AC243830.2 | 14.0483449 | 0.443697211 | 0.41478318 | 1.0697088 | 0.28475041 | 0.48087533 |
| AC243830.3 | 0 | NA | NA | NA | NA | NA |
| AC243830.1 | 1.85556298 | -0.781010426 | 1.06864615 | -0.730841 | 0.46487628 | 0.65205955 |
| LHX1-DT | 604.9894 | 0.738136394 | 0.39040814 | 1.89067878 | 0.05866724 | 0.16287786 |
| LHX1 | 956.587859 | 1.102050901 | 0.37507846 | 2.93818765 | 0.00330137 | 0.01974822 |
| AC243773.1 | 159.623689 | 0.904063176 | 0.35688266 | 2.53322245 | 0.01130192 | 0.04939205 |
| AATF | 2345.06499 | -0.326359815 | 0.17489738 | -1.8660074 | 0.06204033 | 0.16932348 |
| AC244093.5 | 2.50234655 | -0.037811194 | 0.84115418 | -0.0449516 | 0.96414592 | 0.98324765 |
| ACACA | 1425.47711 | -0.08053037 | 0.17855572 | -0.4510098 | 0.65198252 | 0.79555685 |
| C17orf78 | 5.33239808 | 0.964142923 | 0.5622822 | 1.7146958 | 0.08640101 | 0.21401416 |
| TADA2A | 758.716888 | -0.164994059 | 0.12955947 | -1.2735006 | 0.20284046 | 0.38486853 |
| AC243654.1 | 0.92855652 | -1.081947483 | 1.61422812 | -0.6702569 | 0.50269406 | NA |
| DUSP14 | 880.37689 | -0.611171628 | 0.2974517 | -2.054692 | 0.03990876 | 0.12363652 |
| SYNRG | 2378.41429 | 0.058952009 | 0.11459451 | 0.51444006 | 0.60694436 | 0.76327544 |
| DDX52 | 2323.7025 | -1.045839843 | 0.31982847 | -3.2700023 | 0.00107547 | 0.00830212 |
| AC243585.2 | 343.998545 | -1.576686436 | 0.3643631 | -4.3272396 | 1.51E-05 | 0.00028533 |
| AC243571.2 | 65.5170381 | -0.203718669 | 0.42894299 | -0.4749318 | 0.63483558 | 0.78341547 |
| AC243585.1 | 41.9057588 | -3.969191833 | 0.61254381 | -6.4798497 | 9.18E-11 | 1.34E-08 |
| HNF1B | 8164.10585 | 0.112281174 | 0.22484508 | 0.49937127 | 0.61751786 | 0.77138541 |
| AC243571.1 | 22.7181066 | 2.125820357 | 0.39301829 | 5.40896034 | 6.34E-08 | 3.23E-06 |
| TBC1D3K | 1.65004542 | 0.471577161 | 1.06762252 | 0.44170777 | 0.65870069 | 0.79950001 |
| TBC1D3L | 437.88771 | -0.745381915 | 0.34795735 | -2.1421646 | 0.03218024 | 0.10607411 |
| TBC1D3D | 0.16964538 | 1.509808164 | 3.2763376 | 0.46082191 | 0.64492638 | NA |
| TBC1D3C | 0.1905444 | -0.340597073 | 3.29738699 | -0.103293 | 0.91773041 | NA |
| TBC1D3E | 3.67311129 | 0.689330323 | 0.75763116 | 0.90984949 | 0.36290189 | 0.56012425 |
| TBC1D3 | 0.10721287 | -0.642905668 | 3.30558948 | -0.1944905 | 0.84579185 | NA |
| MRPL45 | 1894.96236 | 0.158041934 | 0.18429023 | 0.85757086 | 0.39112947 | 0.58676096 |
| GPR179 | 5.2721193 | 0.187432451 | 0.68452062 | 0.27381564 | 0.7842263 | 0.88211398 |
| SOCS7 | 932.517262 | 0.393304652 | 0.19458532 | 2.02124523 | 0.04325439 | 0.13113568 |
| ARHGAP23 | 1999.70291 | 0.003013091 | 0.28775329 | 0.01047109 | 0.99164543 | 0.99605449 |
| SRCIN1 | 265.922398 | -1.732232698 | 0.38552806 | -4.493143 | 7.02E-06 | 0.0001544 |
| AC006449.1 | 0 | NA | NA | NA | NA | NA |
| EPOP | 190.095503 | -1.873474271 | 0.26777311 | -6.9964988 | 2.62E-12 | 6.35E-10 |
| AC006449.6 | 270.159319 | -0.057799686 | 0.23104265 | -0.2501689 | 0.80245673 | 0.89289776 |
| AC006449.2 | 69.9665761 | -0.2761775 | 0.26579442 | -1.0390643 | 0.29877482 | 0.49563286 |
| MLLT6 | 16595.1935 | -0.147983758 | 0.20786634 | -0.7119179 | 0.47651564 | 0.66179458 |
| AC006449.3 | 939.85545 | 0.621134972 | 0.26959528 | 2.30395344 | 0.02122526 | 0.07831398 |
| CISD3 | 3666.47323 | 0.397413489 | 0.13075031 | 3.03948411 | 0.00236984 | 0.01537333 |
| PCGF2 | 4854.14858 | 0.056916357 | 0.13467469 | 0.42262103 | 0.67257178 | 0.80897677 |
| AC006449.5 | 92.4768384 | 0.727478453 | 0.26353807 | 2.76043026 | 0.00577253 | 0.02996488 |
| PSMB3 | 4458.55119 | -0.630035899 | 0.16150018 | -3.9011468 | 9.57E-05 | 0.00122198 |
| PIP4K2B | 3602.56052 | 0.321284757 | 0.11996759 | 2.67809619 | 0.0074042 | 0.03608365 |
| CWC25 | 2158.24492 | -0.571558396 | 0.26261425 | -2.176418 | 0.02952402 | 0.09976273 |
| C17orf98 | 1.80673589 | 1.985556774 | 1.13449729 | 1.75016441 | 0.08008995 | 0.20261727 |
| RPL23 | 22273.7801 | -0.381180065 | 0.19587301 | -1.9460572 | 0.05164787 | 0.14920046 |
| LASP1 | 12965.0064 | -0.22110891 | 0.19906155 | -1.1107565 | 0.26667317 | 0.46076343 |
| AC006441.1 | 0.95076225 | 0.670388025 | 1.22921955 | 0.54537696 | 0.58549428 | NA |
| AC006441.3 | 2527.55266 | -0.37417379 | 0.19935826 | -1.8768914 | 0.06053298 | 0.16652683 |
| LINC00672 | 70.0693207 | 0.048740635 | 0.31980001 | 0.15240974 | 0.87886378 | 0.93705294 |
| FBXO47 | 0.3348092 | 0.669724679 | 2.23196307 | 0.30006082 | 0.76413076 | NA |
| AC006441.4 | 87.5553029 | 2.858412911 | 0.77507672 | 3.68790962 | 0.0002261 | 0.00241391 |
| LINC02079 | 0.13028434 | 1.049816488 | 3.28477129 | 0.31960109 | 0.74927075 | NA |
| PLXDC1 | 534.813923 | 0.69825847 | 0.53703434 | 1.30021195 | 0.19352834 | 0.37310874 |
| ARL5C | 3.89591227 | 1.894188631 | 0.92268258 | 2.05291469 | 0.04008085 | 0.1239803 |
| AC004408.1 | 68.1730628 | -0.772995411 | 0.38631537 | -2.000944 | 0.04539843 | 0.13571664 |
| CACNB1 | 536.887003 | -0.704636821 | 0.28884521 | -2.4394963 | 0.01470775 | 0.06001023 |
| RPL19 | 35190.2794 | -0.271956368 | 0.13908082 | -1.9553837 | 0.0505378 | 0.14677112 |

| | | | | | | |
|---|---|---|---|---|---|---|
| STAC2 | 2027.59946 | -1.979802758 | 0.39793319 | -4.9752139 | 6.52E-07 | 2.26E-05 |
| FBXL20 | 1525.25543 | -0.244112891 | 0.14228674 | -1.7156404 | 0.08622787 | 0.21368565 |
| AC005288.1 | 397.815842 | 0.22373956 | 0.18348835 | 1.21936655 | 0.2227051 | 0.40870857 |
| MED1 | 937.234508 | 0.001050257 | 0.16879366 | 0.00622214 | 0.99503549 | 0.99770729 |
| CDK12 | 1684.55326 | -0.128034131 | 0.17904056 | -0.7151124 | 0.47453958 | 0.66007262 |
| AC009283.1 | 324.16853 | -0.200550861 | 0.28107868 | -0.7135043 | 0.47553373 | 0.66099965 |
| NEUROD2 | 3.11705301 | 0.164671887 | 0.66080521 | 0.24919883 | 0.80320698 | 0.89318485 |
| AC087491.1 | 12.0722127 | -1.060618244 | 0.63495551 | -1.670382 | 0.09484381 | 0.22913618 |
| PPP1R1B | 1137.54321 | -1.870938699 | 0.55317978 | -3.3821531 | 0.0007192 | 0.00603799 |
| STARD3 | 3407.23559 | -0.582726316 | 0.11093242 | -5.2529846 | 1.50E-07 | 6.69E-06 |
| TCAP | 270.055689 | -0.693330643 | 0.33483598 | -2.0706575 | 0.03839081 | 0.12019063 |
| PNMT | 62.2919055 | -0.183892951 | 0.47171204 | -0.3898415 | 0.69665372 | 0.82558262 |
| PGAP3 | 2680.48254 | 0.601801446 | 0.25742465 | 2.33777707 | 0.01939882 | 0.073277 |
| ERBB2 | 16629.399 | 0.524450753 | 0.24827924 | 2.11234236 | 0.03465709 | 0.11187509 |
| MIEN1 | 2895.98537 | -0.342958364 | 0.12285748 | -2.791514 | 0.00524621 | 0.0278672 |
| GRB7 | 7124.82729 | -0.805111134 | 0.22160037 | -3.6331669 | 0.00027996 | 0.00287822 |
| IKZF3 | 202.54307 | -0.366412404 | 0.61790842 | -0.5929882 | 0.55318904 | 0.72290971 |
| ZPBP2 | 0.37767972 | -1.733137851 | 2.80413468 | -0.6180651 | 0.53653241 | NA |
| GSDMB | 814.93265 | -0.60167705 | 0.37950025 | -1.5854457 | 0.1128651 | 0.2586022 |
| ORMDL3 | 4013.13196 | 0.094741569 | 0.13025873 | 0.72733376 | 0.46702152 | 0.65413276 |
| AC090844.2 | 16.4709192 | 1.196616026 | 0.51262656 | 2.3342841 | 0.01958085 | 0.07374279 |
| LRRC3C | 2.00215653 | 0.422839508 | 1.04338103 | 0.40525896 | 0.68528717 | 0.81810054 |
| GSDMA | 15.9084344 | -0.01402586 | 0.78115521 | -0.0179553 | 0.98567453 | 0.99368571 |
| PSMD3 | 5790.58591 | -0.408665983 | 0.2135813 | -1.9133977 | 0.05569715 | 0.15705025 |
| AC090844.3 | 1.50097916 | -2.626411095 | 1.7961866 | -1.4622151 | 0.14368229 | 0.30526119 |
| CSF3 | 150.443593 | -2.409526925 | 1.27333676 | -1.8922935 | NA | NA |
| MED24 | 4324.85069 | -0.105234639 | 0.10819609 | -0.9726289 | 0.33073778 | 0.52867582 |
| THRA | 4959.3388 | 0.094641515 | 0.14537697 | 0.65100763 | 0.51504156 | 0.6927644 |
| NR1D1 | 3268.86891 | -1.171735137 | 0.37301634 | -3.1412434 | 0.00168232 | 0.01180176 |
| MSL1 | 5548.40803 | -0.657420235 | 0.19510857 | -3.3695099 | 0.00075302 | 0.00625743 |
| CASC3 | 4747.46983 | 0.110633442 | 0.10394212 | 1.06437551 | 0.28715862 | 0.48339437 |
| RAPGEFL1 | 504.606927 | -0.945164109 | 0.32878844 | -2.8746878 | 0.00404427 | 0.02294974 |
| WIPF2 | 3155.2567 | 0.002741381 | 0.21010682 | 0.01304756 | 0.98958985 | 0.99519138 |
| CDC6 | 72.1037801 | -1.168255064 | 0.53390967 | -2.1881137 | 0.02866132 | 0.09767369 |
| RARA | 3103.94958 | -0.07705831 | 0.24759417 | -0.3112283 | 0.75562708 | 0.86427612 |
| RARA-AS1 | 396.623588 | -0.027446108 | 0.28250271 | -0.0971534 | 0.92260455 | 0.96215621 |
| AC080112.1 | 359.924672 | -0.060475939 | 0.22596909 | -0.2676293 | 0.78898471 | 0.88537924 |
| GJD3 | 66.5264419 | -0.112054504 | 0.26821296 | -0.4177818 | 0.67610662 | 0.81149917 |
| TOP2A | 113.399923 | -1.41283718 | 0.72442733 | -1.9502814 | 0.05114258 | 0.14805466 |
| IGFBP4 | 43284.7871 | -0.326999602 | 0.22786231 | -1.4350754 | 0.1512656 | 0.3161758 |
| TNS4 | 102.808669 | 2.728824594 | 0.61787358 | 4.41647724 | 1.00E-05 | 0.00020397 |
| AC004585.1 | 12.0033801 | 0.830101679 | 1.08138332 | 0.76762944 | 0.44270736 | 0.63391579 |
| CCR7 | 73.7519528 | -0.125420506 | 0.68742047 | -0.1824509 | 0.85522886 | 0.92410315 |
| SMARCE1 | 4807.71293 | -0.069490246 | 0.13203531 | -0.5263005 | 0.59867944 | 0.75677743 |
| AC073508.2 | 4475.34031 | -0.098283103 | 0.12900906 | -0.761831 | 0.44616086 | 0.63654503 |
| AC073508.3 | 250.905549 | 0.20901033 | 0.19820925 | 1.05449333 | 0.29165711 | 0.48785272 |
| KRT222 | 60.0120126 | -0.811536652 | 0.60418607 | -1.3431899 | 0.17921048 | 0.3548671 |
| KRT24 | 6.67044125 | -0.422053864 | 0.65152714 | -0.6477917 | 0.5171197 | 0.69438193 |
| KRT25 | 0.3042876 | -0.699687286 | 2.92313922 | -0.2393616 | 0.8108252 | NA |
| KRT26 | 0.12088456 | 0.02007176 | 3.29738704 | 0.00608717 | 0.99514317 | NA |
| KRT27 | 1.02395481 | 0.13095292 | 1.8935493 | 0.06915739 | 0.94486434 | 0.9737607 |
| KRT28 | 0.09997307 | 0.547239394 | 3.29377202 | 0.16614368 | 0.86804388 | NA |
| AC090283.1 | 0.03392908 | 0.380740593 | 3.29738709 | 0.11546736 | 0.90807469 | NA |
| KRT10 | 724.90209 | -0.720732248 | 0.28441459 | -2.5340903 | 0.01127397 | 0.04927806 |
| TMEM99 | 554.513014 | 0.026885719 | 0.23210976 | 0.11583192 | 0.90778575 | 0.95386955 |
| AC004231.3 | 0.37273028 | -0.542388995 | 2.81714319 | -0.1925316 | 0.84732584 | NA |
| KRT12 | 1.33083244 | 0.353459079 | 0.97722961 | 0.36169502 | 0.71757995 | 0.83969751 |
| KRT20 | 0.76383051 | -0.836873674 | 2.30032012 | -0.3638075 | 0.71600177 | NA |
| AC004231.1 | 14.3821224 | -0.249980554 | 0.6181766 | -0.4043837 | 0.68593057 | 0.81838699 |
| KRT23 | 36.6469888 | -0.613101399 | 0.61300944 | -1.00015 | 0.31723792 | 0.51448733 |
| KRT39 | 0.92084072 | 0.868763272 | 1.71848026 | 0.50554161 | 0.61317848 | NA |
| KRT40 | 1.86191279 | 2.680559545 | 1.16225568 | 2.30634239 | 0.02109151 | 0.07791271 |
| KRTAP3-3 | 0 | NA | NA | NA | NA | NA |
| KRTAP3-2 | 0.26427878 | 1.012543026 | 2.9068939 | 0.34832473 | 0.72759633 | NA |
| KRTAP3-1 | 0 | NA | NA | NA | NA | NA |
| KRTAP1-5 | 4.96512053 | 0.253025035 | 1.03766793 | 0.24384008 | 0.80735467 | 0.89590613 |
| KRTAP1-4 | 0 | NA | NA | NA | NA | NA |
| KRTAP1-3 | 0.06785815 | 0.795868262 | 3.28898349 | 0.24198001 | 0.80879565 | NA |
| KRTAP1-1 | 0.07830766 | -0.642905668 | 3.30558948 | -0.1944905 | 0.84579185 | NA |
| KRTAP2-1 | 0 | NA | NA | NA | NA | NA |
| KRTAP2-2 | 0 | NA | NA | NA | NA | NA |
| KRTAP2-3 | 2.06217228 | -2.495211371 | 1.51200411 | -1.6502676 | 0.09888822 | 0.23577598 |
| KRTAP2-4 | 0 | NA | NA | NA | NA | NA |
| KRTAP4-7 | 0.03392908 | 0.380740593 | 3.29738709 | 0.11546736 | 0.90807469 | NA |
| KRTAP4-8 | 0.10721287 | -0.642905668 | 3.30558948 | -0.1944905 | 0.84579185 | NA |
| KRTAP4-16 | 0 | NA | NA | NA | NA | NA |
| KRTAP4-9 | 0 | NA | NA | NA | NA | NA |
| KRTAP4-11 | 0 | NA | NA | NA | NA | NA |
| KRTAP4-12 | 0.19632833 | 1.506623996 | 3.27645372 | 0.45983375 | 0.64563556 | NA |
| KRTAP4-6 | 0.08757577 | 0.547239394 | 3.29377202 | 0.16614368 | 0.86804388 | NA |
| KRTAP4-5 | 0 | NA | NA | NA | NA | NA |
| KRTAP4-4 | 0 | NA | NA | NA | NA | NA |
| KRTAP4-3 | 0 | NA | NA | NA | NA | NA |
| KRTAP4-2 | 0.21442573 | -1.003574173 | 3.30558948 | -0.3035992 | 0.7614333 | NA |
| KRTAP4-1 | 0.42885147 | -1.859940617 | 3.29773397 | -0.5640057 | 0.57275028 | NA |
| KRTAP9-1 | 0 | NA | NA | NA | NA | NA |
| KRTAP9-2 | 0 | NA | NA | NA | NA | NA |
| KRTAP9-3 | 0 | NA | NA | NA | NA | NA |
| KRTAP9-8 | 0 | NA | NA | NA | NA | NA |
| KRTAP9-4 | 0 | NA | NA | NA | NA | NA |
| KRTAP9-9 | 0 | NA | NA | NA | NA | NA |
| KRTAP9-6 | 0 | NA | NA | NA | NA | NA |
| KRTAP9-7 | 0 | NA | NA | NA | NA | NA |
| KRTAP29-1 | 0.11786115 | 0.02007176 | 3.29738704 | 0.00608717 | 0.99514317 | NA |
| KRTAP16-1 | 0.17423948 | 0.18657164 | 3.29377195 | 0.05664376 | 0.95482897 | NA |
| KRTAP17-1 | 1.66682449 | 0.605543093 | 1.94605666 | 0.31116416 | 0.75567582 | 0.86429439 |
| KRT33A | 0.10721287 | -0.642905668 | 3.30558948 | -0.1944905 | 0.84579185 | NA |
| KRT33B | 0.62735043 | -2.538011199 | 1.81763176 | -1.3963286 | 0.16261557 | NA |
| KRT34 | 0.39352299 | -1.054987527 | 1.92936694 | -0.546805 | 0.58451269 | NA |
| KRT31 | 2.14214304 | 3.232294763 | 1.8939218 | 1.70666749 | 0.08788387 | 0.21666185 |
| AC003958.2 | 1.8816881 | 3.935095783 | 1.20525788 | 3.26494092 | 0.00109487 | 0.0084125 |
| KRT37 | 2.13245885 | 2.838894052 | 1.07643028 | 2.63732275 | 0.00835633 | 0.03932549 |
| KRT38 | 0.09997307 | 0.547239394 | 3.29377202 | 0.16614368 | 0.86804388 | NA |
| KRT32 | 0.75684125 | -2.042173781 | 2.31470739 | -0.88226 | 0.37763621 | NA |
| KRT35 | 0.254706 | 0.078718067 | 3.03476391 | 0.02593878 | 0.97930617 | NA |
| KRT36 | 0.82507018 | -1.011086348 | 1.41461382 | -0.7147437 | 0.47476741 | NA |
| KRT13 | 9.6791866 | 0.99005759 | 1.23627001 | 0.80084252 | 0.42322282 | 0.6157794 |
| AC019349.1 | 14.4159618 | -0.825631211 | 2.42883404 | -0.339929 | 0.73390997 | 0.8501319 |
| KRT15 | 55.6337257 | -1.77297689 | 0.7943577 | -2.2319629 | 0.02561742 | 0.0898791 |
| KRT19 | 10929.3234 | -1.588734986 | 0.53065141 | -2.9939334 | 0.00275406 | 0.01722186 |
| LINC00974 | 0.17361919 | -1.003574173 | 3.30558948 | -0.3035992 | 0.7614333 | NA |
| KRT9 | 0.17088756 | -1.003574173 | 3.30558948 | -0.3035992 | 0.7614333 | NA |
| KRT14 | 22.9354383 | 4.426275797 | 1.19728075 | 3.69694058 | NA | NA |

| | | | | | | |
|---|---|---|---|---|---|---|
| KRT16 | 23.5208053 | -0.231634813 | 1.05916516 | -0.2186956 | 0.82688714 | 0.90767239 |
| KRT17 | 220.525598 | -1.958992581 | 0.66588148 | -2.9419538 | 0.00326149 | 0.01956891 |
| EIF1 | 26792.2589 | -0.748232998 | 0.20611309 | -3.6302061 | 0.00028319 | 0.00289806 |
| GAST | 11.4691411 | -5.528309573 | 1.28213183 | -4.3118106 | 1.62E-05 | 0.00030028 |
| HAP1 | 615.288225 | -2.920196494 | 0.39466758 | -7.3991295 | 1.37E-13 | 5.16E-11 |
| JUP | 19768.7134 | 0.051171036 | 0.16003882 | 0.3197414 | 0.74916437 | 0.85990531 |
| P3H4 | 1029.00359 | -0.224820459 | 0.24157924 | -0.9306282 | 0.3520459 | 0.54970692 |
| FKBP10 | 2738.60323 | -1.069477426 | 0.35425255 | -3.0189689 | 0.00253637 | 0.01617117 |
| NT5C3B | 2578.13448 | -0.535669452 | 0.1724867 | -3.1055697 | 0.00189913 | 0.01294727 |
| KLHL10 | 49.6582254 | -0.328999339 | 0.29555886 | -1.1131432 | 0.26564693 | 0.45980371 |
| KLHL11 | 10.7127493 | 0.362186816 | 0.60156163 | 0.60207766 | 0.54712244 | 0.71822077 |
| ACLY | 9225.6682 | -0.004794442 | 0.21035883 | -0.0227917 | 0.98181641 | 0.99174514 |
| AC125257.2 | 1440.54748 | -0.419550275 | 0.2069385 | -2.0274153 | 0.04261996 | 0.12971933 |
| TTC25 | 212.079036 | 0.477758018 | 0.26814533 | 1.78171299 | 0.07479605 | 0.19327504 |
| CNP | 9401.55547 | -0.027168811 | 0.25921106 | -0.1048135 | 0.91652382 | 0.95937676 |
| DNAJC7 | 4907.4322 | -0.252697864 | 0.08344833 | -3.0281957 | 0.00246019 | 0.01579623 |
| NKIRAS2 | 2156.70126 | -0.178413884 | 0.10965931 | -1.6269834 | 0.10374062 | 0.24368787 |
| ZNF385C | 192.972274 | -0.218265843 | 0.29143884 | -0.748925 | 0.45390239 | 0.64358565 |
| C17orf113 | 248.483764 | 0.417727727 | 0.41090312 | 1.01660881 | 0.30933956 | 0.50650632 |
| DHX58 | 1131.45749 | 0.161115517 | 0.25041019 | 0.64340641 | 0.51996043 | 0.69660908 |
| KAT2A | 6000.00843 | 0.258569389 | 0.26383234 | 0.9800519 | 0.3270605 | 0.52489687 |
| AC099811.2 | 9414.85802 | 0.099355519 | 0.13974809 | 0.71096154 | 0.47710807 | 0.66217578 |
| HSPB9 | 24.6162789 | 0.351081182 | 0.52697495 | 0.66621987 | 0.50527058 | 0.68458023 |
| RAB5C | 12809.4428 | -0.065580958 | 0.1326577 | -0.4943622 | 0.62105038 | 0.77363364 |
| AC099811.3 | 35.7316442 | 0.62633054 | 0.38474363 | 1.62791658 | 0.10354257 | 0.24335274 |
| KCNH4 | 16.6003427 | -0.169600395 | 0.49310248 | -0.3439455 | 0.73088725 | 0.84796941 |
| HCRT | 1.29571631 | -1.686319758 | 1.66553069 | -1.0124819 | 0.31130767 | 0.50840301 |
| GHDC | 3998.26125 | 0.72606799 | 0.16625654 | 4.36715443 | 1.26E-05 | 0.00024647 |
| STAT5B | 4629.90039 | 0.051888292 | 0.14760058 | 0.35154531 | 0.72517928 | 0.84416976 |
| AC099811.4 | 0.94070409 | 2.769652433 | 1.69372199 | 1.63524619 | 0.10199745 | NA |
| STAT5A | 2402.49326 | -0.265239942 | 0.12153063 | -2.1824946 | 0.02907305 | 0.0987196 |
| STAT3 | 14649.1236 | -0.842453541 | 0.20983861 | -4.0147689 | 5.95E-05 | 0.00084589 |
| CAVIN1 | 6965.06533 | 0.206124687 | 0.38091057 | 0.5411367 | 0.58841336 | 0.74908617 |
| ATP6V0A1 | 8945.46498 | -0.131130874 | 0.26379702 | -0.4970901 | 0.61912555 | 0.77236923 |
| NAGLU | 5596.45637 | 0.384092 | 0.28277431 | 1.35829877 | 0.1743689 | 0.34829853 |
| HSD17B1 | 1127.5804 | -0.194551204 | 0.14898275 | -1.305864 | 0.19159882 | 0.37079485 |
| AC067852.2 | 1019.31286 | -0.267512261 | 0.14906813 | -1.7945638 | 0.07272323 | 0.18966586 |
| COASY | 7659.58051 | -0.288896493 | 0.16213359 | -1.7818423 | 0.07477495 | 0.19324739 |
| MLX | 3624.68255 | -0.143425756 | 0.17707889 | -0.809954 | 0.41796662 | 0.61076531 |
| PSMC3IP | 1042.87063 | -0.410489062 | 0.14247704 | -2.8810891 | 0.00396304 | 0.02261529 |
| RETREG3 | 3764.60905 | -0.319735042 | 0.10756935 | -2.9723618 | 0.00295518 | 0.01813126 |
| TUBG1 | 2365.85385 | -0.105943372 | 0.24865531 | -0.4260652 | 0.67006034 | 0.80733565 |
| TUBG2 | 2574.18622 | -0.050360305 | 0.24173323 | -0.2083301 | 0.83497124 | 0.91248204 |
| PLEKHH3 | 2071.04954 | -0.206373712 | 0.29136804 | -0.7082922 | 0.47876382 | 0.66367039 |
| CCR10 | 108.527161 | -0.026577165 | 0.24045195 | -0.11053 | 0.91198902 | 0.95679931 |
| AC100793.2 | 43.3084953 | -0.499947877 | 0.35195132 | -1.4205029 | 0.15546132 | 0.32175096 |
| CNTNAP1 | 877.486523 | -0.289740374 | 0.30477657 | -0.9506648 | 0.34177454 | 0.53967632 |
| AC100793.3 | 218.484848 | -0.80765066 | 0.38683987 | -2.0878165 | 0.03681439 | 0.11661207 |
| EZH1 | 3991.32793 | 0.038748003 | 0.12703889 | 0.30500897 | 0.76035932 | 0.86715659 |
| AC100793.4 | 10.4245014 | 0.475058609 | 0.46946249 | 1.01192026 | 0.31157618 | 0.50871332 |
| RAMP2-AS1 | 262.025308 | 0.633190305 | 0.32205513 | 1.96609291 | 0.04928788 | 0.14426256 |
| RAMP2 | 900.904188 | 0.439995097 | 0.38014824 | 1.15743032 | 0.24709659 | 0.43860643 |
| VPS25 | 3543.89365 | -0.62274787 | 0.13636098 | -4.5669066 | 4.95E-06 | 0.00011654 |
| WNK4 | 5610.9674 | 1.846343657 | 0.40435952 | 4.56609414 | 4.97E-06 | 0.00011667 |
| COA3 | 6736.35 | 0.578560522 | 0.20873084 | 2.7718018 | 0.0055747 | 0.02926729 |
| CNTD1 | 2207.46864 | -0.323761953 | 0.12191371 | -2.6556648 | 0.00791523 | 0.03787416 |
| BECN1 | 4534.37839 | -0.104439013 | 0.10827012 | -0.9646153 | 0.33473754 | 0.53285059 |
| PSME3 | 5130.71594 | -0.359858485 | 0.18007655 | -1.9983639 | 0.04567722 | 0.13638002 |
| AOC2 | 184.481542 | -1.281808159 | 0.45093082 | -2.8425827 | 0.00447496 | 0.0247805 |
| AOC3 | 2574.9321 | 0.877088349 | 0.37428937 | 2.3433429 | 0.01911181 | 0.07253575 |
| LINC00671 | 3259.50551 | 2.637457394 | 0.83887391 | 3.14404507 | 0.0016663 | 0.01171116 |
| G6PC | 4854.14422 | 3.020083311 | 1.25134658 | 2.41346671 | 0.01580157 | 0.06323986 |
| PTGES3L-AAI | 1433.53908 | 0.065165991 | 0.12714887 | 0.51251725 | 0.60828905 | 0.76445609 |
| AARSD1 | 1525.78973 | 0.067237397 | 0.12133433 | 0.5541498 | 0.57947633 | 0.74277814 |
| PTGES3L | 39.4510406 | 0.016024236 | 0.31669901 | 0.05059768 | 0.95964611 | 0.98050555 |
| RUNDC1 | 1370.3658 | -0.061821903 | 0.15202854 | -0.4066467 | 0.68426749 | 0.81757775 |
| RPL27 | 14135.9014 | -0.491807077 | 0.20161487 | -2.4393393 | 0.01471415 | 0.06001733 |
| IFI35 | 1605.16664 | -0.361981906 | 0.20012784 | -1.8087534 | 0.07048932 | 0.18515822 |
| VAT1 | 13178.2283 | -0.624786064 | 0.29221895 | -2.1380751 | 0.03251065 | 0.10677944 |
| RND2 | 130.484926 | 0.238828036 | 0.29426215 | 0.81161658 | 0.41701168 | 0.61000709 |
| BRCA1 | 156.874263 | 0.249892246 | 0.30611597 | 0.81633195 | 0.41431031 | 0.6079779 |
| AC060780.1 | 880.917401 | 0.2655839 | 0.26868845 | 0.98844556 | 0.32293449 | 0.52066432 |
| NBR1 | 7920.88028 | 0.60916356 | 0.20757003 | 2.9347376 | 0.0033383 | 0.019894 |
| TMEM106A | 1418.93674 | 2.096208942 | 0.4338339 | 4.8318238 | 1.35E-06 | 4.15E-05 |
| LINC00854 | 599.922364 | 2.336809255 | 0.39284807 | 5.94837908 | 2.71E-09 | 2.31E-07 |
| LINC00910 | 136.383639 | 0.602396698 | 0.34853645 | 1.72836069 | 0.08392358 | 0.20953107 |
| ARL4D | 2109.50476 | -0.85005584 | 0.19739765 | -4.3063118 | 1.66E-05 | 0.00030587 |
| MIR2117HG | 2.963756 | -0.50664703 | 1.34190536 | -0.3775579 | 0.70575901 | 0.83232849 |
| DHX8 | 1715.3186 | -0.02311099 | 0.1248532 | -0.1851053 | 0.85314643 | 0.92319951 |
| ETV4 | 436.515467 | -1.38869264 | 0.40798653 | -3.4037708 | 0.00066462 | 0.00564996 |
| AC068675.1 | 5.42345679 | 0.021233598 | 0.70219044 | 0.03023909 | 0.97587638 | 0.98881307 |
| MEOX1 | 16.2880086 | 0.602475682 | 0.84135861 | 0.71607478 | 0.47394517 | 0.65980014 |
| AC004149.1 | 2.33832258 | 0.63730504 | 0.96913941 | 0.65759893 | 0.51079588 | 0.68919809 |
| SOST | 345.447518 | 2.721158889 | 1.27160438 | 2.13994143 | 0.0323595 | 0.10636156 |
| DUSP3 | 8658.42817 | -0.052684911 | 0.23887083 | -0.2205582 | 0.82543648 | 0.90675212 |
| CFAP97D1 | 2.5336341 | 0.170389467 | 0.80822162 | 0.21082023 | 0.83302755 | 0.91136194 |
| AC003098.1 | 0.19624763 | 0.18657164 | 3.29377195 | 0.05664376 | 0.95482897 | NA |
| MPP3 | 442.133817 | -0.542202585 | 0.22872856 | -2.3705067 | 0.01776372 | 0.068705 |
| CD300LG | 118.065341 | 1.29990918 | 0.59865291 | 2.1713904 | 0.02990167 | 0.10065192 |
| MPP2 | 227.631573 | 0.543319012 | 0.38208371 | 1.42198947 | 0.15502931 | 0.32116543 |
| AC007993.3 | 13.5727956 | 2.137456013 | 0.71690678 | 2.98149784 | 0.00286842 | 0.01773096 |
| FAM215A | 9.79784168 | 2.217409181 | 0.59532674 | 3.72469271 | 0.00019555 | 0.00214556 |
| AC007993.2 | 38.2795604 | 3.321740019 | 0.73313827 | 4.53085064 | 5.87E-06 | 0.0001339 |
| LINC01976 | 29.5656499 | 2.211232521 | 0.74664139 | 2.96157236 | 0.00306073 | 0.01864864 |
| PPY | 0.40866785 | 0.309801681 | 2.21387465 | 0.13993641 | 0.88871023 | NA |
| PYY | 73.9527129 | -1.420845916 | 0.4817422 | -2.9493906 | 0.00318401 | 0.01922657 |
| NAGS | 1035.45953 | 0.326355593 | 0.25072683 | 1.3016381 | 0.19304013 | 0.37251871 |
| TMEM101 | 4172.94572 | -0.887198991 | 0.29574338 | -2.9998947 | 0.00270073 | 0.01695673 |
| LSM12 | 675.535299 | -0.292607045 | 0.14934143 | -1.959316 | 0.0500758 | 0.14578259 |
| G6PC3 | 3774.41168 | -0.181678576 | 0.11636828 | -1.5612379 | 0.11846763 | 0.26753223 |
| HDAC5 | 4258.37011 | 0.039681406 | 0.14441662 | 0.27477035 | 0.78349268 | 0.8815797 |
| AC023855.1 | 1.14810213 | -0.443093318 | 1.2079407 | -0.3668171 | 0.71375544 | 0.83745188 |
| C17orf53 | 63.6402406 | -0.317594231 | 0.36046978 | -0.8810565 | 0.37828726 | 0.57427972 |
| ASB16 | 454.194728 | 0.248931396 | 0.26802885 | 0.9287485 | 0.35301944 | 0.55058469 |
| ASB16-AS1 | 686.456487 | 0.018641867 | 0.24819219 | 0.07511061 | 0.9401267 | 0.97140164 |
| TMUB2 | 3234.66644 | -0.216424812 | 0.115516 | -1.8735483 | 0.06099271 | 0.16736033 |
| ATXN7L3 | 4033.40351 | -0.485224371 | 0.21607234 | -2.2456571 | 0.02472598 | 0.08756703 |
| AC004596.1 | 195.259823 | 0.854301929 | 0.15549886 | 5.49394331 | 3.93E-08 | 2.18E-06 |
| UBTF | 4846.14474 | 0.399911644 | 0.15920325 | 2.51195661 | 0.01200639 | 0.05154809 |
| AC003102.1 | 414.048849 | -0.435015299 | 0.23806169 | -1.8273217 | 0.06765142 | 0.17972676 |
| SLC4A1 | 2414.36412 | 2.928686086 | 0.59947064 | 4.88545372 | 1.03E-06 | 3.34E-05 |
| AC003043.2 | 2.93282999 | 0.158584086 | 1.03050622 | 0.1538895 | 0.87769686 | 0.93635676 |
| RUNDC3A-AS | 72.3678489 | -0.007750256 | 0.39658823 | -0.0195423 | 0.98440847 | 0.99326005 |

| | | | | | | |
|---|---|---|---|---|---|---|
| RUNDC3A | 149.965273 | -0.695819987 | 0.35865081 | -1.9401043 | 0.05236701 | 0.15041042 |
| SLC25A39 | 12458.0336 | 0.031694441 | 0.19390085 | 0.16345694 | 0.87015867 | 0.9323196 |
| AC003043.1 | 0.57492173 | 1.169201638 | 1.84494807 | 0.63373146 | 0.52625609 | NA |
| GRN | 21815.9131 | -0.389950154 | 0.21664353 | -1.7999621 | 0.07186662 | 0.18791499 |
| FAM171A2 | 103.130909 | 0.207092338 | 0.41297268 | 0.50146741 | 0.61604221 | 0.77034489 |
| ITGA2B | 132.658649 | -0.127781595 | 0.42041806 | -0.3039394 | 0.76117409 | 0.86772267 |
| GPATCH8 | 2151.14236 | 0.024725136 | 0.11186155 | 0.22103338 | 0.82506644 | 0.90654176 |
| FZD2 | 164.277097 | 0.075288971 | 0.20600888 | 0.36546468 | 0.71476457 | 0.83791991 |
| LINC01180 | 0.12059279 | 0.02007176 | 3.29738704 | 0.00608717 | 0.99514317 | NA |
| MEIOC | 447.705435 | 0.163598138 | 0.23393285 | 0.69933801 | 0.48434082 | 0.668712 |
| CCDC43 | 924.732836 | -0.012740191 | 0.17623218 | -0.0722921 | 0.94236947 | 0.97270099 |
| AC091152.2 | 37.120607 | 0.121365282 | 0.33777719 | 0.35930573 | 0.71936639 | 0.84022198 |
| DBF4B | 517.478856 | 0.212851786 | 0.3506983 | 0.60693704 | 0.54389269 | 0.71604845 |
| ADAM11 | 65.6810657 | 0.30658522 | 0.27501777 | 1.11478331 | 0.26494329 | 0.45897167 |
| AC005180.2 | 6.91393116 | 1.413830042 | 0.73576319 | 1.92158301 | 0.05465825 | 0.15504927 |
| AC005180.1 | 3.82075605 | 1.420640193 | 0.9083852 | 1.56391826 | 0.11783677 | 0.26654276 |
| GJC1 | 257.23127 | -0.306186595 | 0.55276731 | -0.5539159 | 0.57963641 | 0.74281468 |
| HIGD1B | 84.5843282 | -0.036605711 | 0.48801033 | -0.0750101 | 0.94020666 | 0.97144619 |
| EFTUD2 | 3828.17166 | -0.203393168 | 0.14086292 | -1.4439085 | 0.14876472 | 0.31285794 |
| CCDC103 | 379.786732 | 0.829295579 | 0.2275903 | 3.64380902 | 0.00026863 | 0.00278016 |
| FAM187A | 312.157005 | 0.888746185 | 0.28913177 | 3.07384474 | 0.00211319 | 0.01403297 |
| GFAP | 223.245054 | 0.640448765 | 0.28083549 | 2.28051224 | 0.02257733 | 0.08199478 |
| KIF18B | 43.7915734 | -2.016517058 | 0.67722559 | -2.977615 | 0.00290501 | 0.01789001 |
| AC015936.1 | 0 | NA | NA | NA | NA | NA |
| C1QL1 | 205.894505 | -0.967793401 | 0.53938817 | -1.7942429 | 0.0727744 | 0.18969885 |
| AC015936.2 | 0.16325399 | -1.003574173 | 3.30558948 | -0.3035992 | 0.7614333 | NA |
| DCAKD | 1719.62108 | 0.042112142 | 0.19424381 | 0.21680043 | 0.82836388 | 0.908726 |
| NMT1 | 5870.16466 | -0.043024566 | 0.11838181 | -0.363439 | 0.71627696 | 0.83854488 |
| PLCD3 | 1585.17158 | -0.751029989 | 0.24243682 | -3.097838 | 0.00194938 | 0.0132317 |
| ACBD4 | 2135.61807 | 0.977496193 | 0.34630118 | 2.8226765 | 0.00476246 | 0.02593146 |
| AC142472.1 | 124.210145 | 0.241037365 | 0.2416284 | 0.99755396 | 0.31849569 | 0.51576587 |
| HEXIM1 | 4311.54221 | 0.167911262 | 0.15838756 | 1.06012911 | 0.28908587 | 0.4852447 |
| AC138150.1 | 630.956902 | 0.243014749 | 0.16243718 | 1.49605376 | 0.13463965 | 0.29204964 |
| HEXIM2 | 670.802954 | 1.154569297 | 0.25290687 | 4.56519547 | 4.99E-06 | 0.00011687 |
| AC138150.2 | 519.44351 | 1.293269407 | 0.23596851 | 5.48068631 | 4.24E-08 | 2.31E-06 |
| AC008105.3 | 188.033929 | 0.00185482 | 0.62090605 | 0.00298728 | 0.9976165 | 0.99867691 |
| FMNL1 | 1273.46461 | -0.295380916 | 0.45455625 | -0.6498226 | 0.51580683 | 0.69342133 |
| AC008105.1 | 225.636355 | 0.203983575 | 0.40702781 | 0.5011539 | 0.61626282 | 0.77049015 |
| MAP3K14-AS | 614.581806 | -0.360886964 | 0.25837274 | -1.3967687 | 0.16248314 | 0.33138509 |
| SPATA32 | 14.1080617 | 0.39516697 | 0.33429431 | 1.18209303 | 0.23716879 | 0.42639107 |
| MAP3K14 | 2184.41437 | -0.407187074 | 0.27120621 | -1.5013929 | 0.13325398 | 0.28987852 |
| AC003070.2 | 2.27161344 | -0.784742588 | 1.00230796 | -0.7829356 | 0.43366493 | 0.62562198 |
| ARHGAP27 | 3927.56676 | -0.333861449 | 0.19740406 | -1.6912593 | 0.0907873 | 0.22204556 |
| AC003070.1 | 357.291282 | 0.076269441 | 0.20957915 | 0.36391712 | 0.7159199 | 0.83832645 |
| PLEKHM1 | 3188.04759 | -1.195284249 | 0.47382902 | -2.5226067 | 0.01164886 | 0.05048159 |
| AC091132.2 | 22.0769054 | -1.324444544 | 0.42159548 | -3.1415056 | 0.00168082 | 0.01179748 |
| AC091132.1 | 3.37492334 | -0.522306748 | 0.82784197 | -0.6309257 | 0.52808913 | 0.7026799 |
| AC091132.5 | 19.0173775 | 0.128991769 | 0.49731429 | 0.25937676 | 0.79534456 | 0.8889978 |
| LINC02210-C | 105.315579 | 0.533579598 | 0.35703931 | 1.49445616 | 0.13505643 | 0.29242691 |
| AC217774.1 | 0 | NA | NA | NA | NA | NA |
| AC217774.2 | 0 | NA | NA | NA | NA | NA |
| CRHR1 | 11.9168346 | 3.040292862 | 0.82865109 | 3.66896621 | 0.00024353 | 0.00256674 |
| MAPT-AS1 | 114.015352 | 2.468335663 | 0.53335096 | 4.62797644 | 3.69E-06 | 9.24E-05 |
| SPPL2C | 0.27059475 | 1.100737003 | 2.99309129 | 0.36775925 | 0.71305276 | NA |
| MAPT | 2619.22723 | 1.939562165 | 0.44542724 | 4.3543861 | 1.33E-05 | 0.00025798 |
| CR936218.2 | 9.0693921 | 3.492549957 | 1.51833376 | 2.30025179 | 0.02143396 | 0.07890737 |
| STH | 0.20710523 | -1.003574173 | 3.30558948 | -0.3035992 | 0.7614333 | NA |
| KANSL1 | 2734.6267 | -0.006814296 | 0.14252141 | -0.0478124 | 0.96186572 | 0.98182237 |
| CR936218.1 | 5.13246048 | 1.440623055 | 0.58407982 | 2.46648318 | 0.01364471 | 0.05671676 |
| KANSL1-AS1 | 181.979173 | 1.248482857 | 0.52496682 | 2.37821288 | 0.01739678 | 0.06772468 |
| ARL17B | 69.5697235 | -0.145527927 | 0.44446726 | -0.327421 | 0.74334946 | 0.85602539 |
| LRRC37A | 99.3426043 | 0.063045268 | 0.34557869 | 0.1824339 | 0.85524223 | 0.92410315 |
| LRRC37A2 | 444.346221 | 0.2400928 | 0.32171109 | 0.74629942 | 0.45548656 | 0.64489561 |
| ARL17A | 165.280724 | 0.176584338 | 0.29735258 | 0.59385507 | 0.55260905 | 0.72258908 |
| FAM215B | 4.82125376 | 1.798929935 | 0.61078969 | 2.94525262 | 0.00322691 | 0.01942777 |
| NSF | 1808.05421 | -0.048768287 | 0.11080316 | -0.4401345 | 0.65983973 | 0.79999826 |
| WNT3 | 186.612963 | 0.094017912 | 0.37035025 | 0.25386215 | 0.79960206 | 0.89126326 |
| WNT9B | 143.705663 | 2.500365175 | 0.79993011 | 3.12572953 | 0.00177365 | 0.0123017 |
| LINC01974 | 0.56764698 | 1.708771729 | 1.69205599 | 1.00987895 | 0.31255329 | NA |
| AC005670.1 | 8.4949869 | 1.132522605 | 0.55804423 | 2.02944955 | 0.04241252 | 0.12937046 |
| GOSR2 | 3509.34787 | 0.073573734 | 0.19038005 | 0.38645717 | 0.69915812 | 0.8273188 |
| AC005670.2 | 1144.33823 | 0.319255723 | 0.24624499 | 1.29649632 | 0.19480455 | 0.37480173 |
| RPRML | 3.08439608 | -1.44416194 | 0.92251128 | -1.5654681 | 0.1174732 | 0.26601409 |
| CDC27 | 2269.70208 | -0.528490207 | 0.18580342 | -2.8443513 | 0.00445019 | 0.02469002 |
| AC002558.2 | 49.6715158 | 0.350269154 | 0.36579039 | 0.95756795 | 0.33828067 | 0.53628886 |
| MYL4 | 24.9928401 | -1.104982006 | 0.6104602 | -1.8100803 | 0.07028333 | 0.1847828 |
| ITGB3 | 1613.51052 | -1.247144 | 0.28611904 | -4.3588291 | 1.31E-05 | 0.00025317 |
| AC068234.1 | 870.99104 | -0.935525788 | 0.21815564 | -4.288341 | 1.80E-05 | 0.0003272 |
| AC068234.3 | 8.35891155 | -1.161905491 | 0.71336491 | -1.6287674 | 0.10336227 | 0.24301562 |
| AC068234.2 | 190.303724 | -1.877448438 | 0.32861821 | -5.7131601 | 1.11E-08 | 7.58E-07 |
| EFCAB13 | 492.178369 | -0.883430936 | 0.28778125 | -3.0698002 | 0.00214202 | 0.01420293 |
| AC040934.1 | 37.8629374 | -1.263359276 | 0.30589518 | -4.1300399 | 3.63E-05 | 0.00057341 |
| NPEPPS | 6228.54361 | -0.157798843 | 0.12450022 | -1.2674584 | 0.20499144 | 0.38741664 |
| AC025682.1 | 1417.29337 | -0.220984507 | 0.1436776 | -1.5380581 | 0.1240344 | 0.27575981 |
| KPNB1 | 7786.121 | -0.302770145 | 0.16375293 | -1.8489449 | 0.06446576 | 0.17378472 |
| AC015674.1 | 1301.91921 | -0.488758085 | 0.17321777 | -2.8216393 | 0.00477789 | 0.02597256 |
| TBKBP1 | 2144.33697 | -0.389839128 | 0.25752758 | -1.5137762 | 0.1300826 | 0.28521479 |
| TBX21 | 85.3872392 | -0.806436876 | 0.5686055 | -1.4182713 | 0.15611156 | 0.32262241 |
| OSBPL7 | 1159.33631 | 0.05823771 | 0.22498884 | 0.25884711 | 0.79575321 | 0.88920418 |
| MRPL10 | 2420.27708 | -0.151288773 | 0.1160193 | -1.3039966 | 0.19223475 | 0.37150497 |
| LRRC46 | 234.606999 | 0.324068869 | 0.35750497 | 0.90647374 | 0.36468516 | 0.56182657 |
| SCRN2 | 6498.82546 | 1.102672753 | 0.42687229 | 2.58314438 | 0.00979043 | 0.04436737 |
| SP6 | 136.046822 | -1.014222377 | 0.26865107 | -3.7752404 | 0.00015985 | 0.00184156 |
| AC018521.3 | 0.17656334 | 0.18657164 | 3.29377195 | 0.05664376 | 0.95482897 | NA |
| AC018521.7 | 7.29898896 | 0.857263101 | 0.69608168 | 1.23155532 | 0.21811524 | 0.40325414 |
| SP2 | 1392.19976 | 0.299816333 | 0.18228198 | 1.6447941 | 0.10001228 | 0.23748976 |
| SP2-AS1 | 59.1474996 | -0.030395443 | 0.25521471 | -0.1190975 | 0.90519809 | 0.95244089 |
| AC018521.6 | 24.1947179 | 0.434365092 | 0.48412341 | 0.89721976 | 0.36960166 | 0.56638748 |
| PNPO | 4554.81944 | 0.395434397 | 0.40349956 | 0.98001196 | 0.32708022 | 0.52489687 |
| AC018521.2 | 1494.07775 | 0.458411276 | 0.40471163 | 1.13268621 | 0.25734605 | 0.4499697 |
| PRR15L | 1834.59303 | -0.460750682 | 0.51001071 | -0.9034137 | 0.36630635 | 0.56331382 |
| CDK5RAP3 | 10582.5547 | -0.179924069 | 0.15071289 | -1.1938201 | 0.23254836 | 0.42059697 |
| AC018521.4 | 1947.03523 | -0.250969901 | 0.16461173 | -1.5246174 | 0.12735456 | 0.28093858 |
| COPZ2 | 518.718447 | 0.114962467 | 0.23415489 | 0.49096761 | 0.62344935 | 0.77548865 |
| NFE2L1 | 24145.9703 | 0.639117872 | 0.20066453 | 3.18500671 | 0.00144751 | 0.01051825 |
| AC004477.2 | 5749.64611 | 0.153778668 | 0.15625709 | 0.98413879 | 0.32504728 | 0.52264317 |
| AC004477.3 | 22.8756769 | 0.479902256 | 0.33601032 | 1.42823666 | 0.15322375 | 0.31877794 |
| CBX1 | 2813.8518 | -0.070218853 | 0.14334325 | -0.4898651 | 0.62422938 | 0.77611992 |
| SNX11 | 1523.78142 | -0.315816431 | 0.16353063 | -1.9312372 | 0.05345372 | 0.1527332 |
| SKAP1 | 790.178262 | -0.233940372 | 0.25393568 | -0.9212584 | 0.35691554 | 0.5544217 |
| SKAP1-AS1 | 15.9491056 | -0.291944439 | 0.57265137 | -0.5098118 | 0.6101833 | 0.76610644 |
| AC090627.1 | 1.27582072 | -1.622801406 | 1.22063524 | -1.3294728 | 0.18369202 | 0.36076253 |
| AC036222.1 | 0.20452486 | -0.339987873 | 3.13600186 | -0.1084144 | 0.91366695 | NA |

| | | | | | | |
|---|---|---|---|---|---|---|
| HOXB1 | 12.9296047 | 1.206813656 | 0.48443929 | 2.4911556 | 0.01273283 | 0.05391215 |
| HOXB2 | 2294.93267 | -0.630687989 | 0.28937813 | -2.1794597 | 0.02929753 | 0.09921355 |
| HOXB-AS1 | 1739.04714 | -0.262924139 | 0.25768958 | -1.0203134 | 0.30757983 | 0.5044676 |
| HOXB3 | 4348.46461 | -0.164863538 | 0.2467647 | -0.6681002 | 0.50406965 | 0.68375733 |
| HOXB-AS3 | 3211.80621 | -0.457056365 | 0.21753133 | -2.1011059 | 0.03563167 | 0.11405767 |
| HOXB-AS2 | 15.2463653 | 1.492567302 | 0.55358182 | 2.69620001 | 0.00701355 | 0.03467981 |
| HOXB4 | 1063.31516 | -0.397237406 | 0.18281936 | -2.1728411 | 0.02979228 | 0.10042471 |
| HOXB5 | 1269.17688 | -0.38272487 | 0.30224574 | -1.2662705 | 0.20541625 | 0.38799703 |
| HOXB6 | 3313.0547 | -0.520911831 | 0.23910937 | -2.1785504 | 0.02936508 | 0.09935299 |
| HOXB7 | 1336.24831 | -0.704639691 | 0.17660678 | -3.9898791 | 6.61E-05 | 0.00091895 |
| HOXB8 | 1531.8355 | -0.591151659 | 0.26371597 | -2.2416225 | 0.02498578 | 0.08829737 |
| HOXB9 | 1749.34652 | -0.628150099 | 0.2832971 | -2.2172839 | 0.0266037 | 0.09247818 |
| HOXB-AS4 | 10.6885228 | -0.593113167 | 0.46909436 | -1.2643792 | 0.20609394 | 0.38874802 |
| AC103702.2 | 100.83507 | -1.339100185 | 0.454711 | -2.9449478 | 0.00323009 | 0.01943804 |
| LINC02086 | 231.477014 | -0.935936966 | 0.56188015 | -1.6657235 | 0.0957685 | 0.2306526 |
| PRAC1 | 0.07830766 | -0.642905668 | 3.30558948 | -0.1944905 | 0.84579185 | NA |
| PRAC2 | 0 | NA | NA | NA | NA | NA |
| HOXB13 | 2.48422778 | 0.384766033 | 1.21748121 | 0.31603447 | 0.75197634 | 0.86163275 |
| AC091179.1 | 0.07830766 | -0.642905668 | 3.30558948 | -0.1944905 | 0.84579185 | NA |
| TTLL6 | 196.91121 | 1.337443903 | 0.36192225 | 3.69539016 | 0.00021955 | 0.00235346 |
| CALCOCO2 | 5658.85161 | -0.224789332 | 0.15574946 | -1.4432752 | 0.14894296 | 0.31303205 |
| ATP5MC1 | 4149.0371 | 0.258406142 | 0.23473952 | 1.10082078 | 0.27097466 | 0.46554185 |
| UBE2Z | 6042.27166 | -0.733639369 | 0.18305904 | -4.0076653 | 6.13E-05 | 0.0008652 |
| SNF8 | 3420.5803 | -0.188634821 | 0.13147063 | -1.4348058 | 0.15134245 | 0.31631136 |
| AC091133.1 | 1836.4372 | -0.145723398 | 0.14279063 | -1.020539 | 0.30747293 | 0.50442087 |
| GIP | 0.35900909 | -1.731401174 | 3.30177887 | -0.5243844 | 0.60001124 | NA |
| AC091133.2 | 0 | NA | NA | NA | NA | NA |
| IGF2BP1 | 199.103254 | 1.911252573 | 0.56852502 | 3.36177387 | 0.00077444 | 0.00640384 |
| AC091133.3 | 3.16797765 | 2.174039172 | 1.00620036 | 2.1606424 | 0.03072297 | 0.10263045 |
| B4GALNT2 | 488.692759 | 2.148579917 | 0.54289581 | 3.95762848 | 7.57E-05 | 0.00101688 |
| GNGT2 | 93.174663 | 0.113630418 | 0.32025647 | 0.35481069 | 0.72273142 | 0.8426594 |
| ABI3 | 607.854693 | 0.296306964 | 0.29711749 | 0.99727203 | 0.31863249 | 0.51592402 |
| PHOSPHO1 | 48.0293124 | 0.659512133 | 0.52580087 | 1.2543002 | 0.20973291 | 0.39291396 |
| ZNF652 | 1943.95851 | -0.210915232 | 0.12060099 | -1.7488682 | 0.08031381 | 0.20297913 |
| AC091180.2 | 3.17586016 | 0.359539294 | 0.65114231 | 0.55216699 | 0.58083395 | 0.74363214 |
| AC091180.5 | 2.63117086 | -0.336303768 | 0.97213489 | -0.3459435 | 0.72938516 | 0.84712119 |
| AC091180.4 | 3.26942594 | -0.141488438 | 0.97597316 | -0.1449716 | 0.88473326 | 0.9403634 |
| PHB | 11264.3691 | 0.446049368 | 0.15245814 | 2.92571701 | 0.00343663 | 0.02031929 |
| AC091180.3 | 6658.39493 | 0.413178821 | 0.15663036 | 2.63792293 | 0.00834155 | 0.03927978 |
| LINC02075 | 0 | NA | NA | NA | NA | NA |
| NGFR | 776.811959 | -0.305496834 | 0.49690679 | -0.6147971 | 0.53868875 | 0.71200553 |
| AC006487.1 | 44.2186867 | 0.043901092 | 0.54802566 | 0.08010773 | 0.93615157 | 0.96942144 |
| AC006487.2 | 0.58654082 | -1.405697279 | 2.25360301 | -0.6237555 | 0.53278817 | NA |
| NXPH3 | 108.271886 | -0.595907684 | 0.38987258 | -1.5284678 | 0.12639643 | 0.2796203 |
| SPOP | 2414.46406 | -0.133496541 | 0.11530364 | -1.1577826 | 0.24695278 | 0.43841519 |
| SLC35B1 | 2582.60944 | -0.094623616 | 0.16550833 | -0.5717151 | 0.56751498 | 0.73345276 |
| AC015795.1 | 144.559021 | 0.816297019 | 0.19590976 | 4.16669907 | 3.09E-05 | 0.0005012 |
| FAM117A | 1080.19012 | 0.574380232 | 0.17579505 | 3.26732875 | 0.00108568 | 0.00835561 |
| KAT7 | 2310.91253 | 0.1422247 | 0.09147711 | 1.55475729 | 0.12000389 | 0.26991662 |
| AC027801.4 | 20.3150609 | 0.064959436 | 0.35109957 | 0.18501713 | 0.8532156 | 0.92319951 |
| TAC4 | 29.3644276 | 0.00893151 | 0.31176299 | 0.0286484 | 0.97714501 | 0.98949019 |
| FLJ45513 | 140.389334 | -0.655221063 | 0.32094787 | -2.0415187 | 0.0411993 | 0.12663778 |
| AC027801.1 | 0.83234466 | -1.387325378 | 1.36737774 | -1.0145882 | 0.31030215 | NA |
| AC027801.3 | 1.55641816 | -0.336272677 | 1.12807627 | -0.2980939 | 0.76563148 | 0.87069614 |
| DLX4 | 15.6359011 | -1.5893769 | 0.45587794 | -3.4864089 | 0.00048955 | 0.00445539 |
| DLX3 | 25.4780469 | -0.088197623 | 0.55115477 | -0.1600233 | 0.87286269 | 0.93369257 |
| AC009720.2 | 0 | NA | NA | NA | NA | NA |
| AC009720.1 | 162.131188 | -1.3417141 | 0.38156641 | -3.5163318 | 0.00043755 | 0.00409159 |
| ITGA3 | 22871.8646 | -1.600336437 | 0.28721909 | -5.5718317 | 2.52E-08 | 1.53E-06 |
| AC002401.4 | 59.8539113 | -3.350835677 | 0.69177069 | -4.8438532 | 1.27E-06 | 3.96E-05 |
| PDK2 | 6328.7399 | 1.333491939 | 0.34965932 | 3.81368909 | 0.00013691 | 0.00162259 |
| AC002401.3 | 17.2278341 | 1.732156353 | 0.38449879 | 4.50497216 | 6.64E-06 | 0.00014797 |
| AC002401.2 | 11.4395955 | 2.016384909 | 0.45791419 | 4.40341214 | 1.07E-05 | 0.000215 |
| SAMD14 | 3704.25496 | 1.017768573 | 0.29751186 | 3.42093447 | 0.00062406 | 0.00537796 |
| PPP1R9B | 4856.78065 | -0.463328361 | 0.20522696 | -2.2576388 | 0.02396819 | 0.08554775 |
| AC002401.1 | 1033.61331 | 0.535348388 | 0.23953847 | 2.23491615 | 0.02542285 | 0.08943445 |
| AC015909.2 | 36.1657074 | 4.491078727 | 0.5752004 | 7.80785043 | 5.82E-15 | 3.20E-12 |
| AC015909.3 | 7.64466077 | 3.763295424 | 0.74609891 | 5.04396317 | 4.56E-07 | 1.68E-05 |
| SGCA | 150.428734 | 1.400423669 | 0.39155129 | 3.57660336 | 0.00034809 | 0.00341356 |
| COL1A1 | 9768.85631 | -0.606299313 | 0.47888552 | -1.2660632 | 0.20549047 | 0.38808163 |
| AC015909.1 | 1.91538061 | -1.141434826 | 1.04633478 | -1.0908887 | 0.27532184 | 0.47058835 |
| AC015909.4 | 1.35735859 | -0.155216339 | 1.20764144 | -0.1285285 | 0.89773075 | 0.94798814 |
| LINC01969 | 1.48145084 | -1.207499516 | 1.2802465 | -0.9431774 | 0.34559019 | 0.54326859 |
| TMEM92 | 41.746819 | -2.937139395 | 0.76140025 | -3.8575498 | 0.00011453 | 0.00140865 |
| TMEM92-AS | 7.15972564 | -1.469723136 | 0.6900902 | -2.1297551 | 0.03319184 | 0.1083932 |
| XYLT2 | 2143.01907 | -0.37938512 | 0.23666246 | -1.6030642 | 0.10892048 | 0.25230439 |
| MRPL27 | 2377.57068 | -0.094512067 | 0.14765754 | -0.6400761 | 0.52212309 | 0.69805242 |
| EME1 | 666.684402 | -1.226392137 | 0.26890973 | -4.5606091 | 5.10E-06 | 0.00011861 |
| LRRC59 | 4907.75772 | -0.838075483 | 0.28631278 | -2.9271326 | 0.00342103 | 0.02025882 |
| AC004707.1 | 428.676812 | -0.57712133 | 0.35857651 | -1.6094789 | 0.10751166 | 0.24980901 |
| ACSF2 | 11941.803 | 1.769349391 | 0.47215293 | 3.74740744 | 0.00017867 | 0.00200246 |
| CHAD | 371.367286 | 2.521965179 | 0.39741659 | 6.34589804 | 2.21E-10 | 2.84E-08 |
| AC021491.4 | 139.10528 | 4.168346185 | 0.54837952 | 7.60120694 | 2.93E-14 | 1.27E-11 |
| RSAD1 | 4178.06014 | 0.826896789 | 0.17193054 | 4.80948159 | 1.51E-06 | 4.57E-05 |
| AC021491.1 | 7.96758807 | 1.313830313 | 0.7667174 | 1.71357833 | 0.0866062 | 0.21440156 |
| MYCBPAP | 72.6250616 | 0.698319726 | 0.39473909 | 1.76906656 | 0.07688277 | 0.19694954 |
| EPN3 | 844.174454 | 0.619499721 | 0.47385289 | 1.3073672 | 0.19108803 | 0.37016024 |
| SPATA20 | 6454.92997 | 0.135308939 | 0.25949739 | 0.52142698 | 0.60206936 | 0.75935379 |
| CACNA1G-AS | 4.39132556 | 0.022025265 | 0.70665044 | 0.03116854 | 0.97513513 | 0.9885938 |
| CACNA1G | 7.23948788 | -0.575029141 | 0.53869835 | -1.0674418 | 0.28577238 | 0.48176817 |
| AC021491.3 | 0 | NA | NA | NA | NA | NA |
| AC004590.1 | 0.49285683 | 1.480049846 | 1.78522166 | 0.82905663 | 0.40707237 | NA |
| ABCC3 | 7520.01071 | -1.378949564 | 0.40198749 | -3.4303295 | 0.00060285 | 0.00523621 |
| ANKRD40 | 2595.21989 | -0.120857377 | 0.20870488 | -0.5790826 | 0.56253341 | 0.72998405 |
| AC005921.2 | 962.690354 | -0.541940371 | 0.2212445 | -2.4495089 | 0.01430512 | 0.05882143 |
| LUC7L3 | 8492.0941 | -0.039945804 | 0.14103992 | -0.2832234 | 0.7770056 | 0.8778389 |
| LINC00483 | 0.86742083 | 2.176849779 | 1.55828107 | 1.3969558 | 0.16242686 | NA |
| WFIKKN2 | 13.7307377 | 0.495575467 | 0.45186263 | 1.09673923 | 0.2727554 | 0.46762862 |
| AC091062.1 | 4.32773948 | 0.498650864 | 0.83260067 | 0.59890759 | 0.54923451 | 0.7197393 |
| TOB1 | 4295.28798 | 0.025090862 | 0.25704701 | 0.09761196 | 0.92224043 | 0.96203347 |
| AC005920.4 | 89.468133 | 0.634743341 | 0.61703813 | 1.02869387 | 0.30362355 | 0.5005347 |
| AC005920.1 | 9.52570487 | 0.698192367 | 0.48759792 | 1.43190185 | 0.15217191 | 0.31725782 |
| AC005920.3 | 10.0115739 | 1.012333648 | 0.66066507 | 1.53229479 | 0.12544971 | 0.27808352 |
| SPAG9 | 6680.29718 | -0.330566501 | 0.34721745 | -0.9520446 | 0.34107436 | 0.53911952 |
| AC005920.2 | 0.93136976 | -0.719252726 | 1.60387246 | -0.4484476 | 0.65383021 | NA |
| NME1 | 2064.88651 | -0.87519722 | 0.25022355 | -3.4976613 | 0.00046936 | 0.00432156 |
| NME1-NME2 | 13288.1243 | -0.744792361 | 0.23088758 | -3.2257792 | 0.0012563 | 0.00939022 |
| NME2 | 11699.085 | -0.736602643 | 0.23265046 | -3.1661345 | 0.00154479 | 0.0110546 |
| MBTD1 | 1132.86496 | 0.01789068 | 0.15374663 | 0.1163647 | 0.90736351 | 0.95380594 |
| UTP18 | 1667.66246 | -0.429833066 | 0.22205054 | -1.9357443 | 0.05289903 | 0.15155882 |
| LINC02071 | 0.35900206 | 1.594990198 | 2.13672577 | 0.74646462 | 0.45538679 | NA |
| AC005823.2 | 0.13028434 | 1.049816488 | 3.28477129 | 0.31960109 | 0.74927075 | NA |
| AC005823.1 | 0 | NA | NA | NA | NA | NA |

| | | | | | | |
|---|---|---|---|---|---|---|
| LINC02072 | 0.32445269 | 0.328540111 | 2.26427136 | 0.1450975 | 0.88463389 | NA |
| LINC02073 | 1.29979441 | -1.031430437 | 1.41422336 | -0.7293264 | 0.46580203 | 0.65297961 |
| CA10 | 443.160969 | 0.772976564 | 0.33241831 | 2.32531282 | 0.02005524 | 0.07506837 |
| LINC01982 | 0.51005714 | 0.983574107 | 2.78338876 | 0.35337288 | 0.72380891 | NA |
| LINC02089 | 0.16325399 | -1.003574173 | 3.30558948 | -0.3035992 | 0.7614333 | NA |
| C17orf112 | 0 | NA | NA | NA | NA | NA |
| KIF2B | 0.19289513 | -1.003574173 | 3.30558948 | -0.3035992 | 0.7614333 | NA |
| TOM1L1 | 4059.12801 | 0.387587604 | 0.14176681 | 2.73397996 | 0.00625739 | 0.03180556 |
| COX11 | 4610.64463 | 0.291913577 | 0.1086983 | 2.68553938 | 0.00724128 | 0.03552654 |
| AC007485.1 | 18.9622175 | 1.048987135 | 0.3442816 | 3.04688703 | 0.00231225 | 0.01510751 |
| STXBP4 | 1025.86457 | 0.156929804 | 0.25601038 | 0.61298219 | 0.53988812 | 0.71287615 |
| HLF | 1486.57178 | -0.016240607 | 0.33209654 | -0.0489033 | 0.96099638 | 0.98131485 |
| AC007638.2 | 39.4028565 | -0.999764743 | 0.74451449 | -1.3428412 | 0.17932339 | 0.35493191 |
| AC007638.1 | 0.08695548 | -0.642905668 | 3.30558948 | -0.1944905 | 0.84579185 | NA |
| MMD | 843.087263 | -0.8991552 | 0.26745235 | -3.3619267 | 0.00077401 | 0.00640357 |
| SMIM36 | 0.08666371 | -0.642905668 | 3.30558948 | -0.1944905 | 0.84579185 | NA |
| TMEM100 | 82.1675764 | -0.756913141 | 0.26288588 | -2.8792461 | 0.00398627 | 0.02268407 |
| PCTP | 1709.63426 | 0.388732168 | 0.29928108 | 1.29888655 | 0.19398287 | 0.37378516 |
| AC090618.1 | 16.8069488 | 0.694598477 | 0.66711785 | 1.04119305 | 0.29778596 | 0.4946218 |
| ANKFN1 | 8.50377561 | 0.504837767 | 0.89198099 | 0.56597368 | 0.57141167 | 0.73635956 |
| NOG | 23.0427556 | 0.361919072 | 0.51031931 | 0.70920121 | 0.47819962 | 0.66311492 |
| C17orf67 | 88.1517105 | -0.496527031 | 0.41215351 | -1.2047138 | 0.2283138 | 0.41547408 |
| DGKE | 467.916159 | 0.097226692 | 0.20463897 | 0.47511328 | 0.63470624 | 0.78341547 |
| TRIM25 | 2685.13378 | -0.788738817 | 0.39547511 | -1.9944083 | 0.04610746 | 0.13727601 |
| AC015912.1 | 344.861504 | -0.796006406 | 0.43042178 | -1.8493637 | 0.06440532 | 0.17365728 |
| AC015912.3 | 23.3409476 | -1.966965341 | 0.52188512 | -3.7689623 | 0.00016393 | 0.00187103 |
| COIL | 792.355371 | -0.031021101 | 0.17826956 | -0.1740123 | 0.86185577 | 0.92801702 |
| AC004584.1 | 9.19098631 | 0.061464557 | 0.43711483 | 0.14061421 | 0.88817472 | 0.94217674 |
| SCPEP1 | 5760.01048 | 0.481187758 | 0.16185405 | 2.97297328 | 0.0029493 | 0.0181036 |
| AC007114.1 | 127.242778 | 0.926292076 | 0.37427469 | 2.47489908 | 0.01332738 | 0.05573154 |
| AKAP1 | 3577.20847 | 0.616609647 | 0.16570712 | 3.72108128 | 0.00019837 | 0.00216517 |
| AC007114.2 | 1.30976541 | 2.367686023 | 1.33317694 | 1.77597283 | 0.07573739 | 0.1951148 |
| MSI2 | 5543.40974 | 0.086851204 | 0.1511203 | 0.57471568 | 0.5654836 | 0.73214268 |
| AC007431.2 | 0 | NA | NA | NA | NA | NA |
| AC007431.1 | 0.79178462 | 1.290327478 | 1.72256027 | 0.74907537 | 0.45381178 | NA |
| CCDC182 | 1.05665784 | 0.566029 | 1.57326726 | 0.3597793 | 0.71901218 | 0.84018041 |
| AC015845.2 | 72.3767864 | 1.3414968 | 0.47633582 | 2.81628366 | 0.00485828 | 0.02630111 |
| MRPS23 | 2374.32507 | -0.273004703 | 0.2160704 | -1.2634989 | 0.20640995 | 0.38909985 |
| CUEDC1 | 4692.70876 | -0.425730542 | 0.23233408 | -1.8324068 | 0.06689084 | 0.17829895 |
| AC015845.1 | 0.49742502 | -1.138387487 | 2.20165784 | -0.5170592 | 0.60511482 | NA |
| AC015813.4 | 8.61570503 | -0.478789884 | 0.48521003 | -0.9867683 | 0.32375623 | 0.5215653 |
| VEZF1 | 3059.57665 | -0.109393785 | 0.17187657 | -0.6364671 | 0.524472 | 0.69951978 |
| AC015813.2 | 2884.45855 | -0.231388434 | 0.12469249 | -1.8556726 | 0.06350024 | 0.1719658 |
| AC015813.1 | 1724.03485 | -0.210973592 | 0.20328498 | -1.0378218 | 0.29935301 | 0.49609973 |
| SRSF1 | 9428.46518 | -0.126648977 | 0.11911697 | -1.063232 | 0.28767674 | 0.4838718 |
| AC015813.3 | 0.87713412 | -3.028928908 | 2.72453808 | -1.111722 | 0.2662577 | NA |
| DYNLL2 | 8500.52066 | 0.561012 | 0.17182208 | 3.26507506 | 0.00109435 | 0.00841097 |
| OR4D1 | 5.76075284 | 0.692226577 | 0.69055629 | 1.00241875 | 0.31614139 | 0.51330256 |
| OR4D2 | 0 | NA | NA | NA | NA | NA |
| EPX | 36.2757234 | 1.159880278 | 0.40992635 | 2.82948459 | 0.0046623 | 0.02551479 |
| AC005962.1 | 27.4344259 | 0.748013308 | 0.27767104 | 2.69388309 | 0.00706249 | 0.03485644 |
| MKS1 | 955.194284 | 0.256166128 | 0.21837531 | 1.17305444 | 0.24077397 | 0.43089309 |
| LPO | 98.020584 | 0.656919357 | 0.36116417 | 1.81889404 | 0.0689276 | 0.18210808 |
| MPO | 120.930141 | -4.176266548 | 0.64185233 | -6.5065847 | 7.69E-11 | 1.15E-08 |
| TSPOAP1 | 2845.36843 | 0.893884192 | 0.37079208 | 2.4107424 | 0.01592009 | 0.06360799 |
| TSPOAP1-AS | 1428.12229 | 0.154314118 | 0.11861746 | 1.30093931 | 0.19327923 | 0.37281642 |
| AC004687.1 | 65.6600361 | -0.916954707 | 0.75438381 | -1.2155016 | 0.22417485 | 0.41020684 |
| SUPT4H1 | 2712.75711 | 0.053589863 | 0.12237203 | 0.43792575 | 0.6614401 | 0.80079774 |
| RNF43 | 762.730167 | 0.77399497 | 0.31958534 | 2.42187257 | 0.01544076 | 0.06216381 |
| HSF5 | 3.60979502 | 1.1260526 | 0.97083614 | 1.15987915 | 0.24609801 | 0.4374228 |
| MTMR4 | 2399.30571 | 0.992467307 | 0.27790061 | 3.57130303 | 0.00035521 | 0.00347565 |
| SEPT4-AS1 | 326.367846 | 0.701324548 | 0.35004312 | 2.00353761 | 0.04511962 | 0.13503623 |
| 4-Sep | 1270.0459 | 0.707987991 | 0.45536663 | 1.55476475 | 0.12000211 | 0.26991662 |
| C17orf47 | 63.6777367 | 0.164345336 | 0.43713683 | 0.37595857 | 0.7069477 | 0.83325953 |
| TEX14 | 62.4065676 | 0.288162411 | 0.36584128 | 0.78767056 | 0.43088943 | 0.62295052 |
| AC011195.2 | 0.66559539 | 1.408116591 | 1.84722966 | 0.76228561 | 0.44588956 | NA |
| RAD51C | 506.294914 | 0.084103849 | 0.14063461 | 0.59803095 | 0.54981928 | 0.72018351 |
| AC025521.1 | 0.76351963 | -1.209641812 | 1.4818667 | -0.816296 | 0.41433088 | NA |
| PPM1E | 203.801754 | 0.361310181 | 0.4909346 | 0.73596398 | 0.46175262 | 0.65013703 |
| TRIM37 | 1126.91953 | -0.27393342 | 0.21412974 | -1.2792871 | 0.20079597 | 0.38239256 |
| AC099850.1 | 27.158077 | 0.511561094 | 0.40775844 | 1.254569 | 0.20963526 | 0.39278683 |
| SKA2 | 915.164704 | 0.212527158 | 0.19993079 | 1.06300363 | 0.2877803 | 0.48397746 |
| PRR11 | 26.9847198 | 0.138530487 | 0.42453803 | 0.32630878 | 0.74419074 | 0.85654401 |
| AC099850.2 | 153.657366 | -0.388220117 | 0.17763021 | -2.1855523 | 0.02884838 | 0.0981336 |
| AC099850.3 | 7.48685957 | 0.027366914 | 0.52093038 | 0.05253469 | 0.95810266 | 0.97995373 |
| SMG8 | 592.010103 | 0.196154986 | 0.21177665 | 0.92623518 | 0.35432378 | 0.5519564 |
| GDPD1 | 139.358094 | 0.255302372 | 0.23856689 | 1.07015007 | 0.28455176 | 0.48069379 |
| YPEL2 | 2537.08914 | -0.107533859 | 0.20281635 | -0.5302031 | 0.59597113 | 0.75484247 |
| AC091059.1 | 276.035173 | -0.424487871 | 0.16678873 | -2.5450632 | 0.0109258 | 0.04812344 |
| LINC01476 | 0.20696692 | -0.339951137 | 3.12703692 | -0.1087135 | 0.91342973 | NA |
| DHX40 | 2886.59272 | 0.074763459 | 0.10433822 | 0.71654912 | 0.47365235 | 0.65956659 |
| AC091271.1 | 99.6691826 | 0.882658946 | 0.36369242 | 2.42693799 | 0.01522686 | 0.06150923 |
| CLTC | 16098.9824 | 0.07112668 | 0.1643 | 0.43290737 | 0.66508206 | 0.80354687 |
| PTRH2 | 1273.64155 | -0.247652979 | 0.35751033 | -0.6927156 | 0.48848805 | 0.67195774 |
| VMP1 | 21330.5948 | -1.666009558 | 0.33599753 | -4.9583983 | 7.11E-07 | 2.44E-05 |
| TUBD1 | 414.500664 | -0.316809542 | 0.2138611 | -1.4813799 | 0.13850536 | 0.297668 |
| RPS6KB1 | 1651.44978 | -0.231506356 | 0.14726543 | -1.5720346 | 0.11594253 | 0.26375126 |
| AC005702.1 | 219.14495 | -0.180776238 | 0.15499652 | -1.1663245 | 0.24348327 | 0.43409194 |
| RNFT1 | 595.078692 | 0.23745556 | 0.14252532 | 1.66605879 | 0.0957017 | 0.23057597 |
| RNFT1-DT | 46.041959 | 0.774928472 | 0.33540972 | 2.31039362 | 0.02086637 | 0.07727092 |
| HEATR6 | 480.571981 | 0.000773059 | 0.14007623 | 0.00551885 | 0.99559662 | 0.99794294 |
| AC025048.4 | 23.1305588 | -0.825999282 | 0.49985933 | -1.6524635 | 0.09844011 | 0.23506888 |
| AC025048.1 | 1.545753 | 1.461362022 | 1.06733218 | 1.36917265 | 0.17094531 | 0.34319208 |
| CA4 | 1578.32959 | 0.643882236 | 0.33606807 | 1.91592801 | 0.05537426 | 0.15646496 |
| USP32 | 1392.22179 | -0.352440429 | 0.22149169 | -1.5912128 | 0.11156168 | 0.25650505 |
| C17orf64 | 4.59348755 | -2.035142225 | 0.79518625 | -2.5593277 | 0.01048748 | 0.04668062 |
| APPBP2 | 2309.76809 | -0.151841598 | 0.11883394 | -1.2777628 | 0.20133306 | 0.38307233 |
| AC011921.1 | 32.7383209 | 0.32570017 | 0.33004125 | 0.98684685 | 0.32371772 | 0.5215653 |
| LINC01999 | 0.55668469 | -0.673748778 | 1.86577853 | -0.3611087 | 0.71801822 | NA |
| PPM1D | 885.855744 | -0.154516301 | 0.15094511 | -1.0236589 | 0.30599641 | 0.50296861 |
| BCAS3 | 1245.43142 | 0.574996891 | 0.21399076 | 2.68701742 | 0.00720932 | 0.03542198 |
| AC005884.1 | 100.631689 | 0.890985965 | 0.22835638 | 3.90173442 | 9.55E-05 | 0.0012202 |
| AC005884.2 | 3.685111 | 0.183463868 | 0.59543053 | 0.30811969 | 0.75799126 | 0.86618156 |
| AC005856.1 | 9.65369717 | 0.731937624 | 0.64979716 | 1.1264094 | 0.25999227 | 0.45292965 |
| AC005746.3 | 2.76768381 | -0.236105148 | 1.17321461 | -0.2012463 | 0.84050596 | 0.91507243 |
| AC005746.2 | 34.9612036 | 0.890669304 | 0.43772923 | 2.03474944 | 0.04187609 | 0.12808601 |
| TBX2-AS1 | 486.691287 | 0.250560709 | 0.25510365 | 0.98219178 | 0.32600538 | 0.52368188 |
| TBX2 | 5755.64664 | 0.258990899 | 0.20939051 | 1.23687983 | 0.2161317 | 0.40082938 |
| C17orf82 | 33.7605206 | -0.081491031 | 0.38244665 | -0.2130782 | 0.83126599 | 0.91040231 |
| TBX4 | 2.02382378 | -0.053576648 | 0.93101411 | -0.0575465 | 0.95410983 | 0.9780936 |
| AC005901.1 | 0 | NA | NA | NA | NA | NA |
| NACA2 | 6.88359405 | -0.617785219 | 0.49010449 | -1.2605174 | 0.20748279 | 0.39040822 |
| BRIP1 | 28.3488567 | -0.013671058 | 0.52441322 | -0.0260692 | 0.9792021 | 0.99058424 |

| | | | | | | |
|---|---|---|---|---|---|---|
| INTS2 | 530.04027 | 0.065183481 | 0.13308382 | 0.48979268 | 0.62428061 | 0.77611992 |
| MED13 | 2763.27285 | -0.339243587 | 0.18488494 | -1.8348903 | 0.06652195 | 0.17763886 |
| EFCAB3 | 1.46285294 | -1.495391902 | 1.49418818 | -1.0008056 | 0.3169208 | 0.51422603 |
| METTL2A | 637.35978 | -0.138880316 | 0.17072838 | -0.8134577 | 0.4159557 | 0.60920587 |
| TLK2 | 1569.0413 | -0.378040969 | 0.16790674 | -2.2514937 | 0.02435429 | 0.0865492 |
| AC080038.2 | 5.11591399 | -0.10223315 | 0.53775545 | -0.1901109 | 0.84922226 | 0.92065413 |
| AC080038.1 | 97.9925095 | -0.335934853 | 0.46299803 | -0.7255643 | 0.4681059 | 0.65488634 |
| MRC2 | 3344.58859 | 0.319177524 | 0.2908081 | 1.09755377 | 0.27239939 | 0.46738056 |
| AC005821.1 | 16.7046039 | 1.482663882 | 0.52908543 | 2.80231473 | 0.00507373 | 0.02715535 |
| 10-Mar | 69.789101 | 0.486134186 | 0.36861315 | 1.31881942 | 0.18722949 | 0.3651806 |
| AC068512.1 | 0.09635527 | 0.547239394 | 3.29377202 | 0.16614368 | 0.86804388 | NA |
| AC005972.4 | 2.86511383 | -0.66610427 | 0.74303696 | -0.8964618 | 0.37000616 | 0.56680971 |
| AC005972.1 | 0.55222564 | -0.67756755 | 1.51994368 | -0.4457846 | 0.65575282 | NA |
| TANC2 | 1312.82816 | -0.819176601 | 0.18899363 | -4.3344139 | 1.46E-05 | 0.00027746 |
| AC006270.1 | 1.26329751 | -2.311891701 | 1.42571624 | -1.6215651 | 0.1048965 | 0.24546284 |
| AC015923.1 | 21.1110834 | -1.587072354 | 0.48574798 | -3.2672752 | 0.00108588 | 0.00835561 |
| AC005828.2 | 0.09644756 | -0.642905668 | 3.30558948 | -0.1944905 | 0.84579185 | NA |
| AC005828.1 | 0.53706932 | -0.04116872 | 1.92120927 | -0.0214285 | 0.9829038 | NA |
| AC005828.3 | 0.97888437 | -1.682556268 | 1.33790822 | -1.2576022 | 0.20853567 | NA |
| AC005828.4 | 1.01784029 | 1.597321525 | 1.23332374 | 1.29513564 | 0.19527344 | 0.37542729 |
| CYB561 | 3416.33019 | 0.198207123 | 0.12048855 | 1.64502874 | 0.09996388 | 0.23748358 |
| AC005828.5 | 2.5153776 | -0.568399193 | 0.88282262 | -0.643843 | 0.51967722 | 0.69637118 |
| ACE | 2552.89883 | 1.596005072 | 0.48623332 | 3.28238523 | 0.00102933 | 0.00803059 |
| AC113554.1 | 1369.22289 | 1.511392183 | 0.48405027 | 3.12238679 | 0.00179391 | 0.01240635 |
| KCNH6 | 1305.33867 | 1.969372479 | 0.62518374 | 3.15006992 | 0.00163231 | 0.01154927 |
| DCAF7 | 4450.08477 | 0.065977233 | 0.09278698 | 0.7110613 | 0.47704625 | 0.66212483 |
| TACO1 | 1725.13867 | 0.482407073 | 0.28856189 | 1.67176293 | 0.09457108 | 0.22870867 |
| MAP3K3 | 3404.93277 | -0.295377219 | 0.15879184 | -1.8601536 | 0.0628638 | 0.17079315 |
| STRADA | 2208.32305 | -0.026850027 | 0.15748189 | -0.170496 | 0.86462011 | 0.92966493 |
| LIMD2 | 1336.21413 | -0.590878666 | 0.31830084 | -1.8563528 | 0.06340328 | 0.17179132 |
| AC046185.1 | 2955.44097 | -0.132639053 | 0.09781078 | -1.3560781 | 0.17507433 | 0.3490896 |
| CCDC47 | 5947.86339 | -0.148430971 | 0.15636946 | -0.9492325 | 0.34250239 | 0.54018364 |
| DDX42 | 5100.61784 | 0.060745258 | 0.11718176 | 0.51838494 | 0.60418972 | 0.76129983 |
| FTSJ3 | 5538.46326 | 0.07502825 | 0.13771351 | 0.54481403 | 0.58588143 | 0.74726448 |
| PSMC5 | 8179.99827 | -0.120967848 | 0.11626389 | -1.0404594 | 0.29812653 | 0.49509394 |
| SMARCD2 | 3327.98586 | -0.519871481 | 0.16905694 | -3.0751266 | 0.00210413 | 0.01399821 |
| CSH2 | 0.12059279 | 0.02007176 | 3.29738704 | 0.00608717 | 0.99514317 | NA |
| GH2 | 0 | NA | NA | NA | NA | NA |
| CSH1 | 0.03392908 | 0.380740593 | 3.29738709 | 0.11546736 | 0.90807469 | NA |
| CSHL1 | 0.17332742 | -1.003574173 | 3.30558948 | -0.3035992 | 0.7614333 | NA |
| GH1 | 2.29348862 | 0.19050647 | 1.01204996 | 0.18823821 | 0.85068992 | 0.92161226 |
| CD79B | 205.134431 | -0.278744944 | 0.48861083 | -0.5704846 | 0.56834908 | 0.73404883 |
| SCN4A | 81.4254697 | 0.131373376 | 0.58092184 | 0.22614639 | 0.82108757 | 0.90408983 |
| AC127029.2 | 0.09989236 | -0.642905668 | 3.30558948 | -0.1944905 | 0.84579185 | NA |
| PRR29-AS1 | 18.7845414 | -0.985406019 | 0.44153177 | -2.2317896 | 0.02562887 | 0.08990732 |
| PRR29 | 1363.23208 | -0.965716233 | 0.26999427 | -3.5768027 | 0.00034782 | 0.00341223 |
| ICAM2 | 1685.96853 | -0.838522948 | 0.27156913 | -3.0876961 | 0.00201715 | 0.01358328 |
| ERN1 | 1391.88209 | -0.813429269 | 0.16849764 | -4.8275409 | 1.38E-06 | 4.23E-05 |
| SNHG25 | 204.999317 | -1.205045343 | 0.25422496 | -4.7400749 | 2.14E-06 | 5.99E-05 |
| TEX2 | 2722.28673 | -0.127062972 | 0.2204073 | -0.5764917 | 0.5642829 | 0.73132527 |
| PECAM1 | 9758.82873 | -0.788011848 | 0.37119242 | -2.1229201 | 0.03376055 | 0.10976604 |
| MILR1 | 109.665887 | -0.237462906 | 0.49219991 | -0.4824522 | 0.62948478 | 0.77986909 |
| POLG2 | 633.760819 | -0.259983395 | 0.14585968 | -1.7824213 | 0.07468055 | 0.19311679 |
| DDX5 | 40687.7547 | -0.083120317 | 0.10489148 | -0.7924411 | 0.42810354 | 0.62048424 |
| CEP95 | 2598.373 | -0.194457878 | 0.13545266 | -1.435615 | 0.15111192 | 0.31595474 |
| SMURF2 | 1369.30006 | 0.110346589 | 0.10543205 | 1.04661336 | 0.29527794 | 0.49212571 |
| LRRC37A3 | 997.264171 | 0.645910847 | 0.31513478 | 2.04963363 | 0.0404002 | 0.12469021 |
| AC103810.2 | 34.4639554 | 0.353234683 | 0.63397998 | 0.55717009 | 0.57741122 | 0.7411933 |
| AC103810.5 | 6.05357536 | 0.392546782 | 0.83805167 | 0.46840403 | 0.63949569 | 0.78694765 |
| AC037487.1 | 17.6793526 | 1.035588719 | 0.65857511 | 1.57246867 | 0.1158419 | 0.26356781 |
| GNA13 | 3426.40141 | -1.00102011 | 0.36152069 | -2.7689151 | 0.00562433 | 0.02943928 |
| AC037487.2 | 1.51701303 | -2.367484682 | 1.18549177 | -1.9970486 | 0.0458199 | 0.13667723 |
| RGS9 | 439.379732 | -1.000264939 | 0.39787241 | -2.5140344 | 0.01193588 | 0.05136245 |
| LINC02563 | 0 | NA | NA | NA | NA | NA |
| AXIN2 | 337.082915 | -0.019056902 | 0.29508653 | -0.0645807 | 0.94850783 | 0.97514677 |
| AC004805.1 | 171.236386 | -0.256957084 | 0.24232687 | -1.0603739 | 0.28897455 | 0.4851505 |
| CEP112 | 570.554721 | -0.38345972 | 0.25425861 | -1.5081484 | 0.13151655 | 0.28733152 |
| APOH | 341.588924 | 3.220542503 | 0.88378075 | 3.64405145 | 0.00026838 | 0.0027787 |
| PRKCA | 2459.54635 | 0.441061786 | 0.23641943 | 1.86559027 | 0.06209871 | 0.16939516 |
| PRKCA-AS1 | 0.72352837 | 1.641204679 | 1.99162122 | 0.82405463 | 0.40990852 | NA |
| CACNG5 | 1.34394504 | 2.35218895 | 1.60403901 | 1.4664163 | 0.14253491 | 0.30375348 |
| CACNG4 | 20.0877829 | 0.964022642 | 0.94191959 | 1.02346597 | 0.30608758 | 0.5030244 |
| AC005544.1 | 0.19718306 | -0.340099831 | 3.16382928 | -0.1074963 | 0.91439529 | NA |
| AC005544.2 | 4.27874001 | 0.98230235 | 0.94006945 | 1.0449253 | 0.2960575 | 0.49280217 |
| CACNG1 | 0.99728948 | -0.284029808 | 1.59245961 | -0.1783592 | 0.8584409 | NA |
| HELZ | 2081.59932 | -0.080376914 | 0.13965091 | -0.575556 | 0.56491535 | 0.73174269 |
| AC007448.2 | 0 | NA | NA | NA | NA | NA |
| AC007448.3 | 73.4903932 | 2.09481682 | 0.38843999 | 5.39289693 | 6.93E-08 | 3.48E-06 |
| PSMD12 | 2804.61233 | -0.658026914 | 0.27781758 | -2.3685575 | 0.01785761 | 0.06897204 |
| PITPNC1 | 1571.42663 | -0.59623656 | 0.28196035 | -2.1146114 | 0.03446308 | 0.11142325 |
| NOL11 | 1885.59593 | 0.080096947 | 0.18207534 | 0.43991101 | 0.66000156 | 0.79999826 |
| BPTF | 2214.89574 | 0.140165609 | 0.14715852 | 0.9524804 | 0.3408534 | 0.53888521 |
| AC134407.1 | 6.05213274 | 0.498142339 | 0.5573232 | 0.89381232 | 0.37142232 | 0.56829264 |
| C17orf58 | 446.722479 | -0.46348213 | 0.24548723 | -1.8880091 | 0.05902472 | 0.16363351 |
| KPNA2 | 1818.79485 | -0.952153591 | 0.34989839 | -2.7212289 | 0.00650397 | 0.03271214 |
| AC145343.1 | 8.67889312 | -0.877650842 | 0.61737728 | -1.4215794 | 0.15514838 | 0.32131178 |
| AC005332.7 | 1652.19169 | -0.001652202 | 0.12758887 | -0.0129494 | 0.98966815 | 0.99519138 |
| AC005332.4 | 121.908864 | 0.072146928 | 0.18498852 | 0.39000759 | 0.69653093 | 0.82550655 |
| AC005332.6 | 1039.47416 | -0.485223858 | 0.20464031 | -2.3711059 | 0.01773495 | 0.0686274 |
| AC005332.8 | 14.3229888 | -0.212828718 | 0.50526813 | -0.4212194 | 0.67359491 | 0.80982482 |
| AC005332.3 | 30.9814226 | 0.391993161 | 0.32078903 | 1.22196561 | 0.22172063 | 0.40769633 |
| AC005332.5 | 174.565815 | -0.666788088 | 0.29080954 | -2.2928687 | 0.02185557 | 0.08012391 |
| AC005332.1 | 21.3269145 | 0.11828648 | 0.37589051 | 0.31468334 | 0.7530021 | 0.86232098 |
| AMZ2 | 2386.94511 | 0.241420363 | 0.20097578 | 1.20124107 | 0.2296577 | 0.41717115 |
| ARSG | 1366.3098 | -0.14487319 | 0.12652153 | -1.1450477 | 0.25218939 | 0.44444524 |
| SLC16A6 | 102.413694 | -1.020496489 | 0.58278593 | -1.7510658 | 0.07993458 | 0.2023301 |
| WIPI1 | 2512.04097 | -0.137316347 | 0.14254618 | -0.9633113 | 0.33539131 | 0.53359089 |
| PRKAR1A | 13881.0842 | 0.190622604 | 0.12621137 | 1.51034416 | 0.13095563 | 0.28653335 |
| FAM20A | 4736.75429 | -1.400899776 | 0.55373166 | -2.5299254 | 0.01140868 | 0.04971854 |
| AC079210.1 | 33.8973571 | -1.475785511 | 0.61855008 | -2.3858788 | 0.01703837 | 0.06677095 |
| LINC01482 | 13.6063208 | -0.672446793 | 0.65894237 | -1.0204941 | 0.30749419 | 0.50442438 |
| AC011591.1 | 0.42563236 | 0.330626757 | 1.85358267 | 0.17837173 | 0.85843105 | NA |
| ABCA8 | 1239.00621 | 1.079676625 | 0.39563659 | 2.72896048 | 0.00635343 | 0.03216192 |
| ABCA9-AS1 | 21.1237919 | 1.713484888 | 0.45481233 | 3.76745481 | 0.00016492 | 0.00187941 |
| ABCA9 | 527.116982 | 0.274903761 | 0.26334021 | 1.04391109 | 0.29652653 | 0.49336895 |
| ABCA6 | 418.289838 | 0.187465042 | 0.2926954 | 0.64047826 | 0.52186172 | 0.69787989 |
| ABCA10 | 386.831381 | 0.44600443 | 0.26469719 | 1.68496092 | 0.09199611 | 0.2241672 |
| ABCA5 | 1967.00158 | 0.369768066 | 0.17839046 | 2.07280175 | 0.03819073 | 0.11966369 |
| MAP2K6 | 474.349524 | 1.146511796 | 0.48877413 | 2.34568838 | 0.01899198 | 0.07227861 |
| AC002546.1 | 6.41604299 | -2.988376611 | 1.19881883 | -2.4927675 | 0.01267518 | 0.0537408 |
| LINC01483 | 0.22871771 | 0.689158768 | 3.02668931 | 0.22769392 | 0.8198842 | NA |
| LINC01497 | 3.88152266 | 0.300198946 | 1.39316231 | 0.21548024 | 0.82939292 | 0.90934819 |
| AC005208.1 | 44.206404 | 2.051027229 | 0.76217598 | 2.69101532 | 0.00712349 | 0.03507871 |

| | | | | | | |
|---|---|---|---|---|---|---|
| LINC01028 | 1.95492525 | -0.360138239 | 1.41805684 | -0.253966 | 0.79952182 | 0.8912141 |
| KCNJ16 | 14410.2554 | 0.797039222 | 0.28025842 | 2.84394388 | 0.00445589 | 0.0247101 |
| KCNJ2-AS1 | 43.23045 | -0.694430829 | 0.42744401 | -1.6246124 | 0.10424518 | 0.24447977 |
| KCNJ2 | 132.191607 | -0.436457198 | 0.2506584 | -1.741243 | 0.08164099 | 0.20531714 |
| AC011990.1 | 0.09997307 | 0.547239394 | 3.29377202 | 0.16614368 | 0.86804388 | NA |
| AC007423.1 | 0.34665484 | -1.746384021 | 3.30130528 | -0.528998 | 0.59680681 | NA |
| CASC17 | 0 | NA | NA | NA | NA | NA |
| AC118653.1 | 0 | NA | NA | NA | NA | NA |
| AC005144.1 | 0 | NA | NA | NA | NA | NA |
| ROCR | 0.47599435 | 2.179672787 | 2.70380839 | 0.80614913 | 0.42015686 | NA |
| LINC01152 | 9.18444969 | 0.382411657 | 0.66623386 | 0.57399013 | 0.56597448 | 0.73257484 |
| LINC02097 | 75.0767107 | 0.941757124 | 0.42211173 | 2.23106123 | 0.02567707 | 0.0900314 |
| SOX9 | 3136.35459 | -1.208018427 | 0.42458165 | -2.845197 | 0.0044384 | 0.02464011 |
| LINC00511 | 154.327089 | -1.914077471 | 0.38917981 | -4.9182343 | 8.73E-07 | 2.89E-05 |
| LINC02003 | 0.97661342 | -2.289033809 | 1.40745579 | -1.6263629 | 0.10387249 | NA |
| AC007639.1 | 14.6825076 | -1.577825829 | 0.61091143 | -2.5827407 | 0.0098019 | 0.04441168 |
| AC080037.1 | 2.58185599 | -1.037974414 | 1.40423775 | -0.7391728 | 0.45980205 | 0.64846401 |
| SLC39A11 | 1460.22717 | -0.186820037 | 0.31238657 | -0.5980412 | 0.54981245 | 0.72018351 |
| AC011120.1 | 0.67679946 | 0.702830114 | 1.62838532 | 0.43161167 | 0.66602367 | NA |
| SSTR2 | 1108.73379 | -0.414835367 | 0.47848104 | -0.8669839 | 0.38595082 | 0.58163922 |
| COG1 | 3078.45608 | -0.123478067 | 0.0815513 | -1.5141153 | 0.12999661 | 0.28509735 |
| AC097641.2 | 267.269254 | -0.049906682 | 0.13481401 | -0.3701891 | 0.71124158 | 0.83591569 |
| FAM104A | 1828.40673 | -0.436740278 | 0.13606025 | -3.2099036 | 0.00132779 | 0.00984091 |
| C17orf80 | 1073.28613 | 0.177929053 | 0.13138152 | 1.35429281 | 0.17564298 | 0.34990597 |
| CPSF4L | 198.734462 | -0.269129815 | 0.17185812 | -1.566 | 0.11734863 | 0.26580341 |
| CDC42EP4 | 3903.2456 | -0.844950422 | 0.18887931 | -4.4734937 | 7.70E-06 | 0.00016552 |
| SDK2 | 123.720511 | 0.256871436 | 0.31914525 | 0.80487314 | 0.42089289 | 0.61354889 |
| AC124804.1 | 0.26173473 | 0.079077975 | 3.01575203 | 0.02622164 | 0.97908055 | NA |
| AC032019.1 | 0.07830766 | -0.642905668 | 3.30558948 | -0.1944905 | 0.84579185 | NA |
| AC032019.2 | 0.03392908 | 0.380740593 | 3.29738709 | 0.11546736 | 0.90807469 | NA |
| AC125421.1 | 0.08757577 | 0.547239394 | 3.29377202 | 0.16614368 | 0.86804388 | NA |
| LINC00469 | 0.75049007 | -2.780432901 | 3.26469898 | -0.8516659 | 0.39439953 | NA |
| LINC02092 | 0 | NA | NA | NA | NA | NA |
| AC125421.2 | 0 | NA | NA | NA | NA | NA |
| AC137735.1 | 0 | NA | NA | NA | NA | NA |
| LINC02074 | 0 | NA | NA | NA | NA | NA |
| RPL38 | 8253.2911 | -0.55299028 | 0.1430151 | -3.8666567 | 0.00011034 | 0.00136922 |
| AC100786.1 | 48.675953 | 0.21255902 | 0.26189075 | 0.81163242 | 0.41700258 | 0.61000709 |
| TTYH2 | 305.832365 | 0.405661584 | 0.32302095 | 1.25583676 | 0.20917516 | 0.39235447 |
| AC100786.2 | 57.9460879 | -0.318550566 | 0.3926162 | -0.8113536 | 0.41716263 | 0.61016022 |
| DNAI2 | 7.33095732 | -0.238167889 | 0.52052586 | -0.4575525 | 0.647274 | 0.79230448 |
| AC103809.1 | 7.4544862 | -0.746952687 | 0.67598 | -1.1049923 | 0.26916293 | 0.4633046 |
| KIF19 | 49.4914057 | -1.319514902 | 0.49783213 | -2.6505218 | 0.00803675 | 0.03824588 |
| BTBD17 | 0.40883395 | -1.945586661 | 3.29439637 | -0.5905746 | 0.55480552 | NA |
| GPR142 | 0.67119008 | -0.441349399 | 1.93307284 | -0.2283149 | 0.81940141 | NA |
| GPRC5C | 13638.9332 | -0.683809919 | 0.1331398 | -5.1360293 | 2.81E-07 | 1.13E-05 |
| CD300A | 371.656442 | -1.257790796 | 0.5376192 | -2.3395571 | 0.01930662 | 0.07304385 |
| CD300LB | 32.9206496 | 0.808943404 | 0.68991605 | 1.1725244 | 0.24098658 | 0.43106879 |
| CD300C | 79.1865835 | -0.757527511 | 0.5515148 | -1.3735398 | 0.16958461 | 0.3413747 |
| CD300LD | 0.09635527 | 0.547239394 | 3.29377202 | 0.16614368 | 0.86804388 | NA |
| C17orf77 | 0.16588343 | 0.18657164 | 3.29377195 | 0.05664376 | 0.95482897 | NA |
| AC064805.1 | 17.1386279 | -0.660569609 | 0.98662515 | -0.6695244 | 0.50316102 | 0.68332771 |
| CD300E | 260.992653 | -0.75196232 | 0.83099101 | -0.9048983 | 0.36551929 | 0.56260407 |
| RAB37 | 319.032062 | 0.490849251 | 0.4148299 | 1.18325429 | 0.23670838 | 0.42601653 |
| AC064805.2 | 0.18382444 | -1.003574173 | 3.30558948 | -0.3035992 | 0.7614333 | NA |
| CD300LF | 68.2472772 | -0.867121172 | 0.55463965 | -1.5633956 | 0.11795958 | 0.2666611 |
| AC016888.1 | 1784.41543 | 0.442428243 | 0.43937482 | 1.00694948 | 0.31395905 | 0.51093903 |
| SLC9A3R1 | 14524.7172 | 0.242043077 | 0.37203242 | 0.65059673 | 0.51530684 | 0.69294449 |
| NAT9 | 2627.11398 | -0.575292713 | 0.20511675 | -2.8047086 | 0.00503621 | 0.02700751 |
| TMEM104 | 1743.54415 | -0.469147437 | 0.20213115 | -2.3210051 | 0.02028657 | 0.07566575 |
| GRIN2C | 98.1094635 | 0.097338355 | 0.4816953 | 0.20207454 | 0.83985845 | 0.91484357 |
| FDXR | 828.959907 | -0.084058702 | 0.29322094 | -0.2866736 | 0.77436225 | 0.87594083 |
| FADS6 | 146.073196 | 1.276838991 | 0.68619856 | 1.86074274 | 0.06278052 | 0.17068999 |
| USH1G | 30.454633 | 1.255118998 | 0.74969247 | 1.67417847 | 0.09409553 | 0.22793488 |
| OTOP2 | 0.93832423 | 2.256139081 | 1.7868621 | 1.2626263 | 0.20672351 | NA |
| OTOP3 | 0 | NA | NA | NA | NA | NA |
| HID1 | 1790.41467 | -0.26810458 | 0.24070582 | -1.1138267 | 0.26535352 | 0.45945899 |
| HID1-AS1 | 12.8698041 | 0.323111069 | 0.64419248 | 0.50157535 | 0.61596626 | 0.77032298 |
| CDR2L | 1920.7241 | -1.027071464 | 0.31273892 | -3.2841179 | 0.00102302 | 0.00798611 |
| MRPL58 | 1150.16137 | 0.165127522 | 0.1718555 | 0.96085097 | 0.3366271 | 0.53462158 |
| KCTD2 | 6489.90886 | -0.221974012 | 0.13026148 | -1.7040649 | 0.08836896 | 0.21764169 |
| ATP5PD | 7401.55512 | -0.074769034 | 0.14282078 | -0.5235165 | 0.60061487 | 0.75824464 |
| SLC16A5 | 5409.51829 | -1.562941965 | 0.39630458 | -3.9437899 | 8.02E-05 | 0.00106334 |
| ARMC7 | 2020.79312 | -1.130620837 | 0.30277473 | -3.7341982 | 0.00018831 | 0.0020795 |
| NT5C | 1834.87857 | -0.41977484 | 0.13797768 | -3.0423387 | 0.00234748 | 0.01529334 |
| JPT1 | 4043.93759 | -0.891202734 | 0.15342873 | -5.8085778 | 6.30E-09 | 4.67E-07 |
| SUMO2 | 6624.50909 | -0.328341219 | 0.14349194 | -2.2882206 | 0.02212468 | 0.08079597 |
| NUP85 | 1971.00484 | -0.488920116 | 0.16044084 | -3.0473545 | 0.00230865 | 0.01508777 |
| GGA3 | 2830.29062 | -0.146130024 | 0.13947803 | -1.047692 | 0.29478052 | 0.49164268 |
| MRPS7 | 3397.88702 | 0.065101137 | 0.12476797 | 0.52177764 | 0.60182516 | 0.75915472 |
| MIF4GD | 1305.85741 | -0.293780297 | 0.14160132 | -2.0747003 | 0.03801432 | 0.11925262 |
| AC022211.2 | 348.928887 | -0.324977144 | 0.16140327 | -2.0134483 | 0.04406749 | 0.1329892 |
| SLC25A19 | 590.028332 | -0.28908653 | 0.15637739 | -1.8486466 | 0.06450886 | 0.17386535 |
| GRB2 | 6586.02965 | -0.077073286 | 0.09554637 | -0.8066584 | 0.41986331 | 0.61261525 |
| AC011933.4 | 0.23034971 | 0.68915899 | 3.0220972 | 0.22803998 | 0.81961516 | NA |
| AC011933.3 | 3.50275408 | 1.06930585 | 0.76862926 | 1.39118546 | 0.16416919 | 0.33375543 |
| TMEM94 | 5615.48653 | 0.390948176 | 0.08359473 | 4.67670867 | 2.92E-06 | 7.66E-05 |
| CASKIN2 | 5290.77578 | 0.215290159 | 0.27053702 | 0.79578818 | 0.42615516 | 0.61861233 |
| TSEN54 | 2406.56795 | -0.369265627 | 0.14622054 | -2.5254019 | 0.0115566 | 0.05018074 |
| LLGL2 | 12744.2625 | -0.01877154 | 0.19896167 | -0.0943475 | 0.9248331 | 0.96356574 |
| MYO15B | 15904.0177 | 0.58625347 | 0.1873075 | 3.12989849 | 0.00174867 | 0.01217327 |
| RECQL5 | 3972.15334 | 0.337511902 | 0.28909703 | 1.16746927 | 0.24302091 | 0.43359009 |
| SMIM5 | 1876.55026 | 1.063668761 | 0.34139835 | 3.11562364 | 0.00183556 | 0.01260844 |
| SMIM6 | 315.380354 | 0.319350048 | 0.30534156 | 1.04587811 | 0.29561731 | 0.49250875 |
| SAP30BP | 3512.62575 | -0.383596003 | 0.17460211 | -2.1969724 | 0.02802242 | 0.09616003 |
| AC087749.2 | 0.73530046 | 0.414241332 | 1.8701001 | 0.22150757 | 0.82469724 | NA |
| AC087749.1 | 0.10721287 | -0.642905668 | 3.30558948 | -0.1944905 | 0.84579185 | NA |
| ITGB4 | 2220.33208 | -0.498515705 | 0.34543571 | -1.4431505 | 0.14897809 | 0.31303205 |
| GALK1 | 1878.58537 | 0.613044931 | 0.26576557 | 2.3067131 | 0.02107082 | 0.07785309 |
| H3F3B | 20568.1931 | -0.455316674 | 0.20004038 | -2.2761238 | 0.0228386 | 0.08273355 |
| UNK | 2344.63823 | 0.061152875 | 0.18517602 | 0.33024186 | 0.74121722 | 0.8546523 |
| AC087289.1 | 294.820049 | 0.297521263 | 0.23288629 | 1.27753879 | 0.2014121 | 0.38317848 |
| UNC13D | 805.680637 | -0.152714896 | 0.27975386 | -0.5458902 | 0.5851414 | 0.74685098 |
| WBP2 | 12441.5691 | -0.650865753 | 0.11808048 | -5.5120519 | 3.55E-08 | 2.03E-06 |
| TRIM47 | 4506.32354 | -0.830073458 | 0.20114855 | -4.1266688 | 3.68E-05 | 0.00057875 |
| AC087289.5 | 1218.80783 | -0.666815291 | 0.20110349 | -3.3157817 | 0.00091387 | 0.00731819 |
| TRIM65 | 1775.558 | 0.06352206 | 0.13687606 | 0.46408452 | 0.64258719 | 0.78843403 |
| AC087289.2 | 974.069568 | -0.222277399 | 0.09016199 | -2.4653115 | 0.01368942 | 0.05683986 |
| MRPL38 | 4351.58057 | -0.170671113 | 0.08603227 | -1.9838034 | 0.04727776 | 0.13978187 |
| AC087289.3 | 4542.9443 | -0.131850877 | 0.08040201 | -1.6398954 | 0.10102693 | 0.23918837 |
| FBF1 | 870.773412 | -0.126667792 | 0.31178213 | -0.4062702 | 0.68454407 | 0.81778936 |
| ACOX1 | 5594.77024 | 0.699661146 | 0.30517534 | 2.29265299 | 0.02186799 | 0.08014127 |
| AC087289.4 | 397.647899 | 0.96738108 | 0.33822719 | 2.86015174 | 0.00423438 | 0.02375254 |

| | | | | | | |
|---|---|---|---|---|---|---|
| TEN1 | 1177.59258 | 0.039099275 | 0.24576368 | 0.15909298 | 0.87359562 | 0.93417196 |
| CDK3 | 1277.64734 | -0.092983421 | 0.16263172 | -0.5717422 | 0.56749663 | 0.73345276 |
| EVPL | 3692.01762 | 0.290490419 | 0.25841435 | 1.12412648 | 0.26095937 | 0.45414028 |
| SRP68 | 4033.77316 | -0.202198772 | 0.136303 | -1.4834506 | 0.13795472 | 0.29675336 |
| ZACN | 5536.45301 | -0.117124262 | 0.11748692 | -0.9969132 | 0.31880665 | 0.5160893 |
| GALR2 | 9.85233679 | -1.187643176 | 0.61190781 | -1.9408858 | 0.05227213 | 0.15023609 |
| EXOC7 | 8429.14346 | 0.13312323 | 0.1148309 | 1.15929795 | 0.24633476 | 0.43769955 |
| FOXJ1 | 528.950859 | -1.364075488 | 0.51608918 | -2.6431003 | 0.00821507 | 0.03882282 |
| RNF157-AS1 | 470.814378 | -0.100044261 | 0.34242856 | -0.292161 | 0.77016355 | 0.87291382 |
| RNF157 | 1090.40213 | 0.1011149 | 0.39854683 | 0.25370896 | 0.79972041 | 0.8913389 |
| UBALD2 | 2530.08059 | -1.18102575 | 0.24385515 | -4.8431446 | 1.28E-06 | 3.97E-05 |
| QRICH2 | 293.602576 | 0.348182247 | 0.24369142 | 1.42878337 | 0.15306651 | 0.31857242 |
| PRPSAP1 | 2278.0267 | 0.103882036 | 0.18340343 | 0.56641273 | 0.57111324 | 0.73615487 |
| SPHK1 | 580.493816 | -2.111649894 | 0.50395851 | -4.1901265 | 2.79E-05 | 0.00046268 |
| UBE2O | 1867.86967 | -0.481477516 | 0.19239651 | -2.5025273 | 0.01233101 | 0.05265039 |
| AANAT | 28.3774064 | -0.80856645 | 0.57769414 | -1.3996446 | 0.16161979 | 0.3303041 |
| RHBDF2 | 1982.084 | -1.266516453 | 0.27313745 | -4.6369198 | 3.54E-06 | 8.94E-05 |
| CYGB | 672.034016 | 0.407188365 | 0.44629299 | 0.91237904 | 0.36156922 | 0.55904898 |
| PRCD | 274.458556 | 0.289341726 | 0.37958129 | 0.76226551 | 0.44590156 | 0.63631279 |
| AC015802.4 | 79.6029097 | 0.826581677 | 0.32461187 | 2.5463692 | 0.010885 | 0.04799155 |
| AC015802.6 | 2767.48644 | -0.295455539 | 0.2190267 | -1.3489476 | 0.1773538 | 0.35243554 |
| ST6GALNAC2 | 889.521213 | 0.900122468 | 0.42384931 | 2.12368513 | 0.03369648 | 0.1096289 |
| AC015802.1 | 174.59759 | 1.057166057 | 0.44186837 | 2.39249089 | 0.01673444 | 0.06591296 |
| ST6GALNAC1 | 7.01588398 | 1.572376551 | 0.78037404 | 2.01490114 | 0.04391501 | 0.13277216 |
| AC005837.1 | 40.8121272 | 1.055594919 | 0.37941925 | 2.78213324 | 0.00540029 | 0.02852104 |
| MXRA7 | 4524.69295 | -0.008271473 | 0.33934524 | -0.0243748 | 0.98055365 | 0.99131724 |
| AC005837.3 | 4.00446958 | 0.446246617 | 0.81876213 | 0.54502596 | 0.58573566 | 0.74722314 |
| JMJD6 | 2068.52628 | -1.056836113 | 0.24400437 | -4.3312179 | 1.48E-05 | 0.00028091 |
| METTL23 | 1401.50325 | -0.349490084 | 0.13221112 | -2.6434242 | 0.00820721 | 0.03879721 |
| AC005837.2 | 763.791059 | -0.161599904 | 0.12836462 | -1.2589131 | 0.20806171 | 0.39128421 |
| SRSF2 | 10805.5073 | -0.216177631 | 0.18042251 | -1.1981744 | 0.23084914 | 0.41855692 |
| MFSD11 | 5100.41364 | -0.015860841 | 0.19855353 | -0.0798819 | 0.93633115 | 0.96951778 |
| LINC02080 | 10.4448135 | 1.420603931 | 0.68887396 | 2.06221167 | 0.03918759 | 0.12196422 |
| LINC00868 | 0.46822477 | -0.83393584 | 2.29250231 | -0.3637666 | 0.71603228 | NA |
| MGAT5B | 45.9291129 | 0.534201086 | 0.41581325 | 1.28471396 | 0.19889226 | 0.37994704 |
| AC016168.1 | 0.08393208 | -0.642905668 | 3.30558948 | -0.1944905 | 0.84579185 | NA |
| SEC14L1 | 8163.68116 | -0.556895179 | 0.24970301 | -2.2302302 | 0.02573217 | 0.09019769 |
| AC068594.1 | 7.83976118 | -0.949844095 | 0.51472993 | -1.8453252 | 0.06499025 | 0.17478766 |
| AC068594.2 | 1.05790926 | 0.277157583 | 1.25585899 | 0.22069164 | 0.82533254 | 0.90673291 |
| 9-Sep | 11042.3537 | -0.521821275 | 0.15700532 | -3.3235898 | 0.00088867 | 0.00715547 |
| AC111182.1 | 18.3281918 | 0.544518703 | 0.50873244 | 1.07034397 | 0.28446451 | 0.48060335 |
| AC111170.1 | 2.27718934 | 0.205849355 | 0.83398887 | 0.24682506 | 0.80504361 | 0.89438161 |
| AC021683.3 | 3.63879886 | 1.208144448 | 1.20966409 | 0.99874375 | 0.31791884 | 0.51510153 |
| AC021683.2 | 5.70096405 | 0.548839328 | 0.75024815 | 0.73154372 | 0.46444711 | 0.65187727 |
| AC021683.1 | 8.30082438 | 0.806889952 | 0.65525567 | 1.23141238 | 0.21816867 | 0.40327406 |
| LINC01987 | 7.09122893 | 2.108287561 | 1.0217965 | 2.06331452 | 0.03908276 | 0.12173544 |
| LINC01973 | 0.41931504 | -0.032078219 | 1.8788046 | -0.0170737 | 0.98637779 | NA |
| TNRC6C | 789.699357 | 0.061039805 | 0.12311976 | 0.49577586 | 0.62005256 | 0.77286756 |
| TMC6 | 2504.2554 | -0.609714085 | 0.27543804 | -2.2136161 | 0.0268552 | 0.09307026 |
| TMC8 | 849.190192 | -0.052310519 | 0.57358663 | -0.091199 | 0.92733448 | 0.96473672 |
| C17orf99 | 4.57844881 | 0.32333716 | 0.8097251 | 0.3993172 | 0.6896595 | 0.82118262 |
| SYNGR2 | 7403.82379 | -0.665952971 | 0.14639603 | -4.5489823 | 5.39E-06 | 0.00012447 |
| TK1 | 948.577485 | 0.483610715 | 0.27148375 | 1.78136153 | 0.07485341 | 0.19336642 |
| AFMID | 6580.90531 | 1.159635782 | 0.30768597 | 3.76889395 | 0.00016397 | 0.00187103 |
| BIRC5 | 70.6391728 | -1.277304304 | 0.58057724 | -2.2000592 | 0.02780269 | 0.09561255 |
| TMEM235 | 1.44149986 | -1.997089666 | 1.39472867 | -1.431884 | 0.15217702 | 0.31725782 |
| AC087645.4 | 6.99043678 | -1.345113571 | 0.98749956 | -1.3621409 | 0.17315341 | 0.34678025 |
| LINC01993 | 32.8059318 | -3.962728953 | 0.67048721 | -5.9102231 | 3.42E-09 | 2.77E-07 |
| AC087645.2 | 11.7541858 | -1.451169804 | 0.51227806 | -2.8327776 | 0.00461455 | 0.02537232 |
| SOCS3 | 8893.13603 | -2.361163336 | 0.46387819 | -5.0900503 | 3.58E-07 | 1.38E-05 |
| AC061992.1 | 133.406397 | -3.552524681 | 0.50369859 | -7.0528779 | 1.75E-12 | 4.36E-10 |
| PGS1 | 2125.47214 | -0.800867584 | 0.17281621 | -4.6342155 | 3.58E-06 | 9.03E-05 |
| DNAH17 | 938.750559 | -0.982943241 | 0.19063277 | -5.1562134 | 2.52E-07 | 1.03E-05 |
| LINC02081 | 3.25118327 | -0.842791766 | 0.77175769 | -1.0920419 | 0.27481467 | 0.47005661 |
| CYTH1 | 3807.76489 | -0.392667242 | 0.19829853 | -1.9801823 | 0.04768305 | 0.14074345 |
| USP36 | 2951.67848 | -0.322854316 | 0.29419131 | -1.0974298 | 0.27245354 | 0.46741265 |
| TIMP2 | 10175.4509 | 0.260215214 | 0.26607984 | 0.97795916 | 0.32809452 | 0.52575672 |
| AC022966.1 | 1575.31209 | 0.738151455 | 0.26757507 | 2.75867053 | 0.0058037 | 0.03007287 |
| CEP295NL | 651.179175 | 0.667495116 | 0.28943912 | 2.30616759 | 0.02110127 | 0.07792848 |
| AC100788.1 | 0.26272731 | 1.990198932 | 3.242756 | 0.61373687 | 0.53938922 | NA |
| LGALS3BP | 19328.674 | -0.574819817 | 0.50895151 | -1.1294196 | 0.25872087 | 0.45149026 |
| CANT1 | 2219.35635 | -0.224581428 | 0.14214435 | -1.5799532 | 0.11411758 | 0.26063403 |
| C1QTNF1-AS | 6.78289007 | -0.270262476 | 0.57919315 | -0.4666189 | 0.64077255 | 0.78737988 |
| C1QTNF1 | 2467.37616 | -1.357800723 | 0.49223501 | -2.7584399 | 0.0058078 | 0.03007981 |
| ENGASE | 3989.41225 | 0.048365756 | 0.25185476 | 0.19203828 | 0.84771222 | 0.92003832 |
| RBFOX3 | 28.4678737 | -0.880071167 | 0.60291014 | -1.4597054 | 0.14437107 | 0.3062311 |
| AC021534.1 | 0 | NA | NA | NA | NA | NA |
| LINC02078 | 1.87013085 | 1.673929158 | 1.14869859 | 1.45723967 | 0.14505023 | 0.30727624 |
| ENPP7 | 418.899282 | 1.86483838 | 0.58825045 | 3.17014359 | 0.00152364 | 0.01095672 |
| CBX2 | 150.515992 | 0.060941656 | 0.48611054 | 0.12536584 | 0.90023393 | 0.94951124 |
| CBX8 | 457.513749 | -0.517003302 | 0.27677598 | -1.8679486 | 0.06176924 | 0.16884574 |
| AC100791.2 | 0.62858447 | 0.742415417 | 2.17447011 | 0.3414236 | 0.73278471 | NA |
| LINC01977 | 9.98574993 | 0.6602851 | 0.62414537 | 1.05790274 | 0.29009979 | 0.48620385 |
| CBX4 | 1146.86144 | -0.704811223 | 0.17535408 | -4.0193603 | 5.84E-05 | 0.00083407 |
| AC100791.1 | 0 | NA | NA | NA | NA | NA |
| LINC01979 | 27.3685217 | -0.516670967 | 0.43071242 | -1.199573 | 0.23030523 | 0.41789624 |
| LINC01978 | 18.8901936 | -0.510045722 | 0.41570973 | -1.2269276 | 0.21984982 | 0.40558554 |
| TBC1D16 | 5339.26996 | 0.103998943 | 0.29834879 | 0.34858174 | 0.72740333 | 0.84604913 |
| AC100791.3 | 54.3927468 | 0.702176919 | 0.27139036 | 2.58733183 | 0.00967224 | 0.04394503 |
| AC116025.1 | 14.4830216 | 0.891625749 | 0.6649786 | 1.34083376 | 0.17997443 | 0.3558199 |
| CCDC40 | 951.692798 | -0.159592956 | 0.27022238 | -0.5905986 | 0.55478942 | 0.72430887 |
| GAA | 8796.91574 | 0.155206536 | 0.21312529 | 0.72824081 | 0.46646619 | 0.65366741 |
| EIF4A3 | 3413.49377 | -0.640807061 | 0.17532313 | -3.6550059 | 0.00025718 | 0.00267595 |
| AC087741.2 | 14.3005204 | -0.484873425 | 0.39379812 | -1.2312741 | 0.21822036 | 0.40332906 |
| CARD14 | 832.442177 | -0.0372859 | 0.31320996 | -0.1190444 | 0.90524016 | 0.95244715 |
| AC087741.1 | 1528.79089 | -0.048501168 | 0.33864604 | -0.1432208 | 0.88611578 | 0.94110344 |
| SGSH | 3524.14952 | -0.315679185 | 0.16209971 | -1.9474383 | 0.05148221 | 0.14882546 |
| SLC26A11 | 2851.58904 | -0.541670475 | 0.22637882 | -2.3927613 | 0.01672212 | 0.06588409 |
| RNF213 | 14843.965 | -1.079021765 | 0.43568849 | -2.47659 | 0.01326442 | 0.05552988 |
| AC124319.3 | 37.578082 | 0.472171389 | 0.3794024 | 1.24451343 | 0.21331066 | 0.39731383 |
| AC124319.2 | 233.368098 | 0.64493101 | 0.25550285 | 2.52416368 | 0.01159739 | 0.05033478 |
| ENDOV | 1541.48933 | 0.22988535 | 0.164297 | 1.39920603 | 0.16175121 | 0.33046283 |
| AC120024.1 | 10.7718998 | 0.841683579 | 0.58300928 | 1.44368814 | 0.14882672 | 0.31286357 |
| NPTX1 | 46.6801532 | -0.109403643 | 0.62250754 | -0.1757467 | 0.86049295 | 0.92734057 |
| RPTOR | 1309.02195 | 0.223593513 | 0.10948103 | 2.04230372 | 0.04112141 | 0.12642785 |
| AC016245.1 | 0.5723124 | -1.642189738 | 1.87352841 | -0.8765225 | 0.38074607 | NA |
| AC127496.3 | 44.4196201 | 0.722928117 | 0.25438538 | 2.84186185 | 0.00448509 | 0.02482159 |
| AC127496.4 | 64.1534278 | 0.375859163 | 0.19526176 | 1.92489896 | 0.054242 | 0.1542496 |
| AC127496.1 | 26.9510147 | -0.386460762 | 0.38238725 | -1.0106529 | 0.31218261 | 0.50910117 |
| CHMP6 | 1389.97769 | -0.035951157 | 0.16874193 | -0.2130541 | 0.83128478 | 0.91040231 |
| AC127496.6 | 660.695861 | -0.131039864 | 0.1809778 | -0.7240659 | 0.46902525 | 0.65569182 |
| AC127496.5 | 7.1563715 | -0.23409964 | 0.56984883 | -0.4108101 | 0.68121181 | 0.81527767 |
| AC127496.2 | 1.11108341 | 1.605540427 | 1.23118573 | 1.3040603 | 0.19221302 | 0.37149351 |

| | | | | | | |
|---|---|---|---|---|---|---|
| BAIAP2-DT | 1247.08672 | 0.208283432 | 0.25009834 | 0.83280614 | 0.40495407 | 0.59972246 |
| BAIAP2 | 2535.14352 | 0.700540472 | 0.29306741 | 2.39037318 | 0.01683126 | 0.06622506 |
| AATK | 464.187935 | 0.817070868 | 0.43862195 | 1.86281344 | 0.06248852 | 0.17007145 |
| AC115099.1 | 13.5942105 | 2.623667811 | 0.5842635 | 4.49055576 | 7.10E-06 | 0.00015572 |
| PVALEF | 1.48132725 | -0.37821458 | 1.18849512 | -0.3182298 | 0.75031062 | 0.86062162 |
| CEP131 | 913.665455 | -0.076325807 | 0.24205319 | -0.3153266 | 0.7525137 | 0.86206078 |
| AC027601.2 | 90.9259534 | 0.340717211 | 0.23843964 | 1.42894534 | 0.15301994 | 0.31857242 |
| TEPSIN | 3149.69101 | -0.217827721 | 0.18733776 | -1.162754 | 0.24492934 | 0.43599208 |
| AC027601.1 | 2097.30879 | -0.282754964 | 0.18532138 | -1.5257547 | 0.12707097 | 0.28057117 |
| NDUFAF8 | 842.721827 | -0.285081764 | 0.20313 | -1.4034449 | 0.16048422 | 0.32874767 |
| SLC38A10 | 8793.39606 | 0.334204896 | 0.18195737 | 1.83672086 | 0.06625111 | 0.1772387 |
| AC027601.4 | 689.250335 | 0.393516023 | 0.18479285 | 2.12949811 | 0.03321307 | 0.1084349 |
| LINC00482 | 228.419452 | 0.671307981 | 0.41082133 | 1.6340631 | 0.1022456 | 0.24116427 |
| TMEM105 | 115.177684 | 1.354302351 | 0.89914405 | 1.50621288 | 0.13201253 | 0.28815531 |
| AC027601.3 | 22.4651275 | -0.112185477 | 0.41615639 | -0.2695753 | 0.78748702 | 0.88426302 |
| AC110285.6 | 19.6519774 | -0.438248187 | 0.38356568 | -1.1425636 | 0.25321982 | 0.44575478 |
| AC110285.3 | 0.55472719 | -1.604272687 | 1.65867855 | -0.9671993 | 0.33344443 | NA |
| AC110285.1 | 16.2842514 | 0.377492448 | 0.37487272 | 1.00698832 | 0.31394039 | 0.51093903 |
| AC110285.5 | 2.73560406 | -1.113984968 | 0.9551044 | -1.1663489 | 0.24347341 | 0.43409194 |
| AC110285.2 | 1061.32972 | -0.065309079 | 0.5603568 | -0.1165491 | 0.90721738 | 0.95376639 |
| BAHCC1 | 1986.3807 | 0.611069665 | 0.22523225 | 2.71306469 | 0.00666641 | 0.03333843 |
| AC110285.4 | 130.008883 | 0.850112147 | 0.23468175 | 3.62240413 | 0.00029188 | 0.00297174 |
| LINC01971 | 2.84267679 | -0.24774835 | 1.00489523 | -0.2465415 | 0.80526311 | 0.8944712 |
| ACTG1 | 75857.7931 | 0.011265854 | 0.26032255 | 0.04327652 | 0.96548111 | 0.98376108 |
| AC139149.1 | 20.3137755 | 0.706111653 | 0.38306238 | 1.84333333 | 0.06528036 | 0.17534325 |
| FSCN2 | 46.1439003 | 0.425324015 | 0.37413184 | 1.13682924 | 0.25560968 | 0.44823751 |
| FAAP100 | 2921.45194 | -0.276341521 | 0.14291925 | -1.93355 | 0.05316848 | 0.15213234 |
| NPLOC4 | 8215.91287 | 0.014632251 | 0.20556536 | 0.07118053 | 0.94325408 | 0.97315146 |
| TSPAN10 | 113.037121 | 0.787235185 | 0.32204362 | 2.44449862 | 0.01450536 | 0.05941386 |
| PDE6G | 109.062925 | -0.259331648 | 0.29594015 | -0.8762976 | 0.38086826 | 0.57691402 |
| OXLD1 | 1161.17412 | -0.299452221 | 0.08795643 | -3.4045518 | 0.00066273 | 0.00563928 |
| CCDC137 | 1760.51805 | -0.69076331 | 0.20553777 | -3.360761 | 0.00077728 | 0.00642536 |
| ARL16 | 2111.28119 | 0.321677771 | 0.15462503 | 2.08037324 | 0.03749131 | 0.11801909 |
| HGS | 9249.49493 | -0.740294719 | 0.26813111 | -2.7609431 | 0.00576347 | 0.02993959 |
| AC139530.3 | 61.0574123 | 0.264209388 | 0.31617316 | 0.83564773 | 0.40335311 | 0.59829123 |
| AC139530.1 | 3099.30598 | -0.974455567 | 0.24263342 | -4.0161639 | 5.92E-05 | 0.00084226 |
| MRPL12 | 3081.21592 | -0.146068595 | 0.1754064 | -0.8327438 | 0.40498921 | 0.59973974 |
| AC139530.2 | 6535.98076 | 0.823452484 | 0.31485588 | 2.61533148 | 0.00891409 | 0.04133366 |
| SLC25A10 | 4435.74192 | 1.44702076 | 0.48191682 | 3.00263594 | 0.00267652 | 0.01684083 |
| GCGR | 2992.82305 | 0.922547715 | 0.46487845 | 1.98449231 | 0.04720098 | 0.13963316 |
| MCRIP1 | 3604.73838 | -0.43079045 | 0.19277311 | -2.234702 | 0.02543692 | 0.08943613 |
| PPP1R27 | 8.21531822 | -0.612787726 | 0.60338533 | -1.0155827 | 0.30982813 | 0.50697377 |
| P4HB | 30564.0002 | -0.587131272 | 0.24177595 | -2.4284106 | 0.01516517 | 0.06132578 |
| AC145207.3 | 12988.8566 | -0.805486117 | 0.25302081 | -3.1834777 | 0.00145517 | 0.01056127 |
| ARHGDIA | 18449.9949 | -0.839596167 | 0.25652677 | -3.2729378 | 0.00106436 | 0.00822842 |
| AC145207.2 | 9941.56378 | -0.977490048 | 0.25896851 | -3.7745517 | 0.0001603 | 0.00184424 |
| AC145207.5 | 163.906544 | 0.190379364 | 0.34459591 | 0.55247135 | 0.58062546 | 0.74350374 |
| AC145207.6 | 106.838598 | 0.175439193 | 0.33626976 | 0.52172159 | 0.60186419 | 0.75916764 |
| ALYREF | 1335.44267 | -0.588229659 | 0.24448577 | -2.4059873 | 0.01612883 | 0.06424512 |
| ANAPC11 | 2273.47349 | -0.180332984 | 0.13706316 | -1.3156926 | 0.18827724 | 0.36635705 |
| NPB | 219.11547 | 0.99508165 | 0.39814675 | 2.49928361 | 0.01244447 | 0.05300607 |
| PCYT2 | 3748.71328 | 1.073405815 | 0.36432008 | 2.94632623 | 0.00321573 | 0.01938259 |
| SIRT7 | 2326.76166 | -0.589981177 | 0.28312977 | -2.0837836 | 0.03717985 | 0.11733279 |
| MAFG | 3317.87059 | -1.113024439 | 0.3443775 | -3.2319894 | 0.00122932 | 0.00923037 |
| AC145207.8 | 101.714734 | -0.132379972 | 0.44829925 | -0.2952938 | 0.76776948 | 0.87183611 |
| MAFG-DT | 121.780128 | -0.336416186 | 0.23565561 | -1.4275755 | 0.15341407 | 0.31902287 |
| PYCR1 | 283.554416 | -0.9288167 | 0.26993217 | -3.4409263 | 0.00057973 | 0.0050941 |
| AC145207.4 | 114.434465 | -1.205379773 | 0.30728442 | -3.9226843 | 8.76E-05 | 0.00114146 |
| MYADML2 | 8.50283354 | 0.240953632 | 0.61371165 | 0.39261701 | 0.69460238 | 0.82407273 |
| AC145207.1 | 1.30868014 | -0.31551294 | 1.10822352 | -0.2847015 | 0.77587281 | 0.87702628 |
| NOTUM | 34.7617059 | 2.113969979 | 0.48557524 | 4.35353739 | 1.34E-05 | 0.00025879 |
| AC145207.7 | 0.08757577 | 0.547239394 | 3.29377202 | 0.16614368 | 0.86804388 | NA |
| ASPSCR1 | 2942.49108 | 0.311977068 | 0.16681567 | 1.8701904 | 0.06145738 | 0.1681676 |
| CENPX | 1232.09432 | -0.145724029 | 0.21925752 | -0.664625 | 0.50629038 | 0.68556876 |
| LRRC45 | 857.845774 | 0.412227579 | 0.19254056 | 2.1409909 | 0.03227477 | 0.10622638 |
| RAC3 | 516.179001 | -0.111899564 | 0.27705234 | -0.4038932 | 0.68629123 | 0.81857592 |
| DCXR | 7471.72218 | 1.517749068 | 0.41740222 | 3.63617871 | 0.00027671 | 0.0028555 |
| AC137723.1 | 14.746057 | 2.101056065 | 0.86752715 | 2.42189085 | 0.01543999 | 0.06216381 |
| RFNG | 3698.81116 | -0.168167845 | 0.11621125 | -1.4470875 | 0.14787243 | 0.3113825 |
| GPS1 | 6808.89322 | 0.427857847 | 0.11352164 | 3.76895401 | 0.00016393 | 0.00187103 |
| DUS1L | 5883.66422 | -0.171560342 | 0.08913488 | -1.9247273 | 0.05426348 | 0.15425001 |
| FASN | 6758.48745 | -0.951471051 | 0.35203179 | -2.7027986 | 0.00687584 | 0.03412041 |
| CCDC57 | 2963.91644 | -0.00671693 | 0.20709112 | -0.0324347 | 0.97412542 | 0.98807744 |
| AC129510.1 | 296.73454 | 0.176237125 | 0.26026646 | 0.67714112 | 0.49831642 | 0.67987397 |
| AC129510.2 | 1.92298322 | 0.188342609 | 0.92436594 | 0.2037533 | 0.83854629 | 0.91450415 |
| AC132872.2 | 62.1427097 | -0.201832974 | 0.43973363 | -0.4589892 | 0.64624195 | 0.79140854 |
| SLC16A3 | 6584.50621 | -2.961957505 | 0.41330089 | -7.1665887 | 7.69E-13 | 2.16E-10 |
| CSNK1D | 14203.4601 | -0.871075015 | 0.22058203 | -3.9489846 | 7.85E-05 | 0.00104684 |
| AC132872.3 | 1596.05492 | -0.859668058 | 0.21651593 | -3.970461 | 7.17E-05 | 0.00097305 |
| LINC01970 | 13.7987605 | 0.169537426 | 0.57911166 | 0.29275429 | 0.76970997 | 0.87270139 |
| AC132872.1 | 401.923672 | -0.67240287 | 0.24348503 | -2.7615778 | 0.00575228 | 0.02990097 |
| CD7 | 424.305737 | -2.243643095 | 0.65008245 | -3.4513208 | 0.00055785 | 0.0049497 |
| SECTM1 | 4678.4567 | -3.193761665 | 0.56599541 | -5.6427342 | 1.67E-08 | 1.07E-06 |
| TEX19 | 1.53448005 | -2.405337311 | 1.46828589 | -1.6381941 | 0.10138121 | 0.23980115 |
| UTS2R | 4.54579586 | 0.606630002 | 0.6410241 | 0.94634508 | 0.34397261 | 0.54155816 |
| AC132938.1 | 40.2001718 | -0.305713865 | 0.40214251 | -0.7602128 | 0.44712742 | 0.63733102 |
| OGFOD3 | 1845.97967 | -0.07345141 | 0.10789918 | -0.6807411 | 0.4960353 | 0.67784893 |
| HEXDC | 3255.03513 | -0.318785031 | 0.31262689 | -1.019698 | 0.30787169 | 0.50476104 |
| CYBC1 | 4650.61054 | -0.891702197 | 0.20262192 | -4.400818 | 1.08E-05 | 0.00021709 |
| AC132938.3 | 80.2484051 | 0.80405117 | 0.34396318 | 2.33760826 | 0.01940758 | 0.07329654 |
| NARF | 3546.20945 | -0.958838201 | 0.20485118 | -4.6806574 | 2.86E-06 | 7.56E-05 |
| NARF-AS1 | 6.98420044 | 0.613387735 | 0.59038445 | 1.03896324 | 0.29882183 | 0.49563286 |
| FOXK2 | 2908.30906 | -0.317616024 | 0.15040749 | -2.1117035 | 0.03471189 | 0.11196974 |
| AC124283.2 | 20.8124427 | -0.441662597 | 0.42994431 | -1.0272554 | 0.30430023 | 0.5012257 |
| AC124283.1 | 635.369521 | -0.400207201 | 0.26315535 | -1.5208021 | 0.12830952 | 0.28245446 |
| WDR45B | 6679.6817 | -0.416737839 | 0.18677496 | -2.2312297 | 0.02566592 | 0.09001333 |
| RAB40B | 2429.73382 | -0.168379512 | 0.26923031 | -0.6254107 | 0.53170155 | 0.70599237 |
| AC024361.2 | 8.2250308 | 1.491310904 | 0.68836221 | 2.16646247 | 0.03027586 | 0.1014453 |
| FN3KRP | 2174.88665 | -0.010118728 | 0.18662873 | -0.0542185 | 0.95676109 | 0.97934128 |
| AC024361.3 | 34.7333004 | -0.391820685 | 0.35858207 | -1.0926946 | 0.27452792 | 0.46977945 |
| FN3K | 4786.10082 | 1.42084858 | 0.32996751 | 4.30602566 | 1.66E-05 | 0.00030587 |
| TBCD | 4453.38631 | 0.236438788 | 0.10377043 | 2.27847947 | 0.02269803 | 0.08232997 |
| AC068014.1 | 0.12059279 | 0.02007176 | 3.29738704 | 0.00608717 | 0.99514317 | NA |
| ZNF750 | 40.8828717 | 0.351889872 | 0.49830761 | 0.70616997 | 0.48008244 | 0.66476444 |
| AC087222.1 | 4.01370275 | -0.109212561 | 0.95991712 | -0.1137729 | 0.90941782 | 0.9548873 |
| B3GNTL1 | 886.613982 | -0.172574623 | 0.19317232 | -0.8933714 | 0.37165833 | 0.56836599 |
| AC130371.1 | 1.19703676 | 0.840013115 | 1.42721866 | 0.58856651 | 0.5561521 | 0.72530816 |
| METRNL | 2906.75688 | -0.526413794 | 0.20487188 | -2.569478 | 0.01018519 | 0.04568521 |
| AC130371.2 | 91.3666976 | 0.801132888 | 0.23280116 | 3.44127528 | 0.00057898 | 0.00508923 |
| AC144831.1 | 407.235704 | 0.132913932 | 0.32958276 | 0.40327938 | 0.68674271 | 0.81894837 |
| AC139099.2 | 1.76430458 | -1.287052274 | 1.11314387 | -1.1562317 | 0.24758638 | 0.43915066 |
| AC139099.3 | 7.04811545 | -1.666112001 | 0.79164654 | -2.104616 | 0.03532474 | 0.11344185 |
| AC139099.1 | 8.44988564 | -1.174204017 | 0.76988212 | -1.5251738 | 0.12721576 | 0.28079682 |

| | | | | | | |
|---|---|---|---|---|---|---|
| LINC02564 | 0 | NA | NA | NA | NA | NA |
| AP005530.1 | 0 | NA | NA | NA | NA | NA |
| TUBB8P12 | 13.8318412 | 1.556367522 | 0.68425766 | 2.27453429 | 0.02293388 | 0.08298113 |
| AP001005.3 | 1.83237497 | -1.552186294 | 1.43474496 | -1.0818552 | 0.27931687 | 0.47492119 |
| USP14 | 3292.1676 | -0.431787668 | 0.2093366 | -2.0626477 | 0.03914611 | 0.12188959 |
| THOC1 | 1672.06969 | -0.239792807 | 0.12934013 | -1.8539706 | 0.06374336 | 0.17243664 |
| AP000845.1 | 18.9151796 | 0.844502599 | 0.3557324 | 2.37398277 | 0.01759737 | 0.06829179 |
| AP000915.1 | 16.6524885 | -0.704471215 | 0.76684981 | -0.9186561 | 0.3582755 | 0.55594475 |
| COLEC12 | 434.283798 | -0.13747378 | 0.586961 | -0.2342128 | 0.81481978 | 0.9001968 |
| LINC01925 | 0.42885147 | -1.859940617 | 3.29773397 | -0.5640057 | 0.57275028 | NA |
| CETN1 | 0 | NA | NA | NA | NA | NA |
| CLUL1 | 210.389026 | 1.222853904 | 0.4495087 | 2.72042321 | 0.00651984 | 0.03274199 |
| AP001178.3 | 29.8870768 | 0.72299187 | 0.63579442 | 1.13714724 | 0.25547674 | 0.44814219 |
| TYMS | 1776.13435 | -0.021147676 | 0.21079449 | -0.1003237 | 0.92008737 | 0.96097528 |
| ENOSF1 | 6531.72109 | 0.306055209 | 0.21122277 | 1.44896885 | 0.14734628 | 0.31069259 |
| AP001178.2 | 213.433445 | 0.716434556 | 0.31303004 | 2.28870867 | 0.02209628 | 0.08072583 |
| YES1 | 3540.9399 | -0.469438604 | 0.23638365 | -1.9859182 | 0.0470424 | 0.13931171 |
| AP001020.3 | 102.712684 | 0.569092305 | 0.42986475 | 1.32388688 | 0.18554062 | 0.36286362 |
| AP001020.1 | 2.14917748 | 1.16411271 | 1.06407891 | 1.09400976 | 0.2739507 | 0.46900473 |
| AP000894.4 | 51.0672295 | 0.355097121 | 0.25511154 | 1.3919289 | 0.16394393 | 0.33341303 |
| AP000894.3 | 1.18591806 | 1.328198304 | 1.81171471 | 0.7331167 | 0.46348726 | 0.65128914 |
| ADCYAP1 | 49.045494 | 0.100808391 | 0.56117947 | 0.17963663 | 0.85743784 | 0.9254517 |
| LINC01904 | 0 | NA | NA | NA | NA | NA |
| AP005328.2 | 0.32913742 | -0.556401113 | 2.30333799 | -0.2415629 | 0.80911884 | NA |
| AP005328.1 | 1.31178958 | 0.563932045 | 1.07521997 | 0.52448063 | 0.59994429 | 0.75775321 |
| LINC00470 | 13.8933556 | 0.893258885 | 1.01795946 | 0.87749947 | 0.38021541 | 0.576223 |
| AC008109.1 | 0 | NA | NA | NA | NA | NA |
| AC019183.1 | 0.03392908 | 0.380740593 | 3.29738709 | 0.11546736 | 0.90807469 | NA |
| AP005057.1 | 0.13390214 | 1.04981649 | 3.28477129 | 0.31960109 | 0.74927075 | NA |
| AP005230.1 | 7.0317697 | 1.330681667 | 0.9933771 | 1.33955339 | 0.18039058 | 0.35632208 |
| METTL4 | 569.194677 | -0.165203632 | 0.14453082 | -1.1430339 | 0.2530245 | 0.44550402 |
| NDC80 | 57.0048055 | -0.691186969 | 0.58044321 | -1.1907917 | 0.23373536 | 0.42225125 |
| SMCHD1 | 2066.37273 | -0.540718611 | 0.24450114 | -2.2115178 | 0.027 | 0.09348609 |
| AP001011.1 | 1.80520668 | 0.158882592 | 0.9095876 | 0.17467542 | 0.86133469 | 0.92783061 |
| EMILIN2 | 538.539639 | -0.979680637 | 0.38308441 | -2.5573492 | 0.01054733 | 0.0468624 |
| LPIN2 | 4272.72012 | 0.012624456 | 0.17282434 | 0.07304791 | 0.941768 | 0.97233647 |
| AP000919.4 | 54.4182478 | 1.328317437 | 0.34079088 | 3.89774938 | 9.71E-05 | 0.00123566 |
| AP000919.2 | 22.6958842 | 2.058689298 | 0.49234998 | 4.18135349 | 2.90E-05 | 0.00047789 |
| AP000919.3 | 0.74270343 | -0.672320491 | 1.90639207 | -0.3526664 | 0.72433853 | NA |
| MYOM1 | 213.001173 | 0.966428304 | 0.42731466 | 2.26163153 | 0.02372018 | 0.08495673 |
| AP005329.2 | 83.1890446 | 0.123586322 | 0.22775157 | 0.54263654 | 0.58738008 | 0.74833531 |
| AP005329.3 | 69.254695 | 0.018766977 | 0.2381323 | 0.07880903 | 0.93718452 | 0.96995784 |
| MYL12A | 14320.9021 | -0.713009164 | 0.15123041 | -4.7147207 | 2.42E-06 | 6.64E-05 |
| AP005329.1 | 6316.97156 | -0.699972399 | 0.14927831 | -4.6890428 | 2.74E-06 | 7.32E-05 |
| MYL12B | 22020.0035 | -0.547810585 | 0.15007118 | -3.6503384 | 0.0002619 | 0.00271683 |
| AP001025.1 | 0.55717991 | -1.259246797 | 2.75015651 | -0.4578819 | 0.64703732 | NA |
| LINC01895 | 0.08757577 | 0.547239394 | 3.29377202 | 0.16614368 | 0.86804388 | NA |
| IGLJCOR18 | 0.3143988 | -0.174096114 | 3.29377188 | -0.0528562 | 0.95784652 | NA |
| TGIF1 | 3918.10985 | -0.994372976 | 0.2516938 | -3.950725 | 7.79E-05 | 0.00104189 |
| GAPLINC | 6.50762512 | -3.216390325 | 1.03147983 | -3.118229 | 0.00181941 | 0.01251707 |
| DLGAP1 | 297.370811 | -1.733119403 | 0.62527058 | -2.771791 | 0.00557488 | 0.02926729 |
| DLGAP1-AS1 | 452.868607 | -0.458916714 | 0.22981569 | -1.9968903 | 0.04583711 | 0.13668715 |
| DLGAP1-AS2 | 309.76592 | -1.10783415 | 0.36460815 | -3.038424 | 0.00237819 | 0.01541994 |
| AP002478.1 | 1.60458751 | -1.646640417 | 1.44217134 | -1.1417786 | 0.25354605 | 0.44618013 |
| DLGAP1-AS3 | 71.0735835 | -4.028837701 | 0.88061133 | -4.5750464 | 4.76E-06 | 0.00011329 |
| DLGAP1-AS4 | 0.20543692 | 0.689155168 | 3.10695182 | 0.2218107 | 0.82446125 | NA |
| DLGAP1-AS5 | 0.75282881 | -1.207960769 | 1.61930487 | -0.7459749 | 0.45568259 | NA |
| AP005203.1 | 0 | NA | NA | NA | NA | NA |
| LINC01892 | 0 | NA | NA | NA | NA | NA |
| AKAIN1 | 1.11937315 | 2.629528506 | 1.12717766 | 2.332843 | 0.01965639 | 0.07395333 |
| AP005380.1 | 0.65990388 | 2.749511938 | 1.49665432 | 1.83710553 | 0.06619432 | NA |
| AP001496.2 | 340.740101 | 0.532887924 | 0.28640136 | 1.86063338 | 0.06279597 | 0.1707144 |
| LINC00526 | 695.163853 | 0.545048996 | 0.2666418 | 2.04412433 | 0.04094127 | 0.12606506 |
| LINC00667 | 1744.10166 | 0.383054789 | 0.12963722 | 2.95482107 | 0.0031285 | 0.01896085 |
| ZBTB14 | 693.085939 | 0.726796095 | 0.21041505 | 3.45410695 | 0.00055212 | 0.00490545 |
| AP001496.1 | 0.3283321 | 0.732866073 | 3.27839706 | 0.22354402 | 0.82311212 | NA |
| AP005671.1 | 0.65327515 | 1.348859166 | 1.70960789 | 0.78898745 | 0.43011934 | NA |
| EPB41L3 | 4383.68314 | 0.899687803 | 0.25333393 | 3.55139084 | 0.0003832 | 0.00368796 |
| AP005059.2 | 0.18340304 | -1.003574173 | 3.30558948 | -0.3035992 | 0.7614333 | NA |
| AP005059.1 | 1.35909986 | 2.002315994 | 1.23444025 | 1.62204368 | 0.104794 | 0.24527624 |
| MIR3976HG | 0 | NA | NA | NA | NA | NA |
| TMEM200C | 153.502351 | 0.760514076 | 0.45363908 | 1.67647389 | 0.09364541 | 0.22705309 |
| AP005433.1 | 14.5810695 | 0.121939231 | 0.53293948 | 0.22880502 | 0.81902046 | 0.90287738 |
| L3MBTL4 | 762.298723 | 0.528737433 | 0.29312751 | 1.80377961 | 0.07126586 | 0.18673299 |
| AP001021.1 | 79.5195131 | 0.625016066 | 0.29699918 | 2.10443704 | 0.03534034 | 0.11344185 |
| AP001021.2 | 6.28927476 | 0.016048947 | 0.66777117 | 0.0240336 | 0.98082581 | 0.99132745 |
| L3MBTL4-AS | 4.33840468 | 0.238414736 | 0.72254667 | 0.32996448 | 0.7414268 | 0.85474446 |
| AP005202.2 | 0 | NA | NA | NA | NA | NA |
| LINC01387 | 0.55802599 | 0.202810051 | 2.19942235 | 0.0922106 | 0.92653072 | NA |
| AP005205.1 | 0.09635527 | 0.547239394 | 3.29377202 | 0.16614368 | 0.86804388 | NA |
| AP005205.2 | 13.8781825 | 1.422483388 | 0.50981203 | 2.79021148 | 0.00526736 | 0.02796102 |
| ARHGAP28 | 484.892561 | 0.981808574 | 0.32320649 | 3.03771307 | 0.00238381 | 0.01543359 |
| AP005210.1 | 7.86366998 | 0.927502205 | 0.59409657 | 1.5611977 | 0.11847711 | 0.26753223 |
| LINC00668 | 5.25802617 | -0.504923734 | 1.05451806 | -0.4788194 | 0.6320671 | 0.78149288 |
| LAMA1 | 1642.48011 | -1.014183954 | 0.30669673 | -3.3067974 | 0.00094369 | 0.00750909 |
| AP002409.1 | 251.453449 | -0.811669744 | 0.32052532 | -2.5323109 | 0.01133135 | 0.04949605 |
| AP005062.1 | 5.83374139 | -0.254020621 | 0.63156421 | -0.4022087 | 0.68753044 | 0.81947556 |
| LRRC30 | 0.03392908 | 0.380740593 | 3.29738709 | 0.11546736 | 0.90807469 | NA |
| PTPRM | 5121.45758 | 0.072813082 | 0.18914532 | 0.38495841 | 0.70026823 | 0.82811216 |
| AP000897.2 | 4.52440574 | 1.434858776 | 0.77649471 | 1.84786678 | 0.06462162 | 0.17406252 |
| AP001094.3 | 1.0231634 | 0.971523371 | 1.32843585 | 0.73132878 | 0.46457835 | 0.65193497 |
| AP001094.1 | 213.461164 | -0.510280747 | 0.27257343 | -1.8720854 | 0.06119479 | 0.16769267 |
| RAB12 | 2249.81873 | -0.237352818 | 0.120269 | -1.9735163 | 0.04843677 | 0.14239447 |
| AP001793.2 | 555.853513 | 0.008918137 | 0.11977278 | 0.0744588 | 0.94064532 | 0.97174713 |
| GACAT2 | 2.85395864 | 0.781521685 | 0.87380105 | 0.89439316 | 0.37111157 | 0.56790925 |
| MTCL1 | 472.519716 | -0.117108779 | 0.30857372 | -0.3795164 | 0.70430445 | 0.83113346 |
| NDUFV2 | 5896.99331 | -0.070278291 | 0.12263836 | -0.5730531 | 0.56660876 | 0.73291904 |
| AP005899.1 | 27.8829587 | -1.891092501 | 0.66574262 | -2.840576 | 0.00450321 | 0.02488469 |
| NDUFV2-AS1 | 145.271968 | -0.135302988 | 0.16402352 | -0.8248999 | 0.40942841 | 0.60393954 |
| ANKRD12 | 1856.7053 | -0.339580443 | 0.21911448 | -1.5497855 | 0.121193 | 0.27171055 |
| AP001033.1 | 18.4470146 | -0.073081186 | 0.36057645 | -0.2026788 | 0.83938613 | 0.91472899 |
| TWSG1 | 2514.36874 | 0.148896176 | 0.22455674 | 0.66306705 | 0.50728761 | 0.6862905 |
| AP005432.2 | 128.416454 | 1.661730561 | 0.74921484 | 2.21796269 | 0.02655738 | 0.09236587 |
| RALBP1 | 3913.57822 | 0.208244128 | 0.17047672 | 1.22153998 | 0.22188164 | 0.40780891 |
| PPP4R1 | 3092.38084 | -0.394756517 | 0.20728515 | -1.9044129 | 0.05685643 | 0.15931399 |
| AP001381.1 | 2.28575782 | -1.069481355 | 0.96729007 | -1.105647 | 0.26887934 | 0.46308852 |
| PPP4R1-AS1 | 9.57171246 | -0.387826474 | 0.58743375 | -0.6602046 | 0.50912253 | 0.68799637 |
| RAB31 | 1938.02258 | -0.563327876 | 0.34380377 | -1.6385157 | 0.10131416 | 0.23971776 |
| TXNDC2 | 3.59590152 | -2.65501522 | 1.23825332 | -2.1441616 | 0.03201994 | 0.10571752 |
| AC006238.1 | 12.3161675 | 0.24465508 | 0.44277099 | 0.55255445 | 0.58056854 | 0.74350374 |
| VAPA | 7365.39291 | -0.319717762 | 0.14161871 | -2.2575955 | 0.02397089 | 0.08554775 |
| AP005271.1 | 0.11223674 | 0.02007176 | 3.29738704 | 0.00608717 | 0.99514317 | NA |
| AP005209.1 | 0 | NA | NA | NA | NA | NA |

| | | | | | | |
|---|---|---|---|---|---|---|
| AP006219.1 | 0 | NA | NA | NA | NA | NA |
| LINC01254 | 0.17332742 | -1.003574173 | 3.30558948 | -0.3035992 | 0.7614333 | NA |
| APCDD1 | 522.65234 | -0.200125489 | 0.59153012 | -0.3383183 | 0.7351233 | 0.85108669 |
| NAPG | 1963.39478 | -0.292819722 | 0.1641249 | -1.7841273 | 0.07440297 | 0.19265578 |
| AP001099.1 | 10.2650214 | -0.203404463 | 0.73864727 | -0.2753743 | 0.7830287 | 0.88125233 |
| LINC01887 | 0.50299062 | -2.236486034 | 2.71055215 | -0.8251035 | 0.40931284 | NA |
| AP001180.2 | 0 | NA | NA | NA | NA | NA |
| AP001180.1 | 2.59819704 | 1.093829168 | 1.1179216 | 0.9784489 | 0.32785235 | 0.52559222 |
| PIEZO2 | 167.338096 | 0.783632817 | 0.25064071 | 3.12651849 | 0.00176889 | 0.01228813 |
| AP001180.4 | 3.08359079 | 2.083187045 | 0.86334764 | 2.41291799 | 0.01582538 | 0.06330129 |
| AP005120.1 | 0.08666371 | -0.642905668 | 3.30558948 | -0.1944905 | 0.84579185 | NA |
| AP005229.1 | 9.34844263 | 5.273841744 | 0.82756577 | 6.37271613 | 1.86E-10 | 2.45E-08 |
| AP005229.2 | 2.77250839 | 4.989387047 | 1.09010392 | 4.57698293 | 4.72E-06 | 0.00011255 |
| AP001109.1 | 0.42873441 | 1.24856177 | 2.13264495 | 0.58545224 | 0.55824367 | NA |
| LINC01928 | 38.8728685 | 2.959282995 | 1.52424527 | 1.94147429 | 0.05220077 | 0.15017389 |
| LINC01255 | 4.95963221 | 3.256561873 | 1.29584204 | 2.51308552 | 0.01196803 | 0.05148401 |
| SLC35G4 | 0 | NA | NA | NA | NA | NA |
| AP001120.2 | 0.81037102 | -1.948440464 | 1.8881648 | -1.0319229 | 0.30210825 | NA |
| AP001120.1 | 0 | NA | NA | NA | NA | NA |
| GNAL | 1665.84898 | 0.120569416 | 0.13523362 | 0.89156393 | 0.37262671 | 0.56887433 |
| AP005137.1 | 0 | NA | NA | NA | NA | NA |
| CHMP1B | 4499.28178 | -0.538036018 | 0.23719567 | -2.2683214 | 0.02330963 | 0.0839032 |
| CHMP1B-AS1 | 1837.02872 | -0.65803472 | 0.26868968 | -2.449051 | 0.01432332 | 0.05887854 |
| AP005137.2 | 2.57388124 | -0.46327594 | 1.647017 | -0.2812818 | 0.77849425 | 0.87861887 |
| MPPE1 | 2622.39462 | 0.29555178 | 0.15547498 | 1.90096034 | 0.05730721 | 0.16023657 |
| AP001269.4 | 12.8467104 | 0.702799507 | 0.44803074 | 1.56864127 | 0.11673155 | 0.26504314 |
| AP001269.2 | 2.33459372 | -0.957577894 | 1.17872954 | -0.8123813 | 0.41657285 | 0.6097994 |
| IMPA2 | 9878.30196 | 0.57321652 | 0.3073676 | 1.86492174 | 0.06219238 | 0.16952851 |
| AP001269.1 | 9.34726844 | 1.656178344 | 0.63143599 | 2.62287605 | 0.0087191 | 0.0406152 |
| AP001542.3 | 179.598039 | 0.674113003 | 0.37391295 | 1.80286083 | 0.07141007 | 0.1869808 |
| AP002414.5 | 0 | NA | NA | NA | NA | NA |
| ANKRD62 | 0.06785815 | 0.795868262 | 3.28898349 | 0.24198001 | 0.80879565 | NA |
| AP005264.4 | 0 | NA | NA | NA | NA | NA |
| AP005264.3 | 3.33119806 | 0.15131014 | 0.81501769 | 0.18565258 | 0.85271721 | 0.92285671 |
| CIDEA | 3.50216043 | 1.264967966 | 1.43303298 | 0.88272077 | NA | NA |
| AP005264.1 | 2.33274183 | 1.95612797 | 0.97369041 | 2.0089835 | 0.04453888 | 0.1338298 |
| TUBB6 | 6118.78159 | -0.563969891 | 0.21596536 | -2.6113905 | 0.00901749 | 0.04172675 |
| AFG3L2 | 6942.80165 | 0.061437655 | 0.23011158 | 0.26699071 | 0.78947631 | 0.88557334 |
| PRELID3A | 97.4244595 | -0.39811219 | 0.23797156 | -1.6729402 | 0.09433907 | 0.22833606 |
| AP001029.1 | 9.0198791 | -0.28776032 | 0.61965224 | -0.46439 | 0.64236834 | 0.78831222 |
| AP001029.2 | 1227.59139 | -0.512776971 | 0.21871137 | -2.3445374 | 0.0190507 | 0.07240804 |
| SPIRE1 | 1931.05173 | -0.264257971 | 0.25656481 | -1.0299852 | 0.30301694 | 0.4998366 |
| PSMG2 | 2596.26967 | -0.362261927 | 0.10269516 | -3.5275464 | 0.00041943 | 0.00395862 |
| CEP76 | 262.937177 | 0.004603517 | 0.16019101 | 0.02873767 | 0.97707381 | 0.98945608 |
| AP005482.2 | 0.15318056 | -1.003574173 | 3.30558948 | -0.3035992 | 0.7614333 | NA |
| AP005482.1 | 11.671608 | -1.633622271 | 0.56841687 | -2.8739863 | 0.00405327 | 0.02299089 |
| PTPN2 | 1975.43488 | -0.765967693 | 0.17524753 | -4.370776 | 1.24E-05 | 0.00024369 |
| SEH1L | 1702.31329 | -0.434913007 | 0.22563241 | -1.927529 | 0.05391374 | 0.15364804 |
| AP002449.1 | 297.825103 | -0.444518714 | 0.20514097 | -2.1668938 | 0.03024295 | 0.10141057 |
| CEP192 | 993.53727 | 0.167148297 | 0.33452436 | 0.49965956 | 0.61731481 | 0.77126014 |
| AP001198.2 | 0.17391096 | -1.003574173 | 3.30558948 | -0.3035992 | 0.7614333 | NA |
| AP001198.1 | 0.31330332 | 1.411662933 | 2.86567349 | 0.49261123 | 0.62228731 | NA |
| LDLRAD4 | 681.98977 | 0.231099861 | 0.37429029 | 0.61743482 | 0.53694796 | 0.71066642 |
| AP002505.1 | 0 | NA | NA | NA | NA | NA |
| AP002439.1 | 0 | NA | NA | NA | NA | NA |
| LDLRAD4-AS | 3.58972842 | 0.015648263 | 1.27353353 | 0.01228728 | 0.99019642 | 0.99533983 |
| AP005131.5 | 0.2908067 | 0.328535201 | 2.31399726 | 0.14197735 | 0.8870979 | NA |
| AP005131.2 | 2.68191651 | 0.308557656 | 1.10839265 | 0.27838299 | 0.78071838 | 0.88005505 |
| AP005131.1 | 0.97544246 | 1.43069942 | 1.48926789 | 0.96067298 | 0.33671662 | NA |
| AP005131.3 | 0.50914098 | -0.047757538 | 1.86378376 | -0.025624 | 0.97955727 | NA |
| AP001010.1 | 89.3896566 | -0.312019164 | 0.46311683 | -0.6737375 | 0.50047823 | 0.68155397 |
| FAM210A | 1214.99285 | -0.573671556 | 0.24551177 | -2.3366356 | 0.01945814 | 0.07340646 |
| RNMT | 2000.51634 | -0.440059586 | 0.19602794 | -2.244882 | 0.02477571 | 0.08772939 |
| MC5R | 0.27075863 | -1.364244818 | 2.94219035 | -0.4636834 | 0.64287459 | NA |
| MC2R | 0.23017671 | 0.689158974 | 3.02240241 | 0.22801695 | 0.81963307 | NA |
| ZNF519 | 111.573436 | 0.703321693 | 0.31129296 | 2.25935626 | 0.02386123 | 0.08533949 |
| AC006557.1 | 6.84691699 | 0.001889383 | 0.51336737 | 0.00368037 | 0.99706349 | 0.99850237 |
| AC006557.3 | 5.5472132 | 1.041916506 | 0.55287988 | 1.88452601 | 0.05949386 | 0.16459701 |
| POTEC | 0 | NA | NA | NA | NA | NA |
| ANKRD30B | 4.46601317 | -0.378361379 | 0.77036621 | -0.4911448 | 0.62332402 | 0.7754059 |
| AP006565.1 | 0.51749002 | -0.40522677 | 1.8879926 | -0.2146337 | 0.83005295 | NA |
| LINC01906 | 0 | NA | NA | NA | NA | NA |
| AP005121.1 | 0.34823675 | 1.594838211 | 2.1504546 | 0.7416284 | 0.45831251 | NA |
| LINC01443 | 8.67749511 | 1.129113531 | 1.04205069 | 1.08354952 | 0.27856458 | 0.47419456 |
| LINC01444 | 0.43225995 | -0.032125397 | 2.20891676 | -0.0145435 | 0.98839637 | NA |
| AP005121.2 | 0.7843359 | -0.921693167 | 2.19723068 | -0.4194795 | 0.67486575 | NA |
| AP005242.3 | 0 | NA | NA | NA | NA | NA |
| AP005242.1 | 0.26780428 | 2.026828142 | 3.24034724 | 0.6254972 | 0.53164478 | NA |
| AP005901.2 | 0.09635527 | 0.547239394 | 3.29377202 | 0.16614368 | 0.86804388 | NA |
| ROCK1 | 2425.86828 | -0.02766202 | 0.13242253 | -0.2088921 | 0.83453247 | 0.91222937 |
| AC022809.1 | 0.2839041 | 2.085287735 | 2.91926552 | 0.71431931 | 0.47502975 | NA |
| GREB1L | 69.0385228 | -0.415517782 | 0.66734342 | -0.6226446 | 0.53351809 | 0.70726282 |
| AC015878.1 | 0.10178723 | 1.087385526 | 3.28420134 | 0.33109588 | 0.74057207 | NA |
| ESCO1 | 864.441917 | -0.369408626 | 0.18896175 | -1.9549386 | 0.05059032 | 0.14682662 |
| SNRPD1 | 1272.73746 | -0.649303441 | 0.19986829 | -3.2486565 | 0.00115951 | 0.00881418 |
| ABHD3 | 1350.46317 | -0.528222359 | 0.32594982 | -1.6205634 | 0.10511131 | 0.24587822 |
| AC106037.2 | 3.10006136 | -1.024537587 | 0.75426133 | -1.3583324 | 0.17435824 | 0.34829853 |
| MIB1 | 1831.67509 | 0.049020376 | 0.22467984 | 0.21817879 | 0.82728981 | 0.90800101 |
| MIR133A1HG | 3.09204002 | -0.4181957 | 0.79401253 | -0.5266865 | 0.59841129 | 0.75673073 |
| AC103987.2 | 0.09635527 | 0.547239394 | 3.29377202 | 0.16614368 | 0.86804388 | NA |
| LINC01900 | 0.19271054 | 1.506623853 | 3.27645372 | 0.4598337 | 0.64563559 | NA |
| AC091043.1 | 0.25134551 | -1.364244527 | 2.99415014 | -0.4556366 | 0.64865129 | NA |
| GATA6-AS1 | 105.169515 | -0.569544374 | 0.44574421 | -1.2777381 | 0.20134177 | 0.38307233 |
| GATA6 | 370.476454 | -0.268664516 | 0.41095714 | -0.6537531 | 0.51327087 | 0.69122889 |
| AC091588.1 | 1.94389434 | -0.218858841 | 1.02960964 | -0.2125649 | 0.83166637 | 0.91050001 |
| AC091588.3 | 115.857464 | -0.043324569 | 0.40341327 | -0.107395 | 0.91447562 | 0.95815011 |
| AC091588.2 | 0.34145402 | -1.700893945 | 2.27276716 | -0.7483802 | 0.45423086 | NA |
| AC027449.1 | 5.45100825 | -0.032803003 | 0.83627608 | -0.0392251 | 0.96871093 | 0.98550634 |
| CTAGE1 | 2.06721094 | -0.637565146 | 0.99661796 | -0.6397287 | 0.52234897 | 0.69824821 |
| AC090912.1 | 19.3073678 | -0.707872948 | 0.50160813 | -1.4112071 | 0.15818356 | 0.32536292 |
| AC090912.2 | 13.0761988 | -1.59378738 | 0.68849731 | -2.3148781 | 0.0206196 | 0.07660452 |
| RBBP8 | 3592.91532 | -1.447515911 | 0.23532994 | -6.1510061 | 7.70E-10 | 8.29E-08 |
| CABLES1 | 2246.83981 | -0.298781733 | 0.31559845 | -0.9467148 | 0.34378412 | 0.54151789 |
| TMEM241 | 243.331008 | -0.292917838 | 0.14616445 | -2.0040293 | 0.04506693 | 0.13492448 |
| AC011731.1 | 370.989299 | -0.319559463 | 0.32160674 | -0.9936342 | 0.32040097 | 0.51786538 |
| RIOK3 | 4310.13382 | -0.231425606 | 0.20760306 | -1.1147505 | 0.26495737 | 0.45897167 |
| RMC1 | 2839.50274 | -1.053439718 | 0.28390452 | -3.7105422 | 0.00020682 | 0.00224065 |
| NPC1 | 5178.26954 | -1.617812335 | 0.44234422 | -3.6573606 | 0.00025483 | 0.00265751 |
| ANKRD29 | 319.745632 | 0.163086594 | 0.33846433 | 0.48184279 | 0.62991763 | 0.78011215 |
| LAMA3 | 528.322005 | -1.171038795 | 0.33578871 | -3.4874276 | 0.00048769 | 0.00444228 |
| AC010754.1 | 3.2790953 | -0.81440663 | 0.79299501 | -1.027001 | 0.30442002 | 0.50134623 |
| TTC39C | 369.708227 | -0.920675974 | 0.28899328 | -3.1858041 | 0.00144352 | 0.01050088 |
| TTC39C-AS1 | 12.2376781 | -0.826492035 | 0.46315504 | -1.7844824 | 0.07434531 | 0.19253563 |

| | | | | | | |
|---|---|---|---|---|---|---|
| AC090772.2 | 0.18746814 | 0.18657164 | 3.29377195 | 0.05664376 | 0.95482897 | NA |
| AC090772.1 | 3.62593684 | -0.504535202 | 0.6486635 | -0.7778073 | 0.43668262 | 0.62877147 |
| CABYR | 112.010546 | 0.113273201 | 0.28526699 | 0.39707785 | 0.69131006 | 0.82241065 |
| AC090772.3 | 316.642221 | 0.228305681 | 0.1989065 | 1.14780403 | 0.25104948 | 0.44338455 |
| OSBPL1A | 2417.46679 | 0.914795131 | 0.22356238 | 4.09190095 | 4.28E-05 | 0.00065481 |
| AC023983.1 | 1.3879204 | 0.115965116 | 1.13132429 | 0.10250387 | 0.91835674 | 0.96027072 |
| AC023983.2 | 3.51797101 | -1.161769813 | 0.84294221 | -1.3782319 | 0.16813171 | 0.33945399 |
| IMPACT | 1239.75359 | 0.890007775 | 0.19542273 | 4.55426944 | 5.26E-06 | 0.00012213 |
| HRH4 | 7.08233985 | -1.031408641 | 0.57319999 | -1.7993871 | 0.07195748 | 0.18808549 |
| AC007922.1 | 0.47345575 | -0.376019507 | 1.88982039 | -0.198971 | 0.84228541 | NA |
| AC007922.3 | 0 | NA | NA | NA | NA | NA |
| LINC01915 | 3.52441104 | 0.49373606 | 0.7748163 | 0.63722983 | 0.52397515 | 0.69914611 |
| AC022884.1 | 0 | NA | NA | NA | NA | NA |
| AC018697.1 | 0.14114194 | 0.02007176 | 3.29738704 | 0.00608717 | 0.99514317 | NA |
| LINC01894 | 0.42235908 | 0.310428605 | 2.19911269 | 0.14116084 | 0.88774288 | NA |
| AC104961.1 | 0 | NA | NA | NA | NA | NA |
| ZNF521 | 319.07712 | 0.406090577 | 0.31278372 | 1.29831111 | 0.19418045 | 0.37398031 |
| AC110603.1 | 0.10721287 | -0.642905668 | 3.30558948 | -0.1944905 | 0.84579185 | NA |
| SS18 | 4525.40376 | -0.20105718 | 0.147814 | -1.3602039 | 0.17376542 | 0.3476363 |
| PSMA8 | 1.45970573 | -1.047033326 | 1.15338399 | -0.9077925 | 0.36398787 | 0.56116501 |
| TAF4B | 300.317412 | 0.052834146 | 0.2666774 | 0.19812008 | 0.84295112 | 0.91653078 |
| LINC01543 | 70.5023175 | 1.669841588 | 0.90428202 | 1.84659381 | 0.06480603 | 0.17445233 |
| KCTD1 | 2072.61951 | -0.29101985 | 0.25554676 | -1.1388125 | 0.25478136 | 0.44753011 |
| AQP4-AS1 | 46.5005338 | 1.738630069 | 0.53422518 | 3.2544892 | 0.00113596 | 0.00867769 |
| PCAT18 | 33.3862345 | 0.926178371 | 0.39621412 | 2.33757035 | 0.01940955 | 0.07329654 |
| AC012588.1 | 16.4175462 | 0.461312499 | 0.45300489 | 1.01833888 | 0.30851694 | 0.50550464 |
| AC018371.1 | 0 | NA | NA | NA | NA | NA |
| AQP4 | 137.207079 | 0.379537018 | 0.65227376 | 0.58186768 | 0.56065581 | 0.72862652 |
| CHST9 | 184.141571 | -0.10419429 | 0.390287 | -0.2669684 | 0.7894935 | 0.88557334 |
| LINC01908 | 0 | NA | NA | NA | NA | NA |
| AC090403.1 | 0.27937425 | 1.145957498 | 3.27645355 | 0.34975545 | 0.72652223 | NA |
| AC021382.1 | 0 | NA | NA | NA | NA | NA |
| AC068408.1 | 0.12150485 | 1.04981648 | 3.28477129 | 0.31960109 | 0.74927075 | NA |
| CDH2 | 1984.49135 | 0.125332581 | 0.32937742 | 0.38051358 | 0.70356422 | 0.83059446 |
| AC110015.1 | 235.049438 | 0.069203797 | 0.3426024 | 0.20199449 | 0.83992103 | 0.91485823 |
| AC023932.1 | 0 | NA | NA | NA | NA | NA |
| AC074237.1 | 0.09989236 | -0.642905668 | 3.30558948 | -0.1944905 | 0.84579185 | NA |
| AC117569.1 | 0 | NA | NA | NA | NA | NA |
| AC090506.1 | 0 | NA | NA | NA | NA | NA |
| AC016382.1 | 0 | NA | NA | NA | NA | NA |
| DSC3 | 5.00625991 | 0.11495089 | 0.99619193 | 0.1153903 | 0.90813576 | 0.95414792 |
| DSC2 | 1247.10761 | -0.746606267 | 0.18685856 | -3.995569 | 6.45E-05 | 0.00090095 |
| DSCAS | 7.90738929 | 0.329762623 | 0.77657712 | 0.42463603 | 0.67110203 | 0.80785905 |
| DSC1 | 6.37245673 | 1.368802092 | 0.61538563 | 2.22429972 | 0.02612829 | 0.09125866 |
| AC012417.1 | 2.21020899 | 0.347757805 | 0.95836179 | 0.36286694 | 0.71670429 | 0.83885895 |
| DSG1 | 1.0122414 | 1.25553616 | 2.07223022 | 0.60588642 | 0.54459017 | 0.71643964 |
| DSG1-AS1 | 0 | NA | NA | NA | NA | NA |
| DSG4 | 0.38013148 | 0.707871477 | 2.29477746 | 0.30847064 | 0.75772424 | NA |
| DSG3 | 0.7981915 | 3.098668702 | 2.44480278 | 1.2674514 | 0.20499393 | NA |
| AC021549.1 | 2.5857539 | 0.591776316 | 1.18316503 | 0.5001638 | 0.61695975 | 0.77098041 |
| DSG2 | 2881.7792 | 0.087175474 | 0.26728763 | 0.32614855 | 0.74431196 | 0.85661007 |
| DSG2-AS1 | 93.9261045 | 0.831917897 | 0.33011407 | 2.52009223 | 0.01173241 | 0.05075191 |
| TTR | 60.5241339 | 1.748218526 | 0.65262085 | 2.67876597 | 0.0073894 | 0.03604163 |
| B4GALT6 | 339.340597 | 0.05595116 | 0.26945084 | 0.20764886 | 0.83550315 | 0.91276071 |
| AC017100.1 | 42.5920117 | 0.153929264 | 0.47081188 | 0.3269443 | 0.74371001 | 0.85622495 |
| SLC25A52 | 3.71256229 | 0.715300584 | 0.73827244 | 0.96888431 | 0.33260291 | 0.53055511 |
| TRAPPC8 | 2068.47458 | 0.098042011 | 0.14387569 | 0.68143558 | 0.49559592 | 0.67769982 |
| AC022960.1 | 0.42658574 | 2.681251852 | 2.60604169 | 1.02885992 | 0.3035455 | NA |
| AC009831.1 | 57.9632572 | 0.202948742 | 0.31157667 | 0.65136052 | 0.51481379 | 0.69256857 |
| RNF125 | 214.117173 | 0.535148634 | 0.36729967 | 1.45698097 | 0.14512163 | 0.30735701 |
| AC011825.4 | 53.7326757 | 0.035676465 | 0.33529079 | 0.10640455 | 0.91526138 | 0.95863503 |
| RNF138 | 1084.52672 | -0.330653667 | 0.16227184 | -2.0376528 | 0.04158466 | 0.12745091 |
| AC011825.2 | 7.71222103 | 0.683536162 | 0.50896904 | 1.34298181 | 0.17927787 | 0.35488621 |
| GAREM1 | 2083.63305 | -0.22421415 | 0.18060483 | -1.2414626 | 0.2144349 | 0.39857957 |
| MEP1B | 2.49840384 | -0.028461388 | 0.80653837 | -0.0352883 | 0.97184983 | 0.98670705 |
| AC015563.2 | 125.561234 | -0.077650457 | 0.27795138 | -0.279367 | 0.77996316 | 0.87952228 |
| AC021224.1 | 62.0183834 | -0.303978105 | 0.26926329 | -1.1289252 | 0.2589294 | 0.45167971 |
| AC025887.2 | 18.6495677 | 0.293419744 | 0.37104303 | 0.79079707 | 0.42906242 | 0.62112297 |
| AC025887.1 | 0.33384827 | 2.331878155 | 3.22240111 | 0.72364615 | 0.469283 | NA |
| KLHL14 | 542.130581 | 0.516570925 | 0.34824097 | 1.48337205 | 0.13797559 | 0.29677131 |
| AC012123.1 | 43.7929358 | 1.026374325 | 0.42759225 | 2.40035763 | 0.01637906 | 0.0648912 |
| AC090371.2 | 30.990822 | 0.447455567 | 0.38605172 | 1.15905601 | 0.24643336 | 0.4377238 |
| CCDC178 | 166.516186 | 0.608289668 | 0.3148627 | 1.93192038 | 0.05336933 | 0.15255819 |
| AC090371.1 | 0 | NA | NA | NA | NA | NA |
| AC091198.1 | 0.58325627 | 1.258892647 | 2.10084029 | 0.59923291 | 0.54901758 | NA |
| ASXL3 | 186.616899 | 0.621705645 | 0.24186677 | 2.57044671 | 0.01015675 | 0.04559155 |
| NOL4 | 54.6880894 | 2.662083494 | 0.6307567 | 4.22046011 | 2.44E-05 | 0.00041639 |
| AC104985.1 | 0.38440338 | -0.032130535 | 2.25801197 | -0.0142296 | 0.98864683 | NA |
| AC104985.2 | 0 | NA | NA | NA | NA | NA |
| DTNA | 395.123074 | -0.383644443 | 0.30501159 | -1.2578028 | 0.20846308 | 0.39156431 |
| AC022601.1 | 8.4899322 | 0.064860392 | 0.534391 | 0.12137254 | 0.90339597 | 0.95141786 |
| AC068506.1 | 4.40376764 | -3.241226007 | 1.4211486 | -2.2807087 | 0.02256569 | 0.0819638 |
| MAPRE2 | 1673.3025 | 0.406358967 | 0.17066197 | 2.38107513 | 0.01726219 | 0.06738286 |
| AC015967.1 | 3.93432666 | 0.961523307 | 0.64314576 | 1.49503171 | 0.13490617 | 0.29233919 |
| ZNF397 | 1495.73328 | 0.054009287 | 0.11915337 | 0.45327536 | 0.65035047 | 0.79448448 |
| ZSCAN30 | 1237.61831 | 0.134806533 | 0.21899989 | 0.61555524 | 0.5381881 | 0.71162915 |
| AC011815.2 | 33.0122583 | 0.430972419 | 0.24679178 | 1.7462997 | 0.08075889 | 0.20381011 |
| AC011815.1 | 91.2079877 | -0.095681901 | 0.33252548 | -0.2877431 | 0.77354342 | 0.8753901 |
| AC116447.1 | 14.509632 | -0.478147459 | 0.34049033 | -1.4042909 | 0.16023225 | 0.32835907 |
| ZNF24 | 3349.02913 | -0.062094024 | 0.11520184 | -0.539002 | 0.58988547 | 0.74983272 |
| ZNF396 | 315.503153 | 0.223204763 | 0.19077907 | 1.16996464 | 0.2420152 | 0.43243941 |
| AC007998.4 | 1.28696276 | -0.886786714 | 1.64348957 | -0.5395755 | 0.58948981 | 0.74961886 |
| AC007998.3 | 26.068574 | -0.77360497 | 0.52553939 | -1.4720209 | 0.14101523 | 0.30134292 |
| AC007998.2 | 196.381559 | 0.10947484 | 0.19552779 | 0.55989401 | 0.57555174 | 0.73970755 |
| INO80C | 724.864568 | 0.297336159 | 0.20234908 | 1.46942183 | 0.14171841 | 0.30242894 |
| GALNT1 | 1980.30788 | -0.380710188 | 0.19591927 | -1.9431993 | 0.05199208 | 0.14983948 |
| AC090229.1 | 15.1290515 | -1.305450426 | 0.67976666 | -1.920439 | 0.05480247 | 0.15541114 |
| AC118757.1 | 3.97284316 | 1.401598492 | 0.84328245 | 1.66207479 | 0.09649778 | 0.23183732 |
| C18orf21 | 556.86365 | -0.353074102 | 0.21040196 | -1.6780932 | 0.0933289 | 0.22641058 |
| RPRD1A | 2804.64605 | 0.027296101 | 0.09817159 | 0.27804481 | 0.78097796 | 0.88016021 |
| SLC39A6 | 3120.25853 | -0.191634198 | 0.16429797 | -1.166382 | 0.24346001 | 0.43409194 |
| ELP2 | 3079.03665 | -0.126989897 | 0.08099211 | -1.5679292 | 0.11689766 | 0.26521478 |
| AC023043.1 | 111.240286 | -0.657348681 | 0.23620973 | -2.7829027 | 0.0053875 | 0.02846489 |
| MOCOS | 331.349191 | -1.160049324 | 0.34646271 | -3.3482661 | 0.00081319 | 0.0066637 |
| FHOD3 | 1963.08458 | 1.187567976 | 0.22790621 | 5.21077509 | 1.88E-07 | 8.12E-06 |
| TPGS2 | 3562.88852 | -0.059318011 | 0.15469393 | -0.383454 | 0.70138315 | 0.82894736 |
| KIAA1328 | 344.208077 | 0.087500371 | 0.17808865 | 0.49133043 | 0.62319276 | 0.77531577 |
| AC016493.1 | 0.17059578 | -1.003574173 | 3.30558948 | -0.3035992 | 0.7614333 | NA |
| AC015961.1 | 6.17402516 | -0.635290834 | 0.53219535 | -1.1937174 | 0.23258854 | 0.42064055 |
| CELF4 | 123.521235 | -0.485841707 | 0.25958876 | -1.8715822 | 0.06126442 | 0.16783117 |
| AC015961.2 | 59.418962 | -0.169749464 | 0.24905241 | -0.6815813 | 0.49550374 | 0.67768444 |
| AC090386.1 | 0.16786415 | -1.003574173 | 3.30558948 | -0.3035992 | 0.7614333 | NA |
| AC100781.1 | 1.50013516 | 4.029569231 | 1.37893869 | 2.92222509 | 0.0034754 | 0.02047506 |

| | | | | | | |
|---|---|---|---|---|---|---|
| AC016205.1 | 1.66514764 | -0.003618732 | 1.18341512 | -0.0030579 | 0.99756018 | 0.99867691 |
| AC011139.1 | 0.48285747 | 1.386171748 | 2.20506611 | 0.62863047 | 0.52959101 | NA |
| LINC01901 | 0 | NA | NA | NA | NA | NA |
| LINC01902 | 0.03392908 | 0.380740593 | 3.29738709 | 0.11546736 | 0.90807469 | NA |
| LINC01477 | 0 | NA | NA | NA | NA | NA |
| AC021504.1 | 9.3390061 | -1.005414162 | 0.73951372 | -1.3595612 | 0.17396882 | 0.34788772 |
| PIK3C3 | 2007.83685 | -0.052254863 | 0.11863342 | -0.4404734 | 0.6595943 | 0.79999826 |
| LINC00907 | 37.2856022 | 1.323150593 | 0.40001064 | 3.30778846 | 0.00094036 | 0.00749386 |
| RIT2 | 0.12059279 | 0.02007176 | 3.29738704 | 0.00608717 | 0.99514317 | NA |
| SYT4 | 2.33017946 | 0.823063801 | 1.10507218 | 0.74480547 | 0.45638933 | 0.64586168 |
| LINC01478 | 0.39844907 | -0.543376921 | 2.76931447 | -0.1962135 | 0.84444305 | NA |
| AC120049.1 | 179.433396 | 0.27437831 | 0.28547669 | 0.96112333 | 0.33649016 | 0.53445331 |
| SETBP1 | 1442.26071 | 0.103601797 | 0.19063546 | 0.54345502 | 0.58681655 | 0.74778302 |
| AC090376.1 | 0.69534609 | 0.253650548 | 2.72222702 | 0.09317759 | 0.92576248 | NA |
| SLC14A2 | 78.3398193 | 2.362674874 | 0.82391846 | 2.86760764 | 0.00413588 | 0.023349 |
| SLC14A2-AS1 | 20.7162479 | 0.048142846 | 0.4703989 | 0.10234472 | 0.91848306 | 0.96032664 |
| AC023421.2 | 13.4409892 | 1.517872018 | 1.1798998 | 1.28644146 | 0.19828904 | 0.37923626 |
| SLC14A1 | 928.263617 | 1.828748379 | 0.86358934 | 2.11761343 | 0.03420782 | 0.11074278 |
| SIGLEC15 | 20.1506556 | 1.011806335 | 0.61734314 | 1.63896911 | 0.10121969 | 0.23953722 |
| EPG5 | 1762.64486 | 0.302701017 | 0.15634172 | 1.93614993 | 0.05284934 | 0.15144937 |
| PSTPIP2 | 450.391057 | -1.221100162 | 0.38995048 | -3.1314237 | 0.00173961 | 0.01212331 |
| ATP5F1A | 41117.3631 | 0.589837167 | 0.17033331 | 3.46284094 | 0.0005345 | 0.00478117 |
| HAUS1 | 514.067831 | -0.603571533 | 0.16164629 | -3.7339028 | 0.00018854 | 0.0020802 |
| C18orf25 | 1291.11118 | -0.16088958 | 0.14309334 | -1.124368 | 0.26085693 | 0.45404603 |
| RNF165 | 123.152419 | 0.816222639 | 0.51128013 | 1.59642943 | 0.11039295 | 0.25449125 |
| LOXHD1 | 72.601632 | -1.132905826 | 0.91078182 | -1.2438828 | 0.2135427 | 0.39755147 |
| ST8SIA5 | 299.786113 | -0.592777725 | 0.3245549 | -1.8264329 | 0.06778509 | 0.18000934 |
| AC090241.3 | 21.3908629 | 0.685227053 | 0.39533759 | 1.73327069 | 0.08304757 | 0.20799358 |
| PIAS2 | 1773.04858 | -0.116676981 | 0.15108581 | -0.7722564 | 0.43996261 | 0.63125382 |
| KATNAL2 | 527.140124 | 0.493128886 | 0.18612558 | 2.64944174 | 0.00806249 | 0.03832843 |
| ELOA3D | 0 | NA | NA | NA | NA | NA |
| ELOA3C | 0 | NA | NA | NA | NA | NA |
| ELOA3B | 0.03392908 | 0.380740593 | 3.29738709 | 0.11546736 | 0.90807469 | NA |
| ELOA3 | 0 | NA | NA | NA | NA | NA |
| ELOA2 | 1.16544491 | 0.160413077 | 1.18158486 | 0.13576094 | 0.89201027 | 0.94406449 |
| AC012254.1 | 101.0197 | 0.195365291 | 0.20449407 | 0.9553592 | 0.33939608 | 0.53756229 |
| AC012254.3 | 822.734974 | 0.304532818 | 0.2313707 | 1.31621166 | 0.18810301 | 0.36613054 |
| HDHD2 | 2383.5914 | 0.584210442 | 0.17112192 | 3.41400114 | 0.00064016 | 0.00548979 |
| AC012254.2 | 3334.29569 | 0.297161736 | 0.17698071 | 1.6790628 | 0.0931398 | 0.22605583 |
| IER3IP1 | 3873.72907 | -0.096108424 | 0.21489428 | -0.4472358 | 0.65470479 | 0.7974395 |
| SKOR2 | 0.09644756 | -0.642905668 | 3.30558948 | -0.1944905 | 0.84579185 | NA |
| MIR4527HG | 0 | NA | NA | NA | NA | NA |
| AC102797.1 | 0 | NA | NA | NA | NA | NA |
| SMAD2 | 3621.38684 | -0.235442749 | 0.11323527 | -2.0792351 | 0.03759575 | 0.11829135 |
| AC120349.1 | 4.82536096 | -1.42465095 | 0.9388569 | -1.5174314 | 0.1291578 | 0.28377696 |
| ZBTB7C | 251.482488 | 0.069056601 | 0.40045239 | 0.17244647 | 0.86308654 | 0.9286976 |
| CTIF | 3013.45691 | 0.177826917 | 0.18831477 | 0.94430682 | 0.34501288 | 0.54267758 |
| AC048380.1 | 0.20859201 | 0.689155664 | 3.09576263 | 0.22261257 | 0.82383706 | NA |
| AC093567.1 | 4.58876356 | 0.166390945 | 0.81024715 | 0.20535826 | 0.83729223 | 0.91386632 |
| SMAD7 | 2215.77316 | 0.315454549 | 0.28418383 | 1.11003692 | 0.26698312 | 0.4611392 |
| AC114684.1 | 2.94540394 | 2.278258683 | 0.7144655 | 3.18875954 | 0.00142885 | 0.01042287 |
| AC016866.1 | 11.218796 | 1.168162237 | 0.54549276 | 2.14148075 | 0.03223529 | 0.10621574 |
| DYM | 1850.60557 | 0.050149934 | 0.13343825 | 0.37582877 | 0.7070442 | 0.83325953 |
| AC016866.2 | 0 | NA | NA | NA | NA | NA |
| AC044840.1 | 1.68617539 | 0.630756796 | 1.07793278 | 0.58515411 | 0.5584441 | 0.7270388 |
| C18orf32 | 3774.93259 | -0.168053714 | 0.15426675 | -1.0893709 | 0.27599035 | 0.47133715 |
| RPL17-C18or | 23824.2349 | -0.512480304 | 0.20595686 | -2.4882895 | 0.01283592 | 0.054192 |
| AC100778.3 | 450.81524 | -0.800122307 | 0.38038792 | -2.1034377 | 0.03542752 | 0.11359725 |
| RPL17 | 21023.4168 | -0.577201049 | 0.22186232 | -2.6016181 | 0.00927851 | 0.04256759 |
| AC091170.1 | 0 | NA | NA | NA | NA | NA |
| LIPG | 2367.95282 | -2.41893476 | 0.5009563 | -4.8286343 | 1.37E-06 | 4.22E-05 |
| AC090227.2 | 0.36691442 | -1.731312147 | 2.25778977 | -0.7668173 | 0.44319018 | NA |
| ACAA2 | 17147.0011 | 0.596547652 | 0.34665886 | 1.7208493 | 0.08527817 | 0.21186992 |
| AC090227.1 | 5788.76195 | 0.552099793 | 0.31313734 | 1.76312345 | 0.07787967 | 0.19867404 |
| SNHG22 | 511.526264 | -0.26787385 | 0.25618722 | -1.0456175 | 0.29573766 | 0.492647 |
| MYO5B | 3624.68328 | -0.188455323 | 0.25863456 | -0.7286548 | 0.46621283 | 0.65341655 |
| CFAP53 | 167.130871 | 0.206588609 | 0.26668355 | 0.77465824 | 0.43854163 | 0.63025154 |
| MBD1 | 4201.30506 | -0.136719891 | 0.11741164 | -1.1644492 | 0.24424203 | 0.43529753 |
| CXXC1 | 4339.79418 | -0.11160383 | 0.15739241 | -0.7090801 | 0.47827475 | 0.66313188 |
| AC105227.1 | 0.8352341 | -0.277373463 | 1.83129912 | -0.1514627 | 0.87961075 | NA |
| SKA1 | 14.2351169 | -1.220279236 | 0.5893734 | -2.0704688 | 0.03840846 | 0.12021811 |
| MAPK4 | 431.249113 | 1.694897963 | 0.40267636 | 4.20908237 | 2.56E-05 | 0.00043288 |
| MRO | 1242.16499 | 4.842860814 | 0.62185862 | 7.78772004 | 6.82E-15 | 3.60E-12 |
| ME2 | 1647.20992 | -0.190285798 | 0.11803103 | -1.6121676 | 0.10692548 | 0.24875385 |
| ELAC1 | 450.546868 | 0.306744885 | 0.25209918 | 1.21676272 | 0.22369451 | 0.40978002 |
| AC091551.1 | 509.881115 | 0.699080415 | 0.25387099 | 2.75368377 | 0.00589287 | 0.03039212 |
| SMAD4 | 4060.69379 | -0.052317437 | 0.13055914 | -0.4007183 | 0.68862753 | 0.82026867 |
| MEX3C | 1501.64833 | -0.448213528 | 0.16997075 | -2.6370038 | 0.00836419 | 0.03935123 |
| LINC01630 | 0.46593032 | 0.396282279 | 2.79403232 | 0.14183167 | 0.88721297 | NA |
| AC016383.1 | 0.57567565 | 2.512699208 | 2.58327049 | 0.97268142 | 0.33071166 | NA |
| DCC | 24.269983 | 2.931477012 | 0.55209714 | 5.30971235 | 1.10E-07 | 5.19E-06 |
| AC103949.1 | 5.73313411 | 1.591719284 | 0.61135357 | 2.60359859 | 0.00922507 | 0.04237331 |
| LINC01919 | 0.90296326 | -1.467565106 | 1.67755235 | -0.8748252 | 0.38166901 | NA |
| LINC01917 | 0.86636268 | 0.587464233 | 1.48231121 | 0.39631639 | 0.69187163 | NA |
| MBD2 | 4433.96302 | -0.747046129 | 0.16986538 | -4.3978716 | 1.09E-05 | 0.00021908 |
| AC093462.1 | 58.6404479 | 0.346736172 | 0.32752438 | 1.05865759 | 0.28975575 | 0.48593234 |
| POLI | 1750.07809 | 0.157992924 | 0.165994 | 0.95179898 | 0.34119894 | 0.53921947 |
| STARD6 | 2.05603796 | 0.470746025 | 0.89440019 | 0.52632595 | 0.59866176 | 0.75677743 |
| C18orf54 | 103.609727 | -0.099823466 | 0.2269747 | -0.4398 | 0.660082 | 0.79999826 |
| DYNAP | 0.87011363 | -2.254440526 | 1.62372636 | -1.3884362 | 0.16500425 | NA |
| RAB27B | 44.7058876 | -0.756492272 | 0.44169814 | -1.7126907 | 0.08676947 | 0.21462441 |
| AC007673.1 | 0 | NA | NA | NA | NA | NA |
| AC098848.1 | 2.77410323 | -2.099297119 | 1.05329788 | -1.9930707 | 0.04625371 | 0.13760269 |
| CCDC68 | 279.339774 | 0.278783183 | 0.32305435 | 0.86296063 | 0.38815911 | 0.58379945 |
| LINC01929 | 3.16057188 | 0.171163966 | 0.92239496 | 0.18556472 | 0.85278612 | 0.92285671 |
| TCF4 | 1325.35363 | -0.224719793 | 0.2710607 | -0.8290387 | 0.40708254 | 0.60169085 |
| TCF4-AS1 | 0.16325399 | -1.003574173 | 3.30558948 | -0.3035992 | 0.7614333 | NA |
| TCF4-AS2 | 0 | NA | NA | NA | NA | NA |
| AC022031.2 | 0.31713219 | 0.328539074 | 2.27448921 | 0.14444521 | 0.88514892 | NA |
| LINC01415 | 3.34731038 | -0.34387414 | 1.31417973 | -0.2616645 | 0.79358014 | 0.88835379 |
| AC022031.1 | 0 | NA | NA | NA | NA | NA |
| LINC01416 | 1.35338977 | 0.861371557 | 1.31598795 | 0.65454365 | 0.51276161 | 0.69088659 |
| AC009271.1 | 2.55133196 | 0.423197281 | 0.74379789 | 0.56896811 | 0.56937778 | 0.73485181 |
| AC006305.1 | 45.2942899 | 2.701362534 | 0.56384766 | 4.79094392 | 1.66E-06 | 4.90E-05 |
| LINC01905 | 2.15930974 | 0.534620731 | 0.827729 | 0.6458886 | 0.5183515 | 0.69530036 |
| AC006305.2 | 2.34875835 | 3.914440351 | 1.37490673 | 2.84705884 | 0.00441252 | 0.02452744 |
| LINC01539 | 10.660946 | 2.376317901 | 0.76104543 | 3.12243898 | 0.00179359 | 0.01240635 |
| AC006305.3 | 1.84366962 | 3.073028325 | 1.02114628 | 3.00939092 | 0.00261772 | 0.01655574 |
| TXNL1 | 3271.11694 | -0.109234938 | 0.18010809 | -0.6064966 | 0.54418505 | 0.71620188 |
| WDR7 | 754.423861 | 0.356464495 | 0.08898193 | 4.00603243 | 6.17E-05 | 0.00086836 |
| AC012301.1 | 0 | NA | NA | NA | NA | NA |
| LINC-ROR | 0.54946107 | 1.135929236 | 1.5373479 | 0.73888886 | 0.45997449 | NA |
| AC100775.1 | 0 | NA | NA | NA | NA | NA |

| | | | | | | |
|---|---|---|---|---|---|---|
| BOD1L2 | 0.2874412 | 1.099444791 | 2.9528429 | 0.37233433 | 0.70964394 | NA |
| LINC02565 | 0 | NA | NA | NA | NA | NA |
| ST8SIA3 | 0 | NA | NA | NA | NA | NA |
| AC090340.1 | 0.10721287 | -0.642905668 | 3.30558948 | -0.1944905 | 0.84579185 | NA |
| ONECUT2 | 15.0417389 | -0.307087222 | 0.54751812 | -0.5608713 | 0.57488525 | 0.73915805 |
| FECH | 2580.40986 | 1.370442496 | 0.23681017 | 5.78709297 | 7.16E-09 | 5.20E-07 |
| AC100847.1 | 29.225861 | -0.29688859 | 0.52138031 | -0.5694281 | 0.56906567 | 0.73454422 |
| NARS | 11017.8634 | -0.119713637 | 0.1961297 | -0.61038 | 0.54161014 | 0.71436299 |
| AC027097.1 | 166.136507 | -0.144136983 | 0.22739983 | -0.6338483 | 0.52617986 | 0.70109458 |
| AC027097.2 | 157.740255 | -0.206605429 | 0.28226916 | -0.7319448 | 0.46420229 | 0.65178001 |
| ATP8B1 | 1124.95512 | -0.102761858 | 0.24038785 | -0.4274836 | 0.66902715 | 0.80665017 |
| LINC01897 | 0.96762627 | 3.884437175 | 2.17063935 | 1.78953596 | 0.07352854 | NA |
| NEDD4L | 6455.95311 | 0.192156237 | 0.38156426 | 0.50360125 | 0.61454161 | 0.7694536 |
| AC107896.1 | 12.7953696 | 1.37308636 | 0.65723902 | 2.08917352 | 0.03669211 | 0.1163645 |
| AC090236.1 | 524.196117 | 1.613183577 | 0.3211045 | 5.02385851 | 5.06E-07 | 1.83E-05 |
| AC105105.2 | 0.09989236 | -0.642905668 | 3.30558948 | -0.1944905 | 0.84579185 | NA |
| MIR122HG | 7.34055714 | 0.867164033 | 0.65156996 | 1.33088401 | 0.18322718 | 0.35998366 |
| AC105105.4 | 0 | NA | NA | NA | NA | NA |
| ALPK2 | 1518.47025 | 1.331659107 | 0.37158821 | 3.58369581 | 0.00033877 | 0.00333954 |
| AC105105.1 | 211.119726 | 2.182740591 | 0.47779 | 4.56840998 | 4.91E-06 | 0.00011593 |
| AC104971.2 | 0.2710182 | 1.145957498 | 3.27645355 | 0.34975545 | 0.72652223 | NA |
| AC104971.3 | 0.7910714 | -0.031165896 | 1.62829006 | -0.0191403 | 0.98472921 | NA |
| MALT1 | 2139.32288 | -0.02944172 | 0.21457836 | -0.1372073 | 0.89086693 | 0.9433845 |
| AC104365.1 | 33.4371093 | -0.148994445 | 0.34353837 | -0.4337054 | 0.66450238 | 0.80299367 |
| AC104365.3 | 163.163179 | -0.021415781 | 0.25745349 | -0.0831831 | 0.93370594 | 0.96806738 |
| AC104365.2 | 10.1817831 | 0.377978057 | 0.69250125 | 0.54581571 | 0.58519263 | 0.74685098 |
| LINC01926 | 0.80802063 | -0.848410485 | 2.75745626 | -0.3076787 | 0.75832685 | NA |
| ZNF532 | 1838.95053 | -0.247942289 | 0.18992057 | -1.3055052 | 0.19172087 | 0.3709347 |
| AC040963.1 | 0 | NA | NA | NA | NA | NA |
| SEC11C | 2580.01303 | -0.217008089 | 0.24658966 | -0.8800372 | 0.37883913 | 0.57489716 |
| GRP | 0.39658274 | 0.231398012 | 3.28103693 | 0.07052588 | 0.9437751 | NA |
| RAX | 1.13089107 | -3.379746138 | 2.14160408 | -1.5781377 | 0.11453398 | 0.26149446 |
| CPLX4 | 1.34322908 | -1.123131271 | 1.26671 | -0.8866522 | 0.37526615 | 0.57129724 |
| LMAN1 | 6519.2215 | 0.109367087 | 0.16426468 | 0.66579793 | 0.50554027 | 0.68476972 |
| AC016229.2 | 0.86063168 | -1.711102137 | 1.48193982 | -1.1546367 | 0.24823922 | NA |
| CCBE1 | 308.350511 | 0.541126241 | 0.31872093 | 1.69780578 | 0.08954443 | 0.21981814 |
| AC016229.1 | 0.14616582 | 0.435199075 | 3.28898345 | 0.13232024 | 0.894731 | NA |
| AC090213.1 | 0.18692855 | 0.18657164 | 3.29377195 | 0.05664376 | 0.95482897 | NA |
| AC010776.1 | 0.08695548 | -0.642905668 | 3.30558948 | -0.1944905 | 0.84579185 | NA |
| AC010776.2 | 21.6928284 | 0.547563755 | 0.54459078 | 1.00545911 | 0.31467583 | 0.51173087 |
| AC010776.3 | 2.81786381 | 0.557659228 | 0.76840975 | 0.72573159 | 0.46800333 | 0.65481233 |
| PMAIP1 | 226.923643 | -2.096353962 | 0.69691945 | -3.0080291 | 0.00262948 | 0.01660028 |
| AC090377.1 | 0.46205694 | 0.345619465 | 2.28268788 | 0.15140899 | 0.8796531 | NA |
| AC090771.2 | 0.10721287 | -0.642905668 | 3.30558948 | -0.1944905 | 0.84579185 | NA |
| AC090621.1 | 0 | NA | NA | NA | NA | NA |
| MC4R | 1.17936132 | -1.541479405 | 1.89082058 | -0.8152436 | 0.41493288 | 0.60849924 |
| AC113137.1 | 0.19809512 | 0.689153972 | 3.13478683 | 0.21984078 | 0.82599516 | NA |
| CDH20 | 21.2922558 | 0.933602326 | 0.32408284 | 2.88075207 | 0.00396728 | 0.02262969 |
| AC090409.2 | 6.44308778 | 1.662069112 | 0.54728145 | 3.03695494 | 0.00238981 | 0.0154586 |
| AC090409.1 | 58.5540703 | 0.272137929 | 0.30613596 | 0.88894467 | 0.37403282 | 0.57018461 |
| AC105094.2 | 28.0311946 | 2.034047742 | 0.32448247 | 6.26859064 | 3.64E-10 | 4.31E-08 |
| AC105094.1 | 0.36054175 | 2.45020282 | 2.65903065 | 0.92146468 | 0.35680787 | NA |
| LINC01544 | 37.8361414 | 2.927733445 | 0.62254584 | 4.70283995 | 2.57E-06 | 6.94E-05 |
| RNF152 | 6505.20635 | 1.394296313 | 0.44519764 | 3.13185914 | 0.00173703 | 0.01211251 |
| PIGN | 979.437405 | 0.107314312 | 0.16180936 | 0.66321447 | 0.5071932 | 0.68619798 |
| KIAA1468 | 1713.5502 | 0.109865877 | 0.13075522 | 0.84024085 | 0.40077336 | 0.59557022 |
| AC027514.2 | 0.84933887 | 0.022012476 | 1.55911001 | 0.01411862 | 0.98873535 | NA |
| TNFRSF11A | 89.5659461 | -0.136690627 | 0.40847205 | -0.3346389 | 0.73789752 | 0.85283548 |
| AC100843.1 | 0 | NA | NA | NA | NA | NA |
| ZCCHC2 | 1152.31283 | -0.688223885 | 0.22752765 | -3.0247923 | 0.00248804 | 0.01593626 |
| AC064801.1 | 9.04122936 | 0.729561242 | 0.44301965 | 1.64679207 | 0.09960079 | 0.23681119 |
| PHLPP1 | 685.68618 | 0.174307659 | 0.17530499 | 0.99431086 | 0.32007155 | 0.5175173 |
| BCL2 | 1249.16642 | 0.698208886 | 0.17442723 | 4.00286637 | 6.26E-05 | 0.00087778 |
| AC022726.2 | 0.55006129 | 0.959826378 | 2.20276589 | 0.4357369 | 0.66302763 | NA |
| KDSR | 3959.93532 | -0.761135697 | 0.27023782 | -2.8165403 | 0.0048544 | 0.0262945 |
| AC036176.1 | 84.9308058 | 0.482143417 | 0.31598489 | 1.52584329 | 0.1270489 | 0.28057117 |
| VPS4B | 2479.68418 | -0.109504098 | 0.18502875 | -0.5918221 | 0.55396974 | 0.72353552 |
| SERPINB5 | 0.716155 | 1.740189115 | 2.59969438 | 0.66938219 | 0.50325171 | NA |
| SERPINB12 | 0.03392908 | 0.380740593 | 3.29738709 | 0.11546736 | 0.90807469 | NA |
| SERPINB13 | 0 | NA | NA | NA | NA | NA |
| SERPINB4 | 8.09296035 | -3.912492254 | 1.69570641 | -2.3072934 | 0.02103847 | 0.07777715 |
| SERPINB11 | 4.08598144 | -0.39194568 | 1.65498657 | -0.2368271 | 0.81279092 | 0.89895498 |
| SERPINB3 | 3.34125696 | 0.235150351 | 1.43540141 | 0.16382201 | 0.86987126 | 0.93220007 |
| SERPINB7 | 93.8383644 | -1.249254016 | 0.95038217 | -1.3144754 | 0.18868626 | 0.36687944 |
| SERPINB2 | 6.37875643 | -1.285073631 | 0.70941761 | -1.8114487 | 0.07007142 | 0.18437271 |
| SERPINB10 | 7.86847849 | -1.936123442 | 0.90122 | -2.1483361 | 0.03168707 | 0.10493381 |
| HMSD | 176.698489 | -2.218759983 | 0.60503674 | -3.6671492 | 0.00024527 | 0.00258297 |
| AC009802.1 | 54.5499488 | -0.095175175 | 0.42308394 | -0.2249558 | 0.82201368 | 0.90490334 |
| SERPINB8 | 971.097568 | -2.287581252 | 0.49999606 | -4.5751986 | 4.76E-06 | 0.00011329 |
| LINC00305 | 2.63198685 | -1.875243408 | 1.7246823 | -1.0872979 | 0.27690519 | 0.47229668 |
| LINC01924 | 3.83944927 | -2.55090383 | 1.08252837 | -2.3564314 | 0.01845148 | 0.07077315 |
| AC100848.1 | 0.27637501 | -1.364242679 | 3.30558948 | -0.4127078 | 0.67982068 | NA |
| LINC01538 | 0.8597873 | 2.137400233 | 1.61833961 | 1.32073652 | 0.18658923 | NA |
| AC007948.1 | 0 | NA | NA | NA | NA | NA |
| AC007631.1 | 0 | NA | NA | NA | NA | NA |
| LINC01916 | 0.2838234 | 1.099696404 | 2.96053436 | 0.371452 | 0.7103009 | NA |
| AC090358.1 | 0 | NA | NA | NA | NA | NA |
| CDH7 | 1.00378435 | 2.41311495 | 1.19408887 | 2.02088388 | 0.04329179 | NA |
| AC023394.1 | 0 | NA | NA | NA | NA | NA |
| CDH19 | 50.5666337 | 0.332884938 | 0.37315571 | 0.89208051 | 0.37234978 | 0.56878538 |
| AC110597.1 | 43.8373383 | 1.136322708 | 0.54627452 | 2.08013125 | 0.0375135 | 0.11807482 |
| DSEL | 574.455224 | 0.95291243 | 0.27758808 | 3.43282903 | 0.00059732 | 0.00520642 |
| AC110597.3 | 190.410431 | 0.701204417 | 0.27066217 | 2.59069975 | 0.0095781 | 0.04363008 |
| AC114689.3 | 4.69546198 | 0.758203671 | 0.72448753 | 1.04653792 | 0.29531275 | 0.49212571 |
| AC022655.1 | 0 | NA | NA | NA | NA | NA |
| LINC01903 | 0 | NA | NA | NA | NA | NA |
| AC100844.1 | 0 | NA | NA | NA | NA | NA |
| AC068112.1 | 0 | NA | NA | NA | NA | NA |
| LINC01912 | 0 | NA | NA | NA | NA | NA |
| AC005909.1 | 0 | NA | NA | NA | NA | NA |
| TMX3 | 2566.43186 | 0.041973951 | 0.13615706 | 0.30827598 | 0.75787234 | 0.86613044 |
| CCDC102B | 226.121358 | 0.371927261 | 0.31556749 | 1.17859814 | 0.23855823 | 0.42808987 |
| AC022035.1 | 9.28192916 | 0.999925957 | 0.4676894 | 2.13801289 | 0.0325157 | 0.10677944 |
| AC096708.2 | 0.11786115 | 0.02007176 | 3.29738704 | 0.00608717 | 0.99514317 | NA |
| AC090337.1 | 17.1780567 | -0.231105265 | 0.51236323 | -0.4510575 | 0.65194812 | 0.79555169 |
| AC026585.1 | 1.49151811 | 1.154516384 | 1.05490474 | 1.09442715 | 0.27376768 | 0.46881316 |
| DOK6 | 422.265678 | 0.778872628 | 0.26833721 | 2.90258893 | 0.00370092 | 0.02149632 |
| AC119868.2 | 7.04911996 | 0.198726878 | 0.5994365 | 0.33152282 | 0.74024961 | 0.85420604 |
| AC068254.1 | 0.08757577 | 0.547239394 | 3.29377202 | 0.16614368 | 0.86804388 | NA |
| CD226 | 162.128455 | 0.914704464 | 0.39168237 | 2.33532201 | 0.01952661 | 0.07362262 |
| RTTN | 535.325963 | 0.910131047 | 0.23744123 | 3.83307927 | 0.00012655 | 0.00152379 |
| SOCS6 | 1170.44974 | -0.138037268 | 0.19251128 | -0.7170347 | 0.47335269 | 0.65935822 |
| LINC01909 | 24.1610537 | 0.046163647 | 0.42004869 | 0.1099007 | 0.91248812 | 0.95698027 |

| | | | | | | |
|---|---|---|---|---|---|---|
| LIVAR | 12.6496581 | -1.313364798 | 0.67251854 | -1.952905 | 0.05083086 | 0.14733014 |
| LINC01910 | 0.37494809 | -1.722775267 | 2.8042968 | -0.6143341 | 0.53899455 | NA |
| GTSCR1 | 0.24300969 | 1.904561295 | 3.24861966 | 0.58626786 | 0.55769553 | NA |
| AC090415.2 | 0 | NA | NA | NA | NA | NA |
| AC091691.2 | 0 | NA | NA | NA | NA | NA |
| LINC01541 | 12.7379298 | -1.392357179 | 0.89609904 | -1.5537983 | 0.12023253 | 0.27029257 |
| LINC01899 | 0.28871938 | -0.695500688 | 2.34733066 | -0.2962943 | 0.76700534 | NA |
| AC027458.1 | 29.9577047 | 4.092587358 | 0.72911322 | 5.61310267 | 1.99E-08 | 1.24E-06 |
| AC069114.1 | 0 | NA | NA | NA | NA | NA |
| CBLN2 | 66.7194466 | 3.061561885 | 0.8767295 | 3.49202561 | NA | NA |
| NETO1 | 10.2406009 | -0.883204994 | 0.93846621 | -0.9411154 | 0.34664573 | 0.54425278 |
| AC091138.1 | 1.36208388 | -2.790285997 | 1.37512118 | -2.0291201 | 0.04244606 | 0.12941411 |
| AC079070.1 | 0 | NA | NA | NA | NA | NA |
| AC090125.1 | 0.06785815 | 0.795868262 | 3.28898349 | 0.24198001 | 0.80879565 | NA |
| FBXO15 | 146.627222 | 0.912137839 | 0.43692368 | 2.08763656 | 0.03683063 | 0.11662149 |
| TIMM21 | 1314.26263 | 0.318328229 | 0.16759699 | 1.89936719 | 0.05751621 | 0.16071872 |
| AC090398.1 | 0.20179323 | -0.340597073 | 3.29738699 | -0.103293 | 0.91773041 | NA |
| CYB5A | 15260.2546 | 0.72119054 | 0.31089101 | 2.31975362 | 0.02035421 | 0.07585366 |
| C18orf63 | 2.29162628 | 2.367163182 | 1.04697315 | 2.26095882 | 0.02376181 | 0.08504775 |
| LINC01922 | 0 | NA | NA | NA | NA | NA |
| FAM69C | 3.50831776 | 1.624510294 | 0.84796121 | 1.91578373 | 0.05539263 | 0.15646856 |
| CNDP2 | 37772.3233 | -0.00637552 | 0.33666416 | -0.0189373 | 0.9848911 | 0.99347171 |
| CNDP1 | 165.825213 | 2.07379485 | 0.47930585 | 4.3266629 | 1.51E-05 | 0.00028576 |
| AC009704.1 | 9.23750172 | 2.224956121 | 0.51532643 | 4.31756645 | 1.58E-05 | 0.00029483 |
| LINC00909 | 614.761292 | 0.359690887 | 0.23065299 | 1.559446 | 0.11889086 | 0.26814505 |
| ZNF407 | 511.300376 | 0.131882396 | 0.17923285 | 0.735816 | 0.46184269 | 0.65014385 |
| AC015819.2 | 4.39779759 | -0.559118254 | 0.62118225 | -0.9000873 | 0.36807379 | 0.56489965 |
| AC015819.1 | 6.73533263 | -0.176563619 | 0.65407653 | -0.2699434 | 0.78720383 | 0.88414299 |
| ZADH2 | 2363.5384 | 0.324105009 | 0.12943906 | 2.5039197 | 0.01228259 | 0.05247763 |
| TSHZ1 | 1719.55041 | 0.611894791 | 0.20163622 | 3.03464725 | 0.00240817 | 0.01554162 |
| AC116003.3 | 1.62380559 | -1.178389525 | 1.08902736 | -1.0820569 | 0.27922726 | 0.47486664 |
| AC116003.1 | 0.71419994 | 2.829398736 | 1.66625136 | 1.69806237 | 0.08949599 | NA |
| SMIM21 | 2.04787039 | 1.191275217 | 0.91478563 | 1.30224522 | 0.19283257 | 0.37233616 |
| AC116003.2 | 0.20908062 | 1.675937956 | 3.26598797 | 0.51314885 | 0.60784721 | NA |
| LINC01898 | 0 | NA | NA | NA | NA | NA |
| AC021506.1 | 0.43926382 | 2.777935361 | 3.20192849 | 0.86758195 | 0.38562323 | NA |
| AC090457.1 | 0 | NA | NA | NA | NA | NA |
| AC103808.6 | 0.70096999 | -0.056787716 | 1.80918309 | -0.0313886 | 0.97495964 | NA |
| LINC01893 | 0.07830766 | -0.642905668 | 3.30558948 | -0.1944905 | 0.84579185 | NA |
| AC103808.2 | 0 | NA | NA | NA | NA | NA |
| AC103808.4 | 0 | NA | NA | NA | NA | NA |
| AC103808.3 | 0.11051935 | 0.02007176 | 3.29738704 | 0.00608717 | 0.99514317 | NA |
| AC103808.1 | 0.03392908 | 0.380740593 | 3.29738709 | 0.11546736 | 0.90807469 | NA |
| AC103808.5 | 0.11051935 | 0.02007176 | 3.29738704 | 0.00608717 | 0.99514317 | NA |
| ZNF516 | 1498.06064 | -0.322970265 | 0.20951124 | -1.5415415 | 0.12318506 | 0.2746124 |
| AC009716.1 | 2.72587421 | -0.035940271 | 0.78457503 | -0.0458086 | 0.96346282 | 0.98291994 |
| AC018413.1 | 68.5542262 | -0.111685739 | 0.22511126 | -0.4961357 | 0.61979866 | 0.77271053 |
| C18orf65 | 12.3685713 | -1.510826677 | 0.49801537 | -3.0336949 | 0.00241579 | 0.01557601 |
| LINC00908 | 34.5964457 | -0.412688222 | 0.40780025 | -1.0119862 | 0.31154466 | 0.50869552 |
| LINC00683 | 5.06798201 | 0.606833935 | 0.82151592 | 0.73867581 | 0.46010388 | 0.64871621 |
| LINC01927 | 3.08333453 | 0.428612992 | 0.76320768 | 0.56159418 | 0.57439255 | 0.73879287 |
| LINC01879 | 1.13220054 | 0.399661679 | 1.91489929 | 0.20871159 | 0.83467339 | 0.91223215 |
| ZNF236-DT | 57.2879271 | 0.341128719 | 0.21361503 | 1.59693222 | 0.11028082 | 0.25431615 |
| AC027575.2 | 23.7867372 | -0.210959392 | 0.36100325 | -0.5843698 | 0.55897155 | 0.72754589 |
| ZNF236 | 509.527181 | 0.28204895 | 0.1824695 | 1.54573204 | 0.1221693 | 0.27307983 |
| AC093330.1 | 76.3023693 | -0.96467665 | 0.36091447 | -2.6728678 | 0.00752059 | 0.03652937 |
| MBP | 3072.04177 | -0.459747495 | 0.30435914 | -1.5105427 | 0.13090499 | 0.28647009 |
| AC018529.1 | 3.61237799 | 1.561541592 | 0.67358415 | 2.31825761 | 0.02043532 | 0.07610218 |
| AC018529.2 | 58.1807091 | -0.915711508 | 0.59711613 | -1.5335568 | 0.12513872 | 0.27748974 |
| AC100863.1 | 0.34809709 | 1.665718184 | 2.84122255 | 0.58626811 | 0.55769536 | NA |
| GALR1 | 21.4464396 | -0.065854643 | 0.92706595 | -0.0710356 | 0.94336946 | 0.97315146 |
| AC124254.1 | 2.10538252 | -1.900956879 | 1.34977485 | -1.4083511 | 0.15902713 | 0.32654925 |
| AC123786.1 | 0.18380315 | -1.003574173 | 3.30558948 | -0.3035992 | 0.7614333 | NA |
| LINC01029 | 0.10721287 | -0.642905668 | 3.30558948 | -0.1944905 | 0.84579185 | NA |
| AC134978.1 | 0 | NA | NA | NA | NA | NA |
| AC107892.1 | 0 | NA | NA | NA | NA | NA |
| AC087399.1 | 0 | NA | NA | NA | NA | NA |
| AC012572.1 | 0 | NA | NA | NA | NA | NA |
| AC044873.1 | 0 | NA | NA | NA | NA | NA |
| AC091027.2 | 0 | NA | NA | NA | NA | NA |
| AC091027.1 | 0 | NA | NA | NA | NA | NA |
| LINC01896 | 201.614568 | 1.301216133 | 0.83678564 | 1.55501729 | 0.11994195 | 0.26984637 |
| SALL3 | 162.689001 | 1.224243581 | 0.73649379 | 1.66225921 | 0.09646081 | 0.2317934 |
| ATP9B | 1082.16273 | 0.549232236 | 0.15359193 | 3.57591849 | 0.000349 | 0.00342124 |
| AC104423.1 | 1.61531625 | -0.937172862 | 1.00526214 | -0.9322671 | 0.35119848 | 0.54887121 |
| AC023090.1 | 0.48046688 | 0.652012895 | 1.65316753 | 0.39440219 | 0.69328413 | NA |
| NFATC1 | 654.856885 | -0.109091093 | 0.30624795 | -0.3562182 | 0.72167717 | 0.84180481 |
| AC018445.1 | 0.55328291 | -1.272652786 | 1.88832486 | -0.6739586 | 0.50033763 | NA |
| AC068473.4 | 2.93107628 | 1.934438519 | 0.86585212 | 2.23414424 | 0.02547358 | 0.08950545 |
| AC068473.1 | 3.14398419 | 0.950733713 | 0.74193033 | 1.2814326 | 0.20004176 | 0.38143769 |
| AC068473.5 | 384.041437 | 0.370294421 | 0.26868107 | 1.3781932 | 0.16814365 | 0.33945399 |
| CTDP1 | 1816.11647 | -0.429358443 | 0.1634969 | -2.6260953 | 0.00863706 | 0.0403114 |
| AC068473.2 | 0.17656334 | 0.18657164 | 3.29377195 | 0.05664376 | 0.95482897 | NA |
| AC021594.1 | 0.27637501 | -1.364242679 | 3.30558948 | -0.4127078 | 0.67982068 | NA |
| AC021594.2 | 1.61433816 | 0.38070008 | 1.07324702 | 0.35471804 | 0.72280083 | 0.84270307 |
| KCNG2 | 25.1908787 | 0.361981626 | 0.75990174 | 0.4763532 | 0.63382277 | 0.78295381 |
| AC114341.1 | 8.58886018 | -1.282920634 | 0.64814925 | -1.9793599 | 0.0477755 | 0.14092572 |
| PQLC1 | 5316.36831 | -0.656102914 | 0.28415685 | -2.3089463 | 0.02094656 | 0.07750257 |
| HSBP1L1 | 2242.46011 | -0.872100579 | 0.22470149 | -3.8811518 | 0.00010396 | 0.0013061 |
| TXNL4A | 2286.9632 | -0.363981706 | 0.19747733 | -1.8431569 | 0.06530611 | 0.17536884 |
| AC090360.1 | 695.702667 | 0.04497386 | 0.10546436 | 0.42643655 | 0.66978977 | 0.80715607 |
| RBFA | 893.726848 | 0.015059934 | 0.09173995 | 0.16415895 | 0.86960602 | 0.93213734 |
| RBFADN | 143.304645 | 0.526332582 | 0.37078063 | 1.41952557 | 0.15574585 | 0.32211906 |
| ADNP2 | 1321.33802 | -0.045126323 | 0.19377765 | -0.2328768 | 0.81585706 | 0.90085213 |
| PARD6G-AS1 | 168.943857 | -0.289420109 | 0.35642899 | -0.8119994 | 0.416792 | 0.60986792 |
| PARD6G | 457.558353 | 0.538521415 | 0.26049995 | 2.06726109 | 0.03870956 | 0.12087283 |
| AC139100.1 | 67.3720705 | 0.615547981 | 0.25993389 | 2.3680944 | 0.01787997 | 0.06902282 |
| DEFB125 | 0 | NA | NA | NA | NA | NA |
| DEFB126 | 0.2890658 | 2.115835529 | 3.23473121 | 0.65409933 | 0.51304781 | NA |
| DEFB127 | 0.57381375 | 1.933146219 | 2.01795501 | 0.9579729 | 0.33807642 | NA |
| DEFB128 | 0 | NA | NA | NA | NA | NA |
| DEFB129 | 0 | NA | NA | NA | NA | NA |
| DEFB132 | 3.46410006 | 2.258016535 | 1.3748718 | 1.64234697 | 0.10051812 | 0.23843473 |
| AL034548.1 | 0 | NA | NA | NA | NA | NA |
| C20orf96 | 979.612232 | -0.166696255 | 0.21802663 | -0.7645683 | 0.44452864 | 0.63537357 |
| ZCCHC3 | 1369.98194 | -0.390180587 | 0.123399 | -3.161943 | 0.0015672 | 0.01117856 |
| NRSN2-AS1 | 225.779979 | 0.171574995 | 0.23094046 | 0.74294038 | 0.45751778 | 0.64676468 |
| SOX12 | 2136.73991 | -0.507972052 | 0.23250016 | -2.1848245 | 0.02890172 | 0.09823902 |
| NRSN2 | 2975.46863 | -0.731442722 | 0.20805192 | -3.5156739 | 0.00043864 | 0.00409884 |
| TRIB3 | 823.754882 | -2.673584342 | 0.47178992 | -5.6668959 | 1.45E-08 | 9.49E-07 |
| RBCK1 | 7608.93424 | -0.844624253 | 0.26475397 | -3.1902231 | 0.00142163 | 0.01037884 |
| TBC1D20 | 3141.21914 | -0.310281036 | 0.2652819 | -1.1696276 | 0.24215087 | 0.43253528 |
| CSNK2A1 | 5547.38059 | -0.310613354 | 0.22644657 | -1.371685 | 0.17016154 | 0.34211343 |

| | | | | | | |
|---|---|---|---|---|---|---|
| TCF15 | 41.2775112 | -0.426967016 | 0.52340905 | -0.8157425 | 0.41464741 | 0.60825557 |
| SRXN1 | 6480.20848 | -2.47400661 | 0.67560507 | -3.6619124 | 0.00025034 | 0.0026228 |
| AL121758.1 | 6469.97525 | -2.472791802 | 0.67599196 | -3.6580196 | 0.00025417 | 0.00265341 |
| SCRT2 | 0.17361919 | -1.003574173 | 3.30558948 | -0.3035992 | 0.7614333 | NA |
| SLC52A3 | 1379.67814 | -1.107372177 | 0.41064303 | -2.6966783 | 0.00700349 | 0.03464303 |
| FAM110A | 488.287108 | -0.600662471 | 0.27455678 | -2.1877532 | 0.02868759 | 0.09775057 |
| ANGPT4 | 48.863108 | 0.05972632 | 0.50109337 | 0.119192 | 0.90512325 | 0.95240014 |
| RSPO4 | 76.8501999 | 0.799612638 | 0.52616789 | 1.51969105 | 0.12858864 | 0.28280921 |
| PSMF1 | 5558.36896 | -0.107337842 | 0.13073087 | -0.8210596 | 0.41161229 | 0.60570403 |
| AL031665.2 | 8.85280585 | 0.143528875 | 0.44423548 | 0.32309187 | 0.74662566 | 0.85807447 |
| TMEM74B | 180.191942 | -0.502191937 | 0.37315931 | -1.3457843 | 0.17837209 | 0.35379256 |
| C20orf202 | 16.0873702 | 0.361702898 | 0.28816599 | 1.25518941 | 0.20941001 | 0.39256003 |
| RAD21L1 | 12.3992067 | 0.969201858 | 0.41756954 | 2.32105499 | 0.02028388 | 0.07566575 |
| SNPH | 453.579752 | 0.125763538 | 0.24791926 | 0.50727618 | 0.61196105 | 0.76758665 |
| SDCBP2 | 827.953886 | -2.468848574 | 0.34217854 | -7.2150889 | 5.39E-13 | 1.60E-10 |
| AL136531.2 | 2009.65685 | -0.889135057 | 0.23693064 | -3.7527229 | 0.00017492 | 0.00197126 |
| SDCBP2-AS1 | 286.216984 | 0.823667924 | 0.27049509 | 3.04503833 | 0.00232651 | 0.01518188 |
| AL136531.1 | 39.7106627 | 0.569591024 | 0.33680568 | 1.69115624 | 0.09080697 | 0.2220493 |
| FKBP1A | 10737.5155 | -0.952374584 | 0.21284977 | -4.474398 | 7.66E-06 | 0.00016509 |
| NSFL1C | 3935.49265 | -0.391807959 | 0.16245344 | -2.4118169 | 0.01587325 | 0.06344945 |
| SIRPB2 | 186.797962 | -0.746871547 | 0.36815545 | -2.0286853 | 0.04249035 | 0.12946065 |
| SIRPD | 4.67130826 | -1.331175907 | 0.76188382 | -1.7472164 | 0.08059981 | 0.20350603 |
| AL049634.2 | 56.3346511 | -0.805329577 | 0.6584958 | -1.2229836 | 0.22133589 | 0.40744344 |
| SIRPB1 | 132.60236 | -1.229273631 | 0.44036061 | -2.7915158 | 0.00524618 | 0.0278672 |
| SIRPG | 47.0248338 | 0.144226359 | 0.86011618 | 0.16768242 | 0.86683313 | 0.9308678 |
| SIRPG-AS1 | 0.38280371 | -0.542723645 | 2.80066761 | -0.1937837 | 0.84634527 | NA |
| AL121760.1 | 3.58715269 | -1.481870823 | 0.78747113 | -1.8818097 | 0.05986185 | 0.16529051 |
| AL117335.1 | 39.2701095 | -0.152917158 | 0.31537324 | -0.4848768 | 0.62776376 | 0.77854162 |
| SIRPA | 4385.31671 | -0.088975005 | 0.06318231 | -1.4082266 | 0.159064 | 0.32659952 |
| PDYN-AS1 | 8.08963374 | 0.317390151 | 0.65782367 | 0.48248515 | 0.62946135 | 0.77986909 |
| PDYN | 0.64918787 | 0.038902697 | 1.63123053 | 0.02384868 | 0.98097331 | NA |
| STK35 | 1825.12672 | -0.427769217 | 0.13880423 | -3.0818168 | 0.00205741 | 0.01377665 |
| AL121899.1 | 7.91020365 | -0.921139734 | 0.46521037 | -1.9800499 | 0.04769792 | 0.14075585 |
| TGM3 | 4.82634429 | -1.541438917 | 0.67232584 | -2.2926962 | 0.0218655 | 0.08014127 |
| TGM6 | 0 | NA | NA | NA | NA | NA |
| SNRPB | 5675.64564 | -1.073531786 | 0.26364523 | -4.0718801 | 4.66E-05 | 0.00070192 |
| AL049650.1 | 1888.94527 | -0.941871284 | 0.26669268 | -3.5316727 | 0.00041294 | 0.00391558 |
| ZNF343 | 789.937964 | -0.058322504 | 0.11852146 | -0.4920839 | 0.62266003 | 0.77487235 |
| TMC2 | 11.0375247 | 0.308160375 | 0.63999214 | 0.4815065 | 0.63015656 | 0.7801882 |
| NOP56 | 4194.55153 | -0.683211827 | 0.29258711 | -2.3350715 | 0.01953968 | 0.0736614 |
| IDH3B | 5022.9705 | 0.290460828 | 0.13249124 | 2.19230216 | 0.0283577 | 0.09700252 |
| AL049712.1 | 3.42668735 | 0.488091018 | 0.7003031 | 0.6969711 | 0.48582088 | 0.66986126 |
| EBF4 | 1107.32777 | -0.788879739 | 0.27624673 | -2.855707 | 0.00429411 | 0.02402116 |
| CPXM1 | 828.490735 | -0.851332206 | 0.65243591 | -1.3048518 | 0.1919433 | 0.37121416 |
| AL035460.1 | 2.64173087 | -3.82199955 | 1.27678957 | -2.9934452 | 0.00275847 | 0.01724126 |
| C20orf141 | 1.81091627 | -3.231098398 | 1.36938478 | -2.3595256 | 0.01829832 | 0.07032908 |
| TMEM239 | 1.12104644 | -2.515742983 | 1.41162229 | -1.7821644 | 0.07472243 | 0.19317646 |
| PCED1A | 3857.11621 | -0.080612998 | 0.2290589 | -0.3519313 | 0.72488978 | 0.84398179 |
| VPS16 | 2843.87873 | -0.143226444 | 0.10808332 | -1.3251485 | 0.18512192 | 0.36251782 |
| PTPRA | 4561.62636 | -0.009758598 | 0.08915309 | -0.1094589 | 0.91283852 | 0.95719548 |
| GNRH2 | 19.9621592 | 0.466128282 | 0.4622637 | 1.00836013 | 0.3132816 | 0.51020479 |
| MRPS26 | 2209.25154 | 0.031738082 | 0.17358354 | 0.18284039 | 0.85492325 | 0.92401312 |
| OXT | 47.850633 | -1.194456368 | 0.95497153 | -1.2507769 | 0.21101587 | 0.39438912 |
| AVP | 12.274039 | 0.805398084 | 1.19366265 | 0.6747284 | 0.49984834 | 0.68118867 |
| UBOX5-AS1 | 177.836088 | 0.363422534 | 0.33760748 | 1.07646469 | 0.28171948 | 0.47759195 |
| UBOX5 | 814.399165 | 0.409184887 | 0.25674607 | 1.59373376 | 0.11099568 | 0.25542858 |
| FASTKD5 | 1647.90511 | -0.169588997 | 0.34096418 | -0.4973807 | 0.61892063 | 0.77231656 |
| LZTS3 | 5651.20264 | 0.244233842 | 0.24394557 | 1.00118171 | 0.31673897 | 0.51402589 |
| DDRGK1 | 4179.1574 | -0.801854059 | 0.1442744 | -5.5578401 | 2.73E-08 | 1.62E-06 |
| ITPA | 2541.65724 | -0.669721277 | 0.1316739 | -5.0862113 | 3.65E-07 | 1.40E-05 |
| SLC4A11 | 1287.78366 | 1.308391362 | 0.5976414 | 2.18925824 | 0.02857808 | 0.0974908 |
| AL109976.1 | 90.5102568 | 0.034932537 | 0.61756475 | 0.05656498 | 0.95489173 | 0.97841508 |
| C20orf194 | 2175.99368 | 0.401176025 | 0.15548367 | 2.58018106 | 0.00987485 | 0.04466542 |
| ATRN | 3691.62426 | 0.44662034 | 0.13906575 | 3.21157676 | 0.00132009 | 0.00980031 |
| GFRA4 | 23.9148295 | -1.03739509 | 1.71205278 | -0.6059364 | 0.54455699 | 0.71643964 |
| ADAM33 | 1222.9583 | -1.695297736 | 0.74002061 | -2.2908791 | 0.02197041 | 0.08041078 |
| SIGLEC1 | 1140.28014 | -1.338540704 | 0.64848625 | -2.0641004 | 0.0390082 | 0.12156066 |
| HSPA12B | 832.095281 | 0.457862607 | 0.27254003 | 1.67998298 | 0.09296063 | 0.22576641 |
| C20orf27 | 1075.26912 | -0.21464756 | 0.18539651 | -1.1577757 | 0.2469556 | 0.43841519 |
| SPEF1 | 177.522259 | 0.122232916 | 0.41474833 | 0.29471587 | 0.76821094 | 0.87196142 |
| CENPB | 4748.43733 | -0.572629789 | 0.2126348 | -2.6930202 | 0.0070808 | 0.03492717 |
| CDC25B | 4215.9422 | -1.13870251 | 0.26730108 | -4.2599997 | 2.04E-05 | 0.00036107 |
| LINC01730 | 44.1530843 | -1.097383339 | 0.6185664 | -1.7740752 | 0.07605071 | 0.19565435 |
| AP5S1 | 845.714527 | 0.085956267 | 0.23804321 | 0.36109523 | 0.71802826 | 0.83985952 |
| MAVS | 6031.8898 | -0.00167066 | 0.15095105 | -0.0110676 | 0.99116955 | 0.99583588 |
| AL353194.1 | 52.5045312 | -0.131354509 | 0.30235502 | -0.434438 | 0.66397041 | 0.80260972 |
| PANK2 | 1840.20104 | -0.413778036 | 0.17606633 | -2.3501258 | 0.01876707 | 0.07166052 |
| AL031670.1 | 324.943603 | -0.708096333 | 0.21946409 | -3.2264793 | 0.00125323 | 0.00937444 |
| RNF24 | 2380.93531 | -0.835546167 | 0.29404908 | -2.8415194 | 0.00448991 | 0.02484162 |
| AL356414.1 | 0.9458202 | -3.150342106 | 2.66000524 | -1.1843368 | 0.23627976 | NA |
| SMOX | 1403.98261 | -2.76716954 | 0.28522121 | -9.7018366 | 2.96E-22 | 9.76E-19 |
| LINC01433 | 4.9829052 | -2.718139017 | 0.76807625 | -3.5388922 | 0.00040181 | 0.00383209 |
| ADRA1D | 22.7128669 | 0.196392909 | 0.53933877 | 0.36413646 | 0.71575611 | 0.83823283 |
| AL139350.1 | 0 | NA | NA | NA | NA | NA |
| AL121781.1 | 0.06785815 | 0.795868262 | 3.28898349 | 0.24198001 | 0.80879565 | NA |
| PRNP | 8413.9129 | 0.411392086 | 0.29774049 | 1.38171362 | 0.16705965 | 0.33806792 |
| PRND | 2.23171932 | 1.371885397 | 1.33660169 | 1.02639807 | 0.30470399 | 0.50168857 |
| PRNT | 0 | NA | NA | NA | NA | NA |
| AL133396.2 | 0.25725949 | -1.364242679 | 3.30558948 | -0.4127078 | 0.67982068 | NA |
| RASSF2 | 528.870416 | -0.419300747 | 0.35469154 | -1.1821561 | 0.23714379 | 0.42639107 |
| SLC23A2 | 1463.59111 | 0.287420633 | 0.17755356 | 1.61878271 | 0.10549402 | 0.24646739 |
| AL121890.3 | 2.14297001 | 0.696044328 | 0.8552276 | 0.81387029 | 0.41571927 | 0.60908136 |
| AL121890.2 | 3.79003946 | -0.29483795 | 0.66630242 | -0.4424987 | 0.65812837 | 0.79943711 |
| AL121890.5 | 8.64189901 | -1.070960609 | 0.70511829 | -1.5188382 | 0.12880322 | 0.28311586 |
| AL121890.4 | 4.18836018 | 0.35435695 | 0.5924269 | 0.5981446 | 0.54974346 | 0.72018351 |
| TMEM230 | 5753.52417 | 0.190028385 | 0.13402908 | 1.4178146 | 0.1562449 | 0.32284736 |
| PCNA | 1917.09128 | -0.067067237 | 0.15609776 | -0.4296489 | 0.66745104 | 0.80541271 |
| CDS2 | 4449.35706 | -0.315103908 | 0.17463366 | -1.804371 | 0.07117317 | 0.18656425 |
| PROKR2 | 0 | NA | NA | NA | NA | NA |
| AL121757.2 | 0 | NA | NA | NA | NA | NA |
| LINC00658 | 0.86459369 | -3.017575565 | 1.55594777 | -1.939381 | 0.05245496 | NA |
| AL121757.1 | 1.96976687 | 1.276989346 | 0.99869795 | 1.27865422 | 0.20101885 | 0.38273408 |
| LINC00654 | 54.3786394 | -0.006942924 | 0.32525353 | -0.0213462 | 0.9829695 | 0.9924157 |
| AL109935.2 | 2.00661709 | -0.544325609 | 0.93418099 | -0.5826768 | 0.56011088 | 0.72838393 |
| LINC01729 | 0.34627297 | -0.541481398 | 2.86332064 | -0.1891096 | 0.85000692 | NA |
| AL109935.1 | 0 | NA | NA | NA | NA | NA |
| GPCPD1 | 2376.7453 | -0.070194834 | 0.3328699 | -0.2108777 | 0.83298272 | 0.91136194 |
| SHLD1 | 160.32133 | 0.119168942 | 0.20455719 | 0.5825703 | 0.56018261 | 0.72838393 |
| CHGB | 755.397614 | 1.380355982 | 0.61251697 | 2.25357998 | 0.0242226 | 0.08622436 |
| TRMT6 | 908.242463 | -0.150603172 | 0.23841568 | -0.6316832 | 0.52759392 | 0.70223445 |
| MCM8 | 479.948547 | 0.103990275 | 0.2483721 | 0.41868743 | 0.67544459 | 0.81122383 |
| MCM8-AS1 | 37.5971628 | 0.283959404 | 0.26921049 | 1.05478582 | 0.29152329 | 0.48774871 |
| AL035461.2 | 43.9149658 | -0.312712531 | 0.31614637 | -0.9891385 | 0.3225954 | 0.5203279 |

| | | | | | | |
|---|---|---|---|---|---|---|
| CRLS1 | 4261.48335 | -0.36163172 | 0.23123195 | -1.5639349 | 0.11783286 | 0.26654276 |
| LRRN4 | 666.453374 | -2.170016797 | 0.48672406 | -4.4584128 | 8.26E-06 | 0.00017573 |
| AL118505.1 | 46.0863692 | 2.127525893 | 0.4172532 | 5.09888458 | 3.42E-07 | 1.33E-05 |
| FERMT1 | 314.560393 | 1.849277394 | 0.39368936 | 4.6973009 | 2.64E-06 | 7.08E-05 |
| CASC20 | 0 | NA | NA | NA | NA | NA |
| LINC01713 | 0.44665835 | 2.760059159 | 3.20263664 | 0.8618084 | 0.38879296 | NA |
| BMP2 | 465.953888 | -0.33115065 | 0.33188866 | -0.9977763 | 0.31838783 | 0.51565454 |
| AL096799.1 | 3.53721363 | 1.782920861 | 0.79821894 | 2.23362386 | 0.02550783 | 0.08958998 |
| LINC01428 | 5.50060768 | 2.759105387 | 0.64357543 | 4.28715153 | 1.81E-05 | 0.00032845 |
| AL080248.1 | 16.3438059 | 3.838140164 | 0.92974401 | 4.12816876 | 3.66E-05 | 0.00057624 |
| LINC01751 | 0.2873605 | -0.174096114 | 3.29377188 | -0.0528562 | 0.95784652 | NA |
| LINC01706 | 0.28750021 | -0.17981576 | 2.91671556 | -0.0616501 | 0.95084149 | NA |
| HAO1 | 29.5214877 | 3.481791348 | 1.06353875 | 3.2737795 | 0.00106119 | 0.00820636 |
| TMX4 | 3722.32905 | -0.249416123 | 0.18772422 | -1.3286305 | 0.1839699 | 0.36101253 |
| AL021396.1 | 1.35837554 | -0.30356819 | 1.1989055 | -0.2532044 | 0.80011024 | 0.89152759 |
| PLCB1 | 954.738175 | 0.136485792 | 0.40054464 | 0.34075051 | 0.73329141 | 0.84963726 |
| PLCB4 | 503.964456 | -0.491022178 | 0.26298894 | -1.867083 | 0.06189 | 0.16905317 |
| LAMP5-AS1 | 4.80130295 | 0.439758581 | 0.67179232 | 0.65460496 | 0.51272213 | 0.69087801 |
| LAMP5 | 49.3089198 | 0.142437157 | 0.47485925 | 0.29995658 | 0.76421028 | 0.86949238 |
| PAK5 | 12.5103633 | 0.806327993 | 0.5790611 | 1.39247481 | 0.16377867 | 0.33328303 |
| AL353612.1 | 0 | NA | NA | NA | NA | NA |
| PARAL1 | 4.03984386 | -3.858557616 | 1.55897033 | -2.475068 | 0.01332108 | 0.05571402 |
| ANKEF1 | 403.488574 | -0.096643593 | 0.18757288 | -0.5152322 | 0.60639075 | 0.76308872 |
| SNAP25-AS1 | 47.8520277 | 0.128331071 | 0.28108753 | 0.45655199 | 0.6479931 | 0.79277053 |
| AL354824.1 | 1.20485031 | -2.634142212 | 1.42053911 | -1.8543257 | 0.06369257 | 0.17233733 |
| AL354824.2 | 0.3262184 | -1.662021924 | 3.30365031 | -0.5030865 | 0.61490345 | NA |
| SNAP25 | 144.686348 | -0.916700435 | 0.58891214 | -1.5565997 | 0.11956557 | 0.26920628 |
| MKKS | 2986.02939 | 0.047929789 | 0.2020146 | 0.23725903 | 0.81245585 | 0.89882785 |
| AL034430.1 | 1007.5817 | 0.014280918 | 0.27716532 | 0.05152491 | 0.95890725 | 0.98020639 |
| SLX4IP | 578.737425 | 0.255083041 | 0.23067551 | 1.10580895 | 0.26880922 | 0.46299798 |
| JAG1 | 3678.3258 | 0.450593618 | 0.19264117 | 2.33903076 | 0.01933384 | 0.07310487 |
| AL050403.2 | 20.4373367 | 1.705605194 | 0.41647514 | 4.09533495 | 4.22E-05 | 0.00064781 |
| LINC01752 | 8.53942185 | -0.265978766 | 0.60580653 | -0.439049 | 0.66062602 | 0.80027291 |
| AL135937.1 | 7.54485364 | -0.205449273 | 0.63866054 | -0.3216878 | 0.74768926 | 0.85884766 |
| AL050403.1 | 1.60763492 | 1.666578059 | 1.0632019 | 1.56750854 | 0.11699587 | 0.26531355 |
| AL158042.1 | 2.88022556 | 1.154759208 | 1.19304957 | 0.96790548 | 0.33309158 | 0.5310535 |
| C20orf187 | 0.35809632 | 1.762373218 | 3.23429066 | 0.54490255 | 0.58582054 | NA |
| AL049649.1 | 0.39990806 | -0.559317536 | 2.22736297 | -0.251112 | 0.80172753 | NA |
| AL109837.2 | 0 | NA | NA | NA | NA | NA |
| AL161938.1 | 8.77477737 | -3.634478132 | 0.84866501 | -4.2825827 | 1.85E-05 | 0.00033426 |
| LINC00687 | 0 | NA | NA | NA | NA | NA |
| BTBD3 | 1791.49158 | -0.220261027 | 0.17718844 | -1.2430891 | 0.213835 | 0.39785711 |
| AL035448.1 | 41.8421519 | 0.909188917 | 0.32731764 | 2.77769607 | 0.00547458 | 0.02883833 |
| AL109838.1 | 0.26997446 | -0.179050932 | 2.95977615 | -0.0604948 | 0.95176159 | NA |
| AL136460.1 | 0 | NA | NA | NA | NA | NA |
| LINC01722 | 1.08949526 | -0.608045629 | 1.48050876 | -0.4107005 | 0.68129219 | 0.81532449 |
| AL078623.1 | 0 | NA | NA | NA | NA | NA |
| LINC01723 | 0 | NA | NA | NA | NA | NA |
| SPTLC3 | 2116.65185 | -0.110023588 | 0.28299644 | -0.3887808 | 0.69743829 | 0.82609925 |
| ISM1 | 215.499505 | 0.664965059 | 0.47097432 | 1.4118924 | 0.15798165 | 0.32512868 |
| ISM1-AS1 | 2.98720394 | 0.298188606 | 0.74144646 | 0.40217146 | 0.68755784 | 0.81947556 |
| AL050320.1 | 2.58411157 | 0.025957632 | 0.8902085 | 0.02915905 | 0.97673774 | 0.98926772 |
| AL121782.1 | 10.3168914 | -0.675145304 | 0.44100718 | -1.5309168 | 0.12578995 | 0.27858243 |
| TASP1 | 645.629835 | -0.189172678 | 0.12601048 | -1.5012456 | 0.13329204 | 0.28991815 |
| ESF1 | 625.262755 | -0.158188525 | 0.22662436 | -0.6980208 | 0.48516417 | 0.66944583 |
| NDUFAF5 | 960.628534 | 0.030990064 | 0.14371136 | 0.21564102 | 0.82926758 | 0.90929435 |
| SEL1L2 | 4.7636821 | 0.838461803 | 1.10108056 | 0.76148997 | 0.44636447 | 0.63673218 |
| MACROD2 | 282.825043 | 0.349775208 | 0.21831482 | 1.60215972 | 0.1091203 | 0.25247891 |
| FLRT3 | 2961.10388 | 0.49539323 | 0.24339155 | 2.03537564 | 0.04181309 | 0.12797054 |
| MACROD2-A | 0.97243294 | 0.696665995 | 1.13953363 | 0.61136063 | 0.54096086 | NA |
| AL138808.1 | 0 | NA | NA | NA | NA | NA |
| AL121584.1 | 0.09989236 | -0.642905668 | 3.30558948 | -0.1944905 | 0.84579185 | NA |
| KIF16B | 1513.73698 | 0.445403841 | 0.20413591 | 2.1818985 | 0.02911702 | 0.09883073 |
| AL049794.1 | 3.39163695 | 1.567946374 | 0.76230277 | 2.05685515 | 0.03970016 | 0.12311104 |
| AL049794.2 | 0.13028434 | 1.049816488 | 3.28477129 | 0.31960109 | 0.74927075 | NA |
| SNRPB2 | 1956.36334 | -0.276003944 | 0.17947284 | -1.5378592 | 0.12408306 | 0.27582148 |
| OTOR | 0.51180205 | 0.620910188 | 1.63637214 | 0.37944314 | 0.70435882 | NA |
| AL121892.1 | 0 | NA | NA | NA | NA | NA |
| PCSK2 | 19.3852603 | 1.439202033 | 0.78154738 | 1.84147765 | 0.06555159 | 0.17586687 |
| BFSP1 | 105.613551 | 0.16103993 | 0.51661155 | 0.31172344 | 0.75525071 | 0.86403303 |
| DSTN | 19235.7931 | -0.177651914 | 0.24842618 | -0.7151095 | 0.4745414 | 0.66007262 |
| RRBP1 | 16716.5867 | -0.479452688 | 0.14804407 | -3.2385807 | 0.00120126 | 0.00905244 |
| BANF2 | 0.88547665 | 2.186667307 | 1.54459574 | 1.41568907 | 0.15686656 | NA |
| AL035045.1 | 0.20687462 | 0.68915539 | 3.1020265 | 0.22216296 | 0.82418703 | NA |
| SNX5 | 7478.28299 | 0.557081249 | 0.23327657 | 2.38807204 | 0.01693702 | 0.06649237 |
| OVOL2 | 726.906125 | 0.706880429 | 0.27261567 | 2.59295601 | 0.00951549 | 0.04341238 |
| MGME1 | 1160.98901 | 0.081077429 | 0.20665275 | 0.39233656 | 0.69480955 | 0.82425506 |
| AL160411.1 | 0.26533299 | 0.373589558 | 3.00844626 | 0.12418023 | 0.90117257 | NA |
| PET117 | 806.665132 | -0.063496761 | 0.23749725 | -0.2673579 | 0.78919362 | 0.88546295 |
| KAT14 | 1500.31659 | 0.30852696 | 0.18146749 | 1.70017759 | 0.08909753 | 0.21898609 |
| ZNF133 | 1452.89553 | -0.050454551 | 0.23431397 | -0.2153288 | 0.82951096 | 0.90939253 |
| AL049646.2 | 83.1045185 | 0.308822382 | 0.22320241 | 1.38359789 | 0.16648162 | 0.33726052 |
| AL049646.1 | 0.84375421 | 2.579829762 | 1.31552576 | 1.96106366 | 0.0498716 | NA |
| LINC00851 | 0.19994613 | 1.506623855 | 3.27645372 | 0.45983371 | 0.64563559 | NA |
| DZANK1 | 469.79057 | 0.085426227 | 0.22093995 | 0.38664907 | 0.69901603 | 0.82722491 |
| POLR3F | 642.507906 | 0.042328677 | 0.17787439 | 0.23796948 | 0.81190476 | 0.89845484 |
| RBBP9 | 1494.69643 | 0.686989526 | 0.15512309 | 4.42867363 | 9.48E-06 | 0.00019564 |
| SEC23B | 4249.81112 | -0.224339466 | 0.21063604 | -1.0650574 | 0.28684996 | 0.48312151 |
| SMIM26 | 1544.94121 | -0.336552714 | 0.15673879 | -2.1472203 | 0.03177575 | 0.10506914 |
| AL121900.2 | 2153.07142 | -0.301651515 | 0.18214773 | -1.6560817 | 0.09770528 | 0.23388577 |
| DTD1 | 1771.14205 | -0.3557017 | 0.21449752 | -1.6583022 | 0.09725649 | 0.23298089 |
| AL121900.1 | 0.64757368 | 1.367331889 | 1.8108778 | 0.7550658 | 0.4502095 | NA |
| LINC00652 | 33.3515935 | 1.357513502 | 0.35982345 | 3.77272104 | 0.00016148 | 0.00185218 |
| AL035563.1 | 151.787699 | 0.73881001 | 0.39204523 | 1.88450197 | 0.05949711 | 0.16459701 |
| SCP2D1 | 0.09644756 | -0.642905668 | 3.30558948 | -0.1944905 | 0.84579185 | NA |
| AL135936.1 | 0.4780663 | 1.539239534 | 2.14658453 | 0.71706449 | 0.47333431 | NA |
| SLC24A3 | 192.437502 | -0.245109778 | 0.33739593 | -0.7264752 | 0.46754751 | 0.65455658 |
| AL136090.1 | 1.28343364 | -0.215510263 | 1.32943127 | -0.1621071 | 0.87122151 | 0.93306711 |
| AL049647.1 | 1.08227961 | -0.599719734 | 1.86780633 | -0.3210824 | 0.74814795 | 0.85922484 |
| AL121761.2 | 42.5612907 | 0.408178753 | 0.39514615 | 1.03298173 | 0.30161245 | 0.49840333 |
| AL121761.1 | 51.0148981 | 0.727007573 | 0.21283538 | 3.41582098 | 0.0006359 | 0.00546568 |
| AL132821.1 | 8.1221165 | -0.379132056 | 0.59119946 | -0.641293 | 0.52133236 | 0.69769389 |
| RIN2 | 2552.06605 | 0.190223859 | 0.13787649 | 1.37966859 | 0.16768871 | 0.33887281 |
| NAA20 | 3041.08762 | -0.334369118 | 0.14964164 | -2.2344658 | 0.02545244 | 0.08946693 |
| CRNKL1 | 1233.16639 | -0.123948515 | 0.13160592 | -0.9418156 | 0.34628704 | 0.54387071 |
| CFAP61 | 53.0687289 | 0.485125093 | 0.58463706 | 0.82978848 | 0.40665839 | 0.60140134 |
| AL035454.1 | 0.26523621 | -0.178786168 | 2.97513023 | -0.0600936 | 0.95208112 | NA |
| AL049648.1 | 0.41057065 | 2.632352683 | 2.66025702 | 0.98951066 | 0.32241336 | NA |
| AL121721.1 | 1.64768231 | 0.081082053 | 0.90357958 | 0.08973427 | 0.92849838 | 0.96528691 |
| INSM1 | 1.32934553 | 2.935124162 | 1.77795196 | 1.6508456 | 0.09877011 | 0.23562219 |
| RALGAPA2 | 3241.64187 | -0.58957121 | 0.20215235 | -2.9164698 | 0.00354017 | 0.02075462 |
| AL133465.1 | 0 | NA | NA | NA | NA | NA |
| LINC00237 | 0.96292594 | 2.463724201 | 1.43487257 | 1.71703345 | 0.08597306 | NA |

| | | | | | | |
|---|---|---|---|---|---|---|
| AL121759.2 | 0.06785815 | 0.795868262 | 3.28898349 | 0.24198001 | 0.80879565 | NA |
| KIZ | 1464.08315 | -0.081953438 | 0.29475238 | -0.2780416 | 0.78098039 | 0.88016021 |
| AL121759.1 | 84.5005288 | 0.471300818 | 0.34237763 | 1.37655262 | 0.16865061 | 0.34026905 |
| KIZ-AS1 | 101.710338 | 0.398012666 | 0.30491064 | 1.305342 | 0.19177642 | 0.37100317 |
| AL117332.1 | 132.93476 | 0.250434351 | 0.28090705 | 0.89152035 | 0.37265008 | 0.56887707 |
| XRN2 | 3915.07249 | -0.187167566 | 0.16493937 | -1.1347658 | 0.25647344 | 0.44921834 |
| NKX2-4 | 0 | NA | NA | NA | NA | NA |
| AL158013.1 | 0 | NA | NA | NA | NA | NA |
| AL133325.3 | 4.64627762 | 1.328282519 | 0.72474202 | 1.83276598 | 0.06683738 | 0.17824241 |
| NKX2-2 | 0.42730237 | -0.899974433 | 2.24832821 | -0.4002861 | 0.68894584 | NA |
| NKX2-2-AS1 | 0.42730237 | -0.899974433 | 2.24832821 | -0.4002861 | 0.68894584 | NA |
| AL133325.2 | 0 | NA | NA | NA | NA | NA |
| LINC01727 | 6.13692874 | 4.129576492 | 0.89254563 | 4.6267399 | 3.71E-06 | 9.28E-05 |
| LINC01726 | 2.65197351 | 3.303455338 | 1.09196648 | 3.02523512 | 0.0024844 | 0.01592442 |
| PAX1 | 0.27359226 | -0.179157115 | 2.95368447 | -0.0606555 | 0.9516336 | NA |
| AL109807.1 | 0 | NA | NA | NA | NA | NA |
| LINC01432 | 0.09989236 | -0.642905668 | 3.30558948 | -0.1944905 | 0.84579185 | NA |
| AL035258.1 | 0 | NA | NA | NA | NA | NA |
| LINC01427 | 0 | NA | NA | NA | NA | NA |
| AL133464.1 | 0.03392908 | 0.380740593 | 3.29738709 | 0.11546736 | 0.90807469 | NA |
| LINC00261 | 0.06785815 | 0.795868262 | 3.28898349 | 0.24198001 | 0.80879565 | NA |
| AL121722.1 | 1.02520602 | 1.061700337 | 1.57748058 | 0.67303544 | 0.50092474 | 0.68184512 |
| FOXA2 | 12.287447 | -0.028408711 | 0.78616141 | -0.036136 | 0.97117393 | 0.9865502 |
| LINC01384 | 0 | NA | NA | NA | NA | NA |
| LINC01747 | 1.90785222 | -0.308192796 | 0.9158893 | -0.3364957 | 0.73649711 | 0.85207763 |
| AL158175.1 | 0.20481663 | -0.33998347 | 3.13492333 | -0.1084503 | 0.91363848 | NA |
| AL049651.1 | 0 | NA | NA | NA | NA | NA |
| SSTR4 | 0 | NA | NA | NA | NA | NA |
| THBD | 1959.16525 | -0.415088112 | 0.42991002 | -0.9655232 | 0.33428281 | 0.53240549 |
| CD93 | 4295.31287 | -0.851201664 | 0.40485977 | -2.1024605 | 0.03551296 | 0.11377435 |
| LINC00656 | 0.90161697 | 0.537916224 | 1.31101196 | 0.41030611 | 0.68158141 | NA |
| AL118508.2 | 0.95935475 | 1.745567136 | 1.48434527 | 1.17598457 | 0.23960104 | NA |
| AL118508.1 | 54.9636189 | 1.359813588 | 0.40399926 | 3.36588135 | 0.000763 | 0.00632312 |
| AL390037.1 | 2.82039872 | -1.196837759 | 0.77167809 | -1.5509547 | 0.12091254 | 0.27130065 |
| AL096677.1 | 3.41866386 | 0.092274317 | 0.73938689 | 0.12479842 | 0.90068314 | 0.94969868 |
| NXT1 | 585.388197 | -0.959599015 | 0.24070515 | -3.986616 | 6.70E-05 | 0.00092987 |
| LINC01431 | 62.7846943 | 1.174722407 | 0.37770391 | 3.11016744 | 0.00186981 | 0.01278708 |
| GZF1 | 1089.1256 | 0.187307407 | 0.20423402 | 0.91712146 | 0.359079 | 0.5567329 |
| NAPB | 1220.64626 | -0.158248939 | 0.21748156 | -0.727643 | 0.46683214 | 0.65400646 |
| CSTL1 | 0.2927759 | 1.128512364 | 2.95401749 | 0.3820263 | 0.70244186 | NA |
| CST11 | 2.10658992 | -2.460447251 | 1.55672406 | -1.5805288 | 0.11398581 | 0.26039437 |
| AL109954.2 | 0.08757577 | 0.547239394 | 3.29377202 | 0.16614368 | 0.86804388 | NA |
| AL109954.1 | 0 | NA | NA | NA | NA | NA |
| AL109954.3 | 0.09997307 | 0.547239394 | 3.29377202 | 0.16614368 | 0.86804388 | NA |
| CST8 | 0.51197563 | 0.685232603 | 1.85604875 | 0.3691889 | 0.71198693 | NA |
| CST9L | 0.10721287 | -0.642905668 | 3.30558948 | -0.1944905 | 0.84579185 | NA |
| CST9 | 11.1804081 | 3.442137386 | 1.52836185 | 2.25217437 | 0.02431126 | 0.08643492 |
| CST3 | 17496.8383 | -0.456654513 | 0.17857322 | -2.5572396 | 0.01055065 | 0.0468624 |
| AL121894.2 | 11.8696683 | -0.778755489 | 0.52735719 | -1.4767135 | 0.13975245 | 0.29961562 |
| AL121894.1 | 0.28216687 | 0.374943671 | 2.96645449 | 0.12639455 | 0.89941962 | NA |
| CST4 | 0.59397492 | -2.477332114 | 2.15590242 | -1.1490929 | 0.25051769 | NA |
| CST1 | 1.49351659 | -1.956280016 | 1.64880275 | -1.1864852 | 0.23543075 | 0.42432567 |
| AL591074.1 | 0 | NA | NA | NA | NA | NA |
| CST2 | 1.48405796 | -1.287402697 | 1.38805449 | -0.9274871 | 0.35367366 | 0.55123674 |
| CST5 | 7.52670248 | 2.480335249 | 0.50194155 | 4.9414822 | 7.75E-07 | 2.62E-05 |
| GGTLC1 | 7.62834503 | 0.321807985 | 0.53488534 | 0.6016392 | 0.54741433 | 0.71842513 |
| AL110503.1 | 1.44484696 | 2.675247051 | 1.69866941 | 1.57490742 | 0.11527782 | 0.26271507 |
| LINC01721 | 0.6548473 | -0.975306014 | 1.94033235 | -0.5026489 | 0.61521111 | NA |
| AL158090.1 | 0.29052362 | -0.179944783 | 2.90963392 | -0.0618445 | 0.95068669 | NA |
| SYNDIG1 | 383.98267 | 0.958833459 | 0.34677263 | 2.76502056 | 0.00569192 | 0.02967496 |
| AL157413.1 | 0.36621596 | -0.278376804 | 2.86145791 | -0.097285 | 0.9225001 | NA |
| AL049594.1 | 0.15318056 | -1.003574173 | 3.30558948 | -0.3035992 | 0.7614333 | NA |
| AL035661.1 | 685.997221 | -0.009798511 | 0.28540454 | -0.034332 | 0.9726124 | 0.98717482 |
| CST7 | 238.275714 | -1.463105463 | 0.6406577 | -2.2837554 | 0.02238591 | 0.08151291 |
| APMAP | 5545.19478 | -0.474534684 | 0.18578884 | -2.5541613 | 0.01064439 | 0.04715146 |
| ACSS1 | 7077.99806 | 0.372082666 | 0.16057293 | 2.31721911 | 0.0204918 | 0.07628017 |
| AL080312.2 | 1.63694033 | -0.503988311 | 0.99203323 | -0.5080357 | 0.6114283 | 0.76723122 |
| VSX1 | 3.08641709 | -3.2933351 | 1.1566579 | -2.8472854 | 0.00440938 | 0.024522 |
| AL035252.2 | 1.34833514 | -0.939593089 | 1.65918976 | -0.5662963 | 0.57119235 | 0.73622084 |
| ENTPD6 | 6387.56291 | -0.761883114 | 0.18878408 | -4.0357381 | 5.44E-05 | 0.0007952 |
| AL035252.3 | 126.516536 | -0.218916445 | 0.19135258 | -1.1440475 | 0.25260393 | 0.44493792 |
| AL121772.1 | 28.1850436 | -1.402280375 | 0.51761688 | -2.7091087 | 0.00674642 | 0.0336301 |
| PYGB | 8345.00387 | -0.606976952 | 0.25316117 | -2.397591 | 0.01650328 | 0.06524611 |
| AL121772.2 | 0.61850471 | -0.319816017 | 1.48967928 | -0.2146878 | 0.83001071 | NA |
| AL121772.3 | 12.2329244 | 0.349294749 | 0.35125389 | 0.99442245 | 0.32001724 | 0.51746718 |
| ABHD12 | 3803.52616 | -0.003264098 | 0.16204287 | -0.0201434 | 0.98392896 | 0.99305611 |
| GINS1 | 146.506253 | 0.24002323 | 0.29137245 | 0.82376775 | 0.41007154 | 0.6044565 |
| NINL | 2981.4363 | 0.722494352 | 0.15705295 | 4.6003233 | 4.22E-06 | 0.0001029 |
| NANP | 493.819465 | -0.201748917 | 0.28085501 | -0.7183383 | 0.47254871 | 0.65875468 |
| ZNF337-AS1 | 1651.7602 | -0.032859853 | 0.21580855 | -0.1522639 | 0.87897879 | 0.93705294 |
| ZNF337 | 1671.51419 | -0.031726364 | 0.21482961 | -0.1476815 | 0.88259411 | 0.93902736 |
| AL031673.1 | 29.0607001 | 0.665991812 | 0.31310258 | 2.12707229 | 0.03341408 | 0.10895625 |
| AL390198.1 | 20.3332845 | 0.600961777 | 0.40400915 | 1.48749545 | 0.136884 | 0.29530784 |
| FAM182B | 39.4722912 | 1.093202027 | 0.71905366 | 1.52033442 | 0.12842695 | 0.28259501 |
| LINC01733 | 0.08757577 | 0.547239394 | 3.29377202 | 0.16614368 | 0.86804388 | NA |
| AL078587.1 | 19.1524442 | -0.940127174 | 0.47479972 | -1.98005 | 0.04769791 | 0.14075585 |
| AL391119.1 | 0 | NA | NA | NA | NA | NA |
| AL121904.1 | 0 | NA | NA | NA | NA | NA |
| LINC01597 | 175.841368 | -0.059431368 | 0.60173456 | -0.0987668 | 0.92132347 | 0.96158077 |
| LINC01598 | 0.30399583 | -0.695256667 | 2.32395324 | -0.2991698 | 0.76481048 | NA |
| DEFB115 | 0 | NA | NA | NA | NA | NA |
| DEFB116 | 0 | NA | NA | NA | NA | NA |
| DEFB118 | 0 | NA | NA | NA | NA | NA |
| DEFB119 | 0.20357445 | 1.728110593 | 3.26179266 | 0.52980394 | 0.59624787 | NA |
| DEFB121 | 0 | NA | NA | NA | NA | NA |
| DEFB123 | 0 | NA | NA | NA | NA | NA |
| DEFB124 | 31.6414295 | 1.053335852 | 0.57856859 | 1.82058941 | 0.06866929 | 0.18160759 |
| REM1 | 91.6784124 | 0.988627508 | 0.64060914 | 1.54326164 | 0.12276731 | 0.27398222 |
| LINC00028 | 2.97581145 | -0.030055811 | 0.68524799 | -0.0438612 | 0.96501503 | 0.98359669 |
| HM13 | 8047.60222 | -0.903613824 | 0.19893254 | -4.5423128 | 5.56E-06 | 0.00012792 |
| HM13-AS1 | 67.7482062 | -0.665788149 | 0.30106562 | -2.2114387 | 0.02700548 | 0.09349276 |
| AL110115.2 | 4.86343158 | -0.842184558 | 0.88517034 | -0.9514378 | 0.34138216 | 0.53937972 |
| ID1 | 3787.71403 | -0.2863204 | 0.40022907 | -0.7153913 | 0.47436728 | 0.66007262 |
| COX4I2 | 92.3131343 | 0.000351257 | 0.67816444 | 0.00051795 | 0.99958673 | 0.99979217 |
| BCL2L1 | 10398.2216 | -0.279810261 | 0.24831634 | -1.1268298 | 0.25981444 | 0.45270958 |
| AL117381.1 | 3.48356043 | -0.239799011 | 0.69238394 | -0.3463382 | 0.72908856 | 0.84692592 |
| ABALON | 3154.30983 | -0.006076763 | 0.2749992 | -0.0220974 | 0.98237027 | 0.99215259 |
| TPX2 | 88.8619721 | -1.201964161 | 0.55966995 | -2.1476303 | 0.03174313 | 0.10505031 |
| MYLK2 | 12.3438544 | 0.971315241 | 1.03830161 | 0.93548467 | 0.34953858 | 0.54721682 |
| FOXS1 | 122.67691 | 1.638421433 | 0.61323596 | 2.67176346 | 0.00754538 | 0.03661541 |
| DUSP15 | 1515.804 | 0.220662326 | 0.43366865 | 0.50882701 | 0.61087348 | 0.76671746 |
| TTLL9 | 156.06182 | -0.293291347 | 0.3986749 | -0.7356654 | 0.46193433 | 0.65014988 |
| PDRG1 | 1210.39567 | -0.547795549 | 0.20038088 | -2.7337716 | 0.00626135 | 0.03181814 |

| | | | | | | |
|---|---|---|---|---|---|---|
| XKR7 | 2.40465057 | -0.172570818 | 0.95428181 | -0.1808384 | 0.8564944 | 0.92488775 |
| CCM2L | 448.571672 | 0.553714501 | 0.36826251 | 1.50358639 | 0.1326879 | 0.28905244 |
| AL031658.1 | 246.928387 | 0.59864894 | 0.37500662 | 1.59636898 | 0.11040644 | 0.25449125 |
| HCK | 530.540781 | -0.638995361 | 0.45630863 | -1.4003578 | 0.16140619 | 0.32999545 |
| TM9SF4 | 5076.40718 | -0.081324428 | 0.13854008 | -0.5870101 | 0.55719692 | 0.7260962 |
| PLAGL2 | 878.289253 | -0.158967182 | 0.16047561 | -0.9906003 | 0.3218808 | 0.51955663 |
| POFUT1 | 2769.88283 | 0.302807828 | 0.14150344 | 2.13993263 | 0.03236021 | 0.10636156 |
| KIF3B | 4308.62079 | 0.301221751 | 0.13593478 | 2.21592842 | 0.0266964 | 0.09273697 |
| AL121583.1 | 36.2931107 | 0.411249615 | 0.33775571 | 1.21759484 | 0.22337798 | 0.40943055 |
| ASXL1 | 4569.55929 | -0.401611585 | 0.18343425 | -2.1894035 | 0.02856753 | 0.09748843 |
| NOL4L | 2359.44875 | -0.038446949 | 0.15378191 | -0.2500096 | 0.80257995 | 0.89299719 |
| AL034550.1 | 110.081152 | -0.133894695 | 0.22115102 | -0.6054446 | 0.54488361 | 0.71677397 |
| AL034550.2 | 5.78781989 | 0.293219196 | 0.75854795 | 0.38655328 | 0.69908695 | 0.82727172 |
| AL133343.2 | 0.11786115 | 0.02007176 | 3.29738704 | 0.00608717 | 0.99514317 | NA |
| NOL4L-DT | 0.08757577 | 0.547239394 | 3.29377202 | 0.16614368 | 0.86804388 | NA |
| AL133343.1 | 0.07659028 | -0.642905668 | 3.30558948 | -0.1944905 | 0.84579185 | NA |
| C20orf203 | 18.4894406 | 0.169372932 | 0.67960037 | 0.24922431 | 0.80318728 | 0.89318485 |
| FO393400.1 | 1053.46506 | -0.000884241 | 0.15144778 | -0.0058386 | 0.99534151 | 0.99776308 |
| COMMD7 | 2710.43377 | -0.126473244 | 0.14939796 | -0.8465527 | 0.39724451 | 0.59212807 |
| DNMT3B | 334.816397 | -0.032767264 | 0.24590305 | -0.1332528 | 0.89399347 | 0.9455183 |
| MAPRE1 | 4815.66604 | -0.331849944 | 0.20279382 | -1.6363908 | 0.10175781 | 0.24039429 |
| AL035071.2 | 2.40739614 | 1.659259854 | 0.85491986 | 1.94083672 | 0.05227809 | 0.15023683 |
| AL035071.1 | 823.028304 | -0.561734819 | 0.31165297 | -1.8024369 | 0.0714767 | 0.18711808 |
| EFCAB8 | 11.104629 | -0.314162939 | 0.49013545 | -0.6409717 | 0.52154109 | 0.69776958 |
| SUN5 | 0 | NA | NA | NA | NA | NA |
| BPIFB2 | 0.17332742 | -1.003574173 | 3.30558948 | -0.3035992 | 0.7614333 | NA |
| BPIFB6 | 0 | NA | NA | NA | NA | NA |
| BPIFB3 | 0 | NA | NA | NA | NA | NA |
| BPIFB4 | 0.42525427 | -1.993748192 | 2.78949482 | -0.7147345 | 0.47477311 | NA |
| BPIFA2 | 21.5883871 | -2.138811121 | 0.62109283 | -3.4436255 | 0.00057397 | 0.00505193 |
| AL121901.1 | 0 | NA | NA | NA | NA | NA |
| BPIFA3 | 0 | NA | NA | NA | NA | NA |
| BPIFA1 | 0 | NA | NA | NA | NA | NA |
| BPIFB1 | 3.04174739 | -1.327795824 | 0.99164415 | -1.3389842 | 0.18057582 | 0.35652774 |
| CDK5RAP1 | 1673.82966 | -0.125319495 | 0.10964489 | -1.1429579 | 0.25305608 | 0.44552603 |
| SNTA1 | 1810.46325 | 0.955744057 | 0.22461787 | 4.25497784 | 2.09E-05 | 0.00036706 |
| CBFA2T2 | 1706.63292 | -0.172879192 | 0.13191447 | -1.31054 | 0.19001323 | 0.36870117 |
| AL121906.2 | 9.80727526 | 0.496262853 | 0.40507475 | 1.22511425 | 0.22053218 | 0.40650311 |
| NECAB3 | 1446.87437 | 0.163030408 | 0.1983188 | 0.82206228 | 0.41104145 | 0.60533693 |
| C20orf144 | 31.6819231 | 0.481711769 | 0.30807858 | 1.56360032 | 0.11791146 | 0.26660915 |
| ACTL10 | 70.0686053 | -1.063148187 | 0.28781404 | -3.6938719 | 0.00022087 | 0.00236564 |
| AL121906.1 | 0.40130254 | 1.921760995 | 2.75588123 | 0.69733085 | 0.48559577 | NA |
| E2F1 | 86.7539678 | -0.875285295 | 0.37282234 | -2.3477276 | 0.01888833 | 0.07196724 |
| PXMP4 | 559.212326 | 1.08502261 | 0.23650637 | 4.5877099 | 4.48E-06 | 0.00010812 |
| AL050349.1 | 0 | NA | NA | NA | NA | NA |
| ZNF341 | 395.909377 | -0.360121044 | 0.20967356 | -1.717532 | 0.08588202 | 0.21302877 |
| ZNF341-AS1 | 30.2199063 | -0.34075044 | 0.45161903 | -0.7545086 | 0.45054389 | 0.64040763 |
| CHMP4B | 7434.96732 | -0.848964212 | 0.21493356 | -3.9498914 | 7.82E-05 | 0.00104465 |
| AL034549.1 | 130.303519 | 0.766022032 | 0.26576038 | 2.88237865 | 0.00394685 | 0.02254244 |
| RALY | 6642.89786 | -0.424938071 | 0.10857032 | -3.9139434 | 9.08E-05 | 0.00117316 |
| AL031668.1 | 2.02026874 | 1.213458806 | 0.97560565 | 1.24380051 | 0.21357299 | 0.39755147 |
| EIF2S2 | 4495.15889 | -1.025954317 | 0.23093731 | -4.4425664 | 8.89E-06 | 0.00018574 |
| ASIP | 20.0201356 | -1.026624703 | 0.60426293 | -1.6989702 | 0.0893248 | 0.21944242 |
| AHCY | 10835.5096 | 1.363956066 | 0.28692606 | 4.75368489 | 2.00E-06 | 5.65E-05 |
| ITCH | 2562.47815 | -0.083474483 | 0.15926555 | -0.5241214 | 0.60019411 | 0.75785859 |
| ITCH-AS1 | 0 | NA | NA | NA | NA | NA |
| DYNLRB1 | 5483.02555 | -0.283267952 | 0.11551663 | -2.4521833 | 0.01419923 | 0.05849612 |
| MAP1LC3A | 1904.32459 | -0.739657391 | 0.26371771 | -2.8047316 | 0.00503585 | 0.02700751 |
| PIGU | 1114.77898 | -0.314166569 | 0.16939174 | -1.8546747 | 0.06364269 | 0.17225267 |
| NCOA6 | 2399.23827 | -0.195250478 | 0.16204665 | -1.2049029 | 0.2282408 | 0.4153699 |
| TP53INP2 | 4379.06816 | 0.23572152 | 0.25377369 | 0.9288651 | 0.352959 | 0.55054589 |
| GGT7 | 2938.37481 | 0.062937166 | 0.23805868 | 0.26437669 | 0.79148967 | 0.88679396 |
| ACSS2 | 6448.10699 | 0.837411582 | 0.3176943 | 2.63590367 | 0.00839136 | 0.03943683 |
| GSS | 4468.04304 | 0.363604845 | 0.26959616 | 1.34870188 | 0.17743274 | 0.35252117 |
| MYH7B | 433.586936 | 0.237957548 | 0.21554775 | 1.10396677 | 0.26960755 | 0.46391851 |
| TRPC4AP | 5815.4196 | -0.206164175 | 0.13695176 | -1.505378 | 0.13222691 | 0.28840559 |
| EDEM2 | 2053.38399 | -0.361025821 | 0.11817041 | -3.0551289 | 0.00224964 | 0.0147945 |
| PROCR | 1002.99853 | -1.422871248 | 0.32966747 | -4.3160803 | 1.59E-05 | 0.00029599 |
| AL356652.1 | 325.78982 | -1.297984761 | 0.36354545 | -3.5703507 | 0.0003565 | 0.00348314 |
| MMP24OS | 5281.93837 | -0.118908661 | 0.1814999 | -0.6551445 | 0.51237473 | 0.69051575 |
| MMP24 | 3699.34809 | 0.070786152 | 0.31032843 | 0.22810077 | 0.8195679 | 0.9031808 |
| AL121753.1 | 4217.85536 | -0.467633917 | 0.31730323 | -1.4737761 | 0.14054189 | 0.30069693 |
| EIF6 | 7844.58234 | -0.441223642 | 0.31588712 | -1.3967763 | 0.16248085 | 0.33138509 |
| FAM83C-AS1 | 9.97839165 | 0.814683241 | 0.64842179 | 1.25640941 | 0.20896757 | 0.39217671 |
| FAM83C | 17.7460322 | -1.731845655 | 1.06975026 | -1.6189252 | 0.10546336 | 0.24643941 |
| UQCC1 | 3076.01137 | 0.098916842 | 0.11022802 | 0.89738386 | 0.36951412 | 0.56635208 |
| GDF5OS | 28.3160695 | -2.892249997 | 0.63307853 | -4.5685485 | 4.91E-06 | 0.00011593 |
| GDF5 | 55.8368416 | -3.080824639 | 0.59326325 | -5.1930145 | 2.07E-07 | 8.77E-06 |
| CEP250 | 1408.34517 | -0.561878445 | 0.30858975 | -1.8207942 | 0.06863814 | 0.18157983 |
| FO393401.1 | 15.7614528 | -0.561377754 | 0.46945899 | -1.1957972 | 0.2317757 | 0.4197736 |
| C20orf173 | 0.73521835 | -1.088818078 | 1.6980088 | -0.6412323 | 0.52137177 | NA |
| ERGIC3 | 9418.06724 | -0.171766084 | 0.10315047 | -1.6651993 | 0.095873 | 0.23084194 |
| SPAG4 | 582.445297 | -1.943427814 | 0.43272607 | -4.4911272 | 7.08E-06 | 0.00015556 |
| CPNE1 | 5608.09614 | -0.26846774 | 0.15330212 | -1.7512331 | 0.07990577 | 0.20227805 |
| AL109827.1 | 1897.19432 | 0.336500535 | 0.11679318 | 2.88116602 | 0.00396207 | 0.02261467 |
| RBM12 | 2651.57056 | -0.153100124 | 0.23861023 | -0.6416327 | 0.5211117 | 0.69754901 |
| NFS1 | 2986.46448 | 0.554859794 | 0.22045978 | 2.51683006 | 0.01184159 | 0.05109012 |
| ROMO1 | 2586.06163 | -0.68024639 | 0.14676712 | -4.634869 | 3.57E-06 | 9.01E-05 |
| RBM39 | 13819.3206 | -0.159514468 | 0.11095295 | -1.4376767 | 0.15052581 | 0.31515408 |
| PHF20 | 1292.92832 | -0.322750699 | 0.10992238 | -2.9361692 | 0.00332293 | 0.01984382 |
| SCAND1 | 2717.23027 | -0.464984914 | 0.18239785 | -2.5492895 | 0.01079427 | 0.04767919 |
| CNBD2 | 205.006853 | 0.477286705 | 0.2288007 | 2.08603692 | 0.03697528 | 0.11693909 |
| NORAD | 12813.6282 | 0.038452779 | 0.17122573 | 0.2245736 | 0.82231101 | 0.90506036 |
| AL035420.3 | 7.6358174 | 0.663450916 | 0.43502323 | 1.52509309 | 0.12723588 | 0.28080466 |
| AL035420.1 | 2.96070335 | 0.606931691 | 0.7937152 | 0.76467187 | 0.44446696 | 0.63537357 |
| AL035420.2 | 1.73168361 | 2.52692459 | 1.00157904 | 2.52294076 | 0.0116378 | 0.05047513 |
| EPB41L1 | 6628.37126 | 0.513023858 | 0.23613077 | 2.17262602 | 0.02980848 | 0.10045361 |
| AL121895.1 | 38.772535 | 1.033989494 | 0.57834147 | 1.78785293 | 0.07379974 | 0.19159137 |
| AL121895.2 | 21.3004165 | -0.232803038 | 0.35052135 | -0.6641622 | 0.50658649 | 0.68569375 |
| AAR2 | 2002.01638 | 0.025223274 | 0.15431897 | 0.16344895 | 0.87016496 | 0.9323196 |
| DLGAP4 | 4884.78994 | -0.45493181 | 0.21791351 | -2.0876715 | 0.03682748 | 0.11662149 |
| DLGAP4-AS1 | 1450.56263 | -0.454993616 | 0.19375737 | -2.3482648 | 0.01886111 | 0.0719122 |
| MYL9 | 11517.367 | -0.212274539 | 0.4870988 | -0.4357936 | 0.66298649 | 0.80186001 |
| TGIF2 | 1411.7347 | -0.114410351 | 0.17691844 | -0.6466841 | 0.51783639 | 0.69488619 |
| TGIF2-RAB5I | 5250.7182 | -1.498846409 | 0.38695691 | -3.8734194 | 0.00010732 | 0.0013406 |
| RAB5IF | 5504.99115 | -1.561290223 | 0.38646538 | -4.0399226 | 5.35E-05 | 0.0007855 |
| SLA2 | 1510.89518 | -1.627583111 | 0.36629506 | -4.4433663 | 8.86E-06 | 0.0001852 |
| NDRG3 | 1873.00301 | 0.202970912 | 0.21917641 | 0.92606184 | 0.35441384 | 0.55199607 |
| DSN1 | 452.498578 | 0.035114792 | 0.11556487 | 0.30385352 | 0.76123949 | 0.8677512 |
| SOGA1 | 2108.15224 | 0.15172812 | 0.22561226 | 0.67251716 | 0.50125452 | 0.68208275 |
| TLDC2 | 554.882 | -0.127803516 | 0.39093464 | -0.3269179 | 0.74373001 | 0.85622495 |
| SAMHD1 | 2914.03033 | -0.54845125 | 0.51706424 | -1.0607023 | 0.2888252 | 0.48502327 |
| RBL1 | 197.972903 | 0.240760795 | 0.16103689 | 1.4950661 | 0.13489719 | 0.29233919 |

| | | | | | | |
|---|---|---|---|---|---|---|
| AL136172.1 | 2.7037526 | 2.267665548 | 0.83345861 | 2.72078965 | 0.00651262 | 0.0327286 |
| MROH8 | 851.20687 | 0.334055421 | 0.20691441 | 1.61446187 | 0.10642728 | 0.24790102 |
| RPN2 | 13566.1988 | -0.138247264 | 0.1520273 | -0.9093581 | 0.36316111 | 0.56029027 |
| GHRH | 0.41986265 | -0.032126622 | 2.22020843 | -0.0144701 | 0.98845494 | NA |
| MANBAL | 2827.53455 | -0.29441808 | 0.15840576 | -1.8586324 | 0.06307927 | 0.17122095 |
| SRC | 4749.12067 | -0.378894663 | 0.12700384 | -2.9833324 | 0.00285128 | 0.01763328 |
| BLCAP | 5405.55674 | 0.157210099 | 0.09910516 | 1.58629587 | 0.11267221 | 0.25832775 |
| NNAT | 44.5057571 | 0.136762563 | 0.50240199 | 0.2722174 | 0.78545487 | 0.88299633 |
| LINC01746 | 0 | NA | NA | NA | NA | NA |
| LINC00489 | 0 | NA | NA | NA | NA | NA |
| AL162293.1 | 0 | NA | NA | NA | NA | NA |
| CTNNBL1 | 1643.54568 | -0.143842437 | 0.15236026 | -0.9440942 | 0.34512152 | 0.54275134 |
| VSTM2L | 127.534994 | -0.320488627 | 0.5531837 | -0.579353 | 0.56235102 | 0.72983653 |
| TTI1 | 1141.07734 | 0.123727506 | 0.16680097 | 0.74176731 | 0.45822833 | 0.64717911 |
| RPRD1B | 1919.79848 | 0.145077435 | 0.15825565 | 0.91672829 | 0.35928504 | 0.55692137 |
| AL031651.1 | 1.18262062 | -0.300491854 | 1.11826476 | -0.2687126 | 0.78815084 | 0.88474469 |
| TGM2 | 29097.5385 | -2.416413644 | 0.38210234 | -6.3239959 | 2.55E-10 | 3.25E-08 |
| KIAA1755 | 535.295684 | 0.785458666 | 0.64824511 | 1.21166925 | 0.22563901 | 0.41217102 |
| AL359555.3 | 29.8526055 | 0.730612061 | 0.59765374 | 1.22246714 | 0.22153102 | 0.40759381 |
| AL359555.4 | 29.8455853 | 1.034044505 | 0.57219681 | 1.80714833 | 0.07073915 | 0.18566652 |
| AL359555.2 | 0.92914288 | 0.574726364 | 1.29233777 | 0.44471838 | 0.65652328 | NA |
| AL359555.1 | 39.3789553 | 0.526975547 | 0.54692545 | 0.96352354 | 0.33528486 | 0.53355028 |
| BPI | 149.917016 | 2.108143722 | 0.55259305 | 3.81500225 | 0.00013618 | 0.00161616 |
| LBP | 2116.09985 | -3.280231471 | 0.68725731 | -4.7729306 | 1.82E-06 | 5.26E-05 |
| AL391095.2 | 83.276314 | -2.799668332 | 0.77403607 | -3.616974 | 0.00029807 | 0.00301382 |
| AL391095.1 | 29.8306873 | -0.83023916 | 0.50951735 | -1.629462 | 0.10321525 | 0.24279984 |
| AL391095.3 | 3.85827316 | -0.356556406 | 0.7006218 | -0.5089142 | 0.61081234 | 0.76667721 |
| RALGAPB | 2848.16112 | -0.099908203 | 0.09382029 | -1.0648891 | 0.28692612 | 0.48321891 |
| ADIG | 5.1147752 | 0.784397845 | 0.60511257 | 1.29628417 | 0.1948776 | 0.37491226 |
| ARHGAP40 | 596.582174 | -1.784294919 | 0.53074439 | -3.3618724 | 0.00077416 | 0.00640357 |
| SLC32A1 | 0.28236441 | 1.093535322 | 3.27839713 | 0.33355792 | 0.73871318 | NA |
| ACTR5 | 584.955367 | -0.449285493 | 0.19624281 | -2.2894368 | 0.02205399 | 0.08063834 |
| PPP1R16B | 2605.88716 | 1.41157859 | 0.63289683 | 2.23034547 | 0.02572452 | 0.09018286 |
| FAM83D | 196.952321 | 1.737933058 | 0.5386563 | 3.22642295 | 0.00125348 | 0.00937444 |
| AL023803.2 | 2.11416596 | 0.844074789 | 0.84206527 | 1.00238641 | 0.316157 | 0.51330256 |
| AL023803.1 | 4.44705656 | 0.665037814 | 0.72617307 | 0.91581173 | 0.35976565 | 0.55742181 |
| DHX35 | 826.993297 | 0.506064536 | 0.1370571 | 3.6923627 | 0.00022218 | 0.00237683 |
| LINC01734 | 0.08695548 | -0.642905668 | 3.30558948 | -0.1944905 | 0.84579185 | NA |
| AL118523.1 | 4.12892687 | -0.056709901 | 0.83751815 | -0.0677118 | 0.94601502 | 0.97409658 |
| LINC01370 | 0 | NA | NA | NA | NA | NA |
| AL133419.1 | 0 | NA | NA | NA | NA | NA |
| AL009050.1 | 0 | NA | NA | NA | NA | NA |
| MAFB | 1304.03278 | -0.156448726 | 0.20031153 | -0.7810271 | 0.43478657 | 0.62679433 |
| AL035665.1 | 3.77176689 | -0.467865724 | 1.09379337 | -0.427746 | 0.66883607 | 0.8065673 |
| TOP1 | 3814.82352 | -0.134111347 | 0.15704203 | -0.8539838 | 0.39311398 | 0.58859957 |
| PLCG1-AS1 | 312.832979 | 0.555657654 | 0.26701661 | 2.08098533 | 0.03743525 | 0.11788482 |
| PLCG1 | 5127.32026 | -0.222179825 | 0.19896679 | -1.1166679 | 0.26413635 | 0.45800076 |
| ZHX3 | 4804.23834 | 0.079418497 | 0.26827375 | 0.29603529 | 0.76720313 | 0.87152635 |
| LPIN3 | 7797.16665 | -0.684399567 | 0.45225363 | -1.5133092 | 0.13020114 | 0.2854035 |
| EMILIN3 | 2345.61283 | -1.192345499 | 0.44414673 | -2.6845757 | 0.00726219 | 0.03558593 |
| CHD6 | 1656.39407 | 0.170902567 | 0.19741751 | 0.86569104 | 0.38665962 | 0.58230806 |
| AL049812.2 | 0.08666371 | -0.642905668 | 3.30558948 | -0.1944905 | 0.84579185 | NA |
| AL049812.3 | 0.03392908 | 0.380740593 | 3.29738709 | 0.11546736 | 0.90807469 | NA |
| PTPRT | 26.2035904 | 1.058140865 | 0.72756813 | 1.45435296 | 0.14584848 | 0.30843912 |
| AL031656.1 | 0.09644756 | -0.642905668 | 3.30558948 | -0.1944905 | 0.84579185 | NA |
| AL031676.1 | 0 | NA | NA | NA | NA | NA |
| AL021395.1 | 0.1357163 | 1.313406857 | 3.28103699 | 0.40030236 | 0.68893383 | NA |
| SRSF6 | 10075.2242 | 0.351722478 | 0.15892931 | 2.21307496 | 0.02689248 | 0.09317496 |
| L3MBTL1 | 1121.3193 | -0.347867044 | 0.15672337 | -2.2196245 | 0.02644427 | 0.09206962 |
| Z98752.3 | 783.233914 | 0.852333785 | 0.25622306 | 3.32653033 | 0.00087934 | 0.00709991 |
| Z98752.1 | 136.809244 | -0.801769382 | 0.22912902 | -3.4992049 | 0.00046665 | 0.00430413 |
| SGK2 | 4157.12368 | 0.834639043 | 0.29600242 | 2.81970341 | 0.00480681 | 0.02609431 |
| Z98752.4 | 4.10359281 | -0.10933082 | 0.55663287 | -0.1964146 | 0.84428567 | 0.9176413 |
| IFT52 | 1175.15654 | -0.059159827 | 0.16060685 | -0.3683518 | 0.71261092 | 0.8366674 |
| MYBL2 | 65.5977429 | -2.100623103 | 0.70709141 | -2.9707943 | 0.00297031 | 0.01819568 |
| GTSF1L | 0.54137467 | -0.124622999 | 1.71349343 | -0.0727304 | 0.94202069 | NA |
| LINC01728 | 0 | NA | NA | NA | NA | NA |
| TOX2 | 184.966556 | -0.415624745 | 0.37838024 | -1.0984314 | 0.27201614 | 0.4668445 |
| JPH2 | 277.467795 | -0.271914235 | 0.52838295 | -0.5146158 | 0.60682149 | 0.76327544 |
| OSER1 | 2455.82883 | -0.86531059 | 0.29539744 | -2.9293097 | 0.00339716 | 0.0201446 |
| OSER1-DT | 366.434523 | 0.217959212 | 0.30304372 | 0.71923356 | 0.47199702 | 0.65821986 |
| GDAP1L1 | 6.95083524 | 0.283325491 | 0.5157288 | 0.54936915 | 0.58275214 | 0.74511123 |
| FITM2 | 1199.14226 | 0.3646576 | 0.30149951 | 1.2094799 | 0.22647853 | 0.4133196 |
| R3HDML | 87.6206387 | 5.683365681 | 1.42853543 | 3.97845623 | 6.94E-05 | 0.00094872 |
| AL117382.1 | 91.737557 | 4.898167329 | 1.57642521 | 3.10713588 | 0.0018891 | 0.0129089 |
| HNF4A | 5796.3998 | 2.280531655 | 0.74173303 | 3.07459904 | 0.00210786 | 0.01401165 |
| HNF4A-AS1 | 106.116624 | 4.13931478 | 1.33305337 | 3.10513807 | 0.0019019 | 0.01296282 |
| AL117382.2 | 6.93557698 | 3.67170264 | 1.13324703 | 3.23998435 | 0.00119536 | 0.00902501 |
| LINC01620 | 3.06050848 | 1.027786705 | 0.71191432 | 1.44369439 | 0.14882496 | 0.31286357 |
| TTPAL | 732.963536 | -0.171285375 | 0.17059256 | -1.0040612 | 0.31534911 | 0.51249574 |
| SERINC3 | 6017.44204 | -0.10690447 | 0.10574577 | -1.0109574 | 0.31203681 | 0.50905783 |
| PKIG | 3425.50346 | -0.188169449 | 0.18925233 | -0.9942781 | 0.32008749 | 0.5175173 |
| ADA | 327.64815 | -1.066073884 | 0.39692861 | -2.6858076 | 0.00723547 | 0.03552401 |
| AL139352.1 | 9.20031066 | -0.466980883 | 0.69790676 | -0.6691164 | 0.5034212 | 0.68350956 |
| LINC01260 | 7.78137198 | -0.387132354 | 0.71518373 | -0.5413048 | 0.58829754 | 0.74904598 |
| KCNK15-AS1 | 24.538638 | -2.091394667 | 0.55369019 | -3.7771929 | 0.00015861 | 0.00183119 |
| WISP2 | 55.766577 | 0.687024051 | 0.9970995 | 0.68902256 | NA | NA |
| KCNK15 | 145.897409 | -2.727561507 | 0.49750153 | -5.4825188 | 4.19E-08 | 2.29E-06 |
| RIMS4 | 33.0704178 | -0.540805201 | 0.65837822 | -0.8214203 | 0.41140693 | 0.60563841 |
| YWHAB | 11770.6099 | -0.125966101 | 0.16494472 | -0.7636868 | 0.44505393 | 0.63561916 |
| PABPC1L | 3339.57018 | 0.097296129 | 0.31526875 | 0.3086133 | 0.7576157 | 0.865984 |
| TOMM34 | 2400.698 | -0.208297999 | 0.16501804 | -1.2622741 | 0.20685018 | 0.38947799 |
| STK4-AS1 | 13.5209095 | 1.489338973 | 0.55585303 | 2.67937546 | 0.00737596 | 0.03599672 |
| STK4 | 1135.04998 | -0.373644665 | 0.1659264 | -2.2518699 | 0.0243305 | 0.08648001 |
| KCNS1 | 33.8307877 | -1.478172261 | 0.48922602 | -3.0214506 | 0.00251567 | 0.01607028 |
| WFDC5 | 9.47480421 | 0.632674623 | 0.92185192 | 0.6863083 | 0.49251871 | 0.67538838 |
| WFDC12 | 7.6694211 | -1.039267215 | 1.05463439 | -0.9854289 | 0.32441344 | 0.52192497 |
| PI3 | 24.9808704 | -1.579535686 | 1.08181527 | -1.4600789 | 0.14426838 | 0.30606253 |
| SEMG1 | 5.57498764 | -2.941020047 | 1.13650933 | -2.5877659 | 0.00966006 | 0.04391194 |
| SEMG2 | 72.2908443 | -1.781006109 | 0.66328821 | -2.6851165 | 0.00725045 | 0.03554392 |
| SLPI | 36715.1927 | -6.196339295 | 0.7214929 | -8.5882194 | 8.83E-18 | 1.01E-14 |
| MATN4 | 18.257639 | 0.300019957 | 0.37270602 | 0.80497749 | 0.42083267 | 0.61352042 |
| RBPJL | 5.64883331 | -0.01446676 | 0.60088247 | -0.0240759 | 0.9807921 | 0.99132745 |
| SDC4 | 22509.5999 | -0.573732559 | 0.13687957 | -4.1915134 | 2.77E-05 | 0.00046102 |
| SYS1 | 2875.8562 | -0.230649841 | 0.19835805 | -1.1627955 | 0.24491249 | 0.43599208 |
| SYS1-DBNDD | 2135.40408 | -0.672564809 | 0.21591773 | -3.1149124 | 0.00184 | 0.0126356 |
| TP53TG5 | 131.730933 | -0.055712037 | 0.22234133 | -0.2505699 | 0.80214668 | 0.89266577 |
| DBNDD2 | 1847.25824 | -0.860600656 | 0.25897905 | -3.3230513 | 0.00089039 | 0.00716343 |
| PIGT | 5492.61974 | -0.107988717 | 0.17050138 | -0.6333598 | 0.52649873 | 0.7013777 |
| AL031663.3 | 50.3088415 | -0.446311249 | 0.43750148 | -1.0201365 | 0.30766371 | 0.50448277 |
| WFDC2 | 29298.0046 | -3.316046637 | 0.41594096 | -7.9723926 | 1.56E-15 | 1.11E-12 |
| AL031663.2 | 1.33355982 | -0.975184312 | 1.70206164 | -0.572943 | 0.56668328 | 0.73295263 |
| SPINT3 | 0.35141535 | 0.016916985 | 2.88972606 | 0.00585418 | 0.99532906 | NA |

| | | | | | | | |
|---|---|---|---|---|---|---|---|
| WFDC6 | 1.76585741 | | -1.063185407 | 1.06601059 | -0.9973498 | 0.31859477 | 0.51589463 |
| EPPIN-WFDC | 0.91792312 | | -0.765986067 | 1.28699988 | -0.5951718 | 0.55172862 | NA |
| EPPIN | 4.30797214 | | -2.22842182 | 0.8233518 | -2.7065245 | 0.00679916 | 0.03381625 |
| WFDC8 | 0.9328846 | | -0.38567528 | 1.4959797 | -0.2578078 | 0.79655522 | NA |
| WFDC9 | 0 | NA | | NA | NA | NA | NA |
| WFDC10A | 0.07659028 | | -0.642905668 | 3.30558948 | -0.1944905 | 0.84579185 | NA |
| WFDC11 | 0.52109599 | | -0.05792738 | 1.93551946 | -0.0299286 | 0.976124 | NA |
| WFDC10B | 2.13834437 | | -1.237143416 | 1.12914311 | -1.095648 | 0.27323284 | 0.46819503 |
| WFDC13 | 0.99110201 | | 0.454633601 | 1.48591975 | 0.30596107 | 0.75963428 | NA |
| SPINT4 | 0 | NA | | NA | NA | NA | NA |
| WFDC3 | 25.8743554 | | 0.589686363 | 0.46956289 | 1.25581977 | 0.20918132 | 0.39235447 |
| DNTTIP1 | 1104.48414 | | -0.382060012 | 0.09508445 | -4.0181123 | 5.87E-05 | 0.00083623 |
| UBE2C | 63.1535025 | | -2.40570291 | 0.66271234 | -3.6300862 | 0.00028333 | 0.00289806 |
| TNNC2 | 25.352521 | | 1.040246316 | 0.43433617 | 2.39502574 | 0.0166192 | 0.06559621 |
| SNX21 | 2150.46491 | | 0.167978775 | 0.20472904 | 0.82049316 | 0.41193503 | 0.60590845 |
| ACOT8 | 1926.8892 | | 0.270159934 | 0.17900769 | 1.50920858 | 0.13124549 | 0.28690571 |
| ZSWIM3 | 189.409171 | | 0.481006791 | 0.21084391 | 2.28134072 | 0.02252829 | 0.08186704 |
| ZSWIM1 | 820.116706 | | 0.153668486 | 0.16641285 | 0.9234172 | 0.35578983 | 0.55319445 |
| SPATA25 | 232.978973 | | 1.157818581 | 0.2902495 | 3.98904587 | 6.63E-05 | 0.00092169 |
| NEURL2 | 483.309317 | | -0.038226029 | 0.31112472 | -0.122864 | 0.90221479 | 0.95066747 |
| CTSA | 27723.0531 | | -0.058068828 | 0.2414642 | -0.2404863 | 0.8099533 | 0.89730154 |
| AL008726.1 | 6085.38041 | | 0.292345828 | 0.29689336 | 0.98468295 | 0.32477983 | 0.52234961 |
| PLTP | 17455.4061 | | -1.489780726 | 0.41618908 | -3.5795767 | 0.00034415 | 0.00338251 |
| PCIF1 | 2372.54468 | | 0.071501526 | 0.16798702 | 0.42563722 | 0.67037222 | 0.80748317 |
| ZNF335 | 2013.98332 | | -0.047215415 | 0.25089638 | -0.1881869 | 0.85073013 | 0.92161226 |
| MMP9 | 405.483877 | | -1.277208036 | 0.80523687 | -1.5861271 | 0.11271048 | 0.2583377 |
| SLC12A5-AS1 | 150.624503 | | -1.473800371 | 0.73384439 | -2.0083282 | 0.04460843 | 0.13396245 |
| SLC12A5 | 19.4003759 | | -1.193404974 | 0.47997766 | -2.4863761 | 0.01290515 | 0.0544407 |
| NCOA5 | 2111.60868 | | -0.170481524 | 0.28438332 | -0.5994779 | 0.54885422 | 0.71945547 |
| CD40 | 1367.24614 | | -0.896078472 | 0.37665279 | -2.379057 | 0.01735699 | 0.0676516 |
| AL031687.1 | 0 | NA | | NA | NA | NA | NA |
| CDH22 | 25.4094485 | | 0.331332562 | 0.56758146 | 0.58376213 | 0.55938036 | 0.72772184 |
| SLC35C2 | 3424.33448 | | -0.336284269 | 0.12537008 | -2.6823327 | 0.00731107 | 0.03578191 |
| AL133227.1 | 328.400826 | | -0.537890519 | 0.15136195 | -3.5536706 | 0.00037989 | 0.00366416 |
| ELMO2 | 2521.48771 | | -0.252516491 | 0.12594887 | -2.0049127 | 0.04497238 | 0.1347778 |
| ZNF334 | 540.018339 | | 0.305257913 | 0.40206579 | 0.7592238 | 0.44771869 | 0.63783587 |
| OCSTAMP | 10.6834859 | | -0.097584939 | 0.560468 | -0.1741133 | 0.86177641 | 0.92800735 |
| SLC13A3 | 44287.9153 | | 1.040203284 | 0.43447702 | 2.39415027 | 0.01665892 | 0.06571365 |
| TP53RK | 798.493977 | | -0.530486606 | 0.25226519 | -2.1028927 | 0.03547515 | 0.11369468 |
| AL031055.1 | 9.88602971 | | -0.884256815 | 0.53627119 | -1.6488986 | 0.09916841 | 0.23616643 |
| SLC2A10 | 400.949034 | | 0.106153325 | 0.27020176 | 0.39286688 | 0.6944178 | 0.8239763 |
| EYA2 | 467.703985 | | 0.937712957 | 0.46680353 | 2.00879576 | 0.0445588 | 0.13385914 |
| AL354766.2 | 0.79299602 | | -1.657317883 | 1.83793421 | -0.9017286 | 0.36720104 | NA |
| ZMYND8 | 2324.54508 | | -0.348336117 | 0.22628743 | -1.5393525 | 0.12371826 | 0.27528905 |
| AL031666.2 | 9.66473461 | | -0.305883899 | 0.41804552 | -0.7317 | 0.4643517 | 0.65181276 |
| AL031666.1 | 25.9118359 | | -0.310310331 | 0.39261706 | -0.7903638 | 0.42931531 | 0.62135263 |
| LINC01754 | 0.52131791 | | -1.233797142 | 1.87967573 | -0.6563883 | 0.51157432 | NA |
| NCOA3 | 1938.78644 | | -0.192126306 | 0.1558601 | -1.2326843 | 0.21769356 | 0.40265881 |
| SULF2 | 6808.79964 | | -0.689939127 | 0.26492955 | -2.6042362 | 0.00920793 | 0.0423395 |
| AL354813.1 | 646.921936 | | 0.01720941 | 0.30729019 | 0.05600377 | 0.9553388 | 0.97856915 |
| AL357558.2 | 0 | NA | | NA | NA | NA | NA |
| AL357558.1 | 0.08695548 | | -0.642905668 | 3.30558948 | -0.1944905 | 0.84579185 | NA |
| LINC01522 | 0.43853044 | | -0.653330851 | 2.29601579 | -0.2845498 | 0.77598907 | NA |
| LINC01523 | 0 | NA | | NA | NA | NA | NA |
| AL139351.1 | 2.90838619 | | -1.834331363 | 0.87911734 | -2.0865603 | 0.0369279 | 0.11680324 |
| LINC00494 | 110.432133 | | 1.193298151 | 0.60398146 | 1.97571982 | 0.04818652 | 0.14180097 |
| AL137078.1 | 0.09997307 | | 0.547239394 | 3.29377202 | 0.16614368 | 0.86804388 | NA |
| AL137078.2 | 0.09635527 | | 0.547239394 | 3.29377202 | 0.16614368 | 0.86804388 | NA |
| AL049541.1 | 0 | NA | | NA | NA | NA | NA |
| AL049541.2 | 0 | NA | | NA | NA | NA | NA |
| PREX1 | 1721.54588 | | -0.047091221 | 0.27248165 | -0.1728235 | 0.8627902 | 0.92851314 |
| AL133342.1 | 5.24878677 | | -0.11233777 | 0.60744213 | -0.1849358 | 0.85327943 | 0.92322961 |
| ARFGEF2 | 2874.41051 | | 0.046889524 | 0.15831878 | 0.29617159 | 0.76709905 | 0.87152635 |
| CSE1L-AS1 | 2.10156251 | | 1.95439676 | 1.10281763 | 1.77218491 | 0.07636387 | 0.1961526 |
| CSE1L | 2523.48291 | | 0.005125893 | 0.12419661 | 0.04127241 | 0.96707873 | 0.98447599 |
| STAU1 | 7411.36901 | | 0.362521753 | 0.21526175 | 1.68409739 | 0.09216285 | 0.22442832 |
| DDX27 | 2624.65465 | | -0.322137046 | 0.11266797 | -2.8591715 | 0.00424749 | 0.02380581 |
| ZNFX1 | 3924.44656 | | -0.862862948 | 0.41009802 | -2.1040407 | 0.03537489 | 0.11351131 |
| ZFAS1 | 2242.63788 | | -1.073039451 | 0.23517971 | -4.5626362 | 5.05E-06 | 0.00011784 |
| KCNB1 | 74.8752177 | | 0.669208219 | 0.28930857 | 2.31312961 | 0.02071551 | 0.07688503 |
| PTGIS | 825.369582 | | -0.07325001 | 0.47040072 | -0.1557183 | 0.87625506 | 0.93555133 |
| B4GALT5 | 6315.94801 | | -0.888240336 | 0.28724684 | -3.0922545 | 0.00198642 | 0.01342096 |
| SLC9A8 | 1895.08652 | | 0.114942539 | 0.18065381 | 0.63625858 | 0.5246079 | 0.69953025 |
| SPATA2 | 1000.18011 | | -0.131320974 | 0.16084574 | -0.8164405 | 0.41424826 | 0.60794725 |
| RNF114 | 5752.39337 | | -0.405933909 | 0.19753845 | -2.0549615 | 0.03988272 | 0.12358489 |
| SNAI1 | 448.393484 | | -0.927107888 | 0.44681027 | -2.0749476 | 0.0379914 | 0.11920908 |
| TRERNA1 | 0.08757577 | | 0.547239394 | 3.29377202 | 0.16614368 | 0.86804388 | NA |
| UBE2V1 | 6548.82871 | | -0.494131158 | 0.13612464 | -3.6299905 | 0.00028343 | 0.00289806 |
| TMEM189-U | 6893.60642 | | -0.508756781 | 0.13800022 | -3.6866375 | 0.00022724 | 0.00242208 |
| TMEM189 | 1585.72041 | | -0.825285872 | 0.16418052 | -5.0266978 | 4.99E-07 | 1.80E-05 |
| AL161937.2 | 5.76004907 | | -0.97369147 | 1.11621705 | -0.8723137 | 0.38303724 | 0.57895142 |
| AL161937.1 | 0.95459271 | | -1.157876486 | 1.42974732 | -0.8098469 | 0.41802815 | NA |
| LINC01273 | 57.6452242 | | -0.235519871 | 0.46444252 | -0.5071023 | 0.61208304 | 0.76768751 |
| CEBPB-AS1 | 243.649952 | | -0.92925003 | 0.27918343 | -3.328457 | 0.00087328 | 0.0070618 |
| CEBPB | 3061.90648 | | -2.525494759 | 0.34113531 | -7.4032053 | 1.33E-13 | 5.08E-11 |
| SMIM25 | 66.7312634 | | -1.007839156 | 0.61426089 | -1.6407347 | 0.1008525 | 0.23892285 |
| LINC01270 | 31.6343733 | | -0.748580756 | 0.42767243 | -1.7503601 | 0.08005619 | 0.20256015 |
| LINC01271 | 20.3982208 | | -0.80371189 | 0.41363154 | -1.9430624 | 0.05200862 | 0.14987075 |
| PTPN1 | 9279.66469 | | -1.83488338 | 0.32535383 | -5.6396551 | 1.70E-08 | 1.08E-06 |
| AL133230.1 | 466.218307 | | -1.394575956 | 0.44826248 | -3.11107 | 0.00186411 | 0.01275467 |
| RIPOR3 | 3664.51678 | | 0.526899254 | 0.25201876 | 2.09071445 | 0.03655367 | 0.11609216 |
| AL353653.1 | 16.9380872 | | -0.709190089 | 0.52602435 | -1.3482077 | 0.17759158 | 0.35275879 |
| PARD6B | 2836.35574 | | 0.430071864 | 0.28152566 | 1.5276471 | 0.12660018 | 0.27990666 |
| BCAS4 | 557.732384 | | -0.541986216 | 0.22645404 | -2.3933608 | 0.01669482 | 0.06581586 |
| ADNP | 3548.33634 | | 0.048667405 | 0.12694407 | 0.38337675 | 0.70144044 | 0.82897792 |
| ADNP-AS1 | 91.0580314 | | 0.722601195 | 0.31602213 | 2.28655249 | 0.02222196 | 0.08106142 |
| DPM1 | 1709.76703 | | -0.26212342 | 0.11986027 | -2.1869083 | 0.02874922 | 0.09788469 |
| MOCS3 | 395.972786 | | 0.052985004 | 0.19697341 | 0.26899573 | 0.78793297 | 0.88457544 |
| AL050404.1 | 4.65892708 | | 0.080463 | 0.64952745 | 0.12387929 | 0.90141085 | 0.95006982 |
| KCNG1 | 19.142425 | | 0.380448342 | 0.37509343 | 1.01427622 | 0.31045097 | 0.50741242 |
| NFATC2 | 349.682225 | | 0.15133804 | 0.3389053 | 0.44654964 | 0.65520026 | 0.79767456 |
| ATP9A | 2236.20563 | | -0.236773878 | 0.20851833 | -1.1355063 | 0.25616324 | 0.44898802 |
| SALL4 | 33.3949667 | | -0.380380798 | 0.47516373 | -0.8005257 | 0.42340626 | 0.61597832 |
| LINC01429 | 0.82795587 | | 1.293824276 | 1.85259822 | 0.69838363 | 0.48493731 | NA |
| ZFP64 | 692.675941 | | -0.114376619 | 0.15143781 | -0.7552712 | 0.45008627 | 0.63999865 |
| LINC01524 | 0.08757577 | | 0.547239394 | 3.29377202 | 0.16614368 | 0.86804388 | NA |
| AL109610.1 | 0 | NA | | NA | NA | NA | NA |
| AL049765.1 | 0 | NA | | NA | NA | NA | NA |
| AL031674.1 | 0.53470053 | | 1.674908073 | 1.70067672 | 0.984848 | 0.32469874 | NA |
| AL121902.1 | 0.08666371 | | -0.642905668 | 3.30558948 | -0.1944905 | 0.84579185 | NA |
| TSHZ2 | 220.721803 | | -0.241995148 | 0.32249847 | -0.7503761 | 0.45302821 | 0.64266256 |
| AL391097.1 | 0.08695548 | | -0.642905668 | 3.30558948 | -0.1944905 | 0.84579185 | NA |
| AL391097.2 | 0.14616582 | | 0.435199075 | 3.28898345 | 0.13232024 | 0.894731 | NA |

| | | | | | | |
|---|---|---|---|---|---|---|
| AL109930.1 | 11.3644162 | -0.026278965 | 0.44225748 | -0.0594201 | 0.95261754 | 0.97717271 |
| AL354993.1 | 1.85931143 | -0.693796457 | 1.1145926 | -0.6224664 | 0.53363522 | 0.70738254 |
| AL354993.2 | 0.74929024 | -1.191133711 | 1.89392983 | -0.6289218 | 0.52940027 | NA |
| AL157838.1 | 121.203682 | 0.041738176 | 0.28498041 | 0.14645981 | 0.88355842 | 0.93960703 |
| ZNF217 | 1915.38383 | -0.633490752 | 0.16559577 | -3.825525 | 0.00013049 | 0.0015606 |
| AC005808.1 | 0.12059279 | 0.02007176 | 3.29738704 | 0.00608717 | 0.99514317 | NA |
| AC006076.1 | 0.09989236 | -0.642905668 | 3.30558948 | -0.1944905 | 0.84579185 | NA |
| BCAS1 | 185.87354 | -0.760066555 | 0.72649235 | -1.0462141 | 0.29546218 | 0.4922814 |
| AC005220.1 | 15.4205721 | 1.730072449 | 0.82218347 | 2.10424135 | 0.0353574 | 0.11348279 |
| CYP24A1 | 7888.00758 | -1.620633128 | 0.60273122 | -2.6888156 | 0.0071706 | 0.03526433 |
| PFDN4 | 626.507038 | -0.780614789 | 0.1651497 | -4.7267103 | 2.28E-06 | 6.31E-05 |
| DOK5 | 301.050529 | -0.578281201 | 0.41964251 | -1.3780329 | 0.16819312 | 0.33950753 |
| LINC01440 | 0.03392908 | 0.380740593 | 3.29738709 | 0.11546736 | 0.90807469 | NA |
| LINC01441 | 0 | NA | NA | NA | NA | NA |
| AL160413.1 | 0 | NA | NA | NA | NA | NA |
| CBLN4 | 33.9920213 | 3.237431022 | 0.71974998 | 4.49799389 | 6.86E-06 | 0.00015211 |
| MC3R | 0 | NA | NA | NA | NA | NA |
| AL139824.1 | 0.15543392 | 1.368031615 | 3.28033652 | 0.41704002 | 0.67664913 | NA |
| FAM210B | 2307.67719 | 0.393555021 | 0.12920941 | 3.04586954 | 0.00232008 | 0.01515498 |
| AURKA | 204.97217 | -0.406055292 | 0.19920069 | -2.0384231 | 0.04150764 | 0.1272888 |
| CSTF1 | 1272.46275 | -0.165182098 | 0.13096993 | -1.2612216 | 0.20722903 | 0.39002423 |
| CASS4 | 77.6128596 | 0.741304558 | 0.41480444 | 1.78711818 | 0.07391839 | 0.19179651 |
| RTF2 | 4520.1543 | 0.135162002 | 0.11037566 | 1.2245635 | 0.22073973 | 0.40665828 |
| GCNT7 | 13.997809 | 0.282958612 | 0.4159197 | 0.68032029 | 0.49630168 | 0.67814266 |
| FAM209A | 1532.73067 | 0.124334254 | 0.11089717 | 1.12116705 | 0.26221676 | 0.45590135 |
| FAM209B | 15.152405 | -0.709060826 | 0.44767428 | -1.5838766 | 0.1132218 | 0.25926179 |
| LINC01716 | 0.08393208 | -0.642905668 | 3.30558948 | -0.1944905 | 0.84579185 | NA |
| TFAP2C | 96.4661077 | -0.839329964 | 0.6920487 | -1.2128192 | 0.22519895 | 0.41151291 |
| AL133232.1 | 0 | NA | NA | NA | NA | NA |
| AL157414.2 | 0 | NA | NA | NA | NA | NA |
| AL157414.1 | 0.07830766 | -0.642905668 | 3.30558948 | -0.1944905 | 0.84579185 | NA |
| BMP7 | 426.791461 | 1.19509278 | 0.35053475 | 3.40934186 | 0.0006512 | 0.00556196 |
| BMP7-AS1 | 0 | NA | NA | NA | NA | NA |
| AL122058.1 | 3.31170081 | 1.233770244 | 0.87421682 | 1.41128633 | 0.1581602 | 0.32535179 |
| SPO11 | 0 | NA | NA | NA | NA | NA |
| RAE1 | 1510.76784 | -0.32124807 | 0.17632908 | -1.8218667 | 0.06847521 | 0.18125325 |
| MTRNR2L3 | 0.49616683 | 2.203310989 | 2.68475322 | 0.82067542 | 0.41183117 | NA |
| AL109955.1 | 229.923058 | 0.128617627 | 0.23521257 | 0.54681443 | 0.58450623 | 0.74648301 |
| RBM38 | 1640.19119 | -0.10420471 | 0.20164175 | -0.5167814 | 0.60530875 | 0.76218921 |
| CTCFL | 6.47907023 | 4.752930954 | 1.17398217 | 4.04855465 | 5.15E-05 | 0.0007609 |
| PCK1 | 73972.5391 | 1.061553924 | 0.87012856 | 1.21999664 | 0.22246615 | 0.4084406 |
| AL035541.1 | 0.43394103 | -0.91819521 | 2.24922326 | -0.4082277 | 0.68310652 | NA |
| ZBP1 | 87.7746452 | -0.520648316 | 0.76767519 | -0.6782143 | 0.49763581 | 0.67936717 |
| PMEPA1 | 1244.25843 | 0.124893258 | 0.36364439 | 0.34344888 | 0.73126079 | 0.8482162 |
| NKILA | 75.4635085 | -0.107569877 | 0.32442497 | -0.3315709 | 0.74021331 | 0.85420604 |
| AL354984.1 | 0.10178723 | 1.087385526 | 3.28420134 | 0.33109588 | 0.74057207 | NA |
| LINC01742 | 0 | NA | NA | NA | NA | NA |
| AL354984.2 | 0 | NA | NA | NA | NA | NA |
| C20orf85 | 1.31399613 | -3.604420336 | 2.66920023 | -1.3503747 | 0.17689584 | 0.3518702 |
| ANKRD60 | 0 | NA | NA | NA | NA | NA |
| RAB22A | 2517.48975 | -0.405768039 | 0.25022555 | -1.6216091 | 0.10488707 | 0.24546254 |
| VAPB | 4267.85764 | -0.008235052 | 0.18344849 | -0.0448903 | 0.96419477 | 0.98324824 |
| APCDD1L | 247.824843 | 1.185982802 | 0.32646496 | 3.63280277 | 0.00028036 | 0.00288117 |
| AL050327.1 | 2.38871618 | 2.160778311 | 0.85377768 | 2.53084422 | 0.01137884 | 0.04964592 |
| LINC01711 | 10.5897685 | 1.567739238 | 0.57102134 | 2.74550026 | 0.00604187 | 0.03094517 |
| STX16 | 5542.46446 | -0.106515193 | 0.288938 | -0.3686438 | 0.71239327 | 0.8365236 |
| STX16-NPEPI | 5497.71472 | 0.160203737 | 0.19164803 | 0.83592685 | 0.40319605 | 0.59815921 |
| NPEPL1 | 5655.875 | -0.048776916 | 0.2009146 | -0.2427744 | 0.80818019 | 0.89624087 |
| AL132655.3 | 0 | NA | NA | NA | NA | NA |
| AL132655.2 | 10.785426 | -1.132787426 | 0.67837006 | -1.6698665 | 0.09494578 | 0.22927708 |
| GNAS-AS1 | 65.5361138 | 1.515534814 | 0.39441948 | 3.84244416 | 0.00012182 | 0.00148099 |
| AL132655.1 | 0.35323289 | 0.764092886 | 2.89597703 | 0.26384632 | 0.79189834 | NA |
| GNAS | 37276.5661 | -0.13600674 | 0.15972409 | -0.8515105 | 0.39448584 | 0.58975305 |
| AL121917.1 | 25.6456246 | -0.133802441 | 0.41815481 | -0.319983 | 0.7489812 | 0.8597841 |
| AL121917.2 | 11.6924419 | 0.595109235 | 0.56575437 | 1.05188624 | 0.29285174 | 0.48929231 |
| NELFCD | 4252.15574 | 0.151549301 | 0.10063715 | 1.50589818 | 0.13209331 | 0.2882334 |
| CTSZ | 10253.8306 | -0.736123465 | 0.17639374 | -4.1731835 | 3.00E-05 | 0.00048954 |
| TUBB1 | 95.1988209 | 0.586658142 | 0.33473808 | 1.75258862 | 0.07967265 | 0.20186085 |
| ATP5F1E | 10338.4468 | -0.359471691 | 0.17775259 | -2.0223148 | 0.04314384 | 0.13098124 |
| PRELID3B | 2630.73803 | -0.26864566 | 0.18134334 | -1.48142 | 0.13849468 | 0.297668 |
| ZNF831 | 50.9608094 | -0.079142863 | 0.68639218 | -0.1153027 | 0.90820521 | 0.95415822 |
| EDN3 | 4.76046048 | 1.646647313 | 0.76900386 | 2.14127316 | 0.03225202 | 0.10622638 |
| AL035250.1 | 0 | NA | NA | NA | NA | NA |
| AL035250.2 | 0 | NA | NA | NA | NA | NA |
| AL389889.1 | 0 | NA | NA | NA | NA | NA |
| PHACTR3 | 145.44376 | -1.118233293 | 0.50720272 | -2.2047068 | 0.02747467 | 0.09473158 |
| AL121908.1 | 0 | NA | NA | NA | NA | NA |
| SYCP2 | 208.702712 | 0.253429721 | 0.23271801 | 1.08899919 | 0.27615424 | 0.47149878 |
| FAM217B | 464.737041 | 0.487424455 | 0.2288461 | 2.12992246 | 0.03317802 | 0.10838288 |
| PPP1R3D | 477.677421 | -0.143872737 | 0.16493609 | -0.8722938 | 0.3830481 | 0.57895142 |
| CDH26 | 44.2994722 | 0.044275816 | 0.31795836 | 0.13925036 | 0.88925231 | 0.94277263 |
| C20orf197 | 11.5308923 | 0.529718683 | 0.56582479 | 0.93618854 | 0.34917612 | 0.54677995 |
| AL132822.1 | 2.8185347 | -2.001618593 | 1.31415185 | -1.5231258 | 0.12772724 | 0.28152876 |
| MIR646HG | 22.8964127 | 0.419118981 | 0.48912537 | 0.85687434 | 0.39151434 | 0.5872047 |
| AL117372.1 | 0.20687462 | 0.68915539 | 3.1020265 | 0.22216296 | 0.82418703 | NA |
| AL139348.1 | 0 | NA | NA | NA | NA | NA |
| AL121910.1 | 0 | NA | NA | NA | NA | NA |
| LINC01718 | 0 | NA | NA | NA | NA | NA |
| CDH4 | 118.134226 | 0.023887139 | 0.30368587 | 0.07865739 | 0.93730514 | 0.96999796 |
| BX640515.1 | 0 | NA | NA | NA | NA | NA |
| AL160412.1 | 0.14444843 | 0.435199075 | 3.28898345 | 0.13232024 | 0.894731 | NA |
| AL162457.1 | 0.5593837 | 3.091191619 | 3.19080836 | 0.96878009 | 0.33265492 | NA |
| AL162457.2 | 0.44476003 | 1.401704438 | 3.23429052 | 0.43338854 | 0.66473253 | NA |
| TAF4 | 1141.51904 | -0.013779078 | 0.14948918 | -0.0921744 | 0.92655946 | 0.96437374 |
| AL137077.2 | 90.7167811 | 0.007247657 | 0.18165004 | 0.03989901 | 0.96817364 | 0.98514449 |
| LSM14B | 2234.11937 | 0.155550086 | 0.110085 | 1.41299987 | 0.15765576 | 0.3247955 |
| PSMA7 | 6796.53522 | -0.542451836 | 0.18971803 | -2.8592529 | 0.0042464 | 0.02380476 |
| SS18L1 | 1774.63118 | 0.125240837 | 0.16299725 | 0.76836166 | 0.44227235 | 0.63338963 |
| MTG2 | 1971.42351 | -0.017002021 | 0.13261584 | -0.1282051 | 0.8979867 | 0.94810655 |
| HRH3 | 1.41835733 | -0.924499461 | 1.31105299 | -0.705158 | 0.48071194 | 0.66542234 |
| OSBPL2 | 3376.14384 | -0.082880564 | 0.11638383 | -0.7121313 | 0.4763835 | 0.66176225 |
| ADRM1 | 5660.94355 | -0.584726102 | 0.22820678 | -2.5622643 | 0.01039921 | 0.04642412 |
| AL354836.1 | 1685.45775 | -0.495426588 | 0.20594876 | -2.4055818 | 0.01614673 | 0.06428951 |
| LAMA5 | 28782.1631 | -0.613061407 | 0.20880251 | -2.9360826 | 0.00332386 | 0.01984382 |
| LAMA5-AS1 | 241.869206 | 0.383609928 | 0.34004596 | 1.128112 | 0.25927262 | 0.45207293 |
| AL121832.2 | 91.710044 | 0.047533926 | 0.20259547 | 0.23462482 | 0.81449994 | 0.8999602 |
| RPS21 | 8386.45749 | -0.543188108 | 0.18478199 | -2.9396161 | 0.00328619 | 0.01968579 |
| CABLES2 | 395.14613 | -0.230516168 | 0.21325629 | -1.0809349 | 0.27972607 | 0.47534443 |
| RBBP8NL | 52.3587833 | 1.803065684 | 0.73923891 | 2.43908384 | 0.01472455 | 0.06002772 |
| AL121832.1 | 3.74101579 | 0.204398513 | 0.95839667 | 0.21327131 | 0.83111535 | 0.91040231 |
| GATA5 | 179.45718 | 0.07165779 | 0.39492892 | 0.18144478 | 0.85601847 | 0.92460093 |
| AL499627.2 | 1.0862574 | 0.883740585 | 1.55007908 | 0.57012613 | 0.56859216 | 0.7341972 |
| AL499627.1 | 0.10721287 | -0.642905668 | 3.30558948 | -0.1944905 | 0.84579185 | NA |

| | | | | | | |
|---|---|---|---|---|---|---|
| MIR1-1HG-A | 27.1327908 | 1.438112664 | 0.67120311 | 2.14258941 | 0.03214608 | 0.10598802 |
| MIR1-1HG | 2.61559993 | 1.423468687 | 1.13859342 | 1.2501993 | 0.21122675 | 0.39461916 |
| BX640514.1 | 0.30338168 | -0.695266322 | 2.32486381 | -0.2990568 | 0.76489671 | NA |
| BX640514.2 | 23.7954589 | -3.152681752 | 0.67429713 | -4.6755082 | 2.93E-06 | 7.70E-05 |
| AL450469.2 | 12.6788326 | 0.32469663 | 1.21507508 | 0.26722351 | 0.78929707 | 0.88550367 |
| AL450469.1 | 14.3084512 | 0.39540349 | 1.06365727 | 0.37173957 | 0.71008676 | 0.83522851 |
| AL357033.1 | 16.2833805 | -0.733384615 | 0.54834246 | -1.3374573 | 0.18107342 | 0.35729617 |
| SLCO4A1 | 9343.03317 | -3.427441993 | 0.32836598 | -10.437872 | 1.67E-25 | 1.10E-21 |
| AL357033.4 | 112.345861 | 0.021244987 | 0.37912366 | 0.05603709 | 0.95531226 | 0.97856915 |
| AL357033.3 | 26.1402446 | 0.212568397 | 0.3774834 | 0.56311985 | 0.57335328 | 0.73792405 |
| SLCO4A1-AS | 1712.82492 | -3.235758784 | 0.33489803 | -9.6619223 | 4.38E-22 | 1.28E-18 |
| LINC00686 | 3.50730889 | -5.027121931 | 1.74183471 | -2.8861073 | 0.00390039 | 0.02235456 |
| NTSR1 | 156.966021 | 1.470847065 | 0.32815285 | 4.48220116 | 7.39E-06 | 0.00016064 |
| AL357033.2 | 6.05975598 | 1.116388537 | 0.63943348 | 1.74590254 | 0.08082789 | 0.20389238 |
| LINC00659 | 68.3458232 | 0.506197271 | 0.24417684 | 2.07307653 | 0.03816516 | 0.11961198 |
| MRGBP | 1360.22485 | -0.860449268 | 0.19537435 | -4.4041055 | 1.06E-05 | 0.00021448 |
| OGFR-AS1 | 112.626107 | -0.786712498 | 0.33820056 | -2.3261715 | 0.0200094 | 0.07495 |
| OGFR | 5927.1378 | -1.095242046 | 0.30213796 | -3.6249733 | 0.00028899 | 0.00294577 |
| COL9A3 | 276.567944 | 0.466407436 | 0.34817792 | 1.33956639 | 0.18038636 | 0.35632208 |
| TCFL5 | 575.861437 | 0.092316201 | 0.13721408 | 0.67278957 | 0.50108117 | 0.6819342 |
| DIDO1 | 3608.07102 | 0.038918062 | 0.12210201 | 0.31873399 | 0.74992824 | 0.86037014 |
| AL117379.1 | 105.236205 | -0.443233075 | 0.46206374 | -0.9592466 | 0.33743453 | 0.53545244 |
| GID8 | 2645.31547 | -0.148954681 | 0.11361408 | -1.3110583 | 0.18983806 | 0.36838837 |
| SLC17A9 | 1067.808 | -0.785221231 | 0.45033306 | -1.7436455 | 0.08122092 | 0.20454547 |
| BHLHE23 | 0.08757577 | 0.547239394 | 3.29377202 | 0.16614368 | 0.86804388 | NA |
| LINC01749 | 0.08666371 | -0.642905668 | 3.30558948 | -0.1944905 | 0.84579185 | NA |
| LINC00029 | 0.16052235 | -1.003574173 | 3.30558948 | -0.3035992 | 0.7614333 | NA |
| LINC01056 | 0.08393208 | -0.642905668 | 3.30558948 | -0.1944905 | 0.84579185 | NA |
| HAR1B | 1.7926149 | -0.077288793 | 1.10437357 | -0.0699843 | 0.94420617 | 0.97336656 |
| HAR1A | 8.39912795 | 0.630613093 | 0.54499998 | 1.15708829 | 0.24723629 | 0.43879533 |
| AL096828.2 | 0.28377621 | -0.179599913 | 2.92867844 | -0.0613246 | 0.95110073 | NA |
| AL096828.1 | 0.13037664 | 0.02007176 | 3.29738704 | 0.00608717 | 0.99514317 | NA |
| YTHDF1 | 2975.23997 | -0.478706208 | 0.19524739 | -2.4517931 | 0.01421464 | 0.05855045 |
| AL096828.3 | 9.93700791 | 0.404512187 | 0.53510485 | 0.7559494 | 0.44967953 | 0.63973076 |
| BIRC7 | 113.741134 | -2.444837033 | 0.37237506 | -6.565523 | 5.19E-11 | 8.39E-09 |
| NKAIN4 | 3546.43351 | -0.469625109 | 0.67084833 | -0.7000466 | 0.48389819 | 0.66841062 |
| AL121827.2 | 455.583682 | -1.67113336 | 0.73588577 | -2.2709141 | 0.02315218 | 0.08362183 |
| ARFGAP1 | 8179.51103 | -0.712497343 | 0.1792894 | -3.974007 | 7.07E-05 | 0.00096164 |
| COL20A1 | 12.4809759 | 1.392295964 | 0.70484291 | 1.97532803 | 0.04823093 | 0.14191584 |
| CHRNA4 | 562.31314 | 3.086316776 | 0.81770094 | 3.77438331 | 0.0001604 | 0.00184468 |
| AL121827.1 | 117.580061 | 5.006784571 | 0.74233155 | 6.7446744 | 1.53E-11 | 2.91E-09 |
| KCNQ2 | 77.0002294 | -3.610538346 | 0.94179699 | -3.8336694 | 0.00012625 | 0.00152083 |
| AL353658.2 | 0.08695548 | -0.642905668 | 3.30558948 | -0.1944905 | 0.84579185 | NA |
| AL353658.1 | 0.17332742 | -1.003574173 | 3.30558948 | -0.3035992 | 0.7614333 | NA |
| EEF1A2 | 1113.36752 | -4.27392377 | 0.64593675 | -6.6166289 | 3.67E-11 | 6.29E-09 |
| AL121829.1 | 0 | NA | NA | NA | NA | NA |
| PPDPF | 9362.96679 | -0.57734434 | 0.24559346 | -2.3508131 | 0.01873244 | 0.07159049 |
| PTK6 | 94.6508739 | -1.242380623 | 0.46152466 | -2.6919052 | 0.00710451 | 0.03501141 |
| SRMS | 30.8505714 | 0.606219828 | 0.39440926 | 1.53703242 | 0.12428537 | 0.27620135 |
| AL121829.2 | 5.35849325 | 0.604087524 | 0.9563156 | 0.63168218 | 0.52759457 | 0.70223445 |
| FNDC11 | 56.8186877 | -1.321491898 | 0.5095843 | -2.5932743 | 0.00950669 | 0.04338723 |
| HELZ2 | 1381.32593 | -0.350813794 | 0.23702569 | -1.4800665 | 0.13885549 | 0.29821044 |
| GMEB2 | 1358.52473 | -0.558969353 | 0.13735123 | -4.0696348 | 4.71E-05 | 0.00070751 |
| MHENCR | 634.418835 | -0.342929819 | 0.22124079 | -1.5500298 | 0.12113436 | 0.27162525 |
| STMN3 | 2145.56905 | -1.331295326 | 0.25699303 | -5.1802779 | 2.22E-07 | 9.26E-06 |
| RTEL1 | 2809.64678 | -0.183517226 | 0.22009771 | -0.8337989 | 0.40439433 | 0.59919529 |
| RTEL1-TNFR: | 4523.74747 | -0.681304076 | 0.23657132 | -2.8799099 | 0.00397789 | 0.02264614 |
| TNFRSF6B | 1465.15349 | -2.265720966 | 0.314598 | -7.201956 | 5.94E-13 | 1.72E-10 |
| ARFRP1 | 3319.88886 | -0.258830798 | 0.16707864 | -1.5491555 | 0.12134434 | 0.27192025 |
| ZGPAT | 4473.2362 | 0.862776994 | 0.21950785 | 3.93050639 | 8.48E-05 | 0.0011132 |
| AL121845.3 | 6851.24445 | 1.331337478 | 0.28412065 | 4.68581733 | 2.79E-06 | 7.42E-05 |
| AL121845.4 | 25.2267237 | 2.718853481 | 0.55896297 | 4.86410306 | 1.15E-06 | 3.65E-05 |
| LIME1 | 3525.93348 | 1.665926461 | 0.35237342 | 4.72773024 | 2.27E-06 | 6.29E-05 |
| AL121845.2 | 2511.74897 | 2.036111904 | 0.47622444 | 4.27553001 | 1.91E-05 | 0.0003409 |
| SLC2A4RG | 6716.69767 | 1.165381789 | 0.31841078 | 3.65999476 | 0.00025222 | 0.00263892 |
| ZBTB46 | 1662.49257 | 0.517697187 | 0.17956319 | 2.88309188 | 0.00393793 | 0.02250121 |
| AL121845.1 | 400.224847 | 0.226406455 | 0.23502482 | 0.96332997 | 0.33538196 | 0.53359089 |
| ZBTB46-AS1 | 1.76654703 | -0.42630692 | 1.15129306 | -0.3702853 | 0.71116991 | 0.83587657 |
| AL118506.1 | 627.092089 | -0.046178937 | 0.22966695 | -0.2010691 | 0.84064451 | 0.91507929 |
| ABHD16B | 293.355058 | -0.314137063 | 0.2278145 | -1.378916 | 0.16792065 | 0.33918557 |
| TPD52L2 | 6685.65509 | -0.506264509 | 0.15487024 | -3.2689593 | 0.00107944 | 0.00831817 |
| DNAJC5 | 4952.44343 | -0.288772863 | 0.15135794 | -1.9078806 | 0.05640665 | 0.15840719 |
| UCKL1 | 4077.25166 | -0.071694415 | 0.14889054 | -0.4815243 | 0.63014391 | 0.7801882 |
| UCKL1-AS1 | 553.577406 | 0.309961894 | 0.13514913 | 2.29348046 | 0.02182036 | 0.08003935 |
| ZNF512B | 4012.20703 | 0.503975051 | 0.16985682 | 2.96705807 | 0.00300664 | 0.01838283 |
| SAMD10 | 865.790278 | -0.641333579 | 0.23172351 | -2.7676673 | 0.00564591 | 0.02951091 |
| PRPF6 | 4608.47128 | 0.186626498 | 0.14261811 | 1.30857505 | 0.19067834 | 0.36950233 |
| C20orf204 | 112.972303 | -0.36740501 | 0.24049081 | -1.5277299 | 0.1265796 | 0.27988463 |
| SOX18 | 755.522024 | 0.016939027 | 0.4799515 | 0.03529321 | 0.97184594 | 0.98670705 |
| TCEA2 | 2290.6361 | 0.112676833 | 0.18306865 | 0.6154895 | 0.5382315 | 0.71165086 |
| AL355803.1 | 38.5353821 | 1.34643003 | 0.44741481 | 3.00935506 | 0.00261803 | 0.01655574 |
| RGS19 | 482.16673 | -0.836277469 | 0.31162123 | -2.6836345 | 0.00728267 | 0.03565614 |
| OPRL1 | 69.7461999 | -0.032628883 | 0.42564293 | -0.0766579 | 0.93889571 | 0.97092863 |
| LKAAEAR1 | 0.69851406 | -1.122250899 | 1.84521898 | -0.6081939 | 0.54305889 | NA |
| MYT1 | 13.4803072 | 0.521213156 | 0.68922529 | 0.75623045 | 0.44951104 | 0.63959456 |
| NPBWR2 | 0 | NA | NA | NA | NA | NA |
| PCMTD2 | 3756.98954 | 0.231661026 | 0.1659893 | 1.39563832 | 0.16282344 | 0.33188415 |
| LINC00266-1 | 5.19099299 | 1.087649374 | 0.6863411 | 1.58470676 | 0.11303298 | 0.25894184 |
| AC008993.4 | 0.19387658 | -1.003574173 | 3.30558948 | -0.3035992 | 0.7614333 | NA |
| FAM138F | 0 | NA | NA | NA | NA | NA |
| OR4F17 | 0 | NA | NA | NA | NA | NA |
| AC092192.1 | 0 | NA | NA | NA | NA | NA |
| AC016588.2 | 7.21366995 | -0.064799218 | 0.50192922 | -0.1291003 | 0.89727828 | 0.94754828 |
| PLPP2 | 1195.5001 | -3.262575209 | 0.45466617 | -7.1757598 | 7.19E-13 | 2.04E-10 |
| MIER2 | 1217.05173 | -0.595761948 | 0.2322153 | -2.5655586 | 0.01030098 | 0.04606357 |
| AC016588.1 | 285.411833 | -0.966083058 | 0.24744637 | -3.9042119 | 9.45E-05 | 0.00121247 |
| THEG | 0.32793579 | -1.669260191 | 2.86721581 | -0.5821885 | 0.56043969 | NA |
| AC010641.1 | 2.21509262 | 3.794149668 | 1.59870298 | 2.3732674 | 0.01763149 | 0.06838668 |
| C2CD4C | 220.723644 | 1.023337842 | 0.65158556 | 1.57053488 | 0.11629073 | 0.26433811 |
| SHC2 | 2189.52837 | -0.969367292 | 0.26206277 | -3.698989 | 0.00021646 | 0.00232792 |
| ODF3L2 | 14.0367266 | -0.945640247 | 0.47298718 | -1.9992936 | 0.0455766 | 0.13612582 |
| MADCAM1 | 71.470083 | 0.51585399 | 0.27544695 | 1.87278887 | 0.06109755 | 0.16751569 |
| AC005775.1 | 380.87884 | -0.52881075 | 0.31847479 | -1.6604477 | 0.09682441 | 0.23226233 |
| TPGS1 | 605.915649 | -0.584518673 | 0.29792089 | -1.9619929 | 0.04976331 | 0.1452256 |
| CDC34 | 2430.83387 | -0.783596165 | 0.15749506 | -4.9753698 | 6.51E-07 | 2.26E-05 |
| GZMM | 49.9833151 | -0.620437499 | 0.59926232 | -1.0353354 | 0.3005123 | 0.49734515 |
| AC009005.1 | 34.2706252 | -0.23886806 | 0.26163319 | -0.9129884 | 0.36124866 | 0.55878832 |
| BSG | 37348.8256 | 0.113546873 | 0.13494437 | 0.84143469 | 0.40010446 | 0.59484436 |
| HCN2 | 99.2380866 | 0.395770253 | 0.38624405 | 1.02466368 | 0.3055219 | 0.50259592 |
| POLRMT | 4484.94626 | -0.147225572 | 0.07871593 | -1.8703403 | 0.06143658 | 0.16814558 |
| AC004449.1 | 11.3850069 | -0.244998448 | 0.63465946 | -0.3860314 | 0.69947345 | 0.82758053 |
| FGF22 | 26.7479509 | 0.830596249 | 0.47574616 | 1.74588116 | 0.0808316 | 0.20389238 |
| RNF126 | 2239.70907 | -0.74110017 | 0.16395008 | -4.5202794 | 6.18E-06 | 0.00013955 |

| | | | | | | |
|---|---|---|---|---|---|---|
| AC004156.1 | 29.6981955 | 2.632496854 | 0.50468297 | 5.21613969 | 1.83E-07 | 7.95E-06 |
| FSTL3 | 3659.59771 | -1.361034046 | 0.41428747 | -3.2852407 | 0.00101895 | 0.00795906 |
| PRSS57 | 5.24336686 | -0.090769191 | 0.61876646 | -0.1466938 | 0.88337373 | 0.9394864 |
| PALM | 4386.24792 | 1.238522393 | 0.36118657 | 3.42903782 | 0.00060573 | 0.00525427 |
| MISP | 1617.14277 | -0.037330538 | 0.22752832 | -0.1640698 | 0.86967616 | 0.93214735 |
| AC006273.1 | 28.956627 | -0.14371369 | 0.3747285 | -0.3835142 | 0.70133857 | 0.82893183 |
| LINC01836 | 157.380384 | 1.509502701 | 0.36309109 | 4.15736646 | 3.22E-05 | 0.00051955 |
| PTBP1 | 13466.3863 | -0.34362207 | 0.19031693 | -1.8055255 | 0.07099249 | 0.18622024 |
| PLPPR3 | 14.4017159 | -2.073073985 | 0.90213218 | -2.2979714 | 0.02156341 | 0.07927095 |
| AZU1 | 63.8806832 | -4.301541419 | 0.78437152 | -5.4840612 | 4.16E-08 | 2.27E-06 |
| PRTN3 | 101.997922 | -5.281644952 | 0.95626613 | -5.5231957 | 3.33E-08 | 1.92E-06 |
| ELANE | 156.354114 | -4.278184648 | 0.91230319 | -4.689433 | 2.74E-06 | 7.31E-05 |
| CFD | 798.72885 | -1.535216854 | 0.77178354 | -1.9891806 | 0.04668127 | 0.13849967 |
| MED16 | 3472.88268 | -0.038895413 | 0.19315486 | -0.2013691 | 0.84041 | 0.91505077 |
| R3HDM4 | 4014.66326 | -1.179175268 | 0.21402618 | -5.5094908 | 3.60E-08 | 2.05E-06 |
| KISS1R | 18.3915113 | -1.777295306 | 0.76916242 | -2.3106892 | 0.02085003 | 0.07722124 |
| ARID3A | 1287.88232 | -0.576356508 | 0.21901433 | -2.6315927 | 0.00849857 | 0.03982713 |
| AC005379.1 | 4.60751315 | -0.089495039 | 0.65309861 | -0.1370314 | 0.89100595 | 0.94349383 |
| WDR18 | 2171.48603 | 0.264268508 | 0.18553449 | 1.42436321 | 0.15434136 | 0.32026925 |
| AC004528.1 | 2.14205857 | -0.972504268 | 1.40213233 | -0.6935895 | 0.48793968 | 0.67152554 |
| GRIN3B | 1015.2039 | -0.514909357 | 0.14334074 | -3.5922053 | 0.00032789 | 0.00325585 |
| TMEM259 | 13772.6992 | -0.460839086 | 0.10935098 | -4.2143115 | 2.51E-05 | 0.00042542 |
| AC004528.2 | 4650.87784 | -0.564259267 | 0.11537336 | -4.8907241 | 1.00E-06 | 3.26E-05 |
| CNN2 | 4820.84557 | -0.229704951 | 0.28225591 | -0.813818 | 0.4157492 | 0.60908136 |
| ABCA7 | 2090.57033 | -0.446172081 | 0.29870863 | -1.4936699 | 0.13526192 | 0.29272523 |
| ARHGAP45 | 1921.84554 | -0.264551952 | 0.27723149 | -0.9542637 | 0.33995017 | 0.53809861 |
| POLR2E | 7409.4702 | -0.049257387 | 0.18428531 | -0.2672887 | 0.78924685 | 0.885485 |
| GPX4 | 15856.409 | -0.055053338 | 0.22790894 | -0.2415585 | 0.8091223 | 0.89687074 |
| SBNO2 | 11595.8901 | -1.998665124 | 0.27237876 | -7.3378158 | 2.17E-13 | 7.44E-11 |
| STK11 | 4432.098 | -0.387015343 | 0.12608479 | -3.0694848 | 0.00214428 | 0.01421436 |
| CBARP | 472.116214 | -0.331050434 | 0.34770647 | -0.9520974 | 0.34104757 | 0.53910949 |
| AC004221.1 | 6.2557779 | 0.882727361 | 0.59025661 | 1.49549763 | 0.13478462 | 0.29218265 |
| ATP5F1D | 5285.83239 | -0.082691177 | 0.13114457 | -0.6305345 | 0.52834492 | 0.70294932 |
| MIDN | 8240.05974 | -1.005577884 | 0.2556845 | -3.9328856 | 8.39E-05 | 0.00110554 |
| CIRBP | 19882.8434 | 0.325091537 | 0.25208899 | 1.28959039 | 0.19719291 | 0.37796025 |
| CIRBP-AS1 | 134.963815 | -0.236182026 | 0.34755383 | -0.6795552 | 0.49678612 | 0.67865838 |
| C19orf24 | 1411.61823 | -0.330273057 | 0.22496233 | -1.468126 | 0.14206999 | 0.30303208 |
| EFNA2 | 15.3372563 | -1.560299944 | 0.73175819 | -2.1322617 | 0.03298534 | 0.10797334 |
| AC005330.1 | 2.38600597 | 1.014167997 | 0.90991805 | 1.1145707 | 0.26503443 | 0.45904486 |
| MUM1 | 3411.65724 | -0.017411316 | 0.14430561 | -0.1206558 | 0.90396363 | 0.95167363 |
| AC005329.2 | 786.976097 | -0.148340438 | 0.17779518 | -0.8343333 | 0.40409319 | 0.59897135 |
| NDUFS7 | 5003.0677 | 0.158134888 | 0.12456389 | 1.26950825 | 0.20425985 | 0.38661587 |
| AC005329.3 | 2171.92448 | 0.296185182 | 0.12873948 | 2.30065535 | 0.02141112 | 0.07884531 |
| AC005329.1 | 2693.57376 | 0.13315989 | 0.14077499 | 0.94590589 | 0.34419659 | 0.54181376 |
| GAMT | 2154.06961 | 0.641354727 | 0.37982229 | 1.68856528 | 0.09130277 | 0.22288144 |
| DAZAP1 | 7296.01294 | -0.565724315 | 0.14719007 | -3.8434951 | 0.00012129 | 0.0014767 |
| RPS15 | 11165.9485 | -0.437552043 | 0.12291559 | -3.5597766 | 0.00037117 | 0.00359911 |
| AC027307.3 | 12.6479604 | 0.510610687 | 0.64846394 | 0.7874157 | 0.43103856 | 0.62306129 |
| APC2 | 368.242324 | -0.177236148 | 0.22049187 | -0.8038217 | 0.42149995 | 0.61411977 |
| AC027307.2 | 297.930089 | -0.462775674 | 0.26578268 | -1.7411807 | 0.08165191 | 0.20531714 |
| C19orf25 | 1973.52385 | -0.157587775 | 0.11618167 | -1.356391 | 0.17497478 | 0.34897028 |
| PCSK4 | 614.427892 | -0.031215577 | 0.34053153 | -0.0916672 | 0.92696244 | 0.96452668 |
| REEP6 | 1459.18007 | 1.145188735 | 0.46257939 | 2.47565881 | 0.01329906 | 0.05563957 |
| ADAMTSL5 | 132.500457 | -0.111133954 | 0.44077215 | -0.2521347 | 0.80093695 | 0.89207245 |
| AC027307.1 | 21.8758935 | 0.16296902 | 0.51706362 | 0.31518176 | 0.75262365 | 0.86206078 |
| PLK5 | 5.83286258 | 0.265140599 | 0.73447254 | 0.36099457 | 0.71810351 | 0.83985952 |
| MEX3D | 531.594512 | -0.765821873 | 0.26618767 | -2.8769998 | 0.00401476 | 0.02281172 |
| MBD3 | 5508.50703 | -0.402479177 | 0.15311946 | -2.6285305 | 0.00857547 | 0.04008972 |
| AC005943.1 | 5996.63996 | -0.131799743 | 0.14626592 | -0.9010967 | 0.36753688 | 0.56438321 |
| UQCR11 | 5340.12046 | -0.094850919 | 0.15448656 | -0.6139752 | 0.5392317 | 0.71236611 |
| TCF3 | 2694.64404 | -0.267959214 | 0.18785328 | -1.4264282 | 0.15374479 | 0.31942832 |
| AC005256.1 | 0.08666371 | -0.642905668 | 3.30558948 | -0.1944905 | 0.84579185 | NA |
| ONECUT3 | 0.72115045 | -1.056708752 | 1.65617666 | -0.6380411 | 0.52344694 | NA |
| ATP8B3 | 110.98284 | -1.517541302 | 0.3459116 | -4.3870783 | 1.15E-05 | 0.00022864 |
| REXO1 | 3235.44819 | -0.261205237 | 0.14507394 | -1.8004973 | 0.07178215 | 0.18778723 |
| AC012615.2 | 488.539446 | -0.576514827 | 0.14710764 | -3.919 | 8.89E-05 | 0.00115448 |
| AC012615.3 | 435.686043 | -1.087559096 | 0.22761195 | -4.7781283 | 1.77E-06 | 5.16E-05 |
| KLF16 | 1374.12047 | -1.016772468 | 0.21300561 | -4.773454 | 1.81E-06 | 5.25E-05 |
| AC012615.4 | 10.5705073 | -0.095692784 | 0.57405625 | -0.1666958 | 0.86760939 | 0.93109788 |
| AC012615.1 | 326.555531 | -0.292968767 | 0.21122036 | -1.387029 | 0.16543292 | 0.33583496 |
| AC012615.5 | 253.713211 | -0.26226371 | 0.21579168 | -1.215356 | 0.22423034 | 0.41025145 |
| ABHD17A | 4328.82556 | -0.286251845 | 0.12021044 | -2.381256 | 0.01725371 | 0.06737139 |
| SCAMP4 | 6456.25207 | -0.494555086 | 0.17752893 | -2.7857719 | 0.00534004 | 0.0282765 |
| ADAT3 | 669.585466 | -0.608038579 | 0.18365579 | -3.3107509 | 0.00093046 | 0.00743297 |
| CSNK1G2 | 4710.28906 | -0.520718267 | 0.15921399 | -3.270556 | 0.00107336 | 0.00828831 |
| CSNK1G2-AS | 8.47179386 | -0.519830682 | 0.79060771 | -0.6575077 | 0.5108545 | 0.68920615 |
| BTBD2 | 6011.43709 | 0.13503891 | 0.13103209 | 1.03057891 | 0.30273833 | 0.49963761 |
| MKNK2 | 8799.14018 | -0.407263904 | 0.19982122 | -2.0381414 | 0.04153579 | 0.12736034 |
| MOB3A | 2100.06316 | -0.258930094 | 0.15301513 | -1.6921862 | 0.09061048 | 0.22169218 |
| IZUMO4 | 311.147435 | 0.069057998 | 0.27149912 | 0.25435809 | 0.79921892 | 0.89108294 |
| AP3D1 | 10996.746 | -0.141239391 | 0.18694108 | -0.7555289 | 0.4499317 | 0.63993921 |
| DOT1L | 5439.84893 | -0.0487033 | 0.33360695 | -0.1459901 | 0.88392923 | 0.93980604 |
| AC004490.1 | 234.377221 | 0.427880775 | 0.46722581 | 0.91579011 | 0.35977699 | 0.55742181 |
| PLEKHJ1 | 5082.81229 | -0.223993988 | 0.14554724 | -1.5389779 | 0.12380968 | 0.27542002 |
| SF3A2 | 2523.36958 | -0.572939807 | 0.19452179 | -2.945376 | 0.00322562 | 0.01942446 |
| AMH | 108.153932 | -0.998529092 | 0.51916936 | -1.9233205 | 0.05443981 | 0.15457355 |
| JSRP1 | 5968.38366 | -0.279916132 | 0.13698549 | -2.0433998 | 0.04101287 | 0.12621187 |
| OAZ1 | 22888.6148 | -0.421639374 | 0.1102889 | -3.8230444 | 0.00013181 | 0.00157282 |
| AC005258.1 | 7690.29598 | -0.144176034 | 0.09239914 | -1.5603612 | 0.11867455 | 0.26786339 |
| PEAK3 | 24.3788516 | -0.402170963 | 0.44176531 | -0.9103724 | 0.36262613 | 0.55991227 |
| LINGO3 | 17.34415 | 0.011757398 | 0.50488783 | 0.02328715 | 0.98142122 | 0.9915968 |
| LSM7 | 1782.25166 | -0.858837743 | 0.20469282 | -4.1957394 | 2.72E-05 | 0.0004548 |
| SPPL2B | 7509.99121 | 0.006223484 | 0.19756446 | 0.03150103 | 0.97486997 | 0.98843899 |
| TMPRSS9 | 216.124003 | 0.41770543 | 0.31660722 | 1.31931747 | 0.187063 | 0.36501786 |
| TIMM13 | 3111.85673 | 0.03103734 | 0.14237658 | 0.2179947 | 0.82743324 | 0.90808285 |
| LMNB2 | 1998.61527 | -0.470800042 | 0.21322821 | -2.2079632 | 0.02724684 | 0.0941772 |
| LINC01775 | 0.35475782 | 0.922720074 | 2.78597936 | 0.33120133 | 0.74049241 | NA |
| GADD45B | 12662.6706 | -1.298341124 | 0.42961378 | -3.0221124 | 0.00251017 | 0.0160462 |
| GNG7 | 1436.52808 | 0.967863431 | 0.29035525 | 3.3333767 | 0.00085799 | 0.00697015 |
| AC092068.1 | 3.85704255 | -2.042657591 | 0.77080916 | -2.6500173 | 0.00804877 | 0.0382839 |
| DIRAS1 | 430.750337 | 0.421833603 | 0.45553153 | 0.92602503 | 0.35443297 | 0.55199607 |
| AC006538.2 | 486.83123 | -0.017125254 | 0.17864309 | -0.095863 | 0.92362942 | 0.96284423 |
| SLC39A3 | 1034.08721 | -0.216823984 | 0.17086984 | -1.2689424 | 0.20446162 | 0.38677567 |
| SGTA | 5099.92528 | -0.399580858 | 0.15515825 | -2.5753118 | 0.01001498 | 0.04511621 |
| THOP1 | 2030.72727 | -0.238051281 | 0.14539762 | -1.6372433 | 0.10157964 | 0.24015197 |
| ZNF554 | 461.167399 | 0.244103875 | 0.15335391 | 1.59176814 | 0.11143681 | 0.25626482 |
| ZNF555 | 313.015487 | 0.042776022 | 0.17700492 | 0.24166572 | 0.80903919 | 0.89681632 |
| AC006130.1 | 12.0704542 | 0.658795468 | 0.63429575 | 1.03862506 | 0.29897914 | 0.49579957 |
| ZNF556 | 41.1655173 | 0.614614695 | 0.50553201 | 1.21577799 | 0.2240695 | 0.41012089 |
| ZNF57 | 421.350945 | -0.436887824 | 0.1871363 | -2.3345969 | 0.01956449 | 0.07372329 |
| AC119403.1 | 111.053486 | -0.131834451 | 0.18201256 | -0.7243151 | 0.46887229 | 0.6556107 |
| ZNF77 | 247.900079 | -0.330958878 | 0.27083083 | -1.2220133 | 0.22170259 | 0.40769633 |
| TLE6 | 246.648932 | -0.884405787 | 0.49662437 | -1.7808345 | 0.0749395 | 0.19353015 |

| | | | | | | |
|---|---|---|---|---|---|---|
| TLE2 | 3502.58973 | -0.606010201 | 0.33414789 | -1.8135987 | 0.06973952 | 0.18373776 |
| AC005944.1 | 9773.04978 | -0.611095817 | 0.17418977 | -3.5082188 | 0.00045112 | 0.0041843 |
| AES | 20640.2627 | -0.544924132 | 0.17754942 | -3.0691405 | 0.00214676 | 0.01422718 |
| GNA11 | 6844.58833 | -0.18088054 | 0.11279226 | -1.6036609 | 0.10878882 | 0.25208797 |
| AC005262.2 | 839.617286 | 0.251489277 | 0.16538067 | 1.52066908 | 0.1283429 | 0.28245721 |
| KF456478.1 | 1371.90693 | -0.027425927 | 0.12089984 | -0.2268483 | 0.82054168 | 0.90366551 |
| GNA15 | 270.801499 | -1.373203675 | 0.45513443 | -3.0171387 | 0.00255173 | 0.01624557 |
| AC005264.1 | 51.1858402 | -1.054664847 | 0.35314087 | -2.9865273 | 0.00282166 | 0.0175042 |
| S1PR4 | 98.3990983 | -0.758469665 | 0.43913042 | -1.7272082 | 0.08413028 | 0.2099674 |
| NCLN | 3831.13831 | -0.399582052 | 0.13807586 | -2.8939312 | 0.00380451 | 0.0219722 |
| CELF5 | 21.8991517 | -0.751777879 | 0.66889359 | -1.1239125 | 0.26105014 | 0.45423241 |
| AC010649.1 | 2.57152419 | -1.470814369 | 1.21615878 | -1.2093934 | 0.22651175 | 0.4133196 |
| NFIC | 6933.75215 | -0.020086755 | 0.19775456 | -0.1015742 | 0.91909468 | 0.96062334 |
| SMIM24 | 9634.73518 | 1.54393659 | 0.3937867 | 3.92074337 | 8.83E-05 | 0.00114842 |
| AC005551.1 | 9.2278899 | 0.250296962 | 0.46376427 | 0.5397073 | 0.5893989 | 0.74961886 |
| DOHH | 948.412988 | -0.34297832 | 0.20673641 | -1.6590127 | 0.09711324 | 0.23272229 |
| FZR1 | 3569.83604 | -0.093888563 | 0.15466964 | -0.6070265 | 0.54383335 | 0.71604108 |
| MFSD12 | 9759.6981 | -0.004358131 | 0.24281035 | -0.0179487 | 0.98567977 | 0.99368571 |
| C19orf71 | 1062.5726 | 0.724227042 | 0.22376448 | 3.23655942 | 0.0012098 | 0.00910458 |
| AC005786.3 | 1838.87375 | 0.035013653 | 0.28048026 | 0.12483464 | 0.90065446 | 0.94969868 |
| AC005786.2 | 0.26052132 | 1.101547414 | 3.01931676 | 0.36483334 | 0.71523582 | NA |
| HMG20B | 4481.09987 | -0.140656121 | 0.12813929 | -1.0976814 | 0.27234361 | 0.46731527 |
| GIPC3 | 411.461451 | 0.512671941 | 0.39149716 | 1.30951639 | 0.19035949 | 0.36904718 |
| TBXA2R | 290.382321 | 0.321950846 | 0.42102668 | 0.76468039 | 0.44446189 | 0.63537357 |
| CACTIN-AS1 | 886.80449 | -0.192408849 | 0.16796925 | -1.1455004 | 0.25200191 | 0.44427498 |
| CACTIN | 2033.03446 | -0.168305284 | 0.16420969 | -1.0249412 | 0.30539092 | 0.5024118 |
| PIP5K1C | 3913.47976 | -0.187190062 | 0.16228093 | -1.1534939 | 0.24870771 | 0.44046175 |
| AC004637.1 | 2.55056352 | -0.309395558 | 0.81339808 | -0.3803741 | 0.70366774 | 0.83067948 |
| TJP3 | 1599.60102 | -0.42692375 | 0.24540842 | -1.7396459 | 0.08192122 | 0.20580902 |
| APBA3 | 1522.59832 | -0.319794 | 0.11941971 | -2.6778996 | 0.00740854 | 0.03609816 |
| AC005954.2 | 171.037945 | -0.208497082 | 0.15387759 | -1.3549542 | 0.17543216 | 0.34961805 |
| MRPL54 | 1685.03874 | 0.013527535 | 0.19493176 | 0.06939626 | 0.94467421 | 0.9736968 |
| RAX2 | 0.07659028 | -0.642905668 | 3.30558948 | -0.1944905 | 0.84579185 | NA |
| MATK | 121.666499 | -0.347957778 | 0.52015608 | -0.6689488 | 0.50352815 | 0.68355405 |
| ZFR2 | 13.9726028 | -0.562302406 | 0.6073301 | -0.9258596 | 0.35451895 | 0.55209736 |
| ATCAY | 6.08698166 | 0.329204057 | 1.12886408 | 0.29162418 | 0.77057399 | 0.87330394 |
| NMRK2 | 98.3575208 | -1.136286316 | 0.86528495 | -1.3131932 | 0.18911785 | 0.36739633 |
| DAPK3 | 2546.60943 | -0.77014266 | 0.21147634 | -3.6417439 | 0.0002708 | 0.00280036 |
| EEF2 | 86663.6587 | 0.151857479 | 0.24213913 | 0.62714968 | 0.53056113 | 0.70495094 |
| PIAS4 | 1277.59956 | -0.237983604 | 0.15495121 | -1.5358615 | 0.12457234 | 0.27665257 |
| ZBTB7A | 2900.41971 | -0.382062393 | 0.18419476 | -2.0742305 | 0.03805791 | 0.11936097 |
| MAP2K2 | 5312.03994 | -0.37018083 | 0.11869672 | -3.1187114 | 0.00181644 | 0.01249986 |
| CREB3L3 | 159.210527 | 3.203519334 | 0.96236214 | 3.32880855 | 0.00087218 | 0.00705722 |
| SIRT6 | 1410.67729 | -0.760293274 | 0.18476239 | -4.1149787 | 3.87E-05 | 0.00060635 |
| ANKRD24 | 355.292416 | 1.364887903 | 0.49243288 | 2.77172373 | 0.00557603 | 0.02926752 |
| EBI3 | 146.792262 | -0.897862966 | 0.55940326 | -1.6050371 | 0.10848565 | 0.25154019 |
| YJU2 | 981.242537 | -0.01111912 | 0.29280113 | -0.037975 | 0.96970762 | 0.98601026 |
| SHD | 5.09685748 | -0.487604604 | 1.04717311 | -0.465639 | 0.64147395 | 0.78758115 |
| TMIGD2 | 5.87861011 | -0.933491609 | 0.84885497 | -1.0997068 | 0.27145988 | 0.46613251 |
| FSD1 | 15.6657483 | 0.303306878 | 0.50821633 | 0.59680664 | 0.55063649 | 0.72096744 |
| STAP2 | 2183.36338 | -0.545060437 | 0.33739255 | -1.6155082 | 0.10620069 | 0.24754814 |
| AC104521.1 | 3.63122727 | 1.484685926 | 0.64424573 | 2.30453358 | 0.02119271 | 0.07820483 |
| MPND | 1267.53937 | 0.764321295 | 0.25128124 | 3.04169658 | 0.00235249 | 0.01531354 |
| AC007292.3 | 210.454759 | 0.780612482 | 0.2949579 | 2.64652168 | 0.00813243 | 0.03859136 |
| AC007292.2 | 596.395772 | 0.788591059 | 0.23112673 | 3.41194221 | 0.00064502 | 0.00552065 |
| SH3GL1 | 5262.56653 | -0.981181954 | 0.17663324 | -5.5549111 | 2.78E-08 | 1.63E-06 |
| AC007292.1 | 894.309123 | -0.720996561 | 0.17281385 | -4.1720993 | 3.02E-05 | 0.00049117 |
| CHAF1A | 321.661186 | -0.039996782 | 0.26131899 | -0.1530573 | 0.87835308 | 0.93677394 |
| AC011498.3 | 35.1214787 | -0.090696989 | 0.34738723 | -0.2610833 | 0.7940283 | 0.88844572 |
| UBXN6 | 9993.48955 | -0.111686216 | 0.12556776 | -0.8894498 | 0.37376141 | 0.57001263 |
| AC011498.2 | 124.705611 | -0.351257875 | 0.26632302 | -1.3189167 | 0.18719697 | 0.36514417 |
| AC011498.6 | 841.36373 | 0.009245917 | 0.17030926 | 0.05428898 | 0.95670493 | 0.9793218 |
| AC011498.1 | 10.633431 | 0.483723597 | 0.44421246 | 1.08894648 | 0.27617749 | 0.47149878 |
| HDGFL2 | 2855.33362 | -0.180379758 | 0.17062969 | -1.0571417 | 0.29044694 | 0.48652584 |
| PLIN4 | 2400.70844 | -0.854741891 | 0.38559129 | -2.2167043 | 0.0266433 | 0.09259144 |
| PLIN5 | 4014.22819 | -0.711884659 | 0.44822555 | -1.5882286 | 0.11223465 | 0.25767262 |
| AC011498.4 | 1366.21853 | -0.768999036 | 0.40883374 | -1.8809579 | 0.05997765 | 0.16545775 |
| LRG1 | 3829.66781 | -4.601243612 | 0.56005677 | -8.2156736 | 2.11E-16 | 1.74E-13 |
| SEMA6B | 1400.32695 | -1.692890705 | 0.44414848 | -3.8115422 | 0.0001381 | 0.00163381 |
| TNFAIP8L1 | 1526.53753 | 0.563078965 | 0.24654334 | 2.2838944 | 0.02237774 | 0.0814944 |
| MYDGF | 2933.86688 | -0.893634578 | 0.19539447 | -4.5734896 | 4.80E-06 | 0.00011375 |
| AC005339.1 | 459.202189 | 0.15485127 | 0.24257909 | 0.63835374 | 0.52324343 | 0.69873547 |
| DPP9 | 3277.4261 | -0.309584509 | 0.16164555 | -1.9152059 | 0.05546626 | 0.15661689 |
| DPP9-AS1 | 677.092375 | -0.128735251 | 0.18255878 | -0.7051715 | 0.4807035 | 0.66542234 |
| MIR7-3HG | 0.3035528 | -1.364245242 | 2.86464618 | -0.4762352 | 0.63390685 | NA |
| AC005523.1 | 3.04257969 | 0.77339181 | 0.76822338 | 1.00672778 | 0.31406561 | 0.51094247 |
| FEM1A | 1707.22046 | 0.226150981 | 0.18224366 | 1.24092649 | 0.21463291 | 0.39877895 |
| AC005523.2 | 1583.93878 | 0.260615324 | 0.1833819 | 1.42116164 | 0.15526977 | 0.32148743 |
| TICAM1 | 2004.36431 | -1.096007043 | 0.20619921 | -5.3152825 | 1.06E-07 | 5.05E-06 |
| AC027319.1 | 784.131588 | -1.582831471 | 0.25687884 | -6.1617822 | 7.19E-10 | 7.87E-08 |
| PLIN3 | 3563.95031 | -1.306048487 | 0.23811254 | -5.4850051 | 4.13E-08 | 2.27E-06 |
| ARRDC5 | 12.2691843 | -0.198190788 | 0.75745784 | -0.2616526 | 0.79358931 | 0.88835379 |
| UHRF1 | 55.8798146 | -1.190255819 | 0.4686694 | -2.5396491 | 0.01109637 | 0.04867932 |
| KDM4B | 2570.215 | -0.048367733 | 0.16069361 | -0.3009935 | 0.76341945 | 0.8690784 |
| PTPRS | 2740.68956 | 0.491241358 | 0.2871919 | 1.71049863 | 0.08717371 | 0.2152407 |
| AC022517.1 | 1.00670367 | -3.251743952 | 1.54000705 | -2.1115124 | 0.0347283 | NA |
| AC005790.1 | 6.90911836 | -0.595880358 | 0.68379787 | -0.8714276 | 0.3835207 | 0.57930123 |
| ZNRF4 | 0 | NA | NA | NA | NA | NA |
| TINCR | 67.4207866 | 1.047531367 | 0.29707944 | 3.52609853 | 0.00042173 | 0.00397367 |
| SAFB2 | 5777.79119 | -0.014050307 | 0.13901637 | -0.1010694 | 0.91949533 | 0.96074349 |
| SAFB | 5346.8815 | -0.366111988 | 0.13135354 | -2.7872258 | 0.00531614 | 0.02817848 |
| RPL36 | 10327.7809 | -0.636524617 | 0.09128142 | -6.9732111 | 3.10E-12 | 7.43E-10 |
| C19orf70 | 2578.24909 | -0.086828777 | 0.14202439 | -0.6113652 | 0.54095781 | 0.71385965 |
| HSD11B1L | 654.270728 | -0.080257738 | 0.16213107 | -0.4950176 | 0.62058769 | 0.77335179 |
| LONP1 | 7560.05724 | -0.309366384 | 0.23803455 | -1.2996701 | 0.19371407 | 0.37338207 |
| CATSPERD | 3.35427513 | -3.13239197 | 1.31669695 | -2.3789772 | 0.01736075 | 0.0676516 |
| PRR22 | 322.950521 | 0.20328752 | 0.31999749 | 0.63527849 | 0.5252468 | 0.70002821 |
| AC011499.1 | 467.601678 | -0.227583088 | 0.18947176 | -1.2011452 | 0.22969489 | 0.41719056 |
| DUS3L | 1954.15805 | -0.223195493 | 0.1474063 | -1.5141516 | 0.12998738 | 0.28509735 |
| NRTN | 178.546159 | 1.636661031 | 0.33982741 | 4.81615366 | 1.46E-06 | 4.44E-05 |
| FUT6 | 3773.51934 | 2.159047265 | 0.51559481 | 4.18748836 | 2.82E-05 | 0.00046779 |
| FUT3 | 997.528438 | 3.061718757 | 0.50643903 | 6.04558211 | 1.49E-09 | 1.43E-07 |
| AC024592.2 | 77.1249404 | 3.167417172 | 0.56612134 | 5.59494393 | 2.21E-08 | 1.36E-06 |
| FUT5 | 3.68616499 | 0.816908641 | 0.68710901 | 1.1889069 | 0.23447631 | 0.42322754 |
| AC024592.3 | 4831.07549 | 0.030039099 | 0.08653149 | 0.34714645 | 0.7284813 | 0.84642348 |
| NDUFA11 | 6127.02691 | 0.019992198 | 0.09613308 | 0.20796379 | 0.83525724 | 0.9126433 |
| AC024592.1 | 101.206388 | 0.374248758 | 0.28237494 | 1.32536109 | 0.18505142 | 0.36240667 |
| AC104532.1 | 876.63637 | -0.113470057 | 0.38349308 | -0.2958855 | 0.76731749 | 0.87152635 |
| VMAC | 313.837605 | 0.905021094 | 0.19953522 | 4.53564579 | 5.74E-06 | 0.00013134 |
| AC104532.2 | 211.423837 | 0.510481933 | 0.22117032 | 2.30809419 | 0.0209939 | 0.07762455 |
| CAPS | 2453.9543 | -0.061181829 | 0.36965289 | -0.1655116 | 0.86854134 | 0.93156219 |
| RANBP3 | 3137.66649 | -0.205160578 | 0.12453634 | -1.6473953 | 0.09947683 | 0.23663096 |
| AC011444.1 | 20.9748185 | 0.173718687 | 0.39063893 | 0.44470398 | 0.65653369 | 0.7984501 |

| | | | | | | |
|---|---|---|---|---|---|---|
| RFX2 | 445.221169 | -0.181107233 | 0.29509125 | -0.613733 | 0.53939179 | 0.71247053 |
| AC011444.2 | 0 | NA | NA | NA | NA | NA |
| AC011444.3 | 3.01231947 | 0.649495861 | 0.67566869 | 0.96126382 | 0.33641953 | 0.53442067 |
| ACSBG2 | 58.6111131 | 2.794080517 | 0.57646527 | 4.8469191 | 1.25E-06 | 3.91E-05 |
| MLLT1 | 5137.93297 | -0.2184639 | 0.18275021 | -1.1954235 | 0.2319216 | 0.4198911 |
| AC011471.2 | 0.14616582 | 0.435199075 | 3.28898345 | 0.13232024 | 0.894731 | NA |
| ACER1 | 3.73468295 | 0.445026596 | 0.85513515 | 0.52041668 | 0.60277319 | 0.76009606 |
| AC011491.3 | 15.2172277 | 0.124139279 | 0.46785037 | 0.2653397 | 0.79074778 | 0.886377 |
| CLPP | 2265.47385 | -0.256109959 | 0.13425117 | -1.9076926 | 0.05643096 | 0.15844169 |
| ALKBH7 | 1789.14772 | 0.313366485 | 0.16544579 | 1.89407346 | 0.05821526 | 0.16207099 |
| PSPN | 274.456676 | 0.02714199 | 0.20920291 | 0.12974002 | 0.89677212 | 0.94715114 |
| GTF2F1 | 4272.6742 | -0.115556485 | 0.10934491 | -1.0568072 | 0.29059959 | 0.48667034 |
| KHSRP | 7522.21957 | -0.143497633 | 0.15752701 | -0.9109398 | 0.36232707 | 0.55963011 |
| SLC25A41 | 37.6064955 | -0.642255084 | 0.62725564 | -1.0239128 | 0.30587646 | 0.50286548 |
| SLC25A23 | 5709.61709 | 0.171926794 | 0.17269499 | 0.99555173 | 0.319468 | 0.51688118 |
| CRB3 | 1703.54583 | -0.042260252 | 0.19346703 | -0.2184365 | 0.82708907 | 0.90785626 |
| DENND1C | 2913.84154 | 0.271334488 | 0.18693254 | 1.45151021 | 0.14663784 | 0.30961048 |
| AC010503.4 | 1252.28896 | -0.277942257 | 0.17031433 | -1.631937 | 0.10269276 | 0.24180802 |
| TUBB4A | 185.953668 | 0.672326342 | 0.54179112 | 1.24093275 | 0.21463059 | 0.39877895 |
| AC010503.1 | 85.7452532 | 0.439214808 | 0.54847033 | 0.80079958 | 0.42324768 | 0.61578159 |
| AC010503.2 | 61.0307601 | 0.452820996 | 0.5481376 | 0.82610826 | 0.40874268 | 0.60316422 |
| TNFSF9 | 113.348475 | -3.19242586 | 0.57957872 | -5.5081833 | 3.63E-08 | 2.05E-06 |
| CD70 | 17.2797551 | -0.852819187 | 0.64041551 | -1.3316654 | 0.18297016 | 0.35970684 |
| TNFSF14 | 189.468965 | -2.757412741 | 0.57554971 | -4.7909203 | 1.66E-06 | 4.90E-05 |
| C3 | 23629.3869 | -3.156354654 | 0.61389967 | -5.1414829 | 2.73E-07 | 1.10E-05 |
| GPR108 | 5663.46109 | -0.312604796 | 0.11322573 | -2.760899 | 0.00576425 | 0.02993959 |
| TRIP10 | 8148.99778 | -0.821890008 | 0.22066328 | -3.7246342 | 0.0001956 | 0.00214556 |
| AC008760.1 | 713.529937 | -1.026825544 | 0.24326048 | -4.2210948 | 2.43E-05 | 0.00041576 |
| SH2D3A | 1921.88314 | -1.235080013 | 0.2589207 | -4.7701091 | 1.84E-06 | 5.31E-05 |
| VAV1 | 284.69825 | -0.403948489 | 0.47573141 | -0.8491104 | 0.39581986 | 0.59110713 |
| ADGRE1 | 43.4236489 | -1.214309474 | 0.60699551 | -2.0005246 | 0.04544364 | 0.13582102 |
| AC025278.1 | 6.74005291 | -0.533225744 | 1.03480595 | -0.5152906 | 0.60634999 | 0.76308872 |
| MBD3L2B | 0 | NA | NA | NA | NA | NA |
| MBD3L5 | 0 | NA | NA | NA | NA | NA |
| MBD3L4 | 0 | NA | NA | NA | NA | NA |
| MBD3L2 | 0 | NA | NA | NA | NA | NA |
| MBD3L3 | 0.22977084 | -1.364242679 | 3.30558948 | -0.4127078 | 0.67982068 | NA |
| ZNF557 | 383.435583 | -0.093039363 | 0.194067 | -0.4794188 | 0.63164075 | 0.78108538 |
| INSR | 7883.3794 | 0.28316174 | 0.23202957 | 1.22036918 | 0.22232496 | 0.40835196 |
| AC119396.1 | 29.8436285 | 1.43833491 | 0.39576329 | 3.63433131 | 0.0002787 | 0.00286973 |
| AC008878.3 | 3747.32772 | 0.226211451 | 0.1790935 | 1.26309132 | 0.20655636 | 0.38923048 |
| AC119396.2 | 6.40927677 | 0.297679632 | 0.51918712 | 0.57335712 | 0.56640291 | 0.73284279 |
| ARHGEF18 | 3992.5443 | 0.220745472 | 0.18280361 | 1.20755534 | 0.22721835 | 0.41410932 |
| PEX11G | 361.683523 | 1.122671264 | 0.39697449 | 2.82806906 | 0.00468297 | 0.02558921 |
| TEX45 | 16.5050534 | 0.228993436 | 0.36728973 | 0.62346812 | 0.53297695 | 0.70689184 |
| AC008878.1 | 823.185795 | 0.125059671 | 0.15681248 | 0.79751097 | 0.42515433 | 0.61749944 |
| ZNF358 | 2954.79645 | -0.118606131 | 0.15425836 | -0.7688798 | 0.44196468 | 0.63319617 |
| AC008878.4 | 682.482798 | -0.189440437 | 0.22368838 | -0.8468944 | 0.39705399 | 0.59197794 |
| MCOLN1 | 1888.57625 | -0.477726464 | 0.29888424 | -1.5983662 | 0.1099615 | 0.25384617 |
| AC008878.2 | 926.518975 | -0.21292445 | 0.23595037 | -0.902412 | 0.36683803 | 0.56387888 |
| PNPLA6 | 6677.31972 | -0.363791424 | 0.24877154 | -1.4623515 | 0.14364492 | 0.30523098 |
| CAMSAP3 | 4146.79874 | -0.167987736 | 0.16812754 | -0.9991684 | 0.3177131 | 0.51494128 |
| XAB2 | 4204.54294 | -0.105616362 | 0.13401291 | -0.7881059 | 0.43063477 | 0.62264716 |
| PET100 | 1650.16792 | -0.278893793 | 0.15888847 | -1.7552802 | 0.07921139 | 0.20107062 |
| AC008763.2 | 4999.63662 | -0.241948403 | 0.09204761 | -2.6285138 | 0.00857589 | 0.04008972 |
| PCP2 | 495.502593 | 0.167463754 | 0.42417207 | 0.39480146 | 0.69298941 | 0.82334449 |
| AC008763.1 | 261.825301 | 0.832650834 | 0.34395481 | 2.42081464 | 0.01548577 | 0.06227317 |
| STXBP2 | 7154.24899 | -0.427498248 | 0.12352284 | -3.4608843 | 0.0005384 | 0.00480302 |
| RETN | 39.1905357 | -4.002524709 | 0.93864752 | -4.2641403 | 2.01E-05 | 0.00035515 |
| MCEMP1 | 70.4156283 | -2.999791401 | 0.76837821 | -3.9040558 | 9.46E-05 | 0.00121252 |
| AC008763.3 | 1500.73781 | -0.605723475 | 0.19250223 | -3.146579 | 0.00165193 | 0.01165052 |
| TRAPPC5 | 1528.49911 | -0.586926725 | 0.18850448 | -3.1135957 | 0.00184823 | 0.01267891 |
| FCER2 | 32.0103753 | 1.584116442 | 0.85675857 | 1.8489648 | NA | NA |
| CLEC4G | 32.8559041 | -2.206575845 | 0.78405749 | -2.8143036 | 0.0048883 | 0.02643112 |
| CD209 | 370.184922 | -1.516221658 | 0.560622 | -2.7045347 | 0.00684001 | 0.03398284 |
| CLEC4M | 9.79901951 | 0.787861094 | 0.74167162 | 1.06227753 | 0.28810971 | 0.48426636 |
| EVI5L | 1720.75279 | 0.205887687 | 0.13464504 | 1.5291145 | 0.12623606 | 0.27933582 |
| PRR36 | 572.764225 | -1.459754339 | 0.38727614 | -3.7692855 | 0.00016372 | 0.00187053 |
| AC010336.3 | 151.81061 | -1.327965877 | 0.42042321 | -3.1586408 | 0.00158507 | 0.01127549 |
| LRRC8E | 341.908359 | -1.128811379 | 0.23933982 | -4.7163542 | 2.40E-06 | 6.60E-05 |
| AC010336.1 | 329.236137 | -1.141732066 | 0.24318641 | -4.6948842 | 2.67E-06 | 7.16E-05 |
| MAP2K7 | 3414.18586 | -0.417291202 | 0.20051177 | -2.0811307 | 0.03742195 | 0.11788273 |
| AC010336.4 | 554.599867 | -0.364368414 | 0.21567128 | -1.689462 | 0.09113094 | 0.2225732 |
| TGFBR3L | 142.873423 | 1.918120727 | 0.5518596 | 3.47574043 | 0.00050945 | 0.00460182 |
| AC010336.2 | 53.3760608 | 1.806998803 | 0.44496818 | 4.06096181 | 4.89E-05 | 0.00073099 |
| SNAPC2 | 965.907694 | -0.428451228 | 0.12100584 | -3.5407484 | 0.00039899 | 0.00381489 |
| CTXN1 | 518.788178 | -1.918590696 | 0.34902172 | -5.4970525 | 3.86E-08 | 2.14E-06 |
| TIMM44 | 2609.33884 | -0.28068169 | 0.192485 | -1.4582003 | 0.14478533 | 0.30684991 |
| ELAVL1 | 3540.79145 | -0.284326284 | 0.13121945 | -2.1667999 | 0.03025012 | 0.10141057 |
| AC010336.6 | 108.366353 | 0.07804052 | 0.30690748 | 0.25428028 | 0.79927903 | 0.89108294 |
| AC008946.1 | 24.6395321 | 0.764435598 | 0.37005656 | 2.06572639 | 0.03885432 | 0.12122438 |
| CCL25 | 2.27539003 | -0.484397879 | 0.99567878 | -0.4865002 | 0.6266126 | 0.77783715 |
| FBN3 | 263.025254 | 0.959599765 | 0.60394414 | 1.58888828 | 0.11208562 | 0.25742011 |
| CERS4 | 3067.82755 | 0.217142095 | 0.27341772 | 0.7941771 | 0.42709234 | 0.61942718 |
| AC022146.2 | 3.91727141 | 1.072351841 | 0.80244821 | 1.33635022 | 0.18143483 | 0.35776837 |
| CD320 | 1713.07242 | 0.111214135 | 0.18224904 | 0.61023167 | 0.54170835 | 0.71442107 |
| AC010323.1 | 2133.27161 | -0.022171646 | 0.16105483 | -0.1376652 | 0.89050502 | 0.94322853 |
| NDUFA7 | 2049.67657 | -0.016952491 | 0.15640362 | -0.1083894 | 0.91368683 | 0.95775549 |
| RPS28 | 13730.4611 | -0.4675861 | 0.11584076 | -4.0364557 | 5.43E-05 | 0.0007941 |
| KANK3 | 1520.38418 | -0.415148827 | 0.23613856 | -1.758073 | 0.07873509 | 0.20022821 |
| ANGPTL4 | 9637.20607 | -3.374937308 | 0.66718613 | -5.0584644 | 4.23E-07 | 1.58E-05 |
| RAB11B-AS1 | 1229.31355 | -0.09787115 | 0.157511 | -0.6213607 | 0.53436231 | 0.70808591 |
| RAB11B | 8056.36842 | -0.369067536 | 0.1727756 | -2.1361092 | 0.03267052 | 0.10722109 |
| 2-Mar | 1782.58955 | -0.314950043 | 0.1443987 | -2.1811141 | 0.02917498 | 0.09897648 |
| AC136469.1 | 1.94074797 | -0.312810047 | 0.84480157 | -0.3702764 | 0.71117659 | 0.83587657 |
| HNRNPM | 9071.05792 | -0.176669711 | 0.19715753 | -0.896084 | 0.37020789 | 0.56696095 |
| PRAM1 | 147.702647 | -0.477299595 | 0.41073678 | -1.1620571 | 0.24521227 | 0.43623069 |
| ZNF414 | 630.047018 | -0.144327573 | 0.18656565 | -0.7736021 | 0.43916615 | 0.63069445 |
| MYO1F | 948.699341 | -0.373670744 | 0.46061317 | -0.8112463 | 0.41722422 | 0.61021645 |
| AC092316.1 | 2.82247727 | 0.531952173 | 0.9066784 | 0.58670436 | 0.55740228 | 0.72629087 |
| ADAMTS10 | 1131.0166 | -0.08844084 | 0.43337792 | -0.2040733 | 0.83829626 | 0.91448609 |
| AC130469.1 | 13.4380801 | -0.461487336 | 0.54686893 | -0.8438719 | 0.39874098 | 0.59358697 |
| LINC01862 | 0 | NA | NA | NA | NA | NA |
| ACTL9 | 0 | NA | NA | NA | NA | NA |
| OR2Z1 | 0.10721287 | -0.642905668 | 3.30558948 | -0.1944905 | 0.84579185 | NA |
| ZNF558 | 1004.26132 | -0.288927838 | 0.11085598 | -2.6063352 | 0.00915169 | 0.0421758 |
| AC008734.1 | 42.7064237 | 0.640212987 | 0.28725038 | 2.22876291 | 0.02582968 | 0.0904914 |
| MBD3L1 | 0.78821284 | 1.804036369 | 1.4146525 | 1.27525055 | 0.20222058 | NA |
| MUC16 | 3.62285434 | -0.617794681 | 1.13445125 | -0.5445758 | 0.58604531 | 0.74736507 |
| OR1M1 | 0 | NA | NA | NA | NA | NA |
| OR7G2 | 0.09635527 | 0.547239394 | 3.29377202 | 0.16614368 | 0.86804388 | NA |
| OR7G1 | 0 | NA | NA | NA | NA | NA |
| OR7G3 | 0 | NA | NA | NA | NA | NA |

| | | | | | | |
|---|---|---|---|---|---|---|
| ZNF317 | 1371.66364 | -0.042040915 | 0.1848382 | -0.2274471 | 0.82007609 | 0.90346785 |
| OR7D2 | 15.3589294 | 2.377432739 | 0.92736038 | 2.56365572 | NA | NA |
| OR7D4 | 0 | NA | NA | NA | NA | NA |
| OR7E24 | 0.09989236 | -0.642905668 | 3.30558948 | -0.1944905 | 0.84579185 | NA |
| AC011451.1 | 49.8154436 | -0.234717627 | 0.19566984 | -1.1995596 | 0.23031043 | 0.41789624 |
| ZNF699 | 22.2023578 | 0.337914393 | 0.34806665 | 0.97083242 | 0.33163173 | 0.52955848 |
| ZNF559 | 816.787266 | 0.24274909 | 0.08961784 | 2.7087139 | 0.00675446 | 0.03365105 |
| ZNF559-ZNF | 612.219666 | 0.437595325 | 0.22690746 | 1.92851892 | 0.05379061 | 0.15338003 |
| ZNF177 | 502.271784 | 0.365341488 | 0.26601732 | 1.37337482 | 0.16963585 | 0.34140975 |
| ZNF266 | 1913.96377 | -0.478309108 | 0.14027381 | -3.4098248 | 0.00065005 | 0.00555467 |
| ZNF560 | 2.55620891 | 0.91391308 | 0.98127788 | 0.93134992 | 0.35167258 | 0.54931916 |
| AC008567.2 | 1.45505834 | 0.460982015 | 1.02408134 | 0.45014199 | 0.65260806 | 0.79590266 |
| ZNF426 | 581.6731 | -0.104165007 | 0.11236263 | -0.9270432 | 0.35390411 | 0.55148822 |
| ZNF426-DT | 53.5119025 | -0.213102563 | 0.37225138 | -0.5724695 | 0.56700394 | 0.73311566 |
| ZNF121 | 1453.3466 | -0.69852668 | 0.15580162 | -4.4834365 | 7.35E-06 | 0.00016034 |
| ZNF561 | 1186.20638 | 0.334373736 | 0.15715693 | 2.12764237 | 0.03336675 | 0.10881537 |
| ZNF562 | 1228.61289 | -0.278281114 | 0.14934928 | -1.8632907 | 0.06242138 | 0.16999144 |
| AC008759.3 | 14.0526524 | -2.850290141 | 0.94656026 | -3.0112083 | 0.0026021 | 0.01648533 |
| AC008759.2 | 26.8403497 | 0.000227684 | 0.50534258 | 0.00045055 | 0.99964051 | 0.99979217 |
| ZNF846 | 308.233504 | 0.085390716 | 0.23394195 | 0.36500814 | 0.71510533 | 0.83801965 |
| AC008752.1 | 2.02505874 | -0.582815101 | 0.96477051 | -0.6040971 | 0.54577906 | 0.71715108 |
| FBXL12 | 1256.8849 | -0.170149145 | 0.10251375 | -1.659769 | 0.09696092 | 0.23248405 |
| UBL5 | 3964.22248 | -0.334731825 | 0.12390427 | -2.7015357 | 0.00690201 | 0.0342245 |
| AC008752.2 | 3.81222459 | 0.825008698 | 0.76380047 | 1.08013641 | 0.28008144 | 0.47564225 |
| PIN1 | 2534.00574 | -0.133907734 | 0.1491939 | -0.8975416 | 0.36942998 | 0.56630227 |
| OLFM2 | 268.864059 | 0.002156113 | 0.52981386 | 0.00406957 | 0.99675296 | 0.99841889 |
| COL5A3 | 804.348312 | 0.444131123 | 0.50561218 | 0.87840273 | 0.3797252 | 0.57573766 |
| AC008742.1 | 43.3115631 | 0.836069149 | 0.62778599 | 1.33177414 | 0.18293442 | 0.35970684 |
| RDH8 | 427.696392 | 3.442915245 | 0.95566313 | 3.60264528 | 0.000315 | 0.00314637 |
| C19orf66 | 4825.1075 | -1.19875204 | 0.53107787 | -2.2572058 | 0.02399522 | 0.08559839 |
| AC020931.1 | 1123.86316 | -1.094334276 | 0.5567506 | -1.9655736 | 0.04934789 | 0.14434327 |
| ANGPTL6 | 1428.45823 | -1.130180523 | 0.50399635 | -2.2424379 | 0.02493308 | 0.08817023 |
| PPAN-P2RY1 | 1830.05094 | -0.825224823 | 0.24102117 | -3.4238686 | 0.00061736 | 0.00532894 |
| PPAN | 1508.10565 | -0.726006416 | 0.22811005 | -3.1827024 | 0.00145908 | 0.01058191 |
| P2RY11 | 1975.20824 | -0.674296352 | 0.20146655 | -3.3469395 | 0.00081709 | 0.00668944 |
| EIF3G | 7091.46446 | -0.405897967 | 0.11775262 | -3.4470399 | 0.00056676 | 0.00500858 |
| DNMT1 | 1693.15629 | -0.165551638 | 0.20904483 | -0.7919432 | 0.42839379 | 0.62076844 |
| S1PR2 | 281.781988 | -0.663649082 | 0.3054989 | -2.1723452 | 0.02982963 | 0.10048639 |
| MRPL4 | 2967.12464 | -0.265400464 | 0.10192297 | -2.6039319 | 0.00921611 | 0.0423395 |
| AC011511.2 | 1327.53281 | -0.740179277 | 0.29698974 | -2.4922722 | 0.01269287 | 0.05377748 |
| AC011511.3 | 721.83 | -0.404902041 | 0.09090194 | -4.4542729 | 8.42E-06 | 0.00017801 |
| ICAM1 | 8371.18279 | -1.900043179 | 0.53241819 | -3.5687045 | 0.00035875 | 0.00349985 |
| AC011511.5 | 3176.29374 | -2.303311802 | 0.54232814 | -4.2470815 | 2.17E-05 | 0.00037796 |
| ICAM4 | 20.8869615 | -0.668668845 | 0.54456282 | -1.2279003 | 0.2194844 | 0.40499641 |
| ICAM5 | 189.168532 | -1.468990877 | 0.40399989 | -3.636117 | 0.00027678 | 0.0028555 |
| ZGLP1 | 331.899776 | -0.319549205 | 0.33824079 | -0.9447388 | 0.34479223 | 0.54239522 |
| AC011511.1 | 1303.45113 | -0.297675227 | 0.16644495 | -1.7884305 | 0.07370658 | 0.19141645 |
| FDX2 | 1067.8953 | -0.30769408 | 0.15639324 | -1.9674385 | 0.04913267 | 0.14395874 |
| AC011511.4 | 2468.87264 | -0.401766247 | 0.16134258 | -2.4901439 | 0.01276914 | 0.0539965 |
| RAVER1 | 3496.01846 | -0.250734397 | 0.17155542 | -1.4615358 | 0.14386845 | 0.30558285 |
| AC114271.1 | 909.078607 | 0.155534188 | 0.32010875 | 0.48587921 | 0.62705281 | 0.77806242 |
| ICAM3 | 768.6892 | 0.151294253 | 0.37162222 | 0.40711842 | 0.68392101 | 0.81748139 |
| TYK2 | 6548.64496 | 0.084932281 | 0.13215358 | 0.64267863 | 0.52043265 | 0.69699386 |
| CDC37 | 10034.1282 | -0.575802242 | 0.16410003 | -3.5088491 | 0.00045005 | 0.00418323 |
| PDE4A | 1065.31109 | -0.372078607 | 0.32496615 | -1.1449765 | 0.25221889 | 0.44445342 |
| KEAP1 | 3628.9899 | -0.174307991 | 0.22416205 | -0.7775981 | 0.43680597 | 0.62891432 |
| S1PR5 | 45.6533228 | -0.570107318 | 0.59692331 | -0.9550763 | 0.3395391 | 0.53765964 |
| ATG4D | 2907.81486 | -0.230386492 | 0.10998686 | -2.0946728 | 0.03620009 | 0.11526758 |
| KRI1 | 3026.77243 | -0.451114806 | 0.13866146 | -3.2533538 | 0.00114051 | 0.00870747 |
| CDKN2D | 556.947039 | -1.080952248 | 0.15693683 | -6.8878176 | 5.67E-12 | 1.26E-09 |
| AP1M2 | 3575.17338 | 0.153600778 | 0.26477233 | 0.58012399 | 0.56183101 | 0.72961113 |
| SLC44A2 | 10516.4156 | 0.246245838 | 0.23044042 | 1.06858789 | 0.28525541 | 0.48132743 |
| ILF3-DT | 1269.99571 | 0.327548216 | 0.24508067 | 1.33649145 | 0.18138869 | 0.3577309 |
| ILF3 | 11483.8149 | -0.111191787 | 0.18766003 | -0.5925172 | 0.55350433 | 0.7230174 |
| QTRT1 | 3953.31183 | -0.045246228 | 0.20290995 | -0.2229867 | 0.82354583 | 0.90564302 |
| DNM2 | 16507.2106 | 0.001152126 | 0.1344942 | 0.00856637 | 0.99316511 | 0.99686464 |
| TMED1 | 2664.72926 | -0.364216626 | 0.17706845 | -2.0569256 | 0.03969338 | 0.12311104 |
| C19orf38 | 97.0226633 | -0.642164608 | 0.39319476 | -1.6331973 | 0.10242751 | 0.241399 |
| CARM1 | 4156.95213 | -0.419524856 | 0.14724814 | -2.8491012 | 0.00438429 | 0.0244375 |
| YIPF2 | 3527.26769 | -0.370372 | 0.13115725 | -2.8238775 | 0.00474465 | 0.02585051 |
| TIMM29 | 844.826046 | -0.199848996 | 0.1481848 | -1.348647 | 0.17745036 | 0.35252117 |
| SMARCA4 | 5348.02007 | 0.216007164 | 0.11262137 | 1.91799443 | 0.05511172 | 0.15594978 |
| AC011442.1 | 1.48842102 | 2.083451789 | 1.23263105 | 1.6902477 | 0.09098058 | 0.22230891 |
| LDLR | 2544.49731 | -1.268695275 | 0.50944895 | -2.4903286 | 0.01276251 | 0.05397982 |
| SPC24 | 222.970531 | -1.170304881 | 0.45846128 | -2.5526799 | 0.01068977 | 0.04730479 |
| KANK2 | 4463.24445 | 0.390602974 | 0.26868294 | 1.45376918 | 0.14601031 | 0.30864063 |
| DOCK6 | 4039.48689 | -0.261284382 | 0.17201148 | -1.5189939 | 0.12876404 | 0.28305333 |
| AC011472.1 | 151.534106 | -0.373823519 | 0.24630541 | -1.5177235 | 0.12908411 | 0.2836387 |
| AC011472.4 | 18.6584122 | 0.776349624 | 0.41031027 | 1.89210381 | 0.05847715 | 0.16254269 |
| ANGPTL8 | 383.324015 | -0.932361096 | 0.51923875 | -1.7956308 | 0.07255325 | 0.18940989 |
| TSPAN16 | 3.72366981 | -0.338377857 | 0.85304639 | -0.3966699 | 0.69161087 | 0.82241065 |
| RAB3D | 2930.45951 | -0.53132685 | 0.18611952 | -2.8547616 | 0.00430691 | 0.02407235 |
| TMEM205 | 6179.43838 | -0.206195485 | 0.10982394 | -1.8775094 | 0.06044831 | 0.16639061 |
| CCDC159 | 5699.5621 | 0.01862559 | 0.11130832 | 0.16733331 | 0.86710779 | 0.93098661 |
| PLPPR2 | 1980.97068 | -0.750712413 | 0.22329162 | -3.3620268 | 0.00077373 | 0.00640357 |
| AC024575.1 | 93.3281442 | 0.335409962 | 0.2683232 | 1.25002223 | 0.21129143 | 0.39468406 |
| SWSAP1 | 328.784985 | 0.106065588 | 0.21760464 | 0.48742338 | 0.62595833 | 0.77729476 |
| EPOR | 3481.39567 | -0.099369312 | 0.26724697 | -0.3718258 | 0.71002257 | 0.83521649 |
| RGL3 | 8256.72699 | -0.189282541 | 0.23606277 | -0.8018314 | 0.42265049 | 0.61515031 |
| CCDC151 | 818.115689 | 0.231766592 | 0.12566431 | 1.84433102 | 0.06513491 | 0.17505174 |
| PRKCSH | 13641.2331 | -0.121262967 | 0.09085126 | -1.3347417 | 0.18196091 | 0.35845729 |
| ELAVL3 | 19.9093093 | -1.861323621 | 0.78068849 | -2.3842079 | 0.01711593 | 0.06697539 |
| AC008481.3 | 101.971288 | -0.125811686 | 0.19395948 | -0.6486493 | 0.51656506 | 0.69389362 |
| ZNF653 | 471.130584 | -0.40223559 | 0.17673365 | -2.2759423 | 0.02284946 | 0.08274379 |
| ECSIT | 2546.09286 | 0.228833135 | 0.19790963 | 1.15625061 | 0.24757866 | 0.43915066 |
| CNN1 | 2001.32904 | -0.511522919 | 0.8826337 | -0.5795416 | 0.56222381 | 0.72982024 |
| ELOF1 | 2217.6078 | -0.488535035 | 0.18244785 | -2.6776695 | 0.00741363 | 0.03611629 |
| ZNF627 | 1458.36535 | 0.59781071 | 0.186502 | 3.20538501 | 0.00134882 | 0.00994965 |
| ACP5 | 5174.72145 | 1.180320016 | 0.42074719 | 2.80529509 | 0.00502705 | 0.02698766 |
| AC008543.1 | 103.782755 | 0.039753723 | 0.27796671 | 0.14301613 | 0.88627744 | 0.94110344 |
| ZNF823 | 656.010509 | -0.441692101 | 0.22778517 | -1.9390732 | 0.05249243 | 0.15067219 |
| ZNF441 | 540.091836 | 0.440054576 | 0.2324334 | 1.89325019 | 0.05832461 | 0.1622555 |
| AC008543.5 | 3.71727409 | 0.678289248 | 0.63373185 | 1.07030954 | 0.28448 | 0.48060335 |
| ZNF491 | 110.115255 | 0.637053463 | 0.21901412 | 2.90873234 | 0.00362897 | 0.02117169 |
| ZNF440 | 833.994663 | -0.364417791 | 0.23760576 | -1.5337077 | 0.12510158 | 0.27747452 |
| AC008543.4 | 5.05348926 | -0.533544277 | 0.54922094 | -0.9714565 | 0.33132098 | 0.5292225 |
| ZNF439 | 612.8775 | 0.081417664 | 0.14237502 | 0.57185356 | 0.56742119 | 0.73343938 |
| ZNF69 | 505.71444 | 0.490850777 | 0.18026396 | 2.72295567 | 0.00647007 | 0.03259138 |
| ZNF700 | 1717.67305 | -0.355887446 | 0.17093272 | -2.0820323 | 0.03733952 | 0.11770161 |
| AC008770.2 | 779.791861 | -0.296260842 | 0.18876678 | -1.5694544 | 0.1165421 | 0.26472695 |
| ZNF763 | 406.765939 | 0.562327544 | 0.30422155 | 1.84841455 | 0.0645424 | 0.17391172 |
| ZNF433 | 840.308685 | -0.075234384 | 0.19646277 | -0.3829447 | 0.70176074 | 0.82913351 |
| AC008770.1 | 145.635167 | 0.028942788 | 0.18016026 | 0.16065023 | 0.87236889 | 0.93346728 |

| | | | | | | |
|---|---|---|---|---|---|---|
| ZNF878 | 53.0292569 | -0.470804903 | 0.41252673 | -1.1412713 | 0.25375705 | 0.44642527 |
| ZNF844 | 1079.08393 | 0.514353008 | 0.2071391 | 2.48312849 | 0.01302341 | 0.05481147 |
| ZNF788P | 329.309854 | 0.504186488 | 0.15843164 | 3.18235984 | 0.0014608 | 0.01059152 |
| ZNF20 | 389.643791 | 0.039593647 | 0.13038357 | 0.30367052 | 0.76137892 | 0.8677512 |
| ZNF625-ZNF | 516.240524 | -0.008725674 | 0.15197116 | -0.0574166 | 0.9542133 | 0.9780936 |
| ZNF625 | 247.209368 | -0.309603577 | 0.24786754 | -1.2490687 | 0.21163996 | 0.39519514 |
| ZNF136 | 537.862619 | 0.319004865 | 0.15618899 | 2.04242865 | 0.04110903 | 0.12640451 |
| ZNF44 | 2321.79916 | -0.014898788 | 0.20261499 | -0.0735325 | 0.94138238 | 0.97208964 |
| ZNF563 | 329.799033 | -0.162112153 | 0.19264441 | -0.8415098 | 0.40006242 | 0.59481541 |
| ZNF442 | 242.148192 | 0.008105609 | 0.26237483 | 0.03089324 | 0.97535468 | 0.98862634 |
| AC008758.4 | 74.514099 | -0.170202508 | 0.23013156 | -0.7395879 | 0.45955011 | 0.64821269 |
| ZNF799 | 421.966194 | -0.431051776 | 0.15189933 | -2.8377464 | 0.00454333 | 0.02503815 |
| AC008758.5 | 79.4608936 | 0.00954187 | 0.26257685 | 0.03633934 | 0.97101178 | 0.9865502 |
| ZNF443 | 321.216517 | -0.002896231 | 0.16198314 | -0.0178798 | 0.98573472 | 0.99368571 |
| AC008758.6 | 31.3109288 | 0.581659047 | 0.26157437 | 2.22368513 | 0.02616964 | 0.09139098 |
| AC008758.2 | 13.6463063 | 0.990979675 | 0.42482281 | 2.33268944 | 0.01966445 | 0.07396257 |
| ZNF709 | 279.570593 | 0.317203559 | 0.1697727 | 1.86840146 | 0.06170613 | 0.16869073 |
| AC008758.1 | 288.549324 | 0.387114692 | 0.13420968 | 2.88440212 | 0.00392158 | 0.0224321 |
| ZNF564 | 845.750094 | 0.073840532 | 0.17332212 | 0.42603063 | 0.67008553 | 0.80733565 |
| AC010422.6 | 1033.13 | 0.098547857 | 0.1760764 | 0.55968805 | 0.57569223 | 0.73977897 |
| AC010422.1 | 7.28766987 | 1.560220578 | 0.78116359 | 1.99730325 | 0.04579225 | 0.13661519 |
| ZNF490 | 168.44226 | 0.241071631 | 0.17643369 | 1.36635825 | 0.17182655 | 0.34456776 |
| ZNF791 | 1199.4177 | -0.174869261 | 0.12438356 | -1.4058873 | 0.15975759 | 0.32767428 |
| AC010422.3 | 1493.7908 | -0.484833492 | 0.17781836 | -2.7265661 | 0.00639971 | 0.03231721 |
| MAN2B1 | 4176.8582 | -0.113849749 | 0.1516419 | -0.7507803 | 0.45278489 | 0.64248481 |
| AC010422.5 | 2610.80679 | 0.018942939 | 0.11609353 | 0.16316964 | 0.87038487 | 0.93249822 |
| WDR83 | 4170.08045 | -0.114222169 | 0.09279499 | -1.2309088 | 0.21835697 | 0.40345245 |
| WDR83OS | 4984.31784 | -0.169944119 | 0.09732709 | -1.7461132 | 0.08079129 | 0.20383337 |
| DHPS | 3368.91078 | -0.239679806 | 0.09538835 | -2.5126738 | 0.01198201 | 0.05148699 |
| AC010422.2 | 9.44222959 | -0.477574877 | 0.50082843 | -0.9535698 | 0.34030144 | 0.53834957 |
| GNG14 | 5.08156274 | 1.104425465 | 0.62239511 | 1.77447644 | 0.07598438 | 0.1955982 |
| FBXW9 | 746.945819 | -0.018066085 | 0.13215468 | -0.1367041 | 0.89126469 | 0.94362126 |
| AC018761.4 | 209.492349 | -0.232147117 | 0.19441083 | -1.1941059 | 0.23243654 | 0.42056756 |
| TNPO2 | 4878.78665 | 0.07737507 | 0.1129534 | 0.68501763 | 0.49333279 | 0.67597744 |
| AC018761.1 | 0.09997307 | 0.547239394 | 3.29377202 | 0.16614368 | 0.86804388 | NA |
| TRIR | 5235.49645 | -0.561548238 | 0.14065735 | -3.9923135 | 6.54E-05 | 0.00091065 |
| ASNA1 | 4227.08167 | 0.070699217 | 0.13007242 | 0.54353735 | 0.58675988 | 0.74777006 |
| BEST2 | 1.88402831 | -2.213829714 | 1.19910014 | -1.8462426 | 0.06485699 | 0.17455388 |
| HOOK2 | 7579.79029 | -0.212519121 | 0.20660601 | -1.0286202 | 0.30365818 | 0.50056049 |
| AC018761.3 | 2.54495786 | 0.253357366 | 0.91941858 | 0.27556259 | 0.78288405 | 0.88119568 |
| JUNB | 9247.01819 | -0.445898257 | 0.42661854 | -1.0451919 | 0.29593427 | 0.49275663 |
| PRDX2 | 11806.2812 | 0.195832278 | 0.18935738 | 1.03419405 | 0.30104545 | 0.49768474 |
| AC018761.2 | 4752.16515 | 0.210953994 | 0.18758426 | 1.12458261 | 0.26076595 | 0.45391762 |
| THSD8 | 297.978238 | 0.080094444 | 0.28887202 | 0.27726619 | 0.78157572 | 0.88036187 |
| RNASEH2A | 299.09455 | 0.067078967 | 0.29061483 | 0.23081742 | 0.81745665 | 0.9018631 |
| RTBDN | 1.14554669 | -2.516520547 | 1.61562706 | -1.5576123 | 0.11932519 | 0.26882573 |
| AC020934.1 | 1.13645471 | -2.522962299 | 1.45839513 | -1.729958 | 0.08363778 | 0.209035 |
| MAST1 | 14.9670496 | -0.643593819 | 0.33648519 | -1.9126958 | 0.055787 | 0.15721954 |
| DNASE2 | 1830.38294 | -0.120227693 | 0.17018287 | -0.7064618 | 0.47990101 | 0.66458301 |
| AC020934.2 | 372.509941 | 0.09948459 | 0.19638374 | 0.50658261 | 0.61244771 | 0.76781621 |
| KLF1 | 7.93827659 | -4.607380609 | 1.39410099 | -3.3049116 | 0.00095006 | 0.00754841 |
| GCDH | 2065.13718 | 1.305657001 | 0.37028126 | 3.52612225 | 0.00042169 | 0.00397367 |
| SYCE2 | 116.87673 | 1.044527974 | 0.27951047 | 3.73699056 | 0.00018624 | 0.00206433 |
| FARSA | 2993.96101 | -0.256656368 | 0.12996324 | -1.9748382 | 0.04828651 | 0.14204768 |
| FARSA-AS1 | 280.364584 | 0.049366717 | 0.16759485 | 0.29455987 | 0.76833012 | 0.87199765 |
| CALR | 32956.102 | -0.204271663 | 0.25020757 | -0.8164088 | 0.41426637 | 0.60794725 |
| AC092069.1 | 7195.14865 | -0.53528106 | 0.28263867 | -1.8938706 | 0.05824219 | 0.16207752 |
| RAD23A | 5970.71561 | -0.441627695 | 0.15062405 | -2.9319866 | 0.00336801 | 0.0200213 |
| GADD45GIP1 | 3933.7589 | -0.606068401 | 0.11926551 | -5.0816739 | 3.74E-07 | 1.43E-05 |
| DAND5 | 3.96097667 | -0.142370092 | 0.66742542 | -0.2133124 | 0.83108333 | 0.91040231 |
| NFIX | 1766.74438 | 0.271865285 | 0.25193917 | 1.07909099 | 0.28054717 | 0.47627817 |
| AC138474.1 | 0.16354576 | -1.003574173 | 3.30558948 | -0.3035992 | 0.7614333 | NA |
| AC007787.1 | 0.60368709 | -0.787256271 | 1.88937612 | -0.4166753 | 0.67691596 | NA |
| LYL1 | 308.889655 | -0.092657195 | 0.30149776 | -0.307323 | 0.75859754 | 0.86635037 |
| TRMT1 | 2825.00135 | -0.592222671 | 0.13223432 | -4.4785854 | 7.51E-06 | 0.00016241 |
| NACC1 | 3847.39736 | -0.443642575 | 0.16748705 | -2.6488172 | 0.0080774 | 0.03837165 |
| AC005546.1 | 1.11230043 | 0.117836931 | 1.19832992 | 0.0983343 | 0.92166684 | 0.96174883 |
| AC011446.1 | 696.984064 | -0.641346569 | 0.19148772 | -3.3492831 | 0.00081021 | 0.00664672 |
| STX10 | 2168.96728 | -0.424183124 | 0.16961001 | -2.5009321 | 0.01238669 | 0.05282826 |
| IER2 | 5218.29988 | 0.159064436 | 0.24701677 | 0.64394184 | 0.51961315 | 0.6963207 |
| AC011446.2 | 2772.36399 | 0.336142094 | 0.2821281 | 1.19145201 | 0.23347619 | 0.42186975 |
| CACNA1A | 61.1328448 | -0.326217648 | 0.3953851 | -0.8250631 | 0.40933578 | 0.60383668 |
| CCDC130 | 3098.55595 | -0.434008276 | 0.11234721 | -3.8630979 | 0.00011196 | 0.00138347 |
| MRI1 | 2858.43102 | 0.408887893 | 0.20797771 | 1.96601786 | 0.04929655 | 0.14426256 |
| AC008686.1 | 1628.5974 | 0.312425033 | 0.21478268 | 1.45460999 | 0.14577726 | 0.30837048 |
| C19orf53 | 2720.14126 | -0.255612509 | 0.1074759 | -2.378324 | 0.01739154 | 0.06772468 |
| ZSWIM4 | 1341.34829 | -1.217837286 | 0.27886571 | -4.3671101 | 1.26E-05 | 0.00024647 |
| AC020916.2 | 0.86387931 | -0.627547426 | 1.46402921 | -0.4286441 | 0.66818227 | NA |
| AC020916.1 | 1421.86056 | 0.290998352 | 0.39807588 | 0.73101227 | 0.46477166 | 0.65205955 |
| NANOS3 | 10.351125 | -0.227578297 | 0.45046932 | -0.5052027 | 0.6134165 | 0.76841076 |
| C19orf57 | 215.399669 | -0.016143507 | 0.1349872 | -0.1195929 | 0.90480567 | 0.9522378 |
| CC2D1A | 4129.05095 | -0.51190155 | 0.15188551 | -3.370312 | 0.00075083 | 0.00624773 |
| PODNL1 | 61.6636319 | -0.186470924 | 0.29777036 | -0.6262239 | 0.53116808 | 0.70547921 |
| DCAF15 | 1816.92983 | -0.463037995 | 0.18719599 | -2.4735466 | 0.01337794 | 0.05587206 |
| RFX1 | 1472.52081 | -0.057838858 | 0.14192571 | -0.4075291 | 0.68361943 | 0.81725652 |
| AC022098.4 | 5.45035414 | -0.553841171 | 0.77726126 | -0.7125547 | 0.47612134 | 0.66150263 |
| RLN3 | 1.56733686 | -1.735635625 | 1.17268809 | -1.4800488 | 0.13886022 | 0.29821044 |
| AC022098.2 | 2.61238969 | -3.207643527 | 1.26227842 | -2.5411537 | 0.01104873 | 0.04851224 |
| IL27RA | 580.74212 | -0.494179433 | 0.40699424 | -1.2142173 | 0.22466477 | 0.41084674 |
| PALM3 | 3834.93001 | 0.273836339 | 0.439743 | 0.62271904 | 0.53346917 | 0.70723352 |
| MISP3 | 1832.67718 | 0.096598669 | 0.23020503 | 0.41962015 | 0.67476297 | 0.81067472 |
| C19orf67 | 10.4760269 | 0.146947721 | 0.5641421 | 0.26047998 | 0.79449356 | 0.88857752 |
| SAMD1 | 1823.93734 | -0.998951391 | 0.28300992 | -3.5297399 | 0.00041597 | 0.00393298 |
| PRKACA | 4241.11178 | -0.201306323 | 0.17342857 | -1.1607448 | 0.24574568 | 0.43700274 |
| AC022098.3 | 2.42364729 | -0.328496176 | 0.81917974 | -0.4010062 | 0.68841554 | 0.8201273 |
| ASF1B | 92.8626532 | -1.693895506 | 0.47766688 | -3.5461858 | 0.00039085 | 0.00374926 |
| AC022098.1 | 392.563082 | 0.097560788 | 0.17287927 | 0.564329 | 0.57253025 | 0.73719144 |
| ADGRL1 | 2240.69355 | -0.257883869 | 0.31452221 | -0.8199226 | 0.41226024 | 0.60616749 |
| AC011509.2 | 2.03423993 | 0.621899517 | 1.01180725 | 0.61464228 | 0.53879099 | 0.71207851 |
| AC011509.1 | 1.09676173 | -0.431764315 | 1.85130435 | -0.2332217 | 0.81558927 | 0.90059416 |
| LINC01841 | 2.24867019 | 0.090507166 | 1.05636765 | 0.08567771 | 0.93172262 | 0.96704554 |
| LINC01842 | 1.68489468 | -0.791063658 | 1.26057468 | -0.6275421 | 0.53030396 | 0.70472412 |
| ADGRE5 | 3156.88905 | -0.464612965 | 0.42423705 | -1.095173 | 0.27344085 | 0.46843129 |
| AC008569.1 | 595.630604 | -0.516472177 | 0.40394448 | -1.2785722 | 0.20104775 | 0.38276147 |
| DDX39A | 3612.12054 | -1.140149938 | 0.39388096 | -2.8946562 | 0.00379574 | 0.02193983 |
| PKN1 | 6035.43378 | -0.604339934 | 0.15632411 | -3.865942 | 0.00011066 | 0.00137124 |
| PTGER1 | 382.355737 | 0.281517358 | 0.49367854 | 0.57024427 | 0.56851203 | 0.73412969 |
| GIPC1 | 5556.03127 | -0.502261493 | 0.18070438 | -2.7794649 | 0.00544485 | 0.02871615 |
| DNAJB1 | 8738.13796 | -0.142180329 | 0.21421557 | -0.6637255 | 0.50686604 | 0.68594288 |
| TECR | 6203.81836 | 0.334191528 | 0.0891084 | 3.7503929 | 0.00017656 | 0.00198458 |
| NDUFB7 | 4651.20338 | 0.242045272 | 0.13226886 | 1.82994901 | 0.06725756 | 0.17900504 |
| CLEC17A | 19.935615 | 0.850713547 | 0.88960039 | 0.95628729 | 0.33892711 | 0.53701303 |
| ADGRE3 | 8.95928281 | 0.838028324 | 0.77417862 | 1.08247412 | 0.27904191 | 0.47470067 |

| | | | | | | |
|---|---|---|---|---|---|---|
| ZNF333 | 832.1445 | -0.15109362 | 0.27662614 | -0.5462015 | 0.58492741 | 0.74673623 |
| ADGRE2 | 327.612452 | -1.063234309 | 0.30634121 | -3.4707518 | 0.000519 | 0.00466784 |
| OR7C1 | 18.5207598 | -0.410744885 | 0.4272924 | -0.9612735 | 0.33641464 | 0.53442067 |
| OR7A5 | 4.18852197 | -0.198521732 | 0.98979047 | -0.2005695 | 0.84103525 | 0.91512686 |
| OR7A10 | 0.08393208 | -0.642905668 | 3.30558948 | -0.1944905 | 0.84579185 | NA |
| OR7A17 | 0.06785815 | 0.795868262 | 3.28898349 | 0.24198001 | 0.80879565 | NA |
| OR7C2 | 0 | NA | NA | NA | NA | NA |
| SLC1A6 | 3.46112474 | 0.134817155 | 0.76703187 | 0.17576474 | 0.86047877 | 0.92734057 |
| CCDC105 | 0.40065839 | -1.928496579 | 2.79860735 | -0.6890915 | 0.49076568 | NA |
| CASP14 | 48.9106856 | -4.254554815 | 1.08334358 | -3.9272442 | 8.59E-05 | 0.0011256 |
| OR1I1 | 0 | NA | NA | NA | NA | NA |
| SYDE1 | 1764.6318 | 0.074896367 | 0.24833453 | 0.30159465 | 0.76296109 | 0.86877586 |
| ILVBL | 6536.52406 | 0.549614626 | 0.1503214 | 3.65626326 | 0.00025592 | 0.00266637 |
| AC003956.1 | 32.3809153 | 1.425302397 | 0.42956948 | 3.31797871 | 0.00090671 | 0.0072697 |
| NOTCH3 | 6347.75042 | 0.364307964 | 0.40759797 | 0.89379239 | 0.37143298 | 0.56829264 |
| AC004257.1 | 0.19981251 | 0.689154255 | 3.12815507 | 0.22030693 | 0.82563212 | NA |
| EPHX3 | 158.526968 | -0.790584439 | 0.32421632 | -2.4384474 | 0.0147505 | 0.06009754 |
| BRD4 | 5322.50542 | -0.410172038 | 0.17691029 | -2.3185313 | 0.02042046 | 0.07608276 |
| AC005776.2 | 25.171293 | -0.949593881 | 0.32758034 | -2.8988122 | 0.00374579 | 0.0217139 |
| AKAP8 | 2594.38914 | -0.242690116 | 0.1432506 | -1.6941648 | 0.09023398 | 0.22109924 |
| AKAP8L | 8623.01005 | -0.599656333 | 0.26566848 | -2.2571603 | 0.02399806 | 0.08559839 |
| WIZ | 3854.34243 | -0.225973437 | 0.17963156 | -1.257983 | 0.20839792 | 0.39152061 |
| RASAL3 | 679.050765 | -0.643142883 | 0.53290061 | -1.2068721 | 0.2274814 | 0.41438826 |
| PGLYRP2 | 2.12519474 | 0.670649336 | 1.18604923 | 0.56544814 | 0.57176899 | 0.73660547 |
| CYP4F22 | 7.96153845 | -0.442896961 | 0.78323749 | -0.5654696 | 0.57175442 | 0.73660547 |
| AC093072.1 | 0 | NA | NA | NA | NA | NA |
| CYP4F8 | 0.59501419 | 1.087743134 | 2.14528202 | 0.50703969 | 0.61212697 | NA |
| CYP4F3 | 4404.84197 | 1.244997687 | 0.71398814 | 1.74372321 | 0.08120737 | 0.20454547 |
| CYP4F12 | 410.445039 | 1.469228527 | 0.60507616 | 2.42817124 | 0.01517518 | 0.06134745 |
| OR10H2 | 0.19978473 | -1.003574173 | 3.30558948 | -0.3035992 | 0.7614333 | NA |
| OR10H3 | 0.09997307 | 0.547239394 | 3.29377202 | 0.16614368 | 0.86804388 | NA |
| AC004597.1 | 12.7329519 | -0.195809658 | 1.2580292 | -0.1556479 | 0.87631053 | 0.93555133 |
| OR10H5 | 0.09644756 | -0.642905668 | 3.30558948 | -0.1944905 | 0.84579185 | NA |
| OR10H1 | 0.55995999 | -2.369025895 | 2.70745101 | -0.8750023 | 0.38157265 | NA |
| LINC01764 | 12.5955693 | -1.507077099 | 0.76650234 | -1.9661742 | 0.0492785 | 0.14425776 |
| UCA1 | 37.1025128 | -1.071400668 | 0.67756789 | -1.5812448 | 0.11382209 | 0.26012213 |
| CYP4F2 | 2265.56546 | 4.114922384 | 0.85124489 | 4.83400541 | 1.34E-06 | 4.12E-05 |
| CYP4F11 | 970.70613 | -0.118300818 | 0.78469203 | -0.1507608 | 0.88016439 | 0.9378468 |
| OR10H4 | 0 | NA | NA | NA | NA | NA |
| LINC01855 | 0 | NA | NA | NA | NA | NA |
| TPM4 | 13212.6245 | -0.323361096 | 0.31832243 | -1.0158288 | 0.30971092 | 0.50692551 |
| AC008894.1 | 0.09635527 | 0.547239394 | 3.29377202 | 0.16614368 | 0.86804388 | NA |
| RAB8A | 3238.66419 | -0.279105217 | 0.13850249 | -2.0151639 | 0.04388747 | 0.13270413 |
| AC008894.2 | 2512.12044 | -0.324461565 | 0.14314117 | -2.2667243 | 0.02340707 | 0.08412765 |
| HSH2D | 536.658359 | -1.791260125 | 0.77403256 | -2.3141922 | 0.02065718 | 0.07671704 |
| CIB3 | 0.41019645 | -1.082170709 | 2.85743156 | -0.3787215 | 0.7048947 | NA |
| FAM32A | 3619.53422 | -0.275799328 | 0.20076027 | -1.3737744 | 0.16951172 | 0.3413242 |
| AC020911.1 | 14.1055297 | 1.050511747 | 0.43578119 | 2.41064044 | 0.01592454 | 0.06361324 |
| AP1M1 | 4207.60275 | 0.174186181 | 0.18528971 | 0.94007476 | 0.34717921 | 0.54472033 |
| AC020911.2 | 0.97047018 | 0.982887082 | 1.26552331 | 0.77666454 | 0.43735671 | NA |
| KLF2 | 1575.47547 | 0.482469158 | 0.35551795 | 1.35708805 | 0.17475323 | 0.34864121 |
| AC020917.3 | 0.08393208 | -0.642905668 | 3.30558948 | -0.1944905 | 0.84579185 | NA |
| EPS15L1 | 1998.35097 | 0.072278689 | 0.16412671 | 0.44038347 | 0.65965939 | 0.79999826 |
| AC020917.2 | 10.1702725 | -0.751264983 | 0.57907111 | -1.2973622 | 0.19450659 | 0.37448581 |
| CALR3 | 2.75643577 | 0.511957906 | 0.79538267 | 0.64366239 | 0.51979439 | 0.69649278 |
| AC008764.1 | 804.556914 | 0.231096401 | 0.22898776 | 1.00920853 | 0.31287464 | 0.50973089 |
| C19orf44 | 2129.03762 | -0.054137384 | 0.12135462 | -0.446109 | 0.65551854 | 0.79798836 |
| CHERP | 3471.9995 | 0.029908357 | 0.15698805 | 0.1905136 | 0.84890669 | 0.92057694 |
| AC008764.6 | 89.9569864 | 0.003439064 | 0.24271445 | 0.01416918 | 0.98869501 | 0.99493137 |
| SLC35E1 | 2840.93387 | -0.30106632 | 0.18987568 | -1.5855971 | 0.11283074 | 0.25854594 |
| AC008764.3 | 518.995275 | -0.501417266 | 0.20476361 | -2.4487616 | 0.01433483 | 0.0589075 |
| AC008764.5 | 0.7110371 | -1.742783491 | 1.66163778 | -1.0488348 | 0.29425417 | NA |
| AC008764.8 | 293.35255 | -0.382322874 | 0.17107473 | -2.2348296 | 0.02542854 | 0.08943613 |
| MED26 | 614.536741 | -0.396665953 | 0.19102047 | -2.0765625 | 0.03784196 | 0.11889579 |
| AC008764.4 | 1299.01287 | 0.213126001 | 0.12190237 | 1.74833349 | 0.0804063 | 0.20307577 |
| AC008764.2 | 70.9587584 | 1.068149655 | 0.27802464 | 3.84192449 | 0.00012207 | 0.00148241 |
| AC024075.3 | 233.698609 | -0.222640471 | 0.20171404 | -1.103743 | 0.26970462 | 0.4639947 |
| SMIM7 | 2661.5898 | 0.129092553 | 0.10598466 | 1.21803057 | 0.22321236 | 0.40924076 |
| TMEM38A | 687.947245 | 0.06189658 | 0.34634279 | 0.17871479 | 0.85816165 | 0.92582473 |
| NWD1 | 44.7475623 | 1.461152611 | 0.78541482 | 1.86035783 | 0.06283492 | 0.17074092 |
| SIN3B | 4805.093 | -0.528584339 | 0.19426428 | -2.7209549 | 0.00650936 | 0.03272053 |
| F2RL3 | 760.300305 | -3.014194399 | 0.41587716 | -7.2477997 | 4.24E-13 | 1.29E-10 |
| CPAMD8 | 2123.60962 | -0.447912317 | 0.33633651 | -1.3317386 | 0.1829461 | 0.35970684 |
| AC020908.2 | 23.5570966 | 0.683937884 | 0.42647633 | 1.60369482 | 0.10878134 | 0.25208797 |
| HAUS8 | 249.441114 | -0.553137899 | 0.181067 | -3.0548796 | 0.00225151 | 0.01480236 |
| AC020908.1 | 3.82868319 | -0.344770911 | 0.75508423 | -0.4565993 | 0.6479591 | 0.79277053 |
| MYO9B | 3507.40755 | -0.538885417 | 0.20790466 | -2.5919833 | 0.00954244 | 0.04350878 |
| AC020908.3 | 0.51312181 | -1.195921559 | 2.20571861 | -0.5421914 | 0.58768669 | NA |
| AC020913.1 | 34.9903377 | 1.760443899 | 0.49014779 | 3.59165937 | 0.00032858 | 0.00325983 |
| AC020913.3 | 0.37634252 | 0.953311038 | 2.17901597 | 0.43749612 | 0.66175159 | NA |
| AC020913.2 | 318.864948 | -0.552845574 | 0.2124591 | -2.6021271 | 0.00926475 | 0.04252594 |
| USE1 | 991.940239 | 0.148098809 | 0.1952738 | 0.75841619 | 0.44820187 | 0.63828284 |
| OCEL1 | 1747.51162 | 0.281672918 | 0.18241753 | 1.54411102 | 0.12256144 | 0.27373127 |
| NR2F6 | 2932.11002 | -0.345343751 | 0.20725744 | -1.6662551 | 0.09566262 | 0.23053237 |
| AC010646.1 | 535.807306 | -0.126714363 | 0.22030141 | -0.5751863 | 0.56516528 | 0.73195857 |
| USHBP1 | 703.33693 | 0.448711687 | 0.30024408 | 1.4944897 | 0.13504767 | 0.29242691 |
| BABAM1 | 3326.76578 | -0.146897992 | 0.0824425 | -1.7818235 | 0.07477802 | 0.19324739 |
| AC010463.1 | 2371.03446 | -0.029844134 | 0.08852513 | -0.3371261 | 0.73602181 | 0.85182775 |
| ANKLE1 | 82.0912516 | -0.672718614 | 0.36278478 | -1.8543187 | 0.06369358 | 0.17233733 |
| ABHD8 | 981.660126 | -0.185928375 | 0.23295449 | -0.7981318 | 0.42479403 | 0.61714608 |
| MRPL34 | 3045.86969 | 0.208373914 | 0.22020428 | 0.94627549 | 0.34400809 | 0.5415817 |
| DDA1 | 2101.51227 | -0.41707288 | 0.12316649 | -3.3862528 | 0.00070854 | 0.00596749 |
| ANO8 | 1814.60647 | -0.128982914 | 0.31825348 | -0.4052836 | 0.68526906 | 0.81810054 |
| GTPBP3 | 1309.57161 | -0.099512801 | 0.15637334 | -0.6363796 | 0.52452905 | 0.69951978 |
| PLVAP | 11231.9654 | -0.510225679 | 0.47659785 | -1.0705581 | 0.28436818 | 0.48053758 |
| AC010463.2 | 2.89372155 | -0.229928807 | 1.26109332 | -0.182325 | 0.8553277 | 0.92411978 |
| CCDC194 | 10.6041301 | 0.449351304 | 0.51095343 | 0.87943691 | 0.37916442 | 0.5751591 |
| BST2 | 4790.66811 | -1.169782342 | 0.51913565 | -2.2533269 | 0.02423854 | 0.08625781 |
| BISPR | 201.094414 | 0.136078948 | 0.25289303 | 0.53808897 | 0.59051563 | 0.75023593 |
| MVB12A | 1949.47516 | -0.030892004 | 0.2102047 | -0.1469615 | 0.88316238 | 0.93933739 |
| AC010319.4 | 115.526603 | -0.061513534 | 0.38115471 | -0.1613873 | 0.87178836 | 0.93317628 |
| AC010319.3 | 14.9700284 | -0.433701092 | 0.80886159 | -0.536187 | 0.5918293 | 0.75136193 |
| TMEM221 | 27.1195619 | -0.244075463 | 0.26268976 | -0.9291396 | 0.35281674 | 0.5504868 |
| AC010319.2 | 6.89119069 | 0.40870186 | 0.65542271 | 0.62356988 | 0.5329101 | 0.70688425 |
| AC010319.5 | 0.61127651 | -1.172707601 | 1.67562871 | -0.6998612 | 0.48401396 | NA |
| NXNL1 | 1.38445528 | 2.057116181 | 1.22325942 | 1.68166797 | 0.09263324 | 0.22521167 |
| SLC27A1 | 1798.87988 | 0.240874206 | 0.22853377 | 1.05399829 | 0.2918837 | 0.48807693 |
| AC010618.3 | 549.700971 | 0.015783939 | 0.13054548 | 0.12090758 | 0.90376424 | 0.95167363 |
| PGLS | 2355.42486 | -0.319725264 | 0.08270113 | -3.8660324 | 0.00011062 | 0.00137124 |
| AC010618.2 | 14.6534971 | -0.183175918 | 0.33959731 | -0.5393915 | 0.58961671 | 0.74967828 |
| FAM129C | 76.2865703 | 1.568861784 | 0.79685752 | 1.9688109 | NA | NA |
| COLGALT1 | 3864.51216 | -0.845777663 | 0.20776993 | -4.0707413 | 4.69E-05 | 0.00070496 |
| UNC13A | 67.5530205 | -0.69637893 | 0.57017179 | -1.2213493 | 0.22195379 | 0.40784067 |

| | | | | | | |
|---|---|---|---|---|---|---|
| AC008761.2 | 0 | NA | NA | NA | NA | NA |
| MAP1S | 1887.81821 | -0.77639801 | 0.20112936 | -3.8601922 | 0.0001133 | 0.00139738 |
| AC008761.1 | 513.607842 | -0.962926823 | 0.19440173 | -4.9532832 | 7.30E-07 | 2.49E-05 |
| FCHO1 | 134.102911 | 0.007717604 | 0.47632306 | 0.01620246 | 0.98707288 | 0.99410333 |
| B3GNT3 | 979.606699 | -1.70148143 | 0.27088373 | -6.2812241 | 3.36E-10 | 4.03E-08 |
| INSL3 | 20.1803702 | -1.660750229 | 0.47493618 | -3.4967861 | 0.0004709 | 0.00433018 |
| JAK3 | 1201.83569 | -1.163932087 | 0.46815159 | -2.4862291 | 0.01291048 | 0.05444173 |
| RPL18A | 14427.4082 | -0.717064324 | 0.16640524 | -4.3091451 | 1.64E-05 | 0.00030285 |
| SLC5A5 | 3.39708154 | -0.16064665 | 1.17731063 | -0.1364522 | 0.89146379 | 0.94377583 |
| CCDC124 | 2469.41274 | -0.544311802 | 0.16400443 | -3.3188847 | 0.00090378 | 0.00725057 |
| KCNN1 | 10.8306844 | 0.166034399 | 0.45681027 | 0.36346468 | 0.71625779 | 0.83854488 |
| ARRDC2 | 2792.22176 | -1.281756408 | 0.28753473 | -4.4577447 | 8.28E-06 | 0.00017614 |
| IL12RB1 | 125.835823 | -0.432083588 | 0.47898115 | -0.9020889 | 0.36700964 | 0.56395482 |
| MAST3 | 1943.95409 | 0.770629002 | 0.31721148 | 2.42938557 | 0.01512444 | 0.06124561 |
| AC007192.2 | 483.314448 | 0.341279508 | 0.29775926 | 1.14615918 | 0.2517293 | 0.44399047 |
| PIK3R2 | 2768.14732 | 0.123234195 | 0.125022 | 0.98570008 | 0.32428031 | 0.52186901 |
| AC007192.1 | 9041.68606 | -0.547311017 | 0.219483 | -2.4936374 | 0.01264417 | 0.05363387 |
| IFI30 | 6344.82401 | -0.883705752 | 0.31214755 | -2.8310514 | 0.00463953 | 0.02545024 |
| MPV17L2 | 825.422237 | -0.386928392 | 0.19560925 | -1.978068 | 0.04792104 | 0.14119304 |
| RAB3A | 303.83551 | 0.621513013 | 0.37353542 | 1.66386635 | 0.09613914 | 0.23124958 |
| AC005759.1 | 151.990787 | 0.205669616 | 0.38985989 | 0.52754752 | 0.59781343 | 0.75633738 |
| AC008397.2 | 404.750432 | -0.00813708 | 0.41832392 | -0.0194516 | 0.98448083 | 0.99328299 |
| PDE4C | 362.450539 | 0.19640708 | 0.44607082 | 0.44030471 | 0.65971643 | 0.79999826 |
| AC008397.1 | 32.3298971 | -2.199316367 | 0.90724611 | -2.4241673 | 0.01534353 | 0.06183845 |
| IQCN | 533.22862 | 0.977699441 | 0.45087021 | 2.16847205 | 0.03012279 | 0.10111226 |
| JUND | 7318.26975 | -0.670029155 | 0.26965868 | -2.4847305 | 0.01296495 | 0.0545969 |
| LSM4 | 3980.69853 | -0.122683611 | 0.19973783 | -0.6142232 | 0.53906784 | 0.71225669 |
| PGPEP1 | 8105.08015 | 0.924762471 | 0.3204627 | 2.88571013 | 0.00390532 | 0.02237306 |
| GDF15 | 15208.1126 | -2.042148504 | 0.412787 | -4.947221 | 7.53E-07 | 2.56E-05 |
| LRRC25 | 285.268481 | -0.642995351 | 0.44385487 | -1.4486613 | 0.14743221 | 0.31075551 |
| SSBP4 | 2854.52335 | -0.929746755 | 0.27064662 | -3.4352794 | 0.00059194 | 0.00516701 |
| ISYNA1 | 4502.63345 | 0.487392626 | 0.31530147 | 1.54579876 | 0.12215318 | 0.27307983 |
| AC010335.1 | 597.882276 | -0.888969785 | 0.20790933 | -4.2757571 | 1.90E-05 | 0.00034079 |
| ELL | 2633.70961 | -0.733649095 | 0.17134588 | -4.2816851 | 1.85E-05 | 0.00033458 |
| AC005387.1 | 92.3309299 | -1.076481452 | 0.25326365 | -4.2504381 | 2.13E-05 | 0.00037284 |
| FKBP8 | 13400.9553 | -0.133565824 | 0.14188282 | -0.9413812 | 0.34650953 | 0.54415533 |
| AC005387.2 | 44.8321647 | -0.52335746 | 0.27914863 | -1.8748344 | 0.06081551 | 0.16705259 |
| KXD1 | 3469.8975 | -0.57282333 | 0.09997209 | -5.7298328 | 1.01E-08 | 6.98E-07 |
| AC005253.1 | 500.836156 | -0.072876523 | 0.1228731 | -0.593104 | 0.55311158 | 0.72290971 |
| AC005253.2 | 1461.72864 | -0.650145964 | 0.1180721 | -5.5063469 | 3.66E-08 | 2.06E-06 |
| UBA52 | 13316.1936 | -0.457347993 | 0.1901996 | -2.4045687 | 0.01619156 | 0.0643689 |
| CRLF1 | 642.031269 | -1.537095253 | 0.31363658 | -4.9008801 | 9.54E-07 | 3.13E-05 |
| REX1BD | 1589.56397 | -0.14938661 | 0.23012412 | -0.6491567 | 0.51623711 | 0.69363383 |
| TMEM59L | 209.37913 | -2.017623706 | 0.68854487 | -2.9302719 | 0.00338665 | 0.02010574 |
| KLHL26 | 800.418653 | 0.263283364 | 0.20919244 | 1.25857019 | 0.20818562 | 0.39140559 |
| CRTC1 | 2306.27979 | 0.184913338 | 0.20658281 | 0.89510516 | 0.37073088 | 0.56749134 |
| COMP | 342.924741 | -1.866902939 | 0.62829774 | -2.9713666 | 0.00296478 | 0.01816899 |
| UPF1 | 7047.08438 | 0.267723239 | 0.1360767 | 1.96744361 | 0.04913209 | 0.14395874 |
| GDF1 | 26.8209653 | -1.119018292 | 0.3717948 | -3.0097739 | 0.00261442 | 0.0165488 |
| CERS1 | 45.1527957 | -1.185757402 | 0.36580323 | -3.241517 | 0.00118895 | 0.00898482 |
| AC005197.1 | 2.17016971 | -0.555516417 | 0.90465936 | -0.6140614 | 0.53917474 | 0.71232654 |
| COPE | 6868.54479 | -0.331605179 | 0.14263509 | -2.32485 | 0.02007998 | 0.07513966 |
| AC002985.1 | 5357.87587 | -0.405655593 | 0.2139902 | -1.8956737 | 0.05800321 | 0.16166839 |
| DDX49 | 1836.10625 | -0.121721652 | 0.1851819 | -0.6573086 | 0.51098253 | 0.68923833 |
| HOMER3 | 3590.21324 | -0.606981475 | 0.35398502 | -1.7147095 | 0.0863985 | 0.21401416 |
| HOMER3-AS | 99.9974862 | -0.069953374 | 0.43631472 | -0.1603278 | 0.87262287 | 0.93366644 |
| SUGP2 | 6067.99369 | -0.110548704 | 0.12868983 | -0.8590322 | 0.39032275 | 0.5858507 |
| ARMC6 | 1563.16207 | -0.063167243 | 0.20687002 | -0.3053475 | 0.7601015 | 0.86700095 |
| SLC25A42 | 4791.52139 | 1.127065895 | 0.52617576 | 2.14199511 | 0.03219388 | 0.1061058 |
| TMEM161A | 1977.97201 | -0.0061688 | 0.17482602 | -0.0352854 | 0.97185219 | 0.98670705 |
| BORCS8-MEI | 1110.18544 | -0.073028415 | 0.11705429 | -0.623885 | 0.53270312 | 0.7068228 |
| MEF2B | 456.813865 | -0.386140625 | 0.14933067 | -2.5858092 | 0.00971507 | 0.04410163 |
| BORCS8 | 1905.28528 | 0.154629066 | 0.19950033 | 0.77508175 | 0.43829136 | 0.63008807 |
| RFXANK | 2271.592 | -0.488596044 | 0.11410466 | -4.2819991 | 1.85E-05 | 0.00033454 |
| NR2C2AP | 1221.56576 | -0.762951477 | 0.12896101 | -5.9161406 | 3.30E-09 | 2.69E-07 |
| NCAN | 1.41186781 | 0.622940012 | 1.27177037 | 0.48982114 | 0.62426047 | 0.77611992 |
| HAPLN4 | 69.3813582 | 0.530278036 | 0.71096615 | 0.74585553 | 0.45575469 | 0.64517131 |
| TM6SF2 | 511.568206 | 0.256459353 | 0.51842368 | 0.49469066 | 0.62081849 | 0.77350682 |
| SUGP1 | 1904.85295 | -0.105288791 | 0.10643853 | -0.9891981 | 0.32256624 | 0.5203279 |
| MAU2 | 3831.41281 | 0.16108732 | 0.15224949 | 1.05804833 | 0.29003341 | 0.48615433 |
| GATAD2A | 4756.26889 | -0.586828933 | 0.20278406 | -2.8938612 | 0.00380536 | 0.02197229 |
| AC092067.1 | 0.13382144 | 0.02007176 | 3.29738704 | 0.00608717 | 0.99514317 | NA |
| TSSK6 | 92.2662602 | 0.777949964 | 0.2811791 | 2.76674181 | 0.00566196 | 0.02956551 |
| NDUFA13 | 7932.64865 | -0.183712355 | 0.09003205 | -2.0405217 | 0.04129839 | 0.12689799 |
| AC011448.1 | 8282.25404 | -0.180908979 | 0.09085642 | -1.9911525 | 0.04646413 | 0.13808847 |
| YJEFN3 | 1158.75034 | -0.180471596 | 0.28820051 | -0.6262015 | 0.53118277 | 0.70547921 |
| CILP2 | 13.9181677 | -0.720730635 | 0.46748168 | -1.5417302 | 0.12313917 | 0.2746124 |
| PBX4 | 197.556076 | -0.143507913 | 0.32783211 | -0.4377482 | 0.66156883 | 0.80084282 |
| AC002306.1 | 0.68972249 | -1.898132174 | 1.68116726 | -1.1290561 | 0.25887416 | NA |
| LPAR2 | 510.411798 | -0.996162831 | 0.26689995 | -3.7323455 | 0.00018971 | 0.00209136 |
| GMIP | 819.797886 | -0.879523646 | 0.29229883 | -3.0089879 | 0.0026212 | 0.01656385 |
| ATP13A1 | 5304.38747 | -0.305195875 | 0.13645628 | -2.2365835 | 0.02531357 | 0.08918088 |
| ZNF101 | 612.746283 | -0.273345142 | 0.16900772 | -1.617353 | 0.10580211 | 0.24690281 |
| ZNF14 | 328.9705 | 0.435661166 | 0.1109811 | 3.92554379 | 8.65E-05 | 0.00113133 |
| LINC00663 | 334.234067 | 0.531127284 | 0.25992198 | 2.04341042 | 0.04101183 | 0.12621187 |
| AC011477.4 | 96.5671625 | 0.837393549 | 0.24474854 | 3.42144452 | 0.00062289 | 0.00536964 |
| ZNF506 | 1128.46423 | 0.027929668 | 0.14079529 | 0.19837075 | 0.842755 | 0.91639311 |
| ZNF253 | 404.791512 | 0.377439805 | 0.18555896 | 2.03406945 | 0.04194459 | 0.12824103 |
| AC011477.3 | 73.4530252 | 0.573849867 | 0.24077954 | 2.38329994 | 0.01715821 | 0.06706138 |
| ZNF93 | 165.116478 | -0.332634232 | 0.21822231 | -1.5242907 | 0.12743611 | 0.28104794 |
| ZNF682 | 319.457248 | 0.353619959 | 0.27773189 | 1.27324218 | 0.20293212 | 0.38495415 |
| ZNF90 | 225.55154 | 0.142401615 | 0.22288863 | 0.63889133 | 0.52289363 | 0.69862624 |
| AC011447.3 | 109.446523 | 0.214578034 | 0.19923522 | 1.07700852 | 0.28147646 | 0.47739479 |
| ZNF486 | 786.758931 | -0.301845052 | 0.33004701 | -0.9145517 | 0.36042703 | 0.5580355 |
| AC008554.1 | 37.8840868 | 1.11295908 | 0.57509966 | 1.93524559 | 0.05296017 | 0.15163517 |
| ZNF737 | 297.052579 | 0.812055795 | 0.21270339 | 3.81778492 | 0.00013466 | 0.00159949 |
| AC010636.2 | 136.863284 | 0.614342343 | 0.2822088 | 2.17690708 | 0.0294875 | 0.09969044 |
| AC010636.1 | 0.78020709 | 2.482453895 | 1.31516086 | 1.88756673 | 0.05908414 | NA |
| ZNF626 | 548.084644 | 0.195706776 | 0.20153389 | 0.97108618 | 0.33150536 | 0.52948495 |
| AC010329.1 | 148.512377 | 0.242748409 | 0.3006493 | 0.80741386 | 0.41942808 | 0.61235293 |
| ZNF66 | 33.8362225 | -0.040773221 | 0.44442565 | -0.0917436 | 0.92690173 | 0.96450157 |
| ZNF85 | 170.036826 | 0.225482747 | 0.28280594 | 0.79730555 | 0.4252736 | 0.61763864 |
| AC008739.1 | 0.10178723 | 1.087385526 | 3.28420134 | 0.33109588 | 0.74057207 | NA |
| ZNF430 | 277.632933 | 0.058901897 | 0.19956669 | 0.29514895 | 0.7678801 | 0.87183611 |
| ZNF714 | 111.178915 | -0.023062852 | 0.36725553 | -0.0627978 | 0.94992749 | 0.97598332 |
| ZNF431 | 928.050061 | -0.018607377 | 0.16921065 | -0.1099658 | 0.91243653 | 0.95696422 |
| ZNF708 | 257.120469 | 0.240458098 | 0.17275055 | 1.39193821 | 0.16394111 | 0.33341303 |
| ZNF738 | 249.518707 | 0.054042335 | 0.31095647 | 0.17379389 | 0.86202745 | 0.92808258 |
| ZNF493 | 866.29023 | 0.612163637 | 0.15027545 | 4.07361036 | 4.63E-05 | 0.00069792 |
| AC010615.4 | 29.6024942 | 1.31017166 | 0.53401607 | 2.45343115 | 0.01415006 | 0.05837565 |
| LINC00664 | 9.11226405 | -0.014057002 | 0.49985588 | -0.0281221 | 0.97756476 | 0.98965006 |
| ZNF429 | 323.915218 | 0.681616731 | 0.23022709 | 2.96062786 | 0.00307013 | 0.01869297 |
| AC123912.1 | 15.1594671 | 0.394131643 | 0.45272695 | 0.87057252 | 0.38398761 | 0.57960808 |

| | | | | | | |
|---|---|---|---|---|---|---|
| AC123912.4 | 88.2461576 | 0.580588093 | 0.49054008 | 1.18356912 | 0.23658367 | 0.42590875 |
| AC123912.2 | 0.85691534 | -1.004330772 | 1.87762123 | -0.5348953 | 0.59272226 | NA |
| ZNF100 | 362.087811 | -0.267957349 | 0.29771778 | -0.9000381 | 0.36809998 | 0.56490696 |
| AC092364.1 | 0.03392908 | 0.380740593 | 3.29738709 | 0.11546736 | 0.90807469 | NA |
| ZNF43 | 643.016241 | 0.090891565 | 0.24168569 | 0.37607343 | 0.70686231 | 0.83325953 |
| ZNF208 | 103.106585 | -0.309646821 | 0.34653609 | -0.8935486 | 0.37156348 | 0.56836599 |
| AC003973.2 | 0.16964538 | 1.509808164 | 3.2763376 | 0.46082191 | 0.64492638 | NA |
| AC003973.1 | 0 | NA | NA | NA | NA | NA |
| ZNF257 | 67.1583498 | 0.341074023 | 0.33856298 | 1.00741677 | 0.31373454 | 0.51080113 |
| ZNF676 | 101.845502 | 0.172559217 | 0.34181933 | 0.50482581 | 0.61368119 | 0.76859525 |
| AC073539.1 | 0 | NA | NA | NA | NA | NA |
| AC073539.3 | 0 | NA | NA | NA | NA | NA |
| ZNF729 | 1.05475524 | -0.666181637 | 1.57592546 | -0.4227241 | 0.67249659 | 0.80897677 |
| ZNF98 | 54.8204697 | 1.225811022 | 0.33774876 | 3.62935763 | 0.00028413 | 0.00290292 |
| AC011516.1 | 0.06785815 | 0.795868262 | 3.28898349 | 0.24198001 | 0.80879565 | NA |
| AC025811.1 | 16.0006917 | 1.090180618 | 0.47062839 | 2.31643618 | 0.02053447 | 0.07639586 |
| LINC01233 | 6.22237292 | -0.198706995 | 0.50204613 | -0.3957943 | 0.69225678 | 0.82291897 |
| LINC01785 | 5.60655596 | 1.015381473 | 0.76967498 | 1.31923409 | 0.18709086 | 0.36504522 |
| ZNF492 | 6.45269947 | 0.991363849 | 0.49882891 | 1.98738252 | 0.04688002 | 0.1389799 |
| AC024563.1 | 4.70486869 | 0.449692704 | 0.66293172 | 0.6783394 | 0.49755652 | 0.6792941 |
| ZNF99 | 22.7377241 | 0.403271251 | 0.3942895 | 1.0227796 | 0.30641207 | 0.50327538 |
| ZNF723 | 2.69674149 | -1.690583738 | 1.50262914 | -1.1250838 | 0.26055352 | 0.45363767 |
| AC022145.1 | 0 | NA | NA | NA | NA | NA |
| ZNF728 | 7.56390779 | 0.87552062 | 0.53998478 | 1.62138017 | 0.10493613 | 0.24553379 |
| LINC01859 | 0.09635527 | 0.547239394 | 3.29377202 | 0.16614368 | 0.86804388 | NA |
| LINC01858 | 0.33649131 | -0.541134579 | 2.88157096 | -0.1877915 | 0.85104009 | NA |
| AC074135.2 | 0 | NA | NA | NA | NA | NA |
| AC074135.1 | 1.32379353 | 0.521583283 | 1.15516028 | 0.4515246 | 0.6516115 | 0.79524711 |
| ZNF730 | 6.85419686 | -0.458417816 | 0.68262146 | -0.6715549 | 0.50186707 | 0.68231773 |
| ZNF724 | 14.2768982 | 0.3667807 | 0.39883232 | 0.91963635 | 0.35776282 | 0.55541979 |
| AC092329.3 | 126.307272 | 0.90996493 | 0.2288669 | 3.97595684 | 7.01E-05 | 0.0009563 |
| AC092329.1 | 155.314434 | 0.957764958 | 0.19785041 | 4.84085398 | 1.29E-06 | 4.01E-05 |
| ZNF91 | 1202.72594 | 0.213802425 | 0.16030995 | 1.33368154 | 0.18230825 | 0.35900743 |
| AC010300.1 | 58.3049029 | 0.746519407 | 0.39518565 | 1.88903472 | 0.05888718 | 0.1633064 |
| LINC01224 | 15.5820095 | 1.319825225 | 0.69608708 | 1.89606338 | 0.05795166 | 0.16157596 |
| ZNF675 | 275.814187 | 0.362333846 | 0.20387383 | 1.77724546 | 0.07552786 | 0.19467546 |
| ZNF681 | 106.531088 | -0.01288022 | 0.24637468 | -0.052279 | 0.95830639 | 0.98008608 |
| AC139769.2 | 37.8655921 | 0.011752958 | 0.7544534 | 0.01557811 | 0.98757097 | 0.99440414 |
| ZNF726 | 129.28364 | 0.402999496 | 0.36110963 | 1.11600318 | 0.26442077 | 0.45837338 |
| AC011503.1 | 40.8109745 | 0.475327682 | 0.39226936 | 1.21173799 | 0.22561269 | 0.41215149 |
| ZNF254 | 901.694816 | 0.011397525 | 0.1131734 | 0.10070851 | 0.91978185 | 0.96074349 |
| AC092279.1 | 217.851804 | 0.063223422 | 0.45838738 | 0.13792575 | 0.8902991 | 0.94316 |
| AC073534.2 | 11.0357046 | -0.774214172 | 0.39184685 | -1.9758081 | 0.04817652 | 0.14178736 |
| ERVK-28 | 15.3048181 | -0.080729 | 1.31764005 | -0.0612679 | 0.95114588 | 0.97670237 |
| LINC00662 | 402.095846 | -0.107850903 | 0.1451677 | -0.7429401 | 0.45751795 | 0.64676468 |
| AC006504.1 | 63.7894231 | 0.255365228 | 0.60950208 | 0.41897351 | 0.67523549 | 0.81102072 |
| AC006504.5 | 430.175981 | -0.315944293 | 0.20563492 | -1.5364331 | 0.1244322 | 0.27641117 |
| AC006504.2 | 0.03392908 | 0.380740593 | 3.29738709 | 0.11546736 | 0.90807469 | NA |
| AC010511.1 | 0.06785815 | 0.795868262 | 3.28898349 | 0.24198001 | 0.80879565 | NA |
| AC020905.1 | 2.40988951 | -2.04382254 | 1.57365925 | -1.2987707 | 0.19402263 | 0.37380449 |
| AC005580.1 | 0 | NA | NA | NA | NA | NA |
| AC093074.1 | 0 | NA | NA | NA | NA | NA |
| AC005307.1 | 2.20975875 | 0.464099273 | 1.23278576 | 0.37646385 | 0.70657208 | 0.83306384 |
| AC005381.1 | 2.17063661 | -0.696001386 | 1.32211401 | -0.5264307 | 0.598589 | 0.75677743 |
| AC005394.1 | 0 | NA | NA | NA | NA | NA |
| AC005616.1 | 9.79210687 | -0.633237074 | 1.25122319 | -0.5060944 | 0.61279037 | 0.76799695 |
| AC079466.2 | 0.46224828 | 2.133126024 | 2.56992267 | 0.8300351 | 0.40651894 | NA |
| AC079466.1 | 5.94047255 | -3.740863612 | 1.61225415 | -2.3202692 | 0.02032632 | 0.07577286 |
| AC007795.1 | 0 | NA | NA | NA | NA | NA |
| AC008991.1 | 0.583105 | 1.11533409 | 1.52072245 | 0.73342383 | 0.46329997 | NA |
| AC008991.2 | 1.68954937 | 0.454881138 | 1.16877382 | 0.38919518 | 0.69713177 | 0.82584747 |
| AC011524.1 | 0.08757577 | 0.547239394 | 3.29377202 | 0.16614368 | 0.86804388 | NA |
| AC011524.2 | 0 | NA | NA | NA | NA | NA |
| LINC00906 | 2.11884577 | -1.023931935 | 1.64013604 | -0.624297 | 0.53243258 | 0.70667642 |
| LINC01532 | 75.7463144 | 0.20228726 | 0.36461401 | 0.55479838 | 0.57903258 | 0.74245887 |
| UQCRFS1 | 7911.7911 | 0.022152877 | 0.19607087 | 0.11298403 | 0.91004321 | 0.95536604 |
| AC007786.1 | 12.7009317 | 0.793706352 | 0.64449168 | 1.23152304 | 0.21812731 | 0.40325414 |
| AC007786.2 | 2.35750359 | 1.014626003 | 0.87851531 | 1.15493263 | 0.24811801 | 0.43979781 |
| AC007786.3 | 0.40830231 | -1.802811494 | 3.29956038 | -0.5463793 | 0.58480524 | NA |
| AC011474.3 | 0 | NA | NA | NA | NA | NA |
| AC011474.2 | 0.09644756 | -0.642905668 | 3.30558948 | -0.1944905 | 0.84579185 | NA |
| VSTM2B | 1.51090804 | 0.129460704 | 1.21630525 | 0.10643768 | 0.9152351 | 0.95863503 |
| POP4 | 1471.23381 | -0.036087137 | 0.18189552 | -0.1983949 | 0.84273613 | 0.91639311 |
| PLEKHF1 | 454.273275 | -1.639849966 | 0.17304322 | -9.4765341 | 2.63E-21 | 5.78E-18 |
| C19orf12 | 2138.84716 | -0.104003794 | 0.23907548 | -0.4350249 | 0.66354433 | 0.80246109 |
| CCNE1 | 89.2363257 | -0.759853228 | 0.24100814 | -3.1528115 | 0.00161706 | 0.01145672 |
| URI1 | 2795.89351 | -0.094523361 | 0.13941992 | -0.677976 | 0.49778689 | 0.67953825 |
| AC008507.2 | 0.2444215 | 1.011631552 | 2.96273484 | 0.34145194 | 0.73276338 | NA |
| AC005597.1 | 0.03392908 | 0.380740593 | 3.29738709 | 0.11546736 | 0.90807469 | NA |
| ZNF536 | 12.9968694 | 0.403778022 | 0.47468418 | 0.85062455 | 0.39497795 | 0.59031787 |
| AC011478.1 | 0 | NA | NA | NA | NA | NA |
| AC011507.1 | 0.06785815 | 0.795868262 | 3.28898349 | 0.24198001 | 0.80879565 | NA |
| LINC01834 | 0 | NA | NA | NA | NA | NA |
| AC020897.1 | 0 | NA | NA | NA | NA | NA |
| AC020912.1 | 3.69808368 | -0.369227177 | 0.9738546 | -0.3791399 | 0.70458395 | 0.83125502 |
| LINC01791 | 0.17332742 | -1.003574173 | 3.30558948 | -0.3035992 | 0.7614333 | NA |
| TSHZ3 | 396.845414 | -0.421941057 | 0.3320399 | -1.2707541 | 0.20381615 | 0.38607951 |
| AC025809.1 | 1.27993721 | 1.936132638 | 1.19020649 | 1.62671995 | 0.10379659 | 0.24379764 |
| AC008992.1 | 1.44649839 | -0.328484239 | 1.6168406 | -0.2031643 | 0.83900664 | 0.91458001 |
| AC008794.2 | 0 | NA | NA | NA | NA | NA |
| AC008794.1 | 0 | NA | NA | NA | NA | NA |
| AC011525.1 | 0 | NA | NA | NA | NA | NA |
| AC011525.2 | 0 | NA | NA | NA | NA | NA |
| LINC01837 | 0.13390214 | 1.04981649 | 3.28477129 | 0.31960109 | 0.74927075 | NA |
| AC011456.1 | 0 | NA | NA | NA | NA | NA |
| LINC01533 | 0 | NA | NA | NA | NA | NA |
| LINC01782 | 0.09989236 | -0.642905668 | 3.30558948 | -0.1944905 | 0.84579185 | NA |
| AC011518.1 | 0 | NA | NA | NA | NA | NA |
| ZNF507 | 806.740333 | -0.010319017 | 0.14208126 | -0.0726276 | 0.94210248 | 0.97252858 |
| AC007773.1 | 25.487828 | 0.157032626 | 0.50193609 | 0.31285383 | 0.75439172 | 0.86329626 |
| DPY19L3 | 1409.51428 | -0.437927779 | 0.13778869 | -3.1782563 | 0.00148164 | 0.01070726 |
| PDCD5 | 2456.88441 | -0.830218964 | 0.12128335 | -6.8452838 | 7.63E-12 | 1.58E-09 |
| ANKRD27 | 1574.68785 | -0.276824175 | 0.14774974 | -1.8736018 | 0.06098534 | 0.16736033 |
| AC008474.1 | 0 | NA | NA | NA | NA | NA |
| RGS9BP | 19.5254352 | 1.603508867 | 0.58064275 | 2.76161009 | 0.00575171 | 0.02990097 |
| AC008736.1 | 43.0665214 | -0.145575014 | 0.27651423 | -0.5264648 | 0.59856529 | 0.75677743 |
| NUDT19 | 858.703418 | -0.055858729 | 0.13129553 | -0.4254427 | 0.67051397 | 0.80755667 |
| AC008736.2 | 4.76205092 | -0.977396768 | 0.71897451 | -1.3594317 | 0.17400982 | 0.34788772 |
| TDRD12 | 15.6811547 | 1.275496666 | 0.52379861 | 2.43508981 | 0.01488809 | 0.06052089 |
| SLC7A9 | 3126.5689 | 1.933234431 | 0.58703249 | 3.29323241 | 0.00099043 | 0.0077777 |
| AC008805.2 | 510.635518 | 1.653646939 | 0.52722548 | 3.13650798 | 0.00170973 | 0.01194989 |
| CEP89 | 1320.28768 | -0.250897101 | 0.12904824 | -1.9442118 | 0.05186992 | 0.14959794 |
| FAAP24 | 62.2893 | -0.072253404 | 0.38395792 | -0.1881805 | 0.85073514 | 0.92161226 |

| | | | | | | |
|---|---|---|---|---|---|---|
| RHPN2 | 3100.96121 | -0.485699792 | 0.21754885 | -2.232601 | 0.02557527 | 0.08977897 |
| GPATCH1 | 505.577437 | 0.191034319 | 0.18019075 | 1.06017831 | 0.28906349 | 0.4852447 |
| WDR88 | 52.3980166 | 0.926844508 | 0.39247625 | 2.36153019 | 0.01819969 | 0.06999064 |
| LRP3 | 1299.49614 | -0.149490965 | 0.3140475 | -0.4760139 | 0.6340645 | 0.78307797 |
| SLC7A10 | 50.6300577 | 0.59302102 | 0.65436996 | 0.90624732 | 0.36480496 | 0.56191044 |
| AC008738.4 | 0.43418788 | 1.269895288 | 2.13171719 | 0.59571471 | 0.55136583 | NA |
| AC008738.6 | 0.54620538 | -2.361017783 | 3.27766189 | -0.720336 | 0.47131816 | NA |
| CEBPA | 557.009641 | 0.051146741 | 0.48703095 | 0.10501743 | 0.91636197 | 0.95924994 |
| AC008738.3 | 318.890812 | -0.043095498 | 0.48879492 | -0.0881668 | 0.92974408 | 0.96597439 |
| AC008738.5 | 91.7169304 | 0.421603481 | 0.52019293 | 0.81047522 | 0.4176671 | 0.61042406 |
| CEBPA-DT | 57.7516328 | 0.3856093 | 0.45514522 | 0.84722256 | 0.39687108 | 0.59183908 |
| AC008738.2 | 4.03976473 | 0.693294863 | 0.61862809 | 1.12069735 | 0.26241671 | 0.45603853 |
| CEBPG | 1757.12642 | -0.531271741 | 0.30275204 | -1.7548081 | 0.07929213 | 0.20115808 |
| PEPD | 21032.0205 | 1.558534268 | 0.35259168 | 4.42022414 | 9.86E-06 | 0.00020093 |
| AC010485.1 | 0 | NA | NA | NA | NA | NA |
| CHST8 | 7.31807903 | -0.695832952 | 0.67234191 | -1.0349391 | 0.30069734 | 0.4974536 |
| KCTD15 | 1491.72211 | 0.305380439 | 0.30945718 | 0.98682615 | 0.32372787 | 0.5215653 |
| AC008556.1 | 111.609781 | 0.169098828 | 0.40693117 | 0.41554652 | 0.67774187 | 0.81269852 |
| LSM14A | 5739.95427 | 0.011099914 | 0.1541933 | 0.071987 | 0.94261225 | 0.97275716 |
| KIAA0355 | 1763.92524 | -0.091719214 | 0.15449139 | -0.593685 | 0.55272284 | 0.72263885 |
| AC010504.1 | 36.2200368 | 0.265683776 | 0.21714907 | 1.22350872 | 0.22113761 | 0.40727747 |
| GPI | 23327.5006 | -0.474607661 | 0.18554364 | -2.5579301 | 0.01052973 | 0.04680076 |
| AC092073.1 | 5059.83615 | -0.35216906 | 0.18896451 | -1.8636783 | 0.06236689 | 0.16986313 |
| PDCD2L | 208.78297 | -0.273845551 | 0.17157901 | -1.5960318 | 0.1104817 | 0.25460127 |
| AC008747.1 | 22.1802187 | -0.2369063 | 0.31119497 | -0.7612793 | 0.44649024 | 0.63687714 |
| UBA2 | 3388.376 | -0.287049172 | 0.13584592 | -2.1130496 | 0.03459652 | 0.11173426 |
| WTIP | 1132.65672 | 0.065132208 | 0.30413051 | 0.21415874 | 0.83042327 | 0.91019001 |
| SCGB1B2P | 64.1532323 | -0.090676128 | 0.30751609 | -0.2948663 | 0.76809603 | 0.87195714 |
| SCGB2B2 | 47.4209185 | 0.337645708 | 0.36496128 | 0.92515487 | 0.35488536 | 0.55237438 |
| ZNF302 | 1927.90137 | 0.003743493 | 0.1391286 | 0.02690671 | 0.97853414 | 0.99032643 |
| AC020910.4 | 74.9084634 | -0.316268306 | 0.25202565 | -1.2549052 | 0.20951317 | 0.39264177 |
| ZNF181 | 709.573454 | 0.425460778 | 0.1066146 | 3.99064254 | 6.59E-05 | 0.00091648 |
| ZNF599 | 362.254308 | 0.431941243 | 0.18593229 | 2.32311041 | 0.02017322 | 0.07537091 |
| LINC01801 | 4.70754184 | 0.258572402 | 0.69618574 | 0.37141295 | 0.71032998 | 0.83536599 |
| AC008555.1 | 24.7293058 | 1.086207555 | 0.27930255 | 3.88899978 | 0.00010066 | 0.00127246 |
| AC008555.2 | 23.0049981 | 0.352602558 | 0.36113639 | 0.97636949 | 0.32888139 | 0.5266656 |
| AC008555.5 | 27.8229555 | -0.209340281 | 0.49067284 | -0.4266392 | 0.66964212 | 0.80710223 |
| LINC00904 | 0.6478667 | -0.390430305 | 1.92273924 | -0.2030594 | 0.83908859 | NA |
| AC008555.3 | 0.6478667 | -0.390430305 | 1.92273924 | -0.2030594 | 0.83908859 | NA |
| AC008555.6 | 0.47645233 | -1.374229568 | 2.79589636 | -0.4915166 | 0.62306109 | NA |
| LINC01838 | 0.87125766 | -0.263450169 | 1.56155467 | -0.1687102 | 0.86602461 | NA |
| ZNF30-AS1 | 14.052113 | 1.186627251 | 0.47788016 | 2.48310634 | 0.01302422 | 0.05481147 |
| ZNF30 | 205.802828 | 0.737429023 | 0.16206051 | 4.55033129 | 5.36E-06 | 0.0001239 |
| ZNF792 | 224.04927 | -0.110370056 | 0.27282537 | -0.4045447 | 0.68581222 | 0.81835792 |
| GRAMD1A | 4368.77055 | -1.009370587 | 0.22650848 | -4.4562154 | 8.34E-06 | 0.00017683 |
| AC020907.3 | 2.16867786 | -1.210353658 | 0.96668762 | -1.2520629 | 0.21054696 | 0.3939632 |
| AC020907.4 | 125.234351 | -0.872186122 | 0.23038684 | -3.7857463 | 0.00015325 | 0.0017826 |
| SCN1B | 537.07052 | -0.355912489 | 0.19356342 | -1.8387383 | 0.06595368 | 0.17666047 |
| HPN | 21330.6344 | 0.429230387 | 0.32077715 | 1.33809528 | 0.18086538 | 0.35699252 |
| HPN-AS1 | 2954.66051 | 0.637451648 | 0.32953195 | 1.9344153 | 0.05306209 | 0.151861 |
| AC020907.1 | 0.33376757 | 1.448875046 | 2.83086872 | 0.51181287 | 0.60878198 | NA |
| FXYD3 | 234.950754 | -1.118187602 | 0.39790805 | -2.8101658 | 0.0049516 | 0.0266858 |
| LGI4 | 526.246298 | -0.751164204 | 0.72882909 | -1.0306452 | 0.30270723 | 0.49961756 |
| FXYD1 | 322.328755 | 0.440099338 | 0.50452125 | 0.87231081 | 0.38303884 | 0.57895142 |
| FXYD7 | 9.97123819 | -0.65616985 | 0.67929047 | -0.9659636 | 0.33406241 | 0.53218303 |
| FXYD5 | 1062.95366 | -0.801433108 | 0.39633366 | -2.0221172 | 0.04316425 | 0.13100136 |
| FAM187B | 1.85794977 | -0.039373178 | 0.85283146 | -0.0461676 | 0.96317667 | 0.9827801 |
| LSR | 4535.43944 | -1.212237407 | 0.22289295 | -5.438653 | 5.37E-08 | 2.81E-06 |
| AC002128.2 | 7.99822312 | -0.539157223 | 0.46509544 | -1.15924 | 0.24635838 | 0.43769955 |
| USF2 | 5125.17324 | -0.186093314 | 0.14904879 | -1.2485396 | 0.21183352 | 0.3954386 |
| HAMP | 25.5107021 | -1.505375803 | 1.02874417 | -1.4633141 | 0.14338146 | 0.30496235 |
| MAG | 19.6895874 | 1.115307629 | 0.93940904 | 1.18724388 | 0.23513144 | 0.42393108 |
| CD22 | 172.622837 | 0.719733748 | 0.77247989 | 0.93171843 | 0.35148206 | 0.54914481 |
| FFAR1 | 1.05663528 | 0.605588043 | 1.85375377 | 0.32668203 | 0.74390839 | 0.85633247 |
| FFAR3 | 3.65876337 | -2.744338235 | 1.50902607 | -1.8186155 | 0.06897011 | 0.18216467 |
| GPR42 | 0.8434561 | -0.962144366 | 1.57832522 | -0.6095983 | 0.54212794 | NA |
| LINC01531 | 1.78462128 | -1.562706913 | 1.11374351 | -1.403112 | 0.16058344 | 0.32892537 |
| AC002511.2 | 31.2628338 | -1.738052268 | 0.63053353 | -2.7564787 | 0.00584274 | 0.03020449 |
| AC002511.1 | 12.5069064 | -0.532295015 | 0.5658592 | -0.9406846 | 0.34686653 | 0.54448958 |
| FFAR2 | 47.947861 | -0.646203653 | 0.61027507 | -1.0588728 | 0.28965773 | 0.48588757 |
| KRTDAP | 11.4398439 | -1.439873145 | 0.53167402 | -2.7081879 | 0.00676517 | 0.03368387 |
| DMKN | 2780.11963 | -0.946796731 | 0.24629202 | -3.8442039 | 0.00012094 | 0.00147516 |
| SBSN | 1.49687582 | -1.883586584 | 1.5121619 | -1.2456249 | 0.21290212 | 0.39676223 |
| GAPDHS | 267.497271 | 0.900076243 | 0.31164416 | 2.88815375 | 0.0038751 | 0.02224347 |
| TMEM147-A! | 1819.11192 | 0.122305818 | 0.09408544 | 1.29994413 | 0.19362012 | 0.37322825 |
| TMEM147 | 3971.38116 | -0.276273299 | 0.16012935 | -1.7253133 | 0.08447103 | 0.21057866 |
| ATP4A | 0.96692489 | -0.698252042 | 1.32931797 | -0.5252709 | 0.59939489 | NA |
| AD000090.1 | 0.43519223 | 1.239711137 | 1.77459859 | 0.69858679 | 0.4848103 | NA |
| PMIS2 | 0 | NA | NA | NA | NA | NA |
| LINC01766 | 0.09989236 | -0.642905668 | 3.30558948 | -0.1944905 | 0.84579185 | NA |
| AC002115.1 | 0 | NA | NA | NA | NA | NA |
| HAUS5 | 877.867543 | 0.001819172 | 0.28466265 | 0.00639062 | 0.99490106 | 0.99770729 |
| RBM42 | 3328.65547 | -0.300997752 | 0.14847204 | -2.0273026 | 0.04263147 | 0.1297262 |
| ETV2 | 56.24503 | 0.325420282 | 0.25625426 | 1.26991169 | 0.20411609 | 0.38645471 |
| COX6B1 | 7412.61562 | -0.051851509 | 0.1569793 | -0.3303079 | 0.7411673 | 0.85463211 |
| UPK1A | 11.4814653 | -0.912432031 | 0.75036785 | -1.2159796 | 0.22399268 | 0.41004421 |
| UPK1A-AS1 | 5.12006786 | -0.412847666 | 0.8839366 | -0.4670557 | 0.64046 | 0.78721591 |
| ZBTB32 | 40.9811811 | -0.908190936 | 0.566821 | -1.6022535 | 0.10909956 | 0.25245309 |
| KMT2B | 5810.38764 | -0.240396842 | 0.19805574 | -1.2137837 | 0.22483032 | 0.41104619 |
| IGFLR1 | 1141.10614 | -0.964008461 | 0.29837961 | -3.2308121 | 0.00123439 | 0.00925535 |
| AD000671.2 | 2295.19176 | -0.491539795 | 0.24833423 | -1.9793478 | 0.04777687 | 0.14092572 |
| U2AF1L4 | 1626.58287 | -0.228649177 | 0.23367403 | -0.9784963 | 0.32782891 | 0.5255866 |
| PSENEN | 3131.14444 | -0.665652801 | 0.14585333 | -4.5638505 | 5.02E-06 | 0.00011751 |
| AD000671.1 | 2983.19201 | -0.669931414 | 0.14301593 | -4.6843131 | 2.81E-06 | 7.44E-05 |
| AD000671.3 | 609.39369 | -0.563301371 | 0.15135636 | -3.7216894 | 0.00019789 | 0.00216086 |
| LIN37 | 789.997422 | -0.545152058 | 0.16859181 | -3.2335619 | 0.00122257 | 0.00918494 |
| AC002398.2 | 1463.27918 | -1.160780794 | 0.45672795 | -2.5415147 | 0.01103733 | 0.0484878 |
| HSPB6 | 1748.56733 | -1.157681043 | 0.46523975 | -2.4883537 | 0.0128336 | 0.05419089 |
| PROSER3 | 1195.36214 | -0.33048207 | 0.1841783 | -1.7943594 | 0.07275582 | 0.18969885 |
| AC002398.1 | 266.27312 | -0.835291593 | 0.20178031 | -4.139609 | 3.48E-05 | 0.00055332 |
| ARHGAP33 | 1391.71989 | 0.359070354 | 0.41503088 | 0.86516539 | 0.38694802 | 0.5826093 |
| PRODH2 | 8920.53081 | 2.858059845 | 0.6129383 | 4.66288342 | 3.12E-06 | 8.06E-05 |
| NPHS1 | 2875.47146 | 1.044135605 | 0.87929713 | 1.18746618 | 0.23504379 | 0.42389952 |
| KIRREL2 | 84.3288033 | 0.077282817 | 0.5947048 | 0.12995156 | 0.89660475 | 0.94706458 |
| APLP1 | 232.454576 | 0.318915233 | 0.35860858 | 0.88931289 | 0.37383495 | 0.57001824 |
| NFKBID | 769.114489 | -0.304009223 | 0.3544253 | -0.8577526 | 0.39102909 | 0.58667712 |
| HCST | 284.897979 | -0.699750017 | 0.48132574 | -1.4537972 | 0.14600253 | 0.30864063 |
| TYROBP | 1024.81872 | -0.817943115 | 0.47089253 | -1.7370059 | 0.08238612 | 0.20671 |
| LRFN3 | 734.174947 | -0.336109298 | 0.21523997 | -1.5615562 | 0.11839258 | 0.26743395 |
| AF038458.3 | 1.28676285 | -0.039009644 | 1.05900511 | -0.0368361 | 0.97061567 | 0.98639399 |
| AF038458.2 | 4.21962089 | -0.851697523 | 0.81873188 | -1.0402643 | 0.29821714 | 0.49518204 |
| SDHAF1 | 709.919558 | -0.384313408 | 0.13034557 | -2.9484195 | 0.00319403 | 0.01927384 |
| SYNE4 | 881.834835 | -0.227380682 | 0.18120174 | -1.2548482 | 0.20953386 | 0.39265263 |

| | | | | | | |
|---|---|---|---|---|---|---|
| ALKBH6 | 852.689377 | -0.498845531 | 0.13435509 | -3.712889 | 0.00020491 | 0.00222522 |
| AC002116.2 | 170.217476 | 0.155079566 | 0.24365413 | 0.63647419 | 0.5244674 | 0.69951978 |
| CLIP3 | 965.005282 | -0.072402136 | 0.23878762 | -0.3032072 | 0.76173193 | 0.86791967 |
| THAP8 | 388.309756 | 0.28240739 | 0.2207714 | 1.27918468 | 0.20083203 | 0.38243361 |
| WDR62 | 138.476122 | -0.687068471 | 0.39657764 | -1.7324942 | 0.08318561 | 0.20814161 |
| OVOL3 | 12.7970212 | -0.25652748 | 0.41599894 | -0.6166542 | 0.53746286 | 0.71113301 |
| POLR2I | 1285.22268 | -0.279023801 | 0.11673894 | -2.3901519 | 0.0168414 | 0.06625509 |
| TBCB | 3986.01442 | -0.370861282 | 0.13574229 | -2.7320982 | 0.00629324 | 0.03192011 |
| CAPNS1 | 20574.6411 | -0.382190413 | 0.14066086 | -2.7171057 | 0.00658556 | 0.03299661 |
| AD001527.1 | 12.7209985 | -0.42534574 | 0.3914584 | -1.0865669 | 0.27722826 | 0.47266066 |
| COX7A1 | 907.000537 | 0.346870033 | 0.3709521 | 0.9350804 | 0.34974687 | 0.54738114 |
| ZNF565 | 181.556933 | -0.088649213 | 0.1881901 | -0.471062 | 0.63759643 | 0.78556361 |
| ZNF146 | 3068.03147 | -0.124879289 | 0.16103254 | -0.775491 | 0.43804957 | 0.63000386 |
| AC012617.1 | 161.296124 | 0.278881618 | 0.27050007 | 1.03098538 | 0.30254767 | 0.49944799 |
| LINC00665 | 1433.12143 | 0.235026719 | 0.17649364 | 1.33164413 | 0.18297716 | 0.35970684 |
| ZFP14 | 644.415671 | 0.446670142 | 0.19178443 | 2.32902188 | 0.01985791 | 0.07454135 |
| AC092296.1 | 26.6142563 | 2.92856064 | 0.64771283 | 4.52138744 | 6.14E-06 | 0.00013929 |
| AC092296.3 | 9.26436018 | 3.128680169 | 0.63514824 | 4.92590542 | 8.40E-07 | 2.80E-05 |
| ZFP82 | 242.805808 | -0.085361694 | 0.20597567 | -0.4144261 | 0.67856207 | 0.81342312 |
| ZNF566 | 471.235718 | 0.160593267 | 0.16754541 | 0.95850591 | 0.33780771 | 0.53581891 |
| AC092295.2 | 150.00197 | 0.557533729 | 0.22698676 | 2.45623894 | 0.01403998 | 0.05801225 |
| ZNF260 | 822.974558 | 0.097117388 | 0.10714264 | 0.90643076 | 0.3647079 | 0.56182657 |
| AC092295.1 | 3.88004717 | 1.452126598 | 0.70219642 | 2.06797778 | 0.03864211 | 0.12074802 |
| ZNF529 | 961.997139 | -0.032154583 | 0.17000446 | -0.1891396 | 0.84998337 | 0.92117629 |
| ZNF529-AS1 | 200.02129 | -0.337700422 | 0.20896229 | -1.6160831 | 0.10607636 | 0.24734579 |
| ZNF382 | 277.163049 | 0.012386415 | 0.19880906 | 0.06230307 | 0.95032149 | 0.9761964 |
| ZNF461 | 304.010899 | 0.006236093 | 0.15301898 | 0.04075373 | 0.96749223 | 0.98467589 |
| AC074138.1 | 11.1299552 | 0.53885574 | 0.53273353 | 1.01149207 | 0.31178097 | 0.50882933 |
| LINC01534 | 52.3398599 | 0.167581756 | 0.25111997 | 0.66733743 | 0.50455662 | 0.68396433 |
| ZNF567 | 331.513541 | -0.142189447 | 0.19260315 | -0.7382509 | 0.460362 | 0.64890395 |
| ZNF850 | 86.937814 | -0.125860264 | 0.22197807 | -0.5669941 | 0.57071817 | 0.73576038 |
| AC020928.2 | 23.9456965 | 0.332799933 | 0.31754394 | 1.04804372 | 0.29461847 | 0.49140348 |
| AC020928.3 | 16.7927061 | 0.227833415 | 0.36729424 | 0.62030217 | 0.53505887 | 0.70879884 |
| AC020928.1 | 204.136676 | 0.162676442 | 0.14833327 | 1.09669556 | 0.2727745 | 0.46762862 |
| ZNF790 | 360.305125 | 0.571907318 | 0.14378339 | 3.97756166 | 6.96E-05 | 0.0009518 |
| ZNF345 | 332.075621 | 0.473189491 | 0.20236844 | 2.33825733 | 0.0193739 | 0.07321436 |
| ZNF829 | 138.223022 | 0.037464528 | 0.2696422 | 0.13894164 | 0.88949627 | 0.94296756 |
| ZNF568 | 392.518535 | -0.00351707 | 0.1797238 | -0.0195693 | 0.98438695 | 0.99326005 |
| ZNF420 | 339.458138 | 0.059822539 | 0.16983894 | 0.352231 | 0.72466503 | 0.84389181 |
| AC010632.1 | 14.2747643 | 0.785704795 | 0.39559028 | 1.98615797 | 0.04701579 | 0.13927278 |
| AC012309.1 | 253.401857 | -0.495017945 | 0.15012412 | -3.2973911 | 0.00097588 | 0.00769783 |
| ZNF585A | 567.004975 | -0.076234082 | 0.08427357 | -0.9046025 | 0.36567598 | 0.5626605 |
| AC012309.2 | 48.5990312 | 0.223520619 | 0.24605127 | 0.90843107 | 0.36365052 | 0.56082017 |
| ZNF585B | 1039.96372 | -0.00271956 | 0.14297392 | -0.0190214 | 0.98482406 | 0.99347171 |
| ZNF383 | 598.622694 | 0.086185801 | 0.12286624 | 0.7014604 | 0.48301574 | 0.6675992 |
| LINC01535 | 71.2778351 | 0.652627388 | 0.43635451 | 1.49563572 | 0.13474861 | 0.29218265 |
| AC016590.1 | 125.769979 | -0.211103959 | 0.37858341 | -0.5576155 | 0.577107 | 0.74101922 |
| HKR1 | 3224.254 | 0.104331546 | 0.098497 | 1.0592358 | 0.28949241 | 0.48571068 |
| AC016590.3 | 54.9280055 | 0.50586401 | 0.35497485 | 1.42506998 | 0.15413697 | 0.31999622 |
| ZNF527 | 302.470763 | 0.273842592 | 0.11766741 | 2.3272594 | 0.01995146 | 0.07479042 |
| ZNF569 | 286.65956 | 0.517711399 | 0.17654764 | 2.93241761 | 0.00336334 | 0.02000255 |
| ZNF570 | 339.37858 | 0.037695902 | 0.14632272 | 0.25762165 | 0.79669892 | 0.88968199 |
| ZNF793-AS1 | 137.06236 | 0.21410969 | 0.17048248 | 1.25590438 | 0.20915064 | 0.39235447 |
| AC022148.2 | 3.23495975 | -0.633428233 | 0.71239212 | -0.8891567 | 0.37391886 | 0.57005495 |
| ZNF793 | 500.496095 | 0.546783518 | 0.18310765 | 2.98613142 | 0.00282531 | 0.01751786 |
| AC022148.1 | 18.7229229 | 0.721464908 | 0.52595717 | 1.37171799 | 0.17015125 | 0.34211343 |
| ZNF571-AS1 | 89.9223108 | 0.727735573 | 0.30013083 | 2.42472781 | 0.01531987 | 0.0617714 |
| ZNF540 | 462.276396 | 0.424784881 | 0.30765782 | 1.38070564 | 0.16736949 | 0.33859098 |
| ZNF571 | 313.372579 | 0.103005049 | 0.13299077 | 0.77452779 | 0.43861874 | 0.63025154 |
| ZFP30 | 459.011447 | 0.15151034 | 0.1272025 | 1.19109562 | 0.23361605 | 0.42206462 |
| ZNF781 | 115.162859 | 0.409968364 | 0.35819131 | 1.14455141 | 0.25239503 | 0.44468878 |
| AC093227.2 | 33.6210914 | 0.325360727 | 0.31982633 | 1.01730437 | 0.30900866 | 0.50605877 |
| ZNF607 | 475.600439 | -0.180284233 | 0.17415929 | -1.0351686 | 0.30059017 | 0.49739112 |
| AC093227.1 | 194.783976 | 0.308937503 | 0.22422794 | 1.37778326 | 0.16827022 | 0.33960551 |
| ZNF573 | 60.3027076 | 0.149734579 | 0.19926769 | 0.75142426 | 0.45239737 | 0.64210768 |
| WDR87 | 1.10608456 | -0.491219677 | 1.54193291 | -0.318574 | 0.7500496 | 0.86045777 |
| SIPA1L3 | 2538.87898 | 0.302982162 | 0.15258236 | 1.98569586 | 0.04706711 | 0.13931525 |
| AC011465.1 | 1.50827859 | -0.568955441 | 1.06164271 | -0.5359199 | 0.59201392 | 0.75152395 |
| DPF1 | 17.1488003 | -1.365818603 | 0.57488396 | -2.3758161 | 0.01751019 | 0.06803353 |
| SPINT2 | 16446.7942 | -0.699964465 | 0.13924339 | -5.0269136 | 4.98E-07 | 1.80E-05 |
| PPP1R14A | 717.233638 | -0.336146403 | 0.55559669 | -0.6050187 | 0.54516655 | 0.71686656 |
| AC011479.1 | 6943.86657 | -0.642512642 | 0.14588905 | -4.4041185 | 1.06E-05 | 0.00021448 |
| C19orf33 | 497.711782 | -0.797042687 | 0.45178513 | -1.7642074 | 0.07769706 | 0.19839949 |
| YIF1B | 1238.27917 | -0.172697015 | 0.19342252 | -0.8928486 | 0.37193828 | 0.56855459 |
| KCNK6 | 611.227613 | -0.818559509 | 0.27586936 | -2.9671998 | 0.00300526 | 0.01838016 |
| CATSPERG | 206.934391 | 0.160970904 | 0.29801161 | 0.54014977 | 0.58909375 | 0.74948842 |
| AC005625.1 | 19.7840577 | 0.265203698 | 0.39344001 | 0.67406388 | 0.5002707 | 0.68137691 |
| PSMD8 | 6989.1963 | -0.298681902 | 0.15913816 | -1.8768716 | 0.06053569 | 0.16652683 |
| GGN | 79.1988651 | -1.976056656 | 0.30042046 | -6.5776367 | 4.78E-11 | 7.83E-09 |
| AC005789.1 | 23.9078289 | -2.362572186 | 0.39782004 | -5.9387963 | 2.87E-09 | 2.40E-07 |
| SPRED3 | 393.27382 | -3.186231444 | 0.42086343 | -7.5707016 | 3.71E-14 | 1.55E-11 |
| FAM98C | 1008.19867 | -0.008255039 | 0.18884674 | -0.0437129 | 0.96513326 | 0.98359669 |
| RASGRP4 | 150.668065 | -0.131136571 | 0.31464398 | -0.4167776 | 0.67684107 | 0.81209758 |
| RYR1 | 175.853825 | 0.800902923 | 0.45394041 | 1.76433492 | 0.07767561 | 0.19836392 |
| AC067969.2 | 1.50573163 | 2.171065866 | 1.1838867 | 1.83384599 | 0.06667686 | 0.17796243 |
| AC067969.1 | 0.40355352 | 1.17450744 | 2.71653232 | 0.43235541 | 0.66548312 | NA |
| MAP4K1 | 289.15956 | -0.008494327 | 0.532356 | -0.0159561 | 0.98726941 | 0.99420661 |
| AC008649.1 | 94.5617644 | -0.097725101 | 0.56655903 | -0.1724888 | 0.86305325 | 0.9286976 |
| EIF3K | 8668.99706 | 0.167835725 | 0.15020799 | 1.11735548 | 0.26384236 | 0.45761781 |
| ACTN4 | 34913.7548 | 0.287294226 | 0.1741075 | 1.65009676 | 0.09892315 | 0.23581661 |
| AC008649.2 | 0.35467906 | 0.670828026 | 2.2054785 | 0.30416439 | 0.76100265 | NA |
| CAPN12 | 5553.80833 | -0.028565507 | 0.26727622 | -0.1068764 | 0.91488708 | 0.95850505 |
| AC022144.1 | 259.231327 | 1.179952788 | 0.3515883 | 3.35606385 | 0.0007906 | 0.00650693 |
| LGALS7 | 2.12697884 | 1.152613456 | 1.25138881 | 0.92106741 | 0.35701523 | 0.55454389 |
| LGALS7B | 5.3118132 | -0.37579583 | 0.70655013 | -0.5318743 | 0.59481308 | 0.75384347 |
| LGALS4 | 203.352086 | 0.725394251 | 0.64621028 | 1.12253591 | 0.26163465 | 0.45509932 |
| ECH1 | 12983.0961 | 0.507726972 | 0.27762185 | 1.82884371 | 0.06742302 | 0.17928256 |
| AC008982.1 | 11423.7561 | 0.247967563 | 0.19421998 | 1.27673559 | 0.20169561 | 0.3835795 |
| AC104534.1 | 17.3383252 | 1.064130846 | 0.46888914 | 2.26947214 | 0.02323963 | 0.08377188 |
| HNRNPL | 10943.3866 | -0.09927157 | 0.13960157 | -0.7111064 | 0.4770183 | 0.66212483 |
| RINL | 500.367405 | -0.180653757 | 0.23332727 | -0.7742505 | 0.43878267 | 0.63037483 |
| AC011455.1 | 21.6835638 | -0.101823568 | 0.45042764 | -0.2260598 | 0.82115494 | 0.90408983 |
| SIRT2 | 3812.40775 | 0.209509138 | 0.16392884 | 1.27804928 | 0.20123205 | 0.38300175 |
| NFKBIB | 2136.75565 | -0.54502118 | 0.2390694 | -2.2797613 | 0.02262185 | 0.08213384 |
| CCER2 | 41.2324382 | 1.084247276 | 0.28205744 | 3.84406551 | 0.00012101 | 0.00147531 |
| SARS2 | 1580.02337 | 0.290083878 | 0.15962004 | 1.81733998 | 0.06916507 | 0.18255259 |
| AC011455.2 | 2821.38172 | 0.328074232 | 0.17109064 | 1.91754633 | 0.05516856 | 0.15605013 |
| MRPS12 | 1830.95999 | -0.115581298 | 0.13803717 | -0.8373201 | 0.40241266 | 0.59743396 |
| ERVK9-11 | 377.995471 | 1.032963401 | 0.40803119 | 2.53157948 | 0.01135501 | 0.04957203 |
| FBXO17 | 4338.35051 | 0.114432109 | 0.29979124 | 0.38170598 | 0.70267947 | 0.82987743 |
| FBXO27 | 416.863142 | 0.076575911 | 0.23820309 | 0.3214732 | 0.74785182 | 0.85899697 |
| AC010605.1 | 72.5311379 | 0.344382453 | 0.30965261 | 1.11215743 | 0.26607047 | 0.46020452 |
| ACP7 | 5.53446249 | -1.413057699 | 1.38673342 | -1.0189829 | 0.30821107 | 0.50512901 |

| | | | | | | |
|---|---|---|---|---|---|---|
| PAK4 | 4029.12325 | 0.113348296 | 0.1710654 | 0.6626021 | 0.50758543 | 0.68651729 |
| AC011443.1 | 0.43640398 | 1.429601034 | 2.73025564 | 0.52361435 | 0.6005468 | NA |
| NCCRP1 | 77.0642742 | 0.182736474 | 0.50026628 | 0.36527841 | 0.71490359 | 0.83794149 |
| SYCN | 0 | NA | NA | NA | NA | NA |
| IFNL3 | 0.40065839 | -1.928496579 | 2.79860735 | -0.6890915 | 0.49076568 | NA |
| IFNL2 | 0.30636111 | -1.6406616 | 3.30406282 | -0.4965588 | 0.61950019 | NA |
| IFNL1 | 0.2926836 | 2.132206094 | 3.2337338 | 0.65936352 | 0.50966236 | NA |
| LRFN1 | 65.6263204 | -0.143728287 | 0.38735966 | -0.3710461 | 0.7106032 | 0.83553781 |
| GMFG | 672.85846 | -0.583190609 | 0.27342478 | -2.1329106 | 0.03293206 | 0.10782449 |
| AC011445.2 | 15.5781946 | -0.094053467 | 0.38691646 | -0.2430847 | 0.80793979 | 0.89620864 |
| SAMD4B | 5506.03495 | -0.305502512 | 0.13876432 | -2.2015926 | 0.0276941 | 0.09532611 |
| PAF1 | 3480.42876 | -0.237232852 | 0.15432714 | -1.5372076 | 0.12424247 | 0.27615256 |
| MED29 | 3541.44953 | 0.379248664 | 0.16047466 | 2.36329318 | 0.01811333 | 0.06976028 |
| ZFP36 | 19388.8795 | -0.519972139 | 0.47695795 | -1.0901844 | 0.27563191 | 0.47093517 |
| PLEKHG2 | 2048.64755 | -0.372094184 | 0.34800615 | -1.0692173 | 0.28497177 | 0.48109992 |
| RPS16 | 21244.8544 | -0.544611767 | 0.21625742 | -2.5183495 | 0.01179063 | 0.05092021 |
| SUPT5H | 8662.62607 | -0.127636106 | 0.14520983 | -0.8789771 | 0.37941372 | 0.57543796 |
| TIMM50 | 2549.47513 | -0.073668447 | 0.1467272 | -0.5020776 | 0.6156129 | 0.77018414 |
| DLL3 | 2.04909944 | -0.923169587 | 1.11245736 | -0.8298472 | 0.4066252 | 0.60140134 |
| SELENOV | 2.64730995 | -0.783872328 | 1.38043116 | -0.567846 | 0.57013953 | 0.73533109 |
| EID2B | 176.55738 | -0.047746719 | 0.15073995 | -0.3167489 | 0.75143412 | 0.86135902 |
| AC011500.3 | 57.4914493 | 0.268087699 | 0.19260342 | 1.39191558 | 0.16394796 | 0.33341303 |
| EID2 | 726.318002 | -0.279155699 | 0.17925077 | -1.5573472 | 0.11938807 | 0.26889847 |
| LGALS13 | 0 | NA | NA | NA | NA | NA |
| LGALS16 | 0 | NA | NA | NA | NA | NA |
| AC005515.2 | 1.06066293 | 1.196133808 | 1.56602582 | 0.7638021 | 0.44498519 | 0.6355554 |
| LGALS14 | 0.07659028 | -0.642905668 | 3.30558948 | -0.1944905 | 0.84579185 | NA |
| CLC | 13.9548603 | -1.738371632 | 0.86686064 | -2.0053646 | 0.04492408 | 0.13468032 |
| LEUTX | 0 | NA | NA | NA | NA | NA |
| AC005393.1 | 37.7951357 | -0.492532819 | 0.33092504 | -1.4883516 | 0.13665818 | 0.29498449 |
| DYRK1B | 1667.17643 | -0.090831211 | 0.17982977 | -0.5050955 | 0.61349174 | 0.76843981 |
| FBL | 3187.56731 | -0.556318071 | 0.17140678 | -3.2456014 | 0.00117203 | 0.00889392 |
| FCGBP | 2342.44395 | -0.05726759 | 0.61418993 | -0.0932408 | 0.92571222 | 0.9640363 |
| PSMC4 | 5080.53489 | -0.213523234 | 0.20494037 | -1.0418798 | 0.2974674 | 0.49434178 |
| ZNF546 | 415.988032 | 0.446606283 | 0.22744344 | 1.96359273 | 0.04957734 | 0.14481109 |
| ZNF780B | 987.728162 | 0.356082697 | 0.2882386 | 1.23537476 | 0.21669106 | 0.40152788 |
| ZNF780A | 629.447088 | 0.186543699 | 0.15965969 | 1.16838322 | 0.24265222 | 0.43307879 |
| AC005614.1 | 39.1563331 | -0.47543063 | 0.32539486 | -1.4610883 | 0.1439912 | 0.30574507 |
| MAP3K10 | 1677.9272 | -0.585561954 | 0.18372707 | -3.1871295 | 0.00143692 | 0.01046154 |
| TTC9B | 31.7021048 | -0.980961733 | 0.60467354 | -1.6222998 | 0.10473918 | 0.24522527 |
| CNTD2 | 143.85496 | 0.152679336 | 0.57462459 | 0.26570275 | 0.79046814 | 0.88620494 |
| AKT2 | 10974.2394 | -0.100394775 | 0.11269528 | -0.8908517 | 0.37300876 | 0.56921038 |
| C19orf47 | 590.100553 | -0.415999593 | 0.22662004 | -1.8356699 | 0.06640649 | 0.17742654 |
| PLD3 | 9592.22353 | -0.125409715 | 0.12412348 | -1.0103625 | 0.31232163 | 0.50918254 |
| HIPK4 | 12.4249441 | -0.181907472 | 0.52743221 | -0.3448926 | 0.73017511 | 0.84751607 |
| PRX | 891.901377 | 0.720964529 | 0.37134206 | 1.94151057 | 0.05219638 | 0.15017389 |
| SERTAD1 | 1389.81058 | -1.080130228 | 0.3196788 | -3.3787984 | 0.00072803 | 0.00610157 |
| AC010271.2 | 4.11203144 | 1.439862571 | 0.68681319 | 2.0964399 | 0.03604319 | 0.11495632 |
| SERTAD3 | 1489.07892 | -0.66631912 | 0.12412195 | -5.3682616 | 7.95E-08 | 3.93E-06 |
| AC010271.1 | 64.1365467 | -0.118361613 | 0.20810582 | -0.5687569 | 0.56952116 | 0.73496491 |
| BLVRB | 4314.96421 | -0.301084087 | 0.26029474 | -1.1567045 | 0.24739313 | 0.43894019 |
| SPTBN4 | 204.776896 | 1.103619156 | 0.43681967 | 2.52648688 | 0.01152097 | 0.050092 |
| SHKBP1 | 3177.76562 | -0.37861385 | 0.13470935 | -2.8105982 | 0.00494495 | 0.02666633 |
| LTBP4 | 6890.39306 | 0.931223915 | 0.42237431 | 2.20473614 | 0.02747262 | 0.09473158 |
| NUMBL | 1189.5274 | -0.745352484 | 0.32404016 | -2.3001856 | 0.02143771 | 0.07891016 |
| COQ8B | 2461.87311 | -0.318494343 | 0.15466971 | -2.0591901 | 0.03947603 | 0.12262363 |
| ITPKC | 2810.2791 | -1.375309002 | 0.22175449 | -6.2019444 | 5.58E-10 | 6.21E-08 |
| C19orf54 | 1473.44473 | 0.085337841 | 0.12071198 | 0.70695418 | 0.47959495 | 0.66422893 |
| SNRPA | 2229.31264 | -0.63201287 | 0.11831457 | -5.3418007 | 9.20E-08 | 4.47E-06 |
| MIA-RAB4B | 1746.28239 | -0.692149836 | 0.19385895 | -3.5703785 | 0.00035647 | 0.00348314 |
| MIA | 41.9486633 | -1.298474592 | 0.48351888 | -2.6854682 | 0.00724283 | 0.03552703 |
| RAB4B | 1882.83952 | -0.631861056 | 0.19690371 | -3.208985 | 0.00133204 | 0.0098628 |
| RAB4B-EGLN | 5923.17276 | -0.575956039 | 0.17198099 | -3.3489518 | 0.00081118 | 0.00665069 |
| AC008537.4 | 808.550262 | -0.80097531 | 0.19984773 | -4.0079279 | 6.13E-05 | 0.00086516 |
| EGLN2 | 5136.59141 | -0.504832684 | 0.22011324 | -2.2935135 | 0.02181846 | 0.08003935 |
| AC008537.1 | 515.122239 | -0.417224392 | 0.24869385 | -1.6776627 | 0.09341297 | 0.22657284 |
| AC008537.2 | 22.462615 | -0.057803545 | 0.333939 | -0.1730961 | 0.86257587 | 0.9284133 |
| CYP2A6 | 2.55834329 | -1.104235262 | 1.05182751 | -1.0498254 | 0.29379838 | 0.49041437 |
| CYP2A7 | 0.85664727 | 0.987189458 | 1.45401276 | 0.6789414 | 0.49717499 | NA |
| CYP2B6 | 842.689166 | 2.314191931 | 0.73810829 | 3.13530138 | 0.00171678 | 0.01199242 |
| CYP2A13 | 0 | NA | NA | NA | NA | NA |
| CYP2F1 | 0.36177836 | 0.005377682 | 3.28420126 | 0.00163744 | 0.99869351 | NA |
| CYP2S1 | 138.731468 | -0.42580366 | 0.82011462 | -0.5192002 | 0.60362115 | 0.7608379 |
| AXL | 1674.05056 | -0.579507873 | 0.33573478 | -1.7260883 | 0.08433153 | 0.21039001 |
| AC011510.1 | 40.5523963 | 0.553763352 | 0.4209764 | 1.31542611 | 0.18836674 | 0.36647712 |
| HNRNPUL1 | 8660.40402 | 0.005423623 | 0.16409681 | 0.03305136 | 0.97363363 | 0.98779582 |
| TGFB1 | 3983.08705 | -0.43522677 | 0.19533163 | -2.2281428 | 0.02587099 | 0.09061205 |
| CCDC97 | 1705.12113 | -0.186501267 | 0.19416463 | -0.9605316 | 0.33678774 | 0.53481661 |
| AC011462.1 | 3972.73468 | 0.39801183 | 0.2318977 | 1.71632505 | 0.08610257 | 0.21343816 |
| TMEM91 | 923.454426 | -0.564118906 | 0.25466541 | -2.2151376 | 0.02675062 | 0.09282984 |
| B9D2 | 430.922455 | -0.41491601 | 0.23640318 | -1.7551202 | 0.07923875 | 0.20110931 |
| EXOSC5 | 584.948264 | -0.239327155 | 0.1918206 | -1.2476614 | 0.21215508 | 0.3958767 |
| BCKDHA | 3673.52576 | 0.465568673 | 0.23792394 | 1.95679623 | 0.05037143 | 0.14640768 |
| AC011462.2 | 115.617439 | -0.958966096 | 0.37322889 | -2.569378 | 0.01018813 | 0.04569063 |
| B3GNT8 | 276.6797 | -0.780886763 | 0.35781435 | -2.1823797 | 0.02908152 | 0.09873565 |
| DMAC2 | 2976.96871 | -0.034302796 | 0.09291166 | -0.369198 | 0.71198017 | 0.83626196 |
| ERICH4 | 245.680389 | 2.302356045 | 0.96605957 | 2.3832444 | 0.0171608 | 0.06706138 |
| PCAT19 | 547.607162 | -1.039757312 | 0.31874821 | -3.2620021 | 0.00110628 | 0.00847304 |
| AC243960.3 | 8.1127629 | -0.426330044 | 0.51997691 | -0.8199019 | 0.41227205 | 0.60616749 |
| LINC01480 | 67.2071589 | -0.926245385 | 0.48653725 | -1.9037502 | 0.05694273 | 0.15948612 |
| AC243960.1 | 49.2207155 | 0.146543865 | 0.55171787 | 0.26561377 | 0.79053668 | 0.88621571 |
| CEACAM21 | 56.0214937 | -0.03148378 | 0.51799701 | -0.0607798 | 0.95153454 | 0.97675915 |
| CEACAM4 | 36.801848 | -1.167370706 | 0.56882539 | -2.0522479 | 0.04014558 | 0.12412646 |
| CEACAM7 | 10.7999247 | -1.087211693 | 0.97276886 | -1.1176465 | 0.26371801 | 0.45750525 |
| CEACAM5 | 1.00595665 | -3.219099046 | 2.16801341 | -1.4848151 | 0.13759281 | NA |
| AC243967.1 | 0.76692141 | -2.121491724 | 1.95057117 | -1.0876259 | 0.27676029 | NA |
| CEACAM6 | 10.5637322 | -2.826677747 | 1.22214882 | -2.3128752 | 0.0207295 | 0.0769261 |
| AC243967.2 | 0.24327904 | -1.364242679 | 3.30558948 | -0.4127078 | 0.67982068 | NA |
| CEACAM3 | 21.8740084 | -0.755015246 | 0.56837374 | -1.3283781 | 0.18405322 | 0.36107052 |
| AC243967.3 | 0 | NA | NA | NA | NA | NA |
| LYPD4 | 0 | NA | NA | NA | NA | NA |
| DMRTC2 | 0.03392908 | 0.380740593 | 3.29738709 | 0.11546736 | 0.90807469 | NA |
| RPS19 | 19374.8078 | -1.033605625 | 0.13991918 | -7.3871616 | 1.50E-13 | 5.49E-11 |
| CD79A | 165.066516 | 0.205407351 | 0.87469212 | 0.23483389 | 0.81433766 | 0.89985267 |
| ARHGEF1 | 8182.90123 | -0.387795571 | 0.20385446 | -1.9023159 | 0.05712987 | 0.1598424 |
| AC010616.1 | 18.3817164 | -0.938682682 | 0.57448289 | -1.6339611 | 0.10226702 | 0.24117165 |
| RABAC1 | 4037.7311 | -0.564956443 | 0.18629963 | -3.0325152 | 0.00242525 | 0.01562517 |
| AC010616.2 | 3681.29336 | -0.520871077 | 0.17987881 | -2.8956778 | 0.00378341 | 0.02190245 |
| ATP1A3 | 16.5575274 | 0.291544346 | 0.55814017 | 0.52234969 | 0.60142688 | 0.75890644 |
| GRIK5 | 1138.60115 | -0.331468785 | 0.34011576 | -0.9745764 | 0.32977043 | 0.52764069 |
| ZNF574 | 871.34634 | -0.048080986 | 0.20666647 | -0.2326502 | 0.81603308 | 0.90097104 |
| POU2F2 | 205.9262 | 0.045011781 | 0.43500867 | 0.1034733 | 0.91758734 | 0.96005997 |
| AC010247.2 | 7.03342352 | 0.248648816 | 0.66125331 | 0.37602657 | 0.70689714 | 0.83325953 |

| | | | | | | |
|---|---|---|---|---|---|---|
| DEDD2 | 1980.90928 | -0.425117291 | 0.2208871 | -1.9245908 | 0.05428057 | 0.15425001 |
| ZNF526 | 487.010518 | 0.003397114 | 0.18139221 | 0.01872801 | 0.98505809 | 0.99347171 |
| GSK3A | 4217.15963 | -0.117051846 | 0.16123981 | -0.7259488 | 0.46787016 | 0.65469547 |
| AC006486.1 | 1509.47604 | 0.079105507 | 0.18982815 | 0.41672168 | 0.676882 | 0.81209758 |
| ERF | 4690.14147 | 0.007968098 | 0.18918839 | 0.04211727 | 0.96640521 | 0.98410057 |
| CIC | 8326.18895 | -0.458191936 | 0.17903477 | -2.5592344 | 0.0104903 | 0.04668062 |
| PAFAH1B3 | 736.848226 | -0.355513146 | 0.22104476 | -1.608331 | 0.1077627 | 0.25019391 |
| PRR19 | 319.853297 | -0.427590199 | 0.23378535 | -1.8289863 | 0.06740166 | 0.17927504 |
| TMEM145 | 13.5514308 | 0.073031346 | 0.5295559 | 0.13791055 | 0.89031111 | 0.94316 |
| MEGF8 | 4335.41014 | 0.560877667 | 0.14530628 | 3.85996845 | 0.0001134 | 0.00139738 |
| CNFN | 26.3244261 | -0.577214454 | 0.48228132 | -1.1968418 | 0.23136821 | 0.41926744 |
| LIPE-AS1 | 760.055998 | -0.098475089 | 0.21065419 | -0.4674727 | 0.6401617 | 0.78714964 |
| LIPE | 488.910194 | -0.076457799 | 0.36178295 | -0.2113361 | 0.83262501 | 0.9111315 |
| CXCL17 | 1.20033411 | -0.644050226 | 2.35033759 | -0.2740246 | 0.78406575 | 0.88203634 |
| CEACAM1 | 3749.19932 | -0.025798249 | 0.28957892 | -0.0890888 | 0.92901131 | 0.96558201 |
| CEACAM8 | 14.3280542 | -3.729084428 | 1.17672688 | -3.1690314 | 0.00152948 | 0.01099013 |
| PSG3 | 0.91727073 | -1.505673425 | 1.75117377 | -0.8598081 | 0.38989482 | NA |
| PSG8 | 0.90498791 | -3.044527586 | 2.25746983 | -1.348646 | 0.17745071 | NA |
| PSG8-AS1 | 0.41332618 | -1.067756906 | 2.26711902 | -0.4709752 | 0.63765842 | NA |
| PSG1 | 0.55086675 | -0.131025866 | 1.71311303 | -0.0764841 | 0.93903398 | NA |
| PSG6 | 0 | NA | NA | NA | NA | NA |
| PSG7 | 0.10721287 | -0.642905668 | 3.30558948 | -0.1944905 | 0.84579185 | NA |
| AC004784.1 | 9.8604228 | -2.406247856 | 1.19053401 | -2.02115 | 0.04326424 | 0.13115046 |
| PSG11 | 0.61999641 | -2.505340608 | 3.2727771 | -0.7655091 | 0.44396846 | NA |
| PSG2 | 0.10721287 | -0.642905668 | 3.30558948 | -0.1944905 | 0.84579185 | NA |
| AC005392.1 | 0.21442573 | -1.003574173 | 3.30558948 | -0.3035992 | 0.7614333 | NA |
| PSG5 | 53.5316528 | -2.458829252 | 1.12341038 | -2.1887187 | 0.02861729 | 0.09757414 |
| PSG4 | 81.3531995 | -4.189641335 | 1.45875968 | -2.8720573 | 0.00407809 | 0.02307968 |
| PSG9 | 3.82273982 | -1.674733924 | 1.04495475 | -1.6026856 | 0.10900409 | 0.25236504 |
| AC005392.2 | 4.67492028 | -0.674937335 | 0.82667815 | -0.8164451 | 0.4142456 | 0.60794725 |
| CD177 | 113.625438 | -3.758509263 | 0.68195153 | -5.5114023 | 3.56E-08 | 2.03E-06 |
| AC005392.3 | 21.9770687 | -5.394676083 | 1.08869012 | -4.9551989 | 7.23E-07 | 2.47E-05 |
| TEX101 | 0.92308008 | -2.132188601 | 1.89599959 | -1.1245723 | 0.26077032 | NA |
| LYPD3 | 99.9635935 | 0.20436514 | 0.48671975 | 0.41988257 | 0.67457124 | 0.81055056 |
| PHLDB3 | 690.585959 | -0.297700491 | 0.21635861 | -1.3759586 | 0.16883445 | 0.34041279 |
| ETHE1 | 1887.89196 | -0.884918568 | 0.20193114 | -4.3822789 | 1.17E-05 | 0.00023321 |
| ZNF575 | 1066.08972 | -0.663482008 | 0.18975394 | -3.4965387 | 0.00047134 | 0.00433071 |
| XRCC1 | 1668.02583 | -0.013930268 | 0.15459908 | -0.0901058 | 0.92820317 | 0.96520831 |
| L34079.2 | 182.793034 | 0.218871831 | 0.15790725 | 1.38607841 | 0.16572296 | 0.33621972 |
| L34079.1 | 327.092014 | 0.236152189 | 0.14356889 | 1.64487021 | 0.09999658 | 0.23748976 |
| L34079.3 | 231.081404 | 0.232634704 | 0.13644673 | 1.70494895 | 0.08820395 | 0.21733676 |
| PINLYP | 378.696585 | -0.054511368 | 0.17717284 | -0.3076734 | 0.75833088 | 0.86631656 |
| IRGQ | 2056.07833 | -0.19742828 | 0.17162556 | -1.1503431 | 0.25000259 | 0.4420982 |
| ZNF576 | 601.35313 | 0.008816922 | 0.12778712 | 0.06899696 | 0.94499204 | 0.9737607 |
| SRRM5 | 1071.63703 | 0.000438867 | 0.11890633 | 0.00369086 | 0.99705512 | 0.99850237 |
| ZNF428 | 1626.76931 | -0.161519673 | 0.10988768 | -1.4698616 | 0.14159925 | 0.30224804 |
| CADM4 | 2788.94329 | 0.091136747 | 0.1823371 | 0.49982559 | 0.61719789 | 0.77116837 |
| PLAUR | 997.163771 | -1.499334885 | 0.49303572 | -3.0410269 | 0.00235773 | 0.0153325 |
| IRGC | 0.53080094 | 0.990423151 | 2.22169618 | 0.44579595 | 0.65574465 | NA |
| SMG9 | 1756.5726 | -0.340008763 | 0.1520803 | -2.2357187 | 0.0253702 | 0.08929689 |
| KCNN4 | 119.762866 | 0.466323007 | 0.60907917 | 0.76561969 | 0.44390263 | 0.63486887 |
| AC115522.1 | 9.9318175 | -0.701641037 | 0.63841107 | -1.0990427 | 0.27174943 | 0.46653857 |
| LYPD5 | 84.6807017 | -1.182650477 | 0.48661418 | -2.4303658 | 0.01508359 | 0.06113653 |
| ZNF283 | 234.577354 | -0.238909692 | 0.17721844 | -1.3481085 | 0.17762348 | 0.35275879 |
| ZNF404 | 256.282656 | 0.049172083 | 0.42430739 | 0.11588788 | 0.9077414 | 0.95386955 |
| AC006213.2 | 46.7847559 | 0.59385599 | 0.33607791 | 1.76701882 | 0.07722508 | 0.19763444 |
| AC006213.3 | 6.22624142 | 0.542477753 | 0.56319795 | 0.96320975 | 0.33544227 | 0.53363977 |
| ZNF45 | 540.880018 | 0.36888526 | 0.16478647 | 2.23856521 | 0.02518422 | 0.08883198 |
| ZNF221 | 11.8478992 | -0.023150838 | 0.33974328 | -0.0681422 | 0.94567248 | 0.97395466 |
| ZNF155 | 406.411155 | 0.01185435 | 0.14234901 | 0.08327666 | 0.93363156 | 0.96806638 |
| AC006213.1 | 324.868879 | -0.114884202 | 0.1672239 | -0.6870083 | 0.49207751 | 0.6750144 |
| ZNF230 | 276.638661 | -0.115258936 | 0.10444492 | -1.103538 | 0.26979361 | 0.46402671 |
| AC067968.2 | 0.09635527 | 0.547239394 | 3.29377202 | 0.16614368 | 0.86804388 | NA |
| ZNF222 | 189.006532 | -0.147213012 | 0.16912893 | -0.8704189 | 0.38407154 | 0.57963524 |
| AC067968.1 | 350.376868 | 0.110070225 | 0.11106734 | 0.99102245 | 0.32167462 | 0.51938279 |
| ZNF223 | 265.195052 | 0.138485729 | 0.14880824 | 0.93063211 | 0.3520439 | 0.54970692 |
| ZNF284 | 273.217048 | -0.397741515 | 0.19771878 | -2.0116527 | 0.04425656 | 0.13327002 |
| ZNF224 | 1451.84032 | -0.123855861 | 0.14465333 | -0.8562255 | 0.39187309 | 0.58760908 |
| AC021092.1 | 774.711431 | 0.01814312 | 0.16020775 | 0.11324745 | 0.90983438 | 0.95518483 |
| ZNF225 | 307.307174 | 0.166029518 | 0.16347251 | 1.01564179 | 0.3098 | 0.50697377 |
| ZNF234 | 461.815923 | 0.046259002 | 0.13980877 | 0.33087339 | 0.74074012 | 0.85451319 |
| ZNF226 | 1101.93964 | -0.051986167 | 0.19979154 | -0.260202 | 0.79470793 | 0.8886252 |
| ZNF227 | 699.552729 | -0.017628169 | 0.08587032 | -0.2052883 | 0.83734691 | 0.91386632 |
| ZNF235 | 582.478693 | 0.066133402 | 0.10370412 | 0.63771237 | 0.52366093 | 0.69892989 |
| ZNF233 | 101.190964 | 0.859622568 | 0.27935975 | 3.07711679 | 0.00209013 | 0.01392542 |
| ZNF112 | 290.965143 | 0.480499183 | 0.16089353 | 2.98644194 | 0.00282244 | 0.0175042 |
| AC245748.1 | 26.0061695 | 1.532542607 | 0.41279873 | 3.71256617 | 0.00020517 | 0.00222554 |
| ZNF285 | 167.187056 | 0.381049065 | 0.20234608 | 1.88315518 | 0.05967934 | 0.16496271 |
| ZNF229 | 344.179442 | 0.070427427 | 0.23266273 | 0.3027018 | 0.76211713 | 0.86815969 |
| ZNF180 | 379.013507 | 0.543646235 | 0.17418996 | 3.12099634 | 0.0018024 | 0.01244224 |
| CEACAM20 | 1.08330373 | -0.044021303 | 1.24297018 | -0.0354162 | 0.97174785 | 0.98670705 |
| AC245748.3 | 0.3046877 | -0.695245806 | 2.32293025 | -0.2992969 | 0.76471353 | NA |
| IGSF23 | 19.5574391 | -0.061629088 | 0.56721852 | -0.1086514 | 0.91347899 | 0.95764812 |
| AC243964.2 | 129.216839 | 0.010773733 | 0.6503199 | 0.01656682 | 0.98678219 | 0.99400325 |
| PVR | 3780.01645 | -1.530596692 | 0.46734724 | -3.2750738 | 0.00105634 | 0.00818325 |
| CEACAM19 | 1668.2176 | -1.122404685 | 0.37744022 | -2.9737284 | 0.00294205 | 0.01807174 |
| CEACAM16 | 4.28879519 | 0.581261953 | 0.83830286 | 0.69337942 | 0.48807147 | 0.6715885 |
| BCL3 | 4878.34781 | -1.393928869 | 0.44527163 | -3.1305136 | 0.00174501 | 0.01215102 |
| CBLC | 705.458243 | -0.576524949 | 0.33945881 | -1.698365 | 0.0894389 | 0.21959999 |
| AC092306.1 | 36.8592722 | -0.992375633 | 0.37104808 | -2.6745203 | 0.00748362 | 0.03639675 |
| BCAM | 46063.2904 | -0.537177347 | 0.3026632 | -1.7748353 | 0.07592508 | 0.19548373 |
| NECTIN2 | 6965.74433 | -0.817474519 | 0.19964078 | -4.0947272 | 4.23E-05 | 0.00064876 |
| AC011481.2 | 392.061581 | -0.418159799 | 0.23027513 | -1.8159138 | 0.06938359 | 0.18300112 |
| TOMM40 | 2920.16144 | -0.522820143 | 0.17650364 | -2.9620927 | 0.00305556 | 0.01862146 |
| APOE | 52964.3168 | 0.931227057 | 0.43709246 | 2.1305036 | 0.03313006 | 0.10833158 |
| APOC1 | 172.29804 | -1.384411044 | 0.39239945 | -3.5280657 | 0.00041861 | 0.00395369 |
| APOC4-APOC | 318.803419 | 1.314695341 | 0.69548808 | 1.89032045 | 0.05871512 | 0.16292935 |
| APOC4 | 1.22294774 | 1.284651254 | 1.06095312 | 1.21084639 | 0.22595428 | 0.41266115 |
| APOC2 | 320.29307 | 1.312698295 | 0.69679524 | 1.88390824 | 0.05957739 | 0.16478453 |
| AC011481.1 | 35.9172373 | 0.438755569 | 0.36245903 | 1.210497 | 0.22608824 | 0.41281666 |
| CLPTM1 | 9891.02645 | 0.458004381 | 0.19941978 | 2.29668477 | 0.02163676 | 0.07949858 |
| RELB | 1683.36152 | -1.68392303 | 0.34133847 | -4.933294 | 8.09E-07 | 2.70E-05 |
| CLASRP | 4118.65366 | -0.775254263 | 0.20456906 | -3.7896946 | 0.00015083 | 0.00176072 |
| ZNF296 | 111.946764 | -0.473036251 | 0.35582423 | -1.3294099 | 0.18371277 | 0.36077642 |
| GEMIN7-AS1 | 461.022585 | -0.581401307 | 0.18240335 | -3.1874486 | 0.00143534 | 0.01045578 |
| GEMIN7 | 803.465548 | -0.646512618 | 0.19039966 | -3.3955555 | 0.00068489 | 0.00579239 |
| MARK4 | 2026.35239 | -0.36865389 | 0.15184059 | -2.4279008 | 0.0151865 | 0.06137439 |
| PPP1R37 | 3470.03806 | -0.471631204 | 0.15297658 | -3.0830288 | 0.00204905 | 0.0137385 |
| AC005757.1 | 3.3610015 | -0.213097437 | 0.7549689 | -0.2822599 | 0.77774424 | 0.87833472 |
| NKPD1 | 8.96602512 | -0.104209115 | 0.490595 | -0.2124137 | 0.83178427 | 0.91058416 |
| TRAPPC6A | 1289.60054 | 0.48195325 | 0.19477884 | 2.47436149 | 0.01334746 | 0.05579607 |
| BLOC1S3 | 665.905845 | -0.550532573 | 0.28031834 | -1.9639549 | 0.04953532 | 0.14473095 |
| AC005779.2 | 33.2008425 | -0.254780374 | 0.33995163 | -0.7494607 | 0.45357959 | 0.64323172 |

| | | | | | | |
|---|---|---|---|---|---|---|
| AC005779.1 | 9.70388202 | 0.212113421 | 0.46324251 | 0.45788851 | 0.64703254 | 0.79208245 |
| EXOC3L2 | 1835.39434 | 0.595754202 | 0.42105518 | 1.41490766 | 0.15709557 | 0.32404648 |
| CKM | 65.1866621 | 4.782063787 | 0.87738789 | 5.4503417 | 5.03E-08 | 2.67E-06 |
| KLC3 | 702.546002 | -0.275815644 | 0.25852222 | -1.0668934 | 0.28601999 | 0.48203152 |
| ERCC2 | 1818.07334 | 0.412824173 | 0.20267766 | 2.03685093 | 0.04166498 | 0.12763773 |
| PPP1R13L | 1913.659 | -1.025754394 | 0.25454394 | -4.0297733 | 5.58E-05 | 0.00080804 |
| CD3EAP | 1191.54244 | -0.121112772 | 0.30457336 | -0.3976473 | 0.6908902 | 0.82203558 |
| ERCC1 | 3281.16868 | -0.125295535 | 0.24779374 | -0.5056445 | 0.61310625 | 0.76817248 |
| FOSB | 1412.07367 | 3.433029654 | 0.95569706 | 3.5921735 | 0.00032793 | 0.00325585 |
| RTN2 | 277.42667 | -0.585521118 | 0.2660699 | -2.200629 | 0.0277623 | 0.09551101 |
| PPM1N | 128.754336 | -0.432063368 | 0.30165611 | -1.4323044 | 0.15205673 | 0.31712559 |
| VASP | 4361.32054 | -0.995549999 | 0.2691592 | -3.6987404 | 0.00021667 | 0.00232925 |
| OPA3 | 2402.97215 | 0.17587205 | 0.14966799 | 1.17508126 | 0.23996221 | 0.42987795 |
| GPR4 | 1194.11818 | -1.392000983 | 0.38151818 | -3.6485836 | 0.00026369 | 0.00273437 |
| EML2 | 3051.08132 | -0.470397426 | 0.27513112 | -1.7097209 | 0.0873175 | 0.21553519 |
| GIPR | 192.088007 | -1.074532596 | 0.3774313 | -2.8469621 | 0.00441386 | 0.02452973 |
| SNRPD2 | 3500.16909 | -0.577461631 | 0.11487585 | -5.0268322 | 4.99E-07 | 1.80E-05 |
| QPCTL | 649.905966 | -0.655157143 | 0.13451223 | -4.870614 | 1.11E-06 | 3.56E-05 |
| FBXO46 | 966.746576 | -0.284117443 | 0.19336384 | -1.4693411 | 0.14174029 | 0.30245116 |
| BHMG1 | 0.57796482 | 0.906832076 | 1.64519643 | 0.55119988 | 0.58149666 | NA |
| SIX5 | 1382.43981 | -0.13135798 | 0.23057861 | -0.5696885 | 0.568889 | 0.73440069 |
| DM1-AS | 1433.85304 | 0.160941325 | 0.14138437 | 1.13832474 | 0.25498491 | 0.44769638 |
| DMPK | 5173.01828 | -0.204028765 | 0.16758338 | -1.2174761 | 0.22342312 | 0.40948483 |
| AC011530.1 | 1231.56004 | -0.228415439 | 0.2005522 | -1.1389326 | 0.25473127 | 0.44753011 |
| DMWD | 2328.27303 | -0.190092765 | 0.19492615 | -0.975204 | 0.32945907 | 0.52723848 |
| RSPH6A | 2.58517549 | 1.144345987 | 0.80981155 | 1.41310159 | 0.15762585 | 0.32475926 |
| SYMPK | 6264.41243 | -0.299057497 | 0.14267479 | -2.096078 | 0.03607527 | 0.11503081 |
| AC092301.1 | 241.07916 | 0.095566462 | 0.18556776 | 0.51499495 | 0.60655655 | 0.76313157 |
| FOXA3 | 177.939567 | -2.157942877 | 0.46405289 | -4.6502089 | 3.32E-06 | 8.48E-05 |
| IRF2BP1 | 1468.87366 | -0.010452936 | 0.14989928 | -0.0697331 | 0.94440613 | 0.97345853 |
| MYPOP | 745.574003 | -0.275195493 | 0.1889865 | -1.4561648 | 0.14534706 | 0.30768258 |
| NANOS2 | 0.34095445 | 0.669946673 | 2.22654746 | 0.30089036 | 0.7634981 | NA |
| NOVA2 | 576.439461 | -0.107458585 | 0.35369843 | -0.3038141 | 0.76126949 | 0.8677512 |
| CCDC61 | 722.190356 | -0.107582898 | 0.25605113 | -0.4201618 | 0.67436728 | 0.81034701 |
| PGLYRP1 | 17.0509452 | -1.629001606 | 0.59747623 | -2.726471 | 0.00640156 | 0.03232033 |
| AC007785.3 | 5.2290635 | -0.757921898 | 0.596058 | -1.2715573 | 0.20353046 | 0.3857431 |
| IGFL4 | 5.93051934 | 0.631426689 | 0.53982784 | 1.1696816 | 0.24212913 | 0.43253528 |
| AC007785.1 | 0.30866185 | 1.514719681 | 2.81598423 | 0.53790063 | 0.59064566 | NA |
| IGFL3 | 0.32679975 | -1.692381882 | 2.87486798 | -0.5886816 | 0.55607488 | NA |
| IGFL2 | 1.4983923 | 0.911157566 | 1.62111271 | 0.56205689 | 0.57407726 | 0.73863954 |
| AC006262.2 | 0.16052235 | -1.003574173 | 3.30558948 | -0.3035992 | 0.7614333 | NA |
| IGFL2-AS1 | 0.7888124 | 1.244592639 | 1.46297058 | 0.85072978 | 0.39491948 | NA |
| AC006262.3 | 0.61211833 | 2.637039344 | 3.19388444 | 0.82565271 | 0.40900113 | NA |
| IGFL1 | 1.39905968 | 3.033042331 | 2.53306684 | 1.19737951 | 0.23115867 | 0.41900289 |
| HIF3A | 1028.36032 | -0.309420864 | 0.55524527 | -0.5572688 | 0.57734379 | 0.7411933 |
| AC007193.2 | 41.9777959 | 0.131966486 | 0.66732325 | 0.19775496 | 0.84323678 | 0.91676578 |
| PPP5C | 3993.19527 | -0.14865715 | 0.12862115 | -1.1557753 | 0.24777307 | 0.43933404 |
| AC007193.1 | 123.186358 | 0.167111981 | 0.26054347 | 0.64139769 | 0.52126434 | 0.6976825 |
| AC007193.3 | 366.311394 | -0.551372791 | 0.17187746 | -3.2079413 | 0.00133689 | 0.00989162 |
| CCDC8 | 1220.16287 | 0.730957299 | 0.30189402 | 2.42123808 | 0.01546774 | 0.06223442 |
| PNMA8C | 6.67248963 | 0.985359088 | 0.58216031 | 1.69259063 | 0.09053342 | 0.22158587 |
| PNMA8A | 892.599384 | 0.675270548 | 0.38971664 | 1.73272189 | 0.08314511 | 0.20809952 |
| PPP5D1 | 40.2825782 | 0.57474756 | 0.29551573 | 1.94489665 | 0.05178742 | 0.14945107 |
| PNMA8B | 122.154759 | 0.734645944 | 0.29976126 | 2.45077013 | 0.0142551 | 0.05864382 |
| AC011484.1 | 49.4154174 | 1.018533788 | 0.28568485 | 3.5652356 | 0.00036353 | 0.00353737 |
| CALM3 | 12309.5052 | -0.043304891 | 0.20345456 | -0.212848 | 0.83144554 | 0.91040231 |
| AC093503.2 | 2575.98855 | -0.067420022 | 0.20765408 | -0.3246747 | 0.7454273 | 0.85738944 |
| PTGIR | 992.820516 | -1.049850106 | 0.4413127 | -2.3789256 | 0.01736318 | 0.0676516 |
| GNG8 | 27.402865 | -0.9339129 | 0.53481853 | -1.7462239 | 0.08077205 | 0.20382383 |
| DACT3 | 529.078047 | -0.160317528 | 0.53709556 | -0.2984898 | 0.76532939 | 0.87045645 |
| DACT3-AS1 | 12.9047606 | -0.166681694 | 0.35140023 | -0.4743358 | 0.6352605 | 0.78363752 |
| PRKD2 | 3671.96358 | -0.811518771 | 0.27185537 | -2.9851121 | 0.00283474 | 0.01756395 |
| STRN4 | 5923.61766 | -0.461349386 | 0.17530438 | -2.6317049 | 0.00849576 | 0.03982567 |
| AC008635.1 | 572.624258 | -0.160581515 | 0.17587954 | -0.9130199 | 0.36123209 | 0.55878832 |
| FKRP | 1988.528 | -0.257819674 | 0.15120486 | -1.7051018 | 0.08817545 | 0.21728684 |
| SLC1A5 | 6459.41384 | -0.344866108 | 0.29640085 | -1.1635126 | 0.24462161 | 0.43567957 |
| AC008622.2 | 1277.96093 | 0.033380717 | 0.36528602 | 0.09138241 | 0.92718874 | 0.96468601 |
| AP2S1 | 2751.74979 | -0.63296557 | 0.14940637 | -4.2365367 | 2.27E-05 | 0.0003938 |
| ARHGAP35 | 5413.01797 | 0.345371804 | 0.18412168 | 1.87578026 | 0.06068546 | 0.16679962 |
| NPAS1 | 85.3296572 | -0.413821635 | 0.40178477 | -1.0299585 | 0.30302949 | 0.4998366 |
| TMEM160 | 469.553891 | -0.443298696 | 0.32859796 | -1.349061 | 0.17731737 | 0.35240147 |
| ZC3H4 | 1778.91548 | 0.130609529 | 0.17717507 | 0.73717782 | 0.46101422 | 0.64951355 |
| SAE1 | 2501.95331 | -0.368706338 | 0.18214777 | -2.0242155 | 0.04294799 | 0.13049181 |
| BBC3 | 540.828872 | -1.833796669 | 0.25670466 | -7.143605 | 9.09E-13 | 2.45E-10 |
| CCDC9 | 2361.26983 | -0.73705414 | 0.18521787 | -3.97939 | 6.91E-05 | 0.00094802 |
| INAFM1 | 725.136006 | -1.236251375 | 0.23817231 | -5.1905757 | 2.10E-07 | 8.85E-06 |
| C5AR1 | 1380.02914 | -1.16676422 | 0.37333779 | -3.125224 | 0.0017767 | 0.01231314 |
| C5AR2 | 191.729703 | 1.064366829 | 0.41998862 | 2.53427542 | 0.01126801 | 0.04926836 |
| DHX34 | 1758.80091 | -0.576705621 | 0.20775968 | -2.7758303 | 0.00550609 | 0.0289754 |
| MEIS3 | 294.50079 | -0.685485361 | 0.44465352 | -1.5416169 | 0.12316672 | 0.2746124 |
| SLC8A2 | 18.2905884 | -3.149393511 | 0.8556691 | -3.6806208 | 0.00023267 | 0.00246501 |
| KPTN | 579.535212 | -0.036958626 | 0.21839476 | -0.1692285 | 0.86561689 | 0.93016778 |
| NAPA-AS1 | 2089.19804 | -0.01920315 | 0.18629276 | -0.1030805 | 0.91789909 | 0.9601219 |
| NAPA | 10472.1089 | -0.378470589 | 0.19364255 | -1.9544805 | 0.05064441 | 0.14691895 |
| ZNF541 | 91.9185796 | 1.897473365 | 0.59692944 | 3.17872302 | 0.00147925 | 0.01069296 |
| AC010519.1 | 3.53479147 | -1.216236691 | 0.9306543 | -1.306862 | 0.1912596 | 0.37036838 |
| AC010331.1 | 36.2963491 | -0.310126267 | 0.31198954 | -0.9940278 | 0.32020934 | 0.5176191 |
| BICRA | 1248.17694 | -0.482993063 | 0.19088261 | -2.5303147 | 0.01139603 | 0.04968803 |
| BICRA-AS1 | 2.19362309 | 2.941480081 | 1.28691036 | 2.2856915 | 0.02227231 | 0.08123387 |
| EHD2 | 3013.32372 | -0.343328325 | 0.32282585 | -1.0635094 | 0.287551 | 0.48380077 |
| NOP53 | 9787.50965 | -0.135830746 | 0.18144363 | -0.7486113 | 0.45409153 | 0.64372316 |
| NOP53-AS1 | 89.7675258 | -0.425858507 | 0.21641078 | -1.9678248 | 0.0490882 | 0.14389238 |
| SELENOW | 6348.47998 | -0.166118898 | 0.20358245 | -0.8159785 | 0.41451244 | 0.60817298 |
| TPRX1 | 0 | NA | NA | NA | NA | NA |
| CRX | 0.08757577 | 0.547239394 | 3.29377202 | 0.16614368 | 0.86804388 | NA |
| LINC01595 | 0 | NA | NA | NA | NA | NA |
| SULT2A1 | 0.09989236 | -0.642905668 | 3.30558948 | -0.1944905 | 0.84579185 | NA |
| BSPH1 | 0.08757577 | 0.547239394 | 3.29377202 | 0.16614368 | 0.86804388 | NA |
| ELSPBP1 | 0.06785815 | 0.795868262 | 3.28898349 | 0.24198001 | 0.80879565 | NA |
| CABP5 | 0.49128398 | 0.427800043 | 3.27633741 | 0.13057265 | 0.89611338 | NA |
| PLA2G4C | 1194.42905 | -1.156667889 | 0.3475856 | -3.3277209 | 0.0008756 | 0.00707614 |
| PLA2G4C-AS | 272.951536 | -1.545418427 | 0.3522894 | -4.3867866 | 1.15E-05 | 0.00022877 |
| LIG1 | 972.496888 | 0.318587082 | 0.25657227 | 1.24170504 | 0.21434541 | 0.3984694 |
| AC011466.3 | 203.157319 | 0.349360745 | 0.26945671 | 1.29653756 | 0.19479035 | 0.37480173 |
| AC011466.2 | 0.44483423 | 1.303178837 | 2.75648929 | 0.4727676 | 0.63637899 | NA |
| ZSWIM9 | 679.414975 | -0.185073699 | 0.16701942 | -1.1080969 | 0.26781997 | 0.46200891 |
| CARD8 | 1795.83917 | 0.017492479 | 0.21304466 | 0.0821071 | 0.93456155 | 0.96864979 |
| AC011466.1 | 202.806077 | -0.211605605 | 0.22038084 | -0.9601815 | 0.33696387 | 0.53499533 |
| CARD8-AS1 | 193.787317 | -0.460609033 | 0.17259004 | -2.6688043 | 0.00761218 | 0.03680476 |
| AC008392.1 | 0.40324896 | 0.952179483 | 2.20423338 | 0.43197762 | 0.66575768 | NA |
| ZNF114 | 20.0762918 | 0.530286063 | 0.4647692 | 1.14096645 | 0.25388389 | 0.44653626 |
| CCDC114 | 116.481235 | -0.289760285 | 0.44776362 | -0.6471278 | 0.51754924 | 0.69466488 |
| EMP3 | 1243.86272 | -0.49645461 | 0.30228562 | -1.6423362 | 0.10052035 | 0.23843473 |

| | | | | | | |
|---|---|---|---|---|---|---|
| TMEM143 | 834.02827 | 1.106188287 | 0.28574037 | 3.87130555 | 0.00010825 | 0.00134972 |
| SYNGR4 | 1.2613422 | 1.95629498 | 1.07756959 | 1.81546973 | 0.06945175 | 0.18316259 |
| KDELR1 | 10248.1553 | -0.419454785 | 0.10242643 | -4.0951812 | 4.22E-05 | 0.00064787 |
| GRIN2D | 32.6453228 | -0.067719532 | 0.35244242 | -0.1921435 | 0.84762977 | 0.92002458 |
| GRWD1 | 1866.93495 | -0.230953999 | 0.21564344 | -1.0709994 | 0.28416969 | 0.48023294 |
| KCNJ14 | 167.129158 | -0.859504307 | 0.3838477 | -2.2391805 | 0.02514417 | 0.08875007 |
| AC008403.2 | 483.868996 | -0.359952762 | 0.1778603 | -2.0237949 | 0.04299126 | 0.13060824 |
| AC008403.1 | 1130.82488 | -0.235606735 | 0.18335921 | -1.2849463 | 0.19881104 | 0.37984692 |
| CYTH2 | 6286.44355 | -0.60091608 | 0.14627999 | -4.1079855 | 3.99E-05 | 0.00062241 |
| LMTK3 | 503.383163 | -0.518854956 | 0.42036695 | -1.2342906 | 0.21709465 | 0.40199326 |
| AC008403.3 | 9.58187179 | 0.358749103 | 0.56340002 | 0.63675734 | 0.52428292 | 0.69941521 |
| SULT2B1 | 283.908563 | -1.710718698 | 0.69925888 | -2.446474 | 0.01442612 | 0.05919962 |
| FAM83E | 85.699304 | -2.314891844 | 0.78503965 | -2.9487579 | 0.00319054 | 0.01926115 |
| SPACA4 | 5.41028375 | -2.89151419 | 1.05432915 | -2.7425157 | 0.00609705 | 0.03119506 |
| RPL18 | 17185.4857 | -0.562022087 | 0.16347025 | -3.4380696 | 0.00058588 | 0.00513105 |
| AC022154.1 | 114.914588 | 0.38375203 | 0.27980324 | 1.37150672 | 0.17021705 | 0.34219004 |
| SPHK2 | 3201.60426 | 1.179287476 | 0.27115885 | 4.34906499 | 1.37E-05 | 0.00026297 |
| DBP | 1082.08081 | 0.907225625 | 0.27974699 | 3.24302195 | 0.00118269 | 0.00894652 |
| CA11 | 833.458553 | -0.298959239 | 0.21408717 | -1.3964369 | 0.16258296 | 0.33152742 |
| NTN5 | 54.707472 | 0.2723287 | 0.45491436 | 0.5986373 | 0.54941478 | 0.71992001 |
| FUT2 | 91.6030878 | -0.633710133 | 0.54426867 | -1.1643333 | 0.24428897 | 0.43534851 |
| MAMSTR | 98.5254927 | 0.00165692 | 0.38051787 | 0.00435438 | 0.99652572 | 0.99824844 |
| RASIP1 | 1269.58408 | -0.44048954 | 0.27291636 | -1.6140093 | 0.10652541 | 0.24810767 |
| IZUMO1 | 28.1248145 | -0.380578733 | 0.44857609 | -0.8484151 | 0.39620683 | 0.5912411 |
| FUT1 | 248.807607 | 0.149099371 | 0.27814534 | 0.53604842 | 0.59192509 | 0.75144736 |
| FGF21 | 0 | NA | NA | NA | NA | NA |
| BCAT2 | 3181.59476 | 0.37015194 | 0.16172918 | 2.2887146 | 0.02209594 | 0.08072583 |
| HSD17B14 | 2408.4917 | 1.089996684 | 0.35286446 | 3.08899541 | 0.00200835 | 0.01354477 |
| PLEKHA4 | 2515.63138 | -1.661934844 | 0.48323186 | -3.4392079 | 0.00058342 | 0.00511461 |
| PPP1R15A | 5751.26148 | -1.18571326 | 0.33812819 | -3.5066974 | 0.0004537 | 0.00420682 |
| TULP2 | 10.9806046 | 0.127230475 | 0.45418386 | 0.28012989 | 0.77937785 | 0.87912542 |
| NUCB1 | 25483.0901 | 0.006881142 | 0.12491006 | 0.05508878 | 0.95606774 | 0.97900515 |
| NUCB1-AS1 | 4578.97398 | 0.236488847 | 0.15293716 | 1.54631381 | 0.1220288 | 0.2729125 |
| DHDH | 779.703037 | 2.186255903 | 0.54479129 | 4.01301555 | 5.99E-05 | 0.00084899 |
| BAX | 3073.47896 | -1.273726893 | 0.17391391 | -7.323893 | 2.41E-13 | 8.04E-11 |
| AC026803.2 | 37.3980116 | 0.438370375 | 0.36848176 | 1.18966642 | 0.23417753 | 0.42276042 |
| FTL | 324066.144 | -0.320608505 | 0.46268103 | -0.6929364 | 0.4883495 | 0.67184654 |
| GYS1 | 3607.71409 | -0.171398436 | 0.21281241 | -0.8053968 | 0.42059074 | 0.61330335 |
| RUVBL2 | 3110.96805 | -0.185372723 | 0.09893311 | -1.8737178 | 0.06096933 | 0.16736033 |
| LHB | 12.4569277 | -0.713050235 | 0.59340456 | -1.2016258 | 0.22950854 | 0.41695764 |
| AC008687.4 | 19.9861014 | -0.934534198 | 0.365722 | -2.5553131 | 0.01060923 | 0.04706687 |
| CGB3 | 0.43299836 | -2.018433049 | 1.88025654 | -1.0734881 | 0.28305217 | NA |
| AC008687.1 | 0.07659028 | -0.642905668 | 3.30558948 | -0.1944905 | 0.84579185 | NA |
| CGB2 | 0 | NA | NA | NA | NA | NA |
| CGB1 | 0.07659028 | -0.642905668 | 3.30558948 | -0.1944905 | 0.84579185 | NA |
| CGB5 | 0.3283865 | -1.696004896 | 3.30291516 | -0.5134873 | 0.60761051 | NA |
| CGB8 | 0.07659028 | -0.642905668 | 3.30558948 | -0.1944905 | 0.84579185 | NA |
| CGB7 | 14.0904876 | -0.966095866 | 0.64860963 | -1.4894874 | 0.13635907 | 0.29451952 |
| AC008687.8 | 12.7432246 | -1.178634529 | 0.76244466 | -1.5458624 | 0.1221378 | 0.27307983 |
| AC008687.2 | 0.74612838 | 1.287824808 | 1.46847562 | 0.87698072 | 0.38049711 | NA |
| NTF4 | 8.61810402 | -1.313533788 | 1.10901678 | -1.1844129 | 0.23624965 | 0.425573 |
| AC008687.3 | 2.51961395 | -3.863269062 | 1.67458058 | -2.307007 | 0.02105443 | 0.07782279 |
| KCNA7 | 0.425059 | 0.231398012 | 3.28103693 | 0.07052588 | 0.9437751 | NA |
| SNRNP70 | 22816.5774 | 0.026600056 | 0.18853233 | 0.14109016 | 0.88779872 | 0.94201397 |
| LIN7B | 265.59612 | -0.087339518 | 0.28755137 | -0.3037354 | 0.76132952 | 0.8677512 |
| C19orf73 | 56.3622038 | -0.017219224 | 0.28308857 | -0.0608263 | 0.95149756 | 0.97675915 |
| PPFIA3 | 433.596883 | -0.146490526 | 0.20805183 | -0.7041059 | 0.48136681 | 0.66598336 |
| HRC | 304.923048 | 0.751562746 | 0.4463938 | 1.68363168 | 0.09225287 | 0.22454387 |
| TRPM4 | 5149.57099 | -0.094672136 | 0.35362609 | -0.2677182 | 0.78891625 | 0.88537504 |
| SLC6A16 | 520.638763 | 0.369651959 | 0.28524829 | 1.29589544 | 0.19501151 | 0.37506043 |
| AC011450.1 | 214.143584 | -0.079641371 | 0.31283938 | -0.2545759 | 0.79905066 | 0.89108294 |
| CD37 | 1466.78279 | -0.429538204 | 0.3452723 | -1.2440564 | 0.21347882 | 0.39750293 |
| TEAD2 | 2376.57662 | -1.007441547 | 0.33855335 | -2.9757247 | 0.00292297 | 0.01797965 |
| AC010524.1 | 7.85154364 | -0.636863048 | 0.69024161 | -0.9226668 | 0.35618085 | 0.55354131 |
| DKKL1 | 16.2921628 | -0.016638324 | 0.41449685 | -0.040141 | 0.9679807 | 0.98501394 |
| AC010643.1 | 16.6899316 | -0.068562306 | 0.55279373 | -0.1240287 | 0.90129253 | 0.95003733 |
| CCDC155 | 2.50351557 | -0.817089167 | 0.81089236 | -1.007642 | 0.31362637 | 0.51067166 |
| PTH2 | 0 | NA | NA | NA | NA | NA |
| GFY | 0.08666371 | -0.642905668 | 3.30558948 | -0.1944905 | 0.84579185 | NA |
| SLC17A7 | 49.5537614 | 0.636049786 | 0.41189694 | 1.54419644 | 0.12254075 | 0.27373127 |
| PIH1D1 | 3528.62836 | -0.175007781 | 0.11700587 | -1.4957179 | 0.13472718 | 0.29216741 |
| ALDH16A1 | 2794.84063 | 0.065724061 | 0.17239534 | 0.38124035 | 0.70302491 | 0.83010643 |
| AC010619.1 | 910.084962 | 0.089506794 | 0.16731193 | 0.53496958 | 0.59267089 | 0.75199593 |
| FLT3LG | 407.182806 | -0.341749496 | 0.41330315 | -0.8268737 | 0.40830866 | 0.60278813 |
| RPL13A | 55764.2204 | -0.546685398 | 0.18476171 | -2.9588674 | 0.00308772 | 0.01876974 |
| RPS11 | 28040.2709 | -0.436646814 | 0.1530518 | -2.8529348 | 0.00433175 | 0.02418553 |
| FCGRT | 10990.1004 | 0.611054888 | 0.14775967 | 4.13546458 | 3.54E-05 | 0.0005627 |
| RCN3 | 1687.07091 | -0.236359244 | 0.28342976 | -0.8339253 | 0.40432308 | 0.59912338 |
| NOSIP | 2155.11972 | -0.285925814 | 0.12768934 | -2.2392302 | 0.02514094 | 0.08875007 |
| PRRG2 | 866.557113 | -0.219191703 | 0.17390568 | -1.2604057 | 0.20752306 | 0.39041044 |
| PRR12 | 2343.4122 | 0.294537567 | 0.19938809 | 1.47720741 | 0.13962005 | 0.29945349 |
| AC011495.2 | 97.2675563 | -0.225192465 | 0.24696629 | -0.9118348 | 0.36185567 | 0.55919679 |
| RRAS | 2239.06033 | -0.833404403 | 0.31326082 | -2.6604169 | 0.0078044 | 0.03753455 |
| SCAF1 | 4171.98822 | -0.45916265 | 0.12404557 | -3.7015642 | 0.00021427 | 0.00231006 |
| IRF3 | 6081.40044 | -0.673429383 | 0.18606488 | -3.6193256 | 0.00029537 | 0.00299296 |
| BCL2L12 | 1241.00137 | -0.446262105 | 0.16511556 | -2.7027259 | 0.00687734 | 0.03412145 |
| PRMT1 | 5442.71683 | -0.465724384 | 0.16378488 | -2.8435127 | 0.00446192 | 0.02472791 |
| ADM5 | 173.170656 | -0.721082747 | 0.29912976 | -2.4106018 | 0.01592623 | 0.06361324 |
| AC011495.3 | 57.106156 | -0.675525115 | 0.29466793 | -2.2924962 | 0.02187703 | 0.08015801 |
| CPT1C | 218.047275 | -0.738329502 | 0.4226899 | -1.7467403 | 0.08068239 | 0.20363654 |
| TSKS | 11.1747642 | 0.059429086 | 0.54702797 | 0.10863994 | 0.91348808 | 0.95764812 |
| AP2A1 | 5662.61226 | -0.154031432 | 0.13067077 | -1.178775 | 0.2384878 | 0.42801021 |
| FUZ | 2120.77011 | 0.399001908 | 0.210559 | 1.89496486 | 0.05809706 | 0.16187863 |
| AC006942.1 | 443.35913 | 0.46657451 | 0.27487009 | 1.69743645 | 0.08961418 | 0.21990879 |
| MED25 | 4295.37025 | -0.386773313 | 0.23804656 | -1.6247801 | 0.10420942 | 0.24444081 |
| PTOV1-AS1 | 1017.09745 | -0.368148952 | 0.22829388 | -1.6126098 | 0.1068293 | 0.24863608 |
| PTOV1 | 11677.8107 | -0.354276194 | 0.14847385 | -2.3861185 | 0.01702726 | 0.06674827 |
| AC018766.1 | 916.600956 | 0.096626908 | 0.16157214 | 0.59804188 | 0.54981199 | 0.72018351 |
| PTOV1-AS2 | 4611.66375 | -0.326295172 | 0.1445994 | -2.2565458 | 0.02403647 | 0.08570061 |
| PNKP | 7134.76743 | -0.373872231 | 0.16894342 | -2.2130026 | 0.02689747 | 0.09318001 |
| AKT1S1 | 3305.19516 | -0.318298453 | 0.20670924 | -1.5398366 | 0.1236002 | 0.27514242 |
| TBC1D17 | 4413.75967 | 0.102053368 | 0.11154253 | 0.91492784 | 0.36022951 | 0.55789348 |
| IL4I1 | 1197.14073 | -1.225476776 | 0.36401029 | -3.3665992 | 0.00076101 | 0.00631066 |
| NUP62 | 3119.98189 | -0.343814532 | 0.15159424 | -2.2679921 | 0.02332969 | 0.08394105 |
| AC011452.1 | 423.584497 | 0.297307756 | 0.19970583 | 1.48872845 | 0.13655889 | 0.29487545 |
| ATF5 | 2279.46104 | -0.818923162 | 0.31488597 | -2.6006975 | 0.00930344 | 0.04264416 |
| SIGLEC11 | 99.6322271 | 1.045054157 | 0.6354944 | 1.64447422 | 0.10007828 | 0.23760371 |
| VRK3 | 3767.21593 | -0.455997699 | 0.24836021 | -1.8360337 | 0.06635268 | 0.17734848 |
| ZNF473 | 534.456883 | -0.451057702 | 0.20276584 | -2.2245251 | 0.02611314 | 0.09122992 |
| AC010624.3 | 63.6828236 | -0.224653122 | 0.22075422 | -1.0176617 | 0.30883874 | 0.50587474 |
| AC010624.2 | 2.16891826 | -0.416708746 | 0.80235884 | -0.5193546 | 0.60351349 | 0.76073856 |
| IZUMO2 | 0.28111514 | -0.695625285 | 2.3595665 | -0.2948106 | 0.76813855 | NA |
| MYH14 | 3838.64547 | 0.176124233 | 0.24226947 | 0.7269766 | 0.46724029 | 0.65433492 |

| | | | | | | |
|---|---|---|---|---|---|---|
| AC008655.1 | 606.060707 | -0.426543166 | 0.25632397 | -1.6640783 | 0.09609678 | 0.23121017 |
| KCNC3 | 4433.9706 | -0.317839632 | 0.2937379 | -1.0820518 | 0.27922951 | 0.47486664 |
| NR1H2 | 4889.75294 | -0.493692449 | 0.11557797 | -4.2715098 | 1.94E-05 | 0.0003464 |
| NAPSA | 14307.2501 | -0.849621689 | 0.48330074 | -1.7579565 | 0.07875491 | 0.20022821 |
| AC008655.2 | 22.1916514 | 0.205064457 | 0.43827198 | 0.46789315 | 0.639861 | 0.78705286 |
| POLD1 | 1322.54144 | -0.859195845 | 0.23727975 | -3.6210247 | 0.00029344 | 0.00298533 |
| AC020909.1 | 410.17682 | -1.037507676 | 0.28388126 | -3.654724 | 0.00025746 | 0.00267712 |
| SPIB | 58.5567642 | 0.414495721 | 0.92966867 | 0.44585317 | NA | NA |
| MYBPC2 | 132.437872 | -2.796572371 | 0.53715301 | -5.2062863 | 1.93E-07 | 8.30E-06 |
| FAM71E1 | 1066.5101 | -0.123674788 | 0.18932529 | -0.6532397 | 0.51360178 | 0.69160389 |
| EMC10 | 10866.0591 | -0.157014759 | 0.14646991 | -1.0719933 | 0.28372304 | 0.47972407 |
| AC020909.3 | 3308.48587 | -0.148974395 | 0.16016615 | -0.9301241 | 0.35230683 | 0.54985388 |
| AC020909.2 | 86.698863 | -0.532915118 | 0.40472211 | -1.3167433 | 0.18792469 | 0.36610329 |
| JOSD2 | 865.463717 | -0.276536778 | 0.22078573 | -1.252512 | 0.21038336 | 0.3937409 |
| ASPDH | 1268.92586 | 2.76133971 | 0.49563676 | 5.57129723 | 2.53E-08 | 1.53E-06 |
| LRRC4B | 110.52876 | 0.311545708 | 0.50697504 | 0.61451882 | 0.53887254 | 0.71212457 |
| SYT3 | 40.9463876 | -0.243869506 | 0.34082362 | -0.7155299 | 0.47428165 | 0.66003563 |
| C19orf81 | 11.0209773 | -1.010393176 | 0.71280272 | -1.4174934 | 0.15633871 | 0.32301589 |
| SHANK1 | 40.8193422 | 0.598688152 | 0.33860132 | 1.76812115 | 0.07704065 | 0.19723903 |
| CLEC11A | 614.22625 | -0.071878448 | 0.24789005 | -0.289961 | 0.77184607 | 0.87421716 |
| GPR32 | 0 | NA | NA | NA | NA | NA |
| AC010325.1 | 2.31906909 | -0.158176215 | 0.86422143 | -0.1830274 | 0.85477651 | 0.92400911 |
| ACP4 | 3.64028758 | 0.006967425 | 0.64611844 | 0.01078351 | 0.99139617 | 0.99589033 |
| AC010325.2 | 0.20366043 | -1.003574173 | 3.30558948 | -0.3035992 | 0.7614333 | NA |
| C19orf48 | 1451.80124 | -0.623487096 | 0.33157329 | -1.88039 | 0.06005494 | 0.16558436 |
| LINC01869 | 9.92746141 | -0.027889961 | 0.58844127 | -0.0473963 | 0.96219735 | 0.98204684 |
| KLK1 | 1633.70753 | 2.151824139 | 1.09398583 | 1.96695797 | 0.04918806 | 0.14407298 |
| KLK15 | 16.3533948 | -0.208984443 | 0.82830804 | -0.2523028 | 0.80080702 | 0.89197577 |
| AC011523.1 | 4.05460558 | 0.033135122 | 1.37639584 | 0.02407383 | 0.98079372 | 0.99132745 |
| KLK3 | 22.6927414 | 1.936368754 | 0.72892319 | 2.65647846 | 0.00789615 | 0.03781719 |
| KLK2 | 0.75914191 | 1.787607163 | 1.71353643 | 1.04322682 | 0.29684326 | NA |
| KLK4 | 4.34371629 | 0.516568965 | 1.07188469 | 0.48192587 | 0.62985861 | 0.78011215 |
| KLK5 | 17.5836183 | 1.452347722 | 0.83131339 | 1.74705201 | 0.08062832 | 0.20353903 |
| AC011483.2 | 244.00981 | 1.101752609 | 0.71976511 | 1.53071133 | 0.12584075 | 0.27862533 |
| KLK6 | 441.754584 | 1.081414513 | 0.74237189 | 1.45670186 | 0.1451987 | 0.30741787 |
| AC011483.1 | 0.35696888 | 0.705516065 | 2.30677947 | 0.30584461 | 0.75972296 | NA |
| KLK7 | 327.006282 | 1.172178779 | 0.81277483 | 1.44219375 | 0.14924772 | 0.31340282 |
| KLK8 | 6.3613594 | 0.008139356 | 0.48663044 | 0.01672595 | 0.98665525 | 0.99400325 |
| AC011473.4 | 4.85178621 | -0.318663914 | 0.59886036 | -0.5321172 | 0.59464481 | 0.75370069 |
| KLK9 | 0.11051935 | 0.02007176 | 3.29738704 | 0.00608717 | 0.99514317 | NA |
| KLK10 | 12.7888356 | 0.008524586 | 0.91921732 | 0.00927374 | 0.99260073 | 0.99656886 |
| AC011473.2 | 6.15956593 | -0.593142783 | 0.9982442 | -0.5941861 | 0.55238768 | 0.72236365 |
| KLK11 | 4.06338115 | 0.69461985 | 1.65987474 | 0.41847727 | NA | NA |
| KLK12 | 0.24075633 | -0.174096114 | 3.29377188 | -0.0528562 | 0.95784652 | NA |
| AC011473.3 | 0.07659028 | -0.642905668 | 3.30558948 | -0.1944905 | 0.84579185 | NA |
| AC011473.1 | 0 | NA | NA | NA | NA | NA |
| KLK13 | 1.41581612 | 0.416710167 | 1.28567168 | 0.32411865 | 0.74584821 | 0.85747993 |
| KLK14 | 38.0945849 | -0.520843973 | 0.27257094 | -1.9108566 | 0.05602301 | 0.15771611 |
| CTU1 | 249.525023 | -0.855953541 | 0.24906004 | -3.4367357 | 0.00058877 | 0.00514783 |
| SIGLEC9 | 162.112544 | -1.203094102 | 0.4379388 | -2.747174 | 0.00601112 | 0.03082118 |
| SIGLEC7 | 105.085 | -1.144210175 | 0.58314144 | -1.9621486 | 0.04974518 | 0.14518877 |
| AC063977.6 | 51.0408787 | -1.524400079 | 0.5549434 | -2.7469469 | 0.00601529 | 0.03083653 |
| CD33 | 174.105904 | -0.260756735 | 0.40657463 | -0.6413502 | 0.52129516 | 0.69768834 |
| SIGLECL1 | 0.91784769 | -0.671694336 | 1.29411117 | -0.5190391 | 0.60373345 | NA |
| AC063977.3 | 0 | NA | NA | NA | NA | NA |
| LINC01872 | 0.19416835 | -1.003574173 | 3.30558948 | -0.3035992 | 0.7614333 | NA |
| IGLON5 | 20.4651947 | -2.258424403 | 0.47679105 | -4.7367172 | 2.17E-06 | 6.07E-05 |
| VSIG10L | 314.725739 | -1.165886533 | 0.40379319 | -2.8873358 | 0.00388519 | 0.02229168 |
| AC008750.3 | 5.71741036 | -0.061690669 | 0.76846987 | -0.0802773 | 0.93601673 | 0.96939601 |
| ETFB | 6164.20102 | 0.647028819 | 0.29135685 | 2.22074346 | 0.02636834 | 0.09192422 |
| AC008750.4 | 3457.90776 | 0.700943128 | 0.30311936 | 2.3124327 | 0.02075385 | 0.07697314 |
| AC008750.8 | 4524.2033 | 0.698220505 | 0.31188673 | 2.23869897 | 0.02517551 | 0.08882502 |
| CLDND2 | 83.5571703 | -0.35961259 | 0.24522702 | -1.4664477 | 0.14252637 | 0.30375348 |
| NKG7 | 283.075955 | -1.405509553 | 0.53169215 | -2.6434649 | 0.00820623 | 0.03879721 |
| LIM2 | 0.87098173 | -0.992811905 | 1.42599501 | -0.696224 | 0.48628858 | NA |
| C19orf84 | 2.45980484 | -0.804382583 | 1.05153162 | -0.7649628 | 0.44429369 | 0.63522146 |
| AC008750.7 | 0.46663567 | -1.340588746 | 1.68656684 | -0.7948625 | 0.42669349 | NA |
| SIGLEC10 | 312.402897 | -1.430212832 | 0.5989544 | -2.3878493 | 0.01694729 | 0.06651086 |
| AC008750.1 | 42.5556367 | -1.030979352 | 0.67352112 | -1.5307306 | 0.125836 | 0.27862533 |
| AC008750.2 | 70.9337078 | -0.753851274 | 0.54820192 | -1.3751343 | 0.16908981 | 0.34073806 |
| SIGLEC8 | 23.4089562 | -2.194770632 | 1.20721388 | -1.8180462 | 0.06905707 | 0.18233014 |
| CEACAM18 | 0.08393208 | -0.642905668 | 3.30558948 | -0.1944905 | 0.84579185 | NA |
| SIGLEC12 | 22.7862467 | -0.816297493 | 0.70817577 | -1.1526764 | 0.24904323 | 0.44078927 |
| SIGLEC6 | 5.81179079 | 1.953459834 | 0.96244212 | 2.0296907 | 0.04238799 | 0.12931181 |
| AC020914.1 | 2.74505053 | 2.409695775 | 1.31311865 | 1.83509371 | 0.06649181 | 0.17757635 |
| ZNF175 | 682.373729 | 0.206825008 | 0.16475459 | 1.25535204 | 0.20935099 | 0.3925052 |
| AC018755.1 | 7.39273998 | -0.022884247 | 0.42272679 | -0.0541348 | 0.95682774 | 0.97937148 |
| SIGLEC5 | 71.1592176 | -1.166565333 | 0.51069103 | -2.2842879 | 0.02235462 | 0.0814552 |
| AC018755.2 | 52.6742709 | -1.373848671 | 0.54118289 | -2.5386033 | 0.01112959 | 0.04877637 |
| SIGLEC14 | 136.112778 | -1.220174758 | 0.64417616 | -1.8941632 | 0.05820335 | 0.1620588 |
| SPACA6P-AS | 107.202883 | 0.115544302 | 0.26306263 | 0.43922735 | 0.66049681 | 0.80020679 |
| SPACA6 | 1169.31589 | -0.490787995 | 0.25867492 | -1.8973157 | 0.05778628 | 0.16120007 |
| HAS1 | 11.843735 | 0.085659572 | 0.82641783 | 0.10365165 | 0.9174458 | 0.9600034 |
| FPR1 | 560.059616 | -1.665260309 | 0.51540381 | -3.2309817 | 0.00123366 | 0.00925507 |
| FPR2 | 85.314667 | -2.151667613 | 0.62976035 | -3.4166451 | 0.00063398 | 0.00545093 |
| FPR3 | 386.340797 | -0.022099536 | 0.55406231 | -0.0398864 | 0.96818372 | 0.98514449 |
| ZNF577 | 1150.15957 | 0.182088283 | 0.16405623 | 1.10991385 | 0.26703615 | 0.4611392 |
| AC006272.1 | 7.0100091 | 0.573467135 | 0.74148491 | 0.77340365 | 0.43928353 | 0.63082866 |
| ZNF649-AS1 | 52.3349462 | 0.510568098 | 0.26467898 | 1.92900886 | 0.05372976 | 0.15330706 |
| ZNF649 | 432.385199 | 0.244639991 | 0.19521996 | 1.25315052 | 0.21015093 | 0.3934588 |
| ZNF613 | 386.748634 | -0.254047093 | 0.15368139 | -1.6530765 | 0.0983153 | 0.23485592 |
| ZNF350-AS1 | 125.713958 | -0.487125751 | 0.54256562 | -0.897819 | 0.36928203 | 0.56616088 |
| ZNF350 | 600.966543 | 0.015039347 | 0.21230968 | 0.07083684 | 0.94352761 | 0.97315146 |
| ZNF615 | 518.923308 | 0.204760776 | 0.16718463 | 1.22475836 | 0.22066628 | 0.40657154 |
| ZNF614 | 414.640852 | 0.079518484 | 0.10941618 | 0.72675249 | 0.46737759 | 0.65445768 |
| ZNF432 | 797.898336 | -0.03490995 | 0.17517755 | -0.1992832 | 0.84204119 | 0.91603243 |
| AC011468.5 | 86.5746479 | 0.373569232 | 0.28488138 | 1.311315 | 0.18975136 | 0.36824723 |
| ZNF841 | 600.069313 | -0.224368443 | 0.20716466 | -1.083044 | 0.2787889 | 0.47442329 |
| ZNF616 | 416.463793 | 0.180970006 | 0.15642015 | 1.15694815 | 0.24729354 | 0.4388674 |
| AC011468.3 | 4.3149413 | -0.638808742 | 0.56542124 | -1.1297926 | 0.25856362 | 0.4513652 |
| ZNF836 | 344.46578 | -0.33645715 | 0.16357978 | -2.0568382 | 0.03970179 | 0.12311104 |
| AC010320.1 | 4.27308408 | -0.516811631 | 0.67969818 | -0.7603546 | 0.44704266 | 0.63728592 |
| PPP2R1A | 18373.2728 | -0.362866823 | 0.13171271 | -2.7549872 | 0.00586944 | 0.03032469 |
| AC010320.2 | 2834.28055 | -0.445780473 | 0.14011162 | -3.1816095 | 0.00146459 | 0.01060546 |
| ZNF766 | 729.071918 | -0.186528499 | 0.17701469 | -1.0537459 | 0.29199927 | 0.48820826 |
| AC010320.3 | 10.295053 | -0.077999019 | 0.41853425 | -0.1863623 | 0.85216063 | 0.92251225 |
| ZNF480 | 798.966981 | -0.159345103 | 0.25318598 | -0.6293599 | 0.52911347 | 0.70368783 |
| AC010320.4 | 43.9870613 | -0.489796324 | 0.27001956 | -1.813929 | 0.06968865 | 0.18367289 |
| ZNF610 | 196.253989 | 0.112223748 | 0.28638065 | 0.39186917 | 0.69515488 | 0.82446857 |
| ZNF880 | 402.291649 | 0.223121019 | 0.21086726 | 1.05811122 | 0.29000474 | 0.48615433 |
| ZNF528 | 802.104624 | 0.24025938 | 0.2191745 | 1.09620132 | 0.27299068 | 0.46784716 |
| ZNF534 | 4.33688861 | 0.535878669 | 0.94651563 | 0.56615934 | 0.57128546 | 0.73630487 |
| ZNF578 | 43.6350552 | 0.483363596 | 0.45537237 | 1.06146888 | 0.28847687 | 0.48465437 |

| | | | | | | |
|---|---|---|---|---|---|---|
| AC022150.3 | 4.98842625 | 2.103893805 | 0.6591145 | 3.19200047 | 0.00141291 | 0.01032663 |
| ZNF808 | 557.828685 | 0.1037538 | 0.17299118 | 0.59976353 | 0.54866384 | 0.71927743 |
| ZNF701 | 354.933392 | -0.063489456 | 0.16761181 | -0.3787887 | 0.70484479 | 0.83144403 |
| AC022150.1 | 33.9731608 | 0.332521891 | 0.36599159 | 0.90855063 | 0.36358738 | 0.56082017 |
| ZNF83 | 4249.9907 | 0.171768037 | 0.20717718 | 0.82908762 | 0.40705483 | 0.60168362 |
| ZNF611 | 541.513115 | -0.043271223 | 0.33404379 | -0.1295376 | 0.89693229 | 0.94725878 |
| ZNF600 | 580.937355 | -0.654907197 | 0.27475977 | -2.3835629 | 0.01714595 | 0.06705306 |
| ZNF28 | 490.454193 | 0.09696765 | 0.10869899 | 0.89207497 | 0.37235275 | 0.56878538 |
| ZNF468 | 2114.10496 | -0.210739461 | 0.23173604 | -0.9093944 | 0.36314198 | 0.56029027 |
| ZNF320 | 1463.19237 | -0.206882192 | 0.2824233 | -0.7325252 | 0.46384806 | 0.65155396 |
| AC010487.3 | 215.56095 | 0.689426108 | 0.31180497 | 2.21108122 | 0.02703021 | 0.09356612 |
| ZNF888 | 644.882467 | -0.27705187 | 0.32408344 | -0.8548782 | 0.39261858 | 0.58819298 |
| AC010487.2 | 16.4346242 | 0.37802025 | 0.42467933 | 0.89013103 | 0.37339553 | 0.56961935 |
| ZNF816-ZNF | 197.002761 | 0.182641887 | 0.31564632 | 0.57862828 | 0.56284002 | 0.73016634 |
| ZNF816 | 300.138549 | 0.003797717 | 0.23026602 | 0.01649274 | 0.9868413 | 0.99400325 |
| AC010328.1 | 3.8826263 | 0.676054845 | 0.80263546 | 0.84229377 | 0.39962354 | 0.59436394 |
| ERVV-1 | 22.4841712 | -2.580280952 | 1.04419524 | -2.4710714 | 0.01347089 | 0.05619799 |
| ERVV-2 | 10.7134475 | -2.419369834 | 1.00766284 | -2.4009716 | 0.01635161 | 0.06482513 |
| ZNF160 | 2289.74777 | -0.533121352 | 0.26167082 | -2.0373741 | 0.04161256 | 0.12750678 |
| ZNF415 | 407.714137 | 0.11331871 | 0.21992932 | 0.51525059 | 0.60637792 | 0.76308872 |
| ZNF347 | 509.395718 | -0.008831334 | 0.14968012 | -0.0590014 | 0.95295101 | 0.97725695 |
| ZNF665 | 139.23674 | 0.070159299 | 0.34551896 | 0.20305485 | 0.83909216 | 0.91463543 |
| AC092070.1 | 9.91001732 | -0.614113299 | 0.53057457 | -1.1574495 | 0.24708874 | 0.43860643 |
| ZNF677 | 467.602761 | -0.000282975 | 0.18153002 | -0.0015588 | 0.99875623 | 0.99943844 |
| AC092070.3 | 30.8327156 | 0.567925527 | 0.33543463 | 1.69310343 | 0.09043578 | 0.22142561 |
| VN1R2 | 0.84781283 | 0.748290611 | 1.86258261 | 0.40174895 | 0.68786879 | NA |
| VN1R4 | 0 | NA | NA | NA | NA | NA |
| AC092070.4 | 0.08695548 | -0.642905668 | 3.30558948 | -0.1944905 | 0.84579185 | NA |
| BIRC8 | 0 | NA | NA | NA | NA | NA |
| ZNF845 | 315.763007 | -0.190067831 | 0.14664587 | -1.2961008 | 0.19494077 | 0.37497907 |
| ZNF525 | 390.765374 | -0.470924494 | 0.17348305 | -2.7145274 | 0.00663704 | 0.03321669 |
| ZNF765 | 432.639011 | -0.124372623 | 0.1494044 | -0.8324562 | 0.40515148 | 0.59987898 |
| ZNF761 | 735.673904 | -0.204405968 | 0.1912216 | -1.0689481 | 0.28509304 | 0.48111504 |
| ZNF813 | 175.819816 | -0.057446899 | 0.16501868 | -0.3481236 | 0.72774736 | 0.84611931 |
| ZNF331 | 1345.1871 | 0.682747868 | 0.44330566 | 1.54012893 | 0.12352893 | 0.27505344 |
| DPRX | 0.58016444 | -2.443332344 | 2.69078733 | -0.9080362 | 0.36385909 | NA |
| AC011453.1 | 0.03392908 | 0.380740593 | 3.29738709 | 0.11546736 | 0.90807469 | NA |
| AC008753.3 | 0 | NA | NA | NA | NA | NA |
| AC008753.2 | 0 | NA | NA | NA | NA | NA |
| NLRP12 | 13.7994725 | -0.243292935 | 0.47084809 | -0.5167122 | 0.60535711 | 0.76218921 |
| AC008440.1 | 4.13399461 | 0.63018094 | 0.71957212 | 0.87577176 | 0.38115413 | 0.5772144 |
| AC008440.3 | 2.10627702 | 0.553787772 | 0.76635876 | 0.72262209 | 0.46991209 | 0.65657722 |
| MYADM | 3238.02515 | 0.398346935 | 0.49866339 | 0.79882932 | NA | NA |
| AC008440.2 | 1094.35899 | 0.185419314 | 0.48920728 | 0.37901994 | NA | NA |
| PRKCG | 3.37399635 | -0.482481292 | 0.8138385 | -0.5928465 | 0.55328389 | 0.72292619 |
| CACNG7 | 2.22413018 | -3.682697665 | 2.20286272 | -1.6717781 | 0.09456809 | 0.22870867 |
| CACNG8 | 10.7831696 | -0.855600276 | 1.11007554 | -0.7707586 | 0.44085003 | 0.63214873 |
| CACNG6 | 2.23727124 | 0.956668404 | 1.25476161 | 0.76243041 | 0.44580316 | 0.63620681 |
| VSTM1 | 7.66129435 | -1.601301721 | 0.83390186 | -1.920252 | 0.05482607 | 0.15542502 |
| TARM1 | 0.54824982 | -1.077555328 | 1.87394903 | -0.5750185 | 0.5652788 | NA |
| OSCAR | 130.496471 | -0.894711012 | 0.60453079 | -1.480009 | 0.13887084 | 0.29821044 |
| NDUFA3 | 2767.39165 | -0.026336882 | 0.13019677 | -0.2022852 | 0.83969375 | 0.91484357 |
| TFPT | 842.978968 | -0.633051652 | 0.16220894 | -3.9026927 | 9.51E-05 | 0.00121715 |
| PRPF31 | 3699.75909 | -0.163639932 | 0.11435334 | -1.4310027 | 0.15242945 | 0.3176266 |
| AC245052.4 | 1691.53755 | 0.165703303 | 0.12502206 | 1.32539253 | 0.18504099 | 0.36240667 |
| CNOT3 | 3330.31545 | -0.232741839 | 0.14193984 | -1.6397218 | 0.10106303 | 0.2392309 |
| LENG1 | 944.232634 | -0.44103321 | 0.13979265 | -3.1549099 | 0.00160548 | 0.01138994 |
| TMC4 | 4955.51111 | -0.221516843 | 0.26778733 | -0.8272118 | 0.40811702 | 0.60264172 |
| MBOAT7 | 5037.70197 | -0.844567997 | 0.17555658 | -4.8108023 | 1.50E-06 | 4.55E-05 |
| TSEN34 | 2551.71968 | -0.305457475 | 0.12491079 | -2.445405 | 0.01446896 | 0.05930678 |
| AC245052.1 | 0.62395047 | 0.245263593 | 1.81447599 | 0.13517048 | 0.89247709 | NA |
| RPS9 | 24333.6757 | -0.546610673 | 0.14781595 | -3.6979139 | 0.00021738 | 0.00233589 |
| LILRB3 | 462.103649 | -1.321721809 | 0.47931456 | -2.7575248 | 0.00582408 | 0.03012573 |
| AC245052.7 | 0.62589349 | -2.532823687 | 1.62394398 | -1.5596743 | 0.11883687 | NA |
| LILRA6 | 216.016086 | -1.146957891 | 0.4907667 | -2.3370736 | 0.01943536 | 0.0733415 |
| LILRB5 | 704.652122 | -0.963940094 | 0.35548056 | -2.7116535 | 0.00669485 | 0.03344256 |
| LILRB2 | 601.061697 | -0.825477723 | 0.50649731 | -1.6297771 | 0.10314861 | 0.24268637 |
| LILRA5 | 226.255294 | -2.42745191 | 0.56259024 | -4.3147778 | 1.60E-05 | 0.00029711 |
| LILRA4 | 20.4628589 | 0.006339345 | 1.03114753 | 0.00614785 | 0.99509475 | 0.99770729 |
| LAIR1 | 846.290384 | -0.805849118 | 0.49016962 | -1.6440209 | 0.10017188 | 0.23776169 |
| TTYH1 | 107.107398 | -0.4219615 | 0.35667228 | -1.183051 | 0.23678894 | 0.42601653 |
| AC245884.1 | 15.9808331 | -0.56323973 | 0.50680579 | -1.1113522 | 0.26641678 | 0.46053165 |
| AC245884.1( | 4.24056691 | -0.151535175 | 0.59583709 | -0.2543232 | 0.7992459 | 0.89108294 |
| AC245884.8 | 135.417964 | -0.321649021 | 0.42568674 | -0.7556003 | 0.44988889 | 0.63993921 |
| LENG8-AS1 | 303.972098 | 0.118592719 | 0.21932543 | 0.54071576 | 0.58870351 | 0.74924212 |
| LENG8 | 29968.7398 | -0.008703785 | 0.12704562 | -0.0685091 | 0.94538035 | 0.97392975 |
| LENG9 | 1270.07546 | -0.797729363 | 0.12362317 | -6.4529115 | 1.10E-10 | 1.56E-08 |
| CDC42EP5 | 327.944725 | -1.080792866 | 0.36107855 | -2.9932347 | 0.00276037 | 0.01724906 |
| LAIR2 | 3.3624995 | -0.350349406 | 1.16728686 | -0.3001399 | 0.76407042 | 0.86937077 |
| KIR3DX1 | 10.1247458 | -3.247318241 | 1.034281 | -3.1396866 | 0.00169129 | 0.01184574 |
| LILRA2 | 98.0597391 | -0.383605377 | 0.52367026 | -0.7325323 | 0.46384374 | 0.65155396 |
| AC245036.5 | 0.81495308 | 0.777232145 | 1.35965086 | 0.57164098 | 0.56756523 | NA |
| LILRA1 | 66.6228423 | -0.18958293 | 0.7711365 | -0.2458487 | 0.80579933 | 0.89480475 |
| LILRB1 | 357.319973 | -0.433845456 | 0.48585132 | -0.8929593 | 0.37187896 | 0.56852271 |
| LILRB1-AS1 | 29.2388191 | -0.172765825 | 0.54798835 | -0.3152728 | 0.75255453 | 0.86206078 |
| LILRB4 | 388.293579 | -0.957261949 | 0.58299818 | -1.6419639 | 0.10059749 | 0.23850898 |
| KIR3DL3 | 0.19114494 | -1.003574173 | 3.30558948 | -0.3035992 | 0.7614333 | NA |
| KIR2DL3 | 4.09133881 | -0.96672786 | 0.88666759 | -1.0902934 | 0.2755839 | 0.47090048 |
| KIR2DL1 | 5.99279491 | -1.693386918 | 1.09188021 | -1.5508907 | 0.12092787 | 0.27130065 |
| KIR2DL4 | 19.9939192 | -1.796204002 | 0.80287849 | -2.2372053 | 0.02527293 | 0.08907338 |
| KIR3DL1 | 9.85488664 | -1.928561328 | 1.07220879 | -1.7986808 | 0.07206919 | 0.1882596 |
| KIR3DL2 | 7.65180953 | -0.980349936 | 1.09577752 | -0.8946615 | 0.37096808 | 0.56778851 |
| FCAR | 55.1834122 | -1.909807608 | 0.62401483 | -3.0605164 | 0.00220956 | 0.01458108 |
| AC245128.3 | 29.3885534 | -2.48781958 | 0.7046012 | -3.5308194 | 0.00041427 | 0.003924 |
| NCR1 | 7.47294324 | -0.282942496 | 0.84321483 | -0.3355521 | 0.73720866 | 0.8524157 |
| NLRP7 | 3.47133511 | -0.365243275 | 1.01153183 | -0.3610794 | 0.71804012 | 0.83985952 |
| NLRP2 | 320.475451 | -1.37205275 | 0.41571575 | -3.3004589 | 0.00096527 | 0.0076393 |
| AC011476.3 | 177.334323 | 0.429052822 | 0.29047566 | 1.47706978 | 0.13965693 | 0.29948448 |
| GP6 | 31.6946534 | -0.284210422 | 0.49568918 | -0.5733642 | 0.56639814 | 0.73284279 |
| RDH13 | 1689.69141 | -0.01200195 | 0.18723038 | -0.0641026 | 0.94888855 | 0.97533389 |
| EPS8L1 | 2997.79307 | -0.629467921 | 0.55649924 | -1.1311209 | 0.25800421 | 0.45051502 |
| PPP1R12C | 8108.12757 | -0.422608365 | 0.17576912 | -2.404338 | 0.01620179 | 0.06437456 |
| TNNT1 | 13.6469098 | 2.502881494 | 1.13723843 | 2.20084147 | 0.02774725 | 0.09547168 |
| TNNI3 | 7.01268666 | -1.13802357 | 0.86917924 | -1.3093083 | 0.19042995 | 0.36915664 |
| AC010327.1 | 111.991718 | 0.113909316 | 0.37136328 | 0.30673284 | 0.75904673 | 0.86658803 |
| DNAAF3 | 256.730122 | 0.053085256 | 0.33620922 | 0.15789352 | 0.87454071 | 0.93476707 |
| AC010327.3 | 155.019834 | 0.078619263 | 0.35144607 | 0.22370221 | 0.82298903 | 0.90546648 |
| SYT5 | 4.20765282 | -0.543563297 | 1.51642685 | -0.3584501 | 0.72000653 | 0.84070901 |
| PTPRH | 31.1092753 | -1.845996864 | 0.6464122 | -2.8557581 | 0.00429342 | 0.02402116 |
| AC010327.4 | 6.92770085 | -0.96472506 | 0.58772649 | -1.6414524 | 0.10070354 | 0.23865843 |
| TMEM86B | 514.686687 | -0.287401793 | 0.19051698 | -1.5085363 | 0.13141731 | 0.28718609 |
| PPP6R1 | 6323.75035 | -0.424625189 | 0.20826137 | -2.0389051 | 0.04145949 | 0.12718554 |
| HSPBP1 | 1938.63188 | -0.142411231 | 0.15870413 | -0.8973379 | 0.36953865 | 0.56635675 |

| | | | | | | | |
|---|---|---|---|---|---|---|---|
| BRSK1 | 257.133865 | | -0.86225886 | 0.420736 | -2.0494059 | 0.04042244 | 0.12474426 |
| AC020922.2 | 43.2605854 | | -0.664881152 | 0.40755723 | -1.6313811 | 0.10280994 | 0.24201911 |
| TMEM150B | 279.858214 | | 1.068152008 | 0.43619363 | 2.44880238 | 0.01433321 | 0.0589075 |
| AC020922.4 | 0.81902425 | | -0.473503816 | 1.40703184 | -0.3365267 | 0.7364737 | NA |
| KMT5C | 1075.83206 | | -0.400569101 | 0.2689712 | -1.4892639 | 0.13641789 | 0.29462241 |
| COX6B2 | 62.9383291 | | -1.654621774 | 0.46039975 | -3.5938807 | 0.00032579 | 0.00323824 |
| AC020922.1 | 22.9561917 | | -1.174190609 | 0.61230564 | -1.9176544 | 0.05515485 | 0.15603833 |
| FAM71E2 | 3.55678606 | | -1.574312244 | 0.86769751 | -1.8143561 | 0.06962292 | 0.18354059 |
| IL11 | 219.802537 | | -1.98386665 | 0.77898787 | -2.5467234 | 0.01087396 | 0.04795161 |
| TMEM190 | 20.332783 | | -1.271222408 | 0.5198961 | -2.445147 | 0.01447931 | 0.05933455 |
| AC020922.3 | 15.8292855 | | -1.243534678 | 0.51376262 | -2.420446 | 0.01550148 | 0.06232262 |
| TMEM238 | 108.177387 | | -0.929156256 | 0.32123658 | -2.892436 | 0.00382267 | 0.02202881 |
| RPL28 | 19018.6223 | | -0.942568821 | 0.16198708 | -5.81879 | 5.93E-09 | 4.43E-07 |
| UBE2S | 573.554007 | | -0.702452657 | 0.23014122 | -3.0522678 | 0.00227119 | 0.01489587 |
| SHISA7 | 2.62388032 | | -0.386368023 | 1.30442566 | -0.2961978 | 0.76707902 | 0.87152635 |
| ISOC2 | 4300.91602 | | 0.806503906 | 0.19996385 | 4.03324861 | 5.50E-05 | 0.00080057 |
| C19orf85 | 0.45646629 | | 1.404392505 | 2.11508813 | 0.6639877 | 0.50669818 | NA |
| AC008735.3 | 0.87774955 | | 1.581958661 | 1.59519191 | 0.99170429 | 0.3213418 | NA |
| ZNF628 | 454.024194 | | -0.563738761 | 0.21021356 | -2.6817431 | 0.00732397 | 0.03582508 |
| NAT14 | 984.723604 | | -0.522618732 | 0.3623899 | -1.4421449 | 0.14926151 | 0.31340282 |
| SSC5D | 502.641854 | | -0.48630966 | 0.45860197 | -1.0604177 | 0.2889546 | 0.4851505 |
| SBK2 | 1.6334436 | | -0.747072488 | 1.11834653 | -0.6680152 | 0.50412389 | 0.68375733 |
| SBK3 | 8.87659807 | | -2.626722016 | 0.70555904 | -3.7228947 | 0.00019695 | 0.00215382 |
| ZNF579 | 923.183174 | | -0.767544353 | 0.27443946 | -2.7967711 | 0.00516161 | 0.02754559 |
| FIZ1 | 1225.62428 | | 0.057461055 | 0.14590494 | 0.39382529 | 0.69371003 | 0.82376229 |
| ZNF524 | 754.287938 | | -0.128455546 | 0.11620727 | -1.1054002 | 0.26898621 | 0.46314344 |
| ZNF865 | 1166.29217 | | -0.542968858 | 0.19344816 | -2.8067925 | 0.00500375 | 0.02689539 |
| AC008735.4 | 39.9856265 | | 0.220410754 | 0.24086521 | 0.91507925 | 0.36015003 | 0.55780314 |
| ZNF784 | 522.262664 | | 0.240522339 | 0.2031537 | 1.18394268 | 0.23643575 | 0.42572966 |
| ZNF580 | 2232.33784 | | -0.216072836 | 0.15412363 | -1.4019449 | 0.16093171 | 0.32946417 |
| ZNF581 | 1064.42649 | | -0.47897725 | 0.11038342 | -4.3392139 | 1.43E-05 | 0.00027285 |
| CCDC106 | 2151.29438 | | 0.031329429 | 0.10215519 | 0.30668465 | 0.75908341 | 0.86658803 |
| U2AF2 | 9463.10043 | | -0.313069898 | 0.15540529 | -2.0145382 | 0.04395306 | 0.13282248 |
| AC008735.2 | 654.787495 | | -0.596873419 | 0.1557861 | -3.8313651 | 0.00012743 | 0.00153136 |
| EPN1 | 10122.3532 | | 0.012591838 | 0.09980048 | 0.12617011 | 0.89959727 | 0.94902072 |
| AC010525.1 | 15.5689736 | | -1.232884485 | 0.44781233 | -2.7531276 | 0.00590289 | 0.03042 |
| NLRP9 | 106.836408 | | 1.175184722 | 0.31040359 | 3.78598948 | 0.0001531 | 0.00178164 |
| RFPL4A | 4.91438483 | | 3.072629 | 1.16347448 | 2.64090795 | 0.00826842 | 0.039019 |
| RFPL4AL1 | 2.18865806 | | 4.2039118 | 1.27291566 | 3.30258472 | 0.00095798 | 0.0075953 |
| NLRP11 | 16.2298153 | | 1.213086337 | 0.6118591 | 1.98262366 | 0.04740948 | 0.14005709 |
| NLRP4 | 2.52201811 | | 1.249806517 | 2.30473745 | 0.54227718 | 0.58762758 | 0.74858643 |
| NLRP13 | 0 | NA | | NA | NA | NA | NA |
| NLRP8 | 0 | NA | | NA | NA | NA | NA |
| NLRP5 | 0.07830766 | | -0.642905668 | 3.30558948 | -0.1944905 | 0.84579185 | NA |
| LINC01864 | 0.70180933 | | -0.71893621 | 2.23361983 | -0.3218704 | 0.74755084 | NA |
| ZNF787 | 1341.0701 | | -0.729731564 | 0.21600536 | -3.378303 | 0.00072935 | 0.00610544 |
| ZNF444 | 2397.05895 | | -0.175410332 | 0.15905838 | -1.1028047 | 0.27011198 | 0.4644531 |
| GALP | 0 | NA | | NA | NA | NA | NA |
| ZSCAN5B | 3.67214212 | | -1.036917269 | 0.67322927 | -1.5402142 | 0.12350816 | 0.27503041 |
| ZSCAN5C | 3.78131757 | | -0.874812864 | 0.77523567 | -1.1284476 | 0.25913091 | 0.45187688 |
| ZSCAN5A | 344.060895 | | 0.000716009 | 0.25303761 | 0.00282965 | 0.99774227 | 0.99872698 |
| EDDM13 | 16.3186269 | | 0.1711186 | 0.32149514 | 0.53225875 | 0.5945468 | 0.75365087 |
| AC006116.8 | 49.2632711 | | 0.539975945 | 0.2379438 | 2.26934237 | 0.02324751 | 0.08378255 |
| AC006116.9 | 3.26121075 | | 0.70638501 | 0.74541368 | 0.9476416 | 0.34331194 | 0.54097017 |
| ZNF582 | 207.263799 | | 0.103119013 | 0.23473846 | 0.43929322 | 0.66044908 | 0.80018573 |
| AC006116.4 | 4.64396535 | | -0.187576899 | 0.60737524 | -0.308832 | 0.75744934 | 0.86583463 |
| ZNF582-AS1 | 323.157057 | | 0.25165727 | 0.1823843 | 1.37981873 | 0.16764246 | 0.33885726 |
| ZNF583 | 347.431571 | | 0.24525 | 0.18778288 | 1.30602961 | 0.19154249 | 0.37072406 |
| ZNF667 | 444.808854 | | -0.180157054 | 0.23845983 | -0.7555027 | 0.4499474 | 0.63993921 |
| ZNF667-AS1 | 1115.7008 | | -0.09370557 | 0.14666935 | -0.6388899 | 0.52289455 | 0.69862624 |
| ZNF471 | 581.668336 | | 0.12577144 | 0.16237936 | 0.77455311 | 0.43860378 | 0.63025154 |
| AC005498.1 | 3.47738178 | | -0.32978329 | 0.74966886 | -0.4399053 | 0.66000571 | 0.79999826 |
| AC005498.2 | 17.5579627 | | 1.046307591 | 0.38059811 | 2.74911396 | 0.00597566 | 0.03068708 |
| ZFP28 | 495.382902 | | 0.209732562 | 0.20105832 | 1.04314291 | 0.29688211 | 0.49371137 |
| AC005498.3 | 80.9792952 | | 0.324203707 | 0.35962368 | 0.90150823 | 0.36731816 | 0.56423253 |
| ZNF470 | 466.624544 | | -0.20865384 | 0.21118736 | -0.9880034 | 0.32315096 | 0.52085294 |
| ZNF71 | 404.462919 | | -0.489647635 | 0.2308648 | -2.1209281 | 0.03392785 | 0.11017978 |
| SMIM17 | 15.9449525 | | 1.470514088 | 0.48138141 | 3.05477955 | 0.00225226 | 0.01480244 |
| ZNF835 | 91.7222326 | | 1.174135143 | 0.26414216 | 4.44508793 | 8.79E-06 | 0.00018402 |
| AC006115.1 | 4.29838451 | | -0.012356844 | 0.77737355 | -0.0158956 | 0.98731765 | 0.99421723 |
| AC007228.2 | 51.2627948 | | -0.559238479 | 0.38437001 | -1.4549483 | 0.14568358 | 0.30825717 |
| AC006115.3 | 3.31730804 | | 0.585161746 | 0.75495362 | 0.77509629 | 0.43828277 | 0.63008807 |
| ZIM2-AS1 | 7.7132654 | | 0.499329372 | 0.75160149 | 0.66435389 | 0.50646384 | 0.68566368 |
| ZIM2 | 5.06360364 | | 1.000334241 | 0.59819427 | 1.67225647 | 0.09447376 | 0.22851523 |
| PEG3 | 424.962668 | | 0.873484286 | 0.17592902 | 4.9649813 | 6.87E-07 | 2.36E-05 |
| MIMT1 | 2.09746013 | | -0.228795203 | 0.99010728 | -0.2310812 | 0.8172517 | 0.9017286 |
| USP29 | 0 | NA | | NA | NA | NA | NA |
| ZIM3 | 0.08757577 | | 0.547239394 | 3.29377202 | 0.16614368 | 0.86804388 | NA |
| DUXA | 0.43324851 | | 0.607574816 | 2.1891089 | 0.27754435 | 0.78136215 | NA |
| ZNF264 | 1377.50075 | | -0.030366651 | 0.14024575 | -0.2165246 | 0.82857888 | 0.90880809 |
| AURKC | 157.45254 | | -0.565626161 | 0.26407777 | -2.1418924 | 0.03220214 | 0.10611978 |
| ZNF805 | 51.6666087 | | 0.002258979 | 0.27597884 | 0.00818533 | 0.99346912 | 0.99694729 |
| AC005261.3 | 317.462341 | | -0.982222826 | 0.18913915 | -5.1931229 | 2.07E-07 | 8.77E-06 |
| ZNF460-AS1 | 67.9659411 | | -0.002074331 | 0.45626495 | -0.0045463 | 0.99637257 | 0.99823988 |
| ZNF460 | 71.7756669 | | 0.042381415 | 0.30026074 | 0.14114871 | 0.88775247 | 0.94201397 |
| AC005261.1 | 2737.89192 | | -0.494989107 | 0.18267324 | -2.7096968 | 0.00673447 | 0.03358957 |
| ZNF543 | 140.863093 | | 0.457475304 | 0.22710182 | 2.01440612 | 0.04396691 | 0.13282248 |
| AC005261.2 | 15.6739736 | | 1.666025296 | 0.41458368 | 4.01855009 | 5.86E-05 | 0.00083596 |
| ZNF304 | 610.805061 | | 0.281482589 | 0.16664656 | 1.68909932 | 0.0912004 | 0.22270159 |
| ZNF547 | 395.696133 | | -0.213048025 | 0.13744108 | -1.5501044 | 0.12111646 | 0.27160821 |
| TRAPPC2B | 345.962191 | | -0.659896916 | 0.22458481 | -2.9382972 | 0.00330021 | 0.01974731 |
| AC003002.1 | 244.529916 | | -0.222621545 | 0.18469736 | -1.2053315 | 0.22807536 | 0.41516082 |
| ZNF548 | 687.014124 | | 0.20768515 | 0.1419504 | 1.46308251 | 0.1434448 | 0.30499973 |
| AC003002.3 | 107.269886 | | 0.783610218 | 0.27293072 | 2.87109569 | 0.00409052 | 0.02313252 |
| AC003002.2 | 170.7951 | | 0.533315815 | 0.44324167 | 1.20321677 | 0.22889245 | 0.41621113 |
| ZNF17 | 291.064711 | | 0.155730185 | 0.19324026 | 0.80588892 | 0.4203069 | 0.61305896 |
| ZNF749 | 259.151863 | | 0.336989366 | 0.43222795 | 0.77965658 | 0.43559304 | 0.62768243 |
| AC004076.1 | 81.3454223 | | 0.7302674 | 0.24975891 | 2.92388923 | 0.00345688 | 0.02039782 |
| AC004076.2 | 37.2151227 | | -0.896733014 | 0.37619728 | -2.3836776 | 0.01714061 | 0.06704211 |
| VN1R1 | 122.133722 | | 0.441462439 | 0.2290988 | 1.92695218 | 0.05398559 | 0.15380294 |
| ZNF772 | 342.407575 | | 0.662157164 | 0.24158854 | 2.74084676 | 0.00612811 | 0.03131748 |
| AC003005.2 | 1.9015274 | | 2.148653414 | 1.48213973 | 1.44969693 | 0.14714305 | 0.31038815 |
| ZNF419 | 639.057057 | | -0.151375901 | 0.13895773 | -1.0893665 | 0.2759923 | 0.47133715 |
| AC003005.1 | 73.2716569 | | 0.051441741 | 0.27511341 | 0.18698376 | 0.85167336 | 0.92224978 |
| ZNF773 | 280.888114 | | -0.086660187 | 0.17994834 | -0.4815837 | 0.63010172 | 0.7801882 |
| ZNF549 | 413.856014 | | 0.262938005 | 0.17285469 | 1.52115053 | 0.12822207 | 0.28235623 |
| ZNF550 | 520.08443 | | 0.242129005 | 0.13239084 | 1.82889543 | 0.06741527 | 0.17928003 |
| ZNF416 | 174.278615 | | 0.266820458 | 0.18064705 | 1.47702636 | 0.13966857 | 0.29948448 |
| ZIK1 | 225.453445 | | 0.401501981 | 0.2210583 | 1.81627188 | 0.06932868 | 0.18289288 |
| ZNF530 | 118.917108 | | 0.244772946 | 0.25302951 | 0.96736915 | 0.33335952 | 0.53132008 |
| ZNF134 | 772.954753 | | 0.219346376 | 0.13361429 | 1.64163858 | 0.10066493 | 0.23858837 |
| AC003682.1 | 0.45519172 | | 1.248707418 | 2.10725499 | 0.59257538 | 0.55346536 | NA |
| ZNF211 | 878.423363 | | 0.099881115 | 0.17580863 | 0.56812406 | 0.56995073 | 0.73523142 |

| | | | | | | |
|---|---|---|---|---|---|---|
| ZSCAN4 | 8.07198037 | -0.408802807 | 0.72229072 | -0.565981 | 0.57140671 | 0.73635956 |
| ZNF551 | 281.788472 | 0.43070858 | 0.21156973 | 2.03577596 | 0.04177286 | 0.12790876 |
| AC003006.1 | 94.250303 | 1.163990049 | 0.34666258 | 3.3577032 | 0.00078593 | 0.0064806 |
| ZNF154 | 385.104213 | 0.192704971 | 0.28919204 | 0.66635641 | 0.50518332 | 0.68452998 |
| ZNF671 | 633.601342 | 0.28538613 | 0.16901903 | 1.68848523 | 0.09131812 | 0.22288144 |
| ZNF776 | 922.736232 | 0.462805119 | 0.14020604 | 3.30089293 | 0.00096378 | 0.00762978 |
| ZNF586 | 392.788666 | 0.280781026 | 0.18603206 | 1.50931528 | 0.13121823 | 0.2868699 |
| ZNF552 | 555.013283 | 0.491493809 | 0.15608366 | 3.14891262 | 0.00163879 | 0.01158578 |
| AC010522.1 | 58.2819823 | 0.633330981 | 0.27884277 | 2.27128352 | 0.02312982 | 0.08356396 |
| ZNF587B | 710.696589 | -0.109595895 | 0.28736172 | -0.3813865 | 0.70291644 | 0.83001552 |
| ZNF814 | 1504.42905 | -0.000743005 | 0.12984617 | -0.0057222 | 0.99543437 | 0.99781824 |
| ZNF587 | 1076.7302 | 0.061302979 | 0.18337463 | 0.33430457 | 0.73814974 | 0.85298022 |
| AC010326.2 | 277.139722 | 0.513195944 | 0.10656292 | 4.81589592 | 1.47E-06 | 4.44E-05 |
| ZNF417 | 340.508757 | 0.319735175 | 0.17761049 | 1.80020431 | 0.07182838 | 0.18785227 |
| ZNF418 | 345.891332 | 0.37383201 | 0.21198563 | 1.7634781 | 0.07781989 | 0.19861696 |
| ZNF256 | 231.430067 | -0.394193877 | 0.1741495 | -2.2635373 | 0.02360259 | 0.08464576 |
| C19orf18 | 75.4077428 | 0.331065894 | 0.26264835 | 1.26049106 | 0.20749227 | 0.39040822 |
| ZNF606 | 795.449291 | 0.463223935 | 0.18228964 | 2.54114244 | 0.01104909 | 0.04851224 |
| ZSCAN1 | 115.139255 | 0.333462684 | 0.31063159 | 1.07349894 | 0.28304731 | 0.4789542 |
| ZNF135 | 3735.52498 | -0.193194596 | 0.20313926 | -0.9510451 | 0.34158148 | 0.53956532 |
| ZSCAN18 | 5853.57347 | -0.178614083 | 0.20370834 | -0.8768128 | 0.38058831 | 0.57668871 |
| ZNF329 | 957.448071 | 0.1118073 | 0.1406263 | 0.79506677 | 0.42657467 | 0.61888061 |
| ZNF274 | 1881.17009 | -0.241165472 | 0.14183066 | -1.7003762 | 0.08906019 | 0.21892876 |
| ZNF544 | 1297.81046 | 0.109977871 | 0.10746886 | 1.02334639 | 0.3061441 | 0.50305458 |
| AC020915.5 | 1967.7002 | -0.373256729 | 0.17742336 | -2.1037632 | 0.03539911 | 0.11353376 |
| AC020915.3 | 221.55627 | -0.015241336 | 0.19686797 | -0.0774191 | 0.93829016 | 0.97060692 |
| ZNF8 | 780.486971 | -0.097528494 | 0.13400273 | -0.7278097 | 0.46673006 | 0.65393293 |
| AC020915.1 | 1770.29509 | -0.462533908 | 0.18807606 | -2.4592918 | 0.01392114 | 0.05763864 |
| ERVK3-1 | 1994.80696 | -0.387738781 | 0.19046157 | -2.0357849 | 0.04177196 | 0.12790876 |
| AC010642.2 | 1734.60701 | -0.401598737 | 0.21892932 | -1.8343762 | 0.06659817 | 0.1777884 |
| ZSCAN22 | 292.394805 | -0.408282069 | 0.24990706 | -1.6337356 | 0.10231438 | 0.24120317 |
| A1BG | 163.382858 | -0.502823705 | 0.34046996 | -1.4768519 | 0.13971535 | 0.29956043 |
| A1BG-AS1 | 299.422985 | -0.441621196 | 0.29786073 | -1.4826432 | 0.13816923 | 0.29713933 |
| ZNF497 | 413.45831 | 0.058905335 | 0.23335777 | 0.25242499 | 0.80071258 | 0.89197325 |
| AC012313.2 | 93.3730267 | 0.59755463 | 0.26260132 | 2.27552029 | 0.02287474 | 0.08280121 |
| AC012313.6 | 57.2841828 | 0.796745263 | 0.23277365 | 3.42283275 | 0.00061972 | 0.00534579 |
| ZNF837 | 208.650406 | -0.083615606 | 0.1739677 | -0.4806387 | 0.63077333 | 0.78054869 |
| RPS5 | 15563.8427 | -0.493560976 | 0.20992357 | -2.351146 | 0.01871569 | 0.07153684 |
| RNF225 | 0.12088456 | 0.02007176 | 3.29738704 | 0.00608717 | 0.99514317 | NA |
| LINC02560 | 4.45621188 | 0.924873843 | 0.67330384 | 1.37363519 | 0.16955497 | 0.34136238 |
| ZNF584 | 463.521968 | 0.082381266 | 0.13567363 | 0.60720175 | 0.54371703 | 0.71596016 |
| AC012313.1 | 940.748042 | -0.183370893 | 0.27931532 | -0.6565014 | 0.51150159 | 0.68965625 |
| AC012313.5 | 135.019294 | 0.454170223 | 0.21905791 | 2.07328837 | 0.03814545 | 0.11959064 |
| ZNF132 | 340.500784 | 0.464321934 | 0.11414314 | 4.06789181 | 4.74E-05 | 0.00071242 |
| AC012313.8 | 32.035795 | 0.140050454 | 0.34619406 | 0.4045432 | 0.68581332 | 0.81835792 |
| ZNF324B | 297.660978 | 0.088976859 | 0.2287481 | 0.3889731 | 0.69729604 | 0.82600497 |
| ZNF324 | 541.877315 | 0.049166411 | 0.20853711 | 0.23576816 | 0.81361258 | 0.89931524 |
| ZNF446 | 1211.23645 | 0.085190202 | 0.12656898 | 0.67307329 | 0.50090066 | 0.68184512 |
| AC012313.7 | 27.9861008 | 0.382081213 | 0.27304177 | 1.39935077 | 0.16170782 | 0.33042381 |
| SLC27A5 | 1109.91425 | 0.166423424 | 0.1261202 | 1.31956197 | 0.18698131 | 0.36493946 |
| AC012313.9 | 24.8120797 | 1.410355481 | 0.40222316 | 3.50640047 | 0.00045421 | 0.00421004 |
| ZBTB45 | 743.328455 | -0.253608664 | 0.0965389 | -2.62701 | 0.00861388 | 0.04023168 |
| TRIM28 | 11463.58 | -0.515458708 | 0.16892256 | -3.0514498 | 0.00227739 | 0.01492045 |
| CHMP2A | 5619.45231 | -0.124452172 | 0.1192367 | -1.0437405 | 0.29660546 | 0.49341406 |
| UBE2M | 3060.2866 | -0.664034231 | 0.14629112 | -4.5391289 | 5.65E-06 | 0.00012941 |
| MZF1-AS1 | 615.948216 | 0.338165619 | 0.18518644 | 1.82608196 | 0.06783792 | 0.18004722 |
| MZF1 | 2625.31398 | -0.098451749 | 0.13378339 | -0.7359041 | 0.46178904 | 0.65014385 |
| AC016629.2 | 24.3967067 | 0.001120165 | 0.3356004 | 0.00333779 | 0.99733683 | 0.9986244 |
| SRY | 0.46945845 | -0.393244958 | 2.27744958 | -0.172669 | 0.86291161 | NA |
| RPS4Y1 | 6259.56111 | 0.18287018 | 1.92276797 | 0.09510777 | 0.92422922 | 0.96312684 |
| AC006157.1 | 13.9252101 | 0.527124465 | 1.27046191 | 0.41490773 | 0.67820945 | 0.81313471 |
| ZFY | 401.005506 | 0.646550479 | 1.79648301 | 0.35989791 | 0.71892348 | 0.84015122 |
| ZFY-AS1 | 122.399943 | 1.277632703 | 1.8670474 | 0.68430652 | 0.49378163 | 0.67645185 |
| LINC00278 | 57.9636875 | 1.181180923 | 1.5367078 | 0.7686438 | 0.4421048 | 0.63331779 |
| TGIF2LY | 0 | NA | NA | NA | NA | NA |
| AC010737.1 | 5.27599309 | 0.240280477 | 1.39834883 | 0.17183157 | 0.86356994 | 0.92899031 |
| AC010722.1 | 0.41141449 | 1.217900152 | 3.24620513 | 0.37517658 | 0.70752914 | NA |
| PCDH11Y | 9.59027747 | -0.334614013 | 1.21334256 | -0.2757787 | 0.78271806 | 0.88107472 |
| TSPY2 | 0 | NA | NA | NA | NA | NA |
| LINC00280 | 0 | NA | NA | NA | NA | NA |
| TTTY1B | 0 | NA | NA | NA | NA | NA |
| TTTY2B | 0 | NA | NA | NA | NA | NA |
| TTTY21B | 0 | NA | NA | NA | NA | NA |
| TTTY7 | 0 | NA | NA | NA | NA | NA |
| TTTY8B | 0 | NA | NA | NA | NA | NA |
| AMELY | 2.1119025 | -0.053589129 | 1.23304805 | -0.0434607 | 0.96533429 | 0.9836495 |
| TBL1Y | 38.1316945 | 1.625235495 | 1.64053751 | 0.99067256 | 0.32184549 | 0.51955663 |
| TTTY16 | 0.08695548 | -0.642905668 | 3.30558948 | -0.1944905 | 0.84579185 | NA |
| TTTY12 | 0 | NA | NA | NA | NA | NA |
| LINC00279 | 0 | NA | NA | NA | NA | NA |
| TTTY18 | 0 | NA | NA | NA | NA | NA |
| TTTY19 | 0 | NA | NA | NA | NA | NA |
| TTTY11 | 0 | NA | NA | NA | NA | NA |
| TTTY20 | 0 | NA | NA | NA | NA | NA |
| TSPY4 | 0 | NA | NA | NA | NA | NA |
| TSPY8 | 0 | NA | NA | NA | NA | NA |
| TSPY3 | 0 | NA | NA | NA | NA | NA |
| FAM197Y5 | 0 | 0 | 0 | 0 | 1 | NA |
| TSPY1 | 0.17361919 | -1.003574173 | 3.30558948 | -0.3035992 | 0.7614333 | NA |
| TSPY10 | 0 | NA | NA | NA | NA | NA |
| TTTY8 | 0 | NA | NA | NA | NA | NA |
| TTTY7B | 0 | NA | NA | NA | NA | NA |
| TTTY21 | 0 | NA | NA | NA | NA | NA |
| TTTY2 | 0 | NA | NA | NA | NA | NA |
| TTTY1 | 0 | NA | NA | NA | NA | NA |
| TTTY22 | 0 | NA | NA | NA | NA | NA |
| AC010891.1 | 0 | NA | NA | NA | NA | NA |
| TTTY23 | 0 | NA | NA | NA | NA | NA |
| TTTY15 | 132.486313 | 0.345033225 | 1.62087286 | 0.21286878 | 0.83142932 | 0.91040231 |
| USP9Y | 1349.73326 | 0.770952743 | 1.7651923 | 0.43675284 | 0.66229061 | 0.80131238 |
| DDX3Y | 2488.81442 | 0.082433704 | 1.98524927 | 0.0415231 | 0.96687888 | 0.98431054 |
| UTY | 586.332454 | 0.605602367 | 1.82686176 | 0.33149874 | 0.74026779 | 0.85420604 |
| TMSB4Y | 88.146419 | 0.68445839 | 1.86538217 | 0.36692663 | 0.71367375 | 0.83745188 |
| VCY | 0 | NA | NA | NA | NA | NA |
| VCY1B | 0 | NA | NA | NA | NA | NA |
| AC010723.1 | 0.08695548 | -0.642905668 | 3.30558948 | -0.1944905 | 0.84579185 | NA |
| NLGN4Y | 272.446793 | -0.395689526 | 2.0318238 | -0.194746 | 0.84559181 | 0.91858498 |
| NLGN4Y-AS1 | 2.77710428 | -0.794797558 | 1.32285461 | -0.60082 | 0.54795986 | 0.71876822 |
| FAM41AY1 | 0 | NA | NA | NA | NA | NA |
| FAM224B | 4.86563906 | 0.790846528 | 1.32678299 | 0.59606321 | 0.551133 | 0.72140262 |
| CDY2B | 0 | NA | NA | NA | NA | NA |
| CDY2A | 0 | NA | NA | NA | NA | NA |
| FAM224A | 0 | NA | NA | NA | NA | NA |

| | | | | | | |
|---|---|---|---|---|---|---|
| FAM41AY2 | 0 | NA | NA | NA | NA | NA |
| AC022486.1 | 0 | NA | NA | NA | NA | NA |
| HSFY1 | 4.96864909 | 5.458785155 | 1.38517402 | 3.94086597 | 8.12E-05 | 0.0010753 |
| TTTY9B | 8.59949472 | 4.618952765 | 1.3533017 | 3.41309906 | 0.00064229 | 0.00550262 |
| HSFY2 | 1.32596058 | -0.454079032 | 1.68241113 | -0.2698978 | 0.78723889 | 0.88414299 |
| TTTY14 | 280.395281 | 0.731236757 | 1.77834215 | 0.41119014 | 0.68093312 | 0.81511673 |
| AC010889.1 | 78.5774238 | 0.734101153 | 1.52731139 | 0.4806493 | 0.63076577 | 0.78054869 |
| KDM5D | 2024.91408 | 0.638643372 | 1.92800673 | 0.33124541 | 0.74045913 | 0.85426378 |
| TTTY10 | 10.5621791 | 0.306866701 | 1.11232765 | 0.27587798 | 0.78264179 | 0.88106052 |
| EIF1AY | 833.868554 | 0.348290974 | 1.87468356 | 0.18578654 | 0.85261216 | 0.92285671 |
| RPS4Y2 | 0.57874814 | 0.246295587 | 1.82386569 | 0.13504042 | 0.89257992 | NA |
| AC007876.1 | 0 | NA | NA | NA | NA | NA |
| PRORY | 0.17391096 | -1.003574173 | 3.30558948 | -0.3035992 | 0.7614333 | NA |
| RBMY1B | 1.23360586 | 4.228143416 | 2.96789561 | 1.42462673 | 0.15426513 | 0.32016145 |
| RBMY1A1 | 1.23360586 | 4.228143416 | 2.96789561 | 1.42462673 | 0.15426513 | 0.32016145 |
| TTTY13 | 0 | NA | NA | NA | NA | NA |
| RBMY1D | 0.03392908 | 0.380740593 | 3.29738709 | 0.11546736 | 0.90807469 | NA |
| RBMY1E | 0.33023047 | 2.310906901 | 2.77720112 | 0.83209922 | 0.40535294 | NA |
| PRY2 | 0 | NA | NA | NA | NA | NA |
| AC007359.1 | 0 | NA | NA | NA | NA | NA |
| TTTY6B | 0 | NA | NA | NA | NA | NA |
| RBMY1F | 0 | NA | NA | NA | NA | NA |
| TTTY5 | 0 | NA | NA | NA | NA | NA |
| RBMY1J | 0 | NA | NA | NA | NA | NA |
| AC008175.1 | 0 | NA | NA | NA | NA | NA |
| TTTY6 | 0 | NA | NA | NA | NA | NA |
| PRY | 0 | NA | NA | NA | NA | NA |
| TTTY17A | 0 | NA | NA | NA | NA | NA |
| TTTY4 | 0 | NA | NA | NA | NA | NA |
| BPY2 | 0 | NA | NA | NA | NA | NA |
| DAZ1 | 0 | NA | NA | NA | NA | NA |
| DAZ2 | 0 | NA | NA | NA | NA | NA |
| TTTY3B | 0 | NA | NA | NA | NA | NA |
| CDY1B | 0 | NA | NA | NA | NA | NA |
| TTTY17B | 0 | NA | NA | NA | NA | NA |
| TTTY4B | 0.16783122 | 1.368031398 | 3.28033652 | 0.41703996 | 0.67664918 | NA |
| BPY2B | 0 | NA | NA | NA | NA | NA |
| DAZ3 | 0 | NA | NA | NA | NA | NA |
| DAZ4 | 0 | NA | NA | NA | NA | NA |
| BPY2C | 0.13390214 | 1.04981649 | 3.28477129 | 0.31960109 | 0.74927075 | NA |
| TTTY4C | 0.90826414 | -0.083944647 | 1.96494561 | -0.0427211 | 0.96592386 | NA |
| TTTY17C | 0 | NA | NA | NA | NA | NA |
| LINC00266-4 | 0 | NA | NA | NA | NA | NA |
| CDY1 | 0 | NA | NA | NA | NA | NA |
| TTTY3 | 0 | NA | NA | NA | NA | NA |
| CU104787.1 | 0.57813161 | 3.129191697 | 3.18961246 | 0.98105702 | 0.32656462 | NA |
| AP000542.3 | 0 | NA | NA | NA | NA | NA |
| AP000542.2 | 0 | NA | NA | NA | NA | NA |
| OR11H1 | 0 | NA | NA | NA | NA | NA |
| AP000534.1 | 0 | NA | NA | NA | NA | NA |
| AP000532.2 | 0 | NA | NA | NA | NA | NA |
| POTEH | 0 | NA | NA | NA | NA | NA |
| POTEH-AS1 | 0 | NA | NA | NA | NA | NA |
| AP000527.1 | 0.08757577 | 0.547239394 | 3.29377202 | 0.16614368 | 0.86804388 | NA |
| AC145543.1 | 0 | NA | NA | NA | NA | NA |
| AP000547.2 | 0.03392908 | 0.380740593 | 3.29738709 | 0.11546736 | 0.90807469 | NA |
| CCT8L2 | 0.95770952 | -2.188613114 | 1.50001437 | -1.4590614 | 0.1445482 | NA |
| AP000547.3 | 580.333913 | -0.341744068 | 0.45485404 | -0.7513269 | 0.45245595 | 0.64215627 |
| AP000365.1 | 0 | NA | NA | NA | NA | NA |
| LINC01665 | 4.98782096 | 0.058254509 | 0.87267543 | 0.06675392 | 0.94677761 | 0.97438741 |
| XKR3 | 1.05037305 | -2.374404039 | 1.8592415 | -1.2770821 | 0.20157326 | 0.3834234 |
| AC007064.2 | 0.81593223 | 3.111148283 | 1.9351527 | 1.60770169 | 0.10790053 | NA |
| IGKV1OR22-[illegible] | 0 | NA | NA | NA | NA | NA |
| IGKV2OR22-[illegible] | 0.18301898 | 0.18657164 | 3.29377195 | 0.05664376 | 0.95482897 | NA |
| IGKV2OR22-[illegible] | 0 | NA | NA | NA | NA | NA |
| IGKV3OR22-[illegible] | 0 | NA | NA | NA | NA | NA |
| IGKV1OR22-[illegible] | 0 | NA | NA | NA | NA | NA |
| GAB4 | 0.09989236 | -0.642905668 | 3.30558948 | -0.1944905 | 0.84579185 | NA |
| AC006946.2 | 35.8685522 | -0.174401394 | 0.43724271 | -0.3988663 | 0.6899917 | 0.82144836 |
| AC006946.3 | 16.2506422 | -0.367728356 | 0.42892364 | -0.8573283 | 0.3912635 | 0.58692863 |
| IL17RA | 1605.72518 | -0.613338795 | 0.15971558 | -3.8401939 | 0.00012294 | 0.0014867 |
| TMEM121B | 43.8692231 | -0.259155464 | 0.43981698 | -0.5892348 | 0.55570379 | 0.72497446 |
| LINC01664 | 1.98562585 | 1.776908313 | 0.92747242 | 1.91586108 | 0.05538278 | 0.15646496 |
| HDHD5 | 2437.4554 | -0.145615384 | 0.1503394 | -0.9685777 | 0.33275595 | 0.53071092 |
| HDHD5-AS1 | 35.2128902 | 0.400571744 | 0.32809377 | 1.22090627 | 0.22212151 | 0.40806355 |
| ADA2 | 1905.34942 | 0.862867541 | 0.41368913 | 2.08578731 | 0.03699789 | 0.11699658 |
| CECR3 | 3.94980314 | 1.223533592 | 0.59504684 | 2.0561971 | 0.03976352 | 0.12325897 |
| CECR2 | 300.571755 | 0.126938354 | 0.21672692 | 0.58570645 | 0.5580728 | 0.72665737 |
| SLC25A18 | 236.927896 | -0.558133396 | 0.50351039 | -1.1084844 | 0.26765268 | 0.46182041 |
| AC007666.1 | 61.5390403 | -0.940846426 | 0.58161508 | -1.6176445 | 0.10573924 | 0.24677792 |
| ATP6V1E1 | 9917.62645 | -0.306630544 | 0.22393274 | -1.3692975 | 0.1709063 | 0.34315983 |
| BCL2L13 | 4983.50863 | -0.192296044 | 0.32094822 | -0.5991497 | 0.54907303 | 0.71963498 |
| BID | 1033.27978 | -0.605088019 | 0.15712854 | -3.8509109 | 0.00011768 | 0.00144335 |
| LINC00528 | 39.8188115 | -0.669307064 | 0.51836057 | -1.2911998 | 0.19663441 | 0.37721878 |
| MICAL3 | 3973.11184 | 0.181507443 | 0.214676 | 0.84549481 | 0.39783463 | 0.59270617 |
| LINC01634 | 0.91576902 | 0.078882909 | 1.55667071 | 0.05067411 | 0.9595852 | NA |
| AC016027.1 | 268.954042 | 0.310028937 | 0.25052454 | 1.23751924 | 0.21589438 | 0.40049525 |
| PEX26 | 3865.48934 | -0.115974664 | 0.17372109 | -0.6675912 | 0.50439459 | 0.68392997 |
| TUBA8 | 165.896156 | -0.36360398 | 0.20714135 | -1.7553423 | 0.07920077 | 0.20107062 |
| AC008079.2 | 202.508509 | -0.276818138 | 0.194681 | -1.4219063 | 0.15505347 | 0.32116565 |
| USP18 | 204.927831 | 0.524038927 | 0.2391135 | 2.1915907 | 0.02840907 | 0.09715307 |
| AC008079.1 | 0.18710963 | -0.340597073 | 3.29738699 | -0.103293 | 0.91773041 | NA |
| LINC01660 | 0 | NA | NA | NA | NA | NA |
| AC011718.1 | 0 | NA | NA | NA | NA | NA |
| FAM230A | 0.08695548 | -0.642905668 | 3.30558948 | -0.1944905 | 0.84579185 | NA |
| AC023490.1 | 0 | NA | NA | NA | NA | NA |
| AC023490.3 | 14.9538772 | -2.730931982 | 1.01506752 | -2.6903944 | 0.00713676 | 0.03513749 |
| GGTLC3 | 980.239464 | -2.714432022 | 0.75130503 | -3.612956 | 0.00030273 | 0.00305624 |
| TMEM191B | 47.9890373 | -0.752647839 | 0.68180034 | -1.1039124 | 0.26963115 | 0.46392884 |
| RIMBP3 | 2.9690584 | 1.578454983 | 0.82557471 | 1.91194687 | 0.05588301 | 0.157406 |
| LINC01662 | 0.09635527 | 0.547239394 | 3.29377202 | 0.16614368 | 0.86804388 | NA |
| LINC01663 | 0 | NA | NA | NA | NA | NA |
| DGCR6 | 117.546552 | 1.386828424 | 1.00208001 | 1.38394979 | 0.16637383 | 0.33717168 |
| AC007326.4 | 21.7017465 | -0.550467804 | 0.51001294 | -1.0793212 | 0.28044455 | 0.47616526 |
| PRODH | 122.071781 | 1.252171654 | 1.04022153 | 1.2037548 | 0.22868437 | 0.41594749 |
| AC007326.2 | 0.12150485 | 1.04981648 | 3.28477129 | 0.31960109 | 0.74927075 | NA |
| DGCR9 | 96.7905001 | 1.457576153 | 0.37262846 | 3.91160714 | 9.17E-05 | 0.00118167 |
| DGCR2 | 6402.0237 | 0.025023939 | 0.10526852 | 0.23771531 | 0.81210191 | 0.89858708 |
| AC004471.1 | 16.3219529 | -0.971038262 | 0.39127027 | -2.4817583 | 0.01307359 | 0.05493158 |
| AC004471.2 | 8.52915145 | 0.149513959 | 0.4645899 | 0.32181922 | 0.74758965 | 0.85884766 |
| ESS2 | 940.570286 | -0.135366779 | 0.15780958 | -0.8577856 | 0.39101088 | 0.58667712 |
| TSSK2 | 58.4380732 | -0.030105624 | 0.30705116 | -0.0980476 | 0.92189451 | 0.96183415 |
| GSC2 | 0 | NA | NA | NA | NA | NA |

| | | | | | | |
|---|---|---|---|---|---|---|
| LINC01311 | 89.0794492 | -0.211049055 | 0.27796262 | -0.7592713 | 0.44769026 | 0.63783587 |
| SLC25A1 | 6409.62187 | 0.118392223 | 0.24641916 | 0.48045056 | 0.63090705 | 0.78062237 |
| CLTCL1 | 489.033588 | -0.246237541 | 0.21945156 | -1.1220587 | 0.26183747 | 0.45539205 |
| HIRA | 2878.67243 | 0.21049144 | 0.17449551 | 1.20628568 | 0.22770736 | 0.41471393 |
| C22orf39 | 3012.74691 | 0.497419126 | 0.11243591 | 4.42402354 | 9.69E-06 | 0.0001985 |
| MRPL40 | 1404.39126 | 0.358858381 | 0.136692 | 2.62530641 | 0.0086571 | 0.04039062 |
| AC000068.1 | 52.9850267 | 0.327402814 | 0.30615484 | 1.06940272 | 0.28488824 | 0.48101568 |
| UFD1 | 2819.98361 | -0.600255244 | 0.28036102 | -2.1410082 | 0.03227337 | 0.10622638 |
| AC000068.3 | 30.4777679 | 1.302560894 | 0.31763277 | 4.1008391 | 4.12E-05 | 0.00063556 |
| AC000068.2 | 739.540489 | -0.580511155 | 0.29435698 | -1.9721331 | 0.04859441 | 0.14276232 |
| CDC45 | 30.9011544 | -1.168986409 | 0.56050917 | -2.0855795 | 0.03701673 | 0.11700003 |
| CLDN5 | 1722.65574 | -1.649498261 | 0.42261158 | -3.9031071 | 9.50E-05 | 0.00121565 |
| LINC00895 | 0.3184437 | -1.669100907 | 2.89273496 | -0.5769975 | 0.56394113 | NA |
| AC000067.1 | 1.00895934 | -1.697321353 | 1.36638926 | -1.2421946 | 0.21416477 | 0.39823808 |
| 5-Sep | 688.09916 | 0.184677906 | 0.42907407 | 0.43041031 | 0.66689719 | 0.80500224 |
| AC000093.1 | 738.166514 | 0.155964106 | 0.42830108 | 0.36414596 | 0.71574902 | 0.83823283 |
| GP1BB | 228.35938 | -0.061211453 | 0.42900323 | -0.142683 | 0.88654057 | 0.94122211 |
| TBX1 | 59.8912955 | -1.334589983 | 0.56034142 | -2.3817443 | 0.01723086 | 0.06730525 |
| GNB1L | 275.834632 | -0.348188573 | 0.17507782 | -1.9887646 | 0.04672719 | 0.13860473 |
| RTL10 | 957.925509 | -0.013354474 | 0.14897722 | -0.089641 | 0.92857247 | 0.96531469 |
| TXNRD2 | 2885.7555 | 0.25431782 | 0.12002837 | 2.11881419 | 0.03410617 | 0.11050249 |
| COMT | 5479.56983 | -0.219791264 | 0.224715 | -0.978089 | 0.32803031 | 0.52568577 |
| ARVCF | 2673.49734 | 0.131477132 | 0.19073363 | 0.68932327 | 0.49061986 | 0.67369536 |
| TANGO2 | 2011.4998 | -0.167391677 | 0.16271849 | -1.0287195 | 0.30361152 | 0.5005347 |
| AC006547.1 | 746.298496 | -0.428572171 | 0.18451871 | -2.3226488 | 0.02019803 | 0.07541776 |
| AC006547.3 | 1023.94 | -0.219427416 | 0.16537278 | -1.3268654 | 0.18455322 | 0.36172651 |
| DGCR8 | 2102.88673 | -0.010641983 | 0.19331464 | -0.0550501 | 0.95609858 | 0.97900515 |
| AC006547.2 | 701.612831 | -0.428908436 | 0.1872795 | -2.2902049 | 0.02200944 | 0.08053128 |
| TRMT2A | 2591.80401 | -0.1190318 | 0.12149286 | -0.9797432 | 0.32721289 | 0.52502871 |
| RANBP1 | 2297.63703 | -0.275832217 | 0.11707882 | -2.3559531 | 0.01847525 | 0.07083968 |
| ZDHHC8 | 4235.22796 | 0.354925818 | 0.28372222 | 1.25096236 | 0.21094821 | 0.39435023 |
| CCDC188 | 84.5782281 | 0.57588737 | 0.42107018 | 1.3676755 | 0.17141367 | 0.34388234 |
| AC007663.2 | 13.753021 | 1.848967904 | 0.79917782 | 2.3135876 | 0.02069035 | 0.07681327 |
| LINC00896 | 12.5326548 | -1.018229251 | 0.38661329 | -2.6337151 | 0.00844563 | 0.03964952 |
| RTN4R | 79.3454353 | -0.195092017 | 0.21449406 | -0.9095451 | 0.36306247 | 0.56024093 |
| DGCR6L | 2052.84645 | 0.540929746 | 0.20745307 | 2.60748008 | 0.00912114 | 0.04207179 |
| AC007663.3 | 13.9580143 | -0.217430967 | 0.42828121 | -0.5076827 | 0.61167587 | 0.76749276 |
| AC007731.1 | 0 | NA | NA | NA | NA | NA |
| AC007731.3 | 0 | NA | NA | NA | NA | NA |
| USP41 | 1.52903285 | -1.212768019 | 2.40325866 | -0.5046348 | 0.61381534 | 0.76872676 |
| ZNF74 | 514.821494 | 0.331300854 | 0.1425959 | 2.32335472 | 0.0201601 | 0.07534325 |
| SCARF2 | 965.412999 | -0.343426485 | 0.334461 | -1.0268058 | 0.30451193 | 0.50146628 |
| AC007731.5 | 576.158367 | 0.421320183 | 0.15188504 | 2.77394122 | 0.00553817 | 0.02912094 |
| KLHL22 | 1626.81779 | 0.488403909 | 0.11404674 | 4.28248904 | 1.85E-05 | 0.00033426 |
| MED15 | 7346.67868 | -0.66756389 | 0.18163009 | -3.6754036 | 0.00023747 | 0.00251291 |
| AC007731.4 | 0.17423948 | 0.18657164 | 3.29377195 | 0.05664376 | 0.95482897 | NA |
| PI4KA | 5923.3503 | 0.166816021 | 0.11031111 | 1.51223224 | 0.13047479 | 0.2857658 |
| SERPIND1 | 4.44020343 | 0.231175684 | 0.98783876 | 0.23402168 | 0.81496815 | 0.900323 |
| SNAP29 | 1593.8761 | 0.005474256 | 0.16618225 | 0.03294128 | 0.97372141 | 0.98784418 |
| AC007308.1 | 219.881895 | -0.216438944 | 0.22420115 | -0.9653784 | 0.33435532 | 0.53248882 |
| CRKL | 3028.68996 | -0.186154414 | 0.18746565 | -0.9930054 | 0.3207073 | 0.51819065 |
| LINC01637 | 50.8279759 | 0.224274206 | 0.4965088 | 0.45170238 | 0.65148341 | 0.7952054 |
| AIFM3 | 159.025005 | 0.263279721 | 0.37458979 | 0.70284809 | 0.48215042 | 0.66664779 |
| AC002470.1 | 19.8532897 | 0.579192807 | 0.37155273 | 1.55884416 | 0.11903327 | 0.26835148 |
| LZTR1 | 4989.63975 | 0.069795273 | 0.13625837 | 0.5122274 | 0.60849187 | 0.7646381 |
| THAP7 | 1360.57418 | 0.071202772 | 0.14462894 | 0.49231345 | 0.62249778 | 0.77481557 |
| THAP7-AS1 | 278.801705 | 0.588940999 | 0.30232956 | 1.94801001 | 0.05141377 | 0.1486439 |
| P2RX6 | 81.36867 | 0.94404746 | 0.33523498 | 2.81607687 | 0.0048614 | 0.02630901 |
| SLC7A4 | 32.0869292 | 0.667892282 | 0.6447935 | 1.03582353 | 0.30028448 | 0.49704988 |
| AC002472.1 | 4.30844092 | 1.203375385 | 0.78579954 | 1.5314025 | 0.12566995 | 0.27842664 |
| LRRC74B | 10.3004183 | -2.677412532 | 1.2650107 | -2.1165137 | 0.03430114 | 0.11101393 |
| AP000550.1 | 6.05920309 | -0.650584392 | 0.80606474 | -0.8071118 | 0.41960205 | 0.61243738 |
| AP000550.2 | 0 | NA | NA | NA | NA | NA |
| GGT2 | 2412.96004 | -2.380349785 | 0.64740131 | -3.6767763 | 0.0002362 | 0.00250044 |
| AP000552.1 | 0 | NA | NA | NA | NA | NA |
| AP000552.3 | 0 | NA | NA | NA | NA | NA |
| LINC01651 | 0 | NA | NA | NA | NA | NA |
| RIMBP3B | 1.40790751 | 0.224385102 | 1.17628848 | 0.19075686 | 0.84871609 | 0.9204838 |
| HIC2 | 494.221972 | 0.26412301 | 0.20702995 | 1.27577197 | 0.20203614 | 0.38389487 |
| TMEM191C | 62.5096769 | 0.210922774 | 0.30883335 | 0.68296632 | 0.49462813 | 0.67694119 |
| RIMBP3C | 0.35475976 | 1.663862788 | 2.74343737 | 0.60648834 | 0.54419052 | NA |
| UBE2L3 | 4512.54317 | -0.330695167 | 0.17413201 | -1.8991061 | 0.05755052 | 0.16078052 |
| YDJC | 1155.04887 | -0.59880749 | 0.20164382 | -2.9696298 | 0.00298159 | 0.01824944 |
| CCDC116 | 32.1828 | -0.270186451 | 0.30779416 | -0.8778154 | 0.38004391 | 0.5759962 |
| AP000553.3 | 3.00079657 | -0.291159696 | 0.67486808 | -0.431432 | 0.66615428 | 0.80436322 |
| SDF2L1 | 1070.59124 | -0.622682257 | 0.34424575 | -1.8088306 | 0.07047732 | 0.18515822 |
| PPIL2 | 3852.59671 | 0.397130661 | 0.19808905 | 2.00480877 | 0.04498349 | 0.13478182 |
| YPEL1 | 1164.95718 | 0.160589036 | 0.20451616 | 0.78521442 | 0.43232786 | 0.62437624 |
| MAPK1 | 5729.62167 | 0.04038621 | 0.12695182 | 0.31812233 | 0.75039215 | 0.86064026 |
| PPM1F | 2799.01103 | -0.134676381 | 0.14547829 | -0.925749 | 0.35457645 | 0.55215431 |
| AC245452.1 | 2100.1651 | -0.202719055 | 0.25302319 | -0.8011876 | 0.42302302 | 0.61559059 |
| TOP3B | 1642.68008 | -0.373566611 | 0.28509694 | -1.3103143 | 0.19008952 | 0.36882206 |
| IGLVI-70 | 3.19273647 | -0.98068955 | 1.10623067 | -0.8865145 | 0.37534034 | 0.57129724 |
| IGLV4-69 | 70.7439899 | -0.764693109 | 1.13934864 | -0.6711669 | 0.5021142 | 0.68251297 |
| IGLVI-68 | 0.07830766 | -0.642905668 | 3.30558948 | -0.1944905 | 0.84579185 | NA |
| AC245452.5 | 0 | NA | NA | NA | NA | NA |
| IGLV10-54 | 30.6995576 | 0.934327697 | 1.86580841 | 0.50076293 | 0.61653797 | 0.7706427 |
| IGLV10-67 | 0.08695548 | -0.642905668 | 3.30558948 | -0.1944905 | 0.84579185 | NA |
| IGLVIV-66-1 | 0 | NA | NA | NA | NA | NA |
| IGLVV-66 | 0 | NA | NA | NA | NA | NA |
| IGLVIV-65 | 0 | NA | NA | NA | NA | NA |
| IGLVIV-64 | 0.09989236 | -0.642905668 | 3.30558948 | -0.1944905 | 0.84579185 | NA |
| IGLVI-63 | 0.27150681 | 2.033044511 | 3.2399442 | 0.62749368 | 0.53033568 | NA |
| IGLV1-62 | 0.08757577 | 0.547239394 | 3.29377202 | 0.16614368 | 0.86804388 | NA |
| IGLV8-61 | 30.4178741 | 1.517430123 | 0.80498632 | 1.8850384 | 0.05942465 | 0.16444833 |
| IGLV4-60 | 25.4036055 | 1.262885475 | 1.14966647 | 1.09847987 | 0.271995 | 0.4668386 |
| IGLVIV-59 | 0.11223674 | 0.02007176 | 3.29738704 | 0.00608717 | 0.99514317 | NA |
| IGLVV-58 | 0 | NA | NA | NA | NA | NA |
| IGLV6-57 | 122.232668 | -1.689071186 | 0.79156069 | -2.1338493 | 0.03285512 | 0.1076394 |
| IGLVI-56 | 0.08757577 | 0.547239394 | 3.29377202 | 0.16614368 | 0.86804388 | NA |
| IGLV11-55 | 0.17820003 | -1.003574173 | 3.30558948 | -0.3035992 | 0.7614333 | NA |
| IGLVIV-53 | 0 | NA | NA | NA | NA | NA |
| VPREB1 | 0.03392908 | 0.380740593 | 3.29738709 | 0.11546736 | 0.90807469 | NA |
| AC245060.6 | 221.031051 | 0.524092006 | 0.35539735 | 1.47466491 | 0.14030265 | 0.30042883 |
| AC245060.5 | 103.914774 | 0.662476362 | 0.40164668 | 1.64940083 | 0.09906554 | 0.23602605 |
| AC245060.2 | 39.8652689 | 0.576638783 | 0.40250146 | 1.43263775 | 0.15196139 | 0.31705213 |
| IGLV5-52 | 148.708129 | 0.3119775 | 0.28202422 | 1.10620818 | 0.26863642 | 0.46282128 |
| IGLV1-51 | 334.820975 | -0.26095611 | 0.73299796 | -0.3560121 | 0.72183154 | 0.84194522 |
| IGLV1-50 | 7.17220458 | -0.606198676 | 0.5485598 | -1.1050731 | 0.26912792 | 0.46327458 |
| IGLV9-49 | 19.4847576 | -0.026831619 | 1.06349798 | -0.0252296 | 0.97987183 | 0.99099448 |
| IGLV5-48 | 1.08192206 | -0.104206995 | 1.58850249 | -0.0656008 | 0.94769567 | 0.9747917 |
| IGLV1-47 | 157.144556 | -0.433528754 | 0.9808324 | -0.4420008 | 0.65848859 | 0.79946078 |
| IGLV7-46 | 87.2935431 | 0.268453925 | 1.37861771 | 0.19472688 | 0.84560677 | 0.91858498 |

| | | | | | | | |
|---|---|---|---|---|---|---|---|
| IGLV5-45 | 36.6624039 | | -0.210079179 | 1.25775234 | -0.1670275 | 0.86734844 | 0.93105 |
| IGLV1-44 | 272.7935 | | 1.338950833 | 0.99011051 | 1.35232464 | 0.17627149 | 0.35091945 |
| IGLV7-43 | 49.7781384 | | -0.24145305 | 1.10706453 | -0.2181021 | 0.8273496 | 0.90802884 |
| IGLVI-42 | 0 | NA | | NA | NA | NA | NA |
| IGLVVII-41-1 | 0 | NA | | NA | NA | NA | NA |
| IGLV1-41 | 3.67578179 | | 0.590586778 | 1.51860977 | 0.38889963 | 0.6973504 | 0.82603225 |
| IGLV1-40 | 495.01709 | | -0.982301623 | 0.64968546 | -1.5119649 | 0.13054279 | 0.28586725 |
| IGLVI-38 | 0.10721287 | | -0.642905668 | 3.30558948 | -0.1944905 | 0.84579185 | NA |
| IGLV5-37 | 3.26135698 | | -0.590450312 | 1.66704912 | -0.3541889 | 0.72319731 | 0.84309077 |
| IGLV1-36 | 19.1478134 | | -1.1514438 | 1.19186517 | -0.9660856 | 0.33400134 | 0.53211789 |
| IGLV7-35 | 0.34990677 | | 0.725448958 | 2.88852975 | 0.25114817 | 0.80169956 | NA |
| ZNF280B | 68.8925251 | | -0.489790216 | 0.36302906 | -1.3491763 | 0.17728033 | 0.35236922 |
| ZNF280A | 0.06785815 | | 0.795868262 | 3.28898349 | 0.24198001 | 0.80879565 | NA |
| PRAME | 164.075624 | | -1.132670125 | 0.55081513 | -2.0563526 | 0.03974854 | 0.12322702 |
| AC246793.1 | 0.09989236 | | -0.642905668 | 3.30558948 | -0.1944905 | 0.84579185 | NA |
| IGLV2-34 | 3.09909089 | | 0.183508238 | 0.97476707 | 0.18825855 | 0.85067397 | 0.92161226 |
| IGLV2-33 | 4.59094588 | | 1.007060631 | 1.85967009 | 0.5415265 | NA | NA |
| IGLV3-32 | 0.17515154 | | 1.45420457 | 3.2783972 | 0.44357181 | 0.65735219 | NA |
| IGLV3-31 | 0.60755803 | | -1.447352567 | 3.26883318 | -0.4427735 | 0.6579296 | NA |
| IGLV3-30 | 0.08757577 | | 0.547239394 | 3.29377202 | 0.16614368 | 0.86804388 | NA |
| GGTLC2 | 45.4246646 | | -1.165482798 | 0.46135348 | -2.5262252 | 0.01152955 | 0.05010454 |
| IGLV3-29 | 0.80070735 | | -1.346243862 | 1.66271087 | -0.809668 | 0.41813099 | NA |
| IGLV2-28 | 41.0874045 | | -1.797914684 | 1.12504913 | -1.5980766 | 0.11002593 | 0.25395044 |
| IGLV3-27 | 31.9167512 | | -0.664578904 | 1.59353363 | -0.4170473 | NA | NA |
| IGLV3-26 | 0.19271054 | | 1.506623853 | 3.27645372 | 0.4598337 | 0.64563559 | NA |
| IGLVVI-25-1 | 1.21476342 | | 0.41905056 | 1.18857995 | 0.35256405 | 0.72441529 | 0.84387647 |
| IGLV3-25 | 221.450591 | | -0.869404393 | 0.94025282 | -0.9246496 | 0.3551482 | 0.55265302 |
| AC244250.1 | 12.7519249 | | -0.446885441 | 0.62135563 | -0.7192104 | 0.47201128 | 0.65821986 |
| IGLV3-24 | 0.6107044 | | 1.033829647 | 3.23473075 | 0.319603 | 0.7492693 | NA |
| IGLV2-23 | 543.051583 | | -1.377886615 | 0.94346623 | -1.4604514 | 0.14416604 | 0.3059685 |
| IGLVVI-22-1 | 0 | NA | | NA | NA | NA | NA |
| IGLV3-22 | 0.98984791 | | -2.256757885 | 2.21227573 | -1.020107 | 0.30767773 | NA |
| IGLV3-21 | 340.440769 | | -0.600044822 | 1.21569398 | -0.4935821 | 0.62160132 | 0.77399315 |
| IGLVI-20 | 0 | NA | | NA | NA | NA | NA |
| IGLV3-19 | 427.954798 | | -0.767983646 | 0.99332326 | -0.7731457 | 0.43943613 | 0.63087603 |
| IGLV2-18 | 20.9630851 | | 0.697678273 | 0.97657945 | 0.71441016 | 0.47497359 | 0.66032548 |
| IGLV3-17 | 0.17656334 | | 0.18657164 | 3.29377195 | 0.05664376 | 0.95482897 | NA |
| IGLV3-16 | 6.93100056 | | -1.664338894 | 1.0181442 | -1.634679 | 0.10211637 | 0.24094565 |
| IGLV3-15 | 0 | NA | | NA | NA | NA | NA |
| IGLV2-14 | 1522.76829 | | -2.223293244 | 1.2028642 | -1.8483327 | 0.06455423 | 0.17391652 |
| IGLV3-13 | 0.8389282 | | 1.37991782 | 2.22344085 | 0.62062268 | 0.53484791 | NA |
| IGLV3-12 | 6.90925937 | | -0.818020936 | 1.22558528 | -0.6674533 | 0.50448263 | 0.68396433 |
| IGLV2-11 | 242.332651 | | 1.081901432 | 1.00028625 | 1.08159182 | 0.27943394 | 0.47503049 |
| IGLV3-10 | 118.932721 | | -1.140321925 | 1.2566634 | -0.9074203 | 0.36418456 | 0.56134831 |
| IGLV3-9 | 35.634558 | | -0.322986301 | 1.11455103 | -0.2897905 | 0.77197652 | 0.87421716 |
| IGLV2-8 | 342.446164 | | 0.376537004 | 0.95483978 | 0.39434575 | 0.69332579 | 0.82352701 |
| IGLV3-7 | 0 | NA | | NA | NA | NA | NA |
| IGLV3-6 | 1.14766324 | | -1.504875642 | 1.90171272 | -0.7913265 | 0.4287535 | 0.62111903 |
| IGLV2-5 | 10.2948245 | | -0.670100169 | 1.17693504 | -0.5693604 | 0.56911161 | 0.73454422 |
| IGLV3-4 | 0 | NA | | NA | NA | NA | NA |
| IGLV4-3 | 4.68863554 | | 0.335633434 | 1.28758833 | 0.26066828 | 0.79434833 | 0.88856221 |
| IGLV3-2 | 0.27507702 | | -1.364242679 | 3.30558948 | -0.4127078 | 0.67982068 | NA |
| IGLV3-1 | 374.971451 | | -0.775580264 | 0.95623433 | -0.8110776 | 0.4173211 | 0.6102566 |
| IGLL5 | 5248.09329 | | -1.807393669 | 1.15855539 | -1.5600408 | 0.11875024 | 0.26797223 |
| IGLJ1 | 1492.7891 | | -1.508757637 | 1.19576936 | -1.2617464 | 0.20704007 | 0.38975205 |
| IGLC1 | 5098.69102 | | -1.837383803 | 1.16505424 | -1.5770801 | 0.11477711 | 0.26193616 |
| IGLJ2 | 665.785119 | | -1.26904902 | 0.85452992 | -1.4850844 | 0.13752148 | 0.2961811 |
| IGLC2 | 3057.53958 | | -1.450052543 | 0.83918584 | -1.7279278 | 0.08400117 | 0.20968486 |
| IGLJ3 | 317.661356 | | -0.901419898 | 0.85304505 | -1.0567084 | 0.29064467 | 0.48670756 |
| IGLC3 | 1767.1437 | | -1.453113017 | 0.87276598 | -1.6649515 | 0.09592244 | 0.2309179 |
| IGLJ4 | 0 | NA | | NA | NA | NA | NA |
| IGLC4 | 0.15318056 | | -1.003574173 | 3.30558948 | -0.3035992 | 0.7614333 | NA |
| IGLJ5 | 0.21442573 | | -1.003574173 | 3.30558948 | -0.3035992 | 0.7614333 | NA |
| IGLC5 | 4.67057375 | | -2.656448769 | 1.4549681 | -1.825778 | 0.06788371 | 0.18010799 |
| IGLJ6 | 9.6673564 | | -1.163272255 | 1.22055183 | -0.9530708 | 0.34055422 | 0.53859897 |
| IGLC6 | 12.5990205 | | -2.568359481 | 1.07564716 | -2.3877342 | 0.0169526 | 0.06651392 |
| IGLJ7 | 22.0401786 | | -0.94327367 | 2.1502504 | -0.4386808 | 0.66089281 | 0.80046589 |
| IGLC7 | 88.163543 | | -1.680414596 | 1.61909751 | -1.0378712 | 0.29933005 | 0.49609609 |
| RSPH14 | 289.156873 | | 0.607159909 | 0.33735194 | 1.7997819 | 0.07189508 | 0.18795215 |
| GNAZ | 348.631327 | | -0.483019517 | 0.33687303 | -1.4338326 | 0.15162005 | 0.31666567 |
| RAB36 | 1304.44024 | | 0.3444525 | 0.30825311 | 1.11743398 | 0.26380881 | 0.45761337 |
| BCR | 6214.31582 | | -0.007034825 | 0.18676596 | -0.0376665 | 0.96995356 | 0.98614626 |
| LINC02556 | 0.81195717 | | -1.967625742 | 1.47898654 | -1.3303879 | 0.18339051 | NA |
| LINC01659 | 32.7302176 | | 1.288686948 | 0.51373808 | 2.50845131 | 0.01212617 | 0.05192709 |
| LINC01658 | 11.459901 | | 1.618428962 | 0.60387936 | 2.68005344 | 0.00736104 | 0.03595367 |
| LINC02557 | 0.47392499 | | 0.867512093 | 1.80825914 | 0.47974987 | 0.63140526 | NA |
| PCAT14 | 36.8872351 | | 1.584712864 | 0.77696862 | 2.03960988 | 0.0413892 | 0.12704736 |
| AP000345.1 | 4.73620136 | | 2.056559639 | 0.85388723 | 2.4084675 | 0.01601965 | 0.06390896 |
| IGLL1 | 13.6175669 | | 1.268993381 | 0.58907887 | 2.15419945 | 0.03122453 | 0.10390302 |
| AP000345.2 | 268.182518 | | 1.168951309 | 0.58318306 | 2.00443289 | 0.04502371 | 0.13487166 |
| DRICH1 | 145.608645 | | 1.279460518 | 0.53026024 | 2.41289166 | 0.01582652 | 0.06330129 |
| RGL4 | 1466.67831 | | 0.695780414 | 0.30743859 | 2.26315247 | 0.02362629 | 0.08470773 |
| ZNF70 | 218.184868 | | 0.45119281 | 0.14716814 | 3.06583209 | 0.00217065 | 0.01436749 |
| VPREB3 | 36.6002601 | | -0.338329259 | 0.58722223 | -0.576152 | 0.56451245 | 0.73150827 |
| C22orf15 | 578.957579 | | -0.186143696 | 0.23947586 | -0.7772963 | 0.43698398 | 0.62903366 |
| CHCHD10 | 4161.71907 | | -0.136591864 | 0.21199286 | -0.644323 | 0.51936603 | 0.69627261 |
| MMP11 | 290.18235 | | 0.353943637 | 0.28284502 | 1.25136952 | 0.21079969 | 0.39421232 |
| SMARCB1 | 3853.68835 | | -0.097704248 | 0.1330734 | -0.7342132 | 0.46281882 | 0.65098743 |
| DERL3 | 334.992533 | | -0.258239296 | 0.40473415 | -0.6380467 | 0.52344327 | 0.69882538 |
| AP000350.5 | 95.5535316 | | 1.608937532 | 0.37219189 | 4.32287096 | 1.54E-05 | 0.00029009 |
| SLC2A11 | 2513.29648 | | 1.394728874 | 0.25845404 | 5.39642897 | 6.80E-08 | 3.43E-06 |
| AP000350.4 | 11626.8382 | | -0.604685509 | 0.15552834 | -3.8879442 | 0.0001011 | 0.00127651 |
| AP000350.7 | 260.098876 | | 1.218262083 | 0.31499525 | 3.86755703 | 0.00010993 | 0.00136552 |
| MIF | 12356.5199 | | -0.67517927 | 0.14944529 | -4.5179027 | 6.25E-06 | 0.00014052 |
| MIF-AS1 | 12347.4171 | | -0.671638424 | 0.14982424 | -4.4828421 | 7.37E-06 | 0.00016052 |
| AP000350.6 | 37.6750435 | | 1.175691641 | 0.91761303 | 1.28124994 | 0.20010589 | 0.38149164 |
| GSTT2B | 297.435888 | | -0.038059109 | 0.49341538 | -0.077134 | 0.93851693 | 0.97068919 |
| DDTL | 2788.11332 | | 0.044381653 | 0.15919103 | 0.27879493 | 0.78040221 | 0.8797765 |
| AC253536.6 | 77.485762 | | -0.097642519 | 0.40445714 | -0.2414162 | 0.80923253 | 0.89691753 |
| DDT | 2512.32725 | | 0.067056716 | 0.18009855 | 0.37233346 | 0.70964459 | 0.83500637 |
| GSTT4 | 0 | NA | | NA | NA | NA | NA |
| CABIN1 | 4955.02119 | | -0.029169244 | 0.17898634 | -0.1629691 | 0.87054276 | 0.9326107 |
| SUSD2 | 6008.64603 | | 2.345811284 | 0.50298992 | 4.66373417 | 3.11E-06 | 8.04E-05 |
| GGT5 | 4537.15186 | | 0.246149656 | 0.22015387 | 1.11808009 | 0.26353279 | 0.45731506 |
| SPECC1L | 2727.33315 | | -0.189521843 | 0.17751929 | -1.0676126 | 0.28569528 | 0.48173058 |
| SPECC1L-AD( | 1643.25998 | | -0.259924434 | 0.30161289 | -0.8617816 | 0.38880772 | 0.58437501 |
| ADORA2A | 1253.62278 | | -0.570949382 | 0.33135015 | -1.7230998 | 0.08487048 | 0.21113533 |
| ADORA2A-AS | 885.928327 | | 1.03024567 | 0.34502605 | 2.9859939 | 0.00282658 | 0.01752162 |
| UPB1 | 2783.51159 | | 2.42496167 | 0.58731803 | 4.12887318 | 3.65E-05 | 0.00057529 |
| AP000355.1 | 32.0328626 | | 2.966287229 | 0.64010245 | 4.63408198 | 3.59E-06 | 9.03E-05 |
| GUCD1 | 9329.71801 | | -0.242593307 | 0.26246395 | -0.9242919 | 0.35533435 | 0.55283748 |
| SNRPD3 | 5319.15799 | | 0.194777985 | 0.16403029 | 1.18745136 | 0.23504963 | 0.42389952 |
| GGT1 | 16749.0348 | | 0.322489865 | 0.36135819 | 0.89243824 | 0.37215808 | 0.56863396 |

| | | | | | | | |
|---|---|---|---|---|---|---|---|
| LRRC75B | 2810.29281 | | 0.519765935 | 0.38793096 | 1.33984135 | 0.18029693 | 0.35621444 |
| AP000356.2 | 624.392228 | | -0.056776483 | 0.39110592 | -0.1451691 | 0.88457738 | 0.94023563 |
| AP000357.2 | 25.1312618 | | 0.338950549 | 0.7499789 | 0.45194678 | 0.65130732 | 0.795097 |
| PIWIL3 | 0.34832787 | | 0.328499903 | 2.80900699 | 0.11694521 | 0.90690347 | NA |
| SGSM1 | 383.108809 | | 0.536724956 | 0.18892493 | 2.84094294 | 0.00449804 | 0.0248665 |
| TMEM211 | 6.45484808 | | 0.119900776 | 0.54548453 | 0.21980601 | 0.82602224 | 0.90710059 |
| KIAA1671 | 2707.32643 | | -0.232084907 | 0.210318 | -1.1034952 | 0.26981216 | 0.46402835 |
| AL022323.3 | 1.19360397 | | 0.938432025 | 1.0668197 | 0.87965382 | 0.37904687 | 0.57504694 |
| AL022323.5 | 0.82493203 | | -1.38231192 | 2.33564702 | -0.5918325 | 0.55396273 | NA |
| AL022323.2 | 9.2722259 | | 0.645881536 | 0.44711936 | 1.44453942 | 0.1485873 | 0.31253885 |
| AL022323.4 | 11.5489827 | | -0.913532277 | 0.77357081 | -1.1809291 | 0.23763089 | 0.42700692 |
| AL022323.1 | 27.1215371 | | 0.200857748 | 0.39526985 | 0.50815348 | 0.61134571 | 0.76720061 |
| CRYBB3 | 78.4765685 | | 0.737970739 | 0.42822101 | 1.72334079 | 0.08482692 | 0.21104687 |
| Z99916.1 | 11.5397464 | | -0.843511234 | 0.65464184 | -1.288508 | 0.19756919 | 0.37834197 |
| CRYBB2 | 32.4877141 | | -0.821214266 | 0.38364255 | -2.1405714 | 0.03230862 | 0.10628952 |
| Z99916.3 | 0 | NA | | NA | NA | NA | NA |
| AL022332.1 | 0.86256179 | | 2.134750198 | 1.63376961 | 1.3066409 | 0.1913347 | NA |
| LRP5L | 1105.85265 | | -0.348582201 | 0.22350131 | -1.5596428 | 0.11884432 | 0.26810889 |
| AL022324.3 | 2.26210079 | | 2.37097601 | 0.96367494 | 2.46034831 | 0.01388022 | 0.05750534 |
| AL008721.1 | 1.08983716 | | 3.539235281 | 1.54667055 | 2.28829293 | 0.02212047 | 0.08079179 |
| IGLVIVOR22- | 0.93956989 | | 2.784639098 | 1.2634676 | 2.20396557 | 0.02752677 | NA |
| AL022329.1 | 0.72080354 | | -1.1350728 | 1.89541206 | -0.5988528 | 0.54927106 | NA |
| GRK3 | 534.951185 | | -0.133276403 | 0.19773322 | -0.6740213 | 0.50029777 | 0.68137859 |
| AL022329.2 | 0 | NA | | NA | NA | NA | NA |
| MYO18B | 22.9797801 | | -0.550614722 | 0.71386221 | -0.7713179 | 0.44051851 | 0.63181078 |
| AL022329.3 | 0 | NA | | NA | NA | NA | NA |
| Z98949.1 | 0.78559729 | | -0.506269246 | 1.63218951 | -0.310178 | 0.75642561 | NA |
| Z98949.3 | 0.09635527 | | 0.547239394 | 3.29377202 | 0.16614368 | 0.86804388 | NA |
| LINC02559 | 0 | NA | | NA | NA | NA | NA |
| AL022337.1 | 0 | NA | | NA | NA | NA | NA |
| SEZ6L | 2.51420302 | | 0.212301628 | 1.01691968 | 0.20876932 | 0.83462832 | 0.91223215 |
| AL080273.1 | 0 | NA | | NA | NA | NA | NA |
| ASPHD2 | 461.534261 | | -0.47672205 | 0.39027031 | -1.2215176 | 0.22189011 | 0.40780891 |
| HPS4 | 2398.87641 | | 0.177134992 | 0.16496284 | 1.07378725 | 0.28291804 | 0.47888496 |
| SRRD | 1817.01339 | | -0.19078943 | 0.18480167 | -1.032401 | 0.3018843 | 0.49870254 |
| TFIP11 | 2573.69911 | | -0.09128649 | 0.22447627 | -0.4066643 | 0.68425455 | 0.81757775 |
| Z95115.1 | 225.967972 | | 0.495255835 | 0.25245409 | 1.96176591 | 0.04978974 | 0.14524035 |
| TPST2 | 2499.7407 | | -0.010819273 | 0.14044899 | -0.0770335 | 0.93859692 | 0.97073384 |
| CRYBB1 | 17.9994928 | | -0.423111025 | 0.53576618 | -0.7897307 | 0.42968503 | 0.62180135 |
| CRYBA4 | 1.10425996 | | 0.091462878 | 1.20418306 | 0.0759543 | 0.93945546 | 0.97110795 |
| MIAT | 298.690632 | | -0.906173554 | 0.51499764 | -1.7595684 | 0.07848102 | 0.19976298 |
| Z99774.1 | 26.3670923 | | -0.802804708 | 0.75572137 | -1.0623025 | 0.28809837 | 0.48426636 |
| MIATNB | 127.057018 | | -0.180100507 | 0.22424551 | -0.8031399 | 0.42189391 | 0.61446342 |
| LINC01422 | 44.680365 | | 0.299366528 | 0.31197425 | 0.95958729 | 0.33726297 | 0.53527683 |
| Z97353.2 | 9.50894376 | | -0.136458686 | 0.42419411 | -0.3216893 | 0.74768812 | 0.85884766 |
| AL008638.3 | 0.18746814 | | 0.18657164 | 3.29377195 | 0.05664376 | 0.95482897 | NA |
| AL008638.5 | 0 | NA | | NA | NA | NA | NA |
| AL008638.4 | 0 | NA | | NA | NA | NA | NA |
| AL008638.2 | 0.08393208 | | -0.642905668 | 3.30558948 | -0.1944905 | 0.84579185 | NA |
| AL008638.6 | 0.72957619 | | 2.979574833 | 1.49343981 | 1.99510876 | 0.04603102 | NA |
| AL008638.1 | 0.08666371 | | -0.642905668 | 3.30558948 | -0.1944905 | 0.84579185 | NA |
| AL021153.1 | 0.25315362 | | -0.174096114 | 3.29377188 | -0.0528562 | 0.95784652 | NA |
| LINC01638 | 1.41052295 | | 1.30639131 | 1.23677844 | 1.05628565 | 0.29083773 | 0.48690769 |
| AL020994.3 | 0.24118557 | | 0.074529887 | 3.28898342 | 0.02266046 | 0.98192111 | NA |
| AL020994.1 | 2.39537141 | | 2.516633948 | 1.16698895 | 2.15651909 | 0.03104316 | 0.10341182 |
| LINC02554 | 12.535201 | | 3.645392877 | 0.91696045 | 3.97551811 | 7.02E-05 | 0.00095704 |
| AL020994.2 | 0.09989236 | | -0.642905668 | 3.30558948 | -0.1944905 | 0.84579185 | NA |
| AL049536.1 | 0 | NA | | NA | NA | NA | NA |
| AL050402.1 | 0.03392908 | | 0.380740593 | 3.29738709 | 0.11546736 | 0.90807469 | NA |
| AL133456.1 | 0 | NA | | NA | NA | NA | NA |
| AL121885.2 | 1.34364237 | | -1.793587102 | 1.74127498 | -1.0300424 | 0.30299009 | 0.49983414 |
| AL121885.3 | 0 | NA | | NA | NA | NA | NA |
| AL121885.1 | 0 | NA | | NA | NA | NA | NA |
| MN1 | 192.804225 | | 0.395385978 | 0.35624008 | 1.10988628 | 0.26704803 | 0.4611392 |
| PITPNB | 3127.9807 | | -0.576801085 | 0.16804792 | -3.4323607 | 0.00059835 | 0.00520743 |
| TTC28 | 1551.71536 | | 0.595725823 | 0.16168055 | 3.68458562 | 0.00022908 | 0.00243577 |
| CHEK2 | 588.327293 | | -0.62293591 | 0.19928841 | -3.125801 | 0.00177321 | 0.0123017 |
| HSCB | 644.416237 | | -0.266225385 | 0.18498961 | -1.439137 | 0.15011172 | 0.3145264 |
| CCDC117 | 1292.04975 | | -0.321089029 | 0.1613569 | -1.9899306 | 0.04659858 | 0.13834774 |
| XBP1 | 8447.89165 | | -1.082513676 | 0.19967324 | -5.4214259 | 5.91E-08 | 3.04E-06 |
| Z93930.2 | 78.4264511 | | 0.207660907 | 0.23842769 | 0.87095969 | 0.38377617 | 0.57948708 |
| ZNRF3 | 1695.2584 | | 0.244130181 | 0.1254215 | 1.94647788 | 0.05159736 | 0.14910931 |
| ZNRF3-AS1 | 8.16635748 | | -1.18353111 | 0.65742612 | -1.8002496 | 0.07182123 | 0.1878522 |
| C22orf31 | 4.90674408 | | 1.416400602 | 0.68007438 | 2.08271424 | 0.03727728 | 0.11758395 |
| KREMEN1 | 2192.54696 | | 0.35915197 | 0.31665294 | 1.13421327 | 0.2567051 | 0.44944698 |
| AL021393.1 | 0.61490306 | | 2.097524412 | 1.62641825 | 1.28965868 | 0.19716919 | NA |
| EMID1 | 819.014219 | | 0.402094157 | 0.19577648 | 2.05384303 | 0.03999089 | 0.12382171 |
| RHBDD3 | 1618.64166 | | -0.640036181 | 0.21539262 | -2.9714861 | 0.00296362 | 0.01816614 |
| AL031186.1 | 122.366145 | | -0.612325465 | 0.35866082 | -1.7072549 | 0.08777468 | 0.21648228 |
| EWSR1 | 12994.5572 | | -0.083528193 | 0.18063201 | -0.4624219 | 0.64377882 | 0.78950376 |
| GAS2L1 | 1664.73845 | | -0.143543966 | 0.23659961 | -0.6066957 | 0.54405287 | 0.71615206 |
| RASL10A | 77.4199768 | | 0.002597804 | 0.25652529 | 0.01012689 | 0.99192005 | 0.99613883 |
| AP1B1 | 7198.2636 | | 0.069158657 | 0.14564498 | 0.47484407 | 0.63489812 | 0.78341572 |
| RFPL1S | 15.3116271 | | 0.573649638 | 0.38273498 | 1.49881685 | 0.13392115 | 0.29089792 |
| RFPL1 | 10.8140401 | | -0.626954179 | 0.75142621 | -0.8343523 | 0.40408249 | 0.59897135 |
| AC000035.1 | 3.8646061 | | 1.027900501 | 1.07133605 | 0.95945666 | 0.33732875 | 0.535349 |
| NEFH | 111.246012 | | 1.044922017 | 0.39749684 | 2.62875554 | 0.00856979 | 0.04007808 |
| THOC5 | 2983.90671 | | 0.256587648 | 0.19430792 | 1.32052079 | 0.1866612 | 0.36450245 |
| NIPSNAP1 | 6195.08643 | | 0.962590793 | 0.37781212 | 2.54780286 | 0.01084037 | 0.04783477 |
| NF2 | 2361.17927 | | -0.236089505 | 0.24735408 | -0.9544597 | 0.339851 | 0.5380566 |
| AC004882.1 | 0.57090649 | | -1.297975994 | 2.21792693 | -0.5852204 | 0.55839956 | NA |
| AC004882.2 | 0.07830766 | | -0.642905668 | 3.30558948 | -0.1944905 | 0.84579185 | NA |
| CABP7 | 424.08365 | | -0.163808127 | 0.14446842 | -1.1338681 | 0.2568499 | 0.4495342 |
| ZMAT5 | 801.44419 | | -0.045807067 | 0.13700968 | -0.3343345 | 0.73812714 | 0.85298022 |
| UQCR10 | 3823.10421 | | -0.104932874 | 0.14809497 | -0.7085512 | 0.47860301 | 0.66355212 |
| ASCC2 | 2617.18464 | | 0.054698227 | 0.11369623 | 0.48109094 | 0.63045187 | 0.78040725 |
| MTMR3 | 3091.00541 | | -0.074133718 | 0.17476075 | -0.4242012 | 0.67141909 | 0.80813006 |
| AC003681.1 | 2602.27344 | | -0.084460119 | 0.20002957 | -0.4222382 | 0.67285119 | 0.80918894 |
| HORMAD2-A | 48.3251602 | | 0.836697396 | 0.28986277 | 2.88652939 | 0.00389516 | 0.02233396 |
| HORMAD2 | 12.4602521 | | 0.529422011 | 0.47029632 | 1.12572009 | 0.26028402 | 0.45334805 |
| AC002378.1 | 0.21421642 | | 0.68915655 | 3.07571484 | 0.22406386 | 0.82270761 | NA |
| LIF-AS1 | 28.8388053 | | -3.696464075 | 0.85516344 | -4.3225235 | 1.54E-05 | 0.00029034 |
| LIF | 1986.31053 | | -1.789399394 | 0.59066909 | -3.0294448 | 0.00245004 | 0.01574451 |
| AC004264.1 | 97.3053112 | | -1.152401245 | 0.60789928 | -1.8957108 | 0.05799829 | 0.16166839 |
| OSM | 104.379909 | | 0.050200319 | 0.71124324 | 0.07058108 | 0.94373116 | 0.97315146 |
| CASTOR1 | 865.440553 | | 0.676526182 | 0.35557319 | 1.9026355 | 0.05708812 | 0.15977646 |
| AC004997.1 | 2173.19972 | | 0.206831372 | 0.22920371 | 0.90239102 | 0.36684919 | 0.56387888 |
| TBC1D10A | 2564.5214 | | -0.134660951 | 0.18245117 | -0.7380657 | 0.46047453 | 0.64896122 |
| SF3A1 | 4750.16503 | | 0.299491184 | 0.1184855 | 2.52766104 | 0.01148252 | 0.04997424 |
| CCDC157 | 823.535265 | | -0.100575282 | 0.17269756 | -0.5823781 | 0.56031201 | 0.72842771 |
| RNF215 | 1644.33004 | | -0.300203364 | 0.14090382 | -2.1305552 | 0.03312581 | 0.10833158 |
| SEC14L2 | 973.824514 | | -0.527645616 | 0.21190682 | -2.4899889 | 0.01277471 | 0.0540114 |
| AC004832.3 | 2308.76217 | | -1.077170587 | 0.21985244 | -4.8995163 | 9.61E-07 | 3.14E-05 |

| | | | | | | |
|---|---|---|---|---|---|---|
| MTFP1 | 2063.29973 | -1.225066467 | 0.24543063 | -4.9914978 | 5.99E-07 | 2.12E-05 |
| AC004832.6 | 7.18726926 | -0.703535976 | 0.57862908 | -1.215867 | 0.22403561 | 0.41009433 |
| SEC14L3 | 1.03848096 | -0.918013094 | 1.41244797 | -0.6499447 | 0.51572793 | 0.69339702 |
| AC004832.1 | 13.0733603 | 0.29066068 | 0.47028726 | 0.61804923 | 0.53654289 | 0.71034421 |
| SEC14L4 | 46.9444964 | 0.113065444 | 0.471239 | 0.23993227 | 0.81038276 | 0.89747017 |
| SEC14L6 | 1190.9404 | -0.680405274 | 0.48155214 | -1.4129421 | 0.15767276 | 0.32479992 |
| GAL3ST1 | 1853.73223 | -0.055710312 | 0.64463569 | -0.0864214 | 0.93113144 | 0.96688001 |
| PES1 | 3753.70588 | -0.417656415 | 0.2158009 | -1.9353785 | 0.05294387 | 0.15163517 |
| TCN2 | 11932.2417 | 0.247392534 | 0.3565102 | 0.69392834 | 0.48772715 | 0.67150655 |
| SLC35E4 | 294.405634 | -0.625251643 | 0.20083651 | -3.1132369 | 0.00185047 | 0.01268773 |
| DUSP18 | 251.142266 | -0.247355125 | 0.12514908 | -1.9764838 | 0.04810001 | 0.14162543 |
| OSBP2 | 577.265171 | 0.261710275 | 0.16696199 | 1.56748417 | 0.11700156 | 0.26531355 |
| MORC2-AS1 | 643.565788 | -0.493977413 | 0.21493705 | -2.2982423 | 0.021548 | 0.07922767 |
| MORC2 | 1964.61916 | -0.332675751 | 0.13785405 | -2.4132462 | 0.01581114 | 0.06325894 |
| AC004542.2 | 1322.32318 | -0.095266388 | 0.1429562 | -0.6664027 | 0.50515377 | 0.68452748 |
| AC004542.1 | 1339.38369 | -0.001723489 | 0.17153714 | -0.0100473 | 0.99198353 | 0.99613883 |
| SMTN | 7563.80686 | -0.240469893 | 0.29653328 | -0.8109373 | 0.41740169 | 0.61027293 |
| AC005005.3 | 485.673015 | 0.02658813 | 0.34954004 | 0.07606605 | 0.93936655 | 0.97110795 |
| SELENOM | 3211.88196 | -0.813222797 | 0.26419854 | -3.0780745 | 0.00208343 | 0.01388777 |
| INPP5J | 2889.97869 | 1.17911684 | 0.42414573 | 2.77998043 | 0.00543622 | 0.02869357 |
| PLA2G3 | 36.9194558 | -0.141710944 | 0.83820282 | -0.1690652 | 0.86574534 | 0.93026791 |
| RNF185 | 3930.98491 | 0.181457936 | 0.21153194 | 0.85782761 | 0.39098766 | 0.58667712 |
| RNF185-AS1 | 786.700141 | 0.421196505 | 0.2478086 | 1.69968476 | 0.08919024 | 0.21915637 |
| LIMK2 | 3665.9344 | -0.868250911 | 0.17289437 | -5.0218575 | 5.12E-07 | 1.84E-05 |
| PIK3IP1 | 2852.63732 | -0.537468515 | 0.33822435 | -1.5890888 | 0.11204035 | 0.25733856 |
| PIK3IP1-AS1 | 28.3113811 | -1.020576469 | 0.33573566 | -3.0398214 | 0.00236718 | 0.01536591 |
| PATZ1 | 2690.89062 | 0.061533542 | 0.24438772 | 0.25178656 | 0.80120605 | 0.89225911 |
| LINC01521 | 50.8363675 | 1.328278793 | 0.33733883 | 3.93752121 | 8.23E-05 | 0.00108767 |
| DRG1 | 2094.00803 | -0.243478892 | 0.15732397 | -1.5476275 | 0.12171201 | 0.27250347 |
| EIF4ENIF1 | 2103.07982 | -0.235174898 | 0.15686608 | -1.4992081 | 0.13381964 | 0.29074928 |
| AL096701.3 | 387.526388 | -0.08364889 | 0.1934523 | -0.4324006 | 0.66545029 | 0.8037781 |
| SFI1 | 2227.62568 | -0.383626192 | 0.19639259 | -1.9533639 | 0.05077649 | 0.14722113 |
| PISD | 4744.64412 | -0.786596997 | 0.23228243 | -3.3863818 | 0.00070821 | 0.00596659 |
| PRR14L | 1465.09291 | -0.203818343 | 0.11741256 | -1.735916 | 0.08257869 | 0.20715372 |
| DEPDC5 | 779.844867 | 0.097977234 | 0.12554468 | 0.78041726 | 0.43514531 | 0.62724162 |
| Z82190.2 | 947.665246 | -0.682990095 | 0.28121619 | -2.4287012 | 0.01515302 | 0.06131449 |
| YWHAH | 6469.84794 | -1.106951781 | 0.27532659 | -4.0205044 | 5.81E-05 | 0.00083138 |
| AL008719.1 | 0.12150485 | 1.04981648 | 3.28477129 | 0.31960109 | 0.74927075 | NA |
| LINC02558 | 0.65141345 | 0.529861436 | 2.17028904 | 0.24414326 | 0.80711987 | NA |
| SLC5A1 | 637.098046 | 2.196768054 | 0.56956787 | 3.85690308 | 0.00011483 | 0.00141172 |
| Z83839.2 | 0.09997307 | 0.547239394 | 3.29377202 | 0.16614368 | 0.86804388 | NA |
| C22orf42 | 0.10721287 | -0.642905668 | 3.30558948 | -0.1944905 | 0.84579185 | NA |
| AL008723.2 | 0 | NA | NA | NA | NA | NA |
| RFPL2 | 5.52015233 | 0.324659011 | 0.65918215 | 0.4925179 | 0.62235327 | 0.77470996 |
| IGLCOR22-1 | 0.08393208 | -0.642905668 | 3.30558948 | -0.1944905 | 0.84579185 | NA |
| SLC5A4-AS1 | 1.05832845 | 2.449194262 | 1.77968965 | 1.37619178 | 0.16876227 | 0.3404008 |
| SLC5A4 | 24.825141 | 0.845103604 | 0.47359119 | 1.78445802 | 0.07434927 | 0.19253563 |
| AL008723.1 | 1.56246759 | 0.384006104 | 1.12000298 | 0.34286168 | 0.73170252 | 0.84861659 |
| AL008723.3 | 3.25368653 | 0.649967448 | 1.33294009 | 0.4876194 | 0.62581945 | 0.77729476 |
| AL021937.3 | 0.47483003 | 0.610803665 | 1.65035311 | 0.37010483 | 0.71130438 | NA |
| RFPL3 | 65.2290546 | -0.950978035 | 0.21153918 | -4.4955172 | 6.94E-06 | 0.00015313 |
| IGLCOR22-2 | 0.18746814 | 0.18657164 | 3.29377195 | 0.05664376 | 0.95482897 | NA |
| RFPL3S | 91.0057038 | -1.053432637 | 0.21140608 | -4.9829817 | 6.26E-07 | 2.20E-05 |
| AL021937.1 | 18.3961296 | -0.65815815 | 0.43108025 | -1.5267648 | 0.12681951 | 0.28022712 |
| IGLVIVOR22- | 0.91109601 | 0.524128122 | 1.14771471 | 0.45667109 | 0.64790748 | NA |
| AL021937.4 | 5.21114146 | -0.369582412 | 0.64394804 | -0.573932 | 0.56601379 | 0.73258976 |
| RTCB | 5945.8895 | -0.054402695 | 0.25002279 | -0.2175909 | 0.82774784 | 0.90828919 |
| BPIFC | 0.49625174 | 2.179055541 | 2.67996626 | 0.81309066 | 0.41616608 | NA |
| FBXO7 | 7561.5057 | 0.233272287 | 0.15972364 | 1.46047441 | 0.14415974 | 0.3059685 |
| SYN3 | 23.704486 | 1.527525984 | 0.42201928 | 3.61956441 | 0.0002951 | 0.00299278 |
| Z82246.1 | 4.2895803 | 0.727794406 | 0.68678433 | 1.05971319 | 0.2892751 | 0.48553144 |
| Z73495.1 | 0 | NA | NA | NA | NA | NA |
| TIMP3 | 28804.6066 | 0.498108505 | 0.27077829 | 1.83954371 | 0.06583526 | 0.17644225 |
| Z82198.1 | 0.82466458 | -0.925200807 | 2.00661386 | -0.4610757 | 0.64474433 | NA |
| LINC01640 | 0.17507102 | 0.435199075 | 3.28898345 | 0.13232024 | 0.894731 | NA |
| LARGE1 | 929.449988 | -0.092051007 | 0.20972571 | -0.4389114 | 0.66072574 | 0.80033538 |
| Z82198.2 | 5.40856086 | -0.374835131 | 0.69789405 | -0.5370946 | 0.59120227 | 0.75081891 |
| Z82173.1 | 0.66438071 | 0.637826107 | 2.24435582 | 0.28419117 | 0.77626388 | NA |
| Z73429.1 | 7.76275338 | -0.521540461 | 0.5095683 | -1.0234947 | 0.306074 | 0.5030244 |
| LINC01643 | 0 | NA | NA | NA | NA | NA |
| Z68323.1 | 0 | NA | NA | NA | NA | NA |
| AL021877.2 | 0 | NA | NA | NA | NA | NA |
| Z82196.1 | 0.91599345 | -0.273660179 | 1.54117557 | -0.1775659 | 0.85906393 | NA |
| Z82196.2 | 0.6794669 | -1.606857967 | 1.91518132 | -0.8390109 | 0.4014632 | NA |
| ISX-AS1 | 1.95117476 | -0.428979405 | 1.05686786 | -0.4058969 | 0.68481834 | 0.81786188 |
| ISX | 8.50959338 | 1.117458331 | 0.91109606 | 1.22649892 | 0.22001098 | 0.40574093 |
| Z99755.2 | 0 | NA | NA | NA | NA | NA |
| HMGXB4 | 684.504431 | -0.123574588 | 0.15712021 | -0.7864971 | 0.43157633 | 0.62363681 |
| AL008635.1 | 12.1004546 | -0.367753068 | 0.52438338 | -0.7013057 | 0.48311224 | 0.66769758 |
| TOM1 | 5386.82855 | -0.310208198 | 0.23039177 | -1.3464378 | 0.17816135 | 0.35361395 |
| Z82244.2 | 0.599622 | -1.690059302 | 1.88208524 | -0.8979717 | 0.36920063 | NA |
| Z82244.1 | 0 | NA | NA | NA | NA | NA |
| HMOX1 | 29377.8524 | -2.575837382 | 0.70320403 | -3.6630015 | 0.00024928 | 0.00261577 |
| MCM5 | 1173.38456 | 0.103136511 | 0.27088928 | 0.38073309 | 0.70340132 | 0.83051369 |
| AL022334.2 | 2.68290421 | -1.089097938 | 0.98149734 | -1.109629 | 0.26715892 | 0.46124006 |
| RASD2 | 146.7241 | -1.089700036 | 0.35600285 | -3.0609306 | 0.0022065 | 0.01456457 |
| MB | 17.7877055 | 4.374010894 | 1.05289307 | 4.15427838 | NA | NA |
| APOL6 | 3119.72483 | -1.027652114 | 0.40966801 | -2.5084998 | 0.0121245 | 0.05192709 |
| AL049748.1 | 0 | NA | NA | NA | NA | NA |
| APOL5 | 1.81719174 | 0.333634374 | 1.02257971 | 0.32626735 | 0.74422209 | 0.85654401 |
| RBFOX2 | 3101.24916 | 0.024402814 | 0.12276878 | 0.19877051 | 0.84244227 | 0.91630899 |
| Z82217.1 | 7.42786885 | -0.531143576 | 0.65410808 | -0.8120119 | 0.41678478 | 0.60986792 |
| Z95114.4 | 0.18552053 | -1.003574173 | 3.30558948 | -0.3035992 | 0.7614333 | NA |
| Z95114.1 | 0.75788396 | -1.201187973 | 1.65033092 | -0.7278467 | 0.46670741 | NA |
| Z95114.2 | 0 | NA | NA | NA | NA | NA |
| APOL3 | 1785.15495 | -0.221636835 | 0.33129965 | -0.6689921 | 0.50350055 | 0.68355176 |
| APOL4 | 1145.1259 | -3.004159499 | 0.80182304 | -3.7466615 | 0.0001792 | 0.00200513 |
| APOL2 | 5228.76128 | -1.580916137 | 0.41395553 | -3.8190482 | 0.00013397 | 0.00159244 |
| APOL1 | 5451.21867 | -1.480135988 | 0.40886861 | -3.6200774 | 0.00029451 | 0.00299124 |
| MYH9 | 39111.6536 | 0.004487847 | 0.26295218 | 0.01706716 | 0.98638304 | 0.99391317 |
| FO393418.1 | 6.84893419 | -0.276877434 | 0.75473569 | -0.3668535 | 0.71372829 | 0.83745188 |
| AL022313.2 | 2.42735443 | -0.193216343 | 0.77845919 | -0.2482036 | 0.80397691 | 0.89372191 |
| TXN2 | 4711.29323 | 0.437102761 | 0.14670138 | 2.97954084 | 0.00288681 | 0.01780288 |
| FOXRED2 | 931.721668 | 1.075995771 | 0.34886639 | 3.08426321 | 0.00204057 | 0.01369904 |
| EIF3D | 7499.11987 | -0.357059121 | 0.15561239 | -2.2945417 | 0.0217594 | 0.07987131 |
| AL022313.3 | 0 | NA | NA | NA | NA | NA |
| CACNG2 | 0.22085762 | 0.689157565 | 3.05316003 | 0.22571944 | 0.82141964 | NA |
| AL049749.1 | 0.08666371 | -0.642905668 | 3.30558948 | -0.1944905 | 0.84579185 | NA |
| IFT27 | 1369.56053 | 0.136914352 | 0.18486675 | 0.74061101 | 0.45892933 | 0.64778312 |
| PVALB | 586.782492 | 1.421969997 | 0.57612472 | 2.46816347 | 0.01358083 | 0.05654033 |
| Z82185.1 | 15.9475933 | 1.430633326 | 0.55570869 | 2.57443035 | 0.01004053 | 0.045196 |
| NCF4-AS1 | 63.8668469 | -0.758218136 | 0.46361713 | -1.6354403 | 0.10195678 | 0.24069833 |
| NCF4 | 584.321611 | -0.856313976 | 0.40354725 | -2.1219671 | 0.0338405 | 0.10993794 |

| | | | | | | |
|---|---|---|---|---|---|---|
| CSF2RB | 485.317647 | -0.20602164 | 0.43971521 | -0.4685343 | 0.63940258 | 0.7868698 |
| Z82180.1 | 0 | NA | NA | NA | NA | NA |
| TEX33 | 0.6547064 | -1.568467247 | 1.66127599 | -0.9441341 | 0.34510115 | NA |
| TST | 4932.4134 | 0.582044995 | 0.26483877 | 2.19773333 | 0.02796811 | 0.09605616 |
| MPST | 5504.75059 | -0.008967134 | 0.24756308 | -0.0362216 | 0.97110565 | 0.9865502 |
| KCTD17 | 533.196894 | -1.024683022 | 0.37686047 | -2.7189984 | 0.00654799 | 0.03287084 |
| TMPRSS6 | 30.211171 | -0.926227844 | 0.32370096 | -2.8613688 | 0.00421816 | 0.02368673 |
| AL022314.1 | 0 | NA | NA | NA | NA | NA |
| IL2RB | 228.350737 | 0.02928742 | 0.62915181 | 0.04655064 | 0.96287137 | 0.98262067 |
| Z82188.2 | 110.769847 | 0.121964078 | 0.27644655 | 0.44118502 | 0.65907906 | 0.79973839 |
| C1QTNF6 | 834.92236 | -0.152919315 | 0.25515015 | -0.5993307 | 0.54895239 | 0.71954839 |
| SSTR3 | 10.5393493 | -0.575698338 | 0.82706527 | -0.6960736 | 0.48638271 | 0.67045724 |
| RAC2 | 776.686281 | -0.356479031 | 0.51035948 | -0.6984862 | 0.48487322 | 0.6691144 |
| CYTH4 | 883.653652 | 0.16640544 | 0.38169279 | 0.43596695 | 0.66286071 | 0.80178143 |
| ELFN2 | 171.529621 | -0.37688295 | 0.64665343 | -0.5828206 | 0.56001407 | 0.72832763 |
| Z94160.2 | 19.3106661 | -0.39587445 | 0.61472205 | -0.6439893 | 0.51958234 | 0.69631663 |
| MFNG | 707.151987 | -0.532763895 | 0.34252955 | -1.5553808 | 0.11985539 | 0.26972812 |
| CARD10 | 4064.8576 | -0.18369323 | 0.21782725 | -0.8432977 | 0.39906195 | 0.59390477 |
| AL022315.1 | 0.6629899 | 0.758483214 | 1.44704306 | 0.52416078 | 0.60016672 | NA |
| CDC42EP1 | 4373.03028 | -1.02398213 | 0.24287488 | -4.2160892 | 2.49E-05 | 0.00042302 |
| LGALS2 | 2074.71793 | 0.98786368 | 0.36978623 | 2.67144529 | 0.00755254 | 0.03661711 |
| GGA1 | 4862.42413 | -0.225677774 | 0.14397428 | -1.5674867 | 0.11700097 | 0.26531355 |
| SH3BP1 | 506.842393 | -0.527464394 | 0.38891663 | -1.3562403 | 0.17502273 | 0.34903951 |
| Z83844.3 | 1398.51533 | 0.362510893 | 0.25008621 | 1.44954374 | 0.1471858 | 0.31045204 |
| Z83844.2 | 220.930847 | -0.367642881 | 0.36964091 | -0.9945947 | 0.31993344 | 0.51736339 |
| PDXP | 1082.63097 | 0.572163675 | 0.30954694 | 1.84839067 | 0.06454585 | 0.17391172 |
| LGALS1 | 3447.98018 | -0.845321025 | 0.33461636 | -2.5262394 | 0.01152909 | 0.05010454 |
| NOL12 | 1073.26149 | -0.127213228 | 0.16135516 | -0.7884051 | 0.43045981 | 0.62251837 |
| Z83844.1 | 689.846718 | -0.128083386 | 0.19720689 | -0.6494874 | 0.5160234 | 0.69349777 |
| TRIOBP | 5037.1675 | -0.427481127 | 0.11927913 | -3.5838718 | 0.00033854 | 0.00333854 |
| H1F0 | 14332.4184 | -0.727178943 | 0.37903397 | -1.918506 | 0.05504688 | 0.15579612 |
| GCAT | 1615.41998 | 0.56180579 | 0.36783156 | 1.52734526 | 0.12667518 | 0.28002552 |
| GALR3 | 1.5125841 | -0.552846742 | 1.04919678 | -0.5269238 | 0.59824651 | 0.75659797 |
| ANKRD54 | 1828.84265 | -0.110864855 | 0.13501812 | -0.8211109 | 0.41158308 | 0.60570403 |
| EIF3L | 10800.7169 | 0.222065226 | 0.24653441 | 0.90074738 | 0.36772265 | 0.56452502 |
| MICALL1 | 1605.95336 | -0.753572619 | 0.34258492 | -2.1996667 | 0.02783055 | 0.09569588 |
| C22orf23 | 348.777006 | -0.070871124 | 0.23209688 | -0.3053515 | 0.76009847 | 0.86700095 |
| AL031587.3 | 22.9265802 | -0.150553676 | 0.28434899 | -0.529468 | 0.59648087 | 0.75523121 |
| POLR2F | 1248.01558 | -0.095594045 | 0.08601232 | -1.1113995 | 0.26639643 | 0.46053165 |
| SOX10 | 59.3104927 | -0.447066416 | 0.72380871 | -0.6176582 | 0.53680064 | 0.71053163 |
| AL031587.1 | 0.88859709 | 3.246042879 | 1.94261893 | 1.67096224 | 0.09472914 | NA |
| AL031587.4 | 0.3439232 | -1.739004615 | 3.30153799 | -0.5267256 | 0.59838415 | NA |
| PICK1 | 2705.61363 | -0.320972536 | 0.13018188 | -2.4655699 | 0.01367955 | 0.05681677 |
| AL031587.2 | 339.15359 | -0.363742947 | 0.15483517 | -2.3492269 | 0.01881244 | 0.07177186 |
| SLC16A8 | 74.9832117 | 0.96223373 | 0.42721493 | 2.25234109 | 0.02430073 | 0.08640913 |
| BAIAP2L2 | 3293.25975 | 0.065483731 | 0.31761761 | 0.2061716 | 0.83665687 | 0.91334003 |
| AL022322.1 | 190.843108 | 0.181338109 | 0.32823901 | 0.55245751 | 0.58063494 | 0.74350374 |
| PLA2G6 | 3060.68615 | 0.401506614 | 0.2805048 | 1.43137163 | 0.15232373 | 0.31751267 |
| AL022322.2 | 460.986461 | 0.898070713 | 0.25866832 | 3.47190068 | 0.00051679 | 0.00465109 |
| MAFF | 3027.40048 | -2.059185528 | 0.44566192 | -4.6205104 | 3.83E-06 | 9.51E-05 |
| TMEM184B | 4440.986 | -0.341527879 | 0.18305635 | -1.8656981 | 0.06208362 | 0.16939516 |
| AL020993.1 | 143.891759 | 0.224034639 | 0.18297135 | 1.22442467 | 0.22079207 | 0.40672629 |
| CSNK1E | 6132.07366 | -0.751437738 | 0.19039442 | -3.9467425 | 7.92E-05 | 0.00105246 |
| TPTEP2-CSNI | 2581.26769 | -0.534709841 | 0.18297133 | -2.9223695 | 0.00347379 | 0.02047014 |
| Z97056.1 | 11.5164746 | -1.013873757 | 0.62365005 | -1.6257094 | 0.10401149 | 0.24416075 |
| KCNJ4 | 24.9785801 | -0.50508382 | 0.54391278 | -0.9286118 | 0.35309031 | 0.55065299 |
| KDELR3 | 614.120075 | -0.364226917 | 0.2426813 | -1.5008446 | 0.13339576 | 0.29009123 |
| DDX17 | 33178.0568 | 0.154635648 | 0.16118198 | 0.95938544 | 0.33736461 | 0.5353737 |
| DMC1 | 18.4063281 | 0.773971679 | 0.51692843 | 1.49725115 | 0.13432792 | 0.29154134 |
| FAM227A | 328.181047 | -0.324202608 | 0.36314633 | -0.8927602 | 0.37198558 | 0.56855459 |
| CBY1 | 1772.33461 | -0.102133805 | 0.21280583 | -0.4799389 | 0.63127081 | 0.7807979 |
| AL021707.4 | 17.9287599 | 1.143513806 | 0.42798636 | 2.67184636 | 0.00754352 | 0.03661376 |
| AL021707.2 | 1006.9853 | 0.131751943 | 0.20000919 | 0.65872946 | 0.51006951 | 0.68871137 |
| TOMM22 | 2522.92627 | -0.011317978 | 0.14465725 | -0.07824 | 0.93763717 | 0.97023592 |
| JOSD1 | 2976.3191 | -0.680846487 | 0.21813874 | -3.1211626 | 0.00180139 | 0.01243847 |
| GTPBP1 | 6611.9742 | -0.167658268 | 0.173379 | -0.9670045 | 0.3335418 | 0.53154634 |
| AL021707.5 | 2342.84662 | -0.031408311 | 0.21008674 | -0.1495016 | 0.88115782 | 0.93836253 |
| SUN2 | 11662.2567 | 0.084179729 | 0.18474725 | 0.45564809 | 0.64864306 | 0.7931703 |
| AL021707.3 | 2160.0751 | 0.140811233 | 0.21334288 | 0.66002312 | 0.50923899 | 0.68801272 |
| AL021707.1 | 575.414984 | 0.456270009 | 0.21250029 | 2.14715003 | 0.03178134 | 0.10507446 |
| AL021707.6 | 186.432369 | -0.788751112 | 0.25031962 | -3.150976 | 0.00162726 | 0.01151659 |
| AL021707.7 | 8.15977164 | -0.695977336 | 0.68435345 | -1.0169852 | 0.30916047 | 0.5062445 |
| DNAL4 | 1502.28433 | -0.205986583 | 0.16499755 | -1.2484221 | 0.21187653 | 0.39546886 |
| NPTXR | 524.789198 | -0.926439238 | 0.27867512 | -3.3244418 | 0.00088596 | 0.00714019 |
| CBX6 | 6594.7935 | -0.432447471 | 0.24009213 | -1.8011731 | 0.0716756 | 0.18753166 |
| AL022318.1 | 1.43832448 | -0.049868456 | 1.04026716 | -0.0479381 | 0.96176556 | 0.98175813 |
| APOBEC3A | 119.524188 | -0.72427839 | 0.57278616 | -1.264483 | 0.2060567 | 0.38870557 |
| APOBEC3B | 89.8923114 | -1.378203362 | 0.42541401 | -3.2396755 | 0.00119666 | 0.00903144 |
| APOBEC3B-A | 43.5963319 | -1.694481926 | 0.44680864 | -3.7924108 | 0.00014919 | 0.0017462 |
| APOBEC3C | 583.104417 | -0.299609315 | 0.254869 | -1.1755424 | 0.23977778 | 0.42969352 |
| AL022318.4 | 216.669218 | -0.536729374 | 0.41383175 | -1.2969748 | 0.19463984 | 0.37461884 |
| APOBEC3D | 191.777273 | -0.690142094 | 0.47069119 | -1.4662312 | 0.14258532 | 0.30378955 |
| APOBEC3F | 230.891095 | -0.594410136 | 0.46317476 | -1.2833388 | 0.19937341 | 0.38053539 |
| APOBEC3G | 472.591073 | -0.692896775 | 0.59949642 | -1.155798 | 0.24776379 | 0.43933404 |
| APOBEC3H | 45.843385 | -1.209418533 | 0.7466098 | -1.6198803 | 0.10525798 | 0.24609035 |
| CBX7 | 5349.04444 | 0.551429836 | 0.20660453 | 2.66901137 | 0.00760749 | 0.03679839 |
| AL031846.2 | 1346.64527 | 1.021822834 | 0.18771916 | 5.44335935 | 5.23E-08 | 2.76E-06 |
| PDGFB | 1564.65566 | 0.444336157 | 0.3115191 | 1.42635284 | 0.15376652 | 0.31942832 |
| AL031590.1 | 20.2901959 | 0.600736152 | 0.41172842 | 1.45905922 | 0.14454881 | 0.30653415 |
| AL022326.1 | 1.71943026 | 1.265156412 | 1.07079687 | 1.18150926 | 0.23740048 | 0.42665103 |
| RPL3 | 72073.261 | -0.219006886 | 0.24060412 | -0.9102375 | 0.36269729 | 0.55993956 |
| SYNGR1 | 2151.85191 | -0.014990961 | 0.34960237 | -0.04288 | 0.96579716 | 0.98387257 |
| TAB1 | 2017.85023 | 0.193216882 | 0.1538328 | 1.25601878 | 0.20910916 | 0.39233074 |
| MGAT3 | 680.318954 | 0.992673917 | 0.45768124 | 2.16891985 | 0.03008877 | 0.10103665 |
| MGAT3-AS1 | 2.55951841 | 2.598406422 | 1.06225209 | 2.44612974 | 0.0144399 | 0.05922322 |
| MIEF1 | 2324.59894 | 0.270871525 | 0.21208311 | 1.27719518 | 0.20153335 | 0.3833815 |
| AL022312.1 | 70.2395684 | 1.456389414 | 0.3389225 | 4.29711636 | 1.73E-05 | 0.00031665 |
| ATF4 | 18831.5826 | -1.11542727 | 0.29018084 | -3.8439039 | 0.00012109 | 0.0014756 |
| RPS19BP1 | 3966.87431 | -0.362000334 | 0.20703515 | -1.748497 | 0.08037801 | 0.20304322 |
| CACNA1I | 19.5812051 | -1.001484737 | 0.82685961 | -1.2111908 | 0.2258223 | 0.41244867 |
| ENTHD1 | 0.75459872 | 0.42664798 | 1.33416866 | 0.31978564 | 0.74913083 | NA |
| GRAP2 | 101.051824 | -0.004808872 | 0.64704512 | -0.0074321 | 0.99407014 | 0.99728027 |
| Z82206.1 | 5.58270248 | -0.426702136 | 0.58018981 | -0.7354527 | 0.46206386 | 0.65029749 |
| FAM83F | 1201.5553 | -0.247434629 | 0.2841172 | -0.8708893 | 0.3838146 | 0.57948708 |
| Z83847.1 | 50.3024514 | -0.930973196 | 0.44497811 | -2.0921775 | 0.03642264 | 0.11577447 |
| TNRC6B | 1807.5693 | 0.138007352 | 0.1219769 | 1.13142207 | 0.25787749 | 0.45043581 |
| ADSL | 2427.28371 | -0.084357279 | 0.14725744 | -0.5728558 | 0.56674233 | 0.73296071 |
| AL022238.4 | 7672.41949 | -0.19924313 | 0.11991917 | -1.6614785 | 0.09661738 | 0.23199785 |
| SGSM3 | 5520.48725 | -0.245163282 | 0.17749136 | -1.3812688 | 0.16719632 | 0.33831852 |
| AL022238.2 | 1.35295234 | -1.074578315 | 1.39817233 | -0.7685593 | 0.44215499 | 0.63331779 |
| MKL1 | 2245.15565 | -0.544608262 | 0.1822546 | -2.988173 | 0.00280651 | 0.01744639 |
| AL022238.3 | 8.635155 | 0.747103594 | 0.47153127 | 1.58442003 | 0.11309817 | 0.25906868 |
| AL031594.1 | 2.52904588 | -1.260834544 | 0.81859266 | -1.5402466 | 0.12350026 | 0.27503041 |

| | | | | | | |
|---|---|---|---|---|---|---|
| MCHR1 | 14.4940691 | -0.926020865 | 1.00086073 | -0.9252245 | 0.35484914 | 0.55235062 |
| SLC25A17 | 1519.67984 | -0.155033481 | 0.12146462 | -1.2763674 | 0.20182567 | 0.38371614 |
| ST13 | 16388.6521 | 0.555615778 | 0.19292356 | 2.87997895 | 0.00397702 | 0.02264614 |
| XPNPEP3 | 1400.898 | 0.284820664 | 0.14870684 | 1.91531649 | 0.05545215 | 0.15659384 |
| DNAJB7 | 88.5975634 | -0.008375115 | 0.42025482 | -0.0199287 | 0.98410028 | 0.99314982 |
| RBX1 | 2176.85502 | -0.570870764 | 0.12408001 | -4.6008279 | 4.21E-06 | 0.00010275 |
| EP300 | 3088.4305 | 0.195087959 | 0.21621094 | 0.90230385 | 0.36689548 | 0.56387888 |
| AL035658.1 | 14.2597329 | 1.558598246 | 0.45027831 | 3.46141086 | 0.00053735 | 0.00479525 |
| EP300-AS1 | 8.84787737 | 0.73575922 | 0.59204292 | 1.24274642 | 0.21396129 | 0.39802193 |
| L3MBTL2 | 1784.91665 | 0.172335596 | 0.14039005 | 1.22754852 | 0.2196165 | 0.40518345 |
| AL035681.1 | 257.503287 | 0.863783115 | 0.22891353 | 3.77340352 | 0.00016104 | 0.00184952 |
| CHADL | 270.201846 | 0.364819028 | 0.22186581 | 1.64432289 | 0.10010952 | 0.23765647 |
| RANGAP1 | 5817.28257 | -0.699634895 | 0.17857414 | -3.917896 | 8.93E-05 | 0.0011575 |
| ZC3H7B | 8934.44914 | 0.592570953 | 0.12966402 | 4.57004921 | 4.88E-06 | 0.00011532 |
| TEF | 2061.42943 | 1.083583161 | 0.40229501 | 2.69350386 | 0.00707053 | 0.03488306 |
| AL008582.1 | 29.6134978 | -0.107769688 | 0.41939433 | -0.2569651 | 0.79720575 | 0.88994647 |
| TOB2 | 4086.2344 | 0.421612649 | 0.2503002 | 1.68442794 | 0.092099 | 0.22435571 |
| PHF5A | 1042.86211 | -0.384560463 | 0.1861304 | -2.0660808 | 0.03882085 | 0.12117729 |
| ACO2 | 20022.8351 | 0.974776601 | 0.22732235 | 4.28808078 | 1.80E-05 | 0.00032731 |
| POLR3H | 11682.8879 | 0.604579621 | 0.15455773 | 3.91167511 | 9.17E-05 | 0.00118167 |
| CSDC2 | 6274.48135 | 0.210381166 | 0.37080571 | 0.56736225 | 0.57046809 | 0.7356469 |
| PMM1 | 3848.29865 | 0.496359182 | 0.19638222 | 2.52751591 | 0.01148726 | 0.04998665 |
| DESI1 | 3226.21488 | -0.617729171 | 0.20409299 | -3.0267045 | 0.00247236 | 0.01585728 |
| XRCC6 | 11815.8487 | -0.311240774 | 0.15236737 | -2.0426997 | 0.04108218 | 0.12635142 |
| SNU13 | 5095.68521 | -0.312817009 | 0.10020538 | -3.1217585 | 0.00179774 | 0.01242483 |
| C22orf46 | 745.165909 | 0.671007819 | 0.21286939 | 3.15220433 | 0.00162043 | 0.01147749 |
| MEI1 | 113.933861 | -0.325062844 | 0.4700351 | -0.6915714 | 0.48920651 | 0.67257742 |
| CCDC134 | 136.919124 | -0.05738526 | 0.24684788 | -0.2324722 | 0.81617131 | 0.90104821 |
| AL021453.1 | 385.448858 | 0.167374413 | 0.16284067 | 1.02784158 | 0.30402435 | 0.50088085 |
| SREBF2 | 8860.15373 | -0.065279543 | 0.21318675 | -0.3062083 | 0.75944609 | 0.86676448 |
| SHISA8 | 17.0361648 | 0.231418083 | 0.70433877 | 0.32856076 | 0.74248771 | 0.85544394 |
| TNFRSF13C | 117.236652 | -0.041945241 | 0.70008183 | -0.0599148 | 0.95222352 | 0.97716209 |
| CENPM | 48.2739114 | -1.152698694 | 0.41547621 | -2.7744036 | 0.0055303 | 0.02908537 |
| 3-Sep | 51.1206599 | -0.270606905 | 0.49723853 | -0.5442195 | 0.58629044 | 0.74760536 |
| Z99716.1 | 1.29613003 | 1.48107632 | 1.24441997 | 1.19017402 | 0.233978 | 0.42254485 |
| WBP2NL | 132.72227 | 0.008722724 | 0.28473411 | 0.03063463 | 0.97556092 | 0.98866399 |
| NAGA | 2285.64991 | 0.150245237 | 0.1588994 | 0.94553686 | 0.34438486 | 0.54203477 |
| PHETA2 | 544.119952 | -0.599655757 | 0.20734816 | -2.8920236 | 0.00382769 | 0.02205293 |
| SMDT1 | 2348.66667 | 0.212080885 | 0.14566652 | 1.45593433 | 0.14541077 | 0.30774335 |
| NDUFA6 | 4168.61842 | -0.179114029 | 0.18572138 | -0.9644233 | 0.33483373 | 0.53296103 |
| CYP2D6 | 311.644618 | 0.476296244 | 0.25608626 | 1.85990549 | 0.0628989 | 0.17087091 |
| AC254562.2 | 46.0033254 | 0.744947989 | 0.2695673 | 2.76349535 | 0.00571859 | 0.0297846 |
| AC254562.3 | 84.3009666 | 0.725045583 | 0.28823165 | 2.51549604 | 0.0118865 | 0.05120299 |
| TCF20 | 2187.74623 | -0.877461483 | 0.29727909 | -2.9516421 | 0.00316089 | 0.01912198 |
| OGFRP1 | 35.598433 | -1.393804871 | 0.58664635 | -2.375886 | 0.01750687 | 0.06803067 |
| Z83851.1 | 64.5860609 | -1.484418973 | 0.46559505 | -3.188219 | 0.00143152 | 0.01043612 |
| Z83851.2 | 28.0360668 | -1.547416224 | 0.47942546 | -3.2276472 | 0.00124813 | 0.00934501 |
| LINC01315 | 124.727644 | -0.967587179 | 0.42857244 | -2.2576981 | 0.02396449 | 0.08554775 |
| NFAM1 | 389.328784 | -0.863902503 | 0.43550396 | -1.9836846 | 0.04729101 | 0.13980538 |
| AL022316.1 | 33.3910715 | -1.009203388 | 0.53527196 | -1.8854031 | 0.05937544 | 0.16436433 |
| RRP7A | 2816.07906 | -0.081664339 | 0.24864443 | -0.3284382 | 0.74258034 | 0.85547591 |
| SERHL2 | 469.29538 | -0.204654655 | 0.23504718 | -0.8706961 | 0.38392013 | 0.57957258 |
| POLDIP3 | 4928.28313 | -0.11391008 | 0.10000229 | -1.1390747 | 0.25467198 | 0.44747469 |
| Z93241.1 | 10.9600345 | -1.702328675 | 0.69100314 | -2.4635614 | 0.01375643 | 0.05707318 |
| CYB5R3 | 21561.0687 | -0.552474452 | 0.16224658 | -3.4051532 | 0.00066127 | 0.00563414 |
| ATP5MGL | 2.76517982 | -0.599756795 | 0.71824613 | -0.8350296 | 0.40370103 | 0.59870627 |
| A4GALT | 5328.04911 | -1.236110688 | 0.24904998 | -4.9633037 | 6.93E-07 | 2.38E-05 |
| ARFGAP3 | 5745.49277 | -1.066565247 | 0.17825816 | -5.9832618 | 2.19E-09 | 1.92E-07 |
| PACSIN2 | 8744.06825 | -0.403870786 | 0.29683607 | -1.3605853 | 0.17364479 | 0.34750043 |
| AL022476.1 | 76.7296452 | 0.152108474 | 0.28476275 | 0.5341586 | 0.59323181 | 0.7524542 |
| TTLL1 | 295.22428 | 0.223943466 | 0.24977712 | 0.89657319 | 0.36994671 | 0.56678449 |
| BIK | 144.469649 | -0.776770971 | 0.27360746 | -2.8389978 | 0.00452555 | 0.02496103 |
| MCAT | 1291.91343 | 0.54487091 | 0.16411439 | 3.3200679 | 0.00089996 | 0.0072243 |
| TSPO | 3578.58306 | -0.798557892 | 0.15643257 | -5.1048059 | 3.31E-07 | 1.29E-05 |
| TTLL12 | 1834.78017 | -0.267988188 | 0.2036085 | -1.3161935 | 0.18810909 | 0.36613054 |
| SCUBE1 | 822.036002 | -0.065428554 | 0.41496652 | -0.1576719 | 0.87471535 | 0.93484007 |
| Z82214.2 | 9.46794232 | 0.11696561 | 0.7495598 | 0.15604574 | 0.87599697 | 0.93550556 |
| Z82214.1 | 38.4089251 | 0.512714937 | 0.4905797 | 1.04512057 | 0.29596725 | 0.49277045 |
| Z99756.1 | 0.86516707 | -0.040801046 | 1.60100444 | -0.0254847 | 0.97966839 | NA |
| LINC01639 | 0.08757577 | 0.547239394 | 3.29377202 | 0.16614368 | 0.86804388 | NA |
| MPPED1 | 2.88209602 | 2.447085772 | 1.52678944 | 1.60276572 | 0.10898639 | 0.25234622 |
| EFCAB6-AS1 | 22.2862631 | -0.016097039 | 0.38282881 | -0.0420476 | 0.96646074 | 0.98410057 |
| EFCAB6 | 196.458329 | 0.216928812 | 0.26258332 | 0.82613324 | 0.40872851 | 0.60316422 |
| Z97055.2 | 24.4528273 | 0.800537442 | 0.36989373 | 2.16423632 | 0.03044622 | 0.10188664 |
| SULT4A1 | 37.9240002 | -0.707897809 | 0.65429942 | -1.0819172 | 0.27928932 | 0.47492119 |
| PNPLA5 | 0.09635527 | 0.547239394 | 3.29377202 | 0.16614368 | 0.86804388 | NA |
| PNPLA3 | 519.483361 | 1.305634699 | 0.30738735 | 4.2475225 | 2.16E-05 | 0.00037747 |
| SAMM50 | 2519.76302 | 0.554650695 | 0.19156601 | 2.89535028 | 0.00378736 | 0.02191149 |
| PARVB | 1164.87843 | -0.201761383 | 0.25551348 | -0.7896311 | 0.42974325 | 0.62180135 |
| PARVG | 822.966656 | -0.47630206 | 0.50905954 | -0.935651 | 0.34945292 | 0.54711592 |
| AL031595.2 | 8.76874364 | 0.121915612 | 0.89623272 | 0.1360312 | NA | NA |
| SHISAL1 | 31.5438251 | 0.351971777 | 0.54033654 | 0.65139362 | 0.51479243 | 0.69256857 |
| Z85994.1 | 0.26504122 | 0.373565666 | 3.00920475 | 0.12414099 | 0.90120364 | NA |
| LINC01656 | 0 | NA | NA | NA | NA | NA |
| RTL6 | 1776.56741 | 0.236826688 | 0.1847316 | 1.28200423 | 0.19984116 | 0.38117169 |
| LINC00207 | 0.03392908 | 0.380740593 | 3.29738709 | 0.11546736 | 0.90807469 | NA |
| LINC00229 | 0 | NA | NA | NA | NA | NA |
| PRR5 | 1862.03942 | 0.737028108 | 0.21254819 | 3.46758124 | 0.00052516 | 0.00471201 |
| PRR5-ARHGA | 2771.64839 | -0.060300338 | 0.16487034 | -0.365744 | 0.71455611 | 0.83779488 |
| ARHGAP8 | 2171.79527 | -0.396940797 | 0.22013468 | -1.8031725 | 0.07136113 | 0.18690833 |
| PHF21B | 230.541079 | 0.741386919 | 0.25613364 | 2.89453163 | 0.00379725 | 0.02193983 |
| NUP50-DT | 630.228468 | -1.012478821 | 0.23357963 | -4.3346195 | 1.46E-05 | 0.0002774 |
| Z82243.1 | 12.0140831 | -0.736585752 | 0.42165877 | -1.7468764 | 0.08065878 | 0.20359645 |
| NUP50 | 2173.81661 | -0.503497523 | 0.19672816 | -2.5593566 | 0.01048661 | 0.04668062 |
| KIAA0930 | 3791.25563 | -1.351156502 | 0.32214995 | -4.194185 | 2.74E-05 | 0.00045728 |
| AL008718.2 | 4.34603745 | -1.339914655 | 0.86949129 | -1.5410329 | 0.12330879 | 0.27477513 |
| UPK3A | 2.26464194 | -0.351403299 | 1.01209747 | -0.347203 | 0.7284388 | 0.84642348 |
| FAM118A | 1423.83579 | -0.258354887 | 0.5388041 | -0.4794969 | 0.63158519 | 0.78107683 |
| SMC1B | 7.69521135 | -0.37006364 | 0.57464408 | -0.6439876 | 0.5195835 | 0.69631663 |
| RIBC2 | 17.8882718 | -2.205716131 | 0.66685901 | -3.3076199 | 0.00094092 | 0.00749386 |
| AL021391.1 | 1.20233102 | -2.781641274 | 1.83827052 | -1.5131839 | 0.13023296 | 0.2854258 |
| FBLN1 | 6081.32157 | -0.229199136 | 0.42528978 | -0.5389246 | 0.58993886 | 0.74984642 |
| LINC01589 | 15.7801166 | 0.488217524 | 0.86003405 | 0.56767232 | 0.57025749 | 0.73541129 |
| ATXN10 | 4634.74042 | -0.066647025 | 0.16344301 | -0.4077692 | 0.68344314 | 0.8171569 |
| BX324167.2 | 6.28177045 | 0.062357767 | 0.47898615 | 0.13018699 | 0.89641848 | 0.94698268 |
| BX324167.1 | 4.05028757 | 0.782861509 | 0.64761105 | 1.2088452 | 0.22672232 | 0.41360152 |
| WNT7B | 211.778443 | 1.876109579 | 0.91978394 | 2.03972857 | 0.04137737 | 0.12704736 |
| BX537318.1 | 94.549846 | 0.398758778 | 0.44484672 | 0.89639591 | 0.37004135 | 0.5668307 |
| PRR34 | 197.553693 | 0.467352086 | 0.26140705 | 1.78783272 | 0.073803 | 0.19159137 |
| AL121672.3 | 6.11446567 | 0.322103191 | 0.56647077 | 0.56861396 | 0.56961815 | 0.73500743 |
| PRR34-AS1 | 246.255805 | 0.280917469 | 0.25276851 | 1.11136259 | 0.26641231 | 0.46053165 |
| MIRLET7BHC | 393.384346 | 0.025379125 | 0.31241122 | 0.08123628 | 0.93525405 | 0.96907653 |
| AL121672.1 | 35.4597003 | 0.535253281 | 0.34565815 | 1.54850475 | 0.12150082 | 0.27209992 |

| | | | | | | | |
|---|---|---|---|---|---|---|---|
| AL121672.2 | 23.7216464 | | 0.81627831 | 0.39556939 | 2.06355278 | 0.03906014 | 0.12167992 |
| PPARA | 4057.11181 | | 0.843018138 | 0.33819672 | 2.49268574 | 0.0126781 | 0.0537408 |
| CDPF1 | 481.82568 | | -0.147485761 | 0.12639192 | -1.1668923 | 0.24325387 | 0.43388829 |
| PKDREJ | 15.6393686 | | 1.087540687 | 0.55030879 | 1.97623715 | 0.04812792 | 0.14167717 |
| TTC38 | 6912.70269 | | 1.147189495 | 0.28647882 | 4.00444792 | 6.22E-05 | 0.00087239 |
| GTSE1-DT | 37.3263116 | | 0.423817879 | 0.40138042 | 1.05590072 | 0.29101358 | 0.48702342 |
| GTSE1 | 49.8787134 | | -0.891612288 | 0.39703627 | -2.2456696 | 0.02472518 | 0.08756703 |
| TRMU | 3402.67367 | | -0.223254112 | 0.09887413 | -2.2579628 | 0.02394798 | 0.08553487 |
| CELSR1 | 4593.96616 | | 1.194868715 | 0.22622061 | 5.28187391 | 1.28E-07 | 5.89E-06 |
| AL021392.1 | 95.7488977 | | 0.696534083 | 0.28979735 | 2.40352121 | 0.01623803 | 0.06447775 |
| GRAMD4 | 2374.67232 | | -0.20046516 | 0.22628947 | -0.8858793 | 0.37568256 | 0.57155421 |
| CERK | 3063.49831 | | 0.184974796 | 0.2510476 | 0.73681165 | 0.4612369 | 0.64968843 |
| AL118516.1 | 241.272183 | | -0.594727597 | 0.28663552 | -2.0748566 | 0.03799983 | 0.11922135 |
| TBC1D22A | 1859.48211 | | -0.064789422 | 0.06926672 | -0.9353614 | 0.34960206 | 0.54721946 |
| TBC1D22A-A | 1.38302594 | | 0.616006098 | 1.06606622 | 0.57783099 | 0.56337823 | 0.73057995 |
| Z82186.1 | 1.66071157 | | 1.347379732 | 1.07641008 | 1.25173459 | 0.21066659 | 0.39401929 |
| LINC01644 | 6.4842436 | | -2.784435985 | 0.74904365 | -3.7173214 | 0.00020135 | 0.00219129 |
| LINC00898 | 0 | NA | | NA | NA | NA | NA |
| AL117329.1 | 0 | NA | | NA | NA | NA | NA |
| AL121580.1 | 0.08666371 | | -0.642905668 | 3.30558948 | -0.1944905 | 0.84579185 | NA |
| FP325330.3 | 0 | NA | | NA | NA | NA | NA |
| FP325330.1 | 0 | NA | | NA | NA | NA | NA |
| FP325330.2 | 0 | NA | | NA | NA | NA | NA |
| BX284656.2 | 0 | NA | | NA | NA | NA | NA |
| BX284656.1 | 0.14114194 | | 0.02007176 | 3.29738704 | 0.00608717 | 0.99514317 | NA |
| Z82249.1 | 0 | NA | | NA | NA | NA | NA |
| FAM19A5 | 179.452901 | | -0.044976094 | 0.39637383 | -0.1134689 | 0.90965884 | 0.95506719 |
| LINC01310 | 0.25087713 | | 1.011917048 | 2.94486708 | 0.34362062 | 0.73113162 | NA |
| Z82202.1 | 0 | NA | | NA | NA | NA | NA |
| C22orf34 | 169.89832 | | -1.231136689 | 0.40260233 | -3.0579472 | 0.00222859 | 0.01468096 |
| AL008636.1 | 0.72142399 | | 0.335263749 | 2.22399486 | 0.15074844 | 0.88017416 | NA |
| AL031593.1 | 0.34847896 | | 0.732866073 | 3.27839706 | 0.22354402 | 0.82311212 | NA |
| Z97192.1 | 0.82621481 | | 0.423617125 | 1.5663532 | 0.27044802 | 0.7868156 | NA |
| Z97192.2 | 20.0931798 | | -0.515682243 | 0.38875819 | -1.3264859 | 0.18467882 | 0.36189198 |
| AL023802.1 | 1.99918325 | | -2.772982269 | 2.11723308 | -1.3097199 | 0.19029061 | 0.3689679 |
| BRD1 | 2258.56488 | | -0.123106217 | 0.12925109 | -0.9524579 | 0.34086482 | 0.53888521 |
| Z98885.3 | 8.15851076 | | 0.737945864 | 0.73914824 | 0.9983733 | 0.31809838 | 0.51528068 |
| Z98885.2 | 3.30218935 | | -0.429383864 | 0.71217122 | -0.6029222 | 0.54656042 | 0.71769736 |
| AL117328.2 | 1.01124094 | | -0.835910569 | 1.57631686 | -0.5302935 | 0.59590846 | 0.75484247 |
| ZBED4 | 803.930962 | | -0.463239972 | 0.23313599 | -1.9869947 | 0.04692299 | 0.13909164 |
| ALG12 | 2148.51554 | | 0.027940357 | 0.13257576 | 0.21075011 | 0.83308227 | 0.91136194 |
| AL671710.1 | 1210.43704 | | -0.116199874 | 0.15659874 | -0.742023 | 0.45807339 | 0.64706542 |
| CRELD2 | 1933.99235 | | -0.15018606 | 0.26296766 | -0.5711199 | 0.56791837 | 0.73383024 |
| BX539320.1 | 24.9224829 | | -0.679290787 | 0.5296075 | -1.2826306 | 0.19962153 | 0.3808784 |
| PIM3 | 11633.1526 | | -1.237025788 | 0.26806417 | -4.6146629 | 3.94E-06 | 9.75E-05 |
| IL17REL | 3.78133396 | | 0.737222976 | 0.99042457 | 0.74435045 | 0.45666449 | 0.64604305 |
| TTLL8 | 0.50087417 | | -2.189813856 | 2.73827928 | -0.7997044 | 0.42388211 | NA |
| MLC1 | 16.5372797 | | -0.787461886 | 0.70860885 | -1.1112787 | 0.26644843 | 0.46055617 |
| MOV10L1 | 24.6494362 | | 2.177784812 | 0.57571514 | 3.78274715 | 0.00015511 | 0.00179867 |
| PANX2 | 755.982504 | | -0.868709412 | 0.45479826 | -1.9100984 | 0.05612055 | 0.15787623 |
| TRABD | 3691.02965 | | -0.598232345 | 0.15020517 | -3.9827681 | 6.81E-05 | 0.00093848 |
| AL022328.3 | 82.6483802 | | -0.575703729 | 0.17010517 | -3.3843987 | 0.00071334 | 0.00600027 |
| AL022328.4 | 1014.04658 | | -0.616331378 | 0.11590765 | -5.3174347 | 1.05E-07 | 5.00E-06 |
| SELENOO | 3497.73149 | | -0.088091554 | 0.16497417 | -0.5339718 | 0.59336105 | 0.75254574 |
| AL022328.1 | 54.4190928 | | 0.677628077 | 0.24195377 | 2.800651 | 0.00509996 | 0.02727912 |
| AL022328.2 | 654.880843 | | 0.195041002 | 0.19363929 | 1.00723877 | 0.31382005 | 0.5108293 |
| TUBGCP6 | 5163.80422 | | 0.25374749 | 0.14607884 | 1.73705855 | 0.08237684 | 0.20670636 |
| HDAC10 | 2869.02235 | | -0.137955013 | 0.16207015 | -0.8512056 | 0.39465517 | 0.58990232 |
| MAPK12 | 4614.49553 | | -0.419253099 | 0.14769204 | -2.8386979 | 0.0045298 | 0.02497405 |
| MAPK11 | 1775.59189 | | -0.440505031 | 0.25615259 | -1.7196977 | 0.0854874 | 0.21220962 |
| PLXNB2 | 26991.5066 | | 0.19096425 | 0.18247145 | 1.04654319 | 0.29531032 | 0.49212571 |
| DENND6B | 1890.05087 | | 0.114667302 | 0.26412478 | 0.43414065 | 0.66418631 | 0.80275887 |
| CR559946.1 | 249.025604 | | 0.13665515 | 0.26417299 | 0.51729419 | 0.60495081 | 0.76185839 |
| CR559946.2 | 330.675994 | | 0.214059205 | 0.31432014 | 0.68102288 | 0.49585701 | 0.67772047 |
| PPP6R2 | 7365.46069 | | 0.098898111 | 0.10279036 | 0.96213408 | 0.33598226 | 0.53414434 |
| SBF1 | 9897.51746 | | -0.0294296 | 0.19771697 | -0.1488471 | 0.88167427 | 0.93857409 |
| ADM2 | 2049.2768 | | 0.523677991 | 0.3756607 | 1.39401855 | 0.163312 | 0.33255116 |
| MIOX | 33355.0575 | | 2.906839254 | 0.82908855 | 3.5060661 | 0.00045478 | 0.00421237 |
| LMF2 | 6940.90403 | | -0.395236945 | 0.09623266 | -4.1070978 | 4.01E-05 | 0.00062353 |
| NCAPH2 | 2053.28231 | | -0.123096542 | 0.18595771 | -0.6619599 | 0.50799693 | 0.68682727 |
| SCO2 | 2996.55321 | | -1.704183991 | 0.40833477 | -4.1734971 | 3.00E-05 | 0.00048933 |
| U62317.3 | 1848.38194 | | -1.331479105 | 0.40796054 | -3.2637448 | 0.0010995 | 0.00843741 |
| TYMP | 6644.09715 | | -2.410025099 | 0.5883085 | -4.0965328 | 4.19E-05 | 0.00064598 |
| ODF3B | 3043.78278 | | -2.495731901 | 0.66497141 | -3.7531416 | 0.00017463 | 0.00196881 |
| U62317.2 | 52.3064353 | | -2.413729604 | 0.63943702 | -3.774773 | 0.00016015 | 0.00184341 |
| U62317.5 | 39.8601821 | | -2.178558822 | 0.68087087 | -3.1996652 | 0.00137587 | 0.01012326 |
| KLHDC7B | 420.333564 | | -3.075844307 | 0.73726739 | -4.1719522 | 3.02E-05 | 0.00049117 |
| SYCE3 | 15.1527207 | | -0.293663795 | 0.43716024 | -0.6717532 | 0.50174082 | 0.68226509 |
| CPT1B | 3590.08224 | | -0.358419666 | 0.34362752 | -1.043047 | 0.29692651 | 0.49372822 |
| CHKB-CPT1B | 8457.03002 | | -0.305530364 | 0.19916541 | -1.5340533 | 0.12501653 | 0.27738207 |
| CHKB | 5043.28468 | | -0.257961283 | 0.12122426 | -2.1279674 | 0.03333978 | 0.1087678 |
| CHKB-DT | 24.2883871 | | 0.071147976 | 0.46106281 | 0.15431298 | 0.87736296 | 0.93618992 |
| U62317.4 | 6.34197054 | | 0.954532379 | 0.57916009 | 1.64813217 | 0.09932555 | 0.23639123 |
| MAPK8IP2 | 266.060702 | | -0.310743189 | 0.30233728 | -1.0278031 | 0.30404246 | 0.50088085 |
| ARSA | 4419.70013 | | -0.32357157 | 0.17380827 | -1.861658 | 0.06265131 | 0.17046172 |
| SHANK3 | 7273.05658 | | -0.077562363 | 0.28674377 | -0.2704936 | 0.78678052 | 0.88392136 |
| AC000036.1 | 7.50577666 | | 1.472630711 | 0.75819003 | 1.94229764 | 0.05210108 | 0.15002351 |
| ACR | 75.2868388 | | -0.147998143 | 0.51724655 | -0.2861269 | 0.77478094 | 0.87626409 |
| AC002056.2 | 7.54982322 | | 0.31216179 | 0.51468605 | 0.60650914 | 0.54417672 | 0.71620188 |
| RABL2B | 2031.82333 | | -0.053546901 | 0.15262463 | -0.3508405 | 0.72570802 | 0.84465425 |
| FP565260.4 | 0.2890658 | | 2.115835529 | 3.23473121 | 0.65409933 | 0.51304781 | NA |
| FP565260.3 | 203.061681 | | -0.065505513 | 0.8379932 | -0.0781695 | 0.93769322 | 0.97025584 |
| FP565260.5 | 0.52304738 | | -1.218226069 | 1.8789192 | -0.6483653 | 0.51674869 | NA |
| GATD3B | 947.112416 | | 0.735901313 | 0.61315558 | 1.20018694 | 0.23006675 | 0.41761051 |
| FP565260.2 | 695.17535 | | 0.670025912 | 0.62466173 | 1.072622 | 0.28344075 | 0.47930955 |
| FP565260.1 | 222.132949 | | -0.0589098 | 0.7350294 | -0.0801462 | 0.93612099 | 0.96942144 |
| FP565260.6 | 261.38893 | | -0.064703322 | 0.29522153 | -0.2191687 | 0.82651863 | 0.90741897 |
| AC079801.1 | 0 | NA | | NA | NA | NA | NA |
| LINC01670 | 2.65031055 | | 0.257375496 | 0.85119612 | 0.30236921 | 0.76237063 | 0.86829842 |
| CU633967.1 | 1.17314727 | | 0.142727651 | 1.98349649 | 0.0719576 | 0.94263565 | 0.97275716 |
| FP236315.3 | 0.3794512 | | -1.796052315 | 2.82410681 | -0.6359718 | 0.5247948 | NA |
| FP236315.1 | 0.3794512 | | -1.796052315 | 2.82410681 | -0.6359718 | 0.5247948 | NA |
| H2BFS | 0.43646594 | | -2.023623934 | 2.74692407 | -0.7366872 | 0.46131256 | NA |
| LINC01669 | 0.34544235 | | -0.991105429 | 2.94239498 | -0.3368363 | 0.73624031 | NA |
| CU639417.3 | 0.34607569 | | -1.691981139 | 3.30304428 | -0.512249 | 0.60847675 | NA |
| CU639417.4 | 0.36416359 | | -1.034575391 | 2.30309408 | -0.4492111 | 0.65327939 | NA |
| SIK1B | 899.126507 | | -1.663450798 | 0.49246226 | -3.3778239 | 0.00073062 | 0.00611244 |
| CU633906.1 | 2.08799445 | | 1.476234401 | 1.11612681 | 1.32264039 | 0.185955 | 0.36347441 |
| CU633906.2 | 0.79827416 | | 1.323414439 | 1.37695913 | 0.96111381 | 0.33649495 | NA |
| CBSL | 16.0074866 | | 1.60532981 | 0.85014645 | 1.88829797 | 0.05898596 | 0.16354324 |
| U2AF1L5 | 128.441642 | | -1.39962694 | 1.05456255 | -1.3272109 | 0.18443892 | 0.3615831 |
| FP236240.1 | 0 | NA | | NA | NA | NA | NA |
| CRYAA2 | 153.593735 | | 1.886934691 | 2.40907988 | 0.78325949 | 0.43347474 | 0.62557806 |

| | | | | | | |
|---|---|---|---|---|---|---|
| CU638689.4 | 14.6474557 | 1.975973569 | 1.18404541 | 1.66883259 | 0.09515056 | 0.22960471 |
| CU638689.1 | 0.80990235 | 0.813093425 | 1.50211465 | 0.54129918 | 0.58830139 | NA |
| CU638689.5 | 1.77139986 | 0.04056375 | 1.15680357 | 0.03506537 | 0.97202761 | 0.98680917 |
| FP475955.3 | 0 | NA | NA | NA | NA | NA |
| FP475955.1 | 0.26086644 | -1.364242679 | 3.30558948 | -0.4127078 | 0.67982068 | NA |
| FP475955.2 | 0 | NA | NA | NA | NA | NA |
| CU634019.6 | 4.03729799 | 0.590870423 | 0.65736815 | 0.89884249 | 0.36873656 | 0.56552188 |
| CU634019.4 | 0.99382325 | 0.468024021 | 1.30679446 | 0.35814662 | 0.72023359 | NA |
| CU634019.2 | 0 | NA | NA | NA | NA | NA |
| CU634019.1 | 15.5642753 | 0.072243738 | 0.65479906 | 0.11032963 | 0.91214796 | 0.95681217 |
| CU633904.1 | 15.82358 | -0.026129597 | 0.60547729 | -0.0431554 | 0.96557768 | 0.98382147 |
| FP236241.1 | 0 | NA | NA | NA | NA | NA |
| SMIM11B | 0.79593423 | -1.338233866 | 1.66047866 | -0.8059326 | 0.42028172 | NA |
| CU633980.1 | 0 | NA | NA | NA | NA | NA |
| SMIM34B | 0.69242499 | 2.205682227 | 2.44171745 | 0.90333229 | 0.36634956 | NA |
| KCNE1B | 34.8356202 | 1.448309047 | 0.63113928 | 2.29475346 | 0.02174726 | 0.07984895 |
| FP671120.1 | 2.88364085 | -1.72934153 | 0.90693528 | -1.9067971 | 0.05654687 | 0.15875024 |
| FP671120.3 | 0.19890045 | -0.340073434 | 3.15719848 | -0.1077137 | 0.91422283 | NA |
| FP671120.4 | 3.48989003 | -0.911871593 | 0.73222346 | -1.2453461 | 0.21300457 | 0.39688173 |
| FP236383.1 | 3.47371877 | -2.029278754 | 0.90438614 | -2.2438189 | 0.02484405 | 0.08790255 |
| FP236383.2 | 0 | NA | NA | NA | NA | NA |
| FP236383.3 | 0 | NA | NA | NA | NA | NA |
| LINC01666 | 5.38772616 | -0.022119489 | 1.0004176 | -0.0221103 | 0.98236001 | 0.99215259 |
| CR392039.3 | 0.90122978 | 2.118634933 | 1.4297207 | 1.48185232 | 0.13837959 | NA |
| CR381653.1 | 16.2783288 | 0.387919808 | 0.31749992 | 1.221795 | 0.22178516 | 0.40772969 |
| LINC01667 | 0.25999113 | -1.364242679 | 3.30558948 | -0.4127078 | 0.67982068 | NA |
| AC018692.1 | 0.07659028 | -0.642905668 | 3.30558948 | -0.1944905 | 0.84579185 | NA |
| CR382287.2 | 0.06785815 | 0.795868262 | 3.28898349 | 0.24198001 | 0.80879565 | NA |
| AP003900.1 | 0 | NA | NA | NA | NA | NA |
| TPTE | 1.72902365 | 2.508293066 | 1.15019148 | 2.18076129 | 0.02920108 | 0.09903954 |
| IGHV1OR21- | 0 | NA | NA | NA | NA | NA |
| AP001464.1 | 0 | NA | NA | NA | NA | NA |
| AJ239318.1 | 0.09635527 | 0.547239394 | 3.29377202 | 0.16614368 | 0.86804388 | NA |
| LINC01674 | 0 | NA | NA | NA | NA | NA |
| POTED | 0 | NA | NA | NA | NA | NA |
| AL050303.1 | 0 | NA | NA | NA | NA | NA |
| AP001347.1 | 6.69599943 | 0.189234414 | 0.89182188 | 0.21218858 | 0.83195992 | 0.91069523 |
| LIPI | 0.87685212 | 1.077218979 | 1.54581475 | 0.69686163 | 0.4858894 | NA |
| RBM11 | 187.199309 | -1.208822741 | 0.34133519 | -3.5414537 | 0.00039793 | 0.00380609 |
| HSPA13 | 1485.11105 | -0.342638176 | 0.30797021 | -1.1125692 | 0.26589349 | 0.45998893 |
| SAMSN1 | 390.962046 | -1.118402317 | 0.44760032 | -2.4986629 | 0.01246628 | 0.0530507 |
| SAMSN1-AS: | 1.59937187 | 0.472123715 | 1.32993806 | 0.35499677 | 0.72259201 | 0.84253411 |
| AF127936.2 | 9.18132471 | -0.938900292 | 0.79315899 | -1.1837479 | 0.23651287 | 0.42581037 |
| AF127936.1 | 6.63225489 | -0.323726746 | 1.13565479 | -0.2850574 | 0.7756002 | 0.8768389 |
| AF127577.4 | 174.644338 | 0.191125627 | 0.24803436 | 0.77056109 | 0.44096713 | 0.63221913 |
| LINC02246 | 69.4466815 | 0.199043668 | 0.239363 | 0.83155571 | 0.40565977 | 0.60046299 |
| AF127577.3 | 0.08695548 | -0.642905668 | 3.30558948 | -0.1944905 | 0.84579185 | NA |
| NRIP1 | 2387.54915 | -0.191241934 | 0.36485184 | -0.5241633 | 0.60016499 | 0.75785859 |
| AF127577.1 | 795.436937 | -0.584448768 | 0.30200239 | -1.9352455 | 0.05296018 | 0.15163517 |
| AF127577.2 | 2.71072266 | -1.230172097 | 1.24766644 | -0.9859784 | 0.32414374 | 0.52170839 |
| AF127577.5 | 3.64063703 | 0.806618918 | 0.77886156 | 1.03563837 | 0.30037089 | 0.4971617 |
| AJ009632.2 | 37.0734124 | 3.162874118 | 0.7262513 | 4.35506843 | 1.33E-05 | 0.00025736 |
| AJ009632.1 | 0 | NA | NA | NA | NA | NA |
| USP25 | 2325.31074 | -0.084685302 | 0.16108282 | -0.5257252 | 0.59907913 | 0.7570307 |
| MIR99AHG | 460.894644 | 0.803829832 | 0.29677039 | 2.70859177 | 0.00675694 | 0.03365708 |
| AP001172.1 | 1.18105251 | 1.385652675 | 1.22316323 | 1.13284363 | 0.25727992 | 0.44995832 |
| AP001172.2 | 0.45987722 | 1.392414782 | 2.1094431 | 0.66008644 | 0.50919836 | NA |
| AP000962.1 | 0.5067096 | 1.026046611 | 2.67300335 | 0.38385534 | 0.70108567 | NA |
| AF130359.1 | 0 | NA | NA | NA | NA | NA |
| AF212831.1 | 0 | NA | NA | NA | NA | NA |
| LINC01549 | 0.08666371 | -0.642905668 | 3.30558948 | -0.1944905 | 0.84579185 | NA |
| CXADR | 1183.4157 | -0.719930939 | 0.19949552 | -3.6087575 | 0.00030767 | 0.00309349 |
| BTG3 | 1250.02723 | -0.874684304 | 0.23286257 | -3.7562254 | 0.0001725 | 0.00194972 |
| AP000432.1 | 0.63124536 | 1.376819086 | 2.02363687 | 0.68036865 | 0.49627107 | NA |
| C21orf91-OT | 4.05568636 | 0.781209467 | 0.57218303 | 1.36531394 | 0.1721544 | 0.34506776 |
| C21orf91 | 536.86897 | -0.158086923 | 0.11867643 | -1.3320836 | 0.18283272 | 0.35969104 |
| AL109761.1 | 38.3218794 | -0.547842777 | 0.33416166 | -1.6394543 | 0.10111867 | 0.23934112 |
| CHODL-AS1 | 0.18301898 | 0.18657164 | 3.29377195 | 0.05664376 | 0.95482897 | NA |
| CHODL | 194.944695 | 0.837055661 | 0.66317783 | 1.26218884 | 0.20688084 | 0.38950791 |
| AF130417.1 | 0.19981251 | 0.689154255 | 3.12815507 | 0.22030693 | 0.82563212 | NA |
| TMPRSS15 | 2.75406676 | -0.036834212 | 1.51797452 | -0.0242654 | 0.98064094 | 0.99131724 |
| MIR548XHG | 0 | NA | NA | NA | NA | NA |
| AF240627.1 | 0 | NA | NA | NA | NA | NA |
| AL157359.2 | 0 | NA | NA | NA | NA | NA |
| AL157359.1 | 0 | NA | NA | NA | NA | NA |
| AP000431.2 | 0 | NA | NA | NA | NA | NA |
| AP000431.1 | 0.03392908 | 0.380740593 | 3.29738709 | 0.11546736 | 0.90807469 | NA |
| AP000855.1 | 0 | NA | NA | NA | NA | NA |
| LINC01683 | 0 | NA | NA | NA | NA | NA |
| LINC02573 | 0 | NA | NA | NA | NA | NA |
| LINC00320 | 0.13028434 | 1.049816488 | 3.28477129 | 0.31960109 | 0.74927075 | NA |
| NCAM2 | 31.1204922 | -0.256822432 | 0.35890766 | -0.7155669 | 0.47425883 | 0.66003563 |
| AP001136.1 | 0 | NA | NA | NA | NA | NA |
| AP001117.1 | 0 | NA | NA | NA | NA | NA |
| AF241725.1 | 0 | NA | NA | NA | NA | NA |
| LINC00317 | 0 | NA | NA | NA | NA | NA |
| LINC01425 | 0 | NA | NA | NA | NA | NA |
| AP000472.1 | 0.60270578 | 1.312733831 | 1.72367707 | 0.76158919 | 0.44630523 | NA |
| LINC01687 | 3.36173482 | 1.915040373 | 1.34158181 | 1.42744957 | 0.15345035 | 0.319048 |
| LINC00308 | 0 | NA | NA | NA | NA | NA |
| AP000561.1 | 0 | NA | NA | NA | NA | NA |
| AP000959.1 | 0 | NA | NA | NA | NA | NA |
| AP001116.1 | 0 | NA | NA | NA | NA | NA |
| AP000459.2 | 0 | NA | NA | NA | NA | NA |
| AP000459.1 | 0 | NA | NA | NA | NA | NA |
| AP000474.1 | 0 | NA | NA | NA | NA | NA |
| AP000477.2 | 0 | NA | NA | NA | NA | NA |
| AP000477.1 | 0 | NA | NA | NA | NA | NA |
| AP000470.1 | 0 | NA | NA | NA | NA | NA |
| LINC01689 | 0 | NA | NA | NA | NA | NA |
| LINC01684 | 6.7363624 | 0.151065489 | 0.59416935 | 0.25424652 | 0.79930512 | 0.89108294 |
| LINC01692 | 0 | NA | NA | NA | NA | NA |
| AP000402.1 | 0 | NA | NA | NA | NA | NA |
| AP000146.1 | 0.274636 | 1.131168378 | 2.99720785 | 0.37740739 | 0.70587088 | NA |
| AP000235.1 | 0 | NA | NA | NA | NA | NA |
| AP000233.2 | 0 | NA | NA | NA | NA | NA |
| AP000233.1 | 0 | NA | NA | NA | NA | NA |
| AP001341.1 | 0 | NA | NA | NA | NA | NA |
| LINC00158 | 4.84803004 | 0.08455687 | 0.82160226 | 0.10291704 | 0.91802882 | 0.9601219 |
| AP000221.1 | 0.7073576 | 0.412963534 | 1.77734722 | 0.23234826 | 0.81626753 | NA |
| MIR155HG | 103.987981 | -1.600685387 | 0.61155365 | -2.6174079 | 0.00886004 | 0.04114091 |
| AP000223.1 | 10.6643943 | 0.885033035 | 0.52031772 | 1.70094732 | 0.08895288 | 0.21875291 |
| MRPL39 | 1230.68315 | -0.050442213 | 0.13701251 | -0.3681577 | 0.71275563 | 0.83676279 |

| | | | | | | |
|---|---|---|---|---|---|---|
| JAM2 | 697.526537 | 0.165197111 | 0.4947462 | 0.33390274 | 0.73845295 | 0.85320463 |
| ATP5PF | 5602.25115 | -0.013990718 | 0.16679582 | -0.0838793 | 0.93315239 | 0.96776463 |
| GABPA | 3096.55883 | 0.170191051 | 0.1452194 | 1.17195812 | 0.24121387 | 0.43135832 |
| APP | 70886.6254 | -0.097271411 | 0.19605728 | -0.4961377 | 0.61979726 | 0.77271053 |
| AP001439.1 | 4.06677593 | 0.995763993 | 0.69790292 | 1.42679442 | 0.15363916 | 0.31931468 |
| AP000229.1 | 42.4991785 | 1.575900111 | 0.63185527 | 2.49408401 | 0.01262827 | 0.05359848 |
| AP001595.1 | 0 | NA | NA | NA | NA | NA |
| AP001596.1 | 7.05032092 | 0.285418933 | 0.51916644 | 0.54976384 | 0.58248137 | 0.74490659 |
| CYYR1-AS1 | 5.84273645 | 0.507801974 | 0.5589017 | 0.90857117 | 0.36357653 | 0.56082017 |
| AP001596.2 | 0.97693152 | 1.235265672 | 1.27311118 | 0.97027321 | 0.33191033 | NA |
| CYYR1 | 1136.09119 | 0.135073007 | 0.30711262 | 0.43981587 | 0.66007047 | 0.79999826 |
| ADAMTS1 | 26061.9493 | -2.777241343 | 0.3502116 | -7.9301808 | 2.19E-15 | 1.46E-12 |
| AP001599.1 | 0.2835572 | -0.699921197 | 2.97030165 | -0.2356398 | 0.81371221 | NA |
| ADAMTS5 | 579.839311 | 0.061640719 | 0.34977996 | 0.17622713 | 0.86011551 | 0.92705166 |
| AP001605.1 | 0.15452186 | 0.435199075 | 3.28898345 | 0.13232024 | 0.894731 | NA |
| AP001604.1 | 0.08757577 | 0.547239394 | 3.29377202 | 0.16614368 | 0.86804388 | NA |
| LINC01673 | 0.09635527 | 0.547239394 | 3.29377202 | 0.16614368 | 0.86804388 | NA |
| LINC00113 | 35.869416 | 2.278039145 | 0.52436148 | 4.34440593 | 1.40E-05 | 0.00026763 |
| AJ006995.1 | 0 | NA | NA | NA | NA | NA |
| LINC00314 | 0.55184605 | 1.043467669 | 2.25216518 | 0.46331756 | 0.64313676 | NA |
| LINC01695 | 6.61621123 | 0.177149698 | 0.45010392 | 0.3935751 | 0.69389477 | 0.82376229 |
| AF165147.1 | 19.8712113 | -0.252465149 | 0.49830082 | -0.5066521 | 0.61239896 | 0.76779926 |
| LINC00161 | 1.40342711 | -0.862311079 | 1.10215735 | -0.7823847 | 0.43398851 | 0.62592658 |
| N6AMT1 | 674.485813 | 0.45975617 | 0.17557223 | 2.61861547 | 0.00882874 | 0.04102043 |
| LTN1 | 1289.04532 | 0.015321419 | 0.1105779 | 0.13855769 | 0.88979968 | 0.94296756 |
| RWDD2B | 865.320312 | 0.454457549 | 0.17026324 | 2.66914657 | 0.00760443 | 0.03679619 |
| AF129075.2 | 77.7767936 | 0.533556177 | 0.37511543 | 1.42237865 | 0.15491636 | 0.32103295 |
| USP16 | 2102.33931 | -0.060851761 | 0.22210372 | -0.273979 | 0.78410074 | 0.88203808 |
| CCT8 | 6578.24145 | -0.303795739 | 0.14597891 | -2.0810933 | 0.03742536 | 0.11788273 |
| MAP3K7CL | 428.811988 | -0.740401128 | 0.44658343 | -1.6579234 | 0.09733292 | 0.23312164 |
| AF124730.1 | 6.12786713 | -2.605865742 | 1.204315 | -2.1637742 | 0.03048168 | 0.10197944 |
| BACH1 | 1834.83319 | -0.768884578 | 0.29964861 | -2.5659541 | 0.01028924 | 0.04603451 |
| BACH1-AS1 | 20.1890451 | -0.924373648 | 0.55822635 | -1.6559119 | 0.09773967 | 0.23392585 |
| BACH1-IT2 | 25.6909323 | -1.452902774 | 0.68983017 | -2.1061746 | 0.03518919 | 0.11306676 |
| GRIK1 | 35.8312347 | -0.892914862 | 0.39095286 | -2.2839451 | 0.02237476 | 0.0814944 |
| AP000238.1 | 24.1580758 | -0.974092805 | 0.39005955 | -2.4972925 | 0.01251457 | 0.0532098 |
| GRIK1-AS1 | 24.3954474 | -0.084502774 | 0.33719688 | -0.2506037 | 0.80212055 | 0.89266577 |
| AF096876.1 | 0.08666371 | -0.642905668 | 3.30558948 | -0.1944905 | 0.84579185 | NA |
| CLDN17 | 0.86524097 | -2.035913084 | 2.30424273 | -0.8835497 | 0.37693932 | NA |
| LINC00307 | 1.06836131 | 2.496272223 | 1.16090108 | 2.15028848 | 0.0315324 | 0.10458872 |
| CLDN8 | 2167.55383 | 1.326744713 | 0.42058685 | 3.15450832 | 0.00160769 | 0.0113995 |
| KRTAP24-1 | 0 | NA | NA | NA | NA | NA |
| KRTAP25-1 | 0 | NA | NA | NA | NA | NA |
| KRTAP26-1 | 0 | NA | NA | NA | NA | NA |
| KRTAP27-1 | 0 | NA | NA | NA | NA | NA |
| KRTAP23-1 | 0 | NA | NA | NA | NA | NA |
| KRTAP13-2 | 0 | NA | NA | NA | NA | NA |
| KRTAP13-1 | 0 | NA | NA | NA | NA | NA |
| KRTAP13-3 | 0 | NA | NA | NA | NA | NA |
| KRTAP13-4 | 0 | NA | NA | NA | NA | NA |
| KRTAP15-1 | 0 | NA | NA | NA | NA | NA |
| KRTAP19-1 | 0 | NA | NA | NA | NA | NA |
| KRTAP19-2 | 0 | NA | NA | NA | NA | NA |
| KRTAP19-3 | 0 | NA | NA | NA | NA | NA |
| KRTAP19-4 | 0 | NA | NA | NA | NA | NA |
| KRTAP19-5 | 0 | NA | NA | NA | NA | NA |
| KRTAP19-6 | 0 | NA | NA | NA | NA | NA |
| KRTAP19-7 | 0 | NA | NA | NA | NA | NA |
| KRTAP22-2 | 0 | NA | NA | NA | NA | NA |
| KRTAP6-3 | 0 | NA | NA | NA | NA | NA |
| KRTAP6-2 | 0 | NA | NA | NA | NA | NA |
| KRTAP22-1 | 0 | NA | NA | NA | NA | NA |
| KRTAP6-1 | 0 | NA | NA | NA | NA | NA |
| KRTAP20-1 | 0 | NA | NA | NA | NA | NA |
| KRTAP20-4 | 0 | NA | NA | NA | NA | NA |
| KRTAP20-2 | 0 | NA | NA | NA | NA | NA |
| KRTAP20-3 | 0 | NA | NA | NA | NA | NA |
| KRTAP21-3 | 0 | NA | NA | NA | NA | NA |
| KRTAP21-2 | 0.03392908 | 0.380740593 | 3.29738709 | 0.11546736 | 0.90807469 | NA |
| KRTAP21-1 | 0 | NA | NA | NA | NA | NA |
| KRTAP8-1 | 0 | NA | NA | NA | NA | NA |
| KRTAP7-1 | 0 | NA | NA | NA | NA | NA |
| KRTAP11-1 | 0 | NA | NA | NA | NA | NA |
| KRTAP19-8 | 0 | NA | NA | NA | NA | NA |
| TIAM1 | 381.99096 | -0.264819501 | 0.51014063 | -0.5191108 | 0.60368349 | 0.76088011 |
| AP000251.1 | 2.33236388 | -0.044868827 | 0.85014109 | -0.0527781 | 0.95790872 | 0.97983137 |
| AP000253.1 | 5.72994658 | 1.066963843 | 0.60480958 | 1.76413185 | 0.07770978 | 0.19841276 |
| SOD1 | 15659.443 | 0.009767205 | 0.21256217 | 0.04594988 | 0.9633502 | 0.98289258 |
| AP000254.1 | 237.982804 | -0.052874777 | 0.12606692 | -0.4194183 | 0.67491043 | 0.81077801 |
| SCAF4 | 1688.75148 | -0.184576352 | 0.15876443 | -1.16258 | 0.24499996 | 0.43602949 |
| AP000255.1 | 6.40463681 | 0.826626762 | 0.60834039 | 1.35882275 | 0.17420276 | 0.34816787 |
| HUNK | 887.23087 | -0.641491989 | 0.25254335 | -2.5401262 | 0.01108125 | 0.04862104 |
| HUNK-AS1 | 0.79833283 | -0.51811309 | 2.27063683 | -0.2281796 | 0.81950659 | NA |
| LINC00159 | 0.61428875 | -0.850138969 | 2.23608006 | -0.3801917 | 0.70380316 | NA |
| AP000265.1 | 3.15804961 | -0.279206499 | 1.22901446 | -0.2271792 | 0.82028441 | 0.90356178 |
| MIS18A | 301.357898 | -0.1106066 | 0.23598645 | -0.468699 | 0.63928484 | 0.78676162 |
| MIS18A-AS1 | 13.2675495 | 1.133170307 | 0.43513896 | 2.6041573 | 0.00921005 | 0.0423395 |
| MRAP | 272.970805 | -0.358502321 | 0.17035424 | -2.104452 | 0.03533904 | 0.11344185 |
| AP000266.1 | 3.53432035 | -1.19525453 | 1.44004106 | -0.8300142 | 0.40653075 | 0.60140134 |
| URB1 | 1059.55567 | -0.069883245 | 0.13980723 | -0.4998543 | 0.61717768 | 0.77116837 |
| URB1-AS1 | 296.181853 | 0.416674353 | 0.23059688 | 1.80693841 | 0.07077188 | 0.18573393 |
| EVA1C | 877.981976 | -1.299126011 | 0.34982368 | -3.7136594 | 0.00020428 | 0.00222051 |
| TCP10L | 110.310077 | -0.55926611 | 0.31769175 | -1.7604049 | 0.07833918 | 0.19949823 |
| AP000275.2 | 1864.11774 | -0.132430338 | 0.09213804 | -1.4373036 | 0.15063176 | 0.31532485 |
| CFAP298 | 2389.00985 | -0.208663759 | 0.11210779 | -1.8612779 | 0.06270494 | 0.17053725 |
| SYNJ1 | 1047.33632 | 0.313803407 | 0.10953493 | 2.86487066 | 0.0041718 | 0.02352154 |
| PAXBP1-AS1 | 446.33314 | -0.120895166 | 0.13859319 | -0.8723024 | 0.38304344 | 0.57895142 |
| PAXBP1 | 2363.51014 | -0.340437998 | 0.15366608 | -2.2154401 | 0.02672987 | 0.09279445 |
| C21orf62-AS | 63.6680829 | 0.871668011 | 0.43205939 | 2.01747269 | 0.04364621 | 0.13209578 |
| C21orf62 | 98.9292438 | 1.276240884 | 0.52423732 | 2.43447162 | 0.01491355 | 0.06057487 |
| AP000281.2 | 1.92957843 | 0.620570128 | 0.95024446 | 0.65306366 | 0.51371524 | 0.69166243 |
| AP000282.1 | 14.9415864 | 1.780336502 | 0.46766734 | 3.80684376 | 0.00014075 | 0.00166068 |
| LINC01690 | 0.32670497 | 1.41436792 | 2.2468522 | 0.62948863 | 0.52902921 | NA |
| OLIG2 | 6.121091 | 2.057340267 | 0.73606577 | 2.79504952 | 0.00518918 | 0.02765498 |
| LINC00945 | 0.55868667 | 0.254522886 | 1.81366453 | 0.14033625 | 0.88839432 | NA |
| OLIG1 | 141.690185 | 1.143504337 | 0.38043222 | 3.00580311 | 0.0026488 | 0.01669431 |
| AP000289.1 | 0.37585234 | 0.248264669 | 2.84952959 | 0.08712479 | 0.93057232 | NA |
| AP000290.1 | 0.10721287 | -0.642905668 | 3.30558948 | -0.1944905 | 0.84579185 | NA |
| LINC01548 | 0 | NA | NA | NA | NA | NA |
| IFNAR2 | 1867.79345 | -1.555188778 | 0.25661965 | -6.0602871 | 1.36E-09 | 1.34E-07 |
| AP000295.1 | 2380.40221 | -0.800318827 | 0.19693064 | -4.0639629 | 4.82E-05 | 0.00072246 |
| IL10RB-DT | 1010.93084 | -1.791690171 | 0.29222317 | -6.1312392 | 8.72E-10 | 9.23E-08 |
| IL10RB | 1597.46676 | -0.567083488 | 0.18060529 | -3.1399052 | 0.00169003 | 0.01184574 |

| | | | | | | |
|---|---|---|---|---|---|---|
| IFNAR1 | 3541.3 | -0.307689991 | 0.19184147 | -1.6038764 | 0.10874131 | 0.25202218 |
| IFNGR2 | 4614.18489 | -0.903105841 | 0.17588928 | -5.1345132 | 2.83E-07 | 1.13E-05 |
| TMEM50B | 6207.65439 | -0.098741964 | 0.11377199 | -0.8678935 | 0.38545265 | 0.58128711 |
| AP000302.1 | 12.4803161 | 0.806761997 | 0.6747108 | 1.19571525 | 0.2318077 | 0.41977537 |
| DNAJC28 | 149.765047 | 1.144242435 | 0.18193025 | 6.28945672 | 3.19E-10 | 3.87E-08 |
| GART | 2296.49871 | -0.276833608 | 0.13553599 | -2.0425099 | 0.04110098 | 0.1263945 |
| SON | 9747.40368 | -0.192676265 | 0.16971237 | -1.1353107 | 0.25624518 | 0.44905696 |
| DONSON | 2936.80942 | -0.202027806 | 0.16353489 | -1.2353805 | 0.21668893 | 0.40152788 |
| AP000311.1 | 7561.70219 | 0.212642968 | 0.17259948 | 1.23200236 | 0.2179482 | 0.40306431 |
| CRYZL1 | 1162.36878 | 0.035954938 | 0.11443202 | 0.31420348 | 0.7533665 | 0.86258845 |
| ITSN1 | 1954.02262 | -0.096141088 | 0.13553479 | -0.7093462 | 0.47810968 | 0.66307733 |
| ATP5PO | 8863.7123 | 0.168502775 | 0.1667582 | 1.01046173 | 0.31227413 | 0.50918254 |
| LINC00649 | 1756.7266 | 0.321151827 | 0.17298609 | 1.85651823 | 0.06337973 | 0.17178779 |
| AP000569.1 | 20.8745335 | -0.116113327 | 0.60753867 | -0.1911209 | 0.84843089 | 0.92025018 |
| MRPS6 | 7525.66417 | 0.233982381 | 0.35228256 | 0.66418951 | 0.50656902 | 0.68569375 |
| SLC5A3 | 8628.52813 | 0.181679337 | 0.39541015 | 0.4594706 | 0.64589626 | 0.7910956 |
| LINC00310 | 43.0248538 | 1.328274454 | 0.39842885 | 3.3337808 | 0.00085674 | 0.00696218 |
| AP000317.1 | 27.4995668 | 0.490508757 | 0.32549444 | 1.50696509 | 0.13181961 | 0.28787437 |
| KCNE2 | 44.1856784 | 0.691995174 | 0.36981007 | 1.87121775 | 0.06131491 | 0.16791713 |
| AP000320.1 | 25.231929 | 0.813109091 | 0.36152896 | 2.24908428 | 0.02450713 | 0.08695548 |
| SMIM11A | 0.71354642 | 0.466070235 | 1.38012277 | 0.33770201 | 0.73558776 | NA |
| C21orf140 | 0 | NA | NA | NA | NA | NA |
| AP000322.2 | 0 | NA | NA | NA | NA | NA |
| SMIM34A | 0 | NA | NA | NA | NA | NA |
| AP000322.1 | 0 | NA | NA | NA | NA | NA |
| KCNE1 | 90.5479986 | 0.678146224 | 0.6266056 | 1.0822537 | 0.27913981 | 0.47477534 |
| RCAN1 | 5752.32742 | 0.243961971 | 0.22070095 | 1.10539611 | 0.26898799 | 0.46314344 |
| CLIC6 | 554.560655 | -1.222677185 | 0.33946544 | -3.6017723 | 0.00031606 | 0.00315457 |
| LINC00160 | 0.17088756 | -1.003574173 | 3.30558948 | -0.3035992 | 0.7614333 | NA |
| LINC01426 | 68.5812033 | -2.64485076 | 0.70435054 | -3.7550206 | 0.00017333 | 0.00195745 |
| RUNX1 | 1210.86285 | -1.690240048 | 0.43505907 | -3.8850817 | 0.0001023 | 0.00128893 |
| AF015262.1 | 0 | NA | NA | NA | NA | NA |
| AF015720.1 | 0.22507402 | -0.339686077 | 3.0646501 | -0.1108401 | 0.91174316 | NA |
| LINC01436 | 33.1362088 | -1.162577437 | 0.72943526 | -1.5938048 | 0.11097976 | 0.25541422 |
| SETD4 | 1512.77976 | -0.155150346 | 0.1312358 | -1.1822258 | 0.23711613 | 0.42637248 |
| AP000688.4 | 29.7438417 | -0.092420563 | 0.2495867 | -0.3702944 | 0.71116313 | 0.83587657 |
| CBR1 | 9186.41729 | -0.357612563 | 0.20099504 | -1.7792108 | 0.0752052 | 0.19412306 |
| AP000688.3 | 0.38542107 | 2.537492904 | 3.21220332 | 0.78995401 | 0.42955462 | NA |
| AP000688.2 | 260.105892 | 0.18854767 | 0.32371676 | 0.58244643 | 0.56026602 | 0.72840383 |
| CBR3 | 153.62933 | -0.30721434 | 0.2328578 | -1.3193217 | 0.1870616 | 0.36501786 |
| DOPEY2 | 598.685318 | -0.001845278 | 0.16895452 | -0.0109217 | 0.99128588 | 0.99583588 |
| AP000692.2 | 16.1284474 | 0.830718535 | 0.45294052 | 1.83405655 | 0.0666456 | 0.177897 |
| MORC3 | 1380.56094 | -0.058616949 | 0.14821573 | -0.395484 | 0.69248574 | 0.82304273 |
| CHAF1B | 172.619946 | -0.844745807 | 0.30239365 | -2.7935303 | 0.00521362 | 0.02772394 |
| AP000695.1 | 33.0491904 | -0.926983997 | 0.53180366 | -1.7430944 | 0.08131712 | 0.20470954 |
| AP000695.2 | 92.7658505 | -1.211322219 | 0.37105089 | -3.2645717 | 0.0010963 | 0.00842102 |
| CLDN14 | 506.888911 | -0.581929888 | 0.41808065 | -1.3919082 | 0.1639502 | 0.33341303 |
| AP000696.1 | 223.57936 | 0.698287909 | 0.45342192 | 1.54004003 | 0.1235506 | 0.27507846 |
| AP000697.1 | 54.6200345 | 0.282468286 | 0.69831247 | 0.40450128 | 0.68584414 | 0.81835792 |
| SIM2 | 2947.66025 | 0.334652559 | 0.4513634 | 0.74142599 | 0.45843519 | 0.64733258 |
| HLCS | 1160.53679 | 0.097601728 | 0.20068451 | 0.4863441 | 0.62672321 | 0.77787307 |
| AP000704.1 | 31.8736528 | -0.087147364 | 0.34204572 | -0.2547828 | 0.79889087 | 0.89107318 |
| RIPPLY3 | 29.0703518 | -0.58065882 | 0.41013604 | -1.4157712 | 0.15684249 | 0.32372702 |
| PIGP | 700.179923 | 0.601304284 | 0.19771148 | 3.04132199 | 0.00235542 | 0.01532126 |
| TTC3 | 5153.12381 | 0.24056618 | 0.10487171 | 2.29390919 | 0.02179572 | 0.0799712 |
| TTC3-AS1 | 150.723898 | 0.932109141 | 0.23186055 | 4.02012827 | 5.82E-05 | 0.00083181 |
| DSCR9 | 8.40113547 | -0.265342924 | 0.56503893 | -0.4696011 | 0.63864001 | 0.78618782 |
| AP001432.1 | 5.66740007 | 0.008742696 | 0.72033792 | 0.01213694 | 0.99031636 | 0.99533983 |
| VPS26C | 3833.47624 | -0.015337259 | 0.23067439 | -0.0664888 | 0.94698869 | 0.9744446 |
| AP001412.1 | 37.7528262 | 0.72677964 | 0.26874743 | 2.70432215 | 0.00684439 | 0.03398354 |
| AP001437.1 | 20.7104339 | -0.09840444 | 0.37627907 | -0.2615198 | 0.79369166 | 0.88839293 |
| DYRK1A | 3322.02679 | -0.182953231 | 0.17238993 | -1.0612756 | 0.28856465 | 0.48477097 |
| KCNJ6 | 20.4914601 | 0.604711624 | 0.51284762 | 1.17912533 | 0.23834827 | 0.42788399 |
| DSCR4 | 0.26431171 | -1.364242679 | 3.30558948 | -0.4127078 | 0.67982068 | NA |
| DSCR8 | 4.76727759 | -0.910690638 | 0.85257148 | -1.0681692 | 0.28544417 | 0.4815227 |
| KCNJ15 | 15399.7934 | 1.5904044 | 0.41723043 | 3.81181309 | 0.00013795 | 0.00163284 |
| AP001434.1 | 15.2462661 | -1.375672167 | 0.63223911 | -2.1758732 | 0.02956474 | 0.09983637 |
| LINC01423 | 6.13215214 | -0.694761941 | 0.91029618 | -0.7632262 | 0.44532848 | 0.63583908 |
| ERG | 1494.70843 | -0.299668914 | 0.35038434 | -0.8552577 | 0.39240849 | 0.58811555 |
| LINC00114 | 1.61459908 | 1.237933065 | 1.2977799 | 0.95388522 | 0.34014174 | 0.53822609 |
| ETS2 | 10031.3183 | -1.255926874 | 0.27171742 | -4.6221802 | 3.80E-06 | 9.45E-05 |
| AP001042.1 | 1.0987798 | -1.39828703 | 1.39791152 | -1.0002686 | 0.31718053 | 0.51448733 |
| AP001043.1 | 36.4060621 | -1.412502174 | 0.57076124 | -2.4747689 | 0.01333224 | 0.05574301 |
| AP001043.2 | 0.08393208 | -0.642905668 | 3.30558948 | -0.1944905 | 0.84579185 | NA |
| LINC01700 | 6.28180162 | -1.265561539 | 0.83971621 | -1.5071301 | 0.13177732 | 0.2878297 |
| AF064858.1 | 41.9387361 | -1.921578757 | 0.42042526 | -4.5705597 | 4.86E-06 | 0.00011514 |
| AF064858.3 | 18.6793362 | -1.749614837 | 0.52271818 | -3.3471475 | 0.00081648 | 0.0066865 |
| AF064858.2 | 4.2614584 | -0.720314048 | 0.79352025 | -0.907745 | 0.36401296 | 0.56116501 |
| PSMG1 | 1157.09884 | -0.333164563 | 0.24686105 | -1.3496036 | 0.17714316 | 0.35216954 |
| BRWD1 | 2620.34338 | 0.104341853 | 0.15073582 | 0.69221671 | 0.48880125 | 0.6722483 |
| AF129408.1 | 141.335764 | -0.354358744 | 0.26771598 | -1.3236369 | 0.18562368 | 0.36296132 |
| BRWD1-AS2 | 38.9324478 | 0.627407698 | 0.46422785 | 1.3515081 | 0.17653273 | 0.35133343 |
| BRWD1-AS1 | 2.44460358 | 1.441130172 | 0.89443694 | 1.61121496 | 0.10713288 | 0.24913947 |
| HMGN1 | 6498.28658 | -0.327944144 | 0.13005671 | -2.5215473 | 0.011684 | 0.05060059 |
| WRB | 1863.52669 | -0.009268242 | 0.16816572 | -0.0551137 | 0.95604785 | 0.97900515 |
| LCA5L | 243.583188 | 0.532586332 | 0.18120342 | 2.93916274 | 0.003291 | 0.01970566 |
| SH3BGR | 362.899081 | 0.577425784 | 0.28183364 | 2.0488178 | 0.04047993 | 0.12489246 |
| B3GALT5 | 64.3190223 | 0.122178333 | 0.60604824 | 0.20159836 | 0.84023072 | 0.91500658 |
| B3GALT5-AS | 5.63521362 | 0.260071508 | 0.7253701 | 0.3585363 | 0.71994201 | 0.84070812 |
| AF064860.2 | 0.54293039 | 0.238350753 | 1.8737225 | 0.12720707 | 0.89877651 | NA |
| IGSF5 | 49.3560245 | 0.674567629 | 0.66306831 | 1.01734259 | 0.30899048 | 0.50605877 |
| PCP4 | 748.038191 | 0.804368861 | 0.58912842 | 1.36535402 | 0.17214181 | 0.34506776 |
| DSCAM | 7.66246557 | -0.519393987 | 0.75338651 | -0.6894124 | 0.4905638 | 0.67369536 |
| DSCAM-AS1 | 0 | NA | NA | NA | NA | NA |
| LINC00323 | 87.8014421 | 1.747134995 | 0.60434181 | 2.89097158 | 0.00384053 | 0.02210918 |
| BACE2 | 4903.89988 | -0.444082451 | 0.43979754 | -1.0097429 | 0.31261847 | 0.50941908 |
| PLAC4 | 27.8375429 | 0.231477135 | 0.48482141 | 0.47744825 | 0.63304296 | 0.78218269 |
| BACE2-IT1 | 3.10217187 | 0.463686808 | 0.76956429 | 0.60253161 | 0.54682033 | 0.7179314 |
| FAM3B | 855.964798 | -0.630835572 | 0.27434025 | -2.2994642 | 0.02147859 | 0.07902756 |
| MX2 | 423.25801 | 0.002517517 | 0.39341777 | 0.00639909 | 0.9948943 | 0.99770729 |
| MX1 | 2191.33252 | -0.682684851 | 0.39997853 | -1.7068037 | 0.08785854 | 0.21666185 |
| AP001610.1 | 361.217246 | -0.31499951 | 0.41651344 | -0.7562769 | 0.44948317 | 0.63959456 |
| TMPRSS2 | 4540.01687 | 0.166680063 | 0.21783316 | 0.76517306 | 0.4441685 | 0.63510611 |
| AP001610.2 | 0 | NA | NA | NA | NA | NA |
| AP006748.1 | 0.11786115 | 0.02007176 | 3.29738704 | 0.00608717 | 0.99514317 | NA |
| LINC00111 | 0.34115864 | 0.653678352 | 2.80571377 | 0.23298113 | 0.81577606 | NA |
| LINC00479 | 41.0073665 | 0.594494825 | 0.58824786 | 1.01061962 | 0.31219852 | 0.50910117 |
| LINC00112 | 2.93910665 | 0.804546239 | 1.11779042 | 0.71976484 | 0.4716698 | 0.65803638 |
| RIPK4 | 2798.63272 | 0.114729179 | 0.32445252 | 0.35360853 | 0.72363228 | 0.84337413 |
| AP001615.1 | 626.763713 | -0.222251037 | 0.34637234 | -0.6416535 | 0.52109816 | 0.69754901 |
| PRDM15 | 592.367781 | -0.758793928 | 0.31297644 | -2.4244442 | 0.01533184 | 0.06180295 |
| AP001619.2 | 4.52632246 | -0.819299621 | 0.64846298 | -1.2634486 | 0.20642802 | 0.38909985 |
| C2CD2 | 1641.52816 | -0.857819472 | 0.20549294 | -4.1744474 | 2.99E-05 | 0.00048865 |

| | | | | | | |
|---|---|---|---|---|---|---|
| ZBTB21 | 1027.2779 | -0.585481986 | 0.23129016 | -2.5313744 | 0.01136165 | 0.04958734 |
| ZNF295-AS1 | 16.5450471 | -0.822600837 | 0.5185906 | -1.586224 | 0.11268851 | 0.25832775 |
| UMODL1 | 18.6382018 | 1.690576401 | 0.68112442 | 2.48203757 | 0.01306335 | 0.0548973 |
| ABCG1 | 1484.40249 | -1.168512046 | 0.37130956 | -3.1470023 | 0.00164954 | 0.01164301 |
| TFF3 | 131.893862 | -0.751429886 | 0.42559434 | -1.7656012 | 0.07746278 | 0.19802062 |
| TFF2 | 0.57136233 | 3.153855986 | 3.18885255 | 0.98902534 | 0.32265074 | NA |
| TFF1 | 1.16792841 | 3.173399334 | 1.57061685 | 2.02047962 | 0.04333366 | 0.13131564 |
| TMPRSS3 | 387.166854 | -0.556657184 | 0.4165909 | -1.3362202 | 0.18147731 | 0.35782537 |
| UBASH3A | 58.5267725 | -0.079920567 | 0.77627392 | -0.1029541 | 0.91799942 | 0.9601219 |
| RSPH1 | 370.769375 | -0.73663428 | 0.38270553 | -1.924807 | 0.05425351 | 0.1542496 |
| SLC37A1 | 1073.2691 | -0.794880976 | 0.17866168 | -4.4490849 | 8.62E-06 | 0.00018149 |
| AP001625.2 | 22.0674199 | -0.849792 | 0.43751391 | -1.94232 | 0.05209838 | 0.15002351 |
| LINC01671 | 909.574427 | -0.441673067 | 0.46565094 | -0.9485068 | 0.34287153 | 0.54049181 |
| AP001626.1 | 87.5930711 | -0.044416378 | 0.4296694 | -0.1033734 | 0.91766664 | 0.96005997 |
| PDE9A | 1813.99473 | -0.247681849 | 0.30344208 | -0.816241 | 0.41436234 | 0.60798658 |
| AP001627.1 | 3.67445812 | 1.206506696 | 0.74135258 | 1.6274398 | 0.10364372 | 0.24354705 |
| LINC01668 | 27.7163538 | 0.922025002 | 0.49982571 | 1.84469301 | 0.06508221 | 0.17496359 |
| AP001628.2 | 10.5921258 | 0.888607614 | 0.54831729 | 1.62060842 | 0.10510165 | 0.24587743 |
| AP001628.1 | 17.9447552 | -0.802644467 | 0.47950264 | -1.6739104 | 0.09414821 | 0.22800861 |
| WDR4 | 382.808041 | -0.654043309 | 0.19179277 | -3.4101563 | 0.00064926 | 0.00554972 |
| NDUFV3 | 2287.30228 | 0.238264665 | 0.14901217 | 1.59896107 | 0.10982925 | 0.25360748 |
| ERVH48-1 | 4.49704645 | 0.726635053 | 0.59286769 | 1.22562769 | 0.22033881 | 0.40617508 |
| AP001630.1 | 6.97339647 | 0.793801329 | 0.78173249 | 1.01543858 | 0.30989681 | 0.50700949 |
| PKNOX1 | 1048.25445 | -0.387992146 | 0.15590691 | -2.4886141 | 0.01282421 | 0.05416759 |
| CBS | 67.4204429 | -1.649100538 | 0.70818899 | -2.3286165 | 0.01987939 | 0.07460077 |
| U2AF1 | 129.320844 | 0.461098017 | 0.91703507 | 0.50281394 | 0.61509509 | 0.76990046 |
| FRGCA | 2.83437749 | -0.677574981 | 0.8401991 | -0.8064457 | 0.41998589 | 0.6127263 |
| AP001631.1 | 0.4820466 | 2.87204955 | 1.8895089 | 1.5199979 | 0.1285115 | NA |
| CRYAA | 303.670362 | 0.449909903 | 1.47270461 | 0.30549908 | 0.75998607 | 0.86700095 |
| LINC00322 | 1.46557676 | -0.159620564 | 1.48703058 | -0.1073418 | 0.91451781 | 0.95815625 |
| LINC01679 | 119.859666 | -1.182156258 | 0.58684431 | -2.0144291 | 0.0439645 | 0.13282248 |
| AP001046.1 | 4.92826889 | 0.632010212 | 0.60462075 | 1.04530023 | 0.29588423 | 0.49270442 |
| SIK1 | 135.886322 | 2.409341807 | 0.68162388 | 3.53470865 | NA | NA |
| LINC00319 | 0.55143073 | -2.3414483 | 2.17279077 | -1.0776225 | 0.28120224 | NA |
| LINC00313 | 6.3150224 | -1.002503583 | 0.6441847 | -1.5562363 | 0.11965192 | 0.26937772 |
| AP001048.1 | 0.07659028 | -0.642905668 | 3.30558948 | -0.1944905 | 0.84579185 | NA |
| HSF2BP | 14.5667791 | 0.712970049 | 0.35322232 | 2.01847392 | 0.04354193 | 0.1318407 |
| RRP1B | 1022.03722 | -0.088624416 | 0.17011899 | -0.5209555 | 0.60239781 | 0.75969537 |
| PDXK | 9485.98448 | 0.317497002 | 0.15921387 | 1.99415414 | 0.04613521 | 0.13732764 |
| CSTB | 5593.34839 | -0.888946798 | 0.15949006 | -5.5736816 | 2.49E-08 | 1.52E-06 |
| RRP1 | 1663.58073 | -0.615979587 | 0.19804742 | -3.1102632 | 0.00186921 | 0.01278625 |
| AATBC | 34.7041561 | 1.259650232 | 0.4288914 | 2.93699114 | 0.00331414 | 0.01980371 |
| AGPAT3 | 9037.44846 | 1.025433515 | 0.25500391 | 4.0212462 | 5.79E-05 | 0.00083012 |
| TRAPPC10 | 685.09579 | 0.456650871 | 0.29196014 | 1.56408633 | 0.1177973 | 0.2664992 |
| PWP2 | 129.506723 | -0.390018564 | 1.31924827 | -0.295637 | 0.76750733 | 0.87170163 |
| GATD3A | 469.993228 | 0.275315118 | 1.49569025 | 0.18407228 | 0.85395675 | 0.9235189 |
| LINC01678 | 8.57922881 | -0.723776959 | 0.8872566 | -0.8157471 | 0.41464482 | 0.60825557 |
| AP001056.2 | 0.81670994 | 0.57171562 | 1.37706774 | 0.41516884 | 0.67801831 | NA |
| AP001056.1 | 3.59688884 | -0.803396662 | 1.04536177 | -0.7685346 | 0.44216966 | 0.63331779 |
| ICOSLG | 270.848066 | -0.825393553 | 0.79987463 | -1.0319036 | 0.30211726 | 0.49897235 |
| AP001059.3 | 0.07830766 | -0.642905668 | 3.30558948 | -0.1944905 | 0.84579185 | NA |
| AP001059.2 | 0.08666371 | -0.642905668 | 3.30558948 | -0.1944905 | 0.84579185 | NA |
| DNMT3L | 54.9272487 | 2.866551681 | 0.49673747 | 5.77075795 | 7.89E-09 | 5.62E-07 |
| AP001059.1 | 34.00508 | 2.705889565 | 0.4664146 | 5.80146835 | 6.57E-09 | 4.86E-07 |
| AIRE | 96.7920276 | -0.581840701 | 0.45471228 | -1.2795799 | 0.20069291 | 0.38222757 |
| PFKL | 24528.9046 | 0.028169151 | 0.13815821 | 0.20389054 | 0.83843904 | 0.91449053 |
| C21orf2 | 4142.49418 | 0.024463762 | 0.1765436 | 0.13857066 | 0.88978943 | 0.94296756 |
| AP001062.1 | 2380.76743 | -0.008507978 | 0.19291541 | -0.0441021 | 0.96482301 | 0.98354657 |
| AP001062.2 | 10.9760197 | 1.046813876 | 0.39557448 | 2.64631298 | 0.00813745 | 0.03859434 |
| AP001063.1 | 0.65793607 | -1.363233117 | 2.25704359 | -0.6039906 | 0.54584989 | NA |
| TRPM2 | 301.6077 | -0.866050644 | 0.40141906 | -2.1574727 | 0.03096886 | 0.10321647 |
| TRPM2-AS | 13.3066484 | -0.917082419 | 0.46828704 | -1.9583767 | 0.05018583 | 0.14602601 |
| LRRC3-DT | 1.66059958 | -1.487921164 | 1.20789864 | -1.2318262 | 0.21801402 | 0.4031295 |
| LRRC3 | 133.656892 | -0.52157941 | 0.38835762 | -1.3430389 | 0.17925937 | 0.3548851 |
| AP001065.1 | 13.4318531 | -0.475558466 | 0.45714082 | -1.0402888 | 0.29820576 | 0.49518204 |
| LINC02575 | 4.08793261 | 0.489733591 | 0.89806875 | 0.5453186 | 0.58553441 | 0.74707483 |
| AP001065.2 | 0.16416605 | 0.18657164 | 3.29377195 | 0.05664376 | 0.95482897 | NA |
| TSPEAR | 15.0391548 | 0.889346583 | 0.67644759 | 1.31473095 | 0.18860034 | 0.36682335 |
| TSPEAR-AS1 | 15.9941677 | 0.660070685 | 0.37985632 | 1.73768513 | 0.08222631 | 0.20648797 |
| TSPEAR-AS2 | 22.7511123 | -0.299504209 | 0.8746864 | -0.3424132 | 0.73203992 | 0.84893323 |
| KRTAP10-1 | 0.08393208 | -0.642905668 | 3.30558948 | -0.1944905 | 0.84579185 | NA |
| KRTAP10-2 | 0.08757577 | 0.547239394 | 3.29377202 | 0.16614368 | 0.86804388 | NA |
| KRTAP10-3 | 0 | NA | NA | NA | NA | NA |
| KRTAP10-4 | 0.13028434 | 1.049816488 | 3.28477129 | 0.31960109 | 0.74927075 | NA |
| KRTAP10-5 | 0 | NA | NA | NA | NA | NA |
| KRTAP10-6 | 0 | NA | NA | NA | NA | NA |
| KRTAP10-7 | 0.21773222 | -0.339792043 | 3.08911945 | -0.1099964 | 0.91241222 | NA |
| KRTAP10-8 | 0 | NA | NA | NA | NA | NA |
| KRTAP10-9 | 0 | NA | NA | NA | NA | NA |
| KRTAP10-10 | 0.08757577 | 0.547239394 | 3.29377202 | 0.16614368 | 0.86804388 | NA |
| KRTAP10-11 | 0 | NA | NA | NA | NA | NA |
| KRTAP12-4 | 0 | NA | NA | NA | NA | NA |
| KRTAP12-3 | 0 | NA | NA | NA | NA | NA |
| KRTAP12-2 | 0 | NA | NA | NA | NA | NA |
| KRTAP12-1 | 0.10721287 | -0.642905668 | 3.30558948 | -0.1944905 | 0.84579185 | NA |
| KRTAP10-12 | 0 | NA | NA | NA | NA | NA |
| UBE2G2 | 5427.91727 | -0.207221146 | 0.11832102 | -1.7513468 | 0.07988619 | 0.20226586 |
| LINC01424 | 18.6508883 | 0.659403551 | 0.4265835 | 1.54577836 | 0.1221581 | 0.27307983 |
| SUMO3 | 4535.50934 | -0.336557066 | 0.20674854 | -1.627857 | 0.10355521 | 0.24336074 |
| PTTG1IP | 19653.7757 | -0.59759809 | 0.15447242 | -3.8686394 | 0.00010944 | 0.00136134 |
| ITGB2 | 2123.72045 | -0.131498836 | 0.4944262 | -0.2659625 | 0.79026808 | 0.88610295 |
| ITGB2-AS1 | 265.450581 | 0.005070242 | 0.56664083 | 0.00894789 | 0.99286071 | 0.9966781 |
| LINC01547 | 138.922412 | 0.76628581 | 0.29123081 | 2.63119759 | 0.00850846 | 0.03985221 |
| AL844908.1 | 21.2743343 | -0.255232959 | 0.64311507 | -0.3968698 | 0.69146346 | 0.82241065 |
| FAM207A | 804.595347 | -0.452602108 | 0.14875214 | -3.0426594 | 0.00234498 | 0.01528727 |
| AP001505.1 | 61.8942129 | -0.848326128 | 0.29036934 | -2.9215416 | 0.00348304 | 0.02049624 |
| LINC00163 | 3.75170428 | 0.421763909 | 0.71614807 | 0.58893395 | 0.55590558 | 0.72516598 |
| LINC00165 | 1.71640715 | -2.180366031 | 1.76411769 | -1.2359527 | 0.21647615 | 0.40135527 |
| PICSAR | 2.01333701 | -2.044484057 | 1.3317297 | -1.5352095 | 0.12473238 | 0.27692419 |
| ADARB1 | 1563.2669 | -0.009849012 | 0.25830639 | -0.0381292 | 0.96958468 | 0.98592327 |
| AL133499.1 | 0.18684785 | -1.003574173 | 3.30558948 | -0.3035992 | 0.7614333 | NA |
| LINC00334 | 833.212334 | -0.69747628 | 0.19432594 | -3.5892083 | 0.00033168 | 0.00328322 |
| POFUT2 | 3639.98448 | -0.253745216 | 0.19418279 | -1.3067338 | 0.19130312 | 0.37038645 |
| LINC00315 | 12.1102054 | -0.079005493 | 0.38058589 | -0.2075891 | 0.83554979 | 0.9127696 |
| BX322557.1 | 0.69300389 | 0.169758644 | 1.5654203 | 0.10844285 | 0.91364441 | NA |
| LINC00316 | 3.1514721 | -1.400806272 | 0.82974604 | -1.688235 | 0.09136614 | 0.22294115 |
| BX322559.1 | 0.16052235 | -1.003574173 | 3.30558948 | -0.3035992 | 0.7614333 | NA |
| BX322562.1 | 369.441018 | -1.51808174 | 0.34077591 | -4.4547801 | 8.40E-06 | 0.00017773 |
| COL18A1 | 41751.158 | -0.168287901 | 0.18117514 | -0.9288686 | 0.35295718 | 0.55054589 |
| COL18A1-AS: | 3.96807577 | 1.807816546 | 0.67964109 | 2.6599577 | 0.00781505 | 0.03755837 |
| COL18A1-AS | 202.943359 | 4.360594638 | 0.64515655 | 6.75897131 | 1.39E-11 | 2.73E-09 |
| SLC19A1 | 19026.0563 | -0.30935396 | 0.16062532 | -1.9259352 | 0.05411246 | 0.15399801 |
| LINC01694 | 13.1083854 | -1.450456365 | 0.55121878 | -2.6313624 | 0.00850433 | 0.03983997 |

| | | | | | | | |
|---|---|---|---|---|---|---|---|
| PCBP3 | 203.513085 | | -0.731322618 | 0.55545127 | -1.3166279 | 0.1879634 | 0.36613054 |
| AL133492.1 | 0.29525572 | | 0.328535869 | 2.3070254 | 0.1424067 | 0.88675877 | NA |
| AL592528.1 | 3.30136181 | | -4.942583934 | 1.39649933 | -3.5392669 | 0.00040124 | 0.00382942 |
| AJ011932.1 | 13.7092662 | | -0.45387846 | 0.46517964 | -0.9757058 | 0.32921029 | 0.52700027 |
| COL6A1 | 14598.9313 | | 0.659910925 | 0.24078198 | 2.74069899 | 0.00613086 | 0.03131943 |
| AP001476.1 | 11.2659999 | | -0.741921657 | 0.80575027 | -0.9207836 | 0.3571634 | 0.5547087 |
| AP001476.3 | 0.14114194 | | 0.02007176 | 3.29738704 | 0.00608717 | 0.99514317 | NA |
| AP001476.2 | 0.15179023 | | 0.435199075 | 3.28898345 | 0.13232024 | 0.894731 | NA |
| AP001471.1 | 2.25859191 | | 0.288767407 | 0.87373724 | 0.33049685 | 0.74102457 | 0.8546067 |
| COL6A2 | 10583.7915 | | -0.016120365 | 0.42956427 | -0.0375272 | 0.97006461 | 0.98622114 |
| FTCD | 10514.5714 | | 1.93966189 | 0.45095769 | 4.30120593 | 1.70E-05 | 0.00031194 |
| FTCD-AS1 | 1186.53935 | | 2.718967356 | 0.46580538 | 5.83713166 | 5.31E-09 | 4.07E-07 |
| SPATC1L | 352.913908 | | -0.593515605 | 0.33208599 | -1.7872347 | 0.07389957 | 0.19178541 |
| AP001468.1 | 64.8444745 | | -0.819981153 | 0.32158338 | -2.5498244 | 0.01077772 | 0.04763802 |
| LSS | 3863.31897 | | 0.085147139 | 0.19352066 | 0.43998993 | 0.6599444 | 0.79999826 |
| AP001469.1 | 114.163493 | | 1.038230601 | 0.33562178 | 3.09345421 | 0.00197841 | 0.0133771 |
| MCM3AP-AS | 2967.56374 | | -0.354482361 | 0.10895937 | -3.2533446 | 0.00114055 | 0.00870747 |
| MCM3AP | 3950.02125 | | -0.195645926 | 0.11234159 | -1.7415271 | 0.08159123 | 0.205282 |
| AP001469.2 | 8.18982935 | | 0.595317503 | 0.45438274 | 1.31016752 | 0.19013917 | 0.36885418 |
| AP001469.3 | 339.907491 | | 0.098805586 | 0.14946119 | 0.66107854 | 0.50856195 | 0.68737973 |
| YBEY | 407.979116 | | 0.061711931 | 0.35337549 | 0.17463557 | 0.86136599 | 0.92783061 |
| C21orf58 | 161.62403 | | 0.292186242 | 0.32055536 | 0.9115001 | 0.36203193 | 0.55933625 |
| PCNT | 1349.25848 | | 0.243078294 | 0.13699996 | 1.77429457 | 0.07601444 | 0.1956183 |
| DIP2A | 1813.53666 | | -0.131612954 | 0.12565726 | -1.0473963 | 0.29491683 | 0.4916937 |
| S100B | 70.8569837 | | -0.81212768 | 0.77710884 | -1.045063 | 0.29599386 | 0.49277045 |
| PRMT2 | 4440.36717 | | -0.24488526 | 0.12891326 | -1.8996126 | 0.05748398 | 0.16065005 |
| MT-ND1 | 505233.886 | | 0.370345539 | 0.43295088 | 0.85539852 | 0.39233056 | 0.58809442 |
| MT-ND2 | 496615.392 | | 0.521283794 | 0.41147581 | 1.26686378 | 0.205204 | 0.38772928 |
| MT-CO1 | 1955686.38 | | 0.744947933 | 0.43478911 | 1.71335461 | 0.08664732 | 0.21446175 |
| MT-CO2 | 668460.722 | | 0.596914028 | 0.27784889 | 2.14834055 | 0.03168671 | 0.10493381 |
| MT-ATP8 | 335347.557 | | 0.577840769 | 0.35872784 | 1.61080549 | 0.10722212 | 0.24928824 |
| MT-ATP6 | 830827.33 | | 0.453403995 | 0.36237504 | 1.25120094 | 0.21086118 | 0.39427138 |
| MT-CO3 | 1273295.68 | | 0.539639177 | 0.29448419 | 1.83248948 | 0.06687853 | 0.17828415 |
| MT-ND3 | 147904.553 | | 0.694025009 | 0.30590917 | 2.26872901 | 0.02328481 | 0.08383922 |
| MT-ND4L | 295869.196 | | 0.803474953 | 0.34921302 | 2.30081613 | 0.02140203 | 0.07883384 |
| MT-ND4 | 1382672.78 | | 0.632612159 | 0.31637244 | 1.99958049 | 0.04554558 | 0.13606184 |
| MT-ND5 | 246161.681 | | 0.875005505 | 0.39650713 | 2.20678379 | 0.02732917 | 0.09437711 |
| MT-ND6 | 69666.489 | | 0.770268785 | 0.46882933 | 1.64296203 | 0.10039079 | 0.23819553 |
| MT-CYB | 699405.162 | | 0.740472907 | 0.33959327 | 2.18046992 | 0.02922265 | 0.09906778 |
| AC133551.1 | 0 | NA | | NA | NA | NA | NA |
| AC136612.1 | 0.17423948 | | 0.18657164 | 3.29377195 | 0.05664376 | 0.95482897 | NA |
| AC136616.1 | 4.16127953 | | -1.436539592 | 0.78342176 | -1.8336733 | 0.0667025 | 0.17797682 |
| AC136616.3 | 0 | NA | | NA | NA | NA | NA |
| AC136616.2 | 0.53693986 | | 2.40429994 | 2.65056046 | 0.90709115 | 0.3643586 | NA |
| AC141272.1 | 2.91040811 | | 0.89587643 | 1.52866318 | 0.5860522 | 0.55784043 | 0.72650396 |
| AC136352.3 | 0 | NA | | NA | NA | NA | NA |
| AC136352.2 | 0 | NA | | NA | NA | NA | NA |
| AC171558.3 | 0 | NA | | NA | NA | NA | NA |
| AC171558.1 | 0 | NA | | NA | NA | NA | NA |
| BX004987.1 | 0.67918581 | | 2.143723346 | 2.08770139 | 1.02683428 | 0.30449851 | NA |
| AC145212.1 | 681.616145 | | 0.02604698 | 0.38474956 | 0.06769853 | 0.94602562 | 0.97409658 |
| MAFIP | 637.234131 | | 0.022576259 | 0.38064567 | 0.05931043 | 0.95270485 | 0.97717271 |
| AC011841.1 | 0.07659028 | | -0.642905668 | 3.30558948 | -0.1944905 | 0.84579185 | NA |
| AC011043.1 | 647.727255 | | 0.99565859 | 0.55348261 | 1.79889768 | 0.07203487 | 0.18824293 |
| AC011043.2 | 1.17252721 | | 2.742995457 | 1.41406864 | 1.93978947 | 0.05240528 | 0.15047117 |
| AL592183.1 | 308.603609 | | 0.153492346 | 0.36387301 | 0.42182944 | 0.67314952 | 0.80936316 |
| AC007325.1 | 0 | NA | | NA | NA | NA | NA |
| AC007325.4 | 69.7525336 | | -0.736731032 | 0.41441994 | -1.7777403 | 0.07544651 | 0.19451746 |
| AC007325.2 | 653.187683 | | 0.579043717 | 0.46073292 | 1.25678823 | 0.20883033 | 0.39203074 |
| BX072566.1 | 0 | NA | | NA | NA | NA | NA |
| AL354822.1 | 49.4715878 | | 0.446323881 | 0.48175094 | 0.92646189 | 0.354206 | 0.55180552 |
| AC023491.2 | 0.18393104 | | 1.469068379 | 3.27783996 | 0.44818185 | 0.65402196 | NA |
| AC004556.1 | 119.482583 | | -0.150616277 | 1.69855079 | -0.0886734 | 0.92934147 | 0.96578264 |
| AC233755.2 | 8.53788786 | | -1.289843851 | 1.19992166 | -1.07494 | 0.28240157 | 0.47837923 |
| AC233755.1 | 50.3533158 | | 0.061401153 | 1.55716414 | 0.03943139 | NA | NA |
| AC240274.1 | 327.993044 | | 0.539355001 | 0.37811117 | 1.42644558 | 0.15373976 | 0.31942832 |
| AC213203.1 | 19.3164201 | | 1.549573506 | 0.60721609 | 2.55193092 | 0.01071278 | 0.04739864 |
| FAM231C | 0 | NA | | NA | NA | NA | NA |
| SCoV1_ORF1 | 0 | NA | | NA | NA | NA | NA |
| SCoV1_ORF1 | 0 | NA | | NA | NA | NA | NA |
| SCoV1_S | 0 | NA | | NA | NA | NA | NA |
| SCoV1_ORF3 | 0 | NA | | NA | NA | NA | NA |
| SCoV1_ORF3 | 0 | NA | | NA | NA | NA | NA |
| SCoV1_E | 0 | NA | | NA | NA | NA | NA |
| SCoV1_M | 0 | NA | | NA | NA | NA | NA |
| SCoV1_ORF6 | 0 | NA | | NA | NA | NA | NA |
| SCoV1_ORF7 | 0 | NA | | NA | NA | NA | NA |
| SCoV1_ORF7 | 0 | NA | | NA | NA | NA | NA |
| SCoV1_ORF8 | 0 | NA | | NA | NA | NA | NA |
| SCoV1_ORF8 | 0 | NA | | NA | NA | NA | NA |
| SCoV1_N | 0 | NA | | NA | NA | NA | NA |
| SCoV1_ORF1 | 0 | NA | | NA | NA | NA | NA |
| SCoV1_UTR3 | 0 | NA | | NA | NA | NA | NA |
| SCoV2_orf1a | 0.03392908 | | 0.380740593 | 3.29738709 | 0.11546736 | 0.90807469 | NA |
| SCoV2_ORF1 | 0 | NA | | NA | NA | NA | NA |
| SCoV2_ORF3 | 0 | NA | | NA | NA | NA | NA |
| SCoV2_E | 0 | NA | | NA | NA | NA | NA |
| SCoV2_M | 0 | NA | | NA | NA | NA | NA |
| SCoV2_ORF7 | 0 | NA | | NA | NA | NA | NA |
| SCoV2_ORF8 | 0 | NA | | NA | NA | NA | NA |
| SCoV2_N | 0 | NA | | NA | NA | NA | NA |
| SCoV2_UTR3 | 0 | NA | | NA | NA | NA | NA |
